# Exploring Ruthenium-based organometallic inhibitors against Plasmodium Calcium Dependent Kinase 2 (PfCDPK2): a combined ensemble docking, QM paramterization and molecular dynamics study

**DOI:** 10.1101/2020.03.31.017541

**Authors:** Dhaval Patel, Mohd Athar, Prakash C. Jha

## Abstract

Recent advances in the metal-organic framework (MOF) have accelerated the discovery of novel metal-based anticancer, antibacterial and antimalarial compounds. This is substantiated by many serendipitously discovered metals (Ru, Rh, and Ir) based inhibitors that established the importance of metal inserted into the known organic scaffold. Conversely, it is possible to design novel bioactive compounds by mimicking hypervalent carbon atoms by transition metals. This process can be facilitated by computational drug discovery by treating metal center using optimized parameters that can be used for molecular docking and molecular dynamics simulations. Further, the method can be plugged with high computational power and refined algorithms to interpret chemical phenomena with atomic-level insights. In the present work, we have demonstrated an approach for parameterizing three organometallic ligands (FLL, E52, and staurosporine) using MCPB.py. In particular, we report that E52 and FLL have a better shape complimentary and affinity compared to staurosporine identified inhibitor (staurosporine) against Calcium-dependent protein kinases 2 (CDPK2). This study also revealed that a flexible approach (ensemble) outperforms for the given target with dynamic movements. The calculated MMPBSA energies for staurosporine, FLL and E52 were −66.461 ± 2.192, −67.182 ± 1.971 and −91.339 ± 2.745 kcal/mol respectively.

## Introduction

The biochemistry of half of all known enzymes requires metal ion for proper functioning (Waldron, Rutherford, Ford, & Robinson, 2009). Recent progress in structural biology have provided new avenues with therapeutically and pharmacologically important targets for structure-based drug design (SBDD). This opens up a new landscape for organometallic chemistry which has recently delivered promiscuous organometallic compounds with unique properties (Babak & Ang, 2018; Jürgens & Casini, 2017; Parveen, Arjmand, & Tabassum, 2019). Interestingly, over the past decade, the growth of medicinal organometallic compounds has increased rapidly with approximately 670 reports as evidenced from the published literature (Ndagi, Mhlongo, & Soliman, 2017; Ravera, Moreno-Viguri, Paucar, Pérez-Silanes, & Gabano, 2018). While much attention has been given to anticancer agents, there are numerous reports that organometallics have great potential for antimalarial and antibacterial agents (Jaouen & Metzler-Nolte, 2013). The idea is to introduce a pharmaceutically relevant metal scaffold that can increase the efficacy of a natural product or organic drug without displaying any metal-related cytotoxicity. Such insertion of metal to organic scaffold present new opportunities for the design of bioactive compounds by rendering access to enlarged chemical space that may not be easily accessible with purely organic scaffolds (D. S. Williams et al., 2005). Recent efforts in similar direction have shown that the kinetically inert metal center (Ru, Rh, and Ir) could complement with organic fragment and serve as “hyper-valent carbon” which is not involved in any direct interactions but plays important role in building 3D space (Debreczeni et al., 2006; Meggers, 2009). Earlier, a plethora of literature has claimed the antimalarial properties of transition metal-based compounds (Adams et al., 2015, 2013; Barbosa et al., 2014; Chellan et al., 2014; Glans et al., 2012). One of the potential successes is the antimalarial activity of ferroquine, a chloroquine-derived iron-based organometallic compound which has proceeded to Phase IIB clinical trials (Biot, Castro, Botté, & Navarro, 2012; Dubar et al., 2011, 2013; Held et al., 2015).

Various metals have been introduced in designing metal-organic framework in drug development, in which ruthenium is often considered as the most promising transition metal from the pharmaceutical viewpoint because of its safe and druggable properties (Maschke, Alborzinia, Lieb, Wölfl, & Metzler-Nolte, 2014; Meier et al., 2013). The ruthenium (Ru-based) compounds have also demonstrated promising biological activities such as antibacterial (F. Li, Collins, & Keene, 2015; Mu et al., 2018), antileishmanial (Iniguez et al., 2016; Marcusso Orsini et al., 2016), anticancer (Su, Li, & Li, 2018; Thota, Rodrigues, Crans, & Barreiro, 2018) and antiplasmodial (Ekengard et al., 2015; Rylands et al., 2019). Notably, reports revealed that Ru complexation improves the antiplasmodial activity in comparison to free ligands e.g. Ru-pyridyl ester (Chellan et al., 2014) or ether complexes (Chellan et al., 2013), Ru-lapachol complexes (Barbosa et al., 2014) and Ru-cotrimazole complex (Iniguez et al., 2016). Conversely, the substitution of Fe by Ru in ferroquine derivatives can increase the antiplasmodial activity against the K1 resistance strain (Beagley et al., 2003). Apart from experimental methods in the discovery of organometallic compounds as potent inhibitors, it is also necessary for parallel advancements in computational modeling which will support in unravelling untapped pharmaceutical potential in metal-ligand interactions. In this work, we undertook Calcium-dependent protein kinases 2 (CDPK2) from *P. falciparum* as a case study to showcase the possibilities of computational modeling and simulations of metal-containing inhibitors. *P. falciparum* CDPKs (PfCDPKs) are important mediators of transduction and signalling pathways that regulate calcium ions for physiological processes (Miller, Ackerman, Su, & Wellems, 2013). CDPK2 is a 59 kDa protein exhibiting calcium-dependent kinase activity in an in vitro phosphorylation assay (Färber, Graeser, Franklin, & Kappes, 1997) and lacks considerable homology with human counterpart CDPK as revealed by our BLAST search. The unique structural features of PfCDPK2 compared to human CDPK counterpart additionally serve an opportunity for exploring structure-based drug design strategy.

Recent advances in computational chemistry have achieved reasonable achievements in structure-guided drug discovery that enable the prediction/prioritization of ligand/inhibitors binding in virtual screens (Ferreira, dos Santos, Oliva, & Andricopulo, 2015; Śledź & Caflisch, 2018). Most molecular docking programs consider the flexibility of ligand, though many of them treat the macromolecule as a rigid structure. In practical scenarios where the receptor is derived from an X-ray structure of the apo-kinase or ligand-bound kinases, the rigid-receptor docking often fails to estimate accurate bound conformation. To overcome this limitation, docking followed with Molecular Dynamics (MD) simulations offer major improvement by considering structural flexibility of the drug-target complexes with respect to explicit solvent (De Vivo & Cavalli, 2017; Gioia et al., 2017). MD based studies can also estimate the kinetics and thermodynamics of the ligand-binding event. These computational methods have not been well applied to inorganics as compared to organic compounds due to their atomic diversity/atom types and varied electronic structure (d-orbitals) (Bernhardt & Comba, 1992).

Historically, standard docking approaches either neglect or partially consider the flexibility of target and thereby relies on one receptor conformation. Alternatively, it is reasonable to take multiple distinct conformations from apoprotein and protein-ligand complex. The representative protein conformations can be randomly selected or identified from the clustering schemes. This approach is known as *‘ensemble docking’* where multiple independent target conformations (extracted through clustering algorithms) are used to achieve more realistic complex structure (Amaro et al., 2018; Nichols, Baron, & McCammon, 2012). In this work, we applied this approach to predict the binding mode of PfCDPK2 with Ruthenium-based compounds (structural scaffold of known kinase inhibitor-staurosporine). Parameterization of metal center (includes the bonded model, nonbonded model, cationic dummy atom model, etc.) by quantum mechanical (QM) parameters enables the reliable computational modeling of these chemical entities. Routinely, available force fields contain libraries and parameter files for 20 standard amino acids and 5 standard nucleic acids only but not for metal sites (Case et al., 2005; Van Der Spoel et al., 2005). Transition metals, in particular, have *d* or *f* as outermost orbitals that have varied valencies and complicated shapes. As a result, it can be led to the possibility of multiple oxidation states/spin state, unconventional chemical bonding and secondary orbital interactions (Sridharan & Sridharan, 2016). To overcome these challenges, density functional theory (DFT) is likely to be required to model these systems with reasonable accuracy. For the first time, our approach has addressed the transitions metal parameterization and force constant derivation using metal center parameter builder tool (MCPB.py) (P. Li & Merz, 2016) and its downstream integration in to MD simulation packages GROMACS (Abraham et al., 2015; Van Der Spoel et al., 2005). We believed that our current hybrid (QM/MD) integrated approach provides a connecting bridge between quantum mechanics (QM) and Molecular Dynamics Simulations (MDS) tools that can overcome the challenges associated with metal ion modeling.

## Materials and Methods

The flowchart in scheme 1 demonstrates the complete hybrid approach used for the current study. Briefly, the protein conformations for ensemble generations were sampled with all-atom molecular dynamics simulations with explicit solvent. The diverse poses generated in the entire MDS trajectory were clustered to yield dominant conformation. The ligands STU (Staurosporine), FLL (Octahedral Ru-Pyridocarbazole) and E52 (Methylated Ruthenium Pyridocarbazole) used in the current study were then docked into each of the protein conformations to generate different poses.

**Scheme 1.**
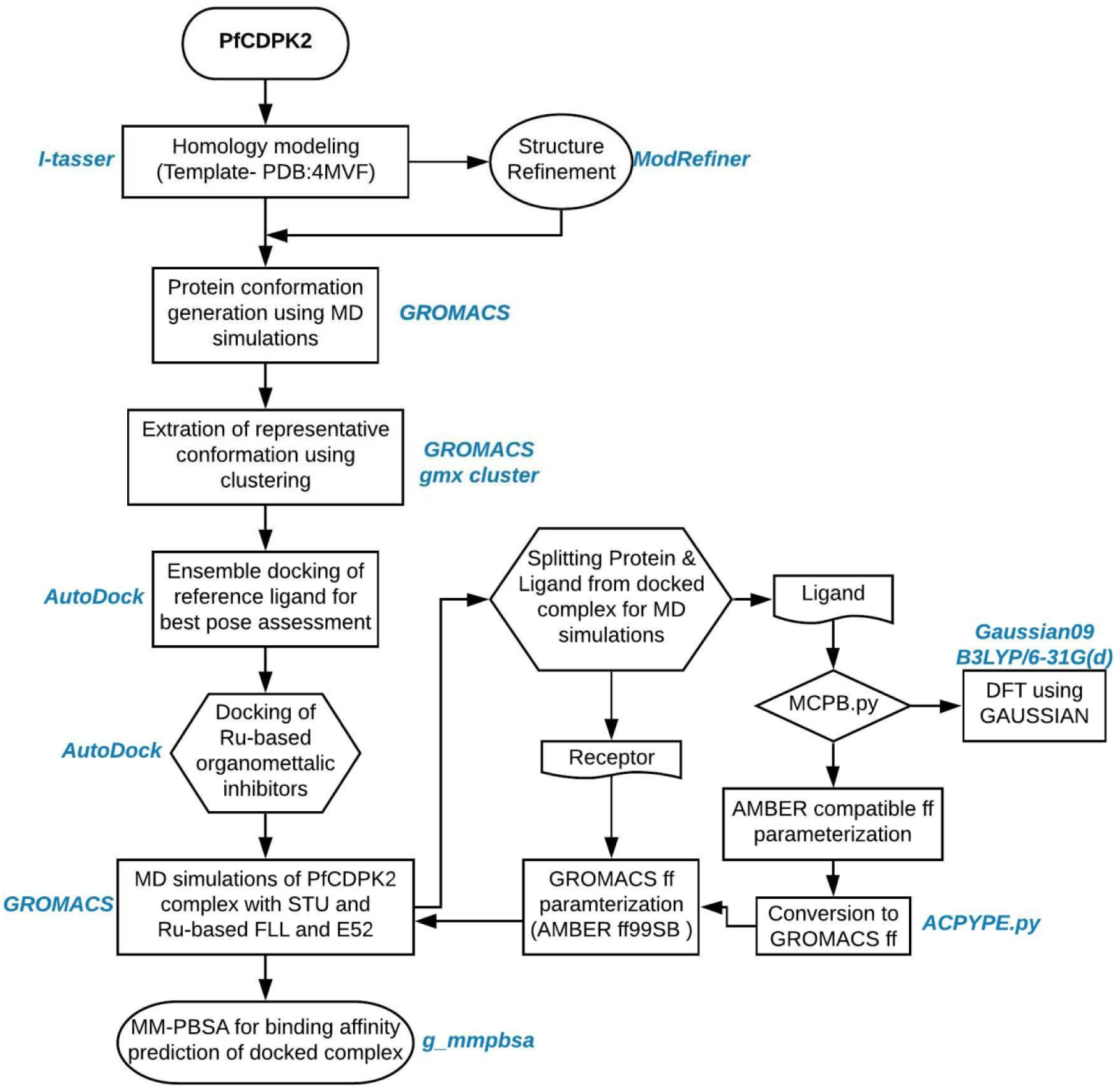
Flow chart for the hybrid methodology of the current study.

Metal parameterization for the compounds was then performed for the selected best pose using MCPB.py. Parameters and force constants were generated followed by topology and detailed input for MDS runs. The AMBER compatible generated parameters and force constant were later converted into GROMACS compatible formats for MDS. Thereafter, the binding affinity was calculated for each protein-ligand complexes using molecular mechanics generalized Born and surface area continuum solvation (MM/GBSA) methods.

### Generation of protein conformation

The protein ensemble conformations of PfCDPK2 were constructed based on the crystallographic structure of the PfCDPK2 complex with inhibitor staurosporine (PDB: 4MVF). The initial structure of PfCDPK2 deposited in the PDB had missing residues mainly due to lack of densities or molecular replacement models while solving the structures. To gratify the missing residues in docking and simulation, the protein sequence of PfCDPK2 was used to generate homology model using I-TASSER (Yang et al., 2015) and PDB: 4MVF as a template structure. The output model quality was assessed with Ramachandran statistics and compared with the original deposited structure. The control docking was performed for staurosporine (STU) with the model to confirm and preserve its native pose as found in PDB: 4MVF using AutoDock 4.2 (Morris et al., 2009b) as mentioned below. The ensembles of conformations were generated by performing two 50 ns molecular dynamics simulations (MDS): one for apo-PfCDPK2 and other for PfCDPK2 complexed with STU (PfCDPK2_STU). The generated model and the docked model with STU respective to crystal structure were used as the starting conformations. For each MDS, the Amber99SB force-field (Lindorff-Larsen et al., 2010) was used with GROMACS ver. 2016.4 (Van Der Spoel et al., 2005) as reported in our earlier studies (P. Patel, Parmar, Vyas, Patel, & Das, 2017). For inhibitor STU topology and parameterization, ACPYPE (Sousa da Silva & Vranken, 2012) tool was used followed by file format conversion into GROMACS compatible force-field format. The MDS systems were prepared as accordingly to earlier studies (Manhas, Patel, Lone, & Jha, 2019; Palak Patel et al., 2018). Briefly, the MDS systems were solvated with Three-site model (TIP3P) model for water in a dodecahedron box maintaining a distance of 1 nm from all the directions of the protein and periodic boundaries. The system was neutralized by adding an equal number of counterions. Subsequently, it was subjected to energy minimization using the steepest descent algorithm in order to remove any steric clashes and bad contacts prior to actual MD run. After the completion of energy minimization, the systems were equilibrated with position restraint simulations of 1 ns carried out under NVT (the constant Number of particles, Volume, and Temperature) and NPT (the constant Number of particles, Pressure, and Temperature) conditions. Finally, the MDS were subjected to 50 ns of production dynamics with default parameters. The trajectories were visualized using VMD (Humphrey, Dalke, & Schulten, 1996) and Chimera (Pettersen et al., 2004). The gmx rms, gmx rmsf, and gmx hbond tools were used for trajectories analysis and GRACE (xmgrace) was used for generation and visualization of plots (http://plasma-gate.weizmann.ac.il/Grace).

### Clustering of conformations and essential dynamics

The entire MDS trajectories were subjected to rmsd based clustering via ‘*gmx cluster’* tool that explores the conformational landscape among the ensemble of protein structures. The GROMOS algorithm as described by Daura and co-workers (Daura et al., 1999), was used to determine the dominant conformation with 0.2 mm Cα RMSD cutoff. The collective motion and dynamics of Cα backbone atoms were also assessed during the entire simulations, as computed by Principal Component Analysis (PCA) using gmx covar and gmx anaeig tools.

### Ensemble Docking with Ru-based organometallic inhibitors

For molecular docking, AutoDock 4.2 program (Morris et al., 2009a) was used to generate and rank a large set of receptor-ligand complex conformations based on their relative stability in conformational space. In biological systems, the majority of interactions are driven by complementing flexible movement of ligand and protein. AutoDock explicitly considers ligand flexibility, while the protein flexibility was taken into account implicitly in our MD simulations as mentioned above. AutoDock calculates the approximate binding free energy based on the evaluation of a single receptor-ligand complex which assumes binding free energy ΔG = ΔG_VdW_ + ΔGhbond + ΔGelec + ΔGconform + ΔGtor + ΔGsol, where the first four terms are molecular mechanics terms, namely, dispersion/repulsion, hydrogen bonding, electrostatics and deviations from the covalent geometry, the fifth term models the restriction of internal rotors, global rotation and translation, and the last term accounts for desolvation and the hydrophobic effect. The grid maps for docking were determined by STU re-docking trials into the generated model for PfCDPK2 till near-native pose was observed as seen in the crystal structure. The grid maps used for each protein conformation were 40×40×40 dimension points with grid spacing 0.375 Å and grid coordinates of 32.24, 84.719 and 21.838 for x, y and z dimensions. The Lamarckian genetic algorithm (LGA) was used for scoring the conformational space of ligand. For each docking run, the individual population in GA was set to 150, the maximum number of energy evaluations and generations were set to 2500000 and 27000 respectively. A total of 10 GA runs was performed for each protein conformation and the entire ensemble docking was achieved using in-house scripts developed for automation. This choice for the set of parameters is adequate enough to reach the convergence of the docking results as performed in our earlier study (B. Patel et al., 2018). The PfCDPK2 conformations generated for ensemble approach were redocked with STU to select the best pose for further docking. Two more organometallic inhibitors FLL (Octahedral Ru-Pyridocarbazole) and E52 (Methylated Ruthenium Pyridocarbazole) were subsequently docked with the selected conformations for further study. A separate control docking was also performed for PAK1 kinase (PDB:3FXZ) with its natural ligand FLL to confirm our docking protocol and parameters. FLL is a very well-known inhibitor of PAK1, GSK3 α and PIM1 (Maksimoska et al., 2008). However, E52 is a potent inhibitor against PI3K lipid kinase inhibitor as well as show 50% inhibition against PIM1 and GSK3α (Xie *et al.*, 2008). For docking of organometallic compounds, the parameter file in Autodock was incorporated with ruthenium metal van der Waals and other needed parameters (Rii-2.96; epsii-0.056; vol-12.00; solpar-0.00110) which were obtained from the Autodock website (“AutoDock - ADL: Parameters for docking with metal ions in receptor,” n.d.). For the Ru atom charge, we applied the native charge of +2 as reported in other complex/receptor interaction studies (Adeniyi & Ajibade, 2013).

### Metal center parameterization, Density Functional Theory calculations, and force-field generation

MD simulation is a well-established tool with a plethora of force-field parameters available for proteins as well as organic molecules. Parameterization of force field for metalloproteins or organometallic inhibitors may seem to be a daunting task due to challenges related to the transferability of ion parameters and development of polarizable non-bonded models (P. Li & Merz, 2016). There always remains the chances of biases from factors like choice of QM methods, basis sets and charge models. To assist in parameterization, Kenneth M. Merz Jr. group have developed MCPB code (P. Li & Merz, 2016) to facilitate framework development (Seminario, Z-matrix and empirical), basis sets (DFT theory); bonded model & non-bonded models for metal ions and charge models. In our current work, the AMBER ff99SB force-field parameters were used to model all standard amino acid residues, while Ru metal parameters were calculated using the python-based metal center parameter builder-MCPB.py program. Briefly, the metal site force-field parameters and RESP charges were derived from QM calculations at B3LYP/6-31G(d) level of theory using Gaussian 09 (Frisch *et al.*, 2009). In particular, based on the experimental results, +2 oxidation state of Ru in FLL (octahedral geometry) and E52 (pseudo tetrahedral geometry) ligands with singlet complex-multiplicity were modeled for the calculation of RESP charges. Frequency calculations were also performed at the same level to approximate the force constants. In particular, the Seminario method based on the Hessian matrix was used to derive the Ru-related force field parameters for AMBER. For Cp ring Ru based molecule, ten harmonic restraints connecting each carbon atom of the Cp ring to the Ru center was used, as described earlier (de Hatten, Cournia, Huc, Smith, & Metzler-Nolte, 2007).

### MD simulations of PfCDPK2 complex with STU and Ru-based FLL and E52

For MDS, the AMBER ff99SB force-field parameter was used to model all standard amino acid residues, while Ru metal parameters were calculated using the metal center parameter builder as mentioned above. The standard output of MCPB.py (P. Li & Merz, 2016) was used to generate prmtop and inpcrd amber files using tleap program with the default setup. The amber parameter and coordinate files were converted to gromacs compatible file format using ACPYPE (Sousa da Silva & Vranken, 2012). Remaining steps for MDS is similar as mentioned earlier in the ensemble conformation generation section. Three independent 50 ns production MD run were generated each for STU, FLL, and E52.

### Binding free-energy calculations

The Poisson–Boltzmann or generalized Born and surface area continuum solvation (MM-PBSA and MM-GBSA) methods (Genheden & Ryde, 2015; Sun et al., 2018; Wang et al., 2016) are used to estimate the free energy of binding in the receptor-ligand complexes. The analysis of free energy and energy contribution by individual residues were used to quantitatively estimate the ligand affinity for PfCDPK2 receptor. The g_mmpbsa tool (Kumari, Kumar, Lynn, & Lynn, 2014) with default parameters was used for molecular mechanics potential energy (electrostatic + Van der Waals interactions) and solvation free energy (polar + non-polar solvation energies) calculations. The last stable 30 ns trajectories which were identified based on the RMSD plot, consisting of 120 frames in the timeframe of 250 ps were used to estimate binding free energy using the g_mmpbsa tool. The frames were selected covering a wide range of trajectory in order to cover different conformational space of the receptor-ligand complexes for better structure-function correlation.

## Results and Discussion

### Ensemble conformation generation

The aim of the procedure was to explore the structural basis for the interaction between known Ru-based inhibitors having staurosporine backbone and Plasmodium calcium-dependent kinase 2 (PfCDPK2). Docking molecules to apoprotein or protein complexes with other ligand have reasonably advanced but it merely considers protein flexibility. Especially proteins from kinase family, which shows higher flexibility in variable loops; the docking interpretation in static structure might result in loss of accurate information (Cozzini et al., 2008). We, therefore, used ensemble conformation for docking with known inhibitor staurosporine to explore the sampling of protein-ligand conformational space. Prior to ensemble generation, the full-length model of PfCDPK2 was generated using PDB: 4MVF as a template structure using I-TASSER. The Ramachandran statistics for crystal structure was compared with the output model for testing model significance. The final output model showed significant statistics with 96.5% residues in the favored region, 2.7% residues in allowed region and 0.9% residues as an outlier. The known inhibitor STU was re-docked on the final output model to get near-native conformation with reference to the crystal structure (Fig. 2a). The generated model and the docked model with STU respective to crystal structure were used to serve as the starting conformations for 50 ns MDS. The RMSD values for Cα backbones from its starting to the final position were calculated for the entire MDS trajectory as shown in the plot of RMSD (nm) vs Time (ns) in Fig. 2b. The RMSD plot indicated that the apoprotein MDS reached equilibrium after 20 ns while the PfCDPK2_STU reached equilibrium around 10 ns. The mean RMSD values for apoprotein and PfCDPK2_STU were 0.57 ± 0.10 and 0.35 ± 0.04 respectively. The decrease in stability of PfCDPK2_STU compared to apoprotein suggested intrinsic flexibility of loop region in the native apo structure which forms induced-fit movement to accommodate the ligand in the active site in a later case.

**Figure 1:**
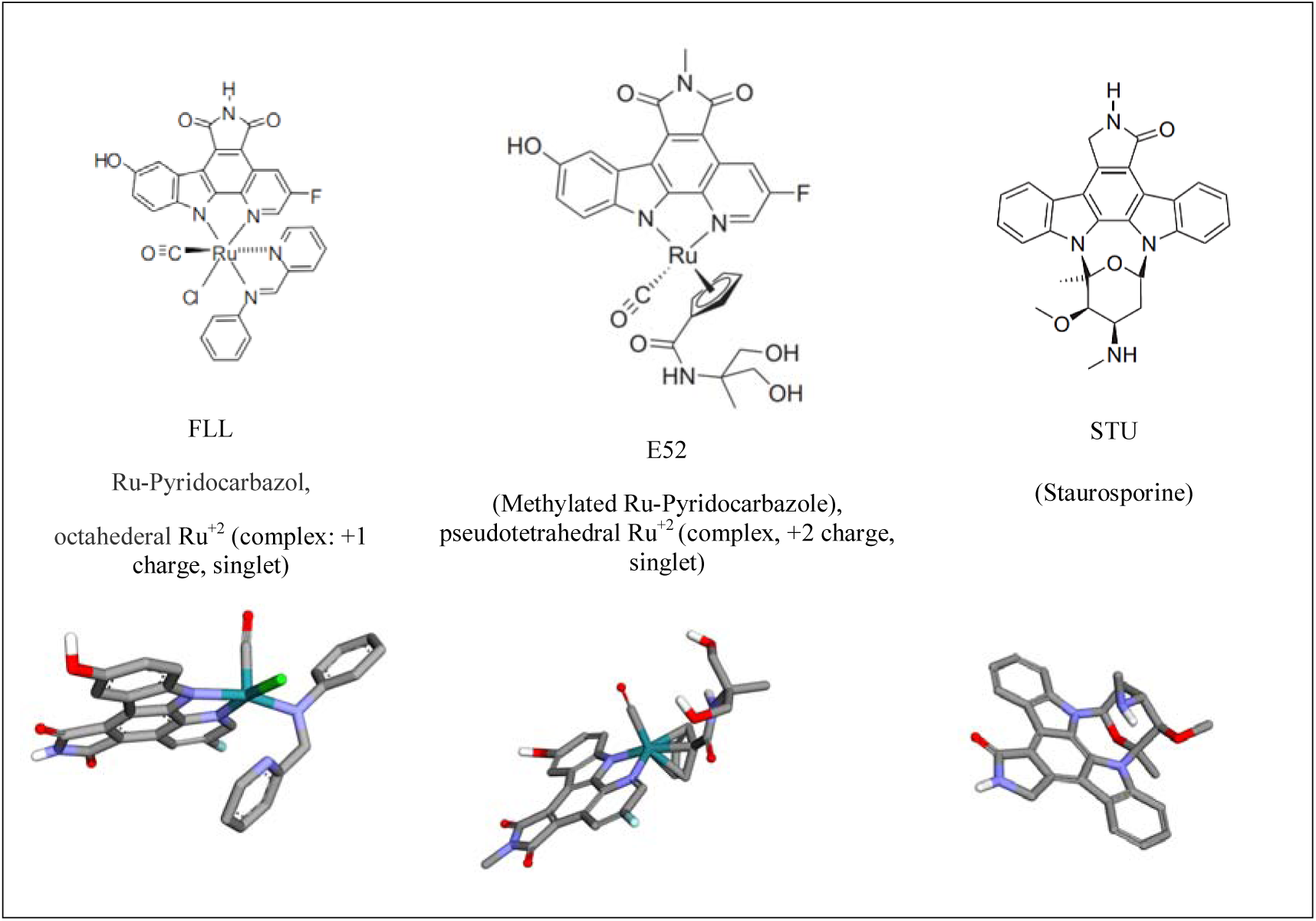
Chemical structures of the modeled Ru-complexes and staurosporine.

**Figure 2.**
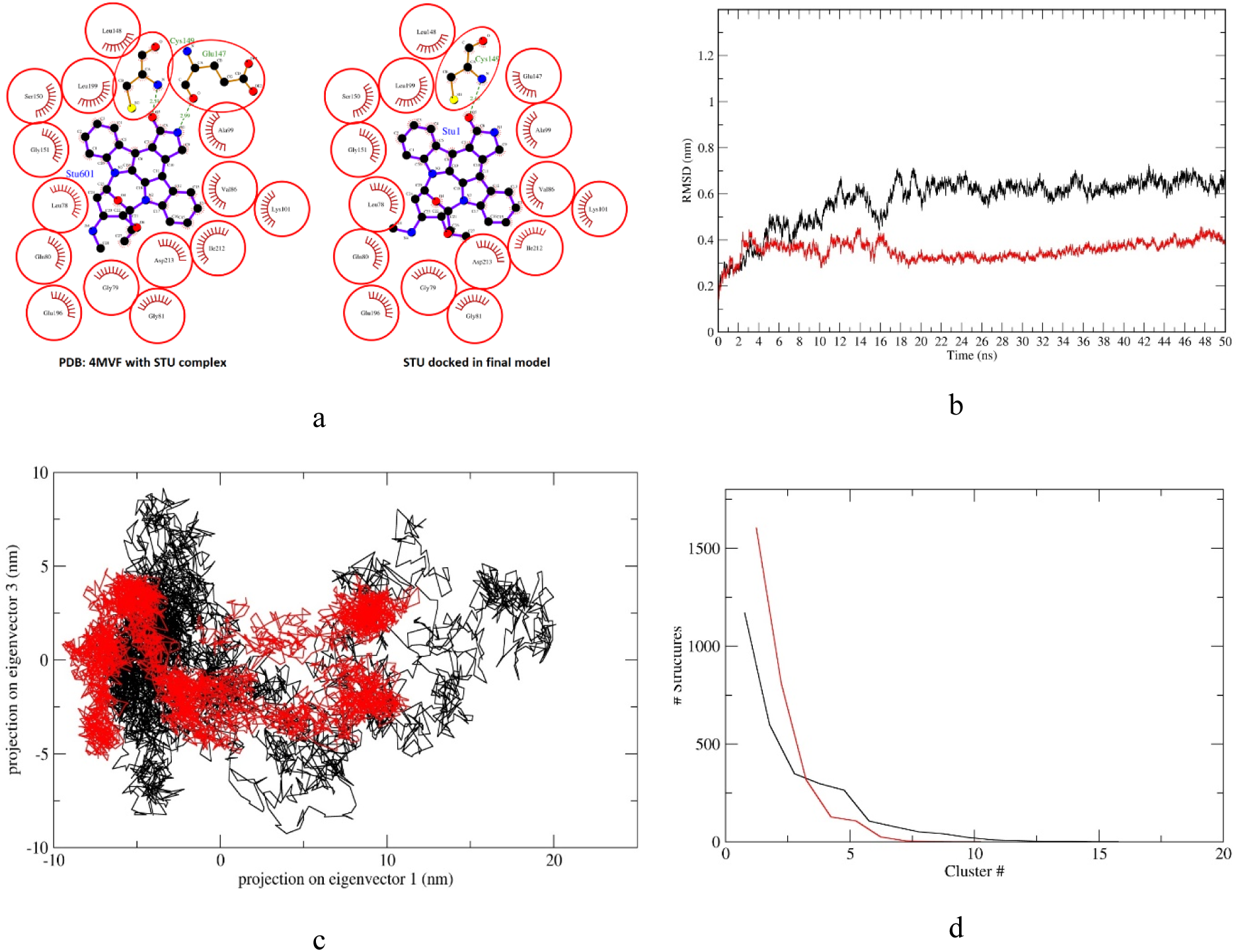
Molecular docking and Dynamics simulation analysis for ensemble conformation generation of PfCDPK2 and Staurosporine (STU) in complex with protein (PfCDPK2_STU). (A) Post docking and 2D comparison of docked STU with PDB:4MVF crystal structure. (B) Computed Cα backbone RMSD (nm) values for 50 ns MDS. (C) PCA 2D scatter plot projecting the motion of the protein in phase space for the two principle components, PC1 and PC3 (D) The plot representing eigenvalues calculated from the covariance matrix of backbone fluctuations vs. the respective eigenvector indices for first 20 eigenvectors from 1000 eigenvectors. In plot (B), (C) and (D) the colour representation is PfCDPK2 (black) and PfCDPK2_STU (red).

### Essential Dynamics and Clustering of MDS trajectories

The conformational space and transitions in the apo and complex structure were inspected by Prinicpal Component Analysis (PCA) analysis. The PCA is a statistical computation which decreases the complexity of the MDS trajectories by extracting only collective motion of Cα atoms while preserving most of the variation. It calculates the covariance matrix of positional fluctuations for backbone atoms which may decipher the dynamics and coherted motions of PfCDPK2 in absence/presence of ligands. The plot which is shown in Fig. 2c is the 2D projection of the trajectories for two major principal components PC1 and PC2 for PfCDPK2 and PfCDPK2_STU which represents different conformations in 2D space. The 2D projection of PfCDPK2 has more variation compare to PfCDPK2_STU due to the stabilization effect on the dynamics motion due to the ligand.

Representative structures for the conformational space traversed by the MDS based ensemble method, GROMOS (Daura et al., 1999) based algorithm in clustering was applied on PfCDPK2 and PfCDPK2_STU MDS. The method basically creates representative RMSD based clusters from the trajectory frames. It counts the number of the neighboring structure using a 0.2 nm cutoff, and then form a cluster set with the largest numbers of neighbor structures followed by its elimination from the pool of clusters. The process is repeated for the remaining frames to identify other clusters with decreasing numbers of neighbor structures and each centroid of the cluster are used as a representative structure. These centroid structure members from each cluster are representative structures of distinct frames. The RMSD values in the clusters ranges from 0.0676 – 0.534 nm (average RMSD 0.267) and 0.0633 - 0.5 nm (average RMSD 0.248) for PfCDPK2 and PfCDPK2_STU respectively. A total of 16 clusters with 310 transitions were found in PfCDPK2 MDS, while the PfCDPK2_STU MDS has 10 clusters with 247 transitions (Fig. 3d). The representative structures from each cluster from PfCDPK2 and the PfCDPK2_STU MDS are shown in supplementary figure (S2, S3) along with the total numbers of structures in each cluster.

**Figure 3.**
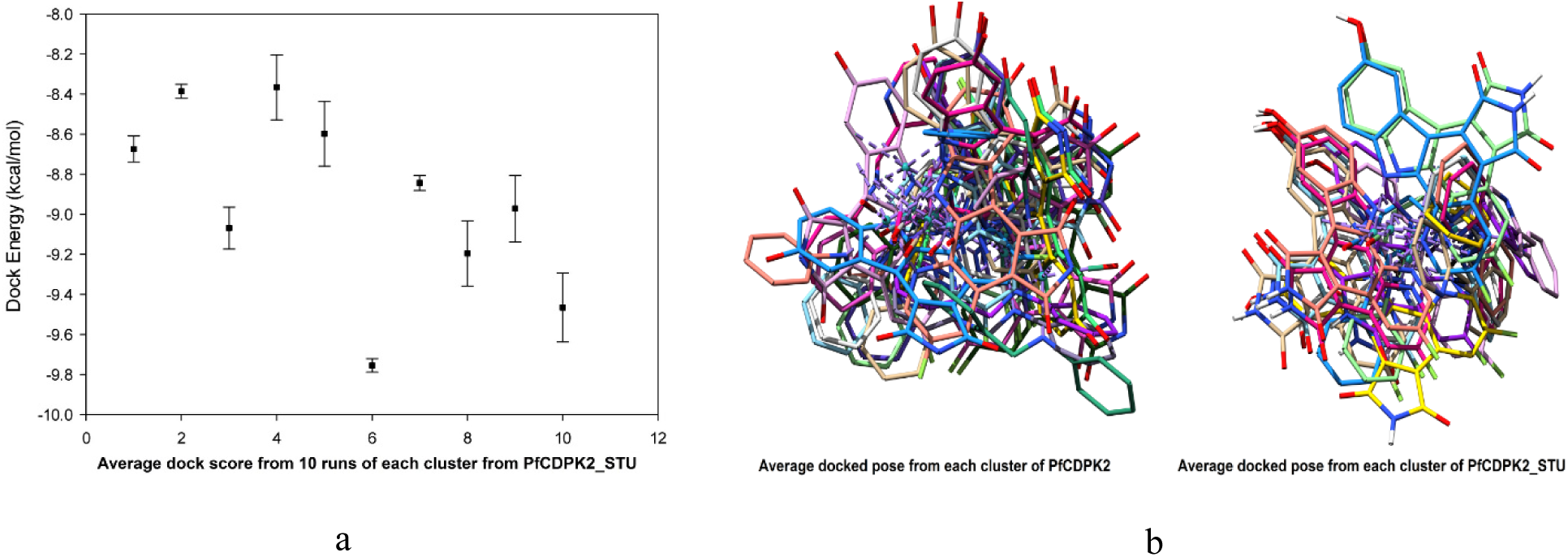
Docking results for each cluster generated from the PfCDPK2 and PfCDPK2_STU MDS. (A) Average docked energy values (kcal/mol) for 10 runs for each cluster generated from the PfCDPK2 MDS. (B) Average docked energy values (kcal/mol) for 10 runs for each cluster from the PfCDPK2_STU MDS. (C) Superposition of average poses of STU for all the clusters from PfCDPK2 MDS (D) Superposition of average poses of STU for all the clusters from PfCDPK2_STU MDS. The standard deviation is denoted by a bar in plot A & B.

**Figure 3.**
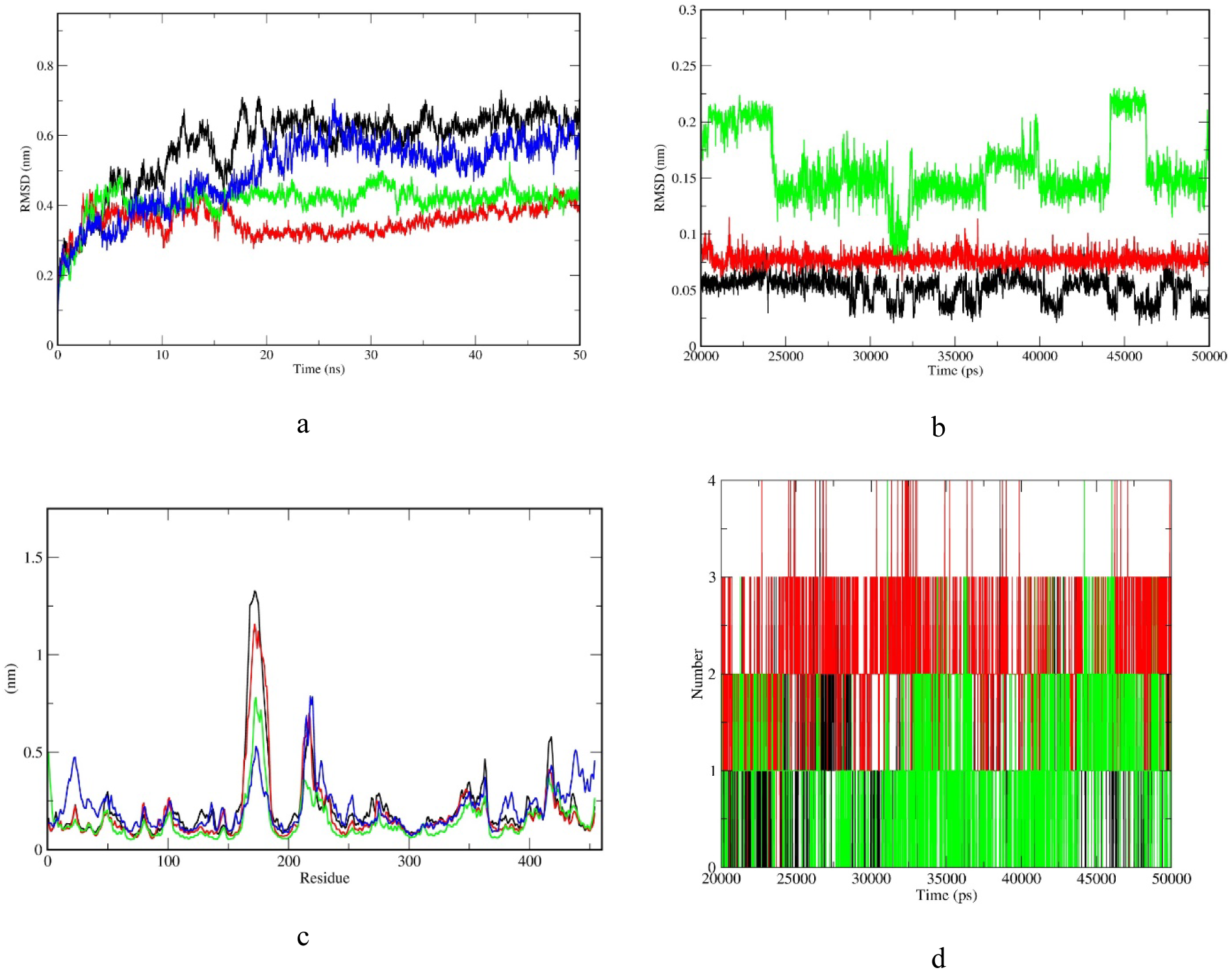
Molecular Dynamics Analysis of PfCDPK2 and PfCDPK2_inhibitor complexes, computing the deviation (nm) vs. function of time (50 ns) for entire MDS. (A) RMSD of the protein Cα backbone atoms. (B) RMSD of the inhibitors PfCDPK2_STU (black), PfCDPK2_FLL (red), PfCDPK2_E52 (green) (C) Computed Cα backbone RMS fluctuations for each residue. (D) Computation of H-bond formation for protein-inhibitor complexes for entire MDS. In (A), (C) & (D) plots, the colour representation is PfCDPK2 (black), PfCDPK2_STU (red), PfCDPK2_FLL (green), PfCDPK2_E52 (blue).

### Ensemble Docking

The overall interest in predicting and comparing the interactions between the Ru-based compounds and PfCDPK2 was to envisage the dynamic residue-wise structural stabilization of the protein-ligand complexes. The docking protocol to non-native protein conformations (apoprotein or protein complexed with the ligands) is still a challenging problem and to overcome this and take flexibility into account, we employed the ensemble-based docking protocol. The current protocol was first tested using PfCDPK2, whose crystal structure has been solved with known inhibitor staurosporine (STU) (PDB ID: 4MVF). The reliability of our docking protocol was determined by ensemble generations using MDS for apo PkCDPK2 structure and PfCDPK2_STU structure with bound STU. The clusters were then generated from these MDS trajectories representing the representative structures for which docking was performed using FLL a known Ru-based inhibitor for PIM1, GSK-3β and CDK2/cyclin A kinase. The ensemble-based approach is already validated as a suitable approach for this family of kinases (Liu, Agrawal, & Radhakrishnan, 2013). In each docking set, the representative clusters from both the MDS were chosen to generate 10 poses of complex docked structure for each cluster representative. A total of 160 poses from 16 clusters and 100 poses from 100 clusters were generated by docking for PkCDPK2 and PfCDPK2_STU. For each cluster from both the MDS, average values for docked energy were calculated for 10 runs with standard deviation and are plotted in Fig. 3a, 3b. The average dock energy for 10 runs of all the clusters from apo PfCDPK2 and PfCDPK2_STU MDS is −9.88 ± 0.42 and −8.93 ± 0.44 kcal/mol respectively. The overall dock energy is significant to indicate a strong interaction between ligand and the protein active site. Fig. 3c, 3d shows the average pose of STU for all the clusters from both the MDS superposed with each other. All the poses were manually inspected for the best pose with respect to the known inhibitor and docked energy, of which one pose was selected from the apo PfCDPK2 ensembles. The basis for selecting the conformation for further MD studies and binding free energy estimation was near-native pose and dock energy. Overall, the poses, as well as docked energy in each cluster in apo conformations, had less variability indicating convergence. The best conformation pose was from the cluster#13 of the apo clusters and the average docked pose with dock energy −10.62 kcal/mol and 3.042 RMSD value was selected for docking with FLL and E52, Ru-based known kinase inhibitor-based on staurosporine scaffold. Our docking protocol and parametrization of grid as well as Ru metal was validated by control docking of PIM1 kinase (PDB:3FXZ) with its co-crystal ligand FLL. The comparison of ligand pose after docking and co-crystal conformation were in agreement (Fig. S4). The same protocol and parameters used in control docking were used for docking of PfCDPK2 with two organometallic compounds FLL and E52, Ru-based known kinase inhibitor-based on staurosporine scaffold. The both docked complex were further subjected to detailed molecular dynamics studies and binding-free energy estimation.

### Molecular Dynamics Simulations of PfCDPK2 with Ru-based inhibitors

A total of four MDS were carried out to study the conformational dynamics and stability of protein-inhibitor complexes. The four MDS consists of one for apoprotein (PfCDPK2), one for PfCDPK complexed with staurosporine (PfCDPK2_STU) and two for protein in complex with two Ru-based known kinase inhibitor having staurosporine scaffold (PfCDPK2_FLL and PfCDPK2_E52). The MDS of protein-inhibitor complex post-molecular docking have been conclusive for deciphering the molecular interaction of protein target on scales where dynamics of individual atoms are investigated. In the current study, we analyzed the RMSD, RMSF, H-bonds as well as PCA of the protein-inhibitor complexes and compared with apo-protein. All the analysis for protein-inhibitor complexes was compared vis-à-vis with PfCDPK2_STU crystal structure MDS as a reference inhibitor (staurosporine). Additionally, the binding free energy of all the complexes for the last stable 30 ns equilibrated trajectory was performed.

For evaluating the dynamic stability of protein–inhibitor complexes and understanding the binding interactions of metal-based compounds, RMSD values for Cα backbone were calculated for the entire 50 ns simulations. RMSD is a means to measure the structural variation between the Cα backbones from its initial conformation to its final position during the entire simulation trajectory. Smaller RMSD values indicate higher stability of the simulation. The RMSD values are shown in the plot of RMSD (nm) vs. Time (ns) in Fig. 4a for PfCDPK2, PfCDPK2_STU, PfCDPK2_FLL, and PfCDPK2_E52. The RMSD plot for Cα backbone values indicated that most of the MDS reached equilibrium within 20 ns. The mean RMSD values for PfCDPK2, PfCDPK2_STU, PfCDPK2_FLL, and PfCDPK2_E52 were 0.573 ± 0.108, 0.354 ± 0.040, 0.412 ± 0.047 and 0.495 ± 0.103 nm respectively. An overall comparison of Ca backbone RMSD showed that the PfCDPK2_STU, PfCDPK2_FLL, and PfCDPK2_E52 were much more stable during the entire simulations in compared to apo-protein PfCDPK2. The RMSD analysis was also carried out with inhibitors to assess the overall stability during the entire course of simulations as shown in Fig. 4b. The mean RMSD values for STU, FLL and E52 were 0.051, 0.077 and 0.15 nm respectively Our MDS results suggested that the Cα backbone RMSD variation of PfCDPK2_FLL is comparable to reference ligand (PfCDPK2_STU).

For analyzing each residue fluctuations during the entire MDS, the Root Mean Squared Fluctuations (RMSF) plot was generated. RMSF values of each protein-inhibitor complexes, as well as the apo-protein and reference inhibitor, were calculated for 50 ns trajectory and overlay on each other to have a comparison of flexible residues in absence and presence of inhibitor binding (Fig 4c). The average RMSF value for PfCDPK2, PfCDPK2_STU, PfCDPK2_FLL, and PfCDPK2_E52 were 0.214 ± 0.19, 0.187 ± 0.18, 0.145 ± 0.10 and 0.211 ± 0.11 respectively. The RMSF analysis of the Cα backbone of protein reveals the residues from 160-185, to be one of the highly flexible regions in the apo-protein as well as the reference inhibitor. Whereas in the case of PfCDPK2_FLL and PfCDPK2_E52 complexes, RMSF values of these flexible residues were decreased substantially. All the protein-inhibitor complexes show less RMS fluctuations compared to apo-protein, which indicates that the binding of inhibitors leads to decrease in flexibility and hence, these compounds have the potential to inhibit the catalytic activity of PfCDPK2. Also, the PfCDPK2_FLL showed very lesser RMSF values compared to PfCDPK2_STU indicating the importance of Ru metal in the ligand moiety.

The essential dynamics of PfCDPK2 and PfCDPK complexes were assessed by calculating Principal Component Analysis (PCA). PCA decreases the complexity of trajectory data by extracting only the collective motion of Cα atoms that are significantly important to investigate the protein-inhibitor complexes stability. The positional fluctuations of the backbone are computed by generating a covariance matrix that reveals the dynamics of PfCDPK2 as well as the difference in concerted motions during ligand binding. Figure 5a shows a plot of eigenvalues in decreasing order vs. the respective eigenvector indices for PfCDPK2, PfCDPK2_STU, PfCDPK2_FLL, and PfCDPK2_E52. The top 15 eigenvectors accounted for 93.7%, 71.8%, 37.3% and 65.6% of the motions observed for 50 ns trajectory for the PfCDPK2, PfCDPK2_STU, PfCDPK2_FLL, and PfCDPK2_E52 respectively. The PCA plot summarized that the values for the first few eigenvalues of PfCDPK2 were much higher compared to the PfCDPK2 complexes indicating that the inhibitor binding leads to a conformational change in protein dynamics. These different correlated motions in absence and presence of inhibitor may be due to the stabilization effect by complex generation. As similar to RMSD and RMSF, the first few eigenvalues PfCDPK2_FLL complex have considerable lesser values even compared to PfCDPK2_STU indicating a better replacement for staurosporine. The plot in Fig. 5b shows the 2D projection of the trajectories for two major principal components PC1 and PC3 for PfCDPK2 and PfCDPK2 complexes. It is evident from the 2D plot that the PfCDPK2 complexes showed higher stability and occupies lesser phase space along with well-defined clusters compare to PfCDPK2. To visualize the essential dynamics, 50 structures were extracted from each MDS projecting the extremely selected eigenvectors. The extreme motion of PfCDPK2 complexes was less deviating when compare to PfCDPK2 indicating a more stable complex formation, especially the PfCDPK2_FLL, showing the least dynamics and stable conformational space.

**Figure 5.**
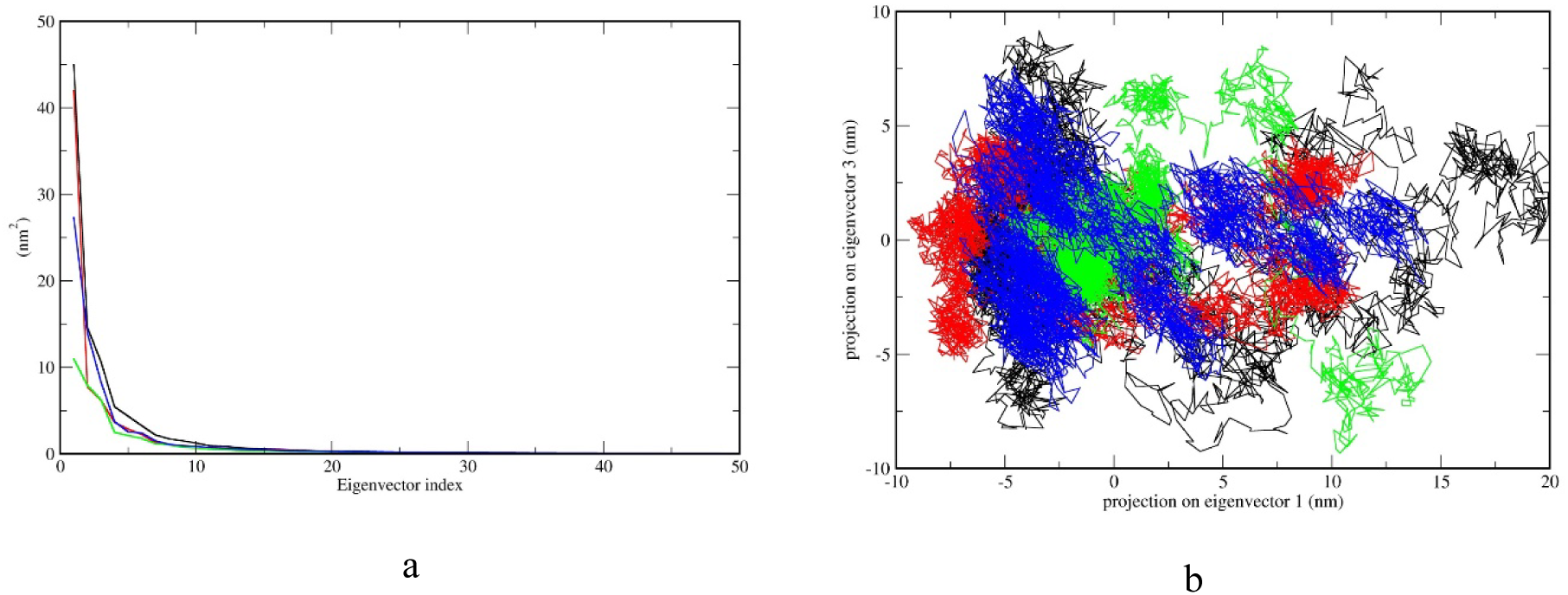
Principle Component Analysis of PfCDPK2 and PfCDPK2_inhibitor complexes for 50 ns MDS. (A) The plot representing eigenvalues calculated from the covariance matrix of backbone fluctuations vs. the respective eigenvector indices for first 20 eigenvectors from 1000 eigenvectors. (B) PCA 2D scatter plot projecting the motion of the protein in phase space for the two principle components, PC1 and PC3. In both the plots, the colour representation is PfCDPK2 (black), PfCDPK2_STU (red), PfCDPK2_FLL (green), PfCDPK2_E52 (blue).

### Molecular interaction and Binding free-energy estimation

The Hydrogen bond formation (H-bond) is the key determinants of specificity and molecular interactions between protein and ligand molecule. The average H-bonds formed between protein and ligand were calculated from stable trajectories 20 to 50 ns of MDS and are plotted in Fig. 4d. The final structures of PfCDPK2_STU, PfCDPK2_E52, and PfCDPK2_FLL formed two H-bonds after simulations and the percentage existence for each H-bond in all three complexes are mentioned in Supplementary file S4. In PfCDPK2_STU complex the residues involved in H-bond interactions throughout the course on entire simulations were GLY20, GLN21, LYS29, CYS90, GLY92, GLU94 and ASP97. For PfCDPK2_FLL complex, the residues involved in H-bond interactions were TYR24, GLY25, CYS26, LYS42, GLU43, ASN138, ARG54, ILE153, and ASP154. While in PfCDPK2_E52 complex, the residues involved in H-bond interactions were LEU19, GLY20, GLN21, GLY22, THR23, CYS26, TYR39, LYS42, GLU43, GLU44, ARG48, LYS50, LYS135, GLU137, ASN138, and ASP154. The residues (GLY, GLN, CYS, ASP and GLU) involved in the active site chemistry of PfCDPK2_STU were also observed to form H-bond in MDS of Ru-based FLL and E52. The 2D protein-ligand plots of post-simulation complexes are shown in Fig. 6. Overall, the molecular interaction pattern from our MDS study revealed that the Ru-based inhibitor has the important interaction with the residues involved in binding with the natural substrate and may have the capability to inhibit the activity of PfCDPK2.

**Figure 6.**
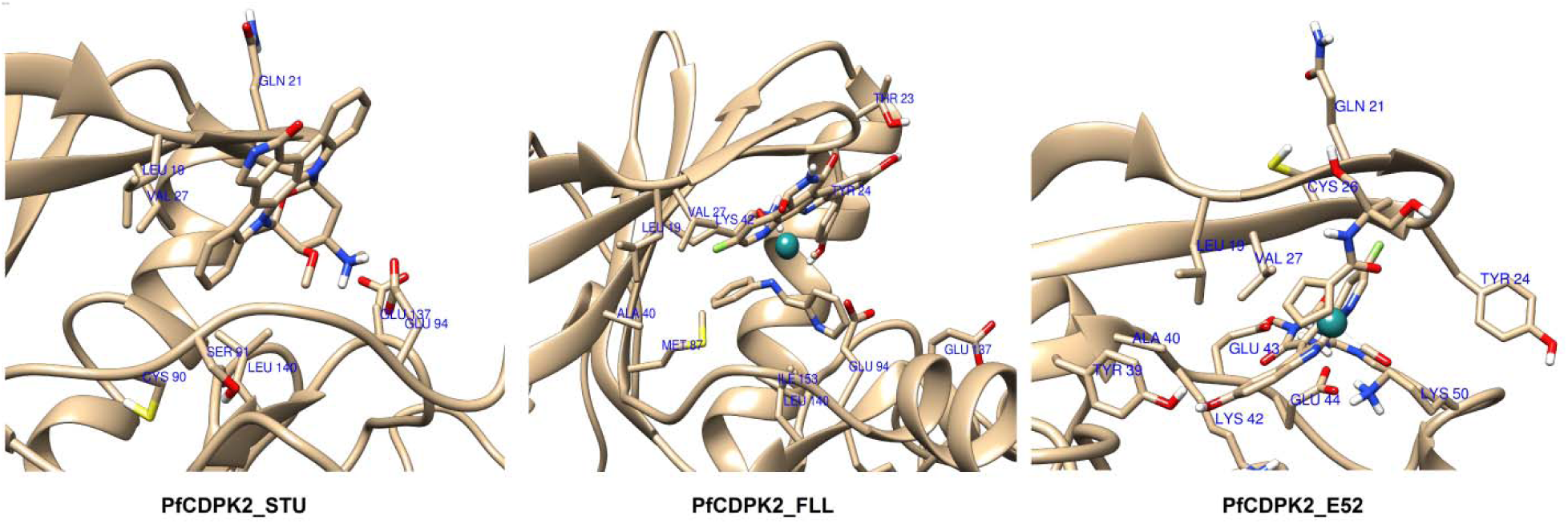
3D protein-ligand plots of post-simulation for PfCDPK2_inhibitor complexes. Th interacting amino acids are labelled with three-letter code and corresponding residue number. 3D plot for PfCDPK2_STU complex, PfCDPK2_FLL complex and PfCDPK2_E52 complex respectively.

The binding free energy between protein-ligand complexes which is often referred to as ΔG was calculated using the MM-PBSA method for the last 30 ns stable trajectories. A total of 120 frames at every 250ps from 30 ns trajectories were taken for the calculations of binding free energy (ΔG) which is an estimation of the non-bonded interaction energies. The ΔG values for PfCDPK2_STU, PfCDPK2_FLL, and PfCDPK2_E52 were negative (Fig. 7a) and for PfCDPK2_FLL the values were similar to known reference inhibitor and for PfCDPK2_E52 values were better compared to known reference inhibitor PfCDPK2_STU. The individual component for binding energy, the Van der Waals, the electrostatic interactions and non-polar solvation energy had contributed negatively to the overall interaction energy as shown in Fig. 7b. The only exception was the polar solvation energy which contributed positively. To gain more insights into key residues involved in ligand binding, the residue-wise energy decomposition plot was generated which shows the total binding energy contribution for each residue for all three MDS (Fig. 7c).

**Figure 7.**
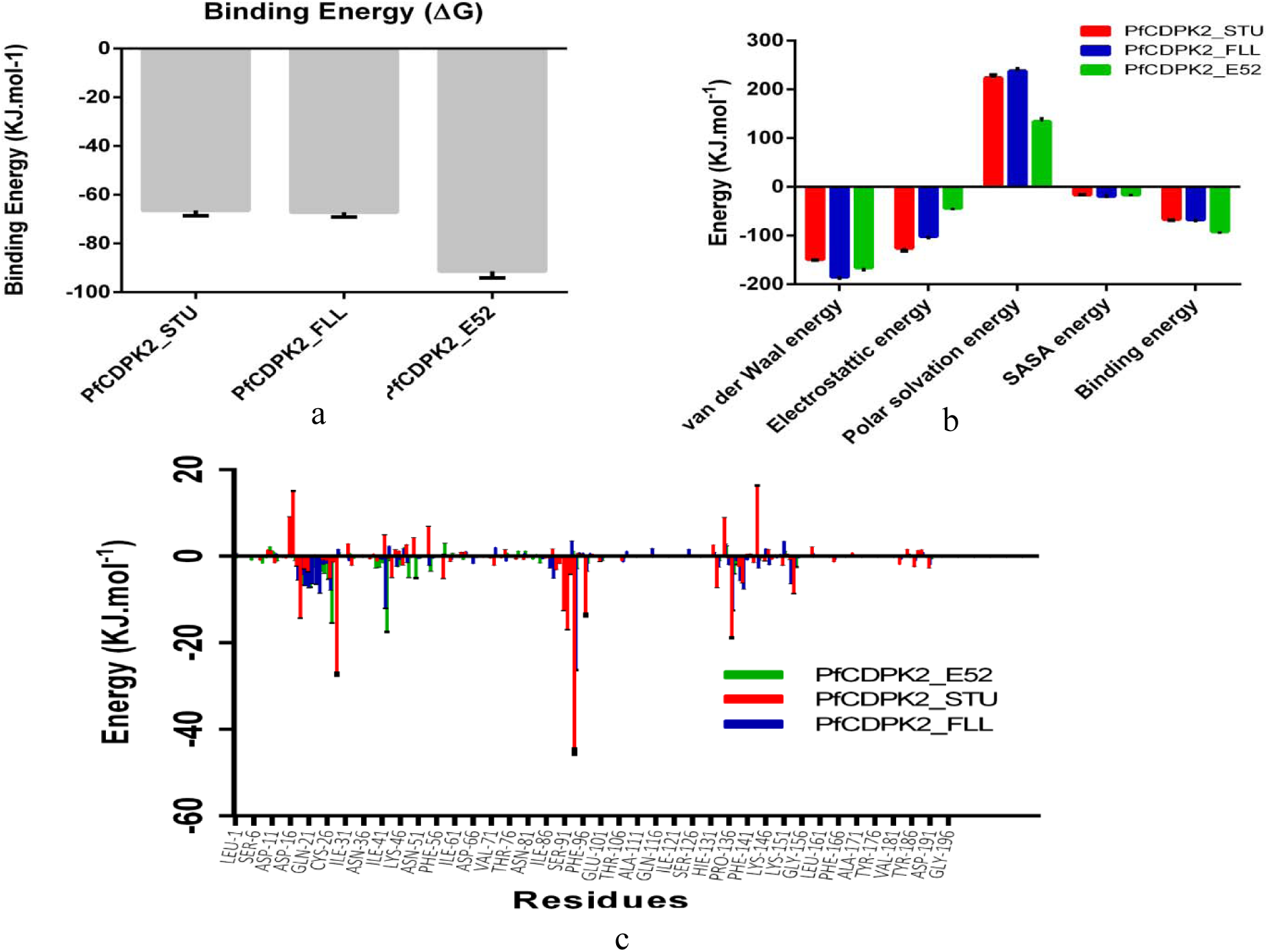
MM-PBSA Calculation of binding free energy. (A) The total binding free energy for all the PfCDPK2_inhibitor complexes calculated for last 30 ns stable trajectory for a total of 120 frames, each at 250 ps interval. (B) Representative contributions of each energy component for binding free energy for PfCDPK2 interactions with inhibitors. (C) The contribution of important binding residues of PfCDPK2 with three inhibitors to the total binding free energy. The (-ve) values indicate stable complex formation for PfCDPK2_inhibitor complexes, while the (+ve) values indicate a destabilizing effect. In all the plots, the colour representation is PfCDPK2_STU (red), PfCDPK2_FLL (blue), PfCDPK2_E52 (green).

## Conclusion and Discussion

Modeling the dynamic behavior of bounded metal-ligand complex using MD simulation requires a comprehensive understanding of the metal-ligand bonding, thermodynamics and coordination chemistry within the receptor binding sites (Riccardi, Genna, & De Vivo, 2018). This would undeniably accelerate the opportunities for novel design and discovery of metalloenzyme-inhibitors (inhibitors for enzymes containing metals) and metallodrugs (organometallic inhibitors with metals). However, there are several methodological challenges associated with the transition metals such as ion parametrization, multiple oxidation states, flexible metal-coordination and schemes for non-bonded models. These challenges can be compensated by utilizing free energy sampling based on DFT geometry optimizations (of metal coordination in protein-ligand complexes) (Gaspari et al., 2016). MD simulations consider structural flexibility and thereby reasonably approximate ligand recognition and binding (Grubmüller, Heymann, & Tavan, 1996). It is worth to mention that unconventional transition metal-centre complexes are difficult to simulate using default molecular mechanics (MM) parameters (Brooks et al., 2009). This issue has been previously addressed by Bosnich and co-workers that introduced the idea of additional five-fold torsion angle parameters to model metal-containing complex viz. ferrocene (Bosnich, 1994; D. J. Williams et al., 1995). Further, this topic was well-reviewed and persuaded by Li and Merz (P. Li & Merz, 2017). As an extension to the practical utility of this idea, we report the ensemble docking and dynamics of Ruthenium-based organometallic inhibitor with staurosporine moiety which can be used as kinase inhibitor against malarial CDPKs. Ruthenium is considered as most promising transition metal for drug development and pharmaceutical values (Abid, Shamsi, & Azam, 2016; Dragutan, Dragutan, & Demonceau, 2017; Meier-Menches, Gerner, Berger, Hartinger, & Keppler, 2018; Meier et al., 2013). In the context of ruthenium-based kinase inhibitor, Meggers and co-workers have pioneered the development of potent and selective kinase inhibitors through the use of octahedral pyridocarbazole complexes mimicking the pan-kinase inhibitor staurosporine (Debreczeni et al., 2006).

Through this study, we observed that apo PfCDPK2 clusters were reliable for molecular docking approach as they were distinct conformation covering flexibility aspects as well as the docking score which was high and clustered together with fewer deviations. Two known inhibitors, FLL, and E52 of protein kinases based on staurosporine moiety were used for molecular docking evaluation. The complexes predicted from docking were further subjected to MD simulations and free energy sampling, which takes into account the structural flexibility of the drug target in order to investigate ligand recognition and binding. Overall, the binding pose, MD simulations and free energy of these complexes suggested that FLL and E52 are two promising drug targets comparable to known STU inhibitor. We also showed that FLL is a more potent drug target against PfCDPK2. Earlier attempt to use these molecules against protein kinase also showed the importance of ensemble docking but the results lacked in dynamic information of these molecules (Liu et al., 2013). Here, we have demonstrated a hybrid-methodology which includes ensemble docking approach, DFT-derived transition-metal-parameterization to prepare the system for MD simulations and free binding energy estimation.

In conclusion, this study along with previous reports establishes a simplistic modeling approach and underpins the attributes needed for modeling metal-ligand complexes for developing bioactive metal compounds. We believe that our approach involving ensemble docking, DFT calculations for metal-ligand parameterization using MCPB.py in AMBER (Case et al., 2005) and MD simulations can be further adapted to explore the potential of transitions metals in drug discovery and overcoming the challenges associated with transition metal parameterization.

## ACKNOWLEDGEMENT

Prakash C Jha acknowledges Science and Engineering Research Board (SERB), Department of Science and Technology (DST) for a project grant, EMR/2016/003025. Dhaval Patel acknowledges the Department of Biotechnology for DISC grant, Govt. of India for the resources and financial assistance. Mohd Athar acknowledges generous support from the DST in the form of SRF INSPIRE Fellowship (IF150167).

## SUPPLEMENTARY FIGURES

**Figure S2.**
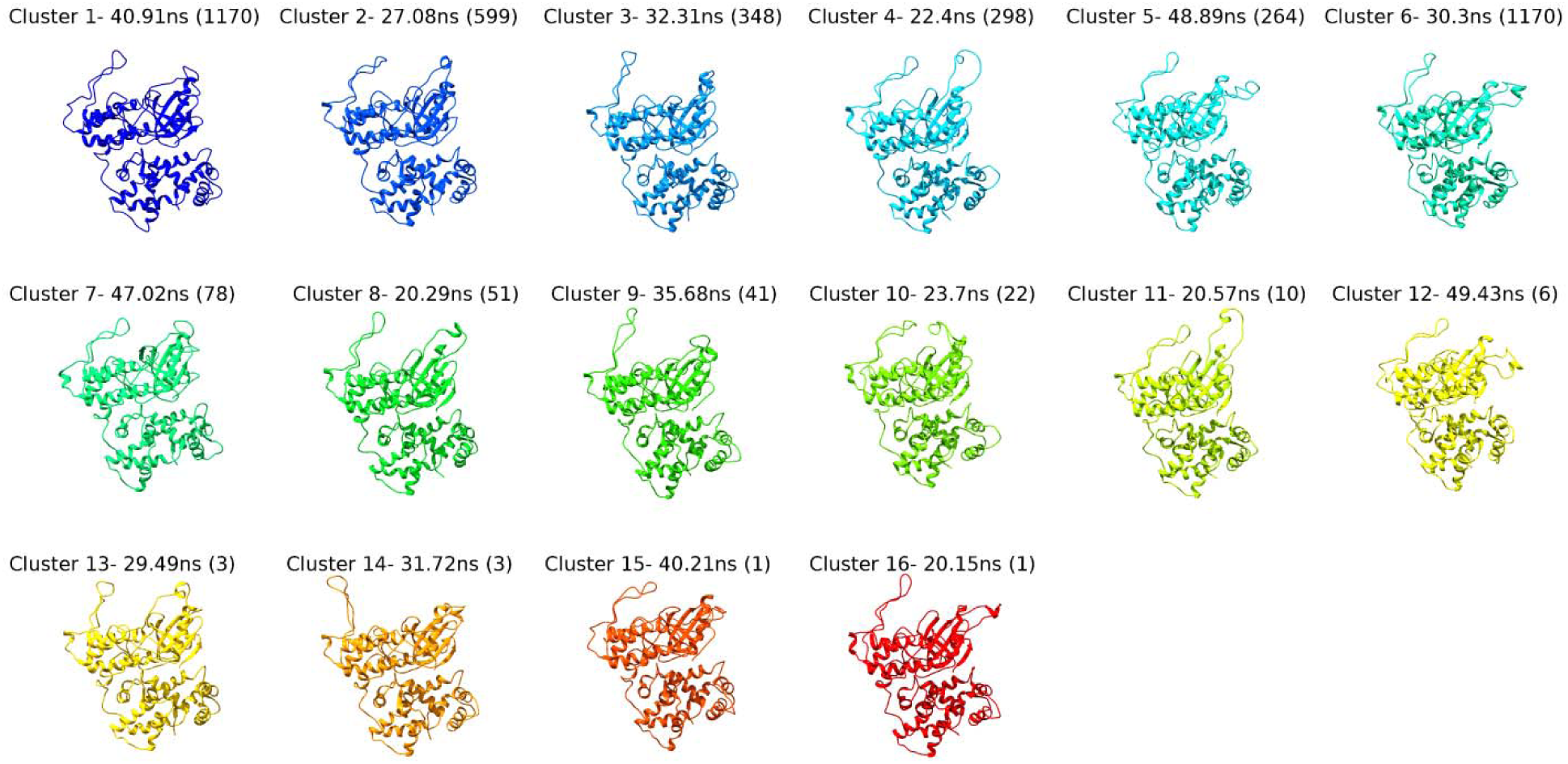
The representative structures from each cluster from PfCDPK2 MDS is shown with the time point (ns) and along with the total numbers of structures in each cluster indicated in parenthesis.

**Figure S3.**
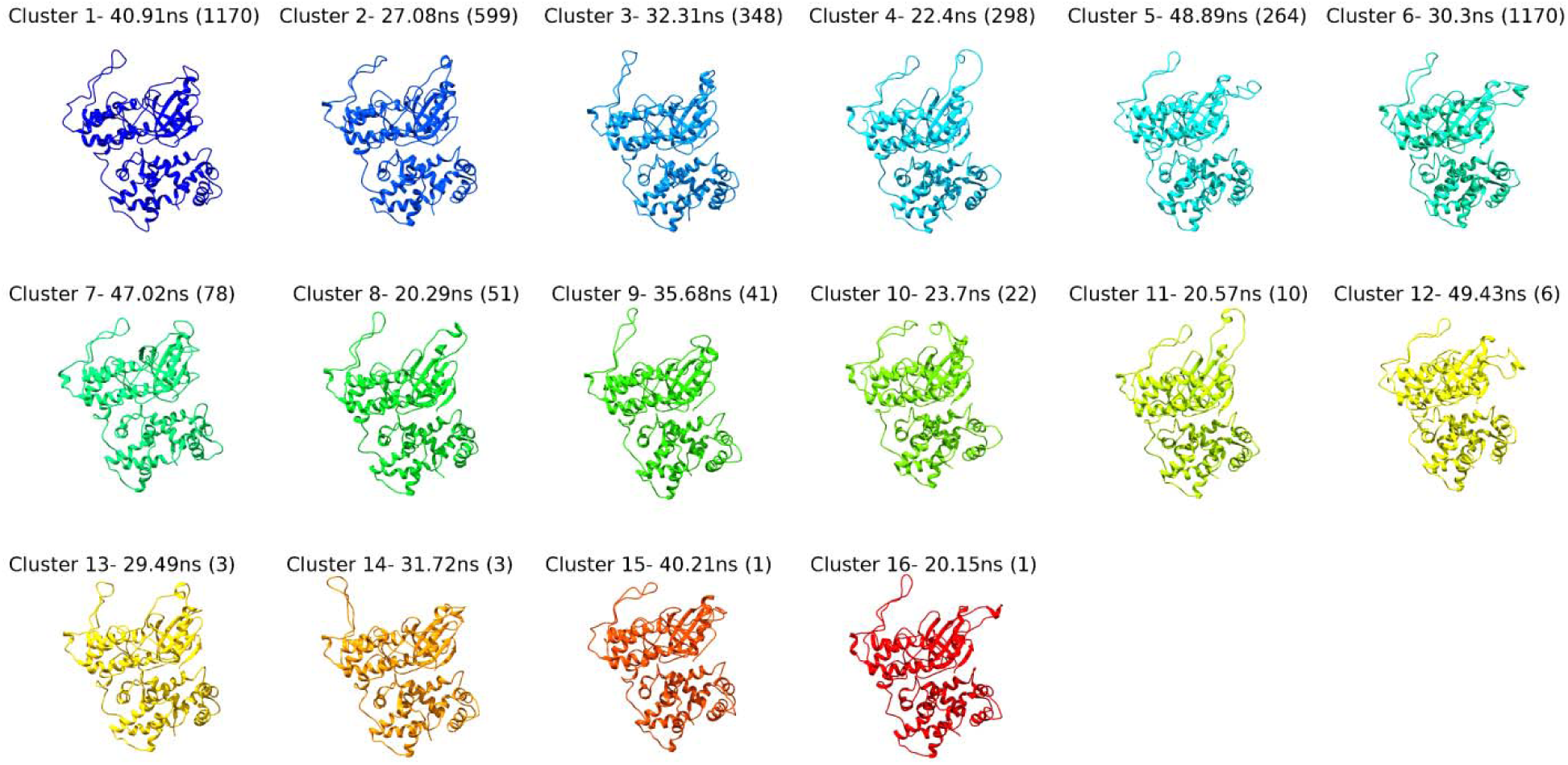
The representative structures from each cluster from the PfCDPK2_STU MDS is shown with the time point (ns) and along with the total numbers of structures in each cluster indicated in parenthesis.

**Figure S4.**
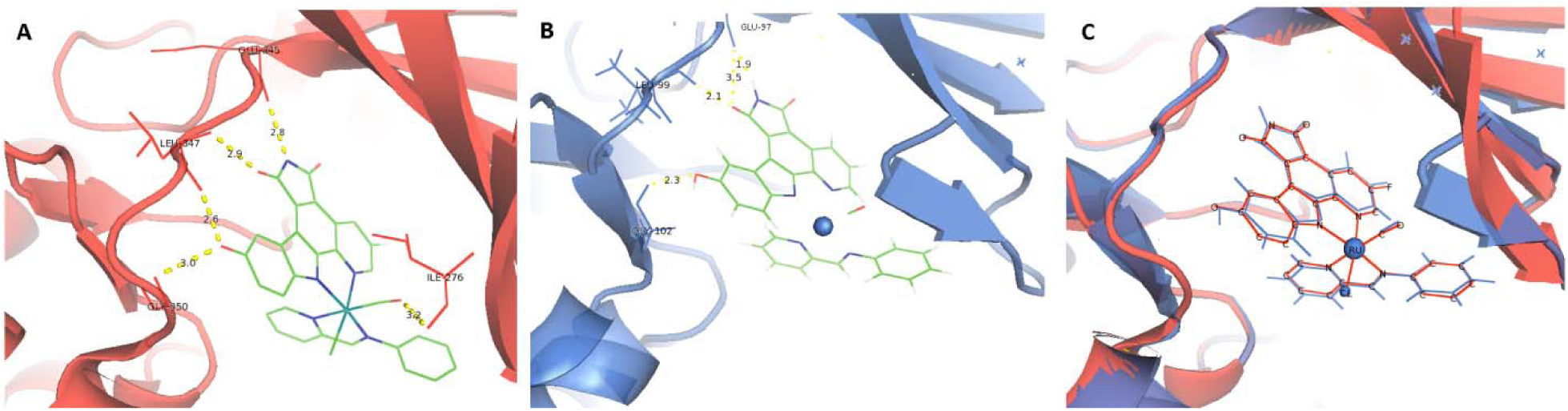
Interaction of Ru-based FLL organometallic inhibitor and PIM1 Kinase (PDB: 3FXZ). (A) 3D interactions of FLL with PIM1 Kinase (PDB: 3FXZ) with interacting labeled with residues name and hydrogen bond length in Å.(B) 3D interactions of FLL after control docking with PIM1 Kinase (PDB: 3FXZ) with interacting labeled with residues name and hydrogen bond length in Å. (C) Superpose 3D images of FLL co-crystal interaction (red) and docked FLL interaction (blue) with PIM1 Kinase (PDB: 3FXZ).

