## Supplementary material for "Exploring Ruthenium-based organometallic inhibitors against Plasmodium Calcium Dependent Kinase 2 (PfCDPK2): a combined ensemble docking, QM paramterization and molecular dynamics study": Table S2

**Title: Topology file for 4MVF with FLL ligand prepared after docking and MCPB.py calculation using Gaussian**

```
; 4MVF_dry_GMX.top created by acpype (Rev: 0) on Mon Nov 12 11:59:34
2018

[ defaults ]
; nbfunc        comb-rule       gen-pairs       fudgeLJ fudgeQQ
1               2               yes             0.5     0.8333

[ atomtypes ]
;name   bond_type     mass     charge   ptype   sigma         epsilon
Amb
 N3       N3          0.00000  0.00000   A     3.25000e-01   7.11280e-
01 ; 1.82  0.1700
 H        H           0.00000  0.00000   A     1.06908e-01   6.56888e-
02 ; 0.60  0.0157
 CX       CX          0.00000  0.00000   A     3.39967e-01   4.57730e-
01 ; 1.91  0.1094
 HP       HP          0.00000  0.00000   A     1.95998e-01   6.56888e-
02 ; 1.10  0.0157
 A2C      A2C         0.00000  0.00000   A     3.39967e-01   4.57730e-
01 ; 1.91  0.1094
 HC       HC          0.00000  0.00000   A     2.64953e-01   6.56888e-
02 ; 1.49  0.0157
 A3C      A3C         0.00000  0.00000   A     3.39967e-01   4.57730e-
01 ; 1.91  0.1094
 CT       CT          0.00000  0.00000   A     3.39967e-01   4.57730e-
01 ; 1.91  0.1094
 C        C           0.00000  0.00000   A     3.39967e-01   3.59824e-
01 ; 1.91  0.0860
 O        O           0.00000  0.00000   A     2.95992e-01   8.78640e-
01 ; 1.66  0.2100
 N        N           0.00000  0.00000   A     3.25000e-01   7.11280e-
01 ; 1.82  0.1700
 H1       H1          0.00000  0.00000   A     2.47135e-01   6.56888e-
02 ; 1.39  0.0157
 SH       SH          0.00000  0.00000   A     3.56359e-01
1.04600e+00 ; 2.00  0.2500
 HS       HS          0.00000  0.00000   A     1.06908e-01   6.56888e-
02 ; 0.60  0.0157
 CC       CC          0.00000  0.00000   A     3.39967e-01   3.59824e-
01 ; 1.91  0.0860
 NB       NB          0.00000  0.00000   A     3.25000e-01   7.11280e-
01 ; 1.82  0.1700
 CR       CR          0.00000  0.00000   A     3.39967e-01   3.59824e-
01 ; 1.91  0.0860
 H5       H5          0.00000  0.00000   A     2.42146e-01   6.27600e-
02 ; 1.36  0.0150
 NA       NA          0.00000  0.00000   A     3.25000e-01   7.11280e-
01 ; 1.82  0.1700
```

```
 CW       CW          0.00000  0.00000   A     3.39967e-01   3.59824e-
01 ; 1.91  0.0860
 H4       H4          0.00000  0.00000   A     2.51055e-01   6.27600e-
02 ; 1.41  0.0150
 OH       OH          0.00000  0.00000   A     3.06647e-01   8.80314e-
01 ; 1.72  0.2104
 HO       HO          0.00000  0.00000   A     0.00000e+00
0.00000e+00 ; 0.00  0.0000
 C8       C8          0.00000  0.00000   A     3.39967e-01   4.57730e-
01 ; 1.91  0.1094
 CO       CO          0.00000  0.00000   A     3.39967e-01   3.59824e-
01 ; 1.91  0.0860
 O2       O2          0.00000  0.00000   A     2.95992e-01   8.78640e-
01 ; 1.66  0.2100
 CA       CA          0.00000  0.00000   A     3.39967e-01   3.59824e-
01 ; 1.91  0.0860
 HA       HA          0.00000  0.00000   A     2.59964e-01   6.27600e-
02 ; 1.46  0.0150
 N2       N2          0.00000  0.00000   A     3.25000e-01   7.11280e-
01 ; 1.82  0.1700
 S        S           0.00000  0.00000   A     3.56359e-01
1.04600e+00 ; 2.00  0.2500
 C*       C*          0.00000  0.00000   A     3.39967e-01   3.59824e-
01 ; 1.91  0.0860
 CN       CN          0.00000  0.00000   A     3.39967e-01   3.59824e-
01 ; 1.91  0.0860
 CB       CB          0.00000  0.00000   A     3.39967e-01   3.59824e-
01 ; 1.91  0.0860
 f        f           0.00000  0.00000   A     3.11815e-01   2.55224e-
01 ; 1.75  0.0610
 c2       c2          0.00000  0.00000   A     3.39967e-01   3.59824e-
01 ; 1.91  0.0860
 Y1       Y1          0.00000  0.00000   A     3.25000e-01   7.11280e-
01 ; 1.82  0.1700
 Y2       Y2          0.00000  0.00000   A     3.25000e-01   7.11280e-
01 ; 1.82  0.1700
 Y5       Y5          0.00000  0.00000   A     3.39967e-01   8.78640e-
01 ; 1.91  0.2100
 Y3       Y3          0.00000  0.00000   A     3.47094e-01
1.10876e+00 ; 1.95  0.2650
 hn       hn          0.00000  0.00000   A     1.06908e-01   6.56888e-
02 ; 0.60  0.0157
 cp       cp          0.00000  0.00000   A     3.39967e-01   3.59824e-
01 ; 1.91  0.0860
 cq       cq          0.00000  0.00000   A     3.39967e-01   3.59824e-
01 ; 1.91  0.0860
 Y4       Y4          0.00000  0.00000   A     3.25000e-01   7.11280e-
01 ; 1.82  0.1700
 M1       M1          0.00000  0.00000   A     2.63973e-01   2.34304e-
01 ; 1.48  0.0560

[ moleculetype ]
```

```
;name            nrexcl
 4MVF_dry         3

[ atoms ]
;   nr  type  resi  res  atom  cgnr     charge      mass       ; qtot
bond_type
     1   N3      1   LEU      N    1     0.101000     14.01000 ; qtot
0.101
     2    H      1   LEU     H1    2     0.214800      1.00800 ; qtot
0.316
     3    H      1   LEU     H2    3     0.214800      1.00800 ; qtot
0.531
     4    H      1   LEU     H3    4     0.214800      1.00800 ; qtot
0.745
     5   CX      1   LEU     CA    5     0.010400     12.01000 ; qtot
0.756
     6   HP      1   LEU     HA    6     0.105300      1.00800 ; qtot
0.861
     7  A2C      1   LEU     CB    7    -0.024400     12.01000 ; qtot
0.837
     8   HC      1   LEU    HB2    8     0.025600      1.00800 ; qtot
0.862
     9   HC      1   LEU    HB3    9     0.025600      1.00800 ; qtot
0.888
    10  A3C      1   LEU     CG   10     0.342100     12.01000 ; qtot
1.230
    11   HC      1   LEU     HG   11    -0.038000      1.00800 ; qtot
1.192
    12   CT      1   LEU    CD1   12    -0.410600     12.01000 ; qtot
0.781
    13   HC      1   LEU   HD11   13     0.098000      1.00800 ; qtot
0.879
    14   HC      1   LEU   HD12   14     0.098000      1.00800 ; qtot
0.977
    15   HC      1   LEU   HD13   15     0.098000      1.00800 ; qtot
1.075
    16   CT      1   LEU    CD2   16    -0.410400     12.01000 ; qtot
0.665
    17   HC      1   LEU   HD21   17     0.098000      1.00800 ; qtot
0.763
    18   HC      1   LEU   HD22   18     0.098000      1.00800 ; qtot
0.861
    19   HC      1   LEU   HD23   19     0.098000      1.00800 ; qtot
0.959
    20    C      1   LEU      C   20     0.612301     12.01000 ; qtot
1.571
    21    O      1   LEU      O   21    -0.571301     16.00000 ; qtot
1.000
    22    N      2   ILE      N   22    -0.415700     14.01000 ; qtot
0.584
    23    H      2   ILE      H   23     0.271900      1.00800 ; qtot
0.856
```

```
    24   CX      2   ILE     CA   24    -0.059700     12.01000 ; qtot
0.797
    25   H1      2   ILE     HA   25     0.086900      1.00800 ; qtot
0.883
    26  A3C      2   ILE     CB   26     0.130300     12.01000 ; qtot
1.014
    27   HC      2   ILE     HB   27     0.018700      1.00800 ; qtot
1.032
    28   CT      2   ILE    CG2   28    -0.320400     12.01000 ; qtot
0.712
    29   HC      2   ILE   HG21   29     0.088200      1.00800 ; qtot
0.800
    30   HC      2   ILE   HG22   30     0.088200      1.00800 ; qtot
0.888
    31   HC      2   ILE   HG23   31     0.088200      1.00800 ; qtot
0.977
    32  A2C      2   ILE    CG1   32    -0.043000     12.01000 ; qtot
0.934
    33   HC      2   ILE   HG12   33     0.023600      1.00800 ; qtot
0.957
    34   HC      2   ILE   HG13   34     0.023600      1.00800 ; qtot
0.981
    35   CT      2   ILE    CD1   35    -0.066000     12.01000 ; qtot
0.915
    36   HC      2   ILE   HD11   36     0.018600      1.00800 ; qtot
0.933
    37   HC      2   ILE   HD12   37     0.018600      1.00800 ; qtot
0.952
    38   HC      2   ILE   HD13   38     0.018600      1.00800 ; qtot
0.971
    39    C      2   ILE      C   39     0.597301     12.01000 ; qtot
1.568
    40    O      2   ILE      O   40    -0.567901     16.00000 ; qtot
1.000
    41    N      3   LEU      N   41    -0.415700     14.01000 ; qtot
0.584
    42    H      3   LEU      H   42     0.271900      1.00800 ; qtot
0.856
    43   CX      3   LEU     CA   43    -0.051800     12.01000 ; qtot
0.804
    44   H1      3   LEU     HA   44     0.092200      1.00800 ; qtot
0.897
    45  A2C      3   LEU     CB   45    -0.110200     12.01000 ; qtot
0.786
    46   HC      3   LEU    HB2   46     0.045700      1.00800 ; qtot
0.832
    47   HC      3   LEU    HB3   47     0.045700      1.00800 ; qtot
0.878
    48  A3C      3   LEU     CG   48     0.353100     12.01000 ; qtot
1.231
    49   HC      3   LEU     HG   49    -0.036100      1.00800 ; qtot
1.195
```

| | | | | | | | | |
|---|---|---|---|---|---|---|---|---|
| 50 | CT | 3 | LEU | CD1 | 50 | -0.412100 | 12.01000 | ; qtot 0.783 |
| 51 | HC | 3 | LEU | HD11 | 51 | 0.100000 | 1.00800 | ; qtot 0.883 |
| 52 | HC | 3 | LEU | HD12 | 52 | 0.100000 | 1.00800 | ; qtot 0.983 |
| 53 | HC | 3 | LEU | HD13 | 53 | 0.100000 | 1.00800 | ; qtot 1.083 |
| 54 | CT | 3 | LEU | CD2 | 54 | -0.412100 | 12.01000 | ; qtot 0.671 |
| 55 | HC | 3 | LEU | HD21 | 55 | 0.100000 | 1.00800 | ; qtot 0.771 |
| 56 | HC | 3 | LEU | HD22 | 56 | 0.100000 | 1.00800 | ; qtot 0.871 |
| 57 | HC | 3 | LEU | HD23 | 57 | 0.100000 | 1.00800 | ; qtot 0.971 |
| 58 | C | 3 | LEU | C | 58 | 0.597301 | 12.01000 | ; qtot 1.568 |
| 59 | O | 3 | LEU | O | 59 | -0.567901 | 16.00000 | ; qtot 1.000 |
| 60 | N | 4 | CYS | N | 60 | -0.415700 | 14.01000 | ; qtot 0.584 |
| 61 | H | 4 | CYS | H | 61 | 0.271900 | 1.00800 | ; qtot 0.856 |
| 62 | CX | 4 | CYS | CA | 62 | 0.021300 | 12.01000 | ; qtot 0.878 |
| 63 | H1 | 4 | CYS | HA | 63 | 0.112400 | 1.00800 | ; qtot 0.990 |
| 64 | A2C | 4 | CYS | CB | 64 | -0.123100 | 12.01000 | ; qtot 0.867 |
| 65 | H1 | 4 | CYS | HB2 | 65 | 0.111200 | 1.00800 | ; qtot 0.978 |
| 66 | H1 | 4 | CYS | HB3 | 66 | 0.111200 | 1.00800 | ; qtot 1.089 |
| 67 | SH | 4 | CYS | SG | 67 | -0.311900 | 32.06000 | ; qtot 0.777 |
| 68 | HS | 4 | CYS | HG | 68 | 0.193300 | 1.00800 | ; qtot 0.971 |
| 69 | C | 4 | CYS | C | 69 | 0.597301 | 12.01000 | ; qtot 1.568 |
| 70 | O | 4 | CYS | O | 70 | -0.567901 | 16.00000 | ; qtot 1.000 |
| 71 | N | 5 | HIE | N | 71 | -0.415700 | 14.01000 | ; qtot 0.584 |
| 72 | H | 5 | HIE | H | 72 | 0.271900 | 1.00800 | ; qtot 0.856 |
| 73 | CX | 5 | HIE | CA | 73 | -0.058100 | 12.01000 | ; qtot 0.798 |
| 74 | H1 | 5 | HIE | HA | 74 | 0.136000 | 1.00800 | ; qtot 0.934 |
| 75 | CT | 5 | HIE | CB | 75 | -0.007400 | 12.01000 | ; qtot 0.927 |

```
     76   HC      5   HIE   HB2   76      0.036700      1.00800 ; qtot
0.963
     77   HC      5   HIE   HB3   77      0.036700      1.00800 ; qtot
1.000
     78   CC      5   HIE    CG   78      0.186800     12.01000 ; qtot
1.187
     79   NB      5   HIE   ND1   79     -0.543201     14.01000 ; qtot
0.644
     80   CR      5   HIE   CE1   80      0.163500     12.01000 ; qtot
0.807
     81   H5      5   HIE   HE1   81      0.143500      1.00800 ; qtot
0.951
     82   NA      5   HIE   NE2   82     -0.279500     14.01000 ; qtot
0.671
     83    H      5   HIE   HE2   83      0.333900      1.00800 ; qtot
1.005
     84   CW      5   HIE   CD2   84     -0.220700     12.01000 ; qtot
0.784
     85   H4      5   HIE   HD2   85      0.186200      1.00800 ; qtot
0.971
     86    C      5   HIE     C   86      0.597301     12.01000 ; qtot
1.568
     87    O      5   HIE     O   87     -0.567901     16.00000 ; qtot
1.000
     88    N      6   SER     N   88     -0.415700     14.01000 ; qtot
0.584
     89    H      6   SER     H   89      0.271900      1.00800 ; qtot
0.856
     90   CX      6   SER    CA   90     -0.024900     12.01000 ; qtot
0.831
     91   H1      6   SER    HA   91      0.084300      1.00800 ; qtot
0.916
     92  A2C      6   SER    CB   92      0.211700     12.01000 ; qtot
1.127
     93   H1      6   SER   HB2   93      0.035200      1.00800 ; qtot
1.163
     94   H1      6   SER   HB3   94      0.035200      1.00800 ; qtot
1.198
     95   OH      6   SER    OG   95     -0.654601     16.00000 ; qtot
0.543
     96   HO      6   SER    HG   96      0.427500      1.00800 ; qtot
0.971
     97    C      6   SER     C   97      0.597301     12.01000 ; qtot
1.568
     98    O      6   SER     O   98     -0.567901     16.00000 ; qtot
1.000
     99    N      7   GLY     N   99     -0.415700     14.01000 ; qtot
0.584
    100    H      7   GLY     H  100      0.271900      1.00800 ; qtot
0.856
    101   CX      7   GLY    CA  101     -0.025200     12.01000 ; qtot
0.831
```

```
   102   H1      7   GLY   HA2  102      0.069800      1.00800 ; qtot
0.901
   103   H1      7   GLY   HA3  103      0.069800      1.00800 ; qtot
0.971
   104    C      7   GLY     C  104      0.597301     12.01000 ; qtot
1.568
   105    O      7   GLY     O  105     -0.567901     16.00000 ; qtot
1.000
   106    N      8   LYS     N  106     -0.347900     14.01000 ; qtot
0.652
   107    H      8   LYS     H  107      0.274700      1.00800 ; qtot
0.927
   108   CX      8   LYS    CA  108     -0.240000     12.01000 ; qtot
0.687
   109   H1      8   LYS    HA  109      0.142600      1.00800 ; qtot
0.829
   110   C8      8   LYS    CB  110     -0.009400     12.01000 ; qtot
0.820
   111   HC      8   LYS   HB2  111      0.036200      1.00800 ; qtot
0.856
   112   HC      8   LYS   HB3  112      0.036200      1.00800 ; qtot
0.892
   113   C8      8   LYS    CG  113      0.018700     12.01000 ; qtot
0.911
   114   HC      8   LYS   HG2  114      0.010300      1.00800 ; qtot
0.921
   115   HC      8   LYS   HG3  115      0.010300      1.00800 ; qtot
0.932
   116   C8      8   LYS    CD  116     -0.047900     12.01000 ; qtot
0.884
   117   HC      8   LYS   HD2  117      0.062100      1.00800 ; qtot
0.946
   118   HC      8   LYS   HD3  118      0.062100      1.00800 ; qtot
1.008
   119   C8      8   LYS    CE  119     -0.014300     12.01000 ; qtot
0.994
   120   HP      8   LYS   HE2  120      0.113500      1.00800 ; qtot
1.107
   121   HP      8   LYS   HE3  121      0.113500      1.00800 ; qtot
1.221
   122   N3      8   LYS    NZ  122     -0.385400     14.01000 ; qtot
0.835
   123    H      8   LYS   HZ1  123      0.340000      1.00800 ; qtot
1.175
   124    H      8   LYS   HZ2  124      0.340000      1.00800 ; qtot
1.515
   125    H      8   LYS   HZ3  125      0.340000      1.00800 ; qtot
1.855
   126    C      8   LYS     C  126      0.734101     12.01000 ; qtot
2.589
   127    O      8   LYS     O  127     -0.589401     16.00000 ; qtot
2.000
```

```
   128    N      9   LEU      N  128    -0.415700     14.01000 ; qtot
1.584
   129    H      9   LEU      H  129     0.271900      1.00800 ; qtot
1.856
   130   CX      9   LEU     CA  130    -0.051800     12.01000 ; qtot
1.804
   131   H1      9   LEU     HA  131     0.092200      1.00800 ; qtot
1.897
   132  A2C      9   LEU     CB  132    -0.110200     12.01000 ; qtot
1.786
   133   HC      9   LEU    HB2  133     0.045700      1.00800 ; qtot
1.832
   134   HC      9   LEU    HB3  134     0.045700      1.00800 ; qtot
1.878
   135  A3C      9   LEU     CG  135     0.353100     12.01000 ; qtot
2.231
   136   HC      9   LEU     HG  136    -0.036100      1.00800 ; qtot
2.195
   137   CT      9   LEU    CD1  137    -0.412100     12.01000 ; qtot
1.783
   138   HC      9   LEU   HD11  138     0.100000      1.00800 ; qtot
1.883
   139   HC      9   LEU   HD12  139     0.100000      1.00800 ; qtot
1.983
   140   HC      9   LEU   HD13  140     0.100000      1.00800 ; qtot
2.083
   141   CT      9   LEU    CD2  141    -0.412100     12.01000 ; qtot
1.671
   142   HC      9   LEU   HD21  142     0.100000      1.00800 ; qtot
1.771
   143   HC      9   LEU   HD22  143     0.100000      1.00800 ; qtot
1.871
   144   HC      9   LEU   HD23  144     0.100000      1.00800 ; qtot
1.971
   145    C      9   LEU      C  145     0.597301     12.01000 ; qtot
2.568
   146    O      9   LEU      O  146    -0.567901     16.00000 ; qtot
2.000
   147    N     10   GLU      N  147    -0.516301     14.01000 ; qtot
1.484
   148    H     10   GLU      H  148     0.293600      1.00800 ; qtot
1.777
   149   CX     10   GLU     CA  149     0.039700     12.01000 ; qtot
1.817
   150   H1     10   GLU     HA  150     0.110500      1.00800 ; qtot
1.928
   151  A2C     10   GLU     CB  151     0.056000     12.01000 ; qtot
1.984
   152   HC     10   GLU    HB2  152    -0.017300      1.00800 ; qtot
1.966
   153   HC     10   GLU    HB3  153    -0.017300      1.00800 ; qtot
1.949
```

```
   154  A2C    10   GLU    CG  154     0.013600     12.01000 ; qtot
1.963
   155   HC    10   GLU   HG2  155    -0.042500      1.00800 ; qtot
1.920
   156   HC    10   GLU   HG3  156    -0.042500      1.00800 ; qtot
1.878
   157   CO    10   GLU    CD  157     0.805401     12.01000 ; qtot
2.683
   158   O2    10   GLU   OE1  158    -0.818801     16.00000 ; qtot
1.864
   159   O2    10   GLU   OE2  159    -0.818801     16.00000 ; qtot
1.045
   160    C    10   GLU     C  160     0.536601     12.01000 ; qtot
1.582
   161    O    10   GLU     O  161    -0.581901     16.00000 ; qtot
1.000
   162    N    11   ASP     N  162    -0.516301     14.01000 ; qtot
0.484
   163    H    11   ASP     H  163     0.293600      1.00800 ; qtot
0.777
   164   CX    11   ASP    CA  164     0.038100     12.01000 ; qtot
0.815
   165   H1    11   ASP    HA  165     0.088000      1.00800 ; qtot
0.903
   166  A2C    11   ASP    CB  166    -0.030300     12.01000 ; qtot
0.873
   167   HC    11   ASP   HB2  167    -0.012200      1.00800 ; qtot
0.861
   168   HC    11   ASP   HB3  168    -0.012200      1.00800 ; qtot
0.849
   169   CO    11   ASP    CG  169     0.799401     12.01000 ; qtot
1.648
   170   O2    11   ASP   OD1  170    -0.801401     16.00000 ; qtot
0.847
   171   O2    11   ASP   OD2  171    -0.801401     16.00000 ; qtot
0.045
   172    C    11   ASP     C  172     0.536601     12.01000 ; qtot
0.582
   173    O    11   ASP     O  173    -0.581901     16.00000 ; qtot -
0.000
   174    N    12   LYS     N  174    -0.347900     14.01000 ; qtot -
0.348
   175    H    12   LYS     H  175     0.274700      1.00800 ; qtot -
0.073
   176   CX    12   LYS    CA  176    -0.240000     12.01000 ; qtot -
0.313
   177   H1    12   LYS    HA  177     0.142600      1.00800 ; qtot -
0.171
   178   C8    12   LYS    CB  178    -0.009400     12.01000 ; qtot -
0.180
   179   HC    12   LYS   HB2  179     0.036200      1.00800 ; qtot -
0.144
```

```
   180    HC     12    LYS    HB3  180      0.036200      1.00800 ; qtot -
0.108
   181    C8     12    LYS     CG  181      0.018700     12.01000 ; qtot -
0.089
   182    HC     12    LYS    HG2  182      0.010300      1.00800 ; qtot -
0.079
   183    HC     12    LYS    HG3  183      0.010300      1.00800 ; qtot -
0.068
   184    C8     12    LYS     CD  184     -0.047900     12.01000 ; qtot -
0.116
   185    HC     12    LYS    HD2  185      0.062100      1.00800 ; qtot -
0.054
   186    HC     12    LYS    HD3  186      0.062100      1.00800 ; qtot
0.008
   187    C8     12    LYS     CE  187     -0.014300     12.01000 ; qtot -
0.006
   188    HP     12    LYS    HE2  188      0.113500      1.00800 ; qtot
0.107
   189    HP     12    LYS    HE3  189      0.113500      1.00800 ; qtot
0.221
   190    N3     12    LYS     NZ  190     -0.385400     14.01000 ; qtot -
0.165
   191     H     12    LYS    HZ1  191      0.340000      1.00800 ; qtot
0.175
   192     H     12    LYS    HZ2  192      0.340000      1.00800 ; qtot
0.515
   193     H     12    LYS    HZ3  193      0.340000      1.00800 ; qtot
0.855
   194     C     12    LYS      C  194      0.734101     12.01000 ; qtot
1.589
   195     O     12    LYS      O  195     -0.589401     16.00000 ; qtot
1.000
   196     N     13    TYR      N  196     -0.415700     14.01000 ; qtot
0.584
   197     H     13    TYR      H  197      0.271900      1.00800 ; qtot
0.856
   198    CX     13    TYR     CA  198     -0.001400     12.01000 ; qtot
0.855
   199    H1     13    TYR     HA  199      0.087600      1.00800 ; qtot
0.942
   200    CT     13    TYR     CB  200     -0.015200     12.01000 ; qtot
0.927
   201    HC     13    TYR    HB2  201      0.029500      1.00800 ; qtot
0.957
   202    HC     13    TYR    HB3  202      0.029500      1.00800 ; qtot
0.986
   203    CA     13    TYR     CG  203     -0.001100     12.01000 ; qtot
0.985
   204    CA     13    TYR    CD1  204     -0.190600     12.01000 ; qtot
0.795
   205    HA     13    TYR    HD1  205      0.169900      1.00800 ; qtot
0.964
```

```
   206   CA    13   TYR   CE1  206    -0.234100     12.01000 ; qtot
0.730
   207   HA    13   TYR   HE1  207     0.165600      1.00800 ; qtot
0.896
   208    C    13   TYR    CZ  208     0.322600     12.01000 ; qtot
1.219
   209   OH    13   TYR    OH  209    -0.557901     16.00000 ; qtot
0.661
   210   HO    13   TYR    HH  210     0.399200      1.00800 ; qtot
1.060
   211   CA    13   TYR   CE2  211    -0.234100     12.01000 ; qtot
0.826
   212   HA    13   TYR   HE2  212     0.165600      1.00800 ; qtot
0.991
   213   CA    13   TYR   CD2  213    -0.190600     12.01000 ; qtot
0.801
   214   HA    13   TYR   HD2  214     0.169900      1.00800 ; qtot
0.971
   215    C    13   TYR     C  215     0.597301     12.01000 ; qtot
1.568
   216    O    13   TYR     O  216    -0.567901     16.00000 ; qtot
1.000
   217    N    14   ILE     N  217    -0.415700     14.01000 ; qtot
0.584
   218    H    14   ILE     H  218     0.271900      1.00800 ; qtot
0.856
   219   CX    14   ILE    CA  219    -0.059700     12.01000 ; qtot
0.797
   220   H1    14   ILE    HA  220     0.086900      1.00800 ; qtot
0.883
   221  A3C    14   ILE    CB  221     0.130300     12.01000 ; qtot
1.014
   222   HC    14   ILE    HB  222     0.018700      1.00800 ; qtot
1.032
   223   CT    14   ILE   CG2  223    -0.320400     12.01000 ; qtot
0.712
   224   HC    14   ILE  HG21  224     0.088200      1.00800 ; qtot
0.800
   225   HC    14   ILE  HG22  225     0.088200      1.00800 ; qtot
0.888
   226   HC    14   ILE  HG23  226     0.088200      1.00800 ; qtot
0.977
   227  A2C    14   ILE   CG1  227    -0.043000     12.01000 ; qtot
0.934
   228   HC    14   ILE  HG12  228     0.023600      1.00800 ; qtot
0.957
   229   HC    14   ILE  HG13  229     0.023600      1.00800 ; qtot
0.981
   230   CT    14   ILE   CD1  230    -0.066000     12.01000 ; qtot
0.915
   231   HC    14   ILE  HD11  231     0.018600      1.00800 ; qtot
0.933
```

```
   232   HC    14   ILE  HD12  232     0.018600      1.00800 ; qtot
0.952
   233   HC    14   ILE  HD13  233     0.018600      1.00800 ; qtot
0.971
   234    C    14   ILE     C  234     0.597301     12.01000 ; qtot
1.568
   235    O    14   ILE     O  235    -0.567901     16.00000 ; qtot
1.000
   236    N    15   ILE     N  236    -0.415700     14.01000 ; qtot
0.584
   237    H    15   ILE     H  237     0.271900      1.00800 ; qtot
0.856
   238   CX    15   ILE    CA  238    -0.059700     12.01000 ; qtot
0.797
   239   H1    15   ILE    HA  239     0.086900      1.00800 ; qtot
0.883
   240  A3C    15   ILE    CB  240     0.130300     12.01000 ; qtot
1.014
   241   HC    15   ILE    HB  241     0.018700      1.00800 ; qtot
1.032
   242   CT    15   ILE   CG2  242    -0.320400     12.01000 ; qtot
0.712
   243   HC    15   ILE  HG21  243     0.088200      1.00800 ; qtot
0.800
   244   HC    15   ILE  HG22  244     0.088200      1.00800 ; qtot
0.888
   245   HC    15   ILE  HG23  245     0.088200      1.00800 ; qtot
0.977
   246  A2C    15   ILE   CG1  246    -0.043000     12.01000 ; qtot
0.934
   247   HC    15   ILE  HG12  247     0.023600      1.00800 ; qtot
0.957
   248   HC    15   ILE  HG13  248     0.023600      1.00800 ; qtot
0.981
   249   CT    15   ILE   CD1  249    -0.066000     12.01000 ; qtot
0.915
   250   HC    15   ILE  HD11  250     0.018600      1.00800 ; qtot
0.933
   251   HC    15   ILE  HD12  251     0.018600      1.00800 ; qtot
0.952
   252   HC    15   ILE  HD13  252     0.018600      1.00800 ; qtot
0.971
   253    C    15   ILE     C  253     0.597301     12.01000 ; qtot
1.568
   254    O    15   ILE     O  254    -0.567901     16.00000 ; qtot
1.000
   255    N    16   ASP     N  255    -0.516301     14.01000 ; qtot
0.484
   256    H    16   ASP     H  256     0.293600      1.00800 ; qtot
0.777
   257   CX    16   ASP    CA  257     0.038100     12.01000 ; qtot
0.815
```

```
   258   H1    16   ASP    HA  258     0.088000      1.00800 ; qtot
0.903
   259  A2C    16   ASP    CB  259    -0.030300     12.01000 ; qtot
0.873
   260   HC    16   ASP   HB2  260    -0.012200      1.00800 ; qtot
0.861
   261   HC    16   ASP   HB3  261    -0.012200      1.00800 ; qtot
0.849
   262   CO    16   ASP    CG  262     0.799401     12.01000 ; qtot
1.648
   263   O2    16   ASP   OD1  263    -0.801401     16.00000 ; qtot
0.847
   264   O2    16   ASP   OD2  264    -0.801401     16.00000 ; qtot
0.045
   265    C    16   ASP     C  265     0.536601     12.01000 ; qtot
0.582
   266    O    16   ASP     O  266    -0.581901     16.00000 ; qtot -
0.000
   267    N    17   GLU     N  267    -0.516301     14.01000 ; qtot -
0.516
   268    H    17   GLU     H  268     0.293600      1.00800 ; qtot -
0.223
   269   CX    17   GLU    CA  269     0.039700     12.01000 ; qtot -
0.183
   270   H1    17   GLU    HA  270     0.110500      1.00800 ; qtot -
0.073
   271  A2C    17   GLU    CB  271     0.056000     12.01000 ; qtot -
0.017
   272   HC    17   GLU   HB2  272    -0.017300      1.00800 ; qtot -
0.034
   273   HC    17   GLU   HB3  273    -0.017300      1.00800 ; qtot -
0.051
   274  A2C    17   GLU    CG  274     0.013600     12.01000 ; qtot -
0.038
   275   HC    17   GLU   HG2  275    -0.042500      1.00800 ; qtot -
0.080
   276   HC    17   GLU   HG3  276    -0.042500      1.00800 ; qtot -
0.123
   277   CO    17   GLU    CD  277     0.805401     12.01000 ; qtot
0.683
   278   O2    17   GLU   OE1  278    -0.818801     16.00000 ; qtot -
0.136
   279   O2    17   GLU   OE2  279    -0.818801     16.00000 ; qtot -
0.955
   280    C    17   GLU     C  280     0.536601     12.01000 ; qtot -
0.418
   281    O    17   GLU     O  281    -0.581901     16.00000 ; qtot -
1.000
   282    N    18   LYS     N  282    -0.347900     14.01000 ; qtot -
1.348
   283    H    18   LYS     H  283     0.274700      1.00800 ; qtot -
1.073
```

```
   284   CX    18   LYS    CA  284    -0.240000     12.01000 ; qtot -
1.313
   285   H1    18   LYS    HA  285     0.142600      1.00800 ; qtot -
1.171
   286   C8    18   LYS    CB  286    -0.009400     12.01000 ; qtot -
1.180
   287   HC    18   LYS   HB2  287     0.036200      1.00800 ; qtot -
1.144
   288   HC    18   LYS   HB3  288     0.036200      1.00800 ; qtot -
1.108
   289   C8    18   LYS    CG  289     0.018700     12.01000 ; qtot -
1.089
   290   HC    18   LYS   HG2  290     0.010300      1.00800 ; qtot -
1.079
   291   HC    18   LYS   HG3  291     0.010300      1.00800 ; qtot -
1.068
   292   C8    18   LYS    CD  292    -0.047900     12.01000 ; qtot -
1.116
   293   HC    18   LYS   HD2  293     0.062100      1.00800 ; qtot -
1.054
   294   HC    18   LYS   HD3  294     0.062100      1.00800 ; qtot -
0.992
   295   C8    18   LYS    CE  295    -0.014300     12.01000 ; qtot -
1.006
   296   HP    18   LYS   HE2  296     0.113500      1.00800 ; qtot -
0.893
   297   HP    18   LYS   HE3  297     0.113500      1.00800 ; qtot -
0.779
   298   N3    18   LYS    NZ  298    -0.385400     14.01000 ; qtot -
1.165
   299    H    18   LYS   HZ1  299     0.340000      1.00800 ; qtot -
0.825
   300    H    18   LYS   HZ2  300     0.340000      1.00800 ; qtot -
0.485
   301    H    18   LYS   HZ3  301     0.340000      1.00800 ; qtot -
0.145
   302    C    18   LYS     C  302     0.734101     12.01000 ; qtot
0.589
   303    O    18   LYS     O  303    -0.589401     16.00000 ; qtot -
0.000
   304    N    19   LEU     N  304    -0.415700     14.01000 ; qtot -
0.416
   305    H    19   LEU     H  305     0.271900      1.00800 ; qtot -
0.144
   306   CX    19   LEU    CA  306    -0.051800     12.01000 ; qtot -
0.196
   307   H1    19   LEU    HA  307     0.092200      1.00800 ; qtot -
0.103
   308  A2C    19   LEU    CB  308    -0.110200     12.01000 ; qtot -
0.214
   309   HC    19   LEU   HB2  309     0.045700      1.00800 ; qtot -
0.168
```

```
   310   HC    19   LEU   HB3  310     0.045700      1.00800 ; qtot -
0.122
   311  A3C    19   LEU    CG  311     0.353100     12.01000 ; qtot
0.231
   312   HC    19   LEU    HG  312    -0.036100      1.00800 ; qtot
0.195
   313   CT    19   LEU   CD1  313    -0.412100     12.01000 ; qtot -
0.217
   314   HC    19   LEU  HD11  314     0.100000      1.00800 ; qtot -
0.117
   315   HC    19   LEU  HD12  315     0.100000      1.00800 ; qtot -
0.017
   316   HC    19   LEU  HD13  316     0.100000      1.00800 ; qtot
0.083
   317   CT    19   LEU   CD2  317    -0.412100     12.01000 ; qtot -
0.329
   318   HC    19   LEU  HD21  318     0.100000      1.00800 ; qtot -
0.229
   319   HC    19   LEU  HD22  319     0.100000      1.00800 ; qtot -
0.129
   320   HC    19   LEU  HD23  320     0.100000      1.00800 ; qtot -
0.029
   321    C    19   LEU     C  321     0.597301     12.01000 ; qtot
0.568
   322    O    19   LEU     O  322    -0.567901     16.00000 ; qtot -
0.000
   323    N    20   GLY     N  323    -0.415700     14.01000 ; qtot -
0.416
   324    H    20   GLY     H  324     0.271900      1.00800 ; qtot -
0.144
   325   CX    20   GLY    CA  325    -0.025200     12.01000 ; qtot -
0.169
   326   H1    20   GLY   HA2  326     0.069800      1.00800 ; qtot -
0.099
   327   H1    20   GLY   HA3  327     0.069800      1.00800 ; qtot -
0.029
   328    C    20   GLY     C  328     0.597301     12.01000 ; qtot
0.568
   329    O    20   GLY     O  329    -0.567901     16.00000 ; qtot -
0.000
   330    N    21   GLN     N  330    -0.415700     14.01000 ; qtot -
0.416
   331    H    21   GLN     H  331     0.271900      1.00800 ; qtot -
0.144
   332   CX    21   GLN    CA  332    -0.003100     12.01000 ; qtot -
0.147
   333   H1    21   GLN    HA  333     0.085000      1.00800 ; qtot -
0.062
   334  A2C    21   GLN    CB  334    -0.003600     12.01000 ; qtot -
0.066
   335   HC    21   GLN   HB2  335     0.017100      1.00800 ; qtot -
0.048
```

336 HC 21 GLN HB3 336 0.017100 1.00800 ; qtot -
0.031
337 A2C 21 GLN CG 337 -0.064500 12.01000 ; qtot -
0.096
338 HC 21 GLN HG2 338 0.035200 1.00800 ; qtot -
0.061
339 HC 21 GLN HG3 339 0.035200 1.00800 ; qtot -
0.025
340 C 21 GLN CD 340 0.695101 12.01000 ; qtot
0.670
341 O 21 GLN OE1 341 -0.608601 16.00000 ; qtot
0.061
342 N 21 GLN NE2 342 -0.940701 14.01000 ; qtot -
0.880
343 H 21 GLN HE21 343 0.425100 1.00800 ; qtot -
0.455
344 H 21 GLN HE22 344 0.425100 1.00800 ; qtot -
0.029
345 C 21 GLN C 345 0.597301 12.01000 ; qtot
0.568
346 O 21 GLN O 346 -0.567901 16.00000 ; qtot
0.000
347 N 22 GLY N 347 -0.415700 14.01000 ; qtot -
0.416
348 H 22 GLY H 348 0.271900 1.00800 ; qtot -
0.144
349 CX 22 GLY CA 349 -0.025200 12.01000 ; qtot -
0.169
350 H1 22 GLY HA2 350 0.069800 1.00800 ; qtot -
0.099
351 H1 22 GLY HA3 351 0.069800 1.00800 ; qtot -
0.029
352 C 22 GLY C 352 0.597301 12.01000 ; qtot
0.568
353 O 22 GLY O 353 -0.567901 16.00000 ; qtot
0.000
354 N 23 THR N 354 -0.415700 14.01000 ; qtot -
0.416
355 H 23 THR H 355 0.271900 1.00800 ; qtot -
0.144
356 CX 23 THR CA 356 -0.038900 12.01000 ; qtot -
0.183
357 H1 23 THR HA 357 0.100700 1.00800 ; qtot -
0.082
358 A3C 23 THR CB 358 0.365400 12.01000 ; qtot
0.283
359 H1 23 THR HB 359 0.004300 1.00800 ; qtot
0.288
360 CT 23 THR CG2 360 -0.243800 12.01000 ; qtot
0.044
361 HC 23 THR HG21 361 0.064200 1.00800 ; qtot
0.108

362 HC 23 THR HG22 362 0.064200 1.00800 ; qtot
0.172
363 HC 23 THR HG23 363 0.064200 1.00800 ; qtot
0.237
364 OH 23 THR OG1 364 -0.676101 16.00000 ; qtot -
0.440
365 HO 23 THR HG1 365 0.410200 1.00800 ; qtot -
0.029
366 C 23 THR C 366 0.597301 12.01000 ; qtot
0.568
367 O 23 THR O 367 -0.567901 16.00000 ; qtot
0.000
368 N 24 TYR N 368 -0.415700 14.01000 ; qtot -
0.416
369 H 24 TYR H 369 0.271900 1.00800 ; qtot -
0.144
370 CX 24 TYR CA 370 -0.001400 12.01000 ; qtot -
0.145
371 H1 24 TYR HA 371 0.087600 1.00800 ; qtot -
0.058
372 CT 24 TYR CB 372 -0.015200 12.01000 ; qtot -
0.073
373 HC 24 TYR HB2 373 0.029500 1.00800 ; qtot -
0.043
374 HC 24 TYR HB3 374 0.029500 1.00800 ; qtot -
0.014
375 CA 24 TYR CG 375 -0.001100 12.01000 ; qtot -
0.015
376 CA 24 TYR CD1 376 -0.190600 12.01000 ; qtot -
0.206
377 HA 24 TYR HD1 377 0.169900 1.00800 ; qtot -
0.036
378 CA 24 TYR CE1 378 -0.234100 12.01000 ; qtot -
0.270
379 HA 24 TYR HE1 379 0.165600 1.00800 ; qtot -
0.104
380 C 24 TYR CZ 380 0.322600 12.01000 ; qtot
0.219
381 OH 24 TYR OH 381 -0.557901 16.00000 ; qtot -
0.339
382 HO 24 TYR HH 382 0.399200 1.00800 ; qtot
0.060
383 CA 24 TYR CE2 383 -0.234100 12.01000 ; qtot -
0.174
384 HA 24 TYR HE2 384 0.165600 1.00800 ; qtot -
0.009
385 CA 24 TYR CD2 385 -0.190600 12.01000 ; qtot -
0.199
386 HA 24 TYR HD2 386 0.169900 1.00800 ; qtot -
0.029
387 C 24 TYR C 387 0.597301 12.01000 ; qtot
0.568

388 O 24 TYR O 388 -0.567901 16.00000 ; qtot
0.000
389 N 25 GLY N 389 -0.415700 14.01000 ; qtot -
0.416
390 H 25 GLY H 390 0.271900 1.00800 ; qtot -
0.144
391 CX 25 GLY CA 391 -0.025200 12.01000 ; qtot -
0.169
392 H1 25 GLY HA2 392 0.069800 1.00800 ; qtot -
0.099
393 H1 25 GLY HA3 393 0.069800 1.00800 ; qtot -
0.029
394 C 25 GLY C 394 0.597301 12.01000 ; qtot
0.568
395 O 25 GLY O 395 -0.567901 16.00000 ; qtot
0.000
396 N 26 CYS N 396 -0.415700 14.01000 ; qtot -
0.416
397 H 26 CYS H 397 0.271900 1.00800 ; qtot -
0.144
398 CX 26 CYS CA 398 0.021300 12.01000 ; qtot -
0.123
399 H1 26 CYS HA 399 0.112400 1.00800 ; qtot -
0.010
400 A2C 26 CYS CB 400 -0.123100 12.01000 ; qtot -
0.133
401 H1 26 CYS HB2 401 0.111200 1.00800 ; qtot -
0.022
402 H1 26 CYS HB3 402 0.111200 1.00800 ; qtot
0.089
403 SH 26 CYS SG 403 -0.311900 32.06000 ; qtot -
0.223
404 HS 26 CYS HG 404 0.193300 1.00800 ; qtot -
0.029
405 C 26 CYS C 405 0.597301 12.01000 ; qtot
0.568
406 O 26 CYS O 406 -0.567901 16.00000 ; qtot
0.000
407 N 27 VAL N 407 -0.415700 14.01000 ; qtot -
0.416
408 H 27 VAL H 408 0.271900 1.00800 ; qtot -
0.144
409 CX 27 VAL CA 409 -0.087500 12.01000 ; qtot -
0.231
410 H1 27 VAL HA 410 0.096900 1.00800 ; qtot -
0.134
411 A3C 27 VAL CB 411 0.298500 12.01000 ; qtot
0.164
412 HC 27 VAL HB 412 -0.029700 1.00800 ; qtot
0.134
413 CT 27 VAL CG1 413 -0.319200 12.01000 ; qtot -
0.185

```
   414   HC     27   VAL  HG11  414      0.079100      1.00800 ; qtot -
0.106
   415   HC     27   VAL  HG12  415      0.079100      1.00800 ; qtot -
0.027
   416   HC     27   VAL  HG13  416      0.079100      1.00800 ; qtot
0.053
   417   CT     27   VAL   CG2  417     -0.319200     12.01000 ; qtot -
0.267
   418   HC     27   VAL  HG21  418      0.079100      1.00800 ; qtot -
0.188
   419   HC     27   VAL  HG22  419      0.079100      1.00800 ; qtot -
0.109
   420   HC     27   VAL  HG23  420      0.079100      1.00800 ; qtot -
0.029
   421    C     27   VAL     C  421      0.597301     12.01000 ; qtot
0.568
   422    O     27   VAL     O  422     -0.567901     16.00000 ; qtot
0.000
   423    N     28   TYR     N  423     -0.415700     14.01000 ; qtot -
0.416
   424    H     28   TYR     H  424      0.271900      1.00800 ; qtot -
0.144
   425   CX     28   TYR    CA  425     -0.001400     12.01000 ; qtot -
0.145
   426   H1     28   TYR    HA  426      0.087600      1.00800 ; qtot -
0.058
   427   CT     28   TYR    CB  427     -0.015200     12.01000 ; qtot -
0.073
   428   HC     28   TYR   HB2  428      0.029500      1.00800 ; qtot -
0.043
   429   HC     28   TYR   HB3  429      0.029500      1.00800 ; qtot -
0.014
   430   CA     28   TYR    CG  430     -0.001100     12.01000 ; qtot -
0.015
   431   CA     28   TYR   CD1  431     -0.190600     12.01000 ; qtot -
0.206
   432   HA     28   TYR   HD1  432      0.169900      1.00800 ; qtot -
0.036
   433   CA     28   TYR   CE1  433     -0.234100     12.01000 ; qtot -
0.270
   434   HA     28   TYR   HE1  434      0.165600      1.00800 ; qtot -
0.104
   435    C     28   TYR    CZ  435      0.322600     12.01000 ; qtot
0.219
   436   OH     28   TYR    OH  436     -0.557901     16.00000 ; qtot -
0.339
   437   HO     28   TYR    HH  437      0.399200      1.00800 ; qtot
0.060
   438   CA     28   TYR   CE2  438     -0.234100     12.01000 ; qtot -
0.174
   439   HA     28   TYR   HE2  439      0.165600      1.00800 ; qtot -
0.009
```

```
   440   CA   28   TYR   CD2  440   -0.190600     12.01000 ; qtot -
0.199
   441   HA   28   TYR   HD2  441    0.169900      1.00800 ; qtot -
0.029
   442    C   28   TYR     C  442    0.597301     12.01000 ; qtot
0.568
   443    O   28   TYR     O  443   -0.567901     16.00000 ; qtot
0.000
   444    N   29   LYS     N  444   -0.347900     14.01000 ; qtot -
0.348
   445    H   29   LYS     H  445    0.274700      1.00800 ; qtot -
0.073
   446   CX   29   LYS    CA  446   -0.240000     12.01000 ; qtot -
0.313
   447   H1   29   LYS    HA  447    0.142600      1.00800 ; qtot -
0.171
   448   C8   29   LYS    CB  448   -0.009400     12.01000 ; qtot -
0.180
   449   HC   29   LYS   HB2  449    0.036200      1.00800 ; qtot -
0.144
   450   HC   29   LYS   HB3  450    0.036200      1.00800 ; qtot -
0.108
   451   C8   29   LYS    CG  451    0.018700     12.01000 ; qtot -
0.089
   452   HC   29   LYS   HG2  452    0.010300      1.00800 ; qtot -
0.079
   453   HC   29   LYS   HG3  453    0.010300      1.00800 ; qtot -
0.068
   454   C8   29   LYS    CD  454   -0.047900     12.01000 ; qtot -
0.116
   455   HC   29   LYS   HD2  455    0.062100      1.00800 ; qtot -
0.054
   456   HC   29   LYS   HD3  456    0.062100      1.00800 ; qtot
0.008
   457   C8   29   LYS    CE  457   -0.014300     12.01000 ; qtot -
0.006
   458   HP   29   LYS   HE2  458    0.113500      1.00800 ; qtot
0.107
   459   HP   29   LYS   HE3  459    0.113500      1.00800 ; qtot
0.221
   460   N3   29   LYS    NZ  460   -0.385400     14.01000 ; qtot -
0.165
   461    H   29   LYS   HZ1  461    0.340000      1.00800 ; qtot
0.175
   462    H   29   LYS   HZ2  462    0.340000      1.00800 ; qtot
0.515
   463    H   29   LYS   HZ3  463    0.340000      1.00800 ; qtot
0.855
   464    C   29   LYS     C  464    0.734101     12.01000 ; qtot
1.589
   465    O   29   LYS     O  465   -0.589401     16.00000 ; qtot
1.000
```

466 N 30 GLY N 466 -0.415700 14.01000 ; qtot
0.584
467 H 30 GLY H 467 0.271900 1.00800 ; qtot
0.856
468 CX 30 GLY CA 468 -0.025200 12.01000 ; qtot
0.831
469 H1 30 GLY HA2 469 0.069800 1.00800 ; qtot
0.901
470 H1 30 GLY HA3 470 0.069800 1.00800 ; qtot
0.971
471 C 30 GLY C 471 0.597301 12.01000 ; qtot
1.568
472 O 30 GLY O 472 -0.567901 16.00000 ; qtot
1.000
473 N 31 ILE N 473 -0.415700 14.01000 ; qtot
0.584
474 H 31 ILE H 474 0.271900 1.00800 ; qtot
0.856
475 CX 31 ILE CA 475 -0.059700 12.01000 ; qtot
0.797
476 H1 31 ILE HA 476 0.086900 1.00800 ; qtot
0.883
477 A3C 31 ILE CB 477 0.130300 12.01000 ; qtot
1.014
478 HC 31 ILE HB 478 0.018700 1.00800 ; qtot
1.032
479 CT 31 ILE CG2 479 -0.320400 12.01000 ; qtot
0.712
480 HC 31 ILE HG21 480 0.088200 1.00800 ; qtot
0.800
481 HC 31 ILE HG22 481 0.088200 1.00800 ; qtot
0.888
482 HC 31 ILE HG23 482 0.088200 1.00800 ; qtot
0.977
483 A2C 31 ILE CG1 483 -0.043000 12.01000 ; qtot
0.934
484 HC 31 ILE HG12 484 0.023600 1.00800 ; qtot
0.957
485 HC 31 ILE HG13 485 0.023600 1.00800 ; qtot
0.981
486 CT 31 ILE CD1 486 -0.066000 12.01000 ; qtot
0.915
487 HC 31 ILE HD11 487 0.018600 1.00800 ; qtot
0.933
488 HC 31 ILE HD12 488 0.018600 1.00800 ; qtot
0.952
489 HC 31 ILE HD13 489 0.018600 1.00800 ; qtot
0.971
490 C 31 ILE C 490 0.597301 12.01000 ; qtot
1.568
491 O 31 ILE O 491 -0.567901 16.00000 ; qtot
1.000

492 N 32 ASP N 492 -0.516301 14.01000 ; qtot
0.484
493 H 32 ASP H 493 0.293600 1.00800 ; qtot
0.777
494 CX 32 ASP CA 494 0.038100 12.01000 ; qtot
0.815
495 H1 32 ASP HA 495 0.088000 1.00800 ; qtot
0.903
496 A2C 32 ASP CB 496 -0.030300 12.01000 ; qtot
0.873
497 HC 32 ASP HB2 497 -0.012200 1.00800 ; qtot
0.861
498 HC 32 ASP HB3 498 -0.012200 1.00800 ; qtot
0.849
499 CO 32 ASP CG 499 0.799401 12.01000 ; qtot
1.648
500 O2 32 ASP OD1 500 -0.801401 16.00000 ; qtot
0.847
501 O2 32 ASP OD2 501 -0.801401 16.00000 ; qtot
0.045
502 C 32 ASP C 502 0.536601 12.01000 ; qtot
0.582
503 O 32 ASP O 503 -0.581901 16.00000 ; qtot
0.000
504 N 33 LYS N 504 -0.347900 14.01000 ; qtot -
0.348
505 H 33 LYS H 505 0.274700 1.00800 ; qtot -
0.073
506 CX 33 LYS CA 506 -0.240000 12.01000 ; qtot -
0.313
507 H1 33 LYS HA 507 0.142600 1.00800 ; qtot -
0.171
508 C8 33 LYS CB 508 -0.009400 12.01000 ; qtot -
0.180
509 HC 33 LYS HB2 509 0.036200 1.00800 ; qtot -
0.144
510 HC 33 LYS HB3 510 0.036200 1.00800 ; qtot -
0.108
511 C8 33 LYS CG 511 0.018700 12.01000 ; qtot -
0.089
512 HC 33 LYS HG2 512 0.010300 1.00800 ; qtot -
0.079
513 HC 33 LYS HG3 513 0.010300 1.00800 ; qtot -
0.068
514 C8 33 LYS CD 514 -0.047900 12.01000 ; qtot -
0.116
515 HC 33 LYS HD2 515 0.062100 1.00800 ; qtot -
0.054
516 HC 33 LYS HD3 516 0.062100 1.00800 ; qtot
0.008
517 C8 33 LYS CE 517 -0.014300 12.01000 ; qtot -
0.006

```
   518    HP    33    LYS    HE2  518     0.113500      1.00800 ; qtot
0.107
   519    HP    33    LYS    HE3  519     0.113500      1.00800 ; qtot
0.221
   520    N3    33    LYS     NZ  520    -0.385400     14.01000 ; qtot -
0.165
   521     H    33    LYS    HZ1  521     0.340000      1.00800 ; qtot
0.175
   522     H    33    LYS    HZ2  522     0.340000      1.00800 ; qtot
0.515
   523     H    33    LYS    HZ3  523     0.340000      1.00800 ; qtot
0.855
   524     C    33    LYS      C  524     0.734101     12.01000 ; qtot
1.589
   525     O    33    LYS      O  525    -0.589401     16.00000 ; qtot
1.000
   526     N    34    VAL      N  526    -0.415700     14.01000 ; qtot
0.584
   527     H    34    VAL      H  527     0.271900      1.00800 ; qtot
0.856
   528    CX    34    VAL     CA  528    -0.087500     12.01000 ; qtot
0.769
   529    H1    34    VAL     HA  529     0.096900      1.00800 ; qtot
0.866
   530   A3C    34    VAL     CB  530     0.298500     12.01000 ; qtot
1.164
   531    HC    34    VAL     HB  531    -0.029700      1.00800 ; qtot
1.134
   532    CT    34    VAL    CG1  532    -0.319200     12.01000 ; qtot
0.815
   533    HC    34    VAL   HG11  533     0.079100      1.00800 ; qtot
0.894
   534    HC    34    VAL   HG12  534     0.079100      1.00800 ; qtot
0.973
   535    HC    34    VAL   HG13  535     0.079100      1.00800 ; qtot
1.053
   536    CT    34    VAL    CG2  536    -0.319200     12.01000 ; qtot
0.733
   537    HC    34    VAL   HG21  537     0.079100      1.00800 ; qtot
0.812
   538    HC    34    VAL   HG22  538     0.079100      1.00800 ; qtot
0.892
   539    HC    34    VAL   HG23  539     0.079100      1.00800 ; qtot
0.971
   540     C    34    VAL      C  540     0.597301     12.01000 ; qtot
1.568
   541     O    34    VAL      O  541    -0.567901     16.00000 ; qtot
1.000
   542     N    35    THR      N  542    -0.415700     14.01000 ; qtot
0.584
   543     H    35    THR      H  543     0.271900      1.00800 ; qtot
0.856
```

```
   544   CX    35   THR    CA  544    -0.038900     12.01000 ; qtot
0.817
   545   H1    35   THR    HA  545     0.100700      1.00800 ; qtot
0.918
   546  A3C    35   THR    CB  546     0.365400     12.01000 ; qtot
1.283
   547   H1    35   THR    HB  547     0.004300      1.00800 ; qtot
1.288
   548   CT    35   THR   CG2  548    -0.243800     12.01000 ; qtot
1.044
   549   HC    35   THR  HG21  549     0.064200      1.00800 ; qtot
1.108
   550   HC    35   THR  HG22  550     0.064200      1.00800 ; qtot
1.172
   551   HC    35   THR  HG23  551     0.064200      1.00800 ; qtot
1.237
   552   OH    35   THR   OG1  552    -0.676101     16.00000 ; qtot
0.560
   553   HO    35   THR   HG1  553     0.410200      1.00800 ; qtot
0.971
   554    C    35   THR     C  554     0.597301     12.01000 ; qtot
1.568
   555    O    35   THR     O  555    -0.567901     16.00000 ; qtot
1.000
   556    N    36   ASN     N  556    -0.415700     14.01000 ; qtot
0.584
   557    H    36   ASN     H  557     0.271900      1.00800 ; qtot
0.856
   558   CX    36   ASN    CA  558     0.014300     12.01000 ; qtot
0.871
   559   H1    36   ASN    HA  559     0.104800      1.00800 ; qtot
0.975
   560  A2C    36   ASN    CB  560    -0.204100     12.01000 ; qtot
0.771
   561   HC    36   ASN   HB2  561     0.079700      1.00800 ; qtot
0.851
   562   HC    36   ASN   HB3  562     0.079700      1.00800 ; qtot
0.931
   563    C    36   ASN    CG  563     0.713001     12.01000 ; qtot
1.644
   564    O    36   ASN   OD1  564    -0.593101     16.00000 ; qtot
1.051
   565    N    36   ASN   ND2  565    -0.919101     14.01000 ; qtot
0.131
   566    H    36   ASN  HD21  566     0.419600      1.00800 ; qtot
0.551
   567    H    36   ASN  HD22  567     0.419600      1.00800 ; qtot
0.971
   568    C    36   ASN     C  568     0.597301     12.01000 ; qtot
1.568
   569    O    36   ASN     O  569    -0.567901     16.00000 ; qtot
1.000
```

570 N 37 GLN N 570 -0.415700 14.01000 ; qtot
0.584
571 H 37 GLN H 571 0.271900 1.00800 ; qtot
0.856
572 CX 37 GLN CA 572 -0.003100 12.01000 ; qtot
0.853
573 H1 37 GLN HA 573 0.085000 1.00800 ; qtot
0.938
574 A2C 37 GLN CB 574 -0.003600 12.01000 ; qtot
0.935
575 HC 37 GLN HB2 575 0.017100 1.00800 ; qtot
0.952
576 HC 37 GLN HB3 576 0.017100 1.00800 ; qtot
0.969
577 A2C 37 GLN CG 577 -0.064500 12.01000 ; qtot
0.904
578 HC 37 GLN HG2 578 0.035200 1.00800 ; qtot
0.939
579 HC 37 GLN HG3 579 0.035200 1.00800 ; qtot
0.975
580 C 37 GLN CD 580 0.695101 12.01000 ; qtot
1.670
581 O 37 GLN OE1 581 -0.608601 16.00000 ; qtot
1.061
582 N 37 GLN NE2 582 -0.940701 14.01000 ; qtot
0.120
583 H 37 GLN HE21 583 0.425100 1.00800 ; qtot
0.546
584 H 37 GLN HE22 584 0.425100 1.00800 ; qtot
0.971
585 C 37 GLN C 585 0.597301 12.01000 ; qtot
1.568
586 O 37 GLN O 586 -0.567901 16.00000 ; qtot
1.000
587 N 38 LEU N 587 -0.415700 14.01000 ; qtot
0.584
588 H 38 LEU H 588 0.271900 1.00800 ; qtot
0.856
589 CX 38 LEU CA 589 -0.051800 12.01000 ; qtot
0.804
590 H1 38 LEU HA 590 0.092200 1.00800 ; qtot
0.897
591 A2C 38 LEU CB 591 -0.110200 12.01000 ; qtot
0.786
592 HC 38 LEU HB2 592 0.045700 1.00800 ; qtot
0.832
593 HC 38 LEU HB3 593 0.045700 1.00800 ; qtot
0.878
594 A3C 38 LEU CG 594 0.353100 12.01000 ; qtot
1.231
595 HC 38 LEU HG 595 -0.036100 1.00800 ; qtot
1.195

596 CT 38 LEU CD1 596 -0.412100 12.01000 ; qtot
0.783
597 HC 38 LEU HD11 597 0.100000 1.00800 ; qtot
0.883
598 HC 38 LEU HD12 598 0.100000 1.00800 ; qtot
0.983
599 HC 38 LEU HD13 599 0.100000 1.00800 ; qtot
1.083
600 CT 38 LEU CD2 600 -0.412100 12.01000 ; qtot
0.671
601 HC 38 LEU HD21 601 0.100000 1.00800 ; qtot
0.771
602 HC 38 LEU HD22 602 0.100000 1.00800 ; qtot
0.871
603 HC 38 LEU HD23 603 0.100000 1.00800 ; qtot
0.971
604 C 38 LEU C 604 0.597301 12.01000 ; qtot
1.568
605 O 38 LEU O 605 -0.567901 16.00000 ; qtot
1.000
606 N 39 TYR N 606 -0.415700 14.01000 ; qtot
0.584
607 H 39 TYR H 607 0.271900 1.00800 ; qtot
0.856
608 CX 39 TYR CA 608 -0.001400 12.01000 ; qtot
0.855
609 H1 39 TYR HA 609 0.087600 1.00800 ; qtot
0.942
610 CT 39 TYR CB 610 -0.015200 12.01000 ; qtot
0.927
611 HC 39 TYR HB2 611 0.029500 1.00800 ; qtot
0.957
612 HC 39 TYR HB3 612 0.029500 1.00800 ; qtot
0.986
613 CA 39 TYR CG 613 -0.001100 12.01000 ; qtot
0.985
614 CA 39 TYR CD1 614 -0.190600 12.01000 ; qtot
0.795
615 HA 39 TYR HD1 615 0.169900 1.00800 ; qtot
0.964
616 CA 39 TYR CE1 616 -0.234100 12.01000 ; qtot
0.730
617 HA 39 TYR HE1 617 0.165600 1.00800 ; qtot
0.896
618 C 39 TYR CZ 618 0.322600 12.01000 ; qtot
1.219
619 OH 39 TYR OH 619 -0.557901 16.00000 ; qtot
0.661
620 HO 39 TYR HH 620 0.399200 1.00800 ; qtot
1.060
621 CA 39 TYR CE2 621 -0.234100 12.01000 ; qtot
0.826

```
   622   HA    39   TYR   HE2  622     0.165600      1.00800 ; qtot
0.991
   623   CA    39   TYR   CD2  623    -0.190600     12.01000 ; qtot
0.801
   624   HA    39   TYR   HD2  624     0.169900      1.00800 ; qtot
0.971
   625    C    39   TYR     C  625     0.597301     12.01000 ; qtot
1.568
   626    O    39   TYR     O  626    -0.567901     16.00000 ; qtot
1.000
   627    N    40   ALA     N  627    -0.415700     14.01000 ; qtot
0.584
   628    H    40   ALA     H  628     0.271900      1.00800 ; qtot
0.856
   629   CX    40   ALA    CA  629     0.033700     12.01000 ; qtot
0.890
   630   H1    40   ALA    HA  630     0.082300      1.00800 ; qtot
0.972
   631   CT    40   ALA    CB  631    -0.182500     12.01000 ; qtot
0.790
   632   HC    40   ALA   HB1  632     0.060300      1.00800 ; qtot
0.850
   633   HC    40   ALA   HB2  633     0.060300      1.00800 ; qtot
0.910
   634   HC    40   ALA   HB3  634     0.060300      1.00800 ; qtot
0.971
   635    C    40   ALA     C  635     0.597301     12.01000 ; qtot
1.568
   636    O    40   ALA     O  636    -0.567901     16.00000 ; qtot
1.000
   637    N    41   ILE     N  637    -0.415700     14.01000 ; qtot
0.584
   638    H    41   ILE     H  638     0.271900      1.00800 ; qtot
0.856
   639   CX    41   ILE    CA  639    -0.059700     12.01000 ; qtot
0.797
   640   H1    41   ILE    HA  640     0.086900      1.00800 ; qtot
0.883
   641  A3C    41   ILE    CB  641     0.130300     12.01000 ; qtot
1.014
   642   HC    41   ILE    HB  642     0.018700      1.00800 ; qtot
1.032
   643   CT    41   ILE   CG2  643    -0.320400     12.01000 ; qtot
0.712
   644   HC    41   ILE  HG21  644     0.088200      1.00800 ; qtot
0.800
   645   HC    41   ILE  HG22  645     0.088200      1.00800 ; qtot
0.888
   646   HC    41   ILE  HG23  646     0.088200      1.00800 ; qtot
0.977
   647  A2C    41   ILE   CG1  647    -0.043000     12.01000 ; qtot
0.934
```

```
   648    HC    41   ILE  HG12  648     0.023600      1.00800 ; qtot
0.957
   649    HC    41   ILE  HG13  649     0.023600      1.00800 ; qtot
0.981
   650    CT    41   ILE   CD1  650    -0.066000     12.01000 ; qtot
0.915
   651    HC    41   ILE  HD11  651     0.018600      1.00800 ; qtot
0.933
   652    HC    41   ILE  HD12  652     0.018600      1.00800 ; qtot
0.952
   653    HC    41   ILE  HD13  653     0.018600      1.00800 ; qtot
0.971
   654     C    41   ILE     C  654     0.597301     12.01000 ; qtot
1.568
   655     O    41   ILE     O  655    -0.567901     16.00000 ; qtot
1.000
   656     N    42   LYS     N  656    -0.347900     14.01000 ; qtot
0.652
   657     H    42   LYS     H  657     0.274700      1.00800 ; qtot
0.927
   658    CX    42   LYS    CA  658    -0.240000     12.01000 ; qtot
0.687
   659    H1    42   LYS    HA  659     0.142600      1.00800 ; qtot
0.829
   660    C8    42   LYS    CB  660    -0.009400     12.01000 ; qtot
0.820
   661    HC    42   LYS   HB2  661     0.036200      1.00800 ; qtot
0.856
   662    HC    42   LYS   HB3  662     0.036200      1.00800 ; qtot
0.892
   663    C8    42   LYS    CG  663     0.018700     12.01000 ; qtot
0.911
   664    HC    42   LYS   HG2  664     0.010300      1.00800 ; qtot
0.921
   665    HC    42   LYS   HG3  665     0.010300      1.00800 ; qtot
0.932
   666    C8    42   LYS    CD  666    -0.047900     12.01000 ; qtot
0.884
   667    HC    42   LYS   HD2  667     0.062100      1.00800 ; qtot
0.946
   668    HC    42   LYS   HD3  668     0.062100      1.00800 ; qtot
1.008
   669    C8    42   LYS    CE  669    -0.014300     12.01000 ; qtot
0.994
   670    HP    42   LYS   HE2  670     0.113500      1.00800 ; qtot
1.107
   671    HP    42   LYS   HE3  671     0.113500      1.00800 ; qtot
1.221
   672    N3    42   LYS    NZ  672    -0.385400     14.01000 ; qtot
0.835
   673     H    42   LYS   HZ1  673     0.340000      1.00800 ; qtot
1.175
```

| | | | | | | | | |
|---|---|---|---|---|---|---|---|---|
| 674 | H | 42 | LYS | HZ2 | 674 | 0.340000 | 1.00800 | ; qtot 1.515 |
| 675 | H | 42 | LYS | HZ3 | 675 | 0.340000 | 1.00800 | ; qtot 1.855 |
| 676 | C | 42 | LYS | C | 676 | 0.734101 | 12.01000 | ; qtot 2.589 |
| 677 | O | 42 | LYS | O | 677 | -0.589401 | 16.00000 | ; qtot 2.000 |
| 678 | N | 43 | GLU | N | 678 | -0.516301 | 14.01000 | ; qtot 1.484 |
| 679 | H | 43 | GLU | H | 679 | 0.293600 | 1.00800 | ; qtot 1.777 |
| 680 | CX | 43 | GLU | CA | 680 | 0.039700 | 12.01000 | ; qtot 1.817 |
| 681 | H1 | 43 | GLU | HA | 681 | 0.110500 | 1.00800 | ; qtot 1.928 |
| 682 | A2C | 43 | GLU | CB | 682 | 0.056000 | 12.01000 | ; qtot 1.984 |
| 683 | HC | 43 | GLU | HB2 | 683 | -0.017300 | 1.00800 | ; qtot 1.966 |
| 684 | HC | 43 | GLU | HB3 | 684 | -0.017300 | 1.00800 | ; qtot 1.949 |
| 685 | A2C | 43 | GLU | CG | 685 | 0.013600 | 12.01000 | ; qtot 1.963 |
| 686 | HC | 43 | GLU | HG2 | 686 | -0.042500 | 1.00800 | ; qtot 1.920 |
| 687 | HC | 43 | GLU | HG3 | 687 | -0.042500 | 1.00800 | ; qtot 1.878 |
| 688 | CO | 43 | GLU | CD | 688 | 0.805401 | 12.01000 | ; qtot 2.683 |
| 689 | O2 | 43 | GLU | OE1 | 689 | -0.818801 | 16.00000 | ; qtot 1.864 |
| 690 | O2 | 43 | GLU | OE2 | 690 | -0.818801 | 16.00000 | ; qtot 1.045 |
| 691 | C | 43 | GLU | C | 691 | 0.536601 | 12.01000 | ; qtot 1.582 |
| 692 | O | 43 | GLU | O | 692 | -0.581901 | 16.00000 | ; qtot 1.000 |
| 693 | N | 44 | GLH | N | 693 | -0.415700 | 14.01000 | ; qtot 0.584 |
| 694 | H | 44 | GLH | H | 694 | 0.271900 | 1.00800 | ; qtot 0.856 |
| 695 | CX | 44 | GLH | CA | 695 | 0.014500 | 12.01000 | ; qtot 0.871 |
| 696 | H1 | 44 | GLH | HA | 696 | 0.077900 | 1.00800 | ; qtot 0.949 |
| 697 | A2C | 44 | GLH | CB | 697 | -0.007100 | 12.01000 | ; qtot 0.942 |
| 698 | HC | 44 | GLH | HB2 | 698 | 0.025600 | 1.00800 | ; qtot 0.967 |
| 699 | HC | 44 | GLH | HB3 | 699 | 0.025600 | 1.00800 | ; qtot 0.993 |

| | | | | | | | | |
|---|---|---|---|---|---|---|---|---|
| 700 | A2C | 44 | GLH | CG | 700 | -0.017400 | 12.01000 | ; qtot 0.975 |
| 701 | HC | 44 | GLH | HG2 | 701 | 0.043000 | 1.00800 | ; qtot 1.018 |
| 702 | HC | 44 | GLH | HG3 | 702 | 0.043000 | 1.00800 | ; qtot 1.061 |
| 703 | C | 44 | GLH | CD | 703 | 0.680101 | 12.01000 | ; qtot 1.741 |
| 704 | O | 44 | GLH | OE1 | 704 | -0.583801 | 16.00000 | ; qtot 1.158 |
| 705 | OH | 44 | GLH | OE2 | 705 | -0.651101 | 16.00000 | ; qtot 0.507 |
| 706 | HO | 44 | GLH | HE2 | 706 | 0.464100 | 1.00800 | ; qtot 0.971 |
| 707 | C | 44 | GLH | C | 707 | 0.597301 | 12.01000 | ; qtot 1.568 |
| 708 | O | 44 | GLH | O | 708 | -0.567901 | 16.00000 | ; qtot 1.000 |
| 709 | N | 45 | LYS | N | 709 | -0.347900 | 14.01000 | ; qtot 0.652 |
| 710 | H | 45 | LYS | H | 710 | 0.274700 | 1.00800 | ; qtot 0.927 |
| 711 | CX | 45 | LYS | CA | 711 | -0.240000 | 12.01000 | ; qtot 0.687 |
| 712 | H1 | 45 | LYS | HA | 712 | 0.142600 | 1.00800 | ; qtot 0.829 |
| 713 | C8 | 45 | LYS | CB | 713 | -0.009400 | 12.01000 | ; qtot 0.820 |
| 714 | HC | 45 | LYS | HB2 | 714 | 0.036200 | 1.00800 | ; qtot 0.856 |
| 715 | HC | 45 | LYS | HB3 | 715 | 0.036200 | 1.00800 | ; qtot 0.892 |
| 716 | C8 | 45 | LYS | CG | 716 | 0.018700 | 12.01000 | ; qtot 0.911 |
| 717 | HC | 45 | LYS | HG2 | 717 | 0.010300 | 1.00800 | ; qtot 0.921 |
| 718 | HC | 45 | LYS | HG3 | 718 | 0.010300 | 1.00800 | ; qtot 0.932 |
| 719 | C8 | 45 | LYS | CD | 719 | -0.047900 | 12.01000 | ; qtot 0.884 |
| 720 | HC | 45 | LYS | HD2 | 720 | 0.062100 | 1.00800 | ; qtot 0.946 |
| 721 | HC | 45 | LYS | HD3 | 721 | 0.062100 | 1.00800 | ; qtot 1.008 |
| 722 | C8 | 45 | LYS | CE | 722 | -0.014300 | 12.01000 | ; qtot 0.994 |
| 723 | HP | 45 | LYS | HE2 | 723 | 0.113500 | 1.00800 | ; qtot 1.107 |
| 724 | HP | 45 | LYS | HE3 | 724 | 0.113500 | 1.00800 | ; qtot 1.221 |
| 725 | N3 | 45 | LYS | NZ | 725 | -0.385400 | 14.01000 | ; qtot 0.835 |

```
   726    H    45   LYS   HZ1  726     0.340000      1.00800 ; qtot
1.175
   727    H    45   LYS   HZ2  727     0.340000      1.00800 ; qtot
1.515
   728    H    45   LYS   HZ3  728     0.340000      1.00800 ; qtot
1.855
   729    C    45   LYS     C  729     0.734101     12.01000 ; qtot
2.589
   730    O    45   LYS     O  730    -0.589401     16.00000 ; qtot
2.000
   731    N    46   LYS     N  731    -0.347900     14.01000 ; qtot
1.652
   732    H    46   LYS     H  732     0.274700      1.00800 ; qtot
1.927
   733   CX    46   LYS    CA  733    -0.240000     12.01000 ; qtot
1.687
   734   H1    46   LYS    HA  734     0.142600      1.00800 ; qtot
1.829
   735   C8    46   LYS    CB  735    -0.009400     12.01000 ; qtot
1.820
   736   HC    46   LYS   HB2  736     0.036200      1.00800 ; qtot
1.856
   737   HC    46   LYS   HB3  737     0.036200      1.00800 ; qtot
1.892
   738   C8    46   LYS    CG  738     0.018700     12.01000 ; qtot
1.911
   739   HC    46   LYS   HG2  739     0.010300      1.00800 ; qtot
1.921
   740   HC    46   LYS   HG3  740     0.010300      1.00800 ; qtot
1.932
   741   C8    46   LYS    CD  741    -0.047900     12.01000 ; qtot
1.884
   742   HC    46   LYS   HD2  742     0.062100      1.00800 ; qtot
1.946
   743   HC    46   LYS   HD3  743     0.062100      1.00800 ; qtot
2.008
   744   C8    46   LYS    CE  744    -0.014300     12.01000 ; qtot
1.994
   745   HP    46   LYS   HE2  745     0.113500      1.00800 ; qtot
2.107
   746   HP    46   LYS   HE3  746     0.113500      1.00800 ; qtot
2.221
   747   N3    46   LYS    NZ  747    -0.385400     14.01000 ; qtot
1.835
   748    H    46   LYS   HZ1  748     0.340000      1.00800 ; qtot
2.175
   749    H    46   LYS   HZ2  749     0.340000      1.00800 ; qtot
2.515
   750    H    46   LYS   HZ3  750     0.340000      1.00800 ; qtot
2.855
   751    C    46   LYS     C  751     0.734101     12.01000 ; qtot
3.589
```

752 O 46 LYS O 752 -0.589401 16.00000 ; qtot
3.000
753 N 47 ASP N 753 -0.516301 14.01000 ; qtot
2.484
754 H 47 ASP H 754 0.293600 1.00800 ; qtot
2.777
755 CX 47 ASP CA 755 0.038100 12.01000 ; qtot
2.815
756 H1 47 ASP HA 756 0.088000 1.00800 ; qtot
2.903
757 A2C 47 ASP CB 757 -0.030300 12.01000 ; qtot
2.873
758 HC 47 ASP HB2 758 -0.012200 1.00800 ; qtot
2.861
759 HC 47 ASP HB3 759 -0.012200 1.00800 ; qtot
2.849
760 CO 47 ASP CG 760 0.799401 12.01000 ; qtot
3.648
761 O2 47 ASP OD1 761 -0.801401 16.00000 ; qtot
2.847
762 O2 47 ASP OD2 762 -0.801401 16.00000 ; qtot
2.045
763 C 47 ASP C 763 0.536601 12.01000 ; qtot
2.582
764 O 47 ASP O 764 -0.581901 16.00000 ; qtot
2.000
765 N 48 ARG N 765 -0.347900 14.01000 ; qtot
1.652
766 H 48 ARG H 766 0.274700 1.00800 ; qtot
1.927
767 CX 48 ARG CA 767 -0.263700 12.01000 ; qtot
1.663
768 H1 48 ARG HA 768 0.156000 1.00800 ; qtot
1.819
769 C8 48 ARG CB 769 -0.000700 12.01000 ; qtot
1.818
770 HC 48 ARG HB2 770 0.032700 1.00800 ; qtot
1.851
771 HC 48 ARG HB3 771 0.032700 1.00800 ; qtot
1.884
772 C8 48 ARG CG 772 0.039000 12.01000 ; qtot
1.923
773 HC 48 ARG HG2 773 0.028500 1.00800 ; qtot
1.951
774 HC 48 ARG HG3 774 0.028500 1.00800 ; qtot
1.980
775 C8 48 ARG CD 775 0.048600 12.01000 ; qtot
2.028
776 H1 48 ARG HD2 776 0.068700 1.00800 ; qtot
2.097
777 H1 48 ARG HD3 777 0.068700 1.00800 ; qtot
2.166

| | | | | | | | | | |
|---|---|---|---|---|---|---|---|---|---|
| 778 | N2 | 48 | ARG | NE | 778 | -0.529501 | 14.01000 | ; qtot | 1.636 |
| 779 | H | 48 | ARG | HE | 779 | 0.345600 | 1.00800 | ; qtot | 1.982 |
| 780 | CA | 48 | ARG | CZ | 780 | 0.807601 | 12.01000 | ; qtot | 2.790 |
| 781 | N2 | 48 | ARG | NH1 | 781 | -0.862701 | 14.01000 | ; qtot | 1.927 |
| 782 | H | 48 | ARG | HH11 | 782 | 0.447800 | 1.00800 | ; qtot | 2.375 |
| 783 | H | 48 | ARG | HH12 | 783 | 0.447800 | 1.00800 | ; qtot | 2.822 |
| 784 | N2 | 48 | ARG | NH2 | 784 | -0.862701 | 14.01000 | ; qtot | 1.960 |
| 785 | H | 48 | ARG | HH21 | 785 | 0.447800 | 1.00800 | ; qtot | 2.408 |
| 786 | H | 48 | ARG | HH22 | 786 | 0.447800 | 1.00800 | ; qtot | 2.855 |
| 787 | C | 48 | ARG | C | 787 | 0.734101 | 12.01000 | ; qtot | 3.589 |
| 788 | O | 48 | ARG | O | 788 | -0.589401 | 16.00000 | ; qtot | 3.000 |
| 789 | N | 49 | LEU | N | 789 | -0.415700 | 14.01000 | ; qtot | 2.584 |
| 790 | H | 49 | LEU | H | 790 | 0.271900 | 1.00800 | ; qtot | 2.856 |
| 791 | CX | 49 | LEU | CA | 791 | -0.051800 | 12.01000 | ; qtot | 2.804 |
| 792 | H1 | 49 | LEU | HA | 792 | 0.092200 | 1.00800 | ; qtot | 2.897 |
| 793 | A2C | 49 | LEU | CB | 793 | -0.110200 | 12.01000 | ; qtot | 2.786 |
| 794 | HC | 49 | LEU | HB2 | 794 | 0.045700 | 1.00800 | ; qtot | 2.832 |
| 795 | HC | 49 | LEU | HB3 | 795 | 0.045700 | 1.00800 | ; qtot | 2.878 |
| 796 | A3C | 49 | LEU | CG | 796 | 0.353100 | 12.01000 | ; qtot | 3.231 |
| 797 | HC | 49 | LEU | HG | 797 | -0.036100 | 1.00800 | ; qtot | 3.195 |
| 798 | CT | 49 | LEU | CD1 | 798 | -0.412100 | 12.01000 | ; qtot | 2.783 |
| 799 | HC | 49 | LEU | HD11 | 799 | 0.100000 | 1.00800 | ; qtot | 2.883 |
| 800 | HC | 49 | LEU | HD12 | 800 | 0.100000 | 1.00800 | ; qtot | 2.983 |
| 801 | HC | 49 | LEU | HD13 | 801 | 0.100000 | 1.00800 | ; qtot | 3.083 |
| 802 | CT | 49 | LEU | CD2 | 802 | -0.412100 | 12.01000 | ; qtot | 2.671 |
| 803 | HC | 49 | LEU | HD21 | 803 | 0.100000 | 1.00800 | ; qtot | 2.771 |

```
   804    HC     49   LEU  HD22  804      0.100000       1.00800 ; qtot
2.871
   805    HC     49   LEU  HD23  805      0.100000       1.00800 ; qtot
2.971
   806     C     49   LEU     C  806      0.597301      12.01000 ; qtot
3.568
   807     O     49   LEU     O  807     -0.567901      16.00000 ; qtot
3.000
   808     N     50   LYS     N  808     -0.347900      14.01000 ; qtot
2.652
   809     H     50   LYS     H  809      0.274700       1.00800 ; qtot
2.927
   810    CX     50   LYS    CA  810     -0.240000      12.01000 ; qtot
2.687
   811    H1     50   LYS    HA  811      0.142600       1.00800 ; qtot
2.829
   812    C8     50   LYS    CB  812     -0.009400      12.01000 ; qtot
2.820
   813    HC     50   LYS   HB2  813      0.036200       1.00800 ; qtot
2.856
   814    HC     50   LYS   HB3  814      0.036200       1.00800 ; qtot
2.892
   815    C8     50   LYS    CG  815      0.018700      12.01000 ; qtot
2.911
   816    HC     50   LYS   HG2  816      0.010300       1.00800 ; qtot
2.921
   817    HC     50   LYS   HG3  817      0.010300       1.00800 ; qtot
2.932
   818    C8     50   LYS    CD  818     -0.047900      12.01000 ; qtot
2.884
   819    HC     50   LYS   HD2  819      0.062100       1.00800 ; qtot
2.946
   820    HC     50   LYS   HD3  820      0.062100       1.00800 ; qtot
3.008
   821    C8     50   LYS    CE  821     -0.014300      12.01000 ; qtot
2.994
   822    HP     50   LYS   HE2  822      0.113500       1.00800 ; qtot
3.107
   823    HP     50   LYS   HE3  823      0.113500       1.00800 ; qtot
3.221
   824    N3     50   LYS    NZ  824     -0.385400      14.01000 ; qtot
2.835
   825     H     50   LYS   HZ1  825      0.340000       1.00800 ; qtot
3.175
   826     H     50   LYS   HZ2  826      0.340000       1.00800 ; qtot
3.515
   827     H     50   LYS   HZ3  827      0.340000       1.00800 ; qtot
3.855
   828     C     50   LYS     C  828      0.734101      12.01000 ; qtot
4.589
   829     O     50   LYS     O  829     -0.589401      16.00000 ; qtot
4.000
```

```
   830    N    51   ASN     N  830   -0.415700    14.01000 ; qtot
3.584
   831    H    51   ASN     H  831    0.271900     1.00800 ; qtot
3.856
   832   CX    51   ASN    CA  832    0.014300    12.01000 ; qtot
3.871
   833   H1    51   ASN    HA  833    0.104800     1.00800 ; qtot
3.975
   834  A2C    51   ASN    CB  834   -0.204100    12.01000 ; qtot
3.771
   835   HC    51   ASN   HB2  835    0.079700     1.00800 ; qtot
3.851
   836   HC    51   ASN   HB3  836    0.079700     1.00800 ; qtot
3.931
   837    C    51   ASN    CG  837    0.713001    12.01000 ; qtot
4.644
   838    O    51   ASN   OD1  838   -0.593101    16.00000 ; qtot
4.051
   839    N    51   ASN   ND2  839   -0.919101    14.01000 ; qtot
3.131
   840    H    51   ASN  HD21  840    0.419600     1.00800 ; qtot
3.551
   841    H    51   ASN  HD22  841    0.419600     1.00800 ; qtot
3.971
   842    C    51   ASN     C  842    0.597301    12.01000 ; qtot
4.568
   843    O    51   ASN     O  843   -0.567901    16.00000 ; qtot
4.000
   844    N    52   ILE     N  844   -0.415700    14.01000 ; qtot
3.584
   845    H    52   ILE     H  845    0.271900     1.00800 ; qtot
3.856
   846   CX    52   ILE    CA  846   -0.059700    12.01000 ; qtot
3.797
   847   H1    52   ILE    HA  847    0.086900     1.00800 ; qtot
3.883
   848  A3C    52   ILE    CB  848    0.130300    12.01000 ; qtot
4.014
   849   HC    52   ILE    HB  849    0.018700     1.00800 ; qtot
4.032
   850   CT    52   ILE   CG2  850   -0.320400    12.01000 ; qtot
3.712
   851   HC    52   ILE  HG21  851    0.088200     1.00800 ; qtot
3.800
   852   HC    52   ILE  HG22  852    0.088200     1.00800 ; qtot
3.888
   853   HC    52   ILE  HG23  853    0.088200     1.00800 ; qtot
3.977
   854  A2C    52   ILE   CG1  854   -0.043000    12.01000 ; qtot
3.934
   855   HC    52   ILE  HG12  855    0.023600     1.00800 ; qtot
3.957
```

| | | | | | | | | | |
|---|---|---|---|---|---|---|---|---|---|
| 856 | HC | 52 | ILE | HG13 | 856 | 0.023600 | 1.00800 | ; qtot | 3.981 |
| 857 | CT | 52 | ILE | CD1 | 857 | -0.066000 | 12.01000 | ; qtot | 3.915 |
| 858 | HC | 52 | ILE | HD11 | 858 | 0.018600 | 1.00800 | ; qtot | 3.933 |
| 859 | HC | 52 | ILE | HD12 | 859 | 0.018600 | 1.00800 | ; qtot | 3.952 |
| 860 | HC | 52 | ILE | HD13 | 860 | 0.018600 | 1.00800 | ; qtot | 3.971 |
| 861 | C | 52 | ILE | C | 861 | 0.597301 | 12.01000 | ; qtot | 4.568 |
| 862 | O | 52 | ILE | O | 862 | -0.567901 | 16.00000 | ; qtot | 4.000 |
| 863 | N | 53 | ASN | N | 863 | -0.415700 | 14.01000 | ; qtot | 3.584 |
| 864 | H | 53 | ASN | H | 864 | 0.271900 | 1.00800 | ; qtot | 3.856 |
| 865 | CX | 53 | ASN | CA | 865 | 0.014300 | 12.01000 | ; qtot | 3.871 |
| 866 | H1 | 53 | ASN | HA | 866 | 0.104800 | 1.00800 | ; qtot | 3.975 |
| 867 | A2C | 53 | ASN | CB | 867 | -0.204100 | 12.01000 | ; qtot | 3.771 |
| 868 | HC | 53 | ASN | HB2 | 868 | 0.079700 | 1.00800 | ; qtot | 3.851 |
| 869 | HC | 53 | ASN | HB3 | 869 | 0.079700 | 1.00800 | ; qtot | 3.931 |
| 870 | C | 53 | ASN | CG | 870 | 0.713001 | 12.01000 | ; qtot | 4.644 |
| 871 | O | 53 | ASN | OD1 | 871 | -0.593101 | 16.00000 | ; qtot | 4.051 |
| 872 | N | 53 | ASN | ND2 | 872 | -0.919101 | 14.01000 | ; qtot | 3.131 |
| 873 | H | 53 | ASN | HD21 | 873 | 0.419600 | 1.00800 | ; qtot | 3.551 |
| 874 | H | 53 | ASN | HD22 | 874 | 0.419600 | 1.00800 | ; qtot | 3.971 |
| 875 | C | 53 | ASN | C | 875 | 0.597301 | 12.01000 | ; qtot | 4.568 |
| 876 | O | 53 | ASN | O | 876 | -0.567901 | 16.00000 | ; qtot | 4.000 |
| 877 | N | 54 | ARG | N | 877 | -0.347900 | 14.01000 | ; qtot | 3.652 |
| 878 | H | 54 | ARG | H | 878 | 0.274700 | 1.00800 | ; qtot | 3.927 |
| 879 | CX | 54 | ARG | CA | 879 | -0.263700 | 12.01000 | ; qtot | 3.663 |
| 880 | H1 | 54 | ARG | HA | 880 | 0.156000 | 1.00800 | ; qtot | 3.819 |
| 881 | C8 | 54 | ARG | CB | 881 | -0.000700 | 12.01000 | ; qtot | 3.818 |

```
   882   HC     54   ARG   HB2  882      0.032700      1.00800 ; qtot
3.851
   883   HC     54   ARG   HB3  883      0.032700      1.00800 ; qtot
3.884
   884   C8     54   ARG    CG  884      0.039000     12.01000 ; qtot
3.923
   885   HC     54   ARG   HG2  885      0.028500      1.00800 ; qtot
3.951
   886   HC     54   ARG   HG3  886      0.028500      1.00800 ; qtot
3.980
   887   C8     54   ARG    CD  887      0.048600     12.01000 ; qtot
4.028
   888   H1     54   ARG   HD2  888      0.068700      1.00800 ; qtot
4.097
   889   H1     54   ARG   HD3  889      0.068700      1.00800 ; qtot
4.166
   890   N2     54   ARG    NE  890     -0.529501     14.01000 ; qtot
3.636
   891    H     54   ARG    HE  891      0.345600      1.00800 ; qtot
3.982
   892   CA     54   ARG    CZ  892      0.807601     12.01000 ; qtot
4.790
   893   N2     54   ARG   NH1  893     -0.862701     14.01000 ; qtot
3.927
   894    H     54   ARG  HH11  894      0.447800      1.00800 ; qtot
4.375
   895    H     54   ARG  HH12  895      0.447800      1.00800 ; qtot
4.822
   896   N2     54   ARG   NH2  896     -0.862701     14.01000 ; qtot
3.960
   897    H     54   ARG  HH21  897      0.447800      1.00800 ; qtot
4.408
   898    H     54   ARG  HH22  898      0.447800      1.00800 ; qtot
4.855
   899    C     54   ARG     C  899      0.734101     12.01000 ; qtot
5.589
   900    O     54   ARG     O  900     -0.589401     16.00000 ; qtot
5.000
   901    N     55   PHE     N  901     -0.415700     14.01000 ; qtot
4.584
   902    H     55   PHE     H  902      0.271900      1.00800 ; qtot
4.856
   903   CX     55   PHE    CA  903     -0.002400     12.01000 ; qtot
4.854
   904   H1     55   PHE    HA  904      0.097800      1.00800 ; qtot
4.952
   905   CT     55   PHE    CB  905     -0.034300     12.01000 ; qtot
4.917
   906   HC     55   PHE   HB2  906      0.029500      1.00800 ; qtot
4.947
   907   HC     55   PHE   HB3  907      0.029500      1.00800 ; qtot
4.976
```

908 CA 55 PHE CG 908 0.011800 12.01000 ; qtot
4.988
909 CA 55 PHE CD1 909 -0.125600 12.01000 ; qtot
4.863
910 HA 55 PHE HD1 910 0.133000 1.00800 ; qtot
4.996
911 CA 55 PHE CE1 911 -0.170400 12.01000 ; qtot
4.825
912 HA 55 PHE HE1 912 0.143000 1.00800 ; qtot
4.968
913 CA 55 PHE CZ 913 -0.107200 12.01000 ; qtot
4.861
914 HA 55 PHE HZ 914 0.129700 1.00800 ; qtot
4.991
915 CA 55 PHE CE2 915 -0.170400 12.01000 ; qtot
4.820
916 HA 55 PHE HE2 916 0.143000 1.00800 ; qtot
4.963
917 CA 55 PHE CD2 917 -0.125600 12.01000 ; qtot
4.838
918 HA 55 PHE HD2 918 0.133000 1.00800 ; qtot
4.971
919 C 55 PHE C 919 0.597301 12.01000 ; qtot
5.568
920 O 55 PHE O 920 -0.567901 16.00000 ; qtot
5.000
921 N 56 PHE N 921 -0.415700 14.01000 ; qtot
4.584
922 H 56 PHE H 922 0.271900 1.00800 ; qtot
4.856
923 CX 56 PHE CA 923 -0.002400 12.01000 ; qtot
4.854
924 H1 56 PHE HA 924 0.097800 1.00800 ; qtot
4.952
925 CT 56 PHE CB 925 -0.034300 12.01000 ; qtot
4.917
926 HC 56 PHE HB2 926 0.029500 1.00800 ; qtot
4.947
927 HC 56 PHE HB3 927 0.029500 1.00800 ; qtot
4.976
928 CA 56 PHE CG 928 0.011800 12.01000 ; qtot
4.988
929 CA 56 PHE CD1 929 -0.125600 12.01000 ; qtot
4.863
930 HA 56 PHE HD1 930 0.133000 1.00800 ; qtot
4.996
931 CA 56 PHE CE1 931 -0.170400 12.01000 ; qtot
4.825
932 HA 56 PHE HE1 932 0.143000 1.00800 ; qtot
4.968
933 CA 56 PHE CZ 933 -0.107200 12.01000 ; qtot
4.861

```
   934   HA    56   PHE    HZ  934     0.129700      1.00800 ; qtot
4.991
   935   CA    56   PHE   CE2  935    -0.170400     12.01000 ; qtot
4.820
   936   HA    56   PHE   HE2  936     0.143000      1.00800 ; qtot
4.963
   937   CA    56   PHE   CD2  937    -0.125600     12.01000 ; qtot
4.838
   938   HA    56   PHE   HD2  938     0.133000      1.00800 ; qtot
4.971
   939    C    56   PHE     C  939     0.597301     12.01000 ; qtot
5.568
   940    O    56   PHE     O  940    -0.567901     16.00000 ; qtot
5.000
   941    N    57   GLN     N  941    -0.415700     14.01000 ; qtot
4.584
   942    H    57   GLN     H  942     0.271900      1.00800 ; qtot
4.856
   943   CX    57   GLN    CA  943    -0.003100     12.01000 ; qtot
4.853
   944   H1    57   GLN    HA  944     0.085000      1.00800 ; qtot
4.938
   945  A2C    57   GLN    CB  945    -0.003600     12.01000 ; qtot
4.935
   946   HC    57   GLN   HB2  946     0.017100      1.00800 ; qtot
4.952
   947   HC    57   GLN   HB3  947     0.017100      1.00800 ; qtot
4.969
   948  A2C    57   GLN    CG  948    -0.064500     12.01000 ; qtot
4.904
   949   HC    57   GLN   HG2  949     0.035200      1.00800 ; qtot
4.939
   950   HC    57   GLN   HG3  950     0.035200      1.00800 ; qtot
4.975
   951    C    57   GLN    CD  951     0.695101     12.01000 ; qtot
5.670
   952    O    57   GLN   OE1  952    -0.608601     16.00000 ; qtot
5.061
   953    N    57   GLN   NE2  953    -0.940701     14.01000 ; qtot
4.120
   954    H    57   GLN  HE21  954     0.425100      1.00800 ; qtot
4.546
   955    H    57   GLN  HE22  955     0.425100      1.00800 ; qtot
4.971
   956    C    57   GLN     C  956     0.597301     12.01000 ; qtot
5.568
   957    O    57   GLN     O  957    -0.567901     16.00000 ; qtot
5.000
   958    N    58   GLU     N  958    -0.516301     14.01000 ; qtot
4.484
   959    H    58   GLU     H  959     0.293600      1.00800 ; qtot
4.777
```

```
   960   CX    58   GLU    CA  960     0.039700     12.01000 ; qtot
4.817
   961   H1    58   GLU    HA  961     0.110500      1.00800 ; qtot
4.928
   962  A2C    58   GLU    CB  962     0.056000     12.01000 ; qtot
4.984
   963   HC    58   GLU   HB2  963    -0.017300      1.00800 ; qtot
4.966
   964   HC    58   GLU   HB3  964    -0.017300      1.00800 ; qtot
4.949
   965  A2C    58   GLU    CG  965     0.013600     12.01000 ; qtot
4.963
   966   HC    58   GLU   HG2  966    -0.042500      1.00800 ; qtot
4.920
   967   HC    58   GLU   HG3  967    -0.042500      1.00800 ; qtot
4.878
   968   CO    58   GLU    CD  968     0.805401     12.01000 ; qtot
5.683
   969   O2    58   GLU   OE1  969    -0.818801     16.00000 ; qtot
4.864
   970   O2    58   GLU   OE2  970    -0.818801     16.00000 ; qtot
4.045
   971    C    58   GLU     C  971     0.536601     12.01000 ; qtot
4.582
   972    O    58   GLU     O  972    -0.581901     16.00000 ; qtot
4.000
   973    N    59   ILE     N  973    -0.415700     14.01000 ; qtot
3.584
   974    H    59   ILE     H  974     0.271900      1.00800 ; qtot
3.856
   975   CX    59   ILE    CA  975    -0.059700     12.01000 ; qtot
3.797
   976   H1    59   ILE    HA  976     0.086900      1.00800 ; qtot
3.883
   977  A3C    59   ILE    CB  977     0.130300     12.01000 ; qtot
4.014
   978   HC    59   ILE    HB  978     0.018700      1.00800 ; qtot
4.032
   979   CT    59   ILE   CG2  979    -0.320400     12.01000 ; qtot
3.712
   980   HC    59   ILE  HG21  980     0.088200      1.00800 ; qtot
3.800
   981   HC    59   ILE  HG22  981     0.088200      1.00800 ; qtot
3.888
   982   HC    59   ILE  HG23  982     0.088200      1.00800 ; qtot
3.977
   983  A2C    59   ILE   CG1  983    -0.043000     12.01000 ; qtot
3.934
   984   HC    59   ILE  HG12  984     0.023600      1.00800 ; qtot
3.957
   985   HC    59   ILE  HG13  985     0.023600      1.00800 ; qtot
3.981
```

986 CT 59 ILE CD1 986 -0.066000 12.01000 ; qtot
3.915
987 HC 59 ILE HD11 987 0.018600 1.00800 ; qtot
3.933
988 HC 59 ILE HD12 988 0.018600 1.00800 ; qtot
3.952
989 HC 59 ILE HD13 989 0.018600 1.00800 ; qtot
3.971
990 C 59 ILE C 990 0.597301 12.01000 ; qtot
4.568
991 O 59 ILE O 991 -0.567901 16.00000 ; qtot
4.000
992 N 60 GLU N 992 -0.516301 14.01000 ; qtot
3.484
993 H 60 GLU H 993 0.293600 1.00800 ; qtot
3.777
994 CX 60 GLU CA 994 0.039700 12.01000 ; qtot
3.817
995 H1 60 GLU HA 995 0.110500 1.00800 ; qtot
3.928
996 A2C 60 GLU CB 996 0.056000 12.01000 ; qtot
3.984
997 HC 60 GLU HB2 997 -0.017300 1.00800 ; qtot
3.966
998 HC 60 GLU HB3 998 -0.017300 1.00800 ; qtot
3.949
999 A2C 60 GLU CG 999 0.013600 12.01000 ; qtot
3.963
1000 HC 60 GLU HG2 1000 -0.042500 1.00800 ; qtot
3.920
1001 HC 60 GLU HG3 1001 -0.042500 1.00800 ; qtot
3.878
1002 CO 60 GLU CD 1002 0.805401 12.01000 ; qtot
4.683
1003 O2 60 GLU OE1 1003 -0.818801 16.00000 ; qtot
3.864
1004 O2 60 GLU OE2 1004 -0.818801 16.00000 ; qtot
3.045
1005 C 60 GLU C 1005 0.536601 12.01000 ; qtot
3.582
1006 O 60 GLU O 1006 -0.581901 16.00000 ; qtot
3.000
1007 N 61 ILE N 1007 -0.415700 14.01000 ; qtot
2.584
1008 H 61 ILE H 1008 0.271900 1.00800 ; qtot
2.856
1009 CX 61 ILE CA 1009 -0.059700 12.01000 ; qtot
2.797
1010 H1 61 ILE HA 1010 0.086900 1.00800 ; qtot
2.883
1011 A3C 61 ILE CB 1011 0.130300 12.01000 ; qtot
3.014

1012 HC 61 ILE HB 1012 0.018700 1.00800 ; qtot
3.032
1013 CT 61 ILE CG2 1013 -0.320400 12.01000 ; qtot
2.712
1014 HC 61 ILE HG21 1014 0.088200 1.00800 ; qtot
2.800
1015 HC 61 ILE HG22 1015 0.088200 1.00800 ; qtot
2.888
1016 HC 61 ILE HG23 1016 0.088200 1.00800 ; qtot
2.977
1017 A2C 61 ILE CG1 1017 -0.043000 12.01000 ; qtot
2.934
1018 HC 61 ILE HG12 1018 0.023600 1.00800 ; qtot
2.957
1019 HC 61 ILE HG13 1019 0.023600 1.00800 ; qtot
2.981
1020 CT 61 ILE CD1 1020 -0.066000 12.01000 ; qtot
2.915
1021 HC 61 ILE HD11 1021 0.018600 1.00800 ; qtot
2.933
1022 HC 61 ILE HD12 1022 0.018600 1.00800 ; qtot
2.952
1023 HC 61 ILE HD13 1023 0.018600 1.00800 ; qtot
2.971
1024 C 61 ILE C 1024 0.597301 12.01000 ; qtot
3.568
1025 O 61 ILE O 1025 -0.567901 16.00000 ; qtot
3.000
1026 N 62 MET N 1026 -0.415700 14.01000 ; qtot
2.584
1027 H 62 MET H 1027 0.271900 1.00800 ; qtot
2.856
1028 CX 62 MET CA 1028 -0.023700 12.01000 ; qtot
2.833
1029 H1 62 MET HA 1029 0.088000 1.00800 ; qtot
2.921
1030 A2C 62 MET CB 1030 0.034200 12.01000 ; qtot
2.955
1031 HC 62 MET HB2 1031 0.024100 1.00800 ; qtot
2.979
1032 HC 62 MET HB3 1032 0.024100 1.00800 ; qtot
3.003
1033 A2C 62 MET CG 1033 0.001800 12.01000 ; qtot
3.005
1034 H1 62 MET HG2 1034 0.044000 1.00800 ; qtot
3.049
1035 H1 62 MET HG3 1035 0.044000 1.00800 ; qtot
3.093
1036 S 62 MET SD 1036 -0.273700 32.06000 ; qtot
2.819
1037 CT 62 MET CE 1037 -0.053600 12.01000 ; qtot
2.765

```
  1038   H1     62   MET   HE1 1038     0.068400      1.00800 ; qtot
2.834
  1039   H1     62   MET   HE2 1039     0.068400      1.00800 ; qtot
2.902
  1040   H1     62   MET   HE3 1040     0.068400      1.00800 ; qtot
2.971
  1041    C     62   MET     C 1041     0.597301     12.01000 ; qtot
3.568
  1042    O     62   MET     O 1042    -0.567901     16.00000 ; qtot
3.000
  1043    N     63   LYS     N 1043    -0.347900     14.01000 ; qtot
2.652
  1044    H     63   LYS     H 1044     0.274700      1.00800 ; qtot
2.927
  1045   CX     63   LYS    CA 1045    -0.240000     12.01000 ; qtot
2.687
  1046   H1     63   LYS    HA 1046     0.142600      1.00800 ; qtot
2.829
  1047   C8     63   LYS    CB 1047    -0.009400     12.01000 ; qtot
2.820
  1048   HC     63   LYS   HB2 1048     0.036200      1.00800 ; qtot
2.856
  1049   HC     63   LYS   HB3 1049     0.036200      1.00800 ; qtot
2.892
  1050   C8     63   LYS    CG 1050     0.018700     12.01000 ; qtot
2.911
  1051   HC     63   LYS   HG2 1051     0.010300      1.00800 ; qtot
2.921
  1052   HC     63   LYS   HG3 1052     0.010300      1.00800 ; qtot
2.932
  1053   C8     63   LYS    CD 1053    -0.047900     12.01000 ; qtot
2.884
  1054   HC     63   LYS   HD2 1054     0.062100      1.00800 ; qtot
2.946
  1055   HC     63   LYS   HD3 1055     0.062100      1.00800 ; qtot
3.008
  1056   C8     63   LYS    CE 1056    -0.014300     12.01000 ; qtot
2.994
  1057   HP     63   LYS   HE2 1057     0.113500      1.00800 ; qtot
3.107
  1058   HP     63   LYS   HE3 1058     0.113500      1.00800 ; qtot
3.221
  1059   N3     63   LYS    NZ 1059    -0.385400     14.01000 ; qtot
2.835
  1060    H     63   LYS   HZ1 1060     0.340000      1.00800 ; qtot
3.175
  1061    H     63   LYS   HZ2 1061     0.340000      1.00800 ; qtot
3.515
  1062    H     63   LYS   HZ3 1062     0.340000      1.00800 ; qtot
3.855
  1063    C     63   LYS     C 1063     0.734101     12.01000 ; qtot
4.589
```

```
  1064    O    63   LYS     O 1064    -0.589401     16.00000 ; qtot
4.000
  1065    N    64   LYS     N 1065    -0.347900     14.01000 ; qtot
3.652
  1066    H    64   LYS     H 1066     0.274700      1.00800 ; qtot
3.927
  1067   CX    64   LYS    CA 1067    -0.240000     12.01000 ; qtot
3.687
  1068   H1    64   LYS    HA 1068     0.142600      1.00800 ; qtot
3.829
  1069   C8    64   LYS    CB 1069    -0.009400     12.01000 ; qtot
3.820
  1070   HC    64   LYS   HB2 1070     0.036200      1.00800 ; qtot
3.856
  1071   HC    64   LYS   HB3 1071     0.036200      1.00800 ; qtot
3.892
  1072   C8    64   LYS    CG 1072     0.018700     12.01000 ; qtot
3.911
  1073   HC    64   LYS   HG2 1073     0.010300      1.00800 ; qtot
3.921
  1074   HC    64   LYS   HG3 1074     0.010300      1.00800 ; qtot
3.932
  1075   C8    64   LYS    CD 1075    -0.047900     12.01000 ; qtot
3.884
  1076   HC    64   LYS   HD2 1076     0.062100      1.00800 ; qtot
3.946
  1077   HC    64   LYS   HD3 1077     0.062100      1.00800 ; qtot
4.008
  1078   C8    64   LYS    CE 1078    -0.014300     12.01000 ; qtot
3.994
  1079   HP    64   LYS   HE2 1079     0.113500      1.00800 ; qtot
4.107
  1080   HP    64   LYS   HE3 1080     0.113500      1.00800 ; qtot
4.221
  1081   N3    64   LYS    NZ 1081    -0.385400     14.01000 ; qtot
3.835
  1082    H    64   LYS   HZ1 1082     0.340000      1.00800 ; qtot
4.175
  1083    H    64   LYS   HZ2 1083     0.340000      1.00800 ; qtot
4.515
  1084    H    64   LYS   HZ3 1084     0.340000      1.00800 ; qtot
4.855
  1085    C    64   LYS     C 1085     0.734101     12.01000 ; qtot
5.589
  1086    O    64   LYS     O 1086    -0.589401     16.00000 ; qtot
5.000
  1087    N    65   LEU     N 1087    -0.415700     14.01000 ; qtot
4.584
  1088    H    65   LEU     H 1088     0.271900      1.00800 ; qtot
4.856
  1089   CX    65   LEU    CA 1089    -0.051800     12.01000 ; qtot
4.804
```

1090 H1 65 LEU HA 1090 0.092200 1.00800 ; qtot
4.897
1091 A2C 65 LEU CB 1091 -0.110200 12.01000 ; qtot
4.786
1092 HC 65 LEU HB2 1092 0.045700 1.00800 ; qtot
4.832
1093 HC 65 LEU HB3 1093 0.045700 1.00800 ; qtot
4.878
1094 A3C 65 LEU CG 1094 0.353100 12.01000 ; qtot
5.231
1095 HC 65 LEU HG 1095 -0.036100 1.00800 ; qtot
5.195
1096 CT 65 LEU CD1 1096 -0.412100 12.01000 ; qtot
4.783
1097 HC 65 LEU HD11 1097 0.100000 1.00800 ; qtot
4.883
1098 HC 65 LEU HD12 1098 0.100000 1.00800 ; qtot
4.983
1099 HC 65 LEU HD13 1099 0.100000 1.00800 ; qtot
5.083
1100 CT 65 LEU CD2 1100 -0.412100 12.01000 ; qtot
4.671
1101 HC 65 LEU HD21 1101 0.100000 1.00800 ; qtot
4.771
1102 HC 65 LEU HD22 1102 0.100000 1.00800 ; qtot
4.871
1103 HC 65 LEU HD23 1103 0.100000 1.00800 ; qtot
4.971
1104 C 65 LEU C 1104 0.597301 12.01000 ; qtot
5.568
1105 O 65 LEU O 1105 -0.567901 16.00000 ; qtot
5.000
1106 N 66 ASP N 1106 -0.516301 14.01000 ; qtot
4.484
1107 H 66 ASP H 1107 0.293600 1.00800 ; qtot
4.777
1108 CX 66 ASP CA 1108 0.038100 12.01000 ; qtot
4.815
1109 H1 66 ASP HA 1109 0.088000 1.00800 ; qtot
4.903
1110 A2C 66 ASP CB 1110 -0.030300 12.01000 ; qtot
4.873
1111 HC 66 ASP HB2 1111 -0.012200 1.00800 ; qtot
4.861
1112 HC 66 ASP HB3 1112 -0.012200 1.00800 ; qtot
4.849
1113 CO 66 ASP CG 1113 0.799401 12.01000 ; qtot
5.648
1114 O2 66 ASP OD1 1114 -0.801401 16.00000 ; qtot
4.847
1115 O2 66 ASP OD2 1115 -0.801401 16.00000 ; qtot
4.045

1116 C 66 ASP C 1116 0.536601 12.01000 ; qtot
4.582
1117 O 66 ASP O 1117 -0.581901 16.00000 ; qtot
4.000
1118 N 67 HIE N 1118 -0.415700 14.01000 ; qtot
3.584
1119 H 67 HIE H 1119 0.271900 1.00800 ; qtot
3.856
1120 CX 67 HIE CA 1120 -0.058100 12.01000 ; qtot
3.798
1121 H1 67 HIE HA 1121 0.136000 1.00800 ; qtot
3.934
1122 CT 67 HIE CB 1122 -0.007400 12.01000 ; qtot
3.927
1123 HC 67 HIE HB2 1123 0.036700 1.00800 ; qtot
3.963
1124 HC 67 HIE HB3 1124 0.036700 1.00800 ; qtot
4.000
1125 CC 67 HIE CG 1125 0.186800 12.01000 ; qtot
4.187
1126 NB 67 HIE ND1 1126 -0.543201 14.01000 ; qtot
3.644
1127 CR 67 HIE CE1 1127 0.163500 12.01000 ; qtot
3.807
1128 H5 67 HIE HE1 1128 0.143500 1.00800 ; qtot
3.951
1129 NA 67 HIE NE2 1129 -0.279500 14.01000 ; qtot
3.671
1130 H 67 HIE HE2 1130 0.333900 1.00800 ; qtot
4.005
1131 CW 67 HIE CD2 1131 -0.220700 12.01000 ; qtot
3.784
1132 H4 67 HIE HD2 1132 0.186200 1.00800 ; qtot
3.971
1133 C 67 HIE C 1133 0.597301 12.01000 ; qtot
4.568
1134 O 67 HIE O 1134 -0.567901 16.00000 ; qtot
4.000
1135 N 68 PRO N 1135 -0.254800 14.01000 ; qtot
3.745
1136 CT 68 PRO CD 1136 0.019200 12.01000 ; qtot
3.764
1137 H1 68 PRO HD2 1137 0.039100 1.00800 ; qtot
3.804
1138 H1 68 PRO HD3 1138 0.039100 1.00800 ; qtot
3.843
1139 CT 68 PRO CG 1139 0.018900 12.01000 ; qtot
3.862
1140 HC 68 PRO HG2 1140 0.021300 1.00800 ; qtot
3.883
1141 HC 68 PRO HG3 1141 0.021300 1.00800 ; qtot
3.904

```
  1142   CT     68   PRO    CB 1142    -0.007000     12.01000 ; qtot
3.897
  1143   HC     68   PRO   HB2 1143     0.025300      1.00800 ; qtot
3.922
  1144   HC     68   PRO   HB3 1144     0.025300      1.00800 ; qtot
3.948
  1145   CX     68   PRO    CA 1145    -0.026600     12.01000 ; qtot
3.921
  1146   H1     68   PRO    HA 1146     0.064100      1.00800 ; qtot
3.985
  1147    C     68   PRO     C 1147     0.589601     12.01000 ; qtot
4.575
  1148    O     68   PRO     O 1148    -0.574801     16.00000 ; qtot
4.000
  1149    N     69   ASN     N 1149    -0.415700     14.01000 ; qtot
3.584
  1150    H     69   ASN     H 1150     0.271900      1.00800 ; qtot
3.856
  1151   CX     69   ASN    CA 1151     0.014300     12.01000 ; qtot
3.871
  1152   H1     69   ASN    HA 1152     0.104800      1.00800 ; qtot
3.975
  1153  A2C     69   ASN    CB 1153    -0.204100     12.01000 ; qtot
3.771
  1154   HC     69   ASN   HB2 1154     0.079700      1.00800 ; qtot
3.851
  1155   HC     69   ASN   HB3 1155     0.079700      1.00800 ; qtot
3.931
  1156    C     69   ASN    CG 1156     0.713001     12.01000 ; qtot
4.644
  1157    O     69   ASN   OD1 1157    -0.593101     16.00000 ; qtot
4.051
  1158    N     69   ASN   ND2 1158    -0.919101     14.01000 ; qtot
3.131
  1159    H     69   ASN  HD21 1159     0.419600      1.00800 ; qtot
3.551
  1160    H     69   ASN  HD22 1160     0.419600      1.00800 ; qtot
3.971
  1161    C     69   ASN     C 1161     0.597301     12.01000 ; qtot
4.568
  1162    O     69   ASN     O 1162    -0.567901     16.00000 ; qtot
4.000
  1163    N     70   ILE     N 1163    -0.415700     14.01000 ; qtot
3.584
  1164    H     70   ILE     H 1164     0.271900      1.00800 ; qtot
3.856
  1165   CX     70   ILE    CA 1165    -0.059700     12.01000 ; qtot
3.797
  1166   H1     70   ILE    HA 1166     0.086900      1.00800 ; qtot
3.883
  1167  A3C     70   ILE    CB 1167     0.130300     12.01000 ; qtot
4.014
```

1168 HC 70 ILE HB 1168 0.018700 1.00800 ; qtot
4.032
1169 CT 70 ILE CG2 1169 -0.320400 12.01000 ; qtot
3.712
1170 HC 70 ILE HG21 1170 0.088200 1.00800 ; qtot
3.800
1171 HC 70 ILE HG22 1171 0.088200 1.00800 ; qtot
3.888
1172 HC 70 ILE HG23 1172 0.088200 1.00800 ; qtot
3.977
1173 A2C 70 ILE CG1 1173 -0.043000 12.01000 ; qtot
3.934
1174 HC 70 ILE HG12 1174 0.023600 1.00800 ; qtot
3.957
1175 HC 70 ILE HG13 1175 0.023600 1.00800 ; qtot
3.981
1176 CT 70 ILE CD1 1176 -0.066000 12.01000 ; qtot
3.915
1177 HC 70 ILE HD11 1177 0.018600 1.00800 ; qtot
3.933
1178 HC 70 ILE HD12 1178 0.018600 1.00800 ; qtot
3.952
1179 HC 70 ILE HD13 1179 0.018600 1.00800 ; qtot
3.971
1180 C 70 ILE C 1180 0.597301 12.01000 ; qtot
4.568
1181 O 70 ILE O 1181 -0.567901 16.00000 ; qtot
4.000
1182 N 71 VAL N 1182 -0.415700 14.01000 ; qtot
3.584
1183 H 71 VAL H 1183 0.271900 1.00800 ; qtot
3.856
1184 CX 71 VAL CA 1184 -0.087500 12.01000 ; qtot
3.769
1185 H1 71 VAL HA 1185 0.096900 1.00800 ; qtot
3.866
1186 A3C 71 VAL CB 1186 0.298500 12.01000 ; qtot
4.164
1187 HC 71 VAL HB 1187 -0.029700 1.00800 ; qtot
4.134
1188 CT 71 VAL CG1 1188 -0.319200 12.01000 ; qtot
3.815
1189 HC 71 VAL HG11 1189 0.079100 1.00800 ; qtot
3.894
1190 HC 71 VAL HG12 1190 0.079100 1.00800 ; qtot
3.973
1191 HC 71 VAL HG13 1191 0.079100 1.00800 ; qtot
4.053
1192 CT 71 VAL CG2 1192 -0.319200 12.01000 ; qtot
3.733
1193 HC 71 VAL HG21 1193 0.079100 1.00800 ; qtot
3.812

```
  1194   HC     71   VAL  HG22 1194     0.079100      1.00800 ; qtot
3.892
  1195   HC     71   VAL  HG23 1195     0.079100      1.00800 ; qtot
3.971
  1196    C     71   VAL     C 1196     0.597301     12.01000 ; qtot
4.568
  1197    O     71   VAL     O 1197    -0.567901     16.00000 ; qtot
4.000
  1198    N     72   LYS     N 1198    -0.347900     14.01000 ; qtot
3.652
  1199    H     72   LYS     H 1199     0.274700      1.00800 ; qtot
3.927
  1200   CX     72   LYS    CA 1200    -0.240000     12.01000 ; qtot
3.687
  1201   H1     72   LYS    HA 1201     0.142600      1.00800 ; qtot
3.829
  1202   C8     72   LYS    CB 1202    -0.009400     12.01000 ; qtot
3.820
  1203   HC     72   LYS   HB2 1203     0.036200      1.00800 ; qtot
3.856
  1204   HC     72   LYS   HB3 1204     0.036200      1.00800 ; qtot
3.892
  1205   C8     72   LYS    CG 1205     0.018700     12.01000 ; qtot
3.911
  1206   HC     72   LYS   HG2 1206     0.010300      1.00800 ; qtot
3.921
  1207   HC     72   LYS   HG3 1207     0.010300      1.00800 ; qtot
3.932
  1208   C8     72   LYS    CD 1208    -0.047900     12.01000 ; qtot
3.884
  1209   HC     72   LYS   HD2 1209     0.062100      1.00800 ; qtot
3.946
  1210   HC     72   LYS   HD3 1210     0.062100      1.00800 ; qtot
4.008
  1211   C8     72   LYS    CE 1211    -0.014300     12.01000 ; qtot
3.994
  1212   HP     72   LYS   HE2 1212     0.113500      1.00800 ; qtot
4.107
  1213   HP     72   LYS   HE3 1213     0.113500      1.00800 ; qtot
4.221
  1214   N3     72   LYS    NZ 1214    -0.385400     14.01000 ; qtot
3.835
  1215    H     72   LYS   HZ1 1215     0.340000      1.00800 ; qtot
4.175
  1216    H     72   LYS   HZ2 1216     0.340000      1.00800 ; qtot
4.515
  1217    H     72   LYS   HZ3 1217     0.340000      1.00800 ; qtot
4.855
  1218    C     72   LYS     C 1218     0.734101     12.01000 ; qtot
5.589
  1219    O     72   LYS     O 1219    -0.589401     16.00000 ; qtot
5.000
```

1220 N 73 LEU N 1220 -0.415700 14.01000 ; qtot
4.584
1221 H 73 LEU H 1221 0.271900 1.00800 ; qtot
4.856
1222 CX 73 LEU CA 1222 -0.051800 12.01000 ; qtot
4.804
1223 H1 73 LEU HA 1223 0.092200 1.00800 ; qtot
4.897
1224 A2C 73 LEU CB 1224 -0.110200 12.01000 ; qtot
4.786
1225 HC 73 LEU HB2 1225 0.045700 1.00800 ; qtot
4.832
1226 HC 73 LEU HB3 1226 0.045700 1.00800 ; qtot
4.878
1227 A3C 73 LEU CG 1227 0.353100 12.01000 ; qtot
5.231
1228 HC 73 LEU HG 1228 -0.036100 1.00800 ; qtot
5.195
1229 CT 73 LEU CD1 1229 -0.412100 12.01000 ; qtot
4.783
1230 HC 73 LEU HD11 1230 0.100000 1.00800 ; qtot
4.883
1231 HC 73 LEU HD12 1231 0.100000 1.00800 ; qtot
4.983
1232 HC 73 LEU HD13 1232 0.100000 1.00800 ; qtot
5.083
1233 CT 73 LEU CD2 1233 -0.412100 12.01000 ; qtot
4.671
1234 HC 73 LEU HD21 1234 0.100000 1.00800 ; qtot
4.771
1235 HC 73 LEU HD22 1235 0.100000 1.00800 ; qtot
4.871
1236 HC 73 LEU HD23 1236 0.100000 1.00800 ; qtot
4.971
1237 C 73 LEU C 1237 0.597301 12.01000 ; qtot
5.568
1238 O 73 LEU O 1238 -0.567901 16.00000 ; qtot
5.000
1239 N 74 TYR N 1239 -0.415700 14.01000 ; qtot
4.584
1240 H 74 TYR H 1240 0.271900 1.00800 ; qtot
4.856
1241 CX 74 TYR CA 1241 -0.001400 12.01000 ; qtot
4.855
1242 H1 74 TYR HA 1242 0.087600 1.00800 ; qtot
4.942
1243 CT 74 TYR CB 1243 -0.015200 12.01000 ; qtot
4.927
1244 HC 74 TYR HB2 1244 0.029500 1.00800 ; qtot
4.957
1245 HC 74 TYR HB3 1245 0.029500 1.00800 ; qtot
4.986

1246 CA 74 TYR CG 1246 -0.001100 12.01000 ; qtot
4.985
1247 CA 74 TYR CD1 1247 -0.190600 12.01000 ; qtot
4.795
1248 HA 74 TYR HD1 1248 0.169900 1.00800 ; qtot
4.964
1249 CA 74 TYR CE1 1249 -0.234100 12.01000 ; qtot
4.730
1250 HA 74 TYR HE1 1250 0.165600 1.00800 ; qtot
4.896
1251 C 74 TYR CZ 1251 0.322600 12.01000 ; qtot
5.219
1252 OH 74 TYR OH 1252 -0.557901 16.00000 ; qtot
4.661
1253 HO 74 TYR HH 1253 0.399200 1.00800 ; qtot
5.060
1254 CA 74 TYR CE2 1254 -0.234100 12.01000 ; qtot
4.826
1255 HA 74 TYR HE2 1255 0.165600 1.00800 ; qtot
4.991
1256 CA 74 TYR CD2 1256 -0.190600 12.01000 ; qtot
4.801
1257 HA 74 TYR HD2 1257 0.169900 1.00800 ; qtot
4.971
1258 C 74 TYR C 1258 0.597301 12.01000 ; qtot
5.568
1259 O 74 TYR O 1259 -0.567901 16.00000 ; qtot
5.000
1260 N 75 GLU N 1260 -0.516301 14.01000 ; qtot
4.484
1261 H 75 GLU H 1261 0.293600 1.00800 ; qtot
4.777
1262 CX 75 GLU CA 1262 0.039700 12.01000 ; qtot
4.817
1263 H1 75 GLU HA 1263 0.110500 1.00800 ; qtot
4.928
1264 A2C 75 GLU CB 1264 0.056000 12.01000 ; qtot
4.984
1265 HC 75 GLU HB2 1265 -0.017300 1.00800 ; qtot
4.966
1266 HC 75 GLU HB3 1266 -0.017300 1.00800 ; qtot
4.949
1267 A2C 75 GLU CG 1267 0.013600 12.01000 ; qtot
4.963
1268 HC 75 GLU HG2 1268 -0.042500 1.00800 ; qtot
4.920
1269 HC 75 GLU HG3 1269 -0.042500 1.00800 ; qtot
4.878
1270 CO 75 GLU CD 1270 0.805401 12.01000 ; qtot
5.683
1271 O2 75 GLU OE1 1271 -0.818801 16.00000 ; qtot
4.864

1272 O2 75 GLU OE2 1272 -0.818801 16.00000 ; qtot 4.045
1273 C 75 GLU C 1273 0.536601 12.01000 ; qtot 4.582
1274 O 75 GLU O 1274 -0.581901 16.00000 ; qtot 4.000
1275 N 76 THR N 1275 -0.415700 14.01000 ; qtot 3.584
1276 H 76 THR H 1276 0.271900 1.00800 ; qtot 3.856
1277 CX 76 THR CA 1277 -0.038900 12.01000 ; qtot 3.817
1278 H1 76 THR HA 1278 0.100700 1.00800 ; qtot 3.918
1279 A3C 76 THR CB 1279 0.365400 12.01000 ; qtot 4.283
1280 H1 76 THR HB 1280 0.004300 1.00800 ; qtot 4.288
1281 CT 76 THR CG2 1281 -0.243800 12.01000 ; qtot 4.044
1282 HC 76 THR HG21 1282 0.064200 1.00800 ; qtot 4.108
1283 HC 76 THR HG22 1283 0.064200 1.00800 ; qtot 4.172
1284 HC 76 THR HG23 1284 0.064200 1.00800 ; qtot 4.237
1285 OH 76 THR OG1 1285 -0.676101 16.00000 ; qtot 3.560
1286 HO 76 THR HG1 1286 0.410200 1.00800 ; qtot 3.971
1287 C 76 THR C 1287 0.597301 12.01000 ; qtot 4.568
1288 O 76 THR O 1288 -0.567901 16.00000 ; qtot 4.000
1289 N 77 TYR N 1289 -0.415700 14.01000 ; qtot 3.584
1290 H 77 TYR H 1290 0.271900 1.00800 ; qtot 3.856
1291 CX 77 TYR CA 1291 -0.001400 12.01000 ; qtot 3.855
1292 H1 77 TYR HA 1292 0.087600 1.00800 ; qtot 3.942
1293 CT 77 TYR CB 1293 -0.015200 12.01000 ; qtot 3.927
1294 HC 77 TYR HB2 1294 0.029500 1.00800 ; qtot 3.957
1295 HC 77 TYR HB3 1295 0.029500 1.00800 ; qtot 3.986
1296 CA 77 TYR CG 1296 -0.001100 12.01000 ; qtot 3.985
1297 CA 77 TYR CD1 1297 -0.190600 12.01000 ; qtot 3.795

1298 HA 77 TYR HD1 1298 0.169900 1.00800 ; qtot
3.964
1299 CA 77 TYR CE1 1299 -0.234100 12.01000 ; qtot
3.730
1300 HA 77 TYR HE1 1300 0.165600 1.00800 ; qtot
3.896
1301 C 77 TYR CZ 1301 0.322600 12.01000 ; qtot
4.219
1302 OH 77 TYR OH 1302 -0.557901 16.00000 ; qtot
3.661
1303 HO 77 TYR HH 1303 0.399200 1.00800 ; qtot
4.060
1304 CA 77 TYR CE2 1304 -0.234100 12.01000 ; qtot
3.826
1305 HA 77 TYR HE2 1305 0.165600 1.00800 ; qtot
3.991
1306 CA 77 TYR CD2 1306 -0.190600 12.01000 ; qtot
3.801
1307 HA 77 TYR HD2 1307 0.169900 1.00800 ; qtot
3.971
1308 C 77 TYR C 1308 0.597301 12.01000 ; qtot
4.568
1309 O 77 TYR O 1309 -0.567901 16.00000 ; qtot
4.000
1310 N 78 GLU N 1310 -0.516301 14.01000 ; qtot
3.484
1311 H 78 GLU H 1311 0.293600 1.00800 ; qtot
3.777
1312 CX 78 GLU CA 1312 0.039700 12.01000 ; qtot
3.817
1313 H1 78 GLU HA 1313 0.110500 1.00800 ; qtot
3.928
1314 A2C 78 GLU CB 1314 0.056000 12.01000 ; qtot
3.984
1315 HC 78 GLU HB2 1315 -0.017300 1.00800 ; qtot
3.966
1316 HC 78 GLU HB3 1316 -0.017300 1.00800 ; qtot
3.949
1317 A2C 78 GLU CG 1317 0.013600 12.01000 ; qtot
3.963
1318 HC 78 GLU HG2 1318 -0.042500 1.00800 ; qtot
3.920
1319 HC 78 GLU HG3 1319 -0.042500 1.00800 ; qtot
3.878
1320 CO 78 GLU CD 1320 0.805401 12.01000 ; qtot
4.683
1321 O2 78 GLU OE1 1321 -0.818801 16.00000 ; qtot
3.864
1322 O2 78 GLU OE2 1322 -0.818801 16.00000 ; qtot
3.045
1323 C 78 GLU C 1323 0.536601 12.01000 ; qtot
3.582

1324 O 78 GLU O 1324 -0.581901 16.00000 ; qtot
3.000
1325 N 79 ASN N 1325 -0.415700 14.01000 ; qtot
2.584
1326 H 79 ASN H 1326 0.271900 1.00800 ; qtot
2.856
1327 CX 79 ASN CA 1327 0.014300 12.01000 ; qtot
2.871
1328 H1 79 ASN HA 1328 0.104800 1.00800 ; qtot
2.975
1329 A2C 79 ASN CB 1329 -0.204100 12.01000 ; qtot
2.771
1330 HC 79 ASN HB2 1330 0.079700 1.00800 ; qtot
2.851
1331 HC 79 ASN HB3 1331 0.079700 1.00800 ; qtot
2.931
1332 C 79 ASN CG 1332 0.713001 12.01000 ; qtot
3.644
1333 O 79 ASN OD1 1333 -0.593101 16.00000 ; qtot
3.051
1334 N 79 ASN ND2 1334 -0.919101 14.01000 ; qtot
2.131
1335 H 79 ASN HD21 1335 0.419600 1.00800 ; qtot
2.551
1336 H 79 ASN HD22 1336 0.419600 1.00800 ; qtot
2.971
1337 C 79 ASN C 1337 0.597301 12.01000 ; qtot
3.568
1338 O 79 ASN O 1338 -0.567901 16.00000 ; qtot
3.000
1339 N 80 ASP N 1339 -0.516301 14.01000 ; qtot
2.484
1340 H 80 ASP H 1340 0.293600 1.00800 ; qtot
2.777
1341 CX 80 ASP CA 1341 0.038100 12.01000 ; qtot
2.815
1342 H1 80 ASP HA 1342 0.088000 1.00800 ; qtot
2.903
1343 A2C 80 ASP CB 1343 -0.030300 12.01000 ; qtot
2.873
1344 HC 80 ASP HB2 1344 -0.012200 1.00800 ; qtot
2.861
1345 HC 80 ASP HB3 1345 -0.012200 1.00800 ; qtot
2.849
1346 CO 80 ASP CG 1346 0.799401 12.01000 ; qtot
3.648
1347 O2 80 ASP OD1 1347 -0.801401 16.00000 ; qtot
2.847
1348 O2 80 ASP OD2 1348 -0.801401 16.00000 ; qtot
2.045
1349 C 80 ASP C 1349 0.536601 12.01000 ; qtot
2.582

1350 O 80 ASP O 1350 -0.581901 16.00000 ; qtot
2.000
1351 N 81 ASN N 1351 -0.415700 14.01000 ; qtot
1.584
1352 H 81 ASN H 1352 0.271900 1.00800 ; qtot
1.856
1353 CX 81 ASN CA 1353 0.014300 12.01000 ; qtot
1.871
1354 H1 81 ASN HA 1354 0.104800 1.00800 ; qtot
1.975
1355 A2C 81 ASN CB 1355 -0.204100 12.01000 ; qtot
1.771
1356 HC 81 ASN HB2 1356 0.079700 1.00800 ; qtot
1.851
1357 HC 81 ASN HB3 1357 0.079700 1.00800 ; qtot
1.931
1358 C 81 ASN CG 1358 0.713001 12.01000 ; qtot
2.644
1359 O 81 ASN OD1 1359 -0.593101 16.00000 ; qtot
2.051
1360 N 81 ASN ND2 1360 -0.919101 14.01000 ; qtot
1.131
1361 H 81 ASN HD21 1361 0.419600 1.00800 ; qtot
1.551
1362 H 81 ASN HD22 1362 0.419600 1.00800 ; qtot
1.971
1363 C 81 ASN C 1363 0.597301 12.01000 ; qtot
2.568
1364 O 81 ASN O 1364 -0.567901 16.00000 ; qtot
2.000
1365 N 82 TYR N 1365 -0.415700 14.01000 ; qtot
1.584
1366 H 82 TYR H 1366 0.271900 1.00800 ; qtot
1.856
1367 CX 82 TYR CA 1367 -0.001400 12.01000 ; qtot
1.855
1368 H1 82 TYR HA 1368 0.087600 1.00800 ; qtot
1.942
1369 CT 82 TYR CB 1369 -0.015200 12.01000 ; qtot
1.927
1370 HC 82 TYR HB2 1370 0.029500 1.00800 ; qtot
1.957
1371 HC 82 TYR HB3 1371 0.029500 1.00800 ; qtot
1.986
1372 CA 82 TYR CG 1372 -0.001100 12.01000 ; qtot
1.985
1373 CA 82 TYR CD1 1373 -0.190600 12.01000 ; qtot
1.795
1374 HA 82 TYR HD1 1374 0.169900 1.00800 ; qtot
1.964
1375 CA 82 TYR CE1 1375 -0.234100 12.01000 ; qtot
1.730

1376 HA 82 TYR HE1 1376 0.165600 1.00800 ; qtot 1.896
1377 C 82 TYR CZ 1377 0.322600 12.01000 ; qtot 2.219
1378 OH 82 TYR OH 1378 -0.557901 16.00000 ; qtot 1.661
1379 HO 82 TYR HH 1379 0.399200 1.00800 ; qtot 2.060
1380 CA 82 TYR CE2 1380 -0.234100 12.01000 ; qtot 1.826
1381 HA 82 TYR HE2 1381 0.165600 1.00800 ; qtot 1.991
1382 CA 82 TYR CD2 1382 -0.190600 12.01000 ; qtot 1.801
1383 HA 82 TYR HD2 1383 0.169900 1.00800 ; qtot 1.971
1384 C 82 TYR C 1384 0.597301 12.01000 ; qtot 2.568
1385 O 82 TYR O 1385 -0.567901 16.00000 ; qtot 2.000
1386 N 83 ILE N 1386 -0.415700 14.01000 ; qtot 1.584
1387 H 83 ILE H 1387 0.271900 1.00800 ; qtot 1.856
1388 CX 83 ILE CA 1388 -0.059700 12.01000 ; qtot 1.797
1389 H1 83 ILE HA 1389 0.086900 1.00800 ; qtot 1.883
1390 A3C 83 ILE CB 1390 0.130300 12.01000 ; qtot 2.014
1391 HC 83 ILE HB 1391 0.018700 1.00800 ; qtot 2.032
1392 CT 83 ILE CG2 1392 -0.320400 12.01000 ; qtot 1.712
1393 HC 83 ILE HG21 1393 0.088200 1.00800 ; qtot 1.800
1394 HC 83 ILE HG22 1394 0.088200 1.00800 ; qtot 1.888
1395 HC 83 ILE HG23 1395 0.088200 1.00800 ; qtot 1.977
1396 A2C 83 ILE CG1 1396 -0.043000 12.01000 ; qtot 1.934
1397 HC 83 ILE HG12 1397 0.023600 1.00800 ; qtot 1.957
1398 HC 83 ILE HG13 1398 0.023600 1.00800 ; qtot 1.981
1399 CT 83 ILE CD1 1399 -0.066000 12.01000 ; qtot 1.915
1400 HC 83 ILE HD11 1400 0.018600 1.00800 ; qtot 1.933
1401 HC 83 ILE HD12 1401 0.018600 1.00800 ; qtot 1.952

1402 HC 83 ILE HD13 1402 0.018600 1.00800 ; qtot
1.971
1403 C 83 ILE C 1403 0.597301 12.01000 ; qtot
2.568
1404 O 83 ILE O 1404 -0.567901 16.00000 ; qtot
2.000
1405 N 84 TYR N 1405 -0.415700 14.01000 ; qtot
1.584
1406 H 84 TYR H 1406 0.271900 1.00800 ; qtot
1.856
1407 CX 84 TYR CA 1407 -0.001400 12.01000 ; qtot
1.855
1408 H1 84 TYR HA 1408 0.087600 1.00800 ; qtot
1.942
1409 CT 84 TYR CB 1409 -0.015200 12.01000 ; qtot
1.927
1410 HC 84 TYR HB2 1410 0.029500 1.00800 ; qtot
1.957
1411 HC 84 TYR HB3 1411 0.029500 1.00800 ; qtot
1.986
1412 CA 84 TYR CG 1412 -0.001100 12.01000 ; qtot
1.985
1413 CA 84 TYR CD1 1413 -0.190600 12.01000 ; qtot
1.795
1414 HA 84 TYR HD1 1414 0.169900 1.00800 ; qtot
1.964
1415 CA 84 TYR CE1 1415 -0.234100 12.01000 ; qtot
1.730
1416 HA 84 TYR HE1 1416 0.165600 1.00800 ; qtot
1.896
1417 C 84 TYR CZ 1417 0.322600 12.01000 ; qtot
2.219
1418 OH 84 TYR OH 1418 -0.557901 16.00000 ; qtot
1.661
1419 HO 84 TYR HH 1419 0.399200 1.00800 ; qtot
2.060
1420 CA 84 TYR CE2 1420 -0.234100 12.01000 ; qtot
1.826
1421 HA 84 TYR HE2 1421 0.165600 1.00800 ; qtot
1.991
1422 CA 84 TYR CD2 1422 -0.190600 12.01000 ; qtot
1.801
1423 HA 84 TYR HD2 1423 0.169900 1.00800 ; qtot
1.971
1424 C 84 TYR C 1424 0.597301 12.01000 ; qtot
2.568
1425 O 84 TYR O 1425 -0.567901 16.00000 ; qtot
2.000
1426 N 85 LEU N 1426 -0.415700 14.01000 ; qtot
1.584
1427 H 85 LEU H 1427 0.271900 1.00800 ; qtot
1.856

1428 CX 85 LEU CA 1428 -0.051800 12.01000 ; qtot
1.804
1429 H1 85 LEU HA 1429 0.092200 1.00800 ; qtot
1.897
1430 A2C 85 LEU CB 1430 -0.110200 12.01000 ; qtot
1.786
1431 HC 85 LEU HB2 1431 0.045700 1.00800 ; qtot
1.832
1432 HC 85 LEU HB3 1432 0.045700 1.00800 ; qtot
1.878
1433 A3C 85 LEU CG 1433 0.353100 12.01000 ; qtot
2.231
1434 HC 85 LEU HG 1434 -0.036100 1.00800 ; qtot
2.195
1435 CT 85 LEU CD1 1435 -0.412100 12.01000 ; qtot
1.783
1436 HC 85 LEU HD11 1436 0.100000 1.00800 ; qtot
1.883
1437 HC 85 LEU HD12 1437 0.100000 1.00800 ; qtot
1.983
1438 HC 85 LEU HD13 1438 0.100000 1.00800 ; qtot
2.083
1439 CT 85 LEU CD2 1439 -0.412100 12.01000 ; qtot
1.671
1440 HC 85 LEU HD21 1440 0.100000 1.00800 ; qtot
1.771
1441 HC 85 LEU HD22 1441 0.100000 1.00800 ; qtot
1.871
1442 HC 85 LEU HD23 1442 0.100000 1.00800 ; qtot
1.971
1443 C 85 LEU C 1443 0.597301 12.01000 ; qtot
2.568
1444 O 85 LEU O 1444 -0.567901 16.00000 ; qtot
2.000
1445 N 86 ILE N 1445 -0.415700 14.01000 ; qtot
1.584
1446 H 86 ILE H 1446 0.271900 1.00800 ; qtot
1.856
1447 CX 86 ILE CA 1447 -0.059700 12.01000 ; qtot
1.797
1448 H1 86 ILE HA 1448 0.086900 1.00800 ; qtot
1.883
1449 A3C 86 ILE CB 1449 0.130300 12.01000 ; qtot
2.014
1450 HC 86 ILE HB 1450 0.018700 1.00800 ; qtot
2.032
1451 CT 86 ILE CG2 1451 -0.320400 12.01000 ; qtot
1.712
1452 HC 86 ILE HG21 1452 0.088200 1.00800 ; qtot
1.800
1453 HC 86 ILE HG22 1453 0.088200 1.00800 ; qtot
1.888

1454 HC 86 ILE HG23 1454 0.088200 1.00800 ; qtot
1.977
1455 A2C 86 ILE CG1 1455 -0.043000 12.01000 ; qtot
1.934
1456 HC 86 ILE HG12 1456 0.023600 1.00800 ; qtot
1.957
1457 HC 86 ILE HG13 1457 0.023600 1.00800 ; qtot
1.981
1458 CT 86 ILE CD1 1458 -0.066000 12.01000 ; qtot
1.915
1459 HC 86 ILE HD11 1459 0.018600 1.00800 ; qtot
1.933
1460 HC 86 ILE HD12 1460 0.018600 1.00800 ; qtot
1.952
1461 HC 86 ILE HD13 1461 0.018600 1.00800 ; qtot
1.971
1462 C 86 ILE C 1462 0.597301 12.01000 ; qtot
2.568
1463 O 86 ILE O 1463 -0.567901 16.00000 ; qtot
2.000
1464 N 87 MET N 1464 -0.415700 14.01000 ; qtot
1.584
1465 H 87 MET H 1465 0.271900 1.00800 ; qtot
1.856
1466 CX 87 MET CA 1466 -0.023700 12.01000 ; qtot
1.833
1467 H1 87 MET HA 1467 0.088000 1.00800 ; qtot
1.921
1468 A2C 87 MET CB 1468 0.034200 12.01000 ; qtot
1.955
1469 HC 87 MET HB2 1469 0.024100 1.00800 ; qtot
1.979
1470 HC 87 MET HB3 1470 0.024100 1.00800 ; qtot
2.003
1471 A2C 87 MET CG 1471 0.001800 12.01000 ; qtot
2.005
1472 H1 87 MET HG2 1472 0.044000 1.00800 ; qtot
2.049
1473 H1 87 MET HG3 1473 0.044000 1.00800 ; qtot
2.093
1474 S 87 MET SD 1474 -0.273700 32.06000 ; qtot
1.819
1475 CT 87 MET CE 1475 -0.053600 12.01000 ; qtot
1.765
1476 H1 87 MET HE1 1476 0.068400 1.00800 ; qtot
1.834
1477 H1 87 MET HE2 1477 0.068400 1.00800 ; qtot
1.902
1478 H1 87 MET HE3 1478 0.068400 1.00800 ; qtot
1.971
1479 C 87 MET C 1479 0.597301 12.01000 ; qtot
2.568

1480 O 87 MET O 1480 -0.567901 16.00000 ; qtot
2.000
1481 N 88 GLU N 1481 -0.516301 14.01000 ; qtot
1.484
1482 H 88 GLU H 1482 0.293600 1.00800 ; qtot
1.777
1483 CX 88 GLU CA 1483 0.039700 12.01000 ; qtot
1.817
1484 H1 88 GLU HA 1484 0.110500 1.00800 ; qtot
1.928
1485 A2C 88 GLU CB 1485 0.056000 12.01000 ; qtot
1.984
1486 HC 88 GLU HB2 1486 -0.017300 1.00800 ; qtot
1.966
1487 HC 88 GLU HB3 1487 -0.017300 1.00800 ; qtot
1.949
1488 A2C 88 GLU CG 1488 0.013600 12.01000 ; qtot
1.963
1489 HC 88 GLU HG2 1489 -0.042500 1.00800 ; qtot
1.920
1490 HC 88 GLU HG3 1490 -0.042500 1.00800 ; qtot
1.878
1491 CO 88 GLU CD 1491 0.805401 12.01000 ; qtot
2.683
1492 O2 88 GLU OE1 1492 -0.818801 16.00000 ; qtot
1.864
1493 O2 88 GLU OE2 1493 -0.818801 16.00000 ; qtot
1.045
1494 C 88 GLU C 1494 0.536601 12.01000 ; qtot
1.582
1495 O 88 GLU O 1495 -0.581901 16.00000 ; qtot
1.000
1496 N 89 LEU N 1496 -0.415700 14.01000 ; qtot
0.584
1497 H 89 LEU H 1497 0.271900 1.00800 ; qtot
0.856
1498 CX 89 LEU CA 1498 -0.051800 12.01000 ; qtot
0.804
1499 H1 89 LEU HA 1499 0.092200 1.00800 ; qtot
0.897
1500 A2C 89 LEU CB 1500 -0.110200 12.01000 ; qtot
0.786
1501 HC 89 LEU HB2 1501 0.045700 1.00800 ; qtot
0.832
1502 HC 89 LEU HB3 1502 0.045700 1.00800 ; qtot
0.878
1503 A3C 89 LEU CG 1503 0.353100 12.01000 ; qtot
1.231
1504 HC 89 LEU HG 1504 -0.036100 1.00800 ; qtot
1.195
1505 CT 89 LEU CD1 1505 -0.412100 12.01000 ; qtot
0.783

1506 HC 89 LEU HD11 1506 0.100000 1.00800 ; qtot
0.883
1507 HC 89 LEU HD12 1507 0.100000 1.00800 ; qtot
0.983
1508 HC 89 LEU HD13 1508 0.100000 1.00800 ; qtot
1.083
1509 CT 89 LEU CD2 1509 -0.412100 12.01000 ; qtot
0.671
1510 HC 89 LEU HD21 1510 0.100000 1.00800 ; qtot
0.771
1511 HC 89 LEU HD22 1511 0.100000 1.00800 ; qtot
0.871
1512 HC 89 LEU HD23 1512 0.100000 1.00800 ; qtot
0.971
1513 C 89 LEU C 1513 0.597301 12.01000 ; qtot
1.568
1514 O 89 LEU O 1514 -0.567901 16.00000 ; qtot
1.000
1515 N 90 CYS N 1515 -0.415700 14.01000 ; qtot
0.584
1516 H 90 CYS H 1516 0.271900 1.00800 ; qtot
0.856
1517 CX 90 CYS CA 1517 0.021300 12.01000 ; qtot
0.878
1518 H1 90 CYS HA 1518 0.112400 1.00800 ; qtot
0.990
1519 A2C 90 CYS CB 1519 -0.123100 12.01000 ; qtot
0.867
1520 H1 90 CYS HB2 1520 0.111200 1.00800 ; qtot
0.978
1521 H1 90 CYS HB3 1521 0.111200 1.00800 ; qtot
1.089
1522 SH 90 CYS SG 1522 -0.311900 32.06000 ; qtot
0.777
1523 HS 90 CYS HG 1523 0.193300 1.00800 ; qtot
0.971
1524 C 90 CYS C 1524 0.597301 12.01000 ; qtot
1.568
1525 O 90 CYS O 1525 -0.567901 16.00000 ; qtot
1.000
1526 N 91 SER N 1526 -0.415700 14.01000 ; qtot
0.584
1527 H 91 SER H 1527 0.271900 1.00800 ; qtot
0.856
1528 CX 91 SER CA 1528 -0.024900 12.01000 ; qtot
0.831
1529 H1 91 SER HA 1529 0.084300 1.00800 ; qtot
0.916
1530 A2C 91 SER CB 1530 0.211700 12.01000 ; qtot
1.127
1531 H1 91 SER HB2 1531 0.035200 1.00800 ; qtot
1.163

1532 H1 91 SER HB3 1532 0.035200 1.00800 ; qtot
1.198
1533 OH 91 SER OG 1533 -0.654601 16.00000 ; qtot
0.543
1534 HO 91 SER HG 1534 0.427500 1.00800 ; qtot
0.971
1535 C 91 SER C 1535 0.597301 12.01000 ; qtot
1.568
1536 O 91 SER O 1536 -0.567901 16.00000 ; qtot
1.000
1537 N 92 GLY N 1537 -0.415700 14.01000 ; qtot
0.584
1538 H 92 GLY H 1538 0.271900 1.00800 ; qtot
0.856
1539 CX 92 GLY CA 1539 -0.025200 12.01000 ; qtot
0.831
1540 H1 92 GLY HA2 1540 0.069800 1.00800 ; qtot
0.901
1541 H1 92 GLY HA3 1541 0.069800 1.00800 ; qtot
0.971
1542 C 92 GLY C 1542 0.597301 12.01000 ; qtot
1.568
1543 O 92 GLY O 1543 -0.567901 16.00000 ; qtot
1.000
1544 N 93 ARG N 1544 -0.347900 14.01000 ; qtot
0.652
1545 H 93 ARG H 1545 0.274700 1.00800 ; qtot
0.927
1546 CX 93 ARG CA 1546 -0.263700 12.01000 ; qtot
0.663
1547 H1 93 ARG HA 1547 0.156000 1.00800 ; qtot
0.819
1548 C8 93 ARG CB 1548 -0.000700 12.01000 ; qtot
0.818
1549 HC 93 ARG HB2 1549 0.032700 1.00800 ; qtot
0.851
1550 HC 93 ARG HB3 1550 0.032700 1.00800 ; qtot
0.884
1551 C8 93 ARG CG 1551 0.039000 12.01000 ; qtot
0.923
1552 HC 93 ARG HG2 1552 0.028500 1.00800 ; qtot
0.951
1553 HC 93 ARG HG3 1553 0.028500 1.00800 ; qtot
0.980
1554 C8 93 ARG CD 1554 0.048600 12.01000 ; qtot
1.028
1555 H1 93 ARG HD2 1555 0.068700 1.00800 ; qtot
1.097
1556 H1 93 ARG HD3 1556 0.068700 1.00800 ; qtot
1.166
1557 N2 93 ARG NE 1557 -0.529501 14.01000 ; qtot
0.636

```
  1558    H    93   ARG    HE 1558     0.345600      1.00800 ; qtot
0.982
  1559   CA    93   ARG    CZ 1559     0.807601     12.01000 ; qtot
1.790
  1560   N2    93   ARG   NH1 1560    -0.862701     14.01000 ; qtot
0.927
  1561    H    93   ARG  HH11 1561     0.447800      1.00800 ; qtot
1.375
  1562    H    93   ARG  HH12 1562     0.447800      1.00800 ; qtot
1.822
  1563   N2    93   ARG   NH2 1563    -0.862701     14.01000 ; qtot
0.960
  1564    H    93   ARG  HH21 1564     0.447800      1.00800 ; qtot
1.408
  1565    H    93   ARG  HH22 1565     0.447800      1.00800 ; qtot
1.855
  1566    C    93   ARG     C 1566     0.734101     12.01000 ; qtot
2.589
  1567    O    93   ARG     O 1567    -0.589401     16.00000 ; qtot
2.000
  1568    N    94   GLU     N 1568    -0.516301     14.01000 ; qtot
1.484
  1569    H    94   GLU     H 1569     0.293600      1.00800 ; qtot
1.777
  1570   CX    94   GLU    CA 1570     0.039700     12.01000 ; qtot
1.817
  1571   H1    94   GLU    HA 1571     0.110500      1.00800 ; qtot
1.928
  1572  A2C    94   GLU    CB 1572     0.056000     12.01000 ; qtot
1.984
  1573   HC    94   GLU   HB2 1573    -0.017300      1.00800 ; qtot
1.966
  1574   HC    94   GLU   HB3 1574    -0.017300      1.00800 ; qtot
1.949
  1575  A2C    94   GLU    CG 1575     0.013600     12.01000 ; qtot
1.963
  1576   HC    94   GLU   HG2 1576    -0.042500      1.00800 ; qtot
1.920
  1577   HC    94   GLU   HG3 1577    -0.042500      1.00800 ; qtot
1.878
  1578   CO    94   GLU    CD 1578     0.805401     12.01000 ; qtot
2.683
  1579   O2    94   GLU   OE1 1579    -0.818801     16.00000 ; qtot
1.864
  1580   O2    94   GLU   OE2 1580    -0.818801     16.00000 ; qtot
1.045
  1581    C    94   GLU     C 1581     0.536601     12.01000 ; qtot
1.582
  1582    O    94   GLU     O 1582    -0.581901     16.00000 ; qtot
1.000
  1583    N    95   LEU     N 1583    -0.415700     14.01000 ; qtot
0.584
```

1584 H 95 LEU H 1584 0.271900 1.00800 ; qtot
0.856
1585 CX 95 LEU CA 1585 -0.051800 12.01000 ; qtot
0.804
1586 H1 95 LEU HA 1586 0.092200 1.00800 ; qtot
0.897
1587 A2C 95 LEU CB 1587 -0.110200 12.01000 ; qtot
0.786
1588 HC 95 LEU HB2 1588 0.045700 1.00800 ; qtot
0.832
1589 HC 95 LEU HB3 1589 0.045700 1.00800 ; qtot
0.878
1590 A3C 95 LEU CG 1590 0.353100 12.01000 ; qtot
1.231
1591 HC 95 LEU HG 1591 -0.036100 1.00800 ; qtot
1.195
1592 CT 95 LEU CD1 1592 -0.412100 12.01000 ; qtot
0.783
1593 HC 95 LEU HD11 1593 0.100000 1.00800 ; qtot
0.883
1594 HC 95 LEU HD12 1594 0.100000 1.00800 ; qtot
0.983
1595 HC 95 LEU HD13 1595 0.100000 1.00800 ; qtot
1.083
1596 CT 95 LEU CD2 1596 -0.412100 12.01000 ; qtot
0.671
1597 HC 95 LEU HD21 1597 0.100000 1.00800 ; qtot
0.771
1598 HC 95 LEU HD22 1598 0.100000 1.00800 ; qtot
0.871
1599 HC 95 LEU HD23 1599 0.100000 1.00800 ; qtot
0.971
1600 C 95 LEU C 1600 0.597301 12.01000 ; qtot
1.568
1601 O 95 LEU O 1601 -0.567901 16.00000 ; qtot
1.000
1602 N 96 PHE N 1602 -0.415700 14.01000 ; qtot
0.584
1603 H 96 PHE H 1603 0.271900 1.00800 ; qtot
0.856
1604 CX 96 PHE CA 1604 -0.002400 12.01000 ; qtot
0.854
1605 H1 96 PHE HA 1605 0.097800 1.00800 ; qtot
0.952
1606 CT 96 PHE CB 1606 -0.034300 12.01000 ; qtot
0.917
1607 HC 96 PHE HB2 1607 0.029500 1.00800 ; qtot
0.947
1608 HC 96 PHE HB3 1608 0.029500 1.00800 ; qtot
0.976
1609 CA 96 PHE CG 1609 0.011800 12.01000 ; qtot
0.988

1610 CA 96 PHE CD1 1610 -0.125600 12.01000 ; qtot
0.863
1611 HA 96 PHE HD1 1611 0.133000 1.00800 ; qtot
0.996
1612 CA 96 PHE CE1 1612 -0.170400 12.01000 ; qtot
0.825
1613 HA 96 PHE HE1 1613 0.143000 1.00800 ; qtot
0.968
1614 CA 96 PHE CZ 1614 -0.107200 12.01000 ; qtot
0.861
1615 HA 96 PHE HZ 1615 0.129700 1.00800 ; qtot
0.991
1616 CA 96 PHE CE2 1616 -0.170400 12.01000 ; qtot
0.820
1617 HA 96 PHE HE2 1617 0.143000 1.00800 ; qtot
0.963
1618 CA 96 PHE CD2 1618 -0.125600 12.01000 ; qtot
0.838
1619 HA 96 PHE HD2 1619 0.133000 1.00800 ; qtot
0.971
1620 C 96 PHE C 1620 0.597301 12.01000 ; qtot
1.568
1621 O 96 PHE O 1621 -0.567901 16.00000 ; qtot
1.000
1622 N 97 ASP N 1622 -0.516301 14.01000 ; qtot
0.484
1623 H 97 ASP H 1623 0.293600 1.00800 ; qtot
0.777
1624 CX 97 ASP CA 1624 0.038100 12.01000 ; qtot
0.815
1625 H1 97 ASP HA 1625 0.088000 1.00800 ; qtot
0.903
1626 A2C 97 ASP CB 1626 -0.030300 12.01000 ; qtot
0.873
1627 HC 97 ASP HB2 1627 -0.012200 1.00800 ; qtot
0.861
1628 HC 97 ASP HB3 1628 -0.012200 1.00800 ; qtot
0.849
1629 CO 97 ASP CG 1629 0.799401 12.01000 ; qtot
1.648
1630 O2 97 ASP OD1 1630 -0.801401 16.00000 ; qtot
0.847
1631 O2 97 ASP OD2 1631 -0.801401 16.00000 ; qtot
0.045
1632 C 97 ASP C 1632 0.536601 12.01000 ; qtot
0.582
1633 O 97 ASP O 1633 -0.581901 16.00000 ; qtot
0.000
1634 N 98 SER N 1634 -0.415700 14.01000 ; qtot -
0.416
1635 H 98 SER H 1635 0.271900 1.00800 ; qtot -
0.144

```
  1636   CX   98   SER    CA 1636   -0.024900    12.01000 ; qtot -0.169
  1637   H1   98   SER    HA 1637    0.084300     1.00800 ; qtot -0.084
  1638  A2C   98   SER    CB 1638    0.211700    12.01000 ; qtot 0.127
  1639   H1   98   SER   HB2 1639    0.035200     1.00800 ; qtot 0.163
  1640   H1   98   SER   HB3 1640    0.035200     1.00800 ; qtot 0.198
  1641   OH   98   SER    OG 1641   -0.654601    16.00000 ; qtot -0.457
  1642   HO   98   SER    HG 1642    0.427500     1.00800 ; qtot -0.029
  1643    C   98   SER     C 1643    0.597301    12.01000 ; qtot 0.568
  1644    O   98   SER     O 1644   -0.567901    16.00000 ; qtot 0.000
  1645    N   99   ILE     N 1645   -0.415700    14.01000 ; qtot -0.416
  1646    H   99   ILE     H 1646    0.271900     1.00800 ; qtot -0.144
  1647   CX   99   ILE    CA 1647   -0.059700    12.01000 ; qtot -0.203
  1648   H1   99   ILE    HA 1648    0.086900     1.00800 ; qtot -0.117
  1649  A3C   99   ILE    CB 1649    0.130300    12.01000 ; qtot 0.014
  1650   HC   99   ILE    HB 1650    0.018700     1.00800 ; qtot 0.032
  1651   CT   99   ILE   CG2 1651   -0.320400    12.01000 ; qtot -0.288
  1652   HC   99   ILE  HG21 1652    0.088200     1.00800 ; qtot -0.200
  1653   HC   99   ILE  HG22 1653    0.088200     1.00800 ; qtot -0.112
  1654   HC   99   ILE  HG23 1654    0.088200     1.00800 ; qtot -0.023
  1655  A2C   99   ILE   CG1 1655   -0.043000    12.01000 ; qtot -0.066
  1656   HC   99   ILE  HG12 1656    0.023600     1.00800 ; qtot -0.043
  1657   HC   99   ILE  HG13 1657    0.023600     1.00800 ; qtot -0.019
  1658   CT   99   ILE   CD1 1658   -0.066000    12.01000 ; qtot -0.085
  1659   HC   99   ILE  HD11 1659    0.018600     1.00800 ; qtot -0.067
  1660   HC   99   ILE  HD12 1660    0.018600     1.00800 ; qtot -0.048
  1661   HC   99   ILE  HD13 1661    0.018600     1.00800 ; qtot -0.029
```

```
  1662    C    99   ILE     C 1662     0.597301     12.01000 ; qtot
0.568
  1663    O    99   ILE     O 1663    -0.567901     16.00000 ; qtot
0.000
  1664    N   100   ILE     N 1664    -0.415700     14.01000 ; qtot -
0.416
  1665    H   100   ILE     H 1665     0.271900      1.00800 ; qtot -
0.144
  1666   CX   100   ILE    CA 1666    -0.059700     12.01000 ; qtot -
0.203
  1667   H1   100   ILE    HA 1667     0.086900      1.00800 ; qtot -
0.117
  1668  A3C   100   ILE    CB 1668     0.130300     12.01000 ; qtot
0.014
  1669   HC   100   ILE    HB 1669     0.018700      1.00800 ; qtot
0.032
  1670   CT   100   ILE   CG2 1670    -0.320400     12.01000 ; qtot -
0.288
  1671   HC   100   ILE  HG21 1671     0.088200      1.00800 ; qtot -
0.200
  1672   HC   100   ILE  HG22 1672     0.088200      1.00800 ; qtot -
0.112
  1673   HC   100   ILE  HG23 1673     0.088200      1.00800 ; qtot -
0.023
  1674  A2C   100   ILE   CG1 1674    -0.043000     12.01000 ; qtot -
0.066
  1675   HC   100   ILE  HG12 1675     0.023600      1.00800 ; qtot -
0.043
  1676   HC   100   ILE  HG13 1676     0.023600      1.00800 ; qtot -
0.019
  1677   CT   100   ILE   CD1 1677    -0.066000     12.01000 ; qtot -
0.085
  1678   HC   100   ILE  HD11 1678     0.018600      1.00800 ; qtot -
0.067
  1679   HC   100   ILE  HD12 1679     0.018600      1.00800 ; qtot -
0.048
  1680   HC   100   ILE  HD13 1680     0.018600      1.00800 ; qtot -
0.029
  1681    C   100   ILE     C 1681     0.597301     12.01000 ; qtot
0.568
  1682    O   100   ILE     O 1682    -0.567901     16.00000 ; qtot
0.000
  1683    N   101   GLU     N 1683    -0.516301     14.01000 ; qtot -
0.516
  1684    H   101   GLU     H 1684     0.293600      1.00800 ; qtot -
0.223
  1685   CX   101   GLU    CA 1685     0.039700     12.01000 ; qtot -
0.183
  1686   H1   101   GLU    HA 1686     0.110500      1.00800 ; qtot -
0.072
  1687  A2C   101   GLU    CB 1687     0.056000     12.01000 ; qtot -
0.016
```

```
  1688   HC   101   GLU   HB2 1688    -0.017300      1.00800 ; qtot -
0.034
  1689   HC   101   GLU   HB3 1689    -0.017300      1.00800 ; qtot -
0.051
  1690  A2C   101   GLU    CG 1690     0.013600     12.01000 ; qtot -
0.037
  1691   HC   101   GLU   HG2 1691    -0.042500      1.00800 ; qtot -
0.080
  1692   HC   101   GLU   HG3 1692    -0.042500      1.00800 ; qtot -
0.122
  1693   CO   101   GLU    CD 1693     0.805401     12.01000 ; qtot
0.683
  1694   O2   101   GLU   OE1 1694    -0.818801     16.00000 ; qtot -
0.136
  1695   O2   101   GLU   OE2 1695    -0.818801     16.00000 ; qtot -
0.955
  1696    C   101   GLU     C 1696     0.536601     12.01000 ; qtot -
0.418
  1697    O   101   GLU     O 1697    -0.581901     16.00000 ; qtot -
1.000
  1698    N   102   ASN     N 1698    -0.415700     14.01000 ; qtot -
1.416
  1699    H   102   ASN     H 1699     0.271900      1.00800 ; qtot -
1.144
  1700   CX   102   ASN    CA 1700     0.014300     12.01000 ; qtot -
1.130
  1701   H1   102   ASN    HA 1701     0.104800      1.00800 ; qtot -
1.025
  1702  A2C   102   ASN    CB 1702    -0.204100     12.01000 ; qtot -
1.229
  1703   HC   102   ASN   HB2 1703     0.079700      1.00800 ; qtot -
1.149
  1704   HC   102   ASN   HB3 1704     0.079700      1.00800 ; qtot -
1.069
  1705    C   102   ASN    CG 1705     0.713001     12.01000 ; qtot -
0.356
  1706    O   102   ASN   OD1 1706    -0.593101     16.00000 ; qtot -
0.950
  1707    N   102   ASN   ND2 1707    -0.919101     14.01000 ; qtot -
1.869
  1708    H   102   ASN  HD21 1708     0.419600      1.00800 ; qtot -
1.449
  1709    H   102   ASN  HD22 1709     0.419600      1.00800 ; qtot -
1.029
  1710    C   102   ASN     C 1710     0.597301     12.01000 ; qtot -
0.432
  1711    O   102   ASN     O 1711    -0.567901     16.00000 ; qtot -
1.000
  1712    N   103   GLY     N 1712    -0.415700     14.01000 ; qtot -
1.416
  1713    H   103   GLY     H 1713     0.271900      1.00800 ; qtot -
1.144
```

1714 CX 103 GLY CA 1714 -0.025200 12.01000 ; qtot -
1.169
1715 H1 103 GLY HA2 1715 0.069800 1.00800 ; qtot -
1.099
1716 H1 103 GLY HA3 1716 0.069800 1.00800 ; qtot -
1.029
1717 C 103 GLY C 1717 0.597301 12.01000 ; qtot -
0.432
1718 O 103 GLY O 1718 -0.567901 16.00000 ; qtot -
1.000
1719 N 104 SER N 1719 -0.415700 14.01000 ; qtot -
1.416
1720 H 104 SER H 1720 0.271900 1.00800 ; qtot -
1.144
1721 CX 104 SER CA 1721 -0.024900 12.01000 ; qtot -
1.169
1722 H1 104 SER HA 1722 0.084300 1.00800 ; qtot -
1.084
1723 A2C 104 SER CB 1723 0.211700 12.01000 ; qtot -
0.873
1724 H1 104 SER HB2 1724 0.035200 1.00800 ; qtot -
0.838
1725 H1 104 SER HB3 1725 0.035200 1.00800 ; qtot -
0.802
1726 OH 104 SER OG 1726 -0.654601 16.00000 ; qtot -
1.457
1727 HO 104 SER HG 1727 0.427500 1.00800 ; qtot -
1.029
1728 C 104 SER C 1728 0.597301 12.01000 ; qtot -
0.432
1729 O 104 SER O 1729 -0.567901 16.00000 ; qtot -
1.000
1730 N 105 PHE N 1730 -0.415700 14.01000 ; qtot -
1.416
1731 H 105 PHE H 1731 0.271900 1.00800 ; qtot -
1.144
1732 CX 105 PHE CA 1732 -0.002400 12.01000 ; qtot -
1.146
1733 H1 105 PHE HA 1733 0.097800 1.00800 ; qtot -
1.048
1734 CT 105 PHE CB 1734 -0.034300 12.01000 ; qtot -
1.083
1735 HC 105 PHE HB2 1735 0.029500 1.00800 ; qtot -
1.053
1736 HC 105 PHE HB3 1736 0.029500 1.00800 ; qtot -
1.024
1737 CA 105 PHE CG 1737 0.011800 12.01000 ; qtot -
1.012
1738 CA 105 PHE CD1 1738 -0.125600 12.01000 ; qtot -
1.138
1739 HA 105 PHE HD1 1739 0.133000 1.00800 ; qtot -
1.005

1740 CA 105 PHE CE1 1740 -0.170400 12.01000 ; qtot -
1.175
1741 HA 105 PHE HE1 1741 0.143000 1.00800 ; qtot -
1.032
1742 CA 105 PHE CZ 1742 -0.107200 12.01000 ; qtot -
1.139
1743 HA 105 PHE HZ 1743 0.129700 1.00800 ; qtot -
1.009
1744 CA 105 PHE CE2 1744 -0.170400 12.01000 ; qtot -
1.180
1745 HA 105 PHE HE2 1745 0.143000 1.00800 ; qtot -
1.037
1746 CA 105 PHE CD2 1746 -0.125600 12.01000 ; qtot -
1.162
1747 HA 105 PHE HD2 1747 0.133000 1.00800 ; qtot -
1.029
1748 C 105 PHE C 1748 0.597301 12.01000 ; qtot -
0.432
1749 O 105 PHE O 1749 -0.567901 16.00000 ; qtot -
1.000
1750 N 106 THR N 1750 -0.415700 14.01000 ; qtot -
1.416
1751 H 106 THR H 1751 0.271900 1.00800 ; qtot -
1.144
1752 CX 106 THR CA 1752 -0.038900 12.01000 ; qtot -
1.183
1753 H1 106 THR HA 1753 0.100700 1.00800 ; qtot -
1.082
1754 A3C 106 THR CB 1754 0.365400 12.01000 ; qtot -
0.717
1755 H1 106 THR HB 1755 0.004300 1.00800 ; qtot -
0.712
1756 CT 106 THR CG2 1756 -0.243800 12.01000 ; qtot -
0.956
1757 HC 106 THR HG21 1757 0.064200 1.00800 ; qtot -
0.892
1758 HC 106 THR HG22 1758 0.064200 1.00800 ; qtot -
0.828
1759 HC 106 THR HG23 1759 0.064200 1.00800 ; qtot -
0.764
1760 OH 106 THR OG1 1760 -0.676101 16.00000 ; qtot -
1.440
1761 HO 106 THR HG1 1761 0.410200 1.00800 ; qtot -
1.029
1762 C 106 THR C 1762 0.597301 12.01000 ; qtot -
0.432
1763 O 106 THR O 1763 -0.567901 16.00000 ; qtot -
1.000
1764 N 107 GLU N 1764 -0.516301 14.01000 ; qtot -
1.516
1765 H 107 GLU H 1765 0.293600 1.00800 ; qtot -
1.223

1766 CX 107 GLU CA 1766 0.039700 12.01000 ; qtot -
1.183
1767 H1 107 GLU HA 1767 0.110500 1.00800 ; qtot -
1.073
1768 A2C 107 GLU CB 1768 0.056000 12.01000 ; qtot -
1.017
1769 HC 107 GLU HB2 1769 -0.017300 1.00800 ; qtot -
1.034
1770 HC 107 GLU HB3 1770 -0.017300 1.00800 ; qtot -
1.051
1771 A2C 107 GLU CG 1771 0.013600 12.01000 ; qtot -
1.038
1772 HC 107 GLU HG2 1772 -0.042500 1.00800 ; qtot -
1.080
1773 HC 107 GLU HG3 1773 -0.042500 1.00800 ; qtot -
1.123
1774 CO 107 GLU CD 1774 0.805401 12.01000 ; qtot -
0.317
1775 O2 107 GLU OE1 1775 -0.818801 16.00000 ; qtot -
1.136
1776 O2 107 GLU OE2 1776 -0.818801 16.00000 ; qtot -
1.955
1777 C 107 GLU C 1777 0.536601 12.01000 ; qtot -
1.418
1778 O 107 GLU O 1778 -0.581901 16.00000 ; qtot -
2.000
1779 N 108 LYS N 1779 -0.347900 14.01000 ; qtot -
2.348
1780 H 108 LYS H 1780 0.274700 1.00800 ; qtot -
2.073
1781 CX 108 LYS CA 1781 -0.240000 12.01000 ; qtot -
2.313
1782 H1 108 LYS HA 1782 0.142600 1.00800 ; qtot -
2.171
1783 C8 108 LYS CB 1783 -0.009400 12.01000 ; qtot -
2.180
1784 HC 108 LYS HB2 1784 0.036200 1.00800 ; qtot -
2.144
1785 HC 108 LYS HB3 1785 0.036200 1.00800 ; qtot -
2.108
1786 C8 108 LYS CG 1786 0.018700 12.01000 ; qtot -
2.089
1787 HC 108 LYS HG2 1787 0.010300 1.00800 ; qtot -
2.079
1788 HC 108 LYS HG3 1788 0.010300 1.00800 ; qtot -
2.068
1789 C8 108 LYS CD 1789 -0.047900 12.01000 ; qtot -
2.116
1790 HC 108 LYS HD2 1790 0.062100 1.00800 ; qtot -
2.054
1791 HC 108 LYS HD3 1791 0.062100 1.00800 ; qtot -
1.992

1792 C8 108 LYS CE 1792 -0.014300 12.01000 ; qtot -
2.006
1793 HP 108 LYS HE2 1793 0.113500 1.00800 ; qtot -
1.893
1794 HP 108 LYS HE3 1794 0.113500 1.00800 ; qtot -
1.779
1795 N3 108 LYS NZ 1795 -0.385400 14.01000 ; qtot -
2.165
1796 H 108 LYS HZ1 1796 0.340000 1.00800 ; qtot -
1.825
1797 H 108 LYS HZ2 1797 0.340000 1.00800 ; qtot -
1.485
1798 H 108 LYS HZ3 1798 0.340000 1.00800 ; qtot -
1.145
1799 C 108 LYS C 1799 0.734101 12.01000 ; qtot -
0.411
1800 O 108 LYS O 1800 -0.589401 16.00000 ; qtot -
1.000
1801 N 109 ASN N 1801 -0.415700 14.01000 ; qtot -
1.416
1802 H 109 ASN H 1802 0.271900 1.00800 ; qtot -
1.144
1803 CX 109 ASN CA 1803 0.014300 12.01000 ; qtot -
1.130
1804 H1 109 ASN HA 1804 0.104800 1.00800 ; qtot -
1.025
1805 A2C 109 ASN CB 1805 -0.204100 12.01000 ; qtot -
1.229
1806 HC 109 ASN HB2 1806 0.079700 1.00800 ; qtot -
1.149
1807 HC 109 ASN HB3 1807 0.079700 1.00800 ; qtot -
1.069
1808 C 109 ASN CG 1808 0.713001 12.01000 ; qtot -
0.356
1809 O 109 ASN OD1 1809 -0.593101 16.00000 ; qtot -
0.950
1810 N 109 ASN ND2 1810 -0.919101 14.01000 ; qtot -
1.869
1811 H 109 ASN HD21 1811 0.419600 1.00800 ; qtot -
1.449
1812 H 109 ASN HD22 1812 0.419600 1.00800 ; qtot -
1.029
1813 C 109 ASN C 1813 0.597301 12.01000 ; qtot -
0.432
1814 O 109 ASN O 1814 -0.567901 16.00000 ; qtot -
1.000
1815 N 110 ALA N 1815 -0.415700 14.01000 ; qtot -
1.416
1816 H 110 ALA H 1816 0.271900 1.00800 ; qtot -
1.144
1817 CX 110 ALA CA 1817 0.033700 12.01000 ; qtot -
1.110

1818 H1 110 ALA HA 1818 0.082300 1.00800 ; qtot -
1.028
1819 CT 110 ALA CB 1819 -0.182500 12.01000 ; qtot -
1.210
1820 HC 110 ALA HB1 1820 0.060300 1.00800 ; qtot -
1.150
1821 HC 110 ALA HB2 1821 0.060300 1.00800 ; qtot -
1.090
1822 HC 110 ALA HB3 1822 0.060300 1.00800 ; qtot -
1.029
1823 C 110 ALA C 1823 0.597301 12.01000 ; qtot -
0.432
1824 O 110 ALA O 1824 -0.567901 16.00000 ; qtot -
1.000
1825 N 111 ALA N 1825 -0.415700 14.01000 ; qtot -
1.416
1826 H 111 ALA H 1826 0.271900 1.00800 ; qtot -
1.144
1827 CX 111 ALA CA 1827 0.033700 12.01000 ; qtot -
1.110
1828 H1 111 ALA HA 1828 0.082300 1.00800 ; qtot -
1.028
1829 CT 111 ALA CB 1829 -0.182500 12.01000 ; qtot -
1.210
1830 HC 111 ALA HB1 1830 0.060300 1.00800 ; qtot -
1.150
1831 HC 111 ALA HB2 1831 0.060300 1.00800 ; qtot -
1.090
1832 HC 111 ALA HB3 1832 0.060300 1.00800 ; qtot -
1.029
1833 C 111 ALA C 1833 0.597301 12.01000 ; qtot -
0.432
1834 O 111 ALA O 1834 -0.567901 16.00000 ; qtot -
1.000
1835 N 112 THR N 1835 -0.415700 14.01000 ; qtot -
1.416
1836 H 112 THR H 1836 0.271900 1.00800 ; qtot -
1.144
1837 CX 112 THR CA 1837 -0.038900 12.01000 ; qtot -
1.183
1838 H1 112 THR HA 1838 0.100700 1.00800 ; qtot -
1.082
1839 A3C 112 THR CB 1839 0.365400 12.01000 ; qtot -
0.717
1840 H1 112 THR HB 1840 0.004300 1.00800 ; qtot -
0.712
1841 CT 112 THR CG2 1841 -0.243800 12.01000 ; qtot -
0.956
1842 HC 112 THR HG21 1842 0.064200 1.00800 ; qtot -
0.892
1843 HC 112 THR HG22 1843 0.064200 1.00800 ; qtot -
0.828

```
  1844   HC   112   THR  HG23 1844     0.064200      1.00800 ; qtot -
0.764
  1845   OH   112   THR   OG1 1845    -0.676101     16.00000 ; qtot -
1.440
  1846   HO   112   THR   HG1 1846     0.410200      1.00800 ; qtot -
1.029
  1847    C   112   THR     C 1847     0.597301     12.01000 ; qtot -
0.432
  1848    O   112   THR     O 1848    -0.567901     16.00000 ; qtot -
1.000
  1849    N   113   ILE     N 1849    -0.415700     14.01000 ; qtot -
1.416
  1850    H   113   ILE     H 1850     0.271900      1.00800 ; qtot -
1.144
  1851   CX   113   ILE    CA 1851    -0.059700     12.01000 ; qtot -
1.204
  1852   H1   113   ILE    HA 1852     0.086900      1.00800 ; qtot -
1.117
  1853  A3C   113   ILE    CB 1853     0.130300     12.01000 ; qtot -
0.986
  1854   HC   113   ILE    HB 1854     0.018700      1.00800 ; qtot -
0.968
  1855   CT   113   ILE   CG2 1855    -0.320400     12.01000 ; qtot -
1.288
  1856   HC   113   ILE  HG21 1856     0.088200      1.00800 ; qtot -
1.200
  1857   HC   113   ILE  HG22 1857     0.088200      1.00800 ; qtot -
1.112
  1858   HC   113   ILE  HG23 1858     0.088200      1.00800 ; qtot -
1.023
  1859  A2C   113   ILE   CG1 1859    -0.043000     12.01000 ; qtot -
1.066
  1860   HC   113   ILE  HG12 1860     0.023600      1.00800 ; qtot -
1.043
  1861   HC   113   ILE  HG13 1861     0.023600      1.00800 ; qtot -
1.019
  1862   CT   113   ILE   CD1 1862    -0.066000     12.01000 ; qtot -
1.085
  1863   HC   113   ILE  HD11 1863     0.018600      1.00800 ; qtot -
1.067
  1864   HC   113   ILE  HD12 1864     0.018600      1.00800 ; qtot -
1.048
  1865   HC   113   ILE  HD13 1865     0.018600      1.00800 ; qtot -
1.029
  1866    C   113   ILE     C 1866     0.597301     12.01000 ; qtot -
0.432
  1867    O   113   ILE     O 1867    -0.567901     16.00000 ; qtot -
1.000
  1868    N   114   MET     N 1868    -0.415700     14.01000 ; qtot -
1.416
  1869    H   114   MET     H 1869     0.271900      1.00800 ; qtot -
1.144
```

1870 CX 114 MET CA 1870 -0.023700 12.01000 ; qtot -1.168
1871 H1 114 MET HA 1871 0.088000 1.00800 ; qtot -1.080
1872 A2C 114 MET CB 1872 0.034200 12.01000 ; qtot -1.045
1873 HC 114 MET HB2 1873 0.024100 1.00800 ; qtot -1.021
1874 HC 114 MET HB3 1874 0.024100 1.00800 ; qtot -0.997
1875 A2C 114 MET CG 1875 0.001800 12.01000 ; qtot -0.995
1876 H1 114 MET HG2 1876 0.044000 1.00800 ; qtot -0.951
1877 H1 114 MET HG3 1877 0.044000 1.00800 ; qtot -0.907
1878 S 114 MET SD 1878 -0.273700 32.06000 ; qtot -1.181
1879 CT 114 MET CE 1879 -0.053600 12.01000 ; qtot -1.235
1880 H1 114 MET HE1 1880 0.068400 1.00800 ; qtot -1.166
1881 H1 114 MET HE2 1881 0.068400 1.00800 ; qtot -1.098
1882 H1 114 MET HE3 1882 0.068400 1.00800 ; qtot -1.029
1883 C 114 MET C 1883 0.597301 12.01000 ; qtot -0.432
1884 O 114 MET O 1884 -0.567901 16.00000 ; qtot -1.000
1885 N 115 LYS N 1885 -0.347900 14.01000 ; qtot -1.348
1886 H 115 LYS H 1886 0.274700 1.00800 ; qtot -1.073
1887 CX 115 LYS CA 1887 -0.240000 12.01000 ; qtot -1.313
1888 H1 115 LYS HA 1888 0.142600 1.00800 ; qtot -1.171
1889 C8 115 LYS CB 1889 -0.009400 12.01000 ; qtot -1.180
1890 HC 115 LYS HB2 1890 0.036200 1.00800 ; qtot -1.144
1891 HC 115 LYS HB3 1891 0.036200 1.00800 ; qtot -1.108
1892 C8 115 LYS CG 1892 0.018700 12.01000 ; qtot -1.089
1893 HC 115 LYS HG2 1893 0.010300 1.00800 ; qtot -1.079
1894 HC 115 LYS HG3 1894 0.010300 1.00800 ; qtot -1.068
1895 C8 115 LYS CD 1895 -0.047900 12.01000 ; qtot -1.116

1896 HC 115 LYS HD2 1896 0.062100 1.00800 ; qtot -
1.054
1897 HC 115 LYS HD3 1897 0.062100 1.00800 ; qtot -
0.992
1898 C8 115 LYS CE 1898 -0.014300 12.01000 ; qtot -
1.006
1899 HP 115 LYS HE2 1899 0.113500 1.00800 ; qtot -
0.893
1900 HP 115 LYS HE3 1900 0.113500 1.00800 ; qtot -
0.779
1901 N3 115 LYS NZ 1901 -0.385400 14.01000 ; qtot -
1.165
1902 H 115 LYS HZ1 1902 0.340000 1.00800 ; qtot -
0.825
1903 H 115 LYS HZ2 1903 0.340000 1.00800 ; qtot -
0.485
1904 H 115 LYS HZ3 1904 0.340000 1.00800 ; qtot -
0.145
1905 C 115 LYS C 1905 0.734101 12.01000 ; qtot
0.589
1906 O 115 LYS O 1906 -0.589401 16.00000 ; qtot
0.000
1907 N 116 GLN N 1907 -0.415700 14.01000 ; qtot -
0.416
1908 H 116 GLN H 1908 0.271900 1.00800 ; qtot -
0.144
1909 CX 116 GLN CA 1909 -0.003100 12.01000 ; qtot -
0.147
1910 H1 116 GLN HA 1910 0.085000 1.00800 ; qtot -
0.062
1911 A2C 116 GLN CB 1911 -0.003600 12.01000 ; qtot -
0.065
1912 HC 116 GLN HB2 1912 0.017100 1.00800 ; qtot -
0.048
1913 HC 116 GLN HB3 1913 0.017100 1.00800 ; qtot -
0.031
1914 A2C 116 GLN CG 1914 -0.064500 12.01000 ; qtot -
0.096
1915 HC 116 GLN HG2 1915 0.035200 1.00800 ; qtot -
0.061
1916 HC 116 GLN HG3 1916 0.035200 1.00800 ; qtot -
0.025
1917 C 116 GLN CD 1917 0.695101 12.01000 ; qtot
0.670
1918 O 116 GLN OE1 1918 -0.608601 16.00000 ; qtot
0.061
1919 N 116 GLN NE2 1919 -0.940701 14.01000 ; qtot -
0.880
1920 H 116 GLN HE21 1920 0.425100 1.00800 ; qtot -
0.455
1921 H 116 GLN HE22 1921 0.425100 1.00800 ; qtot -
0.029

```
  1922    C   116   GLN     C 1922     0.597301     12.01000 ; qtot
0.568
  1923    O   116   GLN     O 1923    -0.567901     16.00000 ; qtot
0.000
  1924    N   117   ILE     N 1924    -0.415700     14.01000 ; qtot -
0.416
  1925    H   117   ILE     H 1925     0.271900      1.00800 ; qtot -
0.144
  1926   CX   117   ILE    CA 1926    -0.059700     12.01000 ; qtot -
0.203
  1927   H1   117   ILE    HA 1927     0.086900      1.00800 ; qtot -
0.117
  1928  A3C   117   ILE    CB 1928     0.130300     12.01000 ; qtot
0.014
  1929   HC   117   ILE    HB 1929     0.018700      1.00800 ; qtot
0.032
  1930   CT   117   ILE   CG2 1930    -0.320400     12.01000 ; qtot -
0.288
  1931   HC   117   ILE  HG21 1931     0.088200      1.00800 ; qtot -
0.200
  1932   HC   117   ILE  HG22 1932     0.088200      1.00800 ; qtot -
0.112
  1933   HC   117   ILE  HG23 1933     0.088200      1.00800 ; qtot -
0.023
  1934  A2C   117   ILE   CG1 1934    -0.043000     12.01000 ; qtot -
0.066
  1935   HC   117   ILE  HG12 1935     0.023600      1.00800 ; qtot -
0.043
  1936   HC   117   ILE  HG13 1936     0.023600      1.00800 ; qtot -
0.019
  1937   CT   117   ILE   CD1 1937    -0.066000     12.01000 ; qtot -
0.085
  1938   HC   117   ILE  HD11 1938     0.018600      1.00800 ; qtot -
0.067
  1939   HC   117   ILE  HD12 1939     0.018600      1.00800 ; qtot -
0.048
  1940   HC   117   ILE  HD13 1940     0.018600      1.00800 ; qtot -
0.029
  1941    C   117   ILE     C 1941     0.597301     12.01000 ; qtot
0.568
  1942    O   117   ILE     O 1942    -0.567901     16.00000 ; qtot
0.000
  1943    N   118   PHE     N 1943    -0.415700     14.01000 ; qtot -
0.416
  1944    H   118   PHE     H 1944     0.271900      1.00800 ; qtot -
0.144
  1945   CX   118   PHE    CA 1945    -0.002400     12.01000 ; qtot -
0.146
  1946   H1   118   PHE    HA 1946     0.097800      1.00800 ; qtot -
0.048
  1947   CT   118   PHE    CB 1947    -0.034300     12.01000 ; qtot -
0.083
```

1948 HC 118 PHE HB2 1948 0.029500 1.00800 ; qtot -0.053
1949 HC 118 PHE HB3 1949 0.029500 1.00800 ; qtot -0.024
1950 CA 118 PHE CG 1950 0.011800 12.01000 ; qtot -0.012
1951 CA 118 PHE CD1 1951 -0.125600 12.01000 ; qtot -0.137
1952 HA 118 PHE HD1 1952 0.133000 1.00800 ; qtot -0.004
1953 CA 118 PHE CE1 1953 -0.170400 12.01000 ; qtot -0.175
1954 HA 118 PHE HE1 1954 0.143000 1.00800 ; qtot -0.032
1955 CA 118 PHE CZ 1955 -0.107200 12.01000 ; qtot -0.139
1956 HA 118 PHE HZ 1956 0.129700 1.00800 ; qtot -0.009
1957 CA 118 PHE CE2 1957 -0.170400 12.01000 ; qtot -0.180
1958 HA 118 PHE HE2 1958 0.143000 1.00800 ; qtot -0.037
1959 CA 118 PHE CD2 1959 -0.125600 12.01000 ; qtot -0.162
1960 HA 118 PHE HD2 1960 0.133000 1.00800 ; qtot -0.029
1961 C 118 PHE C 1961 0.597301 12.01000 ; qtot 0.568
1962 O 118 PHE O 1962 -0.567901 16.00000 ; qtot 0.000
1963 N 119 SER N 1963 -0.415700 14.01000 ; qtot -0.416
1964 H 119 SER H 1964 0.271900 1.00800 ; qtot -0.144
1965 CX 119 SER CA 1965 -0.024900 12.01000 ; qtot -0.169
1966 H1 119 SER HA 1966 0.084300 1.00800 ; qtot -0.084
1967 A2C 119 SER CB 1967 0.211700 12.01000 ; qtot 0.127
1968 H1 119 SER HB2 1968 0.035200 1.00800 ; qtot 0.163
1969 H1 119 SER HB3 1969 0.035200 1.00800 ; qtot 0.198
1970 OH 119 SER OG 1970 -0.654601 16.00000 ; qtot -0.457
1971 HO 119 SER HG 1971 0.427500 1.00800 ; qtot -0.029
1972 C 119 SER C 1972 0.597301 12.01000 ; qtot 0.568
1973 O 119 SER O 1973 -0.567901 16.00000 ; qtot 0.000

```
  1974    N   120   ALA     N 1974    -0.415700     14.01000 ; qtot -
0.416
  1975    H   120   ALA     H 1975     0.271900      1.00800 ; qtot -
0.144
  1976   CX   120   ALA    CA 1976     0.033700     12.01000 ; qtot -
0.110
  1977   H1   120   ALA    HA 1977     0.082300      1.00800 ; qtot -
0.028
  1978   CT   120   ALA    CB 1978    -0.182500     12.01000 ; qtot -
0.210
  1979   HC   120   ALA   HB1 1979     0.060300      1.00800 ; qtot -
0.150
  1980   HC   120   ALA   HB2 1980     0.060300      1.00800 ; qtot -
0.090
  1981   HC   120   ALA   HB3 1981     0.060300      1.00800 ; qtot -
0.029
  1982    C   120   ALA     C 1982     0.597301     12.01000 ; qtot
0.568
  1983    O   120   ALA     O 1983    -0.567901     16.00000 ; qtot
0.000
  1984    N   121   ILE     N 1984    -0.415700     14.01000 ; qtot -
0.416
  1985    H   121   ILE     H 1985     0.271900      1.00800 ; qtot -
0.144
  1986   CX   121   ILE    CA 1986    -0.059700     12.01000 ; qtot -
0.203
  1987   H1   121   ILE    HA 1987     0.086900      1.00800 ; qtot -
0.117
  1988  A3C   121   ILE    CB 1988     0.130300     12.01000 ; qtot
0.014
  1989   HC   121   ILE    HB 1989     0.018700      1.00800 ; qtot
0.032
  1990   CT   121   ILE   CG2 1990    -0.320400     12.01000 ; qtot -
0.288
  1991   HC   121   ILE  HG21 1991     0.088200      1.00800 ; qtot -
0.200
  1992   HC   121   ILE  HG22 1992     0.088200      1.00800 ; qtot -
0.112
  1993   HC   121   ILE  HG23 1993     0.088200      1.00800 ; qtot -
0.023
  1994  A2C   121   ILE   CG1 1994    -0.043000     12.01000 ; qtot -
0.066
  1995   HC   121   ILE  HG12 1995     0.023600      1.00800 ; qtot -
0.043
  1996   HC   121   ILE  HG13 1996     0.023600      1.00800 ; qtot -
0.019
  1997   CT   121   ILE   CD1 1997    -0.066000     12.01000 ; qtot -
0.085
  1998   HC   121   ILE  HD11 1998     0.018600      1.00800 ; qtot -
0.067
  1999   HC   121   ILE  HD12 1999     0.018600      1.00800 ; qtot -
0.048
```

```
  2000   HC   121   ILE  HD13 2000     0.018600      1.00800 ; qtot -
0.029
  2001    C   121   ILE     C 2001     0.597301     12.01000 ; qtot
0.568
  2002    O   121   ILE     O 2002    -0.567901     16.00000 ; qtot
0.000
  2003    N   122   PHE     N 2003    -0.415700     14.01000 ; qtot -
0.416
  2004    H   122   PHE     H 2004     0.271900      1.00800 ; qtot -
0.144
  2005   CX   122   PHE    CA 2005    -0.002400     12.01000 ; qtot -
0.146
  2006   H1   122   PHE    HA 2006     0.097800      1.00800 ; qtot -
0.048
  2007   CT   122   PHE    CB 2007    -0.034300     12.01000 ; qtot -
0.083
  2008   HC   122   PHE   HB2 2008     0.029500      1.00800 ; qtot -
0.053
  2009   HC   122   PHE   HB3 2009     0.029500      1.00800 ; qtot -
0.024
  2010   CA   122   PHE    CG 2010     0.011800     12.01000 ; qtot -
0.012
  2011   CA   122   PHE   CD1 2011    -0.125600     12.01000 ; qtot -
0.137
  2012   HA   122   PHE   HD1 2012     0.133000      1.00800 ; qtot -
0.004
  2013   CA   122   PHE   CE1 2013    -0.170400     12.01000 ; qtot -
0.175
  2014   HA   122   PHE   HE1 2014     0.143000      1.00800 ; qtot -
0.032
  2015   CA   122   PHE    CZ 2015    -0.107200     12.01000 ; qtot -
0.139
  2016   HA   122   PHE    HZ 2016     0.129700      1.00800 ; qtot -
0.009
  2017   CA   122   PHE   CE2 2017    -0.170400     12.01000 ; qtot -
0.180
  2018   HA   122   PHE   HE2 2018     0.143000      1.00800 ; qtot -
0.037
  2019   CA   122   PHE   CD2 2019    -0.125600     12.01000 ; qtot -
0.162
  2020   HA   122   PHE   HD2 2020     0.133000      1.00800 ; qtot -
0.029
  2021    C   122   PHE     C 2021     0.597301     12.01000 ; qtot
0.568
  2022    O   122   PHE     O 2022    -0.567901     16.00000 ; qtot
0.000
  2023    N   123   TYR     N 2023    -0.415700     14.01000 ; qtot -
0.416
  2024    H   123   TYR     H 2024     0.271900      1.00800 ; qtot -
0.144
  2025   CX   123   TYR    CA 2025    -0.001400     12.01000 ; qtot -
0.145
```

2026 H1 123 TYR HA 2026 0.087600 1.00800 ; qtot -
0.058
2027 CT 123 TYR CB 2027 -0.015200 12.01000 ; qtot -
0.073
2028 HC 123 TYR HB2 2028 0.029500 1.00800 ; qtot -
0.043
2029 HC 123 TYR HB3 2029 0.029500 1.00800 ; qtot -
0.014
2030 CA 123 TYR CG 2030 -0.001100 12.01000 ; qtot -
0.015
2031 CA 123 TYR CD1 2031 -0.190600 12.01000 ; qtot -
0.205
2032 HA 123 TYR HD1 2032 0.169900 1.00800 ; qtot -
0.036
2033 CA 123 TYR CE1 2033 -0.234100 12.01000 ; qtot -
0.270
2034 HA 123 TYR HE1 2034 0.165600 1.00800 ; qtot -
0.104
2035 C 123 TYR CZ 2035 0.322600 12.01000 ; qtot
0.219
2036 OH 123 TYR OH 2036 -0.557901 16.00000 ; qtot -
0.339
2037 HO 123 TYR HH 2037 0.399200 1.00800 ; qtot
0.060
2038 CA 123 TYR CE2 2038 -0.234100 12.01000 ; qtot -
0.174
2039 HA 123 TYR HE2 2039 0.165600 1.00800 ; qtot -
0.009
2040 CA 123 TYR CD2 2040 -0.190600 12.01000 ; qtot -
0.199
2041 HA 123 TYR HD2 2041 0.169900 1.00800 ; qtot -
0.029
2042 C 123 TYR C 2042 0.597301 12.01000 ; qtot
0.568
2043 O 123 TYR O 2043 -0.567901 16.00000 ; qtot
0.000
2044 N 124 LEU N 2044 -0.415700 14.01000 ; qtot -
0.416
2045 H 124 LEU H 2045 0.271900 1.00800 ; qtot -
0.144
2046 CX 124 LEU CA 2046 -0.051800 12.01000 ; qtot -
0.196
2047 H1 124 LEU HA 2047 0.092200 1.00800 ; qtot -
0.103
2048 A2C 124 LEU CB 2048 -0.110200 12.01000 ; qtot -
0.214
2049 HC 124 LEU HB2 2049 0.045700 1.00800 ; qtot -
0.168
2050 HC 124 LEU HB3 2050 0.045700 1.00800 ; qtot -
0.122
2051 A3C 124 LEU CG 2051 0.353100 12.01000 ; qtot
0.231

2052 HC 124 LEU HG 2052 -0.036100 1.00800 ; qtot
0.195
2053 CT 124 LEU CD1 2053 -0.412100 12.01000 ; qtot -
0.217
2054 HC 124 LEU HD11 2054 0.100000 1.00800 ; qtot -
0.117
2055 HC 124 LEU HD12 2055 0.100000 1.00800 ; qtot -
0.017
2056 HC 124 LEU HD13 2056 0.100000 1.00800 ; qtot
0.083
2057 CT 124 LEU CD2 2057 -0.412100 12.01000 ; qtot -
0.329
2058 HC 124 LEU HD21 2058 0.100000 1.00800 ; qtot -
0.229
2059 HC 124 LEU HD22 2059 0.100000 1.00800 ; qtot -
0.129
2060 HC 124 LEU HD23 2060 0.100000 1.00800 ; qtot -
0.029
2061 C 124 LEU C 2061 0.597301 12.01000 ; qtot
0.568
2062 O 124 LEU O 2062 -0.567901 16.00000 ; qtot
0.000
2063 N 125 HIP N 2063 -0.347900 14.01000 ; qtot -
0.348
2064 H 125 HIP H 2064 0.274700 1.00800 ; qtot -
0.073
2065 CX 125 HIP CA 2065 -0.135400 12.01000 ; qtot -
0.209
2066 H1 125 HIP HA 2066 0.121200 1.00800 ; qtot -
0.087
2067 CT 125 HIP CB 2067 -0.041400 12.01000 ; qtot -
0.129
2068 HC 125 HIP HB2 2068 0.081000 1.00800 ; qtot -
0.048
2069 HC 125 HIP HB3 2069 0.081000 1.00800 ; qtot
0.033
2070 CC 125 HIP CG 2070 -0.001200 12.01000 ; qtot
0.032
2071 NA 125 HIP ND1 2071 -0.151300 14.01000 ; qtot -
0.119
2072 H 125 HIP HD1 2072 0.386600 1.00800 ; qtot
0.267
2073 CR 125 HIP CE1 2073 -0.017000 12.01000 ; qtot
0.250
2074 H5 125 HIP HE1 2074 0.268100 1.00800 ; qtot
0.518
2075 NA 125 HIP NE2 2075 -0.171800 14.01000 ; qtot
0.347
2076 H 125 HIP HE2 2076 0.391100 1.00800 ; qtot
0.738
2077 CW 125 HIP CD2 2077 -0.114100 12.01000 ; qtot
0.624

2078 H4 125 HIP HD2 2078 0.231700 1.00800 ; qtot
0.855
2079 C 125 HIP C 2079 0.734101 12.01000 ; qtot
1.589
2080 O 125 HIP O 2080 -0.589401 16.00000 ; qtot
1.000
2081 N 126 SER N 2081 -0.415700 14.01000 ; qtot
0.584
2082 H 126 SER H 2082 0.271900 1.00800 ; qtot
0.856
2083 CX 126 SER CA 2083 -0.024900 12.01000 ; qtot
0.831
2084 H1 126 SER HA 2084 0.084300 1.00800 ; qtot
0.916
2085 A2C 126 SER CB 2085 0.211700 12.01000 ; qtot
1.127
2086 H1 126 SER HB2 2086 0.035200 1.00800 ; qtot
1.163
2087 H1 126 SER HB3 2087 0.035200 1.00800 ; qtot
1.198
2088 OH 126 SER OG 2088 -0.654601 16.00000 ; qtot
0.543
2089 HO 126 SER HG 2089 0.427500 1.00800 ; qtot
0.971
2090 C 126 SER C 2090 0.597301 12.01000 ; qtot
1.568
2091 O 126 SER O 2091 -0.567901 16.00000 ; qtot
1.000
2092 N 127 LEU N 2092 -0.415700 14.01000 ; qtot
0.584
2093 H 127 LEU H 2093 0.271900 1.00800 ; qtot
0.856
2094 CX 127 LEU CA 2094 -0.051800 12.01000 ; qtot
0.804
2095 H1 127 LEU HA 2095 0.092200 1.00800 ; qtot
0.897
2096 A2C 127 LEU CB 2096 -0.110200 12.01000 ; qtot
0.786
2097 HC 127 LEU HB2 2097 0.045700 1.00800 ; qtot
0.832
2098 HC 127 LEU HB3 2098 0.045700 1.00800 ; qtot
0.878
2099 A3C 127 LEU CG 2099 0.353100 12.01000 ; qtot
1.231
2100 HC 127 LEU HG 2100 -0.036100 1.00800 ; qtot
1.195
2101 CT 127 LEU CD1 2101 -0.412100 12.01000 ; qtot
0.783
2102 HC 127 LEU HD11 2102 0.100000 1.00800 ; qtot
0.883
2103 HC 127 LEU HD12 2103 0.100000 1.00800 ; qtot
0.983

```
  2104   HC   127   LEU  HD13 2104     0.100000      1.00800 ; qtot
1.083
  2105   CT   127   LEU   CD2 2105    -0.412100     12.01000 ; qtot
0.671
  2106   HC   127   LEU  HD21 2106     0.100000      1.00800 ; qtot
0.771
  2107   HC   127   LEU  HD22 2107     0.100000      1.00800 ; qtot
0.871
  2108   HC   127   LEU  HD23 2108     0.100000      1.00800 ; qtot
0.971
  2109    C   127   LEU     C 2109     0.597301     12.01000 ; qtot
1.568
  2110    O   127   LEU     O 2110    -0.567901     16.00000 ; qtot
1.000
  2111    N   128   ASN     N 2111    -0.415700     14.01000 ; qtot
0.584
  2112    H   128   ASN     H 2112     0.271900      1.00800 ; qtot
0.856
  2113   CX   128   ASN    CA 2113     0.014300     12.01000 ; qtot
0.871
  2114   H1   128   ASN    HA 2114     0.104800      1.00800 ; qtot
0.975
  2115  A2C   128   ASN    CB 2115    -0.204100     12.01000 ; qtot
0.771
  2116   HC   128   ASN   HB2 2116     0.079700      1.00800 ; qtot
0.851
  2117   HC   128   ASN   HB3 2117     0.079700      1.00800 ; qtot
0.931
  2118    C   128   ASN    CG 2118     0.713001     12.01000 ; qtot
1.644
  2119    O   128   ASN   OD1 2119    -0.593101     16.00000 ; qtot
1.051
  2120    N   128   ASN   ND2 2120    -0.919101     14.01000 ; qtot
0.131
  2121    H   128   ASN  HD21 2121     0.419600      1.00800 ; qtot
0.551
  2122    H   128   ASN  HD22 2122     0.419600      1.00800 ; qtot
0.971
  2123    C   128   ASN     C 2123     0.597301     12.01000 ; qtot
1.568
  2124    O   128   ASN     O 2124    -0.567901     16.00000 ; qtot
1.000
  2125    N   129   ILE     N 2125    -0.415700     14.01000 ; qtot
0.584
  2126    H   129   ILE     H 2126     0.271900      1.00800 ; qtot
0.856
  2127   CX   129   ILE    CA 2127    -0.059700     12.01000 ; qtot
0.797
  2128   H1   129   ILE    HA 2128     0.086900      1.00800 ; qtot
0.883
  2129  A3C   129   ILE    CB 2129     0.130300     12.01000 ; qtot
1.014
```

2130 HC 129 ILE HB 2130 0.018700 1.00800 ; qtot
1.032
2131 CT 129 ILE CG2 2131 -0.320400 12.01000 ; qtot
0.712
2132 HC 129 ILE HG21 2132 0.088200 1.00800 ; qtot
0.800
2133 HC 129 ILE HG22 2133 0.088200 1.00800 ; qtot
0.888
2134 HC 129 ILE HG23 2134 0.088200 1.00800 ; qtot
0.977
2135 A2C 129 ILE CG1 2135 -0.043000 12.01000 ; qtot
0.934
2136 HC 129 ILE HG12 2136 0.023600 1.00800 ; qtot
0.957
2137 HC 129 ILE HG13 2137 0.023600 1.00800 ; qtot
0.981
2138 CT 129 ILE CD1 2138 -0.066000 12.01000 ; qtot
0.915
2139 HC 129 ILE HD11 2139 0.018600 1.00800 ; qtot
0.933
2140 HC 129 ILE HD12 2140 0.018600 1.00800 ; qtot
0.952
2141 HC 129 ILE HD13 2141 0.018600 1.00800 ; qtot
0.971
2142 C 129 ILE C 2142 0.597301 12.01000 ; qtot
1.568
2143 O 129 ILE O 2143 -0.567901 16.00000 ; qtot
1.000
2144 N 130 VAL N 2144 -0.415700 14.01000 ; qtot
0.584
2145 H 130 VAL H 2145 0.271900 1.00800 ; qtot
0.856
2146 CX 130 VAL CA 2146 -0.087500 12.01000 ; qtot
0.769
2147 H1 130 VAL HA 2147 0.096900 1.00800 ; qtot
0.866
2148 A3C 130 VAL CB 2148 0.298500 12.01000 ; qtot
1.164
2149 HC 130 VAL HB 2149 -0.029700 1.00800 ; qtot
1.134
2150 CT 130 VAL CG1 2150 -0.319200 12.01000 ; qtot
0.815
2151 HC 130 VAL HG11 2151 0.079100 1.00800 ; qtot
0.894
2152 HC 130 VAL HG12 2152 0.079100 1.00800 ; qtot
0.973
2153 HC 130 VAL HG13 2153 0.079100 1.00800 ; qtot
1.053
2154 CT 130 VAL CG2 2154 -0.319200 12.01000 ; qtot
0.733
2155 HC 130 VAL HG21 2155 0.079100 1.00800 ; qtot
0.812

```
  2156   HC   130   VAL  HG22 2156     0.079100      1.00800 ; qtot
0.892
  2157   HC   130   VAL  HG23 2157     0.079100      1.00800 ; qtot
0.971
  2158    C   130   VAL     C 2158     0.597301     12.01000 ; qtot
1.568
  2159    O   130   VAL     O 2159    -0.567901     16.00000 ; qtot
1.000
  2160    N   131   HIE     N 2160    -0.415700     14.01000 ; qtot
0.584
  2161    H   131   HIE     H 2161     0.271900      1.00800 ; qtot
0.856
  2162   CX   131   HIE    CA 2162    -0.058100     12.01000 ; qtot
0.798
  2163   H1   131   HIE    HA 2163     0.136000      1.00800 ; qtot
0.934
  2164   CT   131   HIE    CB 2164    -0.007400     12.01000 ; qtot
0.927
  2165   HC   131   HIE   HB2 2165     0.036700      1.00800 ; qtot
0.963
  2166   HC   131   HIE   HB3 2166     0.036700      1.00800 ; qtot
1.000
  2167   CC   131   HIE    CG 2167     0.186800     12.01000 ; qtot
1.187
  2168   NB   131   HIE   ND1 2168    -0.543201     14.01000 ; qtot
0.644
  2169   CR   131   HIE   CE1 2169     0.163500     12.01000 ; qtot
0.807
  2170   H5   131   HIE   HE1 2170     0.143500      1.00800 ; qtot
0.951
  2171   NA   131   HIE   NE2 2171    -0.279500     14.01000 ; qtot
0.671
  2172    H   131   HIE   HE2 2172     0.333900      1.00800 ; qtot
1.005
  2173   CW   131   HIE   CD2 2173    -0.220700     12.01000 ; qtot
0.784
  2174   H4   131   HIE   HD2 2174     0.186200      1.00800 ; qtot
0.971
  2175    C   131   HIE     C 2175     0.597301     12.01000 ; qtot
1.568
  2176    O   131   HIE     O 2176    -0.567901     16.00000 ; qtot
1.000
  2177    N   132   ARG     N 2177    -0.347900     14.01000 ; qtot
0.652
  2178    H   132   ARG     H 2178     0.274700      1.00800 ; qtot
0.927
  2179   CX   132   ARG    CA 2179    -0.263700     12.01000 ; qtot
0.663
  2180   H1   132   ARG    HA 2180     0.156000      1.00800 ; qtot
0.819
  2181   C8   132   ARG    CB 2181    -0.000700     12.01000 ; qtot
0.818
```

2182 HC 132 ARG HB2 2182 0.032700 1.00800 ; qtot
0.851
2183 HC 132 ARG HB3 2183 0.032700 1.00800 ; qtot
0.884
2184 C8 132 ARG CG 2184 0.039000 12.01000 ; qtot
0.923
2185 HC 132 ARG HG2 2185 0.028500 1.00800 ; qtot
0.951
2186 HC 132 ARG HG3 2186 0.028500 1.00800 ; qtot
0.980
2187 C8 132 ARG CD 2187 0.048600 12.01000 ; qtot
1.028
2188 H1 132 ARG HD2 2188 0.068700 1.00800 ; qtot
1.097
2189 H1 132 ARG HD3 2189 0.068700 1.00800 ; qtot
1.166
2190 N2 132 ARG NE 2190 -0.529501 14.01000 ; qtot
0.636
2191 H 132 ARG HE 2191 0.345600 1.00800 ; qtot
0.982
2192 CA 132 ARG CZ 2192 0.807601 12.01000 ; qtot
1.790
2193 N2 132 ARG NH1 2193 -0.862701 14.01000 ; qtot
0.927
2194 H 132 ARG HH11 2194 0.447800 1.00800 ; qtot
1.375
2195 H 132 ARG HH12 2195 0.447800 1.00800 ; qtot
1.822
2196 N2 132 ARG NH2 2196 -0.862701 14.01000 ; qtot
0.960
2197 H 132 ARG HH21 2197 0.447800 1.00800 ; qtot
1.408
2198 H 132 ARG HH22 2198 0.447800 1.00800 ; qtot
1.855
2199 C 132 ARG C 2199 0.734101 12.01000 ; qtot
2.589
2200 O 132 ARG O 2200 -0.589401 16.00000 ; qtot
2.000
2201 N 133 ASP N 2201 -0.516301 14.01000 ; qtot
1.484
2202 H 133 ASP H 2202 0.293600 1.00800 ; qtot
1.777
2203 CX 133 ASP CA 2203 0.038100 12.01000 ; qtot
1.815
2204 H1 133 ASP HA 2204 0.088000 1.00800 ; qtot
1.903
2205 A2C 133 ASP CB 2205 -0.030300 12.01000 ; qtot
1.873
2206 HC 133 ASP HB2 2206 -0.012200 1.00800 ; qtot
1.861
2207 HC 133 ASP HB3 2207 -0.012200 1.00800 ; qtot
1.849

2208 CO 133 ASP CG 2208 0.799401 12.01000 ; qtot
2.648
2209 O2 133 ASP OD1 2209 -0.801401 16.00000 ; qtot
1.847
2210 O2 133 ASP OD2 2210 -0.801401 16.00000 ; qtot
1.045
2211 C 133 ASP C 2211 0.536601 12.01000 ; qtot
1.582
2212 O 133 ASP O 2212 -0.581901 16.00000 ; qtot
1.000
2213 N 134 LEU N 2213 -0.415700 14.01000 ; qtot
0.584
2214 H 134 LEU H 2214 0.271900 1.00800 ; qtot
0.856
2215 CX 134 LEU CA 2215 -0.051800 12.01000 ; qtot
0.804
2216 H1 134 LEU HA 2216 0.092200 1.00800 ; qtot
0.897
2217 A2C 134 LEU CB 2217 -0.110200 12.01000 ; qtot
0.786
2218 HC 134 LEU HB2 2218 0.045700 1.00800 ; qtot
0.832
2219 HC 134 LEU HB3 2219 0.045700 1.00800 ; qtot
0.878
2220 A3C 134 LEU CG 2220 0.353100 12.01000 ; qtot
1.231
2221 HC 134 LEU HG 2221 -0.036100 1.00800 ; qtot
1.195
2222 CT 134 LEU CD1 2222 -0.412100 12.01000 ; qtot
0.783
2223 HC 134 LEU HD11 2223 0.100000 1.00800 ; qtot
0.883
2224 HC 134 LEU HD12 2224 0.100000 1.00800 ; qtot
0.983
2225 HC 134 LEU HD13 2225 0.100000 1.00800 ; qtot
1.083
2226 CT 134 LEU CD2 2226 -0.412100 12.01000 ; qtot
0.671
2227 HC 134 LEU HD21 2227 0.100000 1.00800 ; qtot
0.771
2228 HC 134 LEU HD22 2228 0.100000 1.00800 ; qtot
0.871
2229 HC 134 LEU HD23 2229 0.100000 1.00800 ; qtot
0.971
2230 C 134 LEU C 2230 0.597301 12.01000 ; qtot
1.568
2231 O 134 LEU O 2231 -0.567901 16.00000 ; qtot
1.000
2232 N 135 LYS N 2232 -0.347900 14.01000 ; qtot
0.652
2233 H 135 LYS H 2233 0.274700 1.00800 ; qtot
0.927

2234 CX 135 LYS CA 2234 -0.240000 12.01000 ; qtot
0.687
2235 H1 135 LYS HA 2235 0.142600 1.00800 ; qtot
0.829
2236 C8 135 LYS CB 2236 -0.009400 12.01000 ; qtot
0.820
2237 HC 135 LYS HB2 2237 0.036200 1.00800 ; qtot
0.856
2238 HC 135 LYS HB3 2238 0.036200 1.00800 ; qtot
0.892
2239 C8 135 LYS CG 2239 0.018700 12.01000 ; qtot
0.911
2240 HC 135 LYS HG2 2240 0.010300 1.00800 ; qtot
0.921
2241 HC 135 LYS HG3 2241 0.010300 1.00800 ; qtot
0.932
2242 C8 135 LYS CD 2242 -0.047900 12.01000 ; qtot
0.884
2243 HC 135 LYS HD2 2243 0.062100 1.00800 ; qtot
0.946
2244 HC 135 LYS HD3 2244 0.062100 1.00800 ; qtot
1.008
2245 C8 135 LYS CE 2245 -0.014300 12.01000 ; qtot
0.994
2246 HP 135 LYS HE2 2246 0.113500 1.00800 ; qtot
1.107
2247 HP 135 LYS HE3 2247 0.113500 1.00800 ; qtot
1.221
2248 N3 135 LYS NZ 2248 -0.385400 14.01000 ; qtot
0.835
2249 H 135 LYS HZ1 2249 0.340000 1.00800 ; qtot
1.175
2250 H 135 LYS HZ2 2250 0.340000 1.00800 ; qtot
1.515
2251 H 135 LYS HZ3 2251 0.340000 1.00800 ; qtot
1.855
2252 C 135 LYS C 2252 0.734101 12.01000 ; qtot
2.589
2253 O 135 LYS O 2253 -0.589401 16.00000 ; qtot
2.000
2254 N 136 PRO N 2254 -0.254800 14.01000 ; qtot
1.745
2255 CT 136 PRO CD 2255 0.019200 12.01000 ; qtot
1.764
2256 H1 136 PRO HD2 2256 0.039100 1.00800 ; qtot
1.804
2257 H1 136 PRO HD3 2257 0.039100 1.00800 ; qtot
1.843
2258 CT 136 PRO CG 2258 0.018900 12.01000 ; qtot
1.862
2259 HC 136 PRO HG2 2259 0.021300 1.00800 ; qtot
1.883

2260 HC 136 PRO HG3 2260 0.021300 1.00800 ; qtot
1.904
2261 CT 136 PRO CB 2261 -0.007000 12.01000 ; qtot
1.897
2262 HC 136 PRO HB2 2262 0.025300 1.00800 ; qtot
1.922
2263 HC 136 PRO HB3 2263 0.025300 1.00800 ; qtot
1.948
2264 CX 136 PRO CA 2264 -0.026600 12.01000 ; qtot
1.921
2265 H1 136 PRO HA 2265 0.064100 1.00800 ; qtot
1.985
2266 C 136 PRO C 2266 0.589601 12.01000 ; qtot
2.575
2267 O 136 PRO O 2267 -0.574801 16.00000 ; qtot
2.000
2268 N 137 GLU N 2268 -0.516301 14.01000 ; qtot
1.484
2269 H 137 GLU H 2269 0.293600 1.00800 ; qtot
1.777
2270 CX 137 GLU CA 2270 0.039700 12.01000 ; qtot
1.817
2271 H1 137 GLU HA 2271 0.110500 1.00800 ; qtot
1.928
2272 A2C 137 GLU CB 2272 0.056000 12.01000 ; qtot
1.984
2273 HC 137 GLU HB2 2273 -0.017300 1.00800 ; qtot
1.966
2274 HC 137 GLU HB3 2274 -0.017300 1.00800 ; qtot
1.949
2275 A2C 137 GLU CG 2275 0.013600 12.01000 ; qtot
1.963
2276 HC 137 GLU HG2 2276 -0.042500 1.00800 ; qtot
1.920
2277 HC 137 GLU HG3 2277 -0.042500 1.00800 ; qtot
1.878
2278 CO 137 GLU CD 2278 0.805401 12.01000 ; qtot
2.683
2279 O2 137 GLU OE1 2279 -0.818801 16.00000 ; qtot
1.864
2280 O2 137 GLU OE2 2280 -0.818801 16.00000 ; qtot
1.045
2281 C 137 GLU C 2281 0.536601 12.01000 ; qtot
1.582
2282 O 137 GLU O 2282 -0.581901 16.00000 ; qtot
1.000
2283 N 138 ASN N 2283 -0.415700 14.01000 ; qtot
0.584
2284 H 138 ASN H 2284 0.271900 1.00800 ; qtot
0.856
2285 CX 138 ASN CA 2285 0.014300 12.01000 ; qtot
0.871

2286 H1 138 ASN HA 2286 0.104800 1.00800 ; qtot
0.975
2287 A2C 138 ASN CB 2287 -0.204100 12.01000 ; qtot
0.771
2288 HC 138 ASN HB2 2288 0.079700 1.00800 ; qtot
0.851
2289 HC 138 ASN HB3 2289 0.079700 1.00800 ; qtot
0.931
2290 C 138 ASN CG 2290 0.713001 12.01000 ; qtot
1.644
2291 O 138 ASN OD1 2291 -0.593101 16.00000 ; qtot
1.051
2292 N 138 ASN ND2 2292 -0.919101 14.01000 ; qtot
0.131
2293 H 138 ASN HD21 2293 0.419600 1.00800 ; qtot
0.551
2294 H 138 ASN HD22 2294 0.419600 1.00800 ; qtot
0.971
2295 C 138 ASN C 2295 0.597301 12.01000 ; qtot
1.568
2296 O 138 ASN O 2296 -0.567901 16.00000 ; qtot
1.000
2297 N 139 PHE N 2297 -0.415700 14.01000 ; qtot
0.584
2298 H 139 PHE H 2298 0.271900 1.00800 ; qtot
0.856
2299 CX 139 PHE CA 2299 -0.002400 12.01000 ; qtot
0.854
2300 H1 139 PHE HA 2300 0.097800 1.00800 ; qtot
0.952
2301 CT 139 PHE CB 2301 -0.034300 12.01000 ; qtot
0.917
2302 HC 139 PHE HB2 2302 0.029500 1.00800 ; qtot
0.947
2303 HC 139 PHE HB3 2303 0.029500 1.00800 ; qtot
0.976
2304 CA 139 PHE CG 2304 0.011800 12.01000 ; qtot
0.988
2305 CA 139 PHE CD1 2305 -0.125600 12.01000 ; qtot
0.863
2306 HA 139 PHE HD1 2306 0.133000 1.00800 ; qtot
0.996
2307 CA 139 PHE CE1 2307 -0.170400 12.01000 ; qtot
0.825
2308 HA 139 PHE HE1 2308 0.143000 1.00800 ; qtot
0.968
2309 CA 139 PHE CZ 2309 -0.107200 12.01000 ; qtot
0.861
2310 HA 139 PHE HZ 2310 0.129700 1.00800 ; qtot
0.991
2311 CA 139 PHE CE2 2311 -0.170400 12.01000 ; qtot
0.820

2312 HA 139 PHE HE2 2312 0.143000 1.00800 ; qtot
0.963
2313 CA 139 PHE CD2 2313 -0.125600 12.01000 ; qtot
0.838
2314 HA 139 PHE HD2 2314 0.133000 1.00800 ; qtot
0.971
2315 C 139 PHE C 2315 0.597301 12.01000 ; qtot
1.568
2316 O 139 PHE O 2316 -0.567901 16.00000 ; qtot
1.000
2317 N 140 LEU N 2317 -0.415700 14.01000 ; qtot
0.584
2318 H 140 LEU H 2318 0.271900 1.00800 ; qtot
0.856
2319 CX 140 LEU CA 2319 -0.051800 12.01000 ; qtot
0.804
2320 H1 140 LEU HA 2320 0.092200 1.00800 ; qtot
0.897
2321 A2C 140 LEU CB 2321 -0.110200 12.01000 ; qtot
0.786
2322 HC 140 LEU HB2 2322 0.045700 1.00800 ; qtot
0.832
2323 HC 140 LEU HB3 2323 0.045700 1.00800 ; qtot
0.878
2324 A3C 140 LEU CG 2324 0.353100 12.01000 ; qtot
1.231
2325 HC 140 LEU HG 2325 -0.036100 1.00800 ; qtot
1.195
2326 CT 140 LEU CD1 2326 -0.412100 12.01000 ; qtot
0.783
2327 HC 140 LEU HD11 2327 0.100000 1.00800 ; qtot
0.883
2328 HC 140 LEU HD12 2328 0.100000 1.00800 ; qtot
0.983
2329 HC 140 LEU HD13 2329 0.100000 1.00800 ; qtot
1.083
2330 CT 140 LEU CD2 2330 -0.412100 12.01000 ; qtot
0.671
2331 HC 140 LEU HD21 2331 0.100000 1.00800 ; qtot
0.771
2332 HC 140 LEU HD22 2332 0.100000 1.00800 ; qtot
0.871
2333 HC 140 LEU HD23 2333 0.100000 1.00800 ; qtot
0.971
2334 C 140 LEU C 2334 0.597301 12.01000 ; qtot
1.568
2335 O 140 LEU O 2335 -0.567901 16.00000 ; qtot
1.000
2336 N 141 PHE N 2336 -0.415700 14.01000 ; qtot
0.584
2337 H 141 PHE H 2337 0.271900 1.00800 ; qtot
0.856

2338 CX 141 PHE CA 2338 -0.002400 12.01000 ; qtot 0.854
2339 H1 141 PHE HA 2339 0.097800 1.00800 ; qtot 0.952
2340 CT 141 PHE CB 2340 -0.034300 12.01000 ; qtot 0.917
2341 HC 141 PHE HB2 2341 0.029500 1.00800 ; qtot 0.947
2342 HC 141 PHE HB3 2342 0.029500 1.00800 ; qtot 0.976
2343 CA 141 PHE CG 2343 0.011800 12.01000 ; qtot 0.988
2344 CA 141 PHE CD1 2344 -0.125600 12.01000 ; qtot 0.863
2345 HA 141 PHE HD1 2345 0.133000 1.00800 ; qtot 0.996
2346 CA 141 PHE CE1 2346 -0.170400 12.01000 ; qtot 0.825
2347 HA 141 PHE HE1 2347 0.143000 1.00800 ; qtot 0.968
2348 CA 141 PHE CZ 2348 -0.107200 12.01000 ; qtot 0.861
2349 HA 141 PHE HZ 2349 0.129700 1.00800 ; qtot 0.991
2350 CA 141 PHE CE2 2350 -0.170400 12.01000 ; qtot 0.820
2351 HA 141 PHE HE2 2351 0.143000 1.00800 ; qtot 0.963
2352 CA 141 PHE CD2 2352 -0.125600 12.01000 ; qtot 0.838
2353 HA 141 PHE HD2 2353 0.133000 1.00800 ; qtot 0.971
2354 C 141 PHE C 2354 0.597301 12.01000 ; qtot 1.568
2355 O 141 PHE O 2355 -0.567901 16.00000 ; qtot 1.000
2356 N 142 GLN N 2356 -0.415700 14.01000 ; qtot 0.584
2357 H 142 GLN H 2357 0.271900 1.00800 ; qtot 0.856
2358 CX 142 GLN CA 2358 -0.003100 12.01000 ; qtot 0.853
2359 H1 142 GLN HA 2359 0.085000 1.00800 ; qtot 0.938
2360 A2C 142 GLN CB 2360 -0.003600 12.01000 ; qtot 0.935
2361 HC 142 GLN HB2 2361 0.017100 1.00800 ; qtot 0.952
2362 HC 142 GLN HB3 2362 0.017100 1.00800 ; qtot 0.969
2363 A2C 142 GLN CG 2363 -0.064500 12.01000 ; qtot 0.904

2364 HC 142 GLN HG2 2364 0.035200 1.00800 ; qtot
0.939
2365 HC 142 GLN HG3 2365 0.035200 1.00800 ; qtot
0.975
2366 C 142 GLN CD 2366 0.695101 12.01000 ; qtot
1.670
2367 O 142 GLN OE1 2367 -0.608601 16.00000 ; qtot
1.061
2368 N 142 GLN NE2 2368 -0.940701 14.01000 ; qtot
0.120
2369 H 142 GLN HE21 2369 0.425100 1.00800 ; qtot
0.546
2370 H 142 GLN HE22 2370 0.425100 1.00800 ; qtot
0.971
2371 C 142 GLN C 2371 0.597301 12.01000 ; qtot
1.568
2372 O 142 GLN O 2372 -0.567901 16.00000 ; qtot
1.000
2373 N 143 SER N 2373 -0.415700 14.01000 ; qtot
0.584
2374 H 143 SER H 2374 0.271900 1.00800 ; qtot
0.856
2375 CX 143 SER CA 2375 -0.024900 12.01000 ; qtot
0.831
2376 H1 143 SER HA 2376 0.084300 1.00800 ; qtot
0.916
2377 A2C 143 SER CB 2377 0.211700 12.01000 ; qtot
1.127
2378 H1 143 SER HB2 2378 0.035200 1.00800 ; qtot
1.163
2379 H1 143 SER HB3 2379 0.035200 1.00800 ; qtot
1.198
2380 OH 143 SER OG 2380 -0.654601 16.00000 ; qtot
0.543
2381 HO 143 SER HG 2381 0.427500 1.00800 ; qtot
0.971
2382 C 143 SER C 2382 0.597301 12.01000 ; qtot
1.568
2383 O 143 SER O 2383 -0.567901 16.00000 ; qtot
1.000
2384 N 144 GLU N 2384 -0.516301 14.01000 ; qtot
0.484
2385 H 144 GLU H 2385 0.293600 1.00800 ; qtot
0.777
2386 CX 144 GLU CA 2386 0.039700 12.01000 ; qtot
0.817
2387 H1 144 GLU HA 2387 0.110500 1.00800 ; qtot
0.928
2388 A2C 144 GLU CB 2388 0.056000 12.01000 ; qtot
0.984
2389 HC 144 GLU HB2 2389 -0.017300 1.00800 ; qtot
0.966

```
  2390   HC   144   GLU   HB3 2390    -0.017300      1.00800 ; qtot
0.949
  2391  A2C   144   GLU    CG 2391     0.013600     12.01000 ; qtot
0.963
  2392   HC   144   GLU   HG2 2392    -0.042500      1.00800 ; qtot
0.920
  2393   HC   144   GLU   HG3 2393    -0.042500      1.00800 ; qtot
0.878
  2394   CO   144   GLU    CD 2394     0.805401     12.01000 ; qtot
1.683
  2395   O2   144   GLU   OE1 2395    -0.818801     16.00000 ; qtot
0.864
  2396   O2   144   GLU   OE2 2396    -0.818801     16.00000 ; qtot
0.045
  2397    C   144   GLU     C 2397     0.536601     12.01000 ; qtot
0.582
  2398    O   144   GLU     O 2398    -0.581901     16.00000 ; qtot
0.000
  2399    N   145   ASN     N 2399    -0.415700     14.01000 ; qtot -
0.416
  2400    H   145   ASN     H 2400     0.271900      1.00800 ; qtot -
0.144
  2401   CX   145   ASN    CA 2401     0.014300     12.01000 ; qtot -
0.129
  2402   H1   145   ASN    HA 2402     0.104800      1.00800 ; qtot -
0.025
  2403  A2C   145   ASN    CB 2403    -0.204100     12.01000 ; qtot -
0.229
  2404   HC   145   ASN   HB2 2404     0.079700      1.00800 ; qtot -
0.149
  2405   HC   145   ASN   HB3 2405     0.079700      1.00800 ; qtot -
0.069
  2406    C   145   ASN    CG 2406     0.713001     12.01000 ; qtot
0.644
  2407    O   145   ASN   OD1 2407    -0.593101     16.00000 ; qtot
0.051
  2408    N   145   ASN   ND2 2408    -0.919101     14.01000 ; qtot -
0.869
  2409    H   145   ASN  HD21 2409     0.419600      1.00800 ; qtot -
0.449
  2410    H   145   ASN  HD22 2410     0.419600      1.00800 ; qtot -
0.029
  2411    C   145   ASN     C 2411     0.597301     12.01000 ; qtot
0.568
  2412    O   145   ASN     O 2412    -0.567901     16.00000 ; qtot
0.000
  2413    N   146   LYS     N 2413    -0.347900     14.01000 ; qtot -
0.348
  2414    H   146   LYS     H 2414     0.274700      1.00800 ; qtot -
0.073
  2415   CX   146   LYS    CA 2415    -0.240000     12.01000 ; qtot -
0.313
```

```
  2416   H1   146   LYS    HA 2416     0.142600       1.00800 ; qtot -
0.171
  2417   C8   146   LYS    CB 2417    -0.009400      12.01000 ; qtot -
0.180
  2418   HC   146   LYS   HB2 2418     0.036200       1.00800 ; qtot -
0.144
  2419   HC   146   LYS   HB3 2419     0.036200       1.00800 ; qtot -
0.108
  2420   C8   146   LYS    CG 2420     0.018700      12.01000 ; qtot -
0.089
  2421   HC   146   LYS   HG2 2421     0.010300       1.00800 ; qtot -
0.079
  2422   HC   146   LYS   HG3 2422     0.010300       1.00800 ; qtot -
0.068
  2423   C8   146   LYS    CD 2423    -0.047900      12.01000 ; qtot -
0.116
  2424   HC   146   LYS   HD2 2424     0.062100       1.00800 ; qtot -
0.054
  2425   HC   146   LYS   HD3 2425     0.062100       1.00800 ; qtot
0.008
  2426   C8   146   LYS    CE 2426    -0.014300      12.01000 ; qtot -
0.006
  2427   HP   146   LYS   HE2 2427     0.113500       1.00800 ; qtot
0.107
  2428   HP   146   LYS   HE3 2428     0.113500       1.00800 ; qtot
0.221
  2429   N3   146   LYS    NZ 2429    -0.385400      14.01000 ; qtot -
0.165
  2430    H   146   LYS   HZ1 2430     0.340000       1.00800 ; qtot
0.175
  2431    H   146   LYS   HZ2 2431     0.340000       1.00800 ; qtot
0.515
  2432    H   146   LYS   HZ3 2432     0.340000       1.00800 ; qtot
0.855
  2433    C   146   LYS     C 2433     0.734101      12.01000 ; qtot
1.589
  2434    O   146   LYS     O 2434    -0.589401      16.00000 ; qtot
1.000
  2435    N   147   ASP     N 2435    -0.516301      14.01000 ; qtot
0.484
  2436    H   147   ASP     H 2436     0.293600       1.00800 ; qtot
0.777
  2437   CX   147   ASP    CA 2437     0.038100      12.01000 ; qtot
0.815
  2438   H1   147   ASP    HA 2438     0.088000       1.00800 ; qtot
0.903
  2439  A2C   147   ASP    CB 2439    -0.030300      12.01000 ; qtot
0.873
  2440   HC   147   ASP   HB2 2440    -0.012200       1.00800 ; qtot
0.861
  2441   HC   147   ASP   HB3 2441    -0.012200       1.00800 ; qtot
0.849
```

2442 CO 147 ASP CG 2442 0.799401 12.01000 ; qtot
1.648
2443 O2 147 ASP OD1 2443 -0.801401 16.00000 ; qtot
0.847
2444 O2 147 ASP OD2 2444 -0.801401 16.00000 ; qtot
0.045
2445 C 147 ASP C 2445 0.536601 12.01000 ; qtot
0.582
2446 O 147 ASP O 2446 -0.581901 16.00000 ; qtot
0.000
2447 N 148 SER N 2447 -0.415700 14.01000 ; qtot -
0.416
2448 H 148 SER H 2448 0.271900 1.00800 ; qtot -
0.144
2449 CX 148 SER CA 2449 -0.024900 12.01000 ; qtot -
0.169
2450 H1 148 SER HA 2450 0.084300 1.00800 ; qtot -
0.084
2451 A2C 148 SER CB 2451 0.211700 12.01000 ; qtot
0.127
2452 H1 148 SER HB2 2452 0.035200 1.00800 ; qtot
0.163
2453 H1 148 SER HB3 2453 0.035200 1.00800 ; qtot
0.198
2454 OH 148 SER OG 2454 -0.654601 16.00000 ; qtot -
0.457
2455 HO 148 SER HG 2455 0.427500 1.00800 ; qtot -
0.029
2456 C 148 SER C 2456 0.597301 12.01000 ; qtot
0.568
2457 O 148 SER O 2457 -0.567901 16.00000 ; qtot
0.000
2458 N 149 LEU N 2458 -0.415700 14.01000 ; qtot -
0.416
2459 H 149 LEU H 2459 0.271900 1.00800 ; qtot -
0.144
2460 CX 149 LEU CA 2460 -0.051800 12.01000 ; qtot -
0.196
2461 H1 149 LEU HA 2461 0.092200 1.00800 ; qtot -
0.103
2462 A2C 149 LEU CB 2462 -0.110200 12.01000 ; qtot -
0.214
2463 HC 149 LEU HB2 2463 0.045700 1.00800 ; qtot -
0.168
2464 HC 149 LEU HB3 2464 0.045700 1.00800 ; qtot -
0.122
2465 A3C 149 LEU CG 2465 0.353100 12.01000 ; qtot
0.231
2466 HC 149 LEU HG 2466 -0.036100 1.00800 ; qtot
0.195
2467 CT 149 LEU CD1 2467 -0.412100 12.01000 ; qtot -
0.217

2468 HC 149 LEU HD11 2468 0.100000 1.00800 ; qtot -
0.117
2469 HC 149 LEU HD12 2469 0.100000 1.00800 ; qtot -
0.017
2470 HC 149 LEU HD13 2470 0.100000 1.00800 ; qtot
0.083
2471 CT 149 LEU CD2 2471 -0.412100 12.01000 ; qtot -
0.329
2472 HC 149 LEU HD21 2472 0.100000 1.00800 ; qtot -
0.229
2473 HC 149 LEU HD22 2473 0.100000 1.00800 ; qtot -
0.129
2474 HC 149 LEU HD23 2474 0.100000 1.00800 ; qtot -
0.029
2475 C 149 LEU C 2475 0.597301 12.01000 ; qtot
0.568
2476 O 149 LEU O 2476 -0.567901 16.00000 ; qtot
0.000
2477 N 150 LEU N 2477 -0.415700 14.01000 ; qtot -
0.416
2478 H 150 LEU H 2478 0.271900 1.00800 ; qtot -
0.144
2479 CX 150 LEU CA 2479 -0.051800 12.01000 ; qtot -
0.196
2480 H1 150 LEU HA 2480 0.092200 1.00800 ; qtot -
0.103
2481 A2C 150 LEU CB 2481 -0.110200 12.01000 ; qtot -
0.214
2482 HC 150 LEU HB2 2482 0.045700 1.00800 ; qtot -
0.168
2483 HC 150 LEU HB3 2483 0.045700 1.00800 ; qtot -
0.122
2484 A3C 150 LEU CG 2484 0.353100 12.01000 ; qtot
0.231
2485 HC 150 LEU HG 2485 -0.036100 1.00800 ; qtot
0.195
2486 CT 150 LEU CD1 2486 -0.412100 12.01000 ; qtot -
0.217
2487 HC 150 LEU HD11 2487 0.100000 1.00800 ; qtot -
0.117
2488 HC 150 LEU HD12 2488 0.100000 1.00800 ; qtot -
0.017
2489 HC 150 LEU HD13 2489 0.100000 1.00800 ; qtot
0.083
2490 CT 150 LEU CD2 2490 -0.412100 12.01000 ; qtot -
0.329
2491 HC 150 LEU HD21 2491 0.100000 1.00800 ; qtot -
0.229
2492 HC 150 LEU HD22 2492 0.100000 1.00800 ; qtot -
0.129
2493 HC 150 LEU HD23 2493 0.100000 1.00800 ; qtot -
0.029

```
  2494    C   150   LEU      C 2494     0.597301     12.01000 ; qtot
0.568
  2495    O   150   LEU      O 2495    -0.567901     16.00000 ; qtot
0.000
  2496    N   151   LYS      N 2496    -0.347900     14.01000 ; qtot -
0.348
  2497    H   151   LYS      H 2497     0.274700      1.00800 ; qtot -
0.073
  2498   CX   151   LYS     CA 2498    -0.240000     12.01000 ; qtot -
0.313
  2499   H1   151   LYS     HA 2499     0.142600      1.00800 ; qtot -
0.171
  2500   C8   151   LYS     CB 2500    -0.009400     12.01000 ; qtot -
0.180
  2501   HC   151   LYS    HB2 2501     0.036200      1.00800 ; qtot -
0.144
  2502   HC   151   LYS    HB3 2502     0.036200      1.00800 ; qtot -
0.108
  2503   C8   151   LYS     CG 2503     0.018700     12.01000 ; qtot -
0.089
  2504   HC   151   LYS    HG2 2504     0.010300      1.00800 ; qtot -
0.079
  2505   HC   151   LYS    HG3 2505     0.010300      1.00800 ; qtot -
0.068
  2506   C8   151   LYS     CD 2506    -0.047900     12.01000 ; qtot -
0.116
  2507   HC   151   LYS    HD2 2507     0.062100      1.00800 ; qtot -
0.054
  2508   HC   151   LYS    HD3 2508     0.062100      1.00800 ; qtot
0.008
  2509   C8   151   LYS     CE 2509    -0.014300     12.01000 ; qtot -
0.006
  2510   HP   151   LYS    HE2 2510     0.113500      1.00800 ; qtot
0.107
  2511   HP   151   LYS    HE3 2511     0.113500      1.00800 ; qtot
0.221
  2512   N3   151   LYS     NZ 2512    -0.385400     14.01000 ; qtot -
0.165
  2513    H   151   LYS    HZ1 2513     0.340000      1.00800 ; qtot
0.175
  2514    H   151   LYS    HZ2 2514     0.340000      1.00800 ; qtot
0.515
  2515    H   151   LYS    HZ3 2515     0.340000      1.00800 ; qtot
0.855
  2516    C   151   LYS      C 2516     0.734101     12.01000 ; qtot
1.589
  2517    O   151   LYS      O 2517    -0.589401     16.00000 ; qtot
1.000
  2518    N   152   ILE      N 2518    -0.415700     14.01000 ; qtot
0.584
  2519    H   152   ILE      H 2519     0.271900      1.00800 ; qtot
0.856
```

2520 CX 152 ILE CA 2520 -0.059700 12.01000 ; qtot
0.797
2521 H1 152 ILE HA 2521 0.086900 1.00800 ; qtot
0.883
2522 A3C 152 ILE CB 2522 0.130300 12.01000 ; qtot
1.014
2523 HC 152 ILE HB 2523 0.018700 1.00800 ; qtot
1.032
2524 CT 152 ILE CG2 2524 -0.320400 12.01000 ; qtot
0.712
2525 HC 152 ILE HG21 2525 0.088200 1.00800 ; qtot
0.800
2526 HC 152 ILE HG22 2526 0.088200 1.00800 ; qtot
0.888
2527 HC 152 ILE HG23 2527 0.088200 1.00800 ; qtot
0.977
2528 A2C 152 ILE CG1 2528 -0.043000 12.01000 ; qtot
0.934
2529 HC 152 ILE HG12 2529 0.023600 1.00800 ; qtot
0.957
2530 HC 152 ILE HG13 2530 0.023600 1.00800 ; qtot
0.981
2531 CT 152 ILE CD1 2531 -0.066000 12.01000 ; qtot
0.915
2532 HC 152 ILE HD11 2532 0.018600 1.00800 ; qtot
0.933
2533 HC 152 ILE HD12 2533 0.018600 1.00800 ; qtot
0.952
2534 HC 152 ILE HD13 2534 0.018600 1.00800 ; qtot
0.971
2535 C 152 ILE C 2535 0.597301 12.01000 ; qtot
1.568
2536 O 152 ILE O 2536 -0.567901 16.00000 ; qtot
1.000
2537 N 153 ILE N 2537 -0.415700 14.01000 ; qtot
0.584
2538 H 153 ILE H 2538 0.271900 1.00800 ; qtot
0.856
2539 CX 153 ILE CA 2539 -0.059700 12.01000 ; qtot
0.797
2540 H1 153 ILE HA 2540 0.086900 1.00800 ; qtot
0.883
2541 A3C 153 ILE CB 2541 0.130300 12.01000 ; qtot
1.014
2542 HC 153 ILE HB 2542 0.018700 1.00800 ; qtot
1.032
2543 CT 153 ILE CG2 2543 -0.320400 12.01000 ; qtot
0.712
2544 HC 153 ILE HG21 2544 0.088200 1.00800 ; qtot
0.800
2545 HC 153 ILE HG22 2545 0.088200 1.00800 ; qtot
0.888

2546 HC 153 ILE HG23 2546 0.088200 1.00800 ; qtot 0.977
2547 A2C 153 ILE CG1 2547 -0.043000 12.01000 ; qtot 0.934
2548 HC 153 ILE HG12 2548 0.023600 1.00800 ; qtot 0.957
2549 HC 153 ILE HG13 2549 0.023600 1.00800 ; qtot 0.981
2550 CT 153 ILE CD1 2550 -0.066000 12.01000 ; qtot 0.915
2551 HC 153 ILE HD11 2551 0.018600 1.00800 ; qtot 0.933
2552 HC 153 ILE HD12 2552 0.018600 1.00800 ; qtot 0.952
2553 HC 153 ILE HD13 2553 0.018600 1.00800 ; qtot 0.971
2554 C 153 ILE C 2554 0.597301 12.01000 ; qtot 1.568
2555 O 153 ILE O 2555 -0.567901 16.00000 ; qtot 1.000
2556 N 154 ASP N 2556 -0.516301 14.01000 ; qtot 0.484
2557 H 154 ASP H 2557 0.293600 1.00800 ; qtot 0.777
2558 CX 154 ASP CA 2558 0.038100 12.01000 ; qtot 0.815
2559 H1 154 ASP HA 2559 0.088000 1.00800 ; qtot 0.903
2560 A2C 154 ASP CB 2560 -0.030300 12.01000 ; qtot 0.873
2561 HC 154 ASP HB2 2561 -0.012200 1.00800 ; qtot 0.861
2562 HC 154 ASP HB3 2562 -0.012200 1.00800 ; qtot 0.849
2563 CO 154 ASP CG 2563 0.799401 12.01000 ; qtot 1.648
2564 O2 154 ASP OD1 2564 -0.801401 16.00000 ; qtot 0.847
2565 O2 154 ASP OD2 2565 -0.801401 16.00000 ; qtot 0.045
2566 C 154 ASP C 2566 0.536601 12.01000 ; qtot 0.582
2567 O 154 ASP O 2567 -0.581901 16.00000 ; qtot 0.000
2568 N 155 PHE N 2568 -0.415700 14.01000 ; qtot -0.416
2569 H 155 PHE H 2569 0.271900 1.00800 ; qtot -0.144
2570 CX 155 PHE CA 2570 -0.002400 12.01000 ; qtot -0.146
2571 H1 155 PHE HA 2571 0.097800 1.00800 ; qtot -0.048

2572 CT 155 PHE CB 2572 -0.034300 12.01000 ; qtot -
0.083
2573 HC 155 PHE HB2 2573 0.029500 1.00800 ; qtot -
0.053
2574 HC 155 PHE HB3 2574 0.029500 1.00800 ; qtot -
0.024
2575 CA 155 PHE CG 2575 0.011800 12.01000 ; qtot -
0.012
2576 CA 155 PHE CD1 2576 -0.125600 12.01000 ; qtot -
0.137
2577 HA 155 PHE HD1 2577 0.133000 1.00800 ; qtot -
0.004
2578 CA 155 PHE CE1 2578 -0.170400 12.01000 ; qtot -
0.175
2579 HA 155 PHE HE1 2579 0.143000 1.00800 ; qtot -
0.032
2580 CA 155 PHE CZ 2580 -0.107200 12.01000 ; qtot -
0.139
2581 HA 155 PHE HZ 2581 0.129700 1.00800 ; qtot -
0.009
2582 CA 155 PHE CE2 2582 -0.170400 12.01000 ; qtot -
0.180
2583 HA 155 PHE HE2 2583 0.143000 1.00800 ; qtot -
0.037
2584 CA 155 PHE CD2 2584 -0.125600 12.01000 ; qtot -
0.162
2585 HA 155 PHE HD2 2585 0.133000 1.00800 ; qtot -
0.029
2586 C 155 PHE C 2586 0.597301 12.01000 ; qtot
0.568
2587 O 155 PHE O 2587 -0.567901 16.00000 ; qtot
0.000
2588 N 156 GLY N 2588 -0.415700 14.01000 ; qtot -
0.416
2589 H 156 GLY H 2589 0.271900 1.00800 ; qtot -
0.144
2590 CX 156 GLY CA 2590 -0.025200 12.01000 ; qtot -
0.169
2591 H1 156 GLY HA2 2591 0.069800 1.00800 ; qtot -
0.099
2592 H1 156 GLY HA3 2592 0.069800 1.00800 ; qtot -
0.029
2593 C 156 GLY C 2593 0.597301 12.01000 ; qtot
0.568
2594 O 156 GLY O 2594 -0.567901 16.00000 ; qtot
0.000
2595 N 157 LEU N 2595 -0.415700 14.01000 ; qtot -
0.416
2596 H 157 LEU H 2596 0.271900 1.00800 ; qtot -
0.144
2597 CX 157 LEU CA 2597 -0.051800 12.01000 ; qtot -
0.196

```
  2598   H1   157   LEU    HA 2598     0.092200      1.00800 ; qtot -
0.103
  2599  A2C   157   LEU    CB 2599    -0.110200     12.01000 ; qtot -
0.214
  2600   HC   157   LEU   HB2 2600     0.045700      1.00800 ; qtot -
0.168
  2601   HC   157   LEU   HB3 2601     0.045700      1.00800 ; qtot -
0.122
  2602  A3C   157   LEU    CG 2602     0.353100     12.01000 ; qtot
0.231
  2603   HC   157   LEU    HG 2603    -0.036100      1.00800 ; qtot
0.195
  2604   CT   157   LEU   CD1 2604    -0.412100     12.01000 ; qtot -
0.217
  2605   HC   157   LEU  HD11 2605     0.100000      1.00800 ; qtot -
0.117
  2606   HC   157   LEU  HD12 2606     0.100000      1.00800 ; qtot -
0.017
  2607   HC   157   LEU  HD13 2607     0.100000      1.00800 ; qtot
0.083
  2608   CT   157   LEU   CD2 2608    -0.412100     12.01000 ; qtot -
0.329
  2609   HC   157   LEU  HD21 2609     0.100000      1.00800 ; qtot -
0.229
  2610   HC   157   LEU  HD22 2610     0.100000      1.00800 ; qtot -
0.129
  2611   HC   157   LEU  HD23 2611     0.100000      1.00800 ; qtot -
0.029
  2612    C   157   LEU     C 2612     0.597301     12.01000 ; qtot
0.568
  2613    O   157   LEU     O 2613    -0.567901     16.00000 ; qtot
0.000
  2614    N   158   SER     N 2614    -0.415700     14.01000 ; qtot -
0.416
  2615    H   158   SER     H 2615     0.271900      1.00800 ; qtot -
0.144
  2616   CX   158   SER    CA 2616    -0.024900     12.01000 ; qtot -
0.169
  2617   H1   158   SER    HA 2617     0.084300      1.00800 ; qtot -
0.084
  2618  A2C   158   SER    CB 2618     0.211700     12.01000 ; qtot
0.127
  2619   H1   158   SER   HB2 2619     0.035200      1.00800 ; qtot
0.163
  2620   H1   158   SER   HB3 2620     0.035200      1.00800 ; qtot
0.198
  2621   OH   158   SER    OG 2621    -0.654601     16.00000 ; qtot -
0.457
  2622   HO   158   SER    HG 2622     0.427500      1.00800 ; qtot -
0.029
  2623    C   158   SER     C 2623     0.597301     12.01000 ; qtot
0.568
```

2624 O 158 SER O 2624 -0.567901 16.00000 ; qtot
0.000
2625 N 159 LYS N 2625 -0.347900 14.01000 ; qtot -
0.348
2626 H 159 LYS H 2626 0.274700 1.00800 ; qtot -
0.073
2627 CX 159 LYS CA 2627 -0.240000 12.01000 ; qtot -
0.313
2628 H1 159 LYS HA 2628 0.142600 1.00800 ; qtot -
0.171
2629 C8 159 LYS CB 2629 -0.009400 12.01000 ; qtot -
0.180
2630 HC 159 LYS HB2 2630 0.036200 1.00800 ; qtot -
0.144
2631 HC 159 LYS HB3 2631 0.036200 1.00800 ; qtot -
0.108
2632 C8 159 LYS CG 2632 0.018700 12.01000 ; qtot -
0.089
2633 HC 159 LYS HG2 2633 0.010300 1.00800 ; qtot -
0.079
2634 HC 159 LYS HG3 2634 0.010300 1.00800 ; qtot -
0.068
2635 C8 159 LYS CD 2635 -0.047900 12.01000 ; qtot -
0.116
2636 HC 159 LYS HD2 2636 0.062100 1.00800 ; qtot -
0.054
2637 HC 159 LYS HD3 2637 0.062100 1.00800 ; qtot
0.008
2638 C8 159 LYS CE 2638 -0.014300 12.01000 ; qtot -
0.006
2639 HP 159 LYS HE2 2639 0.113500 1.00800 ; qtot
0.107
2640 HP 159 LYS HE3 2640 0.113500 1.00800 ; qtot
0.221
2641 N3 159 LYS NZ 2641 -0.385400 14.01000 ; qtot -
0.165
2642 H 159 LYS HZ1 2642 0.340000 1.00800 ; qtot
0.175
2643 H 159 LYS HZ2 2643 0.340000 1.00800 ; qtot
0.515
2644 H 159 LYS HZ3 2644 0.340000 1.00800 ; qtot
0.855
2645 C 159 LYS C 2645 0.734101 12.01000 ; qtot
1.589
2646 O 159 LYS O 2646 -0.589401 16.00000 ; qtot
1.000
2647 N 160 ASN N 2647 -0.415700 14.01000 ; qtot
0.584
2648 H 160 ASN H 2648 0.271900 1.00800 ; qtot
0.856
2649 CX 160 ASN CA 2649 0.014300 12.01000 ; qtot
0.871

2650 H1 160 ASN HA 2650 0.104800 1.00800 ; qtot
0.975
2651 A2C 160 ASN CB 2651 -0.204100 12.01000 ; qtot
0.771
2652 HC 160 ASN HB2 2652 0.079700 1.00800 ; qtot
0.851
2653 HC 160 ASN HB3 2653 0.079700 1.00800 ; qtot
0.931
2654 C 160 ASN CG 2654 0.713001 12.01000 ; qtot
1.644
2655 O 160 ASN OD1 2655 -0.593101 16.00000 ; qtot
1.051
2656 N 160 ASN ND2 2656 -0.919101 14.01000 ; qtot
0.131
2657 H 160 ASN HD21 2657 0.419600 1.00800 ; qtot
0.551
2658 H 160 ASN HD22 2658 0.419600 1.00800 ; qtot
0.971
2659 C 160 ASN C 2659 0.597301 12.01000 ; qtot
1.568
2660 O 160 ASN O 2660 -0.567901 16.00000 ; qtot
1.000
2661 N 161 LEU N 2661 -0.415700 14.01000 ; qtot
0.584
2662 H 161 LEU H 2662 0.271900 1.00800 ; qtot
0.856
2663 CX 161 LEU CA 2663 -0.051800 12.01000 ; qtot
0.804
2664 H1 161 LEU HA 2664 0.092200 1.00800 ; qtot
0.897
2665 A2C 161 LEU CB 2665 -0.110200 12.01000 ; qtot
0.786
2666 HC 161 LEU HB2 2666 0.045700 1.00800 ; qtot
0.832
2667 HC 161 LEU HB3 2667 0.045700 1.00800 ; qtot
0.878
2668 A3C 161 LEU CG 2668 0.353100 12.01000 ; qtot
1.231
2669 HC 161 LEU HG 2669 -0.036100 1.00800 ; qtot
1.195
2670 CT 161 LEU CD1 2670 -0.412100 12.01000 ; qtot
0.783
2671 HC 161 LEU HD11 2671 0.100000 1.00800 ; qtot
0.883
2672 HC 161 LEU HD12 2672 0.100000 1.00800 ; qtot
0.983
2673 HC 161 LEU HD13 2673 0.100000 1.00800 ; qtot
1.083
2674 CT 161 LEU CD2 2674 -0.412100 12.01000 ; qtot
0.671
2675 HC 161 LEU HD21 2675 0.100000 1.00800 ; qtot
0.771

2676 HC 161 LEU HD22 2676 0.100000 1.00800 ; qtot
0.871
2677 HC 161 LEU HD23 2677 0.100000 1.00800 ; qtot
0.971
2678 C 161 LEU C 2678 0.597301 12.01000 ; qtot
1.568
2679 O 161 LEU O 2679 -0.567901 16.00000 ; qtot
1.000
2680 N 162 GLY N 2680 -0.415700 14.01000 ; qtot
0.584
2681 H 162 GLY H 2681 0.271900 1.00800 ; qtot
0.856
2682 CX 162 GLY CA 2682 -0.025200 12.01000 ; qtot
0.831
2683 H1 162 GLY HA2 2683 0.069800 1.00800 ; qtot
0.901
2684 H1 162 GLY HA3 2684 0.069800 1.00800 ; qtot
0.971
2685 C 162 GLY C 2685 0.597301 12.01000 ; qtot
1.568
2686 O 162 GLY O 2686 -0.567901 16.00000 ; qtot
1.000
2687 N 163 THR N 2687 -0.415700 14.01000 ; qtot
0.584
2688 H 163 THR H 2688 0.271900 1.00800 ; qtot
0.856
2689 CX 163 THR CA 2689 -0.038900 12.01000 ; qtot
0.817
2690 H1 163 THR HA 2690 0.100700 1.00800 ; qtot
0.918
2691 A3C 163 THR CB 2691 0.365400 12.01000 ; qtot
1.283
2692 H1 163 THR HB 2692 0.004300 1.00800 ; qtot
1.288
2693 CT 163 THR CG2 2693 -0.243800 12.01000 ; qtot
1.044
2694 HC 163 THR HG21 2694 0.064200 1.00800 ; qtot
1.108
2695 HC 163 THR HG22 2695 0.064200 1.00800 ; qtot
1.172
2696 HC 163 THR HG23 2696 0.064200 1.00800 ; qtot
1.237
2697 OH 163 THR OG1 2697 -0.676101 16.00000 ; qtot
0.560
2698 HO 163 THR HG1 2698 0.410200 1.00800 ; qtot
0.971
2699 C 163 THR C 2699 0.597301 12.01000 ; qtot
1.568
2700 O 163 THR O 2700 -0.567901 16.00000 ; qtot
1.000
2701 N 164 GLY N 2701 -0.415700 14.01000 ; qtot
0.584

2702 H 164 GLY H 2702 0.271900 1.00800 ; qtot
0.856
2703 CX 164 GLY CA 2703 -0.025200 12.01000 ; qtot
0.831
2704 H1 164 GLY HA2 2704 0.069800 1.00800 ; qtot
0.901
2705 H1 164 GLY HA3 2705 0.069800 1.00800 ; qtot
0.971
2706 C 164 GLY C 2706 0.597301 12.01000 ; qtot
1.568
2707 O 164 GLY O 2707 -0.567901 16.00000 ; qtot
1.000
2708 N 165 GLU N 2708 -0.516301 14.01000 ; qtot
0.484
2709 H 165 GLU H 2709 0.293600 1.00800 ; qtot
0.777
2710 CX 165 GLU CA 2710 0.039700 12.01000 ; qtot
0.817
2711 H1 165 GLU HA 2711 0.110500 1.00800 ; qtot
0.928
2712 A2C 165 GLU CB 2712 0.056000 12.01000 ; qtot
0.984
2713 HC 165 GLU HB2 2713 -0.017300 1.00800 ; qtot
0.966
2714 HC 165 GLU HB3 2714 -0.017300 1.00800 ; qtot
0.949
2715 A2C 165 GLU CG 2715 0.013600 12.01000 ; qtot
0.963
2716 HC 165 GLU HG2 2716 -0.042500 1.00800 ; qtot
0.920
2717 HC 165 GLU HG3 2717 -0.042500 1.00800 ; qtot
0.878
2718 CO 165 GLU CD 2718 0.805401 12.01000 ; qtot
1.683
2719 O2 165 GLU OE1 2719 -0.818801 16.00000 ; qtot
0.864
2720 O2 165 GLU OE2 2720 -0.818801 16.00000 ; qtot
0.045
2721 C 165 GLU C 2721 0.536601 12.01000 ; qtot
0.582
2722 O 165 GLU O 2722 -0.581901 16.00000 ; qtot
0.000
2723 N 166 PHE N 2723 -0.415700 14.01000 ; qtot -
0.416
2724 H 166 PHE H 2724 0.271900 1.00800 ; qtot -
0.144
2725 CX 166 PHE CA 2725 -0.002400 12.01000 ; qtot -
0.146
2726 H1 166 PHE HA 2726 0.097800 1.00800 ; qtot -
0.048
2727 CT 166 PHE CB 2727 -0.034300 12.01000 ; qtot -
0.083

```
  2728   HC   166   PHE   HB2 2728     0.029500      1.00800 ; qtot -
0.053
  2729   HC   166   PHE   HB3 2729     0.029500      1.00800 ; qtot -
0.024
  2730   CA   166   PHE    CG 2730     0.011800     12.01000 ; qtot -
0.012
  2731   CA   166   PHE   CD1 2731    -0.125600     12.01000 ; qtot -
0.137
  2732   HA   166   PHE   HD1 2732     0.133000      1.00800 ; qtot -
0.004
  2733   CA   166   PHE   CE1 2733    -0.170400     12.01000 ; qtot -
0.175
  2734   HA   166   PHE   HE1 2734     0.143000      1.00800 ; qtot -
0.032
  2735   CA   166   PHE    CZ 2735    -0.107200     12.01000 ; qtot -
0.139
  2736   HA   166   PHE    HZ 2736     0.129700      1.00800 ; qtot -
0.009
  2737   CA   166   PHE   CE2 2737    -0.170400     12.01000 ; qtot -
0.180
  2738   HA   166   PHE   HE2 2738     0.143000      1.00800 ; qtot -
0.037
  2739   CA   166   PHE   CD2 2739    -0.125600     12.01000 ; qtot -
0.162
  2740   HA   166   PHE   HD2 2740     0.133000      1.00800 ; qtot -
0.029
  2741    C   166   PHE     C 2741     0.597301     12.01000 ; qtot
0.568
  2742    O   166   PHE     O 2742    -0.567901     16.00000 ; qtot
0.000
  2743    N   167   THR     N 2743    -0.415700     14.01000 ; qtot -
0.416
  2744    H   167   THR     H 2744     0.271900      1.00800 ; qtot -
0.144
  2745   CX   167   THR    CA 2745    -0.038900     12.01000 ; qtot -
0.183
  2746   H1   167   THR    HA 2746     0.100700      1.00800 ; qtot -
0.082
  2747  A3C   167   THR    CB 2747     0.365400     12.01000 ; qtot
0.283
  2748   H1   167   THR    HB 2748     0.004300      1.00800 ; qtot
0.288
  2749   CT   167   THR   CG2 2749    -0.243800     12.01000 ; qtot
0.044
  2750   HC   167   THR  HG21 2750     0.064200      1.00800 ; qtot
0.108
  2751   HC   167   THR  HG22 2751     0.064200      1.00800 ; qtot
0.172
  2752   HC   167   THR  HG23 2752     0.064200      1.00800 ; qtot
0.237
  2753   OH   167   THR   OG1 2753    -0.676101     16.00000 ; qtot -
0.440
```

2754 HO 167 THR HG1 2754 0.410200 1.00800 ; qtot -
0.029
2755 C 167 THR C 2755 0.597301 12.01000 ; qtot
0.568
2756 O 167 THR O 2756 -0.567901 16.00000 ; qtot
0.000
2757 N 168 THR N 2757 -0.415700 14.01000 ; qtot -
0.416
2758 H 168 THR H 2758 0.271900 1.00800 ; qtot -
0.144
2759 CX 168 THR CA 2759 -0.038900 12.01000 ; qtot -
0.183
2760 H1 168 THR HA 2760 0.100700 1.00800 ; qtot -
0.082
2761 A3C 168 THR CB 2761 0.365400 12.01000 ; qtot
0.283
2762 H1 168 THR HB 2762 0.004300 1.00800 ; qtot
0.288
2763 CT 168 THR CG2 2763 -0.243800 12.01000 ; qtot
0.044
2764 HC 168 THR HG21 2764 0.064200 1.00800 ; qtot
0.108
2765 HC 168 THR HG22 2765 0.064200 1.00800 ; qtot
0.172
2766 HC 168 THR HG23 2766 0.064200 1.00800 ; qtot
0.237
2767 OH 168 THR OG1 2767 -0.676101 16.00000 ; qtot -
0.440
2768 HO 168 THR HG1 2768 0.410200 1.00800 ; qtot -
0.029
2769 C 168 THR C 2769 0.597301 12.01000 ; qtot
0.568
2770 O 168 THR O 2770 -0.567901 16.00000 ; qtot
0.000
2771 N 169 THR N 2771 -0.415700 14.01000 ; qtot -
0.416
2772 H 169 THR H 2772 0.271900 1.00800 ; qtot -
0.144
2773 CX 169 THR CA 2773 -0.038900 12.01000 ; qtot -
0.183
2774 H1 169 THR HA 2774 0.100700 1.00800 ; qtot -
0.082
2775 A3C 169 THR CB 2775 0.365400 12.01000 ; qtot
0.283
2776 H1 169 THR HB 2776 0.004300 1.00800 ; qtot
0.288
2777 CT 169 THR CG2 2777 -0.243800 12.01000 ; qtot
0.044
2778 HC 169 THR HG21 2778 0.064200 1.00800 ; qtot
0.108
2779 HC 169 THR HG22 2779 0.064200 1.00800 ; qtot
0.172

```
  2780   HC   169   THR  HG23 2780     0.064200      1.00800 ; qtot
0.237
  2781   OH   169   THR   OG1 2781    -0.676101     16.00000 ; qtot -
0.440
  2782   HO   169   THR   HG1 2782     0.410200      1.00800 ; qtot -
0.029
  2783    C   169   THR     C 2783     0.597301     12.01000 ; qtot
0.568
  2784    O   169   THR     O 2784    -0.567901     16.00000 ; qtot
0.000
  2785    N   170   LYS     N 2785    -0.347900     14.01000 ; qtot -
0.348
  2786    H   170   LYS     H 2786     0.274700      1.00800 ; qtot -
0.073
  2787   CX   170   LYS    CA 2787    -0.240000     12.01000 ; qtot -
0.313
  2788   H1   170   LYS    HA 2788     0.142600      1.00800 ; qtot -
0.171
  2789   C8   170   LYS    CB 2789    -0.009400     12.01000 ; qtot -
0.180
  2790   HC   170   LYS   HB2 2790     0.036200      1.00800 ; qtot -
0.144
  2791   HC   170   LYS   HB3 2791     0.036200      1.00800 ; qtot -
0.108
  2792   C8   170   LYS    CG 2792     0.018700     12.01000 ; qtot -
0.089
  2793   HC   170   LYS   HG2 2793     0.010300      1.00800 ; qtot -
0.079
  2794   HC   170   LYS   HG3 2794     0.010300      1.00800 ; qtot -
0.068
  2795   C8   170   LYS    CD 2795    -0.047900     12.01000 ; qtot -
0.116
  2796   HC   170   LYS   HD2 2796     0.062100      1.00800 ; qtot -
0.054
  2797   HC   170   LYS   HD3 2797     0.062100      1.00800 ; qtot
0.008
  2798   C8   170   LYS    CE 2798    -0.014300     12.01000 ; qtot -
0.006
  2799   HP   170   LYS   HE2 2799     0.113500      1.00800 ; qtot
0.107
  2800   HP   170   LYS   HE3 2800     0.113500      1.00800 ; qtot
0.221
  2801   N3   170   LYS    NZ 2801    -0.385400     14.01000 ; qtot -
0.165
  2802    H   170   LYS   HZ1 2802     0.340000      1.00800 ; qtot
0.175
  2803    H   170   LYS   HZ2 2803     0.340000      1.00800 ; qtot
0.515
  2804    H   170   LYS   HZ3 2804     0.340000      1.00800 ; qtot
0.855
  2805    C   170   LYS     C 2805     0.734101     12.01000 ; qtot
1.589
```

2806 O 170 LYS O 2806 -0.589401 16.00000 ; qtot
1.000
2807 N 171 ALA N 2807 -0.415700 14.01000 ; qtot
0.584
2808 H 171 ALA H 2808 0.271900 1.00800 ; qtot
0.856
2809 CX 171 ALA CA 2809 0.033700 12.01000 ; qtot
0.890
2810 H1 171 ALA HA 2810 0.082300 1.00800 ; qtot
0.972
2811 CT 171 ALA CB 2811 -0.182500 12.01000 ; qtot
0.790
2812 HC 171 ALA HB1 2812 0.060300 1.00800 ; qtot
0.850
2813 HC 171 ALA HB2 2813 0.060300 1.00800 ; qtot
0.910
2814 HC 171 ALA HB3 2814 0.060300 1.00800 ; qtot
0.971
2815 C 171 ALA C 2815 0.597301 12.01000 ; qtot
1.568
2816 O 171 ALA O 2816 -0.567901 16.00000 ; qtot
1.000
2817 N 172 GLY N 2817 -0.415700 14.01000 ; qtot
0.584
2818 H 172 GLY H 2818 0.271900 1.00800 ; qtot
0.856
2819 CX 172 GLY CA 2819 -0.025200 12.01000 ; qtot
0.831
2820 H1 172 GLY HA2 2820 0.069800 1.00800 ; qtot
0.901
2821 H1 172 GLY HA3 2821 0.069800 1.00800 ; qtot
0.971
2822 C 172 GLY C 2822 0.597301 12.01000 ; qtot
1.568
2823 O 172 GLY O 2823 -0.567901 16.00000 ; qtot
1.000
2824 N 173 THR N 2824 -0.415700 14.01000 ; qtot
0.584
2825 H 173 THR H 2825 0.271900 1.00800 ; qtot
0.856
2826 CX 173 THR CA 2826 -0.038900 12.01000 ; qtot
0.817
2827 H1 173 THR HA 2827 0.100700 1.00800 ; qtot
0.918
2828 A3C 173 THR CB 2828 0.365400 12.01000 ; qtot
1.283
2829 H1 173 THR HB 2829 0.004300 1.00800 ; qtot
1.288
2830 CT 173 THR CG2 2830 -0.243800 12.01000 ; qtot
1.044
2831 HC 173 THR HG21 2831 0.064200 1.00800 ; qtot
1.108

2832 HC 173 THR HG22 2832 0.064200 1.00800 ; qtot
1.172
2833 HC 173 THR HG23 2833 0.064200 1.00800 ; qtot
1.237
2834 OH 173 THR OG1 2834 -0.676101 16.00000 ; qtot
0.560
2835 HO 173 THR HG1 2835 0.410200 1.00800 ; qtot
0.971
2836 C 173 THR C 2836 0.597301 12.01000 ; qtot
1.568
2837 O 173 THR O 2837 -0.567901 16.00000 ; qtot
1.000
2838 N 174 PRO N 2838 -0.254800 14.01000 ; qtot
0.745
2839 CT 174 PRO CD 2839 0.019200 12.01000 ; qtot
0.764
2840 H1 174 PRO HD2 2840 0.039100 1.00800 ; qtot
0.804
2841 H1 174 PRO HD3 2841 0.039100 1.00800 ; qtot
0.843
2842 CT 174 PRO CG 2842 0.018900 12.01000 ; qtot
0.862
2843 HC 174 PRO HG2 2843 0.021300 1.00800 ; qtot
0.883
2844 HC 174 PRO HG3 2844 0.021300 1.00800 ; qtot
0.904
2845 CT 174 PRO CB 2845 -0.007000 12.01000 ; qtot
0.897
2846 HC 174 PRO HB2 2846 0.025300 1.00800 ; qtot
0.922
2847 HC 174 PRO HB3 2847 0.025300 1.00800 ; qtot
0.948
2848 CX 174 PRO CA 2848 -0.026600 12.01000 ; qtot
0.921
2849 H1 174 PRO HA 2849 0.064100 1.00800 ; qtot
0.985
2850 C 174 PRO C 2850 0.589601 12.01000 ; qtot
1.575
2851 O 174 PRO O 2851 -0.574801 16.00000 ; qtot
1.000
2852 N 175 TYR N 2852 -0.415700 14.01000 ; qtot
0.584
2853 H 175 TYR H 2853 0.271900 1.00800 ; qtot
0.856
2854 CX 175 TYR CA 2854 -0.001400 12.01000 ; qtot
0.855
2855 H1 175 TYR HA 2855 0.087600 1.00800 ; qtot
0.942
2856 CT 175 TYR CB 2856 -0.015200 12.01000 ; qtot
0.927
2857 HC 175 TYR HB2 2857 0.029500 1.00800 ; qtot
0.957

2858 HC 175 TYR HB3 2858 0.029500 1.00800 ; qtot
0.986
2859 CA 175 TYR CG 2859 -0.001100 12.01000 ; qtot
0.985
2860 CA 175 TYR CD1 2860 -0.190600 12.01000 ; qtot
0.795
2861 HA 175 TYR HD1 2861 0.169900 1.00800 ; qtot
0.964
2862 CA 175 TYR CE1 2862 -0.234100 12.01000 ; qtot
0.730
2863 HA 175 TYR HE1 2863 0.165600 1.00800 ; qtot
0.896
2864 C 175 TYR CZ 2864 0.322600 12.01000 ; qtot
1.219
2865 OH 175 TYR OH 2865 -0.557901 16.00000 ; qtot
0.661
2866 HO 175 TYR HH 2866 0.399200 1.00800 ; qtot
1.060
2867 CA 175 TYR CE2 2867 -0.234100 12.01000 ; qtot
0.826
2868 HA 175 TYR HE2 2868 0.165600 1.00800 ; qtot
0.991
2869 CA 175 TYR CD2 2869 -0.190600 12.01000 ; qtot
0.801
2870 HA 175 TYR HD2 2870 0.169900 1.00800 ; qtot
0.971
2871 C 175 TYR C 2871 0.597301 12.01000 ; qtot
1.568
2872 O 175 TYR O 2872 -0.567901 16.00000 ; qtot
1.000
2873 N 176 TYR N 2873 -0.415700 14.01000 ; qtot
0.584
2874 H 176 TYR H 2874 0.271900 1.00800 ; qtot
0.856
2875 CX 176 TYR CA 2875 -0.001400 12.01000 ; qtot
0.855
2876 H1 176 TYR HA 2876 0.087600 1.00800 ; qtot
0.942
2877 CT 176 TYR CB 2877 -0.015200 12.01000 ; qtot
0.927
2878 HC 176 TYR HB2 2878 0.029500 1.00800 ; qtot
0.957
2879 HC 176 TYR HB3 2879 0.029500 1.00800 ; qtot
0.986
2880 CA 176 TYR CG 2880 -0.001100 12.01000 ; qtot
0.985
2881 CA 176 TYR CD1 2881 -0.190600 12.01000 ; qtot
0.795
2882 HA 176 TYR HD1 2882 0.169900 1.00800 ; qtot
0.964
2883 CA 176 TYR CE1 2883 -0.234100 12.01000 ; qtot
0.730

2884 HA 176 TYR HE1 2884 0.165600 1.00800 ; qtot
0.896
2885 C 176 TYR CZ 2885 0.322600 12.01000 ; qtot
1.219
2886 OH 176 TYR OH 2886 -0.557901 16.00000 ; qtot
0.661
2887 HO 176 TYR HH 2887 0.399200 1.00800 ; qtot
1.060
2888 CA 176 TYR CE2 2888 -0.234100 12.01000 ; qtot
0.826
2889 HA 176 TYR HE2 2889 0.165600 1.00800 ; qtot
0.991
2890 CA 176 TYR CD2 2890 -0.190600 12.01000 ; qtot
0.801
2891 HA 176 TYR HD2 2891 0.169900 1.00800 ; qtot
0.971
2892 C 176 TYR C 2892 0.597301 12.01000 ; qtot
1.568
2893 O 176 TYR O 2893 -0.567901 16.00000 ; qtot
1.000
2894 N 177 VAL N 2894 -0.415700 14.01000 ; qtot
0.584
2895 H 177 VAL H 2895 0.271900 1.00800 ; qtot
0.856
2896 CX 177 VAL CA 2896 -0.087500 12.01000 ; qtot
0.769
2897 H1 177 VAL HA 2897 0.096900 1.00800 ; qtot
0.866
2898 A3C 177 VAL CB 2898 0.298500 12.01000 ; qtot
1.164
2899 HC 177 VAL HB 2899 -0.029700 1.00800 ; qtot
1.134
2900 CT 177 VAL CG1 2900 -0.319200 12.01000 ; qtot
0.815
2901 HC 177 VAL HG11 2901 0.079100 1.00800 ; qtot
0.894
2902 HC 177 VAL HG12 2902 0.079100 1.00800 ; qtot
0.973
2903 HC 177 VAL HG13 2903 0.079100 1.00800 ; qtot
1.053
2904 CT 177 VAL CG2 2904 -0.319200 12.01000 ; qtot
0.733
2905 HC 177 VAL HG21 2905 0.079100 1.00800 ; qtot
0.812
2906 HC 177 VAL HG22 2906 0.079100 1.00800 ; qtot
0.892
2907 HC 177 VAL HG23 2907 0.079100 1.00800 ; qtot
0.971
2908 C 177 VAL C 2908 0.597301 12.01000 ; qtot
1.568
2909 O 177 VAL O 2909 -0.567901 16.00000 ; qtot
1.000

2910 N 178 ALA N 2910 -0.415700 14.01000 ; qtot
0.584
2911 H 178 ALA H 2911 0.271900 1.00800 ; qtot
0.856
2912 CX 178 ALA CA 2912 0.033700 12.01000 ; qtot
0.890
2913 H1 178 ALA HA 2913 0.082300 1.00800 ; qtot
0.972
2914 CT 178 ALA CB 2914 -0.182500 12.01000 ; qtot
0.790
2915 HC 178 ALA HB1 2915 0.060300 1.00800 ; qtot
0.850
2916 HC 178 ALA HB2 2916 0.060300 1.00800 ; qtot
0.910
2917 HC 178 ALA HB3 2917 0.060300 1.00800 ; qtot
0.971
2918 C 178 ALA C 2918 0.597301 12.01000 ; qtot
1.568
2919 O 178 ALA O 2919 -0.567901 16.00000 ; qtot
1.000
2920 N 179 PRO N 2920 -0.254800 14.01000 ; qtot
0.745
2921 CT 179 PRO CD 2921 0.019200 12.01000 ; qtot
0.764
2922 H1 179 PRO HD2 2922 0.039100 1.00800 ; qtot
0.804
2923 H1 179 PRO HD3 2923 0.039100 1.00800 ; qtot
0.843
2924 CT 179 PRO CG 2924 0.018900 12.01000 ; qtot
0.862
2925 HC 179 PRO HG2 2925 0.021300 1.00800 ; qtot
0.883
2926 HC 179 PRO HG3 2926 0.021300 1.00800 ; qtot
0.904
2927 CT 179 PRO CB 2927 -0.007000 12.01000 ; qtot
0.897
2928 HC 179 PRO HB2 2928 0.025300 1.00800 ; qtot
0.922
2929 HC 179 PRO HB3 2929 0.025300 1.00800 ; qtot
0.948
2930 CX 179 PRO CA 2930 -0.026600 12.01000 ; qtot
0.921
2931 H1 179 PRO HA 2931 0.064100 1.00800 ; qtot
0.985
2932 C 179 PRO C 2932 0.589601 12.01000 ; qtot
1.575
2933 O 179 PRO O 2933 -0.574801 16.00000 ; qtot
1.000
2934 N 180 GLN N 2934 -0.415700 14.01000 ; qtot
0.584
2935 H 180 GLN H 2935 0.271900 1.00800 ; qtot
0.856

2936 CX 180 GLN CA 2936 -0.003100 12.01000 ; qtot
0.853
2937 H1 180 GLN HA 2937 0.085000 1.00800 ; qtot
0.938
2938 A2C 180 GLN CB 2938 -0.003600 12.01000 ; qtot
0.935
2939 HC 180 GLN HB2 2939 0.017100 1.00800 ; qtot
0.952
2940 HC 180 GLN HB3 2940 0.017100 1.00800 ; qtot
0.969
2941 A2C 180 GLN CG 2941 -0.064500 12.01000 ; qtot
0.904
2942 HC 180 GLN HG2 2942 0.035200 1.00800 ; qtot
0.939
2943 HC 180 GLN HG3 2943 0.035200 1.00800 ; qtot
0.975
2944 C 180 GLN CD 2944 0.695101 12.01000 ; qtot
1.670
2945 O 180 GLN OE1 2945 -0.608601 16.00000 ; qtot
1.061
2946 N 180 GLN NE2 2946 -0.940701 14.01000 ; qtot
0.120
2947 H 180 GLN HE21 2947 0.425100 1.00800 ; qtot
0.546
2948 H 180 GLN HE22 2948 0.425100 1.00800 ; qtot
0.971
2949 C 180 GLN C 2949 0.597301 12.01000 ; qtot
1.568
2950 O 180 GLN O 2950 -0.567901 16.00000 ; qtot
1.000
2951 N 181 VAL N 2951 -0.415700 14.01000 ; qtot
0.584
2952 H 181 VAL H 2952 0.271900 1.00800 ; qtot
0.856
2953 CX 181 VAL CA 2953 -0.087500 12.01000 ; qtot
0.769
2954 H1 181 VAL HA 2954 0.096900 1.00800 ; qtot
0.866
2955 A3C 181 VAL CB 2955 0.298500 12.01000 ; qtot
1.164
2956 HC 181 VAL HB 2956 -0.029700 1.00800 ; qtot
1.134
2957 CT 181 VAL CG1 2957 -0.319200 12.01000 ; qtot
0.815
2958 HC 181 VAL HG11 2958 0.079100 1.00800 ; qtot
0.894
2959 HC 181 VAL HG12 2959 0.079100 1.00800 ; qtot
0.973
2960 HC 181 VAL HG13 2960 0.079100 1.00800 ; qtot
1.053
2961 CT 181 VAL CG2 2961 -0.319200 12.01000 ; qtot
0.733

2962 HC 181 VAL HG21 2962 0.079100 1.00800 ; qtot
0.812
2963 HC 181 VAL HG22 2963 0.079100 1.00800 ; qtot
0.892
2964 HC 181 VAL HG23 2964 0.079100 1.00800 ; qtot
0.971
2965 C 181 VAL C 2965 0.597301 12.01000 ; qtot
1.568
2966 O 181 VAL O 2966 -0.567901 16.00000 ; qtot
1.000
2967 N 182 LEU N 2967 -0.415700 14.01000 ; qtot
0.584
2968 H 182 LEU H 2968 0.271900 1.00800 ; qtot
0.856
2969 CX 182 LEU CA 2969 -0.051800 12.01000 ; qtot
0.804
2970 H1 182 LEU HA 2970 0.092200 1.00800 ; qtot
0.897
2971 A2C 182 LEU CB 2971 -0.110200 12.01000 ; qtot
0.786
2972 HC 182 LEU HB2 2972 0.045700 1.00800 ; qtot
0.832
2973 HC 182 LEU HB3 2973 0.045700 1.00800 ; qtot
0.878
2974 A3C 182 LEU CG 2974 0.353100 12.01000 ; qtot
1.231
2975 HC 182 LEU HG 2975 -0.036100 1.00800 ; qtot
1.195
2976 CT 182 LEU CD1 2976 -0.412100 12.01000 ; qtot
0.783
2977 HC 182 LEU HD11 2977 0.100000 1.00800 ; qtot
0.883
2978 HC 182 LEU HD12 2978 0.100000 1.00800 ; qtot
0.983
2979 HC 182 LEU HD13 2979 0.100000 1.00800 ; qtot
1.083
2980 CT 182 LEU CD2 2980 -0.412100 12.01000 ; qtot
0.671
2981 HC 182 LEU HD21 2981 0.100000 1.00800 ; qtot
0.771
2982 HC 182 LEU HD22 2982 0.100000 1.00800 ; qtot
0.871
2983 HC 182 LEU HD23 2983 0.100000 1.00800 ; qtot
0.971
2984 C 182 LEU C 2984 0.597301 12.01000 ; qtot
1.568
2985 O 182 LEU O 2985 -0.567901 16.00000 ; qtot
1.000
2986 N 183 ASP N 2986 -0.516301 14.01000 ; qtot
0.484
2987 H 183 ASP H 2987 0.293600 1.00800 ; qtot
0.777

```
  2988   CX   183   ASP    CA 2988     0.038100     12.01000 ; qtot
0.815
  2989   H1   183   ASP    HA 2989     0.088000      1.00800 ; qtot
0.903
  2990  A2C   183   ASP    CB 2990    -0.030300     12.01000 ; qtot
0.873
  2991   HC   183   ASP   HB2 2991    -0.012200      1.00800 ; qtot
0.861
  2992   HC   183   ASP   HB3 2992    -0.012200      1.00800 ; qtot
0.849
  2993   CO   183   ASP    CG 2993     0.799401     12.01000 ; qtot
1.648
  2994   O2   183   ASP   OD1 2994    -0.801401     16.00000 ; qtot
0.847
  2995   O2   183   ASP   OD2 2995    -0.801401     16.00000 ; qtot
0.045
  2996    C   183   ASP     C 2996     0.536601     12.01000 ; qtot
0.582
  2997    O   183   ASP     O 2997    -0.581901     16.00000 ; qtot
0.000
  2998    N   184   GLY     N 2998    -0.415700     14.01000 ; qtot -
0.416
  2999    H   184   GLY     H 2999     0.271900      1.00800 ; qtot -
0.144
  3000   CX   184   GLY    CA 3000    -0.025200     12.01000 ; qtot -
0.169
  3001   H1   184   GLY   HA2 3001     0.069800      1.00800 ; qtot -
0.099
  3002   H1   184   GLY   HA3 3002     0.069800      1.00800 ; qtot -
0.029
  3003    C   184   GLY     C 3003     0.597301     12.01000 ; qtot
0.568
  3004    O   184   GLY     O 3004    -0.567901     16.00000 ; qtot
0.000
  3005    N   185   LYS     N 3005    -0.347900     14.01000 ; qtot -
0.348
  3006    H   185   LYS     H 3006     0.274700      1.00800 ; qtot -
0.073
  3007   CX   185   LYS    CA 3007    -0.240000     12.01000 ; qtot -
0.313
  3008   H1   185   LYS    HA 3008     0.142600      1.00800 ; qtot -
0.171
  3009   C8   185   LYS    CB 3009    -0.009400     12.01000 ; qtot -
0.180
  3010   HC   185   LYS   HB2 3010     0.036200      1.00800 ; qtot -
0.144
  3011   HC   185   LYS   HB3 3011     0.036200      1.00800 ; qtot -
0.108
  3012   C8   185   LYS    CG 3012     0.018700     12.01000 ; qtot -
0.089
  3013   HC   185   LYS   HG2 3013     0.010300      1.00800 ; qtot -
0.079
```

```
  3014   HC   185   LYS   HG3 3014     0.010300      1.00800 ; qtot -
0.068
  3015   C8   185   LYS    CD 3015    -0.047900     12.01000 ; qtot -
0.116
  3016   HC   185   LYS   HD2 3016     0.062100      1.00800 ; qtot -
0.054
  3017   HC   185   LYS   HD3 3017     0.062100      1.00800 ; qtot
0.008
  3018   C8   185   LYS    CE 3018    -0.014300     12.01000 ; qtot -
0.006
  3019   HP   185   LYS   HE2 3019     0.113500      1.00800 ; qtot
0.107
  3020   HP   185   LYS   HE3 3020     0.113500      1.00800 ; qtot
0.221
  3021   N3   185   LYS    NZ 3021    -0.385400     14.01000 ; qtot -
0.165
  3022    H   185   LYS   HZ1 3022     0.340000      1.00800 ; qtot
0.175
  3023    H   185   LYS   HZ2 3023     0.340000      1.00800 ; qtot
0.515
  3024    H   185   LYS   HZ3 3024     0.340000      1.00800 ; qtot
0.855
  3025    C   185   LYS     C 3025     0.734101     12.01000 ; qtot
1.589
  3026    O   185   LYS     O 3026    -0.589401     16.00000 ; qtot
1.000
  3027    N   186   TYR     N 3027    -0.415700     14.01000 ; qtot
0.584
  3028    H   186   TYR     H 3028     0.271900      1.00800 ; qtot
0.856
  3029   CX   186   TYR    CA 3029    -0.001400     12.01000 ; qtot
0.855
  3030   H1   186   TYR    HA 3030     0.087600      1.00800 ; qtot
0.942
  3031   CT   186   TYR    CB 3031    -0.015200     12.01000 ; qtot
0.927
  3032   HC   186   TYR   HB2 3032     0.029500      1.00800 ; qtot
0.957
  3033   HC   186   TYR   HB3 3033     0.029500      1.00800 ; qtot
0.986
  3034   CA   186   TYR    CG 3034    -0.001100     12.01000 ; qtot
0.985
  3035   CA   186   TYR   CD1 3035    -0.190600     12.01000 ; qtot
0.795
  3036   HA   186   TYR   HD1 3036     0.169900      1.00800 ; qtot
0.964
  3037   CA   186   TYR   CE1 3037    -0.234100     12.01000 ; qtot
0.730
  3038   HA   186   TYR   HE1 3038     0.165600      1.00800 ; qtot
0.896
  3039    C   186   TYR    CZ 3039     0.322600     12.01000 ; qtot
1.219
```

```
  3040   OH   186   TYR    OH 3040    -0.557901     16.00000 ; qtot
0.661
  3041   HO   186   TYR    HH 3041     0.399200      1.00800 ; qtot
1.060
  3042   CA   186   TYR   CE2 3042    -0.234100     12.01000 ; qtot
0.826
  3043   HA   186   TYR   HE2 3043     0.165600      1.00800 ; qtot
0.991
  3044   CA   186   TYR   CD2 3044    -0.190600     12.01000 ; qtot
0.801
  3045   HA   186   TYR   HD2 3045     0.169900      1.00800 ; qtot
0.971
  3046    C   186   TYR     C 3046     0.597301     12.01000 ; qtot
1.568
  3047    O   186   TYR     O 3047    -0.567901     16.00000 ; qtot
1.000
  3048    N   187   ASP     N 3048    -0.516301     14.01000 ; qtot
0.484
  3049    H   187   ASP     H 3049     0.293600      1.00800 ; qtot
0.777
  3050   CX   187   ASP    CA 3050     0.038100     12.01000 ; qtot
0.815
  3051   H1   187   ASP    HA 3051     0.088000      1.00800 ; qtot
0.903
  3052  A2C   187   ASP    CB 3052    -0.030300     12.01000 ; qtot
0.873
  3053   HC   187   ASP   HB2 3053    -0.012200      1.00800 ; qtot
0.861
  3054   HC   187   ASP   HB3 3054    -0.012200      1.00800 ; qtot
0.849
  3055   CO   187   ASP    CG 3055     0.799401     12.01000 ; qtot
1.648
  3056   O2   187   ASP   OD1 3056    -0.801401     16.00000 ; qtot
0.847
  3057   O2   187   ASP   OD2 3057    -0.801401     16.00000 ; qtot
0.045
  3058    C   187   ASP     C 3058     0.536601     12.01000 ; qtot
0.582
  3059    O   187   ASP     O 3059    -0.581901     16.00000 ; qtot
0.000
  3060    N   188   LYS     N 3060    -0.347900     14.01000 ; qtot -
0.348
  3061    H   188   LYS     H 3061     0.274700      1.00800 ; qtot -
0.073
  3062   CX   188   LYS    CA 3062    -0.240000     12.01000 ; qtot -
0.313
  3063   H1   188   LYS    HA 3063     0.142600      1.00800 ; qtot -
0.171
  3064   C8   188   LYS    CB 3064    -0.009400     12.01000 ; qtot -
0.180
  3065   HC   188   LYS   HB2 3065     0.036200      1.00800 ; qtot -
0.144
```

3066 HC 188 LYS HB3 3066 0.036200 1.00800 ; qtot -
0.108
3067 C8 188 LYS CG 3067 0.018700 12.01000 ; qtot -
0.089
3068 HC 188 LYS HG2 3068 0.010300 1.00800 ; qtot -
0.079
3069 HC 188 LYS HG3 3069 0.010300 1.00800 ; qtot -
0.068
3070 C8 188 LYS CD 3070 -0.047900 12.01000 ; qtot -
0.116
3071 HC 188 LYS HD2 3071 0.062100 1.00800 ; qtot -
0.054
3072 HC 188 LYS HD3 3072 0.062100 1.00800 ; qtot
0.008
3073 C8 188 LYS CE 3073 -0.014300 12.01000 ; qtot -
0.006
3074 HP 188 LYS HE2 3074 0.113500 1.00800 ; qtot
0.107
3075 HP 188 LYS HE3 3075 0.113500 1.00800 ; qtot
0.221
3076 N3 188 LYS NZ 3076 -0.385400 14.01000 ; qtot -
0.165
3077 H 188 LYS HZ1 3077 0.340000 1.00800 ; qtot
0.175
3078 H 188 LYS HZ2 3078 0.340000 1.00800 ; qtot
0.515
3079 H 188 LYS HZ3 3079 0.340000 1.00800 ; qtot
0.855
3080 C 188 LYS C 3080 0.734101 12.01000 ; qtot
1.589
3081 O 188 LYS O 3081 -0.589401 16.00000 ; qtot
1.000
3082 N 189 LYS N 3082 -0.347900 14.01000 ; qtot
0.652
3083 H 189 LYS H 3083 0.274700 1.00800 ; qtot
0.927
3084 CX 189 LYS CA 3084 -0.240000 12.01000 ; qtot
0.687
3085 H1 189 LYS HA 3085 0.142600 1.00800 ; qtot
0.829
3086 C8 189 LYS CB 3086 -0.009400 12.01000 ; qtot
0.820
3087 HC 189 LYS HB2 3087 0.036200 1.00800 ; qtot
0.856
3088 HC 189 LYS HB3 3088 0.036200 1.00800 ; qtot
0.892
3089 C8 189 LYS CG 3089 0.018700 12.01000 ; qtot
0.911
3090 HC 189 LYS HG2 3090 0.010300 1.00800 ; qtot
0.921
3091 HC 189 LYS HG3 3091 0.010300 1.00800 ; qtot
0.932

3092 C8 189 LYS CD 3092 -0.047900 12.01000 ; qtot
0.884
3093 HC 189 LYS HD2 3093 0.062100 1.00800 ; qtot
0.946
3094 HC 189 LYS HD3 3094 0.062100 1.00800 ; qtot
1.008
3095 C8 189 LYS CE 3095 -0.014300 12.01000 ; qtot
0.994
3096 HP 189 LYS HE2 3096 0.113500 1.00800 ; qtot
1.107
3097 HP 189 LYS HE3 3097 0.113500 1.00800 ; qtot
1.221
3098 N3 189 LYS NZ 3098 -0.385400 14.01000 ; qtot
0.835
3099 H 189 LYS HZ1 3099 0.340000 1.00800 ; qtot
1.175
3100 H 189 LYS HZ2 3100 0.340000 1.00800 ; qtot
1.515
3101 H 189 LYS HZ3 3101 0.340000 1.00800 ; qtot
1.855
3102 C 189 LYS C 3102 0.734101 12.01000 ; qtot
2.589
3103 O 189 LYS O 3103 -0.589401 16.00000 ; qtot
2.000
3104 N 190 CYS N 3104 -0.415700 14.01000 ; qtot
1.584
3105 H 190 CYS H 3105 0.271900 1.00800 ; qtot
1.856
3106 CX 190 CYS CA 3106 0.021300 12.01000 ; qtot
1.878
3107 H1 190 CYS HA 3107 0.112400 1.00800 ; qtot
1.990
3108 A2C 190 CYS CB 3108 -0.123100 12.01000 ; qtot
1.867
3109 H1 190 CYS HB2 3109 0.111200 1.00800 ; qtot
1.978
3110 H1 190 CYS HB3 3110 0.111200 1.00800 ; qtot
2.089
3111 SH 190 CYS SG 3111 -0.311900 32.06000 ; qtot
1.777
3112 HS 190 CYS HG 3112 0.193300 1.00800 ; qtot
1.971
3113 C 190 CYS C 3113 0.597301 12.01000 ; qtot
2.568
3114 O 190 CYS O 3114 -0.567901 16.00000 ; qtot
2.000
3115 N 191 ASP N 3115 -0.516301 14.01000 ; qtot
1.484
3116 H 191 ASP H 3116 0.293600 1.00800 ; qtot
1.777
3117 CX 191 ASP CA 3117 0.038100 12.01000 ; qtot
1.815

3118 H1 191 ASP HA 3118 0.088000 1.00800 ; qtot
1.903
3119 A2C 191 ASP CB 3119 -0.030300 12.01000 ; qtot
1.873
3120 HC 191 ASP HB2 3120 -0.012200 1.00800 ; qtot
1.861
3121 HC 191 ASP HB3 3121 -0.012200 1.00800 ; qtot
1.849
3122 CO 191 ASP CG 3122 0.799401 12.01000 ; qtot
2.648
3123 O2 191 ASP OD1 3123 -0.801401 16.00000 ; qtot
1.847
3124 O2 191 ASP OD2 3124 -0.801401 16.00000 ; qtot
1.045
3125 C 191 ASP C 3125 0.536601 12.01000 ; qtot
1.582
3126 O 191 ASP O 3126 -0.581901 16.00000 ; qtot
1.000
3127 N 192 ILE N 3127 -0.415700 14.01000 ; qtot
0.584
3128 H 192 ILE H 3128 0.271900 1.00800 ; qtot
0.856
3129 CX 192 ILE CA 3129 -0.059700 12.01000 ; qtot
0.797
3130 H1 192 ILE HA 3130 0.086900 1.00800 ; qtot
0.883
3131 A3C 192 ILE CB 3131 0.130300 12.01000 ; qtot
1.014
3132 HC 192 ILE HB 3132 0.018700 1.00800 ; qtot
1.032
3133 CT 192 ILE CG2 3133 -0.320400 12.01000 ; qtot
0.712
3134 HC 192 ILE HG21 3134 0.088200 1.00800 ; qtot
0.800
3135 HC 192 ILE HG22 3135 0.088200 1.00800 ; qtot
0.888
3136 HC 192 ILE HG23 3136 0.088200 1.00800 ; qtot
0.977
3137 A2C 192 ILE CG1 3137 -0.043000 12.01000 ; qtot
0.934
3138 HC 192 ILE HG12 3138 0.023600 1.00800 ; qtot
0.957
3139 HC 192 ILE HG13 3139 0.023600 1.00800 ; qtot
0.981
3140 CT 192 ILE CD1 3140 -0.066000 12.01000 ; qtot
0.915
3141 HC 192 ILE HD11 3141 0.018600 1.00800 ; qtot
0.933
3142 HC 192 ILE HD12 3142 0.018600 1.00800 ; qtot
0.952
3143 HC 192 ILE HD13 3143 0.018600 1.00800 ; qtot
0.971

3144 C 192 ILE C 3144 0.597301 12.01000 ; qtot
1.568
3145 O 192 ILE O 3145 -0.567901 16.00000 ; qtot
1.000
3146 N 193 TRP N 3146 -0.415700 14.01000 ; qtot
0.584
3147 H 193 TRP H 3147 0.271900 1.00800 ; qtot
0.856
3148 CX 193 TRP CA 3148 -0.027500 12.01000 ; qtot
0.829
3149 H1 193 TRP HA 3149 0.112300 1.00800 ; qtot
0.941
3150 CT 193 TRP CB 3150 -0.005000 12.01000 ; qtot
0.936
3151 HC 193 TRP HB2 3151 0.033900 1.00800 ; qtot
0.970
3152 HC 193 TRP HB3 3152 0.033900 1.00800 ; qtot
1.004
3153 C* 193 TRP CG 3153 -0.141500 12.01000 ; qtot
0.862
3154 CW 193 TRP CD1 3154 -0.163800 12.01000 ; qtot
0.699
3155 H4 193 TRP HD1 3155 0.206200 1.00800 ; qtot
0.905
3156 NA 193 TRP NE1 3156 -0.341800 14.01000 ; qtot
0.563
3157 H 193 TRP HE1 3157 0.341200 1.00800 ; qtot
0.904
3158 CN 193 TRP CE2 3158 0.138000 12.01000 ; qtot
1.042
3159 CA 193 TRP CZ2 3159 -0.260100 12.01000 ; qtot
0.782
3160 HA 193 TRP HZ2 3160 0.157200 1.00800 ; qtot
0.939
3161 CA 193 TRP CH2 3161 -0.113400 12.01000 ; qtot
0.826
3162 HA 193 TRP HH2 3162 0.141700 1.00800 ; qtot
0.968
3163 CA 193 TRP CZ3 3163 -0.197200 12.01000 ; qtot
0.770
3164 HA 193 TRP HZ3 3164 0.144700 1.00800 ; qtot
0.915
3165 CA 193 TRP CE3 3165 -0.238700 12.01000 ; qtot
0.676
3166 HA 193 TRP HE3 3166 0.170000 1.00800 ; qtot
0.846
3167 CB 193 TRP CD2 3167 0.124300 12.01000 ; qtot
0.971
3168 C 193 TRP C 3168 0.597301 12.01000 ; qtot
1.568
3169 O 193 TRP O 3169 -0.567901 16.00000 ; qtot
1.000

3170 N 194 SER N 3170 -0.415700 14.01000 ; qtot
0.584
3171 H 194 SER H 3171 0.271900 1.00800 ; qtot
0.856
3172 CX 194 SER CA 3172 -0.024900 12.01000 ; qtot
0.831
3173 H1 194 SER HA 3173 0.084300 1.00800 ; qtot
0.916
3174 A2C 194 SER CB 3174 0.211700 12.01000 ; qtot
1.127
3175 H1 194 SER HB2 3175 0.035200 1.00800 ; qtot
1.163
3176 H1 194 SER HB3 3176 0.035200 1.00800 ; qtot
1.198
3177 OH 194 SER OG 3177 -0.654601 16.00000 ; qtot
0.543
3178 HO 194 SER HG 3178 0.427500 1.00800 ; qtot
0.971
3179 C 194 SER C 3179 0.597301 12.01000 ; qtot
1.568
3180 O 194 SER O 3180 -0.567901 16.00000 ; qtot
1.000
3181 N 195 SER N 3181 -0.415700 14.01000 ; qtot
0.584
3182 H 195 SER H 3182 0.271900 1.00800 ; qtot
0.856
3183 CX 195 SER CA 3183 -0.024900 12.01000 ; qtot
0.831
3184 H1 195 SER HA 3184 0.084300 1.00800 ; qtot
0.916
3185 A2C 195 SER CB 3185 0.211700 12.01000 ; qtot
1.127
3186 H1 195 SER HB2 3186 0.035200 1.00800 ; qtot
1.163
3187 H1 195 SER HB3 3187 0.035200 1.00800 ; qtot
1.198
3188 OH 195 SER OG 3188 -0.654601 16.00000 ; qtot
0.543
3189 HO 195 SER HG 3189 0.427500 1.00800 ; qtot
0.971
3190 C 195 SER C 3190 0.597301 12.01000 ; qtot
1.568
3191 O 195 SER O 3191 -0.567901 16.00000 ; qtot
1.000
3192 N 196 GLY N 3192 -0.415700 14.01000 ; qtot
0.584
3193 H 196 GLY H 3193 0.271900 1.00800 ; qtot
0.856
3194 CX 196 GLY CA 3194 -0.025200 12.01000 ; qtot
0.831
3195 H1 196 GLY HA2 3195 0.069800 1.00800 ; qtot
0.901

```
  3196   H1   196   GLY   HA3 3196     0.069800      1.00800 ; qtot
0.971
  3197    C   196   GLY     C 3197     0.597301     12.01000 ; qtot
1.568
  3198    O   196   GLY     O 3198    -0.567901     16.00000 ; qtot
1.000
  3199    N   197   VAL     N 3199    -0.415700     14.01000 ; qtot
0.584
  3200    H   197   VAL     H 3200     0.271900      1.00800 ; qtot
0.856
  3201   CX   197   VAL    CA 3201    -0.087500     12.01000 ; qtot
0.769
  3202   H1   197   VAL    HA 3202     0.096900      1.00800 ; qtot
0.866
  3203  A3C   197   VAL    CB 3203     0.298500     12.01000 ; qtot
1.164
  3204   HC   197   VAL    HB 3204    -0.029700      1.00800 ; qtot
1.134
  3205   CT   197   VAL   CG1 3205    -0.319200     12.01000 ; qtot
0.815
  3206   HC   197   VAL  HG11 3206     0.079100      1.00800 ; qtot
0.894
  3207   HC   197   VAL  HG12 3207     0.079100      1.00800 ; qtot
0.973
  3208   HC   197   VAL  HG13 3208     0.079100      1.00800 ; qtot
1.053
  3209   CT   197   VAL   CG2 3209    -0.319200     12.01000 ; qtot
0.733
  3210   HC   197   VAL  HG21 3210     0.079100      1.00800 ; qtot
0.812
  3211   HC   197   VAL  HG22 3211     0.079100      1.00800 ; qtot
0.892
  3212   HC   197   VAL  HG23 3212     0.079100      1.00800 ; qtot
0.971
  3213    C   197   VAL     C 3213     0.597301     12.01000 ; qtot
1.568
  3214    O   197   VAL     O 3214    -0.567901     16.00000 ; qtot
1.000
  3215    N   198   ILE     N 3215    -0.415700     14.01000 ; qtot
0.584
  3216    H   198   ILE     H 3216     0.271900      1.00800 ; qtot
0.856
  3217   CX   198   ILE    CA 3217    -0.059700     12.01000 ; qtot
0.797
  3218   H1   198   ILE    HA 3218     0.086900      1.00800 ; qtot
0.883
  3219  A3C   198   ILE    CB 3219     0.130300     12.01000 ; qtot
1.014
  3220   HC   198   ILE    HB 3220     0.018700      1.00800 ; qtot
1.032
  3221   CT   198   ILE   CG2 3221    -0.320400     12.01000 ; qtot
0.712
```

3222 HC 198 ILE HG21 3222 0.088200 1.00800 ; qtot
0.800
3223 HC 198 ILE HG22 3223 0.088200 1.00800 ; qtot
0.888
3224 HC 198 ILE HG23 3224 0.088200 1.00800 ; qtot
0.977
3225 A2C 198 ILE CG1 3225 -0.043000 12.01000 ; qtot
0.934
3226 HC 198 ILE HG12 3226 0.023600 1.00800 ; qtot
0.957
3227 HC 198 ILE HG13 3227 0.023600 1.00800 ; qtot
0.981
3228 CT 198 ILE CD1 3228 -0.066000 12.01000 ; qtot
0.915
3229 HC 198 ILE HD11 3229 0.018600 1.00800 ; qtot
0.933
3230 HC 198 ILE HD12 3230 0.018600 1.00800 ; qtot
0.952
3231 HC 198 ILE HD13 3231 0.018600 1.00800 ; qtot
0.971
3232 C 198 ILE C 3232 0.597301 12.01000 ; qtot
1.568
3233 O 198 ILE O 3233 -0.567901 16.00000 ; qtot
1.000
3234 N 199 MET N 3234 -0.415700 14.01000 ; qtot
0.584
3235 H 199 MET H 3235 0.271900 1.00800 ; qtot
0.856
3236 CX 199 MET CA 3236 -0.023700 12.01000 ; qtot
0.833
3237 H1 199 MET HA 3237 0.088000 1.00800 ; qtot
0.921
3238 A2C 199 MET CB 3238 0.034200 12.01000 ; qtot
0.955
3239 HC 199 MET HB2 3239 0.024100 1.00800 ; qtot
0.979
3240 HC 199 MET HB3 3240 0.024100 1.00800 ; qtot
1.003
3241 A2C 199 MET CG 3241 0.001800 12.01000 ; qtot
1.005
3242 H1 199 MET HG2 3242 0.044000 1.00800 ; qtot
1.049
3243 H1 199 MET HG3 3243 0.044000 1.00800 ; qtot
1.093
3244 S 199 MET SD 3244 -0.273700 32.06000 ; qtot
0.819
3245 CT 199 MET CE 3245 -0.053600 12.01000 ; qtot
0.765
3246 H1 199 MET HE1 3246 0.068400 1.00800 ; qtot
0.834
3247 H1 199 MET HE2 3247 0.068400 1.00800 ; qtot
0.902

3248 H1 199 MET HE3 3248 0.068400 1.00800 ; qtot
0.971
3249 C 199 MET C 3249 0.597301 12.01000 ; qtot
1.568
3250 O 199 MET O 3250 -0.567901 16.00000 ; qtot
1.000
3251 N 200 TYR N 3251 -0.415700 14.01000 ; qtot
0.584
3252 H 200 TYR H 3252 0.271900 1.00800 ; qtot
0.856
3253 CX 200 TYR CA 3253 -0.001400 12.01000 ; qtot
0.855
3254 H1 200 TYR HA 3254 0.087600 1.00800 ; qtot
0.942
3255 CT 200 TYR CB 3255 -0.015200 12.01000 ; qtot
0.927
3256 HC 200 TYR HB2 3256 0.029500 1.00800 ; qtot
0.957
3257 HC 200 TYR HB3 3257 0.029500 1.00800 ; qtot
0.986
3258 CA 200 TYR CG 3258 -0.001100 12.01000 ; qtot
0.985
3259 CA 200 TYR CD1 3259 -0.190600 12.01000 ; qtot
0.795
3260 HA 200 TYR HD1 3260 0.169900 1.00800 ; qtot
0.964
3261 CA 200 TYR CE1 3261 -0.234100 12.01000 ; qtot
0.730
3262 HA 200 TYR HE1 3262 0.165600 1.00800 ; qtot
0.896
3263 C 200 TYR CZ 3263 0.322600 12.01000 ; qtot
1.219
3264 OH 200 TYR OH 3264 -0.557901 16.00000 ; qtot
0.661
3265 HO 200 TYR HH 3265 0.399200 1.00800 ; qtot
1.060
3266 CA 200 TYR CE2 3266 -0.234100 12.01000 ; qtot
0.826
3267 HA 200 TYR HE2 3267 0.165600 1.00800 ; qtot
0.991
3268 CA 200 TYR CD2 3268 -0.190600 12.01000 ; qtot
0.801
3269 HA 200 TYR HD2 3269 0.169900 1.00800 ; qtot
0.971
3270 C 200 TYR C 3270 0.597301 12.01000 ; qtot
1.568
3271 O 200 TYR O 3271 -0.567901 16.00000 ; qtot
1.000
3272 N 201 THR N 3272 -0.415700 14.01000 ; qtot
0.584
3273 H 201 THR H 3273 0.271900 1.00800 ; qtot
0.856

3274 CX 201 THR CA 3274 -0.038900 12.01000 ; qtot
0.817
3275 H1 201 THR HA 3275 0.100700 1.00800 ; qtot
0.918
3276 A3C 201 THR CB 3276 0.365400 12.01000 ; qtot
1.283
3277 H1 201 THR HB 3277 0.004300 1.00800 ; qtot
1.288
3278 CT 201 THR CG2 3278 -0.243800 12.01000 ; qtot
1.044
3279 HC 201 THR HG21 3279 0.064200 1.00800 ; qtot
1.108
3280 HC 201 THR HG22 3280 0.064200 1.00800 ; qtot
1.172
3281 HC 201 THR HG23 3281 0.064200 1.00800 ; qtot
1.237
3282 OH 201 THR OG1 3282 -0.676101 16.00000 ; qtot
0.560
3283 HO 201 THR HG1 3283 0.410200 1.00800 ; qtot
0.971
3284 C 201 THR C 3284 0.597301 12.01000 ; qtot
1.568
3285 O 201 THR O 3285 -0.567901 16.00000 ; qtot
1.000
3286 N 202 LEU N 3286 -0.415700 14.01000 ; qtot
0.584
3287 H 202 LEU H 3287 0.271900 1.00800 ; qtot
0.856
3288 CX 202 LEU CA 3288 -0.051800 12.01000 ; qtot
0.804
3289 H1 202 LEU HA 3289 0.092200 1.00800 ; qtot
0.897
3290 A2C 202 LEU CB 3290 -0.110200 12.01000 ; qtot
0.786
3291 HC 202 LEU HB2 3291 0.045700 1.00800 ; qtot
0.832
3292 HC 202 LEU HB3 3292 0.045700 1.00800 ; qtot
0.878
3293 A3C 202 LEU CG 3293 0.353100 12.01000 ; qtot
1.231
3294 HC 202 LEU HG 3294 -0.036100 1.00800 ; qtot
1.195
3295 CT 202 LEU CD1 3295 -0.412100 12.01000 ; qtot
0.783
3296 HC 202 LEU HD11 3296 0.100000 1.00800 ; qtot
0.883
3297 HC 202 LEU HD12 3297 0.100000 1.00800 ; qtot
0.983
3298 HC 202 LEU HD13 3298 0.100000 1.00800 ; qtot
1.083
3299 CT 202 LEU CD2 3299 -0.412100 12.01000 ; qtot
0.671

3300 HC 202 LEU HD21 3300 0.100000 1.00800 ; qtot
0.771
3301 HC 202 LEU HD22 3301 0.100000 1.00800 ; qtot
0.871
3302 HC 202 LEU HD23 3302 0.100000 1.00800 ; qtot
0.971
3303 C 202 LEU C 3303 0.597301 12.01000 ; qtot
1.568
3304 O 202 LEU O 3304 -0.567901 16.00000 ; qtot
1.000
3305 N 203 LEU N 3305 -0.415700 14.01000 ; qtot
0.584
3306 H 203 LEU H 3306 0.271900 1.00800 ; qtot
0.856
3307 CX 203 LEU CA 3307 -0.051800 12.01000 ; qtot
0.804
3308 H1 203 LEU HA 3308 0.092200 1.00800 ; qtot
0.897
3309 A2C 203 LEU CB 3309 -0.110200 12.01000 ; qtot
0.786
3310 HC 203 LEU HB2 3310 0.045700 1.00800 ; qtot
0.832
3311 HC 203 LEU HB3 3311 0.045700 1.00800 ; qtot
0.878
3312 A3C 203 LEU CG 3312 0.353100 12.01000 ; qtot
1.231
3313 HC 203 LEU HG 3313 -0.036100 1.00800 ; qtot
1.195
3314 CT 203 LEU CD1 3314 -0.412100 12.01000 ; qtot
0.783
3315 HC 203 LEU HD11 3315 0.100000 1.00800 ; qtot
0.883
3316 HC 203 LEU HD12 3316 0.100000 1.00800 ; qtot
0.983
3317 HC 203 LEU HD13 3317 0.100000 1.00800 ; qtot
1.083
3318 CT 203 LEU CD2 3318 -0.412100 12.01000 ; qtot
0.671
3319 HC 203 LEU HD21 3319 0.100000 1.00800 ; qtot
0.771
3320 HC 203 LEU HD22 3320 0.100000 1.00800 ; qtot
0.871
3321 HC 203 LEU HD23 3321 0.100000 1.00800 ; qtot
0.971
3322 C 203 LEU C 3322 0.597301 12.01000 ; qtot
1.568
3323 O 203 LEU O 3323 -0.567901 16.00000 ; qtot
1.000
3324 N 204 CYS N 3324 -0.415700 14.01000 ; qtot
0.584
3325 H 204 CYS H 3325 0.271900 1.00800 ; qtot
0.856

3326 CX 204 CYS CA 3326 0.021300 12.01000 ; qtot
0.878
3327 H1 204 CYS HA 3327 0.112400 1.00800 ; qtot
0.990
3328 A2C 204 CYS CB 3328 -0.123100 12.01000 ; qtot
0.867
3329 H1 204 CYS HB2 3329 0.111200 1.00800 ; qtot
0.978
3330 H1 204 CYS HB3 3330 0.111200 1.00800 ; qtot
1.089
3331 SH 204 CYS SG 3331 -0.311900 32.06000 ; qtot
0.777
3332 HS 204 CYS HG 3332 0.193300 1.00800 ; qtot
0.971
3333 C 204 CYS C 3333 0.597301 12.01000 ; qtot
1.568
3334 O 204 CYS O 3334 -0.567901 16.00000 ; qtot
1.000
3335 N 205 GLY N 3335 -0.415700 14.01000 ; qtot
0.584
3336 H 205 GLY H 3336 0.271900 1.00800 ; qtot
0.856
3337 CX 205 GLY CA 3337 -0.025200 12.01000 ; qtot
0.831
3338 H1 205 GLY HA2 3338 0.069800 1.00800 ; qtot
0.901
3339 H1 205 GLY HA3 3339 0.069800 1.00800 ; qtot
0.971
3340 C 205 GLY C 3340 0.597301 12.01000 ; qtot
1.568
3341 O 205 GLY O 3341 -0.567901 16.00000 ; qtot
1.000
3342 N 206 TYR N 3342 -0.415700 14.01000 ; qtot
0.584
3343 H 206 TYR H 3343 0.271900 1.00800 ; qtot
0.856
3344 CX 206 TYR CA 3344 -0.001400 12.01000 ; qtot
0.855
3345 H1 206 TYR HA 3345 0.087600 1.00800 ; qtot
0.942
3346 CT 206 TYR CB 3346 -0.015200 12.01000 ; qtot
0.927
3347 HC 206 TYR HB2 3347 0.029500 1.00800 ; qtot
0.957
3348 HC 206 TYR HB3 3348 0.029500 1.00800 ; qtot
0.986
3349 CA 206 TYR CG 3349 -0.001100 12.01000 ; qtot
0.985
3350 CA 206 TYR CD1 3350 -0.190600 12.01000 ; qtot
0.795
3351 HA 206 TYR HD1 3351 0.169900 1.00800 ; qtot
0.964

3352 CA 206 TYR CE1 3352 -0.234100 12.01000 ; qtot 0.730
3353 HA 206 TYR HE1 3353 0.165600 1.00800 ; qtot 0.896
3354 C 206 TYR CZ 3354 0.322600 12.01000 ; qtot 1.219
3355 OH 206 TYR OH 3355 -0.557901 16.00000 ; qtot 0.661
3356 HO 206 TYR HH 3356 0.399200 1.00800 ; qtot 1.060
3357 CA 206 TYR CE2 3357 -0.234100 12.01000 ; qtot 0.826
3358 HA 206 TYR HE2 3358 0.165600 1.00800 ; qtot 0.991
3359 CA 206 TYR CD2 3359 -0.190600 12.01000 ; qtot 0.801
3360 HA 206 TYR HD2 3360 0.169900 1.00800 ; qtot 0.971
3361 C 206 TYR C 3361 0.597301 12.01000 ; qtot 1.568
3362 O 206 TYR O 3362 -0.567901 16.00000 ; qtot 1.000
3363 N 207 PRO N 3363 -0.254800 14.01000 ; qtot 0.745
3364 CT 207 PRO CD 3364 0.019200 12.01000 ; qtot 0.764
3365 H1 207 PRO HD2 3365 0.039100 1.00800 ; qtot 0.804
3366 H1 207 PRO HD3 3366 0.039100 1.00800 ; qtot 0.843
3367 CT 207 PRO CG 3367 0.018900 12.01000 ; qtot 0.862
3368 HC 207 PRO HG2 3368 0.021300 1.00800 ; qtot 0.883
3369 HC 207 PRO HG3 3369 0.021300 1.00800 ; qtot 0.904
3370 CT 207 PRO CB 3370 -0.007000 12.01000 ; qtot 0.897
3371 HC 207 PRO HB2 3371 0.025300 1.00800 ; qtot 0.922
3372 HC 207 PRO HB3 3372 0.025300 1.00800 ; qtot 0.948
3373 CX 207 PRO CA 3373 -0.026600 12.01000 ; qtot 0.921
3374 H1 207 PRO HA 3374 0.064100 1.00800 ; qtot 0.985
3375 C 207 PRO C 3375 0.589601 12.01000 ; qtot 1.575
3376 O 207 PRO O 3376 -0.574801 16.00000 ; qtot 1.000
3377 N 208 PRO N 3377 -0.254800 14.01000 ; qtot 0.745

3378 CT 208 PRO CD 3378 0.019200 12.01000 ; qtot
0.764
3379 H1 208 PRO HD2 3379 0.039100 1.00800 ; qtot
0.804
3380 H1 208 PRO HD3 3380 0.039100 1.00800 ; qtot
0.843
3381 CT 208 PRO CG 3381 0.018900 12.01000 ; qtot
0.862
3382 HC 208 PRO HG2 3382 0.021300 1.00800 ; qtot
0.883
3383 HC 208 PRO HG3 3383 0.021300 1.00800 ; qtot
0.904
3384 CT 208 PRO CB 3384 -0.007000 12.01000 ; qtot
0.897
3385 HC 208 PRO HB2 3385 0.025300 1.00800 ; qtot
0.922
3386 HC 208 PRO HB3 3386 0.025300 1.00800 ; qtot
0.948
3387 CX 208 PRO CA 3387 -0.026600 12.01000 ; qtot
0.921
3388 H1 208 PRO HA 3388 0.064100 1.00800 ; qtot
0.985
3389 C 208 PRO C 3389 0.589601 12.01000 ; qtot
1.575
3390 O 208 PRO O 3390 -0.574801 16.00000 ; qtot
1.000
3391 N 209 PHE N 3391 -0.415700 14.01000 ; qtot
0.584
3392 H 209 PHE H 3392 0.271900 1.00800 ; qtot
0.856
3393 CX 209 PHE CA 3393 -0.002400 12.01000 ; qtot
0.854
3394 H1 209 PHE HA 3394 0.097800 1.00800 ; qtot
0.952
3395 CT 209 PHE CB 3395 -0.034300 12.01000 ; qtot
0.917
3396 HC 209 PHE HB2 3396 0.029500 1.00800 ; qtot
0.947
3397 HC 209 PHE HB3 3397 0.029500 1.00800 ; qtot
0.976
3398 CA 209 PHE CG 3398 0.011800 12.01000 ; qtot
0.988
3399 CA 209 PHE CD1 3399 -0.125600 12.01000 ; qtot
0.863
3400 HA 209 PHE HD1 3400 0.133000 1.00800 ; qtot
0.996
3401 CA 209 PHE CE1 3401 -0.170400 12.01000 ; qtot
0.825
3402 HA 209 PHE HE1 3402 0.143000 1.00800 ; qtot
0.968
3403 CA 209 PHE CZ 3403 -0.107200 12.01000 ; qtot
0.861

3404 HA 209 PHE HZ 3404 0.129700 1.00800 ; qtot 0.991
3405 CA 209 PHE CE2 3405 -0.170400 12.01000 ; qtot 0.820
3406 HA 209 PHE HE2 3406 0.143000 1.00800 ; qtot 0.963
3407 CA 209 PHE CD2 3407 -0.125600 12.01000 ; qtot 0.838
3408 HA 209 PHE HD2 3408 0.133000 1.00800 ; qtot 0.971
3409 C 209 PHE C 3409 0.597301 12.01000 ; qtot 1.568
3410 O 209 PHE O 3410 -0.567901 16.00000 ; qtot 1.000
3411 N 210 TYR N 3411 -0.415700 14.01000 ; qtot 0.584
3412 H 210 TYR H 3412 0.271900 1.00800 ; qtot 0.856
3413 CX 210 TYR CA 3413 -0.001400 12.01000 ; qtot 0.855
3414 H1 210 TYR HA 3414 0.087600 1.00800 ; qtot 0.942
3415 CT 210 TYR CB 3415 -0.015200 12.01000 ; qtot 0.927
3416 HC 210 TYR HB2 3416 0.029500 1.00800 ; qtot 0.957
3417 HC 210 TYR HB3 3417 0.029500 1.00800 ; qtot 0.986
3418 CA 210 TYR CG 3418 -0.001100 12.01000 ; qtot 0.985
3419 CA 210 TYR CD1 3419 -0.190600 12.01000 ; qtot 0.795
3420 HA 210 TYR HD1 3420 0.169900 1.00800 ; qtot 0.964
3421 CA 210 TYR CE1 3421 -0.234100 12.01000 ; qtot 0.730
3422 HA 210 TYR HE1 3422 0.165600 1.00800 ; qtot 0.896
3423 C 210 TYR CZ 3423 0.322600 12.01000 ; qtot 1.219
3424 OH 210 TYR OH 3424 -0.557901 16.00000 ; qtot 0.661
3425 HO 210 TYR HH 3425 0.399200 1.00800 ; qtot 1.060
3426 CA 210 TYR CE2 3426 -0.234100 12.01000 ; qtot 0.826
3427 HA 210 TYR HE2 3427 0.165600 1.00800 ; qtot 0.991
3428 CA 210 TYR CD2 3428 -0.190600 12.01000 ; qtot 0.801
3429 HA 210 TYR HD2 3429 0.169900 1.00800 ; qtot 0.971

3430 C 210 TYR C 3430 0.597301 12.01000 ; qtot
1.568
3431 O 210 TYR O 3431 -0.567901 16.00000 ; qtot
1.000
3432 N 211 GLY N 3432 -0.415700 14.01000 ; qtot
0.584
3433 H 211 GLY H 3433 0.271900 1.00800 ; qtot
0.856
3434 CX 211 GLY CA 3434 -0.025200 12.01000 ; qtot
0.831
3435 H1 211 GLY HA2 3435 0.069800 1.00800 ; qtot
0.901
3436 H1 211 GLY HA3 3436 0.069800 1.00800 ; qtot
0.971
3437 C 211 GLY C 3437 0.597301 12.01000 ; qtot
1.568
3438 O 211 GLY O 3438 -0.567901 16.00000 ; qtot
1.000
3439 N 212 ASP N 3439 -0.516301 14.01000 ; qtot
0.484
3440 H 212 ASP H 3440 0.293600 1.00800 ; qtot
0.777
3441 CX 212 ASP CA 3441 0.038100 12.01000 ; qtot
0.815
3442 H1 212 ASP HA 3442 0.088000 1.00800 ; qtot
0.903
3443 A2C 212 ASP CB 3443 -0.030300 12.01000 ; qtot
0.873
3444 HC 212 ASP HB2 3444 -0.012200 1.00800 ; qtot
0.861
3445 HC 212 ASP HB3 3445 -0.012200 1.00800 ; qtot
0.849
3446 CO 212 ASP CG 3446 0.799401 12.01000 ; qtot
1.648
3447 O2 212 ASP OD1 3447 -0.801401 16.00000 ; qtot
0.847
3448 O2 212 ASP OD2 3448 -0.801401 16.00000 ; qtot
0.045
3449 C 212 ASP C 3449 0.536601 12.01000 ; qtot
0.582
3450 O 212 ASP O 3450 -0.581901 16.00000 ; qtot
0.000
3451 N 213 THR N 3451 -0.415700 14.01000 ; qtot -
0.416
3452 H 213 THR H 3452 0.271900 1.00800 ; qtot -
0.144
3453 CX 213 THR CA 3453 -0.038900 12.01000 ; qtot -
0.183
3454 H1 213 THR HA 3454 0.100700 1.00800 ; qtot -
0.082
3455 A3C 213 THR CB 3455 0.365400 12.01000 ; qtot
0.283

3456 H1 213 THR HB 3456 0.004300 1.00800 ; qtot
0.288
3457 CT 213 THR CG2 3457 -0.243800 12.01000 ; qtot
0.044
3458 HC 213 THR HG21 3458 0.064200 1.00800 ; qtot
0.108
3459 HC 213 THR HG22 3459 0.064200 1.00800 ; qtot
0.172
3460 HC 213 THR HG23 3460 0.064200 1.00800 ; qtot
0.237
3461 OH 213 THR OG1 3461 -0.676101 16.00000 ; qtot -
0.440
3462 HO 213 THR HG1 3462 0.410200 1.00800 ; qtot -
0.029
3463 C 213 THR C 3463 0.597301 12.01000 ; qtot
0.568
3464 O 213 THR O 3464 -0.567901 16.00000 ; qtot
0.000
3465 N 214 ASP N 3465 -0.516301 14.01000 ; qtot -
0.516
3466 H 214 ASP H 3466 0.293600 1.00800 ; qtot -
0.223
3467 CX 214 ASP CA 3467 0.038100 12.01000 ; qtot -
0.185
3468 H1 214 ASP HA 3468 0.088000 1.00800 ; qtot -
0.097
3469 A2C 214 ASP CB 3469 -0.030300 12.01000 ; qtot -
0.127
3470 HC 214 ASP HB2 3470 -0.012200 1.00800 ; qtot -
0.139
3471 HC 214 ASP HB3 3471 -0.012200 1.00800 ; qtot -
0.151
3472 CO 214 ASP CG 3472 0.799401 12.01000 ; qtot
0.648
3473 O2 214 ASP OD1 3473 -0.801401 16.00000 ; qtot -
0.153
3474 O2 214 ASP OD2 3474 -0.801401 16.00000 ; qtot -
0.955
3475 C 214 ASP C 3475 0.536601 12.01000 ; qtot -
0.418
3476 O 214 ASP O 3476 -0.581901 16.00000 ; qtot -
1.000
3477 N 215 ASN N 3477 -0.415700 14.01000 ; qtot -
1.416
3478 H 215 ASN H 3478 0.271900 1.00800 ; qtot -
1.144
3479 CX 215 ASN CA 3479 0.014300 12.01000 ; qtot -
1.130
3480 H1 215 ASN HA 3480 0.104800 1.00800 ; qtot -
1.025
3481 A2C 215 ASN CB 3481 -0.204100 12.01000 ; qtot -
1.229

3482 HC 215 ASN HB2 3482 0.079700 1.00800 ; qtot -
1.149
3483 HC 215 ASN HB3 3483 0.079700 1.00800 ; qtot -
1.069
3484 C 215 ASN CG 3484 0.713001 12.01000 ; qtot -
0.356
3485 O 215 ASN OD1 3485 -0.593101 16.00000 ; qtot -
0.950
3486 N 215 ASN ND2 3486 -0.919101 14.01000 ; qtot -
1.869
3487 H 215 ASN HD21 3487 0.419600 1.00800 ; qtot -
1.449
3488 H 215 ASN HD22 3488 0.419600 1.00800 ; qtot -
1.029
3489 C 215 ASN C 3489 0.597301 12.01000 ; qtot -
0.432
3490 O 215 ASN O 3490 -0.567901 16.00000 ; qtot -
1.000
3491 N 216 GLU N 3491 -0.516301 14.01000 ; qtot -
1.516
3492 H 216 GLU H 3492 0.293600 1.00800 ; qtot -
1.223
3493 CX 216 GLU CA 3493 0.039700 12.01000 ; qtot -
1.183
3494 H1 216 GLU HA 3494 0.110500 1.00800 ; qtot -
1.073
3495 A2C 216 GLU CB 3495 0.056000 12.01000 ; qtot -
1.017
3496 HC 216 GLU HB2 3496 -0.017300 1.00800 ; qtot -
1.034
3497 HC 216 GLU HB3 3497 -0.017300 1.00800 ; qtot -
1.051
3498 A2C 216 GLU CG 3498 0.013600 12.01000 ; qtot -
1.038
3499 HC 216 GLU HG2 3499 -0.042500 1.00800 ; qtot -
1.080
3500 HC 216 GLU HG3 3500 -0.042500 1.00800 ; qtot -
1.123
3501 CO 216 GLU CD 3501 0.805401 12.01000 ; qtot -
0.317
3502 O2 216 GLU OE1 3502 -0.818801 16.00000 ; qtot -
1.136
3503 O2 216 GLU OE2 3503 -0.818801 16.00000 ; qtot -
1.955
3504 C 216 GLU C 3504 0.536601 12.01000 ; qtot -
1.418
3505 O 216 GLU O 3505 -0.581901 16.00000 ; qtot -
2.000
3506 N 217 VAL N 3506 -0.415700 14.01000 ; qtot -
2.416
3507 H 217 VAL H 3507 0.271900 1.00800 ; qtot -
2.144

3508 CX 217 VAL CA 3508 -0.087500 12.01000 ; qtot -
2.231
3509 H1 217 VAL HA 3509 0.096900 1.00800 ; qtot -
2.134
3510 A3C 217 VAL CB 3510 0.298500 12.01000 ; qtot -
1.836
3511 HC 217 VAL HB 3511 -0.029700 1.00800 ; qtot -
1.866
3512 CT 217 VAL CG1 3512 -0.319200 12.01000 ; qtot -
2.185
3513 HC 217 VAL HG11 3513 0.079100 1.00800 ; qtot -
2.106
3514 HC 217 VAL HG12 3514 0.079100 1.00800 ; qtot -
2.027
3515 HC 217 VAL HG13 3515 0.079100 1.00800 ; qtot -
1.948
3516 CT 217 VAL CG2 3516 -0.319200 12.01000 ; qtot -
2.267
3517 HC 217 VAL HG21 3517 0.079100 1.00800 ; qtot -
2.188
3518 HC 217 VAL HG22 3518 0.079100 1.00800 ; qtot -
2.109
3519 HC 217 VAL HG23 3519 0.079100 1.00800 ; qtot -
2.029
3520 C 217 VAL C 3520 0.597301 12.01000 ; qtot -
1.432
3521 O 217 VAL O 3521 -0.567901 16.00000 ; qtot -
2.000
3522 N 218 LEU N 3522 -0.415700 14.01000 ; qtot -
2.416
3523 H 218 LEU H 3523 0.271900 1.00800 ; qtot -
2.144
3524 CX 218 LEU CA 3524 -0.051800 12.01000 ; qtot -
2.196
3525 H1 218 LEU HA 3525 0.092200 1.00800 ; qtot -
2.103
3526 A2C 218 LEU CB 3526 -0.110200 12.01000 ; qtot -
2.214
3527 HC 218 LEU HB2 3527 0.045700 1.00800 ; qtot -
2.168
3528 HC 218 LEU HB3 3528 0.045700 1.00800 ; qtot -
2.122
3529 A3C 218 LEU CG 3529 0.353100 12.01000 ; qtot -
1.769
3530 HC 218 LEU HG 3530 -0.036100 1.00800 ; qtot -
1.805
3531 CT 218 LEU CD1 3531 -0.412100 12.01000 ; qtot -
2.217
3532 HC 218 LEU HD11 3532 0.100000 1.00800 ; qtot -
2.117
3533 HC 218 LEU HD12 3533 0.100000 1.00800 ; qtot -
2.017

3534 HC 218 LEU HD13 3534 0.100000 1.00800 ; qtot -
1.917
3535 CT 218 LEU CD2 3535 -0.412100 12.01000 ; qtot -
2.329
3536 HC 218 LEU HD21 3536 0.100000 1.00800 ; qtot -
2.229
3537 HC 218 LEU HD22 3537 0.100000 1.00800 ; qtot -
2.129
3538 HC 218 LEU HD23 3538 0.100000 1.00800 ; qtot -
2.029
3539 C 218 LEU C 3539 0.597301 12.01000 ; qtot -
1.432
3540 O 218 LEU O 3540 -0.567901 16.00000 ; qtot -
2.000
3541 N 219 LYS N 3541 -0.347900 14.01000 ; qtot -
2.348
3542 H 219 LYS H 3542 0.274700 1.00800 ; qtot -
2.073
3543 CX 219 LYS CA 3543 -0.240000 12.01000 ; qtot -
2.313
3544 H1 219 LYS HA 3544 0.142600 1.00800 ; qtot -
2.171
3545 C8 219 LYS CB 3545 -0.009400 12.01000 ; qtot -
2.180
3546 HC 219 LYS HB2 3546 0.036200 1.00800 ; qtot -
2.144
3547 HC 219 LYS HB3 3547 0.036200 1.00800 ; qtot -
2.108
3548 C8 219 LYS CG 3548 0.018700 12.01000 ; qtot -
2.089
3549 HC 219 LYS HG2 3549 0.010300 1.00800 ; qtot -
2.079
3550 HC 219 LYS HG3 3550 0.010300 1.00800 ; qtot -
2.068
3551 C8 219 LYS CD 3551 -0.047900 12.01000 ; qtot -
2.116
3552 HC 219 LYS HD2 3552 0.062100 1.00800 ; qtot -
2.054
3553 HC 219 LYS HD3 3553 0.062100 1.00800 ; qtot -
1.992
3554 C8 219 LYS CE 3554 -0.014300 12.01000 ; qtot -
2.006
3555 HP 219 LYS HE2 3555 0.113500 1.00800 ; qtot -
1.893
3556 HP 219 LYS HE3 3556 0.113500 1.00800 ; qtot -
1.779
3557 N3 219 LYS NZ 3557 -0.385400 14.01000 ; qtot -
2.165
3558 H 219 LYS HZ1 3558 0.340000 1.00800 ; qtot -
1.825
3559 H 219 LYS HZ2 3559 0.340000 1.00800 ; qtot -
1.485

3560 H 219 LYS HZ3 3560 0.340000 1.00800 ; qtot -
1.145
3561 C 219 LYS C 3561 0.734101 12.01000 ; qtot -
0.411
3562 O 219 LYS O 3562 -0.589401 16.00000 ; qtot -
1.000
3563 N 220 LYS N 3563 -0.347900 14.01000 ; qtot -
1.348
3564 H 220 LYS H 3564 0.274700 1.00800 ; qtot -
1.073
3565 CX 220 LYS CA 3565 -0.240000 12.01000 ; qtot -
1.313
3566 H1 220 LYS HA 3566 0.142600 1.00800 ; qtot -
1.171
3567 C8 220 LYS CB 3567 -0.009400 12.01000 ; qtot -
1.180
3568 HC 220 LYS HB2 3568 0.036200 1.00800 ; qtot -
1.144
3569 HC 220 LYS HB3 3569 0.036200 1.00800 ; qtot -
1.108
3570 C8 220 LYS CG 3570 0.018700 12.01000 ; qtot -
1.089
3571 HC 220 LYS HG2 3571 0.010300 1.00800 ; qtot -
1.079
3572 HC 220 LYS HG3 3572 0.010300 1.00800 ; qtot -
1.068
3573 C8 220 LYS CD 3573 -0.047900 12.01000 ; qtot -
1.116
3574 HC 220 LYS HD2 3574 0.062100 1.00800 ; qtot -
1.054
3575 HC 220 LYS HD3 3575 0.062100 1.00800 ; qtot -
0.992
3576 C8 220 LYS CE 3576 -0.014300 12.01000 ; qtot -
1.006
3577 HP 220 LYS HE2 3577 0.113500 1.00800 ; qtot -
0.893
3578 HP 220 LYS HE3 3578 0.113500 1.00800 ; qtot -
0.779
3579 N3 220 LYS NZ 3579 -0.385400 14.01000 ; qtot -
1.165
3580 H 220 LYS HZ1 3580 0.340000 1.00800 ; qtot -
0.825
3581 H 220 LYS HZ2 3581 0.340000 1.00800 ; qtot -
0.485
3582 H 220 LYS HZ3 3582 0.340000 1.00800 ; qtot -
0.145
3583 C 220 LYS C 3583 0.734101 12.01000 ; qtot
0.589
3584 O 220 LYS O 3584 -0.589401 16.00000 ; qtot
0.000
3585 N 221 VAL N 3585 -0.415700 14.01000 ; qtot -
0.416

```
  3586    H   221   VAL     H 3586     0.271900      1.00800 ; qtot -
0.144
  3587   CX   221   VAL    CA 3587    -0.087500     12.01000 ; qtot -
0.231
  3588   H1   221   VAL    HA 3588     0.096900      1.00800 ; qtot -
0.134
  3589  A3C   221   VAL    CB 3589     0.298500     12.01000 ; qtot
0.164
  3590   HC   221   VAL    HB 3590    -0.029700      1.00800 ; qtot
0.134
  3591   CT   221   VAL   CG1 3591    -0.319200     12.01000 ; qtot -
0.185
  3592   HC   221   VAL  HG11 3592     0.079100      1.00800 ; qtot -
0.106
  3593   HC   221   VAL  HG12 3593     0.079100      1.00800 ; qtot -
0.027
  3594   HC   221   VAL  HG13 3594     0.079100      1.00800 ; qtot
0.053
  3595   CT   221   VAL   CG2 3595    -0.319200     12.01000 ; qtot -
0.267
  3596   HC   221   VAL  HG21 3596     0.079100      1.00800 ; qtot -
0.188
  3597   HC   221   VAL  HG22 3597     0.079100      1.00800 ; qtot -
0.108
  3598   HC   221   VAL  HG23 3598     0.079100      1.00800 ; qtot -
0.029
  3599    C   221   VAL     C 3599     0.597301     12.01000 ; qtot
0.568
  3600    O   221   VAL     O 3600    -0.567901     16.00000 ; qtot
0.000
  3601    N   222   LYS     N 3601    -0.347900     14.01000 ; qtot -
0.348
  3602    H   222   LYS     H 3602     0.274700      1.00800 ; qtot -
0.073
  3603   CX   222   LYS    CA 3603    -0.240000     12.01000 ; qtot -
0.313
  3604   H1   222   LYS    HA 3604     0.142600      1.00800 ; qtot -
0.171
  3605   C8   222   LYS    CB 3605    -0.009400     12.01000 ; qtot -
0.180
  3606   HC   222   LYS   HB2 3606     0.036200      1.00800 ; qtot -
0.144
  3607   HC   222   LYS   HB3 3607     0.036200      1.00800 ; qtot -
0.108
  3608   C8   222   LYS    CG 3608     0.018700     12.01000 ; qtot -
0.089
  3609   HC   222   LYS   HG2 3609     0.010300      1.00800 ; qtot -
0.079
  3610   HC   222   LYS   HG3 3610     0.010300      1.00800 ; qtot -
0.068
  3611   C8   222   LYS    CD 3611    -0.047900     12.01000 ; qtot -
0.116
```

3612 HC 222 LYS HD2 3612 0.062100 1.00800 ; qtot -
0.054
3613 HC 222 LYS HD3 3613 0.062100 1.00800 ; qtot
0.008
3614 C8 222 LYS CE 3614 -0.014300 12.01000 ; qtot -
0.006
3615 HP 222 LYS HE2 3615 0.113500 1.00800 ; qtot
0.107
3616 HP 222 LYS HE3 3616 0.113500 1.00800 ; qtot
0.221
3617 N3 222 LYS NZ 3617 -0.385400 14.01000 ; qtot -
0.165
3618 H 222 LYS HZ1 3618 0.340000 1.00800 ; qtot
0.175
3619 H 222 LYS HZ2 3619 0.340000 1.00800 ; qtot
0.515
3620 H 222 LYS HZ3 3620 0.340000 1.00800 ; qtot
0.855
3621 C 222 LYS C 3621 0.734101 12.01000 ; qtot
1.589
3622 O 222 LYS O 3622 -0.589401 16.00000 ; qtot
1.000
3623 N 223 LYS N 3623 -0.347900 14.01000 ; qtot
0.652
3624 H 223 LYS H 3624 0.274700 1.00800 ; qtot
0.927
3625 CX 223 LYS CA 3625 -0.240000 12.01000 ; qtot
0.687
3626 H1 223 LYS HA 3626 0.142600 1.00800 ; qtot
0.829
3627 C8 223 LYS CB 3627 -0.009400 12.01000 ; qtot
0.820
3628 HC 223 LYS HB2 3628 0.036200 1.00800 ; qtot
0.856
3629 HC 223 LYS HB3 3629 0.036200 1.00800 ; qtot
0.892
3630 C8 223 LYS CG 3630 0.018700 12.01000 ; qtot
0.911
3631 HC 223 LYS HG2 3631 0.010300 1.00800 ; qtot
0.921
3632 HC 223 LYS HG3 3632 0.010300 1.00800 ; qtot
0.932
3633 C8 223 LYS CD 3633 -0.047900 12.01000 ; qtot
0.884
3634 HC 223 LYS HD2 3634 0.062100 1.00800 ; qtot
0.946
3635 HC 223 LYS HD3 3635 0.062100 1.00800 ; qtot
1.008
3636 C8 223 LYS CE 3636 -0.014300 12.01000 ; qtot
0.994
3637 HP 223 LYS HE2 3637 0.113500 1.00800 ; qtot
1.107

3638 HP 223 LYS HE3 3638 0.113500 1.00800 ; qtot
1.221
3639 N3 223 LYS NZ 3639 -0.385400 14.01000 ; qtot
0.835
3640 H 223 LYS HZ1 3640 0.340000 1.00800 ; qtot
1.175
3641 H 223 LYS HZ2 3641 0.340000 1.00800 ; qtot
1.515
3642 H 223 LYS HZ3 3642 0.340000 1.00800 ; qtot
1.855
3643 C 223 LYS C 3643 0.734101 12.01000 ; qtot
2.589
3644 O 223 LYS O 3644 -0.589401 16.00000 ; qtot
2.000
3645 N 224 GLY N 3645 -0.415700 14.01000 ; qtot
1.584
3646 H 224 GLY H 3646 0.271900 1.00800 ; qtot
1.856
3647 CX 224 GLY CA 3647 -0.025200 12.01000 ; qtot
1.831
3648 H1 224 GLY HA2 3648 0.069800 1.00800 ; qtot
1.901
3649 H1 224 GLY HA3 3649 0.069800 1.00800 ; qtot
1.971
3650 C 224 GLY C 3650 0.597301 12.01000 ; qtot
2.568
3651 O 224 GLY O 3651 -0.567901 16.00000 ; qtot
2.000
3652 N 225 GLU N 3652 -0.516301 14.01000 ; qtot
1.484
3653 H 225 GLU H 3653 0.293600 1.00800 ; qtot
1.777
3654 CX 225 GLU CA 3654 0.039700 12.01000 ; qtot
1.817
3655 H1 225 GLU HA 3655 0.110500 1.00800 ; qtot
1.928
3656 A2C 225 GLU CB 3656 0.056000 12.01000 ; qtot
1.984
3657 HC 225 GLU HB2 3657 -0.017300 1.00800 ; qtot
1.966
3658 HC 225 GLU HB3 3658 -0.017300 1.00800 ; qtot
1.949
3659 A2C 225 GLU CG 3659 0.013600 12.01000 ; qtot
1.963
3660 HC 225 GLU HG2 3660 -0.042500 1.00800 ; qtot
1.920
3661 HC 225 GLU HG3 3661 -0.042500 1.00800 ; qtot
1.878
3662 CO 225 GLU CD 3662 0.805401 12.01000 ; qtot
2.683
3663 O2 225 GLU OE1 3663 -0.818801 16.00000 ; qtot
1.864

3664 O2 225 GLU OE2 3664 -0.818801 16.00000 ; qtot
1.045
3665 C 225 GLU C 3665 0.536601 12.01000 ; qtot
1.582
3666 O 225 GLU O 3666 -0.581901 16.00000 ; qtot
1.000
3667 N 226 PHE N 3667 -0.415700 14.01000 ; qtot
0.584
3668 H 226 PHE H 3668 0.271900 1.00800 ; qtot
0.856
3669 CX 226 PHE CA 3669 -0.002400 12.01000 ; qtot
0.854
3670 H1 226 PHE HA 3670 0.097800 1.00800 ; qtot
0.952
3671 CT 226 PHE CB 3671 -0.034300 12.01000 ; qtot
0.917
3672 HC 226 PHE HB2 3672 0.029500 1.00800 ; qtot
0.947
3673 HC 226 PHE HB3 3673 0.029500 1.00800 ; qtot
0.976
3674 CA 226 PHE CG 3674 0.011800 12.01000 ; qtot
0.988
3675 CA 226 PHE CD1 3675 -0.125600 12.01000 ; qtot
0.863
3676 HA 226 PHE HD1 3676 0.133000 1.00800 ; qtot
0.996
3677 CA 226 PHE CE1 3677 -0.170400 12.01000 ; qtot
0.825
3678 HA 226 PHE HE1 3678 0.143000 1.00800 ; qtot
0.968
3679 CA 226 PHE CZ 3679 -0.107200 12.01000 ; qtot
0.861
3680 HA 226 PHE HZ 3680 0.129700 1.00800 ; qtot
0.991
3681 CA 226 PHE CE2 3681 -0.170400 12.01000 ; qtot
0.820
3682 HA 226 PHE HE2 3682 0.143000 1.00800 ; qtot
0.963
3683 CA 226 PHE CD2 3683 -0.125600 12.01000 ; qtot
0.838
3684 HA 226 PHE HD2 3684 0.133000 1.00800 ; qtot
0.971
3685 C 226 PHE C 3685 0.597301 12.01000 ; qtot
1.568
3686 O 226 PHE O 3686 -0.567901 16.00000 ; qtot
1.000
3687 N 227 CYS N 3687 -0.415700 14.01000 ; qtot
0.584
3688 H 227 CYS H 3688 0.271900 1.00800 ; qtot
0.856
3689 CX 227 CYS CA 3689 0.021300 12.01000 ; qtot
0.878

3690 H1 227 CYS HA 3690 0.112400 1.00800 ; qtot
0.990
3691 A2C 227 CYS CB 3691 -0.123100 12.01000 ; qtot
0.867
3692 H1 227 CYS HB2 3692 0.111200 1.00800 ; qtot
0.978
3693 H1 227 CYS HB3 3693 0.111200 1.00800 ; qtot
1.089
3694 SH 227 CYS SG 3694 -0.311900 32.06000 ; qtot
0.777
3695 HS 227 CYS HG 3695 0.193300 1.00800 ; qtot
0.971
3696 C 227 CYS C 3696 0.597301 12.01000 ; qtot
1.568
3697 O 227 CYS O 3697 -0.567901 16.00000 ; qtot
1.000
3698 N 228 PHE N 3698 -0.415700 14.01000 ; qtot
0.584
3699 H 228 PHE H 3699 0.271900 1.00800 ; qtot
0.856
3700 CX 228 PHE CA 3700 -0.002400 12.01000 ; qtot
0.854
3701 H1 228 PHE HA 3701 0.097800 1.00800 ; qtot
0.952
3702 CT 228 PHE CB 3702 -0.034300 12.01000 ; qtot
0.917
3703 HC 228 PHE HB2 3703 0.029500 1.00800 ; qtot
0.947
3704 HC 228 PHE HB3 3704 0.029500 1.00800 ; qtot
0.976
3705 CA 228 PHE CG 3705 0.011800 12.01000 ; qtot
0.988
3706 CA 228 PHE CD1 3706 -0.125600 12.01000 ; qtot
0.863
3707 HA 228 PHE HD1 3707 0.133000 1.00800 ; qtot
0.996
3708 CA 228 PHE CE1 3708 -0.170400 12.01000 ; qtot
0.825
3709 HA 228 PHE HE1 3709 0.143000 1.00800 ; qtot
0.968
3710 CA 228 PHE CZ 3710 -0.107200 12.01000 ; qtot
0.861
3711 HA 228 PHE HZ 3711 0.129700 1.00800 ; qtot
0.991
3712 CA 228 PHE CE2 3712 -0.170400 12.01000 ; qtot
0.820
3713 HA 228 PHE HE2 3713 0.143000 1.00800 ; qtot
0.963
3714 CA 228 PHE CD2 3714 -0.125600 12.01000 ; qtot
0.838
3715 HA 228 PHE HD2 3715 0.133000 1.00800 ; qtot
0.971

3716 C 228 PHE C 3716 0.597301 12.01000 ; qtot
1.568
3717 O 228 PHE O 3717 -0.567901 16.00000 ; qtot
1.000
3718 N 229 TYR N 3718 -0.415700 14.01000 ; qtot
0.584
3719 H 229 TYR H 3719 0.271900 1.00800 ; qtot
0.856
3720 CX 229 TYR CA 3720 -0.001400 12.01000 ; qtot
0.855
3721 H1 229 TYR HA 3721 0.087600 1.00800 ; qtot
0.942
3722 CT 229 TYR CB 3722 -0.015200 12.01000 ; qtot
0.927
3723 HC 229 TYR HB2 3723 0.029500 1.00800 ; qtot
0.957
3724 HC 229 TYR HB3 3724 0.029500 1.00800 ; qtot
0.986
3725 CA 229 TYR CG 3725 -0.001100 12.01000 ; qtot
0.985
3726 CA 229 TYR CD1 3726 -0.190600 12.01000 ; qtot
0.795
3727 HA 229 TYR HD1 3727 0.169900 1.00800 ; qtot
0.964
3728 CA 229 TYR CE1 3728 -0.234100 12.01000 ; qtot
0.730
3729 HA 229 TYR HE1 3729 0.165600 1.00800 ; qtot
0.896
3730 C 229 TYR CZ 3730 0.322600 12.01000 ; qtot
1.219
3731 OH 229 TYR OH 3731 -0.557901 16.00000 ; qtot
0.661
3732 HO 229 TYR HH 3732 0.399200 1.00800 ; qtot
1.060
3733 CA 229 TYR CE2 3733 -0.234100 12.01000 ; qtot
0.826
3734 HA 229 TYR HE2 3734 0.165600 1.00800 ; qtot
0.991
3735 CA 229 TYR CD2 3735 -0.190600 12.01000 ; qtot
0.801
3736 HA 229 TYR HD2 3736 0.169900 1.00800 ; qtot
0.971
3737 C 229 TYR C 3737 0.597301 12.01000 ; qtot
1.568
3738 O 229 TYR O 3738 -0.567901 16.00000 ; qtot
1.000
3739 N 230 GLU N 3739 -0.516301 14.01000 ; qtot
0.484
3740 H 230 GLU H 3740 0.293600 1.00800 ; qtot
0.777
3741 CX 230 GLU CA 3741 0.039700 12.01000 ; qtot
0.817

3742 H1 230 GLU HA 3742 0.110500 1.00800 ; qtot
0.928
3743 A2C 230 GLU CB 3743 0.056000 12.01000 ; qtot
0.984
3744 HC 230 GLU HB2 3744 -0.017300 1.00800 ; qtot
0.966
3745 HC 230 GLU HB3 3745 -0.017300 1.00800 ; qtot
0.949
3746 A2C 230 GLU CG 3746 0.013600 12.01000 ; qtot
0.963
3747 HC 230 GLU HG2 3747 -0.042500 1.00800 ; qtot
0.920
3748 HC 230 GLU HG3 3748 -0.042500 1.00800 ; qtot
0.878
3749 CO 230 GLU CD 3749 0.805401 12.01000 ; qtot
1.683
3750 O2 230 GLU OE1 3750 -0.818801 16.00000 ; qtot
0.864
3751 O2 230 GLU OE2 3751 -0.818801 16.00000 ; qtot
0.045
3752 C 230 GLU C 3752 0.536601 12.01000 ; qtot
0.582
3753 O 230 GLU O 3753 -0.581901 16.00000 ; qtot
0.000
3754 N 231 ASN N 3754 -0.415700 14.01000 ; qtot -
0.416
3755 H 231 ASN H 3755 0.271900 1.00800 ; qtot -
0.144
3756 CX 231 ASN CA 3756 0.014300 12.01000 ; qtot -
0.129
3757 H1 231 ASN HA 3757 0.104800 1.00800 ; qtot -
0.025
3758 A2C 231 ASN CB 3758 -0.204100 12.01000 ; qtot -
0.229
3759 HC 231 ASN HB2 3759 0.079700 1.00800 ; qtot -
0.149
3760 HC 231 ASN HB3 3760 0.079700 1.00800 ; qtot -
0.069
3761 C 231 ASN CG 3761 0.713001 12.01000 ; qtot
0.644
3762 O 231 ASN OD1 3762 -0.593101 16.00000 ; qtot
0.051
3763 N 231 ASN ND2 3763 -0.919101 14.01000 ; qtot -
0.869
3764 H 231 ASN HD21 3764 0.419600 1.00800 ; qtot -
0.449
3765 H 231 ASN HD22 3765 0.419600 1.00800 ; qtot -
0.029
3766 C 231 ASN C 3766 0.597301 12.01000 ; qtot
0.568
3767 O 231 ASN O 3767 -0.567901 16.00000 ; qtot
0.000

3768 N 232 ASP N 3768 -0.516301 14.01000 ; qtot -
0.516
3769 H 232 ASP H 3769 0.293600 1.00800 ; qtot -
0.223
3770 CX 232 ASP CA 3770 0.038100 12.01000 ; qtot -
0.185
3771 H1 232 ASP HA 3771 0.088000 1.00800 ; qtot -
0.097
3772 A2C 232 ASP CB 3772 -0.030300 12.01000 ; qtot -
0.127
3773 HC 232 ASP HB2 3773 -0.012200 1.00800 ; qtot -
0.139
3774 HC 232 ASP HB3 3774 -0.012200 1.00800 ; qtot -
0.151
3775 CO 232 ASP CG 3775 0.799401 12.01000 ; qtot
0.648
3776 O2 232 ASP OD1 3776 -0.801401 16.00000 ; qtot -
0.153
3777 O2 232 ASP OD2 3777 -0.801401 16.00000 ; qtot -
0.955
3778 C 232 ASP C 3778 0.536601 12.01000 ; qtot -
0.418
3779 O 232 ASP O 3779 -0.581901 16.00000 ; qtot -
1.000
3780 N 233 TRP N 3780 -0.415700 14.01000 ; qtot -
1.416
3781 H 233 TRP H 3781 0.271900 1.00800 ; qtot -
1.144
3782 CX 233 TRP CA 3782 -0.027500 12.01000 ; qtot -
1.171
3783 H1 233 TRP HA 3783 0.112300 1.00800 ; qtot -
1.059
3784 CT 233 TRP CB 3784 -0.005000 12.01000 ; qtot -
1.064
3785 HC 233 TRP HB2 3785 0.033900 1.00800 ; qtot -
1.030
3786 HC 233 TRP HB3 3786 0.033900 1.00800 ; qtot -
0.996
3787 C* 233 TRP CG 3787 -0.141500 12.01000 ; qtot -
1.138
3788 CW 233 TRP CD1 3788 -0.163800 12.01000 ; qtot -
1.302
3789 H4 233 TRP HD1 3789 0.206200 1.00800 ; qtot -
1.095
3790 NA 233 TRP NE1 3790 -0.341800 14.01000 ; qtot -
1.437
3791 H 233 TRP HE1 3791 0.341200 1.00800 ; qtot -
1.096
3792 CN 233 TRP CE2 3792 0.138000 12.01000 ; qtot -
0.958
3793 CA 233 TRP CZ2 3793 -0.260100 12.01000 ; qtot -
1.218

3794 HA 233 TRP HZ2 3794 0.157200 1.00800 ; qtot -
1.061
3795 CA 233 TRP CH2 3795 -0.113400 12.01000 ; qtot -
1.174
3796 HA 233 TRP HH2 3796 0.141700 1.00800 ; qtot -
1.033
3797 CA 233 TRP CZ3 3797 -0.197200 12.01000 ; qtot -
1.230
3798 HA 233 TRP HZ3 3798 0.144700 1.00800 ; qtot -
1.085
3799 CA 233 TRP CE3 3799 -0.238700 12.01000 ; qtot -
1.324
3800 HA 233 TRP HE3 3800 0.170000 1.00800 ; qtot -
1.154
3801 CB 233 TRP CD2 3801 0.124300 12.01000 ; qtot -
1.029
3802 C 233 TRP C 3802 0.597301 12.01000 ; qtot -
0.432
3803 O 233 TRP O 3803 -0.567901 16.00000 ; qtot -
1.000
3804 N 234 GLY N 3804 -0.415700 14.01000 ; qtot -
1.416
3805 H 234 GLY H 3805 0.271900 1.00800 ; qtot -
1.144
3806 CX 234 GLY CA 3806 -0.025200 12.01000 ; qtot -
1.169
3807 H1 234 GLY HA2 3807 0.069800 1.00800 ; qtot -
1.099
3808 H1 234 GLY HA3 3808 0.069800 1.00800 ; qtot -
1.029
3809 C 234 GLY C 3809 0.597301 12.01000 ; qtot -
0.432
3810 O 234 GLY O 3810 -0.567901 16.00000 ; qtot -
1.000
3811 N 235 SER N 3811 -0.415700 14.01000 ; qtot -
1.416
3812 H 235 SER H 3812 0.271900 1.00800 ; qtot -
1.144
3813 CX 235 SER CA 3813 -0.024900 12.01000 ; qtot -
1.169
3814 H1 235 SER HA 3814 0.084300 1.00800 ; qtot -
1.084
3815 A2C 235 SER CB 3815 0.211700 12.01000 ; qtot -
0.873
3816 H1 235 SER HB2 3816 0.035200 1.00800 ; qtot -
0.838
3817 H1 235 SER HB3 3817 0.035200 1.00800 ; qtot -
0.802
3818 OH 235 SER OG 3818 -0.654601 16.00000 ; qtot -
1.457
3819 HO 235 SER HG 3819 0.427500 1.00800 ; qtot -
1.029

3820 C 235 SER C 3820 0.597301 12.01000 ; qtot -
0.432
3821 O 235 SER O 3821 -0.567901 16.00000 ; qtot -
1.000
3822 N 236 ILE N 3822 -0.415700 14.01000 ; qtot -
1.416
3823 H 236 ILE H 3823 0.271900 1.00800 ; qtot -
1.144
3824 CX 236 ILE CA 3824 -0.059700 12.01000 ; qtot -
1.204
3825 H1 236 ILE HA 3825 0.086900 1.00800 ; qtot -
1.117
3826 A3C 236 ILE CB 3826 0.130300 12.01000 ; qtot -
0.986
3827 HC 236 ILE HB 3827 0.018700 1.00800 ; qtot -
0.968
3828 CT 236 ILE CG2 3828 -0.320400 12.01000 ; qtot -
1.288
3829 HC 236 ILE HG21 3829 0.088200 1.00800 ; qtot -
1.200
3830 HC 236 ILE HG22 3830 0.088200 1.00800 ; qtot -
1.112
3831 HC 236 ILE HG23 3831 0.088200 1.00800 ; qtot -
1.023
3832 A2C 236 ILE CG1 3832 -0.043000 12.01000 ; qtot -
1.066
3833 HC 236 ILE HG12 3833 0.023600 1.00800 ; qtot -
1.043
3834 HC 236 ILE HG13 3834 0.023600 1.00800 ; qtot -
1.019
3835 CT 236 ILE CD1 3835 -0.066000 12.01000 ; qtot -
1.085
3836 HC 236 ILE HD11 3836 0.018600 1.00800 ; qtot -
1.067
3837 HC 236 ILE HD12 3837 0.018600 1.00800 ; qtot -
1.048
3838 HC 236 ILE HD13 3838 0.018600 1.00800 ; qtot -
1.029
3839 C 236 ILE C 3839 0.597301 12.01000 ; qtot -
0.432
3840 O 236 ILE O 3840 -0.567901 16.00000 ; qtot -
1.000
3841 N 237 SER N 3841 -0.415700 14.01000 ; qtot -
1.416
3842 H 237 SER H 3842 0.271900 1.00800 ; qtot -
1.144
3843 CX 237 SER CA 3843 -0.024900 12.01000 ; qtot -
1.169
3844 H1 237 SER HA 3844 0.084300 1.00800 ; qtot -
1.084
3845 A2C 237 SER CB 3845 0.211700 12.01000 ; qtot -
0.873

3846 H1 237 SER HB2 3846 0.035200 1.00800 ; qtot -0.838
3847 H1 237 SER HB3 3847 0.035200 1.00800 ; qtot -0.802
3848 OH 237 SER OG 3848 -0.654601 16.00000 ; qtot -1.457
3849 HO 237 SER HG 3849 0.427500 1.00800 ; qtot -1.029
3850 C 237 SER C 3850 0.597301 12.01000 ; qtot -0.432
3851 O 237 SER O 3851 -0.567901 16.00000 ; qtot -1.000
3852 N 238 SER N 3852 -0.415700 14.01000 ; qtot -1.416
3853 H 238 SER H 3853 0.271900 1.00800 ; qtot -1.144
3854 CX 238 SER CA 3854 -0.024900 12.01000 ; qtot -1.169
3855 H1 238 SER HA 3855 0.084300 1.00800 ; qtot -1.084
3856 A2C 238 SER CB 3856 0.211700 12.01000 ; qtot -0.873
3857 H1 238 SER HB2 3857 0.035200 1.00800 ; qtot -0.838
3858 H1 238 SER HB3 3858 0.035200 1.00800 ; qtot -0.802
3859 OH 238 SER OG 3859 -0.654601 16.00000 ; qtot -1.457
3860 HO 238 SER HG 3860 0.427500 1.00800 ; qtot -1.029
3861 C 238 SER C 3861 0.597301 12.01000 ; qtot -0.432
3862 O 238 SER O 3862 -0.567901 16.00000 ; qtot -1.000
3863 N 239 ASP N 3863 -0.516301 14.01000 ; qtot -1.516
3864 H 239 ASP H 3864 0.293600 1.00800 ; qtot -1.223
3865 CX 239 ASP CA 3865 0.038100 12.01000 ; qtot -1.185
3866 H1 239 ASP HA 3866 0.088000 1.00800 ; qtot -1.097
3867 A2C 239 ASP CB 3867 -0.030300 12.01000 ; qtot -1.127
3868 HC 239 ASP HB2 3868 -0.012200 1.00800 ; qtot -1.139
3869 HC 239 ASP HB3 3869 -0.012200 1.00800 ; qtot -1.151
3870 CO 239 ASP CG 3870 0.799401 12.01000 ; qtot -0.352
3871 O2 239 ASP OD1 3871 -0.801401 16.00000 ; qtot -1.153

```
  3872   O2   239   ASP   OD2 3872    -0.801401     16.00000 ; qtot -
1.955
  3873    C   239   ASP     C 3873     0.536601     12.01000 ; qtot -
1.418
  3874    O   239   ASP     O 3874    -0.581901     16.00000 ; qtot -
2.000
  3875    N   240   ALA     N 3875    -0.415700     14.01000 ; qtot -
2.416
  3876    H   240   ALA     H 3876     0.271900      1.00800 ; qtot -
2.144
  3877   CX   240   ALA    CA 3877     0.033700     12.01000 ; qtot -
2.110
  3878   H1   240   ALA    HA 3878     0.082300      1.00800 ; qtot -
2.028
  3879   CT   240   ALA    CB 3879    -0.182500     12.01000 ; qtot -
2.210
  3880   HC   240   ALA   HB1 3880     0.060300      1.00800 ; qtot -
2.150
  3881   HC   240   ALA   HB2 3881     0.060300      1.00800 ; qtot -
2.090
  3882   HC   240   ALA   HB3 3882     0.060300      1.00800 ; qtot -
2.029
  3883    C   240   ALA     C 3883     0.597301     12.01000 ; qtot -
1.432
  3884    O   240   ALA     O 3884    -0.567901     16.00000 ; qtot -
2.000
  3885    N   241   LYS     N 3885    -0.347900     14.01000 ; qtot -
2.348
  3886    H   241   LYS     H 3886     0.274700      1.00800 ; qtot -
2.073
  3887   CX   241   LYS    CA 3887    -0.240000     12.01000 ; qtot -
2.313
  3888   H1   241   LYS    HA 3888     0.142600      1.00800 ; qtot -
2.171
  3889   C8   241   LYS    CB 3889    -0.009400     12.01000 ; qtot -
2.180
  3890   HC   241   LYS   HB2 3890     0.036200      1.00800 ; qtot -
2.144
  3891   HC   241   LYS   HB3 3891     0.036200      1.00800 ; qtot -
2.108
  3892   C8   241   LYS    CG 3892     0.018700     12.01000 ; qtot -
2.089
  3893   HC   241   LYS   HG2 3893     0.010300      1.00800 ; qtot -
2.079
  3894   HC   241   LYS   HG3 3894     0.010300      1.00800 ; qtot -
2.068
  3895   C8   241   LYS    CD 3895    -0.047900     12.01000 ; qtot -
2.116
  3896   HC   241   LYS   HD2 3896     0.062100      1.00800 ; qtot -
2.054
  3897   HC   241   LYS   HD3 3897     0.062100      1.00800 ; qtot -
1.992
```

3898 C8 241 LYS CE 3898 -0.014300 12.01000 ; qtot -
2.006
3899 HP 241 LYS HE2 3899 0.113500 1.00800 ; qtot -
1.893
3900 HP 241 LYS HE3 3900 0.113500 1.00800 ; qtot -
1.779
3901 N3 241 LYS NZ 3901 -0.385400 14.01000 ; qtot -
2.165
3902 H 241 LYS HZ1 3902 0.340000 1.00800 ; qtot -
1.825
3903 H 241 LYS HZ2 3903 0.340000 1.00800 ; qtot -
1.485
3904 H 241 LYS HZ3 3904 0.340000 1.00800 ; qtot -
1.145
3905 C 241 LYS C 3905 0.734101 12.01000 ; qtot -
0.411
3906 O 241 LYS O 3906 -0.589401 16.00000 ; qtot -
1.000
3907 N 242 ASN N 3907 -0.415700 14.01000 ; qtot -
1.416
3908 H 242 ASN H 3908 0.271900 1.00800 ; qtot -
1.144
3909 CX 242 ASN CA 3909 0.014300 12.01000 ; qtot -
1.130
3910 H1 242 ASN HA 3910 0.104800 1.00800 ; qtot -
1.025
3911 A2C 242 ASN CB 3911 -0.204100 12.01000 ; qtot -
1.229
3912 HC 242 ASN HB2 3912 0.079700 1.00800 ; qtot -
1.149
3913 HC 242 ASN HB3 3913 0.079700 1.00800 ; qtot -
1.069
3914 C 242 ASN CG 3914 0.713001 12.01000 ; qtot -
0.356
3915 O 242 ASN OD1 3915 -0.593101 16.00000 ; qtot -
0.950
3916 N 242 ASN ND2 3916 -0.919101 14.01000 ; qtot -
1.869
3917 H 242 ASN HD21 3917 0.419600 1.00800 ; qtot -
1.449
3918 H 242 ASN HD22 3918 0.419600 1.00800 ; qtot -
1.029
3919 C 242 ASN C 3919 0.597301 12.01000 ; qtot -
0.432
3920 O 242 ASN O 3920 -0.567901 16.00000 ; qtot -
1.000
3921 N 243 LEU N 3921 -0.415700 14.01000 ; qtot -
1.416
3922 H 243 LEU H 3922 0.271900 1.00800 ; qtot -
1.144
3923 CX 243 LEU CA 3923 -0.051800 12.01000 ; qtot -
1.196

3924 H1 243 LEU HA 3924 0.092200 1.00800 ; qtot -1.103
3925 A2C 243 LEU CB 3925 -0.110200 12.01000 ; qtot -1.214
3926 HC 243 LEU HB2 3926 0.045700 1.00800 ; qtot -1.168
3927 HC 243 LEU HB3 3927 0.045700 1.00800 ; qtot -1.122
3928 A3C 243 LEU CG 3928 0.353100 12.01000 ; qtot -0.769
3929 HC 243 LEU HG 3929 -0.036100 1.00800 ; qtot -0.805
3930 CT 243 LEU CD1 3930 -0.412100 12.01000 ; qtot -1.217
3931 HC 243 LEU HD11 3931 0.100000 1.00800 ; qtot -1.117
3932 HC 243 LEU HD12 3932 0.100000 1.00800 ; qtot -1.017
3933 HC 243 LEU HD13 3933 0.100000 1.00800 ; qtot -0.917
3934 CT 243 LEU CD2 3934 -0.412100 12.01000 ; qtot -1.329
3935 HC 243 LEU HD21 3935 0.100000 1.00800 ; qtot -1.229
3936 HC 243 LEU HD22 3936 0.100000 1.00800 ; qtot -1.129
3937 HC 243 LEU HD23 3937 0.100000 1.00800 ; qtot -1.029
3938 C 243 LEU C 3938 0.597301 12.01000 ; qtot -0.432
3939 O 243 LEU O 3939 -0.567901 16.00000 ; qtot -1.000
3940 N 244 ILE N 3940 -0.415700 14.01000 ; qtot -1.416
3941 H 244 ILE H 3941 0.271900 1.00800 ; qtot -1.144
3942 CX 244 ILE CA 3942 -0.059700 12.01000 ; qtot -1.204
3943 H1 244 ILE HA 3943 0.086900 1.00800 ; qtot -1.117
3944 A3C 244 ILE CB 3944 0.130300 12.01000 ; qtot -0.986
3945 HC 244 ILE HB 3945 0.018700 1.00800 ; qtot -0.968
3946 CT 244 ILE CG2 3946 -0.320400 12.01000 ; qtot -1.288
3947 HC 244 ILE HG21 3947 0.088200 1.00800 ; qtot -1.200
3948 HC 244 ILE HG22 3948 0.088200 1.00800 ; qtot -1.112
3949 HC 244 ILE HG23 3949 0.088200 1.00800 ; qtot -1.023

3950 A2C 244 ILE CG1 3950 -0.043000 12.01000 ; qtot -
1.066
3951 HC 244 ILE HG12 3951 0.023600 1.00800 ; qtot -
1.043
3952 HC 244 ILE HG13 3952 0.023600 1.00800 ; qtot -
1.019
3953 CT 244 ILE CD1 3953 -0.066000 12.01000 ; qtot -
1.085
3954 HC 244 ILE HD11 3954 0.018600 1.00800 ; qtot -
1.067
3955 HC 244 ILE HD12 3955 0.018600 1.00800 ; qtot -
1.048
3956 HC 244 ILE HD13 3956 0.018600 1.00800 ; qtot -
1.029
3957 C 244 ILE C 3957 0.597301 12.01000 ; qtot -
0.432
3958 O 244 ILE O 3958 -0.567901 16.00000 ; qtot -
1.000
3959 N 245 THR N 3959 -0.415700 14.01000 ; qtot -
1.416
3960 H 245 THR H 3960 0.271900 1.00800 ; qtot -
1.144
3961 CX 245 THR CA 3961 -0.038900 12.01000 ; qtot -
1.183
3962 H1 245 THR HA 3962 0.100700 1.00800 ; qtot -
1.082
3963 A3C 245 THR CB 3963 0.365400 12.01000 ; qtot -
0.717
3964 H1 245 THR HB 3964 0.004300 1.00800 ; qtot -
0.712
3965 CT 245 THR CG2 3965 -0.243800 12.01000 ; qtot -
0.956
3966 HC 245 THR HG21 3966 0.064200 1.00800 ; qtot -
0.892
3967 HC 245 THR HG22 3967 0.064200 1.00800 ; qtot -
0.828
3968 HC 245 THR HG23 3968 0.064200 1.00800 ; qtot -
0.764
3969 OH 245 THR OG1 3969 -0.676101 16.00000 ; qtot -
1.440
3970 HO 245 THR HG1 3970 0.410200 1.00800 ; qtot -
1.029
3971 C 245 THR C 3971 0.597301 12.01000 ; qtot -
0.432
3972 O 245 THR O 3972 -0.567901 16.00000 ; qtot -
1.000
3973 N 246 LYS N 3973 -0.347900 14.01000 ; qtot -
1.348
3974 H 246 LYS H 3974 0.274700 1.00800 ; qtot -
1.073
3975 CX 246 LYS CA 3975 -0.240000 12.01000 ; qtot -
1.313

3976 H1 246 LYS HA 3976 0.142600 1.00800 ; qtot -
1.171
3977 C8 246 LYS CB 3977 -0.009400 12.01000 ; qtot -
1.180
3978 HC 246 LYS HB2 3978 0.036200 1.00800 ; qtot -
1.144
3979 HC 246 LYS HB3 3979 0.036200 1.00800 ; qtot -
1.108
3980 C8 246 LYS CG 3980 0.018700 12.01000 ; qtot -
1.089
3981 HC 246 LYS HG2 3981 0.010300 1.00800 ; qtot -
1.079
3982 HC 246 LYS HG3 3982 0.010300 1.00800 ; qtot -
1.068
3983 C8 246 LYS CD 3983 -0.047900 12.01000 ; qtot -
1.116
3984 HC 246 LYS HD2 3984 0.062100 1.00800 ; qtot -
1.054
3985 HC 246 LYS HD3 3985 0.062100 1.00800 ; qtot -
0.992
3986 C8 246 LYS CE 3986 -0.014300 12.01000 ; qtot -
1.006
3987 HP 246 LYS HE2 3987 0.113500 1.00800 ; qtot -
0.893
3988 HP 246 LYS HE3 3988 0.113500 1.00800 ; qtot -
0.779
3989 N3 246 LYS NZ 3989 -0.385400 14.01000 ; qtot -
1.165
3990 H 246 LYS HZ1 3990 0.340000 1.00800 ; qtot -
0.825
3991 H 246 LYS HZ2 3991 0.340000 1.00800 ; qtot -
0.485
3992 H 246 LYS HZ3 3992 0.340000 1.00800 ; qtot -
0.145
3993 C 246 LYS C 3993 0.734101 12.01000 ; qtot
0.589
3994 O 246 LYS O 3994 -0.589401 16.00000 ; qtot
0.000
3995 N 247 LEU N 3995 -0.415700 14.01000 ; qtot -
0.416
3996 H 247 LEU H 3996 0.271900 1.00800 ; qtot -
0.144
3997 CX 247 LEU CA 3997 -0.051800 12.01000 ; qtot -
0.196
3998 H1 247 LEU HA 3998 0.092200 1.00800 ; qtot -
0.103
3999 A2C 247 LEU CB 3999 -0.110200 12.01000 ; qtot -
0.214
4000 HC 247 LEU HB2 4000 0.045700 1.00800 ; qtot -
0.168
4001 HC 247 LEU HB3 4001 0.045700 1.00800 ; qtot -
0.122

4002 A3C 247 LEU CG 4002 0.353100 12.01000 ; qtot
0.231
4003 HC 247 LEU HG 4003 -0.036100 1.00800 ; qtot
0.195
4004 CT 247 LEU CD1 4004 -0.412100 12.01000 ; qtot -
0.217
4005 HC 247 LEU HD11 4005 0.100000 1.00800 ; qtot -
0.117
4006 HC 247 LEU HD12 4006 0.100000 1.00800 ; qtot -
0.017
4007 HC 247 LEU HD13 4007 0.100000 1.00800 ; qtot
0.083
4008 CT 247 LEU CD2 4008 -0.412100 12.01000 ; qtot -
0.329
4009 HC 247 LEU HD21 4009 0.100000 1.00800 ; qtot -
0.229
4010 HC 247 LEU HD22 4010 0.100000 1.00800 ; qtot -
0.129
4011 HC 247 LEU HD23 4011 0.100000 1.00800 ; qtot -
0.029
4012 C 247 LEU C 4012 0.597301 12.01000 ; qtot
0.568
4013 O 247 LEU O 4013 -0.567901 16.00000 ; qtot
0.000
4014 N 248 LEU N 4014 -0.415700 14.01000 ; qtot -
0.416
4015 H 248 LEU H 4015 0.271900 1.00800 ; qtot -
0.144
4016 CX 248 LEU CA 4016 -0.051800 12.01000 ; qtot -
0.196
4017 H1 248 LEU HA 4017 0.092200 1.00800 ; qtot -
0.103
4018 A2C 248 LEU CB 4018 -0.110200 12.01000 ; qtot -
0.214
4019 HC 248 LEU HB2 4019 0.045700 1.00800 ; qtot -
0.168
4020 HC 248 LEU HB3 4020 0.045700 1.00800 ; qtot -
0.122
4021 A3C 248 LEU CG 4021 0.353100 12.01000 ; qtot
0.231
4022 HC 248 LEU HG 4022 -0.036100 1.00800 ; qtot
0.195
4023 CT 248 LEU CD1 4023 -0.412100 12.01000 ; qtot -
0.217
4024 HC 248 LEU HD11 4024 0.100000 1.00800 ; qtot -
0.117
4025 HC 248 LEU HD12 4025 0.100000 1.00800 ; qtot -
0.017
4026 HC 248 LEU HD13 4026 0.100000 1.00800 ; qtot
0.083
4027 CT 248 LEU CD2 4027 -0.412100 12.01000 ; qtot -
0.329

4028 HC 248 LEU HD21 4028 0.100000 1.00800 ; qtot -0.229
4029 HC 248 LEU HD22 4029 0.100000 1.00800 ; qtot -0.129
4030 HC 248 LEU HD23 4030 0.100000 1.00800 ; qtot -0.029
4031 C 248 LEU C 4031 0.597301 12.01000 ; qtot 0.568
4032 O 248 LEU O 4032 -0.567901 16.00000 ; qtot 0.000
4033 N 249 THR N 4033 -0.415700 14.01000 ; qtot -0.416
4034 H 249 THR H 4034 0.271900 1.00800 ; qtot -0.144
4035 CX 249 THR CA 4035 -0.038900 12.01000 ; qtot -0.183
4036 H1 249 THR HA 4036 0.100700 1.00800 ; qtot -0.082
4037 A3C 249 THR CB 4037 0.365400 12.01000 ; qtot 0.283
4038 H1 249 THR HB 4038 0.004300 1.00800 ; qtot 0.288
4039 CT 249 THR CG2 4039 -0.243800 12.01000 ; qtot 0.044
4040 HC 249 THR HG21 4040 0.064200 1.00800 ; qtot 0.108
4041 HC 249 THR HG22 4041 0.064200 1.00800 ; qtot 0.172
4042 HC 249 THR HG23 4042 0.064200 1.00800 ; qtot 0.237
4043 OH 249 THR OG1 4043 -0.676101 16.00000 ; qtot -0.440
4044 HO 249 THR HG1 4044 0.410200 1.00800 ; qtot -0.029
4045 C 249 THR C 4045 0.597301 12.01000 ; qtot 0.568
4046 O 249 THR O 4046 -0.567901 16.00000 ; qtot 0.000
4047 N 250 TYR N 4047 -0.415700 14.01000 ; qtot -0.416
4048 H 250 TYR H 4048 0.271900 1.00800 ; qtot -0.144
4049 CX 250 TYR CA 4049 -0.001400 12.01000 ; qtot -0.145
4050 H1 250 TYR HA 4050 0.087600 1.00800 ; qtot -0.058
4051 CT 250 TYR CB 4051 -0.015200 12.01000 ; qtot -0.073
4052 HC 250 TYR HB2 4052 0.029500 1.00800 ; qtot -0.043
4053 HC 250 TYR HB3 4053 0.029500 1.00800 ; qtot -0.014

```
  4054   CA   250   TYR    CG 4054    -0.001100     12.01000 ; qtot -
0.015
  4055   CA   250   TYR   CD1 4055    -0.190600     12.01000 ; qtot -
0.205
  4056   HA   250   TYR   HD1 4056     0.169900      1.00800 ; qtot -
0.036
  4057   CA   250   TYR   CE1 4057    -0.234100     12.01000 ; qtot -
0.270
  4058   HA   250   TYR   HE1 4058     0.165600      1.00800 ; qtot -
0.104
  4059    C   250   TYR    CZ 4059     0.322600     12.01000 ; qtot
0.219
  4060   OH   250   TYR    OH 4060    -0.557901     16.00000 ; qtot -
0.339
  4061   HO   250   TYR    HH 4061     0.399200      1.00800 ; qtot
0.060
  4062   CA   250   TYR   CE2 4062    -0.234100     12.01000 ; qtot -
0.174
  4063   HA   250   TYR   HE2 4063     0.165600      1.00800 ; qtot -
0.009
  4064   CA   250   TYR   CD2 4064    -0.190600     12.01000 ; qtot -
0.199
  4065   HA   250   TYR   HD2 4065     0.169900      1.00800 ; qtot -
0.029
  4066    C   250   TYR     C 4066     0.597301     12.01000 ; qtot
0.568
  4067    O   250   TYR     O 4067    -0.567901     16.00000 ; qtot
0.000
  4068    N   251   ASN     N 4068    -0.415700     14.01000 ; qtot -
0.416
  4069    H   251   ASN     H 4069     0.271900      1.00800 ; qtot -
0.144
  4070   CX   251   ASN    CA 4070     0.014300     12.01000 ; qtot -
0.129
  4071   H1   251   ASN    HA 4071     0.104800      1.00800 ; qtot -
0.025
  4072  A2C   251   ASN    CB 4072    -0.204100     12.01000 ; qtot -
0.229
  4073   HC   251   ASN   HB2 4073     0.079700      1.00800 ; qtot -
0.149
  4074   HC   251   ASN   HB3 4074     0.079700      1.00800 ; qtot -
0.069
  4075    C   251   ASN    CG 4075     0.713001     12.01000 ; qtot
0.644
  4076    O   251   ASN   OD1 4076    -0.593101     16.00000 ; qtot
0.051
  4077    N   251   ASN   ND2 4077    -0.919101     14.01000 ; qtot -
0.869
  4078    H   251   ASN  HD21 4078     0.419600      1.00800 ; qtot -
0.449
  4079    H   251   ASN  HD22 4079     0.419600      1.00800 ; qtot -
0.029
```

```
  4080    C   251   ASN     C 4080     0.597301     12.01000 ; qtot
0.568
  4081    O   251   ASN     O 4081    -0.567901     16.00000 ; qtot
0.000
  4082    N   252   PRO     N 4082    -0.254800     14.01000 ; qtot -
0.255
  4083   CT   252   PRO    CD 4083     0.019200     12.01000 ; qtot -
0.236
  4084   H1   252   PRO   HD2 4084     0.039100      1.00800 ; qtot -
0.196
  4085   H1   252   PRO   HD3 4085     0.039100      1.00800 ; qtot -
0.157
  4086   CT   252   PRO    CG 4086     0.018900     12.01000 ; qtot -
0.138
  4087   HC   252   PRO   HG2 4087     0.021300      1.00800 ; qtot -
0.117
  4088   HC   252   PRO   HG3 4088     0.021300      1.00800 ; qtot -
0.096
  4089   CT   252   PRO    CB 4089    -0.007000     12.01000 ; qtot -
0.103
  4090   HC   252   PRO   HB2 4090     0.025300      1.00800 ; qtot -
0.078
  4091   HC   252   PRO   HB3 4091     0.025300      1.00800 ; qtot -
0.052
  4092   CX   252   PRO    CA 4092    -0.026600     12.01000 ; qtot -
0.079
  4093   H1   252   PRO    HA 4093     0.064100      1.00800 ; qtot -
0.015
  4094    C   252   PRO     C 4094     0.589601     12.01000 ; qtot
0.575
  4095    O   252   PRO     O 4095    -0.574801     16.00000 ; qtot
0.000
  4096    N   253   ASN     N 4096    -0.415700     14.01000 ; qtot -
0.416
  4097    H   253   ASN     H 4097     0.271900      1.00800 ; qtot -
0.144
  4098   CX   253   ASN    CA 4098     0.014300     12.01000 ; qtot -
0.129
  4099   H1   253   ASN    HA 4099     0.104800      1.00800 ; qtot -
0.025
  4100  A2C   253   ASN    CB 4100    -0.204100     12.01000 ; qtot -
0.229
  4101   HC   253   ASN   HB2 4101     0.079700      1.00800 ; qtot -
0.149
  4102   HC   253   ASN   HB3 4102     0.079700      1.00800 ; qtot -
0.069
  4103    C   253   ASN    CG 4103     0.713001     12.01000 ; qtot
0.644
  4104    O   253   ASN   OD1 4104    -0.593101     16.00000 ; qtot
0.051
  4105    N   253   ASN   ND2 4105    -0.919101     14.01000 ; qtot -
0.869
```

4106 H 253 ASN HD21 4106 0.419600 1.00800 ; qtot -
0.449
4107 H 253 ASN HD22 4107 0.419600 1.00800 ; qtot -
0.029
4108 C 253 ASN C 4108 0.597301 12.01000 ; qtot
0.568
4109 O 253 ASN O 4109 -0.567901 16.00000 ; qtot
0.000
4110 N 254 GLU N 4110 -0.516301 14.01000 ; qtot -
0.516
4111 H 254 GLU H 4111 0.293600 1.00800 ; qtot -
0.223
4112 CX 254 GLU CA 4112 0.039700 12.01000 ; qtot -
0.183
4113 H1 254 GLU HA 4113 0.110500 1.00800 ; qtot -
0.072
4114 A2C 254 GLU CB 4114 0.056000 12.01000 ; qtot -
0.016
4115 HC 254 GLU HB2 4115 -0.017300 1.00800 ; qtot -
0.034
4116 HC 254 GLU HB3 4116 -0.017300 1.00800 ; qtot -
0.051
4117 A2C 254 GLU CG 4117 0.013600 12.01000 ; qtot -
0.037
4118 HC 254 GLU HG2 4118 -0.042500 1.00800 ; qtot -
0.080
4119 HC 254 GLU HG3 4119 -0.042500 1.00800 ; qtot -
0.122
4120 CO 254 GLU CD 4120 0.805401 12.01000 ; qtot
0.683
4121 O2 254 GLU OE1 4121 -0.818801 16.00000 ; qtot -
0.136
4122 O2 254 GLU OE2 4122 -0.818801 16.00000 ; qtot -
0.955
4123 C 254 GLU C 4123 0.536601 12.01000 ; qtot -
0.418
4124 O 254 GLU O 4124 -0.581901 16.00000 ; qtot -
1.000
4125 N 255 ARG N 4125 -0.347900 14.01000 ; qtot -
1.348
4126 H 255 ARG H 4126 0.274700 1.00800 ; qtot -
1.073
4127 CX 255 ARG CA 4127 -0.263700 12.01000 ; qtot -
1.337
4128 H1 255 ARG HA 4128 0.156000 1.00800 ; qtot -
1.181
4129 C8 255 ARG CB 4129 -0.000700 12.01000 ; qtot -
1.182
4130 HC 255 ARG HB2 4130 0.032700 1.00800 ; qtot -
1.149
4131 HC 255 ARG HB3 4131 0.032700 1.00800 ; qtot -
1.116

4132 C8 255 ARG CG 4132 0.039000 12.01000 ; qtot -
1.077
4133 HC 255 ARG HG2 4133 0.028500 1.00800 ; qtot -
1.049
4134 HC 255 ARG HG3 4134 0.028500 1.00800 ; qtot -
1.020
4135 C8 255 ARG CD 4135 0.048600 12.01000 ; qtot -
0.972
4136 H1 255 ARG HD2 4136 0.068700 1.00800 ; qtot -
0.903
4137 H1 255 ARG HD3 4137 0.068700 1.00800 ; qtot -
0.834
4138 N2 255 ARG NE 4138 -0.529501 14.01000 ; qtot -
1.364
4139 H 255 ARG HE 4139 0.345600 1.00800 ; qtot -
1.018
4140 CA 255 ARG CZ 4140 0.807601 12.01000 ; qtot -
0.210
4141 N2 255 ARG NH1 4141 -0.862701 14.01000 ; qtot -
1.073
4142 H 255 ARG HH11 4142 0.447800 1.00800 ; qtot -
0.625
4143 H 255 ARG HH12 4143 0.447800 1.00800 ; qtot -
0.178
4144 N2 255 ARG NH2 4144 -0.862701 14.01000 ; qtot -
1.040
4145 H 255 ARG HH21 4145 0.447800 1.00800 ; qtot -
0.593
4146 H 255 ARG HH22 4146 0.447800 1.00800 ; qtot -
0.145
4147 C 255 ARG C 4147 0.734101 12.01000 ; qtot
0.589
4148 O 255 ARG O 4148 -0.589401 16.00000 ; qtot
0.000
4149 N 256 CYS N 4149 -0.415700 14.01000 ; qtot -
0.416
4150 H 256 CYS H 4150 0.271900 1.00800 ; qtot -
0.144
4151 CX 256 CYS CA 4151 0.021300 12.01000 ; qtot -
0.122
4152 H1 256 CYS HA 4152 0.112400 1.00800 ; qtot -
0.010
4153 A2C 256 CYS CB 4153 -0.123100 12.01000 ; qtot -
0.133
4154 H1 256 CYS HB2 4154 0.111200 1.00800 ; qtot -
0.022
4155 H1 256 CYS HB3 4155 0.111200 1.00800 ; qtot
0.089
4156 SH 256 CYS SG 4156 -0.311900 32.06000 ; qtot -
0.223
4157 HS 256 CYS HG 4157 0.193300 1.00800 ; qtot -
0.029

4158 C 256 CYS C 4158 0.597301 12.01000 ; qtot
0.568
4159 O 256 CYS O 4159 -0.567901 16.00000 ; qtot
0.000
4160 N 257 THR N 4160 -0.415700 14.01000 ; qtot -
0.416
4161 H 257 THR H 4161 0.271900 1.00800 ; qtot -
0.144
4162 CX 257 THR CA 4162 -0.038900 12.01000 ; qtot -
0.183
4163 H1 257 THR HA 4163 0.100700 1.00800 ; qtot -
0.082
4164 A3C 257 THR CB 4164 0.365400 12.01000 ; qtot
0.283
4165 H1 257 THR HB 4165 0.004300 1.00800 ; qtot
0.288
4166 CT 257 THR CG2 4166 -0.243800 12.01000 ; qtot
0.044
4167 HC 257 THR HG21 4167 0.064200 1.00800 ; qtot
0.108
4168 HC 257 THR HG22 4168 0.064200 1.00800 ; qtot
0.172
4169 HC 257 THR HG23 4169 0.064200 1.00800 ; qtot
0.237
4170 OH 257 THR OG1 4170 -0.676101 16.00000 ; qtot -
0.440
4171 HO 257 THR HG1 4171 0.410200 1.00800 ; qtot -
0.029
4172 C 257 THR C 4172 0.597301 12.01000 ; qtot
0.568
4173 O 257 THR O 4173 -0.567901 16.00000 ; qtot
0.000
4174 N 258 ILE N 4174 -0.415700 14.01000 ; qtot -
0.416
4175 H 258 ILE H 4175 0.271900 1.00800 ; qtot -
0.144
4176 CX 258 ILE CA 4176 -0.059700 12.01000 ; qtot -
0.203
4177 H1 258 ILE HA 4177 0.086900 1.00800 ; qtot -
0.117
4178 A3C 258 ILE CB 4178 0.130300 12.01000 ; qtot
0.014
4179 HC 258 ILE HB 4179 0.018700 1.00800 ; qtot
0.032
4180 CT 258 ILE CG2 4180 -0.320400 12.01000 ; qtot -
0.288
4181 HC 258 ILE HG21 4181 0.088200 1.00800 ; qtot -
0.200
4182 HC 258 ILE HG22 4182 0.088200 1.00800 ; qtot -
0.112
4183 HC 258 ILE HG23 4183 0.088200 1.00800 ; qtot -
0.023

4184 A2C 258 ILE CG1 4184 -0.043000 12.01000 ; qtot -
0.066
4185 HC 258 ILE HG12 4185 0.023600 1.00800 ; qtot -
0.043
4186 HC 258 ILE HG13 4186 0.023600 1.00800 ; qtot -
0.019
4187 CT 258 ILE CD1 4187 -0.066000 12.01000 ; qtot -
0.085
4188 HC 258 ILE HD11 4188 0.018600 1.00800 ; qtot -
0.067
4189 HC 258 ILE HD12 4189 0.018600 1.00800 ; qtot -
0.048
4190 HC 258 ILE HD13 4190 0.018600 1.00800 ; qtot -
0.029
4191 C 258 ILE C 4191 0.597301 12.01000 ; qtot
0.568
4192 O 258 ILE O 4192 -0.567901 16.00000 ; qtot
0.000
4193 N 259 GLU N 4193 -0.516301 14.01000 ; qtot -
0.516
4194 H 259 GLU H 4194 0.293600 1.00800 ; qtot -
0.223
4195 CX 259 GLU CA 4195 0.039700 12.01000 ; qtot -
0.183
4196 H1 259 GLU HA 4196 0.110500 1.00800 ; qtot -
0.072
4197 A2C 259 GLU CB 4197 0.056000 12.01000 ; qtot -
0.016
4198 HC 259 GLU HB2 4198 -0.017300 1.00800 ; qtot -
0.034
4199 HC 259 GLU HB3 4199 -0.017300 1.00800 ; qtot -
0.051
4200 A2C 259 GLU CG 4200 0.013600 12.01000 ; qtot -
0.037
4201 HC 259 GLU HG2 4201 -0.042500 1.00800 ; qtot -
0.080
4202 HC 259 GLU HG3 4202 -0.042500 1.00800 ; qtot -
0.122
4203 CO 259 GLU CD 4203 0.805401 12.01000 ; qtot
0.683
4204 O2 259 GLU OE1 4204 -0.818801 16.00000 ; qtot -
0.136
4205 O2 259 GLU OE2 4205 -0.818801 16.00000 ; qtot -
0.955
4206 C 259 GLU C 4206 0.536601 12.01000 ; qtot -
0.418
4207 O 259 GLU O 4207 -0.581901 16.00000 ; qtot -
1.000
4208 N 260 GLU N 4208 -0.516301 14.01000 ; qtot -
1.516
4209 H 260 GLU H 4209 0.293600 1.00800 ; qtot -
1.223

4210 CX 260 GLU CA 4210 0.039700 12.01000 ; qtot -
1.183
4211 H1 260 GLU HA 4211 0.110500 1.00800 ; qtot -
1.073
4212 A2C 260 GLU CB 4212 0.056000 12.01000 ; qtot -
1.017
4213 HC 260 GLU HB2 4213 -0.017300 1.00800 ; qtot -
1.034
4214 HC 260 GLU HB3 4214 -0.017300 1.00800 ; qtot -
1.051
4215 A2C 260 GLU CG 4215 0.013600 12.01000 ; qtot -
1.038
4216 HC 260 GLU HG2 4216 -0.042500 1.00800 ; qtot -
1.080
4217 HC 260 GLU HG3 4217 -0.042500 1.00800 ; qtot -
1.123
4218 CO 260 GLU CD 4218 0.805401 12.01000 ; qtot -
0.317
4219 O2 260 GLU OE1 4219 -0.818801 16.00000 ; qtot -
1.136
4220 O2 260 GLU OE2 4220 -0.818801 16.00000 ; qtot -
1.955
4221 C 260 GLU C 4221 0.536601 12.01000 ; qtot -
1.418
4222 O 260 GLU O 4222 -0.581901 16.00000 ; qtot -
2.000
4223 N 261 ALA N 4223 -0.415700 14.01000 ; qtot -
2.416
4224 H 261 ALA H 4224 0.271900 1.00800 ; qtot -
2.144
4225 CX 261 ALA CA 4225 0.033700 12.01000 ; qtot -
2.110
4226 H1 261 ALA HA 4226 0.082300 1.00800 ; qtot -
2.028
4227 CT 261 ALA CB 4227 -0.182500 12.01000 ; qtot -
2.210
4228 HC 261 ALA HB1 4228 0.060300 1.00800 ; qtot -
2.150
4229 HC 261 ALA HB2 4229 0.060300 1.00800 ; qtot -
2.090
4230 HC 261 ALA HB3 4230 0.060300 1.00800 ; qtot -
2.029
4231 C 261 ALA C 4231 0.597301 12.01000 ; qtot -
1.432
4232 O 261 ALA O 4232 -0.567901 16.00000 ; qtot -
2.000
4233 N 262 LEU N 4233 -0.415700 14.01000 ; qtot -
2.416
4234 H 262 LEU H 4234 0.271900 1.00800 ; qtot -
2.144
4235 CX 262 LEU CA 4235 -0.051800 12.01000 ; qtot -
2.196

4236 H1 262 LEU HA 4236 0.092200 1.00800 ; qtot -
2.103
4237 A2C 262 LEU CB 4237 -0.110200 12.01000 ; qtot -
2.214
4238 HC 262 LEU HB2 4238 0.045700 1.00800 ; qtot -
2.168
4239 HC 262 LEU HB3 4239 0.045700 1.00800 ; qtot -
2.122
4240 A3C 262 LEU CG 4240 0.353100 12.01000 ; qtot -
1.769
4241 HC 262 LEU HG 4241 -0.036100 1.00800 ; qtot -
1.805
4242 CT 262 LEU CD1 4242 -0.412100 12.01000 ; qtot -
2.217
4243 HC 262 LEU HD11 4243 0.100000 1.00800 ; qtot -
2.117
4244 HC 262 LEU HD12 4244 0.100000 1.00800 ; qtot -
2.017
4245 HC 262 LEU HD13 4245 0.100000 1.00800 ; qtot -
1.917
4246 CT 262 LEU CD2 4246 -0.412100 12.01000 ; qtot -
2.329
4247 HC 262 LEU HD21 4247 0.100000 1.00800 ; qtot -
2.229
4248 HC 262 LEU HD22 4248 0.100000 1.00800 ; qtot -
2.129
4249 HC 262 LEU HD23 4249 0.100000 1.00800 ; qtot -
2.029
4250 C 262 LEU C 4250 0.597301 12.01000 ; qtot -
1.432
4251 O 262 LEU O 4251 -0.567901 16.00000 ; qtot -
2.000
4252 N 263 ASN N 4252 -0.415700 14.01000 ; qtot -
2.416
4253 H 263 ASN H 4253 0.271900 1.00800 ; qtot -
2.144
4254 CX 263 ASN CA 4254 0.014300 12.01000 ; qtot -
2.130
4255 H1 263 ASN HA 4255 0.104800 1.00800 ; qtot -
2.025
4256 A2C 263 ASN CB 4256 -0.204100 12.01000 ; qtot -
2.229
4257 HC 263 ASN HB2 4257 0.079700 1.00800 ; qtot -
2.149
4258 HC 263 ASN HB3 4258 0.079700 1.00800 ; qtot -
2.069
4259 C 263 ASN CG 4259 0.713001 12.01000 ; qtot -
1.356
4260 O 263 ASN OD1 4260 -0.593101 16.00000 ; qtot -
1.950
4261 N 263 ASN ND2 4261 -0.919101 14.01000 ; qtot -
2.869

```
  4262    H   263   ASN  HD21 4262     0.419600      1.00800 ; qtot -
2.449
  4263    H   263   ASN  HD22 4263     0.419600      1.00800 ; qtot -
2.029
  4264    C   263   ASN     C 4264     0.597301     12.01000 ; qtot -
1.432
  4265    O   263   ASN     O 4265    -0.567901     16.00000 ; qtot -
2.000
  4266    N   264   HIE     N 4266    -0.415700     14.01000 ; qtot -
2.416
  4267    H   264   HIE     H 4267     0.271900      1.00800 ; qtot -
2.144
  4268   CX   264   HIE    CA 4268    -0.058100     12.01000 ; qtot -
2.202
  4269   H1   264   HIE    HA 4269     0.136000      1.00800 ; qtot -
2.066
  4270   CT   264   HIE    CB 4270    -0.007400     12.01000 ; qtot -
2.073
  4271   HC   264   HIE   HB2 4271     0.036700      1.00800 ; qtot -
2.037
  4272   HC   264   HIE   HB3 4272     0.036700      1.00800 ; qtot -
2.000
  4273   CC   264   HIE    CG 4273     0.186800     12.01000 ; qtot -
1.813
  4274   NB   264   HIE   ND1 4274    -0.543201     14.01000 ; qtot -
2.356
  4275   CR   264   HIE   CE1 4275     0.163500     12.01000 ; qtot -
2.193
  4276   H5   264   HIE   HE1 4276     0.143500      1.00800 ; qtot -
2.049
  4277   NA   264   HIE   NE2 4277    -0.279500     14.01000 ; qtot -
2.329
  4278    H   264   HIE   HE2 4278     0.333900      1.00800 ; qtot -
1.995
  4279   CW   264   HIE   CD2 4279    -0.220700     12.01000 ; qtot -
2.216
  4280   H4   264   HIE   HD2 4280     0.186200      1.00800 ; qtot -
2.029
  4281    C   264   HIE     C 4281     0.597301     12.01000 ; qtot -
1.432
  4282    O   264   HIE     O 4282    -0.567901     16.00000 ; qtot -
2.000
  4283    N   265   PRO     N 4283    -0.254800     14.01000 ; qtot -
2.255
  4284   CT   265   PRO    CD 4284     0.019200     12.01000 ; qtot -
2.236
  4285   H1   265   PRO   HD2 4285     0.039100      1.00800 ; qtot -
2.197
  4286   H1   265   PRO   HD3 4286     0.039100      1.00800 ; qtot -
2.157
  4287   CT   265   PRO    CG 4287     0.018900     12.01000 ; qtot -
2.139
```

```
  4288   HC   265   PRO   HG2 4288    0.021300      1.00800 ; qtot -
2.117
  4289   HC   265   PRO   HG3 4289    0.021300      1.00800 ; qtot -
2.096
  4290   CT   265   PRO    CB 4290   -0.007000     12.01000 ; qtot -
2.103
  4291   HC   265   PRO   HB2 4291    0.025300      1.00800 ; qtot -
2.078
  4292   HC   265   PRO   HB3 4292    0.025300      1.00800 ; qtot -
2.052
  4293   CX   265   PRO    CA 4293   -0.026600     12.01000 ; qtot -
2.079
  4294   H1   265   PRO    HA 4294    0.064100      1.00800 ; qtot -
2.015
  4295    C   265   PRO     C 4295    0.589601     12.01000 ; qtot -
1.425
  4296    O   265   PRO     O 4296   -0.574801     16.00000 ; qtot -
2.000
  4297    N   266   TRP     N 4297   -0.415700     14.01000 ; qtot -
2.416
  4298    H   266   TRP     H 4298    0.271900      1.00800 ; qtot -
2.144
  4299   CX   266   TRP    CA 4299   -0.027500     12.01000 ; qtot -
2.171
  4300   H1   266   TRP    HA 4300    0.112300      1.00800 ; qtot -
2.059
  4301   CT   266   TRP    CB 4301   -0.005000     12.01000 ; qtot -
2.064
  4302   HC   266   TRP   HB2 4302    0.033900      1.00800 ; qtot -
2.030
  4303   HC   266   TRP   HB3 4303    0.033900      1.00800 ; qtot -
1.996
  4304   C*   266   TRP    CG 4304   -0.141500     12.01000 ; qtot -
2.138
  4305   CW   266   TRP   CD1 4305   -0.163800     12.01000 ; qtot -
2.302
  4306   H4   266   TRP   HD1 4306    0.206200      1.00800 ; qtot -
2.095
  4307   NA   266   TRP   NE1 4307   -0.341800     14.01000 ; qtot -
2.437
  4308    H   266   TRP   HE1 4308    0.341200      1.00800 ; qtot -
2.096
  4309   CN   266   TRP   CE2 4309    0.138000     12.01000 ; qtot -
1.958
  4310   CA   266   TRP   CZ2 4310   -0.260100     12.01000 ; qtot -
2.218
  4311   HA   266   TRP   HZ2 4311    0.157200      1.00800 ; qtot -
2.061
  4312   CA   266   TRP   CH2 4312   -0.113400     12.01000 ; qtot -
2.174
  4313   HA   266   TRP   HH2 4313    0.141700      1.00800 ; qtot -
2.033
```

```
  4314   CA   266   TRP   CZ3 4314    -0.197200     12.01000 ; qtot -
2.230
  4315   HA   266   TRP   HZ3 4315     0.144700      1.00800 ; qtot -
2.085
  4316   CA   266   TRP   CE3 4316    -0.238700     12.01000 ; qtot -
2.324
  4317   HA   266   TRP   HE3 4317     0.170000      1.00800 ; qtot -
2.154
  4318   CB   266   TRP   CD2 4318     0.124300     12.01000 ; qtot -
2.029
  4319    C   266   TRP     C 4319     0.597301     12.01000 ; qtot -
1.432
  4320    O   266   TRP     O 4320    -0.567901     16.00000 ; qtot -
2.000
  4321    N   267   ILE     N 4321    -0.415700     14.01000 ; qtot -
2.416
  4322    H   267   ILE     H 4322     0.271900      1.00800 ; qtot -
2.144
  4323   CX   267   ILE    CA 4323    -0.059700     12.01000 ; qtot -
2.204
  4324   H1   267   ILE    HA 4324     0.086900      1.00800 ; qtot -
2.117
  4325  A3C   267   ILE    CB 4325     0.130300     12.01000 ; qtot -
1.986
  4326   HC   267   ILE    HB 4326     0.018700      1.00800 ; qtot -
1.968
  4327   CT   267   ILE   CG2 4327    -0.320400     12.01000 ; qtot -
2.288
  4328   HC   267   ILE  HG21 4328     0.088200      1.00800 ; qtot -
2.200
  4329   HC   267   ILE  HG22 4329     0.088200      1.00800 ; qtot -
2.112
  4330   HC   267   ILE  HG23 4330     0.088200      1.00800 ; qtot -
2.023
  4331  A2C   267   ILE   CG1 4331    -0.043000     12.01000 ; qtot -
2.066
  4332   HC   267   ILE  HG12 4332     0.023600      1.00800 ; qtot -
2.043
  4333   HC   267   ILE  HG13 4333     0.023600      1.00800 ; qtot -
2.019
  4334   CT   267   ILE   CD1 4334    -0.066000     12.01000 ; qtot -
2.085
  4335   HC   267   ILE  HD11 4335     0.018600      1.00800 ; qtot -
2.067
  4336   HC   267   ILE  HD12 4336     0.018600      1.00800 ; qtot -
2.048
  4337   HC   267   ILE  HD13 4337     0.018600      1.00800 ; qtot -
2.029
  4338    C   267   ILE     C 4338     0.597301     12.01000 ; qtot -
1.432
  4339    O   267   ILE     O 4339    -0.567901     16.00000 ; qtot -
2.000
```

4340 N 268 THR N 4340 -0.415700 14.01000 ; qtot -
2.416
4341 H 268 THR H 4341 0.271900 1.00800 ; qtot -
2.144
4342 CX 268 THR CA 4342 -0.038900 12.01000 ; qtot -
2.183
4343 H1 268 THR HA 4343 0.100700 1.00800 ; qtot -
2.082
4344 A3C 268 THR CB 4344 0.365400 12.01000 ; qtot -
1.717
4345 H1 268 THR HB 4345 0.004300 1.00800 ; qtot -
1.712
4346 CT 268 THR CG2 4346 -0.243800 12.01000 ; qtot -
1.956
4347 HC 268 THR HG21 4347 0.064200 1.00800 ; qtot -
1.892
4348 HC 268 THR HG22 4348 0.064200 1.00800 ; qtot -
1.828
4349 HC 268 THR HG23 4349 0.064200 1.00800 ; qtot -
1.764
4350 OH 268 THR OG1 4350 -0.676101 16.00000 ; qtot -
2.440
4351 HO 268 THR HG1 4351 0.410200 1.00800 ; qtot -
2.029
4352 C 268 THR C 4352 0.597301 12.01000 ; qtot -
1.432
4353 O 268 THR O 4353 -0.567901 16.00000 ; qtot -
2.000
4354 N 269 GLN N 4354 -0.415700 14.01000 ; qtot -
2.416
4355 H 269 GLN H 4355 0.271900 1.00800 ; qtot -
2.144
4356 CX 269 GLN CA 4356 -0.003100 12.01000 ; qtot -
2.147
4357 H1 269 GLN HA 4357 0.085000 1.00800 ; qtot -
2.062
4358 A2C 269 GLN CB 4358 -0.003600 12.01000 ; qtot -
2.066
4359 HC 269 GLN HB2 4359 0.017100 1.00800 ; qtot -
2.048
4360 HC 269 GLN HB3 4360 0.017100 1.00800 ; qtot -
2.031
4361 A2C 269 GLN CG 4361 -0.064500 12.01000 ; qtot -
2.096
4362 HC 269 GLN HG2 4362 0.035200 1.00800 ; qtot -
2.061
4363 HC 269 GLN HG3 4363 0.035200 1.00800 ; qtot -
2.025
4364 C 269 GLN CD 4364 0.695101 12.01000 ; qtot -
1.330
4365 O 269 GLN OE1 4365 -0.608601 16.00000 ; qtot -
1.939

4366 N 269 GLN NE2 4366 -0.940701 14.01000 ; qtot -
2.880
4367 H 269 GLN HE21 4367 0.425100 1.00800 ; qtot -
2.455
4368 H 269 GLN HE22 4368 0.425100 1.00800 ; qtot -
2.029
4369 C 269 GLN C 4369 0.597301 12.01000 ; qtot -
1.432
4370 O 269 GLN O 4370 -0.567901 16.00000 ; qtot -
2.000
4371 N 270 MET N 4371 -0.415700 14.01000 ; qtot -
2.416
4372 H 270 MET H 4372 0.271900 1.00800 ; qtot -
2.144
4373 CX 270 MET CA 4373 -0.023700 12.01000 ; qtot -
2.168
4374 H1 270 MET HA 4374 0.088000 1.00800 ; qtot -
2.080
4375 A2C 270 MET CB 4375 0.034200 12.01000 ; qtot -
2.045
4376 HC 270 MET HB2 4376 0.024100 1.00800 ; qtot -
2.021
4377 HC 270 MET HB3 4377 0.024100 1.00800 ; qtot -
1.997
4378 A2C 270 MET CG 4378 0.001800 12.01000 ; qtot -
1.995
4379 H1 270 MET HG2 4379 0.044000 1.00800 ; qtot -
1.951
4380 H1 270 MET HG3 4380 0.044000 1.00800 ; qtot -
1.907
4381 S 270 MET SD 4381 -0.273700 32.06000 ; qtot -
2.181
4382 CT 270 MET CE 4382 -0.053600 12.01000 ; qtot -
2.235
4383 H1 270 MET HE1 4383 0.068400 1.00800 ; qtot -
2.166
4384 H1 270 MET HE2 4384 0.068400 1.00800 ; qtot -
2.098
4385 H1 270 MET HE3 4385 0.068400 1.00800 ; qtot -
2.029
4386 C 270 MET C 4386 0.597301 12.01000 ; qtot -
1.432
4387 O 270 MET O 4387 -0.567901 16.00000 ; qtot -
2.000
4388 N 271 THR N 4388 -0.415700 14.01000 ; qtot -
2.416
4389 H 271 THR H 4389 0.271900 1.00800 ; qtot -
2.144
4390 CX 271 THR CA 4390 -0.038900 12.01000 ; qtot -
2.183
4391 H1 271 THR HA 4391 0.100700 1.00800 ; qtot -
2.082

4392 A3C 271 THR CB 4392 0.365400 12.01000 ; qtot -1.717
4393 H1 271 THR HB 4393 0.004300 1.00800 ; qtot -1.712
4394 CT 271 THR CG2 4394 -0.243800 12.01000 ; qtot -1.956
4395 HC 271 THR HG21 4395 0.064200 1.00800 ; qtot -1.892
4396 HC 271 THR HG22 4396 0.064200 1.00800 ; qtot -1.828
4397 HC 271 THR HG23 4397 0.064200 1.00800 ; qtot -1.764
4398 OH 271 THR OG1 4398 -0.676101 16.00000 ; qtot -2.440
4399 HO 271 THR HG1 4399 0.410200 1.00800 ; qtot -2.029
4400 C 271 THR C 4400 0.597301 12.01000 ; qtot -1.432
4401 O 271 THR O 4401 -0.567901 16.00000 ; qtot -2.000
4402 N 272 LYS N 4402 -0.347900 14.01000 ; qtot -2.348
4403 H 272 LYS H 4403 0.274700 1.00800 ; qtot -2.073
4404 CX 272 LYS CA 4404 -0.240000 12.01000 ; qtot -2.313
4405 H1 272 LYS HA 4405 0.142600 1.00800 ; qtot -2.171
4406 C8 272 LYS CB 4406 -0.009400 12.01000 ; qtot -2.180
4407 HC 272 LYS HB2 4407 0.036200 1.00800 ; qtot -2.144
4408 HC 272 LYS HB3 4408 0.036200 1.00800 ; qtot -2.108
4409 C8 272 LYS CG 4409 0.018700 12.01000 ; qtot -2.089
4410 HC 272 LYS HG2 4410 0.010300 1.00800 ; qtot -2.079
4411 HC 272 LYS HG3 4411 0.010300 1.00800 ; qtot -2.068
4412 C8 272 LYS CD 4412 -0.047900 12.01000 ; qtot -2.116
4413 HC 272 LYS HD2 4413 0.062100 1.00800 ; qtot -2.054
4414 HC 272 LYS HD3 4414 0.062100 1.00800 ; qtot -1.992
4415 C8 272 LYS CE 4415 -0.014300 12.01000 ; qtot -2.006
4416 HP 272 LYS HE2 4416 0.113500 1.00800 ; qtot -1.893
4417 HP 272 LYS HE3 4417 0.113500 1.00800 ; qtot -1.779

```
  4418   N3   272   LYS    NZ 4418    -0.385400     14.01000 ; qtot -
2.165
  4419    H   272   LYS   HZ1 4419     0.340000      1.00800 ; qtot -
1.825
  4420    H   272   LYS   HZ2 4420     0.340000      1.00800 ; qtot -
1.485
  4421    H   272   LYS   HZ3 4421     0.340000      1.00800 ; qtot -
1.145
  4422    C   272   LYS     C 4422     0.734101     12.01000 ; qtot -
0.411
  4423    O   272   LYS     O 4423    -0.589401     16.00000 ; qtot -
1.000
  4424    N   273   SER     N 4424    -0.415700     14.01000 ; qtot -
1.416
  4425    H   273   SER     H 4425     0.271900      1.00800 ; qtot -
1.144
  4426   CX   273   SER    CA 4426    -0.024900     12.01000 ; qtot -
1.169
  4427   H1   273   SER    HA 4427     0.084300      1.00800 ; qtot -
1.084
  4428  A2C   273   SER    CB 4428     0.211700     12.01000 ; qtot -
0.873
  4429   H1   273   SER   HB2 4429     0.035200      1.00800 ; qtot -
0.838
  4430   H1   273   SER   HB3 4430     0.035200      1.00800 ; qtot -
0.802
  4431   OH   273   SER    OG 4431    -0.654601     16.00000 ; qtot -
1.457
  4432   HO   273   SER    HG 4432     0.427500      1.00800 ; qtot -
1.029
  4433    C   273   SER     C 4433     0.597301     12.01000 ; qtot -
0.432
  4434    O   273   SER     O 4434    -0.567901     16.00000 ; qtot -
1.000
  4435    N   274   HIE     N 4435    -0.415700     14.01000 ; qtot -
1.416
  4436    H   274   HIE     H 4436     0.271900      1.00800 ; qtot -
1.144
  4437   CX   274   HIE    CA 4437    -0.058100     12.01000 ; qtot -
1.202
  4438   H1   274   HIE    HA 4438     0.136000      1.00800 ; qtot -
1.066
  4439   CT   274   HIE    CB 4439    -0.007400     12.01000 ; qtot -
1.073
  4440   HC   274   HIE   HB2 4440     0.036700      1.00800 ; qtot -
1.037
  4441   HC   274   HIE   HB3 4441     0.036700      1.00800 ; qtot -
1.000
  4442   CC   274   HIE    CG 4442     0.186800     12.01000 ; qtot -
0.813
  4443   NB   274   HIE   ND1 4443    -0.543201     14.01000 ; qtot -
1.356
```

4444 CR 274 HIE CE1 4444 0.163500 12.01000 ; qtot -1.193
4445 H5 274 HIE HE1 4445 0.143500 1.00800 ; qtot -1.049
4446 NA 274 HIE NE2 4446 -0.279500 14.01000 ; qtot -1.329
4447 H 274 HIE HE2 4447 0.333900 1.00800 ; qtot -0.995
4448 CW 274 HIE CD2 4448 -0.220700 12.01000 ; qtot -1.216
4449 H4 274 HIE HD2 4449 0.186200 1.00800 ; qtot -1.029
4450 C 274 HIE C 4450 0.597301 12.01000 ; qtot -0.432
4451 O 274 HIE O 4451 -0.567901 16.00000 ; qtot -1.000
4452 N 275 GLU N 4452 -0.516301 14.01000 ; qtot -1.516
4453 H 275 GLU H 4453 0.293600 1.00800 ; qtot -1.223
4454 CX 275 GLU CA 4454 0.039700 12.01000 ; qtot -1.183
4455 H1 275 GLU HA 4455 0.110500 1.00800 ; qtot -1.073
4456 A2C 275 GLU CB 4456 0.056000 12.01000 ; qtot -1.017
4457 HC 275 GLU HB2 4457 -0.017300 1.00800 ; qtot -1.034
4458 HC 275 GLU HB3 4458 -0.017300 1.00800 ; qtot -1.051
4459 A2C 275 GLU CG 4459 0.013600 12.01000 ; qtot -1.038
4460 HC 275 GLU HG2 4460 -0.042500 1.00800 ; qtot -1.080
4461 HC 275 GLU HG3 4461 -0.042500 1.00800 ; qtot -1.123
4462 CO 275 GLU CD 4462 0.805401 12.01000 ; qtot -0.317
4463 O2 275 GLU OE1 4463 -0.818801 16.00000 ; qtot -1.136
4464 O2 275 GLU OE2 4464 -0.818801 16.00000 ; qtot -1.955
4465 C 275 GLU C 4465 0.536601 12.01000 ; qtot -1.418
4466 O 275 GLU O 4466 -0.581901 16.00000 ; qtot -2.000
4467 N 276 HIE N 4467 -0.415700 14.01000 ; qtot -2.416
4468 H 276 HIE H 4468 0.271900 1.00800 ; qtot -2.144
4469 CX 276 HIE CA 4469 -0.058100 12.01000 ; qtot -2.202

4470 H1 276 HIE HA 4470 0.136000 1.00800 ; qtot -
2.066
4471 CT 276 HIE CB 4471 -0.007400 12.01000 ; qtot -
2.073
4472 HC 276 HIE HB2 4472 0.036700 1.00800 ; qtot -
2.037
4473 HC 276 HIE HB3 4473 0.036700 1.00800 ; qtot -
2.000
4474 CC 276 HIE CG 4474 0.186800 12.01000 ; qtot -
1.813
4475 NB 276 HIE ND1 4475 -0.543201 14.01000 ; qtot -
2.356
4476 CR 276 HIE CE1 4476 0.163500 12.01000 ; qtot -
2.193
4477 H5 276 HIE HE1 4477 0.143500 1.00800 ; qtot -
2.049
4478 NA 276 HIE NE2 4478 -0.279500 14.01000 ; qtot -
2.329
4479 H 276 HIE HE2 4479 0.333900 1.00800 ; qtot -
1.995
4480 CW 276 HIE CD2 4480 -0.220700 12.01000 ; qtot -
2.216
4481 H4 276 HIE HD2 4481 0.186200 1.00800 ; qtot -
2.029
4482 C 276 HIE C 4482 0.597301 12.01000 ; qtot -
1.432
4483 O 276 HIE O 4483 -0.567901 16.00000 ; qtot -
2.000
4484 N 277 VAL N 4484 -0.415700 14.01000 ; qtot -
2.416
4485 H 277 VAL H 4485 0.271900 1.00800 ; qtot -
2.144
4486 CX 277 VAL CA 4486 -0.087500 12.01000 ; qtot -
2.231
4487 H1 277 VAL HA 4487 0.096900 1.00800 ; qtot -
2.134
4488 A3C 277 VAL CB 4488 0.298500 12.01000 ; qtot -
1.836
4489 HC 277 VAL HB 4489 -0.029700 1.00800 ; qtot -
1.866
4490 CT 277 VAL CG1 4490 -0.319200 12.01000 ; qtot -
2.185
4491 HC 277 VAL HG11 4491 0.079100 1.00800 ; qtot -
2.106
4492 HC 277 VAL HG12 4492 0.079100 1.00800 ; qtot -
2.027
4493 HC 277 VAL HG13 4493 0.079100 1.00800 ; qtot -
1.948
4494 CT 277 VAL CG2 4494 -0.319200 12.01000 ; qtot -
2.267
4495 HC 277 VAL HG21 4495 0.079100 1.00800 ; qtot -
2.188

```
  4496   HC   277   VAL  HG22 4496     0.079100      1.00800 ; qtot -
2.109
  4497   HC   277   VAL  HG23 4497     0.079100      1.00800 ; qtot -
2.029
  4498    C   277   VAL     C 4498     0.597301     12.01000 ; qtot -
1.432
  4499    O   277   VAL     O 4499    -0.567901     16.00000 ; qtot -
2.000
  4500    N   278   GLU     N 4500    -0.516301     14.01000 ; qtot -
2.516
  4501    H   278   GLU     H 4501     0.293600      1.00800 ; qtot -
2.223
  4502   CX   278   GLU    CA 4502     0.039700     12.01000 ; qtot -
2.183
  4503   H1   278   GLU    HA 4503     0.110500      1.00800 ; qtot -
2.073
  4504  A2C   278   GLU    CB 4504     0.056000     12.01000 ; qtot -
2.017
  4505   HC   278   GLU   HB2 4505    -0.017300      1.00800 ; qtot -
2.034
  4506   HC   278   GLU   HB3 4506    -0.017300      1.00800 ; qtot -
2.051
  4507  A2C   278   GLU    CG 4507     0.013600     12.01000 ; qtot -
2.038
  4508   HC   278   GLU   HG2 4508    -0.042500      1.00800 ; qtot -
2.080
  4509   HC   278   GLU   HG3 4509    -0.042500      1.00800 ; qtot -
2.123
  4510   CO   278   GLU    CD 4510     0.805401     12.01000 ; qtot -
1.317
  4511   O2   278   GLU   OE1 4511    -0.818801     16.00000 ; qtot -
2.136
  4512   O2   278   GLU   OE2 4512    -0.818801     16.00000 ; qtot -
2.955
  4513    C   278   GLU     C 4513     0.536601     12.01000 ; qtot -
2.418
  4514    O   278   GLU     O 4514    -0.581901     16.00000 ; qtot -
3.000
  4515    N   279   LEU     N 4515    -0.415700     14.01000 ; qtot -
3.416
  4516    H   279   LEU     H 4516     0.271900      1.00800 ; qtot -
3.144
  4517   CX   279   LEU    CA 4517    -0.051800     12.01000 ; qtot -
3.196
  4518   H1   279   LEU    HA 4518     0.092200      1.00800 ; qtot -
3.103
  4519  A2C   279   LEU    CB 4519    -0.110200     12.01000 ; qtot -
3.214
  4520   HC   279   LEU   HB2 4520     0.045700      1.00800 ; qtot -
3.168
  4521   HC   279   LEU   HB3 4521     0.045700      1.00800 ; qtot -
3.122
```

4522 A3C 279 LEU CG 4522 0.353100 12.01000 ; qtot -
2.769
4523 HC 279 LEU HG 4523 -0.036100 1.00800 ; qtot -
2.805
4524 CT 279 LEU CD1 4524 -0.412100 12.01000 ; qtot -
3.217
4525 HC 279 LEU HD11 4525 0.100000 1.00800 ; qtot -
3.117
4526 HC 279 LEU HD12 4526 0.100000 1.00800 ; qtot -
3.017
4527 HC 279 LEU HD13 4527 0.100000 1.00800 ; qtot -
2.917
4528 CT 279 LEU CD2 4528 -0.412100 12.01000 ; qtot -
3.329
4529 HC 279 LEU HD21 4529 0.100000 1.00800 ; qtot -
3.229
4530 HC 279 LEU HD22 4530 0.100000 1.00800 ; qtot -
3.129
4531 HC 279 LEU HD23 4531 0.100000 1.00800 ; qtot -
3.029
4532 C 279 LEU C 4532 0.597301 12.01000 ; qtot -
2.432
4533 O 279 LEU O 4533 -0.567901 16.00000 ; qtot -
3.000
4534 N 280 SER N 4534 -0.415700 14.01000 ; qtot -
3.416
4535 H 280 SER H 4535 0.271900 1.00800 ; qtot -
3.144
4536 CX 280 SER CA 4536 -0.024900 12.01000 ; qtot -
3.169
4537 H1 280 SER HA 4537 0.084300 1.00800 ; qtot -
3.084
4538 A2C 280 SER CB 4538 0.211700 12.01000 ; qtot -
2.873
4539 H1 280 SER HB2 4539 0.035200 1.00800 ; qtot -
2.838
4540 H1 280 SER HB3 4540 0.035200 1.00800 ; qtot -
2.802
4541 OH 280 SER OG 4541 -0.654601 16.00000 ; qtot -
3.457
4542 HO 280 SER HG 4542 0.427500 1.00800 ; qtot -
3.029
4543 C 280 SER C 4543 0.597301 12.01000 ; qtot -
2.432
4544 O 280 SER O 4544 -0.567901 16.00000 ; qtot -
3.000
4545 N 281 SER N 4545 -0.415700 14.01000 ; qtot -
3.416
4546 H 281 SER H 4546 0.271900 1.00800 ; qtot -
3.144
4547 CX 281 SER CA 4547 -0.024900 12.01000 ; qtot -
3.169

```
  4548   H1   281   SER    HA 4548     0.084300      1.00800 ; qtot -
3.084
  4549  A2C   281   SER    CB 4549     0.211700     12.01000 ; qtot -
2.873
  4550   H1   281   SER   HB2 4550     0.035200      1.00800 ; qtot -
2.838
  4551   H1   281   SER   HB3 4551     0.035200      1.00800 ; qtot -
2.802
  4552   OH   281   SER    OG 4552    -0.654601     16.00000 ; qtot -
3.457
  4553   HO   281   SER    HG 4553     0.427500      1.00800 ; qtot -
3.029
  4554    C   281   SER     C 4554     0.597301     12.01000 ; qtot -
2.432
  4555    O   281   SER     O 4555    -0.567901     16.00000 ; qtot -
3.000
  4556    N   282   THR     N 4556    -0.415700     14.01000 ; qtot -
3.416
  4557    H   282   THR     H 4557     0.271900      1.00800 ; qtot -
3.144
  4558   CX   282   THR    CA 4558    -0.038900     12.01000 ; qtot -
3.183
  4559   H1   282   THR    HA 4559     0.100700      1.00800 ; qtot -
3.082
  4560  A3C   282   THR    CB 4560     0.365400     12.01000 ; qtot -
2.717
  4561   H1   282   THR    HB 4561     0.004300      1.00800 ; qtot -
2.712
  4562   CT   282   THR   CG2 4562    -0.243800     12.01000 ; qtot -
2.956
  4563   HC   282   THR  HG21 4563     0.064200      1.00800 ; qtot -
2.892
  4564   HC   282   THR  HG22 4564     0.064200      1.00800 ; qtot -
2.828
  4565   HC   282   THR  HG23 4565     0.064200      1.00800 ; qtot -
2.764
  4566   OH   282   THR   OG1 4566    -0.676101     16.00000 ; qtot -
3.440
  4567   HO   282   THR   HG1 4567     0.410200      1.00800 ; qtot -
3.029
  4568    C   282   THR     C 4568     0.597301     12.01000 ; qtot -
2.432
  4569    O   282   THR     O 4569    -0.567901     16.00000 ; qtot -
3.000
  4570    N   283   LEU     N 4570    -0.415700     14.01000 ; qtot -
3.416
  4571    H   283   LEU     H 4571     0.271900      1.00800 ; qtot -
3.144
  4572   CX   283   LEU    CA 4572    -0.051800     12.01000 ; qtot -
3.196
  4573   H1   283   LEU    HA 4573     0.092200      1.00800 ; qtot -
3.103
```

4574 A2C 283 LEU CB 4574 -0.110200 12.01000 ; qtot -
3.214
4575 HC 283 LEU HB2 4575 0.045700 1.00800 ; qtot -
3.168
4576 HC 283 LEU HB3 4576 0.045700 1.00800 ; qtot -
3.122
4577 A3C 283 LEU CG 4577 0.353100 12.01000 ; qtot -
2.769
4578 HC 283 LEU HG 4578 -0.036100 1.00800 ; qtot -
2.805
4579 CT 283 LEU CD1 4579 -0.412100 12.01000 ; qtot -
3.217
4580 HC 283 LEU HD11 4580 0.100000 1.00800 ; qtot -
3.117
4581 HC 283 LEU HD12 4581 0.100000 1.00800 ; qtot -
3.017
4582 HC 283 LEU HD13 4582 0.100000 1.00800 ; qtot -
2.917
4583 CT 283 LEU CD2 4583 -0.412100 12.01000 ; qtot -
3.329
4584 HC 283 LEU HD21 4584 0.100000 1.00800 ; qtot -
3.229
4585 HC 283 LEU HD22 4585 0.100000 1.00800 ; qtot -
3.129
4586 HC 283 LEU HD23 4586 0.100000 1.00800 ; qtot -
3.029
4587 C 283 LEU C 4587 0.597301 12.01000 ; qtot -
2.432
4588 O 283 LEU O 4588 -0.567901 16.00000 ; qtot -
3.000
4589 N 284 LEU N 4589 -0.415700 14.01000 ; qtot -
3.416
4590 H 284 LEU H 4590 0.271900 1.00800 ; qtot -
3.144
4591 CX 284 LEU CA 4591 -0.051800 12.01000 ; qtot -
3.196
4592 H1 284 LEU HA 4592 0.092200 1.00800 ; qtot -
3.103
4593 A2C 284 LEU CB 4593 -0.110200 12.01000 ; qtot -
3.214
4594 HC 284 LEU HB2 4594 0.045700 1.00800 ; qtot -
3.168
4595 HC 284 LEU HB3 4595 0.045700 1.00800 ; qtot -
3.122
4596 A3C 284 LEU CG 4596 0.353100 12.01000 ; qtot -
2.769
4597 HC 284 LEU HG 4597 -0.036100 1.00800 ; qtot -
2.805
4598 CT 284 LEU CD1 4598 -0.412100 12.01000 ; qtot -
3.217
4599 HC 284 LEU HD11 4599 0.100000 1.00800 ; qtot -
3.117

```
  4600   HC   284   LEU  HD12 4600     0.100000      1.00800 ; qtot -
3.017
  4601   HC   284   LEU  HD13 4601     0.100000      1.00800 ; qtot -
2.917
  4602   CT   284   LEU   CD2 4602    -0.412100     12.01000 ; qtot -
3.329
  4603   HC   284   LEU  HD21 4603     0.100000      1.00800 ; qtot -
3.229
  4604   HC   284   LEU  HD22 4604     0.100000      1.00800 ; qtot -
3.129
  4605   HC   284   LEU  HD23 4605     0.100000      1.00800 ; qtot -
3.029
  4606    C   284   LEU     C 4606     0.597301     12.01000 ; qtot -
2.432
  4607    O   284   LEU     O 4607    -0.567901     16.00000 ; qtot -
3.000
  4608    N   285   LYS     N 4608    -0.347900     14.01000 ; qtot -
3.348
  4609    H   285   LYS     H 4609     0.274700      1.00800 ; qtot -
3.073
  4610   CX   285   LYS    CA 4610    -0.240000     12.01000 ; qtot -
3.313
  4611   H1   285   LYS    HA 4611     0.142600      1.00800 ; qtot -
3.171
  4612   C8   285   LYS    CB 4612    -0.009400     12.01000 ; qtot -
3.180
  4613   HC   285   LYS   HB2 4613     0.036200      1.00800 ; qtot -
3.144
  4614   HC   285   LYS   HB3 4614     0.036200      1.00800 ; qtot -
3.108
  4615   C8   285   LYS    CG 4615     0.018700     12.01000 ; qtot -
3.089
  4616   HC   285   LYS   HG2 4616     0.010300      1.00800 ; qtot -
3.079
  4617   HC   285   LYS   HG3 4617     0.010300      1.00800 ; qtot -
3.068
  4618   C8   285   LYS    CD 4618    -0.047900     12.01000 ; qtot -
3.116
  4619   HC   285   LYS   HD2 4619     0.062100      1.00800 ; qtot -
3.054
  4620   HC   285   LYS   HD3 4620     0.062100      1.00800 ; qtot -
2.992
  4621   C8   285   LYS    CE 4621    -0.014300     12.01000 ; qtot -
3.006
  4622   HP   285   LYS   HE2 4622     0.113500      1.00800 ; qtot -
2.893
  4623   HP   285   LYS   HE3 4623     0.113500      1.00800 ; qtot -
2.779
  4624   N3   285   LYS    NZ 4624    -0.385400     14.01000 ; qtot -
3.165
  4625    H   285   LYS   HZ1 4625     0.340000      1.00800 ; qtot -
2.825
```

4626 H 285 LYS HZ2 4626 0.340000 1.00800 ; qtot -
2.485
4627 H 285 LYS HZ3 4627 0.340000 1.00800 ; qtot -
2.145
4628 C 285 LYS C 4628 0.734101 12.01000 ; qtot -
1.411
4629 O 285 LYS O 4629 -0.589401 16.00000 ; qtot -
2.000
4630 N 286 ASN N 4630 -0.415700 14.01000 ; qtot -
2.416
4631 H 286 ASN H 4631 0.271900 1.00800 ; qtot -
2.144
4632 CX 286 ASN CA 4632 0.014300 12.01000 ; qtot -
2.130
4633 H1 286 ASN HA 4633 0.104800 1.00800 ; qtot -
2.025
4634 A2C 286 ASN CB 4634 -0.204100 12.01000 ; qtot -
2.229
4635 HC 286 ASN HB2 4635 0.079700 1.00800 ; qtot -
2.149
4636 HC 286 ASN HB3 4636 0.079700 1.00800 ; qtot -
2.069
4637 C 286 ASN CG 4637 0.713001 12.01000 ; qtot -
1.356
4638 O 286 ASN OD1 4638 -0.593101 16.00000 ; qtot -
1.950
4639 N 286 ASN ND2 4639 -0.919101 14.01000 ; qtot -
2.869
4640 H 286 ASN HD21 4640 0.419600 1.00800 ; qtot -
2.449
4641 H 286 ASN HD22 4641 0.419600 1.00800 ; qtot -
2.029
4642 C 286 ASN C 4642 0.597301 12.01000 ; qtot -
1.432
4643 O 286 ASN O 4643 -0.567901 16.00000 ; qtot -
2.000
4644 N 287 LEU N 4644 -0.415700 14.01000 ; qtot -
2.416
4645 H 287 LEU H 4645 0.271900 1.00800 ; qtot -
2.144
4646 CX 287 LEU CA 4646 -0.051800 12.01000 ; qtot -
2.196
4647 H1 287 LEU HA 4647 0.092200 1.00800 ; qtot -
2.103
4648 A2C 287 LEU CB 4648 -0.110200 12.01000 ; qtot -
2.214
4649 HC 287 LEU HB2 4649 0.045700 1.00800 ; qtot -
2.168
4650 HC 287 LEU HB3 4650 0.045700 1.00800 ; qtot -
2.122
4651 A3C 287 LEU CG 4651 0.353100 12.01000 ; qtot -
1.769

4652 HC 287 LEU HG 4652 -0.036100 1.00800 ; qtot -
1.805
4653 CT 287 LEU CD1 4653 -0.412100 12.01000 ; qtot -
2.217
4654 HC 287 LEU HD11 4654 0.100000 1.00800 ; qtot -
2.117
4655 HC 287 LEU HD12 4655 0.100000 1.00800 ; qtot -
2.017
4656 HC 287 LEU HD13 4656 0.100000 1.00800 ; qtot -
1.917
4657 CT 287 LEU CD2 4657 -0.412100 12.01000 ; qtot -
2.329
4658 HC 287 LEU HD21 4658 0.100000 1.00800 ; qtot -
2.229
4659 HC 287 LEU HD22 4659 0.100000 1.00800 ; qtot -
2.129
4660 HC 287 LEU HD23 4660 0.100000 1.00800 ; qtot -
2.029
4661 C 287 LEU C 4661 0.597301 12.01000 ; qtot -
1.432
4662 O 287 LEU O 4662 -0.567901 16.00000 ; qtot -
2.000
4663 N 288 LYS N 4663 -0.347900 14.01000 ; qtot -
2.348
4664 H 288 LYS H 4664 0.274700 1.00800 ; qtot -
2.073
4665 CX 288 LYS CA 4665 -0.240000 12.01000 ; qtot -
2.313
4666 H1 288 LYS HA 4666 0.142600 1.00800 ; qtot -
2.171
4667 C8 288 LYS CB 4667 -0.009400 12.01000 ; qtot -
2.180
4668 HC 288 LYS HB2 4668 0.036200 1.00800 ; qtot -
2.144
4669 HC 288 LYS HB3 4669 0.036200 1.00800 ; qtot -
2.108
4670 C8 288 LYS CG 4670 0.018700 12.01000 ; qtot -
2.089
4671 HC 288 LYS HG2 4671 0.010300 1.00800 ; qtot -
2.079
4672 HC 288 LYS HG3 4672 0.010300 1.00800 ; qtot -
2.068
4673 C8 288 LYS CD 4673 -0.047900 12.01000 ; qtot -
2.116
4674 HC 288 LYS HD2 4674 0.062100 1.00800 ; qtot -
2.054
4675 HC 288 LYS HD3 4675 0.062100 1.00800 ; qtot -
1.992
4676 C8 288 LYS CE 4676 -0.014300 12.01000 ; qtot -
2.006
4677 HP 288 LYS HE2 4677 0.113500 1.00800 ; qtot -
1.893

```
  4678   HP   288   LYS   HE3 4678     0.113500      1.00800 ; qtot -
1.779
  4679   N3   288   LYS    NZ 4679    -0.385400     14.01000 ; qtot -
2.165
  4680    H   288   LYS   HZ1 4680     0.340000      1.00800 ; qtot -
1.825
  4681    H   288   LYS   HZ2 4681     0.340000      1.00800 ; qtot -
1.485
  4682    H   288   LYS   HZ3 4682     0.340000      1.00800 ; qtot -
1.145
  4683    C   288   LYS     C 4683     0.734101     12.01000 ; qtot -
0.411
  4684    O   288   LYS     O 4684    -0.589401     16.00000 ; qtot -
1.000
  4685    N   289   ASN     N 4685    -0.415700     14.01000 ; qtot -
1.416
  4686    H   289   ASN     H 4686     0.271900      1.00800 ; qtot -
1.144
  4687   CX   289   ASN    CA 4687     0.014300     12.01000 ; qtot -
1.130
  4688   H1   289   ASN    HA 4688     0.104800      1.00800 ; qtot -
1.025
  4689  A2C   289   ASN    CB 4689    -0.204100     12.01000 ; qtot -
1.229
  4690   HC   289   ASN   HB2 4690     0.079700      1.00800 ; qtot -
1.149
  4691   HC   289   ASN   HB3 4691     0.079700      1.00800 ; qtot -
1.069
  4692    C   289   ASN    CG 4692     0.713001     12.01000 ; qtot -
0.356
  4693    O   289   ASN   OD1 4693    -0.593101     16.00000 ; qtot -
0.950
  4694    N   289   ASN   ND2 4694    -0.919101     14.01000 ; qtot -
1.869
  4695    H   289   ASN  HD21 4695     0.419600      1.00800 ; qtot -
1.449
  4696    H   289   ASN  HD22 4696     0.419600      1.00800 ; qtot -
1.029
  4697    C   289   ASN     C 4697     0.597301     12.01000 ; qtot -
0.432
  4698    O   289   ASN     O 4698    -0.567901     16.00000 ; qtot -
1.000
  4699    N   290   PHE     N 4699    -0.415700     14.01000 ; qtot -
1.416
  4700    H   290   PHE     H 4700     0.271900      1.00800 ; qtot -
1.144
  4701   CX   290   PHE    CA 4701    -0.002400     12.01000 ; qtot -
1.146
  4702   H1   290   PHE    HA 4702     0.097800      1.00800 ; qtot -
1.048
  4703   CT   290   PHE    CB 4703    -0.034300     12.01000 ; qtot -
1.083
```

4704 HC 290 PHE HB2 4704 0.029500 1.00800 ; qtot -
1.053
4705 HC 290 PHE HB3 4705 0.029500 1.00800 ; qtot -
1.024
4706 CA 290 PHE CG 4706 0.011800 12.01000 ; qtot -
1.012
4707 CA 290 PHE CD1 4707 -0.125600 12.01000 ; qtot -
1.138
4708 HA 290 PHE HD1 4708 0.133000 1.00800 ; qtot -
1.005
4709 CA 290 PHE CE1 4709 -0.170400 12.01000 ; qtot -
1.175
4710 HA 290 PHE HE1 4710 0.143000 1.00800 ; qtot -
1.032
4711 CA 290 PHE CZ 4711 -0.107200 12.01000 ; qtot -
1.139
4712 HA 290 PHE HZ 4712 0.129700 1.00800 ; qtot -
1.009
4713 CA 290 PHE CE2 4713 -0.170400 12.01000 ; qtot -
1.180
4714 HA 290 PHE HE2 4714 0.143000 1.00800 ; qtot -
1.037
4715 CA 290 PHE CD2 4715 -0.125600 12.01000 ; qtot -
1.162
4716 HA 290 PHE HD2 4716 0.133000 1.00800 ; qtot -
1.029
4717 C 290 PHE C 4717 0.597301 12.01000 ; qtot -
0.432
4718 O 290 PHE O 4718 -0.567901 16.00000 ; qtot -
1.000
4719 N 291 LYS N 4719 -0.347900 14.01000 ; qtot -
1.348
4720 H 291 LYS H 4720 0.274700 1.00800 ; qtot -
1.073
4721 CX 291 LYS CA 4721 -0.240000 12.01000 ; qtot -
1.313
4722 H1 291 LYS HA 4722 0.142600 1.00800 ; qtot -
1.171
4723 C8 291 LYS CB 4723 -0.009400 12.01000 ; qtot -
1.180
4724 HC 291 LYS HB2 4724 0.036200 1.00800 ; qtot -
1.144
4725 HC 291 LYS HB3 4725 0.036200 1.00800 ; qtot -
1.108
4726 C8 291 LYS CG 4726 0.018700 12.01000 ; qtot -
1.089
4727 HC 291 LYS HG2 4727 0.010300 1.00800 ; qtot -
1.079
4728 HC 291 LYS HG3 4728 0.010300 1.00800 ; qtot -
1.068
4729 C8 291 LYS CD 4729 -0.047900 12.01000 ; qtot -
1.116

4730 HC 291 LYS HD2 4730 0.062100 1.00800 ; qtot -
1.054
4731 HC 291 LYS HD3 4731 0.062100 1.00800 ; qtot -
0.992
4732 C8 291 LYS CE 4732 -0.014300 12.01000 ; qtot -
1.006
4733 HP 291 LYS HE2 4733 0.113500 1.00800 ; qtot -
0.893
4734 HP 291 LYS HE3 4734 0.113500 1.00800 ; qtot -
0.779
4735 N3 291 LYS NZ 4735 -0.385400 14.01000 ; qtot -
1.165
4736 H 291 LYS HZ1 4736 0.340000 1.00800 ; qtot -
0.825
4737 H 291 LYS HZ2 4737 0.340000 1.00800 ; qtot -
0.485
4738 H 291 LYS HZ3 4738 0.340000 1.00800 ; qtot -
0.145
4739 C 291 LYS C 4739 0.734101 12.01000 ; qtot
0.589
4740 O 291 LYS O 4740 -0.589401 16.00000 ; qtot
0.000
4741 N 292 LYS N 4741 -0.347900 14.01000 ; qtot -
0.348
4742 H 292 LYS H 4742 0.274700 1.00800 ; qtot -
0.073
4743 CX 292 LYS CA 4743 -0.240000 12.01000 ; qtot -
0.313
4744 H1 292 LYS HA 4744 0.142600 1.00800 ; qtot -
0.171
4745 C8 292 LYS CB 4745 -0.009400 12.01000 ; qtot -
0.180
4746 HC 292 LYS HB2 4746 0.036200 1.00800 ; qtot -
0.144
4747 HC 292 LYS HB3 4747 0.036200 1.00800 ; qtot -
0.108
4748 C8 292 LYS CG 4748 0.018700 12.01000 ; qtot -
0.089
4749 HC 292 LYS HG2 4749 0.010300 1.00800 ; qtot -
0.079
4750 HC 292 LYS HG3 4750 0.010300 1.00800 ; qtot -
0.068
4751 C8 292 LYS CD 4751 -0.047900 12.01000 ; qtot -
0.116
4752 HC 292 LYS HD2 4752 0.062100 1.00800 ; qtot -
0.054
4753 HC 292 LYS HD3 4753 0.062100 1.00800 ; qtot
0.008
4754 C8 292 LYS CE 4754 -0.014300 12.01000 ; qtot -
0.006
4755 HP 292 LYS HE2 4755 0.113500 1.00800 ; qtot
0.107

4756 HP 292 LYS HE3 4756 0.113500 1.00800 ; qtot
0.221
4757 N3 292 LYS NZ 4757 -0.385400 14.01000 ; qtot -
0.165
4758 H 292 LYS HZ1 4758 0.340000 1.00800 ; qtot
0.175
4759 H 292 LYS HZ2 4759 0.340000 1.00800 ; qtot
0.515
4760 H 292 LYS HZ3 4760 0.340000 1.00800 ; qtot
0.855
4761 C 292 LYS C 4761 0.734101 12.01000 ; qtot
1.589
4762 O 292 LYS O 4762 -0.589401 16.00000 ; qtot
1.000
4763 N 293 GLU N 4763 -0.516301 14.01000 ; qtot
0.484
4764 H 293 GLU H 4764 0.293600 1.00800 ; qtot
0.777
4765 CX 293 GLU CA 4765 0.039700 12.01000 ; qtot
0.817
4766 H1 293 GLU HA 4766 0.110500 1.00800 ; qtot
0.928
4767 A2C 293 GLU CB 4767 0.056000 12.01000 ; qtot
0.984
4768 HC 293 GLU HB2 4768 -0.017300 1.00800 ; qtot
0.966
4769 HC 293 GLU HB3 4769 -0.017300 1.00800 ; qtot
0.949
4770 A2C 293 GLU CG 4770 0.013600 12.01000 ; qtot
0.963
4771 HC 293 GLU HG2 4771 -0.042500 1.00800 ; qtot
0.920
4772 HC 293 GLU HG3 4772 -0.042500 1.00800 ; qtot
0.878
4773 CO 293 GLU CD 4773 0.805401 12.01000 ; qtot
1.683
4774 O2 293 GLU OE1 4774 -0.818801 16.00000 ; qtot
0.864
4775 O2 293 GLU OE2 4775 -0.818801 16.00000 ; qtot
0.045
4776 C 293 GLU C 4776 0.536601 12.01000 ; qtot
0.582
4777 O 293 GLU O 4777 -0.581901 16.00000 ; qtot
0.000
4778 N 294 ASN N 4778 -0.415700 14.01000 ; qtot -
0.416
4779 H 294 ASN H 4779 0.271900 1.00800 ; qtot -
0.144
4780 CX 294 ASN CA 4780 0.014300 12.01000 ; qtot -
0.129
4781 H1 294 ASN HA 4781 0.104800 1.00800 ; qtot -
0.025

4782 A2C 294 ASN CB 4782 -0.204100 12.01000 ; qtot -0.229
4783 HC 294 ASN HB2 4783 0.079700 1.00800 ; qtot -0.149
4784 HC 294 ASN HB3 4784 0.079700 1.00800 ; qtot -0.069
4785 C 294 ASN CG 4785 0.713001 12.01000 ; qtot 0.644
4786 O 294 ASN OD1 4786 -0.593101 16.00000 ; qtot 0.051
4787 N 294 ASN ND2 4787 -0.919101 14.01000 ; qtot -0.869
4788 H 294 ASN HD21 4788 0.419600 1.00800 ; qtot -0.449
4789 H 294 ASN HD22 4789 0.419600 1.00800 ; qtot -0.029
4790 C 294 ASN C 4790 0.597301 12.01000 ; qtot 0.568
4791 O 294 ASN O 4791 -0.567901 16.00000 ; qtot 0.000
4792 N 295 GLU N 4792 -0.516301 14.01000 ; qtot -0.516
4793 H 295 GLU H 4793 0.293600 1.00800 ; qtot -0.223
4794 CX 295 GLU CA 4794 0.039700 12.01000 ; qtot -0.183
4795 H1 295 GLU HA 4795 0.110500 1.00800 ; qtot -0.072
4796 A2C 295 GLU CB 4796 0.056000 12.01000 ; qtot -0.016
4797 HC 295 GLU HB2 4797 -0.017300 1.00800 ; qtot -0.034
4798 HC 295 GLU HB3 4798 -0.017300 1.00800 ; qtot -0.051
4799 A2C 295 GLU CG 4799 0.013600 12.01000 ; qtot -0.037
4800 HC 295 GLU HG2 4800 -0.042500 1.00800 ; qtot -0.080
4801 HC 295 GLU HG3 4801 -0.042500 1.00800 ; qtot -0.122
4802 CO 295 GLU CD 4802 0.805401 12.01000 ; qtot 0.683
4803 O2 295 GLU OE1 4803 -0.818801 16.00000 ; qtot -0.136
4804 O2 295 GLU OE2 4804 -0.818801 16.00000 ; qtot -0.955
4805 C 295 GLU C 4805 0.536601 12.01000 ; qtot -0.418
4806 O 295 GLU O 4806 -0.581901 16.00000 ; qtot -1.000
4807 N 296 LEU N 4807 -0.415700 14.01000 ; qtot -1.416

4808 H 296 LEU H 4808 0.271900 1.00800 ; qtot -
1.144
4809 CX 296 LEU CA 4809 -0.051800 12.01000 ; qtot -
1.196
4810 H1 296 LEU HA 4810 0.092200 1.00800 ; qtot -
1.103
4811 A2C 296 LEU CB 4811 -0.110200 12.01000 ; qtot -
1.214
4812 HC 296 LEU HB2 4812 0.045700 1.00800 ; qtot -
1.168
4813 HC 296 LEU HB3 4813 0.045700 1.00800 ; qtot -
1.122
4814 A3C 296 LEU CG 4814 0.353100 12.01000 ; qtot -
0.769
4815 HC 296 LEU HG 4815 -0.036100 1.00800 ; qtot -
0.805
4816 CT 296 LEU CD1 4816 -0.412100 12.01000 ; qtot -
1.217
4817 HC 296 LEU HD11 4817 0.100000 1.00800 ; qtot -
1.117
4818 HC 296 LEU HD12 4818 0.100000 1.00800 ; qtot -
1.017
4819 HC 296 LEU HD13 4819 0.100000 1.00800 ; qtot -
0.917
4820 CT 296 LEU CD2 4820 -0.412100 12.01000 ; qtot -
1.329
4821 HC 296 LEU HD21 4821 0.100000 1.00800 ; qtot -
1.229
4822 HC 296 LEU HD22 4822 0.100000 1.00800 ; qtot -
1.129
4823 HC 296 LEU HD23 4823 0.100000 1.00800 ; qtot -
1.029
4824 C 296 LEU C 4824 0.597301 12.01000 ; qtot -
0.432
4825 O 296 LEU O 4825 -0.567901 16.00000 ; qtot -
1.000
4826 N 297 LYS N 4826 -0.347900 14.01000 ; qtot -
1.348
4827 H 297 LYS H 4827 0.274700 1.00800 ; qtot -
1.073
4828 CX 297 LYS CA 4828 -0.240000 12.01000 ; qtot -
1.313
4829 H1 297 LYS HA 4829 0.142600 1.00800 ; qtot -
1.171
4830 C8 297 LYS CB 4830 -0.009400 12.01000 ; qtot -
1.180
4831 HC 297 LYS HB2 4831 0.036200 1.00800 ; qtot -
1.144
4832 HC 297 LYS HB3 4832 0.036200 1.00800 ; qtot -
1.108
4833 C8 297 LYS CG 4833 0.018700 12.01000 ; qtot -
1.089

4834 HC 297 LYS HG2 4834 0.010300 1.00800 ; qtot -1.079
4835 HC 297 LYS HG3 4835 0.010300 1.00800 ; qtot -1.068
4836 C8 297 LYS CD 4836 -0.047900 12.01000 ; qtot -1.116
4837 HC 297 LYS HD2 4837 0.062100 1.00800 ; qtot -1.054
4838 HC 297 LYS HD3 4838 0.062100 1.00800 ; qtot -0.992
4839 C8 297 LYS CE 4839 -0.014300 12.01000 ; qtot -1.006
4840 HP 297 LYS HE2 4840 0.113500 1.00800 ; qtot -0.893
4841 HP 297 LYS HE3 4841 0.113500 1.00800 ; qtot -0.779
4842 N3 297 LYS NZ 4842 -0.385400 14.01000 ; qtot -1.165
4843 H 297 LYS HZ1 4843 0.340000 1.00800 ; qtot -0.825
4844 H 297 LYS HZ2 4844 0.340000 1.00800 ; qtot -0.485
4845 H 297 LYS HZ3 4845 0.340000 1.00800 ; qtot -0.145
4846 C 297 LYS C 4846 0.734101 12.01000 ; qtot 0.589
4847 O 297 LYS O 4847 -0.589401 16.00000 ; qtot 0.000
4848 N 298 LYS N 4848 -0.347900 14.01000 ; qtot -0.348
4849 H 298 LYS H 4849 0.274700 1.00800 ; qtot -0.073
4850 CX 298 LYS CA 4850 -0.240000 12.01000 ; qtot -0.313
4851 H1 298 LYS HA 4851 0.142600 1.00800 ; qtot -0.171
4852 C8 298 LYS CB 4852 -0.009400 12.01000 ; qtot -0.180
4853 HC 298 LYS HB2 4853 0.036200 1.00800 ; qtot -0.144
4854 HC 298 LYS HB3 4854 0.036200 1.00800 ; qtot -0.108
4855 C8 298 LYS CG 4855 0.018700 12.01000 ; qtot -0.089
4856 HC 298 LYS HG2 4856 0.010300 1.00800 ; qtot -0.079
4857 HC 298 LYS HG3 4857 0.010300 1.00800 ; qtot -0.068
4858 C8 298 LYS CD 4858 -0.047900 12.01000 ; qtot -0.116
4859 HC 298 LYS HD2 4859 0.062100 1.00800 ; qtot -0.054

4860 HC 298 LYS HD3 4860 0.062100 1.00800 ; qtot
0.008
4861 C8 298 LYS CE 4861 -0.014300 12.01000 ; qtot -
0.006
4862 HP 298 LYS HE2 4862 0.113500 1.00800 ; qtot
0.107
4863 HP 298 LYS HE3 4863 0.113500 1.00800 ; qtot
0.221
4864 N3 298 LYS NZ 4864 -0.385400 14.01000 ; qtot -
0.165
4865 H 298 LYS HZ1 4865 0.340000 1.00800 ; qtot
0.175
4866 H 298 LYS HZ2 4866 0.340000 1.00800 ; qtot
0.515
4867 H 298 LYS HZ3 4867 0.340000 1.00800 ; qtot
0.855
4868 C 298 LYS C 4868 0.734101 12.01000 ; qtot
1.589
4869 O 298 LYS O 4869 -0.589401 16.00000 ; qtot
1.000
4870 N 299 ILE N 4870 -0.415700 14.01000 ; qtot
0.584
4871 H 299 ILE H 4871 0.271900 1.00800 ; qtot
0.856
4872 CX 299 ILE CA 4872 -0.059700 12.01000 ; qtot
0.797
4873 H1 299 ILE HA 4873 0.086900 1.00800 ; qtot
0.883
4874 A3C 299 ILE CB 4874 0.130300 12.01000 ; qtot
1.014
4875 HC 299 ILE HB 4875 0.018700 1.00800 ; qtot
1.032
4876 CT 299 ILE CG2 4876 -0.320400 12.01000 ; qtot
0.712
4877 HC 299 ILE HG21 4877 0.088200 1.00800 ; qtot
0.800
4878 HC 299 ILE HG22 4878 0.088200 1.00800 ; qtot
0.888
4879 HC 299 ILE HG23 4879 0.088200 1.00800 ; qtot
0.977
4880 A2C 299 ILE CG1 4880 -0.043000 12.01000 ; qtot
0.934
4881 HC 299 ILE HG12 4881 0.023600 1.00800 ; qtot
0.957
4882 HC 299 ILE HG13 4882 0.023600 1.00800 ; qtot
0.981
4883 CT 299 ILE CD1 4883 -0.066000 12.01000 ; qtot
0.915
4884 HC 299 ILE HD11 4884 0.018600 1.00800 ; qtot
0.933
4885 HC 299 ILE HD12 4885 0.018600 1.00800 ; qtot
0.952

```
  4886   HC   299   ILE  HD13 4886     0.018600      1.00800 ; qtot
0.971
  4887    C   299   ILE     C 4887     0.597301     12.01000 ; qtot
1.568
  4888    O   299   ILE     O 4888    -0.567901     16.00000 ; qtot
1.000
  4889    N   300   ALA     N 4889    -0.415700     14.01000 ; qtot
0.584
  4890    H   300   ALA     H 4890     0.271900      1.00800 ; qtot
0.856
  4891   CX   300   ALA    CA 4891     0.033700     12.01000 ; qtot
0.890
  4892   H1   300   ALA    HA 4892     0.082300      1.00800 ; qtot
0.972
  4893   CT   300   ALA    CB 4893    -0.182500     12.01000 ; qtot
0.790
  4894   HC   300   ALA   HB1 4894     0.060300      1.00800 ; qtot
0.850
  4895   HC   300   ALA   HB2 4895     0.060300      1.00800 ; qtot
0.910
  4896   HC   300   ALA   HB3 4896     0.060300      1.00800 ; qtot
0.971
  4897    C   300   ALA     C 4897     0.597301     12.01000 ; qtot
1.568
  4898    O   300   ALA     O 4898    -0.567901     16.00000 ; qtot
1.000
  4899    N   301   LEU     N 4899    -0.415700     14.01000 ; qtot
0.584
  4900    H   301   LEU     H 4900     0.271900      1.00800 ; qtot
0.856
  4901   CX   301   LEU    CA 4901    -0.051800     12.01000 ; qtot
0.804
  4902   H1   301   LEU    HA 4902     0.092200      1.00800 ; qtot
0.897
  4903  A2C   301   LEU    CB 4903    -0.110200     12.01000 ; qtot
0.786
  4904   HC   301   LEU   HB2 4904     0.045700      1.00800 ; qtot
0.832
  4905   HC   301   LEU   HB3 4905     0.045700      1.00800 ; qtot
0.878
  4906  A3C   301   LEU    CG 4906     0.353100     12.01000 ; qtot
1.231
  4907   HC   301   LEU    HG 4907    -0.036100      1.00800 ; qtot
1.195
  4908   CT   301   LEU   CD1 4908    -0.412100     12.01000 ; qtot
0.783
  4909   HC   301   LEU  HD11 4909     0.100000      1.00800 ; qtot
0.883
  4910   HC   301   LEU  HD12 4910     0.100000      1.00800 ; qtot
0.983
  4911   HC   301   LEU  HD13 4911     0.100000      1.00800 ; qtot
1.083
```

4912 CT 301 LEU CD2 4912 -0.412100 12.01000 ; qtot
0.671
4913 HC 301 LEU HD21 4913 0.100000 1.00800 ; qtot
0.771
4914 HC 301 LEU HD22 4914 0.100000 1.00800 ; qtot
0.871
4915 HC 301 LEU HD23 4915 0.100000 1.00800 ; qtot
0.971
4916 C 301 LEU C 4916 0.597301 12.01000 ; qtot
1.568
4917 O 301 LEU O 4917 -0.567901 16.00000 ; qtot
1.000
4918 N 302 THR N 4918 -0.415700 14.01000 ; qtot
0.584
4919 H 302 THR H 4919 0.271900 1.00800 ; qtot
0.856
4920 CX 302 THR CA 4920 -0.038900 12.01000 ; qtot
0.817
4921 H1 302 THR HA 4921 0.100700 1.00800 ; qtot
0.918
4922 A3C 302 THR CB 4922 0.365400 12.01000 ; qtot
1.283
4923 H1 302 THR HB 4923 0.004300 1.00800 ; qtot
1.288
4924 CT 302 THR CG2 4924 -0.243800 12.01000 ; qtot
1.044
4925 HC 302 THR HG21 4925 0.064200 1.00800 ; qtot
1.108
4926 HC 302 THR HG22 4926 0.064200 1.00800 ; qtot
1.172
4927 HC 302 THR HG23 4927 0.064200 1.00800 ; qtot
1.237
4928 OH 302 THR OG1 4928 -0.676101 16.00000 ; qtot
0.560
4929 HO 302 THR HG1 4929 0.410200 1.00800 ; qtot
0.971
4930 C 302 THR C 4930 0.597301 12.01000 ; qtot
1.568
4931 O 302 THR O 4931 -0.567901 16.00000 ; qtot
1.000
4932 N 303 ILE N 4932 -0.415700 14.01000 ; qtot
0.584
4933 H 303 ILE H 4933 0.271900 1.00800 ; qtot
0.856
4934 CX 303 ILE CA 4934 -0.059700 12.01000 ; qtot
0.797
4935 H1 303 ILE HA 4935 0.086900 1.00800 ; qtot
0.883
4936 A3C 303 ILE CB 4936 0.130300 12.01000 ; qtot
1.014
4937 HC 303 ILE HB 4937 0.018700 1.00800 ; qtot
1.032

4938 CT 303 ILE CG2 4938 -0.320400 12.01000 ; qtot 0.712
4939 HC 303 ILE HG21 4939 0.088200 1.00800 ; qtot 0.800
4940 HC 303 ILE HG22 4940 0.088200 1.00800 ; qtot 0.888
4941 HC 303 ILE HG23 4941 0.088200 1.00800 ; qtot 0.977
4942 A2C 303 ILE CG1 4942 -0.043000 12.01000 ; qtot 0.934
4943 HC 303 ILE HG12 4943 0.023600 1.00800 ; qtot 0.957
4944 HC 303 ILE HG13 4944 0.023600 1.00800 ; qtot 0.981
4945 CT 303 ILE CD1 4945 -0.066000 12.01000 ; qtot 0.915
4946 HC 303 ILE HD11 4946 0.018600 1.00800 ; qtot 0.933
4947 HC 303 ILE HD12 4947 0.018600 1.00800 ; qtot 0.952
4948 HC 303 ILE HD13 4948 0.018600 1.00800 ; qtot 0.971
4949 C 303 ILE C 4949 0.597301 12.01000 ; qtot 1.568
4950 O 303 ILE O 4950 -0.567901 16.00000 ; qtot 1.000
4951 N 304 ILE N 4951 -0.415700 14.01000 ; qtot 0.584
4952 H 304 ILE H 4952 0.271900 1.00800 ; qtot 0.856
4953 CX 304 ILE CA 4953 -0.059700 12.01000 ; qtot 0.797
4954 H1 304 ILE HA 4954 0.086900 1.00800 ; qtot 0.883
4955 A3C 304 ILE CB 4955 0.130300 12.01000 ; qtot 1.014
4956 HC 304 ILE HB 4956 0.018700 1.00800 ; qtot 1.032
4957 CT 304 ILE CG2 4957 -0.320400 12.01000 ; qtot 0.712
4958 HC 304 ILE HG21 4958 0.088200 1.00800 ; qtot 0.800
4959 HC 304 ILE HG22 4959 0.088200 1.00800 ; qtot 0.888
4960 HC 304 ILE HG23 4960 0.088200 1.00800 ; qtot 0.977
4961 A2C 304 ILE CG1 4961 -0.043000 12.01000 ; qtot 0.934
4962 HC 304 ILE HG12 4962 0.023600 1.00800 ; qtot 0.957
4963 HC 304 ILE HG13 4963 0.023600 1.00800 ; qtot 0.981

4964 CT 304 ILE CD1 4964 -0.066000 12.01000 ; qtot
0.915
4965 HC 304 ILE HD11 4965 0.018600 1.00800 ; qtot
0.933
4966 HC 304 ILE HD12 4966 0.018600 1.00800 ; qtot
0.952
4967 HC 304 ILE HD13 4967 0.018600 1.00800 ; qtot
0.971
4968 C 304 ILE C 4968 0.597301 12.01000 ; qtot
1.568
4969 O 304 ILE O 4969 -0.567901 16.00000 ; qtot
1.000
4970 N 305 ALA N 4970 -0.415700 14.01000 ; qtot
0.584
4971 H 305 ALA H 4971 0.271900 1.00800 ; qtot
0.856
4972 CX 305 ALA CA 4972 0.033700 12.01000 ; qtot
0.890
4973 H1 305 ALA HA 4973 0.082300 1.00800 ; qtot
0.972
4974 CT 305 ALA CB 4974 -0.182500 12.01000 ; qtot
0.790
4975 HC 305 ALA HB1 4975 0.060300 1.00800 ; qtot
0.850
4976 HC 305 ALA HB2 4976 0.060300 1.00800 ; qtot
0.910
4977 HC 305 ALA HB3 4977 0.060300 1.00800 ; qtot
0.971
4978 C 305 ALA C 4978 0.597301 12.01000 ; qtot
1.568
4979 O 305 ALA O 4979 -0.567901 16.00000 ; qtot
1.000
4980 N 306 LYS N 4980 -0.347900 14.01000 ; qtot
0.652
4981 H 306 LYS H 4981 0.274700 1.00800 ; qtot
0.927
4982 CX 306 LYS CA 4982 -0.240000 12.01000 ; qtot
0.687
4983 H1 306 LYS HA 4983 0.142600 1.00800 ; qtot
0.829
4984 C8 306 LYS CB 4984 -0.009400 12.01000 ; qtot
0.820
4985 HC 306 LYS HB2 4985 0.036200 1.00800 ; qtot
0.856
4986 HC 306 LYS HB3 4986 0.036200 1.00800 ; qtot
0.892
4987 C8 306 LYS CG 4987 0.018700 12.01000 ; qtot
0.911
4988 HC 306 LYS HG2 4988 0.010300 1.00800 ; qtot
0.921
4989 HC 306 LYS HG3 4989 0.010300 1.00800 ; qtot
0.932

4990 C8 306 LYS CD 4990 -0.047900 12.01000 ; qtot
0.884
4991 HC 306 LYS HD2 4991 0.062100 1.00800 ; qtot
0.946
4992 HC 306 LYS HD3 4992 0.062100 1.00800 ; qtot
1.008
4993 C8 306 LYS CE 4993 -0.014300 12.01000 ; qtot
0.994
4994 HP 306 LYS HE2 4994 0.113500 1.00800 ; qtot
1.107
4995 HP 306 LYS HE3 4995 0.113500 1.00800 ; qtot
1.221
4996 N3 306 LYS NZ 4996 -0.385400 14.01000 ; qtot
0.835
4997 H 306 LYS HZ1 4997 0.340000 1.00800 ; qtot
1.175
4998 H 306 LYS HZ2 4998 0.340000 1.00800 ; qtot
1.515
4999 H 306 LYS HZ3 4999 0.340000 1.00800 ; qtot
1.855
5000 C 306 LYS C 5000 0.734101 12.01000 ; qtot
2.589
5001 O 306 LYS O 5001 -0.589401 16.00000 ; qtot
2.000
5002 N 307 HIP N 5002 -0.347900 14.01000 ; qtot
1.652
5003 H 307 HIP H 5003 0.274700 1.00800 ; qtot
1.927
5004 CX 307 HIP CA 5004 -0.135400 12.01000 ; qtot
1.791
5005 H1 307 HIP HA 5005 0.121200 1.00800 ; qtot
1.913
5006 CT 307 HIP CB 5006 -0.041400 12.01000 ; qtot
1.871
5007 HC 307 HIP HB2 5007 0.081000 1.00800 ; qtot
1.952
5008 HC 307 HIP HB3 5008 0.081000 1.00800 ; qtot
2.033
5009 CC 307 HIP CG 5009 -0.001200 12.01000 ; qtot
2.032
5010 NA 307 HIP ND1 5010 -0.151300 14.01000 ; qtot
1.881
5011 H 307 HIP HD1 5011 0.386600 1.00800 ; qtot
2.267
5012 CR 307 HIP CE1 5012 -0.017000 12.01000 ; qtot
2.250
5013 H5 307 HIP HE1 5013 0.268100 1.00800 ; qtot
2.518
5014 NA 307 HIP NE2 5014 -0.171800 14.01000 ; qtot
2.347
5015 H 307 HIP HE2 5015 0.391100 1.00800 ; qtot
2.738

5016 CW 307 HIP CD2 5016 -0.114100 12.01000 ; qtot 2.624
5017 H4 307 HIP HD2 5017 0.231700 1.00800 ; qtot 2.855
5018 C 307 HIP C 5018 0.734101 12.01000 ; qtot 3.589
5019 O 307 HIP O 5019 -0.589401 16.00000 ; qtot 3.000
5020 N 308 LEU N 5020 -0.415700 14.01000 ; qtot 2.584
5021 H 308 LEU H 5021 0.271900 1.00800 ; qtot 2.856
5022 CX 308 LEU CA 5022 -0.051800 12.01000 ; qtot 2.804
5023 H1 308 LEU HA 5023 0.092200 1.00800 ; qtot 2.897
5024 A2C 308 LEU CB 5024 -0.110200 12.01000 ; qtot 2.786
5025 HC 308 LEU HB2 5025 0.045700 1.00800 ; qtot 2.832
5026 HC 308 LEU HB3 5026 0.045700 1.00800 ; qtot 2.878
5027 A3C 308 LEU CG 5027 0.353100 12.01000 ; qtot 3.231
5028 HC 308 LEU HG 5028 -0.036100 1.00800 ; qtot 3.195
5029 CT 308 LEU CD1 5029 -0.412100 12.01000 ; qtot 2.783
5030 HC 308 LEU HD11 5030 0.100000 1.00800 ; qtot 2.883
5031 HC 308 LEU HD12 5031 0.100000 1.00800 ; qtot 2.983
5032 HC 308 LEU HD13 5032 0.100000 1.00800 ; qtot 3.083
5033 CT 308 LEU CD2 5033 -0.412100 12.01000 ; qtot 2.671
5034 HC 308 LEU HD21 5034 0.100000 1.00800 ; qtot 2.771
5035 HC 308 LEU HD22 5035 0.100000 1.00800 ; qtot 2.871
5036 HC 308 LEU HD23 5036 0.100000 1.00800 ; qtot 2.971
5037 C 308 LEU C 5037 0.597301 12.01000 ; qtot 3.568
5038 O 308 LEU O 5038 -0.567901 16.00000 ; qtot 3.000
5039 N 309 CYS N 5039 -0.415700 14.01000 ; qtot 2.584
5040 H 309 CYS H 5040 0.271900 1.00800 ; qtot 2.856
5041 CX 309 CYS CA 5041 0.021300 12.01000 ; qtot 2.878

5042 H1 309 CYS HA 5042 0.112400 1.00800 ; qtot
2.990
5043 A2C 309 CYS CB 5043 -0.123100 12.01000 ; qtot
2.867
5044 H1 309 CYS HB2 5044 0.111200 1.00800 ; qtot
2.978
5045 H1 309 CYS HB3 5045 0.111200 1.00800 ; qtot
3.089
5046 SH 309 CYS SG 5046 -0.311900 32.06000 ; qtot
2.777
5047 HS 309 CYS HG 5047 0.193300 1.00800 ; qtot
2.971
5048 C 309 CYS C 5048 0.597301 12.01000 ; qtot
3.568
5049 O 309 CYS O 5049 -0.567901 16.00000 ; qtot
3.000
5050 N 310 ASP N 5050 -0.516301 14.01000 ; qtot
2.484
5051 H 310 ASP H 5051 0.293600 1.00800 ; qtot
2.777
5052 CX 310 ASP CA 5052 0.038100 12.01000 ; qtot
2.815
5053 H1 310 ASP HA 5053 0.088000 1.00800 ; qtot
2.903
5054 A2C 310 ASP CB 5054 -0.030300 12.01000 ; qtot
2.873
5055 HC 310 ASP HB2 5055 -0.012200 1.00800 ; qtot
2.861
5056 HC 310 ASP HB3 5056 -0.012200 1.00800 ; qtot
2.849
5057 CO 310 ASP CG 5057 0.799401 12.01000 ; qtot
3.648
5058 O2 310 ASP OD1 5058 -0.801401 16.00000 ; qtot
2.847
5059 O2 310 ASP OD2 5059 -0.801401 16.00000 ; qtot
2.045
5060 C 310 ASP C 5060 0.536601 12.01000 ; qtot
2.582
5061 O 310 ASP O 5061 -0.581901 16.00000 ; qtot
2.000
5062 N 311 VAL N 5062 -0.415700 14.01000 ; qtot
1.584
5063 H 311 VAL H 5063 0.271900 1.00800 ; qtot
1.856
5064 CX 311 VAL CA 5064 -0.087500 12.01000 ; qtot
1.769
5065 H1 311 VAL HA 5065 0.096900 1.00800 ; qtot
1.866
5066 A3C 311 VAL CB 5066 0.298500 12.01000 ; qtot
2.164
5067 HC 311 VAL HB 5067 -0.029700 1.00800 ; qtot
2.134

5068 CT 311 VAL CG1 5068 -0.319200 12.01000 ; qtot
1.815
5069 HC 311 VAL HG11 5069 0.079100 1.00800 ; qtot
1.894
5070 HC 311 VAL HG12 5070 0.079100 1.00800 ; qtot
1.973
5071 HC 311 VAL HG13 5071 0.079100 1.00800 ; qtot
2.053
5072 CT 311 VAL CG2 5072 -0.319200 12.01000 ; qtot
1.733
5073 HC 311 VAL HG21 5073 0.079100 1.00800 ; qtot
1.812
5074 HC 311 VAL HG22 5074 0.079100 1.00800 ; qtot
1.892
5075 HC 311 VAL HG23 5075 0.079100 1.00800 ; qtot
1.971
5076 C 311 VAL C 5076 0.597301 12.01000 ; qtot
2.568
5077 O 311 VAL O 5077 -0.567901 16.00000 ; qtot
2.000
5078 N 312 GLU N 5078 -0.516301 14.01000 ; qtot
1.484
5079 H 312 GLU H 5079 0.293600 1.00800 ; qtot
1.777
5080 CX 312 GLU CA 5080 0.039700 12.01000 ; qtot
1.817
5081 H1 312 GLU HA 5081 0.110500 1.00800 ; qtot
1.928
5082 A2C 312 GLU CB 5082 0.056000 12.01000 ; qtot
1.984
5083 HC 312 GLU HB2 5083 -0.017300 1.00800 ; qtot
1.966
5084 HC 312 GLU HB3 5084 -0.017300 1.00800 ; qtot
1.949
5085 A2C 312 GLU CG 5085 0.013600 12.01000 ; qtot
1.963
5086 HC 312 GLU HG2 5086 -0.042500 1.00800 ; qtot
1.920
5087 HC 312 GLU HG3 5087 -0.042500 1.00800 ; qtot
1.878
5088 CO 312 GLU CD 5088 0.805401 12.01000 ; qtot
2.683
5089 O2 312 GLU OE1 5089 -0.818801 16.00000 ; qtot
1.864
5090 O2 312 GLU OE2 5090 -0.818801 16.00000 ; qtot
1.045
5091 C 312 GLU C 5091 0.536601 12.01000 ; qtot
1.582
5092 O 312 GLU O 5092 -0.581901 16.00000 ; qtot
1.000
5093 N 313 ILE N 5093 -0.415700 14.01000 ; qtot
0.584

5094 H 313 ILE H 5094 0.271900 1.00800 ; qtot
0.856
5095 CX 313 ILE CA 5095 -0.059700 12.01000 ; qtot
0.797
5096 H1 313 ILE HA 5096 0.086900 1.00800 ; qtot
0.883
5097 A3C 313 ILE CB 5097 0.130300 12.01000 ; qtot
1.014
5098 HC 313 ILE HB 5098 0.018700 1.00800 ; qtot
1.032
5099 CT 313 ILE CG2 5099 -0.320400 12.01000 ; qtot
0.712
5100 HC 313 ILE HG21 5100 0.088200 1.00800 ; qtot
0.800
5101 HC 313 ILE HG22 5101 0.088200 1.00800 ; qtot
0.888
5102 HC 313 ILE HG23 5102 0.088200 1.00800 ; qtot
0.977
5103 A2C 313 ILE CG1 5103 -0.043000 12.01000 ; qtot
0.934
5104 HC 313 ILE HG12 5104 0.023600 1.00800 ; qtot
0.957
5105 HC 313 ILE HG13 5105 0.023600 1.00800 ; qtot
0.981
5106 CT 313 ILE CD1 5106 -0.066000 12.01000 ; qtot
0.915
5107 HC 313 ILE HD11 5107 0.018600 1.00800 ; qtot
0.933
5108 HC 313 ILE HD12 5108 0.018600 1.00800 ; qtot
0.952
5109 HC 313 ILE HD13 5109 0.018600 1.00800 ; qtot
0.971
5110 C 313 ILE C 5110 0.597301 12.01000 ; qtot
1.568
5111 O 313 ILE O 5111 -0.567901 16.00000 ; qtot
1.000
5112 N 314 ASN N 5112 -0.415700 14.01000 ; qtot
0.584
5113 H 314 ASN H 5113 0.271900 1.00800 ; qtot
0.856
5114 CX 314 ASN CA 5114 0.014300 12.01000 ; qtot
0.871
5115 H1 314 ASN HA 5115 0.104800 1.00800 ; qtot
0.975
5116 A2C 314 ASN CB 5116 -0.204100 12.01000 ; qtot
0.771
5117 HC 314 ASN HB2 5117 0.079700 1.00800 ; qtot
0.851
5118 HC 314 ASN HB3 5118 0.079700 1.00800 ; qtot
0.931
5119 C 314 ASN CG 5119 0.713001 12.01000 ; qtot
1.644

5120 O 314 ASN OD1 5120 -0.593101 16.00000 ; qtot
1.051
5121 N 314 ASN ND2 5121 -0.919101 14.01000 ; qtot
0.131
5122 H 314 ASN HD21 5122 0.419600 1.00800 ; qtot
0.551
5123 H 314 ASN HD22 5123 0.419600 1.00800 ; qtot
0.971
5124 C 314 ASN C 5124 0.597301 12.01000 ; qtot
1.568
5125 O 314 ASN O 5125 -0.567901 16.00000 ; qtot
1.000
5126 N 315 ASN N 5126 -0.415700 14.01000 ; qtot
0.584
5127 H 315 ASN H 5127 0.271900 1.00800 ; qtot
0.856
5128 CX 315 ASN CA 5128 0.014300 12.01000 ; qtot
0.871
5129 H1 315 ASN HA 5129 0.104800 1.00800 ; qtot
0.975
5130 A2C 315 ASN CB 5130 -0.204100 12.01000 ; qtot
0.771
5131 HC 315 ASN HB2 5131 0.079700 1.00800 ; qtot
0.851
5132 HC 315 ASN HB3 5132 0.079700 1.00800 ; qtot
0.931
5133 C 315 ASN CG 5133 0.713001 12.01000 ; qtot
1.644
5134 O 315 ASN OD1 5134 -0.593101 16.00000 ; qtot
1.051
5135 N 315 ASN ND2 5135 -0.919101 14.01000 ; qtot
0.131
5136 H 315 ASN HD21 5136 0.419600 1.00800 ; qtot
0.551
5137 H 315 ASN HD22 5137 0.419600 1.00800 ; qtot
0.971
5138 C 315 ASN C 5138 0.597301 12.01000 ; qtot
1.568
5139 O 315 ASN O 5139 -0.567901 16.00000 ; qtot
1.000
5140 N 316 LEU N 5140 -0.415700 14.01000 ; qtot
0.584
5141 H 316 LEU H 5141 0.271900 1.00800 ; qtot
0.856
5142 CX 316 LEU CA 5142 -0.051800 12.01000 ; qtot
0.804
5143 H1 316 LEU HA 5143 0.092200 1.00800 ; qtot
0.897
5144 A2C 316 LEU CB 5144 -0.110200 12.01000 ; qtot
0.786
5145 HC 316 LEU HB2 5145 0.045700 1.00800 ; qtot
0.832

5146 HC 316 LEU HB3 5146 0.045700 1.00800 ; qtot
0.878
5147 A3C 316 LEU CG 5147 0.353100 12.01000 ; qtot
1.231
5148 HC 316 LEU HG 5148 -0.036100 1.00800 ; qtot
1.195
5149 CT 316 LEU CD1 5149 -0.412100 12.01000 ; qtot
0.783
5150 HC 316 LEU HD11 5150 0.100000 1.00800 ; qtot
0.883
5151 HC 316 LEU HD12 5151 0.100000 1.00800 ; qtot
0.983
5152 HC 316 LEU HD13 5152 0.100000 1.00800 ; qtot
1.083
5153 CT 316 LEU CD2 5153 -0.412100 12.01000 ; qtot
0.671
5154 HC 316 LEU HD21 5154 0.100000 1.00800 ; qtot
0.771
5155 HC 316 LEU HD22 5155 0.100000 1.00800 ; qtot
0.871
5156 HC 316 LEU HD23 5156 0.100000 1.00800 ; qtot
0.971
5157 C 316 LEU C 5157 0.597301 12.01000 ; qtot
1.568
5158 O 316 LEU O 5158 -0.567901 16.00000 ; qtot
1.000
5159 N 317 ARG N 5159 -0.347900 14.01000 ; qtot
0.652
5160 H 317 ARG H 5160 0.274700 1.00800 ; qtot
0.927
5161 CX 317 ARG CA 5161 -0.263700 12.01000 ; qtot
0.663
5162 H1 317 ARG HA 5162 0.156000 1.00800 ; qtot
0.819
5163 C8 317 ARG CB 5163 -0.000700 12.01000 ; qtot
0.818
5164 HC 317 ARG HB2 5164 0.032700 1.00800 ; qtot
0.851
5165 HC 317 ARG HB3 5165 0.032700 1.00800 ; qtot
0.884
5166 C8 317 ARG CG 5166 0.039000 12.01000 ; qtot
0.923
5167 HC 317 ARG HG2 5167 0.028500 1.00800 ; qtot
0.951
5168 HC 317 ARG HG3 5168 0.028500 1.00800 ; qtot
0.980
5169 C8 317 ARG CD 5169 0.048600 12.01000 ; qtot
1.028
5170 H1 317 ARG HD2 5170 0.068700 1.00800 ; qtot
1.097
5171 H1 317 ARG HD3 5171 0.068700 1.00800 ; qtot
1.166

5172 N2 317 ARG NE 5172 -0.529501 14.01000 ; qtot
0.636
5173 H 317 ARG HE 5173 0.345600 1.00800 ; qtot
0.982
5174 CA 317 ARG CZ 5174 0.807601 12.01000 ; qtot
1.790
5175 N2 317 ARG NH1 5175 -0.862701 14.01000 ; qtot
0.927
5176 H 317 ARG HH11 5176 0.447800 1.00800 ; qtot
1.375
5177 H 317 ARG HH12 5177 0.447800 1.00800 ; qtot
1.822
5178 N2 317 ARG NH2 5178 -0.862701 14.01000 ; qtot
0.960
5179 H 317 ARG HH21 5179 0.447800 1.00800 ; qtot
1.408
5180 H 317 ARG HH22 5180 0.447800 1.00800 ; qtot
1.855
5181 C 317 ARG C 5181 0.734101 12.01000 ; qtot
2.589
5182 O 317 ARG O 5182 -0.589401 16.00000 ; qtot
2.000
5183 N 318 ASN N 5183 -0.415700 14.01000 ; qtot
1.584
5184 H 318 ASN H 5184 0.271900 1.00800 ; qtot
1.856
5185 CX 318 ASN CA 5185 0.014300 12.01000 ; qtot
1.871
5186 H1 318 ASN HA 5186 0.104800 1.00800 ; qtot
1.975
5187 A2C 318 ASN CB 5187 -0.204100 12.01000 ; qtot
1.771
5188 HC 318 ASN HB2 5188 0.079700 1.00800 ; qtot
1.851
5189 HC 318 ASN HB3 5189 0.079700 1.00800 ; qtot
1.931
5190 C 318 ASN CG 5190 0.713001 12.01000 ; qtot
2.644
5191 O 318 ASN OD1 5191 -0.593101 16.00000 ; qtot
2.051
5192 N 318 ASN ND2 5192 -0.919101 14.01000 ; qtot
1.131
5193 H 318 ASN HD21 5193 0.419600 1.00800 ; qtot
1.551
5194 H 318 ASN HD22 5194 0.419600 1.00800 ; qtot
1.971
5195 C 318 ASN C 5195 0.597301 12.01000 ; qtot
2.568
5196 O 318 ASN O 5196 -0.567901 16.00000 ; qtot
2.000
5197 N 319 ILE N 5197 -0.415700 14.01000 ; qtot
1.584

5198 H 319 ILE H 5198 0.271900 1.00800 ; qtot
1.856
5199 CX 319 ILE CA 5199 -0.059700 12.01000 ; qtot
1.797
5200 H1 319 ILE HA 5200 0.086900 1.00800 ; qtot
1.883
5201 A3C 319 ILE CB 5201 0.130300 12.01000 ; qtot
2.014
5202 HC 319 ILE HB 5202 0.018700 1.00800 ; qtot
2.032
5203 CT 319 ILE CG2 5203 -0.320400 12.01000 ; qtot
1.712
5204 HC 319 ILE HG21 5204 0.088200 1.00800 ; qtot
1.800
5205 HC 319 ILE HG22 5205 0.088200 1.00800 ; qtot
1.888
5206 HC 319 ILE HG23 5206 0.088200 1.00800 ; qtot
1.977
5207 A2C 319 ILE CG1 5207 -0.043000 12.01000 ; qtot
1.934
5208 HC 319 ILE HG12 5208 0.023600 1.00800 ; qtot
1.957
5209 HC 319 ILE HG13 5209 0.023600 1.00800 ; qtot
1.981
5210 CT 319 ILE CD1 5210 -0.066000 12.01000 ; qtot
1.915
5211 HC 319 ILE HD11 5211 0.018600 1.00800 ; qtot
1.933
5212 HC 319 ILE HD12 5212 0.018600 1.00800 ; qtot
1.952
5213 HC 319 ILE HD13 5213 0.018600 1.00800 ; qtot
1.971
5214 C 319 ILE C 5214 0.597301 12.01000 ; qtot
2.568
5215 O 319 ILE O 5215 -0.567901 16.00000 ; qtot
2.000
5216 N 320 PHE N 5216 -0.415700 14.01000 ; qtot
1.584
5217 H 320 PHE H 5217 0.271900 1.00800 ; qtot
1.856
5218 CX 320 PHE CA 5218 -0.002400 12.01000 ; qtot
1.854
5219 H1 320 PHE HA 5219 0.097800 1.00800 ; qtot
1.952
5220 CT 320 PHE CB 5220 -0.034300 12.01000 ; qtot
1.917
5221 HC 320 PHE HB2 5221 0.029500 1.00800 ; qtot
1.947
5222 HC 320 PHE HB3 5222 0.029500 1.00800 ; qtot
1.976
5223 CA 320 PHE CG 5223 0.011800 12.01000 ; qtot
1.988

5224 CA 320 PHE CD1 5224 -0.125600 12.01000 ; qtot
1.863
5225 HA 320 PHE HD1 5225 0.133000 1.00800 ; qtot
1.996
5226 CA 320 PHE CE1 5226 -0.170400 12.01000 ; qtot
1.825
5227 HA 320 PHE HE1 5227 0.143000 1.00800 ; qtot
1.968
5228 CA 320 PHE CZ 5228 -0.107200 12.01000 ; qtot
1.861
5229 HA 320 PHE HZ 5229 0.129700 1.00800 ; qtot
1.991
5230 CA 320 PHE CE2 5230 -0.170400 12.01000 ; qtot
1.820
5231 HA 320 PHE HE2 5231 0.143000 1.00800 ; qtot
1.963
5232 CA 320 PHE CD2 5232 -0.125600 12.01000 ; qtot
1.838
5233 HA 320 PHE HD2 5233 0.133000 1.00800 ; qtot
1.971
5234 C 320 PHE C 5234 0.597301 12.01000 ; qtot
2.568
5235 O 320 PHE O 5235 -0.567901 16.00000 ; qtot
2.000
5236 N 321 ILE N 5236 -0.415700 14.01000 ; qtot
1.584
5237 H 321 ILE H 5237 0.271900 1.00800 ; qtot
1.856
5238 CX 321 ILE CA 5238 -0.059700 12.01000 ; qtot
1.797
5239 H1 321 ILE HA 5239 0.086900 1.00800 ; qtot
1.883
5240 A3C 321 ILE CB 5240 0.130300 12.01000 ; qtot
2.014
5241 HC 321 ILE HB 5241 0.018700 1.00800 ; qtot
2.032
5242 CT 321 ILE CG2 5242 -0.320400 12.01000 ; qtot
1.712
5243 HC 321 ILE HG21 5243 0.088200 1.00800 ; qtot
1.800
5244 HC 321 ILE HG22 5244 0.088200 1.00800 ; qtot
1.888
5245 HC 321 ILE HG23 5245 0.088200 1.00800 ; qtot
1.977
5246 A2C 321 ILE CG1 5246 -0.043000 12.01000 ; qtot
1.934
5247 HC 321 ILE HG12 5247 0.023600 1.00800 ; qtot
1.957
5248 HC 321 ILE HG13 5248 0.023600 1.00800 ; qtot
1.981
5249 CT 321 ILE CD1 5249 -0.066000 12.01000 ; qtot
1.915

5250 HC 321 ILE HD11 5250 0.018600 1.00800 ; qtot 1.933
5251 HC 321 ILE HD12 5251 0.018600 1.00800 ; qtot 1.952
5252 HC 321 ILE HD13 5252 0.018600 1.00800 ; qtot 1.971
5253 C 321 ILE C 5253 0.597301 12.01000 ; qtot 2.568
5254 O 321 ILE O 5254 -0.567901 16.00000 ; qtot 2.000
5255 N 322 ALA N 5255 -0.415700 14.01000 ; qtot 1.584
5256 H 322 ALA H 5256 0.271900 1.00800 ; qtot 1.856
5257 CX 322 ALA CA 5257 0.033700 12.01000 ; qtot 1.890
5258 H1 322 ALA HA 5258 0.082300 1.00800 ; qtot 1.972
5259 CT 322 ALA CB 5259 -0.182500 12.01000 ; qtot 1.790
5260 HC 322 ALA HB1 5260 0.060300 1.00800 ; qtot 1.850
5261 HC 322 ALA HB2 5261 0.060300 1.00800 ; qtot 1.910
5262 HC 322 ALA HB3 5262 0.060300 1.00800 ; qtot 1.971
5263 C 322 ALA C 5263 0.597301 12.01000 ; qtot 2.568
5264 O 322 ALA O 5264 -0.567901 16.00000 ; qtot 2.000
5265 N 323 LEU N 5265 -0.415700 14.01000 ; qtot 1.584
5266 H 323 LEU H 5266 0.271900 1.00800 ; qtot 1.856
5267 CX 323 LEU CA 5267 -0.051800 12.01000 ; qtot 1.804
5268 H1 323 LEU HA 5268 0.092200 1.00800 ; qtot 1.897
5269 A2C 323 LEU CB 5269 -0.110200 12.01000 ; qtot 1.786
5270 HC 323 LEU HB2 5270 0.045700 1.00800 ; qtot 1.832
5271 HC 323 LEU HB3 5271 0.045700 1.00800 ; qtot 1.878
5272 A3C 323 LEU CG 5272 0.353100 12.01000 ; qtot 2.231
5273 HC 323 LEU HG 5273 -0.036100 1.00800 ; qtot 2.195
5274 CT 323 LEU CD1 5274 -0.412100 12.01000 ; qtot 1.783
5275 HC 323 LEU HD11 5275 0.100000 1.00800 ; qtot 1.883

5276 HC 323 LEU HD12 5276 0.100000 1.00800 ; qtot 1.983
5277 HC 323 LEU HD13 5277 0.100000 1.00800 ; qtot 2.083
5278 CT 323 LEU CD2 5278 -0.412100 12.01000 ; qtot 1.671
5279 HC 323 LEU HD21 5279 0.100000 1.00800 ; qtot 1.771
5280 HC 323 LEU HD22 5280 0.100000 1.00800 ; qtot 1.871
5281 HC 323 LEU HD23 5281 0.100000 1.00800 ; qtot 1.971
5282 C 323 LEU C 5282 0.597301 12.01000 ; qtot 2.568
5283 O 323 LEU O 5283 -0.567901 16.00000 ; qtot 2.000
5284 N 324 ASP N 5284 -0.516301 14.01000 ; qtot 1.484
5285 H 324 ASP H 5285 0.293600 1.00800 ; qtot 1.777
5286 CX 324 ASP CA 5286 0.038100 12.01000 ; qtot 1.815
5287 H1 324 ASP HA 5287 0.088000 1.00800 ; qtot 1.903
5288 A2C 324 ASP CB 5288 -0.030300 12.01000 ; qtot 1.873
5289 HC 324 ASP HB2 5289 -0.012200 1.00800 ; qtot 1.861
5290 HC 324 ASP HB3 5290 -0.012200 1.00800 ; qtot 1.849
5291 CO 324 ASP CG 5291 0.799401 12.01000 ; qtot 2.648
5292 O2 324 ASP OD1 5292 -0.801401 16.00000 ; qtot 1.847
5293 O2 324 ASP OD2 5293 -0.801401 16.00000 ; qtot 1.045
5294 C 324 ASP C 5294 0.536601 12.01000 ; qtot 1.582
5295 O 324 ASP O 5295 -0.581901 16.00000 ; qtot 1.000
5296 N 325 VAL N 5296 -0.415700 14.01000 ; qtot 0.584
5297 H 325 VAL H 5297 0.271900 1.00800 ; qtot 0.856
5298 CX 325 VAL CA 5298 -0.087500 12.01000 ; qtot 0.769
5299 H1 325 VAL HA 5299 0.096900 1.00800 ; qtot 0.866
5300 A3C 325 VAL CB 5300 0.298500 12.01000 ; qtot 1.164
5301 HC 325 VAL HB 5301 -0.029700 1.00800 ; qtot 1.134

5302 CT 325 VAL CG1 5302 -0.319200 12.01000 ; qtot
0.815
5303 HC 325 VAL HG11 5303 0.079100 1.00800 ; qtot
0.894
5304 HC 325 VAL HG12 5304 0.079100 1.00800 ; qtot
0.973
5305 HC 325 VAL HG13 5305 0.079100 1.00800 ; qtot
1.053
5306 CT 325 VAL CG2 5306 -0.319200 12.01000 ; qtot
0.733
5307 HC 325 VAL HG21 5307 0.079100 1.00800 ; qtot
0.812
5308 HC 325 VAL HG22 5308 0.079100 1.00800 ; qtot
0.892
5309 HC 325 VAL HG23 5309 0.079100 1.00800 ; qtot
0.971
5310 C 325 VAL C 5310 0.597301 12.01000 ; qtot
1.568
5311 O 325 VAL O 5311 -0.567901 16.00000 ; qtot
1.000
5312 N 326 ASP N 5312 -0.516301 14.01000 ; qtot
0.484
5313 H 326 ASP H 5313 0.293600 1.00800 ; qtot
0.777
5314 CX 326 ASP CA 5314 0.038100 12.01000 ; qtot
0.815
5315 H1 326 ASP HA 5315 0.088000 1.00800 ; qtot
0.903
5316 A2C 326 ASP CB 5316 -0.030300 12.01000 ; qtot
0.873
5317 HC 326 ASP HB2 5317 -0.012200 1.00800 ; qtot
0.861
5318 HC 326 ASP HB3 5318 -0.012200 1.00800 ; qtot
0.849
5319 CO 326 ASP CG 5319 0.799401 12.01000 ; qtot
1.648
5320 O2 326 ASP OD1 5320 -0.801401 16.00000 ; qtot
0.847
5321 O2 326 ASP OD2 5321 -0.801401 16.00000 ; qtot
0.045
5322 C 326 ASP C 5322 0.536601 12.01000 ; qtot
0.582
5323 O 326 ASP O 5323 -0.581901 16.00000 ; qtot
0.000
5324 N 327 ASN N 5324 -0.415700 14.01000 ; qtot -
0.416
5325 H 327 ASN H 5325 0.271900 1.00800 ; qtot -
0.144
5326 CX 327 ASN CA 5326 0.014300 12.01000 ; qtot -
0.129
5327 H1 327 ASN HA 5327 0.104800 1.00800 ; qtot -
0.025

5328 A2C 327 ASN CB 5328 -0.204100 12.01000 ; qtot -
0.229
5329 HC 327 ASN HB2 5329 0.079700 1.00800 ; qtot -
0.149
5330 HC 327 ASN HB3 5330 0.079700 1.00800 ; qtot -
0.069
5331 C 327 ASN CG 5331 0.713001 12.01000 ; qtot
0.644
5332 O 327 ASN OD1 5332 -0.593101 16.00000 ; qtot
0.051
5333 N 327 ASN ND2 5333 -0.919101 14.01000 ; qtot -
0.869
5334 H 327 ASN HD21 5334 0.419600 1.00800 ; qtot -
0.449
5335 H 327 ASN HD22 5335 0.419600 1.00800 ; qtot -
0.029
5336 C 327 ASN C 5336 0.597301 12.01000 ; qtot
0.568
5337 O 327 ASN O 5337 -0.567901 16.00000 ; qtot
0.000
5338 N 328 SER N 5338 -0.415700 14.01000 ; qtot -
0.416
5339 H 328 SER H 5339 0.271900 1.00800 ; qtot -
0.144
5340 CX 328 SER CA 5340 -0.024900 12.01000 ; qtot -
0.169
5341 H1 328 SER HA 5341 0.084300 1.00800 ; qtot -
0.084
5342 A2C 328 SER CB 5342 0.211700 12.01000 ; qtot
0.127
5343 H1 328 SER HB2 5343 0.035200 1.00800 ; qtot
0.163
5344 H1 328 SER HB3 5344 0.035200 1.00800 ; qtot
0.198
5345 OH 328 SER OG 5345 -0.654601 16.00000 ; qtot -
0.457
5346 HO 328 SER HG 5346 0.427500 1.00800 ; qtot -
0.029
5347 C 328 SER C 5347 0.597301 12.01000 ; qtot
0.568
5348 O 328 SER O 5348 -0.567901 16.00000 ; qtot
0.000
5349 N 329 GLY N 5349 -0.415700 14.01000 ; qtot -
0.416
5350 H 329 GLY H 5350 0.271900 1.00800 ; qtot -
0.144
5351 CX 329 GLY CA 5351 -0.025200 12.01000 ; qtot -
0.169
5352 H1 329 GLY HA2 5352 0.069800 1.00800 ; qtot -
0.099
5353 H1 329 GLY HA3 5353 0.069800 1.00800 ; qtot -
0.029

5354 C 329 GLY C 5354 0.597301 12.01000 ; qtot
0.568
5355 O 329 GLY O 5355 -0.567901 16.00000 ; qtot
0.000
5356 N 330 THR N 5356 -0.415700 14.01000 ; qtot -
0.416
5357 H 330 THR H 5357 0.271900 1.00800 ; qtot -
0.144
5358 CX 330 THR CA 5358 -0.038900 12.01000 ; qtot -
0.183
5359 H1 330 THR HA 5359 0.100700 1.00800 ; qtot -
0.082
5360 A3C 330 THR CB 5360 0.365400 12.01000 ; qtot
0.283
5361 H1 330 THR HB 5361 0.004300 1.00800 ; qtot
0.288
5362 CT 330 THR CG2 5362 -0.243800 12.01000 ; qtot
0.044
5363 HC 330 THR HG21 5363 0.064200 1.00800 ; qtot
0.108
5364 HC 330 THR HG22 5364 0.064200 1.00800 ; qtot
0.172
5365 HC 330 THR HG23 5365 0.064200 1.00800 ; qtot
0.237
5366 OH 330 THR OG1 5366 -0.676101 16.00000 ; qtot -
0.440
5367 HO 330 THR HG1 5367 0.410200 1.00800 ; qtot -
0.029
5368 C 330 THR C 5368 0.597301 12.01000 ; qtot
0.568
5369 O 330 THR O 5369 -0.567901 16.00000 ; qtot
0.000
5370 N 331 LEU N 5370 -0.415700 14.01000 ; qtot -
0.416
5371 H 331 LEU H 5371 0.271900 1.00800 ; qtot -
0.144
5372 CX 331 LEU CA 5372 -0.051800 12.01000 ; qtot -
0.196
5373 H1 331 LEU HA 5373 0.092200 1.00800 ; qtot -
0.103
5374 A2C 331 LEU CB 5374 -0.110200 12.01000 ; qtot -
0.214
5375 HC 331 LEU HB2 5375 0.045700 1.00800 ; qtot -
0.168
5376 HC 331 LEU HB3 5376 0.045700 1.00800 ; qtot -
0.122
5377 A3C 331 LEU CG 5377 0.353100 12.01000 ; qtot
0.231
5378 HC 331 LEU HG 5378 -0.036100 1.00800 ; qtot
0.195
5379 CT 331 LEU CD1 5379 -0.412100 12.01000 ; qtot -
0.217

5380 HC 331 LEU HD11 5380 0.100000 1.00800 ; qtot -
0.117
5381 HC 331 LEU HD12 5381 0.100000 1.00800 ; qtot -
0.017
5382 HC 331 LEU HD13 5382 0.100000 1.00800 ; qtot
0.083
5383 CT 331 LEU CD2 5383 -0.412100 12.01000 ; qtot -
0.329
5384 HC 331 LEU HD21 5384 0.100000 1.00800 ; qtot -
0.229
5385 HC 331 LEU HD22 5385 0.100000 1.00800 ; qtot -
0.129
5386 HC 331 LEU HD23 5386 0.100000 1.00800 ; qtot -
0.029
5387 C 331 LEU C 5387 0.597301 12.01000 ; qtot
0.568
5388 O 331 LEU O 5388 -0.567901 16.00000 ; qtot
0.000
5389 N 332 SER N 5389 -0.415700 14.01000 ; qtot -
0.416
5390 H 332 SER H 5390 0.271900 1.00800 ; qtot -
0.144
5391 CX 332 SER CA 5391 -0.024900 12.01000 ; qtot -
0.169
5392 H1 332 SER HA 5392 0.084300 1.00800 ; qtot -
0.084
5393 A2C 332 SER CB 5393 0.211700 12.01000 ; qtot
0.127
5394 H1 332 SER HB2 5394 0.035200 1.00800 ; qtot
0.163
5395 H1 332 SER HB3 5395 0.035200 1.00800 ; qtot
0.198
5396 OH 332 SER OG 5396 -0.654601 16.00000 ; qtot -
0.457
5397 HO 332 SER HG 5397 0.427500 1.00800 ; qtot -
0.029
5398 C 332 SER C 5398 0.597301 12.01000 ; qtot
0.568
5399 O 332 SER O 5399 -0.567901 16.00000 ; qtot
0.000
5400 N 333 SER N 5400 -0.415700 14.01000 ; qtot -
0.416
5401 H 333 SER H 5401 0.271900 1.00800 ; qtot -
0.144
5402 CX 333 SER CA 5402 -0.024900 12.01000 ; qtot -
0.169
5403 H1 333 SER HA 5403 0.084300 1.00800 ; qtot -
0.084
5404 A2C 333 SER CB 5404 0.211700 12.01000 ; qtot
0.127
5405 H1 333 SER HB2 5405 0.035200 1.00800 ; qtot
0.163

5406 H1 333 SER HB3 5406 0.035200 1.00800 ; qtot
0.198
5407 OH 333 SER OG 5407 -0.654601 16.00000 ; qtot -
0.457
5408 HO 333 SER HG 5408 0.427500 1.00800 ; qtot -
0.029
5409 C 333 SER C 5409 0.597301 12.01000 ; qtot
0.568
5410 O 333 SER O 5410 -0.567901 16.00000 ; qtot
0.000
5411 N 334 GLN N 5411 -0.415700 14.01000 ; qtot -
0.416
5412 H 334 GLN H 5412 0.271900 1.00800 ; qtot -
0.144
5413 CX 334 GLN CA 5413 -0.003100 12.01000 ; qtot -
0.147
5414 H1 334 GLN HA 5414 0.085000 1.00800 ; qtot -
0.062
5415 A2C 334 GLN CB 5415 -0.003600 12.01000 ; qtot -
0.065
5416 HC 334 GLN HB2 5416 0.017100 1.00800 ; qtot -
0.048
5417 HC 334 GLN HB3 5417 0.017100 1.00800 ; qtot -
0.031
5418 A2C 334 GLN CG 5418 -0.064500 12.01000 ; qtot -
0.096
5419 HC 334 GLN HG2 5419 0.035200 1.00800 ; qtot -
0.061
5420 HC 334 GLN HG3 5420 0.035200 1.00800 ; qtot -
0.025
5421 C 334 GLN CD 5421 0.695101 12.01000 ; qtot
0.670
5422 O 334 GLN OE1 5422 -0.608601 16.00000 ; qtot
0.061
5423 N 334 GLN NE2 5423 -0.940701 14.01000 ; qtot -
0.880
5424 H 334 GLN HE21 5424 0.425100 1.00800 ; qtot -
0.454
5425 H 334 GLN HE22 5425 0.425100 1.00800 ; qtot -
0.029
5426 C 334 GLN C 5426 0.597301 12.01000 ; qtot
0.568
5427 O 334 GLN O 5427 -0.567901 16.00000 ; qtot
0.000
5428 N 335 GLH N 5428 -0.415700 14.01000 ; qtot -
0.416
5429 H 335 GLH H 5429 0.271900 1.00800 ; qtot -
0.144
5430 CX 335 GLH CA 5430 0.014500 12.01000 ; qtot -
0.129
5431 H1 335 GLH HA 5431 0.077900 1.00800 ; qtot -
0.051

5432 A2C 335 GLH CB 5432 -0.007100 12.01000 ; qtot -
0.058
5433 HC 335 GLH HB2 5433 0.025600 1.00800 ; qtot -
0.033
5434 HC 335 GLH HB3 5434 0.025600 1.00800 ; qtot -
0.007
5435 A2C 335 GLH CG 5435 -0.017400 12.01000 ; qtot -
0.025
5436 HC 335 GLH HG2 5436 0.043000 1.00800 ; qtot
0.018
5437 HC 335 GLH HG3 5437 0.043000 1.00800 ; qtot
0.061
5438 C 335 GLH CD 5438 0.680101 12.01000 ; qtot
0.741
5439 O 335 GLH OE1 5439 -0.583801 16.00000 ; qtot
0.158
5440 OH 335 GLH OE2 5440 -0.651101 16.00000 ; qtot -
0.493
5441 HO 335 GLH HE2 5441 0.464100 1.00800 ; qtot -
0.029
5442 C 335 GLH C 5442 0.597301 12.01000 ; qtot
0.568
5443 O 335 GLH O 5443 -0.567901 16.00000 ; qtot
0.000
5444 N 336 ILE N 5444 -0.415700 14.01000 ; qtot -
0.416
5445 H 336 ILE H 5445 0.271900 1.00800 ; qtot -
0.144
5446 CX 336 ILE CA 5446 -0.059700 12.01000 ; qtot -
0.203
5447 H1 336 ILE HA 5447 0.086900 1.00800 ; qtot -
0.117
5448 A3C 336 ILE CB 5448 0.130300 12.01000 ; qtot
0.014
5449 HC 336 ILE HB 5449 0.018700 1.00800 ; qtot
0.032
5450 CT 336 ILE CG2 5450 -0.320400 12.01000 ; qtot -
0.288
5451 HC 336 ILE HG21 5451 0.088200 1.00800 ; qtot -
0.200
5452 HC 336 ILE HG22 5452 0.088200 1.00800 ; qtot -
0.112
5453 HC 336 ILE HG23 5453 0.088200 1.00800 ; qtot -
0.023
5454 A2C 336 ILE CG1 5454 -0.043000 12.01000 ; qtot -
0.066
5455 HC 336 ILE HG12 5455 0.023600 1.00800 ; qtot -
0.043
5456 HC 336 ILE HG13 5456 0.023600 1.00800 ; qtot -
0.019
5457 CT 336 ILE CD1 5457 -0.066000 12.01000 ; qtot -
0.085

```
  5458   HC   336   ILE  HD11 5458     0.018600      1.00800 ; qtot -
0.067
  5459   HC   336   ILE  HD12 5459     0.018600      1.00800 ; qtot -
0.048
  5460   HC   336   ILE  HD13 5460     0.018600      1.00800 ; qtot -
0.029
  5461    C   336   ILE     C 5461     0.597301     12.01000 ; qtot
0.568
  5462    O   336   ILE     O 5462    -0.567901     16.00000 ; qtot
0.000
  5463    N   337   LEU     N 5463    -0.415700     14.01000 ; qtot -
0.416
  5464    H   337   LEU     H 5464     0.271900      1.00800 ; qtot -
0.144
  5465   CX   337   LEU    CA 5465    -0.051800     12.01000 ; qtot -
0.196
  5466   H1   337   LEU    HA 5466     0.092200      1.00800 ; qtot -
0.103
  5467  A2C   337   LEU    CB 5467    -0.110200     12.01000 ; qtot -
0.214
  5468   HC   337   LEU   HB2 5468     0.045700      1.00800 ; qtot -
0.168
  5469   HC   337   LEU   HB3 5469     0.045700      1.00800 ; qtot -
0.122
  5470  A3C   337   LEU    CG 5470     0.353100     12.01000 ; qtot
0.231
  5471   HC   337   LEU    HG 5471    -0.036100      1.00800 ; qtot
0.195
  5472   CT   337   LEU   CD1 5472    -0.412100     12.01000 ; qtot -
0.217
  5473   HC   337   LEU  HD11 5473     0.100000      1.00800 ; qtot -
0.117
  5474   HC   337   LEU  HD12 5474     0.100000      1.00800 ; qtot -
0.017
  5475   HC   337   LEU  HD13 5475     0.100000      1.00800 ; qtot
0.083
  5476   CT   337   LEU   CD2 5476    -0.412100     12.01000 ; qtot -
0.329
  5477   HC   337   LEU  HD21 5477     0.100000      1.00800 ; qtot -
0.229
  5478   HC   337   LEU  HD22 5478     0.100000      1.00800 ; qtot -
0.129
  5479   HC   337   LEU  HD23 5479     0.100000      1.00800 ; qtot -
0.029
  5480    C   337   LEU     C 5480     0.597301     12.01000 ; qtot
0.568
  5481    O   337   LEU     O 5481    -0.567901     16.00000 ; qtot
0.000
  5482    N   338   ASP     N 5482    -0.516301     14.01000 ; qtot -
0.516
  5483    H   338   ASP     H 5483     0.293600      1.00800 ; qtot -
0.223
```

5484 CX 338 ASP CA 5484 0.038100 12.01000 ; qtot -
0.185
5485 H1 338 ASP HA 5485 0.088000 1.00800 ; qtot -
0.097
5486 A2C 338 ASP CB 5486 -0.030300 12.01000 ; qtot -
0.127
5487 HC 338 ASP HB2 5487 -0.012200 1.00800 ; qtot -
0.139
5488 HC 338 ASP HB3 5488 -0.012200 1.00800 ; qtot -
0.151
5489 CO 338 ASP CG 5489 0.799401 12.01000 ; qtot
0.648
5490 O2 338 ASP OD1 5490 -0.801401 16.00000 ; qtot -
0.153
5491 O2 338 ASP OD2 5491 -0.801401 16.00000 ; qtot -
0.955
5492 C 338 ASP C 5492 0.536601 12.01000 ; qtot -
0.418
5493 O 338 ASP O 5493 -0.581901 16.00000 ; qtot -
1.000
5494 N 339 GLY N 5494 -0.415700 14.01000 ; qtot -
1.416
5495 H 339 GLY H 5495 0.271900 1.00800 ; qtot -
1.144
5496 CX 339 GLY CA 5496 -0.025200 12.01000 ; qtot -
1.169
5497 H1 339 GLY HA2 5497 0.069800 1.00800 ; qtot -
1.099
5498 H1 339 GLY HA3 5498 0.069800 1.00800 ; qtot -
1.029
5499 C 339 GLY C 5499 0.597301 12.01000 ; qtot -
0.432
5500 O 339 GLY O 5500 -0.567901 16.00000 ; qtot -
1.000
5501 N 340 LEU N 5501 -0.415700 14.01000 ; qtot -
1.416
5502 H 340 LEU H 5502 0.271900 1.00800 ; qtot -
1.144
5503 CX 340 LEU CA 5503 -0.051800 12.01000 ; qtot -
1.196
5504 H1 340 LEU HA 5504 0.092200 1.00800 ; qtot -
1.103
5505 A2C 340 LEU CB 5505 -0.110200 12.01000 ; qtot -
1.214
5506 HC 340 LEU HB2 5506 0.045700 1.00800 ; qtot -
1.168
5507 HC 340 LEU HB3 5507 0.045700 1.00800 ; qtot -
1.122
5508 A3C 340 LEU CG 5508 0.353100 12.01000 ; qtot -
0.769
5509 HC 340 LEU HG 5509 -0.036100 1.00800 ; qtot -
0.805

```
  5510   CT   340   LEU   CD1 5510    -0.412100     12.01000 ; qtot -
1.217
  5511   HC   340   LEU  HD11 5511     0.100000      1.00800 ; qtot -
1.117
  5512   HC   340   LEU  HD12 5512     0.100000      1.00800 ; qtot -
1.017
  5513   HC   340   LEU  HD13 5513     0.100000      1.00800 ; qtot -
0.917
  5514   CT   340   LEU   CD2 5514    -0.412100     12.01000 ; qtot -
1.329
  5515   HC   340   LEU  HD21 5515     0.100000      1.00800 ; qtot -
1.229
  5516   HC   340   LEU  HD22 5516     0.100000      1.00800 ; qtot -
1.129
  5517   HC   340   LEU  HD23 5517     0.100000      1.00800 ; qtot -
1.029
  5518    C   340   LEU     C 5518     0.597301     12.01000 ; qtot -
0.432
  5519    O   340   LEU     O 5519    -0.567901     16.00000 ; qtot -
1.000
  5520    N   341   LYS     N 5520    -0.347900     14.01000 ; qtot -
1.348
  5521    H   341   LYS     H 5521     0.274700      1.00800 ; qtot -
1.073
  5522   CX   341   LYS    CA 5522    -0.240000     12.01000 ; qtot -
1.313
  5523   H1   341   LYS    HA 5523     0.142600      1.00800 ; qtot -
1.171
  5524   C8   341   LYS    CB 5524    -0.009400     12.01000 ; qtot -
1.180
  5525   HC   341   LYS   HB2 5525     0.036200      1.00800 ; qtot -
1.144
  5526   HC   341   LYS   HB3 5526     0.036200      1.00800 ; qtot -
1.108
  5527   C8   341   LYS    CG 5527     0.018700     12.01000 ; qtot -
1.089
  5528   HC   341   LYS   HG2 5528     0.010300      1.00800 ; qtot -
1.079
  5529   HC   341   LYS   HG3 5529     0.010300      1.00800 ; qtot -
1.068
  5530   C8   341   LYS    CD 5530    -0.047900     12.01000 ; qtot -
1.116
  5531   HC   341   LYS   HD2 5531     0.062100      1.00800 ; qtot -
1.054
  5532   HC   341   LYS   HD3 5532     0.062100      1.00800 ; qtot -
0.992
  5533   C8   341   LYS    CE 5533    -0.014300     12.01000 ; qtot -
1.006
  5534   HP   341   LYS   HE2 5534     0.113500      1.00800 ; qtot -
0.893
  5535   HP   341   LYS   HE3 5535     0.113500      1.00800 ; qtot -
0.779
```

5536 N3 341 LYS NZ 5536 -0.385400 14.01000 ; qtot -
1.165
5537 H 341 LYS HZ1 5537 0.340000 1.00800 ; qtot -
0.825
5538 H 341 LYS HZ2 5538 0.340000 1.00800 ; qtot -
0.485
5539 H 341 LYS HZ3 5539 0.340000 1.00800 ; qtot -
0.145
5540 C 341 LYS C 5540 0.734101 12.01000 ; qtot
0.589
5541 O 341 LYS O 5541 -0.589401 16.00000 ; qtot
0.000
5542 N 342 LYS N 5542 -0.347900 14.01000 ; qtot -
0.348
5543 H 342 LYS H 5543 0.274700 1.00800 ; qtot -
0.073
5544 CX 342 LYS CA 5544 -0.240000 12.01000 ; qtot -
0.313
5545 H1 342 LYS HA 5545 0.142600 1.00800 ; qtot -
0.171
5546 C8 342 LYS CB 5546 -0.009400 12.01000 ; qtot -
0.180
5547 HC 342 LYS HB2 5547 0.036200 1.00800 ; qtot -
0.144
5548 HC 342 LYS HB3 5548 0.036200 1.00800 ; qtot -
0.108
5549 C8 342 LYS CG 5549 0.018700 12.01000 ; qtot -
0.089
5550 HC 342 LYS HG2 5550 0.010300 1.00800 ; qtot -
0.079
5551 HC 342 LYS HG3 5551 0.010300 1.00800 ; qtot -
0.068
5552 C8 342 LYS CD 5552 -0.047900 12.01000 ; qtot -
0.116
5553 HC 342 LYS HD2 5553 0.062100 1.00800 ; qtot -
0.054
5554 HC 342 LYS HD3 5554 0.062100 1.00800 ; qtot
0.008
5555 C8 342 LYS CE 5555 -0.014300 12.01000 ; qtot -
0.006
5556 HP 342 LYS HE2 5556 0.113500 1.00800 ; qtot
0.107
5557 HP 342 LYS HE3 5557 0.113500 1.00800 ; qtot
0.221
5558 N3 342 LYS NZ 5558 -0.385400 14.01000 ; qtot -
0.165
5559 H 342 LYS HZ1 5559 0.340000 1.00800 ; qtot
0.175
5560 H 342 LYS HZ2 5560 0.340000 1.00800 ; qtot
0.515
5561 H 342 LYS HZ3 5561 0.340000 1.00800 ; qtot
0.855

```
  5562    C   342   LYS     C 5562     0.734101     12.01000 ; qtot
1.589
  5563    O   342   LYS     O 5563    -0.589401     16.00000 ; qtot
1.000
  5564    N   343   ILE     N 5564    -0.415700     14.01000 ; qtot
0.584
  5565    H   343   ILE     H 5565     0.271900      1.00800 ; qtot
0.856
  5566   CX   343   ILE    CA 5566    -0.059700     12.01000 ; qtot
0.797
  5567   H1   343   ILE    HA 5567     0.086900      1.00800 ; qtot
0.883
  5568  A3C   343   ILE    CB 5568     0.130300     12.01000 ; qtot
1.014
  5569   HC   343   ILE    HB 5569     0.018700      1.00800 ; qtot
1.032
  5570   CT   343   ILE   CG2 5570    -0.320400     12.01000 ; qtot
0.712
  5571   HC   343   ILE  HG21 5571     0.088200      1.00800 ; qtot
0.800
  5572   HC   343   ILE  HG22 5572     0.088200      1.00800 ; qtot
0.888
  5573   HC   343   ILE  HG23 5573     0.088200      1.00800 ; qtot
0.977
  5574  A2C   343   ILE   CG1 5574    -0.043000     12.01000 ; qtot
0.934
  5575   HC   343   ILE  HG12 5575     0.023600      1.00800 ; qtot
0.957
  5576   HC   343   ILE  HG13 5576     0.023600      1.00800 ; qtot
0.981
  5577   CT   343   ILE   CD1 5577    -0.066000     12.01000 ; qtot
0.915
  5578   HC   343   ILE  HD11 5578     0.018600      1.00800 ; qtot
0.933
  5579   HC   343   ILE  HD12 5579     0.018600      1.00800 ; qtot
0.952
  5580   HC   343   ILE  HD13 5580     0.018600      1.00800 ; qtot
0.971
  5581    C   343   ILE     C 5581     0.597301     12.01000 ; qtot
1.568
  5582    O   343   ILE     O 5582    -0.567901     16.00000 ; qtot
1.000
  5583    N   344   GLY     N 5583    -0.415700     14.01000 ; qtot
0.584
  5584    H   344   GLY     H 5584     0.271900      1.00800 ; qtot
0.856
  5585   CX   344   GLY    CA 5585    -0.025200     12.01000 ; qtot
0.831
  5586   H1   344   GLY   HA2 5586     0.069800      1.00800 ; qtot
0.901
  5587   H1   344   GLY   HA3 5587     0.069800      1.00800 ; qtot
0.971
```

5588 C 344 GLY C 5588 0.597301 12.01000 ; qtot
1.568
5589 O 344 GLY O 5589 -0.567901 16.00000 ; qtot
1.000
5590 N 345 TYR N 5590 -0.415700 14.01000 ; qtot
0.584
5591 H 345 TYR H 5591 0.271900 1.00800 ; qtot
0.856
5592 CX 345 TYR CA 5592 -0.001400 12.01000 ; qtot
0.855
5593 H1 345 TYR HA 5593 0.087600 1.00800 ; qtot
0.942
5594 CT 345 TYR CB 5594 -0.015200 12.01000 ; qtot
0.927
5595 HC 345 TYR HB2 5595 0.029500 1.00800 ; qtot
0.957
5596 HC 345 TYR HB3 5596 0.029500 1.00800 ; qtot
0.986
5597 CA 345 TYR CG 5597 -0.001100 12.01000 ; qtot
0.985
5598 CA 345 TYR CD1 5598 -0.190600 12.01000 ; qtot
0.795
5599 HA 345 TYR HD1 5599 0.169900 1.00800 ; qtot
0.964
5600 CA 345 TYR CE1 5600 -0.234100 12.01000 ; qtot
0.730
5601 HA 345 TYR HE1 5601 0.165600 1.00800 ; qtot
0.896
5602 C 345 TYR CZ 5602 0.322600 12.01000 ; qtot
1.219
5603 OH 345 TYR OH 5603 -0.557901 16.00000 ; qtot
0.661
5604 HO 345 TYR HH 5604 0.399200 1.00800 ; qtot
1.060
5605 CA 345 TYR CE2 5605 -0.234100 12.01000 ; qtot
0.826
5606 HA 345 TYR HE2 5606 0.165600 1.00800 ; qtot
0.991
5607 CA 345 TYR CD2 5607 -0.190600 12.01000 ; qtot
0.801
5608 HA 345 TYR HD2 5608 0.169900 1.00800 ; qtot
0.971
5609 C 345 TYR C 5609 0.597301 12.01000 ; qtot
1.568
5610 O 345 TYR O 5610 -0.567901 16.00000 ; qtot
1.000
5611 N 346 GLN N 5611 -0.415700 14.01000 ; qtot
0.584
5612 H 346 GLN H 5612 0.271900 1.00800 ; qtot
0.856
5613 CX 346 GLN CA 5613 -0.003100 12.01000 ; qtot
0.853

5614 H1 346 GLN HA 5614 0.085000 1.00800 ; qtot
0.938
5615 A2C 346 GLN CB 5615 -0.003600 12.01000 ; qtot
0.935
5616 HC 346 GLN HB2 5616 0.017100 1.00800 ; qtot
0.952
5617 HC 346 GLN HB3 5617 0.017100 1.00800 ; qtot
0.969
5618 A2C 346 GLN CG 5618 -0.064500 12.01000 ; qtot
0.904
5619 HC 346 GLN HG2 5619 0.035200 1.00800 ; qtot
0.939
5620 HC 346 GLN HG3 5620 0.035200 1.00800 ; qtot
0.975
5621 C 346 GLN CD 5621 0.695101 12.01000 ; qtot
1.670
5622 O 346 GLN OE1 5622 -0.608601 16.00000 ; qtot
1.061
5623 N 346 GLN NE2 5623 -0.940701 14.01000 ; qtot
0.120
5624 H 346 GLN HE21 5624 0.425100 1.00800 ; qtot
0.546
5625 H 346 GLN HE22 5625 0.425100 1.00800 ; qtot
0.971
5626 C 346 GLN C 5626 0.597301 12.01000 ; qtot
1.568
5627 O 346 GLN O 5627 -0.567901 16.00000 ; qtot
1.000
5628 N 347 LYS N 5628 -0.347900 14.01000 ; qtot
0.652
5629 H 347 LYS H 5629 0.274700 1.00800 ; qtot
0.927
5630 CX 347 LYS CA 5630 -0.240000 12.01000 ; qtot
0.687
5631 H1 347 LYS HA 5631 0.142600 1.00800 ; qtot
0.829
5632 C8 347 LYS CB 5632 -0.009400 12.01000 ; qtot
0.820
5633 HC 347 LYS HB2 5633 0.036200 1.00800 ; qtot
0.856
5634 HC 347 LYS HB3 5634 0.036200 1.00800 ; qtot
0.892
5635 C8 347 LYS CG 5635 0.018700 12.01000 ; qtot
0.911
5636 HC 347 LYS HG2 5636 0.010300 1.00800 ; qtot
0.921
5637 HC 347 LYS HG3 5637 0.010300 1.00800 ; qtot
0.932
5638 C8 347 LYS CD 5638 -0.047900 12.01000 ; qtot
0.884
5639 HC 347 LYS HD2 5639 0.062100 1.00800 ; qtot
0.946

5640 HC 347 LYS HD3 5640 0.062100 1.00800 ; qtot
1.008
5641 C8 347 LYS CE 5641 -0.014300 12.01000 ; qtot
0.994
5642 HP 347 LYS HE2 5642 0.113500 1.00800 ; qtot
1.107
5643 HP 347 LYS HE3 5643 0.113500 1.00800 ; qtot
1.221
5644 N3 347 LYS NZ 5644 -0.385400 14.01000 ; qtot
0.835
5645 H 347 LYS HZ1 5645 0.340000 1.00800 ; qtot
1.175
5646 H 347 LYS HZ2 5646 0.340000 1.00800 ; qtot
1.515
5647 H 347 LYS HZ3 5647 0.340000 1.00800 ; qtot
1.855
5648 C 347 LYS C 5648 0.734101 12.01000 ; qtot
2.589
5649 O 347 LYS O 5649 -0.589401 16.00000 ; qtot
2.000
5650 N 348 ILE N 5650 -0.415700 14.01000 ; qtot
1.584
5651 H 348 ILE H 5651 0.271900 1.00800 ; qtot
1.856
5652 CX 348 ILE CA 5652 -0.059700 12.01000 ; qtot
1.797
5653 H1 348 ILE HA 5653 0.086900 1.00800 ; qtot
1.883
5654 A3C 348 ILE CB 5654 0.130300 12.01000 ; qtot
2.014
5655 HC 348 ILE HB 5655 0.018700 1.00800 ; qtot
2.032
5656 CT 348 ILE CG2 5656 -0.320400 12.01000 ; qtot
1.712
5657 HC 348 ILE HG21 5657 0.088200 1.00800 ; qtot
1.800
5658 HC 348 ILE HG22 5658 0.088200 1.00800 ; qtot
1.888
5659 HC 348 ILE HG23 5659 0.088200 1.00800 ; qtot
1.977
5660 A2C 348 ILE CG1 5660 -0.043000 12.01000 ; qtot
1.934
5661 HC 348 ILE HG12 5661 0.023600 1.00800 ; qtot
1.957
5662 HC 348 ILE HG13 5662 0.023600 1.00800 ; qtot
1.981
5663 CT 348 ILE CD1 5663 -0.066000 12.01000 ; qtot
1.915
5664 HC 348 ILE HD11 5664 0.018600 1.00800 ; qtot
1.933
5665 HC 348 ILE HD12 5665 0.018600 1.00800 ; qtot
1.952

```
  5666   HC   348   ILE  HD13 5666     0.018600       1.00800 ; qtot
1.971
  5667    C   348   ILE     C 5667     0.597301      12.01000 ; qtot
2.568
  5668    O   348   ILE     O 5668    -0.567901      16.00000 ; qtot
2.000
  5669    N   349   PRO     N 5669    -0.254800      14.01000 ; qtot
1.745
  5670   CT   349   PRO    CD 5670     0.019200      12.01000 ; qtot
1.764
  5671   H1   349   PRO   HD2 5671     0.039100       1.00800 ; qtot
1.804
  5672   H1   349   PRO   HD3 5672     0.039100       1.00800 ; qtot
1.843
  5673   CT   349   PRO    CG 5673     0.018900      12.01000 ; qtot
1.862
  5674   HC   349   PRO   HG2 5674     0.021300       1.00800 ; qtot
1.883
  5675   HC   349   PRO   HG3 5675     0.021300       1.00800 ; qtot
1.904
  5676   CT   349   PRO    CB 5676    -0.007000      12.01000 ; qtot
1.897
  5677   HC   349   PRO   HB2 5677     0.025300       1.00800 ; qtot
1.922
  5678   HC   349   PRO   HB3 5678     0.025300       1.00800 ; qtot
1.948
  5679   CX   349   PRO    CA 5679    -0.026600      12.01000 ; qtot
1.921
  5680   H1   349   PRO    HA 5680     0.064100       1.00800 ; qtot
1.985
  5681    C   349   PRO     C 5681     0.589601      12.01000 ; qtot
2.575
  5682    O   349   PRO     O 5682    -0.574801      16.00000 ; qtot
2.000
  5683    N   350   PRO     N 5683    -0.254800      14.01000 ; qtot
1.745
  5684   CT   350   PRO    CD 5684     0.019200      12.01000 ; qtot
1.764
  5685   H1   350   PRO   HD2 5685     0.039100       1.00800 ; qtot
1.804
  5686   H1   350   PRO   HD3 5686     0.039100       1.00800 ; qtot
1.843
  5687   CT   350   PRO    CG 5687     0.018900      12.01000 ; qtot
1.862
  5688   HC   350   PRO   HG2 5688     0.021300       1.00800 ; qtot
1.883
  5689   HC   350   PRO   HG3 5689     0.021300       1.00800 ; qtot
1.904
  5690   CT   350   PRO    CB 5690    -0.007000      12.01000 ; qtot
1.897
  5691   HC   350   PRO   HB2 5691     0.025300       1.00800 ; qtot
1.922
```

5692 HC 350 PRO HB3 5692 0.025300 1.00800 ; qtot
1.948
5693 CX 350 PRO CA 5693 -0.026600 12.01000 ; qtot
1.921
5694 H1 350 PRO HA 5694 0.064100 1.00800 ; qtot
1.985
5695 C 350 PRO C 5695 0.589601 12.01000 ; qtot
2.575
5696 O 350 PRO O 5696 -0.574801 16.00000 ; qtot
2.000
5697 N 351 ASP N 5697 -0.516301 14.01000 ; qtot
1.484
5698 H 351 ASP H 5698 0.293600 1.00800 ; qtot
1.777
5699 CX 351 ASP CA 5699 0.038100 12.01000 ; qtot
1.815
5700 H1 351 ASP HA 5700 0.088000 1.00800 ; qtot
1.903
5701 A2C 351 ASP CB 5701 -0.030300 12.01000 ; qtot
1.873
5702 HC 351 ASP HB2 5702 -0.012200 1.00800 ; qtot
1.861
5703 HC 351 ASP HB3 5703 -0.012200 1.00800 ; qtot
1.849
5704 CO 351 ASP CG 5704 0.799401 12.01000 ; qtot
2.648
5705 O2 351 ASP OD1 5705 -0.801401 16.00000 ; qtot
1.847
5706 O2 351 ASP OD2 5706 -0.801401 16.00000 ; qtot
1.045
5707 C 351 ASP C 5707 0.536601 12.01000 ; qtot
1.582
5708 O 351 ASP O 5708 -0.581901 16.00000 ; qtot
1.000
5709 N 352 ILE N 5709 -0.415700 14.01000 ; qtot
0.584
5710 H 352 ILE H 5710 0.271900 1.00800 ; qtot
0.856
5711 CX 352 ILE CA 5711 -0.059700 12.01000 ; qtot
0.797
5712 H1 352 ILE HA 5712 0.086900 1.00800 ; qtot
0.883
5713 A3C 352 ILE CB 5713 0.130300 12.01000 ; qtot
1.014
5714 HC 352 ILE HB 5714 0.018700 1.00800 ; qtot
1.032
5715 CT 352 ILE CG2 5715 -0.320400 12.01000 ; qtot
0.712
5716 HC 352 ILE HG21 5716 0.088200 1.00800 ; qtot
0.800
5717 HC 352 ILE HG22 5717 0.088200 1.00800 ; qtot
0.888

5718 HC 352 ILE HG23 5718 0.088200 1.00800 ; qtot
0.977
5719 A2C 352 ILE CG1 5719 -0.043000 12.01000 ; qtot
0.934
5720 HC 352 ILE HG12 5720 0.023600 1.00800 ; qtot
0.957
5721 HC 352 ILE HG13 5721 0.023600 1.00800 ; qtot
0.981
5722 CT 352 ILE CD1 5722 -0.066000 12.01000 ; qtot
0.915
5723 HC 352 ILE HD11 5723 0.018600 1.00800 ; qtot
0.933
5724 HC 352 ILE HD12 5724 0.018600 1.00800 ; qtot
0.952
5725 HC 352 ILE HD13 5725 0.018600 1.00800 ; qtot
0.971
5726 C 352 ILE C 5726 0.597301 12.01000 ; qtot
1.568
5727 O 352 ILE O 5727 -0.567901 16.00000 ; qtot
1.000
5728 N 353 HIE N 5728 -0.415700 14.01000 ; qtot
0.584
5729 H 353 HIE H 5729 0.271900 1.00800 ; qtot
0.856
5730 CX 353 HIE CA 5730 -0.058100 12.01000 ; qtot
0.798
5731 H1 353 HIE HA 5731 0.136000 1.00800 ; qtot
0.934
5732 CT 353 HIE CB 5732 -0.007400 12.01000 ; qtot
0.927
5733 HC 353 HIE HB2 5733 0.036700 1.00800 ; qtot
0.963
5734 HC 353 HIE HB3 5734 0.036700 1.00800 ; qtot
1.000
5735 CC 353 HIE CG 5735 0.186800 12.01000 ; qtot
1.187
5736 NB 353 HIE ND1 5736 -0.543201 14.01000 ; qtot
0.644
5737 CR 353 HIE CE1 5737 0.163500 12.01000 ; qtot
0.807
5738 H5 353 HIE HE1 5738 0.143500 1.00800 ; qtot
0.951
5739 NA 353 HIE NE2 5739 -0.279500 14.01000 ; qtot
0.671
5740 H 353 HIE HE2 5740 0.333900 1.00800 ; qtot
1.005
5741 CW 353 HIE CD2 5741 -0.220700 12.01000 ; qtot
0.784
5742 H4 353 HIE HD2 5742 0.186200 1.00800 ; qtot
0.971
5743 C 353 HIE C 5743 0.597301 12.01000 ; qtot
1.568

5744 O 353 HIE O 5744 -0.567901 16.00000 ; qtot
1.000
5745 N 354 GLN N 5745 -0.415700 14.01000 ; qtot
0.584
5746 H 354 GLN H 5746 0.271900 1.00800 ; qtot
0.856
5747 CX 354 GLN CA 5747 -0.003100 12.01000 ; qtot
0.853
5748 H1 354 GLN HA 5748 0.085000 1.00800 ; qtot
0.938
5749 A2C 354 GLN CB 5749 -0.003600 12.01000 ; qtot
0.935
5750 HC 354 GLN HB2 5750 0.017100 1.00800 ; qtot
0.952
5751 HC 354 GLN HB3 5751 0.017100 1.00800 ; qtot
0.969
5752 A2C 354 GLN CG 5752 -0.064500 12.01000 ; qtot
0.904
5753 HC 354 GLN HG2 5753 0.035200 1.00800 ; qtot
0.939
5754 HC 354 GLN HG3 5754 0.035200 1.00800 ; qtot
0.975
5755 C 354 GLN CD 5755 0.695101 12.01000 ; qtot
1.670
5756 O 354 GLN OE1 5756 -0.608601 16.00000 ; qtot
1.061
5757 N 354 GLN NE2 5757 -0.940701 14.01000 ; qtot
0.120
5758 H 354 GLN HE21 5758 0.425100 1.00800 ; qtot
0.546
5759 H 354 GLN HE22 5759 0.425100 1.00800 ; qtot
0.971
5760 C 354 GLN C 5760 0.597301 12.01000 ; qtot
1.568
5761 O 354 GLN O 5761 -0.567901 16.00000 ; qtot
1.000
5762 N 355 VAL N 5762 -0.415700 14.01000 ; qtot
0.584
5763 H 355 VAL H 5763 0.271900 1.00800 ; qtot
0.856
5764 CX 355 VAL CA 5764 -0.087500 12.01000 ; qtot
0.769
5765 H1 355 VAL HA 5765 0.096900 1.00800 ; qtot
0.866
5766 A3C 355 VAL CB 5766 0.298500 12.01000 ; qtot
1.164
5767 HC 355 VAL HB 5767 -0.029700 1.00800 ; qtot
1.134
5768 CT 355 VAL CG1 5768 -0.319200 12.01000 ; qtot
0.815
5769 HC 355 VAL HG11 5769 0.079100 1.00800 ; qtot
0.894

5770 HC 355 VAL HG12 5770 0.079100 1.00800 ; qtot
0.973
5771 HC 355 VAL HG13 5771 0.079100 1.00800 ; qtot
1.053
5772 CT 355 VAL CG2 5772 -0.319200 12.01000 ; qtot
0.733
5773 HC 355 VAL HG21 5773 0.079100 1.00800 ; qtot
0.812
5774 HC 355 VAL HG22 5774 0.079100 1.00800 ; qtot
0.892
5775 HC 355 VAL HG23 5775 0.079100 1.00800 ; qtot
0.971
5776 C 355 VAL C 5776 0.597301 12.01000 ; qtot
1.568
5777 O 355 VAL O 5777 -0.567901 16.00000 ; qtot
1.000
5778 N 356 LEU N 5778 -0.415700 14.01000 ; qtot
0.584
5779 H 356 LEU H 5779 0.271900 1.00800 ; qtot
0.856
5780 CX 356 LEU CA 5780 -0.051800 12.01000 ; qtot
0.804
5781 H1 356 LEU HA 5781 0.092200 1.00800 ; qtot
0.897
5782 A2C 356 LEU CB 5782 -0.110200 12.01000 ; qtot
0.786
5783 HC 356 LEU HB2 5783 0.045700 1.00800 ; qtot
0.832
5784 HC 356 LEU HB3 5784 0.045700 1.00800 ; qtot
0.878
5785 A3C 356 LEU CG 5785 0.353100 12.01000 ; qtot
1.231
5786 HC 356 LEU HG 5786 -0.036100 1.00800 ; qtot
1.195
5787 CT 356 LEU CD1 5787 -0.412100 12.01000 ; qtot
0.783
5788 HC 356 LEU HD11 5788 0.100000 1.00800 ; qtot
0.883
5789 HC 356 LEU HD12 5789 0.100000 1.00800 ; qtot
0.983
5790 HC 356 LEU HD13 5790 0.100000 1.00800 ; qtot
1.083
5791 CT 356 LEU CD2 5791 -0.412100 12.01000 ; qtot
0.671
5792 HC 356 LEU HD21 5792 0.100000 1.00800 ; qtot
0.771
5793 HC 356 LEU HD22 5793 0.100000 1.00800 ; qtot
0.871
5794 HC 356 LEU HD23 5794 0.100000 1.00800 ; qtot
0.971
5795 C 356 LEU C 5795 0.597301 12.01000 ; qtot
1.568

```
  5796    O   356   LEU     O 5796    -0.567901     16.00000 ; qtot
1.000
  5797    N   357   ARG     N 5797    -0.347900     14.01000 ; qtot
0.652
  5798    H   357   ARG     H 5798     0.274700      1.00800 ; qtot
0.927
  5799   CX   357   ARG    CA 5799    -0.263700     12.01000 ; qtot
0.663
  5800   H1   357   ARG    HA 5800     0.156000      1.00800 ; qtot
0.819
  5801   C8   357   ARG    CB 5801    -0.000700     12.01000 ; qtot
0.818
  5802   HC   357   ARG   HB2 5802     0.032700      1.00800 ; qtot
0.851
  5803   HC   357   ARG   HB3 5803     0.032700      1.00800 ; qtot
0.884
  5804   C8   357   ARG    CG 5804     0.039000     12.01000 ; qtot
0.923
  5805   HC   357   ARG   HG2 5805     0.028500      1.00800 ; qtot
0.951
  5806   HC   357   ARG   HG3 5806     0.028500      1.00800 ; qtot
0.980
  5807   C8   357   ARG    CD 5807     0.048600     12.01000 ; qtot
1.028
  5808   H1   357   ARG   HD2 5808     0.068700      1.00800 ; qtot
1.097
  5809   H1   357   ARG   HD3 5809     0.068700      1.00800 ; qtot
1.166
  5810   N2   357   ARG    NE 5810    -0.529501     14.01000 ; qtot
0.636
  5811    H   357   ARG    HE 5811     0.345600      1.00800 ; qtot
0.982
  5812   CA   357   ARG    CZ 5812     0.807601     12.01000 ; qtot
1.790
  5813   N2   357   ARG   NH1 5813    -0.862701     14.01000 ; qtot
0.927
  5814    H   357   ARG  HH11 5814     0.447800      1.00800 ; qtot
1.375
  5815    H   357   ARG  HH12 5815     0.447800      1.00800 ; qtot
1.822
  5816   N2   357   ARG   NH2 5816    -0.862701     14.01000 ; qtot
0.960
  5817    H   357   ARG  HH21 5817     0.447800      1.00800 ; qtot
1.408
  5818    H   357   ARG  HH22 5818     0.447800      1.00800 ; qtot
1.855
  5819    C   357   ARG     C 5819     0.734101     12.01000 ; qtot
2.589
  5820    O   357   ARG     O 5820    -0.589401     16.00000 ; qtot
2.000
  5821    N   358   ASP     N 5821    -0.516301     14.01000 ; qtot
1.484
```

5822 H 358 ASP H 5822 0.293600 1.00800 ; qtot
1.777
5823 CX 358 ASP CA 5823 0.038100 12.01000 ; qtot
1.815
5824 H1 358 ASP HA 5824 0.088000 1.00800 ; qtot
1.903
5825 A2C 358 ASP CB 5825 -0.030300 12.01000 ; qtot
1.873
5826 HC 358 ASP HB2 5826 -0.012200 1.00800 ; qtot
1.861
5827 HC 358 ASP HB3 5827 -0.012200 1.00800 ; qtot
1.849
5828 CO 358 ASP CG 5828 0.799401 12.01000 ; qtot
2.648
5829 O2 358 ASP OD1 5829 -0.801401 16.00000 ; qtot
1.847
5830 O2 358 ASP OD2 5830 -0.801401 16.00000 ; qtot
1.045
5831 C 358 ASP C 5831 0.536601 12.01000 ; qtot
1.582
5832 O 358 ASP O 5832 -0.581901 16.00000 ; qtot
1.000
5833 N 359 ILE N 5833 -0.415700 14.01000 ; qtot
0.584
5834 H 359 ILE H 5834 0.271900 1.00800 ; qtot
0.856
5835 CX 359 ILE CA 5835 -0.059700 12.01000 ; qtot
0.797
5836 H1 359 ILE HA 5836 0.086900 1.00800 ; qtot
0.883
5837 A3C 359 ILE CB 5837 0.130300 12.01000 ; qtot
1.014
5838 HC 359 ILE HB 5838 0.018700 1.00800 ; qtot
1.032
5839 CT 359 ILE CG2 5839 -0.320400 12.01000 ; qtot
0.712
5840 HC 359 ILE HG21 5840 0.088200 1.00800 ; qtot
0.800
5841 HC 359 ILE HG22 5841 0.088200 1.00800 ; qtot
0.888
5842 HC 359 ILE HG23 5842 0.088200 1.00800 ; qtot
0.977
5843 A2C 359 ILE CG1 5843 -0.043000 12.01000 ; qtot
0.934
5844 HC 359 ILE HG12 5844 0.023600 1.00800 ; qtot
0.957
5845 HC 359 ILE HG13 5845 0.023600 1.00800 ; qtot
0.981
5846 CT 359 ILE CD1 5846 -0.066000 12.01000 ; qtot
0.915
5847 HC 359 ILE HD11 5847 0.018600 1.00800 ; qtot
0.933

5848 HC 359 ILE HD12 5848 0.018600 1.00800 ; qtot
0.952
5849 HC 359 ILE HD13 5849 0.018600 1.00800 ; qtot
0.971
5850 C 359 ILE C 5850 0.597301 12.01000 ; qtot
1.568
5851 O 359 ILE O 5851 -0.567901 16.00000 ; qtot
1.000
5852 N 360 ASP N 5852 -0.516301 14.01000 ; qtot
0.484
5853 H 360 ASP H 5853 0.293600 1.00800 ; qtot
0.777
5854 CX 360 ASP CA 5854 0.038100 12.01000 ; qtot
0.815
5855 H1 360 ASP HA 5855 0.088000 1.00800 ; qtot
0.903
5856 A2C 360 ASP CB 5856 -0.030300 12.01000 ; qtot
0.873
5857 HC 360 ASP HB2 5857 -0.012200 1.00800 ; qtot
0.861
5858 HC 360 ASP HB3 5858 -0.012200 1.00800 ; qtot
0.849
5859 CO 360 ASP CG 5859 0.799401 12.01000 ; qtot
1.648
5860 O2 360 ASP OD1 5860 -0.801401 16.00000 ; qtot
0.847
5861 O2 360 ASP OD2 5861 -0.801401 16.00000 ; qtot
0.045
5862 C 360 ASP C 5862 0.536601 12.01000 ; qtot
0.582
5863 O 360 ASP O 5863 -0.581901 16.00000 ; qtot
0.000
5864 N 361 SER N 5864 -0.415700 14.01000 ; qtot -
0.416
5865 H 361 SER H 5865 0.271900 1.00800 ; qtot -
0.144
5866 CX 361 SER CA 5866 -0.024900 12.01000 ; qtot -
0.169
5867 H1 361 SER HA 5867 0.084300 1.00800 ; qtot -
0.084
5868 A2C 361 SER CB 5868 0.211700 12.01000 ; qtot
0.127
5869 H1 361 SER HB2 5869 0.035200 1.00800 ; qtot
0.163
5870 H1 361 SER HB3 5870 0.035200 1.00800 ; qtot
0.198
5871 OH 361 SER OG 5871 -0.654601 16.00000 ; qtot -
0.457
5872 HO 361 SER HG 5872 0.427500 1.00800 ; qtot -
0.029
5873 C 361 SER C 5873 0.597301 12.01000 ; qtot
0.568

5874 O 361 SER O 5874 -0.567901 16.00000 ; qtot
0.000
5875 N 362 ASN N 5875 -0.415700 14.01000 ; qtot -
0.416
5876 H 362 ASN H 5876 0.271900 1.00800 ; qtot -
0.144
5877 CX 362 ASN CA 5877 0.014300 12.01000 ; qtot -
0.129
5878 H1 362 ASN HA 5878 0.104800 1.00800 ; qtot -
0.025
5879 A2C 362 ASN CB 5879 -0.204100 12.01000 ; qtot -
0.229
5880 HC 362 ASN HB2 5880 0.079700 1.00800 ; qtot -
0.149
5881 HC 362 ASN HB3 5881 0.079700 1.00800 ; qtot -
0.069
5882 C 362 ASN CG 5882 0.713001 12.01000 ; qtot
0.644
5883 O 362 ASN OD1 5883 -0.593101 16.00000 ; qtot
0.051
5884 N 362 ASN ND2 5884 -0.919101 14.01000 ; qtot -
0.869
5885 H 362 ASN HD21 5885 0.419600 1.00800 ; qtot -
0.449
5886 H 362 ASN HD22 5886 0.419600 1.00800 ; qtot -
0.029
5887 C 362 ASN C 5887 0.597301 12.01000 ; qtot
0.568
5888 O 362 ASN O 5888 -0.567901 16.00000 ; qtot
0.000
5889 N 363 ALA N 5889 -0.415700 14.01000 ; qtot -
0.416
5890 H 363 ALA H 5890 0.271900 1.00800 ; qtot -
0.144
5891 CX 363 ALA CA 5891 0.033700 12.01000 ; qtot -
0.110
5892 H1 363 ALA HA 5892 0.082300 1.00800 ; qtot -
0.028
5893 CT 363 ALA CB 5893 -0.182500 12.01000 ; qtot -
0.210
5894 HC 363 ALA HB1 5894 0.060300 1.00800 ; qtot -
0.150
5895 HC 363 ALA HB2 5895 0.060300 1.00800 ; qtot -
0.090
5896 HC 363 ALA HB3 5896 0.060300 1.00800 ; qtot -
0.029
5897 C 363 ALA C 5897 0.597301 12.01000 ; qtot
0.568
5898 O 363 ALA O 5898 -0.567901 16.00000 ; qtot
0.000
5899 N 364 SER N 5899 -0.415700 14.01000 ; qtot -
0.416

5900 H 364 SER H 5900 0.271900 1.00800 ; qtot -
0.144
5901 CX 364 SER CA 5901 -0.024900 12.01000 ; qtot -
0.169
5902 H1 364 SER HA 5902 0.084300 1.00800 ; qtot -
0.084
5903 A2C 364 SER CB 5903 0.211700 12.01000 ; qtot
0.127
5904 H1 364 SER HB2 5904 0.035200 1.00800 ; qtot
0.163
5905 H1 364 SER HB3 5905 0.035200 1.00800 ; qtot
0.198
5906 OH 364 SER OG 5906 -0.654601 16.00000 ; qtot -
0.457
5907 HO 364 SER HG 5907 0.427500 1.00800 ; qtot -
0.029
5908 C 364 SER C 5908 0.597301 12.01000 ; qtot
0.568
5909 O 364 SER O 5909 -0.567901 16.00000 ; qtot
0.000
5910 N 365 GLY N 5910 -0.415700 14.01000 ; qtot -
0.416
5911 H 365 GLY H 5911 0.271900 1.00800 ; qtot -
0.144
5912 CX 365 GLY CA 5912 -0.025200 12.01000 ; qtot -
0.169
5913 H1 365 GLY HA2 5913 0.069800 1.00800 ; qtot -
0.099
5914 H1 365 GLY HA3 5914 0.069800 1.00800 ; qtot -
0.029
5915 C 365 GLY C 5915 0.597301 12.01000 ; qtot
0.568
5916 O 365 GLY O 5916 -0.567901 16.00000 ; qtot
0.000
5917 N 366 GLN N 5917 -0.415700 14.01000 ; qtot -
0.416
5918 H 366 GLN H 5918 0.271900 1.00800 ; qtot -
0.144
5919 CX 366 GLN CA 5919 -0.003100 12.01000 ; qtot -
0.147
5920 H1 366 GLN HA 5920 0.085000 1.00800 ; qtot -
0.062
5921 A2C 366 GLN CB 5921 -0.003600 12.01000 ; qtot -
0.065
5922 HC 366 GLN HB2 5922 0.017100 1.00800 ; qtot -
0.048
5923 HC 366 GLN HB3 5923 0.017100 1.00800 ; qtot -
0.031
5924 A2C 366 GLN CG 5924 -0.064500 12.01000 ; qtot -
0.096
5925 HC 366 GLN HG2 5925 0.035200 1.00800 ; qtot -
0.061

5926 HC 366 GLN HG3 5926 0.035200 1.00800 ; qtot -
0.025
5927 C 366 GLN CD 5927 0.695101 12.01000 ; qtot
0.670
5928 O 366 GLN OE1 5928 -0.608601 16.00000 ; qtot
0.061
5929 N 366 GLN NE2 5929 -0.940701 14.01000 ; qtot -
0.880
5930 H 366 GLN HE21 5930 0.425100 1.00800 ; qtot -
0.454
5931 H 366 GLN HE22 5931 0.425100 1.00800 ; qtot -
0.029
5932 C 366 GLN C 5932 0.597301 12.01000 ; qtot
0.568
5933 O 366 GLN O 5933 -0.567901 16.00000 ; qtot
0.000
5934 N 367 ILE N 5934 -0.415700 14.01000 ; qtot -
0.416
5935 H 367 ILE H 5935 0.271900 1.00800 ; qtot -
0.144
5936 CX 367 ILE CA 5936 -0.059700 12.01000 ; qtot -
0.203
5937 H1 367 ILE HA 5937 0.086900 1.00800 ; qtot -
0.117
5938 A3C 367 ILE CB 5938 0.130300 12.01000 ; qtot
0.014
5939 HC 367 ILE HB 5939 0.018700 1.00800 ; qtot
0.032
5940 CT 367 ILE CG2 5940 -0.320400 12.01000 ; qtot -
0.288
5941 HC 367 ILE HG21 5941 0.088200 1.00800 ; qtot -
0.200
5942 HC 367 ILE HG22 5942 0.088200 1.00800 ; qtot -
0.112
5943 HC 367 ILE HG23 5943 0.088200 1.00800 ; qtot -
0.023
5944 A2C 367 ILE CG1 5944 -0.043000 12.01000 ; qtot -
0.066
5945 HC 367 ILE HG12 5945 0.023600 1.00800 ; qtot -
0.043
5946 HC 367 ILE HG13 5946 0.023600 1.00800 ; qtot -
0.019
5947 CT 367 ILE CD1 5947 -0.066000 12.01000 ; qtot -
0.085
5948 HC 367 ILE HD11 5948 0.018600 1.00800 ; qtot -
0.067
5949 HC 367 ILE HD12 5949 0.018600 1.00800 ; qtot -
0.048
5950 HC 367 ILE HD13 5950 0.018600 1.00800 ; qtot -
0.029
5951 C 367 ILE C 5951 0.597301 12.01000 ; qtot
0.568

5952 O 367 ILE O 5952 -0.567901 16.00000 ; qtot
0.000
5953 N 368 HIP N 5953 -0.347900 14.01000 ; qtot -
0.348
5954 H 368 HIP H 5954 0.274700 1.00800 ; qtot -
0.073
5955 CX 368 HIP CA 5955 -0.135400 12.01000 ; qtot -
0.209
5956 H1 368 HIP HA 5956 0.121200 1.00800 ; qtot -
0.087
5957 CT 368 HIP CB 5957 -0.041400 12.01000 ; qtot -
0.129
5958 HC 368 HIP HB2 5958 0.081000 1.00800 ; qtot -
0.048
5959 HC 368 HIP HB3 5959 0.081000 1.00800 ; qtot
0.033
5960 CC 368 HIP CG 5960 -0.001200 12.01000 ; qtot
0.032
5961 NA 368 HIP ND1 5961 -0.151300 14.01000 ; qtot -
0.119
5962 H 368 HIP HD1 5962 0.386600 1.00800 ; qtot
0.267
5963 CR 368 HIP CE1 5963 -0.017000 12.01000 ; qtot
0.250
5964 H5 368 HIP HE1 5964 0.268100 1.00800 ; qtot
0.518
5965 NA 368 HIP NE2 5965 -0.171800 14.01000 ; qtot
0.347
5966 H 368 HIP HE2 5966 0.391100 1.00800 ; qtot
0.738
5967 CW 368 HIP CD2 5967 -0.114100 12.01000 ; qtot
0.624
5968 H4 368 HIP HD2 5968 0.231700 1.00800 ; qtot
0.855
5969 C 368 HIP C 5969 0.734101 12.01000 ; qtot
1.589
5970 O 368 HIP O 5970 -0.589401 16.00000 ; qtot
1.000
5971 N 369 TYR N 5971 -0.415700 14.01000 ; qtot
0.584
5972 H 369 TYR H 5972 0.271900 1.00800 ; qtot
0.856
5973 CX 369 TYR CA 5973 -0.001400 12.01000 ; qtot
0.855
5974 H1 369 TYR HA 5974 0.087600 1.00800 ; qtot
0.942
5975 CT 369 TYR CB 5975 -0.015200 12.01000 ; qtot
0.927
5976 HC 369 TYR HB2 5976 0.029500 1.00800 ; qtot
0.957
5977 HC 369 TYR HB3 5977 0.029500 1.00800 ; qtot
0.986

5978 CA 369 TYR CG 5978 -0.001100 12.01000 ; qtot
0.985
5979 CA 369 TYR CD1 5979 -0.190600 12.01000 ; qtot
0.795
5980 HA 369 TYR HD1 5980 0.169900 1.00800 ; qtot
0.964
5981 CA 369 TYR CE1 5981 -0.234100 12.01000 ; qtot
0.730
5982 HA 369 TYR HE1 5982 0.165600 1.00800 ; qtot
0.896
5983 C 369 TYR CZ 5983 0.322600 12.01000 ; qtot
1.219
5984 OH 369 TYR OH 5984 -0.557901 16.00000 ; qtot
0.661
5985 HO 369 TYR HH 5985 0.399200 1.00800 ; qtot
1.060
5986 CA 369 TYR CE2 5986 -0.234100 12.01000 ; qtot
0.826
5987 HA 369 TYR HE2 5987 0.165600 1.00800 ; qtot
0.991
5988 CA 369 TYR CD2 5988 -0.190600 12.01000 ; qtot
0.801
5989 HA 369 TYR HD2 5989 0.169900 1.00800 ; qtot
0.971
5990 C 369 TYR C 5990 0.597301 12.01000 ; qtot
1.568
5991 O 369 TYR O 5991 -0.567901 16.00000 ; qtot
1.000
5992 N 370 THR N 5992 -0.415700 14.01000 ; qtot
0.584
5993 H 370 THR H 5993 0.271900 1.00800 ; qtot
0.856
5994 CX 370 THR CA 5994 -0.038900 12.01000 ; qtot
0.817
5995 H1 370 THR HA 5995 0.100700 1.00800 ; qtot
0.918
5996 A3C 370 THR CB 5996 0.365400 12.01000 ; qtot
1.283
5997 H1 370 THR HB 5997 0.004300 1.00800 ; qtot
1.288
5998 CT 370 THR CG2 5998 -0.243800 12.01000 ; qtot
1.044
5999 HC 370 THR HG21 5999 0.064200 1.00800 ; qtot
1.108
6000 HC 370 THR HG22 6000 0.064200 1.00800 ; qtot
1.172
6001 HC 370 THR HG23 6001 0.064200 1.00800 ; qtot
1.237
6002 OH 370 THR OG1 6002 -0.676101 16.00000 ; qtot
0.560
6003 HO 370 THR HG1 6003 0.410200 1.00800 ; qtot
0.971

```
  6004    C   370   THR     C 6004     0.597301     12.01000 ; qtot
1.568
  6005    O   370   THR     O 6005    -0.567901     16.00000 ; qtot
1.000
  6006    N   371   ASP     N 6006    -0.516301     14.01000 ; qtot
0.484
  6007    H   371   ASP     H 6007     0.293600      1.00800 ; qtot
0.777
  6008   CX   371   ASP    CA 6008     0.038100     12.01000 ; qtot
0.815
  6009   H1   371   ASP    HA 6009     0.088000      1.00800 ; qtot
0.903
  6010  A2C   371   ASP    CB 6010    -0.030300     12.01000 ; qtot
0.873
  6011   HC   371   ASP   HB2 6011    -0.012200      1.00800 ; qtot
0.861
  6012   HC   371   ASP   HB3 6012    -0.012200      1.00800 ; qtot
0.849
  6013   CO   371   ASP    CG 6013     0.799401     12.01000 ; qtot
1.648
  6014   O2   371   ASP   OD1 6014    -0.801401     16.00000 ; qtot
0.847
  6015   O2   371   ASP   OD2 6015    -0.801401     16.00000 ; qtot
0.045
  6016    C   371   ASP     C 6016     0.536601     12.01000 ; qtot
0.582
  6017    O   371   ASP     O 6017    -0.581901     16.00000 ; qtot
0.000
  6018    N   372   PHE     N 6018    -0.415700     14.01000 ; qtot -
0.416
  6019    H   372   PHE     H 6019     0.271900      1.00800 ; qtot -
0.144
  6020   CX   372   PHE    CA 6020    -0.002400     12.01000 ; qtot -
0.146
  6021   H1   372   PHE    HA 6021     0.097800      1.00800 ; qtot -
0.048
  6022   CT   372   PHE    CB 6022    -0.034300     12.01000 ; qtot -
0.083
  6023   HC   372   PHE   HB2 6023     0.029500      1.00800 ; qtot -
0.053
  6024   HC   372   PHE   HB3 6024     0.029500      1.00800 ; qtot -
0.024
  6025   CA   372   PHE    CG 6025     0.011800     12.01000 ; qtot -
0.012
  6026   CA   372   PHE   CD1 6026    -0.125600     12.01000 ; qtot -
0.137
  6027   HA   372   PHE   HD1 6027     0.133000      1.00800 ; qtot -
0.004
  6028   CA   372   PHE   CE1 6028    -0.170400     12.01000 ; qtot -
0.175
  6029   HA   372   PHE   HE1 6029     0.143000      1.00800 ; qtot -
0.032
```

6030 CA 372 PHE CZ 6030 -0.107200 12.01000 ; qtot -
0.139
6031 HA 372 PHE HZ 6031 0.129700 1.00800 ; qtot -
0.009
6032 CA 372 PHE CE2 6032 -0.170400 12.01000 ; qtot -
0.180
6033 HA 372 PHE HE2 6033 0.143000 1.00800 ; qtot -
0.037
6034 CA 372 PHE CD2 6034 -0.125600 12.01000 ; qtot -
0.162
6035 HA 372 PHE HD2 6035 0.133000 1.00800 ; qtot -
0.029
6036 C 372 PHE C 6036 0.597301 12.01000 ; qtot
0.568
6037 O 372 PHE O 6037 -0.567901 16.00000 ; qtot
0.000
6038 N 373 LEU N 6038 -0.415700 14.01000 ; qtot -
0.416
6039 H 373 LEU H 6039 0.271900 1.00800 ; qtot -
0.144
6040 CX 373 LEU CA 6040 -0.051800 12.01000 ; qtot -
0.196
6041 H1 373 LEU HA 6041 0.092200 1.00800 ; qtot -
0.103
6042 A2C 373 LEU CB 6042 -0.110200 12.01000 ; qtot -
0.214
6043 HC 373 LEU HB2 6043 0.045700 1.00800 ; qtot -
0.168
6044 HC 373 LEU HB3 6044 0.045700 1.00800 ; qtot -
0.122
6045 A3C 373 LEU CG 6045 0.353100 12.01000 ; qtot
0.231
6046 HC 373 LEU HG 6046 -0.036100 1.00800 ; qtot
0.195
6047 CT 373 LEU CD1 6047 -0.412100 12.01000 ; qtot -
0.217
6048 HC 373 LEU HD11 6048 0.100000 1.00800 ; qtot -
0.117
6049 HC 373 LEU HD12 6049 0.100000 1.00800 ; qtot -
0.017
6050 HC 373 LEU HD13 6050 0.100000 1.00800 ; qtot
0.083
6051 CT 373 LEU CD2 6051 -0.412100 12.01000 ; qtot -
0.329
6052 HC 373 LEU HD21 6052 0.100000 1.00800 ; qtot -
0.229
6053 HC 373 LEU HD22 6053 0.100000 1.00800 ; qtot -
0.129
6054 HC 373 LEU HD23 6054 0.100000 1.00800 ; qtot -
0.029
6055 C 373 LEU C 6055 0.597301 12.01000 ; qtot
0.568

6056 O 373 LEU O 6056 -0.567901 16.00000 ; qtot
0.000
6057 N 374 ALA N 6057 -0.415700 14.01000 ; qtot -
0.416
6058 H 374 ALA H 6058 0.271900 1.00800 ; qtot -
0.144
6059 CX 374 ALA CA 6059 0.033700 12.01000 ; qtot -
0.110
6060 H1 374 ALA HA 6060 0.082300 1.00800 ; qtot -
0.028
6061 CT 374 ALA CB 6061 -0.182500 12.01000 ; qtot -
0.210
6062 HC 374 ALA HB1 6062 0.060300 1.00800 ; qtot -
0.150
6063 HC 374 ALA HB2 6063 0.060300 1.00800 ; qtot -
0.090
6064 HC 374 ALA HB3 6064 0.060300 1.00800 ; qtot -
0.029
6065 C 374 ALA C 6065 0.597301 12.01000 ; qtot
0.568
6066 O 374 ALA O 6066 -0.567901 16.00000 ; qtot
0.000
6067 N 375 ALA N 6067 -0.415700 14.01000 ; qtot -
0.416
6068 H 375 ALA H 6068 0.271900 1.00800 ; qtot -
0.144
6069 CX 375 ALA CA 6069 0.033700 12.01000 ; qtot -
0.110
6070 H1 375 ALA HA 6070 0.082300 1.00800 ; qtot -
0.028
6071 CT 375 ALA CB 6071 -0.182500 12.01000 ; qtot -
0.210
6072 HC 375 ALA HB1 6072 0.060300 1.00800 ; qtot -
0.150
6073 HC 375 ALA HB2 6073 0.060300 1.00800 ; qtot -
0.090
6074 HC 375 ALA HB3 6074 0.060300 1.00800 ; qtot -
0.029
6075 C 375 ALA C 6075 0.597301 12.01000 ; qtot
0.568
6076 O 375 ALA O 6076 -0.567901 16.00000 ; qtot
0.000
6077 N 376 THR N 6077 -0.415700 14.01000 ; qtot -
0.416
6078 H 376 THR H 6078 0.271900 1.00800 ; qtot -
0.144
6079 CX 376 THR CA 6079 -0.038900 12.01000 ; qtot -
0.183
6080 H1 376 THR HA 6080 0.100700 1.00800 ; qtot -
0.082
6081 A3C 376 THR CB 6081 0.365400 12.01000 ; qtot
0.283

6082 H1 376 THR HB 6082 0.004300 1.00800 ; qtot
0.288
6083 CT 376 THR CG2 6083 -0.243800 12.01000 ; qtot
0.044
6084 HC 376 THR HG21 6084 0.064200 1.00800 ; qtot
0.108
6085 HC 376 THR HG22 6085 0.064200 1.00800 ; qtot
0.172
6086 HC 376 THR HG23 6086 0.064200 1.00800 ; qtot
0.237
6087 OH 376 THR OG1 6087 -0.676101 16.00000 ; qtot -
0.440
6088 HO 376 THR HG1 6088 0.410200 1.00800 ; qtot -
0.029
6089 C 376 THR C 6089 0.597301 12.01000 ; qtot
0.568
6090 O 376 THR O 6090 -0.567901 16.00000 ; qtot
0.000
6091 N 377 ILE N 6091 -0.415700 14.01000 ; qtot -
0.416
6092 H 377 ILE H 6092 0.271900 1.00800 ; qtot -
0.144
6093 CX 377 ILE CA 6093 -0.059700 12.01000 ; qtot -
0.203
6094 H1 377 ILE HA 6094 0.086900 1.00800 ; qtot -
0.117
6095 A3C 377 ILE CB 6095 0.130300 12.01000 ; qtot
0.014
6096 HC 377 ILE HB 6096 0.018700 1.00800 ; qtot
0.032
6097 CT 377 ILE CG2 6097 -0.320400 12.01000 ; qtot -
0.288
6098 HC 377 ILE HG21 6098 0.088200 1.00800 ; qtot -
0.200
6099 HC 377 ILE HG22 6099 0.088200 1.00800 ; qtot -
0.112
6100 HC 377 ILE HG23 6100 0.088200 1.00800 ; qtot -
0.023
6101 A2C 377 ILE CG1 6101 -0.043000 12.01000 ; qtot -
0.066
6102 HC 377 ILE HG12 6102 0.023600 1.00800 ; qtot -
0.043
6103 HC 377 ILE HG13 6103 0.023600 1.00800 ; qtot -
0.019
6104 CT 377 ILE CD1 6104 -0.066000 12.01000 ; qtot -
0.085
6105 HC 377 ILE HD11 6105 0.018600 1.00800 ; qtot -
0.067
6106 HC 377 ILE HD12 6106 0.018600 1.00800 ; qtot -
0.048
6107 HC 377 ILE HD13 6107 0.018600 1.00800 ; qtot -
0.029

6108 C 377 ILE C 6108 0.597301 12.01000 ; qtot 0.568
6109 O 377 ILE O 6109 -0.567901 16.00000 ; qtot 0.000
6110 N 378 ASP N 6110 -0.516301 14.01000 ; qtot -0.516
6111 H 378 ASP H 6111 0.293600 1.00800 ; qtot -0.223
6112 CX 378 ASP CA 6112 0.038100 12.01000 ; qtot -0.185
6113 H1 378 ASP HA 6113 0.088000 1.00800 ; qtot -0.097
6114 A2C 378 ASP CB 6114 -0.030300 12.01000 ; qtot -0.127
6115 HC 378 ASP HB2 6115 -0.012200 1.00800 ; qtot -0.139
6116 HC 378 ASP HB3 6116 -0.012200 1.00800 ; qtot -0.151
6117 CO 378 ASP CG 6117 0.799401 12.01000 ; qtot 0.648
6118 O2 378 ASP OD1 6118 -0.801401 16.00000 ; qtot -0.153
6119 O2 378 ASP OD2 6119 -0.801401 16.00000 ; qtot -0.955
6120 C 378 ASP C 6120 0.536601 12.01000 ; qtot -0.418
6121 O 378 ASP O 6121 -0.581901 16.00000 ; qtot -1.000
6122 N 379 LYS N 6122 -0.347900 14.01000 ; qtot -1.348
6123 H 379 LYS H 6123 0.274700 1.00800 ; qtot -1.073
6124 CX 379 LYS CA 6124 -0.240000 12.01000 ; qtot -1.313
6125 H1 379 LYS HA 6125 0.142600 1.00800 ; qtot -1.171
6126 C8 379 LYS CB 6126 -0.009400 12.01000 ; qtot -1.180
6127 HC 379 LYS HB2 6127 0.036200 1.00800 ; qtot -1.144
6128 HC 379 LYS HB3 6128 0.036200 1.00800 ; qtot -1.108
6129 C8 379 LYS CG 6129 0.018700 12.01000 ; qtot -1.089
6130 HC 379 LYS HG2 6130 0.010300 1.00800 ; qtot -1.079
6131 HC 379 LYS HG3 6131 0.010300 1.00800 ; qtot -1.068
6132 C8 379 LYS CD 6132 -0.047900 12.01000 ; qtot -1.116
6133 HC 379 LYS HD2 6133 0.062100 1.00800 ; qtot -1.054

6134 HC 379 LYS HD3 6134 0.062100 1.00800 ; qtot -
0.992
6135 C8 379 LYS CE 6135 -0.014300 12.01000 ; qtot -
1.006
6136 HP 379 LYS HE2 6136 0.113500 1.00800 ; qtot -
0.893
6137 HP 379 LYS HE3 6137 0.113500 1.00800 ; qtot -
0.779
6138 N3 379 LYS NZ 6138 -0.385400 14.01000 ; qtot -
1.165
6139 H 379 LYS HZ1 6139 0.340000 1.00800 ; qtot -
0.825
6140 H 379 LYS HZ2 6140 0.340000 1.00800 ; qtot -
0.485
6141 H 379 LYS HZ3 6141 0.340000 1.00800 ; qtot -
0.145
6142 C 379 LYS C 6142 0.734101 12.01000 ; qtot
0.589
6143 O 379 LYS O 6143 -0.589401 16.00000 ; qtot
0.000
6144 N 380 GLN N 6144 -0.415700 14.01000 ; qtot -
0.416
6145 H 380 GLN H 6145 0.271900 1.00800 ; qtot -
0.144
6146 CX 380 GLN CA 6146 -0.003100 12.01000 ; qtot -
0.147
6147 H1 380 GLN HA 6147 0.085000 1.00800 ; qtot -
0.062
6148 A2C 380 GLN CB 6148 -0.003600 12.01000 ; qtot -
0.065
6149 HC 380 GLN HB2 6149 0.017100 1.00800 ; qtot -
0.048
6150 HC 380 GLN HB3 6150 0.017100 1.00800 ; qtot -
0.031
6151 A2C 380 GLN CG 6151 -0.064500 12.01000 ; qtot -
0.096
6152 HC 380 GLN HG2 6152 0.035200 1.00800 ; qtot -
0.061
6153 HC 380 GLN HG3 6153 0.035200 1.00800 ; qtot -
0.025
6154 C 380 GLN CD 6154 0.695101 12.01000 ; qtot
0.670
6155 O 380 GLN OE1 6155 -0.608601 16.00000 ; qtot
0.061
6156 N 380 GLN NE2 6156 -0.940701 14.01000 ; qtot -
0.880
6157 H 380 GLN HE21 6157 0.425100 1.00800 ; qtot -
0.454
6158 H 380 GLN HE22 6158 0.425100 1.00800 ; qtot -
0.029
6159 C 380 GLN C 6159 0.597301 12.01000 ; qtot
0.568

```
  6160    O   380   GLN     O 6160    -0.567901     16.00000 ; qtot
0.000
  6161    N   381   THR     N 6161    -0.415700     14.01000 ; qtot -
0.416
  6162    H   381   THR     H 6162     0.271900      1.00800 ; qtot -
0.144
  6163   CX   381   THR    CA 6163    -0.038900     12.01000 ; qtot -
0.183
  6164   H1   381   THR    HA 6164     0.100700      1.00800 ; qtot -
0.082
  6165  A3C   381   THR    CB 6165     0.365400     12.01000 ; qtot
0.283
  6166   H1   381   THR    HB 6166     0.004300      1.00800 ; qtot
0.288
  6167   CT   381   THR   CG2 6167    -0.243800     12.01000 ; qtot
0.044
  6168   HC   381   THR  HG21 6168     0.064200      1.00800 ; qtot
0.108
  6169   HC   381   THR  HG22 6169     0.064200      1.00800 ; qtot
0.172
  6170   HC   381   THR  HG23 6170     0.064200      1.00800 ; qtot
0.237
  6171   OH   381   THR   OG1 6171    -0.676101     16.00000 ; qtot -
0.440
  6172   HO   381   THR   HG1 6172     0.410200      1.00800 ; qtot -
0.029
  6173    C   381   THR     C 6173     0.597301     12.01000 ; qtot
0.568
  6174    O   381   THR     O 6174    -0.567901     16.00000 ; qtot
0.000
  6175    N   382   TYR     N 6175    -0.415700     14.01000 ; qtot -
0.416
  6176    H   382   TYR     H 6176     0.271900      1.00800 ; qtot -
0.144
  6177   CX   382   TYR    CA 6177    -0.001400     12.01000 ; qtot -
0.145
  6178   H1   382   TYR    HA 6178     0.087600      1.00800 ; qtot -
0.058
  6179   CT   382   TYR    CB 6179    -0.015200     12.01000 ; qtot -
0.073
  6180   HC   382   TYR   HB2 6180     0.029500      1.00800 ; qtot -
0.043
  6181   HC   382   TYR   HB3 6181     0.029500      1.00800 ; qtot -
0.014
  6182   CA   382   TYR    CG 6182    -0.001100     12.01000 ; qtot -
0.015
  6183   CA   382   TYR   CD1 6183    -0.190600     12.01000 ; qtot -
0.205
  6184   HA   382   TYR   HD1 6184     0.169900      1.00800 ; qtot -
0.036
  6185   CA   382   TYR   CE1 6185    -0.234100     12.01000 ; qtot -
0.270
```

```
  6186   HA   382   TYR   HE1 6186     0.165600      1.00800 ; qtot -
0.104
  6187    C   382   TYR    CZ 6187     0.322600     12.01000 ; qtot
0.219
  6188   OH   382   TYR    OH 6188    -0.557901     16.00000 ; qtot -
0.339
  6189   HO   382   TYR    HH 6189     0.399200      1.00800 ; qtot
0.060
  6190   CA   382   TYR   CE2 6190    -0.234100     12.01000 ; qtot -
0.174
  6191   HA   382   TYR   HE2 6191     0.165600      1.00800 ; qtot -
0.009
  6192   CA   382   TYR   CD2 6192    -0.190600     12.01000 ; qtot -
0.199
  6193   HA   382   TYR   HD2 6193     0.169900      1.00800 ; qtot -
0.029
  6194    C   382   TYR     C 6194     0.597301     12.01000 ; qtot
0.568
  6195    O   382   TYR     O 6195    -0.567901     16.00000 ; qtot
0.000
  6196    N   383   LEU     N 6196    -0.415700     14.01000 ; qtot -
0.416
  6197    H   383   LEU     H 6197     0.271900      1.00800 ; qtot -
0.144
  6198   CX   383   LEU    CA 6198    -0.051800     12.01000 ; qtot -
0.196
  6199   H1   383   LEU    HA 6199     0.092200      1.00800 ; qtot -
0.103
  6200  A2C   383   LEU    CB 6200    -0.110200     12.01000 ; qtot -
0.214
  6201   HC   383   LEU   HB2 6201     0.045700      1.00800 ; qtot -
0.168
  6202   HC   383   LEU   HB3 6202     0.045700      1.00800 ; qtot -
0.122
  6203  A3C   383   LEU    CG 6203     0.353100     12.01000 ; qtot
0.231
  6204   HC   383   LEU    HG 6204    -0.036100      1.00800 ; qtot
0.195
  6205   CT   383   LEU   CD1 6205    -0.412100     12.01000 ; qtot -
0.217
  6206   HC   383   LEU  HD11 6206     0.100000      1.00800 ; qtot -
0.117
  6207   HC   383   LEU  HD12 6207     0.100000      1.00800 ; qtot -
0.017
  6208   HC   383   LEU  HD13 6208     0.100000      1.00800 ; qtot
0.083
  6209   CT   383   LEU   CD2 6209    -0.412100     12.01000 ; qtot -
0.329
  6210   HC   383   LEU  HD21 6210     0.100000      1.00800 ; qtot -
0.229
  6211   HC   383   LEU  HD22 6211     0.100000      1.00800 ; qtot -
0.129
```

6212 HC 383 LEU HD23 6212 0.100000 1.00800 ; qtot -
0.029
6213 C 383 LEU C 6213 0.597301 12.01000 ; qtot
0.568
6214 O 383 LEU O 6214 -0.567901 16.00000 ; qtot
0.000
6215 N 384 LYS N 6215 -0.347900 14.01000 ; qtot -
0.348
6216 H 384 LYS H 6216 0.274700 1.00800 ; qtot -
0.073
6217 CX 384 LYS CA 6217 -0.240000 12.01000 ; qtot -
0.313
6218 H1 384 LYS HA 6218 0.142600 1.00800 ; qtot -
0.171
6219 C8 384 LYS CB 6219 -0.009400 12.01000 ; qtot -
0.180
6220 HC 384 LYS HB2 6220 0.036200 1.00800 ; qtot -
0.144
6221 HC 384 LYS HB3 6221 0.036200 1.00800 ; qtot -
0.108
6222 C8 384 LYS CG 6222 0.018700 12.01000 ; qtot -
0.089
6223 HC 384 LYS HG2 6223 0.010300 1.00800 ; qtot -
0.079
6224 HC 384 LYS HG3 6224 0.010300 1.00800 ; qtot -
0.068
6225 C8 384 LYS CD 6225 -0.047900 12.01000 ; qtot -
0.116
6226 HC 384 LYS HD2 6226 0.062100 1.00800 ; qtot -
0.054
6227 HC 384 LYS HD3 6227 0.062100 1.00800 ; qtot
0.008
6228 C8 384 LYS CE 6228 -0.014300 12.01000 ; qtot -
0.006
6229 HP 384 LYS HE2 6229 0.113500 1.00800 ; qtot
0.107
6230 HP 384 LYS HE3 6230 0.113500 1.00800 ; qtot
0.221
6231 N3 384 LYS NZ 6231 -0.385400 14.01000 ; qtot -
0.165
6232 H 384 LYS HZ1 6232 0.340000 1.00800 ; qtot
0.175
6233 H 384 LYS HZ2 6233 0.340000 1.00800 ; qtot
0.515
6234 H 384 LYS HZ3 6234 0.340000 1.00800 ; qtot
0.855
6235 C 384 LYS C 6235 0.734101 12.01000 ; qtot
1.589
6236 O 384 LYS O 6236 -0.589401 16.00000 ; qtot
1.000
6237 N 385 LYS N 6237 -0.347900 14.01000 ; qtot
0.652

| | | | | | | | | | | |
|---|---|---|---|---|---|---|---|---|---|---|
| 6238 | H | 385 | LYS | H | 6238 | 0.274700 | 1.00800 | ; | qtot | 0.927 |
| 6239 | CX | 385 | LYS | CA | 6239 | -0.240000 | 12.01000 | ; | qtot | 0.687 |
| 6240 | H1 | 385 | LYS | HA | 6240 | 0.142600 | 1.00800 | ; | qtot | 0.829 |
| 6241 | C8 | 385 | LYS | CB | 6241 | -0.009400 | 12.01000 | ; | qtot | 0.820 |
| 6242 | HC | 385 | LYS | HB2 | 6242 | 0.036200 | 1.00800 | ; | qtot | 0.856 |
| 6243 | HC | 385 | LYS | HB3 | 6243 | 0.036200 | 1.00800 | ; | qtot | 0.892 |
| 6244 | C8 | 385 | LYS | CG | 6244 | 0.018700 | 12.01000 | ; | qtot | 0.911 |
| 6245 | HC | 385 | LYS | HG2 | 6245 | 0.010300 | 1.00800 | ; | qtot | 0.921 |
| 6246 | HC | 385 | LYS | HG3 | 6246 | 0.010300 | 1.00800 | ; | qtot | 0.932 |
| 6247 | C8 | 385 | LYS | CD | 6247 | -0.047900 | 12.01000 | ; | qtot | 0.884 |
| 6248 | HC | 385 | LYS | HD2 | 6248 | 0.062100 | 1.00800 | ; | qtot | 0.946 |
| 6249 | HC | 385 | LYS | HD3 | 6249 | 0.062100 | 1.00800 | ; | qtot | 1.008 |
| 6250 | C8 | 385 | LYS | CE | 6250 | -0.014300 | 12.01000 | ; | qtot | 0.994 |
| 6251 | HP | 385 | LYS | HE2 | 6251 | 0.113500 | 1.00800 | ; | qtot | 1.107 |
| 6252 | HP | 385 | LYS | HE3 | 6252 | 0.113500 | 1.00800 | ; | qtot | 1.221 |
| 6253 | N3 | 385 | LYS | NZ | 6253 | -0.385400 | 14.01000 | ; | qtot | 0.835 |
| 6254 | H | 385 | LYS | HZ1 | 6254 | 0.340000 | 1.00800 | ; | qtot | 1.175 |
| 6255 | H | 385 | LYS | HZ2 | 6255 | 0.340000 | 1.00800 | ; | qtot | 1.515 |
| 6256 | H | 385 | LYS | HZ3 | 6256 | 0.340000 | 1.00800 | ; | qtot | 1.855 |
| 6257 | C | 385 | LYS | C | 6257 | 0.734101 | 12.01000 | ; | qtot | 2.589 |
| 6258 | O | 385 | LYS | O | 6258 | -0.589401 | 16.00000 | ; | qtot | 2.000 |
| 6259 | N | 386 | GLU | N | 6259 | -0.516301 | 14.01000 | ; | qtot | 1.484 |
| 6260 | H | 386 | GLU | H | 6260 | 0.293600 | 1.00800 | ; | qtot | 1.777 |
| 6261 | CX | 386 | GLU | CA | 6261 | 0.039700 | 12.01000 | ; | qtot | 1.817 |
| 6262 | H1 | 386 | GLU | HA | 6262 | 0.110500 | 1.00800 | ; | qtot | 1.928 |
| 6263 | A2C | 386 | GLU | CB | 6263 | 0.056000 | 12.01000 | ; | qtot | 1.984 |

6264 HC 386 GLU HB2 6264 -0.017300 1.00800 ; qtot
1.966
6265 HC 386 GLU HB3 6265 -0.017300 1.00800 ; qtot
1.949
6266 A2C 386 GLU CG 6266 0.013600 12.01000 ; qtot
1.963
6267 HC 386 GLU HG2 6267 -0.042500 1.00800 ; qtot
1.920
6268 HC 386 GLU HG3 6268 -0.042500 1.00800 ; qtot
1.878
6269 CO 386 GLU CD 6269 0.805401 12.01000 ; qtot
2.683
6270 O2 386 GLU OE1 6270 -0.818801 16.00000 ; qtot
1.864
6271 O2 386 GLU OE2 6271 -0.818801 16.00000 ; qtot
1.045
6272 C 386 GLU C 6272 0.536601 12.01000 ; qtot
1.582
6273 O 386 GLU O 6273 -0.581901 16.00000 ; qtot
1.000
6274 N 387 VAL N 6274 -0.415700 14.01000 ; qtot
0.584
6275 H 387 VAL H 6275 0.271900 1.00800 ; qtot
0.856
6276 CX 387 VAL CA 6276 -0.087500 12.01000 ; qtot
0.769
6277 H1 387 VAL HA 6277 0.096900 1.00800 ; qtot
0.866
6278 A3C 387 VAL CB 6278 0.298500 12.01000 ; qtot
1.164
6279 HC 387 VAL HB 6279 -0.029700 1.00800 ; qtot
1.134
6280 CT 387 VAL CG1 6280 -0.319200 12.01000 ; qtot
0.815
6281 HC 387 VAL HG11 6281 0.079100 1.00800 ; qtot
0.894
6282 HC 387 VAL HG12 6282 0.079100 1.00800 ; qtot
0.973
6283 HC 387 VAL HG13 6283 0.079100 1.00800 ; qtot
1.053
6284 CT 387 VAL CG2 6284 -0.319200 12.01000 ; qtot
0.733
6285 HC 387 VAL HG21 6285 0.079100 1.00800 ; qtot
0.812
6286 HC 387 VAL HG22 6286 0.079100 1.00800 ; qtot
0.892
6287 HC 387 VAL HG23 6287 0.079100 1.00800 ; qtot
0.971
6288 C 387 VAL C 6288 0.597301 12.01000 ; qtot
1.568
6289 O 387 VAL O 6289 -0.567901 16.00000 ; qtot
1.000

6290 N 388 CYS N 6290 -0.415700 14.01000 ; qtot
0.584
6291 H 388 CYS H 6291 0.271900 1.00800 ; qtot
0.856
6292 CX 388 CYS CA 6292 0.021300 12.01000 ; qtot
0.878
6293 H1 388 CYS HA 6293 0.112400 1.00800 ; qtot
0.990
6294 A2C 388 CYS CB 6294 -0.123100 12.01000 ; qtot
0.867
6295 H1 388 CYS HB2 6295 0.111200 1.00800 ; qtot
0.978
6296 H1 388 CYS HB3 6296 0.111200 1.00800 ; qtot
1.089
6297 SH 388 CYS SG 6297 -0.311900 32.06000 ; qtot
0.777
6298 HS 388 CYS HG 6298 0.193300 1.00800 ; qtot
0.971
6299 C 388 CYS C 6299 0.597301 12.01000 ; qtot
1.568
6300 O 388 CYS O 6300 -0.567901 16.00000 ; qtot
1.000
6301 N 389 LEU N 6301 -0.415700 14.01000 ; qtot
0.584
6302 H 389 LEU H 6302 0.271900 1.00800 ; qtot
0.856
6303 CX 389 LEU CA 6303 -0.051800 12.01000 ; qtot
0.804
6304 H1 389 LEU HA 6304 0.092200 1.00800 ; qtot
0.897
6305 A2C 389 LEU CB 6305 -0.110200 12.01000 ; qtot
0.786
6306 HC 389 LEU HB2 6306 0.045700 1.00800 ; qtot
0.832
6307 HC 389 LEU HB3 6307 0.045700 1.00800 ; qtot
0.878
6308 A3C 389 LEU CG 6308 0.353100 12.01000 ; qtot
1.231
6309 HC 389 LEU HG 6309 -0.036100 1.00800 ; qtot
1.195
6310 CT 389 LEU CD1 6310 -0.412100 12.01000 ; qtot
0.783
6311 HC 389 LEU HD11 6311 0.100000 1.00800 ; qtot
0.883
6312 HC 389 LEU HD12 6312 0.100000 1.00800 ; qtot
0.983
6313 HC 389 LEU HD13 6313 0.100000 1.00800 ; qtot
1.083
6314 CT 389 LEU CD2 6314 -0.412100 12.01000 ; qtot
0.671
6315 HC 389 LEU HD21 6315 0.100000 1.00800 ; qtot
0.771

```
  6316   HC   389   LEU  HD22 6316     0.100000      1.00800 ; qtot
0.871
  6317   HC   389   LEU  HD23 6317     0.100000      1.00800 ; qtot
0.971
  6318    C   389   LEU     C 6318     0.597301     12.01000 ; qtot
1.568
  6319    O   389   LEU     O 6319    -0.567901     16.00000 ; qtot
1.000
  6320    N   390   ILE     N 6320    -0.415700     14.01000 ; qtot
0.584
  6321    H   390   ILE     H 6321     0.271900      1.00800 ; qtot
0.856
  6322   CX   390   ILE    CA 6322    -0.059700     12.01000 ; qtot
0.797
  6323   H1   390   ILE    HA 6323     0.086900      1.00800 ; qtot
0.883
  6324  A3C   390   ILE    CB 6324     0.130300     12.01000 ; qtot
1.014
  6325   HC   390   ILE    HB 6325     0.018700      1.00800 ; qtot
1.032
  6326   CT   390   ILE   CG2 6326    -0.320400     12.01000 ; qtot
0.712
  6327   HC   390   ILE  HG21 6327     0.088200      1.00800 ; qtot
0.800
  6328   HC   390   ILE  HG22 6328     0.088200      1.00800 ; qtot
0.888
  6329   HC   390   ILE  HG23 6329     0.088200      1.00800 ; qtot
0.977
  6330  A2C   390   ILE   CG1 6330    -0.043000     12.01000 ; qtot
0.934
  6331   HC   390   ILE  HG12 6331     0.023600      1.00800 ; qtot
0.957
  6332   HC   390   ILE  HG13 6332     0.023600      1.00800 ; qtot
0.981
  6333   CT   390   ILE   CD1 6333    -0.066000     12.01000 ; qtot
0.915
  6334   HC   390   ILE  HD11 6334     0.018600      1.00800 ; qtot
0.933
  6335   HC   390   ILE  HD12 6335     0.018600      1.00800 ; qtot
0.952
  6336   HC   390   ILE  HD13 6336     0.018600      1.00800 ; qtot
0.971
  6337    C   390   ILE     C 6337     0.597301     12.01000 ; qtot
1.568
  6338    O   390   ILE     O 6338    -0.567901     16.00000 ; qtot
1.000
  6339    N   391   PRO     N 6339    -0.254800     14.01000 ; qtot
0.745
  6340   CT   391   PRO    CD 6340     0.019200     12.01000 ; qtot
0.764
  6341   H1   391   PRO   HD2 6341     0.039100      1.00800 ; qtot
0.804
```

6342 H1 391 PRO HD3 6342 0.039100 1.00800 ; qtot
0.843
6343 CT 391 PRO CG 6343 0.018900 12.01000 ; qtot
0.862
6344 HC 391 PRO HG2 6344 0.021300 1.00800 ; qtot
0.883
6345 HC 391 PRO HG3 6345 0.021300 1.00800 ; qtot
0.904
6346 CT 391 PRO CB 6346 -0.007000 12.01000 ; qtot
0.897
6347 HC 391 PRO HB2 6347 0.025300 1.00800 ; qtot
0.922
6348 HC 391 PRO HB3 6348 0.025300 1.00800 ; qtot
0.948
6349 CX 391 PRO CA 6349 -0.026600 12.01000 ; qtot
0.921
6350 H1 391 PRO HA 6350 0.064100 1.00800 ; qtot
0.985
6351 C 391 PRO C 6351 0.589601 12.01000 ; qtot
1.575
6352 O 391 PRO O 6352 -0.574801 16.00000 ; qtot
1.000
6353 N 392 PHE N 6353 -0.415700 14.01000 ; qtot
0.584
6354 H 392 PHE H 6354 0.271900 1.00800 ; qtot
0.856
6355 CX 392 PHE CA 6355 -0.002400 12.01000 ; qtot
0.854
6356 H1 392 PHE HA 6356 0.097800 1.00800 ; qtot
0.952
6357 CT 392 PHE CB 6357 -0.034300 12.01000 ; qtot
0.917
6358 HC 392 PHE HB2 6358 0.029500 1.00800 ; qtot
0.947
6359 HC 392 PHE HB3 6359 0.029500 1.00800 ; qtot
0.976
6360 CA 392 PHE CG 6360 0.011800 12.01000 ; qtot
0.988
6361 CA 392 PHE CD1 6361 -0.125600 12.01000 ; qtot
0.863
6362 HA 392 PHE HD1 6362 0.133000 1.00800 ; qtot
0.996
6363 CA 392 PHE CE1 6363 -0.170400 12.01000 ; qtot
0.825
6364 HA 392 PHE HE1 6364 0.143000 1.00800 ; qtot
0.968
6365 CA 392 PHE CZ 6365 -0.107200 12.01000 ; qtot
0.861
6366 HA 392 PHE HZ 6366 0.129700 1.00800 ; qtot
0.991
6367 CA 392 PHE CE2 6367 -0.170400 12.01000 ; qtot
0.820

6368 HA 392 PHE HE2 6368 0.143000 1.00800 ; qtot
0.963
6369 CA 392 PHE CD2 6369 -0.125600 12.01000 ; qtot
0.838
6370 HA 392 PHE HD2 6370 0.133000 1.00800 ; qtot
0.971
6371 C 392 PHE C 6371 0.597301 12.01000 ; qtot
1.568
6372 O 392 PHE O 6372 -0.567901 16.00000 ; qtot
1.000
6373 N 393 LYS N 6373 -0.347900 14.01000 ; qtot
0.652
6374 H 393 LYS H 6374 0.274700 1.00800 ; qtot
0.927
6375 CX 393 LYS CA 6375 -0.240000 12.01000 ; qtot
0.687
6376 H1 393 LYS HA 6376 0.142600 1.00800 ; qtot
0.829
6377 C8 393 LYS CB 6377 -0.009400 12.01000 ; qtot
0.820
6378 HC 393 LYS HB2 6378 0.036200 1.00800 ; qtot
0.856
6379 HC 393 LYS HB3 6379 0.036200 1.00800 ; qtot
0.892
6380 C8 393 LYS CG 6380 0.018700 12.01000 ; qtot
0.911
6381 HC 393 LYS HG2 6381 0.010300 1.00800 ; qtot
0.921
6382 HC 393 LYS HG3 6382 0.010300 1.00800 ; qtot
0.932
6383 C8 393 LYS CD 6383 -0.047900 12.01000 ; qtot
0.884
6384 HC 393 LYS HD2 6384 0.062100 1.00800 ; qtot
0.946
6385 HC 393 LYS HD3 6385 0.062100 1.00800 ; qtot
1.008
6386 C8 393 LYS CE 6386 -0.014300 12.01000 ; qtot
0.994
6387 HP 393 LYS HE2 6387 0.113500 1.00800 ; qtot
1.107
6388 HP 393 LYS HE3 6388 0.113500 1.00800 ; qtot
1.221
6389 N3 393 LYS NZ 6389 -0.385400 14.01000 ; qtot
0.835
6390 H 393 LYS HZ1 6390 0.340000 1.00800 ; qtot
1.175
6391 H 393 LYS HZ2 6391 0.340000 1.00800 ; qtot
1.515
6392 H 393 LYS HZ3 6392 0.340000 1.00800 ; qtot
1.855
6393 C 393 LYS C 6393 0.734101 12.01000 ; qtot
2.589

6394 O 393 LYS O 6394 -0.589401 16.00000 ; qtot
2.000
6395 N 394 PHE N 6395 -0.415700 14.01000 ; qtot
1.584
6396 H 394 PHE H 6396 0.271900 1.00800 ; qtot
1.856
6397 CX 394 PHE CA 6397 -0.002400 12.01000 ; qtot
1.854
6398 H1 394 PHE HA 6398 0.097800 1.00800 ; qtot
1.952
6399 CT 394 PHE CB 6399 -0.034300 12.01000 ; qtot
1.917
6400 HC 394 PHE HB2 6400 0.029500 1.00800 ; qtot
1.947
6401 HC 394 PHE HB3 6401 0.029500 1.00800 ; qtot
1.976
6402 CA 394 PHE CG 6402 0.011800 12.01000 ; qtot
1.988
6403 CA 394 PHE CD1 6403 -0.125600 12.01000 ; qtot
1.863
6404 HA 394 PHE HD1 6404 0.133000 1.00800 ; qtot
1.996
6405 CA 394 PHE CE1 6405 -0.170400 12.01000 ; qtot
1.825
6406 HA 394 PHE HE1 6406 0.143000 1.00800 ; qtot
1.968
6407 CA 394 PHE CZ 6407 -0.107200 12.01000 ; qtot
1.861
6408 HA 394 PHE HZ 6408 0.129700 1.00800 ; qtot
1.991
6409 CA 394 PHE CE2 6409 -0.170400 12.01000 ; qtot
1.820
6410 HA 394 PHE HE2 6410 0.143000 1.00800 ; qtot
1.963
6411 CA 394 PHE CD2 6411 -0.125600 12.01000 ; qtot
1.838
6412 HA 394 PHE HD2 6412 0.133000 1.00800 ; qtot
1.971
6413 C 394 PHE C 6413 0.597301 12.01000 ; qtot
2.568
6414 O 394 PHE O 6414 -0.567901 16.00000 ; qtot
2.000
6415 N 395 PHE N 6415 -0.415700 14.01000 ; qtot
1.584
6416 H 395 PHE H 6416 0.271900 1.00800 ; qtot
1.856
6417 CX 395 PHE CA 6417 -0.002400 12.01000 ; qtot
1.854
6418 H1 395 PHE HA 6418 0.097800 1.00800 ; qtot
1.952
6419 CT 395 PHE CB 6419 -0.034300 12.01000 ; qtot
1.917

6420 HC 395 PHE HB2 6420 0.029500 1.00800 ; qtot
1.947
6421 HC 395 PHE HB3 6421 0.029500 1.00800 ; qtot
1.976
6422 CA 395 PHE CG 6422 0.011800 12.01000 ; qtot
1.988
6423 CA 395 PHE CD1 6423 -0.125600 12.01000 ; qtot
1.863
6424 HA 395 PHE HD1 6424 0.133000 1.00800 ; qtot
1.996
6425 CA 395 PHE CE1 6425 -0.170400 12.01000 ; qtot
1.825
6426 HA 395 PHE HE1 6426 0.143000 1.00800 ; qtot
1.968
6427 CA 395 PHE CZ 6427 -0.107200 12.01000 ; qtot
1.861
6428 HA 395 PHE HZ 6428 0.129700 1.00800 ; qtot
1.991
6429 CA 395 PHE CE2 6429 -0.170400 12.01000 ; qtot
1.820
6430 HA 395 PHE HE2 6430 0.143000 1.00800 ; qtot
1.963
6431 CA 395 PHE CD2 6431 -0.125600 12.01000 ; qtot
1.838
6432 HA 395 PHE HD2 6432 0.133000 1.00800 ; qtot
1.971
6433 C 395 PHE C 6433 0.597301 12.01000 ; qtot
2.568
6434 O 395 PHE O 6434 -0.567901 16.00000 ; qtot
2.000
6435 N 396 ASP N 6435 -0.516301 14.01000 ; qtot
1.484
6436 H 396 ASP H 6436 0.293600 1.00800 ; qtot
1.777
6437 CX 396 ASP CA 6437 0.038100 12.01000 ; qtot
1.815
6438 H1 396 ASP HA 6438 0.088000 1.00800 ; qtot
1.903
6439 A2C 396 ASP CB 6439 -0.030300 12.01000 ; qtot
1.873
6440 HC 396 ASP HB2 6440 -0.012200 1.00800 ; qtot
1.861
6441 HC 396 ASP HB3 6441 -0.012200 1.00800 ; qtot
1.849
6442 CO 396 ASP CG 6442 0.799401 12.01000 ; qtot
2.648
6443 O2 396 ASP OD1 6443 -0.801401 16.00000 ; qtot
1.847
6444 O2 396 ASP OD2 6444 -0.801401 16.00000 ; qtot
1.045
6445 C 396 ASP C 6445 0.536601 12.01000 ; qtot
1.582

```
  6446    O   396   ASP     O 6446    -0.581901     16.00000 ; qtot
1.000
  6447    N   397   ILE     N 6447    -0.415700     14.01000 ; qtot
0.584
  6448    H   397   ILE     H 6448     0.271900      1.00800 ; qtot
0.856
  6449   CX   397   ILE    CA 6449    -0.059700     12.01000 ; qtot
0.797
  6450   H1   397   ILE    HA 6450     0.086900      1.00800 ; qtot
0.883
  6451  A3C   397   ILE    CB 6451     0.130300     12.01000 ; qtot
1.014
  6452   HC   397   ILE    HB 6452     0.018700      1.00800 ; qtot
1.032
  6453   CT   397   ILE   CG2 6453    -0.320400     12.01000 ; qtot
0.712
  6454   HC   397   ILE  HG21 6454     0.088200      1.00800 ; qtot
0.800
  6455   HC   397   ILE  HG22 6455     0.088200      1.00800 ; qtot
0.888
  6456   HC   397   ILE  HG23 6456     0.088200      1.00800 ; qtot
0.977
  6457  A2C   397   ILE   CG1 6457    -0.043000     12.01000 ; qtot
0.934
  6458   HC   397   ILE  HG12 6458     0.023600      1.00800 ; qtot
0.957
  6459   HC   397   ILE  HG13 6459     0.023600      1.00800 ; qtot
0.981
  6460   CT   397   ILE   CD1 6460    -0.066000     12.01000 ; qtot
0.915
  6461   HC   397   ILE  HD11 6461     0.018600      1.00800 ; qtot
0.933
  6462   HC   397   ILE  HD12 6462     0.018600      1.00800 ; qtot
0.952
  6463   HC   397   ILE  HD13 6463     0.018600      1.00800 ; qtot
0.971
  6464    C   397   ILE     C 6464     0.597301     12.01000 ; qtot
1.568
  6465    O   397   ILE     O 6465    -0.567901     16.00000 ; qtot
1.000
  6466    N   398   ASP     N 6466    -0.516301     14.01000 ; qtot
0.484
  6467    H   398   ASP     H 6467     0.293600      1.00800 ; qtot
0.777
  6468   CX   398   ASP    CA 6468     0.038100     12.01000 ; qtot
0.815
  6469   H1   398   ASP    HA 6469     0.088000      1.00800 ; qtot
0.903
  6470  A2C   398   ASP    CB 6470    -0.030300     12.01000 ; qtot
0.873
  6471   HC   398   ASP   HB2 6471    -0.012200      1.00800 ; qtot
0.861
```

```
  6472   HC   398   ASP   HB3 6472    -0.012200      1.00800 ; qtot
0.849
  6473   CO   398   ASP    CG 6473     0.799401     12.01000 ; qtot
1.648
  6474   O2   398   ASP   OD1 6474    -0.801401     16.00000 ; qtot
0.847
  6475   O2   398   ASP   OD2 6475    -0.801401     16.00000 ; qtot
0.045
  6476    C   398   ASP     C 6476     0.536601     12.01000 ; qtot
0.582
  6477    O   398   ASP     O 6477    -0.581901     16.00000 ; qtot
0.000
  6478    N   399   GLY     N 6478    -0.415700     14.01000 ; qtot -
0.416
  6479    H   399   GLY     H 6479     0.271900      1.00800 ; qtot -
0.144
  6480   CX   399   GLY    CA 6480    -0.025200     12.01000 ; qtot -
0.169
  6481   H1   399   GLY   HA2 6481     0.069800      1.00800 ; qtot -
0.099
  6482   H1   399   GLY   HA3 6482     0.069800      1.00800 ; qtot -
0.029
  6483    C   399   GLY     C 6483     0.597301     12.01000 ; qtot
0.568
  6484    O   399   GLY     O 6484    -0.567901     16.00000 ; qtot
0.000
  6485    N   400   ASN     N 6485    -0.415700     14.01000 ; qtot -
0.416
  6486    H   400   ASN     H 6486     0.271900      1.00800 ; qtot -
0.144
  6487   CX   400   ASN    CA 6487     0.014300     12.01000 ; qtot -
0.129
  6488   H1   400   ASN    HA 6488     0.104800      1.00800 ; qtot -
0.025
  6489  A2C   400   ASN    CB 6489    -0.204100     12.01000 ; qtot -
0.229
  6490   HC   400   ASN   HB2 6490     0.079700      1.00800 ; qtot -
0.149
  6491   HC   400   ASN   HB3 6491     0.079700      1.00800 ; qtot -
0.069
  6492    C   400   ASN    CG 6492     0.713001     12.01000 ; qtot
0.644
  6493    O   400   ASN   OD1 6493    -0.593101     16.00000 ; qtot
0.051
  6494    N   400   ASN   ND2 6494    -0.919101     14.01000 ; qtot -
0.869
  6495    H   400   ASN  HD21 6495     0.419600      1.00800 ; qtot -
0.449
  6496    H   400   ASN  HD22 6496     0.419600      1.00800 ; qtot -
0.029
  6497    C   400   ASN     C 6497     0.597301     12.01000 ; qtot
0.568
```

6498 O 400 ASN O 6498 -0.567901 16.00000 ; qtot
0.000
6499 N 401 GLY N 6499 -0.415700 14.01000 ; qtot -
0.416
6500 H 401 GLY H 6500 0.271900 1.00800 ; qtot -
0.144
6501 CX 401 GLY CA 6501 -0.025200 12.01000 ; qtot -
0.169
6502 H1 401 GLY HA2 6502 0.069800 1.00800 ; qtot -
0.099
6503 H1 401 GLY HA3 6503 0.069800 1.00800 ; qtot -
0.029
6504 C 401 GLY C 6504 0.597301 12.01000 ; qtot
0.568
6505 O 401 GLY O 6505 -0.567901 16.00000 ; qtot
0.000
6506 N 402 LYS N 6506 -0.347900 14.01000 ; qtot -
0.348
6507 H 402 LYS H 6507 0.274700 1.00800 ; qtot -
0.073
6508 CX 402 LYS CA 6508 -0.240000 12.01000 ; qtot -
0.313
6509 H1 402 LYS HA 6509 0.142600 1.00800 ; qtot -
0.171
6510 C8 402 LYS CB 6510 -0.009400 12.01000 ; qtot -
0.180
6511 HC 402 LYS HB2 6511 0.036200 1.00800 ; qtot -
0.144
6512 HC 402 LYS HB3 6512 0.036200 1.00800 ; qtot -
0.108
6513 C8 402 LYS CG 6513 0.018700 12.01000 ; qtot -
0.089
6514 HC 402 LYS HG2 6514 0.010300 1.00800 ; qtot -
0.079
6515 HC 402 LYS HG3 6515 0.010300 1.00800 ; qtot -
0.068
6516 C8 402 LYS CD 6516 -0.047900 12.01000 ; qtot -
0.116
6517 HC 402 LYS HD2 6517 0.062100 1.00800 ; qtot -
0.054
6518 HC 402 LYS HD3 6518 0.062100 1.00800 ; qtot
0.008
6519 C8 402 LYS CE 6519 -0.014300 12.01000 ; qtot -
0.006
6520 HP 402 LYS HE2 6520 0.113500 1.00800 ; qtot
0.107
6521 HP 402 LYS HE3 6521 0.113500 1.00800 ; qtot
0.221
6522 N3 402 LYS NZ 6522 -0.385400 14.01000 ; qtot -
0.165
6523 H 402 LYS HZ1 6523 0.340000 1.00800 ; qtot
0.175

6524 H 402 LYS HZ2 6524 0.340000 1.00800 ; qtot
0.515
6525 H 402 LYS HZ3 6525 0.340000 1.00800 ; qtot
0.855
6526 C 402 LYS C 6526 0.734101 12.01000 ; qtot
1.589
6527 O 402 LYS O 6527 -0.589401 16.00000 ; qtot
1.000
6528 N 403 ILE N 6528 -0.415700 14.01000 ; qtot
0.584
6529 H 403 ILE H 6529 0.271900 1.00800 ; qtot
0.856
6530 CX 403 ILE CA 6530 -0.059700 12.01000 ; qtot
0.797
6531 H1 403 ILE HA 6531 0.086900 1.00800 ; qtot
0.883
6532 A3C 403 ILE CB 6532 0.130300 12.01000 ; qtot
1.014
6533 HC 403 ILE HB 6533 0.018700 1.00800 ; qtot
1.032
6534 CT 403 ILE CG2 6534 -0.320400 12.01000 ; qtot
0.712
6535 HC 403 ILE HG21 6535 0.088200 1.00800 ; qtot
0.800
6536 HC 403 ILE HG22 6536 0.088200 1.00800 ; qtot
0.888
6537 HC 403 ILE HG23 6537 0.088200 1.00800 ; qtot
0.977
6538 A2C 403 ILE CG1 6538 -0.043000 12.01000 ; qtot
0.934
6539 HC 403 ILE HG12 6539 0.023600 1.00800 ; qtot
0.957
6540 HC 403 ILE HG13 6540 0.023600 1.00800 ; qtot
0.981
6541 CT 403 ILE CD1 6541 -0.066000 12.01000 ; qtot
0.915
6542 HC 403 ILE HD11 6542 0.018600 1.00800 ; qtot
0.933
6543 HC 403 ILE HD12 6543 0.018600 1.00800 ; qtot
0.952
6544 HC 403 ILE HD13 6544 0.018600 1.00800 ; qtot
0.971
6545 C 403 ILE C 6545 0.597301 12.01000 ; qtot
1.568
6546 O 403 ILE O 6546 -0.567901 16.00000 ; qtot
1.000
6547 N 404 SER N 6547 -0.415700 14.01000 ; qtot
0.584
6548 H 404 SER H 6548 0.271900 1.00800 ; qtot
0.856
6549 CX 404 SER CA 6549 -0.024900 12.01000 ; qtot
0.831

```
  6550   H1   404   SER    HA 6550     0.084300      1.00800 ; qtot
0.916
  6551  A2C   404   SER    CB 6551     0.211700     12.01000 ; qtot
1.127
  6552   H1   404   SER   HB2 6552     0.035200      1.00800 ; qtot
1.163
  6553   H1   404   SER   HB3 6553     0.035200      1.00800 ; qtot
1.198
  6554   OH   404   SER    OG 6554    -0.654601     16.00000 ; qtot
0.543
  6555   HO   404   SER    HG 6555     0.427500      1.00800 ; qtot
0.971
  6556    C   404   SER     C 6556     0.597301     12.01000 ; qtot
1.568
  6557    O   404   SER     O 6557    -0.567901     16.00000 ; qtot
1.000
  6558    N   405   VAL     N 6558    -0.415700     14.01000 ; qtot
0.584
  6559    H   405   VAL     H 6559     0.271900      1.00800 ; qtot
0.856
  6560   CX   405   VAL    CA 6560    -0.087500     12.01000 ; qtot
0.769
  6561   H1   405   VAL    HA 6561     0.096900      1.00800 ; qtot
0.866
  6562  A3C   405   VAL    CB 6562     0.298500     12.01000 ; qtot
1.164
  6563   HC   405   VAL    HB 6563    -0.029700      1.00800 ; qtot
1.134
  6564   CT   405   VAL   CG1 6564    -0.319200     12.01000 ; qtot
0.815
  6565   HC   405   VAL  HG11 6565     0.079100      1.00800 ; qtot
0.894
  6566   HC   405   VAL  HG12 6566     0.079100      1.00800 ; qtot
0.973
  6567   HC   405   VAL  HG13 6567     0.079100      1.00800 ; qtot
1.053
  6568   CT   405   VAL   CG2 6568    -0.319200     12.01000 ; qtot
0.733
  6569   HC   405   VAL  HG21 6569     0.079100      1.00800 ; qtot
0.812
  6570   HC   405   VAL  HG22 6570     0.079100      1.00800 ; qtot
0.892
  6571   HC   405   VAL  HG23 6571     0.079100      1.00800 ; qtot
0.971
  6572    C   405   VAL     C 6572     0.597301     12.01000 ; qtot
1.568
  6573    O   405   VAL     O 6573    -0.567901     16.00000 ; qtot
1.000
  6574    N   406   GLU     N 6574    -0.516301     14.01000 ; qtot
0.484
  6575    H   406   GLU     H 6575     0.293600      1.00800 ; qtot
0.777
```

```
  6576   CX   406   GLU    CA 6576     0.039700     12.01000 ; qtot
0.817
  6577   H1   406   GLU    HA 6577     0.110500      1.00800 ; qtot
0.928
  6578  A2C   406   GLU    CB 6578     0.056000     12.01000 ; qtot
0.984
  6579   HC   406   GLU   HB2 6579    -0.017300      1.00800 ; qtot
0.966
  6580   HC   406   GLU   HB3 6580    -0.017300      1.00800 ; qtot
0.949
  6581  A2C   406   GLU    CG 6581     0.013600     12.01000 ; qtot
0.963
  6582   HC   406   GLU   HG2 6582    -0.042500      1.00800 ; qtot
0.920
  6583   HC   406   GLU   HG3 6583    -0.042500      1.00800 ; qtot
0.878
  6584   CO   406   GLU    CD 6584     0.805401     12.01000 ; qtot
1.683
  6585   O2   406   GLU   OE1 6585    -0.818801     16.00000 ; qtot
0.864
  6586   O2   406   GLU   OE2 6586    -0.818801     16.00000 ; qtot
0.045
  6587    C   406   GLU     C 6587     0.536601     12.01000 ; qtot
0.582
  6588    O   406   GLU     O 6588    -0.581901     16.00000 ; qtot
0.000
  6589    N   407   GLH     N 6589    -0.415700     14.01000 ; qtot -
0.416
  6590    H   407   GLH     H 6590     0.271900      1.00800 ; qtot -
0.144
  6591   CX   407   GLH    CA 6591     0.014500     12.01000 ; qtot -
0.129
  6592   H1   407   GLH    HA 6592     0.077900      1.00800 ; qtot -
0.051
  6593  A2C   407   GLH    CB 6593    -0.007100     12.01000 ; qtot -
0.058
  6594   HC   407   GLH   HB2 6594     0.025600      1.00800 ; qtot -
0.033
  6595   HC   407   GLH   HB3 6595     0.025600      1.00800 ; qtot -
0.007
  6596  A2C   407   GLH    CG 6596    -0.017400     12.01000 ; qtot -
0.025
  6597   HC   407   GLH   HG2 6597     0.043000      1.00800 ; qtot
0.018
  6598   HC   407   GLH   HG3 6598     0.043000      1.00800 ; qtot
0.061
  6599    C   407   GLH    CD 6599     0.680101     12.01000 ; qtot
0.741
  6600    O   407   GLH   OE1 6600    -0.583801     16.00000 ; qtot
0.158
  6601   OH   407   GLH   OE2 6601    -0.651101     16.00000 ; qtot -
0.493
```

6602 HO 407 GLH HE2 6602 0.464100 1.00800 ; qtot -
0.029
6603 C 407 GLH C 6603 0.597301 12.01000 ; qtot
0.568
6604 O 407 GLH O 6604 -0.567901 16.00000 ; qtot
0.000
6605 N 408 LEU N 6605 -0.415700 14.01000 ; qtot -
0.416
6606 H 408 LEU H 6606 0.271900 1.00800 ; qtot -
0.144
6607 CX 408 LEU CA 6607 -0.051800 12.01000 ; qtot -
0.196
6608 H1 408 LEU HA 6608 0.092200 1.00800 ; qtot -
0.103
6609 A2C 408 LEU CB 6609 -0.110200 12.01000 ; qtot -
0.214
6610 HC 408 LEU HB2 6610 0.045700 1.00800 ; qtot -
0.168
6611 HC 408 LEU HB3 6611 0.045700 1.00800 ; qtot -
0.122
6612 A3C 408 LEU CG 6612 0.353100 12.01000 ; qtot
0.231
6613 HC 408 LEU HG 6613 -0.036100 1.00800 ; qtot
0.195
6614 CT 408 LEU CD1 6614 -0.412100 12.01000 ; qtot -
0.217
6615 HC 408 LEU HD11 6615 0.100000 1.00800 ; qtot -
0.117
6616 HC 408 LEU HD12 6616 0.100000 1.00800 ; qtot -
0.017
6617 HC 408 LEU HD13 6617 0.100000 1.00800 ; qtot
0.083
6618 CT 408 LEU CD2 6618 -0.412100 12.01000 ; qtot -
0.329
6619 HC 408 LEU HD21 6619 0.100000 1.00800 ; qtot -
0.229
6620 HC 408 LEU HD22 6620 0.100000 1.00800 ; qtot -
0.129
6621 HC 408 LEU HD23 6621 0.100000 1.00800 ; qtot -
0.029
6622 C 408 LEU C 6622 0.597301 12.01000 ; qtot
0.568
6623 O 408 LEU O 6623 -0.567901 16.00000 ; qtot
0.000
6624 N 409 LYS N 6624 -0.347900 14.01000 ; qtot -
0.348
6625 H 409 LYS H 6625 0.274700 1.00800 ; qtot -
0.073
6626 CX 409 LYS CA 6626 -0.240000 12.01000 ; qtot -
0.313
6627 H1 409 LYS HA 6627 0.142600 1.00800 ; qtot -
0.171

6628 C8 409 LYS CB 6628 -0.009400 12.01000 ; qtot -
0.180
6629 HC 409 LYS HB2 6629 0.036200 1.00800 ; qtot -
0.144
6630 HC 409 LYS HB3 6630 0.036200 1.00800 ; qtot -
0.108
6631 C8 409 LYS CG 6631 0.018700 12.01000 ; qtot -
0.089
6632 HC 409 LYS HG2 6632 0.010300 1.00800 ; qtot -
0.079
6633 HC 409 LYS HG3 6633 0.010300 1.00800 ; qtot -
0.068
6634 C8 409 LYS CD 6634 -0.047900 12.01000 ; qtot -
0.116
6635 HC 409 LYS HD2 6635 0.062100 1.00800 ; qtot -
0.054
6636 HC 409 LYS HD3 6636 0.062100 1.00800 ; qtot
0.008
6637 C8 409 LYS CE 6637 -0.014300 12.01000 ; qtot -
0.006
6638 HP 409 LYS HE2 6638 0.113500 1.00800 ; qtot
0.107
6639 HP 409 LYS HE3 6639 0.113500 1.00800 ; qtot
0.221
6640 N3 409 LYS NZ 6640 -0.385400 14.01000 ; qtot -
0.165
6641 H 409 LYS HZ1 6641 0.340000 1.00800 ; qtot
0.175
6642 H 409 LYS HZ2 6642 0.340000 1.00800 ; qtot
0.515
6643 H 409 LYS HZ3 6643 0.340000 1.00800 ; qtot
0.855
6644 C 409 LYS C 6644 0.734101 12.01000 ; qtot
1.589
6645 O 409 LYS O 6645 -0.589401 16.00000 ; qtot
1.000
6646 N 410 ARG N 6646 -0.347900 14.01000 ; qtot
0.652
6647 H 410 ARG H 6647 0.274700 1.00800 ; qtot
0.927
6648 CX 410 ARG CA 6648 -0.263700 12.01000 ; qtot
0.663
6649 H1 410 ARG HA 6649 0.156000 1.00800 ; qtot
0.819
6650 C8 410 ARG CB 6650 -0.000700 12.01000 ; qtot
0.818
6651 HC 410 ARG HB2 6651 0.032700 1.00800 ; qtot
0.851
6652 HC 410 ARG HB3 6652 0.032700 1.00800 ; qtot
0.884
6653 C8 410 ARG CG 6653 0.039000 12.01000 ; qtot
0.923

6654 HC 410 ARG HG2 6654 0.028500 1.00800 ; qtot
0.951
6655 HC 410 ARG HG3 6655 0.028500 1.00800 ; qtot
0.980
6656 C8 410 ARG CD 6656 0.048600 12.01000 ; qtot
1.028
6657 H1 410 ARG HD2 6657 0.068700 1.00800 ; qtot
1.097
6658 H1 410 ARG HD3 6658 0.068700 1.00800 ; qtot
1.166
6659 N2 410 ARG NE 6659 -0.529501 14.01000 ; qtot
0.636
6660 H 410 ARG HE 6660 0.345600 1.00800 ; qtot
0.982
6661 CA 410 ARG CZ 6661 0.807601 12.01000 ; qtot
1.790
6662 N2 410 ARG NH1 6662 -0.862701 14.01000 ; qtot
0.927
6663 H 410 ARG HH11 6663 0.447800 1.00800 ; qtot
1.375
6664 H 410 ARG HH12 6664 0.447800 1.00800 ; qtot
1.822
6665 N2 410 ARG NH2 6665 -0.862701 14.01000 ; qtot
0.960
6666 H 410 ARG HH21 6666 0.447800 1.00800 ; qtot
1.408
6667 H 410 ARG HH22 6667 0.447800 1.00800 ; qtot
1.855
6668 C 410 ARG C 6668 0.734101 12.01000 ; qtot
2.589
6669 O 410 ARG O 6669 -0.589401 16.00000 ; qtot
2.000
6670 N 411 ILE N 6670 -0.415700 14.01000 ; qtot
1.584
6671 H 411 ILE H 6671 0.271900 1.00800 ; qtot
1.856
6672 CX 411 ILE CA 6672 -0.059700 12.01000 ; qtot
1.797
6673 H1 411 ILE HA 6673 0.086900 1.00800 ; qtot
1.883
6674 A3C 411 ILE CB 6674 0.130300 12.01000 ; qtot
2.014
6675 HC 411 ILE HB 6675 0.018700 1.00800 ; qtot
2.032
6676 CT 411 ILE CG2 6676 -0.320400 12.01000 ; qtot
1.712
6677 HC 411 ILE HG21 6677 0.088200 1.00800 ; qtot
1.800
6678 HC 411 ILE HG22 6678 0.088200 1.00800 ; qtot
1.888
6679 HC 411 ILE HG23 6679 0.088200 1.00800 ; qtot
1.977

6680  A2C   411   ILE   CG1 6680    -0.043000     12.01000 ; qtot
1.934
6681   HC   411   ILE  HG12 6681     0.023600      1.00800 ; qtot
1.957
6682   HC   411   ILE  HG13 6682     0.023600      1.00800 ; qtot
1.981
6683   CT   411   ILE   CD1 6683    -0.066000     12.01000 ; qtot
1.915
6684   HC   411   ILE  HD11 6684     0.018600      1.00800 ; qtot
1.933
6685   HC   411   ILE  HD12 6685     0.018600      1.00800 ; qtot
1.952
6686   HC   411   ILE  HD13 6686     0.018600      1.00800 ; qtot
1.971
6687    C   411   ILE     C 6687     0.597301     12.01000 ; qtot
2.568
6688    O   411   ILE     O 6688    -0.567901     16.00000 ; qtot
2.000
6689    N   412   PHE     N 6689    -0.415700     14.01000 ; qtot
1.584
6690    H   412   PHE     H 6690     0.271900      1.00800 ; qtot
1.856
6691   CX   412   PHE    CA 6691    -0.002400     12.01000 ; qtot
1.854
6692   H1   412   PHE    HA 6692     0.097800      1.00800 ; qtot
1.952
6693   CT   412   PHE    CB 6693    -0.034300     12.01000 ; qtot
1.917
6694   HC   412   PHE   HB2 6694     0.029500      1.00800 ; qtot
1.947
6695   HC   412   PHE   HB3 6695     0.029500      1.00800 ; qtot
1.976
6696   CA   412   PHE    CG 6696     0.011800     12.01000 ; qtot
1.988
6697   CA   412   PHE   CD1 6697    -0.125600     12.01000 ; qtot
1.863
6698   HA   412   PHE   HD1 6698     0.133000      1.00800 ; qtot
1.996
6699   CA   412   PHE   CE1 6699    -0.170400     12.01000 ; qtot
1.825
6700   HA   412   PHE   HE1 6700     0.143000      1.00800 ; qtot
1.968
6701   CA   412   PHE    CZ 6701    -0.107200     12.01000 ; qtot
1.861
6702   HA   412   PHE    HZ 6702     0.129700      1.00800 ; qtot
1.991
6703   CA   412   PHE   CE2 6703    -0.170400     12.01000 ; qtot
1.820
6704   HA   412   PHE   HE2 6704     0.143000      1.00800 ; qtot
1.963
6705   CA   412   PHE   CD2 6705    -0.125600     12.01000 ; qtot
1.838

6706 HA 412 PHE HD2 6706 0.133000 1.00800 ; qtot 1.971
6707 C 412 PHE C 6707 0.597301 12.01000 ; qtot 2.568
6708 O 412 PHE O 6708 -0.567901 16.00000 ; qtot 2.000
6709 N 413 GLY N 6709 -0.415700 14.01000 ; qtot 1.584
6710 H 413 GLY H 6710 0.271900 1.00800 ; qtot 1.856
6711 CX 413 GLY CA 6711 -0.025200 12.01000 ; qtot 1.831
6712 H1 413 GLY HA2 6712 0.069800 1.00800 ; qtot 1.901
6713 H1 413 GLY HA3 6713 0.069800 1.00800 ; qtot 1.971
6714 C 413 GLY C 6714 0.597301 12.01000 ; qtot 2.568
6715 O 413 GLY O 6715 -0.567901 16.00000 ; qtot 2.000
6716 N 414 ARG N 6716 -0.347900 14.01000 ; qtot 1.652
6717 H 414 ARG H 6717 0.274700 1.00800 ; qtot 1.927
6718 CX 414 ARG CA 6718 -0.263700 12.01000 ; qtot 1.663
6719 H1 414 ARG HA 6719 0.156000 1.00800 ; qtot 1.819
6720 C8 414 ARG CB 6720 -0.000700 12.01000 ; qtot 1.818
6721 HC 414 ARG HB2 6721 0.032700 1.00800 ; qtot 1.851
6722 HC 414 ARG HB3 6722 0.032700 1.00800 ; qtot 1.884
6723 C8 414 ARG CG 6723 0.039000 12.01000 ; qtot 1.923
6724 HC 414 ARG HG2 6724 0.028500 1.00800 ; qtot 1.951
6725 HC 414 ARG HG3 6725 0.028500 1.00800 ; qtot 1.980
6726 C8 414 ARG CD 6726 0.048600 12.01000 ; qtot 2.028
6727 H1 414 ARG HD2 6727 0.068700 1.00800 ; qtot 2.097
6728 H1 414 ARG HD3 6728 0.068700 1.00800 ; qtot 2.166
6729 N2 414 ARG NE 6729 -0.529501 14.01000 ; qtot 1.636
6730 H 414 ARG HE 6730 0.345600 1.00800 ; qtot 1.982
6731 CA 414 ARG CZ 6731 0.807601 12.01000 ; qtot 2.790

```
  6732   N2   414   ARG   NH1 6732    -0.862701     14.01000 ; qtot
1.927
  6733    H   414   ARG  HH11 6733     0.447800      1.00800 ; qtot
2.375
  6734    H   414   ARG  HH12 6734     0.447800      1.00800 ; qtot
2.822
  6735   N2   414   ARG   NH2 6735    -0.862701     14.01000 ; qtot
1.960
  6736    H   414   ARG  HH21 6736     0.447800      1.00800 ; qtot
2.408
  6737    H   414   ARG  HH22 6737     0.447800      1.00800 ; qtot
2.855
  6738    C   414   ARG     C 6738     0.734101     12.01000 ; qtot
3.589
  6739    O   414   ARG     O 6739    -0.589401     16.00000 ; qtot
3.000
  6740    N   415   ASP     N 6740    -0.516301     14.01000 ; qtot
2.484
  6741    H   415   ASP     H 6741     0.293600      1.00800 ; qtot
2.777
  6742   CX   415   ASP    CA 6742     0.038100     12.01000 ; qtot
2.815
  6743   H1   415   ASP    HA 6743     0.088000      1.00800 ; qtot
2.903
  6744  A2C   415   ASP    CB 6744    -0.030300     12.01000 ; qtot
2.873
  6745   HC   415   ASP   HB2 6745    -0.012200      1.00800 ; qtot
2.861
  6746   HC   415   ASP   HB3 6746    -0.012200      1.00800 ; qtot
2.849
  6747   CO   415   ASP    CG 6747     0.799401     12.01000 ; qtot
3.648
  6748   O2   415   ASP   OD1 6748    -0.801401     16.00000 ; qtot
2.847
  6749   O2   415   ASP   OD2 6749    -0.801401     16.00000 ; qtot
2.045
  6750    C   415   ASP     C 6750     0.536601     12.01000 ; qtot
2.582
  6751    O   415   ASP     O 6751    -0.581901     16.00000 ; qtot
2.000
  6752    N   416   ASP     N 6752    -0.516301     14.01000 ; qtot
1.484
  6753    H   416   ASP     H 6753     0.293600      1.00800 ; qtot
1.777
  6754   CX   416   ASP    CA 6754     0.038100     12.01000 ; qtot
1.815
  6755   H1   416   ASP    HA 6755     0.088000      1.00800 ; qtot
1.903
  6756  A2C   416   ASP    CB 6756    -0.030300     12.01000 ; qtot
1.873
  6757   HC   416   ASP   HB2 6757    -0.012200      1.00800 ; qtot
1.861
```

```
  6758   HC   416   ASP   HB3 6758    -0.012200      1.00800 ; qtot
1.849
  6759   CO   416   ASP    CG 6759     0.799401     12.01000 ; qtot
2.648
  6760   O2   416   ASP   OD1 6760    -0.801401     16.00000 ; qtot
1.847
  6761   O2   416   ASP   OD2 6761    -0.801401     16.00000 ; qtot
1.045
  6762    C   416   ASP     C 6762     0.536601     12.01000 ; qtot
1.582
  6763    O   416   ASP     O 6763    -0.581901     16.00000 ; qtot
1.000
  6764    N   417   ILE     N 6764    -0.415700     14.01000 ; qtot
0.584
  6765    H   417   ILE     H 6765     0.271900      1.00800 ; qtot
0.856
  6766   CX   417   ILE    CA 6766    -0.059700     12.01000 ; qtot
0.797
  6767   H1   417   ILE    HA 6767     0.086900      1.00800 ; qtot
0.883
  6768  A3C   417   ILE    CB 6768     0.130300     12.01000 ; qtot
1.014
  6769   HC   417   ILE    HB 6769     0.018700      1.00800 ; qtot
1.032
  6770   CT   417   ILE   CG2 6770    -0.320400     12.01000 ; qtot
0.712
  6771   HC   417   ILE  HG21 6771     0.088200      1.00800 ; qtot
0.800
  6772   HC   417   ILE  HG22 6772     0.088200      1.00800 ; qtot
0.888
  6773   HC   417   ILE  HG23 6773     0.088200      1.00800 ; qtot
0.977
  6774  A2C   417   ILE   CG1 6774    -0.043000     12.01000 ; qtot
0.934
  6775   HC   417   ILE  HG12 6775     0.023600      1.00800 ; qtot
0.957
  6776   HC   417   ILE  HG13 6776     0.023600      1.00800 ; qtot
0.981
  6777   CT   417   ILE   CD1 6777    -0.066000     12.01000 ; qtot
0.915
  6778   HC   417   ILE  HD11 6778     0.018600      1.00800 ; qtot
0.933
  6779   HC   417   ILE  HD12 6779     0.018600      1.00800 ; qtot
0.952
  6780   HC   417   ILE  HD13 6780     0.018600      1.00800 ; qtot
0.971
  6781    C   417   ILE     C 6781     0.597301     12.01000 ; qtot
1.568
  6782    O   417   ILE     O 6782    -0.567901     16.00000 ; qtot
1.000
  6783    N   418   GLU     N 6783    -0.516301     14.01000 ; qtot
0.484
```

```
  6784    H   418   GLU     H 6784     0.293600      1.00800 ; qtot
0.777
  6785   CX   418   GLU    CA 6785     0.039700     12.01000 ; qtot
0.817
  6786   H1   418   GLU    HA 6786     0.110500      1.00800 ; qtot
0.928
  6787  A2C   418   GLU    CB 6787     0.056000     12.01000 ; qtot
0.984
  6788   HC   418   GLU   HB2 6788    -0.017300      1.00800 ; qtot
0.966
  6789   HC   418   GLU   HB3 6789    -0.017300      1.00800 ; qtot
0.949
  6790  A2C   418   GLU    CG 6790     0.013600     12.01000 ; qtot
0.963
  6791   HC   418   GLU   HG2 6791    -0.042500      1.00800 ; qtot
0.920
  6792   HC   418   GLU   HG3 6792    -0.042500      1.00800 ; qtot
0.878
  6793   CO   418   GLU    CD 6793     0.805401     12.01000 ; qtot
1.683
  6794   O2   418   GLU   OE1 6794    -0.818801     16.00000 ; qtot
0.864
  6795   O2   418   GLU   OE2 6795    -0.818801     16.00000 ; qtot
0.045
  6796    C   418   GLU     C 6796     0.536601     12.01000 ; qtot
0.582
  6797    O   418   GLU     O 6797    -0.581901     16.00000 ; qtot
0.000
  6798    N   419   ASN     N 6798    -0.415700     14.01000 ; qtot -
0.416
  6799    H   419   ASN     H 6799     0.271900      1.00800 ; qtot -
0.144
  6800   CX   419   ASN    CA 6800     0.014300     12.01000 ; qtot -
0.129
  6801   H1   419   ASN    HA 6801     0.104800      1.00800 ; qtot -
0.025
  6802  A2C   419   ASN    CB 6802    -0.204100     12.01000 ; qtot -
0.229
  6803   HC   419   ASN   HB2 6803     0.079700      1.00800 ; qtot -
0.149
  6804   HC   419   ASN   HB3 6804     0.079700      1.00800 ; qtot -
0.069
  6805    C   419   ASN    CG 6805     0.713001     12.01000 ; qtot
0.644
  6806    O   419   ASN   OD1 6806    -0.593101     16.00000 ; qtot
0.051
  6807    N   419   ASN   ND2 6807    -0.919101     14.01000 ; qtot -
0.869
  6808    H   419   ASN  HD21 6808     0.419600      1.00800 ; qtot -
0.449
  6809    H   419   ASN  HD22 6809     0.419600      1.00800 ; qtot -
0.029
```

6810 C 419 ASN C 6810 0.597301 12.01000 ; qtot
0.568
6811 O 419 ASN O 6811 -0.567901 16.00000 ; qtot
0.000
6812 N 420 PRO N 6812 -0.254800 14.01000 ; qtot -
0.255
6813 CT 420 PRO CD 6813 0.019200 12.01000 ; qtot -
0.236
6814 H1 420 PRO HD2 6814 0.039100 1.00800 ; qtot -
0.196
6815 H1 420 PRO HD3 6815 0.039100 1.00800 ; qtot -
0.157
6816 CT 420 PRO CG 6816 0.018900 12.01000 ; qtot -
0.138
6817 HC 420 PRO HG2 6817 0.021300 1.00800 ; qtot -
0.117
6818 HC 420 PRO HG3 6818 0.021300 1.00800 ; qtot -
0.096
6819 CT 420 PRO CB 6819 -0.007000 12.01000 ; qtot -
0.103
6820 HC 420 PRO HB2 6820 0.025300 1.00800 ; qtot -
0.078
6821 HC 420 PRO HB3 6821 0.025300 1.00800 ; qtot -
0.052
6822 CX 420 PRO CA 6822 -0.026600 12.01000 ; qtot -
0.079
6823 H1 420 PRO HA 6823 0.064100 1.00800 ; qtot -
0.015
6824 C 420 PRO C 6824 0.589601 12.01000 ; qtot
0.575
6825 O 420 PRO O 6825 -0.574801 16.00000 ; qtot
0.000
6826 N 421 LEU N 6826 -0.415700 14.01000 ; qtot -
0.416
6827 H 421 LEU H 6827 0.271900 1.00800 ; qtot -
0.144
6828 CX 421 LEU CA 6828 -0.051800 12.01000 ; qtot -
0.196
6829 H1 421 LEU HA 6829 0.092200 1.00800 ; qtot -
0.103
6830 A2C 421 LEU CB 6830 -0.110200 12.01000 ; qtot -
0.214
6831 HC 421 LEU HB2 6831 0.045700 1.00800 ; qtot -
0.168
6832 HC 421 LEU HB3 6832 0.045700 1.00800 ; qtot -
0.122
6833 A3C 421 LEU CG 6833 0.353100 12.01000 ; qtot
0.231
6834 HC 421 LEU HG 6834 -0.036100 1.00800 ; qtot
0.195
6835 CT 421 LEU CD1 6835 -0.412100 12.01000 ; qtot -
0.217

6836 HC 421 LEU HD11 6836 0.100000 1.00800 ; qtot -
0.117
6837 HC 421 LEU HD12 6837 0.100000 1.00800 ; qtot -
0.017
6838 HC 421 LEU HD13 6838 0.100000 1.00800 ; qtot
0.083
6839 CT 421 LEU CD2 6839 -0.412100 12.01000 ; qtot -
0.329
6840 HC 421 LEU HD21 6840 0.100000 1.00800 ; qtot -
0.229
6841 HC 421 LEU HD22 6841 0.100000 1.00800 ; qtot -
0.129
6842 HC 421 LEU HD23 6842 0.100000 1.00800 ; qtot -
0.029
6843 C 421 LEU C 6843 0.597301 12.01000 ; qtot
0.568
6844 O 421 LEU O 6844 -0.567901 16.00000 ; qtot
0.000
6845 N 422 ILE N 6845 -0.415700 14.01000 ; qtot -
0.416
6846 H 422 ILE H 6846 0.271900 1.00800 ; qtot -
0.144
6847 CX 422 ILE CA 6847 -0.059700 12.01000 ; qtot -
0.203
6848 H1 422 ILE HA 6848 0.086900 1.00800 ; qtot -
0.117
6849 A3C 422 ILE CB 6849 0.130300 12.01000 ; qtot
0.014
6850 HC 422 ILE HB 6850 0.018700 1.00800 ; qtot
0.032
6851 CT 422 ILE CG2 6851 -0.320400 12.01000 ; qtot -
0.288
6852 HC 422 ILE HG21 6852 0.088200 1.00800 ; qtot -
0.200
6853 HC 422 ILE HG22 6853 0.088200 1.00800 ; qtot -
0.112
6854 HC 422 ILE HG23 6854 0.088200 1.00800 ; qtot -
0.023
6855 A2C 422 ILE CG1 6855 -0.043000 12.01000 ; qtot -
0.066
6856 HC 422 ILE HG12 6856 0.023600 1.00800 ; qtot -
0.043
6857 HC 422 ILE HG13 6857 0.023600 1.00800 ; qtot -
0.019
6858 CT 422 ILE CD1 6858 -0.066000 12.01000 ; qtot -
0.085
6859 HC 422 ILE HD11 6859 0.018600 1.00800 ; qtot -
0.067
6860 HC 422 ILE HD12 6860 0.018600 1.00800 ; qtot -
0.048
6861 HC 422 ILE HD13 6861 0.018600 1.00800 ; qtot -
0.029

6862 C 422 ILE C 6862 0.597301 12.01000 ; qtot
0.568
6863 O 422 ILE O 6863 -0.567901 16.00000 ; qtot
0.000
6864 N 423 ASP N 6864 -0.516301 14.01000 ; qtot -
0.516
6865 H 423 ASP H 6865 0.293600 1.00800 ; qtot -
0.223
6866 CX 423 ASP CA 6866 0.038100 12.01000 ; qtot -
0.185
6867 H1 423 ASP HA 6867 0.088000 1.00800 ; qtot -
0.097
6868 A2C 423 ASP CB 6868 -0.030300 12.01000 ; qtot -
0.127
6869 HC 423 ASP HB2 6869 -0.012200 1.00800 ; qtot -
0.139
6870 HC 423 ASP HB3 6870 -0.012200 1.00800 ; qtot -
0.151
6871 CO 423 ASP CG 6871 0.799401 12.01000 ; qtot
0.648
6872 O2 423 ASP OD1 6872 -0.801401 16.00000 ; qtot -
0.153
6873 O2 423 ASP OD2 6873 -0.801401 16.00000 ; qtot -
0.955
6874 C 423 ASP C 6874 0.536601 12.01000 ; qtot -
0.418
6875 O 423 ASP O 6875 -0.581901 16.00000 ; qtot -
1.000
6876 N 424 LYS N 6876 -0.347900 14.01000 ; qtot -
1.348
6877 H 424 LYS H 6877 0.274700 1.00800 ; qtot -
1.073
6878 CX 424 LYS CA 6878 -0.240000 12.01000 ; qtot -
1.313
6879 H1 424 LYS HA 6879 0.142600 1.00800 ; qtot -
1.171
6880 C8 424 LYS CB 6880 -0.009400 12.01000 ; qtot -
1.180
6881 HC 424 LYS HB2 6881 0.036200 1.00800 ; qtot -
1.144
6882 HC 424 LYS HB3 6882 0.036200 1.00800 ; qtot -
1.108
6883 C8 424 LYS CG 6883 0.018700 12.01000 ; qtot -
1.089
6884 HC 424 LYS HG2 6884 0.010300 1.00800 ; qtot -
1.079
6885 HC 424 LYS HG3 6885 0.010300 1.00800 ; qtot -
1.068
6886 C8 424 LYS CD 6886 -0.047900 12.01000 ; qtot -
1.116
6887 HC 424 LYS HD2 6887 0.062100 1.00800 ; qtot -
1.054

```
  6888   HC   424   LYS   HD3 6888     0.062100      1.00800 ; qtot -
0.992
  6889   C8   424   LYS    CE 6889    -0.014300     12.01000 ; qtot -
1.006
  6890   HP   424   LYS   HE2 6890     0.113500      1.00800 ; qtot -
0.893
  6891   HP   424   LYS   HE3 6891     0.113500      1.00800 ; qtot -
0.779
  6892   N3   424   LYS    NZ 6892    -0.385400     14.01000 ; qtot -
1.165
  6893    H   424   LYS   HZ1 6893     0.340000      1.00800 ; qtot -
0.825
  6894    H   424   LYS   HZ2 6894     0.340000      1.00800 ; qtot -
0.485
  6895    H   424   LYS   HZ3 6895     0.340000      1.00800 ; qtot -
0.145
  6896    C   424   LYS     C 6896     0.734101     12.01000 ; qtot
0.589
  6897    O   424   LYS     O 6897    -0.589401     16.00000 ; qtot
0.000
  6898    N   425   ALA     N 6898    -0.415700     14.01000 ; qtot -
0.416
  6899    H   425   ALA     H 6899     0.271900      1.00800 ; qtot -
0.144
  6900   CX   425   ALA    CA 6900     0.033700     12.01000 ; qtot -
0.110
  6901   H1   425   ALA    HA 6901     0.082300      1.00800 ; qtot -
0.028
  6902   CT   425   ALA    CB 6902    -0.182500     12.01000 ; qtot -
0.210
  6903   HC   425   ALA   HB1 6903     0.060300      1.00800 ; qtot -
0.150
  6904   HC   425   ALA   HB2 6904     0.060300      1.00800 ; qtot -
0.090
  6905   HC   425   ALA   HB3 6905     0.060300      1.00800 ; qtot -
0.029
  6906    C   425   ALA     C 6906     0.597301     12.01000 ; qtot
0.568
  6907    O   425   ALA     O 6907    -0.567901     16.00000 ; qtot
0.000
  6908    N   426   ILE     N 6908    -0.415700     14.01000 ; qtot -
0.416
  6909    H   426   ILE     H 6909     0.271900      1.00800 ; qtot -
0.144
  6910   CX   426   ILE    CA 6910    -0.059700     12.01000 ; qtot -
0.203
  6911   H1   426   ILE    HA 6911     0.086900      1.00800 ; qtot -
0.117
  6912  A3C   426   ILE    CB 6912     0.130300     12.01000 ; qtot
0.014
  6913   HC   426   ILE    HB 6913     0.018700      1.00800 ; qtot
0.032
```

```
  6914   CT   426   ILE   CG2 6914    -0.320400     12.01000 ; qtot -
0.288
  6915   HC   426   ILE  HG21 6915     0.088200      1.00800 ; qtot -
0.200
  6916   HC   426   ILE  HG22 6916     0.088200      1.00800 ; qtot -
0.112
  6917   HC   426   ILE  HG23 6917     0.088200      1.00800 ; qtot -
0.023
  6918  A2C   426   ILE   CG1 6918    -0.043000     12.01000 ; qtot -
0.066
  6919   HC   426   ILE  HG12 6919     0.023600      1.00800 ; qtot -
0.043
  6920   HC   426   ILE  HG13 6920     0.023600      1.00800 ; qtot -
0.019
  6921   CT   426   ILE   CD1 6921    -0.066000     12.01000 ; qtot -
0.085
  6922   HC   426   ILE  HD11 6922     0.018600      1.00800 ; qtot -
0.067
  6923   HC   426   ILE  HD12 6923     0.018600      1.00800 ; qtot -
0.048
  6924   HC   426   ILE  HD13 6924     0.018600      1.00800 ; qtot -
0.029
  6925    C   426   ILE     C 6925     0.597301     12.01000 ; qtot
0.568
  6926    O   426   ILE     O 6926    -0.567901     16.00000 ; qtot
0.000
  6927    N   427   ASP     N 6927    -0.516301     14.01000 ; qtot -
0.516
  6928    H   427   ASP     H 6928     0.293600      1.00800 ; qtot -
0.223
  6929   CX   427   ASP    CA 6929     0.038100     12.01000 ; qtot -
0.185
  6930   H1   427   ASP    HA 6930     0.088000      1.00800 ; qtot -
0.097
  6931  A2C   427   ASP    CB 6931    -0.030300     12.01000 ; qtot -
0.127
  6932   HC   427   ASP   HB2 6932    -0.012200      1.00800 ; qtot -
0.139
  6933   HC   427   ASP   HB3 6933    -0.012200      1.00800 ; qtot -
0.151
  6934   CO   427   ASP    CG 6934     0.799401     12.01000 ; qtot
0.648
  6935   O2   427   ASP   OD1 6935    -0.801401     16.00000 ; qtot -
0.153
  6936   O2   427   ASP   OD2 6936    -0.801401     16.00000 ; qtot -
0.955
  6937    C   427   ASP     C 6937     0.536601     12.01000 ; qtot -
0.418
  6938    O   427   ASP     O 6938    -0.581901     16.00000 ; qtot -
1.000
  6939    N   428   SER     N 6939    -0.415700     14.01000 ; qtot -
1.416
```

6940 H 428 SER H 6940 0.271900 1.00800 ; qtot -
1.144
6941 CX 428 SER CA 6941 -0.024900 12.01000 ; qtot -
1.169
6942 H1 428 SER HA 6942 0.084300 1.00800 ; qtot -
1.084
6943 A2C 428 SER CB 6943 0.211700 12.01000 ; qtot -
0.873
6944 H1 428 SER HB2 6944 0.035200 1.00800 ; qtot -
0.837
6945 H1 428 SER HB3 6945 0.035200 1.00800 ; qtot -
0.802
6946 OH 428 SER OG 6946 -0.654601 16.00000 ; qtot -
1.457
6947 HO 428 SER HG 6947 0.427500 1.00800 ; qtot -
1.029
6948 C 428 SER C 6948 0.597301 12.01000 ; qtot -
0.432
6949 O 428 SER O 6949 -0.567901 16.00000 ; qtot -
1.000
6950 N 429 LEU N 6950 -0.415700 14.01000 ; qtot -
1.416
6951 H 429 LEU H 6951 0.271900 1.00800 ; qtot -
1.144
6952 CX 429 LEU CA 6952 -0.051800 12.01000 ; qtot -
1.196
6953 H1 429 LEU HA 6953 0.092200 1.00800 ; qtot -
1.103
6954 A2C 429 LEU CB 6954 -0.110200 12.01000 ; qtot -
1.214
6955 HC 429 LEU HB2 6955 0.045700 1.00800 ; qtot -
1.168
6956 HC 429 LEU HB3 6956 0.045700 1.00800 ; qtot -
1.122
6957 A3C 429 LEU CG 6957 0.353100 12.01000 ; qtot -
0.769
6958 HC 429 LEU HG 6958 -0.036100 1.00800 ; qtot -
0.805
6959 CT 429 LEU CD1 6959 -0.412100 12.01000 ; qtot -
1.217
6960 HC 429 LEU HD11 6960 0.100000 1.00800 ; qtot -
1.117
6961 HC 429 LEU HD12 6961 0.100000 1.00800 ; qtot -
1.017
6962 HC 429 LEU HD13 6962 0.100000 1.00800 ; qtot -
0.917
6963 CT 429 LEU CD2 6963 -0.412100 12.01000 ; qtot -
1.329
6964 HC 429 LEU HD21 6964 0.100000 1.00800 ; qtot -
1.229
6965 HC 429 LEU HD22 6965 0.100000 1.00800 ; qtot -
1.129

```
  6966   HC   429   LEU  HD23 6966     0.100000      1.00800 ; qtot -
1.029
  6967    C   429   LEU     C 6967     0.597301     12.01000 ; qtot -
0.432
  6968    O   429   LEU     O 6968    -0.567901     16.00000 ; qtot -
1.000
  6969    N   430   LEU     N 6969    -0.415700     14.01000 ; qtot -
1.416
  6970    H   430   LEU     H 6970     0.271900      1.00800 ; qtot -
1.144
  6971   CX   430   LEU    CA 6971    -0.051800     12.01000 ; qtot -
1.196
  6972   H1   430   LEU    HA 6972     0.092200      1.00800 ; qtot -
1.103
  6973  A2C   430   LEU    CB 6973    -0.110200     12.01000 ; qtot -
1.214
  6974   HC   430   LEU   HB2 6974     0.045700      1.00800 ; qtot -
1.168
  6975   HC   430   LEU   HB3 6975     0.045700      1.00800 ; qtot -
1.122
  6976  A3C   430   LEU    CG 6976     0.353100     12.01000 ; qtot -
0.769
  6977   HC   430   LEU    HG 6977    -0.036100      1.00800 ; qtot -
0.805
  6978   CT   430   LEU   CD1 6978    -0.412100     12.01000 ; qtot -
1.217
  6979   HC   430   LEU  HD11 6979     0.100000      1.00800 ; qtot -
1.117
  6980   HC   430   LEU  HD12 6980     0.100000      1.00800 ; qtot -
1.017
  6981   HC   430   LEU  HD13 6981     0.100000      1.00800 ; qtot -
0.917
  6982   CT   430   LEU   CD2 6982    -0.412100     12.01000 ; qtot -
1.329
  6983   HC   430   LEU  HD21 6983     0.100000      1.00800 ; qtot -
1.229
  6984   HC   430   LEU  HD22 6984     0.100000      1.00800 ; qtot -
1.129
  6985   HC   430   LEU  HD23 6985     0.100000      1.00800 ; qtot -
1.029
  6986    C   430   LEU     C 6986     0.597301     12.01000 ; qtot -
0.432
  6987    O   430   LEU     O 6987    -0.567901     16.00000 ; qtot -
1.000
  6988    N   431   GLN     N 6988    -0.415700     14.01000 ; qtot -
1.416
  6989    H   431   GLN     H 6989     0.271900      1.00800 ; qtot -
1.144
  6990   CX   431   GLN    CA 6990    -0.003100     12.01000 ; qtot -
1.147
  6991   H1   431   GLN    HA 6991     0.085000      1.00800 ; qtot -
1.062
```

6992 A2C 431 GLN CB 6992 -0.003600 12.01000 ; qtot -1.066
6993 HC 431 GLN HB2 6993 0.017100 1.00800 ; qtot -1.048
6994 HC 431 GLN HB3 6994 0.017100 1.00800 ; qtot -1.031
6995 A2C 431 GLN CG 6995 -0.064500 12.01000 ; qtot -1.096
6996 HC 431 GLN HG2 6996 0.035200 1.00800 ; qtot -1.061
6997 HC 431 GLN HG3 6997 0.035200 1.00800 ; qtot -1.025
6998 C 431 GLN CD 6998 0.695101 12.01000 ; qtot -0.330
6999 O 431 GLN OE1 6999 -0.608601 16.00000 ; qtot -0.939
7000 N 431 GLN NE2 7000 -0.940701 14.01000 ; qtot -1.880
7001 H 431 GLN HE21 7001 0.425100 1.00800 ; qtot -1.455
7002 H 431 GLN HE22 7002 0.425100 1.00800 ; qtot -1.029
7003 C 431 GLN C 7003 0.597301 12.01000 ; qtot -0.432
7004 O 431 GLN O 7004 -0.567901 16.00000 ; qtot -1.000
7005 N 432 GLU N 7005 -0.516301 14.01000 ; qtot -1.516
7006 H 432 GLU H 7006 0.293600 1.00800 ; qtot -1.223
7007 CX 432 GLU CA 7007 0.039700 12.01000 ; qtot -1.183
7008 H1 432 GLU HA 7008 0.110500 1.00800 ; qtot -1.073
7009 A2C 432 GLU CB 7009 0.056000 12.01000 ; qtot -1.016
7010 HC 432 GLU HB2 7010 -0.017300 1.00800 ; qtot -1.034
7011 HC 432 GLU HB3 7011 -0.017300 1.00800 ; qtot -1.051
7012 A2C 432 GLU CG 7012 0.013600 12.01000 ; qtot -1.038
7013 HC 432 GLU HG2 7013 -0.042500 1.00800 ; qtot -1.080
7014 HC 432 GLU HG3 7014 -0.042500 1.00800 ; qtot -1.123
7015 CO 432 GLU CD 7015 0.805401 12.01000 ; qtot -0.317
7016 O2 432 GLU OE1 7016 -0.818801 16.00000 ; qtot -1.136
7017 O2 432 GLU OE2 7017 -0.818801 16.00000 ; qtot -1.955

7018 C 432 GLU C 7018 0.536601 12.01000 ; qtot -
1.418
7019 O 432 GLU O 7019 -0.581901 16.00000 ; qtot -
2.000
7020 N 433 VAL N 7020 -0.415700 14.01000 ; qtot -
2.416
7021 H 433 VAL H 7021 0.271900 1.00800 ; qtot -
2.144
7022 CX 433 VAL CA 7022 -0.087500 12.01000 ; qtot -
2.231
7023 H1 433 VAL HA 7023 0.096900 1.00800 ; qtot -
2.134
7024 A3C 433 VAL CB 7024 0.298500 12.01000 ; qtot -
1.836
7025 HC 433 VAL HB 7025 -0.029700 1.00800 ; qtot -
1.866
7026 CT 433 VAL CG1 7026 -0.319200 12.01000 ; qtot -
2.185
7027 HC 433 VAL HG11 7027 0.079100 1.00800 ; qtot -
2.106
7028 HC 433 VAL HG12 7028 0.079100 1.00800 ; qtot -
2.027
7029 HC 433 VAL HG13 7029 0.079100 1.00800 ; qtot -
1.948
7030 CT 433 VAL CG2 7030 -0.319200 12.01000 ; qtot -
2.267
7031 HC 433 VAL HG21 7031 0.079100 1.00800 ; qtot -
2.188
7032 HC 433 VAL HG22 7032 0.079100 1.00800 ; qtot -
2.109
7033 HC 433 VAL HG23 7033 0.079100 1.00800 ; qtot -
2.029
7034 C 433 VAL C 7034 0.597301 12.01000 ; qtot -
1.432
7035 O 433 VAL O 7035 -0.567901 16.00000 ; qtot -
2.000
7036 N 434 ASP N 7036 -0.516301 14.01000 ; qtot -
2.516
7037 H 434 ASP H 7037 0.293600 1.00800 ; qtot -
2.223
7038 CX 434 ASP CA 7038 0.038100 12.01000 ; qtot -
2.185
7039 H1 434 ASP HA 7039 0.088000 1.00800 ; qtot -
2.097
7040 A2C 434 ASP CB 7040 -0.030300 12.01000 ; qtot -
2.127
7041 HC 434 ASP HB2 7041 -0.012200 1.00800 ; qtot -
2.139
7042 HC 434 ASP HB3 7042 -0.012200 1.00800 ; qtot -
2.151
7043 CO 434 ASP CG 7043 0.799401 12.01000 ; qtot -
1.352

7044 O2 434 ASP OD1 7044 -0.801401 16.00000 ; qtot -2.153
7045 O2 434 ASP OD2 7045 -0.801401 16.00000 ; qtot -2.955
7046 C 434 ASP C 7046 0.536601 12.01000 ; qtot -2.418
7047 O 434 ASP O 7047 -0.581901 16.00000 ; qtot -3.000
7048 N 435 LEU N 7048 -0.415700 14.01000 ; qtot -3.416
7049 H 435 LEU H 7049 0.271900 1.00800 ; qtot -3.144
7050 CX 435 LEU CA 7050 -0.051800 12.01000 ; qtot -3.196
7051 H1 435 LEU HA 7051 0.092200 1.00800 ; qtot -3.103
7052 A2C 435 LEU CB 7052 -0.110200 12.01000 ; qtot -3.214
7053 HC 435 LEU HB2 7053 0.045700 1.00800 ; qtot -3.168
7054 HC 435 LEU HB3 7054 0.045700 1.00800 ; qtot -3.122
7055 A3C 435 LEU CG 7055 0.353100 12.01000 ; qtot -2.769
7056 HC 435 LEU HG 7056 -0.036100 1.00800 ; qtot -2.805
7057 CT 435 LEU CD1 7057 -0.412100 12.01000 ; qtot -3.217
7058 HC 435 LEU HD11 7058 0.100000 1.00800 ; qtot -3.117
7059 HC 435 LEU HD12 7059 0.100000 1.00800 ; qtot -3.017
7060 HC 435 LEU HD13 7060 0.100000 1.00800 ; qtot -2.917
7061 CT 435 LEU CD2 7061 -0.412100 12.01000 ; qtot -3.329
7062 HC 435 LEU HD21 7062 0.100000 1.00800 ; qtot -3.229
7063 HC 435 LEU HD22 7063 0.100000 1.00800 ; qtot -3.129
7064 HC 435 LEU HD23 7064 0.100000 1.00800 ; qtot -3.029
7065 C 435 LEU C 7065 0.597301 12.01000 ; qtot -2.432
7066 O 435 LEU O 7066 -0.567901 16.00000 ; qtot -3.000
7067 N 436 ASN N 7067 -0.415700 14.01000 ; qtot -3.416
7068 H 436 ASN H 7068 0.271900 1.00800 ; qtot -3.144
7069 CX 436 ASN CA 7069 0.014300 12.01000 ; qtot -3.130

7070 H1 436 ASN HA 7070 0.104800 1.00800 ; qtot -
3.025
7071 A2C 436 ASN CB 7071 -0.204100 12.01000 ; qtot -
3.229
7072 HC 436 ASN HB2 7072 0.079700 1.00800 ; qtot -
3.149
7073 HC 436 ASN HB3 7073 0.079700 1.00800 ; qtot -
3.069
7074 C 436 ASN CG 7074 0.713001 12.01000 ; qtot -
2.356
7075 O 436 ASN OD1 7075 -0.593101 16.00000 ; qtot -
2.950
7076 N 436 ASN ND2 7076 -0.919101 14.01000 ; qtot -
3.869
7077 H 436 ASN HD21 7077 0.419600 1.00800 ; qtot -
3.449
7078 H 436 ASN HD22 7078 0.419600 1.00800 ; qtot -
3.029
7079 C 436 ASN C 7079 0.597301 12.01000 ; qtot -
2.432
7080 O 436 ASN O 7080 -0.567901 16.00000 ; qtot -
3.000
7081 N 437 GLY N 7081 -0.415700 14.01000 ; qtot -
3.416
7082 H 437 GLY H 7082 0.271900 1.00800 ; qtot -
3.144
7083 CX 437 GLY CA 7083 -0.025200 12.01000 ; qtot -
3.169
7084 H1 437 GLY HA2 7084 0.069800 1.00800 ; qtot -
3.099
7085 H1 437 GLY HA3 7085 0.069800 1.00800 ; qtot -
3.029
7086 C 437 GLY C 7086 0.597301 12.01000 ; qtot -
2.432
7087 O 437 GLY O 7087 -0.567901 16.00000 ; qtot -
3.000
7088 N 438 ASH N 7088 -0.415700 14.01000 ; qtot -
3.416
7089 H 438 ASH H 7089 0.271900 1.00800 ; qtot -
3.144
7090 CX 438 ASH CA 7090 0.034100 12.01000 ; qtot -
3.110
7091 H1 438 ASH HA 7091 0.086400 1.00800 ; qtot -
3.023
7092 A2C 438 ASH CB 7092 -0.031600 12.01000 ; qtot -
3.055
7093 HC 438 ASH HB2 7093 0.048800 1.00800 ; qtot -
3.006
7094 HC 438 ASH HB3 7094 0.048800 1.00800 ; qtot -
2.957
7095 C 438 ASH CG 7095 0.646201 12.01000 ; qtot -
2.311

7096 O 438 ASH OD1 7096 -0.555401 16.00000 ; qtot -2.867
7097 OH 438 ASH OD2 7097 -0.637601 16.00000 ; qtot -3.504
7098 HO 438 ASH HD2 7098 0.474700 1.00800 ; qtot -3.029
7099 C 438 ASH C 7099 0.597301 12.01000 ; qtot -2.432
7100 O 438 ASH O 7100 -0.567901 16.00000 ; qtot -3.000
7101 N 439 GLY N 7101 -0.415700 14.01000 ; qtot -3.416
7102 H 439 GLY H 7102 0.271900 1.00800 ; qtot -3.144
7103 CX 439 GLY CA 7103 -0.025200 12.01000 ; qtot -3.169
7104 H1 439 GLY HA2 7104 0.069800 1.00800 ; qtot -3.099
7105 H1 439 GLY HA3 7105 0.069800 1.00800 ; qtot -3.029
7106 C 439 GLY C 7106 0.597301 12.01000 ; qtot -2.432
7107 O 439 GLY O 7107 -0.567901 16.00000 ; qtot -3.000
7108 N 440 GLU N 7108 -0.516301 14.01000 ; qtot -3.516
7109 H 440 GLU H 7109 0.293600 1.00800 ; qtot -3.223
7110 CX 440 GLU CA 7110 0.039700 12.01000 ; qtot -3.183
7111 H1 440 GLU HA 7111 0.110500 1.00800 ; qtot -3.073
7112 A2C 440 GLU CB 7112 0.056000 12.01000 ; qtot -3.017
7113 HC 440 GLU HB2 7113 -0.017300 1.00800 ; qtot -3.034
7114 HC 440 GLU HB3 7114 -0.017300 1.00800 ; qtot -3.051
7115 A2C 440 GLU CG 7115 0.013600 12.01000 ; qtot -3.038
7116 HC 440 GLU HG2 7116 -0.042500 1.00800 ; qtot -3.080
7117 HC 440 GLU HG3 7117 -0.042500 1.00800 ; qtot -3.123
7118 CO 440 GLU CD 7118 0.805401 12.01000 ; qtot -2.317
7119 O2 440 GLU OE1 7119 -0.818801 16.00000 ; qtot -3.136
7120 O2 440 GLU OE2 7120 -0.818801 16.00000 ; qtot -3.955
7121 C 440 GLU C 7121 0.536601 12.01000 ; qtot -3.418

7122 O 440 GLU O 7122 -0.581901 16.00000 ; qtot -
4.000
7123 N 441 ILE N 7123 -0.415700 14.01000 ; qtot -
4.416
7124 H 441 ILE H 7124 0.271900 1.00800 ; qtot -
4.144
7125 CX 441 ILE CA 7125 -0.059700 12.01000 ; qtot -
4.204
7126 H1 441 ILE HA 7126 0.086900 1.00800 ; qtot -
4.117
7127 A3C 441 ILE CB 7127 0.130300 12.01000 ; qtot -
3.986
7128 HC 441 ILE HB 7128 0.018700 1.00800 ; qtot -
3.968
7129 CT 441 ILE CG2 7129 -0.320400 12.01000 ; qtot -
4.288
7130 HC 441 ILE HG21 7130 0.088200 1.00800 ; qtot -
4.200
7131 HC 441 ILE HG22 7131 0.088200 1.00800 ; qtot -
4.112
7132 HC 441 ILE HG23 7132 0.088200 1.00800 ; qtot -
4.023
7133 A2C 441 ILE CG1 7133 -0.043000 12.01000 ; qtot -
4.066
7134 HC 441 ILE HG12 7134 0.023600 1.00800 ; qtot -
4.043
7135 HC 441 ILE HG13 7135 0.023600 1.00800 ; qtot -
4.019
7136 CT 441 ILE CD1 7136 -0.066000 12.01000 ; qtot -
4.085
7137 HC 441 ILE HD11 7137 0.018600 1.00800 ; qtot -
4.067
7138 HC 441 ILE HD12 7138 0.018600 1.00800 ; qtot -
4.048
7139 HC 441 ILE HD13 7139 0.018600 1.00800 ; qtot -
4.029
7140 C 441 ILE C 7140 0.597301 12.01000 ; qtot -
3.432
7141 O 441 ILE O 7141 -0.567901 16.00000 ; qtot -
4.000
7142 N 442 ASP N 7142 -0.516301 14.01000 ; qtot -
4.516
7143 H 442 ASP H 7143 0.293600 1.00800 ; qtot -
4.223
7144 CX 442 ASP CA 7144 0.038100 12.01000 ; qtot -
4.185
7145 H1 442 ASP HA 7145 0.088000 1.00800 ; qtot -
4.097
7146 A2C 442 ASP CB 7146 -0.030300 12.01000 ; qtot -
4.127
7147 HC 442 ASP HB2 7147 -0.012200 1.00800 ; qtot -
4.139

7148 HC 442 ASP HB3 7148 -0.012200 1.00800 ; qtot -4.151
7149 CO 442 ASP CG 7149 0.799401 12.01000 ; qtot -3.352
7150 O2 442 ASP OD1 7150 -0.801401 16.00000 ; qtot -4.153
7151 O2 442 ASP OD2 7151 -0.801401 16.00000 ; qtot -4.955
7152 C 442 ASP C 7152 0.536601 12.01000 ; qtot -4.418
7153 O 442 ASP O 7153 -0.581901 16.00000 ; qtot -5.000
7154 N 443 PHE N 7154 -0.415700 14.01000 ; qtot -5.416
7155 H 443 PHE H 7155 0.271900 1.00800 ; qtot -5.144
7156 CX 443 PHE CA 7156 -0.002400 12.01000 ; qtot -5.146
7157 H1 443 PHE HA 7157 0.097800 1.00800 ; qtot -5.048
7158 CT 443 PHE CB 7158 -0.034300 12.01000 ; qtot -5.083
7159 HC 443 PHE HB2 7159 0.029500 1.00800 ; qtot -5.053
7160 HC 443 PHE HB3 7160 0.029500 1.00800 ; qtot -5.024
7161 CA 443 PHE CG 7161 0.011800 12.01000 ; qtot -5.012
7162 CA 443 PHE CD1 7162 -0.125600 12.01000 ; qtot -5.138
7163 HA 443 PHE HD1 7163 0.133000 1.00800 ; qtot -5.005
7164 CA 443 PHE CE1 7164 -0.170400 12.01000 ; qtot -5.175
7165 HA 443 PHE HE1 7165 0.143000 1.00800 ; qtot -5.032
7166 CA 443 PHE CZ 7166 -0.107200 12.01000 ; qtot -5.139
7167 HA 443 PHE HZ 7167 0.129700 1.00800 ; qtot -5.009
7168 CA 443 PHE CE2 7168 -0.170400 12.01000 ; qtot -5.180
7169 HA 443 PHE HE2 7169 0.143000 1.00800 ; qtot -5.037
7170 CA 443 PHE CD2 7170 -0.125600 12.01000 ; qtot -5.162
7171 HA 443 PHE HD2 7171 0.133000 1.00800 ; qtot -5.029
7172 C 443 PHE C 7172 0.597301 12.01000 ; qtot -4.432
7173 O 443 PHE O 7173 -0.567901 16.00000 ; qtot -5.000

7174 N 444 HIE N 7174 -0.415700 14.01000 ; qtot -5.416
7175 H 444 HIE H 7175 0.271900 1.00800 ; qtot -5.144
7176 CX 444 HIE CA 7176 -0.058100 12.01000 ; qtot -5.202
7177 H1 444 HIE HA 7177 0.136000 1.00800 ; qtot -5.066
7178 CT 444 HIE CB 7178 -0.007400 12.01000 ; qtot -5.073
7179 HC 444 HIE HB2 7179 0.036700 1.00800 ; qtot -5.037
7180 HC 444 HIE HB3 7180 0.036700 1.00800 ; qtot -5.000
7181 CC 444 HIE CG 7181 0.186800 12.01000 ; qtot -4.813
7182 NB 444 HIE ND1 7182 -0.543201 14.01000 ; qtot -5.356
7183 CR 444 HIE CE1 7183 0.163500 12.01000 ; qtot -5.193
7184 H5 444 HIE HE1 7184 0.143500 1.00800 ; qtot -5.049
7185 NA 444 HIE NE2 7185 -0.279500 14.01000 ; qtot -5.329
7186 H 444 HIE HE2 7186 0.333900 1.00800 ; qtot -4.995
7187 CW 444 HIE CD2 7187 -0.220700 12.01000 ; qtot -5.216
7188 H4 444 HIE HD2 7188 0.186200 1.00800 ; qtot -5.029
7189 C 444 HIE C 7189 0.597301 12.01000 ; qtot -4.432
7190 O 444 HIE O 7190 -0.567901 16.00000 ; qtot -5.000
7191 N 445 GLH N 7191 -0.415700 14.01000 ; qtot -5.416
7192 H 445 GLH H 7192 0.271900 1.00800 ; qtot -5.144
7193 CX 445 GLH CA 7193 0.014500 12.01000 ; qtot -5.129
7194 H1 445 GLH HA 7194 0.077900 1.00800 ; qtot -5.051
7195 A2C 445 GLH CB 7195 -0.007100 12.01000 ; qtot -5.059
7196 HC 445 GLH HB2 7196 0.025600 1.00800 ; qtot -5.033
7197 HC 445 GLH HB3 7197 0.025600 1.00800 ; qtot -5.007
7198 A2C 445 GLH CG 7198 -0.017400 12.01000 ; qtot -5.025
7199 HC 445 GLH HG2 7199 0.043000 1.00800 ; qtot -4.982

7200 HC 445 GLH HG3 7200 0.043000 1.00800 ; qtot -4.939
7201 C 445 GLH CD 7201 0.680101 12.01000 ; qtot -4.259
7202 O 445 GLH OE1 7202 -0.583801 16.00000 ; qtot -4.842
7203 OH 445 GLH OE2 7203 -0.651101 16.00000 ; qtot -5.494
7204 HO 445 GLH HE2 7204 0.464100 1.00800 ; qtot -5.029
7205 C 445 GLH C 7205 0.597301 12.01000 ; qtot -4.432
7206 O 445 GLH O 7206 -0.567901 16.00000 ; qtot -5.000
7207 N 446 PHE N 7207 -0.415700 14.01000 ; qtot -5.416
7208 H 446 PHE H 7208 0.271900 1.00800 ; qtot -5.144
7209 CX 446 PHE CA 7209 -0.002400 12.01000 ; qtot -5.146
7210 H1 446 PHE HA 7210 0.097800 1.00800 ; qtot -5.048
7211 CT 446 PHE CB 7211 -0.034300 12.01000 ; qtot -5.083
7212 HC 446 PHE HB2 7212 0.029500 1.00800 ; qtot -5.053
7213 HC 446 PHE HB3 7213 0.029500 1.00800 ; qtot -5.024
7214 CA 446 PHE CG 7214 0.011800 12.01000 ; qtot -5.012
7215 CA 446 PHE CD1 7215 -0.125600 12.01000 ; qtot -5.138
7216 HA 446 PHE HD1 7216 0.133000 1.00800 ; qtot -5.005
7217 CA 446 PHE CE1 7217 -0.170400 12.01000 ; qtot -5.175
7218 HA 446 PHE HE1 7218 0.143000 1.00800 ; qtot -5.032
7219 CA 446 PHE CZ 7219 -0.107200 12.01000 ; qtot -5.139
7220 HA 446 PHE HZ 7220 0.129700 1.00800 ; qtot -5.009
7221 CA 446 PHE CE2 7221 -0.170400 12.01000 ; qtot -5.180
7222 HA 446 PHE HE2 7222 0.143000 1.00800 ; qtot -5.037
7223 CA 446 PHE CD2 7223 -0.125600 12.01000 ; qtot -5.162
7224 HA 446 PHE HD2 7224 0.133000 1.00800 ; qtot -5.029
7225 C 446 PHE C 7225 0.597301 12.01000 ; qtot -4.432

```
  7226    O   446   PHE      O 7226    -0.567901     16.00000 ; qtot -
5.000
  7227    N   447   MET      N 7227    -0.415700     14.01000 ; qtot -
5.416
  7228    H   447   MET      H 7228     0.271900      1.00800 ; qtot -
5.144
  7229   CX   447   MET     CA 7229    -0.023700     12.01000 ; qtot -
5.168
  7230   H1   447   MET     HA 7230     0.088000      1.00800 ; qtot -
5.080
  7231  A2C   447   MET     CB 7231     0.034200     12.01000 ; qtot -
5.045
  7232   HC   447   MET    HB2 7232     0.024100      1.00800 ; qtot -
5.021
  7233   HC   447   MET    HB3 7233     0.024100      1.00800 ; qtot -
4.997
  7234  A2C   447   MET     CG 7234     0.001800     12.01000 ; qtot -
4.995
  7235   H1   447   MET    HG2 7235     0.044000      1.00800 ; qtot -
4.951
  7236   H1   447   MET    HG3 7236     0.044000      1.00800 ; qtot -
4.907
  7237    S   447   MET     SD 7237    -0.273700     32.06000 ; qtot -
5.181
  7238   CT   447   MET     CE 7238    -0.053600     12.01000 ; qtot -
5.235
  7239   H1   447   MET    HE1 7239     0.068400      1.00800 ; qtot -
5.166
  7240   H1   447   MET    HE2 7240     0.068400      1.00800 ; qtot -
5.098
  7241   H1   447   MET    HE3 7241     0.068400      1.00800 ; qtot -
5.029
  7242    C   447   MET      C 7242     0.597301     12.01000 ; qtot -
4.432
  7243    O   447   MET      O 7243    -0.567901     16.00000 ; qtot -
5.000
  7244    N   448   LEU      N 7244    -0.415700     14.01000 ; qtot -
5.416
  7245    H   448   LEU      H 7245     0.271900      1.00800 ; qtot -
5.144
  7246   CX   448   LEU     CA 7246    -0.051800     12.01000 ; qtot -
5.196
  7247   H1   448   LEU     HA 7247     0.092200      1.00800 ; qtot -
5.103
  7248  A2C   448   LEU     CB 7248    -0.110200     12.01000 ; qtot -
5.214
  7249   HC   448   LEU    HB2 7249     0.045700      1.00800 ; qtot -
5.168
  7250   HC   448   LEU    HB3 7250     0.045700      1.00800 ; qtot -
5.122
  7251  A3C   448   LEU     CG 7251     0.353100     12.01000 ; qtot -
4.769
```

```
  7252   HC   448   LEU    HG 7252    -0.036100      1.00800 ; qtot -
4.805
  7253   CT   448   LEU   CD1 7253    -0.412100     12.01000 ; qtot -
5.217
  7254   HC   448   LEU  HD11 7254     0.100000      1.00800 ; qtot -
5.117
  7255   HC   448   LEU  HD12 7255     0.100000      1.00800 ; qtot -
5.017
  7256   HC   448   LEU  HD13 7256     0.100000      1.00800 ; qtot -
4.917
  7257   CT   448   LEU   CD2 7257    -0.412100     12.01000 ; qtot -
5.329
  7258   HC   448   LEU  HD21 7258     0.100000      1.00800 ; qtot -
5.229
  7259   HC   448   LEU  HD22 7259     0.100000      1.00800 ; qtot -
5.129
  7260   HC   448   LEU  HD23 7260     0.100000      1.00800 ; qtot -
5.029
  7261    C   448   LEU     C 7261     0.597301     12.01000 ; qtot -
4.432
  7262    O   448   LEU     O 7262    -0.567901     16.00000 ; qtot -
5.000
  7263    N   449   MET     N 7263    -0.415700     14.01000 ; qtot -
5.416
  7264    H   449   MET     H 7264     0.271900      1.00800 ; qtot -
5.144
  7265   CX   449   MET    CA 7265    -0.023700     12.01000 ; qtot -
5.168
  7266   H1   449   MET    HA 7266     0.088000      1.00800 ; qtot -
5.080
  7267  A2C   449   MET    CB 7267     0.034200     12.01000 ; qtot -
5.045
  7268   HC   449   MET   HB2 7268     0.024100      1.00800 ; qtot -
5.021
  7269   HC   449   MET   HB3 7269     0.024100      1.00800 ; qtot -
4.997
  7270  A2C   449   MET    CG 7270     0.001800     12.01000 ; qtot -
4.995
  7271   H1   449   MET   HG2 7271     0.044000      1.00800 ; qtot -
4.951
  7272   H1   449   MET   HG3 7272     0.044000      1.00800 ; qtot -
4.907
  7273    S   449   MET    SD 7273    -0.273700     32.06000 ; qtot -
5.181
  7274   CT   449   MET    CE 7274    -0.053600     12.01000 ; qtot -
5.235
  7275   H1   449   MET   HE1 7275     0.068400      1.00800 ; qtot -
5.166
  7276   H1   449   MET   HE2 7276     0.068400      1.00800 ; qtot -
5.098
  7277   H1   449   MET   HE3 7277     0.068400      1.00800 ; qtot -
5.029
```

7278 C 449 MET C 7278 0.597301 12.01000 ; qtot -
4.432
7279 O 449 MET O 7279 -0.567901 16.00000 ; qtot -
5.000
7280 N 450 MET N 7280 -0.415700 14.01000 ; qtot -
5.416
7281 H 450 MET H 7281 0.271900 1.00800 ; qtot -
5.144
7282 CX 450 MET CA 7282 -0.023700 12.01000 ; qtot -
5.168
7283 H1 450 MET HA 7283 0.088000 1.00800 ; qtot -
5.080
7284 A2C 450 MET CB 7284 0.034200 12.01000 ; qtot -
5.045
7285 HC 450 MET HB2 7285 0.024100 1.00800 ; qtot -
5.021
7286 HC 450 MET HB3 7286 0.024100 1.00800 ; qtot -
4.997
7287 A2C 450 MET CG 7287 0.001800 12.01000 ; qtot -
4.995
7288 H1 450 MET HG2 7288 0.044000 1.00800 ; qtot -
4.951
7289 H1 450 MET HG3 7289 0.044000 1.00800 ; qtot -
4.907
7290 S 450 MET SD 7290 -0.273700 32.06000 ; qtot -
5.181
7291 CT 450 MET CE 7291 -0.053600 12.01000 ; qtot -
5.235
7292 H1 450 MET HE1 7292 0.068400 1.00800 ; qtot -
5.166
7293 H1 450 MET HE2 7293 0.068400 1.00800 ; qtot -
5.098
7294 H1 450 MET HE3 7294 0.068400 1.00800 ; qtot -
5.029
7295 C 450 MET C 7295 0.597301 12.01000 ; qtot -
4.432
7296 O 450 MET O 7296 -0.567901 16.00000 ; qtot -
5.000
7297 N 451 SER N 7297 -0.415700 14.01000 ; qtot -
5.416
7298 H 451 SER H 7298 0.271900 1.00800 ; qtot -
5.144
7299 CX 451 SER CA 7299 -0.024900 12.01000 ; qtot -
5.169
7300 H1 451 SER HA 7300 0.084300 1.00800 ; qtot -
5.084
7301 A2C 451 SER CB 7301 0.211700 12.01000 ; qtot -
4.873
7302 H1 451 SER HB2 7302 0.035200 1.00800 ; qtot -
4.838
7303 H1 451 SER HB3 7303 0.035200 1.00800 ; qtot -
4.802

7304 OH 451 SER OG 7304 -0.654601 16.00000 ; qtot -5.457
7305 HO 451 SER HG 7305 0.427500 1.00800 ; qtot -5.029
7306 C 451 SER C 7306 0.597301 12.01000 ; qtot -4.432
7307 O 451 SER O 7307 -0.567901 16.00000 ; qtot -5.000
7308 N 452 LYS N 7308 -0.347900 14.01000 ; qtot -5.348
7309 H 452 LYS H 7309 0.274700 1.00800 ; qtot -5.073
7310 CX 452 LYS CA 7310 -0.240000 12.01000 ; qtot -5.313
7311 H1 452 LYS HA 7311 0.142600 1.00800 ; qtot -5.171
7312 C8 452 LYS CB 7312 -0.009400 12.01000 ; qtot -5.180
7313 HC 452 LYS HB2 7313 0.036200 1.00800 ; qtot -5.144
7314 HC 452 LYS HB3 7314 0.036200 1.00800 ; qtot -5.108
7315 C8 452 LYS CG 7315 0.018700 12.01000 ; qtot -5.089
7316 HC 452 LYS HG2 7316 0.010300 1.00800 ; qtot -5.079
7317 HC 452 LYS HG3 7317 0.010300 1.00800 ; qtot -5.068
7318 C8 452 LYS CD 7318 -0.047900 12.01000 ; qtot -5.116
7319 HC 452 LYS HD2 7319 0.062100 1.00800 ; qtot -5.054
7320 HC 452 LYS HD3 7320 0.062100 1.00800 ; qtot -4.992
7321 C8 452 LYS CE 7321 -0.014300 12.01000 ; qtot -5.006
7322 HP 452 LYS HE2 7322 0.113500 1.00800 ; qtot -4.893
7323 HP 452 LYS HE3 7323 0.113500 1.00800 ; qtot -4.779
7324 N3 452 LYS NZ 7324 -0.385400 14.01000 ; qtot -5.165
7325 H 452 LYS HZ1 7325 0.340000 1.00800 ; qtot -4.825
7326 H 452 LYS HZ2 7326 0.340000 1.00800 ; qtot -4.485
7327 H 452 LYS HZ3 7327 0.340000 1.00800 ; qtot -4.145
7328 C 452 LYS C 7328 0.734101 12.01000 ; qtot -3.411
7329 O 452 LYS O 7329 -0.589401 16.00000 ; qtot -4.000

7330 N 453 LYS N 7330 -0.347900 14.01000 ; qtot -
4.348
7331 H 453 LYS H 7331 0.274700 1.00800 ; qtot -
4.073
7332 CX 453 LYS CA 7332 -0.240000 12.01000 ; qtot -
4.313
7333 H1 453 LYS HA 7333 0.142600 1.00800 ; qtot -
4.171
7334 C8 453 LYS CB 7334 -0.009400 12.01000 ; qtot -
4.180
7335 HC 453 LYS HB2 7335 0.036200 1.00800 ; qtot -
4.144
7336 HC 453 LYS HB3 7336 0.036200 1.00800 ; qtot -
4.108
7337 C8 453 LYS CG 7337 0.018700 12.01000 ; qtot -
4.089
7338 HC 453 LYS HG2 7338 0.010300 1.00800 ; qtot -
4.079
7339 HC 453 LYS HG3 7339 0.010300 1.00800 ; qtot -
4.068
7340 C8 453 LYS CD 7340 -0.047900 12.01000 ; qtot -
4.116
7341 HC 453 LYS HD2 7341 0.062100 1.00800 ; qtot -
4.054
7342 HC 453 LYS HD3 7342 0.062100 1.00800 ; qtot -
3.992
7343 C8 453 LYS CE 7343 -0.014300 12.01000 ; qtot -
4.006
7344 HP 453 LYS HE2 7344 0.113500 1.00800 ; qtot -
3.893
7345 HP 453 LYS HE3 7345 0.113500 1.00800 ; qtot -
3.779
7346 N3 453 LYS NZ 7346 -0.385400 14.01000 ; qtot -
4.165
7347 H 453 LYS HZ1 7347 0.340000 1.00800 ; qtot -
3.825
7348 H 453 LYS HZ2 7348 0.340000 1.00800 ; qtot -
3.485
7349 H 453 LYS HZ3 7349 0.340000 1.00800 ; qtot -
3.145
7350 C 453 LYS C 7350 0.734101 12.01000 ; qtot -
2.411
7351 O 453 LYS O 7351 -0.589401 16.00000 ; qtot -
3.000
7352 N 454 LYS N 7352 -0.348100 14.01000 ; qtot -
3.348
7353 H 454 LYS H 7353 0.276400 1.00800 ; qtot -
3.072
7354 CX 454 LYS CA 7354 -0.290300 12.01000 ; qtot -
3.362
7355 H1 454 LYS HA 7355 0.143800 1.00800 ; qtot -
3.218

7356 C8 454 LYS CB 7356 -0.053800 12.01000 ; qtot -
3.272
7357 HC 454 LYS HB2 7357 0.048200 1.00800 ; qtot -
3.224
7358 HC 454 LYS HB3 7358 0.048200 1.00800 ; qtot -
3.176
7359 C8 454 LYS CG 7359 0.022700 12.01000 ; qtot -
3.153
7360 HC 454 LYS HG2 7360 0.013400 1.00800 ; qtot -
3.140
7361 HC 454 LYS HG3 7361 0.013400 1.00800 ; qtot -
3.126
7362 C8 454 LYS CD 7362 -0.039200 12.01000 ; qtot -
3.165
7363 HC 454 LYS HD2 7363 0.061100 1.00800 ; qtot -
3.104
7364 HC 454 LYS HD3 7364 0.061100 1.00800 ; qtot -
3.043
7365 C8 454 LYS CE 7365 -0.017600 12.01000 ; qtot -
3.061
7366 HP 454 LYS HE2 7366 0.112100 1.00800 ; qtot -
2.949
7367 HP 454 LYS HE3 7367 0.112100 1.00800 ; qtot -
2.837
7368 N3 454 LYS NZ 7368 -0.374100 14.01000 ; qtot -
3.211
7369 H 454 LYS HZ1 7369 0.337400 1.00800 ; qtot -
2.873
7370 H 454 LYS HZ2 7370 0.337400 1.00800 ; qtot -
2.536
7371 H 454 LYS HZ3 7371 0.337400 1.00800 ; qtot -
2.198
7372 C 454 LYS C 7372 0.848801 12.01000 ; qtot -
1.350
7373 O2 454 LYS O 7373 -0.825201 16.00000 ; qtot -
2.175
7374 O2 454 LYS OXT 7374 -0.825201 16.00000 ; qtot -
3.000
7375 f 455 FL1 F2 7375 -0.152091 19.00000 ; qtot -
3.152
7376 c2 455 FL1 C38 7376 0.104019 12.01000 ; qtot -
3.048
7377 c2 455 FL1 C39 7377 -0.071000 12.01000 ; qtot -
3.119
7378 Y1 455 FL1 N23 7378 -0.050516 14.01000 ; qtot -
3.170
7379 Y2 455 FL1 N21 7379 0.013398 14.01000 ; qtot -
3.156
7380 Y5 455 FL1 C13 7380 0.291175 12.01000 ; qtot -
2.865
7381 O 455 FL1 O14 7381 -0.219494 16.00000 ; qtot -
3.085

7382 Y3 455 FL1 CL17 7382 -0.440553 35.45000 ; qtot -3.525
7383 c2 455 FL1 C37 7383 -0.038661 12.01000 ; qtot -3.564
7384 CA 455 FL1 C36 7384 -0.021785 12.01000 ; qtot -3.586
7385 CA 455 FL1 C40 7385 0.087057 12.01000 ; qtot -3.498
7386 CA 455 FL1 C35 7386 -0.252013 12.01000 ; qtot -3.750
7387 c2 455 FL1 C34 7387 0.550463 12.01000 ; qtot -3.200
7388 N 455 FL1 N22 7388 -0.477191 14.01000 ; qtot -3.677
7389 hn 455 FL1 H22 7389 -0.446766 1.00800 ; qtot -4.124
7390 O 455 FL1 O42 7390 -0.045686 16.00000 ; qtot -4.170
7391 CA 455 FL1 C32 7391 0.496379 12.01000 ; qtot -3.673
7392 c2 455 FL1 C33 7392 -0.399186 12.01000 ; qtot -4.072
7393 O 455 FL1 O41 7393 -0.021222 16.00000 ; qtot -4.094
7394 cp 455 FL1 C31 7394 0.009439 12.01000 ; qtot -4.084
7395 CA 455 FL1 C24 7395 -0.081068 12.01000 ; qtot -4.165
7396 cq 455 FL1 C30 7396 -0.018991 12.01000 ; qtot -4.184
7397 CA 455 FL1 C25 7397 -0.094537 12.01000 ; qtot -4.279
7398 CA 455 FL1 C29 7398 0.203475 12.01000 ; qtot -4.075
7399 CA 455 FL1 C28 7399 -0.552211 12.01000 ; qtot -4.628
7400 CA 455 FL1 C27 7400 -0.200637 12.01000 ; qtot -4.828
7401 CA 455 FL1 C26 7401 -0.205457 12.01000 ; qtot -5.034
7402 Y4 455 FL1 N15 7402 0.004397 14.01000 ; qtot -5.029
7403 c2 455 FL1 C7 7403 -0.120053 12.01000 ; qtot -5.149
7404 CA 455 FL1 C18 7404 0.023489 12.01000 ; qtot -5.126
7405 CA 455 FL1 C5 7405 -0.082755 12.01000 ; qtot -5.209
7406 CA 455 FL1 C6 7406 -0.205989 12.01000 ; qtot -5.415
7407 CA 455 FL1 C4 7407 -0.101475 12.01000 ; qtot -5.516

7408 CA 455 FL1 C3 7408 -0.217360 12.01000 ; qtot -
5.733
7409 CA 455 FL1 C19 7409 -0.061505 12.01000 ; qtot -
5.795
7410 c2 455 FL1 C8 7410 0.222378 12.01000 ; qtot -
5.573
7411 c2 455 FL1 C9 7411 -0.141964 12.01000 ; qtot -
5.715
7412 c2 455 FL1 C10 7412 -0.162282 12.01000 ; qtot -
5.877
7413 c2 455 FL1 C12 7413 -0.085409 12.01000 ; qtot -
5.962
7414 c2 455 FL1 C11 7414 0.019314 12.01000 ; qtot -
5.943
7415 N2 455 FL1 N16 7415 -0.190712 14.01000 ; qtot -
6.134
7416 OH 455 FL1 O1 7416 0.345138 16.00000 ; qtot -
5.788
7417 HO 455 FL1 H1 7417 0.156204 1.00800 ; qtot -
5.632
7418 HA 455 FL1 H9 7418 0.116601 1.00800 ; qtot -
5.516
7419 H4 455 FL1 H7A 7419 0.181930 1.00800 ; qtot -
5.334
7420 HA 455 FL1 H6 7420 0.163520 1.00800 ; qtot -
5.170
7421 HA 455 FL1 H5 7421 0.152187 1.00800 ; qtot -
5.018
7422 HA 455 FL1 H4 7422 0.179263 1.00800 ; qtot -
4.839
7423 H4 455 FL1 H39 7423 0.152550 1.00800 ; qtot -
4.686
7424 HA 455 FL1 H37 7424 0.179402 1.00800 ; qtot -
4.507
7425 HA 455 FL1 H3 7425 0.132471 1.00800 ; qtot -
4.374
7426 HA 455 FL1 H29 7426 0.165463 1.00800 ; qtot -
4.209
7427 HA 455 FL1 H27 7427 0.187733 1.00800 ; qtot -
4.021
7428 HA 455 FL1 H26 7428 0.100013 1.00800 ; qtot -
3.921
7429 HA 455 FL1 H19 7429 0.144523 1.00800 ; qtot -
3.777
7430 HA 455 FL1 H12 7430 0.165882 1.00800 ; qtot -
3.611
7431 H4 455 FL1 H11 7431 0.175823 1.00800 ; qtot -
3.435
7432 HA 455 FL1 H10 7432 0.420758 1.00800 ; qtot -
3.014
7433 M1 456 RU1 RU 7433 0.014127 101.07000 ; qtot -
3.000

```
[ bonds ]
;   ai     aj funct   r              k
     1      2   1    1.0100e-01    3.6317e+05 ;      N - H1
     1      3   1    1.0100e-01    3.6317e+05 ;      N - H2
     1      4   1    1.0100e-01    3.6317e+05 ;      N - H3
     1      5   1    1.4710e-01    3.0711e+05 ;      N - CA
     5      6   1    1.0900e-01    2.8451e+05 ;     CA - HA
     5      7   1    1.5260e-01    2.5941e+05 ;     CA - CB
     5     20   1    1.5220e-01    2.6527e+05 ;     CA - C
     7      8   1    1.0900e-01    2.8451e+05 ;     CB - HB2
     7      9   1    1.0900e-01    2.8451e+05 ;     CB - HB3
     7     10   1    1.5260e-01    2.5941e+05 ;     CB - CG
    10     11   1    1.0900e-01    2.8451e+05 ;     CG - HG
    10     12   1    1.5260e-01    2.5941e+05 ;     CG - CD1
    10     16   1    1.5260e-01    2.5941e+05 ;     CG - CD2
    12     13   1    1.0900e-01    2.8451e+05 ;    CD1 - HD11
    12     14   1    1.0900e-01    2.8451e+05 ;    CD1 - HD12
    12     15   1    1.0900e-01    2.8451e+05 ;    CD1 - HD13
    16     17   1    1.0900e-01    2.8451e+05 ;    CD2 - HD21
    16     18   1    1.0900e-01    2.8451e+05 ;    CD2 - HD22
    16     19   1    1.0900e-01    2.8451e+05 ;    CD2 - HD23
    20     21   1    1.2290e-01    4.7698e+05 ;      C - O
    20     22   1    1.3350e-01    4.1003e+05 ;      C - N
    22     23   1    1.0100e-01    3.6317e+05 ;      N - H
    22     24   1    1.4490e-01    2.8200e+05 ;      N - CA
    24     25   1    1.0900e-01    2.8451e+05 ;     CA - HA
    24     26   1    1.5260e-01    2.5941e+05 ;     CA - CB
    24     39   1    1.5220e-01    2.6527e+05 ;     CA - C
    26     27   1    1.0900e-01    2.8451e+05 ;     CB - HB
    26     28   1    1.5260e-01    2.5941e+05 ;     CB - CG2
    26     32   1    1.5260e-01    2.5941e+05 ;     CB - CG1
    28     29   1    1.0900e-01    2.8451e+05 ;    CG2 - HG21
    28     30   1    1.0900e-01    2.8451e+05 ;    CG2 - HG22
    28     31   1    1.0900e-01    2.8451e+05 ;    CG2 - HG23
    32     33   1    1.0900e-01    2.8451e+05 ;    CG1 - HG12
    32     34   1    1.0900e-01    2.8451e+05 ;    CG1 - HG13
    32     35   1    1.5260e-01    2.5941e+05 ;    CG1 - CD1
    35     36   1    1.0900e-01    2.8451e+05 ;    CD1 - HD11
    35     37   1    1.0900e-01    2.8451e+05 ;    CD1 - HD12
    35     38   1    1.0900e-01    2.8451e+05 ;    CD1 - HD13
    39     40   1    1.2290e-01    4.7698e+05 ;      C - O
    39     41   1    1.3350e-01    4.1003e+05 ;      C - N
    41     42   1    1.0100e-01    3.6317e+05 ;      N - H
    41     43   1    1.4490e-01    2.8200e+05 ;      N - CA
    43     44   1    1.0900e-01    2.8451e+05 ;     CA - HA
    43     45   1    1.5260e-01    2.5941e+05 ;     CA - CB
    43     58   1    1.5220e-01    2.6527e+05 ;     CA - C
    45     46   1    1.0900e-01    2.8451e+05 ;     CB - HB2
    45     47   1    1.0900e-01    2.8451e+05 ;     CB - HB3
    45     48   1    1.5260e-01    2.5941e+05 ;     CB - CG
    48     49   1    1.0900e-01    2.8451e+05 ;     CG - HG
```

| | | | | | | |
|---|---|---|---|---|---|---|
| 48 | 50 | 1 | 1.5260e-01 | 2.5941e+05 | ; | CG - CD1 |
| 48 | 54 | 1 | 1.5260e-01 | 2.5941e+05 | ; | CG - CD2 |
| 50 | 51 | 1 | 1.0900e-01 | 2.8451e+05 | ; | CD1 - HD11 |
| 50 | 52 | 1 | 1.0900e-01 | 2.8451e+05 | ; | CD1 - HD12 |
| 50 | 53 | 1 | 1.0900e-01 | 2.8451e+05 | ; | CD1 - HD13 |
| 54 | 55 | 1 | 1.0900e-01 | 2.8451e+05 | ; | CD2 - HD21 |
| 54 | 56 | 1 | 1.0900e-01 | 2.8451e+05 | ; | CD2 - HD22 |
| 54 | 57 | 1 | 1.0900e-01 | 2.8451e+05 | ; | CD2 - HD23 |
| 58 | 59 | 1 | 1.2290e-01 | 4.7698e+05 | ; | C - O |
| 58 | 60 | 1 | 1.3350e-01 | 4.1003e+05 | ; | C - N |
| 60 | 61 | 1 | 1.0100e-01 | 3.6317e+05 | ; | N - H |
| 60 | 62 | 1 | 1.4490e-01 | 2.8200e+05 | ; | N - CA |
| 62 | 63 | 1 | 1.0900e-01 | 2.8451e+05 | ; | CA - HA |
| 62 | 64 | 1 | 1.5260e-01 | 2.5941e+05 | ; | CA - CB |
| 62 | 69 | 1 | 1.5220e-01 | 2.6527e+05 | ; | CA - C |
| 64 | 65 | 1 | 1.0900e-01 | 2.8451e+05 | ; | CB - HB2 |
| 64 | 66 | 1 | 1.0900e-01 | 2.8451e+05 | ; | CB - HB3 |
| 64 | 67 | 1 | 1.8100e-01 | 1.9832e+05 | ; | CB - SG |
| 67 | 68 | 1 | 1.3360e-01 | 2.2928e+05 | ; | SG - HG |
| 69 | 70 | 1 | 1.2290e-01 | 4.7698e+05 | ; | C - O |
| 69 | 71 | 1 | 1.3350e-01 | 4.1003e+05 | ; | C - N |
| 71 | 72 | 1 | 1.0100e-01 | 3.6317e+05 | ; | N - H |
| 71 | 73 | 1 | 1.4490e-01 | 2.8200e+05 | ; | N - CA |
| 73 | 74 | 1 | 1.0900e-01 | 2.8451e+05 | ; | CA - HA |
| 73 | 75 | 1 | 1.5260e-01 | 2.5941e+05 | ; | CA - CB |
| 73 | 86 | 1 | 1.5220e-01 | 2.6527e+05 | ; | CA - C |
| 75 | 76 | 1 | 1.0900e-01 | 2.8451e+05 | ; | CB - HB2 |
| 75 | 77 | 1 | 1.0900e-01 | 2.8451e+05 | ; | CB - HB3 |
| 75 | 78 | 1 | 1.5040e-01 | 2.6527e+05 | ; | CB - CG |
| 78 | 79 | 1 | 1.3940e-01 | 3.4309e+05 | ; | CG - ND1 |
| 78 | 84 | 1 | 1.3710e-01 | 4.3346e+05 | ; | CG - CD2 |
| 79 | 80 | 1 | 1.3350e-01 | 4.0836e+05 | ; | ND1 - CE1 |
| 80 | 81 | 1 | 1.0800e-01 | 3.0711e+05 | ; | CE1 - HE1 |
| 80 | 82 | 1 | 1.3430e-01 | 3.9915e+05 | ; | CE1 - NE2 |
| 82 | 83 | 1 | 1.0100e-01 | 3.6317e+05 | ; | NE2 - HE2 |
| 82 | 84 | 1 | 1.3810e-01 | 3.5731e+05 | ; | NE2 - CD2 |
| 84 | 85 | 1 | 1.0800e-01 | 3.0711e+05 | ; | CD2 - HD2 |
| 86 | 87 | 1 | 1.2290e-01 | 4.7698e+05 | ; | C - O |
| 86 | 88 | 1 | 1.3350e-01 | 4.1003e+05 | ; | C - N |
| 88 | 89 | 1 | 1.0100e-01 | 3.6317e+05 | ; | N - H |
| 88 | 90 | 1 | 1.4490e-01 | 2.8200e+05 | ; | N - CA |
| 90 | 91 | 1 | 1.0900e-01 | 2.8451e+05 | ; | CA - HA |
| 90 | 92 | 1 | 1.5260e-01 | 2.5941e+05 | ; | CA - CB |
| 90 | 97 | 1 | 1.5220e-01 | 2.6527e+05 | ; | CA - C |
| 92 | 93 | 1 | 1.0900e-01 | 2.8451e+05 | ; | CB - HB2 |
| 92 | 94 | 1 | 1.0900e-01 | 2.8451e+05 | ; | CB - HB3 |
| 92 | 95 | 1 | 1.4100e-01 | 2.6778e+05 | ; | CB - OG |
| 95 | 96 | 1 | 9.6000e-02 | 4.6275e+05 | ; | OG - HG |
| 97 | 98 | 1 | 1.2290e-01 | 4.7698e+05 | ; | C - O |
| 97 | 99 | 1 | 1.3350e-01 | 4.1003e+05 | ; | C - N |
| 99 | 100 | 1 | 1.0100e-01 | 3.6317e+05 | ; | N - H |
| 99 | 101 | 1 | 1.4490e-01 | 2.8200e+05 | ; | N - CA |

| | | | | | | |
|---|---|---|---|---|---|---|
| 101 | 102 | 1 | 1.0900e-01 | 2.8451e+05 | ; | CA - HA2 |
| 101 | 103 | 1 | 1.0900e-01 | 2.8451e+05 | ; | CA - HA3 |
| 101 | 104 | 1 | 1.5220e-01 | 2.6527e+05 | ; | CA - C |
| 104 | 105 | 1 | 1.2290e-01 | 4.7698e+05 | ; | C - O |
| 104 | 106 | 1 | 1.3350e-01 | 4.1003e+05 | ; | C - N |
| 106 | 107 | 1 | 1.0100e-01 | 3.6317e+05 | ; | N - H |
| 106 | 108 | 1 | 1.4490e-01 | 2.8200e+05 | ; | N - CA |
| 108 | 109 | 1 | 1.0900e-01 | 2.8451e+05 | ; | CA - HA |
| 108 | 110 | 1 | 1.5260e-01 | 2.5941e+05 | ; | CA - CB |
| 108 | 126 | 1 | 1.5220e-01 | 2.6527e+05 | ; | CA - C |
| 110 | 111 | 1 | 1.0900e-01 | 2.8451e+05 | ; | CB - HB2 |
| 110 | 112 | 1 | 1.0900e-01 | 2.8451e+05 | ; | CB - HB3 |
| 110 | 113 | 1 | 1.5260e-01 | 2.5941e+05 | ; | CB - CG |
| 113 | 114 | 1 | 1.0900e-01 | 2.8451e+05 | ; | CG - HG2 |
| 113 | 115 | 1 | 1.0900e-01 | 2.8451e+05 | ; | CG - HG3 |
| 113 | 116 | 1 | 1.5260e-01 | 2.5941e+05 | ; | CG - CD |
| 116 | 117 | 1 | 1.0900e-01 | 2.8451e+05 | ; | CD - HD2 |
| 116 | 118 | 1 | 1.0900e-01 | 2.8451e+05 | ; | CD - HD3 |
| 116 | 119 | 1 | 1.5260e-01 | 2.5941e+05 | ; | CD - CE |
| 119 | 120 | 1 | 1.0900e-01 | 2.8451e+05 | ; | CE - HE2 |
| 119 | 121 | 1 | 1.0900e-01 | 2.8451e+05 | ; | CE - HE3 |
| 119 | 122 | 1 | 1.4710e-01 | 3.0711e+05 | ; | CE - NZ |
| 122 | 123 | 1 | 1.0100e-01 | 3.6317e+05 | ; | NZ - HZ1 |
| 122 | 124 | 1 | 1.0100e-01 | 3.6317e+05 | ; | NZ - HZ2 |
| 122 | 125 | 1 | 1.0100e-01 | 3.6317e+05 | ; | NZ - HZ3 |
| 126 | 127 | 1 | 1.2290e-01 | 4.7698e+05 | ; | C - O |
| 126 | 128 | 1 | 1.3350e-01 | 4.1003e+05 | ; | C - N |
| 128 | 129 | 1 | 1.0100e-01 | 3.6317e+05 | ; | N - H |
| 128 | 130 | 1 | 1.4490e-01 | 2.8200e+05 | ; | N - CA |
| 130 | 131 | 1 | 1.0900e-01 | 2.8451e+05 | ; | CA - HA |
| 130 | 132 | 1 | 1.5260e-01 | 2.5941e+05 | ; | CA - CB |
| 130 | 145 | 1 | 1.5220e-01 | 2.6527e+05 | ; | CA - C |
| 132 | 133 | 1 | 1.0900e-01 | 2.8451e+05 | ; | CB - HB2 |
| 132 | 134 | 1 | 1.0900e-01 | 2.8451e+05 | ; | CB - HB3 |
| 132 | 135 | 1 | 1.5260e-01 | 2.5941e+05 | ; | CB - CG |
| 135 | 136 | 1 | 1.0900e-01 | 2.8451e+05 | ; | CG - HG |
| 135 | 137 | 1 | 1.5260e-01 | 2.5941e+05 | ; | CG - CD1 |
| 135 | 141 | 1 | 1.5260e-01 | 2.5941e+05 | ; | CG - CD2 |
| 137 | 138 | 1 | 1.0900e-01 | 2.8451e+05 | ; | CD1 - HD11 |
| 137 | 139 | 1 | 1.0900e-01 | 2.8451e+05 | ; | CD1 - HD12 |
| 137 | 140 | 1 | 1.0900e-01 | 2.8451e+05 | ; | CD1 - HD13 |
| 141 | 142 | 1 | 1.0900e-01 | 2.8451e+05 | ; | CD2 - HD21 |
| 141 | 143 | 1 | 1.0900e-01 | 2.8451e+05 | ; | CD2 - HD22 |
| 141 | 144 | 1 | 1.0900e-01 | 2.8451e+05 | ; | CD2 - HD23 |
| 145 | 146 | 1 | 1.2290e-01 | 4.7698e+05 | ; | C - O |
| 145 | 147 | 1 | 1.3350e-01 | 4.1003e+05 | ; | C - N |
| 147 | 148 | 1 | 1.0100e-01 | 3.6317e+05 | ; | N - H |
| 147 | 149 | 1 | 1.4490e-01 | 2.8200e+05 | ; | N - CA |
| 149 | 150 | 1 | 1.0900e-01 | 2.8451e+05 | ; | CA - HA |
| 149 | 151 | 1 | 1.5260e-01 | 2.5941e+05 | ; | CA - CB |
| 149 | 160 | 1 | 1.5220e-01 | 2.6527e+05 | ; | CA - C |
| 151 | 152 | 1 | 1.0900e-01 | 2.8451e+05 | ; | CB - HB2 |

| | | | | | | |
|---|---|---|---|---|---|---|
| 151 | 153 | 1 | 1.0900e-01 | 2.8451e+05 | ; | CB - HB3 |
| 151 | 154 | 1 | 1.5260e-01 | 2.5941e+05 | ; | CB - CG |
| 154 | 155 | 1 | 1.0900e-01 | 2.8451e+05 | ; | CG - HG2 |
| 154 | 156 | 1 | 1.0900e-01 | 2.8451e+05 | ; | CG - HG3 |
| 154 | 157 | 1 | 1.5220e-01 | 2.6527e+05 | ; | CG - CD |
| 157 | 158 | 1 | 1.2500e-01 | 5.4894e+05 | ; | CD - OE1 |
| 157 | 159 | 1 | 1.2500e-01 | 5.4894e+05 | ; | CD - OE2 |
| 160 | 161 | 1 | 1.2290e-01 | 4.7698e+05 | ; | C - O |
| 160 | 162 | 1 | 1.3350e-01 | 4.1003e+05 | ; | C - N |
| 162 | 163 | 1 | 1.0100e-01 | 3.6317e+05 | ; | N - H |
| 162 | 164 | 1 | 1.4490e-01 | 2.8200e+05 | ; | N - CA |
| 164 | 165 | 1 | 1.0900e-01 | 2.8451e+05 | ; | CA - HA |
| 164 | 166 | 1 | 1.5260e-01 | 2.5941e+05 | ; | CA - CB |
| 164 | 172 | 1 | 1.5220e-01 | 2.6527e+05 | ; | CA - C |
| 166 | 167 | 1 | 1.0900e-01 | 2.8451e+05 | ; | CB - HB2 |
| 166 | 168 | 1 | 1.0900e-01 | 2.8451e+05 | ; | CB - HB3 |
| 166 | 169 | 1 | 1.5220e-01 | 2.6527e+05 | ; | CB - CG |
| 169 | 170 | 1 | 1.2500e-01 | 5.4894e+05 | ; | CG - OD1 |
| 169 | 171 | 1 | 1.2500e-01 | 5.4894e+05 | ; | CG - OD2 |
| 172 | 173 | 1 | 1.2290e-01 | 4.7698e+05 | ; | C - O |
| 172 | 174 | 1 | 1.3350e-01 | 4.1003e+05 | ; | C - N |
| 174 | 175 | 1 | 1.0100e-01 | 3.6317e+05 | ; | N - H |
| 174 | 176 | 1 | 1.4490e-01 | 2.8200e+05 | ; | N - CA |
| 176 | 177 | 1 | 1.0900e-01 | 2.8451e+05 | ; | CA - HA |
| 176 | 178 | 1 | 1.5260e-01 | 2.5941e+05 | ; | CA - CB |
| 176 | 194 | 1 | 1.5220e-01 | 2.6527e+05 | ; | CA - C |
| 178 | 179 | 1 | 1.0900e-01 | 2.8451e+05 | ; | CB - HB2 |
| 178 | 180 | 1 | 1.0900e-01 | 2.8451e+05 | ; | CB - HB3 |
| 178 | 181 | 1 | 1.5260e-01 | 2.5941e+05 | ; | CB - CG |
| 181 | 182 | 1 | 1.0900e-01 | 2.8451e+05 | ; | CG - HG2 |
| 181 | 183 | 1 | 1.0900e-01 | 2.8451e+05 | ; | CG - HG3 |
| 181 | 184 | 1 | 1.5260e-01 | 2.5941e+05 | ; | CG - CD |
| 184 | 185 | 1 | 1.0900e-01 | 2.8451e+05 | ; | CD - HD2 |
| 184 | 186 | 1 | 1.0900e-01 | 2.8451e+05 | ; | CD - HD3 |
| 184 | 187 | 1 | 1.5260e-01 | 2.5941e+05 | ; | CD - CE |
| 187 | 188 | 1 | 1.0900e-01 | 2.8451e+05 | ; | CE - HE2 |
| 187 | 189 | 1 | 1.0900e-01 | 2.8451e+05 | ; | CE - HE3 |
| 187 | 190 | 1 | 1.4710e-01 | 3.0711e+05 | ; | CE - NZ |
| 190 | 191 | 1 | 1.0100e-01 | 3.6317e+05 | ; | NZ - HZ1 |
| 190 | 192 | 1 | 1.0100e-01 | 3.6317e+05 | ; | NZ - HZ2 |
| 190 | 193 | 1 | 1.0100e-01 | 3.6317e+05 | ; | NZ - HZ3 |
| 194 | 195 | 1 | 1.2290e-01 | 4.7698e+05 | ; | C - O |
| 194 | 196 | 1 | 1.3350e-01 | 4.1003e+05 | ; | C - N |
| 196 | 197 | 1 | 1.0100e-01 | 3.6317e+05 | ; | N - H |
| 196 | 198 | 1 | 1.4490e-01 | 2.8200e+05 | ; | N - CA |
| 198 | 199 | 1 | 1.0900e-01 | 2.8451e+05 | ; | CA - HA |
| 198 | 200 | 1 | 1.5260e-01 | 2.5941e+05 | ; | CA - CB |
| 198 | 215 | 1 | 1.5220e-01 | 2.6527e+05 | ; | CA - C |
| 200 | 201 | 1 | 1.0900e-01 | 2.8451e+05 | ; | CB - HB2 |
| 200 | 202 | 1 | 1.0900e-01 | 2.8451e+05 | ; | CB - HB3 |
| 200 | 203 | 1 | 1.5100e-01 | 2.6527e+05 | ; | CB - CG |
| 203 | 204 | 1 | 1.4000e-01 | 3.9246e+05 | ; | CG - CD1 |

| | | | | | | |
|---|---|---|---|---|---|---|
| 203 | 213 | 1 | 1.4000e-01 | 3.9246e+05 | ; | CG - CD2 |
| 204 | 205 | 1 | 1.0800e-01 | 3.0711e+05 | ; | CD1 - HD1 |
| 204 | 206 | 1 | 1.4000e-01 | 3.9246e+05 | ; | CD1 - CE1 |
| 206 | 207 | 1 | 1.0800e-01 | 3.0711e+05 | ; | CE1 - HE1 |
| 206 | 208 | 1 | 1.4090e-01 | 3.9246e+05 | ; | CE1 - CZ |
| 208 | 209 | 1 | 1.3640e-01 | 3.7656e+05 | ; | CZ - OH |
| 208 | 211 | 1 | 1.4090e-01 | 3.9246e+05 | ; | CZ - CE2 |
| 209 | 210 | 1 | 9.6000e-02 | 4.6275e+05 | ; | OH - HH |
| 211 | 212 | 1 | 1.0800e-01 | 3.0711e+05 | ; | CE2 - HE2 |
| 211 | 213 | 1 | 1.4000e-01 | 3.9246e+05 | ; | CE2 - CD2 |
| 213 | 214 | 1 | 1.0800e-01 | 3.0711e+05 | ; | CD2 - HD2 |
| 215 | 216 | 1 | 1.2290e-01 | 4.7698e+05 | ; | C - O |
| 215 | 217 | 1 | 1.3350e-01 | 4.1003e+05 | ; | C - N |
| 217 | 218 | 1 | 1.0100e-01 | 3.6317e+05 | ; | N - H |
| 217 | 219 | 1 | 1.4490e-01 | 2.8200e+05 | ; | N - CA |
| 219 | 220 | 1 | 1.0900e-01 | 2.8451e+05 | ; | CA - HA |
| 219 | 221 | 1 | 1.5260e-01 | 2.5941e+05 | ; | CA - CB |
| 219 | 234 | 1 | 1.5220e-01 | 2.6527e+05 | ; | CA - C |
| 221 | 222 | 1 | 1.0900e-01 | 2.8451e+05 | ; | CB - HB |
| 221 | 223 | 1 | 1.5260e-01 | 2.5941e+05 | ; | CB - CG2 |
| 221 | 227 | 1 | 1.5260e-01 | 2.5941e+05 | ; | CB - CG1 |
| 223 | 224 | 1 | 1.0900e-01 | 2.8451e+05 | ; | CG2 - HG21 |
| 223 | 225 | 1 | 1.0900e-01 | 2.8451e+05 | ; | CG2 - HG22 |
| 223 | 226 | 1 | 1.0900e-01 | 2.8451e+05 | ; | CG2 - HG23 |
| 227 | 228 | 1 | 1.0900e-01 | 2.8451e+05 | ; | CG1 - HG12 |
| 227 | 229 | 1 | 1.0900e-01 | 2.8451e+05 | ; | CG1 - HG13 |
| 227 | 230 | 1 | 1.5260e-01 | 2.5941e+05 | ; | CG1 - CD1 |
| 230 | 231 | 1 | 1.0900e-01 | 2.8451e+05 | ; | CD1 - HD11 |
| 230 | 232 | 1 | 1.0900e-01 | 2.8451e+05 | ; | CD1 - HD12 |
| 230 | 233 | 1 | 1.0900e-01 | 2.8451e+05 | ; | CD1 - HD13 |
| 234 | 235 | 1 | 1.2290e-01 | 4.7698e+05 | ; | C - O |
| 234 | 236 | 1 | 1.3350e-01 | 4.1003e+05 | ; | C - N |
| 236 | 237 | 1 | 1.0100e-01 | 3.6317e+05 | ; | N - H |
| 236 | 238 | 1 | 1.4490e-01 | 2.8200e+05 | ; | N - CA |
| 238 | 239 | 1 | 1.0900e-01 | 2.8451e+05 | ; | CA - HA |
| 238 | 240 | 1 | 1.5260e-01 | 2.5941e+05 | ; | CA - CB |
| 238 | 253 | 1 | 1.5220e-01 | 2.6527e+05 | ; | CA - C |
| 240 | 241 | 1 | 1.0900e-01 | 2.8451e+05 | ; | CB - HB |
| 240 | 242 | 1 | 1.5260e-01 | 2.5941e+05 | ; | CB - CG2 |
| 240 | 246 | 1 | 1.5260e-01 | 2.5941e+05 | ; | CB - CG1 |
| 242 | 243 | 1 | 1.0900e-01 | 2.8451e+05 | ; | CG2 - HG21 |
| 242 | 244 | 1 | 1.0900e-01 | 2.8451e+05 | ; | CG2 - HG22 |
| 242 | 245 | 1 | 1.0900e-01 | 2.8451e+05 | ; | CG2 - HG23 |
| 246 | 247 | 1 | 1.0900e-01 | 2.8451e+05 | ; | CG1 - HG12 |
| 246 | 248 | 1 | 1.0900e-01 | 2.8451e+05 | ; | CG1 - HG13 |
| 246 | 249 | 1 | 1.5260e-01 | 2.5941e+05 | ; | CG1 - CD1 |
| 249 | 250 | 1 | 1.0900e-01 | 2.8451e+05 | ; | CD1 - HD11 |
| 249 | 251 | 1 | 1.0900e-01 | 2.8451e+05 | ; | CD1 - HD12 |
| 249 | 252 | 1 | 1.0900e-01 | 2.8451e+05 | ; | CD1 - HD13 |
| 253 | 254 | 1 | 1.2290e-01 | 4.7698e+05 | ; | C - O |
| 253 | 255 | 1 | 1.3350e-01 | 4.1003e+05 | ; | C - N |
| 255 | 256 | 1 | 1.0100e-01 | 3.6317e+05 | ; | N - H |

| | | | | | | |
|---|---|---|---|---|---|---|
| 255 | 257 | 1 | 1.4490e-01 | 2.8200e+05 | ; | N - CA |
| 257 | 258 | 1 | 1.0900e-01 | 2.8451e+05 | ; | CA - HA |
| 257 | 259 | 1 | 1.5260e-01 | 2.5941e+05 | ; | CA - CB |
| 257 | 265 | 1 | 1.5220e-01 | 2.6527e+05 | ; | CA - C |
| 259 | 260 | 1 | 1.0900e-01 | 2.8451e+05 | ; | CB - HB2 |
| 259 | 261 | 1 | 1.0900e-01 | 2.8451e+05 | ; | CB - HB3 |
| 259 | 262 | 1 | 1.5220e-01 | 2.6527e+05 | ; | CB - CG |
| 262 | 263 | 1 | 1.2500e-01 | 5.4894e+05 | ; | CG - OD1 |
| 262 | 264 | 1 | 1.2500e-01 | 5.4894e+05 | ; | CG - OD2 |
| 265 | 266 | 1 | 1.2290e-01 | 4.7698e+05 | ; | C - O |
| 265 | 267 | 1 | 1.3350e-01 | 4.1003e+05 | ; | C - N |
| 267 | 268 | 1 | 1.0100e-01 | 3.6317e+05 | ; | N - H |
| 267 | 269 | 1 | 1.4490e-01 | 2.8200e+05 | ; | N - CA |
| 269 | 270 | 1 | 1.0900e-01 | 2.8451e+05 | ; | CA - HA |
| 269 | 271 | 1 | 1.5260e-01 | 2.5941e+05 | ; | CA - CB |
| 269 | 280 | 1 | 1.5220e-01 | 2.6527e+05 | ; | CA - C |
| 271 | 272 | 1 | 1.0900e-01 | 2.8451e+05 | ; | CB - HB2 |
| 271 | 273 | 1 | 1.0900e-01 | 2.8451e+05 | ; | CB - HB3 |
| 271 | 274 | 1 | 1.5260e-01 | 2.5941e+05 | ; | CB - CG |
| 274 | 275 | 1 | 1.0900e-01 | 2.8451e+05 | ; | CG - HG2 |
| 274 | 276 | 1 | 1.0900e-01 | 2.8451e+05 | ; | CG - HG3 |
| 274 | 277 | 1 | 1.5220e-01 | 2.6527e+05 | ; | CG - CD |
| 277 | 278 | 1 | 1.2500e-01 | 5.4894e+05 | ; | CD - OE1 |
| 277 | 279 | 1 | 1.2500e-01 | 5.4894e+05 | ; | CD - OE2 |
| 280 | 281 | 1 | 1.2290e-01 | 4.7698e+05 | ; | C - O |
| 280 | 282 | 1 | 1.3350e-01 | 4.1003e+05 | ; | C - N |
| 282 | 283 | 1 | 1.0100e-01 | 3.6317e+05 | ; | N - H |
| 282 | 284 | 1 | 1.4490e-01 | 2.8200e+05 | ; | N - CA |
| 284 | 285 | 1 | 1.0900e-01 | 2.8451e+05 | ; | CA - HA |
| 284 | 286 | 1 | 1.5260e-01 | 2.5941e+05 | ; | CA - CB |
| 284 | 302 | 1 | 1.5220e-01 | 2.6527e+05 | ; | CA - C |
| 286 | 287 | 1 | 1.0900e-01 | 2.8451e+05 | ; | CB - HB2 |
| 286 | 288 | 1 | 1.0900e-01 | 2.8451e+05 | ; | CB - HB3 |
| 286 | 289 | 1 | 1.5260e-01 | 2.5941e+05 | ; | CB - CG |
| 289 | 290 | 1 | 1.0900e-01 | 2.8451e+05 | ; | CG - HG2 |
| 289 | 291 | 1 | 1.0900e-01 | 2.8451e+05 | ; | CG - HG3 |
| 289 | 292 | 1 | 1.5260e-01 | 2.5941e+05 | ; | CG - CD |
| 292 | 293 | 1 | 1.0900e-01 | 2.8451e+05 | ; | CD - HD2 |
| 292 | 294 | 1 | 1.0900e-01 | 2.8451e+05 | ; | CD - HD3 |
| 292 | 295 | 1 | 1.5260e-01 | 2.5941e+05 | ; | CD - CE |
| 295 | 296 | 1 | 1.0900e-01 | 2.8451e+05 | ; | CE - HE2 |
| 295 | 297 | 1 | 1.0900e-01 | 2.8451e+05 | ; | CE - HE3 |
| 295 | 298 | 1 | 1.4710e-01 | 3.0711e+05 | ; | CE - NZ |
| 298 | 299 | 1 | 1.0100e-01 | 3.6317e+05 | ; | NZ - HZ1 |
| 298 | 300 | 1 | 1.0100e-01 | 3.6317e+05 | ; | NZ - HZ2 |
| 298 | 301 | 1 | 1.0100e-01 | 3.6317e+05 | ; | NZ - HZ3 |
| 302 | 303 | 1 | 1.2290e-01 | 4.7698e+05 | ; | C - O |
| 302 | 304 | 1 | 1.3350e-01 | 4.1003e+05 | ; | C - N |
| 304 | 305 | 1 | 1.0100e-01 | 3.6317e+05 | ; | N - H |
| 304 | 306 | 1 | 1.4490e-01 | 2.8200e+05 | ; | N - CA |
| 306 | 307 | 1 | 1.0900e-01 | 2.8451e+05 | ; | CA - HA |
| 306 | 308 | 1 | 1.5260e-01 | 2.5941e+05 | ; | CA - CB |

| | | | | | |
|---|---|---|---|---|---|
| 306 | 321 | 1 | 1.5220e-01 | 2.6527e+05 ; | CA - C |
| 308 | 309 | 1 | 1.0900e-01 | 2.8451e+05 ; | CB - HB2 |
| 308 | 310 | 1 | 1.0900e-01 | 2.8451e+05 ; | CB - HB3 |
| 308 | 311 | 1 | 1.5260e-01 | 2.5941e+05 ; | CB - CG |
| 311 | 312 | 1 | 1.0900e-01 | 2.8451e+05 ; | CG - HG |
| 311 | 313 | 1 | 1.5260e-01 | 2.5941e+05 ; | CG - CD1 |
| 311 | 317 | 1 | 1.5260e-01 | 2.5941e+05 ; | CG - CD2 |
| 313 | 314 | 1 | 1.0900e-01 | 2.8451e+05 ; | CD1 - HD11 |
| 313 | 315 | 1 | 1.0900e-01 | 2.8451e+05 ; | CD1 - HD12 |
| 313 | 316 | 1 | 1.0900e-01 | 2.8451e+05 ; | CD1 - HD13 |
| 317 | 318 | 1 | 1.0900e-01 | 2.8451e+05 ; | CD2 - HD21 |
| 317 | 319 | 1 | 1.0900e-01 | 2.8451e+05 ; | CD2 - HD22 |
| 317 | 320 | 1 | 1.0900e-01 | 2.8451e+05 ; | CD2 - HD23 |
| 321 | 322 | 1 | 1.2290e-01 | 4.7698e+05 ; | C - O |
| 321 | 323 | 1 | 1.3350e-01 | 4.1003e+05 ; | C - N |
| 323 | 324 | 1 | 1.0100e-01 | 3.6317e+05 ; | N - H |
| 323 | 325 | 1 | 1.4490e-01 | 2.8200e+05 ; | N - CA |
| 325 | 326 | 1 | 1.0900e-01 | 2.8451e+05 ; | CA - HA2 |
| 325 | 327 | 1 | 1.0900e-01 | 2.8451e+05 ; | CA - HA3 |
| 325 | 328 | 1 | 1.5220e-01 | 2.6527e+05 ; | CA - C |
| 328 | 329 | 1 | 1.2290e-01 | 4.7698e+05 ; | C - O |
| 328 | 330 | 1 | 1.3350e-01 | 4.1003e+05 ; | C - N |
| 330 | 331 | 1 | 1.0100e-01 | 3.6317e+05 ; | N - H |
| 330 | 332 | 1 | 1.4490e-01 | 2.8200e+05 ; | N - CA |
| 332 | 333 | 1 | 1.0900e-01 | 2.8451e+05 ; | CA - HA |
| 332 | 334 | 1 | 1.5260e-01 | 2.5941e+05 ; | CA - CB |
| 332 | 345 | 1 | 1.5220e-01 | 2.6527e+05 ; | CA - C |
| 334 | 335 | 1 | 1.0900e-01 | 2.8451e+05 ; | CB - HB2 |
| 334 | 336 | 1 | 1.0900e-01 | 2.8451e+05 ; | CB - HB3 |
| 334 | 337 | 1 | 1.5260e-01 | 2.5941e+05 ; | CB - CG |
| 337 | 338 | 1 | 1.0900e-01 | 2.8451e+05 ; | CG - HG2 |
| 337 | 339 | 1 | 1.0900e-01 | 2.8451e+05 ; | CG - HG3 |
| 337 | 340 | 1 | 1.5220e-01 | 2.6527e+05 ; | CG - CD |
| 340 | 341 | 1 | 1.2290e-01 | 4.7698e+05 ; | CD - OE1 |
| 340 | 342 | 1 | 1.3350e-01 | 4.1003e+05 ; | CD - NE2 |
| 342 | 343 | 1 | 1.0100e-01 | 3.6317e+05 ; | NE2 - HE21 |
| 342 | 344 | 1 | 1.0100e-01 | 3.6317e+05 ; | NE2 - HE22 |
| 345 | 346 | 1 | 1.2290e-01 | 4.7698e+05 ; | C - O |
| 345 | 347 | 1 | 1.3350e-01 | 4.1003e+05 ; | C - N |
| 347 | 348 | 1 | 1.0100e-01 | 3.6317e+05 ; | N - H |
| 347 | 349 | 1 | 1.4490e-01 | 2.8200e+05 ; | N - CA |
| 349 | 350 | 1 | 1.0900e-01 | 2.8451e+05 ; | CA - HA2 |
| 349 | 351 | 1 | 1.0900e-01 | 2.8451e+05 ; | CA - HA3 |
| 349 | 352 | 1 | 1.5220e-01 | 2.6527e+05 ; | CA - C |
| 352 | 353 | 1 | 1.2290e-01 | 4.7698e+05 ; | C - O |
| 352 | 354 | 1 | 1.3350e-01 | 4.1003e+05 ; | C - N |
| 354 | 355 | 1 | 1.0100e-01 | 3.6317e+05 ; | N - H |
| 354 | 356 | 1 | 1.4490e-01 | 2.8200e+05 ; | N - CA |
| 356 | 357 | 1 | 1.0900e-01 | 2.8451e+05 ; | CA - HA |
| 356 | 358 | 1 | 1.5260e-01 | 2.5941e+05 ; | CA - CB |
| 356 | 366 | 1 | 1.5220e-01 | 2.6527e+05 ; | CA - C |
| 358 | 359 | 1 | 1.0900e-01 | 2.8451e+05 ; | CB - HB |

| | | | | | | |
|---|---|---|---|---|---|---|
| 358 | 360 | 1 | 1.5260e-01 | 2.5941e+05 | ; | CB - CG2 |
| 358 | 364 | 1 | 1.4100e-01 | 2.6778e+05 | ; | CB - OG1 |
| 360 | 361 | 1 | 1.0900e-01 | 2.8451e+05 | ; | CG2 - HG21 |
| 360 | 362 | 1 | 1.0900e-01 | 2.8451e+05 | ; | CG2 - HG22 |
| 360 | 363 | 1 | 1.0900e-01 | 2.8451e+05 | ; | CG2 - HG23 |
| 364 | 365 | 1 | 9.6000e-02 | 4.6275e+05 | ; | OG1 - HG1 |
| 366 | 367 | 1 | 1.2290e-01 | 4.7698e+05 | ; | C - O |
| 366 | 368 | 1 | 1.3350e-01 | 4.1003e+05 | ; | C - N |
| 368 | 369 | 1 | 1.0100e-01 | 3.6317e+05 | ; | N - H |
| 368 | 370 | 1 | 1.4490e-01 | 2.8200e+05 | ; | N - CA |
| 370 | 371 | 1 | 1.0900e-01 | 2.8451e+05 | ; | CA - HA |
| 370 | 372 | 1 | 1.5260e-01 | 2.5941e+05 | ; | CA - CB |
| 370 | 387 | 1 | 1.5220e-01 | 2.6527e+05 | ; | CA - C |
| 372 | 373 | 1 | 1.0900e-01 | 2.8451e+05 | ; | CB - HB2 |
| 372 | 374 | 1 | 1.0900e-01 | 2.8451e+05 | ; | CB - HB3 |
| 372 | 375 | 1 | 1.5100e-01 | 2.6527e+05 | ; | CB - CG |
| 375 | 376 | 1 | 1.4000e-01 | 3.9246e+05 | ; | CG - CD1 |
| 375 | 385 | 1 | 1.4000e-01 | 3.9246e+05 | ; | CG - CD2 |
| 376 | 377 | 1 | 1.0800e-01 | 3.0711e+05 | ; | CD1 - HD1 |
| 376 | 378 | 1 | 1.4000e-01 | 3.9246e+05 | ; | CD1 - CE1 |
| 378 | 379 | 1 | 1.0800e-01 | 3.0711e+05 | ; | CE1 - HE1 |
| 378 | 380 | 1 | 1.4090e-01 | 3.9246e+05 | ; | CE1 - CZ |
| 380 | 381 | 1 | 1.3640e-01 | 3.7656e+05 | ; | CZ - OH |
| 380 | 383 | 1 | 1.4090e-01 | 3.9246e+05 | ; | CZ - CE2 |
| 381 | 382 | 1 | 9.6000e-02 | 4.6275e+05 | ; | OH - HH |
| 383 | 384 | 1 | 1.0800e-01 | 3.0711e+05 | ; | CE2 - HE2 |
| 383 | 385 | 1 | 1.4000e-01 | 3.9246e+05 | ; | CE2 - CD2 |
| 385 | 386 | 1 | 1.0800e-01 | 3.0711e+05 | ; | CD2 - HD2 |
| 387 | 388 | 1 | 1.2290e-01 | 4.7698e+05 | ; | C - O |
| 387 | 389 | 1 | 1.3350e-01 | 4.1003e+05 | ; | C - N |
| 389 | 390 | 1 | 1.0100e-01 | 3.6317e+05 | ; | N - H |
| 389 | 391 | 1 | 1.4490e-01 | 2.8200e+05 | ; | N - CA |
| 391 | 392 | 1 | 1.0900e-01 | 2.8451e+05 | ; | CA - HA2 |
| 391 | 393 | 1 | 1.0900e-01 | 2.8451e+05 | ; | CA - HA3 |
| 391 | 394 | 1 | 1.5220e-01 | 2.6527e+05 | ; | CA - C |
| 394 | 395 | 1 | 1.2290e-01 | 4.7698e+05 | ; | C - O |
| 394 | 396 | 1 | 1.3350e-01 | 4.1003e+05 | ; | C - N |
| 396 | 397 | 1 | 1.0100e-01 | 3.6317e+05 | ; | N - H |
| 396 | 398 | 1 | 1.4490e-01 | 2.8200e+05 | ; | N - CA |
| 398 | 399 | 1 | 1.0900e-01 | 2.8451e+05 | ; | CA - HA |
| 398 | 400 | 1 | 1.5260e-01 | 2.5941e+05 | ; | CA - CB |
| 398 | 405 | 1 | 1.5220e-01 | 2.6527e+05 | ; | CA - C |
| 400 | 401 | 1 | 1.0900e-01 | 2.8451e+05 | ; | CB - HB2 |
| 400 | 402 | 1 | 1.0900e-01 | 2.8451e+05 | ; | CB - HB3 |
| 400 | 403 | 1 | 1.8100e-01 | 1.9832e+05 | ; | CB - SG |
| 403 | 404 | 1 | 1.3360e-01 | 2.2928e+05 | ; | SG - HG |
| 405 | 406 | 1 | 1.2290e-01 | 4.7698e+05 | ; | C - O |
| 405 | 407 | 1 | 1.3350e-01 | 4.1003e+05 | ; | C - N |
| 407 | 408 | 1 | 1.0100e-01 | 3.6317e+05 | ; | N - H |
| 407 | 409 | 1 | 1.4490e-01 | 2.8200e+05 | ; | N - CA |
| 409 | 410 | 1 | 1.0900e-01 | 2.8451e+05 | ; | CA - HA |
| 409 | 411 | 1 | 1.5260e-01 | 2.5941e+05 | ; | CA - CB |

| | | | | | |
|---|---|---|---|---|---|
| 409 | 421 | 1 | 1.5220e-01 | 2.6527e+05 ; | CA - C |
| 411 | 412 | 1 | 1.0900e-01 | 2.8451e+05 ; | CB - HB |
| 411 | 413 | 1 | 1.5260e-01 | 2.5941e+05 ; | CB - CG1 |
| 411 | 417 | 1 | 1.5260e-01 | 2.5941e+05 ; | CB - CG2 |
| 413 | 414 | 1 | 1.0900e-01 | 2.8451e+05 ; | CG1 - HG11 |
| 413 | 415 | 1 | 1.0900e-01 | 2.8451e+05 ; | CG1 - HG12 |
| 413 | 416 | 1 | 1.0900e-01 | 2.8451e+05 ; | CG1 - HG13 |
| 417 | 418 | 1 | 1.0900e-01 | 2.8451e+05 ; | CG2 - HG21 |
| 417 | 419 | 1 | 1.0900e-01 | 2.8451e+05 ; | CG2 - HG22 |
| 417 | 420 | 1 | 1.0900e-01 | 2.8451e+05 ; | CG2 - HG23 |
| 421 | 422 | 1 | 1.2290e-01 | 4.7698e+05 ; | C - O |
| 421 | 423 | 1 | 1.3350e-01 | 4.1003e+05 ; | C - N |
| 423 | 424 | 1 | 1.0100e-01 | 3.6317e+05 ; | N - H |
| 423 | 425 | 1 | 1.4490e-01 | 2.8200e+05 ; | N - CA |
| 425 | 426 | 1 | 1.0900e-01 | 2.8451e+05 ; | CA - HA |
| 425 | 427 | 1 | 1.5260e-01 | 2.5941e+05 ; | CA - CB |
| 425 | 442 | 1 | 1.5220e-01 | 2.6527e+05 ; | CA - C |
| 427 | 428 | 1 | 1.0900e-01 | 2.8451e+05 ; | CB - HB2 |
| 427 | 429 | 1 | 1.0900e-01 | 2.8451e+05 ; | CB - HB3 |
| 427 | 430 | 1 | 1.5100e-01 | 2.6527e+05 ; | CB - CG |
| 430 | 431 | 1 | 1.4000e-01 | 3.9246e+05 ; | CG - CD1 |
| 430 | 440 | 1 | 1.4000e-01 | 3.9246e+05 ; | CG - CD2 |
| 431 | 432 | 1 | 1.0800e-01 | 3.0711e+05 ; | CD1 - HD1 |
| 431 | 433 | 1 | 1.4000e-01 | 3.9246e+05 ; | CD1 - CE1 |
| 433 | 434 | 1 | 1.0800e-01 | 3.0711e+05 ; | CE1 - HE1 |
| 433 | 435 | 1 | 1.4090e-01 | 3.9246e+05 ; | CE1 - CZ |
| 435 | 436 | 1 | 1.3640e-01 | 3.7656e+05 ; | CZ - OH |
| 435 | 438 | 1 | 1.4090e-01 | 3.9246e+05 ; | CZ - CE2 |
| 436 | 437 | 1 | 9.6000e-02 | 4.6275e+05 ; | OH - HH |
| 438 | 439 | 1 | 1.0800e-01 | 3.0711e+05 ; | CE2 - HE2 |
| 438 | 440 | 1 | 1.4000e-01 | 3.9246e+05 ; | CE2 - CD2 |
| 440 | 441 | 1 | 1.0800e-01 | 3.0711e+05 ; | CD2 - HD2 |
| 442 | 443 | 1 | 1.2290e-01 | 4.7698e+05 ; | C - O |
| 442 | 444 | 1 | 1.3350e-01 | 4.1003e+05 ; | C - N |
| 444 | 445 | 1 | 1.0100e-01 | 3.6317e+05 ; | N - H |
| 444 | 446 | 1 | 1.4490e-01 | 2.8200e+05 ; | N - CA |
| 446 | 447 | 1 | 1.0900e-01 | 2.8451e+05 ; | CA - HA |
| 446 | 448 | 1 | 1.5260e-01 | 2.5941e+05 ; | CA - CB |
| 446 | 464 | 1 | 1.5220e-01 | 2.6527e+05 ; | CA - C |
| 448 | 449 | 1 | 1.0900e-01 | 2.8451e+05 ; | CB - HB2 |
| 448 | 450 | 1 | 1.0900e-01 | 2.8451e+05 ; | CB - HB3 |
| 448 | 451 | 1 | 1.5260e-01 | 2.5941e+05 ; | CB - CG |
| 451 | 452 | 1 | 1.0900e-01 | 2.8451e+05 ; | CG - HG2 |
| 451 | 453 | 1 | 1.0900e-01 | 2.8451e+05 ; | CG - HG3 |
| 451 | 454 | 1 | 1.5260e-01 | 2.5941e+05 ; | CG - CD |
| 454 | 455 | 1 | 1.0900e-01 | 2.8451e+05 ; | CD - HD2 |
| 454 | 456 | 1 | 1.0900e-01 | 2.8451e+05 ; | CD - HD3 |
| 454 | 457 | 1 | 1.5260e-01 | 2.5941e+05 ; | CD - CE |
| 457 | 458 | 1 | 1.0900e-01 | 2.8451e+05 ; | CE - HE2 |
| 457 | 459 | 1 | 1.0900e-01 | 2.8451e+05 ; | CE - HE3 |
| 457 | 460 | 1 | 1.4710e-01 | 3.0711e+05 ; | CE - NZ |
| 460 | 461 | 1 | 1.0100e-01 | 3.6317e+05 ; | NZ - HZ1 |

| | | | | | | |
|---|---|---|---|---|---|---|
| 460 | 462 | 1 | 1.0100e-01 | 3.6317e+05 | ; | NZ - HZ2 |
| 460 | 463 | 1 | 1.0100e-01 | 3.6317e+05 | ; | NZ - HZ3 |
| 464 | 465 | 1 | 1.2290e-01 | 4.7698e+05 | ; | C - O |
| 464 | 466 | 1 | 1.3350e-01 | 4.1003e+05 | ; | C - N |
| 466 | 467 | 1 | 1.0100e-01 | 3.6317e+05 | ; | N - H |
| 466 | 468 | 1 | 1.4490e-01 | 2.8200e+05 | ; | N - CA |
| 468 | 469 | 1 | 1.0900e-01 | 2.8451e+05 | ; | CA - HA2 |
| 468 | 470 | 1 | 1.0900e-01 | 2.8451e+05 | ; | CA - HA3 |
| 468 | 471 | 1 | 1.5220e-01 | 2.6527e+05 | ; | CA - C |
| 471 | 472 | 1 | 1.2290e-01 | 4.7698e+05 | ; | C - O |
| 471 | 473 | 1 | 1.3350e-01 | 4.1003e+05 | ; | C - N |
| 473 | 474 | 1 | 1.0100e-01 | 3.6317e+05 | ; | N - H |
| 473 | 475 | 1 | 1.4490e-01 | 2.8200e+05 | ; | N - CA |
| 475 | 476 | 1 | 1.0900e-01 | 2.8451e+05 | ; | CA - HA |
| 475 | 477 | 1 | 1.5260e-01 | 2.5941e+05 | ; | CA - CB |
| 475 | 490 | 1 | 1.5220e-01 | 2.6527e+05 | ; | CA - C |
| 477 | 478 | 1 | 1.0900e-01 | 2.8451e+05 | ; | CB - HB |
| 477 | 479 | 1 | 1.5260e-01 | 2.5941e+05 | ; | CB - CG2 |
| 477 | 483 | 1 | 1.5260e-01 | 2.5941e+05 | ; | CB - CG1 |
| 479 | 480 | 1 | 1.0900e-01 | 2.8451e+05 | ; | CG2 - HG21 |
| 479 | 481 | 1 | 1.0900e-01 | 2.8451e+05 | ; | CG2 - HG22 |
| 479 | 482 | 1 | 1.0900e-01 | 2.8451e+05 | ; | CG2 - HG23 |
| 483 | 484 | 1 | 1.0900e-01 | 2.8451e+05 | ; | CG1 - HG12 |
| 483 | 485 | 1 | 1.0900e-01 | 2.8451e+05 | ; | CG1 - HG13 |
| 483 | 486 | 1 | 1.5260e-01 | 2.5941e+05 | ; | CG1 - CD1 |
| 486 | 487 | 1 | 1.0900e-01 | 2.8451e+05 | ; | CD1 - HD11 |
| 486 | 488 | 1 | 1.0900e-01 | 2.8451e+05 | ; | CD1 - HD12 |
| 486 | 489 | 1 | 1.0900e-01 | 2.8451e+05 | ; | CD1 - HD13 |
| 490 | 491 | 1 | 1.2290e-01 | 4.7698e+05 | ; | C - O |
| 490 | 492 | 1 | 1.3350e-01 | 4.1003e+05 | ; | C - N |
| 492 | 493 | 1 | 1.0100e-01 | 3.6317e+05 | ; | N - H |
| 492 | 494 | 1 | 1.4490e-01 | 2.8200e+05 | ; | N - CA |
| 494 | 495 | 1 | 1.0900e-01 | 2.8451e+05 | ; | CA - HA |
| 494 | 496 | 1 | 1.5260e-01 | 2.5941e+05 | ; | CA - CB |
| 494 | 502 | 1 | 1.5220e-01 | 2.6527e+05 | ; | CA - C |
| 496 | 497 | 1 | 1.0900e-01 | 2.8451e+05 | ; | CB - HB2 |
| 496 | 498 | 1 | 1.0900e-01 | 2.8451e+05 | ; | CB - HB3 |
| 496 | 499 | 1 | 1.5220e-01 | 2.6527e+05 | ; | CB - CG |
| 499 | 500 | 1 | 1.2500e-01 | 5.4894e+05 | ; | CG - OD1 |
| 499 | 501 | 1 | 1.2500e-01 | 5.4894e+05 | ; | CG - OD2 |
| 502 | 503 | 1 | 1.2290e-01 | 4.7698e+05 | ; | C - O |
| 502 | 504 | 1 | 1.3350e-01 | 4.1003e+05 | ; | C - N |
| 504 | 505 | 1 | 1.0100e-01 | 3.6317e+05 | ; | N - H |
| 504 | 506 | 1 | 1.4490e-01 | 2.8200e+05 | ; | N - CA |
| 506 | 507 | 1 | 1.0900e-01 | 2.8451e+05 | ; | CA - HA |
| 506 | 508 | 1 | 1.5260e-01 | 2.5941e+05 | ; | CA - CB |
| 506 | 524 | 1 | 1.5220e-01 | 2.6527e+05 | ; | CA - C |
| 508 | 509 | 1 | 1.0900e-01 | 2.8451e+05 | ; | CB - HB2 |
| 508 | 510 | 1 | 1.0900e-01 | 2.8451e+05 | ; | CB - HB3 |
| 508 | 511 | 1 | 1.5260e-01 | 2.5941e+05 | ; | CB - CG |
| 511 | 512 | 1 | 1.0900e-01 | 2.8451e+05 | ; | CG - HG2 |
| 511 | 513 | 1 | 1.0900e-01 | 2.8451e+05 | ; | CG - HG3 |

| | | | | | | |
|---|---|---|---|---|---|---|
| 511 | 514 | 1 | 1.5260e-01 | 2.5941e+05 | ; | CG - CD |
| 514 | 515 | 1 | 1.0900e-01 | 2.8451e+05 | ; | CD - HD2 |
| 514 | 516 | 1 | 1.0900e-01 | 2.8451e+05 | ; | CD - HD3 |
| 514 | 517 | 1 | 1.5260e-01 | 2.5941e+05 | ; | CD - CE |
| 517 | 518 | 1 | 1.0900e-01 | 2.8451e+05 | ; | CE - HE2 |
| 517 | 519 | 1 | 1.0900e-01 | 2.8451e+05 | ; | CE - HE3 |
| 517 | 520 | 1 | 1.4710e-01 | 3.0711e+05 | ; | CE - NZ |
| 520 | 521 | 1 | 1.0100e-01 | 3.6317e+05 | ; | NZ - HZ1 |
| 520 | 522 | 1 | 1.0100e-01 | 3.6317e+05 | ; | NZ - HZ2 |
| 520 | 523 | 1 | 1.0100e-01 | 3.6317e+05 | ; | NZ - HZ3 |
| 524 | 525 | 1 | 1.2290e-01 | 4.7698e+05 | ; | C - O |
| 524 | 526 | 1 | 1.3350e-01 | 4.1003e+05 | ; | C - N |
| 526 | 527 | 1 | 1.0100e-01 | 3.6317e+05 | ; | N - H |
| 526 | 528 | 1 | 1.4490e-01 | 2.8200e+05 | ; | N - CA |
| 528 | 529 | 1 | 1.0900e-01 | 2.8451e+05 | ; | CA - HA |
| 528 | 530 | 1 | 1.5260e-01 | 2.5941e+05 | ; | CA - CB |
| 528 | 540 | 1 | 1.5220e-01 | 2.6527e+05 | ; | CA - C |
| 530 | 531 | 1 | 1.0900e-01 | 2.8451e+05 | ; | CB - HB |
| 530 | 532 | 1 | 1.5260e-01 | 2.5941e+05 | ; | CB - CG1 |
| 530 | 536 | 1 | 1.5260e-01 | 2.5941e+05 | ; | CB - CG2 |
| 532 | 533 | 1 | 1.0900e-01 | 2.8451e+05 | ; | CG1 - HG11 |
| 532 | 534 | 1 | 1.0900e-01 | 2.8451e+05 | ; | CG1 - HG12 |
| 532 | 535 | 1 | 1.0900e-01 | 2.8451e+05 | ; | CG1 - HG13 |
| 536 | 537 | 1 | 1.0900e-01 | 2.8451e+05 | ; | CG2 - HG21 |
| 536 | 538 | 1 | 1.0900e-01 | 2.8451e+05 | ; | CG2 - HG22 |
| 536 | 539 | 1 | 1.0900e-01 | 2.8451e+05 | ; | CG2 - HG23 |
| 540 | 541 | 1 | 1.2290e-01 | 4.7698e+05 | ; | C - O |
| 540 | 542 | 1 | 1.3350e-01 | 4.1003e+05 | ; | C - N |
| 542 | 543 | 1 | 1.0100e-01 | 3.6317e+05 | ; | N - H |
| 542 | 544 | 1 | 1.4490e-01 | 2.8200e+05 | ; | N - CA |
| 544 | 545 | 1 | 1.0900e-01 | 2.8451e+05 | ; | CA - HA |
| 544 | 546 | 1 | 1.5260e-01 | 2.5941e+05 | ; | CA - CB |
| 544 | 554 | 1 | 1.5220e-01 | 2.6527e+05 | ; | CA - C |
| 546 | 547 | 1 | 1.0900e-01 | 2.8451e+05 | ; | CB - HB |
| 546 | 548 | 1 | 1.5260e-01 | 2.5941e+05 | ; | CB - CG2 |
| 546 | 552 | 1 | 1.4100e-01 | 2.6778e+05 | ; | CB - OG1 |
| 548 | 549 | 1 | 1.0900e-01 | 2.8451e+05 | ; | CG2 - HG21 |
| 548 | 550 | 1 | 1.0900e-01 | 2.8451e+05 | ; | CG2 - HG22 |
| 548 | 551 | 1 | 1.0900e-01 | 2.8451e+05 | ; | CG2 - HG23 |
| 552 | 553 | 1 | 9.6000e-02 | 4.6275e+05 | ; | OG1 - HG1 |
| 554 | 555 | 1 | 1.2290e-01 | 4.7698e+05 | ; | C - O |
| 554 | 556 | 1 | 1.3350e-01 | 4.1003e+05 | ; | C - N |
| 556 | 557 | 1 | 1.0100e-01 | 3.6317e+05 | ; | N - H |
| 556 | 558 | 1 | 1.4490e-01 | 2.8200e+05 | ; | N - CA |
| 558 | 559 | 1 | 1.0900e-01 | 2.8451e+05 | ; | CA - HA |
| 558 | 560 | 1 | 1.5260e-01 | 2.5941e+05 | ; | CA - CB |
| 558 | 568 | 1 | 1.5220e-01 | 2.6527e+05 | ; | CA - C |
| 560 | 561 | 1 | 1.0900e-01 | 2.8451e+05 | ; | CB - HB2 |
| 560 | 562 | 1 | 1.0900e-01 | 2.8451e+05 | ; | CB - HB3 |
| 560 | 563 | 1 | 1.5220e-01 | 2.6527e+05 | ; | CB - CG |
| 563 | 564 | 1 | 1.2290e-01 | 4.7698e+05 | ; | CG - OD1 |
| 563 | 565 | 1 | 1.3350e-01 | 4.1003e+05 | ; | CG - ND2 |

| | | | | | | |
|---|---|---|---|---|---|---|
| 565 | 566 | 1 | 1.0100e-01 | 3.6317e+05 | ; | ND2 - HD21 |
| 565 | 567 | 1 | 1.0100e-01 | 3.6317e+05 | ; | ND2 - HD22 |
| 568 | 569 | 1 | 1.2290e-01 | 4.7698e+05 | ; | C - O |
| 568 | 570 | 1 | 1.3350e-01 | 4.1003e+05 | ; | C - N |
| 570 | 571 | 1 | 1.0100e-01 | 3.6317e+05 | ; | N - H |
| 570 | 572 | 1 | 1.4490e-01 | 2.8200e+05 | ; | N - CA |
| 572 | 573 | 1 | 1.0900e-01 | 2.8451e+05 | ; | CA - HA |
| 572 | 574 | 1 | 1.5260e-01 | 2.5941e+05 | ; | CA - CB |
| 572 | 585 | 1 | 1.5220e-01 | 2.6527e+05 | ; | CA - C |
| 574 | 575 | 1 | 1.0900e-01 | 2.8451e+05 | ; | CB - HB2 |
| 574 | 576 | 1 | 1.0900e-01 | 2.8451e+05 | ; | CB - HB3 |
| 574 | 577 | 1 | 1.5260e-01 | 2.5941e+05 | ; | CB - CG |
| 577 | 578 | 1 | 1.0900e-01 | 2.8451e+05 | ; | CG - HG2 |
| 577 | 579 | 1 | 1.0900e-01 | 2.8451e+05 | ; | CG - HG3 |
| 577 | 580 | 1 | 1.5220e-01 | 2.6527e+05 | ; | CG - CD |
| 580 | 581 | 1 | 1.2290e-01 | 4.7698e+05 | ; | CD - OE1 |
| 580 | 582 | 1 | 1.3350e-01 | 4.1003e+05 | ; | CD - NE2 |
| 582 | 583 | 1 | 1.0100e-01 | 3.6317e+05 | ; | NE2 - HE21 |
| 582 | 584 | 1 | 1.0100e-01 | 3.6317e+05 | ; | NE2 - HE22 |
| 585 | 586 | 1 | 1.2290e-01 | 4.7698e+05 | ; | C - O |
| 585 | 587 | 1 | 1.3350e-01 | 4.1003e+05 | ; | C - N |
| 587 | 588 | 1 | 1.0100e-01 | 3.6317e+05 | ; | N - H |
| 587 | 589 | 1 | 1.4490e-01 | 2.8200e+05 | ; | N - CA |
| 589 | 590 | 1 | 1.0900e-01 | 2.8451e+05 | ; | CA - HA |
| 589 | 591 | 1 | 1.5260e-01 | 2.5941e+05 | ; | CA - CB |
| 589 | 604 | 1 | 1.5220e-01 | 2.6527e+05 | ; | CA - C |
| 591 | 592 | 1 | 1.0900e-01 | 2.8451e+05 | ; | CB - HB2 |
| 591 | 593 | 1 | 1.0900e-01 | 2.8451e+05 | ; | CB - HB3 |
| 591 | 594 | 1 | 1.5260e-01 | 2.5941e+05 | ; | CB - CG |
| 594 | 595 | 1 | 1.0900e-01 | 2.8451e+05 | ; | CG - HG |
| 594 | 596 | 1 | 1.5260e-01 | 2.5941e+05 | ; | CG - CD1 |
| 594 | 600 | 1 | 1.5260e-01 | 2.5941e+05 | ; | CG - CD2 |
| 596 | 597 | 1 | 1.0900e-01 | 2.8451e+05 | ; | CD1 - HD11 |
| 596 | 598 | 1 | 1.0900e-01 | 2.8451e+05 | ; | CD1 - HD12 |
| 596 | 599 | 1 | 1.0900e-01 | 2.8451e+05 | ; | CD1 - HD13 |
| 600 | 601 | 1 | 1.0900e-01 | 2.8451e+05 | ; | CD2 - HD21 |
| 600 | 602 | 1 | 1.0900e-01 | 2.8451e+05 | ; | CD2 - HD22 |
| 600 | 603 | 1 | 1.0900e-01 | 2.8451e+05 | ; | CD2 - HD23 |
| 604 | 605 | 1 | 1.2290e-01 | 4.7698e+05 | ; | C - O |
| 604 | 606 | 1 | 1.3350e-01 | 4.1003e+05 | ; | C - N |
| 606 | 607 | 1 | 1.0100e-01 | 3.6317e+05 | ; | N - H |
| 606 | 608 | 1 | 1.4490e-01 | 2.8200e+05 | ; | N - CA |
| 608 | 609 | 1 | 1.0900e-01 | 2.8451e+05 | ; | CA - HA |
| 608 | 610 | 1 | 1.5260e-01 | 2.5941e+05 | ; | CA - CB |
| 608 | 625 | 1 | 1.5220e-01 | 2.6527e+05 | ; | CA - C |
| 610 | 611 | 1 | 1.0900e-01 | 2.8451e+05 | ; | CB - HB2 |
| 610 | 612 | 1 | 1.0900e-01 | 2.8451e+05 | ; | CB - HB3 |
| 610 | 613 | 1 | 1.5100e-01 | 2.6527e+05 | ; | CB - CG |
| 613 | 614 | 1 | 1.4000e-01 | 3.9246e+05 | ; | CG - CD1 |
| 613 | 623 | 1 | 1.4000e-01 | 3.9246e+05 | ; | CG - CD2 |
| 614 | 615 | 1 | 1.0800e-01 | 3.0711e+05 | ; | CD1 - HD1 |
| 614 | 616 | 1 | 1.4000e-01 | 3.9246e+05 | ; | CD1 - CE1 |

| | | | | | | |
|---|---|---|---|---|---|---|
| 616 | 617 | 1 | 1.0800e-01 | 3.0711e+05 | ; | CE1 - HE1 |
| 616 | 618 | 1 | 1.4090e-01 | 3.9246e+05 | ; | CE1 - CZ |
| 618 | 619 | 1 | 1.3640e-01 | 3.7656e+05 | ; | CZ - OH |
| 618 | 621 | 1 | 1.4090e-01 | 3.9246e+05 | ; | CZ - CE2 |
| 619 | 620 | 1 | 9.6000e-02 | 4.6275e+05 | ; | OH - HH |
| 621 | 622 | 1 | 1.0800e-01 | 3.0711e+05 | ; | CE2 - HE2 |
| 621 | 623 | 1 | 1.4000e-01 | 3.9246e+05 | ; | CE2 - CD2 |
| 623 | 624 | 1 | 1.0800e-01 | 3.0711e+05 | ; | CD2 - HD2 |
| 625 | 626 | 1 | 1.2290e-01 | 4.7698e+05 | ; | C - O |
| 625 | 627 | 1 | 1.3350e-01 | 4.1003e+05 | ; | C - N |
| 627 | 628 | 1 | 1.0100e-01 | 3.6317e+05 | ; | N - H |
| 627 | 629 | 1 | 1.4490e-01 | 2.8200e+05 | ; | N - CA |
| 629 | 630 | 1 | 1.0900e-01 | 2.8451e+05 | ; | CA - HA |
| 629 | 631 | 1 | 1.5260e-01 | 2.5941e+05 | ; | CA - CB |
| 629 | 635 | 1 | 1.5220e-01 | 2.6527e+05 | ; | CA - C |
| 631 | 632 | 1 | 1.0900e-01 | 2.8451e+05 | ; | CB - HB1 |
| 631 | 633 | 1 | 1.0900e-01 | 2.8451e+05 | ; | CB - HB2 |
| 631 | 634 | 1 | 1.0900e-01 | 2.8451e+05 | ; | CB - HB3 |
| 635 | 636 | 1 | 1.2290e-01 | 4.7698e+05 | ; | C - O |
| 635 | 637 | 1 | 1.3350e-01 | 4.1003e+05 | ; | C - N |
| 637 | 638 | 1 | 1.0100e-01 | 3.6317e+05 | ; | N - H |
| 637 | 639 | 1 | 1.4490e-01 | 2.8200e+05 | ; | N - CA |
| 639 | 640 | 1 | 1.0900e-01 | 2.8451e+05 | ; | CA - HA |
| 639 | 641 | 1 | 1.5260e-01 | 2.5941e+05 | ; | CA - CB |
| 639 | 654 | 1 | 1.5220e-01 | 2.6527e+05 | ; | CA - C |
| 641 | 642 | 1 | 1.0900e-01 | 2.8451e+05 | ; | CB - HB |
| 641 | 643 | 1 | 1.5260e-01 | 2.5941e+05 | ; | CB - CG2 |
| 641 | 647 | 1 | 1.5260e-01 | 2.5941e+05 | ; | CB - CG1 |
| 643 | 644 | 1 | 1.0900e-01 | 2.8451e+05 | ; | CG2 - HG21 |
| 643 | 645 | 1 | 1.0900e-01 | 2.8451e+05 | ; | CG2 - HG22 |
| 643 | 646 | 1 | 1.0900e-01 | 2.8451e+05 | ; | CG2 - HG23 |
| 647 | 648 | 1 | 1.0900e-01 | 2.8451e+05 | ; | CG1 - HG12 |
| 647 | 649 | 1 | 1.0900e-01 | 2.8451e+05 | ; | CG1 - HG13 |
| 647 | 650 | 1 | 1.5260e-01 | 2.5941e+05 | ; | CG1 - CD1 |
| 650 | 651 | 1 | 1.0900e-01 | 2.8451e+05 | ; | CD1 - HD11 |
| 650 | 652 | 1 | 1.0900e-01 | 2.8451e+05 | ; | CD1 - HD12 |
| 650 | 653 | 1 | 1.0900e-01 | 2.8451e+05 | ; | CD1 - HD13 |
| 654 | 655 | 1 | 1.2290e-01 | 4.7698e+05 | ; | C - O |
| 654 | 656 | 1 | 1.3350e-01 | 4.1003e+05 | ; | C - N |
| 656 | 657 | 1 | 1.0100e-01 | 3.6317e+05 | ; | N - H |
| 656 | 658 | 1 | 1.4490e-01 | 2.8200e+05 | ; | N - CA |
| 658 | 659 | 1 | 1.0900e-01 | 2.8451e+05 | ; | CA - HA |
| 658 | 660 | 1 | 1.5260e-01 | 2.5941e+05 | ; | CA - CB |
| 658 | 676 | 1 | 1.5220e-01 | 2.6527e+05 | ; | CA - C |
| 660 | 661 | 1 | 1.0900e-01 | 2.8451e+05 | ; | CB - HB2 |
| 660 | 662 | 1 | 1.0900e-01 | 2.8451e+05 | ; | CB - HB3 |
| 660 | 663 | 1 | 1.5260e-01 | 2.5941e+05 | ; | CB - CG |
| 663 | 664 | 1 | 1.0900e-01 | 2.8451e+05 | ; | CG - HG2 |
| 663 | 665 | 1 | 1.0900e-01 | 2.8451e+05 | ; | CG - HG3 |
| 663 | 666 | 1 | 1.5260e-01 | 2.5941e+05 | ; | CG - CD |
| 666 | 667 | 1 | 1.0900e-01 | 2.8451e+05 | ; | CD - HD2 |
| 666 | 668 | 1 | 1.0900e-01 | 2.8451e+05 | ; | CD - HD3 |

| | | | | | | |
|---|---|---|---|---|---|---|
| 666 | 669 | 1 | 1.5260e-01 | 2.5941e+05 | ; | CD - CE |
| 669 | 670 | 1 | 1.0900e-01 | 2.8451e+05 | ; | CE - HE2 |
| 669 | 671 | 1 | 1.0900e-01 | 2.8451e+05 | ; | CE - HE3 |
| 669 | 672 | 1 | 1.4710e-01 | 3.0711e+05 | ; | CE - NZ |
| 672 | 673 | 1 | 1.0100e-01 | 3.6317e+05 | ; | NZ - HZ1 |
| 672 | 674 | 1 | 1.0100e-01 | 3.6317e+05 | ; | NZ - HZ2 |
| 672 | 675 | 1 | 1.0100e-01 | 3.6317e+05 | ; | NZ - HZ3 |
| 676 | 677 | 1 | 1.2290e-01 | 4.7698e+05 | ; | C - O |
| 676 | 678 | 1 | 1.3350e-01 | 4.1003e+05 | ; | C - N |
| 678 | 679 | 1 | 1.0100e-01 | 3.6317e+05 | ; | N - H |
| 678 | 680 | 1 | 1.4490e-01 | 2.8200e+05 | ; | N - CA |
| 680 | 681 | 1 | 1.0900e-01 | 2.8451e+05 | ; | CA - HA |
| 680 | 682 | 1 | 1.5260e-01 | 2.5941e+05 | ; | CA - CB |
| 680 | 691 | 1 | 1.5220e-01 | 2.6527e+05 | ; | CA - C |
| 682 | 683 | 1 | 1.0900e-01 | 2.8451e+05 | ; | CB - HB2 |
| 682 | 684 | 1 | 1.0900e-01 | 2.8451e+05 | ; | CB - HB3 |
| 682 | 685 | 1 | 1.5260e-01 | 2.5941e+05 | ; | CB - CG |
| 685 | 686 | 1 | 1.0900e-01 | 2.8451e+05 | ; | CG - HG2 |
| 685 | 687 | 1 | 1.0900e-01 | 2.8451e+05 | ; | CG - HG3 |
| 685 | 688 | 1 | 1.5220e-01 | 2.6527e+05 | ; | CG - CD |
| 688 | 689 | 1 | 1.2500e-01 | 5.4894e+05 | ; | CD - OE1 |
| 688 | 690 | 1 | 1.2500e-01 | 5.4894e+05 | ; | CD - OE2 |
| 691 | 692 | 1 | 1.2290e-01 | 4.7698e+05 | ; | C - O |
| 691 | 693 | 1 | 1.3350e-01 | 4.1003e+05 | ; | C - N |
| 693 | 694 | 1 | 1.0100e-01 | 3.6317e+05 | ; | N - H |
| 693 | 695 | 1 | 1.4490e-01 | 2.8200e+05 | ; | N - CA |
| 695 | 696 | 1 | 1.0900e-01 | 2.8451e+05 | ; | CA - HA |
| 695 | 697 | 1 | 1.5260e-01 | 2.5941e+05 | ; | CA - CB |
| 695 | 707 | 1 | 1.5220e-01 | 2.6527e+05 | ; | CA - C |
| 697 | 698 | 1 | 1.0900e-01 | 2.8451e+05 | ; | CB - HB2 |
| 697 | 699 | 1 | 1.0900e-01 | 2.8451e+05 | ; | CB - HB3 |
| 697 | 700 | 1 | 1.5260e-01 | 2.5941e+05 | ; | CB - CG |
| 700 | 701 | 1 | 1.0900e-01 | 2.8451e+05 | ; | CG - HG2 |
| 700 | 702 | 1 | 1.0900e-01 | 2.8451e+05 | ; | CG - HG3 |
| 700 | 703 | 1 | 1.5220e-01 | 2.6527e+05 | ; | CG - CD |
| 703 | 704 | 1 | 1.2290e-01 | 4.7698e+05 | ; | CD - OE1 |
| 703 | 705 | 1 | 1.3640e-01 | 3.7656e+05 | ; | CD - OE2 |
| 705 | 706 | 1 | 9.6000e-02 | 4.6275e+05 | ; | OE2 - HE2 |
| 707 | 708 | 1 | 1.2290e-01 | 4.7698e+05 | ; | C - O |
| 707 | 709 | 1 | 1.3350e-01 | 4.1003e+05 | ; | C - N |
| 709 | 710 | 1 | 1.0100e-01 | 3.6317e+05 | ; | N - H |
| 709 | 711 | 1 | 1.4490e-01 | 2.8200e+05 | ; | N - CA |
| 711 | 712 | 1 | 1.0900e-01 | 2.8451e+05 | ; | CA - HA |
| 711 | 713 | 1 | 1.5260e-01 | 2.5941e+05 | ; | CA - CB |
| 711 | 729 | 1 | 1.5220e-01 | 2.6527e+05 | ; | CA - C |
| 713 | 714 | 1 | 1.0900e-01 | 2.8451e+05 | ; | CB - HB2 |
| 713 | 715 | 1 | 1.0900e-01 | 2.8451e+05 | ; | CB - HB3 |
| 713 | 716 | 1 | 1.5260e-01 | 2.5941e+05 | ; | CB - CG |
| 716 | 717 | 1 | 1.0900e-01 | 2.8451e+05 | ; | CG - HG2 |
| 716 | 718 | 1 | 1.0900e-01 | 2.8451e+05 | ; | CG - HG3 |
| 716 | 719 | 1 | 1.5260e-01 | 2.5941e+05 | ; | CG - CD |
| 719 | 720 | 1 | 1.0900e-01 | 2.8451e+05 | ; | CD - HD2 |

| | | | | | | |
|---|---|---|---|---|---|---|
| 719 | 721 | 1 | 1.0900e-01 | 2.8451e+05 | ; | CD - HD3 |
| 719 | 722 | 1 | 1.5260e-01 | 2.5941e+05 | ; | CD - CE |
| 722 | 723 | 1 | 1.0900e-01 | 2.8451e+05 | ; | CE - HE2 |
| 722 | 724 | 1 | 1.0900e-01 | 2.8451e+05 | ; | CE - HE3 |
| 722 | 725 | 1 | 1.4710e-01 | 3.0711e+05 | ; | CE - NZ |
| 725 | 726 | 1 | 1.0100e-01 | 3.6317e+05 | ; | NZ - HZ1 |
| 725 | 727 | 1 | 1.0100e-01 | 3.6317e+05 | ; | NZ - HZ2 |
| 725 | 728 | 1 | 1.0100e-01 | 3.6317e+05 | ; | NZ - HZ3 |
| 729 | 730 | 1 | 1.2290e-01 | 4.7698e+05 | ; | C - O |
| 729 | 731 | 1 | 1.3350e-01 | 4.1003e+05 | ; | C - N |
| 731 | 732 | 1 | 1.0100e-01 | 3.6317e+05 | ; | N - H |
| 731 | 733 | 1 | 1.4490e-01 | 2.8200e+05 | ; | N - CA |
| 733 | 734 | 1 | 1.0900e-01 | 2.8451e+05 | ; | CA - HA |
| 733 | 735 | 1 | 1.5260e-01 | 2.5941e+05 | ; | CA - CB |
| 733 | 751 | 1 | 1.5220e-01 | 2.6527e+05 | ; | CA - C |
| 735 | 736 | 1 | 1.0900e-01 | 2.8451e+05 | ; | CB - HB2 |
| 735 | 737 | 1 | 1.0900e-01 | 2.8451e+05 | ; | CB - HB3 |
| 735 | 738 | 1 | 1.5260e-01 | 2.5941e+05 | ; | CB - CG |
| 738 | 739 | 1 | 1.0900e-01 | 2.8451e+05 | ; | CG - HG2 |
| 738 | 740 | 1 | 1.0900e-01 | 2.8451e+05 | ; | CG - HG3 |
| 738 | 741 | 1 | 1.5260e-01 | 2.5941e+05 | ; | CG - CD |
| 741 | 742 | 1 | 1.0900e-01 | 2.8451e+05 | ; | CD - HD2 |
| 741 | 743 | 1 | 1.0900e-01 | 2.8451e+05 | ; | CD - HD3 |
| 741 | 744 | 1 | 1.5260e-01 | 2.5941e+05 | ; | CD - CE |
| 744 | 745 | 1 | 1.0900e-01 | 2.8451e+05 | ; | CE - HE2 |
| 744 | 746 | 1 | 1.0900e-01 | 2.8451e+05 | ; | CE - HE3 |
| 744 | 747 | 1 | 1.4710e-01 | 3.0711e+05 | ; | CE - NZ |
| 747 | 748 | 1 | 1.0100e-01 | 3.6317e+05 | ; | NZ - HZ1 |
| 747 | 749 | 1 | 1.0100e-01 | 3.6317e+05 | ; | NZ - HZ2 |
| 747 | 750 | 1 | 1.0100e-01 | 3.6317e+05 | ; | NZ - HZ3 |
| 751 | 752 | 1 | 1.2290e-01 | 4.7698e+05 | ; | C - O |
| 751 | 753 | 1 | 1.3350e-01 | 4.1003e+05 | ; | C - N |
| 753 | 754 | 1 | 1.0100e-01 | 3.6317e+05 | ; | N - H |
| 753 | 755 | 1 | 1.4490e-01 | 2.8200e+05 | ; | N - CA |
| 755 | 756 | 1 | 1.0900e-01 | 2.8451e+05 | ; | CA - HA |
| 755 | 757 | 1 | 1.5260e-01 | 2.5941e+05 | ; | CA - CB |
| 755 | 763 | 1 | 1.5220e-01 | 2.6527e+05 | ; | CA - C |
| 757 | 758 | 1 | 1.0900e-01 | 2.8451e+05 | ; | CB - HB2 |
| 757 | 759 | 1 | 1.0900e-01 | 2.8451e+05 | ; | CB - HB3 |
| 757 | 760 | 1 | 1.5220e-01 | 2.6527e+05 | ; | CB - CG |
| 760 | 761 | 1 | 1.2500e-01 | 5.4894e+05 | ; | CG - OD1 |
| 760 | 762 | 1 | 1.2500e-01 | 5.4894e+05 | ; | CG - OD2 |
| 763 | 764 | 1 | 1.2290e-01 | 4.7698e+05 | ; | C - O |
| 763 | 765 | 1 | 1.3350e-01 | 4.1003e+05 | ; | C - N |
| 765 | 766 | 1 | 1.0100e-01 | 3.6317e+05 | ; | N - H |
| 765 | 767 | 1 | 1.4490e-01 | 2.8200e+05 | ; | N - CA |
| 767 | 768 | 1 | 1.0900e-01 | 2.8451e+05 | ; | CA - HA |
| 767 | 769 | 1 | 1.5260e-01 | 2.5941e+05 | ; | CA - CB |
| 767 | 787 | 1 | 1.5220e-01 | 2.6527e+05 | ; | CA - C |
| 769 | 770 | 1 | 1.0900e-01 | 2.8451e+05 | ; | CB - HB2 |
| 769 | 771 | 1 | 1.0900e-01 | 2.8451e+05 | ; | CB - HB3 |
| 769 | 772 | 1 | 1.5260e-01 | 2.5941e+05 | ; | CB - CG |

| | | | | | | |
|---|---|---|---|---|---|---|
| 772 | 773 | 1 | 1.0900e-01 | 2.8451e+05 | ; | CG - HG2 |
| 772 | 774 | 1 | 1.0900e-01 | 2.8451e+05 | ; | CG - HG3 |
| 772 | 775 | 1 | 1.5260e-01 | 2.5941e+05 | ; | CG - CD |
| 775 | 776 | 1 | 1.0900e-01 | 2.8451e+05 | ; | CD - HD2 |
| 775 | 777 | 1 | 1.0900e-01 | 2.8451e+05 | ; | CD - HD3 |
| 775 | 778 | 1 | 1.4630e-01 | 2.8200e+05 | ; | CD - NE |
| 778 | 779 | 1 | 1.0100e-01 | 3.6317e+05 | ; | NE - HE |
| 778 | 780 | 1 | 1.3400e-01 | 4.0250e+05 | ; | NE - CZ |
| 780 | 781 | 1 | 1.3400e-01 | 4.0250e+05 | ; | CZ - NH1 |
| 780 | 784 | 1 | 1.3400e-01 | 4.0250e+05 | ; | CZ - NH2 |
| 781 | 782 | 1 | 1.0100e-01 | 3.6317e+05 | ; | NH1 - HH11 |
| 781 | 783 | 1 | 1.0100e-01 | 3.6317e+05 | ; | NH1 - HH12 |
| 784 | 785 | 1 | 1.0100e-01 | 3.6317e+05 | ; | NH2 - HH21 |
| 784 | 786 | 1 | 1.0100e-01 | 3.6317e+05 | ; | NH2 - HH22 |
| 787 | 788 | 1 | 1.2290e-01 | 4.7698e+05 | ; | C - O |
| 787 | 789 | 1 | 1.3350e-01 | 4.1003e+05 | ; | C - N |
| 789 | 790 | 1 | 1.0100e-01 | 3.6317e+05 | ; | N - H |
| 789 | 791 | 1 | 1.4490e-01 | 2.8200e+05 | ; | N - CA |
| 791 | 792 | 1 | 1.0900e-01 | 2.8451e+05 | ; | CA - HA |
| 791 | 793 | 1 | 1.5260e-01 | 2.5941e+05 | ; | CA - CB |
| 791 | 806 | 1 | 1.5220e-01 | 2.6527e+05 | ; | CA - C |
| 793 | 794 | 1 | 1.0900e-01 | 2.8451e+05 | ; | CB - HB2 |
| 793 | 795 | 1 | 1.0900e-01 | 2.8451e+05 | ; | CB - HB3 |
| 793 | 796 | 1 | 1.5260e-01 | 2.5941e+05 | ; | CB - CG |
| 796 | 797 | 1 | 1.0900e-01 | 2.8451e+05 | ; | CG - HG |
| 796 | 798 | 1 | 1.5260e-01 | 2.5941e+05 | ; | CG - CD1 |
| 796 | 802 | 1 | 1.5260e-01 | 2.5941e+05 | ; | CG - CD2 |
| 798 | 799 | 1 | 1.0900e-01 | 2.8451e+05 | ; | CD1 - HD11 |
| 798 | 800 | 1 | 1.0900e-01 | 2.8451e+05 | ; | CD1 - HD12 |
| 798 | 801 | 1 | 1.0900e-01 | 2.8451e+05 | ; | CD1 - HD13 |
| 802 | 803 | 1 | 1.0900e-01 | 2.8451e+05 | ; | CD2 - HD21 |
| 802 | 804 | 1 | 1.0900e-01 | 2.8451e+05 | ; | CD2 - HD22 |
| 802 | 805 | 1 | 1.0900e-01 | 2.8451e+05 | ; | CD2 - HD23 |
| 806 | 807 | 1 | 1.2290e-01 | 4.7698e+05 | ; | C - O |
| 806 | 808 | 1 | 1.3350e-01 | 4.1003e+05 | ; | C - N |
| 808 | 809 | 1 | 1.0100e-01 | 3.6317e+05 | ; | N - H |
| 808 | 810 | 1 | 1.4490e-01 | 2.8200e+05 | ; | N - CA |
| 810 | 811 | 1 | 1.0900e-01 | 2.8451e+05 | ; | CA - HA |
| 810 | 812 | 1 | 1.5260e-01 | 2.5941e+05 | ; | CA - CB |
| 810 | 828 | 1 | 1.5220e-01 | 2.6527e+05 | ; | CA - C |
| 812 | 813 | 1 | 1.0900e-01 | 2.8451e+05 | ; | CB - HB2 |
| 812 | 814 | 1 | 1.0900e-01 | 2.8451e+05 | ; | CB - HB3 |
| 812 | 815 | 1 | 1.5260e-01 | 2.5941e+05 | ; | CB - CG |
| 815 | 816 | 1 | 1.0900e-01 | 2.8451e+05 | ; | CG - HG2 |
| 815 | 817 | 1 | 1.0900e-01 | 2.8451e+05 | ; | CG - HG3 |
| 815 | 818 | 1 | 1.5260e-01 | 2.5941e+05 | ; | CG - CD |
| 818 | 819 | 1 | 1.0900e-01 | 2.8451e+05 | ; | CD - HD2 |
| 818 | 820 | 1 | 1.0900e-01 | 2.8451e+05 | ; | CD - HD3 |
| 818 | 821 | 1 | 1.5260e-01 | 2.5941e+05 | ; | CD - CE |
| 821 | 822 | 1 | 1.0900e-01 | 2.8451e+05 | ; | CE - HE2 |
| 821 | 823 | 1 | 1.0900e-01 | 2.8451e+05 | ; | CE - HE3 |
| 821 | 824 | 1 | 1.4710e-01 | 3.0711e+05 | ; | CE - NZ |

| | | | | | | |
|---|---|---|---|---|---|---|
| 824 | 825 | 1 | 1.0100e-01 | 3.6317e+05 | ; | NZ - HZ1 |
| 824 | 826 | 1 | 1.0100e-01 | 3.6317e+05 | ; | NZ - HZ2 |
| 824 | 827 | 1 | 1.0100e-01 | 3.6317e+05 | ; | NZ - HZ3 |
| 828 | 829 | 1 | 1.2290e-01 | 4.7698e+05 | ; | C - O |
| 828 | 830 | 1 | 1.3350e-01 | 4.1003e+05 | ; | C - N |
| 830 | 831 | 1 | 1.0100e-01 | 3.6317e+05 | ; | N - H |
| 830 | 832 | 1 | 1.4490e-01 | 2.8200e+05 | ; | N - CA |
| 832 | 833 | 1 | 1.0900e-01 | 2.8451e+05 | ; | CA - HA |
| 832 | 834 | 1 | 1.5260e-01 | 2.5941e+05 | ; | CA - CB |
| 832 | 842 | 1 | 1.5220e-01 | 2.6527e+05 | ; | CA - C |
| 834 | 835 | 1 | 1.0900e-01 | 2.8451e+05 | ; | CB - HB2 |
| 834 | 836 | 1 | 1.0900e-01 | 2.8451e+05 | ; | CB - HB3 |
| 834 | 837 | 1 | 1.5220e-01 | 2.6527e+05 | ; | CB - CG |
| 837 | 838 | 1 | 1.2290e-01 | 4.7698e+05 | ; | CG - OD1 |
| 837 | 839 | 1 | 1.3350e-01 | 4.1003e+05 | ; | CG - ND2 |
| 839 | 840 | 1 | 1.0100e-01 | 3.6317e+05 | ; | ND2 - HD21 |
| 839 | 841 | 1 | 1.0100e-01 | 3.6317e+05 | ; | ND2 - HD22 |
| 842 | 843 | 1 | 1.2290e-01 | 4.7698e+05 | ; | C - O |
| 842 | 844 | 1 | 1.3350e-01 | 4.1003e+05 | ; | C - N |
| 844 | 845 | 1 | 1.0100e-01 | 3.6317e+05 | ; | N - H |
| 844 | 846 | 1 | 1.4490e-01 | 2.8200e+05 | ; | N - CA |
| 846 | 847 | 1 | 1.0900e-01 | 2.8451e+05 | ; | CA - HA |
| 846 | 848 | 1 | 1.5260e-01 | 2.5941e+05 | ; | CA - CB |
| 846 | 861 | 1 | 1.5220e-01 | 2.6527e+05 | ; | CA - C |
| 848 | 849 | 1 | 1.0900e-01 | 2.8451e+05 | ; | CB - HB |
| 848 | 850 | 1 | 1.5260e-01 | 2.5941e+05 | ; | CB - CG2 |
| 848 | 854 | 1 | 1.5260e-01 | 2.5941e+05 | ; | CB - CG1 |
| 850 | 851 | 1 | 1.0900e-01 | 2.8451e+05 | ; | CG2 - HG21 |
| 850 | 852 | 1 | 1.0900e-01 | 2.8451e+05 | ; | CG2 - HG22 |
| 850 | 853 | 1 | 1.0900e-01 | 2.8451e+05 | ; | CG2 - HG23 |
| 854 | 855 | 1 | 1.0900e-01 | 2.8451e+05 | ; | CG1 - HG12 |
| 854 | 856 | 1 | 1.0900e-01 | 2.8451e+05 | ; | CG1 - HG13 |
| 854 | 857 | 1 | 1.5260e-01 | 2.5941e+05 | ; | CG1 - CD1 |
| 857 | 858 | 1 | 1.0900e-01 | 2.8451e+05 | ; | CD1 - HD11 |
| 857 | 859 | 1 | 1.0900e-01 | 2.8451e+05 | ; | CD1 - HD12 |
| 857 | 860 | 1 | 1.0900e-01 | 2.8451e+05 | ; | CD1 - HD13 |
| 861 | 862 | 1 | 1.2290e-01 | 4.7698e+05 | ; | C - O |
| 861 | 863 | 1 | 1.3350e-01 | 4.1003e+05 | ; | C - N |
| 863 | 864 | 1 | 1.0100e-01 | 3.6317e+05 | ; | N - H |
| 863 | 865 | 1 | 1.4490e-01 | 2.8200e+05 | ; | N - CA |
| 865 | 866 | 1 | 1.0900e-01 | 2.8451e+05 | ; | CA - HA |
| 865 | 867 | 1 | 1.5260e-01 | 2.5941e+05 | ; | CA - CB |
| 865 | 875 | 1 | 1.5220e-01 | 2.6527e+05 | ; | CA - C |
| 867 | 868 | 1 | 1.0900e-01 | 2.8451e+05 | ; | CB - HB2 |
| 867 | 869 | 1 | 1.0900e-01 | 2.8451e+05 | ; | CB - HB3 |
| 867 | 870 | 1 | 1.5220e-01 | 2.6527e+05 | ; | CB - CG |
| 870 | 871 | 1 | 1.2290e-01 | 4.7698e+05 | ; | CG - OD1 |
| 870 | 872 | 1 | 1.3350e-01 | 4.1003e+05 | ; | CG - ND2 |
| 872 | 873 | 1 | 1.0100e-01 | 3.6317e+05 | ; | ND2 - HD21 |
| 872 | 874 | 1 | 1.0100e-01 | 3.6317e+05 | ; | ND2 - HD22 |
| 875 | 876 | 1 | 1.2290e-01 | 4.7698e+05 | ; | C - O |
| 875 | 877 | 1 | 1.3350e-01 | 4.1003e+05 | ; | C - N |

| | | | | | | |
|---|---|---|---|---|---|---|
| 877 | 878 | 1 | 1.0100e-01 | 3.6317e+05 | ; | N - H |
| 877 | 879 | 1 | 1.4490e-01 | 2.8200e+05 | ; | N - CA |
| 879 | 880 | 1 | 1.0900e-01 | 2.8451e+05 | ; | CA - HA |
| 879 | 881 | 1 | 1.5260e-01 | 2.5941e+05 | ; | CA - CB |
| 879 | 899 | 1 | 1.5220e-01 | 2.6527e+05 | ; | CA - C |
| 881 | 882 | 1 | 1.0900e-01 | 2.8451e+05 | ; | CB - HB2 |
| 881 | 883 | 1 | 1.0900e-01 | 2.8451e+05 | ; | CB - HB3 |
| 881 | 884 | 1 | 1.5260e-01 | 2.5941e+05 | ; | CB - CG |
| 884 | 885 | 1 | 1.0900e-01 | 2.8451e+05 | ; | CG - HG2 |
| 884 | 886 | 1 | 1.0900e-01 | 2.8451e+05 | ; | CG - HG3 |
| 884 | 887 | 1 | 1.5260e-01 | 2.5941e+05 | ; | CG - CD |
| 887 | 888 | 1 | 1.0900e-01 | 2.8451e+05 | ; | CD - HD2 |
| 887 | 889 | 1 | 1.0900e-01 | 2.8451e+05 | ; | CD - HD3 |
| 887 | 890 | 1 | 1.4630e-01 | 2.8200e+05 | ; | CD - NE |
| 890 | 891 | 1 | 1.0100e-01 | 3.6317e+05 | ; | NE - HE |
| 890 | 892 | 1 | 1.3400e-01 | 4.0250e+05 | ; | NE - CZ |
| 892 | 893 | 1 | 1.3400e-01 | 4.0250e+05 | ; | CZ - NH1 |
| 892 | 896 | 1 | 1.3400e-01 | 4.0250e+05 | ; | CZ - NH2 |
| 893 | 894 | 1 | 1.0100e-01 | 3.6317e+05 | ; | NH1 - HH11 |
| 893 | 895 | 1 | 1.0100e-01 | 3.6317e+05 | ; | NH1 - HH12 |
| 896 | 897 | 1 | 1.0100e-01 | 3.6317e+05 | ; | NH2 - HH21 |
| 896 | 898 | 1 | 1.0100e-01 | 3.6317e+05 | ; | NH2 - HH22 |
| 899 | 900 | 1 | 1.2290e-01 | 4.7698e+05 | ; | C - O |
| 899 | 901 | 1 | 1.3350e-01 | 4.1003e+05 | ; | C - N |
| 901 | 902 | 1 | 1.0100e-01 | 3.6317e+05 | ; | N - H |
| 901 | 903 | 1 | 1.4490e-01 | 2.8200e+05 | ; | N - CA |
| 903 | 904 | 1 | 1.0900e-01 | 2.8451e+05 | ; | CA - HA |
| 903 | 905 | 1 | 1.5260e-01 | 2.5941e+05 | ; | CA - CB |
| 903 | 919 | 1 | 1.5220e-01 | 2.6527e+05 | ; | CA - C |
| 905 | 906 | 1 | 1.0900e-01 | 2.8451e+05 | ; | CB - HB2 |
| 905 | 907 | 1 | 1.0900e-01 | 2.8451e+05 | ; | CB - HB3 |
| 905 | 908 | 1 | 1.5100e-01 | 2.6527e+05 | ; | CB - CG |
| 908 | 909 | 1 | 1.4000e-01 | 3.9246e+05 | ; | CG - CD1 |
| 908 | 917 | 1 | 1.4000e-01 | 3.9246e+05 | ; | CG - CD2 |
| 909 | 910 | 1 | 1.0800e-01 | 3.0711e+05 | ; | CD1 - HD1 |
| 909 | 911 | 1 | 1.4000e-01 | 3.9246e+05 | ; | CD1 - CE1 |
| 911 | 912 | 1 | 1.0800e-01 | 3.0711e+05 | ; | CE1 - HE1 |
| 911 | 913 | 1 | 1.4000e-01 | 3.9246e+05 | ; | CE1 - CZ |
| 913 | 914 | 1 | 1.0800e-01 | 3.0711e+05 | ; | CZ - HZ |
| 913 | 915 | 1 | 1.4000e-01 | 3.9246e+05 | ; | CZ - CE2 |
| 915 | 916 | 1 | 1.0800e-01 | 3.0711e+05 | ; | CE2 - HE2 |
| 915 | 917 | 1 | 1.4000e-01 | 3.9246e+05 | ; | CE2 - CD2 |
| 917 | 918 | 1 | 1.0800e-01 | 3.0711e+05 | ; | CD2 - HD2 |
| 919 | 920 | 1 | 1.2290e-01 | 4.7698e+05 | ; | C - O |
| 919 | 921 | 1 | 1.3350e-01 | 4.1003e+05 | ; | C - N |
| 921 | 922 | 1 | 1.0100e-01 | 3.6317e+05 | ; | N - H |
| 921 | 923 | 1 | 1.4490e-01 | 2.8200e+05 | ; | N - CA |
| 923 | 924 | 1 | 1.0900e-01 | 2.8451e+05 | ; | CA - HA |
| 923 | 925 | 1 | 1.5260e-01 | 2.5941e+05 | ; | CA - CB |
| 923 | 939 | 1 | 1.5220e-01 | 2.6527e+05 | ; | CA - C |
| 925 | 926 | 1 | 1.0900e-01 | 2.8451e+05 | ; | CB - HB2 |
| 925 | 927 | 1 | 1.0900e-01 | 2.8451e+05 | ; | CB - HB3 |

| | | | | | | |
|---|---|---|---|---|---|---|
| 925 | 928 | 1 | 1.5100e-01 | 2.6527e+05 | ; | CB - CG |
| 928 | 929 | 1 | 1.4000e-01 | 3.9246e+05 | ; | CG - CD1 |
| 928 | 937 | 1 | 1.4000e-01 | 3.9246e+05 | ; | CG - CD2 |
| 929 | 930 | 1 | 1.0800e-01 | 3.0711e+05 | ; | CD1 - HD1 |
| 929 | 931 | 1 | 1.4000e-01 | 3.9246e+05 | ; | CD1 - CE1 |
| 931 | 932 | 1 | 1.0800e-01 | 3.0711e+05 | ; | CE1 - HE1 |
| 931 | 933 | 1 | 1.4000e-01 | 3.9246e+05 | ; | CE1 - CZ |
| 933 | 934 | 1 | 1.0800e-01 | 3.0711e+05 | ; | CZ - HZ |
| 933 | 935 | 1 | 1.4000e-01 | 3.9246e+05 | ; | CZ - CE2 |
| 935 | 936 | 1 | 1.0800e-01 | 3.0711e+05 | ; | CE2 - HE2 |
| 935 | 937 | 1 | 1.4000e-01 | 3.9246e+05 | ; | CE2 - CD2 |
| 937 | 938 | 1 | 1.0800e-01 | 3.0711e+05 | ; | CD2 - HD2 |
| 939 | 940 | 1 | 1.2290e-01 | 4.7698e+05 | ; | C - O |
| 939 | 941 | 1 | 1.3350e-01 | 4.1003e+05 | ; | C - N |
| 941 | 942 | 1 | 1.0100e-01 | 3.6317e+05 | ; | N - H |
| 941 | 943 | 1 | 1.4490e-01 | 2.8200e+05 | ; | N - CA |
| 943 | 944 | 1 | 1.0900e-01 | 2.8451e+05 | ; | CA - HA |
| 943 | 945 | 1 | 1.5260e-01 | 2.5941e+05 | ; | CA - CB |
| 943 | 956 | 1 | 1.5220e-01 | 2.6527e+05 | ; | CA - C |
| 945 | 946 | 1 | 1.0900e-01 | 2.8451e+05 | ; | CB - HB2 |
| 945 | 947 | 1 | 1.0900e-01 | 2.8451e+05 | ; | CB - HB3 |
| 945 | 948 | 1 | 1.5260e-01 | 2.5941e+05 | ; | CB - CG |
| 948 | 949 | 1 | 1.0900e-01 | 2.8451e+05 | ; | CG - HG2 |
| 948 | 950 | 1 | 1.0900e-01 | 2.8451e+05 | ; | CG - HG3 |
| 948 | 951 | 1 | 1.5220e-01 | 2.6527e+05 | ; | CG - CD |
| 951 | 952 | 1 | 1.2290e-01 | 4.7698e+05 | ; | CD - OE1 |
| 951 | 953 | 1 | 1.3350e-01 | 4.1003e+05 | ; | CD - NE2 |
| 953 | 954 | 1 | 1.0100e-01 | 3.6317e+05 | ; | NE2 - HE21 |
| 953 | 955 | 1 | 1.0100e-01 | 3.6317e+05 | ; | NE2 - HE22 |
| 956 | 957 | 1 | 1.2290e-01 | 4.7698e+05 | ; | C - O |
| 956 | 958 | 1 | 1.3350e-01 | 4.1003e+05 | ; | C - N |
| 958 | 959 | 1 | 1.0100e-01 | 3.6317e+05 | ; | N - H |
| 958 | 960 | 1 | 1.4490e-01 | 2.8200e+05 | ; | N - CA |
| 960 | 961 | 1 | 1.0900e-01 | 2.8451e+05 | ; | CA - HA |
| 960 | 962 | 1 | 1.5260e-01 | 2.5941e+05 | ; | CA - CB |
| 960 | 971 | 1 | 1.5220e-01 | 2.6527e+05 | ; | CA - C |
| 962 | 963 | 1 | 1.0900e-01 | 2.8451e+05 | ; | CB - HB2 |
| 962 | 964 | 1 | 1.0900e-01 | 2.8451e+05 | ; | CB - HB3 |
| 962 | 965 | 1 | 1.5260e-01 | 2.5941e+05 | ; | CB - CG |
| 965 | 966 | 1 | 1.0900e-01 | 2.8451e+05 | ; | CG - HG2 |
| 965 | 967 | 1 | 1.0900e-01 | 2.8451e+05 | ; | CG - HG3 |
| 965 | 968 | 1 | 1.5220e-01 | 2.6527e+05 | ; | CG - CD |
| 968 | 969 | 1 | 1.2500e-01 | 5.4894e+05 | ; | CD - OE1 |
| 968 | 970 | 1 | 1.2500e-01 | 5.4894e+05 | ; | CD - OE2 |
| 971 | 972 | 1 | 1.2290e-01 | 4.7698e+05 | ; | C - O |
| 971 | 973 | 1 | 1.3350e-01 | 4.1003e+05 | ; | C - N |
| 973 | 974 | 1 | 1.0100e-01 | 3.6317e+05 | ; | N - H |
| 973 | 975 | 1 | 1.4490e-01 | 2.8200e+05 | ; | N - CA |
| 975 | 976 | 1 | 1.0900e-01 | 2.8451e+05 | ; | CA - HA |
| 975 | 977 | 1 | 1.5260e-01 | 2.5941e+05 | ; | CA - CB |
| 975 | 990 | 1 | 1.5220e-01 | 2.6527e+05 | ; | CA - C |
| 977 | 978 | 1 | 1.0900e-01 | 2.8451e+05 | ; | CB - HB |

| | | | | | | |
|---|---|---|---|---|---|---|
| 977 | 979 | 1 | 1.5260e-01 | 2.5941e+05 | ; | CB - CG2 |
| 977 | 983 | 1 | 1.5260e-01 | 2.5941e+05 | ; | CB - CG1 |
| 979 | 980 | 1 | 1.0900e-01 | 2.8451e+05 | ; | CG2 - HG21 |
| 979 | 981 | 1 | 1.0900e-01 | 2.8451e+05 | ; | CG2 - HG22 |
| 979 | 982 | 1 | 1.0900e-01 | 2.8451e+05 | ; | CG2 - HG23 |
| 983 | 984 | 1 | 1.0900e-01 | 2.8451e+05 | ; | CG1 - HG12 |
| 983 | 985 | 1 | 1.0900e-01 | 2.8451e+05 | ; | CG1 - HG13 |
| 983 | 986 | 1 | 1.5260e-01 | 2.5941e+05 | ; | CG1 - CD1 |
| 986 | 987 | 1 | 1.0900e-01 | 2.8451e+05 | ; | CD1 - HD11 |
| 986 | 988 | 1 | 1.0900e-01 | 2.8451e+05 | ; | CD1 - HD12 |
| 986 | 989 | 1 | 1.0900e-01 | 2.8451e+05 | ; | CD1 - HD13 |
| 990 | 991 | 1 | 1.2290e-01 | 4.7698e+05 | ; | C - O |
| 990 | 992 | 1 | 1.3350e-01 | 4.1003e+05 | ; | C - N |
| 992 | 993 | 1 | 1.0100e-01 | 3.6317e+05 | ; | N - H |
| 992 | 994 | 1 | 1.4490e-01 | 2.8200e+05 | ; | N - CA |
| 994 | 995 | 1 | 1.0900e-01 | 2.8451e+05 | ; | CA - HA |
| 994 | 996 | 1 | 1.5260e-01 | 2.5941e+05 | ; | CA - CB |
| 994 | 1005 | 1 | 1.5220e-01 | 2.6527e+05 | ; | CA - C |
| 996 | 997 | 1 | 1.0900e-01 | 2.8451e+05 | ; | CB - HB2 |
| 996 | 998 | 1 | 1.0900e-01 | 2.8451e+05 | ; | CB - HB3 |
| 996 | 999 | 1 | 1.5260e-01 | 2.5941e+05 | ; | CB - CG |
| 999 | 1000 | 1 | 1.0900e-01 | 2.8451e+05 | ; | CG - HG2 |
| 999 | 1001 | 1 | 1.0900e-01 | 2.8451e+05 | ; | CG - HG3 |
| 999 | 1002 | 1 | 1.5220e-01 | 2.6527e+05 | ; | CG - CD |
| 1002 | 1003 | 1 | 1.2500e-01 | 5.4894e+05 | ; | CD - OE1 |
| 1002 | 1004 | 1 | 1.2500e-01 | 5.4894e+05 | ; | CD - OE2 |
| 1005 | 1006 | 1 | 1.2290e-01 | 4.7698e+05 | ; | C - O |
| 1005 | 1007 | 1 | 1.3350e-01 | 4.1003e+05 | ; | C - N |
| 1007 | 1008 | 1 | 1.0100e-01 | 3.6317e+05 | ; | N - H |
| 1007 | 1009 | 1 | 1.4490e-01 | 2.8200e+05 | ; | N - CA |
| 1009 | 1010 | 1 | 1.0900e-01 | 2.8451e+05 | ; | CA - HA |
| 1009 | 1011 | 1 | 1.5260e-01 | 2.5941e+05 | ; | CA - CB |
| 1009 | 1024 | 1 | 1.5220e-01 | 2.6527e+05 | ; | CA - C |
| 1011 | 1012 | 1 | 1.0900e-01 | 2.8451e+05 | ; | CB - HB |
| 1011 | 1013 | 1 | 1.5260e-01 | 2.5941e+05 | ; | CB - CG2 |
| 1011 | 1017 | 1 | 1.5260e-01 | 2.5941e+05 | ; | CB - CG1 |
| 1013 | 1014 | 1 | 1.0900e-01 | 2.8451e+05 | ; | CG2 - HG21 |
| 1013 | 1015 | 1 | 1.0900e-01 | 2.8451e+05 | ; | CG2 - HG22 |
| 1013 | 1016 | 1 | 1.0900e-01 | 2.8451e+05 | ; | CG2 - HG23 |
| 1017 | 1018 | 1 | 1.0900e-01 | 2.8451e+05 | ; | CG1 - HG12 |
| 1017 | 1019 | 1 | 1.0900e-01 | 2.8451e+05 | ; | CG1 - HG13 |
| 1017 | 1020 | 1 | 1.5260e-01 | 2.5941e+05 | ; | CG1 - CD1 |
| 1020 | 1021 | 1 | 1.0900e-01 | 2.8451e+05 | ; | CD1 - HD11 |
| 1020 | 1022 | 1 | 1.0900e-01 | 2.8451e+05 | ; | CD1 - HD12 |
| 1020 | 1023 | 1 | 1.0900e-01 | 2.8451e+05 | ; | CD1 - HD13 |
| 1024 | 1025 | 1 | 1.2290e-01 | 4.7698e+05 | ; | C - O |
| 1024 | 1026 | 1 | 1.3350e-01 | 4.1003e+05 | ; | C - N |
| 1026 | 1027 | 1 | 1.0100e-01 | 3.6317e+05 | ; | N - H |
| 1026 | 1028 | 1 | 1.4490e-01 | 2.8200e+05 | ; | N - CA |
| 1028 | 1029 | 1 | 1.0900e-01 | 2.8451e+05 | ; | CA - HA |
| 1028 | 1030 | 1 | 1.5260e-01 | 2.5941e+05 | ; | CA - CB |
| 1028 | 1041 | 1 | 1.5220e-01 | 2.6527e+05 | ; | CA - C |

| | | | | | | |
|---|---|---|---|---|---|---|
| 1030 | 1031 | 1 | 1.0900e-01 | 2.8451e+05 | ; | CB - HB2 |
| 1030 | 1032 | 1 | 1.0900e-01 | 2.8451e+05 | ; | CB - HB3 |
| 1030 | 1033 | 1 | 1.5260e-01 | 2.5941e+05 | ; | CB - CG |
| 1033 | 1034 | 1 | 1.0900e-01 | 2.8451e+05 | ; | CG - HG2 |
| 1033 | 1035 | 1 | 1.0900e-01 | 2.8451e+05 | ; | CG - HG3 |
| 1033 | 1036 | 1 | 1.8100e-01 | 1.8995e+05 | ; | CG - SD |
| 1036 | 1037 | 1 | 1.8100e-01 | 1.8995e+05 | ; | SD - CE |
| 1037 | 1038 | 1 | 1.0900e-01 | 2.8451e+05 | ; | CE - HE1 |
| 1037 | 1039 | 1 | 1.0900e-01 | 2.8451e+05 | ; | CE - HE2 |
| 1037 | 1040 | 1 | 1.0900e-01 | 2.8451e+05 | ; | CE - HE3 |
| 1041 | 1042 | 1 | 1.2290e-01 | 4.7698e+05 | ; | C - O |
| 1041 | 1043 | 1 | 1.3350e-01 | 4.1003e+05 | ; | C - N |
| 1043 | 1044 | 1 | 1.0100e-01 | 3.6317e+05 | ; | N - H |
| 1043 | 1045 | 1 | 1.4490e-01 | 2.8200e+05 | ; | N - CA |
| 1045 | 1046 | 1 | 1.0900e-01 | 2.8451e+05 | ; | CA - HA |
| 1045 | 1047 | 1 | 1.5260e-01 | 2.5941e+05 | ; | CA - CB |
| 1045 | 1063 | 1 | 1.5220e-01 | 2.6527e+05 | ; | CA - C |
| 1047 | 1048 | 1 | 1.0900e-01 | 2.8451e+05 | ; | CB - HB2 |
| 1047 | 1049 | 1 | 1.0900e-01 | 2.8451e+05 | ; | CB - HB3 |
| 1047 | 1050 | 1 | 1.5260e-01 | 2.5941e+05 | ; | CB - CG |
| 1050 | 1051 | 1 | 1.0900e-01 | 2.8451e+05 | ; | CG - HG2 |
| 1050 | 1052 | 1 | 1.0900e-01 | 2.8451e+05 | ; | CG - HG3 |
| 1050 | 1053 | 1 | 1.5260e-01 | 2.5941e+05 | ; | CG - CD |
| 1053 | 1054 | 1 | 1.0900e-01 | 2.8451e+05 | ; | CD - HD2 |
| 1053 | 1055 | 1 | 1.0900e-01 | 2.8451e+05 | ; | CD - HD3 |
| 1053 | 1056 | 1 | 1.5260e-01 | 2.5941e+05 | ; | CD - CE |
| 1056 | 1057 | 1 | 1.0900e-01 | 2.8451e+05 | ; | CE - HE2 |
| 1056 | 1058 | 1 | 1.0900e-01 | 2.8451e+05 | ; | CE - HE3 |
| 1056 | 1059 | 1 | 1.4710e-01 | 3.0711e+05 | ; | CE - NZ |
| 1059 | 1060 | 1 | 1.0100e-01 | 3.6317e+05 | ; | NZ - HZ1 |
| 1059 | 1061 | 1 | 1.0100e-01 | 3.6317e+05 | ; | NZ - HZ2 |
| 1059 | 1062 | 1 | 1.0100e-01 | 3.6317e+05 | ; | NZ - HZ3 |
| 1063 | 1064 | 1 | 1.2290e-01 | 4.7698e+05 | ; | C - O |
| 1063 | 1065 | 1 | 1.3350e-01 | 4.1003e+05 | ; | C - N |
| 1065 | 1066 | 1 | 1.0100e-01 | 3.6317e+05 | ; | N - H |
| 1065 | 1067 | 1 | 1.4490e-01 | 2.8200e+05 | ; | N - CA |
| 1067 | 1068 | 1 | 1.0900e-01 | 2.8451e+05 | ; | CA - HA |
| 1067 | 1069 | 1 | 1.5260e-01 | 2.5941e+05 | ; | CA - CB |
| 1067 | 1085 | 1 | 1.5220e-01 | 2.6527e+05 | ; | CA - C |
| 1069 | 1070 | 1 | 1.0900e-01 | 2.8451e+05 | ; | CB - HB2 |
| 1069 | 1071 | 1 | 1.0900e-01 | 2.8451e+05 | ; | CB - HB3 |
| 1069 | 1072 | 1 | 1.5260e-01 | 2.5941e+05 | ; | CB - CG |
| 1072 | 1073 | 1 | 1.0900e-01 | 2.8451e+05 | ; | CG - HG2 |
| 1072 | 1074 | 1 | 1.0900e-01 | 2.8451e+05 | ; | CG - HG3 |
| 1072 | 1075 | 1 | 1.5260e-01 | 2.5941e+05 | ; | CG - CD |
| 1075 | 1076 | 1 | 1.0900e-01 | 2.8451e+05 | ; | CD - HD2 |
| 1075 | 1077 | 1 | 1.0900e-01 | 2.8451e+05 | ; | CD - HD3 |
| 1075 | 1078 | 1 | 1.5260e-01 | 2.5941e+05 | ; | CD - CE |
| 1078 | 1079 | 1 | 1.0900e-01 | 2.8451e+05 | ; | CE - HE2 |
| 1078 | 1080 | 1 | 1.0900e-01 | 2.8451e+05 | ; | CE - HE3 |
| 1078 | 1081 | 1 | 1.4710e-01 | 3.0711e+05 | ; | CE - NZ |
| 1081 | 1082 | 1 | 1.0100e-01 | 3.6317e+05 | ; | NZ - HZ1 |

| | | | | | | |
|---|---|---|---|---|---|---|
| 1081 | 1083 | 1 | 1.0100e-01 | 3.6317e+05 | ; | NZ - HZ2 |
| 1081 | 1084 | 1 | 1.0100e-01 | 3.6317e+05 | ; | NZ - HZ3 |
| 1085 | 1086 | 1 | 1.2290e-01 | 4.7698e+05 | ; | C - O |
| 1085 | 1087 | 1 | 1.3350e-01 | 4.1003e+05 | ; | C - N |
| 1087 | 1088 | 1 | 1.0100e-01 | 3.6317e+05 | ; | N - H |
| 1087 | 1089 | 1 | 1.4490e-01 | 2.8200e+05 | ; | N - CA |
| 1089 | 1090 | 1 | 1.0900e-01 | 2.8451e+05 | ; | CA - HA |
| 1089 | 1091 | 1 | 1.5260e-01 | 2.5941e+05 | ; | CA - CB |
| 1089 | 1104 | 1 | 1.5220e-01 | 2.6527e+05 | ; | CA - C |
| 1091 | 1092 | 1 | 1.0900e-01 | 2.8451e+05 | ; | CB - HB2 |
| 1091 | 1093 | 1 | 1.0900e-01 | 2.8451e+05 | ; | CB - HB3 |
| 1091 | 1094 | 1 | 1.5260e-01 | 2.5941e+05 | ; | CB - CG |
| 1094 | 1095 | 1 | 1.0900e-01 | 2.8451e+05 | ; | CG - HG |
| 1094 | 1096 | 1 | 1.5260e-01 | 2.5941e+05 | ; | CG - CD1 |
| 1094 | 1100 | 1 | 1.5260e-01 | 2.5941e+05 | ; | CG - CD2 |
| 1096 | 1097 | 1 | 1.0900e-01 | 2.8451e+05 | ; | CD1 - HD11 |
| 1096 | 1098 | 1 | 1.0900e-01 | 2.8451e+05 | ; | CD1 - HD12 |
| 1096 | 1099 | 1 | 1.0900e-01 | 2.8451e+05 | ; | CD1 - HD13 |
| 1100 | 1101 | 1 | 1.0900e-01 | 2.8451e+05 | ; | CD2 - HD21 |
| 1100 | 1102 | 1 | 1.0900e-01 | 2.8451e+05 | ; | CD2 - HD22 |
| 1100 | 1103 | 1 | 1.0900e-01 | 2.8451e+05 | ; | CD2 - HD23 |
| 1104 | 1105 | 1 | 1.2290e-01 | 4.7698e+05 | ; | C - O |
| 1104 | 1106 | 1 | 1.3350e-01 | 4.1003e+05 | ; | C - N |
| 1106 | 1107 | 1 | 1.0100e-01 | 3.6317e+05 | ; | N - H |
| 1106 | 1108 | 1 | 1.4490e-01 | 2.8200e+05 | ; | N - CA |
| 1108 | 1109 | 1 | 1.0900e-01 | 2.8451e+05 | ; | CA - HA |
| 1108 | 1110 | 1 | 1.5260e-01 | 2.5941e+05 | ; | CA - CB |
| 1108 | 1116 | 1 | 1.5220e-01 | 2.6527e+05 | ; | CA - C |
| 1110 | 1111 | 1 | 1.0900e-01 | 2.8451e+05 | ; | CB - HB2 |
| 1110 | 1112 | 1 | 1.0900e-01 | 2.8451e+05 | ; | CB - HB3 |
| 1110 | 1113 | 1 | 1.5220e-01 | 2.6527e+05 | ; | CB - CG |
| 1113 | 1114 | 1 | 1.2500e-01 | 5.4894e+05 | ; | CG - OD1 |
| 1113 | 1115 | 1 | 1.2500e-01 | 5.4894e+05 | ; | CG - OD2 |
| 1116 | 1117 | 1 | 1.2290e-01 | 4.7698e+05 | ; | C - O |
| 1116 | 1118 | 1 | 1.3350e-01 | 4.1003e+05 | ; | C - N |
| 1118 | 1119 | 1 | 1.0100e-01 | 3.6317e+05 | ; | N - H |
| 1118 | 1120 | 1 | 1.4490e-01 | 2.8200e+05 | ; | N - CA |
| 1120 | 1121 | 1 | 1.0900e-01 | 2.8451e+05 | ; | CA - HA |
| 1120 | 1122 | 1 | 1.5260e-01 | 2.5941e+05 | ; | CA - CB |
| 1120 | 1133 | 1 | 1.5220e-01 | 2.6527e+05 | ; | CA - C |
| 1122 | 1123 | 1 | 1.0900e-01 | 2.8451e+05 | ; | CB - HB2 |
| 1122 | 1124 | 1 | 1.0900e-01 | 2.8451e+05 | ; | CB - HB3 |
| 1122 | 1125 | 1 | 1.5040e-01 | 2.6527e+05 | ; | CB - CG |
| 1125 | 1126 | 1 | 1.3940e-01 | 3.4309e+05 | ; | CG - ND1 |
| 1125 | 1131 | 1 | 1.3710e-01 | 4.3346e+05 | ; | CG - CD2 |
| 1126 | 1127 | 1 | 1.3350e-01 | 4.0836e+05 | ; | ND1 - CE1 |
| 1127 | 1128 | 1 | 1.0800e-01 | 3.0711e+05 | ; | CE1 - HE1 |
| 1127 | 1129 | 1 | 1.3430e-01 | 3.9915e+05 | ; | CE1 - NE2 |
| 1129 | 1130 | 1 | 1.0100e-01 | 3.6317e+05 | ; | NE2 - HE2 |
| 1129 | 1131 | 1 | 1.3810e-01 | 3.5731e+05 | ; | NE2 - CD2 |
| 1131 | 1132 | 1 | 1.0800e-01 | 3.0711e+05 | ; | CD2 - HD2 |
| 1133 | 1134 | 1 | 1.2290e-01 | 4.7698e+05 | ; | C - O |

| | | | | | | |
|---|---|---|---|---|---|---|
| 1133 | 1135 | 1 | 1.3350e-01 | 4.1003e+05 | ; | C - N |
| 1135 | 1136 | 1 | 1.4490e-01 | 2.8200e+05 | ; | N - CD |
| 1135 | 1145 | 1 | 1.4490e-01 | 2.8200e+05 | ; | N - CA |
| 1136 | 1137 | 1 | 1.0900e-01 | 2.8451e+05 | ; | CD - HD2 |
| 1136 | 1138 | 1 | 1.0900e-01 | 2.8451e+05 | ; | CD - HD3 |
| 1136 | 1139 | 1 | 1.5260e-01 | 2.5941e+05 | ; | CD - CG |
| 1139 | 1140 | 1 | 1.0900e-01 | 2.8451e+05 | ; | CG - HG2 |
| 1139 | 1141 | 1 | 1.0900e-01 | 2.8451e+05 | ; | CG - HG3 |
| 1139 | 1142 | 1 | 1.5260e-01 | 2.5941e+05 | ; | CG - CB |
| 1142 | 1143 | 1 | 1.0900e-01 | 2.8451e+05 | ; | CB - HB2 |
| 1142 | 1144 | 1 | 1.0900e-01 | 2.8451e+05 | ; | CB - HB3 |
| 1142 | 1145 | 1 | 1.5260e-01 | 2.5941e+05 | ; | CB - CA |
| 1145 | 1146 | 1 | 1.0900e-01 | 2.8451e+05 | ; | CA - HA |
| 1145 | 1147 | 1 | 1.5220e-01 | 2.6527e+05 | ; | CA - C |
| 1147 | 1148 | 1 | 1.2290e-01 | 4.7698e+05 | ; | C - O |
| 1147 | 1149 | 1 | 1.3350e-01 | 4.1003e+05 | ; | C - N |
| 1149 | 1150 | 1 | 1.0100e-01 | 3.6317e+05 | ; | N - H |
| 1149 | 1151 | 1 | 1.4490e-01 | 2.8200e+05 | ; | N - CA |
| 1151 | 1152 | 1 | 1.0900e-01 | 2.8451e+05 | ; | CA - HA |
| 1151 | 1153 | 1 | 1.5260e-01 | 2.5941e+05 | ; | CA - CB |
| 1151 | 1161 | 1 | 1.5220e-01 | 2.6527e+05 | ; | CA - C |
| 1153 | 1154 | 1 | 1.0900e-01 | 2.8451e+05 | ; | CB - HB2 |
| 1153 | 1155 | 1 | 1.0900e-01 | 2.8451e+05 | ; | CB - HB3 |
| 1153 | 1156 | 1 | 1.5220e-01 | 2.6527e+05 | ; | CB - CG |
| 1156 | 1157 | 1 | 1.2290e-01 | 4.7698e+05 | ; | CG - OD1 |
| 1156 | 1158 | 1 | 1.3350e-01 | 4.1003e+05 | ; | CG - ND2 |
| 1158 | 1159 | 1 | 1.0100e-01 | 3.6317e+05 | ; | ND2 - HD21 |
| 1158 | 1160 | 1 | 1.0100e-01 | 3.6317e+05 | ; | ND2 - HD22 |
| 1161 | 1162 | 1 | 1.2290e-01 | 4.7698e+05 | ; | C - O |
| 1161 | 1163 | 1 | 1.3350e-01 | 4.1003e+05 | ; | C - N |
| 1163 | 1164 | 1 | 1.0100e-01 | 3.6317e+05 | ; | N - H |
| 1163 | 1165 | 1 | 1.4490e-01 | 2.8200e+05 | ; | N - CA |
| 1165 | 1166 | 1 | 1.0900e-01 | 2.8451e+05 | ; | CA - HA |
| 1165 | 1167 | 1 | 1.5260e-01 | 2.5941e+05 | ; | CA - CB |
| 1165 | 1180 | 1 | 1.5220e-01 | 2.6527e+05 | ; | CA - C |
| 1167 | 1168 | 1 | 1.0900e-01 | 2.8451e+05 | ; | CB - HB |
| 1167 | 1169 | 1 | 1.5260e-01 | 2.5941e+05 | ; | CB - CG2 |
| 1167 | 1173 | 1 | 1.5260e-01 | 2.5941e+05 | ; | CB - CG1 |
| 1169 | 1170 | 1 | 1.0900e-01 | 2.8451e+05 | ; | CG2 - HG21 |
| 1169 | 1171 | 1 | 1.0900e-01 | 2.8451e+05 | ; | CG2 - HG22 |
| 1169 | 1172 | 1 | 1.0900e-01 | 2.8451e+05 | ; | CG2 - HG23 |
| 1173 | 1174 | 1 | 1.0900e-01 | 2.8451e+05 | ; | CG1 - HG12 |
| 1173 | 1175 | 1 | 1.0900e-01 | 2.8451e+05 | ; | CG1 - HG13 |
| 1173 | 1176 | 1 | 1.5260e-01 | 2.5941e+05 | ; | CG1 - CD1 |
| 1176 | 1177 | 1 | 1.0900e-01 | 2.8451e+05 | ; | CD1 - HD11 |
| 1176 | 1178 | 1 | 1.0900e-01 | 2.8451e+05 | ; | CD1 - HD12 |
| 1176 | 1179 | 1 | 1.0900e-01 | 2.8451e+05 | ; | CD1 - HD13 |
| 1180 | 1181 | 1 | 1.2290e-01 | 4.7698e+05 | ; | C - O |
| 1180 | 1182 | 1 | 1.3350e-01 | 4.1003e+05 | ; | C - N |
| 1182 | 1183 | 1 | 1.0100e-01 | 3.6317e+05 | ; | N - H |
| 1182 | 1184 | 1 | 1.4490e-01 | 2.8200e+05 | ; | N - CA |
| 1184 | 1185 | 1 | 1.0900e-01 | 2.8451e+05 | ; | CA - HA |

| | | | | | | |
|---|---|---|---|---|---|---|
| 1184 | 1186 | 1 | 1.5260e-01 | 2.5941e+05 | ; | CA - CB |
| 1184 | 1196 | 1 | 1.5220e-01 | 2.6527e+05 | ; | CA - C |
| 1186 | 1187 | 1 | 1.0900e-01 | 2.8451e+05 | ; | CB - HB |
| 1186 | 1188 | 1 | 1.5260e-01 | 2.5941e+05 | ; | CB - CG1 |
| 1186 | 1192 | 1 | 1.5260e-01 | 2.5941e+05 | ; | CB - CG2 |
| 1188 | 1189 | 1 | 1.0900e-01 | 2.8451e+05 | ; | CG1 - HG11 |
| 1188 | 1190 | 1 | 1.0900e-01 | 2.8451e+05 | ; | CG1 - HG12 |
| 1188 | 1191 | 1 | 1.0900e-01 | 2.8451e+05 | ; | CG1 - HG13 |
| 1192 | 1193 | 1 | 1.0900e-01 | 2.8451e+05 | ; | CG2 - HG21 |
| 1192 | 1194 | 1 | 1.0900e-01 | 2.8451e+05 | ; | CG2 - HG22 |
| 1192 | 1195 | 1 | 1.0900e-01 | 2.8451e+05 | ; | CG2 - HG23 |
| 1196 | 1197 | 1 | 1.2290e-01 | 4.7698e+05 | ; | C - O |
| 1196 | 1198 | 1 | 1.3350e-01 | 4.1003e+05 | ; | C - N |
| 1198 | 1199 | 1 | 1.0100e-01 | 3.6317e+05 | ; | N - H |
| 1198 | 1200 | 1 | 1.4490e-01 | 2.8200e+05 | ; | N - CA |
| 1200 | 1201 | 1 | 1.0900e-01 | 2.8451e+05 | ; | CA - HA |
| 1200 | 1202 | 1 | 1.5260e-01 | 2.5941e+05 | ; | CA - CB |
| 1200 | 1218 | 1 | 1.5220e-01 | 2.6527e+05 | ; | CA - C |
| 1202 | 1203 | 1 | 1.0900e-01 | 2.8451e+05 | ; | CB - HB2 |
| 1202 | 1204 | 1 | 1.0900e-01 | 2.8451e+05 | ; | CB - HB3 |
| 1202 | 1205 | 1 | 1.5260e-01 | 2.5941e+05 | ; | CB - CG |
| 1205 | 1206 | 1 | 1.0900e-01 | 2.8451e+05 | ; | CG - HG2 |
| 1205 | 1207 | 1 | 1.0900e-01 | 2.8451e+05 | ; | CG - HG3 |
| 1205 | 1208 | 1 | 1.5260e-01 | 2.5941e+05 | ; | CG - CD |
| 1208 | 1209 | 1 | 1.0900e-01 | 2.8451e+05 | ; | CD - HD2 |
| 1208 | 1210 | 1 | 1.0900e-01 | 2.8451e+05 | ; | CD - HD3 |
| 1208 | 1211 | 1 | 1.5260e-01 | 2.5941e+05 | ; | CD - CE |
| 1211 | 1212 | 1 | 1.0900e-01 | 2.8451e+05 | ; | CE - HE2 |
| 1211 | 1213 | 1 | 1.0900e-01 | 2.8451e+05 | ; | CE - HE3 |
| 1211 | 1214 | 1 | 1.4710e-01 | 3.0711e+05 | ; | CE - NZ |
| 1214 | 1215 | 1 | 1.0100e-01 | 3.6317e+05 | ; | NZ - HZ1 |
| 1214 | 1216 | 1 | 1.0100e-01 | 3.6317e+05 | ; | NZ - HZ2 |
| 1214 | 1217 | 1 | 1.0100e-01 | 3.6317e+05 | ; | NZ - HZ3 |
| 1218 | 1219 | 1 | 1.2290e-01 | 4.7698e+05 | ; | C - O |
| 1218 | 1220 | 1 | 1.3350e-01 | 4.1003e+05 | ; | C - N |
| 1220 | 1221 | 1 | 1.0100e-01 | 3.6317e+05 | ; | N - H |
| 1220 | 1222 | 1 | 1.4490e-01 | 2.8200e+05 | ; | N - CA |
| 1222 | 1223 | 1 | 1.0900e-01 | 2.8451e+05 | ; | CA - HA |
| 1222 | 1224 | 1 | 1.5260e-01 | 2.5941e+05 | ; | CA - CB |
| 1222 | 1237 | 1 | 1.5220e-01 | 2.6527e+05 | ; | CA - C |
| 1224 | 1225 | 1 | 1.0900e-01 | 2.8451e+05 | ; | CB - HB2 |
| 1224 | 1226 | 1 | 1.0900e-01 | 2.8451e+05 | ; | CB - HB3 |
| 1224 | 1227 | 1 | 1.5260e-01 | 2.5941e+05 | ; | CB - CG |
| 1227 | 1228 | 1 | 1.0900e-01 | 2.8451e+05 | ; | CG - HG |
| 1227 | 1229 | 1 | 1.5260e-01 | 2.5941e+05 | ; | CG - CD1 |
| 1227 | 1233 | 1 | 1.5260e-01 | 2.5941e+05 | ; | CG - CD2 |
| 1229 | 1230 | 1 | 1.0900e-01 | 2.8451e+05 | ; | CD1 - HD11 |
| 1229 | 1231 | 1 | 1.0900e-01 | 2.8451e+05 | ; | CD1 - HD12 |
| 1229 | 1232 | 1 | 1.0900e-01 | 2.8451e+05 | ; | CD1 - HD13 |
| 1233 | 1234 | 1 | 1.0900e-01 | 2.8451e+05 | ; | CD2 - HD21 |
| 1233 | 1235 | 1 | 1.0900e-01 | 2.8451e+05 | ; | CD2 - HD22 |
| 1233 | 1236 | 1 | 1.0900e-01 | 2.8451e+05 | ; | CD2 - HD23 |

| | | | | | | |
|---|---|---|---|---|---|---|
| 1237 | 1238 | 1 | 1.2290e-01 | 4.7698e+05 | ; | C - O |
| 1237 | 1239 | 1 | 1.3350e-01 | 4.1003e+05 | ; | C - N |
| 1239 | 1240 | 1 | 1.0100e-01 | 3.6317e+05 | ; | N - H |
| 1239 | 1241 | 1 | 1.4490e-01 | 2.8200e+05 | ; | N - CA |
| 1241 | 1242 | 1 | 1.0900e-01 | 2.8451e+05 | ; | CA - HA |
| 1241 | 1243 | 1 | 1.5260e-01 | 2.5941e+05 | ; | CA - CB |
| 1241 | 1258 | 1 | 1.5220e-01 | 2.6527e+05 | ; | CA - C |
| 1243 | 1244 | 1 | 1.0900e-01 | 2.8451e+05 | ; | CB - HB2 |
| 1243 | 1245 | 1 | 1.0900e-01 | 2.8451e+05 | ; | CB - HB3 |
| 1243 | 1246 | 1 | 1.5100e-01 | 2.6527e+05 | ; | CB - CG |
| 1246 | 1247 | 1 | 1.4000e-01 | 3.9246e+05 | ; | CG - CD1 |
| 1246 | 1256 | 1 | 1.4000e-01 | 3.9246e+05 | ; | CG - CD2 |
| 1247 | 1248 | 1 | 1.0800e-01 | 3.0711e+05 | ; | CD1 - HD1 |
| 1247 | 1249 | 1 | 1.4000e-01 | 3.9246e+05 | ; | CD1 - CE1 |
| 1249 | 1250 | 1 | 1.0800e-01 | 3.0711e+05 | ; | CE1 - HE1 |
| 1249 | 1251 | 1 | 1.4090e-01 | 3.9246e+05 | ; | CE1 - CZ |
| 1251 | 1252 | 1 | 1.3640e-01 | 3.7656e+05 | ; | CZ - OH |
| 1251 | 1254 | 1 | 1.4090e-01 | 3.9246e+05 | ; | CZ - CE2 |
| 1252 | 1253 | 1 | 9.6000e-02 | 4.6275e+05 | ; | OH - HH |
| 1254 | 1255 | 1 | 1.0800e-01 | 3.0711e+05 | ; | CE2 - HE2 |
| 1254 | 1256 | 1 | 1.4000e-01 | 3.9246e+05 | ; | CE2 - CD2 |
| 1256 | 1257 | 1 | 1.0800e-01 | 3.0711e+05 | ; | CD2 - HD2 |
| 1258 | 1259 | 1 | 1.2290e-01 | 4.7698e+05 | ; | C - O |
| 1258 | 1260 | 1 | 1.3350e-01 | 4.1003e+05 | ; | C - N |
| 1260 | 1261 | 1 | 1.0100e-01 | 3.6317e+05 | ; | N - H |
| 1260 | 1262 | 1 | 1.4490e-01 | 2.8200e+05 | ; | N - CA |
| 1262 | 1263 | 1 | 1.0900e-01 | 2.8451e+05 | ; | CA - HA |
| 1262 | 1264 | 1 | 1.5260e-01 | 2.5941e+05 | ; | CA - CB |
| 1262 | 1273 | 1 | 1.5220e-01 | 2.6527e+05 | ; | CA - C |
| 1264 | 1265 | 1 | 1.0900e-01 | 2.8451e+05 | ; | CB - HB2 |
| 1264 | 1266 | 1 | 1.0900e-01 | 2.8451e+05 | ; | CB - HB3 |
| 1264 | 1267 | 1 | 1.5260e-01 | 2.5941e+05 | ; | CB - CG |
| 1267 | 1268 | 1 | 1.0900e-01 | 2.8451e+05 | ; | CG - HG2 |
| 1267 | 1269 | 1 | 1.0900e-01 | 2.8451e+05 | ; | CG - HG3 |
| 1267 | 1270 | 1 | 1.5220e-01 | 2.6527e+05 | ; | CG - CD |
| 1270 | 1271 | 1 | 1.2500e-01 | 5.4894e+05 | ; | CD - OE1 |
| 1270 | 1272 | 1 | 1.2500e-01 | 5.4894e+05 | ; | CD - OE2 |
| 1273 | 1274 | 1 | 1.2290e-01 | 4.7698e+05 | ; | C - O |
| 1273 | 1275 | 1 | 1.3350e-01 | 4.1003e+05 | ; | C - N |
| 1275 | 1276 | 1 | 1.0100e-01 | 3.6317e+05 | ; | N - H |
| 1275 | 1277 | 1 | 1.4490e-01 | 2.8200e+05 | ; | N - CA |
| 1277 | 1278 | 1 | 1.0900e-01 | 2.8451e+05 | ; | CA - HA |
| 1277 | 1279 | 1 | 1.5260e-01 | 2.5941e+05 | ; | CA - CB |
| 1277 | 1287 | 1 | 1.5220e-01 | 2.6527e+05 | ; | CA - C |
| 1279 | 1280 | 1 | 1.0900e-01 | 2.8451e+05 | ; | CB - HB |
| 1279 | 1281 | 1 | 1.5260e-01 | 2.5941e+05 | ; | CB - CG2 |
| 1279 | 1285 | 1 | 1.4100e-01 | 2.6778e+05 | ; | CB - OG1 |
| 1281 | 1282 | 1 | 1.0900e-01 | 2.8451e+05 | ; | CG2 - HG21 |
| 1281 | 1283 | 1 | 1.0900e-01 | 2.8451e+05 | ; | CG2 - HG22 |
| 1281 | 1284 | 1 | 1.0900e-01 | 2.8451e+05 | ; | CG2 - HG23 |
| 1285 | 1286 | 1 | 9.6000e-02 | 4.6275e+05 | ; | OG1 - HG1 |
| 1287 | 1288 | 1 | 1.2290e-01 | 4.7698e+05 | ; | C - O |

| | | | | | |
|---|---|---|---|---|---|
| 1287 | 1289 | 1 | 1.3350e-01 | 4.1003e+05 ; | C - N |
| 1289 | 1290 | 1 | 1.0100e-01 | 3.6317e+05 ; | N - H |
| 1289 | 1291 | 1 | 1.4490e-01 | 2.8200e+05 ; | N - CA |
| 1291 | 1292 | 1 | 1.0900e-01 | 2.8451e+05 ; | CA - HA |
| 1291 | 1293 | 1 | 1.5260e-01 | 2.5941e+05 ; | CA - CB |
| 1291 | 1308 | 1 | 1.5220e-01 | 2.6527e+05 ; | CA - C |
| 1293 | 1294 | 1 | 1.0900e-01 | 2.8451e+05 ; | CB - HB2 |
| 1293 | 1295 | 1 | 1.0900e-01 | 2.8451e+05 ; | CB - HB3 |
| 1293 | 1296 | 1 | 1.5100e-01 | 2.6527e+05 ; | CB - CG |
| 1296 | 1297 | 1 | 1.4000e-01 | 3.9246e+05 ; | CG - CD1 |
| 1296 | 1306 | 1 | 1.4000e-01 | 3.9246e+05 ; | CG - CD2 |
| 1297 | 1298 | 1 | 1.0800e-01 | 3.0711e+05 ; | CD1 - HD1 |
| 1297 | 1299 | 1 | 1.4000e-01 | 3.9246e+05 ; | CD1 - CE1 |
| 1299 | 1300 | 1 | 1.0800e-01 | 3.0711e+05 ; | CE1 - HE1 |
| 1299 | 1301 | 1 | 1.4090e-01 | 3.9246e+05 ; | CE1 - CZ |
| 1301 | 1302 | 1 | 1.3640e-01 | 3.7656e+05 ; | CZ - OH |
| 1301 | 1304 | 1 | 1.4090e-01 | 3.9246e+05 ; | CZ - CE2 |
| 1302 | 1303 | 1 | 9.6000e-02 | 4.6275e+05 ; | OH - HH |
| 1304 | 1305 | 1 | 1.0800e-01 | 3.0711e+05 ; | CE2 - HE2 |
| 1304 | 1306 | 1 | 1.4000e-01 | 3.9246e+05 ; | CE2 - CD2 |
| 1306 | 1307 | 1 | 1.0800e-01 | 3.0711e+05 ; | CD2 - HD2 |
| 1308 | 1309 | 1 | 1.2290e-01 | 4.7698e+05 ; | C - O |
| 1308 | 1310 | 1 | 1.3350e-01 | 4.1003e+05 ; | C - N |
| 1310 | 1311 | 1 | 1.0100e-01 | 3.6317e+05 ; | N - H |
| 1310 | 1312 | 1 | 1.4490e-01 | 2.8200e+05 ; | N - CA |
| 1312 | 1313 | 1 | 1.0900e-01 | 2.8451e+05 ; | CA - HA |
| 1312 | 1314 | 1 | 1.5260e-01 | 2.5941e+05 ; | CA - CB |
| 1312 | 1323 | 1 | 1.5220e-01 | 2.6527e+05 ; | CA - C |
| 1314 | 1315 | 1 | 1.0900e-01 | 2.8451e+05 ; | CB - HB2 |
| 1314 | 1316 | 1 | 1.0900e-01 | 2.8451e+05 ; | CB - HB3 |
| 1314 | 1317 | 1 | 1.5260e-01 | 2.5941e+05 ; | CB - CG |
| 1317 | 1318 | 1 | 1.0900e-01 | 2.8451e+05 ; | CG - HG2 |
| 1317 | 1319 | 1 | 1.0900e-01 | 2.8451e+05 ; | CG - HG3 |
| 1317 | 1320 | 1 | 1.5220e-01 | 2.6527e+05 ; | CG - CD |
| 1320 | 1321 | 1 | 1.2500e-01 | 5.4894e+05 ; | CD - OE1 |
| 1320 | 1322 | 1 | 1.2500e-01 | 5.4894e+05 ; | CD - OE2 |
| 1323 | 1324 | 1 | 1.2290e-01 | 4.7698e+05 ; | C - O |
| 1323 | 1325 | 1 | 1.3350e-01 | 4.1003e+05 ; | C - N |
| 1325 | 1326 | 1 | 1.0100e-01 | 3.6317e+05 ; | N - H |
| 1325 | 1327 | 1 | 1.4490e-01 | 2.8200e+05 ; | N - CA |
| 1327 | 1328 | 1 | 1.0900e-01 | 2.8451e+05 ; | CA - HA |
| 1327 | 1329 | 1 | 1.5260e-01 | 2.5941e+05 ; | CA - CB |
| 1327 | 1337 | 1 | 1.5220e-01 | 2.6527e+05 ; | CA - C |
| 1329 | 1330 | 1 | 1.0900e-01 | 2.8451e+05 ; | CB - HB2 |
| 1329 | 1331 | 1 | 1.0900e-01 | 2.8451e+05 ; | CB - HB3 |
| 1329 | 1332 | 1 | 1.5220e-01 | 2.6527e+05 ; | CB - CG |
| 1332 | 1333 | 1 | 1.2290e-01 | 4.7698e+05 ; | CG - OD1 |
| 1332 | 1334 | 1 | 1.3350e-01 | 4.1003e+05 ; | CG - ND2 |
| 1334 | 1335 | 1 | 1.0100e-01 | 3.6317e+05 ; | ND2 - HD21 |
| 1334 | 1336 | 1 | 1.0100e-01 | 3.6317e+05 ; | ND2 - HD22 |
| 1337 | 1338 | 1 | 1.2290e-01 | 4.7698e+05 ; | C - O |
| 1337 | 1339 | 1 | 1.3350e-01 | 4.1003e+05 ; | C - N |

| | | | | | | | | |
|---|---|---|---|---|---|---|---|---|
| 1339 | 1340 | 1 | 1.0100e-01 | 3.6317e+05 | ; | N | - | H |
| 1339 | 1341 | 1 | 1.4490e-01 | 2.8200e+05 | ; | N | - | CA |
| 1341 | 1342 | 1 | 1.0900e-01 | 2.8451e+05 | ; | CA | - | HA |
| 1341 | 1343 | 1 | 1.5260e-01 | 2.5941e+05 | ; | CA | - | CB |
| 1341 | 1349 | 1 | 1.5220e-01 | 2.6527e+05 | ; | CA | - | C |
| 1343 | 1344 | 1 | 1.0900e-01 | 2.8451e+05 | ; | CB | - | HB2 |
| 1343 | 1345 | 1 | 1.0900e-01 | 2.8451e+05 | ; | CB | - | HB3 |
| 1343 | 1346 | 1 | 1.5220e-01 | 2.6527e+05 | ; | CB | - | CG |
| 1346 | 1347 | 1 | 1.2500e-01 | 5.4894e+05 | ; | CG | - | OD1 |
| 1346 | 1348 | 1 | 1.2500e-01 | 5.4894e+05 | ; | CG | - | OD2 |
| 1349 | 1350 | 1 | 1.2290e-01 | 4.7698e+05 | ; | C | - | O |
| 1349 | 1351 | 1 | 1.3350e-01 | 4.1003e+05 | ; | C | - | N |
| 1351 | 1352 | 1 | 1.0100e-01 | 3.6317e+05 | ; | N | - | H |
| 1351 | 1353 | 1 | 1.4490e-01 | 2.8200e+05 | ; | N | - | CA |
| 1353 | 1354 | 1 | 1.0900e-01 | 2.8451e+05 | ; | CA | - | HA |
| 1353 | 1355 | 1 | 1.5260e-01 | 2.5941e+05 | ; | CA | - | CB |
| 1353 | 1363 | 1 | 1.5220e-01 | 2.6527e+05 | ; | CA | - | C |
| 1355 | 1356 | 1 | 1.0900e-01 | 2.8451e+05 | ; | CB | - | HB2 |
| 1355 | 1357 | 1 | 1.0900e-01 | 2.8451e+05 | ; | CB | - | HB3 |
| 1355 | 1358 | 1 | 1.5220e-01 | 2.6527e+05 | ; | CB | - | CG |
| 1358 | 1359 | 1 | 1.2290e-01 | 4.7698e+05 | ; | CG | - | OD1 |
| 1358 | 1360 | 1 | 1.3350e-01 | 4.1003e+05 | ; | CG | - | ND2 |
| 1360 | 1361 | 1 | 1.0100e-01 | 3.6317e+05 | ; | ND2 | - | HD21 |
| 1360 | 1362 | 1 | 1.0100e-01 | 3.6317e+05 | ; | ND2 | - | HD22 |
| 1363 | 1364 | 1 | 1.2290e-01 | 4.7698e+05 | ; | C | - | O |
| 1363 | 1365 | 1 | 1.3350e-01 | 4.1003e+05 | ; | C | - | N |
| 1365 | 1366 | 1 | 1.0100e-01 | 3.6317e+05 | ; | N | - | H |
| 1365 | 1367 | 1 | 1.4490e-01 | 2.8200e+05 | ; | N | - | CA |
| 1367 | 1368 | 1 | 1.0900e-01 | 2.8451e+05 | ; | CA | - | HA |
| 1367 | 1369 | 1 | 1.5260e-01 | 2.5941e+05 | ; | CA | - | CB |
| 1367 | 1384 | 1 | 1.5220e-01 | 2.6527e+05 | ; | CA | - | C |
| 1369 | 1370 | 1 | 1.0900e-01 | 2.8451e+05 | ; | CB | - | HB2 |
| 1369 | 1371 | 1 | 1.0900e-01 | 2.8451e+05 | ; | CB | - | HB3 |
| 1369 | 1372 | 1 | 1.5100e-01 | 2.6527e+05 | ; | CB | - | CG |
| 1372 | 1373 | 1 | 1.4000e-01 | 3.9246e+05 | ; | CG | - | CD1 |
| 1372 | 1382 | 1 | 1.4000e-01 | 3.9246e+05 | ; | CG | - | CD2 |
| 1373 | 1374 | 1 | 1.0800e-01 | 3.0711e+05 | ; | CD1 | - | HD1 |
| 1373 | 1375 | 1 | 1.4000e-01 | 3.9246e+05 | ; | CD1 | - | CE1 |
| 1375 | 1376 | 1 | 1.0800e-01 | 3.0711e+05 | ; | CE1 | - | HE1 |
| 1375 | 1377 | 1 | 1.4090e-01 | 3.9246e+05 | ; | CE1 | - | CZ |
| 1377 | 1378 | 1 | 1.3640e-01 | 3.7656e+05 | ; | CZ | - | OH |
| 1377 | 1380 | 1 | 1.4090e-01 | 3.9246e+05 | ; | CZ | - | CE2 |
| 1378 | 1379 | 1 | 9.6000e-02 | 4.6275e+05 | ; | OH | - | HH |
| 1380 | 1381 | 1 | 1.0800e-01 | 3.0711e+05 | ; | CE2 | - | HE2 |
| 1380 | 1382 | 1 | 1.4000e-01 | 3.9246e+05 | ; | CE2 | - | CD2 |
| 1382 | 1383 | 1 | 1.0800e-01 | 3.0711e+05 | ; | CD2 | - | HD2 |
| 1384 | 1385 | 1 | 1.2290e-01 | 4.7698e+05 | ; | C | - | O |
| 1384 | 1386 | 1 | 1.3350e-01 | 4.1003e+05 | ; | C | - | N |
| 1386 | 1387 | 1 | 1.0100e-01 | 3.6317e+05 | ; | N | - | H |
| 1386 | 1388 | 1 | 1.4490e-01 | 2.8200e+05 | ; | N | - | CA |
| 1388 | 1389 | 1 | 1.0900e-01 | 2.8451e+05 | ; | CA | - | HA |
| 1388 | 1390 | 1 | 1.5260e-01 | 2.5941e+05 | ; | CA | - | CB |

```
1388   1403   1    1.5220e-01    2.6527e+05 ;      CA - C
1390   1391   1    1.0900e-01    2.8451e+05 ;      CB - HB
1390   1392   1    1.5260e-01    2.5941e+05 ;      CB - CG2
1390   1396   1    1.5260e-01    2.5941e+05 ;      CB - CG1
1392   1393   1    1.0900e-01    2.8451e+05 ;     CG2 - HG21
1392   1394   1    1.0900e-01    2.8451e+05 ;     CG2 - HG22
1392   1395   1    1.0900e-01    2.8451e+05 ;     CG2 - HG23
1396   1397   1    1.0900e-01    2.8451e+05 ;     CG1 - HG12
1396   1398   1    1.0900e-01    2.8451e+05 ;     CG1 - HG13
1396   1399   1    1.5260e-01    2.5941e+05 ;     CG1 - CD1
1399   1400   1    1.0900e-01    2.8451e+05 ;     CD1 - HD11
1399   1401   1    1.0900e-01    2.8451e+05 ;     CD1 - HD12
1399   1402   1    1.0900e-01    2.8451e+05 ;     CD1 - HD13
1403   1404   1    1.2290e-01    4.7698e+05 ;       C - O
1403   1405   1    1.3350e-01    4.1003e+05 ;       C - N
1405   1406   1    1.0100e-01    3.6317e+05 ;       N - H
1405   1407   1    1.4490e-01    2.8200e+05 ;       N - CA
1407   1408   1    1.0900e-01    2.8451e+05 ;      CA - HA
1407   1409   1    1.5260e-01    2.5941e+05 ;      CA - CB
1407   1424   1    1.5220e-01    2.6527e+05 ;      CA - C
1409   1410   1    1.0900e-01    2.8451e+05 ;      CB - HB2
1409   1411   1    1.0900e-01    2.8451e+05 ;      CB - HB3
1409   1412   1    1.5100e-01    2.6527e+05 ;      CB - CG
1412   1413   1    1.4000e-01    3.9246e+05 ;      CG - CD1
1412   1422   1    1.4000e-01    3.9246e+05 ;      CG - CD2
1413   1414   1    1.0800e-01    3.0711e+05 ;     CD1 - HD1
1413   1415   1    1.4000e-01    3.9246e+05 ;     CD1 - CE1
1415   1416   1    1.0800e-01    3.0711e+05 ;     CE1 - HE1
1415   1417   1    1.4090e-01    3.9246e+05 ;     CE1 - CZ
1417   1418   1    1.3640e-01    3.7656e+05 ;      CZ - OH
1417   1420   1    1.4090e-01    3.9246e+05 ;      CZ - CE2
1418   1419   1    9.6000e-02    4.6275e+05 ;      OH - HH
1420   1421   1    1.0800e-01    3.0711e+05 ;     CE2 - HE2
1420   1422   1    1.4000e-01    3.9246e+05 ;     CE2 - CD2
1422   1423   1    1.0800e-01    3.0711e+05 ;     CD2 - HD2
1424   1425   1    1.2290e-01    4.7698e+05 ;       C - O
1424   1426   1    1.3350e-01    4.1003e+05 ;       C - N
1426   1427   1    1.0100e-01    3.6317e+05 ;       N - H
1426   1428   1    1.4490e-01    2.8200e+05 ;       N - CA
1428   1429   1    1.0900e-01    2.8451e+05 ;      CA - HA
1428   1430   1    1.5260e-01    2.5941e+05 ;      CA - CB
1428   1443   1    1.5220e-01    2.6527e+05 ;      CA - C
1430   1431   1    1.0900e-01    2.8451e+05 ;      CB - HB2
1430   1432   1    1.0900e-01    2.8451e+05 ;      CB - HB3
1430   1433   1    1.5260e-01    2.5941e+05 ;      CB - CG
1433   1434   1    1.0900e-01    2.8451e+05 ;      CG - HG
1433   1435   1    1.5260e-01    2.5941e+05 ;      CG - CD1
1433   1439   1    1.5260e-01    2.5941e+05 ;      CG - CD2
1435   1436   1    1.0900e-01    2.8451e+05 ;     CD1 - HD11
1435   1437   1    1.0900e-01    2.8451e+05 ;     CD1 - HD12
1435   1438   1    1.0900e-01    2.8451e+05 ;     CD1 - HD13
1439   1440   1    1.0900e-01    2.8451e+05 ;     CD2 - HD21
```

| | | | | | | |
|---|---|---|---|---|---|---|
| 1439 | 1441 | 1 | 1.0900e-01 | 2.8451e+05 | ; | CD2 - HD22 |
| 1439 | 1442 | 1 | 1.0900e-01 | 2.8451e+05 | ; | CD2 - HD23 |
| 1443 | 1444 | 1 | 1.2290e-01 | 4.7698e+05 | ; | C - O |
| 1443 | 1445 | 1 | 1.3350e-01 | 4.1003e+05 | ; | C - N |
| 1445 | 1446 | 1 | 1.0100e-01 | 3.6317e+05 | ; | N - H |
| 1445 | 1447 | 1 | 1.4490e-01 | 2.8200e+05 | ; | N - CA |
| 1447 | 1448 | 1 | 1.0900e-01 | 2.8451e+05 | ; | CA - HA |
| 1447 | 1449 | 1 | 1.5260e-01 | 2.5941e+05 | ; | CA - CB |
| 1447 | 1462 | 1 | 1.5220e-01 | 2.6527e+05 | ; | CA - C |
| 1449 | 1450 | 1 | 1.0900e-01 | 2.8451e+05 | ; | CB - HB |
| 1449 | 1451 | 1 | 1.5260e-01 | 2.5941e+05 | ; | CB - CG2 |
| 1449 | 1455 | 1 | 1.5260e-01 | 2.5941e+05 | ; | CB - CG1 |
| 1451 | 1452 | 1 | 1.0900e-01 | 2.8451e+05 | ; | CG2 - HG21 |
| 1451 | 1453 | 1 | 1.0900e-01 | 2.8451e+05 | ; | CG2 - HG22 |
| 1451 | 1454 | 1 | 1.0900e-01 | 2.8451e+05 | ; | CG2 - HG23 |
| 1455 | 1456 | 1 | 1.0900e-01 | 2.8451e+05 | ; | CG1 - HG12 |
| 1455 | 1457 | 1 | 1.0900e-01 | 2.8451e+05 | ; | CG1 - HG13 |
| 1455 | 1458 | 1 | 1.5260e-01 | 2.5941e+05 | ; | CG1 - CD1 |
| 1458 | 1459 | 1 | 1.0900e-01 | 2.8451e+05 | ; | CD1 - HD11 |
| 1458 | 1460 | 1 | 1.0900e-01 | 2.8451e+05 | ; | CD1 - HD12 |
| 1458 | 1461 | 1 | 1.0900e-01 | 2.8451e+05 | ; | CD1 - HD13 |
| 1462 | 1463 | 1 | 1.2290e-01 | 4.7698e+05 | ; | C - O |
| 1462 | 1464 | 1 | 1.3350e-01 | 4.1003e+05 | ; | C - N |
| 1464 | 1465 | 1 | 1.0100e-01 | 3.6317e+05 | ; | N - H |
| 1464 | 1466 | 1 | 1.4490e-01 | 2.8200e+05 | ; | N - CA |
| 1466 | 1467 | 1 | 1.0900e-01 | 2.8451e+05 | ; | CA - HA |
| 1466 | 1468 | 1 | 1.5260e-01 | 2.5941e+05 | ; | CA - CB |
| 1466 | 1479 | 1 | 1.5220e-01 | 2.6527e+05 | ; | CA - C |
| 1468 | 1469 | 1 | 1.0900e-01 | 2.8451e+05 | ; | CB - HB2 |
| 1468 | 1470 | 1 | 1.0900e-01 | 2.8451e+05 | ; | CB - HB3 |
| 1468 | 1471 | 1 | 1.5260e-01 | 2.5941e+05 | ; | CB - CG |
| 1471 | 1472 | 1 | 1.0900e-01 | 2.8451e+05 | ; | CG - HG2 |
| 1471 | 1473 | 1 | 1.0900e-01 | 2.8451e+05 | ; | CG - HG3 |
| 1471 | 1474 | 1 | 1.8100e-01 | 1.8995e+05 | ; | CG - SD |
| 1474 | 1475 | 1 | 1.8100e-01 | 1.8995e+05 | ; | SD - CE |
| 1475 | 1476 | 1 | 1.0900e-01 | 2.8451e+05 | ; | CE - HE1 |
| 1475 | 1477 | 1 | 1.0900e-01 | 2.8451e+05 | ; | CE - HE2 |
| 1475 | 1478 | 1 | 1.0900e-01 | 2.8451e+05 | ; | CE - HE3 |
| 1479 | 1480 | 1 | 1.2290e-01 | 4.7698e+05 | ; | C - O |
| 1479 | 1481 | 1 | 1.3350e-01 | 4.1003e+05 | ; | C - N |
| 1481 | 1482 | 1 | 1.0100e-01 | 3.6317e+05 | ; | N - H |
| 1481 | 1483 | 1 | 1.4490e-01 | 2.8200e+05 | ; | N - CA |
| 1483 | 1484 | 1 | 1.0900e-01 | 2.8451e+05 | ; | CA - HA |
| 1483 | 1485 | 1 | 1.5260e-01 | 2.5941e+05 | ; | CA - CB |
| 1483 | 1494 | 1 | 1.5220e-01 | 2.6527e+05 | ; | CA - C |
| 1485 | 1486 | 1 | 1.0900e-01 | 2.8451e+05 | ; | CB - HB2 |
| 1485 | 1487 | 1 | 1.0900e-01 | 2.8451e+05 | ; | CB - HB3 |
| 1485 | 1488 | 1 | 1.5260e-01 | 2.5941e+05 | ; | CB - CG |
| 1488 | 1489 | 1 | 1.0900e-01 | 2.8451e+05 | ; | CG - HG2 |
| 1488 | 1490 | 1 | 1.0900e-01 | 2.8451e+05 | ; | CG - HG3 |
| 1488 | 1491 | 1 | 1.5220e-01 | 2.6527e+05 | ; | CG - CD |
| 1491 | 1492 | 1 | 1.2500e-01 | 5.4894e+05 | ; | CD - OE1 |

| | | | | | | | | |
|---|---|---|---|---|---|---|---|---|
| 1491 | 1493 | 1 | 1.2500e-01 | 5.4894e+05 | ; | CD | - | OE2 |
| 1494 | 1495 | 1 | 1.2290e-01 | 4.7698e+05 | ; | C | - | O |
| 1494 | 1496 | 1 | 1.3350e-01 | 4.1003e+05 | ; | C | - | N |
| 1496 | 1497 | 1 | 1.0100e-01 | 3.6317e+05 | ; | N | - | H |
| 1496 | 1498 | 1 | 1.4490e-01 | 2.8200e+05 | ; | N | - | CA |
| 1498 | 1499 | 1 | 1.0900e-01 | 2.8451e+05 | ; | CA | - | HA |
| 1498 | 1500 | 1 | 1.5260e-01 | 2.5941e+05 | ; | CA | - | CB |
| 1498 | 1513 | 1 | 1.5220e-01 | 2.6527e+05 | ; | CA | - | C |
| 1500 | 1501 | 1 | 1.0900e-01 | 2.8451e+05 | ; | CB | - | HB2 |
| 1500 | 1502 | 1 | 1.0900e-01 | 2.8451e+05 | ; | CB | - | HB3 |
| 1500 | 1503 | 1 | 1.5260e-01 | 2.5941e+05 | ; | CB | - | CG |
| 1503 | 1504 | 1 | 1.0900e-01 | 2.8451e+05 | ; | CG | - | HG |
| 1503 | 1505 | 1 | 1.5260e-01 | 2.5941e+05 | ; | CG | - | CD1 |
| 1503 | 1509 | 1 | 1.5260e-01 | 2.5941e+05 | ; | CG | - | CD2 |
| 1505 | 1506 | 1 | 1.0900e-01 | 2.8451e+05 | ; | CD1 | - | HD11 |
| 1505 | 1507 | 1 | 1.0900e-01 | 2.8451e+05 | ; | CD1 | - | HD12 |
| 1505 | 1508 | 1 | 1.0900e-01 | 2.8451e+05 | ; | CD1 | - | HD13 |
| 1509 | 1510 | 1 | 1.0900e-01 | 2.8451e+05 | ; | CD2 | - | HD21 |
| 1509 | 1511 | 1 | 1.0900e-01 | 2.8451e+05 | ; | CD2 | - | HD22 |
| 1509 | 1512 | 1 | 1.0900e-01 | 2.8451e+05 | ; | CD2 | - | HD23 |
| 1513 | 1514 | 1 | 1.2290e-01 | 4.7698e+05 | ; | C | - | O |
| 1513 | 1515 | 1 | 1.3350e-01 | 4.1003e+05 | ; | C | - | N |
| 1515 | 1516 | 1 | 1.0100e-01 | 3.6317e+05 | ; | N | - | H |
| 1515 | 1517 | 1 | 1.4490e-01 | 2.8200e+05 | ; | N | - | CA |
| 1517 | 1518 | 1 | 1.0900e-01 | 2.8451e+05 | ; | CA | - | HA |
| 1517 | 1519 | 1 | 1.5260e-01 | 2.5941e+05 | ; | CA | - | CB |
| 1517 | 1524 | 1 | 1.5220e-01 | 2.6527e+05 | ; | CA | - | C |
| 1519 | 1520 | 1 | 1.0900e-01 | 2.8451e+05 | ; | CB | - | HB2 |
| 1519 | 1521 | 1 | 1.0900e-01 | 2.8451e+05 | ; | CB | - | HB3 |
| 1519 | 1522 | 1 | 1.8100e-01 | 1.9832e+05 | ; | CB | - | SG |
| 1522 | 1523 | 1 | 1.3360e-01 | 2.2928e+05 | ; | SG | - | HG |
| 1524 | 1525 | 1 | 1.2290e-01 | 4.7698e+05 | ; | C | - | O |
| 1524 | 1526 | 1 | 1.3350e-01 | 4.1003e+05 | ; | C | - | N |
| 1526 | 1527 | 1 | 1.0100e-01 | 3.6317e+05 | ; | N | - | H |
| 1526 | 1528 | 1 | 1.4490e-01 | 2.8200e+05 | ; | N | - | CA |
| 1528 | 1529 | 1 | 1.0900e-01 | 2.8451e+05 | ; | CA | - | HA |
| 1528 | 1530 | 1 | 1.5260e-01 | 2.5941e+05 | ; | CA | - | CB |
| 1528 | 1535 | 1 | 1.5220e-01 | 2.6527e+05 | ; | CA | - | C |
| 1530 | 1531 | 1 | 1.0900e-01 | 2.8451e+05 | ; | CB | - | HB2 |
| 1530 | 1532 | 1 | 1.0900e-01 | 2.8451e+05 | ; | CB | - | HB3 |
| 1530 | 1533 | 1 | 1.4100e-01 | 2.6778e+05 | ; | CB | - | OG |
| 1533 | 1534 | 1 | 9.6000e-02 | 4.6275e+05 | ; | OG | - | HG |
| 1535 | 1536 | 1 | 1.2290e-01 | 4.7698e+05 | ; | C | - | O |
| 1535 | 1537 | 1 | 1.3350e-01 | 4.1003e+05 | ; | C | - | N |
| 1537 | 1538 | 1 | 1.0100e-01 | 3.6317e+05 | ; | N | - | H |
| 1537 | 1539 | 1 | 1.4490e-01 | 2.8200e+05 | ; | N | - | CA |
| 1539 | 1540 | 1 | 1.0900e-01 | 2.8451e+05 | ; | CA | - | HA2 |
| 1539 | 1541 | 1 | 1.0900e-01 | 2.8451e+05 | ; | CA | - | HA3 |
| 1539 | 1542 | 1 | 1.5220e-01 | 2.6527e+05 | ; | CA | - | C |
| 1542 | 1543 | 1 | 1.2290e-01 | 4.7698e+05 | ; | C | - | O |
| 1542 | 1544 | 1 | 1.3350e-01 | 4.1003e+05 | ; | C | - | N |
| 1544 | 1545 | 1 | 1.0100e-01 | 3.6317e+05 | ; | N | - | H |

| | | | | | | |
|---|---|---|---|---|---|---|
| 1544 | 1546 | 1 | 1.4490e-01 | 2.8200e+05 | ; | N - CA |
| 1546 | 1547 | 1 | 1.0900e-01 | 2.8451e+05 | ; | CA - HA |
| 1546 | 1548 | 1 | 1.5260e-01 | 2.5941e+05 | ; | CA - CB |
| 1546 | 1566 | 1 | 1.5220e-01 | 2.6527e+05 | ; | CA - C |
| 1548 | 1549 | 1 | 1.0900e-01 | 2.8451e+05 | ; | CB - HB2 |
| 1548 | 1550 | 1 | 1.0900e-01 | 2.8451e+05 | ; | CB - HB3 |
| 1548 | 1551 | 1 | 1.5260e-01 | 2.5941e+05 | ; | CB - CG |
| 1551 | 1552 | 1 | 1.0900e-01 | 2.8451e+05 | ; | CG - HG2 |
| 1551 | 1553 | 1 | 1.0900e-01 | 2.8451e+05 | ; | CG - HG3 |
| 1551 | 1554 | 1 | 1.5260e-01 | 2.5941e+05 | ; | CG - CD |
| 1554 | 1555 | 1 | 1.0900e-01 | 2.8451e+05 | ; | CD - HD2 |
| 1554 | 1556 | 1 | 1.0900e-01 | 2.8451e+05 | ; | CD - HD3 |
| 1554 | 1557 | 1 | 1.4630e-01 | 2.8200e+05 | ; | CD - NE |
| 1557 | 1558 | 1 | 1.0100e-01 | 3.6317e+05 | ; | NE - HE |
| 1557 | 1559 | 1 | 1.3400e-01 | 4.0250e+05 | ; | NE - CZ |
| 1559 | 1560 | 1 | 1.3400e-01 | 4.0250e+05 | ; | CZ - NH1 |
| 1559 | 1563 | 1 | 1.3400e-01 | 4.0250e+05 | ; | CZ - NH2 |
| 1560 | 1561 | 1 | 1.0100e-01 | 3.6317e+05 | ; | NH1 - HH11 |
| 1560 | 1562 | 1 | 1.0100e-01 | 3.6317e+05 | ; | NH1 - HH12 |
| 1563 | 1564 | 1 | 1.0100e-01 | 3.6317e+05 | ; | NH2 - HH21 |
| 1563 | 1565 | 1 | 1.0100e-01 | 3.6317e+05 | ; | NH2 - HH22 |
| 1566 | 1567 | 1 | 1.2290e-01 | 4.7698e+05 | ; | C - O |
| 1566 | 1568 | 1 | 1.3350e-01 | 4.1003e+05 | ; | C - N |
| 1568 | 1569 | 1 | 1.0100e-01 | 3.6317e+05 | ; | N - H |
| 1568 | 1570 | 1 | 1.4490e-01 | 2.8200e+05 | ; | N - CA |
| 1570 | 1571 | 1 | 1.0900e-01 | 2.8451e+05 | ; | CA - HA |
| 1570 | 1572 | 1 | 1.5260e-01 | 2.5941e+05 | ; | CA - CB |
| 1570 | 1581 | 1 | 1.5220e-01 | 2.6527e+05 | ; | CA - C |
| 1572 | 1573 | 1 | 1.0900e-01 | 2.8451e+05 | ; | CB - HB2 |
| 1572 | 1574 | 1 | 1.0900e-01 | 2.8451e+05 | ; | CB - HB3 |
| 1572 | 1575 | 1 | 1.5260e-01 | 2.5941e+05 | ; | CB - CG |
| 1575 | 1576 | 1 | 1.0900e-01 | 2.8451e+05 | ; | CG - HG2 |
| 1575 | 1577 | 1 | 1.0900e-01 | 2.8451e+05 | ; | CG - HG3 |
| 1575 | 1578 | 1 | 1.5220e-01 | 2.6527e+05 | ; | CG - CD |
| 1578 | 1579 | 1 | 1.2500e-01 | 5.4894e+05 | ; | CD - OE1 |
| 1578 | 1580 | 1 | 1.2500e-01 | 5.4894e+05 | ; | CD - OE2 |
| 1581 | 1582 | 1 | 1.2290e-01 | 4.7698e+05 | ; | C - O |
| 1581 | 1583 | 1 | 1.3350e-01 | 4.1003e+05 | ; | C - N |
| 1583 | 1584 | 1 | 1.0100e-01 | 3.6317e+05 | ; | N - H |
| 1583 | 1585 | 1 | 1.4490e-01 | 2.8200e+05 | ; | N - CA |
| 1585 | 1586 | 1 | 1.0900e-01 | 2.8451e+05 | ; | CA - HA |
| 1585 | 1587 | 1 | 1.5260e-01 | 2.5941e+05 | ; | CA - CB |
| 1585 | 1600 | 1 | 1.5220e-01 | 2.6527e+05 | ; | CA - C |
| 1587 | 1588 | 1 | 1.0900e-01 | 2.8451e+05 | ; | CB - HB2 |
| 1587 | 1589 | 1 | 1.0900e-01 | 2.8451e+05 | ; | CB - HB3 |
| 1587 | 1590 | 1 | 1.5260e-01 | 2.5941e+05 | ; | CB - CG |
| 1590 | 1591 | 1 | 1.0900e-01 | 2.8451e+05 | ; | CG - HG |
| 1590 | 1592 | 1 | 1.5260e-01 | 2.5941e+05 | ; | CG - CD1 |
| 1590 | 1596 | 1 | 1.5260e-01 | 2.5941e+05 | ; | CG - CD2 |
| 1592 | 1593 | 1 | 1.0900e-01 | 2.8451e+05 | ; | CD1 - HD11 |
| 1592 | 1594 | 1 | 1.0900e-01 | 2.8451e+05 | ; | CD1 - HD12 |
| 1592 | 1595 | 1 | 1.0900e-01 | 2.8451e+05 | ; | CD1 - HD13 |

| | | | | | |
|---|---|---|---|---|---|
| 1596 | 1597 | 1 | 1.0900e-01 | 2.8451e+05 ; | CD2 - HD21 |
| 1596 | 1598 | 1 | 1.0900e-01 | 2.8451e+05 ; | CD2 - HD22 |
| 1596 | 1599 | 1 | 1.0900e-01 | 2.8451e+05 ; | CD2 - HD23 |
| 1600 | 1601 | 1 | 1.2290e-01 | 4.7698e+05 ; | C - O |
| 1600 | 1602 | 1 | 1.3350e-01 | 4.1003e+05 ; | C - N |
| 1602 | 1603 | 1 | 1.0100e-01 | 3.6317e+05 ; | N - H |
| 1602 | 1604 | 1 | 1.4490e-01 | 2.8200e+05 ; | N - CA |
| 1604 | 1605 | 1 | 1.0900e-01 | 2.8451e+05 ; | CA - HA |
| 1604 | 1606 | 1 | 1.5260e-01 | 2.5941e+05 ; | CA - CB |
| 1604 | 1620 | 1 | 1.5220e-01 | 2.6527e+05 ; | CA - C |
| 1606 | 1607 | 1 | 1.0900e-01 | 2.8451e+05 ; | CB - HB2 |
| 1606 | 1608 | 1 | 1.0900e-01 | 2.8451e+05 ; | CB - HB3 |
| 1606 | 1609 | 1 | 1.5100e-01 | 2.6527e+05 ; | CB - CG |
| 1609 | 1610 | 1 | 1.4000e-01 | 3.9246e+05 ; | CG - CD1 |
| 1609 | 1618 | 1 | 1.4000e-01 | 3.9246e+05 ; | CG - CD2 |
| 1610 | 1611 | 1 | 1.0800e-01 | 3.0711e+05 ; | CD1 - HD1 |
| 1610 | 1612 | 1 | 1.4000e-01 | 3.9246e+05 ; | CD1 - CE1 |
| 1612 | 1613 | 1 | 1.0800e-01 | 3.0711e+05 ; | CE1 - HE1 |
| 1612 | 1614 | 1 | 1.4000e-01 | 3.9246e+05 ; | CE1 - CZ |
| 1614 | 1615 | 1 | 1.0800e-01 | 3.0711e+05 ; | CZ - HZ |
| 1614 | 1616 | 1 | 1.4000e-01 | 3.9246e+05 ; | CZ - CE2 |
| 1616 | 1617 | 1 | 1.0800e-01 | 3.0711e+05 ; | CE2 - HE2 |
| 1616 | 1618 | 1 | 1.4000e-01 | 3.9246e+05 ; | CE2 - CD2 |
| 1618 | 1619 | 1 | 1.0800e-01 | 3.0711e+05 ; | CD2 - HD2 |
| 1620 | 1621 | 1 | 1.2290e-01 | 4.7698e+05 ; | C - O |
| 1620 | 1622 | 1 | 1.3350e-01 | 4.1003e+05 ; | C - N |
| 1622 | 1623 | 1 | 1.0100e-01 | 3.6317e+05 ; | N - H |
| 1622 | 1624 | 1 | 1.4490e-01 | 2.8200e+05 ; | N - CA |
| 1624 | 1625 | 1 | 1.0900e-01 | 2.8451e+05 ; | CA - HA |
| 1624 | 1626 | 1 | 1.5260e-01 | 2.5941e+05 ; | CA - CB |
| 1624 | 1632 | 1 | 1.5220e-01 | 2.6527e+05 ; | CA - C |
| 1626 | 1627 | 1 | 1.0900e-01 | 2.8451e+05 ; | CB - HB2 |
| 1626 | 1628 | 1 | 1.0900e-01 | 2.8451e+05 ; | CB - HB3 |
| 1626 | 1629 | 1 | 1.5220e-01 | 2.6527e+05 ; | CB - CG |
| 1629 | 1630 | 1 | 1.2500e-01 | 5.4894e+05 ; | CG - OD1 |
| 1629 | 1631 | 1 | 1.2500e-01 | 5.4894e+05 ; | CG - OD2 |
| 1632 | 1633 | 1 | 1.2290e-01 | 4.7698e+05 ; | C - O |
| 1632 | 1634 | 1 | 1.3350e-01 | 4.1003e+05 ; | C - N |
| 1634 | 1635 | 1 | 1.0100e-01 | 3.6317e+05 ; | N - H |
| 1634 | 1636 | 1 | 1.4490e-01 | 2.8200e+05 ; | N - CA |
| 1636 | 1637 | 1 | 1.0900e-01 | 2.8451e+05 ; | CA - HA |
| 1636 | 1638 | 1 | 1.5260e-01 | 2.5941e+05 ; | CA - CB |
| 1636 | 1643 | 1 | 1.5220e-01 | 2.6527e+05 ; | CA - C |
| 1638 | 1639 | 1 | 1.0900e-01 | 2.8451e+05 ; | CB - HB2 |
| 1638 | 1640 | 1 | 1.0900e-01 | 2.8451e+05 ; | CB - HB3 |
| 1638 | 1641 | 1 | 1.4100e-01 | 2.6778e+05 ; | CB - OG |
| 1641 | 1642 | 1 | 9.6000e-02 | 4.6275e+05 ; | OG - HG |
| 1643 | 1644 | 1 | 1.2290e-01 | 4.7698e+05 ; | C - O |
| 1643 | 1645 | 1 | 1.3350e-01 | 4.1003e+05 ; | C - N |
| 1645 | 1646 | 1 | 1.0100e-01 | 3.6317e+05 ; | N - H |
| 1645 | 1647 | 1 | 1.4490e-01 | 2.8200e+05 ; | N - CA |
| 1647 | 1648 | 1 | 1.0900e-01 | 2.8451e+05 ; | CA - HA |

| | | | | | | |
|---|---|---|---|---|---|---|
| 1647 | 1649 | 1 | 1.5260e-01 | 2.5941e+05 | ; | CA - CB |
| 1647 | 1662 | 1 | 1.5220e-01 | 2.6527e+05 | ; | CA - C |
| 1649 | 1650 | 1 | 1.0900e-01 | 2.8451e+05 | ; | CB - HB |
| 1649 | 1651 | 1 | 1.5260e-01 | 2.5941e+05 | ; | CB - CG2 |
| 1649 | 1655 | 1 | 1.5260e-01 | 2.5941e+05 | ; | CB - CG1 |
| 1651 | 1652 | 1 | 1.0900e-01 | 2.8451e+05 | ; | CG2 - HG21 |
| 1651 | 1653 | 1 | 1.0900e-01 | 2.8451e+05 | ; | CG2 - HG22 |
| 1651 | 1654 | 1 | 1.0900e-01 | 2.8451e+05 | ; | CG2 - HG23 |
| 1655 | 1656 | 1 | 1.0900e-01 | 2.8451e+05 | ; | CG1 - HG12 |
| 1655 | 1657 | 1 | 1.0900e-01 | 2.8451e+05 | ; | CG1 - HG13 |
| 1655 | 1658 | 1 | 1.5260e-01 | 2.5941e+05 | ; | CG1 - CD1 |
| 1658 | 1659 | 1 | 1.0900e-01 | 2.8451e+05 | ; | CD1 - HD11 |
| 1658 | 1660 | 1 | 1.0900e-01 | 2.8451e+05 | ; | CD1 - HD12 |
| 1658 | 1661 | 1 | 1.0900e-01 | 2.8451e+05 | ; | CD1 - HD13 |
| 1662 | 1663 | 1 | 1.2290e-01 | 4.7698e+05 | ; | C - O |
| 1662 | 1664 | 1 | 1.3350e-01 | 4.1003e+05 | ; | C - N |
| 1664 | 1665 | 1 | 1.0100e-01 | 3.6317e+05 | ; | N - H |
| 1664 | 1666 | 1 | 1.4490e-01 | 2.8200e+05 | ; | N - CA |
| 1666 | 1667 | 1 | 1.0900e-01 | 2.8451e+05 | ; | CA - HA |
| 1666 | 1668 | 1 | 1.5260e-01 | 2.5941e+05 | ; | CA - CB |
| 1666 | 1681 | 1 | 1.5220e-01 | 2.6527e+05 | ; | CA - C |
| 1668 | 1669 | 1 | 1.0900e-01 | 2.8451e+05 | ; | CB - HB |
| 1668 | 1670 | 1 | 1.5260e-01 | 2.5941e+05 | ; | CB - CG2 |
| 1668 | 1674 | 1 | 1.5260e-01 | 2.5941e+05 | ; | CB - CG1 |
| 1670 | 1671 | 1 | 1.0900e-01 | 2.8451e+05 | ; | CG2 - HG21 |
| 1670 | 1672 | 1 | 1.0900e-01 | 2.8451e+05 | ; | CG2 - HG22 |
| 1670 | 1673 | 1 | 1.0900e-01 | 2.8451e+05 | ; | CG2 - HG23 |
| 1674 | 1675 | 1 | 1.0900e-01 | 2.8451e+05 | ; | CG1 - HG12 |
| 1674 | 1676 | 1 | 1.0900e-01 | 2.8451e+05 | ; | CG1 - HG13 |
| 1674 | 1677 | 1 | 1.5260e-01 | 2.5941e+05 | ; | CG1 - CD1 |
| 1677 | 1678 | 1 | 1.0900e-01 | 2.8451e+05 | ; | CD1 - HD11 |
| 1677 | 1679 | 1 | 1.0900e-01 | 2.8451e+05 | ; | CD1 - HD12 |
| 1677 | 1680 | 1 | 1.0900e-01 | 2.8451e+05 | ; | CD1 - HD13 |
| 1681 | 1682 | 1 | 1.2290e-01 | 4.7698e+05 | ; | C - O |
| 1681 | 1683 | 1 | 1.3350e-01 | 4.1003e+05 | ; | C - N |
| 1683 | 1684 | 1 | 1.0100e-01 | 3.6317e+05 | ; | N - H |
| 1683 | 1685 | 1 | 1.4490e-01 | 2.8200e+05 | ; | N - CA |
| 1685 | 1686 | 1 | 1.0900e-01 | 2.8451e+05 | ; | CA - HA |
| 1685 | 1687 | 1 | 1.5260e-01 | 2.5941e+05 | ; | CA - CB |
| 1685 | 1696 | 1 | 1.5220e-01 | 2.6527e+05 | ; | CA - C |
| 1687 | 1688 | 1 | 1.0900e-01 | 2.8451e+05 | ; | CB - HB2 |
| 1687 | 1689 | 1 | 1.0900e-01 | 2.8451e+05 | ; | CB - HB3 |
| 1687 | 1690 | 1 | 1.5260e-01 | 2.5941e+05 | ; | CB - CG |
| 1690 | 1691 | 1 | 1.0900e-01 | 2.8451e+05 | ; | CG - HG2 |
| 1690 | 1692 | 1 | 1.0900e-01 | 2.8451e+05 | ; | CG - HG3 |
| 1690 | 1693 | 1 | 1.5220e-01 | 2.6527e+05 | ; | CG - CD |
| 1693 | 1694 | 1 | 1.2500e-01 | 5.4894e+05 | ; | CD - OE1 |
| 1693 | 1695 | 1 | 1.2500e-01 | 5.4894e+05 | ; | CD - OE2 |
| 1696 | 1697 | 1 | 1.2290e-01 | 4.7698e+05 | ; | C - O |
| 1696 | 1698 | 1 | 1.3350e-01 | 4.1003e+05 | ; | C - N |
| 1698 | 1699 | 1 | 1.0100e-01 | 3.6317e+05 | ; | N - H |
| 1698 | 1700 | 1 | 1.4490e-01 | 2.8200e+05 | ; | N - CA |

```
1700   1701   1    1.0900e-01    2.8451e+05 ;     CA - HA
1700   1702   1    1.5260e-01    2.5941e+05 ;     CA - CB
1700   1710   1    1.5220e-01    2.6527e+05 ;     CA - C
1702   1703   1    1.0900e-01    2.8451e+05 ;     CB - HB2
1702   1704   1    1.0900e-01    2.8451e+05 ;     CB - HB3
1702   1705   1    1.5220e-01    2.6527e+05 ;     CB - CG
1705   1706   1    1.2290e-01    4.7698e+05 ;     CG - OD1
1705   1707   1    1.3350e-01    4.1003e+05 ;     CG - ND2
1707   1708   1    1.0100e-01    3.6317e+05 ;    ND2 - HD21
1707   1709   1    1.0100e-01    3.6317e+05 ;    ND2 - HD22
1710   1711   1    1.2290e-01    4.7698e+05 ;      C - O
1710   1712   1    1.3350e-01    4.1003e+05 ;      C - N
1712   1713   1    1.0100e-01    3.6317e+05 ;      N - H
1712   1714   1    1.4490e-01    2.8200e+05 ;      N - CA
1714   1715   1    1.0900e-01    2.8451e+05 ;     CA - HA2
1714   1716   1    1.0900e-01    2.8451e+05 ;     CA - HA3
1714   1717   1    1.5220e-01    2.6527e+05 ;     CA - C
1717   1718   1    1.2290e-01    4.7698e+05 ;      C - O
1717   1719   1    1.3350e-01    4.1003e+05 ;      C - N
1719   1720   1    1.0100e-01    3.6317e+05 ;      N - H
1719   1721   1    1.4490e-01    2.8200e+05 ;      N - CA
1721   1722   1    1.0900e-01    2.8451e+05 ;     CA - HA
1721   1723   1    1.5260e-01    2.5941e+05 ;     CA - CB
1721   1728   1    1.5220e-01    2.6527e+05 ;     CA - C
1723   1724   1    1.0900e-01    2.8451e+05 ;     CB - HB2
1723   1725   1    1.0900e-01    2.8451e+05 ;     CB - HB3
1723   1726   1    1.4100e-01    2.6778e+05 ;     CB - OG
1726   1727   1    9.6000e-02    4.6275e+05 ;     OG - HG
1728   1729   1    1.2290e-01    4.7698e+05 ;      C - O
1728   1730   1    1.3350e-01    4.1003e+05 ;      C - N
1730   1731   1    1.0100e-01    3.6317e+05 ;      N - H
1730   1732   1    1.4490e-01    2.8200e+05 ;      N - CA
1732   1733   1    1.0900e-01    2.8451e+05 ;     CA - HA
1732   1734   1    1.5260e-01    2.5941e+05 ;     CA - CB
1732   1748   1    1.5220e-01    2.6527e+05 ;     CA - C
1734   1735   1    1.0900e-01    2.8451e+05 ;     CB - HB2
1734   1736   1    1.0900e-01    2.8451e+05 ;     CB - HB3
1734   1737   1    1.5100e-01    2.6527e+05 ;     CB - CG
1737   1738   1    1.4000e-01    3.9246e+05 ;     CG - CD1
1737   1746   1    1.4000e-01    3.9246e+05 ;     CG - CD2
1738   1739   1    1.0800e-01    3.0711e+05 ;    CD1 - HD1
1738   1740   1    1.4000e-01    3.9246e+05 ;    CD1 - CE1
1740   1741   1    1.0800e-01    3.0711e+05 ;    CE1 - HE1
1740   1742   1    1.4000e-01    3.9246e+05 ;    CE1 - CZ
1742   1743   1    1.0800e-01    3.0711e+05 ;     CZ - HZ
1742   1744   1    1.4000e-01    3.9246e+05 ;     CZ - CE2
1744   1745   1    1.0800e-01    3.0711e+05 ;    CE2 - HE2
1744   1746   1    1.4000e-01    3.9246e+05 ;    CE2 - CD2
1746   1747   1    1.0800e-01    3.0711e+05 ;    CD2 - HD2
1748   1749   1    1.2290e-01    4.7698e+05 ;      C - O
1748   1750   1    1.3350e-01    4.1003e+05 ;      C - N
1750   1751   1    1.0100e-01    3.6317e+05 ;      N - H
```

| | | | | | | |
|---|---|---|---|---|---|---|
| 1750 | 1752 | 1 | 1.4490e-01 | 2.8200e+05 | ; | N - CA |
| 1752 | 1753 | 1 | 1.0900e-01 | 2.8451e+05 | ; | CA - HA |
| 1752 | 1754 | 1 | 1.5260e-01 | 2.5941e+05 | ; | CA - CB |
| 1752 | 1762 | 1 | 1.5220e-01 | 2.6527e+05 | ; | CA - C |
| 1754 | 1755 | 1 | 1.0900e-01 | 2.8451e+05 | ; | CB - HB |
| 1754 | 1756 | 1 | 1.5260e-01 | 2.5941e+05 | ; | CB - CG2 |
| 1754 | 1760 | 1 | 1.4100e-01 | 2.6778e+05 | ; | CB - OG1 |
| 1756 | 1757 | 1 | 1.0900e-01 | 2.8451e+05 | ; | CG2 - HG21 |
| 1756 | 1758 | 1 | 1.0900e-01 | 2.8451e+05 | ; | CG2 - HG22 |
| 1756 | 1759 | 1 | 1.0900e-01 | 2.8451e+05 | ; | CG2 - HG23 |
| 1760 | 1761 | 1 | 9.6000e-02 | 4.6275e+05 | ; | OG1 - HG1 |
| 1762 | 1763 | 1 | 1.2290e-01 | 4.7698e+05 | ; | C - O |
| 1762 | 1764 | 1 | 1.3350e-01 | 4.1003e+05 | ; | C - N |
| 1764 | 1765 | 1 | 1.0100e-01 | 3.6317e+05 | ; | N - H |
| 1764 | 1766 | 1 | 1.4490e-01 | 2.8200e+05 | ; | N - CA |
| 1766 | 1767 | 1 | 1.0900e-01 | 2.8451e+05 | ; | CA - HA |
| 1766 | 1768 | 1 | 1.5260e-01 | 2.5941e+05 | ; | CA - CB |
| 1766 | 1777 | 1 | 1.5220e-01 | 2.6527e+05 | ; | CA - C |
| 1768 | 1769 | 1 | 1.0900e-01 | 2.8451e+05 | ; | CB - HB2 |
| 1768 | 1770 | 1 | 1.0900e-01 | 2.8451e+05 | ; | CB - HB3 |
| 1768 | 1771 | 1 | 1.5260e-01 | 2.5941e+05 | ; | CB - CG |
| 1771 | 1772 | 1 | 1.0900e-01 | 2.8451e+05 | ; | CG - HG2 |
| 1771 | 1773 | 1 | 1.0900e-01 | 2.8451e+05 | ; | CG - HG3 |
| 1771 | 1774 | 1 | 1.5220e-01 | 2.6527e+05 | ; | CG - CD |
| 1774 | 1775 | 1 | 1.2500e-01 | 5.4894e+05 | ; | CD - OE1 |
| 1774 | 1776 | 1 | 1.2500e-01 | 5.4894e+05 | ; | CD - OE2 |
| 1777 | 1778 | 1 | 1.2290e-01 | 4.7698e+05 | ; | C - O |
| 1777 | 1779 | 1 | 1.3350e-01 | 4.1003e+05 | ; | C - N |
| 1779 | 1780 | 1 | 1.0100e-01 | 3.6317e+05 | ; | N - H |
| 1779 | 1781 | 1 | 1.4490e-01 | 2.8200e+05 | ; | N - CA |
| 1781 | 1782 | 1 | 1.0900e-01 | 2.8451e+05 | ; | CA - HA |
| 1781 | 1783 | 1 | 1.5260e-01 | 2.5941e+05 | ; | CA - CB |
| 1781 | 1799 | 1 | 1.5220e-01 | 2.6527e+05 | ; | CA - C |
| 1783 | 1784 | 1 | 1.0900e-01 | 2.8451e+05 | ; | CB - HB2 |
| 1783 | 1785 | 1 | 1.0900e-01 | 2.8451e+05 | ; | CB - HB3 |
| 1783 | 1786 | 1 | 1.5260e-01 | 2.5941e+05 | ; | CB - CG |
| 1786 | 1787 | 1 | 1.0900e-01 | 2.8451e+05 | ; | CG - HG2 |
| 1786 | 1788 | 1 | 1.0900e-01 | 2.8451e+05 | ; | CG - HG3 |
| 1786 | 1789 | 1 | 1.5260e-01 | 2.5941e+05 | ; | CG - CD |
| 1789 | 1790 | 1 | 1.0900e-01 | 2.8451e+05 | ; | CD - HD2 |
| 1789 | 1791 | 1 | 1.0900e-01 | 2.8451e+05 | ; | CD - HD3 |
| 1789 | 1792 | 1 | 1.5260e-01 | 2.5941e+05 | ; | CD - CE |
| 1792 | 1793 | 1 | 1.0900e-01 | 2.8451e+05 | ; | CE - HE2 |
| 1792 | 1794 | 1 | 1.0900e-01 | 2.8451e+05 | ; | CE - HE3 |
| 1792 | 1795 | 1 | 1.4710e-01 | 3.0711e+05 | ; | CE - NZ |
| 1795 | 1796 | 1 | 1.0100e-01 | 3.6317e+05 | ; | NZ - HZ1 |
| 1795 | 1797 | 1 | 1.0100e-01 | 3.6317e+05 | ; | NZ - HZ2 |
| 1795 | 1798 | 1 | 1.0100e-01 | 3.6317e+05 | ; | NZ - HZ3 |
| 1799 | 1800 | 1 | 1.2290e-01 | 4.7698e+05 | ; | C - O |
| 1799 | 1801 | 1 | 1.3350e-01 | 4.1003e+05 | ; | C - N |
| 1801 | 1802 | 1 | 1.0100e-01 | 3.6317e+05 | ; | N - H |
| 1801 | 1803 | 1 | 1.4490e-01 | 2.8200e+05 | ; | N - CA |

| | | | | | | |
|---|---|---|---|---|---|---|
| 1803 | 1804 | 1 | 1.0900e-01 | 2.8451e+05 | ; | CA - HA |
| 1803 | 1805 | 1 | 1.5260e-01 | 2.5941e+05 | ; | CA - CB |
| 1803 | 1813 | 1 | 1.5220e-01 | 2.6527e+05 | ; | CA - C |
| 1805 | 1806 | 1 | 1.0900e-01 | 2.8451e+05 | ; | CB - HB2 |
| 1805 | 1807 | 1 | 1.0900e-01 | 2.8451e+05 | ; | CB - HB3 |
| 1805 | 1808 | 1 | 1.5220e-01 | 2.6527e+05 | ; | CB - CG |
| 1808 | 1809 | 1 | 1.2290e-01 | 4.7698e+05 | ; | CG - OD1 |
| 1808 | 1810 | 1 | 1.3350e-01 | 4.1003e+05 | ; | CG - ND2 |
| 1810 | 1811 | 1 | 1.0100e-01 | 3.6317e+05 | ; | ND2 - HD21 |
| 1810 | 1812 | 1 | 1.0100e-01 | 3.6317e+05 | ; | ND2 - HD22 |
| 1813 | 1814 | 1 | 1.2290e-01 | 4.7698e+05 | ; | C - O |
| 1813 | 1815 | 1 | 1.3350e-01 | 4.1003e+05 | ; | C - N |
| 1815 | 1816 | 1 | 1.0100e-01 | 3.6317e+05 | ; | N - H |
| 1815 | 1817 | 1 | 1.4490e-01 | 2.8200e+05 | ; | N - CA |
| 1817 | 1818 | 1 | 1.0900e-01 | 2.8451e+05 | ; | CA - HA |
| 1817 | 1819 | 1 | 1.5260e-01 | 2.5941e+05 | ; | CA - CB |
| 1817 | 1823 | 1 | 1.5220e-01 | 2.6527e+05 | ; | CA - C |
| 1819 | 1820 | 1 | 1.0900e-01 | 2.8451e+05 | ; | CB - HB1 |
| 1819 | 1821 | 1 | 1.0900e-01 | 2.8451e+05 | ; | CB - HB2 |
| 1819 | 1822 | 1 | 1.0900e-01 | 2.8451e+05 | ; | CB - HB3 |
| 1823 | 1824 | 1 | 1.2290e-01 | 4.7698e+05 | ; | C - O |
| 1823 | 1825 | 1 | 1.3350e-01 | 4.1003e+05 | ; | C - N |
| 1825 | 1826 | 1 | 1.0100e-01 | 3.6317e+05 | ; | N - H |
| 1825 | 1827 | 1 | 1.4490e-01 | 2.8200e+05 | ; | N - CA |
| 1827 | 1828 | 1 | 1.0900e-01 | 2.8451e+05 | ; | CA - HA |
| 1827 | 1829 | 1 | 1.5260e-01 | 2.5941e+05 | ; | CA - CB |
| 1827 | 1833 | 1 | 1.5220e-01 | 2.6527e+05 | ; | CA - C |
| 1829 | 1830 | 1 | 1.0900e-01 | 2.8451e+05 | ; | CB - HB1 |
| 1829 | 1831 | 1 | 1.0900e-01 | 2.8451e+05 | ; | CB - HB2 |
| 1829 | 1832 | 1 | 1.0900e-01 | 2.8451e+05 | ; | CB - HB3 |
| 1833 | 1834 | 1 | 1.2290e-01 | 4.7698e+05 | ; | C - O |
| 1833 | 1835 | 1 | 1.3350e-01 | 4.1003e+05 | ; | C - N |
| 1835 | 1836 | 1 | 1.0100e-01 | 3.6317e+05 | ; | N - H |
| 1835 | 1837 | 1 | 1.4490e-01 | 2.8200e+05 | ; | N - CA |
| 1837 | 1838 | 1 | 1.0900e-01 | 2.8451e+05 | ; | CA - HA |
| 1837 | 1839 | 1 | 1.5260e-01 | 2.5941e+05 | ; | CA - CB |
| 1837 | 1847 | 1 | 1.5220e-01 | 2.6527e+05 | ; | CA - C |
| 1839 | 1840 | 1 | 1.0900e-01 | 2.8451e+05 | ; | CB - HB |
| 1839 | 1841 | 1 | 1.5260e-01 | 2.5941e+05 | ; | CB - CG2 |
| 1839 | 1845 | 1 | 1.4100e-01 | 2.6778e+05 | ; | CB - OG1 |
| 1841 | 1842 | 1 | 1.0900e-01 | 2.8451e+05 | ; | CG2 - HG21 |
| 1841 | 1843 | 1 | 1.0900e-01 | 2.8451e+05 | ; | CG2 - HG22 |
| 1841 | 1844 | 1 | 1.0900e-01 | 2.8451e+05 | ; | CG2 - HG23 |
| 1845 | 1846 | 1 | 9.6000e-02 | 4.6275e+05 | ; | OG1 - HG1 |
| 1847 | 1848 | 1 | 1.2290e-01 | 4.7698e+05 | ; | C - O |
| 1847 | 1849 | 1 | 1.3350e-01 | 4.1003e+05 | ; | C - N |
| 1849 | 1850 | 1 | 1.0100e-01 | 3.6317e+05 | ; | N - H |
| 1849 | 1851 | 1 | 1.4490e-01 | 2.8200e+05 | ; | N - CA |
| 1851 | 1852 | 1 | 1.0900e-01 | 2.8451e+05 | ; | CA - HA |
| 1851 | 1853 | 1 | 1.5260e-01 | 2.5941e+05 | ; | CA - CB |
| 1851 | 1866 | 1 | 1.5220e-01 | 2.6527e+05 | ; | CA - C |
| 1853 | 1854 | 1 | 1.0900e-01 | 2.8451e+05 | ; | CB - HB |

| | | | | | |
|---|---|---|---|---|---|
| 1853 | 1855 | 1 | 1.5260e-01 | 2.5941e+05 ; | CB - CG2 |
| 1853 | 1859 | 1 | 1.5260e-01 | 2.5941e+05 ; | CB - CG1 |
| 1855 | 1856 | 1 | 1.0900e-01 | 2.8451e+05 ; | CG2 - HG21 |
| 1855 | 1857 | 1 | 1.0900e-01 | 2.8451e+05 ; | CG2 - HG22 |
| 1855 | 1858 | 1 | 1.0900e-01 | 2.8451e+05 ; | CG2 - HG23 |
| 1859 | 1860 | 1 | 1.0900e-01 | 2.8451e+05 ; | CG1 - HG12 |
| 1859 | 1861 | 1 | 1.0900e-01 | 2.8451e+05 ; | CG1 - HG13 |
| 1859 | 1862 | 1 | 1.5260e-01 | 2.5941e+05 ; | CG1 - CD1 |
| 1862 | 1863 | 1 | 1.0900e-01 | 2.8451e+05 ; | CD1 - HD11 |
| 1862 | 1864 | 1 | 1.0900e-01 | 2.8451e+05 ; | CD1 - HD12 |
| 1862 | 1865 | 1 | 1.0900e-01 | 2.8451e+05 ; | CD1 - HD13 |
| 1866 | 1867 | 1 | 1.2290e-01 | 4.7698e+05 ; | C - O |
| 1866 | 1868 | 1 | 1.3350e-01 | 4.1003e+05 ; | C - N |
| 1868 | 1869 | 1 | 1.0100e-01 | 3.6317e+05 ; | N - H |
| 1868 | 1870 | 1 | 1.4490e-01 | 2.8200e+05 ; | N - CA |
| 1870 | 1871 | 1 | 1.0900e-01 | 2.8451e+05 ; | CA - HA |
| 1870 | 1872 | 1 | 1.5260e-01 | 2.5941e+05 ; | CA - CB |
| 1870 | 1883 | 1 | 1.5220e-01 | 2.6527e+05 ; | CA - C |
| 1872 | 1873 | 1 | 1.0900e-01 | 2.8451e+05 ; | CB - HB2 |
| 1872 | 1874 | 1 | 1.0900e-01 | 2.8451e+05 ; | CB - HB3 |
| 1872 | 1875 | 1 | 1.5260e-01 | 2.5941e+05 ; | CB - CG |
| 1875 | 1876 | 1 | 1.0900e-01 | 2.8451e+05 ; | CG - HG2 |
| 1875 | 1877 | 1 | 1.0900e-01 | 2.8451e+05 ; | CG - HG3 |
| 1875 | 1878 | 1 | 1.8100e-01 | 1.8995e+05 ; | CG - SD |
| 1878 | 1879 | 1 | 1.8100e-01 | 1.8995e+05 ; | SD - CE |
| 1879 | 1880 | 1 | 1.0900e-01 | 2.8451e+05 ; | CE - HE1 |
| 1879 | 1881 | 1 | 1.0900e-01 | 2.8451e+05 ; | CE - HE2 |
| 1879 | 1882 | 1 | 1.0900e-01 | 2.8451e+05 ; | CE - HE3 |
| 1883 | 1884 | 1 | 1.2290e-01 | 4.7698e+05 ; | C - O |
| 1883 | 1885 | 1 | 1.3350e-01 | 4.1003e+05 ; | C - N |
| 1885 | 1886 | 1 | 1.0100e-01 | 3.6317e+05 ; | N - H |
| 1885 | 1887 | 1 | 1.4490e-01 | 2.8200e+05 ; | N - CA |
| 1887 | 1888 | 1 | 1.0900e-01 | 2.8451e+05 ; | CA - HA |
| 1887 | 1889 | 1 | 1.5260e-01 | 2.5941e+05 ; | CA - CB |
| 1887 | 1905 | 1 | 1.5220e-01 | 2.6527e+05 ; | CA - C |
| 1889 | 1890 | 1 | 1.0900e-01 | 2.8451e+05 ; | CB - HB2 |
| 1889 | 1891 | 1 | 1.0900e-01 | 2.8451e+05 ; | CB - HB3 |
| 1889 | 1892 | 1 | 1.5260e-01 | 2.5941e+05 ; | CB - CG |
| 1892 | 1893 | 1 | 1.0900e-01 | 2.8451e+05 ; | CG - HG2 |
| 1892 | 1894 | 1 | 1.0900e-01 | 2.8451e+05 ; | CG - HG3 |
| 1892 | 1895 | 1 | 1.5260e-01 | 2.5941e+05 ; | CG - CD |
| 1895 | 1896 | 1 | 1.0900e-01 | 2.8451e+05 ; | CD - HD2 |
| 1895 | 1897 | 1 | 1.0900e-01 | 2.8451e+05 ; | CD - HD3 |
| 1895 | 1898 | 1 | 1.5260e-01 | 2.5941e+05 ; | CD - CE |
| 1898 | 1899 | 1 | 1.0900e-01 | 2.8451e+05 ; | CE - HE2 |
| 1898 | 1900 | 1 | 1.0900e-01 | 2.8451e+05 ; | CE - HE3 |
| 1898 | 1901 | 1 | 1.4710e-01 | 3.0711e+05 ; | CE - NZ |
| 1901 | 1902 | 1 | 1.0100e-01 | 3.6317e+05 ; | NZ - HZ1 |
| 1901 | 1903 | 1 | 1.0100e-01 | 3.6317e+05 ; | NZ - HZ2 |
| 1901 | 1904 | 1 | 1.0100e-01 | 3.6317e+05 ; | NZ - HZ3 |
| 1905 | 1906 | 1 | 1.2290e-01 | 4.7698e+05 ; | C - O |
| 1905 | 1907 | 1 | 1.3350e-01 | 4.1003e+05 ; | C - N |

| | | | | | | |
|---|---|---|---|---|---|---|
| 1907 | 1908 | 1 | 1.0100e-01 | 3.6317e+05 | ; | N - H |
| 1907 | 1909 | 1 | 1.4490e-01 | 2.8200e+05 | ; | N - CA |
| 1909 | 1910 | 1 | 1.0900e-01 | 2.8451e+05 | ; | CA - HA |
| 1909 | 1911 | 1 | 1.5260e-01 | 2.5941e+05 | ; | CA - CB |
| 1909 | 1922 | 1 | 1.5220e-01 | 2.6527e+05 | ; | CA - C |
| 1911 | 1912 | 1 | 1.0900e-01 | 2.8451e+05 | ; | CB - HB2 |
| 1911 | 1913 | 1 | 1.0900e-01 | 2.8451e+05 | ; | CB - HB3 |
| 1911 | 1914 | 1 | 1.5260e-01 | 2.5941e+05 | ; | CB - CG |
| 1914 | 1915 | 1 | 1.0900e-01 | 2.8451e+05 | ; | CG - HG2 |
| 1914 | 1916 | 1 | 1.0900e-01 | 2.8451e+05 | ; | CG - HG3 |
| 1914 | 1917 | 1 | 1.5220e-01 | 2.6527e+05 | ; | CG - CD |
| 1917 | 1918 | 1 | 1.2290e-01 | 4.7698e+05 | ; | CD - OE1 |
| 1917 | 1919 | 1 | 1.3350e-01 | 4.1003e+05 | ; | CD - NE2 |
| 1919 | 1920 | 1 | 1.0100e-01 | 3.6317e+05 | ; | NE2 - HE21 |
| 1919 | 1921 | 1 | 1.0100e-01 | 3.6317e+05 | ; | NE2 - HE22 |
| 1922 | 1923 | 1 | 1.2290e-01 | 4.7698e+05 | ; | C - O |
| 1922 | 1924 | 1 | 1.3350e-01 | 4.1003e+05 | ; | C - N |
| 1924 | 1925 | 1 | 1.0100e-01 | 3.6317e+05 | ; | N - H |
| 1924 | 1926 | 1 | 1.4490e-01 | 2.8200e+05 | ; | N - CA |
| 1926 | 1927 | 1 | 1.0900e-01 | 2.8451e+05 | ; | CA - HA |
| 1926 | 1928 | 1 | 1.5260e-01 | 2.5941e+05 | ; | CA - CB |
| 1926 | 1941 | 1 | 1.5220e-01 | 2.6527e+05 | ; | CA - C |
| 1928 | 1929 | 1 | 1.0900e-01 | 2.8451e+05 | ; | CB - HB |
| 1928 | 1930 | 1 | 1.5260e-01 | 2.5941e+05 | ; | CB - CG2 |
| 1928 | 1934 | 1 | 1.5260e-01 | 2.5941e+05 | ; | CB - CG1 |
| 1930 | 1931 | 1 | 1.0900e-01 | 2.8451e+05 | ; | CG2 - HG21 |
| 1930 | 1932 | 1 | 1.0900e-01 | 2.8451e+05 | ; | CG2 - HG22 |
| 1930 | 1933 | 1 | 1.0900e-01 | 2.8451e+05 | ; | CG2 - HG23 |
| 1934 | 1935 | 1 | 1.0900e-01 | 2.8451e+05 | ; | CG1 - HG12 |
| 1934 | 1936 | 1 | 1.0900e-01 | 2.8451e+05 | ; | CG1 - HG13 |
| 1934 | 1937 | 1 | 1.5260e-01 | 2.5941e+05 | ; | CG1 - CD1 |
| 1937 | 1938 | 1 | 1.0900e-01 | 2.8451e+05 | ; | CD1 - HD11 |
| 1937 | 1939 | 1 | 1.0900e-01 | 2.8451e+05 | ; | CD1 - HD12 |
| 1937 | 1940 | 1 | 1.0900e-01 | 2.8451e+05 | ; | CD1 - HD13 |
| 1941 | 1942 | 1 | 1.2290e-01 | 4.7698e+05 | ; | C - O |
| 1941 | 1943 | 1 | 1.3350e-01 | 4.1003e+05 | ; | C - N |
| 1943 | 1944 | 1 | 1.0100e-01 | 3.6317e+05 | ; | N - H |
| 1943 | 1945 | 1 | 1.4490e-01 | 2.8200e+05 | ; | N - CA |
| 1945 | 1946 | 1 | 1.0900e-01 | 2.8451e+05 | ; | CA - HA |
| 1945 | 1947 | 1 | 1.5260e-01 | 2.5941e+05 | ; | CA - CB |
| 1945 | 1961 | 1 | 1.5220e-01 | 2.6527e+05 | ; | CA - C |
| 1947 | 1948 | 1 | 1.0900e-01 | 2.8451e+05 | ; | CB - HB2 |
| 1947 | 1949 | 1 | 1.0900e-01 | 2.8451e+05 | ; | CB - HB3 |
| 1947 | 1950 | 1 | 1.5100e-01 | 2.6527e+05 | ; | CB - CG |
| 1950 | 1951 | 1 | 1.4000e-01 | 3.9246e+05 | ; | CG - CD1 |
| 1950 | 1959 | 1 | 1.4000e-01 | 3.9246e+05 | ; | CG - CD2 |
| 1951 | 1952 | 1 | 1.0800e-01 | 3.0711e+05 | ; | CD1 - HD1 |
| 1951 | 1953 | 1 | 1.4000e-01 | 3.9246e+05 | ; | CD1 - CE1 |
| 1953 | 1954 | 1 | 1.0800e-01 | 3.0711e+05 | ; | CE1 - HE1 |
| 1953 | 1955 | 1 | 1.4000e-01 | 3.9246e+05 | ; | CE1 - CZ |
| 1955 | 1956 | 1 | 1.0800e-01 | 3.0711e+05 | ; | CZ - HZ |
| 1955 | 1957 | 1 | 1.4000e-01 | 3.9246e+05 | ; | CZ - CE2 |

| | | | | | | |
|---|---|---|---|---|---|---|
| 1957 | 1958 | 1 | 1.0800e-01 | 3.0711e+05 | ; | CE2 - HE2 |
| 1957 | 1959 | 1 | 1.4000e-01 | 3.9246e+05 | ; | CE2 - CD2 |
| 1959 | 1960 | 1 | 1.0800e-01 | 3.0711e+05 | ; | CD2 - HD2 |
| 1961 | 1962 | 1 | 1.2290e-01 | 4.7698e+05 | ; | C - O |
| 1961 | 1963 | 1 | 1.3350e-01 | 4.1003e+05 | ; | C - N |
| 1963 | 1964 | 1 | 1.0100e-01 | 3.6317e+05 | ; | N - H |
| 1963 | 1965 | 1 | 1.4490e-01 | 2.8200e+05 | ; | N - CA |
| 1965 | 1966 | 1 | 1.0900e-01 | 2.8451e+05 | ; | CA - HA |
| 1965 | 1967 | 1 | 1.5260e-01 | 2.5941e+05 | ; | CA - CB |
| 1965 | 1972 | 1 | 1.5220e-01 | 2.6527e+05 | ; | CA - C |
| 1967 | 1968 | 1 | 1.0900e-01 | 2.8451e+05 | ; | CB - HB2 |
| 1967 | 1969 | 1 | 1.0900e-01 | 2.8451e+05 | ; | CB - HB3 |
| 1967 | 1970 | 1 | 1.4100e-01 | 2.6778e+05 | ; | CB - OG |
| 1970 | 1971 | 1 | 9.6000e-02 | 4.6275e+05 | ; | OG - HG |
| 1972 | 1973 | 1 | 1.2290e-01 | 4.7698e+05 | ; | C - O |
| 1972 | 1974 | 1 | 1.3350e-01 | 4.1003e+05 | ; | C - N |
| 1974 | 1975 | 1 | 1.0100e-01 | 3.6317e+05 | ; | N - H |
| 1974 | 1976 | 1 | 1.4490e-01 | 2.8200e+05 | ; | N - CA |
| 1976 | 1977 | 1 | 1.0900e-01 | 2.8451e+05 | ; | CA - HA |
| 1976 | 1978 | 1 | 1.5260e-01 | 2.5941e+05 | ; | CA - CB |
| 1976 | 1982 | 1 | 1.5220e-01 | 2.6527e+05 | ; | CA - C |
| 1978 | 1979 | 1 | 1.0900e-01 | 2.8451e+05 | ; | CB - HB1 |
| 1978 | 1980 | 1 | 1.0900e-01 | 2.8451e+05 | ; | CB - HB2 |
| 1978 | 1981 | 1 | 1.0900e-01 | 2.8451e+05 | ; | CB - HB3 |
| 1982 | 1983 | 1 | 1.2290e-01 | 4.7698e+05 | ; | C - O |
| 1982 | 1984 | 1 | 1.3350e-01 | 4.1003e+05 | ; | C - N |
| 1984 | 1985 | 1 | 1.0100e-01 | 3.6317e+05 | ; | N - H |
| 1984 | 1986 | 1 | 1.4490e-01 | 2.8200e+05 | ; | N - CA |
| 1986 | 1987 | 1 | 1.0900e-01 | 2.8451e+05 | ; | CA - HA |
| 1986 | 1988 | 1 | 1.5260e-01 | 2.5941e+05 | ; | CA - CB |
| 1986 | 2001 | 1 | 1.5220e-01 | 2.6527e+05 | ; | CA - C |
| 1988 | 1989 | 1 | 1.0900e-01 | 2.8451e+05 | ; | CB - HB |
| 1988 | 1990 | 1 | 1.5260e-01 | 2.5941e+05 | ; | CB - CG2 |
| 1988 | 1994 | 1 | 1.5260e-01 | 2.5941e+05 | ; | CB - CG1 |
| 1990 | 1991 | 1 | 1.0900e-01 | 2.8451e+05 | ; | CG2 - HG21 |
| 1990 | 1992 | 1 | 1.0900e-01 | 2.8451e+05 | ; | CG2 - HG22 |
| 1990 | 1993 | 1 | 1.0900e-01 | 2.8451e+05 | ; | CG2 - HG23 |
| 1994 | 1995 | 1 | 1.0900e-01 | 2.8451e+05 | ; | CG1 - HG12 |
| 1994 | 1996 | 1 | 1.0900e-01 | 2.8451e+05 | ; | CG1 - HG13 |
| 1994 | 1997 | 1 | 1.5260e-01 | 2.5941e+05 | ; | CG1 - CD1 |
| 1997 | 1998 | 1 | 1.0900e-01 | 2.8451e+05 | ; | CD1 - HD11 |
| 1997 | 1999 | 1 | 1.0900e-01 | 2.8451e+05 | ; | CD1 - HD12 |
| 1997 | 2000 | 1 | 1.0900e-01 | 2.8451e+05 | ; | CD1 - HD13 |
| 2001 | 2002 | 1 | 1.2290e-01 | 4.7698e+05 | ; | C - O |
| 2001 | 2003 | 1 | 1.3350e-01 | 4.1003e+05 | ; | C - N |
| 2003 | 2004 | 1 | 1.0100e-01 | 3.6317e+05 | ; | N - H |
| 2003 | 2005 | 1 | 1.4490e-01 | 2.8200e+05 | ; | N - CA |
| 2005 | 2006 | 1 | 1.0900e-01 | 2.8451e+05 | ; | CA - HA |
| 2005 | 2007 | 1 | 1.5260e-01 | 2.5941e+05 | ; | CA - CB |
| 2005 | 2021 | 1 | 1.5220e-01 | 2.6527e+05 | ; | CA - C |
| 2007 | 2008 | 1 | 1.0900e-01 | 2.8451e+05 | ; | CB - HB2 |
| 2007 | 2009 | 1 | 1.0900e-01 | 2.8451e+05 | ; | CB - HB3 |

| | | | | | | |
|---|---|---|---|---|---|---|
| 2007 | 2010 | 1 | 1.5100e-01 | 2.6527e+05 | ; | CB - CG |
| 2010 | 2011 | 1 | 1.4000e-01 | 3.9246e+05 | ; | CG - CD1 |
| 2010 | 2019 | 1 | 1.4000e-01 | 3.9246e+05 | ; | CG - CD2 |
| 2011 | 2012 | 1 | 1.0800e-01 | 3.0711e+05 | ; | CD1 - HD1 |
| 2011 | 2013 | 1 | 1.4000e-01 | 3.9246e+05 | ; | CD1 - CE1 |
| 2013 | 2014 | 1 | 1.0800e-01 | 3.0711e+05 | ; | CE1 - HE1 |
| 2013 | 2015 | 1 | 1.4000e-01 | 3.9246e+05 | ; | CE1 - CZ |
| 2015 | 2016 | 1 | 1.0800e-01 | 3.0711e+05 | ; | CZ - HZ |
| 2015 | 2017 | 1 | 1.4000e-01 | 3.9246e+05 | ; | CZ - CE2 |
| 2017 | 2018 | 1 | 1.0800e-01 | 3.0711e+05 | ; | CE2 - HE2 |
| 2017 | 2019 | 1 | 1.4000e-01 | 3.9246e+05 | ; | CE2 - CD2 |
| 2019 | 2020 | 1 | 1.0800e-01 | 3.0711e+05 | ; | CD2 - HD2 |
| 2021 | 2022 | 1 | 1.2290e-01 | 4.7698e+05 | ; | C - O |
| 2021 | 2023 | 1 | 1.3350e-01 | 4.1003e+05 | ; | C - N |
| 2023 | 2024 | 1 | 1.0100e-01 | 3.6317e+05 | ; | N - H |
| 2023 | 2025 | 1 | 1.4490e-01 | 2.8200e+05 | ; | N - CA |
| 2025 | 2026 | 1 | 1.0900e-01 | 2.8451e+05 | ; | CA - HA |
| 2025 | 2027 | 1 | 1.5260e-01 | 2.5941e+05 | ; | CA - CB |
| 2025 | 2042 | 1 | 1.5220e-01 | 2.6527e+05 | ; | CA - C |
| 2027 | 2028 | 1 | 1.0900e-01 | 2.8451e+05 | ; | CB - HB2 |
| 2027 | 2029 | 1 | 1.0900e-01 | 2.8451e+05 | ; | CB - HB3 |
| 2027 | 2030 | 1 | 1.5100e-01 | 2.6527e+05 | ; | CB - CG |
| 2030 | 2031 | 1 | 1.4000e-01 | 3.9246e+05 | ; | CG - CD1 |
| 2030 | 2040 | 1 | 1.4000e-01 | 3.9246e+05 | ; | CG - CD2 |
| 2031 | 2032 | 1 | 1.0800e-01 | 3.0711e+05 | ; | CD1 - HD1 |
| 2031 | 2033 | 1 | 1.4000e-01 | 3.9246e+05 | ; | CD1 - CE1 |
| 2033 | 2034 | 1 | 1.0800e-01 | 3.0711e+05 | ; | CE1 - HE1 |
| 2033 | 2035 | 1 | 1.4090e-01 | 3.9246e+05 | ; | CE1 - CZ |
| 2035 | 2036 | 1 | 1.3640e-01 | 3.7656e+05 | ; | CZ - OH |
| 2035 | 2038 | 1 | 1.4090e-01 | 3.9246e+05 | ; | CZ - CE2 |
| 2036 | 2037 | 1 | 9.6000e-02 | 4.6275e+05 | ; | OH - HH |
| 2038 | 2039 | 1 | 1.0800e-01 | 3.0711e+05 | ; | CE2 - HE2 |
| 2038 | 2040 | 1 | 1.4000e-01 | 3.9246e+05 | ; | CE2 - CD2 |
| 2040 | 2041 | 1 | 1.0800e-01 | 3.0711e+05 | ; | CD2 - HD2 |
| 2042 | 2043 | 1 | 1.2290e-01 | 4.7698e+05 | ; | C - O |
| 2042 | 2044 | 1 | 1.3350e-01 | 4.1003e+05 | ; | C - N |
| 2044 | 2045 | 1 | 1.0100e-01 | 3.6317e+05 | ; | N - H |
| 2044 | 2046 | 1 | 1.4490e-01 | 2.8200e+05 | ; | N - CA |
| 2046 | 2047 | 1 | 1.0900e-01 | 2.8451e+05 | ; | CA - HA |
| 2046 | 2048 | 1 | 1.5260e-01 | 2.5941e+05 | ; | CA - CB |
| 2046 | 2061 | 1 | 1.5220e-01 | 2.6527e+05 | ; | CA - C |
| 2048 | 2049 | 1 | 1.0900e-01 | 2.8451e+05 | ; | CB - HB2 |
| 2048 | 2050 | 1 | 1.0900e-01 | 2.8451e+05 | ; | CB - HB3 |
| 2048 | 2051 | 1 | 1.5260e-01 | 2.5941e+05 | ; | CB - CG |
| 2051 | 2052 | 1 | 1.0900e-01 | 2.8451e+05 | ; | CG - HG |
| 2051 | 2053 | 1 | 1.5260e-01 | 2.5941e+05 | ; | CG - CD1 |
| 2051 | 2057 | 1 | 1.5260e-01 | 2.5941e+05 | ; | CG - CD2 |
| 2053 | 2054 | 1 | 1.0900e-01 | 2.8451e+05 | ; | CD1 - HD11 |
| 2053 | 2055 | 1 | 1.0900e-01 | 2.8451e+05 | ; | CD1 - HD12 |
| 2053 | 2056 | 1 | 1.0900e-01 | 2.8451e+05 | ; | CD1 - HD13 |
| 2057 | 2058 | 1 | 1.0900e-01 | 2.8451e+05 | ; | CD2 - HD21 |
| 2057 | 2059 | 1 | 1.0900e-01 | 2.8451e+05 | ; | CD2 - HD22 |

| | | | | | |
|---|---|---|---|---|---|
| 2057 | 2060 | 1 | 1.0900e-01 | 2.8451e+05 ; | CD2 - HD23 |
| 2061 | 2062 | 1 | 1.2290e-01 | 4.7698e+05 ; | C - O |
| 2061 | 2063 | 1 | 1.3350e-01 | 4.1003e+05 ; | C - N |
| 2063 | 2064 | 1 | 1.0100e-01 | 3.6317e+05 ; | N - H |
| 2063 | 2065 | 1 | 1.4490e-01 | 2.8200e+05 ; | N - CA |
| 2065 | 2066 | 1 | 1.0900e-01 | 2.8451e+05 ; | CA - HA |
| 2065 | 2067 | 1 | 1.5260e-01 | 2.5941e+05 ; | CA - CB |
| 2065 | 2079 | 1 | 1.5220e-01 | 2.6527e+05 ; | CA - C |
| 2067 | 2068 | 1 | 1.0900e-01 | 2.8451e+05 ; | CB - HB2 |
| 2067 | 2069 | 1 | 1.0900e-01 | 2.8451e+05 ; | CB - HB3 |
| 2067 | 2070 | 1 | 1.5040e-01 | 2.6527e+05 ; | CB - CG |
| 2070 | 2071 | 1 | 1.3850e-01 | 3.5313e+05 ; | CG - ND1 |
| 2070 | 2077 | 1 | 1.3710e-01 | 4.3346e+05 ; | CG - CD2 |
| 2071 | 2072 | 1 | 1.0100e-01 | 3.6317e+05 ; | ND1 - HD1 |
| 2071 | 2073 | 1 | 1.3430e-01 | 3.9915e+05 ; | ND1 - CE1 |
| 2073 | 2074 | 1 | 1.0800e-01 | 3.0711e+05 ; | CE1 - HE1 |
| 2073 | 2075 | 1 | 1.3430e-01 | 3.9915e+05 ; | CE1 - NE2 |
| 2075 | 2076 | 1 | 1.0100e-01 | 3.6317e+05 ; | NE2 - HE2 |
| 2075 | 2077 | 1 | 1.3810e-01 | 3.5731e+05 ; | NE2 - CD2 |
| 2077 | 2078 | 1 | 1.0800e-01 | 3.0711e+05 ; | CD2 - HD2 |
| 2079 | 2080 | 1 | 1.2290e-01 | 4.7698e+05 ; | C - O |
| 2079 | 2081 | 1 | 1.3350e-01 | 4.1003e+05 ; | C - N |
| 2081 | 2082 | 1 | 1.0100e-01 | 3.6317e+05 ; | N - H |
| 2081 | 2083 | 1 | 1.4490e-01 | 2.8200e+05 ; | N - CA |
| 2083 | 2084 | 1 | 1.0900e-01 | 2.8451e+05 ; | CA - HA |
| 2083 | 2085 | 1 | 1.5260e-01 | 2.5941e+05 ; | CA - CB |
| 2083 | 2090 | 1 | 1.5220e-01 | 2.6527e+05 ; | CA - C |
| 2085 | 2086 | 1 | 1.0900e-01 | 2.8451e+05 ; | CB - HB2 |
| 2085 | 2087 | 1 | 1.0900e-01 | 2.8451e+05 ; | CB - HB3 |
| 2085 | 2088 | 1 | 1.4100e-01 | 2.6778e+05 ; | CB - OG |
| 2088 | 2089 | 1 | 9.6000e-02 | 4.6275e+05 ; | OG - HG |
| 2090 | 2091 | 1 | 1.2290e-01 | 4.7698e+05 ; | C - O |
| 2090 | 2092 | 1 | 1.3350e-01 | 4.1003e+05 ; | C - N |
| 2092 | 2093 | 1 | 1.0100e-01 | 3.6317e+05 ; | N - H |
| 2092 | 2094 | 1 | 1.4490e-01 | 2.8200e+05 ; | N - CA |
| 2094 | 2095 | 1 | 1.0900e-01 | 2.8451e+05 ; | CA - HA |
| 2094 | 2096 | 1 | 1.5260e-01 | 2.5941e+05 ; | CA - CB |
| 2094 | 2109 | 1 | 1.5220e-01 | 2.6527e+05 ; | CA - C |
| 2096 | 2097 | 1 | 1.0900e-01 | 2.8451e+05 ; | CB - HB2 |
| 2096 | 2098 | 1 | 1.0900e-01 | 2.8451e+05 ; | CB - HB3 |
| 2096 | 2099 | 1 | 1.5260e-01 | 2.5941e+05 ; | CB - CG |
| 2099 | 2100 | 1 | 1.0900e-01 | 2.8451e+05 ; | CG - HG |
| 2099 | 2101 | 1 | 1.5260e-01 | 2.5941e+05 ; | CG - CD1 |
| 2099 | 2105 | 1 | 1.5260e-01 | 2.5941e+05 ; | CG - CD2 |
| 2101 | 2102 | 1 | 1.0900e-01 | 2.8451e+05 ; | CD1 - HD11 |
| 2101 | 2103 | 1 | 1.0900e-01 | 2.8451e+05 ; | CD1 - HD12 |
| 2101 | 2104 | 1 | 1.0900e-01 | 2.8451e+05 ; | CD1 - HD13 |
| 2105 | 2106 | 1 | 1.0900e-01 | 2.8451e+05 ; | CD2 - HD21 |
| 2105 | 2107 | 1 | 1.0900e-01 | 2.8451e+05 ; | CD2 - HD22 |
| 2105 | 2108 | 1 | 1.0900e-01 | 2.8451e+05 ; | CD2 - HD23 |
| 2109 | 2110 | 1 | 1.2290e-01 | 4.7698e+05 ; | C - O |
| 2109 | 2111 | 1 | 1.3350e-01 | 4.1003e+05 ; | C - N |

| | | | | | |
|---|---|---|---|---|---|
| 2111 | 2112 | 1 | 1.0100e-01 | 3.6317e+05 ; | N - H |
| 2111 | 2113 | 1 | 1.4490e-01 | 2.8200e+05 ; | N - CA |
| 2113 | 2114 | 1 | 1.0900e-01 | 2.8451e+05 ; | CA - HA |
| 2113 | 2115 | 1 | 1.5260e-01 | 2.5941e+05 ; | CA - CB |
| 2113 | 2123 | 1 | 1.5220e-01 | 2.6527e+05 ; | CA - C |
| 2115 | 2116 | 1 | 1.0900e-01 | 2.8451e+05 ; | CB - HB2 |
| 2115 | 2117 | 1 | 1.0900e-01 | 2.8451e+05 ; | CB - HB3 |
| 2115 | 2118 | 1 | 1.5220e-01 | 2.6527e+05 ; | CB - CG |
| 2118 | 2119 | 1 | 1.2290e-01 | 4.7698e+05 ; | CG - OD1 |
| 2118 | 2120 | 1 | 1.3350e-01 | 4.1003e+05 ; | CG - ND2 |
| 2120 | 2121 | 1 | 1.0100e-01 | 3.6317e+05 ; | ND2 - HD21 |
| 2120 | 2122 | 1 | 1.0100e-01 | 3.6317e+05 ; | ND2 - HD22 |
| 2123 | 2124 | 1 | 1.2290e-01 | 4.7698e+05 ; | C - O |
| 2123 | 2125 | 1 | 1.3350e-01 | 4.1003e+05 ; | C - N |
| 2125 | 2126 | 1 | 1.0100e-01 | 3.6317e+05 ; | N - H |
| 2125 | 2127 | 1 | 1.4490e-01 | 2.8200e+05 ; | N - CA |
| 2127 | 2128 | 1 | 1.0900e-01 | 2.8451e+05 ; | CA - HA |
| 2127 | 2129 | 1 | 1.5260e-01 | 2.5941e+05 ; | CA - CB |
| 2127 | 2142 | 1 | 1.5220e-01 | 2.6527e+05 ; | CA - C |
| 2129 | 2130 | 1 | 1.0900e-01 | 2.8451e+05 ; | CB - HB |
| 2129 | 2131 | 1 | 1.5260e-01 | 2.5941e+05 ; | CB - CG2 |
| 2129 | 2135 | 1 | 1.5260e-01 | 2.5941e+05 ; | CB - CG1 |
| 2131 | 2132 | 1 | 1.0900e-01 | 2.8451e+05 ; | CG2 - HG21 |
| 2131 | 2133 | 1 | 1.0900e-01 | 2.8451e+05 ; | CG2 - HG22 |
| 2131 | 2134 | 1 | 1.0900e-01 | 2.8451e+05 ; | CG2 - HG23 |
| 2135 | 2136 | 1 | 1.0900e-01 | 2.8451e+05 ; | CG1 - HG12 |
| 2135 | 2137 | 1 | 1.0900e-01 | 2.8451e+05 ; | CG1 - HG13 |
| 2135 | 2138 | 1 | 1.5260e-01 | 2.5941e+05 ; | CG1 - CD1 |
| 2138 | 2139 | 1 | 1.0900e-01 | 2.8451e+05 ; | CD1 - HD11 |
| 2138 | 2140 | 1 | 1.0900e-01 | 2.8451e+05 ; | CD1 - HD12 |
| 2138 | 2141 | 1 | 1.0900e-01 | 2.8451e+05 ; | CD1 - HD13 |
| 2142 | 2143 | 1 | 1.2290e-01 | 4.7698e+05 ; | C - O |
| 2142 | 2144 | 1 | 1.3350e-01 | 4.1003e+05 ; | C - N |
| 2144 | 2145 | 1 | 1.0100e-01 | 3.6317e+05 ; | N - H |
| 2144 | 2146 | 1 | 1.4490e-01 | 2.8200e+05 ; | N - CA |
| 2146 | 2147 | 1 | 1.0900e-01 | 2.8451e+05 ; | CA - HA |
| 2146 | 2148 | 1 | 1.5260e-01 | 2.5941e+05 ; | CA - CB |
| 2146 | 2158 | 1 | 1.5220e-01 | 2.6527e+05 ; | CA - C |
| 2148 | 2149 | 1 | 1.0900e-01 | 2.8451e+05 ; | CB - HB |
| 2148 | 2150 | 1 | 1.5260e-01 | 2.5941e+05 ; | CB - CG1 |
| 2148 | 2154 | 1 | 1.5260e-01 | 2.5941e+05 ; | CB - CG2 |
| 2150 | 2151 | 1 | 1.0900e-01 | 2.8451e+05 ; | CG1 - HG11 |
| 2150 | 2152 | 1 | 1.0900e-01 | 2.8451e+05 ; | CG1 - HG12 |
| 2150 | 2153 | 1 | 1.0900e-01 | 2.8451e+05 ; | CG1 - HG13 |
| 2154 | 2155 | 1 | 1.0900e-01 | 2.8451e+05 ; | CG2 - HG21 |
| 2154 | 2156 | 1 | 1.0900e-01 | 2.8451e+05 ; | CG2 - HG22 |
| 2154 | 2157 | 1 | 1.0900e-01 | 2.8451e+05 ; | CG2 - HG23 |
| 2158 | 2159 | 1 | 1.2290e-01 | 4.7698e+05 ; | C - O |
| 2158 | 2160 | 1 | 1.3350e-01 | 4.1003e+05 ; | C - N |
| 2160 | 2161 | 1 | 1.0100e-01 | 3.6317e+05 ; | N - H |
| 2160 | 2162 | 1 | 1.4490e-01 | 2.8200e+05 ; | N - CA |
| 2162 | 2163 | 1 | 1.0900e-01 | 2.8451e+05 ; | CA - HA |

| | | | | | |
|---|---|---|---|---|---|
| 2162 | 2164 | 1 | 1.5260e-01 | 2.5941e+05 ; | CA - CB |
| 2162 | 2175 | 1 | 1.5220e-01 | 2.6527e+05 ; | CA - C |
| 2164 | 2165 | 1 | 1.0900e-01 | 2.8451e+05 ; | CB - HB2 |
| 2164 | 2166 | 1 | 1.0900e-01 | 2.8451e+05 ; | CB - HB3 |
| 2164 | 2167 | 1 | 1.5040e-01 | 2.6527e+05 ; | CB - CG |
| 2167 | 2168 | 1 | 1.3940e-01 | 3.4309e+05 ; | CG - ND1 |
| 2167 | 2173 | 1 | 1.3710e-01 | 4.3346e+05 ; | CG - CD2 |
| 2168 | 2169 | 1 | 1.3350e-01 | 4.0836e+05 ; | ND1 - CE1 |
| 2169 | 2170 | 1 | 1.0800e-01 | 3.0711e+05 ; | CE1 - HE1 |
| 2169 | 2171 | 1 | 1.3430e-01 | 3.9915e+05 ; | CE1 - NE2 |
| 2171 | 2172 | 1 | 1.0100e-01 | 3.6317e+05 ; | NE2 - HE2 |
| 2171 | 2173 | 1 | 1.3810e-01 | 3.5731e+05 ; | NE2 - CD2 |
| 2173 | 2174 | 1 | 1.0800e-01 | 3.0711e+05 ; | CD2 - HD2 |
| 2175 | 2176 | 1 | 1.2290e-01 | 4.7698e+05 ; | C - O |
| 2175 | 2177 | 1 | 1.3350e-01 | 4.1003e+05 ; | C - N |
| 2177 | 2178 | 1 | 1.0100e-01 | 3.6317e+05 ; | N - H |
| 2177 | 2179 | 1 | 1.4490e-01 | 2.8200e+05 ; | N - CA |
| 2179 | 2180 | 1 | 1.0900e-01 | 2.8451e+05 ; | CA - HA |
| 2179 | 2181 | 1 | 1.5260e-01 | 2.5941e+05 ; | CA - CB |
| 2179 | 2199 | 1 | 1.5220e-01 | 2.6527e+05 ; | CA - C |
| 2181 | 2182 | 1 | 1.0900e-01 | 2.8451e+05 ; | CB - HB2 |
| 2181 | 2183 | 1 | 1.0900e-01 | 2.8451e+05 ; | CB - HB3 |
| 2181 | 2184 | 1 | 1.5260e-01 | 2.5941e+05 ; | CB - CG |
| 2184 | 2185 | 1 | 1.0900e-01 | 2.8451e+05 ; | CG - HG2 |
| 2184 | 2186 | 1 | 1.0900e-01 | 2.8451e+05 ; | CG - HG3 |
| 2184 | 2187 | 1 | 1.5260e-01 | 2.5941e+05 ; | CG - CD |
| 2187 | 2188 | 1 | 1.0900e-01 | 2.8451e+05 ; | CD - HD2 |
| 2187 | 2189 | 1 | 1.0900e-01 | 2.8451e+05 ; | CD - HD3 |
| 2187 | 2190 | 1 | 1.4630e-01 | 2.8200e+05 ; | CD - NE |
| 2190 | 2191 | 1 | 1.0100e-01 | 3.6317e+05 ; | NE - HE |
| 2190 | 2192 | 1 | 1.3400e-01 | 4.0250e+05 ; | NE - CZ |
| 2192 | 2193 | 1 | 1.3400e-01 | 4.0250e+05 ; | CZ - NH1 |
| 2192 | 2196 | 1 | 1.3400e-01 | 4.0250e+05 ; | CZ - NH2 |
| 2193 | 2194 | 1 | 1.0100e-01 | 3.6317e+05 ; | NH1 - HH11 |
| 2193 | 2195 | 1 | 1.0100e-01 | 3.6317e+05 ; | NH1 - HH12 |
| 2196 | 2197 | 1 | 1.0100e-01 | 3.6317e+05 ; | NH2 - HH21 |
| 2196 | 2198 | 1 | 1.0100e-01 | 3.6317e+05 ; | NH2 - HH22 |
| 2199 | 2200 | 1 | 1.2290e-01 | 4.7698e+05 ; | C - O |
| 2199 | 2201 | 1 | 1.3350e-01 | 4.1003e+05 ; | C - N |
| 2201 | 2202 | 1 | 1.0100e-01 | 3.6317e+05 ; | N - H |
| 2201 | 2203 | 1 | 1.4490e-01 | 2.8200e+05 ; | N - CA |
| 2203 | 2204 | 1 | 1.0900e-01 | 2.8451e+05 ; | CA - HA |
| 2203 | 2205 | 1 | 1.5260e-01 | 2.5941e+05 ; | CA - CB |
| 2203 | 2211 | 1 | 1.5220e-01 | 2.6527e+05 ; | CA - C |
| 2205 | 2206 | 1 | 1.0900e-01 | 2.8451e+05 ; | CB - HB2 |
| 2205 | 2207 | 1 | 1.0900e-01 | 2.8451e+05 ; | CB - HB3 |
| 2205 | 2208 | 1 | 1.5220e-01 | 2.6527e+05 ; | CB - CG |
| 2208 | 2209 | 1 | 1.2500e-01 | 5.4894e+05 ; | CG - OD1 |
| 2208 | 2210 | 1 | 1.2500e-01 | 5.4894e+05 ; | CG - OD2 |
| 2211 | 2212 | 1 | 1.2290e-01 | 4.7698e+05 ; | C - O |
| 2211 | 2213 | 1 | 1.3350e-01 | 4.1003e+05 ; | C - N |
| 2213 | 2214 | 1 | 1.0100e-01 | 3.6317e+05 ; | N - H |

| | | | | | | |
|---|---|---|---|---|---|---|
| 2213 | 2215 | 1 | 1.4490e-01 | 2.8200e+05 | ; | N - CA |
| 2215 | 2216 | 1 | 1.0900e-01 | 2.8451e+05 | ; | CA - HA |
| 2215 | 2217 | 1 | 1.5260e-01 | 2.5941e+05 | ; | CA - CB |
| 2215 | 2230 | 1 | 1.5220e-01 | 2.6527e+05 | ; | CA - C |
| 2217 | 2218 | 1 | 1.0900e-01 | 2.8451e+05 | ; | CB - HB2 |
| 2217 | 2219 | 1 | 1.0900e-01 | 2.8451e+05 | ; | CB - HB3 |
| 2217 | 2220 | 1 | 1.5260e-01 | 2.5941e+05 | ; | CB - CG |
| 2220 | 2221 | 1 | 1.0900e-01 | 2.8451e+05 | ; | CG - HG |
| 2220 | 2222 | 1 | 1.5260e-01 | 2.5941e+05 | ; | CG - CD1 |
| 2220 | 2226 | 1 | 1.5260e-01 | 2.5941e+05 | ; | CG - CD2 |
| 2222 | 2223 | 1 | 1.0900e-01 | 2.8451e+05 | ; | CD1 - HD11 |
| 2222 | 2224 | 1 | 1.0900e-01 | 2.8451e+05 | ; | CD1 - HD12 |
| 2222 | 2225 | 1 | 1.0900e-01 | 2.8451e+05 | ; | CD1 - HD13 |
| 2226 | 2227 | 1 | 1.0900e-01 | 2.8451e+05 | ; | CD2 - HD21 |
| 2226 | 2228 | 1 | 1.0900e-01 | 2.8451e+05 | ; | CD2 - HD22 |
| 2226 | 2229 | 1 | 1.0900e-01 | 2.8451e+05 | ; | CD2 - HD23 |
| 2230 | 2231 | 1 | 1.2290e-01 | 4.7698e+05 | ; | C - O |
| 2230 | 2232 | 1 | 1.3350e-01 | 4.1003e+05 | ; | C - N |
| 2232 | 2233 | 1 | 1.0100e-01 | 3.6317e+05 | ; | N - H |
| 2232 | 2234 | 1 | 1.4490e-01 | 2.8200e+05 | ; | N - CA |
| 2234 | 2235 | 1 | 1.0900e-01 | 2.8451e+05 | ; | CA - HA |
| 2234 | 2236 | 1 | 1.5260e-01 | 2.5941e+05 | ; | CA - CB |
| 2234 | 2252 | 1 | 1.5220e-01 | 2.6527e+05 | ; | CA - C |
| 2236 | 2237 | 1 | 1.0900e-01 | 2.8451e+05 | ; | CB - HB2 |
| 2236 | 2238 | 1 | 1.0900e-01 | 2.8451e+05 | ; | CB - HB3 |
| 2236 | 2239 | 1 | 1.5260e-01 | 2.5941e+05 | ; | CB - CG |
| 2239 | 2240 | 1 | 1.0900e-01 | 2.8451e+05 | ; | CG - HG2 |
| 2239 | 2241 | 1 | 1.0900e-01 | 2.8451e+05 | ; | CG - HG3 |
| 2239 | 2242 | 1 | 1.5260e-01 | 2.5941e+05 | ; | CG - CD |
| 2242 | 2243 | 1 | 1.0900e-01 | 2.8451e+05 | ; | CD - HD2 |
| 2242 | 2244 | 1 | 1.0900e-01 | 2.8451e+05 | ; | CD - HD3 |
| 2242 | 2245 | 1 | 1.5260e-01 | 2.5941e+05 | ; | CD - CE |
| 2245 | 2246 | 1 | 1.0900e-01 | 2.8451e+05 | ; | CE - HE2 |
| 2245 | 2247 | 1 | 1.0900e-01 | 2.8451e+05 | ; | CE - HE3 |
| 2245 | 2248 | 1 | 1.4710e-01 | 3.0711e+05 | ; | CE - NZ |
| 2248 | 2249 | 1 | 1.0100e-01 | 3.6317e+05 | ; | NZ - HZ1 |
| 2248 | 2250 | 1 | 1.0100e-01 | 3.6317e+05 | ; | NZ - HZ2 |
| 2248 | 2251 | 1 | 1.0100e-01 | 3.6317e+05 | ; | NZ - HZ3 |
| 2252 | 2253 | 1 | 1.2290e-01 | 4.7698e+05 | ; | C - O |
| 2252 | 2254 | 1 | 1.3350e-01 | 4.1003e+05 | ; | C - N |
| 2254 | 2255 | 1 | 1.4490e-01 | 2.8200e+05 | ; | N - CD |
| 2254 | 2264 | 1 | 1.4490e-01 | 2.8200e+05 | ; | N - CA |
| 2255 | 2256 | 1 | 1.0900e-01 | 2.8451e+05 | ; | CD - HD2 |
| 2255 | 2257 | 1 | 1.0900e-01 | 2.8451e+05 | ; | CD - HD3 |
| 2255 | 2258 | 1 | 1.5260e-01 | 2.5941e+05 | ; | CD - CG |
| 2258 | 2259 | 1 | 1.0900e-01 | 2.8451e+05 | ; | CG - HG2 |
| 2258 | 2260 | 1 | 1.0900e-01 | 2.8451e+05 | ; | CG - HG3 |
| 2258 | 2261 | 1 | 1.5260e-01 | 2.5941e+05 | ; | CG - CB |
| 2261 | 2262 | 1 | 1.0900e-01 | 2.8451e+05 | ; | CB - HB2 |
| 2261 | 2263 | 1 | 1.0900e-01 | 2.8451e+05 | ; | CB - HB3 |
| 2261 | 2264 | 1 | 1.5260e-01 | 2.5941e+05 | ; | CB - CA |
| 2264 | 2265 | 1 | 1.0900e-01 | 2.8451e+05 | ; | CA - HA |

| | | | | | | |
|---|---|---|---|---|---|---|
| 2264 | 2266 | 1 | 1.5220e-01 | 2.6527e+05 | ; | CA - C |
| 2266 | 2267 | 1 | 1.2290e-01 | 4.7698e+05 | ; | C - O |
| 2266 | 2268 | 1 | 1.3350e-01 | 4.1003e+05 | ; | C - N |
| 2268 | 2269 | 1 | 1.0100e-01 | 3.6317e+05 | ; | N - H |
| 2268 | 2270 | 1 | 1.4490e-01 | 2.8200e+05 | ; | N - CA |
| 2270 | 2271 | 1 | 1.0900e-01 | 2.8451e+05 | ; | CA - HA |
| 2270 | 2272 | 1 | 1.5260e-01 | 2.5941e+05 | ; | CA - CB |
| 2270 | 2281 | 1 | 1.5220e-01 | 2.6527e+05 | ; | CA - C |
| 2272 | 2273 | 1 | 1.0900e-01 | 2.8451e+05 | ; | CB - HB2 |
| 2272 | 2274 | 1 | 1.0900e-01 | 2.8451e+05 | ; | CB - HB3 |
| 2272 | 2275 | 1 | 1.5260e-01 | 2.5941e+05 | ; | CB - CG |
| 2275 | 2276 | 1 | 1.0900e-01 | 2.8451e+05 | ; | CG - HG2 |
| 2275 | 2277 | 1 | 1.0900e-01 | 2.8451e+05 | ; | CG - HG3 |
| 2275 | 2278 | 1 | 1.5220e-01 | 2.6527e+05 | ; | CG - CD |
| 2278 | 2279 | 1 | 1.2500e-01 | 5.4894e+05 | ; | CD - OE1 |
| 2278 | 2280 | 1 | 1.2500e-01 | 5.4894e+05 | ; | CD - OE2 |
| 2281 | 2282 | 1 | 1.2290e-01 | 4.7698e+05 | ; | C - O |
| 2281 | 2283 | 1 | 1.3350e-01 | 4.1003e+05 | ; | C - N |
| 2283 | 2284 | 1 | 1.0100e-01 | 3.6317e+05 | ; | N - H |
| 2283 | 2285 | 1 | 1.4490e-01 | 2.8200e+05 | ; | N - CA |
| 2285 | 2286 | 1 | 1.0900e-01 | 2.8451e+05 | ; | CA - HA |
| 2285 | 2287 | 1 | 1.5260e-01 | 2.5941e+05 | ; | CA - CB |
| 2285 | 2295 | 1 | 1.5220e-01 | 2.6527e+05 | ; | CA - C |
| 2287 | 2288 | 1 | 1.0900e-01 | 2.8451e+05 | ; | CB - HB2 |
| 2287 | 2289 | 1 | 1.0900e-01 | 2.8451e+05 | ; | CB - HB3 |
| 2287 | 2290 | 1 | 1.5220e-01 | 2.6527e+05 | ; | CB - CG |
| 2290 | 2291 | 1 | 1.2290e-01 | 4.7698e+05 | ; | CG - OD1 |
| 2290 | 2292 | 1 | 1.3350e-01 | 4.1003e+05 | ; | CG - ND2 |
| 2292 | 2293 | 1 | 1.0100e-01 | 3.6317e+05 | ; | ND2 - HD21 |
| 2292 | 2294 | 1 | 1.0100e-01 | 3.6317e+05 | ; | ND2 - HD22 |
| 2295 | 2296 | 1 | 1.2290e-01 | 4.7698e+05 | ; | C - O |
| 2295 | 2297 | 1 | 1.3350e-01 | 4.1003e+05 | ; | C - N |
| 2297 | 2298 | 1 | 1.0100e-01 | 3.6317e+05 | ; | N - H |
| 2297 | 2299 | 1 | 1.4490e-01 | 2.8200e+05 | ; | N - CA |
| 2299 | 2300 | 1 | 1.0900e-01 | 2.8451e+05 | ; | CA - HA |
| 2299 | 2301 | 1 | 1.5260e-01 | 2.5941e+05 | ; | CA - CB |
| 2299 | 2315 | 1 | 1.5220e-01 | 2.6527e+05 | ; | CA - C |
| 2301 | 2302 | 1 | 1.0900e-01 | 2.8451e+05 | ; | CB - HB2 |
| 2301 | 2303 | 1 | 1.0900e-01 | 2.8451e+05 | ; | CB - HB3 |
| 2301 | 2304 | 1 | 1.5100e-01 | 2.6527e+05 | ; | CB - CG |
| 2304 | 2305 | 1 | 1.4000e-01 | 3.9246e+05 | ; | CG - CD1 |
| 2304 | 2313 | 1 | 1.4000e-01 | 3.9246e+05 | ; | CG - CD2 |
| 2305 | 2306 | 1 | 1.0800e-01 | 3.0711e+05 | ; | CD1 - HD1 |
| 2305 | 2307 | 1 | 1.4000e-01 | 3.9246e+05 | ; | CD1 - CE1 |
| 2307 | 2308 | 1 | 1.0800e-01 | 3.0711e+05 | ; | CE1 - HE1 |
| 2307 | 2309 | 1 | 1.4000e-01 | 3.9246e+05 | ; | CE1 - CZ |
| 2309 | 2310 | 1 | 1.0800e-01 | 3.0711e+05 | ; | CZ - HZ |
| 2309 | 2311 | 1 | 1.4000e-01 | 3.9246e+05 | ; | CZ - CE2 |
| 2311 | 2312 | 1 | 1.0800e-01 | 3.0711e+05 | ; | CE2 - HE2 |
| 2311 | 2313 | 1 | 1.4000e-01 | 3.9246e+05 | ; | CE2 - CD2 |
| 2313 | 2314 | 1 | 1.0800e-01 | 3.0711e+05 | ; | CD2 - HD2 |
| 2315 | 2316 | 1 | 1.2290e-01 | 4.7698e+05 | ; | C - O |

| | | | | | |
|---|---|---|---|---|---|
| 2315 | 2317 | 1 | 1.3350e-01 | 4.1003e+05 ; | C - N |
| 2317 | 2318 | 1 | 1.0100e-01 | 3.6317e+05 ; | N - H |
| 2317 | 2319 | 1 | 1.4490e-01 | 2.8200e+05 ; | N - CA |
| 2319 | 2320 | 1 | 1.0900e-01 | 2.8451e+05 ; | CA - HA |
| 2319 | 2321 | 1 | 1.5260e-01 | 2.5941e+05 ; | CA - CB |
| 2319 | 2334 | 1 | 1.5220e-01 | 2.6527e+05 ; | CA - C |
| 2321 | 2322 | 1 | 1.0900e-01 | 2.8451e+05 ; | CB - HB2 |
| 2321 | 2323 | 1 | 1.0900e-01 | 2.8451e+05 ; | CB - HB3 |
| 2321 | 2324 | 1 | 1.5260e-01 | 2.5941e+05 ; | CB - CG |
| 2324 | 2325 | 1 | 1.0900e-01 | 2.8451e+05 ; | CG - HG |
| 2324 | 2326 | 1 | 1.5260e-01 | 2.5941e+05 ; | CG - CD1 |
| 2324 | 2330 | 1 | 1.5260e-01 | 2.5941e+05 ; | CG - CD2 |
| 2326 | 2327 | 1 | 1.0900e-01 | 2.8451e+05 ; | CD1 - HD11 |
| 2326 | 2328 | 1 | 1.0900e-01 | 2.8451e+05 ; | CD1 - HD12 |
| 2326 | 2329 | 1 | 1.0900e-01 | 2.8451e+05 ; | CD1 - HD13 |
| 2330 | 2331 | 1 | 1.0900e-01 | 2.8451e+05 ; | CD2 - HD21 |
| 2330 | 2332 | 1 | 1.0900e-01 | 2.8451e+05 ; | CD2 - HD22 |
| 2330 | 2333 | 1 | 1.0900e-01 | 2.8451e+05 ; | CD2 - HD23 |
| 2334 | 2335 | 1 | 1.2290e-01 | 4.7698e+05 ; | C - O |
| 2334 | 2336 | 1 | 1.3350e-01 | 4.1003e+05 ; | C - N |
| 2336 | 2337 | 1 | 1.0100e-01 | 3.6317e+05 ; | N - H |
| 2336 | 2338 | 1 | 1.4490e-01 | 2.8200e+05 ; | N - CA |
| 2338 | 2339 | 1 | 1.0900e-01 | 2.8451e+05 ; | CA - HA |
| 2338 | 2340 | 1 | 1.5260e-01 | 2.5941e+05 ; | CA - CB |
| 2338 | 2354 | 1 | 1.5220e-01 | 2.6527e+05 ; | CA - C |
| 2340 | 2341 | 1 | 1.0900e-01 | 2.8451e+05 ; | CB - HB2 |
| 2340 | 2342 | 1 | 1.0900e-01 | 2.8451e+05 ; | CB - HB3 |
| 2340 | 2343 | 1 | 1.5100e-01 | 2.6527e+05 ; | CB - CG |
| 2343 | 2344 | 1 | 1.4000e-01 | 3.9246e+05 ; | CG - CD1 |
| 2343 | 2352 | 1 | 1.4000e-01 | 3.9246e+05 ; | CG - CD2 |
| 2344 | 2345 | 1 | 1.0800e-01 | 3.0711e+05 ; | CD1 - HD1 |
| 2344 | 2346 | 1 | 1.4000e-01 | 3.9246e+05 ; | CD1 - CE1 |
| 2346 | 2347 | 1 | 1.0800e-01 | 3.0711e+05 ; | CE1 - HE1 |
| 2346 | 2348 | 1 | 1.4000e-01 | 3.9246e+05 ; | CE1 - CZ |
| 2348 | 2349 | 1 | 1.0800e-01 | 3.0711e+05 ; | CZ - HZ |
| 2348 | 2350 | 1 | 1.4000e-01 | 3.9246e+05 ; | CZ - CE2 |
| 2350 | 2351 | 1 | 1.0800e-01 | 3.0711e+05 ; | CE2 - HE2 |
| 2350 | 2352 | 1 | 1.4000e-01 | 3.9246e+05 ; | CE2 - CD2 |
| 2352 | 2353 | 1 | 1.0800e-01 | 3.0711e+05 ; | CD2 - HD2 |
| 2354 | 2355 | 1 | 1.2290e-01 | 4.7698e+05 ; | C - O |
| 2354 | 2356 | 1 | 1.3350e-01 | 4.1003e+05 ; | C - N |
| 2356 | 2357 | 1 | 1.0100e-01 | 3.6317e+05 ; | N - H |
| 2356 | 2358 | 1 | 1.4490e-01 | 2.8200e+05 ; | N - CA |
| 2358 | 2359 | 1 | 1.0900e-01 | 2.8451e+05 ; | CA - HA |
| 2358 | 2360 | 1 | 1.5260e-01 | 2.5941e+05 ; | CA - CB |
| 2358 | 2371 | 1 | 1.5220e-01 | 2.6527e+05 ; | CA - C |
| 2360 | 2361 | 1 | 1.0900e-01 | 2.8451e+05 ; | CB - HB2 |
| 2360 | 2362 | 1 | 1.0900e-01 | 2.8451e+05 ; | CB - HB3 |
| 2360 | 2363 | 1 | 1.5260e-01 | 2.5941e+05 ; | CB - CG |
| 2363 | 2364 | 1 | 1.0900e-01 | 2.8451e+05 ; | CG - HG2 |
| 2363 | 2365 | 1 | 1.0900e-01 | 2.8451e+05 ; | CG - HG3 |
| 2363 | 2366 | 1 | 1.5220e-01 | 2.6527e+05 ; | CG - CD |

| | | | | | | |
|---|---|---|---|---|---|---|
| 2366 | 2367 | 1 | 1.2290e-01 | 4.7698e+05 | ; | CD - OE1 |
| 2366 | 2368 | 1 | 1.3350e-01 | 4.1003e+05 | ; | CD - NE2 |
| 2368 | 2369 | 1 | 1.0100e-01 | 3.6317e+05 | ; | NE2 - HE21 |
| 2368 | 2370 | 1 | 1.0100e-01 | 3.6317e+05 | ; | NE2 - HE22 |
| 2371 | 2372 | 1 | 1.2290e-01 | 4.7698e+05 | ; | C - O |
| 2371 | 2373 | 1 | 1.3350e-01 | 4.1003e+05 | ; | C - N |
| 2373 | 2374 | 1 | 1.0100e-01 | 3.6317e+05 | ; | N - H |
| 2373 | 2375 | 1 | 1.4490e-01 | 2.8200e+05 | ; | N - CA |
| 2375 | 2376 | 1 | 1.0900e-01 | 2.8451e+05 | ; | CA - HA |
| 2375 | 2377 | 1 | 1.5260e-01 | 2.5941e+05 | ; | CA - CB |
| 2375 | 2382 | 1 | 1.5220e-01 | 2.6527e+05 | ; | CA - C |
| 2377 | 2378 | 1 | 1.0900e-01 | 2.8451e+05 | ; | CB - HB2 |
| 2377 | 2379 | 1 | 1.0900e-01 | 2.8451e+05 | ; | CB - HB3 |
| 2377 | 2380 | 1 | 1.4100e-01 | 2.6778e+05 | ; | CB - OG |
| 2380 | 2381 | 1 | 9.6000e-02 | 4.6275e+05 | ; | OG - HG |
| 2382 | 2383 | 1 | 1.2290e-01 | 4.7698e+05 | ; | C - O |
| 2382 | 2384 | 1 | 1.3350e-01 | 4.1003e+05 | ; | C - N |
| 2384 | 2385 | 1 | 1.0100e-01 | 3.6317e+05 | ; | N - H |
| 2384 | 2386 | 1 | 1.4490e-01 | 2.8200e+05 | ; | N - CA |
| 2386 | 2387 | 1 | 1.0900e-01 | 2.8451e+05 | ; | CA - HA |
| 2386 | 2388 | 1 | 1.5260e-01 | 2.5941e+05 | ; | CA - CB |
| 2386 | 2397 | 1 | 1.5220e-01 | 2.6527e+05 | ; | CA - C |
| 2388 | 2389 | 1 | 1.0900e-01 | 2.8451e+05 | ; | CB - HB2 |
| 2388 | 2390 | 1 | 1.0900e-01 | 2.8451e+05 | ; | CB - HB3 |
| 2388 | 2391 | 1 | 1.5260e-01 | 2.5941e+05 | ; | CB - CG |
| 2391 | 2392 | 1 | 1.0900e-01 | 2.8451e+05 | ; | CG - HG2 |
| 2391 | 2393 | 1 | 1.0900e-01 | 2.8451e+05 | ; | CG - HG3 |
| 2391 | 2394 | 1 | 1.5220e-01 | 2.6527e+05 | ; | CG - CD |
| 2394 | 2395 | 1 | 1.2500e-01 | 5.4894e+05 | ; | CD - OE1 |
| 2394 | 2396 | 1 | 1.2500e-01 | 5.4894e+05 | ; | CD - OE2 |
| 2397 | 2398 | 1 | 1.2290e-01 | 4.7698e+05 | ; | C - O |
| 2397 | 2399 | 1 | 1.3350e-01 | 4.1003e+05 | ; | C - N |
| 2399 | 2400 | 1 | 1.0100e-01 | 3.6317e+05 | ; | N - H |
| 2399 | 2401 | 1 | 1.4490e-01 | 2.8200e+05 | ; | N - CA |
| 2401 | 2402 | 1 | 1.0900e-01 | 2.8451e+05 | ; | CA - HA |
| 2401 | 2403 | 1 | 1.5260e-01 | 2.5941e+05 | ; | CA - CB |
| 2401 | 2411 | 1 | 1.5220e-01 | 2.6527e+05 | ; | CA - C |
| 2403 | 2404 | 1 | 1.0900e-01 | 2.8451e+05 | ; | CB - HB2 |
| 2403 | 2405 | 1 | 1.0900e-01 | 2.8451e+05 | ; | CB - HB3 |
| 2403 | 2406 | 1 | 1.5220e-01 | 2.6527e+05 | ; | CB - CG |
| 2406 | 2407 | 1 | 1.2290e-01 | 4.7698e+05 | ; | CG - OD1 |
| 2406 | 2408 | 1 | 1.3350e-01 | 4.1003e+05 | ; | CG - ND2 |
| 2408 | 2409 | 1 | 1.0100e-01 | 3.6317e+05 | ; | ND2 - HD21 |
| 2408 | 2410 | 1 | 1.0100e-01 | 3.6317e+05 | ; | ND2 - HD22 |
| 2411 | 2412 | 1 | 1.2290e-01 | 4.7698e+05 | ; | C - O |
| 2411 | 2413 | 1 | 1.3350e-01 | 4.1003e+05 | ; | C - N |
| 2413 | 2414 | 1 | 1.0100e-01 | 3.6317e+05 | ; | N - H |
| 2413 | 2415 | 1 | 1.4490e-01 | 2.8200e+05 | ; | N - CA |
| 2415 | 2416 | 1 | 1.0900e-01 | 2.8451e+05 | ; | CA - HA |
| 2415 | 2417 | 1 | 1.5260e-01 | 2.5941e+05 | ; | CA - CB |
| 2415 | 2433 | 1 | 1.5220e-01 | 2.6527e+05 | ; | CA - C |
| 2417 | 2418 | 1 | 1.0900e-01 | 2.8451e+05 | ; | CB - HB2 |

| | | | | | |
|---|---|---|---|---|---|
| 2417 | 2419 | 1 | 1.0900e-01 | 2.8451e+05 ; | CB - HB3 |
| 2417 | 2420 | 1 | 1.5260e-01 | 2.5941e+05 ; | CB - CG |
| 2420 | 2421 | 1 | 1.0900e-01 | 2.8451e+05 ; | CG - HG2 |
| 2420 | 2422 | 1 | 1.0900e-01 | 2.8451e+05 ; | CG - HG3 |
| 2420 | 2423 | 1 | 1.5260e-01 | 2.5941e+05 ; | CG - CD |
| 2423 | 2424 | 1 | 1.0900e-01 | 2.8451e+05 ; | CD - HD2 |
| 2423 | 2425 | 1 | 1.0900e-01 | 2.8451e+05 ; | CD - HD3 |
| 2423 | 2426 | 1 | 1.5260e-01 | 2.5941e+05 ; | CD - CE |
| 2426 | 2427 | 1 | 1.0900e-01 | 2.8451e+05 ; | CE - HE2 |
| 2426 | 2428 | 1 | 1.0900e-01 | 2.8451e+05 ; | CE - HE3 |
| 2426 | 2429 | 1 | 1.4710e-01 | 3.0711e+05 ; | CE - NZ |
| 2429 | 2430 | 1 | 1.0100e-01 | 3.6317e+05 ; | NZ - HZ1 |
| 2429 | 2431 | 1 | 1.0100e-01 | 3.6317e+05 ; | NZ - HZ2 |
| 2429 | 2432 | 1 | 1.0100e-01 | 3.6317e+05 ; | NZ - HZ3 |
| 2433 | 2434 | 1 | 1.2290e-01 | 4.7698e+05 ; | C - O |
| 2433 | 2435 | 1 | 1.3350e-01 | 4.1003e+05 ; | C - N |
| 2435 | 2436 | 1 | 1.0100e-01 | 3.6317e+05 ; | N - H |
| 2435 | 2437 | 1 | 1.4490e-01 | 2.8200e+05 ; | N - CA |
| 2437 | 2438 | 1 | 1.0900e-01 | 2.8451e+05 ; | CA - HA |
| 2437 | 2439 | 1 | 1.5260e-01 | 2.5941e+05 ; | CA - CB |
| 2437 | 2445 | 1 | 1.5220e-01 | 2.6527e+05 ; | CA - C |
| 2439 | 2440 | 1 | 1.0900e-01 | 2.8451e+05 ; | CB - HB2 |
| 2439 | 2441 | 1 | 1.0900e-01 | 2.8451e+05 ; | CB - HB3 |
| 2439 | 2442 | 1 | 1.5220e-01 | 2.6527e+05 ; | CB - CG |
| 2442 | 2443 | 1 | 1.2500e-01 | 5.4894e+05 ; | CG - OD1 |
| 2442 | 2444 | 1 | 1.2500e-01 | 5.4894e+05 ; | CG - OD2 |
| 2445 | 2446 | 1 | 1.2290e-01 | 4.7698e+05 ; | C - O |
| 2445 | 2447 | 1 | 1.3350e-01 | 4.1003e+05 ; | C - N |
| 2447 | 2448 | 1 | 1.0100e-01 | 3.6317e+05 ; | N - H |
| 2447 | 2449 | 1 | 1.4490e-01 | 2.8200e+05 ; | N - CA |
| 2449 | 2450 | 1 | 1.0900e-01 | 2.8451e+05 ; | CA - HA |
| 2449 | 2451 | 1 | 1.5260e-01 | 2.5941e+05 ; | CA - CB |
| 2449 | 2456 | 1 | 1.5220e-01 | 2.6527e+05 ; | CA - C |
| 2451 | 2452 | 1 | 1.0900e-01 | 2.8451e+05 ; | CB - HB2 |
| 2451 | 2453 | 1 | 1.0900e-01 | 2.8451e+05 ; | CB - HB3 |
| 2451 | 2454 | 1 | 1.4100e-01 | 2.6778e+05 ; | CB - OG |
| 2454 | 2455 | 1 | 9.6000e-02 | 4.6275e+05 ; | OG - HG |
| 2456 | 2457 | 1 | 1.2290e-01 | 4.7698e+05 ; | C - O |
| 2456 | 2458 | 1 | 1.3350e-01 | 4.1003e+05 ; | C - N |
| 2458 | 2459 | 1 | 1.0100e-01 | 3.6317e+05 ; | N - H |
| 2458 | 2460 | 1 | 1.4490e-01 | 2.8200e+05 ; | N - CA |
| 2460 | 2461 | 1 | 1.0900e-01 | 2.8451e+05 ; | CA - HA |
| 2460 | 2462 | 1 | 1.5260e-01 | 2.5941e+05 ; | CA - CB |
| 2460 | 2475 | 1 | 1.5220e-01 | 2.6527e+05 ; | CA - C |
| 2462 | 2463 | 1 | 1.0900e-01 | 2.8451e+05 ; | CB - HB2 |
| 2462 | 2464 | 1 | 1.0900e-01 | 2.8451e+05 ; | CB - HB3 |
| 2462 | 2465 | 1 | 1.5260e-01 | 2.5941e+05 ; | CB - CG |
| 2465 | 2466 | 1 | 1.0900e-01 | 2.8451e+05 ; | CG - HG |
| 2465 | 2467 | 1 | 1.5260e-01 | 2.5941e+05 ; | CG - CD1 |
| 2465 | 2471 | 1 | 1.5260e-01 | 2.5941e+05 ; | CG - CD2 |
| 2467 | 2468 | 1 | 1.0900e-01 | 2.8451e+05 ; | CD1 - HD11 |
| 2467 | 2469 | 1 | 1.0900e-01 | 2.8451e+05 ; | CD1 - HD12 |

| | | | | | | |
|---|---|---|---|---|---|---|
| 2467 | 2470 | 1 | 1.0900e-01 | 2.8451e+05 | ; | CD1 - HD13 |
| 2471 | 2472 | 1 | 1.0900e-01 | 2.8451e+05 | ; | CD2 - HD21 |
| 2471 | 2473 | 1 | 1.0900e-01 | 2.8451e+05 | ; | CD2 - HD22 |
| 2471 | 2474 | 1 | 1.0900e-01 | 2.8451e+05 | ; | CD2 - HD23 |
| 2475 | 2476 | 1 | 1.2290e-01 | 4.7698e+05 | ; | C - O |
| 2475 | 2477 | 1 | 1.3350e-01 | 4.1003e+05 | ; | C - N |
| 2477 | 2478 | 1 | 1.0100e-01 | 3.6317e+05 | ; | N - H |
| 2477 | 2479 | 1 | 1.4490e-01 | 2.8200e+05 | ; | N - CA |
| 2479 | 2480 | 1 | 1.0900e-01 | 2.8451e+05 | ; | CA - HA |
| 2479 | 2481 | 1 | 1.5260e-01 | 2.5941e+05 | ; | CA - CB |
| 2479 | 2494 | 1 | 1.5220e-01 | 2.6527e+05 | ; | CA - C |
| 2481 | 2482 | 1 | 1.0900e-01 | 2.8451e+05 | ; | CB - HB2 |
| 2481 | 2483 | 1 | 1.0900e-01 | 2.8451e+05 | ; | CB - HB3 |
| 2481 | 2484 | 1 | 1.5260e-01 | 2.5941e+05 | ; | CB - CG |
| 2484 | 2485 | 1 | 1.0900e-01 | 2.8451e+05 | ; | CG - HG |
| 2484 | 2486 | 1 | 1.5260e-01 | 2.5941e+05 | ; | CG - CD1 |
| 2484 | 2490 | 1 | 1.5260e-01 | 2.5941e+05 | ; | CG - CD2 |
| 2486 | 2487 | 1 | 1.0900e-01 | 2.8451e+05 | ; | CD1 - HD11 |
| 2486 | 2488 | 1 | 1.0900e-01 | 2.8451e+05 | ; | CD1 - HD12 |
| 2486 | 2489 | 1 | 1.0900e-01 | 2.8451e+05 | ; | CD1 - HD13 |
| 2490 | 2491 | 1 | 1.0900e-01 | 2.8451e+05 | ; | CD2 - HD21 |
| 2490 | 2492 | 1 | 1.0900e-01 | 2.8451e+05 | ; | CD2 - HD22 |
| 2490 | 2493 | 1 | 1.0900e-01 | 2.8451e+05 | ; | CD2 - HD23 |
| 2494 | 2495 | 1 | 1.2290e-01 | 4.7698e+05 | ; | C - O |
| 2494 | 2496 | 1 | 1.3350e-01 | 4.1003e+05 | ; | C - N |
| 2496 | 2497 | 1 | 1.0100e-01 | 3.6317e+05 | ; | N - H |
| 2496 | 2498 | 1 | 1.4490e-01 | 2.8200e+05 | ; | N - CA |
| 2498 | 2499 | 1 | 1.0900e-01 | 2.8451e+05 | ; | CA - HA |
| 2498 | 2500 | 1 | 1.5260e-01 | 2.5941e+05 | ; | CA - CB |
| 2498 | 2516 | 1 | 1.5220e-01 | 2.6527e+05 | ; | CA - C |
| 2500 | 2501 | 1 | 1.0900e-01 | 2.8451e+05 | ; | CB - HB2 |
| 2500 | 2502 | 1 | 1.0900e-01 | 2.8451e+05 | ; | CB - HB3 |
| 2500 | 2503 | 1 | 1.5260e-01 | 2.5941e+05 | ; | CB - CG |
| 2503 | 2504 | 1 | 1.0900e-01 | 2.8451e+05 | ; | CG - HG2 |
| 2503 | 2505 | 1 | 1.0900e-01 | 2.8451e+05 | ; | CG - HG3 |
| 2503 | 2506 | 1 | 1.5260e-01 | 2.5941e+05 | ; | CG - CD |
| 2506 | 2507 | 1 | 1.0900e-01 | 2.8451e+05 | ; | CD - HD2 |
| 2506 | 2508 | 1 | 1.0900e-01 | 2.8451e+05 | ; | CD - HD3 |
| 2506 | 2509 | 1 | 1.5260e-01 | 2.5941e+05 | ; | CD - CE |
| 2509 | 2510 | 1 | 1.0900e-01 | 2.8451e+05 | ; | CE - HE2 |
| 2509 | 2511 | 1 | 1.0900e-01 | 2.8451e+05 | ; | CE - HE3 |
| 2509 | 2512 | 1 | 1.4710e-01 | 3.0711e+05 | ; | CE - NZ |
| 2512 | 2513 | 1 | 1.0100e-01 | 3.6317e+05 | ; | NZ - HZ1 |
| 2512 | 2514 | 1 | 1.0100e-01 | 3.6317e+05 | ; | NZ - HZ2 |
| 2512 | 2515 | 1 | 1.0100e-01 | 3.6317e+05 | ; | NZ - HZ3 |
| 2516 | 2517 | 1 | 1.2290e-01 | 4.7698e+05 | ; | C - O |
| 2516 | 2518 | 1 | 1.3350e-01 | 4.1003e+05 | ; | C - N |
| 2518 | 2519 | 1 | 1.0100e-01 | 3.6317e+05 | ; | N - H |
| 2518 | 2520 | 1 | 1.4490e-01 | 2.8200e+05 | ; | N - CA |
| 2520 | 2521 | 1 | 1.0900e-01 | 2.8451e+05 | ; | CA - HA |
| 2520 | 2522 | 1 | 1.5260e-01 | 2.5941e+05 | ; | CA - CB |
| 2520 | 2535 | 1 | 1.5220e-01 | 2.6527e+05 | ; | CA - C |

| | | | | | | |
|---|---|---|---|---|---|---|
| 2522 | 2523 | 1 | 1.0900e-01 | 2.8451e+05 | ; | CB - HB |
| 2522 | 2524 | 1 | 1.5260e-01 | 2.5941e+05 | ; | CB - CG2 |
| 2522 | 2528 | 1 | 1.5260e-01 | 2.5941e+05 | ; | CB - CG1 |
| 2524 | 2525 | 1 | 1.0900e-01 | 2.8451e+05 | ; | CG2 - HG21 |
| 2524 | 2526 | 1 | 1.0900e-01 | 2.8451e+05 | ; | CG2 - HG22 |
| 2524 | 2527 | 1 | 1.0900e-01 | 2.8451e+05 | ; | CG2 - HG23 |
| 2528 | 2529 | 1 | 1.0900e-01 | 2.8451e+05 | ; | CG1 - HG12 |
| 2528 | 2530 | 1 | 1.0900e-01 | 2.8451e+05 | ; | CG1 - HG13 |
| 2528 | 2531 | 1 | 1.5260e-01 | 2.5941e+05 | ; | CG1 - CD1 |
| 2531 | 2532 | 1 | 1.0900e-01 | 2.8451e+05 | ; | CD1 - HD11 |
| 2531 | 2533 | 1 | 1.0900e-01 | 2.8451e+05 | ; | CD1 - HD12 |
| 2531 | 2534 | 1 | 1.0900e-01 | 2.8451e+05 | ; | CD1 - HD13 |
| 2535 | 2536 | 1 | 1.2290e-01 | 4.7698e+05 | ; | C - O |
| 2535 | 2537 | 1 | 1.3350e-01 | 4.1003e+05 | ; | C - N |
| 2537 | 2538 | 1 | 1.0100e-01 | 3.6317e+05 | ; | N - H |
| 2537 | 2539 | 1 | 1.4490e-01 | 2.8200e+05 | ; | N - CA |
| 2539 | 2540 | 1 | 1.0900e-01 | 2.8451e+05 | ; | CA - HA |
| 2539 | 2541 | 1 | 1.5260e-01 | 2.5941e+05 | ; | CA - CB |
| 2539 | 2554 | 1 | 1.5220e-01 | 2.6527e+05 | ; | CA - C |
| 2541 | 2542 | 1 | 1.0900e-01 | 2.8451e+05 | ; | CB - HB |
| 2541 | 2543 | 1 | 1.5260e-01 | 2.5941e+05 | ; | CB - CG2 |
| 2541 | 2547 | 1 | 1.5260e-01 | 2.5941e+05 | ; | CB - CG1 |
| 2543 | 2544 | 1 | 1.0900e-01 | 2.8451e+05 | ; | CG2 - HG21 |
| 2543 | 2545 | 1 | 1.0900e-01 | 2.8451e+05 | ; | CG2 - HG22 |
| 2543 | 2546 | 1 | 1.0900e-01 | 2.8451e+05 | ; | CG2 - HG23 |
| 2547 | 2548 | 1 | 1.0900e-01 | 2.8451e+05 | ; | CG1 - HG12 |
| 2547 | 2549 | 1 | 1.0900e-01 | 2.8451e+05 | ; | CG1 - HG13 |
| 2547 | 2550 | 1 | 1.5260e-01 | 2.5941e+05 | ; | CG1 - CD1 |
| 2550 | 2551 | 1 | 1.0900e-01 | 2.8451e+05 | ; | CD1 - HD11 |
| 2550 | 2552 | 1 | 1.0900e-01 | 2.8451e+05 | ; | CD1 - HD12 |
| 2550 | 2553 | 1 | 1.0900e-01 | 2.8451e+05 | ; | CD1 - HD13 |
| 2554 | 2555 | 1 | 1.2290e-01 | 4.7698e+05 | ; | C - O |
| 2554 | 2556 | 1 | 1.3350e-01 | 4.1003e+05 | ; | C - N |
| 2556 | 2557 | 1 | 1.0100e-01 | 3.6317e+05 | ; | N - H |
| 2556 | 2558 | 1 | 1.4490e-01 | 2.8200e+05 | ; | N - CA |
| 2558 | 2559 | 1 | 1.0900e-01 | 2.8451e+05 | ; | CA - HA |
| 2558 | 2560 | 1 | 1.5260e-01 | 2.5941e+05 | ; | CA - CB |
| 2558 | 2566 | 1 | 1.5220e-01 | 2.6527e+05 | ; | CA - C |
| 2560 | 2561 | 1 | 1.0900e-01 | 2.8451e+05 | ; | CB - HB2 |
| 2560 | 2562 | 1 | 1.0900e-01 | 2.8451e+05 | ; | CB - HB3 |
| 2560 | 2563 | 1 | 1.5220e-01 | 2.6527e+05 | ; | CB - CG |
| 2563 | 2564 | 1 | 1.2500e-01 | 5.4894e+05 | ; | CG - OD1 |
| 2563 | 2565 | 1 | 1.2500e-01 | 5.4894e+05 | ; | CG - OD2 |
| 2566 | 2567 | 1 | 1.2290e-01 | 4.7698e+05 | ; | C - O |
| 2566 | 2568 | 1 | 1.3350e-01 | 4.1003e+05 | ; | C - N |
| 2568 | 2569 | 1 | 1.0100e-01 | 3.6317e+05 | ; | N - H |
| 2568 | 2570 | 1 | 1.4490e-01 | 2.8200e+05 | ; | N - CA |
| 2570 | 2571 | 1 | 1.0900e-01 | 2.8451e+05 | ; | CA - HA |
| 2570 | 2572 | 1 | 1.5260e-01 | 2.5941e+05 | ; | CA - CB |
| 2570 | 2586 | 1 | 1.5220e-01 | 2.6527e+05 | ; | CA - C |
| 2572 | 2573 | 1 | 1.0900e-01 | 2.8451e+05 | ; | CB - HB2 |
| 2572 | 2574 | 1 | 1.0900e-01 | 2.8451e+05 | ; | CB - HB3 |

| | | | | | | |
|---|---|---|---|---|---|---|
| 2572 | 2575 | 1 | 1.5100e-01 | 2.6527e+05 | ; | CB - CG |
| 2575 | 2576 | 1 | 1.4000e-01 | 3.9246e+05 | ; | CG - CD1 |
| 2575 | 2584 | 1 | 1.4000e-01 | 3.9246e+05 | ; | CG - CD2 |
| 2576 | 2577 | 1 | 1.0800e-01 | 3.0711e+05 | ; | CD1 - HD1 |
| 2576 | 2578 | 1 | 1.4000e-01 | 3.9246e+05 | ; | CD1 - CE1 |
| 2578 | 2579 | 1 | 1.0800e-01 | 3.0711e+05 | ; | CE1 - HE1 |
| 2578 | 2580 | 1 | 1.4000e-01 | 3.9246e+05 | ; | CE1 - CZ |
| 2580 | 2581 | 1 | 1.0800e-01 | 3.0711e+05 | ; | CZ - HZ |
| 2580 | 2582 | 1 | 1.4000e-01 | 3.9246e+05 | ; | CZ - CE2 |
| 2582 | 2583 | 1 | 1.0800e-01 | 3.0711e+05 | ; | CE2 - HE2 |
| 2582 | 2584 | 1 | 1.4000e-01 | 3.9246e+05 | ; | CE2 - CD2 |
| 2584 | 2585 | 1 | 1.0800e-01 | 3.0711e+05 | ; | CD2 - HD2 |
| 2586 | 2587 | 1 | 1.2290e-01 | 4.7698e+05 | ; | C - O |
| 2586 | 2588 | 1 | 1.3350e-01 | 4.1003e+05 | ; | C - N |
| 2588 | 2589 | 1 | 1.0100e-01 | 3.6317e+05 | ; | N - H |
| 2588 | 2590 | 1 | 1.4490e-01 | 2.8200e+05 | ; | N - CA |
| 2590 | 2591 | 1 | 1.0900e-01 | 2.8451e+05 | ; | CA - HA2 |
| 2590 | 2592 | 1 | 1.0900e-01 | 2.8451e+05 | ; | CA - HA3 |
| 2590 | 2593 | 1 | 1.5220e-01 | 2.6527e+05 | ; | CA - C |
| 2593 | 2594 | 1 | 1.2290e-01 | 4.7698e+05 | ; | C - O |
| 2593 | 2595 | 1 | 1.3350e-01 | 4.1003e+05 | ; | C - N |
| 2595 | 2596 | 1 | 1.0100e-01 | 3.6317e+05 | ; | N - H |
| 2595 | 2597 | 1 | 1.4490e-01 | 2.8200e+05 | ; | N - CA |
| 2597 | 2598 | 1 | 1.0900e-01 | 2.8451e+05 | ; | CA - HA |
| 2597 | 2599 | 1 | 1.5260e-01 | 2.5941e+05 | ; | CA - CB |
| 2597 | 2612 | 1 | 1.5220e-01 | 2.6527e+05 | ; | CA - C |
| 2599 | 2600 | 1 | 1.0900e-01 | 2.8451e+05 | ; | CB - HB2 |
| 2599 | 2601 | 1 | 1.0900e-01 | 2.8451e+05 | ; | CB - HB3 |
| 2599 | 2602 | 1 | 1.5260e-01 | 2.5941e+05 | ; | CB - CG |
| 2602 | 2603 | 1 | 1.0900e-01 | 2.8451e+05 | ; | CG - HG |
| 2602 | 2604 | 1 | 1.5260e-01 | 2.5941e+05 | ; | CG - CD1 |
| 2602 | 2608 | 1 | 1.5260e-01 | 2.5941e+05 | ; | CG - CD2 |
| 2604 | 2605 | 1 | 1.0900e-01 | 2.8451e+05 | ; | CD1 - HD11 |
| 2604 | 2606 | 1 | 1.0900e-01 | 2.8451e+05 | ; | CD1 - HD12 |
| 2604 | 2607 | 1 | 1.0900e-01 | 2.8451e+05 | ; | CD1 - HD13 |
| 2608 | 2609 | 1 | 1.0900e-01 | 2.8451e+05 | ; | CD2 - HD21 |
| 2608 | 2610 | 1 | 1.0900e-01 | 2.8451e+05 | ; | CD2 - HD22 |
| 2608 | 2611 | 1 | 1.0900e-01 | 2.8451e+05 | ; | CD2 - HD23 |
| 2612 | 2613 | 1 | 1.2290e-01 | 4.7698e+05 | ; | C - O |
| 2612 | 2614 | 1 | 1.3350e-01 | 4.1003e+05 | ; | C - N |
| 2614 | 2615 | 1 | 1.0100e-01 | 3.6317e+05 | ; | N - H |
| 2614 | 2616 | 1 | 1.4490e-01 | 2.8200e+05 | ; | N - CA |
| 2616 | 2617 | 1 | 1.0900e-01 | 2.8451e+05 | ; | CA - HA |
| 2616 | 2618 | 1 | 1.5260e-01 | 2.5941e+05 | ; | CA - CB |
| 2616 | 2623 | 1 | 1.5220e-01 | 2.6527e+05 | ; | CA - C |
| 2618 | 2619 | 1 | 1.0900e-01 | 2.8451e+05 | ; | CB - HB2 |
| 2618 | 2620 | 1 | 1.0900e-01 | 2.8451e+05 | ; | CB - HB3 |
| 2618 | 2621 | 1 | 1.4100e-01 | 2.6778e+05 | ; | CB - OG |
| 2621 | 2622 | 1 | 9.6000e-02 | 4.6275e+05 | ; | OG - HG |
| 2623 | 2624 | 1 | 1.2290e-01 | 4.7698e+05 | ; | C - O |
| 2623 | 2625 | 1 | 1.3350e-01 | 4.1003e+05 | ; | C - N |
| 2625 | 2626 | 1 | 1.0100e-01 | 3.6317e+05 | ; | N - H |

| | | | | | | |
|---|---|---|---|---|---|---|
| 2625 | 2627 | 1 | 1.4490e-01 | 2.8200e+05 | ; | N - CA |
| 2627 | 2628 | 1 | 1.0900e-01 | 2.8451e+05 | ; | CA - HA |
| 2627 | 2629 | 1 | 1.5260e-01 | 2.5941e+05 | ; | CA - CB |
| 2627 | 2645 | 1 | 1.5220e-01 | 2.6527e+05 | ; | CA - C |
| 2629 | 2630 | 1 | 1.0900e-01 | 2.8451e+05 | ; | CB - HB2 |
| 2629 | 2631 | 1 | 1.0900e-01 | 2.8451e+05 | ; | CB - HB3 |
| 2629 | 2632 | 1 | 1.5260e-01 | 2.5941e+05 | ; | CB - CG |
| 2632 | 2633 | 1 | 1.0900e-01 | 2.8451e+05 | ; | CG - HG2 |
| 2632 | 2634 | 1 | 1.0900e-01 | 2.8451e+05 | ; | CG - HG3 |
| 2632 | 2635 | 1 | 1.5260e-01 | 2.5941e+05 | ; | CG - CD |
| 2635 | 2636 | 1 | 1.0900e-01 | 2.8451e+05 | ; | CD - HD2 |
| 2635 | 2637 | 1 | 1.0900e-01 | 2.8451e+05 | ; | CD - HD3 |
| 2635 | 2638 | 1 | 1.5260e-01 | 2.5941e+05 | ; | CD - CE |
| 2638 | 2639 | 1 | 1.0900e-01 | 2.8451e+05 | ; | CE - HE2 |
| 2638 | 2640 | 1 | 1.0900e-01 | 2.8451e+05 | ; | CE - HE3 |
| 2638 | 2641 | 1 | 1.4710e-01 | 3.0711e+05 | ; | CE - NZ |
| 2641 | 2642 | 1 | 1.0100e-01 | 3.6317e+05 | ; | NZ - HZ1 |
| 2641 | 2643 | 1 | 1.0100e-01 | 3.6317e+05 | ; | NZ - HZ2 |
| 2641 | 2644 | 1 | 1.0100e-01 | 3.6317e+05 | ; | NZ - HZ3 |
| 2645 | 2646 | 1 | 1.2290e-01 | 4.7698e+05 | ; | C - O |
| 2645 | 2647 | 1 | 1.3350e-01 | 4.1003e+05 | ; | C - N |
| 2647 | 2648 | 1 | 1.0100e-01 | 3.6317e+05 | ; | N - H |
| 2647 | 2649 | 1 | 1.4490e-01 | 2.8200e+05 | ; | N - CA |
| 2649 | 2650 | 1 | 1.0900e-01 | 2.8451e+05 | ; | CA - HA |
| 2649 | 2651 | 1 | 1.5260e-01 | 2.5941e+05 | ; | CA - CB |
| 2649 | 2659 | 1 | 1.5220e-01 | 2.6527e+05 | ; | CA - C |
| 2651 | 2652 | 1 | 1.0900e-01 | 2.8451e+05 | ; | CB - HB2 |
| 2651 | 2653 | 1 | 1.0900e-01 | 2.8451e+05 | ; | CB - HB3 |
| 2651 | 2654 | 1 | 1.5220e-01 | 2.6527e+05 | ; | CB - CG |
| 2654 | 2655 | 1 | 1.2290e-01 | 4.7698e+05 | ; | CG - OD1 |
| 2654 | 2656 | 1 | 1.3350e-01 | 4.1003e+05 | ; | CG - ND2 |
| 2656 | 2657 | 1 | 1.0100e-01 | 3.6317e+05 | ; | ND2 - HD21 |
| 2656 | 2658 | 1 | 1.0100e-01 | 3.6317e+05 | ; | ND2 - HD22 |
| 2659 | 2660 | 1 | 1.2290e-01 | 4.7698e+05 | ; | C - O |
| 2659 | 2661 | 1 | 1.3350e-01 | 4.1003e+05 | ; | C - N |
| 2661 | 2662 | 1 | 1.0100e-01 | 3.6317e+05 | ; | N - H |
| 2661 | 2663 | 1 | 1.4490e-01 | 2.8200e+05 | ; | N - CA |
| 2663 | 2664 | 1 | 1.0900e-01 | 2.8451e+05 | ; | CA - HA |
| 2663 | 2665 | 1 | 1.5260e-01 | 2.5941e+05 | ; | CA - CB |
| 2663 | 2678 | 1 | 1.5220e-01 | 2.6527e+05 | ; | CA - C |
| 2665 | 2666 | 1 | 1.0900e-01 | 2.8451e+05 | ; | CB - HB2 |
| 2665 | 2667 | 1 | 1.0900e-01 | 2.8451e+05 | ; | CB - HB3 |
| 2665 | 2668 | 1 | 1.5260e-01 | 2.5941e+05 | ; | CB - CG |
| 2668 | 2669 | 1 | 1.0900e-01 | 2.8451e+05 | ; | CG - HG |
| 2668 | 2670 | 1 | 1.5260e-01 | 2.5941e+05 | ; | CG - CD1 |
| 2668 | 2674 | 1 | 1.5260e-01 | 2.5941e+05 | ; | CG - CD2 |
| 2670 | 2671 | 1 | 1.0900e-01 | 2.8451e+05 | ; | CD1 - HD11 |
| 2670 | 2672 | 1 | 1.0900e-01 | 2.8451e+05 | ; | CD1 - HD12 |
| 2670 | 2673 | 1 | 1.0900e-01 | 2.8451e+05 | ; | CD1 - HD13 |
| 2674 | 2675 | 1 | 1.0900e-01 | 2.8451e+05 | ; | CD2 - HD21 |
| 2674 | 2676 | 1 | 1.0900e-01 | 2.8451e+05 | ; | CD2 - HD22 |
| 2674 | 2677 | 1 | 1.0900e-01 | 2.8451e+05 | ; | CD2 - HD23 |

| | | | | | | | | |
|---|---|---|---|---|---|---|---|---|
| 2678 | 2679 | 1 | 1.2290e-01 | 4.7698e+05 | ; | C | - | O |
| 2678 | 2680 | 1 | 1.3350e-01 | 4.1003e+05 | ; | C | - | N |
| 2680 | 2681 | 1 | 1.0100e-01 | 3.6317e+05 | ; | N | - | H |
| 2680 | 2682 | 1 | 1.4490e-01 | 2.8200e+05 | ; | N | - | CA |
| 2682 | 2683 | 1 | 1.0900e-01 | 2.8451e+05 | ; | CA | - | HA2 |
| 2682 | 2684 | 1 | 1.0900e-01 | 2.8451e+05 | ; | CA | - | HA3 |
| 2682 | 2685 | 1 | 1.5220e-01 | 2.6527e+05 | ; | CA | - | C |
| 2685 | 2686 | 1 | 1.2290e-01 | 4.7698e+05 | ; | C | - | O |
| 2685 | 2687 | 1 | 1.3350e-01 | 4.1003e+05 | ; | C | - | N |
| 2687 | 2688 | 1 | 1.0100e-01 | 3.6317e+05 | ; | N | - | H |
| 2687 | 2689 | 1 | 1.4490e-01 | 2.8200e+05 | ; | N | - | CA |
| 2689 | 2690 | 1 | 1.0900e-01 | 2.8451e+05 | ; | CA | - | HA |
| 2689 | 2691 | 1 | 1.5260e-01 | 2.5941e+05 | ; | CA | - | CB |
| 2689 | 2699 | 1 | 1.5220e-01 | 2.6527e+05 | ; | CA | - | C |
| 2691 | 2692 | 1 | 1.0900e-01 | 2.8451e+05 | ; | CB | - | HB |
| 2691 | 2693 | 1 | 1.5260e-01 | 2.5941e+05 | ; | CB | - | CG2 |
| 2691 | 2697 | 1 | 1.4100e-01 | 2.6778e+05 | ; | CB | - | OG1 |
| 2693 | 2694 | 1 | 1.0900e-01 | 2.8451e+05 | ; | CG2 | - | HG21 |
| 2693 | 2695 | 1 | 1.0900e-01 | 2.8451e+05 | ; | CG2 | - | HG22 |
| 2693 | 2696 | 1 | 1.0900e-01 | 2.8451e+05 | ; | CG2 | - | HG23 |
| 2697 | 2698 | 1 | 9.6000e-02 | 4.6275e+05 | ; | OG1 | - | HG1 |
| 2699 | 2700 | 1 | 1.2290e-01 | 4.7698e+05 | ; | C | - | O |
| 2699 | 2701 | 1 | 1.3350e-01 | 4.1003e+05 | ; | C | - | N |
| 2701 | 2702 | 1 | 1.0100e-01 | 3.6317e+05 | ; | N | - | H |
| 2701 | 2703 | 1 | 1.4490e-01 | 2.8200e+05 | ; | N | - | CA |
| 2703 | 2704 | 1 | 1.0900e-01 | 2.8451e+05 | ; | CA | - | HA2 |
| 2703 | 2705 | 1 | 1.0900e-01 | 2.8451e+05 | ; | CA | - | HA3 |
| 2703 | 2706 | 1 | 1.5220e-01 | 2.6527e+05 | ; | CA | - | C |
| 2706 | 2707 | 1 | 1.2290e-01 | 4.7698e+05 | ; | C | - | O |
| 2706 | 2708 | 1 | 1.3350e-01 | 4.1003e+05 | ; | C | - | N |
| 2708 | 2709 | 1 | 1.0100e-01 | 3.6317e+05 | ; | N | - | H |
| 2708 | 2710 | 1 | 1.4490e-01 | 2.8200e+05 | ; | N | - | CA |
| 2710 | 2711 | 1 | 1.0900e-01 | 2.8451e+05 | ; | CA | - | HA |
| 2710 | 2712 | 1 | 1.5260e-01 | 2.5941e+05 | ; | CA | - | CB |
| 2710 | 2721 | 1 | 1.5220e-01 | 2.6527e+05 | ; | CA | - | C |
| 2712 | 2713 | 1 | 1.0900e-01 | 2.8451e+05 | ; | CB | - | HB2 |
| 2712 | 2714 | 1 | 1.0900e-01 | 2.8451e+05 | ; | CB | - | HB3 |
| 2712 | 2715 | 1 | 1.5260e-01 | 2.5941e+05 | ; | CB | - | CG |
| 2715 | 2716 | 1 | 1.0900e-01 | 2.8451e+05 | ; | CG | - | HG2 |
| 2715 | 2717 | 1 | 1.0900e-01 | 2.8451e+05 | ; | CG | - | HG3 |
| 2715 | 2718 | 1 | 1.5220e-01 | 2.6527e+05 | ; | CG | - | CD |
| 2718 | 2719 | 1 | 1.2500e-01 | 5.4894e+05 | ; | CD | - | OE1 |
| 2718 | 2720 | 1 | 1.2500e-01 | 5.4894e+05 | ; | CD | - | OE2 |
| 2721 | 2722 | 1 | 1.2290e-01 | 4.7698e+05 | ; | C | - | O |
| 2721 | 2723 | 1 | 1.3350e-01 | 4.1003e+05 | ; | C | - | N |
| 2723 | 2724 | 1 | 1.0100e-01 | 3.6317e+05 | ; | N | - | H |
| 2723 | 2725 | 1 | 1.4490e-01 | 2.8200e+05 | ; | N | - | CA |
| 2725 | 2726 | 1 | 1.0900e-01 | 2.8451e+05 | ; | CA | - | HA |
| 2725 | 2727 | 1 | 1.5260e-01 | 2.5941e+05 | ; | CA | - | CB |
| 2725 | 2741 | 1 | 1.5220e-01 | 2.6527e+05 | ; | CA | - | C |
| 2727 | 2728 | 1 | 1.0900e-01 | 2.8451e+05 | ; | CB | - | HB2 |
| 2727 | 2729 | 1 | 1.0900e-01 | 2.8451e+05 | ; | CB | - | HB3 |

| | | | | | |
|---|---|---|---|---|---|
| 2727 | 2730 | 1 | 1.5100e-01 | 2.6527e+05 ; | CB - CG |
| 2730 | 2731 | 1 | 1.4000e-01 | 3.9246e+05 ; | CG - CD1 |
| 2730 | 2739 | 1 | 1.4000e-01 | 3.9246e+05 ; | CG - CD2 |
| 2731 | 2732 | 1 | 1.0800e-01 | 3.0711e+05 ; | CD1 - HD1 |
| 2731 | 2733 | 1 | 1.4000e-01 | 3.9246e+05 ; | CD1 - CE1 |
| 2733 | 2734 | 1 | 1.0800e-01 | 3.0711e+05 ; | CE1 - HE1 |
| 2733 | 2735 | 1 | 1.4000e-01 | 3.9246e+05 ; | CE1 - CZ |
| 2735 | 2736 | 1 | 1.0800e-01 | 3.0711e+05 ; | CZ - HZ |
| 2735 | 2737 | 1 | 1.4000e-01 | 3.9246e+05 ; | CZ - CE2 |
| 2737 | 2738 | 1 | 1.0800e-01 | 3.0711e+05 ; | CE2 - HE2 |
| 2737 | 2739 | 1 | 1.4000e-01 | 3.9246e+05 ; | CE2 - CD2 |
| 2739 | 2740 | 1 | 1.0800e-01 | 3.0711e+05 ; | CD2 - HD2 |
| 2741 | 2742 | 1 | 1.2290e-01 | 4.7698e+05 ; | C - O |
| 2741 | 2743 | 1 | 1.3350e-01 | 4.1003e+05 ; | C - N |
| 2743 | 2744 | 1 | 1.0100e-01 | 3.6317e+05 ; | N - H |
| 2743 | 2745 | 1 | 1.4490e-01 | 2.8200e+05 ; | N - CA |
| 2745 | 2746 | 1 | 1.0900e-01 | 2.8451e+05 ; | CA - HA |
| 2745 | 2747 | 1 | 1.5260e-01 | 2.5941e+05 ; | CA - CB |
| 2745 | 2755 | 1 | 1.5220e-01 | 2.6527e+05 ; | CA - C |
| 2747 | 2748 | 1 | 1.0900e-01 | 2.8451e+05 ; | CB - HB |
| 2747 | 2749 | 1 | 1.5260e-01 | 2.5941e+05 ; | CB - CG2 |
| 2747 | 2753 | 1 | 1.4100e-01 | 2.6778e+05 ; | CB - OG1 |
| 2749 | 2750 | 1 | 1.0900e-01 | 2.8451e+05 ; | CG2 - HG21 |
| 2749 | 2751 | 1 | 1.0900e-01 | 2.8451e+05 ; | CG2 - HG22 |
| 2749 | 2752 | 1 | 1.0900e-01 | 2.8451e+05 ; | CG2 - HG23 |
| 2753 | 2754 | 1 | 9.6000e-02 | 4.6275e+05 ; | OG1 - HG1 |
| 2755 | 2756 | 1 | 1.2290e-01 | 4.7698e+05 ; | C - O |
| 2755 | 2757 | 1 | 1.3350e-01 | 4.1003e+05 ; | C - N |
| 2757 | 2758 | 1 | 1.0100e-01 | 3.6317e+05 ; | N - H |
| 2757 | 2759 | 1 | 1.4490e-01 | 2.8200e+05 ; | N - CA |
| 2759 | 2760 | 1 | 1.0900e-01 | 2.8451e+05 ; | CA - HA |
| 2759 | 2761 | 1 | 1.5260e-01 | 2.5941e+05 ; | CA - CB |
| 2759 | 2769 | 1 | 1.5220e-01 | 2.6527e+05 ; | CA - C |
| 2761 | 2762 | 1 | 1.0900e-01 | 2.8451e+05 ; | CB - HB |
| 2761 | 2763 | 1 | 1.5260e-01 | 2.5941e+05 ; | CB - CG2 |
| 2761 | 2767 | 1 | 1.4100e-01 | 2.6778e+05 ; | CB - OG1 |
| 2763 | 2764 | 1 | 1.0900e-01 | 2.8451e+05 ; | CG2 - HG21 |
| 2763 | 2765 | 1 | 1.0900e-01 | 2.8451e+05 ; | CG2 - HG22 |
| 2763 | 2766 | 1 | 1.0900e-01 | 2.8451e+05 ; | CG2 - HG23 |
| 2767 | 2768 | 1 | 9.6000e-02 | 4.6275e+05 ; | OG1 - HG1 |
| 2769 | 2770 | 1 | 1.2290e-01 | 4.7698e+05 ; | C - O |
| 2769 | 2771 | 1 | 1.3350e-01 | 4.1003e+05 ; | C - N |
| 2771 | 2772 | 1 | 1.0100e-01 | 3.6317e+05 ; | N - H |
| 2771 | 2773 | 1 | 1.4490e-01 | 2.8200e+05 ; | N - CA |
| 2773 | 2774 | 1 | 1.0900e-01 | 2.8451e+05 ; | CA - HA |
| 2773 | 2775 | 1 | 1.5260e-01 | 2.5941e+05 ; | CA - CB |
| 2773 | 2783 | 1 | 1.5220e-01 | 2.6527e+05 ; | CA - C |
| 2775 | 2776 | 1 | 1.0900e-01 | 2.8451e+05 ; | CB - HB |
| 2775 | 2777 | 1 | 1.5260e-01 | 2.5941e+05 ; | CB - CG2 |
| 2775 | 2781 | 1 | 1.4100e-01 | 2.6778e+05 ; | CB - OG1 |
| 2777 | 2778 | 1 | 1.0900e-01 | 2.8451e+05 ; | CG2 - HG21 |
| 2777 | 2779 | 1 | 1.0900e-01 | 2.8451e+05 ; | CG2 - HG22 |

| | | | | | | |
|---|---|---|---|---|---|---|
| 2777 | 2780 | 1 | 1.0900e-01 | 2.8451e+05 | ; | CG2 - HG23 |
| 2781 | 2782 | 1 | 9.6000e-02 | 4.6275e+05 | ; | OG1 - HG1 |
| 2783 | 2784 | 1 | 1.2290e-01 | 4.7698e+05 | ; | C - O |
| 2783 | 2785 | 1 | 1.3350e-01 | 4.1003e+05 | ; | C - N |
| 2785 | 2786 | 1 | 1.0100e-01 | 3.6317e+05 | ; | N - H |
| 2785 | 2787 | 1 | 1.4490e-01 | 2.8200e+05 | ; | N - CA |
| 2787 | 2788 | 1 | 1.0900e-01 | 2.8451e+05 | ; | CA - HA |
| 2787 | 2789 | 1 | 1.5260e-01 | 2.5941e+05 | ; | CA - CB |
| 2787 | 2805 | 1 | 1.5220e-01 | 2.6527e+05 | ; | CA - C |
| 2789 | 2790 | 1 | 1.0900e-01 | 2.8451e+05 | ; | CB - HB2 |
| 2789 | 2791 | 1 | 1.0900e-01 | 2.8451e+05 | ; | CB - HB3 |
| 2789 | 2792 | 1 | 1.5260e-01 | 2.5941e+05 | ; | CB - CG |
| 2792 | 2793 | 1 | 1.0900e-01 | 2.8451e+05 | ; | CG - HG2 |
| 2792 | 2794 | 1 | 1.0900e-01 | 2.8451e+05 | ; | CG - HG3 |
| 2792 | 2795 | 1 | 1.5260e-01 | 2.5941e+05 | ; | CG - CD |
| 2795 | 2796 | 1 | 1.0900e-01 | 2.8451e+05 | ; | CD - HD2 |
| 2795 | 2797 | 1 | 1.0900e-01 | 2.8451e+05 | ; | CD - HD3 |
| 2795 | 2798 | 1 | 1.5260e-01 | 2.5941e+05 | ; | CD - CE |
| 2798 | 2799 | 1 | 1.0900e-01 | 2.8451e+05 | ; | CE - HE2 |
| 2798 | 2800 | 1 | 1.0900e-01 | 2.8451e+05 | ; | CE - HE3 |
| 2798 | 2801 | 1 | 1.4710e-01 | 3.0711e+05 | ; | CE - NZ |
| 2801 | 2802 | 1 | 1.0100e-01 | 3.6317e+05 | ; | NZ - HZ1 |
| 2801 | 2803 | 1 | 1.0100e-01 | 3.6317e+05 | ; | NZ - HZ2 |
| 2801 | 2804 | 1 | 1.0100e-01 | 3.6317e+05 | ; | NZ - HZ3 |
| 2805 | 2806 | 1 | 1.2290e-01 | 4.7698e+05 | ; | C - O |
| 2805 | 2807 | 1 | 1.3350e-01 | 4.1003e+05 | ; | C - N |
| 2807 | 2808 | 1 | 1.0100e-01 | 3.6317e+05 | ; | N - H |
| 2807 | 2809 | 1 | 1.4490e-01 | 2.8200e+05 | ; | N - CA |
| 2809 | 2810 | 1 | 1.0900e-01 | 2.8451e+05 | ; | CA - HA |
| 2809 | 2811 | 1 | 1.5260e-01 | 2.5941e+05 | ; | CA - CB |
| 2809 | 2815 | 1 | 1.5220e-01 | 2.6527e+05 | ; | CA - C |
| 2811 | 2812 | 1 | 1.0900e-01 | 2.8451e+05 | ; | CB - HB1 |
| 2811 | 2813 | 1 | 1.0900e-01 | 2.8451e+05 | ; | CB - HB2 |
| 2811 | 2814 | 1 | 1.0900e-01 | 2.8451e+05 | ; | CB - HB3 |
| 2815 | 2816 | 1 | 1.2290e-01 | 4.7698e+05 | ; | C - O |
| 2815 | 2817 | 1 | 1.3350e-01 | 4.1003e+05 | ; | C - N |
| 2817 | 2818 | 1 | 1.0100e-01 | 3.6317e+05 | ; | N - H |
| 2817 | 2819 | 1 | 1.4490e-01 | 2.8200e+05 | ; | N - CA |
| 2819 | 2820 | 1 | 1.0900e-01 | 2.8451e+05 | ; | CA - HA2 |
| 2819 | 2821 | 1 | 1.0900e-01 | 2.8451e+05 | ; | CA - HA3 |
| 2819 | 2822 | 1 | 1.5220e-01 | 2.6527e+05 | ; | CA - C |
| 2822 | 2823 | 1 | 1.2290e-01 | 4.7698e+05 | ; | C - O |
| 2822 | 2824 | 1 | 1.3350e-01 | 4.1003e+05 | ; | C - N |
| 2824 | 2825 | 1 | 1.0100e-01 | 3.6317e+05 | ; | N - H |
| 2824 | 2826 | 1 | 1.4490e-01 | 2.8200e+05 | ; | N - CA |
| 2826 | 2827 | 1 | 1.0900e-01 | 2.8451e+05 | ; | CA - HA |
| 2826 | 2828 | 1 | 1.5260e-01 | 2.5941e+05 | ; | CA - CB |
| 2826 | 2836 | 1 | 1.5220e-01 | 2.6527e+05 | ; | CA - C |
| 2828 | 2829 | 1 | 1.0900e-01 | 2.8451e+05 | ; | CB - HB |
| 2828 | 2830 | 1 | 1.5260e-01 | 2.5941e+05 | ; | CB - CG2 |
| 2828 | 2834 | 1 | 1.4100e-01 | 2.6778e+05 | ; | CB - OG1 |
| 2830 | 2831 | 1 | 1.0900e-01 | 2.8451e+05 | ; | CG2 - HG21 |

| | | | | | | | | |
|---|---|---|---|---|---|---|---|---|
| 2830 | 2832 | 1 | 1.0900e-01 | 2.8451e+05 | ; | CG2 | - | HG22 |
| 2830 | 2833 | 1 | 1.0900e-01 | 2.8451e+05 | ; | CG2 | - | HG23 |
| 2834 | 2835 | 1 | 9.6000e-02 | 4.6275e+05 | ; | OG1 | - | HG1 |
| 2836 | 2837 | 1 | 1.2290e-01 | 4.7698e+05 | ; | C | - | O |
| 2836 | 2838 | 1 | 1.3350e-01 | 4.1003e+05 | ; | C | - | N |
| 2838 | 2839 | 1 | 1.4490e-01 | 2.8200e+05 | ; | N | - | CD |
| 2838 | 2848 | 1 | 1.4490e-01 | 2.8200e+05 | ; | N | - | CA |
| 2839 | 2840 | 1 | 1.0900e-01 | 2.8451e+05 | ; | CD | - | HD2 |
| 2839 | 2841 | 1 | 1.0900e-01 | 2.8451e+05 | ; | CD | - | HD3 |
| 2839 | 2842 | 1 | 1.5260e-01 | 2.5941e+05 | ; | CD | - | CG |
| 2842 | 2843 | 1 | 1.0900e-01 | 2.8451e+05 | ; | CG | - | HG2 |
| 2842 | 2844 | 1 | 1.0900e-01 | 2.8451e+05 | ; | CG | - | HG3 |
| 2842 | 2845 | 1 | 1.5260e-01 | 2.5941e+05 | ; | CG | - | CB |
| 2845 | 2846 | 1 | 1.0900e-01 | 2.8451e+05 | ; | CB | - | HB2 |
| 2845 | 2847 | 1 | 1.0900e-01 | 2.8451e+05 | ; | CB | - | HB3 |
| 2845 | 2848 | 1 | 1.5260e-01 | 2.5941e+05 | ; | CB | - | CA |
| 2848 | 2849 | 1 | 1.0900e-01 | 2.8451e+05 | ; | CA | - | HA |
| 2848 | 2850 | 1 | 1.5220e-01 | 2.6527e+05 | ; | CA | - | C |
| 2850 | 2851 | 1 | 1.2290e-01 | 4.7698e+05 | ; | C | - | O |
| 2850 | 2852 | 1 | 1.3350e-01 | 4.1003e+05 | ; | C | - | N |
| 2852 | 2853 | 1 | 1.0100e-01 | 3.6317e+05 | ; | N | - | H |
| 2852 | 2854 | 1 | 1.4490e-01 | 2.8200e+05 | ; | N | - | CA |
| 2854 | 2855 | 1 | 1.0900e-01 | 2.8451e+05 | ; | CA | - | HA |
| 2854 | 2856 | 1 | 1.5260e-01 | 2.5941e+05 | ; | CA | - | CB |
| 2854 | 2871 | 1 | 1.5220e-01 | 2.6527e+05 | ; | CA | - | C |
| 2856 | 2857 | 1 | 1.0900e-01 | 2.8451e+05 | ; | CB | - | HB2 |
| 2856 | 2858 | 1 | 1.0900e-01 | 2.8451e+05 | ; | CB | - | HB3 |
| 2856 | 2859 | 1 | 1.5100e-01 | 2.6527e+05 | ; | CB | - | CG |
| 2859 | 2860 | 1 | 1.4000e-01 | 3.9246e+05 | ; | CG | - | CD1 |
| 2859 | 2869 | 1 | 1.4000e-01 | 3.9246e+05 | ; | CG | - | CD2 |
| 2860 | 2861 | 1 | 1.0800e-01 | 3.0711e+05 | ; | CD1 | - | HD1 |
| 2860 | 2862 | 1 | 1.4000e-01 | 3.9246e+05 | ; | CD1 | - | CE1 |
| 2862 | 2863 | 1 | 1.0800e-01 | 3.0711e+05 | ; | CE1 | - | HE1 |
| 2862 | 2864 | 1 | 1.4090e-01 | 3.9246e+05 | ; | CE1 | - | CZ |
| 2864 | 2865 | 1 | 1.3640e-01 | 3.7656e+05 | ; | CZ | - | OH |
| 2864 | 2867 | 1 | 1.4090e-01 | 3.9246e+05 | ; | CZ | - | CE2 |
| 2865 | 2866 | 1 | 9.6000e-02 | 4.6275e+05 | ; | OH | - | HH |
| 2867 | 2868 | 1 | 1.0800e-01 | 3.0711e+05 | ; | CE2 | - | HE2 |
| 2867 | 2869 | 1 | 1.4000e-01 | 3.9246e+05 | ; | CE2 | - | CD2 |
| 2869 | 2870 | 1 | 1.0800e-01 | 3.0711e+05 | ; | CD2 | - | HD2 |
| 2871 | 2872 | 1 | 1.2290e-01 | 4.7698e+05 | ; | C | - | O |
| 2871 | 2873 | 1 | 1.3350e-01 | 4.1003e+05 | ; | C | - | N |
| 2873 | 2874 | 1 | 1.0100e-01 | 3.6317e+05 | ; | N | - | H |
| 2873 | 2875 | 1 | 1.4490e-01 | 2.8200e+05 | ; | N | - | CA |
| 2875 | 2876 | 1 | 1.0900e-01 | 2.8451e+05 | ; | CA | - | HA |
| 2875 | 2877 | 1 | 1.5260e-01 | 2.5941e+05 | ; | CA | - | CB |
| 2875 | 2892 | 1 | 1.5220e-01 | 2.6527e+05 | ; | CA | - | C |
| 2877 | 2878 | 1 | 1.0900e-01 | 2.8451e+05 | ; | CB | - | HB2 |
| 2877 | 2879 | 1 | 1.0900e-01 | 2.8451e+05 | ; | CB | - | HB3 |
| 2877 | 2880 | 1 | 1.5100e-01 | 2.6527e+05 | ; | CB | - | CG |
| 2880 | 2881 | 1 | 1.4000e-01 | 3.9246e+05 | ; | CG | - | CD1 |
| 2880 | 2890 | 1 | 1.4000e-01 | 3.9246e+05 | ; | CG | - | CD2 |

| | | | | | | |
|---|---|---|---|---|---|---|
| 2881 | 2882 | 1 | 1.0800e-01 | 3.0711e+05 | ; | CD1 - HD1 |
| 2881 | 2883 | 1 | 1.4000e-01 | 3.9246e+05 | ; | CD1 - CE1 |
| 2883 | 2884 | 1 | 1.0800e-01 | 3.0711e+05 | ; | CE1 - HE1 |
| 2883 | 2885 | 1 | 1.4090e-01 | 3.9246e+05 | ; | CE1 - CZ |
| 2885 | 2886 | 1 | 1.3640e-01 | 3.7656e+05 | ; | CZ - OH |
| 2885 | 2888 | 1 | 1.4090e-01 | 3.9246e+05 | ; | CZ - CE2 |
| 2886 | 2887 | 1 | 9.6000e-02 | 4.6275e+05 | ; | OH - HH |
| 2888 | 2889 | 1 | 1.0800e-01 | 3.0711e+05 | ; | CE2 - HE2 |
| 2888 | 2890 | 1 | 1.4000e-01 | 3.9246e+05 | ; | CE2 - CD2 |
| 2890 | 2891 | 1 | 1.0800e-01 | 3.0711e+05 | ; | CD2 - HD2 |
| 2892 | 2893 | 1 | 1.2290e-01 | 4.7698e+05 | ; | C - O |
| 2892 | 2894 | 1 | 1.3350e-01 | 4.1003e+05 | ; | C - N |
| 2894 | 2895 | 1 | 1.0100e-01 | 3.6317e+05 | ; | N - H |
| 2894 | 2896 | 1 | 1.4490e-01 | 2.8200e+05 | ; | N - CA |
| 2896 | 2897 | 1 | 1.0900e-01 | 2.8451e+05 | ; | CA - HA |
| 2896 | 2898 | 1 | 1.5260e-01 | 2.5941e+05 | ; | CA - CB |
| 2896 | 2908 | 1 | 1.5220e-01 | 2.6527e+05 | ; | CA - C |
| 2898 | 2899 | 1 | 1.0900e-01 | 2.8451e+05 | ; | CB - HB |
| 2898 | 2900 | 1 | 1.5260e-01 | 2.5941e+05 | ; | CB - CG1 |
| 2898 | 2904 | 1 | 1.5260e-01 | 2.5941e+05 | ; | CB - CG2 |
| 2900 | 2901 | 1 | 1.0900e-01 | 2.8451e+05 | ; | CG1 - HG11 |
| 2900 | 2902 | 1 | 1.0900e-01 | 2.8451e+05 | ; | CG1 - HG12 |
| 2900 | 2903 | 1 | 1.0900e-01 | 2.8451e+05 | ; | CG1 - HG13 |
| 2904 | 2905 | 1 | 1.0900e-01 | 2.8451e+05 | ; | CG2 - HG21 |
| 2904 | 2906 | 1 | 1.0900e-01 | 2.8451e+05 | ; | CG2 - HG22 |
| 2904 | 2907 | 1 | 1.0900e-01 | 2.8451e+05 | ; | CG2 - HG23 |
| 2908 | 2909 | 1 | 1.2290e-01 | 4.7698e+05 | ; | C - O |
| 2908 | 2910 | 1 | 1.3350e-01 | 4.1003e+05 | ; | C - N |
| 2910 | 2911 | 1 | 1.0100e-01 | 3.6317e+05 | ; | N - H |
| 2910 | 2912 | 1 | 1.4490e-01 | 2.8200e+05 | ; | N - CA |
| 2912 | 2913 | 1 | 1.0900e-01 | 2.8451e+05 | ; | CA - HA |
| 2912 | 2914 | 1 | 1.5260e-01 | 2.5941e+05 | ; | CA - CB |
| 2912 | 2918 | 1 | 1.5220e-01 | 2.6527e+05 | ; | CA - C |
| 2914 | 2915 | 1 | 1.0900e-01 | 2.8451e+05 | ; | CB - HB1 |
| 2914 | 2916 | 1 | 1.0900e-01 | 2.8451e+05 | ; | CB - HB2 |
| 2914 | 2917 | 1 | 1.0900e-01 | 2.8451e+05 | ; | CB - HB3 |
| 2918 | 2919 | 1 | 1.2290e-01 | 4.7698e+05 | ; | C - O |
| 2918 | 2920 | 1 | 1.3350e-01 | 4.1003e+05 | ; | C - N |
| 2920 | 2921 | 1 | 1.4490e-01 | 2.8200e+05 | ; | N - CD |
| 2920 | 2930 | 1 | 1.4490e-01 | 2.8200e+05 | ; | N - CA |
| 2921 | 2922 | 1 | 1.0900e-01 | 2.8451e+05 | ; | CD - HD2 |
| 2921 | 2923 | 1 | 1.0900e-01 | 2.8451e+05 | ; | CD - HD3 |
| 2921 | 2924 | 1 | 1.5260e-01 | 2.5941e+05 | ; | CD - CG |
| 2924 | 2925 | 1 | 1.0900e-01 | 2.8451e+05 | ; | CG - HG2 |
| 2924 | 2926 | 1 | 1.0900e-01 | 2.8451e+05 | ; | CG - HG3 |
| 2924 | 2927 | 1 | 1.5260e-01 | 2.5941e+05 | ; | CG - CB |
| 2927 | 2928 | 1 | 1.0900e-01 | 2.8451e+05 | ; | CB - HB2 |
| 2927 | 2929 | 1 | 1.0900e-01 | 2.8451e+05 | ; | CB - HB3 |
| 2927 | 2930 | 1 | 1.5260e-01 | 2.5941e+05 | ; | CB - CA |
| 2930 | 2931 | 1 | 1.0900e-01 | 2.8451e+05 | ; | CA - HA |
| 2930 | 2932 | 1 | 1.5220e-01 | 2.6527e+05 | ; | CA - C |
| 2932 | 2933 | 1 | 1.2290e-01 | 4.7698e+05 | ; | C - O |

| | | | | | |
|---|---|---|---|---|---|
| 2932 | 2934 | 1 | 1.3350e-01 | 4.1003e+05 ; | C - N |
| 2934 | 2935 | 1 | 1.0100e-01 | 3.6317e+05 ; | N - H |
| 2934 | 2936 | 1 | 1.4490e-01 | 2.8200e+05 ; | N - CA |
| 2936 | 2937 | 1 | 1.0900e-01 | 2.8451e+05 ; | CA - HA |
| 2936 | 2938 | 1 | 1.5260e-01 | 2.5941e+05 ; | CA - CB |
| 2936 | 2949 | 1 | 1.5220e-01 | 2.6527e+05 ; | CA - C |
| 2938 | 2939 | 1 | 1.0900e-01 | 2.8451e+05 ; | CB - HB2 |
| 2938 | 2940 | 1 | 1.0900e-01 | 2.8451e+05 ; | CB - HB3 |
| 2938 | 2941 | 1 | 1.5260e-01 | 2.5941e+05 ; | CB - CG |
| 2941 | 2942 | 1 | 1.0900e-01 | 2.8451e+05 ; | CG - HG2 |
| 2941 | 2943 | 1 | 1.0900e-01 | 2.8451e+05 ; | CG - HG3 |
| 2941 | 2944 | 1 | 1.5220e-01 | 2.6527e+05 ; | CG - CD |
| 2944 | 2945 | 1 | 1.2290e-01 | 4.7698e+05 ; | CD - OE1 |
| 2944 | 2946 | 1 | 1.3350e-01 | 4.1003e+05 ; | CD - NE2 |
| 2946 | 2947 | 1 | 1.0100e-01 | 3.6317e+05 ; | NE2 - HE21 |
| 2946 | 2948 | 1 | 1.0100e-01 | 3.6317e+05 ; | NE2 - HE22 |
| 2949 | 2950 | 1 | 1.2290e-01 | 4.7698e+05 ; | C - O |
| 2949 | 2951 | 1 | 1.3350e-01 | 4.1003e+05 ; | C - N |
| 2951 | 2952 | 1 | 1.0100e-01 | 3.6317e+05 ; | N - H |
| 2951 | 2953 | 1 | 1.4490e-01 | 2.8200e+05 ; | N - CA |
| 2953 | 2954 | 1 | 1.0900e-01 | 2.8451e+05 ; | CA - HA |
| 2953 | 2955 | 1 | 1.5260e-01 | 2.5941e+05 ; | CA - CB |
| 2953 | 2965 | 1 | 1.5220e-01 | 2.6527e+05 ; | CA - C |
| 2955 | 2956 | 1 | 1.0900e-01 | 2.8451e+05 ; | CB - HB |
| 2955 | 2957 | 1 | 1.5260e-01 | 2.5941e+05 ; | CB - CG1 |
| 2955 | 2961 | 1 | 1.5260e-01 | 2.5941e+05 ; | CB - CG2 |
| 2957 | 2958 | 1 | 1.0900e-01 | 2.8451e+05 ; | CG1 - HG11 |
| 2957 | 2959 | 1 | 1.0900e-01 | 2.8451e+05 ; | CG1 - HG12 |
| 2957 | 2960 | 1 | 1.0900e-01 | 2.8451e+05 ; | CG1 - HG13 |
| 2961 | 2962 | 1 | 1.0900e-01 | 2.8451e+05 ; | CG2 - HG21 |
| 2961 | 2963 | 1 | 1.0900e-01 | 2.8451e+05 ; | CG2 - HG22 |
| 2961 | 2964 | 1 | 1.0900e-01 | 2.8451e+05 ; | CG2 - HG23 |
| 2965 | 2966 | 1 | 1.2290e-01 | 4.7698e+05 ; | C - O |
| 2965 | 2967 | 1 | 1.3350e-01 | 4.1003e+05 ; | C - N |
| 2967 | 2968 | 1 | 1.0100e-01 | 3.6317e+05 ; | N - H |
| 2967 | 2969 | 1 | 1.4490e-01 | 2.8200e+05 ; | N - CA |
| 2969 | 2970 | 1 | 1.0900e-01 | 2.8451e+05 ; | CA - HA |
| 2969 | 2971 | 1 | 1.5260e-01 | 2.5941e+05 ; | CA - CB |
| 2969 | 2984 | 1 | 1.5220e-01 | 2.6527e+05 ; | CA - C |
| 2971 | 2972 | 1 | 1.0900e-01 | 2.8451e+05 ; | CB - HB2 |
| 2971 | 2973 | 1 | 1.0900e-01 | 2.8451e+05 ; | CB - HB3 |
| 2971 | 2974 | 1 | 1.5260e-01 | 2.5941e+05 ; | CB - CG |
| 2974 | 2975 | 1 | 1.0900e-01 | 2.8451e+05 ; | CG - HG |
| 2974 | 2976 | 1 | 1.5260e-01 | 2.5941e+05 ; | CG - CD1 |
| 2974 | 2980 | 1 | 1.5260e-01 | 2.5941e+05 ; | CG - CD2 |
| 2976 | 2977 | 1 | 1.0900e-01 | 2.8451e+05 ; | CD1 - HD11 |
| 2976 | 2978 | 1 | 1.0900e-01 | 2.8451e+05 ; | CD1 - HD12 |
| 2976 | 2979 | 1 | 1.0900e-01 | 2.8451e+05 ; | CD1 - HD13 |
| 2980 | 2981 | 1 | 1.0900e-01 | 2.8451e+05 ; | CD2 - HD21 |
| 2980 | 2982 | 1 | 1.0900e-01 | 2.8451e+05 ; | CD2 - HD22 |
| 2980 | 2983 | 1 | 1.0900e-01 | 2.8451e+05 ; | CD2 - HD23 |
| 2984 | 2985 | 1 | 1.2290e-01 | 4.7698e+05 ; | C - O |

| | | | | | | |
|---|---|---|---|---|---|---|
| 2984 | 2986 | 1 | 1.3350e-01 | 4.1003e+05 | ; | C - N |
| 2986 | 2987 | 1 | 1.0100e-01 | 3.6317e+05 | ; | N - H |
| 2986 | 2988 | 1 | 1.4490e-01 | 2.8200e+05 | ; | N - CA |
| 2988 | 2989 | 1 | 1.0900e-01 | 2.8451e+05 | ; | CA - HA |
| 2988 | 2990 | 1 | 1.5260e-01 | 2.5941e+05 | ; | CA - CB |
| 2988 | 2996 | 1 | 1.5220e-01 | 2.6527e+05 | ; | CA - C |
| 2990 | 2991 | 1 | 1.0900e-01 | 2.8451e+05 | ; | CB - HB2 |
| 2990 | 2992 | 1 | 1.0900e-01 | 2.8451e+05 | ; | CB - HB3 |
| 2990 | 2993 | 1 | 1.5220e-01 | 2.6527e+05 | ; | CB - CG |
| 2993 | 2994 | 1 | 1.2500e-01 | 5.4894e+05 | ; | CG - OD1 |
| 2993 | 2995 | 1 | 1.2500e-01 | 5.4894e+05 | ; | CG - OD2 |
| 2996 | 2997 | 1 | 1.2290e-01 | 4.7698e+05 | ; | C - O |
| 2996 | 2998 | 1 | 1.3350e-01 | 4.1003e+05 | ; | C - N |
| 2998 | 2999 | 1 | 1.0100e-01 | 3.6317e+05 | ; | N - H |
| 2998 | 3000 | 1 | 1.4490e-01 | 2.8200e+05 | ; | N - CA |
| 3000 | 3001 | 1 | 1.0900e-01 | 2.8451e+05 | ; | CA - HA2 |
| 3000 | 3002 | 1 | 1.0900e-01 | 2.8451e+05 | ; | CA - HA3 |
| 3000 | 3003 | 1 | 1.5220e-01 | 2.6527e+05 | ; | CA - C |
| 3003 | 3004 | 1 | 1.2290e-01 | 4.7698e+05 | ; | C - O |
| 3003 | 3005 | 1 | 1.3350e-01 | 4.1003e+05 | ; | C - N |
| 3005 | 3006 | 1 | 1.0100e-01 | 3.6317e+05 | ; | N - H |
| 3005 | 3007 | 1 | 1.4490e-01 | 2.8200e+05 | ; | N - CA |
| 3007 | 3008 | 1 | 1.0900e-01 | 2.8451e+05 | ; | CA - HA |
| 3007 | 3009 | 1 | 1.5260e-01 | 2.5941e+05 | ; | CA - CB |
| 3007 | 3025 | 1 | 1.5220e-01 | 2.6527e+05 | ; | CA - C |
| 3009 | 3010 | 1 | 1.0900e-01 | 2.8451e+05 | ; | CB - HB2 |
| 3009 | 3011 | 1 | 1.0900e-01 | 2.8451e+05 | ; | CB - HB3 |
| 3009 | 3012 | 1 | 1.5260e-01 | 2.5941e+05 | ; | CB - CG |
| 3012 | 3013 | 1 | 1.0900e-01 | 2.8451e+05 | ; | CG - HG2 |
| 3012 | 3014 | 1 | 1.0900e-01 | 2.8451e+05 | ; | CG - HG3 |
| 3012 | 3015 | 1 | 1.5260e-01 | 2.5941e+05 | ; | CG - CD |
| 3015 | 3016 | 1 | 1.0900e-01 | 2.8451e+05 | ; | CD - HD2 |
| 3015 | 3017 | 1 | 1.0900e-01 | 2.8451e+05 | ; | CD - HD3 |
| 3015 | 3018 | 1 | 1.5260e-01 | 2.5941e+05 | ; | CD - CE |
| 3018 | 3019 | 1 | 1.0900e-01 | 2.8451e+05 | ; | CE - HE2 |
| 3018 | 3020 | 1 | 1.0900e-01 | 2.8451e+05 | ; | CE - HE3 |
| 3018 | 3021 | 1 | 1.4710e-01 | 3.0711e+05 | ; | CE - NZ |
| 3021 | 3022 | 1 | 1.0100e-01 | 3.6317e+05 | ; | NZ - HZ1 |
| 3021 | 3023 | 1 | 1.0100e-01 | 3.6317e+05 | ; | NZ - HZ2 |
| 3021 | 3024 | 1 | 1.0100e-01 | 3.6317e+05 | ; | NZ - HZ3 |
| 3025 | 3026 | 1 | 1.2290e-01 | 4.7698e+05 | ; | C - O |
| 3025 | 3027 | 1 | 1.3350e-01 | 4.1003e+05 | ; | C - N |
| 3027 | 3028 | 1 | 1.0100e-01 | 3.6317e+05 | ; | N - H |
| 3027 | 3029 | 1 | 1.4490e-01 | 2.8200e+05 | ; | N - CA |
| 3029 | 3030 | 1 | 1.0900e-01 | 2.8451e+05 | ; | CA - HA |
| 3029 | 3031 | 1 | 1.5260e-01 | 2.5941e+05 | ; | CA - CB |
| 3029 | 3046 | 1 | 1.5220e-01 | 2.6527e+05 | ; | CA - C |
| 3031 | 3032 | 1 | 1.0900e-01 | 2.8451e+05 | ; | CB - HB2 |
| 3031 | 3033 | 1 | 1.0900e-01 | 2.8451e+05 | ; | CB - HB3 |
| 3031 | 3034 | 1 | 1.5100e-01 | 2.6527e+05 | ; | CB - CG |
| 3034 | 3035 | 1 | 1.4000e-01 | 3.9246e+05 | ; | CG - CD1 |
| 3034 | 3044 | 1 | 1.4000e-01 | 3.9246e+05 | ; | CG - CD2 |

| | | | | | | |
|---|---|---|---|---|---|---|
| 3035 | 3036 | 1 | 1.0800e-01 | 3.0711e+05 | ; | CD1 - HD1 |
| 3035 | 3037 | 1 | 1.4000e-01 | 3.9246e+05 | ; | CD1 - CE1 |
| 3037 | 3038 | 1 | 1.0800e-01 | 3.0711e+05 | ; | CE1 - HE1 |
| 3037 | 3039 | 1 | 1.4090e-01 | 3.9246e+05 | ; | CE1 - CZ |
| 3039 | 3040 | 1 | 1.3640e-01 | 3.7656e+05 | ; | CZ - OH |
| 3039 | 3042 | 1 | 1.4090e-01 | 3.9246e+05 | ; | CZ - CE2 |
| 3040 | 3041 | 1 | 9.6000e-02 | 4.6275e+05 | ; | OH - HH |
| 3042 | 3043 | 1 | 1.0800e-01 | 3.0711e+05 | ; | CE2 - HE2 |
| 3042 | 3044 | 1 | 1.4000e-01 | 3.9246e+05 | ; | CE2 - CD2 |
| 3044 | 3045 | 1 | 1.0800e-01 | 3.0711e+05 | ; | CD2 - HD2 |
| 3046 | 3047 | 1 | 1.2290e-01 | 4.7698e+05 | ; | C - O |
| 3046 | 3048 | 1 | 1.3350e-01 | 4.1003e+05 | ; | C - N |
| 3048 | 3049 | 1 | 1.0100e-01 | 3.6317e+05 | ; | N - H |
| 3048 | 3050 | 1 | 1.4490e-01 | 2.8200e+05 | ; | N - CA |
| 3050 | 3051 | 1 | 1.0900e-01 | 2.8451e+05 | ; | CA - HA |
| 3050 | 3052 | 1 | 1.5260e-01 | 2.5941e+05 | ; | CA - CB |
| 3050 | 3058 | 1 | 1.5220e-01 | 2.6527e+05 | ; | CA - C |
| 3052 | 3053 | 1 | 1.0900e-01 | 2.8451e+05 | ; | CB - HB2 |
| 3052 | 3054 | 1 | 1.0900e-01 | 2.8451e+05 | ; | CB - HB3 |
| 3052 | 3055 | 1 | 1.5220e-01 | 2.6527e+05 | ; | CB - CG |
| 3055 | 3056 | 1 | 1.2500e-01 | 5.4894e+05 | ; | CG - OD1 |
| 3055 | 3057 | 1 | 1.2500e-01 | 5.4894e+05 | ; | CG - OD2 |
| 3058 | 3059 | 1 | 1.2290e-01 | 4.7698e+05 | ; | C - O |
| 3058 | 3060 | 1 | 1.3350e-01 | 4.1003e+05 | ; | C - N |
| 3060 | 3061 | 1 | 1.0100e-01 | 3.6317e+05 | ; | N - H |
| 3060 | 3062 | 1 | 1.4490e-01 | 2.8200e+05 | ; | N - CA |
| 3062 | 3063 | 1 | 1.0900e-01 | 2.8451e+05 | ; | CA - HA |
| 3062 | 3064 | 1 | 1.5260e-01 | 2.5941e+05 | ; | CA - CB |
| 3062 | 3080 | 1 | 1.5220e-01 | 2.6527e+05 | ; | CA - C |
| 3064 | 3065 | 1 | 1.0900e-01 | 2.8451e+05 | ; | CB - HB2 |
| 3064 | 3066 | 1 | 1.0900e-01 | 2.8451e+05 | ; | CB - HB3 |
| 3064 | 3067 | 1 | 1.5260e-01 | 2.5941e+05 | ; | CB - CG |
| 3067 | 3068 | 1 | 1.0900e-01 | 2.8451e+05 | ; | CG - HG2 |
| 3067 | 3069 | 1 | 1.0900e-01 | 2.8451e+05 | ; | CG - HG3 |
| 3067 | 3070 | 1 | 1.5260e-01 | 2.5941e+05 | ; | CG - CD |
| 3070 | 3071 | 1 | 1.0900e-01 | 2.8451e+05 | ; | CD - HD2 |
| 3070 | 3072 | 1 | 1.0900e-01 | 2.8451e+05 | ; | CD - HD3 |
| 3070 | 3073 | 1 | 1.5260e-01 | 2.5941e+05 | ; | CD - CE |
| 3073 | 3074 | 1 | 1.0900e-01 | 2.8451e+05 | ; | CE - HE2 |
| 3073 | 3075 | 1 | 1.0900e-01 | 2.8451e+05 | ; | CE - HE3 |
| 3073 | 3076 | 1 | 1.4710e-01 | 3.0711e+05 | ; | CE - NZ |
| 3076 | 3077 | 1 | 1.0100e-01 | 3.6317e+05 | ; | NZ - HZ1 |
| 3076 | 3078 | 1 | 1.0100e-01 | 3.6317e+05 | ; | NZ - HZ2 |
| 3076 | 3079 | 1 | 1.0100e-01 | 3.6317e+05 | ; | NZ - HZ3 |
| 3080 | 3081 | 1 | 1.2290e-01 | 4.7698e+05 | ; | C - O |
| 3080 | 3082 | 1 | 1.3350e-01 | 4.1003e+05 | ; | C - N |
| 3082 | 3083 | 1 | 1.0100e-01 | 3.6317e+05 | ; | N - H |
| 3082 | 3084 | 1 | 1.4490e-01 | 2.8200e+05 | ; | N - CA |
| 3084 | 3085 | 1 | 1.0900e-01 | 2.8451e+05 | ; | CA - HA |
| 3084 | 3086 | 1 | 1.5260e-01 | 2.5941e+05 | ; | CA - CB |
| 3084 | 3102 | 1 | 1.5220e-01 | 2.6527e+05 | ; | CA - C |
| 3086 | 3087 | 1 | 1.0900e-01 | 2.8451e+05 | ; | CB - HB2 |

| | | | | | | |
|---|---|---|---|---|---|---|
| 3086 | 3088 | 1 | 1.0900e-01 | 2.8451e+05 | ; | CB - HB3 |
| 3086 | 3089 | 1 | 1.5260e-01 | 2.5941e+05 | ; | CB - CG |
| 3089 | 3090 | 1 | 1.0900e-01 | 2.8451e+05 | ; | CG - HG2 |
| 3089 | 3091 | 1 | 1.0900e-01 | 2.8451e+05 | ; | CG - HG3 |
| 3089 | 3092 | 1 | 1.5260e-01 | 2.5941e+05 | ; | CG - CD |
| 3092 | 3093 | 1 | 1.0900e-01 | 2.8451e+05 | ; | CD - HD2 |
| 3092 | 3094 | 1 | 1.0900e-01 | 2.8451e+05 | ; | CD - HD3 |
| 3092 | 3095 | 1 | 1.5260e-01 | 2.5941e+05 | ; | CD - CE |
| 3095 | 3096 | 1 | 1.0900e-01 | 2.8451e+05 | ; | CE - HE2 |
| 3095 | 3097 | 1 | 1.0900e-01 | 2.8451e+05 | ; | CE - HE3 |
| 3095 | 3098 | 1 | 1.4710e-01 | 3.0711e+05 | ; | CE - NZ |
| 3098 | 3099 | 1 | 1.0100e-01 | 3.6317e+05 | ; | NZ - HZ1 |
| 3098 | 3100 | 1 | 1.0100e-01 | 3.6317e+05 | ; | NZ - HZ2 |
| 3098 | 3101 | 1 | 1.0100e-01 | 3.6317e+05 | ; | NZ - HZ3 |
| 3102 | 3103 | 1 | 1.2290e-01 | 4.7698e+05 | ; | C - O |
| 3102 | 3104 | 1 | 1.3350e-01 | 4.1003e+05 | ; | C - N |
| 3104 | 3105 | 1 | 1.0100e-01 | 3.6317e+05 | ; | N - H |
| 3104 | 3106 | 1 | 1.4490e-01 | 2.8200e+05 | ; | N - CA |
| 3106 | 3107 | 1 | 1.0900e-01 | 2.8451e+05 | ; | CA - HA |
| 3106 | 3108 | 1 | 1.5260e-01 | 2.5941e+05 | ; | CA - CB |
| 3106 | 3113 | 1 | 1.5220e-01 | 2.6527e+05 | ; | CA - C |
| 3108 | 3109 | 1 | 1.0900e-01 | 2.8451e+05 | ; | CB - HB2 |
| 3108 | 3110 | 1 | 1.0900e-01 | 2.8451e+05 | ; | CB - HB3 |
| 3108 | 3111 | 1 | 1.8100e-01 | 1.9832e+05 | ; | CB - SG |
| 3111 | 3112 | 1 | 1.3360e-01 | 2.2928e+05 | ; | SG - HG |
| 3113 | 3114 | 1 | 1.2290e-01 | 4.7698e+05 | ; | C - O |
| 3113 | 3115 | 1 | 1.3350e-01 | 4.1003e+05 | ; | C - N |
| 3115 | 3116 | 1 | 1.0100e-01 | 3.6317e+05 | ; | N - H |
| 3115 | 3117 | 1 | 1.4490e-01 | 2.8200e+05 | ; | N - CA |
| 3117 | 3118 | 1 | 1.0900e-01 | 2.8451e+05 | ; | CA - HA |
| 3117 | 3119 | 1 | 1.5260e-01 | 2.5941e+05 | ; | CA - CB |
| 3117 | 3125 | 1 | 1.5220e-01 | 2.6527e+05 | ; | CA - C |
| 3119 | 3120 | 1 | 1.0900e-01 | 2.8451e+05 | ; | CB - HB2 |
| 3119 | 3121 | 1 | 1.0900e-01 | 2.8451e+05 | ; | CB - HB3 |
| 3119 | 3122 | 1 | 1.5220e-01 | 2.6527e+05 | ; | CB - CG |
| 3122 | 3123 | 1 | 1.2500e-01 | 5.4894e+05 | ; | CG - OD1 |
| 3122 | 3124 | 1 | 1.2500e-01 | 5.4894e+05 | ; | CG - OD2 |
| 3125 | 3126 | 1 | 1.2290e-01 | 4.7698e+05 | ; | C - O |
| 3125 | 3127 | 1 | 1.3350e-01 | 4.1003e+05 | ; | C - N |
| 3127 | 3128 | 1 | 1.0100e-01 | 3.6317e+05 | ; | N - H |
| 3127 | 3129 | 1 | 1.4490e-01 | 2.8200e+05 | ; | N - CA |
| 3129 | 3130 | 1 | 1.0900e-01 | 2.8451e+05 | ; | CA - HA |
| 3129 | 3131 | 1 | 1.5260e-01 | 2.5941e+05 | ; | CA - CB |
| 3129 | 3144 | 1 | 1.5220e-01 | 2.6527e+05 | ; | CA - C |
| 3131 | 3132 | 1 | 1.0900e-01 | 2.8451e+05 | ; | CB - HB |
| 3131 | 3133 | 1 | 1.5260e-01 | 2.5941e+05 | ; | CB - CG2 |
| 3131 | 3137 | 1 | 1.5260e-01 | 2.5941e+05 | ; | CB - CG1 |
| 3133 | 3134 | 1 | 1.0900e-01 | 2.8451e+05 | ; | CG2 - HG21 |
| 3133 | 3135 | 1 | 1.0900e-01 | 2.8451e+05 | ; | CG2 - HG22 |
| 3133 | 3136 | 1 | 1.0900e-01 | 2.8451e+05 | ; | CG2 - HG23 |
| 3137 | 3138 | 1 | 1.0900e-01 | 2.8451e+05 | ; | CG1 - HG12 |
| 3137 | 3139 | 1 | 1.0900e-01 | 2.8451e+05 | ; | CG1 - HG13 |

| | | | | | | |
|---|---|---|---|---|---|---|
| 3137 | 3140 | 1 | 1.5260e-01 | 2.5941e+05 | ; | CG1 - CD1 |
| 3140 | 3141 | 1 | 1.0900e-01 | 2.8451e+05 | ; | CD1 - HD11 |
| 3140 | 3142 | 1 | 1.0900e-01 | 2.8451e+05 | ; | CD1 - HD12 |
| 3140 | 3143 | 1 | 1.0900e-01 | 2.8451e+05 | ; | CD1 - HD13 |
| 3144 | 3145 | 1 | 1.2290e-01 | 4.7698e+05 | ; | C - O |
| 3144 | 3146 | 1 | 1.3350e-01 | 4.1003e+05 | ; | C - N |
| 3146 | 3147 | 1 | 1.0100e-01 | 3.6317e+05 | ; | N - H |
| 3146 | 3148 | 1 | 1.4490e-01 | 2.8200e+05 | ; | N - CA |
| 3148 | 3149 | 1 | 1.0900e-01 | 2.8451e+05 | ; | CA - HA |
| 3148 | 3150 | 1 | 1.5260e-01 | 2.5941e+05 | ; | CA - CB |
| 3148 | 3168 | 1 | 1.5220e-01 | 2.6527e+05 | ; | CA - C |
| 3150 | 3151 | 1 | 1.0900e-01 | 2.8451e+05 | ; | CB - HB2 |
| 3150 | 3152 | 1 | 1.0900e-01 | 2.8451e+05 | ; | CB - HB3 |
| 3150 | 3153 | 1 | 1.4950e-01 | 2.6527e+05 | ; | CB - CG |
| 3153 | 3154 | 1 | 1.3520e-01 | 4.5689e+05 | ; | CG - CD1 |
| 3153 | 3167 | 1 | 1.4590e-01 | 3.2468e+05 | ; | CG - CD2 |
| 3154 | 3155 | 1 | 1.0800e-01 | 3.0711e+05 | ; | CD1 - HD1 |
| 3154 | 3156 | 1 | 1.3810e-01 | 3.5731e+05 | ; | CD1 - NE1 |
| 3156 | 3157 | 1 | 1.0100e-01 | 3.6317e+05 | ; | NE1 - HE1 |
| 3156 | 3158 | 1 | 1.3800e-01 | 3.5815e+05 | ; | NE1 - CE2 |
| 3158 | 3159 | 1 | 1.4000e-01 | 3.9246e+05 | ; | CE2 - CZ2 |
| 3158 | 3167 | 1 | 1.4190e-01 | 3.7405e+05 | ; | CE2 - CD2 |
| 3159 | 3160 | 1 | 1.0800e-01 | 3.0711e+05 | ; | CZ2 - HZ2 |
| 3159 | 3161 | 1 | 1.4000e-01 | 3.9246e+05 | ; | CZ2 - CH2 |
| 3161 | 3162 | 1 | 1.0800e-01 | 3.0711e+05 | ; | CH2 - HH2 |
| 3161 | 3163 | 1 | 1.4000e-01 | 3.9246e+05 | ; | CH2 - CZ3 |
| 3163 | 3164 | 1 | 1.0800e-01 | 3.0711e+05 | ; | CZ3 - HZ3 |
| 3163 | 3165 | 1 | 1.4000e-01 | 3.9246e+05 | ; | CZ3 - CE3 |
| 3165 | 3166 | 1 | 1.0800e-01 | 3.0711e+05 | ; | CE3 - HE3 |
| 3165 | 3167 | 1 | 1.4040e-01 | 3.9246e+05 | ; | CE3 - CD2 |
| 3168 | 3169 | 1 | 1.2290e-01 | 4.7698e+05 | ; | C - O |
| 3168 | 3170 | 1 | 1.3350e-01 | 4.1003e+05 | ; | C - N |
| 3170 | 3171 | 1 | 1.0100e-01 | 3.6317e+05 | ; | N - H |
| 3170 | 3172 | 1 | 1.4490e-01 | 2.8200e+05 | ; | N - CA |
| 3172 | 3173 | 1 | 1.0900e-01 | 2.8451e+05 | ; | CA - HA |
| 3172 | 3174 | 1 | 1.5260e-01 | 2.5941e+05 | ; | CA - CB |
| 3172 | 3179 | 1 | 1.5220e-01 | 2.6527e+05 | ; | CA - C |
| 3174 | 3175 | 1 | 1.0900e-01 | 2.8451e+05 | ; | CB - HB2 |
| 3174 | 3176 | 1 | 1.0900e-01 | 2.8451e+05 | ; | CB - HB3 |
| 3174 | 3177 | 1 | 1.4100e-01 | 2.6778e+05 | ; | CB - OG |
| 3177 | 3178 | 1 | 9.6000e-02 | 4.6275e+05 | ; | OG - HG |
| 3179 | 3180 | 1 | 1.2290e-01 | 4.7698e+05 | ; | C - O |
| 3179 | 3181 | 1 | 1.3350e-01 | 4.1003e+05 | ; | C - N |
| 3181 | 3182 | 1 | 1.0100e-01 | 3.6317e+05 | ; | N - H |
| 3181 | 3183 | 1 | 1.4490e-01 | 2.8200e+05 | ; | N - CA |
| 3183 | 3184 | 1 | 1.0900e-01 | 2.8451e+05 | ; | CA - HA |
| 3183 | 3185 | 1 | 1.5260e-01 | 2.5941e+05 | ; | CA - CB |
| 3183 | 3190 | 1 | 1.5220e-01 | 2.6527e+05 | ; | CA - C |
| 3185 | 3186 | 1 | 1.0900e-01 | 2.8451e+05 | ; | CB - HB2 |
| 3185 | 3187 | 1 | 1.0900e-01 | 2.8451e+05 | ; | CB - HB3 |
| 3185 | 3188 | 1 | 1.4100e-01 | 2.6778e+05 | ; | CB - OG |
| 3188 | 3189 | 1 | 9.6000e-02 | 4.6275e+05 | ; | OG - HG |

| | | | | | | |
|---|---|---|---|---|---|---|
| 3190 | 3191 | 1 | 1.2290e-01 | 4.7698e+05 | ; | C - O |
| 3190 | 3192 | 1 | 1.3350e-01 | 4.1003e+05 | ; | C - N |
| 3192 | 3193 | 1 | 1.0100e-01 | 3.6317e+05 | ; | N - H |
| 3192 | 3194 | 1 | 1.4490e-01 | 2.8200e+05 | ; | N - CA |
| 3194 | 3195 | 1 | 1.0900e-01 | 2.8451e+05 | ; | CA - HA2 |
| 3194 | 3196 | 1 | 1.0900e-01 | 2.8451e+05 | ; | CA - HA3 |
| 3194 | 3197 | 1 | 1.5220e-01 | 2.6527e+05 | ; | CA - C |
| 3197 | 3198 | 1 | 1.2290e-01 | 4.7698e+05 | ; | C - O |
| 3197 | 3199 | 1 | 1.3350e-01 | 4.1003e+05 | ; | C - N |
| 3199 | 3200 | 1 | 1.0100e-01 | 3.6317e+05 | ; | N - H |
| 3199 | 3201 | 1 | 1.4490e-01 | 2.8200e+05 | ; | N - CA |
| 3201 | 3202 | 1 | 1.0900e-01 | 2.8451e+05 | ; | CA - HA |
| 3201 | 3203 | 1 | 1.5260e-01 | 2.5941e+05 | ; | CA - CB |
| 3201 | 3213 | 1 | 1.5220e-01 | 2.6527e+05 | ; | CA - C |
| 3203 | 3204 | 1 | 1.0900e-01 | 2.8451e+05 | ; | CB - HB |
| 3203 | 3205 | 1 | 1.5260e-01 | 2.5941e+05 | ; | CB - CG1 |
| 3203 | 3209 | 1 | 1.5260e-01 | 2.5941e+05 | ; | CB - CG2 |
| 3205 | 3206 | 1 | 1.0900e-01 | 2.8451e+05 | ; | CG1 - HG11 |
| 3205 | 3207 | 1 | 1.0900e-01 | 2.8451e+05 | ; | CG1 - HG12 |
| 3205 | 3208 | 1 | 1.0900e-01 | 2.8451e+05 | ; | CG1 - HG13 |
| 3209 | 3210 | 1 | 1.0900e-01 | 2.8451e+05 | ; | CG2 - HG21 |
| 3209 | 3211 | 1 | 1.0900e-01 | 2.8451e+05 | ; | CG2 - HG22 |
| 3209 | 3212 | 1 | 1.0900e-01 | 2.8451e+05 | ; | CG2 - HG23 |
| 3213 | 3214 | 1 | 1.2290e-01 | 4.7698e+05 | ; | C - O |
| 3213 | 3215 | 1 | 1.3350e-01 | 4.1003e+05 | ; | C - N |
| 3215 | 3216 | 1 | 1.0100e-01 | 3.6317e+05 | ; | N - H |
| 3215 | 3217 | 1 | 1.4490e-01 | 2.8200e+05 | ; | N - CA |
| 3217 | 3218 | 1 | 1.0900e-01 | 2.8451e+05 | ; | CA - HA |
| 3217 | 3219 | 1 | 1.5260e-01 | 2.5941e+05 | ; | CA - CB |
| 3217 | 3232 | 1 | 1.5220e-01 | 2.6527e+05 | ; | CA - C |
| 3219 | 3220 | 1 | 1.0900e-01 | 2.8451e+05 | ; | CB - HB |
| 3219 | 3221 | 1 | 1.5260e-01 | 2.5941e+05 | ; | CB - CG2 |
| 3219 | 3225 | 1 | 1.5260e-01 | 2.5941e+05 | ; | CB - CG1 |
| 3221 | 3222 | 1 | 1.0900e-01 | 2.8451e+05 | ; | CG2 - HG21 |
| 3221 | 3223 | 1 | 1.0900e-01 | 2.8451e+05 | ; | CG2 - HG22 |
| 3221 | 3224 | 1 | 1.0900e-01 | 2.8451e+05 | ; | CG2 - HG23 |
| 3225 | 3226 | 1 | 1.0900e-01 | 2.8451e+05 | ; | CG1 - HG12 |
| 3225 | 3227 | 1 | 1.0900e-01 | 2.8451e+05 | ; | CG1 - HG13 |
| 3225 | 3228 | 1 | 1.5260e-01 | 2.5941e+05 | ; | CG1 - CD1 |
| 3228 | 3229 | 1 | 1.0900e-01 | 2.8451e+05 | ; | CD1 - HD11 |
| 3228 | 3230 | 1 | 1.0900e-01 | 2.8451e+05 | ; | CD1 - HD12 |
| 3228 | 3231 | 1 | 1.0900e-01 | 2.8451e+05 | ; | CD1 - HD13 |
| 3232 | 3233 | 1 | 1.2290e-01 | 4.7698e+05 | ; | C - O |
| 3232 | 3234 | 1 | 1.3350e-01 | 4.1003e+05 | ; | C - N |
| 3234 | 3235 | 1 | 1.0100e-01 | 3.6317e+05 | ; | N - H |
| 3234 | 3236 | 1 | 1.4490e-01 | 2.8200e+05 | ; | N - CA |
| 3236 | 3237 | 1 | 1.0900e-01 | 2.8451e+05 | ; | CA - HA |
| 3236 | 3238 | 1 | 1.5260e-01 | 2.5941e+05 | ; | CA - CB |
| 3236 | 3249 | 1 | 1.5220e-01 | 2.6527e+05 | ; | CA - C |
| 3238 | 3239 | 1 | 1.0900e-01 | 2.8451e+05 | ; | CB - HB2 |
| 3238 | 3240 | 1 | 1.0900e-01 | 2.8451e+05 | ; | CB - HB3 |
| 3238 | 3241 | 1 | 1.5260e-01 | 2.5941e+05 | ; | CB - CG |

| | | | | | | |
|---|---|---|---|---|---|---|
| 3241 | 3242 | 1 | 1.0900e-01 | 2.8451e+05 | ; | CG - HG2 |
| 3241 | 3243 | 1 | 1.0900e-01 | 2.8451e+05 | ; | CG - HG3 |
| 3241 | 3244 | 1 | 1.8100e-01 | 1.8995e+05 | ; | CG - SD |
| 3244 | 3245 | 1 | 1.8100e-01 | 1.8995e+05 | ; | SD - CE |
| 3245 | 3246 | 1 | 1.0900e-01 | 2.8451e+05 | ; | CE - HE1 |
| 3245 | 3247 | 1 | 1.0900e-01 | 2.8451e+05 | ; | CE - HE2 |
| 3245 | 3248 | 1 | 1.0900e-01 | 2.8451e+05 | ; | CE - HE3 |
| 3249 | 3250 | 1 | 1.2290e-01 | 4.7698e+05 | ; | C - O |
| 3249 | 3251 | 1 | 1.3350e-01 | 4.1003e+05 | ; | C - N |
| 3251 | 3252 | 1 | 1.0100e-01 | 3.6317e+05 | ; | N - H |
| 3251 | 3253 | 1 | 1.4490e-01 | 2.8200e+05 | ; | N - CA |
| 3253 | 3254 | 1 | 1.0900e-01 | 2.8451e+05 | ; | CA - HA |
| 3253 | 3255 | 1 | 1.5260e-01 | 2.5941e+05 | ; | CA - CB |
| 3253 | 3270 | 1 | 1.5220e-01 | 2.6527e+05 | ; | CA - C |
| 3255 | 3256 | 1 | 1.0900e-01 | 2.8451e+05 | ; | CB - HB2 |
| 3255 | 3257 | 1 | 1.0900e-01 | 2.8451e+05 | ; | CB - HB3 |
| 3255 | 3258 | 1 | 1.5100e-01 | 2.6527e+05 | ; | CB - CG |
| 3258 | 3259 | 1 | 1.4000e-01 | 3.9246e+05 | ; | CG - CD1 |
| 3258 | 3268 | 1 | 1.4000e-01 | 3.9246e+05 | ; | CG - CD2 |
| 3259 | 3260 | 1 | 1.0800e-01 | 3.0711e+05 | ; | CD1 - HD1 |
| 3259 | 3261 | 1 | 1.4000e-01 | 3.9246e+05 | ; | CD1 - CE1 |
| 3261 | 3262 | 1 | 1.0800e-01 | 3.0711e+05 | ; | CE1 - HE1 |
| 3261 | 3263 | 1 | 1.4090e-01 | 3.9246e+05 | ; | CE1 - CZ |
| 3263 | 3264 | 1 | 1.3640e-01 | 3.7656e+05 | ; | CZ - OH |
| 3263 | 3266 | 1 | 1.4090e-01 | 3.9246e+05 | ; | CZ - CE2 |
| 3264 | 3265 | 1 | 9.6000e-02 | 4.6275e+05 | ; | OH - HH |
| 3266 | 3267 | 1 | 1.0800e-01 | 3.0711e+05 | ; | CE2 - HE2 |
| 3266 | 3268 | 1 | 1.4000e-01 | 3.9246e+05 | ; | CE2 - CD2 |
| 3268 | 3269 | 1 | 1.0800e-01 | 3.0711e+05 | ; | CD2 - HD2 |
| 3270 | 3271 | 1 | 1.2290e-01 | 4.7698e+05 | ; | C - O |
| 3270 | 3272 | 1 | 1.3350e-01 | 4.1003e+05 | ; | C - N |
| 3272 | 3273 | 1 | 1.0100e-01 | 3.6317e+05 | ; | N - H |
| 3272 | 3274 | 1 | 1.4490e-01 | 2.8200e+05 | ; | N - CA |
| 3274 | 3275 | 1 | 1.0900e-01 | 2.8451e+05 | ; | CA - HA |
| 3274 | 3276 | 1 | 1.5260e-01 | 2.5941e+05 | ; | CA - CB |
| 3274 | 3284 | 1 | 1.5220e-01 | 2.6527e+05 | ; | CA - C |
| 3276 | 3277 | 1 | 1.0900e-01 | 2.8451e+05 | ; | CB - HB |
| 3276 | 3278 | 1 | 1.5260e-01 | 2.5941e+05 | ; | CB - CG2 |
| 3276 | 3282 | 1 | 1.4100e-01 | 2.6778e+05 | ; | CB - OG1 |
| 3278 | 3279 | 1 | 1.0900e-01 | 2.8451e+05 | ; | CG2 - HG21 |
| 3278 | 3280 | 1 | 1.0900e-01 | 2.8451e+05 | ; | CG2 - HG22 |
| 3278 | 3281 | 1 | 1.0900e-01 | 2.8451e+05 | ; | CG2 - HG23 |
| 3282 | 3283 | 1 | 9.6000e-02 | 4.6275e+05 | ; | OG1 - HG1 |
| 3284 | 3285 | 1 | 1.2290e-01 | 4.7698e+05 | ; | C - O |
| 3284 | 3286 | 1 | 1.3350e-01 | 4.1003e+05 | ; | C - N |
| 3286 | 3287 | 1 | 1.0100e-01 | 3.6317e+05 | ; | N - H |
| 3286 | 3288 | 1 | 1.4490e-01 | 2.8200e+05 | ; | N - CA |
| 3288 | 3289 | 1 | 1.0900e-01 | 2.8451e+05 | ; | CA - HA |
| 3288 | 3290 | 1 | 1.5260e-01 | 2.5941e+05 | ; | CA - CB |
| 3288 | 3303 | 1 | 1.5220e-01 | 2.6527e+05 | ; | CA - C |
| 3290 | 3291 | 1 | 1.0900e-01 | 2.8451e+05 | ; | CB - HB2 |
| 3290 | 3292 | 1 | 1.0900e-01 | 2.8451e+05 | ; | CB - HB3 |

| | | | | | | |
|---|---|---|---|---|---|---|
| 3290 | 3293 | 1 | 1.5260e-01 | 2.5941e+05 | ; | CB - CG |
| 3293 | 3294 | 1 | 1.0900e-01 | 2.8451e+05 | ; | CG - HG |
| 3293 | 3295 | 1 | 1.5260e-01 | 2.5941e+05 | ; | CG - CD1 |
| 3293 | 3299 | 1 | 1.5260e-01 | 2.5941e+05 | ; | CG - CD2 |
| 3295 | 3296 | 1 | 1.0900e-01 | 2.8451e+05 | ; | CD1 - HD11 |
| 3295 | 3297 | 1 | 1.0900e-01 | 2.8451e+05 | ; | CD1 - HD12 |
| 3295 | 3298 | 1 | 1.0900e-01 | 2.8451e+05 | ; | CD1 - HD13 |
| 3299 | 3300 | 1 | 1.0900e-01 | 2.8451e+05 | ; | CD2 - HD21 |
| 3299 | 3301 | 1 | 1.0900e-01 | 2.8451e+05 | ; | CD2 - HD22 |
| 3299 | 3302 | 1 | 1.0900e-01 | 2.8451e+05 | ; | CD2 - HD23 |
| 3303 | 3304 | 1 | 1.2290e-01 | 4.7698e+05 | ; | C - O |
| 3303 | 3305 | 1 | 1.3350e-01 | 4.1003e+05 | ; | C - N |
| 3305 | 3306 | 1 | 1.0100e-01 | 3.6317e+05 | ; | N - H |
| 3305 | 3307 | 1 | 1.4490e-01 | 2.8200e+05 | ; | N - CA |
| 3307 | 3308 | 1 | 1.0900e-01 | 2.8451e+05 | ; | CA - HA |
| 3307 | 3309 | 1 | 1.5260e-01 | 2.5941e+05 | ; | CA - CB |
| 3307 | 3322 | 1 | 1.5220e-01 | 2.6527e+05 | ; | CA - C |
| 3309 | 3310 | 1 | 1.0900e-01 | 2.8451e+05 | ; | CB - HB2 |
| 3309 | 3311 | 1 | 1.0900e-01 | 2.8451e+05 | ; | CB - HB3 |
| 3309 | 3312 | 1 | 1.5260e-01 | 2.5941e+05 | ; | CB - CG |
| 3312 | 3313 | 1 | 1.0900e-01 | 2.8451e+05 | ; | CG - HG |
| 3312 | 3314 | 1 | 1.5260e-01 | 2.5941e+05 | ; | CG - CD1 |
| 3312 | 3318 | 1 | 1.5260e-01 | 2.5941e+05 | ; | CG - CD2 |
| 3314 | 3315 | 1 | 1.0900e-01 | 2.8451e+05 | ; | CD1 - HD11 |
| 3314 | 3316 | 1 | 1.0900e-01 | 2.8451e+05 | ; | CD1 - HD12 |
| 3314 | 3317 | 1 | 1.0900e-01 | 2.8451e+05 | ; | CD1 - HD13 |
| 3318 | 3319 | 1 | 1.0900e-01 | 2.8451e+05 | ; | CD2 - HD21 |
| 3318 | 3320 | 1 | 1.0900e-01 | 2.8451e+05 | ; | CD2 - HD22 |
| 3318 | 3321 | 1 | 1.0900e-01 | 2.8451e+05 | ; | CD2 - HD23 |
| 3322 | 3323 | 1 | 1.2290e-01 | 4.7698e+05 | ; | C - O |
| 3322 | 3324 | 1 | 1.3350e-01 | 4.1003e+05 | ; | C - N |
| 3324 | 3325 | 1 | 1.0100e-01 | 3.6317e+05 | ; | N - H |
| 3324 | 3326 | 1 | 1.4490e-01 | 2.8200e+05 | ; | N - CA |
| 3326 | 3327 | 1 | 1.0900e-01 | 2.8451e+05 | ; | CA - HA |
| 3326 | 3328 | 1 | 1.5260e-01 | 2.5941e+05 | ; | CA - CB |
| 3326 | 3333 | 1 | 1.5220e-01 | 2.6527e+05 | ; | CA - C |
| 3328 | 3329 | 1 | 1.0900e-01 | 2.8451e+05 | ; | CB - HB2 |
| 3328 | 3330 | 1 | 1.0900e-01 | 2.8451e+05 | ; | CB - HB3 |
| 3328 | 3331 | 1 | 1.8100e-01 | 1.9832e+05 | ; | CB - SG |
| 3331 | 3332 | 1 | 1.3360e-01 | 2.2928e+05 | ; | SG - HG |
| 3333 | 3334 | 1 | 1.2290e-01 | 4.7698e+05 | ; | C - O |
| 3333 | 3335 | 1 | 1.3350e-01 | 4.1003e+05 | ; | C - N |
| 3335 | 3336 | 1 | 1.0100e-01 | 3.6317e+05 | ; | N - H |
| 3335 | 3337 | 1 | 1.4490e-01 | 2.8200e+05 | ; | N - CA |
| 3337 | 3338 | 1 | 1.0900e-01 | 2.8451e+05 | ; | CA - HA2 |
| 3337 | 3339 | 1 | 1.0900e-01 | 2.8451e+05 | ; | CA - HA3 |
| 3337 | 3340 | 1 | 1.5220e-01 | 2.6527e+05 | ; | CA - C |
| 3340 | 3341 | 1 | 1.2290e-01 | 4.7698e+05 | ; | C - O |
| 3340 | 3342 | 1 | 1.3350e-01 | 4.1003e+05 | ; | C - N |
| 3342 | 3343 | 1 | 1.0100e-01 | 3.6317e+05 | ; | N - H |
| 3342 | 3344 | 1 | 1.4490e-01 | 2.8200e+05 | ; | N - CA |
| 3344 | 3345 | 1 | 1.0900e-01 | 2.8451e+05 | ; | CA - HA |

| | | | | | | |
|---|---|---|---|---|---|---|
| 3344 | 3346 | 1 | 1.5260e-01 | 2.5941e+05 | ; | CA - CB |
| 3344 | 3361 | 1 | 1.5220e-01 | 2.6527e+05 | ; | CA - C |
| 3346 | 3347 | 1 | 1.0900e-01 | 2.8451e+05 | ; | CB - HB2 |
| 3346 | 3348 | 1 | 1.0900e-01 | 2.8451e+05 | ; | CB - HB3 |
| 3346 | 3349 | 1 | 1.5100e-01 | 2.6527e+05 | ; | CB - CG |
| 3349 | 3350 | 1 | 1.4000e-01 | 3.9246e+05 | ; | CG - CD1 |
| 3349 | 3359 | 1 | 1.4000e-01 | 3.9246e+05 | ; | CG - CD2 |
| 3350 | 3351 | 1 | 1.0800e-01 | 3.0711e+05 | ; | CD1 - HD1 |
| 3350 | 3352 | 1 | 1.4000e-01 | 3.9246e+05 | ; | CD1 - CE1 |
| 3352 | 3353 | 1 | 1.0800e-01 | 3.0711e+05 | ; | CE1 - HE1 |
| 3352 | 3354 | 1 | 1.4090e-01 | 3.9246e+05 | ; | CE1 - CZ |
| 3354 | 3355 | 1 | 1.3640e-01 | 3.7656e+05 | ; | CZ - OH |
| 3354 | 3357 | 1 | 1.4090e-01 | 3.9246e+05 | ; | CZ - CE2 |
| 3355 | 3356 | 1 | 9.6000e-02 | 4.6275e+05 | ; | OH - HH |
| 3357 | 3358 | 1 | 1.0800e-01 | 3.0711e+05 | ; | CE2 - HE2 |
| 3357 | 3359 | 1 | 1.4000e-01 | 3.9246e+05 | ; | CE2 - CD2 |
| 3359 | 3360 | 1 | 1.0800e-01 | 3.0711e+05 | ; | CD2 - HD2 |
| 3361 | 3362 | 1 | 1.2290e-01 | 4.7698e+05 | ; | C - O |
| 3361 | 3363 | 1 | 1.3350e-01 | 4.1003e+05 | ; | C - N |
| 3363 | 3364 | 1 | 1.4490e-01 | 2.8200e+05 | ; | N - CD |
| 3363 | 3373 | 1 | 1.4490e-01 | 2.8200e+05 | ; | N - CA |
| 3364 | 3365 | 1 | 1.0900e-01 | 2.8451e+05 | ; | CD - HD2 |
| 3364 | 3366 | 1 | 1.0900e-01 | 2.8451e+05 | ; | CD - HD3 |
| 3364 | 3367 | 1 | 1.5260e-01 | 2.5941e+05 | ; | CD - CG |
| 3367 | 3368 | 1 | 1.0900e-01 | 2.8451e+05 | ; | CG - HG2 |
| 3367 | 3369 | 1 | 1.0900e-01 | 2.8451e+05 | ; | CG - HG3 |
| 3367 | 3370 | 1 | 1.5260e-01 | 2.5941e+05 | ; | CG - CB |
| 3370 | 3371 | 1 | 1.0900e-01 | 2.8451e+05 | ; | CB - HB2 |
| 3370 | 3372 | 1 | 1.0900e-01 | 2.8451e+05 | ; | CB - HB3 |
| 3370 | 3373 | 1 | 1.5260e-01 | 2.5941e+05 | ; | CB - CA |
| 3373 | 3374 | 1 | 1.0900e-01 | 2.8451e+05 | ; | CA - HA |
| 3373 | 3375 | 1 | 1.5220e-01 | 2.6527e+05 | ; | CA - C |
| 3375 | 3376 | 1 | 1.2290e-01 | 4.7698e+05 | ; | C - O |
| 3375 | 3377 | 1 | 1.3350e-01 | 4.1003e+05 | ; | C - N |
| 3377 | 3378 | 1 | 1.4490e-01 | 2.8200e+05 | ; | N - CD |
| 3377 | 3387 | 1 | 1.4490e-01 | 2.8200e+05 | ; | N - CA |
| 3378 | 3379 | 1 | 1.0900e-01 | 2.8451e+05 | ; | CD - HD2 |
| 3378 | 3380 | 1 | 1.0900e-01 | 2.8451e+05 | ; | CD - HD3 |
| 3378 | 3381 | 1 | 1.5260e-01 | 2.5941e+05 | ; | CD - CG |
| 3381 | 3382 | 1 | 1.0900e-01 | 2.8451e+05 | ; | CG - HG2 |
| 3381 | 3383 | 1 | 1.0900e-01 | 2.8451e+05 | ; | CG - HG3 |
| 3381 | 3384 | 1 | 1.5260e-01 | 2.5941e+05 | ; | CG - CB |
| 3384 | 3385 | 1 | 1.0900e-01 | 2.8451e+05 | ; | CB - HB2 |
| 3384 | 3386 | 1 | 1.0900e-01 | 2.8451e+05 | ; | CB - HB3 |
| 3384 | 3387 | 1 | 1.5260e-01 | 2.5941e+05 | ; | CB - CA |
| 3387 | 3388 | 1 | 1.0900e-01 | 2.8451e+05 | ; | CA - HA |
| 3387 | 3389 | 1 | 1.5220e-01 | 2.6527e+05 | ; | CA - C |
| 3389 | 3390 | 1 | 1.2290e-01 | 4.7698e+05 | ; | C - O |
| 3389 | 3391 | 1 | 1.3350e-01 | 4.1003e+05 | ; | C - N |
| 3391 | 3392 | 1 | 1.0100e-01 | 3.6317e+05 | ; | N - H |
| 3391 | 3393 | 1 | 1.4490e-01 | 2.8200e+05 | ; | N - CA |
| 3393 | 3394 | 1 | 1.0900e-01 | 2.8451e+05 | ; | CA - HA |

| | | | | | | |
|---|---|---|---|---|---|---|
| 3393 | 3395 | 1 | 1.5260e-01 | 2.5941e+05 | ; | CA - CB |
| 3393 | 3409 | 1 | 1.5220e-01 | 2.6527e+05 | ; | CA - C |
| 3395 | 3396 | 1 | 1.0900e-01 | 2.8451e+05 | ; | CB - HB2 |
| 3395 | 3397 | 1 | 1.0900e-01 | 2.8451e+05 | ; | CB - HB3 |
| 3395 | 3398 | 1 | 1.5100e-01 | 2.6527e+05 | ; | CB - CG |
| 3398 | 3399 | 1 | 1.4000e-01 | 3.9246e+05 | ; | CG - CD1 |
| 3398 | 3407 | 1 | 1.4000e-01 | 3.9246e+05 | ; | CG - CD2 |
| 3399 | 3400 | 1 | 1.0800e-01 | 3.0711e+05 | ; | CD1 - HD1 |
| 3399 | 3401 | 1 | 1.4000e-01 | 3.9246e+05 | ; | CD1 - CE1 |
| 3401 | 3402 | 1 | 1.0800e-01 | 3.0711e+05 | ; | CE1 - HE1 |
| 3401 | 3403 | 1 | 1.4000e-01 | 3.9246e+05 | ; | CE1 - CZ |
| 3403 | 3404 | 1 | 1.0800e-01 | 3.0711e+05 | ; | CZ - HZ |
| 3403 | 3405 | 1 | 1.4000e-01 | 3.9246e+05 | ; | CZ - CE2 |
| 3405 | 3406 | 1 | 1.0800e-01 | 3.0711e+05 | ; | CE2 - HE2 |
| 3405 | 3407 | 1 | 1.4000e-01 | 3.9246e+05 | ; | CE2 - CD2 |
| 3407 | 3408 | 1 | 1.0800e-01 | 3.0711e+05 | ; | CD2 - HD2 |
| 3409 | 3410 | 1 | 1.2290e-01 | 4.7698e+05 | ; | C - O |
| 3409 | 3411 | 1 | 1.3350e-01 | 4.1003e+05 | ; | C - N |
| 3411 | 3412 | 1 | 1.0100e-01 | 3.6317e+05 | ; | N - H |
| 3411 | 3413 | 1 | 1.4490e-01 | 2.8200e+05 | ; | N - CA |
| 3413 | 3414 | 1 | 1.0900e-01 | 2.8451e+05 | ; | CA - HA |
| 3413 | 3415 | 1 | 1.5260e-01 | 2.5941e+05 | ; | CA - CB |
| 3413 | 3430 | 1 | 1.5220e-01 | 2.6527e+05 | ; | CA - C |
| 3415 | 3416 | 1 | 1.0900e-01 | 2.8451e+05 | ; | CB - HB2 |
| 3415 | 3417 | 1 | 1.0900e-01 | 2.8451e+05 | ; | CB - HB3 |
| 3415 | 3418 | 1 | 1.5100e-01 | 2.6527e+05 | ; | CB - CG |
| 3418 | 3419 | 1 | 1.4000e-01 | 3.9246e+05 | ; | CG - CD1 |
| 3418 | 3428 | 1 | 1.4000e-01 | 3.9246e+05 | ; | CG - CD2 |
| 3419 | 3420 | 1 | 1.0800e-01 | 3.0711e+05 | ; | CD1 - HD1 |
| 3419 | 3421 | 1 | 1.4000e-01 | 3.9246e+05 | ; | CD1 - CE1 |
| 3421 | 3422 | 1 | 1.0800e-01 | 3.0711e+05 | ; | CE1 - HE1 |
| 3421 | 3423 | 1 | 1.4090e-01 | 3.9246e+05 | ; | CE1 - CZ |
| 3423 | 3424 | 1 | 1.3640e-01 | 3.7656e+05 | ; | CZ - OH |
| 3423 | 3426 | 1 | 1.4090e-01 | 3.9246e+05 | ; | CZ - CE2 |
| 3424 | 3425 | 1 | 9.6000e-02 | 4.6275e+05 | ; | OH - HH |
| 3426 | 3427 | 1 | 1.0800e-01 | 3.0711e+05 | ; | CE2 - HE2 |
| 3426 | 3428 | 1 | 1.4000e-01 | 3.9246e+05 | ; | CE2 - CD2 |
| 3428 | 3429 | 1 | 1.0800e-01 | 3.0711e+05 | ; | CD2 - HD2 |
| 3430 | 3431 | 1 | 1.2290e-01 | 4.7698e+05 | ; | C - O |
| 3430 | 3432 | 1 | 1.3350e-01 | 4.1003e+05 | ; | C - N |
| 3432 | 3433 | 1 | 1.0100e-01 | 3.6317e+05 | ; | N - H |
| 3432 | 3434 | 1 | 1.4490e-01 | 2.8200e+05 | ; | N - CA |
| 3434 | 3435 | 1 | 1.0900e-01 | 2.8451e+05 | ; | CA - HA2 |
| 3434 | 3436 | 1 | 1.0900e-01 | 2.8451e+05 | ; | CA - HA3 |
| 3434 | 3437 | 1 | 1.5220e-01 | 2.6527e+05 | ; | CA - C |
| 3437 | 3438 | 1 | 1.2290e-01 | 4.7698e+05 | ; | C - O |
| 3437 | 3439 | 1 | 1.3350e-01 | 4.1003e+05 | ; | C - N |
| 3439 | 3440 | 1 | 1.0100e-01 | 3.6317e+05 | ; | N - H |
| 3439 | 3441 | 1 | 1.4490e-01 | 2.8200e+05 | ; | N - CA |
| 3441 | 3442 | 1 | 1.0900e-01 | 2.8451e+05 | ; | CA - HA |
| 3441 | 3443 | 1 | 1.5260e-01 | 2.5941e+05 | ; | CA - CB |
| 3441 | 3449 | 1 | 1.5220e-01 | 2.6527e+05 | ; | CA - C |

| | | | | | | | | |
|---|---|---|---|---|---|---|---|---|
| 3443 | 3444 | 1 | 1.0900e-01 | 2.8451e+05 | ; | CB | - | HB2 |
| 3443 | 3445 | 1 | 1.0900e-01 | 2.8451e+05 | ; | CB | - | HB3 |
| 3443 | 3446 | 1 | 1.5220e-01 | 2.6527e+05 | ; | CB | - | CG |
| 3446 | 3447 | 1 | 1.2500e-01 | 5.4894e+05 | ; | CG | - | OD1 |
| 3446 | 3448 | 1 | 1.2500e-01 | 5.4894e+05 | ; | CG | - | OD2 |
| 3449 | 3450 | 1 | 1.2290e-01 | 4.7698e+05 | ; | C | - | O |
| 3449 | 3451 | 1 | 1.3350e-01 | 4.1003e+05 | ; | C | - | N |
| 3451 | 3452 | 1 | 1.0100e-01 | 3.6317e+05 | ; | N | - | H |
| 3451 | 3453 | 1 | 1.4490e-01 | 2.8200e+05 | ; | N | - | CA |
| 3453 | 3454 | 1 | 1.0900e-01 | 2.8451e+05 | ; | CA | - | HA |
| 3453 | 3455 | 1 | 1.5260e-01 | 2.5941e+05 | ; | CA | - | CB |
| 3453 | 3463 | 1 | 1.5220e-01 | 2.6527e+05 | ; | CA | - | C |
| 3455 | 3456 | 1 | 1.0900e-01 | 2.8451e+05 | ; | CB | - | HB |
| 3455 | 3457 | 1 | 1.5260e-01 | 2.5941e+05 | ; | CB | - | CG2 |
| 3455 | 3461 | 1 | 1.4100e-01 | 2.6778e+05 | ; | CB | - | OG1 |
| 3457 | 3458 | 1 | 1.0900e-01 | 2.8451e+05 | ; | CG2 | - | HG21 |
| 3457 | 3459 | 1 | 1.0900e-01 | 2.8451e+05 | ; | CG2 | - | HG22 |
| 3457 | 3460 | 1 | 1.0900e-01 | 2.8451e+05 | ; | CG2 | - | HG23 |
| 3461 | 3462 | 1 | 9.6000e-02 | 4.6275e+05 | ; | OG1 | - | HG1 |
| 3463 | 3464 | 1 | 1.2290e-01 | 4.7698e+05 | ; | C | - | O |
| 3463 | 3465 | 1 | 1.3350e-01 | 4.1003e+05 | ; | C | - | N |
| 3465 | 3466 | 1 | 1.0100e-01 | 3.6317e+05 | ; | N | - | H |
| 3465 | 3467 | 1 | 1.4490e-01 | 2.8200e+05 | ; | N | - | CA |
| 3467 | 3468 | 1 | 1.0900e-01 | 2.8451e+05 | ; | CA | - | HA |
| 3467 | 3469 | 1 | 1.5260e-01 | 2.5941e+05 | ; | CA | - | CB |
| 3467 | 3475 | 1 | 1.5220e-01 | 2.6527e+05 | ; | CA | - | C |
| 3469 | 3470 | 1 | 1.0900e-01 | 2.8451e+05 | ; | CB | - | HB2 |
| 3469 | 3471 | 1 | 1.0900e-01 | 2.8451e+05 | ; | CB | - | HB3 |
| 3469 | 3472 | 1 | 1.5220e-01 | 2.6527e+05 | ; | CB | - | CG |
| 3472 | 3473 | 1 | 1.2500e-01 | 5.4894e+05 | ; | CG | - | OD1 |
| 3472 | 3474 | 1 | 1.2500e-01 | 5.4894e+05 | ; | CG | - | OD2 |
| 3475 | 3476 | 1 | 1.2290e-01 | 4.7698e+05 | ; | C | - | O |
| 3475 | 3477 | 1 | 1.3350e-01 | 4.1003e+05 | ; | C | - | N |
| 3477 | 3478 | 1 | 1.0100e-01 | 3.6317e+05 | ; | N | - | H |
| 3477 | 3479 | 1 | 1.4490e-01 | 2.8200e+05 | ; | N | - | CA |
| 3479 | 3480 | 1 | 1.0900e-01 | 2.8451e+05 | ; | CA | - | HA |
| 3479 | 3481 | 1 | 1.5260e-01 | 2.5941e+05 | ; | CA | - | CB |
| 3479 | 3489 | 1 | 1.5220e-01 | 2.6527e+05 | ; | CA | - | C |
| 3481 | 3482 | 1 | 1.0900e-01 | 2.8451e+05 | ; | CB | - | HB2 |
| 3481 | 3483 | 1 | 1.0900e-01 | 2.8451e+05 | ; | CB | - | HB3 |
| 3481 | 3484 | 1 | 1.5220e-01 | 2.6527e+05 | ; | CB | - | CG |
| 3484 | 3485 | 1 | 1.2290e-01 | 4.7698e+05 | ; | CG | - | OD1 |
| 3484 | 3486 | 1 | 1.3350e-01 | 4.1003e+05 | ; | CG | - | ND2 |
| 3486 | 3487 | 1 | 1.0100e-01 | 3.6317e+05 | ; | ND2 | - | HD21 |
| 3486 | 3488 | 1 | 1.0100e-01 | 3.6317e+05 | ; | ND2 | - | HD22 |
| 3489 | 3490 | 1 | 1.2290e-01 | 4.7698e+05 | ; | C | - | O |
| 3489 | 3491 | 1 | 1.3350e-01 | 4.1003e+05 | ; | C | - | N |
| 3491 | 3492 | 1 | 1.0100e-01 | 3.6317e+05 | ; | N | - | H |
| 3491 | 3493 | 1 | 1.4490e-01 | 2.8200e+05 | ; | N | - | CA |
| 3493 | 3494 | 1 | 1.0900e-01 | 2.8451e+05 | ; | CA | - | HA |
| 3493 | 3495 | 1 | 1.5260e-01 | 2.5941e+05 | ; | CA | - | CB |
| 3493 | 3504 | 1 | 1.5220e-01 | 2.6527e+05 | ; | CA | - | C |

| | | | | | | |
|---|---|---|---|---|---|---|
| 3495 | 3496 | 1 | 1.0900e-01 | 2.8451e+05 | ; | CB - HB2 |
| 3495 | 3497 | 1 | 1.0900e-01 | 2.8451e+05 | ; | CB - HB3 |
| 3495 | 3498 | 1 | 1.5260e-01 | 2.5941e+05 | ; | CB - CG |
| 3498 | 3499 | 1 | 1.0900e-01 | 2.8451e+05 | ; | CG - HG2 |
| 3498 | 3500 | 1 | 1.0900e-01 | 2.8451e+05 | ; | CG - HG3 |
| 3498 | 3501 | 1 | 1.5220e-01 | 2.6527e+05 | ; | CG - CD |
| 3501 | 3502 | 1 | 1.2500e-01 | 5.4894e+05 | ; | CD - OE1 |
| 3501 | 3503 | 1 | 1.2500e-01 | 5.4894e+05 | ; | CD - OE2 |
| 3504 | 3505 | 1 | 1.2290e-01 | 4.7698e+05 | ; | C - O |
| 3504 | 3506 | 1 | 1.3350e-01 | 4.1003e+05 | ; | C - N |
| 3506 | 3507 | 1 | 1.0100e-01 | 3.6317e+05 | ; | N - H |
| 3506 | 3508 | 1 | 1.4490e-01 | 2.8200e+05 | ; | N - CA |
| 3508 | 3509 | 1 | 1.0900e-01 | 2.8451e+05 | ; | CA - HA |
| 3508 | 3510 | 1 | 1.5260e-01 | 2.5941e+05 | ; | CA - CB |
| 3508 | 3520 | 1 | 1.5220e-01 | 2.6527e+05 | ; | CA - C |
| 3510 | 3511 | 1 | 1.0900e-01 | 2.8451e+05 | ; | CB - HB |
| 3510 | 3512 | 1 | 1.5260e-01 | 2.5941e+05 | ; | CB - CG1 |
| 3510 | 3516 | 1 | 1.5260e-01 | 2.5941e+05 | ; | CB - CG2 |
| 3512 | 3513 | 1 | 1.0900e-01 | 2.8451e+05 | ; | CG1 - HG11 |
| 3512 | 3514 | 1 | 1.0900e-01 | 2.8451e+05 | ; | CG1 - HG12 |
| 3512 | 3515 | 1 | 1.0900e-01 | 2.8451e+05 | ; | CG1 - HG13 |
| 3516 | 3517 | 1 | 1.0900e-01 | 2.8451e+05 | ; | CG2 - HG21 |
| 3516 | 3518 | 1 | 1.0900e-01 | 2.8451e+05 | ; | CG2 - HG22 |
| 3516 | 3519 | 1 | 1.0900e-01 | 2.8451e+05 | ; | CG2 - HG23 |
| 3520 | 3521 | 1 | 1.2290e-01 | 4.7698e+05 | ; | C - O |
| 3520 | 3522 | 1 | 1.3350e-01 | 4.1003e+05 | ; | C - N |
| 3522 | 3523 | 1 | 1.0100e-01 | 3.6317e+05 | ; | N - H |
| 3522 | 3524 | 1 | 1.4490e-01 | 2.8200e+05 | ; | N - CA |
| 3524 | 3525 | 1 | 1.0900e-01 | 2.8451e+05 | ; | CA - HA |
| 3524 | 3526 | 1 | 1.5260e-01 | 2.5941e+05 | ; | CA - CB |
| 3524 | 3539 | 1 | 1.5220e-01 | 2.6527e+05 | ; | CA - C |
| 3526 | 3527 | 1 | 1.0900e-01 | 2.8451e+05 | ; | CB - HB2 |
| 3526 | 3528 | 1 | 1.0900e-01 | 2.8451e+05 | ; | CB - HB3 |
| 3526 | 3529 | 1 | 1.5260e-01 | 2.5941e+05 | ; | CB - CG |
| 3529 | 3530 | 1 | 1.0900e-01 | 2.8451e+05 | ; | CG - HG |
| 3529 | 3531 | 1 | 1.5260e-01 | 2.5941e+05 | ; | CG - CD1 |
| 3529 | 3535 | 1 | 1.5260e-01 | 2.5941e+05 | ; | CG - CD2 |
| 3531 | 3532 | 1 | 1.0900e-01 | 2.8451e+05 | ; | CD1 - HD11 |
| 3531 | 3533 | 1 | 1.0900e-01 | 2.8451e+05 | ; | CD1 - HD12 |
| 3531 | 3534 | 1 | 1.0900e-01 | 2.8451e+05 | ; | CD1 - HD13 |
| 3535 | 3536 | 1 | 1.0900e-01 | 2.8451e+05 | ; | CD2 - HD21 |
| 3535 | 3537 | 1 | 1.0900e-01 | 2.8451e+05 | ; | CD2 - HD22 |
| 3535 | 3538 | 1 | 1.0900e-01 | 2.8451e+05 | ; | CD2 - HD23 |
| 3539 | 3540 | 1 | 1.2290e-01 | 4.7698e+05 | ; | C - O |
| 3539 | 3541 | 1 | 1.3350e-01 | 4.1003e+05 | ; | C - N |
| 3541 | 3542 | 1 | 1.0100e-01 | 3.6317e+05 | ; | N - H |
| 3541 | 3543 | 1 | 1.4490e-01 | 2.8200e+05 | ; | N - CA |
| 3543 | 3544 | 1 | 1.0900e-01 | 2.8451e+05 | ; | CA - HA |
| 3543 | 3545 | 1 | 1.5260e-01 | 2.5941e+05 | ; | CA - CB |
| 3543 | 3561 | 1 | 1.5220e-01 | 2.6527e+05 | ; | CA - C |
| 3545 | 3546 | 1 | 1.0900e-01 | 2.8451e+05 | ; | CB - HB2 |
| 3545 | 3547 | 1 | 1.0900e-01 | 2.8451e+05 | ; | CB - HB3 |

| | | | | | | |
|---|---|---|---|---|---|---|
| 3545 | 3548 | 1 | 1.5260e-01 | 2.5941e+05 | ; | CB - CG |
| 3548 | 3549 | 1 | 1.0900e-01 | 2.8451e+05 | ; | CG - HG2 |
| 3548 | 3550 | 1 | 1.0900e-01 | 2.8451e+05 | ; | CG - HG3 |
| 3548 | 3551 | 1 | 1.5260e-01 | 2.5941e+05 | ; | CG - CD |
| 3551 | 3552 | 1 | 1.0900e-01 | 2.8451e+05 | ; | CD - HD2 |
| 3551 | 3553 | 1 | 1.0900e-01 | 2.8451e+05 | ; | CD - HD3 |
| 3551 | 3554 | 1 | 1.5260e-01 | 2.5941e+05 | ; | CD - CE |
| 3554 | 3555 | 1 | 1.0900e-01 | 2.8451e+05 | ; | CE - HE2 |
| 3554 | 3556 | 1 | 1.0900e-01 | 2.8451e+05 | ; | CE - HE3 |
| 3554 | 3557 | 1 | 1.4710e-01 | 3.0711e+05 | ; | CE - NZ |
| 3557 | 3558 | 1 | 1.0100e-01 | 3.6317e+05 | ; | NZ - HZ1 |
| 3557 | 3559 | 1 | 1.0100e-01 | 3.6317e+05 | ; | NZ - HZ2 |
| 3557 | 3560 | 1 | 1.0100e-01 | 3.6317e+05 | ; | NZ - HZ3 |
| 3561 | 3562 | 1 | 1.2290e-01 | 4.7698e+05 | ; | C - O |
| 3561 | 3563 | 1 | 1.3350e-01 | 4.1003e+05 | ; | C - N |
| 3563 | 3564 | 1 | 1.0100e-01 | 3.6317e+05 | ; | N - H |
| 3563 | 3565 | 1 | 1.4490e-01 | 2.8200e+05 | ; | N - CA |
| 3565 | 3566 | 1 | 1.0900e-01 | 2.8451e+05 | ; | CA - HA |
| 3565 | 3567 | 1 | 1.5260e-01 | 2.5941e+05 | ; | CA - CB |
| 3565 | 3583 | 1 | 1.5220e-01 | 2.6527e+05 | ; | CA - C |
| 3567 | 3568 | 1 | 1.0900e-01 | 2.8451e+05 | ; | CB - HB2 |
| 3567 | 3569 | 1 | 1.0900e-01 | 2.8451e+05 | ; | CB - HB3 |
| 3567 | 3570 | 1 | 1.5260e-01 | 2.5941e+05 | ; | CB - CG |
| 3570 | 3571 | 1 | 1.0900e-01 | 2.8451e+05 | ; | CG - HG2 |
| 3570 | 3572 | 1 | 1.0900e-01 | 2.8451e+05 | ; | CG - HG3 |
| 3570 | 3573 | 1 | 1.5260e-01 | 2.5941e+05 | ; | CG - CD |
| 3573 | 3574 | 1 | 1.0900e-01 | 2.8451e+05 | ; | CD - HD2 |
| 3573 | 3575 | 1 | 1.0900e-01 | 2.8451e+05 | ; | CD - HD3 |
| 3573 | 3576 | 1 | 1.5260e-01 | 2.5941e+05 | ; | CD - CE |
| 3576 | 3577 | 1 | 1.0900e-01 | 2.8451e+05 | ; | CE - HE2 |
| 3576 | 3578 | 1 | 1.0900e-01 | 2.8451e+05 | ; | CE - HE3 |
| 3576 | 3579 | 1 | 1.4710e-01 | 3.0711e+05 | ; | CE - NZ |
| 3579 | 3580 | 1 | 1.0100e-01 | 3.6317e+05 | ; | NZ - HZ1 |
| 3579 | 3581 | 1 | 1.0100e-01 | 3.6317e+05 | ; | NZ - HZ2 |
| 3579 | 3582 | 1 | 1.0100e-01 | 3.6317e+05 | ; | NZ - HZ3 |
| 3583 | 3584 | 1 | 1.2290e-01 | 4.7698e+05 | ; | C - O |
| 3583 | 3585 | 1 | 1.3350e-01 | 4.1003e+05 | ; | C - N |
| 3585 | 3586 | 1 | 1.0100e-01 | 3.6317e+05 | ; | N - H |
| 3585 | 3587 | 1 | 1.4490e-01 | 2.8200e+05 | ; | N - CA |
| 3587 | 3588 | 1 | 1.0900e-01 | 2.8451e+05 | ; | CA - HA |
| 3587 | 3589 | 1 | 1.5260e-01 | 2.5941e+05 | ; | CA - CB |
| 3587 | 3599 | 1 | 1.5220e-01 | 2.6527e+05 | ; | CA - C |
| 3589 | 3590 | 1 | 1.0900e-01 | 2.8451e+05 | ; | CB - HB |
| 3589 | 3591 | 1 | 1.5260e-01 | 2.5941e+05 | ; | CB - CG1 |
| 3589 | 3595 | 1 | 1.5260e-01 | 2.5941e+05 | ; | CB - CG2 |
| 3591 | 3592 | 1 | 1.0900e-01 | 2.8451e+05 | ; | CG1 - HG11 |
| 3591 | 3593 | 1 | 1.0900e-01 | 2.8451e+05 | ; | CG1 - HG12 |
| 3591 | 3594 | 1 | 1.0900e-01 | 2.8451e+05 | ; | CG1 - HG13 |
| 3595 | 3596 | 1 | 1.0900e-01 | 2.8451e+05 | ; | CG2 - HG21 |
| 3595 | 3597 | 1 | 1.0900e-01 | 2.8451e+05 | ; | CG2 - HG22 |
| 3595 | 3598 | 1 | 1.0900e-01 | 2.8451e+05 | ; | CG2 - HG23 |
| 3599 | 3600 | 1 | 1.2290e-01 | 4.7698e+05 | ; | C - O |

| | | | | | | | | |
|---|---|---|---|---|---|---|---|---|
| 3599 | 3601 | 1 | 1.3350e-01 | 4.1003e+05 | ; | C | - | N |
| 3601 | 3602 | 1 | 1.0100e-01 | 3.6317e+05 | ; | N | - | H |
| 3601 | 3603 | 1 | 1.4490e-01 | 2.8200e+05 | ; | N | - | CA |
| 3603 | 3604 | 1 | 1.0900e-01 | 2.8451e+05 | ; | CA | - | HA |
| 3603 | 3605 | 1 | 1.5260e-01 | 2.5941e+05 | ; | CA | - | CB |
| 3603 | 3621 | 1 | 1.5220e-01 | 2.6527e+05 | ; | CA | - | C |
| 3605 | 3606 | 1 | 1.0900e-01 | 2.8451e+05 | ; | CB | - | HB2 |
| 3605 | 3607 | 1 | 1.0900e-01 | 2.8451e+05 | ; | CB | - | HB3 |
| 3605 | 3608 | 1 | 1.5260e-01 | 2.5941e+05 | ; | CB | - | CG |
| 3608 | 3609 | 1 | 1.0900e-01 | 2.8451e+05 | ; | CG | - | HG2 |
| 3608 | 3610 | 1 | 1.0900e-01 | 2.8451e+05 | ; | CG | - | HG3 |
| 3608 | 3611 | 1 | 1.5260e-01 | 2.5941e+05 | ; | CG | - | CD |
| 3611 | 3612 | 1 | 1.0900e-01 | 2.8451e+05 | ; | CD | - | HD2 |
| 3611 | 3613 | 1 | 1.0900e-01 | 2.8451e+05 | ; | CD | - | HD3 |
| 3611 | 3614 | 1 | 1.5260e-01 | 2.5941e+05 | ; | CD | - | CE |
| 3614 | 3615 | 1 | 1.0900e-01 | 2.8451e+05 | ; | CE | - | HE2 |
| 3614 | 3616 | 1 | 1.0900e-01 | 2.8451e+05 | ; | CE | - | HE3 |
| 3614 | 3617 | 1 | 1.4710e-01 | 3.0711e+05 | ; | CE | - | NZ |
| 3617 | 3618 | 1 | 1.0100e-01 | 3.6317e+05 | ; | NZ | - | HZ1 |
| 3617 | 3619 | 1 | 1.0100e-01 | 3.6317e+05 | ; | NZ | - | HZ2 |
| 3617 | 3620 | 1 | 1.0100e-01 | 3.6317e+05 | ; | NZ | - | HZ3 |
| 3621 | 3622 | 1 | 1.2290e-01 | 4.7698e+05 | ; | C | - | O |
| 3621 | 3623 | 1 | 1.3350e-01 | 4.1003e+05 | ; | C | - | N |
| 3623 | 3624 | 1 | 1.0100e-01 | 3.6317e+05 | ; | N | - | H |
| 3623 | 3625 | 1 | 1.4490e-01 | 2.8200e+05 | ; | N | - | CA |
| 3625 | 3626 | 1 | 1.0900e-01 | 2.8451e+05 | ; | CA | - | HA |
| 3625 | 3627 | 1 | 1.5260e-01 | 2.5941e+05 | ; | CA | - | CB |
| 3625 | 3643 | 1 | 1.5220e-01 | 2.6527e+05 | ; | CA | - | C |
| 3627 | 3628 | 1 | 1.0900e-01 | 2.8451e+05 | ; | CB | - | HB2 |
| 3627 | 3629 | 1 | 1.0900e-01 | 2.8451e+05 | ; | CB | - | HB3 |
| 3627 | 3630 | 1 | 1.5260e-01 | 2.5941e+05 | ; | CB | - | CG |
| 3630 | 3631 | 1 | 1.0900e-01 | 2.8451e+05 | ; | CG | - | HG2 |
| 3630 | 3632 | 1 | 1.0900e-01 | 2.8451e+05 | ; | CG | - | HG3 |
| 3630 | 3633 | 1 | 1.5260e-01 | 2.5941e+05 | ; | CG | - | CD |
| 3633 | 3634 | 1 | 1.0900e-01 | 2.8451e+05 | ; | CD | - | HD2 |
| 3633 | 3635 | 1 | 1.0900e-01 | 2.8451e+05 | ; | CD | - | HD3 |
| 3633 | 3636 | 1 | 1.5260e-01 | 2.5941e+05 | ; | CD | - | CE |
| 3636 | 3637 | 1 | 1.0900e-01 | 2.8451e+05 | ; | CE | - | HE2 |
| 3636 | 3638 | 1 | 1.0900e-01 | 2.8451e+05 | ; | CE | - | HE3 |
| 3636 | 3639 | 1 | 1.4710e-01 | 3.0711e+05 | ; | CE | - | NZ |
| 3639 | 3640 | 1 | 1.0100e-01 | 3.6317e+05 | ; | NZ | - | HZ1 |
| 3639 | 3641 | 1 | 1.0100e-01 | 3.6317e+05 | ; | NZ | - | HZ2 |
| 3639 | 3642 | 1 | 1.0100e-01 | 3.6317e+05 | ; | NZ | - | HZ3 |
| 3643 | 3644 | 1 | 1.2290e-01 | 4.7698e+05 | ; | C | - | O |
| 3643 | 3645 | 1 | 1.3350e-01 | 4.1003e+05 | ; | C | - | N |
| 3645 | 3646 | 1 | 1.0100e-01 | 3.6317e+05 | ; | N | - | H |
| 3645 | 3647 | 1 | 1.4490e-01 | 2.8200e+05 | ; | N | - | CA |
| 3647 | 3648 | 1 | 1.0900e-01 | 2.8451e+05 | ; | CA | - | HA2 |
| 3647 | 3649 | 1 | 1.0900e-01 | 2.8451e+05 | ; | CA | - | HA3 |
| 3647 | 3650 | 1 | 1.5220e-01 | 2.6527e+05 | ; | CA | - | C |
| 3650 | 3651 | 1 | 1.2290e-01 | 4.7698e+05 | ; | C | - | O |
| 3650 | 3652 | 1 | 1.3350e-01 | 4.1003e+05 | ; | C | - | N |

| | | | | | | | | |
|---|---|---|---|---|---|---|---|---|
| 3652 | 3653 | 1 | 1.0100e-01 | 3.6317e+05 | ; | N | - | H |
| 3652 | 3654 | 1 | 1.4490e-01 | 2.8200e+05 | ; | N | - | CA |
| 3654 | 3655 | 1 | 1.0900e-01 | 2.8451e+05 | ; | CA | - | HA |
| 3654 | 3656 | 1 | 1.5260e-01 | 2.5941e+05 | ; | CA | - | CB |
| 3654 | 3665 | 1 | 1.5220e-01 | 2.6527e+05 | ; | CA | - | C |
| 3656 | 3657 | 1 | 1.0900e-01 | 2.8451e+05 | ; | CB | - | HB2 |
| 3656 | 3658 | 1 | 1.0900e-01 | 2.8451e+05 | ; | CB | - | HB3 |
| 3656 | 3659 | 1 | 1.5260e-01 | 2.5941e+05 | ; | CB | - | CG |
| 3659 | 3660 | 1 | 1.0900e-01 | 2.8451e+05 | ; | CG | - | HG2 |
| 3659 | 3661 | 1 | 1.0900e-01 | 2.8451e+05 | ; | CG | - | HG3 |
| 3659 | 3662 | 1 | 1.5220e-01 | 2.6527e+05 | ; | CG | - | CD |
| 3662 | 3663 | 1 | 1.2500e-01 | 5.4894e+05 | ; | CD | - | OE1 |
| 3662 | 3664 | 1 | 1.2500e-01 | 5.4894e+05 | ; | CD | - | OE2 |
| 3665 | 3666 | 1 | 1.2290e-01 | 4.7698e+05 | ; | C | - | O |
| 3665 | 3667 | 1 | 1.3350e-01 | 4.1003e+05 | ; | C | - | N |
| 3667 | 3668 | 1 | 1.0100e-01 | 3.6317e+05 | ; | N | - | H |
| 3667 | 3669 | 1 | 1.4490e-01 | 2.8200e+05 | ; | N | - | CA |
| 3669 | 3670 | 1 | 1.0900e-01 | 2.8451e+05 | ; | CA | - | HA |
| 3669 | 3671 | 1 | 1.5260e-01 | 2.5941e+05 | ; | CA | - | CB |
| 3669 | 3685 | 1 | 1.5220e-01 | 2.6527e+05 | ; | CA | - | C |
| 3671 | 3672 | 1 | 1.0900e-01 | 2.8451e+05 | ; | CB | - | HB2 |
| 3671 | 3673 | 1 | 1.0900e-01 | 2.8451e+05 | ; | CB | - | HB3 |
| 3671 | 3674 | 1 | 1.5100e-01 | 2.6527e+05 | ; | CB | - | CG |
| 3674 | 3675 | 1 | 1.4000e-01 | 3.9246e+05 | ; | CG | - | CD1 |
| 3674 | 3683 | 1 | 1.4000e-01 | 3.9246e+05 | ; | CG | - | CD2 |
| 3675 | 3676 | 1 | 1.0800e-01 | 3.0711e+05 | ; | CD1 | - | HD1 |
| 3675 | 3677 | 1 | 1.4000e-01 | 3.9246e+05 | ; | CD1 | - | CE1 |
| 3677 | 3678 | 1 | 1.0800e-01 | 3.0711e+05 | ; | CE1 | - | HE1 |
| 3677 | 3679 | 1 | 1.4000e-01 | 3.9246e+05 | ; | CE1 | - | CZ |
| 3679 | 3680 | 1 | 1.0800e-01 | 3.0711e+05 | ; | CZ | - | HZ |
| 3679 | 3681 | 1 | 1.4000e-01 | 3.9246e+05 | ; | CZ | - | CE2 |
| 3681 | 3682 | 1 | 1.0800e-01 | 3.0711e+05 | ; | CE2 | - | HE2 |
| 3681 | 3683 | 1 | 1.4000e-01 | 3.9246e+05 | ; | CE2 | - | CD2 |
| 3683 | 3684 | 1 | 1.0800e-01 | 3.0711e+05 | ; | CD2 | - | HD2 |
| 3685 | 3686 | 1 | 1.2290e-01 | 4.7698e+05 | ; | C | - | O |
| 3685 | 3687 | 1 | 1.3350e-01 | 4.1003e+05 | ; | C | - | N |
| 3687 | 3688 | 1 | 1.0100e-01 | 3.6317e+05 | ; | N | - | H |
| 3687 | 3689 | 1 | 1.4490e-01 | 2.8200e+05 | ; | N | - | CA |
| 3689 | 3690 | 1 | 1.0900e-01 | 2.8451e+05 | ; | CA | - | HA |
| 3689 | 3691 | 1 | 1.5260e-01 | 2.5941e+05 | ; | CA | - | CB |
| 3689 | 3696 | 1 | 1.5220e-01 | 2.6527e+05 | ; | CA | - | C |
| 3691 | 3692 | 1 | 1.0900e-01 | 2.8451e+05 | ; | CB | - | HB2 |
| 3691 | 3693 | 1 | 1.0900e-01 | 2.8451e+05 | ; | CB | - | HB3 |
| 3691 | 3694 | 1 | 1.8100e-01 | 1.9832e+05 | ; | CB | - | SG |
| 3694 | 3695 | 1 | 1.3360e-01 | 2.2928e+05 | ; | SG | - | HG |
| 3696 | 3697 | 1 | 1.2290e-01 | 4.7698e+05 | ; | C | - | O |
| 3696 | 3698 | 1 | 1.3350e-01 | 4.1003e+05 | ; | C | - | N |
| 3698 | 3699 | 1 | 1.0100e-01 | 3.6317e+05 | ; | N | - | H |
| 3698 | 3700 | 1 | 1.4490e-01 | 2.8200e+05 | ; | N | - | CA |
| 3700 | 3701 | 1 | 1.0900e-01 | 2.8451e+05 | ; | CA | - | HA |
| 3700 | 3702 | 1 | 1.5260e-01 | 2.5941e+05 | ; | CA | - | CB |
| 3700 | 3716 | 1 | 1.5220e-01 | 2.6527e+05 | ; | CA | - | C |

| | | | | | |
|---|---|---|---|---|---|
| 3702 | 3703 | 1 | 1.0900e-01 | 2.8451e+05 ; | CB - HB2 |
| 3702 | 3704 | 1 | 1.0900e-01 | 2.8451e+05 ; | CB - HB3 |
| 3702 | 3705 | 1 | 1.5100e-01 | 2.6527e+05 ; | CB - CG |
| 3705 | 3706 | 1 | 1.4000e-01 | 3.9246e+05 ; | CG - CD1 |
| 3705 | 3714 | 1 | 1.4000e-01 | 3.9246e+05 ; | CG - CD2 |
| 3706 | 3707 | 1 | 1.0800e-01 | 3.0711e+05 ; | CD1 - HD1 |
| 3706 | 3708 | 1 | 1.4000e-01 | 3.9246e+05 ; | CD1 - CE1 |
| 3708 | 3709 | 1 | 1.0800e-01 | 3.0711e+05 ; | CE1 - HE1 |
| 3708 | 3710 | 1 | 1.4000e-01 | 3.9246e+05 ; | CE1 - CZ |
| 3710 | 3711 | 1 | 1.0800e-01 | 3.0711e+05 ; | CZ - HZ |
| 3710 | 3712 | 1 | 1.4000e-01 | 3.9246e+05 ; | CZ - CE2 |
| 3712 | 3713 | 1 | 1.0800e-01 | 3.0711e+05 ; | CE2 - HE2 |
| 3712 | 3714 | 1 | 1.4000e-01 | 3.9246e+05 ; | CE2 - CD2 |
| 3714 | 3715 | 1 | 1.0800e-01 | 3.0711e+05 ; | CD2 - HD2 |
| 3716 | 3717 | 1 | 1.2290e-01 | 4.7698e+05 ; | C - O |
| 3716 | 3718 | 1 | 1.3350e-01 | 4.1003e+05 ; | C - N |
| 3718 | 3719 | 1 | 1.0100e-01 | 3.6317e+05 ; | N - H |
| 3718 | 3720 | 1 | 1.4490e-01 | 2.8200e+05 ; | N - CA |
| 3720 | 3721 | 1 | 1.0900e-01 | 2.8451e+05 ; | CA - HA |
| 3720 | 3722 | 1 | 1.5260e-01 | 2.5941e+05 ; | CA - CB |
| 3720 | 3737 | 1 | 1.5220e-01 | 2.6527e+05 ; | CA - C |
| 3722 | 3723 | 1 | 1.0900e-01 | 2.8451e+05 ; | CB - HB2 |
| 3722 | 3724 | 1 | 1.0900e-01 | 2.8451e+05 ; | CB - HB3 |
| 3722 | 3725 | 1 | 1.5100e-01 | 2.6527e+05 ; | CB - CG |
| 3725 | 3726 | 1 | 1.4000e-01 | 3.9246e+05 ; | CG - CD1 |
| 3725 | 3735 | 1 | 1.4000e-01 | 3.9246e+05 ; | CG - CD2 |
| 3726 | 3727 | 1 | 1.0800e-01 | 3.0711e+05 ; | CD1 - HD1 |
| 3726 | 3728 | 1 | 1.4000e-01 | 3.9246e+05 ; | CD1 - CE1 |
| 3728 | 3729 | 1 | 1.0800e-01 | 3.0711e+05 ; | CE1 - HE1 |
| 3728 | 3730 | 1 | 1.4090e-01 | 3.9246e+05 ; | CE1 - CZ |
| 3730 | 3731 | 1 | 1.3640e-01 | 3.7656e+05 ; | CZ - OH |
| 3730 | 3733 | 1 | 1.4090e-01 | 3.9246e+05 ; | CZ - CE2 |
| 3731 | 3732 | 1 | 9.6000e-02 | 4.6275e+05 ; | OH - HH |
| 3733 | 3734 | 1 | 1.0800e-01 | 3.0711e+05 ; | CE2 - HE2 |
| 3733 | 3735 | 1 | 1.4000e-01 | 3.9246e+05 ; | CE2 - CD2 |
| 3735 | 3736 | 1 | 1.0800e-01 | 3.0711e+05 ; | CD2 - HD2 |
| 3737 | 3738 | 1 | 1.2290e-01 | 4.7698e+05 ; | C - O |
| 3737 | 3739 | 1 | 1.3350e-01 | 4.1003e+05 ; | C - N |
| 3739 | 3740 | 1 | 1.0100e-01 | 3.6317e+05 ; | N - H |
| 3739 | 3741 | 1 | 1.4490e-01 | 2.8200e+05 ; | N - CA |
| 3741 | 3742 | 1 | 1.0900e-01 | 2.8451e+05 ; | CA - HA |
| 3741 | 3743 | 1 | 1.5260e-01 | 2.5941e+05 ; | CA - CB |
| 3741 | 3752 | 1 | 1.5220e-01 | 2.6527e+05 ; | CA - C |
| 3743 | 3744 | 1 | 1.0900e-01 | 2.8451e+05 ; | CB - HB2 |
| 3743 | 3745 | 1 | 1.0900e-01 | 2.8451e+05 ; | CB - HB3 |
| 3743 | 3746 | 1 | 1.5260e-01 | 2.5941e+05 ; | CB - CG |
| 3746 | 3747 | 1 | 1.0900e-01 | 2.8451e+05 ; | CG - HG2 |
| 3746 | 3748 | 1 | 1.0900e-01 | 2.8451e+05 ; | CG - HG3 |
| 3746 | 3749 | 1 | 1.5220e-01 | 2.6527e+05 ; | CG - CD |
| 3749 | 3750 | 1 | 1.2500e-01 | 5.4894e+05 ; | CD - OE1 |
| 3749 | 3751 | 1 | 1.2500e-01 | 5.4894e+05 ; | CD - OE2 |
| 3752 | 3753 | 1 | 1.2290e-01 | 4.7698e+05 ; | C - O |

| | | | | | | |
|---|---|---|---|---|---|---|
| 3752 | 3754 | 1 | 1.3350e-01 | 4.1003e+05 | ; | C - N |
| 3754 | 3755 | 1 | 1.0100e-01 | 3.6317e+05 | ; | N - H |
| 3754 | 3756 | 1 | 1.4490e-01 | 2.8200e+05 | ; | N - CA |
| 3756 | 3757 | 1 | 1.0900e-01 | 2.8451e+05 | ; | CA - HA |
| 3756 | 3758 | 1 | 1.5260e-01 | 2.5941e+05 | ; | CA - CB |
| 3756 | 3766 | 1 | 1.5220e-01 | 2.6527e+05 | ; | CA - C |
| 3758 | 3759 | 1 | 1.0900e-01 | 2.8451e+05 | ; | CB - HB2 |
| 3758 | 3760 | 1 | 1.0900e-01 | 2.8451e+05 | ; | CB - HB3 |
| 3758 | 3761 | 1 | 1.5220e-01 | 2.6527e+05 | ; | CB - CG |
| 3761 | 3762 | 1 | 1.2290e-01 | 4.7698e+05 | ; | CG - OD1 |
| 3761 | 3763 | 1 | 1.3350e-01 | 4.1003e+05 | ; | CG - ND2 |
| 3763 | 3764 | 1 | 1.0100e-01 | 3.6317e+05 | ; | ND2 - HD21 |
| 3763 | 3765 | 1 | 1.0100e-01 | 3.6317e+05 | ; | ND2 - HD22 |
| 3766 | 3767 | 1 | 1.2290e-01 | 4.7698e+05 | ; | C - O |
| 3766 | 3768 | 1 | 1.3350e-01 | 4.1003e+05 | ; | C - N |
| 3768 | 3769 | 1 | 1.0100e-01 | 3.6317e+05 | ; | N - H |
| 3768 | 3770 | 1 | 1.4490e-01 | 2.8200e+05 | ; | N - CA |
| 3770 | 3771 | 1 | 1.0900e-01 | 2.8451e+05 | ; | CA - HA |
| 3770 | 3772 | 1 | 1.5260e-01 | 2.5941e+05 | ; | CA - CB |
| 3770 | 3778 | 1 | 1.5220e-01 | 2.6527e+05 | ; | CA - C |
| 3772 | 3773 | 1 | 1.0900e-01 | 2.8451e+05 | ; | CB - HB2 |
| 3772 | 3774 | 1 | 1.0900e-01 | 2.8451e+05 | ; | CB - HB3 |
| 3772 | 3775 | 1 | 1.5220e-01 | 2.6527e+05 | ; | CB - CG |
| 3775 | 3776 | 1 | 1.2500e-01 | 5.4894e+05 | ; | CG - OD1 |
| 3775 | 3777 | 1 | 1.2500e-01 | 5.4894e+05 | ; | CG - OD2 |
| 3778 | 3779 | 1 | 1.2290e-01 | 4.7698e+05 | ; | C - O |
| 3778 | 3780 | 1 | 1.3350e-01 | 4.1003e+05 | ; | C - N |
| 3780 | 3781 | 1 | 1.0100e-01 | 3.6317e+05 | ; | N - H |
| 3780 | 3782 | 1 | 1.4490e-01 | 2.8200e+05 | ; | N - CA |
| 3782 | 3783 | 1 | 1.0900e-01 | 2.8451e+05 | ; | CA - HA |
| 3782 | 3784 | 1 | 1.5260e-01 | 2.5941e+05 | ; | CA - CB |
| 3782 | 3802 | 1 | 1.5220e-01 | 2.6527e+05 | ; | CA - C |
| 3784 | 3785 | 1 | 1.0900e-01 | 2.8451e+05 | ; | CB - HB2 |
| 3784 | 3786 | 1 | 1.0900e-01 | 2.8451e+05 | ; | CB - HB3 |
| 3784 | 3787 | 1 | 1.4950e-01 | 2.6527e+05 | ; | CB - CG |
| 3787 | 3788 | 1 | 1.3520e-01 | 4.5689e+05 | ; | CG - CD1 |
| 3787 | 3801 | 1 | 1.4590e-01 | 3.2468e+05 | ; | CG - CD2 |
| 3788 | 3789 | 1 | 1.0800e-01 | 3.0711e+05 | ; | CD1 - HD1 |
| 3788 | 3790 | 1 | 1.3810e-01 | 3.5731e+05 | ; | CD1 - NE1 |
| 3790 | 3791 | 1 | 1.0100e-01 | 3.6317e+05 | ; | NE1 - HE1 |
| 3790 | 3792 | 1 | 1.3800e-01 | 3.5815e+05 | ; | NE1 - CE2 |
| 3792 | 3793 | 1 | 1.4000e-01 | 3.9246e+05 | ; | CE2 - CZ2 |
| 3792 | 3801 | 1 | 1.4190e-01 | 3.7405e+05 | ; | CE2 - CD2 |
| 3793 | 3794 | 1 | 1.0800e-01 | 3.0711e+05 | ; | CZ2 - HZ2 |
| 3793 | 3795 | 1 | 1.4000e-01 | 3.9246e+05 | ; | CZ2 - CH2 |
| 3795 | 3796 | 1 | 1.0800e-01 | 3.0711e+05 | ; | CH2 - HH2 |
| 3795 | 3797 | 1 | 1.4000e-01 | 3.9246e+05 | ; | CH2 - CZ3 |
| 3797 | 3798 | 1 | 1.0800e-01 | 3.0711e+05 | ; | CZ3 - HZ3 |
| 3797 | 3799 | 1 | 1.4000e-01 | 3.9246e+05 | ; | CZ3 - CE3 |
| 3799 | 3800 | 1 | 1.0800e-01 | 3.0711e+05 | ; | CE3 - HE3 |
| 3799 | 3801 | 1 | 1.4040e-01 | 3.9246e+05 | ; | CE3 - CD2 |
| 3802 | 3803 | 1 | 1.2290e-01 | 4.7698e+05 | ; | C - O |

| | | | | | | |
|---|---|---|---|---|---|---|
| 3802 | 3804 | 1 | 1.3350e-01 | 4.1003e+05 | ; | C - N |
| 3804 | 3805 | 1 | 1.0100e-01 | 3.6317e+05 | ; | N - H |
| 3804 | 3806 | 1 | 1.4490e-01 | 2.8200e+05 | ; | N - CA |
| 3806 | 3807 | 1 | 1.0900e-01 | 2.8451e+05 | ; | CA - HA2 |
| 3806 | 3808 | 1 | 1.0900e-01 | 2.8451e+05 | ; | CA - HA3 |
| 3806 | 3809 | 1 | 1.5220e-01 | 2.6527e+05 | ; | CA - C |
| 3809 | 3810 | 1 | 1.2290e-01 | 4.7698e+05 | ; | C - O |
| 3809 | 3811 | 1 | 1.3350e-01 | 4.1003e+05 | ; | C - N |
| 3811 | 3812 | 1 | 1.0100e-01 | 3.6317e+05 | ; | N - H |
| 3811 | 3813 | 1 | 1.4490e-01 | 2.8200e+05 | ; | N - CA |
| 3813 | 3814 | 1 | 1.0900e-01 | 2.8451e+05 | ; | CA - HA |
| 3813 | 3815 | 1 | 1.5260e-01 | 2.5941e+05 | ; | CA - CB |
| 3813 | 3820 | 1 | 1.5220e-01 | 2.6527e+05 | ; | CA - C |
| 3815 | 3816 | 1 | 1.0900e-01 | 2.8451e+05 | ; | CB - HB2 |
| 3815 | 3817 | 1 | 1.0900e-01 | 2.8451e+05 | ; | CB - HB3 |
| 3815 | 3818 | 1 | 1.4100e-01 | 2.6778e+05 | ; | CB - OG |
| 3818 | 3819 | 1 | 9.6000e-02 | 4.6275e+05 | ; | OG - HG |
| 3820 | 3821 | 1 | 1.2290e-01 | 4.7698e+05 | ; | C - O |
| 3820 | 3822 | 1 | 1.3350e-01 | 4.1003e+05 | ; | C - N |
| 3822 | 3823 | 1 | 1.0100e-01 | 3.6317e+05 | ; | N - H |
| 3822 | 3824 | 1 | 1.4490e-01 | 2.8200e+05 | ; | N - CA |
| 3824 | 3825 | 1 | 1.0900e-01 | 2.8451e+05 | ; | CA - HA |
| 3824 | 3826 | 1 | 1.5260e-01 | 2.5941e+05 | ; | CA - CB |
| 3824 | 3839 | 1 | 1.5220e-01 | 2.6527e+05 | ; | CA - C |
| 3826 | 3827 | 1 | 1.0900e-01 | 2.8451e+05 | ; | CB - HB |
| 3826 | 3828 | 1 | 1.5260e-01 | 2.5941e+05 | ; | CB - CG2 |
| 3826 | 3832 | 1 | 1.5260e-01 | 2.5941e+05 | ; | CB - CG1 |
| 3828 | 3829 | 1 | 1.0900e-01 | 2.8451e+05 | ; | CG2 - HG21 |
| 3828 | 3830 | 1 | 1.0900e-01 | 2.8451e+05 | ; | CG2 - HG22 |
| 3828 | 3831 | 1 | 1.0900e-01 | 2.8451e+05 | ; | CG2 - HG23 |
| 3832 | 3833 | 1 | 1.0900e-01 | 2.8451e+05 | ; | CG1 - HG12 |
| 3832 | 3834 | 1 | 1.0900e-01 | 2.8451e+05 | ; | CG1 - HG13 |
| 3832 | 3835 | 1 | 1.5260e-01 | 2.5941e+05 | ; | CG1 - CD1 |
| 3835 | 3836 | 1 | 1.0900e-01 | 2.8451e+05 | ; | CD1 - HD11 |
| 3835 | 3837 | 1 | 1.0900e-01 | 2.8451e+05 | ; | CD1 - HD12 |
| 3835 | 3838 | 1 | 1.0900e-01 | 2.8451e+05 | ; | CD1 - HD13 |
| 3839 | 3840 | 1 | 1.2290e-01 | 4.7698e+05 | ; | C - O |
| 3839 | 3841 | 1 | 1.3350e-01 | 4.1003e+05 | ; | C - N |
| 3841 | 3842 | 1 | 1.0100e-01 | 3.6317e+05 | ; | N - H |
| 3841 | 3843 | 1 | 1.4490e-01 | 2.8200e+05 | ; | N - CA |
| 3843 | 3844 | 1 | 1.0900e-01 | 2.8451e+05 | ; | CA - HA |
| 3843 | 3845 | 1 | 1.5260e-01 | 2.5941e+05 | ; | CA - CB |
| 3843 | 3850 | 1 | 1.5220e-01 | 2.6527e+05 | ; | CA - C |
| 3845 | 3846 | 1 | 1.0900e-01 | 2.8451e+05 | ; | CB - HB2 |
| 3845 | 3847 | 1 | 1.0900e-01 | 2.8451e+05 | ; | CB - HB3 |
| 3845 | 3848 | 1 | 1.4100e-01 | 2.6778e+05 | ; | CB - OG |
| 3848 | 3849 | 1 | 9.6000e-02 | 4.6275e+05 | ; | OG - HG |
| 3850 | 3851 | 1 | 1.2290e-01 | 4.7698e+05 | ; | C - O |
| 3850 | 3852 | 1 | 1.3350e-01 | 4.1003e+05 | ; | C - N |
| 3852 | 3853 | 1 | 1.0100e-01 | 3.6317e+05 | ; | N - H |
| 3852 | 3854 | 1 | 1.4490e-01 | 2.8200e+05 | ; | N - CA |
| 3854 | 3855 | 1 | 1.0900e-01 | 2.8451e+05 | ; | CA - HA |

| | | | | | | |
|---|---|---|---|---|---|---|
| 3854 | 3856 | 1 | 1.5260e-01 | 2.5941e+05 | ; | CA - CB |
| 3854 | 3861 | 1 | 1.5220e-01 | 2.6527e+05 | ; | CA - C |
| 3856 | 3857 | 1 | 1.0900e-01 | 2.8451e+05 | ; | CB - HB2 |
| 3856 | 3858 | 1 | 1.0900e-01 | 2.8451e+05 | ; | CB - HB3 |
| 3856 | 3859 | 1 | 1.4100e-01 | 2.6778e+05 | ; | CB - OG |
| 3859 | 3860 | 1 | 9.6000e-02 | 4.6275e+05 | ; | OG - HG |
| 3861 | 3862 | 1 | 1.2290e-01 | 4.7698e+05 | ; | C - O |
| 3861 | 3863 | 1 | 1.3350e-01 | 4.1003e+05 | ; | C - N |
| 3863 | 3864 | 1 | 1.0100e-01 | 3.6317e+05 | ; | N - H |
| 3863 | 3865 | 1 | 1.4490e-01 | 2.8200e+05 | ; | N - CA |
| 3865 | 3866 | 1 | 1.0900e-01 | 2.8451e+05 | ; | CA - HA |
| 3865 | 3867 | 1 | 1.5260e-01 | 2.5941e+05 | ; | CA - CB |
| 3865 | 3873 | 1 | 1.5220e-01 | 2.6527e+05 | ; | CA - C |
| 3867 | 3868 | 1 | 1.0900e-01 | 2.8451e+05 | ; | CB - HB2 |
| 3867 | 3869 | 1 | 1.0900e-01 | 2.8451e+05 | ; | CB - HB3 |
| 3867 | 3870 | 1 | 1.5220e-01 | 2.6527e+05 | ; | CB - CG |
| 3870 | 3871 | 1 | 1.2500e-01 | 5.4894e+05 | ; | CG - OD1 |
| 3870 | 3872 | 1 | 1.2500e-01 | 5.4894e+05 | ; | CG - OD2 |
| 3873 | 3874 | 1 | 1.2290e-01 | 4.7698e+05 | ; | C - O |
| 3873 | 3875 | 1 | 1.3350e-01 | 4.1003e+05 | ; | C - N |
| 3875 | 3876 | 1 | 1.0100e-01 | 3.6317e+05 | ; | N - H |
| 3875 | 3877 | 1 | 1.4490e-01 | 2.8200e+05 | ; | N - CA |
| 3877 | 3878 | 1 | 1.0900e-01 | 2.8451e+05 | ; | CA - HA |
| 3877 | 3879 | 1 | 1.5260e-01 | 2.5941e+05 | ; | CA - CB |
| 3877 | 3883 | 1 | 1.5220e-01 | 2.6527e+05 | ; | CA - C |
| 3879 | 3880 | 1 | 1.0900e-01 | 2.8451e+05 | ; | CB - HB1 |
| 3879 | 3881 | 1 | 1.0900e-01 | 2.8451e+05 | ; | CB - HB2 |
| 3879 | 3882 | 1 | 1.0900e-01 | 2.8451e+05 | ; | CB - HB3 |
| 3883 | 3884 | 1 | 1.2290e-01 | 4.7698e+05 | ; | C - O |
| 3883 | 3885 | 1 | 1.3350e-01 | 4.1003e+05 | ; | C - N |
| 3885 | 3886 | 1 | 1.0100e-01 | 3.6317e+05 | ; | N - H |
| 3885 | 3887 | 1 | 1.4490e-01 | 2.8200e+05 | ; | N - CA |
| 3887 | 3888 | 1 | 1.0900e-01 | 2.8451e+05 | ; | CA - HA |
| 3887 | 3889 | 1 | 1.5260e-01 | 2.5941e+05 | ; | CA - CB |
| 3887 | 3905 | 1 | 1.5220e-01 | 2.6527e+05 | ; | CA - C |
| 3889 | 3890 | 1 | 1.0900e-01 | 2.8451e+05 | ; | CB - HB2 |
| 3889 | 3891 | 1 | 1.0900e-01 | 2.8451e+05 | ; | CB - HB3 |
| 3889 | 3892 | 1 | 1.5260e-01 | 2.5941e+05 | ; | CB - CG |
| 3892 | 3893 | 1 | 1.0900e-01 | 2.8451e+05 | ; | CG - HG2 |
| 3892 | 3894 | 1 | 1.0900e-01 | 2.8451e+05 | ; | CG - HG3 |
| 3892 | 3895 | 1 | 1.5260e-01 | 2.5941e+05 | ; | CG - CD |
| 3895 | 3896 | 1 | 1.0900e-01 | 2.8451e+05 | ; | CD - HD2 |
| 3895 | 3897 | 1 | 1.0900e-01 | 2.8451e+05 | ; | CD - HD3 |
| 3895 | 3898 | 1 | 1.5260e-01 | 2.5941e+05 | ; | CD - CE |
| 3898 | 3899 | 1 | 1.0900e-01 | 2.8451e+05 | ; | CE - HE2 |
| 3898 | 3900 | 1 | 1.0900e-01 | 2.8451e+05 | ; | CE - HE3 |
| 3898 | 3901 | 1 | 1.4710e-01 | 3.0711e+05 | ; | CE - NZ |
| 3901 | 3902 | 1 | 1.0100e-01 | 3.6317e+05 | ; | NZ - HZ1 |
| 3901 | 3903 | 1 | 1.0100e-01 | 3.6317e+05 | ; | NZ - HZ2 |
| 3901 | 3904 | 1 | 1.0100e-01 | 3.6317e+05 | ; | NZ - HZ3 |
| 3905 | 3906 | 1 | 1.2290e-01 | 4.7698e+05 | ; | C - O |
| 3905 | 3907 | 1 | 1.3350e-01 | 4.1003e+05 | ; | C - N |

| | | | | | | |
|---|---|---|---|---|---|---|
| 3907 | 3908 | 1 | 1.0100e-01 | 3.6317e+05 | ; | N - H |
| 3907 | 3909 | 1 | 1.4490e-01 | 2.8200e+05 | ; | N - CA |
| 3909 | 3910 | 1 | 1.0900e-01 | 2.8451e+05 | ; | CA - HA |
| 3909 | 3911 | 1 | 1.5260e-01 | 2.5941e+05 | ; | CA - CB |
| 3909 | 3919 | 1 | 1.5220e-01 | 2.6527e+05 | ; | CA - C |
| 3911 | 3912 | 1 | 1.0900e-01 | 2.8451e+05 | ; | CB - HB2 |
| 3911 | 3913 | 1 | 1.0900e-01 | 2.8451e+05 | ; | CB - HB3 |
| 3911 | 3914 | 1 | 1.5220e-01 | 2.6527e+05 | ; | CB - CG |
| 3914 | 3915 | 1 | 1.2290e-01 | 4.7698e+05 | ; | CG - OD1 |
| 3914 | 3916 | 1 | 1.3350e-01 | 4.1003e+05 | ; | CG - ND2 |
| 3916 | 3917 | 1 | 1.0100e-01 | 3.6317e+05 | ; | ND2 - HD21 |
| 3916 | 3918 | 1 | 1.0100e-01 | 3.6317e+05 | ; | ND2 - HD22 |
| 3919 | 3920 | 1 | 1.2290e-01 | 4.7698e+05 | ; | C - O |
| 3919 | 3921 | 1 | 1.3350e-01 | 4.1003e+05 | ; | C - N |
| 3921 | 3922 | 1 | 1.0100e-01 | 3.6317e+05 | ; | N - H |
| 3921 | 3923 | 1 | 1.4490e-01 | 2.8200e+05 | ; | N - CA |
| 3923 | 3924 | 1 | 1.0900e-01 | 2.8451e+05 | ; | CA - HA |
| 3923 | 3925 | 1 | 1.5260e-01 | 2.5941e+05 | ; | CA - CB |
| 3923 | 3938 | 1 | 1.5220e-01 | 2.6527e+05 | ; | CA - C |
| 3925 | 3926 | 1 | 1.0900e-01 | 2.8451e+05 | ; | CB - HB2 |
| 3925 | 3927 | 1 | 1.0900e-01 | 2.8451e+05 | ; | CB - HB3 |
| 3925 | 3928 | 1 | 1.5260e-01 | 2.5941e+05 | ; | CB - CG |
| 3928 | 3929 | 1 | 1.0900e-01 | 2.8451e+05 | ; | CG - HG |
| 3928 | 3930 | 1 | 1.5260e-01 | 2.5941e+05 | ; | CG - CD1 |
| 3928 | 3934 | 1 | 1.5260e-01 | 2.5941e+05 | ; | CG - CD2 |
| 3930 | 3931 | 1 | 1.0900e-01 | 2.8451e+05 | ; | CD1 - HD11 |
| 3930 | 3932 | 1 | 1.0900e-01 | 2.8451e+05 | ; | CD1 - HD12 |
| 3930 | 3933 | 1 | 1.0900e-01 | 2.8451e+05 | ; | CD1 - HD13 |
| 3934 | 3935 | 1 | 1.0900e-01 | 2.8451e+05 | ; | CD2 - HD21 |
| 3934 | 3936 | 1 | 1.0900e-01 | 2.8451e+05 | ; | CD2 - HD22 |
| 3934 | 3937 | 1 | 1.0900e-01 | 2.8451e+05 | ; | CD2 - HD23 |
| 3938 | 3939 | 1 | 1.2290e-01 | 4.7698e+05 | ; | C - O |
| 3938 | 3940 | 1 | 1.3350e-01 | 4.1003e+05 | ; | C - N |
| 3940 | 3941 | 1 | 1.0100e-01 | 3.6317e+05 | ; | N - H |
| 3940 | 3942 | 1 | 1.4490e-01 | 2.8200e+05 | ; | N - CA |
| 3942 | 3943 | 1 | 1.0900e-01 | 2.8451e+05 | ; | CA - HA |
| 3942 | 3944 | 1 | 1.5260e-01 | 2.5941e+05 | ; | CA - CB |
| 3942 | 3957 | 1 | 1.5220e-01 | 2.6527e+05 | ; | CA - C |
| 3944 | 3945 | 1 | 1.0900e-01 | 2.8451e+05 | ; | CB - HB |
| 3944 | 3946 | 1 | 1.5260e-01 | 2.5941e+05 | ; | CB - CG2 |
| 3944 | 3950 | 1 | 1.5260e-01 | 2.5941e+05 | ; | CB - CG1 |
| 3946 | 3947 | 1 | 1.0900e-01 | 2.8451e+05 | ; | CG2 - HG21 |
| 3946 | 3948 | 1 | 1.0900e-01 | 2.8451e+05 | ; | CG2 - HG22 |
| 3946 | 3949 | 1 | 1.0900e-01 | 2.8451e+05 | ; | CG2 - HG23 |
| 3950 | 3951 | 1 | 1.0900e-01 | 2.8451e+05 | ; | CG1 - HG12 |
| 3950 | 3952 | 1 | 1.0900e-01 | 2.8451e+05 | ; | CG1 - HG13 |
| 3950 | 3953 | 1 | 1.5260e-01 | 2.5941e+05 | ; | CG1 - CD1 |
| 3953 | 3954 | 1 | 1.0900e-01 | 2.8451e+05 | ; | CD1 - HD11 |
| 3953 | 3955 | 1 | 1.0900e-01 | 2.8451e+05 | ; | CD1 - HD12 |
| 3953 | 3956 | 1 | 1.0900e-01 | 2.8451e+05 | ; | CD1 - HD13 |
| 3957 | 3958 | 1 | 1.2290e-01 | 4.7698e+05 | ; | C - O |
| 3957 | 3959 | 1 | 1.3350e-01 | 4.1003e+05 | ; | C - N |

| | | | | | | |
|---|---|---|---|---|---|---|
| 3959 | 3960 | 1 | 1.0100e-01 | 3.6317e+05 | ; | N - H |
| 3959 | 3961 | 1 | 1.4490e-01 | 2.8200e+05 | ; | N - CA |
| 3961 | 3962 | 1 | 1.0900e-01 | 2.8451e+05 | ; | CA - HA |
| 3961 | 3963 | 1 | 1.5260e-01 | 2.5941e+05 | ; | CA - CB |
| 3961 | 3971 | 1 | 1.5220e-01 | 2.6527e+05 | ; | CA - C |
| 3963 | 3964 | 1 | 1.0900e-01 | 2.8451e+05 | ; | CB - HB |
| 3963 | 3965 | 1 | 1.5260e-01 | 2.5941e+05 | ; | CB - CG2 |
| 3963 | 3969 | 1 | 1.4100e-01 | 2.6778e+05 | ; | CB - OG1 |
| 3965 | 3966 | 1 | 1.0900e-01 | 2.8451e+05 | ; | CG2 - HG21 |
| 3965 | 3967 | 1 | 1.0900e-01 | 2.8451e+05 | ; | CG2 - HG22 |
| 3965 | 3968 | 1 | 1.0900e-01 | 2.8451e+05 | ; | CG2 - HG23 |
| 3969 | 3970 | 1 | 9.6000e-02 | 4.6275e+05 | ; | OG1 - HG1 |
| 3971 | 3972 | 1 | 1.2290e-01 | 4.7698e+05 | ; | C - O |
| 3971 | 3973 | 1 | 1.3350e-01 | 4.1003e+05 | ; | C - N |
| 3973 | 3974 | 1 | 1.0100e-01 | 3.6317e+05 | ; | N - H |
| 3973 | 3975 | 1 | 1.4490e-01 | 2.8200e+05 | ; | N - CA |
| 3975 | 3976 | 1 | 1.0900e-01 | 2.8451e+05 | ; | CA - HA |
| 3975 | 3977 | 1 | 1.5260e-01 | 2.5941e+05 | ; | CA - CB |
| 3975 | 3993 | 1 | 1.5220e-01 | 2.6527e+05 | ; | CA - C |
| 3977 | 3978 | 1 | 1.0900e-01 | 2.8451e+05 | ; | CB - HB2 |
| 3977 | 3979 | 1 | 1.0900e-01 | 2.8451e+05 | ; | CB - HB3 |
| 3977 | 3980 | 1 | 1.5260e-01 | 2.5941e+05 | ; | CB - CG |
| 3980 | 3981 | 1 | 1.0900e-01 | 2.8451e+05 | ; | CG - HG2 |
| 3980 | 3982 | 1 | 1.0900e-01 | 2.8451e+05 | ; | CG - HG3 |
| 3980 | 3983 | 1 | 1.5260e-01 | 2.5941e+05 | ; | CG - CD |
| 3983 | 3984 | 1 | 1.0900e-01 | 2.8451e+05 | ; | CD - HD2 |
| 3983 | 3985 | 1 | 1.0900e-01 | 2.8451e+05 | ; | CD - HD3 |
| 3983 | 3986 | 1 | 1.5260e-01 | 2.5941e+05 | ; | CD - CE |
| 3986 | 3987 | 1 | 1.0900e-01 | 2.8451e+05 | ; | CE - HE2 |
| 3986 | 3988 | 1 | 1.0900e-01 | 2.8451e+05 | ; | CE - HE3 |
| 3986 | 3989 | 1 | 1.4710e-01 | 3.0711e+05 | ; | CE - NZ |
| 3989 | 3990 | 1 | 1.0100e-01 | 3.6317e+05 | ; | NZ - HZ1 |
| 3989 | 3991 | 1 | 1.0100e-01 | 3.6317e+05 | ; | NZ - HZ2 |
| 3989 | 3992 | 1 | 1.0100e-01 | 3.6317e+05 | ; | NZ - HZ3 |
| 3993 | 3994 | 1 | 1.2290e-01 | 4.7698e+05 | ; | C - O |
| 3993 | 3995 | 1 | 1.3350e-01 | 4.1003e+05 | ; | C - N |
| 3995 | 3996 | 1 | 1.0100e-01 | 3.6317e+05 | ; | N - H |
| 3995 | 3997 | 1 | 1.4490e-01 | 2.8200e+05 | ; | N - CA |
| 3997 | 3998 | 1 | 1.0900e-01 | 2.8451e+05 | ; | CA - HA |
| 3997 | 3999 | 1 | 1.5260e-01 | 2.5941e+05 | ; | CA - CB |
| 3997 | 4012 | 1 | 1.5220e-01 | 2.6527e+05 | ; | CA - C |
| 3999 | 4000 | 1 | 1.0900e-01 | 2.8451e+05 | ; | CB - HB2 |
| 3999 | 4001 | 1 | 1.0900e-01 | 2.8451e+05 | ; | CB - HB3 |
| 3999 | 4002 | 1 | 1.5260e-01 | 2.5941e+05 | ; | CB - CG |
| 4002 | 4003 | 1 | 1.0900e-01 | 2.8451e+05 | ; | CG - HG |
| 4002 | 4004 | 1 | 1.5260e-01 | 2.5941e+05 | ; | CG - CD1 |
| 4002 | 4008 | 1 | 1.5260e-01 | 2.5941e+05 | ; | CG - CD2 |
| 4004 | 4005 | 1 | 1.0900e-01 | 2.8451e+05 | ; | CD1 - HD11 |
| 4004 | 4006 | 1 | 1.0900e-01 | 2.8451e+05 | ; | CD1 - HD12 |
| 4004 | 4007 | 1 | 1.0900e-01 | 2.8451e+05 | ; | CD1 - HD13 |
| 4008 | 4009 | 1 | 1.0900e-01 | 2.8451e+05 | ; | CD2 - HD21 |
| 4008 | 4010 | 1 | 1.0900e-01 | 2.8451e+05 | ; | CD2 - HD22 |

| | | | | | | |
|---|---|---|---|---|---|---|
| 4008 | 4011 | 1 | 1.0900e-01 | 2.8451e+05 | ; | CD2 - HD23 |
| 4012 | 4013 | 1 | 1.2290e-01 | 4.7698e+05 | ; | C - O |
| 4012 | 4014 | 1 | 1.3350e-01 | 4.1003e+05 | ; | C - N |
| 4014 | 4015 | 1 | 1.0100e-01 | 3.6317e+05 | ; | N - H |
| 4014 | 4016 | 1 | 1.4490e-01 | 2.8200e+05 | ; | N - CA |
| 4016 | 4017 | 1 | 1.0900e-01 | 2.8451e+05 | ; | CA - HA |
| 4016 | 4018 | 1 | 1.5260e-01 | 2.5941e+05 | ; | CA - CB |
| 4016 | 4031 | 1 | 1.5220e-01 | 2.6527e+05 | ; | CA - C |
| 4018 | 4019 | 1 | 1.0900e-01 | 2.8451e+05 | ; | CB - HB2 |
| 4018 | 4020 | 1 | 1.0900e-01 | 2.8451e+05 | ; | CB - HB3 |
| 4018 | 4021 | 1 | 1.5260e-01 | 2.5941e+05 | ; | CB - CG |
| 4021 | 4022 | 1 | 1.0900e-01 | 2.8451e+05 | ; | CG - HG |
| 4021 | 4023 | 1 | 1.5260e-01 | 2.5941e+05 | ; | CG - CD1 |
| 4021 | 4027 | 1 | 1.5260e-01 | 2.5941e+05 | ; | CG - CD2 |
| 4023 | 4024 | 1 | 1.0900e-01 | 2.8451e+05 | ; | CD1 - HD11 |
| 4023 | 4025 | 1 | 1.0900e-01 | 2.8451e+05 | ; | CD1 - HD12 |
| 4023 | 4026 | 1 | 1.0900e-01 | 2.8451e+05 | ; | CD1 - HD13 |
| 4027 | 4028 | 1 | 1.0900e-01 | 2.8451e+05 | ; | CD2 - HD21 |
| 4027 | 4029 | 1 | 1.0900e-01 | 2.8451e+05 | ; | CD2 - HD22 |
| 4027 | 4030 | 1 | 1.0900e-01 | 2.8451e+05 | ; | CD2 - HD23 |
| 4031 | 4032 | 1 | 1.2290e-01 | 4.7698e+05 | ; | C - O |
| 4031 | 4033 | 1 | 1.3350e-01 | 4.1003e+05 | ; | C - N |
| 4033 | 4034 | 1 | 1.0100e-01 | 3.6317e+05 | ; | N - H |
| 4033 | 4035 | 1 | 1.4490e-01 | 2.8200e+05 | ; | N - CA |
| 4035 | 4036 | 1 | 1.0900e-01 | 2.8451e+05 | ; | CA - HA |
| 4035 | 4037 | 1 | 1.5260e-01 | 2.5941e+05 | ; | CA - CB |
| 4035 | 4045 | 1 | 1.5220e-01 | 2.6527e+05 | ; | CA - C |
| 4037 | 4038 | 1 | 1.0900e-01 | 2.8451e+05 | ; | CB - HB |
| 4037 | 4039 | 1 | 1.5260e-01 | 2.5941e+05 | ; | CB - CG2 |
| 4037 | 4043 | 1 | 1.4100e-01 | 2.6778e+05 | ; | CB - OG1 |
| 4039 | 4040 | 1 | 1.0900e-01 | 2.8451e+05 | ; | CG2 - HG21 |
| 4039 | 4041 | 1 | 1.0900e-01 | 2.8451e+05 | ; | CG2 - HG22 |
| 4039 | 4042 | 1 | 1.0900e-01 | 2.8451e+05 | ; | CG2 - HG23 |
| 4043 | 4044 | 1 | 9.6000e-02 | 4.6275e+05 | ; | OG1 - HG1 |
| 4045 | 4046 | 1 | 1.2290e-01 | 4.7698e+05 | ; | C - O |
| 4045 | 4047 | 1 | 1.3350e-01 | 4.1003e+05 | ; | C - N |
| 4047 | 4048 | 1 | 1.0100e-01 | 3.6317e+05 | ; | N - H |
| 4047 | 4049 | 1 | 1.4490e-01 | 2.8200e+05 | ; | N - CA |
| 4049 | 4050 | 1 | 1.0900e-01 | 2.8451e+05 | ; | CA - HA |
| 4049 | 4051 | 1 | 1.5260e-01 | 2.5941e+05 | ; | CA - CB |
| 4049 | 4066 | 1 | 1.5220e-01 | 2.6527e+05 | ; | CA - C |
| 4051 | 4052 | 1 | 1.0900e-01 | 2.8451e+05 | ; | CB - HB2 |
| 4051 | 4053 | 1 | 1.0900e-01 | 2.8451e+05 | ; | CB - HB3 |
| 4051 | 4054 | 1 | 1.5100e-01 | 2.6527e+05 | ; | CB - CG |
| 4054 | 4055 | 1 | 1.4000e-01 | 3.9246e+05 | ; | CG - CD1 |
| 4054 | 4064 | 1 | 1.4000e-01 | 3.9246e+05 | ; | CG - CD2 |
| 4055 | 4056 | 1 | 1.0800e-01 | 3.0711e+05 | ; | CD1 - HD1 |
| 4055 | 4057 | 1 | 1.4000e-01 | 3.9246e+05 | ; | CD1 - CE1 |
| 4057 | 4058 | 1 | 1.0800e-01 | 3.0711e+05 | ; | CE1 - HE1 |
| 4057 | 4059 | 1 | 1.4090e-01 | 3.9246e+05 | ; | CE1 - CZ |
| 4059 | 4060 | 1 | 1.3640e-01 | 3.7656e+05 | ; | CZ - OH |
| 4059 | 4062 | 1 | 1.4090e-01 | 3.9246e+05 | ; | CZ - CE2 |

| | | | | | | |
|---|---|---|---|---|---|---|
| 4060 | 4061 | 1 | 9.6000e-02 | 4.6275e+05 | ; | OH - HH |
| 4062 | 4063 | 1 | 1.0800e-01 | 3.0711e+05 | ; | CE2 - HE2 |
| 4062 | 4064 | 1 | 1.4000e-01 | 3.9246e+05 | ; | CE2 - CD2 |
| 4064 | 4065 | 1 | 1.0800e-01 | 3.0711e+05 | ; | CD2 - HD2 |
| 4066 | 4067 | 1 | 1.2290e-01 | 4.7698e+05 | ; | C - O |
| 4066 | 4068 | 1 | 1.3350e-01 | 4.1003e+05 | ; | C - N |
| 4068 | 4069 | 1 | 1.0100e-01 | 3.6317e+05 | ; | N - H |
| 4068 | 4070 | 1 | 1.4490e-01 | 2.8200e+05 | ; | N - CA |
| 4070 | 4071 | 1 | 1.0900e-01 | 2.8451e+05 | ; | CA - HA |
| 4070 | 4072 | 1 | 1.5260e-01 | 2.5941e+05 | ; | CA - CB |
| 4070 | 4080 | 1 | 1.5220e-01 | 2.6527e+05 | ; | CA - C |
| 4072 | 4073 | 1 | 1.0900e-01 | 2.8451e+05 | ; | CB - HB2 |
| 4072 | 4074 | 1 | 1.0900e-01 | 2.8451e+05 | ; | CB - HB3 |
| 4072 | 4075 | 1 | 1.5220e-01 | 2.6527e+05 | ; | CB - CG |
| 4075 | 4076 | 1 | 1.2290e-01 | 4.7698e+05 | ; | CG - OD1 |
| 4075 | 4077 | 1 | 1.3350e-01 | 4.1003e+05 | ; | CG - ND2 |
| 4077 | 4078 | 1 | 1.0100e-01 | 3.6317e+05 | ; | ND2 - HD21 |
| 4077 | 4079 | 1 | 1.0100e-01 | 3.6317e+05 | ; | ND2 - HD22 |
| 4080 | 4081 | 1 | 1.2290e-01 | 4.7698e+05 | ; | C - O |
| 4080 | 4082 | 1 | 1.3350e-01 | 4.1003e+05 | ; | C - N |
| 4082 | 4083 | 1 | 1.4490e-01 | 2.8200e+05 | ; | N - CD |
| 4082 | 4092 | 1 | 1.4490e-01 | 2.8200e+05 | ; | N - CA |
| 4083 | 4084 | 1 | 1.0900e-01 | 2.8451e+05 | ; | CD - HD2 |
| 4083 | 4085 | 1 | 1.0900e-01 | 2.8451e+05 | ; | CD - HD3 |
| 4083 | 4086 | 1 | 1.5260e-01 | 2.5941e+05 | ; | CD - CG |
| 4086 | 4087 | 1 | 1.0900e-01 | 2.8451e+05 | ; | CG - HG2 |
| 4086 | 4088 | 1 | 1.0900e-01 | 2.8451e+05 | ; | CG - HG3 |
| 4086 | 4089 | 1 | 1.5260e-01 | 2.5941e+05 | ; | CG - CB |
| 4089 | 4090 | 1 | 1.0900e-01 | 2.8451e+05 | ; | CB - HB2 |
| 4089 | 4091 | 1 | 1.0900e-01 | 2.8451e+05 | ; | CB - HB3 |
| 4089 | 4092 | 1 | 1.5260e-01 | 2.5941e+05 | ; | CB - CA |
| 4092 | 4093 | 1 | 1.0900e-01 | 2.8451e+05 | ; | CA - HA |
| 4092 | 4094 | 1 | 1.5220e-01 | 2.6527e+05 | ; | CA - C |
| 4094 | 4095 | 1 | 1.2290e-01 | 4.7698e+05 | ; | C - O |
| 4094 | 4096 | 1 | 1.3350e-01 | 4.1003e+05 | ; | C - N |
| 4096 | 4097 | 1 | 1.0100e-01 | 3.6317e+05 | ; | N - H |
| 4096 | 4098 | 1 | 1.4490e-01 | 2.8200e+05 | ; | N - CA |
| 4098 | 4099 | 1 | 1.0900e-01 | 2.8451e+05 | ; | CA - HA |
| 4098 | 4100 | 1 | 1.5260e-01 | 2.5941e+05 | ; | CA - CB |
| 4098 | 4108 | 1 | 1.5220e-01 | 2.6527e+05 | ; | CA - C |
| 4100 | 4101 | 1 | 1.0900e-01 | 2.8451e+05 | ; | CB - HB2 |
| 4100 | 4102 | 1 | 1.0900e-01 | 2.8451e+05 | ; | CB - HB3 |
| 4100 | 4103 | 1 | 1.5220e-01 | 2.6527e+05 | ; | CB - CG |
| 4103 | 4104 | 1 | 1.2290e-01 | 4.7698e+05 | ; | CG - OD1 |
| 4103 | 4105 | 1 | 1.3350e-01 | 4.1003e+05 | ; | CG - ND2 |
| 4105 | 4106 | 1 | 1.0100e-01 | 3.6317e+05 | ; | ND2 - HD21 |
| 4105 | 4107 | 1 | 1.0100e-01 | 3.6317e+05 | ; | ND2 - HD22 |
| 4108 | 4109 | 1 | 1.2290e-01 | 4.7698e+05 | ; | C - O |
| 4108 | 4110 | 1 | 1.3350e-01 | 4.1003e+05 | ; | C - N |
| 4110 | 4111 | 1 | 1.0100e-01 | 3.6317e+05 | ; | N - H |
| 4110 | 4112 | 1 | 1.4490e-01 | 2.8200e+05 | ; | N - CA |
| 4112 | 4113 | 1 | 1.0900e-01 | 2.8451e+05 | ; | CA - HA |

| | | | | | | |
|---|---|---|---|---|---|---|
| 4112 | 4114 | 1 | 1.5260e-01 | 2.5941e+05 | ; | CA - CB |
| 4112 | 4123 | 1 | 1.5220e-01 | 2.6527e+05 | ; | CA - C |
| 4114 | 4115 | 1 | 1.0900e-01 | 2.8451e+05 | ; | CB - HB2 |
| 4114 | 4116 | 1 | 1.0900e-01 | 2.8451e+05 | ; | CB - HB3 |
| 4114 | 4117 | 1 | 1.5260e-01 | 2.5941e+05 | ; | CB - CG |
| 4117 | 4118 | 1 | 1.0900e-01 | 2.8451e+05 | ; | CG - HG2 |
| 4117 | 4119 | 1 | 1.0900e-01 | 2.8451e+05 | ; | CG - HG3 |
| 4117 | 4120 | 1 | 1.5220e-01 | 2.6527e+05 | ; | CG - CD |
| 4120 | 4121 | 1 | 1.2500e-01 | 5.4894e+05 | ; | CD - OE1 |
| 4120 | 4122 | 1 | 1.2500e-01 | 5.4894e+05 | ; | CD - OE2 |
| 4123 | 4124 | 1 | 1.2290e-01 | 4.7698e+05 | ; | C - O |
| 4123 | 4125 | 1 | 1.3350e-01 | 4.1003e+05 | ; | C - N |
| 4125 | 4126 | 1 | 1.0100e-01 | 3.6317e+05 | ; | N - H |
| 4125 | 4127 | 1 | 1.4490e-01 | 2.8200e+05 | ; | N - CA |
| 4127 | 4128 | 1 | 1.0900e-01 | 2.8451e+05 | ; | CA - HA |
| 4127 | 4129 | 1 | 1.5260e-01 | 2.5941e+05 | ; | CA - CB |
| 4127 | 4147 | 1 | 1.5220e-01 | 2.6527e+05 | ; | CA - C |
| 4129 | 4130 | 1 | 1.0900e-01 | 2.8451e+05 | ; | CB - HB2 |
| 4129 | 4131 | 1 | 1.0900e-01 | 2.8451e+05 | ; | CB - HB3 |
| 4129 | 4132 | 1 | 1.5260e-01 | 2.5941e+05 | ; | CB - CG |
| 4132 | 4133 | 1 | 1.0900e-01 | 2.8451e+05 | ; | CG - HG2 |
| 4132 | 4134 | 1 | 1.0900e-01 | 2.8451e+05 | ; | CG - HG3 |
| 4132 | 4135 | 1 | 1.5260e-01 | 2.5941e+05 | ; | CG - CD |
| 4135 | 4136 | 1 | 1.0900e-01 | 2.8451e+05 | ; | CD - HD2 |
| 4135 | 4137 | 1 | 1.0900e-01 | 2.8451e+05 | ; | CD - HD3 |
| 4135 | 4138 | 1 | 1.4630e-01 | 2.8200e+05 | ; | CD - NE |
| 4138 | 4139 | 1 | 1.0100e-01 | 3.6317e+05 | ; | NE - HE |
| 4138 | 4140 | 1 | 1.3400e-01 | 4.0250e+05 | ; | NE - CZ |
| 4140 | 4141 | 1 | 1.3400e-01 | 4.0250e+05 | ; | CZ - NH1 |
| 4140 | 4144 | 1 | 1.3400e-01 | 4.0250e+05 | ; | CZ - NH2 |
| 4141 | 4142 | 1 | 1.0100e-01 | 3.6317e+05 | ; | NH1 - HH11 |
| 4141 | 4143 | 1 | 1.0100e-01 | 3.6317e+05 | ; | NH1 - HH12 |
| 4144 | 4145 | 1 | 1.0100e-01 | 3.6317e+05 | ; | NH2 - HH21 |
| 4144 | 4146 | 1 | 1.0100e-01 | 3.6317e+05 | ; | NH2 - HH22 |
| 4147 | 4148 | 1 | 1.2290e-01 | 4.7698e+05 | ; | C - O |
| 4147 | 4149 | 1 | 1.3350e-01 | 4.1003e+05 | ; | C - N |
| 4149 | 4150 | 1 | 1.0100e-01 | 3.6317e+05 | ; | N - H |
| 4149 | 4151 | 1 | 1.4490e-01 | 2.8200e+05 | ; | N - CA |
| 4151 | 4152 | 1 | 1.0900e-01 | 2.8451e+05 | ; | CA - HA |
| 4151 | 4153 | 1 | 1.5260e-01 | 2.5941e+05 | ; | CA - CB |
| 4151 | 4158 | 1 | 1.5220e-01 | 2.6527e+05 | ; | CA - C |
| 4153 | 4154 | 1 | 1.0900e-01 | 2.8451e+05 | ; | CB - HB2 |
| 4153 | 4155 | 1 | 1.0900e-01 | 2.8451e+05 | ; | CB - HB3 |
| 4153 | 4156 | 1 | 1.8100e-01 | 1.9832e+05 | ; | CB - SG |
| 4156 | 4157 | 1 | 1.3360e-01 | 2.2928e+05 | ; | SG - HG |
| 4158 | 4159 | 1 | 1.2290e-01 | 4.7698e+05 | ; | C - O |
| 4158 | 4160 | 1 | 1.3350e-01 | 4.1003e+05 | ; | C - N |
| 4160 | 4161 | 1 | 1.0100e-01 | 3.6317e+05 | ; | N - H |
| 4160 | 4162 | 1 | 1.4490e-01 | 2.8200e+05 | ; | N - CA |
| 4162 | 4163 | 1 | 1.0900e-01 | 2.8451e+05 | ; | CA - HA |
| 4162 | 4164 | 1 | 1.5260e-01 | 2.5941e+05 | ; | CA - CB |
| 4162 | 4172 | 1 | 1.5220e-01 | 2.6527e+05 | ; | CA - C |

| | | | | | | |
|---|---|---|---|---|---|---|
| 4164 | 4165 | 1 | 1.0900e-01 | 2.8451e+05 | ; | CB - HB |
| 4164 | 4166 | 1 | 1.5260e-01 | 2.5941e+05 | ; | CB - CG2 |
| 4164 | 4170 | 1 | 1.4100e-01 | 2.6778e+05 | ; | CB - OG1 |
| 4166 | 4167 | 1 | 1.0900e-01 | 2.8451e+05 | ; | CG2 - HG21 |
| 4166 | 4168 | 1 | 1.0900e-01 | 2.8451e+05 | ; | CG2 - HG22 |
| 4166 | 4169 | 1 | 1.0900e-01 | 2.8451e+05 | ; | CG2 - HG23 |
| 4170 | 4171 | 1 | 9.6000e-02 | 4.6275e+05 | ; | OG1 - HG1 |
| 4172 | 4173 | 1 | 1.2290e-01 | 4.7698e+05 | ; | C - O |
| 4172 | 4174 | 1 | 1.3350e-01 | 4.1003e+05 | ; | C - N |
| 4174 | 4175 | 1 | 1.0100e-01 | 3.6317e+05 | ; | N - H |
| 4174 | 4176 | 1 | 1.4490e-01 | 2.8200e+05 | ; | N - CA |
| 4176 | 4177 | 1 | 1.0900e-01 | 2.8451e+05 | ; | CA - HA |
| 4176 | 4178 | 1 | 1.5260e-01 | 2.5941e+05 | ; | CA - CB |
| 4176 | 4191 | 1 | 1.5220e-01 | 2.6527e+05 | ; | CA - C |
| 4178 | 4179 | 1 | 1.0900e-01 | 2.8451e+05 | ; | CB - HB |
| 4178 | 4180 | 1 | 1.5260e-01 | 2.5941e+05 | ; | CB - CG2 |
| 4178 | 4184 | 1 | 1.5260e-01 | 2.5941e+05 | ; | CB - CG1 |
| 4180 | 4181 | 1 | 1.0900e-01 | 2.8451e+05 | ; | CG2 - HG21 |
| 4180 | 4182 | 1 | 1.0900e-01 | 2.8451e+05 | ; | CG2 - HG22 |
| 4180 | 4183 | 1 | 1.0900e-01 | 2.8451e+05 | ; | CG2 - HG23 |
| 4184 | 4185 | 1 | 1.0900e-01 | 2.8451e+05 | ; | CG1 - HG12 |
| 4184 | 4186 | 1 | 1.0900e-01 | 2.8451e+05 | ; | CG1 - HG13 |
| 4184 | 4187 | 1 | 1.5260e-01 | 2.5941e+05 | ; | CG1 - CD1 |
| 4187 | 4188 | 1 | 1.0900e-01 | 2.8451e+05 | ; | CD1 - HD11 |
| 4187 | 4189 | 1 | 1.0900e-01 | 2.8451e+05 | ; | CD1 - HD12 |
| 4187 | 4190 | 1 | 1.0900e-01 | 2.8451e+05 | ; | CD1 - HD13 |
| 4191 | 4192 | 1 | 1.2290e-01 | 4.7698e+05 | ; | C - O |
| 4191 | 4193 | 1 | 1.3350e-01 | 4.1003e+05 | ; | C - N |
| 4193 | 4194 | 1 | 1.0100e-01 | 3.6317e+05 | ; | N - H |
| 4193 | 4195 | 1 | 1.4490e-01 | 2.8200e+05 | ; | N - CA |
| 4195 | 4196 | 1 | 1.0900e-01 | 2.8451e+05 | ; | CA - HA |
| 4195 | 4197 | 1 | 1.5260e-01 | 2.5941e+05 | ; | CA - CB |
| 4195 | 4206 | 1 | 1.5220e-01 | 2.6527e+05 | ; | CA - C |
| 4197 | 4198 | 1 | 1.0900e-01 | 2.8451e+05 | ; | CB - HB2 |
| 4197 | 4199 | 1 | 1.0900e-01 | 2.8451e+05 | ; | CB - HB3 |
| 4197 | 4200 | 1 | 1.5260e-01 | 2.5941e+05 | ; | CB - CG |
| 4200 | 4201 | 1 | 1.0900e-01 | 2.8451e+05 | ; | CG - HG2 |
| 4200 | 4202 | 1 | 1.0900e-01 | 2.8451e+05 | ; | CG - HG3 |
| 4200 | 4203 | 1 | 1.5220e-01 | 2.6527e+05 | ; | CG - CD |
| 4203 | 4204 | 1 | 1.2500e-01 | 5.4894e+05 | ; | CD - OE1 |
| 4203 | 4205 | 1 | 1.2500e-01 | 5.4894e+05 | ; | CD - OE2 |
| 4206 | 4207 | 1 | 1.2290e-01 | 4.7698e+05 | ; | C - O |
| 4206 | 4208 | 1 | 1.3350e-01 | 4.1003e+05 | ; | C - N |
| 4208 | 4209 | 1 | 1.0100e-01 | 3.6317e+05 | ; | N - H |
| 4208 | 4210 | 1 | 1.4490e-01 | 2.8200e+05 | ; | N - CA |
| 4210 | 4211 | 1 | 1.0900e-01 | 2.8451e+05 | ; | CA - HA |
| 4210 | 4212 | 1 | 1.5260e-01 | 2.5941e+05 | ; | CA - CB |
| 4210 | 4221 | 1 | 1.5220e-01 | 2.6527e+05 | ; | CA - C |
| 4212 | 4213 | 1 | 1.0900e-01 | 2.8451e+05 | ; | CB - HB2 |
| 4212 | 4214 | 1 | 1.0900e-01 | 2.8451e+05 | ; | CB - HB3 |
| 4212 | 4215 | 1 | 1.5260e-01 | 2.5941e+05 | ; | CB - CG |
| 4215 | 4216 | 1 | 1.0900e-01 | 2.8451e+05 | ; | CG - HG2 |

| | | | | | | |
|---|---|---|---|---|---|---|
| 4215 | 4217 | 1 | 1.0900e-01 | 2.8451e+05 | ; | CG - HG3 |
| 4215 | 4218 | 1 | 1.5220e-01 | 2.6527e+05 | ; | CG - CD |
| 4218 | 4219 | 1 | 1.2500e-01 | 5.4894e+05 | ; | CD - OE1 |
| 4218 | 4220 | 1 | 1.2500e-01 | 5.4894e+05 | ; | CD - OE2 |
| 4221 | 4222 | 1 | 1.2290e-01 | 4.7698e+05 | ; | C - O |
| 4221 | 4223 | 1 | 1.3350e-01 | 4.1003e+05 | ; | C - N |
| 4223 | 4224 | 1 | 1.0100e-01 | 3.6317e+05 | ; | N - H |
| 4223 | 4225 | 1 | 1.4490e-01 | 2.8200e+05 | ; | N - CA |
| 4225 | 4226 | 1 | 1.0900e-01 | 2.8451e+05 | ; | CA - HA |
| 4225 | 4227 | 1 | 1.5260e-01 | 2.5941e+05 | ; | CA - CB |
| 4225 | 4231 | 1 | 1.5220e-01 | 2.6527e+05 | ; | CA - C |
| 4227 | 4228 | 1 | 1.0900e-01 | 2.8451e+05 | ; | CB - HB1 |
| 4227 | 4229 | 1 | 1.0900e-01 | 2.8451e+05 | ; | CB - HB2 |
| 4227 | 4230 | 1 | 1.0900e-01 | 2.8451e+05 | ; | CB - HB3 |
| 4231 | 4232 | 1 | 1.2290e-01 | 4.7698e+05 | ; | C - O |
| 4231 | 4233 | 1 | 1.3350e-01 | 4.1003e+05 | ; | C - N |
| 4233 | 4234 | 1 | 1.0100e-01 | 3.6317e+05 | ; | N - H |
| 4233 | 4235 | 1 | 1.4490e-01 | 2.8200e+05 | ; | N - CA |
| 4235 | 4236 | 1 | 1.0900e-01 | 2.8451e+05 | ; | CA - HA |
| 4235 | 4237 | 1 | 1.5260e-01 | 2.5941e+05 | ; | CA - CB |
| 4235 | 4250 | 1 | 1.5220e-01 | 2.6527e+05 | ; | CA - C |
| 4237 | 4238 | 1 | 1.0900e-01 | 2.8451e+05 | ; | CB - HB2 |
| 4237 | 4239 | 1 | 1.0900e-01 | 2.8451e+05 | ; | CB - HB3 |
| 4237 | 4240 | 1 | 1.5260e-01 | 2.5941e+05 | ; | CB - CG |
| 4240 | 4241 | 1 | 1.0900e-01 | 2.8451e+05 | ; | CG - HG |
| 4240 | 4242 | 1 | 1.5260e-01 | 2.5941e+05 | ; | CG - CD1 |
| 4240 | 4246 | 1 | 1.5260e-01 | 2.5941e+05 | ; | CG - CD2 |
| 4242 | 4243 | 1 | 1.0900e-01 | 2.8451e+05 | ; | CD1 - HD11 |
| 4242 | 4244 | 1 | 1.0900e-01 | 2.8451e+05 | ; | CD1 - HD12 |
| 4242 | 4245 | 1 | 1.0900e-01 | 2.8451e+05 | ; | CD1 - HD13 |
| 4246 | 4247 | 1 | 1.0900e-01 | 2.8451e+05 | ; | CD2 - HD21 |
| 4246 | 4248 | 1 | 1.0900e-01 | 2.8451e+05 | ; | CD2 - HD22 |
| 4246 | 4249 | 1 | 1.0900e-01 | 2.8451e+05 | ; | CD2 - HD23 |
| 4250 | 4251 | 1 | 1.2290e-01 | 4.7698e+05 | ; | C - O |
| 4250 | 4252 | 1 | 1.3350e-01 | 4.1003e+05 | ; | C - N |
| 4252 | 4253 | 1 | 1.0100e-01 | 3.6317e+05 | ; | N - H |
| 4252 | 4254 | 1 | 1.4490e-01 | 2.8200e+05 | ; | N - CA |
| 4254 | 4255 | 1 | 1.0900e-01 | 2.8451e+05 | ; | CA - HA |
| 4254 | 4256 | 1 | 1.5260e-01 | 2.5941e+05 | ; | CA - CB |
| 4254 | 4264 | 1 | 1.5220e-01 | 2.6527e+05 | ; | CA - C |
| 4256 | 4257 | 1 | 1.0900e-01 | 2.8451e+05 | ; | CB - HB2 |
| 4256 | 4258 | 1 | 1.0900e-01 | 2.8451e+05 | ; | CB - HB3 |
| 4256 | 4259 | 1 | 1.5220e-01 | 2.6527e+05 | ; | CB - CG |
| 4259 | 4260 | 1 | 1.2290e-01 | 4.7698e+05 | ; | CG - OD1 |
| 4259 | 4261 | 1 | 1.3350e-01 | 4.1003e+05 | ; | CG - ND2 |
| 4261 | 4262 | 1 | 1.0100e-01 | 3.6317e+05 | ; | ND2 - HD21 |
| 4261 | 4263 | 1 | 1.0100e-01 | 3.6317e+05 | ; | ND2 - HD22 |
| 4264 | 4265 | 1 | 1.2290e-01 | 4.7698e+05 | ; | C - O |
| 4264 | 4266 | 1 | 1.3350e-01 | 4.1003e+05 | ; | C - N |
| 4266 | 4267 | 1 | 1.0100e-01 | 3.6317e+05 | ; | N - H |
| 4266 | 4268 | 1 | 1.4490e-01 | 2.8200e+05 | ; | N - CA |
| 4268 | 4269 | 1 | 1.0900e-01 | 2.8451e+05 | ; | CA - HA |

| | | | | | | |
|---|---|---|---|---|---|---|
| 4268 | 4270 | 1 | 1.5260e-01 | 2.5941e+05 | ; | CA - CB |
| 4268 | 4281 | 1 | 1.5220e-01 | 2.6527e+05 | ; | CA - C |
| 4270 | 4271 | 1 | 1.0900e-01 | 2.8451e+05 | ; | CB - HB2 |
| 4270 | 4272 | 1 | 1.0900e-01 | 2.8451e+05 | ; | CB - HB3 |
| 4270 | 4273 | 1 | 1.5040e-01 | 2.6527e+05 | ; | CB - CG |
| 4273 | 4274 | 1 | 1.3940e-01 | 3.4309e+05 | ; | CG - ND1 |
| 4273 | 4279 | 1 | 1.3710e-01 | 4.3346e+05 | ; | CG - CD2 |
| 4274 | 4275 | 1 | 1.3350e-01 | 4.0836e+05 | ; | ND1 - CE1 |
| 4275 | 4276 | 1 | 1.0800e-01 | 3.0711e+05 | ; | CE1 - HE1 |
| 4275 | 4277 | 1 | 1.3430e-01 | 3.9915e+05 | ; | CE1 - NE2 |
| 4277 | 4278 | 1 | 1.0100e-01 | 3.6317e+05 | ; | NE2 - HE2 |
| 4277 | 4279 | 1 | 1.3810e-01 | 3.5731e+05 | ; | NE2 - CD2 |
| 4279 | 4280 | 1 | 1.0800e-01 | 3.0711e+05 | ; | CD2 - HD2 |
| 4281 | 4282 | 1 | 1.2290e-01 | 4.7698e+05 | ; | C - O |
| 4281 | 4283 | 1 | 1.3350e-01 | 4.1003e+05 | ; | C - N |
| 4283 | 4284 | 1 | 1.4490e-01 | 2.8200e+05 | ; | N - CD |
| 4283 | 4293 | 1 | 1.4490e-01 | 2.8200e+05 | ; | N - CA |
| 4284 | 4285 | 1 | 1.0900e-01 | 2.8451e+05 | ; | CD - HD2 |
| 4284 | 4286 | 1 | 1.0900e-01 | 2.8451e+05 | ; | CD - HD3 |
| 4284 | 4287 | 1 | 1.5260e-01 | 2.5941e+05 | ; | CD - CG |
| 4287 | 4288 | 1 | 1.0900e-01 | 2.8451e+05 | ; | CG - HG2 |
| 4287 | 4289 | 1 | 1.0900e-01 | 2.8451e+05 | ; | CG - HG3 |
| 4287 | 4290 | 1 | 1.5260e-01 | 2.5941e+05 | ; | CG - CB |
| 4290 | 4291 | 1 | 1.0900e-01 | 2.8451e+05 | ; | CB - HB2 |
| 4290 | 4292 | 1 | 1.0900e-01 | 2.8451e+05 | ; | CB - HB3 |
| 4290 | 4293 | 1 | 1.5260e-01 | 2.5941e+05 | ; | CB - CA |
| 4293 | 4294 | 1 | 1.0900e-01 | 2.8451e+05 | ; | CA - HA |
| 4293 | 4295 | 1 | 1.5220e-01 | 2.6527e+05 | ; | CA - C |
| 4295 | 4296 | 1 | 1.2290e-01 | 4.7698e+05 | ; | C - O |
| 4295 | 4297 | 1 | 1.3350e-01 | 4.1003e+05 | ; | C - N |
| 4297 | 4298 | 1 | 1.0100e-01 | 3.6317e+05 | ; | N - H |
| 4297 | 4299 | 1 | 1.4490e-01 | 2.8200e+05 | ; | N - CA |
| 4299 | 4300 | 1 | 1.0900e-01 | 2.8451e+05 | ; | CA - HA |
| 4299 | 4301 | 1 | 1.5260e-01 | 2.5941e+05 | ; | CA - CB |
| 4299 | 4319 | 1 | 1.5220e-01 | 2.6527e+05 | ; | CA - C |
| 4301 | 4302 | 1 | 1.0900e-01 | 2.8451e+05 | ; | CB - HB2 |
| 4301 | 4303 | 1 | 1.0900e-01 | 2.8451e+05 | ; | CB - HB3 |
| 4301 | 4304 | 1 | 1.4950e-01 | 2.6527e+05 | ; | CB - CG |
| 4304 | 4305 | 1 | 1.3520e-01 | 4.5689e+05 | ; | CG - CD1 |
| 4304 | 4318 | 1 | 1.4590e-01 | 3.2468e+05 | ; | CG - CD2 |
| 4305 | 4306 | 1 | 1.0800e-01 | 3.0711e+05 | ; | CD1 - HD1 |
| 4305 | 4307 | 1 | 1.3810e-01 | 3.5731e+05 | ; | CD1 - NE1 |
| 4307 | 4308 | 1 | 1.0100e-01 | 3.6317e+05 | ; | NE1 - HE1 |
| 4307 | 4309 | 1 | 1.3800e-01 | 3.5815e+05 | ; | NE1 - CE2 |
| 4309 | 4310 | 1 | 1.4000e-01 | 3.9246e+05 | ; | CE2 - CZ2 |
| 4309 | 4318 | 1 | 1.4190e-01 | 3.7405e+05 | ; | CE2 - CD2 |
| 4310 | 4311 | 1 | 1.0800e-01 | 3.0711e+05 | ; | CZ2 - HZ2 |
| 4310 | 4312 | 1 | 1.4000e-01 | 3.9246e+05 | ; | CZ2 - CH2 |
| 4312 | 4313 | 1 | 1.0800e-01 | 3.0711e+05 | ; | CH2 - HH2 |
| 4312 | 4314 | 1 | 1.4000e-01 | 3.9246e+05 | ; | CH2 - CZ3 |
| 4314 | 4315 | 1 | 1.0800e-01 | 3.0711e+05 | ; | CZ3 - HZ3 |
| 4314 | 4316 | 1 | 1.4000e-01 | 3.9246e+05 | ; | CZ3 - CE3 |

| | | | | | | |
|---|---|---|---|---|---|---|
| 4316 | 4317 | 1 | 1.0800e-01 | 3.0711e+05 | ; | CE3 - HE3 |
| 4316 | 4318 | 1 | 1.4040e-01 | 3.9246e+05 | ; | CE3 - CD2 |
| 4319 | 4320 | 1 | 1.2290e-01 | 4.7698e+05 | ; | C - O |
| 4319 | 4321 | 1 | 1.3350e-01 | 4.1003e+05 | ; | C - N |
| 4321 | 4322 | 1 | 1.0100e-01 | 3.6317e+05 | ; | N - H |
| 4321 | 4323 | 1 | 1.4490e-01 | 2.8200e+05 | ; | N - CA |
| 4323 | 4324 | 1 | 1.0900e-01 | 2.8451e+05 | ; | CA - HA |
| 4323 | 4325 | 1 | 1.5260e-01 | 2.5941e+05 | ; | CA - CB |
| 4323 | 4338 | 1 | 1.5220e-01 | 2.6527e+05 | ; | CA - C |
| 4325 | 4326 | 1 | 1.0900e-01 | 2.8451e+05 | ; | CB - HB |
| 4325 | 4327 | 1 | 1.5260e-01 | 2.5941e+05 | ; | CB - CG2 |
| 4325 | 4331 | 1 | 1.5260e-01 | 2.5941e+05 | ; | CB - CG1 |
| 4327 | 4328 | 1 | 1.0900e-01 | 2.8451e+05 | ; | CG2 - HG21 |
| 4327 | 4329 | 1 | 1.0900e-01 | 2.8451e+05 | ; | CG2 - HG22 |
| 4327 | 4330 | 1 | 1.0900e-01 | 2.8451e+05 | ; | CG2 - HG23 |
| 4331 | 4332 | 1 | 1.0900e-01 | 2.8451e+05 | ; | CG1 - HG12 |
| 4331 | 4333 | 1 | 1.0900e-01 | 2.8451e+05 | ; | CG1 - HG13 |
| 4331 | 4334 | 1 | 1.5260e-01 | 2.5941e+05 | ; | CG1 - CD1 |
| 4334 | 4335 | 1 | 1.0900e-01 | 2.8451e+05 | ; | CD1 - HD11 |
| 4334 | 4336 | 1 | 1.0900e-01 | 2.8451e+05 | ; | CD1 - HD12 |
| 4334 | 4337 | 1 | 1.0900e-01 | 2.8451e+05 | ; | CD1 - HD13 |
| 4338 | 4339 | 1 | 1.2290e-01 | 4.7698e+05 | ; | C - O |
| 4338 | 4340 | 1 | 1.3350e-01 | 4.1003e+05 | ; | C - N |
| 4340 | 4341 | 1 | 1.0100e-01 | 3.6317e+05 | ; | N - H |
| 4340 | 4342 | 1 | 1.4490e-01 | 2.8200e+05 | ; | N - CA |
| 4342 | 4343 | 1 | 1.0900e-01 | 2.8451e+05 | ; | CA - HA |
| 4342 | 4344 | 1 | 1.5260e-01 | 2.5941e+05 | ; | CA - CB |
| 4342 | 4352 | 1 | 1.5220e-01 | 2.6527e+05 | ; | CA - C |
| 4344 | 4345 | 1 | 1.0900e-01 | 2.8451e+05 | ; | CB - HB |
| 4344 | 4346 | 1 | 1.5260e-01 | 2.5941e+05 | ; | CB - CG2 |
| 4344 | 4350 | 1 | 1.4100e-01 | 2.6778e+05 | ; | CB - OG1 |
| 4346 | 4347 | 1 | 1.0900e-01 | 2.8451e+05 | ; | CG2 - HG21 |
| 4346 | 4348 | 1 | 1.0900e-01 | 2.8451e+05 | ; | CG2 - HG22 |
| 4346 | 4349 | 1 | 1.0900e-01 | 2.8451e+05 | ; | CG2 - HG23 |
| 4350 | 4351 | 1 | 9.6000e-02 | 4.6275e+05 | ; | OG1 - HG1 |
| 4352 | 4353 | 1 | 1.2290e-01 | 4.7698e+05 | ; | C - O |
| 4352 | 4354 | 1 | 1.3350e-01 | 4.1003e+05 | ; | C - N |
| 4354 | 4355 | 1 | 1.0100e-01 | 3.6317e+05 | ; | N - H |
| 4354 | 4356 | 1 | 1.4490e-01 | 2.8200e+05 | ; | N - CA |
| 4356 | 4357 | 1 | 1.0900e-01 | 2.8451e+05 | ; | CA - HA |
| 4356 | 4358 | 1 | 1.5260e-01 | 2.5941e+05 | ; | CA - CB |
| 4356 | 4369 | 1 | 1.5220e-01 | 2.6527e+05 | ; | CA - C |
| 4358 | 4359 | 1 | 1.0900e-01 | 2.8451e+05 | ; | CB - HB2 |
| 4358 | 4360 | 1 | 1.0900e-01 | 2.8451e+05 | ; | CB - HB3 |
| 4358 | 4361 | 1 | 1.5260e-01 | 2.5941e+05 | ; | CB - CG |
| 4361 | 4362 | 1 | 1.0900e-01 | 2.8451e+05 | ; | CG - HG2 |
| 4361 | 4363 | 1 | 1.0900e-01 | 2.8451e+05 | ; | CG - HG3 |
| 4361 | 4364 | 1 | 1.5220e-01 | 2.6527e+05 | ; | CG - CD |
| 4364 | 4365 | 1 | 1.2290e-01 | 4.7698e+05 | ; | CD - OE1 |
| 4364 | 4366 | 1 | 1.3350e-01 | 4.1003e+05 | ; | CD - NE2 |
| 4366 | 4367 | 1 | 1.0100e-01 | 3.6317e+05 | ; | NE2 - HE21 |
| 4366 | 4368 | 1 | 1.0100e-01 | 3.6317e+05 | ; | NE2 - HE22 |

| | | | | | | |
|---|---|---|---|---|---|---|
| 4369 | 4370 | 1 | 1.2290e-01 | 4.7698e+05 | ; | C - O |
| 4369 | 4371 | 1 | 1.3350e-01 | 4.1003e+05 | ; | C - N |
| 4371 | 4372 | 1 | 1.0100e-01 | 3.6317e+05 | ; | N - H |
| 4371 | 4373 | 1 | 1.4490e-01 | 2.8200e+05 | ; | N - CA |
| 4373 | 4374 | 1 | 1.0900e-01 | 2.8451e+05 | ; | CA - HA |
| 4373 | 4375 | 1 | 1.5260e-01 | 2.5941e+05 | ; | CA - CB |
| 4373 | 4386 | 1 | 1.5220e-01 | 2.6527e+05 | ; | CA - C |
| 4375 | 4376 | 1 | 1.0900e-01 | 2.8451e+05 | ; | CB - HB2 |
| 4375 | 4377 | 1 | 1.0900e-01 | 2.8451e+05 | ; | CB - HB3 |
| 4375 | 4378 | 1 | 1.5260e-01 | 2.5941e+05 | ; | CB - CG |
| 4378 | 4379 | 1 | 1.0900e-01 | 2.8451e+05 | ; | CG - HG2 |
| 4378 | 4380 | 1 | 1.0900e-01 | 2.8451e+05 | ; | CG - HG3 |
| 4378 | 4381 | 1 | 1.8100e-01 | 1.8995e+05 | ; | CG - SD |
| 4381 | 4382 | 1 | 1.8100e-01 | 1.8995e+05 | ; | SD - CE |
| 4382 | 4383 | 1 | 1.0900e-01 | 2.8451e+05 | ; | CE - HE1 |
| 4382 | 4384 | 1 | 1.0900e-01 | 2.8451e+05 | ; | CE - HE2 |
| 4382 | 4385 | 1 | 1.0900e-01 | 2.8451e+05 | ; | CE - HE3 |
| 4386 | 4387 | 1 | 1.2290e-01 | 4.7698e+05 | ; | C - O |
| 4386 | 4388 | 1 | 1.3350e-01 | 4.1003e+05 | ; | C - N |
| 4388 | 4389 | 1 | 1.0100e-01 | 3.6317e+05 | ; | N - H |
| 4388 | 4390 | 1 | 1.4490e-01 | 2.8200e+05 | ; | N - CA |
| 4390 | 4391 | 1 | 1.0900e-01 | 2.8451e+05 | ; | CA - HA |
| 4390 | 4392 | 1 | 1.5260e-01 | 2.5941e+05 | ; | CA - CB |
| 4390 | 4400 | 1 | 1.5220e-01 | 2.6527e+05 | ; | CA - C |
| 4392 | 4393 | 1 | 1.0900e-01 | 2.8451e+05 | ; | CB - HB |
| 4392 | 4394 | 1 | 1.5260e-01 | 2.5941e+05 | ; | CB - CG2 |
| 4392 | 4398 | 1 | 1.4100e-01 | 2.6778e+05 | ; | CB - OG1 |
| 4394 | 4395 | 1 | 1.0900e-01 | 2.8451e+05 | ; | CG2 - HG21 |
| 4394 | 4396 | 1 | 1.0900e-01 | 2.8451e+05 | ; | CG2 - HG22 |
| 4394 | 4397 | 1 | 1.0900e-01 | 2.8451e+05 | ; | CG2 - HG23 |
| 4398 | 4399 | 1 | 9.6000e-02 | 4.6275e+05 | ; | OG1 - HG1 |
| 4400 | 4401 | 1 | 1.2290e-01 | 4.7698e+05 | ; | C - O |
| 4400 | 4402 | 1 | 1.3350e-01 | 4.1003e+05 | ; | C - N |
| 4402 | 4403 | 1 | 1.0100e-01 | 3.6317e+05 | ; | N - H |
| 4402 | 4404 | 1 | 1.4490e-01 | 2.8200e+05 | ; | N - CA |
| 4404 | 4405 | 1 | 1.0900e-01 | 2.8451e+05 | ; | CA - HA |
| 4404 | 4406 | 1 | 1.5260e-01 | 2.5941e+05 | ; | CA - CB |
| 4404 | 4422 | 1 | 1.5220e-01 | 2.6527e+05 | ; | CA - C |
| 4406 | 4407 | 1 | 1.0900e-01 | 2.8451e+05 | ; | CB - HB2 |
| 4406 | 4408 | 1 | 1.0900e-01 | 2.8451e+05 | ; | CB - HB3 |
| 4406 | 4409 | 1 | 1.5260e-01 | 2.5941e+05 | ; | CB - CG |
| 4409 | 4410 | 1 | 1.0900e-01 | 2.8451e+05 | ; | CG - HG2 |
| 4409 | 4411 | 1 | 1.0900e-01 | 2.8451e+05 | ; | CG - HG3 |
| 4409 | 4412 | 1 | 1.5260e-01 | 2.5941e+05 | ; | CG - CD |
| 4412 | 4413 | 1 | 1.0900e-01 | 2.8451e+05 | ; | CD - HD2 |
| 4412 | 4414 | 1 | 1.0900e-01 | 2.8451e+05 | ; | CD - HD3 |
| 4412 | 4415 | 1 | 1.5260e-01 | 2.5941e+05 | ; | CD - CE |
| 4415 | 4416 | 1 | 1.0900e-01 | 2.8451e+05 | ; | CE - HE2 |
| 4415 | 4417 | 1 | 1.0900e-01 | 2.8451e+05 | ; | CE - HE3 |
| 4415 | 4418 | 1 | 1.4710e-01 | 3.0711e+05 | ; | CE - NZ |
| 4418 | 4419 | 1 | 1.0100e-01 | 3.6317e+05 | ; | NZ - HZ1 |
| 4418 | 4420 | 1 | 1.0100e-01 | 3.6317e+05 | ; | NZ - HZ2 |

| | | | | | | |
|---|---|---|---|---|---|---|
| 4418 | 4421 | 1 | 1.0100e-01 | 3.6317e+05 | ; | NZ - HZ3 |
| 4422 | 4423 | 1 | 1.2290e-01 | 4.7698e+05 | ; | C - O |
| 4422 | 4424 | 1 | 1.3350e-01 | 4.1003e+05 | ; | C - N |
| 4424 | 4425 | 1 | 1.0100e-01 | 3.6317e+05 | ; | N - H |
| 4424 | 4426 | 1 | 1.4490e-01 | 2.8200e+05 | ; | N - CA |
| 4426 | 4427 | 1 | 1.0900e-01 | 2.8451e+05 | ; | CA - HA |
| 4426 | 4428 | 1 | 1.5260e-01 | 2.5941e+05 | ; | CA - CB |
| 4426 | 4433 | 1 | 1.5220e-01 | 2.6527e+05 | ; | CA - C |
| 4428 | 4429 | 1 | 1.0900e-01 | 2.8451e+05 | ; | CB - HB2 |
| 4428 | 4430 | 1 | 1.0900e-01 | 2.8451e+05 | ; | CB - HB3 |
| 4428 | 4431 | 1 | 1.4100e-01 | 2.6778e+05 | ; | CB - OG |
| 4431 | 4432 | 1 | 9.6000e-02 | 4.6275e+05 | ; | OG - HG |
| 4433 | 4434 | 1 | 1.2290e-01 | 4.7698e+05 | ; | C - O |
| 4433 | 4435 | 1 | 1.3350e-01 | 4.1003e+05 | ; | C - N |
| 4435 | 4436 | 1 | 1.0100e-01 | 3.6317e+05 | ; | N - H |
| 4435 | 4437 | 1 | 1.4490e-01 | 2.8200e+05 | ; | N - CA |
| 4437 | 4438 | 1 | 1.0900e-01 | 2.8451e+05 | ; | CA - HA |
| 4437 | 4439 | 1 | 1.5260e-01 | 2.5941e+05 | ; | CA - CB |
| 4437 | 4450 | 1 | 1.5220e-01 | 2.6527e+05 | ; | CA - C |
| 4439 | 4440 | 1 | 1.0900e-01 | 2.8451e+05 | ; | CB - HB2 |
| 4439 | 4441 | 1 | 1.0900e-01 | 2.8451e+05 | ; | CB - HB3 |
| 4439 | 4442 | 1 | 1.5040e-01 | 2.6527e+05 | ; | CB - CG |
| 4442 | 4443 | 1 | 1.3940e-01 | 3.4309e+05 | ; | CG - ND1 |
| 4442 | 4448 | 1 | 1.3710e-01 | 4.3346e+05 | ; | CG - CD2 |
| 4443 | 4444 | 1 | 1.3350e-01 | 4.0836e+05 | ; | ND1 - CE1 |
| 4444 | 4445 | 1 | 1.0800e-01 | 3.0711e+05 | ; | CE1 - HE1 |
| 4444 | 4446 | 1 | 1.3430e-01 | 3.9915e+05 | ; | CE1 - NE2 |
| 4446 | 4447 | 1 | 1.0100e-01 | 3.6317e+05 | ; | NE2 - HE2 |
| 4446 | 4448 | 1 | 1.3810e-01 | 3.5731e+05 | ; | NE2 - CD2 |
| 4448 | 4449 | 1 | 1.0800e-01 | 3.0711e+05 | ; | CD2 - HD2 |
| 4450 | 4451 | 1 | 1.2290e-01 | 4.7698e+05 | ; | C - O |
| 4450 | 4452 | 1 | 1.3350e-01 | 4.1003e+05 | ; | C - N |
| 4452 | 4453 | 1 | 1.0100e-01 | 3.6317e+05 | ; | N - H |
| 4452 | 4454 | 1 | 1.4490e-01 | 2.8200e+05 | ; | N - CA |
| 4454 | 4455 | 1 | 1.0900e-01 | 2.8451e+05 | ; | CA - HA |
| 4454 | 4456 | 1 | 1.5260e-01 | 2.5941e+05 | ; | CA - CB |
| 4454 | 4465 | 1 | 1.5220e-01 | 2.6527e+05 | ; | CA - C |
| 4456 | 4457 | 1 | 1.0900e-01 | 2.8451e+05 | ; | CB - HB2 |
| 4456 | 4458 | 1 | 1.0900e-01 | 2.8451e+05 | ; | CB - HB3 |
| 4456 | 4459 | 1 | 1.5260e-01 | 2.5941e+05 | ; | CB - CG |
| 4459 | 4460 | 1 | 1.0900e-01 | 2.8451e+05 | ; | CG - HG2 |
| 4459 | 4461 | 1 | 1.0900e-01 | 2.8451e+05 | ; | CG - HG3 |
| 4459 | 4462 | 1 | 1.5220e-01 | 2.6527e+05 | ; | CG - CD |
| 4462 | 4463 | 1 | 1.2500e-01 | 5.4894e+05 | ; | CD - OE1 |
| 4462 | 4464 | 1 | 1.2500e-01 | 5.4894e+05 | ; | CD - OE2 |
| 4465 | 4466 | 1 | 1.2290e-01 | 4.7698e+05 | ; | C - O |
| 4465 | 4467 | 1 | 1.3350e-01 | 4.1003e+05 | ; | C - N |
| 4467 | 4468 | 1 | 1.0100e-01 | 3.6317e+05 | ; | N - H |
| 4467 | 4469 | 1 | 1.4490e-01 | 2.8200e+05 | ; | N - CA |
| 4469 | 4470 | 1 | 1.0900e-01 | 2.8451e+05 | ; | CA - HA |
| 4469 | 4471 | 1 | 1.5260e-01 | 2.5941e+05 | ; | CA - CB |
| 4469 | 4482 | 1 | 1.5220e-01 | 2.6527e+05 | ; | CA - C |

| | | | | | | | | |
|---|---|---|---|---|---|---|---|---|
| 4471 | 4472 | 1 | 1.0900e-01 | 2.8451e+05 | ; | CB | - | HB2 |
| 4471 | 4473 | 1 | 1.0900e-01 | 2.8451e+05 | ; | CB | - | HB3 |
| 4471 | 4474 | 1 | 1.5040e-01 | 2.6527e+05 | ; | CB | - | CG |
| 4474 | 4475 | 1 | 1.3940e-01 | 3.4309e+05 | ; | CG | - | ND1 |
| 4474 | 4480 | 1 | 1.3710e-01 | 4.3346e+05 | ; | CG | - | CD2 |
| 4475 | 4476 | 1 | 1.3350e-01 | 4.0836e+05 | ; | ND1 | - | CE1 |
| 4476 | 4477 | 1 | 1.0800e-01 | 3.0711e+05 | ; | CE1 | - | HE1 |
| 4476 | 4478 | 1 | 1.3430e-01 | 3.9915e+05 | ; | CE1 | - | NE2 |
| 4478 | 4479 | 1 | 1.0100e-01 | 3.6317e+05 | ; | NE2 | - | HE2 |
| 4478 | 4480 | 1 | 1.3810e-01 | 3.5731e+05 | ; | NE2 | - | CD2 |
| 4480 | 4481 | 1 | 1.0800e-01 | 3.0711e+05 | ; | CD2 | - | HD2 |
| 4482 | 4483 | 1 | 1.2290e-01 | 4.7698e+05 | ; | C | - | O |
| 4482 | 4484 | 1 | 1.3350e-01 | 4.1003e+05 | ; | C | - | N |
| 4484 | 4485 | 1 | 1.0100e-01 | 3.6317e+05 | ; | N | - | H |
| 4484 | 4486 | 1 | 1.4490e-01 | 2.8200e+05 | ; | N | - | CA |
| 4486 | 4487 | 1 | 1.0900e-01 | 2.8451e+05 | ; | CA | - | HA |
| 4486 | 4488 | 1 | 1.5260e-01 | 2.5941e+05 | ; | CA | - | CB |
| 4486 | 4498 | 1 | 1.5220e-01 | 2.6527e+05 | ; | CA | - | C |
| 4488 | 4489 | 1 | 1.0900e-01 | 2.8451e+05 | ; | CB | - | HB |
| 4488 | 4490 | 1 | 1.5260e-01 | 2.5941e+05 | ; | CB | - | CG1 |
| 4488 | 4494 | 1 | 1.5260e-01 | 2.5941e+05 | ; | CB | - | CG2 |
| 4490 | 4491 | 1 | 1.0900e-01 | 2.8451e+05 | ; | CG1 | - | HG11 |
| 4490 | 4492 | 1 | 1.0900e-01 | 2.8451e+05 | ; | CG1 | - | HG12 |
| 4490 | 4493 | 1 | 1.0900e-01 | 2.8451e+05 | ; | CG1 | - | HG13 |
| 4494 | 4495 | 1 | 1.0900e-01 | 2.8451e+05 | ; | CG2 | - | HG21 |
| 4494 | 4496 | 1 | 1.0900e-01 | 2.8451e+05 | ; | CG2 | - | HG22 |
| 4494 | 4497 | 1 | 1.0900e-01 | 2.8451e+05 | ; | CG2 | - | HG23 |
| 4498 | 4499 | 1 | 1.2290e-01 | 4.7698e+05 | ; | C | - | O |
| 4498 | 4500 | 1 | 1.3350e-01 | 4.1003e+05 | ; | C | - | N |
| 4500 | 4501 | 1 | 1.0100e-01 | 3.6317e+05 | ; | N | - | H |
| 4500 | 4502 | 1 | 1.4490e-01 | 2.8200e+05 | ; | N | - | CA |
| 4502 | 4503 | 1 | 1.0900e-01 | 2.8451e+05 | ; | CA | - | HA |
| 4502 | 4504 | 1 | 1.5260e-01 | 2.5941e+05 | ; | CA | - | CB |
| 4502 | 4513 | 1 | 1.5220e-01 | 2.6527e+05 | ; | CA | - | C |
| 4504 | 4505 | 1 | 1.0900e-01 | 2.8451e+05 | ; | CB | - | HB2 |
| 4504 | 4506 | 1 | 1.0900e-01 | 2.8451e+05 | ; | CB | - | HB3 |
| 4504 | 4507 | 1 | 1.5260e-01 | 2.5941e+05 | ; | CB | - | CG |
| 4507 | 4508 | 1 | 1.0900e-01 | 2.8451e+05 | ; | CG | - | HG2 |
| 4507 | 4509 | 1 | 1.0900e-01 | 2.8451e+05 | ; | CG | - | HG3 |
| 4507 | 4510 | 1 | 1.5220e-01 | 2.6527e+05 | ; | CG | - | CD |
| 4510 | 4511 | 1 | 1.2500e-01 | 5.4894e+05 | ; | CD | - | OE1 |
| 4510 | 4512 | 1 | 1.2500e-01 | 5.4894e+05 | ; | CD | - | OE2 |
| 4513 | 4514 | 1 | 1.2290e-01 | 4.7698e+05 | ; | C | - | O |
| 4513 | 4515 | 1 | 1.3350e-01 | 4.1003e+05 | ; | C | - | N |
| 4515 | 4516 | 1 | 1.0100e-01 | 3.6317e+05 | ; | N | - | H |
| 4515 | 4517 | 1 | 1.4490e-01 | 2.8200e+05 | ; | N | - | CA |
| 4517 | 4518 | 1 | 1.0900e-01 | 2.8451e+05 | ; | CA | - | HA |
| 4517 | 4519 | 1 | 1.5260e-01 | 2.5941e+05 | ; | CA | - | CB |
| 4517 | 4532 | 1 | 1.5220e-01 | 2.6527e+05 | ; | CA | - | C |
| 4519 | 4520 | 1 | 1.0900e-01 | 2.8451e+05 | ; | CB | - | HB2 |
| 4519 | 4521 | 1 | 1.0900e-01 | 2.8451e+05 | ; | CB | - | HB3 |
| 4519 | 4522 | 1 | 1.5260e-01 | 2.5941e+05 | ; | CB | - | CG |

| | | | | | |
|---|---|---|---|---|---|
| 4522 | 4523 | 1 | 1.0900e-01 | 2.8451e+05 ; | CG - HG |
| 4522 | 4524 | 1 | 1.5260e-01 | 2.5941e+05 ; | CG - CD1 |
| 4522 | 4528 | 1 | 1.5260e-01 | 2.5941e+05 ; | CG - CD2 |
| 4524 | 4525 | 1 | 1.0900e-01 | 2.8451e+05 ; | CD1 - HD11 |
| 4524 | 4526 | 1 | 1.0900e-01 | 2.8451e+05 ; | CD1 - HD12 |
| 4524 | 4527 | 1 | 1.0900e-01 | 2.8451e+05 ; | CD1 - HD13 |
| 4528 | 4529 | 1 | 1.0900e-01 | 2.8451e+05 ; | CD2 - HD21 |
| 4528 | 4530 | 1 | 1.0900e-01 | 2.8451e+05 ; | CD2 - HD22 |
| 4528 | 4531 | 1 | 1.0900e-01 | 2.8451e+05 ; | CD2 - HD23 |
| 4532 | 4533 | 1 | 1.2290e-01 | 4.7698e+05 ; | C - O |
| 4532 | 4534 | 1 | 1.3350e-01 | 4.1003e+05 ; | C - N |
| 4534 | 4535 | 1 | 1.0100e-01 | 3.6317e+05 ; | N - H |
| 4534 | 4536 | 1 | 1.4490e-01 | 2.8200e+05 ; | N - CA |
| 4536 | 4537 | 1 | 1.0900e-01 | 2.8451e+05 ; | CA - HA |
| 4536 | 4538 | 1 | 1.5260e-01 | 2.5941e+05 ; | CA - CB |
| 4536 | 4543 | 1 | 1.5220e-01 | 2.6527e+05 ; | CA - C |
| 4538 | 4539 | 1 | 1.0900e-01 | 2.8451e+05 ; | CB - HB2 |
| 4538 | 4540 | 1 | 1.0900e-01 | 2.8451e+05 ; | CB - HB3 |
| 4538 | 4541 | 1 | 1.4100e-01 | 2.6778e+05 ; | CB - OG |
| 4541 | 4542 | 1 | 9.6000e-02 | 4.6275e+05 ; | OG - HG |
| 4543 | 4544 | 1 | 1.2290e-01 | 4.7698e+05 ; | C - O |
| 4543 | 4545 | 1 | 1.3350e-01 | 4.1003e+05 ; | C - N |
| 4545 | 4546 | 1 | 1.0100e-01 | 3.6317e+05 ; | N - H |
| 4545 | 4547 | 1 | 1.4490e-01 | 2.8200e+05 ; | N - CA |
| 4547 | 4548 | 1 | 1.0900e-01 | 2.8451e+05 ; | CA - HA |
| 4547 | 4549 | 1 | 1.5260e-01 | 2.5941e+05 ; | CA - CB |
| 4547 | 4554 | 1 | 1.5220e-01 | 2.6527e+05 ; | CA - C |
| 4549 | 4550 | 1 | 1.0900e-01 | 2.8451e+05 ; | CB - HB2 |
| 4549 | 4551 | 1 | 1.0900e-01 | 2.8451e+05 ; | CB - HB3 |
| 4549 | 4552 | 1 | 1.4100e-01 | 2.6778e+05 ; | CB - OG |
| 4552 | 4553 | 1 | 9.6000e-02 | 4.6275e+05 ; | OG - HG |
| 4554 | 4555 | 1 | 1.2290e-01 | 4.7698e+05 ; | C - O |
| 4554 | 4556 | 1 | 1.3350e-01 | 4.1003e+05 ; | C - N |
| 4556 | 4557 | 1 | 1.0100e-01 | 3.6317e+05 ; | N - H |
| 4556 | 4558 | 1 | 1.4490e-01 | 2.8200e+05 ; | N - CA |
| 4558 | 4559 | 1 | 1.0900e-01 | 2.8451e+05 ; | CA - HA |
| 4558 | 4560 | 1 | 1.5260e-01 | 2.5941e+05 ; | CA - CB |
| 4558 | 4568 | 1 | 1.5220e-01 | 2.6527e+05 ; | CA - C |
| 4560 | 4561 | 1 | 1.0900e-01 | 2.8451e+05 ; | CB - HB |
| 4560 | 4562 | 1 | 1.5260e-01 | 2.5941e+05 ; | CB - CG2 |
| 4560 | 4566 | 1 | 1.4100e-01 | 2.6778e+05 ; | CB - OG1 |
| 4562 | 4563 | 1 | 1.0900e-01 | 2.8451e+05 ; | CG2 - HG21 |
| 4562 | 4564 | 1 | 1.0900e-01 | 2.8451e+05 ; | CG2 - HG22 |
| 4562 | 4565 | 1 | 1.0900e-01 | 2.8451e+05 ; | CG2 - HG23 |
| 4566 | 4567 | 1 | 9.6000e-02 | 4.6275e+05 ; | OG1 - HG1 |
| 4568 | 4569 | 1 | 1.2290e-01 | 4.7698e+05 ; | C - O |
| 4568 | 4570 | 1 | 1.3350e-01 | 4.1003e+05 ; | C - N |
| 4570 | 4571 | 1 | 1.0100e-01 | 3.6317e+05 ; | N - H |
| 4570 | 4572 | 1 | 1.4490e-01 | 2.8200e+05 ; | N - CA |
| 4572 | 4573 | 1 | 1.0900e-01 | 2.8451e+05 ; | CA - HA |
| 4572 | 4574 | 1 | 1.5260e-01 | 2.5941e+05 ; | CA - CB |
| 4572 | 4587 | 1 | 1.5220e-01 | 2.6527e+05 ; | CA - C |

| | | | | | | |
|---|---|---|---|---|---|---|
| 4574 | 4575 | 1 | 1.0900e-01 | 2.8451e+05 | ; | CB - HB2 |
| 4574 | 4576 | 1 | 1.0900e-01 | 2.8451e+05 | ; | CB - HB3 |
| 4574 | 4577 | 1 | 1.5260e-01 | 2.5941e+05 | ; | CB - CG |
| 4577 | 4578 | 1 | 1.0900e-01 | 2.8451e+05 | ; | CG - HG |
| 4577 | 4579 | 1 | 1.5260e-01 | 2.5941e+05 | ; | CG - CD1 |
| 4577 | 4583 | 1 | 1.5260e-01 | 2.5941e+05 | ; | CG - CD2 |
| 4579 | 4580 | 1 | 1.0900e-01 | 2.8451e+05 | ; | CD1 - HD11 |
| 4579 | 4581 | 1 | 1.0900e-01 | 2.8451e+05 | ; | CD1 - HD12 |
| 4579 | 4582 | 1 | 1.0900e-01 | 2.8451e+05 | ; | CD1 - HD13 |
| 4583 | 4584 | 1 | 1.0900e-01 | 2.8451e+05 | ; | CD2 - HD21 |
| 4583 | 4585 | 1 | 1.0900e-01 | 2.8451e+05 | ; | CD2 - HD22 |
| 4583 | 4586 | 1 | 1.0900e-01 | 2.8451e+05 | ; | CD2 - HD23 |
| 4587 | 4588 | 1 | 1.2290e-01 | 4.7698e+05 | ; | C - O |
| 4587 | 4589 | 1 | 1.3350e-01 | 4.1003e+05 | ; | C - N |
| 4589 | 4590 | 1 | 1.0100e-01 | 3.6317e+05 | ; | N - H |
| 4589 | 4591 | 1 | 1.4490e-01 | 2.8200e+05 | ; | N - CA |
| 4591 | 4592 | 1 | 1.0900e-01 | 2.8451e+05 | ; | CA - HA |
| 4591 | 4593 | 1 | 1.5260e-01 | 2.5941e+05 | ; | CA - CB |
| 4591 | 4606 | 1 | 1.5220e-01 | 2.6527e+05 | ; | CA - C |
| 4593 | 4594 | 1 | 1.0900e-01 | 2.8451e+05 | ; | CB - HB2 |
| 4593 | 4595 | 1 | 1.0900e-01 | 2.8451e+05 | ; | CB - HB3 |
| 4593 | 4596 | 1 | 1.5260e-01 | 2.5941e+05 | ; | CB - CG |
| 4596 | 4597 | 1 | 1.0900e-01 | 2.8451e+05 | ; | CG - HG |
| 4596 | 4598 | 1 | 1.5260e-01 | 2.5941e+05 | ; | CG - CD1 |
| 4596 | 4602 | 1 | 1.5260e-01 | 2.5941e+05 | ; | CG - CD2 |
| 4598 | 4599 | 1 | 1.0900e-01 | 2.8451e+05 | ; | CD1 - HD11 |
| 4598 | 4600 | 1 | 1.0900e-01 | 2.8451e+05 | ; | CD1 - HD12 |
| 4598 | 4601 | 1 | 1.0900e-01 | 2.8451e+05 | ; | CD1 - HD13 |
| 4602 | 4603 | 1 | 1.0900e-01 | 2.8451e+05 | ; | CD2 - HD21 |
| 4602 | 4604 | 1 | 1.0900e-01 | 2.8451e+05 | ; | CD2 - HD22 |
| 4602 | 4605 | 1 | 1.0900e-01 | 2.8451e+05 | ; | CD2 - HD23 |
| 4606 | 4607 | 1 | 1.2290e-01 | 4.7698e+05 | ; | C - O |
| 4606 | 4608 | 1 | 1.3350e-01 | 4.1003e+05 | ; | C - N |
| 4608 | 4609 | 1 | 1.0100e-01 | 3.6317e+05 | ; | N - H |
| 4608 | 4610 | 1 | 1.4490e-01 | 2.8200e+05 | ; | N - CA |
| 4610 | 4611 | 1 | 1.0900e-01 | 2.8451e+05 | ; | CA - HA |
| 4610 | 4612 | 1 | 1.5260e-01 | 2.5941e+05 | ; | CA - CB |
| 4610 | 4628 | 1 | 1.5220e-01 | 2.6527e+05 | ; | CA - C |
| 4612 | 4613 | 1 | 1.0900e-01 | 2.8451e+05 | ; | CB - HB2 |
| 4612 | 4614 | 1 | 1.0900e-01 | 2.8451e+05 | ; | CB - HB3 |
| 4612 | 4615 | 1 | 1.5260e-01 | 2.5941e+05 | ; | CB - CG |
| 4615 | 4616 | 1 | 1.0900e-01 | 2.8451e+05 | ; | CG - HG2 |
| 4615 | 4617 | 1 | 1.0900e-01 | 2.8451e+05 | ; | CG - HG3 |
| 4615 | 4618 | 1 | 1.5260e-01 | 2.5941e+05 | ; | CG - CD |
| 4618 | 4619 | 1 | 1.0900e-01 | 2.8451e+05 | ; | CD - HD2 |
| 4618 | 4620 | 1 | 1.0900e-01 | 2.8451e+05 | ; | CD - HD3 |
| 4618 | 4621 | 1 | 1.5260e-01 | 2.5941e+05 | ; | CD - CE |
| 4621 | 4622 | 1 | 1.0900e-01 | 2.8451e+05 | ; | CE - HE2 |
| 4621 | 4623 | 1 | 1.0900e-01 | 2.8451e+05 | ; | CE - HE3 |
| 4621 | 4624 | 1 | 1.4710e-01 | 3.0711e+05 | ; | CE - NZ |
| 4624 | 4625 | 1 | 1.0100e-01 | 3.6317e+05 | ; | NZ - HZ1 |
| 4624 | 4626 | 1 | 1.0100e-01 | 3.6317e+05 | ; | NZ - HZ2 |

| | | | | | | |
|---|---|---|---|---|---|---|
| 4624 | 4627 | 1 | 1.0100e-01 | 3.6317e+05 | ; | NZ - HZ3 |
| 4628 | 4629 | 1 | 1.2290e-01 | 4.7698e+05 | ; | C - O |
| 4628 | 4630 | 1 | 1.3350e-01 | 4.1003e+05 | ; | C - N |
| 4630 | 4631 | 1 | 1.0100e-01 | 3.6317e+05 | ; | N - H |
| 4630 | 4632 | 1 | 1.4490e-01 | 2.8200e+05 | ; | N - CA |
| 4632 | 4633 | 1 | 1.0900e-01 | 2.8451e+05 | ; | CA - HA |
| 4632 | 4634 | 1 | 1.5260e-01 | 2.5941e+05 | ; | CA - CB |
| 4632 | 4642 | 1 | 1.5220e-01 | 2.6527e+05 | ; | CA - C |
| 4634 | 4635 | 1 | 1.0900e-01 | 2.8451e+05 | ; | CB - HB2 |
| 4634 | 4636 | 1 | 1.0900e-01 | 2.8451e+05 | ; | CB - HB3 |
| 4634 | 4637 | 1 | 1.5220e-01 | 2.6527e+05 | ; | CB - CG |
| 4637 | 4638 | 1 | 1.2290e-01 | 4.7698e+05 | ; | CG - OD1 |
| 4637 | 4639 | 1 | 1.3350e-01 | 4.1003e+05 | ; | CG - ND2 |
| 4639 | 4640 | 1 | 1.0100e-01 | 3.6317e+05 | ; | ND2 - HD21 |
| 4639 | 4641 | 1 | 1.0100e-01 | 3.6317e+05 | ; | ND2 - HD22 |
| 4642 | 4643 | 1 | 1.2290e-01 | 4.7698e+05 | ; | C - O |
| 4642 | 4644 | 1 | 1.3350e-01 | 4.1003e+05 | ; | C - N |
| 4644 | 4645 | 1 | 1.0100e-01 | 3.6317e+05 | ; | N - H |
| 4644 | 4646 | 1 | 1.4490e-01 | 2.8200e+05 | ; | N - CA |
| 4646 | 4647 | 1 | 1.0900e-01 | 2.8451e+05 | ; | CA - HA |
| 4646 | 4648 | 1 | 1.5260e-01 | 2.5941e+05 | ; | CA - CB |
| 4646 | 4661 | 1 | 1.5220e-01 | 2.6527e+05 | ; | CA - C |
| 4648 | 4649 | 1 | 1.0900e-01 | 2.8451e+05 | ; | CB - HB2 |
| 4648 | 4650 | 1 | 1.0900e-01 | 2.8451e+05 | ; | CB - HB3 |
| 4648 | 4651 | 1 | 1.5260e-01 | 2.5941e+05 | ; | CB - CG |
| 4651 | 4652 | 1 | 1.0900e-01 | 2.8451e+05 | ; | CG - HG |
| 4651 | 4653 | 1 | 1.5260e-01 | 2.5941e+05 | ; | CG - CD1 |
| 4651 | 4657 | 1 | 1.5260e-01 | 2.5941e+05 | ; | CG - CD2 |
| 4653 | 4654 | 1 | 1.0900e-01 | 2.8451e+05 | ; | CD1 - HD11 |
| 4653 | 4655 | 1 | 1.0900e-01 | 2.8451e+05 | ; | CD1 - HD12 |
| 4653 | 4656 | 1 | 1.0900e-01 | 2.8451e+05 | ; | CD1 - HD13 |
| 4657 | 4658 | 1 | 1.0900e-01 | 2.8451e+05 | ; | CD2 - HD21 |
| 4657 | 4659 | 1 | 1.0900e-01 | 2.8451e+05 | ; | CD2 - HD22 |
| 4657 | 4660 | 1 | 1.0900e-01 | 2.8451e+05 | ; | CD2 - HD23 |
| 4661 | 4662 | 1 | 1.2290e-01 | 4.7698e+05 | ; | C - O |
| 4661 | 4663 | 1 | 1.3350e-01 | 4.1003e+05 | ; | C - N |
| 4663 | 4664 | 1 | 1.0100e-01 | 3.6317e+05 | ; | N - H |
| 4663 | 4665 | 1 | 1.4490e-01 | 2.8200e+05 | ; | N - CA |
| 4665 | 4666 | 1 | 1.0900e-01 | 2.8451e+05 | ; | CA - HA |
| 4665 | 4667 | 1 | 1.5260e-01 | 2.5941e+05 | ; | CA - CB |
| 4665 | 4683 | 1 | 1.5220e-01 | 2.6527e+05 | ; | CA - C |
| 4667 | 4668 | 1 | 1.0900e-01 | 2.8451e+05 | ; | CB - HB2 |
| 4667 | 4669 | 1 | 1.0900e-01 | 2.8451e+05 | ; | CB - HB3 |
| 4667 | 4670 | 1 | 1.5260e-01 | 2.5941e+05 | ; | CB - CG |
| 4670 | 4671 | 1 | 1.0900e-01 | 2.8451e+05 | ; | CG - HG2 |
| 4670 | 4672 | 1 | 1.0900e-01 | 2.8451e+05 | ; | CG - HG3 |
| 4670 | 4673 | 1 | 1.5260e-01 | 2.5941e+05 | ; | CG - CD |
| 4673 | 4674 | 1 | 1.0900e-01 | 2.8451e+05 | ; | CD - HD2 |
| 4673 | 4675 | 1 | 1.0900e-01 | 2.8451e+05 | ; | CD - HD3 |
| 4673 | 4676 | 1 | 1.5260e-01 | 2.5941e+05 | ; | CD - CE |
| 4676 | 4677 | 1 | 1.0900e-01 | 2.8451e+05 | ; | CE - HE2 |
| 4676 | 4678 | 1 | 1.0900e-01 | 2.8451e+05 | ; | CE - HE3 |

| | | | | | | |
|---|---|---|---|---|---|---|
| 4676 | 4679 | 1 | 1.4710e-01 | 3.0711e+05 | ; | CE - NZ |
| 4679 | 4680 | 1 | 1.0100e-01 | 3.6317e+05 | ; | NZ - HZ1 |
| 4679 | 4681 | 1 | 1.0100e-01 | 3.6317e+05 | ; | NZ - HZ2 |
| 4679 | 4682 | 1 | 1.0100e-01 | 3.6317e+05 | ; | NZ - HZ3 |
| 4683 | 4684 | 1 | 1.2290e-01 | 4.7698e+05 | ; | C - O |
| 4683 | 4685 | 1 | 1.3350e-01 | 4.1003e+05 | ; | C - N |
| 4685 | 4686 | 1 | 1.0100e-01 | 3.6317e+05 | ; | N - H |
| 4685 | 4687 | 1 | 1.4490e-01 | 2.8200e+05 | ; | N - CA |
| 4687 | 4688 | 1 | 1.0900e-01 | 2.8451e+05 | ; | CA - HA |
| 4687 | 4689 | 1 | 1.5260e-01 | 2.5941e+05 | ; | CA - CB |
| 4687 | 4697 | 1 | 1.5220e-01 | 2.6527e+05 | ; | CA - C |
| 4689 | 4690 | 1 | 1.0900e-01 | 2.8451e+05 | ; | CB - HB2 |
| 4689 | 4691 | 1 | 1.0900e-01 | 2.8451e+05 | ; | CB - HB3 |
| 4689 | 4692 | 1 | 1.5220e-01 | 2.6527e+05 | ; | CB - CG |
| 4692 | 4693 | 1 | 1.2290e-01 | 4.7698e+05 | ; | CG - OD1 |
| 4692 | 4694 | 1 | 1.3350e-01 | 4.1003e+05 | ; | CG - ND2 |
| 4694 | 4695 | 1 | 1.0100e-01 | 3.6317e+05 | ; | ND2 - HD21 |
| 4694 | 4696 | 1 | 1.0100e-01 | 3.6317e+05 | ; | ND2 - HD22 |
| 4697 | 4698 | 1 | 1.2290e-01 | 4.7698e+05 | ; | C - O |
| 4697 | 4699 | 1 | 1.3350e-01 | 4.1003e+05 | ; | C - N |
| 4699 | 4700 | 1 | 1.0100e-01 | 3.6317e+05 | ; | N - H |
| 4699 | 4701 | 1 | 1.4490e-01 | 2.8200e+05 | ; | N - CA |
| 4701 | 4702 | 1 | 1.0900e-01 | 2.8451e+05 | ; | CA - HA |
| 4701 | 4703 | 1 | 1.5260e-01 | 2.5941e+05 | ; | CA - CB |
| 4701 | 4717 | 1 | 1.5220e-01 | 2.6527e+05 | ; | CA - C |
| 4703 | 4704 | 1 | 1.0900e-01 | 2.8451e+05 | ; | CB - HB2 |
| 4703 | 4705 | 1 | 1.0900e-01 | 2.8451e+05 | ; | CB - HB3 |
| 4703 | 4706 | 1 | 1.5100e-01 | 2.6527e+05 | ; | CB - CG |
| 4706 | 4707 | 1 | 1.4000e-01 | 3.9246e+05 | ; | CG - CD1 |
| 4706 | 4715 | 1 | 1.4000e-01 | 3.9246e+05 | ; | CG - CD2 |
| 4707 | 4708 | 1 | 1.0800e-01 | 3.0711e+05 | ; | CD1 - HD1 |
| 4707 | 4709 | 1 | 1.4000e-01 | 3.9246e+05 | ; | CD1 - CE1 |
| 4709 | 4710 | 1 | 1.0800e-01 | 3.0711e+05 | ; | CE1 - HE1 |
| 4709 | 4711 | 1 | 1.4000e-01 | 3.9246e+05 | ; | CE1 - CZ |
| 4711 | 4712 | 1 | 1.0800e-01 | 3.0711e+05 | ; | CZ - HZ |
| 4711 | 4713 | 1 | 1.4000e-01 | 3.9246e+05 | ; | CZ - CE2 |
| 4713 | 4714 | 1 | 1.0800e-01 | 3.0711e+05 | ; | CE2 - HE2 |
| 4713 | 4715 | 1 | 1.4000e-01 | 3.9246e+05 | ; | CE2 - CD2 |
| 4715 | 4716 | 1 | 1.0800e-01 | 3.0711e+05 | ; | CD2 - HD2 |
| 4717 | 4718 | 1 | 1.2290e-01 | 4.7698e+05 | ; | C - O |
| 4717 | 4719 | 1 | 1.3350e-01 | 4.1003e+05 | ; | C - N |
| 4719 | 4720 | 1 | 1.0100e-01 | 3.6317e+05 | ; | N - H |
| 4719 | 4721 | 1 | 1.4490e-01 | 2.8200e+05 | ; | N - CA |
| 4721 | 4722 | 1 | 1.0900e-01 | 2.8451e+05 | ; | CA - HA |
| 4721 | 4723 | 1 | 1.5260e-01 | 2.5941e+05 | ; | CA - CB |
| 4721 | 4739 | 1 | 1.5220e-01 | 2.6527e+05 | ; | CA - C |
| 4723 | 4724 | 1 | 1.0900e-01 | 2.8451e+05 | ; | CB - HB2 |
| 4723 | 4725 | 1 | 1.0900e-01 | 2.8451e+05 | ; | CB - HB3 |
| 4723 | 4726 | 1 | 1.5260e-01 | 2.5941e+05 | ; | CB - CG |
| 4726 | 4727 | 1 | 1.0900e-01 | 2.8451e+05 | ; | CG - HG2 |
| 4726 | 4728 | 1 | 1.0900e-01 | 2.8451e+05 | ; | CG - HG3 |
| 4726 | 4729 | 1 | 1.5260e-01 | 2.5941e+05 | ; | CG - CD |

| | | | | | |
|---|---|---|---|---|---|
| 4729 | 4730 | 1 | 1.0900e-01 | 2.8451e+05 ; | CD - HD2 |
| 4729 | 4731 | 1 | 1.0900e-01 | 2.8451e+05 ; | CD - HD3 |
| 4729 | 4732 | 1 | 1.5260e-01 | 2.5941e+05 ; | CD - CE |
| 4732 | 4733 | 1 | 1.0900e-01 | 2.8451e+05 ; | CE - HE2 |
| 4732 | 4734 | 1 | 1.0900e-01 | 2.8451e+05 ; | CE - HE3 |
| 4732 | 4735 | 1 | 1.4710e-01 | 3.0711e+05 ; | CE - NZ |
| 4735 | 4736 | 1 | 1.0100e-01 | 3.6317e+05 ; | NZ - HZ1 |
| 4735 | 4737 | 1 | 1.0100e-01 | 3.6317e+05 ; | NZ - HZ2 |
| 4735 | 4738 | 1 | 1.0100e-01 | 3.6317e+05 ; | NZ - HZ3 |
| 4739 | 4740 | 1 | 1.2290e-01 | 4.7698e+05 ; | C - O |
| 4739 | 4741 | 1 | 1.3350e-01 | 4.1003e+05 ; | C - N |
| 4741 | 4742 | 1 | 1.0100e-01 | 3.6317e+05 ; | N - H |
| 4741 | 4743 | 1 | 1.4490e-01 | 2.8200e+05 ; | N - CA |
| 4743 | 4744 | 1 | 1.0900e-01 | 2.8451e+05 ; | CA - HA |
| 4743 | 4745 | 1 | 1.5260e-01 | 2.5941e+05 ; | CA - CB |
| 4743 | 4761 | 1 | 1.5220e-01 | 2.6527e+05 ; | CA - C |
| 4745 | 4746 | 1 | 1.0900e-01 | 2.8451e+05 ; | CB - HB2 |
| 4745 | 4747 | 1 | 1.0900e-01 | 2.8451e+05 ; | CB - HB3 |
| 4745 | 4748 | 1 | 1.5260e-01 | 2.5941e+05 ; | CB - CG |
| 4748 | 4749 | 1 | 1.0900e-01 | 2.8451e+05 ; | CG - HG2 |
| 4748 | 4750 | 1 | 1.0900e-01 | 2.8451e+05 ; | CG - HG3 |
| 4748 | 4751 | 1 | 1.5260e-01 | 2.5941e+05 ; | CG - CD |
| 4751 | 4752 | 1 | 1.0900e-01 | 2.8451e+05 ; | CD - HD2 |
| 4751 | 4753 | 1 | 1.0900e-01 | 2.8451e+05 ; | CD - HD3 |
| 4751 | 4754 | 1 | 1.5260e-01 | 2.5941e+05 ; | CD - CE |
| 4754 | 4755 | 1 | 1.0900e-01 | 2.8451e+05 ; | CE - HE2 |
| 4754 | 4756 | 1 | 1.0900e-01 | 2.8451e+05 ; | CE - HE3 |
| 4754 | 4757 | 1 | 1.4710e-01 | 3.0711e+05 ; | CE - NZ |
| 4757 | 4758 | 1 | 1.0100e-01 | 3.6317e+05 ; | NZ - HZ1 |
| 4757 | 4759 | 1 | 1.0100e-01 | 3.6317e+05 ; | NZ - HZ2 |
| 4757 | 4760 | 1 | 1.0100e-01 | 3.6317e+05 ; | NZ - HZ3 |
| 4761 | 4762 | 1 | 1.2290e-01 | 4.7698e+05 ; | C - O |
| 4761 | 4763 | 1 | 1.3350e-01 | 4.1003e+05 ; | C - N |
| 4763 | 4764 | 1 | 1.0100e-01 | 3.6317e+05 ; | N - H |
| 4763 | 4765 | 1 | 1.4490e-01 | 2.8200e+05 ; | N - CA |
| 4765 | 4766 | 1 | 1.0900e-01 | 2.8451e+05 ; | CA - HA |
| 4765 | 4767 | 1 | 1.5260e-01 | 2.5941e+05 ; | CA - CB |
| 4765 | 4776 | 1 | 1.5220e-01 | 2.6527e+05 ; | CA - C |
| 4767 | 4768 | 1 | 1.0900e-01 | 2.8451e+05 ; | CB - HB2 |
| 4767 | 4769 | 1 | 1.0900e-01 | 2.8451e+05 ; | CB - HB3 |
| 4767 | 4770 | 1 | 1.5260e-01 | 2.5941e+05 ; | CB - CG |
| 4770 | 4771 | 1 | 1.0900e-01 | 2.8451e+05 ; | CG - HG2 |
| 4770 | 4772 | 1 | 1.0900e-01 | 2.8451e+05 ; | CG - HG3 |
| 4770 | 4773 | 1 | 1.5220e-01 | 2.6527e+05 ; | CG - CD |
| 4773 | 4774 | 1 | 1.2500e-01 | 5.4894e+05 ; | CD - OE1 |
| 4773 | 4775 | 1 | 1.2500e-01 | 5.4894e+05 ; | CD - OE2 |
| 4776 | 4777 | 1 | 1.2290e-01 | 4.7698e+05 ; | C - O |
| 4776 | 4778 | 1 | 1.3350e-01 | 4.1003e+05 ; | C - N |
| 4778 | 4779 | 1 | 1.0100e-01 | 3.6317e+05 ; | N - H |
| 4778 | 4780 | 1 | 1.4490e-01 | 2.8200e+05 ; | N - CA |
| 4780 | 4781 | 1 | 1.0900e-01 | 2.8451e+05 ; | CA - HA |
| 4780 | 4782 | 1 | 1.5260e-01 | 2.5941e+05 ; | CA - CB |

| | | | | | |
|---|---|---|---|---|---|
| 4780 | 4790 | 1 | 1.5220e-01 | 2.6527e+05 ; | CA - C |
| 4782 | 4783 | 1 | 1.0900e-01 | 2.8451e+05 ; | CB - HB2 |
| 4782 | 4784 | 1 | 1.0900e-01 | 2.8451e+05 ; | CB - HB3 |
| 4782 | 4785 | 1 | 1.5220e-01 | 2.6527e+05 ; | CB - CG |
| 4785 | 4786 | 1 | 1.2290e-01 | 4.7698e+05 ; | CG - OD1 |
| 4785 | 4787 | 1 | 1.3350e-01 | 4.1003e+05 ; | CG - ND2 |
| 4787 | 4788 | 1 | 1.0100e-01 | 3.6317e+05 ; | ND2 - HD21 |
| 4787 | 4789 | 1 | 1.0100e-01 | 3.6317e+05 ; | ND2 - HD22 |
| 4790 | 4791 | 1 | 1.2290e-01 | 4.7698e+05 ; | C - O |
| 4790 | 4792 | 1 | 1.3350e-01 | 4.1003e+05 ; | C - N |
| 4792 | 4793 | 1 | 1.0100e-01 | 3.6317e+05 ; | N - H |
| 4792 | 4794 | 1 | 1.4490e-01 | 2.8200e+05 ; | N - CA |
| 4794 | 4795 | 1 | 1.0900e-01 | 2.8451e+05 ; | CA - HA |
| 4794 | 4796 | 1 | 1.5260e-01 | 2.5941e+05 ; | CA - CB |
| 4794 | 4805 | 1 | 1.5220e-01 | 2.6527e+05 ; | CA - C |
| 4796 | 4797 | 1 | 1.0900e-01 | 2.8451e+05 ; | CB - HB2 |
| 4796 | 4798 | 1 | 1.0900e-01 | 2.8451e+05 ; | CB - HB3 |
| 4796 | 4799 | 1 | 1.5260e-01 | 2.5941e+05 ; | CB - CG |
| 4799 | 4800 | 1 | 1.0900e-01 | 2.8451e+05 ; | CG - HG2 |
| 4799 | 4801 | 1 | 1.0900e-01 | 2.8451e+05 ; | CG - HG3 |
| 4799 | 4802 | 1 | 1.5220e-01 | 2.6527e+05 ; | CG - CD |
| 4802 | 4803 | 1 | 1.2500e-01 | 5.4894e+05 ; | CD - OE1 |
| 4802 | 4804 | 1 | 1.2500e-01 | 5.4894e+05 ; | CD - OE2 |
| 4805 | 4806 | 1 | 1.2290e-01 | 4.7698e+05 ; | C - O |
| 4805 | 4807 | 1 | 1.3350e-01 | 4.1003e+05 ; | C - N |
| 4807 | 4808 | 1 | 1.0100e-01 | 3.6317e+05 ; | N - H |
| 4807 | 4809 | 1 | 1.4490e-01 | 2.8200e+05 ; | N - CA |
| 4809 | 4810 | 1 | 1.0900e-01 | 2.8451e+05 ; | CA - HA |
| 4809 | 4811 | 1 | 1.5260e-01 | 2.5941e+05 ; | CA - CB |
| 4809 | 4824 | 1 | 1.5220e-01 | 2.6527e+05 ; | CA - C |
| 4811 | 4812 | 1 | 1.0900e-01 | 2.8451e+05 ; | CB - HB2 |
| 4811 | 4813 | 1 | 1.0900e-01 | 2.8451e+05 ; | CB - HB3 |
| 4811 | 4814 | 1 | 1.5260e-01 | 2.5941e+05 ; | CB - CG |
| 4814 | 4815 | 1 | 1.0900e-01 | 2.8451e+05 ; | CG - HG |
| 4814 | 4816 | 1 | 1.5260e-01 | 2.5941e+05 ; | CG - CD1 |
| 4814 | 4820 | 1 | 1.5260e-01 | 2.5941e+05 ; | CG - CD2 |
| 4816 | 4817 | 1 | 1.0900e-01 | 2.8451e+05 ; | CD1 - HD11 |
| 4816 | 4818 | 1 | 1.0900e-01 | 2.8451e+05 ; | CD1 - HD12 |
| 4816 | 4819 | 1 | 1.0900e-01 | 2.8451e+05 ; | CD1 - HD13 |
| 4820 | 4821 | 1 | 1.0900e-01 | 2.8451e+05 ; | CD2 - HD21 |
| 4820 | 4822 | 1 | 1.0900e-01 | 2.8451e+05 ; | CD2 - HD22 |
| 4820 | 4823 | 1 | 1.0900e-01 | 2.8451e+05 ; | CD2 - HD23 |
| 4824 | 4825 | 1 | 1.2290e-01 | 4.7698e+05 ; | C - O |
| 4824 | 4826 | 1 | 1.3350e-01 | 4.1003e+05 ; | C - N |
| 4826 | 4827 | 1 | 1.0100e-01 | 3.6317e+05 ; | N - H |
| 4826 | 4828 | 1 | 1.4490e-01 | 2.8200e+05 ; | N - CA |
| 4828 | 4829 | 1 | 1.0900e-01 | 2.8451e+05 ; | CA - HA |
| 4828 | 4830 | 1 | 1.5260e-01 | 2.5941e+05 ; | CA - CB |
| 4828 | 4846 | 1 | 1.5220e-01 | 2.6527e+05 ; | CA - C |
| 4830 | 4831 | 1 | 1.0900e-01 | 2.8451e+05 ; | CB - HB2 |
| 4830 | 4832 | 1 | 1.0900e-01 | 2.8451e+05 ; | CB - HB3 |
| 4830 | 4833 | 1 | 1.5260e-01 | 2.5941e+05 ; | CB - CG |

| | | | | | |
|---|---|---|---|---|---|
| 4833 | 4834 | 1 | 1.0900e-01 | 2.8451e+05 ; | CG - HG2 |
| 4833 | 4835 | 1 | 1.0900e-01 | 2.8451e+05 ; | CG - HG3 |
| 4833 | 4836 | 1 | 1.5260e-01 | 2.5941e+05 ; | CG - CD |
| 4836 | 4837 | 1 | 1.0900e-01 | 2.8451e+05 ; | CD - HD2 |
| 4836 | 4838 | 1 | 1.0900e-01 | 2.8451e+05 ; | CD - HD3 |
| 4836 | 4839 | 1 | 1.5260e-01 | 2.5941e+05 ; | CD - CE |
| 4839 | 4840 | 1 | 1.0900e-01 | 2.8451e+05 ; | CE - HE2 |
| 4839 | 4841 | 1 | 1.0900e-01 | 2.8451e+05 ; | CE - HE3 |
| 4839 | 4842 | 1 | 1.4710e-01 | 3.0711e+05 ; | CE - NZ |
| 4842 | 4843 | 1 | 1.0100e-01 | 3.6317e+05 ; | NZ - HZ1 |
| 4842 | 4844 | 1 | 1.0100e-01 | 3.6317e+05 ; | NZ - HZ2 |
| 4842 | 4845 | 1 | 1.0100e-01 | 3.6317e+05 ; | NZ - HZ3 |
| 4846 | 4847 | 1 | 1.2290e-01 | 4.7698e+05 ; | C - O |
| 4846 | 4848 | 1 | 1.3350e-01 | 4.1003e+05 ; | C - N |
| 4848 | 4849 | 1 | 1.0100e-01 | 3.6317e+05 ; | N - H |
| 4848 | 4850 | 1 | 1.4490e-01 | 2.8200e+05 ; | N - CA |
| 4850 | 4851 | 1 | 1.0900e-01 | 2.8451e+05 ; | CA - HA |
| 4850 | 4852 | 1 | 1.5260e-01 | 2.5941e+05 ; | CA - CB |
| 4850 | 4868 | 1 | 1.5220e-01 | 2.6527e+05 ; | CA - C |
| 4852 | 4853 | 1 | 1.0900e-01 | 2.8451e+05 ; | CB - HB2 |
| 4852 | 4854 | 1 | 1.0900e-01 | 2.8451e+05 ; | CB - HB3 |
| 4852 | 4855 | 1 | 1.5260e-01 | 2.5941e+05 ; | CB - CG |
| 4855 | 4856 | 1 | 1.0900e-01 | 2.8451e+05 ; | CG - HG2 |
| 4855 | 4857 | 1 | 1.0900e-01 | 2.8451e+05 ; | CG - HG3 |
| 4855 | 4858 | 1 | 1.5260e-01 | 2.5941e+05 ; | CG - CD |
| 4858 | 4859 | 1 | 1.0900e-01 | 2.8451e+05 ; | CD - HD2 |
| 4858 | 4860 | 1 | 1.0900e-01 | 2.8451e+05 ; | CD - HD3 |
| 4858 | 4861 | 1 | 1.5260e-01 | 2.5941e+05 ; | CD - CE |
| 4861 | 4862 | 1 | 1.0900e-01 | 2.8451e+05 ; | CE - HE2 |
| 4861 | 4863 | 1 | 1.0900e-01 | 2.8451e+05 ; | CE - HE3 |
| 4861 | 4864 | 1 | 1.4710e-01 | 3.0711e+05 ; | CE - NZ |
| 4864 | 4865 | 1 | 1.0100e-01 | 3.6317e+05 ; | NZ - HZ1 |
| 4864 | 4866 | 1 | 1.0100e-01 | 3.6317e+05 ; | NZ - HZ2 |
| 4864 | 4867 | 1 | 1.0100e-01 | 3.6317e+05 ; | NZ - HZ3 |
| 4868 | 4869 | 1 | 1.2290e-01 | 4.7698e+05 ; | C - O |
| 4868 | 4870 | 1 | 1.3350e-01 | 4.1003e+05 ; | C - N |
| 4870 | 4871 | 1 | 1.0100e-01 | 3.6317e+05 ; | N - H |
| 4870 | 4872 | 1 | 1.4490e-01 | 2.8200e+05 ; | N - CA |
| 4872 | 4873 | 1 | 1.0900e-01 | 2.8451e+05 ; | CA - HA |
| 4872 | 4874 | 1 | 1.5260e-01 | 2.5941e+05 ; | CA - CB |
| 4872 | 4887 | 1 | 1.5220e-01 | 2.6527e+05 ; | CA - C |
| 4874 | 4875 | 1 | 1.0900e-01 | 2.8451e+05 ; | CB - HB |
| 4874 | 4876 | 1 | 1.5260e-01 | 2.5941e+05 ; | CB - CG2 |
| 4874 | 4880 | 1 | 1.5260e-01 | 2.5941e+05 ; | CB - CG1 |
| 4876 | 4877 | 1 | 1.0900e-01 | 2.8451e+05 ; | CG2 - HG21 |
| 4876 | 4878 | 1 | 1.0900e-01 | 2.8451e+05 ; | CG2 - HG22 |
| 4876 | 4879 | 1 | 1.0900e-01 | 2.8451e+05 ; | CG2 - HG23 |
| 4880 | 4881 | 1 | 1.0900e-01 | 2.8451e+05 ; | CG1 - HG12 |
| 4880 | 4882 | 1 | 1.0900e-01 | 2.8451e+05 ; | CG1 - HG13 |
| 4880 | 4883 | 1 | 1.5260e-01 | 2.5941e+05 ; | CG1 - CD1 |
| 4883 | 4884 | 1 | 1.0900e-01 | 2.8451e+05 ; | CD1 - HD11 |
| 4883 | 4885 | 1 | 1.0900e-01 | 2.8451e+05 ; | CD1 - HD12 |

| | | | | | | | | |
|---|---|---|---|---|---|---|---|---|
| 4883 | 4886 | 1 | 1.0900e-01 | 2.8451e+05 | ; | CD1 | - | HD13 |
| 4887 | 4888 | 1 | 1.2290e-01 | 4.7698e+05 | ; | C | - | O |
| 4887 | 4889 | 1 | 1.3350e-01 | 4.1003e+05 | ; | C | - | N |
| 4889 | 4890 | 1 | 1.0100e-01 | 3.6317e+05 | ; | N | - | H |
| 4889 | 4891 | 1 | 1.4490e-01 | 2.8200e+05 | ; | N | - | CA |
| 4891 | 4892 | 1 | 1.0900e-01 | 2.8451e+05 | ; | CA | - | HA |
| 4891 | 4893 | 1 | 1.5260e-01 | 2.5941e+05 | ; | CA | - | CB |
| 4891 | 4897 | 1 | 1.5220e-01 | 2.6527e+05 | ; | CA | - | C |
| 4893 | 4894 | 1 | 1.0900e-01 | 2.8451e+05 | ; | CB | - | HB1 |
| 4893 | 4895 | 1 | 1.0900e-01 | 2.8451e+05 | ; | CB | - | HB2 |
| 4893 | 4896 | 1 | 1.0900e-01 | 2.8451e+05 | ; | CB | - | HB3 |
| 4897 | 4898 | 1 | 1.2290e-01 | 4.7698e+05 | ; | C | - | O |
| 4897 | 4899 | 1 | 1.3350e-01 | 4.1003e+05 | ; | C | - | N |
| 4899 | 4900 | 1 | 1.0100e-01 | 3.6317e+05 | ; | N | - | H |
| 4899 | 4901 | 1 | 1.4490e-01 | 2.8200e+05 | ; | N | - | CA |
| 4901 | 4902 | 1 | 1.0900e-01 | 2.8451e+05 | ; | CA | - | HA |
| 4901 | 4903 | 1 | 1.5260e-01 | 2.5941e+05 | ; | CA | - | CB |
| 4901 | 4916 | 1 | 1.5220e-01 | 2.6527e+05 | ; | CA | - | C |
| 4903 | 4904 | 1 | 1.0900e-01 | 2.8451e+05 | ; | CB | - | HB2 |
| 4903 | 4905 | 1 | 1.0900e-01 | 2.8451e+05 | ; | CB | - | HB3 |
| 4903 | 4906 | 1 | 1.5260e-01 | 2.5941e+05 | ; | CB | - | CG |
| 4906 | 4907 | 1 | 1.0900e-01 | 2.8451e+05 | ; | CG | - | HG |
| 4906 | 4908 | 1 | 1.5260e-01 | 2.5941e+05 | ; | CG | - | CD1 |
| 4906 | 4912 | 1 | 1.5260e-01 | 2.5941e+05 | ; | CG | - | CD2 |
| 4908 | 4909 | 1 | 1.0900e-01 | 2.8451e+05 | ; | CD1 | - | HD11 |
| 4908 | 4910 | 1 | 1.0900e-01 | 2.8451e+05 | ; | CD1 | - | HD12 |
| 4908 | 4911 | 1 | 1.0900e-01 | 2.8451e+05 | ; | CD1 | - | HD13 |
| 4912 | 4913 | 1 | 1.0900e-01 | 2.8451e+05 | ; | CD2 | - | HD21 |
| 4912 | 4914 | 1 | 1.0900e-01 | 2.8451e+05 | ; | CD2 | - | HD22 |
| 4912 | 4915 | 1 | 1.0900e-01 | 2.8451e+05 | ; | CD2 | - | HD23 |
| 4916 | 4917 | 1 | 1.2290e-01 | 4.7698e+05 | ; | C | - | O |
| 4916 | 4918 | 1 | 1.3350e-01 | 4.1003e+05 | ; | C | - | N |
| 4918 | 4919 | 1 | 1.0100e-01 | 3.6317e+05 | ; | N | - | H |
| 4918 | 4920 | 1 | 1.4490e-01 | 2.8200e+05 | ; | N | - | CA |
| 4920 | 4921 | 1 | 1.0900e-01 | 2.8451e+05 | ; | CA | - | HA |
| 4920 | 4922 | 1 | 1.5260e-01 | 2.5941e+05 | ; | CA | - | CB |
| 4920 | 4930 | 1 | 1.5220e-01 | 2.6527e+05 | ; | CA | - | C |
| 4922 | 4923 | 1 | 1.0900e-01 | 2.8451e+05 | ; | CB | - | HB |
| 4922 | 4924 | 1 | 1.5260e-01 | 2.5941e+05 | ; | CB | - | CG2 |
| 4922 | 4928 | 1 | 1.4100e-01 | 2.6778e+05 | ; | CB | - | OG1 |
| 4924 | 4925 | 1 | 1.0900e-01 | 2.8451e+05 | ; | CG2 | - | HG21 |
| 4924 | 4926 | 1 | 1.0900e-01 | 2.8451e+05 | ; | CG2 | - | HG22 |
| 4924 | 4927 | 1 | 1.0900e-01 | 2.8451e+05 | ; | CG2 | - | HG23 |
| 4928 | 4929 | 1 | 9.6000e-02 | 4.6275e+05 | ; | OG1 | - | HG1 |
| 4930 | 4931 | 1 | 1.2290e-01 | 4.7698e+05 | ; | C | - | O |
| 4930 | 4932 | 1 | 1.3350e-01 | 4.1003e+05 | ; | C | - | N |
| 4932 | 4933 | 1 | 1.0100e-01 | 3.6317e+05 | ; | N | - | H |
| 4932 | 4934 | 1 | 1.4490e-01 | 2.8200e+05 | ; | N | - | CA |
| 4934 | 4935 | 1 | 1.0900e-01 | 2.8451e+05 | ; | CA | - | HA |
| 4934 | 4936 | 1 | 1.5260e-01 | 2.5941e+05 | ; | CA | - | CB |
| 4934 | 4949 | 1 | 1.5220e-01 | 2.6527e+05 | ; | CA | - | C |
| 4936 | 4937 | 1 | 1.0900e-01 | 2.8451e+05 | ; | CB | - | HB |

| | | | | | | |
|---|---|---|---|---|---|---|
| 4936 | 4938 | 1 | 1.5260e-01 | 2.5941e+05 | ; | CB - CG2 |
| 4936 | 4942 | 1 | 1.5260e-01 | 2.5941e+05 | ; | CB - CG1 |
| 4938 | 4939 | 1 | 1.0900e-01 | 2.8451e+05 | ; | CG2 - HG21 |
| 4938 | 4940 | 1 | 1.0900e-01 | 2.8451e+05 | ; | CG2 - HG22 |
| 4938 | 4941 | 1 | 1.0900e-01 | 2.8451e+05 | ; | CG2 - HG23 |
| 4942 | 4943 | 1 | 1.0900e-01 | 2.8451e+05 | ; | CG1 - HG12 |
| 4942 | 4944 | 1 | 1.0900e-01 | 2.8451e+05 | ; | CG1 - HG13 |
| 4942 | 4945 | 1 | 1.5260e-01 | 2.5941e+05 | ; | CG1 - CD1 |
| 4945 | 4946 | 1 | 1.0900e-01 | 2.8451e+05 | ; | CD1 - HD11 |
| 4945 | 4947 | 1 | 1.0900e-01 | 2.8451e+05 | ; | CD1 - HD12 |
| 4945 | 4948 | 1 | 1.0900e-01 | 2.8451e+05 | ; | CD1 - HD13 |
| 4949 | 4950 | 1 | 1.2290e-01 | 4.7698e+05 | ; | C - O |
| 4949 | 4951 | 1 | 1.3350e-01 | 4.1003e+05 | ; | C - N |
| 4951 | 4952 | 1 | 1.0100e-01 | 3.6317e+05 | ; | N - H |
| 4951 | 4953 | 1 | 1.4490e-01 | 2.8200e+05 | ; | N - CA |
| 4953 | 4954 | 1 | 1.0900e-01 | 2.8451e+05 | ; | CA - HA |
| 4953 | 4955 | 1 | 1.5260e-01 | 2.5941e+05 | ; | CA - CB |
| 4953 | 4968 | 1 | 1.5220e-01 | 2.6527e+05 | ; | CA - C |
| 4955 | 4956 | 1 | 1.0900e-01 | 2.8451e+05 | ; | CB - HB |
| 4955 | 4957 | 1 | 1.5260e-01 | 2.5941e+05 | ; | CB - CG2 |
| 4955 | 4961 | 1 | 1.5260e-01 | 2.5941e+05 | ; | CB - CG1 |
| 4957 | 4958 | 1 | 1.0900e-01 | 2.8451e+05 | ; | CG2 - HG21 |
| 4957 | 4959 | 1 | 1.0900e-01 | 2.8451e+05 | ; | CG2 - HG22 |
| 4957 | 4960 | 1 | 1.0900e-01 | 2.8451e+05 | ; | CG2 - HG23 |
| 4961 | 4962 | 1 | 1.0900e-01 | 2.8451e+05 | ; | CG1 - HG12 |
| 4961 | 4963 | 1 | 1.0900e-01 | 2.8451e+05 | ; | CG1 - HG13 |
| 4961 | 4964 | 1 | 1.5260e-01 | 2.5941e+05 | ; | CG1 - CD1 |
| 4964 | 4965 | 1 | 1.0900e-01 | 2.8451e+05 | ; | CD1 - HD11 |
| 4964 | 4966 | 1 | 1.0900e-01 | 2.8451e+05 | ; | CD1 - HD12 |
| 4964 | 4967 | 1 | 1.0900e-01 | 2.8451e+05 | ; | CD1 - HD13 |
| 4968 | 4969 | 1 | 1.2290e-01 | 4.7698e+05 | ; | C - O |
| 4968 | 4970 | 1 | 1.3350e-01 | 4.1003e+05 | ; | C - N |
| 4970 | 4971 | 1 | 1.0100e-01 | 3.6317e+05 | ; | N - H |
| 4970 | 4972 | 1 | 1.4490e-01 | 2.8200e+05 | ; | N - CA |
| 4972 | 4973 | 1 | 1.0900e-01 | 2.8451e+05 | ; | CA - HA |
| 4972 | 4974 | 1 | 1.5260e-01 | 2.5941e+05 | ; | CA - CB |
| 4972 | 4978 | 1 | 1.5220e-01 | 2.6527e+05 | ; | CA - C |
| 4974 | 4975 | 1 | 1.0900e-01 | 2.8451e+05 | ; | CB - HB1 |
| 4974 | 4976 | 1 | 1.0900e-01 | 2.8451e+05 | ; | CB - HB2 |
| 4974 | 4977 | 1 | 1.0900e-01 | 2.8451e+05 | ; | CB - HB3 |
| 4978 | 4979 | 1 | 1.2290e-01 | 4.7698e+05 | ; | C - O |
| 4978 | 4980 | 1 | 1.3350e-01 | 4.1003e+05 | ; | C - N |
| 4980 | 4981 | 1 | 1.0100e-01 | 3.6317e+05 | ; | N - H |
| 4980 | 4982 | 1 | 1.4490e-01 | 2.8200e+05 | ; | N - CA |
| 4982 | 4983 | 1 | 1.0900e-01 | 2.8451e+05 | ; | CA - HA |
| 4982 | 4984 | 1 | 1.5260e-01 | 2.5941e+05 | ; | CA - CB |
| 4982 | 5000 | 1 | 1.5220e-01 | 2.6527e+05 | ; | CA - C |
| 4984 | 4985 | 1 | 1.0900e-01 | 2.8451e+05 | ; | CB - HB2 |
| 4984 | 4986 | 1 | 1.0900e-01 | 2.8451e+05 | ; | CB - HB3 |
| 4984 | 4987 | 1 | 1.5260e-01 | 2.5941e+05 | ; | CB - CG |
| 4987 | 4988 | 1 | 1.0900e-01 | 2.8451e+05 | ; | CG - HG2 |
| 4987 | 4989 | 1 | 1.0900e-01 | 2.8451e+05 | ; | CG - HG3 |

| | | | | | | |
|---|---|---|---|---|---|---|
| 4987 | 4990 | 1 | 1.5260e-01 | 2.5941e+05 | ; | CG - CD |
| 4990 | 4991 | 1 | 1.0900e-01 | 2.8451e+05 | ; | CD - HD2 |
| 4990 | 4992 | 1 | 1.0900e-01 | 2.8451e+05 | ; | CD - HD3 |
| 4990 | 4993 | 1 | 1.5260e-01 | 2.5941e+05 | ; | CD - CE |
| 4993 | 4994 | 1 | 1.0900e-01 | 2.8451e+05 | ; | CE - HE2 |
| 4993 | 4995 | 1 | 1.0900e-01 | 2.8451e+05 | ; | CE - HE3 |
| 4993 | 4996 | 1 | 1.4710e-01 | 3.0711e+05 | ; | CE - NZ |
| 4996 | 4997 | 1 | 1.0100e-01 | 3.6317e+05 | ; | NZ - HZ1 |
| 4996 | 4998 | 1 | 1.0100e-01 | 3.6317e+05 | ; | NZ - HZ2 |
| 4996 | 4999 | 1 | 1.0100e-01 | 3.6317e+05 | ; | NZ - HZ3 |
| 5000 | 5001 | 1 | 1.2290e-01 | 4.7698e+05 | ; | C - O |
| 5000 | 5002 | 1 | 1.3350e-01 | 4.1003e+05 | ; | C - N |
| 5002 | 5003 | 1 | 1.0100e-01 | 3.6317e+05 | ; | N - H |
| 5002 | 5004 | 1 | 1.4490e-01 | 2.8200e+05 | ; | N - CA |
| 5004 | 5005 | 1 | 1.0900e-01 | 2.8451e+05 | ; | CA - HA |
| 5004 | 5006 | 1 | 1.5260e-01 | 2.5941e+05 | ; | CA - CB |
| 5004 | 5018 | 1 | 1.5220e-01 | 2.6527e+05 | ; | CA - C |
| 5006 | 5007 | 1 | 1.0900e-01 | 2.8451e+05 | ; | CB - HB2 |
| 5006 | 5008 | 1 | 1.0900e-01 | 2.8451e+05 | ; | CB - HB3 |
| 5006 | 5009 | 1 | 1.5040e-01 | 2.6527e+05 | ; | CB - CG |
| 5009 | 5010 | 1 | 1.3850e-01 | 3.5313e+05 | ; | CG - ND1 |
| 5009 | 5016 | 1 | 1.3710e-01 | 4.3346e+05 | ; | CG - CD2 |
| 5010 | 5011 | 1 | 1.0100e-01 | 3.6317e+05 | ; | ND1 - HD1 |
| 5010 | 5012 | 1 | 1.3430e-01 | 3.9915e+05 | ; | ND1 - CE1 |
| 5012 | 5013 | 1 | 1.0800e-01 | 3.0711e+05 | ; | CE1 - HE1 |
| 5012 | 5014 | 1 | 1.3430e-01 | 3.9915e+05 | ; | CE1 - NE2 |
| 5014 | 5015 | 1 | 1.0100e-01 | 3.6317e+05 | ; | NE2 - HE2 |
| 5014 | 5016 | 1 | 1.3810e-01 | 3.5731e+05 | ; | NE2 - CD2 |
| 5016 | 5017 | 1 | 1.0800e-01 | 3.0711e+05 | ; | CD2 - HD2 |
| 5018 | 5019 | 1 | 1.2290e-01 | 4.7698e+05 | ; | C - O |
| 5018 | 5020 | 1 | 1.3350e-01 | 4.1003e+05 | ; | C - N |
| 5020 | 5021 | 1 | 1.0100e-01 | 3.6317e+05 | ; | N - H |
| 5020 | 5022 | 1 | 1.4490e-01 | 2.8200e+05 | ; | N - CA |
| 5022 | 5023 | 1 | 1.0900e-01 | 2.8451e+05 | ; | CA - HA |
| 5022 | 5024 | 1 | 1.5260e-01 | 2.5941e+05 | ; | CA - CB |
| 5022 | 5037 | 1 | 1.5220e-01 | 2.6527e+05 | ; | CA - C |
| 5024 | 5025 | 1 | 1.0900e-01 | 2.8451e+05 | ; | CB - HB2 |
| 5024 | 5026 | 1 | 1.0900e-01 | 2.8451e+05 | ; | CB - HB3 |
| 5024 | 5027 | 1 | 1.5260e-01 | 2.5941e+05 | ; | CB - CG |
| 5027 | 5028 | 1 | 1.0900e-01 | 2.8451e+05 | ; | CG - HG |
| 5027 | 5029 | 1 | 1.5260e-01 | 2.5941e+05 | ; | CG - CD1 |
| 5027 | 5033 | 1 | 1.5260e-01 | 2.5941e+05 | ; | CG - CD2 |
| 5029 | 5030 | 1 | 1.0900e-01 | 2.8451e+05 | ; | CD1 - HD11 |
| 5029 | 5031 | 1 | 1.0900e-01 | 2.8451e+05 | ; | CD1 - HD12 |
| 5029 | 5032 | 1 | 1.0900e-01 | 2.8451e+05 | ; | CD1 - HD13 |
| 5033 | 5034 | 1 | 1.0900e-01 | 2.8451e+05 | ; | CD2 - HD21 |
| 5033 | 5035 | 1 | 1.0900e-01 | 2.8451e+05 | ; | CD2 - HD22 |
| 5033 | 5036 | 1 | 1.0900e-01 | 2.8451e+05 | ; | CD2 - HD23 |
| 5037 | 5038 | 1 | 1.2290e-01 | 4.7698e+05 | ; | C - O |
| 5037 | 5039 | 1 | 1.3350e-01 | 4.1003e+05 | ; | C - N |
| 5039 | 5040 | 1 | 1.0100e-01 | 3.6317e+05 | ; | N - H |
| 5039 | 5041 | 1 | 1.4490e-01 | 2.8200e+05 | ; | N - CA |

| | | | | | | |
|---|---|---|---|---|---|---|
| 5041 | 5042 | 1 | 1.0900e-01 | 2.8451e+05 | ; | CA - HA |
| 5041 | 5043 | 1 | 1.5260e-01 | 2.5941e+05 | ; | CA - CB |
| 5041 | 5048 | 1 | 1.5220e-01 | 2.6527e+05 | ; | CA - C |
| 5043 | 5044 | 1 | 1.0900e-01 | 2.8451e+05 | ; | CB - HB2 |
| 5043 | 5045 | 1 | 1.0900e-01 | 2.8451e+05 | ; | CB - HB3 |
| 5043 | 5046 | 1 | 1.8100e-01 | 1.9832e+05 | ; | CB - SG |
| 5046 | 5047 | 1 | 1.3360e-01 | 2.2928e+05 | ; | SG - HG |
| 5048 | 5049 | 1 | 1.2290e-01 | 4.7698e+05 | ; | C - O |
| 5048 | 5050 | 1 | 1.3350e-01 | 4.1003e+05 | ; | C - N |
| 5050 | 5051 | 1 | 1.0100e-01 | 3.6317e+05 | ; | N - H |
| 5050 | 5052 | 1 | 1.4490e-01 | 2.8200e+05 | ; | N - CA |
| 5052 | 5053 | 1 | 1.0900e-01 | 2.8451e+05 | ; | CA - HA |
| 5052 | 5054 | 1 | 1.5260e-01 | 2.5941e+05 | ; | CA - CB |
| 5052 | 5060 | 1 | 1.5220e-01 | 2.6527e+05 | ; | CA - C |
| 5054 | 5055 | 1 | 1.0900e-01 | 2.8451e+05 | ; | CB - HB2 |
| 5054 | 5056 | 1 | 1.0900e-01 | 2.8451e+05 | ; | CB - HB3 |
| 5054 | 5057 | 1 | 1.5220e-01 | 2.6527e+05 | ; | CB - CG |
| 5057 | 5058 | 1 | 1.2500e-01 | 5.4894e+05 | ; | CG - OD1 |
| 5057 | 5059 | 1 | 1.2500e-01 | 5.4894e+05 | ; | CG - OD2 |
| 5060 | 5061 | 1 | 1.2290e-01 | 4.7698e+05 | ; | C - O |
| 5060 | 5062 | 1 | 1.3350e-01 | 4.1003e+05 | ; | C - N |
| 5062 | 5063 | 1 | 1.0100e-01 | 3.6317e+05 | ; | N - H |
| 5062 | 5064 | 1 | 1.4490e-01 | 2.8200e+05 | ; | N - CA |
| 5064 | 5065 | 1 | 1.0900e-01 | 2.8451e+05 | ; | CA - HA |
| 5064 | 5066 | 1 | 1.5260e-01 | 2.5941e+05 | ; | CA - CB |
| 5064 | 5076 | 1 | 1.5220e-01 | 2.6527e+05 | ; | CA - C |
| 5066 | 5067 | 1 | 1.0900e-01 | 2.8451e+05 | ; | CB - HB |
| 5066 | 5068 | 1 | 1.5260e-01 | 2.5941e+05 | ; | CB - CG1 |
| 5066 | 5072 | 1 | 1.5260e-01 | 2.5941e+05 | ; | CB - CG2 |
| 5068 | 5069 | 1 | 1.0900e-01 | 2.8451e+05 | ; | CG1 - HG11 |
| 5068 | 5070 | 1 | 1.0900e-01 | 2.8451e+05 | ; | CG1 - HG12 |
| 5068 | 5071 | 1 | 1.0900e-01 | 2.8451e+05 | ; | CG1 - HG13 |
| 5072 | 5073 | 1 | 1.0900e-01 | 2.8451e+05 | ; | CG2 - HG21 |
| 5072 | 5074 | 1 | 1.0900e-01 | 2.8451e+05 | ; | CG2 - HG22 |
| 5072 | 5075 | 1 | 1.0900e-01 | 2.8451e+05 | ; | CG2 - HG23 |
| 5076 | 5077 | 1 | 1.2290e-01 | 4.7698e+05 | ; | C - O |
| 5076 | 5078 | 1 | 1.3350e-01 | 4.1003e+05 | ; | C - N |
| 5078 | 5079 | 1 | 1.0100e-01 | 3.6317e+05 | ; | N - H |
| 5078 | 5080 | 1 | 1.4490e-01 | 2.8200e+05 | ; | N - CA |
| 5080 | 5081 | 1 | 1.0900e-01 | 2.8451e+05 | ; | CA - HA |
| 5080 | 5082 | 1 | 1.5260e-01 | 2.5941e+05 | ; | CA - CB |
| 5080 | 5091 | 1 | 1.5220e-01 | 2.6527e+05 | ; | CA - C |
| 5082 | 5083 | 1 | 1.0900e-01 | 2.8451e+05 | ; | CB - HB2 |
| 5082 | 5084 | 1 | 1.0900e-01 | 2.8451e+05 | ; | CB - HB3 |
| 5082 | 5085 | 1 | 1.5260e-01 | 2.5941e+05 | ; | CB - CG |
| 5085 | 5086 | 1 | 1.0900e-01 | 2.8451e+05 | ; | CG - HG2 |
| 5085 | 5087 | 1 | 1.0900e-01 | 2.8451e+05 | ; | CG - HG3 |
| 5085 | 5088 | 1 | 1.5220e-01 | 2.6527e+05 | ; | CG - CD |
| 5088 | 5089 | 1 | 1.2500e-01 | 5.4894e+05 | ; | CD - OE1 |
| 5088 | 5090 | 1 | 1.2500e-01 | 5.4894e+05 | ; | CD - OE2 |
| 5091 | 5092 | 1 | 1.2290e-01 | 4.7698e+05 | ; | C - O |
| 5091 | 5093 | 1 | 1.3350e-01 | 4.1003e+05 | ; | C - N |

| | | | | | | |
|---|---|---|---|---|---|---|
| 5093 | 5094 | 1 | 1.0100e-01 | 3.6317e+05 | ; | N - H |
| 5093 | 5095 | 1 | 1.4490e-01 | 2.8200e+05 | ; | N - CA |
| 5095 | 5096 | 1 | 1.0900e-01 | 2.8451e+05 | ; | CA - HA |
| 5095 | 5097 | 1 | 1.5260e-01 | 2.5941e+05 | ; | CA - CB |
| 5095 | 5110 | 1 | 1.5220e-01 | 2.6527e+05 | ; | CA - C |
| 5097 | 5098 | 1 | 1.0900e-01 | 2.8451e+05 | ; | CB - HB |
| 5097 | 5099 | 1 | 1.5260e-01 | 2.5941e+05 | ; | CB - CG2 |
| 5097 | 5103 | 1 | 1.5260e-01 | 2.5941e+05 | ; | CB - CG1 |
| 5099 | 5100 | 1 | 1.0900e-01 | 2.8451e+05 | ; | CG2 - HG21 |
| 5099 | 5101 | 1 | 1.0900e-01 | 2.8451e+05 | ; | CG2 - HG22 |
| 5099 | 5102 | 1 | 1.0900e-01 | 2.8451e+05 | ; | CG2 - HG23 |
| 5103 | 5104 | 1 | 1.0900e-01 | 2.8451e+05 | ; | CG1 - HG12 |
| 5103 | 5105 | 1 | 1.0900e-01 | 2.8451e+05 | ; | CG1 - HG13 |
| 5103 | 5106 | 1 | 1.5260e-01 | 2.5941e+05 | ; | CG1 - CD1 |
| 5106 | 5107 | 1 | 1.0900e-01 | 2.8451e+05 | ; | CD1 - HD11 |
| 5106 | 5108 | 1 | 1.0900e-01 | 2.8451e+05 | ; | CD1 - HD12 |
| 5106 | 5109 | 1 | 1.0900e-01 | 2.8451e+05 | ; | CD1 - HD13 |
| 5110 | 5111 | 1 | 1.2290e-01 | 4.7698e+05 | ; | C - O |
| 5110 | 5112 | 1 | 1.3350e-01 | 4.1003e+05 | ; | C - N |
| 5112 | 5113 | 1 | 1.0100e-01 | 3.6317e+05 | ; | N - H |
| 5112 | 5114 | 1 | 1.4490e-01 | 2.8200e+05 | ; | N - CA |
| 5114 | 5115 | 1 | 1.0900e-01 | 2.8451e+05 | ; | CA - HA |
| 5114 | 5116 | 1 | 1.5260e-01 | 2.5941e+05 | ; | CA - CB |
| 5114 | 5124 | 1 | 1.5220e-01 | 2.6527e+05 | ; | CA - C |
| 5116 | 5117 | 1 | 1.0900e-01 | 2.8451e+05 | ; | CB - HB2 |
| 5116 | 5118 | 1 | 1.0900e-01 | 2.8451e+05 | ; | CB - HB3 |
| 5116 | 5119 | 1 | 1.5220e-01 | 2.6527e+05 | ; | CB - CG |
| 5119 | 5120 | 1 | 1.2290e-01 | 4.7698e+05 | ; | CG - OD1 |
| 5119 | 5121 | 1 | 1.3350e-01 | 4.1003e+05 | ; | CG - ND2 |
| 5121 | 5122 | 1 | 1.0100e-01 | 3.6317e+05 | ; | ND2 - HD21 |
| 5121 | 5123 | 1 | 1.0100e-01 | 3.6317e+05 | ; | ND2 - HD22 |
| 5124 | 5125 | 1 | 1.2290e-01 | 4.7698e+05 | ; | C - O |
| 5124 | 5126 | 1 | 1.3350e-01 | 4.1003e+05 | ; | C - N |
| 5126 | 5127 | 1 | 1.0100e-01 | 3.6317e+05 | ; | N - H |
| 5126 | 5128 | 1 | 1.4490e-01 | 2.8200e+05 | ; | N - CA |
| 5128 | 5129 | 1 | 1.0900e-01 | 2.8451e+05 | ; | CA - HA |
| 5128 | 5130 | 1 | 1.5260e-01 | 2.5941e+05 | ; | CA - CB |
| 5128 | 5138 | 1 | 1.5220e-01 | 2.6527e+05 | ; | CA - C |
| 5130 | 5131 | 1 | 1.0900e-01 | 2.8451e+05 | ; | CB - HB2 |
| 5130 | 5132 | 1 | 1.0900e-01 | 2.8451e+05 | ; | CB - HB3 |
| 5130 | 5133 | 1 | 1.5220e-01 | 2.6527e+05 | ; | CB - CG |
| 5133 | 5134 | 1 | 1.2290e-01 | 4.7698e+05 | ; | CG - OD1 |
| 5133 | 5135 | 1 | 1.3350e-01 | 4.1003e+05 | ; | CG - ND2 |
| 5135 | 5136 | 1 | 1.0100e-01 | 3.6317e+05 | ; | ND2 - HD21 |
| 5135 | 5137 | 1 | 1.0100e-01 | 3.6317e+05 | ; | ND2 - HD22 |
| 5138 | 5139 | 1 | 1.2290e-01 | 4.7698e+05 | ; | C - O |
| 5138 | 5140 | 1 | 1.3350e-01 | 4.1003e+05 | ; | C - N |
| 5140 | 5141 | 1 | 1.0100e-01 | 3.6317e+05 | ; | N - H |
| 5140 | 5142 | 1 | 1.4490e-01 | 2.8200e+05 | ; | N - CA |
| 5142 | 5143 | 1 | 1.0900e-01 | 2.8451e+05 | ; | CA - HA |
| 5142 | 5144 | 1 | 1.5260e-01 | 2.5941e+05 | ; | CA - CB |
| 5142 | 5157 | 1 | 1.5220e-01 | 2.6527e+05 | ; | CA - C |

| | | | | | |
|---|---|---|---|---|---|
| 5144 | 5145 | 1 | 1.0900e-01 | 2.8451e+05 ; | CB - HB2 |
| 5144 | 5146 | 1 | 1.0900e-01 | 2.8451e+05 ; | CB - HB3 |
| 5144 | 5147 | 1 | 1.5260e-01 | 2.5941e+05 ; | CB - CG |
| 5147 | 5148 | 1 | 1.0900e-01 | 2.8451e+05 ; | CG - HG |
| 5147 | 5149 | 1 | 1.5260e-01 | 2.5941e+05 ; | CG - CD1 |
| 5147 | 5153 | 1 | 1.5260e-01 | 2.5941e+05 ; | CG - CD2 |
| 5149 | 5150 | 1 | 1.0900e-01 | 2.8451e+05 ; | CD1 - HD11 |
| 5149 | 5151 | 1 | 1.0900e-01 | 2.8451e+05 ; | CD1 - HD12 |
| 5149 | 5152 | 1 | 1.0900e-01 | 2.8451e+05 ; | CD1 - HD13 |
| 5153 | 5154 | 1 | 1.0900e-01 | 2.8451e+05 ; | CD2 - HD21 |
| 5153 | 5155 | 1 | 1.0900e-01 | 2.8451e+05 ; | CD2 - HD22 |
| 5153 | 5156 | 1 | 1.0900e-01 | 2.8451e+05 ; | CD2 - HD23 |
| 5157 | 5158 | 1 | 1.2290e-01 | 4.7698e+05 ; | C - O |
| 5157 | 5159 | 1 | 1.3350e-01 | 4.1003e+05 ; | C - N |
| 5159 | 5160 | 1 | 1.0100e-01 | 3.6317e+05 ; | N - H |
| 5159 | 5161 | 1 | 1.4490e-01 | 2.8200e+05 ; | N - CA |
| 5161 | 5162 | 1 | 1.0900e-01 | 2.8451e+05 ; | CA - HA |
| 5161 | 5163 | 1 | 1.5260e-01 | 2.5941e+05 ; | CA - CB |
| 5161 | 5181 | 1 | 1.5220e-01 | 2.6527e+05 ; | CA - C |
| 5163 | 5164 | 1 | 1.0900e-01 | 2.8451e+05 ; | CB - HB2 |
| 5163 | 5165 | 1 | 1.0900e-01 | 2.8451e+05 ; | CB - HB3 |
| 5163 | 5166 | 1 | 1.5260e-01 | 2.5941e+05 ; | CB - CG |
| 5166 | 5167 | 1 | 1.0900e-01 | 2.8451e+05 ; | CG - HG2 |
| 5166 | 5168 | 1 | 1.0900e-01 | 2.8451e+05 ; | CG - HG3 |
| 5166 | 5169 | 1 | 1.5260e-01 | 2.5941e+05 ; | CG - CD |
| 5169 | 5170 | 1 | 1.0900e-01 | 2.8451e+05 ; | CD - HD2 |
| 5169 | 5171 | 1 | 1.0900e-01 | 2.8451e+05 ; | CD - HD3 |
| 5169 | 5172 | 1 | 1.4630e-01 | 2.8200e+05 ; | CD - NE |
| 5172 | 5173 | 1 | 1.0100e-01 | 3.6317e+05 ; | NE - HE |
| 5172 | 5174 | 1 | 1.3400e-01 | 4.0250e+05 ; | NE - CZ |
| 5174 | 5175 | 1 | 1.3400e-01 | 4.0250e+05 ; | CZ - NH1 |
| 5174 | 5178 | 1 | 1.3400e-01 | 4.0250e+05 ; | CZ - NH2 |
| 5175 | 5176 | 1 | 1.0100e-01 | 3.6317e+05 ; | NH1 - HH11 |
| 5175 | 5177 | 1 | 1.0100e-01 | 3.6317e+05 ; | NH1 - HH12 |
| 5178 | 5179 | 1 | 1.0100e-01 | 3.6317e+05 ; | NH2 - HH21 |
| 5178 | 5180 | 1 | 1.0100e-01 | 3.6317e+05 ; | NH2 - HH22 |
| 5181 | 5182 | 1 | 1.2290e-01 | 4.7698e+05 ; | C - O |
| 5181 | 5183 | 1 | 1.3350e-01 | 4.1003e+05 ; | C - N |
| 5183 | 5184 | 1 | 1.0100e-01 | 3.6317e+05 ; | N - H |
| 5183 | 5185 | 1 | 1.4490e-01 | 2.8200e+05 ; | N - CA |
| 5185 | 5186 | 1 | 1.0900e-01 | 2.8451e+05 ; | CA - HA |
| 5185 | 5187 | 1 | 1.5260e-01 | 2.5941e+05 ; | CA - CB |
| 5185 | 5195 | 1 | 1.5220e-01 | 2.6527e+05 ; | CA - C |
| 5187 | 5188 | 1 | 1.0900e-01 | 2.8451e+05 ; | CB - HB2 |
| 5187 | 5189 | 1 | 1.0900e-01 | 2.8451e+05 ; | CB - HB3 |
| 5187 | 5190 | 1 | 1.5220e-01 | 2.6527e+05 ; | CB - CG |
| 5190 | 5191 | 1 | 1.2290e-01 | 4.7698e+05 ; | CG - OD1 |
| 5190 | 5192 | 1 | 1.3350e-01 | 4.1003e+05 ; | CG - ND2 |
| 5192 | 5193 | 1 | 1.0100e-01 | 3.6317e+05 ; | ND2 - HD21 |
| 5192 | 5194 | 1 | 1.0100e-01 | 3.6317e+05 ; | ND2 - HD22 |
| 5195 | 5196 | 1 | 1.2290e-01 | 4.7698e+05 ; | C - O |
| 5195 | 5197 | 1 | 1.3350e-01 | 4.1003e+05 ; | C - N |

| | | | | | | |
|---|---|---|---|---|---|---|
| 5197 | 5198 | 1 | 1.0100e-01 | 3.6317e+05 | ; | N - H |
| 5197 | 5199 | 1 | 1.4490e-01 | 2.8200e+05 | ; | N - CA |
| 5199 | 5200 | 1 | 1.0900e-01 | 2.8451e+05 | ; | CA - HA |
| 5199 | 5201 | 1 | 1.5260e-01 | 2.5941e+05 | ; | CA - CB |
| 5199 | 5214 | 1 | 1.5220e-01 | 2.6527e+05 | ; | CA - C |
| 5201 | 5202 | 1 | 1.0900e-01 | 2.8451e+05 | ; | CB - HB |
| 5201 | 5203 | 1 | 1.5260e-01 | 2.5941e+05 | ; | CB - CG2 |
| 5201 | 5207 | 1 | 1.5260e-01 | 2.5941e+05 | ; | CB - CG1 |
| 5203 | 5204 | 1 | 1.0900e-01 | 2.8451e+05 | ; | CG2 - HG21 |
| 5203 | 5205 | 1 | 1.0900e-01 | 2.8451e+05 | ; | CG2 - HG22 |
| 5203 | 5206 | 1 | 1.0900e-01 | 2.8451e+05 | ; | CG2 - HG23 |
| 5207 | 5208 | 1 | 1.0900e-01 | 2.8451e+05 | ; | CG1 - HG12 |
| 5207 | 5209 | 1 | 1.0900e-01 | 2.8451e+05 | ; | CG1 - HG13 |
| 5207 | 5210 | 1 | 1.5260e-01 | 2.5941e+05 | ; | CG1 - CD1 |
| 5210 | 5211 | 1 | 1.0900e-01 | 2.8451e+05 | ; | CD1 - HD11 |
| 5210 | 5212 | 1 | 1.0900e-01 | 2.8451e+05 | ; | CD1 - HD12 |
| 5210 | 5213 | 1 | 1.0900e-01 | 2.8451e+05 | ; | CD1 - HD13 |
| 5214 | 5215 | 1 | 1.2290e-01 | 4.7698e+05 | ; | C - O |
| 5214 | 5216 | 1 | 1.3350e-01 | 4.1003e+05 | ; | C - N |
| 5216 | 5217 | 1 | 1.0100e-01 | 3.6317e+05 | ; | N - H |
| 5216 | 5218 | 1 | 1.4490e-01 | 2.8200e+05 | ; | N - CA |
| 5218 | 5219 | 1 | 1.0900e-01 | 2.8451e+05 | ; | CA - HA |
| 5218 | 5220 | 1 | 1.5260e-01 | 2.5941e+05 | ; | CA - CB |
| 5218 | 5234 | 1 | 1.5220e-01 | 2.6527e+05 | ; | CA - C |
| 5220 | 5221 | 1 | 1.0900e-01 | 2.8451e+05 | ; | CB - HB2 |
| 5220 | 5222 | 1 | 1.0900e-01 | 2.8451e+05 | ; | CB - HB3 |
| 5220 | 5223 | 1 | 1.5100e-01 | 2.6527e+05 | ; | CB - CG |
| 5223 | 5224 | 1 | 1.4000e-01 | 3.9246e+05 | ; | CG - CD1 |
| 5223 | 5232 | 1 | 1.4000e-01 | 3.9246e+05 | ; | CG - CD2 |
| 5224 | 5225 | 1 | 1.0800e-01 | 3.0711e+05 | ; | CD1 - HD1 |
| 5224 | 5226 | 1 | 1.4000e-01 | 3.9246e+05 | ; | CD1 - CE1 |
| 5226 | 5227 | 1 | 1.0800e-01 | 3.0711e+05 | ; | CE1 - HE1 |
| 5226 | 5228 | 1 | 1.4000e-01 | 3.9246e+05 | ; | CE1 - CZ |
| 5228 | 5229 | 1 | 1.0800e-01 | 3.0711e+05 | ; | CZ - HZ |
| 5228 | 5230 | 1 | 1.4000e-01 | 3.9246e+05 | ; | CZ - CE2 |
| 5230 | 5231 | 1 | 1.0800e-01 | 3.0711e+05 | ; | CE2 - HE2 |
| 5230 | 5232 | 1 | 1.4000e-01 | 3.9246e+05 | ; | CE2 - CD2 |
| 5232 | 5233 | 1 | 1.0800e-01 | 3.0711e+05 | ; | CD2 - HD2 |
| 5234 | 5235 | 1 | 1.2290e-01 | 4.7698e+05 | ; | C - O |
| 5234 | 5236 | 1 | 1.3350e-01 | 4.1003e+05 | ; | C - N |
| 5236 | 5237 | 1 | 1.0100e-01 | 3.6317e+05 | ; | N - H |
| 5236 | 5238 | 1 | 1.4490e-01 | 2.8200e+05 | ; | N - CA |
| 5238 | 5239 | 1 | 1.0900e-01 | 2.8451e+05 | ; | CA - HA |
| 5238 | 5240 | 1 | 1.5260e-01 | 2.5941e+05 | ; | CA - CB |
| 5238 | 5253 | 1 | 1.5220e-01 | 2.6527e+05 | ; | CA - C |
| 5240 | 5241 | 1 | 1.0900e-01 | 2.8451e+05 | ; | CB - HB |
| 5240 | 5242 | 1 | 1.5260e-01 | 2.5941e+05 | ; | CB - CG2 |
| 5240 | 5246 | 1 | 1.5260e-01 | 2.5941e+05 | ; | CB - CG1 |
| 5242 | 5243 | 1 | 1.0900e-01 | 2.8451e+05 | ; | CG2 - HG21 |
| 5242 | 5244 | 1 | 1.0900e-01 | 2.8451e+05 | ; | CG2 - HG22 |
| 5242 | 5245 | 1 | 1.0900e-01 | 2.8451e+05 | ; | CG2 - HG23 |
| 5246 | 5247 | 1 | 1.0900e-01 | 2.8451e+05 | ; | CG1 - HG12 |

| | | | | | | |
|---|---|---|---|---|---|---|
| 5246 | 5248 | 1 | 1.0900e-01 | 2.8451e+05 | ; | CG1 - HG13 |
| 5246 | 5249 | 1 | 1.5260e-01 | 2.5941e+05 | ; | CG1 - CD1 |
| 5249 | 5250 | 1 | 1.0900e-01 | 2.8451e+05 | ; | CD1 - HD11 |
| 5249 | 5251 | 1 | 1.0900e-01 | 2.8451e+05 | ; | CD1 - HD12 |
| 5249 | 5252 | 1 | 1.0900e-01 | 2.8451e+05 | ; | CD1 - HD13 |
| 5253 | 5254 | 1 | 1.2290e-01 | 4.7698e+05 | ; | C - O |
| 5253 | 5255 | 1 | 1.3350e-01 | 4.1003e+05 | ; | C - N |
| 5255 | 5256 | 1 | 1.0100e-01 | 3.6317e+05 | ; | N - H |
| 5255 | 5257 | 1 | 1.4490e-01 | 2.8200e+05 | ; | N - CA |
| 5257 | 5258 | 1 | 1.0900e-01 | 2.8451e+05 | ; | CA - HA |
| 5257 | 5259 | 1 | 1.5260e-01 | 2.5941e+05 | ; | CA - CB |
| 5257 | 5263 | 1 | 1.5220e-01 | 2.6527e+05 | ; | CA - C |
| 5259 | 5260 | 1 | 1.0900e-01 | 2.8451e+05 | ; | CB - HB1 |
| 5259 | 5261 | 1 | 1.0900e-01 | 2.8451e+05 | ; | CB - HB2 |
| 5259 | 5262 | 1 | 1.0900e-01 | 2.8451e+05 | ; | CB - HB3 |
| 5263 | 5264 | 1 | 1.2290e-01 | 4.7698e+05 | ; | C - O |
| 5263 | 5265 | 1 | 1.3350e-01 | 4.1003e+05 | ; | C - N |
| 5265 | 5266 | 1 | 1.0100e-01 | 3.6317e+05 | ; | N - H |
| 5265 | 5267 | 1 | 1.4490e-01 | 2.8200e+05 | ; | N - CA |
| 5267 | 5268 | 1 | 1.0900e-01 | 2.8451e+05 | ; | CA - HA |
| 5267 | 5269 | 1 | 1.5260e-01 | 2.5941e+05 | ; | CA - CB |
| 5267 | 5282 | 1 | 1.5220e-01 | 2.6527e+05 | ; | CA - C |
| 5269 | 5270 | 1 | 1.0900e-01 | 2.8451e+05 | ; | CB - HB2 |
| 5269 | 5271 | 1 | 1.0900e-01 | 2.8451e+05 | ; | CB - HB3 |
| 5269 | 5272 | 1 | 1.5260e-01 | 2.5941e+05 | ; | CB - CG |
| 5272 | 5273 | 1 | 1.0900e-01 | 2.8451e+05 | ; | CG - HG |
| 5272 | 5274 | 1 | 1.5260e-01 | 2.5941e+05 | ; | CG - CD1 |
| 5272 | 5278 | 1 | 1.5260e-01 | 2.5941e+05 | ; | CG - CD2 |
| 5274 | 5275 | 1 | 1.0900e-01 | 2.8451e+05 | ; | CD1 - HD11 |
| 5274 | 5276 | 1 | 1.0900e-01 | 2.8451e+05 | ; | CD1 - HD12 |
| 5274 | 5277 | 1 | 1.0900e-01 | 2.8451e+05 | ; | CD1 - HD13 |
| 5278 | 5279 | 1 | 1.0900e-01 | 2.8451e+05 | ; | CD2 - HD21 |
| 5278 | 5280 | 1 | 1.0900e-01 | 2.8451e+05 | ; | CD2 - HD22 |
| 5278 | 5281 | 1 | 1.0900e-01 | 2.8451e+05 | ; | CD2 - HD23 |
| 5282 | 5283 | 1 | 1.2290e-01 | 4.7698e+05 | ; | C - O |
| 5282 | 5284 | 1 | 1.3350e-01 | 4.1003e+05 | ; | C - N |
| 5284 | 5285 | 1 | 1.0100e-01 | 3.6317e+05 | ; | N - H |
| 5284 | 5286 | 1 | 1.4490e-01 | 2.8200e+05 | ; | N - CA |
| 5286 | 5287 | 1 | 1.0900e-01 | 2.8451e+05 | ; | CA - HA |
| 5286 | 5288 | 1 | 1.5260e-01 | 2.5941e+05 | ; | CA - CB |
| 5286 | 5294 | 1 | 1.5220e-01 | 2.6527e+05 | ; | CA - C |
| 5288 | 5289 | 1 | 1.0900e-01 | 2.8451e+05 | ; | CB - HB2 |
| 5288 | 5290 | 1 | 1.0900e-01 | 2.8451e+05 | ; | CB - HB3 |
| 5288 | 5291 | 1 | 1.5220e-01 | 2.6527e+05 | ; | CB - CG |
| 5291 | 5292 | 1 | 1.2500e-01 | 5.4894e+05 | ; | CG - OD1 |
| 5291 | 5293 | 1 | 1.2500e-01 | 5.4894e+05 | ; | CG - OD2 |
| 5294 | 5295 | 1 | 1.2290e-01 | 4.7698e+05 | ; | C - O |
| 5294 | 5296 | 1 | 1.3350e-01 | 4.1003e+05 | ; | C - N |
| 5296 | 5297 | 1 | 1.0100e-01 | 3.6317e+05 | ; | N - H |
| 5296 | 5298 | 1 | 1.4490e-01 | 2.8200e+05 | ; | N - CA |
| 5298 | 5299 | 1 | 1.0900e-01 | 2.8451e+05 | ; | CA - HA |
| 5298 | 5300 | 1 | 1.5260e-01 | 2.5941e+05 | ; | CA - CB |

| | | | | | | |
|---|---|---|---|---|---|---|
| 5298 | 5310 | 1 | 1.5220e-01 | 2.6527e+05 | ; | CA - C |
| 5300 | 5301 | 1 | 1.0900e-01 | 2.8451e+05 | ; | CB - HB |
| 5300 | 5302 | 1 | 1.5260e-01 | 2.5941e+05 | ; | CB - CG1 |
| 5300 | 5306 | 1 | 1.5260e-01 | 2.5941e+05 | ; | CB - CG2 |
| 5302 | 5303 | 1 | 1.0900e-01 | 2.8451e+05 | ; | CG1 - HG11 |
| 5302 | 5304 | 1 | 1.0900e-01 | 2.8451e+05 | ; | CG1 - HG12 |
| 5302 | 5305 | 1 | 1.0900e-01 | 2.8451e+05 | ; | CG1 - HG13 |
| 5306 | 5307 | 1 | 1.0900e-01 | 2.8451e+05 | ; | CG2 - HG21 |
| 5306 | 5308 | 1 | 1.0900e-01 | 2.8451e+05 | ; | CG2 - HG22 |
| 5306 | 5309 | 1 | 1.0900e-01 | 2.8451e+05 | ; | CG2 - HG23 |
| 5310 | 5311 | 1 | 1.2290e-01 | 4.7698e+05 | ; | C - O |
| 5310 | 5312 | 1 | 1.3350e-01 | 4.1003e+05 | ; | C - N |
| 5312 | 5313 | 1 | 1.0100e-01 | 3.6317e+05 | ; | N - H |
| 5312 | 5314 | 1 | 1.4490e-01 | 2.8200e+05 | ; | N - CA |
| 5314 | 5315 | 1 | 1.0900e-01 | 2.8451e+05 | ; | CA - HA |
| 5314 | 5316 | 1 | 1.5260e-01 | 2.5941e+05 | ; | CA - CB |
| 5314 | 5322 | 1 | 1.5220e-01 | 2.6527e+05 | ; | CA - C |
| 5316 | 5317 | 1 | 1.0900e-01 | 2.8451e+05 | ; | CB - HB2 |
| 5316 | 5318 | 1 | 1.0900e-01 | 2.8451e+05 | ; | CB - HB3 |
| 5316 | 5319 | 1 | 1.5220e-01 | 2.6527e+05 | ; | CB - CG |
| 5319 | 5320 | 1 | 1.2500e-01 | 5.4894e+05 | ; | CG - OD1 |
| 5319 | 5321 | 1 | 1.2500e-01 | 5.4894e+05 | ; | CG - OD2 |
| 5322 | 5323 | 1 | 1.2290e-01 | 4.7698e+05 | ; | C - O |
| 5322 | 5324 | 1 | 1.3350e-01 | 4.1003e+05 | ; | C - N |
| 5324 | 5325 | 1 | 1.0100e-01 | 3.6317e+05 | ; | N - H |
| 5324 | 5326 | 1 | 1.4490e-01 | 2.8200e+05 | ; | N - CA |
| 5326 | 5327 | 1 | 1.0900e-01 | 2.8451e+05 | ; | CA - HA |
| 5326 | 5328 | 1 | 1.5260e-01 | 2.5941e+05 | ; | CA - CB |
| 5326 | 5336 | 1 | 1.5220e-01 | 2.6527e+05 | ; | CA - C |
| 5328 | 5329 | 1 | 1.0900e-01 | 2.8451e+05 | ; | CB - HB2 |
| 5328 | 5330 | 1 | 1.0900e-01 | 2.8451e+05 | ; | CB - HB3 |
| 5328 | 5331 | 1 | 1.5220e-01 | 2.6527e+05 | ; | CB - CG |
| 5331 | 5332 | 1 | 1.2290e-01 | 4.7698e+05 | ; | CG - OD1 |
| 5331 | 5333 | 1 | 1.3350e-01 | 4.1003e+05 | ; | CG - ND2 |
| 5333 | 5334 | 1 | 1.0100e-01 | 3.6317e+05 | ; | ND2 - HD21 |
| 5333 | 5335 | 1 | 1.0100e-01 | 3.6317e+05 | ; | ND2 - HD22 |
| 5336 | 5337 | 1 | 1.2290e-01 | 4.7698e+05 | ; | C - O |
| 5336 | 5338 | 1 | 1.3350e-01 | 4.1003e+05 | ; | C - N |
| 5338 | 5339 | 1 | 1.0100e-01 | 3.6317e+05 | ; | N - H |
| 5338 | 5340 | 1 | 1.4490e-01 | 2.8200e+05 | ; | N - CA |
| 5340 | 5341 | 1 | 1.0900e-01 | 2.8451e+05 | ; | CA - HA |
| 5340 | 5342 | 1 | 1.5260e-01 | 2.5941e+05 | ; | CA - CB |
| 5340 | 5347 | 1 | 1.5220e-01 | 2.6527e+05 | ; | CA - C |
| 5342 | 5343 | 1 | 1.0900e-01 | 2.8451e+05 | ; | CB - HB2 |
| 5342 | 5344 | 1 | 1.0900e-01 | 2.8451e+05 | ; | CB - HB3 |
| 5342 | 5345 | 1 | 1.4100e-01 | 2.6778e+05 | ; | CB - OG |
| 5345 | 5346 | 1 | 9.6000e-02 | 4.6275e+05 | ; | OG - HG |
| 5347 | 5348 | 1 | 1.2290e-01 | 4.7698e+05 | ; | C - O |
| 5347 | 5349 | 1 | 1.3350e-01 | 4.1003e+05 | ; | C - N |
| 5349 | 5350 | 1 | 1.0100e-01 | 3.6317e+05 | ; | N - H |
| 5349 | 5351 | 1 | 1.4490e-01 | 2.8200e+05 | ; | N - CA |
| 5351 | 5352 | 1 | 1.0900e-01 | 2.8451e+05 | ; | CA - HA2 |

| | | | | | | |
|---|---|---|---|---|---|---|
| 5351 | 5353 | 1 | 1.0900e-01 | 2.8451e+05 | ; | CA - HA3 |
| 5351 | 5354 | 1 | 1.5220e-01 | 2.6527e+05 | ; | CA - C |
| 5354 | 5355 | 1 | 1.2290e-01 | 4.7698e+05 | ; | C - O |
| 5354 | 5356 | 1 | 1.3350e-01 | 4.1003e+05 | ; | C - N |
| 5356 | 5357 | 1 | 1.0100e-01 | 3.6317e+05 | ; | N - H |
| 5356 | 5358 | 1 | 1.4490e-01 | 2.8200e+05 | ; | N - CA |
| 5358 | 5359 | 1 | 1.0900e-01 | 2.8451e+05 | ; | CA - HA |
| 5358 | 5360 | 1 | 1.5260e-01 | 2.5941e+05 | ; | CA - CB |
| 5358 | 5368 | 1 | 1.5220e-01 | 2.6527e+05 | ; | CA - C |
| 5360 | 5361 | 1 | 1.0900e-01 | 2.8451e+05 | ; | CB - HB |
| 5360 | 5362 | 1 | 1.5260e-01 | 2.5941e+05 | ; | CB - CG2 |
| 5360 | 5366 | 1 | 1.4100e-01 | 2.6778e+05 | ; | CB - OG1 |
| 5362 | 5363 | 1 | 1.0900e-01 | 2.8451e+05 | ; | CG2 - HG21 |
| 5362 | 5364 | 1 | 1.0900e-01 | 2.8451e+05 | ; | CG2 - HG22 |
| 5362 | 5365 | 1 | 1.0900e-01 | 2.8451e+05 | ; | CG2 - HG23 |
| 5366 | 5367 | 1 | 9.6000e-02 | 4.6275e+05 | ; | OG1 - HG1 |
| 5368 | 5369 | 1 | 1.2290e-01 | 4.7698e+05 | ; | C - O |
| 5368 | 5370 | 1 | 1.3350e-01 | 4.1003e+05 | ; | C - N |
| 5370 | 5371 | 1 | 1.0100e-01 | 3.6317e+05 | ; | N - H |
| 5370 | 5372 | 1 | 1.4490e-01 | 2.8200e+05 | ; | N - CA |
| 5372 | 5373 | 1 | 1.0900e-01 | 2.8451e+05 | ; | CA - HA |
| 5372 | 5374 | 1 | 1.5260e-01 | 2.5941e+05 | ; | CA - CB |
| 5372 | 5387 | 1 | 1.5220e-01 | 2.6527e+05 | ; | CA - C |
| 5374 | 5375 | 1 | 1.0900e-01 | 2.8451e+05 | ; | CB - HB2 |
| 5374 | 5376 | 1 | 1.0900e-01 | 2.8451e+05 | ; | CB - HB3 |
| 5374 | 5377 | 1 | 1.5260e-01 | 2.5941e+05 | ; | CB - CG |
| 5377 | 5378 | 1 | 1.0900e-01 | 2.8451e+05 | ; | CG - HG |
| 5377 | 5379 | 1 | 1.5260e-01 | 2.5941e+05 | ; | CG - CD1 |
| 5377 | 5383 | 1 | 1.5260e-01 | 2.5941e+05 | ; | CG - CD2 |
| 5379 | 5380 | 1 | 1.0900e-01 | 2.8451e+05 | ; | CD1 - HD11 |
| 5379 | 5381 | 1 | 1.0900e-01 | 2.8451e+05 | ; | CD1 - HD12 |
| 5379 | 5382 | 1 | 1.0900e-01 | 2.8451e+05 | ; | CD1 - HD13 |
| 5383 | 5384 | 1 | 1.0900e-01 | 2.8451e+05 | ; | CD2 - HD21 |
| 5383 | 5385 | 1 | 1.0900e-01 | 2.8451e+05 | ; | CD2 - HD22 |
| 5383 | 5386 | 1 | 1.0900e-01 | 2.8451e+05 | ; | CD2 - HD23 |
| 5387 | 5388 | 1 | 1.2290e-01 | 4.7698e+05 | ; | C - O |
| 5387 | 5389 | 1 | 1.3350e-01 | 4.1003e+05 | ; | C - N |
| 5389 | 5390 | 1 | 1.0100e-01 | 3.6317e+05 | ; | N - H |
| 5389 | 5391 | 1 | 1.4490e-01 | 2.8200e+05 | ; | N - CA |
| 5391 | 5392 | 1 | 1.0900e-01 | 2.8451e+05 | ; | CA - HA |
| 5391 | 5393 | 1 | 1.5260e-01 | 2.5941e+05 | ; | CA - CB |
| 5391 | 5398 | 1 | 1.5220e-01 | 2.6527e+05 | ; | CA - C |
| 5393 | 5394 | 1 | 1.0900e-01 | 2.8451e+05 | ; | CB - HB2 |
| 5393 | 5395 | 1 | 1.0900e-01 | 2.8451e+05 | ; | CB - HB3 |
| 5393 | 5396 | 1 | 1.4100e-01 | 2.6778e+05 | ; | CB - OG |
| 5396 | 5397 | 1 | 9.6000e-02 | 4.6275e+05 | ; | OG - HG |
| 5398 | 5399 | 1 | 1.2290e-01 | 4.7698e+05 | ; | C - O |
| 5398 | 5400 | 1 | 1.3350e-01 | 4.1003e+05 | ; | C - N |
| 5400 | 5401 | 1 | 1.0100e-01 | 3.6317e+05 | ; | N - H |
| 5400 | 5402 | 1 | 1.4490e-01 | 2.8200e+05 | ; | N - CA |
| 5402 | 5403 | 1 | 1.0900e-01 | 2.8451e+05 | ; | CA - HA |
| 5402 | 5404 | 1 | 1.5260e-01 | 2.5941e+05 | ; | CA - CB |

| | | | | | |
|---|---|---|---|---|---|
| 5402 | 5409 | 1 | 1.5220e-01 | 2.6527e+05 ; | CA - C |
| 5404 | 5405 | 1 | 1.0900e-01 | 2.8451e+05 ; | CB - HB2 |
| 5404 | 5406 | 1 | 1.0900e-01 | 2.8451e+05 ; | CB - HB3 |
| 5404 | 5407 | 1 | 1.4100e-01 | 2.6778e+05 ; | CB - OG |
| 5407 | 5408 | 1 | 9.6000e-02 | 4.6275e+05 ; | OG - HG |
| 5409 | 5410 | 1 | 1.2290e-01 | 4.7698e+05 ; | C - O |
| 5409 | 5411 | 1 | 1.3350e-01 | 4.1003e+05 ; | C - N |
| 5411 | 5412 | 1 | 1.0100e-01 | 3.6317e+05 ; | N - H |
| 5411 | 5413 | 1 | 1.4490e-01 | 2.8200e+05 ; | N - CA |
| 5413 | 5414 | 1 | 1.0900e-01 | 2.8451e+05 ; | CA - HA |
| 5413 | 5415 | 1 | 1.5260e-01 | 2.5941e+05 ; | CA - CB |
| 5413 | 5426 | 1 | 1.5220e-01 | 2.6527e+05 ; | CA - C |
| 5415 | 5416 | 1 | 1.0900e-01 | 2.8451e+05 ; | CB - HB2 |
| 5415 | 5417 | 1 | 1.0900e-01 | 2.8451e+05 ; | CB - HB3 |
| 5415 | 5418 | 1 | 1.5260e-01 | 2.5941e+05 ; | CB - CG |
| 5418 | 5419 | 1 | 1.0900e-01 | 2.8451e+05 ; | CG - HG2 |
| 5418 | 5420 | 1 | 1.0900e-01 | 2.8451e+05 ; | CG - HG3 |
| 5418 | 5421 | 1 | 1.5220e-01 | 2.6527e+05 ; | CG - CD |
| 5421 | 5422 | 1 | 1.2290e-01 | 4.7698e+05 ; | CD - OE1 |
| 5421 | 5423 | 1 | 1.3350e-01 | 4.1003e+05 ; | CD - NE2 |
| 5423 | 5424 | 1 | 1.0100e-01 | 3.6317e+05 ; | NE2 - HE21 |
| 5423 | 5425 | 1 | 1.0100e-01 | 3.6317e+05 ; | NE2 - HE22 |
| 5426 | 5427 | 1 | 1.2290e-01 | 4.7698e+05 ; | C - O |
| 5426 | 5428 | 1 | 1.3350e-01 | 4.1003e+05 ; | C - N |
| 5428 | 5429 | 1 | 1.0100e-01 | 3.6317e+05 ; | N - H |
| 5428 | 5430 | 1 | 1.4490e-01 | 2.8200e+05 ; | N - CA |
| 5430 | 5431 | 1 | 1.0900e-01 | 2.8451e+05 ; | CA - HA |
| 5430 | 5432 | 1 | 1.5260e-01 | 2.5941e+05 ; | CA - CB |
| 5430 | 5442 | 1 | 1.5220e-01 | 2.6527e+05 ; | CA - C |
| 5432 | 5433 | 1 | 1.0900e-01 | 2.8451e+05 ; | CB - HB2 |
| 5432 | 5434 | 1 | 1.0900e-01 | 2.8451e+05 ; | CB - HB3 |
| 5432 | 5435 | 1 | 1.5260e-01 | 2.5941e+05 ; | CB - CG |
| 5435 | 5436 | 1 | 1.0900e-01 | 2.8451e+05 ; | CG - HG2 |
| 5435 | 5437 | 1 | 1.0900e-01 | 2.8451e+05 ; | CG - HG3 |
| 5435 | 5438 | 1 | 1.5220e-01 | 2.6527e+05 ; | CG - CD |
| 5438 | 5439 | 1 | 1.2290e-01 | 4.7698e+05 ; | CD - OE1 |
| 5438 | 5440 | 1 | 1.3640e-01 | 3.7656e+05 ; | CD - OE2 |
| 5440 | 5441 | 1 | 9.6000e-02 | 4.6275e+05 ; | OE2 - HE2 |
| 5442 | 5443 | 1 | 1.2290e-01 | 4.7698e+05 ; | C - O |
| 5442 | 5444 | 1 | 1.3350e-01 | 4.1003e+05 ; | C - N |
| 5444 | 5445 | 1 | 1.0100e-01 | 3.6317e+05 ; | N - H |
| 5444 | 5446 | 1 | 1.4490e-01 | 2.8200e+05 ; | N - CA |
| 5446 | 5447 | 1 | 1.0900e-01 | 2.8451e+05 ; | CA - HA |
| 5446 | 5448 | 1 | 1.5260e-01 | 2.5941e+05 ; | CA - CB |
| 5446 | 5461 | 1 | 1.5220e-01 | 2.6527e+05 ; | CA - C |
| 5448 | 5449 | 1 | 1.0900e-01 | 2.8451e+05 ; | CB - HB |
| 5448 | 5450 | 1 | 1.5260e-01 | 2.5941e+05 ; | CB - CG2 |
| 5448 | 5454 | 1 | 1.5260e-01 | 2.5941e+05 ; | CB - CG1 |
| 5450 | 5451 | 1 | 1.0900e-01 | 2.8451e+05 ; | CG2 - HG21 |
| 5450 | 5452 | 1 | 1.0900e-01 | 2.8451e+05 ; | CG2 - HG22 |
| 5450 | 5453 | 1 | 1.0900e-01 | 2.8451e+05 ; | CG2 - HG23 |
| 5454 | 5455 | 1 | 1.0900e-01 | 2.8451e+05 ; | CG1 - HG12 |

| | | | | | | |
|---|---|---|---|---|---|---|
| 5454 | 5456 | 1 | 1.0900e-01 | 2.8451e+05 | ; | CG1 - HG13 |
| 5454 | 5457 | 1 | 1.5260e-01 | 2.5941e+05 | ; | CG1 - CD1 |
| 5457 | 5458 | 1 | 1.0900e-01 | 2.8451e+05 | ; | CD1 - HD11 |
| 5457 | 5459 | 1 | 1.0900e-01 | 2.8451e+05 | ; | CD1 - HD12 |
| 5457 | 5460 | 1 | 1.0900e-01 | 2.8451e+05 | ; | CD1 - HD13 |
| 5461 | 5462 | 1 | 1.2290e-01 | 4.7698e+05 | ; | C - O |
| 5461 | 5463 | 1 | 1.3350e-01 | 4.1003e+05 | ; | C - N |
| 5463 | 5464 | 1 | 1.0100e-01 | 3.6317e+05 | ; | N - H |
| 5463 | 5465 | 1 | 1.4490e-01 | 2.8200e+05 | ; | N - CA |
| 5465 | 5466 | 1 | 1.0900e-01 | 2.8451e+05 | ; | CA - HA |
| 5465 | 5467 | 1 | 1.5260e-01 | 2.5941e+05 | ; | CA - CB |
| 5465 | 5480 | 1 | 1.5220e-01 | 2.6527e+05 | ; | CA - C |
| 5467 | 5468 | 1 | 1.0900e-01 | 2.8451e+05 | ; | CB - HB2 |
| 5467 | 5469 | 1 | 1.0900e-01 | 2.8451e+05 | ; | CB - HB3 |
| 5467 | 5470 | 1 | 1.5260e-01 | 2.5941e+05 | ; | CB - CG |
| 5470 | 5471 | 1 | 1.0900e-01 | 2.8451e+05 | ; | CG - HG |
| 5470 | 5472 | 1 | 1.5260e-01 | 2.5941e+05 | ; | CG - CD1 |
| 5470 | 5476 | 1 | 1.5260e-01 | 2.5941e+05 | ; | CG - CD2 |
| 5472 | 5473 | 1 | 1.0900e-01 | 2.8451e+05 | ; | CD1 - HD11 |
| 5472 | 5474 | 1 | 1.0900e-01 | 2.8451e+05 | ; | CD1 - HD12 |
| 5472 | 5475 | 1 | 1.0900e-01 | 2.8451e+05 | ; | CD1 - HD13 |
| 5476 | 5477 | 1 | 1.0900e-01 | 2.8451e+05 | ; | CD2 - HD21 |
| 5476 | 5478 | 1 | 1.0900e-01 | 2.8451e+05 | ; | CD2 - HD22 |
| 5476 | 5479 | 1 | 1.0900e-01 | 2.8451e+05 | ; | CD2 - HD23 |
| 5480 | 5481 | 1 | 1.2290e-01 | 4.7698e+05 | ; | C - O |
| 5480 | 5482 | 1 | 1.3350e-01 | 4.1003e+05 | ; | C - N |
| 5482 | 5483 | 1 | 1.0100e-01 | 3.6317e+05 | ; | N - H |
| 5482 | 5484 | 1 | 1.4490e-01 | 2.8200e+05 | ; | N - CA |
| 5484 | 5485 | 1 | 1.0900e-01 | 2.8451e+05 | ; | CA - HA |
| 5484 | 5486 | 1 | 1.5260e-01 | 2.5941e+05 | ; | CA - CB |
| 5484 | 5492 | 1 | 1.5220e-01 | 2.6527e+05 | ; | CA - C |
| 5486 | 5487 | 1 | 1.0900e-01 | 2.8451e+05 | ; | CB - HB2 |
| 5486 | 5488 | 1 | 1.0900e-01 | 2.8451e+05 | ; | CB - HB3 |
| 5486 | 5489 | 1 | 1.5220e-01 | 2.6527e+05 | ; | CB - CG |
| 5489 | 5490 | 1 | 1.2500e-01 | 5.4894e+05 | ; | CG - OD1 |
| 5489 | 5491 | 1 | 1.2500e-01 | 5.4894e+05 | ; | CG - OD2 |
| 5492 | 5493 | 1 | 1.2290e-01 | 4.7698e+05 | ; | C - O |
| 5492 | 5494 | 1 | 1.3350e-01 | 4.1003e+05 | ; | C - N |
| 5494 | 5495 | 1 | 1.0100e-01 | 3.6317e+05 | ; | N - H |
| 5494 | 5496 | 1 | 1.4490e-01 | 2.8200e+05 | ; | N - CA |
| 5496 | 5497 | 1 | 1.0900e-01 | 2.8451e+05 | ; | CA - HA2 |
| 5496 | 5498 | 1 | 1.0900e-01 | 2.8451e+05 | ; | CA - HA3 |
| 5496 | 5499 | 1 | 1.5220e-01 | 2.6527e+05 | ; | CA - C |
| 5499 | 5500 | 1 | 1.2290e-01 | 4.7698e+05 | ; | C - O |
| 5499 | 5501 | 1 | 1.3350e-01 | 4.1003e+05 | ; | C - N |
| 5501 | 5502 | 1 | 1.0100e-01 | 3.6317e+05 | ; | N - H |
| 5501 | 5503 | 1 | 1.4490e-01 | 2.8200e+05 | ; | N - CA |
| 5503 | 5504 | 1 | 1.0900e-01 | 2.8451e+05 | ; | CA - HA |
| 5503 | 5505 | 1 | 1.5260e-01 | 2.5941e+05 | ; | CA - CB |
| 5503 | 5518 | 1 | 1.5220e-01 | 2.6527e+05 | ; | CA - C |
| 5505 | 5506 | 1 | 1.0900e-01 | 2.8451e+05 | ; | CB - HB2 |
| 5505 | 5507 | 1 | 1.0900e-01 | 2.8451e+05 | ; | CB - HB3 |

| | | | | | | | |
|---|---|---|---|---|---|---|---|
| 5505 | 5508 | 1 | 1.5260e-01 | 2.5941e+05 ; | CB | - | CG |
| 5508 | 5509 | 1 | 1.0900e-01 | 2.8451e+05 ; | CG | - | HG |
| 5508 | 5510 | 1 | 1.5260e-01 | 2.5941e+05 ; | CG | - | CD1 |
| 5508 | 5514 | 1 | 1.5260e-01 | 2.5941e+05 ; | CG | - | CD2 |
| 5510 | 5511 | 1 | 1.0900e-01 | 2.8451e+05 ; | CD1 | - | HD11 |
| 5510 | 5512 | 1 | 1.0900e-01 | 2.8451e+05 ; | CD1 | - | HD12 |
| 5510 | 5513 | 1 | 1.0900e-01 | 2.8451e+05 ; | CD1 | - | HD13 |
| 5514 | 5515 | 1 | 1.0900e-01 | 2.8451e+05 ; | CD2 | - | HD21 |
| 5514 | 5516 | 1 | 1.0900e-01 | 2.8451e+05 ; | CD2 | - | HD22 |
| 5514 | 5517 | 1 | 1.0900e-01 | 2.8451e+05 ; | CD2 | - | HD23 |
| 5518 | 5519 | 1 | 1.2290e-01 | 4.7698e+05 ; | C | - | O |
| 5518 | 5520 | 1 | 1.3350e-01 | 4.1003e+05 ; | C | - | N |
| 5520 | 5521 | 1 | 1.0100e-01 | 3.6317e+05 ; | N | - | H |
| 5520 | 5522 | 1 | 1.4490e-01 | 2.8200e+05 ; | N | - | CA |
| 5522 | 5523 | 1 | 1.0900e-01 | 2.8451e+05 ; | CA | - | HA |
| 5522 | 5524 | 1 | 1.5260e-01 | 2.5941e+05 ; | CA | - | CB |
| 5522 | 5540 | 1 | 1.5220e-01 | 2.6527e+05 ; | CA | - | C |
| 5524 | 5525 | 1 | 1.0900e-01 | 2.8451e+05 ; | CB | - | HB2 |
| 5524 | 5526 | 1 | 1.0900e-01 | 2.8451e+05 ; | CB | - | HB3 |
| 5524 | 5527 | 1 | 1.5260e-01 | 2.5941e+05 ; | CB | - | CG |
| 5527 | 5528 | 1 | 1.0900e-01 | 2.8451e+05 ; | CG | - | HG2 |
| 5527 | 5529 | 1 | 1.0900e-01 | 2.8451e+05 ; | CG | - | HG3 |
| 5527 | 5530 | 1 | 1.5260e-01 | 2.5941e+05 ; | CG | - | CD |
| 5530 | 5531 | 1 | 1.0900e-01 | 2.8451e+05 ; | CD | - | HD2 |
| 5530 | 5532 | 1 | 1.0900e-01 | 2.8451e+05 ; | CD | - | HD3 |
| 5530 | 5533 | 1 | 1.5260e-01 | 2.5941e+05 ; | CD | - | CE |
| 5533 | 5534 | 1 | 1.0900e-01 | 2.8451e+05 ; | CE | - | HE2 |
| 5533 | 5535 | 1 | 1.0900e-01 | 2.8451e+05 ; | CE | - | HE3 |
| 5533 | 5536 | 1 | 1.4710e-01 | 3.0711e+05 ; | CE | - | NZ |
| 5536 | 5537 | 1 | 1.0100e-01 | 3.6317e+05 ; | NZ | - | HZ1 |
| 5536 | 5538 | 1 | 1.0100e-01 | 3.6317e+05 ; | NZ | - | HZ2 |
| 5536 | 5539 | 1 | 1.0100e-01 | 3.6317e+05 ; | NZ | - | HZ3 |
| 5540 | 5541 | 1 | 1.2290e-01 | 4.7698e+05 ; | C | - | O |
| 5540 | 5542 | 1 | 1.3350e-01 | 4.1003e+05 ; | C | - | N |
| 5542 | 5543 | 1 | 1.0100e-01 | 3.6317e+05 ; | N | - | H |
| 5542 | 5544 | 1 | 1.4490e-01 | 2.8200e+05 ; | N | - | CA |
| 5544 | 5545 | 1 | 1.0900e-01 | 2.8451e+05 ; | CA | - | HA |
| 5544 | 5546 | 1 | 1.5260e-01 | 2.5941e+05 ; | CA | - | CB |
| 5544 | 5562 | 1 | 1.5220e-01 | 2.6527e+05 ; | CA | - | C |
| 5546 | 5547 | 1 | 1.0900e-01 | 2.8451e+05 ; | CB | - | HB2 |
| 5546 | 5548 | 1 | 1.0900e-01 | 2.8451e+05 ; | CB | - | HB3 |
| 5546 | 5549 | 1 | 1.5260e-01 | 2.5941e+05 ; | CB | - | CG |
| 5549 | 5550 | 1 | 1.0900e-01 | 2.8451e+05 ; | CG | - | HG2 |
| 5549 | 5551 | 1 | 1.0900e-01 | 2.8451e+05 ; | CG | - | HG3 |
| 5549 | 5552 | 1 | 1.5260e-01 | 2.5941e+05 ; | CG | - | CD |
| 5552 | 5553 | 1 | 1.0900e-01 | 2.8451e+05 ; | CD | - | HD2 |
| 5552 | 5554 | 1 | 1.0900e-01 | 2.8451e+05 ; | CD | - | HD3 |
| 5552 | 5555 | 1 | 1.5260e-01 | 2.5941e+05 ; | CD | - | CE |
| 5555 | 5556 | 1 | 1.0900e-01 | 2.8451e+05 ; | CE | - | HE2 |
| 5555 | 5557 | 1 | 1.0900e-01 | 2.8451e+05 ; | CE | - | HE3 |
| 5555 | 5558 | 1 | 1.4710e-01 | 3.0711e+05 ; | CE | - | NZ |
| 5558 | 5559 | 1 | 1.0100e-01 | 3.6317e+05 ; | NZ | - | HZ1 |

| | | | | | | |
|---|---|---|---|---|---|---|
| 5558 | 5560 | 1 | 1.0100e-01 | 3.6317e+05 | ; | NZ - HZ2 |
| 5558 | 5561 | 1 | 1.0100e-01 | 3.6317e+05 | ; | NZ - HZ3 |
| 5562 | 5563 | 1 | 1.2290e-01 | 4.7698e+05 | ; | C - O |
| 5562 | 5564 | 1 | 1.3350e-01 | 4.1003e+05 | ; | C - N |
| 5564 | 5565 | 1 | 1.0100e-01 | 3.6317e+05 | ; | N - H |
| 5564 | 5566 | 1 | 1.4490e-01 | 2.8200e+05 | ; | N - CA |
| 5566 | 5567 | 1 | 1.0900e-01 | 2.8451e+05 | ; | CA - HA |
| 5566 | 5568 | 1 | 1.5260e-01 | 2.5941e+05 | ; | CA - CB |
| 5566 | 5581 | 1 | 1.5220e-01 | 2.6527e+05 | ; | CA - C |
| 5568 | 5569 | 1 | 1.0900e-01 | 2.8451e+05 | ; | CB - HB |
| 5568 | 5570 | 1 | 1.5260e-01 | 2.5941e+05 | ; | CB - CG2 |
| 5568 | 5574 | 1 | 1.5260e-01 | 2.5941e+05 | ; | CB - CG1 |
| 5570 | 5571 | 1 | 1.0900e-01 | 2.8451e+05 | ; | CG2 - HG21 |
| 5570 | 5572 | 1 | 1.0900e-01 | 2.8451e+05 | ; | CG2 - HG22 |
| 5570 | 5573 | 1 | 1.0900e-01 | 2.8451e+05 | ; | CG2 - HG23 |
| 5574 | 5575 | 1 | 1.0900e-01 | 2.8451e+05 | ; | CG1 - HG12 |
| 5574 | 5576 | 1 | 1.0900e-01 | 2.8451e+05 | ; | CG1 - HG13 |
| 5574 | 5577 | 1 | 1.5260e-01 | 2.5941e+05 | ; | CG1 - CD1 |
| 5577 | 5578 | 1 | 1.0900e-01 | 2.8451e+05 | ; | CD1 - HD11 |
| 5577 | 5579 | 1 | 1.0900e-01 | 2.8451e+05 | ; | CD1 - HD12 |
| 5577 | 5580 | 1 | 1.0900e-01 | 2.8451e+05 | ; | CD1 - HD13 |
| 5581 | 5582 | 1 | 1.2290e-01 | 4.7698e+05 | ; | C - O |
| 5581 | 5583 | 1 | 1.3350e-01 | 4.1003e+05 | ; | C - N |
| 5583 | 5584 | 1 | 1.0100e-01 | 3.6317e+05 | ; | N - H |
| 5583 | 5585 | 1 | 1.4490e-01 | 2.8200e+05 | ; | N - CA |
| 5585 | 5586 | 1 | 1.0900e-01 | 2.8451e+05 | ; | CA - HA2 |
| 5585 | 5587 | 1 | 1.0900e-01 | 2.8451e+05 | ; | CA - HA3 |
| 5585 | 5588 | 1 | 1.5220e-01 | 2.6527e+05 | ; | CA - C |
| 5588 | 5589 | 1 | 1.2290e-01 | 4.7698e+05 | ; | C - O |
| 5588 | 5590 | 1 | 1.3350e-01 | 4.1003e+05 | ; | C - N |
| 5590 | 5591 | 1 | 1.0100e-01 | 3.6317e+05 | ; | N - H |
| 5590 | 5592 | 1 | 1.4490e-01 | 2.8200e+05 | ; | N - CA |
| 5592 | 5593 | 1 | 1.0900e-01 | 2.8451e+05 | ; | CA - HA |
| 5592 | 5594 | 1 | 1.5260e-01 | 2.5941e+05 | ; | CA - CB |
| 5592 | 5609 | 1 | 1.5220e-01 | 2.6527e+05 | ; | CA - C |
| 5594 | 5595 | 1 | 1.0900e-01 | 2.8451e+05 | ; | CB - HB2 |
| 5594 | 5596 | 1 | 1.0900e-01 | 2.8451e+05 | ; | CB - HB3 |
| 5594 | 5597 | 1 | 1.5100e-01 | 2.6527e+05 | ; | CB - CG |
| 5597 | 5598 | 1 | 1.4000e-01 | 3.9246e+05 | ; | CG - CD1 |
| 5597 | 5607 | 1 | 1.4000e-01 | 3.9246e+05 | ; | CG - CD2 |
| 5598 | 5599 | 1 | 1.0800e-01 | 3.0711e+05 | ; | CD1 - HD1 |
| 5598 | 5600 | 1 | 1.4000e-01 | 3.9246e+05 | ; | CD1 - CE1 |
| 5600 | 5601 | 1 | 1.0800e-01 | 3.0711e+05 | ; | CE1 - HE1 |
| 5600 | 5602 | 1 | 1.4090e-01 | 3.9246e+05 | ; | CE1 - CZ |
| 5602 | 5603 | 1 | 1.3640e-01 | 3.7656e+05 | ; | CZ - OH |
| 5602 | 5605 | 1 | 1.4090e-01 | 3.9246e+05 | ; | CZ - CE2 |
| 5603 | 5604 | 1 | 9.6000e-02 | 4.6275e+05 | ; | OH - HH |
| 5605 | 5606 | 1 | 1.0800e-01 | 3.0711e+05 | ; | CE2 - HE2 |
| 5605 | 5607 | 1 | 1.4000e-01 | 3.9246e+05 | ; | CE2 - CD2 |
| 5607 | 5608 | 1 | 1.0800e-01 | 3.0711e+05 | ; | CD2 - HD2 |
| 5609 | 5610 | 1 | 1.2290e-01 | 4.7698e+05 | ; | C - O |
| 5609 | 5611 | 1 | 1.3350e-01 | 4.1003e+05 | ; | C - N |

| | | | | | |
|---|---|---|---|---|---|
| 5611 | 5612 | 1 | 1.0100e-01 | 3.6317e+05 ; | N - H |
| 5611 | 5613 | 1 | 1.4490e-01 | 2.8200e+05 ; | N - CA |
| 5613 | 5614 | 1 | 1.0900e-01 | 2.8451e+05 ; | CA - HA |
| 5613 | 5615 | 1 | 1.5260e-01 | 2.5941e+05 ; | CA - CB |
| 5613 | 5626 | 1 | 1.5220e-01 | 2.6527e+05 ; | CA - C |
| 5615 | 5616 | 1 | 1.0900e-01 | 2.8451e+05 ; | CB - HB2 |
| 5615 | 5617 | 1 | 1.0900e-01 | 2.8451e+05 ; | CB - HB3 |
| 5615 | 5618 | 1 | 1.5260e-01 | 2.5941e+05 ; | CB - CG |
| 5618 | 5619 | 1 | 1.0900e-01 | 2.8451e+05 ; | CG - HG2 |
| 5618 | 5620 | 1 | 1.0900e-01 | 2.8451e+05 ; | CG - HG3 |
| 5618 | 5621 | 1 | 1.5220e-01 | 2.6527e+05 ; | CG - CD |
| 5621 | 5622 | 1 | 1.2290e-01 | 4.7698e+05 ; | CD - OE1 |
| 5621 | 5623 | 1 | 1.3350e-01 | 4.1003e+05 ; | CD - NE2 |
| 5623 | 5624 | 1 | 1.0100e-01 | 3.6317e+05 ; | NE2 - HE21 |
| 5623 | 5625 | 1 | 1.0100e-01 | 3.6317e+05 ; | NE2 - HE22 |
| 5626 | 5627 | 1 | 1.2290e-01 | 4.7698e+05 ; | C - O |
| 5626 | 5628 | 1 | 1.3350e-01 | 4.1003e+05 ; | C - N |
| 5628 | 5629 | 1 | 1.0100e-01 | 3.6317e+05 ; | N - H |
| 5628 | 5630 | 1 | 1.4490e-01 | 2.8200e+05 ; | N - CA |
| 5630 | 5631 | 1 | 1.0900e-01 | 2.8451e+05 ; | CA - HA |
| 5630 | 5632 | 1 | 1.5260e-01 | 2.5941e+05 ; | CA - CB |
| 5630 | 5648 | 1 | 1.5220e-01 | 2.6527e+05 ; | CA - C |
| 5632 | 5633 | 1 | 1.0900e-01 | 2.8451e+05 ; | CB - HB2 |
| 5632 | 5634 | 1 | 1.0900e-01 | 2.8451e+05 ; | CB - HB3 |
| 5632 | 5635 | 1 | 1.5260e-01 | 2.5941e+05 ; | CB - CG |
| 5635 | 5636 | 1 | 1.0900e-01 | 2.8451e+05 ; | CG - HG2 |
| 5635 | 5637 | 1 | 1.0900e-01 | 2.8451e+05 ; | CG - HG3 |
| 5635 | 5638 | 1 | 1.5260e-01 | 2.5941e+05 ; | CG - CD |
| 5638 | 5639 | 1 | 1.0900e-01 | 2.8451e+05 ; | CD - HD2 |
| 5638 | 5640 | 1 | 1.0900e-01 | 2.8451e+05 ; | CD - HD3 |
| 5638 | 5641 | 1 | 1.5260e-01 | 2.5941e+05 ; | CD - CE |
| 5641 | 5642 | 1 | 1.0900e-01 | 2.8451e+05 ; | CE - HE2 |
| 5641 | 5643 | 1 | 1.0900e-01 | 2.8451e+05 ; | CE - HE3 |
| 5641 | 5644 | 1 | 1.4710e-01 | 3.0711e+05 ; | CE - NZ |
| 5644 | 5645 | 1 | 1.0100e-01 | 3.6317e+05 ; | NZ - HZ1 |
| 5644 | 5646 | 1 | 1.0100e-01 | 3.6317e+05 ; | NZ - HZ2 |
| 5644 | 5647 | 1 | 1.0100e-01 | 3.6317e+05 ; | NZ - HZ3 |
| 5648 | 5649 | 1 | 1.2290e-01 | 4.7698e+05 ; | C - O |
| 5648 | 5650 | 1 | 1.3350e-01 | 4.1003e+05 ; | C - N |
| 5650 | 5651 | 1 | 1.0100e-01 | 3.6317e+05 ; | N - H |
| 5650 | 5652 | 1 | 1.4490e-01 | 2.8200e+05 ; | N - CA |
| 5652 | 5653 | 1 | 1.0900e-01 | 2.8451e+05 ; | CA - HA |
| 5652 | 5654 | 1 | 1.5260e-01 | 2.5941e+05 ; | CA - CB |
| 5652 | 5667 | 1 | 1.5220e-01 | 2.6527e+05 ; | CA - C |
| 5654 | 5655 | 1 | 1.0900e-01 | 2.8451e+05 ; | CB - HB |
| 5654 | 5656 | 1 | 1.5260e-01 | 2.5941e+05 ; | CB - CG2 |
| 5654 | 5660 | 1 | 1.5260e-01 | 2.5941e+05 ; | CB - CG1 |
| 5656 | 5657 | 1 | 1.0900e-01 | 2.8451e+05 ; | CG2 - HG21 |
| 5656 | 5658 | 1 | 1.0900e-01 | 2.8451e+05 ; | CG2 - HG22 |
| 5656 | 5659 | 1 | 1.0900e-01 | 2.8451e+05 ; | CG2 - HG23 |
| 5660 | 5661 | 1 | 1.0900e-01 | 2.8451e+05 ; | CG1 - HG12 |
| 5660 | 5662 | 1 | 1.0900e-01 | 2.8451e+05 ; | CG1 - HG13 |

| | | | | | | |
|---|---|---|---|---|---|---|
| 5660 | 5663 | 1 | 1.5260e-01 | 2.5941e+05 | ; | CG1 - CD1 |
| 5663 | 5664 | 1 | 1.0900e-01 | 2.8451e+05 | ; | CD1 - HD11 |
| 5663 | 5665 | 1 | 1.0900e-01 | 2.8451e+05 | ; | CD1 - HD12 |
| 5663 | 5666 | 1 | 1.0900e-01 | 2.8451e+05 | ; | CD1 - HD13 |
| 5667 | 5668 | 1 | 1.2290e-01 | 4.7698e+05 | ; | C - O |
| 5667 | 5669 | 1 | 1.3350e-01 | 4.1003e+05 | ; | C - N |
| 5669 | 5670 | 1 | 1.4490e-01 | 2.8200e+05 | ; | N - CD |
| 5669 | 5679 | 1 | 1.4490e-01 | 2.8200e+05 | ; | N - CA |
| 5670 | 5671 | 1 | 1.0900e-01 | 2.8451e+05 | ; | CD - HD2 |
| 5670 | 5672 | 1 | 1.0900e-01 | 2.8451e+05 | ; | CD - HD3 |
| 5670 | 5673 | 1 | 1.5260e-01 | 2.5941e+05 | ; | CD - CG |
| 5673 | 5674 | 1 | 1.0900e-01 | 2.8451e+05 | ; | CG - HG2 |
| 5673 | 5675 | 1 | 1.0900e-01 | 2.8451e+05 | ; | CG - HG3 |
| 5673 | 5676 | 1 | 1.5260e-01 | 2.5941e+05 | ; | CG - CB |
| 5676 | 5677 | 1 | 1.0900e-01 | 2.8451e+05 | ; | CB - HB2 |
| 5676 | 5678 | 1 | 1.0900e-01 | 2.8451e+05 | ; | CB - HB3 |
| 5676 | 5679 | 1 | 1.5260e-01 | 2.5941e+05 | ; | CB - CA |
| 5679 | 5680 | 1 | 1.0900e-01 | 2.8451e+05 | ; | CA - HA |
| 5679 | 5681 | 1 | 1.5220e-01 | 2.6527e+05 | ; | CA - C |
| 5681 | 5682 | 1 | 1.2290e-01 | 4.7698e+05 | ; | C - O |
| 5681 | 5683 | 1 | 1.3350e-01 | 4.1003e+05 | ; | C - N |
| 5683 | 5684 | 1 | 1.4490e-01 | 2.8200e+05 | ; | N - CD |
| 5683 | 5693 | 1 | 1.4490e-01 | 2.8200e+05 | ; | N - CA |
| 5684 | 5685 | 1 | 1.0900e-01 | 2.8451e+05 | ; | CD - HD2 |
| 5684 | 5686 | 1 | 1.0900e-01 | 2.8451e+05 | ; | CD - HD3 |
| 5684 | 5687 | 1 | 1.5260e-01 | 2.5941e+05 | ; | CD - CG |
| 5687 | 5688 | 1 | 1.0900e-01 | 2.8451e+05 | ; | CG - HG2 |
| 5687 | 5689 | 1 | 1.0900e-01 | 2.8451e+05 | ; | CG - HG3 |
| 5687 | 5690 | 1 | 1.5260e-01 | 2.5941e+05 | ; | CG - CB |
| 5690 | 5691 | 1 | 1.0900e-01 | 2.8451e+05 | ; | CB - HB2 |
| 5690 | 5692 | 1 | 1.0900e-01 | 2.8451e+05 | ; | CB - HB3 |
| 5690 | 5693 | 1 | 1.5260e-01 | 2.5941e+05 | ; | CB - CA |
| 5693 | 5694 | 1 | 1.0900e-01 | 2.8451e+05 | ; | CA - HA |
| 5693 | 5695 | 1 | 1.5220e-01 | 2.6527e+05 | ; | CA - C |
| 5695 | 5696 | 1 | 1.2290e-01 | 4.7698e+05 | ; | C - O |
| 5695 | 5697 | 1 | 1.3350e-01 | 4.1003e+05 | ; | C - N |
| 5697 | 5698 | 1 | 1.0100e-01 | 3.6317e+05 | ; | N - H |
| 5697 | 5699 | 1 | 1.4490e-01 | 2.8200e+05 | ; | N - CA |
| 5699 | 5700 | 1 | 1.0900e-01 | 2.8451e+05 | ; | CA - HA |
| 5699 | 5701 | 1 | 1.5260e-01 | 2.5941e+05 | ; | CA - CB |
| 5699 | 5707 | 1 | 1.5220e-01 | 2.6527e+05 | ; | CA - C |
| 5701 | 5702 | 1 | 1.0900e-01 | 2.8451e+05 | ; | CB - HB2 |
| 5701 | 5703 | 1 | 1.0900e-01 | 2.8451e+05 | ; | CB - HB3 |
| 5701 | 5704 | 1 | 1.5220e-01 | 2.6527e+05 | ; | CB - CG |
| 5704 | 5705 | 1 | 1.2500e-01 | 5.4894e+05 | ; | CG - OD1 |
| 5704 | 5706 | 1 | 1.2500e-01 | 5.4894e+05 | ; | CG - OD2 |
| 5707 | 5708 | 1 | 1.2290e-01 | 4.7698e+05 | ; | C - O |
| 5707 | 5709 | 1 | 1.3350e-01 | 4.1003e+05 | ; | C - N |
| 5709 | 5710 | 1 | 1.0100e-01 | 3.6317e+05 | ; | N - H |
| 5709 | 5711 | 1 | 1.4490e-01 | 2.8200e+05 | ; | N - CA |
| 5711 | 5712 | 1 | 1.0900e-01 | 2.8451e+05 | ; | CA - HA |
| 5711 | 5713 | 1 | 1.5260e-01 | 2.5941e+05 | ; | CA - CB |

| | | | | | | |
|---|---|---|---|---|---|---|
| 5711 | 5726 | 1 | 1.5220e-01 | 2.6527e+05 | ; | CA - C |
| 5713 | 5714 | 1 | 1.0900e-01 | 2.8451e+05 | ; | CB - HB |
| 5713 | 5715 | 1 | 1.5260e-01 | 2.5941e+05 | ; | CB - CG2 |
| 5713 | 5719 | 1 | 1.5260e-01 | 2.5941e+05 | ; | CB - CG1 |
| 5715 | 5716 | 1 | 1.0900e-01 | 2.8451e+05 | ; | CG2 - HG21 |
| 5715 | 5717 | 1 | 1.0900e-01 | 2.8451e+05 | ; | CG2 - HG22 |
| 5715 | 5718 | 1 | 1.0900e-01 | 2.8451e+05 | ; | CG2 - HG23 |
| 5719 | 5720 | 1 | 1.0900e-01 | 2.8451e+05 | ; | CG1 - HG12 |
| 5719 | 5721 | 1 | 1.0900e-01 | 2.8451e+05 | ; | CG1 - HG13 |
| 5719 | 5722 | 1 | 1.5260e-01 | 2.5941e+05 | ; | CG1 - CD1 |
| 5722 | 5723 | 1 | 1.0900e-01 | 2.8451e+05 | ; | CD1 - HD11 |
| 5722 | 5724 | 1 | 1.0900e-01 | 2.8451e+05 | ; | CD1 - HD12 |
| 5722 | 5725 | 1 | 1.0900e-01 | 2.8451e+05 | ; | CD1 - HD13 |
| 5726 | 5727 | 1 | 1.2290e-01 | 4.7698e+05 | ; | C - O |
| 5726 | 5728 | 1 | 1.3350e-01 | 4.1003e+05 | ; | C - N |
| 5728 | 5729 | 1 | 1.0100e-01 | 3.6317e+05 | ; | N - H |
| 5728 | 5730 | 1 | 1.4490e-01 | 2.8200e+05 | ; | N - CA |
| 5730 | 5731 | 1 | 1.0900e-01 | 2.8451e+05 | ; | CA - HA |
| 5730 | 5732 | 1 | 1.5260e-01 | 2.5941e+05 | ; | CA - CB |
| 5730 | 5743 | 1 | 1.5220e-01 | 2.6527e+05 | ; | CA - C |
| 5732 | 5733 | 1 | 1.0900e-01 | 2.8451e+05 | ; | CB - HB2 |
| 5732 | 5734 | 1 | 1.0900e-01 | 2.8451e+05 | ; | CB - HB3 |
| 5732 | 5735 | 1 | 1.5040e-01 | 2.6527e+05 | ; | CB - CG |
| 5735 | 5736 | 1 | 1.3940e-01 | 3.4309e+05 | ; | CG - ND1 |
| 5735 | 5741 | 1 | 1.3710e-01 | 4.3346e+05 | ; | CG - CD2 |
| 5736 | 5737 | 1 | 1.3350e-01 | 4.0836e+05 | ; | ND1 - CE1 |
| 5737 | 5738 | 1 | 1.0800e-01 | 3.0711e+05 | ; | CE1 - HE1 |
| 5737 | 5739 | 1 | 1.3430e-01 | 3.9915e+05 | ; | CE1 - NE2 |
| 5739 | 5740 | 1 | 1.0100e-01 | 3.6317e+05 | ; | NE2 - HE2 |
| 5739 | 5741 | 1 | 1.3810e-01 | 3.5731e+05 | ; | NE2 - CD2 |
| 5741 | 5742 | 1 | 1.0800e-01 | 3.0711e+05 | ; | CD2 - HD2 |
| 5743 | 5744 | 1 | 1.2290e-01 | 4.7698e+05 | ; | C - O |
| 5743 | 5745 | 1 | 1.3350e-01 | 4.1003e+05 | ; | C - N |
| 5745 | 5746 | 1 | 1.0100e-01 | 3.6317e+05 | ; | N - H |
| 5745 | 5747 | 1 | 1.4490e-01 | 2.8200e+05 | ; | N - CA |
| 5747 | 5748 | 1 | 1.0900e-01 | 2.8451e+05 | ; | CA - HA |
| 5747 | 5749 | 1 | 1.5260e-01 | 2.5941e+05 | ; | CA - CB |
| 5747 | 5760 | 1 | 1.5220e-01 | 2.6527e+05 | ; | CA - C |
| 5749 | 5750 | 1 | 1.0900e-01 | 2.8451e+05 | ; | CB - HB2 |
| 5749 | 5751 | 1 | 1.0900e-01 | 2.8451e+05 | ; | CB - HB3 |
| 5749 | 5752 | 1 | 1.5260e-01 | 2.5941e+05 | ; | CB - CG |
| 5752 | 5753 | 1 | 1.0900e-01 | 2.8451e+05 | ; | CG - HG2 |
| 5752 | 5754 | 1 | 1.0900e-01 | 2.8451e+05 | ; | CG - HG3 |
| 5752 | 5755 | 1 | 1.5220e-01 | 2.6527e+05 | ; | CG - CD |
| 5755 | 5756 | 1 | 1.2290e-01 | 4.7698e+05 | ; | CD - OE1 |
| 5755 | 5757 | 1 | 1.3350e-01 | 4.1003e+05 | ; | CD - NE2 |
| 5757 | 5758 | 1 | 1.0100e-01 | 3.6317e+05 | ; | NE2 - HE21 |
| 5757 | 5759 | 1 | 1.0100e-01 | 3.6317e+05 | ; | NE2 - HE22 |
| 5760 | 5761 | 1 | 1.2290e-01 | 4.7698e+05 | ; | C - O |
| 5760 | 5762 | 1 | 1.3350e-01 | 4.1003e+05 | ; | C - N |
| 5762 | 5763 | 1 | 1.0100e-01 | 3.6317e+05 | ; | N - H |
| 5762 | 5764 | 1 | 1.4490e-01 | 2.8200e+05 | ; | N - CA |

| | | | | | | | | |
|---|---|---|---|---|---|---|---|---|
| 5764 | 5765 | 1 | 1.0900e-01 | 2.8451e+05 | ; | CA | - | HA |
| 5764 | 5766 | 1 | 1.5260e-01 | 2.5941e+05 | ; | CA | - | CB |
| 5764 | 5776 | 1 | 1.5220e-01 | 2.6527e+05 | ; | CA | - | C |
| 5766 | 5767 | 1 | 1.0900e-01 | 2.8451e+05 | ; | CB | - | HB |
| 5766 | 5768 | 1 | 1.5260e-01 | 2.5941e+05 | ; | CB | - | CG1 |
| 5766 | 5772 | 1 | 1.5260e-01 | 2.5941e+05 | ; | CB | - | CG2 |
| 5768 | 5769 | 1 | 1.0900e-01 | 2.8451e+05 | ; | CG1 | - | HG11 |
| 5768 | 5770 | 1 | 1.0900e-01 | 2.8451e+05 | ; | CG1 | - | HG12 |
| 5768 | 5771 | 1 | 1.0900e-01 | 2.8451e+05 | ; | CG1 | - | HG13 |
| 5772 | 5773 | 1 | 1.0900e-01 | 2.8451e+05 | ; | CG2 | - | HG21 |
| 5772 | 5774 | 1 | 1.0900e-01 | 2.8451e+05 | ; | CG2 | - | HG22 |
| 5772 | 5775 | 1 | 1.0900e-01 | 2.8451e+05 | ; | CG2 | - | HG23 |
| 5776 | 5777 | 1 | 1.2290e-01 | 4.7698e+05 | ; | C | - | O |
| 5776 | 5778 | 1 | 1.3350e-01 | 4.1003e+05 | ; | C | - | N |
| 5778 | 5779 | 1 | 1.0100e-01 | 3.6317e+05 | ; | N | - | H |
| 5778 | 5780 | 1 | 1.4490e-01 | 2.8200e+05 | ; | N | - | CA |
| 5780 | 5781 | 1 | 1.0900e-01 | 2.8451e+05 | ; | CA | - | HA |
| 5780 | 5782 | 1 | 1.5260e-01 | 2.5941e+05 | ; | CA | - | CB |
| 5780 | 5795 | 1 | 1.5220e-01 | 2.6527e+05 | ; | CA | - | C |
| 5782 | 5783 | 1 | 1.0900e-01 | 2.8451e+05 | ; | CB | - | HB2 |
| 5782 | 5784 | 1 | 1.0900e-01 | 2.8451e+05 | ; | CB | - | HB3 |
| 5782 | 5785 | 1 | 1.5260e-01 | 2.5941e+05 | ; | CB | - | CG |
| 5785 | 5786 | 1 | 1.0900e-01 | 2.8451e+05 | ; | CG | - | HG |
| 5785 | 5787 | 1 | 1.5260e-01 | 2.5941e+05 | ; | CG | - | CD1 |
| 5785 | 5791 | 1 | 1.5260e-01 | 2.5941e+05 | ; | CG | - | CD2 |
| 5787 | 5788 | 1 | 1.0900e-01 | 2.8451e+05 | ; | CD1 | - | HD11 |
| 5787 | 5789 | 1 | 1.0900e-01 | 2.8451e+05 | ; | CD1 | - | HD12 |
| 5787 | 5790 | 1 | 1.0900e-01 | 2.8451e+05 | ; | CD1 | - | HD13 |
| 5791 | 5792 | 1 | 1.0900e-01 | 2.8451e+05 | ; | CD2 | - | HD21 |
| 5791 | 5793 | 1 | 1.0900e-01 | 2.8451e+05 | ; | CD2 | - | HD22 |
| 5791 | 5794 | 1 | 1.0900e-01 | 2.8451e+05 | ; | CD2 | - | HD23 |
| 5795 | 5796 | 1 | 1.2290e-01 | 4.7698e+05 | ; | C | - | O |
| 5795 | 5797 | 1 | 1.3350e-01 | 4.1003e+05 | ; | C | - | N |
| 5797 | 5798 | 1 | 1.0100e-01 | 3.6317e+05 | ; | N | - | H |
| 5797 | 5799 | 1 | 1.4490e-01 | 2.8200e+05 | ; | N | - | CA |
| 5799 | 5800 | 1 | 1.0900e-01 | 2.8451e+05 | ; | CA | - | HA |
| 5799 | 5801 | 1 | 1.5260e-01 | 2.5941e+05 | ; | CA | - | CB |
| 5799 | 5819 | 1 | 1.5220e-01 | 2.6527e+05 | ; | CA | - | C |
| 5801 | 5802 | 1 | 1.0900e-01 | 2.8451e+05 | ; | CB | - | HB2 |
| 5801 | 5803 | 1 | 1.0900e-01 | 2.8451e+05 | ; | CB | - | HB3 |
| 5801 | 5804 | 1 | 1.5260e-01 | 2.5941e+05 | ; | CB | - | CG |
| 5804 | 5805 | 1 | 1.0900e-01 | 2.8451e+05 | ; | CG | - | HG2 |
| 5804 | 5806 | 1 | 1.0900e-01 | 2.8451e+05 | ; | CG | - | HG3 |
| 5804 | 5807 | 1 | 1.5260e-01 | 2.5941e+05 | ; | CG | - | CD |
| 5807 | 5808 | 1 | 1.0900e-01 | 2.8451e+05 | ; | CD | - | HD2 |
| 5807 | 5809 | 1 | 1.0900e-01 | 2.8451e+05 | ; | CD | - | HD3 |
| 5807 | 5810 | 1 | 1.4630e-01 | 2.8200e+05 | ; | CD | - | NE |
| 5810 | 5811 | 1 | 1.0100e-01 | 3.6317e+05 | ; | NE | - | HE |
| 5810 | 5812 | 1 | 1.3400e-01 | 4.0250e+05 | ; | NE | - | CZ |
| 5812 | 5813 | 1 | 1.3400e-01 | 4.0250e+05 | ; | CZ | - | NH1 |
| 5812 | 5816 | 1 | 1.3400e-01 | 4.0250e+05 | ; | CZ | - | NH2 |
| 5813 | 5814 | 1 | 1.0100e-01 | 3.6317e+05 | ; | NH1 | - | HH11 |

| | | | | | | |
|---|---|---|---|---|---|---|
| 5813 | 5815 | 1 | 1.0100e-01 | 3.6317e+05 | ; | NH1 - HH12 |
| 5816 | 5817 | 1 | 1.0100e-01 | 3.6317e+05 | ; | NH2 - HH21 |
| 5816 | 5818 | 1 | 1.0100e-01 | 3.6317e+05 | ; | NH2 - HH22 |
| 5819 | 5820 | 1 | 1.2290e-01 | 4.7698e+05 | ; | C - O |
| 5819 | 5821 | 1 | 1.3350e-01 | 4.1003e+05 | ; | C - N |
| 5821 | 5822 | 1 | 1.0100e-01 | 3.6317e+05 | ; | N - H |
| 5821 | 5823 | 1 | 1.4490e-01 | 2.8200e+05 | ; | N - CA |
| 5823 | 5824 | 1 | 1.0900e-01 | 2.8451e+05 | ; | CA - HA |
| 5823 | 5825 | 1 | 1.5260e-01 | 2.5941e+05 | ; | CA - CB |
| 5823 | 5831 | 1 | 1.5220e-01 | 2.6527e+05 | ; | CA - C |
| 5825 | 5826 | 1 | 1.0900e-01 | 2.8451e+05 | ; | CB - HB2 |
| 5825 | 5827 | 1 | 1.0900e-01 | 2.8451e+05 | ; | CB - HB3 |
| 5825 | 5828 | 1 | 1.5220e-01 | 2.6527e+05 | ; | CB - CG |
| 5828 | 5829 | 1 | 1.2500e-01 | 5.4894e+05 | ; | CG - OD1 |
| 5828 | 5830 | 1 | 1.2500e-01 | 5.4894e+05 | ; | CG - OD2 |
| 5831 | 5832 | 1 | 1.2290e-01 | 4.7698e+05 | ; | C - O |
| 5831 | 5833 | 1 | 1.3350e-01 | 4.1003e+05 | ; | C - N |
| 5833 | 5834 | 1 | 1.0100e-01 | 3.6317e+05 | ; | N - H |
| 5833 | 5835 | 1 | 1.4490e-01 | 2.8200e+05 | ; | N - CA |
| 5835 | 5836 | 1 | 1.0900e-01 | 2.8451e+05 | ; | CA - HA |
| 5835 | 5837 | 1 | 1.5260e-01 | 2.5941e+05 | ; | CA - CB |
| 5835 | 5850 | 1 | 1.5220e-01 | 2.6527e+05 | ; | CA - C |
| 5837 | 5838 | 1 | 1.0900e-01 | 2.8451e+05 | ; | CB - HB |
| 5837 | 5839 | 1 | 1.5260e-01 | 2.5941e+05 | ; | CB - CG2 |
| 5837 | 5843 | 1 | 1.5260e-01 | 2.5941e+05 | ; | CB - CG1 |
| 5839 | 5840 | 1 | 1.0900e-01 | 2.8451e+05 | ; | CG2 - HG21 |
| 5839 | 5841 | 1 | 1.0900e-01 | 2.8451e+05 | ; | CG2 - HG22 |
| 5839 | 5842 | 1 | 1.0900e-01 | 2.8451e+05 | ; | CG2 - HG23 |
| 5843 | 5844 | 1 | 1.0900e-01 | 2.8451e+05 | ; | CG1 - HG12 |
| 5843 | 5845 | 1 | 1.0900e-01 | 2.8451e+05 | ; | CG1 - HG13 |
| 5843 | 5846 | 1 | 1.5260e-01 | 2.5941e+05 | ; | CG1 - CD1 |
| 5846 | 5847 | 1 | 1.0900e-01 | 2.8451e+05 | ; | CD1 - HD11 |
| 5846 | 5848 | 1 | 1.0900e-01 | 2.8451e+05 | ; | CD1 - HD12 |
| 5846 | 5849 | 1 | 1.0900e-01 | 2.8451e+05 | ; | CD1 - HD13 |
| 5850 | 5851 | 1 | 1.2290e-01 | 4.7698e+05 | ; | C - O |
| 5850 | 5852 | 1 | 1.3350e-01 | 4.1003e+05 | ; | C - N |
| 5852 | 5853 | 1 | 1.0100e-01 | 3.6317e+05 | ; | N - H |
| 5852 | 5854 | 1 | 1.4490e-01 | 2.8200e+05 | ; | N - CA |
| 5854 | 5855 | 1 | 1.0900e-01 | 2.8451e+05 | ; | CA - HA |
| 5854 | 5856 | 1 | 1.5260e-01 | 2.5941e+05 | ; | CA - CB |
| 5854 | 5862 | 1 | 1.5220e-01 | 2.6527e+05 | ; | CA - C |
| 5856 | 5857 | 1 | 1.0900e-01 | 2.8451e+05 | ; | CB - HB2 |
| 5856 | 5858 | 1 | 1.0900e-01 | 2.8451e+05 | ; | CB - HB3 |
| 5856 | 5859 | 1 | 1.5220e-01 | 2.6527e+05 | ; | CB - CG |
| 5859 | 5860 | 1 | 1.2500e-01 | 5.4894e+05 | ; | CG - OD1 |
| 5859 | 5861 | 1 | 1.2500e-01 | 5.4894e+05 | ; | CG - OD2 |
| 5862 | 5863 | 1 | 1.2290e-01 | 4.7698e+05 | ; | C - O |
| 5862 | 5864 | 1 | 1.3350e-01 | 4.1003e+05 | ; | C - N |
| 5864 | 5865 | 1 | 1.0100e-01 | 3.6317e+05 | ; | N - H |
| 5864 | 5866 | 1 | 1.4490e-01 | 2.8200e+05 | ; | N - CA |
| 5866 | 5867 | 1 | 1.0900e-01 | 2.8451e+05 | ; | CA - HA |
| 5866 | 5868 | 1 | 1.5260e-01 | 2.5941e+05 | ; | CA - CB |

| | | | | | | |
|---|---|---|---|---|---|---|
| 5866 | 5873 | 1 | 1.5220e-01 | 2.6527e+05 | ; | CA - C |
| 5868 | 5869 | 1 | 1.0900e-01 | 2.8451e+05 | ; | CB - HB2 |
| 5868 | 5870 | 1 | 1.0900e-01 | 2.8451e+05 | ; | CB - HB3 |
| 5868 | 5871 | 1 | 1.4100e-01 | 2.6778e+05 | ; | CB - OG |
| 5871 | 5872 | 1 | 9.6000e-02 | 4.6275e+05 | ; | OG - HG |
| 5873 | 5874 | 1 | 1.2290e-01 | 4.7698e+05 | ; | C - O |
| 5873 | 5875 | 1 | 1.3350e-01 | 4.1003e+05 | ; | C - N |
| 5875 | 5876 | 1 | 1.0100e-01 | 3.6317e+05 | ; | N - H |
| 5875 | 5877 | 1 | 1.4490e-01 | 2.8200e+05 | ; | N - CA |
| 5877 | 5878 | 1 | 1.0900e-01 | 2.8451e+05 | ; | CA - HA |
| 5877 | 5879 | 1 | 1.5260e-01 | 2.5941e+05 | ; | CA - CB |
| 5877 | 5887 | 1 | 1.5220e-01 | 2.6527e+05 | ; | CA - C |
| 5879 | 5880 | 1 | 1.0900e-01 | 2.8451e+05 | ; | CB - HB2 |
| 5879 | 5881 | 1 | 1.0900e-01 | 2.8451e+05 | ; | CB - HB3 |
| 5879 | 5882 | 1 | 1.5220e-01 | 2.6527e+05 | ; | CB - CG |
| 5882 | 5883 | 1 | 1.2290e-01 | 4.7698e+05 | ; | CG - OD1 |
| 5882 | 5884 | 1 | 1.3350e-01 | 4.1003e+05 | ; | CG - ND2 |
| 5884 | 5885 | 1 | 1.0100e-01 | 3.6317e+05 | ; | ND2 - HD21 |
| 5884 | 5886 | 1 | 1.0100e-01 | 3.6317e+05 | ; | ND2 - HD22 |
| 5887 | 5888 | 1 | 1.2290e-01 | 4.7698e+05 | ; | C - O |
| 5887 | 5889 | 1 | 1.3350e-01 | 4.1003e+05 | ; | C - N |
| 5889 | 5890 | 1 | 1.0100e-01 | 3.6317e+05 | ; | N - H |
| 5889 | 5891 | 1 | 1.4490e-01 | 2.8200e+05 | ; | N - CA |
| 5891 | 5892 | 1 | 1.0900e-01 | 2.8451e+05 | ; | CA - HA |
| 5891 | 5893 | 1 | 1.5260e-01 | 2.5941e+05 | ; | CA - CB |
| 5891 | 5897 | 1 | 1.5220e-01 | 2.6527e+05 | ; | CA - C |
| 5893 | 5894 | 1 | 1.0900e-01 | 2.8451e+05 | ; | CB - HB1 |
| 5893 | 5895 | 1 | 1.0900e-01 | 2.8451e+05 | ; | CB - HB2 |
| 5893 | 5896 | 1 | 1.0900e-01 | 2.8451e+05 | ; | CB - HB3 |
| 5897 | 5898 | 1 | 1.2290e-01 | 4.7698e+05 | ; | C - O |
| 5897 | 5899 | 1 | 1.3350e-01 | 4.1003e+05 | ; | C - N |
| 5899 | 5900 | 1 | 1.0100e-01 | 3.6317e+05 | ; | N - H |
| 5899 | 5901 | 1 | 1.4490e-01 | 2.8200e+05 | ; | N - CA |
| 5901 | 5902 | 1 | 1.0900e-01 | 2.8451e+05 | ; | CA - HA |
| 5901 | 5903 | 1 | 1.5260e-01 | 2.5941e+05 | ; | CA - CB |
| 5901 | 5908 | 1 | 1.5220e-01 | 2.6527e+05 | ; | CA - C |
| 5903 | 5904 | 1 | 1.0900e-01 | 2.8451e+05 | ; | CB - HB2 |
| 5903 | 5905 | 1 | 1.0900e-01 | 2.8451e+05 | ; | CB - HB3 |
| 5903 | 5906 | 1 | 1.4100e-01 | 2.6778e+05 | ; | CB - OG |
| 5906 | 5907 | 1 | 9.6000e-02 | 4.6275e+05 | ; | OG - HG |
| 5908 | 5909 | 1 | 1.2290e-01 | 4.7698e+05 | ; | C - O |
| 5908 | 5910 | 1 | 1.3350e-01 | 4.1003e+05 | ; | C - N |
| 5910 | 5911 | 1 | 1.0100e-01 | 3.6317e+05 | ; | N - H |
| 5910 | 5912 | 1 | 1.4490e-01 | 2.8200e+05 | ; | N - CA |
| 5912 | 5913 | 1 | 1.0900e-01 | 2.8451e+05 | ; | CA - HA2 |
| 5912 | 5914 | 1 | 1.0900e-01 | 2.8451e+05 | ; | CA - HA3 |
| 5912 | 5915 | 1 | 1.5220e-01 | 2.6527e+05 | ; | CA - C |
| 5915 | 5916 | 1 | 1.2290e-01 | 4.7698e+05 | ; | C - O |
| 5915 | 5917 | 1 | 1.3350e-01 | 4.1003e+05 | ; | C - N |
| 5917 | 5918 | 1 | 1.0100e-01 | 3.6317e+05 | ; | N - H |
| 5917 | 5919 | 1 | 1.4490e-01 | 2.8200e+05 | ; | N - CA |
| 5919 | 5920 | 1 | 1.0900e-01 | 2.8451e+05 | ; | CA - HA |

| | | | | | |
|---|---|---|---|---|---|
| 5919 | 5921 | 1 | 1.5260e-01 | 2.5941e+05 ; | CA - CB |
| 5919 | 5932 | 1 | 1.5220e-01 | 2.6527e+05 ; | CA - C |
| 5921 | 5922 | 1 | 1.0900e-01 | 2.8451e+05 ; | CB - HB2 |
| 5921 | 5923 | 1 | 1.0900e-01 | 2.8451e+05 ; | CB - HB3 |
| 5921 | 5924 | 1 | 1.5260e-01 | 2.5941e+05 ; | CB - CG |
| 5924 | 5925 | 1 | 1.0900e-01 | 2.8451e+05 ; | CG - HG2 |
| 5924 | 5926 | 1 | 1.0900e-01 | 2.8451e+05 ; | CG - HG3 |
| 5924 | 5927 | 1 | 1.5220e-01 | 2.6527e+05 ; | CG - CD |
| 5927 | 5928 | 1 | 1.2290e-01 | 4.7698e+05 ; | CD - OE1 |
| 5927 | 5929 | 1 | 1.3350e-01 | 4.1003e+05 ; | CD - NE2 |
| 5929 | 5930 | 1 | 1.0100e-01 | 3.6317e+05 ; | NE2 - HE21 |
| 5929 | 5931 | 1 | 1.0100e-01 | 3.6317e+05 ; | NE2 - HE22 |
| 5932 | 5933 | 1 | 1.2290e-01 | 4.7698e+05 ; | C - O |
| 5932 | 5934 | 1 | 1.3350e-01 | 4.1003e+05 ; | C - N |
| 5934 | 5935 | 1 | 1.0100e-01 | 3.6317e+05 ; | N - H |
| 5934 | 5936 | 1 | 1.4490e-01 | 2.8200e+05 ; | N - CA |
| 5936 | 5937 | 1 | 1.0900e-01 | 2.8451e+05 ; | CA - HA |
| 5936 | 5938 | 1 | 1.5260e-01 | 2.5941e+05 ; | CA - CB |
| 5936 | 5951 | 1 | 1.5220e-01 | 2.6527e+05 ; | CA - C |
| 5938 | 5939 | 1 | 1.0900e-01 | 2.8451e+05 ; | CB - HB |
| 5938 | 5940 | 1 | 1.5260e-01 | 2.5941e+05 ; | CB - CG2 |
| 5938 | 5944 | 1 | 1.5260e-01 | 2.5941e+05 ; | CB - CG1 |
| 5940 | 5941 | 1 | 1.0900e-01 | 2.8451e+05 ; | CG2 - HG21 |
| 5940 | 5942 | 1 | 1.0900e-01 | 2.8451e+05 ; | CG2 - HG22 |
| 5940 | 5943 | 1 | 1.0900e-01 | 2.8451e+05 ; | CG2 - HG23 |
| 5944 | 5945 | 1 | 1.0900e-01 | 2.8451e+05 ; | CG1 - HG12 |
| 5944 | 5946 | 1 | 1.0900e-01 | 2.8451e+05 ; | CG1 - HG13 |
| 5944 | 5947 | 1 | 1.5260e-01 | 2.5941e+05 ; | CG1 - CD1 |
| 5947 | 5948 | 1 | 1.0900e-01 | 2.8451e+05 ; | CD1 - HD11 |
| 5947 | 5949 | 1 | 1.0900e-01 | 2.8451e+05 ; | CD1 - HD12 |
| 5947 | 5950 | 1 | 1.0900e-01 | 2.8451e+05 ; | CD1 - HD13 |
| 5951 | 5952 | 1 | 1.2290e-01 | 4.7698e+05 ; | C - O |
| 5951 | 5953 | 1 | 1.3350e-01 | 4.1003e+05 ; | C - N |
| 5953 | 5954 | 1 | 1.0100e-01 | 3.6317e+05 ; | N - H |
| 5953 | 5955 | 1 | 1.4490e-01 | 2.8200e+05 ; | N - CA |
| 5955 | 5956 | 1 | 1.0900e-01 | 2.8451e+05 ; | CA - HA |
| 5955 | 5957 | 1 | 1.5260e-01 | 2.5941e+05 ; | CA - CB |
| 5955 | 5969 | 1 | 1.5220e-01 | 2.6527e+05 ; | CA - C |
| 5957 | 5958 | 1 | 1.0900e-01 | 2.8451e+05 ; | CB - HB2 |
| 5957 | 5959 | 1 | 1.0900e-01 | 2.8451e+05 ; | CB - HB3 |
| 5957 | 5960 | 1 | 1.5040e-01 | 2.6527e+05 ; | CB - CG |
| 5960 | 5961 | 1 | 1.3850e-01 | 3.5313e+05 ; | CG - ND1 |
| 5960 | 5967 | 1 | 1.3710e-01 | 4.3346e+05 ; | CG - CD2 |
| 5961 | 5962 | 1 | 1.0100e-01 | 3.6317e+05 ; | ND1 - HD1 |
| 5961 | 5963 | 1 | 1.3430e-01 | 3.9915e+05 ; | ND1 - CE1 |
| 5963 | 5964 | 1 | 1.0800e-01 | 3.0711e+05 ; | CE1 - HE1 |
| 5963 | 5965 | 1 | 1.3430e-01 | 3.9915e+05 ; | CE1 - NE2 |
| 5965 | 5966 | 1 | 1.0100e-01 | 3.6317e+05 ; | NE2 - HE2 |
| 5965 | 5967 | 1 | 1.3810e-01 | 3.5731e+05 ; | NE2 - CD2 |
| 5967 | 5968 | 1 | 1.0800e-01 | 3.0711e+05 ; | CD2 - HD2 |
| 5969 | 5970 | 1 | 1.2290e-01 | 4.7698e+05 ; | C - O |
| 5969 | 5971 | 1 | 1.3350e-01 | 4.1003e+05 ; | C - N |

| | | | | | | |
|---|---|---|---|---|---|---|
| 5971 | 5972 | 1 | 1.0100e-01 | 3.6317e+05 | ; | N - H |
| 5971 | 5973 | 1 | 1.4490e-01 | 2.8200e+05 | ; | N - CA |
| 5973 | 5974 | 1 | 1.0900e-01 | 2.8451e+05 | ; | CA - HA |
| 5973 | 5975 | 1 | 1.5260e-01 | 2.5941e+05 | ; | CA - CB |
| 5973 | 5990 | 1 | 1.5220e-01 | 2.6527e+05 | ; | CA - C |
| 5975 | 5976 | 1 | 1.0900e-01 | 2.8451e+05 | ; | CB - HB2 |
| 5975 | 5977 | 1 | 1.0900e-01 | 2.8451e+05 | ; | CB - HB3 |
| 5975 | 5978 | 1 | 1.5100e-01 | 2.6527e+05 | ; | CB - CG |
| 5978 | 5979 | 1 | 1.4000e-01 | 3.9246e+05 | ; | CG - CD1 |
| 5978 | 5988 | 1 | 1.4000e-01 | 3.9246e+05 | ; | CG - CD2 |
| 5979 | 5980 | 1 | 1.0800e-01 | 3.0711e+05 | ; | CD1 - HD1 |
| 5979 | 5981 | 1 | 1.4000e-01 | 3.9246e+05 | ; | CD1 - CE1 |
| 5981 | 5982 | 1 | 1.0800e-01 | 3.0711e+05 | ; | CE1 - HE1 |
| 5981 | 5983 | 1 | 1.4090e-01 | 3.9246e+05 | ; | CE1 - CZ |
| 5983 | 5984 | 1 | 1.3640e-01 | 3.7656e+05 | ; | CZ - OH |
| 5983 | 5986 | 1 | 1.4090e-01 | 3.9246e+05 | ; | CZ - CE2 |
| 5984 | 5985 | 1 | 9.6000e-02 | 4.6275e+05 | ; | OH - HH |
| 5986 | 5987 | 1 | 1.0800e-01 | 3.0711e+05 | ; | CE2 - HE2 |
| 5986 | 5988 | 1 | 1.4000e-01 | 3.9246e+05 | ; | CE2 - CD2 |
| 5988 | 5989 | 1 | 1.0800e-01 | 3.0711e+05 | ; | CD2 - HD2 |
| 5990 | 5991 | 1 | 1.2290e-01 | 4.7698e+05 | ; | C - O |
| 5990 | 5992 | 1 | 1.3350e-01 | 4.1003e+05 | ; | C - N |
| 5992 | 5993 | 1 | 1.0100e-01 | 3.6317e+05 | ; | N - H |
| 5992 | 5994 | 1 | 1.4490e-01 | 2.8200e+05 | ; | N - CA |
| 5994 | 5995 | 1 | 1.0900e-01 | 2.8451e+05 | ; | CA - HA |
| 5994 | 5996 | 1 | 1.5260e-01 | 2.5941e+05 | ; | CA - CB |
| 5994 | 6004 | 1 | 1.5220e-01 | 2.6527e+05 | ; | CA - C |
| 5996 | 5997 | 1 | 1.0900e-01 | 2.8451e+05 | ; | CB - HB |
| 5996 | 5998 | 1 | 1.5260e-01 | 2.5941e+05 | ; | CB - CG2 |
| 5996 | 6002 | 1 | 1.4100e-01 | 2.6778e+05 | ; | CB - OG1 |
| 5998 | 5999 | 1 | 1.0900e-01 | 2.8451e+05 | ; | CG2 - HG21 |
| 5998 | 6000 | 1 | 1.0900e-01 | 2.8451e+05 | ; | CG2 - HG22 |
| 5998 | 6001 | 1 | 1.0900e-01 | 2.8451e+05 | ; | CG2 - HG23 |
| 6002 | 6003 | 1 | 9.6000e-02 | 4.6275e+05 | ; | OG1 - HG1 |
| 6004 | 6005 | 1 | 1.2290e-01 | 4.7698e+05 | ; | C - O |
| 6004 | 6006 | 1 | 1.3350e-01 | 4.1003e+05 | ; | C - N |
| 6006 | 6007 | 1 | 1.0100e-01 | 3.6317e+05 | ; | N - H |
| 6006 | 6008 | 1 | 1.4490e-01 | 2.8200e+05 | ; | N - CA |
| 6008 | 6009 | 1 | 1.0900e-01 | 2.8451e+05 | ; | CA - HA |
| 6008 | 6010 | 1 | 1.5260e-01 | 2.5941e+05 | ; | CA - CB |
| 6008 | 6016 | 1 | 1.5220e-01 | 2.6527e+05 | ; | CA - C |
| 6010 | 6011 | 1 | 1.0900e-01 | 2.8451e+05 | ; | CB - HB2 |
| 6010 | 6012 | 1 | 1.0900e-01 | 2.8451e+05 | ; | CB - HB3 |
| 6010 | 6013 | 1 | 1.5220e-01 | 2.6527e+05 | ; | CB - CG |
| 6013 | 6014 | 1 | 1.2500e-01 | 5.4894e+05 | ; | CG - OD1 |
| 6013 | 6015 | 1 | 1.2500e-01 | 5.4894e+05 | ; | CG - OD2 |
| 6016 | 6017 | 1 | 1.2290e-01 | 4.7698e+05 | ; | C - O |
| 6016 | 6018 | 1 | 1.3350e-01 | 4.1003e+05 | ; | C - N |
| 6018 | 6019 | 1 | 1.0100e-01 | 3.6317e+05 | ; | N - H |
| 6018 | 6020 | 1 | 1.4490e-01 | 2.8200e+05 | ; | N - CA |
| 6020 | 6021 | 1 | 1.0900e-01 | 2.8451e+05 | ; | CA - HA |
| 6020 | 6022 | 1 | 1.5260e-01 | 2.5941e+05 | ; | CA - CB |

| | | | | | |
|---|---|---|---|---|---|
| 6020 | 6036 | 1 | 1.5220e-01 | 2.6527e+05 ; | CA - C |
| 6022 | 6023 | 1 | 1.0900e-01 | 2.8451e+05 ; | CB - HB2 |
| 6022 | 6024 | 1 | 1.0900e-01 | 2.8451e+05 ; | CB - HB3 |
| 6022 | 6025 | 1 | 1.5100e-01 | 2.6527e+05 ; | CB - CG |
| 6025 | 6026 | 1 | 1.4000e-01 | 3.9246e+05 ; | CG - CD1 |
| 6025 | 6034 | 1 | 1.4000e-01 | 3.9246e+05 ; | CG - CD2 |
| 6026 | 6027 | 1 | 1.0800e-01 | 3.0711e+05 ; | CD1 - HD1 |
| 6026 | 6028 | 1 | 1.4000e-01 | 3.9246e+05 ; | CD1 - CE1 |
| 6028 | 6029 | 1 | 1.0800e-01 | 3.0711e+05 ; | CE1 - HE1 |
| 6028 | 6030 | 1 | 1.4000e-01 | 3.9246e+05 ; | CE1 - CZ |
| 6030 | 6031 | 1 | 1.0800e-01 | 3.0711e+05 ; | CZ - HZ |
| 6030 | 6032 | 1 | 1.4000e-01 | 3.9246e+05 ; | CZ - CE2 |
| 6032 | 6033 | 1 | 1.0800e-01 | 3.0711e+05 ; | CE2 - HE2 |
| 6032 | 6034 | 1 | 1.4000e-01 | 3.9246e+05 ; | CE2 - CD2 |
| 6034 | 6035 | 1 | 1.0800e-01 | 3.0711e+05 ; | CD2 - HD2 |
| 6036 | 6037 | 1 | 1.2290e-01 | 4.7698e+05 ; | C - O |
| 6036 | 6038 | 1 | 1.3350e-01 | 4.1003e+05 ; | C - N |
| 6038 | 6039 | 1 | 1.0100e-01 | 3.6317e+05 ; | N - H |
| 6038 | 6040 | 1 | 1.4490e-01 | 2.8200e+05 ; | N - CA |
| 6040 | 6041 | 1 | 1.0900e-01 | 2.8451e+05 ; | CA - HA |
| 6040 | 6042 | 1 | 1.5260e-01 | 2.5941e+05 ; | CA - CB |
| 6040 | 6055 | 1 | 1.5220e-01 | 2.6527e+05 ; | CA - C |
| 6042 | 6043 | 1 | 1.0900e-01 | 2.8451e+05 ; | CB - HB2 |
| 6042 | 6044 | 1 | 1.0900e-01 | 2.8451e+05 ; | CB - HB3 |
| 6042 | 6045 | 1 | 1.5260e-01 | 2.5941e+05 ; | CB - CG |
| 6045 | 6046 | 1 | 1.0900e-01 | 2.8451e+05 ; | CG - HG |
| 6045 | 6047 | 1 | 1.5260e-01 | 2.5941e+05 ; | CG - CD1 |
| 6045 | 6051 | 1 | 1.5260e-01 | 2.5941e+05 ; | CG - CD2 |
| 6047 | 6048 | 1 | 1.0900e-01 | 2.8451e+05 ; | CD1 - HD11 |
| 6047 | 6049 | 1 | 1.0900e-01 | 2.8451e+05 ; | CD1 - HD12 |
| 6047 | 6050 | 1 | 1.0900e-01 | 2.8451e+05 ; | CD1 - HD13 |
| 6051 | 6052 | 1 | 1.0900e-01 | 2.8451e+05 ; | CD2 - HD21 |
| 6051 | 6053 | 1 | 1.0900e-01 | 2.8451e+05 ; | CD2 - HD22 |
| 6051 | 6054 | 1 | 1.0900e-01 | 2.8451e+05 ; | CD2 - HD23 |
| 6055 | 6056 | 1 | 1.2290e-01 | 4.7698e+05 ; | C - O |
| 6055 | 6057 | 1 | 1.3350e-01 | 4.1003e+05 ; | C - N |
| 6057 | 6058 | 1 | 1.0100e-01 | 3.6317e+05 ; | N - H |
| 6057 | 6059 | 1 | 1.4490e-01 | 2.8200e+05 ; | N - CA |
| 6059 | 6060 | 1 | 1.0900e-01 | 2.8451e+05 ; | CA - HA |
| 6059 | 6061 | 1 | 1.5260e-01 | 2.5941e+05 ; | CA - CB |
| 6059 | 6065 | 1 | 1.5220e-01 | 2.6527e+05 ; | CA - C |
| 6061 | 6062 | 1 | 1.0900e-01 | 2.8451e+05 ; | CB - HB1 |
| 6061 | 6063 | 1 | 1.0900e-01 | 2.8451e+05 ; | CB - HB2 |
| 6061 | 6064 | 1 | 1.0900e-01 | 2.8451e+05 ; | CB - HB3 |
| 6065 | 6066 | 1 | 1.2290e-01 | 4.7698e+05 ; | C - O |
| 6065 | 6067 | 1 | 1.3350e-01 | 4.1003e+05 ; | C - N |
| 6067 | 6068 | 1 | 1.0100e-01 | 3.6317e+05 ; | N - H |
| 6067 | 6069 | 1 | 1.4490e-01 | 2.8200e+05 ; | N - CA |
| 6069 | 6070 | 1 | 1.0900e-01 | 2.8451e+05 ; | CA - HA |
| 6069 | 6071 | 1 | 1.5260e-01 | 2.5941e+05 ; | CA - CB |
| 6069 | 6075 | 1 | 1.5220e-01 | 2.6527e+05 ; | CA - C |
| 6071 | 6072 | 1 | 1.0900e-01 | 2.8451e+05 ; | CB - HB1 |

| | | | | | | |
|---|---|---|---|---|---|---|
| 6071 | 6073 | 1 | 1.0900e-01 | 2.8451e+05 | ; | CB - HB2 |
| 6071 | 6074 | 1 | 1.0900e-01 | 2.8451e+05 | ; | CB - HB3 |
| 6075 | 6076 | 1 | 1.2290e-01 | 4.7698e+05 | ; | C - O |
| 6075 | 6077 | 1 | 1.3350e-01 | 4.1003e+05 | ; | C - N |
| 6077 | 6078 | 1 | 1.0100e-01 | 3.6317e+05 | ; | N - H |
| 6077 | 6079 | 1 | 1.4490e-01 | 2.8200e+05 | ; | N - CA |
| 6079 | 6080 | 1 | 1.0900e-01 | 2.8451e+05 | ; | CA - HA |
| 6079 | 6081 | 1 | 1.5260e-01 | 2.5941e+05 | ; | CA - CB |
| 6079 | 6089 | 1 | 1.5220e-01 | 2.6527e+05 | ; | CA - C |
| 6081 | 6082 | 1 | 1.0900e-01 | 2.8451e+05 | ; | CB - HB |
| 6081 | 6083 | 1 | 1.5260e-01 | 2.5941e+05 | ; | CB - CG2 |
| 6081 | 6087 | 1 | 1.4100e-01 | 2.6778e+05 | ; | CB - OG1 |
| 6083 | 6084 | 1 | 1.0900e-01 | 2.8451e+05 | ; | CG2 - HG21 |
| 6083 | 6085 | 1 | 1.0900e-01 | 2.8451e+05 | ; | CG2 - HG22 |
| 6083 | 6086 | 1 | 1.0900e-01 | 2.8451e+05 | ; | CG2 - HG23 |
| 6087 | 6088 | 1 | 9.6000e-02 | 4.6275e+05 | ; | OG1 - HG1 |
| 6089 | 6090 | 1 | 1.2290e-01 | 4.7698e+05 | ; | C - O |
| 6089 | 6091 | 1 | 1.3350e-01 | 4.1003e+05 | ; | C - N |
| 6091 | 6092 | 1 | 1.0100e-01 | 3.6317e+05 | ; | N - H |
| 6091 | 6093 | 1 | 1.4490e-01 | 2.8200e+05 | ; | N - CA |
| 6093 | 6094 | 1 | 1.0900e-01 | 2.8451e+05 | ; | CA - HA |
| 6093 | 6095 | 1 | 1.5260e-01 | 2.5941e+05 | ; | CA - CB |
| 6093 | 6108 | 1 | 1.5220e-01 | 2.6527e+05 | ; | CA - C |
| 6095 | 6096 | 1 | 1.0900e-01 | 2.8451e+05 | ; | CB - HB |
| 6095 | 6097 | 1 | 1.5260e-01 | 2.5941e+05 | ; | CB - CG2 |
| 6095 | 6101 | 1 | 1.5260e-01 | 2.5941e+05 | ; | CB - CG1 |
| 6097 | 6098 | 1 | 1.0900e-01 | 2.8451e+05 | ; | CG2 - HG21 |
| 6097 | 6099 | 1 | 1.0900e-01 | 2.8451e+05 | ; | CG2 - HG22 |
| 6097 | 6100 | 1 | 1.0900e-01 | 2.8451e+05 | ; | CG2 - HG23 |
| 6101 | 6102 | 1 | 1.0900e-01 | 2.8451e+05 | ; | CG1 - HG12 |
| 6101 | 6103 | 1 | 1.0900e-01 | 2.8451e+05 | ; | CG1 - HG13 |
| 6101 | 6104 | 1 | 1.5260e-01 | 2.5941e+05 | ; | CG1 - CD1 |
| 6104 | 6105 | 1 | 1.0900e-01 | 2.8451e+05 | ; | CD1 - HD11 |
| 6104 | 6106 | 1 | 1.0900e-01 | 2.8451e+05 | ; | CD1 - HD12 |
| 6104 | 6107 | 1 | 1.0900e-01 | 2.8451e+05 | ; | CD1 - HD13 |
| 6108 | 6109 | 1 | 1.2290e-01 | 4.7698e+05 | ; | C - O |
| 6108 | 6110 | 1 | 1.3350e-01 | 4.1003e+05 | ; | C - N |
| 6110 | 6111 | 1 | 1.0100e-01 | 3.6317e+05 | ; | N - H |
| 6110 | 6112 | 1 | 1.4490e-01 | 2.8200e+05 | ; | N - CA |
| 6112 | 6113 | 1 | 1.0900e-01 | 2.8451e+05 | ; | CA - HA |
| 6112 | 6114 | 1 | 1.5260e-01 | 2.5941e+05 | ; | CA - CB |
| 6112 | 6120 | 1 | 1.5220e-01 | 2.6527e+05 | ; | CA - C |
| 6114 | 6115 | 1 | 1.0900e-01 | 2.8451e+05 | ; | CB - HB2 |
| 6114 | 6116 | 1 | 1.0900e-01 | 2.8451e+05 | ; | CB - HB3 |
| 6114 | 6117 | 1 | 1.5220e-01 | 2.6527e+05 | ; | CB - CG |
| 6117 | 6118 | 1 | 1.2500e-01 | 5.4894e+05 | ; | CG - OD1 |
| 6117 | 6119 | 1 | 1.2500e-01 | 5.4894e+05 | ; | CG - OD2 |
| 6120 | 6121 | 1 | 1.2290e-01 | 4.7698e+05 | ; | C - O |
| 6120 | 6122 | 1 | 1.3350e-01 | 4.1003e+05 | ; | C - N |
| 6122 | 6123 | 1 | 1.0100e-01 | 3.6317e+05 | ; | N - H |
| 6122 | 6124 | 1 | 1.4490e-01 | 2.8200e+05 | ; | N - CA |
| 6124 | 6125 | 1 | 1.0900e-01 | 2.8451e+05 | ; | CA - HA |

| | | | | | | |
|---|---|---|---|---|---|---|
| 6124 | 6126 | 1 | 1.5260e-01 | 2.5941e+05 | ; | CA - CB |
| 6124 | 6142 | 1 | 1.5220e-01 | 2.6527e+05 | ; | CA - C |
| 6126 | 6127 | 1 | 1.0900e-01 | 2.8451e+05 | ; | CB - HB2 |
| 6126 | 6128 | 1 | 1.0900e-01 | 2.8451e+05 | ; | CB - HB3 |
| 6126 | 6129 | 1 | 1.5260e-01 | 2.5941e+05 | ; | CB - CG |
| 6129 | 6130 | 1 | 1.0900e-01 | 2.8451e+05 | ; | CG - HG2 |
| 6129 | 6131 | 1 | 1.0900e-01 | 2.8451e+05 | ; | CG - HG3 |
| 6129 | 6132 | 1 | 1.5260e-01 | 2.5941e+05 | ; | CG - CD |
| 6132 | 6133 | 1 | 1.0900e-01 | 2.8451e+05 | ; | CD - HD2 |
| 6132 | 6134 | 1 | 1.0900e-01 | 2.8451e+05 | ; | CD - HD3 |
| 6132 | 6135 | 1 | 1.5260e-01 | 2.5941e+05 | ; | CD - CE |
| 6135 | 6136 | 1 | 1.0900e-01 | 2.8451e+05 | ; | CE - HE2 |
| 6135 | 6137 | 1 | 1.0900e-01 | 2.8451e+05 | ; | CE - HE3 |
| 6135 | 6138 | 1 | 1.4710e-01 | 3.0711e+05 | ; | CE - NZ |
| 6138 | 6139 | 1 | 1.0100e-01 | 3.6317e+05 | ; | NZ - HZ1 |
| 6138 | 6140 | 1 | 1.0100e-01 | 3.6317e+05 | ; | NZ - HZ2 |
| 6138 | 6141 | 1 | 1.0100e-01 | 3.6317e+05 | ; | NZ - HZ3 |
| 6142 | 6143 | 1 | 1.2290e-01 | 4.7698e+05 | ; | C - O |
| 6142 | 6144 | 1 | 1.3350e-01 | 4.1003e+05 | ; | C - N |
| 6144 | 6145 | 1 | 1.0100e-01 | 3.6317e+05 | ; | N - H |
| 6144 | 6146 | 1 | 1.4490e-01 | 2.8200e+05 | ; | N - CA |
| 6146 | 6147 | 1 | 1.0900e-01 | 2.8451e+05 | ; | CA - HA |
| 6146 | 6148 | 1 | 1.5260e-01 | 2.5941e+05 | ; | CA - CB |
| 6146 | 6159 | 1 | 1.5220e-01 | 2.6527e+05 | ; | CA - C |
| 6148 | 6149 | 1 | 1.0900e-01 | 2.8451e+05 | ; | CB - HB2 |
| 6148 | 6150 | 1 | 1.0900e-01 | 2.8451e+05 | ; | CB - HB3 |
| 6148 | 6151 | 1 | 1.5260e-01 | 2.5941e+05 | ; | CB - CG |
| 6151 | 6152 | 1 | 1.0900e-01 | 2.8451e+05 | ; | CG - HG2 |
| 6151 | 6153 | 1 | 1.0900e-01 | 2.8451e+05 | ; | CG - HG3 |
| 6151 | 6154 | 1 | 1.5220e-01 | 2.6527e+05 | ; | CG - CD |
| 6154 | 6155 | 1 | 1.2290e-01 | 4.7698e+05 | ; | CD - OE1 |
| 6154 | 6156 | 1 | 1.3350e-01 | 4.1003e+05 | ; | CD - NE2 |
| 6156 | 6157 | 1 | 1.0100e-01 | 3.6317e+05 | ; | NE2 - HE21 |
| 6156 | 6158 | 1 | 1.0100e-01 | 3.6317e+05 | ; | NE2 - HE22 |
| 6159 | 6160 | 1 | 1.2290e-01 | 4.7698e+05 | ; | C - O |
| 6159 | 6161 | 1 | 1.3350e-01 | 4.1003e+05 | ; | C - N |
| 6161 | 6162 | 1 | 1.0100e-01 | 3.6317e+05 | ; | N - H |
| 6161 | 6163 | 1 | 1.4490e-01 | 2.8200e+05 | ; | N - CA |
| 6163 | 6164 | 1 | 1.0900e-01 | 2.8451e+05 | ; | CA - HA |
| 6163 | 6165 | 1 | 1.5260e-01 | 2.5941e+05 | ; | CA - CB |
| 6163 | 6173 | 1 | 1.5220e-01 | 2.6527e+05 | ; | CA - C |
| 6165 | 6166 | 1 | 1.0900e-01 | 2.8451e+05 | ; | CB - HB |
| 6165 | 6167 | 1 | 1.5260e-01 | 2.5941e+05 | ; | CB - CG2 |
| 6165 | 6171 | 1 | 1.4100e-01 | 2.6778e+05 | ; | CB - OG1 |
| 6167 | 6168 | 1 | 1.0900e-01 | 2.8451e+05 | ; | CG2 - HG21 |
| 6167 | 6169 | 1 | 1.0900e-01 | 2.8451e+05 | ; | CG2 - HG22 |
| 6167 | 6170 | 1 | 1.0900e-01 | 2.8451e+05 | ; | CG2 - HG23 |
| 6171 | 6172 | 1 | 9.6000e-02 | 4.6275e+05 | ; | OG1 - HG1 |
| 6173 | 6174 | 1 | 1.2290e-01 | 4.7698e+05 | ; | C - O |
| 6173 | 6175 | 1 | 1.3350e-01 | 4.1003e+05 | ; | C - N |
| 6175 | 6176 | 1 | 1.0100e-01 | 3.6317e+05 | ; | N - H |
| 6175 | 6177 | 1 | 1.4490e-01 | 2.8200e+05 | ; | N - CA |

| | | | | | | |
|---|---|---|---|---|---|---|
| 6177 | 6178 | 1 | 1.0900e-01 | 2.8451e+05 | ; | CA - HA |
| 6177 | 6179 | 1 | 1.5260e-01 | 2.5941e+05 | ; | CA - CB |
| 6177 | 6194 | 1 | 1.5220e-01 | 2.6527e+05 | ; | CA - C |
| 6179 | 6180 | 1 | 1.0900e-01 | 2.8451e+05 | ; | CB - HB2 |
| 6179 | 6181 | 1 | 1.0900e-01 | 2.8451e+05 | ; | CB - HB3 |
| 6179 | 6182 | 1 | 1.5100e-01 | 2.6527e+05 | ; | CB - CG |
| 6182 | 6183 | 1 | 1.4000e-01 | 3.9246e+05 | ; | CG - CD1 |
| 6182 | 6192 | 1 | 1.4000e-01 | 3.9246e+05 | ; | CG - CD2 |
| 6183 | 6184 | 1 | 1.0800e-01 | 3.0711e+05 | ; | CD1 - HD1 |
| 6183 | 6185 | 1 | 1.4000e-01 | 3.9246e+05 | ; | CD1 - CE1 |
| 6185 | 6186 | 1 | 1.0800e-01 | 3.0711e+05 | ; | CE1 - HE1 |
| 6185 | 6187 | 1 | 1.4090e-01 | 3.9246e+05 | ; | CE1 - CZ |
| 6187 | 6188 | 1 | 1.3640e-01 | 3.7656e+05 | ; | CZ - OH |
| 6187 | 6190 | 1 | 1.4090e-01 | 3.9246e+05 | ; | CZ - CE2 |
| 6188 | 6189 | 1 | 9.6000e-02 | 4.6275e+05 | ; | OH - HH |
| 6190 | 6191 | 1 | 1.0800e-01 | 3.0711e+05 | ; | CE2 - HE2 |
| 6190 | 6192 | 1 | 1.4000e-01 | 3.9246e+05 | ; | CE2 - CD2 |
| 6192 | 6193 | 1 | 1.0800e-01 | 3.0711e+05 | ; | CD2 - HD2 |
| 6194 | 6195 | 1 | 1.2290e-01 | 4.7698e+05 | ; | C - O |
| 6194 | 6196 | 1 | 1.3350e-01 | 4.1003e+05 | ; | C - N |
| 6196 | 6197 | 1 | 1.0100e-01 | 3.6317e+05 | ; | N - H |
| 6196 | 6198 | 1 | 1.4490e-01 | 2.8200e+05 | ; | N - CA |
| 6198 | 6199 | 1 | 1.0900e-01 | 2.8451e+05 | ; | CA - HA |
| 6198 | 6200 | 1 | 1.5260e-01 | 2.5941e+05 | ; | CA - CB |
| 6198 | 6213 | 1 | 1.5220e-01 | 2.6527e+05 | ; | CA - C |
| 6200 | 6201 | 1 | 1.0900e-01 | 2.8451e+05 | ; | CB - HB2 |
| 6200 | 6202 | 1 | 1.0900e-01 | 2.8451e+05 | ; | CB - HB3 |
| 6200 | 6203 | 1 | 1.5260e-01 | 2.5941e+05 | ; | CB - CG |
| 6203 | 6204 | 1 | 1.0900e-01 | 2.8451e+05 | ; | CG - HG |
| 6203 | 6205 | 1 | 1.5260e-01 | 2.5941e+05 | ; | CG - CD1 |
| 6203 | 6209 | 1 | 1.5260e-01 | 2.5941e+05 | ; | CG - CD2 |
| 6205 | 6206 | 1 | 1.0900e-01 | 2.8451e+05 | ; | CD1 - HD11 |
| 6205 | 6207 | 1 | 1.0900e-01 | 2.8451e+05 | ; | CD1 - HD12 |
| 6205 | 6208 | 1 | 1.0900e-01 | 2.8451e+05 | ; | CD1 - HD13 |
| 6209 | 6210 | 1 | 1.0900e-01 | 2.8451e+05 | ; | CD2 - HD21 |
| 6209 | 6211 | 1 | 1.0900e-01 | 2.8451e+05 | ; | CD2 - HD22 |
| 6209 | 6212 | 1 | 1.0900e-01 | 2.8451e+05 | ; | CD2 - HD23 |
| 6213 | 6214 | 1 | 1.2290e-01 | 4.7698e+05 | ; | C - O |
| 6213 | 6215 | 1 | 1.3350e-01 | 4.1003e+05 | ; | C - N |
| 6215 | 6216 | 1 | 1.0100e-01 | 3.6317e+05 | ; | N - H |
| 6215 | 6217 | 1 | 1.4490e-01 | 2.8200e+05 | ; | N - CA |
| 6217 | 6218 | 1 | 1.0900e-01 | 2.8451e+05 | ; | CA - HA |
| 6217 | 6219 | 1 | 1.5260e-01 | 2.5941e+05 | ; | CA - CB |
| 6217 | 6235 | 1 | 1.5220e-01 | 2.6527e+05 | ; | CA - C |
| 6219 | 6220 | 1 | 1.0900e-01 | 2.8451e+05 | ; | CB - HB2 |
| 6219 | 6221 | 1 | 1.0900e-01 | 2.8451e+05 | ; | CB - HB3 |
| 6219 | 6222 | 1 | 1.5260e-01 | 2.5941e+05 | ; | CB - CG |
| 6222 | 6223 | 1 | 1.0900e-01 | 2.8451e+05 | ; | CG - HG2 |
| 6222 | 6224 | 1 | 1.0900e-01 | 2.8451e+05 | ; | CG - HG3 |
| 6222 | 6225 | 1 | 1.5260e-01 | 2.5941e+05 | ; | CG - CD |
| 6225 | 6226 | 1 | 1.0900e-01 | 2.8451e+05 | ; | CD - HD2 |
| 6225 | 6227 | 1 | 1.0900e-01 | 2.8451e+05 | ; | CD - HD3 |

| | | | | | | |
|---|---|---|---|---|---|---|
| 6225 | 6228 | 1 | 1.5260e-01 | 2.5941e+05 | ; | CD - CE |
| 6228 | 6229 | 1 | 1.0900e-01 | 2.8451e+05 | ; | CE - HE2 |
| 6228 | 6230 | 1 | 1.0900e-01 | 2.8451e+05 | ; | CE - HE3 |
| 6228 | 6231 | 1 | 1.4710e-01 | 3.0711e+05 | ; | CE - NZ |
| 6231 | 6232 | 1 | 1.0100e-01 | 3.6317e+05 | ; | NZ - HZ1 |
| 6231 | 6233 | 1 | 1.0100e-01 | 3.6317e+05 | ; | NZ - HZ2 |
| 6231 | 6234 | 1 | 1.0100e-01 | 3.6317e+05 | ; | NZ - HZ3 |
| 6235 | 6236 | 1 | 1.2290e-01 | 4.7698e+05 | ; | C - O |
| 6235 | 6237 | 1 | 1.3350e-01 | 4.1003e+05 | ; | C - N |
| 6237 | 6238 | 1 | 1.0100e-01 | 3.6317e+05 | ; | N - H |
| 6237 | 6239 | 1 | 1.4490e-01 | 2.8200e+05 | ; | N - CA |
| 6239 | 6240 | 1 | 1.0900e-01 | 2.8451e+05 | ; | CA - HA |
| 6239 | 6241 | 1 | 1.5260e-01 | 2.5941e+05 | ; | CA - CB |
| 6239 | 6257 | 1 | 1.5220e-01 | 2.6527e+05 | ; | CA - C |
| 6241 | 6242 | 1 | 1.0900e-01 | 2.8451e+05 | ; | CB - HB2 |
| 6241 | 6243 | 1 | 1.0900e-01 | 2.8451e+05 | ; | CB - HB3 |
| 6241 | 6244 | 1 | 1.5260e-01 | 2.5941e+05 | ; | CB - CG |
| 6244 | 6245 | 1 | 1.0900e-01 | 2.8451e+05 | ; | CG - HG2 |
| 6244 | 6246 | 1 | 1.0900e-01 | 2.8451e+05 | ; | CG - HG3 |
| 6244 | 6247 | 1 | 1.5260e-01 | 2.5941e+05 | ; | CG - CD |
| 6247 | 6248 | 1 | 1.0900e-01 | 2.8451e+05 | ; | CD - HD2 |
| 6247 | 6249 | 1 | 1.0900e-01 | 2.8451e+05 | ; | CD - HD3 |
| 6247 | 6250 | 1 | 1.5260e-01 | 2.5941e+05 | ; | CD - CE |
| 6250 | 6251 | 1 | 1.0900e-01 | 2.8451e+05 | ; | CE - HE2 |
| 6250 | 6252 | 1 | 1.0900e-01 | 2.8451e+05 | ; | CE - HE3 |
| 6250 | 6253 | 1 | 1.4710e-01 | 3.0711e+05 | ; | CE - NZ |
| 6253 | 6254 | 1 | 1.0100e-01 | 3.6317e+05 | ; | NZ - HZ1 |
| 6253 | 6255 | 1 | 1.0100e-01 | 3.6317e+05 | ; | NZ - HZ2 |
| 6253 | 6256 | 1 | 1.0100e-01 | 3.6317e+05 | ; | NZ - HZ3 |
| 6257 | 6258 | 1 | 1.2290e-01 | 4.7698e+05 | ; | C - O |
| 6257 | 6259 | 1 | 1.3350e-01 | 4.1003e+05 | ; | C - N |
| 6259 | 6260 | 1 | 1.0100e-01 | 3.6317e+05 | ; | N - H |
| 6259 | 6261 | 1 | 1.4490e-01 | 2.8200e+05 | ; | N - CA |
| 6261 | 6262 | 1 | 1.0900e-01 | 2.8451e+05 | ; | CA - HA |
| 6261 | 6263 | 1 | 1.5260e-01 | 2.5941e+05 | ; | CA - CB |
| 6261 | 6272 | 1 | 1.5220e-01 | 2.6527e+05 | ; | CA - C |
| 6263 | 6264 | 1 | 1.0900e-01 | 2.8451e+05 | ; | CB - HB2 |
| 6263 | 6265 | 1 | 1.0900e-01 | 2.8451e+05 | ; | CB - HB3 |
| 6263 | 6266 | 1 | 1.5260e-01 | 2.5941e+05 | ; | CB - CG |
| 6266 | 6267 | 1 | 1.0900e-01 | 2.8451e+05 | ; | CG - HG2 |
| 6266 | 6268 | 1 | 1.0900e-01 | 2.8451e+05 | ; | CG - HG3 |
| 6266 | 6269 | 1 | 1.5220e-01 | 2.6527e+05 | ; | CG - CD |
| 6269 | 6270 | 1 | 1.2500e-01 | 5.4894e+05 | ; | CD - OE1 |
| 6269 | 6271 | 1 | 1.2500e-01 | 5.4894e+05 | ; | CD - OE2 |
| 6272 | 6273 | 1 | 1.2290e-01 | 4.7698e+05 | ; | C - O |
| 6272 | 6274 | 1 | 1.3350e-01 | 4.1003e+05 | ; | C - N |
| 6274 | 6275 | 1 | 1.0100e-01 | 3.6317e+05 | ; | N - H |
| 6274 | 6276 | 1 | 1.4490e-01 | 2.8200e+05 | ; | N - CA |
| 6276 | 6277 | 1 | 1.0900e-01 | 2.8451e+05 | ; | CA - HA |
| 6276 | 6278 | 1 | 1.5260e-01 | 2.5941e+05 | ; | CA - CB |
| 6276 | 6288 | 1 | 1.5220e-01 | 2.6527e+05 | ; | CA - C |
| 6278 | 6279 | 1 | 1.0900e-01 | 2.8451e+05 | ; | CB - HB |

| | | | | | |
|---|---|---|---|---|---|
| 6278 | 6280 | 1 | 1.5260e-01 | 2.5941e+05 ; | CB - CG1 |
| 6278 | 6284 | 1 | 1.5260e-01 | 2.5941e+05 ; | CB - CG2 |
| 6280 | 6281 | 1 | 1.0900e-01 | 2.8451e+05 ; | CG1 - HG11 |
| 6280 | 6282 | 1 | 1.0900e-01 | 2.8451e+05 ; | CG1 - HG12 |
| 6280 | 6283 | 1 | 1.0900e-01 | 2.8451e+05 ; | CG1 - HG13 |
| 6284 | 6285 | 1 | 1.0900e-01 | 2.8451e+05 ; | CG2 - HG21 |
| 6284 | 6286 | 1 | 1.0900e-01 | 2.8451e+05 ; | CG2 - HG22 |
| 6284 | 6287 | 1 | 1.0900e-01 | 2.8451e+05 ; | CG2 - HG23 |
| 6288 | 6289 | 1 | 1.2290e-01 | 4.7698e+05 ; | C - O |
| 6288 | 6290 | 1 | 1.3350e-01 | 4.1003e+05 ; | C - N |
| 6290 | 6291 | 1 | 1.0100e-01 | 3.6317e+05 ; | N - H |
| 6290 | 6292 | 1 | 1.4490e-01 | 2.8200e+05 ; | N - CA |
| 6292 | 6293 | 1 | 1.0900e-01 | 2.8451e+05 ; | CA - HA |
| 6292 | 6294 | 1 | 1.5260e-01 | 2.5941e+05 ; | CA - CB |
| 6292 | 6299 | 1 | 1.5220e-01 | 2.6527e+05 ; | CA - C |
| 6294 | 6295 | 1 | 1.0900e-01 | 2.8451e+05 ; | CB - HB2 |
| 6294 | 6296 | 1 | 1.0900e-01 | 2.8451e+05 ; | CB - HB3 |
| 6294 | 6297 | 1 | 1.8100e-01 | 1.9832e+05 ; | CB - SG |
| 6297 | 6298 | 1 | 1.3360e-01 | 2.2928e+05 ; | SG - HG |
| 6299 | 6300 | 1 | 1.2290e-01 | 4.7698e+05 ; | C - O |
| 6299 | 6301 | 1 | 1.3350e-01 | 4.1003e+05 ; | C - N |
| 6301 | 6302 | 1 | 1.0100e-01 | 3.6317e+05 ; | N - H |
| 6301 | 6303 | 1 | 1.4490e-01 | 2.8200e+05 ; | N - CA |
| 6303 | 6304 | 1 | 1.0900e-01 | 2.8451e+05 ; | CA - HA |
| 6303 | 6305 | 1 | 1.5260e-01 | 2.5941e+05 ; | CA - CB |
| 6303 | 6318 | 1 | 1.5220e-01 | 2.6527e+05 ; | CA - C |
| 6305 | 6306 | 1 | 1.0900e-01 | 2.8451e+05 ; | CB - HB2 |
| 6305 | 6307 | 1 | 1.0900e-01 | 2.8451e+05 ; | CB - HB3 |
| 6305 | 6308 | 1 | 1.5260e-01 | 2.5941e+05 ; | CB - CG |
| 6308 | 6309 | 1 | 1.0900e-01 | 2.8451e+05 ; | CG - HG |
| 6308 | 6310 | 1 | 1.5260e-01 | 2.5941e+05 ; | CG - CD1 |
| 6308 | 6314 | 1 | 1.5260e-01 | 2.5941e+05 ; | CG - CD2 |
| 6310 | 6311 | 1 | 1.0900e-01 | 2.8451e+05 ; | CD1 - HD11 |
| 6310 | 6312 | 1 | 1.0900e-01 | 2.8451e+05 ; | CD1 - HD12 |
| 6310 | 6313 | 1 | 1.0900e-01 | 2.8451e+05 ; | CD1 - HD13 |
| 6314 | 6315 | 1 | 1.0900e-01 | 2.8451e+05 ; | CD2 - HD21 |
| 6314 | 6316 | 1 | 1.0900e-01 | 2.8451e+05 ; | CD2 - HD22 |
| 6314 | 6317 | 1 | 1.0900e-01 | 2.8451e+05 ; | CD2 - HD23 |
| 6318 | 6319 | 1 | 1.2290e-01 | 4.7698e+05 ; | C - O |
| 6318 | 6320 | 1 | 1.3350e-01 | 4.1003e+05 ; | C - N |
| 6320 | 6321 | 1 | 1.0100e-01 | 3.6317e+05 ; | N - H |
| 6320 | 6322 | 1 | 1.4490e-01 | 2.8200e+05 ; | N - CA |
| 6322 | 6323 | 1 | 1.0900e-01 | 2.8451e+05 ; | CA - HA |
| 6322 | 6324 | 1 | 1.5260e-01 | 2.5941e+05 ; | CA - CB |
| 6322 | 6337 | 1 | 1.5220e-01 | 2.6527e+05 ; | CA - C |
| 6324 | 6325 | 1 | 1.0900e-01 | 2.8451e+05 ; | CB - HB |
| 6324 | 6326 | 1 | 1.5260e-01 | 2.5941e+05 ; | CB - CG2 |
| 6324 | 6330 | 1 | 1.5260e-01 | 2.5941e+05 ; | CB - CG1 |
| 6326 | 6327 | 1 | 1.0900e-01 | 2.8451e+05 ; | CG2 - HG21 |
| 6326 | 6328 | 1 | 1.0900e-01 | 2.8451e+05 ; | CG2 - HG22 |
| 6326 | 6329 | 1 | 1.0900e-01 | 2.8451e+05 ; | CG2 - HG23 |
| 6330 | 6331 | 1 | 1.0900e-01 | 2.8451e+05 ; | CG1 - HG12 |

| | | | | | | |
|---|---|---|---|---|---|---|
| 6330 | 6332 | 1 | 1.0900e-01 | 2.8451e+05 | ; | CG1 - HG13 |
| 6330 | 6333 | 1 | 1.5260e-01 | 2.5941e+05 | ; | CG1 - CD1 |
| 6333 | 6334 | 1 | 1.0900e-01 | 2.8451e+05 | ; | CD1 - HD11 |
| 6333 | 6335 | 1 | 1.0900e-01 | 2.8451e+05 | ; | CD1 - HD12 |
| 6333 | 6336 | 1 | 1.0900e-01 | 2.8451e+05 | ; | CD1 - HD13 |
| 6337 | 6338 | 1 | 1.2290e-01 | 4.7698e+05 | ; | C - O |
| 6337 | 6339 | 1 | 1.3350e-01 | 4.1003e+05 | ; | C - N |
| 6339 | 6340 | 1 | 1.4490e-01 | 2.8200e+05 | ; | N - CD |
| 6339 | 6349 | 1 | 1.4490e-01 | 2.8200e+05 | ; | N - CA |
| 6340 | 6341 | 1 | 1.0900e-01 | 2.8451e+05 | ; | CD - HD2 |
| 6340 | 6342 | 1 | 1.0900e-01 | 2.8451e+05 | ; | CD - HD3 |
| 6340 | 6343 | 1 | 1.5260e-01 | 2.5941e+05 | ; | CD - CG |
| 6343 | 6344 | 1 | 1.0900e-01 | 2.8451e+05 | ; | CG - HG2 |
| 6343 | 6345 | 1 | 1.0900e-01 | 2.8451e+05 | ; | CG - HG3 |
| 6343 | 6346 | 1 | 1.5260e-01 | 2.5941e+05 | ; | CG - CB |
| 6346 | 6347 | 1 | 1.0900e-01 | 2.8451e+05 | ; | CB - HB2 |
| 6346 | 6348 | 1 | 1.0900e-01 | 2.8451e+05 | ; | CB - HB3 |
| 6346 | 6349 | 1 | 1.5260e-01 | 2.5941e+05 | ; | CB - CA |
| 6349 | 6350 | 1 | 1.0900e-01 | 2.8451e+05 | ; | CA - HA |
| 6349 | 6351 | 1 | 1.5220e-01 | 2.6527e+05 | ; | CA - C |
| 6351 | 6352 | 1 | 1.2290e-01 | 4.7698e+05 | ; | C - O |
| 6351 | 6353 | 1 | 1.3350e-01 | 4.1003e+05 | ; | C - N |
| 6353 | 6354 | 1 | 1.0100e-01 | 3.6317e+05 | ; | N - H |
| 6353 | 6355 | 1 | 1.4490e-01 | 2.8200e+05 | ; | N - CA |
| 6355 | 6356 | 1 | 1.0900e-01 | 2.8451e+05 | ; | CA - HA |
| 6355 | 6357 | 1 | 1.5260e-01 | 2.5941e+05 | ; | CA - CB |
| 6355 | 6371 | 1 | 1.5220e-01 | 2.6527e+05 | ; | CA - C |
| 6357 | 6358 | 1 | 1.0900e-01 | 2.8451e+05 | ; | CB - HB2 |
| 6357 | 6359 | 1 | 1.0900e-01 | 2.8451e+05 | ; | CB - HB3 |
| 6357 | 6360 | 1 | 1.5100e-01 | 2.6527e+05 | ; | CB - CG |
| 6360 | 6361 | 1 | 1.4000e-01 | 3.9246e+05 | ; | CG - CD1 |
| 6360 | 6369 | 1 | 1.4000e-01 | 3.9246e+05 | ; | CG - CD2 |
| 6361 | 6362 | 1 | 1.0800e-01 | 3.0711e+05 | ; | CD1 - HD1 |
| 6361 | 6363 | 1 | 1.4000e-01 | 3.9246e+05 | ; | CD1 - CE1 |
| 6363 | 6364 | 1 | 1.0800e-01 | 3.0711e+05 | ; | CE1 - HE1 |
| 6363 | 6365 | 1 | 1.4000e-01 | 3.9246e+05 | ; | CE1 - CZ |
| 6365 | 6366 | 1 | 1.0800e-01 | 3.0711e+05 | ; | CZ - HZ |
| 6365 | 6367 | 1 | 1.4000e-01 | 3.9246e+05 | ; | CZ - CE2 |
| 6367 | 6368 | 1 | 1.0800e-01 | 3.0711e+05 | ; | CE2 - HE2 |
| 6367 | 6369 | 1 | 1.4000e-01 | 3.9246e+05 | ; | CE2 - CD2 |
| 6369 | 6370 | 1 | 1.0800e-01 | 3.0711e+05 | ; | CD2 - HD2 |
| 6371 | 6372 | 1 | 1.2290e-01 | 4.7698e+05 | ; | C - O |
| 6371 | 6373 | 1 | 1.3350e-01 | 4.1003e+05 | ; | C - N |
| 6373 | 6374 | 1 | 1.0100e-01 | 3.6317e+05 | ; | N - H |
| 6373 | 6375 | 1 | 1.4490e-01 | 2.8200e+05 | ; | N - CA |
| 6375 | 6376 | 1 | 1.0900e-01 | 2.8451e+05 | ; | CA - HA |
| 6375 | 6377 | 1 | 1.5260e-01 | 2.5941e+05 | ; | CA - CB |
| 6375 | 6393 | 1 | 1.5220e-01 | 2.6527e+05 | ; | CA - C |
| 6377 | 6378 | 1 | 1.0900e-01 | 2.8451e+05 | ; | CB - HB2 |
| 6377 | 6379 | 1 | 1.0900e-01 | 2.8451e+05 | ; | CB - HB3 |
| 6377 | 6380 | 1 | 1.5260e-01 | 2.5941e+05 | ; | CB - CG |
| 6380 | 6381 | 1 | 1.0900e-01 | 2.8451e+05 | ; | CG - HG2 |

| | | | | | | |
|---|---|---|---|---|---|---|
| 6380 | 6382 | 1 | 1.0900e-01 | 2.8451e+05 | ; | CG - HG3 |
| 6380 | 6383 | 1 | 1.5260e-01 | 2.5941e+05 | ; | CG - CD |
| 6383 | 6384 | 1 | 1.0900e-01 | 2.8451e+05 | ; | CD - HD2 |
| 6383 | 6385 | 1 | 1.0900e-01 | 2.8451e+05 | ; | CD - HD3 |
| 6383 | 6386 | 1 | 1.5260e-01 | 2.5941e+05 | ; | CD - CE |
| 6386 | 6387 | 1 | 1.0900e-01 | 2.8451e+05 | ; | CE - HE2 |
| 6386 | 6388 | 1 | 1.0900e-01 | 2.8451e+05 | ; | CE - HE3 |
| 6386 | 6389 | 1 | 1.4710e-01 | 3.0711e+05 | ; | CE - NZ |
| 6389 | 6390 | 1 | 1.0100e-01 | 3.6317e+05 | ; | NZ - HZ1 |
| 6389 | 6391 | 1 | 1.0100e-01 | 3.6317e+05 | ; | NZ - HZ2 |
| 6389 | 6392 | 1 | 1.0100e-01 | 3.6317e+05 | ; | NZ - HZ3 |
| 6393 | 6394 | 1 | 1.2290e-01 | 4.7698e+05 | ; | C - O |
| 6393 | 6395 | 1 | 1.3350e-01 | 4.1003e+05 | ; | C - N |
| 6395 | 6396 | 1 | 1.0100e-01 | 3.6317e+05 | ; | N - H |
| 6395 | 6397 | 1 | 1.4490e-01 | 2.8200e+05 | ; | N - CA |
| 6397 | 6398 | 1 | 1.0900e-01 | 2.8451e+05 | ; | CA - HA |
| 6397 | 6399 | 1 | 1.5260e-01 | 2.5941e+05 | ; | CA - CB |
| 6397 | 6413 | 1 | 1.5220e-01 | 2.6527e+05 | ; | CA - C |
| 6399 | 6400 | 1 | 1.0900e-01 | 2.8451e+05 | ; | CB - HB2 |
| 6399 | 6401 | 1 | 1.0900e-01 | 2.8451e+05 | ; | CB - HB3 |
| 6399 | 6402 | 1 | 1.5100e-01 | 2.6527e+05 | ; | CB - CG |
| 6402 | 6403 | 1 | 1.4000e-01 | 3.9246e+05 | ; | CG - CD1 |
| 6402 | 6411 | 1 | 1.4000e-01 | 3.9246e+05 | ; | CG - CD2 |
| 6403 | 6404 | 1 | 1.0800e-01 | 3.0711e+05 | ; | CD1 - HD1 |
| 6403 | 6405 | 1 | 1.4000e-01 | 3.9246e+05 | ; | CD1 - CE1 |
| 6405 | 6406 | 1 | 1.0800e-01 | 3.0711e+05 | ; | CE1 - HE1 |
| 6405 | 6407 | 1 | 1.4000e-01 | 3.9246e+05 | ; | CE1 - CZ |
| 6407 | 6408 | 1 | 1.0800e-01 | 3.0711e+05 | ; | CZ - HZ |
| 6407 | 6409 | 1 | 1.4000e-01 | 3.9246e+05 | ; | CZ - CE2 |
| 6409 | 6410 | 1 | 1.0800e-01 | 3.0711e+05 | ; | CE2 - HE2 |
| 6409 | 6411 | 1 | 1.4000e-01 | 3.9246e+05 | ; | CE2 - CD2 |
| 6411 | 6412 | 1 | 1.0800e-01 | 3.0711e+05 | ; | CD2 - HD2 |
| 6413 | 6414 | 1 | 1.2290e-01 | 4.7698e+05 | ; | C - O |
| 6413 | 6415 | 1 | 1.3350e-01 | 4.1003e+05 | ; | C - N |
| 6415 | 6416 | 1 | 1.0100e-01 | 3.6317e+05 | ; | N - H |
| 6415 | 6417 | 1 | 1.4490e-01 | 2.8200e+05 | ; | N - CA |
| 6417 | 6418 | 1 | 1.0900e-01 | 2.8451e+05 | ; | CA - HA |
| 6417 | 6419 | 1 | 1.5260e-01 | 2.5941e+05 | ; | CA - CB |
| 6417 | 6433 | 1 | 1.5220e-01 | 2.6527e+05 | ; | CA - C |
| 6419 | 6420 | 1 | 1.0900e-01 | 2.8451e+05 | ; | CB - HB2 |
| 6419 | 6421 | 1 | 1.0900e-01 | 2.8451e+05 | ; | CB - HB3 |
| 6419 | 6422 | 1 | 1.5100e-01 | 2.6527e+05 | ; | CB - CG |
| 6422 | 6423 | 1 | 1.4000e-01 | 3.9246e+05 | ; | CG - CD1 |
| 6422 | 6431 | 1 | 1.4000e-01 | 3.9246e+05 | ; | CG - CD2 |
| 6423 | 6424 | 1 | 1.0800e-01 | 3.0711e+05 | ; | CD1 - HD1 |
| 6423 | 6425 | 1 | 1.4000e-01 | 3.9246e+05 | ; | CD1 - CE1 |
| 6425 | 6426 | 1 | 1.0800e-01 | 3.0711e+05 | ; | CE1 - HE1 |
| 6425 | 6427 | 1 | 1.4000e-01 | 3.9246e+05 | ; | CE1 - CZ |
| 6427 | 6428 | 1 | 1.0800e-01 | 3.0711e+05 | ; | CZ - HZ |
| 6427 | 6429 | 1 | 1.4000e-01 | 3.9246e+05 | ; | CZ - CE2 |
| 6429 | 6430 | 1 | 1.0800e-01 | 3.0711e+05 | ; | CE2 - HE2 |
| 6429 | 6431 | 1 | 1.4000e-01 | 3.9246e+05 | ; | CE2 - CD2 |

| | | | | | | |
|---|---|---|---|---|---|---|
| 6431 | 6432 | 1 | 1.0800e-01 | 3.0711e+05 | ; | CD2 - HD2 |
| 6433 | 6434 | 1 | 1.2290e-01 | 4.7698e+05 | ; | C - O |
| 6433 | 6435 | 1 | 1.3350e-01 | 4.1003e+05 | ; | C - N |
| 6435 | 6436 | 1 | 1.0100e-01 | 3.6317e+05 | ; | N - H |
| 6435 | 6437 | 1 | 1.4490e-01 | 2.8200e+05 | ; | N - CA |
| 6437 | 6438 | 1 | 1.0900e-01 | 2.8451e+05 | ; | CA - HA |
| 6437 | 6439 | 1 | 1.5260e-01 | 2.5941e+05 | ; | CA - CB |
| 6437 | 6445 | 1 | 1.5220e-01 | 2.6527e+05 | ; | CA - C |
| 6439 | 6440 | 1 | 1.0900e-01 | 2.8451e+05 | ; | CB - HB2 |
| 6439 | 6441 | 1 | 1.0900e-01 | 2.8451e+05 | ; | CB - HB3 |
| 6439 | 6442 | 1 | 1.5220e-01 | 2.6527e+05 | ; | CB - CG |
| 6442 | 6443 | 1 | 1.2500e-01 | 5.4894e+05 | ; | CG - OD1 |
| 6442 | 6444 | 1 | 1.2500e-01 | 5.4894e+05 | ; | CG - OD2 |
| 6445 | 6446 | 1 | 1.2290e-01 | 4.7698e+05 | ; | C - O |
| 6445 | 6447 | 1 | 1.3350e-01 | 4.1003e+05 | ; | C - N |
| 6447 | 6448 | 1 | 1.0100e-01 | 3.6317e+05 | ; | N - H |
| 6447 | 6449 | 1 | 1.4490e-01 | 2.8200e+05 | ; | N - CA |
| 6449 | 6450 | 1 | 1.0900e-01 | 2.8451e+05 | ; | CA - HA |
| 6449 | 6451 | 1 | 1.5260e-01 | 2.5941e+05 | ; | CA - CB |
| 6449 | 6464 | 1 | 1.5220e-01 | 2.6527e+05 | ; | CA - C |
| 6451 | 6452 | 1 | 1.0900e-01 | 2.8451e+05 | ; | CB - HB |
| 6451 | 6453 | 1 | 1.5260e-01 | 2.5941e+05 | ; | CB - CG2 |
| 6451 | 6457 | 1 | 1.5260e-01 | 2.5941e+05 | ; | CB - CG1 |
| 6453 | 6454 | 1 | 1.0900e-01 | 2.8451e+05 | ; | CG2 - HG21 |
| 6453 | 6455 | 1 | 1.0900e-01 | 2.8451e+05 | ; | CG2 - HG22 |
| 6453 | 6456 | 1 | 1.0900e-01 | 2.8451e+05 | ; | CG2 - HG23 |
| 6457 | 6458 | 1 | 1.0900e-01 | 2.8451e+05 | ; | CG1 - HG12 |
| 6457 | 6459 | 1 | 1.0900e-01 | 2.8451e+05 | ; | CG1 - HG13 |
| 6457 | 6460 | 1 | 1.5260e-01 | 2.5941e+05 | ; | CG1 - CD1 |
| 6460 | 6461 | 1 | 1.0900e-01 | 2.8451e+05 | ; | CD1 - HD11 |
| 6460 | 6462 | 1 | 1.0900e-01 | 2.8451e+05 | ; | CD1 - HD12 |
| 6460 | 6463 | 1 | 1.0900e-01 | 2.8451e+05 | ; | CD1 - HD13 |
| 6464 | 6465 | 1 | 1.2290e-01 | 4.7698e+05 | ; | C - O |
| 6464 | 6466 | 1 | 1.3350e-01 | 4.1003e+05 | ; | C - N |
| 6466 | 6467 | 1 | 1.0100e-01 | 3.6317e+05 | ; | N - H |
| 6466 | 6468 | 1 | 1.4490e-01 | 2.8200e+05 | ; | N - CA |
| 6468 | 6469 | 1 | 1.0900e-01 | 2.8451e+05 | ; | CA - HA |
| 6468 | 6470 | 1 | 1.5260e-01 | 2.5941e+05 | ; | CA - CB |
| 6468 | 6476 | 1 | 1.5220e-01 | 2.6527e+05 | ; | CA - C |
| 6470 | 6471 | 1 | 1.0900e-01 | 2.8451e+05 | ; | CB - HB2 |
| 6470 | 6472 | 1 | 1.0900e-01 | 2.8451e+05 | ; | CB - HB3 |
| 6470 | 6473 | 1 | 1.5220e-01 | 2.6527e+05 | ; | CB - CG |
| 6473 | 6474 | 1 | 1.2500e-01 | 5.4894e+05 | ; | CG - OD1 |
| 6473 | 6475 | 1 | 1.2500e-01 | 5.4894e+05 | ; | CG - OD2 |
| 6476 | 6477 | 1 | 1.2290e-01 | 4.7698e+05 | ; | C - O |
| 6476 | 6478 | 1 | 1.3350e-01 | 4.1003e+05 | ; | C - N |
| 6478 | 6479 | 1 | 1.0100e-01 | 3.6317e+05 | ; | N - H |
| 6478 | 6480 | 1 | 1.4490e-01 | 2.8200e+05 | ; | N - CA |
| 6480 | 6481 | 1 | 1.0900e-01 | 2.8451e+05 | ; | CA - HA2 |
| 6480 | 6482 | 1 | 1.0900e-01 | 2.8451e+05 | ; | CA - HA3 |
| 6480 | 6483 | 1 | 1.5220e-01 | 2.6527e+05 | ; | CA - C |
| 6483 | 6484 | 1 | 1.2290e-01 | 4.7698e+05 | ; | C - O |

| | | | | | | |
|---|---|---|---|---|---|---|
| 6483 | 6485 | 1 | 1.3350e-01 | 4.1003e+05 | ; | C - N |
| 6485 | 6486 | 1 | 1.0100e-01 | 3.6317e+05 | ; | N - H |
| 6485 | 6487 | 1 | 1.4490e-01 | 2.8200e+05 | ; | N - CA |
| 6487 | 6488 | 1 | 1.0900e-01 | 2.8451e+05 | ; | CA - HA |
| 6487 | 6489 | 1 | 1.5260e-01 | 2.5941e+05 | ; | CA - CB |
| 6487 | 6497 | 1 | 1.5220e-01 | 2.6527e+05 | ; | CA - C |
| 6489 | 6490 | 1 | 1.0900e-01 | 2.8451e+05 | ; | CB - HB2 |
| 6489 | 6491 | 1 | 1.0900e-01 | 2.8451e+05 | ; | CB - HB3 |
| 6489 | 6492 | 1 | 1.5220e-01 | 2.6527e+05 | ; | CB - CG |
| 6492 | 6493 | 1 | 1.2290e-01 | 4.7698e+05 | ; | CG - OD1 |
| 6492 | 6494 | 1 | 1.3350e-01 | 4.1003e+05 | ; | CG - ND2 |
| 6494 | 6495 | 1 | 1.0100e-01 | 3.6317e+05 | ; | ND2 - HD21 |
| 6494 | 6496 | 1 | 1.0100e-01 | 3.6317e+05 | ; | ND2 - HD22 |
| 6497 | 6498 | 1 | 1.2290e-01 | 4.7698e+05 | ; | C - O |
| 6497 | 6499 | 1 | 1.3350e-01 | 4.1003e+05 | ; | C - N |
| 6499 | 6500 | 1 | 1.0100e-01 | 3.6317e+05 | ; | N - H |
| 6499 | 6501 | 1 | 1.4490e-01 | 2.8200e+05 | ; | N - CA |
| 6501 | 6502 | 1 | 1.0900e-01 | 2.8451e+05 | ; | CA - HA2 |
| 6501 | 6503 | 1 | 1.0900e-01 | 2.8451e+05 | ; | CA - HA3 |
| 6501 | 6504 | 1 | 1.5220e-01 | 2.6527e+05 | ; | CA - C |
| 6504 | 6505 | 1 | 1.2290e-01 | 4.7698e+05 | ; | C - O |
| 6504 | 6506 | 1 | 1.3350e-01 | 4.1003e+05 | ; | C - N |
| 6506 | 6507 | 1 | 1.0100e-01 | 3.6317e+05 | ; | N - H |
| 6506 | 6508 | 1 | 1.4490e-01 | 2.8200e+05 | ; | N - CA |
| 6508 | 6509 | 1 | 1.0900e-01 | 2.8451e+05 | ; | CA - HA |
| 6508 | 6510 | 1 | 1.5260e-01 | 2.5941e+05 | ; | CA - CB |
| 6508 | 6526 | 1 | 1.5220e-01 | 2.6527e+05 | ; | CA - C |
| 6510 | 6511 | 1 | 1.0900e-01 | 2.8451e+05 | ; | CB - HB2 |
| 6510 | 6512 | 1 | 1.0900e-01 | 2.8451e+05 | ; | CB - HB3 |
| 6510 | 6513 | 1 | 1.5260e-01 | 2.5941e+05 | ; | CB - CG |
| 6513 | 6514 | 1 | 1.0900e-01 | 2.8451e+05 | ; | CG - HG2 |
| 6513 | 6515 | 1 | 1.0900e-01 | 2.8451e+05 | ; | CG - HG3 |
| 6513 | 6516 | 1 | 1.5260e-01 | 2.5941e+05 | ; | CG - CD |
| 6516 | 6517 | 1 | 1.0900e-01 | 2.8451e+05 | ; | CD - HD2 |
| 6516 | 6518 | 1 | 1.0900e-01 | 2.8451e+05 | ; | CD - HD3 |
| 6516 | 6519 | 1 | 1.5260e-01 | 2.5941e+05 | ; | CD - CE |
| 6519 | 6520 | 1 | 1.0900e-01 | 2.8451e+05 | ; | CE - HE2 |
| 6519 | 6521 | 1 | 1.0900e-01 | 2.8451e+05 | ; | CE - HE3 |
| 6519 | 6522 | 1 | 1.4710e-01 | 3.0711e+05 | ; | CE - NZ |
| 6522 | 6523 | 1 | 1.0100e-01 | 3.6317e+05 | ; | NZ - HZ1 |
| 6522 | 6524 | 1 | 1.0100e-01 | 3.6317e+05 | ; | NZ - HZ2 |
| 6522 | 6525 | 1 | 1.0100e-01 | 3.6317e+05 | ; | NZ - HZ3 |
| 6526 | 6527 | 1 | 1.2290e-01 | 4.7698e+05 | ; | C - O |
| 6526 | 6528 | 1 | 1.3350e-01 | 4.1003e+05 | ; | C - N |
| 6528 | 6529 | 1 | 1.0100e-01 | 3.6317e+05 | ; | N - H |
| 6528 | 6530 | 1 | 1.4490e-01 | 2.8200e+05 | ; | N - CA |
| 6530 | 6531 | 1 | 1.0900e-01 | 2.8451e+05 | ; | CA - HA |
| 6530 | 6532 | 1 | 1.5260e-01 | 2.5941e+05 | ; | CA - CB |
| 6530 | 6545 | 1 | 1.5220e-01 | 2.6527e+05 | ; | CA - C |
| 6532 | 6533 | 1 | 1.0900e-01 | 2.8451e+05 | ; | CB - HB |
| 6532 | 6534 | 1 | 1.5260e-01 | 2.5941e+05 | ; | CB - CG2 |
| 6532 | 6538 | 1 | 1.5260e-01 | 2.5941e+05 | ; | CB - CG1 |

| | | | | | | |
|---|---|---|---|---|---|---|
| 6534 | 6535 | 1 | 1.0900e-01 | 2.8451e+05 | ; | CG2 - HG21 |
| 6534 | 6536 | 1 | 1.0900e-01 | 2.8451e+05 | ; | CG2 - HG22 |
| 6534 | 6537 | 1 | 1.0900e-01 | 2.8451e+05 | ; | CG2 - HG23 |
| 6538 | 6539 | 1 | 1.0900e-01 | 2.8451e+05 | ; | CG1 - HG12 |
| 6538 | 6540 | 1 | 1.0900e-01 | 2.8451e+05 | ; | CG1 - HG13 |
| 6538 | 6541 | 1 | 1.5260e-01 | 2.5941e+05 | ; | CG1 - CD1 |
| 6541 | 6542 | 1 | 1.0900e-01 | 2.8451e+05 | ; | CD1 - HD11 |
| 6541 | 6543 | 1 | 1.0900e-01 | 2.8451e+05 | ; | CD1 - HD12 |
| 6541 | 6544 | 1 | 1.0900e-01 | 2.8451e+05 | ; | CD1 - HD13 |
| 6545 | 6546 | 1 | 1.2290e-01 | 4.7698e+05 | ; | C - O |
| 6545 | 6547 | 1 | 1.3350e-01 | 4.1003e+05 | ; | C - N |
| 6547 | 6548 | 1 | 1.0100e-01 | 3.6317e+05 | ; | N - H |
| 6547 | 6549 | 1 | 1.4490e-01 | 2.8200e+05 | ; | N - CA |
| 6549 | 6550 | 1 | 1.0900e-01 | 2.8451e+05 | ; | CA - HA |
| 6549 | 6551 | 1 | 1.5260e-01 | 2.5941e+05 | ; | CA - CB |
| 6549 | 6556 | 1 | 1.5220e-01 | 2.6527e+05 | ; | CA - C |
| 6551 | 6552 | 1 | 1.0900e-01 | 2.8451e+05 | ; | CB - HB2 |
| 6551 | 6553 | 1 | 1.0900e-01 | 2.8451e+05 | ; | CB - HB3 |
| 6551 | 6554 | 1 | 1.4100e-01 | 2.6778e+05 | ; | CB - OG |
| 6554 | 6555 | 1 | 9.6000e-02 | 4.6275e+05 | ; | OG - HG |
| 6556 | 6557 | 1 | 1.2290e-01 | 4.7698e+05 | ; | C - O |
| 6556 | 6558 | 1 | 1.3350e-01 | 4.1003e+05 | ; | C - N |
| 6558 | 6559 | 1 | 1.0100e-01 | 3.6317e+05 | ; | N - H |
| 6558 | 6560 | 1 | 1.4490e-01 | 2.8200e+05 | ; | N - CA |
| 6560 | 6561 | 1 | 1.0900e-01 | 2.8451e+05 | ; | CA - HA |
| 6560 | 6562 | 1 | 1.5260e-01 | 2.5941e+05 | ; | CA - CB |
| 6560 | 6572 | 1 | 1.5220e-01 | 2.6527e+05 | ; | CA - C |
| 6562 | 6563 | 1 | 1.0900e-01 | 2.8451e+05 | ; | CB - HB |
| 6562 | 6564 | 1 | 1.5260e-01 | 2.5941e+05 | ; | CB - CG1 |
| 6562 | 6568 | 1 | 1.5260e-01 | 2.5941e+05 | ; | CB - CG2 |
| 6564 | 6565 | 1 | 1.0900e-01 | 2.8451e+05 | ; | CG1 - HG11 |
| 6564 | 6566 | 1 | 1.0900e-01 | 2.8451e+05 | ; | CG1 - HG12 |
| 6564 | 6567 | 1 | 1.0900e-01 | 2.8451e+05 | ; | CG1 - HG13 |
| 6568 | 6569 | 1 | 1.0900e-01 | 2.8451e+05 | ; | CG2 - HG21 |
| 6568 | 6570 | 1 | 1.0900e-01 | 2.8451e+05 | ; | CG2 - HG22 |
| 6568 | 6571 | 1 | 1.0900e-01 | 2.8451e+05 | ; | CG2 - HG23 |
| 6572 | 6573 | 1 | 1.2290e-01 | 4.7698e+05 | ; | C - O |
| 6572 | 6574 | 1 | 1.3350e-01 | 4.1003e+05 | ; | C - N |
| 6574 | 6575 | 1 | 1.0100e-01 | 3.6317e+05 | ; | N - H |
| 6574 | 6576 | 1 | 1.4490e-01 | 2.8200e+05 | ; | N - CA |
| 6576 | 6577 | 1 | 1.0900e-01 | 2.8451e+05 | ; | CA - HA |
| 6576 | 6578 | 1 | 1.5260e-01 | 2.5941e+05 | ; | CA - CB |
| 6576 | 6587 | 1 | 1.5220e-01 | 2.6527e+05 | ; | CA - C |
| 6578 | 6579 | 1 | 1.0900e-01 | 2.8451e+05 | ; | CB - HB2 |
| 6578 | 6580 | 1 | 1.0900e-01 | 2.8451e+05 | ; | CB - HB3 |
| 6578 | 6581 | 1 | 1.5260e-01 | 2.5941e+05 | ; | CB - CG |
| 6581 | 6582 | 1 | 1.0900e-01 | 2.8451e+05 | ; | CG - HG2 |
| 6581 | 6583 | 1 | 1.0900e-01 | 2.8451e+05 | ; | CG - HG3 |
| 6581 | 6584 | 1 | 1.5220e-01 | 2.6527e+05 | ; | CG - CD |
| 6584 | 6585 | 1 | 1.2500e-01 | 5.4894e+05 | ; | CD - OE1 |
| 6584 | 6586 | 1 | 1.2500e-01 | 5.4894e+05 | ; | CD - OE2 |
| 6587 | 6588 | 1 | 1.2290e-01 | 4.7698e+05 | ; | C - O |

| | | | | | | | | |
|---|---|---|---|---|---|---|---|---|
| 6587 | 6589 | 1 | 1.3350e-01 | 4.1003e+05 | ; | C | - | N |
| 6589 | 6590 | 1 | 1.0100e-01 | 3.6317e+05 | ; | N | - | H |
| 6589 | 6591 | 1 | 1.4490e-01 | 2.8200e+05 | ; | N | - | CA |
| 6591 | 6592 | 1 | 1.0900e-01 | 2.8451e+05 | ; | CA | - | HA |
| 6591 | 6593 | 1 | 1.5260e-01 | 2.5941e+05 | ; | CA | - | CB |
| 6591 | 6603 | 1 | 1.5220e-01 | 2.6527e+05 | ; | CA | - | C |
| 6593 | 6594 | 1 | 1.0900e-01 | 2.8451e+05 | ; | CB | - | HB2 |
| 6593 | 6595 | 1 | 1.0900e-01 | 2.8451e+05 | ; | CB | - | HB3 |
| 6593 | 6596 | 1 | 1.5260e-01 | 2.5941e+05 | ; | CB | - | CG |
| 6596 | 6597 | 1 | 1.0900e-01 | 2.8451e+05 | ; | CG | - | HG2 |
| 6596 | 6598 | 1 | 1.0900e-01 | 2.8451e+05 | ; | CG | - | HG3 |
| 6596 | 6599 | 1 | 1.5220e-01 | 2.6527e+05 | ; | CG | - | CD |
| 6599 | 6600 | 1 | 1.2290e-01 | 4.7698e+05 | ; | CD | - | OE1 |
| 6599 | 6601 | 1 | 1.3640e-01 | 3.7656e+05 | ; | CD | - | OE2 |
| 6601 | 6602 | 1 | 9.6000e-02 | 4.6275e+05 | ; | OE2 | - | HE2 |
| 6603 | 6604 | 1 | 1.2290e-01 | 4.7698e+05 | ; | C | - | O |
| 6603 | 6605 | 1 | 1.3350e-01 | 4.1003e+05 | ; | C | - | N |
| 6605 | 6606 | 1 | 1.0100e-01 | 3.6317e+05 | ; | N | - | H |
| 6605 | 6607 | 1 | 1.4490e-01 | 2.8200e+05 | ; | N | - | CA |
| 6607 | 6608 | 1 | 1.0900e-01 | 2.8451e+05 | ; | CA | - | HA |
| 6607 | 6609 | 1 | 1.5260e-01 | 2.5941e+05 | ; | CA | - | CB |
| 6607 | 6622 | 1 | 1.5220e-01 | 2.6527e+05 | ; | CA | - | C |
| 6609 | 6610 | 1 | 1.0900e-01 | 2.8451e+05 | ; | CB | - | HB2 |
| 6609 | 6611 | 1 | 1.0900e-01 | 2.8451e+05 | ; | CB | - | HB3 |
| 6609 | 6612 | 1 | 1.5260e-01 | 2.5941e+05 | ; | CB | - | CG |
| 6612 | 6613 | 1 | 1.0900e-01 | 2.8451e+05 | ; | CG | - | HG |
| 6612 | 6614 | 1 | 1.5260e-01 | 2.5941e+05 | ; | CG | - | CD1 |
| 6612 | 6618 | 1 | 1.5260e-01 | 2.5941e+05 | ; | CG | - | CD2 |
| 6614 | 6615 | 1 | 1.0900e-01 | 2.8451e+05 | ; | CD1 | - | HD11 |
| 6614 | 6616 | 1 | 1.0900e-01 | 2.8451e+05 | ; | CD1 | - | HD12 |
| 6614 | 6617 | 1 | 1.0900e-01 | 2.8451e+05 | ; | CD1 | - | HD13 |
| 6618 | 6619 | 1 | 1.0900e-01 | 2.8451e+05 | ; | CD2 | - | HD21 |
| 6618 | 6620 | 1 | 1.0900e-01 | 2.8451e+05 | ; | CD2 | - | HD22 |
| 6618 | 6621 | 1 | 1.0900e-01 | 2.8451e+05 | ; | CD2 | - | HD23 |
| 6622 | 6623 | 1 | 1.2290e-01 | 4.7698e+05 | ; | C | - | O |
| 6622 | 6624 | 1 | 1.3350e-01 | 4.1003e+05 | ; | C | - | N |
| 6624 | 6625 | 1 | 1.0100e-01 | 3.6317e+05 | ; | N | - | H |
| 6624 | 6626 | 1 | 1.4490e-01 | 2.8200e+05 | ; | N | - | CA |
| 6626 | 6627 | 1 | 1.0900e-01 | 2.8451e+05 | ; | CA | - | HA |
| 6626 | 6628 | 1 | 1.5260e-01 | 2.5941e+05 | ; | CA | - | CB |
| 6626 | 6644 | 1 | 1.5220e-01 | 2.6527e+05 | ; | CA | - | C |
| 6628 | 6629 | 1 | 1.0900e-01 | 2.8451e+05 | ; | CB | - | HB2 |
| 6628 | 6630 | 1 | 1.0900e-01 | 2.8451e+05 | ; | CB | - | HB3 |
| 6628 | 6631 | 1 | 1.5260e-01 | 2.5941e+05 | ; | CB | - | CG |
| 6631 | 6632 | 1 | 1.0900e-01 | 2.8451e+05 | ; | CG | - | HG2 |
| 6631 | 6633 | 1 | 1.0900e-01 | 2.8451e+05 | ; | CG | - | HG3 |
| 6631 | 6634 | 1 | 1.5260e-01 | 2.5941e+05 | ; | CG | - | CD |
| 6634 | 6635 | 1 | 1.0900e-01 | 2.8451e+05 | ; | CD | - | HD2 |
| 6634 | 6636 | 1 | 1.0900e-01 | 2.8451e+05 | ; | CD | - | HD3 |
| 6634 | 6637 | 1 | 1.5260e-01 | 2.5941e+05 | ; | CD | - | CE |
| 6637 | 6638 | 1 | 1.0900e-01 | 2.8451e+05 | ; | CE | - | HE2 |
| 6637 | 6639 | 1 | 1.0900e-01 | 2.8451e+05 | ; | CE | - | HE3 |

| | | | | | | |
|---|---|---|---|---|---|---|
| 6637 | 6640 | 1 | 1.4710e-01 | 3.0711e+05 | ; | CE - NZ |
| 6640 | 6641 | 1 | 1.0100e-01 | 3.6317e+05 | ; | NZ - HZ1 |
| 6640 | 6642 | 1 | 1.0100e-01 | 3.6317e+05 | ; | NZ - HZ2 |
| 6640 | 6643 | 1 | 1.0100e-01 | 3.6317e+05 | ; | NZ - HZ3 |
| 6644 | 6645 | 1 | 1.2290e-01 | 4.7698e+05 | ; | C - O |
| 6644 | 6646 | 1 | 1.3350e-01 | 4.1003e+05 | ; | C - N |
| 6646 | 6647 | 1 | 1.0100e-01 | 3.6317e+05 | ; | N - H |
| 6646 | 6648 | 1 | 1.4490e-01 | 2.8200e+05 | ; | N - CA |
| 6648 | 6649 | 1 | 1.0900e-01 | 2.8451e+05 | ; | CA - HA |
| 6648 | 6650 | 1 | 1.5260e-01 | 2.5941e+05 | ; | CA - CB |
| 6648 | 6668 | 1 | 1.5220e-01 | 2.6527e+05 | ; | CA - C |
| 6650 | 6651 | 1 | 1.0900e-01 | 2.8451e+05 | ; | CB - HB2 |
| 6650 | 6652 | 1 | 1.0900e-01 | 2.8451e+05 | ; | CB - HB3 |
| 6650 | 6653 | 1 | 1.5260e-01 | 2.5941e+05 | ; | CB - CG |
| 6653 | 6654 | 1 | 1.0900e-01 | 2.8451e+05 | ; | CG - HG2 |
| 6653 | 6655 | 1 | 1.0900e-01 | 2.8451e+05 | ; | CG - HG3 |
| 6653 | 6656 | 1 | 1.5260e-01 | 2.5941e+05 | ; | CG - CD |
| 6656 | 6657 | 1 | 1.0900e-01 | 2.8451e+05 | ; | CD - HD2 |
| 6656 | 6658 | 1 | 1.0900e-01 | 2.8451e+05 | ; | CD - HD3 |
| 6656 | 6659 | 1 | 1.4630e-01 | 2.8200e+05 | ; | CD - NE |
| 6659 | 6660 | 1 | 1.0100e-01 | 3.6317e+05 | ; | NE - HE |
| 6659 | 6661 | 1 | 1.3400e-01 | 4.0250e+05 | ; | NE - CZ |
| 6661 | 6662 | 1 | 1.3400e-01 | 4.0250e+05 | ; | CZ - NH1 |
| 6661 | 6665 | 1 | 1.3400e-01 | 4.0250e+05 | ; | CZ - NH2 |
| 6662 | 6663 | 1 | 1.0100e-01 | 3.6317e+05 | ; | NH1 - HH11 |
| 6662 | 6664 | 1 | 1.0100e-01 | 3.6317e+05 | ; | NH1 - HH12 |
| 6665 | 6666 | 1 | 1.0100e-01 | 3.6317e+05 | ; | NH2 - HH21 |
| 6665 | 6667 | 1 | 1.0100e-01 | 3.6317e+05 | ; | NH2 - HH22 |
| 6668 | 6669 | 1 | 1.2290e-01 | 4.7698e+05 | ; | C - O |
| 6668 | 6670 | 1 | 1.3350e-01 | 4.1003e+05 | ; | C - N |
| 6670 | 6671 | 1 | 1.0100e-01 | 3.6317e+05 | ; | N - H |
| 6670 | 6672 | 1 | 1.4490e-01 | 2.8200e+05 | ; | N - CA |
| 6672 | 6673 | 1 | 1.0900e-01 | 2.8451e+05 | ; | CA - HA |
| 6672 | 6674 | 1 | 1.5260e-01 | 2.5941e+05 | ; | CA - CB |
| 6672 | 6687 | 1 | 1.5220e-01 | 2.6527e+05 | ; | CA - C |
| 6674 | 6675 | 1 | 1.0900e-01 | 2.8451e+05 | ; | CB - HB |
| 6674 | 6676 | 1 | 1.5260e-01 | 2.5941e+05 | ; | CB - CG2 |
| 6674 | 6680 | 1 | 1.5260e-01 | 2.5941e+05 | ; | CB - CG1 |
| 6676 | 6677 | 1 | 1.0900e-01 | 2.8451e+05 | ; | CG2 - HG21 |
| 6676 | 6678 | 1 | 1.0900e-01 | 2.8451e+05 | ; | CG2 - HG22 |
| 6676 | 6679 | 1 | 1.0900e-01 | 2.8451e+05 | ; | CG2 - HG23 |
| 6680 | 6681 | 1 | 1.0900e-01 | 2.8451e+05 | ; | CG1 - HG12 |
| 6680 | 6682 | 1 | 1.0900e-01 | 2.8451e+05 | ; | CG1 - HG13 |
| 6680 | 6683 | 1 | 1.5260e-01 | 2.5941e+05 | ; | CG1 - CD1 |
| 6683 | 6684 | 1 | 1.0900e-01 | 2.8451e+05 | ; | CD1 - HD11 |
| 6683 | 6685 | 1 | 1.0900e-01 | 2.8451e+05 | ; | CD1 - HD12 |
| 6683 | 6686 | 1 | 1.0900e-01 | 2.8451e+05 | ; | CD1 - HD13 |
| 6687 | 6688 | 1 | 1.2290e-01 | 4.7698e+05 | ; | C - O |
| 6687 | 6689 | 1 | 1.3350e-01 | 4.1003e+05 | ; | C - N |
| 6689 | 6690 | 1 | 1.0100e-01 | 3.6317e+05 | ; | N - H |
| 6689 | 6691 | 1 | 1.4490e-01 | 2.8200e+05 | ; | N - CA |
| 6691 | 6692 | 1 | 1.0900e-01 | 2.8451e+05 | ; | CA - HA |

| | | | | | | |
|---|---|---|---|---|---|---|
| 6691 | 6693 | 1 | 1.5260e-01 | 2.5941e+05 | ; | CA - CB |
| 6691 | 6707 | 1 | 1.5220e-01 | 2.6527e+05 | ; | CA - C |
| 6693 | 6694 | 1 | 1.0900e-01 | 2.8451e+05 | ; | CB - HB2 |
| 6693 | 6695 | 1 | 1.0900e-01 | 2.8451e+05 | ; | CB - HB3 |
| 6693 | 6696 | 1 | 1.5100e-01 | 2.6527e+05 | ; | CB - CG |
| 6696 | 6697 | 1 | 1.4000e-01 | 3.9246e+05 | ; | CG - CD1 |
| 6696 | 6705 | 1 | 1.4000e-01 | 3.9246e+05 | ; | CG - CD2 |
| 6697 | 6698 | 1 | 1.0800e-01 | 3.0711e+05 | ; | CD1 - HD1 |
| 6697 | 6699 | 1 | 1.4000e-01 | 3.9246e+05 | ; | CD1 - CE1 |
| 6699 | 6700 | 1 | 1.0800e-01 | 3.0711e+05 | ; | CE1 - HE1 |
| 6699 | 6701 | 1 | 1.4000e-01 | 3.9246e+05 | ; | CE1 - CZ |
| 6701 | 6702 | 1 | 1.0800e-01 | 3.0711e+05 | ; | CZ - HZ |
| 6701 | 6703 | 1 | 1.4000e-01 | 3.9246e+05 | ; | CZ - CE2 |
| 6703 | 6704 | 1 | 1.0800e-01 | 3.0711e+05 | ; | CE2 - HE2 |
| 6703 | 6705 | 1 | 1.4000e-01 | 3.9246e+05 | ; | CE2 - CD2 |
| 6705 | 6706 | 1 | 1.0800e-01 | 3.0711e+05 | ; | CD2 - HD2 |
| 6707 | 6708 | 1 | 1.2290e-01 | 4.7698e+05 | ; | C - O |
| 6707 | 6709 | 1 | 1.3350e-01 | 4.1003e+05 | ; | C - N |
| 6709 | 6710 | 1 | 1.0100e-01 | 3.6317e+05 | ; | N - H |
| 6709 | 6711 | 1 | 1.4490e-01 | 2.8200e+05 | ; | N - CA |
| 6711 | 6712 | 1 | 1.0900e-01 | 2.8451e+05 | ; | CA - HA2 |
| 6711 | 6713 | 1 | 1.0900e-01 | 2.8451e+05 | ; | CA - HA3 |
| 6711 | 6714 | 1 | 1.5220e-01 | 2.6527e+05 | ; | CA - C |
| 6714 | 6715 | 1 | 1.2290e-01 | 4.7698e+05 | ; | C - O |
| 6714 | 6716 | 1 | 1.3350e-01 | 4.1003e+05 | ; | C - N |
| 6716 | 6717 | 1 | 1.0100e-01 | 3.6317e+05 | ; | N - H |
| 6716 | 6718 | 1 | 1.4490e-01 | 2.8200e+05 | ; | N - CA |
| 6718 | 6719 | 1 | 1.0900e-01 | 2.8451e+05 | ; | CA - HA |
| 6718 | 6720 | 1 | 1.5260e-01 | 2.5941e+05 | ; | CA - CB |
| 6718 | 6738 | 1 | 1.5220e-01 | 2.6527e+05 | ; | CA - C |
| 6720 | 6721 | 1 | 1.0900e-01 | 2.8451e+05 | ; | CB - HB2 |
| 6720 | 6722 | 1 | 1.0900e-01 | 2.8451e+05 | ; | CB - HB3 |
| 6720 | 6723 | 1 | 1.5260e-01 | 2.5941e+05 | ; | CB - CG |
| 6723 | 6724 | 1 | 1.0900e-01 | 2.8451e+05 | ; | CG - HG2 |
| 6723 | 6725 | 1 | 1.0900e-01 | 2.8451e+05 | ; | CG - HG3 |
| 6723 | 6726 | 1 | 1.5260e-01 | 2.5941e+05 | ; | CG - CD |
| 6726 | 6727 | 1 | 1.0900e-01 | 2.8451e+05 | ; | CD - HD2 |
| 6726 | 6728 | 1 | 1.0900e-01 | 2.8451e+05 | ; | CD - HD3 |
| 6726 | 6729 | 1 | 1.4630e-01 | 2.8200e+05 | ; | CD - NE |
| 6729 | 6730 | 1 | 1.0100e-01 | 3.6317e+05 | ; | NE - HE |
| 6729 | 6731 | 1 | 1.3400e-01 | 4.0250e+05 | ; | NE - CZ |
| 6731 | 6732 | 1 | 1.3400e-01 | 4.0250e+05 | ; | CZ - NH1 |
| 6731 | 6735 | 1 | 1.3400e-01 | 4.0250e+05 | ; | CZ - NH2 |
| 6732 | 6733 | 1 | 1.0100e-01 | 3.6317e+05 | ; | NH1 - HH11 |
| 6732 | 6734 | 1 | 1.0100e-01 | 3.6317e+05 | ; | NH1 - HH12 |
| 6735 | 6736 | 1 | 1.0100e-01 | 3.6317e+05 | ; | NH2 - HH21 |
| 6735 | 6737 | 1 | 1.0100e-01 | 3.6317e+05 | ; | NH2 - HH22 |
| 6738 | 6739 | 1 | 1.2290e-01 | 4.7698e+05 | ; | C - O |
| 6738 | 6740 | 1 | 1.3350e-01 | 4.1003e+05 | ; | C - N |
| 6740 | 6741 | 1 | 1.0100e-01 | 3.6317e+05 | ; | N - H |
| 6740 | 6742 | 1 | 1.4490e-01 | 2.8200e+05 | ; | N - CA |
| 6742 | 6743 | 1 | 1.0900e-01 | 2.8451e+05 | ; | CA - HA |

| | | | | | | |
|---|---|---|---|---|---|---|
| 6742 | 6744 | 1 | 1.5260e-01 | 2.5941e+05 | ; | CA - CB |
| 6742 | 6750 | 1 | 1.5220e-01 | 2.6527e+05 | ; | CA - C |
| 6744 | 6745 | 1 | 1.0900e-01 | 2.8451e+05 | ; | CB - HB2 |
| 6744 | 6746 | 1 | 1.0900e-01 | 2.8451e+05 | ; | CB - HB3 |
| 6744 | 6747 | 1 | 1.5220e-01 | 2.6527e+05 | ; | CB - CG |
| 6747 | 6748 | 1 | 1.2500e-01 | 5.4894e+05 | ; | CG - OD1 |
| 6747 | 6749 | 1 | 1.2500e-01 | 5.4894e+05 | ; | CG - OD2 |
| 6750 | 6751 | 1 | 1.2290e-01 | 4.7698e+05 | ; | C - O |
| 6750 | 6752 | 1 | 1.3350e-01 | 4.1003e+05 | ; | C - N |
| 6752 | 6753 | 1 | 1.0100e-01 | 3.6317e+05 | ; | N - H |
| 6752 | 6754 | 1 | 1.4490e-01 | 2.8200e+05 | ; | N - CA |
| 6754 | 6755 | 1 | 1.0900e-01 | 2.8451e+05 | ; | CA - HA |
| 6754 | 6756 | 1 | 1.5260e-01 | 2.5941e+05 | ; | CA - CB |
| 6754 | 6762 | 1 | 1.5220e-01 | 2.6527e+05 | ; | CA - C |
| 6756 | 6757 | 1 | 1.0900e-01 | 2.8451e+05 | ; | CB - HB2 |
| 6756 | 6758 | 1 | 1.0900e-01 | 2.8451e+05 | ; | CB - HB3 |
| 6756 | 6759 | 1 | 1.5220e-01 | 2.6527e+05 | ; | CB - CG |
| 6759 | 6760 | 1 | 1.2500e-01 | 5.4894e+05 | ; | CG - OD1 |
| 6759 | 6761 | 1 | 1.2500e-01 | 5.4894e+05 | ; | CG - OD2 |
| 6762 | 6763 | 1 | 1.2290e-01 | 4.7698e+05 | ; | C - O |
| 6762 | 6764 | 1 | 1.3350e-01 | 4.1003e+05 | ; | C - N |
| 6764 | 6765 | 1 | 1.0100e-01 | 3.6317e+05 | ; | N - H |
| 6764 | 6766 | 1 | 1.4490e-01 | 2.8200e+05 | ; | N - CA |
| 6766 | 6767 | 1 | 1.0900e-01 | 2.8451e+05 | ; | CA - HA |
| 6766 | 6768 | 1 | 1.5260e-01 | 2.5941e+05 | ; | CA - CB |
| 6766 | 6781 | 1 | 1.5220e-01 | 2.6527e+05 | ; | CA - C |
| 6768 | 6769 | 1 | 1.0900e-01 | 2.8451e+05 | ; | CB - HB |
| 6768 | 6770 | 1 | 1.5260e-01 | 2.5941e+05 | ; | CB - CG2 |
| 6768 | 6774 | 1 | 1.5260e-01 | 2.5941e+05 | ; | CB - CG1 |
| 6770 | 6771 | 1 | 1.0900e-01 | 2.8451e+05 | ; | CG2 - HG21 |
| 6770 | 6772 | 1 | 1.0900e-01 | 2.8451e+05 | ; | CG2 - HG22 |
| 6770 | 6773 | 1 | 1.0900e-01 | 2.8451e+05 | ; | CG2 - HG23 |
| 6774 | 6775 | 1 | 1.0900e-01 | 2.8451e+05 | ; | CG1 - HG12 |
| 6774 | 6776 | 1 | 1.0900e-01 | 2.8451e+05 | ; | CG1 - HG13 |
| 6774 | 6777 | 1 | 1.5260e-01 | 2.5941e+05 | ; | CG1 - CD1 |
| 6777 | 6778 | 1 | 1.0900e-01 | 2.8451e+05 | ; | CD1 - HD11 |
| 6777 | 6779 | 1 | 1.0900e-01 | 2.8451e+05 | ; | CD1 - HD12 |
| 6777 | 6780 | 1 | 1.0900e-01 | 2.8451e+05 | ; | CD1 - HD13 |
| 6781 | 6782 | 1 | 1.2290e-01 | 4.7698e+05 | ; | C - O |
| 6781 | 6783 | 1 | 1.3350e-01 | 4.1003e+05 | ; | C - N |
| 6783 | 6784 | 1 | 1.0100e-01 | 3.6317e+05 | ; | N - H |
| 6783 | 6785 | 1 | 1.4490e-01 | 2.8200e+05 | ; | N - CA |
| 6785 | 6786 | 1 | 1.0900e-01 | 2.8451e+05 | ; | CA - HA |
| 6785 | 6787 | 1 | 1.5260e-01 | 2.5941e+05 | ; | CA - CB |
| 6785 | 6796 | 1 | 1.5220e-01 | 2.6527e+05 | ; | CA - C |
| 6787 | 6788 | 1 | 1.0900e-01 | 2.8451e+05 | ; | CB - HB2 |
| 6787 | 6789 | 1 | 1.0900e-01 | 2.8451e+05 | ; | CB - HB3 |
| 6787 | 6790 | 1 | 1.5260e-01 | 2.5941e+05 | ; | CB - CG |
| 6790 | 6791 | 1 | 1.0900e-01 | 2.8451e+05 | ; | CG - HG2 |
| 6790 | 6792 | 1 | 1.0900e-01 | 2.8451e+05 | ; | CG - HG3 |
| 6790 | 6793 | 1 | 1.5220e-01 | 2.6527e+05 | ; | CG - CD |
| 6793 | 6794 | 1 | 1.2500e-01 | 5.4894e+05 | ; | CD - OE1 |

| | | | | | | |
|---|---|---|---|---|---|---|
| 6793 | 6795 | 1 | 1.2500e-01 | 5.4894e+05 | ; | CD - OE2 |
| 6796 | 6797 | 1 | 1.2290e-01 | 4.7698e+05 | ; | C - O |
| 6796 | 6798 | 1 | 1.3350e-01 | 4.1003e+05 | ; | C - N |
| 6798 | 6799 | 1 | 1.0100e-01 | 3.6317e+05 | ; | N - H |
| 6798 | 6800 | 1 | 1.4490e-01 | 2.8200e+05 | ; | N - CA |
| 6800 | 6801 | 1 | 1.0900e-01 | 2.8451e+05 | ; | CA - HA |
| 6800 | 6802 | 1 | 1.5260e-01 | 2.5941e+05 | ; | CA - CB |
| 6800 | 6810 | 1 | 1.5220e-01 | 2.6527e+05 | ; | CA - C |
| 6802 | 6803 | 1 | 1.0900e-01 | 2.8451e+05 | ; | CB - HB2 |
| 6802 | 6804 | 1 | 1.0900e-01 | 2.8451e+05 | ; | CB - HB3 |
| 6802 | 6805 | 1 | 1.5220e-01 | 2.6527e+05 | ; | CB - CG |
| 6805 | 6806 | 1 | 1.2290e-01 | 4.7698e+05 | ; | CG - OD1 |
| 6805 | 6807 | 1 | 1.3350e-01 | 4.1003e+05 | ; | CG - ND2 |
| 6807 | 6808 | 1 | 1.0100e-01 | 3.6317e+05 | ; | ND2 - HD21 |
| 6807 | 6809 | 1 | 1.0100e-01 | 3.6317e+05 | ; | ND2 - HD22 |
| 6810 | 6811 | 1 | 1.2290e-01 | 4.7698e+05 | ; | C - O |
| 6810 | 6812 | 1 | 1.3350e-01 | 4.1003e+05 | ; | C - N |
| 6812 | 6813 | 1 | 1.4490e-01 | 2.8200e+05 | ; | N - CD |
| 6812 | 6822 | 1 | 1.4490e-01 | 2.8200e+05 | ; | N - CA |
| 6813 | 6814 | 1 | 1.0900e-01 | 2.8451e+05 | ; | CD - HD2 |
| 6813 | 6815 | 1 | 1.0900e-01 | 2.8451e+05 | ; | CD - HD3 |
| 6813 | 6816 | 1 | 1.5260e-01 | 2.5941e+05 | ; | CD - CG |
| 6816 | 6817 | 1 | 1.0900e-01 | 2.8451e+05 | ; | CG - HG2 |
| 6816 | 6818 | 1 | 1.0900e-01 | 2.8451e+05 | ; | CG - HG3 |
| 6816 | 6819 | 1 | 1.5260e-01 | 2.5941e+05 | ; | CG - CB |
| 6819 | 6820 | 1 | 1.0900e-01 | 2.8451e+05 | ; | CB - HB2 |
| 6819 | 6821 | 1 | 1.0900e-01 | 2.8451e+05 | ; | CB - HB3 |
| 6819 | 6822 | 1 | 1.5260e-01 | 2.5941e+05 | ; | CB - CA |
| 6822 | 6823 | 1 | 1.0900e-01 | 2.8451e+05 | ; | CA - HA |
| 6822 | 6824 | 1 | 1.5220e-01 | 2.6527e+05 | ; | CA - C |
| 6824 | 6825 | 1 | 1.2290e-01 | 4.7698e+05 | ; | C - O |
| 6824 | 6826 | 1 | 1.3350e-01 | 4.1003e+05 | ; | C - N |
| 6826 | 6827 | 1 | 1.0100e-01 | 3.6317e+05 | ; | N - H |
| 6826 | 6828 | 1 | 1.4490e-01 | 2.8200e+05 | ; | N - CA |
| 6828 | 6829 | 1 | 1.0900e-01 | 2.8451e+05 | ; | CA - HA |
| 6828 | 6830 | 1 | 1.5260e-01 | 2.5941e+05 | ; | CA - CB |
| 6828 | 6843 | 1 | 1.5220e-01 | 2.6527e+05 | ; | CA - C |
| 6830 | 6831 | 1 | 1.0900e-01 | 2.8451e+05 | ; | CB - HB2 |
| 6830 | 6832 | 1 | 1.0900e-01 | 2.8451e+05 | ; | CB - HB3 |
| 6830 | 6833 | 1 | 1.5260e-01 | 2.5941e+05 | ; | CB - CG |
| 6833 | 6834 | 1 | 1.0900e-01 | 2.8451e+05 | ; | CG - HG |
| 6833 | 6835 | 1 | 1.5260e-01 | 2.5941e+05 | ; | CG - CD1 |
| 6833 | 6839 | 1 | 1.5260e-01 | 2.5941e+05 | ; | CG - CD2 |
| 6835 | 6836 | 1 | 1.0900e-01 | 2.8451e+05 | ; | CD1 - HD11 |
| 6835 | 6837 | 1 | 1.0900e-01 | 2.8451e+05 | ; | CD1 - HD12 |
| 6835 | 6838 | 1 | 1.0900e-01 | 2.8451e+05 | ; | CD1 - HD13 |
| 6839 | 6840 | 1 | 1.0900e-01 | 2.8451e+05 | ; | CD2 - HD21 |
| 6839 | 6841 | 1 | 1.0900e-01 | 2.8451e+05 | ; | CD2 - HD22 |
| 6839 | 6842 | 1 | 1.0900e-01 | 2.8451e+05 | ; | CD2 - HD23 |
| 6843 | 6844 | 1 | 1.2290e-01 | 4.7698e+05 | ; | C - O |
| 6843 | 6845 | 1 | 1.3350e-01 | 4.1003e+05 | ; | C - N |
| 6845 | 6846 | 1 | 1.0100e-01 | 3.6317e+05 | ; | N - H |

| | | | | | | |
|---|---|---|---|---|---|---|
| 6845 | 6847 | 1 | 1.4490e-01 | 2.8200e+05 | ; | N - CA |
| 6847 | 6848 | 1 | 1.0900e-01 | 2.8451e+05 | ; | CA - HA |
| 6847 | 6849 | 1 | 1.5260e-01 | 2.5941e+05 | ; | CA - CB |
| 6847 | 6862 | 1 | 1.5220e-01 | 2.6527e+05 | ; | CA - C |
| 6849 | 6850 | 1 | 1.0900e-01 | 2.8451e+05 | ; | CB - HB |
| 6849 | 6851 | 1 | 1.5260e-01 | 2.5941e+05 | ; | CB - CG2 |
| 6849 | 6855 | 1 | 1.5260e-01 | 2.5941e+05 | ; | CB - CG1 |
| 6851 | 6852 | 1 | 1.0900e-01 | 2.8451e+05 | ; | CG2 - HG21 |
| 6851 | 6853 | 1 | 1.0900e-01 | 2.8451e+05 | ; | CG2 - HG22 |
| 6851 | 6854 | 1 | 1.0900e-01 | 2.8451e+05 | ; | CG2 - HG23 |
| 6855 | 6856 | 1 | 1.0900e-01 | 2.8451e+05 | ; | CG1 - HG12 |
| 6855 | 6857 | 1 | 1.0900e-01 | 2.8451e+05 | ; | CG1 - HG13 |
| 6855 | 6858 | 1 | 1.5260e-01 | 2.5941e+05 | ; | CG1 - CD1 |
| 6858 | 6859 | 1 | 1.0900e-01 | 2.8451e+05 | ; | CD1 - HD11 |
| 6858 | 6860 | 1 | 1.0900e-01 | 2.8451e+05 | ; | CD1 - HD12 |
| 6858 | 6861 | 1 | 1.0900e-01 | 2.8451e+05 | ; | CD1 - HD13 |
| 6862 | 6863 | 1 | 1.2290e-01 | 4.7698e+05 | ; | C - O |
| 6862 | 6864 | 1 | 1.3350e-01 | 4.1003e+05 | ; | C - N |
| 6864 | 6865 | 1 | 1.0100e-01 | 3.6317e+05 | ; | N - H |
| 6864 | 6866 | 1 | 1.4490e-01 | 2.8200e+05 | ; | N - CA |
| 6866 | 6867 | 1 | 1.0900e-01 | 2.8451e+05 | ; | CA - HA |
| 6866 | 6868 | 1 | 1.5260e-01 | 2.5941e+05 | ; | CA - CB |
| 6866 | 6874 | 1 | 1.5220e-01 | 2.6527e+05 | ; | CA - C |
| 6868 | 6869 | 1 | 1.0900e-01 | 2.8451e+05 | ; | CB - HB2 |
| 6868 | 6870 | 1 | 1.0900e-01 | 2.8451e+05 | ; | CB - HB3 |
| 6868 | 6871 | 1 | 1.5220e-01 | 2.6527e+05 | ; | CB - CG |
| 6871 | 6872 | 1 | 1.2500e-01 | 5.4894e+05 | ; | CG - OD1 |
| 6871 | 6873 | 1 | 1.2500e-01 | 5.4894e+05 | ; | CG - OD2 |
| 6874 | 6875 | 1 | 1.2290e-01 | 4.7698e+05 | ; | C - O |
| 6874 | 6876 | 1 | 1.3350e-01 | 4.1003e+05 | ; | C - N |
| 6876 | 6877 | 1 | 1.0100e-01 | 3.6317e+05 | ; | N - H |
| 6876 | 6878 | 1 | 1.4490e-01 | 2.8200e+05 | ; | N - CA |
| 6878 | 6879 | 1 | 1.0900e-01 | 2.8451e+05 | ; | CA - HA |
| 6878 | 6880 | 1 | 1.5260e-01 | 2.5941e+05 | ; | CA - CB |
| 6878 | 6896 | 1 | 1.5220e-01 | 2.6527e+05 | ; | CA - C |
| 6880 | 6881 | 1 | 1.0900e-01 | 2.8451e+05 | ; | CB - HB2 |
| 6880 | 6882 | 1 | 1.0900e-01 | 2.8451e+05 | ; | CB - HB3 |
| 6880 | 6883 | 1 | 1.5260e-01 | 2.5941e+05 | ; | CB - CG |
| 6883 | 6884 | 1 | 1.0900e-01 | 2.8451e+05 | ; | CG - HG2 |
| 6883 | 6885 | 1 | 1.0900e-01 | 2.8451e+05 | ; | CG - HG3 |
| 6883 | 6886 | 1 | 1.5260e-01 | 2.5941e+05 | ; | CG - CD |
| 6886 | 6887 | 1 | 1.0900e-01 | 2.8451e+05 | ; | CD - HD2 |
| 6886 | 6888 | 1 | 1.0900e-01 | 2.8451e+05 | ; | CD - HD3 |
| 6886 | 6889 | 1 | 1.5260e-01 | 2.5941e+05 | ; | CD - CE |
| 6889 | 6890 | 1 | 1.0900e-01 | 2.8451e+05 | ; | CE - HE2 |
| 6889 | 6891 | 1 | 1.0900e-01 | 2.8451e+05 | ; | CE - HE3 |
| 6889 | 6892 | 1 | 1.4710e-01 | 3.0711e+05 | ; | CE - NZ |
| 6892 | 6893 | 1 | 1.0100e-01 | 3.6317e+05 | ; | NZ - HZ1 |
| 6892 | 6894 | 1 | 1.0100e-01 | 3.6317e+05 | ; | NZ - HZ2 |
| 6892 | 6895 | 1 | 1.0100e-01 | 3.6317e+05 | ; | NZ - HZ3 |
| 6896 | 6897 | 1 | 1.2290e-01 | 4.7698e+05 | ; | C - O |
| 6896 | 6898 | 1 | 1.3350e-01 | 4.1003e+05 | ; | C - N |

| | | | | | | |
|---|---|---|---|---|---|---|
| 6898 | 6899 | 1 | 1.0100e-01 | 3.6317e+05 | ; | N - H |
| 6898 | 6900 | 1 | 1.4490e-01 | 2.8200e+05 | ; | N - CA |
| 6900 | 6901 | 1 | 1.0900e-01 | 2.8451e+05 | ; | CA - HA |
| 6900 | 6902 | 1 | 1.5260e-01 | 2.5941e+05 | ; | CA - CB |
| 6900 | 6906 | 1 | 1.5220e-01 | 2.6527e+05 | ; | CA - C |
| 6902 | 6903 | 1 | 1.0900e-01 | 2.8451e+05 | ; | CB - HB1 |
| 6902 | 6904 | 1 | 1.0900e-01 | 2.8451e+05 | ; | CB - HB2 |
| 6902 | 6905 | 1 | 1.0900e-01 | 2.8451e+05 | ; | CB - HB3 |
| 6906 | 6907 | 1 | 1.2290e-01 | 4.7698e+05 | ; | C - O |
| 6906 | 6908 | 1 | 1.3350e-01 | 4.1003e+05 | ; | C - N |
| 6908 | 6909 | 1 | 1.0100e-01 | 3.6317e+05 | ; | N - H |
| 6908 | 6910 | 1 | 1.4490e-01 | 2.8200e+05 | ; | N - CA |
| 6910 | 6911 | 1 | 1.0900e-01 | 2.8451e+05 | ; | CA - HA |
| 6910 | 6912 | 1 | 1.5260e-01 | 2.5941e+05 | ; | CA - CB |
| 6910 | 6925 | 1 | 1.5220e-01 | 2.6527e+05 | ; | CA - C |
| 6912 | 6913 | 1 | 1.0900e-01 | 2.8451e+05 | ; | CB - HB |
| 6912 | 6914 | 1 | 1.5260e-01 | 2.5941e+05 | ; | CB - CG2 |
| 6912 | 6918 | 1 | 1.5260e-01 | 2.5941e+05 | ; | CB - CG1 |
| 6914 | 6915 | 1 | 1.0900e-01 | 2.8451e+05 | ; | CG2 - HG21 |
| 6914 | 6916 | 1 | 1.0900e-01 | 2.8451e+05 | ; | CG2 - HG22 |
| 6914 | 6917 | 1 | 1.0900e-01 | 2.8451e+05 | ; | CG2 - HG23 |
| 6918 | 6919 | 1 | 1.0900e-01 | 2.8451e+05 | ; | CG1 - HG12 |
| 6918 | 6920 | 1 | 1.0900e-01 | 2.8451e+05 | ; | CG1 - HG13 |
| 6918 | 6921 | 1 | 1.5260e-01 | 2.5941e+05 | ; | CG1 - CD1 |
| 6921 | 6922 | 1 | 1.0900e-01 | 2.8451e+05 | ; | CD1 - HD11 |
| 6921 | 6923 | 1 | 1.0900e-01 | 2.8451e+05 | ; | CD1 - HD12 |
| 6921 | 6924 | 1 | 1.0900e-01 | 2.8451e+05 | ; | CD1 - HD13 |
| 6925 | 6926 | 1 | 1.2290e-01 | 4.7698e+05 | ; | C - O |
| 6925 | 6927 | 1 | 1.3350e-01 | 4.1003e+05 | ; | C - N |
| 6927 | 6928 | 1 | 1.0100e-01 | 3.6317e+05 | ; | N - H |
| 6927 | 6929 | 1 | 1.4490e-01 | 2.8200e+05 | ; | N - CA |
| 6929 | 6930 | 1 | 1.0900e-01 | 2.8451e+05 | ; | CA - HA |
| 6929 | 6931 | 1 | 1.5260e-01 | 2.5941e+05 | ; | CA - CB |
| 6929 | 6937 | 1 | 1.5220e-01 | 2.6527e+05 | ; | CA - C |
| 6931 | 6932 | 1 | 1.0900e-01 | 2.8451e+05 | ; | CB - HB2 |
| 6931 | 6933 | 1 | 1.0900e-01 | 2.8451e+05 | ; | CB - HB3 |
| 6931 | 6934 | 1 | 1.5220e-01 | 2.6527e+05 | ; | CB - CG |
| 6934 | 6935 | 1 | 1.2500e-01 | 5.4894e+05 | ; | CG - OD1 |
| 6934 | 6936 | 1 | 1.2500e-01 | 5.4894e+05 | ; | CG - OD2 |
| 6937 | 6938 | 1 | 1.2290e-01 | 4.7698e+05 | ; | C - O |
| 6937 | 6939 | 1 | 1.3350e-01 | 4.1003e+05 | ; | C - N |
| 6939 | 6940 | 1 | 1.0100e-01 | 3.6317e+05 | ; | N - H |
| 6939 | 6941 | 1 | 1.4490e-01 | 2.8200e+05 | ; | N - CA |
| 6941 | 6942 | 1 | 1.0900e-01 | 2.8451e+05 | ; | CA - HA |
| 6941 | 6943 | 1 | 1.5260e-01 | 2.5941e+05 | ; | CA - CB |
| 6941 | 6948 | 1 | 1.5220e-01 | 2.6527e+05 | ; | CA - C |
| 6943 | 6944 | 1 | 1.0900e-01 | 2.8451e+05 | ; | CB - HB2 |
| 6943 | 6945 | 1 | 1.0900e-01 | 2.8451e+05 | ; | CB - HB3 |
| 6943 | 6946 | 1 | 1.4100e-01 | 2.6778e+05 | ; | CB - OG |
| 6946 | 6947 | 1 | 9.6000e-02 | 4.6275e+05 | ; | OG - HG |
| 6948 | 6949 | 1 | 1.2290e-01 | 4.7698e+05 | ; | C - O |
| 6948 | 6950 | 1 | 1.3350e-01 | 4.1003e+05 | ; | C - N |

| | | | | | | |
|---|---|---|---|---|---|---|
| 6950 | 6951 | 1 | 1.0100e-01 | 3.6317e+05 | ; | N - H |
| 6950 | 6952 | 1 | 1.4490e-01 | 2.8200e+05 | ; | N - CA |
| 6952 | 6953 | 1 | 1.0900e-01 | 2.8451e+05 | ; | CA - HA |
| 6952 | 6954 | 1 | 1.5260e-01 | 2.5941e+05 | ; | CA - CB |
| 6952 | 6967 | 1 | 1.5220e-01 | 2.6527e+05 | ; | CA - C |
| 6954 | 6955 | 1 | 1.0900e-01 | 2.8451e+05 | ; | CB - HB2 |
| 6954 | 6956 | 1 | 1.0900e-01 | 2.8451e+05 | ; | CB - HB3 |
| 6954 | 6957 | 1 | 1.5260e-01 | 2.5941e+05 | ; | CB - CG |
| 6957 | 6958 | 1 | 1.0900e-01 | 2.8451e+05 | ; | CG - HG |
| 6957 | 6959 | 1 | 1.5260e-01 | 2.5941e+05 | ; | CG - CD1 |
| 6957 | 6963 | 1 | 1.5260e-01 | 2.5941e+05 | ; | CG - CD2 |
| 6959 | 6960 | 1 | 1.0900e-01 | 2.8451e+05 | ; | CD1 - HD11 |
| 6959 | 6961 | 1 | 1.0900e-01 | 2.8451e+05 | ; | CD1 - HD12 |
| 6959 | 6962 | 1 | 1.0900e-01 | 2.8451e+05 | ; | CD1 - HD13 |
| 6963 | 6964 | 1 | 1.0900e-01 | 2.8451e+05 | ; | CD2 - HD21 |
| 6963 | 6965 | 1 | 1.0900e-01 | 2.8451e+05 | ; | CD2 - HD22 |
| 6963 | 6966 | 1 | 1.0900e-01 | 2.8451e+05 | ; | CD2 - HD23 |
| 6967 | 6968 | 1 | 1.2290e-01 | 4.7698e+05 | ; | C - O |
| 6967 | 6969 | 1 | 1.3350e-01 | 4.1003e+05 | ; | C - N |
| 6969 | 6970 | 1 | 1.0100e-01 | 3.6317e+05 | ; | N - H |
| 6969 | 6971 | 1 | 1.4490e-01 | 2.8200e+05 | ; | N - CA |
| 6971 | 6972 | 1 | 1.0900e-01 | 2.8451e+05 | ; | CA - HA |
| 6971 | 6973 | 1 | 1.5260e-01 | 2.5941e+05 | ; | CA - CB |
| 6971 | 6986 | 1 | 1.5220e-01 | 2.6527e+05 | ; | CA - C |
| 6973 | 6974 | 1 | 1.0900e-01 | 2.8451e+05 | ; | CB - HB2 |
| 6973 | 6975 | 1 | 1.0900e-01 | 2.8451e+05 | ; | CB - HB3 |
| 6973 | 6976 | 1 | 1.5260e-01 | 2.5941e+05 | ; | CB - CG |
| 6976 | 6977 | 1 | 1.0900e-01 | 2.8451e+05 | ; | CG - HG |
| 6976 | 6978 | 1 | 1.5260e-01 | 2.5941e+05 | ; | CG - CD1 |
| 6976 | 6982 | 1 | 1.5260e-01 | 2.5941e+05 | ; | CG - CD2 |
| 6978 | 6979 | 1 | 1.0900e-01 | 2.8451e+05 | ; | CD1 - HD11 |
| 6978 | 6980 | 1 | 1.0900e-01 | 2.8451e+05 | ; | CD1 - HD12 |
| 6978 | 6981 | 1 | 1.0900e-01 | 2.8451e+05 | ; | CD1 - HD13 |
| 6982 | 6983 | 1 | 1.0900e-01 | 2.8451e+05 | ; | CD2 - HD21 |
| 6982 | 6984 | 1 | 1.0900e-01 | 2.8451e+05 | ; | CD2 - HD22 |
| 6982 | 6985 | 1 | 1.0900e-01 | 2.8451e+05 | ; | CD2 - HD23 |
| 6986 | 6987 | 1 | 1.2290e-01 | 4.7698e+05 | ; | C - O |
| 6986 | 6988 | 1 | 1.3350e-01 | 4.1003e+05 | ; | C - N |
| 6988 | 6989 | 1 | 1.0100e-01 | 3.6317e+05 | ; | N - H |
| 6988 | 6990 | 1 | 1.4490e-01 | 2.8200e+05 | ; | N - CA |
| 6990 | 6991 | 1 | 1.0900e-01 | 2.8451e+05 | ; | CA - HA |
| 6990 | 6992 | 1 | 1.5260e-01 | 2.5941e+05 | ; | CA - CB |
| 6990 | 7003 | 1 | 1.5220e-01 | 2.6527e+05 | ; | CA - C |
| 6992 | 6993 | 1 | 1.0900e-01 | 2.8451e+05 | ; | CB - HB2 |
| 6992 | 6994 | 1 | 1.0900e-01 | 2.8451e+05 | ; | CB - HB3 |
| 6992 | 6995 | 1 | 1.5260e-01 | 2.5941e+05 | ; | CB - CG |
| 6995 | 6996 | 1 | 1.0900e-01 | 2.8451e+05 | ; | CG - HG2 |
| 6995 | 6997 | 1 | 1.0900e-01 | 2.8451e+05 | ; | CG - HG3 |
| 6995 | 6998 | 1 | 1.5220e-01 | 2.6527e+05 | ; | CG - CD |
| 6998 | 6999 | 1 | 1.2290e-01 | 4.7698e+05 | ; | CD - OE1 |
| 6998 | 7000 | 1 | 1.3350e-01 | 4.1003e+05 | ; | CD - NE2 |
| 7000 | 7001 | 1 | 1.0100e-01 | 3.6317e+05 | ; | NE2 - HE21 |

| | | | | | | |
|---|---|---|---|---|---|---|
| 7000 | 7002 | 1 | 1.0100e-01 | 3.6317e+05 | ; | NE2 - HE22 |
| 7003 | 7004 | 1 | 1.2290e-01 | 4.7698e+05 | ; | C - O |
| 7003 | 7005 | 1 | 1.3350e-01 | 4.1003e+05 | ; | C - N |
| 7005 | 7006 | 1 | 1.0100e-01 | 3.6317e+05 | ; | N - H |
| 7005 | 7007 | 1 | 1.4490e-01 | 2.8200e+05 | ; | N - CA |
| 7007 | 7008 | 1 | 1.0900e-01 | 2.8451e+05 | ; | CA - HA |
| 7007 | 7009 | 1 | 1.5260e-01 | 2.5941e+05 | ; | CA - CB |
| 7007 | 7018 | 1 | 1.5220e-01 | 2.6527e+05 | ; | CA - C |
| 7009 | 7010 | 1 | 1.0900e-01 | 2.8451e+05 | ; | CB - HB2 |
| 7009 | 7011 | 1 | 1.0900e-01 | 2.8451e+05 | ; | CB - HB3 |
| 7009 | 7012 | 1 | 1.5260e-01 | 2.5941e+05 | ; | CB - CG |
| 7012 | 7013 | 1 | 1.0900e-01 | 2.8451e+05 | ; | CG - HG2 |
| 7012 | 7014 | 1 | 1.0900e-01 | 2.8451e+05 | ; | CG - HG3 |
| 7012 | 7015 | 1 | 1.5220e-01 | 2.6527e+05 | ; | CG - CD |
| 7015 | 7016 | 1 | 1.2500e-01 | 5.4894e+05 | ; | CD - OE1 |
| 7015 | 7017 | 1 | 1.2500e-01 | 5.4894e+05 | ; | CD - OE2 |
| 7018 | 7019 | 1 | 1.2290e-01 | 4.7698e+05 | ; | C - O |
| 7018 | 7020 | 1 | 1.3350e-01 | 4.1003e+05 | ; | C - N |
| 7020 | 7021 | 1 | 1.0100e-01 | 3.6317e+05 | ; | N - H |
| 7020 | 7022 | 1 | 1.4490e-01 | 2.8200e+05 | ; | N - CA |
| 7022 | 7023 | 1 | 1.0900e-01 | 2.8451e+05 | ; | CA - HA |
| 7022 | 7024 | 1 | 1.5260e-01 | 2.5941e+05 | ; | CA - CB |
| 7022 | 7034 | 1 | 1.5220e-01 | 2.6527e+05 | ; | CA - C |
| 7024 | 7025 | 1 | 1.0900e-01 | 2.8451e+05 | ; | CB - HB |
| 7024 | 7026 | 1 | 1.5260e-01 | 2.5941e+05 | ; | CB - CG1 |
| 7024 | 7030 | 1 | 1.5260e-01 | 2.5941e+05 | ; | CB - CG2 |
| 7026 | 7027 | 1 | 1.0900e-01 | 2.8451e+05 | ; | CG1 - HG11 |
| 7026 | 7028 | 1 | 1.0900e-01 | 2.8451e+05 | ; | CG1 - HG12 |
| 7026 | 7029 | 1 | 1.0900e-01 | 2.8451e+05 | ; | CG1 - HG13 |
| 7030 | 7031 | 1 | 1.0900e-01 | 2.8451e+05 | ; | CG2 - HG21 |
| 7030 | 7032 | 1 | 1.0900e-01 | 2.8451e+05 | ; | CG2 - HG22 |
| 7030 | 7033 | 1 | 1.0900e-01 | 2.8451e+05 | ; | CG2 - HG23 |
| 7034 | 7035 | 1 | 1.2290e-01 | 4.7698e+05 | ; | C - O |
| 7034 | 7036 | 1 | 1.3350e-01 | 4.1003e+05 | ; | C - N |
| 7036 | 7037 | 1 | 1.0100e-01 | 3.6317e+05 | ; | N - H |
| 7036 | 7038 | 1 | 1.4490e-01 | 2.8200e+05 | ; | N - CA |
| 7038 | 7039 | 1 | 1.0900e-01 | 2.8451e+05 | ; | CA - HA |
| 7038 | 7040 | 1 | 1.5260e-01 | 2.5941e+05 | ; | CA - CB |
| 7038 | 7046 | 1 | 1.5220e-01 | 2.6527e+05 | ; | CA - C |
| 7040 | 7041 | 1 | 1.0900e-01 | 2.8451e+05 | ; | CB - HB2 |
| 7040 | 7042 | 1 | 1.0900e-01 | 2.8451e+05 | ; | CB - HB3 |
| 7040 | 7043 | 1 | 1.5220e-01 | 2.6527e+05 | ; | CB - CG |
| 7043 | 7044 | 1 | 1.2500e-01 | 5.4894e+05 | ; | CG - OD1 |
| 7043 | 7045 | 1 | 1.2500e-01 | 5.4894e+05 | ; | CG - OD2 |
| 7046 | 7047 | 1 | 1.2290e-01 | 4.7698e+05 | ; | C - O |
| 7046 | 7048 | 1 | 1.3350e-01 | 4.1003e+05 | ; | C - N |
| 7048 | 7049 | 1 | 1.0100e-01 | 3.6317e+05 | ; | N - H |
| 7048 | 7050 | 1 | 1.4490e-01 | 2.8200e+05 | ; | N - CA |
| 7050 | 7051 | 1 | 1.0900e-01 | 2.8451e+05 | ; | CA - HA |
| 7050 | 7052 | 1 | 1.5260e-01 | 2.5941e+05 | ; | CA - CB |
| 7050 | 7065 | 1 | 1.5220e-01 | 2.6527e+05 | ; | CA - C |
| 7052 | 7053 | 1 | 1.0900e-01 | 2.8451e+05 | ; | CB - HB2 |

| | | | | | |
|---|---|---|---|---|---|
| 7052 | 7054 | 1 | 1.0900e-01 | 2.8451e+05 ; | CB - HB3 |
| 7052 | 7055 | 1 | 1.5260e-01 | 2.5941e+05 ; | CB - CG |
| 7055 | 7056 | 1 | 1.0900e-01 | 2.8451e+05 ; | CG - HG |
| 7055 | 7057 | 1 | 1.5260e-01 | 2.5941e+05 ; | CG - CD1 |
| 7055 | 7061 | 1 | 1.5260e-01 | 2.5941e+05 ; | CG - CD2 |
| 7057 | 7058 | 1 | 1.0900e-01 | 2.8451e+05 ; | CD1 - HD11 |
| 7057 | 7059 | 1 | 1.0900e-01 | 2.8451e+05 ; | CD1 - HD12 |
| 7057 | 7060 | 1 | 1.0900e-01 | 2.8451e+05 ; | CD1 - HD13 |
| 7061 | 7062 | 1 | 1.0900e-01 | 2.8451e+05 ; | CD2 - HD21 |
| 7061 | 7063 | 1 | 1.0900e-01 | 2.8451e+05 ; | CD2 - HD22 |
| 7061 | 7064 | 1 | 1.0900e-01 | 2.8451e+05 ; | CD2 - HD23 |
| 7065 | 7066 | 1 | 1.2290e-01 | 4.7698e+05 ; | C - O |
| 7065 | 7067 | 1 | 1.3350e-01 | 4.1003e+05 ; | C - N |
| 7067 | 7068 | 1 | 1.0100e-01 | 3.6317e+05 ; | N - H |
| 7067 | 7069 | 1 | 1.4490e-01 | 2.8200e+05 ; | N - CA |
| 7069 | 7070 | 1 | 1.0900e-01 | 2.8451e+05 ; | CA - HA |
| 7069 | 7071 | 1 | 1.5260e-01 | 2.5941e+05 ; | CA - CB |
| 7069 | 7079 | 1 | 1.5220e-01 | 2.6527e+05 ; | CA - C |
| 7071 | 7072 | 1 | 1.0900e-01 | 2.8451e+05 ; | CB - HB2 |
| 7071 | 7073 | 1 | 1.0900e-01 | 2.8451e+05 ; | CB - HB3 |
| 7071 | 7074 | 1 | 1.5220e-01 | 2.6527e+05 ; | CB - CG |
| 7074 | 7075 | 1 | 1.2290e-01 | 4.7698e+05 ; | CG - OD1 |
| 7074 | 7076 | 1 | 1.3350e-01 | 4.1003e+05 ; | CG - ND2 |
| 7076 | 7077 | 1 | 1.0100e-01 | 3.6317e+05 ; | ND2 - HD21 |
| 7076 | 7078 | 1 | 1.0100e-01 | 3.6317e+05 ; | ND2 - HD22 |
| 7079 | 7080 | 1 | 1.2290e-01 | 4.7698e+05 ; | C - O |
| 7079 | 7081 | 1 | 1.3350e-01 | 4.1003e+05 ; | C - N |
| 7081 | 7082 | 1 | 1.0100e-01 | 3.6317e+05 ; | N - H |
| 7081 | 7083 | 1 | 1.4490e-01 | 2.8200e+05 ; | N - CA |
| 7083 | 7084 | 1 | 1.0900e-01 | 2.8451e+05 ; | CA - HA2 |
| 7083 | 7085 | 1 | 1.0900e-01 | 2.8451e+05 ; | CA - HA3 |
| 7083 | 7086 | 1 | 1.5220e-01 | 2.6527e+05 ; | CA - C |
| 7086 | 7087 | 1 | 1.2290e-01 | 4.7698e+05 ; | C - O |
| 7086 | 7088 | 1 | 1.3350e-01 | 4.1003e+05 ; | C - N |
| 7088 | 7089 | 1 | 1.0100e-01 | 3.6317e+05 ; | N - H |
| 7088 | 7090 | 1 | 1.4490e-01 | 2.8200e+05 ; | N - CA |
| 7090 | 7091 | 1 | 1.0900e-01 | 2.8451e+05 ; | CA - HA |
| 7090 | 7092 | 1 | 1.5260e-01 | 2.5941e+05 ; | CA - CB |
| 7090 | 7099 | 1 | 1.5220e-01 | 2.6527e+05 ; | CA - C |
| 7092 | 7093 | 1 | 1.0900e-01 | 2.8451e+05 ; | CB - HB2 |
| 7092 | 7094 | 1 | 1.0900e-01 | 2.8451e+05 ; | CB - HB3 |
| 7092 | 7095 | 1 | 1.5220e-01 | 2.6527e+05 ; | CB - CG |
| 7095 | 7096 | 1 | 1.2290e-01 | 4.7698e+05 ; | CG - OD1 |
| 7095 | 7097 | 1 | 1.3640e-01 | 3.7656e+05 ; | CG - OD2 |
| 7097 | 7098 | 1 | 9.6000e-02 | 4.6275e+05 ; | OD2 - HD2 |
| 7099 | 7100 | 1 | 1.2290e-01 | 4.7698e+05 ; | C - O |
| 7099 | 7101 | 1 | 1.3350e-01 | 4.1003e+05 ; | C - N |
| 7101 | 7102 | 1 | 1.0100e-01 | 3.6317e+05 ; | N - H |
| 7101 | 7103 | 1 | 1.4490e-01 | 2.8200e+05 ; | N - CA |
| 7103 | 7104 | 1 | 1.0900e-01 | 2.8451e+05 ; | CA - HA2 |
| 7103 | 7105 | 1 | 1.0900e-01 | 2.8451e+05 ; | CA - HA3 |
| 7103 | 7106 | 1 | 1.5220e-01 | 2.6527e+05 ; | CA - C |

| | | | | | | | | |
|---|---|---|---|---|---|---|---|---|
| 7106 | 7107 | 1 | 1.2290e-01 | 4.7698e+05 | ; | C | - | O |
| 7106 | 7108 | 1 | 1.3350e-01 | 4.1003e+05 | ; | C | - | N |
| 7108 | 7109 | 1 | 1.0100e-01 | 3.6317e+05 | ; | N | - | H |
| 7108 | 7110 | 1 | 1.4490e-01 | 2.8200e+05 | ; | N | - | CA |
| 7110 | 7111 | 1 | 1.0900e-01 | 2.8451e+05 | ; | CA | - | HA |
| 7110 | 7112 | 1 | 1.5260e-01 | 2.5941e+05 | ; | CA | - | CB |
| 7110 | 7121 | 1 | 1.5220e-01 | 2.6527e+05 | ; | CA | - | C |
| 7112 | 7113 | 1 | 1.0900e-01 | 2.8451e+05 | ; | CB | - | HB2 |
| 7112 | 7114 | 1 | 1.0900e-01 | 2.8451e+05 | ; | CB | - | HB3 |
| 7112 | 7115 | 1 | 1.5260e-01 | 2.5941e+05 | ; | CB | - | CG |
| 7115 | 7116 | 1 | 1.0900e-01 | 2.8451e+05 | ; | CG | - | HG2 |
| 7115 | 7117 | 1 | 1.0900e-01 | 2.8451e+05 | ; | CG | - | HG3 |
| 7115 | 7118 | 1 | 1.5220e-01 | 2.6527e+05 | ; | CG | - | CD |
| 7118 | 7119 | 1 | 1.2500e-01 | 5.4894e+05 | ; | CD | - | OE1 |
| 7118 | 7120 | 1 | 1.2500e-01 | 5.4894e+05 | ; | CD | - | OE2 |
| 7121 | 7122 | 1 | 1.2290e-01 | 4.7698e+05 | ; | C | - | O |
| 7121 | 7123 | 1 | 1.3350e-01 | 4.1003e+05 | ; | C | - | N |
| 7123 | 7124 | 1 | 1.0100e-01 | 3.6317e+05 | ; | N | - | H |
| 7123 | 7125 | 1 | 1.4490e-01 | 2.8200e+05 | ; | N | - | CA |
| 7125 | 7126 | 1 | 1.0900e-01 | 2.8451e+05 | ; | CA | - | HA |
| 7125 | 7127 | 1 | 1.5260e-01 | 2.5941e+05 | ; | CA | - | CB |
| 7125 | 7140 | 1 | 1.5220e-01 | 2.6527e+05 | ; | CA | - | C |
| 7127 | 7128 | 1 | 1.0900e-01 | 2.8451e+05 | ; | CB | - | HB |
| 7127 | 7129 | 1 | 1.5260e-01 | 2.5941e+05 | ; | CB | - | CG2 |
| 7127 | 7133 | 1 | 1.5260e-01 | 2.5941e+05 | ; | CB | - | CG1 |
| 7129 | 7130 | 1 | 1.0900e-01 | 2.8451e+05 | ; | CG2 | - | HG21 |
| 7129 | 7131 | 1 | 1.0900e-01 | 2.8451e+05 | ; | CG2 | - | HG22 |
| 7129 | 7132 | 1 | 1.0900e-01 | 2.8451e+05 | ; | CG2 | - | HG23 |
| 7133 | 7134 | 1 | 1.0900e-01 | 2.8451e+05 | ; | CG1 | - | HG12 |
| 7133 | 7135 | 1 | 1.0900e-01 | 2.8451e+05 | ; | CG1 | - | HG13 |
| 7133 | 7136 | 1 | 1.5260e-01 | 2.5941e+05 | ; | CG1 | - | CD1 |
| 7136 | 7137 | 1 | 1.0900e-01 | 2.8451e+05 | ; | CD1 | - | HD11 |
| 7136 | 7138 | 1 | 1.0900e-01 | 2.8451e+05 | ; | CD1 | - | HD12 |
| 7136 | 7139 | 1 | 1.0900e-01 | 2.8451e+05 | ; | CD1 | - | HD13 |
| 7140 | 7141 | 1 | 1.2290e-01 | 4.7698e+05 | ; | C | - | O |
| 7140 | 7142 | 1 | 1.3350e-01 | 4.1003e+05 | ; | C | - | N |
| 7142 | 7143 | 1 | 1.0100e-01 | 3.6317e+05 | ; | N | - | H |
| 7142 | 7144 | 1 | 1.4490e-01 | 2.8200e+05 | ; | N | - | CA |
| 7144 | 7145 | 1 | 1.0900e-01 | 2.8451e+05 | ; | CA | - | HA |
| 7144 | 7146 | 1 | 1.5260e-01 | 2.5941e+05 | ; | CA | - | CB |
| 7144 | 7152 | 1 | 1.5220e-01 | 2.6527e+05 | ; | CA | - | C |
| 7146 | 7147 | 1 | 1.0900e-01 | 2.8451e+05 | ; | CB | - | HB2 |
| 7146 | 7148 | 1 | 1.0900e-01 | 2.8451e+05 | ; | CB | - | HB3 |
| 7146 | 7149 | 1 | 1.5220e-01 | 2.6527e+05 | ; | CB | - | CG |
| 7149 | 7150 | 1 | 1.2500e-01 | 5.4894e+05 | ; | CG | - | OD1 |
| 7149 | 7151 | 1 | 1.2500e-01 | 5.4894e+05 | ; | CG | - | OD2 |
| 7152 | 7153 | 1 | 1.2290e-01 | 4.7698e+05 | ; | C | - | O |
| 7152 | 7154 | 1 | 1.3350e-01 | 4.1003e+05 | ; | C | - | N |
| 7154 | 7155 | 1 | 1.0100e-01 | 3.6317e+05 | ; | N | - | H |
| 7154 | 7156 | 1 | 1.4490e-01 | 2.8200e+05 | ; | N | - | CA |
| 7156 | 7157 | 1 | 1.0900e-01 | 2.8451e+05 | ; | CA | - | HA |
| 7156 | 7158 | 1 | 1.5260e-01 | 2.5941e+05 | ; | CA | - | CB |

| | | | | | | |
|---|---|---|---|---|---|---|
| 7156 | 7172 | 1 | 1.5220e-01 | 2.6527e+05 | ; | CA - C |
| 7158 | 7159 | 1 | 1.0900e-01 | 2.8451e+05 | ; | CB - HB2 |
| 7158 | 7160 | 1 | 1.0900e-01 | 2.8451e+05 | ; | CB - HB3 |
| 7158 | 7161 | 1 | 1.5100e-01 | 2.6527e+05 | ; | CB - CG |
| 7161 | 7162 | 1 | 1.4000e-01 | 3.9246e+05 | ; | CG - CD1 |
| 7161 | 7170 | 1 | 1.4000e-01 | 3.9246e+05 | ; | CG - CD2 |
| 7162 | 7163 | 1 | 1.0800e-01 | 3.0711e+05 | ; | CD1 - HD1 |
| 7162 | 7164 | 1 | 1.4000e-01 | 3.9246e+05 | ; | CD1 - CE1 |
| 7164 | 7165 | 1 | 1.0800e-01 | 3.0711e+05 | ; | CE1 - HE1 |
| 7164 | 7166 | 1 | 1.4000e-01 | 3.9246e+05 | ; | CE1 - CZ |
| 7166 | 7167 | 1 | 1.0800e-01 | 3.0711e+05 | ; | CZ - HZ |
| 7166 | 7168 | 1 | 1.4000e-01 | 3.9246e+05 | ; | CZ - CE2 |
| 7168 | 7169 | 1 | 1.0800e-01 | 3.0711e+05 | ; | CE2 - HE2 |
| 7168 | 7170 | 1 | 1.4000e-01 | 3.9246e+05 | ; | CE2 - CD2 |
| 7170 | 7171 | 1 | 1.0800e-01 | 3.0711e+05 | ; | CD2 - HD2 |
| 7172 | 7173 | 1 | 1.2290e-01 | 4.7698e+05 | ; | C - O |
| 7172 | 7174 | 1 | 1.3350e-01 | 4.1003e+05 | ; | C - N |
| 7174 | 7175 | 1 | 1.0100e-01 | 3.6317e+05 | ; | N - H |
| 7174 | 7176 | 1 | 1.4490e-01 | 2.8200e+05 | ; | N - CA |
| 7176 | 7177 | 1 | 1.0900e-01 | 2.8451e+05 | ; | CA - HA |
| 7176 | 7178 | 1 | 1.5260e-01 | 2.5941e+05 | ; | CA - CB |
| 7176 | 7189 | 1 | 1.5220e-01 | 2.6527e+05 | ; | CA - C |
| 7178 | 7179 | 1 | 1.0900e-01 | 2.8451e+05 | ; | CB - HB2 |
| 7178 | 7180 | 1 | 1.0900e-01 | 2.8451e+05 | ; | CB - HB3 |
| 7178 | 7181 | 1 | 1.5040e-01 | 2.6527e+05 | ; | CB - CG |
| 7181 | 7182 | 1 | 1.3940e-01 | 3.4309e+05 | ; | CG - ND1 |
| 7181 | 7187 | 1 | 1.3710e-01 | 4.3346e+05 | ; | CG - CD2 |
| 7182 | 7183 | 1 | 1.3350e-01 | 4.0836e+05 | ; | ND1 - CE1 |
| 7183 | 7184 | 1 | 1.0800e-01 | 3.0711e+05 | ; | CE1 - HE1 |
| 7183 | 7185 | 1 | 1.3430e-01 | 3.9915e+05 | ; | CE1 - NE2 |
| 7185 | 7186 | 1 | 1.0100e-01 | 3.6317e+05 | ; | NE2 - HE2 |
| 7185 | 7187 | 1 | 1.3810e-01 | 3.5731e+05 | ; | NE2 - CD2 |
| 7187 | 7188 | 1 | 1.0800e-01 | 3.0711e+05 | ; | CD2 - HD2 |
| 7189 | 7190 | 1 | 1.2290e-01 | 4.7698e+05 | ; | C - O |
| 7189 | 7191 | 1 | 1.3350e-01 | 4.1003e+05 | ; | C - N |
| 7191 | 7192 | 1 | 1.0100e-01 | 3.6317e+05 | ; | N - H |
| 7191 | 7193 | 1 | 1.4490e-01 | 2.8200e+05 | ; | N - CA |
| 7193 | 7194 | 1 | 1.0900e-01 | 2.8451e+05 | ; | CA - HA |
| 7193 | 7195 | 1 | 1.5260e-01 | 2.5941e+05 | ; | CA - CB |
| 7193 | 7205 | 1 | 1.5220e-01 | 2.6527e+05 | ; | CA - C |
| 7195 | 7196 | 1 | 1.0900e-01 | 2.8451e+05 | ; | CB - HB2 |
| 7195 | 7197 | 1 | 1.0900e-01 | 2.8451e+05 | ; | CB - HB3 |
| 7195 | 7198 | 1 | 1.5260e-01 | 2.5941e+05 | ; | CB - CG |
| 7198 | 7199 | 1 | 1.0900e-01 | 2.8451e+05 | ; | CG - HG2 |
| 7198 | 7200 | 1 | 1.0900e-01 | 2.8451e+05 | ; | CG - HG3 |
| 7198 | 7201 | 1 | 1.5220e-01 | 2.6527e+05 | ; | CG - CD |
| 7201 | 7202 | 1 | 1.2290e-01 | 4.7698e+05 | ; | CD - OE1 |
| 7201 | 7203 | 1 | 1.3640e-01 | 3.7656e+05 | ; | CD - OE2 |
| 7203 | 7204 | 1 | 9.6000e-02 | 4.6275e+05 | ; | OE2 - HE2 |
| 7205 | 7206 | 1 | 1.2290e-01 | 4.7698e+05 | ; | C - O |
| 7205 | 7207 | 1 | 1.3350e-01 | 4.1003e+05 | ; | C - N |
| 7207 | 7208 | 1 | 1.0100e-01 | 3.6317e+05 | ; | N - H |

| | | | | | | | | |
|---|---|---|---|---|---|---|---|---|
| 7207 | 7209 | 1 | 1.4490e-01 | 2.8200e+05 | ; | N | - | CA |
| 7209 | 7210 | 1 | 1.0900e-01 | 2.8451e+05 | ; | CA | - | HA |
| 7209 | 7211 | 1 | 1.5260e-01 | 2.5941e+05 | ; | CA | - | CB |
| 7209 | 7225 | 1 | 1.5220e-01 | 2.6527e+05 | ; | CA | - | C |
| 7211 | 7212 | 1 | 1.0900e-01 | 2.8451e+05 | ; | CB | - | HB2 |
| 7211 | 7213 | 1 | 1.0900e-01 | 2.8451e+05 | ; | CB | - | HB3 |
| 7211 | 7214 | 1 | 1.5100e-01 | 2.6527e+05 | ; | CB | - | CG |
| 7214 | 7215 | 1 | 1.4000e-01 | 3.9246e+05 | ; | CG | - | CD1 |
| 7214 | 7223 | 1 | 1.4000e-01 | 3.9246e+05 | ; | CG | - | CD2 |
| 7215 | 7216 | 1 | 1.0800e-01 | 3.0711e+05 | ; | CD1 | - | HD1 |
| 7215 | 7217 | 1 | 1.4000e-01 | 3.9246e+05 | ; | CD1 | - | CE1 |
| 7217 | 7218 | 1 | 1.0800e-01 | 3.0711e+05 | ; | CE1 | - | HE1 |
| 7217 | 7219 | 1 | 1.4000e-01 | 3.9246e+05 | ; | CE1 | - | CZ |
| 7219 | 7220 | 1 | 1.0800e-01 | 3.0711e+05 | ; | CZ | - | HZ |
| 7219 | 7221 | 1 | 1.4000e-01 | 3.9246e+05 | ; | CZ | - | CE2 |
| 7221 | 7222 | 1 | 1.0800e-01 | 3.0711e+05 | ; | CE2 | - | HE2 |
| 7221 | 7223 | 1 | 1.4000e-01 | 3.9246e+05 | ; | CE2 | - | CD2 |
| 7223 | 7224 | 1 | 1.0800e-01 | 3.0711e+05 | ; | CD2 | - | HD2 |
| 7225 | 7226 | 1 | 1.2290e-01 | 4.7698e+05 | ; | C | - | O |
| 7225 | 7227 | 1 | 1.3350e-01 | 4.1003e+05 | ; | C | - | N |
| 7227 | 7228 | 1 | 1.0100e-01 | 3.6317e+05 | ; | N | - | H |
| 7227 | 7229 | 1 | 1.4490e-01 | 2.8200e+05 | ; | N | - | CA |
| 7229 | 7230 | 1 | 1.0900e-01 | 2.8451e+05 | ; | CA | - | HA |
| 7229 | 7231 | 1 | 1.5260e-01 | 2.5941e+05 | ; | CA | - | CB |
| 7229 | 7242 | 1 | 1.5220e-01 | 2.6527e+05 | ; | CA | - | C |
| 7231 | 7232 | 1 | 1.0900e-01 | 2.8451e+05 | ; | CB | - | HB2 |
| 7231 | 7233 | 1 | 1.0900e-01 | 2.8451e+05 | ; | CB | - | HB3 |
| 7231 | 7234 | 1 | 1.5260e-01 | 2.5941e+05 | ; | CB | - | CG |
| 7234 | 7235 | 1 | 1.0900e-01 | 2.8451e+05 | ; | CG | - | HG2 |
| 7234 | 7236 | 1 | 1.0900e-01 | 2.8451e+05 | ; | CG | - | HG3 |
| 7234 | 7237 | 1 | 1.8100e-01 | 1.8995e+05 | ; | CG | - | SD |
| 7237 | 7238 | 1 | 1.8100e-01 | 1.8995e+05 | ; | SD | - | CE |
| 7238 | 7239 | 1 | 1.0900e-01 | 2.8451e+05 | ; | CE | - | HE1 |
| 7238 | 7240 | 1 | 1.0900e-01 | 2.8451e+05 | ; | CE | - | HE2 |
| 7238 | 7241 | 1 | 1.0900e-01 | 2.8451e+05 | ; | CE | - | HE3 |
| 7242 | 7243 | 1 | 1.2290e-01 | 4.7698e+05 | ; | C | - | O |
| 7242 | 7244 | 1 | 1.3350e-01 | 4.1003e+05 | ; | C | - | N |
| 7244 | 7245 | 1 | 1.0100e-01 | 3.6317e+05 | ; | N | - | H |
| 7244 | 7246 | 1 | 1.4490e-01 | 2.8200e+05 | ; | N | - | CA |
| 7246 | 7247 | 1 | 1.0900e-01 | 2.8451e+05 | ; | CA | - | HA |
| 7246 | 7248 | 1 | 1.5260e-01 | 2.5941e+05 | ; | CA | - | CB |
| 7246 | 7261 | 1 | 1.5220e-01 | 2.6527e+05 | ; | CA | - | C |
| 7248 | 7249 | 1 | 1.0900e-01 | 2.8451e+05 | ; | CB | - | HB2 |
| 7248 | 7250 | 1 | 1.0900e-01 | 2.8451e+05 | ; | CB | - | HB3 |
| 7248 | 7251 | 1 | 1.5260e-01 | 2.5941e+05 | ; | CB | - | CG |
| 7251 | 7252 | 1 | 1.0900e-01 | 2.8451e+05 | ; | CG | - | HG |
| 7251 | 7253 | 1 | 1.5260e-01 | 2.5941e+05 | ; | CG | - | CD1 |
| 7251 | 7257 | 1 | 1.5260e-01 | 2.5941e+05 | ; | CG | - | CD2 |
| 7253 | 7254 | 1 | 1.0900e-01 | 2.8451e+05 | ; | CD1 | - | HD11 |
| 7253 | 7255 | 1 | 1.0900e-01 | 2.8451e+05 | ; | CD1 | - | HD12 |
| 7253 | 7256 | 1 | 1.0900e-01 | 2.8451e+05 | ; | CD1 | - | HD13 |
| 7257 | 7258 | 1 | 1.0900e-01 | 2.8451e+05 | ; | CD2 | - | HD21 |

| | | | | | | |
|---|---|---|---|---|---|---|
| 7257 | 7259 | 1 | 1.0900e-01 | 2.8451e+05 | ; | CD2 - HD22 |
| 7257 | 7260 | 1 | 1.0900e-01 | 2.8451e+05 | ; | CD2 - HD23 |
| 7261 | 7262 | 1 | 1.2290e-01 | 4.7698e+05 | ; | C - O |
| 7261 | 7263 | 1 | 1.3350e-01 | 4.1003e+05 | ; | C - N |
| 7263 | 7264 | 1 | 1.0100e-01 | 3.6317e+05 | ; | N - H |
| 7263 | 7265 | 1 | 1.4490e-01 | 2.8200e+05 | ; | N - CA |
| 7265 | 7266 | 1 | 1.0900e-01 | 2.8451e+05 | ; | CA - HA |
| 7265 | 7267 | 1 | 1.5260e-01 | 2.5941e+05 | ; | CA - CB |
| 7265 | 7278 | 1 | 1.5220e-01 | 2.6527e+05 | ; | CA - C |
| 7267 | 7268 | 1 | 1.0900e-01 | 2.8451e+05 | ; | CB - HB2 |
| 7267 | 7269 | 1 | 1.0900e-01 | 2.8451e+05 | ; | CB - HB3 |
| 7267 | 7270 | 1 | 1.5260e-01 | 2.5941e+05 | ; | CB - CG |
| 7270 | 7271 | 1 | 1.0900e-01 | 2.8451e+05 | ; | CG - HG2 |
| 7270 | 7272 | 1 | 1.0900e-01 | 2.8451e+05 | ; | CG - HG3 |
| 7270 | 7273 | 1 | 1.8100e-01 | 1.8995e+05 | ; | CG - SD |
| 7273 | 7274 | 1 | 1.8100e-01 | 1.8995e+05 | ; | SD - CE |
| 7274 | 7275 | 1 | 1.0900e-01 | 2.8451e+05 | ; | CE - HE1 |
| 7274 | 7276 | 1 | 1.0900e-01 | 2.8451e+05 | ; | CE - HE2 |
| 7274 | 7277 | 1 | 1.0900e-01 | 2.8451e+05 | ; | CE - HE3 |
| 7278 | 7279 | 1 | 1.2290e-01 | 4.7698e+05 | ; | C - O |
| 7278 | 7280 | 1 | 1.3350e-01 | 4.1003e+05 | ; | C - N |
| 7280 | 7281 | 1 | 1.0100e-01 | 3.6317e+05 | ; | N - H |
| 7280 | 7282 | 1 | 1.4490e-01 | 2.8200e+05 | ; | N - CA |
| 7282 | 7283 | 1 | 1.0900e-01 | 2.8451e+05 | ; | CA - HA |
| 7282 | 7284 | 1 | 1.5260e-01 | 2.5941e+05 | ; | CA - CB |
| 7282 | 7295 | 1 | 1.5220e-01 | 2.6527e+05 | ; | CA - C |
| 7284 | 7285 | 1 | 1.0900e-01 | 2.8451e+05 | ; | CB - HB2 |
| 7284 | 7286 | 1 | 1.0900e-01 | 2.8451e+05 | ; | CB - HB3 |
| 7284 | 7287 | 1 | 1.5260e-01 | 2.5941e+05 | ; | CB - CG |
| 7287 | 7288 | 1 | 1.0900e-01 | 2.8451e+05 | ; | CG - HG2 |
| 7287 | 7289 | 1 | 1.0900e-01 | 2.8451e+05 | ; | CG - HG3 |
| 7287 | 7290 | 1 | 1.8100e-01 | 1.8995e+05 | ; | CG - SD |
| 7290 | 7291 | 1 | 1.8100e-01 | 1.8995e+05 | ; | SD - CE |
| 7291 | 7292 | 1 | 1.0900e-01 | 2.8451e+05 | ; | CE - HE1 |
| 7291 | 7293 | 1 | 1.0900e-01 | 2.8451e+05 | ; | CE - HE2 |
| 7291 | 7294 | 1 | 1.0900e-01 | 2.8451e+05 | ; | CE - HE3 |
| 7295 | 7296 | 1 | 1.2290e-01 | 4.7698e+05 | ; | C - O |
| 7295 | 7297 | 1 | 1.3350e-01 | 4.1003e+05 | ; | C - N |
| 7297 | 7298 | 1 | 1.0100e-01 | 3.6317e+05 | ; | N - H |
| 7297 | 7299 | 1 | 1.4490e-01 | 2.8200e+05 | ; | N - CA |
| 7299 | 7300 | 1 | 1.0900e-01 | 2.8451e+05 | ; | CA - HA |
| 7299 | 7301 | 1 | 1.5260e-01 | 2.5941e+05 | ; | CA - CB |
| 7299 | 7306 | 1 | 1.5220e-01 | 2.6527e+05 | ; | CA - C |
| 7301 | 7302 | 1 | 1.0900e-01 | 2.8451e+05 | ; | CB - HB2 |
| 7301 | 7303 | 1 | 1.0900e-01 | 2.8451e+05 | ; | CB - HB3 |
| 7301 | 7304 | 1 | 1.4100e-01 | 2.6778e+05 | ; | CB - OG |
| 7304 | 7305 | 1 | 9.6000e-02 | 4.6275e+05 | ; | OG - HG |
| 7306 | 7307 | 1 | 1.2290e-01 | 4.7698e+05 | ; | C - O |
| 7306 | 7308 | 1 | 1.3350e-01 | 4.1003e+05 | ; | C - N |
| 7308 | 7309 | 1 | 1.0100e-01 | 3.6317e+05 | ; | N - H |
| 7308 | 7310 | 1 | 1.4490e-01 | 2.8200e+05 | ; | N - CA |
| 7310 | 7311 | 1 | 1.0900e-01 | 2.8451e+05 | ; | CA - HA |

| | | | | | |
|---|---|---|---|---|---|
| 7310 | 7312 | 1 | 1.5260e-01 | 2.5941e+05 ; | CA - CB |
| 7310 | 7328 | 1 | 1.5220e-01 | 2.6527e+05 ; | CA - C |
| 7312 | 7313 | 1 | 1.0900e-01 | 2.8451e+05 ; | CB - HB2 |
| 7312 | 7314 | 1 | 1.0900e-01 | 2.8451e+05 ; | CB - HB3 |
| 7312 | 7315 | 1 | 1.5260e-01 | 2.5941e+05 ; | CB - CG |
| 7315 | 7316 | 1 | 1.0900e-01 | 2.8451e+05 ; | CG - HG2 |
| 7315 | 7317 | 1 | 1.0900e-01 | 2.8451e+05 ; | CG - HG3 |
| 7315 | 7318 | 1 | 1.5260e-01 | 2.5941e+05 ; | CG - CD |
| 7318 | 7319 | 1 | 1.0900e-01 | 2.8451e+05 ; | CD - HD2 |
| 7318 | 7320 | 1 | 1.0900e-01 | 2.8451e+05 ; | CD - HD3 |
| 7318 | 7321 | 1 | 1.5260e-01 | 2.5941e+05 ; | CD - CE |
| 7321 | 7322 | 1 | 1.0900e-01 | 2.8451e+05 ; | CE - HE2 |
| 7321 | 7323 | 1 | 1.0900e-01 | 2.8451e+05 ; | CE - HE3 |
| 7321 | 7324 | 1 | 1.4710e-01 | 3.0711e+05 ; | CE - NZ |
| 7324 | 7325 | 1 | 1.0100e-01 | 3.6317e+05 ; | NZ - HZ1 |
| 7324 | 7326 | 1 | 1.0100e-01 | 3.6317e+05 ; | NZ - HZ2 |
| 7324 | 7327 | 1 | 1.0100e-01 | 3.6317e+05 ; | NZ - HZ3 |
| 7328 | 7329 | 1 | 1.2290e-01 | 4.7698e+05 ; | C - O |
| 7328 | 7330 | 1 | 1.3350e-01 | 4.1003e+05 ; | C - N |
| 7330 | 7331 | 1 | 1.0100e-01 | 3.6317e+05 ; | N - H |
| 7330 | 7332 | 1 | 1.4490e-01 | 2.8200e+05 ; | N - CA |
| 7332 | 7333 | 1 | 1.0900e-01 | 2.8451e+05 ; | CA - HA |
| 7332 | 7334 | 1 | 1.5260e-01 | 2.5941e+05 ; | CA - CB |
| 7332 | 7350 | 1 | 1.5220e-01 | 2.6527e+05 ; | CA - C |
| 7334 | 7335 | 1 | 1.0900e-01 | 2.8451e+05 ; | CB - HB2 |
| 7334 | 7336 | 1 | 1.0900e-01 | 2.8451e+05 ; | CB - HB3 |
| 7334 | 7337 | 1 | 1.5260e-01 | 2.5941e+05 ; | CB - CG |
| 7337 | 7338 | 1 | 1.0900e-01 | 2.8451e+05 ; | CG - HG2 |
| 7337 | 7339 | 1 | 1.0900e-01 | 2.8451e+05 ; | CG - HG3 |
| 7337 | 7340 | 1 | 1.5260e-01 | 2.5941e+05 ; | CG - CD |
| 7340 | 7341 | 1 | 1.0900e-01 | 2.8451e+05 ; | CD - HD2 |
| 7340 | 7342 | 1 | 1.0900e-01 | 2.8451e+05 ; | CD - HD3 |
| 7340 | 7343 | 1 | 1.5260e-01 | 2.5941e+05 ; | CD - CE |
| 7343 | 7344 | 1 | 1.0900e-01 | 2.8451e+05 ; | CE - HE2 |
| 7343 | 7345 | 1 | 1.0900e-01 | 2.8451e+05 ; | CE - HE3 |
| 7343 | 7346 | 1 | 1.4710e-01 | 3.0711e+05 ; | CE - NZ |
| 7346 | 7347 | 1 | 1.0100e-01 | 3.6317e+05 ; | NZ - HZ1 |
| 7346 | 7348 | 1 | 1.0100e-01 | 3.6317e+05 ; | NZ - HZ2 |
| 7346 | 7349 | 1 | 1.0100e-01 | 3.6317e+05 ; | NZ - HZ3 |
| 7350 | 7351 | 1 | 1.2290e-01 | 4.7698e+05 ; | C - O |
| 7350 | 7352 | 1 | 1.3350e-01 | 4.1003e+05 ; | C - N |
| 7352 | 7353 | 1 | 1.0100e-01 | 3.6317e+05 ; | N - H |
| 7352 | 7354 | 1 | 1.4490e-01 | 2.8200e+05 ; | N - CA |
| 7354 | 7355 | 1 | 1.0900e-01 | 2.8451e+05 ; | CA - HA |
| 7354 | 7356 | 1 | 1.5260e-01 | 2.5941e+05 ; | CA - CB |
| 7354 | 7372 | 1 | 1.5220e-01 | 2.6527e+05 ; | CA - C |
| 7356 | 7357 | 1 | 1.0900e-01 | 2.8451e+05 ; | CB - HB2 |
| 7356 | 7358 | 1 | 1.0900e-01 | 2.8451e+05 ; | CB - HB3 |
| 7356 | 7359 | 1 | 1.5260e-01 | 2.5941e+05 ; | CB - CG |
| 7359 | 7360 | 1 | 1.0900e-01 | 2.8451e+05 ; | CG - HG2 |
| 7359 | 7361 | 1 | 1.0900e-01 | 2.8451e+05 ; | CG - HG3 |
| 7359 | 7362 | 1 | 1.5260e-01 | 2.5941e+05 ; | CG - CD |

| | | | | | | | | |
|---|---|---|---|---|---|---|---|---|
| 7362 | 7363 | 1 | 1.0900e-01 | 2.8451e+05 | ; | CD | - | HD2 |
| 7362 | 7364 | 1 | 1.0900e-01 | 2.8451e+05 | ; | CD | - | HD3 |
| 7362 | 7365 | 1 | 1.5260e-01 | 2.5941e+05 | ; | CD | - | CE |
| 7365 | 7366 | 1 | 1.0900e-01 | 2.8451e+05 | ; | CE | - | HE2 |
| 7365 | 7367 | 1 | 1.0900e-01 | 2.8451e+05 | ; | CE | - | HE3 |
| 7365 | 7368 | 1 | 1.4710e-01 | 3.0711e+05 | ; | CE | - | NZ |
| 7368 | 7369 | 1 | 1.0100e-01 | 3.6317e+05 | ; | NZ | - | HZ1 |
| 7368 | 7370 | 1 | 1.0100e-01 | 3.6317e+05 | ; | NZ | - | HZ2 |
| 7368 | 7371 | 1 | 1.0100e-01 | 3.6317e+05 | ; | NZ | - | HZ3 |
| 7372 | 7373 | 1 | 1.2500e-01 | 5.4894e+05 | ; | C | - | O |
| 7372 | 7374 | 1 | 1.2500e-01 | 5.4894e+05 | ; | C | - | OXT |
| 7375 | 7376 | 1 | 1.3385e-01 | 3.1012e+05 | ; | F2 | - | C38 |
| 7376 | 7377 | 1 | 1.3343e-01 | 4.7647e+05 | ; | C38 | - | C39 |
| 7376 | 7383 | 1 | 1.3343e-01 | 4.7647e+05 | ; | C38 | - | C37 |
| 7377 | 7378 | 1 | 1.2817e-01 | 4.9714e+05 | ; | C39 | - | N23 |
| 7377 | 7423 | 1 | 1.0868e-01 | 2.8836e+05 | ; | C39 | - | H39 |
| 7378 | 7385 | 1 | 1.3030e-01 | 4.6158e+05 | ; | N23 | - | C40 |
| 7378 | 7433 | 1 | 2.1594e-01 | 5.0710e+04 | ; | N23 | - | RU |
| 7379 | 7395 | 1 | 1.3030e-01 | 4.6158e+05 | ; | N21 | - | C24 |
| 7379 | 7397 | 1 | 1.3030e-01 | 4.6158e+05 | ; | N21 | - | C25 |
| 7379 | 7433 | 1 | 2.1162e-01 | 8.2090e+04 | ; | N21 | - | RU |
| 7380 | 7381 | 1 | 1.1724e-01 | 6.3438e+05 | ; | C13 | - | O14 |
| 7380 | 7433 | 1 | 1.9055e-01 | 1.3422e+05 | ; | C13 | - | RU |
| 7382 | 7433 | 1 | 2.4338e-01 | 7.3304e+04 | ; | CL17 | - | RU |
| 7383 | 7384 | 1 | 1.3846e-01 | 4.0342e+05 | ; | C37 | - | C36 |
| 7383 | 7424 | 1 | 1.0879e-01 | 2.8711e+05 | ; | C37 | - | H37 |
| 7384 | 7385 | 1 | 1.3984e-01 | 3.8585e+05 | ; | C36 | - | C40 |
| 7384 | 7386 | 1 | 1.3984e-01 | 3.8585e+05 | ; | C36 | - | C35 |
| 7385 | 7395 | 1 | 1.3984e-01 | 3.8585e+05 | ; | C40 | - | C24 |
| 7386 | 7387 | 1 | 1.3846e-01 | 4.0342e+05 | ; | C35 | - | C34 |
| 7386 | 7391 | 1 | 1.3984e-01 | 3.8585e+05 | ; | C35 | - | C32 |
| 7387 | 7388 | 1 | 1.3994e-01 | 3.3480e+05 | ; | C34 | - | N22 |
| 7387 | 7390 | 1 | 1.2247e-01 | 5.2124e+05 | ; | C34 | - | O42 |
| 7388 | 7389 | 1 | 1.0129e-01 | 3.3740e+05 | ; | N22 | - | H22 |
| 7388 | 7392 | 1 | 1.3994e-01 | 3.3480e+05 | ; | N22 | - | C33 |
| 7391 | 7392 | 1 | 1.3846e-01 | 4.0342e+05 | ; | C32 | - | C33 |
| 7391 | 7394 | 1 | 1.4058e-01 | 3.7673e+05 | ; | C32 | - | C31 |
| 7392 | 7393 | 1 | 1.2247e-01 | 5.2124e+05 | ; | C33 | - | O41 |
| 7394 | 7395 | 1 | 1.4058e-01 | 3.7673e+05 | ; | C31 | - | C24 |
| 7394 | 7396 | 1 | 1.4542e-01 | 3.2351e+05 | ; | C31 | - | C30 |
| 7396 | 7397 | 1 | 1.4058e-01 | 3.7673e+05 | ; | C30 | - | C25 |
| 7396 | 7398 | 1 | 1.4058e-01 | 3.7673e+05 | ; | C30 | - | C29 |
| 7397 | 7401 | 1 | 1.3984e-01 | 3.8585e+05 | ; | C25 | - | C26 |
| 7398 | 7399 | 1 | 1.3984e-01 | 3.8585e+05 | ; | C29 | - | C28 |
| 7398 | 7426 | 1 | 1.0860e-01 | 2.8937e+05 | ; | C29 | - | H29 |
| 7399 | 7400 | 1 | 1.3984e-01 | 3.8585e+05 | ; | C28 | - | C27 |
| 7399 | 7416 | 1 | 1.3637e-01 | 3.2133e+05 | ; | C28 | - | O1 |
| 7400 | 7401 | 1 | 1.3984e-01 | 3.8585e+05 | ; | C27 | - | C26 |
| 7400 | 7427 | 1 | 1.0860e-01 | 2.8937e+05 | ; | C27 | - | H27 |
| 7401 | 7428 | 1 | 1.0860e-01 | 2.8937e+05 | ; | C26 | - | H26 |
| 7402 | 7403 | 1 | 1.2817e-01 | 4.9714e+05 | ; | N15 | - | C7 |
| 7402 | 7404 | 1 | 1.3030e-01 | 4.6158e+05 | ; | N15 | - | C18 |

```
  7402   7433   1    2.0700e-01    6.9873e+04 ;    N15 - RU
  7403   7410   1    1.3343e-01    4.7647e+05 ;     C7 - C8
  7403   7419   1    1.0868e-01    2.8836e+05 ;     C7 - H7A
  7404   7405   1    1.3984e-01    3.8585e+05 ;    C18 - C5
  7404   7409   1    1.3984e-01    3.8585e+05 ;    C18 - C19
  7405   7406   1    1.3984e-01    3.8585e+05 ;     C5 - C6
  7405   7421   1    1.0860e-01    2.8937e+05 ;     C5 - H5
  7406   7407   1    1.3984e-01    3.8585e+05 ;     C6 - C4
  7406   7420   1    1.0860e-01    2.8937e+05 ;     C6 - H6
  7407   7408   1    1.3984e-01    3.8585e+05 ;     C4 - C3
  7407   7422   1    1.0860e-01    2.8937e+05 ;     C4 - H4
  7408   7409   1    1.3984e-01    3.8585e+05 ;     C3 - C19
  7408   7425   1    1.0860e-01    2.8937e+05 ;     C3 - H3
  7409   7429   1    1.0860e-01    2.8937e+05 ;    C19 - H19
  7410   7411   1    1.3343e-01    4.7647e+05 ;     C8 - C9
  7410   7415   1    1.2817e-01    4.9714e+05 ;     C8 - N16
  7411   7412   1    1.3343e-01    4.7647e+05 ;     C9 - C10
  7411   7418   1    1.0879e-01    2.8711e+05 ;     C9 - H9
  7412   7413   1    1.3343e-01    4.7647e+05 ;    C10 - C12
  7412   7432   1    1.0879e-01    2.8711e+05 ;    C10 - H10
  7413   7414   1    1.3343e-01    4.7647e+05 ;    C12 - C11
  7413   7430   1    1.0879e-01    2.8711e+05 ;    C12 - H12
  7414   7415   1    1.2817e-01    4.9714e+05 ;    C11 - N16
  7414   7431   1    1.0868e-01    2.8836e+05 ;    C11 - H11
  7416   7417   1    9.7300e-02    3.1079e+05 ;     O1 - H1

[ pairs ]
;   ai     aj    funct
     1      8      1 ;      N - HB2
     1      9      1 ;      N - HB3
     1     10      1 ;      N - CG
     1     21      1 ;      N - O
     1     22      1 ;      N - N
     2      6      1 ;     H1 - HA
     2      7      1 ;     H1 - CB
     2     20      1 ;     H1 - C
     3      6      1 ;     H2 - HA
     3      7      1 ;     H2 - CB
     3     20      1 ;     H2 - C
     4      6      1 ;     H3 - HA
     4      7      1 ;     H3 - CB
     4     20      1 ;     H3 - C
     5     11      1 ;     CA - HG
     5     12      1 ;     CA - CD1
     5     16      1 ;     CA - CD2
     5     23      1 ;     CA - H
     5     24      1 ;     CA - CA
     6      8      1 ;     HA - HB2
     6      9      1 ;     HA - HB3
     6     10      1 ;     HA - CG
     6     21      1 ;     HA - O
     6     22      1 ;     HA - N
```

| | | | | |
|---|---|---|---|---|
| 7 | 13 | 1 | ; | CB - HD11 |
| 7 | 14 | 1 | ; | CB - HD12 |
| 7 | 15 | 1 | ; | CB - HD13 |
| 7 | 17 | 1 | ; | CB - HD21 |
| 7 | 18 | 1 | ; | CB - HD22 |
| 7 | 19 | 1 | ; | CB - HD23 |
| 7 | 21 | 1 | ; | CB - O |
| 7 | 22 | 1 | ; | CB - N |
| 8 | 11 | 1 | ; | HB2 - HG |
| 8 | 12 | 1 | ; | HB2 - CD1 |
| 8 | 16 | 1 | ; | HB2 - CD2 |
| 8 | 20 | 1 | ; | HB2 - C |
| 9 | 11 | 1 | ; | HB3 - HG |
| 9 | 12 | 1 | ; | HB3 - CD1 |
| 9 | 16 | 1 | ; | HB3 - CD2 |
| 9 | 20 | 1 | ; | HB3 - C |
| 10 | 20 | 1 | ; | CG - C |
| 11 | 13 | 1 | ; | HG - HD11 |
| 11 | 14 | 1 | ; | HG - HD12 |
| 11 | 15 | 1 | ; | HG - HD13 |
| 11 | 17 | 1 | ; | HG - HD21 |
| 11 | 18 | 1 | ; | HG - HD22 |
| 11 | 19 | 1 | ; | HG - HD23 |
| 12 | 17 | 1 | ; | CD1 - HD21 |
| 12 | 18 | 1 | ; | CD1 - HD22 |
| 12 | 19 | 1 | ; | CD1 - HD23 |
| 13 | 16 | 1 | ; | HD11 - CD2 |
| 14 | 16 | 1 | ; | HD12 - CD2 |
| 15 | 16 | 1 | ; | HD13 - CD2 |
| 20 | 25 | 1 | ; | C - HA |
| 20 | 26 | 1 | ; | C - CB |
| 20 | 39 | 1 | ; | C - C |
| 21 | 23 | 1 | ; | O - H |
| 21 | 24 | 1 | ; | O - CA |
| 22 | 27 | 1 | ; | N - HB |
| 22 | 28 | 1 | ; | N - CG2 |
| 22 | 32 | 1 | ; | N - CG1 |
| 22 | 40 | 1 | ; | N - O |
| 22 | 41 | 1 | ; | N - N |
| 23 | 25 | 1 | ; | H - HA |
| 23 | 26 | 1 | ; | H - CB |
| 23 | 39 | 1 | ; | H - C |
| 24 | 29 | 1 | ; | CA - HG21 |
| 24 | 30 | 1 | ; | CA - HG22 |
| 24 | 31 | 1 | ; | CA - HG23 |
| 24 | 33 | 1 | ; | CA - HG12 |
| 24 | 34 | 1 | ; | CA - HG13 |
| 24 | 35 | 1 | ; | CA - CD1 |
| 24 | 42 | 1 | ; | CA - H |
| 24 | 43 | 1 | ; | CA - CA |
| 25 | 27 | 1 | ; | HA - HB |
| 25 | 28 | 1 | ; | HA - CG2 |

```
25      32       1 ;      HA - CG1
25      40       1 ;      HA - O
25      41       1 ;      HA - N
26      36       1 ;      CB - HD11
26      37       1 ;      CB - HD12
26      38       1 ;      CB - HD13
26      40       1 ;      CB - O
26      41       1 ;      CB - N
27      29       1 ;      HB - HG21
27      30       1 ;      HB - HG22
27      31       1 ;      HB - HG23
27      33       1 ;      HB - HG12
27      34       1 ;      HB - HG13
27      35       1 ;      HB - CD1
27      39       1 ;      HB - C
28      33       1 ;     CG2 - HG12
28      34       1 ;     CG2 - HG13
28      35       1 ;     CG2 - CD1
28      39       1 ;     CG2 - C
29      32       1 ;    HG21 - CG1
30      32       1 ;    HG22 - CG1
31      32       1 ;    HG23 - CG1
32      39       1 ;     CG1 - C
33      36       1 ;    HG12 - HD11
33      37       1 ;    HG12 - HD12
33      38       1 ;    HG12 - HD13
34      36       1 ;    HG13 - HD11
34      37       1 ;    HG13 - HD12
34      38       1 ;    HG13 - HD13
39      44       1 ;       C - HA
39      45       1 ;       C - CB
39      58       1 ;       C - C
40      42       1 ;       O - H
40      43       1 ;       O - CA
41      46       1 ;       N - HB2
41      47       1 ;       N - HB3
41      48       1 ;       N - CG
41      59       1 ;       N - O
41      60       1 ;       N - N
42      44       1 ;       H - HA
42      45       1 ;       H - CB
42      58       1 ;       H - C
43      49       1 ;      CA - HG
43      50       1 ;      CA - CD1
43      54       1 ;      CA - CD2
43      61       1 ;      CA - H
43      62       1 ;      CA - CA
44      46       1 ;      HA - HB2
44      47       1 ;      HA - HB3
44      48       1 ;      HA - CG
44      59       1 ;      HA - O
44      60       1 ;      HA - N
```

| | | | | |
|---|---|---|---|---|
| 45 | 51 | 1 | ; | CB - HD11 |
| 45 | 52 | 1 | ; | CB - HD12 |
| 45 | 53 | 1 | ; | CB - HD13 |
| 45 | 55 | 1 | ; | CB - HD21 |
| 45 | 56 | 1 | ; | CB - HD22 |
| 45 | 57 | 1 | ; | CB - HD23 |
| 45 | 59 | 1 | ; | CB - O |
| 45 | 60 | 1 | ; | CB - N |
| 46 | 49 | 1 | ; | HB2 - HG |
| 46 | 50 | 1 | ; | HB2 - CD1 |
| 46 | 54 | 1 | ; | HB2 - CD2 |
| 46 | 58 | 1 | ; | HB2 - C |
| 47 | 49 | 1 | ; | HB3 - HG |
| 47 | 50 | 1 | ; | HB3 - CD1 |
| 47 | 54 | 1 | ; | HB3 - CD2 |
| 47 | 58 | 1 | ; | HB3 - C |
| 48 | 58 | 1 | ; | CG - C |
| 49 | 51 | 1 | ; | HG - HD11 |
| 49 | 52 | 1 | ; | HG - HD12 |
| 49 | 53 | 1 | ; | HG - HD13 |
| 49 | 55 | 1 | ; | HG - HD21 |
| 49 | 56 | 1 | ; | HG - HD22 |
| 49 | 57 | 1 | ; | HG - HD23 |
| 50 | 55 | 1 | ; | CD1 - HD21 |
| 50 | 56 | 1 | ; | CD1 - HD22 |
| 50 | 57 | 1 | ; | CD1 - HD23 |
| 51 | 54 | 1 | ; | HD11 - CD2 |
| 52 | 54 | 1 | ; | HD12 - CD2 |
| 53 | 54 | 1 | ; | HD13 - CD2 |
| 58 | 63 | 1 | ; | C - HA |
| 58 | 64 | 1 | ; | C - CB |
| 58 | 69 | 1 | ; | C - C |
| 59 | 61 | 1 | ; | O - H |
| 59 | 62 | 1 | ; | O - CA |
| 60 | 65 | 1 | ; | N - HB2 |
| 60 | 66 | 1 | ; | N - HB3 |
| 60 | 67 | 1 | ; | N - SG |
| 60 | 70 | 1 | ; | N - O |
| 60 | 71 | 1 | ; | N - N |
| 61 | 63 | 1 | ; | H - HA |
| 61 | 64 | 1 | ; | H - CB |
| 61 | 69 | 1 | ; | H - C |
| 62 | 68 | 1 | ; | CA - HG |
| 62 | 72 | 1 | ; | CA - H |
| 62 | 73 | 1 | ; | CA - CA |
| 63 | 65 | 1 | ; | HA - HB2 |
| 63 | 66 | 1 | ; | HA - HB3 |
| 63 | 67 | 1 | ; | HA - SG |
| 63 | 70 | 1 | ; | HA - O |
| 63 | 71 | 1 | ; | HA - N |
| 64 | 70 | 1 | ; | CB - O |
| 64 | 71 | 1 | ; | CB - N |

```
   65      68       1 ;     HB2 - HG
   65      69       1 ;     HB2 - C
   66      68       1 ;     HB3 - HG
   66      69       1 ;     HB3 - C
   67      69       1 ;      SG - C
   69      74       1 ;       C - HA
   69      75       1 ;       C - CB
   69      86       1 ;       C - C
   70      72       1 ;       O - H
   70      73       1 ;       O - CA
   71      76       1 ;       N - HB2
   71      77       1 ;       N - HB3
   71      78       1 ;       N - CG
   71      87       1 ;       N - O
   71      88       1 ;       N - N
   72      74       1 ;       H - HA
   72      75       1 ;       H - CB
   72      86       1 ;       H - C
   73      79       1 ;      CA - ND1
   73      84       1 ;      CA - CD2
   73      89       1 ;      CA - H
   73      90       1 ;      CA - CA
   74      76       1 ;      HA - HB2
   74      77       1 ;      HA - HB3
   74      78       1 ;      HA - CG
   74      87       1 ;      HA - O
   74      88       1 ;      HA - N
   75      80       1 ;      CB - CE1
   75      82       1 ;      CB - NE2
   75      85       1 ;      CB - HD2
   75      87       1 ;      CB - O
   75      88       1 ;      CB - N
   76      79       1 ;     HB2 - ND1
   76      84       1 ;     HB2 - CD2
   76      86       1 ;     HB2 - C
   77      79       1 ;     HB3 - ND1
   77      84       1 ;     HB3 - CD2
   77      86       1 ;     HB3 - C
   78      81       1 ;      CG - HE1
   78      83       1 ;      CG - HE2
   78      86       1 ;      CG - C
   79      83       1 ;     ND1 - HE2
   79      85       1 ;     ND1 - HD2
   80      85       1 ;     CE1 - HD2
   81      83       1 ;     HE1 - HE2
   81      84       1 ;     HE1 - CD2
   83      85       1 ;     HE2 - HD2
   86      91       1 ;       C - HA
   86      92       1 ;       C - CB
   86      97       1 ;       C - C
   87      89       1 ;       O - H
   87      90       1 ;       O - CA
```

| | | | | |
|---|---|---|---|---|
| 88 | 93 | 1 ; | N - | HB2 |
| 88 | 94 | 1 ; | N - | HB3 |
| 88 | 95 | 1 ; | N - | OG |
| 88 | 98 | 1 ; | N - | O |
| 88 | 99 | 1 ; | N - | N |
| 89 | 91 | 1 ; | H - | HA |
| 89 | 92 | 1 ; | H - | CB |
| 89 | 97 | 1 ; | H - | C |
| 90 | 96 | 1 ; | CA - | HG |
| 90 | 100 | 1 ; | CA - | H |
| 90 | 101 | 1 ; | CA - | CA |
| 91 | 93 | 1 ; | HA - | HB2 |
| 91 | 94 | 1 ; | HA - | HB3 |
| 91 | 95 | 1 ; | HA - | OG |
| 91 | 98 | 1 ; | HA - | O |
| 91 | 99 | 1 ; | HA - | N |
| 92 | 98 | 1 ; | CB - | O |
| 92 | 99 | 1 ; | CB - | N |
| 93 | 96 | 1 ; | HB2 - | HG |
| 93 | 97 | 1 ; | HB2 - | C |
| 94 | 96 | 1 ; | HB3 - | HG |
| 94 | 97 | 1 ; | HB3 - | C |
| 95 | 97 | 1 ; | OG - | C |
| 97 | 102 | 1 ; | C - | HA2 |
| 97 | 103 | 1 ; | C - | HA3 |
| 97 | 104 | 1 ; | C - | C |
| 98 | 100 | 1 ; | O - | H |
| 98 | 101 | 1 ; | O - | CA |
| 99 | 105 | 1 ; | N - | O |
| 99 | 106 | 1 ; | N - | N |
| 100 | 102 | 1 ; | H - | HA2 |
| 100 | 103 | 1 ; | H - | HA3 |
| 100 | 104 | 1 ; | H - | C |
| 101 | 107 | 1 ; | CA - | H |
| 101 | 108 | 1 ; | CA - | CA |
| 102 | 105 | 1 ; | HA2 - | O |
| 102 | 106 | 1 ; | HA2 - | N |
| 103 | 105 | 1 ; | HA3 - | O |
| 103 | 106 | 1 ; | HA3 - | N |
| 104 | 109 | 1 ; | C - | HA |
| 104 | 110 | 1 ; | C - | CB |
| 104 | 126 | 1 ; | C - | C |
| 105 | 107 | 1 ; | O - | H |
| 105 | 108 | 1 ; | O - | CA |
| 106 | 111 | 1 ; | N - | HB2 |
| 106 | 112 | 1 ; | N - | HB3 |
| 106 | 113 | 1 ; | N - | CG |
| 106 | 127 | 1 ; | N - | O |
| 106 | 128 | 1 ; | N - | N |
| 107 | 109 | 1 ; | H - | HA |
| 107 | 110 | 1 ; | H - | CB |
| 107 | 126 | 1 ; | H - | C |

```
  108    114       1 ;      CA - HG2
  108    115       1 ;      CA - HG3
  108    116       1 ;      CA - CD
  108    129       1 ;      CA - H
  108    130       1 ;      CA - CA
  109    111       1 ;      HA - HB2
  109    112       1 ;      HA - HB3
  109    113       1 ;      HA - CG
  109    127       1 ;      HA - O
  109    128       1 ;      HA - N
  110    117       1 ;      CB - HD2
  110    118       1 ;      CB - HD3
  110    119       1 ;      CB - CE
  110    127       1 ;      CB - O
  110    128       1 ;      CB - N
  111    114       1 ;     HB2 - HG2
  111    115       1 ;     HB2 - HG3
  111    116       1 ;     HB2 - CD
  111    126       1 ;     HB2 - C
  112    114       1 ;     HB3 - HG2
  112    115       1 ;     HB3 - HG3
  112    116       1 ;     HB3 - CD
  112    126       1 ;     HB3 - C
  113    120       1 ;      CG - HE2
  113    121       1 ;      CG - HE3
  113    122       1 ;      CG - NZ
  113    126       1 ;      CG - C
  114    117       1 ;     HG2 - HD2
  114    118       1 ;     HG2 - HD3
  114    119       1 ;     HG2 - CE
  115    117       1 ;     HG3 - HD2
  115    118       1 ;     HG3 - HD3
  115    119       1 ;     HG3 - CE
  116    123       1 ;      CD - HZ1
  116    124       1 ;      CD - HZ2
  116    125       1 ;      CD - HZ3
  117    120       1 ;     HD2 - HE2
  117    121       1 ;     HD2 - HE3
  117    122       1 ;     HD2 - NZ
  118    120       1 ;     HD3 - HE2
  118    121       1 ;     HD3 - HE3
  118    122       1 ;     HD3 - NZ
  120    123       1 ;     HE2 - HZ1
  120    124       1 ;     HE2 - HZ2
  120    125       1 ;     HE2 - HZ3
  121    123       1 ;     HE3 - HZ1
  121    124       1 ;     HE3 - HZ2
  121    125       1 ;     HE3 - HZ3
  126    131       1 ;       C - HA
  126    132       1 ;       C - CB
  126    145       1 ;       C - C
  127    129       1 ;       O - H
```

| | | | |
|---|---|---|---|
| 127 | 130 | 1 ; | O - CA |
| 128 | 133 | 1 ; | N - HB2 |
| 128 | 134 | 1 ; | N - HB3 |
| 128 | 135 | 1 ; | N - CG |
| 128 | 146 | 1 ; | N - O |
| 128 | 147 | 1 ; | N - N |
| 129 | 131 | 1 ; | H - HA |
| 129 | 132 | 1 ; | H - CB |
| 129 | 145 | 1 ; | H - C |
| 130 | 136 | 1 ; | CA - HG |
| 130 | 137 | 1 ; | CA - CD1 |
| 130 | 141 | 1 ; | CA - CD2 |
| 130 | 148 | 1 ; | CA - H |
| 130 | 149 | 1 ; | CA - CA |
| 131 | 133 | 1 ; | HA - HB2 |
| 131 | 134 | 1 ; | HA - HB3 |
| 131 | 135 | 1 ; | HA - CG |
| 131 | 146 | 1 ; | HA - O |
| 131 | 147 | 1 ; | HA - N |
| 132 | 138 | 1 ; | CB - HD11 |
| 132 | 139 | 1 ; | CB - HD12 |
| 132 | 140 | 1 ; | CB - HD13 |
| 132 | 142 | 1 ; | CB - HD21 |
| 132 | 143 | 1 ; | CB - HD22 |
| 132 | 144 | 1 ; | CB - HD23 |
| 132 | 146 | 1 ; | CB - O |
| 132 | 147 | 1 ; | CB - N |
| 133 | 136 | 1 ; | HB2 - HG |
| 133 | 137 | 1 ; | HB2 - CD1 |
| 133 | 141 | 1 ; | HB2 - CD2 |
| 133 | 145 | 1 ; | HB2 - C |
| 134 | 136 | 1 ; | HB3 - HG |
| 134 | 137 | 1 ; | HB3 - CD1 |
| 134 | 141 | 1 ; | HB3 - CD2 |
| 134 | 145 | 1 ; | HB3 - C |
| 135 | 145 | 1 ; | CG - C |
| 136 | 138 | 1 ; | HG - HD11 |
| 136 | 139 | 1 ; | HG - HD12 |
| 136 | 140 | 1 ; | HG - HD13 |
| 136 | 142 | 1 ; | HG - HD21 |
| 136 | 143 | 1 ; | HG - HD22 |
| 136 | 144 | 1 ; | HG - HD23 |
| 137 | 142 | 1 ; | CD1 - HD21 |
| 137 | 143 | 1 ; | CD1 - HD22 |
| 137 | 144 | 1 ; | CD1 - HD23 |
| 138 | 141 | 1 ; | HD11 - CD2 |
| 139 | 141 | 1 ; | HD12 - CD2 |
| 140 | 141 | 1 ; | HD13 - CD2 |
| 145 | 150 | 1 ; | C - HA |
| 145 | 151 | 1 ; | C - CB |
| 145 | 160 | 1 ; | C - C |
| 146 | 148 | 1 ; | O - H |

| | | | | |
|---|---|---|---|---|
| 146 | 149 | 1 ; | O - | CA |
| 147 | 152 | 1 ; | N - | HB2 |
| 147 | 153 | 1 ; | N - | HB3 |
| 147 | 154 | 1 ; | N - | CG |
| 147 | 161 | 1 ; | N - | O |
| 147 | 162 | 1 ; | N - | N |
| 148 | 150 | 1 ; | H - | HA |
| 148 | 151 | 1 ; | H - | CB |
| 148 | 160 | 1 ; | H - | C |
| 149 | 155 | 1 ; | CA - | HG2 |
| 149 | 156 | 1 ; | CA - | HG3 |
| 149 | 157 | 1 ; | CA - | CD |
| 149 | 163 | 1 ; | CA - | H |
| 149 | 164 | 1 ; | CA - | CA |
| 150 | 152 | 1 ; | HA - | HB2 |
| 150 | 153 | 1 ; | HA - | HB3 |
| 150 | 154 | 1 ; | HA - | CG |
| 150 | 161 | 1 ; | HA - | O |
| 150 | 162 | 1 ; | HA - | N |
| 151 | 158 | 1 ; | CB - | OE1 |
| 151 | 159 | 1 ; | CB - | OE2 |
| 151 | 161 | 1 ; | CB - | O |
| 151 | 162 | 1 ; | CB - | N |
| 152 | 155 | 1 ; | HB2 - | HG2 |
| 152 | 156 | 1 ; | HB2 - | HG3 |
| 152 | 157 | 1 ; | HB2 - | CD |
| 152 | 160 | 1 ; | HB2 - | C |
| 153 | 155 | 1 ; | HB3 - | HG2 |
| 153 | 156 | 1 ; | HB3 - | HG3 |
| 153 | 157 | 1 ; | HB3 - | CD |
| 153 | 160 | 1 ; | HB3 - | C |
| 154 | 160 | 1 ; | CG - | C |
| 155 | 158 | 1 ; | HG2 - | OE1 |
| 155 | 159 | 1 ; | HG2 - | OE2 |
| 156 | 158 | 1 ; | HG3 - | OE1 |
| 156 | 159 | 1 ; | HG3 - | OE2 |
| 160 | 165 | 1 ; | C - | HA |
| 160 | 166 | 1 ; | C - | CB |
| 160 | 172 | 1 ; | C - | C |
| 161 | 163 | 1 ; | O - | H |
| 161 | 164 | 1 ; | O - | CA |
| 162 | 167 | 1 ; | N - | HB2 |
| 162 | 168 | 1 ; | N - | HB3 |
| 162 | 169 | 1 ; | N - | CG |
| 162 | 173 | 1 ; | N - | O |
| 162 | 174 | 1 ; | N - | N |
| 163 | 165 | 1 ; | H - | HA |
| 163 | 166 | 1 ; | H - | CB |
| 163 | 172 | 1 ; | H - | C |
| 164 | 170 | 1 ; | CA - | OD1 |
| 164 | 171 | 1 ; | CA - | OD2 |
| 164 | 175 | 1 ; | CA - | H |

```
164    176     1 ;     CA - CA
165    167     1 ;     HA - HB2
165    168     1 ;     HA - HB3
165    169     1 ;     HA - CG
165    173     1 ;     HA - O
165    174     1 ;     HA - N
166    173     1 ;     CB - O
166    174     1 ;     CB - N
167    170     1 ;    HB2 - OD1
167    171     1 ;    HB2 - OD2
167    172     1 ;    HB2 - C
168    170     1 ;    HB3 - OD1
168    171     1 ;    HB3 - OD2
168    172     1 ;    HB3 - C
169    172     1 ;     CG - C
172    177     1 ;      C - HA
172    178     1 ;      C - CB
172    194     1 ;      C - C
173    175     1 ;      O - H
173    176     1 ;      O - CA
174    179     1 ;      N - HB2
174    180     1 ;      N - HB3
174    181     1 ;      N - CG
174    195     1 ;      N - O
174    196     1 ;      N - N
175    177     1 ;      H - HA
175    178     1 ;      H - CB
175    194     1 ;      H - C
176    182     1 ;     CA - HG2
176    183     1 ;     CA - HG3
176    184     1 ;     CA - CD
176    197     1 ;     CA - H
176    198     1 ;     CA - CA
177    179     1 ;     HA - HB2
177    180     1 ;     HA - HB3
177    181     1 ;     HA - CG
177    195     1 ;     HA - O
177    196     1 ;     HA - N
178    185     1 ;     CB - HD2
178    186     1 ;     CB - HD3
178    187     1 ;     CB - CE
178    195     1 ;     CB - O
178    196     1 ;     CB - N
179    182     1 ;    HB2 - HG2
179    183     1 ;    HB2 - HG3
179    184     1 ;    HB2 - CD
179    194     1 ;    HB2 - C
180    182     1 ;    HB3 - HG2
180    183     1 ;    HB3 - HG3
180    184     1 ;    HB3 - CD
180    194     1 ;    HB3 - C
181    188     1 ;     CG - HE2
```

| | | | |
|---|---|---|---|
| 181 | 189 | 1 ; | CG - HE3 |
| 181 | 190 | 1 ; | CG - NZ |
| 181 | 194 | 1 ; | CG - C |
| 182 | 185 | 1 ; | HG2 - HD2 |
| 182 | 186 | 1 ; | HG2 - HD3 |
| 182 | 187 | 1 ; | HG2 - CE |
| 183 | 185 | 1 ; | HG3 - HD2 |
| 183 | 186 | 1 ; | HG3 - HD3 |
| 183 | 187 | 1 ; | HG3 - CE |
| 184 | 191 | 1 ; | CD - HZ1 |
| 184 | 192 | 1 ; | CD - HZ2 |
| 184 | 193 | 1 ; | CD - HZ3 |
| 185 | 188 | 1 ; | HD2 - HE2 |
| 185 | 189 | 1 ; | HD2 - HE3 |
| 185 | 190 | 1 ; | HD2 - NZ |
| 186 | 188 | 1 ; | HD3 - HE2 |
| 186 | 189 | 1 ; | HD3 - HE3 |
| 186 | 190 | 1 ; | HD3 - NZ |
| 188 | 191 | 1 ; | HE2 - HZ1 |
| 188 | 192 | 1 ; | HE2 - HZ2 |
| 188 | 193 | 1 ; | HE2 - HZ3 |
| 189 | 191 | 1 ; | HE3 - HZ1 |
| 189 | 192 | 1 ; | HE3 - HZ2 |
| 189 | 193 | 1 ; | HE3 - HZ3 |
| 194 | 199 | 1 ; | C - HA |
| 194 | 200 | 1 ; | C - CB |
| 194 | 215 | 1 ; | C - C |
| 195 | 197 | 1 ; | O - H |
| 195 | 198 | 1 ; | O - CA |
| 196 | 201 | 1 ; | N - HB2 |
| 196 | 202 | 1 ; | N - HB3 |
| 196 | 203 | 1 ; | N - CG |
| 196 | 216 | 1 ; | N - O |
| 196 | 217 | 1 ; | N - N |
| 197 | 199 | 1 ; | H - HA |
| 197 | 200 | 1 ; | H - CB |
| 197 | 215 | 1 ; | H - C |
| 198 | 204 | 1 ; | CA - CD1 |
| 198 | 213 | 1 ; | CA - CD2 |
| 198 | 218 | 1 ; | CA - H |
| 198 | 219 | 1 ; | CA - CA |
| 199 | 201 | 1 ; | HA - HB2 |
| 199 | 202 | 1 ; | HA - HB3 |
| 199 | 203 | 1 ; | HA - CG |
| 199 | 216 | 1 ; | HA - O |
| 199 | 217 | 1 ; | HA - N |
| 200 | 205 | 1 ; | CB - HD1 |
| 200 | 206 | 1 ; | CB - CE1 |
| 200 | 211 | 1 ; | CB - CE2 |
| 200 | 214 | 1 ; | CB - HD2 |
| 200 | 216 | 1 ; | CB - O |
| 200 | 217 | 1 ; | CB - N |

```
 201    204       1 ;     HB2 - CD1
 201    213       1 ;     HB2 - CD2
 201    215       1 ;     HB2 - C
 202    204       1 ;     HB3 - CD1
 202    213       1 ;     HB3 - CD2
 202    215       1 ;     HB3 - C
 203    207       1 ;      CG - HE1
 203    208       1 ;      CG - CZ
 203    212       1 ;      CG - HE2
 203    215       1 ;      CG - C
 204    209       1 ;     CD1 - OH
 204    211       1 ;     CD1 - CE2
 204    214       1 ;     CD1 - HD2
 205    207       1 ;     HD1 - HE1
 205    208       1 ;     HD1 - CZ
 205    213       1 ;     HD1 - CD2
 206    210       1 ;     CE1 - HH
 206    212       1 ;     CE1 - HE2
 206    213       1 ;     CE1 - CD2
 207    209       1 ;     HE1 - OH
 207    211       1 ;     HE1 - CE2
 208    214       1 ;      CZ - HD2
 209    212       1 ;      OH - HE2
 209    213       1 ;      OH - CD2
 210    211       1 ;      HH - CE2
 212    214       1 ;     HE2 - HD2
 215    220       1 ;       C - HA
 215    221       1 ;       C - CB
 215    234       1 ;       C - C
 216    218       1 ;       O - H
 216    219       1 ;       O - CA
 217    222       1 ;       N - HB
 217    223       1 ;       N - CG2
 217    227       1 ;       N - CG1
 217    235       1 ;       N - O
 217    236       1 ;       N - N
 218    220       1 ;       H - HA
 218    221       1 ;       H - CB
 218    234       1 ;       H - C
 219    224       1 ;      CA - HG21
 219    225       1 ;      CA - HG22
 219    226       1 ;      CA - HG23
 219    228       1 ;      CA - HG12
 219    229       1 ;      CA - HG13
 219    230       1 ;      CA - CD1
 219    237       1 ;      CA - H
 219    238       1 ;      CA - CA
 220    222       1 ;      HA - HB
 220    223       1 ;      HA - CG2
 220    227       1 ;      HA - CG1
 220    235       1 ;      HA - O
 220    236       1 ;      HA - N
```

| | | | | |
|---|---|---|---|---|
| 221 | 231 | 1 | ; | CB - HD11 |
| 221 | 232 | 1 | ; | CB - HD12 |
| 221 | 233 | 1 | ; | CB - HD13 |
| 221 | 235 | 1 | ; | CB - O |
| 221 | 236 | 1 | ; | CB - N |
| 222 | 224 | 1 | ; | HB - HG21 |
| 222 | 225 | 1 | ; | HB - HG22 |
| 222 | 226 | 1 | ; | HB - HG23 |
| 222 | 228 | 1 | ; | HB - HG12 |
| 222 | 229 | 1 | ; | HB - HG13 |
| 222 | 230 | 1 | ; | HB - CD1 |
| 222 | 234 | 1 | ; | HB - C |
| 223 | 228 | 1 | ; | CG2 - HG12 |
| 223 | 229 | 1 | ; | CG2 - HG13 |
| 223 | 230 | 1 | ; | CG2 - CD1 |
| 223 | 234 | 1 | ; | CG2 - C |
| 224 | 227 | 1 | ; | HG21 - CG1 |
| 225 | 227 | 1 | ; | HG22 - CG1 |
| 226 | 227 | 1 | ; | HG23 - CG1 |
| 227 | 234 | 1 | ; | CG1 - C |
| 228 | 231 | 1 | ; | HG12 - HD11 |
| 228 | 232 | 1 | ; | HG12 - HD12 |
| 228 | 233 | 1 | ; | HG12 - HD13 |
| 229 | 231 | 1 | ; | HG13 - HD11 |
| 229 | 232 | 1 | ; | HG13 - HD12 |
| 229 | 233 | 1 | ; | HG13 - HD13 |
| 234 | 239 | 1 | ; | C - HA |
| 234 | 240 | 1 | ; | C - CB |
| 234 | 253 | 1 | ; | C - C |
| 235 | 237 | 1 | ; | O - H |
| 235 | 238 | 1 | ; | O - CA |
| 236 | 241 | 1 | ; | N - HB |
| 236 | 242 | 1 | ; | N - CG2 |
| 236 | 246 | 1 | ; | N - CG1 |
| 236 | 254 | 1 | ; | N - O |
| 236 | 255 | 1 | ; | N - N |
| 237 | 239 | 1 | ; | H - HA |
| 237 | 240 | 1 | ; | H - CB |
| 237 | 253 | 1 | ; | H - C |
| 238 | 243 | 1 | ; | CA - HG21 |
| 238 | 244 | 1 | ; | CA - HG22 |
| 238 | 245 | 1 | ; | CA - HG23 |
| 238 | 247 | 1 | ; | CA - HG12 |
| 238 | 248 | 1 | ; | CA - HG13 |
| 238 | 249 | 1 | ; | CA - CD1 |
| 238 | 256 | 1 | ; | CA - H |
| 238 | 257 | 1 | ; | CA - CA |
| 239 | 241 | 1 | ; | HA - HB |
| 239 | 242 | 1 | ; | HA - CG2 |
| 239 | 246 | 1 | ; | HA - CG1 |
| 239 | 254 | 1 | ; | HA - O |
| 239 | 255 | 1 | ; | HA - N |

```
  240   250     1 ;    CB - HD11
  240   251     1 ;    CB - HD12
  240   252     1 ;    CB - HD13
  240   254     1 ;    CB - O
  240   255     1 ;    CB - N
  241   243     1 ;    HB - HG21
  241   244     1 ;    HB - HG22
  241   245     1 ;    HB - HG23
  241   247     1 ;    HB - HG12
  241   248     1 ;    HB - HG13
  241   249     1 ;    HB - CD1
  241   253     1 ;    HB - C
  242   247     1 ;   CG2 - HG12
  242   248     1 ;   CG2 - HG13
  242   249     1 ;   CG2 - CD1
  242   253     1 ;   CG2 - C
  243   246     1 ;  HG21 - CG1
  244   246     1 ;  HG22 - CG1
  245   246     1 ;  HG23 - CG1
  246   253     1 ;   CG1 - C
  247   250     1 ;  HG12 - HD11
  247   251     1 ;  HG12 - HD12
  247   252     1 ;  HG12 - HD13
  248   250     1 ;  HG13 - HD11
  248   251     1 ;  HG13 - HD12
  248   252     1 ;  HG13 - HD13
  253   258     1 ;     C - HA
  253   259     1 ;     C - CB
  253   265     1 ;     C - C
  254   256     1 ;     O - H
  254   257     1 ;     O - CA
  255   260     1 ;     N - HB2
  255   261     1 ;     N - HB3
  255   262     1 ;     N - CG
  255   266     1 ;     N - O
  255   267     1 ;     N - N
  256   258     1 ;     H - HA
  256   259     1 ;     H - CB
  256   265     1 ;     H - C
  257   263     1 ;    CA - OD1
  257   264     1 ;    CA - OD2
  257   268     1 ;    CA - H
  257   269     1 ;    CA - CA
  258   260     1 ;    HA - HB2
  258   261     1 ;    HA - HB3
  258   262     1 ;    HA - CG
  258   266     1 ;    HA - O
  258   267     1 ;    HA - N
  259   266     1 ;    CB - O
  259   267     1 ;    CB - N
  260   263     1 ;   HB2 - OD1
  260   264     1 ;   HB2 - OD2
```

| | | | |
|---|---|---|---|
| 260 | 265 | 1 ; | HB2 - C |
| 261 | 263 | 1 ; | HB3 - OD1 |
| 261 | 264 | 1 ; | HB3 - OD2 |
| 261 | 265 | 1 ; | HB3 - C |
| 262 | 265 | 1 ; | CG - C |
| 265 | 270 | 1 ; | C - HA |
| 265 | 271 | 1 ; | C - CB |
| 265 | 280 | 1 ; | C - C |
| 266 | 268 | 1 ; | O - H |
| 266 | 269 | 1 ; | O - CA |
| 267 | 272 | 1 ; | N - HB2 |
| 267 | 273 | 1 ; | N - HB3 |
| 267 | 274 | 1 ; | N - CG |
| 267 | 281 | 1 ; | N - O |
| 267 | 282 | 1 ; | N - N |
| 268 | 270 | 1 ; | H - HA |
| 268 | 271 | 1 ; | H - CB |
| 268 | 280 | 1 ; | H - C |
| 269 | 275 | 1 ; | CA - HG2 |
| 269 | 276 | 1 ; | CA - HG3 |
| 269 | 277 | 1 ; | CA - CD |
| 269 | 283 | 1 ; | CA - H |
| 269 | 284 | 1 ; | CA - CA |
| 270 | 272 | 1 ; | HA - HB2 |
| 270 | 273 | 1 ; | HA - HB3 |
| 270 | 274 | 1 ; | HA - CG |
| 270 | 281 | 1 ; | HA - O |
| 270 | 282 | 1 ; | HA - N |
| 271 | 278 | 1 ; | CB - OE1 |
| 271 | 279 | 1 ; | CB - OE2 |
| 271 | 281 | 1 ; | CB - O |
| 271 | 282 | 1 ; | CB - N |
| 272 | 275 | 1 ; | HB2 - HG2 |
| 272 | 276 | 1 ; | HB2 - HG3 |
| 272 | 277 | 1 ; | HB2 - CD |
| 272 | 280 | 1 ; | HB2 - C |
| 273 | 275 | 1 ; | HB3 - HG2 |
| 273 | 276 | 1 ; | HB3 - HG3 |
| 273 | 277 | 1 ; | HB3 - CD |
| 273 | 280 | 1 ; | HB3 - C |
| 274 | 280 | 1 ; | CG - C |
| 275 | 278 | 1 ; | HG2 - OE1 |
| 275 | 279 | 1 ; | HG2 - OE2 |
| 276 | 278 | 1 ; | HG3 - OE1 |
| 276 | 279 | 1 ; | HG3 - OE2 |
| 280 | 285 | 1 ; | C - HA |
| 280 | 286 | 1 ; | C - CB |
| 280 | 302 | 1 ; | C - C |
| 281 | 283 | 1 ; | O - H |
| 281 | 284 | 1 ; | O - CA |
| 282 | 287 | 1 ; | N - HB2 |
| 282 | 288 | 1 ; | N - HB3 |

| | | | |
|---|---|---|---|
| 282 | 289 | 1 ; | N - CG |
| 282 | 303 | 1 ; | N - O |
| 282 | 304 | 1 ; | N - N |
| 283 | 285 | 1 ; | H - HA |
| 283 | 286 | 1 ; | H - CB |
| 283 | 302 | 1 ; | H - C |
| 284 | 290 | 1 ; | CA - HG2 |
| 284 | 291 | 1 ; | CA - HG3 |
| 284 | 292 | 1 ; | CA - CD |
| 284 | 305 | 1 ; | CA - H |
| 284 | 306 | 1 ; | CA - CA |
| 285 | 287 | 1 ; | HA - HB2 |
| 285 | 288 | 1 ; | HA - HB3 |
| 285 | 289 | 1 ; | HA - CG |
| 285 | 303 | 1 ; | HA - O |
| 285 | 304 | 1 ; | HA - N |
| 286 | 293 | 1 ; | CB - HD2 |
| 286 | 294 | 1 ; | CB - HD3 |
| 286 | 295 | 1 ; | CB - CE |
| 286 | 303 | 1 ; | CB - O |
| 286 | 304 | 1 ; | CB - N |
| 287 | 290 | 1 ; | HB2 - HG2 |
| 287 | 291 | 1 ; | HB2 - HG3 |
| 287 | 292 | 1 ; | HB2 - CD |
| 287 | 302 | 1 ; | HB2 - C |
| 288 | 290 | 1 ; | HB3 - HG2 |
| 288 | 291 | 1 ; | HB3 - HG3 |
| 288 | 292 | 1 ; | HB3 - CD |
| 288 | 302 | 1 ; | HB3 - C |
| 289 | 296 | 1 ; | CG - HE2 |
| 289 | 297 | 1 ; | CG - HE3 |
| 289 | 298 | 1 ; | CG - NZ |
| 289 | 302 | 1 ; | CG - C |
| 290 | 293 | 1 ; | HG2 - HD2 |
| 290 | 294 | 1 ; | HG2 - HD3 |
| 290 | 295 | 1 ; | HG2 - CE |
| 291 | 293 | 1 ; | HG3 - HD2 |
| 291 | 294 | 1 ; | HG3 - HD3 |
| 291 | 295 | 1 ; | HG3 - CE |
| 292 | 299 | 1 ; | CD - HZ1 |
| 292 | 300 | 1 ; | CD - HZ2 |
| 292 | 301 | 1 ; | CD - HZ3 |
| 293 | 296 | 1 ; | HD2 - HE2 |
| 293 | 297 | 1 ; | HD2 - HE3 |
| 293 | 298 | 1 ; | HD2 - NZ |
| 294 | 296 | 1 ; | HD3 - HE2 |
| 294 | 297 | 1 ; | HD3 - HE3 |
| 294 | 298 | 1 ; | HD3 - NZ |
| 296 | 299 | 1 ; | HE2 - HZ1 |
| 296 | 300 | 1 ; | HE2 - HZ2 |
| 296 | 301 | 1 ; | HE2 - HZ3 |
| 297 | 299 | 1 ; | HE3 - HZ1 |

```
297    300      1 ;    HE3 - HZ2
297    301      1 ;    HE3 - HZ3
302    307      1 ;      C - HA
302    308      1 ;      C - CB
302    321      1 ;      C - C
303    305      1 ;      O - H
303    306      1 ;      O - CA
304    309      1 ;      N - HB2
304    310      1 ;      N - HB3
304    311      1 ;      N - CG
304    322      1 ;      N - O
304    323      1 ;      N - N
305    307      1 ;      H - HA
305    308      1 ;      H - CB
305    321      1 ;      H - C
306    312      1 ;     CA - HG
306    313      1 ;     CA - CD1
306    317      1 ;     CA - CD2
306    324      1 ;     CA - H
306    325      1 ;     CA - CA
307    309      1 ;     HA - HB2
307    310      1 ;     HA - HB3
307    311      1 ;     HA - CG
307    322      1 ;     HA - O
307    323      1 ;     HA - N
308    314      1 ;     CB - HD11
308    315      1 ;     CB - HD12
308    316      1 ;     CB - HD13
308    318      1 ;     CB - HD21
308    319      1 ;     CB - HD22
308    320      1 ;     CB - HD23
308    322      1 ;     CB - O
308    323      1 ;     CB - N
309    312      1 ;    HB2 - HG
309    313      1 ;    HB2 - CD1
309    317      1 ;    HB2 - CD2
309    321      1 ;    HB2 - C
310    312      1 ;    HB3 - HG
310    313      1 ;    HB3 - CD1
310    317      1 ;    HB3 - CD2
310    321      1 ;    HB3 - C
311    321      1 ;     CG - C
312    314      1 ;     HG - HD11
312    315      1 ;     HG - HD12
312    316      1 ;     HG - HD13
312    318      1 ;     HG - HD21
312    319      1 ;     HG - HD22
312    320      1 ;     HG - HD23
313    318      1 ;    CD1 - HD21
313    319      1 ;    CD1 - HD22
313    320      1 ;    CD1 - HD23
314    317      1 ;   HD11 - CD2
```

```
   315     317       1 ;   HD12 - CD2
   316     317       1 ;   HD13 - CD2
   321     326       1 ;      C - HA2
   321     327       1 ;      C - HA3
   321     328       1 ;      C - C
   322     324       1 ;      O - H
   322     325       1 ;      O - CA
   323     329       1 ;      N - O
   323     330       1 ;      N - N
   324     326       1 ;      H - HA2
   324     327       1 ;      H - HA3
   324     328       1 ;      H - C
   325     331       1 ;     CA - H
   325     332       1 ;     CA - CA
   326     329       1 ;    HA2 - O
   326     330       1 ;    HA2 - N
   327     329       1 ;    HA3 - O
   327     330       1 ;    HA3 - N
   328     333       1 ;      C - HA
   328     334       1 ;      C - CB
   328     345       1 ;      C - C
   329     331       1 ;      O - H
   329     332       1 ;      O - CA
   330     335       1 ;      N - HB2
   330     336       1 ;      N - HB3
   330     337       1 ;      N - CG
   330     346       1 ;      N - O
   330     347       1 ;      N - N
   331     333       1 ;      H - HA
   331     334       1 ;      H - CB
   331     345       1 ;      H - C
   332     338       1 ;     CA - HG2
   332     339       1 ;     CA - HG3
   332     340       1 ;     CA - CD
   332     348       1 ;     CA - H
   332     349       1 ;     CA - CA
   333     335       1 ;     HA - HB2
   333     336       1 ;     HA - HB3
   333     337       1 ;     HA - CG
   333     346       1 ;     HA - O
   333     347       1 ;     HA - N
   334     341       1 ;     CB - OE1
   334     342       1 ;     CB - NE2
   334     346       1 ;     CB - O
   334     347       1 ;     CB - N
   335     338       1 ;    HB2 - HG2
   335     339       1 ;    HB2 - HG3
   335     340       1 ;    HB2 - CD
   335     345       1 ;    HB2 - C
   336     338       1 ;    HB3 - HG2
   336     339       1 ;    HB3 - HG3
   336     340       1 ;    HB3 - CD
```

```
336    345      1 ;    HB3 - C
337    343      1 ;     CG - HE21
337    344      1 ;     CG - HE22
337    345      1 ;     CG - C
338    341      1 ;    HG2 - OE1
338    342      1 ;    HG2 - NE2
339    341      1 ;    HG3 - OE1
339    342      1 ;    HG3 - NE2
341    343      1 ;    OE1 - HE21
341    344      1 ;    OE1 - HE22
345    350      1 ;      C - HA2
345    351      1 ;      C - HA3
345    352      1 ;      C - C
346    348      1 ;      O - H
346    349      1 ;      O - CA
347    353      1 ;      N - O
347    354      1 ;      N - N
348    350      1 ;      H - HA2
348    351      1 ;      H - HA3
348    352      1 ;      H - C
349    355      1 ;     CA - H
349    356      1 ;     CA - CA
350    353      1 ;    HA2 - O
350    354      1 ;    HA2 - N
351    353      1 ;    HA3 - O
351    354      1 ;    HA3 - N
352    357      1 ;      C - HA
352    358      1 ;      C - CB
352    366      1 ;      C - C
353    355      1 ;      O - H
353    356      1 ;      O - CA
354    359      1 ;      N - HB
354    360      1 ;      N - CG2
354    364      1 ;      N - OG1
354    367      1 ;      N - O
354    368      1 ;      N - N
355    357      1 ;      H - HA
355    358      1 ;      H - CB
355    366      1 ;      H - C
356    361      1 ;     CA - HG21
356    362      1 ;     CA - HG22
356    363      1 ;     CA - HG23
356    365      1 ;     CA - HG1
356    369      1 ;     CA - H
356    370      1 ;     CA - CA
357    359      1 ;     HA - HB
357    360      1 ;     HA - CG2
357    364      1 ;     HA - OG1
357    367      1 ;     HA - O
357    368      1 ;     HA - N
358    367      1 ;     CB - O
358    368      1 ;     CB - N
```

```
359    361       1 ;      HB - HG21
359    362       1 ;      HB - HG22
359    363       1 ;      HB - HG23
359    365       1 ;      HB - HG1
359    366       1 ;      HB - C
360    365       1 ;     CG2 - HG1
360    366       1 ;     CG2 - C
361    364       1 ;    HG21 - OG1
362    364       1 ;    HG22 - OG1
363    364       1 ;    HG23 - OG1
364    366       1 ;     OG1 - C
366    371       1 ;       C - HA
366    372       1 ;       C - CB
366    387       1 ;       C - C
367    369       1 ;       O - H
367    370       1 ;       O - CA
368    373       1 ;       N - HB2
368    374       1 ;       N - HB3
368    375       1 ;       N - CG
368    388       1 ;       N - O
368    389       1 ;       N - N
369    371       1 ;       H - HA
369    372       1 ;       H - CB
369    387       1 ;       H - C
370    376       1 ;      CA - CD1
370    385       1 ;      CA - CD2
370    390       1 ;      CA - H
370    391       1 ;      CA - CA
371    373       1 ;      HA - HB2
371    374       1 ;      HA - HB3
371    375       1 ;      HA - CG
371    388       1 ;      HA - O
371    389       1 ;      HA - N
372    377       1 ;      CB - HD1
372    378       1 ;      CB - CE1
372    383       1 ;      CB - CE2
372    386       1 ;      CB - HD2
372    388       1 ;      CB - O
372    389       1 ;      CB - N
373    376       1 ;     HB2 - CD1
373    385       1 ;     HB2 - CD2
373    387       1 ;     HB2 - C
374    376       1 ;     HB3 - CD1
374    385       1 ;     HB3 - CD2
374    387       1 ;     HB3 - C
375    379       1 ;      CG - HE1
375    380       1 ;      CG - CZ
375    384       1 ;      CG - HE2
375    387       1 ;      CG - C
376    381       1 ;     CD1 - OH
376    383       1 ;     CD1 - CE2
376    386       1 ;     CD1 - HD2
```

| | | | | | |
|---|---|---|---|---|---|
| 377 | 379 | 1 ; | HD1 | - | HE1 |
| 377 | 380 | 1 ; | HD1 | - | CZ |
| 377 | 385 | 1 ; | HD1 | - | CD2 |
| 378 | 382 | 1 ; | CE1 | - | HH |
| 378 | 384 | 1 ; | CE1 | - | HE2 |
| 378 | 385 | 1 ; | CE1 | - | CD2 |
| 379 | 381 | 1 ; | HE1 | - | OH |
| 379 | 383 | 1 ; | HE1 | - | CE2 |
| 380 | 386 | 1 ; | CZ | - | HD2 |
| 381 | 384 | 1 ; | OH | - | HE2 |
| 381 | 385 | 1 ; | OH | - | CD2 |
| 382 | 383 | 1 ; | HH | - | CE2 |
| 384 | 386 | 1 ; | HE2 | - | HD2 |
| 387 | 392 | 1 ; | C | - | HA2 |
| 387 | 393 | 1 ; | C | - | HA3 |
| 387 | 394 | 1 ; | C | - | C |
| 388 | 390 | 1 ; | O | - | H |
| 388 | 391 | 1 ; | O | - | CA |
| 389 | 395 | 1 ; | N | - | O |
| 389 | 396 | 1 ; | N | - | N |
| 390 | 392 | 1 ; | H | - | HA2 |
| 390 | 393 | 1 ; | H | - | HA3 |
| 390 | 394 | 1 ; | H | - | C |
| 391 | 397 | 1 ; | CA | - | H |
| 391 | 398 | 1 ; | CA | - | CA |
| 392 | 395 | 1 ; | HA2 | - | O |
| 392 | 396 | 1 ; | HA2 | - | N |
| 393 | 395 | 1 ; | HA3 | - | O |
| 393 | 396 | 1 ; | HA3 | - | N |
| 394 | 399 | 1 ; | C | - | HA |
| 394 | 400 | 1 ; | C | - | CB |
| 394 | 405 | 1 ; | C | - | C |
| 395 | 397 | 1 ; | O | - | H |
| 395 | 398 | 1 ; | O | - | CA |
| 396 | 401 | 1 ; | N | - | HB2 |
| 396 | 402 | 1 ; | N | - | HB3 |
| 396 | 403 | 1 ; | N | - | SG |
| 396 | 406 | 1 ; | N | - | O |
| 396 | 407 | 1 ; | N | - | N |
| 397 | 399 | 1 ; | H | - | HA |
| 397 | 400 | 1 ; | H | - | CB |
| 397 | 405 | 1 ; | H | - | C |
| 398 | 404 | 1 ; | CA | - | HG |
| 398 | 408 | 1 ; | CA | - | H |
| 398 | 409 | 1 ; | CA | - | CA |
| 399 | 401 | 1 ; | HA | - | HB2 |
| 399 | 402 | 1 ; | HA | - | HB3 |
| 399 | 403 | 1 ; | HA | - | SG |
| 399 | 406 | 1 ; | HA | - | O |
| 399 | 407 | 1 ; | HA | - | N |
| 400 | 406 | 1 ; | CB | - | O |
| 400 | 407 | 1 ; | CB | - | N |

```
401    404      1 ;    HB2 - HG
401    405      1 ;    HB2 - C
402    404      1 ;    HB3 - HG
402    405      1 ;    HB3 - C
403    405      1 ;     SG - C
405    410      1 ;      C - HA
405    411      1 ;      C - CB
405    421      1 ;      C - C
406    408      1 ;      O - H
406    409      1 ;      O - CA
407    412      1 ;      N - HB
407    413      1 ;      N - CG1
407    417      1 ;      N - CG2
407    422      1 ;      N - O
407    423      1 ;      N - N
408    410      1 ;      H - HA
408    411      1 ;      H - CB
408    421      1 ;      H - C
409    414      1 ;     CA - HG11
409    415      1 ;     CA - HG12
409    416      1 ;     CA - HG13
409    418      1 ;     CA - HG21
409    419      1 ;     CA - HG22
409    420      1 ;     CA - HG23
409    424      1 ;     CA - H
409    425      1 ;     CA - CA
410    412      1 ;     HA - HB
410    413      1 ;     HA - CG1
410    417      1 ;     HA - CG2
410    422      1 ;     HA - O
410    423      1 ;     HA - N
411    422      1 ;     CB - O
411    423      1 ;     CB - N
412    414      1 ;     HB - HG11
412    415      1 ;     HB - HG12
412    416      1 ;     HB - HG13
412    418      1 ;     HB - HG21
412    419      1 ;     HB - HG22
412    420      1 ;     HB - HG23
412    421      1 ;     HB - C
413    418      1 ;    CG1 - HG21
413    419      1 ;    CG1 - HG22
413    420      1 ;    CG1 - HG23
413    421      1 ;    CG1 - C
414    417      1 ;   HG11 - CG2
415    417      1 ;   HG12 - CG2
416    417      1 ;   HG13 - CG2
417    421      1 ;    CG2 - C
421    426      1 ;      C - HA
421    427      1 ;      C - CB
421    442      1 ;      C - C
422    424      1 ;      O - H
```

| | | | | |
|---|---|---|---|---|
| 422 | 425 | 1 ; | O - | CA |
| 423 | 428 | 1 ; | N - | HB2 |
| 423 | 429 | 1 ; | N - | HB3 |
| 423 | 430 | 1 ; | N - | CG |
| 423 | 443 | 1 ; | N - | O |
| 423 | 444 | 1 ; | N - | N |
| 424 | 426 | 1 ; | H - | HA |
| 424 | 427 | 1 ; | H - | CB |
| 424 | 442 | 1 ; | H - | C |
| 425 | 431 | 1 ; | CA - | CD1 |
| 425 | 440 | 1 ; | CA - | CD2 |
| 425 | 445 | 1 ; | CA - | H |
| 425 | 446 | 1 ; | CA - | CA |
| 426 | 428 | 1 ; | HA - | HB2 |
| 426 | 429 | 1 ; | HA - | HB3 |
| 426 | 430 | 1 ; | HA - | CG |
| 426 | 443 | 1 ; | HA - | O |
| 426 | 444 | 1 ; | HA - | N |
| 427 | 432 | 1 ; | CB - | HD1 |
| 427 | 433 | 1 ; | CB - | CE1 |
| 427 | 438 | 1 ; | CB - | CE2 |
| 427 | 441 | 1 ; | CB - | HD2 |
| 427 | 443 | 1 ; | CB - | O |
| 427 | 444 | 1 ; | CB - | N |
| 428 | 431 | 1 ; | HB2 - | CD1 |
| 428 | 440 | 1 ; | HB2 - | CD2 |
| 428 | 442 | 1 ; | HB2 - | C |
| 429 | 431 | 1 ; | HB3 - | CD1 |
| 429 | 440 | 1 ; | HB3 - | CD2 |
| 429 | 442 | 1 ; | HB3 - | C |
| 430 | 434 | 1 ; | CG - | HE1 |
| 430 | 435 | 1 ; | CG - | CZ |
| 430 | 439 | 1 ; | CG - | HE2 |
| 430 | 442 | 1 ; | CG - | C |
| 431 | 436 | 1 ; | CD1 - | OH |
| 431 | 438 | 1 ; | CD1 - | CE2 |
| 431 | 441 | 1 ; | CD1 - | HD2 |
| 432 | 434 | 1 ; | HD1 - | HE1 |
| 432 | 435 | 1 ; | HD1 - | CZ |
| 432 | 440 | 1 ; | HD1 - | CD2 |
| 433 | 437 | 1 ; | CE1 - | HH |
| 433 | 439 | 1 ; | CE1 - | HE2 |
| 433 | 440 | 1 ; | CE1 - | CD2 |
| 434 | 436 | 1 ; | HE1 - | OH |
| 434 | 438 | 1 ; | HE1 - | CE2 |
| 435 | 441 | 1 ; | CZ - | HD2 |
| 436 | 439 | 1 ; | OH - | HE2 |
| 436 | 440 | 1 ; | OH - | CD2 |
| 437 | 438 | 1 ; | HH - | CE2 |
| 439 | 441 | 1 ; | HE2 - | HD2 |
| 442 | 447 | 1 ; | C - | HA |
| 442 | 448 | 1 ; | C - | CB |

```
442    464       1 ;       C - C
443    445       1 ;       O - H
443    446       1 ;       O - CA
444    449       1 ;       N - HB2
444    450       1 ;       N - HB3
444    451       1 ;       N - CG
444    465       1 ;       N - O
444    466       1 ;       N - N
445    447       1 ;       H - HA
445    448       1 ;       H - CB
445    464       1 ;       H - C
446    452       1 ;      CA - HG2
446    453       1 ;      CA - HG3
446    454       1 ;      CA - CD
446    467       1 ;      CA - H
446    468       1 ;      CA - CA
447    449       1 ;      HA - HB2
447    450       1 ;      HA - HB3
447    451       1 ;      HA - CG
447    465       1 ;      HA - O
447    466       1 ;      HA - N
448    455       1 ;      CB - HD2
448    456       1 ;      CB - HD3
448    457       1 ;      CB - CE
448    465       1 ;      CB - O
448    466       1 ;      CB - N
449    452       1 ;     HB2 - HG2
449    453       1 ;     HB2 - HG3
449    454       1 ;     HB2 - CD
449    464       1 ;     HB2 - C
450    452       1 ;     HB3 - HG2
450    453       1 ;     HB3 - HG3
450    454       1 ;     HB3 - CD
450    464       1 ;     HB3 - C
451    458       1 ;      CG - HE2
451    459       1 ;      CG - HE3
451    460       1 ;      CG - NZ
451    464       1 ;      CG - C
452    455       1 ;     HG2 - HD2
452    456       1 ;     HG2 - HD3
452    457       1 ;     HG2 - CE
453    455       1 ;     HG3 - HD2
453    456       1 ;     HG3 - HD3
453    457       1 ;     HG3 - CE
454    461       1 ;      CD - HZ1
454    462       1 ;      CD - HZ2
454    463       1 ;      CD - HZ3
455    458       1 ;     HD2 - HE2
455    459       1 ;     HD2 - HE3
455    460       1 ;     HD2 - NZ
456    458       1 ;     HD3 - HE2
456    459       1 ;     HD3 - HE3
```

| | | | | |
|---|---|---|---|---|
| 456 | 460 | 1 | ; | HD3 - NZ |
| 458 | 461 | 1 | ; | HE2 - HZ1 |
| 458 | 462 | 1 | ; | HE2 - HZ2 |
| 458 | 463 | 1 | ; | HE2 - HZ3 |
| 459 | 461 | 1 | ; | HE3 - HZ1 |
| 459 | 462 | 1 | ; | HE3 - HZ2 |
| 459 | 463 | 1 | ; | HE3 - HZ3 |
| 464 | 469 | 1 | ; | C - HA2 |
| 464 | 470 | 1 | ; | C - HA3 |
| 464 | 471 | 1 | ; | C - C |
| 465 | 467 | 1 | ; | O - H |
| 465 | 468 | 1 | ; | O - CA |
| 466 | 472 | 1 | ; | N - O |
| 466 | 473 | 1 | ; | N - N |
| 467 | 469 | 1 | ; | H - HA2 |
| 467 | 470 | 1 | ; | H - HA3 |
| 467 | 471 | 1 | ; | H - C |
| 468 | 474 | 1 | ; | CA - H |
| 468 | 475 | 1 | ; | CA - CA |
| 469 | 472 | 1 | ; | HA2 - O |
| 469 | 473 | 1 | ; | HA2 - N |
| 470 | 472 | 1 | ; | HA3 - O |
| 470 | 473 | 1 | ; | HA3 - N |
| 471 | 476 | 1 | ; | C - HA |
| 471 | 477 | 1 | ; | C - CB |
| 471 | 490 | 1 | ; | C - C |
| 472 | 474 | 1 | ; | O - H |
| 472 | 475 | 1 | ; | O - CA |
| 473 | 478 | 1 | ; | N - HB |
| 473 | 479 | 1 | ; | N - CG2 |
| 473 | 483 | 1 | ; | N - CG1 |
| 473 | 491 | 1 | ; | N - O |
| 473 | 492 | 1 | ; | N - N |
| 474 | 476 | 1 | ; | H - HA |
| 474 | 477 | 1 | ; | H - CB |
| 474 | 490 | 1 | ; | H - C |
| 475 | 480 | 1 | ; | CA - HG21 |
| 475 | 481 | 1 | ; | CA - HG22 |
| 475 | 482 | 1 | ; | CA - HG23 |
| 475 | 484 | 1 | ; | CA - HG12 |
| 475 | 485 | 1 | ; | CA - HG13 |
| 475 | 486 | 1 | ; | CA - CD1 |
| 475 | 493 | 1 | ; | CA - H |
| 475 | 494 | 1 | ; | CA - CA |
| 476 | 478 | 1 | ; | HA - HB |
| 476 | 479 | 1 | ; | HA - CG2 |
| 476 | 483 | 1 | ; | HA - CG1 |
| 476 | 491 | 1 | ; | HA - O |
| 476 | 492 | 1 | ; | HA - N |
| 477 | 487 | 1 | ; | CB - HD11 |
| 477 | 488 | 1 | ; | CB - HD12 |
| 477 | 489 | 1 | ; | CB - HD13 |

| | | | |
|---|---|---|---|
| 477 | 491 | 1 ; | CB - O |
| 477 | 492 | 1 ; | CB - N |
| 478 | 480 | 1 ; | HB - HG21 |
| 478 | 481 | 1 ; | HB - HG22 |
| 478 | 482 | 1 ; | HB - HG23 |
| 478 | 484 | 1 ; | HB - HG12 |
| 478 | 485 | 1 ; | HB - HG13 |
| 478 | 486 | 1 ; | HB - CD1 |
| 478 | 490 | 1 ; | HB - C |
| 479 | 484 | 1 ; | CG2 - HG12 |
| 479 | 485 | 1 ; | CG2 - HG13 |
| 479 | 486 | 1 ; | CG2 - CD1 |
| 479 | 490 | 1 ; | CG2 - C |
| 480 | 483 | 1 ; | HG21 - CG1 |
| 481 | 483 | 1 ; | HG22 - CG1 |
| 482 | 483 | 1 ; | HG23 - CG1 |
| 483 | 490 | 1 ; | CG1 - C |
| 484 | 487 | 1 ; | HG12 - HD11 |
| 484 | 488 | 1 ; | HG12 - HD12 |
| 484 | 489 | 1 ; | HG12 - HD13 |
| 485 | 487 | 1 ; | HG13 - HD11 |
| 485 | 488 | 1 ; | HG13 - HD12 |
| 485 | 489 | 1 ; | HG13 - HD13 |
| 490 | 495 | 1 ; | C - HA |
| 490 | 496 | 1 ; | C - CB |
| 490 | 502 | 1 ; | C - C |
| 491 | 493 | 1 ; | O - H |
| 491 | 494 | 1 ; | O - CA |
| 492 | 497 | 1 ; | N - HB2 |
| 492 | 498 | 1 ; | N - HB3 |
| 492 | 499 | 1 ; | N - CG |
| 492 | 503 | 1 ; | N - O |
| 492 | 504 | 1 ; | N - N |
| 493 | 495 | 1 ; | H - HA |
| 493 | 496 | 1 ; | H - CB |
| 493 | 502 | 1 ; | H - C |
| 494 | 500 | 1 ; | CA - OD1 |
| 494 | 501 | 1 ; | CA - OD2 |
| 494 | 505 | 1 ; | CA - H |
| 494 | 506 | 1 ; | CA - CA |
| 495 | 497 | 1 ; | HA - HB2 |
| 495 | 498 | 1 ; | HA - HB3 |
| 495 | 499 | 1 ; | HA - CG |
| 495 | 503 | 1 ; | HA - O |
| 495 | 504 | 1 ; | HA - N |
| 496 | 503 | 1 ; | CB - O |
| 496 | 504 | 1 ; | CB - N |
| 497 | 500 | 1 ; | HB2 - OD1 |
| 497 | 501 | 1 ; | HB2 - OD2 |
| 497 | 502 | 1 ; | HB2 - C |
| 498 | 500 | 1 ; | HB3 - OD1 |
| 498 | 501 | 1 ; | HB3 - OD2 |

```
498    502       1 ;    HB3 - C
499    502       1 ;     CG - C
502    507       1 ;      C - HA
502    508       1 ;      C - CB
502    524       1 ;      C - C
503    505       1 ;      O - H
503    506       1 ;      O - CA
504    509       1 ;      N - HB2
504    510       1 ;      N - HB3
504    511       1 ;      N - CG
504    525       1 ;      N - O
504    526       1 ;      N - N
505    507       1 ;      H - HA
505    508       1 ;      H - CB
505    524       1 ;      H - C
506    512       1 ;     CA - HG2
506    513       1 ;     CA - HG3
506    514       1 ;     CA - CD
506    527       1 ;     CA - H
506    528       1 ;     CA - CA
507    509       1 ;     HA - HB2
507    510       1 ;     HA - HB3
507    511       1 ;     HA - CG
507    525       1 ;     HA - O
507    526       1 ;     HA - N
508    515       1 ;     CB - HD2
508    516       1 ;     CB - HD3
508    517       1 ;     CB - CE
508    525       1 ;     CB - O
508    526       1 ;     CB - N
509    512       1 ;    HB2 - HG2
509    513       1 ;    HB2 - HG3
509    514       1 ;    HB2 - CD
509    524       1 ;    HB2 - C
510    512       1 ;    HB3 - HG2
510    513       1 ;    HB3 - HG3
510    514       1 ;    HB3 - CD
510    524       1 ;    HB3 - C
511    518       1 ;     CG - HE2
511    519       1 ;     CG - HE3
511    520       1 ;     CG - NZ
511    524       1 ;     CG - C
512    515       1 ;    HG2 - HD2
512    516       1 ;    HG2 - HD3
512    517       1 ;    HG2 - CE
513    515       1 ;    HG3 - HD2
513    516       1 ;    HG3 - HD3
513    517       1 ;    HG3 - CE
514    521       1 ;     CD - HZ1
514    522       1 ;     CD - HZ2
514    523       1 ;     CD - HZ3
515    518       1 ;    HD2 - HE2
```

```
515    519       1 ;     HD2 - HE3
515    520       1 ;     HD2 - NZ
516    518       1 ;     HD3 - HE2
516    519       1 ;     HD3 - HE3
516    520       1 ;     HD3 - NZ
518    521       1 ;     HE2 - HZ1
518    522       1 ;     HE2 - HZ2
518    523       1 ;     HE2 - HZ3
519    521       1 ;     HE3 - HZ1
519    522       1 ;     HE3 - HZ2
519    523       1 ;     HE3 - HZ3
524    529       1 ;       C - HA
524    530       1 ;       C - CB
524    540       1 ;       C - C
525    527       1 ;       O - H
525    528       1 ;       O - CA
526    531       1 ;       N - HB
526    532       1 ;       N - CG1
526    536       1 ;       N - CG2
526    541       1 ;       N - O
526    542       1 ;       N - N
527    529       1 ;       H - HA
527    530       1 ;       H - CB
527    540       1 ;       H - C
528    533       1 ;      CA - HG11
528    534       1 ;      CA - HG12
528    535       1 ;      CA - HG13
528    537       1 ;      CA - HG21
528    538       1 ;      CA - HG22
528    539       1 ;      CA - HG23
528    543       1 ;      CA - H
528    544       1 ;      CA - CA
529    531       1 ;      HA - HB
529    532       1 ;      HA - CG1
529    536       1 ;      HA - CG2
529    541       1 ;      HA - O
529    542       1 ;      HA - N
530    541       1 ;      CB - O
530    542       1 ;      CB - N
531    533       1 ;      HB - HG11
531    534       1 ;      HB - HG12
531    535       1 ;      HB - HG13
531    537       1 ;      HB - HG21
531    538       1 ;      HB - HG22
531    539       1 ;      HB - HG23
531    540       1 ;      HB - C
532    537       1 ;     CG1 - HG21
532    538       1 ;     CG1 - HG22
532    539       1 ;     CG1 - HG23
532    540       1 ;     CG1 - C
533    536       1 ;    HG11 - CG2
534    536       1 ;    HG12 - CG2
```

```
535    536       1 ;   HG13 - CG2
536    540       1 ;    CG2 - C
540    545       1 ;      C - HA
540    546       1 ;      C - CB
540    554       1 ;      C - C
541    543       1 ;      O - H
541    544       1 ;      O - CA
542    547       1 ;      N - HB
542    548       1 ;      N - CG2
542    552       1 ;      N - OG1
542    555       1 ;      N - O
542    556       1 ;      N - N
543    545       1 ;      H - HA
543    546       1 ;      H - CB
543    554       1 ;      H - C
544    549       1 ;     CA - HG21
544    550       1 ;     CA - HG22
544    551       1 ;     CA - HG23
544    553       1 ;     CA - HG1
544    557       1 ;     CA - H
544    558       1 ;     CA - CA
545    547       1 ;     HA - HB
545    548       1 ;     HA - CG2
545    552       1 ;     HA - OG1
545    555       1 ;     HA - O
545    556       1 ;     HA - N
546    555       1 ;     CB - O
546    556       1 ;     CB - N
547    549       1 ;     HB - HG21
547    550       1 ;     HB - HG22
547    551       1 ;     HB - HG23
547    553       1 ;     HB - HG1
547    554       1 ;     HB - C
548    553       1 ;    CG2 - HG1
548    554       1 ;    CG2 - C
549    552       1 ;   HG21 - OG1
550    552       1 ;   HG22 - OG1
551    552       1 ;   HG23 - OG1
552    554       1 ;    OG1 - C
554    559       1 ;      C - HA
554    560       1 ;      C - CB
554    568       1 ;      C - C
555    557       1 ;      O - H
555    558       1 ;      O - CA
556    561       1 ;      N - HB2
556    562       1 ;      N - HB3
556    563       1 ;      N - CG
556    569       1 ;      N - O
556    570       1 ;      N - N
557    559       1 ;      H - HA
557    560       1 ;      H - CB
557    568       1 ;      H - C
```

| | | | | |
|---|---|---|---|---|
| 558 | 564 | 1 | ; | CA - OD1 |
| 558 | 565 | 1 | ; | CA - ND2 |
| 558 | 571 | 1 | ; | CA - H |
| 558 | 572 | 1 | ; | CA - CA |
| 559 | 561 | 1 | ; | HA - HB2 |
| 559 | 562 | 1 | ; | HA - HB3 |
| 559 | 563 | 1 | ; | HA - CG |
| 559 | 569 | 1 | ; | HA - O |
| 559 | 570 | 1 | ; | HA - N |
| 560 | 566 | 1 | ; | CB - HD21 |
| 560 | 567 | 1 | ; | CB - HD22 |
| 560 | 569 | 1 | ; | CB - O |
| 560 | 570 | 1 | ; | CB - N |
| 561 | 564 | 1 | ; | HB2 - OD1 |
| 561 | 565 | 1 | ; | HB2 - ND2 |
| 561 | 568 | 1 | ; | HB2 - C |
| 562 | 564 | 1 | ; | HB3 - OD1 |
| 562 | 565 | 1 | ; | HB3 - ND2 |
| 562 | 568 | 1 | ; | HB3 - C |
| 563 | 568 | 1 | ; | CG - C |
| 564 | 566 | 1 | ; | OD1 - HD21 |
| 564 | 567 | 1 | ; | OD1 - HD22 |
| 568 | 573 | 1 | ; | C - HA |
| 568 | 574 | 1 | ; | C - CB |
| 568 | 585 | 1 | ; | C - C |
| 569 | 571 | 1 | ; | O - H |
| 569 | 572 | 1 | ; | O - CA |
| 570 | 575 | 1 | ; | N - HB2 |
| 570 | 576 | 1 | ; | N - HB3 |
| 570 | 577 | 1 | ; | N - CG |
| 570 | 586 | 1 | ; | N - O |
| 570 | 587 | 1 | ; | N - N |
| 571 | 573 | 1 | ; | H - HA |
| 571 | 574 | 1 | ; | H - CB |
| 571 | 585 | 1 | ; | H - C |
| 572 | 578 | 1 | ; | CA - HG2 |
| 572 | 579 | 1 | ; | CA - HG3 |
| 572 | 580 | 1 | ; | CA - CD |
| 572 | 588 | 1 | ; | CA - H |
| 572 | 589 | 1 | ; | CA - CA |
| 573 | 575 | 1 | ; | HA - HB2 |
| 573 | 576 | 1 | ; | HA - HB3 |
| 573 | 577 | 1 | ; | HA - CG |
| 573 | 586 | 1 | ; | HA - O |
| 573 | 587 | 1 | ; | HA - N |
| 574 | 581 | 1 | ; | CB - OE1 |
| 574 | 582 | 1 | ; | CB - NE2 |
| 574 | 586 | 1 | ; | CB - O |
| 574 | 587 | 1 | ; | CB - N |
| 575 | 578 | 1 | ; | HB2 - HG2 |
| 575 | 579 | 1 | ; | HB2 - HG3 |
| 575 | 580 | 1 | ; | HB2 - CD |

| | | | | | |
|---|---|---|---|---|---|
| 575 | 585 | 1 ; | HB2 | - | C |
| 576 | 578 | 1 ; | HB3 | - | HG2 |
| 576 | 579 | 1 ; | HB3 | - | HG3 |
| 576 | 580 | 1 ; | HB3 | - | CD |
| 576 | 585 | 1 ; | HB3 | - | C |
| 577 | 583 | 1 ; | CG | - | HE21 |
| 577 | 584 | 1 ; | CG | - | HE22 |
| 577 | 585 | 1 ; | CG | - | C |
| 578 | 581 | 1 ; | HG2 | - | OE1 |
| 578 | 582 | 1 ; | HG2 | - | NE2 |
| 579 | 581 | 1 ; | HG3 | - | OE1 |
| 579 | 582 | 1 ; | HG3 | - | NE2 |
| 581 | 583 | 1 ; | OE1 | - | HE21 |
| 581 | 584 | 1 ; | OE1 | - | HE22 |
| 585 | 590 | 1 ; | C | - | HA |
| 585 | 591 | 1 ; | C | - | CB |
| 585 | 604 | 1 ; | C | - | C |
| 586 | 588 | 1 ; | O | - | H |
| 586 | 589 | 1 ; | O | - | CA |
| 587 | 592 | 1 ; | N | - | HB2 |
| 587 | 593 | 1 ; | N | - | HB3 |
| 587 | 594 | 1 ; | N | - | CG |
| 587 | 605 | 1 ; | N | - | O |
| 587 | 606 | 1 ; | N | - | N |
| 588 | 590 | 1 ; | H | - | HA |
| 588 | 591 | 1 ; | H | - | CB |
| 588 | 604 | 1 ; | H | - | C |
| 589 | 595 | 1 ; | CA | - | HG |
| 589 | 596 | 1 ; | CA | - | CD1 |
| 589 | 600 | 1 ; | CA | - | CD2 |
| 589 | 607 | 1 ; | CA | - | H |
| 589 | 608 | 1 ; | CA | - | CA |
| 590 | 592 | 1 ; | HA | - | HB2 |
| 590 | 593 | 1 ; | HA | - | HB3 |
| 590 | 594 | 1 ; | HA | - | CG |
| 590 | 605 | 1 ; | HA | - | O |
| 590 | 606 | 1 ; | HA | - | N |
| 591 | 597 | 1 ; | CB | - | HD11 |
| 591 | 598 | 1 ; | CB | - | HD12 |
| 591 | 599 | 1 ; | CB | - | HD13 |
| 591 | 601 | 1 ; | CB | - | HD21 |
| 591 | 602 | 1 ; | CB | - | HD22 |
| 591 | 603 | 1 ; | CB | - | HD23 |
| 591 | 605 | 1 ; | CB | - | O |
| 591 | 606 | 1 ; | CB | - | N |
| 592 | 595 | 1 ; | HB2 | - | HG |
| 592 | 596 | 1 ; | HB2 | - | CD1 |
| 592 | 600 | 1 ; | HB2 | - | CD2 |
| 592 | 604 | 1 ; | HB2 | - | C |
| 593 | 595 | 1 ; | HB3 | - | HG |
| 593 | 596 | 1 ; | HB3 | - | CD1 |
| 593 | 600 | 1 ; | HB3 | - | CD2 |

| | | | | | | |
|---|---|---|---|---|---|---|
| 593 | 604 | 1 | ; | HB3 | - | C |
| 594 | 604 | 1 | ; | CG | - | C |
| 595 | 597 | 1 | ; | HG | - | HD11 |
| 595 | 598 | 1 | ; | HG | - | HD12 |
| 595 | 599 | 1 | ; | HG | - | HD13 |
| 595 | 601 | 1 | ; | HG | - | HD21 |
| 595 | 602 | 1 | ; | HG | - | HD22 |
| 595 | 603 | 1 | ; | HG | - | HD23 |
| 596 | 601 | 1 | ; | CD1 | - | HD21 |
| 596 | 602 | 1 | ; | CD1 | - | HD22 |
| 596 | 603 | 1 | ; | CD1 | - | HD23 |
| 597 | 600 | 1 | ; | HD11 | - | CD2 |
| 598 | 600 | 1 | ; | HD12 | - | CD2 |
| 599 | 600 | 1 | ; | HD13 | - | CD2 |
| 604 | 609 | 1 | ; | C | - | HA |
| 604 | 610 | 1 | ; | C | - | CB |
| 604 | 625 | 1 | ; | C | - | C |
| 605 | 607 | 1 | ; | O | - | H |
| 605 | 608 | 1 | ; | O | - | CA |
| 606 | 611 | 1 | ; | N | - | HB2 |
| 606 | 612 | 1 | ; | N | - | HB3 |
| 606 | 613 | 1 | ; | N | - | CG |
| 606 | 626 | 1 | ; | N | - | O |
| 606 | 627 | 1 | ; | N | - | N |
| 607 | 609 | 1 | ; | H | - | HA |
| 607 | 610 | 1 | ; | H | - | CB |
| 607 | 625 | 1 | ; | H | - | C |
| 608 | 614 | 1 | ; | CA | - | CD1 |
| 608 | 623 | 1 | ; | CA | - | CD2 |
| 608 | 628 | 1 | ; | CA | - | H |
| 608 | 629 | 1 | ; | CA | - | CA |
| 609 | 611 | 1 | ; | HA | - | HB2 |
| 609 | 612 | 1 | ; | HA | - | HB3 |
| 609 | 613 | 1 | ; | HA | - | CG |
| 609 | 626 | 1 | ; | HA | - | O |
| 609 | 627 | 1 | ; | HA | - | N |
| 610 | 615 | 1 | ; | CB | - | HD1 |
| 610 | 616 | 1 | ; | CB | - | CE1 |
| 610 | 621 | 1 | ; | CB | - | CE2 |
| 610 | 624 | 1 | ; | CB | - | HD2 |
| 610 | 626 | 1 | ; | CB | - | O |
| 610 | 627 | 1 | ; | CB | - | N |
| 611 | 614 | 1 | ; | HB2 | - | CD1 |
| 611 | 623 | 1 | ; | HB2 | - | CD2 |
| 611 | 625 | 1 | ; | HB2 | - | C |
| 612 | 614 | 1 | ; | HB3 | - | CD1 |
| 612 | 623 | 1 | ; | HB3 | - | CD2 |
| 612 | 625 | 1 | ; | HB3 | - | C |
| 613 | 617 | 1 | ; | CG | - | HE1 |
| 613 | 618 | 1 | ; | CG | - | CZ |
| 613 | 622 | 1 | ; | CG | - | HE2 |
| 613 | 625 | 1 | ; | CG | - | C |

| | | | |
|---|---|---|---|
| 614 | 619 | 1 ; | CD1 - OH |
| 614 | 621 | 1 ; | CD1 - CE2 |
| 614 | 624 | 1 ; | CD1 - HD2 |
| 615 | 617 | 1 ; | HD1 - HE1 |
| 615 | 618 | 1 ; | HD1 - CZ |
| 615 | 623 | 1 ; | HD1 - CD2 |
| 616 | 620 | 1 ; | CE1 - HH |
| 616 | 622 | 1 ; | CE1 - HE2 |
| 616 | 623 | 1 ; | CE1 - CD2 |
| 617 | 619 | 1 ; | HE1 - OH |
| 617 | 621 | 1 ; | HE1 - CE2 |
| 618 | 624 | 1 ; | CZ - HD2 |
| 619 | 622 | 1 ; | OH - HE2 |
| 619 | 623 | 1 ; | OH - CD2 |
| 620 | 621 | 1 ; | HH - CE2 |
| 622 | 624 | 1 ; | HE2 - HD2 |
| 625 | 630 | 1 ; | C - HA |
| 625 | 631 | 1 ; | C - CB |
| 625 | 635 | 1 ; | C - C |
| 626 | 628 | 1 ; | O - H |
| 626 | 629 | 1 ; | O - CA |
| 627 | 632 | 1 ; | N - HB1 |
| 627 | 633 | 1 ; | N - HB2 |
| 627 | 634 | 1 ; | N - HB3 |
| 627 | 636 | 1 ; | N - O |
| 627 | 637 | 1 ; | N - N |
| 628 | 630 | 1 ; | H - HA |
| 628 | 631 | 1 ; | H - CB |
| 628 | 635 | 1 ; | H - C |
| 629 | 638 | 1 ; | CA - H |
| 629 | 639 | 1 ; | CA - CA |
| 630 | 632 | 1 ; | HA - HB1 |
| 630 | 633 | 1 ; | HA - HB2 |
| 630 | 634 | 1 ; | HA - HB3 |
| 630 | 636 | 1 ; | HA - O |
| 630 | 637 | 1 ; | HA - N |
| 631 | 636 | 1 ; | CB - O |
| 631 | 637 | 1 ; | CB - N |
| 632 | 635 | 1 ; | HB1 - C |
| 633 | 635 | 1 ; | HB2 - C |
| 634 | 635 | 1 ; | HB3 - C |
| 635 | 640 | 1 ; | C - HA |
| 635 | 641 | 1 ; | C - CB |
| 635 | 654 | 1 ; | C - C |
| 636 | 638 | 1 ; | O - H |
| 636 | 639 | 1 ; | O - CA |
| 637 | 642 | 1 ; | N - HB |
| 637 | 643 | 1 ; | N - CG2 |
| 637 | 647 | 1 ; | N - CG1 |
| 637 | 655 | 1 ; | N - O |
| 637 | 656 | 1 ; | N - N |
| 638 | 640 | 1 ; | H - HA |

```
638    641       1 ;       H - CB
638    654       1 ;       H - C
639    644       1 ;      CA - HG21
639    645       1 ;      CA - HG22
639    646       1 ;      CA - HG23
639    648       1 ;      CA - HG12
639    649       1 ;      CA - HG13
639    650       1 ;      CA - CD1
639    657       1 ;      CA - H
639    658       1 ;      CA - CA
640    642       1 ;      HA - HB
640    643       1 ;      HA - CG2
640    647       1 ;      HA - CG1
640    655       1 ;      HA - O
640    656       1 ;      HA - N
641    651       1 ;      CB - HD11
641    652       1 ;      CB - HD12
641    653       1 ;      CB - HD13
641    655       1 ;      CB - O
641    656       1 ;      CB - N
642    644       1 ;      HB - HG21
642    645       1 ;      HB - HG22
642    646       1 ;      HB - HG23
642    648       1 ;      HB - HG12
642    649       1 ;      HB - HG13
642    650       1 ;      HB - CD1
642    654       1 ;      HB - C
643    648       1 ;     CG2 - HG12
643    649       1 ;     CG2 - HG13
643    650       1 ;     CG2 - CD1
643    654       1 ;     CG2 - C
644    647       1 ;    HG21 - CG1
645    647       1 ;    HG22 - CG1
646    647       1 ;    HG23 - CG1
647    654       1 ;     CG1 - C
648    651       1 ;    HG12 - HD11
648    652       1 ;    HG12 - HD12
648    653       1 ;    HG12 - HD13
649    651       1 ;    HG13 - HD11
649    652       1 ;    HG13 - HD12
649    653       1 ;    HG13 - HD13
654    659       1 ;       C - HA
654    660       1 ;       C - CB
654    676       1 ;       C - C
655    657       1 ;       O - H
655    658       1 ;       O - CA
656    661       1 ;       N - HB2
656    662       1 ;       N - HB3
656    663       1 ;       N - CG
656    677       1 ;       N - O
656    678       1 ;       N - N
657    659       1 ;       H - HA
```

```
657    660      1 ;      H - CB
657    676      1 ;      H - C
658    664      1 ;     CA - HG2
658    665      1 ;     CA - HG3
658    666      1 ;     CA - CD
658    679      1 ;     CA - H
658    680      1 ;     CA - CA
659    661      1 ;     HA - HB2
659    662      1 ;     HA - HB3
659    663      1 ;     HA - CG
659    677      1 ;     HA - O
659    678      1 ;     HA - N
660    667      1 ;     CB - HD2
660    668      1 ;     CB - HD3
660    669      1 ;     CB - CE
660    677      1 ;     CB - O
660    678      1 ;     CB - N
661    664      1 ;    HB2 - HG2
661    665      1 ;    HB2 - HG3
661    666      1 ;    HB2 - CD
661    676      1 ;    HB2 - C
662    664      1 ;    HB3 - HG2
662    665      1 ;    HB3 - HG3
662    666      1 ;    HB3 - CD
662    676      1 ;    HB3 - C
663    670      1 ;     CG - HE2
663    671      1 ;     CG - HE3
663    672      1 ;     CG - NZ
663    676      1 ;     CG - C
664    667      1 ;    HG2 - HD2
664    668      1 ;    HG2 - HD3
664    669      1 ;    HG2 - CE
665    667      1 ;    HG3 - HD2
665    668      1 ;    HG3 - HD3
665    669      1 ;    HG3 - CE
666    673      1 ;     CD - HZ1
666    674      1 ;     CD - HZ2
666    675      1 ;     CD - HZ3
667    670      1 ;    HD2 - HE2
667    671      1 ;    HD2 - HE3
667    672      1 ;    HD2 - NZ
668    670      1 ;    HD3 - HE2
668    671      1 ;    HD3 - HE3
668    672      1 ;    HD3 - NZ
670    673      1 ;    HE2 - HZ1
670    674      1 ;    HE2 - HZ2
670    675      1 ;    HE2 - HZ3
671    673      1 ;    HE3 - HZ1
671    674      1 ;    HE3 - HZ2
671    675      1 ;    HE3 - HZ3
676    681      1 ;      C - HA
676    682      1 ;      C - CB
```

| | | | | | |
|---|---|---|---|---|---|
| 676 | 691 | 1 ; | C | - | C |
| 677 | 679 | 1 ; | O | - | H |
| 677 | 680 | 1 ; | O | - | CA |
| 678 | 683 | 1 ; | N | - | HB2 |
| 678 | 684 | 1 ; | N | - | HB3 |
| 678 | 685 | 1 ; | N | - | CG |
| 678 | 692 | 1 ; | N | - | O |
| 678 | 693 | 1 ; | N | - | N |
| 679 | 681 | 1 ; | H | - | HA |
| 679 | 682 | 1 ; | H | - | CB |
| 679 | 691 | 1 ; | H | - | C |
| 680 | 686 | 1 ; | CA | - | HG2 |
| 680 | 687 | 1 ; | CA | - | HG3 |
| 680 | 688 | 1 ; | CA | - | CD |
| 680 | 694 | 1 ; | CA | - | H |
| 680 | 695 | 1 ; | CA | - | CA |
| 681 | 683 | 1 ; | HA | - | HB2 |
| 681 | 684 | 1 ; | HA | - | HB3 |
| 681 | 685 | 1 ; | HA | - | CG |
| 681 | 692 | 1 ; | HA | - | O |
| 681 | 693 | 1 ; | HA | - | N |
| 682 | 689 | 1 ; | CB | - | OE1 |
| 682 | 690 | 1 ; | CB | - | OE2 |
| 682 | 692 | 1 ; | CB | - | O |
| 682 | 693 | 1 ; | CB | - | N |
| 683 | 686 | 1 ; | HB2 | - | HG2 |
| 683 | 687 | 1 ; | HB2 | - | HG3 |
| 683 | 688 | 1 ; | HB2 | - | CD |
| 683 | 691 | 1 ; | HB2 | - | C |
| 684 | 686 | 1 ; | HB3 | - | HG2 |
| 684 | 687 | 1 ; | HB3 | - | HG3 |
| 684 | 688 | 1 ; | HB3 | - | CD |
| 684 | 691 | 1 ; | HB3 | - | C |
| 685 | 691 | 1 ; | CG | - | C |
| 686 | 689 | 1 ; | HG2 | - | OE1 |
| 686 | 690 | 1 ; | HG2 | - | OE2 |
| 687 | 689 | 1 ; | HG3 | - | OE1 |
| 687 | 690 | 1 ; | HG3 | - | OE2 |
| 691 | 696 | 1 ; | C | - | HA |
| 691 | 697 | 1 ; | C | - | CB |
| 691 | 707 | 1 ; | C | - | C |
| 692 | 694 | 1 ; | O | - | H |
| 692 | 695 | 1 ; | O | - | CA |
| 693 | 698 | 1 ; | N | - | HB2 |
| 693 | 699 | 1 ; | N | - | HB3 |
| 693 | 700 | 1 ; | N | - | CG |
| 693 | 708 | 1 ; | N | - | O |
| 693 | 709 | 1 ; | N | - | N |
| 694 | 696 | 1 ; | H | - | HA |
| 694 | 697 | 1 ; | H | - | CB |
| 694 | 707 | 1 ; | H | - | C |
| 695 | 701 | 1 ; | CA | - | HG2 |

| | | | |
|---|---|---|---|
| 695 | 702 | 1 ; | CA - HG3 |
| 695 | 703 | 1 ; | CA - CD |
| 695 | 710 | 1 ; | CA - H |
| 695 | 711 | 1 ; | CA - CA |
| 696 | 698 | 1 ; | HA - HB2 |
| 696 | 699 | 1 ; | HA - HB3 |
| 696 | 700 | 1 ; | HA - CG |
| 696 | 708 | 1 ; | HA - O |
| 696 | 709 | 1 ; | HA - N |
| 697 | 704 | 1 ; | CB - OE1 |
| 697 | 705 | 1 ; | CB - OE2 |
| 697 | 708 | 1 ; | CB - O |
| 697 | 709 | 1 ; | CB - N |
| 698 | 701 | 1 ; | HB2 - HG2 |
| 698 | 702 | 1 ; | HB2 - HG3 |
| 698 | 703 | 1 ; | HB2 - CD |
| 698 | 707 | 1 ; | HB2 - C |
| 699 | 701 | 1 ; | HB3 - HG2 |
| 699 | 702 | 1 ; | HB3 - HG3 |
| 699 | 703 | 1 ; | HB3 - CD |
| 699 | 707 | 1 ; | HB3 - C |
| 700 | 706 | 1 ; | CG - HE2 |
| 700 | 707 | 1 ; | CG - C |
| 701 | 704 | 1 ; | HG2 - OE1 |
| 701 | 705 | 1 ; | HG2 - OE2 |
| 702 | 704 | 1 ; | HG3 - OE1 |
| 702 | 705 | 1 ; | HG3 - OE2 |
| 704 | 706 | 1 ; | OE1 - HE2 |
| 707 | 712 | 1 ; | C - HA |
| 707 | 713 | 1 ; | C - CB |
| 707 | 729 | 1 ; | C - C |
| 708 | 710 | 1 ; | O - H |
| 708 | 711 | 1 ; | O - CA |
| 709 | 714 | 1 ; | N - HB2 |
| 709 | 715 | 1 ; | N - HB3 |
| 709 | 716 | 1 ; | N - CG |
| 709 | 730 | 1 ; | N - O |
| 709 | 731 | 1 ; | N - N |
| 710 | 712 | 1 ; | H - HA |
| 710 | 713 | 1 ; | H - CB |
| 710 | 729 | 1 ; | H - C |
| 711 | 717 | 1 ; | CA - HG2 |
| 711 | 718 | 1 ; | CA - HG3 |
| 711 | 719 | 1 ; | CA - CD |
| 711 | 732 | 1 ; | CA - H |
| 711 | 733 | 1 ; | CA - CA |
| 712 | 714 | 1 ; | HA - HB2 |
| 712 | 715 | 1 ; | HA - HB3 |
| 712 | 716 | 1 ; | HA - CG |
| 712 | 730 | 1 ; | HA - O |
| 712 | 731 | 1 ; | HA - N |
| 713 | 720 | 1 ; | CB - HD2 |

```
713    721      1 ;     CB - HD3
713    722      1 ;     CB - CE
713    730      1 ;     CB - O
713    731      1 ;     CB - N
714    717      1 ;    HB2 - HG2
714    718      1 ;    HB2 - HG3
714    719      1 ;    HB2 - CD
714    729      1 ;    HB2 - C
715    717      1 ;    HB3 - HG2
715    718      1 ;    HB3 - HG3
715    719      1 ;    HB3 - CD
715    729      1 ;    HB3 - C
716    723      1 ;     CG - HE2
716    724      1 ;     CG - HE3
716    725      1 ;     CG - NZ
716    729      1 ;     CG - C
717    720      1 ;    HG2 - HD2
717    721      1 ;    HG2 - HD3
717    722      1 ;    HG2 - CE
718    720      1 ;    HG3 - HD2
718    721      1 ;    HG3 - HD3
718    722      1 ;    HG3 - CE
719    726      1 ;     CD - HZ1
719    727      1 ;     CD - HZ2
719    728      1 ;     CD - HZ3
720    723      1 ;    HD2 - HE2
720    724      1 ;    HD2 - HE3
720    725      1 ;    HD2 - NZ
721    723      1 ;    HD3 - HE2
721    724      1 ;    HD3 - HE3
721    725      1 ;    HD3 - NZ
723    726      1 ;    HE2 - HZ1
723    727      1 ;    HE2 - HZ2
723    728      1 ;    HE2 - HZ3
724    726      1 ;    HE3 - HZ1
724    727      1 ;    HE3 - HZ2
724    728      1 ;    HE3 - HZ3
729    734      1 ;      C - HA
729    735      1 ;      C - CB
729    751      1 ;      C - C
730    732      1 ;      O - H
730    733      1 ;      O - CA
731    736      1 ;      N - HB2
731    737      1 ;      N - HB3
731    738      1 ;      N - CG
731    752      1 ;      N - O
731    753      1 ;      N - N
732    734      1 ;      H - HA
732    735      1 ;      H - CB
732    751      1 ;      H - C
733    739      1 ;     CA - HG2
733    740      1 ;     CA - HG3
```

```
733    741      1 ;      CA - CD
733    754      1 ;      CA - H
733    755      1 ;      CA - CA
734    736      1 ;      HA - HB2
734    737      1 ;      HA - HB3
734    738      1 ;      HA - CG
734    752      1 ;      HA - O
734    753      1 ;      HA - N
735    742      1 ;      CB - HD2
735    743      1 ;      CB - HD3
735    744      1 ;      CB - CE
735    752      1 ;      CB - O
735    753      1 ;      CB - N
736    739      1 ;     HB2 - HG2
736    740      1 ;     HB2 - HG3
736    741      1 ;     HB2 - CD
736    751      1 ;     HB2 - C
737    739      1 ;     HB3 - HG2
737    740      1 ;     HB3 - HG3
737    741      1 ;     HB3 - CD
737    751      1 ;     HB3 - C
738    745      1 ;      CG - HE2
738    746      1 ;      CG - HE3
738    747      1 ;      CG - NZ
738    751      1 ;      CG - C
739    742      1 ;     HG2 - HD2
739    743      1 ;     HG2 - HD3
739    744      1 ;     HG2 - CE
740    742      1 ;     HG3 - HD2
740    743      1 ;     HG3 - HD3
740    744      1 ;     HG3 - CE
741    748      1 ;      CD - HZ1
741    749      1 ;      CD - HZ2
741    750      1 ;      CD - HZ3
742    745      1 ;     HD2 - HE2
742    746      1 ;     HD2 - HE3
742    747      1 ;     HD2 - NZ
743    745      1 ;     HD3 - HE2
743    746      1 ;     HD3 - HE3
743    747      1 ;     HD3 - NZ
745    748      1 ;     HE2 - HZ1
745    749      1 ;     HE2 - HZ2
745    750      1 ;     HE2 - HZ3
746    748      1 ;     HE3 - HZ1
746    749      1 ;     HE3 - HZ2
746    750      1 ;     HE3 - HZ3
751    756      1 ;       C - HA
751    757      1 ;       C - CB
751    763      1 ;       C - C
752    754      1 ;       O - H
752    755      1 ;       O - CA
753    758      1 ;       N - HB2
```

| | | | | |
|---|---|---|---|---|
| 753 | 759 | 1 | ; | N - HB3 |
| 753 | 760 | 1 | ; | N - CG |
| 753 | 764 | 1 | ; | N - O |
| 753 | 765 | 1 | ; | N - N |
| 754 | 756 | 1 | ; | H - HA |
| 754 | 757 | 1 | ; | H - CB |
| 754 | 763 | 1 | ; | H - C |
| 755 | 761 | 1 | ; | CA - OD1 |
| 755 | 762 | 1 | ; | CA - OD2 |
| 755 | 766 | 1 | ; | CA - H |
| 755 | 767 | 1 | ; | CA - CA |
| 756 | 758 | 1 | ; | HA - HB2 |
| 756 | 759 | 1 | ; | HA - HB3 |
| 756 | 760 | 1 | ; | HA - CG |
| 756 | 764 | 1 | ; | HA - O |
| 756 | 765 | 1 | ; | HA - N |
| 757 | 764 | 1 | ; | CB - O |
| 757 | 765 | 1 | ; | CB - N |
| 758 | 761 | 1 | ; | HB2 - OD1 |
| 758 | 762 | 1 | ; | HB2 - OD2 |
| 758 | 763 | 1 | ; | HB2 - C |
| 759 | 761 | 1 | ; | HB3 - OD1 |
| 759 | 762 | 1 | ; | HB3 - OD2 |
| 759 | 763 | 1 | ; | HB3 - C |
| 760 | 763 | 1 | ; | CG - C |
| 763 | 768 | 1 | ; | C - HA |
| 763 | 769 | 1 | ; | C - CB |
| 763 | 787 | 1 | ; | C - C |
| 764 | 766 | 1 | ; | O - H |
| 764 | 767 | 1 | ; | O - CA |
| 765 | 770 | 1 | ; | N - HB2 |
| 765 | 771 | 1 | ; | N - HB3 |
| 765 | 772 | 1 | ; | N - CG |
| 765 | 788 | 1 | ; | N - O |
| 765 | 789 | 1 | ; | N - N |
| 766 | 768 | 1 | ; | H - HA |
| 766 | 769 | 1 | ; | H - CB |
| 766 | 787 | 1 | ; | H - C |
| 767 | 773 | 1 | ; | CA - HG2 |
| 767 | 774 | 1 | ; | CA - HG3 |
| 767 | 775 | 1 | ; | CA - CD |
| 767 | 790 | 1 | ; | CA - H |
| 767 | 791 | 1 | ; | CA - CA |
| 768 | 770 | 1 | ; | HA - HB2 |
| 768 | 771 | 1 | ; | HA - HB3 |
| 768 | 772 | 1 | ; | HA - CG |
| 768 | 788 | 1 | ; | HA - O |
| 768 | 789 | 1 | ; | HA - N |
| 769 | 776 | 1 | ; | CB - HD2 |
| 769 | 777 | 1 | ; | CB - HD3 |
| 769 | 778 | 1 | ; | CB - NE |
| 769 | 788 | 1 | ; | CB - O |

| | | | | | |
|---|---|---|---|---|---|
| 769 | 789 | 1 ; | CB | - | N |
| 770 | 773 | 1 ; | HB2 | - | HG2 |
| 770 | 774 | 1 ; | HB2 | - | HG3 |
| 770 | 775 | 1 ; | HB2 | - | CD |
| 770 | 787 | 1 ; | HB2 | - | C |
| 771 | 773 | 1 ; | HB3 | - | HG2 |
| 771 | 774 | 1 ; | HB3 | - | HG3 |
| 771 | 775 | 1 ; | HB3 | - | CD |
| 771 | 787 | 1 ; | HB3 | - | C |
| 772 | 779 | 1 ; | CG | - | HE |
| 772 | 780 | 1 ; | CG | - | CZ |
| 772 | 787 | 1 ; | CG | - | C |
| 773 | 776 | 1 ; | HG2 | - | HD2 |
| 773 | 777 | 1 ; | HG2 | - | HD3 |
| 773 | 778 | 1 ; | HG2 | - | NE |
| 774 | 776 | 1 ; | HG3 | - | HD2 |
| 774 | 777 | 1 ; | HG3 | - | HD3 |
| 774 | 778 | 1 ; | HG3 | - | NE |
| 775 | 781 | 1 ; | CD | - | NH1 |
| 775 | 784 | 1 ; | CD | - | NH2 |
| 776 | 779 | 1 ; | HD2 | - | HE |
| 776 | 780 | 1 ; | HD2 | - | CZ |
| 777 | 779 | 1 ; | HD3 | - | HE |
| 777 | 780 | 1 ; | HD3 | - | CZ |
| 778 | 782 | 1 ; | NE | - | HH11 |
| 778 | 783 | 1 ; | NE | - | HH12 |
| 778 | 785 | 1 ; | NE | - | HH21 |
| 778 | 786 | 1 ; | NE | - | HH22 |
| 779 | 781 | 1 ; | HE | - | NH1 |
| 779 | 784 | 1 ; | HE | - | NH2 |
| 781 | 785 | 1 ; | NH1 | - | HH21 |
| 781 | 786 | 1 ; | NH1 | - | HH22 |
| 782 | 784 | 1 ; | HH11 | - | NH2 |
| 783 | 784 | 1 ; | HH12 | - | NH2 |
| 787 | 792 | 1 ; | C | - | HA |
| 787 | 793 | 1 ; | C | - | CB |
| 787 | 806 | 1 ; | C | - | C |
| 788 | 790 | 1 ; | O | - | H |
| 788 | 791 | 1 ; | O | - | CA |
| 789 | 794 | 1 ; | N | - | HB2 |
| 789 | 795 | 1 ; | N | - | HB3 |
| 789 | 796 | 1 ; | N | - | CG |
| 789 | 807 | 1 ; | N | - | O |
| 789 | 808 | 1 ; | N | - | N |
| 790 | 792 | 1 ; | H | - | HA |
| 790 | 793 | 1 ; | H | - | CB |
| 790 | 806 | 1 ; | H | - | C |
| 791 | 797 | 1 ; | CA | - | HG |
| 791 | 798 | 1 ; | CA | - | CD1 |
| 791 | 802 | 1 ; | CA | - | CD2 |
| 791 | 809 | 1 ; | CA | - | H |
| 791 | 810 | 1 ; | CA | - | CA |

| | | | | |
|---|---|---|---|---|
| 792 | 794 | 1 | ; | HA - HB2 |
| 792 | 795 | 1 | ; | HA - HB3 |
| 792 | 796 | 1 | ; | HA - CG |
| 792 | 807 | 1 | ; | HA - O |
| 792 | 808 | 1 | ; | HA - N |
| 793 | 799 | 1 | ; | CB - HD11 |
| 793 | 800 | 1 | ; | CB - HD12 |
| 793 | 801 | 1 | ; | CB - HD13 |
| 793 | 803 | 1 | ; | CB - HD21 |
| 793 | 804 | 1 | ; | CB - HD22 |
| 793 | 805 | 1 | ; | CB - HD23 |
| 793 | 807 | 1 | ; | CB - O |
| 793 | 808 | 1 | ; | CB - N |
| 794 | 797 | 1 | ; | HB2 - HG |
| 794 | 798 | 1 | ; | HB2 - CD1 |
| 794 | 802 | 1 | ; | HB2 - CD2 |
| 794 | 806 | 1 | ; | HB2 - C |
| 795 | 797 | 1 | ; | HB3 - HG |
| 795 | 798 | 1 | ; | HB3 - CD1 |
| 795 | 802 | 1 | ; | HB3 - CD2 |
| 795 | 806 | 1 | ; | HB3 - C |
| 796 | 806 | 1 | ; | CG - C |
| 797 | 799 | 1 | ; | HG - HD11 |
| 797 | 800 | 1 | ; | HG - HD12 |
| 797 | 801 | 1 | ; | HG - HD13 |
| 797 | 803 | 1 | ; | HG - HD21 |
| 797 | 804 | 1 | ; | HG - HD22 |
| 797 | 805 | 1 | ; | HG - HD23 |
| 798 | 803 | 1 | ; | CD1 - HD21 |
| 798 | 804 | 1 | ; | CD1 - HD22 |
| 798 | 805 | 1 | ; | CD1 - HD23 |
| 799 | 802 | 1 | ; | HD11 - CD2 |
| 800 | 802 | 1 | ; | HD12 - CD2 |
| 801 | 802 | 1 | ; | HD13 - CD2 |
| 806 | 811 | 1 | ; | C - HA |
| 806 | 812 | 1 | ; | C - CB |
| 806 | 828 | 1 | ; | C - C |
| 807 | 809 | 1 | ; | O - H |
| 807 | 810 | 1 | ; | O - CA |
| 808 | 813 | 1 | ; | N - HB2 |
| 808 | 814 | 1 | ; | N - HB3 |
| 808 | 815 | 1 | ; | N - CG |
| 808 | 829 | 1 | ; | N - O |
| 808 | 830 | 1 | ; | N - N |
| 809 | 811 | 1 | ; | H - HA |
| 809 | 812 | 1 | ; | H - CB |
| 809 | 828 | 1 | ; | H - C |
| 810 | 816 | 1 | ; | CA - HG2 |
| 810 | 817 | 1 | ; | CA - HG3 |
| 810 | 818 | 1 | ; | CA - CD |
| 810 | 831 | 1 | ; | CA - H |
| 810 | 832 | 1 | ; | CA - CA |

| | | | | |
|---|---|---|---|---|
| 811 | 813 | 1 | ; | HA - HB2 |
| 811 | 814 | 1 | ; | HA - HB3 |
| 811 | 815 | 1 | ; | HA - CG |
| 811 | 829 | 1 | ; | HA - O |
| 811 | 830 | 1 | ; | HA - N |
| 812 | 819 | 1 | ; | CB - HD2 |
| 812 | 820 | 1 | ; | CB - HD3 |
| 812 | 821 | 1 | ; | CB - CE |
| 812 | 829 | 1 | ; | CB - O |
| 812 | 830 | 1 | ; | CB - N |
| 813 | 816 | 1 | ; | HB2 - HG2 |
| 813 | 817 | 1 | ; | HB2 - HG3 |
| 813 | 818 | 1 | ; | HB2 - CD |
| 813 | 828 | 1 | ; | HB2 - C |
| 814 | 816 | 1 | ; | HB3 - HG2 |
| 814 | 817 | 1 | ; | HB3 - HG3 |
| 814 | 818 | 1 | ; | HB3 - CD |
| 814 | 828 | 1 | ; | HB3 - C |
| 815 | 822 | 1 | ; | CG - HE2 |
| 815 | 823 | 1 | ; | CG - HE3 |
| 815 | 824 | 1 | ; | CG - NZ |
| 815 | 828 | 1 | ; | CG - C |
| 816 | 819 | 1 | ; | HG2 - HD2 |
| 816 | 820 | 1 | ; | HG2 - HD3 |
| 816 | 821 | 1 | ; | HG2 - CE |
| 817 | 819 | 1 | ; | HG3 - HD2 |
| 817 | 820 | 1 | ; | HG3 - HD3 |
| 817 | 821 | 1 | ; | HG3 - CE |
| 818 | 825 | 1 | ; | CD - HZ1 |
| 818 | 826 | 1 | ; | CD - HZ2 |
| 818 | 827 | 1 | ; | CD - HZ3 |
| 819 | 822 | 1 | ; | HD2 - HE2 |
| 819 | 823 | 1 | ; | HD2 - HE3 |
| 819 | 824 | 1 | ; | HD2 - NZ |
| 820 | 822 | 1 | ; | HD3 - HE2 |
| 820 | 823 | 1 | ; | HD3 - HE3 |
| 820 | 824 | 1 | ; | HD3 - NZ |
| 822 | 825 | 1 | ; | HE2 - HZ1 |
| 822 | 826 | 1 | ; | HE2 - HZ2 |
| 822 | 827 | 1 | ; | HE2 - HZ3 |
| 823 | 825 | 1 | ; | HE3 - HZ1 |
| 823 | 826 | 1 | ; | HE3 - HZ2 |
| 823 | 827 | 1 | ; | HE3 - HZ3 |
| 828 | 833 | 1 | ; | C - HA |
| 828 | 834 | 1 | ; | C - CB |
| 828 | 842 | 1 | ; | C - C |
| 829 | 831 | 1 | ; | O - H |
| 829 | 832 | 1 | ; | O - CA |
| 830 | 835 | 1 | ; | N - HB2 |
| 830 | 836 | 1 | ; | N - HB3 |
| 830 | 837 | 1 | ; | N - CG |
| 830 | 843 | 1 | ; | N - O |

```
830    844       1 ;       N - N
831    833       1 ;       H - HA
831    834       1 ;       H - CB
831    842       1 ;       H - C
832    838       1 ;      CA - OD1
832    839       1 ;      CA - ND2
832    845       1 ;      CA - H
832    846       1 ;      CA - CA
833    835       1 ;      HA - HB2
833    836       1 ;      HA - HB3
833    837       1 ;      HA - CG
833    843       1 ;      HA - O
833    844       1 ;      HA - N
834    840       1 ;      CB - HD21
834    841       1 ;      CB - HD22
834    843       1 ;      CB - O
834    844       1 ;      CB - N
835    838       1 ;     HB2 - OD1
835    839       1 ;     HB2 - ND2
835    842       1 ;     HB2 - C
836    838       1 ;     HB3 - OD1
836    839       1 ;     HB3 - ND2
836    842       1 ;     HB3 - C
837    842       1 ;      CG - C
838    840       1 ;     OD1 - HD21
838    841       1 ;     OD1 - HD22
842    847       1 ;       C - HA
842    848       1 ;       C - CB
842    861       1 ;       C - C
843    845       1 ;       O - H
843    846       1 ;       O - CA
844    849       1 ;       N - HB
844    850       1 ;       N - CG2
844    854       1 ;       N - CG1
844    862       1 ;       N - O
844    863       1 ;       N - N
845    847       1 ;       H - HA
845    848       1 ;       H - CB
845    861       1 ;       H - C
846    851       1 ;      CA - HG21
846    852       1 ;      CA - HG22
846    853       1 ;      CA - HG23
846    855       1 ;      CA - HG12
846    856       1 ;      CA - HG13
846    857       1 ;      CA - CD1
846    864       1 ;      CA - H
846    865       1 ;      CA - CA
847    849       1 ;      HA - HB
847    850       1 ;      HA - CG2
847    854       1 ;      HA - CG1
847    862       1 ;      HA - O
847    863       1 ;      HA - N
```

```
848    858       1 ;      CB - HD11
848    859       1 ;      CB - HD12
848    860       1 ;      CB - HD13
848    862       1 ;      CB - O
848    863       1 ;      CB - N
849    851       1 ;      HB - HG21
849    852       1 ;      HB - HG22
849    853       1 ;      HB - HG23
849    855       1 ;      HB - HG12
849    856       1 ;      HB - HG13
849    857       1 ;      HB - CD1
849    861       1 ;      HB - C
850    855       1 ;     CG2 - HG12
850    856       1 ;     CG2 - HG13
850    857       1 ;     CG2 - CD1
850    861       1 ;     CG2 - C
851    854       1 ;    HG21 - CG1
852    854       1 ;    HG22 - CG1
853    854       1 ;    HG23 - CG1
854    861       1 ;     CG1 - C
855    858       1 ;    HG12 - HD11
855    859       1 ;    HG12 - HD12
855    860       1 ;    HG12 - HD13
856    858       1 ;    HG13 - HD11
856    859       1 ;    HG13 - HD12
856    860       1 ;    HG13 - HD13
861    866       1 ;       C - HA
861    867       1 ;       C - CB
861    875       1 ;       C - C
862    864       1 ;       O - H
862    865       1 ;       O - CA
863    868       1 ;       N - HB2
863    869       1 ;       N - HB3
863    870       1 ;       N - CG
863    876       1 ;       N - O
863    877       1 ;       N - N
864    866       1 ;       H - HA
864    867       1 ;       H - CB
864    875       1 ;       H - C
865    871       1 ;      CA - OD1
865    872       1 ;      CA - ND2
865    878       1 ;      CA - H
865    879       1 ;      CA - CA
866    868       1 ;      HA - HB2
866    869       1 ;      HA - HB3
866    870       1 ;      HA - CG
866    876       1 ;      HA - O
866    877       1 ;      HA - N
867    873       1 ;      CB - HD21
867    874       1 ;      CB - HD22
867    876       1 ;      CB - O
867    877       1 ;      CB - N
```

| | | | | |
|---|---|---|---|---|
| 868 | 871 | 1 | ; | HB2 - OD1 |
| 868 | 872 | 1 | ; | HB2 - ND2 |
| 868 | 875 | 1 | ; | HB2 - C |
| 869 | 871 | 1 | ; | HB3 - OD1 |
| 869 | 872 | 1 | ; | HB3 - ND2 |
| 869 | 875 | 1 | ; | HB3 - C |
| 870 | 875 | 1 | ; | CG - C |
| 871 | 873 | 1 | ; | OD1 - HD21 |
| 871 | 874 | 1 | ; | OD1 - HD22 |
| 875 | 880 | 1 | ; | C - HA |
| 875 | 881 | 1 | ; | C - CB |
| 875 | 899 | 1 | ; | C - C |
| 876 | 878 | 1 | ; | O - H |
| 876 | 879 | 1 | ; | O - CA |
| 877 | 882 | 1 | ; | N - HB2 |
| 877 | 883 | 1 | ; | N - HB3 |
| 877 | 884 | 1 | ; | N - CG |
| 877 | 900 | 1 | ; | N - O |
| 877 | 901 | 1 | ; | N - N |
| 878 | 880 | 1 | ; | H - HA |
| 878 | 881 | 1 | ; | H - CB |
| 878 | 899 | 1 | ; | H - C |
| 879 | 885 | 1 | ; | CA - HG2 |
| 879 | 886 | 1 | ; | CA - HG3 |
| 879 | 887 | 1 | ; | CA - CD |
| 879 | 902 | 1 | ; | CA - H |
| 879 | 903 | 1 | ; | CA - CA |
| 880 | 882 | 1 | ; | HA - HB2 |
| 880 | 883 | 1 | ; | HA - HB3 |
| 880 | 884 | 1 | ; | HA - CG |
| 880 | 900 | 1 | ; | HA - O |
| 880 | 901 | 1 | ; | HA - N |
| 881 | 888 | 1 | ; | CB - HD2 |
| 881 | 889 | 1 | ; | CB - HD3 |
| 881 | 890 | 1 | ; | CB - NE |
| 881 | 900 | 1 | ; | CB - O |
| 881 | 901 | 1 | ; | CB - N |
| 882 | 885 | 1 | ; | HB2 - HG2 |
| 882 | 886 | 1 | ; | HB2 - HG3 |
| 882 | 887 | 1 | ; | HB2 - CD |
| 882 | 899 | 1 | ; | HB2 - C |
| 883 | 885 | 1 | ; | HB3 - HG2 |
| 883 | 886 | 1 | ; | HB3 - HG3 |
| 883 | 887 | 1 | ; | HB3 - CD |
| 883 | 899 | 1 | ; | HB3 - C |
| 884 | 891 | 1 | ; | CG - HE |
| 884 | 892 | 1 | ; | CG - CZ |
| 884 | 899 | 1 | ; | CG - C |
| 885 | 888 | 1 | ; | HG2 - HD2 |
| 885 | 889 | 1 | ; | HG2 - HD3 |
| 885 | 890 | 1 | ; | HG2 - NE |
| 886 | 888 | 1 | ; | HG3 - HD2 |

| | | | |
|---|---|---|---|
| 886 | 889 | 1 ; | HG3 - HD3 |
| 886 | 890 | 1 ; | HG3 - NE |
| 887 | 893 | 1 ; | CD - NH1 |
| 887 | 896 | 1 ; | CD - NH2 |
| 888 | 891 | 1 ; | HD2 - HE |
| 888 | 892 | 1 ; | HD2 - CZ |
| 889 | 891 | 1 ; | HD3 - HE |
| 889 | 892 | 1 ; | HD3 - CZ |
| 890 | 894 | 1 ; | NE - HH11 |
| 890 | 895 | 1 ; | NE - HH12 |
| 890 | 897 | 1 ; | NE - HH21 |
| 890 | 898 | 1 ; | NE - HH22 |
| 891 | 893 | 1 ; | HE - NH1 |
| 891 | 896 | 1 ; | HE - NH2 |
| 893 | 897 | 1 ; | NH1 - HH21 |
| 893 | 898 | 1 ; | NH1 - HH22 |
| 894 | 896 | 1 ; | HH11 - NH2 |
| 895 | 896 | 1 ; | HH12 - NH2 |
| 899 | 904 | 1 ; | C - HA |
| 899 | 905 | 1 ; | C - CB |
| 899 | 919 | 1 ; | C - C |
| 900 | 902 | 1 ; | O - H |
| 900 | 903 | 1 ; | O - CA |
| 901 | 906 | 1 ; | N - HB2 |
| 901 | 907 | 1 ; | N - HB3 |
| 901 | 908 | 1 ; | N - CG |
| 901 | 920 | 1 ; | N - O |
| 901 | 921 | 1 ; | N - N |
| 902 | 904 | 1 ; | H - HA |
| 902 | 905 | 1 ; | H - CB |
| 902 | 919 | 1 ; | H - C |
| 903 | 909 | 1 ; | CA - CD1 |
| 903 | 917 | 1 ; | CA - CD2 |
| 903 | 922 | 1 ; | CA - H |
| 903 | 923 | 1 ; | CA - CA |
| 904 | 906 | 1 ; | HA - HB2 |
| 904 | 907 | 1 ; | HA - HB3 |
| 904 | 908 | 1 ; | HA - CG |
| 904 | 920 | 1 ; | HA - O |
| 904 | 921 | 1 ; | HA - N |
| 905 | 910 | 1 ; | CB - HD1 |
| 905 | 911 | 1 ; | CB - CE1 |
| 905 | 915 | 1 ; | CB - CE2 |
| 905 | 918 | 1 ; | CB - HD2 |
| 905 | 920 | 1 ; | CB - O |
| 905 | 921 | 1 ; | CB - N |
| 906 | 909 | 1 ; | HB2 - CD1 |
| 906 | 917 | 1 ; | HB2 - CD2 |
| 906 | 919 | 1 ; | HB2 - C |
| 907 | 909 | 1 ; | HB3 - CD1 |
| 907 | 917 | 1 ; | HB3 - CD2 |
| 907 | 919 | 1 ; | HB3 - C |

| | | | | |
|---|---|---|---|---|
| 908 | 912 | 1 | ; | CG - HE1 |
| 908 | 913 | 1 | ; | CG - CZ |
| 908 | 916 | 1 | ; | CG - HE2 |
| 908 | 919 | 1 | ; | CG - C |
| 909 | 914 | 1 | ; | CD1 - HZ |
| 909 | 915 | 1 | ; | CD1 - CE2 |
| 909 | 918 | 1 | ; | CD1 - HD2 |
| 910 | 912 | 1 | ; | HD1 - HE1 |
| 910 | 913 | 1 | ; | HD1 - CZ |
| 910 | 917 | 1 | ; | HD1 - CD2 |
| 911 | 916 | 1 | ; | CE1 - HE2 |
| 911 | 917 | 1 | ; | CE1 - CD2 |
| 912 | 914 | 1 | ; | HE1 - HZ |
| 912 | 915 | 1 | ; | HE1 - CE2 |
| 913 | 918 | 1 | ; | CZ - HD2 |
| 914 | 916 | 1 | ; | HZ - HE2 |
| 914 | 917 | 1 | ; | HZ - CD2 |
| 916 | 918 | 1 | ; | HE2 - HD2 |
| 919 | 924 | 1 | ; | C - HA |
| 919 | 925 | 1 | ; | C - CB |
| 919 | 939 | 1 | ; | C - C |
| 920 | 922 | 1 | ; | O - H |
| 920 | 923 | 1 | ; | O - CA |
| 921 | 926 | 1 | ; | N - HB2 |
| 921 | 927 | 1 | ; | N - HB3 |
| 921 | 928 | 1 | ; | N - CG |
| 921 | 940 | 1 | ; | N - O |
| 921 | 941 | 1 | ; | N - N |
| 922 | 924 | 1 | ; | H - HA |
| 922 | 925 | 1 | ; | H - CB |
| 922 | 939 | 1 | ; | H - C |
| 923 | 929 | 1 | ; | CA - CD1 |
| 923 | 937 | 1 | ; | CA - CD2 |
| 923 | 942 | 1 | ; | CA - H |
| 923 | 943 | 1 | ; | CA - CA |
| 924 | 926 | 1 | ; | HA - HB2 |
| 924 | 927 | 1 | ; | HA - HB3 |
| 924 | 928 | 1 | ; | HA - CG |
| 924 | 940 | 1 | ; | HA - O |
| 924 | 941 | 1 | ; | HA - N |
| 925 | 930 | 1 | ; | CB - HD1 |
| 925 | 931 | 1 | ; | CB - CE1 |
| 925 | 935 | 1 | ; | CB - CE2 |
| 925 | 938 | 1 | ; | CB - HD2 |
| 925 | 940 | 1 | ; | CB - O |
| 925 | 941 | 1 | ; | CB - N |
| 926 | 929 | 1 | ; | HB2 - CD1 |
| 926 | 937 | 1 | ; | HB2 - CD2 |
| 926 | 939 | 1 | ; | HB2 - C |
| 927 | 929 | 1 | ; | HB3 - CD1 |
| 927 | 937 | 1 | ; | HB3 - CD2 |
| 927 | 939 | 1 | ; | HB3 - C |

| | | | | | | |
|---|---|---|---|---|---|---|
| 928 | 932 | 1 | ; | CG | - | HE1 |
| 928 | 933 | 1 | ; | CG | - | CZ |
| 928 | 936 | 1 | ; | CG | - | HE2 |
| 928 | 939 | 1 | ; | CG | - | C |
| 929 | 934 | 1 | ; | CD1 | - | HZ |
| 929 | 935 | 1 | ; | CD1 | - | CE2 |
| 929 | 938 | 1 | ; | CD1 | - | HD2 |
| 930 | 932 | 1 | ; | HD1 | - | HE1 |
| 930 | 933 | 1 | ; | HD1 | - | CZ |
| 930 | 937 | 1 | ; | HD1 | - | CD2 |
| 931 | 936 | 1 | ; | CE1 | - | HE2 |
| 931 | 937 | 1 | ; | CE1 | - | CD2 |
| 932 | 934 | 1 | ; | HE1 | - | HZ |
| 932 | 935 | 1 | ; | HE1 | - | CE2 |
| 933 | 938 | 1 | ; | CZ | - | HD2 |
| 934 | 936 | 1 | ; | HZ | - | HE2 |
| 934 | 937 | 1 | ; | HZ | - | CD2 |
| 936 | 938 | 1 | ; | HE2 | - | HD2 |
| 939 | 944 | 1 | ; | C | - | HA |
| 939 | 945 | 1 | ; | C | - | CB |
| 939 | 956 | 1 | ; | C | - | C |
| 940 | 942 | 1 | ; | O | - | H |
| 940 | 943 | 1 | ; | O | - | CA |
| 941 | 946 | 1 | ; | N | - | HB2 |
| 941 | 947 | 1 | ; | N | - | HB3 |
| 941 | 948 | 1 | ; | N | - | CG |
| 941 | 957 | 1 | ; | N | - | O |
| 941 | 958 | 1 | ; | N | - | N |
| 942 | 944 | 1 | ; | H | - | HA |
| 942 | 945 | 1 | ; | H | - | CB |
| 942 | 956 | 1 | ; | H | - | C |
| 943 | 949 | 1 | ; | CA | - | HG2 |
| 943 | 950 | 1 | ; | CA | - | HG3 |
| 943 | 951 | 1 | ; | CA | - | CD |
| 943 | 959 | 1 | ; | CA | - | H |
| 943 | 960 | 1 | ; | CA | - | CA |
| 944 | 946 | 1 | ; | HA | - | HB2 |
| 944 | 947 | 1 | ; | HA | - | HB3 |
| 944 | 948 | 1 | ; | HA | - | CG |
| 944 | 957 | 1 | ; | HA | - | O |
| 944 | 958 | 1 | ; | HA | - | N |
| 945 | 952 | 1 | ; | CB | - | OE1 |
| 945 | 953 | 1 | ; | CB | - | NE2 |
| 945 | 957 | 1 | ; | CB | - | O |
| 945 | 958 | 1 | ; | CB | - | N |
| 946 | 949 | 1 | ; | HB2 | - | HG2 |
| 946 | 950 | 1 | ; | HB2 | - | HG3 |
| 946 | 951 | 1 | ; | HB2 | - | CD |
| 946 | 956 | 1 | ; | HB2 | - | C |
| 947 | 949 | 1 | ; | HB3 | - | HG2 |
| 947 | 950 | 1 | ; | HB3 | - | HG3 |
| 947 | 951 | 1 | ; | HB3 | - | CD |

```
947    956     1 ;   HB3 - C
948    954     1 ;    CG - HE21
948    955     1 ;    CG - HE22
948    956     1 ;    CG - C
949    952     1 ;   HG2 - OE1
949    953     1 ;   HG2 - NE2
950    952     1 ;   HG3 - OE1
950    953     1 ;   HG3 - NE2
952    954     1 ;   OE1 - HE21
952    955     1 ;   OE1 - HE22
956    961     1 ;     C - HA
956    962     1 ;     C - CB
956    971     1 ;     C - C
957    959     1 ;     O - H
957    960     1 ;     O - CA
958    963     1 ;     N - HB2
958    964     1 ;     N - HB3
958    965     1 ;     N - CG
958    972     1 ;     N - O
958    973     1 ;     N - N
959    961     1 ;     H - HA
959    962     1 ;     H - CB
959    971     1 ;     H - C
960    966     1 ;    CA - HG2
960    967     1 ;    CA - HG3
960    968     1 ;    CA - CD
960    974     1 ;    CA - H
960    975     1 ;    CA - CA
961    963     1 ;    HA - HB2
961    964     1 ;    HA - HB3
961    965     1 ;    HA - CG
961    972     1 ;    HA - O
961    973     1 ;    HA - N
962    969     1 ;    CB - OE1
962    970     1 ;    CB - OE2
962    972     1 ;    CB - O
962    973     1 ;    CB - N
963    966     1 ;   HB2 - HG2
963    967     1 ;   HB2 - HG3
963    968     1 ;   HB2 - CD
963    971     1 ;   HB2 - C
964    966     1 ;   HB3 - HG2
964    967     1 ;   HB3 - HG3
964    968     1 ;   HB3 - CD
964    971     1 ;   HB3 - C
965    971     1 ;    CG - C
966    969     1 ;   HG2 - OE1
966    970     1 ;   HG2 - OE2
967    969     1 ;   HG3 - OE1
967    970     1 ;   HG3 - OE2
971    976     1 ;     C - HA
971    977     1 ;     C - CB
```

```
971     990       1 ;       C - C
972     974       1 ;       O - H
972     975       1 ;       O - CA
973     978       1 ;       N - HB
973     979       1 ;       N - CG2
973     983       1 ;       N - CG1
973     991       1 ;       N - O
973     992       1 ;       N - N
974     976       1 ;       H - HA
974     977       1 ;       H - CB
974     990       1 ;       H - C
975     980       1 ;      CA - HG21
975     981       1 ;      CA - HG22
975     982       1 ;      CA - HG23
975     984       1 ;      CA - HG12
975     985       1 ;      CA - HG13
975     986       1 ;      CA - CD1
975     993       1 ;      CA - H
975     994       1 ;      CA - CA
976     978       1 ;      HA - HB
976     979       1 ;      HA - CG2
976     983       1 ;      HA - CG1
976     991       1 ;      HA - O
976     992       1 ;      HA - N
977     987       1 ;      CB - HD11
977     988       1 ;      CB - HD12
977     989       1 ;      CB - HD13
977     991       1 ;      CB - O
977     992       1 ;      CB - N
978     980       1 ;      HB - HG21
978     981       1 ;      HB - HG22
978     982       1 ;      HB - HG23
978     984       1 ;      HB - HG12
978     985       1 ;      HB - HG13
978     986       1 ;      HB - CD1
978     990       1 ;      HB - C
979     984       1 ;     CG2 - HG12
979     985       1 ;     CG2 - HG13
979     986       1 ;     CG2 - CD1
979     990       1 ;     CG2 - C
980     983       1 ;    HG21 - CG1
981     983       1 ;    HG22 - CG1
982     983       1 ;    HG23 - CG1
983     990       1 ;     CG1 - C
984     987       1 ;    HG12 - HD11
984     988       1 ;    HG12 - HD12
984     989       1 ;    HG12 - HD13
985     987       1 ;    HG13 - HD11
985     988       1 ;    HG13 - HD12
985     989       1 ;    HG13 - HD13
990     995       1 ;       C - HA
990     996       1 ;       C - CB
```

```
  990   1005       1 ;       C - C
  991    993       1 ;       O - H
  991    994       1 ;       O - CA
  992    997       1 ;       N - HB2
  992    998       1 ;       N - HB3
  992    999       1 ;       N - CG
  992   1006       1 ;       N - O
  992   1007       1 ;       N - N
  993    995       1 ;       H - HA
  993    996       1 ;       H - CB
  993   1005       1 ;       H - C
  994   1000       1 ;      CA - HG2
  994   1001       1 ;      CA - HG3
  994   1002       1 ;      CA - CD
  994   1008       1 ;      CA - H
  994   1009       1 ;      CA - CA
  995    997       1 ;      HA - HB2
  995    998       1 ;      HA - HB3
  995    999       1 ;      HA - CG
  995   1006       1 ;      HA - O
  995   1007       1 ;      HA - N
  996   1003       1 ;      CB - OE1
  996   1004       1 ;      CB - OE2
  996   1006       1 ;      CB - O
  996   1007       1 ;      CB - N
  997   1000       1 ;     HB2 - HG2
  997   1001       1 ;     HB2 - HG3
  997   1002       1 ;     HB2 - CD
  997   1005       1 ;     HB2 - C
  998   1000       1 ;     HB3 - HG2
  998   1001       1 ;     HB3 - HG3
  998   1002       1 ;     HB3 - CD
  998   1005       1 ;     HB3 - C
  999   1005       1 ;      CG - C
 1000   1003       1 ;     HG2 - OE1
 1000   1004       1 ;     HG2 - OE2
 1001   1003       1 ;     HG3 - OE1
 1001   1004       1 ;     HG3 - OE2
 1005   1010       1 ;       C - HA
 1005   1011       1 ;       C - CB
 1005   1024       1 ;       C - C
 1006   1008       1 ;       O - H
 1006   1009       1 ;       O - CA
 1007   1012       1 ;       N - HB
 1007   1013       1 ;       N - CG2
 1007   1017       1 ;       N - CG1
 1007   1025       1 ;       N - O
 1007   1026       1 ;       N - N
 1008   1010       1 ;       H - HA
 1008   1011       1 ;       H - CB
 1008   1024       1 ;       H - C
 1009   1014       1 ;      CA - HG21
```

```
1009   1015      1 ;     CA - HG22
1009   1016      1 ;     CA - HG23
1009   1018      1 ;     CA - HG12
1009   1019      1 ;     CA - HG13
1009   1020      1 ;     CA - CD1
1009   1027      1 ;     CA - H
1009   1028      1 ;     CA - CA
1010   1012      1 ;     HA - HB
1010   1013      1 ;     HA - CG2
1010   1017      1 ;     HA - CG1
1010   1025      1 ;     HA - O
1010   1026      1 ;     HA - N
1011   1021      1 ;     CB - HD11
1011   1022      1 ;     CB - HD12
1011   1023      1 ;     CB - HD13
1011   1025      1 ;     CB - O
1011   1026      1 ;     CB - N
1012   1014      1 ;     HB - HG21
1012   1015      1 ;     HB - HG22
1012   1016      1 ;     HB - HG23
1012   1018      1 ;     HB - HG12
1012   1019      1 ;     HB - HG13
1012   1020      1 ;     HB - CD1
1012   1024      1 ;     HB - C
1013   1018      1 ;    CG2 - HG12
1013   1019      1 ;    CG2 - HG13
1013   1020      1 ;    CG2 - CD1
1013   1024      1 ;    CG2 - C
1014   1017      1 ;   HG21 - CG1
1015   1017      1 ;   HG22 - CG1
1016   1017      1 ;   HG23 - CG1
1017   1024      1 ;    CG1 - C
1018   1021      1 ;   HG12 - HD11
1018   1022      1 ;   HG12 - HD12
1018   1023      1 ;   HG12 - HD13
1019   1021      1 ;   HG13 - HD11
1019   1022      1 ;   HG13 - HD12
1019   1023      1 ;   HG13 - HD13
1024   1029      1 ;      C - HA
1024   1030      1 ;      C - CB
1024   1041      1 ;      C - C
1025   1027      1 ;      O - H
1025   1028      1 ;      O - CA
1026   1031      1 ;      N - HB2
1026   1032      1 ;      N - HB3
1026   1033      1 ;      N - CG
1026   1042      1 ;      N - O
1026   1043      1 ;      N - N
1027   1029      1 ;      H - HA
1027   1030      1 ;      H - CB
1027   1041      1 ;      H - C
1028   1034      1 ;     CA - HG2
```

```
1028   1035      1 ;     CA - HG3
1028   1036      1 ;     CA - SD
1028   1044      1 ;     CA - H
1028   1045      1 ;     CA - CA
1029   1031      1 ;     HA - HB2
1029   1032      1 ;     HA - HB3
1029   1033      1 ;     HA - CG
1029   1042      1 ;     HA - O
1029   1043      1 ;     HA - N
1030   1037      1 ;     CB - CE
1030   1042      1 ;     CB - O
1030   1043      1 ;     CB - N
1031   1034      1 ;    HB2 - HG2
1031   1035      1 ;    HB2 - HG3
1031   1036      1 ;    HB2 - SD
1031   1041      1 ;    HB2 - C
1032   1034      1 ;    HB3 - HG2
1032   1035      1 ;    HB3 - HG3
1032   1036      1 ;    HB3 - SD
1032   1041      1 ;    HB3 - C
1033   1038      1 ;     CG - HE1
1033   1039      1 ;     CG - HE2
1033   1040      1 ;     CG - HE3
1033   1041      1 ;     CG - C
1034   1037      1 ;    HG2 - CE
1035   1037      1 ;    HG3 - CE
1041   1046      1 ;      C - HA
1041   1047      1 ;      C - CB
1041   1063      1 ;      C - C
1042   1044      1 ;      O - H
1042   1045      1 ;      O - CA
1043   1048      1 ;      N - HB2
1043   1049      1 ;      N - HB3
1043   1050      1 ;      N - CG
1043   1064      1 ;      N - O
1043   1065      1 ;      N - N
1044   1046      1 ;      H - HA
1044   1047      1 ;      H - CB
1044   1063      1 ;      H - C
1045   1051      1 ;     CA - HG2
1045   1052      1 ;     CA - HG3
1045   1053      1 ;     CA - CD
1045   1066      1 ;     CA - H
1045   1067      1 ;     CA - CA
1046   1048      1 ;     HA - HB2
1046   1049      1 ;     HA - HB3
1046   1050      1 ;     HA - CG
1046   1064      1 ;     HA - O
1046   1065      1 ;     HA - N
1047   1054      1 ;     CB - HD2
1047   1055      1 ;     CB - HD3
1047   1056      1 ;     CB - CE
```

```
1047   1064       1 ;     CB - O
1047   1065       1 ;     CB - N
1048   1051       1 ;    HB2 - HG2
1048   1052       1 ;    HB2 - HG3
1048   1053       1 ;    HB2 - CD
1048   1063       1 ;    HB2 - C
1049   1051       1 ;    HB3 - HG2
1049   1052       1 ;    HB3 - HG3
1049   1053       1 ;    HB3 - CD
1049   1063       1 ;    HB3 - C
1050   1057       1 ;     CG - HE2
1050   1058       1 ;     CG - HE3
1050   1059       1 ;     CG - NZ
1050   1063       1 ;     CG - C
1051   1054       1 ;    HG2 - HD2
1051   1055       1 ;    HG2 - HD3
1051   1056       1 ;    HG2 - CE
1052   1054       1 ;    HG3 - HD2
1052   1055       1 ;    HG3 - HD3
1052   1056       1 ;    HG3 - CE
1053   1060       1 ;     CD - HZ1
1053   1061       1 ;     CD - HZ2
1053   1062       1 ;     CD - HZ3
1054   1057       1 ;    HD2 - HE2
1054   1058       1 ;    HD2 - HE3
1054   1059       1 ;    HD2 - NZ
1055   1057       1 ;    HD3 - HE2
1055   1058       1 ;    HD3 - HE3
1055   1059       1 ;    HD3 - NZ
1057   1060       1 ;    HE2 - HZ1
1057   1061       1 ;    HE2 - HZ2
1057   1062       1 ;    HE2 - HZ3
1058   1060       1 ;    HE3 - HZ1
1058   1061       1 ;    HE3 - HZ2
1058   1062       1 ;    HE3 - HZ3
1063   1068       1 ;      C - HA
1063   1069       1 ;      C - CB
1063   1085       1 ;      C - C
1064   1066       1 ;      O - H
1064   1067       1 ;      O - CA
1065   1070       1 ;      N - HB2
1065   1071       1 ;      N - HB3
1065   1072       1 ;      N - CG
1065   1086       1 ;      N - O
1065   1087       1 ;      N - N
1066   1068       1 ;      H - HA
1066   1069       1 ;      H - CB
1066   1085       1 ;      H - C
1067   1073       1 ;     CA - HG2
1067   1074       1 ;     CA - HG3
1067   1075       1 ;     CA - CD
1067   1088       1 ;     CA - H
```

```
1067   1089      1 ;     CA - CA
1068   1070      1 ;     HA - HB2
1068   1071      1 ;     HA - HB3
1068   1072      1 ;     HA - CG
1068   1086      1 ;     HA - O
1068   1087      1 ;     HA - N
1069   1076      1 ;     CB - HD2
1069   1077      1 ;     CB - HD3
1069   1078      1 ;     CB - CE
1069   1086      1 ;     CB - O
1069   1087      1 ;     CB - N
1070   1073      1 ;    HB2 - HG2
1070   1074      1 ;    HB2 - HG3
1070   1075      1 ;    HB2 - CD
1070   1085      1 ;    HB2 - C
1071   1073      1 ;    HB3 - HG2
1071   1074      1 ;    HB3 - HG3
1071   1075      1 ;    HB3 - CD
1071   1085      1 ;    HB3 - C
1072   1079      1 ;     CG - HE2
1072   1080      1 ;     CG - HE3
1072   1081      1 ;     CG - NZ
1072   1085      1 ;     CG - C
1073   1076      1 ;    HG2 - HD2
1073   1077      1 ;    HG2 - HD3
1073   1078      1 ;    HG2 - CE
1074   1076      1 ;    HG3 - HD2
1074   1077      1 ;    HG3 - HD3
1074   1078      1 ;    HG3 - CE
1075   1082      1 ;     CD - HZ1
1075   1083      1 ;     CD - HZ2
1075   1084      1 ;     CD - HZ3
1076   1079      1 ;    HD2 - HE2
1076   1080      1 ;    HD2 - HE3
1076   1081      1 ;    HD2 - NZ
1077   1079      1 ;    HD3 - HE2
1077   1080      1 ;    HD3 - HE3
1077   1081      1 ;    HD3 - NZ
1079   1082      1 ;    HE2 - HZ1
1079   1083      1 ;    HE2 - HZ2
1079   1084      1 ;    HE2 - HZ3
1080   1082      1 ;    HE3 - HZ1
1080   1083      1 ;    HE3 - HZ2
1080   1084      1 ;    HE3 - HZ3
1085   1090      1 ;      C - HA
1085   1091      1 ;      C - CB
1085   1104      1 ;      C - C
1086   1088      1 ;      O - H
1086   1089      1 ;      O - CA
1087   1092      1 ;      N - HB2
1087   1093      1 ;      N - HB3
1087   1094      1 ;      N - CG
```

```
1087   1105       1 ;       N - O
1087   1106       1 ;       N - N
1088   1090       1 ;       H - HA
1088   1091       1 ;       H - CB
1088   1104       1 ;       H - C
1089   1095       1 ;      CA - HG
1089   1096       1 ;      CA - CD1
1089   1100       1 ;      CA - CD2
1089   1107       1 ;      CA - H
1089   1108       1 ;      CA - CA
1090   1092       1 ;      HA - HB2
1090   1093       1 ;      HA - HB3
1090   1094       1 ;      HA - CG
1090   1105       1 ;      HA - O
1090   1106       1 ;      HA - N
1091   1097       1 ;      CB - HD11
1091   1098       1 ;      CB - HD12
1091   1099       1 ;      CB - HD13
1091   1101       1 ;      CB - HD21
1091   1102       1 ;      CB - HD22
1091   1103       1 ;      CB - HD23
1091   1105       1 ;      CB - O
1091   1106       1 ;      CB - N
1092   1095       1 ;     HB2 - HG
1092   1096       1 ;     HB2 - CD1
1092   1100       1 ;     HB2 - CD2
1092   1104       1 ;     HB2 - C
1093   1095       1 ;     HB3 - HG
1093   1096       1 ;     HB3 - CD1
1093   1100       1 ;     HB3 - CD2
1093   1104       1 ;     HB3 - C
1094   1104       1 ;      CG - C
1095   1097       1 ;      HG - HD11
1095   1098       1 ;      HG - HD12
1095   1099       1 ;      HG - HD13
1095   1101       1 ;      HG - HD21
1095   1102       1 ;      HG - HD22
1095   1103       1 ;      HG - HD23
1096   1101       1 ;     CD1 - HD21
1096   1102       1 ;     CD1 - HD22
1096   1103       1 ;     CD1 - HD23
1097   1100       1 ;    HD11 - CD2
1098   1100       1 ;    HD12 - CD2
1099   1100       1 ;    HD13 - CD2
1104   1109       1 ;       C - HA
1104   1110       1 ;       C - CB
1104   1116       1 ;       C - C
1105   1107       1 ;       O - H
1105   1108       1 ;       O - CA
1106   1111       1 ;       N - HB2
1106   1112       1 ;       N - HB3
1106   1113       1 ;       N - CG
```

| | | | | | |
|---|---|---|---|---|---|
| 1106 | 1117 | 1 ; | N | - | O |
| 1106 | 1118 | 1 ; | N | - | N |
| 1107 | 1109 | 1 ; | H | - | HA |
| 1107 | 1110 | 1 ; | H | - | CB |
| 1107 | 1116 | 1 ; | H | - | C |
| 1108 | 1114 | 1 ; | CA | - | OD1 |
| 1108 | 1115 | 1 ; | CA | - | OD2 |
| 1108 | 1119 | 1 ; | CA | - | H |
| 1108 | 1120 | 1 ; | CA | - | CA |
| 1109 | 1111 | 1 ; | HA | - | HB2 |
| 1109 | 1112 | 1 ; | HA | - | HB3 |
| 1109 | 1113 | 1 ; | HA | - | CG |
| 1109 | 1117 | 1 ; | HA | - | O |
| 1109 | 1118 | 1 ; | HA | - | N |
| 1110 | 1117 | 1 ; | CB | - | O |
| 1110 | 1118 | 1 ; | CB | - | N |
| 1111 | 1114 | 1 ; | HB2 | - | OD1 |
| 1111 | 1115 | 1 ; | HB2 | - | OD2 |
| 1111 | 1116 | 1 ; | HB2 | - | C |
| 1112 | 1114 | 1 ; | HB3 | - | OD1 |
| 1112 | 1115 | 1 ; | HB3 | - | OD2 |
| 1112 | 1116 | 1 ; | HB3 | - | C |
| 1113 | 1116 | 1 ; | CG | - | C |
| 1116 | 1121 | 1 ; | C | - | HA |
| 1116 | 1122 | 1 ; | C | - | CB |
| 1116 | 1133 | 1 ; | C | - | C |
| 1117 | 1119 | 1 ; | O | - | H |
| 1117 | 1120 | 1 ; | O | - | CA |
| 1118 | 1123 | 1 ; | N | - | HB2 |
| 1118 | 1124 | 1 ; | N | - | HB3 |
| 1118 | 1125 | 1 ; | N | - | CG |
| 1118 | 1134 | 1 ; | N | - | O |
| 1118 | 1135 | 1 ; | N | - | N |
| 1119 | 1121 | 1 ; | H | - | HA |
| 1119 | 1122 | 1 ; | H | - | CB |
| 1119 | 1133 | 1 ; | H | - | C |
| 1120 | 1126 | 1 ; | CA | - | ND1 |
| 1120 | 1131 | 1 ; | CA | - | CD2 |
| 1120 | 1136 | 1 ; | CA | - | CD |
| 1120 | 1145 | 1 ; | CA | - | CA |
| 1121 | 1123 | 1 ; | HA | - | HB2 |
| 1121 | 1124 | 1 ; | HA | - | HB3 |
| 1121 | 1125 | 1 ; | HA | - | CG |
| 1121 | 1134 | 1 ; | HA | - | O |
| 1121 | 1135 | 1 ; | HA | - | N |
| 1122 | 1127 | 1 ; | CB | - | CE1 |
| 1122 | 1129 | 1 ; | CB | - | NE2 |
| 1122 | 1132 | 1 ; | CB | - | HD2 |
| 1122 | 1134 | 1 ; | CB | - | O |
| 1122 | 1135 | 1 ; | CB | - | N |
| 1123 | 1126 | 1 ; | HB2 | - | ND1 |
| 1123 | 1131 | 1 ; | HB2 | - | CD2 |

| | | | | |
|---|---|---|---|---|
| 1123 | 1133 | 1 | ; | HB2 - C |
| 1124 | 1126 | 1 | ; | HB3 - ND1 |
| 1124 | 1131 | 1 | ; | HB3 - CD2 |
| 1124 | 1133 | 1 | ; | HB3 - C |
| 1125 | 1128 | 1 | ; | CG - HE1 |
| 1125 | 1130 | 1 | ; | CG - HE2 |
| 1125 | 1133 | 1 | ; | CG - C |
| 1126 | 1130 | 1 | ; | ND1 - HE2 |
| 1126 | 1132 | 1 | ; | ND1 - HD2 |
| 1127 | 1132 | 1 | ; | CE1 - HD2 |
| 1128 | 1130 | 1 | ; | HE1 - HE2 |
| 1128 | 1131 | 1 | ; | HE1 - CD2 |
| 1130 | 1132 | 1 | ; | HE2 - HD2 |
| 1133 | 1137 | 1 | ; | C - HD2 |
| 1133 | 1138 | 1 | ; | C - HD3 |
| 1133 | 1139 | 1 | ; | C - CG |
| 1133 | 1142 | 1 | ; | C - CB |
| 1133 | 1146 | 1 | ; | C - HA |
| 1133 | 1147 | 1 | ; | C - C |
| 1134 | 1136 | 1 | ; | O - CD |
| 1134 | 1145 | 1 | ; | O - CA |
| 1135 | 1140 | 1 | ; | N - HG2 |
| 1135 | 1141 | 1 | ; | N - HG3 |
| 1135 | 1143 | 1 | ; | N - HB2 |
| 1135 | 1144 | 1 | ; | N - HB3 |
| 1135 | 1148 | 1 | ; | N - O |
| 1135 | 1149 | 1 | ; | N - N |
| 1136 | 1143 | 1 | ; | CD - HB2 |
| 1136 | 1144 | 1 | ; | CD - HB3 |
| 1136 | 1146 | 1 | ; | CD - HA |
| 1136 | 1147 | 1 | ; | CD - C |
| 1137 | 1140 | 1 | ; | HD2 - HG2 |
| 1137 | 1141 | 1 | ; | HD2 - HG3 |
| 1137 | 1142 | 1 | ; | HD2 - CB |
| 1137 | 1145 | 1 | ; | HD2 - CA |
| 1138 | 1140 | 1 | ; | HD3 - HG2 |
| 1138 | 1141 | 1 | ; | HD3 - HG3 |
| 1138 | 1142 | 1 | ; | HD3 - CB |
| 1138 | 1145 | 1 | ; | HD3 - CA |
| 1139 | 1146 | 1 | ; | CG - HA |
| 1139 | 1147 | 1 | ; | CG - C |
| 1140 | 1143 | 1 | ; | HG2 - HB2 |
| 1140 | 1144 | 1 | ; | HG2 - HB3 |
| 1140 | 1145 | 1 | ; | HG2 - CA |
| 1141 | 1143 | 1 | ; | HG3 - HB2 |
| 1141 | 1144 | 1 | ; | HG3 - HB3 |
| 1141 | 1145 | 1 | ; | HG3 - CA |
| 1142 | 1148 | 1 | ; | CB - O |
| 1142 | 1149 | 1 | ; | CB - N |
| 1143 | 1146 | 1 | ; | HB2 - HA |
| 1143 | 1147 | 1 | ; | HB2 - C |
| 1144 | 1146 | 1 | ; | HB3 - HA |

```
1144   1147       1 ;     HB3 - C
1145   1150       1 ;      CA - H
1145   1151       1 ;      CA - CA
1146   1148       1 ;      HA - O
1146   1149       1 ;      HA - N
1147   1152       1 ;       C - HA
1147   1153       1 ;       C - CB
1147   1161       1 ;       C - C
1148   1150       1 ;       O - H
1148   1151       1 ;       O - CA
1149   1154       1 ;       N - HB2
1149   1155       1 ;       N - HB3
1149   1156       1 ;       N - CG
1149   1162       1 ;       N - O
1149   1163       1 ;       N - N
1150   1152       1 ;       H - HA
1150   1153       1 ;       H - CB
1150   1161       1 ;       H - C
1151   1157       1 ;      CA - OD1
1151   1158       1 ;      CA - ND2
1151   1164       1 ;      CA - H
1151   1165       1 ;      CA - CA
1152   1154       1 ;      HA - HB2
1152   1155       1 ;      HA - HB3
1152   1156       1 ;      HA - CG
1152   1162       1 ;      HA - O
1152   1163       1 ;      HA - N
1153   1159       1 ;      CB - HD21
1153   1160       1 ;      CB - HD22
1153   1162       1 ;      CB - O
1153   1163       1 ;      CB - N
1154   1157       1 ;     HB2 - OD1
1154   1158       1 ;     HB2 - ND2
1154   1161       1 ;     HB2 - C
1155   1157       1 ;     HB3 - OD1
1155   1158       1 ;     HB3 - ND2
1155   1161       1 ;     HB3 - C
1156   1161       1 ;      CG - C
1157   1159       1 ;     OD1 - HD21
1157   1160       1 ;     OD1 - HD22
1161   1166       1 ;       C - HA
1161   1167       1 ;       C - CB
1161   1180       1 ;       C - C
1162   1164       1 ;       O - H
1162   1165       1 ;       O - CA
1163   1168       1 ;       N - HB
1163   1169       1 ;       N - CG2
1163   1173       1 ;       N - CG1
1163   1181       1 ;       N - O
1163   1182       1 ;       N - N
1164   1166       1 ;       H - HA
1164   1167       1 ;       H - CB
```

```
 1164   1180       1 ;       H - C
 1165   1170       1 ;      CA - HG21
 1165   1171       1 ;      CA - HG22
 1165   1172       1 ;      CA - HG23
 1165   1174       1 ;      CA - HG12
 1165   1175       1 ;      CA - HG13
 1165   1176       1 ;      CA - CD1
 1165   1183       1 ;      CA - H
 1165   1184       1 ;      CA - CA
 1166   1168       1 ;      HA - HB
 1166   1169       1 ;      HA - CG2
 1166   1173       1 ;      HA - CG1
 1166   1181       1 ;      HA - O
 1166   1182       1 ;      HA - N
 1167   1177       1 ;      CB - HD11
 1167   1178       1 ;      CB - HD12
 1167   1179       1 ;      CB - HD13
 1167   1181       1 ;      CB - O
 1167   1182       1 ;      CB - N
 1168   1170       1 ;      HB - HG21
 1168   1171       1 ;      HB - HG22
 1168   1172       1 ;      HB - HG23
 1168   1174       1 ;      HB - HG12
 1168   1175       1 ;      HB - HG13
 1168   1176       1 ;      HB - CD1
 1168   1180       1 ;      HB - C
 1169   1174       1 ;     CG2 - HG12
 1169   1175       1 ;     CG2 - HG13
 1169   1176       1 ;     CG2 - CD1
 1169   1180       1 ;     CG2 - C
 1170   1173       1 ;    HG21 - CG1
 1171   1173       1 ;    HG22 - CG1
 1172   1173       1 ;    HG23 - CG1
 1173   1180       1 ;     CG1 - C
 1174   1177       1 ;    HG12 - HD11
 1174   1178       1 ;    HG12 - HD12
 1174   1179       1 ;    HG12 - HD13
 1175   1177       1 ;    HG13 - HD11
 1175   1178       1 ;    HG13 - HD12
 1175   1179       1 ;    HG13 - HD13
 1180   1185       1 ;       C - HA
 1180   1186       1 ;       C - CB
 1180   1196       1 ;       C - C
 1181   1183       1 ;       O - H
 1181   1184       1 ;       O - CA
 1182   1187       1 ;       N - HB
 1182   1188       1 ;       N - CG1
 1182   1192       1 ;       N - CG2
 1182   1197       1 ;       N - O
 1182   1198       1 ;       N - N
 1183   1185       1 ;       H - HA
 1183   1186       1 ;       H - CB
```

| | | | |
|---|---|---|---|
| 1183 | 1196 | 1 | ; H - C |
| 1184 | 1189 | 1 | ; CA - HG11 |
| 1184 | 1190 | 1 | ; CA - HG12 |
| 1184 | 1191 | 1 | ; CA - HG13 |
| 1184 | 1193 | 1 | ; CA - HG21 |
| 1184 | 1194 | 1 | ; CA - HG22 |
| 1184 | 1195 | 1 | ; CA - HG23 |
| 1184 | 1199 | 1 | ; CA - H |
| 1184 | 1200 | 1 | ; CA - CA |
| 1185 | 1187 | 1 | ; HA - HB |
| 1185 | 1188 | 1 | ; HA - CG1 |
| 1185 | 1192 | 1 | ; HA - CG2 |
| 1185 | 1197 | 1 | ; HA - O |
| 1185 | 1198 | 1 | ; HA - N |
| 1186 | 1197 | 1 | ; CB - O |
| 1186 | 1198 | 1 | ; CB - N |
| 1187 | 1189 | 1 | ; HB - HG11 |
| 1187 | 1190 | 1 | ; HB - HG12 |
| 1187 | 1191 | 1 | ; HB - HG13 |
| 1187 | 1193 | 1 | ; HB - HG21 |
| 1187 | 1194 | 1 | ; HB - HG22 |
| 1187 | 1195 | 1 | ; HB - HG23 |
| 1187 | 1196 | 1 | ; HB - C |
| 1188 | 1193 | 1 | ; CG1 - HG21 |
| 1188 | 1194 | 1 | ; CG1 - HG22 |
| 1188 | 1195 | 1 | ; CG1 - HG23 |
| 1188 | 1196 | 1 | ; CG1 - C |
| 1189 | 1192 | 1 | ; HG11 - CG2 |
| 1190 | 1192 | 1 | ; HG12 - CG2 |
| 1191 | 1192 | 1 | ; HG13 - CG2 |
| 1192 | 1196 | 1 | ; CG2 - C |
| 1196 | 1201 | 1 | ; C - HA |
| 1196 | 1202 | 1 | ; C - CB |
| 1196 | 1218 | 1 | ; C - C |
| 1197 | 1199 | 1 | ; O - H |
| 1197 | 1200 | 1 | ; O - CA |
| 1198 | 1203 | 1 | ; N - HB2 |
| 1198 | 1204 | 1 | ; N - HB3 |
| 1198 | 1205 | 1 | ; N - CG |
| 1198 | 1219 | 1 | ; N - O |
| 1198 | 1220 | 1 | ; N - N |
| 1199 | 1201 | 1 | ; H - HA |
| 1199 | 1202 | 1 | ; H - CB |
| 1199 | 1218 | 1 | ; H - C |
| 1200 | 1206 | 1 | ; CA - HG2 |
| 1200 | 1207 | 1 | ; CA - HG3 |
| 1200 | 1208 | 1 | ; CA - CD |
| 1200 | 1221 | 1 | ; CA - H |
| 1200 | 1222 | 1 | ; CA - CA |
| 1201 | 1203 | 1 | ; HA - HB2 |
| 1201 | 1204 | 1 | ; HA - HB3 |
| 1201 | 1205 | 1 | ; HA - CG |

| | | | | | |
|---|---|---|---|---|---|
| 1201 | 1219 | 1 ; | HA | - | O |
| 1201 | 1220 | 1 ; | HA | - | N |
| 1202 | 1209 | 1 ; | CB | - | HD2 |
| 1202 | 1210 | 1 ; | CB | - | HD3 |
| 1202 | 1211 | 1 ; | CB | - | CE |
| 1202 | 1219 | 1 ; | CB | - | O |
| 1202 | 1220 | 1 ; | CB | - | N |
| 1203 | 1206 | 1 ; | HB2 | - | HG2 |
| 1203 | 1207 | 1 ; | HB2 | - | HG3 |
| 1203 | 1208 | 1 ; | HB2 | - | CD |
| 1203 | 1218 | 1 ; | HB2 | - | C |
| 1204 | 1206 | 1 ; | HB3 | - | HG2 |
| 1204 | 1207 | 1 ; | HB3 | - | HG3 |
| 1204 | 1208 | 1 ; | HB3 | - | CD |
| 1204 | 1218 | 1 ; | HB3 | - | C |
| 1205 | 1212 | 1 ; | CG | - | HE2 |
| 1205 | 1213 | 1 ; | CG | - | HE3 |
| 1205 | 1214 | 1 ; | CG | - | NZ |
| 1205 | 1218 | 1 ; | CG | - | C |
| 1206 | 1209 | 1 ; | HG2 | - | HD2 |
| 1206 | 1210 | 1 ; | HG2 | - | HD3 |
| 1206 | 1211 | 1 ; | HG2 | - | CE |
| 1207 | 1209 | 1 ; | HG3 | - | HD2 |
| 1207 | 1210 | 1 ; | HG3 | - | HD3 |
| 1207 | 1211 | 1 ; | HG3 | - | CE |
| 1208 | 1215 | 1 ; | CD | - | HZ1 |
| 1208 | 1216 | 1 ; | CD | - | HZ2 |
| 1208 | 1217 | 1 ; | CD | - | HZ3 |
| 1209 | 1212 | 1 ; | HD2 | - | HE2 |
| 1209 | 1213 | 1 ; | HD2 | - | HE3 |
| 1209 | 1214 | 1 ; | HD2 | - | NZ |
| 1210 | 1212 | 1 ; | HD3 | - | HE2 |
| 1210 | 1213 | 1 ; | HD3 | - | HE3 |
| 1210 | 1214 | 1 ; | HD3 | - | NZ |
| 1212 | 1215 | 1 ; | HE2 | - | HZ1 |
| 1212 | 1216 | 1 ; | HE2 | - | HZ2 |
| 1212 | 1217 | 1 ; | HE2 | - | HZ3 |
| 1213 | 1215 | 1 ; | HE3 | - | HZ1 |
| 1213 | 1216 | 1 ; | HE3 | - | HZ2 |
| 1213 | 1217 | 1 ; | HE3 | - | HZ3 |
| 1218 | 1223 | 1 ; | C | - | HA |
| 1218 | 1224 | 1 ; | C | - | CB |
| 1218 | 1237 | 1 ; | C | - | C |
| 1219 | 1221 | 1 ; | O | - | H |
| 1219 | 1222 | 1 ; | O | - | CA |
| 1220 | 1225 | 1 ; | N | - | HB2 |
| 1220 | 1226 | 1 ; | N | - | HB3 |
| 1220 | 1227 | 1 ; | N | - | CG |
| 1220 | 1238 | 1 ; | N | - | O |
| 1220 | 1239 | 1 ; | N | - | N |
| 1221 | 1223 | 1 ; | H | - | HA |
| 1221 | 1224 | 1 ; | H | - | CB |

```
1221   1237      1 ;     H - C
1222   1228      1 ;    CA - HG
1222   1229      1 ;    CA - CD1
1222   1233      1 ;    CA - CD2
1222   1240      1 ;    CA - H
1222   1241      1 ;    CA - CA
1223   1225      1 ;    HA - HB2
1223   1226      1 ;    HA - HB3
1223   1227      1 ;    HA - CG
1223   1238      1 ;    HA - O
1223   1239      1 ;    HA - N
1224   1230      1 ;    CB - HD11
1224   1231      1 ;    CB - HD12
1224   1232      1 ;    CB - HD13
1224   1234      1 ;    CB - HD21
1224   1235      1 ;    CB - HD22
1224   1236      1 ;    CB - HD23
1224   1238      1 ;    CB - O
1224   1239      1 ;    CB - N
1225   1228      1 ;   HB2 - HG
1225   1229      1 ;   HB2 - CD1
1225   1233      1 ;   HB2 - CD2
1225   1237      1 ;   HB2 - C
1226   1228      1 ;   HB3 - HG
1226   1229      1 ;   HB3 - CD1
1226   1233      1 ;   HB3 - CD2
1226   1237      1 ;   HB3 - C
1227   1237      1 ;    CG - C
1228   1230      1 ;    HG - HD11
1228   1231      1 ;    HG - HD12
1228   1232      1 ;    HG - HD13
1228   1234      1 ;    HG - HD21
1228   1235      1 ;    HG - HD22
1228   1236      1 ;    HG - HD23
1229   1234      1 ;   CD1 - HD21
1229   1235      1 ;   CD1 - HD22
1229   1236      1 ;   CD1 - HD23
1230   1233      1 ;  HD11 - CD2
1231   1233      1 ;  HD12 - CD2
1232   1233      1 ;  HD13 - CD2
1237   1242      1 ;     C - HA
1237   1243      1 ;     C - CB
1237   1258      1 ;     C - C
1238   1240      1 ;     O - H
1238   1241      1 ;     O - CA
1239   1244      1 ;     N - HB2
1239   1245      1 ;     N - HB3
1239   1246      1 ;     N - CG
1239   1259      1 ;     N - O
1239   1260      1 ;     N - N
1240   1242      1 ;     H - HA
1240   1243      1 ;     H - CB
```

| | | | | | | |
|---|---|---|---|---|---|---|
| 1240 | 1258 | 1 | ; | H | - | C |
| 1241 | 1247 | 1 | ; | CA | - | CD1 |
| 1241 | 1256 | 1 | ; | CA | - | CD2 |
| 1241 | 1261 | 1 | ; | CA | - | H |
| 1241 | 1262 | 1 | ; | CA | - | CA |
| 1242 | 1244 | 1 | ; | HA | - | HB2 |
| 1242 | 1245 | 1 | ; | HA | - | HB3 |
| 1242 | 1246 | 1 | ; | HA | - | CG |
| 1242 | 1259 | 1 | ; | HA | - | O |
| 1242 | 1260 | 1 | ; | HA | - | N |
| 1243 | 1248 | 1 | ; | CB | - | HD1 |
| 1243 | 1249 | 1 | ; | CB | - | CE1 |
| 1243 | 1254 | 1 | ; | CB | - | CE2 |
| 1243 | 1257 | 1 | ; | CB | - | HD2 |
| 1243 | 1259 | 1 | ; | CB | - | O |
| 1243 | 1260 | 1 | ; | CB | - | N |
| 1244 | 1247 | 1 | ; | HB2 | - | CD1 |
| 1244 | 1256 | 1 | ; | HB2 | - | CD2 |
| 1244 | 1258 | 1 | ; | HB2 | - | C |
| 1245 | 1247 | 1 | ; | HB3 | - | CD1 |
| 1245 | 1256 | 1 | ; | HB3 | - | CD2 |
| 1245 | 1258 | 1 | ; | HB3 | - | C |
| 1246 | 1250 | 1 | ; | CG | - | HE1 |
| 1246 | 1251 | 1 | ; | CG | - | CZ |
| 1246 | 1255 | 1 | ; | CG | - | HE2 |
| 1246 | 1258 | 1 | ; | CG | - | C |
| 1247 | 1252 | 1 | ; | CD1 | - | OH |
| 1247 | 1254 | 1 | ; | CD1 | - | CE2 |
| 1247 | 1257 | 1 | ; | CD1 | - | HD2 |
| 1248 | 1250 | 1 | ; | HD1 | - | HE1 |
| 1248 | 1251 | 1 | ; | HD1 | - | CZ |
| 1248 | 1256 | 1 | ; | HD1 | - | CD2 |
| 1249 | 1253 | 1 | ; | CE1 | - | HH |
| 1249 | 1255 | 1 | ; | CE1 | - | HE2 |
| 1249 | 1256 | 1 | ; | CE1 | - | CD2 |
| 1250 | 1252 | 1 | ; | HE1 | - | OH |
| 1250 | 1254 | 1 | ; | HE1 | - | CE2 |
| 1251 | 1257 | 1 | ; | CZ | - | HD2 |
| 1252 | 1255 | 1 | ; | OH | - | HE2 |
| 1252 | 1256 | 1 | ; | OH | - | CD2 |
| 1253 | 1254 | 1 | ; | HH | - | CE2 |
| 1255 | 1257 | 1 | ; | HE2 | - | HD2 |
| 1258 | 1263 | 1 | ; | C | - | HA |
| 1258 | 1264 | 1 | ; | C | - | CB |
| 1258 | 1273 | 1 | ; | C | - | C |
| 1259 | 1261 | 1 | ; | O | - | H |
| 1259 | 1262 | 1 | ; | O | - | CA |
| 1260 | 1265 | 1 | ; | N | - | HB2 |
| 1260 | 1266 | 1 | ; | N | - | HB3 |
| 1260 | 1267 | 1 | ; | N | - | CG |
| 1260 | 1274 | 1 | ; | N | - | O |
| 1260 | 1275 | 1 | ; | N | - | N |

```
1261   1263       1 ;       H - HA
1261   1264       1 ;       H - CB
1261   1273       1 ;       H - C
1262   1268       1 ;      CA - HG2
1262   1269       1 ;      CA - HG3
1262   1270       1 ;      CA - CD
1262   1276       1 ;      CA - H
1262   1277       1 ;      CA - CA
1263   1265       1 ;      HA - HB2
1263   1266       1 ;      HA - HB3
1263   1267       1 ;      HA - CG
1263   1274       1 ;      HA - O
1263   1275       1 ;      HA - N
1264   1271       1 ;      CB - OE1
1264   1272       1 ;      CB - OE2
1264   1274       1 ;      CB - O
1264   1275       1 ;      CB - N
1265   1268       1 ;     HB2 - HG2
1265   1269       1 ;     HB2 - HG3
1265   1270       1 ;     HB2 - CD
1265   1273       1 ;     HB2 - C
1266   1268       1 ;     HB3 - HG2
1266   1269       1 ;     HB3 - HG3
1266   1270       1 ;     HB3 - CD
1266   1273       1 ;     HB3 - C
1267   1273       1 ;      CG - C
1268   1271       1 ;     HG2 - OE1
1268   1272       1 ;     HG2 - OE2
1269   1271       1 ;     HG3 - OE1
1269   1272       1 ;     HG3 - OE2
1273   1278       1 ;       C - HA
1273   1279       1 ;       C - CB
1273   1287       1 ;       C - C
1274   1276       1 ;       O - H
1274   1277       1 ;       O - CA
1275   1280       1 ;       N - HB
1275   1281       1 ;       N - CG2
1275   1285       1 ;       N - OG1
1275   1288       1 ;       N - O
1275   1289       1 ;       N - N
1276   1278       1 ;       H - HA
1276   1279       1 ;       H - CB
1276   1287       1 ;       H - C
1277   1282       1 ;      CA - HG21
1277   1283       1 ;      CA - HG22
1277   1284       1 ;      CA - HG23
1277   1286       1 ;      CA - HG1
1277   1290       1 ;      CA - H
1277   1291       1 ;      CA - CA
1278   1280       1 ;      HA - HB
1278   1281       1 ;      HA - CG2
1278   1285       1 ;      HA - OG1
```

```
1278   1288      1 ;      HA - O
1278   1289      1 ;      HA - N
1279   1288      1 ;      CB - O
1279   1289      1 ;      CB - N
1280   1282      1 ;      HB - HG21
1280   1283      1 ;      HB - HG22
1280   1284      1 ;      HB - HG23
1280   1286      1 ;      HB - HG1
1280   1287      1 ;      HB - C
1281   1286      1 ;     CG2 - HG1
1281   1287      1 ;     CG2 - C
1282   1285      1 ;    HG21 - OG1
1283   1285      1 ;    HG22 - OG1
1284   1285      1 ;    HG23 - OG1
1285   1287      1 ;     OG1 - C
1287   1292      1 ;       C - HA
1287   1293      1 ;       C - CB
1287   1308      1 ;       C - C
1288   1290      1 ;       O - H
1288   1291      1 ;       O - CA
1289   1294      1 ;       N - HB2
1289   1295      1 ;       N - HB3
1289   1296      1 ;       N - CG
1289   1309      1 ;       N - O
1289   1310      1 ;       N - N
1290   1292      1 ;       H - HA
1290   1293      1 ;       H - CB
1290   1308      1 ;       H - C
1291   1297      1 ;      CA - CD1
1291   1306      1 ;      CA - CD2
1291   1311      1 ;      CA - H
1291   1312      1 ;      CA - CA
1292   1294      1 ;      HA - HB2
1292   1295      1 ;      HA - HB3
1292   1296      1 ;      HA - CG
1292   1309      1 ;      HA - O
1292   1310      1 ;      HA - N
1293   1298      1 ;      CB - HD1
1293   1299      1 ;      CB - CE1
1293   1304      1 ;      CB - CE2
1293   1307      1 ;      CB - HD2
1293   1309      1 ;      CB - O
1293   1310      1 ;      CB - N
1294   1297      1 ;     HB2 - CD1
1294   1306      1 ;     HB2 - CD2
1294   1308      1 ;     HB2 - C
1295   1297      1 ;     HB3 - CD1
1295   1306      1 ;     HB3 - CD2
1295   1308      1 ;     HB3 - C
1296   1300      1 ;      CG - HE1
1296   1301      1 ;      CG - CZ
1296   1305      1 ;      CG - HE2
```

```
1296   1308       1 ;      CG - C
1297   1302       1 ;     CD1 - OH
1297   1304       1 ;     CD1 - CE2
1297   1307       1 ;     CD1 - HD2
1298   1300       1 ;     HD1 - HE1
1298   1301       1 ;     HD1 - CZ
1298   1306       1 ;     HD1 - CD2
1299   1303       1 ;     CE1 - HH
1299   1305       1 ;     CE1 - HE2
1299   1306       1 ;     CE1 - CD2
1300   1302       1 ;     HE1 - OH
1300   1304       1 ;     HE1 - CE2
1301   1307       1 ;      CZ - HD2
1302   1305       1 ;      OH - HE2
1302   1306       1 ;      OH - CD2
1303   1304       1 ;      HH - CE2
1305   1307       1 ;     HE2 - HD2
1308   1313       1 ;       C - HA
1308   1314       1 ;       C - CB
1308   1323       1 ;       C - C
1309   1311       1 ;       O - H
1309   1312       1 ;       O - CA
1310   1315       1 ;       N - HB2
1310   1316       1 ;       N - HB3
1310   1317       1 ;       N - CG
1310   1324       1 ;       N - O
1310   1325       1 ;       N - N
1311   1313       1 ;       H - HA
1311   1314       1 ;       H - CB
1311   1323       1 ;       H - C
1312   1318       1 ;      CA - HG2
1312   1319       1 ;      CA - HG3
1312   1320       1 ;      CA - CD
1312   1326       1 ;      CA - H
1312   1327       1 ;      CA - CA
1313   1315       1 ;      HA - HB2
1313   1316       1 ;      HA - HB3
1313   1317       1 ;      HA - CG
1313   1324       1 ;      HA - O
1313   1325       1 ;      HA - N
1314   1321       1 ;      CB - OE1
1314   1322       1 ;      CB - OE2
1314   1324       1 ;      CB - O
1314   1325       1 ;      CB - N
1315   1318       1 ;     HB2 - HG2
1315   1319       1 ;     HB2 - HG3
1315   1320       1 ;     HB2 - CD
1315   1323       1 ;     HB2 - C
1316   1318       1 ;     HB3 - HG2
1316   1319       1 ;     HB3 - HG3
1316   1320       1 ;     HB3 - CD
1316   1323       1 ;     HB3 - C
```

| | | | | | |
|---|---|---|---|---|---|
| 1317 | 1323 | 1 ; | CG | - | C |
| 1318 | 1321 | 1 ; | HG2 | - | OE1 |
| 1318 | 1322 | 1 ; | HG2 | - | OE2 |
| 1319 | 1321 | 1 ; | HG3 | - | OE1 |
| 1319 | 1322 | 1 ; | HG3 | - | OE2 |
| 1323 | 1328 | 1 ; | C | - | HA |
| 1323 | 1329 | 1 ; | C | - | CB |
| 1323 | 1337 | 1 ; | C | - | C |
| 1324 | 1326 | 1 ; | O | - | H |
| 1324 | 1327 | 1 ; | O | - | CA |
| 1325 | 1330 | 1 ; | N | - | HB2 |
| 1325 | 1331 | 1 ; | N | - | HB3 |
| 1325 | 1332 | 1 ; | N | - | CG |
| 1325 | 1338 | 1 ; | N | - | O |
| 1325 | 1339 | 1 ; | N | - | N |
| 1326 | 1328 | 1 ; | H | - | HA |
| 1326 | 1329 | 1 ; | H | - | CB |
| 1326 | 1337 | 1 ; | H | - | C |
| 1327 | 1333 | 1 ; | CA | - | OD1 |
| 1327 | 1334 | 1 ; | CA | - | ND2 |
| 1327 | 1340 | 1 ; | CA | - | H |
| 1327 | 1341 | 1 ; | CA | - | CA |
| 1328 | 1330 | 1 ; | HA | - | HB2 |
| 1328 | 1331 | 1 ; | HA | - | HB3 |
| 1328 | 1332 | 1 ; | HA | - | CG |
| 1328 | 1338 | 1 ; | HA | - | O |
| 1328 | 1339 | 1 ; | HA | - | N |
| 1329 | 1335 | 1 ; | CB | - | HD21 |
| 1329 | 1336 | 1 ; | CB | - | HD22 |
| 1329 | 1338 | 1 ; | CB | - | O |
| 1329 | 1339 | 1 ; | CB | - | N |
| 1330 | 1333 | 1 ; | HB2 | - | OD1 |
| 1330 | 1334 | 1 ; | HB2 | - | ND2 |
| 1330 | 1337 | 1 ; | HB2 | - | C |
| 1331 | 1333 | 1 ; | HB3 | - | OD1 |
| 1331 | 1334 | 1 ; | HB3 | - | ND2 |
| 1331 | 1337 | 1 ; | HB3 | - | C |
| 1332 | 1337 | 1 ; | CG | - | C |
| 1333 | 1335 | 1 ; | OD1 | - | HD21 |
| 1333 | 1336 | 1 ; | OD1 | - | HD22 |
| 1337 | 1342 | 1 ; | C | - | HA |
| 1337 | 1343 | 1 ; | C | - | CB |
| 1337 | 1349 | 1 ; | C | - | C |
| 1338 | 1340 | 1 ; | O | - | H |
| 1338 | 1341 | 1 ; | O | - | CA |
| 1339 | 1344 | 1 ; | N | - | HB2 |
| 1339 | 1345 | 1 ; | N | - | HB3 |
| 1339 | 1346 | 1 ; | N | - | CG |
| 1339 | 1350 | 1 ; | N | - | O |
| 1339 | 1351 | 1 ; | N | - | N |
| 1340 | 1342 | 1 ; | H | - | HA |
| 1340 | 1343 | 1 ; | H | - | CB |

```
1340   1349     1 ;    H - C
1341   1347     1 ;   CA - OD1
1341   1348     1 ;   CA - OD2
1341   1352     1 ;   CA - H
1341   1353     1 ;   CA - CA
1342   1344     1 ;   HA - HB2
1342   1345     1 ;   HA - HB3
1342   1346     1 ;   HA - CG
1342   1350     1 ;   HA - O
1342   1351     1 ;   HA - N
1343   1350     1 ;   CB - O
1343   1351     1 ;   CB - N
1344   1347     1 ;  HB2 - OD1
1344   1348     1 ;  HB2 - OD2
1344   1349     1 ;  HB2 - C
1345   1347     1 ;  HB3 - OD1
1345   1348     1 ;  HB3 - OD2
1345   1349     1 ;  HB3 - C
1346   1349     1 ;   CG - C
1349   1354     1 ;    C - HA
1349   1355     1 ;    C - CB
1349   1363     1 ;    C - C
1350   1352     1 ;    O - H
1350   1353     1 ;    O - CA
1351   1356     1 ;    N - HB2
1351   1357     1 ;    N - HB3
1351   1358     1 ;    N - CG
1351   1364     1 ;    N - O
1351   1365     1 ;    N - N
1352   1354     1 ;    H - HA
1352   1355     1 ;    H - CB
1352   1363     1 ;    H - C
1353   1359     1 ;   CA - OD1
1353   1360     1 ;   CA - ND2
1353   1366     1 ;   CA - H
1353   1367     1 ;   CA - CA
1354   1356     1 ;   HA - HB2
1354   1357     1 ;   HA - HB3
1354   1358     1 ;   HA - CG
1354   1364     1 ;   HA - O
1354   1365     1 ;   HA - N
1355   1361     1 ;   CB - HD21
1355   1362     1 ;   CB - HD22
1355   1364     1 ;   CB - O
1355   1365     1 ;   CB - N
1356   1359     1 ;  HB2 - OD1
1356   1360     1 ;  HB2 - ND2
1356   1363     1 ;  HB2 - C
1357   1359     1 ;  HB3 - OD1
1357   1360     1 ;  HB3 - ND2
1357   1363     1 ;  HB3 - C
1358   1363     1 ;   CG - C
```

| | | | | | |
|---|---|---|---|---|---|
| 1359 | 1361 | 1 ; | OD1 | - | HD21 |
| 1359 | 1362 | 1 ; | OD1 | - | HD22 |
| 1363 | 1368 | 1 ; | C | - | HA |
| 1363 | 1369 | 1 ; | C | - | CB |
| 1363 | 1384 | 1 ; | C | - | C |
| 1364 | 1366 | 1 ; | O | - | H |
| 1364 | 1367 | 1 ; | O | - | CA |
| 1365 | 1370 | 1 ; | N | - | HB2 |
| 1365 | 1371 | 1 ; | N | - | HB3 |
| 1365 | 1372 | 1 ; | N | - | CG |
| 1365 | 1385 | 1 ; | N | - | O |
| 1365 | 1386 | 1 ; | N | - | N |
| 1366 | 1368 | 1 ; | H | - | HA |
| 1366 | 1369 | 1 ; | H | - | CB |
| 1366 | 1384 | 1 ; | H | - | C |
| 1367 | 1373 | 1 ; | CA | - | CD1 |
| 1367 | 1382 | 1 ; | CA | - | CD2 |
| 1367 | 1387 | 1 ; | CA | - | H |
| 1367 | 1388 | 1 ; | CA | - | CA |
| 1368 | 1370 | 1 ; | HA | - | HB2 |
| 1368 | 1371 | 1 ; | HA | - | HB3 |
| 1368 | 1372 | 1 ; | HA | - | CG |
| 1368 | 1385 | 1 ; | HA | - | O |
| 1368 | 1386 | 1 ; | HA | - | N |
| 1369 | 1374 | 1 ; | CB | - | HD1 |
| 1369 | 1375 | 1 ; | CB | - | CE1 |
| 1369 | 1380 | 1 ; | CB | - | CE2 |
| 1369 | 1383 | 1 ; | CB | - | HD2 |
| 1369 | 1385 | 1 ; | CB | - | O |
| 1369 | 1386 | 1 ; | CB | - | N |
| 1370 | 1373 | 1 ; | HB2 | - | CD1 |
| 1370 | 1382 | 1 ; | HB2 | - | CD2 |
| 1370 | 1384 | 1 ; | HB2 | - | C |
| 1371 | 1373 | 1 ; | HB3 | - | CD1 |
| 1371 | 1382 | 1 ; | HB3 | - | CD2 |
| 1371 | 1384 | 1 ; | HB3 | - | C |
| 1372 | 1376 | 1 ; | CG | - | HE1 |
| 1372 | 1377 | 1 ; | CG | - | CZ |
| 1372 | 1381 | 1 ; | CG | - | HE2 |
| 1372 | 1384 | 1 ; | CG | - | C |
| 1373 | 1378 | 1 ; | CD1 | - | OH |
| 1373 | 1380 | 1 ; | CD1 | - | CE2 |
| 1373 | 1383 | 1 ; | CD1 | - | HD2 |
| 1374 | 1376 | 1 ; | HD1 | - | HE1 |
| 1374 | 1377 | 1 ; | HD1 | - | CZ |
| 1374 | 1382 | 1 ; | HD1 | - | CD2 |
| 1375 | 1379 | 1 ; | CE1 | - | HH |
| 1375 | 1381 | 1 ; | CE1 | - | HE2 |
| 1375 | 1382 | 1 ; | CE1 | - | CD2 |
| 1376 | 1378 | 1 ; | HE1 | - | OH |
| 1376 | 1380 | 1 ; | HE1 | - | CE2 |
| 1377 | 1383 | 1 ; | CZ | - | HD2 |

```
  1378   1381       1 ;      OH - HE2
  1378   1382       1 ;      OH - CD2
  1379   1380       1 ;      HH - CE2
  1381   1383       1 ;     HE2 - HD2
  1384   1389       1 ;       C - HA
  1384   1390       1 ;       C - CB
  1384   1403       1 ;       C - C
  1385   1387       1 ;       O - H
  1385   1388       1 ;       O - CA
  1386   1391       1 ;       N - HB
  1386   1392       1 ;       N - CG2
  1386   1396       1 ;       N - CG1
  1386   1404       1 ;       N - O
  1386   1405       1 ;       N - N
  1387   1389       1 ;       H - HA
  1387   1390       1 ;       H - CB
  1387   1403       1 ;       H - C
  1388   1393       1 ;      CA - HG21
  1388   1394       1 ;      CA - HG22
  1388   1395       1 ;      CA - HG23
  1388   1397       1 ;      CA - HG12
  1388   1398       1 ;      CA - HG13
  1388   1399       1 ;      CA - CD1
  1388   1406       1 ;      CA - H
  1388   1407       1 ;      CA - CA
  1389   1391       1 ;      HA - HB
  1389   1392       1 ;      HA - CG2
  1389   1396       1 ;      HA - CG1
  1389   1404       1 ;      HA - O
  1389   1405       1 ;      HA - N
  1390   1400       1 ;      CB - HD11
  1390   1401       1 ;      CB - HD12
  1390   1402       1 ;      CB - HD13
  1390   1404       1 ;      CB - O
  1390   1405       1 ;      CB - N
  1391   1393       1 ;      HB - HG21
  1391   1394       1 ;      HB - HG22
  1391   1395       1 ;      HB - HG23
  1391   1397       1 ;      HB - HG12
  1391   1398       1 ;      HB - HG13
  1391   1399       1 ;      HB - CD1
  1391   1403       1 ;      HB - C
  1392   1397       1 ;     CG2 - HG12
  1392   1398       1 ;     CG2 - HG13
  1392   1399       1 ;     CG2 - CD1
  1392   1403       1 ;     CG2 - C
  1393   1396       1 ;    HG21 - CG1
  1394   1396       1 ;    HG22 - CG1
  1395   1396       1 ;    HG23 - CG1
  1396   1403       1 ;     CG1 - C
  1397   1400       1 ;    HG12 - HD11
  1397   1401       1 ;    HG12 - HD12
```

```
1397   1402       1 ;   HG12 - HD13
1398   1400       1 ;   HG13 - HD11
1398   1401       1 ;   HG13 - HD12
1398   1402       1 ;   HG13 - HD13
1403   1408       1 ;      C - HA
1403   1409       1 ;      C - CB
1403   1424       1 ;      C - C
1404   1406       1 ;      O - H
1404   1407       1 ;      O - CA
1405   1410       1 ;      N - HB2
1405   1411       1 ;      N - HB3
1405   1412       1 ;      N - CG
1405   1425       1 ;      N - O
1405   1426       1 ;      N - N
1406   1408       1 ;      H - HA
1406   1409       1 ;      H - CB
1406   1424       1 ;      H - C
1407   1413       1 ;     CA - CD1
1407   1422       1 ;     CA - CD2
1407   1427       1 ;     CA - H
1407   1428       1 ;     CA - CA
1408   1410       1 ;     HA - HB2
1408   1411       1 ;     HA - HB3
1408   1412       1 ;     HA - CG
1408   1425       1 ;     HA - O
1408   1426       1 ;     HA - N
1409   1414       1 ;     CB - HD1
1409   1415       1 ;     CB - CE1
1409   1420       1 ;     CB - CE2
1409   1423       1 ;     CB - HD2
1409   1425       1 ;     CB - O
1409   1426       1 ;     CB - N
1410   1413       1 ;    HB2 - CD1
1410   1422       1 ;    HB2 - CD2
1410   1424       1 ;    HB2 - C
1411   1413       1 ;    HB3 - CD1
1411   1422       1 ;    HB3 - CD2
1411   1424       1 ;    HB3 - C
1412   1416       1 ;     CG - HE1
1412   1417       1 ;     CG - CZ
1412   1421       1 ;     CG - HE2
1412   1424       1 ;     CG - C
1413   1418       1 ;    CD1 - OH
1413   1420       1 ;    CD1 - CE2
1413   1423       1 ;    CD1 - HD2
1414   1416       1 ;    HD1 - HE1
1414   1417       1 ;    HD1 - CZ
1414   1422       1 ;    HD1 - CD2
1415   1419       1 ;    CE1 - HH
1415   1421       1 ;    CE1 - HE2
1415   1422       1 ;    CE1 - CD2
1416   1418       1 ;    HE1 - OH
```

| | | | | | |
|---|---|---|---|---|---|
| 1416 | 1420 | 1 ; | HE1 | - | CE2 |
| 1417 | 1423 | 1 ; | CZ | - | HD2 |
| 1418 | 1421 | 1 ; | OH | - | HE2 |
| 1418 | 1422 | 1 ; | OH | - | CD2 |
| 1419 | 1420 | 1 ; | HH | - | CE2 |
| 1421 | 1423 | 1 ; | HE2 | - | HD2 |
| 1424 | 1429 | 1 ; | C | - | HA |
| 1424 | 1430 | 1 ; | C | - | CB |
| 1424 | 1443 | 1 ; | C | - | C |
| 1425 | 1427 | 1 ; | O | - | H |
| 1425 | 1428 | 1 ; | O | - | CA |
| 1426 | 1431 | 1 ; | N | - | HB2 |
| 1426 | 1432 | 1 ; | N | - | HB3 |
| 1426 | 1433 | 1 ; | N | - | CG |
| 1426 | 1444 | 1 ; | N | - | O |
| 1426 | 1445 | 1 ; | N | - | N |
| 1427 | 1429 | 1 ; | H | - | HA |
| 1427 | 1430 | 1 ; | H | - | CB |
| 1427 | 1443 | 1 ; | H | - | C |
| 1428 | 1434 | 1 ; | CA | - | HG |
| 1428 | 1435 | 1 ; | CA | - | CD1 |
| 1428 | 1439 | 1 ; | CA | - | CD2 |
| 1428 | 1446 | 1 ; | CA | - | H |
| 1428 | 1447 | 1 ; | CA | - | CA |
| 1429 | 1431 | 1 ; | HA | - | HB2 |
| 1429 | 1432 | 1 ; | HA | - | HB3 |
| 1429 | 1433 | 1 ; | HA | - | CG |
| 1429 | 1444 | 1 ; | HA | - | O |
| 1429 | 1445 | 1 ; | HA | - | N |
| 1430 | 1436 | 1 ; | CB | - | HD11 |
| 1430 | 1437 | 1 ; | CB | - | HD12 |
| 1430 | 1438 | 1 ; | CB | - | HD13 |
| 1430 | 1440 | 1 ; | CB | - | HD21 |
| 1430 | 1441 | 1 ; | CB | - | HD22 |
| 1430 | 1442 | 1 ; | CB | - | HD23 |
| 1430 | 1444 | 1 ; | CB | - | O |
| 1430 | 1445 | 1 ; | CB | - | N |
| 1431 | 1434 | 1 ; | HB2 | - | HG |
| 1431 | 1435 | 1 ; | HB2 | - | CD1 |
| 1431 | 1439 | 1 ; | HB2 | - | CD2 |
| 1431 | 1443 | 1 ; | HB2 | - | C |
| 1432 | 1434 | 1 ; | HB3 | - | HG |
| 1432 | 1435 | 1 ; | HB3 | - | CD1 |
| 1432 | 1439 | 1 ; | HB3 | - | CD2 |
| 1432 | 1443 | 1 ; | HB3 | - | C |
| 1433 | 1443 | 1 ; | CG | - | C |
| 1434 | 1436 | 1 ; | HG | - | HD11 |
| 1434 | 1437 | 1 ; | HG | - | HD12 |
| 1434 | 1438 | 1 ; | HG | - | HD13 |
| 1434 | 1440 | 1 ; | HG | - | HD21 |
| 1434 | 1441 | 1 ; | HG | - | HD22 |
| 1434 | 1442 | 1 ; | HG | - | HD23 |

```
1435   1440       1 ;     CD1 - HD21
1435   1441       1 ;     CD1 - HD22
1435   1442       1 ;     CD1 - HD23
1436   1439       1 ;    HD11 - CD2
1437   1439       1 ;    HD12 - CD2
1438   1439       1 ;    HD13 - CD2
1443   1448       1 ;       C - HA
1443   1449       1 ;       C - CB
1443   1462       1 ;       C - C
1444   1446       1 ;       O - H
1444   1447       1 ;       O - CA
1445   1450       1 ;       N - HB
1445   1451       1 ;       N - CG2
1445   1455       1 ;       N - CG1
1445   1463       1 ;       N - O
1445   1464       1 ;       N - N
1446   1448       1 ;       H - HA
1446   1449       1 ;       H - CB
1446   1462       1 ;       H - C
1447   1452       1 ;      CA - HG21
1447   1453       1 ;      CA - HG22
1447   1454       1 ;      CA - HG23
1447   1456       1 ;      CA - HG12
1447   1457       1 ;      CA - HG13
1447   1458       1 ;      CA - CD1
1447   1465       1 ;      CA - H
1447   1466       1 ;      CA - CA
1448   1450       1 ;      HA - HB
1448   1451       1 ;      HA - CG2
1448   1455       1 ;      HA - CG1
1448   1463       1 ;      HA - O
1448   1464       1 ;      HA - N
1449   1459       1 ;      CB - HD11
1449   1460       1 ;      CB - HD12
1449   1461       1 ;      CB - HD13
1449   1463       1 ;      CB - O
1449   1464       1 ;      CB - N
1450   1452       1 ;      HB - HG21
1450   1453       1 ;      HB - HG22
1450   1454       1 ;      HB - HG23
1450   1456       1 ;      HB - HG12
1450   1457       1 ;      HB - HG13
1450   1458       1 ;      HB - CD1
1450   1462       1 ;      HB - C
1451   1456       1 ;     CG2 - HG12
1451   1457       1 ;     CG2 - HG13
1451   1458       1 ;     CG2 - CD1
1451   1462       1 ;     CG2 - C
1452   1455       1 ;    HG21 - CG1
1453   1455       1 ;    HG22 - CG1
1454   1455       1 ;    HG23 - CG1
1455   1462       1 ;     CG1 - C
```

```
 1456   1459      1 ;   HG12 - HD11
 1456   1460      1 ;   HG12 - HD12
 1456   1461      1 ;   HG12 - HD13
 1457   1459      1 ;   HG13 - HD11
 1457   1460      1 ;   HG13 - HD12
 1457   1461      1 ;   HG13 - HD13
 1462   1467      1 ;      C - HA
 1462   1468      1 ;      C - CB
 1462   1479      1 ;      C - C
 1463   1465      1 ;      O - H
 1463   1466      1 ;      O - CA
 1464   1469      1 ;      N - HB2
 1464   1470      1 ;      N - HB3
 1464   1471      1 ;      N - CG
 1464   1480      1 ;      N - O
 1464   1481      1 ;      N - N
 1465   1467      1 ;      H - HA
 1465   1468      1 ;      H - CB
 1465   1479      1 ;      H - C
 1466   1472      1 ;     CA - HG2
 1466   1473      1 ;     CA - HG3
 1466   1474      1 ;     CA - SD
 1466   1482      1 ;     CA - H
 1466   1483      1 ;     CA - CA
 1467   1469      1 ;     HA - HB2
 1467   1470      1 ;     HA - HB3
 1467   1471      1 ;     HA - CG
 1467   1480      1 ;     HA - O
 1467   1481      1 ;     HA - N
 1468   1475      1 ;     CB - CE
 1468   1480      1 ;     CB - O
 1468   1481      1 ;     CB - N
 1469   1472      1 ;    HB2 - HG2
 1469   1473      1 ;    HB2 - HG3
 1469   1474      1 ;    HB2 - SD
 1469   1479      1 ;    HB2 - C
 1470   1472      1 ;    HB3 - HG2
 1470   1473      1 ;    HB3 - HG3
 1470   1474      1 ;    HB3 - SD
 1470   1479      1 ;    HB3 - C
 1471   1476      1 ;     CG - HE1
 1471   1477      1 ;     CG - HE2
 1471   1478      1 ;     CG - HE3
 1471   1479      1 ;     CG - C
 1472   1475      1 ;    HG2 - CE
 1473   1475      1 ;    HG3 - CE
 1479   1484      1 ;      C - HA
 1479   1485      1 ;      C - CB
 1479   1494      1 ;      C - C
 1480   1482      1 ;      O - H
 1480   1483      1 ;      O - CA
 1481   1486      1 ;      N - HB2
```

| | | | | | | |
|---|---|---|---|---|---|---|
| 1481 | 1487 | 1 | ; | N | - | HB3 |
| 1481 | 1488 | 1 | ; | N | - | CG |
| 1481 | 1495 | 1 | ; | N | - | O |
| 1481 | 1496 | 1 | ; | N | - | N |
| 1482 | 1484 | 1 | ; | H | - | HA |
| 1482 | 1485 | 1 | ; | H | - | CB |
| 1482 | 1494 | 1 | ; | H | - | C |
| 1483 | 1489 | 1 | ; | CA | - | HG2 |
| 1483 | 1490 | 1 | ; | CA | - | HG3 |
| 1483 | 1491 | 1 | ; | CA | - | CD |
| 1483 | 1497 | 1 | ; | CA | - | H |
| 1483 | 1498 | 1 | ; | CA | - | CA |
| 1484 | 1486 | 1 | ; | HA | - | HB2 |
| 1484 | 1487 | 1 | ; | HA | - | HB3 |
| 1484 | 1488 | 1 | ; | HA | - | CG |
| 1484 | 1495 | 1 | ; | HA | - | O |
| 1484 | 1496 | 1 | ; | HA | - | N |
| 1485 | 1492 | 1 | ; | CB | - | OE1 |
| 1485 | 1493 | 1 | ; | CB | - | OE2 |
| 1485 | 1495 | 1 | ; | CB | - | O |
| 1485 | 1496 | 1 | ; | CB | - | N |
| 1486 | 1489 | 1 | ; | HB2 | - | HG2 |
| 1486 | 1490 | 1 | ; | HB2 | - | HG3 |
| 1486 | 1491 | 1 | ; | HB2 | - | CD |
| 1486 | 1494 | 1 | ; | HB2 | - | C |
| 1487 | 1489 | 1 | ; | HB3 | - | HG2 |
| 1487 | 1490 | 1 | ; | HB3 | - | HG3 |
| 1487 | 1491 | 1 | ; | HB3 | - | CD |
| 1487 | 1494 | 1 | ; | HB3 | - | C |
| 1488 | 1494 | 1 | ; | CG | - | C |
| 1489 | 1492 | 1 | ; | HG2 | - | OE1 |
| 1489 | 1493 | 1 | ; | HG2 | - | OE2 |
| 1490 | 1492 | 1 | ; | HG3 | - | OE1 |
| 1490 | 1493 | 1 | ; | HG3 | - | OE2 |
| 1494 | 1499 | 1 | ; | C | - | HA |
| 1494 | 1500 | 1 | ; | C | - | CB |
| 1494 | 1513 | 1 | ; | C | - | C |
| 1495 | 1497 | 1 | ; | O | - | H |
| 1495 | 1498 | 1 | ; | O | - | CA |
| 1496 | 1501 | 1 | ; | N | - | HB2 |
| 1496 | 1502 | 1 | ; | N | - | HB3 |
| 1496 | 1503 | 1 | ; | N | - | CG |
| 1496 | 1514 | 1 | ; | N | - | O |
| 1496 | 1515 | 1 | ; | N | - | N |
| 1497 | 1499 | 1 | ; | H | - | HA |
| 1497 | 1500 | 1 | ; | H | - | CB |
| 1497 | 1513 | 1 | ; | H | - | C |
| 1498 | 1504 | 1 | ; | CA | - | HG |
| 1498 | 1505 | 1 | ; | CA | - | CD1 |
| 1498 | 1509 | 1 | ; | CA | - | CD2 |
| 1498 | 1516 | 1 | ; | CA | - | H |
| 1498 | 1517 | 1 | ; | CA | - | CA |

```
1499   1501      1 ;      HA - HB2
1499   1502      1 ;      HA - HB3
1499   1503      1 ;      HA - CG
1499   1514      1 ;      HA - O
1499   1515      1 ;      HA - N
1500   1506      1 ;      CB - HD11
1500   1507      1 ;      CB - HD12
1500   1508      1 ;      CB - HD13
1500   1510      1 ;      CB - HD21
1500   1511      1 ;      CB - HD22
1500   1512      1 ;      CB - HD23
1500   1514      1 ;      CB - O
1500   1515      1 ;      CB - N
1501   1504      1 ;     HB2 - HG
1501   1505      1 ;     HB2 - CD1
1501   1509      1 ;     HB2 - CD2
1501   1513      1 ;     HB2 - C
1502   1504      1 ;     HB3 - HG
1502   1505      1 ;     HB3 - CD1
1502   1509      1 ;     HB3 - CD2
1502   1513      1 ;     HB3 - C
1503   1513      1 ;      CG - C
1504   1506      1 ;      HG - HD11
1504   1507      1 ;      HG - HD12
1504   1508      1 ;      HG - HD13
1504   1510      1 ;      HG - HD21
1504   1511      1 ;      HG - HD22
1504   1512      1 ;      HG - HD23
1505   1510      1 ;     CD1 - HD21
1505   1511      1 ;     CD1 - HD22
1505   1512      1 ;     CD1 - HD23
1506   1509      1 ;    HD11 - CD2
1507   1509      1 ;    HD12 - CD2
1508   1509      1 ;    HD13 - CD2
1513   1518      1 ;       C - HA
1513   1519      1 ;       C - CB
1513   1524      1 ;       C - C
1514   1516      1 ;       O - H
1514   1517      1 ;       O - CA
1515   1520      1 ;       N - HB2
1515   1521      1 ;       N - HB3
1515   1522      1 ;       N - SG
1515   1525      1 ;       N - O
1515   1526      1 ;       N - N
1516   1518      1 ;       H - HA
1516   1519      1 ;       H - CB
1516   1524      1 ;       H - C
1517   1523      1 ;      CA - HG
1517   1527      1 ;      CA - H
1517   1528      1 ;      CA - CA
1518   1520      1 ;      HA - HB2
1518   1521      1 ;      HA - HB3
```

```
1518   1522       1 ;      HA - SG
1518   1525       1 ;      HA - O
1518   1526       1 ;      HA - N
1519   1525       1 ;      CB - O
1519   1526       1 ;      CB - N
1520   1523       1 ;     HB2 - HG
1520   1524       1 ;     HB2 - C
1521   1523       1 ;     HB3 - HG
1521   1524       1 ;     HB3 - C
1522   1524       1 ;      SG - C
1524   1529       1 ;       C - HA
1524   1530       1 ;       C - CB
1524   1535       1 ;       C - C
1525   1527       1 ;       O - H
1525   1528       1 ;       O - CA
1526   1531       1 ;       N - HB2
1526   1532       1 ;       N - HB3
1526   1533       1 ;       N - OG
1526   1536       1 ;       N - O
1526   1537       1 ;       N - N
1527   1529       1 ;       H - HA
1527   1530       1 ;       H - CB
1527   1535       1 ;       H - C
1528   1534       1 ;      CA - HG
1528   1538       1 ;      CA - H
1528   1539       1 ;      CA - CA
1529   1531       1 ;      HA - HB2
1529   1532       1 ;      HA - HB3
1529   1533       1 ;      HA - OG
1529   1536       1 ;      HA - O
1529   1537       1 ;      HA - N
1530   1536       1 ;      CB - O
1530   1537       1 ;      CB - N
1531   1534       1 ;     HB2 - HG
1531   1535       1 ;     HB2 - C
1532   1534       1 ;     HB3 - HG
1532   1535       1 ;     HB3 - C
1533   1535       1 ;      OG - C
1535   1540       1 ;       C - HA2
1535   1541       1 ;       C - HA3
1535   1542       1 ;       C - C
1536   1538       1 ;       O - H
1536   1539       1 ;       O - CA
1537   1543       1 ;       N - O
1537   1544       1 ;       N - N
1538   1540       1 ;       H - HA2
1538   1541       1 ;       H - HA3
1538   1542       1 ;       H - C
1539   1545       1 ;      CA - H
1539   1546       1 ;      CA - CA
1540   1543       1 ;     HA2 - O
1540   1544       1 ;     HA2 - N
```

```
1541   1543      1 ;    HA3 - O
1541   1544      1 ;    HA3 - N
1542   1547      1 ;      C - HA
1542   1548      1 ;      C - CB
1542   1566      1 ;      C - C
1543   1545      1 ;      O - H
1543   1546      1 ;      O - CA
1544   1549      1 ;      N - HB2
1544   1550      1 ;      N - HB3
1544   1551      1 ;      N - CG
1544   1567      1 ;      N - O
1544   1568      1 ;      N - N
1545   1547      1 ;      H - HA
1545   1548      1 ;      H - CB
1545   1566      1 ;      H - C
1546   1552      1 ;     CA - HG2
1546   1553      1 ;     CA - HG3
1546   1554      1 ;     CA - CD
1546   1569      1 ;     CA - H
1546   1570      1 ;     CA - CA
1547   1549      1 ;     HA - HB2
1547   1550      1 ;     HA - HB3
1547   1551      1 ;     HA - CG
1547   1567      1 ;     HA - O
1547   1568      1 ;     HA - N
1548   1555      1 ;     CB - HD2
1548   1556      1 ;     CB - HD3
1548   1557      1 ;     CB - NE
1548   1567      1 ;     CB - O
1548   1568      1 ;     CB - N
1549   1552      1 ;    HB2 - HG2
1549   1553      1 ;    HB2 - HG3
1549   1554      1 ;    HB2 - CD
1549   1566      1 ;    HB2 - C
1550   1552      1 ;    HB3 - HG2
1550   1553      1 ;    HB3 - HG3
1550   1554      1 ;    HB3 - CD
1550   1566      1 ;    HB3 - C
1551   1558      1 ;     CG - HE
1551   1559      1 ;     CG - CZ
1551   1566      1 ;     CG - C
1552   1555      1 ;    HG2 - HD2
1552   1556      1 ;    HG2 - HD3
1552   1557      1 ;    HG2 - NE
1553   1555      1 ;    HG3 - HD2
1553   1556      1 ;    HG3 - HD3
1553   1557      1 ;    HG3 - NE
1554   1560      1 ;     CD - NH1
1554   1563      1 ;     CD - NH2
1555   1558      1 ;    HD2 - HE
1555   1559      1 ;    HD2 - CZ
1556   1558      1 ;    HD3 - HE
```

```
1556   1559      1 ;    HD3 - CZ
1557   1561      1 ;     NE - HH11
1557   1562      1 ;     NE - HH12
1557   1564      1 ;     NE - HH21
1557   1565      1 ;     NE - HH22
1558   1560      1 ;     HE - NH1
1558   1563      1 ;     HE - NH2
1560   1564      1 ;    NH1 - HH21
1560   1565      1 ;    NH1 - HH22
1561   1563      1 ;   HH11 - NH2
1562   1563      1 ;   HH12 - NH2
1566   1571      1 ;      C - HA
1566   1572      1 ;      C - CB
1566   1581      1 ;      C - C
1567   1569      1 ;      O - H
1567   1570      1 ;      O - CA
1568   1573      1 ;      N - HB2
1568   1574      1 ;      N - HB3
1568   1575      1 ;      N - CG
1568   1582      1 ;      N - O
1568   1583      1 ;      N - N
1569   1571      1 ;      H - HA
1569   1572      1 ;      H - CB
1569   1581      1 ;      H - C
1570   1576      1 ;     CA - HG2
1570   1577      1 ;     CA - HG3
1570   1578      1 ;     CA - CD
1570   1584      1 ;     CA - H
1570   1585      1 ;     CA - CA
1571   1573      1 ;     HA - HB2
1571   1574      1 ;     HA - HB3
1571   1575      1 ;     HA - CG
1571   1582      1 ;     HA - O
1571   1583      1 ;     HA - N
1572   1579      1 ;     CB - OE1
1572   1580      1 ;     CB - OE2
1572   1582      1 ;     CB - O
1572   1583      1 ;     CB - N
1573   1576      1 ;    HB2 - HG2
1573   1577      1 ;    HB2 - HG3
1573   1578      1 ;    HB2 - CD
1573   1581      1 ;    HB2 - C
1574   1576      1 ;    HB3 - HG2
1574   1577      1 ;    HB3 - HG3
1574   1578      1 ;    HB3 - CD
1574   1581      1 ;    HB3 - C
1575   1581      1 ;     CG - C
1576   1579      1 ;    HG2 - OE1
1576   1580      1 ;    HG2 - OE2
1577   1579      1 ;    HG3 - OE1
1577   1580      1 ;    HG3 - OE2
1581   1586      1 ;      C - HA
```

```
1581   1587       1 ;       C - CB
1581   1600       1 ;       C - C
1582   1584       1 ;       O - H
1582   1585       1 ;       O - CA
1583   1588       1 ;       N - HB2
1583   1589       1 ;       N - HB3
1583   1590       1 ;       N - CG
1583   1601       1 ;       N - O
1583   1602       1 ;       N - N
1584   1586       1 ;       H - HA
1584   1587       1 ;       H - CB
1584   1600       1 ;       H - C
1585   1591       1 ;      CA - HG
1585   1592       1 ;      CA - CD1
1585   1596       1 ;      CA - CD2
1585   1603       1 ;      CA - H
1585   1604       1 ;      CA - CA
1586   1588       1 ;      HA - HB2
1586   1589       1 ;      HA - HB3
1586   1590       1 ;      HA - CG
1586   1601       1 ;      HA - O
1586   1602       1 ;      HA - N
1587   1593       1 ;      CB - HD11
1587   1594       1 ;      CB - HD12
1587   1595       1 ;      CB - HD13
1587   1597       1 ;      CB - HD21
1587   1598       1 ;      CB - HD22
1587   1599       1 ;      CB - HD23
1587   1601       1 ;      CB - O
1587   1602       1 ;      CB - N
1588   1591       1 ;     HB2 - HG
1588   1592       1 ;     HB2 - CD1
1588   1596       1 ;     HB2 - CD2
1588   1600       1 ;     HB2 - C
1589   1591       1 ;     HB3 - HG
1589   1592       1 ;     HB3 - CD1
1589   1596       1 ;     HB3 - CD2
1589   1600       1 ;     HB3 - C
1590   1600       1 ;      CG - C
1591   1593       1 ;      HG - HD11
1591   1594       1 ;      HG - HD12
1591   1595       1 ;      HG - HD13
1591   1597       1 ;      HG - HD21
1591   1598       1 ;      HG - HD22
1591   1599       1 ;      HG - HD23
1592   1597       1 ;     CD1 - HD21
1592   1598       1 ;     CD1 - HD22
1592   1599       1 ;     CD1 - HD23
1593   1596       1 ;    HD11 - CD2
1594   1596       1 ;    HD12 - CD2
1595   1596       1 ;    HD13 - CD2
1600   1605       1 ;       C - HA
```

| | | | | | |
|---|---|---|---|---|---|
| 1600 | 1606 | 1 ; | C | - | CB |
| 1600 | 1620 | 1 ; | C | - | C |
| 1601 | 1603 | 1 ; | O | - | H |
| 1601 | 1604 | 1 ; | O | - | CA |
| 1602 | 1607 | 1 ; | N | - | HB2 |
| 1602 | 1608 | 1 ; | N | - | HB3 |
| 1602 | 1609 | 1 ; | N | - | CG |
| 1602 | 1621 | 1 ; | N | - | O |
| 1602 | 1622 | 1 ; | N | - | N |
| 1603 | 1605 | 1 ; | H | - | HA |
| 1603 | 1606 | 1 ; | H | - | CB |
| 1603 | 1620 | 1 ; | H | - | C |
| 1604 | 1610 | 1 ; | CA | - | CD1 |
| 1604 | 1618 | 1 ; | CA | - | CD2 |
| 1604 | 1623 | 1 ; | CA | - | H |
| 1604 | 1624 | 1 ; | CA | - | CA |
| 1605 | 1607 | 1 ; | HA | - | HB2 |
| 1605 | 1608 | 1 ; | HA | - | HB3 |
| 1605 | 1609 | 1 ; | HA | - | CG |
| 1605 | 1621 | 1 ; | HA | - | O |
| 1605 | 1622 | 1 ; | HA | - | N |
| 1606 | 1611 | 1 ; | CB | - | HD1 |
| 1606 | 1612 | 1 ; | CB | - | CE1 |
| 1606 | 1616 | 1 ; | CB | - | CE2 |
| 1606 | 1619 | 1 ; | CB | - | HD2 |
| 1606 | 1621 | 1 ; | CB | - | O |
| 1606 | 1622 | 1 ; | CB | - | N |
| 1607 | 1610 | 1 ; | HB2 | - | CD1 |
| 1607 | 1618 | 1 ; | HB2 | - | CD2 |
| 1607 | 1620 | 1 ; | HB2 | - | C |
| 1608 | 1610 | 1 ; | HB3 | - | CD1 |
| 1608 | 1618 | 1 ; | HB3 | - | CD2 |
| 1608 | 1620 | 1 ; | HB3 | - | C |
| 1609 | 1613 | 1 ; | CG | - | HE1 |
| 1609 | 1614 | 1 ; | CG | - | CZ |
| 1609 | 1617 | 1 ; | CG | - | HE2 |
| 1609 | 1620 | 1 ; | CG | - | C |
| 1610 | 1615 | 1 ; | CD1 | - | HZ |
| 1610 | 1616 | 1 ; | CD1 | - | CE2 |
| 1610 | 1619 | 1 ; | CD1 | - | HD2 |
| 1611 | 1613 | 1 ; | HD1 | - | HE1 |
| 1611 | 1614 | 1 ; | HD1 | - | CZ |
| 1611 | 1618 | 1 ; | HD1 | - | CD2 |
| 1612 | 1617 | 1 ; | CE1 | - | HE2 |
| 1612 | 1618 | 1 ; | CE1 | - | CD2 |
| 1613 | 1615 | 1 ; | HE1 | - | HZ |
| 1613 | 1616 | 1 ; | HE1 | - | CE2 |
| 1614 | 1619 | 1 ; | CZ | - | HD2 |
| 1615 | 1617 | 1 ; | HZ | - | HE2 |
| 1615 | 1618 | 1 ; | HZ | - | CD2 |
| 1617 | 1619 | 1 ; | HE2 | - | HD2 |
| 1620 | 1625 | 1 ; | C | - | HA |

| | | | | | | |
|---|---|---|---|---|---|---|
| 1620 | 1626 | 1 | ; | C | - | CB |
| 1620 | 1632 | 1 | ; | C | - | C |
| 1621 | 1623 | 1 | ; | O | - | H |
| 1621 | 1624 | 1 | ; | O | - | CA |
| 1622 | 1627 | 1 | ; | N | - | HB2 |
| 1622 | 1628 | 1 | ; | N | - | HB3 |
| 1622 | 1629 | 1 | ; | N | - | CG |
| 1622 | 1633 | 1 | ; | N | - | O |
| 1622 | 1634 | 1 | ; | N | - | N |
| 1623 | 1625 | 1 | ; | H | - | HA |
| 1623 | 1626 | 1 | ; | H | - | CB |
| 1623 | 1632 | 1 | ; | H | - | C |
| 1624 | 1630 | 1 | ; | CA | - | OD1 |
| 1624 | 1631 | 1 | ; | CA | - | OD2 |
| 1624 | 1635 | 1 | ; | CA | - | H |
| 1624 | 1636 | 1 | ; | CA | - | CA |
| 1625 | 1627 | 1 | ; | HA | - | HB2 |
| 1625 | 1628 | 1 | ; | HA | - | HB3 |
| 1625 | 1629 | 1 | ; | HA | - | CG |
| 1625 | 1633 | 1 | ; | HA | - | O |
| 1625 | 1634 | 1 | ; | HA | - | N |
| 1626 | 1633 | 1 | ; | CB | - | O |
| 1626 | 1634 | 1 | ; | CB | - | N |
| 1627 | 1630 | 1 | ; | HB2 | - | OD1 |
| 1627 | 1631 | 1 | ; | HB2 | - | OD2 |
| 1627 | 1632 | 1 | ; | HB2 | - | C |
| 1628 | 1630 | 1 | ; | HB3 | - | OD1 |
| 1628 | 1631 | 1 | ; | HB3 | - | OD2 |
| 1628 | 1632 | 1 | ; | HB3 | - | C |
| 1629 | 1632 | 1 | ; | CG | - | C |
| 1632 | 1637 | 1 | ; | C | - | HA |
| 1632 | 1638 | 1 | ; | C | - | CB |
| 1632 | 1643 | 1 | ; | C | - | C |
| 1633 | 1635 | 1 | ; | O | - | H |
| 1633 | 1636 | 1 | ; | O | - | CA |
| 1634 | 1639 | 1 | ; | N | - | HB2 |
| 1634 | 1640 | 1 | ; | N | - | HB3 |
| 1634 | 1641 | 1 | ; | N | - | OG |
| 1634 | 1644 | 1 | ; | N | - | O |
| 1634 | 1645 | 1 | ; | N | - | N |
| 1635 | 1637 | 1 | ; | H | - | HA |
| 1635 | 1638 | 1 | ; | H | - | CB |
| 1635 | 1643 | 1 | ; | H | - | C |
| 1636 | 1642 | 1 | ; | CA | - | HG |
| 1636 | 1646 | 1 | ; | CA | - | H |
| 1636 | 1647 | 1 | ; | CA | - | CA |
| 1637 | 1639 | 1 | ; | HA | - | HB2 |
| 1637 | 1640 | 1 | ; | HA | - | HB3 |
| 1637 | 1641 | 1 | ; | HA | - | OG |
| 1637 | 1644 | 1 | ; | HA | - | O |
| 1637 | 1645 | 1 | ; | HA | - | N |
| 1638 | 1644 | 1 | ; | CB | - | O |

| | | | |
|---|---|---|---|
| 1638 | 1645 | 1 ; | CB - N |
| 1639 | 1642 | 1 ; | HB2 - HG |
| 1639 | 1643 | 1 ; | HB2 - C |
| 1640 | 1642 | 1 ; | HB3 - HG |
| 1640 | 1643 | 1 ; | HB3 - C |
| 1641 | 1643 | 1 ; | OG - C |
| 1643 | 1648 | 1 ; | C - HA |
| 1643 | 1649 | 1 ; | C - CB |
| 1643 | 1662 | 1 ; | C - C |
| 1644 | 1646 | 1 ; | O - H |
| 1644 | 1647 | 1 ; | O - CA |
| 1645 | 1650 | 1 ; | N - HB |
| 1645 | 1651 | 1 ; | N - CG2 |
| 1645 | 1655 | 1 ; | N - CG1 |
| 1645 | 1663 | 1 ; | N - O |
| 1645 | 1664 | 1 ; | N - N |
| 1646 | 1648 | 1 ; | H - HA |
| 1646 | 1649 | 1 ; | H - CB |
| 1646 | 1662 | 1 ; | H - C |
| 1647 | 1652 | 1 ; | CA - HG21 |
| 1647 | 1653 | 1 ; | CA - HG22 |
| 1647 | 1654 | 1 ; | CA - HG23 |
| 1647 | 1656 | 1 ; | CA - HG12 |
| 1647 | 1657 | 1 ; | CA - HG13 |
| 1647 | 1658 | 1 ; | CA - CD1 |
| 1647 | 1665 | 1 ; | CA - H |
| 1647 | 1666 | 1 ; | CA - CA |
| 1648 | 1650 | 1 ; | HA - HB |
| 1648 | 1651 | 1 ; | HA - CG2 |
| 1648 | 1655 | 1 ; | HA - CG1 |
| 1648 | 1663 | 1 ; | HA - O |
| 1648 | 1664 | 1 ; | HA - N |
| 1649 | 1659 | 1 ; | CB - HD11 |
| 1649 | 1660 | 1 ; | CB - HD12 |
| 1649 | 1661 | 1 ; | CB - HD13 |
| 1649 | 1663 | 1 ; | CB - O |
| 1649 | 1664 | 1 ; | CB - N |
| 1650 | 1652 | 1 ; | HB - HG21 |
| 1650 | 1653 | 1 ; | HB - HG22 |
| 1650 | 1654 | 1 ; | HB - HG23 |
| 1650 | 1656 | 1 ; | HB - HG12 |
| 1650 | 1657 | 1 ; | HB - HG13 |
| 1650 | 1658 | 1 ; | HB - CD1 |
| 1650 | 1662 | 1 ; | HB - C |
| 1651 | 1656 | 1 ; | CG2 - HG12 |
| 1651 | 1657 | 1 ; | CG2 - HG13 |
| 1651 | 1658 | 1 ; | CG2 - CD1 |
| 1651 | 1662 | 1 ; | CG2 - C |
| 1652 | 1655 | 1 ; | HG21 - CG1 |
| 1653 | 1655 | 1 ; | HG22 - CG1 |
| 1654 | 1655 | 1 ; | HG23 - CG1 |
| 1655 | 1662 | 1 ; | CG1 - C |

```
1656   1659      1 ;   HG12 - HD11
1656   1660      1 ;   HG12 - HD12
1656   1661      1 ;   HG12 - HD13
1657   1659      1 ;   HG13 - HD11
1657   1660      1 ;   HG13 - HD12
1657   1661      1 ;   HG13 - HD13
1662   1667      1 ;      C - HA
1662   1668      1 ;      C - CB
1662   1681      1 ;      C - C
1663   1665      1 ;      O - H
1663   1666      1 ;      O - CA
1664   1669      1 ;      N - HB
1664   1670      1 ;      N - CG2
1664   1674      1 ;      N - CG1
1664   1682      1 ;      N - O
1664   1683      1 ;      N - N
1665   1667      1 ;      H - HA
1665   1668      1 ;      H - CB
1665   1681      1 ;      H - C
1666   1671      1 ;     CA - HG21
1666   1672      1 ;     CA - HG22
1666   1673      1 ;     CA - HG23
1666   1675      1 ;     CA - HG12
1666   1676      1 ;     CA - HG13
1666   1677      1 ;     CA - CD1
1666   1684      1 ;     CA - H
1666   1685      1 ;     CA - CA
1667   1669      1 ;     HA - HB
1667   1670      1 ;     HA - CG2
1667   1674      1 ;     HA - CG1
1667   1682      1 ;     HA - O
1667   1683      1 ;     HA - N
1668   1678      1 ;     CB - HD11
1668   1679      1 ;     CB - HD12
1668   1680      1 ;     CB - HD13
1668   1682      1 ;     CB - O
1668   1683      1 ;     CB - N
1669   1671      1 ;     HB - HG21
1669   1672      1 ;     HB - HG22
1669   1673      1 ;     HB - HG23
1669   1675      1 ;     HB - HG12
1669   1676      1 ;     HB - HG13
1669   1677      1 ;     HB - CD1
1669   1681      1 ;     HB - C
1670   1675      1 ;    CG2 - HG12
1670   1676      1 ;    CG2 - HG13
1670   1677      1 ;    CG2 - CD1
1670   1681      1 ;    CG2 - C
1671   1674      1 ;   HG21 - CG1
1672   1674      1 ;   HG22 - CG1
1673   1674      1 ;   HG23 - CG1
1674   1681      1 ;    CG1 - C
```

| | | | | | |
|---|---|---|---|---|---|
| 1675 | 1678 | 1 ; | HG12 | - | HD11 |
| 1675 | 1679 | 1 ; | HG12 | - | HD12 |
| 1675 | 1680 | 1 ; | HG12 | - | HD13 |
| 1676 | 1678 | 1 ; | HG13 | - | HD11 |
| 1676 | 1679 | 1 ; | HG13 | - | HD12 |
| 1676 | 1680 | 1 ; | HG13 | - | HD13 |
| 1681 | 1686 | 1 ; | C | - | HA |
| 1681 | 1687 | 1 ; | C | - | CB |
| 1681 | 1696 | 1 ; | C | - | C |
| 1682 | 1684 | 1 ; | O | - | H |
| 1682 | 1685 | 1 ; | O | - | CA |
| 1683 | 1688 | 1 ; | N | - | HB2 |
| 1683 | 1689 | 1 ; | N | - | HB3 |
| 1683 | 1690 | 1 ; | N | - | CG |
| 1683 | 1697 | 1 ; | N | - | O |
| 1683 | 1698 | 1 ; | N | - | N |
| 1684 | 1686 | 1 ; | H | - | HA |
| 1684 | 1687 | 1 ; | H | - | CB |
| 1684 | 1696 | 1 ; | H | - | C |
| 1685 | 1691 | 1 ; | CA | - | HG2 |
| 1685 | 1692 | 1 ; | CA | - | HG3 |
| 1685 | 1693 | 1 ; | CA | - | CD |
| 1685 | 1699 | 1 ; | CA | - | H |
| 1685 | 1700 | 1 ; | CA | - | CA |
| 1686 | 1688 | 1 ; | HA | - | HB2 |
| 1686 | 1689 | 1 ; | HA | - | HB3 |
| 1686 | 1690 | 1 ; | HA | - | CG |
| 1686 | 1697 | 1 ; | HA | - | O |
| 1686 | 1698 | 1 ; | HA | - | N |
| 1687 | 1694 | 1 ; | CB | - | OE1 |
| 1687 | 1695 | 1 ; | CB | - | OE2 |
| 1687 | 1697 | 1 ; | CB | - | O |
| 1687 | 1698 | 1 ; | CB | - | N |
| 1688 | 1691 | 1 ; | HB2 | - | HG2 |
| 1688 | 1692 | 1 ; | HB2 | - | HG3 |
| 1688 | 1693 | 1 ; | HB2 | - | CD |
| 1688 | 1696 | 1 ; | HB2 | - | C |
| 1689 | 1691 | 1 ; | HB3 | - | HG2 |
| 1689 | 1692 | 1 ; | HB3 | - | HG3 |
| 1689 | 1693 | 1 ; | HB3 | - | CD |
| 1689 | 1696 | 1 ; | HB3 | - | C |
| 1690 | 1696 | 1 ; | CG | - | C |
| 1691 | 1694 | 1 ; | HG2 | - | OE1 |
| 1691 | 1695 | 1 ; | HG2 | - | OE2 |
| 1692 | 1694 | 1 ; | HG3 | - | OE1 |
| 1692 | 1695 | 1 ; | HG3 | - | OE2 |
| 1696 | 1701 | 1 ; | C | - | HA |
| 1696 | 1702 | 1 ; | C | - | CB |
| 1696 | 1710 | 1 ; | C | - | C |
| 1697 | 1699 | 1 ; | O | - | H |
| 1697 | 1700 | 1 ; | O | - | CA |
| 1698 | 1703 | 1 ; | N | - | HB2 |

| | | | | | |
|---|---|---|---|---|---|
| 1698 | 1704 | 1 ; | N | - | HB3 |
| 1698 | 1705 | 1 ; | N | - | CG |
| 1698 | 1711 | 1 ; | N | - | O |
| 1698 | 1712 | 1 ; | N | - | N |
| 1699 | 1701 | 1 ; | H | - | HA |
| 1699 | 1702 | 1 ; | H | - | CB |
| 1699 | 1710 | 1 ; | H | - | C |
| 1700 | 1706 | 1 ; | CA | - | OD1 |
| 1700 | 1707 | 1 ; | CA | - | ND2 |
| 1700 | 1713 | 1 ; | CA | - | H |
| 1700 | 1714 | 1 ; | CA | - | CA |
| 1701 | 1703 | 1 ; | HA | - | HB2 |
| 1701 | 1704 | 1 ; | HA | - | HB3 |
| 1701 | 1705 | 1 ; | HA | - | CG |
| 1701 | 1711 | 1 ; | HA | - | O |
| 1701 | 1712 | 1 ; | HA | - | N |
| 1702 | 1708 | 1 ; | CB | - | HD21 |
| 1702 | 1709 | 1 ; | CB | - | HD22 |
| 1702 | 1711 | 1 ; | CB | - | O |
| 1702 | 1712 | 1 ; | CB | - | N |
| 1703 | 1706 | 1 ; | HB2 | - | OD1 |
| 1703 | 1707 | 1 ; | HB2 | - | ND2 |
| 1703 | 1710 | 1 ; | HB2 | - | C |
| 1704 | 1706 | 1 ; | HB3 | - | OD1 |
| 1704 | 1707 | 1 ; | HB3 | - | ND2 |
| 1704 | 1710 | 1 ; | HB3 | - | C |
| 1705 | 1710 | 1 ; | CG | - | C |
| 1706 | 1708 | 1 ; | OD1 | - | HD21 |
| 1706 | 1709 | 1 ; | OD1 | - | HD22 |
| 1710 | 1715 | 1 ; | C | - | HA2 |
| 1710 | 1716 | 1 ; | C | - | HA3 |
| 1710 | 1717 | 1 ; | C | - | C |
| 1711 | 1713 | 1 ; | O | - | H |
| 1711 | 1714 | 1 ; | O | - | CA |
| 1712 | 1718 | 1 ; | N | - | O |
| 1712 | 1719 | 1 ; | N | - | N |
| 1713 | 1715 | 1 ; | H | - | HA2 |
| 1713 | 1716 | 1 ; | H | - | HA3 |
| 1713 | 1717 | 1 ; | H | - | C |
| 1714 | 1720 | 1 ; | CA | - | H |
| 1714 | 1721 | 1 ; | CA | - | CA |
| 1715 | 1718 | 1 ; | HA2 | - | O |
| 1715 | 1719 | 1 ; | HA2 | - | N |
| 1716 | 1718 | 1 ; | HA3 | - | O |
| 1716 | 1719 | 1 ; | HA3 | - | N |
| 1717 | 1722 | 1 ; | C | - | HA |
| 1717 | 1723 | 1 ; | C | - | CB |
| 1717 | 1728 | 1 ; | C | - | C |
| 1718 | 1720 | 1 ; | O | - | H |
| 1718 | 1721 | 1 ; | O | - | CA |
| 1719 | 1724 | 1 ; | N | - | HB2 |
| 1719 | 1725 | 1 ; | N | - | HB3 |

| | | | |
|---|---|---|---|
| 1719 | 1726 | 1 ; | N - OG |
| 1719 | 1729 | 1 ; | N - O |
| 1719 | 1730 | 1 ; | N - N |
| 1720 | 1722 | 1 ; | H - HA |
| 1720 | 1723 | 1 ; | H - CB |
| 1720 | 1728 | 1 ; | H - C |
| 1721 | 1727 | 1 ; | CA - HG |
| 1721 | 1731 | 1 ; | CA - H |
| 1721 | 1732 | 1 ; | CA - CA |
| 1722 | 1724 | 1 ; | HA - HB2 |
| 1722 | 1725 | 1 ; | HA - HB3 |
| 1722 | 1726 | 1 ; | HA - OG |
| 1722 | 1729 | 1 ; | HA - O |
| 1722 | 1730 | 1 ; | HA - N |
| 1723 | 1729 | 1 ; | CB - O |
| 1723 | 1730 | 1 ; | CB - N |
| 1724 | 1727 | 1 ; | HB2 - HG |
| 1724 | 1728 | 1 ; | HB2 - C |
| 1725 | 1727 | 1 ; | HB3 - HG |
| 1725 | 1728 | 1 ; | HB3 - C |
| 1726 | 1728 | 1 ; | OG - C |
| 1728 | 1733 | 1 ; | C - HA |
| 1728 | 1734 | 1 ; | C - CB |
| 1728 | 1748 | 1 ; | C - C |
| 1729 | 1731 | 1 ; | O - H |
| 1729 | 1732 | 1 ; | O - CA |
| 1730 | 1735 | 1 ; | N - HB2 |
| 1730 | 1736 | 1 ; | N - HB3 |
| 1730 | 1737 | 1 ; | N - CG |
| 1730 | 1749 | 1 ; | N - O |
| 1730 | 1750 | 1 ; | N - N |
| 1731 | 1733 | 1 ; | H - HA |
| 1731 | 1734 | 1 ; | H - CB |
| 1731 | 1748 | 1 ; | H - C |
| 1732 | 1738 | 1 ; | CA - CD1 |
| 1732 | 1746 | 1 ; | CA - CD2 |
| 1732 | 1751 | 1 ; | CA - H |
| 1732 | 1752 | 1 ; | CA - CA |
| 1733 | 1735 | 1 ; | HA - HB2 |
| 1733 | 1736 | 1 ; | HA - HB3 |
| 1733 | 1737 | 1 ; | HA - CG |
| 1733 | 1749 | 1 ; | HA - O |
| 1733 | 1750 | 1 ; | HA - N |
| 1734 | 1739 | 1 ; | CB - HD1 |
| 1734 | 1740 | 1 ; | CB - CE1 |
| 1734 | 1744 | 1 ; | CB - CE2 |
| 1734 | 1747 | 1 ; | CB - HD2 |
| 1734 | 1749 | 1 ; | CB - O |
| 1734 | 1750 | 1 ; | CB - N |
| 1735 | 1738 | 1 ; | HB2 - CD1 |
| 1735 | 1746 | 1 ; | HB2 - CD2 |
| 1735 | 1748 | 1 ; | HB2 - C |

| | | | |
|---|---|---|---|
| 1736 | 1738 | 1 ; | HB3 - CD1 |
| 1736 | 1746 | 1 ; | HB3 - CD2 |
| 1736 | 1748 | 1 ; | HB3 - C |
| 1737 | 1741 | 1 ; | CG - HE1 |
| 1737 | 1742 | 1 ; | CG - CZ |
| 1737 | 1745 | 1 ; | CG - HE2 |
| 1737 | 1748 | 1 ; | CG - C |
| 1738 | 1743 | 1 ; | CD1 - HZ |
| 1738 | 1744 | 1 ; | CD1 - CE2 |
| 1738 | 1747 | 1 ; | CD1 - HD2 |
| 1739 | 1741 | 1 ; | HD1 - HE1 |
| 1739 | 1742 | 1 ; | HD1 - CZ |
| 1739 | 1746 | 1 ; | HD1 - CD2 |
| 1740 | 1745 | 1 ; | CE1 - HE2 |
| 1740 | 1746 | 1 ; | CE1 - CD2 |
| 1741 | 1743 | 1 ; | HE1 - HZ |
| 1741 | 1744 | 1 ; | HE1 - CE2 |
| 1742 | 1747 | 1 ; | CZ - HD2 |
| 1743 | 1745 | 1 ; | HZ - HE2 |
| 1743 | 1746 | 1 ; | HZ - CD2 |
| 1745 | 1747 | 1 ; | HE2 - HD2 |
| 1748 | 1753 | 1 ; | C - HA |
| 1748 | 1754 | 1 ; | C - CB |
| 1748 | 1762 | 1 ; | C - C |
| 1749 | 1751 | 1 ; | O - H |
| 1749 | 1752 | 1 ; | O - CA |
| 1750 | 1755 | 1 ; | N - HB |
| 1750 | 1756 | 1 ; | N - CG2 |
| 1750 | 1760 | 1 ; | N - OG1 |
| 1750 | 1763 | 1 ; | N - O |
| 1750 | 1764 | 1 ; | N - N |
| 1751 | 1753 | 1 ; | H - HA |
| 1751 | 1754 | 1 ; | H - CB |
| 1751 | 1762 | 1 ; | H - C |
| 1752 | 1757 | 1 ; | CA - HG21 |
| 1752 | 1758 | 1 ; | CA - HG22 |
| 1752 | 1759 | 1 ; | CA - HG23 |
| 1752 | 1761 | 1 ; | CA - HG1 |
| 1752 | 1765 | 1 ; | CA - H |
| 1752 | 1766 | 1 ; | CA - CA |
| 1753 | 1755 | 1 ; | HA - HB |
| 1753 | 1756 | 1 ; | HA - CG2 |
| 1753 | 1760 | 1 ; | HA - OG1 |
| 1753 | 1763 | 1 ; | HA - O |
| 1753 | 1764 | 1 ; | HA - N |
| 1754 | 1763 | 1 ; | CB - O |
| 1754 | 1764 | 1 ; | CB - N |
| 1755 | 1757 | 1 ; | HB - HG21 |
| 1755 | 1758 | 1 ; | HB - HG22 |
| 1755 | 1759 | 1 ; | HB - HG23 |
| 1755 | 1761 | 1 ; | HB - HG1 |
| 1755 | 1762 | 1 ; | HB - C |

```
1756   1761       1 ;    CG2 - HG1
1756   1762       1 ;    CG2 - C
1757   1760       1 ;   HG21 - OG1
1758   1760       1 ;   HG22 - OG1
1759   1760       1 ;   HG23 - OG1
1760   1762       1 ;    OG1 - C
1762   1767       1 ;      C - HA
1762   1768       1 ;      C - CB
1762   1777       1 ;      C - C
1763   1765       1 ;      O - H
1763   1766       1 ;      O - CA
1764   1769       1 ;      N - HB2
1764   1770       1 ;      N - HB3
1764   1771       1 ;      N - CG
1764   1778       1 ;      N - O
1764   1779       1 ;      N - N
1765   1767       1 ;      H - HA
1765   1768       1 ;      H - CB
1765   1777       1 ;      H - C
1766   1772       1 ;     CA - HG2
1766   1773       1 ;     CA - HG3
1766   1774       1 ;     CA - CD
1766   1780       1 ;     CA - H
1766   1781       1 ;     CA - CA
1767   1769       1 ;     HA - HB2
1767   1770       1 ;     HA - HB3
1767   1771       1 ;     HA - CG
1767   1778       1 ;     HA - O
1767   1779       1 ;     HA - N
1768   1775       1 ;     CB - OE1
1768   1776       1 ;     CB - OE2
1768   1778       1 ;     CB - O
1768   1779       1 ;     CB - N
1769   1772       1 ;    HB2 - HG2
1769   1773       1 ;    HB2 - HG3
1769   1774       1 ;    HB2 - CD
1769   1777       1 ;    HB2 - C
1770   1772       1 ;    HB3 - HG2
1770   1773       1 ;    HB3 - HG3
1770   1774       1 ;    HB3 - CD
1770   1777       1 ;    HB3 - C
1771   1777       1 ;     CG - C
1772   1775       1 ;    HG2 - OE1
1772   1776       1 ;    HG2 - OE2
1773   1775       1 ;    HG3 - OE1
1773   1776       1 ;    HG3 - OE2
1777   1782       1 ;      C - HA
1777   1783       1 ;      C - CB
1777   1799       1 ;      C - C
1778   1780       1 ;      O - H
1778   1781       1 ;      O - CA
1779   1784       1 ;      N - HB2
```

| | | | | | |
|---|---|---|---|---|---|
| 1779 | 1785 | 1 ; | N | - | HB3 |
| 1779 | 1786 | 1 ; | N | - | CG |
| 1779 | 1800 | 1 ; | N | - | O |
| 1779 | 1801 | 1 ; | N | - | N |
| 1780 | 1782 | 1 ; | H | - | HA |
| 1780 | 1783 | 1 ; | H | - | CB |
| 1780 | 1799 | 1 ; | H | - | C |
| 1781 | 1787 | 1 ; | CA | - | HG2 |
| 1781 | 1788 | 1 ; | CA | - | HG3 |
| 1781 | 1789 | 1 ; | CA | - | CD |
| 1781 | 1802 | 1 ; | CA | - | H |
| 1781 | 1803 | 1 ; | CA | - | CA |
| 1782 | 1784 | 1 ; | HA | - | HB2 |
| 1782 | 1785 | 1 ; | HA | - | HB3 |
| 1782 | 1786 | 1 ; | HA | - | CG |
| 1782 | 1800 | 1 ; | HA | - | O |
| 1782 | 1801 | 1 ; | HA | - | N |
| 1783 | 1790 | 1 ; | CB | - | HD2 |
| 1783 | 1791 | 1 ; | CB | - | HD3 |
| 1783 | 1792 | 1 ; | CB | - | CE |
| 1783 | 1800 | 1 ; | CB | - | O |
| 1783 | 1801 | 1 ; | CB | - | N |
| 1784 | 1787 | 1 ; | HB2 | - | HG2 |
| 1784 | 1788 | 1 ; | HB2 | - | HG3 |
| 1784 | 1789 | 1 ; | HB2 | - | CD |
| 1784 | 1799 | 1 ; | HB2 | - | C |
| 1785 | 1787 | 1 ; | HB3 | - | HG2 |
| 1785 | 1788 | 1 ; | HB3 | - | HG3 |
| 1785 | 1789 | 1 ; | HB3 | - | CD |
| 1785 | 1799 | 1 ; | HB3 | - | C |
| 1786 | 1793 | 1 ; | CG | - | HE2 |
| 1786 | 1794 | 1 ; | CG | - | HE3 |
| 1786 | 1795 | 1 ; | CG | - | NZ |
| 1786 | 1799 | 1 ; | CG | - | C |
| 1787 | 1790 | 1 ; | HG2 | - | HD2 |
| 1787 | 1791 | 1 ; | HG2 | - | HD3 |
| 1787 | 1792 | 1 ; | HG2 | - | CE |
| 1788 | 1790 | 1 ; | HG3 | - | HD2 |
| 1788 | 1791 | 1 ; | HG3 | - | HD3 |
| 1788 | 1792 | 1 ; | HG3 | - | CE |
| 1789 | 1796 | 1 ; | CD | - | HZ1 |
| 1789 | 1797 | 1 ; | CD | - | HZ2 |
| 1789 | 1798 | 1 ; | CD | - | HZ3 |
| 1790 | 1793 | 1 ; | HD2 | - | HE2 |
| 1790 | 1794 | 1 ; | HD2 | - | HE3 |
| 1790 | 1795 | 1 ; | HD2 | - | NZ |
| 1791 | 1793 | 1 ; | HD3 | - | HE2 |
| 1791 | 1794 | 1 ; | HD3 | - | HE3 |
| 1791 | 1795 | 1 ; | HD3 | - | NZ |
| 1793 | 1796 | 1 ; | HE2 | - | HZ1 |
| 1793 | 1797 | 1 ; | HE2 | - | HZ2 |
| 1793 | 1798 | 1 ; | HE2 | - | HZ3 |

```
1794   1796       1 ;    HE3 - HZ1
1794   1797       1 ;    HE3 - HZ2
1794   1798       1 ;    HE3 - HZ3
1799   1804       1 ;      C - HA
1799   1805       1 ;      C - CB
1799   1813       1 ;      C - C
1800   1802       1 ;      O - H
1800   1803       1 ;      O - CA
1801   1806       1 ;      N - HB2
1801   1807       1 ;      N - HB3
1801   1808       1 ;      N - CG
1801   1814       1 ;      N - O
1801   1815       1 ;      N - N
1802   1804       1 ;      H - HA
1802   1805       1 ;      H - CB
1802   1813       1 ;      H - C
1803   1809       1 ;     CA - OD1
1803   1810       1 ;     CA - ND2
1803   1816       1 ;     CA - H
1803   1817       1 ;     CA - CA
1804   1806       1 ;     HA - HB2
1804   1807       1 ;     HA - HB3
1804   1808       1 ;     HA - CG
1804   1814       1 ;     HA - O
1804   1815       1 ;     HA - N
1805   1811       1 ;     CB - HD21
1805   1812       1 ;     CB - HD22
1805   1814       1 ;     CB - O
1805   1815       1 ;     CB - N
1806   1809       1 ;    HB2 - OD1
1806   1810       1 ;    HB2 - ND2
1806   1813       1 ;    HB2 - C
1807   1809       1 ;    HB3 - OD1
1807   1810       1 ;    HB3 - ND2
1807   1813       1 ;    HB3 - C
1808   1813       1 ;     CG - C
1809   1811       1 ;    OD1 - HD21
1809   1812       1 ;    OD1 - HD22
1813   1818       1 ;      C - HA
1813   1819       1 ;      C - CB
1813   1823       1 ;      C - C
1814   1816       1 ;      O - H
1814   1817       1 ;      O - CA
1815   1820       1 ;      N - HB1
1815   1821       1 ;      N - HB2
1815   1822       1 ;      N - HB3
1815   1824       1 ;      N - O
1815   1825       1 ;      N - N
1816   1818       1 ;      H - HA
1816   1819       1 ;      H - CB
1816   1823       1 ;      H - C
1817   1826       1 ;     CA - H
```

| | | | |
|---|---|---|---|
| 1817 | 1827 | 1 ; | CA - CA |
| 1818 | 1820 | 1 ; | HA - HB1 |
| 1818 | 1821 | 1 ; | HA - HB2 |
| 1818 | 1822 | 1 ; | HA - HB3 |
| 1818 | 1824 | 1 ; | HA - O |
| 1818 | 1825 | 1 ; | HA - N |
| 1819 | 1824 | 1 ; | CB - O |
| 1819 | 1825 | 1 ; | CB - N |
| 1820 | 1823 | 1 ; | HB1 - C |
| 1821 | 1823 | 1 ; | HB2 - C |
| 1822 | 1823 | 1 ; | HB3 - C |
| 1823 | 1828 | 1 ; | C - HA |
| 1823 | 1829 | 1 ; | C - CB |
| 1823 | 1833 | 1 ; | C - C |
| 1824 | 1826 | 1 ; | O - H |
| 1824 | 1827 | 1 ; | O - CA |
| 1825 | 1830 | 1 ; | N - HB1 |
| 1825 | 1831 | 1 ; | N - HB2 |
| 1825 | 1832 | 1 ; | N - HB3 |
| 1825 | 1834 | 1 ; | N - O |
| 1825 | 1835 | 1 ; | N - N |
| 1826 | 1828 | 1 ; | H - HA |
| 1826 | 1829 | 1 ; | H - CB |
| 1826 | 1833 | 1 ; | H - C |
| 1827 | 1836 | 1 ; | CA - H |
| 1827 | 1837 | 1 ; | CA - CA |
| 1828 | 1830 | 1 ; | HA - HB1 |
| 1828 | 1831 | 1 ; | HA - HB2 |
| 1828 | 1832 | 1 ; | HA - HB3 |
| 1828 | 1834 | 1 ; | HA - O |
| 1828 | 1835 | 1 ; | HA - N |
| 1829 | 1834 | 1 ; | CB - O |
| 1829 | 1835 | 1 ; | CB - N |
| 1830 | 1833 | 1 ; | HB1 - C |
| 1831 | 1833 | 1 ; | HB2 - C |
| 1832 | 1833 | 1 ; | HB3 - C |
| 1833 | 1838 | 1 ; | C - HA |
| 1833 | 1839 | 1 ; | C - CB |
| 1833 | 1847 | 1 ; | C - C |
| 1834 | 1836 | 1 ; | O - H |
| 1834 | 1837 | 1 ; | O - CA |
| 1835 | 1840 | 1 ; | N - HB |
| 1835 | 1841 | 1 ; | N - CG2 |
| 1835 | 1845 | 1 ; | N - OG1 |
| 1835 | 1848 | 1 ; | N - O |
| 1835 | 1849 | 1 ; | N - N |
| 1836 | 1838 | 1 ; | H - HA |
| 1836 | 1839 | 1 ; | H - CB |
| 1836 | 1847 | 1 ; | H - C |
| 1837 | 1842 | 1 ; | CA - HG21 |
| 1837 | 1843 | 1 ; | CA - HG22 |
| 1837 | 1844 | 1 ; | CA - HG23 |

```
1837   1846       1 ;      CA - HG1
1837   1850       1 ;      CA - H
1837   1851       1 ;      CA - CA
1838   1840       1 ;      HA - HB
1838   1841       1 ;      HA - CG2
1838   1845       1 ;      HA - OG1
1838   1848       1 ;      HA - O
1838   1849       1 ;      HA - N
1839   1848       1 ;      CB - O
1839   1849       1 ;      CB - N
1840   1842       1 ;      HB - HG21
1840   1843       1 ;      HB - HG22
1840   1844       1 ;      HB - HG23
1840   1846       1 ;      HB - HG1
1840   1847       1 ;      HB - C
1841   1846       1 ;     CG2 - HG1
1841   1847       1 ;     CG2 - C
1842   1845       1 ;    HG21 - OG1
1843   1845       1 ;    HG22 - OG1
1844   1845       1 ;    HG23 - OG1
1845   1847       1 ;     OG1 - C
1847   1852       1 ;       C - HA
1847   1853       1 ;       C - CB
1847   1866       1 ;       C - C
1848   1850       1 ;       O - H
1848   1851       1 ;       O - CA
1849   1854       1 ;       N - HB
1849   1855       1 ;       N - CG2
1849   1859       1 ;       N - CG1
1849   1867       1 ;       N - O
1849   1868       1 ;       N - N
1850   1852       1 ;       H - HA
1850   1853       1 ;       H - CB
1850   1866       1 ;       H - C
1851   1856       1 ;      CA - HG21
1851   1857       1 ;      CA - HG22
1851   1858       1 ;      CA - HG23
1851   1860       1 ;      CA - HG12
1851   1861       1 ;      CA - HG13
1851   1862       1 ;      CA - CD1
1851   1869       1 ;      CA - H
1851   1870       1 ;      CA - CA
1852   1854       1 ;      HA - HB
1852   1855       1 ;      HA - CG2
1852   1859       1 ;      HA - CG1
1852   1867       1 ;      HA - O
1852   1868       1 ;      HA - N
1853   1863       1 ;      CB - HD11
1853   1864       1 ;      CB - HD12
1853   1865       1 ;      CB - HD13
1853   1867       1 ;      CB - O
1853   1868       1 ;      CB - N
```

| | | | | |
|---|---|---|---|---|
| 1854 | 1856 | 1 | ; | HB - HG21 |
| 1854 | 1857 | 1 | ; | HB - HG22 |
| 1854 | 1858 | 1 | ; | HB - HG23 |
| 1854 | 1860 | 1 | ; | HB - HG12 |
| 1854 | 1861 | 1 | ; | HB - HG13 |
| 1854 | 1862 | 1 | ; | HB - CD1 |
| 1854 | 1866 | 1 | ; | HB - C |
| 1855 | 1860 | 1 | ; | CG2 - HG12 |
| 1855 | 1861 | 1 | ; | CG2 - HG13 |
| 1855 | 1862 | 1 | ; | CG2 - CD1 |
| 1855 | 1866 | 1 | ; | CG2 - C |
| 1856 | 1859 | 1 | ; | HG21 - CG1 |
| 1857 | 1859 | 1 | ; | HG22 - CG1 |
| 1858 | 1859 | 1 | ; | HG23 - CG1 |
| 1859 | 1866 | 1 | ; | CG1 - C |
| 1860 | 1863 | 1 | ; | HG12 - HD11 |
| 1860 | 1864 | 1 | ; | HG12 - HD12 |
| 1860 | 1865 | 1 | ; | HG12 - HD13 |
| 1861 | 1863 | 1 | ; | HG13 - HD11 |
| 1861 | 1864 | 1 | ; | HG13 - HD12 |
| 1861 | 1865 | 1 | ; | HG13 - HD13 |
| 1866 | 1871 | 1 | ; | C - HA |
| 1866 | 1872 | 1 | ; | C - CB |
| 1866 | 1883 | 1 | ; | C - C |
| 1867 | 1869 | 1 | ; | O - H |
| 1867 | 1870 | 1 | ; | O - CA |
| 1868 | 1873 | 1 | ; | N - HB2 |
| 1868 | 1874 | 1 | ; | N - HB3 |
| 1868 | 1875 | 1 | ; | N - CG |
| 1868 | 1884 | 1 | ; | N - O |
| 1868 | 1885 | 1 | ; | N - N |
| 1869 | 1871 | 1 | ; | H - HA |
| 1869 | 1872 | 1 | ; | H - CB |
| 1869 | 1883 | 1 | ; | H - C |
| 1870 | 1876 | 1 | ; | CA - HG2 |
| 1870 | 1877 | 1 | ; | CA - HG3 |
| 1870 | 1878 | 1 | ; | CA - SD |
| 1870 | 1886 | 1 | ; | CA - H |
| 1870 | 1887 | 1 | ; | CA - CA |
| 1871 | 1873 | 1 | ; | HA - HB2 |
| 1871 | 1874 | 1 | ; | HA - HB3 |
| 1871 | 1875 | 1 | ; | HA - CG |
| 1871 | 1884 | 1 | ; | HA - O |
| 1871 | 1885 | 1 | ; | HA - N |
| 1872 | 1879 | 1 | ; | CB - CE |
| 1872 | 1884 | 1 | ; | CB - O |
| 1872 | 1885 | 1 | ; | CB - N |
| 1873 | 1876 | 1 | ; | HB2 - HG2 |
| 1873 | 1877 | 1 | ; | HB2 - HG3 |
| 1873 | 1878 | 1 | ; | HB2 - SD |
| 1873 | 1883 | 1 | ; | HB2 - C |
| 1874 | 1876 | 1 | ; | HB3 - HG2 |

| | | | | |
|---|---|---|---|---|
| 1874 | 1877 | 1 | ; | HB3 - HG3 |
| 1874 | 1878 | 1 | ; | HB3 - SD |
| 1874 | 1883 | 1 | ; | HB3 - C |
| 1875 | 1880 | 1 | ; | CG - HE1 |
| 1875 | 1881 | 1 | ; | CG - HE2 |
| 1875 | 1882 | 1 | ; | CG - HE3 |
| 1875 | 1883 | 1 | ; | CG - C |
| 1876 | 1879 | 1 | ; | HG2 - CE |
| 1877 | 1879 | 1 | ; | HG3 - CE |
| 1883 | 1888 | 1 | ; | C - HA |
| 1883 | 1889 | 1 | ; | C - CB |
| 1883 | 1905 | 1 | ; | C - C |
| 1884 | 1886 | 1 | ; | O - H |
| 1884 | 1887 | 1 | ; | O - CA |
| 1885 | 1890 | 1 | ; | N - HB2 |
| 1885 | 1891 | 1 | ; | N - HB3 |
| 1885 | 1892 | 1 | ; | N - CG |
| 1885 | 1906 | 1 | ; | N - O |
| 1885 | 1907 | 1 | ; | N - N |
| 1886 | 1888 | 1 | ; | H - HA |
| 1886 | 1889 | 1 | ; | H - CB |
| 1886 | 1905 | 1 | ; | H - C |
| 1887 | 1893 | 1 | ; | CA - HG2 |
| 1887 | 1894 | 1 | ; | CA - HG3 |
| 1887 | 1895 | 1 | ; | CA - CD |
| 1887 | 1908 | 1 | ; | CA - H |
| 1887 | 1909 | 1 | ; | CA - CA |
| 1888 | 1890 | 1 | ; | HA - HB2 |
| 1888 | 1891 | 1 | ; | HA - HB3 |
| 1888 | 1892 | 1 | ; | HA - CG |
| 1888 | 1906 | 1 | ; | HA - O |
| 1888 | 1907 | 1 | ; | HA - N |
| 1889 | 1896 | 1 | ; | CB - HD2 |
| 1889 | 1897 | 1 | ; | CB - HD3 |
| 1889 | 1898 | 1 | ; | CB - CE |
| 1889 | 1906 | 1 | ; | CB - O |
| 1889 | 1907 | 1 | ; | CB - N |
| 1890 | 1893 | 1 | ; | HB2 - HG2 |
| 1890 | 1894 | 1 | ; | HB2 - HG3 |
| 1890 | 1895 | 1 | ; | HB2 - CD |
| 1890 | 1905 | 1 | ; | HB2 - C |
| 1891 | 1893 | 1 | ; | HB3 - HG2 |
| 1891 | 1894 | 1 | ; | HB3 - HG3 |
| 1891 | 1895 | 1 | ; | HB3 - CD |
| 1891 | 1905 | 1 | ; | HB3 - C |
| 1892 | 1899 | 1 | ; | CG - HE2 |
| 1892 | 1900 | 1 | ; | CG - HE3 |
| 1892 | 1901 | 1 | ; | CG - NZ |
| 1892 | 1905 | 1 | ; | CG - C |
| 1893 | 1896 | 1 | ; | HG2 - HD2 |
| 1893 | 1897 | 1 | ; | HG2 - HD3 |
| 1893 | 1898 | 1 | ; | HG2 - CE |

| | | | | |
|---|---|---|---|---|
| 1894 | 1896 | 1 | ; | HG3 - HD2 |
| 1894 | 1897 | 1 | ; | HG3 - HD3 |
| 1894 | 1898 | 1 | ; | HG3 - CE |
| 1895 | 1902 | 1 | ; | CD - HZ1 |
| 1895 | 1903 | 1 | ; | CD - HZ2 |
| 1895 | 1904 | 1 | ; | CD - HZ3 |
| 1896 | 1899 | 1 | ; | HD2 - HE2 |
| 1896 | 1900 | 1 | ; | HD2 - HE3 |
| 1896 | 1901 | 1 | ; | HD2 - NZ |
| 1897 | 1899 | 1 | ; | HD3 - HE2 |
| 1897 | 1900 | 1 | ; | HD3 - HE3 |
| 1897 | 1901 | 1 | ; | HD3 - NZ |
| 1899 | 1902 | 1 | ; | HE2 - HZ1 |
| 1899 | 1903 | 1 | ; | HE2 - HZ2 |
| 1899 | 1904 | 1 | ; | HE2 - HZ3 |
| 1900 | 1902 | 1 | ; | HE3 - HZ1 |
| 1900 | 1903 | 1 | ; | HE3 - HZ2 |
| 1900 | 1904 | 1 | ; | HE3 - HZ3 |
| 1905 | 1910 | 1 | ; | C - HA |
| 1905 | 1911 | 1 | ; | C - CB |
| 1905 | 1922 | 1 | ; | C - C |
| 1906 | 1908 | 1 | ; | O - H |
| 1906 | 1909 | 1 | ; | O - CA |
| 1907 | 1912 | 1 | ; | N - HB2 |
| 1907 | 1913 | 1 | ; | N - HB3 |
| 1907 | 1914 | 1 | ; | N - CG |
| 1907 | 1923 | 1 | ; | N - O |
| 1907 | 1924 | 1 | ; | N - N |
| 1908 | 1910 | 1 | ; | H - HA |
| 1908 | 1911 | 1 | ; | H - CB |
| 1908 | 1922 | 1 | ; | H - C |
| 1909 | 1915 | 1 | ; | CA - HG2 |
| 1909 | 1916 | 1 | ; | CA - HG3 |
| 1909 | 1917 | 1 | ; | CA - CD |
| 1909 | 1925 | 1 | ; | CA - H |
| 1909 | 1926 | 1 | ; | CA - CA |
| 1910 | 1912 | 1 | ; | HA - HB2 |
| 1910 | 1913 | 1 | ; | HA - HB3 |
| 1910 | 1914 | 1 | ; | HA - CG |
| 1910 | 1923 | 1 | ; | HA - O |
| 1910 | 1924 | 1 | ; | HA - N |
| 1911 | 1918 | 1 | ; | CB - OE1 |
| 1911 | 1919 | 1 | ; | CB - NE2 |
| 1911 | 1923 | 1 | ; | CB - O |
| 1911 | 1924 | 1 | ; | CB - N |
| 1912 | 1915 | 1 | ; | HB2 - HG2 |
| 1912 | 1916 | 1 | ; | HB2 - HG3 |
| 1912 | 1917 | 1 | ; | HB2 - CD |
| 1912 | 1922 | 1 | ; | HB2 - C |
| 1913 | 1915 | 1 | ; | HB3 - HG2 |
| 1913 | 1916 | 1 | ; | HB3 - HG3 |
| 1913 | 1917 | 1 | ; | HB3 - CD |

```
1913   1922      1 ;    HB3 - C
1914   1920      1 ;     CG - HE21
1914   1921      1 ;     CG - HE22
1914   1922      1 ;     CG - C
1915   1918      1 ;    HG2 - OE1
1915   1919      1 ;    HG2 - NE2
1916   1918      1 ;    HG3 - OE1
1916   1919      1 ;    HG3 - NE2
1918   1920      1 ;    OE1 - HE21
1918   1921      1 ;    OE1 - HE22
1922   1927      1 ;      C - HA
1922   1928      1 ;      C - CB
1922   1941      1 ;      C - C
1923   1925      1 ;      O - H
1923   1926      1 ;      O - CA
1924   1929      1 ;      N - HB
1924   1930      1 ;      N - CG2
1924   1934      1 ;      N - CG1
1924   1942      1 ;      N - O
1924   1943      1 ;      N - N
1925   1927      1 ;      H - HA
1925   1928      1 ;      H - CB
1925   1941      1 ;      H - C
1926   1931      1 ;     CA - HG21
1926   1932      1 ;     CA - HG22
1926   1933      1 ;     CA - HG23
1926   1935      1 ;     CA - HG12
1926   1936      1 ;     CA - HG13
1926   1937      1 ;     CA - CD1
1926   1944      1 ;     CA - H
1926   1945      1 ;     CA - CA
1927   1929      1 ;     HA - HB
1927   1930      1 ;     HA - CG2
1927   1934      1 ;     HA - CG1
1927   1942      1 ;     HA - O
1927   1943      1 ;     HA - N
1928   1938      1 ;     CB - HD11
1928   1939      1 ;     CB - HD12
1928   1940      1 ;     CB - HD13
1928   1942      1 ;     CB - O
1928   1943      1 ;     CB - N
1929   1931      1 ;     HB - HG21
1929   1932      1 ;     HB - HG22
1929   1933      1 ;     HB - HG23
1929   1935      1 ;     HB - HG12
1929   1936      1 ;     HB - HG13
1929   1937      1 ;     HB - CD1
1929   1941      1 ;     HB - C
1930   1935      1 ;    CG2 - HG12
1930   1936      1 ;    CG2 - HG13
1930   1937      1 ;    CG2 - CD1
1930   1941      1 ;    CG2 - C
```

```
 1931   1934       1 ;   HG21 - CG1
 1932   1934       1 ;   HG22 - CG1
 1933   1934       1 ;   HG23 - CG1
 1934   1941       1 ;    CG1 - C
 1935   1938       1 ;   HG12 - HD11
 1935   1939       1 ;   HG12 - HD12
 1935   1940       1 ;   HG12 - HD13
 1936   1938       1 ;   HG13 - HD11
 1936   1939       1 ;   HG13 - HD12
 1936   1940       1 ;   HG13 - HD13
 1941   1946       1 ;      C - HA
 1941   1947       1 ;      C - CB
 1941   1961       1 ;      C - C
 1942   1944       1 ;      O - H
 1942   1945       1 ;      O - CA
 1943   1948       1 ;      N - HB2
 1943   1949       1 ;      N - HB3
 1943   1950       1 ;      N - CG
 1943   1962       1 ;      N - O
 1943   1963       1 ;      N - N
 1944   1946       1 ;      H - HA
 1944   1947       1 ;      H - CB
 1944   1961       1 ;      H - C
 1945   1951       1 ;     CA - CD1
 1945   1959       1 ;     CA - CD2
 1945   1964       1 ;     CA - H
 1945   1965       1 ;     CA - CA
 1946   1948       1 ;     HA - HB2
 1946   1949       1 ;     HA - HB3
 1946   1950       1 ;     HA - CG
 1946   1962       1 ;     HA - O
 1946   1963       1 ;     HA - N
 1947   1952       1 ;     CB - HD1
 1947   1953       1 ;     CB - CE1
 1947   1957       1 ;     CB - CE2
 1947   1960       1 ;     CB - HD2
 1947   1962       1 ;     CB - O
 1947   1963       1 ;     CB - N
 1948   1951       1 ;    HB2 - CD1
 1948   1959       1 ;    HB2 - CD2
 1948   1961       1 ;    HB2 - C
 1949   1951       1 ;    HB3 - CD1
 1949   1959       1 ;    HB3 - CD2
 1949   1961       1 ;    HB3 - C
 1950   1954       1 ;     CG - HE1
 1950   1955       1 ;     CG - CZ
 1950   1958       1 ;     CG - HE2
 1950   1961       1 ;     CG - C
 1951   1956       1 ;    CD1 - HZ
 1951   1957       1 ;    CD1 - CE2
 1951   1960       1 ;    CD1 - HD2
 1952   1954       1 ;    HD1 - HE1
```

```
1952   1955      1 ;    HD1 - CZ
1952   1959      1 ;    HD1 - CD2
1953   1958      1 ;    CE1 - HE2
1953   1959      1 ;    CE1 - CD2
1954   1956      1 ;    HE1 - HZ
1954   1957      1 ;    HE1 - CE2
1955   1960      1 ;     CZ - HD2
1956   1958      1 ;     HZ - HE2
1956   1959      1 ;     HZ - CD2
1958   1960      1 ;    HE2 - HD2
1961   1966      1 ;      C - HA
1961   1967      1 ;      C - CB
1961   1972      1 ;      C - C
1962   1964      1 ;      O - H
1962   1965      1 ;      O - CA
1963   1968      1 ;      N - HB2
1963   1969      1 ;      N - HB3
1963   1970      1 ;      N - OG
1963   1973      1 ;      N - O
1963   1974      1 ;      N - N
1964   1966      1 ;      H - HA
1964   1967      1 ;      H - CB
1964   1972      1 ;      H - C
1965   1971      1 ;     CA - HG
1965   1975      1 ;     CA - H
1965   1976      1 ;     CA - CA
1966   1968      1 ;     HA - HB2
1966   1969      1 ;     HA - HB3
1966   1970      1 ;     HA - OG
1966   1973      1 ;     HA - O
1966   1974      1 ;     HA - N
1967   1973      1 ;     CB - O
1967   1974      1 ;     CB - N
1968   1971      1 ;    HB2 - HG
1968   1972      1 ;    HB2 - C
1969   1971      1 ;    HB3 - HG
1969   1972      1 ;    HB3 - C
1970   1972      1 ;     OG - C
1972   1977      1 ;      C - HA
1972   1978      1 ;      C - CB
1972   1982      1 ;      C - C
1973   1975      1 ;      O - H
1973   1976      1 ;      O - CA
1974   1979      1 ;      N - HB1
1974   1980      1 ;      N - HB2
1974   1981      1 ;      N - HB3
1974   1983      1 ;      N - O
1974   1984      1 ;      N - N
1975   1977      1 ;      H - HA
1975   1978      1 ;      H - CB
1975   1982      1 ;      H - C
1976   1985      1 ;     CA - H
```

```
1976   1986     1 ;    CA - CA
1977   1979     1 ;    HA - HB1
1977   1980     1 ;    HA - HB2
1977   1981     1 ;    HA - HB3
1977   1983     1 ;    HA - O
1977   1984     1 ;    HA - N
1978   1983     1 ;    CB - O
1978   1984     1 ;    CB - N
1979   1982     1 ;   HB1 - C
1980   1982     1 ;   HB2 - C
1981   1982     1 ;   HB3 - C
1982   1987     1 ;     C - HA
1982   1988     1 ;     C - CB
1982   2001     1 ;     C - C
1983   1985     1 ;     O - H
1983   1986     1 ;     O - CA
1984   1989     1 ;     N - HB
1984   1990     1 ;     N - CG2
1984   1994     1 ;     N - CG1
1984   2002     1 ;     N - O
1984   2003     1 ;     N - N
1985   1987     1 ;     H - HA
1985   1988     1 ;     H - CB
1985   2001     1 ;     H - C
1986   1991     1 ;    CA - HG21
1986   1992     1 ;    CA - HG22
1986   1993     1 ;    CA - HG23
1986   1995     1 ;    CA - HG12
1986   1996     1 ;    CA - HG13
1986   1997     1 ;    CA - CD1
1986   2004     1 ;    CA - H
1986   2005     1 ;    CA - CA
1987   1989     1 ;    HA - HB
1987   1990     1 ;    HA - CG2
1987   1994     1 ;    HA - CG1
1987   2002     1 ;    HA - O
1987   2003     1 ;    HA - N
1988   1998     1 ;    CB - HD11
1988   1999     1 ;    CB - HD12
1988   2000     1 ;    CB - HD13
1988   2002     1 ;    CB - O
1988   2003     1 ;    CB - N
1989   1991     1 ;    HB - HG21
1989   1992     1 ;    HB - HG22
1989   1993     1 ;    HB - HG23
1989   1995     1 ;    HB - HG12
1989   1996     1 ;    HB - HG13
1989   1997     1 ;    HB - CD1
1989   2001     1 ;    HB - C
1990   1995     1 ;   CG2 - HG12
1990   1996     1 ;   CG2 - HG13
1990   1997     1 ;   CG2 - CD1
```

```
1990   2001      1 ;    CG2 - C
1991   1994      1 ;   HG21 - CG1
1992   1994      1 ;   HG22 - CG1
1993   1994      1 ;   HG23 - CG1
1994   2001      1 ;    CG1 - C
1995   1998      1 ;   HG12 - HD11
1995   1999      1 ;   HG12 - HD12
1995   2000      1 ;   HG12 - HD13
1996   1998      1 ;   HG13 - HD11
1996   1999      1 ;   HG13 - HD12
1996   2000      1 ;   HG13 - HD13
2001   2006      1 ;      C - HA
2001   2007      1 ;      C - CB
2001   2021      1 ;      C - C
2002   2004      1 ;      O - H
2002   2005      1 ;      O - CA
2003   2008      1 ;      N - HB2
2003   2009      1 ;      N - HB3
2003   2010      1 ;      N - CG
2003   2022      1 ;      N - O
2003   2023      1 ;      N - N
2004   2006      1 ;      H - HA
2004   2007      1 ;      H - CB
2004   2021      1 ;      H - C
2005   2011      1 ;     CA - CD1
2005   2019      1 ;     CA - CD2
2005   2024      1 ;     CA - H
2005   2025      1 ;     CA - CA
2006   2008      1 ;     HA - HB2
2006   2009      1 ;     HA - HB3
2006   2010      1 ;     HA - CG
2006   2022      1 ;     HA - O
2006   2023      1 ;     HA - N
2007   2012      1 ;     CB - HD1
2007   2013      1 ;     CB - CE1
2007   2017      1 ;     CB - CE2
2007   2020      1 ;     CB - HD2
2007   2022      1 ;     CB - O
2007   2023      1 ;     CB - N
2008   2011      1 ;    HB2 - CD1
2008   2019      1 ;    HB2 - CD2
2008   2021      1 ;    HB2 - C
2009   2011      1 ;    HB3 - CD1
2009   2019      1 ;    HB3 - CD2
2009   2021      1 ;    HB3 - C
2010   2014      1 ;     CG - HE1
2010   2015      1 ;     CG - CZ
2010   2018      1 ;     CG - HE2
2010   2021      1 ;     CG - C
2011   2016      1 ;    CD1 - HZ
2011   2017      1 ;    CD1 - CE2
2011   2020      1 ;    CD1 - HD2
```

```
2012   2014      1 ;    HD1 - HE1
2012   2015      1 ;    HD1 - CZ
2012   2019      1 ;    HD1 - CD2
2013   2018      1 ;    CE1 - HE2
2013   2019      1 ;    CE1 - CD2
2014   2016      1 ;    HE1 - HZ
2014   2017      1 ;    HE1 - CE2
2015   2020      1 ;     CZ - HD2
2016   2018      1 ;     HZ - HE2
2016   2019      1 ;     HZ - CD2
2018   2020      1 ;    HE2 - HD2
2021   2026      1 ;      C - HA
2021   2027      1 ;      C - CB
2021   2042      1 ;      C - C
2022   2024      1 ;      O - H
2022   2025      1 ;      O - CA
2023   2028      1 ;      N - HB2
2023   2029      1 ;      N - HB3
2023   2030      1 ;      N - CG
2023   2043      1 ;      N - O
2023   2044      1 ;      N - N
2024   2026      1 ;      H - HA
2024   2027      1 ;      H - CB
2024   2042      1 ;      H - C
2025   2031      1 ;     CA - CD1
2025   2040      1 ;     CA - CD2
2025   2045      1 ;     CA - H
2025   2046      1 ;     CA - CA
2026   2028      1 ;     HA - HB2
2026   2029      1 ;     HA - HB3
2026   2030      1 ;     HA - CG
2026   2043      1 ;     HA - O
2026   2044      1 ;     HA - N
2027   2032      1 ;     CB - HD1
2027   2033      1 ;     CB - CE1
2027   2038      1 ;     CB - CE2
2027   2041      1 ;     CB - HD2
2027   2043      1 ;     CB - O
2027   2044      1 ;     CB - N
2028   2031      1 ;    HB2 - CD1
2028   2040      1 ;    HB2 - CD2
2028   2042      1 ;    HB2 - C
2029   2031      1 ;    HB3 - CD1
2029   2040      1 ;    HB3 - CD2
2029   2042      1 ;    HB3 - C
2030   2034      1 ;     CG - HE1
2030   2035      1 ;     CG - CZ
2030   2039      1 ;     CG - HE2
2030   2042      1 ;     CG - C
2031   2036      1 ;    CD1 - OH
2031   2038      1 ;    CD1 - CE2
2031   2041      1 ;    CD1 - HD2
```

```
2032   2034       1 ;    HD1 - HE1
2032   2035       1 ;    HD1 - CZ
2032   2040       1 ;    HD1 - CD2
2033   2037       1 ;    CE1 - HH
2033   2039       1 ;    CE1 - HE2
2033   2040       1 ;    CE1 - CD2
2034   2036       1 ;    HE1 - OH
2034   2038       1 ;    HE1 - CE2
2035   2041       1 ;     CZ - HD2
2036   2039       1 ;     OH - HE2
2036   2040       1 ;     OH - CD2
2037   2038       1 ;     HH - CE2
2039   2041       1 ;    HE2 - HD2
2042   2047       1 ;      C - HA
2042   2048       1 ;      C - CB
2042   2061       1 ;      C - C
2043   2045       1 ;      O - H
2043   2046       1 ;      O - CA
2044   2049       1 ;      N - HB2
2044   2050       1 ;      N - HB3
2044   2051       1 ;      N - CG
2044   2062       1 ;      N - O
2044   2063       1 ;      N - N
2045   2047       1 ;      H - HA
2045   2048       1 ;      H - CB
2045   2061       1 ;      H - C
2046   2052       1 ;     CA - HG
2046   2053       1 ;     CA - CD1
2046   2057       1 ;     CA - CD2
2046   2064       1 ;     CA - H
2046   2065       1 ;     CA - CA
2047   2049       1 ;     HA - HB2
2047   2050       1 ;     HA - HB3
2047   2051       1 ;     HA - CG
2047   2062       1 ;     HA - O
2047   2063       1 ;     HA - N
2048   2054       1 ;     CB - HD11
2048   2055       1 ;     CB - HD12
2048   2056       1 ;     CB - HD13
2048   2058       1 ;     CB - HD21
2048   2059       1 ;     CB - HD22
2048   2060       1 ;     CB - HD23
2048   2062       1 ;     CB - O
2048   2063       1 ;     CB - N
2049   2052       1 ;    HB2 - HG
2049   2053       1 ;    HB2 - CD1
2049   2057       1 ;    HB2 - CD2
2049   2061       1 ;    HB2 - C
2050   2052       1 ;    HB3 - HG
2050   2053       1 ;    HB3 - CD1
2050   2057       1 ;    HB3 - CD2
2050   2061       1 ;    HB3 - C
```

```
2051   2061       1 ;      CG - C
2052   2054       1 ;      HG - HD11
2052   2055       1 ;      HG - HD12
2052   2056       1 ;      HG - HD13
2052   2058       1 ;      HG - HD21
2052   2059       1 ;      HG - HD22
2052   2060       1 ;      HG - HD23
2053   2058       1 ;     CD1 - HD21
2053   2059       1 ;     CD1 - HD22
2053   2060       1 ;     CD1 - HD23
2054   2057       1 ;    HD11 - CD2
2055   2057       1 ;    HD12 - CD2
2056   2057       1 ;    HD13 - CD2
2061   2066       1 ;       C - HA
2061   2067       1 ;       C - CB
2061   2079       1 ;       C - C
2062   2064       1 ;       O - H
2062   2065       1 ;       O - CA
2063   2068       1 ;       N - HB2
2063   2069       1 ;       N - HB3
2063   2070       1 ;       N - CG
2063   2080       1 ;       N - O
2063   2081       1 ;       N - N
2064   2066       1 ;       H - HA
2064   2067       1 ;       H - CB
2064   2079       1 ;       H - C
2065   2071       1 ;      CA - ND1
2065   2077       1 ;      CA - CD2
2065   2082       1 ;      CA - H
2065   2083       1 ;      CA - CA
2066   2068       1 ;      HA - HB2
2066   2069       1 ;      HA - HB3
2066   2070       1 ;      HA - CG
2066   2080       1 ;      HA - O
2066   2081       1 ;      HA - N
2067   2072       1 ;      CB - HD1
2067   2073       1 ;      CB - CE1
2067   2075       1 ;      CB - NE2
2067   2078       1 ;      CB - HD2
2067   2080       1 ;      CB - O
2067   2081       1 ;      CB - N
2068   2071       1 ;     HB2 - ND1
2068   2077       1 ;     HB2 - CD2
2068   2079       1 ;     HB2 - C
2069   2071       1 ;     HB3 - ND1
2069   2077       1 ;     HB3 - CD2
2069   2079       1 ;     HB3 - C
2070   2074       1 ;      CG - HE1
2070   2076       1 ;      CG - HE2
2070   2079       1 ;      CG - C
2071   2076       1 ;     ND1 - HE2
2071   2078       1 ;     ND1 - HD2
```

| | | | | |
|---|---|---|---|---|
| 2072 | 2074 | 1 | ; | HD1 - HE1 |
| 2072 | 2075 | 1 | ; | HD1 - NE2 |
| 2072 | 2077 | 1 | ; | HD1 - CD2 |
| 2073 | 2078 | 1 | ; | CE1 - HD2 |
| 2074 | 2076 | 1 | ; | HE1 - HE2 |
| 2074 | 2077 | 1 | ; | HE1 - CD2 |
| 2076 | 2078 | 1 | ; | HE2 - HD2 |
| 2079 | 2084 | 1 | ; | C - HA |
| 2079 | 2085 | 1 | ; | C - CB |
| 2079 | 2090 | 1 | ; | C - C |
| 2080 | 2082 | 1 | ; | O - H |
| 2080 | 2083 | 1 | ; | O - CA |
| 2081 | 2086 | 1 | ; | N - HB2 |
| 2081 | 2087 | 1 | ; | N - HB3 |
| 2081 | 2088 | 1 | ; | N - OG |
| 2081 | 2091 | 1 | ; | N - O |
| 2081 | 2092 | 1 | ; | N - N |
| 2082 | 2084 | 1 | ; | H - HA |
| 2082 | 2085 | 1 | ; | H - CB |
| 2082 | 2090 | 1 | ; | H - C |
| 2083 | 2089 | 1 | ; | CA - HG |
| 2083 | 2093 | 1 | ; | CA - H |
| 2083 | 2094 | 1 | ; | CA - CA |
| 2084 | 2086 | 1 | ; | HA - HB2 |
| 2084 | 2087 | 1 | ; | HA - HB3 |
| 2084 | 2088 | 1 | ; | HA - OG |
| 2084 | 2091 | 1 | ; | HA - O |
| 2084 | 2092 | 1 | ; | HA - N |
| 2085 | 2091 | 1 | ; | CB - O |
| 2085 | 2092 | 1 | ; | CB - N |
| 2086 | 2089 | 1 | ; | HB2 - HG |
| 2086 | 2090 | 1 | ; | HB2 - C |
| 2087 | 2089 | 1 | ; | HB3 - HG |
| 2087 | 2090 | 1 | ; | HB3 - C |
| 2088 | 2090 | 1 | ; | OG - C |
| 2090 | 2095 | 1 | ; | C - HA |
| 2090 | 2096 | 1 | ; | C - CB |
| 2090 | 2109 | 1 | ; | C - C |
| 2091 | 2093 | 1 | ; | O - H |
| 2091 | 2094 | 1 | ; | O - CA |
| 2092 | 2097 | 1 | ; | N - HB2 |
| 2092 | 2098 | 1 | ; | N - HB3 |
| 2092 | 2099 | 1 | ; | N - CG |
| 2092 | 2110 | 1 | ; | N - O |
| 2092 | 2111 | 1 | ; | N - N |
| 2093 | 2095 | 1 | ; | H - HA |
| 2093 | 2096 | 1 | ; | H - CB |
| 2093 | 2109 | 1 | ; | H - C |
| 2094 | 2100 | 1 | ; | CA - HG |
| 2094 | 2101 | 1 | ; | CA - CD1 |
| 2094 | 2105 | 1 | ; | CA - CD2 |
| 2094 | 2112 | 1 | ; | CA - H |

| | | | |
|---|---|---|---|
| 2094 | 2113 | 1 ; | CA - CA |
| 2095 | 2097 | 1 ; | HA - HB2 |
| 2095 | 2098 | 1 ; | HA - HB3 |
| 2095 | 2099 | 1 ; | HA - CG |
| 2095 | 2110 | 1 ; | HA - O |
| 2095 | 2111 | 1 ; | HA - N |
| 2096 | 2102 | 1 ; | CB - HD11 |
| 2096 | 2103 | 1 ; | CB - HD12 |
| 2096 | 2104 | 1 ; | CB - HD13 |
| 2096 | 2106 | 1 ; | CB - HD21 |
| 2096 | 2107 | 1 ; | CB - HD22 |
| 2096 | 2108 | 1 ; | CB - HD23 |
| 2096 | 2110 | 1 ; | CB - O |
| 2096 | 2111 | 1 ; | CB - N |
| 2097 | 2100 | 1 ; | HB2 - HG |
| 2097 | 2101 | 1 ; | HB2 - CD1 |
| 2097 | 2105 | 1 ; | HB2 - CD2 |
| 2097 | 2109 | 1 ; | HB2 - C |
| 2098 | 2100 | 1 ; | HB3 - HG |
| 2098 | 2101 | 1 ; | HB3 - CD1 |
| 2098 | 2105 | 1 ; | HB3 - CD2 |
| 2098 | 2109 | 1 ; | HB3 - C |
| 2099 | 2109 | 1 ; | CG - C |
| 2100 | 2102 | 1 ; | HG - HD11 |
| 2100 | 2103 | 1 ; | HG - HD12 |
| 2100 | 2104 | 1 ; | HG - HD13 |
| 2100 | 2106 | 1 ; | HG - HD21 |
| 2100 | 2107 | 1 ; | HG - HD22 |
| 2100 | 2108 | 1 ; | HG - HD23 |
| 2101 | 2106 | 1 ; | CD1 - HD21 |
| 2101 | 2107 | 1 ; | CD1 - HD22 |
| 2101 | 2108 | 1 ; | CD1 - HD23 |
| 2102 | 2105 | 1 ; | HD11 - CD2 |
| 2103 | 2105 | 1 ; | HD12 - CD2 |
| 2104 | 2105 | 1 ; | HD13 - CD2 |
| 2109 | 2114 | 1 ; | C - HA |
| 2109 | 2115 | 1 ; | C - CB |
| 2109 | 2123 | 1 ; | C - C |
| 2110 | 2112 | 1 ; | O - H |
| 2110 | 2113 | 1 ; | O - CA |
| 2111 | 2116 | 1 ; | N - HB2 |
| 2111 | 2117 | 1 ; | N - HB3 |
| 2111 | 2118 | 1 ; | N - CG |
| 2111 | 2124 | 1 ; | N - O |
| 2111 | 2125 | 1 ; | N - N |
| 2112 | 2114 | 1 ; | H - HA |
| 2112 | 2115 | 1 ; | H - CB |
| 2112 | 2123 | 1 ; | H - C |
| 2113 | 2119 | 1 ; | CA - OD1 |
| 2113 | 2120 | 1 ; | CA - ND2 |
| 2113 | 2126 | 1 ; | CA - H |
| 2113 | 2127 | 1 ; | CA - CA |

```
2114   2116       1 ;      HA - HB2
2114   2117       1 ;      HA - HB3
2114   2118       1 ;      HA - CG
2114   2124       1 ;      HA - O
2114   2125       1 ;      HA - N
2115   2121       1 ;      CB - HD21
2115   2122       1 ;      CB - HD22
2115   2124       1 ;      CB - O
2115   2125       1 ;      CB - N
2116   2119       1 ;     HB2 - OD1
2116   2120       1 ;     HB2 - ND2
2116   2123       1 ;     HB2 - C
2117   2119       1 ;     HB3 - OD1
2117   2120       1 ;     HB3 - ND2
2117   2123       1 ;     HB3 - C
2118   2123       1 ;      CG - C
2119   2121       1 ;     OD1 - HD21
2119   2122       1 ;     OD1 - HD22
2123   2128       1 ;       C - HA
2123   2129       1 ;       C - CB
2123   2142       1 ;       C - C
2124   2126       1 ;       O - H
2124   2127       1 ;       O - CA
2125   2130       1 ;       N - HB
2125   2131       1 ;       N - CG2
2125   2135       1 ;       N - CG1
2125   2143       1 ;       N - O
2125   2144       1 ;       N - N
2126   2128       1 ;       H - HA
2126   2129       1 ;       H - CB
2126   2142       1 ;       H - C
2127   2132       1 ;      CA - HG21
2127   2133       1 ;      CA - HG22
2127   2134       1 ;      CA - HG23
2127   2136       1 ;      CA - HG12
2127   2137       1 ;      CA - HG13
2127   2138       1 ;      CA - CD1
2127   2145       1 ;      CA - H
2127   2146       1 ;      CA - CA
2128   2130       1 ;      HA - HB
2128   2131       1 ;      HA - CG2
2128   2135       1 ;      HA - CG1
2128   2143       1 ;      HA - O
2128   2144       1 ;      HA - N
2129   2139       1 ;      CB - HD11
2129   2140       1 ;      CB - HD12
2129   2141       1 ;      CB - HD13
2129   2143       1 ;      CB - O
2129   2144       1 ;      CB - N
2130   2132       1 ;      HB - HG21
2130   2133       1 ;      HB - HG22
2130   2134       1 ;      HB - HG23
```

| | | | |
|---|---|---|---|
| 2130 | 2136 | 1 ; | HB - HG12 |
| 2130 | 2137 | 1 ; | HB - HG13 |
| 2130 | 2138 | 1 ; | HB - CD1 |
| 2130 | 2142 | 1 ; | HB - C |
| 2131 | 2136 | 1 ; | CG2 - HG12 |
| 2131 | 2137 | 1 ; | CG2 - HG13 |
| 2131 | 2138 | 1 ; | CG2 - CD1 |
| 2131 | 2142 | 1 ; | CG2 - C |
| 2132 | 2135 | 1 ; | HG21 - CG1 |
| 2133 | 2135 | 1 ; | HG22 - CG1 |
| 2134 | 2135 | 1 ; | HG23 - CG1 |
| 2135 | 2142 | 1 ; | CG1 - C |
| 2136 | 2139 | 1 ; | HG12 - HD11 |
| 2136 | 2140 | 1 ; | HG12 - HD12 |
| 2136 | 2141 | 1 ; | HG12 - HD13 |
| 2137 | 2139 | 1 ; | HG13 - HD11 |
| 2137 | 2140 | 1 ; | HG13 - HD12 |
| 2137 | 2141 | 1 ; | HG13 - HD13 |
| 2142 | 2147 | 1 ; | C - HA |
| 2142 | 2148 | 1 ; | C - CB |
| 2142 | 2158 | 1 ; | C - C |
| 2143 | 2145 | 1 ; | O - H |
| 2143 | 2146 | 1 ; | O - CA |
| 2144 | 2149 | 1 ; | N - HB |
| 2144 | 2150 | 1 ; | N - CG1 |
| 2144 | 2154 | 1 ; | N - CG2 |
| 2144 | 2159 | 1 ; | N - O |
| 2144 | 2160 | 1 ; | N - N |
| 2145 | 2147 | 1 ; | H - HA |
| 2145 | 2148 | 1 ; | H - CB |
| 2145 | 2158 | 1 ; | H - C |
| 2146 | 2151 | 1 ; | CA - HG11 |
| 2146 | 2152 | 1 ; | CA - HG12 |
| 2146 | 2153 | 1 ; | CA - HG13 |
| 2146 | 2155 | 1 ; | CA - HG21 |
| 2146 | 2156 | 1 ; | CA - HG22 |
| 2146 | 2157 | 1 ; | CA - HG23 |
| 2146 | 2161 | 1 ; | CA - H |
| 2146 | 2162 | 1 ; | CA - CA |
| 2147 | 2149 | 1 ; | HA - HB |
| 2147 | 2150 | 1 ; | HA - CG1 |
| 2147 | 2154 | 1 ; | HA - CG2 |
| 2147 | 2159 | 1 ; | HA - O |
| 2147 | 2160 | 1 ; | HA - N |
| 2148 | 2159 | 1 ; | CB - O |
| 2148 | 2160 | 1 ; | CB - N |
| 2149 | 2151 | 1 ; | HB - HG11 |
| 2149 | 2152 | 1 ; | HB - HG12 |
| 2149 | 2153 | 1 ; | HB - HG13 |
| 2149 | 2155 | 1 ; | HB - HG21 |
| 2149 | 2156 | 1 ; | HB - HG22 |
| 2149 | 2157 | 1 ; | HB - HG23 |

```
2149   2158      1 ;     HB - C
2150   2155      1 ;    CG1 - HG21
2150   2156      1 ;    CG1 - HG22
2150   2157      1 ;    CG1 - HG23
2150   2158      1 ;    CG1 - C
2151   2154      1 ;   HG11 - CG2
2152   2154      1 ;   HG12 - CG2
2153   2154      1 ;   HG13 - CG2
2154   2158      1 ;    CG2 - C
2158   2163      1 ;      C - HA
2158   2164      1 ;      C - CB
2158   2175      1 ;      C - C
2159   2161      1 ;      O - H
2159   2162      1 ;      O - CA
2160   2165      1 ;      N - HB2
2160   2166      1 ;      N - HB3
2160   2167      1 ;      N - CG
2160   2176      1 ;      N - O
2160   2177      1 ;      N - N
2161   2163      1 ;      H - HA
2161   2164      1 ;      H - CB
2161   2175      1 ;      H - C
2162   2168      1 ;     CA - ND1
2162   2173      1 ;     CA - CD2
2162   2178      1 ;     CA - H
2162   2179      1 ;     CA - CA
2163   2165      1 ;     HA - HB2
2163   2166      1 ;     HA - HB3
2163   2167      1 ;     HA - CG
2163   2176      1 ;     HA - O
2163   2177      1 ;     HA - N
2164   2169      1 ;     CB - CE1
2164   2171      1 ;     CB - NE2
2164   2174      1 ;     CB - HD2
2164   2176      1 ;     CB - O
2164   2177      1 ;     CB - N
2165   2168      1 ;    HB2 - ND1
2165   2173      1 ;    HB2 - CD2
2165   2175      1 ;    HB2 - C
2166   2168      1 ;    HB3 - ND1
2166   2173      1 ;    HB3 - CD2
2166   2175      1 ;    HB3 - C
2167   2170      1 ;     CG - HE1
2167   2172      1 ;     CG - HE2
2167   2175      1 ;     CG - C
2168   2172      1 ;    ND1 - HE2
2168   2174      1 ;    ND1 - HD2
2169   2174      1 ;    CE1 - HD2
2170   2172      1 ;    HE1 - HE2
2170   2173      1 ;    HE1 - CD2
2172   2174      1 ;    HE2 - HD2
2175   2180      1 ;      C - HA
```

```
2175   2181       1 ;       C - CB
2175   2199       1 ;       C - C
2176   2178       1 ;       O - H
2176   2179       1 ;       O - CA
2177   2182       1 ;       N - HB2
2177   2183       1 ;       N - HB3
2177   2184       1 ;       N - CG
2177   2200       1 ;       N - O
2177   2201       1 ;       N - N
2178   2180       1 ;       H - HA
2178   2181       1 ;       H - CB
2178   2199       1 ;       H - C
2179   2185       1 ;      CA - HG2
2179   2186       1 ;      CA - HG3
2179   2187       1 ;      CA - CD
2179   2202       1 ;      CA - H
2179   2203       1 ;      CA - CA
2180   2182       1 ;      HA - HB2
2180   2183       1 ;      HA - HB3
2180   2184       1 ;      HA - CG
2180   2200       1 ;      HA - O
2180   2201       1 ;      HA - N
2181   2188       1 ;      CB - HD2
2181   2189       1 ;      CB - HD3
2181   2190       1 ;      CB - NE
2181   2200       1 ;      CB - O
2181   2201       1 ;      CB - N
2182   2185       1 ;     HB2 - HG2
2182   2186       1 ;     HB2 - HG3
2182   2187       1 ;     HB2 - CD
2182   2199       1 ;     HB2 - C
2183   2185       1 ;     HB3 - HG2
2183   2186       1 ;     HB3 - HG3
2183   2187       1 ;     HB3 - CD
2183   2199       1 ;     HB3 - C
2184   2191       1 ;      CG - HE
2184   2192       1 ;      CG - CZ
2184   2199       1 ;      CG - C
2185   2188       1 ;     HG2 - HD2
2185   2189       1 ;     HG2 - HD3
2185   2190       1 ;     HG2 - NE
2186   2188       1 ;     HG3 - HD2
2186   2189       1 ;     HG3 - HD3
2186   2190       1 ;     HG3 - NE
2187   2193       1 ;      CD - NH1
2187   2196       1 ;      CD - NH2
2188   2191       1 ;     HD2 - HE
2188   2192       1 ;     HD2 - CZ
2189   2191       1 ;     HD3 - HE
2189   2192       1 ;     HD3 - CZ
2190   2194       1 ;      NE - HH11
2190   2195       1 ;      NE - HH12
```

| | | | | | |
|---|---|---|---|---|---|
| 2190 | 2197 | 1 ; | NE | - | HH21 |
| 2190 | 2198 | 1 ; | NE | - | HH22 |
| 2191 | 2193 | 1 ; | HE | - | NH1 |
| 2191 | 2196 | 1 ; | HE | - | NH2 |
| 2193 | 2197 | 1 ; | NH1 | - | HH21 |
| 2193 | 2198 | 1 ; | NH1 | - | HH22 |
| 2194 | 2196 | 1 ; | HH11 | - | NH2 |
| 2195 | 2196 | 1 ; | HH12 | - | NH2 |
| 2199 | 2204 | 1 ; | C | - | HA |
| 2199 | 2205 | 1 ; | C | - | CB |
| 2199 | 2211 | 1 ; | C | - | C |
| 2200 | 2202 | 1 ; | O | - | H |
| 2200 | 2203 | 1 ; | O | - | CA |
| 2201 | 2206 | 1 ; | N | - | HB2 |
| 2201 | 2207 | 1 ; | N | - | HB3 |
| 2201 | 2208 | 1 ; | N | - | CG |
| 2201 | 2212 | 1 ; | N | - | O |
| 2201 | 2213 | 1 ; | N | - | N |
| 2202 | 2204 | 1 ; | H | - | HA |
| 2202 | 2205 | 1 ; | H | - | CB |
| 2202 | 2211 | 1 ; | H | - | C |
| 2203 | 2209 | 1 ; | CA | - | OD1 |
| 2203 | 2210 | 1 ; | CA | - | OD2 |
| 2203 | 2214 | 1 ; | CA | - | H |
| 2203 | 2215 | 1 ; | CA | - | CA |
| 2204 | 2206 | 1 ; | HA | - | HB2 |
| 2204 | 2207 | 1 ; | HA | - | HB3 |
| 2204 | 2208 | 1 ; | HA | - | CG |
| 2204 | 2212 | 1 ; | HA | - | O |
| 2204 | 2213 | 1 ; | HA | - | N |
| 2205 | 2212 | 1 ; | CB | - | O |
| 2205 | 2213 | 1 ; | CB | - | N |
| 2206 | 2209 | 1 ; | HB2 | - | OD1 |
| 2206 | 2210 | 1 ; | HB2 | - | OD2 |
| 2206 | 2211 | 1 ; | HB2 | - | C |
| 2207 | 2209 | 1 ; | HB3 | - | OD1 |
| 2207 | 2210 | 1 ; | HB3 | - | OD2 |
| 2207 | 2211 | 1 ; | HB3 | - | C |
| 2208 | 2211 | 1 ; | CG | - | C |
| 2211 | 2216 | 1 ; | C | - | HA |
| 2211 | 2217 | 1 ; | C | - | CB |
| 2211 | 2230 | 1 ; | C | - | C |
| 2212 | 2214 | 1 ; | O | - | H |
| 2212 | 2215 | 1 ; | O | - | CA |
| 2213 | 2218 | 1 ; | N | - | HB2 |
| 2213 | 2219 | 1 ; | N | - | HB3 |
| 2213 | 2220 | 1 ; | N | - | CG |
| 2213 | 2231 | 1 ; | N | - | O |
| 2213 | 2232 | 1 ; | N | - | N |
| 2214 | 2216 | 1 ; | H | - | HA |
| 2214 | 2217 | 1 ; | H | - | CB |
| 2214 | 2230 | 1 ; | H | - | C |

| | | | |
|---|---|---|---|
| 2215 | 2221 | 1 ; | CA - HG |
| 2215 | 2222 | 1 ; | CA - CD1 |
| 2215 | 2226 | 1 ; | CA - CD2 |
| 2215 | 2233 | 1 ; | CA - H |
| 2215 | 2234 | 1 ; | CA - CA |
| 2216 | 2218 | 1 ; | HA - HB2 |
| 2216 | 2219 | 1 ; | HA - HB3 |
| 2216 | 2220 | 1 ; | HA - CG |
| 2216 | 2231 | 1 ; | HA - O |
| 2216 | 2232 | 1 ; | HA - N |
| 2217 | 2223 | 1 ; | CB - HD11 |
| 2217 | 2224 | 1 ; | CB - HD12 |
| 2217 | 2225 | 1 ; | CB - HD13 |
| 2217 | 2227 | 1 ; | CB - HD21 |
| 2217 | 2228 | 1 ; | CB - HD22 |
| 2217 | 2229 | 1 ; | CB - HD23 |
| 2217 | 2231 | 1 ; | CB - O |
| 2217 | 2232 | 1 ; | CB - N |
| 2218 | 2221 | 1 ; | HB2 - HG |
| 2218 | 2222 | 1 ; | HB2 - CD1 |
| 2218 | 2226 | 1 ; | HB2 - CD2 |
| 2218 | 2230 | 1 ; | HB2 - C |
| 2219 | 2221 | 1 ; | HB3 - HG |
| 2219 | 2222 | 1 ; | HB3 - CD1 |
| 2219 | 2226 | 1 ; | HB3 - CD2 |
| 2219 | 2230 | 1 ; | HB3 - C |
| 2220 | 2230 | 1 ; | CG - C |
| 2221 | 2223 | 1 ; | HG - HD11 |
| 2221 | 2224 | 1 ; | HG - HD12 |
| 2221 | 2225 | 1 ; | HG - HD13 |
| 2221 | 2227 | 1 ; | HG - HD21 |
| 2221 | 2228 | 1 ; | HG - HD22 |
| 2221 | 2229 | 1 ; | HG - HD23 |
| 2222 | 2227 | 1 ; | CD1 - HD21 |
| 2222 | 2228 | 1 ; | CD1 - HD22 |
| 2222 | 2229 | 1 ; | CD1 - HD23 |
| 2223 | 2226 | 1 ; | HD11 - CD2 |
| 2224 | 2226 | 1 ; | HD12 - CD2 |
| 2225 | 2226 | 1 ; | HD13 - CD2 |
| 2230 | 2235 | 1 ; | C - HA |
| 2230 | 2236 | 1 ; | C - CB |
| 2230 | 2252 | 1 ; | C - C |
| 2231 | 2233 | 1 ; | O - H |
| 2231 | 2234 | 1 ; | O - CA |
| 2232 | 2237 | 1 ; | N - HB2 |
| 2232 | 2238 | 1 ; | N - HB3 |
| 2232 | 2239 | 1 ; | N - CG |
| 2232 | 2253 | 1 ; | N - O |
| 2232 | 2254 | 1 ; | N - N |
| 2233 | 2235 | 1 ; | H - HA |
| 2233 | 2236 | 1 ; | H - CB |
| 2233 | 2252 | 1 ; | H - C |

| | | | | |
|---|---|---|---|---|
| 2234 | 2240 | 1 | ; | CA - HG2 |
| 2234 | 2241 | 1 | ; | CA - HG3 |
| 2234 | 2242 | 1 | ; | CA - CD |
| 2234 | 2255 | 1 | ; | CA - CD |
| 2234 | 2264 | 1 | ; | CA - CA |
| 2235 | 2237 | 1 | ; | HA - HB2 |
| 2235 | 2238 | 1 | ; | HA - HB3 |
| 2235 | 2239 | 1 | ; | HA - CG |
| 2235 | 2253 | 1 | ; | HA - O |
| 2235 | 2254 | 1 | ; | HA - N |
| 2236 | 2243 | 1 | ; | CB - HD2 |
| 2236 | 2244 | 1 | ; | CB - HD3 |
| 2236 | 2245 | 1 | ; | CB - CE |
| 2236 | 2253 | 1 | ; | CB - O |
| 2236 | 2254 | 1 | ; | CB - N |
| 2237 | 2240 | 1 | ; | HB2 - HG2 |
| 2237 | 2241 | 1 | ; | HB2 - HG3 |
| 2237 | 2242 | 1 | ; | HB2 - CD |
| 2237 | 2252 | 1 | ; | HB2 - C |
| 2238 | 2240 | 1 | ; | HB3 - HG2 |
| 2238 | 2241 | 1 | ; | HB3 - HG3 |
| 2238 | 2242 | 1 | ; | HB3 - CD |
| 2238 | 2252 | 1 | ; | HB3 - C |
| 2239 | 2246 | 1 | ; | CG - HE2 |
| 2239 | 2247 | 1 | ; | CG - HE3 |
| 2239 | 2248 | 1 | ; | CG - NZ |
| 2239 | 2252 | 1 | ; | CG - C |
| 2240 | 2243 | 1 | ; | HG2 - HD2 |
| 2240 | 2244 | 1 | ; | HG2 - HD3 |
| 2240 | 2245 | 1 | ; | HG2 - CE |
| 2241 | 2243 | 1 | ; | HG3 - HD2 |
| 2241 | 2244 | 1 | ; | HG3 - HD3 |
| 2241 | 2245 | 1 | ; | HG3 - CE |
| 2242 | 2249 | 1 | ; | CD - HZ1 |
| 2242 | 2250 | 1 | ; | CD - HZ2 |
| 2242 | 2251 | 1 | ; | CD - HZ3 |
| 2243 | 2246 | 1 | ; | HD2 - HE2 |
| 2243 | 2247 | 1 | ; | HD2 - HE3 |
| 2243 | 2248 | 1 | ; | HD2 - NZ |
| 2244 | 2246 | 1 | ; | HD3 - HE2 |
| 2244 | 2247 | 1 | ; | HD3 - HE3 |
| 2244 | 2248 | 1 | ; | HD3 - NZ |
| 2246 | 2249 | 1 | ; | HE2 - HZ1 |
| 2246 | 2250 | 1 | ; | HE2 - HZ2 |
| 2246 | 2251 | 1 | ; | HE2 - HZ3 |
| 2247 | 2249 | 1 | ; | HE3 - HZ1 |
| 2247 | 2250 | 1 | ; | HE3 - HZ2 |
| 2247 | 2251 | 1 | ; | HE3 - HZ3 |
| 2252 | 2256 | 1 | ; | C - HD2 |
| 2252 | 2257 | 1 | ; | C - HD3 |
| 2252 | 2258 | 1 | ; | C - CG |
| 2252 | 2261 | 1 | ; | C - CB |

| | | | | | |
|---|---|---|---|---|---|
| 2252 | 2265 | 1 ; | C | - | HA |
| 2252 | 2266 | 1 ; | C | - | C |
| 2253 | 2255 | 1 ; | O | - | CD |
| 2253 | 2264 | 1 ; | O | - | CA |
| 2254 | 2259 | 1 ; | N | - | HG2 |
| 2254 | 2260 | 1 ; | N | - | HG3 |
| 2254 | 2262 | 1 ; | N | - | HB2 |
| 2254 | 2263 | 1 ; | N | - | HB3 |
| 2254 | 2267 | 1 ; | N | - | O |
| 2254 | 2268 | 1 ; | N | - | N |
| 2255 | 2262 | 1 ; | CD | - | HB2 |
| 2255 | 2263 | 1 ; | CD | - | HB3 |
| 2255 | 2265 | 1 ; | CD | - | HA |
| 2255 | 2266 | 1 ; | CD | - | C |
| 2256 | 2259 | 1 ; | HD2 | - | HG2 |
| 2256 | 2260 | 1 ; | HD2 | - | HG3 |
| 2256 | 2261 | 1 ; | HD2 | - | CB |
| 2256 | 2264 | 1 ; | HD2 | - | CA |
| 2257 | 2259 | 1 ; | HD3 | - | HG2 |
| 2257 | 2260 | 1 ; | HD3 | - | HG3 |
| 2257 | 2261 | 1 ; | HD3 | - | CB |
| 2257 | 2264 | 1 ; | HD3 | - | CA |
| 2258 | 2265 | 1 ; | CG | - | HA |
| 2258 | 2266 | 1 ; | CG | - | C |
| 2259 | 2262 | 1 ; | HG2 | - | HB2 |
| 2259 | 2263 | 1 ; | HG2 | - | HB3 |
| 2259 | 2264 | 1 ; | HG2 | - | CA |
| 2260 | 2262 | 1 ; | HG3 | - | HB2 |
| 2260 | 2263 | 1 ; | HG3 | - | HB3 |
| 2260 | 2264 | 1 ; | HG3 | - | CA |
| 2261 | 2267 | 1 ; | CB | - | O |
| 2261 | 2268 | 1 ; | CB | - | N |
| 2262 | 2265 | 1 ; | HB2 | - | HA |
| 2262 | 2266 | 1 ; | HB2 | - | C |
| 2263 | 2265 | 1 ; | HB3 | - | HA |
| 2263 | 2266 | 1 ; | HB3 | - | C |
| 2264 | 2269 | 1 ; | CA | - | H |
| 2264 | 2270 | 1 ; | CA | - | CA |
| 2265 | 2267 | 1 ; | HA | - | O |
| 2265 | 2268 | 1 ; | HA | - | N |
| 2266 | 2271 | 1 ; | C | - | HA |
| 2266 | 2272 | 1 ; | C | - | CB |
| 2266 | 2281 | 1 ; | C | - | C |
| 2267 | 2269 | 1 ; | O | - | H |
| 2267 | 2270 | 1 ; | O | - | CA |
| 2268 | 2273 | 1 ; | N | - | HB2 |
| 2268 | 2274 | 1 ; | N | - | HB3 |
| 2268 | 2275 | 1 ; | N | - | CG |
| 2268 | 2282 | 1 ; | N | - | O |
| 2268 | 2283 | 1 ; | N | - | N |
| 2269 | 2271 | 1 ; | H | - | HA |
| 2269 | 2272 | 1 ; | H | - | CB |

| | | | | |
|---|---|---|---|---|
| 2269 | 2281 | 1 ; | H - | C |
| 2270 | 2276 | 1 ; | CA - | HG2 |
| 2270 | 2277 | 1 ; | CA - | HG3 |
| 2270 | 2278 | 1 ; | CA - | CD |
| 2270 | 2284 | 1 ; | CA - | H |
| 2270 | 2285 | 1 ; | CA - | CA |
| 2271 | 2273 | 1 ; | HA - | HB2 |
| 2271 | 2274 | 1 ; | HA - | HB3 |
| 2271 | 2275 | 1 ; | HA - | CG |
| 2271 | 2282 | 1 ; | HA - | O |
| 2271 | 2283 | 1 ; | HA - | N |
| 2272 | 2279 | 1 ; | CB - | OE1 |
| 2272 | 2280 | 1 ; | CB - | OE2 |
| 2272 | 2282 | 1 ; | CB - | O |
| 2272 | 2283 | 1 ; | CB - | N |
| 2273 | 2276 | 1 ; | HB2 - | HG2 |
| 2273 | 2277 | 1 ; | HB2 - | HG3 |
| 2273 | 2278 | 1 ; | HB2 - | CD |
| 2273 | 2281 | 1 ; | HB2 - | C |
| 2274 | 2276 | 1 ; | HB3 - | HG2 |
| 2274 | 2277 | 1 ; | HB3 - | HG3 |
| 2274 | 2278 | 1 ; | HB3 - | CD |
| 2274 | 2281 | 1 ; | HB3 - | C |
| 2275 | 2281 | 1 ; | CG - | C |
| 2276 | 2279 | 1 ; | HG2 - | OE1 |
| 2276 | 2280 | 1 ; | HG2 - | OE2 |
| 2277 | 2279 | 1 ; | HG3 - | OE1 |
| 2277 | 2280 | 1 ; | HG3 - | OE2 |
| 2281 | 2286 | 1 ; | C - | HA |
| 2281 | 2287 | 1 ; | C - | CB |
| 2281 | 2295 | 1 ; | C - | C |
| 2282 | 2284 | 1 ; | O - | H |
| 2282 | 2285 | 1 ; | O - | CA |
| 2283 | 2288 | 1 ; | N - | HB2 |
| 2283 | 2289 | 1 ; | N - | HB3 |
| 2283 | 2290 | 1 ; | N - | CG |
| 2283 | 2296 | 1 ; | N - | O |
| 2283 | 2297 | 1 ; | N - | N |
| 2284 | 2286 | 1 ; | H - | HA |
| 2284 | 2287 | 1 ; | H - | CB |
| 2284 | 2295 | 1 ; | H - | C |
| 2285 | 2291 | 1 ; | CA - | OD1 |
| 2285 | 2292 | 1 ; | CA - | ND2 |
| 2285 | 2298 | 1 ; | CA - | H |
| 2285 | 2299 | 1 ; | CA - | CA |
| 2286 | 2288 | 1 ; | HA - | HB2 |
| 2286 | 2289 | 1 ; | HA - | HB3 |
| 2286 | 2290 | 1 ; | HA - | CG |
| 2286 | 2296 | 1 ; | HA - | O |
| 2286 | 2297 | 1 ; | HA - | N |
| 2287 | 2293 | 1 ; | CB - | HD21 |
| 2287 | 2294 | 1 ; | CB - | HD22 |

| | | | |
|---|---|---|---|
| 2287 | 2296 | 1 ; | CB - O |
| 2287 | 2297 | 1 ; | CB - N |
| 2288 | 2291 | 1 ; | HB2 - OD1 |
| 2288 | 2292 | 1 ; | HB2 - ND2 |
| 2288 | 2295 | 1 ; | HB2 - C |
| 2289 | 2291 | 1 ; | HB3 - OD1 |
| 2289 | 2292 | 1 ; | HB3 - ND2 |
| 2289 | 2295 | 1 ; | HB3 - C |
| 2290 | 2295 | 1 ; | CG - C |
| 2291 | 2293 | 1 ; | OD1 - HD21 |
| 2291 | 2294 | 1 ; | OD1 - HD22 |
| 2295 | 2300 | 1 ; | C - HA |
| 2295 | 2301 | 1 ; | C - CB |
| 2295 | 2315 | 1 ; | C - C |
| 2296 | 2298 | 1 ; | O - H |
| 2296 | 2299 | 1 ; | O - CA |
| 2297 | 2302 | 1 ; | N - HB2 |
| 2297 | 2303 | 1 ; | N - HB3 |
| 2297 | 2304 | 1 ; | N - CG |
| 2297 | 2316 | 1 ; | N - O |
| 2297 | 2317 | 1 ; | N - N |
| 2298 | 2300 | 1 ; | H - HA |
| 2298 | 2301 | 1 ; | H - CB |
| 2298 | 2315 | 1 ; | H - C |
| 2299 | 2305 | 1 ; | CA - CD1 |
| 2299 | 2313 | 1 ; | CA - CD2 |
| 2299 | 2318 | 1 ; | CA - H |
| 2299 | 2319 | 1 ; | CA - CA |
| 2300 | 2302 | 1 ; | HA - HB2 |
| 2300 | 2303 | 1 ; | HA - HB3 |
| 2300 | 2304 | 1 ; | HA - CG |
| 2300 | 2316 | 1 ; | HA - O |
| 2300 | 2317 | 1 ; | HA - N |
| 2301 | 2306 | 1 ; | CB - HD1 |
| 2301 | 2307 | 1 ; | CB - CE1 |
| 2301 | 2311 | 1 ; | CB - CE2 |
| 2301 | 2314 | 1 ; | CB - HD2 |
| 2301 | 2316 | 1 ; | CB - O |
| 2301 | 2317 | 1 ; | CB - N |
| 2302 | 2305 | 1 ; | HB2 - CD1 |
| 2302 | 2313 | 1 ; | HB2 - CD2 |
| 2302 | 2315 | 1 ; | HB2 - C |
| 2303 | 2305 | 1 ; | HB3 - CD1 |
| 2303 | 2313 | 1 ; | HB3 - CD2 |
| 2303 | 2315 | 1 ; | HB3 - C |
| 2304 | 2308 | 1 ; | CG - HE1 |
| 2304 | 2309 | 1 ; | CG - CZ |
| 2304 | 2312 | 1 ; | CG - HE2 |
| 2304 | 2315 | 1 ; | CG - C |
| 2305 | 2310 | 1 ; | CD1 - HZ |
| 2305 | 2311 | 1 ; | CD1 - CE2 |
| 2305 | 2314 | 1 ; | CD1 - HD2 |

```
2306   2308       1 ;    HD1 - HE1
2306   2309       1 ;    HD1 - CZ
2306   2313       1 ;    HD1 - CD2
2307   2312       1 ;    CE1 - HE2
2307   2313       1 ;    CE1 - CD2
2308   2310       1 ;    HE1 - HZ
2308   2311       1 ;    HE1 - CE2
2309   2314       1 ;     CZ - HD2
2310   2312       1 ;     HZ - HE2
2310   2313       1 ;     HZ - CD2
2312   2314       1 ;    HE2 - HD2
2315   2320       1 ;      C - HA
2315   2321       1 ;      C - CB
2315   2334       1 ;      C - C
2316   2318       1 ;      O - H
2316   2319       1 ;      O - CA
2317   2322       1 ;      N - HB2
2317   2323       1 ;      N - HB3
2317   2324       1 ;      N - CG
2317   2335       1 ;      N - O
2317   2336       1 ;      N - N
2318   2320       1 ;      H - HA
2318   2321       1 ;      H - CB
2318   2334       1 ;      H - C
2319   2325       1 ;     CA - HG
2319   2326       1 ;     CA - CD1
2319   2330       1 ;     CA - CD2
2319   2337       1 ;     CA - H
2319   2338       1 ;     CA - CA
2320   2322       1 ;     HA - HB2
2320   2323       1 ;     HA - HB3
2320   2324       1 ;     HA - CG
2320   2335       1 ;     HA - O
2320   2336       1 ;     HA - N
2321   2327       1 ;     CB - HD11
2321   2328       1 ;     CB - HD12
2321   2329       1 ;     CB - HD13
2321   2331       1 ;     CB - HD21
2321   2332       1 ;     CB - HD22
2321   2333       1 ;     CB - HD23
2321   2335       1 ;     CB - O
2321   2336       1 ;     CB - N
2322   2325       1 ;    HB2 - HG
2322   2326       1 ;    HB2 - CD1
2322   2330       1 ;    HB2 - CD2
2322   2334       1 ;    HB2 - C
2323   2325       1 ;    HB3 - HG
2323   2326       1 ;    HB3 - CD1
2323   2330       1 ;    HB3 - CD2
2323   2334       1 ;    HB3 - C
2324   2334       1 ;     CG - C
2325   2327       1 ;     HG - HD11
```

| | | | | | | |
|---|---|---|---|---|---|---|
| 2325 | 2328 | 1 | ; | HG | - | HD12 |
| 2325 | 2329 | 1 | ; | HG | - | HD13 |
| 2325 | 2331 | 1 | ; | HG | - | HD21 |
| 2325 | 2332 | 1 | ; | HG | - | HD22 |
| 2325 | 2333 | 1 | ; | HG | - | HD23 |
| 2326 | 2331 | 1 | ; | CD1 | - | HD21 |
| 2326 | 2332 | 1 | ; | CD1 | - | HD22 |
| 2326 | 2333 | 1 | ; | CD1 | - | HD23 |
| 2327 | 2330 | 1 | ; | HD11 | - | CD2 |
| 2328 | 2330 | 1 | ; | HD12 | - | CD2 |
| 2329 | 2330 | 1 | ; | HD13 | - | CD2 |
| 2334 | 2339 | 1 | ; | C | - | HA |
| 2334 | 2340 | 1 | ; | C | - | CB |
| 2334 | 2354 | 1 | ; | C | - | C |
| 2335 | 2337 | 1 | ; | O | - | H |
| 2335 | 2338 | 1 | ; | O | - | CA |
| 2336 | 2341 | 1 | ; | N | - | HB2 |
| 2336 | 2342 | 1 | ; | N | - | HB3 |
| 2336 | 2343 | 1 | ; | N | - | CG |
| 2336 | 2355 | 1 | ; | N | - | O |
| 2336 | 2356 | 1 | ; | N | - | N |
| 2337 | 2339 | 1 | ; | H | - | HA |
| 2337 | 2340 | 1 | ; | H | - | CB |
| 2337 | 2354 | 1 | ; | H | - | C |
| 2338 | 2344 | 1 | ; | CA | - | CD1 |
| 2338 | 2352 | 1 | ; | CA | - | CD2 |
| 2338 | 2357 | 1 | ; | CA | - | H |
| 2338 | 2358 | 1 | ; | CA | - | CA |
| 2339 | 2341 | 1 | ; | HA | - | HB2 |
| 2339 | 2342 | 1 | ; | HA | - | HB3 |
| 2339 | 2343 | 1 | ; | HA | - | CG |
| 2339 | 2355 | 1 | ; | HA | - | O |
| 2339 | 2356 | 1 | ; | HA | - | N |
| 2340 | 2345 | 1 | ; | CB | - | HD1 |
| 2340 | 2346 | 1 | ; | CB | - | CE1 |
| 2340 | 2350 | 1 | ; | CB | - | CE2 |
| 2340 | 2353 | 1 | ; | CB | - | HD2 |
| 2340 | 2355 | 1 | ; | CB | - | O |
| 2340 | 2356 | 1 | ; | CB | - | N |
| 2341 | 2344 | 1 | ; | HB2 | - | CD1 |
| 2341 | 2352 | 1 | ; | HB2 | - | CD2 |
| 2341 | 2354 | 1 | ; | HB2 | - | C |
| 2342 | 2344 | 1 | ; | HB3 | - | CD1 |
| 2342 | 2352 | 1 | ; | HB3 | - | CD2 |
| 2342 | 2354 | 1 | ; | HB3 | - | C |
| 2343 | 2347 | 1 | ; | CG | - | HE1 |
| 2343 | 2348 | 1 | ; | CG | - | CZ |
| 2343 | 2351 | 1 | ; | CG | - | HE2 |
| 2343 | 2354 | 1 | ; | CG | - | C |
| 2344 | 2349 | 1 | ; | CD1 | - | HZ |
| 2344 | 2350 | 1 | ; | CD1 | - | CE2 |
| 2344 | 2353 | 1 | ; | CD1 | - | HD2 |

| | | | | | | |
|---|---|---|---|---|---|---|
| 2345 | 2347 | 1 | ; | HD1 | - | HE1 |
| 2345 | 2348 | 1 | ; | HD1 | - | CZ |
| 2345 | 2352 | 1 | ; | HD1 | - | CD2 |
| 2346 | 2351 | 1 | ; | CE1 | - | HE2 |
| 2346 | 2352 | 1 | ; | CE1 | - | CD2 |
| 2347 | 2349 | 1 | ; | HE1 | - | HZ |
| 2347 | 2350 | 1 | ; | HE1 | - | CE2 |
| 2348 | 2353 | 1 | ; | CZ | - | HD2 |
| 2349 | 2351 | 1 | ; | HZ | - | HE2 |
| 2349 | 2352 | 1 | ; | HZ | - | CD2 |
| 2351 | 2353 | 1 | ; | HE2 | - | HD2 |
| 2354 | 2359 | 1 | ; | C | - | HA |
| 2354 | 2360 | 1 | ; | C | - | CB |
| 2354 | 2371 | 1 | ; | C | - | C |
| 2355 | 2357 | 1 | ; | O | - | H |
| 2355 | 2358 | 1 | ; | O | - | CA |
| 2356 | 2361 | 1 | ; | N | - | HB2 |
| 2356 | 2362 | 1 | ; | N | - | HB3 |
| 2356 | 2363 | 1 | ; | N | - | CG |
| 2356 | 2372 | 1 | ; | N | - | O |
| 2356 | 2373 | 1 | ; | N | - | N |
| 2357 | 2359 | 1 | ; | H | - | HA |
| 2357 | 2360 | 1 | ; | H | - | CB |
| 2357 | 2371 | 1 | ; | H | - | C |
| 2358 | 2364 | 1 | ; | CA | - | HG2 |
| 2358 | 2365 | 1 | ; | CA | - | HG3 |
| 2358 | 2366 | 1 | ; | CA | - | CD |
| 2358 | 2374 | 1 | ; | CA | - | H |
| 2358 | 2375 | 1 | ; | CA | - | CA |
| 2359 | 2361 | 1 | ; | HA | - | HB2 |
| 2359 | 2362 | 1 | ; | HA | - | HB3 |
| 2359 | 2363 | 1 | ; | HA | - | CG |
| 2359 | 2372 | 1 | ; | HA | - | O |
| 2359 | 2373 | 1 | ; | HA | - | N |
| 2360 | 2367 | 1 | ; | CB | - | OE1 |
| 2360 | 2368 | 1 | ; | CB | - | NE2 |
| 2360 | 2372 | 1 | ; | CB | - | O |
| 2360 | 2373 | 1 | ; | CB | - | N |
| 2361 | 2364 | 1 | ; | HB2 | - | HG2 |
| 2361 | 2365 | 1 | ; | HB2 | - | HG3 |
| 2361 | 2366 | 1 | ; | HB2 | - | CD |
| 2361 | 2371 | 1 | ; | HB2 | - | C |
| 2362 | 2364 | 1 | ; | HB3 | - | HG2 |
| 2362 | 2365 | 1 | ; | HB3 | - | HG3 |
| 2362 | 2366 | 1 | ; | HB3 | - | CD |
| 2362 | 2371 | 1 | ; | HB3 | - | C |
| 2363 | 2369 | 1 | ; | CG | - | HE21 |
| 2363 | 2370 | 1 | ; | CG | - | HE22 |
| 2363 | 2371 | 1 | ; | CG | - | C |
| 2364 | 2367 | 1 | ; | HG2 | - | OE1 |
| 2364 | 2368 | 1 | ; | HG2 | - | NE2 |
| 2365 | 2367 | 1 | ; | HG3 | - | OE1 |

| | | | |
|---|---|---|---|
| 2365 | 2368 | 1 ; | HG3 - NE2 |
| 2367 | 2369 | 1 ; | OE1 - HE21 |
| 2367 | 2370 | 1 ; | OE1 - HE22 |
| 2371 | 2376 | 1 ; | C - HA |
| 2371 | 2377 | 1 ; | C - CB |
| 2371 | 2382 | 1 ; | C - C |
| 2372 | 2374 | 1 ; | O - H |
| 2372 | 2375 | 1 ; | O - CA |
| 2373 | 2378 | 1 ; | N - HB2 |
| 2373 | 2379 | 1 ; | N - HB3 |
| 2373 | 2380 | 1 ; | N - OG |
| 2373 | 2383 | 1 ; | N - O |
| 2373 | 2384 | 1 ; | N - N |
| 2374 | 2376 | 1 ; | H - HA |
| 2374 | 2377 | 1 ; | H - CB |
| 2374 | 2382 | 1 ; | H - C |
| 2375 | 2381 | 1 ; | CA - HG |
| 2375 | 2385 | 1 ; | CA - H |
| 2375 | 2386 | 1 ; | CA - CA |
| 2376 | 2378 | 1 ; | HA - HB2 |
| 2376 | 2379 | 1 ; | HA - HB3 |
| 2376 | 2380 | 1 ; | HA - OG |
| 2376 | 2383 | 1 ; | HA - O |
| 2376 | 2384 | 1 ; | HA - N |
| 2377 | 2383 | 1 ; | CB - O |
| 2377 | 2384 | 1 ; | CB - N |
| 2378 | 2381 | 1 ; | HB2 - HG |
| 2378 | 2382 | 1 ; | HB2 - C |
| 2379 | 2381 | 1 ; | HB3 - HG |
| 2379 | 2382 | 1 ; | HB3 - C |
| 2380 | 2382 | 1 ; | OG - C |
| 2382 | 2387 | 1 ; | C - HA |
| 2382 | 2388 | 1 ; | C - CB |
| 2382 | 2397 | 1 ; | C - C |
| 2383 | 2385 | 1 ; | O - H |
| 2383 | 2386 | 1 ; | O - CA |
| 2384 | 2389 | 1 ; | N - HB2 |
| 2384 | 2390 | 1 ; | N - HB3 |
| 2384 | 2391 | 1 ; | N - CG |
| 2384 | 2398 | 1 ; | N - O |
| 2384 | 2399 | 1 ; | N - N |
| 2385 | 2387 | 1 ; | H - HA |
| 2385 | 2388 | 1 ; | H - CB |
| 2385 | 2397 | 1 ; | H - C |
| 2386 | 2392 | 1 ; | CA - HG2 |
| 2386 | 2393 | 1 ; | CA - HG3 |
| 2386 | 2394 | 1 ; | CA - CD |
| 2386 | 2400 | 1 ; | CA - H |
| 2386 | 2401 | 1 ; | CA - CA |
| 2387 | 2389 | 1 ; | HA - HB2 |
| 2387 | 2390 | 1 ; | HA - HB3 |
| 2387 | 2391 | 1 ; | HA - CG |

| | | | |
|---|---|---|---|
| 2387 | 2398 | 1 ; | HA - O |
| 2387 | 2399 | 1 ; | HA - N |
| 2388 | 2395 | 1 ; | CB - OE1 |
| 2388 | 2396 | 1 ; | CB - OE2 |
| 2388 | 2398 | 1 ; | CB - O |
| 2388 | 2399 | 1 ; | CB - N |
| 2389 | 2392 | 1 ; | HB2 - HG2 |
| 2389 | 2393 | 1 ; | HB2 - HG3 |
| 2389 | 2394 | 1 ; | HB2 - CD |
| 2389 | 2397 | 1 ; | HB2 - C |
| 2390 | 2392 | 1 ; | HB3 - HG2 |
| 2390 | 2393 | 1 ; | HB3 - HG3 |
| 2390 | 2394 | 1 ; | HB3 - CD |
| 2390 | 2397 | 1 ; | HB3 - C |
| 2391 | 2397 | 1 ; | CG - C |
| 2392 | 2395 | 1 ; | HG2 - OE1 |
| 2392 | 2396 | 1 ; | HG2 - OE2 |
| 2393 | 2395 | 1 ; | HG3 - OE1 |
| 2393 | 2396 | 1 ; | HG3 - OE2 |
| 2397 | 2402 | 1 ; | C - HA |
| 2397 | 2403 | 1 ; | C - CB |
| 2397 | 2411 | 1 ; | C - C |
| 2398 | 2400 | 1 ; | O - H |
| 2398 | 2401 | 1 ; | O - CA |
| 2399 | 2404 | 1 ; | N - HB2 |
| 2399 | 2405 | 1 ; | N - HB3 |
| 2399 | 2406 | 1 ; | N - CG |
| 2399 | 2412 | 1 ; | N - O |
| 2399 | 2413 | 1 ; | N - N |
| 2400 | 2402 | 1 ; | H - HA |
| 2400 | 2403 | 1 ; | H - CB |
| 2400 | 2411 | 1 ; | H - C |
| 2401 | 2407 | 1 ; | CA - OD1 |
| 2401 | 2408 | 1 ; | CA - ND2 |
| 2401 | 2414 | 1 ; | CA - H |
| 2401 | 2415 | 1 ; | CA - CA |
| 2402 | 2404 | 1 ; | HA - HB2 |
| 2402 | 2405 | 1 ; | HA - HB3 |
| 2402 | 2406 | 1 ; | HA - CG |
| 2402 | 2412 | 1 ; | HA - O |
| 2402 | 2413 | 1 ; | HA - N |
| 2403 | 2409 | 1 ; | CB - HD21 |
| 2403 | 2410 | 1 ; | CB - HD22 |
| 2403 | 2412 | 1 ; | CB - O |
| 2403 | 2413 | 1 ; | CB - N |
| 2404 | 2407 | 1 ; | HB2 - OD1 |
| 2404 | 2408 | 1 ; | HB2 - ND2 |
| 2404 | 2411 | 1 ; | HB2 - C |
| 2405 | 2407 | 1 ; | HB3 - OD1 |
| 2405 | 2408 | 1 ; | HB3 - ND2 |
| 2405 | 2411 | 1 ; | HB3 - C |
| 2406 | 2411 | 1 ; | CG - C |

```
2407   2409       1 ;    OD1 - HD21
2407   2410       1 ;    OD1 - HD22
2411   2416       1 ;      C - HA
2411   2417       1 ;      C - CB
2411   2433       1 ;      C - C
2412   2414       1 ;      O - H
2412   2415       1 ;      O - CA
2413   2418       1 ;      N - HB2
2413   2419       1 ;      N - HB3
2413   2420       1 ;      N - CG
2413   2434       1 ;      N - O
2413   2435       1 ;      N - N
2414   2416       1 ;      H - HA
2414   2417       1 ;      H - CB
2414   2433       1 ;      H - C
2415   2421       1 ;     CA - HG2
2415   2422       1 ;     CA - HG3
2415   2423       1 ;     CA - CD
2415   2436       1 ;     CA - H
2415   2437       1 ;     CA - CA
2416   2418       1 ;     HA - HB2
2416   2419       1 ;     HA - HB3
2416   2420       1 ;     HA - CG
2416   2434       1 ;     HA - O
2416   2435       1 ;     HA - N
2417   2424       1 ;     CB - HD2
2417   2425       1 ;     CB - HD3
2417   2426       1 ;     CB - CE
2417   2434       1 ;     CB - O
2417   2435       1 ;     CB - N
2418   2421       1 ;    HB2 - HG2
2418   2422       1 ;    HB2 - HG3
2418   2423       1 ;    HB2 - CD
2418   2433       1 ;    HB2 - C
2419   2421       1 ;    HB3 - HG2
2419   2422       1 ;    HB3 - HG3
2419   2423       1 ;    HB3 - CD
2419   2433       1 ;    HB3 - C
2420   2427       1 ;     CG - HE2
2420   2428       1 ;     CG - HE3
2420   2429       1 ;     CG - NZ
2420   2433       1 ;     CG - C
2421   2424       1 ;    HG2 - HD2
2421   2425       1 ;    HG2 - HD3
2421   2426       1 ;    HG2 - CE
2422   2424       1 ;    HG3 - HD2
2422   2425       1 ;    HG3 - HD3
2422   2426       1 ;    HG3 - CE
2423   2430       1 ;     CD - HZ1
2423   2431       1 ;     CD - HZ2
2423   2432       1 ;     CD - HZ3
2424   2427       1 ;    HD2 - HE2
```

| | | | |
|---|---|---|---|
| 2424 | 2428 | 1 ; | HD2 - HE3 |
| 2424 | 2429 | 1 ; | HD2 - NZ |
| 2425 | 2427 | 1 ; | HD3 - HE2 |
| 2425 | 2428 | 1 ; | HD3 - HE3 |
| 2425 | 2429 | 1 ; | HD3 - NZ |
| 2427 | 2430 | 1 ; | HE2 - HZ1 |
| 2427 | 2431 | 1 ; | HE2 - HZ2 |
| 2427 | 2432 | 1 ; | HE2 - HZ3 |
| 2428 | 2430 | 1 ; | HE3 - HZ1 |
| 2428 | 2431 | 1 ; | HE3 - HZ2 |
| 2428 | 2432 | 1 ; | HE3 - HZ3 |
| 2433 | 2438 | 1 ; | C - HA |
| 2433 | 2439 | 1 ; | C - CB |
| 2433 | 2445 | 1 ; | C - C |
| 2434 | 2436 | 1 ; | O - H |
| 2434 | 2437 | 1 ; | O - CA |
| 2435 | 2440 | 1 ; | N - HB2 |
| 2435 | 2441 | 1 ; | N - HB3 |
| 2435 | 2442 | 1 ; | N - CG |
| 2435 | 2446 | 1 ; | N - O |
| 2435 | 2447 | 1 ; | N - N |
| 2436 | 2438 | 1 ; | H - HA |
| 2436 | 2439 | 1 ; | H - CB |
| 2436 | 2445 | 1 ; | H - C |
| 2437 | 2443 | 1 ; | CA - OD1 |
| 2437 | 2444 | 1 ; | CA - OD2 |
| 2437 | 2448 | 1 ; | CA - H |
| 2437 | 2449 | 1 ; | CA - CA |
| 2438 | 2440 | 1 ; | HA - HB2 |
| 2438 | 2441 | 1 ; | HA - HB3 |
| 2438 | 2442 | 1 ; | HA - CG |
| 2438 | 2446 | 1 ; | HA - O |
| 2438 | 2447 | 1 ; | HA - N |
| 2439 | 2446 | 1 ; | CB - O |
| 2439 | 2447 | 1 ; | CB - N |
| 2440 | 2443 | 1 ; | HB2 - OD1 |
| 2440 | 2444 | 1 ; | HB2 - OD2 |
| 2440 | 2445 | 1 ; | HB2 - C |
| 2441 | 2443 | 1 ; | HB3 - OD1 |
| 2441 | 2444 | 1 ; | HB3 - OD2 |
| 2441 | 2445 | 1 ; | HB3 - C |
| 2442 | 2445 | 1 ; | CG - C |
| 2445 | 2450 | 1 ; | C - HA |
| 2445 | 2451 | 1 ; | C - CB |
| 2445 | 2456 | 1 ; | C - C |
| 2446 | 2448 | 1 ; | O - H |
| 2446 | 2449 | 1 ; | O - CA |
| 2447 | 2452 | 1 ; | N - HB2 |
| 2447 | 2453 | 1 ; | N - HB3 |
| 2447 | 2454 | 1 ; | N - OG |
| 2447 | 2457 | 1 ; | N - O |
| 2447 | 2458 | 1 ; | N - N |

```
2448   2450       1 ;       H - HA
2448   2451       1 ;       H - CB
2448   2456       1 ;       H - C
2449   2455       1 ;      CA - HG
2449   2459       1 ;      CA - H
2449   2460       1 ;      CA - CA
2450   2452       1 ;      HA - HB2
2450   2453       1 ;      HA - HB3
2450   2454       1 ;      HA - OG
2450   2457       1 ;      HA - O
2450   2458       1 ;      HA - N
2451   2457       1 ;      CB - O
2451   2458       1 ;      CB - N
2452   2455       1 ;     HB2 - HG
2452   2456       1 ;     HB2 - C
2453   2455       1 ;     HB3 - HG
2453   2456       1 ;     HB3 - C
2454   2456       1 ;      OG - C
2456   2461       1 ;       C - HA
2456   2462       1 ;       C - CB
2456   2475       1 ;       C - C
2457   2459       1 ;       O - H
2457   2460       1 ;       O - CA
2458   2463       1 ;       N - HB2
2458   2464       1 ;       N - HB3
2458   2465       1 ;       N - CG
2458   2476       1 ;       N - O
2458   2477       1 ;       N - N
2459   2461       1 ;       H - HA
2459   2462       1 ;       H - CB
2459   2475       1 ;       H - C
2460   2466       1 ;      CA - HG
2460   2467       1 ;      CA - CD1
2460   2471       1 ;      CA - CD2
2460   2478       1 ;      CA - H
2460   2479       1 ;      CA - CA
2461   2463       1 ;      HA - HB2
2461   2464       1 ;      HA - HB3
2461   2465       1 ;      HA - CG
2461   2476       1 ;      HA - O
2461   2477       1 ;      HA - N
2462   2468       1 ;      CB - HD11
2462   2469       1 ;      CB - HD12
2462   2470       1 ;      CB - HD13
2462   2472       1 ;      CB - HD21
2462   2473       1 ;      CB - HD22
2462   2474       1 ;      CB - HD23
2462   2476       1 ;      CB - O
2462   2477       1 ;      CB - N
2463   2466       1 ;     HB2 - HG
2463   2467       1 ;     HB2 - CD1
2463   2471       1 ;     HB2 - CD2
```

```
2463   2475       1 ;     HB2 - C
2464   2466       1 ;     HB3 - HG
2464   2467       1 ;     HB3 - CD1
2464   2471       1 ;     HB3 - CD2
2464   2475       1 ;     HB3 - C
2465   2475       1 ;      CG - C
2466   2468       1 ;      HG - HD11
2466   2469       1 ;      HG - HD12
2466   2470       1 ;      HG - HD13
2466   2472       1 ;      HG - HD21
2466   2473       1 ;      HG - HD22
2466   2474       1 ;      HG - HD23
2467   2472       1 ;     CD1 - HD21
2467   2473       1 ;     CD1 - HD22
2467   2474       1 ;     CD1 - HD23
2468   2471       1 ;    HD11 - CD2
2469   2471       1 ;    HD12 - CD2
2470   2471       1 ;    HD13 - CD2
2475   2480       1 ;       C - HA
2475   2481       1 ;       C - CB
2475   2494       1 ;       C - C
2476   2478       1 ;       O - H
2476   2479       1 ;       O - CA
2477   2482       1 ;       N - HB2
2477   2483       1 ;       N - HB3
2477   2484       1 ;       N - CG
2477   2495       1 ;       N - O
2477   2496       1 ;       N - N
2478   2480       1 ;       H - HA
2478   2481       1 ;       H - CB
2478   2494       1 ;       H - C
2479   2485       1 ;      CA - HG
2479   2486       1 ;      CA - CD1
2479   2490       1 ;      CA - CD2
2479   2497       1 ;      CA - H
2479   2498       1 ;      CA - CA
2480   2482       1 ;      HA - HB2
2480   2483       1 ;      HA - HB3
2480   2484       1 ;      HA - CG
2480   2495       1 ;      HA - O
2480   2496       1 ;      HA - N
2481   2487       1 ;      CB - HD11
2481   2488       1 ;      CB - HD12
2481   2489       1 ;      CB - HD13
2481   2491       1 ;      CB - HD21
2481   2492       1 ;      CB - HD22
2481   2493       1 ;      CB - HD23
2481   2495       1 ;      CB - O
2481   2496       1 ;      CB - N
2482   2485       1 ;     HB2 - HG
2482   2486       1 ;     HB2 - CD1
2482   2490       1 ;     HB2 - CD2
```

| | | | | | |
|---|---|---|---|---|---|
| 2482 | 2494 | 1 ; | HB2 | - | C |
| 2483 | 2485 | 1 ; | HB3 | - | HG |
| 2483 | 2486 | 1 ; | HB3 | - | CD1 |
| 2483 | 2490 | 1 ; | HB3 | - | CD2 |
| 2483 | 2494 | 1 ; | HB3 | - | C |
| 2484 | 2494 | 1 ; | CG | - | C |
| 2485 | 2487 | 1 ; | HG | - | HD11 |
| 2485 | 2488 | 1 ; | HG | - | HD12 |
| 2485 | 2489 | 1 ; | HG | - | HD13 |
| 2485 | 2491 | 1 ; | HG | - | HD21 |
| 2485 | 2492 | 1 ; | HG | - | HD22 |
| 2485 | 2493 | 1 ; | HG | - | HD23 |
| 2486 | 2491 | 1 ; | CD1 | - | HD21 |
| 2486 | 2492 | 1 ; | CD1 | - | HD22 |
| 2486 | 2493 | 1 ; | CD1 | - | HD23 |
| 2487 | 2490 | 1 ; | HD11 | - | CD2 |
| 2488 | 2490 | 1 ; | HD12 | - | CD2 |
| 2489 | 2490 | 1 ; | HD13 | - | CD2 |
| 2494 | 2499 | 1 ; | C | - | HA |
| 2494 | 2500 | 1 ; | C | - | CB |
| 2494 | 2516 | 1 ; | C | - | C |
| 2495 | 2497 | 1 ; | O | - | H |
| 2495 | 2498 | 1 ; | O | - | CA |
| 2496 | 2501 | 1 ; | N | - | HB2 |
| 2496 | 2502 | 1 ; | N | - | HB3 |
| 2496 | 2503 | 1 ; | N | - | CG |
| 2496 | 2517 | 1 ; | N | - | O |
| 2496 | 2518 | 1 ; | N | - | N |
| 2497 | 2499 | 1 ; | H | - | HA |
| 2497 | 2500 | 1 ; | H | - | CB |
| 2497 | 2516 | 1 ; | H | - | C |
| 2498 | 2504 | 1 ; | CA | - | HG2 |
| 2498 | 2505 | 1 ; | CA | - | HG3 |
| 2498 | 2506 | 1 ; | CA | - | CD |
| 2498 | 2519 | 1 ; | CA | - | H |
| 2498 | 2520 | 1 ; | CA | - | CA |
| 2499 | 2501 | 1 ; | HA | - | HB2 |
| 2499 | 2502 | 1 ; | HA | - | HB3 |
| 2499 | 2503 | 1 ; | HA | - | CG |
| 2499 | 2517 | 1 ; | HA | - | O |
| 2499 | 2518 | 1 ; | HA | - | N |
| 2500 | 2507 | 1 ; | CB | - | HD2 |
| 2500 | 2508 | 1 ; | CB | - | HD3 |
| 2500 | 2509 | 1 ; | CB | - | CE |
| 2500 | 2517 | 1 ; | CB | - | O |
| 2500 | 2518 | 1 ; | CB | - | N |
| 2501 | 2504 | 1 ; | HB2 | - | HG2 |
| 2501 | 2505 | 1 ; | HB2 | - | HG3 |
| 2501 | 2506 | 1 ; | HB2 | - | CD |
| 2501 | 2516 | 1 ; | HB2 | - | C |
| 2502 | 2504 | 1 ; | HB3 | - | HG2 |
| 2502 | 2505 | 1 ; | HB3 | - | HG3 |

| | | | | |
|---|---|---|---|---|
| 2502 | 2506 | 1 | ; | HB3 - CD |
| 2502 | 2516 | 1 | ; | HB3 - C |
| 2503 | 2510 | 1 | ; | CG - HE2 |
| 2503 | 2511 | 1 | ; | CG - HE3 |
| 2503 | 2512 | 1 | ; | CG - NZ |
| 2503 | 2516 | 1 | ; | CG - C |
| 2504 | 2507 | 1 | ; | HG2 - HD2 |
| 2504 | 2508 | 1 | ; | HG2 - HD3 |
| 2504 | 2509 | 1 | ; | HG2 - CE |
| 2505 | 2507 | 1 | ; | HG3 - HD2 |
| 2505 | 2508 | 1 | ; | HG3 - HD3 |
| 2505 | 2509 | 1 | ; | HG3 - CE |
| 2506 | 2513 | 1 | ; | CD - HZ1 |
| 2506 | 2514 | 1 | ; | CD - HZ2 |
| 2506 | 2515 | 1 | ; | CD - HZ3 |
| 2507 | 2510 | 1 | ; | HD2 - HE2 |
| 2507 | 2511 | 1 | ; | HD2 - HE3 |
| 2507 | 2512 | 1 | ; | HD2 - NZ |
| 2508 | 2510 | 1 | ; | HD3 - HE2 |
| 2508 | 2511 | 1 | ; | HD3 - HE3 |
| 2508 | 2512 | 1 | ; | HD3 - NZ |
| 2510 | 2513 | 1 | ; | HE2 - HZ1 |
| 2510 | 2514 | 1 | ; | HE2 - HZ2 |
| 2510 | 2515 | 1 | ; | HE2 - HZ3 |
| 2511 | 2513 | 1 | ; | HE3 - HZ1 |
| 2511 | 2514 | 1 | ; | HE3 - HZ2 |
| 2511 | 2515 | 1 | ; | HE3 - HZ3 |
| 2516 | 2521 | 1 | ; | C - HA |
| 2516 | 2522 | 1 | ; | C - CB |
| 2516 | 2535 | 1 | ; | C - C |
| 2517 | 2519 | 1 | ; | O - H |
| 2517 | 2520 | 1 | ; | O - CA |
| 2518 | 2523 | 1 | ; | N - HB |
| 2518 | 2524 | 1 | ; | N - CG2 |
| 2518 | 2528 | 1 | ; | N - CG1 |
| 2518 | 2536 | 1 | ; | N - O |
| 2518 | 2537 | 1 | ; | N - N |
| 2519 | 2521 | 1 | ; | H - HA |
| 2519 | 2522 | 1 | ; | H - CB |
| 2519 | 2535 | 1 | ; | H - C |
| 2520 | 2525 | 1 | ; | CA - HG21 |
| 2520 | 2526 | 1 | ; | CA - HG22 |
| 2520 | 2527 | 1 | ; | CA - HG23 |
| 2520 | 2529 | 1 | ; | CA - HG12 |
| 2520 | 2530 | 1 | ; | CA - HG13 |
| 2520 | 2531 | 1 | ; | CA - CD1 |
| 2520 | 2538 | 1 | ; | CA - H |
| 2520 | 2539 | 1 | ; | CA - CA |
| 2521 | 2523 | 1 | ; | HA - HB |
| 2521 | 2524 | 1 | ; | HA - CG2 |
| 2521 | 2528 | 1 | ; | HA - CG1 |
| 2521 | 2536 | 1 | ; | HA - O |

| | | | | |
|---|---|---|---|---|
| 2521 | 2537 | 1 | ; | HA - N |
| 2522 | 2532 | 1 | ; | CB - HD11 |
| 2522 | 2533 | 1 | ; | CB - HD12 |
| 2522 | 2534 | 1 | ; | CB - HD13 |
| 2522 | 2536 | 1 | ; | CB - O |
| 2522 | 2537 | 1 | ; | CB - N |
| 2523 | 2525 | 1 | ; | HB - HG21 |
| 2523 | 2526 | 1 | ; | HB - HG22 |
| 2523 | 2527 | 1 | ; | HB - HG23 |
| 2523 | 2529 | 1 | ; | HB - HG12 |
| 2523 | 2530 | 1 | ; | HB - HG13 |
| 2523 | 2531 | 1 | ; | HB - CD1 |
| 2523 | 2535 | 1 | ; | HB - C |
| 2524 | 2529 | 1 | ; | CG2 - HG12 |
| 2524 | 2530 | 1 | ; | CG2 - HG13 |
| 2524 | 2531 | 1 | ; | CG2 - CD1 |
| 2524 | 2535 | 1 | ; | CG2 - C |
| 2525 | 2528 | 1 | ; | HG21 - CG1 |
| 2526 | 2528 | 1 | ; | HG22 - CG1 |
| 2527 | 2528 | 1 | ; | HG23 - CG1 |
| 2528 | 2535 | 1 | ; | CG1 - C |
| 2529 | 2532 | 1 | ; | HG12 - HD11 |
| 2529 | 2533 | 1 | ; | HG12 - HD12 |
| 2529 | 2534 | 1 | ; | HG12 - HD13 |
| 2530 | 2532 | 1 | ; | HG13 - HD11 |
| 2530 | 2533 | 1 | ; | HG13 - HD12 |
| 2530 | 2534 | 1 | ; | HG13 - HD13 |
| 2535 | 2540 | 1 | ; | C - HA |
| 2535 | 2541 | 1 | ; | C - CB |
| 2535 | 2554 | 1 | ; | C - C |
| 2536 | 2538 | 1 | ; | O - H |
| 2536 | 2539 | 1 | ; | O - CA |
| 2537 | 2542 | 1 | ; | N - HB |
| 2537 | 2543 | 1 | ; | N - CG2 |
| 2537 | 2547 | 1 | ; | N - CG1 |
| 2537 | 2555 | 1 | ; | N - O |
| 2537 | 2556 | 1 | ; | N - N |
| 2538 | 2540 | 1 | ; | H - HA |
| 2538 | 2541 | 1 | ; | H - CB |
| 2538 | 2554 | 1 | ; | H - C |
| 2539 | 2544 | 1 | ; | CA - HG21 |
| 2539 | 2545 | 1 | ; | CA - HG22 |
| 2539 | 2546 | 1 | ; | CA - HG23 |
| 2539 | 2548 | 1 | ; | CA - HG12 |
| 2539 | 2549 | 1 | ; | CA - HG13 |
| 2539 | 2550 | 1 | ; | CA - CD1 |
| 2539 | 2557 | 1 | ; | CA - H |
| 2539 | 2558 | 1 | ; | CA - CA |
| 2540 | 2542 | 1 | ; | HA - HB |
| 2540 | 2543 | 1 | ; | HA - CG2 |
| 2540 | 2547 | 1 | ; | HA - CG1 |
| 2540 | 2555 | 1 | ; | HA - O |

```
2540   2556       1 ;      HA - N
2541   2551       1 ;      CB - HD11
2541   2552       1 ;      CB - HD12
2541   2553       1 ;      CB - HD13
2541   2555       1 ;      CB - O
2541   2556       1 ;      CB - N
2542   2544       1 ;      HB - HG21
2542   2545       1 ;      HB - HG22
2542   2546       1 ;      HB - HG23
2542   2548       1 ;      HB - HG12
2542   2549       1 ;      HB - HG13
2542   2550       1 ;      HB - CD1
2542   2554       1 ;      HB - C
2543   2548       1 ;     CG2 - HG12
2543   2549       1 ;     CG2 - HG13
2543   2550       1 ;     CG2 - CD1
2543   2554       1 ;     CG2 - C
2544   2547       1 ;    HG21 - CG1
2545   2547       1 ;    HG22 - CG1
2546   2547       1 ;    HG23 - CG1
2547   2554       1 ;     CG1 - C
2548   2551       1 ;    HG12 - HD11
2548   2552       1 ;    HG12 - HD12
2548   2553       1 ;    HG12 - HD13
2549   2551       1 ;    HG13 - HD11
2549   2552       1 ;    HG13 - HD12
2549   2553       1 ;    HG13 - HD13
2554   2559       1 ;       C - HA
2554   2560       1 ;       C - CB
2554   2566       1 ;       C - C
2555   2557       1 ;       O - H
2555   2558       1 ;       O - CA
2556   2561       1 ;       N - HB2
2556   2562       1 ;       N - HB3
2556   2563       1 ;       N - CG
2556   2567       1 ;       N - O
2556   2568       1 ;       N - N
2557   2559       1 ;       H - HA
2557   2560       1 ;       H - CB
2557   2566       1 ;       H - C
2558   2564       1 ;      CA - OD1
2558   2565       1 ;      CA - OD2
2558   2569       1 ;      CA - H
2558   2570       1 ;      CA - CA
2559   2561       1 ;      HA - HB2
2559   2562       1 ;      HA - HB3
2559   2563       1 ;      HA - CG
2559   2567       1 ;      HA - O
2559   2568       1 ;      HA - N
2560   2567       1 ;      CB - O
2560   2568       1 ;      CB - N
2561   2564       1 ;     HB2 - OD1
```

```
2561   2565       1 ;     HB2 - OD2
2561   2566       1 ;     HB2 - C
2562   2564       1 ;     HB3 - OD1
2562   2565       1 ;     HB3 - OD2
2562   2566       1 ;     HB3 - C
2563   2566       1 ;      CG - C
2566   2571       1 ;       C - HA
2566   2572       1 ;       C - CB
2566   2586       1 ;       C - C
2567   2569       1 ;       O - H
2567   2570       1 ;       O - CA
2568   2573       1 ;       N - HB2
2568   2574       1 ;       N - HB3
2568   2575       1 ;       N - CG
2568   2587       1 ;       N - O
2568   2588       1 ;       N - N
2569   2571       1 ;       H - HA
2569   2572       1 ;       H - CB
2569   2586       1 ;       H - C
2570   2576       1 ;      CA - CD1
2570   2584       1 ;      CA - CD2
2570   2589       1 ;      CA - H
2570   2590       1 ;      CA - CA
2571   2573       1 ;      HA - HB2
2571   2574       1 ;      HA - HB3
2571   2575       1 ;      HA - CG
2571   2587       1 ;      HA - O
2571   2588       1 ;      HA - N
2572   2577       1 ;      CB - HD1
2572   2578       1 ;      CB - CE1
2572   2582       1 ;      CB - CE2
2572   2585       1 ;      CB - HD2
2572   2587       1 ;      CB - O
2572   2588       1 ;      CB - N
2573   2576       1 ;     HB2 - CD1
2573   2584       1 ;     HB2 - CD2
2573   2586       1 ;     HB2 - C
2574   2576       1 ;     HB3 - CD1
2574   2584       1 ;     HB3 - CD2
2574   2586       1 ;     HB3 - C
2575   2579       1 ;      CG - HE1
2575   2580       1 ;      CG - CZ
2575   2583       1 ;      CG - HE2
2575   2586       1 ;      CG - C
2576   2581       1 ;     CD1 - HZ
2576   2582       1 ;     CD1 - CE2
2576   2585       1 ;     CD1 - HD2
2577   2579       1 ;     HD1 - HE1
2577   2580       1 ;     HD1 - CZ
2577   2584       1 ;     HD1 - CD2
2578   2583       1 ;     CE1 - HE2
2578   2584       1 ;     CE1 - CD2
```

| | | | | | | |
|---|---|---|---|---|---|---|
| 2579 | 2581 | 1 | ; | HE1 | - | HZ |
| 2579 | 2582 | 1 | ; | HE1 | - | CE2 |
| 2580 | 2585 | 1 | ; | CZ | - | HD2 |
| 2581 | 2583 | 1 | ; | HZ | - | HE2 |
| 2581 | 2584 | 1 | ; | HZ | - | CD2 |
| 2583 | 2585 | 1 | ; | HE2 | - | HD2 |
| 2586 | 2591 | 1 | ; | C | - | HA2 |
| 2586 | 2592 | 1 | ; | C | - | HA3 |
| 2586 | 2593 | 1 | ; | C | - | C |
| 2587 | 2589 | 1 | ; | O | - | H |
| 2587 | 2590 | 1 | ; | O | - | CA |
| 2588 | 2594 | 1 | ; | N | - | O |
| 2588 | 2595 | 1 | ; | N | - | N |
| 2589 | 2591 | 1 | ; | H | - | HA2 |
| 2589 | 2592 | 1 | ; | H | - | HA3 |
| 2589 | 2593 | 1 | ; | H | - | C |
| 2590 | 2596 | 1 | ; | CA | - | H |
| 2590 | 2597 | 1 | ; | CA | - | CA |
| 2591 | 2594 | 1 | ; | HA2 | - | O |
| 2591 | 2595 | 1 | ; | HA2 | - | N |
| 2592 | 2594 | 1 | ; | HA3 | - | O |
| 2592 | 2595 | 1 | ; | HA3 | - | N |
| 2593 | 2598 | 1 | ; | C | - | HA |
| 2593 | 2599 | 1 | ; | C | - | CB |
| 2593 | 2612 | 1 | ; | C | - | C |
| 2594 | 2596 | 1 | ; | O | - | H |
| 2594 | 2597 | 1 | ; | O | - | CA |
| 2595 | 2600 | 1 | ; | N | - | HB2 |
| 2595 | 2601 | 1 | ; | N | - | HB3 |
| 2595 | 2602 | 1 | ; | N | - | CG |
| 2595 | 2613 | 1 | ; | N | - | O |
| 2595 | 2614 | 1 | ; | N | - | N |
| 2596 | 2598 | 1 | ; | H | - | HA |
| 2596 | 2599 | 1 | ; | H | - | CB |
| 2596 | 2612 | 1 | ; | H | - | C |
| 2597 | 2603 | 1 | ; | CA | - | HG |
| 2597 | 2604 | 1 | ; | CA | - | CD1 |
| 2597 | 2608 | 1 | ; | CA | - | CD2 |
| 2597 | 2615 | 1 | ; | CA | - | H |
| 2597 | 2616 | 1 | ; | CA | - | CA |
| 2598 | 2600 | 1 | ; | HA | - | HB2 |
| 2598 | 2601 | 1 | ; | HA | - | HB3 |
| 2598 | 2602 | 1 | ; | HA | - | CG |
| 2598 | 2613 | 1 | ; | HA | - | O |
| 2598 | 2614 | 1 | ; | HA | - | N |
| 2599 | 2605 | 1 | ; | CB | - | HD11 |
| 2599 | 2606 | 1 | ; | CB | - | HD12 |
| 2599 | 2607 | 1 | ; | CB | - | HD13 |
| 2599 | 2609 | 1 | ; | CB | - | HD21 |
| 2599 | 2610 | 1 | ; | CB | - | HD22 |
| 2599 | 2611 | 1 | ; | CB | - | HD23 |
| 2599 | 2613 | 1 | ; | CB | - | O |

| | | | | | |
|---|---|---|---|---|---|
| 2599 | 2614 | 1 ; | CB | - | N |
| 2600 | 2603 | 1 ; | HB2 | - | HG |
| 2600 | 2604 | 1 ; | HB2 | - | CD1 |
| 2600 | 2608 | 1 ; | HB2 | - | CD2 |
| 2600 | 2612 | 1 ; | HB2 | - | C |
| 2601 | 2603 | 1 ; | HB3 | - | HG |
| 2601 | 2604 | 1 ; | HB3 | - | CD1 |
| 2601 | 2608 | 1 ; | HB3 | - | CD2 |
| 2601 | 2612 | 1 ; | HB3 | - | C |
| 2602 | 2612 | 1 ; | CG | - | C |
| 2603 | 2605 | 1 ; | HG | - | HD11 |
| 2603 | 2606 | 1 ; | HG | - | HD12 |
| 2603 | 2607 | 1 ; | HG | - | HD13 |
| 2603 | 2609 | 1 ; | HG | - | HD21 |
| 2603 | 2610 | 1 ; | HG | - | HD22 |
| 2603 | 2611 | 1 ; | HG | - | HD23 |
| 2604 | 2609 | 1 ; | CD1 | - | HD21 |
| 2604 | 2610 | 1 ; | CD1 | - | HD22 |
| 2604 | 2611 | 1 ; | CD1 | - | HD23 |
| 2605 | 2608 | 1 ; | HD11 | - | CD2 |
| 2606 | 2608 | 1 ; | HD12 | - | CD2 |
| 2607 | 2608 | 1 ; | HD13 | - | CD2 |
| 2612 | 2617 | 1 ; | C | - | HA |
| 2612 | 2618 | 1 ; | C | - | CB |
| 2612 | 2623 | 1 ; | C | - | C |
| 2613 | 2615 | 1 ; | O | - | H |
| 2613 | 2616 | 1 ; | O | - | CA |
| 2614 | 2619 | 1 ; | N | - | HB2 |
| 2614 | 2620 | 1 ; | N | - | HB3 |
| 2614 | 2621 | 1 ; | N | - | OG |
| 2614 | 2624 | 1 ; | N | - | O |
| 2614 | 2625 | 1 ; | N | - | N |
| 2615 | 2617 | 1 ; | H | - | HA |
| 2615 | 2618 | 1 ; | H | - | CB |
| 2615 | 2623 | 1 ; | H | - | C |
| 2616 | 2622 | 1 ; | CA | - | HG |
| 2616 | 2626 | 1 ; | CA | - | H |
| 2616 | 2627 | 1 ; | CA | - | CA |
| 2617 | 2619 | 1 ; | HA | - | HB2 |
| 2617 | 2620 | 1 ; | HA | - | HB3 |
| 2617 | 2621 | 1 ; | HA | - | OG |
| 2617 | 2624 | 1 ; | HA | - | O |
| 2617 | 2625 | 1 ; | HA | - | N |
| 2618 | 2624 | 1 ; | CB | - | O |
| 2618 | 2625 | 1 ; | CB | - | N |
| 2619 | 2622 | 1 ; | HB2 | - | HG |
| 2619 | 2623 | 1 ; | HB2 | - | C |
| 2620 | 2622 | 1 ; | HB3 | - | HG |
| 2620 | 2623 | 1 ; | HB3 | - | C |
| 2621 | 2623 | 1 ; | OG | - | C |
| 2623 | 2628 | 1 ; | C | - | HA |
| 2623 | 2629 | 1 ; | C | - | CB |

| | | | | | |
|---|---|---|---|---|---|
| 2623 | 2645 | 1 ; | C | - | C |
| 2624 | 2626 | 1 ; | O | - | H |
| 2624 | 2627 | 1 ; | O | - | CA |
| 2625 | 2630 | 1 ; | N | - | HB2 |
| 2625 | 2631 | 1 ; | N | - | HB3 |
| 2625 | 2632 | 1 ; | N | - | CG |
| 2625 | 2646 | 1 ; | N | - | O |
| 2625 | 2647 | 1 ; | N | - | N |
| 2626 | 2628 | 1 ; | H | - | HA |
| 2626 | 2629 | 1 ; | H | - | CB |
| 2626 | 2645 | 1 ; | H | - | C |
| 2627 | 2633 | 1 ; | CA | - | HG2 |
| 2627 | 2634 | 1 ; | CA | - | HG3 |
| 2627 | 2635 | 1 ; | CA | - | CD |
| 2627 | 2648 | 1 ; | CA | - | H |
| 2627 | 2649 | 1 ; | CA | - | CA |
| 2628 | 2630 | 1 ; | HA | - | HB2 |
| 2628 | 2631 | 1 ; | HA | - | HB3 |
| 2628 | 2632 | 1 ; | HA | - | CG |
| 2628 | 2646 | 1 ; | HA | - | O |
| 2628 | 2647 | 1 ; | HA | - | N |
| 2629 | 2636 | 1 ; | CB | - | HD2 |
| 2629 | 2637 | 1 ; | CB | - | HD3 |
| 2629 | 2638 | 1 ; | CB | - | CE |
| 2629 | 2646 | 1 ; | CB | - | O |
| 2629 | 2647 | 1 ; | CB | - | N |
| 2630 | 2633 | 1 ; | HB2 | - | HG2 |
| 2630 | 2634 | 1 ; | HB2 | - | HG3 |
| 2630 | 2635 | 1 ; | HB2 | - | CD |
| 2630 | 2645 | 1 ; | HB2 | - | C |
| 2631 | 2633 | 1 ; | HB3 | - | HG2 |
| 2631 | 2634 | 1 ; | HB3 | - | HG3 |
| 2631 | 2635 | 1 ; | HB3 | - | CD |
| 2631 | 2645 | 1 ; | HB3 | - | C |
| 2632 | 2639 | 1 ; | CG | - | HE2 |
| 2632 | 2640 | 1 ; | CG | - | HE3 |
| 2632 | 2641 | 1 ; | CG | - | NZ |
| 2632 | 2645 | 1 ; | CG | - | C |
| 2633 | 2636 | 1 ; | HG2 | - | HD2 |
| 2633 | 2637 | 1 ; | HG2 | - | HD3 |
| 2633 | 2638 | 1 ; | HG2 | - | CE |
| 2634 | 2636 | 1 ; | HG3 | - | HD2 |
| 2634 | 2637 | 1 ; | HG3 | - | HD3 |
| 2634 | 2638 | 1 ; | HG3 | - | CE |
| 2635 | 2642 | 1 ; | CD | - | HZ1 |
| 2635 | 2643 | 1 ; | CD | - | HZ2 |
| 2635 | 2644 | 1 ; | CD | - | HZ3 |
| 2636 | 2639 | 1 ; | HD2 | - | HE2 |
| 2636 | 2640 | 1 ; | HD2 | - | HE3 |
| 2636 | 2641 | 1 ; | HD2 | - | NZ |
| 2637 | 2639 | 1 ; | HD3 | - | HE2 |
| 2637 | 2640 | 1 ; | HD3 | - | HE3 |

```
2637   2641       1 ;    HD3 - NZ
2639   2642       1 ;    HE2 - HZ1
2639   2643       1 ;    HE2 - HZ2
2639   2644       1 ;    HE2 - HZ3
2640   2642       1 ;    HE3 - HZ1
2640   2643       1 ;    HE3 - HZ2
2640   2644       1 ;    HE3 - HZ3
2645   2650       1 ;      C - HA
2645   2651       1 ;      C - CB
2645   2659       1 ;      C - C
2646   2648       1 ;      O - H
2646   2649       1 ;      O - CA
2647   2652       1 ;      N - HB2
2647   2653       1 ;      N - HB3
2647   2654       1 ;      N - CG
2647   2660       1 ;      N - O
2647   2661       1 ;      N - N
2648   2650       1 ;      H - HA
2648   2651       1 ;      H - CB
2648   2659       1 ;      H - C
2649   2655       1 ;     CA - OD1
2649   2656       1 ;     CA - ND2
2649   2662       1 ;     CA - H
2649   2663       1 ;     CA - CA
2650   2652       1 ;     HA - HB2
2650   2653       1 ;     HA - HB3
2650   2654       1 ;     HA - CG
2650   2660       1 ;     HA - O
2650   2661       1 ;     HA - N
2651   2657       1 ;     CB - HD21
2651   2658       1 ;     CB - HD22
2651   2660       1 ;     CB - O
2651   2661       1 ;     CB - N
2652   2655       1 ;    HB2 - OD1
2652   2656       1 ;    HB2 - ND2
2652   2659       1 ;    HB2 - C
2653   2655       1 ;    HB3 - OD1
2653   2656       1 ;    HB3 - ND2
2653   2659       1 ;    HB3 - C
2654   2659       1 ;     CG - C
2655   2657       1 ;    OD1 - HD21
2655   2658       1 ;    OD1 - HD22
2659   2664       1 ;      C - HA
2659   2665       1 ;      C - CB
2659   2678       1 ;      C - C
2660   2662       1 ;      O - H
2660   2663       1 ;      O - CA
2661   2666       1 ;      N - HB2
2661   2667       1 ;      N - HB3
2661   2668       1 ;      N - CG
2661   2679       1 ;      N - O
2661   2680       1 ;      N - N
```

```
2662   2664       1 ;       H - HA
2662   2665       1 ;       H - CB
2662   2678       1 ;       H - C
2663   2669       1 ;      CA - HG
2663   2670       1 ;      CA - CD1
2663   2674       1 ;      CA - CD2
2663   2681       1 ;      CA - H
2663   2682       1 ;      CA - CA
2664   2666       1 ;      HA - HB2
2664   2667       1 ;      HA - HB3
2664   2668       1 ;      HA - CG
2664   2679       1 ;      HA - O
2664   2680       1 ;      HA - N
2665   2671       1 ;      CB - HD11
2665   2672       1 ;      CB - HD12
2665   2673       1 ;      CB - HD13
2665   2675       1 ;      CB - HD21
2665   2676       1 ;      CB - HD22
2665   2677       1 ;      CB - HD23
2665   2679       1 ;      CB - O
2665   2680       1 ;      CB - N
2666   2669       1 ;     HB2 - HG
2666   2670       1 ;     HB2 - CD1
2666   2674       1 ;     HB2 - CD2
2666   2678       1 ;     HB2 - C
2667   2669       1 ;     HB3 - HG
2667   2670       1 ;     HB3 - CD1
2667   2674       1 ;     HB3 - CD2
2667   2678       1 ;     HB3 - C
2668   2678       1 ;      CG - C
2669   2671       1 ;      HG - HD11
2669   2672       1 ;      HG - HD12
2669   2673       1 ;      HG - HD13
2669   2675       1 ;      HG - HD21
2669   2676       1 ;      HG - HD22
2669   2677       1 ;      HG - HD23
2670   2675       1 ;     CD1 - HD21
2670   2676       1 ;     CD1 - HD22
2670   2677       1 ;     CD1 - HD23
2671   2674       1 ;    HD11 - CD2
2672   2674       1 ;    HD12 - CD2
2673   2674       1 ;    HD13 - CD2
2678   2683       1 ;       C - HA2
2678   2684       1 ;       C - HA3
2678   2685       1 ;       C - C
2679   2681       1 ;       O - H
2679   2682       1 ;       O - CA
2680   2686       1 ;       N - O
2680   2687       1 ;       N - N
2681   2683       1 ;       H - HA2
2681   2684       1 ;       H - HA3
2681   2685       1 ;       H - C
```

| | | | | | | |
|---|---|---|---|---|---|---|
| 2682 | 2688 | 1 | ; | CA | - | H |
| 2682 | 2689 | 1 | ; | CA | - | CA |
| 2683 | 2686 | 1 | ; | HA2 | - | O |
| 2683 | 2687 | 1 | ; | HA2 | - | N |
| 2684 | 2686 | 1 | ; | HA3 | - | O |
| 2684 | 2687 | 1 | ; | HA3 | - | N |
| 2685 | 2690 | 1 | ; | C | - | HA |
| 2685 | 2691 | 1 | ; | C | - | CB |
| 2685 | 2699 | 1 | ; | C | - | C |
| 2686 | 2688 | 1 | ; | O | - | H |
| 2686 | 2689 | 1 | ; | O | - | CA |
| 2687 | 2692 | 1 | ; | N | - | HB |
| 2687 | 2693 | 1 | ; | N | - | CG2 |
| 2687 | 2697 | 1 | ; | N | - | OG1 |
| 2687 | 2700 | 1 | ; | N | - | O |
| 2687 | 2701 | 1 | ; | N | - | N |
| 2688 | 2690 | 1 | ; | H | - | HA |
| 2688 | 2691 | 1 | ; | H | - | CB |
| 2688 | 2699 | 1 | ; | H | - | C |
| 2689 | 2694 | 1 | ; | CA | - | HG21 |
| 2689 | 2695 | 1 | ; | CA | - | HG22 |
| 2689 | 2696 | 1 | ; | CA | - | HG23 |
| 2689 | 2698 | 1 | ; | CA | - | HG1 |
| 2689 | 2702 | 1 | ; | CA | - | H |
| 2689 | 2703 | 1 | ; | CA | - | CA |
| 2690 | 2692 | 1 | ; | HA | - | HB |
| 2690 | 2693 | 1 | ; | HA | - | CG2 |
| 2690 | 2697 | 1 | ; | HA | - | OG1 |
| 2690 | 2700 | 1 | ; | HA | - | O |
| 2690 | 2701 | 1 | ; | HA | - | N |
| 2691 | 2700 | 1 | ; | CB | - | O |
| 2691 | 2701 | 1 | ; | CB | - | N |
| 2692 | 2694 | 1 | ; | HB | - | HG21 |
| 2692 | 2695 | 1 | ; | HB | - | HG22 |
| 2692 | 2696 | 1 | ; | HB | - | HG23 |
| 2692 | 2698 | 1 | ; | HB | - | HG1 |
| 2692 | 2699 | 1 | ; | HB | - | C |
| 2693 | 2698 | 1 | ; | CG2 | - | HG1 |
| 2693 | 2699 | 1 | ; | CG2 | - | C |
| 2694 | 2697 | 1 | ; | HG21 | - | OG1 |
| 2695 | 2697 | 1 | ; | HG22 | - | OG1 |
| 2696 | 2697 | 1 | ; | HG23 | - | OG1 |
| 2697 | 2699 | 1 | ; | OG1 | - | C |
| 2699 | 2704 | 1 | ; | C | - | HA2 |
| 2699 | 2705 | 1 | ; | C | - | HA3 |
| 2699 | 2706 | 1 | ; | C | - | C |
| 2700 | 2702 | 1 | ; | O | - | H |
| 2700 | 2703 | 1 | ; | O | - | CA |
| 2701 | 2707 | 1 | ; | N | - | O |
| 2701 | 2708 | 1 | ; | N | - | N |
| 2702 | 2704 | 1 | ; | H | - | HA2 |
| 2702 | 2705 | 1 | ; | H | - | HA3 |

| | | | | | | |
|---|---|---|---|---|---|---|
| 2702 | 2706 | 1 | ; | H | - | C |
| 2703 | 2709 | 1 | ; | CA | - | H |
| 2703 | 2710 | 1 | ; | CA | - | CA |
| 2704 | 2707 | 1 | ; | HA2 | - | O |
| 2704 | 2708 | 1 | ; | HA2 | - | N |
| 2705 | 2707 | 1 | ; | HA3 | - | O |
| 2705 | 2708 | 1 | ; | HA3 | - | N |
| 2706 | 2711 | 1 | ; | C | - | HA |
| 2706 | 2712 | 1 | ; | C | - | CB |
| 2706 | 2721 | 1 | ; | C | - | C |
| 2707 | 2709 | 1 | ; | O | - | H |
| 2707 | 2710 | 1 | ; | O | - | CA |
| 2708 | 2713 | 1 | ; | N | - | HB2 |
| 2708 | 2714 | 1 | ; | N | - | HB3 |
| 2708 | 2715 | 1 | ; | N | - | CG |
| 2708 | 2722 | 1 | ; | N | - | O |
| 2708 | 2723 | 1 | ; | N | - | N |
| 2709 | 2711 | 1 | ; | H | - | HA |
| 2709 | 2712 | 1 | ; | H | - | CB |
| 2709 | 2721 | 1 | ; | H | - | C |
| 2710 | 2716 | 1 | ; | CA | - | HG2 |
| 2710 | 2717 | 1 | ; | CA | - | HG3 |
| 2710 | 2718 | 1 | ; | CA | - | CD |
| 2710 | 2724 | 1 | ; | CA | - | H |
| 2710 | 2725 | 1 | ; | CA | - | CA |
| 2711 | 2713 | 1 | ; | HA | - | HB2 |
| 2711 | 2714 | 1 | ; | HA | - | HB3 |
| 2711 | 2715 | 1 | ; | HA | - | CG |
| 2711 | 2722 | 1 | ; | HA | - | O |
| 2711 | 2723 | 1 | ; | HA | - | N |
| 2712 | 2719 | 1 | ; | CB | - | OE1 |
| 2712 | 2720 | 1 | ; | CB | - | OE2 |
| 2712 | 2722 | 1 | ; | CB | - | O |
| 2712 | 2723 | 1 | ; | CB | - | N |
| 2713 | 2716 | 1 | ; | HB2 | - | HG2 |
| 2713 | 2717 | 1 | ; | HB2 | - | HG3 |
| 2713 | 2718 | 1 | ; | HB2 | - | CD |
| 2713 | 2721 | 1 | ; | HB2 | - | C |
| 2714 | 2716 | 1 | ; | HB3 | - | HG2 |
| 2714 | 2717 | 1 | ; | HB3 | - | HG3 |
| 2714 | 2718 | 1 | ; | HB3 | - | CD |
| 2714 | 2721 | 1 | ; | HB3 | - | C |
| 2715 | 2721 | 1 | ; | CG | - | C |
| 2716 | 2719 | 1 | ; | HG2 | - | OE1 |
| 2716 | 2720 | 1 | ; | HG2 | - | OE2 |
| 2717 | 2719 | 1 | ; | HG3 | - | OE1 |
| 2717 | 2720 | 1 | ; | HG3 | - | OE2 |
| 2721 | 2726 | 1 | ; | C | - | HA |
| 2721 | 2727 | 1 | ; | C | - | CB |
| 2721 | 2741 | 1 | ; | C | - | C |
| 2722 | 2724 | 1 | ; | O | - | H |
| 2722 | 2725 | 1 | ; | O | - | CA |

| | | | | | | |
|---|---|---|---|---|---|---|
| 2723 | 2728 | 1 | ; | N | - | HB2 |
| 2723 | 2729 | 1 | ; | N | - | HB3 |
| 2723 | 2730 | 1 | ; | N | - | CG |
| 2723 | 2742 | 1 | ; | N | - | O |
| 2723 | 2743 | 1 | ; | N | - | N |
| 2724 | 2726 | 1 | ; | H | - | HA |
| 2724 | 2727 | 1 | ; | H | - | CB |
| 2724 | 2741 | 1 | ; | H | - | C |
| 2725 | 2731 | 1 | ; | CA | - | CD1 |
| 2725 | 2739 | 1 | ; | CA | - | CD2 |
| 2725 | 2744 | 1 | ; | CA | - | H |
| 2725 | 2745 | 1 | ; | CA | - | CA |
| 2726 | 2728 | 1 | ; | HA | - | HB2 |
| 2726 | 2729 | 1 | ; | HA | - | HB3 |
| 2726 | 2730 | 1 | ; | HA | - | CG |
| 2726 | 2742 | 1 | ; | HA | - | O |
| 2726 | 2743 | 1 | ; | HA | - | N |
| 2727 | 2732 | 1 | ; | CB | - | HD1 |
| 2727 | 2733 | 1 | ; | CB | - | CE1 |
| 2727 | 2737 | 1 | ; | CB | - | CE2 |
| 2727 | 2740 | 1 | ; | CB | - | HD2 |
| 2727 | 2742 | 1 | ; | CB | - | O |
| 2727 | 2743 | 1 | ; | CB | - | N |
| 2728 | 2731 | 1 | ; | HB2 | - | CD1 |
| 2728 | 2739 | 1 | ; | HB2 | - | CD2 |
| 2728 | 2741 | 1 | ; | HB2 | - | C |
| 2729 | 2731 | 1 | ; | HB3 | - | CD1 |
| 2729 | 2739 | 1 | ; | HB3 | - | CD2 |
| 2729 | 2741 | 1 | ; | HB3 | - | C |
| 2730 | 2734 | 1 | ; | CG | - | HE1 |
| 2730 | 2735 | 1 | ; | CG | - | CZ |
| 2730 | 2738 | 1 | ; | CG | - | HE2 |
| 2730 | 2741 | 1 | ; | CG | - | C |
| 2731 | 2736 | 1 | ; | CD1 | - | HZ |
| 2731 | 2737 | 1 | ; | CD1 | - | CE2 |
| 2731 | 2740 | 1 | ; | CD1 | - | HD2 |
| 2732 | 2734 | 1 | ; | HD1 | - | HE1 |
| 2732 | 2735 | 1 | ; | HD1 | - | CZ |
| 2732 | 2739 | 1 | ; | HD1 | - | CD2 |
| 2733 | 2738 | 1 | ; | CE1 | - | HE2 |
| 2733 | 2739 | 1 | ; | CE1 | - | CD2 |
| 2734 | 2736 | 1 | ; | HE1 | - | HZ |
| 2734 | 2737 | 1 | ; | HE1 | - | CE2 |
| 2735 | 2740 | 1 | ; | CZ | - | HD2 |
| 2736 | 2738 | 1 | ; | HZ | - | HE2 |
| 2736 | 2739 | 1 | ; | HZ | - | CD2 |
| 2738 | 2740 | 1 | ; | HE2 | - | HD2 |
| 2741 | 2746 | 1 | ; | C | - | HA |
| 2741 | 2747 | 1 | ; | C | - | CB |
| 2741 | 2755 | 1 | ; | C | - | C |
| 2742 | 2744 | 1 | ; | O | - | H |
| 2742 | 2745 | 1 | ; | O | - | CA |

| | | | |
|---|---|---|---|
| 2743 | 2748 | 1 ; | N - HB |
| 2743 | 2749 | 1 ; | N - CG2 |
| 2743 | 2753 | 1 ; | N - OG1 |
| 2743 | 2756 | 1 ; | N - O |
| 2743 | 2757 | 1 ; | N - N |
| 2744 | 2746 | 1 ; | H - HA |
| 2744 | 2747 | 1 ; | H - CB |
| 2744 | 2755 | 1 ; | H - C |
| 2745 | 2750 | 1 ; | CA - HG21 |
| 2745 | 2751 | 1 ; | CA - HG22 |
| 2745 | 2752 | 1 ; | CA - HG23 |
| 2745 | 2754 | 1 ; | CA - HG1 |
| 2745 | 2758 | 1 ; | CA - H |
| 2745 | 2759 | 1 ; | CA - CA |
| 2746 | 2748 | 1 ; | HA - HB |
| 2746 | 2749 | 1 ; | HA - CG2 |
| 2746 | 2753 | 1 ; | HA - OG1 |
| 2746 | 2756 | 1 ; | HA - O |
| 2746 | 2757 | 1 ; | HA - N |
| 2747 | 2756 | 1 ; | CB - O |
| 2747 | 2757 | 1 ; | CB - N |
| 2748 | 2750 | 1 ; | HB - HG21 |
| 2748 | 2751 | 1 ; | HB - HG22 |
| 2748 | 2752 | 1 ; | HB - HG23 |
| 2748 | 2754 | 1 ; | HB - HG1 |
| 2748 | 2755 | 1 ; | HB - C |
| 2749 | 2754 | 1 ; | CG2 - HG1 |
| 2749 | 2755 | 1 ; | CG2 - C |
| 2750 | 2753 | 1 ; | HG21 - OG1 |
| 2751 | 2753 | 1 ; | HG22 - OG1 |
| 2752 | 2753 | 1 ; | HG23 - OG1 |
| 2753 | 2755 | 1 ; | OG1 - C |
| 2755 | 2760 | 1 ; | C - HA |
| 2755 | 2761 | 1 ; | C - CB |
| 2755 | 2769 | 1 ; | C - C |
| 2756 | 2758 | 1 ; | O - H |
| 2756 | 2759 | 1 ; | O - CA |
| 2757 | 2762 | 1 ; | N - HB |
| 2757 | 2763 | 1 ; | N - CG2 |
| 2757 | 2767 | 1 ; | N - OG1 |
| 2757 | 2770 | 1 ; | N - O |
| 2757 | 2771 | 1 ; | N - N |
| 2758 | 2760 | 1 ; | H - HA |
| 2758 | 2761 | 1 ; | H - CB |
| 2758 | 2769 | 1 ; | H - C |
| 2759 | 2764 | 1 ; | CA - HG21 |
| 2759 | 2765 | 1 ; | CA - HG22 |
| 2759 | 2766 | 1 ; | CA - HG23 |
| 2759 | 2768 | 1 ; | CA - HG1 |
| 2759 | 2772 | 1 ; | CA - H |
| 2759 | 2773 | 1 ; | CA - CA |
| 2760 | 2762 | 1 ; | HA - HB |

| | | | | | |
|---|---|---|---|---|---|
| 2760 | 2763 | 1 ; | HA | - | CG2 |
| 2760 | 2767 | 1 ; | HA | - | OG1 |
| 2760 | 2770 | 1 ; | HA | - | O |
| 2760 | 2771 | 1 ; | HA | - | N |
| 2761 | 2770 | 1 ; | CB | - | O |
| 2761 | 2771 | 1 ; | CB | - | N |
| 2762 | 2764 | 1 ; | HB | - | HG21 |
| 2762 | 2765 | 1 ; | HB | - | HG22 |
| 2762 | 2766 | 1 ; | HB | - | HG23 |
| 2762 | 2768 | 1 ; | HB | - | HG1 |
| 2762 | 2769 | 1 ; | HB | - | C |
| 2763 | 2768 | 1 ; | CG2 | - | HG1 |
| 2763 | 2769 | 1 ; | CG2 | - | C |
| 2764 | 2767 | 1 ; | HG21 | - | OG1 |
| 2765 | 2767 | 1 ; | HG22 | - | OG1 |
| 2766 | 2767 | 1 ; | HG23 | - | OG1 |
| 2767 | 2769 | 1 ; | OG1 | - | C |
| 2769 | 2774 | 1 ; | C | - | HA |
| 2769 | 2775 | 1 ; | C | - | CB |
| 2769 | 2783 | 1 ; | C | - | C |
| 2770 | 2772 | 1 ; | O | - | H |
| 2770 | 2773 | 1 ; | O | - | CA |
| 2771 | 2776 | 1 ; | N | - | HB |
| 2771 | 2777 | 1 ; | N | - | CG2 |
| 2771 | 2781 | 1 ; | N | - | OG1 |
| 2771 | 2784 | 1 ; | N | - | O |
| 2771 | 2785 | 1 ; | N | - | N |
| 2772 | 2774 | 1 ; | H | - | HA |
| 2772 | 2775 | 1 ; | H | - | CB |
| 2772 | 2783 | 1 ; | H | - | C |
| 2773 | 2778 | 1 ; | CA | - | HG21 |
| 2773 | 2779 | 1 ; | CA | - | HG22 |
| 2773 | 2780 | 1 ; | CA | - | HG23 |
| 2773 | 2782 | 1 ; | CA | - | HG1 |
| 2773 | 2786 | 1 ; | CA | - | H |
| 2773 | 2787 | 1 ; | CA | - | CA |
| 2774 | 2776 | 1 ; | HA | - | HB |
| 2774 | 2777 | 1 ; | HA | - | CG2 |
| 2774 | 2781 | 1 ; | HA | - | OG1 |
| 2774 | 2784 | 1 ; | HA | - | O |
| 2774 | 2785 | 1 ; | HA | - | N |
| 2775 | 2784 | 1 ; | CB | - | O |
| 2775 | 2785 | 1 ; | CB | - | N |
| 2776 | 2778 | 1 ; | HB | - | HG21 |
| 2776 | 2779 | 1 ; | HB | - | HG22 |
| 2776 | 2780 | 1 ; | HB | - | HG23 |
| 2776 | 2782 | 1 ; | HB | - | HG1 |
| 2776 | 2783 | 1 ; | HB | - | C |
| 2777 | 2782 | 1 ; | CG2 | - | HG1 |
| 2777 | 2783 | 1 ; | CG2 | - | C |
| 2778 | 2781 | 1 ; | HG21 | - | OG1 |
| 2779 | 2781 | 1 ; | HG22 | - | OG1 |

| | | | |
|---|---|---|---|
| 2780 | 2781 | 1 ; | HG23 - OG1 |
| 2781 | 2783 | 1 ; | OG1 - C |
| 2783 | 2788 | 1 ; | C - HA |
| 2783 | 2789 | 1 ; | C - CB |
| 2783 | 2805 | 1 ; | C - C |
| 2784 | 2786 | 1 ; | O - H |
| 2784 | 2787 | 1 ; | O - CA |
| 2785 | 2790 | 1 ; | N - HB2 |
| 2785 | 2791 | 1 ; | N - HB3 |
| 2785 | 2792 | 1 ; | N - CG |
| 2785 | 2806 | 1 ; | N - O |
| 2785 | 2807 | 1 ; | N - N |
| 2786 | 2788 | 1 ; | H - HA |
| 2786 | 2789 | 1 ; | H - CB |
| 2786 | 2805 | 1 ; | H - C |
| 2787 | 2793 | 1 ; | CA - HG2 |
| 2787 | 2794 | 1 ; | CA - HG3 |
| 2787 | 2795 | 1 ; | CA - CD |
| 2787 | 2808 | 1 ; | CA - H |
| 2787 | 2809 | 1 ; | CA - CA |
| 2788 | 2790 | 1 ; | HA - HB2 |
| 2788 | 2791 | 1 ; | HA - HB3 |
| 2788 | 2792 | 1 ; | HA - CG |
| 2788 | 2806 | 1 ; | HA - O |
| 2788 | 2807 | 1 ; | HA - N |
| 2789 | 2796 | 1 ; | CB - HD2 |
| 2789 | 2797 | 1 ; | CB - HD3 |
| 2789 | 2798 | 1 ; | CB - CE |
| 2789 | 2806 | 1 ; | CB - O |
| 2789 | 2807 | 1 ; | CB - N |
| 2790 | 2793 | 1 ; | HB2 - HG2 |
| 2790 | 2794 | 1 ; | HB2 - HG3 |
| 2790 | 2795 | 1 ; | HB2 - CD |
| 2790 | 2805 | 1 ; | HB2 - C |
| 2791 | 2793 | 1 ; | HB3 - HG2 |
| 2791 | 2794 | 1 ; | HB3 - HG3 |
| 2791 | 2795 | 1 ; | HB3 - CD |
| 2791 | 2805 | 1 ; | HB3 - C |
| 2792 | 2799 | 1 ; | CG - HE2 |
| 2792 | 2800 | 1 ; | CG - HE3 |
| 2792 | 2801 | 1 ; | CG - NZ |
| 2792 | 2805 | 1 ; | CG - C |
| 2793 | 2796 | 1 ; | HG2 - HD2 |
| 2793 | 2797 | 1 ; | HG2 - HD3 |
| 2793 | 2798 | 1 ; | HG2 - CE |
| 2794 | 2796 | 1 ; | HG3 - HD2 |
| 2794 | 2797 | 1 ; | HG3 - HD3 |
| 2794 | 2798 | 1 ; | HG3 - CE |
| 2795 | 2802 | 1 ; | CD - HZ1 |
| 2795 | 2803 | 1 ; | CD - HZ2 |
| 2795 | 2804 | 1 ; | CD - HZ3 |
| 2796 | 2799 | 1 ; | HD2 - HE2 |

```
2796   2800      1 ;    HD2 - HE3
2796   2801      1 ;    HD2 - NZ
2797   2799      1 ;    HD3 - HE2
2797   2800      1 ;    HD3 - HE3
2797   2801      1 ;    HD3 - NZ
2799   2802      1 ;    HE2 - HZ1
2799   2803      1 ;    HE2 - HZ2
2799   2804      1 ;    HE2 - HZ3
2800   2802      1 ;    HE3 - HZ1
2800   2803      1 ;    HE3 - HZ2
2800   2804      1 ;    HE3 - HZ3
2805   2810      1 ;      C - HA
2805   2811      1 ;      C - CB
2805   2815      1 ;      C - C
2806   2808      1 ;      O - H
2806   2809      1 ;      O - CA
2807   2812      1 ;      N - HB1
2807   2813      1 ;      N - HB2
2807   2814      1 ;      N - HB3
2807   2816      1 ;      N - O
2807   2817      1 ;      N - N
2808   2810      1 ;      H - HA
2808   2811      1 ;      H - CB
2808   2815      1 ;      H - C
2809   2818      1 ;     CA - H
2809   2819      1 ;     CA - CA
2810   2812      1 ;     HA - HB1
2810   2813      1 ;     HA - HB2
2810   2814      1 ;     HA - HB3
2810   2816      1 ;     HA - O
2810   2817      1 ;     HA - N
2811   2816      1 ;     CB - O
2811   2817      1 ;     CB - N
2812   2815      1 ;    HB1 - C
2813   2815      1 ;    HB2 - C
2814   2815      1 ;    HB3 - C
2815   2820      1 ;      C - HA2
2815   2821      1 ;      C - HA3
2815   2822      1 ;      C - C
2816   2818      1 ;      O - H
2816   2819      1 ;      O - CA
2817   2823      1 ;      N - O
2817   2824      1 ;      N - N
2818   2820      1 ;      H - HA2
2818   2821      1 ;      H - HA3
2818   2822      1 ;      H - C
2819   2825      1 ;     CA - H
2819   2826      1 ;     CA - CA
2820   2823      1 ;    HA2 - O
2820   2824      1 ;    HA2 - N
2821   2823      1 ;    HA3 - O
2821   2824      1 ;    HA3 - N
```

| | | | | | | |
|---|---|---|---|---|---|---|
| 2822 | 2827 | 1 | ; | C | - | HA |
| 2822 | 2828 | 1 | ; | C | - | CB |
| 2822 | 2836 | 1 | ; | C | - | C |
| 2823 | 2825 | 1 | ; | O | - | H |
| 2823 | 2826 | 1 | ; | O | - | CA |
| 2824 | 2829 | 1 | ; | N | - | HB |
| 2824 | 2830 | 1 | ; | N | - | CG2 |
| 2824 | 2834 | 1 | ; | N | - | OG1 |
| 2824 | 2837 | 1 | ; | N | - | O |
| 2824 | 2838 | 1 | ; | N | - | N |
| 2825 | 2827 | 1 | ; | H | - | HA |
| 2825 | 2828 | 1 | ; | H | - | CB |
| 2825 | 2836 | 1 | ; | H | - | C |
| 2826 | 2831 | 1 | ; | CA | - | HG21 |
| 2826 | 2832 | 1 | ; | CA | - | HG22 |
| 2826 | 2833 | 1 | ; | CA | - | HG23 |
| 2826 | 2835 | 1 | ; | CA | - | HG1 |
| 2826 | 2839 | 1 | ; | CA | - | CD |
| 2826 | 2848 | 1 | ; | CA | - | CA |
| 2827 | 2829 | 1 | ; | HA | - | HB |
| 2827 | 2830 | 1 | ; | HA | - | CG2 |
| 2827 | 2834 | 1 | ; | HA | - | OG1 |
| 2827 | 2837 | 1 | ; | HA | - | O |
| 2827 | 2838 | 1 | ; | HA | - | N |
| 2828 | 2837 | 1 | ; | CB | - | O |
| 2828 | 2838 | 1 | ; | CB | - | N |
| 2829 | 2831 | 1 | ; | HB | - | HG21 |
| 2829 | 2832 | 1 | ; | HB | - | HG22 |
| 2829 | 2833 | 1 | ; | HB | - | HG23 |
| 2829 | 2835 | 1 | ; | HB | - | HG1 |
| 2829 | 2836 | 1 | ; | HB | - | C |
| 2830 | 2835 | 1 | ; | CG2 | - | HG1 |
| 2830 | 2836 | 1 | ; | CG2 | - | C |
| 2831 | 2834 | 1 | ; | HG21 | - | OG1 |
| 2832 | 2834 | 1 | ; | HG22 | - | OG1 |
| 2833 | 2834 | 1 | ; | HG23 | - | OG1 |
| 2834 | 2836 | 1 | ; | OG1 | - | C |
| 2836 | 2840 | 1 | ; | C | - | HD2 |
| 2836 | 2841 | 1 | ; | C | - | HD3 |
| 2836 | 2842 | 1 | ; | C | - | CG |
| 2836 | 2845 | 1 | ; | C | - | CB |
| 2836 | 2849 | 1 | ; | C | - | HA |
| 2836 | 2850 | 1 | ; | C | - | C |
| 2837 | 2839 | 1 | ; | O | - | CD |
| 2837 | 2848 | 1 | ; | O | - | CA |
| 2838 | 2843 | 1 | ; | N | - | HG2 |
| 2838 | 2844 | 1 | ; | N | - | HG3 |
| 2838 | 2846 | 1 | ; | N | - | HB2 |
| 2838 | 2847 | 1 | ; | N | - | HB3 |
| 2838 | 2851 | 1 | ; | N | - | O |
| 2838 | 2852 | 1 | ; | N | - | N |
| 2839 | 2846 | 1 | ; | CD | - | HB2 |

| | | | | | |
|---|---|---|---|---|---|
| 2839 | 2847 | 1 ; | CD | - | HB3 |
| 2839 | 2849 | 1 ; | CD | - | HA |
| 2839 | 2850 | 1 ; | CD | - | C |
| 2840 | 2843 | 1 ; | HD2 | - | HG2 |
| 2840 | 2844 | 1 ; | HD2 | - | HG3 |
| 2840 | 2845 | 1 ; | HD2 | - | CB |
| 2840 | 2848 | 1 ; | HD2 | - | CA |
| 2841 | 2843 | 1 ; | HD3 | - | HG2 |
| 2841 | 2844 | 1 ; | HD3 | - | HG3 |
| 2841 | 2845 | 1 ; | HD3 | - | CB |
| 2841 | 2848 | 1 ; | HD3 | - | CA |
| 2842 | 2849 | 1 ; | CG | - | HA |
| 2842 | 2850 | 1 ; | CG | - | C |
| 2843 | 2846 | 1 ; | HG2 | - | HB2 |
| 2843 | 2847 | 1 ; | HG2 | - | HB3 |
| 2843 | 2848 | 1 ; | HG2 | - | CA |
| 2844 | 2846 | 1 ; | HG3 | - | HB2 |
| 2844 | 2847 | 1 ; | HG3 | - | HB3 |
| 2844 | 2848 | 1 ; | HG3 | - | CA |
| 2845 | 2851 | 1 ; | CB | - | O |
| 2845 | 2852 | 1 ; | CB | - | N |
| 2846 | 2849 | 1 ; | HB2 | - | HA |
| 2846 | 2850 | 1 ; | HB2 | - | C |
| 2847 | 2849 | 1 ; | HB3 | - | HA |
| 2847 | 2850 | 1 ; | HB3 | - | C |
| 2848 | 2853 | 1 ; | CA | - | H |
| 2848 | 2854 | 1 ; | CA | - | CA |
| 2849 | 2851 | 1 ; | HA | - | O |
| 2849 | 2852 | 1 ; | HA | - | N |
| 2850 | 2855 | 1 ; | C | - | HA |
| 2850 | 2856 | 1 ; | C | - | CB |
| 2850 | 2871 | 1 ; | C | - | C |
| 2851 | 2853 | 1 ; | O | - | H |
| 2851 | 2854 | 1 ; | O | - | CA |
| 2852 | 2857 | 1 ; | N | - | HB2 |
| 2852 | 2858 | 1 ; | N | - | HB3 |
| 2852 | 2859 | 1 ; | N | - | CG |
| 2852 | 2872 | 1 ; | N | - | O |
| 2852 | 2873 | 1 ; | N | - | N |
| 2853 | 2855 | 1 ; | H | - | HA |
| 2853 | 2856 | 1 ; | H | - | CB |
| 2853 | 2871 | 1 ; | H | - | C |
| 2854 | 2860 | 1 ; | CA | - | CD1 |
| 2854 | 2869 | 1 ; | CA | - | CD2 |
| 2854 | 2874 | 1 ; | CA | - | H |
| 2854 | 2875 | 1 ; | CA | - | CA |
| 2855 | 2857 | 1 ; | HA | - | HB2 |
| 2855 | 2858 | 1 ; | HA | - | HB3 |
| 2855 | 2859 | 1 ; | HA | - | CG |
| 2855 | 2872 | 1 ; | HA | - | O |
| 2855 | 2873 | 1 ; | HA | - | N |
| 2856 | 2861 | 1 ; | CB | - | HD1 |

| | | | | |
|---|---|---|---|---|
| 2856 | 2862 | 1 ; | CB - | CE1 |
| 2856 | 2867 | 1 ; | CB - | CE2 |
| 2856 | 2870 | 1 ; | CB - | HD2 |
| 2856 | 2872 | 1 ; | CB - | O |
| 2856 | 2873 | 1 ; | CB - | N |
| 2857 | 2860 | 1 ; | HB2 - | CD1 |
| 2857 | 2869 | 1 ; | HB2 - | CD2 |
| 2857 | 2871 | 1 ; | HB2 - | C |
| 2858 | 2860 | 1 ; | HB3 - | CD1 |
| 2858 | 2869 | 1 ; | HB3 - | CD2 |
| 2858 | 2871 | 1 ; | HB3 - | C |
| 2859 | 2863 | 1 ; | CG - | HE1 |
| 2859 | 2864 | 1 ; | CG - | CZ |
| 2859 | 2868 | 1 ; | CG - | HE2 |
| 2859 | 2871 | 1 ; | CG - | C |
| 2860 | 2865 | 1 ; | CD1 - | OH |
| 2860 | 2867 | 1 ; | CD1 - | CE2 |
| 2860 | 2870 | 1 ; | CD1 - | HD2 |
| 2861 | 2863 | 1 ; | HD1 - | HE1 |
| 2861 | 2864 | 1 ; | HD1 - | CZ |
| 2861 | 2869 | 1 ; | HD1 - | CD2 |
| 2862 | 2866 | 1 ; | CE1 - | HH |
| 2862 | 2868 | 1 ; | CE1 - | HE2 |
| 2862 | 2869 | 1 ; | CE1 - | CD2 |
| 2863 | 2865 | 1 ; | HE1 - | OH |
| 2863 | 2867 | 1 ; | HE1 - | CE2 |
| 2864 | 2870 | 1 ; | CZ - | HD2 |
| 2865 | 2868 | 1 ; | OH - | HE2 |
| 2865 | 2869 | 1 ; | OH - | CD2 |
| 2866 | 2867 | 1 ; | HH - | CE2 |
| 2868 | 2870 | 1 ; | HE2 - | HD2 |
| 2871 | 2876 | 1 ; | C - | HA |
| 2871 | 2877 | 1 ; | C - | CB |
| 2871 | 2892 | 1 ; | C - | C |
| 2872 | 2874 | 1 ; | O - | H |
| 2872 | 2875 | 1 ; | O - | CA |
| 2873 | 2878 | 1 ; | N - | HB2 |
| 2873 | 2879 | 1 ; | N - | HB3 |
| 2873 | 2880 | 1 ; | N - | CG |
| 2873 | 2893 | 1 ; | N - | O |
| 2873 | 2894 | 1 ; | N - | N |
| 2874 | 2876 | 1 ; | H - | HA |
| 2874 | 2877 | 1 ; | H - | CB |
| 2874 | 2892 | 1 ; | H - | C |
| 2875 | 2881 | 1 ; | CA - | CD1 |
| 2875 | 2890 | 1 ; | CA - | CD2 |
| 2875 | 2895 | 1 ; | CA - | H |
| 2875 | 2896 | 1 ; | CA - | CA |
| 2876 | 2878 | 1 ; | HA - | HB2 |
| 2876 | 2879 | 1 ; | HA - | HB3 |
| 2876 | 2880 | 1 ; | HA - | CG |
| 2876 | 2893 | 1 ; | HA - | O |

| | | | | | |
|---|---|---|---|---|---|
| 2876 | 2894 | 1 ; | HA | - | N |
| 2877 | 2882 | 1 ; | CB | - | HD1 |
| 2877 | 2883 | 1 ; | CB | - | CE1 |
| 2877 | 2888 | 1 ; | CB | - | CE2 |
| 2877 | 2891 | 1 ; | CB | - | HD2 |
| 2877 | 2893 | 1 ; | CB | - | O |
| 2877 | 2894 | 1 ; | CB | - | N |
| 2878 | 2881 | 1 ; | HB2 | - | CD1 |
| 2878 | 2890 | 1 ; | HB2 | - | CD2 |
| 2878 | 2892 | 1 ; | HB2 | - | C |
| 2879 | 2881 | 1 ; | HB3 | - | CD1 |
| 2879 | 2890 | 1 ; | HB3 | - | CD2 |
| 2879 | 2892 | 1 ; | HB3 | - | C |
| 2880 | 2884 | 1 ; | CG | - | HE1 |
| 2880 | 2885 | 1 ; | CG | - | CZ |
| 2880 | 2889 | 1 ; | CG | - | HE2 |
| 2880 | 2892 | 1 ; | CG | - | C |
| 2881 | 2886 | 1 ; | CD1 | - | OH |
| 2881 | 2888 | 1 ; | CD1 | - | CE2 |
| 2881 | 2891 | 1 ; | CD1 | - | HD2 |
| 2882 | 2884 | 1 ; | HD1 | - | HE1 |
| 2882 | 2885 | 1 ; | HD1 | - | CZ |
| 2882 | 2890 | 1 ; | HD1 | - | CD2 |
| 2883 | 2887 | 1 ; | CE1 | - | HH |
| 2883 | 2889 | 1 ; | CE1 | - | HE2 |
| 2883 | 2890 | 1 ; | CE1 | - | CD2 |
| 2884 | 2886 | 1 ; | HE1 | - | OH |
| 2884 | 2888 | 1 ; | HE1 | - | CE2 |
| 2885 | 2891 | 1 ; | CZ | - | HD2 |
| 2886 | 2889 | 1 ; | OH | - | HE2 |
| 2886 | 2890 | 1 ; | OH | - | CD2 |
| 2887 | 2888 | 1 ; | HH | - | CE2 |
| 2889 | 2891 | 1 ; | HE2 | - | HD2 |
| 2892 | 2897 | 1 ; | C | - | HA |
| 2892 | 2898 | 1 ; | C | - | CB |
| 2892 | 2908 | 1 ; | C | - | C |
| 2893 | 2895 | 1 ; | O | - | H |
| 2893 | 2896 | 1 ; | O | - | CA |
| 2894 | 2899 | 1 ; | N | - | HB |
| 2894 | 2900 | 1 ; | N | - | CG1 |
| 2894 | 2904 | 1 ; | N | - | CG2 |
| 2894 | 2909 | 1 ; | N | - | O |
| 2894 | 2910 | 1 ; | N | - | N |
| 2895 | 2897 | 1 ; | H | - | HA |
| 2895 | 2898 | 1 ; | H | - | CB |
| 2895 | 2908 | 1 ; | H | - | C |
| 2896 | 2901 | 1 ; | CA | - | HG11 |
| 2896 | 2902 | 1 ; | CA | - | HG12 |
| 2896 | 2903 | 1 ; | CA | - | HG13 |
| 2896 | 2905 | 1 ; | CA | - | HG21 |
| 2896 | 2906 | 1 ; | CA | - | HG22 |
| 2896 | 2907 | 1 ; | CA | - | HG23 |

| | | | | | | |
|---|---|---|---|---|---|---|
| 2896 | 2911 | 1 | ; | CA | - | H |
| 2896 | 2912 | 1 | ; | CA | - | CA |
| 2897 | 2899 | 1 | ; | HA | - | HB |
| 2897 | 2900 | 1 | ; | HA | - | CG1 |
| 2897 | 2904 | 1 | ; | HA | - | CG2 |
| 2897 | 2909 | 1 | ; | HA | - | O |
| 2897 | 2910 | 1 | ; | HA | - | N |
| 2898 | 2909 | 1 | ; | CB | - | O |
| 2898 | 2910 | 1 | ; | CB | - | N |
| 2899 | 2901 | 1 | ; | HB | - | HG11 |
| 2899 | 2902 | 1 | ; | HB | - | HG12 |
| 2899 | 2903 | 1 | ; | HB | - | HG13 |
| 2899 | 2905 | 1 | ; | HB | - | HG21 |
| 2899 | 2906 | 1 | ; | HB | - | HG22 |
| 2899 | 2907 | 1 | ; | HB | - | HG23 |
| 2899 | 2908 | 1 | ; | HB | - | C |
| 2900 | 2905 | 1 | ; | CG1 | - | HG21 |
| 2900 | 2906 | 1 | ; | CG1 | - | HG22 |
| 2900 | 2907 | 1 | ; | CG1 | - | HG23 |
| 2900 | 2908 | 1 | ; | CG1 | - | C |
| 2901 | 2904 | 1 | ; | HG11 | - | CG2 |
| 2902 | 2904 | 1 | ; | HG12 | - | CG2 |
| 2903 | 2904 | 1 | ; | HG13 | - | CG2 |
| 2904 | 2908 | 1 | ; | CG2 | - | C |
| 2908 | 2913 | 1 | ; | C | - | HA |
| 2908 | 2914 | 1 | ; | C | - | CB |
| 2908 | 2918 | 1 | ; | C | - | C |
| 2909 | 2911 | 1 | ; | O | - | H |
| 2909 | 2912 | 1 | ; | O | - | CA |
| 2910 | 2915 | 1 | ; | N | - | HB1 |
| 2910 | 2916 | 1 | ; | N | - | HB2 |
| 2910 | 2917 | 1 | ; | N | - | HB3 |
| 2910 | 2919 | 1 | ; | N | - | O |
| 2910 | 2920 | 1 | ; | N | - | N |
| 2911 | 2913 | 1 | ; | H | - | HA |
| 2911 | 2914 | 1 | ; | H | - | CB |
| 2911 | 2918 | 1 | ; | H | - | C |
| 2912 | 2921 | 1 | ; | CA | - | CD |
| 2912 | 2930 | 1 | ; | CA | - | CA |
| 2913 | 2915 | 1 | ; | HA | - | HB1 |
| 2913 | 2916 | 1 | ; | HA | - | HB2 |
| 2913 | 2917 | 1 | ; | HA | - | HB3 |
| 2913 | 2919 | 1 | ; | HA | - | O |
| 2913 | 2920 | 1 | ; | HA | - | N |
| 2914 | 2919 | 1 | ; | CB | - | O |
| 2914 | 2920 | 1 | ; | CB | - | N |
| 2915 | 2918 | 1 | ; | HB1 | - | C |
| 2916 | 2918 | 1 | ; | HB2 | - | C |
| 2917 | 2918 | 1 | ; | HB3 | - | C |
| 2918 | 2922 | 1 | ; | C | - | HD2 |
| 2918 | 2923 | 1 | ; | C | - | HD3 |
| 2918 | 2924 | 1 | ; | C | - | CG |

| | | | | | |
|---|---|---|---|---|---|
| 2918 | 2927 | 1 ; | C | - | CB |
| 2918 | 2931 | 1 ; | C | - | HA |
| 2918 | 2932 | 1 ; | C | - | C |
| 2919 | 2921 | 1 ; | O | - | CD |
| 2919 | 2930 | 1 ; | O | - | CA |
| 2920 | 2925 | 1 ; | N | - | HG2 |
| 2920 | 2926 | 1 ; | N | - | HG3 |
| 2920 | 2928 | 1 ; | N | - | HB2 |
| 2920 | 2929 | 1 ; | N | - | HB3 |
| 2920 | 2933 | 1 ; | N | - | O |
| 2920 | 2934 | 1 ; | N | - | N |
| 2921 | 2928 | 1 ; | CD | - | HB2 |
| 2921 | 2929 | 1 ; | CD | - | HB3 |
| 2921 | 2931 | 1 ; | CD | - | HA |
| 2921 | 2932 | 1 ; | CD | - | C |
| 2922 | 2925 | 1 ; | HD2 | - | HG2 |
| 2922 | 2926 | 1 ; | HD2 | - | HG3 |
| 2922 | 2927 | 1 ; | HD2 | - | CB |
| 2922 | 2930 | 1 ; | HD2 | - | CA |
| 2923 | 2925 | 1 ; | HD3 | - | HG2 |
| 2923 | 2926 | 1 ; | HD3 | - | HG3 |
| 2923 | 2927 | 1 ; | HD3 | - | CB |
| 2923 | 2930 | 1 ; | HD3 | - | CA |
| 2924 | 2931 | 1 ; | CG | - | HA |
| 2924 | 2932 | 1 ; | CG | - | C |
| 2925 | 2928 | 1 ; | HG2 | - | HB2 |
| 2925 | 2929 | 1 ; | HG2 | - | HB3 |
| 2925 | 2930 | 1 ; | HG2 | - | CA |
| 2926 | 2928 | 1 ; | HG3 | - | HB2 |
| 2926 | 2929 | 1 ; | HG3 | - | HB3 |
| 2926 | 2930 | 1 ; | HG3 | - | CA |
| 2927 | 2933 | 1 ; | CB | - | O |
| 2927 | 2934 | 1 ; | CB | - | N |
| 2928 | 2931 | 1 ; | HB2 | - | HA |
| 2928 | 2932 | 1 ; | HB2 | - | C |
| 2929 | 2931 | 1 ; | HB3 | - | HA |
| 2929 | 2932 | 1 ; | HB3 | - | C |
| 2930 | 2935 | 1 ; | CA | - | H |
| 2930 | 2936 | 1 ; | CA | - | CA |
| 2931 | 2933 | 1 ; | HA | - | O |
| 2931 | 2934 | 1 ; | HA | - | N |
| 2932 | 2937 | 1 ; | C | - | HA |
| 2932 | 2938 | 1 ; | C | - | CB |
| 2932 | 2949 | 1 ; | C | - | C |
| 2933 | 2935 | 1 ; | O | - | H |
| 2933 | 2936 | 1 ; | O | - | CA |
| 2934 | 2939 | 1 ; | N | - | HB2 |
| 2934 | 2940 | 1 ; | N | - | HB3 |
| 2934 | 2941 | 1 ; | N | - | CG |
| 2934 | 2950 | 1 ; | N | - | O |
| 2934 | 2951 | 1 ; | N | - | N |
| 2935 | 2937 | 1 ; | H | - | HA |

```
2935   2938       1 ;       H - CB
2935   2949       1 ;       H - C
2936   2942       1 ;      CA - HG2
2936   2943       1 ;      CA - HG3
2936   2944       1 ;      CA - CD
2936   2952       1 ;      CA - H
2936   2953       1 ;      CA - CA
2937   2939       1 ;      HA - HB2
2937   2940       1 ;      HA - HB3
2937   2941       1 ;      HA - CG
2937   2950       1 ;      HA - O
2937   2951       1 ;      HA - N
2938   2945       1 ;      CB - OE1
2938   2946       1 ;      CB - NE2
2938   2950       1 ;      CB - O
2938   2951       1 ;      CB - N
2939   2942       1 ;     HB2 - HG2
2939   2943       1 ;     HB2 - HG3
2939   2944       1 ;     HB2 - CD
2939   2949       1 ;     HB2 - C
2940   2942       1 ;     HB3 - HG2
2940   2943       1 ;     HB3 - HG3
2940   2944       1 ;     HB3 - CD
2940   2949       1 ;     HB3 - C
2941   2947       1 ;      CG - HE21
2941   2948       1 ;      CG - HE22
2941   2949       1 ;      CG - C
2942   2945       1 ;     HG2 - OE1
2942   2946       1 ;     HG2 - NE2
2943   2945       1 ;     HG3 - OE1
2943   2946       1 ;     HG3 - NE2
2945   2947       1 ;     OE1 - HE21
2945   2948       1 ;     OE1 - HE22
2949   2954       1 ;       C - HA
2949   2955       1 ;       C - CB
2949   2965       1 ;       C - C
2950   2952       1 ;       O - H
2950   2953       1 ;       O - CA
2951   2956       1 ;       N - HB
2951   2957       1 ;       N - CG1
2951   2961       1 ;       N - CG2
2951   2966       1 ;       N - O
2951   2967       1 ;       N - N
2952   2954       1 ;       H - HA
2952   2955       1 ;       H - CB
2952   2965       1 ;       H - C
2953   2958       1 ;      CA - HG11
2953   2959       1 ;      CA - HG12
2953   2960       1 ;      CA - HG13
2953   2962       1 ;      CA - HG21
2953   2963       1 ;      CA - HG22
2953   2964       1 ;      CA - HG23
```

```
2953   2968      1 ;      CA - H
2953   2969      1 ;      CA - CA
2954   2956      1 ;      HA - HB
2954   2957      1 ;      HA - CG1
2954   2961      1 ;      HA - CG2
2954   2966      1 ;      HA - O
2954   2967      1 ;      HA - N
2955   2966      1 ;      CB - O
2955   2967      1 ;      CB - N
2956   2958      1 ;      HB - HG11
2956   2959      1 ;      HB - HG12
2956   2960      1 ;      HB - HG13
2956   2962      1 ;      HB - HG21
2956   2963      1 ;      HB - HG22
2956   2964      1 ;      HB - HG23
2956   2965      1 ;      HB - C
2957   2962      1 ;     CG1 - HG21
2957   2963      1 ;     CG1 - HG22
2957   2964      1 ;     CG1 - HG23
2957   2965      1 ;     CG1 - C
2958   2961      1 ;    HG11 - CG2
2959   2961      1 ;    HG12 - CG2
2960   2961      1 ;    HG13 - CG2
2961   2965      1 ;     CG2 - C
2965   2970      1 ;       C - HA
2965   2971      1 ;       C - CB
2965   2984      1 ;       C - C
2966   2968      1 ;       O - H
2966   2969      1 ;       O - CA
2967   2972      1 ;       N - HB2
2967   2973      1 ;       N - HB3
2967   2974      1 ;       N - CG
2967   2985      1 ;       N - O
2967   2986      1 ;       N - N
2968   2970      1 ;       H - HA
2968   2971      1 ;       H - CB
2968   2984      1 ;       H - C
2969   2975      1 ;      CA - HG
2969   2976      1 ;      CA - CD1
2969   2980      1 ;      CA - CD2
2969   2987      1 ;      CA - H
2969   2988      1 ;      CA - CA
2970   2972      1 ;      HA - HB2
2970   2973      1 ;      HA - HB3
2970   2974      1 ;      HA - CG
2970   2985      1 ;      HA - O
2970   2986      1 ;      HA - N
2971   2977      1 ;      CB - HD11
2971   2978      1 ;      CB - HD12
2971   2979      1 ;      CB - HD13
2971   2981      1 ;      CB - HD21
2971   2982      1 ;      CB - HD22
```

```
2971   2983       1 ;      CB - HD23
2971   2985       1 ;      CB - O
2971   2986       1 ;      CB - N
2972   2975       1 ;     HB2 - HG
2972   2976       1 ;     HB2 - CD1
2972   2980       1 ;     HB2 - CD2
2972   2984       1 ;     HB2 - C
2973   2975       1 ;     HB3 - HG
2973   2976       1 ;     HB3 - CD1
2973   2980       1 ;     HB3 - CD2
2973   2984       1 ;     HB3 - C
2974   2984       1 ;      CG - C
2975   2977       1 ;      HG - HD11
2975   2978       1 ;      HG - HD12
2975   2979       1 ;      HG - HD13
2975   2981       1 ;      HG - HD21
2975   2982       1 ;      HG - HD22
2975   2983       1 ;      HG - HD23
2976   2981       1 ;     CD1 - HD21
2976   2982       1 ;     CD1 - HD22
2976   2983       1 ;     CD1 - HD23
2977   2980       1 ;    HD11 - CD2
2978   2980       1 ;    HD12 - CD2
2979   2980       1 ;    HD13 - CD2
2984   2989       1 ;       C - HA
2984   2990       1 ;       C - CB
2984   2996       1 ;       C - C
2985   2987       1 ;       O - H
2985   2988       1 ;       O - CA
2986   2991       1 ;       N - HB2
2986   2992       1 ;       N - HB3
2986   2993       1 ;       N - CG
2986   2997       1 ;       N - O
2986   2998       1 ;       N - N
2987   2989       1 ;       H - HA
2987   2990       1 ;       H - CB
2987   2996       1 ;       H - C
2988   2994       1 ;      CA - OD1
2988   2995       1 ;      CA - OD2
2988   2999       1 ;      CA - H
2988   3000       1 ;      CA - CA
2989   2991       1 ;      HA - HB2
2989   2992       1 ;      HA - HB3
2989   2993       1 ;      HA - CG
2989   2997       1 ;      HA - O
2989   2998       1 ;      HA - N
2990   2997       1 ;      CB - O
2990   2998       1 ;      CB - N
2991   2994       1 ;     HB2 - OD1
2991   2995       1 ;     HB2 - OD2
2991   2996       1 ;     HB2 - C
2992   2994       1 ;     HB3 - OD1
```

| | | | |
|---|---|---|---|
| 2992 | 2995 | 1 ; | HB3 - OD2 |
| 2992 | 2996 | 1 ; | HB3 - C |
| 2993 | 2996 | 1 ; | CG - C |
| 2996 | 3001 | 1 ; | C - HA2 |
| 2996 | 3002 | 1 ; | C - HA3 |
| 2996 | 3003 | 1 ; | C - C |
| 2997 | 2999 | 1 ; | O - H |
| 2997 | 3000 | 1 ; | O - CA |
| 2998 | 3004 | 1 ; | N - O |
| 2998 | 3005 | 1 ; | N - N |
| 2999 | 3001 | 1 ; | H - HA2 |
| 2999 | 3002 | 1 ; | H - HA3 |
| 2999 | 3003 | 1 ; | H - C |
| 3000 | 3006 | 1 ; | CA - H |
| 3000 | 3007 | 1 ; | CA - CA |
| 3001 | 3004 | 1 ; | HA2 - O |
| 3001 | 3005 | 1 ; | HA2 - N |
| 3002 | 3004 | 1 ; | HA3 - O |
| 3002 | 3005 | 1 ; | HA3 - N |
| 3003 | 3008 | 1 ; | C - HA |
| 3003 | 3009 | 1 ; | C - CB |
| 3003 | 3025 | 1 ; | C - C |
| 3004 | 3006 | 1 ; | O - H |
| 3004 | 3007 | 1 ; | O - CA |
| 3005 | 3010 | 1 ; | N - HB2 |
| 3005 | 3011 | 1 ; | N - HB3 |
| 3005 | 3012 | 1 ; | N - CG |
| 3005 | 3026 | 1 ; | N - O |
| 3005 | 3027 | 1 ; | N - N |
| 3006 | 3008 | 1 ; | H - HA |
| 3006 | 3009 | 1 ; | H - CB |
| 3006 | 3025 | 1 ; | H - C |
| 3007 | 3013 | 1 ; | CA - HG2 |
| 3007 | 3014 | 1 ; | CA - HG3 |
| 3007 | 3015 | 1 ; | CA - CD |
| 3007 | 3028 | 1 ; | CA - H |
| 3007 | 3029 | 1 ; | CA - CA |
| 3008 | 3010 | 1 ; | HA - HB2 |
| 3008 | 3011 | 1 ; | HA - HB3 |
| 3008 | 3012 | 1 ; | HA - CG |
| 3008 | 3026 | 1 ; | HA - O |
| 3008 | 3027 | 1 ; | HA - N |
| 3009 | 3016 | 1 ; | CB - HD2 |
| 3009 | 3017 | 1 ; | CB - HD3 |
| 3009 | 3018 | 1 ; | CB - CE |
| 3009 | 3026 | 1 ; | CB - O |
| 3009 | 3027 | 1 ; | CB - N |
| 3010 | 3013 | 1 ; | HB2 - HG2 |
| 3010 | 3014 | 1 ; | HB2 - HG3 |
| 3010 | 3015 | 1 ; | HB2 - CD |
| 3010 | 3025 | 1 ; | HB2 - C |
| 3011 | 3013 | 1 ; | HB3 - HG2 |

```
3011   3014       1 ;     HB3 - HG3
3011   3015       1 ;     HB3 - CD
3011   3025       1 ;     HB3 - C
3012   3019       1 ;      CG - HE2
3012   3020       1 ;      CG - HE3
3012   3021       1 ;      CG - NZ
3012   3025       1 ;      CG - C
3013   3016       1 ;     HG2 - HD2
3013   3017       1 ;     HG2 - HD3
3013   3018       1 ;     HG2 - CE
3014   3016       1 ;     HG3 - HD2
3014   3017       1 ;     HG3 - HD3
3014   3018       1 ;     HG3 - CE
3015   3022       1 ;      CD - HZ1
3015   3023       1 ;      CD - HZ2
3015   3024       1 ;      CD - HZ3
3016   3019       1 ;     HD2 - HE2
3016   3020       1 ;     HD2 - HE3
3016   3021       1 ;     HD2 - NZ
3017   3019       1 ;     HD3 - HE2
3017   3020       1 ;     HD3 - HE3
3017   3021       1 ;     HD3 - NZ
3019   3022       1 ;     HE2 - HZ1
3019   3023       1 ;     HE2 - HZ2
3019   3024       1 ;     HE2 - HZ3
3020   3022       1 ;     HE3 - HZ1
3020   3023       1 ;     HE3 - HZ2
3020   3024       1 ;     HE3 - HZ3
3025   3030       1 ;       C - HA
3025   3031       1 ;       C - CB
3025   3046       1 ;       C - C
3026   3028       1 ;       O - H
3026   3029       1 ;       O - CA
3027   3032       1 ;       N - HB2
3027   3033       1 ;       N - HB3
3027   3034       1 ;       N - CG
3027   3047       1 ;       N - O
3027   3048       1 ;       N - N
3028   3030       1 ;       H - HA
3028   3031       1 ;       H - CB
3028   3046       1 ;       H - C
3029   3035       1 ;      CA - CD1
3029   3044       1 ;      CA - CD2
3029   3049       1 ;      CA - H
3029   3050       1 ;      CA - CA
3030   3032       1 ;      HA - HB2
3030   3033       1 ;      HA - HB3
3030   3034       1 ;      HA - CG
3030   3047       1 ;      HA - O
3030   3048       1 ;      HA - N
3031   3036       1 ;      CB - HD1
3031   3037       1 ;      CB - CE1
```

| | | | | |
|---|---|---|---|---|
| 3031 | 3042 | 1 | ; | CB - CE2 |
| 3031 | 3045 | 1 | ; | CB - HD2 |
| 3031 | 3047 | 1 | ; | CB - O |
| 3031 | 3048 | 1 | ; | CB - N |
| 3032 | 3035 | 1 | ; | HB2 - CD1 |
| 3032 | 3044 | 1 | ; | HB2 - CD2 |
| 3032 | 3046 | 1 | ; | HB2 - C |
| 3033 | 3035 | 1 | ; | HB3 - CD1 |
| 3033 | 3044 | 1 | ; | HB3 - CD2 |
| 3033 | 3046 | 1 | ; | HB3 - C |
| 3034 | 3038 | 1 | ; | CG - HE1 |
| 3034 | 3039 | 1 | ; | CG - CZ |
| 3034 | 3043 | 1 | ; | CG - HE2 |
| 3034 | 3046 | 1 | ; | CG - C |
| 3035 | 3040 | 1 | ; | CD1 - OH |
| 3035 | 3042 | 1 | ; | CD1 - CE2 |
| 3035 | 3045 | 1 | ; | CD1 - HD2 |
| 3036 | 3038 | 1 | ; | HD1 - HE1 |
| 3036 | 3039 | 1 | ; | HD1 - CZ |
| 3036 | 3044 | 1 | ; | HD1 - CD2 |
| 3037 | 3041 | 1 | ; | CE1 - HH |
| 3037 | 3043 | 1 | ; | CE1 - HE2 |
| 3037 | 3044 | 1 | ; | CE1 - CD2 |
| 3038 | 3040 | 1 | ; | HE1 - OH |
| 3038 | 3042 | 1 | ; | HE1 - CE2 |
| 3039 | 3045 | 1 | ; | CZ - HD2 |
| 3040 | 3043 | 1 | ; | OH - HE2 |
| 3040 | 3044 | 1 | ; | OH - CD2 |
| 3041 | 3042 | 1 | ; | HH - CE2 |
| 3043 | 3045 | 1 | ; | HE2 - HD2 |
| 3046 | 3051 | 1 | ; | C - HA |
| 3046 | 3052 | 1 | ; | C - CB |
| 3046 | 3058 | 1 | ; | C - C |
| 3047 | 3049 | 1 | ; | O - H |
| 3047 | 3050 | 1 | ; | O - CA |
| 3048 | 3053 | 1 | ; | N - HB2 |
| 3048 | 3054 | 1 | ; | N - HB3 |
| 3048 | 3055 | 1 | ; | N - CG |
| 3048 | 3059 | 1 | ; | N - O |
| 3048 | 3060 | 1 | ; | N - N |
| 3049 | 3051 | 1 | ; | H - HA |
| 3049 | 3052 | 1 | ; | H - CB |
| 3049 | 3058 | 1 | ; | H - C |
| 3050 | 3056 | 1 | ; | CA - OD1 |
| 3050 | 3057 | 1 | ; | CA - OD2 |
| 3050 | 3061 | 1 | ; | CA - H |
| 3050 | 3062 | 1 | ; | CA - CA |
| 3051 | 3053 | 1 | ; | HA - HB2 |
| 3051 | 3054 | 1 | ; | HA - HB3 |
| 3051 | 3055 | 1 | ; | HA - CG |
| 3051 | 3059 | 1 | ; | HA - O |
| 3051 | 3060 | 1 | ; | HA - N |

| | | | | | |
|---|---|---|---|---|---|
| 3052 | 3059 | 1 ; | CB | - | O |
| 3052 | 3060 | 1 ; | CB | - | N |
| 3053 | 3056 | 1 ; | HB2 | - | OD1 |
| 3053 | 3057 | 1 ; | HB2 | - | OD2 |
| 3053 | 3058 | 1 ; | HB2 | - | C |
| 3054 | 3056 | 1 ; | HB3 | - | OD1 |
| 3054 | 3057 | 1 ; | HB3 | - | OD2 |
| 3054 | 3058 | 1 ; | HB3 | - | C |
| 3055 | 3058 | 1 ; | CG | - | C |
| 3058 | 3063 | 1 ; | C | - | HA |
| 3058 | 3064 | 1 ; | C | - | CB |
| 3058 | 3080 | 1 ; | C | - | C |
| 3059 | 3061 | 1 ; | O | - | H |
| 3059 | 3062 | 1 ; | O | - | CA |
| 3060 | 3065 | 1 ; | N | - | HB2 |
| 3060 | 3066 | 1 ; | N | - | HB3 |
| 3060 | 3067 | 1 ; | N | - | CG |
| 3060 | 3081 | 1 ; | N | - | O |
| 3060 | 3082 | 1 ; | N | - | N |
| 3061 | 3063 | 1 ; | H | - | HA |
| 3061 | 3064 | 1 ; | H | - | CB |
| 3061 | 3080 | 1 ; | H | - | C |
| 3062 | 3068 | 1 ; | CA | - | HG2 |
| 3062 | 3069 | 1 ; | CA | - | HG3 |
| 3062 | 3070 | 1 ; | CA | - | CD |
| 3062 | 3083 | 1 ; | CA | - | H |
| 3062 | 3084 | 1 ; | CA | - | CA |
| 3063 | 3065 | 1 ; | HA | - | HB2 |
| 3063 | 3066 | 1 ; | HA | - | HB3 |
| 3063 | 3067 | 1 ; | HA | - | CG |
| 3063 | 3081 | 1 ; | HA | - | O |
| 3063 | 3082 | 1 ; | HA | - | N |
| 3064 | 3071 | 1 ; | CB | - | HD2 |
| 3064 | 3072 | 1 ; | CB | - | HD3 |
| 3064 | 3073 | 1 ; | CB | - | CE |
| 3064 | 3081 | 1 ; | CB | - | O |
| 3064 | 3082 | 1 ; | CB | - | N |
| 3065 | 3068 | 1 ; | HB2 | - | HG2 |
| 3065 | 3069 | 1 ; | HB2 | - | HG3 |
| 3065 | 3070 | 1 ; | HB2 | - | CD |
| 3065 | 3080 | 1 ; | HB2 | - | C |
| 3066 | 3068 | 1 ; | HB3 | - | HG2 |
| 3066 | 3069 | 1 ; | HB3 | - | HG3 |
| 3066 | 3070 | 1 ; | HB3 | - | CD |
| 3066 | 3080 | 1 ; | HB3 | - | C |
| 3067 | 3074 | 1 ; | CG | - | HE2 |
| 3067 | 3075 | 1 ; | CG | - | HE3 |
| 3067 | 3076 | 1 ; | CG | - | NZ |
| 3067 | 3080 | 1 ; | CG | - | C |
| 3068 | 3071 | 1 ; | HG2 | - | HD2 |
| 3068 | 3072 | 1 ; | HG2 | - | HD3 |
| 3068 | 3073 | 1 ; | HG2 | - | CE |

| | | | | |
|---|---|---|---|---|
| 3069 | 3071 | 1 ; | HG3 - | HD2 |
| 3069 | 3072 | 1 ; | HG3 - | HD3 |
| 3069 | 3073 | 1 ; | HG3 - | CE |
| 3070 | 3077 | 1 ; | CD - | HZ1 |
| 3070 | 3078 | 1 ; | CD - | HZ2 |
| 3070 | 3079 | 1 ; | CD - | HZ3 |
| 3071 | 3074 | 1 ; | HD2 - | HE2 |
| 3071 | 3075 | 1 ; | HD2 - | HE3 |
| 3071 | 3076 | 1 ; | HD2 - | NZ |
| 3072 | 3074 | 1 ; | HD3 - | HE2 |
| 3072 | 3075 | 1 ; | HD3 - | HE3 |
| 3072 | 3076 | 1 ; | HD3 - | NZ |
| 3074 | 3077 | 1 ; | HE2 - | HZ1 |
| 3074 | 3078 | 1 ; | HE2 - | HZ2 |
| 3074 | 3079 | 1 ; | HE2 - | HZ3 |
| 3075 | 3077 | 1 ; | HE3 - | HZ1 |
| 3075 | 3078 | 1 ; | HE3 - | HZ2 |
| 3075 | 3079 | 1 ; | HE3 - | HZ3 |
| 3080 | 3085 | 1 ; | C - | HA |
| 3080 | 3086 | 1 ; | C - | CB |
| 3080 | 3102 | 1 ; | C - | C |
| 3081 | 3083 | 1 ; | O - | H |
| 3081 | 3084 | 1 ; | O - | CA |
| 3082 | 3087 | 1 ; | N - | HB2 |
| 3082 | 3088 | 1 ; | N - | HB3 |
| 3082 | 3089 | 1 ; | N - | CG |
| 3082 | 3103 | 1 ; | N - | O |
| 3082 | 3104 | 1 ; | N - | N |
| 3083 | 3085 | 1 ; | H - | HA |
| 3083 | 3086 | 1 ; | H - | CB |
| 3083 | 3102 | 1 ; | H - | C |
| 3084 | 3090 | 1 ; | CA - | HG2 |
| 3084 | 3091 | 1 ; | CA - | HG3 |
| 3084 | 3092 | 1 ; | CA - | CD |
| 3084 | 3105 | 1 ; | CA - | H |
| 3084 | 3106 | 1 ; | CA - | CA |
| 3085 | 3087 | 1 ; | HA - | HB2 |
| 3085 | 3088 | 1 ; | HA - | HB3 |
| 3085 | 3089 | 1 ; | HA - | CG |
| 3085 | 3103 | 1 ; | HA - | O |
| 3085 | 3104 | 1 ; | HA - | N |
| 3086 | 3093 | 1 ; | CB - | HD2 |
| 3086 | 3094 | 1 ; | CB - | HD3 |
| 3086 | 3095 | 1 ; | CB - | CE |
| 3086 | 3103 | 1 ; | CB - | O |
| 3086 | 3104 | 1 ; | CB - | N |
| 3087 | 3090 | 1 ; | HB2 - | HG2 |
| 3087 | 3091 | 1 ; | HB2 - | HG3 |
| 3087 | 3092 | 1 ; | HB2 - | CD |
| 3087 | 3102 | 1 ; | HB2 - | C |
| 3088 | 3090 | 1 ; | HB3 - | HG2 |
| 3088 | 3091 | 1 ; | HB3 - | HG3 |

| | | | | | |
|---|---|---|---|---|---|
| 3088 | 3092 | 1 ; | HB3 | - | CD |
| 3088 | 3102 | 1 ; | HB3 | - | C |
| 3089 | 3096 | 1 ; | CG | - | HE2 |
| 3089 | 3097 | 1 ; | CG | - | HE3 |
| 3089 | 3098 | 1 ; | CG | - | NZ |
| 3089 | 3102 | 1 ; | CG | - | C |
| 3090 | 3093 | 1 ; | HG2 | - | HD2 |
| 3090 | 3094 | 1 ; | HG2 | - | HD3 |
| 3090 | 3095 | 1 ; | HG2 | - | CE |
| 3091 | 3093 | 1 ; | HG3 | - | HD2 |
| 3091 | 3094 | 1 ; | HG3 | - | HD3 |
| 3091 | 3095 | 1 ; | HG3 | - | CE |
| 3092 | 3099 | 1 ; | CD | - | HZ1 |
| 3092 | 3100 | 1 ; | CD | - | HZ2 |
| 3092 | 3101 | 1 ; | CD | - | HZ3 |
| 3093 | 3096 | 1 ; | HD2 | - | HE2 |
| 3093 | 3097 | 1 ; | HD2 | - | HE3 |
| 3093 | 3098 | 1 ; | HD2 | - | NZ |
| 3094 | 3096 | 1 ; | HD3 | - | HE2 |
| 3094 | 3097 | 1 ; | HD3 | - | HE3 |
| 3094 | 3098 | 1 ; | HD3 | - | NZ |
| 3096 | 3099 | 1 ; | HE2 | - | HZ1 |
| 3096 | 3100 | 1 ; | HE2 | - | HZ2 |
| 3096 | 3101 | 1 ; | HE2 | - | HZ3 |
| 3097 | 3099 | 1 ; | HE3 | - | HZ1 |
| 3097 | 3100 | 1 ; | HE3 | - | HZ2 |
| 3097 | 3101 | 1 ; | HE3 | - | HZ3 |
| 3102 | 3107 | 1 ; | C | - | HA |
| 3102 | 3108 | 1 ; | C | - | CB |
| 3102 | 3113 | 1 ; | C | - | C |
| 3103 | 3105 | 1 ; | O | - | H |
| 3103 | 3106 | 1 ; | O | - | CA |
| 3104 | 3109 | 1 ; | N | - | HB2 |
| 3104 | 3110 | 1 ; | N | - | HB3 |
| 3104 | 3111 | 1 ; | N | - | SG |
| 3104 | 3114 | 1 ; | N | - | O |
| 3104 | 3115 | 1 ; | N | - | N |
| 3105 | 3107 | 1 ; | H | - | HA |
| 3105 | 3108 | 1 ; | H | - | CB |
| 3105 | 3113 | 1 ; | H | - | C |
| 3106 | 3112 | 1 ; | CA | - | HG |
| 3106 | 3116 | 1 ; | CA | - | H |
| 3106 | 3117 | 1 ; | CA | - | CA |
| 3107 | 3109 | 1 ; | HA | - | HB2 |
| 3107 | 3110 | 1 ; | HA | - | HB3 |
| 3107 | 3111 | 1 ; | HA | - | SG |
| 3107 | 3114 | 1 ; | HA | - | O |
| 3107 | 3115 | 1 ; | HA | - | N |
| 3108 | 3114 | 1 ; | CB | - | O |
| 3108 | 3115 | 1 ; | CB | - | N |
| 3109 | 3112 | 1 ; | HB2 | - | HG |
| 3109 | 3113 | 1 ; | HB2 | - | C |

3110 3112 1 ; HB3 - HG
3110 3113 1 ; HB3 - C
3111 3113 1 ; SG - C
3113 3118 1 ; C - HA
3113 3119 1 ; C - CB
3113 3125 1 ; C - C
3114 3116 1 ; O - H
3114 3117 1 ; O - CA
3115 3120 1 ; N - HB2
3115 3121 1 ; N - HB3
3115 3122 1 ; N - CG
3115 3126 1 ; N - O
3115 3127 1 ; N - N
3116 3118 1 ; H - HA
3116 3119 1 ; H - CB
3116 3125 1 ; H - C
3117 3123 1 ; CA - OD1
3117 3124 1 ; CA - OD2
3117 3128 1 ; CA - H
3117 3129 1 ; CA - CA
3118 3120 1 ; HA - HB2
3118 3121 1 ; HA - HB3
3118 3122 1 ; HA - CG
3118 3126 1 ; HA - O
3118 3127 1 ; HA - N
3119 3126 1 ; CB - O
3119 3127 1 ; CB - N
3120 3123 1 ; HB2 - OD1
3120 3124 1 ; HB2 - OD2
3120 3125 1 ; HB2 - C
3121 3123 1 ; HB3 - OD1
3121 3124 1 ; HB3 - OD2
3121 3125 1 ; HB3 - C
3122 3125 1 ; CG - C
3125 3130 1 ; C - HA
3125 3131 1 ; C - CB
3125 3144 1 ; C - C
3126 3128 1 ; O - H
3126 3129 1 ; O - CA
3127 3132 1 ; N - HB
3127 3133 1 ; N - CG2
3127 3137 1 ; N - CG1
3127 3145 1 ; N - O
3127 3146 1 ; N - N
3128 3130 1 ; H - HA
3128 3131 1 ; H - CB
3128 3144 1 ; H - C
3129 3134 1 ; CA - HG21
3129 3135 1 ; CA - HG22
3129 3136 1 ; CA - HG23
3129 3138 1 ; CA - HG12
3129 3139 1 ; CA - HG13

| | | | |
|---|---|---|---|
| 3129 | 3140 | 1 ; | CA - CD1 |
| 3129 | 3147 | 1 ; | CA - H |
| 3129 | 3148 | 1 ; | CA - CA |
| 3130 | 3132 | 1 ; | HA - HB |
| 3130 | 3133 | 1 ; | HA - CG2 |
| 3130 | 3137 | 1 ; | HA - CG1 |
| 3130 | 3145 | 1 ; | HA - O |
| 3130 | 3146 | 1 ; | HA - N |
| 3131 | 3141 | 1 ; | CB - HD11 |
| 3131 | 3142 | 1 ; | CB - HD12 |
| 3131 | 3143 | 1 ; | CB - HD13 |
| 3131 | 3145 | 1 ; | CB - O |
| 3131 | 3146 | 1 ; | CB - N |
| 3132 | 3134 | 1 ; | HB - HG21 |
| 3132 | 3135 | 1 ; | HB - HG22 |
| 3132 | 3136 | 1 ; | HB - HG23 |
| 3132 | 3138 | 1 ; | HB - HG12 |
| 3132 | 3139 | 1 ; | HB - HG13 |
| 3132 | 3140 | 1 ; | HB - CD1 |
| 3132 | 3144 | 1 ; | HB - C |
| 3133 | 3138 | 1 ; | CG2 - HG12 |
| 3133 | 3139 | 1 ; | CG2 - HG13 |
| 3133 | 3140 | 1 ; | CG2 - CD1 |
| 3133 | 3144 | 1 ; | CG2 - C |
| 3134 | 3137 | 1 ; | HG21 - CG1 |
| 3135 | 3137 | 1 ; | HG22 - CG1 |
| 3136 | 3137 | 1 ; | HG23 - CG1 |
| 3137 | 3144 | 1 ; | CG1 - C |
| 3138 | 3141 | 1 ; | HG12 - HD11 |
| 3138 | 3142 | 1 ; | HG12 - HD12 |
| 3138 | 3143 | 1 ; | HG12 - HD13 |
| 3139 | 3141 | 1 ; | HG13 - HD11 |
| 3139 | 3142 | 1 ; | HG13 - HD12 |
| 3139 | 3143 | 1 ; | HG13 - HD13 |
| 3144 | 3149 | 1 ; | C - HA |
| 3144 | 3150 | 1 ; | C - CB |
| 3144 | 3168 | 1 ; | C - C |
| 3145 | 3147 | 1 ; | O - H |
| 3145 | 3148 | 1 ; | O - CA |
| 3146 | 3151 | 1 ; | N - HB2 |
| 3146 | 3152 | 1 ; | N - HB3 |
| 3146 | 3153 | 1 ; | N - CG |
| 3146 | 3169 | 1 ; | N - O |
| 3146 | 3170 | 1 ; | N - N |
| 3147 | 3149 | 1 ; | H - HA |
| 3147 | 3150 | 1 ; | H - CB |
| 3147 | 3168 | 1 ; | H - C |
| 3148 | 3154 | 1 ; | CA - CD1 |
| 3148 | 3167 | 1 ; | CA - CD2 |
| 3148 | 3171 | 1 ; | CA - H |
| 3148 | 3172 | 1 ; | CA - CA |
| 3149 | 3151 | 1 ; | HA - HB2 |

| | | | |
|---|---|---|---|
| 3149 | 3152 | 1 ; | HA - HB3 |
| 3149 | 3153 | 1 ; | HA - CG |
| 3149 | 3169 | 1 ; | HA - O |
| 3149 | 3170 | 1 ; | HA - N |
| 3150 | 3155 | 1 ; | CB - HD1 |
| 3150 | 3156 | 1 ; | CB - NE1 |
| 3150 | 3158 | 1 ; | CB - CE2 |
| 3150 | 3165 | 1 ; | CB - CE3 |
| 3150 | 3169 | 1 ; | CB - O |
| 3150 | 3170 | 1 ; | CB - N |
| 3151 | 3154 | 1 ; | HB2 - CD1 |
| 3151 | 3167 | 1 ; | HB2 - CD2 |
| 3151 | 3168 | 1 ; | HB2 - C |
| 3152 | 3154 | 1 ; | HB3 - CD1 |
| 3152 | 3167 | 1 ; | HB3 - CD2 |
| 3152 | 3168 | 1 ; | HB3 - C |
| 3153 | 3157 | 1 ; | CG - HE1 |
| 3153 | 3159 | 1 ; | CG - CZ2 |
| 3153 | 3163 | 1 ; | CG - CZ3 |
| 3153 | 3166 | 1 ; | CG - HE3 |
| 3153 | 3168 | 1 ; | CG - C |
| 3154 | 3159 | 1 ; | CD1 - CZ2 |
| 3154 | 3165 | 1 ; | CD1 - CE3 |
| 3155 | 3157 | 1 ; | HD1 - HE1 |
| 3155 | 3158 | 1 ; | HD1 - CE2 |
| 3155 | 3167 | 1 ; | HD1 - CD2 |
| 3156 | 3160 | 1 ; | NE1 - HZ2 |
| 3156 | 3161 | 1 ; | NE1 - CH2 |
| 3156 | 3165 | 1 ; | NE1 - CE3 |
| 3157 | 3159 | 1 ; | HE1 - CZ2 |
| 3157 | 3167 | 1 ; | HE1 - CD2 |
| 3158 | 3162 | 1 ; | CE2 - HH2 |
| 3158 | 3163 | 1 ; | CE2 - CZ3 |
| 3158 | 3166 | 1 ; | CE2 - HE3 |
| 3159 | 3164 | 1 ; | CZ2 - HZ3 |
| 3159 | 3165 | 1 ; | CZ2 - CE3 |
| 3160 | 3162 | 1 ; | HZ2 - HH2 |
| 3160 | 3163 | 1 ; | HZ2 - CZ3 |
| 3160 | 3167 | 1 ; | HZ2 - CD2 |
| 3161 | 3166 | 1 ; | CH2 - HE3 |
| 3161 | 3167 | 1 ; | CH2 - CD2 |
| 3162 | 3164 | 1 ; | HH2 - HZ3 |
| 3162 | 3165 | 1 ; | HH2 - CE3 |
| 3164 | 3166 | 1 ; | HZ3 - HE3 |
| 3164 | 3167 | 1 ; | HZ3 - CD2 |
| 3168 | 3173 | 1 ; | C - HA |
| 3168 | 3174 | 1 ; | C - CB |
| 3168 | 3179 | 1 ; | C - C |
| 3169 | 3171 | 1 ; | O - H |
| 3169 | 3172 | 1 ; | O - CA |
| 3170 | 3175 | 1 ; | N - HB2 |
| 3170 | 3176 | 1 ; | N - HB3 |

| | | | | |
|---|---|---|---|---|
| 3170 | 3177 | 1 ; | N | - OG |
| 3170 | 3180 | 1 ; | N | - O |
| 3170 | 3181 | 1 ; | N | - N |
| 3171 | 3173 | 1 ; | H | - HA |
| 3171 | 3174 | 1 ; | H | - CB |
| 3171 | 3179 | 1 ; | H | - C |
| 3172 | 3178 | 1 ; | CA | - HG |
| 3172 | 3182 | 1 ; | CA | - H |
| 3172 | 3183 | 1 ; | CA | - CA |
| 3173 | 3175 | 1 ; | HA | - HB2 |
| 3173 | 3176 | 1 ; | HA | - HB3 |
| 3173 | 3177 | 1 ; | HA | - OG |
| 3173 | 3180 | 1 ; | HA | - O |
| 3173 | 3181 | 1 ; | HA | - N |
| 3174 | 3180 | 1 ; | CB | - O |
| 3174 | 3181 | 1 ; | CB | - N |
| 3175 | 3178 | 1 ; | HB2 | - HG |
| 3175 | 3179 | 1 ; | HB2 | - C |
| 3176 | 3178 | 1 ; | HB3 | - HG |
| 3176 | 3179 | 1 ; | HB3 | - C |
| 3177 | 3179 | 1 ; | OG | - C |
| 3179 | 3184 | 1 ; | C | - HA |
| 3179 | 3185 | 1 ; | C | - CB |
| 3179 | 3190 | 1 ; | C | - C |
| 3180 | 3182 | 1 ; | O | - H |
| 3180 | 3183 | 1 ; | O | - CA |
| 3181 | 3186 | 1 ; | N | - HB2 |
| 3181 | 3187 | 1 ; | N | - HB3 |
| 3181 | 3188 | 1 ; | N | - OG |
| 3181 | 3191 | 1 ; | N | - O |
| 3181 | 3192 | 1 ; | N | - N |
| 3182 | 3184 | 1 ; | H | - HA |
| 3182 | 3185 | 1 ; | H | - CB |
| 3182 | 3190 | 1 ; | H | - C |
| 3183 | 3189 | 1 ; | CA | - HG |
| 3183 | 3193 | 1 ; | CA | - H |
| 3183 | 3194 | 1 ; | CA | - CA |
| 3184 | 3186 | 1 ; | HA | - HB2 |
| 3184 | 3187 | 1 ; | HA | - HB3 |
| 3184 | 3188 | 1 ; | HA | - OG |
| 3184 | 3191 | 1 ; | HA | - O |
| 3184 | 3192 | 1 ; | HA | - N |
| 3185 | 3191 | 1 ; | CB | - O |
| 3185 | 3192 | 1 ; | CB | - N |
| 3186 | 3189 | 1 ; | HB2 | - HG |
| 3186 | 3190 | 1 ; | HB2 | - C |
| 3187 | 3189 | 1 ; | HB3 | - HG |
| 3187 | 3190 | 1 ; | HB3 | - C |
| 3188 | 3190 | 1 ; | OG | - C |
| 3190 | 3195 | 1 ; | C | - HA2 |
| 3190 | 3196 | 1 ; | C | - HA3 |
| 3190 | 3197 | 1 ; | C | - C |

| | | | | |
|---|---|---|---|---|
| 3191 | 3193 | 1 | ; | O - H |
| 3191 | 3194 | 1 | ; | O - CA |
| 3192 | 3198 | 1 | ; | N - O |
| 3192 | 3199 | 1 | ; | N - N |
| 3193 | 3195 | 1 | ; | H - HA2 |
| 3193 | 3196 | 1 | ; | H - HA3 |
| 3193 | 3197 | 1 | ; | H - C |
| 3194 | 3200 | 1 | ; | CA - H |
| 3194 | 3201 | 1 | ; | CA - CA |
| 3195 | 3198 | 1 | ; | HA2 - O |
| 3195 | 3199 | 1 | ; | HA2 - N |
| 3196 | 3198 | 1 | ; | HA3 - O |
| 3196 | 3199 | 1 | ; | HA3 - N |
| 3197 | 3202 | 1 | ; | C - HA |
| 3197 | 3203 | 1 | ; | C - CB |
| 3197 | 3213 | 1 | ; | C - C |
| 3198 | 3200 | 1 | ; | O - H |
| 3198 | 3201 | 1 | ; | O - CA |
| 3199 | 3204 | 1 | ; | N - HB |
| 3199 | 3205 | 1 | ; | N - CG1 |
| 3199 | 3209 | 1 | ; | N - CG2 |
| 3199 | 3214 | 1 | ; | N - O |
| 3199 | 3215 | 1 | ; | N - N |
| 3200 | 3202 | 1 | ; | H - HA |
| 3200 | 3203 | 1 | ; | H - CB |
| 3200 | 3213 | 1 | ; | H - C |
| 3201 | 3206 | 1 | ; | CA - HG11 |
| 3201 | 3207 | 1 | ; | CA - HG12 |
| 3201 | 3208 | 1 | ; | CA - HG13 |
| 3201 | 3210 | 1 | ; | CA - HG21 |
| 3201 | 3211 | 1 | ; | CA - HG22 |
| 3201 | 3212 | 1 | ; | CA - HG23 |
| 3201 | 3216 | 1 | ; | CA - H |
| 3201 | 3217 | 1 | ; | CA - CA |
| 3202 | 3204 | 1 | ; | HA - HB |
| 3202 | 3205 | 1 | ; | HA - CG1 |
| 3202 | 3209 | 1 | ; | HA - CG2 |
| 3202 | 3214 | 1 | ; | HA - O |
| 3202 | 3215 | 1 | ; | HA - N |
| 3203 | 3214 | 1 | ; | CB - O |
| 3203 | 3215 | 1 | ; | CB - N |
| 3204 | 3206 | 1 | ; | HB - HG11 |
| 3204 | 3207 | 1 | ; | HB - HG12 |
| 3204 | 3208 | 1 | ; | HB - HG13 |
| 3204 | 3210 | 1 | ; | HB - HG21 |
| 3204 | 3211 | 1 | ; | HB - HG22 |
| 3204 | 3212 | 1 | ; | HB - HG23 |
| 3204 | 3213 | 1 | ; | HB - C |
| 3205 | 3210 | 1 | ; | CG1 - HG21 |
| 3205 | 3211 | 1 | ; | CG1 - HG22 |
| 3205 | 3212 | 1 | ; | CG1 - HG23 |
| 3205 | 3213 | 1 | ; | CG1 - C |

```
3206   3209      1 ;   HG11 - CG2
3207   3209      1 ;   HG12 - CG2
3208   3209      1 ;   HG13 - CG2
3209   3213      1 ;    CG2 - C
3213   3218      1 ;      C - HA
3213   3219      1 ;      C - CB
3213   3232      1 ;      C - C
3214   3216      1 ;      O - H
3214   3217      1 ;      O - CA
3215   3220      1 ;      N - HB
3215   3221      1 ;      N - CG2
3215   3225      1 ;      N - CG1
3215   3233      1 ;      N - O
3215   3234      1 ;      N - N
3216   3218      1 ;      H - HA
3216   3219      1 ;      H - CB
3216   3232      1 ;      H - C
3217   3222      1 ;     CA - HG21
3217   3223      1 ;     CA - HG22
3217   3224      1 ;     CA - HG23
3217   3226      1 ;     CA - HG12
3217   3227      1 ;     CA - HG13
3217   3228      1 ;     CA - CD1
3217   3235      1 ;     CA - H
3217   3236      1 ;     CA - CA
3218   3220      1 ;     HA - HB
3218   3221      1 ;     HA - CG2
3218   3225      1 ;     HA - CG1
3218   3233      1 ;     HA - O
3218   3234      1 ;     HA - N
3219   3229      1 ;     CB - HD11
3219   3230      1 ;     CB - HD12
3219   3231      1 ;     CB - HD13
3219   3233      1 ;     CB - O
3219   3234      1 ;     CB - N
3220   3222      1 ;     HB - HG21
3220   3223      1 ;     HB - HG22
3220   3224      1 ;     HB - HG23
3220   3226      1 ;     HB - HG12
3220   3227      1 ;     HB - HG13
3220   3228      1 ;     HB - CD1
3220   3232      1 ;     HB - C
3221   3226      1 ;    CG2 - HG12
3221   3227      1 ;    CG2 - HG13
3221   3228      1 ;    CG2 - CD1
3221   3232      1 ;    CG2 - C
3222   3225      1 ;   HG21 - CG1
3223   3225      1 ;   HG22 - CG1
3224   3225      1 ;   HG23 - CG1
3225   3232      1 ;    CG1 - C
3226   3229      1 ;   HG12 - HD11
3226   3230      1 ;   HG12 - HD12
```

```
3226   3231       1 ;   HG12 - HD13
3227   3229       1 ;   HG13 - HD11
3227   3230       1 ;   HG13 - HD12
3227   3231       1 ;   HG13 - HD13
3232   3237       1 ;      C - HA
3232   3238       1 ;      C - CB
3232   3249       1 ;      C - C
3233   3235       1 ;      O - H
3233   3236       1 ;      O - CA
3234   3239       1 ;      N - HB2
3234   3240       1 ;      N - HB3
3234   3241       1 ;      N - CG
3234   3250       1 ;      N - O
3234   3251       1 ;      N - N
3235   3237       1 ;      H - HA
3235   3238       1 ;      H - CB
3235   3249       1 ;      H - C
3236   3242       1 ;     CA - HG2
3236   3243       1 ;     CA - HG3
3236   3244       1 ;     CA - SD
3236   3252       1 ;     CA - H
3236   3253       1 ;     CA - CA
3237   3239       1 ;     HA - HB2
3237   3240       1 ;     HA - HB3
3237   3241       1 ;     HA - CG
3237   3250       1 ;     HA - O
3237   3251       1 ;     HA - N
3238   3245       1 ;     CB - CE
3238   3250       1 ;     CB - O
3238   3251       1 ;     CB - N
3239   3242       1 ;    HB2 - HG2
3239   3243       1 ;    HB2 - HG3
3239   3244       1 ;    HB2 - SD
3239   3249       1 ;    HB2 - C
3240   3242       1 ;    HB3 - HG2
3240   3243       1 ;    HB3 - HG3
3240   3244       1 ;    HB3 - SD
3240   3249       1 ;    HB3 - C
3241   3246       1 ;     CG - HE1
3241   3247       1 ;     CG - HE2
3241   3248       1 ;     CG - HE3
3241   3249       1 ;     CG - C
3242   3245       1 ;    HG2 - CE
3243   3245       1 ;    HG3 - CE
3249   3254       1 ;      C - HA
3249   3255       1 ;      C - CB
3249   3270       1 ;      C - C
3250   3252       1 ;      O - H
3250   3253       1 ;      O - CA
3251   3256       1 ;      N - HB2
3251   3257       1 ;      N - HB3
3251   3258       1 ;      N - CG
```

| | | | | | |
|---|---|---|---|---|---|
| 3251 | 3271 | 1 ; | N | - | O |
| 3251 | 3272 | 1 ; | N | - | N |
| 3252 | 3254 | 1 ; | H | - | HA |
| 3252 | 3255 | 1 ; | H | - | CB |
| 3252 | 3270 | 1 ; | H | - | C |
| 3253 | 3259 | 1 ; | CA | - | CD1 |
| 3253 | 3268 | 1 ; | CA | - | CD2 |
| 3253 | 3273 | 1 ; | CA | - | H |
| 3253 | 3274 | 1 ; | CA | - | CA |
| 3254 | 3256 | 1 ; | HA | - | HB2 |
| 3254 | 3257 | 1 ; | HA | - | HB3 |
| 3254 | 3258 | 1 ; | HA | - | CG |
| 3254 | 3271 | 1 ; | HA | - | O |
| 3254 | 3272 | 1 ; | HA | - | N |
| 3255 | 3260 | 1 ; | CB | - | HD1 |
| 3255 | 3261 | 1 ; | CB | - | CE1 |
| 3255 | 3266 | 1 ; | CB | - | CE2 |
| 3255 | 3269 | 1 ; | CB | - | HD2 |
| 3255 | 3271 | 1 ; | CB | - | O |
| 3255 | 3272 | 1 ; | CB | - | N |
| 3256 | 3259 | 1 ; | HB2 | - | CD1 |
| 3256 | 3268 | 1 ; | HB2 | - | CD2 |
| 3256 | 3270 | 1 ; | HB2 | - | C |
| 3257 | 3259 | 1 ; | HB3 | - | CD1 |
| 3257 | 3268 | 1 ; | HB3 | - | CD2 |
| 3257 | 3270 | 1 ; | HB3 | - | C |
| 3258 | 3262 | 1 ; | CG | - | HE1 |
| 3258 | 3263 | 1 ; | CG | - | CZ |
| 3258 | 3267 | 1 ; | CG | - | HE2 |
| 3258 | 3270 | 1 ; | CG | - | C |
| 3259 | 3264 | 1 ; | CD1 | - | OH |
| 3259 | 3266 | 1 ; | CD1 | - | CE2 |
| 3259 | 3269 | 1 ; | CD1 | - | HD2 |
| 3260 | 3262 | 1 ; | HD1 | - | HE1 |
| 3260 | 3263 | 1 ; | HD1 | - | CZ |
| 3260 | 3268 | 1 ; | HD1 | - | CD2 |
| 3261 | 3265 | 1 ; | CE1 | - | HH |
| 3261 | 3267 | 1 ; | CE1 | - | HE2 |
| 3261 | 3268 | 1 ; | CE1 | - | CD2 |
| 3262 | 3264 | 1 ; | HE1 | - | OH |
| 3262 | 3266 | 1 ; | HE1 | - | CE2 |
| 3263 | 3269 | 1 ; | CZ | - | HD2 |
| 3264 | 3267 | 1 ; | OH | - | HE2 |
| 3264 | 3268 | 1 ; | OH | - | CD2 |
| 3265 | 3266 | 1 ; | HH | - | CE2 |
| 3267 | 3269 | 1 ; | HE2 | - | HD2 |
| 3270 | 3275 | 1 ; | C | - | HA |
| 3270 | 3276 | 1 ; | C | - | CB |
| 3270 | 3284 | 1 ; | C | - | C |
| 3271 | 3273 | 1 ; | O | - | H |
| 3271 | 3274 | 1 ; | O | - | CA |
| 3272 | 3277 | 1 ; | N | - | HB |

```
3272   3278     1 ;      N - CG2
3272   3282     1 ;      N - OG1
3272   3285     1 ;      N - O
3272   3286     1 ;      N - N
3273   3275     1 ;      H - HA
3273   3276     1 ;      H - CB
3273   3284     1 ;      H - C
3274   3279     1 ;     CA - HG21
3274   3280     1 ;     CA - HG22
3274   3281     1 ;     CA - HG23
3274   3283     1 ;     CA - HG1
3274   3287     1 ;     CA - H
3274   3288     1 ;     CA - CA
3275   3277     1 ;     HA - HB
3275   3278     1 ;     HA - CG2
3275   3282     1 ;     HA - OG1
3275   3285     1 ;     HA - O
3275   3286     1 ;     HA - N
3276   3285     1 ;     CB - O
3276   3286     1 ;     CB - N
3277   3279     1 ;     HB - HG21
3277   3280     1 ;     HB - HG22
3277   3281     1 ;     HB - HG23
3277   3283     1 ;     HB - HG1
3277   3284     1 ;     HB - C
3278   3283     1 ;    CG2 - HG1
3278   3284     1 ;    CG2 - C
3279   3282     1 ;   HG21 - OG1
3280   3282     1 ;   HG22 - OG1
3281   3282     1 ;   HG23 - OG1
3282   3284     1 ;    OG1 - C
3284   3289     1 ;      C - HA
3284   3290     1 ;      C - CB
3284   3303     1 ;      C - C
3285   3287     1 ;      O - H
3285   3288     1 ;      O - CA
3286   3291     1 ;      N - HB2
3286   3292     1 ;      N - HB3
3286   3293     1 ;      N - CG
3286   3304     1 ;      N - O
3286   3305     1 ;      N - N
3287   3289     1 ;      H - HA
3287   3290     1 ;      H - CB
3287   3303     1 ;      H - C
3288   3294     1 ;     CA - HG
3288   3295     1 ;     CA - CD1
3288   3299     1 ;     CA - CD2
3288   3306     1 ;     CA - H
3288   3307     1 ;     CA - CA
3289   3291     1 ;     HA - HB2
3289   3292     1 ;     HA - HB3
3289   3293     1 ;     HA - CG
```

| | | | | | | |
|---|---|---|---|---|---|---|
| 3289 | 3304 | 1 | ; | HA | - | O |
| 3289 | 3305 | 1 | ; | HA | - | N |
| 3290 | 3296 | 1 | ; | CB | - | HD11 |
| 3290 | 3297 | 1 | ; | CB | - | HD12 |
| 3290 | 3298 | 1 | ; | CB | - | HD13 |
| 3290 | 3300 | 1 | ; | CB | - | HD21 |
| 3290 | 3301 | 1 | ; | CB | - | HD22 |
| 3290 | 3302 | 1 | ; | CB | - | HD23 |
| 3290 | 3304 | 1 | ; | CB | - | O |
| 3290 | 3305 | 1 | ; | CB | - | N |
| 3291 | 3294 | 1 | ; | HB2 | - | HG |
| 3291 | 3295 | 1 | ; | HB2 | - | CD1 |
| 3291 | 3299 | 1 | ; | HB2 | - | CD2 |
| 3291 | 3303 | 1 | ; | HB2 | - | C |
| 3292 | 3294 | 1 | ; | HB3 | - | HG |
| 3292 | 3295 | 1 | ; | HB3 | - | CD1 |
| 3292 | 3299 | 1 | ; | HB3 | - | CD2 |
| 3292 | 3303 | 1 | ; | HB3 | - | C |
| 3293 | 3303 | 1 | ; | CG | - | C |
| 3294 | 3296 | 1 | ; | HG | - | HD11 |
| 3294 | 3297 | 1 | ; | HG | - | HD12 |
| 3294 | 3298 | 1 | ; | HG | - | HD13 |
| 3294 | 3300 | 1 | ; | HG | - | HD21 |
| 3294 | 3301 | 1 | ; | HG | - | HD22 |
| 3294 | 3302 | 1 | ; | HG | - | HD23 |
| 3295 | 3300 | 1 | ; | CD1 | - | HD21 |
| 3295 | 3301 | 1 | ; | CD1 | - | HD22 |
| 3295 | 3302 | 1 | ; | CD1 | - | HD23 |
| 3296 | 3299 | 1 | ; | HD11 | - | CD2 |
| 3297 | 3299 | 1 | ; | HD12 | - | CD2 |
| 3298 | 3299 | 1 | ; | HD13 | - | CD2 |
| 3303 | 3308 | 1 | ; | C | - | HA |
| 3303 | 3309 | 1 | ; | C | - | CB |
| 3303 | 3322 | 1 | ; | C | - | C |
| 3304 | 3306 | 1 | ; | O | - | H |
| 3304 | 3307 | 1 | ; | O | - | CA |
| 3305 | 3310 | 1 | ; | N | - | HB2 |
| 3305 | 3311 | 1 | ; | N | - | HB3 |
| 3305 | 3312 | 1 | ; | N | - | CG |
| 3305 | 3323 | 1 | ; | N | - | O |
| 3305 | 3324 | 1 | ; | N | - | N |
| 3306 | 3308 | 1 | ; | H | - | HA |
| 3306 | 3309 | 1 | ; | H | - | CB |
| 3306 | 3322 | 1 | ; | H | - | C |
| 3307 | 3313 | 1 | ; | CA | - | HG |
| 3307 | 3314 | 1 | ; | CA | - | CD1 |
| 3307 | 3318 | 1 | ; | CA | - | CD2 |
| 3307 | 3325 | 1 | ; | CA | - | H |
| 3307 | 3326 | 1 | ; | CA | - | CA |
| 3308 | 3310 | 1 | ; | HA | - | HB2 |
| 3308 | 3311 | 1 | ; | HA | - | HB3 |
| 3308 | 3312 | 1 | ; | HA | - | CG |

| | | | | | | |
|---|---|---|---|---|---|---|
| 3308 | 3323 | 1 | ; | HA | - | O |
| 3308 | 3324 | 1 | ; | HA | - | N |
| 3309 | 3315 | 1 | ; | CB | - | HD11 |
| 3309 | 3316 | 1 | ; | CB | - | HD12 |
| 3309 | 3317 | 1 | ; | CB | - | HD13 |
| 3309 | 3319 | 1 | ; | CB | - | HD21 |
| 3309 | 3320 | 1 | ; | CB | - | HD22 |
| 3309 | 3321 | 1 | ; | CB | - | HD23 |
| 3309 | 3323 | 1 | ; | CB | - | O |
| 3309 | 3324 | 1 | ; | CB | - | N |
| 3310 | 3313 | 1 | ; | HB2 | - | HG |
| 3310 | 3314 | 1 | ; | HB2 | - | CD1 |
| 3310 | 3318 | 1 | ; | HB2 | - | CD2 |
| 3310 | 3322 | 1 | ; | HB2 | - | C |
| 3311 | 3313 | 1 | ; | HB3 | - | HG |
| 3311 | 3314 | 1 | ; | HB3 | - | CD1 |
| 3311 | 3318 | 1 | ; | HB3 | - | CD2 |
| 3311 | 3322 | 1 | ; | HB3 | - | C |
| 3312 | 3322 | 1 | ; | CG | - | C |
| 3313 | 3315 | 1 | ; | HG | - | HD11 |
| 3313 | 3316 | 1 | ; | HG | - | HD12 |
| 3313 | 3317 | 1 | ; | HG | - | HD13 |
| 3313 | 3319 | 1 | ; | HG | - | HD21 |
| 3313 | 3320 | 1 | ; | HG | - | HD22 |
| 3313 | 3321 | 1 | ; | HG | - | HD23 |
| 3314 | 3319 | 1 | ; | CD1 | - | HD21 |
| 3314 | 3320 | 1 | ; | CD1 | - | HD22 |
| 3314 | 3321 | 1 | ; | CD1 | - | HD23 |
| 3315 | 3318 | 1 | ; | HD11 | - | CD2 |
| 3316 | 3318 | 1 | ; | HD12 | - | CD2 |
| 3317 | 3318 | 1 | ; | HD13 | - | CD2 |
| 3322 | 3327 | 1 | ; | C | - | HA |
| 3322 | 3328 | 1 | ; | C | - | CB |
| 3322 | 3333 | 1 | ; | C | - | C |
| 3323 | 3325 | 1 | ; | O | - | H |
| 3323 | 3326 | 1 | ; | O | - | CA |
| 3324 | 3329 | 1 | ; | N | - | HB2 |
| 3324 | 3330 | 1 | ; | N | - | HB3 |
| 3324 | 3331 | 1 | ; | N | - | SG |
| 3324 | 3334 | 1 | ; | N | - | O |
| 3324 | 3335 | 1 | ; | N | - | N |
| 3325 | 3327 | 1 | ; | H | - | HA |
| 3325 | 3328 | 1 | ; | H | - | CB |
| 3325 | 3333 | 1 | ; | H | - | C |
| 3326 | 3332 | 1 | ; | CA | - | HG |
| 3326 | 3336 | 1 | ; | CA | - | H |
| 3326 | 3337 | 1 | ; | CA | - | CA |
| 3327 | 3329 | 1 | ; | HA | - | HB2 |
| 3327 | 3330 | 1 | ; | HA | - | HB3 |
| 3327 | 3331 | 1 | ; | HA | - | SG |
| 3327 | 3334 | 1 | ; | HA | - | O |
| 3327 | 3335 | 1 | ; | HA | - | N |

```
3328   3334       1 ;      CB - O
3328   3335       1 ;      CB - N
3329   3332       1 ;     HB2 - HG
3329   3333       1 ;     HB2 - C
3330   3332       1 ;     HB3 - HG
3330   3333       1 ;     HB3 - C
3331   3333       1 ;      SG - C
3333   3338       1 ;       C - HA2
3333   3339       1 ;       C - HA3
3333   3340       1 ;       C - C
3334   3336       1 ;       O - H
3334   3337       1 ;       O - CA
3335   3341       1 ;       N - O
3335   3342       1 ;       N - N
3336   3338       1 ;       H - HA2
3336   3339       1 ;       H - HA3
3336   3340       1 ;       H - C
3337   3343       1 ;      CA - H
3337   3344       1 ;      CA - CA
3338   3341       1 ;     HA2 - O
3338   3342       1 ;     HA2 - N
3339   3341       1 ;     HA3 - O
3339   3342       1 ;     HA3 - N
3340   3345       1 ;       C - HA
3340   3346       1 ;       C - CB
3340   3361       1 ;       C - C
3341   3343       1 ;       O - H
3341   3344       1 ;       O - CA
3342   3347       1 ;       N - HB2
3342   3348       1 ;       N - HB3
3342   3349       1 ;       N - CG
3342   3362       1 ;       N - O
3342   3363       1 ;       N - N
3343   3345       1 ;       H - HA
3343   3346       1 ;       H - CB
3343   3361       1 ;       H - C
3344   3350       1 ;      CA - CD1
3344   3359       1 ;      CA - CD2
3344   3364       1 ;      CA - CD
3344   3373       1 ;      CA - CA
3345   3347       1 ;      HA - HB2
3345   3348       1 ;      HA - HB3
3345   3349       1 ;      HA - CG
3345   3362       1 ;      HA - O
3345   3363       1 ;      HA - N
3346   3351       1 ;      CB - HD1
3346   3352       1 ;      CB - CE1
3346   3357       1 ;      CB - CE2
3346   3360       1 ;      CB - HD2
3346   3362       1 ;      CB - O
3346   3363       1 ;      CB - N
3347   3350       1 ;     HB2 - CD1
```

```
3347   3359      1 ;    HB2 - CD2
3347   3361      1 ;    HB2 - C
3348   3350      1 ;    HB3 - CD1
3348   3359      1 ;    HB3 - CD2
3348   3361      1 ;    HB3 - C
3349   3353      1 ;     CG - HE1
3349   3354      1 ;     CG - CZ
3349   3358      1 ;     CG - HE2
3349   3361      1 ;     CG - C
3350   3355      1 ;    CD1 - OH
3350   3357      1 ;    CD1 - CE2
3350   3360      1 ;    CD1 - HD2
3351   3353      1 ;    HD1 - HE1
3351   3354      1 ;    HD1 - CZ
3351   3359      1 ;    HD1 - CD2
3352   3356      1 ;    CE1 - HH
3352   3358      1 ;    CE1 - HE2
3352   3359      1 ;    CE1 - CD2
3353   3355      1 ;    HE1 - OH
3353   3357      1 ;    HE1 - CE2
3354   3360      1 ;     CZ - HD2
3355   3358      1 ;     OH - HE2
3355   3359      1 ;     OH - CD2
3356   3357      1 ;     HH - CE2
3358   3360      1 ;    HE2 - HD2
3361   3365      1 ;      C - HD2
3361   3366      1 ;      C - HD3
3361   3367      1 ;      C - CG
3361   3370      1 ;      C - CB
3361   3374      1 ;      C - HA
3361   3375      1 ;      C - C
3362   3364      1 ;      O - CD
3362   3373      1 ;      O - CA
3363   3368      1 ;      N - HG2
3363   3369      1 ;      N - HG3
3363   3371      1 ;      N - HB2
3363   3372      1 ;      N - HB3
3363   3376      1 ;      N - O
3363   3377      1 ;      N - N
3364   3371      1 ;     CD - HB2
3364   3372      1 ;     CD - HB3
3364   3374      1 ;     CD - HA
3364   3375      1 ;     CD - C
3365   3368      1 ;    HD2 - HG2
3365   3369      1 ;    HD2 - HG3
3365   3370      1 ;    HD2 - CB
3365   3373      1 ;    HD2 - CA
3366   3368      1 ;    HD3 - HG2
3366   3369      1 ;    HD3 - HG3
3366   3370      1 ;    HD3 - CB
3366   3373      1 ;    HD3 - CA
3367   3374      1 ;     CG - HA
```

| | | | | | |
|---|---|---|---|---|---|
| 3367 | 3375 | 1 ; | CG | - | C |
| 3368 | 3371 | 1 ; | HG2 | - | HB2 |
| 3368 | 3372 | 1 ; | HG2 | - | HB3 |
| 3368 | 3373 | 1 ; | HG2 | - | CA |
| 3369 | 3371 | 1 ; | HG3 | - | HB2 |
| 3369 | 3372 | 1 ; | HG3 | - | HB3 |
| 3369 | 3373 | 1 ; | HG3 | - | CA |
| 3370 | 3376 | 1 ; | CB | - | O |
| 3370 | 3377 | 1 ; | CB | - | N |
| 3371 | 3374 | 1 ; | HB2 | - | HA |
| 3371 | 3375 | 1 ; | HB2 | - | C |
| 3372 | 3374 | 1 ; | HB3 | - | HA |
| 3372 | 3375 | 1 ; | HB3 | - | C |
| 3373 | 3378 | 1 ; | CA | - | CD |
| 3373 | 3387 | 1 ; | CA | - | CA |
| 3374 | 3376 | 1 ; | HA | - | O |
| 3374 | 3377 | 1 ; | HA | - | N |
| 3375 | 3379 | 1 ; | C | - | HD2 |
| 3375 | 3380 | 1 ; | C | - | HD3 |
| 3375 | 3381 | 1 ; | C | - | CG |
| 3375 | 3384 | 1 ; | C | - | CB |
| 3375 | 3388 | 1 ; | C | - | HA |
| 3375 | 3389 | 1 ; | C | - | C |
| 3376 | 3378 | 1 ; | O | - | CD |
| 3376 | 3387 | 1 ; | O | - | CA |
| 3377 | 3382 | 1 ; | N | - | HG2 |
| 3377 | 3383 | 1 ; | N | - | HG3 |
| 3377 | 3385 | 1 ; | N | - | HB2 |
| 3377 | 3386 | 1 ; | N | - | HB3 |
| 3377 | 3390 | 1 ; | N | - | O |
| 3377 | 3391 | 1 ; | N | - | N |
| 3378 | 3385 | 1 ; | CD | - | HB2 |
| 3378 | 3386 | 1 ; | CD | - | HB3 |
| 3378 | 3388 | 1 ; | CD | - | HA |
| 3378 | 3389 | 1 ; | CD | - | C |
| 3379 | 3382 | 1 ; | HD2 | - | HG2 |
| 3379 | 3383 | 1 ; | HD2 | - | HG3 |
| 3379 | 3384 | 1 ; | HD2 | - | CB |
| 3379 | 3387 | 1 ; | HD2 | - | CA |
| 3380 | 3382 | 1 ; | HD3 | - | HG2 |
| 3380 | 3383 | 1 ; | HD3 | - | HG3 |
| 3380 | 3384 | 1 ; | HD3 | - | CB |
| 3380 | 3387 | 1 ; | HD3 | - | CA |
| 3381 | 3388 | 1 ; | CG | - | HA |
| 3381 | 3389 | 1 ; | CG | - | C |
| 3382 | 3385 | 1 ; | HG2 | - | HB2 |
| 3382 | 3386 | 1 ; | HG2 | - | HB3 |
| 3382 | 3387 | 1 ; | HG2 | - | CA |
| 3383 | 3385 | 1 ; | HG3 | - | HB2 |
| 3383 | 3386 | 1 ; | HG3 | - | HB3 |
| 3383 | 3387 | 1 ; | HG3 | - | CA |
| 3384 | 3390 | 1 ; | CB | - | O |

| | | | | |
|---|---|---|---|---|
| 3384 | 3391 | 1 ; | CB - | N |
| 3385 | 3388 | 1 ; | HB2 - | HA |
| 3385 | 3389 | 1 ; | HB2 - | C |
| 3386 | 3388 | 1 ; | HB3 - | HA |
| 3386 | 3389 | 1 ; | HB3 - | C |
| 3387 | 3392 | 1 ; | CA - | H |
| 3387 | 3393 | 1 ; | CA - | CA |
| 3388 | 3390 | 1 ; | HA - | O |
| 3388 | 3391 | 1 ; | HA - | N |
| 3389 | 3394 | 1 ; | C - | HA |
| 3389 | 3395 | 1 ; | C - | CB |
| 3389 | 3409 | 1 ; | C - | C |
| 3390 | 3392 | 1 ; | O - | H |
| 3390 | 3393 | 1 ; | O - | CA |
| 3391 | 3396 | 1 ; | N - | HB2 |
| 3391 | 3397 | 1 ; | N - | HB3 |
| 3391 | 3398 | 1 ; | N - | CG |
| 3391 | 3410 | 1 ; | N - | O |
| 3391 | 3411 | 1 ; | N - | N |
| 3392 | 3394 | 1 ; | H - | HA |
| 3392 | 3395 | 1 ; | H - | CB |
| 3392 | 3409 | 1 ; | H - | C |
| 3393 | 3399 | 1 ; | CA - | CD1 |
| 3393 | 3407 | 1 ; | CA - | CD2 |
| 3393 | 3412 | 1 ; | CA - | H |
| 3393 | 3413 | 1 ; | CA - | CA |
| 3394 | 3396 | 1 ; | HA - | HB2 |
| 3394 | 3397 | 1 ; | HA - | HB3 |
| 3394 | 3398 | 1 ; | HA - | CG |
| 3394 | 3410 | 1 ; | HA - | O |
| 3394 | 3411 | 1 ; | HA - | N |
| 3395 | 3400 | 1 ; | CB - | HD1 |
| 3395 | 3401 | 1 ; | CB - | CE1 |
| 3395 | 3405 | 1 ; | CB - | CE2 |
| 3395 | 3408 | 1 ; | CB - | HD2 |
| 3395 | 3410 | 1 ; | CB - | O |
| 3395 | 3411 | 1 ; | CB - | N |
| 3396 | 3399 | 1 ; | HB2 - | CD1 |
| 3396 | 3407 | 1 ; | HB2 - | CD2 |
| 3396 | 3409 | 1 ; | HB2 - | C |
| 3397 | 3399 | 1 ; | HB3 - | CD1 |
| 3397 | 3407 | 1 ; | HB3 - | CD2 |
| 3397 | 3409 | 1 ; | HB3 - | C |
| 3398 | 3402 | 1 ; | CG - | HE1 |
| 3398 | 3403 | 1 ; | CG - | CZ |
| 3398 | 3406 | 1 ; | CG - | HE2 |
| 3398 | 3409 | 1 ; | CG - | C |
| 3399 | 3404 | 1 ; | CD1 - | HZ |
| 3399 | 3405 | 1 ; | CD1 - | CE2 |
| 3399 | 3408 | 1 ; | CD1 - | HD2 |
| 3400 | 3402 | 1 ; | HD1 - | HE1 |
| 3400 | 3403 | 1 ; | HD1 - | CZ |

| | | | | |
|---|---|---|---|---|
| 3400 | 3407 | 1 | ; | HD1 - CD2 |
| 3401 | 3406 | 1 | ; | CE1 - HE2 |
| 3401 | 3407 | 1 | ; | CE1 - CD2 |
| 3402 | 3404 | 1 | ; | HE1 - HZ |
| 3402 | 3405 | 1 | ; | HE1 - CE2 |
| 3403 | 3408 | 1 | ; | CZ - HD2 |
| 3404 | 3406 | 1 | ; | HZ - HE2 |
| 3404 | 3407 | 1 | ; | HZ - CD2 |
| 3406 | 3408 | 1 | ; | HE2 - HD2 |
| 3409 | 3414 | 1 | ; | C - HA |
| 3409 | 3415 | 1 | ; | C - CB |
| 3409 | 3430 | 1 | ; | C - C |
| 3410 | 3412 | 1 | ; | O - H |
| 3410 | 3413 | 1 | ; | O - CA |
| 3411 | 3416 | 1 | ; | N - HB2 |
| 3411 | 3417 | 1 | ; | N - HB3 |
| 3411 | 3418 | 1 | ; | N - CG |
| 3411 | 3431 | 1 | ; | N - O |
| 3411 | 3432 | 1 | ; | N - N |
| 3412 | 3414 | 1 | ; | H - HA |
| 3412 | 3415 | 1 | ; | H - CB |
| 3412 | 3430 | 1 | ; | H - C |
| 3413 | 3419 | 1 | ; | CA - CD1 |
| 3413 | 3428 | 1 | ; | CA - CD2 |
| 3413 | 3433 | 1 | ; | CA - H |
| 3413 | 3434 | 1 | ; | CA - CA |
| 3414 | 3416 | 1 | ; | HA - HB2 |
| 3414 | 3417 | 1 | ; | HA - HB3 |
| 3414 | 3418 | 1 | ; | HA - CG |
| 3414 | 3431 | 1 | ; | HA - O |
| 3414 | 3432 | 1 | ; | HA - N |
| 3415 | 3420 | 1 | ; | CB - HD1 |
| 3415 | 3421 | 1 | ; | CB - CE1 |
| 3415 | 3426 | 1 | ; | CB - CE2 |
| 3415 | 3429 | 1 | ; | CB - HD2 |
| 3415 | 3431 | 1 | ; | CB - O |
| 3415 | 3432 | 1 | ; | CB - N |
| 3416 | 3419 | 1 | ; | HB2 - CD1 |
| 3416 | 3428 | 1 | ; | HB2 - CD2 |
| 3416 | 3430 | 1 | ; | HB2 - C |
| 3417 | 3419 | 1 | ; | HB3 - CD1 |
| 3417 | 3428 | 1 | ; | HB3 - CD2 |
| 3417 | 3430 | 1 | ; | HB3 - C |
| 3418 | 3422 | 1 | ; | CG - HE1 |
| 3418 | 3423 | 1 | ; | CG - CZ |
| 3418 | 3427 | 1 | ; | CG - HE2 |
| 3418 | 3430 | 1 | ; | CG - C |
| 3419 | 3424 | 1 | ; | CD1 - OH |
| 3419 | 3426 | 1 | ; | CD1 - CE2 |
| 3419 | 3429 | 1 | ; | CD1 - HD2 |
| 3420 | 3422 | 1 | ; | HD1 - HE1 |
| 3420 | 3423 | 1 | ; | HD1 - CZ |

```
3420   3428       1 ;     HD1 - CD2
3421   3425       1 ;     CE1 - HH
3421   3427       1 ;     CE1 - HE2
3421   3428       1 ;     CE1 - CD2
3422   3424       1 ;     HE1 - OH
3422   3426       1 ;     HE1 - CE2
3423   3429       1 ;      CZ - HD2
3424   3427       1 ;      OH - HE2
3424   3428       1 ;      OH - CD2
3425   3426       1 ;      HH - CE2
3427   3429       1 ;     HE2 - HD2
3430   3435       1 ;       C - HA2
3430   3436       1 ;       C - HA3
3430   3437       1 ;       C - C
3431   3433       1 ;       O - H
3431   3434       1 ;       O - CA
3432   3438       1 ;       N - O
3432   3439       1 ;       N - N
3433   3435       1 ;       H - HA2
3433   3436       1 ;       H - HA3
3433   3437       1 ;       H - C
3434   3440       1 ;      CA - H
3434   3441       1 ;      CA - CA
3435   3438       1 ;     HA2 - O
3435   3439       1 ;     HA2 - N
3436   3438       1 ;     HA3 - O
3436   3439       1 ;     HA3 - N
3437   3442       1 ;       C - HA
3437   3443       1 ;       C - CB
3437   3449       1 ;       C - C
3438   3440       1 ;       O - H
3438   3441       1 ;       O - CA
3439   3444       1 ;       N - HB2
3439   3445       1 ;       N - HB3
3439   3446       1 ;       N - CG
3439   3450       1 ;       N - O
3439   3451       1 ;       N - N
3440   3442       1 ;       H - HA
3440   3443       1 ;       H - CB
3440   3449       1 ;       H - C
3441   3447       1 ;      CA - OD1
3441   3448       1 ;      CA - OD2
3441   3452       1 ;      CA - H
3441   3453       1 ;      CA - CA
3442   3444       1 ;      HA - HB2
3442   3445       1 ;      HA - HB3
3442   3446       1 ;      HA - CG
3442   3450       1 ;      HA - O
3442   3451       1 ;      HA - N
3443   3450       1 ;      CB - O
3443   3451       1 ;      CB - N
3444   3447       1 ;     HB2 - OD1
```

| | | | | | |
|---|---|---|---|---|---|
| 3444 | 3448 | 1 ; | HB2 | - | OD2 |
| 3444 | 3449 | 1 ; | HB2 | - | C |
| 3445 | 3447 | 1 ; | HB3 | - | OD1 |
| 3445 | 3448 | 1 ; | HB3 | - | OD2 |
| 3445 | 3449 | 1 ; | HB3 | - | C |
| 3446 | 3449 | 1 ; | CG | - | C |
| 3449 | 3454 | 1 ; | C | - | HA |
| 3449 | 3455 | 1 ; | C | - | CB |
| 3449 | 3463 | 1 ; | C | - | C |
| 3450 | 3452 | 1 ; | O | - | H |
| 3450 | 3453 | 1 ; | O | - | CA |
| 3451 | 3456 | 1 ; | N | - | HB |
| 3451 | 3457 | 1 ; | N | - | CG2 |
| 3451 | 3461 | 1 ; | N | - | OG1 |
| 3451 | 3464 | 1 ; | N | - | O |
| 3451 | 3465 | 1 ; | N | - | N |
| 3452 | 3454 | 1 ; | H | - | HA |
| 3452 | 3455 | 1 ; | H | - | CB |
| 3452 | 3463 | 1 ; | H | - | C |
| 3453 | 3458 | 1 ; | CA | - | HG21 |
| 3453 | 3459 | 1 ; | CA | - | HG22 |
| 3453 | 3460 | 1 ; | CA | - | HG23 |
| 3453 | 3462 | 1 ; | CA | - | HG1 |
| 3453 | 3466 | 1 ; | CA | - | H |
| 3453 | 3467 | 1 ; | CA | - | CA |
| 3454 | 3456 | 1 ; | HA | - | HB |
| 3454 | 3457 | 1 ; | HA | - | CG2 |
| 3454 | 3461 | 1 ; | HA | - | OG1 |
| 3454 | 3464 | 1 ; | HA | - | O |
| 3454 | 3465 | 1 ; | HA | - | N |
| 3455 | 3464 | 1 ; | CB | - | O |
| 3455 | 3465 | 1 ; | CB | - | N |
| 3456 | 3458 | 1 ; | HB | - | HG21 |
| 3456 | 3459 | 1 ; | HB | - | HG22 |
| 3456 | 3460 | 1 ; | HB | - | HG23 |
| 3456 | 3462 | 1 ; | HB | - | HG1 |
| 3456 | 3463 | 1 ; | HB | - | C |
| 3457 | 3462 | 1 ; | CG2 | - | HG1 |
| 3457 | 3463 | 1 ; | CG2 | - | C |
| 3458 | 3461 | 1 ; | HG21 | - | OG1 |
| 3459 | 3461 | 1 ; | HG22 | - | OG1 |
| 3460 | 3461 | 1 ; | HG23 | - | OG1 |
| 3461 | 3463 | 1 ; | OG1 | - | C |
| 3463 | 3468 | 1 ; | C | - | HA |
| 3463 | 3469 | 1 ; | C | - | CB |
| 3463 | 3475 | 1 ; | C | - | C |
| 3464 | 3466 | 1 ; | O | - | H |
| 3464 | 3467 | 1 ; | O | - | CA |
| 3465 | 3470 | 1 ; | N | - | HB2 |
| 3465 | 3471 | 1 ; | N | - | HB3 |
| 3465 | 3472 | 1 ; | N | - | CG |
| 3465 | 3476 | 1 ; | N | - | O |

| | | | |
|---|---|---|---|
| 3465 | 3477 | 1 ; | N - N |
| 3466 | 3468 | 1 ; | H - HA |
| 3466 | 3469 | 1 ; | H - CB |
| 3466 | 3475 | 1 ; | H - C |
| 3467 | 3473 | 1 ; | CA - OD1 |
| 3467 | 3474 | 1 ; | CA - OD2 |
| 3467 | 3478 | 1 ; | CA - H |
| 3467 | 3479 | 1 ; | CA - CA |
| 3468 | 3470 | 1 ; | HA - HB2 |
| 3468 | 3471 | 1 ; | HA - HB3 |
| 3468 | 3472 | 1 ; | HA - CG |
| 3468 | 3476 | 1 ; | HA - O |
| 3468 | 3477 | 1 ; | HA - N |
| 3469 | 3476 | 1 ; | CB - O |
| 3469 | 3477 | 1 ; | CB - N |
| 3470 | 3473 | 1 ; | HB2 - OD1 |
| 3470 | 3474 | 1 ; | HB2 - OD2 |
| 3470 | 3475 | 1 ; | HB2 - C |
| 3471 | 3473 | 1 ; | HB3 - OD1 |
| 3471 | 3474 | 1 ; | HB3 - OD2 |
| 3471 | 3475 | 1 ; | HB3 - C |
| 3472 | 3475 | 1 ; | CG - C |
| 3475 | 3480 | 1 ; | C - HA |
| 3475 | 3481 | 1 ; | C - CB |
| 3475 | 3489 | 1 ; | C - C |
| 3476 | 3478 | 1 ; | O - H |
| 3476 | 3479 | 1 ; | O - CA |
| 3477 | 3482 | 1 ; | N - HB2 |
| 3477 | 3483 | 1 ; | N - HB3 |
| 3477 | 3484 | 1 ; | N - CG |
| 3477 | 3490 | 1 ; | N - O |
| 3477 | 3491 | 1 ; | N - N |
| 3478 | 3480 | 1 ; | H - HA |
| 3478 | 3481 | 1 ; | H - CB |
| 3478 | 3489 | 1 ; | H - C |
| 3479 | 3485 | 1 ; | CA - OD1 |
| 3479 | 3486 | 1 ; | CA - ND2 |
| 3479 | 3492 | 1 ; | CA - H |
| 3479 | 3493 | 1 ; | CA - CA |
| 3480 | 3482 | 1 ; | HA - HB2 |
| 3480 | 3483 | 1 ; | HA - HB3 |
| 3480 | 3484 | 1 ; | HA - CG |
| 3480 | 3490 | 1 ; | HA - O |
| 3480 | 3491 | 1 ; | HA - N |
| 3481 | 3487 | 1 ; | CB - HD21 |
| 3481 | 3488 | 1 ; | CB - HD22 |
| 3481 | 3490 | 1 ; | CB - O |
| 3481 | 3491 | 1 ; | CB - N |
| 3482 | 3485 | 1 ; | HB2 - OD1 |
| 3482 | 3486 | 1 ; | HB2 - ND2 |
| 3482 | 3489 | 1 ; | HB2 - C |
| 3483 | 3485 | 1 ; | HB3 - OD1 |

```
3483   3486       1 ;    HB3 - ND2
3483   3489       1 ;    HB3 - C
3484   3489       1 ;     CG - C
3485   3487       1 ;    OD1 - HD21
3485   3488       1 ;    OD1 - HD22
3489   3494       1 ;      C - HA
3489   3495       1 ;      C - CB
3489   3504       1 ;      C - C
3490   3492       1 ;      O - H
3490   3493       1 ;      O - CA
3491   3496       1 ;      N - HB2
3491   3497       1 ;      N - HB3
3491   3498       1 ;      N - CG
3491   3505       1 ;      N - O
3491   3506       1 ;      N - N
3492   3494       1 ;      H - HA
3492   3495       1 ;      H - CB
3492   3504       1 ;      H - C
3493   3499       1 ;     CA - HG2
3493   3500       1 ;     CA - HG3
3493   3501       1 ;     CA - CD
3493   3507       1 ;     CA - H
3493   3508       1 ;     CA - CA
3494   3496       1 ;     HA - HB2
3494   3497       1 ;     HA - HB3
3494   3498       1 ;     HA - CG
3494   3505       1 ;     HA - O
3494   3506       1 ;     HA - N
3495   3502       1 ;     CB - OE1
3495   3503       1 ;     CB - OE2
3495   3505       1 ;     CB - O
3495   3506       1 ;     CB - N
3496   3499       1 ;    HB2 - HG2
3496   3500       1 ;    HB2 - HG3
3496   3501       1 ;    HB2 - CD
3496   3504       1 ;    HB2 - C
3497   3499       1 ;    HB3 - HG2
3497   3500       1 ;    HB3 - HG3
3497   3501       1 ;    HB3 - CD
3497   3504       1 ;    HB3 - C
3498   3504       1 ;     CG - C
3499   3502       1 ;    HG2 - OE1
3499   3503       1 ;    HG2 - OE2
3500   3502       1 ;    HG3 - OE1
3500   3503       1 ;    HG3 - OE2
3504   3509       1 ;      C - HA
3504   3510       1 ;      C - CB
3504   3520       1 ;      C - C
3505   3507       1 ;      O - H
3505   3508       1 ;      O - CA
3506   3511       1 ;      N - HB
3506   3512       1 ;      N - CG1
```

| | | | | |
|---|---|---|---|---|
| 3506 | 3516 | 1 | ; | N - CG2 |
| 3506 | 3521 | 1 | ; | N - O |
| 3506 | 3522 | 1 | ; | N - N |
| 3507 | 3509 | 1 | ; | H - HA |
| 3507 | 3510 | 1 | ; | H - CB |
| 3507 | 3520 | 1 | ; | H - C |
| 3508 | 3513 | 1 | ; | CA - HG11 |
| 3508 | 3514 | 1 | ; | CA - HG12 |
| 3508 | 3515 | 1 | ; | CA - HG13 |
| 3508 | 3517 | 1 | ; | CA - HG21 |
| 3508 | 3518 | 1 | ; | CA - HG22 |
| 3508 | 3519 | 1 | ; | CA - HG23 |
| 3508 | 3523 | 1 | ; | CA - H |
| 3508 | 3524 | 1 | ; | CA - CA |
| 3509 | 3511 | 1 | ; | HA - HB |
| 3509 | 3512 | 1 | ; | HA - CG1 |
| 3509 | 3516 | 1 | ; | HA - CG2 |
| 3509 | 3521 | 1 | ; | HA - O |
| 3509 | 3522 | 1 | ; | HA - N |
| 3510 | 3521 | 1 | ; | CB - O |
| 3510 | 3522 | 1 | ; | CB - N |
| 3511 | 3513 | 1 | ; | HB - HG11 |
| 3511 | 3514 | 1 | ; | HB - HG12 |
| 3511 | 3515 | 1 | ; | HB - HG13 |
| 3511 | 3517 | 1 | ; | HB - HG21 |
| 3511 | 3518 | 1 | ; | HB - HG22 |
| 3511 | 3519 | 1 | ; | HB - HG23 |
| 3511 | 3520 | 1 | ; | HB - C |
| 3512 | 3517 | 1 | ; | CG1 - HG21 |
| 3512 | 3518 | 1 | ; | CG1 - HG22 |
| 3512 | 3519 | 1 | ; | CG1 - HG23 |
| 3512 | 3520 | 1 | ; | CG1 - C |
| 3513 | 3516 | 1 | ; | HG11 - CG2 |
| 3514 | 3516 | 1 | ; | HG12 - CG2 |
| 3515 | 3516 | 1 | ; | HG13 - CG2 |
| 3516 | 3520 | 1 | ; | CG2 - C |
| 3520 | 3525 | 1 | ; | C - HA |
| 3520 | 3526 | 1 | ; | C - CB |
| 3520 | 3539 | 1 | ; | C - C |
| 3521 | 3523 | 1 | ; | O - H |
| 3521 | 3524 | 1 | ; | O - CA |
| 3522 | 3527 | 1 | ; | N - HB2 |
| 3522 | 3528 | 1 | ; | N - HB3 |
| 3522 | 3529 | 1 | ; | N - CG |
| 3522 | 3540 | 1 | ; | N - O |
| 3522 | 3541 | 1 | ; | N - N |
| 3523 | 3525 | 1 | ; | H - HA |
| 3523 | 3526 | 1 | ; | H - CB |
| 3523 | 3539 | 1 | ; | H - C |
| 3524 | 3530 | 1 | ; | CA - HG |
| 3524 | 3531 | 1 | ; | CA - CD1 |
| 3524 | 3535 | 1 | ; | CA - CD2 |

```
3524   3542       1 ;      CA - H
3524   3543       1 ;      CA - CA
3525   3527       1 ;      HA - HB2
3525   3528       1 ;      HA - HB3
3525   3529       1 ;      HA - CG
3525   3540       1 ;      HA - O
3525   3541       1 ;      HA - N
3526   3532       1 ;      CB - HD11
3526   3533       1 ;      CB - HD12
3526   3534       1 ;      CB - HD13
3526   3536       1 ;      CB - HD21
3526   3537       1 ;      CB - HD22
3526   3538       1 ;      CB - HD23
3526   3540       1 ;      CB - O
3526   3541       1 ;      CB - N
3527   3530       1 ;     HB2 - HG
3527   3531       1 ;     HB2 - CD1
3527   3535       1 ;     HB2 - CD2
3527   3539       1 ;     HB2 - C
3528   3530       1 ;     HB3 - HG
3528   3531       1 ;     HB3 - CD1
3528   3535       1 ;     HB3 - CD2
3528   3539       1 ;     HB3 - C
3529   3539       1 ;      CG - C
3530   3532       1 ;      HG - HD11
3530   3533       1 ;      HG - HD12
3530   3534       1 ;      HG - HD13
3530   3536       1 ;      HG - HD21
3530   3537       1 ;      HG - HD22
3530   3538       1 ;      HG - HD23
3531   3536       1 ;     CD1 - HD21
3531   3537       1 ;     CD1 - HD22
3531   3538       1 ;     CD1 - HD23
3532   3535       1 ;    HD11 - CD2
3533   3535       1 ;    HD12 - CD2
3534   3535       1 ;    HD13 - CD2
3539   3544       1 ;       C - HA
3539   3545       1 ;       C - CB
3539   3561       1 ;       C - C
3540   3542       1 ;       O - H
3540   3543       1 ;       O - CA
3541   3546       1 ;       N - HB2
3541   3547       1 ;       N - HB3
3541   3548       1 ;       N - CG
3541   3562       1 ;       N - O
3541   3563       1 ;       N - N
3542   3544       1 ;       H - HA
3542   3545       1 ;       H - CB
3542   3561       1 ;       H - C
3543   3549       1 ;      CA - HG2
3543   3550       1 ;      CA - HG3
3543   3551       1 ;      CA - CD
```

| | | | |
|---|---|---|---|
| 3543 | 3564 | 1 ; | CA - H |
| 3543 | 3565 | 1 ; | CA - CA |
| 3544 | 3546 | 1 ; | HA - HB2 |
| 3544 | 3547 | 1 ; | HA - HB3 |
| 3544 | 3548 | 1 ; | HA - CG |
| 3544 | 3562 | 1 ; | HA - O |
| 3544 | 3563 | 1 ; | HA - N |
| 3545 | 3552 | 1 ; | CB - HD2 |
| 3545 | 3553 | 1 ; | CB - HD3 |
| 3545 | 3554 | 1 ; | CB - CE |
| 3545 | 3562 | 1 ; | CB - O |
| 3545 | 3563 | 1 ; | CB - N |
| 3546 | 3549 | 1 ; | HB2 - HG2 |
| 3546 | 3550 | 1 ; | HB2 - HG3 |
| 3546 | 3551 | 1 ; | HB2 - CD |
| 3546 | 3561 | 1 ; | HB2 - C |
| 3547 | 3549 | 1 ; | HB3 - HG2 |
| 3547 | 3550 | 1 ; | HB3 - HG3 |
| 3547 | 3551 | 1 ; | HB3 - CD |
| 3547 | 3561 | 1 ; | HB3 - C |
| 3548 | 3555 | 1 ; | CG - HE2 |
| 3548 | 3556 | 1 ; | CG - HE3 |
| 3548 | 3557 | 1 ; | CG - NZ |
| 3548 | 3561 | 1 ; | CG - C |
| 3549 | 3552 | 1 ; | HG2 - HD2 |
| 3549 | 3553 | 1 ; | HG2 - HD3 |
| 3549 | 3554 | 1 ; | HG2 - CE |
| 3550 | 3552 | 1 ; | HG3 - HD2 |
| 3550 | 3553 | 1 ; | HG3 - HD3 |
| 3550 | 3554 | 1 ; | HG3 - CE |
| 3551 | 3558 | 1 ; | CD - HZ1 |
| 3551 | 3559 | 1 ; | CD - HZ2 |
| 3551 | 3560 | 1 ; | CD - HZ3 |
| 3552 | 3555 | 1 ; | HD2 - HE2 |
| 3552 | 3556 | 1 ; | HD2 - HE3 |
| 3552 | 3557 | 1 ; | HD2 - NZ |
| 3553 | 3555 | 1 ; | HD3 - HE2 |
| 3553 | 3556 | 1 ; | HD3 - HE3 |
| 3553 | 3557 | 1 ; | HD3 - NZ |
| 3555 | 3558 | 1 ; | HE2 - HZ1 |
| 3555 | 3559 | 1 ; | HE2 - HZ2 |
| 3555 | 3560 | 1 ; | HE2 - HZ3 |
| 3556 | 3558 | 1 ; | HE3 - HZ1 |
| 3556 | 3559 | 1 ; | HE3 - HZ2 |
| 3556 | 3560 | 1 ; | HE3 - HZ3 |
| 3561 | 3566 | 1 ; | C - HA |
| 3561 | 3567 | 1 ; | C - CB |
| 3561 | 3583 | 1 ; | C - C |
| 3562 | 3564 | 1 ; | O - H |
| 3562 | 3565 | 1 ; | O - CA |
| 3563 | 3568 | 1 ; | N - HB2 |
| 3563 | 3569 | 1 ; | N - HB3 |

```
3563   3570       1 ;       N - CG
3563   3584       1 ;       N - O
3563   3585       1 ;       N - N
3564   3566       1 ;       H - HA
3564   3567       1 ;       H - CB
3564   3583       1 ;       H - C
3565   3571       1 ;      CA - HG2
3565   3572       1 ;      CA - HG3
3565   3573       1 ;      CA - CD
3565   3586       1 ;      CA - H
3565   3587       1 ;      CA - CA
3566   3568       1 ;      HA - HB2
3566   3569       1 ;      HA - HB3
3566   3570       1 ;      HA - CG
3566   3584       1 ;      HA - O
3566   3585       1 ;      HA - N
3567   3574       1 ;      CB - HD2
3567   3575       1 ;      CB - HD3
3567   3576       1 ;      CB - CE
3567   3584       1 ;      CB - O
3567   3585       1 ;      CB - N
3568   3571       1 ;     HB2 - HG2
3568   3572       1 ;     HB2 - HG3
3568   3573       1 ;     HB2 - CD
3568   3583       1 ;     HB2 - C
3569   3571       1 ;     HB3 - HG2
3569   3572       1 ;     HB3 - HG3
3569   3573       1 ;     HB3 - CD
3569   3583       1 ;     HB3 - C
3570   3577       1 ;      CG - HE2
3570   3578       1 ;      CG - HE3
3570   3579       1 ;      CG - NZ
3570   3583       1 ;      CG - C
3571   3574       1 ;     HG2 - HD2
3571   3575       1 ;     HG2 - HD3
3571   3576       1 ;     HG2 - CE
3572   3574       1 ;     HG3 - HD2
3572   3575       1 ;     HG3 - HD3
3572   3576       1 ;     HG3 - CE
3573   3580       1 ;      CD - HZ1
3573   3581       1 ;      CD - HZ2
3573   3582       1 ;      CD - HZ3
3574   3577       1 ;     HD2 - HE2
3574   3578       1 ;     HD2 - HE3
3574   3579       1 ;     HD2 - NZ
3575   3577       1 ;     HD3 - HE2
3575   3578       1 ;     HD3 - HE3
3575   3579       1 ;     HD3 - NZ
3577   3580       1 ;     HE2 - HZ1
3577   3581       1 ;     HE2 - HZ2
3577   3582       1 ;     HE2 - HZ3
3578   3580       1 ;     HE3 - HZ1
```

```
3578   3581       1 ;     HE3 - HZ2
3578   3582       1 ;     HE3 - HZ3
3583   3588       1 ;       C - HA
3583   3589       1 ;       C - CB
3583   3599       1 ;       C - C
3584   3586       1 ;       O - H
3584   3587       1 ;       O - CA
3585   3590       1 ;       N - HB
3585   3591       1 ;       N - CG1
3585   3595       1 ;       N - CG2
3585   3600       1 ;       N - O
3585   3601       1 ;       N - N
3586   3588       1 ;       H - HA
3586   3589       1 ;       H - CB
3586   3599       1 ;       H - C
3587   3592       1 ;      CA - HG11
3587   3593       1 ;      CA - HG12
3587   3594       1 ;      CA - HG13
3587   3596       1 ;      CA - HG21
3587   3597       1 ;      CA - HG22
3587   3598       1 ;      CA - HG23
3587   3602       1 ;      CA - H
3587   3603       1 ;      CA - CA
3588   3590       1 ;      HA - HB
3588   3591       1 ;      HA - CG1
3588   3595       1 ;      HA - CG2
3588   3600       1 ;      HA - O
3588   3601       1 ;      HA - N
3589   3600       1 ;      CB - O
3589   3601       1 ;      CB - N
3590   3592       1 ;      HB - HG11
3590   3593       1 ;      HB - HG12
3590   3594       1 ;      HB - HG13
3590   3596       1 ;      HB - HG21
3590   3597       1 ;      HB - HG22
3590   3598       1 ;      HB - HG23
3590   3599       1 ;      HB - C
3591   3596       1 ;     CG1 - HG21
3591   3597       1 ;     CG1 - HG22
3591   3598       1 ;     CG1 - HG23
3591   3599       1 ;     CG1 - C
3592   3595       1 ;    HG11 - CG2
3593   3595       1 ;    HG12 - CG2
3594   3595       1 ;    HG13 - CG2
3595   3599       1 ;     CG2 - C
3599   3604       1 ;       C - HA
3599   3605       1 ;       C - CB
3599   3621       1 ;       C - C
3600   3602       1 ;       O - H
3600   3603       1 ;       O - CA
3601   3606       1 ;       N - HB2
3601   3607       1 ;       N - HB3
```

| | | | |
|---|---|---|---|
| 3601 | 3608 | 1 ; | N - CG |
| 3601 | 3622 | 1 ; | N - O |
| 3601 | 3623 | 1 ; | N - N |
| 3602 | 3604 | 1 ; | H - HA |
| 3602 | 3605 | 1 ; | H - CB |
| 3602 | 3621 | 1 ; | H - C |
| 3603 | 3609 | 1 ; | CA - HG2 |
| 3603 | 3610 | 1 ; | CA - HG3 |
| 3603 | 3611 | 1 ; | CA - CD |
| 3603 | 3624 | 1 ; | CA - H |
| 3603 | 3625 | 1 ; | CA - CA |
| 3604 | 3606 | 1 ; | HA - HB2 |
| 3604 | 3607 | 1 ; | HA - HB3 |
| 3604 | 3608 | 1 ; | HA - CG |
| 3604 | 3622 | 1 ; | HA - O |
| 3604 | 3623 | 1 ; | HA - N |
| 3605 | 3612 | 1 ; | CB - HD2 |
| 3605 | 3613 | 1 ; | CB - HD3 |
| 3605 | 3614 | 1 ; | CB - CE |
| 3605 | 3622 | 1 ; | CB - O |
| 3605 | 3623 | 1 ; | CB - N |
| 3606 | 3609 | 1 ; | HB2 - HG2 |
| 3606 | 3610 | 1 ; | HB2 - HG3 |
| 3606 | 3611 | 1 ; | HB2 - CD |
| 3606 | 3621 | 1 ; | HB2 - C |
| 3607 | 3609 | 1 ; | HB3 - HG2 |
| 3607 | 3610 | 1 ; | HB3 - HG3 |
| 3607 | 3611 | 1 ; | HB3 - CD |
| 3607 | 3621 | 1 ; | HB3 - C |
| 3608 | 3615 | 1 ; | CG - HE2 |
| 3608 | 3616 | 1 ; | CG - HE3 |
| 3608 | 3617 | 1 ; | CG - NZ |
| 3608 | 3621 | 1 ; | CG - C |
| 3609 | 3612 | 1 ; | HG2 - HD2 |
| 3609 | 3613 | 1 ; | HG2 - HD3 |
| 3609 | 3614 | 1 ; | HG2 - CE |
| 3610 | 3612 | 1 ; | HG3 - HD2 |
| 3610 | 3613 | 1 ; | HG3 - HD3 |
| 3610 | 3614 | 1 ; | HG3 - CE |
| 3611 | 3618 | 1 ; | CD - HZ1 |
| 3611 | 3619 | 1 ; | CD - HZ2 |
| 3611 | 3620 | 1 ; | CD - HZ3 |
| 3612 | 3615 | 1 ; | HD2 - HE2 |
| 3612 | 3616 | 1 ; | HD2 - HE3 |
| 3612 | 3617 | 1 ; | HD2 - NZ |
| 3613 | 3615 | 1 ; | HD3 - HE2 |
| 3613 | 3616 | 1 ; | HD3 - HE3 |
| 3613 | 3617 | 1 ; | HD3 - NZ |
| 3615 | 3618 | 1 ; | HE2 - HZ1 |
| 3615 | 3619 | 1 ; | HE2 - HZ2 |
| 3615 | 3620 | 1 ; | HE2 - HZ3 |
| 3616 | 3618 | 1 ; | HE3 - HZ1 |

```
3616   3619       1 ;    HE3 - HZ2
3616   3620       1 ;    HE3 - HZ3
3621   3626       1 ;      C - HA
3621   3627       1 ;      C - CB
3621   3643       1 ;      C - C
3622   3624       1 ;      O - H
3622   3625       1 ;      O - CA
3623   3628       1 ;      N - HB2
3623   3629       1 ;      N - HB3
3623   3630       1 ;      N - CG
3623   3644       1 ;      N - O
3623   3645       1 ;      N - N
3624   3626       1 ;      H - HA
3624   3627       1 ;      H - CB
3624   3643       1 ;      H - C
3625   3631       1 ;     CA - HG2
3625   3632       1 ;     CA - HG3
3625   3633       1 ;     CA - CD
3625   3646       1 ;     CA - H
3625   3647       1 ;     CA - CA
3626   3628       1 ;     HA - HB2
3626   3629       1 ;     HA - HB3
3626   3630       1 ;     HA - CG
3626   3644       1 ;     HA - O
3626   3645       1 ;     HA - N
3627   3634       1 ;     CB - HD2
3627   3635       1 ;     CB - HD3
3627   3636       1 ;     CB - CE
3627   3644       1 ;     CB - O
3627   3645       1 ;     CB - N
3628   3631       1 ;    HB2 - HG2
3628   3632       1 ;    HB2 - HG3
3628   3633       1 ;    HB2 - CD
3628   3643       1 ;    HB2 - C
3629   3631       1 ;    HB3 - HG2
3629   3632       1 ;    HB3 - HG3
3629   3633       1 ;    HB3 - CD
3629   3643       1 ;    HB3 - C
3630   3637       1 ;     CG - HE2
3630   3638       1 ;     CG - HE3
3630   3639       1 ;     CG - NZ
3630   3643       1 ;     CG - C
3631   3634       1 ;    HG2 - HD2
3631   3635       1 ;    HG2 - HD3
3631   3636       1 ;    HG2 - CE
3632   3634       1 ;    HG3 - HD2
3632   3635       1 ;    HG3 - HD3
3632   3636       1 ;    HG3 - CE
3633   3640       1 ;     CD - HZ1
3633   3641       1 ;     CD - HZ2
3633   3642       1 ;     CD - HZ3
3634   3637       1 ;    HD2 - HE2
```

```
3634   3638       1 ;    HD2 - HE3
3634   3639       1 ;    HD2 - NZ
3635   3637       1 ;    HD3 - HE2
3635   3638       1 ;    HD3 - HE3
3635   3639       1 ;    HD3 - NZ
3637   3640       1 ;    HE2 - HZ1
3637   3641       1 ;    HE2 - HZ2
3637   3642       1 ;    HE2 - HZ3
3638   3640       1 ;    HE3 - HZ1
3638   3641       1 ;    HE3 - HZ2
3638   3642       1 ;    HE3 - HZ3
3643   3648       1 ;      C - HA2
3643   3649       1 ;      C - HA3
3643   3650       1 ;      C - C
3644   3646       1 ;      O - H
3644   3647       1 ;      O - CA
3645   3651       1 ;      N - O
3645   3652       1 ;      N - N
3646   3648       1 ;      H - HA2
3646   3649       1 ;      H - HA3
3646   3650       1 ;      H - C
3647   3653       1 ;     CA - H
3647   3654       1 ;     CA - CA
3648   3651       1 ;    HA2 - O
3648   3652       1 ;    HA2 - N
3649   3651       1 ;    HA3 - O
3649   3652       1 ;    HA3 - N
3650   3655       1 ;      C - HA
3650   3656       1 ;      C - CB
3650   3665       1 ;      C - C
3651   3653       1 ;      O - H
3651   3654       1 ;      O - CA
3652   3657       1 ;      N - HB2
3652   3658       1 ;      N - HB3
3652   3659       1 ;      N - CG
3652   3666       1 ;      N - O
3652   3667       1 ;      N - N
3653   3655       1 ;      H - HA
3653   3656       1 ;      H - CB
3653   3665       1 ;      H - C
3654   3660       1 ;     CA - HG2
3654   3661       1 ;     CA - HG3
3654   3662       1 ;     CA - CD
3654   3668       1 ;     CA - H
3654   3669       1 ;     CA - CA
3655   3657       1 ;     HA - HB2
3655   3658       1 ;     HA - HB3
3655   3659       1 ;     HA - CG
3655   3666       1 ;     HA - O
3655   3667       1 ;     HA - N
3656   3663       1 ;     CB - OE1
3656   3664       1 ;     CB - OE2
```

| | | | | | | |
|---|---|---|---|---|---|---|
| 3656 | 3666 | 1 | ; | CB | - | O |
| 3656 | 3667 | 1 | ; | CB | - | N |
| 3657 | 3660 | 1 | ; | HB2 | - | HG2 |
| 3657 | 3661 | 1 | ; | HB2 | - | HG3 |
| 3657 | 3662 | 1 | ; | HB2 | - | CD |
| 3657 | 3665 | 1 | ; | HB2 | - | C |
| 3658 | 3660 | 1 | ; | HB3 | - | HG2 |
| 3658 | 3661 | 1 | ; | HB3 | - | HG3 |
| 3658 | 3662 | 1 | ; | HB3 | - | CD |
| 3658 | 3665 | 1 | ; | HB3 | - | C |
| 3659 | 3665 | 1 | ; | CG | - | C |
| 3660 | 3663 | 1 | ; | HG2 | - | OE1 |
| 3660 | 3664 | 1 | ; | HG2 | - | OE2 |
| 3661 | 3663 | 1 | ; | HG3 | - | OE1 |
| 3661 | 3664 | 1 | ; | HG3 | - | OE2 |
| 3665 | 3670 | 1 | ; | C | - | HA |
| 3665 | 3671 | 1 | ; | C | - | CB |
| 3665 | 3685 | 1 | ; | C | - | C |
| 3666 | 3668 | 1 | ; | O | - | H |
| 3666 | 3669 | 1 | ; | O | - | CA |
| 3667 | 3672 | 1 | ; | N | - | HB2 |
| 3667 | 3673 | 1 | ; | N | - | HB3 |
| 3667 | 3674 | 1 | ; | N | - | CG |
| 3667 | 3686 | 1 | ; | N | - | O |
| 3667 | 3687 | 1 | ; | N | - | N |
| 3668 | 3670 | 1 | ; | H | - | HA |
| 3668 | 3671 | 1 | ; | H | - | CB |
| 3668 | 3685 | 1 | ; | H | - | C |
| 3669 | 3675 | 1 | ; | CA | - | CD1 |
| 3669 | 3683 | 1 | ; | CA | - | CD2 |
| 3669 | 3688 | 1 | ; | CA | - | H |
| 3669 | 3689 | 1 | ; | CA | - | CA |
| 3670 | 3672 | 1 | ; | HA | - | HB2 |
| 3670 | 3673 | 1 | ; | HA | - | HB3 |
| 3670 | 3674 | 1 | ; | HA | - | CG |
| 3670 | 3686 | 1 | ; | HA | - | O |
| 3670 | 3687 | 1 | ; | HA | - | N |
| 3671 | 3676 | 1 | ; | CB | - | HD1 |
| 3671 | 3677 | 1 | ; | CB | - | CE1 |
| 3671 | 3681 | 1 | ; | CB | - | CE2 |
| 3671 | 3684 | 1 | ; | CB | - | HD2 |
| 3671 | 3686 | 1 | ; | CB | - | O |
| 3671 | 3687 | 1 | ; | CB | - | N |
| 3672 | 3675 | 1 | ; | HB2 | - | CD1 |
| 3672 | 3683 | 1 | ; | HB2 | - | CD2 |
| 3672 | 3685 | 1 | ; | HB2 | - | C |
| 3673 | 3675 | 1 | ; | HB3 | - | CD1 |
| 3673 | 3683 | 1 | ; | HB3 | - | CD2 |
| 3673 | 3685 | 1 | ; | HB3 | - | C |
| 3674 | 3678 | 1 | ; | CG | - | HE1 |
| 3674 | 3679 | 1 | ; | CG | - | CZ |
| 3674 | 3682 | 1 | ; | CG | - | HE2 |

| | | | | | |
|---|---|---|---|---|---|
| 3674 | 3685 | 1 ; | CG | - | C |
| 3675 | 3680 | 1 ; | CD1 | - | HZ |
| 3675 | 3681 | 1 ; | CD1 | - | CE2 |
| 3675 | 3684 | 1 ; | CD1 | - | HD2 |
| 3676 | 3678 | 1 ; | HD1 | - | HE1 |
| 3676 | 3679 | 1 ; | HD1 | - | CZ |
| 3676 | 3683 | 1 ; | HD1 | - | CD2 |
| 3677 | 3682 | 1 ; | CE1 | - | HE2 |
| 3677 | 3683 | 1 ; | CE1 | - | CD2 |
| 3678 | 3680 | 1 ; | HE1 | - | HZ |
| 3678 | 3681 | 1 ; | HE1 | - | CE2 |
| 3679 | 3684 | 1 ; | CZ | - | HD2 |
| 3680 | 3682 | 1 ; | HZ | - | HE2 |
| 3680 | 3683 | 1 ; | HZ | - | CD2 |
| 3682 | 3684 | 1 ; | HE2 | - | HD2 |
| 3685 | 3690 | 1 ; | C | - | HA |
| 3685 | 3691 | 1 ; | C | - | CB |
| 3685 | 3696 | 1 ; | C | - | C |
| 3686 | 3688 | 1 ; | O | - | H |
| 3686 | 3689 | 1 ; | O | - | CA |
| 3687 | 3692 | 1 ; | N | - | HB2 |
| 3687 | 3693 | 1 ; | N | - | HB3 |
| 3687 | 3694 | 1 ; | N | - | SG |
| 3687 | 3697 | 1 ; | N | - | O |
| 3687 | 3698 | 1 ; | N | - | N |
| 3688 | 3690 | 1 ; | H | - | HA |
| 3688 | 3691 | 1 ; | H | - | CB |
| 3688 | 3696 | 1 ; | H | - | C |
| 3689 | 3695 | 1 ; | CA | - | HG |
| 3689 | 3699 | 1 ; | CA | - | H |
| 3689 | 3700 | 1 ; | CA | - | CA |
| 3690 | 3692 | 1 ; | HA | - | HB2 |
| 3690 | 3693 | 1 ; | HA | - | HB3 |
| 3690 | 3694 | 1 ; | HA | - | SG |
| 3690 | 3697 | 1 ; | HA | - | O |
| 3690 | 3698 | 1 ; | HA | - | N |
| 3691 | 3697 | 1 ; | CB | - | O |
| 3691 | 3698 | 1 ; | CB | - | N |
| 3692 | 3695 | 1 ; | HB2 | - | HG |
| 3692 | 3696 | 1 ; | HB2 | - | C |
| 3693 | 3695 | 1 ; | HB3 | - | HG |
| 3693 | 3696 | 1 ; | HB3 | - | C |
| 3694 | 3696 | 1 ; | SG | - | C |
| 3696 | 3701 | 1 ; | C | - | HA |
| 3696 | 3702 | 1 ; | C | - | CB |
| 3696 | 3716 | 1 ; | C | - | C |
| 3697 | 3699 | 1 ; | O | - | H |
| 3697 | 3700 | 1 ; | O | - | CA |
| 3698 | 3703 | 1 ; | N | - | HB2 |
| 3698 | 3704 | 1 ; | N | - | HB3 |
| 3698 | 3705 | 1 ; | N | - | CG |
| 3698 | 3717 | 1 ; | N | - | O |

| | | | |
|---|---|---|---|
| 3698 | 3718 | 1 ; | N - N |
| 3699 | 3701 | 1 ; | H - HA |
| 3699 | 3702 | 1 ; | H - CB |
| 3699 | 3716 | 1 ; | H - C |
| 3700 | 3706 | 1 ; | CA - CD1 |
| 3700 | 3714 | 1 ; | CA - CD2 |
| 3700 | 3719 | 1 ; | CA - H |
| 3700 | 3720 | 1 ; | CA - CA |
| 3701 | 3703 | 1 ; | HA - HB2 |
| 3701 | 3704 | 1 ; | HA - HB3 |
| 3701 | 3705 | 1 ; | HA - CG |
| 3701 | 3717 | 1 ; | HA - O |
| 3701 | 3718 | 1 ; | HA - N |
| 3702 | 3707 | 1 ; | CB - HD1 |
| 3702 | 3708 | 1 ; | CB - CE1 |
| 3702 | 3712 | 1 ; | CB - CE2 |
| 3702 | 3715 | 1 ; | CB - HD2 |
| 3702 | 3717 | 1 ; | CB - O |
| 3702 | 3718 | 1 ; | CB - N |
| 3703 | 3706 | 1 ; | HB2 - CD1 |
| 3703 | 3714 | 1 ; | HB2 - CD2 |
| 3703 | 3716 | 1 ; | HB2 - C |
| 3704 | 3706 | 1 ; | HB3 - CD1 |
| 3704 | 3714 | 1 ; | HB3 - CD2 |
| 3704 | 3716 | 1 ; | HB3 - C |
| 3705 | 3709 | 1 ; | CG - HE1 |
| 3705 | 3710 | 1 ; | CG - CZ |
| 3705 | 3713 | 1 ; | CG - HE2 |
| 3705 | 3716 | 1 ; | CG - C |
| 3706 | 3711 | 1 ; | CD1 - HZ |
| 3706 | 3712 | 1 ; | CD1 - CE2 |
| 3706 | 3715 | 1 ; | CD1 - HD2 |
| 3707 | 3709 | 1 ; | HD1 - HE1 |
| 3707 | 3710 | 1 ; | HD1 - CZ |
| 3707 | 3714 | 1 ; | HD1 - CD2 |
| 3708 | 3713 | 1 ; | CE1 - HE2 |
| 3708 | 3714 | 1 ; | CE1 - CD2 |
| 3709 | 3711 | 1 ; | HE1 - HZ |
| 3709 | 3712 | 1 ; | HE1 - CE2 |
| 3710 | 3715 | 1 ; | CZ - HD2 |
| 3711 | 3713 | 1 ; | HZ - HE2 |
| 3711 | 3714 | 1 ; | HZ - CD2 |
| 3713 | 3715 | 1 ; | HE2 - HD2 |
| 3716 | 3721 | 1 ; | C - HA |
| 3716 | 3722 | 1 ; | C - CB |
| 3716 | 3737 | 1 ; | C - C |
| 3717 | 3719 | 1 ; | O - H |
| 3717 | 3720 | 1 ; | O - CA |
| 3718 | 3723 | 1 ; | N - HB2 |
| 3718 | 3724 | 1 ; | N - HB3 |
| 3718 | 3725 | 1 ; | N - CG |
| 3718 | 3738 | 1 ; | N - O |

| | | | |
|---|---|---|---|
| 3718 | 3739 | 1 ; | N - N |
| 3719 | 3721 | 1 ; | H - HA |
| 3719 | 3722 | 1 ; | H - CB |
| 3719 | 3737 | 1 ; | H - C |
| 3720 | 3726 | 1 ; | CA - CD1 |
| 3720 | 3735 | 1 ; | CA - CD2 |
| 3720 | 3740 | 1 ; | CA - H |
| 3720 | 3741 | 1 ; | CA - CA |
| 3721 | 3723 | 1 ; | HA - HB2 |
| 3721 | 3724 | 1 ; | HA - HB3 |
| 3721 | 3725 | 1 ; | HA - CG |
| 3721 | 3738 | 1 ; | HA - O |
| 3721 | 3739 | 1 ; | HA - N |
| 3722 | 3727 | 1 ; | CB - HD1 |
| 3722 | 3728 | 1 ; | CB - CE1 |
| 3722 | 3733 | 1 ; | CB - CE2 |
| 3722 | 3736 | 1 ; | CB - HD2 |
| 3722 | 3738 | 1 ; | CB - O |
| 3722 | 3739 | 1 ; | CB - N |
| 3723 | 3726 | 1 ; | HB2 - CD1 |
| 3723 | 3735 | 1 ; | HB2 - CD2 |
| 3723 | 3737 | 1 ; | HB2 - C |
| 3724 | 3726 | 1 ; | HB3 - CD1 |
| 3724 | 3735 | 1 ; | HB3 - CD2 |
| 3724 | 3737 | 1 ; | HB3 - C |
| 3725 | 3729 | 1 ; | CG - HE1 |
| 3725 | 3730 | 1 ; | CG - CZ |
| 3725 | 3734 | 1 ; | CG - HE2 |
| 3725 | 3737 | 1 ; | CG - C |
| 3726 | 3731 | 1 ; | CD1 - OH |
| 3726 | 3733 | 1 ; | CD1 - CE2 |
| 3726 | 3736 | 1 ; | CD1 - HD2 |
| 3727 | 3729 | 1 ; | HD1 - HE1 |
| 3727 | 3730 | 1 ; | HD1 - CZ |
| 3727 | 3735 | 1 ; | HD1 - CD2 |
| 3728 | 3732 | 1 ; | CE1 - HH |
| 3728 | 3734 | 1 ; | CE1 - HE2 |
| 3728 | 3735 | 1 ; | CE1 - CD2 |
| 3729 | 3731 | 1 ; | HE1 - OH |
| 3729 | 3733 | 1 ; | HE1 - CE2 |
| 3730 | 3736 | 1 ; | CZ - HD2 |
| 3731 | 3734 | 1 ; | OH - HE2 |
| 3731 | 3735 | 1 ; | OH - CD2 |
| 3732 | 3733 | 1 ; | HH - CE2 |
| 3734 | 3736 | 1 ; | HE2 - HD2 |
| 3737 | 3742 | 1 ; | C - HA |
| 3737 | 3743 | 1 ; | C - CB |
| 3737 | 3752 | 1 ; | C - C |
| 3738 | 3740 | 1 ; | O - H |
| 3738 | 3741 | 1 ; | O - CA |
| 3739 | 3744 | 1 ; | N - HB2 |
| 3739 | 3745 | 1 ; | N - HB3 |

| | | | |
|---|---|---|---|
| 3739 | 3746 | 1 ; | N - CG |
| 3739 | 3753 | 1 ; | N - O |
| 3739 | 3754 | 1 ; | N - N |
| 3740 | 3742 | 1 ; | H - HA |
| 3740 | 3743 | 1 ; | H - CB |
| 3740 | 3752 | 1 ; | H - C |
| 3741 | 3747 | 1 ; | CA - HG2 |
| 3741 | 3748 | 1 ; | CA - HG3 |
| 3741 | 3749 | 1 ; | CA - CD |
| 3741 | 3755 | 1 ; | CA - H |
| 3741 | 3756 | 1 ; | CA - CA |
| 3742 | 3744 | 1 ; | HA - HB2 |
| 3742 | 3745 | 1 ; | HA - HB3 |
| 3742 | 3746 | 1 ; | HA - CG |
| 3742 | 3753 | 1 ; | HA - O |
| 3742 | 3754 | 1 ; | HA - N |
| 3743 | 3750 | 1 ; | CB - OE1 |
| 3743 | 3751 | 1 ; | CB - OE2 |
| 3743 | 3753 | 1 ; | CB - O |
| 3743 | 3754 | 1 ; | CB - N |
| 3744 | 3747 | 1 ; | HB2 - HG2 |
| 3744 | 3748 | 1 ; | HB2 - HG3 |
| 3744 | 3749 | 1 ; | HB2 - CD |
| 3744 | 3752 | 1 ; | HB2 - C |
| 3745 | 3747 | 1 ; | HB3 - HG2 |
| 3745 | 3748 | 1 ; | HB3 - HG3 |
| 3745 | 3749 | 1 ; | HB3 - CD |
| 3745 | 3752 | 1 ; | HB3 - C |
| 3746 | 3752 | 1 ; | CG - C |
| 3747 | 3750 | 1 ; | HG2 - OE1 |
| 3747 | 3751 | 1 ; | HG2 - OE2 |
| 3748 | 3750 | 1 ; | HG3 - OE1 |
| 3748 | 3751 | 1 ; | HG3 - OE2 |
| 3752 | 3757 | 1 ; | C - HA |
| 3752 | 3758 | 1 ; | C - CB |
| 3752 | 3766 | 1 ; | C - C |
| 3753 | 3755 | 1 ; | O - H |
| 3753 | 3756 | 1 ; | O - CA |
| 3754 | 3759 | 1 ; | N - HB2 |
| 3754 | 3760 | 1 ; | N - HB3 |
| 3754 | 3761 | 1 ; | N - CG |
| 3754 | 3767 | 1 ; | N - O |
| 3754 | 3768 | 1 ; | N - N |
| 3755 | 3757 | 1 ; | H - HA |
| 3755 | 3758 | 1 ; | H - CB |
| 3755 | 3766 | 1 ; | H - C |
| 3756 | 3762 | 1 ; | CA - OD1 |
| 3756 | 3763 | 1 ; | CA - ND2 |
| 3756 | 3769 | 1 ; | CA - H |
| 3756 | 3770 | 1 ; | CA - CA |
| 3757 | 3759 | 1 ; | HA - HB2 |
| 3757 | 3760 | 1 ; | HA - HB3 |

| | | | | | |
|---|---|---|---|---|---|
| 3757 | 3761 | 1 ; | HA | - | CG |
| 3757 | 3767 | 1 ; | HA | - | O |
| 3757 | 3768 | 1 ; | HA | - | N |
| 3758 | 3764 | 1 ; | CB | - | HD21 |
| 3758 | 3765 | 1 ; | CB | - | HD22 |
| 3758 | 3767 | 1 ; | CB | - | O |
| 3758 | 3768 | 1 ; | CB | - | N |
| 3759 | 3762 | 1 ; | HB2 | - | OD1 |
| 3759 | 3763 | 1 ; | HB2 | - | ND2 |
| 3759 | 3766 | 1 ; | HB2 | - | C |
| 3760 | 3762 | 1 ; | HB3 | - | OD1 |
| 3760 | 3763 | 1 ; | HB3 | - | ND2 |
| 3760 | 3766 | 1 ; | HB3 | - | C |
| 3761 | 3766 | 1 ; | CG | - | C |
| 3762 | 3764 | 1 ; | OD1 | - | HD21 |
| 3762 | 3765 | 1 ; | OD1 | - | HD22 |
| 3766 | 3771 | 1 ; | C | - | HA |
| 3766 | 3772 | 1 ; | C | - | CB |
| 3766 | 3778 | 1 ; | C | - | C |
| 3767 | 3769 | 1 ; | O | - | H |
| 3767 | 3770 | 1 ; | O | - | CA |
| 3768 | 3773 | 1 ; | N | - | HB2 |
| 3768 | 3774 | 1 ; | N | - | HB3 |
| 3768 | 3775 | 1 ; | N | - | CG |
| 3768 | 3779 | 1 ; | N | - | O |
| 3768 | 3780 | 1 ; | N | - | N |
| 3769 | 3771 | 1 ; | H | - | HA |
| 3769 | 3772 | 1 ; | H | - | CB |
| 3769 | 3778 | 1 ; | H | - | C |
| 3770 | 3776 | 1 ; | CA | - | OD1 |
| 3770 | 3777 | 1 ; | CA | - | OD2 |
| 3770 | 3781 | 1 ; | CA | - | H |
| 3770 | 3782 | 1 ; | CA | - | CA |
| 3771 | 3773 | 1 ; | HA | - | HB2 |
| 3771 | 3774 | 1 ; | HA | - | HB3 |
| 3771 | 3775 | 1 ; | HA | - | CG |
| 3771 | 3779 | 1 ; | HA | - | O |
| 3771 | 3780 | 1 ; | HA | - | N |
| 3772 | 3779 | 1 ; | CB | - | O |
| 3772 | 3780 | 1 ; | CB | - | N |
| 3773 | 3776 | 1 ; | HB2 | - | OD1 |
| 3773 | 3777 | 1 ; | HB2 | - | OD2 |
| 3773 | 3778 | 1 ; | HB2 | - | C |
| 3774 | 3776 | 1 ; | HB3 | - | OD1 |
| 3774 | 3777 | 1 ; | HB3 | - | OD2 |
| 3774 | 3778 | 1 ; | HB3 | - | C |
| 3775 | 3778 | 1 ; | CG | - | C |
| 3778 | 3783 | 1 ; | C | - | HA |
| 3778 | 3784 | 1 ; | C | - | CB |
| 3778 | 3802 | 1 ; | C | - | C |
| 3779 | 3781 | 1 ; | O | - | H |
| 3779 | 3782 | 1 ; | O | - | CA |

```
3780   3785      1 ;      N - HB2
3780   3786      1 ;      N - HB3
3780   3787      1 ;      N - CG
3780   3803      1 ;      N - O
3780   3804      1 ;      N - N
3781   3783      1 ;      H - HA
3781   3784      1 ;      H - CB
3781   3802      1 ;      H - C
3782   3788      1 ;     CA - CD1
3782   3801      1 ;     CA - CD2
3782   3805      1 ;     CA - H
3782   3806      1 ;     CA - CA
3783   3785      1 ;     HA - HB2
3783   3786      1 ;     HA - HB3
3783   3787      1 ;     HA - CG
3783   3803      1 ;     HA - O
3783   3804      1 ;     HA - N
3784   3789      1 ;     CB - HD1
3784   3790      1 ;     CB - NE1
3784   3792      1 ;     CB - CE2
3784   3799      1 ;     CB - CE3
3784   3803      1 ;     CB - O
3784   3804      1 ;     CB - N
3785   3788      1 ;    HB2 - CD1
3785   3801      1 ;    HB2 - CD2
3785   3802      1 ;    HB2 - C
3786   3788      1 ;    HB3 - CD1
3786   3801      1 ;    HB3 - CD2
3786   3802      1 ;    HB3 - C
3787   3791      1 ;     CG - HE1
3787   3793      1 ;     CG - CZ2
3787   3797      1 ;     CG - CZ3
3787   3800      1 ;     CG - HE3
3787   3802      1 ;     CG - C
3788   3793      1 ;    CD1 - CZ2
3788   3799      1 ;    CD1 - CE3
3789   3791      1 ;    HD1 - HE1
3789   3792      1 ;    HD1 - CE2
3789   3801      1 ;    HD1 - CD2
3790   3794      1 ;    NE1 - HZ2
3790   3795      1 ;    NE1 - CH2
3790   3799      1 ;    NE1 - CE3
3791   3793      1 ;    HE1 - CZ2
3791   3801      1 ;    HE1 - CD2
3792   3796      1 ;    CE2 - HH2
3792   3797      1 ;    CE2 - CZ3
3792   3800      1 ;    CE2 - HE3
3793   3798      1 ;    CZ2 - HZ3
3793   3799      1 ;    CZ2 - CE3
3794   3796      1 ;    HZ2 - HH2
3794   3797      1 ;    HZ2 - CZ3
3794   3801      1 ;    HZ2 - CD2
```

| | | | | | |
|---|---|---|---|---|---|
| 3795 | 3800 | 1 ; | CH2 | - | HE3 |
| 3795 | 3801 | 1 ; | CH2 | - | CD2 |
| 3796 | 3798 | 1 ; | HH2 | - | HZ3 |
| 3796 | 3799 | 1 ; | HH2 | - | CE3 |
| 3798 | 3800 | 1 ; | HZ3 | - | HE3 |
| 3798 | 3801 | 1 ; | HZ3 | - | CD2 |
| 3802 | 3807 | 1 ; | C | - | HA2 |
| 3802 | 3808 | 1 ; | C | - | HA3 |
| 3802 | 3809 | 1 ; | C | - | C |
| 3803 | 3805 | 1 ; | O | - | H |
| 3803 | 3806 | 1 ; | O | - | CA |
| 3804 | 3810 | 1 ; | N | - | O |
| 3804 | 3811 | 1 ; | N | - | N |
| 3805 | 3807 | 1 ; | H | - | HA2 |
| 3805 | 3808 | 1 ; | H | - | HA3 |
| 3805 | 3809 | 1 ; | H | - | C |
| 3806 | 3812 | 1 ; | CA | - | H |
| 3806 | 3813 | 1 ; | CA | - | CA |
| 3807 | 3810 | 1 ; | HA2 | - | O |
| 3807 | 3811 | 1 ; | HA2 | - | N |
| 3808 | 3810 | 1 ; | HA3 | - | O |
| 3808 | 3811 | 1 ; | HA3 | - | N |
| 3809 | 3814 | 1 ; | C | - | HA |
| 3809 | 3815 | 1 ; | C | - | CB |
| 3809 | 3820 | 1 ; | C | - | C |
| 3810 | 3812 | 1 ; | O | - | H |
| 3810 | 3813 | 1 ; | O | - | CA |
| 3811 | 3816 | 1 ; | N | - | HB2 |
| 3811 | 3817 | 1 ; | N | - | HB3 |
| 3811 | 3818 | 1 ; | N | - | OG |
| 3811 | 3821 | 1 ; | N | - | O |
| 3811 | 3822 | 1 ; | N | - | N |
| 3812 | 3814 | 1 ; | H | - | HA |
| 3812 | 3815 | 1 ; | H | - | CB |
| 3812 | 3820 | 1 ; | H | - | C |
| 3813 | 3819 | 1 ; | CA | - | HG |
| 3813 | 3823 | 1 ; | CA | - | H |
| 3813 | 3824 | 1 ; | CA | - | CA |
| 3814 | 3816 | 1 ; | HA | - | HB2 |
| 3814 | 3817 | 1 ; | HA | - | HB3 |
| 3814 | 3818 | 1 ; | HA | - | OG |
| 3814 | 3821 | 1 ; | HA | - | O |
| 3814 | 3822 | 1 ; | HA | - | N |
| 3815 | 3821 | 1 ; | CB | - | O |
| 3815 | 3822 | 1 ; | CB | - | N |
| 3816 | 3819 | 1 ; | HB2 | - | HG |
| 3816 | 3820 | 1 ; | HB2 | - | C |
| 3817 | 3819 | 1 ; | HB3 | - | HG |
| 3817 | 3820 | 1 ; | HB3 | - | C |
| 3818 | 3820 | 1 ; | OG | - | C |
| 3820 | 3825 | 1 ; | C | - | HA |
| 3820 | 3826 | 1 ; | C | - | CB |

```
3820   3839       1 ;       C - C
3821   3823       1 ;       O - H
3821   3824       1 ;       O - CA
3822   3827       1 ;       N - HB
3822   3828       1 ;       N - CG2
3822   3832       1 ;       N - CG1
3822   3840       1 ;       N - O
3822   3841       1 ;       N - N
3823   3825       1 ;       H - HA
3823   3826       1 ;       H - CB
3823   3839       1 ;       H - C
3824   3829       1 ;      CA - HG21
3824   3830       1 ;      CA - HG22
3824   3831       1 ;      CA - HG23
3824   3833       1 ;      CA - HG12
3824   3834       1 ;      CA - HG13
3824   3835       1 ;      CA - CD1
3824   3842       1 ;      CA - H
3824   3843       1 ;      CA - CA
3825   3827       1 ;      HA - HB
3825   3828       1 ;      HA - CG2
3825   3832       1 ;      HA - CG1
3825   3840       1 ;      HA - O
3825   3841       1 ;      HA - N
3826   3836       1 ;      CB - HD11
3826   3837       1 ;      CB - HD12
3826   3838       1 ;      CB - HD13
3826   3840       1 ;      CB - O
3826   3841       1 ;      CB - N
3827   3829       1 ;      HB - HG21
3827   3830       1 ;      HB - HG22
3827   3831       1 ;      HB - HG23
3827   3833       1 ;      HB - HG12
3827   3834       1 ;      HB - HG13
3827   3835       1 ;      HB - CD1
3827   3839       1 ;      HB - C
3828   3833       1 ;     CG2 - HG12
3828   3834       1 ;     CG2 - HG13
3828   3835       1 ;     CG2 - CD1
3828   3839       1 ;     CG2 - C
3829   3832       1 ;    HG21 - CG1
3830   3832       1 ;    HG22 - CG1
3831   3832       1 ;    HG23 - CG1
3832   3839       1 ;     CG1 - C
3833   3836       1 ;    HG12 - HD11
3833   3837       1 ;    HG12 - HD12
3833   3838       1 ;    HG12 - HD13
3834   3836       1 ;    HG13 - HD11
3834   3837       1 ;    HG13 - HD12
3834   3838       1 ;    HG13 - HD13
3839   3844       1 ;       C - HA
3839   3845       1 ;       C - CB
```

| | | | | | |
|---|---|---|---|---|---|
| 3839 | 3850 | 1 ; | C | - | C |
| 3840 | 3842 | 1 ; | O | - | H |
| 3840 | 3843 | 1 ; | O | - | CA |
| 3841 | 3846 | 1 ; | N | - | HB2 |
| 3841 | 3847 | 1 ; | N | - | HB3 |
| 3841 | 3848 | 1 ; | N | - | OG |
| 3841 | 3851 | 1 ; | N | - | O |
| 3841 | 3852 | 1 ; | N | - | N |
| 3842 | 3844 | 1 ; | H | - | HA |
| 3842 | 3845 | 1 ; | H | - | CB |
| 3842 | 3850 | 1 ; | H | - | C |
| 3843 | 3849 | 1 ; | CA | - | HG |
| 3843 | 3853 | 1 ; | CA | - | H |
| 3843 | 3854 | 1 ; | CA | - | CA |
| 3844 | 3846 | 1 ; | HA | - | HB2 |
| 3844 | 3847 | 1 ; | HA | - | HB3 |
| 3844 | 3848 | 1 ; | HA | - | OG |
| 3844 | 3851 | 1 ; | HA | - | O |
| 3844 | 3852 | 1 ; | HA | - | N |
| 3845 | 3851 | 1 ; | CB | - | O |
| 3845 | 3852 | 1 ; | CB | - | N |
| 3846 | 3849 | 1 ; | HB2 | - | HG |
| 3846 | 3850 | 1 ; | HB2 | - | C |
| 3847 | 3849 | 1 ; | HB3 | - | HG |
| 3847 | 3850 | 1 ; | HB3 | - | C |
| 3848 | 3850 | 1 ; | OG | - | C |
| 3850 | 3855 | 1 ; | C | - | HA |
| 3850 | 3856 | 1 ; | C | - | CB |
| 3850 | 3861 | 1 ; | C | - | C |
| 3851 | 3853 | 1 ; | O | - | H |
| 3851 | 3854 | 1 ; | O | - | CA |
| 3852 | 3857 | 1 ; | N | - | HB2 |
| 3852 | 3858 | 1 ; | N | - | HB3 |
| 3852 | 3859 | 1 ; | N | - | OG |
| 3852 | 3862 | 1 ; | N | - | O |
| 3852 | 3863 | 1 ; | N | - | N |
| 3853 | 3855 | 1 ; | H | - | HA |
| 3853 | 3856 | 1 ; | H | - | CB |
| 3853 | 3861 | 1 ; | H | - | C |
| 3854 | 3860 | 1 ; | CA | - | HG |
| 3854 | 3864 | 1 ; | CA | - | H |
| 3854 | 3865 | 1 ; | CA | - | CA |
| 3855 | 3857 | 1 ; | HA | - | HB2 |
| 3855 | 3858 | 1 ; | HA | - | HB3 |
| 3855 | 3859 | 1 ; | HA | - | OG |
| 3855 | 3862 | 1 ; | HA | - | O |
| 3855 | 3863 | 1 ; | HA | - | N |
| 3856 | 3862 | 1 ; | CB | - | O |
| 3856 | 3863 | 1 ; | CB | - | N |
| 3857 | 3860 | 1 ; | HB2 | - | HG |
| 3857 | 3861 | 1 ; | HB2 | - | C |
| 3858 | 3860 | 1 ; | HB3 | - | HG |

| | | | |
|---|---|---|---|
| 3858 | 3861 | 1 ; | HB3 - C |
| 3859 | 3861 | 1 ; | OG - C |
| 3861 | 3866 | 1 ; | C - HA |
| 3861 | 3867 | 1 ; | C - CB |
| 3861 | 3873 | 1 ; | C - C |
| 3862 | 3864 | 1 ; | O - H |
| 3862 | 3865 | 1 ; | O - CA |
| 3863 | 3868 | 1 ; | N - HB2 |
| 3863 | 3869 | 1 ; | N - HB3 |
| 3863 | 3870 | 1 ; | N - CG |
| 3863 | 3874 | 1 ; | N - O |
| 3863 | 3875 | 1 ; | N - N |
| 3864 | 3866 | 1 ; | H - HA |
| 3864 | 3867 | 1 ; | H - CB |
| 3864 | 3873 | 1 ; | H - C |
| 3865 | 3871 | 1 ; | CA - OD1 |
| 3865 | 3872 | 1 ; | CA - OD2 |
| 3865 | 3876 | 1 ; | CA - H |
| 3865 | 3877 | 1 ; | CA - CA |
| 3866 | 3868 | 1 ; | HA - HB2 |
| 3866 | 3869 | 1 ; | HA - HB3 |
| 3866 | 3870 | 1 ; | HA - CG |
| 3866 | 3874 | 1 ; | HA - O |
| 3866 | 3875 | 1 ; | HA - N |
| 3867 | 3874 | 1 ; | CB - O |
| 3867 | 3875 | 1 ; | CB - N |
| 3868 | 3871 | 1 ; | HB2 - OD1 |
| 3868 | 3872 | 1 ; | HB2 - OD2 |
| 3868 | 3873 | 1 ; | HB2 - C |
| 3869 | 3871 | 1 ; | HB3 - OD1 |
| 3869 | 3872 | 1 ; | HB3 - OD2 |
| 3869 | 3873 | 1 ; | HB3 - C |
| 3870 | 3873 | 1 ; | CG - C |
| 3873 | 3878 | 1 ; | C - HA |
| 3873 | 3879 | 1 ; | C - CB |
| 3873 | 3883 | 1 ; | C - C |
| 3874 | 3876 | 1 ; | O - H |
| 3874 | 3877 | 1 ; | O - CA |
| 3875 | 3880 | 1 ; | N - HB1 |
| 3875 | 3881 | 1 ; | N - HB2 |
| 3875 | 3882 | 1 ; | N - HB3 |
| 3875 | 3884 | 1 ; | N - O |
| 3875 | 3885 | 1 ; | N - N |
| 3876 | 3878 | 1 ; | H - HA |
| 3876 | 3879 | 1 ; | H - CB |
| 3876 | 3883 | 1 ; | H - C |
| 3877 | 3886 | 1 ; | CA - H |
| 3877 | 3887 | 1 ; | CA - CA |
| 3878 | 3880 | 1 ; | HA - HB1 |
| 3878 | 3881 | 1 ; | HA - HB2 |
| 3878 | 3882 | 1 ; | HA - HB3 |
| 3878 | 3884 | 1 ; | HA - O |

| | | | | |
|---|---|---|---|---|
| 3878 | 3885 | 1 | ; | HA - N |
| 3879 | 3884 | 1 | ; | CB - O |
| 3879 | 3885 | 1 | ; | CB - N |
| 3880 | 3883 | 1 | ; | HB1 - C |
| 3881 | 3883 | 1 | ; | HB2 - C |
| 3882 | 3883 | 1 | ; | HB3 - C |
| 3883 | 3888 | 1 | ; | C - HA |
| 3883 | 3889 | 1 | ; | C - CB |
| 3883 | 3905 | 1 | ; | C - C |
| 3884 | 3886 | 1 | ; | O - H |
| 3884 | 3887 | 1 | ; | O - CA |
| 3885 | 3890 | 1 | ; | N - HB2 |
| 3885 | 3891 | 1 | ; | N - HB3 |
| 3885 | 3892 | 1 | ; | N - CG |
| 3885 | 3906 | 1 | ; | N - O |
| 3885 | 3907 | 1 | ; | N - N |
| 3886 | 3888 | 1 | ; | H - HA |
| 3886 | 3889 | 1 | ; | H - CB |
| 3886 | 3905 | 1 | ; | H - C |
| 3887 | 3893 | 1 | ; | CA - HG2 |
| 3887 | 3894 | 1 | ; | CA - HG3 |
| 3887 | 3895 | 1 | ; | CA - CD |
| 3887 | 3908 | 1 | ; | CA - H |
| 3887 | 3909 | 1 | ; | CA - CA |
| 3888 | 3890 | 1 | ; | HA - HB2 |
| 3888 | 3891 | 1 | ; | HA - HB3 |
| 3888 | 3892 | 1 | ; | HA - CG |
| 3888 | 3906 | 1 | ; | HA - O |
| 3888 | 3907 | 1 | ; | HA - N |
| 3889 | 3896 | 1 | ; | CB - HD2 |
| 3889 | 3897 | 1 | ; | CB - HD3 |
| 3889 | 3898 | 1 | ; | CB - CE |
| 3889 | 3906 | 1 | ; | CB - O |
| 3889 | 3907 | 1 | ; | CB - N |
| 3890 | 3893 | 1 | ; | HB2 - HG2 |
| 3890 | 3894 | 1 | ; | HB2 - HG3 |
| 3890 | 3895 | 1 | ; | HB2 - CD |
| 3890 | 3905 | 1 | ; | HB2 - C |
| 3891 | 3893 | 1 | ; | HB3 - HG2 |
| 3891 | 3894 | 1 | ; | HB3 - HG3 |
| 3891 | 3895 | 1 | ; | HB3 - CD |
| 3891 | 3905 | 1 | ; | HB3 - C |
| 3892 | 3899 | 1 | ; | CG - HE2 |
| 3892 | 3900 | 1 | ; | CG - HE3 |
| 3892 | 3901 | 1 | ; | CG - NZ |
| 3892 | 3905 | 1 | ; | CG - C |
| 3893 | 3896 | 1 | ; | HG2 - HD2 |
| 3893 | 3897 | 1 | ; | HG2 - HD3 |
| 3893 | 3898 | 1 | ; | HG2 - CE |
| 3894 | 3896 | 1 | ; | HG3 - HD2 |
| 3894 | 3897 | 1 | ; | HG3 - HD3 |
| 3894 | 3898 | 1 | ; | HG3 - CE |

| | | | | | |
|---|---|---|---|---|---|
| 3895 | 3902 | 1 ; | CD | - | HZ1 |
| 3895 | 3903 | 1 ; | CD | - | HZ2 |
| 3895 | 3904 | 1 ; | CD | - | HZ3 |
| 3896 | 3899 | 1 ; | HD2 | - | HE2 |
| 3896 | 3900 | 1 ; | HD2 | - | HE3 |
| 3896 | 3901 | 1 ; | HD2 | - | NZ |
| 3897 | 3899 | 1 ; | HD3 | - | HE2 |
| 3897 | 3900 | 1 ; | HD3 | - | HE3 |
| 3897 | 3901 | 1 ; | HD3 | - | NZ |
| 3899 | 3902 | 1 ; | HE2 | - | HZ1 |
| 3899 | 3903 | 1 ; | HE2 | - | HZ2 |
| 3899 | 3904 | 1 ; | HE2 | - | HZ3 |
| 3900 | 3902 | 1 ; | HE3 | - | HZ1 |
| 3900 | 3903 | 1 ; | HE3 | - | HZ2 |
| 3900 | 3904 | 1 ; | HE3 | - | HZ3 |
| 3905 | 3910 | 1 ; | C | - | HA |
| 3905 | 3911 | 1 ; | C | - | CB |
| 3905 | 3919 | 1 ; | C | - | C |
| 3906 | 3908 | 1 ; | O | - | H |
| 3906 | 3909 | 1 ; | O | - | CA |
| 3907 | 3912 | 1 ; | N | - | HB2 |
| 3907 | 3913 | 1 ; | N | - | HB3 |
| 3907 | 3914 | 1 ; | N | - | CG |
| 3907 | 3920 | 1 ; | N | - | O |
| 3907 | 3921 | 1 ; | N | - | N |
| 3908 | 3910 | 1 ; | H | - | HA |
| 3908 | 3911 | 1 ; | H | - | CB |
| 3908 | 3919 | 1 ; | H | - | C |
| 3909 | 3915 | 1 ; | CA | - | OD1 |
| 3909 | 3916 | 1 ; | CA | - | ND2 |
| 3909 | 3922 | 1 ; | CA | - | H |
| 3909 | 3923 | 1 ; | CA | - | CA |
| 3910 | 3912 | 1 ; | HA | - | HB2 |
| 3910 | 3913 | 1 ; | HA | - | HB3 |
| 3910 | 3914 | 1 ; | HA | - | CG |
| 3910 | 3920 | 1 ; | HA | - | O |
| 3910 | 3921 | 1 ; | HA | - | N |
| 3911 | 3917 | 1 ; | CB | - | HD21 |
| 3911 | 3918 | 1 ; | CB | - | HD22 |
| 3911 | 3920 | 1 ; | CB | - | O |
| 3911 | 3921 | 1 ; | CB | - | N |
| 3912 | 3915 | 1 ; | HB2 | - | OD1 |
| 3912 | 3916 | 1 ; | HB2 | - | ND2 |
| 3912 | 3919 | 1 ; | HB2 | - | C |
| 3913 | 3915 | 1 ; | HB3 | - | OD1 |
| 3913 | 3916 | 1 ; | HB3 | - | ND2 |
| 3913 | 3919 | 1 ; | HB3 | - | C |
| 3914 | 3919 | 1 ; | CG | - | C |
| 3915 | 3917 | 1 ; | OD1 | - | HD21 |
| 3915 | 3918 | 1 ; | OD1 | - | HD22 |
| 3919 | 3924 | 1 ; | C | - | HA |
| 3919 | 3925 | 1 ; | C | - | CB |

```
3919   3938       1 ;       C - C
3920   3922       1 ;       O - H
3920   3923       1 ;       O - CA
3921   3926       1 ;       N - HB2
3921   3927       1 ;       N - HB3
3921   3928       1 ;       N - CG
3921   3939       1 ;       N - O
3921   3940       1 ;       N - N
3922   3924       1 ;       H - HA
3922   3925       1 ;       H - CB
3922   3938       1 ;       H - C
3923   3929       1 ;      CA - HG
3923   3930       1 ;      CA - CD1
3923   3934       1 ;      CA - CD2
3923   3941       1 ;      CA - H
3923   3942       1 ;      CA - CA
3924   3926       1 ;      HA - HB2
3924   3927       1 ;      HA - HB3
3924   3928       1 ;      HA - CG
3924   3939       1 ;      HA - O
3924   3940       1 ;      HA - N
3925   3931       1 ;      CB - HD11
3925   3932       1 ;      CB - HD12
3925   3933       1 ;      CB - HD13
3925   3935       1 ;      CB - HD21
3925   3936       1 ;      CB - HD22
3925   3937       1 ;      CB - HD23
3925   3939       1 ;      CB - O
3925   3940       1 ;      CB - N
3926   3929       1 ;     HB2 - HG
3926   3930       1 ;     HB2 - CD1
3926   3934       1 ;     HB2 - CD2
3926   3938       1 ;     HB2 - C
3927   3929       1 ;     HB3 - HG
3927   3930       1 ;     HB3 - CD1
3927   3934       1 ;     HB3 - CD2
3927   3938       1 ;     HB3 - C
3928   3938       1 ;      CG - C
3929   3931       1 ;      HG - HD11
3929   3932       1 ;      HG - HD12
3929   3933       1 ;      HG - HD13
3929   3935       1 ;      HG - HD21
3929   3936       1 ;      HG - HD22
3929   3937       1 ;      HG - HD23
3930   3935       1 ;     CD1 - HD21
3930   3936       1 ;     CD1 - HD22
3930   3937       1 ;     CD1 - HD23
3931   3934       1 ;    HD11 - CD2
3932   3934       1 ;    HD12 - CD2
3933   3934       1 ;    HD13 - CD2
3938   3943       1 ;       C - HA
3938   3944       1 ;       C - CB
```

```
3938   3957      1 ;      C - C
3939   3941      1 ;      O - H
3939   3942      1 ;      O - CA
3940   3945      1 ;      N - HB
3940   3946      1 ;      N - CG2
3940   3950      1 ;      N - CG1
3940   3958      1 ;      N - O
3940   3959      1 ;      N - N
3941   3943      1 ;      H - HA
3941   3944      1 ;      H - CB
3941   3957      1 ;      H - C
3942   3947      1 ;     CA - HG21
3942   3948      1 ;     CA - HG22
3942   3949      1 ;     CA - HG23
3942   3951      1 ;     CA - HG12
3942   3952      1 ;     CA - HG13
3942   3953      1 ;     CA - CD1
3942   3960      1 ;     CA - H
3942   3961      1 ;     CA - CA
3943   3945      1 ;     HA - HB
3943   3946      1 ;     HA - CG2
3943   3950      1 ;     HA - CG1
3943   3958      1 ;     HA - O
3943   3959      1 ;     HA - N
3944   3954      1 ;     CB - HD11
3944   3955      1 ;     CB - HD12
3944   3956      1 ;     CB - HD13
3944   3958      1 ;     CB - O
3944   3959      1 ;     CB - N
3945   3947      1 ;     HB - HG21
3945   3948      1 ;     HB - HG22
3945   3949      1 ;     HB - HG23
3945   3951      1 ;     HB - HG12
3945   3952      1 ;     HB - HG13
3945   3953      1 ;     HB - CD1
3945   3957      1 ;     HB - C
3946   3951      1 ;    CG2 - HG12
3946   3952      1 ;    CG2 - HG13
3946   3953      1 ;    CG2 - CD1
3946   3957      1 ;    CG2 - C
3947   3950      1 ;   HG21 - CG1
3948   3950      1 ;   HG22 - CG1
3949   3950      1 ;   HG23 - CG1
3950   3957      1 ;    CG1 - C
3951   3954      1 ;   HG12 - HD11
3951   3955      1 ;   HG12 - HD12
3951   3956      1 ;   HG12 - HD13
3952   3954      1 ;   HG13 - HD11
3952   3955      1 ;   HG13 - HD12
3952   3956      1 ;   HG13 - HD13
3957   3962      1 ;      C - HA
3957   3963      1 ;      C - CB
```

| | | | |
|---|---|---|---|
| 3957 | 3971 | 1 ; | C - C |
| 3958 | 3960 | 1 ; | O - H |
| 3958 | 3961 | 1 ; | O - CA |
| 3959 | 3964 | 1 ; | N - HB |
| 3959 | 3965 | 1 ; | N - CG2 |
| 3959 | 3969 | 1 ; | N - OG1 |
| 3959 | 3972 | 1 ; | N - O |
| 3959 | 3973 | 1 ; | N - N |
| 3960 | 3962 | 1 ; | H - HA |
| 3960 | 3963 | 1 ; | H - CB |
| 3960 | 3971 | 1 ; | H - C |
| 3961 | 3966 | 1 ; | CA - HG21 |
| 3961 | 3967 | 1 ; | CA - HG22 |
| 3961 | 3968 | 1 ; | CA - HG23 |
| 3961 | 3970 | 1 ; | CA - HG1 |
| 3961 | 3974 | 1 ; | CA - H |
| 3961 | 3975 | 1 ; | CA - CA |
| 3962 | 3964 | 1 ; | HA - HB |
| 3962 | 3965 | 1 ; | HA - CG2 |
| 3962 | 3969 | 1 ; | HA - OG1 |
| 3962 | 3972 | 1 ; | HA - O |
| 3962 | 3973 | 1 ; | HA - N |
| 3963 | 3972 | 1 ; | CB - O |
| 3963 | 3973 | 1 ; | CB - N |
| 3964 | 3966 | 1 ; | HB - HG21 |
| 3964 | 3967 | 1 ; | HB - HG22 |
| 3964 | 3968 | 1 ; | HB - HG23 |
| 3964 | 3970 | 1 ; | HB - HG1 |
| 3964 | 3971 | 1 ; | HB - C |
| 3965 | 3970 | 1 ; | CG2 - HG1 |
| 3965 | 3971 | 1 ; | CG2 - C |
| 3966 | 3969 | 1 ; | HG21 - OG1 |
| 3967 | 3969 | 1 ; | HG22 - OG1 |
| 3968 | 3969 | 1 ; | HG23 - OG1 |
| 3969 | 3971 | 1 ; | OG1 - C |
| 3971 | 3976 | 1 ; | C - HA |
| 3971 | 3977 | 1 ; | C - CB |
| 3971 | 3993 | 1 ; | C - C |
| 3972 | 3974 | 1 ; | O - H |
| 3972 | 3975 | 1 ; | O - CA |
| 3973 | 3978 | 1 ; | N - HB2 |
| 3973 | 3979 | 1 ; | N - HB3 |
| 3973 | 3980 | 1 ; | N - CG |
| 3973 | 3994 | 1 ; | N - O |
| 3973 | 3995 | 1 ; | N - N |
| 3974 | 3976 | 1 ; | H - HA |
| 3974 | 3977 | 1 ; | H - CB |
| 3974 | 3993 | 1 ; | H - C |
| 3975 | 3981 | 1 ; | CA - HG2 |
| 3975 | 3982 | 1 ; | CA - HG3 |
| 3975 | 3983 | 1 ; | CA - CD |
| 3975 | 3996 | 1 ; | CA - H |

```
3975   3997       1 ;      CA - CA
3976   3978       1 ;      HA - HB2
3976   3979       1 ;      HA - HB3
3976   3980       1 ;      HA - CG
3976   3994       1 ;      HA - O
3976   3995       1 ;      HA - N
3977   3984       1 ;      CB - HD2
3977   3985       1 ;      CB - HD3
3977   3986       1 ;      CB - CE
3977   3994       1 ;      CB - O
3977   3995       1 ;      CB - N
3978   3981       1 ;     HB2 - HG2
3978   3982       1 ;     HB2 - HG3
3978   3983       1 ;     HB2 - CD
3978   3993       1 ;     HB2 - C
3979   3981       1 ;     HB3 - HG2
3979   3982       1 ;     HB3 - HG3
3979   3983       1 ;     HB3 - CD
3979   3993       1 ;     HB3 - C
3980   3987       1 ;      CG - HE2
3980   3988       1 ;      CG - HE3
3980   3989       1 ;      CG - NZ
3980   3993       1 ;      CG - C
3981   3984       1 ;     HG2 - HD2
3981   3985       1 ;     HG2 - HD3
3981   3986       1 ;     HG2 - CE
3982   3984       1 ;     HG3 - HD2
3982   3985       1 ;     HG3 - HD3
3982   3986       1 ;     HG3 - CE
3983   3990       1 ;      CD - HZ1
3983   3991       1 ;      CD - HZ2
3983   3992       1 ;      CD - HZ3
3984   3987       1 ;     HD2 - HE2
3984   3988       1 ;     HD2 - HE3
3984   3989       1 ;     HD2 - NZ
3985   3987       1 ;     HD3 - HE2
3985   3988       1 ;     HD3 - HE3
3985   3989       1 ;     HD3 - NZ
3987   3990       1 ;     HE2 - HZ1
3987   3991       1 ;     HE2 - HZ2
3987   3992       1 ;     HE2 - HZ3
3988   3990       1 ;     HE3 - HZ1
3988   3991       1 ;     HE3 - HZ2
3988   3992       1 ;     HE3 - HZ3
3993   3998       1 ;       C - HA
3993   3999       1 ;       C - CB
3993   4012       1 ;       C - C
3994   3996       1 ;       O - H
3994   3997       1 ;       O - CA
3995   4000       1 ;       N - HB2
3995   4001       1 ;       N - HB3
3995   4002       1 ;       N - CG
```

```
3995   4013       1 ;       N - O
3995   4014       1 ;       N - N
3996   3998       1 ;       H - HA
3996   3999       1 ;       H - CB
3996   4012       1 ;       H - C
3997   4003       1 ;      CA - HG
3997   4004       1 ;      CA - CD1
3997   4008       1 ;      CA - CD2
3997   4015       1 ;      CA - H
3997   4016       1 ;      CA - CA
3998   4000       1 ;      HA - HB2
3998   4001       1 ;      HA - HB3
3998   4002       1 ;      HA - CG
3998   4013       1 ;      HA - O
3998   4014       1 ;      HA - N
3999   4005       1 ;      CB - HD11
3999   4006       1 ;      CB - HD12
3999   4007       1 ;      CB - HD13
3999   4009       1 ;      CB - HD21
3999   4010       1 ;      CB - HD22
3999   4011       1 ;      CB - HD23
3999   4013       1 ;      CB - O
3999   4014       1 ;      CB - N
4000   4003       1 ;     HB2 - HG
4000   4004       1 ;     HB2 - CD1
4000   4008       1 ;     HB2 - CD2
4000   4012       1 ;     HB2 - C
4001   4003       1 ;     HB3 - HG
4001   4004       1 ;     HB3 - CD1
4001   4008       1 ;     HB3 - CD2
4001   4012       1 ;     HB3 - C
4002   4012       1 ;      CG - C
4003   4005       1 ;      HG - HD11
4003   4006       1 ;      HG - HD12
4003   4007       1 ;      HG - HD13
4003   4009       1 ;      HG - HD21
4003   4010       1 ;      HG - HD22
4003   4011       1 ;      HG - HD23
4004   4009       1 ;     CD1 - HD21
4004   4010       1 ;     CD1 - HD22
4004   4011       1 ;     CD1 - HD23
4005   4008       1 ;    HD11 - CD2
4006   4008       1 ;    HD12 - CD2
4007   4008       1 ;    HD13 - CD2
4012   4017       1 ;       C - HA
4012   4018       1 ;       C - CB
4012   4031       1 ;       C - C
4013   4015       1 ;       O - H
4013   4016       1 ;       O - CA
4014   4019       1 ;       N - HB2
4014   4020       1 ;       N - HB3
4014   4021       1 ;       N - CG
```

| | | | | |
|---|---|---|---|---|
| 4014 | 4032 | 1 ; | N - | O |
| 4014 | 4033 | 1 ; | N - | N |
| 4015 | 4017 | 1 ; | H - | HA |
| 4015 | 4018 | 1 ; | H - | CB |
| 4015 | 4031 | 1 ; | H - | C |
| 4016 | 4022 | 1 ; | CA - | HG |
| 4016 | 4023 | 1 ; | CA - | CD1 |
| 4016 | 4027 | 1 ; | CA - | CD2 |
| 4016 | 4034 | 1 ; | CA - | H |
| 4016 | 4035 | 1 ; | CA - | CA |
| 4017 | 4019 | 1 ; | HA - | HB2 |
| 4017 | 4020 | 1 ; | HA - | HB3 |
| 4017 | 4021 | 1 ; | HA - | CG |
| 4017 | 4032 | 1 ; | HA - | O |
| 4017 | 4033 | 1 ; | HA - | N |
| 4018 | 4024 | 1 ; | CB - | HD11 |
| 4018 | 4025 | 1 ; | CB - | HD12 |
| 4018 | 4026 | 1 ; | CB - | HD13 |
| 4018 | 4028 | 1 ; | CB - | HD21 |
| 4018 | 4029 | 1 ; | CB - | HD22 |
| 4018 | 4030 | 1 ; | CB - | HD23 |
| 4018 | 4032 | 1 ; | CB - | O |
| 4018 | 4033 | 1 ; | CB - | N |
| 4019 | 4022 | 1 ; | HB2 - | HG |
| 4019 | 4023 | 1 ; | HB2 - | CD1 |
| 4019 | 4027 | 1 ; | HB2 - | CD2 |
| 4019 | 4031 | 1 ; | HB2 - | C |
| 4020 | 4022 | 1 ; | HB3 - | HG |
| 4020 | 4023 | 1 ; | HB3 - | CD1 |
| 4020 | 4027 | 1 ; | HB3 - | CD2 |
| 4020 | 4031 | 1 ; | HB3 - | C |
| 4021 | 4031 | 1 ; | CG - | C |
| 4022 | 4024 | 1 ; | HG - | HD11 |
| 4022 | 4025 | 1 ; | HG - | HD12 |
| 4022 | 4026 | 1 ; | HG - | HD13 |
| 4022 | 4028 | 1 ; | HG - | HD21 |
| 4022 | 4029 | 1 ; | HG - | HD22 |
| 4022 | 4030 | 1 ; | HG - | HD23 |
| 4023 | 4028 | 1 ; | CD1 - | HD21 |
| 4023 | 4029 | 1 ; | CD1 - | HD22 |
| 4023 | 4030 | 1 ; | CD1 - | HD23 |
| 4024 | 4027 | 1 ; | HD11 - | CD2 |
| 4025 | 4027 | 1 ; | HD12 - | CD2 |
| 4026 | 4027 | 1 ; | HD13 - | CD2 |
| 4031 | 4036 | 1 ; | C - | HA |
| 4031 | 4037 | 1 ; | C - | CB |
| 4031 | 4045 | 1 ; | C - | C |
| 4032 | 4034 | 1 ; | O - | H |
| 4032 | 4035 | 1 ; | O - | CA |
| 4033 | 4038 | 1 ; | N - | HB |
| 4033 | 4039 | 1 ; | N - | CG2 |
| 4033 | 4043 | 1 ; | N - | OG1 |

```
4033   4046      1 ;      N - O
4033   4047      1 ;      N - N
4034   4036      1 ;      H - HA
4034   4037      1 ;      H - CB
4034   4045      1 ;      H - C
4035   4040      1 ;     CA - HG21
4035   4041      1 ;     CA - HG22
4035   4042      1 ;     CA - HG23
4035   4044      1 ;     CA - HG1
4035   4048      1 ;     CA - H
4035   4049      1 ;     CA - CA
4036   4038      1 ;     HA - HB
4036   4039      1 ;     HA - CG2
4036   4043      1 ;     HA - OG1
4036   4046      1 ;     HA - O
4036   4047      1 ;     HA - N
4037   4046      1 ;     CB - O
4037   4047      1 ;     CB - N
4038   4040      1 ;     HB - HG21
4038   4041      1 ;     HB - HG22
4038   4042      1 ;     HB - HG23
4038   4044      1 ;     HB - HG1
4038   4045      1 ;     HB - C
4039   4044      1 ;    CG2 - HG1
4039   4045      1 ;    CG2 - C
4040   4043      1 ;   HG21 - OG1
4041   4043      1 ;   HG22 - OG1
4042   4043      1 ;   HG23 - OG1
4043   4045      1 ;    OG1 - C
4045   4050      1 ;      C - HA
4045   4051      1 ;      C - CB
4045   4066      1 ;      C - C
4046   4048      1 ;      O - H
4046   4049      1 ;      O - CA
4047   4052      1 ;      N - HB2
4047   4053      1 ;      N - HB3
4047   4054      1 ;      N - CG
4047   4067      1 ;      N - O
4047   4068      1 ;      N - N
4048   4050      1 ;      H - HA
4048   4051      1 ;      H - CB
4048   4066      1 ;      H - C
4049   4055      1 ;     CA - CD1
4049   4064      1 ;     CA - CD2
4049   4069      1 ;     CA - H
4049   4070      1 ;     CA - CA
4050   4052      1 ;     HA - HB2
4050   4053      1 ;     HA - HB3
4050   4054      1 ;     HA - CG
4050   4067      1 ;     HA - O
4050   4068      1 ;     HA - N
4051   4056      1 ;     CB - HD1
```

```
4051   4057       1 ;      CB - CE1
4051   4062       1 ;      CB - CE2
4051   4065       1 ;      CB - HD2
4051   4067       1 ;      CB - O
4051   4068       1 ;      CB - N
4052   4055       1 ;     HB2 - CD1
4052   4064       1 ;     HB2 - CD2
4052   4066       1 ;     HB2 - C
4053   4055       1 ;     HB3 - CD1
4053   4064       1 ;     HB3 - CD2
4053   4066       1 ;     HB3 - C
4054   4058       1 ;      CG - HE1
4054   4059       1 ;      CG - CZ
4054   4063       1 ;      CG - HE2
4054   4066       1 ;      CG - C
4055   4060       1 ;     CD1 - OH
4055   4062       1 ;     CD1 - CE2
4055   4065       1 ;     CD1 - HD2
4056   4058       1 ;     HD1 - HE1
4056   4059       1 ;     HD1 - CZ
4056   4064       1 ;     HD1 - CD2
4057   4061       1 ;     CE1 - HH
4057   4063       1 ;     CE1 - HE2
4057   4064       1 ;     CE1 - CD2
4058   4060       1 ;     HE1 - OH
4058   4062       1 ;     HE1 - CE2
4059   4065       1 ;      CZ - HD2
4060   4063       1 ;      OH - HE2
4060   4064       1 ;      OH - CD2
4061   4062       1 ;      HH - CE2
4063   4065       1 ;     HE2 - HD2
4066   4071       1 ;       C - HA
4066   4072       1 ;       C - CB
4066   4080       1 ;       C - C
4067   4069       1 ;       O - H
4067   4070       1 ;       O - CA
4068   4073       1 ;       N - HB2
4068   4074       1 ;       N - HB3
4068   4075       1 ;       N - CG
4068   4081       1 ;       N - O
4068   4082       1 ;       N - N
4069   4071       1 ;       H - HA
4069   4072       1 ;       H - CB
4069   4080       1 ;       H - C
4070   4076       1 ;      CA - OD1
4070   4077       1 ;      CA - ND2
4070   4083       1 ;      CA - CD
4070   4092       1 ;      CA - CA
4071   4073       1 ;      HA - HB2
4071   4074       1 ;      HA - HB3
4071   4075       1 ;      HA - CG
4071   4081       1 ;      HA - O
```

```
4071   4082      1 ;      HA - N
4072   4078      1 ;      CB - HD21
4072   4079      1 ;      CB - HD22
4072   4081      1 ;      CB - O
4072   4082      1 ;      CB - N
4073   4076      1 ;     HB2 - OD1
4073   4077      1 ;     HB2 - ND2
4073   4080      1 ;     HB2 - C
4074   4076      1 ;     HB3 - OD1
4074   4077      1 ;     HB3 - ND2
4074   4080      1 ;     HB3 - C
4075   4080      1 ;      CG - C
4076   4078      1 ;     OD1 - HD21
4076   4079      1 ;     OD1 - HD22
4080   4084      1 ;       C - HD2
4080   4085      1 ;       C - HD3
4080   4086      1 ;       C - CG
4080   4089      1 ;       C - CB
4080   4093      1 ;       C - HA
4080   4094      1 ;       C - C
4081   4083      1 ;       O - CD
4081   4092      1 ;       O - CA
4082   4087      1 ;       N - HG2
4082   4088      1 ;       N - HG3
4082   4090      1 ;       N - HB2
4082   4091      1 ;       N - HB3
4082   4095      1 ;       N - O
4082   4096      1 ;       N - N
4083   4090      1 ;      CD - HB2
4083   4091      1 ;      CD - HB3
4083   4093      1 ;      CD - HA
4083   4094      1 ;      CD - C
4084   4087      1 ;     HD2 - HG2
4084   4088      1 ;     HD2 - HG3
4084   4089      1 ;     HD2 - CB
4084   4092      1 ;     HD2 - CA
4085   4087      1 ;     HD3 - HG2
4085   4088      1 ;     HD3 - HG3
4085   4089      1 ;     HD3 - CB
4085   4092      1 ;     HD3 - CA
4086   4093      1 ;      CG - HA
4086   4094      1 ;      CG - C
4087   4090      1 ;     HG2 - HB2
4087   4091      1 ;     HG2 - HB3
4087   4092      1 ;     HG2 - CA
4088   4090      1 ;     HG3 - HB2
4088   4091      1 ;     HG3 - HB3
4088   4092      1 ;     HG3 - CA
4089   4095      1 ;      CB - O
4089   4096      1 ;      CB - N
4090   4093      1 ;     HB2 - HA
4090   4094      1 ;     HB2 - C
```

```
4091   4093       1 ;    HB3 - HA
4091   4094       1 ;    HB3 - C
4092   4097       1 ;     CA - H
4092   4098       1 ;     CA - CA
4093   4095       1 ;     HA - O
4093   4096       1 ;     HA - N
4094   4099       1 ;      C - HA
4094   4100       1 ;      C - CB
4094   4108       1 ;      C - C
4095   4097       1 ;      O - H
4095   4098       1 ;      O - CA
4096   4101       1 ;      N - HB2
4096   4102       1 ;      N - HB3
4096   4103       1 ;      N - CG
4096   4109       1 ;      N - O
4096   4110       1 ;      N - N
4097   4099       1 ;      H - HA
4097   4100       1 ;      H - CB
4097   4108       1 ;      H - C
4098   4104       1 ;     CA - OD1
4098   4105       1 ;     CA - ND2
4098   4111       1 ;     CA - H
4098   4112       1 ;     CA - CA
4099   4101       1 ;     HA - HB2
4099   4102       1 ;     HA - HB3
4099   4103       1 ;     HA - CG
4099   4109       1 ;     HA - O
4099   4110       1 ;     HA - N
4100   4106       1 ;     CB - HD21
4100   4107       1 ;     CB - HD22
4100   4109       1 ;     CB - O
4100   4110       1 ;     CB - N
4101   4104       1 ;    HB2 - OD1
4101   4105       1 ;    HB2 - ND2
4101   4108       1 ;    HB2 - C
4102   4104       1 ;    HB3 - OD1
4102   4105       1 ;    HB3 - ND2
4102   4108       1 ;    HB3 - C
4103   4108       1 ;     CG - C
4104   4106       1 ;    OD1 - HD21
4104   4107       1 ;    OD1 - HD22
4108   4113       1 ;      C - HA
4108   4114       1 ;      C - CB
4108   4123       1 ;      C - C
4109   4111       1 ;      O - H
4109   4112       1 ;      O - CA
4110   4115       1 ;      N - HB2
4110   4116       1 ;      N - HB3
4110   4117       1 ;      N - CG
4110   4124       1 ;      N - O
4110   4125       1 ;      N - N
4111   4113       1 ;      H - HA
```

| | | | |
|---|---|---|---|
| 4111 | 4114 | 1 ; | H - CB |
| 4111 | 4123 | 1 ; | H - C |
| 4112 | 4118 | 1 ; | CA - HG2 |
| 4112 | 4119 | 1 ; | CA - HG3 |
| 4112 | 4120 | 1 ; | CA - CD |
| 4112 | 4126 | 1 ; | CA - H |
| 4112 | 4127 | 1 ; | CA - CA |
| 4113 | 4115 | 1 ; | HA - HB2 |
| 4113 | 4116 | 1 ; | HA - HB3 |
| 4113 | 4117 | 1 ; | HA - CG |
| 4113 | 4124 | 1 ; | HA - O |
| 4113 | 4125 | 1 ; | HA - N |
| 4114 | 4121 | 1 ; | CB - OE1 |
| 4114 | 4122 | 1 ; | CB - OE2 |
| 4114 | 4124 | 1 ; | CB - O |
| 4114 | 4125 | 1 ; | CB - N |
| 4115 | 4118 | 1 ; | HB2 - HG2 |
| 4115 | 4119 | 1 ; | HB2 - HG3 |
| 4115 | 4120 | 1 ; | HB2 - CD |
| 4115 | 4123 | 1 ; | HB2 - C |
| 4116 | 4118 | 1 ; | HB3 - HG2 |
| 4116 | 4119 | 1 ; | HB3 - HG3 |
| 4116 | 4120 | 1 ; | HB3 - CD |
| 4116 | 4123 | 1 ; | HB3 - C |
| 4117 | 4123 | 1 ; | CG - C |
| 4118 | 4121 | 1 ; | HG2 - OE1 |
| 4118 | 4122 | 1 ; | HG2 - OE2 |
| 4119 | 4121 | 1 ; | HG3 - OE1 |
| 4119 | 4122 | 1 ; | HG3 - OE2 |
| 4123 | 4128 | 1 ; | C - HA |
| 4123 | 4129 | 1 ; | C - CB |
| 4123 | 4147 | 1 ; | C - C |
| 4124 | 4126 | 1 ; | O - H |
| 4124 | 4127 | 1 ; | O - CA |
| 4125 | 4130 | 1 ; | N - HB2 |
| 4125 | 4131 | 1 ; | N - HB3 |
| 4125 | 4132 | 1 ; | N - CG |
| 4125 | 4148 | 1 ; | N - O |
| 4125 | 4149 | 1 ; | N - N |
| 4126 | 4128 | 1 ; | H - HA |
| 4126 | 4129 | 1 ; | H - CB |
| 4126 | 4147 | 1 ; | H - C |
| 4127 | 4133 | 1 ; | CA - HG2 |
| 4127 | 4134 | 1 ; | CA - HG3 |
| 4127 | 4135 | 1 ; | CA - CD |
| 4127 | 4150 | 1 ; | CA - H |
| 4127 | 4151 | 1 ; | CA - CA |
| 4128 | 4130 | 1 ; | HA - HB2 |
| 4128 | 4131 | 1 ; | HA - HB3 |
| 4128 | 4132 | 1 ; | HA - CG |
| 4128 | 4148 | 1 ; | HA - O |
| 4128 | 4149 | 1 ; | HA - N |

| | | | | | |
|---|---|---|---|---|---|
| 4129 | 4136 | 1 ; | CB | - | HD2 |
| 4129 | 4137 | 1 ; | CB | - | HD3 |
| 4129 | 4138 | 1 ; | CB | - | NE |
| 4129 | 4148 | 1 ; | CB | - | O |
| 4129 | 4149 | 1 ; | CB | - | N |
| 4130 | 4133 | 1 ; | HB2 | - | HG2 |
| 4130 | 4134 | 1 ; | HB2 | - | HG3 |
| 4130 | 4135 | 1 ; | HB2 | - | CD |
| 4130 | 4147 | 1 ; | HB2 | - | C |
| 4131 | 4133 | 1 ; | HB3 | - | HG2 |
| 4131 | 4134 | 1 ; | HB3 | - | HG3 |
| 4131 | 4135 | 1 ; | HB3 | - | CD |
| 4131 | 4147 | 1 ; | HB3 | - | C |
| 4132 | 4139 | 1 ; | CG | - | HE |
| 4132 | 4140 | 1 ; | CG | - | CZ |
| 4132 | 4147 | 1 ; | CG | - | C |
| 4133 | 4136 | 1 ; | HG2 | - | HD2 |
| 4133 | 4137 | 1 ; | HG2 | - | HD3 |
| 4133 | 4138 | 1 ; | HG2 | - | NE |
| 4134 | 4136 | 1 ; | HG3 | - | HD2 |
| 4134 | 4137 | 1 ; | HG3 | - | HD3 |
| 4134 | 4138 | 1 ; | HG3 | - | NE |
| 4135 | 4141 | 1 ; | CD | - | NH1 |
| 4135 | 4144 | 1 ; | CD | - | NH2 |
| 4136 | 4139 | 1 ; | HD2 | - | HE |
| 4136 | 4140 | 1 ; | HD2 | - | CZ |
| 4137 | 4139 | 1 ; | HD3 | - | HE |
| 4137 | 4140 | 1 ; | HD3 | - | CZ |
| 4138 | 4142 | 1 ; | NE | - | HH11 |
| 4138 | 4143 | 1 ; | NE | - | HH12 |
| 4138 | 4145 | 1 ; | NE | - | HH21 |
| 4138 | 4146 | 1 ; | NE | - | HH22 |
| 4139 | 4141 | 1 ; | HE | - | NH1 |
| 4139 | 4144 | 1 ; | HE | - | NH2 |
| 4141 | 4145 | 1 ; | NH1 | - | HH21 |
| 4141 | 4146 | 1 ; | NH1 | - | HH22 |
| 4142 | 4144 | 1 ; | HH11 | - | NH2 |
| 4143 | 4144 | 1 ; | HH12 | - | NH2 |
| 4147 | 4152 | 1 ; | C | - | HA |
| 4147 | 4153 | 1 ; | C | - | CB |
| 4147 | 4158 | 1 ; | C | - | C |
| 4148 | 4150 | 1 ; | O | - | H |
| 4148 | 4151 | 1 ; | O | - | CA |
| 4149 | 4154 | 1 ; | N | - | HB2 |
| 4149 | 4155 | 1 ; | N | - | HB3 |
| 4149 | 4156 | 1 ; | N | - | SG |
| 4149 | 4159 | 1 ; | N | - | O |
| 4149 | 4160 | 1 ; | N | - | N |
| 4150 | 4152 | 1 ; | H | - | HA |
| 4150 | 4153 | 1 ; | H | - | CB |
| 4150 | 4158 | 1 ; | H | - | C |
| 4151 | 4157 | 1 ; | CA | - | HG |

| | | | |
|---|---|---|---|
| 4151 | 4161 | 1 ; | CA - H |
| 4151 | 4162 | 1 ; | CA - CA |
| 4152 | 4154 | 1 ; | HA - HB2 |
| 4152 | 4155 | 1 ; | HA - HB3 |
| 4152 | 4156 | 1 ; | HA - SG |
| 4152 | 4159 | 1 ; | HA - O |
| 4152 | 4160 | 1 ; | HA - N |
| 4153 | 4159 | 1 ; | CB - O |
| 4153 | 4160 | 1 ; | CB - N |
| 4154 | 4157 | 1 ; | HB2 - HG |
| 4154 | 4158 | 1 ; | HB2 - C |
| 4155 | 4157 | 1 ; | HB3 - HG |
| 4155 | 4158 | 1 ; | HB3 - C |
| 4156 | 4158 | 1 ; | SG - C |
| 4158 | 4163 | 1 ; | C - HA |
| 4158 | 4164 | 1 ; | C - CB |
| 4158 | 4172 | 1 ; | C - C |
| 4159 | 4161 | 1 ; | O - H |
| 4159 | 4162 | 1 ; | O - CA |
| 4160 | 4165 | 1 ; | N - HB |
| 4160 | 4166 | 1 ; | N - CG2 |
| 4160 | 4170 | 1 ; | N - OG1 |
| 4160 | 4173 | 1 ; | N - O |
| 4160 | 4174 | 1 ; | N - N |
| 4161 | 4163 | 1 ; | H - HA |
| 4161 | 4164 | 1 ; | H - CB |
| 4161 | 4172 | 1 ; | H - C |
| 4162 | 4167 | 1 ; | CA - HG21 |
| 4162 | 4168 | 1 ; | CA - HG22 |
| 4162 | 4169 | 1 ; | CA - HG23 |
| 4162 | 4171 | 1 ; | CA - HG1 |
| 4162 | 4175 | 1 ; | CA - H |
| 4162 | 4176 | 1 ; | CA - CA |
| 4163 | 4165 | 1 ; | HA - HB |
| 4163 | 4166 | 1 ; | HA - CG2 |
| 4163 | 4170 | 1 ; | HA - OG1 |
| 4163 | 4173 | 1 ; | HA - O |
| 4163 | 4174 | 1 ; | HA - N |
| 4164 | 4173 | 1 ; | CB - O |
| 4164 | 4174 | 1 ; | CB - N |
| 4165 | 4167 | 1 ; | HB - HG21 |
| 4165 | 4168 | 1 ; | HB - HG22 |
| 4165 | 4169 | 1 ; | HB - HG23 |
| 4165 | 4171 | 1 ; | HB - HG1 |
| 4165 | 4172 | 1 ; | HB - C |
| 4166 | 4171 | 1 ; | CG2 - HG1 |
| 4166 | 4172 | 1 ; | CG2 - C |
| 4167 | 4170 | 1 ; | HG21 - OG1 |
| 4168 | 4170 | 1 ; | HG22 - OG1 |
| 4169 | 4170 | 1 ; | HG23 - OG1 |
| 4170 | 4172 | 1 ; | OG1 - C |
| 4172 | 4177 | 1 ; | C - HA |

```
4172   4178       1 ;      C - CB
4172   4191       1 ;      C - C
4173   4175       1 ;      O - H
4173   4176       1 ;      O - CA
4174   4179       1 ;      N - HB
4174   4180       1 ;      N - CG2
4174   4184       1 ;      N - CG1
4174   4192       1 ;      N - O
4174   4193       1 ;      N - N
4175   4177       1 ;      H - HA
4175   4178       1 ;      H - CB
4175   4191       1 ;      H - C
4176   4181       1 ;     CA - HG21
4176   4182       1 ;     CA - HG22
4176   4183       1 ;     CA - HG23
4176   4185       1 ;     CA - HG12
4176   4186       1 ;     CA - HG13
4176   4187       1 ;     CA - CD1
4176   4194       1 ;     CA - H
4176   4195       1 ;     CA - CA
4177   4179       1 ;     HA - HB
4177   4180       1 ;     HA - CG2
4177   4184       1 ;     HA - CG1
4177   4192       1 ;     HA - O
4177   4193       1 ;     HA - N
4178   4188       1 ;     CB - HD11
4178   4189       1 ;     CB - HD12
4178   4190       1 ;     CB - HD13
4178   4192       1 ;     CB - O
4178   4193       1 ;     CB - N
4179   4181       1 ;     HB - HG21
4179   4182       1 ;     HB - HG22
4179   4183       1 ;     HB - HG23
4179   4185       1 ;     HB - HG12
4179   4186       1 ;     HB - HG13
4179   4187       1 ;     HB - CD1
4179   4191       1 ;     HB - C
4180   4185       1 ;    CG2 - HG12
4180   4186       1 ;    CG2 - HG13
4180   4187       1 ;    CG2 - CD1
4180   4191       1 ;    CG2 - C
4181   4184       1 ;   HG21 - CG1
4182   4184       1 ;   HG22 - CG1
4183   4184       1 ;   HG23 - CG1
4184   4191       1 ;    CG1 - C
4185   4188       1 ;   HG12 - HD11
4185   4189       1 ;   HG12 - HD12
4185   4190       1 ;   HG12 - HD13
4186   4188       1 ;   HG13 - HD11
4186   4189       1 ;   HG13 - HD12
4186   4190       1 ;   HG13 - HD13
4191   4196       1 ;      C - HA
```

| | | | | | | |
|---|---|---|---|---|---|---|
| 4191 | 4197 | 1 | ; | C | - | CB |
| 4191 | 4206 | 1 | ; | C | - | C |
| 4192 | 4194 | 1 | ; | O | - | H |
| 4192 | 4195 | 1 | ; | O | - | CA |
| 4193 | 4198 | 1 | ; | N | - | HB2 |
| 4193 | 4199 | 1 | ; | N | - | HB3 |
| 4193 | 4200 | 1 | ; | N | - | CG |
| 4193 | 4207 | 1 | ; | N | - | O |
| 4193 | 4208 | 1 | ; | N | - | N |
| 4194 | 4196 | 1 | ; | H | - | HA |
| 4194 | 4197 | 1 | ; | H | - | CB |
| 4194 | 4206 | 1 | ; | H | - | C |
| 4195 | 4201 | 1 | ; | CA | - | HG2 |
| 4195 | 4202 | 1 | ; | CA | - | HG3 |
| 4195 | 4203 | 1 | ; | CA | - | CD |
| 4195 | 4209 | 1 | ; | CA | - | H |
| 4195 | 4210 | 1 | ; | CA | - | CA |
| 4196 | 4198 | 1 | ; | HA | - | HB2 |
| 4196 | 4199 | 1 | ; | HA | - | HB3 |
| 4196 | 4200 | 1 | ; | HA | - | CG |
| 4196 | 4207 | 1 | ; | HA | - | O |
| 4196 | 4208 | 1 | ; | HA | - | N |
| 4197 | 4204 | 1 | ; | CB | - | OE1 |
| 4197 | 4205 | 1 | ; | CB | - | OE2 |
| 4197 | 4207 | 1 | ; | CB | - | O |
| 4197 | 4208 | 1 | ; | CB | - | N |
| 4198 | 4201 | 1 | ; | HB2 | - | HG2 |
| 4198 | 4202 | 1 | ; | HB2 | - | HG3 |
| 4198 | 4203 | 1 | ; | HB2 | - | CD |
| 4198 | 4206 | 1 | ; | HB2 | - | C |
| 4199 | 4201 | 1 | ; | HB3 | - | HG2 |
| 4199 | 4202 | 1 | ; | HB3 | - | HG3 |
| 4199 | 4203 | 1 | ; | HB3 | - | CD |
| 4199 | 4206 | 1 | ; | HB3 | - | C |
| 4200 | 4206 | 1 | ; | CG | - | C |
| 4201 | 4204 | 1 | ; | HG2 | - | OE1 |
| 4201 | 4205 | 1 | ; | HG2 | - | OE2 |
| 4202 | 4204 | 1 | ; | HG3 | - | OE1 |
| 4202 | 4205 | 1 | ; | HG3 | - | OE2 |
| 4206 | 4211 | 1 | ; | C | - | HA |
| 4206 | 4212 | 1 | ; | C | - | CB |
| 4206 | 4221 | 1 | ; | C | - | C |
| 4207 | 4209 | 1 | ; | O | - | H |
| 4207 | 4210 | 1 | ; | O | - | CA |
| 4208 | 4213 | 1 | ; | N | - | HB2 |
| 4208 | 4214 | 1 | ; | N | - | HB3 |
| 4208 | 4215 | 1 | ; | N | - | CG |
| 4208 | 4222 | 1 | ; | N | - | O |
| 4208 | 4223 | 1 | ; | N | - | N |
| 4209 | 4211 | 1 | ; | H | - | HA |
| 4209 | 4212 | 1 | ; | H | - | CB |
| 4209 | 4221 | 1 | ; | H | - | C |

```
4210   4216      1 ;     CA - HG2
4210   4217      1 ;     CA - HG3
4210   4218      1 ;     CA - CD
4210   4224      1 ;     CA - H
4210   4225      1 ;     CA - CA
4211   4213      1 ;     HA - HB2
4211   4214      1 ;     HA - HB3
4211   4215      1 ;     HA - CG
4211   4222      1 ;     HA - O
4211   4223      1 ;     HA - N
4212   4219      1 ;     CB - OE1
4212   4220      1 ;     CB - OE2
4212   4222      1 ;     CB - O
4212   4223      1 ;     CB - N
4213   4216      1 ;    HB2 - HG2
4213   4217      1 ;    HB2 - HG3
4213   4218      1 ;    HB2 - CD
4213   4221      1 ;    HB2 - C
4214   4216      1 ;    HB3 - HG2
4214   4217      1 ;    HB3 - HG3
4214   4218      1 ;    HB3 - CD
4214   4221      1 ;    HB3 - C
4215   4221      1 ;     CG - C
4216   4219      1 ;    HG2 - OE1
4216   4220      1 ;    HG2 - OE2
4217   4219      1 ;    HG3 - OE1
4217   4220      1 ;    HG3 - OE2
4221   4226      1 ;      C - HA
4221   4227      1 ;      C - CB
4221   4231      1 ;      C - C
4222   4224      1 ;      O - H
4222   4225      1 ;      O - CA
4223   4228      1 ;      N - HB1
4223   4229      1 ;      N - HB2
4223   4230      1 ;      N - HB3
4223   4232      1 ;      N - O
4223   4233      1 ;      N - N
4224   4226      1 ;      H - HA
4224   4227      1 ;      H - CB
4224   4231      1 ;      H - C
4225   4234      1 ;     CA - H
4225   4235      1 ;     CA - CA
4226   4228      1 ;     HA - HB1
4226   4229      1 ;     HA - HB2
4226   4230      1 ;     HA - HB3
4226   4232      1 ;     HA - O
4226   4233      1 ;     HA - N
4227   4232      1 ;     CB - O
4227   4233      1 ;     CB - N
4228   4231      1 ;    HB1 - C
4229   4231      1 ;    HB2 - C
4230   4231      1 ;    HB3 - C
```

| | | | |
|---|---|---|---|
| 4231 | 4236 | 1 ; | C - HA |
| 4231 | 4237 | 1 ; | C - CB |
| 4231 | 4250 | 1 ; | C - C |
| 4232 | 4234 | 1 ; | O - H |
| 4232 | 4235 | 1 ; | O - CA |
| 4233 | 4238 | 1 ; | N - HB2 |
| 4233 | 4239 | 1 ; | N - HB3 |
| 4233 | 4240 | 1 ; | N - CG |
| 4233 | 4251 | 1 ; | N - O |
| 4233 | 4252 | 1 ; | N - N |
| 4234 | 4236 | 1 ; | H - HA |
| 4234 | 4237 | 1 ; | H - CB |
| 4234 | 4250 | 1 ; | H - C |
| 4235 | 4241 | 1 ; | CA - HG |
| 4235 | 4242 | 1 ; | CA - CD1 |
| 4235 | 4246 | 1 ; | CA - CD2 |
| 4235 | 4253 | 1 ; | CA - H |
| 4235 | 4254 | 1 ; | CA - CA |
| 4236 | 4238 | 1 ; | HA - HB2 |
| 4236 | 4239 | 1 ; | HA - HB3 |
| 4236 | 4240 | 1 ; | HA - CG |
| 4236 | 4251 | 1 ; | HA - O |
| 4236 | 4252 | 1 ; | HA - N |
| 4237 | 4243 | 1 ; | CB - HD11 |
| 4237 | 4244 | 1 ; | CB - HD12 |
| 4237 | 4245 | 1 ; | CB - HD13 |
| 4237 | 4247 | 1 ; | CB - HD21 |
| 4237 | 4248 | 1 ; | CB - HD22 |
| 4237 | 4249 | 1 ; | CB - HD23 |
| 4237 | 4251 | 1 ; | CB - O |
| 4237 | 4252 | 1 ; | CB - N |
| 4238 | 4241 | 1 ; | HB2 - HG |
| 4238 | 4242 | 1 ; | HB2 - CD1 |
| 4238 | 4246 | 1 ; | HB2 - CD2 |
| 4238 | 4250 | 1 ; | HB2 - C |
| 4239 | 4241 | 1 ; | HB3 - HG |
| 4239 | 4242 | 1 ; | HB3 - CD1 |
| 4239 | 4246 | 1 ; | HB3 - CD2 |
| 4239 | 4250 | 1 ; | HB3 - C |
| 4240 | 4250 | 1 ; | CG - C |
| 4241 | 4243 | 1 ; | HG - HD11 |
| 4241 | 4244 | 1 ; | HG - HD12 |
| 4241 | 4245 | 1 ; | HG - HD13 |
| 4241 | 4247 | 1 ; | HG - HD21 |
| 4241 | 4248 | 1 ; | HG - HD22 |
| 4241 | 4249 | 1 ; | HG - HD23 |
| 4242 | 4247 | 1 ; | CD1 - HD21 |
| 4242 | 4248 | 1 ; | CD1 - HD22 |
| 4242 | 4249 | 1 ; | CD1 - HD23 |
| 4243 | 4246 | 1 ; | HD11 - CD2 |
| 4244 | 4246 | 1 ; | HD12 - CD2 |
| 4245 | 4246 | 1 ; | HD13 - CD2 |

| | | | |
|---|---|---|---|
| 4250 | 4255 | 1 ; | C - HA |
| 4250 | 4256 | 1 ; | C - CB |
| 4250 | 4264 | 1 ; | C - C |
| 4251 | 4253 | 1 ; | O - H |
| 4251 | 4254 | 1 ; | O - CA |
| 4252 | 4257 | 1 ; | N - HB2 |
| 4252 | 4258 | 1 ; | N - HB3 |
| 4252 | 4259 | 1 ; | N - CG |
| 4252 | 4265 | 1 ; | N - O |
| 4252 | 4266 | 1 ; | N - N |
| 4253 | 4255 | 1 ; | H - HA |
| 4253 | 4256 | 1 ; | H - CB |
| 4253 | 4264 | 1 ; | H - C |
| 4254 | 4260 | 1 ; | CA - OD1 |
| 4254 | 4261 | 1 ; | CA - ND2 |
| 4254 | 4267 | 1 ; | CA - H |
| 4254 | 4268 | 1 ; | CA - CA |
| 4255 | 4257 | 1 ; | HA - HB2 |
| 4255 | 4258 | 1 ; | HA - HB3 |
| 4255 | 4259 | 1 ; | HA - CG |
| 4255 | 4265 | 1 ; | HA - O |
| 4255 | 4266 | 1 ; | HA - N |
| 4256 | 4262 | 1 ; | CB - HD21 |
| 4256 | 4263 | 1 ; | CB - HD22 |
| 4256 | 4265 | 1 ; | CB - O |
| 4256 | 4266 | 1 ; | CB - N |
| 4257 | 4260 | 1 ; | HB2 - OD1 |
| 4257 | 4261 | 1 ; | HB2 - ND2 |
| 4257 | 4264 | 1 ; | HB2 - C |
| 4258 | 4260 | 1 ; | HB3 - OD1 |
| 4258 | 4261 | 1 ; | HB3 - ND2 |
| 4258 | 4264 | 1 ; | HB3 - C |
| 4259 | 4264 | 1 ; | CG - C |
| 4260 | 4262 | 1 ; | OD1 - HD21 |
| 4260 | 4263 | 1 ; | OD1 - HD22 |
| 4264 | 4269 | 1 ; | C - HA |
| 4264 | 4270 | 1 ; | C - CB |
| 4264 | 4281 | 1 ; | C - C |
| 4265 | 4267 | 1 ; | O - H |
| 4265 | 4268 | 1 ; | O - CA |
| 4266 | 4271 | 1 ; | N - HB2 |
| 4266 | 4272 | 1 ; | N - HB3 |
| 4266 | 4273 | 1 ; | N - CG |
| 4266 | 4282 | 1 ; | N - O |
| 4266 | 4283 | 1 ; | N - N |
| 4267 | 4269 | 1 ; | H - HA |
| 4267 | 4270 | 1 ; | H - CB |
| 4267 | 4281 | 1 ; | H - C |
| 4268 | 4274 | 1 ; | CA - ND1 |
| 4268 | 4279 | 1 ; | CA - CD2 |
| 4268 | 4284 | 1 ; | CA - CD |
| 4268 | 4293 | 1 ; | CA - CA |

```
4269   4271       1 ;      HA - HB2
4269   4272       1 ;      HA - HB3
4269   4273       1 ;      HA - CG
4269   4282       1 ;      HA - O
4269   4283       1 ;      HA - N
4270   4275       1 ;      CB - CE1
4270   4277       1 ;      CB - NE2
4270   4280       1 ;      CB - HD2
4270   4282       1 ;      CB - O
4270   4283       1 ;      CB - N
4271   4274       1 ;     HB2 - ND1
4271   4279       1 ;     HB2 - CD2
4271   4281       1 ;     HB2 - C
4272   4274       1 ;     HB3 - ND1
4272   4279       1 ;     HB3 - CD2
4272   4281       1 ;     HB3 - C
4273   4276       1 ;      CG - HE1
4273   4278       1 ;      CG - HE2
4273   4281       1 ;      CG - C
4274   4278       1 ;     ND1 - HE2
4274   4280       1 ;     ND1 - HD2
4275   4280       1 ;     CE1 - HD2
4276   4278       1 ;     HE1 - HE2
4276   4279       1 ;     HE1 - CD2
4278   4280       1 ;     HE2 - HD2
4281   4285       1 ;       C - HD2
4281   4286       1 ;       C - HD3
4281   4287       1 ;       C - CG
4281   4290       1 ;       C - CB
4281   4294       1 ;       C - HA
4281   4295       1 ;       C - C
4282   4284       1 ;       O - CD
4282   4293       1 ;       O - CA
4283   4288       1 ;       N - HG2
4283   4289       1 ;       N - HG3
4283   4291       1 ;       N - HB2
4283   4292       1 ;       N - HB3
4283   4296       1 ;       N - O
4283   4297       1 ;       N - N
4284   4291       1 ;      CD - HB2
4284   4292       1 ;      CD - HB3
4284   4294       1 ;      CD - HA
4284   4295       1 ;      CD - C
4285   4288       1 ;     HD2 - HG2
4285   4289       1 ;     HD2 - HG3
4285   4290       1 ;     HD2 - CB
4285   4293       1 ;     HD2 - CA
4286   4288       1 ;     HD3 - HG2
4286   4289       1 ;     HD3 - HG3
4286   4290       1 ;     HD3 - CB
4286   4293       1 ;     HD3 - CA
4287   4294       1 ;      CG - HA
```

| | | | |
|---|---|---|---|
| 4287 | 4295 | 1 ; | CG - C |
| 4288 | 4291 | 1 ; | HG2 - HB2 |
| 4288 | 4292 | 1 ; | HG2 - HB3 |
| 4288 | 4293 | 1 ; | HG2 - CA |
| 4289 | 4291 | 1 ; | HG3 - HB2 |
| 4289 | 4292 | 1 ; | HG3 - HB3 |
| 4289 | 4293 | 1 ; | HG3 - CA |
| 4290 | 4296 | 1 ; | CB - O |
| 4290 | 4297 | 1 ; | CB - N |
| 4291 | 4294 | 1 ; | HB2 - HA |
| 4291 | 4295 | 1 ; | HB2 - C |
| 4292 | 4294 | 1 ; | HB3 - HA |
| 4292 | 4295 | 1 ; | HB3 - C |
| 4293 | 4298 | 1 ; | CA - H |
| 4293 | 4299 | 1 ; | CA - CA |
| 4294 | 4296 | 1 ; | HA - O |
| 4294 | 4297 | 1 ; | HA - N |
| 4295 | 4300 | 1 ; | C - HA |
| 4295 | 4301 | 1 ; | C - CB |
| 4295 | 4319 | 1 ; | C - C |
| 4296 | 4298 | 1 ; | O - H |
| 4296 | 4299 | 1 ; | O - CA |
| 4297 | 4302 | 1 ; | N - HB2 |
| 4297 | 4303 | 1 ; | N - HB3 |
| 4297 | 4304 | 1 ; | N - CG |
| 4297 | 4320 | 1 ; | N - O |
| 4297 | 4321 | 1 ; | N - N |
| 4298 | 4300 | 1 ; | H - HA |
| 4298 | 4301 | 1 ; | H - CB |
| 4298 | 4319 | 1 ; | H - C |
| 4299 | 4305 | 1 ; | CA - CD1 |
| 4299 | 4318 | 1 ; | CA - CD2 |
| 4299 | 4322 | 1 ; | CA - H |
| 4299 | 4323 | 1 ; | CA - CA |
| 4300 | 4302 | 1 ; | HA - HB2 |
| 4300 | 4303 | 1 ; | HA - HB3 |
| 4300 | 4304 | 1 ; | HA - CG |
| 4300 | 4320 | 1 ; | HA - O |
| 4300 | 4321 | 1 ; | HA - N |
| 4301 | 4306 | 1 ; | CB - HD1 |
| 4301 | 4307 | 1 ; | CB - NE1 |
| 4301 | 4309 | 1 ; | CB - CE2 |
| 4301 | 4316 | 1 ; | CB - CE3 |
| 4301 | 4320 | 1 ; | CB - O |
| 4301 | 4321 | 1 ; | CB - N |
| 4302 | 4305 | 1 ; | HB2 - CD1 |
| 4302 | 4318 | 1 ; | HB2 - CD2 |
| 4302 | 4319 | 1 ; | HB2 - C |
| 4303 | 4305 | 1 ; | HB3 - CD1 |
| 4303 | 4318 | 1 ; | HB3 - CD2 |
| 4303 | 4319 | 1 ; | HB3 - C |
| 4304 | 4308 | 1 ; | CG - HE1 |

```
4304   4310      1 ;     CG - CZ2
4304   4314      1 ;     CG - CZ3
4304   4317      1 ;     CG - HE3
4304   4319      1 ;     CG - C
4305   4310      1 ;    CD1 - CZ2
4305   4316      1 ;    CD1 - CE3
4306   4308      1 ;    HD1 - HE1
4306   4309      1 ;    HD1 - CE2
4306   4318      1 ;    HD1 - CD2
4307   4311      1 ;    NE1 - HZ2
4307   4312      1 ;    NE1 - CH2
4307   4316      1 ;    NE1 - CE3
4308   4310      1 ;    HE1 - CZ2
4308   4318      1 ;    HE1 - CD2
4309   4313      1 ;    CE2 - HH2
4309   4314      1 ;    CE2 - CZ3
4309   4317      1 ;    CE2 - HE3
4310   4315      1 ;    CZ2 - HZ3
4310   4316      1 ;    CZ2 - CE3
4311   4313      1 ;    HZ2 - HH2
4311   4314      1 ;    HZ2 - CZ3
4311   4318      1 ;    HZ2 - CD2
4312   4317      1 ;    CH2 - HE3
4312   4318      1 ;    CH2 - CD2
4313   4315      1 ;    HH2 - HZ3
4313   4316      1 ;    HH2 - CE3
4315   4317      1 ;    HZ3 - HE3
4315   4318      1 ;    HZ3 - CD2
4319   4324      1 ;      C - HA
4319   4325      1 ;      C - CB
4319   4338      1 ;      C - C
4320   4322      1 ;      O - H
4320   4323      1 ;      O - CA
4321   4326      1 ;      N - HB
4321   4327      1 ;      N - CG2
4321   4331      1 ;      N - CG1
4321   4339      1 ;      N - O
4321   4340      1 ;      N - N
4322   4324      1 ;      H - HA
4322   4325      1 ;      H - CB
4322   4338      1 ;      H - C
4323   4328      1 ;     CA - HG21
4323   4329      1 ;     CA - HG22
4323   4330      1 ;     CA - HG23
4323   4332      1 ;     CA - HG12
4323   4333      1 ;     CA - HG13
4323   4334      1 ;     CA - CD1
4323   4341      1 ;     CA - H
4323   4342      1 ;     CA - CA
4324   4326      1 ;     HA - HB
4324   4327      1 ;     HA - CG2
4324   4331      1 ;     HA - CG1
```

```
4324   4339      1 ;      HA - O
4324   4340      1 ;      HA - N
4325   4335      1 ;      CB - HD11
4325   4336      1 ;      CB - HD12
4325   4337      1 ;      CB - HD13
4325   4339      1 ;      CB - O
4325   4340      1 ;      CB - N
4326   4328      1 ;      HB - HG21
4326   4329      1 ;      HB - HG22
4326   4330      1 ;      HB - HG23
4326   4332      1 ;      HB - HG12
4326   4333      1 ;      HB - HG13
4326   4334      1 ;      HB - CD1
4326   4338      1 ;      HB - C
4327   4332      1 ;     CG2 - HG12
4327   4333      1 ;     CG2 - HG13
4327   4334      1 ;     CG2 - CD1
4327   4338      1 ;     CG2 - C
4328   4331      1 ;    HG21 - CG1
4329   4331      1 ;    HG22 - CG1
4330   4331      1 ;    HG23 - CG1
4331   4338      1 ;     CG1 - C
4332   4335      1 ;    HG12 - HD11
4332   4336      1 ;    HG12 - HD12
4332   4337      1 ;    HG12 - HD13
4333   4335      1 ;    HG13 - HD11
4333   4336      1 ;    HG13 - HD12
4333   4337      1 ;    HG13 - HD13
4338   4343      1 ;       C - HA
4338   4344      1 ;       C - CB
4338   4352      1 ;       C - C
4339   4341      1 ;       O - H
4339   4342      1 ;       O - CA
4340   4345      1 ;       N - HB
4340   4346      1 ;       N - CG2
4340   4350      1 ;       N - OG1
4340   4353      1 ;       N - O
4340   4354      1 ;       N - N
4341   4343      1 ;       H - HA
4341   4344      1 ;       H - CB
4341   4352      1 ;       H - C
4342   4347      1 ;      CA - HG21
4342   4348      1 ;      CA - HG22
4342   4349      1 ;      CA - HG23
4342   4351      1 ;      CA - HG1
4342   4355      1 ;      CA - H
4342   4356      1 ;      CA - CA
4343   4345      1 ;      HA - HB
4343   4346      1 ;      HA - CG2
4343   4350      1 ;      HA - OG1
4343   4353      1 ;      HA - O
4343   4354      1 ;      HA - N
```

```
4344   4353      1 ;     CB - O
4344   4354      1 ;     CB - N
4345   4347      1 ;     HB - HG21
4345   4348      1 ;     HB - HG22
4345   4349      1 ;     HB - HG23
4345   4351      1 ;     HB - HG1
4345   4352      1 ;     HB - C
4346   4351      1 ;    CG2 - HG1
4346   4352      1 ;    CG2 - C
4347   4350      1 ;   HG21 - OG1
4348   4350      1 ;   HG22 - OG1
4349   4350      1 ;   HG23 - OG1
4350   4352      1 ;    OG1 - C
4352   4357      1 ;      C - HA
4352   4358      1 ;      C - CB
4352   4369      1 ;      C - C
4353   4355      1 ;      O - H
4353   4356      1 ;      O - CA
4354   4359      1 ;      N - HB2
4354   4360      1 ;      N - HB3
4354   4361      1 ;      N - CG
4354   4370      1 ;      N - O
4354   4371      1 ;      N - N
4355   4357      1 ;      H - HA
4355   4358      1 ;      H - CB
4355   4369      1 ;      H - C
4356   4362      1 ;     CA - HG2
4356   4363      1 ;     CA - HG3
4356   4364      1 ;     CA - CD
4356   4372      1 ;     CA - H
4356   4373      1 ;     CA - CA
4357   4359      1 ;     HA - HB2
4357   4360      1 ;     HA - HB3
4357   4361      1 ;     HA - CG
4357   4370      1 ;     HA - O
4357   4371      1 ;     HA - N
4358   4365      1 ;     CB - OE1
4358   4366      1 ;     CB - NE2
4358   4370      1 ;     CB - O
4358   4371      1 ;     CB - N
4359   4362      1 ;    HB2 - HG2
4359   4363      1 ;    HB2 - HG3
4359   4364      1 ;    HB2 - CD
4359   4369      1 ;    HB2 - C
4360   4362      1 ;    HB3 - HG2
4360   4363      1 ;    HB3 - HG3
4360   4364      1 ;    HB3 - CD
4360   4369      1 ;    HB3 - C
4361   4367      1 ;     CG - HE21
4361   4368      1 ;     CG - HE22
4361   4369      1 ;     CG - C
4362   4365      1 ;    HG2 - OE1
```

```
4362   4366       1 ;     HG2 - NE2
4363   4365       1 ;     HG3 - OE1
4363   4366       1 ;     HG3 - NE2
4365   4367       1 ;     OE1 - HE21
4365   4368       1 ;     OE1 - HE22
4369   4374       1 ;       C - HA
4369   4375       1 ;       C - CB
4369   4386       1 ;       C - C
4370   4372       1 ;       O - H
4370   4373       1 ;       O - CA
4371   4376       1 ;       N - HB2
4371   4377       1 ;       N - HB3
4371   4378       1 ;       N - CG
4371   4387       1 ;       N - O
4371   4388       1 ;       N - N
4372   4374       1 ;       H - HA
4372   4375       1 ;       H - CB
4372   4386       1 ;       H - C
4373   4379       1 ;      CA - HG2
4373   4380       1 ;      CA - HG3
4373   4381       1 ;      CA - SD
4373   4389       1 ;      CA - H
4373   4390       1 ;      CA - CA
4374   4376       1 ;      HA - HB2
4374   4377       1 ;      HA - HB3
4374   4378       1 ;      HA - CG
4374   4387       1 ;      HA - O
4374   4388       1 ;      HA - N
4375   4382       1 ;      CB - CE
4375   4387       1 ;      CB - O
4375   4388       1 ;      CB - N
4376   4379       1 ;     HB2 - HG2
4376   4380       1 ;     HB2 - HG3
4376   4381       1 ;     HB2 - SD
4376   4386       1 ;     HB2 - C
4377   4379       1 ;     HB3 - HG2
4377   4380       1 ;     HB3 - HG3
4377   4381       1 ;     HB3 - SD
4377   4386       1 ;     HB3 - C
4378   4383       1 ;      CG - HE1
4378   4384       1 ;      CG - HE2
4378   4385       1 ;      CG - HE3
4378   4386       1 ;      CG - C
4379   4382       1 ;     HG2 - CE
4380   4382       1 ;     HG3 - CE
4386   4391       1 ;       C - HA
4386   4392       1 ;       C - CB
4386   4400       1 ;       C - C
4387   4389       1 ;       O - H
4387   4390       1 ;       O - CA
4388   4393       1 ;       N - HB
4388   4394       1 ;       N - CG2
```

| | | | | | |
|---|---|---|---|---|---|
| 4388 | 4398 | 1 ; | N | - | OG1 |
| 4388 | 4401 | 1 ; | N | - | O |
| 4388 | 4402 | 1 ; | N | - | N |
| 4389 | 4391 | 1 ; | H | - | HA |
| 4389 | 4392 | 1 ; | H | - | CB |
| 4389 | 4400 | 1 ; | H | - | C |
| 4390 | 4395 | 1 ; | CA | - | HG21 |
| 4390 | 4396 | 1 ; | CA | - | HG22 |
| 4390 | 4397 | 1 ; | CA | - | HG23 |
| 4390 | 4399 | 1 ; | CA | - | HG1 |
| 4390 | 4403 | 1 ; | CA | - | H |
| 4390 | 4404 | 1 ; | CA | - | CA |
| 4391 | 4393 | 1 ; | HA | - | HB |
| 4391 | 4394 | 1 ; | HA | - | CG2 |
| 4391 | 4398 | 1 ; | HA | - | OG1 |
| 4391 | 4401 | 1 ; | HA | - | O |
| 4391 | 4402 | 1 ; | HA | - | N |
| 4392 | 4401 | 1 ; | CB | - | O |
| 4392 | 4402 | 1 ; | CB | - | N |
| 4393 | 4395 | 1 ; | HB | - | HG21 |
| 4393 | 4396 | 1 ; | HB | - | HG22 |
| 4393 | 4397 | 1 ; | HB | - | HG23 |
| 4393 | 4399 | 1 ; | HB | - | HG1 |
| 4393 | 4400 | 1 ; | HB | - | C |
| 4394 | 4399 | 1 ; | CG2 | - | HG1 |
| 4394 | 4400 | 1 ; | CG2 | - | C |
| 4395 | 4398 | 1 ; | HG21 | - | OG1 |
| 4396 | 4398 | 1 ; | HG22 | - | OG1 |
| 4397 | 4398 | 1 ; | HG23 | - | OG1 |
| 4398 | 4400 | 1 ; | OG1 | - | C |
| 4400 | 4405 | 1 ; | C | - | HA |
| 4400 | 4406 | 1 ; | C | - | CB |
| 4400 | 4422 | 1 ; | C | - | C |
| 4401 | 4403 | 1 ; | O | - | H |
| 4401 | 4404 | 1 ; | O | - | CA |
| 4402 | 4407 | 1 ; | N | - | HB2 |
| 4402 | 4408 | 1 ; | N | - | HB3 |
| 4402 | 4409 | 1 ; | N | - | CG |
| 4402 | 4423 | 1 ; | N | - | O |
| 4402 | 4424 | 1 ; | N | - | N |
| 4403 | 4405 | 1 ; | H | - | HA |
| 4403 | 4406 | 1 ; | H | - | CB |
| 4403 | 4422 | 1 ; | H | - | C |
| 4404 | 4410 | 1 ; | CA | - | HG2 |
| 4404 | 4411 | 1 ; | CA | - | HG3 |
| 4404 | 4412 | 1 ; | CA | - | CD |
| 4404 | 4425 | 1 ; | CA | - | H |
| 4404 | 4426 | 1 ; | CA | - | CA |
| 4405 | 4407 | 1 ; | HA | - | HB2 |
| 4405 | 4408 | 1 ; | HA | - | HB3 |
| 4405 | 4409 | 1 ; | HA | - | CG |
| 4405 | 4423 | 1 ; | HA | - | O |

| | | | |
|---|---|---|---|
| 4405 | 4424 | 1 ; | HA - N |
| 4406 | 4413 | 1 ; | CB - HD2 |
| 4406 | 4414 | 1 ; | CB - HD3 |
| 4406 | 4415 | 1 ; | CB - CE |
| 4406 | 4423 | 1 ; | CB - O |
| 4406 | 4424 | 1 ; | CB - N |
| 4407 | 4410 | 1 ; | HB2 - HG2 |
| 4407 | 4411 | 1 ; | HB2 - HG3 |
| 4407 | 4412 | 1 ; | HB2 - CD |
| 4407 | 4422 | 1 ; | HB2 - C |
| 4408 | 4410 | 1 ; | HB3 - HG2 |
| 4408 | 4411 | 1 ; | HB3 - HG3 |
| 4408 | 4412 | 1 ; | HB3 - CD |
| 4408 | 4422 | 1 ; | HB3 - C |
| 4409 | 4416 | 1 ; | CG - HE2 |
| 4409 | 4417 | 1 ; | CG - HE3 |
| 4409 | 4418 | 1 ; | CG - NZ |
| 4409 | 4422 | 1 ; | CG - C |
| 4410 | 4413 | 1 ; | HG2 - HD2 |
| 4410 | 4414 | 1 ; | HG2 - HD3 |
| 4410 | 4415 | 1 ; | HG2 - CE |
| 4411 | 4413 | 1 ; | HG3 - HD2 |
| 4411 | 4414 | 1 ; | HG3 - HD3 |
| 4411 | 4415 | 1 ; | HG3 - CE |
| 4412 | 4419 | 1 ; | CD - HZ1 |
| 4412 | 4420 | 1 ; | CD - HZ2 |
| 4412 | 4421 | 1 ; | CD - HZ3 |
| 4413 | 4416 | 1 ; | HD2 - HE2 |
| 4413 | 4417 | 1 ; | HD2 - HE3 |
| 4413 | 4418 | 1 ; | HD2 - NZ |
| 4414 | 4416 | 1 ; | HD3 - HE2 |
| 4414 | 4417 | 1 ; | HD3 - HE3 |
| 4414 | 4418 | 1 ; | HD3 - NZ |
| 4416 | 4419 | 1 ; | HE2 - HZ1 |
| 4416 | 4420 | 1 ; | HE2 - HZ2 |
| 4416 | 4421 | 1 ; | HE2 - HZ3 |
| 4417 | 4419 | 1 ; | HE3 - HZ1 |
| 4417 | 4420 | 1 ; | HE3 - HZ2 |
| 4417 | 4421 | 1 ; | HE3 - HZ3 |
| 4422 | 4427 | 1 ; | C - HA |
| 4422 | 4428 | 1 ; | C - CB |
| 4422 | 4433 | 1 ; | C - C |
| 4423 | 4425 | 1 ; | O - H |
| 4423 | 4426 | 1 ; | O - CA |
| 4424 | 4429 | 1 ; | N - HB2 |
| 4424 | 4430 | 1 ; | N - HB3 |
| 4424 | 4431 | 1 ; | N - OG |
| 4424 | 4434 | 1 ; | N - O |
| 4424 | 4435 | 1 ; | N - N |
| 4425 | 4427 | 1 ; | H - HA |
| 4425 | 4428 | 1 ; | H - CB |
| 4425 | 4433 | 1 ; | H - C |

```
 4426   4432       1 ;      CA - HG
 4426   4436       1 ;      CA - H
 4426   4437       1 ;      CA - CA
 4427   4429       1 ;      HA - HB2
 4427   4430       1 ;      HA - HB3
 4427   4431       1 ;      HA - OG
 4427   4434       1 ;      HA - O
 4427   4435       1 ;      HA - N
 4428   4434       1 ;      CB - O
 4428   4435       1 ;      CB - N
 4429   4432       1 ;     HB2 - HG
 4429   4433       1 ;     HB2 - C
 4430   4432       1 ;     HB3 - HG
 4430   4433       1 ;     HB3 - C
 4431   4433       1 ;      OG - C
 4433   4438       1 ;       C - HA
 4433   4439       1 ;       C - CB
 4433   4450       1 ;       C - C
 4434   4436       1 ;       O - H
 4434   4437       1 ;       O - CA
 4435   4440       1 ;       N - HB2
 4435   4441       1 ;       N - HB3
 4435   4442       1 ;       N - CG
 4435   4451       1 ;       N - O
 4435   4452       1 ;       N - N
 4436   4438       1 ;       H - HA
 4436   4439       1 ;       H - CB
 4436   4450       1 ;       H - C
 4437   4443       1 ;      CA - ND1
 4437   4448       1 ;      CA - CD2
 4437   4453       1 ;      CA - H
 4437   4454       1 ;      CA - CA
 4438   4440       1 ;      HA - HB2
 4438   4441       1 ;      HA - HB3
 4438   4442       1 ;      HA - CG
 4438   4451       1 ;      HA - O
 4438   4452       1 ;      HA - N
 4439   4444       1 ;      CB - CE1
 4439   4446       1 ;      CB - NE2
 4439   4449       1 ;      CB - HD2
 4439   4451       1 ;      CB - O
 4439   4452       1 ;      CB - N
 4440   4443       1 ;     HB2 - ND1
 4440   4448       1 ;     HB2 - CD2
 4440   4450       1 ;     HB2 - C
 4441   4443       1 ;     HB3 - ND1
 4441   4448       1 ;     HB3 - CD2
 4441   4450       1 ;     HB3 - C
 4442   4445       1 ;      CG - HE1
 4442   4447       1 ;      CG - HE2
 4442   4450       1 ;      CG - C
 4443   4447       1 ;     ND1 - HE2
```

```
4443   4449      1 ;    ND1 - HD2
4444   4449      1 ;    CE1 - HD2
4445   4447      1 ;    HE1 - HE2
4445   4448      1 ;    HE1 - CD2
4447   4449      1 ;    HE2 - HD2
4450   4455      1 ;      C - HA
4450   4456      1 ;      C - CB
4450   4465      1 ;      C - C
4451   4453      1 ;      O - H
4451   4454      1 ;      O - CA
4452   4457      1 ;      N - HB2
4452   4458      1 ;      N - HB3
4452   4459      1 ;      N - CG
4452   4466      1 ;      N - O
4452   4467      1 ;      N - N
4453   4455      1 ;      H - HA
4453   4456      1 ;      H - CB
4453   4465      1 ;      H - C
4454   4460      1 ;     CA - HG2
4454   4461      1 ;     CA - HG3
4454   4462      1 ;     CA - CD
4454   4468      1 ;     CA - H
4454   4469      1 ;     CA - CA
4455   4457      1 ;     HA - HB2
4455   4458      1 ;     HA - HB3
4455   4459      1 ;     HA - CG
4455   4466      1 ;     HA - O
4455   4467      1 ;     HA - N
4456   4463      1 ;     CB - OE1
4456   4464      1 ;     CB - OE2
4456   4466      1 ;     CB - O
4456   4467      1 ;     CB - N
4457   4460      1 ;    HB2 - HG2
4457   4461      1 ;    HB2 - HG3
4457   4462      1 ;    HB2 - CD
4457   4465      1 ;    HB2 - C
4458   4460      1 ;    HB3 - HG2
4458   4461      1 ;    HB3 - HG3
4458   4462      1 ;    HB3 - CD
4458   4465      1 ;    HB3 - C
4459   4465      1 ;     CG - C
4460   4463      1 ;    HG2 - OE1
4460   4464      1 ;    HG2 - OE2
4461   4463      1 ;    HG3 - OE1
4461   4464      1 ;    HG3 - OE2
4465   4470      1 ;      C - HA
4465   4471      1 ;      C - CB
4465   4482      1 ;      C - C
4466   4468      1 ;      O - H
4466   4469      1 ;      O - CA
4467   4472      1 ;      N - HB2
4467   4473      1 ;      N - HB3
```

```
4467   4474       1 ;       N - CG
4467   4483       1 ;       N - O
4467   4484       1 ;       N - N
4468   4470       1 ;       H - HA
4468   4471       1 ;       H - CB
4468   4482       1 ;       H - C
4469   4475       1 ;      CA - ND1
4469   4480       1 ;      CA - CD2
4469   4485       1 ;      CA - H
4469   4486       1 ;      CA - CA
4470   4472       1 ;      HA - HB2
4470   4473       1 ;      HA - HB3
4470   4474       1 ;      HA - CG
4470   4483       1 ;      HA - O
4470   4484       1 ;      HA - N
4471   4476       1 ;      CB - CE1
4471   4478       1 ;      CB - NE2
4471   4481       1 ;      CB - HD2
4471   4483       1 ;      CB - O
4471   4484       1 ;      CB - N
4472   4475       1 ;     HB2 - ND1
4472   4480       1 ;     HB2 - CD2
4472   4482       1 ;     HB2 - C
4473   4475       1 ;     HB3 - ND1
4473   4480       1 ;     HB3 - CD2
4473   4482       1 ;     HB3 - C
4474   4477       1 ;      CG - HE1
4474   4479       1 ;      CG - HE2
4474   4482       1 ;      CG - C
4475   4479       1 ;     ND1 - HE2
4475   4481       1 ;     ND1 - HD2
4476   4481       1 ;     CE1 - HD2
4477   4479       1 ;     HE1 - HE2
4477   4480       1 ;     HE1 - CD2
4479   4481       1 ;     HE2 - HD2
4482   4487       1 ;       C - HA
4482   4488       1 ;       C - CB
4482   4498       1 ;       C - C
4483   4485       1 ;       O - H
4483   4486       1 ;       O - CA
4484   4489       1 ;       N - HB
4484   4490       1 ;       N - CG1
4484   4494       1 ;       N - CG2
4484   4499       1 ;       N - O
4484   4500       1 ;       N - N
4485   4487       1 ;       H - HA
4485   4488       1 ;       H - CB
4485   4498       1 ;       H - C
4486   4491       1 ;      CA - HG11
4486   4492       1 ;      CA - HG12
4486   4493       1 ;      CA - HG13
4486   4495       1 ;      CA - HG21
```

```
4486   4496       1 ;      CA - HG22
4486   4497       1 ;      CA - HG23
4486   4501       1 ;      CA - H
4486   4502       1 ;      CA - CA
4487   4489       1 ;      HA - HB
4487   4490       1 ;      HA - CG1
4487   4494       1 ;      HA - CG2
4487   4499       1 ;      HA - O
4487   4500       1 ;      HA - N
4488   4499       1 ;      CB - O
4488   4500       1 ;      CB - N
4489   4491       1 ;      HB - HG11
4489   4492       1 ;      HB - HG12
4489   4493       1 ;      HB - HG13
4489   4495       1 ;      HB - HG21
4489   4496       1 ;      HB - HG22
4489   4497       1 ;      HB - HG23
4489   4498       1 ;      HB - C
4490   4495       1 ;     CG1 - HG21
4490   4496       1 ;     CG1 - HG22
4490   4497       1 ;     CG1 - HG23
4490   4498       1 ;     CG1 - C
4491   4494       1 ;    HG11 - CG2
4492   4494       1 ;    HG12 - CG2
4493   4494       1 ;    HG13 - CG2
4494   4498       1 ;     CG2 - C
4498   4503       1 ;       C - HA
4498   4504       1 ;       C - CB
4498   4513       1 ;       C - C
4499   4501       1 ;       O - H
4499   4502       1 ;       O - CA
4500   4505       1 ;       N - HB2
4500   4506       1 ;       N - HB3
4500   4507       1 ;       N - CG
4500   4514       1 ;       N - O
4500   4515       1 ;       N - N
4501   4503       1 ;       H - HA
4501   4504       1 ;       H - CB
4501   4513       1 ;       H - C
4502   4508       1 ;      CA - HG2
4502   4509       1 ;      CA - HG3
4502   4510       1 ;      CA - CD
4502   4516       1 ;      CA - H
4502   4517       1 ;      CA - CA
4503   4505       1 ;      HA - HB2
4503   4506       1 ;      HA - HB3
4503   4507       1 ;      HA - CG
4503   4514       1 ;      HA - O
4503   4515       1 ;      HA - N
4504   4511       1 ;      CB - OE1
4504   4512       1 ;      CB - OE2
4504   4514       1 ;      CB - O
```

| | | | |
|---|---|---|---|
| 4504 | 4515 | 1 ; | CB - N |
| 4505 | 4508 | 1 ; | HB2 - HG2 |
| 4505 | 4509 | 1 ; | HB2 - HG3 |
| 4505 | 4510 | 1 ; | HB2 - CD |
| 4505 | 4513 | 1 ; | HB2 - C |
| 4506 | 4508 | 1 ; | HB3 - HG2 |
| 4506 | 4509 | 1 ; | HB3 - HG3 |
| 4506 | 4510 | 1 ; | HB3 - CD |
| 4506 | 4513 | 1 ; | HB3 - C |
| 4507 | 4513 | 1 ; | CG - C |
| 4508 | 4511 | 1 ; | HG2 - OE1 |
| 4508 | 4512 | 1 ; | HG2 - OE2 |
| 4509 | 4511 | 1 ; | HG3 - OE1 |
| 4509 | 4512 | 1 ; | HG3 - OE2 |
| 4513 | 4518 | 1 ; | C - HA |
| 4513 | 4519 | 1 ; | C - CB |
| 4513 | 4532 | 1 ; | C - C |
| 4514 | 4516 | 1 ; | O - H |
| 4514 | 4517 | 1 ; | O - CA |
| 4515 | 4520 | 1 ; | N - HB2 |
| 4515 | 4521 | 1 ; | N - HB3 |
| 4515 | 4522 | 1 ; | N - CG |
| 4515 | 4533 | 1 ; | N - O |
| 4515 | 4534 | 1 ; | N - N |
| 4516 | 4518 | 1 ; | H - HA |
| 4516 | 4519 | 1 ; | H - CB |
| 4516 | 4532 | 1 ; | H - C |
| 4517 | 4523 | 1 ; | CA - HG |
| 4517 | 4524 | 1 ; | CA - CD1 |
| 4517 | 4528 | 1 ; | CA - CD2 |
| 4517 | 4535 | 1 ; | CA - H |
| 4517 | 4536 | 1 ; | CA - CA |
| 4518 | 4520 | 1 ; | HA - HB2 |
| 4518 | 4521 | 1 ; | HA - HB3 |
| 4518 | 4522 | 1 ; | HA - CG |
| 4518 | 4533 | 1 ; | HA - O |
| 4518 | 4534 | 1 ; | HA - N |
| 4519 | 4525 | 1 ; | CB - HD11 |
| 4519 | 4526 | 1 ; | CB - HD12 |
| 4519 | 4527 | 1 ; | CB - HD13 |
| 4519 | 4529 | 1 ; | CB - HD21 |
| 4519 | 4530 | 1 ; | CB - HD22 |
| 4519 | 4531 | 1 ; | CB - HD23 |
| 4519 | 4533 | 1 ; | CB - O |
| 4519 | 4534 | 1 ; | CB - N |
| 4520 | 4523 | 1 ; | HB2 - HG |
| 4520 | 4524 | 1 ; | HB2 - CD1 |
| 4520 | 4528 | 1 ; | HB2 - CD2 |
| 4520 | 4532 | 1 ; | HB2 - C |
| 4521 | 4523 | 1 ; | HB3 - HG |
| 4521 | 4524 | 1 ; | HB3 - CD1 |
| 4521 | 4528 | 1 ; | HB3 - CD2 |

```
4521   4532       1 ;    HB3 - C
4522   4532       1 ;     CG - C
4523   4525       1 ;     HG - HD11
4523   4526       1 ;     HG - HD12
4523   4527       1 ;     HG - HD13
4523   4529       1 ;     HG - HD21
4523   4530       1 ;     HG - HD22
4523   4531       1 ;     HG - HD23
4524   4529       1 ;    CD1 - HD21
4524   4530       1 ;    CD1 - HD22
4524   4531       1 ;    CD1 - HD23
4525   4528       1 ;   HD11 - CD2
4526   4528       1 ;   HD12 - CD2
4527   4528       1 ;   HD13 - CD2
4532   4537       1 ;      C - HA
4532   4538       1 ;      C - CB
4532   4543       1 ;      C - C
4533   4535       1 ;      O - H
4533   4536       1 ;      O - CA
4534   4539       1 ;      N - HB2
4534   4540       1 ;      N - HB3
4534   4541       1 ;      N - OG
4534   4544       1 ;      N - O
4534   4545       1 ;      N - N
4535   4537       1 ;      H - HA
4535   4538       1 ;      H - CB
4535   4543       1 ;      H - C
4536   4542       1 ;     CA - HG
4536   4546       1 ;     CA - H
4536   4547       1 ;     CA - CA
4537   4539       1 ;     HA - HB2
4537   4540       1 ;     HA - HB3
4537   4541       1 ;     HA - OG
4537   4544       1 ;     HA - O
4537   4545       1 ;     HA - N
4538   4544       1 ;     CB - O
4538   4545       1 ;     CB - N
4539   4542       1 ;    HB2 - HG
4539   4543       1 ;    HB2 - C
4540   4542       1 ;    HB3 - HG
4540   4543       1 ;    HB3 - C
4541   4543       1 ;     OG - C
4543   4548       1 ;      C - HA
4543   4549       1 ;      C - CB
4543   4554       1 ;      C - C
4544   4546       1 ;      O - H
4544   4547       1 ;      O - CA
4545   4550       1 ;      N - HB2
4545   4551       1 ;      N - HB3
4545   4552       1 ;      N - OG
4545   4555       1 ;      N - O
4545   4556       1 ;      N - N
```

```
4546   4548      1 ;      H - HA
4546   4549      1 ;      H - CB
4546   4554      1 ;      H - C
4547   4553      1 ;     CA - HG
4547   4557      1 ;     CA - H
4547   4558      1 ;     CA - CA
4548   4550      1 ;     HA - HB2
4548   4551      1 ;     HA - HB3
4548   4552      1 ;     HA - OG
4548   4555      1 ;     HA - O
4548   4556      1 ;     HA - N
4549   4555      1 ;     CB - O
4549   4556      1 ;     CB - N
4550   4553      1 ;    HB2 - HG
4550   4554      1 ;    HB2 - C
4551   4553      1 ;    HB3 - HG
4551   4554      1 ;    HB3 - C
4552   4554      1 ;     OG - C
4554   4559      1 ;      C - HA
4554   4560      1 ;      C - CB
4554   4568      1 ;      C - C
4555   4557      1 ;      O - H
4555   4558      1 ;      O - CA
4556   4561      1 ;      N - HB
4556   4562      1 ;      N - CG2
4556   4566      1 ;      N - OG1
4556   4569      1 ;      N - O
4556   4570      1 ;      N - N
4557   4559      1 ;      H - HA
4557   4560      1 ;      H - CB
4557   4568      1 ;      H - C
4558   4563      1 ;     CA - HG21
4558   4564      1 ;     CA - HG22
4558   4565      1 ;     CA - HG23
4558   4567      1 ;     CA - HG1
4558   4571      1 ;     CA - H
4558   4572      1 ;     CA - CA
4559   4561      1 ;     HA - HB
4559   4562      1 ;     HA - CG2
4559   4566      1 ;     HA - OG1
4559   4569      1 ;     HA - O
4559   4570      1 ;     HA - N
4560   4569      1 ;     CB - O
4560   4570      1 ;     CB - N
4561   4563      1 ;     HB - HG21
4561   4564      1 ;     HB - HG22
4561   4565      1 ;     HB - HG23
4561   4567      1 ;     HB - HG1
4561   4568      1 ;     HB - C
4562   4567      1 ;    CG2 - HG1
4562   4568      1 ;    CG2 - C
4563   4566      1 ;   HG21 - OG1
```

| | | | | |
|---|---|---|---|---|
| 4564 | 4566 | 1 | ; | HG22 - OG1 |
| 4565 | 4566 | 1 | ; | HG23 - OG1 |
| 4566 | 4568 | 1 | ; | OG1 - C |
| 4568 | 4573 | 1 | ; | C - HA |
| 4568 | 4574 | 1 | ; | C - CB |
| 4568 | 4587 | 1 | ; | C - C |
| 4569 | 4571 | 1 | ; | O - H |
| 4569 | 4572 | 1 | ; | O - CA |
| 4570 | 4575 | 1 | ; | N - HB2 |
| 4570 | 4576 | 1 | ; | N - HB3 |
| 4570 | 4577 | 1 | ; | N - CG |
| 4570 | 4588 | 1 | ; | N - O |
| 4570 | 4589 | 1 | ; | N - N |
| 4571 | 4573 | 1 | ; | H - HA |
| 4571 | 4574 | 1 | ; | H - CB |
| 4571 | 4587 | 1 | ; | H - C |
| 4572 | 4578 | 1 | ; | CA - HG |
| 4572 | 4579 | 1 | ; | CA - CD1 |
| 4572 | 4583 | 1 | ; | CA - CD2 |
| 4572 | 4590 | 1 | ; | CA - H |
| 4572 | 4591 | 1 | ; | CA - CA |
| 4573 | 4575 | 1 | ; | HA - HB2 |
| 4573 | 4576 | 1 | ; | HA - HB3 |
| 4573 | 4577 | 1 | ; | HA - CG |
| 4573 | 4588 | 1 | ; | HA - O |
| 4573 | 4589 | 1 | ; | HA - N |
| 4574 | 4580 | 1 | ; | CB - HD11 |
| 4574 | 4581 | 1 | ; | CB - HD12 |
| 4574 | 4582 | 1 | ; | CB - HD13 |
| 4574 | 4584 | 1 | ; | CB - HD21 |
| 4574 | 4585 | 1 | ; | CB - HD22 |
| 4574 | 4586 | 1 | ; | CB - HD23 |
| 4574 | 4588 | 1 | ; | CB - O |
| 4574 | 4589 | 1 | ; | CB - N |
| 4575 | 4578 | 1 | ; | HB2 - HG |
| 4575 | 4579 | 1 | ; | HB2 - CD1 |
| 4575 | 4583 | 1 | ; | HB2 - CD2 |
| 4575 | 4587 | 1 | ; | HB2 - C |
| 4576 | 4578 | 1 | ; | HB3 - HG |
| 4576 | 4579 | 1 | ; | HB3 - CD1 |
| 4576 | 4583 | 1 | ; | HB3 - CD2 |
| 4576 | 4587 | 1 | ; | HB3 - C |
| 4577 | 4587 | 1 | ; | CG - C |
| 4578 | 4580 | 1 | ; | HG - HD11 |
| 4578 | 4581 | 1 | ; | HG - HD12 |
| 4578 | 4582 | 1 | ; | HG - HD13 |
| 4578 | 4584 | 1 | ; | HG - HD21 |
| 4578 | 4585 | 1 | ; | HG - HD22 |
| 4578 | 4586 | 1 | ; | HG - HD23 |
| 4579 | 4584 | 1 | ; | CD1 - HD21 |
| 4579 | 4585 | 1 | ; | CD1 - HD22 |
| 4579 | 4586 | 1 | ; | CD1 - HD23 |

```
4580   4583      1 ;   HD11 - CD2
4581   4583      1 ;   HD12 - CD2
4582   4583      1 ;   HD13 - CD2
4587   4592      1 ;      C - HA
4587   4593      1 ;      C - CB
4587   4606      1 ;      C - C
4588   4590      1 ;      O - H
4588   4591      1 ;      O - CA
4589   4594      1 ;      N - HB2
4589   4595      1 ;      N - HB3
4589   4596      1 ;      N - CG
4589   4607      1 ;      N - O
4589   4608      1 ;      N - N
4590   4592      1 ;      H - HA
4590   4593      1 ;      H - CB
4590   4606      1 ;      H - C
4591   4597      1 ;     CA - HG
4591   4598      1 ;     CA - CD1
4591   4602      1 ;     CA - CD2
4591   4609      1 ;     CA - H
4591   4610      1 ;     CA - CA
4592   4594      1 ;     HA - HB2
4592   4595      1 ;     HA - HB3
4592   4596      1 ;     HA - CG
4592   4607      1 ;     HA - O
4592   4608      1 ;     HA - N
4593   4599      1 ;     CB - HD11
4593   4600      1 ;     CB - HD12
4593   4601      1 ;     CB - HD13
4593   4603      1 ;     CB - HD21
4593   4604      1 ;     CB - HD22
4593   4605      1 ;     CB - HD23
4593   4607      1 ;     CB - O
4593   4608      1 ;     CB - N
4594   4597      1 ;    HB2 - HG
4594   4598      1 ;    HB2 - CD1
4594   4602      1 ;    HB2 - CD2
4594   4606      1 ;    HB2 - C
4595   4597      1 ;    HB3 - HG
4595   4598      1 ;    HB3 - CD1
4595   4602      1 ;    HB3 - CD2
4595   4606      1 ;    HB3 - C
4596   4606      1 ;     CG - C
4597   4599      1 ;     HG - HD11
4597   4600      1 ;     HG - HD12
4597   4601      1 ;     HG - HD13
4597   4603      1 ;     HG - HD21
4597   4604      1 ;     HG - HD22
4597   4605      1 ;     HG - HD23
4598   4603      1 ;    CD1 - HD21
4598   4604      1 ;    CD1 - HD22
4598   4605      1 ;    CD1 - HD23
```

```
4599   4602       1 ;   HD11 - CD2
4600   4602       1 ;   HD12 - CD2
4601   4602       1 ;   HD13 - CD2
4606   4611       1 ;      C - HA
4606   4612       1 ;      C - CB
4606   4628       1 ;      C - C
4607   4609       1 ;      O - H
4607   4610       1 ;      O - CA
4608   4613       1 ;      N - HB2
4608   4614       1 ;      N - HB3
4608   4615       1 ;      N - CG
4608   4629       1 ;      N - O
4608   4630       1 ;      N - N
4609   4611       1 ;      H - HA
4609   4612       1 ;      H - CB
4609   4628       1 ;      H - C
4610   4616       1 ;     CA - HG2
4610   4617       1 ;     CA - HG3
4610   4618       1 ;     CA - CD
4610   4631       1 ;     CA - H
4610   4632       1 ;     CA - CA
4611   4613       1 ;     HA - HB2
4611   4614       1 ;     HA - HB3
4611   4615       1 ;     HA - CG
4611   4629       1 ;     HA - O
4611   4630       1 ;     HA - N
4612   4619       1 ;     CB - HD2
4612   4620       1 ;     CB - HD3
4612   4621       1 ;     CB - CE
4612   4629       1 ;     CB - O
4612   4630       1 ;     CB - N
4613   4616       1 ;    HB2 - HG2
4613   4617       1 ;    HB2 - HG3
4613   4618       1 ;    HB2 - CD
4613   4628       1 ;    HB2 - C
4614   4616       1 ;    HB3 - HG2
4614   4617       1 ;    HB3 - HG3
4614   4618       1 ;    HB3 - CD
4614   4628       1 ;    HB3 - C
4615   4622       1 ;     CG - HE2
4615   4623       1 ;     CG - HE3
4615   4624       1 ;     CG - NZ
4615   4628       1 ;     CG - C
4616   4619       1 ;    HG2 - HD2
4616   4620       1 ;    HG2 - HD3
4616   4621       1 ;    HG2 - CE
4617   4619       1 ;    HG3 - HD2
4617   4620       1 ;    HG3 - HD3
4617   4621       1 ;    HG3 - CE
4618   4625       1 ;     CD - HZ1
4618   4626       1 ;     CD - HZ2
4618   4627       1 ;     CD - HZ3
```

```
4619   4622      1 ;    HD2 - HE2
4619   4623      1 ;    HD2 - HE3
4619   4624      1 ;    HD2 - NZ
4620   4622      1 ;    HD3 - HE2
4620   4623      1 ;    HD3 - HE3
4620   4624      1 ;    HD3 - NZ
4622   4625      1 ;    HE2 - HZ1
4622   4626      1 ;    HE2 - HZ2
4622   4627      1 ;    HE2 - HZ3
4623   4625      1 ;    HE3 - HZ1
4623   4626      1 ;    HE3 - HZ2
4623   4627      1 ;    HE3 - HZ3
4628   4633      1 ;      C - HA
4628   4634      1 ;      C - CB
4628   4642      1 ;      C - C
4629   4631      1 ;      O - H
4629   4632      1 ;      O - CA
4630   4635      1 ;      N - HB2
4630   4636      1 ;      N - HB3
4630   4637      1 ;      N - CG
4630   4643      1 ;      N - O
4630   4644      1 ;      N - N
4631   4633      1 ;      H - HA
4631   4634      1 ;      H - CB
4631   4642      1 ;      H - C
4632   4638      1 ;     CA - OD1
4632   4639      1 ;     CA - ND2
4632   4645      1 ;     CA - H
4632   4646      1 ;     CA - CA
4633   4635      1 ;     HA - HB2
4633   4636      1 ;     HA - HB3
4633   4637      1 ;     HA - CG
4633   4643      1 ;     HA - O
4633   4644      1 ;     HA - N
4634   4640      1 ;     CB - HD21
4634   4641      1 ;     CB - HD22
4634   4643      1 ;     CB - O
4634   4644      1 ;     CB - N
4635   4638      1 ;    HB2 - OD1
4635   4639      1 ;    HB2 - ND2
4635   4642      1 ;    HB2 - C
4636   4638      1 ;    HB3 - OD1
4636   4639      1 ;    HB3 - ND2
4636   4642      1 ;    HB3 - C
4637   4642      1 ;     CG - C
4638   4640      1 ;    OD1 - HD21
4638   4641      1 ;    OD1 - HD22
4642   4647      1 ;      C - HA
4642   4648      1 ;      C - CB
4642   4661      1 ;      C - C
4643   4645      1 ;      O - H
4643   4646      1 ;      O - CA
```

| | | | | | |
|---|---|---|---|---|---|
| 4644 | 4649 | 1 ; | N | - | HB2 |
| 4644 | 4650 | 1 ; | N | - | HB3 |
| 4644 | 4651 | 1 ; | N | - | CG |
| 4644 | 4662 | 1 ; | N | - | O |
| 4644 | 4663 | 1 ; | N | - | N |
| 4645 | 4647 | 1 ; | H | - | HA |
| 4645 | 4648 | 1 ; | H | - | CB |
| 4645 | 4661 | 1 ; | H | - | C |
| 4646 | 4652 | 1 ; | CA | - | HG |
| 4646 | 4653 | 1 ; | CA | - | CD1 |
| 4646 | 4657 | 1 ; | CA | - | CD2 |
| 4646 | 4664 | 1 ; | CA | - | H |
| 4646 | 4665 | 1 ; | CA | - | CA |
| 4647 | 4649 | 1 ; | HA | - | HB2 |
| 4647 | 4650 | 1 ; | HA | - | HB3 |
| 4647 | 4651 | 1 ; | HA | - | CG |
| 4647 | 4662 | 1 ; | HA | - | O |
| 4647 | 4663 | 1 ; | HA | - | N |
| 4648 | 4654 | 1 ; | CB | - | HD11 |
| 4648 | 4655 | 1 ; | CB | - | HD12 |
| 4648 | 4656 | 1 ; | CB | - | HD13 |
| 4648 | 4658 | 1 ; | CB | - | HD21 |
| 4648 | 4659 | 1 ; | CB | - | HD22 |
| 4648 | 4660 | 1 ; | CB | - | HD23 |
| 4648 | 4662 | 1 ; | CB | - | O |
| 4648 | 4663 | 1 ; | CB | - | N |
| 4649 | 4652 | 1 ; | HB2 | - | HG |
| 4649 | 4653 | 1 ; | HB2 | - | CD1 |
| 4649 | 4657 | 1 ; | HB2 | - | CD2 |
| 4649 | 4661 | 1 ; | HB2 | - | C |
| 4650 | 4652 | 1 ; | HB3 | - | HG |
| 4650 | 4653 | 1 ; | HB3 | - | CD1 |
| 4650 | 4657 | 1 ; | HB3 | - | CD2 |
| 4650 | 4661 | 1 ; | HB3 | - | C |
| 4651 | 4661 | 1 ; | CG | - | C |
| 4652 | 4654 | 1 ; | HG | - | HD11 |
| 4652 | 4655 | 1 ; | HG | - | HD12 |
| 4652 | 4656 | 1 ; | HG | - | HD13 |
| 4652 | 4658 | 1 ; | HG | - | HD21 |
| 4652 | 4659 | 1 ; | HG | - | HD22 |
| 4652 | 4660 | 1 ; | HG | - | HD23 |
| 4653 | 4658 | 1 ; | CD1 | - | HD21 |
| 4653 | 4659 | 1 ; | CD1 | - | HD22 |
| 4653 | 4660 | 1 ; | CD1 | - | HD23 |
| 4654 | 4657 | 1 ; | HD11 | - | CD2 |
| 4655 | 4657 | 1 ; | HD12 | - | CD2 |
| 4656 | 4657 | 1 ; | HD13 | - | CD2 |
| 4661 | 4666 | 1 ; | C | - | HA |
| 4661 | 4667 | 1 ; | C | - | CB |
| 4661 | 4683 | 1 ; | C | - | C |
| 4662 | 4664 | 1 ; | O | - | H |
| 4662 | 4665 | 1 ; | O | - | CA |

```
4663   4668       1 ;      N - HB2
4663   4669       1 ;      N - HB3
4663   4670       1 ;      N - CG
4663   4684       1 ;      N - O
4663   4685       1 ;      N - N
4664   4666       1 ;      H - HA
4664   4667       1 ;      H - CB
4664   4683       1 ;      H - C
4665   4671       1 ;     CA - HG2
4665   4672       1 ;     CA - HG3
4665   4673       1 ;     CA - CD
4665   4686       1 ;     CA - H
4665   4687       1 ;     CA - CA
4666   4668       1 ;     HA - HB2
4666   4669       1 ;     HA - HB3
4666   4670       1 ;     HA - CG
4666   4684       1 ;     HA - O
4666   4685       1 ;     HA - N
4667   4674       1 ;     CB - HD2
4667   4675       1 ;     CB - HD3
4667   4676       1 ;     CB - CE
4667   4684       1 ;     CB - O
4667   4685       1 ;     CB - N
4668   4671       1 ;    HB2 - HG2
4668   4672       1 ;    HB2 - HG3
4668   4673       1 ;    HB2 - CD
4668   4683       1 ;    HB2 - C
4669   4671       1 ;    HB3 - HG2
4669   4672       1 ;    HB3 - HG3
4669   4673       1 ;    HB3 - CD
4669   4683       1 ;    HB3 - C
4670   4677       1 ;     CG - HE2
4670   4678       1 ;     CG - HE3
4670   4679       1 ;     CG - NZ
4670   4683       1 ;     CG - C
4671   4674       1 ;    HG2 - HD2
4671   4675       1 ;    HG2 - HD3
4671   4676       1 ;    HG2 - CE
4672   4674       1 ;    HG3 - HD2
4672   4675       1 ;    HG3 - HD3
4672   4676       1 ;    HG3 - CE
4673   4680       1 ;     CD - HZ1
4673   4681       1 ;     CD - HZ2
4673   4682       1 ;     CD - HZ3
4674   4677       1 ;    HD2 - HE2
4674   4678       1 ;    HD2 - HE3
4674   4679       1 ;    HD2 - NZ
4675   4677       1 ;    HD3 - HE2
4675   4678       1 ;    HD3 - HE3
4675   4679       1 ;    HD3 - NZ
4677   4680       1 ;    HE2 - HZ1
4677   4681       1 ;    HE2 - HZ2
```

```
4677   4682       1 ;    HE2 - HZ3
4678   4680       1 ;    HE3 - HZ1
4678   4681       1 ;    HE3 - HZ2
4678   4682       1 ;    HE3 - HZ3
4683   4688       1 ;      C - HA
4683   4689       1 ;      C - CB
4683   4697       1 ;      C - C
4684   4686       1 ;      O - H
4684   4687       1 ;      O - CA
4685   4690       1 ;      N - HB2
4685   4691       1 ;      N - HB3
4685   4692       1 ;      N - CG
4685   4698       1 ;      N - O
4685   4699       1 ;      N - N
4686   4688       1 ;      H - HA
4686   4689       1 ;      H - CB
4686   4697       1 ;      H - C
4687   4693       1 ;     CA - OD1
4687   4694       1 ;     CA - ND2
4687   4700       1 ;     CA - H
4687   4701       1 ;     CA - CA
4688   4690       1 ;     HA - HB2
4688   4691       1 ;     HA - HB3
4688   4692       1 ;     HA - CG
4688   4698       1 ;     HA - O
4688   4699       1 ;     HA - N
4689   4695       1 ;     CB - HD21
4689   4696       1 ;     CB - HD22
4689   4698       1 ;     CB - O
4689   4699       1 ;     CB - N
4690   4693       1 ;    HB2 - OD1
4690   4694       1 ;    HB2 - ND2
4690   4697       1 ;    HB2 - C
4691   4693       1 ;    HB3 - OD1
4691   4694       1 ;    HB3 - ND2
4691   4697       1 ;    HB3 - C
4692   4697       1 ;     CG - C
4693   4695       1 ;    OD1 - HD21
4693   4696       1 ;    OD1 - HD22
4697   4702       1 ;      C - HA
4697   4703       1 ;      C - CB
4697   4717       1 ;      C - C
4698   4700       1 ;      O - H
4698   4701       1 ;      O - CA
4699   4704       1 ;      N - HB2
4699   4705       1 ;      N - HB3
4699   4706       1 ;      N - CG
4699   4718       1 ;      N - O
4699   4719       1 ;      N - N
4700   4702       1 ;      H - HA
4700   4703       1 ;      H - CB
4700   4717       1 ;      H - C
```

```
4701   4707       1 ;      CA - CD1
4701   4715       1 ;      CA - CD2
4701   4720       1 ;      CA - H
4701   4721       1 ;      CA - CA
4702   4704       1 ;      HA - HB2
4702   4705       1 ;      HA - HB3
4702   4706       1 ;      HA - CG
4702   4718       1 ;      HA - O
4702   4719       1 ;      HA - N
4703   4708       1 ;      CB - HD1
4703   4709       1 ;      CB - CE1
4703   4713       1 ;      CB - CE2
4703   4716       1 ;      CB - HD2
4703   4718       1 ;      CB - O
4703   4719       1 ;      CB - N
4704   4707       1 ;     HB2 - CD1
4704   4715       1 ;     HB2 - CD2
4704   4717       1 ;     HB2 - C
4705   4707       1 ;     HB3 - CD1
4705   4715       1 ;     HB3 - CD2
4705   4717       1 ;     HB3 - C
4706   4710       1 ;      CG - HE1
4706   4711       1 ;      CG - CZ
4706   4714       1 ;      CG - HE2
4706   4717       1 ;      CG - C
4707   4712       1 ;     CD1 - HZ
4707   4713       1 ;     CD1 - CE2
4707   4716       1 ;     CD1 - HD2
4708   4710       1 ;     HD1 - HE1
4708   4711       1 ;     HD1 - CZ
4708   4715       1 ;     HD1 - CD2
4709   4714       1 ;     CE1 - HE2
4709   4715       1 ;     CE1 - CD2
4710   4712       1 ;     HE1 - HZ
4710   4713       1 ;     HE1 - CE2
4711   4716       1 ;      CZ - HD2
4712   4714       1 ;      HZ - HE2
4712   4715       1 ;      HZ - CD2
4714   4716       1 ;     HE2 - HD2
4717   4722       1 ;       C - HA
4717   4723       1 ;       C - CB
4717   4739       1 ;       C - C
4718   4720       1 ;       O - H
4718   4721       1 ;       O - CA
4719   4724       1 ;       N - HB2
4719   4725       1 ;       N - HB3
4719   4726       1 ;       N - CG
4719   4740       1 ;       N - O
4719   4741       1 ;       N - N
4720   4722       1 ;       H - HA
4720   4723       1 ;       H - CB
4720   4739       1 ;       H - C
```

```
4721   4727       1 ;      CA - HG2
4721   4728       1 ;      CA - HG3
4721   4729       1 ;      CA - CD
4721   4742       1 ;      CA - H
4721   4743       1 ;      CA - CA
4722   4724       1 ;      HA - HB2
4722   4725       1 ;      HA - HB3
4722   4726       1 ;      HA - CG
4722   4740       1 ;      HA - O
4722   4741       1 ;      HA - N
4723   4730       1 ;      CB - HD2
4723   4731       1 ;      CB - HD3
4723   4732       1 ;      CB - CE
4723   4740       1 ;      CB - O
4723   4741       1 ;      CB - N
4724   4727       1 ;     HB2 - HG2
4724   4728       1 ;     HB2 - HG3
4724   4729       1 ;     HB2 - CD
4724   4739       1 ;     HB2 - C
4725   4727       1 ;     HB3 - HG2
4725   4728       1 ;     HB3 - HG3
4725   4729       1 ;     HB3 - CD
4725   4739       1 ;     HB3 - C
4726   4733       1 ;      CG - HE2
4726   4734       1 ;      CG - HE3
4726   4735       1 ;      CG - NZ
4726   4739       1 ;      CG - C
4727   4730       1 ;     HG2 - HD2
4727   4731       1 ;     HG2 - HD3
4727   4732       1 ;     HG2 - CE
4728   4730       1 ;     HG3 - HD2
4728   4731       1 ;     HG3 - HD3
4728   4732       1 ;     HG3 - CE
4729   4736       1 ;      CD - HZ1
4729   4737       1 ;      CD - HZ2
4729   4738       1 ;      CD - HZ3
4730   4733       1 ;     HD2 - HE2
4730   4734       1 ;     HD2 - HE3
4730   4735       1 ;     HD2 - NZ
4731   4733       1 ;     HD3 - HE2
4731   4734       1 ;     HD3 - HE3
4731   4735       1 ;     HD3 - NZ
4733   4736       1 ;     HE2 - HZ1
4733   4737       1 ;     HE2 - HZ2
4733   4738       1 ;     HE2 - HZ3
4734   4736       1 ;     HE3 - HZ1
4734   4737       1 ;     HE3 - HZ2
4734   4738       1 ;     HE3 - HZ3
4739   4744       1 ;       C - HA
4739   4745       1 ;       C - CB
4739   4761       1 ;       C - C
4740   4742       1 ;       O - H
```

| | | | | |
|---|---|---|---|---|
| 4740 | 4743 | 1 ; | O - | CA |
| 4741 | 4746 | 1 ; | N - | HB2 |
| 4741 | 4747 | 1 ; | N - | HB3 |
| 4741 | 4748 | 1 ; | N - | CG |
| 4741 | 4762 | 1 ; | N - | O |
| 4741 | 4763 | 1 ; | N - | N |
| 4742 | 4744 | 1 ; | H - | HA |
| 4742 | 4745 | 1 ; | H - | CB |
| 4742 | 4761 | 1 ; | H - | C |
| 4743 | 4749 | 1 ; | CA - | HG2 |
| 4743 | 4750 | 1 ; | CA - | HG3 |
| 4743 | 4751 | 1 ; | CA - | CD |
| 4743 | 4764 | 1 ; | CA - | H |
| 4743 | 4765 | 1 ; | CA - | CA |
| 4744 | 4746 | 1 ; | HA - | HB2 |
| 4744 | 4747 | 1 ; | HA - | HB3 |
| 4744 | 4748 | 1 ; | HA - | CG |
| 4744 | 4762 | 1 ; | HA - | O |
| 4744 | 4763 | 1 ; | HA - | N |
| 4745 | 4752 | 1 ; | CB - | HD2 |
| 4745 | 4753 | 1 ; | CB - | HD3 |
| 4745 | 4754 | 1 ; | CB - | CE |
| 4745 | 4762 | 1 ; | CB - | O |
| 4745 | 4763 | 1 ; | CB - | N |
| 4746 | 4749 | 1 ; | HB2 - | HG2 |
| 4746 | 4750 | 1 ; | HB2 - | HG3 |
| 4746 | 4751 | 1 ; | HB2 - | CD |
| 4746 | 4761 | 1 ; | HB2 - | C |
| 4747 | 4749 | 1 ; | HB3 - | HG2 |
| 4747 | 4750 | 1 ; | HB3 - | HG3 |
| 4747 | 4751 | 1 ; | HB3 - | CD |
| 4747 | 4761 | 1 ; | HB3 - | C |
| 4748 | 4755 | 1 ; | CG - | HE2 |
| 4748 | 4756 | 1 ; | CG - | HE3 |
| 4748 | 4757 | 1 ; | CG - | NZ |
| 4748 | 4761 | 1 ; | CG - | C |
| 4749 | 4752 | 1 ; | HG2 - | HD2 |
| 4749 | 4753 | 1 ; | HG2 - | HD3 |
| 4749 | 4754 | 1 ; | HG2 - | CE |
| 4750 | 4752 | 1 ; | HG3 - | HD2 |
| 4750 | 4753 | 1 ; | HG3 - | HD3 |
| 4750 | 4754 | 1 ; | HG3 - | CE |
| 4751 | 4758 | 1 ; | CD - | HZ1 |
| 4751 | 4759 | 1 ; | CD - | HZ2 |
| 4751 | 4760 | 1 ; | CD - | HZ3 |
| 4752 | 4755 | 1 ; | HD2 - | HE2 |
| 4752 | 4756 | 1 ; | HD2 - | HE3 |
| 4752 | 4757 | 1 ; | HD2 - | NZ |
| 4753 | 4755 | 1 ; | HD3 - | HE2 |
| 4753 | 4756 | 1 ; | HD3 - | HE3 |
| 4753 | 4757 | 1 ; | HD3 - | NZ |
| 4755 | 4758 | 1 ; | HE2 - | HZ1 |

| | | | |
|---|---|---|---|
| 4755 | 4759 | 1 ; | HE2 - HZ2 |
| 4755 | 4760 | 1 ; | HE2 - HZ3 |
| 4756 | 4758 | 1 ; | HE3 - HZ1 |
| 4756 | 4759 | 1 ; | HE3 - HZ2 |
| 4756 | 4760 | 1 ; | HE3 - HZ3 |
| 4761 | 4766 | 1 ; | C - HA |
| 4761 | 4767 | 1 ; | C - CB |
| 4761 | 4776 | 1 ; | C - C |
| 4762 | 4764 | 1 ; | O - H |
| 4762 | 4765 | 1 ; | O - CA |
| 4763 | 4768 | 1 ; | N - HB2 |
| 4763 | 4769 | 1 ; | N - HB3 |
| 4763 | 4770 | 1 ; | N - CG |
| 4763 | 4777 | 1 ; | N - O |
| 4763 | 4778 | 1 ; | N - N |
| 4764 | 4766 | 1 ; | H - HA |
| 4764 | 4767 | 1 ; | H - CB |
| 4764 | 4776 | 1 ; | H - C |
| 4765 | 4771 | 1 ; | CA - HG2 |
| 4765 | 4772 | 1 ; | CA - HG3 |
| 4765 | 4773 | 1 ; | CA - CD |
| 4765 | 4779 | 1 ; | CA - H |
| 4765 | 4780 | 1 ; | CA - CA |
| 4766 | 4768 | 1 ; | HA - HB2 |
| 4766 | 4769 | 1 ; | HA - HB3 |
| 4766 | 4770 | 1 ; | HA - CG |
| 4766 | 4777 | 1 ; | HA - O |
| 4766 | 4778 | 1 ; | HA - N |
| 4767 | 4774 | 1 ; | CB - OE1 |
| 4767 | 4775 | 1 ; | CB - OE2 |
| 4767 | 4777 | 1 ; | CB - O |
| 4767 | 4778 | 1 ; | CB - N |
| 4768 | 4771 | 1 ; | HB2 - HG2 |
| 4768 | 4772 | 1 ; | HB2 - HG3 |
| 4768 | 4773 | 1 ; | HB2 - CD |
| 4768 | 4776 | 1 ; | HB2 - C |
| 4769 | 4771 | 1 ; | HB3 - HG2 |
| 4769 | 4772 | 1 ; | HB3 - HG3 |
| 4769 | 4773 | 1 ; | HB3 - CD |
| 4769 | 4776 | 1 ; | HB3 - C |
| 4770 | 4776 | 1 ; | CG - C |
| 4771 | 4774 | 1 ; | HG2 - OE1 |
| 4771 | 4775 | 1 ; | HG2 - OE2 |
| 4772 | 4774 | 1 ; | HG3 - OE1 |
| 4772 | 4775 | 1 ; | HG3 - OE2 |
| 4776 | 4781 | 1 ; | C - HA |
| 4776 | 4782 | 1 ; | C - CB |
| 4776 | 4790 | 1 ; | C - C |
| 4777 | 4779 | 1 ; | O - H |
| 4777 | 4780 | 1 ; | O - CA |
| 4778 | 4783 | 1 ; | N - HB2 |
| 4778 | 4784 | 1 ; | N - HB3 |

| | | | | | |
|---|---|---|---|---|---|
| 4778 | 4785 | 1 ; | N | - | CG |
| 4778 | 4791 | 1 ; | N | - | O |
| 4778 | 4792 | 1 ; | N | - | N |
| 4779 | 4781 | 1 ; | H | - | HA |
| 4779 | 4782 | 1 ; | H | - | CB |
| 4779 | 4790 | 1 ; | H | - | C |
| 4780 | 4786 | 1 ; | CA | - | OD1 |
| 4780 | 4787 | 1 ; | CA | - | ND2 |
| 4780 | 4793 | 1 ; | CA | - | H |
| 4780 | 4794 | 1 ; | CA | - | CA |
| 4781 | 4783 | 1 ; | HA | - | HB2 |
| 4781 | 4784 | 1 ; | HA | - | HB3 |
| 4781 | 4785 | 1 ; | HA | - | CG |
| 4781 | 4791 | 1 ; | HA | - | O |
| 4781 | 4792 | 1 ; | HA | - | N |
| 4782 | 4788 | 1 ; | CB | - | HD21 |
| 4782 | 4789 | 1 ; | CB | - | HD22 |
| 4782 | 4791 | 1 ; | CB | - | O |
| 4782 | 4792 | 1 ; | CB | - | N |
| 4783 | 4786 | 1 ; | HB2 | - | OD1 |
| 4783 | 4787 | 1 ; | HB2 | - | ND2 |
| 4783 | 4790 | 1 ; | HB2 | - | C |
| 4784 | 4786 | 1 ; | HB3 | - | OD1 |
| 4784 | 4787 | 1 ; | HB3 | - | ND2 |
| 4784 | 4790 | 1 ; | HB3 | - | C |
| 4785 | 4790 | 1 ; | CG | - | C |
| 4786 | 4788 | 1 ; | OD1 | - | HD21 |
| 4786 | 4789 | 1 ; | OD1 | - | HD22 |
| 4790 | 4795 | 1 ; | C | - | HA |
| 4790 | 4796 | 1 ; | C | - | CB |
| 4790 | 4805 | 1 ; | C | - | C |
| 4791 | 4793 | 1 ; | O | - | H |
| 4791 | 4794 | 1 ; | O | - | CA |
| 4792 | 4797 | 1 ; | N | - | HB2 |
| 4792 | 4798 | 1 ; | N | - | HB3 |
| 4792 | 4799 | 1 ; | N | - | CG |
| 4792 | 4806 | 1 ; | N | - | O |
| 4792 | 4807 | 1 ; | N | - | N |
| 4793 | 4795 | 1 ; | H | - | HA |
| 4793 | 4796 | 1 ; | H | - | CB |
| 4793 | 4805 | 1 ; | H | - | C |
| 4794 | 4800 | 1 ; | CA | - | HG2 |
| 4794 | 4801 | 1 ; | CA | - | HG3 |
| 4794 | 4802 | 1 ; | CA | - | CD |
| 4794 | 4808 | 1 ; | CA | - | H |
| 4794 | 4809 | 1 ; | CA | - | CA |
| 4795 | 4797 | 1 ; | HA | - | HB2 |
| 4795 | 4798 | 1 ; | HA | - | HB3 |
| 4795 | 4799 | 1 ; | HA | - | CG |
| 4795 | 4806 | 1 ; | HA | - | O |
| 4795 | 4807 | 1 ; | HA | - | N |
| 4796 | 4803 | 1 ; | CB | - | OE1 |

| | | | |
|---|---|---|---|
| 4796 | 4804 | 1 ; | CB - OE2 |
| 4796 | 4806 | 1 ; | CB - O |
| 4796 | 4807 | 1 ; | CB - N |
| 4797 | 4800 | 1 ; | HB2 - HG2 |
| 4797 | 4801 | 1 ; | HB2 - HG3 |
| 4797 | 4802 | 1 ; | HB2 - CD |
| 4797 | 4805 | 1 ; | HB2 - C |
| 4798 | 4800 | 1 ; | HB3 - HG2 |
| 4798 | 4801 | 1 ; | HB3 - HG3 |
| 4798 | 4802 | 1 ; | HB3 - CD |
| 4798 | 4805 | 1 ; | HB3 - C |
| 4799 | 4805 | 1 ; | CG - C |
| 4800 | 4803 | 1 ; | HG2 - OE1 |
| 4800 | 4804 | 1 ; | HG2 - OE2 |
| 4801 | 4803 | 1 ; | HG3 - OE1 |
| 4801 | 4804 | 1 ; | HG3 - OE2 |
| 4805 | 4810 | 1 ; | C - HA |
| 4805 | 4811 | 1 ; | C - CB |
| 4805 | 4824 | 1 ; | C - C |
| 4806 | 4808 | 1 ; | O - H |
| 4806 | 4809 | 1 ; | O - CA |
| 4807 | 4812 | 1 ; | N - HB2 |
| 4807 | 4813 | 1 ; | N - HB3 |
| 4807 | 4814 | 1 ; | N - CG |
| 4807 | 4825 | 1 ; | N - O |
| 4807 | 4826 | 1 ; | N - N |
| 4808 | 4810 | 1 ; | H - HA |
| 4808 | 4811 | 1 ; | H - CB |
| 4808 | 4824 | 1 ; | H - C |
| 4809 | 4815 | 1 ; | CA - HG |
| 4809 | 4816 | 1 ; | CA - CD1 |
| 4809 | 4820 | 1 ; | CA - CD2 |
| 4809 | 4827 | 1 ; | CA - H |
| 4809 | 4828 | 1 ; | CA - CA |
| 4810 | 4812 | 1 ; | HA - HB2 |
| 4810 | 4813 | 1 ; | HA - HB3 |
| 4810 | 4814 | 1 ; | HA - CG |
| 4810 | 4825 | 1 ; | HA - O |
| 4810 | 4826 | 1 ; | HA - N |
| 4811 | 4817 | 1 ; | CB - HD11 |
| 4811 | 4818 | 1 ; | CB - HD12 |
| 4811 | 4819 | 1 ; | CB - HD13 |
| 4811 | 4821 | 1 ; | CB - HD21 |
| 4811 | 4822 | 1 ; | CB - HD22 |
| 4811 | 4823 | 1 ; | CB - HD23 |
| 4811 | 4825 | 1 ; | CB - O |
| 4811 | 4826 | 1 ; | CB - N |
| 4812 | 4815 | 1 ; | HB2 - HG |
| 4812 | 4816 | 1 ; | HB2 - CD1 |
| 4812 | 4820 | 1 ; | HB2 - CD2 |
| 4812 | 4824 | 1 ; | HB2 - C |
| 4813 | 4815 | 1 ; | HB3 - HG |

| | | | | | |
|---|---|---|---|---|---|
| 4813 | 4816 | 1 ; | HB3 | - | CD1 |
| 4813 | 4820 | 1 ; | HB3 | - | CD2 |
| 4813 | 4824 | 1 ; | HB3 | - | C |
| 4814 | 4824 | 1 ; | CG | - | C |
| 4815 | 4817 | 1 ; | HG | - | HD11 |
| 4815 | 4818 | 1 ; | HG | - | HD12 |
| 4815 | 4819 | 1 ; | HG | - | HD13 |
| 4815 | 4821 | 1 ; | HG | - | HD21 |
| 4815 | 4822 | 1 ; | HG | - | HD22 |
| 4815 | 4823 | 1 ; | HG | - | HD23 |
| 4816 | 4821 | 1 ; | CD1 | - | HD21 |
| 4816 | 4822 | 1 ; | CD1 | - | HD22 |
| 4816 | 4823 | 1 ; | CD1 | - | HD23 |
| 4817 | 4820 | 1 ; | HD11 | - | CD2 |
| 4818 | 4820 | 1 ; | HD12 | - | CD2 |
| 4819 | 4820 | 1 ; | HD13 | - | CD2 |
| 4824 | 4829 | 1 ; | C | - | HA |
| 4824 | 4830 | 1 ; | C | - | CB |
| 4824 | 4846 | 1 ; | C | - | C |
| 4825 | 4827 | 1 ; | O | - | H |
| 4825 | 4828 | 1 ; | O | - | CA |
| 4826 | 4831 | 1 ; | N | - | HB2 |
| 4826 | 4832 | 1 ; | N | - | HB3 |
| 4826 | 4833 | 1 ; | N | - | CG |
| 4826 | 4847 | 1 ; | N | - | O |
| 4826 | 4848 | 1 ; | N | - | N |
| 4827 | 4829 | 1 ; | H | - | HA |
| 4827 | 4830 | 1 ; | H | - | CB |
| 4827 | 4846 | 1 ; | H | - | C |
| 4828 | 4834 | 1 ; | CA | - | HG2 |
| 4828 | 4835 | 1 ; | CA | - | HG3 |
| 4828 | 4836 | 1 ; | CA | - | CD |
| 4828 | 4849 | 1 ; | CA | - | H |
| 4828 | 4850 | 1 ; | CA | - | CA |
| 4829 | 4831 | 1 ; | HA | - | HB2 |
| 4829 | 4832 | 1 ; | HA | - | HB3 |
| 4829 | 4833 | 1 ; | HA | - | CG |
| 4829 | 4847 | 1 ; | HA | - | O |
| 4829 | 4848 | 1 ; | HA | - | N |
| 4830 | 4837 | 1 ; | CB | - | HD2 |
| 4830 | 4838 | 1 ; | CB | - | HD3 |
| 4830 | 4839 | 1 ; | CB | - | CE |
| 4830 | 4847 | 1 ; | CB | - | O |
| 4830 | 4848 | 1 ; | CB | - | N |
| 4831 | 4834 | 1 ; | HB2 | - | HG2 |
| 4831 | 4835 | 1 ; | HB2 | - | HG3 |
| 4831 | 4836 | 1 ; | HB2 | - | CD |
| 4831 | 4846 | 1 ; | HB2 | - | C |
| 4832 | 4834 | 1 ; | HB3 | - | HG2 |
| 4832 | 4835 | 1 ; | HB3 | - | HG3 |
| 4832 | 4836 | 1 ; | HB3 | - | CD |
| 4832 | 4846 | 1 ; | HB3 | - | C |

| | | | |
|---|---|---|---|
| 4833 | 4840 | 1 ; | CG - HE2 |
| 4833 | 4841 | 1 ; | CG - HE3 |
| 4833 | 4842 | 1 ; | CG - NZ |
| 4833 | 4846 | 1 ; | CG - C |
| 4834 | 4837 | 1 ; | HG2 - HD2 |
| 4834 | 4838 | 1 ; | HG2 - HD3 |
| 4834 | 4839 | 1 ; | HG2 - CE |
| 4835 | 4837 | 1 ; | HG3 - HD2 |
| 4835 | 4838 | 1 ; | HG3 - HD3 |
| 4835 | 4839 | 1 ; | HG3 - CE |
| 4836 | 4843 | 1 ; | CD - HZ1 |
| 4836 | 4844 | 1 ; | CD - HZ2 |
| 4836 | 4845 | 1 ; | CD - HZ3 |
| 4837 | 4840 | 1 ; | HD2 - HE2 |
| 4837 | 4841 | 1 ; | HD2 - HE3 |
| 4837 | 4842 | 1 ; | HD2 - NZ |
| 4838 | 4840 | 1 ; | HD3 - HE2 |
| 4838 | 4841 | 1 ; | HD3 - HE3 |
| 4838 | 4842 | 1 ; | HD3 - NZ |
| 4840 | 4843 | 1 ; | HE2 - HZ1 |
| 4840 | 4844 | 1 ; | HE2 - HZ2 |
| 4840 | 4845 | 1 ; | HE2 - HZ3 |
| 4841 | 4843 | 1 ; | HE3 - HZ1 |
| 4841 | 4844 | 1 ; | HE3 - HZ2 |
| 4841 | 4845 | 1 ; | HE3 - HZ3 |
| 4846 | 4851 | 1 ; | C - HA |
| 4846 | 4852 | 1 ; | C - CB |
| 4846 | 4868 | 1 ; | C - C |
| 4847 | 4849 | 1 ; | O - H |
| 4847 | 4850 | 1 ; | O - CA |
| 4848 | 4853 | 1 ; | N - HB2 |
| 4848 | 4854 | 1 ; | N - HB3 |
| 4848 | 4855 | 1 ; | N - CG |
| 4848 | 4869 | 1 ; | N - O |
| 4848 | 4870 | 1 ; | N - N |
| 4849 | 4851 | 1 ; | H - HA |
| 4849 | 4852 | 1 ; | H - CB |
| 4849 | 4868 | 1 ; | H - C |
| 4850 | 4856 | 1 ; | CA - HG2 |
| 4850 | 4857 | 1 ; | CA - HG3 |
| 4850 | 4858 | 1 ; | CA - CD |
| 4850 | 4871 | 1 ; | CA - H |
| 4850 | 4872 | 1 ; | CA - CA |
| 4851 | 4853 | 1 ; | HA - HB2 |
| 4851 | 4854 | 1 ; | HA - HB3 |
| 4851 | 4855 | 1 ; | HA - CG |
| 4851 | 4869 | 1 ; | HA - O |
| 4851 | 4870 | 1 ; | HA - N |
| 4852 | 4859 | 1 ; | CB - HD2 |
| 4852 | 4860 | 1 ; | CB - HD3 |
| 4852 | 4861 | 1 ; | CB - CE |
| 4852 | 4869 | 1 ; | CB - O |

| | | | | |
|---|---|---|---|---|
| 4852 | 4870 | 1 | ; | CB - N |
| 4853 | 4856 | 1 | ; | HB2 - HG2 |
| 4853 | 4857 | 1 | ; | HB2 - HG3 |
| 4853 | 4858 | 1 | ; | HB2 - CD |
| 4853 | 4868 | 1 | ; | HB2 - C |
| 4854 | 4856 | 1 | ; | HB3 - HG2 |
| 4854 | 4857 | 1 | ; | HB3 - HG3 |
| 4854 | 4858 | 1 | ; | HB3 - CD |
| 4854 | 4868 | 1 | ; | HB3 - C |
| 4855 | 4862 | 1 | ; | CG - HE2 |
| 4855 | 4863 | 1 | ; | CG - HE3 |
| 4855 | 4864 | 1 | ; | CG - NZ |
| 4855 | 4868 | 1 | ; | CG - C |
| 4856 | 4859 | 1 | ; | HG2 - HD2 |
| 4856 | 4860 | 1 | ; | HG2 - HD3 |
| 4856 | 4861 | 1 | ; | HG2 - CE |
| 4857 | 4859 | 1 | ; | HG3 - HD2 |
| 4857 | 4860 | 1 | ; | HG3 - HD3 |
| 4857 | 4861 | 1 | ; | HG3 - CE |
| 4858 | 4865 | 1 | ; | CD - HZ1 |
| 4858 | 4866 | 1 | ; | CD - HZ2 |
| 4858 | 4867 | 1 | ; | CD - HZ3 |
| 4859 | 4862 | 1 | ; | HD2 - HE2 |
| 4859 | 4863 | 1 | ; | HD2 - HE3 |
| 4859 | 4864 | 1 | ; | HD2 - NZ |
| 4860 | 4862 | 1 | ; | HD3 - HE2 |
| 4860 | 4863 | 1 | ; | HD3 - HE3 |
| 4860 | 4864 | 1 | ; | HD3 - NZ |
| 4862 | 4865 | 1 | ; | HE2 - HZ1 |
| 4862 | 4866 | 1 | ; | HE2 - HZ2 |
| 4862 | 4867 | 1 | ; | HE2 - HZ3 |
| 4863 | 4865 | 1 | ; | HE3 - HZ1 |
| 4863 | 4866 | 1 | ; | HE3 - HZ2 |
| 4863 | 4867 | 1 | ; | HE3 - HZ3 |
| 4868 | 4873 | 1 | ; | C - HA |
| 4868 | 4874 | 1 | ; | C - CB |
| 4868 | 4887 | 1 | ; | C - C |
| 4869 | 4871 | 1 | ; | O - H |
| 4869 | 4872 | 1 | ; | O - CA |
| 4870 | 4875 | 1 | ; | N - HB |
| 4870 | 4876 | 1 | ; | N - CG2 |
| 4870 | 4880 | 1 | ; | N - CG1 |
| 4870 | 4888 | 1 | ; | N - O |
| 4870 | 4889 | 1 | ; | N - N |
| 4871 | 4873 | 1 | ; | H - HA |
| 4871 | 4874 | 1 | ; | H - CB |
| 4871 | 4887 | 1 | ; | H - C |
| 4872 | 4877 | 1 | ; | CA - HG21 |
| 4872 | 4878 | 1 | ; | CA - HG22 |
| 4872 | 4879 | 1 | ; | CA - HG23 |
| 4872 | 4881 | 1 | ; | CA - HG12 |
| 4872 | 4882 | 1 | ; | CA - HG13 |

| | | | | |
|---|---|---|---|---|
| 4872 | 4883 | 1 | ; | CA - CD1 |
| 4872 | 4890 | 1 | ; | CA - H |
| 4872 | 4891 | 1 | ; | CA - CA |
| 4873 | 4875 | 1 | ; | HA - HB |
| 4873 | 4876 | 1 | ; | HA - CG2 |
| 4873 | 4880 | 1 | ; | HA - CG1 |
| 4873 | 4888 | 1 | ; | HA - O |
| 4873 | 4889 | 1 | ; | HA - N |
| 4874 | 4884 | 1 | ; | CB - HD11 |
| 4874 | 4885 | 1 | ; | CB - HD12 |
| 4874 | 4886 | 1 | ; | CB - HD13 |
| 4874 | 4888 | 1 | ; | CB - O |
| 4874 | 4889 | 1 | ; | CB - N |
| 4875 | 4877 | 1 | ; | HB - HG21 |
| 4875 | 4878 | 1 | ; | HB - HG22 |
| 4875 | 4879 | 1 | ; | HB - HG23 |
| 4875 | 4881 | 1 | ; | HB - HG12 |
| 4875 | 4882 | 1 | ; | HB - HG13 |
| 4875 | 4883 | 1 | ; | HB - CD1 |
| 4875 | 4887 | 1 | ; | HB - C |
| 4876 | 4881 | 1 | ; | CG2 - HG12 |
| 4876 | 4882 | 1 | ; | CG2 - HG13 |
| 4876 | 4883 | 1 | ; | CG2 - CD1 |
| 4876 | 4887 | 1 | ; | CG2 - C |
| 4877 | 4880 | 1 | ; | HG21 - CG1 |
| 4878 | 4880 | 1 | ; | HG22 - CG1 |
| 4879 | 4880 | 1 | ; | HG23 - CG1 |
| 4880 | 4887 | 1 | ; | CG1 - C |
| 4881 | 4884 | 1 | ; | HG12 - HD11 |
| 4881 | 4885 | 1 | ; | HG12 - HD12 |
| 4881 | 4886 | 1 | ; | HG12 - HD13 |
| 4882 | 4884 | 1 | ; | HG13 - HD11 |
| 4882 | 4885 | 1 | ; | HG13 - HD12 |
| 4882 | 4886 | 1 | ; | HG13 - HD13 |
| 4887 | 4892 | 1 | ; | C - HA |
| 4887 | 4893 | 1 | ; | C - CB |
| 4887 | 4897 | 1 | ; | C - C |
| 4888 | 4890 | 1 | ; | O - H |
| 4888 | 4891 | 1 | ; | O - CA |
| 4889 | 4894 | 1 | ; | N - HB1 |
| 4889 | 4895 | 1 | ; | N - HB2 |
| 4889 | 4896 | 1 | ; | N - HB3 |
| 4889 | 4898 | 1 | ; | N - O |
| 4889 | 4899 | 1 | ; | N - N |
| 4890 | 4892 | 1 | ; | H - HA |
| 4890 | 4893 | 1 | ; | H - CB |
| 4890 | 4897 | 1 | ; | H - C |
| 4891 | 4900 | 1 | ; | CA - H |
| 4891 | 4901 | 1 | ; | CA - CA |
| 4892 | 4894 | 1 | ; | HA - HB1 |
| 4892 | 4895 | 1 | ; | HA - HB2 |
| 4892 | 4896 | 1 | ; | HA - HB3 |

| | | | |
|---|---|---|---|
| 4892 | 4898 | 1 ; | HA - O |
| 4892 | 4899 | 1 ; | HA - N |
| 4893 | 4898 | 1 ; | CB - O |
| 4893 | 4899 | 1 ; | CB - N |
| 4894 | 4897 | 1 ; | HB1 - C |
| 4895 | 4897 | 1 ; | HB2 - C |
| 4896 | 4897 | 1 ; | HB3 - C |
| 4897 | 4902 | 1 ; | C - HA |
| 4897 | 4903 | 1 ; | C - CB |
| 4897 | 4916 | 1 ; | C - C |
| 4898 | 4900 | 1 ; | O - H |
| 4898 | 4901 | 1 ; | O - CA |
| 4899 | 4904 | 1 ; | N - HB2 |
| 4899 | 4905 | 1 ; | N - HB3 |
| 4899 | 4906 | 1 ; | N - CG |
| 4899 | 4917 | 1 ; | N - O |
| 4899 | 4918 | 1 ; | N - N |
| 4900 | 4902 | 1 ; | H - HA |
| 4900 | 4903 | 1 ; | H - CB |
| 4900 | 4916 | 1 ; | H - C |
| 4901 | 4907 | 1 ; | CA - HG |
| 4901 | 4908 | 1 ; | CA - CD1 |
| 4901 | 4912 | 1 ; | CA - CD2 |
| 4901 | 4919 | 1 ; | CA - H |
| 4901 | 4920 | 1 ; | CA - CA |
| 4902 | 4904 | 1 ; | HA - HB2 |
| 4902 | 4905 | 1 ; | HA - HB3 |
| 4902 | 4906 | 1 ; | HA - CG |
| 4902 | 4917 | 1 ; | HA - O |
| 4902 | 4918 | 1 ; | HA - N |
| 4903 | 4909 | 1 ; | CB - HD11 |
| 4903 | 4910 | 1 ; | CB - HD12 |
| 4903 | 4911 | 1 ; | CB - HD13 |
| 4903 | 4913 | 1 ; | CB - HD21 |
| 4903 | 4914 | 1 ; | CB - HD22 |
| 4903 | 4915 | 1 ; | CB - HD23 |
| 4903 | 4917 | 1 ; | CB - O |
| 4903 | 4918 | 1 ; | CB - N |
| 4904 | 4907 | 1 ; | HB2 - HG |
| 4904 | 4908 | 1 ; | HB2 - CD1 |
| 4904 | 4912 | 1 ; | HB2 - CD2 |
| 4904 | 4916 | 1 ; | HB2 - C |
| 4905 | 4907 | 1 ; | HB3 - HG |
| 4905 | 4908 | 1 ; | HB3 - CD1 |
| 4905 | 4912 | 1 ; | HB3 - CD2 |
| 4905 | 4916 | 1 ; | HB3 - C |
| 4906 | 4916 | 1 ; | CG - C |
| 4907 | 4909 | 1 ; | HG - HD11 |
| 4907 | 4910 | 1 ; | HG - HD12 |
| 4907 | 4911 | 1 ; | HG - HD13 |
| 4907 | 4913 | 1 ; | HG - HD21 |
| 4907 | 4914 | 1 ; | HG - HD22 |

```
4907   4915       1 ;      HG - HD23
4908   4913       1 ;     CD1 - HD21
4908   4914       1 ;     CD1 - HD22
4908   4915       1 ;     CD1 - HD23
4909   4912       1 ;    HD11 - CD2
4910   4912       1 ;    HD12 - CD2
4911   4912       1 ;    HD13 - CD2
4916   4921       1 ;       C - HA
4916   4922       1 ;       C - CB
4916   4930       1 ;       C - C
4917   4919       1 ;       O - H
4917   4920       1 ;       O - CA
4918   4923       1 ;       N - HB
4918   4924       1 ;       N - CG2
4918   4928       1 ;       N - OG1
4918   4931       1 ;       N - O
4918   4932       1 ;       N - N
4919   4921       1 ;       H - HA
4919   4922       1 ;       H - CB
4919   4930       1 ;       H - C
4920   4925       1 ;      CA - HG21
4920   4926       1 ;      CA - HG22
4920   4927       1 ;      CA - HG23
4920   4929       1 ;      CA - HG1
4920   4933       1 ;      CA - H
4920   4934       1 ;      CA - CA
4921   4923       1 ;      HA - HB
4921   4924       1 ;      HA - CG2
4921   4928       1 ;      HA - OG1
4921   4931       1 ;      HA - O
4921   4932       1 ;      HA - N
4922   4931       1 ;      CB - O
4922   4932       1 ;      CB - N
4923   4925       1 ;      HB - HG21
4923   4926       1 ;      HB - HG22
4923   4927       1 ;      HB - HG23
4923   4929       1 ;      HB - HG1
4923   4930       1 ;      HB - C
4924   4929       1 ;     CG2 - HG1
4924   4930       1 ;     CG2 - C
4925   4928       1 ;    HG21 - OG1
4926   4928       1 ;    HG22 - OG1
4927   4928       1 ;    HG23 - OG1
4928   4930       1 ;     OG1 - C
4930   4935       1 ;       C - HA
4930   4936       1 ;       C - CB
4930   4949       1 ;       C - C
4931   4933       1 ;       O - H
4931   4934       1 ;       O - CA
4932   4937       1 ;       N - HB
4932   4938       1 ;       N - CG2
4932   4942       1 ;       N - CG1
```

```
4932   4950      1 ;      N - O
4932   4951      1 ;      N - N
4933   4935      1 ;      H - HA
4933   4936      1 ;      H - CB
4933   4949      1 ;      H - C
4934   4939      1 ;     CA - HG21
4934   4940      1 ;     CA - HG22
4934   4941      1 ;     CA - HG23
4934   4943      1 ;     CA - HG12
4934   4944      1 ;     CA - HG13
4934   4945      1 ;     CA - CD1
4934   4952      1 ;     CA - H
4934   4953      1 ;     CA - CA
4935   4937      1 ;     HA - HB
4935   4938      1 ;     HA - CG2
4935   4942      1 ;     HA - CG1
4935   4950      1 ;     HA - O
4935   4951      1 ;     HA - N
4936   4946      1 ;     CB - HD11
4936   4947      1 ;     CB - HD12
4936   4948      1 ;     CB - HD13
4936   4950      1 ;     CB - O
4936   4951      1 ;     CB - N
4937   4939      1 ;     HB - HG21
4937   4940      1 ;     HB - HG22
4937   4941      1 ;     HB - HG23
4937   4943      1 ;     HB - HG12
4937   4944      1 ;     HB - HG13
4937   4945      1 ;     HB - CD1
4937   4949      1 ;     HB - C
4938   4943      1 ;    CG2 - HG12
4938   4944      1 ;    CG2 - HG13
4938   4945      1 ;    CG2 - CD1
4938   4949      1 ;    CG2 - C
4939   4942      1 ;   HG21 - CG1
4940   4942      1 ;   HG22 - CG1
4941   4942      1 ;   HG23 - CG1
4942   4949      1 ;    CG1 - C
4943   4946      1 ;   HG12 - HD11
4943   4947      1 ;   HG12 - HD12
4943   4948      1 ;   HG12 - HD13
4944   4946      1 ;   HG13 - HD11
4944   4947      1 ;   HG13 - HD12
4944   4948      1 ;   HG13 - HD13
4949   4954      1 ;      C - HA
4949   4955      1 ;      C - CB
4949   4968      1 ;      C - C
4950   4952      1 ;      O - H
4950   4953      1 ;      O - CA
4951   4956      1 ;      N - HB
4951   4957      1 ;      N - CG2
4951   4961      1 ;      N - CG1
```

| | | | | | | |
|---|---|---|---|---|---|---|
| 4951 | 4969 | 1 | ; | N | - | O |
| 4951 | 4970 | 1 | ; | N | - | N |
| 4952 | 4954 | 1 | ; | H | - | HA |
| 4952 | 4955 | 1 | ; | H | - | CB |
| 4952 | 4968 | 1 | ; | H | - | C |
| 4953 | 4958 | 1 | ; | CA | - | HG21 |
| 4953 | 4959 | 1 | ; | CA | - | HG22 |
| 4953 | 4960 | 1 | ; | CA | - | HG23 |
| 4953 | 4962 | 1 | ; | CA | - | HG12 |
| 4953 | 4963 | 1 | ; | CA | - | HG13 |
| 4953 | 4964 | 1 | ; | CA | - | CD1 |
| 4953 | 4971 | 1 | ; | CA | - | H |
| 4953 | 4972 | 1 | ; | CA | - | CA |
| 4954 | 4956 | 1 | ; | HA | - | HB |
| 4954 | 4957 | 1 | ; | HA | - | CG2 |
| 4954 | 4961 | 1 | ; | HA | - | CG1 |
| 4954 | 4969 | 1 | ; | HA | - | O |
| 4954 | 4970 | 1 | ; | HA | - | N |
| 4955 | 4965 | 1 | ; | CB | - | HD11 |
| 4955 | 4966 | 1 | ; | CB | - | HD12 |
| 4955 | 4967 | 1 | ; | CB | - | HD13 |
| 4955 | 4969 | 1 | ; | CB | - | O |
| 4955 | 4970 | 1 | ; | CB | - | N |
| 4956 | 4958 | 1 | ; | HB | - | HG21 |
| 4956 | 4959 | 1 | ; | HB | - | HG22 |
| 4956 | 4960 | 1 | ; | HB | - | HG23 |
| 4956 | 4962 | 1 | ; | HB | - | HG12 |
| 4956 | 4963 | 1 | ; | HB | - | HG13 |
| 4956 | 4964 | 1 | ; | HB | - | CD1 |
| 4956 | 4968 | 1 | ; | HB | - | C |
| 4957 | 4962 | 1 | ; | CG2 | - | HG12 |
| 4957 | 4963 | 1 | ; | CG2 | - | HG13 |
| 4957 | 4964 | 1 | ; | CG2 | - | CD1 |
| 4957 | 4968 | 1 | ; | CG2 | - | C |
| 4958 | 4961 | 1 | ; | HG21 | - | CG1 |
| 4959 | 4961 | 1 | ; | HG22 | - | CG1 |
| 4960 | 4961 | 1 | ; | HG23 | - | CG1 |
| 4961 | 4968 | 1 | ; | CG1 | - | C |
| 4962 | 4965 | 1 | ; | HG12 | - | HD11 |
| 4962 | 4966 | 1 | ; | HG12 | - | HD12 |
| 4962 | 4967 | 1 | ; | HG12 | - | HD13 |
| 4963 | 4965 | 1 | ; | HG13 | - | HD11 |
| 4963 | 4966 | 1 | ; | HG13 | - | HD12 |
| 4963 | 4967 | 1 | ; | HG13 | - | HD13 |
| 4968 | 4973 | 1 | ; | C | - | HA |
| 4968 | 4974 | 1 | ; | C | - | CB |
| 4968 | 4978 | 1 | ; | C | - | C |
| 4969 | 4971 | 1 | ; | O | - | H |
| 4969 | 4972 | 1 | ; | O | - | CA |
| 4970 | 4975 | 1 | ; | N | - | HB1 |
| 4970 | 4976 | 1 | ; | N | - | HB2 |
| 4970 | 4977 | 1 | ; | N | - | HB3 |

| | | | |
|---|---|---|---|
| 4970 | 4979 | 1 ; | N - O |
| 4970 | 4980 | 1 ; | N - N |
| 4971 | 4973 | 1 ; | H - HA |
| 4971 | 4974 | 1 ; | H - CB |
| 4971 | 4978 | 1 ; | H - C |
| 4972 | 4981 | 1 ; | CA - H |
| 4972 | 4982 | 1 ; | CA - CA |
| 4973 | 4975 | 1 ; | HA - HB1 |
| 4973 | 4976 | 1 ; | HA - HB2 |
| 4973 | 4977 | 1 ; | HA - HB3 |
| 4973 | 4979 | 1 ; | HA - O |
| 4973 | 4980 | 1 ; | HA - N |
| 4974 | 4979 | 1 ; | CB - O |
| 4974 | 4980 | 1 ; | CB - N |
| 4975 | 4978 | 1 ; | HB1 - C |
| 4976 | 4978 | 1 ; | HB2 - C |
| 4977 | 4978 | 1 ; | HB3 - C |
| 4978 | 4983 | 1 ; | C - HA |
| 4978 | 4984 | 1 ; | C - CB |
| 4978 | 5000 | 1 ; | C - C |
| 4979 | 4981 | 1 ; | O - H |
| 4979 | 4982 | 1 ; | O - CA |
| 4980 | 4985 | 1 ; | N - HB2 |
| 4980 | 4986 | 1 ; | N - HB3 |
| 4980 | 4987 | 1 ; | N - CG |
| 4980 | 5001 | 1 ; | N - O |
| 4980 | 5002 | 1 ; | N - N |
| 4981 | 4983 | 1 ; | H - HA |
| 4981 | 4984 | 1 ; | H - CB |
| 4981 | 5000 | 1 ; | H - C |
| 4982 | 4988 | 1 ; | CA - HG2 |
| 4982 | 4989 | 1 ; | CA - HG3 |
| 4982 | 4990 | 1 ; | CA - CD |
| 4982 | 5003 | 1 ; | CA - H |
| 4982 | 5004 | 1 ; | CA - CA |
| 4983 | 4985 | 1 ; | HA - HB2 |
| 4983 | 4986 | 1 ; | HA - HB3 |
| 4983 | 4987 | 1 ; | HA - CG |
| 4983 | 5001 | 1 ; | HA - O |
| 4983 | 5002 | 1 ; | HA - N |
| 4984 | 4991 | 1 ; | CB - HD2 |
| 4984 | 4992 | 1 ; | CB - HD3 |
| 4984 | 4993 | 1 ; | CB - CE |
| 4984 | 5001 | 1 ; | CB - O |
| 4984 | 5002 | 1 ; | CB - N |
| 4985 | 4988 | 1 ; | HB2 - HG2 |
| 4985 | 4989 | 1 ; | HB2 - HG3 |
| 4985 | 4990 | 1 ; | HB2 - CD |
| 4985 | 5000 | 1 ; | HB2 - C |
| 4986 | 4988 | 1 ; | HB3 - HG2 |
| 4986 | 4989 | 1 ; | HB3 - HG3 |
| 4986 | 4990 | 1 ; | HB3 - CD |

```
4986   5000       1 ;     HB3 - C
4987   4994       1 ;      CG - HE2
4987   4995       1 ;      CG - HE3
4987   4996       1 ;      CG - NZ
4987   5000       1 ;      CG - C
4988   4991       1 ;     HG2 - HD2
4988   4992       1 ;     HG2 - HD3
4988   4993       1 ;     HG2 - CE
4989   4991       1 ;     HG3 - HD2
4989   4992       1 ;     HG3 - HD3
4989   4993       1 ;     HG3 - CE
4990   4997       1 ;      CD - HZ1
4990   4998       1 ;      CD - HZ2
4990   4999       1 ;      CD - HZ3
4991   4994       1 ;     HD2 - HE2
4991   4995       1 ;     HD2 - HE3
4991   4996       1 ;     HD2 - NZ
4992   4994       1 ;     HD3 - HE2
4992   4995       1 ;     HD3 - HE3
4992   4996       1 ;     HD3 - NZ
4994   4997       1 ;     HE2 - HZ1
4994   4998       1 ;     HE2 - HZ2
4994   4999       1 ;     HE2 - HZ3
4995   4997       1 ;     HE3 - HZ1
4995   4998       1 ;     HE3 - HZ2
4995   4999       1 ;     HE3 - HZ3
5000   5005       1 ;       C - HA
5000   5006       1 ;       C - CB
5000   5018       1 ;       C - C
5001   5003       1 ;       O - H
5001   5004       1 ;       O - CA
5002   5007       1 ;       N - HB2
5002   5008       1 ;       N - HB3
5002   5009       1 ;       N - CG
5002   5019       1 ;       N - O
5002   5020       1 ;       N - N
5003   5005       1 ;       H - HA
5003   5006       1 ;       H - CB
5003   5018       1 ;       H - C
5004   5010       1 ;      CA - ND1
5004   5016       1 ;      CA - CD2
5004   5021       1 ;      CA - H
5004   5022       1 ;      CA - CA
5005   5007       1 ;      HA - HB2
5005   5008       1 ;      HA - HB3
5005   5009       1 ;      HA - CG
5005   5019       1 ;      HA - O
5005   5020       1 ;      HA - N
5006   5011       1 ;      CB - HD1
5006   5012       1 ;      CB - CE1
5006   5014       1 ;      CB - NE2
5006   5017       1 ;      CB - HD2
```

5006 5019 1 ; CB - O
5006 5020 1 ; CB - N
5007 5010 1 ; HB2 - ND1
5007 5016 1 ; HB2 - CD2
5007 5018 1 ; HB2 - C
5008 5010 1 ; HB3 - ND1
5008 5016 1 ; HB3 - CD2
5008 5018 1 ; HB3 - C
5009 5013 1 ; CG - HE1
5009 5015 1 ; CG - HE2
5009 5018 1 ; CG - C
5010 5015 1 ; ND1 - HE2
5010 5017 1 ; ND1 - HD2
5011 5013 1 ; HD1 - HE1
5011 5014 1 ; HD1 - NE2
5011 5016 1 ; HD1 - CD2
5012 5017 1 ; CE1 - HD2
5013 5015 1 ; HE1 - HE2
5013 5016 1 ; HE1 - CD2
5015 5017 1 ; HE2 - HD2
5018 5023 1 ; C - HA
5018 5024 1 ; C - CB
5018 5037 1 ; C - C
5019 5021 1 ; O - H
5019 5022 1 ; O - CA
5020 5025 1 ; N - HB2
5020 5026 1 ; N - HB3
5020 5027 1 ; N - CG
5020 5038 1 ; N - O
5020 5039 1 ; N - N
5021 5023 1 ; H - HA
5021 5024 1 ; H - CB
5021 5037 1 ; H - C
5022 5028 1 ; CA - HG
5022 5029 1 ; CA - CD1
5022 5033 1 ; CA - CD2
5022 5040 1 ; CA - H
5022 5041 1 ; CA - CA
5023 5025 1 ; HA - HB2
5023 5026 1 ; HA - HB3
5023 5027 1 ; HA - CG
5023 5038 1 ; HA - O
5023 5039 1 ; HA - N
5024 5030 1 ; CB - HD11
5024 5031 1 ; CB - HD12
5024 5032 1 ; CB - HD13
5024 5034 1 ; CB - HD21
5024 5035 1 ; CB - HD22
5024 5036 1 ; CB - HD23
5024 5038 1 ; CB - O
5024 5039 1 ; CB - N
5025 5028 1 ; HB2 - HG

```
5025   5029      1 ;    HB2 - CD1
5025   5033      1 ;    HB2 - CD2
5025   5037      1 ;    HB2 - C
5026   5028      1 ;    HB3 - HG
5026   5029      1 ;    HB3 - CD1
5026   5033      1 ;    HB3 - CD2
5026   5037      1 ;    HB3 - C
5027   5037      1 ;     CG - C
5028   5030      1 ;     HG - HD11
5028   5031      1 ;     HG - HD12
5028   5032      1 ;     HG - HD13
5028   5034      1 ;     HG - HD21
5028   5035      1 ;     HG - HD22
5028   5036      1 ;     HG - HD23
5029   5034      1 ;    CD1 - HD21
5029   5035      1 ;    CD1 - HD22
5029   5036      1 ;    CD1 - HD23
5030   5033      1 ;   HD11 - CD2
5031   5033      1 ;   HD12 - CD2
5032   5033      1 ;   HD13 - CD2
5037   5042      1 ;      C - HA
5037   5043      1 ;      C - CB
5037   5048      1 ;      C - C
5038   5040      1 ;      O - H
5038   5041      1 ;      O - CA
5039   5044      1 ;      N - HB2
5039   5045      1 ;      N - HB3
5039   5046      1 ;      N - SG
5039   5049      1 ;      N - O
5039   5050      1 ;      N - N
5040   5042      1 ;      H - HA
5040   5043      1 ;      H - CB
5040   5048      1 ;      H - C
5041   5047      1 ;     CA - HG
5041   5051      1 ;     CA - H
5041   5052      1 ;     CA - CA
5042   5044      1 ;     HA - HB2
5042   5045      1 ;     HA - HB3
5042   5046      1 ;     HA - SG
5042   5049      1 ;     HA - O
5042   5050      1 ;     HA - N
5043   5049      1 ;     CB - O
5043   5050      1 ;     CB - N
5044   5047      1 ;    HB2 - HG
5044   5048      1 ;    HB2 - C
5045   5047      1 ;    HB3 - HG
5045   5048      1 ;    HB3 - C
5046   5048      1 ;     SG - C
5048   5053      1 ;      C - HA
5048   5054      1 ;      C - CB
5048   5060      1 ;      C - C
5049   5051      1 ;      O - H
```

| | | | | |
|---|---|---|---|---|
| 5049 | 5052 | 1 ; | O - | CA |
| 5050 | 5055 | 1 ; | N - | HB2 |
| 5050 | 5056 | 1 ; | N - | HB3 |
| 5050 | 5057 | 1 ; | N - | CG |
| 5050 | 5061 | 1 ; | N - | O |
| 5050 | 5062 | 1 ; | N - | N |
| 5051 | 5053 | 1 ; | H - | HA |
| 5051 | 5054 | 1 ; | H - | CB |
| 5051 | 5060 | 1 ; | H - | C |
| 5052 | 5058 | 1 ; | CA - | OD1 |
| 5052 | 5059 | 1 ; | CA - | OD2 |
| 5052 | 5063 | 1 ; | CA - | H |
| 5052 | 5064 | 1 ; | CA - | CA |
| 5053 | 5055 | 1 ; | HA - | HB2 |
| 5053 | 5056 | 1 ; | HA - | HB3 |
| 5053 | 5057 | 1 ; | HA - | CG |
| 5053 | 5061 | 1 ; | HA - | O |
| 5053 | 5062 | 1 ; | HA - | N |
| 5054 | 5061 | 1 ; | CB - | O |
| 5054 | 5062 | 1 ; | CB - | N |
| 5055 | 5058 | 1 ; | HB2 - | OD1 |
| 5055 | 5059 | 1 ; | HB2 - | OD2 |
| 5055 | 5060 | 1 ; | HB2 - | C |
| 5056 | 5058 | 1 ; | HB3 - | OD1 |
| 5056 | 5059 | 1 ; | HB3 - | OD2 |
| 5056 | 5060 | 1 ; | HB3 - | C |
| 5057 | 5060 | 1 ; | CG - | C |
| 5060 | 5065 | 1 ; | C - | HA |
| 5060 | 5066 | 1 ; | C - | CB |
| 5060 | 5076 | 1 ; | C - | C |
| 5061 | 5063 | 1 ; | O - | H |
| 5061 | 5064 | 1 ; | O - | CA |
| 5062 | 5067 | 1 ; | N - | HB |
| 5062 | 5068 | 1 ; | N - | CG1 |
| 5062 | 5072 | 1 ; | N - | CG2 |
| 5062 | 5077 | 1 ; | N - | O |
| 5062 | 5078 | 1 ; | N - | N |
| 5063 | 5065 | 1 ; | H - | HA |
| 5063 | 5066 | 1 ; | H - | CB |
| 5063 | 5076 | 1 ; | H - | C |
| 5064 | 5069 | 1 ; | CA - | HG11 |
| 5064 | 5070 | 1 ; | CA - | HG12 |
| 5064 | 5071 | 1 ; | CA - | HG13 |
| 5064 | 5073 | 1 ; | CA - | HG21 |
| 5064 | 5074 | 1 ; | CA - | HG22 |
| 5064 | 5075 | 1 ; | CA - | HG23 |
| 5064 | 5079 | 1 ; | CA - | H |
| 5064 | 5080 | 1 ; | CA - | CA |
| 5065 | 5067 | 1 ; | HA - | HB |
| 5065 | 5068 | 1 ; | HA - | CG1 |
| 5065 | 5072 | 1 ; | HA - | CG2 |
| 5065 | 5077 | 1 ; | HA - | O |

| | | | | | | |
|---|---|---|---|---|---|---|
| 5065 | 5078 | 1 | ; | HA | - | N |
| 5066 | 5077 | 1 | ; | CB | - | O |
| 5066 | 5078 | 1 | ; | CB | - | N |
| 5067 | 5069 | 1 | ; | HB | - | HG11 |
| 5067 | 5070 | 1 | ; | HB | - | HG12 |
| 5067 | 5071 | 1 | ; | HB | - | HG13 |
| 5067 | 5073 | 1 | ; | HB | - | HG21 |
| 5067 | 5074 | 1 | ; | HB | - | HG22 |
| 5067 | 5075 | 1 | ; | HB | - | HG23 |
| 5067 | 5076 | 1 | ; | HB | - | C |
| 5068 | 5073 | 1 | ; | CG1 | - | HG21 |
| 5068 | 5074 | 1 | ; | CG1 | - | HG22 |
| 5068 | 5075 | 1 | ; | CG1 | - | HG23 |
| 5068 | 5076 | 1 | ; | CG1 | - | C |
| 5069 | 5072 | 1 | ; | HG11 | - | CG2 |
| 5070 | 5072 | 1 | ; | HG12 | - | CG2 |
| 5071 | 5072 | 1 | ; | HG13 | - | CG2 |
| 5072 | 5076 | 1 | ; | CG2 | - | C |
| 5076 | 5081 | 1 | ; | C | - | HA |
| 5076 | 5082 | 1 | ; | C | - | CB |
| 5076 | 5091 | 1 | ; | C | - | C |
| 5077 | 5079 | 1 | ; | O | - | H |
| 5077 | 5080 | 1 | ; | O | - | CA |
| 5078 | 5083 | 1 | ; | N | - | HB2 |
| 5078 | 5084 | 1 | ; | N | - | HB3 |
| 5078 | 5085 | 1 | ; | N | - | CG |
| 5078 | 5092 | 1 | ; | N | - | O |
| 5078 | 5093 | 1 | ; | N | - | N |
| 5079 | 5081 | 1 | ; | H | - | HA |
| 5079 | 5082 | 1 | ; | H | - | CB |
| 5079 | 5091 | 1 | ; | H | - | C |
| 5080 | 5086 | 1 | ; | CA | - | HG2 |
| 5080 | 5087 | 1 | ; | CA | - | HG3 |
| 5080 | 5088 | 1 | ; | CA | - | CD |
| 5080 | 5094 | 1 | ; | CA | - | H |
| 5080 | 5095 | 1 | ; | CA | - | CA |
| 5081 | 5083 | 1 | ; | HA | - | HB2 |
| 5081 | 5084 | 1 | ; | HA | - | HB3 |
| 5081 | 5085 | 1 | ; | HA | - | CG |
| 5081 | 5092 | 1 | ; | HA | - | O |
| 5081 | 5093 | 1 | ; | HA | - | N |
| 5082 | 5089 | 1 | ; | CB | - | OE1 |
| 5082 | 5090 | 1 | ; | CB | - | OE2 |
| 5082 | 5092 | 1 | ; | CB | - | O |
| 5082 | 5093 | 1 | ; | CB | - | N |
| 5083 | 5086 | 1 | ; | HB2 | - | HG2 |
| 5083 | 5087 | 1 | ; | HB2 | - | HG3 |
| 5083 | 5088 | 1 | ; | HB2 | - | CD |
| 5083 | 5091 | 1 | ; | HB2 | - | C |
| 5084 | 5086 | 1 | ; | HB3 | - | HG2 |
| 5084 | 5087 | 1 | ; | HB3 | - | HG3 |
| 5084 | 5088 | 1 | ; | HB3 | - | CD |

```
5084   5091       1 ;     HB3 - C
5085   5091       1 ;      CG - C
5086   5089       1 ;     HG2 - OE1
5086   5090       1 ;     HG2 - OE2
5087   5089       1 ;     HG3 - OE1
5087   5090       1 ;     HG3 - OE2
5091   5096       1 ;       C - HA
5091   5097       1 ;       C - CB
5091   5110       1 ;       C - C
5092   5094       1 ;       O - H
5092   5095       1 ;       O - CA
5093   5098       1 ;       N - HB
5093   5099       1 ;       N - CG2
5093   5103       1 ;       N - CG1
5093   5111       1 ;       N - O
5093   5112       1 ;       N - N
5094   5096       1 ;       H - HA
5094   5097       1 ;       H - CB
5094   5110       1 ;       H - C
5095   5100       1 ;      CA - HG21
5095   5101       1 ;      CA - HG22
5095   5102       1 ;      CA - HG23
5095   5104       1 ;      CA - HG12
5095   5105       1 ;      CA - HG13
5095   5106       1 ;      CA - CD1
5095   5113       1 ;      CA - H
5095   5114       1 ;      CA - CA
5096   5098       1 ;      HA - HB
5096   5099       1 ;      HA - CG2
5096   5103       1 ;      HA - CG1
5096   5111       1 ;      HA - O
5096   5112       1 ;      HA - N
5097   5107       1 ;      CB - HD11
5097   5108       1 ;      CB - HD12
5097   5109       1 ;      CB - HD13
5097   5111       1 ;      CB - O
5097   5112       1 ;      CB - N
5098   5100       1 ;      HB - HG21
5098   5101       1 ;      HB - HG22
5098   5102       1 ;      HB - HG23
5098   5104       1 ;      HB - HG12
5098   5105       1 ;      HB - HG13
5098   5106       1 ;      HB - CD1
5098   5110       1 ;      HB - C
5099   5104       1 ;     CG2 - HG12
5099   5105       1 ;     CG2 - HG13
5099   5106       1 ;     CG2 - CD1
5099   5110       1 ;     CG2 - C
5100   5103       1 ;    HG21 - CG1
5101   5103       1 ;    HG22 - CG1
5102   5103       1 ;    HG23 - CG1
5103   5110       1 ;     CG1 - C
```

```
5104   5107      1 ;   HG12 - HD11
5104   5108      1 ;   HG12 - HD12
5104   5109      1 ;   HG12 - HD13
5105   5107      1 ;   HG13 - HD11
5105   5108      1 ;   HG13 - HD12
5105   5109      1 ;   HG13 - HD13
5110   5115      1 ;      C - HA
5110   5116      1 ;      C - CB
5110   5124      1 ;      C - C
5111   5113      1 ;      O - H
5111   5114      1 ;      O - CA
5112   5117      1 ;      N - HB2
5112   5118      1 ;      N - HB3
5112   5119      1 ;      N - CG
5112   5125      1 ;      N - O
5112   5126      1 ;      N - N
5113   5115      1 ;      H - HA
5113   5116      1 ;      H - CB
5113   5124      1 ;      H - C
5114   5120      1 ;     CA - OD1
5114   5121      1 ;     CA - ND2
5114   5127      1 ;     CA - H
5114   5128      1 ;     CA - CA
5115   5117      1 ;     HA - HB2
5115   5118      1 ;     HA - HB3
5115   5119      1 ;     HA - CG
5115   5125      1 ;     HA - O
5115   5126      1 ;     HA - N
5116   5122      1 ;     CB - HD21
5116   5123      1 ;     CB - HD22
5116   5125      1 ;     CB - O
5116   5126      1 ;     CB - N
5117   5120      1 ;    HB2 - OD1
5117   5121      1 ;    HB2 - ND2
5117   5124      1 ;    HB2 - C
5118   5120      1 ;    HB3 - OD1
5118   5121      1 ;    HB3 - ND2
5118   5124      1 ;    HB3 - C
5119   5124      1 ;     CG - C
5120   5122      1 ;    OD1 - HD21
5120   5123      1 ;    OD1 - HD22
5124   5129      1 ;      C - HA
5124   5130      1 ;      C - CB
5124   5138      1 ;      C - C
5125   5127      1 ;      O - H
5125   5128      1 ;      O - CA
5126   5131      1 ;      N - HB2
5126   5132      1 ;      N - HB3
5126   5133      1 ;      N - CG
5126   5139      1 ;      N - O
5126   5140      1 ;      N - N
5127   5129      1 ;      H - HA
```

| | | | | | |
|---|---|---|---|---|---|
| 5127 | 5130 | 1 ; | H | - | CB |
| 5127 | 5138 | 1 ; | H | - | C |
| 5128 | 5134 | 1 ; | CA | - | OD1 |
| 5128 | 5135 | 1 ; | CA | - | ND2 |
| 5128 | 5141 | 1 ; | CA | - | H |
| 5128 | 5142 | 1 ; | CA | - | CA |
| 5129 | 5131 | 1 ; | HA | - | HB2 |
| 5129 | 5132 | 1 ; | HA | - | HB3 |
| 5129 | 5133 | 1 ; | HA | - | CG |
| 5129 | 5139 | 1 ; | HA | - | O |
| 5129 | 5140 | 1 ; | HA | - | N |
| 5130 | 5136 | 1 ; | CB | - | HD21 |
| 5130 | 5137 | 1 ; | CB | - | HD22 |
| 5130 | 5139 | 1 ; | CB | - | O |
| 5130 | 5140 | 1 ; | CB | - | N |
| 5131 | 5134 | 1 ; | HB2 | - | OD1 |
| 5131 | 5135 | 1 ; | HB2 | - | ND2 |
| 5131 | 5138 | 1 ; | HB2 | - | C |
| 5132 | 5134 | 1 ; | HB3 | - | OD1 |
| 5132 | 5135 | 1 ; | HB3 | - | ND2 |
| 5132 | 5138 | 1 ; | HB3 | - | C |
| 5133 | 5138 | 1 ; | CG | - | C |
| 5134 | 5136 | 1 ; | OD1 | - | HD21 |
| 5134 | 5137 | 1 ; | OD1 | - | HD22 |
| 5138 | 5143 | 1 ; | C | - | HA |
| 5138 | 5144 | 1 ; | C | - | CB |
| 5138 | 5157 | 1 ; | C | - | C |
| 5139 | 5141 | 1 ; | O | - | H |
| 5139 | 5142 | 1 ; | O | - | CA |
| 5140 | 5145 | 1 ; | N | - | HB2 |
| 5140 | 5146 | 1 ; | N | - | HB3 |
| 5140 | 5147 | 1 ; | N | - | CG |
| 5140 | 5158 | 1 ; | N | - | O |
| 5140 | 5159 | 1 ; | N | - | N |
| 5141 | 5143 | 1 ; | H | - | HA |
| 5141 | 5144 | 1 ; | H | - | CB |
| 5141 | 5157 | 1 ; | H | - | C |
| 5142 | 5148 | 1 ; | CA | - | HG |
| 5142 | 5149 | 1 ; | CA | - | CD1 |
| 5142 | 5153 | 1 ; | CA | - | CD2 |
| 5142 | 5160 | 1 ; | CA | - | H |
| 5142 | 5161 | 1 ; | CA | - | CA |
| 5143 | 5145 | 1 ; | HA | - | HB2 |
| 5143 | 5146 | 1 ; | HA | - | HB3 |
| 5143 | 5147 | 1 ; | HA | - | CG |
| 5143 | 5158 | 1 ; | HA | - | O |
| 5143 | 5159 | 1 ; | HA | - | N |
| 5144 | 5150 | 1 ; | CB | - | HD11 |
| 5144 | 5151 | 1 ; | CB | - | HD12 |
| 5144 | 5152 | 1 ; | CB | - | HD13 |
| 5144 | 5154 | 1 ; | CB | - | HD21 |
| 5144 | 5155 | 1 ; | CB | - | HD22 |

| | | | | | |
|---|---|---|---|---|---|
| 5144 | 5156 | 1 ; | CB | - | HD23 |
| 5144 | 5158 | 1 ; | CB | - | O |
| 5144 | 5159 | 1 ; | CB | - | N |
| 5145 | 5148 | 1 ; | HB2 | - | HG |
| 5145 | 5149 | 1 ; | HB2 | - | CD1 |
| 5145 | 5153 | 1 ; | HB2 | - | CD2 |
| 5145 | 5157 | 1 ; | HB2 | - | C |
| 5146 | 5148 | 1 ; | HB3 | - | HG |
| 5146 | 5149 | 1 ; | HB3 | - | CD1 |
| 5146 | 5153 | 1 ; | HB3 | - | CD2 |
| 5146 | 5157 | 1 ; | HB3 | - | C |
| 5147 | 5157 | 1 ; | CG | - | C |
| 5148 | 5150 | 1 ; | HG | - | HD11 |
| 5148 | 5151 | 1 ; | HG | - | HD12 |
| 5148 | 5152 | 1 ; | HG | - | HD13 |
| 5148 | 5154 | 1 ; | HG | - | HD21 |
| 5148 | 5155 | 1 ; | HG | - | HD22 |
| 5148 | 5156 | 1 ; | HG | - | HD23 |
| 5149 | 5154 | 1 ; | CD1 | - | HD21 |
| 5149 | 5155 | 1 ; | CD1 | - | HD22 |
| 5149 | 5156 | 1 ; | CD1 | - | HD23 |
| 5150 | 5153 | 1 ; | HD11 | - | CD2 |
| 5151 | 5153 | 1 ; | HD12 | - | CD2 |
| 5152 | 5153 | 1 ; | HD13 | - | CD2 |
| 5157 | 5162 | 1 ; | C | - | HA |
| 5157 | 5163 | 1 ; | C | - | CB |
| 5157 | 5181 | 1 ; | C | - | C |
| 5158 | 5160 | 1 ; | O | - | H |
| 5158 | 5161 | 1 ; | O | - | CA |
| 5159 | 5164 | 1 ; | N | - | HB2 |
| 5159 | 5165 | 1 ; | N | - | HB3 |
| 5159 | 5166 | 1 ; | N | - | CG |
| 5159 | 5182 | 1 ; | N | - | O |
| 5159 | 5183 | 1 ; | N | - | N |
| 5160 | 5162 | 1 ; | H | - | HA |
| 5160 | 5163 | 1 ; | H | - | CB |
| 5160 | 5181 | 1 ; | H | - | C |
| 5161 | 5167 | 1 ; | CA | - | HG2 |
| 5161 | 5168 | 1 ; | CA | - | HG3 |
| 5161 | 5169 | 1 ; | CA | - | CD |
| 5161 | 5184 | 1 ; | CA | - | H |
| 5161 | 5185 | 1 ; | CA | - | CA |
| 5162 | 5164 | 1 ; | HA | - | HB2 |
| 5162 | 5165 | 1 ; | HA | - | HB3 |
| 5162 | 5166 | 1 ; | HA | - | CG |
| 5162 | 5182 | 1 ; | HA | - | O |
| 5162 | 5183 | 1 ; | HA | - | N |
| 5163 | 5170 | 1 ; | CB | - | HD2 |
| 5163 | 5171 | 1 ; | CB | - | HD3 |
| 5163 | 5172 | 1 ; | CB | - | NE |
| 5163 | 5182 | 1 ; | CB | - | O |
| 5163 | 5183 | 1 ; | CB | - | N |

| | | | | | | |
|---|---|---|---|---|---|---|
| 5164 | 5167 | 1 | ; | HB2 | - | HG2 |
| 5164 | 5168 | 1 | ; | HB2 | - | HG3 |
| 5164 | 5169 | 1 | ; | HB2 | - | CD |
| 5164 | 5181 | 1 | ; | HB2 | - | C |
| 5165 | 5167 | 1 | ; | HB3 | - | HG2 |
| 5165 | 5168 | 1 | ; | HB3 | - | HG3 |
| 5165 | 5169 | 1 | ; | HB3 | - | CD |
| 5165 | 5181 | 1 | ; | HB3 | - | C |
| 5166 | 5173 | 1 | ; | CG | - | HE |
| 5166 | 5174 | 1 | ; | CG | - | CZ |
| 5166 | 5181 | 1 | ; | CG | - | C |
| 5167 | 5170 | 1 | ; | HG2 | - | HD2 |
| 5167 | 5171 | 1 | ; | HG2 | - | HD3 |
| 5167 | 5172 | 1 | ; | HG2 | - | NE |
| 5168 | 5170 | 1 | ; | HG3 | - | HD2 |
| 5168 | 5171 | 1 | ; | HG3 | - | HD3 |
| 5168 | 5172 | 1 | ; | HG3 | - | NE |
| 5169 | 5175 | 1 | ; | CD | - | NH1 |
| 5169 | 5178 | 1 | ; | CD | - | NH2 |
| 5170 | 5173 | 1 | ; | HD2 | - | HE |
| 5170 | 5174 | 1 | ; | HD2 | - | CZ |
| 5171 | 5173 | 1 | ; | HD3 | - | HE |
| 5171 | 5174 | 1 | ; | HD3 | - | CZ |
| 5172 | 5176 | 1 | ; | NE | - | HH11 |
| 5172 | 5177 | 1 | ; | NE | - | HH12 |
| 5172 | 5179 | 1 | ; | NE | - | HH21 |
| 5172 | 5180 | 1 | ; | NE | - | HH22 |
| 5173 | 5175 | 1 | ; | HE | - | NH1 |
| 5173 | 5178 | 1 | ; | HE | - | NH2 |
| 5175 | 5179 | 1 | ; | NH1 | - | HH21 |
| 5175 | 5180 | 1 | ; | NH1 | - | HH22 |
| 5176 | 5178 | 1 | ; | HH11 | - | NH2 |
| 5177 | 5178 | 1 | ; | HH12 | - | NH2 |
| 5181 | 5186 | 1 | ; | C | - | HA |
| 5181 | 5187 | 1 | ; | C | - | CB |
| 5181 | 5195 | 1 | ; | C | - | C |
| 5182 | 5184 | 1 | ; | O | - | H |
| 5182 | 5185 | 1 | ; | O | - | CA |
| 5183 | 5188 | 1 | ; | N | - | HB2 |
| 5183 | 5189 | 1 | ; | N | - | HB3 |
| 5183 | 5190 | 1 | ; | N | - | CG |
| 5183 | 5196 | 1 | ; | N | - | O |
| 5183 | 5197 | 1 | ; | N | - | N |
| 5184 | 5186 | 1 | ; | H | - | HA |
| 5184 | 5187 | 1 | ; | H | - | CB |
| 5184 | 5195 | 1 | ; | H | - | C |
| 5185 | 5191 | 1 | ; | CA | - | OD1 |
| 5185 | 5192 | 1 | ; | CA | - | ND2 |
| 5185 | 5198 | 1 | ; | CA | - | H |
| 5185 | 5199 | 1 | ; | CA | - | CA |
| 5186 | 5188 | 1 | ; | HA | - | HB2 |
| 5186 | 5189 | 1 | ; | HA | - | HB3 |

| | | | |
|---|---|---|---|
| 5186 | 5190 | 1 ; | HA - CG |
| 5186 | 5196 | 1 ; | HA - O |
| 5186 | 5197 | 1 ; | HA - N |
| 5187 | 5193 | 1 ; | CB - HD21 |
| 5187 | 5194 | 1 ; | CB - HD22 |
| 5187 | 5196 | 1 ; | CB - O |
| 5187 | 5197 | 1 ; | CB - N |
| 5188 | 5191 | 1 ; | HB2 - OD1 |
| 5188 | 5192 | 1 ; | HB2 - ND2 |
| 5188 | 5195 | 1 ; | HB2 - C |
| 5189 | 5191 | 1 ; | HB3 - OD1 |
| 5189 | 5192 | 1 ; | HB3 - ND2 |
| 5189 | 5195 | 1 ; | HB3 - C |
| 5190 | 5195 | 1 ; | CG - C |
| 5191 | 5193 | 1 ; | OD1 - HD21 |
| 5191 | 5194 | 1 ; | OD1 - HD22 |
| 5195 | 5200 | 1 ; | C - HA |
| 5195 | 5201 | 1 ; | C - CB |
| 5195 | 5214 | 1 ; | C - C |
| 5196 | 5198 | 1 ; | O - H |
| 5196 | 5199 | 1 ; | O - CA |
| 5197 | 5202 | 1 ; | N - HB |
| 5197 | 5203 | 1 ; | N - CG2 |
| 5197 | 5207 | 1 ; | N - CG1 |
| 5197 | 5215 | 1 ; | N - O |
| 5197 | 5216 | 1 ; | N - N |
| 5198 | 5200 | 1 ; | H - HA |
| 5198 | 5201 | 1 ; | H - CB |
| 5198 | 5214 | 1 ; | H - C |
| 5199 | 5204 | 1 ; | CA - HG21 |
| 5199 | 5205 | 1 ; | CA - HG22 |
| 5199 | 5206 | 1 ; | CA - HG23 |
| 5199 | 5208 | 1 ; | CA - HG12 |
| 5199 | 5209 | 1 ; | CA - HG13 |
| 5199 | 5210 | 1 ; | CA - CD1 |
| 5199 | 5217 | 1 ; | CA - H |
| 5199 | 5218 | 1 ; | CA - CA |
| 5200 | 5202 | 1 ; | HA - HB |
| 5200 | 5203 | 1 ; | HA - CG2 |
| 5200 | 5207 | 1 ; | HA - CG1 |
| 5200 | 5215 | 1 ; | HA - O |
| 5200 | 5216 | 1 ; | HA - N |
| 5201 | 5211 | 1 ; | CB - HD11 |
| 5201 | 5212 | 1 ; | CB - HD12 |
| 5201 | 5213 | 1 ; | CB - HD13 |
| 5201 | 5215 | 1 ; | CB - O |
| 5201 | 5216 | 1 ; | CB - N |
| 5202 | 5204 | 1 ; | HB - HG21 |
| 5202 | 5205 | 1 ; | HB - HG22 |
| 5202 | 5206 | 1 ; | HB - HG23 |
| 5202 | 5208 | 1 ; | HB - HG12 |
| 5202 | 5209 | 1 ; | HB - HG13 |

| | | | | | |
|---|---|---|---|---|---|
| 5202 | 5210 | 1 ; | HB | - | CD1 |
| 5202 | 5214 | 1 ; | HB | - | C |
| 5203 | 5208 | 1 ; | CG2 | - | HG12 |
| 5203 | 5209 | 1 ; | CG2 | - | HG13 |
| 5203 | 5210 | 1 ; | CG2 | - | CD1 |
| 5203 | 5214 | 1 ; | CG2 | - | C |
| 5204 | 5207 | 1 ; | HG21 | - | CG1 |
| 5205 | 5207 | 1 ; | HG22 | - | CG1 |
| 5206 | 5207 | 1 ; | HG23 | - | CG1 |
| 5207 | 5214 | 1 ; | CG1 | - | C |
| 5208 | 5211 | 1 ; | HG12 | - | HD11 |
| 5208 | 5212 | 1 ; | HG12 | - | HD12 |
| 5208 | 5213 | 1 ; | HG12 | - | HD13 |
| 5209 | 5211 | 1 ; | HG13 | - | HD11 |
| 5209 | 5212 | 1 ; | HG13 | - | HD12 |
| 5209 | 5213 | 1 ; | HG13 | - | HD13 |
| 5214 | 5219 | 1 ; | C | - | HA |
| 5214 | 5220 | 1 ; | C | - | CB |
| 5214 | 5234 | 1 ; | C | - | C |
| 5215 | 5217 | 1 ; | O | - | H |
| 5215 | 5218 | 1 ; | O | - | CA |
| 5216 | 5221 | 1 ; | N | - | HB2 |
| 5216 | 5222 | 1 ; | N | - | HB3 |
| 5216 | 5223 | 1 ; | N | - | CG |
| 5216 | 5235 | 1 ; | N | - | O |
| 5216 | 5236 | 1 ; | N | - | N |
| 5217 | 5219 | 1 ; | H | - | HA |
| 5217 | 5220 | 1 ; | H | - | CB |
| 5217 | 5234 | 1 ; | H | - | C |
| 5218 | 5224 | 1 ; | CA | - | CD1 |
| 5218 | 5232 | 1 ; | CA | - | CD2 |
| 5218 | 5237 | 1 ; | CA | - | H |
| 5218 | 5238 | 1 ; | CA | - | CA |
| 5219 | 5221 | 1 ; | HA | - | HB2 |
| 5219 | 5222 | 1 ; | HA | - | HB3 |
| 5219 | 5223 | 1 ; | HA | - | CG |
| 5219 | 5235 | 1 ; | HA | - | O |
| 5219 | 5236 | 1 ; | HA | - | N |
| 5220 | 5225 | 1 ; | CB | - | HD1 |
| 5220 | 5226 | 1 ; | CB | - | CE1 |
| 5220 | 5230 | 1 ; | CB | - | CE2 |
| 5220 | 5233 | 1 ; | CB | - | HD2 |
| 5220 | 5235 | 1 ; | CB | - | O |
| 5220 | 5236 | 1 ; | CB | - | N |
| 5221 | 5224 | 1 ; | HB2 | - | CD1 |
| 5221 | 5232 | 1 ; | HB2 | - | CD2 |
| 5221 | 5234 | 1 ; | HB2 | - | C |
| 5222 | 5224 | 1 ; | HB3 | - | CD1 |
| 5222 | 5232 | 1 ; | HB3 | - | CD2 |
| 5222 | 5234 | 1 ; | HB3 | - | C |
| 5223 | 5227 | 1 ; | CG | - | HE1 |
| 5223 | 5228 | 1 ; | CG | - | CZ |

```
5223   5231      1 ;     CG - HE2
5223   5234      1 ;     CG - C
5224   5229      1 ;    CD1 - HZ
5224   5230      1 ;    CD1 - CE2
5224   5233      1 ;    CD1 - HD2
5225   5227      1 ;    HD1 - HE1
5225   5228      1 ;    HD1 - CZ
5225   5232      1 ;    HD1 - CD2
5226   5231      1 ;    CE1 - HE2
5226   5232      1 ;    CE1 - CD2
5227   5229      1 ;    HE1 - HZ
5227   5230      1 ;    HE1 - CE2
5228   5233      1 ;     CZ - HD2
5229   5231      1 ;     HZ - HE2
5229   5232      1 ;     HZ - CD2
5231   5233      1 ;    HE2 - HD2
5234   5239      1 ;      C - HA
5234   5240      1 ;      C - CB
5234   5253      1 ;      C - C
5235   5237      1 ;      O - H
5235   5238      1 ;      O - CA
5236   5241      1 ;      N - HB
5236   5242      1 ;      N - CG2
5236   5246      1 ;      N - CG1
5236   5254      1 ;      N - O
5236   5255      1 ;      N - N
5237   5239      1 ;      H - HA
5237   5240      1 ;      H - CB
5237   5253      1 ;      H - C
5238   5243      1 ;     CA - HG21
5238   5244      1 ;     CA - HG22
5238   5245      1 ;     CA - HG23
5238   5247      1 ;     CA - HG12
5238   5248      1 ;     CA - HG13
5238   5249      1 ;     CA - CD1
5238   5256      1 ;     CA - H
5238   5257      1 ;     CA - CA
5239   5241      1 ;     HA - HB
5239   5242      1 ;     HA - CG2
5239   5246      1 ;     HA - CG1
5239   5254      1 ;     HA - O
5239   5255      1 ;     HA - N
5240   5250      1 ;     CB - HD11
5240   5251      1 ;     CB - HD12
5240   5252      1 ;     CB - HD13
5240   5254      1 ;     CB - O
5240   5255      1 ;     CB - N
5241   5243      1 ;     HB - HG21
5241   5244      1 ;     HB - HG22
5241   5245      1 ;     HB - HG23
5241   5247      1 ;     HB - HG12
5241   5248      1 ;     HB - HG13
```

| | | | | | | |
|---|---|---|---|---|---|---|
| 5241 | 5249 | 1 | ; | HB | - | CD1 |
| 5241 | 5253 | 1 | ; | HB | - | C |
| 5242 | 5247 | 1 | ; | CG2 | - | HG12 |
| 5242 | 5248 | 1 | ; | CG2 | - | HG13 |
| 5242 | 5249 | 1 | ; | CG2 | - | CD1 |
| 5242 | 5253 | 1 | ; | CG2 | - | C |
| 5243 | 5246 | 1 | ; | HG21 | - | CG1 |
| 5244 | 5246 | 1 | ; | HG22 | - | CG1 |
| 5245 | 5246 | 1 | ; | HG23 | - | CG1 |
| 5246 | 5253 | 1 | ; | CG1 | - | C |
| 5247 | 5250 | 1 | ; | HG12 | - | HD11 |
| 5247 | 5251 | 1 | ; | HG12 | - | HD12 |
| 5247 | 5252 | 1 | ; | HG12 | - | HD13 |
| 5248 | 5250 | 1 | ; | HG13 | - | HD11 |
| 5248 | 5251 | 1 | ; | HG13 | - | HD12 |
| 5248 | 5252 | 1 | ; | HG13 | - | HD13 |
| 5253 | 5258 | 1 | ; | C | - | HA |
| 5253 | 5259 | 1 | ; | C | - | CB |
| 5253 | 5263 | 1 | ; | C | - | C |
| 5254 | 5256 | 1 | ; | O | - | H |
| 5254 | 5257 | 1 | ; | O | - | CA |
| 5255 | 5260 | 1 | ; | N | - | HB1 |
| 5255 | 5261 | 1 | ; | N | - | HB2 |
| 5255 | 5262 | 1 | ; | N | - | HB3 |
| 5255 | 5264 | 1 | ; | N | - | O |
| 5255 | 5265 | 1 | ; | N | - | N |
| 5256 | 5258 | 1 | ; | H | - | HA |
| 5256 | 5259 | 1 | ; | H | - | CB |
| 5256 | 5263 | 1 | ; | H | - | C |
| 5257 | 5266 | 1 | ; | CA | - | H |
| 5257 | 5267 | 1 | ; | CA | - | CA |
| 5258 | 5260 | 1 | ; | HA | - | HB1 |
| 5258 | 5261 | 1 | ; | HA | - | HB2 |
| 5258 | 5262 | 1 | ; | HA | - | HB3 |
| 5258 | 5264 | 1 | ; | HA | - | O |
| 5258 | 5265 | 1 | ; | HA | - | N |
| 5259 | 5264 | 1 | ; | CB | - | O |
| 5259 | 5265 | 1 | ; | CB | - | N |
| 5260 | 5263 | 1 | ; | HB1 | - | C |
| 5261 | 5263 | 1 | ; | HB2 | - | C |
| 5262 | 5263 | 1 | ; | HB3 | - | C |
| 5263 | 5268 | 1 | ; | C | - | HA |
| 5263 | 5269 | 1 | ; | C | - | CB |
| 5263 | 5282 | 1 | ; | C | - | C |
| 5264 | 5266 | 1 | ; | O | - | H |
| 5264 | 5267 | 1 | ; | O | - | CA |
| 5265 | 5270 | 1 | ; | N | - | HB2 |
| 5265 | 5271 | 1 | ; | N | - | HB3 |
| 5265 | 5272 | 1 | ; | N | - | CG |
| 5265 | 5283 | 1 | ; | N | - | O |
| 5265 | 5284 | 1 | ; | N | - | N |
| 5266 | 5268 | 1 | ; | H | - | HA |

```
5266   5269       1 ;       H - CB
5266   5282       1 ;       H - C
5267   5273       1 ;      CA - HG
5267   5274       1 ;      CA - CD1
5267   5278       1 ;      CA - CD2
5267   5285       1 ;      CA - H
5267   5286       1 ;      CA - CA
5268   5270       1 ;      HA - HB2
5268   5271       1 ;      HA - HB3
5268   5272       1 ;      HA - CG
5268   5283       1 ;      HA - O
5268   5284       1 ;      HA - N
5269   5275       1 ;      CB - HD11
5269   5276       1 ;      CB - HD12
5269   5277       1 ;      CB - HD13
5269   5279       1 ;      CB - HD21
5269   5280       1 ;      CB - HD22
5269   5281       1 ;      CB - HD23
5269   5283       1 ;      CB - O
5269   5284       1 ;      CB - N
5270   5273       1 ;     HB2 - HG
5270   5274       1 ;     HB2 - CD1
5270   5278       1 ;     HB2 - CD2
5270   5282       1 ;     HB2 - C
5271   5273       1 ;     HB3 - HG
5271   5274       1 ;     HB3 - CD1
5271   5278       1 ;     HB3 - CD2
5271   5282       1 ;     HB3 - C
5272   5282       1 ;      CG - C
5273   5275       1 ;      HG - HD11
5273   5276       1 ;      HG - HD12
5273   5277       1 ;      HG - HD13
5273   5279       1 ;      HG - HD21
5273   5280       1 ;      HG - HD22
5273   5281       1 ;      HG - HD23
5274   5279       1 ;     CD1 - HD21
5274   5280       1 ;     CD1 - HD22
5274   5281       1 ;     CD1 - HD23
5275   5278       1 ;    HD11 - CD2
5276   5278       1 ;    HD12 - CD2
5277   5278       1 ;    HD13 - CD2
5282   5287       1 ;       C - HA
5282   5288       1 ;       C - CB
5282   5294       1 ;       C - C
5283   5285       1 ;       O - H
5283   5286       1 ;       O - CA
5284   5289       1 ;       N - HB2
5284   5290       1 ;       N - HB3
5284   5291       1 ;       N - CG
5284   5295       1 ;       N - O
5284   5296       1 ;       N - N
5285   5287       1 ;       H - HA
```

| | | | | | |
|---|---|---|---|---|---|
| 5285 | 5288 | 1 ; | H | - | CB |
| 5285 | 5294 | 1 ; | H | - | C |
| 5286 | 5292 | 1 ; | CA | - | OD1 |
| 5286 | 5293 | 1 ; | CA | - | OD2 |
| 5286 | 5297 | 1 ; | CA | - | H |
| 5286 | 5298 | 1 ; | CA | - | CA |
| 5287 | 5289 | 1 ; | HA | - | HB2 |
| 5287 | 5290 | 1 ; | HA | - | HB3 |
| 5287 | 5291 | 1 ; | HA | - | CG |
| 5287 | 5295 | 1 ; | HA | - | O |
| 5287 | 5296 | 1 ; | HA | - | N |
| 5288 | 5295 | 1 ; | CB | - | O |
| 5288 | 5296 | 1 ; | CB | - | N |
| 5289 | 5292 | 1 ; | HB2 | - | OD1 |
| 5289 | 5293 | 1 ; | HB2 | - | OD2 |
| 5289 | 5294 | 1 ; | HB2 | - | C |
| 5290 | 5292 | 1 ; | HB3 | - | OD1 |
| 5290 | 5293 | 1 ; | HB3 | - | OD2 |
| 5290 | 5294 | 1 ; | HB3 | - | C |
| 5291 | 5294 | 1 ; | CG | - | C |
| 5294 | 5299 | 1 ; | C | - | HA |
| 5294 | 5300 | 1 ; | C | - | CB |
| 5294 | 5310 | 1 ; | C | - | C |
| 5295 | 5297 | 1 ; | O | - | H |
| 5295 | 5298 | 1 ; | O | - | CA |
| 5296 | 5301 | 1 ; | N | - | HB |
| 5296 | 5302 | 1 ; | N | - | CG1 |
| 5296 | 5306 | 1 ; | N | - | CG2 |
| 5296 | 5311 | 1 ; | N | - | O |
| 5296 | 5312 | 1 ; | N | - | N |
| 5297 | 5299 | 1 ; | H | - | HA |
| 5297 | 5300 | 1 ; | H | - | CB |
| 5297 | 5310 | 1 ; | H | - | C |
| 5298 | 5303 | 1 ; | CA | - | HG11 |
| 5298 | 5304 | 1 ; | CA | - | HG12 |
| 5298 | 5305 | 1 ; | CA | - | HG13 |
| 5298 | 5307 | 1 ; | CA | - | HG21 |
| 5298 | 5308 | 1 ; | CA | - | HG22 |
| 5298 | 5309 | 1 ; | CA | - | HG23 |
| 5298 | 5313 | 1 ; | CA | - | H |
| 5298 | 5314 | 1 ; | CA | - | CA |
| 5299 | 5301 | 1 ; | HA | - | HB |
| 5299 | 5302 | 1 ; | HA | - | CG1 |
| 5299 | 5306 | 1 ; | HA | - | CG2 |
| 5299 | 5311 | 1 ; | HA | - | O |
| 5299 | 5312 | 1 ; | HA | - | N |
| 5300 | 5311 | 1 ; | CB | - | O |
| 5300 | 5312 | 1 ; | CB | - | N |
| 5301 | 5303 | 1 ; | HB | - | HG11 |
| 5301 | 5304 | 1 ; | HB | - | HG12 |
| 5301 | 5305 | 1 ; | HB | - | HG13 |
| 5301 | 5307 | 1 ; | HB | - | HG21 |

| | | | | | | |
|---|---|---|---|---|---|---|
| 5301 | 5308 | 1 | ; | HB | - | HG22 |
| 5301 | 5309 | 1 | ; | HB | - | HG23 |
| 5301 | 5310 | 1 | ; | HB | - | C |
| 5302 | 5307 | 1 | ; | CG1 | - | HG21 |
| 5302 | 5308 | 1 | ; | CG1 | - | HG22 |
| 5302 | 5309 | 1 | ; | CG1 | - | HG23 |
| 5302 | 5310 | 1 | ; | CG1 | - | C |
| 5303 | 5306 | 1 | ; | HG11 | - | CG2 |
| 5304 | 5306 | 1 | ; | HG12 | - | CG2 |
| 5305 | 5306 | 1 | ; | HG13 | - | CG2 |
| 5306 | 5310 | 1 | ; | CG2 | - | C |
| 5310 | 5315 | 1 | ; | C | - | HA |
| 5310 | 5316 | 1 | ; | C | - | CB |
| 5310 | 5322 | 1 | ; | C | - | C |
| 5311 | 5313 | 1 | ; | O | - | H |
| 5311 | 5314 | 1 | ; | O | - | CA |
| 5312 | 5317 | 1 | ; | N | - | HB2 |
| 5312 | 5318 | 1 | ; | N | - | HB3 |
| 5312 | 5319 | 1 | ; | N | - | CG |
| 5312 | 5323 | 1 | ; | N | - | O |
| 5312 | 5324 | 1 | ; | N | - | N |
| 5313 | 5315 | 1 | ; | H | - | HA |
| 5313 | 5316 | 1 | ; | H | - | CB |
| 5313 | 5322 | 1 | ; | H | - | C |
| 5314 | 5320 | 1 | ; | CA | - | OD1 |
| 5314 | 5321 | 1 | ; | CA | - | OD2 |
| 5314 | 5325 | 1 | ; | CA | - | H |
| 5314 | 5326 | 1 | ; | CA | - | CA |
| 5315 | 5317 | 1 | ; | HA | - | HB2 |
| 5315 | 5318 | 1 | ; | HA | - | HB3 |
| 5315 | 5319 | 1 | ; | HA | - | CG |
| 5315 | 5323 | 1 | ; | HA | - | O |
| 5315 | 5324 | 1 | ; | HA | - | N |
| 5316 | 5323 | 1 | ; | CB | - | O |
| 5316 | 5324 | 1 | ; | CB | - | N |
| 5317 | 5320 | 1 | ; | HB2 | - | OD1 |
| 5317 | 5321 | 1 | ; | HB2 | - | OD2 |
| 5317 | 5322 | 1 | ; | HB2 | - | C |
| 5318 | 5320 | 1 | ; | HB3 | - | OD1 |
| 5318 | 5321 | 1 | ; | HB3 | - | OD2 |
| 5318 | 5322 | 1 | ; | HB3 | - | C |
| 5319 | 5322 | 1 | ; | CG | - | C |
| 5322 | 5327 | 1 | ; | C | - | HA |
| 5322 | 5328 | 1 | ; | C | - | CB |
| 5322 | 5336 | 1 | ; | C | - | C |
| 5323 | 5325 | 1 | ; | O | - | H |
| 5323 | 5326 | 1 | ; | O | - | CA |
| 5324 | 5329 | 1 | ; | N | - | HB2 |
| 5324 | 5330 | 1 | ; | N | - | HB3 |
| 5324 | 5331 | 1 | ; | N | - | CG |
| 5324 | 5337 | 1 | ; | N | - | O |
| 5324 | 5338 | 1 | ; | N | - | N |

| | | | |
|---|---|---|---|
| 5325 | 5327 | 1 ; | H - HA |
| 5325 | 5328 | 1 ; | H - CB |
| 5325 | 5336 | 1 ; | H - C |
| 5326 | 5332 | 1 ; | CA - OD1 |
| 5326 | 5333 | 1 ; | CA - ND2 |
| 5326 | 5339 | 1 ; | CA - H |
| 5326 | 5340 | 1 ; | CA - CA |
| 5327 | 5329 | 1 ; | HA - HB2 |
| 5327 | 5330 | 1 ; | HA - HB3 |
| 5327 | 5331 | 1 ; | HA - CG |
| 5327 | 5337 | 1 ; | HA - O |
| 5327 | 5338 | 1 ; | HA - N |
| 5328 | 5334 | 1 ; | CB - HD21 |
| 5328 | 5335 | 1 ; | CB - HD22 |
| 5328 | 5337 | 1 ; | CB - O |
| 5328 | 5338 | 1 ; | CB - N |
| 5329 | 5332 | 1 ; | HB2 - OD1 |
| 5329 | 5333 | 1 ; | HB2 - ND2 |
| 5329 | 5336 | 1 ; | HB2 - C |
| 5330 | 5332 | 1 ; | HB3 - OD1 |
| 5330 | 5333 | 1 ; | HB3 - ND2 |
| 5330 | 5336 | 1 ; | HB3 - C |
| 5331 | 5336 | 1 ; | CG - C |
| 5332 | 5334 | 1 ; | OD1 - HD21 |
| 5332 | 5335 | 1 ; | OD1 - HD22 |
| 5336 | 5341 | 1 ; | C - HA |
| 5336 | 5342 | 1 ; | C - CB |
| 5336 | 5347 | 1 ; | C - C |
| 5337 | 5339 | 1 ; | O - H |
| 5337 | 5340 | 1 ; | O - CA |
| 5338 | 5343 | 1 ; | N - HB2 |
| 5338 | 5344 | 1 ; | N - HB3 |
| 5338 | 5345 | 1 ; | N - OG |
| 5338 | 5348 | 1 ; | N - O |
| 5338 | 5349 | 1 ; | N - N |
| 5339 | 5341 | 1 ; | H - HA |
| 5339 | 5342 | 1 ; | H - CB |
| 5339 | 5347 | 1 ; | H - C |
| 5340 | 5346 | 1 ; | CA - HG |
| 5340 | 5350 | 1 ; | CA - H |
| 5340 | 5351 | 1 ; | CA - CA |
| 5341 | 5343 | 1 ; | HA - HB2 |
| 5341 | 5344 | 1 ; | HA - HB3 |
| 5341 | 5345 | 1 ; | HA - OG |
| 5341 | 5348 | 1 ; | HA - O |
| 5341 | 5349 | 1 ; | HA - N |
| 5342 | 5348 | 1 ; | CB - O |
| 5342 | 5349 | 1 ; | CB - N |
| 5343 | 5346 | 1 ; | HB2 - HG |
| 5343 | 5347 | 1 ; | HB2 - C |
| 5344 | 5346 | 1 ; | HB3 - HG |
| 5344 | 5347 | 1 ; | HB3 - C |

```
5345   5347      1 ;     OG - C
5347   5352      1 ;      C - HA2
5347   5353      1 ;      C - HA3
5347   5354      1 ;      C - C
5348   5350      1 ;      O - H
5348   5351      1 ;      O - CA
5349   5355      1 ;      N - O
5349   5356      1 ;      N - N
5350   5352      1 ;      H - HA2
5350   5353      1 ;      H - HA3
5350   5354      1 ;      H - C
5351   5357      1 ;     CA - H
5351   5358      1 ;     CA - CA
5352   5355      1 ;    HA2 - O
5352   5356      1 ;    HA2 - N
5353   5355      1 ;    HA3 - O
5353   5356      1 ;    HA3 - N
5354   5359      1 ;      C - HA
5354   5360      1 ;      C - CB
5354   5368      1 ;      C - C
5355   5357      1 ;      O - H
5355   5358      1 ;      O - CA
5356   5361      1 ;      N - HB
5356   5362      1 ;      N - CG2
5356   5366      1 ;      N - OG1
5356   5369      1 ;      N - O
5356   5370      1 ;      N - N
5357   5359      1 ;      H - HA
5357   5360      1 ;      H - CB
5357   5368      1 ;      H - C
5358   5363      1 ;     CA - HG21
5358   5364      1 ;     CA - HG22
5358   5365      1 ;     CA - HG23
5358   5367      1 ;     CA - HG1
5358   5371      1 ;     CA - H
5358   5372      1 ;     CA - CA
5359   5361      1 ;     HA - HB
5359   5362      1 ;     HA - CG2
5359   5366      1 ;     HA - OG1
5359   5369      1 ;     HA - O
5359   5370      1 ;     HA - N
5360   5369      1 ;     CB - O
5360   5370      1 ;     CB - N
5361   5363      1 ;     HB - HG21
5361   5364      1 ;     HB - HG22
5361   5365      1 ;     HB - HG23
5361   5367      1 ;     HB - HG1
5361   5368      1 ;     HB - C
5362   5367      1 ;    CG2 - HG1
5362   5368      1 ;    CG2 - C
5363   5366      1 ;   HG21 - OG1
5364   5366      1 ;   HG22 - OG1
```

```
5365   5366      1 ;   HG23 - OG1
5366   5368      1 ;    OG1 - C
5368   5373      1 ;      C - HA
5368   5374      1 ;      C - CB
5368   5387      1 ;      C - C
5369   5371      1 ;      O - H
5369   5372      1 ;      O - CA
5370   5375      1 ;      N - HB2
5370   5376      1 ;      N - HB3
5370   5377      1 ;      N - CG
5370   5388      1 ;      N - O
5370   5389      1 ;      N - N
5371   5373      1 ;      H - HA
5371   5374      1 ;      H - CB
5371   5387      1 ;      H - C
5372   5378      1 ;     CA - HG
5372   5379      1 ;     CA - CD1
5372   5383      1 ;     CA - CD2
5372   5390      1 ;     CA - H
5372   5391      1 ;     CA - CA
5373   5375      1 ;     HA - HB2
5373   5376      1 ;     HA - HB3
5373   5377      1 ;     HA - CG
5373   5388      1 ;     HA - O
5373   5389      1 ;     HA - N
5374   5380      1 ;     CB - HD11
5374   5381      1 ;     CB - HD12
5374   5382      1 ;     CB - HD13
5374   5384      1 ;     CB - HD21
5374   5385      1 ;     CB - HD22
5374   5386      1 ;     CB - HD23
5374   5388      1 ;     CB - O
5374   5389      1 ;     CB - N
5375   5378      1 ;    HB2 - HG
5375   5379      1 ;    HB2 - CD1
5375   5383      1 ;    HB2 - CD2
5375   5387      1 ;    HB2 - C
5376   5378      1 ;    HB3 - HG
5376   5379      1 ;    HB3 - CD1
5376   5383      1 ;    HB3 - CD2
5376   5387      1 ;    HB3 - C
5377   5387      1 ;     CG - C
5378   5380      1 ;     HG - HD11
5378   5381      1 ;     HG - HD12
5378   5382      1 ;     HG - HD13
5378   5384      1 ;     HG - HD21
5378   5385      1 ;     HG - HD22
5378   5386      1 ;     HG - HD23
5379   5384      1 ;    CD1 - HD21
5379   5385      1 ;    CD1 - HD22
5379   5386      1 ;    CD1 - HD23
5380   5383      1 ;   HD11 - CD2
```

| | | | | | |
|---|---|---|---|---|---|
| 5381 | 5383 | 1 ; | HD12 | - | CD2 |
| 5382 | 5383 | 1 ; | HD13 | - | CD2 |
| 5387 | 5392 | 1 ; | C | - | HA |
| 5387 | 5393 | 1 ; | C | - | CB |
| 5387 | 5398 | 1 ; | C | - | C |
| 5388 | 5390 | 1 ; | O | - | H |
| 5388 | 5391 | 1 ; | O | - | CA |
| 5389 | 5394 | 1 ; | N | - | HB2 |
| 5389 | 5395 | 1 ; | N | - | HB3 |
| 5389 | 5396 | 1 ; | N | - | OG |
| 5389 | 5399 | 1 ; | N | - | O |
| 5389 | 5400 | 1 ; | N | - | N |
| 5390 | 5392 | 1 ; | H | - | HA |
| 5390 | 5393 | 1 ; | H | - | CB |
| 5390 | 5398 | 1 ; | H | - | C |
| 5391 | 5397 | 1 ; | CA | - | HG |
| 5391 | 5401 | 1 ; | CA | - | H |
| 5391 | 5402 | 1 ; | CA | - | CA |
| 5392 | 5394 | 1 ; | HA | - | HB2 |
| 5392 | 5395 | 1 ; | HA | - | HB3 |
| 5392 | 5396 | 1 ; | HA | - | OG |
| 5392 | 5399 | 1 ; | HA | - | O |
| 5392 | 5400 | 1 ; | HA | - | N |
| 5393 | 5399 | 1 ; | CB | - | O |
| 5393 | 5400 | 1 ; | CB | - | N |
| 5394 | 5397 | 1 ; | HB2 | - | HG |
| 5394 | 5398 | 1 ; | HB2 | - | C |
| 5395 | 5397 | 1 ; | HB3 | - | HG |
| 5395 | 5398 | 1 ; | HB3 | - | C |
| 5396 | 5398 | 1 ; | OG | - | C |
| 5398 | 5403 | 1 ; | C | - | HA |
| 5398 | 5404 | 1 ; | C | - | CB |
| 5398 | 5409 | 1 ; | C | - | C |
| 5399 | 5401 | 1 ; | O | - | H |
| 5399 | 5402 | 1 ; | O | - | CA |
| 5400 | 5405 | 1 ; | N | - | HB2 |
| 5400 | 5406 | 1 ; | N | - | HB3 |
| 5400 | 5407 | 1 ; | N | - | OG |
| 5400 | 5410 | 1 ; | N | - | O |
| 5400 | 5411 | 1 ; | N | - | N |
| 5401 | 5403 | 1 ; | H | - | HA |
| 5401 | 5404 | 1 ; | H | - | CB |
| 5401 | 5409 | 1 ; | H | - | C |
| 5402 | 5408 | 1 ; | CA | - | HG |
| 5402 | 5412 | 1 ; | CA | - | H |
| 5402 | 5413 | 1 ; | CA | - | CA |
| 5403 | 5405 | 1 ; | HA | - | HB2 |
| 5403 | 5406 | 1 ; | HA | - | HB3 |
| 5403 | 5407 | 1 ; | HA | - | OG |
| 5403 | 5410 | 1 ; | HA | - | O |
| 5403 | 5411 | 1 ; | HA | - | N |
| 5404 | 5410 | 1 ; | CB | - | O |

```
5404   5411       1 ;      CB - N
5405   5408       1 ;     HB2 - HG
5405   5409       1 ;     HB2 - C
5406   5408       1 ;     HB3 - HG
5406   5409       1 ;     HB3 - C
5407   5409       1 ;      OG - C
5409   5414       1 ;       C - HA
5409   5415       1 ;       C - CB
5409   5426       1 ;       C - C
5410   5412       1 ;       O - H
5410   5413       1 ;       O - CA
5411   5416       1 ;       N - HB2
5411   5417       1 ;       N - HB3
5411   5418       1 ;       N - CG
5411   5427       1 ;       N - O
5411   5428       1 ;       N - N
5412   5414       1 ;       H - HA
5412   5415       1 ;       H - CB
5412   5426       1 ;       H - C
5413   5419       1 ;      CA - HG2
5413   5420       1 ;      CA - HG3
5413   5421       1 ;      CA - CD
5413   5429       1 ;      CA - H
5413   5430       1 ;      CA - CA
5414   5416       1 ;      HA - HB2
5414   5417       1 ;      HA - HB3
5414   5418       1 ;      HA - CG
5414   5427       1 ;      HA - O
5414   5428       1 ;      HA - N
5415   5422       1 ;      CB - OE1
5415   5423       1 ;      CB - NE2
5415   5427       1 ;      CB - O
5415   5428       1 ;      CB - N
5416   5419       1 ;     HB2 - HG2
5416   5420       1 ;     HB2 - HG3
5416   5421       1 ;     HB2 - CD
5416   5426       1 ;     HB2 - C
5417   5419       1 ;     HB3 - HG2
5417   5420       1 ;     HB3 - HG3
5417   5421       1 ;     HB3 - CD
5417   5426       1 ;     HB3 - C
5418   5424       1 ;      CG - HE21
5418   5425       1 ;      CG - HE22
5418   5426       1 ;      CG - C
5419   5422       1 ;     HG2 - OE1
5419   5423       1 ;     HG2 - NE2
5420   5422       1 ;     HG3 - OE1
5420   5423       1 ;     HG3 - NE2
5422   5424       1 ;     OE1 - HE21
5422   5425       1 ;     OE1 - HE22
5426   5431       1 ;       C - HA
5426   5432       1 ;       C - CB
```

| | | | |
|---|---|---|---|
| 5426 | 5442 | 1 ; | C - C |
| 5427 | 5429 | 1 ; | O - H |
| 5427 | 5430 | 1 ; | O - CA |
| 5428 | 5433 | 1 ; | N - HB2 |
| 5428 | 5434 | 1 ; | N - HB3 |
| 5428 | 5435 | 1 ; | N - CG |
| 5428 | 5443 | 1 ; | N - O |
| 5428 | 5444 | 1 ; | N - N |
| 5429 | 5431 | 1 ; | H - HA |
| 5429 | 5432 | 1 ; | H - CB |
| 5429 | 5442 | 1 ; | H - C |
| 5430 | 5436 | 1 ; | CA - HG2 |
| 5430 | 5437 | 1 ; | CA - HG3 |
| 5430 | 5438 | 1 ; | CA - CD |
| 5430 | 5445 | 1 ; | CA - H |
| 5430 | 5446 | 1 ; | CA - CA |
| 5431 | 5433 | 1 ; | HA - HB2 |
| 5431 | 5434 | 1 ; | HA - HB3 |
| 5431 | 5435 | 1 ; | HA - CG |
| 5431 | 5443 | 1 ; | HA - O |
| 5431 | 5444 | 1 ; | HA - N |
| 5432 | 5439 | 1 ; | CB - OE1 |
| 5432 | 5440 | 1 ; | CB - OE2 |
| 5432 | 5443 | 1 ; | CB - O |
| 5432 | 5444 | 1 ; | CB - N |
| 5433 | 5436 | 1 ; | HB2 - HG2 |
| 5433 | 5437 | 1 ; | HB2 - HG3 |
| 5433 | 5438 | 1 ; | HB2 - CD |
| 5433 | 5442 | 1 ; | HB2 - C |
| 5434 | 5436 | 1 ; | HB3 - HG2 |
| 5434 | 5437 | 1 ; | HB3 - HG3 |
| 5434 | 5438 | 1 ; | HB3 - CD |
| 5434 | 5442 | 1 ; | HB3 - C |
| 5435 | 5441 | 1 ; | CG - HE2 |
| 5435 | 5442 | 1 ; | CG - C |
| 5436 | 5439 | 1 ; | HG2 - OE1 |
| 5436 | 5440 | 1 ; | HG2 - OE2 |
| 5437 | 5439 | 1 ; | HG3 - OE1 |
| 5437 | 5440 | 1 ; | HG3 - OE2 |
| 5439 | 5441 | 1 ; | OE1 - HE2 |
| 5442 | 5447 | 1 ; | C - HA |
| 5442 | 5448 | 1 ; | C - CB |
| 5442 | 5461 | 1 ; | C - C |
| 5443 | 5445 | 1 ; | O - H |
| 5443 | 5446 | 1 ; | O - CA |
| 5444 | 5449 | 1 ; | N - HB |
| 5444 | 5450 | 1 ; | N - CG2 |
| 5444 | 5454 | 1 ; | N - CG1 |
| 5444 | 5462 | 1 ; | N - O |
| 5444 | 5463 | 1 ; | N - N |
| 5445 | 5447 | 1 ; | H - HA |
| 5445 | 5448 | 1 ; | H - CB |

| | | | |
|---|---|---|---|
| 5445 | 5461 | 1 ; | H - C |
| 5446 | 5451 | 1 ; | CA - HG21 |
| 5446 | 5452 | 1 ; | CA - HG22 |
| 5446 | 5453 | 1 ; | CA - HG23 |
| 5446 | 5455 | 1 ; | CA - HG12 |
| 5446 | 5456 | 1 ; | CA - HG13 |
| 5446 | 5457 | 1 ; | CA - CD1 |
| 5446 | 5464 | 1 ; | CA - H |
| 5446 | 5465 | 1 ; | CA - CA |
| 5447 | 5449 | 1 ; | HA - HB |
| 5447 | 5450 | 1 ; | HA - CG2 |
| 5447 | 5454 | 1 ; | HA - CG1 |
| 5447 | 5462 | 1 ; | HA - O |
| 5447 | 5463 | 1 ; | HA - N |
| 5448 | 5458 | 1 ; | CB - HD11 |
| 5448 | 5459 | 1 ; | CB - HD12 |
| 5448 | 5460 | 1 ; | CB - HD13 |
| 5448 | 5462 | 1 ; | CB - O |
| 5448 | 5463 | 1 ; | CB - N |
| 5449 | 5451 | 1 ; | HB - HG21 |
| 5449 | 5452 | 1 ; | HB - HG22 |
| 5449 | 5453 | 1 ; | HB - HG23 |
| 5449 | 5455 | 1 ; | HB - HG12 |
| 5449 | 5456 | 1 ; | HB - HG13 |
| 5449 | 5457 | 1 ; | HB - CD1 |
| 5449 | 5461 | 1 ; | HB - C |
| 5450 | 5455 | 1 ; | CG2 - HG12 |
| 5450 | 5456 | 1 ; | CG2 - HG13 |
| 5450 | 5457 | 1 ; | CG2 - CD1 |
| 5450 | 5461 | 1 ; | CG2 - C |
| 5451 | 5454 | 1 ; | HG21 - CG1 |
| 5452 | 5454 | 1 ; | HG22 - CG1 |
| 5453 | 5454 | 1 ; | HG23 - CG1 |
| 5454 | 5461 | 1 ; | CG1 - C |
| 5455 | 5458 | 1 ; | HG12 - HD11 |
| 5455 | 5459 | 1 ; | HG12 - HD12 |
| 5455 | 5460 | 1 ; | HG12 - HD13 |
| 5456 | 5458 | 1 ; | HG13 - HD11 |
| 5456 | 5459 | 1 ; | HG13 - HD12 |
| 5456 | 5460 | 1 ; | HG13 - HD13 |
| 5461 | 5466 | 1 ; | C - HA |
| 5461 | 5467 | 1 ; | C - CB |
| 5461 | 5480 | 1 ; | C - C |
| 5462 | 5464 | 1 ; | O - H |
| 5462 | 5465 | 1 ; | O - CA |
| 5463 | 5468 | 1 ; | N - HB2 |
| 5463 | 5469 | 1 ; | N - HB3 |
| 5463 | 5470 | 1 ; | N - CG |
| 5463 | 5481 | 1 ; | N - O |
| 5463 | 5482 | 1 ; | N - N |
| 5464 | 5466 | 1 ; | H - HA |
| 5464 | 5467 | 1 ; | H - CB |

| | | | | | |
|---|---|---|---|---|---|
| 5464 | 5480 | 1 ; | H | - | C |
| 5465 | 5471 | 1 ; | CA | - | HG |
| 5465 | 5472 | 1 ; | CA | - | CD1 |
| 5465 | 5476 | 1 ; | CA | - | CD2 |
| 5465 | 5483 | 1 ; | CA | - | H |
| 5465 | 5484 | 1 ; | CA | - | CA |
| 5466 | 5468 | 1 ; | HA | - | HB2 |
| 5466 | 5469 | 1 ; | HA | - | HB3 |
| 5466 | 5470 | 1 ; | HA | - | CG |
| 5466 | 5481 | 1 ; | HA | - | O |
| 5466 | 5482 | 1 ; | HA | - | N |
| 5467 | 5473 | 1 ; | CB | - | HD11 |
| 5467 | 5474 | 1 ; | CB | - | HD12 |
| 5467 | 5475 | 1 ; | CB | - | HD13 |
| 5467 | 5477 | 1 ; | CB | - | HD21 |
| 5467 | 5478 | 1 ; | CB | - | HD22 |
| 5467 | 5479 | 1 ; | CB | - | HD23 |
| 5467 | 5481 | 1 ; | CB | - | O |
| 5467 | 5482 | 1 ; | CB | - | N |
| 5468 | 5471 | 1 ; | HB2 | - | HG |
| 5468 | 5472 | 1 ; | HB2 | - | CD1 |
| 5468 | 5476 | 1 ; | HB2 | - | CD2 |
| 5468 | 5480 | 1 ; | HB2 | - | C |
| 5469 | 5471 | 1 ; | HB3 | - | HG |
| 5469 | 5472 | 1 ; | HB3 | - | CD1 |
| 5469 | 5476 | 1 ; | HB3 | - | CD2 |
| 5469 | 5480 | 1 ; | HB3 | - | C |
| 5470 | 5480 | 1 ; | CG | - | C |
| 5471 | 5473 | 1 ; | HG | - | HD11 |
| 5471 | 5474 | 1 ; | HG | - | HD12 |
| 5471 | 5475 | 1 ; | HG | - | HD13 |
| 5471 | 5477 | 1 ; | HG | - | HD21 |
| 5471 | 5478 | 1 ; | HG | - | HD22 |
| 5471 | 5479 | 1 ; | HG | - | HD23 |
| 5472 | 5477 | 1 ; | CD1 | - | HD21 |
| 5472 | 5478 | 1 ; | CD1 | - | HD22 |
| 5472 | 5479 | 1 ; | CD1 | - | HD23 |
| 5473 | 5476 | 1 ; | HD11 | - | CD2 |
| 5474 | 5476 | 1 ; | HD12 | - | CD2 |
| 5475 | 5476 | 1 ; | HD13 | - | CD2 |
| 5480 | 5485 | 1 ; | C | - | HA |
| 5480 | 5486 | 1 ; | C | - | CB |
| 5480 | 5492 | 1 ; | C | - | C |
| 5481 | 5483 | 1 ; | O | - | H |
| 5481 | 5484 | 1 ; | O | - | CA |
| 5482 | 5487 | 1 ; | N | - | HB2 |
| 5482 | 5488 | 1 ; | N | - | HB3 |
| 5482 | 5489 | 1 ; | N | - | CG |
| 5482 | 5493 | 1 ; | N | - | O |
| 5482 | 5494 | 1 ; | N | - | N |
| 5483 | 5485 | 1 ; | H | - | HA |
| 5483 | 5486 | 1 ; | H | - | CB |

| | | | | | |
|---|---|---|---|---|---|
| 5483 | 5492 | 1 ; | H | - | C |
| 5484 | 5490 | 1 ; | CA | - | OD1 |
| 5484 | 5491 | 1 ; | CA | - | OD2 |
| 5484 | 5495 | 1 ; | CA | - | H |
| 5484 | 5496 | 1 ; | CA | - | CA |
| 5485 | 5487 | 1 ; | HA | - | HB2 |
| 5485 | 5488 | 1 ; | HA | - | HB3 |
| 5485 | 5489 | 1 ; | HA | - | CG |
| 5485 | 5493 | 1 ; | HA | - | O |
| 5485 | 5494 | 1 ; | HA | - | N |
| 5486 | 5493 | 1 ; | CB | - | O |
| 5486 | 5494 | 1 ; | CB | - | N |
| 5487 | 5490 | 1 ; | HB2 | - | OD1 |
| 5487 | 5491 | 1 ; | HB2 | - | OD2 |
| 5487 | 5492 | 1 ; | HB2 | - | C |
| 5488 | 5490 | 1 ; | HB3 | - | OD1 |
| 5488 | 5491 | 1 ; | HB3 | - | OD2 |
| 5488 | 5492 | 1 ; | HB3 | - | C |
| 5489 | 5492 | 1 ; | CG | - | C |
| 5492 | 5497 | 1 ; | C | - | HA2 |
| 5492 | 5498 | 1 ; | C | - | HA3 |
| 5492 | 5499 | 1 ; | C | - | C |
| 5493 | 5495 | 1 ; | O | - | H |
| 5493 | 5496 | 1 ; | O | - | CA |
| 5494 | 5500 | 1 ; | N | - | O |
| 5494 | 5501 | 1 ; | N | - | N |
| 5495 | 5497 | 1 ; | H | - | HA2 |
| 5495 | 5498 | 1 ; | H | - | HA3 |
| 5495 | 5499 | 1 ; | H | - | C |
| 5496 | 5502 | 1 ; | CA | - | H |
| 5496 | 5503 | 1 ; | CA | - | CA |
| 5497 | 5500 | 1 ; | HA2 | - | O |
| 5497 | 5501 | 1 ; | HA2 | - | N |
| 5498 | 5500 | 1 ; | HA3 | - | O |
| 5498 | 5501 | 1 ; | HA3 | - | N |
| 5499 | 5504 | 1 ; | C | - | HA |
| 5499 | 5505 | 1 ; | C | - | CB |
| 5499 | 5518 | 1 ; | C | - | C |
| 5500 | 5502 | 1 ; | O | - | H |
| 5500 | 5503 | 1 ; | O | - | CA |
| 5501 | 5506 | 1 ; | N | - | HB2 |
| 5501 | 5507 | 1 ; | N | - | HB3 |
| 5501 | 5508 | 1 ; | N | - | CG |
| 5501 | 5519 | 1 ; | N | - | O |
| 5501 | 5520 | 1 ; | N | - | N |
| 5502 | 5504 | 1 ; | H | - | HA |
| 5502 | 5505 | 1 ; | H | - | CB |
| 5502 | 5518 | 1 ; | H | - | C |
| 5503 | 5509 | 1 ; | CA | - | HG |
| 5503 | 5510 | 1 ; | CA | - | CD1 |
| 5503 | 5514 | 1 ; | CA | - | CD2 |
| 5503 | 5521 | 1 ; | CA | - | H |

```
5503   5522       1 ;      CA - CA
5504   5506       1 ;      HA - HB2
5504   5507       1 ;      HA - HB3
5504   5508       1 ;      HA - CG
5504   5519       1 ;      HA - O
5504   5520       1 ;      HA - N
5505   5511       1 ;      CB - HD11
5505   5512       1 ;      CB - HD12
5505   5513       1 ;      CB - HD13
5505   5515       1 ;      CB - HD21
5505   5516       1 ;      CB - HD22
5505   5517       1 ;      CB - HD23
5505   5519       1 ;      CB - O
5505   5520       1 ;      CB - N
5506   5509       1 ;     HB2 - HG
5506   5510       1 ;     HB2 - CD1
5506   5514       1 ;     HB2 - CD2
5506   5518       1 ;     HB2 - C
5507   5509       1 ;     HB3 - HG
5507   5510       1 ;     HB3 - CD1
5507   5514       1 ;     HB3 - CD2
5507   5518       1 ;     HB3 - C
5508   5518       1 ;      CG - C
5509   5511       1 ;      HG - HD11
5509   5512       1 ;      HG - HD12
5509   5513       1 ;      HG - HD13
5509   5515       1 ;      HG - HD21
5509   5516       1 ;      HG - HD22
5509   5517       1 ;      HG - HD23
5510   5515       1 ;     CD1 - HD21
5510   5516       1 ;     CD1 - HD22
5510   5517       1 ;     CD1 - HD23
5511   5514       1 ;    HD11 - CD2
5512   5514       1 ;    HD12 - CD2
5513   5514       1 ;    HD13 - CD2
5518   5523       1 ;       C - HA
5518   5524       1 ;       C - CB
5518   5540       1 ;       C - C
5519   5521       1 ;       O - H
5519   5522       1 ;       O - CA
5520   5525       1 ;       N - HB2
5520   5526       1 ;       N - HB3
5520   5527       1 ;       N - CG
5520   5541       1 ;       N - O
5520   5542       1 ;       N - N
5521   5523       1 ;       H - HA
5521   5524       1 ;       H - CB
5521   5540       1 ;       H - C
5522   5528       1 ;      CA - HG2
5522   5529       1 ;      CA - HG3
5522   5530       1 ;      CA - CD
5522   5543       1 ;      CA - H
```

| | | | |
|---|---|---|---|
| 5522 | 5544 | 1 ; | CA - CA |
| 5523 | 5525 | 1 ; | HA - HB2 |
| 5523 | 5526 | 1 ; | HA - HB3 |
| 5523 | 5527 | 1 ; | HA - CG |
| 5523 | 5541 | 1 ; | HA - O |
| 5523 | 5542 | 1 ; | HA - N |
| 5524 | 5531 | 1 ; | CB - HD2 |
| 5524 | 5532 | 1 ; | CB - HD3 |
| 5524 | 5533 | 1 ; | CB - CE |
| 5524 | 5541 | 1 ; | CB - O |
| 5524 | 5542 | 1 ; | CB - N |
| 5525 | 5528 | 1 ; | HB2 - HG2 |
| 5525 | 5529 | 1 ; | HB2 - HG3 |
| 5525 | 5530 | 1 ; | HB2 - CD |
| 5525 | 5540 | 1 ; | HB2 - C |
| 5526 | 5528 | 1 ; | HB3 - HG2 |
| 5526 | 5529 | 1 ; | HB3 - HG3 |
| 5526 | 5530 | 1 ; | HB3 - CD |
| 5526 | 5540 | 1 ; | HB3 - C |
| 5527 | 5534 | 1 ; | CG - HE2 |
| 5527 | 5535 | 1 ; | CG - HE3 |
| 5527 | 5536 | 1 ; | CG - NZ |
| 5527 | 5540 | 1 ; | CG - C |
| 5528 | 5531 | 1 ; | HG2 - HD2 |
| 5528 | 5532 | 1 ; | HG2 - HD3 |
| 5528 | 5533 | 1 ; | HG2 - CE |
| 5529 | 5531 | 1 ; | HG3 - HD2 |
| 5529 | 5532 | 1 ; | HG3 - HD3 |
| 5529 | 5533 | 1 ; | HG3 - CE |
| 5530 | 5537 | 1 ; | CD - HZ1 |
| 5530 | 5538 | 1 ; | CD - HZ2 |
| 5530 | 5539 | 1 ; | CD - HZ3 |
| 5531 | 5534 | 1 ; | HD2 - HE2 |
| 5531 | 5535 | 1 ; | HD2 - HE3 |
| 5531 | 5536 | 1 ; | HD2 - NZ |
| 5532 | 5534 | 1 ; | HD3 - HE2 |
| 5532 | 5535 | 1 ; | HD3 - HE3 |
| 5532 | 5536 | 1 ; | HD3 - NZ |
| 5534 | 5537 | 1 ; | HE2 - HZ1 |
| 5534 | 5538 | 1 ; | HE2 - HZ2 |
| 5534 | 5539 | 1 ; | HE2 - HZ3 |
| 5535 | 5537 | 1 ; | HE3 - HZ1 |
| 5535 | 5538 | 1 ; | HE3 - HZ2 |
| 5535 | 5539 | 1 ; | HE3 - HZ3 |
| 5540 | 5545 | 1 ; | C - HA |
| 5540 | 5546 | 1 ; | C - CB |
| 5540 | 5562 | 1 ; | C - C |
| 5541 | 5543 | 1 ; | O - H |
| 5541 | 5544 | 1 ; | O - CA |
| 5542 | 5547 | 1 ; | N - HB2 |
| 5542 | 5548 | 1 ; | N - HB3 |
| 5542 | 5549 | 1 ; | N - CG |

```
5542   5563       1 ;       N - O
5542   5564       1 ;       N - N
5543   5545       1 ;       H - HA
5543   5546       1 ;       H - CB
5543   5562       1 ;       H - C
5544   5550       1 ;      CA - HG2
5544   5551       1 ;      CA - HG3
5544   5552       1 ;      CA - CD
5544   5565       1 ;      CA - H
5544   5566       1 ;      CA - CA
5545   5547       1 ;      HA - HB2
5545   5548       1 ;      HA - HB3
5545   5549       1 ;      HA - CG
5545   5563       1 ;      HA - O
5545   5564       1 ;      HA - N
5546   5553       1 ;      CB - HD2
5546   5554       1 ;      CB - HD3
5546   5555       1 ;      CB - CE
5546   5563       1 ;      CB - O
5546   5564       1 ;      CB - N
5547   5550       1 ;     HB2 - HG2
5547   5551       1 ;     HB2 - HG3
5547   5552       1 ;     HB2 - CD
5547   5562       1 ;     HB2 - C
5548   5550       1 ;     HB3 - HG2
5548   5551       1 ;     HB3 - HG3
5548   5552       1 ;     HB3 - CD
5548   5562       1 ;     HB3 - C
5549   5556       1 ;      CG - HE2
5549   5557       1 ;      CG - HE3
5549   5558       1 ;      CG - NZ
5549   5562       1 ;      CG - C
5550   5553       1 ;     HG2 - HD2
5550   5554       1 ;     HG2 - HD3
5550   5555       1 ;     HG2 - CE
5551   5553       1 ;     HG3 - HD2
5551   5554       1 ;     HG3 - HD3
5551   5555       1 ;     HG3 - CE
5552   5559       1 ;      CD - HZ1
5552   5560       1 ;      CD - HZ2
5552   5561       1 ;      CD - HZ3
5553   5556       1 ;     HD2 - HE2
5553   5557       1 ;     HD2 - HE3
5553   5558       1 ;     HD2 - NZ
5554   5556       1 ;     HD3 - HE2
5554   5557       1 ;     HD3 - HE3
5554   5558       1 ;     HD3 - NZ
5556   5559       1 ;     HE2 - HZ1
5556   5560       1 ;     HE2 - HZ2
5556   5561       1 ;     HE2 - HZ3
5557   5559       1 ;     HE3 - HZ1
5557   5560       1 ;     HE3 - HZ2
```

```
5557   5561       1 ;    HE3 - HZ3
5562   5567       1 ;      C - HA
5562   5568       1 ;      C - CB
5562   5581       1 ;      C - C
5563   5565       1 ;      O - H
5563   5566       1 ;      O - CA
5564   5569       1 ;      N - HB
5564   5570       1 ;      N - CG2
5564   5574       1 ;      N - CG1
5564   5582       1 ;      N - O
5564   5583       1 ;      N - N
5565   5567       1 ;      H - HA
5565   5568       1 ;      H - CB
5565   5581       1 ;      H - C
5566   5571       1 ;     CA - HG21
5566   5572       1 ;     CA - HG22
5566   5573       1 ;     CA - HG23
5566   5575       1 ;     CA - HG12
5566   5576       1 ;     CA - HG13
5566   5577       1 ;     CA - CD1
5566   5584       1 ;     CA - H
5566   5585       1 ;     CA - CA
5567   5569       1 ;     HA - HB
5567   5570       1 ;     HA - CG2
5567   5574       1 ;     HA - CG1
5567   5582       1 ;     HA - O
5567   5583       1 ;     HA - N
5568   5578       1 ;     CB - HD11
5568   5579       1 ;     CB - HD12
5568   5580       1 ;     CB - HD13
5568   5582       1 ;     CB - O
5568   5583       1 ;     CB - N
5569   5571       1 ;     HB - HG21
5569   5572       1 ;     HB - HG22
5569   5573       1 ;     HB - HG23
5569   5575       1 ;     HB - HG12
5569   5576       1 ;     HB - HG13
5569   5577       1 ;     HB - CD1
5569   5581       1 ;     HB - C
5570   5575       1 ;    CG2 - HG12
5570   5576       1 ;    CG2 - HG13
5570   5577       1 ;    CG2 - CD1
5570   5581       1 ;    CG2 - C
5571   5574       1 ;   HG21 - CG1
5572   5574       1 ;   HG22 - CG1
5573   5574       1 ;   HG23 - CG1
5574   5581       1 ;    CG1 - C
5575   5578       1 ;   HG12 - HD11
5575   5579       1 ;   HG12 - HD12
5575   5580       1 ;   HG12 - HD13
5576   5578       1 ;   HG13 - HD11
5576   5579       1 ;   HG13 - HD12
```

| | | | | | |
|---|---|---|---|---|---|
| 5576 | 5580 | 1 ; | HG13 | - | HD13 |
| 5581 | 5586 | 1 ; | C | - | HA2 |
| 5581 | 5587 | 1 ; | C | - | HA3 |
| 5581 | 5588 | 1 ; | C | - | C |
| 5582 | 5584 | 1 ; | O | - | H |
| 5582 | 5585 | 1 ; | O | - | CA |
| 5583 | 5589 | 1 ; | N | - | O |
| 5583 | 5590 | 1 ; | N | - | N |
| 5584 | 5586 | 1 ; | H | - | HA2 |
| 5584 | 5587 | 1 ; | H | - | HA3 |
| 5584 | 5588 | 1 ; | H | - | C |
| 5585 | 5591 | 1 ; | CA | - | H |
| 5585 | 5592 | 1 ; | CA | - | CA |
| 5586 | 5589 | 1 ; | HA2 | - | O |
| 5586 | 5590 | 1 ; | HA2 | - | N |
| 5587 | 5589 | 1 ; | HA3 | - | O |
| 5587 | 5590 | 1 ; | HA3 | - | N |
| 5588 | 5593 | 1 ; | C | - | HA |
| 5588 | 5594 | 1 ; | C | - | CB |
| 5588 | 5609 | 1 ; | C | - | C |
| 5589 | 5591 | 1 ; | O | - | H |
| 5589 | 5592 | 1 ; | O | - | CA |
| 5590 | 5595 | 1 ; | N | - | HB2 |
| 5590 | 5596 | 1 ; | N | - | HB3 |
| 5590 | 5597 | 1 ; | N | - | CG |
| 5590 | 5610 | 1 ; | N | - | O |
| 5590 | 5611 | 1 ; | N | - | N |
| 5591 | 5593 | 1 ; | H | - | HA |
| 5591 | 5594 | 1 ; | H | - | CB |
| 5591 | 5609 | 1 ; | H | - | C |
| 5592 | 5598 | 1 ; | CA | - | CD1 |
| 5592 | 5607 | 1 ; | CA | - | CD2 |
| 5592 | 5612 | 1 ; | CA | - | H |
| 5592 | 5613 | 1 ; | CA | - | CA |
| 5593 | 5595 | 1 ; | HA | - | HB2 |
| 5593 | 5596 | 1 ; | HA | - | HB3 |
| 5593 | 5597 | 1 ; | HA | - | CG |
| 5593 | 5610 | 1 ; | HA | - | O |
| 5593 | 5611 | 1 ; | HA | - | N |
| 5594 | 5599 | 1 ; | CB | - | HD1 |
| 5594 | 5600 | 1 ; | CB | - | CE1 |
| 5594 | 5605 | 1 ; | CB | - | CE2 |
| 5594 | 5608 | 1 ; | CB | - | HD2 |
| 5594 | 5610 | 1 ; | CB | - | O |
| 5594 | 5611 | 1 ; | CB | - | N |
| 5595 | 5598 | 1 ; | HB2 | - | CD1 |
| 5595 | 5607 | 1 ; | HB2 | - | CD2 |
| 5595 | 5609 | 1 ; | HB2 | - | C |
| 5596 | 5598 | 1 ; | HB3 | - | CD1 |
| 5596 | 5607 | 1 ; | HB3 | - | CD2 |
| 5596 | 5609 | 1 ; | HB3 | - | C |
| 5597 | 5601 | 1 ; | CG | - | HE1 |

| | | | | | |
|---|---|---|---|---|---|
| 5597 | 5602 | 1 ; | CG | - | CZ |
| 5597 | 5606 | 1 ; | CG | - | HE2 |
| 5597 | 5609 | 1 ; | CG | - | C |
| 5598 | 5603 | 1 ; | CD1 | - | OH |
| 5598 | 5605 | 1 ; | CD1 | - | CE2 |
| 5598 | 5608 | 1 ; | CD1 | - | HD2 |
| 5599 | 5601 | 1 ; | HD1 | - | HE1 |
| 5599 | 5602 | 1 ; | HD1 | - | CZ |
| 5599 | 5607 | 1 ; | HD1 | - | CD2 |
| 5600 | 5604 | 1 ; | CE1 | - | HH |
| 5600 | 5606 | 1 ; | CE1 | - | HE2 |
| 5600 | 5607 | 1 ; | CE1 | - | CD2 |
| 5601 | 5603 | 1 ; | HE1 | - | OH |
| 5601 | 5605 | 1 ; | HE1 | - | CE2 |
| 5602 | 5608 | 1 ; | CZ | - | HD2 |
| 5603 | 5606 | 1 ; | OH | - | HE2 |
| 5603 | 5607 | 1 ; | OH | - | CD2 |
| 5604 | 5605 | 1 ; | HH | - | CE2 |
| 5606 | 5608 | 1 ; | HE2 | - | HD2 |
| 5609 | 5614 | 1 ; | C | - | HA |
| 5609 | 5615 | 1 ; | C | - | CB |
| 5609 | 5626 | 1 ; | C | - | C |
| 5610 | 5612 | 1 ; | O | - | H |
| 5610 | 5613 | 1 ; | O | - | CA |
| 5611 | 5616 | 1 ; | N | - | HB2 |
| 5611 | 5617 | 1 ; | N | - | HB3 |
| 5611 | 5618 | 1 ; | N | - | CG |
| 5611 | 5627 | 1 ; | N | - | O |
| 5611 | 5628 | 1 ; | N | - | N |
| 5612 | 5614 | 1 ; | H | - | HA |
| 5612 | 5615 | 1 ; | H | - | CB |
| 5612 | 5626 | 1 ; | H | - | C |
| 5613 | 5619 | 1 ; | CA | - | HG2 |
| 5613 | 5620 | 1 ; | CA | - | HG3 |
| 5613 | 5621 | 1 ; | CA | - | CD |
| 5613 | 5629 | 1 ; | CA | - | H |
| 5613 | 5630 | 1 ; | CA | - | CA |
| 5614 | 5616 | 1 ; | HA | - | HB2 |
| 5614 | 5617 | 1 ; | HA | - | HB3 |
| 5614 | 5618 | 1 ; | HA | - | CG |
| 5614 | 5627 | 1 ; | HA | - | O |
| 5614 | 5628 | 1 ; | HA | - | N |
| 5615 | 5622 | 1 ; | CB | - | OE1 |
| 5615 | 5623 | 1 ; | CB | - | NE2 |
| 5615 | 5627 | 1 ; | CB | - | O |
| 5615 | 5628 | 1 ; | CB | - | N |
| 5616 | 5619 | 1 ; | HB2 | - | HG2 |
| 5616 | 5620 | 1 ; | HB2 | - | HG3 |
| 5616 | 5621 | 1 ; | HB2 | - | CD |
| 5616 | 5626 | 1 ; | HB2 | - | C |
| 5617 | 5619 | 1 ; | HB3 | - | HG2 |
| 5617 | 5620 | 1 ; | HB3 | - | HG3 |

| | | | | | |
|---|---|---|---|---|---|
| 5617 | 5621 | 1 ; | HB3 | - | CD |
| 5617 | 5626 | 1 ; | HB3 | - | C |
| 5618 | 5624 | 1 ; | CG | - | HE21 |
| 5618 | 5625 | 1 ; | CG | - | HE22 |
| 5618 | 5626 | 1 ; | CG | - | C |
| 5619 | 5622 | 1 ; | HG2 | - | OE1 |
| 5619 | 5623 | 1 ; | HG2 | - | NE2 |
| 5620 | 5622 | 1 ; | HG3 | - | OE1 |
| 5620 | 5623 | 1 ; | HG3 | - | NE2 |
| 5622 | 5624 | 1 ; | OE1 | - | HE21 |
| 5622 | 5625 | 1 ; | OE1 | - | HE22 |
| 5626 | 5631 | 1 ; | C | - | HA |
| 5626 | 5632 | 1 ; | C | - | CB |
| 5626 | 5648 | 1 ; | C | - | C |
| 5627 | 5629 | 1 ; | O | - | H |
| 5627 | 5630 | 1 ; | O | - | CA |
| 5628 | 5633 | 1 ; | N | - | HB2 |
| 5628 | 5634 | 1 ; | N | - | HB3 |
| 5628 | 5635 | 1 ; | N | - | CG |
| 5628 | 5649 | 1 ; | N | - | O |
| 5628 | 5650 | 1 ; | N | - | N |
| 5629 | 5631 | 1 ; | H | - | HA |
| 5629 | 5632 | 1 ; | H | - | CB |
| 5629 | 5648 | 1 ; | H | - | C |
| 5630 | 5636 | 1 ; | CA | - | HG2 |
| 5630 | 5637 | 1 ; | CA | - | HG3 |
| 5630 | 5638 | 1 ; | CA | - | CD |
| 5630 | 5651 | 1 ; | CA | - | H |
| 5630 | 5652 | 1 ; | CA | - | CA |
| 5631 | 5633 | 1 ; | HA | - | HB2 |
| 5631 | 5634 | 1 ; | HA | - | HB3 |
| 5631 | 5635 | 1 ; | HA | - | CG |
| 5631 | 5649 | 1 ; | HA | - | O |
| 5631 | 5650 | 1 ; | HA | - | N |
| 5632 | 5639 | 1 ; | CB | - | HD2 |
| 5632 | 5640 | 1 ; | CB | - | HD3 |
| 5632 | 5641 | 1 ; | CB | - | CE |
| 5632 | 5649 | 1 ; | CB | - | O |
| 5632 | 5650 | 1 ; | CB | - | N |
| 5633 | 5636 | 1 ; | HB2 | - | HG2 |
| 5633 | 5637 | 1 ; | HB2 | - | HG3 |
| 5633 | 5638 | 1 ; | HB2 | - | CD |
| 5633 | 5648 | 1 ; | HB2 | - | C |
| 5634 | 5636 | 1 ; | HB3 | - | HG2 |
| 5634 | 5637 | 1 ; | HB3 | - | HG3 |
| 5634 | 5638 | 1 ; | HB3 | - | CD |
| 5634 | 5648 | 1 ; | HB3 | - | C |
| 5635 | 5642 | 1 ; | CG | - | HE2 |
| 5635 | 5643 | 1 ; | CG | - | HE3 |
| 5635 | 5644 | 1 ; | CG | - | NZ |
| 5635 | 5648 | 1 ; | CG | - | C |
| 5636 | 5639 | 1 ; | HG2 | - | HD2 |

| | | | | | |
|---|---|---|---|---|---|
| 5636 | 5640 | 1 ; | HG2 | - | HD3 |
| 5636 | 5641 | 1 ; | HG2 | - | CE |
| 5637 | 5639 | 1 ; | HG3 | - | HD2 |
| 5637 | 5640 | 1 ; | HG3 | - | HD3 |
| 5637 | 5641 | 1 ; | HG3 | - | CE |
| 5638 | 5645 | 1 ; | CD | - | HZ1 |
| 5638 | 5646 | 1 ; | CD | - | HZ2 |
| 5638 | 5647 | 1 ; | CD | - | HZ3 |
| 5639 | 5642 | 1 ; | HD2 | - | HE2 |
| 5639 | 5643 | 1 ; | HD2 | - | HE3 |
| 5639 | 5644 | 1 ; | HD2 | - | NZ |
| 5640 | 5642 | 1 ; | HD3 | - | HE2 |
| 5640 | 5643 | 1 ; | HD3 | - | HE3 |
| 5640 | 5644 | 1 ; | HD3 | - | NZ |
| 5642 | 5645 | 1 ; | HE2 | - | HZ1 |
| 5642 | 5646 | 1 ; | HE2 | - | HZ2 |
| 5642 | 5647 | 1 ; | HE2 | - | HZ3 |
| 5643 | 5645 | 1 ; | HE3 | - | HZ1 |
| 5643 | 5646 | 1 ; | HE3 | - | HZ2 |
| 5643 | 5647 | 1 ; | HE3 | - | HZ3 |
| 5648 | 5653 | 1 ; | C | - | HA |
| 5648 | 5654 | 1 ; | C | - | CB |
| 5648 | 5667 | 1 ; | C | - | C |
| 5649 | 5651 | 1 ; | O | - | H |
| 5649 | 5652 | 1 ; | O | - | CA |
| 5650 | 5655 | 1 ; | N | - | HB |
| 5650 | 5656 | 1 ; | N | - | CG2 |
| 5650 | 5660 | 1 ; | N | - | CG1 |
| 5650 | 5668 | 1 ; | N | - | O |
| 5650 | 5669 | 1 ; | N | - | N |
| 5651 | 5653 | 1 ; | H | - | HA |
| 5651 | 5654 | 1 ; | H | - | CB |
| 5651 | 5667 | 1 ; | H | - | C |
| 5652 | 5657 | 1 ; | CA | - | HG21 |
| 5652 | 5658 | 1 ; | CA | - | HG22 |
| 5652 | 5659 | 1 ; | CA | - | HG23 |
| 5652 | 5661 | 1 ; | CA | - | HG12 |
| 5652 | 5662 | 1 ; | CA | - | HG13 |
| 5652 | 5663 | 1 ; | CA | - | CD1 |
| 5652 | 5670 | 1 ; | CA | - | CD |
| 5652 | 5679 | 1 ; | CA | - | CA |
| 5653 | 5655 | 1 ; | HA | - | HB |
| 5653 | 5656 | 1 ; | HA | - | CG2 |
| 5653 | 5660 | 1 ; | HA | - | CG1 |
| 5653 | 5668 | 1 ; | HA | - | O |
| 5653 | 5669 | 1 ; | HA | - | N |
| 5654 | 5664 | 1 ; | CB | - | HD11 |
| 5654 | 5665 | 1 ; | CB | - | HD12 |
| 5654 | 5666 | 1 ; | CB | - | HD13 |
| 5654 | 5668 | 1 ; | CB | - | O |
| 5654 | 5669 | 1 ; | CB | - | N |
| 5655 | 5657 | 1 ; | HB | - | HG21 |

| | | | | |
|---|---|---|---|---|
| 5655 | 5658 | 1 | ; | HB - HG22 |
| 5655 | 5659 | 1 | ; | HB - HG23 |
| 5655 | 5661 | 1 | ; | HB - HG12 |
| 5655 | 5662 | 1 | ; | HB - HG13 |
| 5655 | 5663 | 1 | ; | HB - CD1 |
| 5655 | 5667 | 1 | ; | HB - C |
| 5656 | 5661 | 1 | ; | CG2 - HG12 |
| 5656 | 5662 | 1 | ; | CG2 - HG13 |
| 5656 | 5663 | 1 | ; | CG2 - CD1 |
| 5656 | 5667 | 1 | ; | CG2 - C |
| 5657 | 5660 | 1 | ; | HG21 - CG1 |
| 5658 | 5660 | 1 | ; | HG22 - CG1 |
| 5659 | 5660 | 1 | ; | HG23 - CG1 |
| 5660 | 5667 | 1 | ; | CG1 - C |
| 5661 | 5664 | 1 | ; | HG12 - HD11 |
| 5661 | 5665 | 1 | ; | HG12 - HD12 |
| 5661 | 5666 | 1 | ; | HG12 - HD13 |
| 5662 | 5664 | 1 | ; | HG13 - HD11 |
| 5662 | 5665 | 1 | ; | HG13 - HD12 |
| 5662 | 5666 | 1 | ; | HG13 - HD13 |
| 5667 | 5671 | 1 | ; | C - HD2 |
| 5667 | 5672 | 1 | ; | C - HD3 |
| 5667 | 5673 | 1 | ; | C - CG |
| 5667 | 5676 | 1 | ; | C - CB |
| 5667 | 5680 | 1 | ; | C - HA |
| 5667 | 5681 | 1 | ; | C - C |
| 5668 | 5670 | 1 | ; | O - CD |
| 5668 | 5679 | 1 | ; | O - CA |
| 5669 | 5674 | 1 | ; | N - HG2 |
| 5669 | 5675 | 1 | ; | N - HG3 |
| 5669 | 5677 | 1 | ; | N - HB2 |
| 5669 | 5678 | 1 | ; | N - HB3 |
| 5669 | 5682 | 1 | ; | N - O |
| 5669 | 5683 | 1 | ; | N - N |
| 5670 | 5677 | 1 | ; | CD - HB2 |
| 5670 | 5678 | 1 | ; | CD - HB3 |
| 5670 | 5680 | 1 | ; | CD - HA |
| 5670 | 5681 | 1 | ; | CD - C |
| 5671 | 5674 | 1 | ; | HD2 - HG2 |
| 5671 | 5675 | 1 | ; | HD2 - HG3 |
| 5671 | 5676 | 1 | ; | HD2 - CB |
| 5671 | 5679 | 1 | ; | HD2 - CA |
| 5672 | 5674 | 1 | ; | HD3 - HG2 |
| 5672 | 5675 | 1 | ; | HD3 - HG3 |
| 5672 | 5676 | 1 | ; | HD3 - CB |
| 5672 | 5679 | 1 | ; | HD3 - CA |
| 5673 | 5680 | 1 | ; | CG - HA |
| 5673 | 5681 | 1 | ; | CG - C |
| 5674 | 5677 | 1 | ; | HG2 - HB2 |
| 5674 | 5678 | 1 | ; | HG2 - HB3 |
| 5674 | 5679 | 1 | ; | HG2 - CA |
| 5675 | 5677 | 1 | ; | HG3 - HB2 |

| | | | |
|---|---|---|---|
| 5675 | 5678 | 1 ; | HG3 - HB3 |
| 5675 | 5679 | 1 ; | HG3 - CA |
| 5676 | 5682 | 1 ; | CB - O |
| 5676 | 5683 | 1 ; | CB - N |
| 5677 | 5680 | 1 ; | HB2 - HA |
| 5677 | 5681 | 1 ; | HB2 - C |
| 5678 | 5680 | 1 ; | HB3 - HA |
| 5678 | 5681 | 1 ; | HB3 - C |
| 5679 | 5684 | 1 ; | CA - CD |
| 5679 | 5693 | 1 ; | CA - CA |
| 5680 | 5682 | 1 ; | HA - O |
| 5680 | 5683 | 1 ; | HA - N |
| 5681 | 5685 | 1 ; | C - HD2 |
| 5681 | 5686 | 1 ; | C - HD3 |
| 5681 | 5687 | 1 ; | C - CG |
| 5681 | 5690 | 1 ; | C - CB |
| 5681 | 5694 | 1 ; | C - HA |
| 5681 | 5695 | 1 ; | C - C |
| 5682 | 5684 | 1 ; | O - CD |
| 5682 | 5693 | 1 ; | O - CA |
| 5683 | 5688 | 1 ; | N - HG2 |
| 5683 | 5689 | 1 ; | N - HG3 |
| 5683 | 5691 | 1 ; | N - HB2 |
| 5683 | 5692 | 1 ; | N - HB3 |
| 5683 | 5696 | 1 ; | N - O |
| 5683 | 5697 | 1 ; | N - N |
| 5684 | 5691 | 1 ; | CD - HB2 |
| 5684 | 5692 | 1 ; | CD - HB3 |
| 5684 | 5694 | 1 ; | CD - HA |
| 5684 | 5695 | 1 ; | CD - C |
| 5685 | 5688 | 1 ; | HD2 - HG2 |
| 5685 | 5689 | 1 ; | HD2 - HG3 |
| 5685 | 5690 | 1 ; | HD2 - CB |
| 5685 | 5693 | 1 ; | HD2 - CA |
| 5686 | 5688 | 1 ; | HD3 - HG2 |
| 5686 | 5689 | 1 ; | HD3 - HG3 |
| 5686 | 5690 | 1 ; | HD3 - CB |
| 5686 | 5693 | 1 ; | HD3 - CA |
| 5687 | 5694 | 1 ; | CG - HA |
| 5687 | 5695 | 1 ; | CG - C |
| 5688 | 5691 | 1 ; | HG2 - HB2 |
| 5688 | 5692 | 1 ; | HG2 - HB3 |
| 5688 | 5693 | 1 ; | HG2 - CA |
| 5689 | 5691 | 1 ; | HG3 - HB2 |
| 5689 | 5692 | 1 ; | HG3 - HB3 |
| 5689 | 5693 | 1 ; | HG3 - CA |
| 5690 | 5696 | 1 ; | CB - O |
| 5690 | 5697 | 1 ; | CB - N |
| 5691 | 5694 | 1 ; | HB2 - HA |
| 5691 | 5695 | 1 ; | HB2 - C |
| 5692 | 5694 | 1 ; | HB3 - HA |
| 5692 | 5695 | 1 ; | HB3 - C |

| | | | |
|---|---|---|---|
| 5693 | 5698 | 1 ; | CA - H |
| 5693 | 5699 | 1 ; | CA - CA |
| 5694 | 5696 | 1 ; | HA - O |
| 5694 | 5697 | 1 ; | HA - N |
| 5695 | 5700 | 1 ; | C - HA |
| 5695 | 5701 | 1 ; | C - CB |
| 5695 | 5707 | 1 ; | C - C |
| 5696 | 5698 | 1 ; | O - H |
| 5696 | 5699 | 1 ; | O - CA |
| 5697 | 5702 | 1 ; | N - HB2 |
| 5697 | 5703 | 1 ; | N - HB3 |
| 5697 | 5704 | 1 ; | N - CG |
| 5697 | 5708 | 1 ; | N - O |
| 5697 | 5709 | 1 ; | N - N |
| 5698 | 5700 | 1 ; | H - HA |
| 5698 | 5701 | 1 ; | H - CB |
| 5698 | 5707 | 1 ; | H - C |
| 5699 | 5705 | 1 ; | CA - OD1 |
| 5699 | 5706 | 1 ; | CA - OD2 |
| 5699 | 5710 | 1 ; | CA - H |
| 5699 | 5711 | 1 ; | CA - CA |
| 5700 | 5702 | 1 ; | HA - HB2 |
| 5700 | 5703 | 1 ; | HA - HB3 |
| 5700 | 5704 | 1 ; | HA - CG |
| 5700 | 5708 | 1 ; | HA - O |
| 5700 | 5709 | 1 ; | HA - N |
| 5701 | 5708 | 1 ; | CB - O |
| 5701 | 5709 | 1 ; | CB - N |
| 5702 | 5705 | 1 ; | HB2 - OD1 |
| 5702 | 5706 | 1 ; | HB2 - OD2 |
| 5702 | 5707 | 1 ; | HB2 - C |
| 5703 | 5705 | 1 ; | HB3 - OD1 |
| 5703 | 5706 | 1 ; | HB3 - OD2 |
| 5703 | 5707 | 1 ; | HB3 - C |
| 5704 | 5707 | 1 ; | CG - C |
| 5707 | 5712 | 1 ; | C - HA |
| 5707 | 5713 | 1 ; | C - CB |
| 5707 | 5726 | 1 ; | C - C |
| 5708 | 5710 | 1 ; | O - H |
| 5708 | 5711 | 1 ; | O - CA |
| 5709 | 5714 | 1 ; | N - HB |
| 5709 | 5715 | 1 ; | N - CG2 |
| 5709 | 5719 | 1 ; | N - CG1 |
| 5709 | 5727 | 1 ; | N - O |
| 5709 | 5728 | 1 ; | N - N |
| 5710 | 5712 | 1 ; | H - HA |
| 5710 | 5713 | 1 ; | H - CB |
| 5710 | 5726 | 1 ; | H - C |
| 5711 | 5716 | 1 ; | CA - HG21 |
| 5711 | 5717 | 1 ; | CA - HG22 |
| 5711 | 5718 | 1 ; | CA - HG23 |
| 5711 | 5720 | 1 ; | CA - HG12 |

| | | | |
|---|---|---|---|
| 5711 | 5721 | 1 ; | CA - HG13 |
| 5711 | 5722 | 1 ; | CA - CD1 |
| 5711 | 5729 | 1 ; | CA - H |
| 5711 | 5730 | 1 ; | CA - CA |
| 5712 | 5714 | 1 ; | HA - HB |
| 5712 | 5715 | 1 ; | HA - CG2 |
| 5712 | 5719 | 1 ; | HA - CG1 |
| 5712 | 5727 | 1 ; | HA - O |
| 5712 | 5728 | 1 ; | HA - N |
| 5713 | 5723 | 1 ; | CB - HD11 |
| 5713 | 5724 | 1 ; | CB - HD12 |
| 5713 | 5725 | 1 ; | CB - HD13 |
| 5713 | 5727 | 1 ; | CB - O |
| 5713 | 5728 | 1 ; | CB - N |
| 5714 | 5716 | 1 ; | HB - HG21 |
| 5714 | 5717 | 1 ; | HB - HG22 |
| 5714 | 5718 | 1 ; | HB - HG23 |
| 5714 | 5720 | 1 ; | HB - HG12 |
| 5714 | 5721 | 1 ; | HB - HG13 |
| 5714 | 5722 | 1 ; | HB - CD1 |
| 5714 | 5726 | 1 ; | HB - C |
| 5715 | 5720 | 1 ; | CG2 - HG12 |
| 5715 | 5721 | 1 ; | CG2 - HG13 |
| 5715 | 5722 | 1 ; | CG2 - CD1 |
| 5715 | 5726 | 1 ; | CG2 - C |
| 5716 | 5719 | 1 ; | HG21 - CG1 |
| 5717 | 5719 | 1 ; | HG22 - CG1 |
| 5718 | 5719 | 1 ; | HG23 - CG1 |
| 5719 | 5726 | 1 ; | CG1 - C |
| 5720 | 5723 | 1 ; | HG12 - HD11 |
| 5720 | 5724 | 1 ; | HG12 - HD12 |
| 5720 | 5725 | 1 ; | HG12 - HD13 |
| 5721 | 5723 | 1 ; | HG13 - HD11 |
| 5721 | 5724 | 1 ; | HG13 - HD12 |
| 5721 | 5725 | 1 ; | HG13 - HD13 |
| 5726 | 5731 | 1 ; | C - HA |
| 5726 | 5732 | 1 ; | C - CB |
| 5726 | 5743 | 1 ; | C - C |
| 5727 | 5729 | 1 ; | O - H |
| 5727 | 5730 | 1 ; | O - CA |
| 5728 | 5733 | 1 ; | N - HB2 |
| 5728 | 5734 | 1 ; | N - HB3 |
| 5728 | 5735 | 1 ; | N - CG |
| 5728 | 5744 | 1 ; | N - O |
| 5728 | 5745 | 1 ; | N - N |
| 5729 | 5731 | 1 ; | H - HA |
| 5729 | 5732 | 1 ; | H - CB |
| 5729 | 5743 | 1 ; | H - C |
| 5730 | 5736 | 1 ; | CA - ND1 |
| 5730 | 5741 | 1 ; | CA - CD2 |
| 5730 | 5746 | 1 ; | CA - H |
| 5730 | 5747 | 1 ; | CA - CA |

| | | | | | | |
|---|---|---|---|---|---|---|
| 5731 | 5733 | 1 | ; | HA | - | HB2 |
| 5731 | 5734 | 1 | ; | HA | - | HB3 |
| 5731 | 5735 | 1 | ; | HA | - | CG |
| 5731 | 5744 | 1 | ; | HA | - | O |
| 5731 | 5745 | 1 | ; | HA | - | N |
| 5732 | 5737 | 1 | ; | CB | - | CE1 |
| 5732 | 5739 | 1 | ; | CB | - | NE2 |
| 5732 | 5742 | 1 | ; | CB | - | HD2 |
| 5732 | 5744 | 1 | ; | CB | - | O |
| 5732 | 5745 | 1 | ; | CB | - | N |
| 5733 | 5736 | 1 | ; | HB2 | - | ND1 |
| 5733 | 5741 | 1 | ; | HB2 | - | CD2 |
| 5733 | 5743 | 1 | ; | HB2 | - | C |
| 5734 | 5736 | 1 | ; | HB3 | - | ND1 |
| 5734 | 5741 | 1 | ; | HB3 | - | CD2 |
| 5734 | 5743 | 1 | ; | HB3 | - | C |
| 5735 | 5738 | 1 | ; | CG | - | HE1 |
| 5735 | 5740 | 1 | ; | CG | - | HE2 |
| 5735 | 5743 | 1 | ; | CG | - | C |
| 5736 | 5740 | 1 | ; | ND1 | - | HE2 |
| 5736 | 5742 | 1 | ; | ND1 | - | HD2 |
| 5737 | 5742 | 1 | ; | CE1 | - | HD2 |
| 5738 | 5740 | 1 | ; | HE1 | - | HE2 |
| 5738 | 5741 | 1 | ; | HE1 | - | CD2 |
| 5740 | 5742 | 1 | ; | HE2 | - | HD2 |
| 5743 | 5748 | 1 | ; | C | - | HA |
| 5743 | 5749 | 1 | ; | C | - | CB |
| 5743 | 5760 | 1 | ; | C | - | C |
| 5744 | 5746 | 1 | ; | O | - | H |
| 5744 | 5747 | 1 | ; | O | - | CA |
| 5745 | 5750 | 1 | ; | N | - | HB2 |
| 5745 | 5751 | 1 | ; | N | - | HB3 |
| 5745 | 5752 | 1 | ; | N | - | CG |
| 5745 | 5761 | 1 | ; | N | - | O |
| 5745 | 5762 | 1 | ; | N | - | N |
| 5746 | 5748 | 1 | ; | H | - | HA |
| 5746 | 5749 | 1 | ; | H | - | CB |
| 5746 | 5760 | 1 | ; | H | - | C |
| 5747 | 5753 | 1 | ; | CA | - | HG2 |
| 5747 | 5754 | 1 | ; | CA | - | HG3 |
| 5747 | 5755 | 1 | ; | CA | - | CD |
| 5747 | 5763 | 1 | ; | CA | - | H |
| 5747 | 5764 | 1 | ; | CA | - | CA |
| 5748 | 5750 | 1 | ; | HA | - | HB2 |
| 5748 | 5751 | 1 | ; | HA | - | HB3 |
| 5748 | 5752 | 1 | ; | HA | - | CG |
| 5748 | 5761 | 1 | ; | HA | - | O |
| 5748 | 5762 | 1 | ; | HA | - | N |
| 5749 | 5756 | 1 | ; | CB | - | OE1 |
| 5749 | 5757 | 1 | ; | CB | - | NE2 |
| 5749 | 5761 | 1 | ; | CB | - | O |
| 5749 | 5762 | 1 | ; | CB | - | N |

| | | | | | | |
|---|---|---|---|---|---|---|
| 5750 | 5753 | 1 | ; | HB2 | - | HG2 |
| 5750 | 5754 | 1 | ; | HB2 | - | HG3 |
| 5750 | 5755 | 1 | ; | HB2 | - | CD |
| 5750 | 5760 | 1 | ; | HB2 | - | C |
| 5751 | 5753 | 1 | ; | HB3 | - | HG2 |
| 5751 | 5754 | 1 | ; | HB3 | - | HG3 |
| 5751 | 5755 | 1 | ; | HB3 | - | CD |
| 5751 | 5760 | 1 | ; | HB3 | - | C |
| 5752 | 5758 | 1 | ; | CG | - | HE21 |
| 5752 | 5759 | 1 | ; | CG | - | HE22 |
| 5752 | 5760 | 1 | ; | CG | - | C |
| 5753 | 5756 | 1 | ; | HG2 | - | OE1 |
| 5753 | 5757 | 1 | ; | HG2 | - | NE2 |
| 5754 | 5756 | 1 | ; | HG3 | - | OE1 |
| 5754 | 5757 | 1 | ; | HG3 | - | NE2 |
| 5756 | 5758 | 1 | ; | OE1 | - | HE21 |
| 5756 | 5759 | 1 | ; | OE1 | - | HE22 |
| 5760 | 5765 | 1 | ; | C | - | HA |
| 5760 | 5766 | 1 | ; | C | - | CB |
| 5760 | 5776 | 1 | ; | C | - | C |
| 5761 | 5763 | 1 | ; | O | - | H |
| 5761 | 5764 | 1 | ; | O | - | CA |
| 5762 | 5767 | 1 | ; | N | - | HB |
| 5762 | 5768 | 1 | ; | N | - | CG1 |
| 5762 | 5772 | 1 | ; | N | - | CG2 |
| 5762 | 5777 | 1 | ; | N | - | O |
| 5762 | 5778 | 1 | ; | N | - | N |
| 5763 | 5765 | 1 | ; | H | - | HA |
| 5763 | 5766 | 1 | ; | H | - | CB |
| 5763 | 5776 | 1 | ; | H | - | C |
| 5764 | 5769 | 1 | ; | CA | - | HG11 |
| 5764 | 5770 | 1 | ; | CA | - | HG12 |
| 5764 | 5771 | 1 | ; | CA | - | HG13 |
| 5764 | 5773 | 1 | ; | CA | - | HG21 |
| 5764 | 5774 | 1 | ; | CA | - | HG22 |
| 5764 | 5775 | 1 | ; | CA | - | HG23 |
| 5764 | 5779 | 1 | ; | CA | - | H |
| 5764 | 5780 | 1 | ; | CA | - | CA |
| 5765 | 5767 | 1 | ; | HA | - | HB |
| 5765 | 5768 | 1 | ; | HA | - | CG1 |
| 5765 | 5772 | 1 | ; | HA | - | CG2 |
| 5765 | 5777 | 1 | ; | HA | - | O |
| 5765 | 5778 | 1 | ; | HA | - | N |
| 5766 | 5777 | 1 | ; | CB | - | O |
| 5766 | 5778 | 1 | ; | CB | - | N |
| 5767 | 5769 | 1 | ; | HB | - | HG11 |
| 5767 | 5770 | 1 | ; | HB | - | HG12 |
| 5767 | 5771 | 1 | ; | HB | - | HG13 |
| 5767 | 5773 | 1 | ; | HB | - | HG21 |
| 5767 | 5774 | 1 | ; | HB | - | HG22 |
| 5767 | 5775 | 1 | ; | HB | - | HG23 |
| 5767 | 5776 | 1 | ; | HB | - | C |

```
 5768   5773       1 ;    CG1 - HG21
 5768   5774       1 ;    CG1 - HG22
 5768   5775       1 ;    CG1 - HG23
 5768   5776       1 ;    CG1 - C
 5769   5772       1 ;   HG11 - CG2
 5770   5772       1 ;   HG12 - CG2
 5771   5772       1 ;   HG13 - CG2
 5772   5776       1 ;    CG2 - C
 5776   5781       1 ;      C - HA
 5776   5782       1 ;      C - CB
 5776   5795       1 ;      C - C
 5777   5779       1 ;      O - H
 5777   5780       1 ;      O - CA
 5778   5783       1 ;      N - HB2
 5778   5784       1 ;      N - HB3
 5778   5785       1 ;      N - CG
 5778   5796       1 ;      N - O
 5778   5797       1 ;      N - N
 5779   5781       1 ;      H - HA
 5779   5782       1 ;      H - CB
 5779   5795       1 ;      H - C
 5780   5786       1 ;     CA - HG
 5780   5787       1 ;     CA - CD1
 5780   5791       1 ;     CA - CD2
 5780   5798       1 ;     CA - H
 5780   5799       1 ;     CA - CA
 5781   5783       1 ;     HA - HB2
 5781   5784       1 ;     HA - HB3
 5781   5785       1 ;     HA - CG
 5781   5796       1 ;     HA - O
 5781   5797       1 ;     HA - N
 5782   5788       1 ;     CB - HD11
 5782   5789       1 ;     CB - HD12
 5782   5790       1 ;     CB - HD13
 5782   5792       1 ;     CB - HD21
 5782   5793       1 ;     CB - HD22
 5782   5794       1 ;     CB - HD23
 5782   5796       1 ;     CB - O
 5782   5797       1 ;     CB - N
 5783   5786       1 ;    HB2 - HG
 5783   5787       1 ;    HB2 - CD1
 5783   5791       1 ;    HB2 - CD2
 5783   5795       1 ;    HB2 - C
 5784   5786       1 ;    HB3 - HG
 5784   5787       1 ;    HB3 - CD1
 5784   5791       1 ;    HB3 - CD2
 5784   5795       1 ;    HB3 - C
 5785   5795       1 ;     CG - C
 5786   5788       1 ;     HG - HD11
 5786   5789       1 ;     HG - HD12
 5786   5790       1 ;     HG - HD13
 5786   5792       1 ;     HG - HD21
```

| | | | |
|---|---|---|---|
| 5786 | 5793 | 1 ; | HG - HD22 |
| 5786 | 5794 | 1 ; | HG - HD23 |
| 5787 | 5792 | 1 ; | CD1 - HD21 |
| 5787 | 5793 | 1 ; | CD1 - HD22 |
| 5787 | 5794 | 1 ; | CD1 - HD23 |
| 5788 | 5791 | 1 ; | HD11 - CD2 |
| 5789 | 5791 | 1 ; | HD12 - CD2 |
| 5790 | 5791 | 1 ; | HD13 - CD2 |
| 5795 | 5800 | 1 ; | C - HA |
| 5795 | 5801 | 1 ; | C - CB |
| 5795 | 5819 | 1 ; | C - C |
| 5796 | 5798 | 1 ; | O - H |
| 5796 | 5799 | 1 ; | O - CA |
| 5797 | 5802 | 1 ; | N - HB2 |
| 5797 | 5803 | 1 ; | N - HB3 |
| 5797 | 5804 | 1 ; | N - CG |
| 5797 | 5820 | 1 ; | N - O |
| 5797 | 5821 | 1 ; | N - N |
| 5798 | 5800 | 1 ; | H - HA |
| 5798 | 5801 | 1 ; | H - CB |
| 5798 | 5819 | 1 ; | H - C |
| 5799 | 5805 | 1 ; | CA - HG2 |
| 5799 | 5806 | 1 ; | CA - HG3 |
| 5799 | 5807 | 1 ; | CA - CD |
| 5799 | 5822 | 1 ; | CA - H |
| 5799 | 5823 | 1 ; | CA - CA |
| 5800 | 5802 | 1 ; | HA - HB2 |
| 5800 | 5803 | 1 ; | HA - HB3 |
| 5800 | 5804 | 1 ; | HA - CG |
| 5800 | 5820 | 1 ; | HA - O |
| 5800 | 5821 | 1 ; | HA - N |
| 5801 | 5808 | 1 ; | CB - HD2 |
| 5801 | 5809 | 1 ; | CB - HD3 |
| 5801 | 5810 | 1 ; | CB - NE |
| 5801 | 5820 | 1 ; | CB - O |
| 5801 | 5821 | 1 ; | CB - N |
| 5802 | 5805 | 1 ; | HB2 - HG2 |
| 5802 | 5806 | 1 ; | HB2 - HG3 |
| 5802 | 5807 | 1 ; | HB2 - CD |
| 5802 | 5819 | 1 ; | HB2 - C |
| 5803 | 5805 | 1 ; | HB3 - HG2 |
| 5803 | 5806 | 1 ; | HB3 - HG3 |
| 5803 | 5807 | 1 ; | HB3 - CD |
| 5803 | 5819 | 1 ; | HB3 - C |
| 5804 | 5811 | 1 ; | CG - HE |
| 5804 | 5812 | 1 ; | CG - CZ |
| 5804 | 5819 | 1 ; | CG - C |
| 5805 | 5808 | 1 ; | HG2 - HD2 |
| 5805 | 5809 | 1 ; | HG2 - HD3 |
| 5805 | 5810 | 1 ; | HG2 - NE |
| 5806 | 5808 | 1 ; | HG3 - HD2 |
| 5806 | 5809 | 1 ; | HG3 - HD3 |

| | | | | | |
|---|---|---|---|---|---|
| 5806 | 5810 | 1 ; | HG3 | - | NE |
| 5807 | 5813 | 1 ; | CD | - | NH1 |
| 5807 | 5816 | 1 ; | CD | - | NH2 |
| 5808 | 5811 | 1 ; | HD2 | - | HE |
| 5808 | 5812 | 1 ; | HD2 | - | CZ |
| 5809 | 5811 | 1 ; | HD3 | - | HE |
| 5809 | 5812 | 1 ; | HD3 | - | CZ |
| 5810 | 5814 | 1 ; | NE | - | HH11 |
| 5810 | 5815 | 1 ; | NE | - | HH12 |
| 5810 | 5817 | 1 ; | NE | - | HH21 |
| 5810 | 5818 | 1 ; | NE | - | HH22 |
| 5811 | 5813 | 1 ; | HE | - | NH1 |
| 5811 | 5816 | 1 ; | HE | - | NH2 |
| 5813 | 5817 | 1 ; | NH1 | - | HH21 |
| 5813 | 5818 | 1 ; | NH1 | - | HH22 |
| 5814 | 5816 | 1 ; | HH11 | - | NH2 |
| 5815 | 5816 | 1 ; | HH12 | - | NH2 |
| 5819 | 5824 | 1 ; | C | - | HA |
| 5819 | 5825 | 1 ; | C | - | CB |
| 5819 | 5831 | 1 ; | C | - | C |
| 5820 | 5822 | 1 ; | O | - | H |
| 5820 | 5823 | 1 ; | O | - | CA |
| 5821 | 5826 | 1 ; | N | - | HB2 |
| 5821 | 5827 | 1 ; | N | - | HB3 |
| 5821 | 5828 | 1 ; | N | - | CG |
| 5821 | 5832 | 1 ; | N | - | O |
| 5821 | 5833 | 1 ; | N | - | N |
| 5822 | 5824 | 1 ; | H | - | HA |
| 5822 | 5825 | 1 ; | H | - | CB |
| 5822 | 5831 | 1 ; | H | - | C |
| 5823 | 5829 | 1 ; | CA | - | OD1 |
| 5823 | 5830 | 1 ; | CA | - | OD2 |
| 5823 | 5834 | 1 ; | CA | - | H |
| 5823 | 5835 | 1 ; | CA | - | CA |
| 5824 | 5826 | 1 ; | HA | - | HB2 |
| 5824 | 5827 | 1 ; | HA | - | HB3 |
| 5824 | 5828 | 1 ; | HA | - | CG |
| 5824 | 5832 | 1 ; | HA | - | O |
| 5824 | 5833 | 1 ; | HA | - | N |
| 5825 | 5832 | 1 ; | CB | - | O |
| 5825 | 5833 | 1 ; | CB | - | N |
| 5826 | 5829 | 1 ; | HB2 | - | OD1 |
| 5826 | 5830 | 1 ; | HB2 | - | OD2 |
| 5826 | 5831 | 1 ; | HB2 | - | C |
| 5827 | 5829 | 1 ; | HB3 | - | OD1 |
| 5827 | 5830 | 1 ; | HB3 | - | OD2 |
| 5827 | 5831 | 1 ; | HB3 | - | C |
| 5828 | 5831 | 1 ; | CG | - | C |
| 5831 | 5836 | 1 ; | C | - | HA |
| 5831 | 5837 | 1 ; | C | - | CB |
| 5831 | 5850 | 1 ; | C | - | C |
| 5832 | 5834 | 1 ; | O | - | H |

| | | | |
|---|---|---|---|
| 5832 | 5835 | 1 ; | O - CA |
| 5833 | 5838 | 1 ; | N - HB |
| 5833 | 5839 | 1 ; | N - CG2 |
| 5833 | 5843 | 1 ; | N - CG1 |
| 5833 | 5851 | 1 ; | N - O |
| 5833 | 5852 | 1 ; | N - N |
| 5834 | 5836 | 1 ; | H - HA |
| 5834 | 5837 | 1 ; | H - CB |
| 5834 | 5850 | 1 ; | H - C |
| 5835 | 5840 | 1 ; | CA - HG21 |
| 5835 | 5841 | 1 ; | CA - HG22 |
| 5835 | 5842 | 1 ; | CA - HG23 |
| 5835 | 5844 | 1 ; | CA - HG12 |
| 5835 | 5845 | 1 ; | CA - HG13 |
| 5835 | 5846 | 1 ; | CA - CD1 |
| 5835 | 5853 | 1 ; | CA - H |
| 5835 | 5854 | 1 ; | CA - CA |
| 5836 | 5838 | 1 ; | HA - HB |
| 5836 | 5839 | 1 ; | HA - CG2 |
| 5836 | 5843 | 1 ; | HA - CG1 |
| 5836 | 5851 | 1 ; | HA - O |
| 5836 | 5852 | 1 ; | HA - N |
| 5837 | 5847 | 1 ; | CB - HD11 |
| 5837 | 5848 | 1 ; | CB - HD12 |
| 5837 | 5849 | 1 ; | CB - HD13 |
| 5837 | 5851 | 1 ; | CB - O |
| 5837 | 5852 | 1 ; | CB - N |
| 5838 | 5840 | 1 ; | HB - HG21 |
| 5838 | 5841 | 1 ; | HB - HG22 |
| 5838 | 5842 | 1 ; | HB - HG23 |
| 5838 | 5844 | 1 ; | HB - HG12 |
| 5838 | 5845 | 1 ; | HB - HG13 |
| 5838 | 5846 | 1 ; | HB - CD1 |
| 5838 | 5850 | 1 ; | HB - C |
| 5839 | 5844 | 1 ; | CG2 - HG12 |
| 5839 | 5845 | 1 ; | CG2 - HG13 |
| 5839 | 5846 | 1 ; | CG2 - CD1 |
| 5839 | 5850 | 1 ; | CG2 - C |
| 5840 | 5843 | 1 ; | HG21 - CG1 |
| 5841 | 5843 | 1 ; | HG22 - CG1 |
| 5842 | 5843 | 1 ; | HG23 - CG1 |
| 5843 | 5850 | 1 ; | CG1 - C |
| 5844 | 5847 | 1 ; | HG12 - HD11 |
| 5844 | 5848 | 1 ; | HG12 - HD12 |
| 5844 | 5849 | 1 ; | HG12 - HD13 |
| 5845 | 5847 | 1 ; | HG13 - HD11 |
| 5845 | 5848 | 1 ; | HG13 - HD12 |
| 5845 | 5849 | 1 ; | HG13 - HD13 |
| 5850 | 5855 | 1 ; | C - HA |
| 5850 | 5856 | 1 ; | C - CB |
| 5850 | 5862 | 1 ; | C - C |
| 5851 | 5853 | 1 ; | O - H |

| | | | | | |
|---|---|---|---|---|---|
| 5851 | 5854 | 1 ; | O | - | CA |
| 5852 | 5857 | 1 ; | N | - | HB2 |
| 5852 | 5858 | 1 ; | N | - | HB3 |
| 5852 | 5859 | 1 ; | N | - | CG |
| 5852 | 5863 | 1 ; | N | - | O |
| 5852 | 5864 | 1 ; | N | - | N |
| 5853 | 5855 | 1 ; | H | - | HA |
| 5853 | 5856 | 1 ; | H | - | CB |
| 5853 | 5862 | 1 ; | H | - | C |
| 5854 | 5860 | 1 ; | CA | - | OD1 |
| 5854 | 5861 | 1 ; | CA | - | OD2 |
| 5854 | 5865 | 1 ; | CA | - | H |
| 5854 | 5866 | 1 ; | CA | - | CA |
| 5855 | 5857 | 1 ; | HA | - | HB2 |
| 5855 | 5858 | 1 ; | HA | - | HB3 |
| 5855 | 5859 | 1 ; | HA | - | CG |
| 5855 | 5863 | 1 ; | HA | - | O |
| 5855 | 5864 | 1 ; | HA | - | N |
| 5856 | 5863 | 1 ; | CB | - | O |
| 5856 | 5864 | 1 ; | CB | - | N |
| 5857 | 5860 | 1 ; | HB2 | - | OD1 |
| 5857 | 5861 | 1 ; | HB2 | - | OD2 |
| 5857 | 5862 | 1 ; | HB2 | - | C |
| 5858 | 5860 | 1 ; | HB3 | - | OD1 |
| 5858 | 5861 | 1 ; | HB3 | - | OD2 |
| 5858 | 5862 | 1 ; | HB3 | - | C |
| 5859 | 5862 | 1 ; | CG | - | C |
| 5862 | 5867 | 1 ; | C | - | HA |
| 5862 | 5868 | 1 ; | C | - | CB |
| 5862 | 5873 | 1 ; | C | - | C |
| 5863 | 5865 | 1 ; | O | - | H |
| 5863 | 5866 | 1 ; | O | - | CA |
| 5864 | 5869 | 1 ; | N | - | HB2 |
| 5864 | 5870 | 1 ; | N | - | HB3 |
| 5864 | 5871 | 1 ; | N | - | OG |
| 5864 | 5874 | 1 ; | N | - | O |
| 5864 | 5875 | 1 ; | N | - | N |
| 5865 | 5867 | 1 ; | H | - | HA |
| 5865 | 5868 | 1 ; | H | - | CB |
| 5865 | 5873 | 1 ; | H | - | C |
| 5866 | 5872 | 1 ; | CA | - | HG |
| 5866 | 5876 | 1 ; | CA | - | H |
| 5866 | 5877 | 1 ; | CA | - | CA |
| 5867 | 5869 | 1 ; | HA | - | HB2 |
| 5867 | 5870 | 1 ; | HA | - | HB3 |
| 5867 | 5871 | 1 ; | HA | - | OG |
| 5867 | 5874 | 1 ; | HA | - | O |
| 5867 | 5875 | 1 ; | HA | - | N |
| 5868 | 5874 | 1 ; | CB | - | O |
| 5868 | 5875 | 1 ; | CB | - | N |
| 5869 | 5872 | 1 ; | HB2 | - | HG |
| 5869 | 5873 | 1 ; | HB2 | - | C |

| | | | | | | |
|---|---|---|---|---|---|---|
| 5870 | 5872 | 1 | ; | HB3 | - | HG |
| 5870 | 5873 | 1 | ; | HB3 | - | C |
| 5871 | 5873 | 1 | ; | OG | - | C |
| 5873 | 5878 | 1 | ; | C | - | HA |
| 5873 | 5879 | 1 | ; | C | - | CB |
| 5873 | 5887 | 1 | ; | C | - | C |
| 5874 | 5876 | 1 | ; | O | - | H |
| 5874 | 5877 | 1 | ; | O | - | CA |
| 5875 | 5880 | 1 | ; | N | - | HB2 |
| 5875 | 5881 | 1 | ; | N | - | HB3 |
| 5875 | 5882 | 1 | ; | N | - | CG |
| 5875 | 5888 | 1 | ; | N | - | O |
| 5875 | 5889 | 1 | ; | N | - | N |
| 5876 | 5878 | 1 | ; | H | - | HA |
| 5876 | 5879 | 1 | ; | H | - | CB |
| 5876 | 5887 | 1 | ; | H | - | C |
| 5877 | 5883 | 1 | ; | CA | - | OD1 |
| 5877 | 5884 | 1 | ; | CA | - | ND2 |
| 5877 | 5890 | 1 | ; | CA | - | H |
| 5877 | 5891 | 1 | ; | CA | - | CA |
| 5878 | 5880 | 1 | ; | HA | - | HB2 |
| 5878 | 5881 | 1 | ; | HA | - | HB3 |
| 5878 | 5882 | 1 | ; | HA | - | CG |
| 5878 | 5888 | 1 | ; | HA | - | O |
| 5878 | 5889 | 1 | ; | HA | - | N |
| 5879 | 5885 | 1 | ; | CB | - | HD21 |
| 5879 | 5886 | 1 | ; | CB | - | HD22 |
| 5879 | 5888 | 1 | ; | CB | - | O |
| 5879 | 5889 | 1 | ; | CB | - | N |
| 5880 | 5883 | 1 | ; | HB2 | - | OD1 |
| 5880 | 5884 | 1 | ; | HB2 | - | ND2 |
| 5880 | 5887 | 1 | ; | HB2 | - | C |
| 5881 | 5883 | 1 | ; | HB3 | - | OD1 |
| 5881 | 5884 | 1 | ; | HB3 | - | ND2 |
| 5881 | 5887 | 1 | ; | HB3 | - | C |
| 5882 | 5887 | 1 | ; | CG | - | C |
| 5883 | 5885 | 1 | ; | OD1 | - | HD21 |
| 5883 | 5886 | 1 | ; | OD1 | - | HD22 |
| 5887 | 5892 | 1 | ; | C | - | HA |
| 5887 | 5893 | 1 | ; | C | - | CB |
| 5887 | 5897 | 1 | ; | C | - | C |
| 5888 | 5890 | 1 | ; | O | - | H |
| 5888 | 5891 | 1 | ; | O | - | CA |
| 5889 | 5894 | 1 | ; | N | - | HB1 |
| 5889 | 5895 | 1 | ; | N | - | HB2 |
| 5889 | 5896 | 1 | ; | N | - | HB3 |
| 5889 | 5898 | 1 | ; | N | - | O |
| 5889 | 5899 | 1 | ; | N | - | N |
| 5890 | 5892 | 1 | ; | H | - | HA |
| 5890 | 5893 | 1 | ; | H | - | CB |
| 5890 | 5897 | 1 | ; | H | - | C |
| 5891 | 5900 | 1 | ; | CA | - | H |

```
5891   5901    1 ;    CA - CA
5892   5894    1 ;    HA - HB1
5892   5895    1 ;    HA - HB2
5892   5896    1 ;    HA - HB3
5892   5898    1 ;    HA - O
5892   5899    1 ;    HA - N
5893   5898    1 ;    CB - O
5893   5899    1 ;    CB - N
5894   5897    1 ;   HB1 - C
5895   5897    1 ;   HB2 - C
5896   5897    1 ;   HB3 - C
5897   5902    1 ;     C - HA
5897   5903    1 ;     C - CB
5897   5908    1 ;     C - C
5898   5900    1 ;     O - H
5898   5901    1 ;     O - CA
5899   5904    1 ;     N - HB2
5899   5905    1 ;     N - HB3
5899   5906    1 ;     N - OG
5899   5909    1 ;     N - O
5899   5910    1 ;     N - N
5900   5902    1 ;     H - HA
5900   5903    1 ;     H - CB
5900   5908    1 ;     H - C
5901   5907    1 ;    CA - HG
5901   5911    1 ;    CA - H
5901   5912    1 ;    CA - CA
5902   5904    1 ;    HA - HB2
5902   5905    1 ;    HA - HB3
5902   5906    1 ;    HA - OG
5902   5909    1 ;    HA - O
5902   5910    1 ;    HA - N
5903   5909    1 ;    CB - O
5903   5910    1 ;    CB - N
5904   5907    1 ;   HB2 - HG
5904   5908    1 ;   HB2 - C
5905   5907    1 ;   HB3 - HG
5905   5908    1 ;   HB3 - C
5906   5908    1 ;    OG - C
5908   5913    1 ;     C - HA2
5908   5914    1 ;     C - HA3
5908   5915    1 ;     C - C
5909   5911    1 ;     O - H
5909   5912    1 ;     O - CA
5910   5916    1 ;     N - O
5910   5917    1 ;     N - N
5911   5913    1 ;     H - HA2
5911   5914    1 ;     H - HA3
5911   5915    1 ;     H - C
5912   5918    1 ;    CA - H
5912   5919    1 ;    CA - CA
5913   5916    1 ;   HA2 - O
```

| | | | |
|---|---|---|---|
| 5913 | 5917 | 1 ; | HA2 - N |
| 5914 | 5916 | 1 ; | HA3 - O |
| 5914 | 5917 | 1 ; | HA3 - N |
| 5915 | 5920 | 1 ; | C - HA |
| 5915 | 5921 | 1 ; | C - CB |
| 5915 | 5932 | 1 ; | C - C |
| 5916 | 5918 | 1 ; | O - H |
| 5916 | 5919 | 1 ; | O - CA |
| 5917 | 5922 | 1 ; | N - HB2 |
| 5917 | 5923 | 1 ; | N - HB3 |
| 5917 | 5924 | 1 ; | N - CG |
| 5917 | 5933 | 1 ; | N - O |
| 5917 | 5934 | 1 ; | N - N |
| 5918 | 5920 | 1 ; | H - HA |
| 5918 | 5921 | 1 ; | H - CB |
| 5918 | 5932 | 1 ; | H - C |
| 5919 | 5925 | 1 ; | CA - HG2 |
| 5919 | 5926 | 1 ; | CA - HG3 |
| 5919 | 5927 | 1 ; | CA - CD |
| 5919 | 5935 | 1 ; | CA - H |
| 5919 | 5936 | 1 ; | CA - CA |
| 5920 | 5922 | 1 ; | HA - HB2 |
| 5920 | 5923 | 1 ; | HA - HB3 |
| 5920 | 5924 | 1 ; | HA - CG |
| 5920 | 5933 | 1 ; | HA - O |
| 5920 | 5934 | 1 ; | HA - N |
| 5921 | 5928 | 1 ; | CB - OE1 |
| 5921 | 5929 | 1 ; | CB - NE2 |
| 5921 | 5933 | 1 ; | CB - O |
| 5921 | 5934 | 1 ; | CB - N |
| 5922 | 5925 | 1 ; | HB2 - HG2 |
| 5922 | 5926 | 1 ; | HB2 - HG3 |
| 5922 | 5927 | 1 ; | HB2 - CD |
| 5922 | 5932 | 1 ; | HB2 - C |
| 5923 | 5925 | 1 ; | HB3 - HG2 |
| 5923 | 5926 | 1 ; | HB3 - HG3 |
| 5923 | 5927 | 1 ; | HB3 - CD |
| 5923 | 5932 | 1 ; | HB3 - C |
| 5924 | 5930 | 1 ; | CG - HE21 |
| 5924 | 5931 | 1 ; | CG - HE22 |
| 5924 | 5932 | 1 ; | CG - C |
| 5925 | 5928 | 1 ; | HG2 - OE1 |
| 5925 | 5929 | 1 ; | HG2 - NE2 |
| 5926 | 5928 | 1 ; | HG3 - OE1 |
| 5926 | 5929 | 1 ; | HG3 - NE2 |
| 5928 | 5930 | 1 ; | OE1 - HE21 |
| 5928 | 5931 | 1 ; | OE1 - HE22 |
| 5932 | 5937 | 1 ; | C - HA |
| 5932 | 5938 | 1 ; | C - CB |
| 5932 | 5951 | 1 ; | C - C |
| 5933 | 5935 | 1 ; | O - H |
| 5933 | 5936 | 1 ; | O - CA |

```
5934   5939      1 ;      N - HB
5934   5940      1 ;      N - CG2
5934   5944      1 ;      N - CG1
5934   5952      1 ;      N - O
5934   5953      1 ;      N - N
5935   5937      1 ;      H - HA
5935   5938      1 ;      H - CB
5935   5951      1 ;      H - C
5936   5941      1 ;     CA - HG21
5936   5942      1 ;     CA - HG22
5936   5943      1 ;     CA - HG23
5936   5945      1 ;     CA - HG12
5936   5946      1 ;     CA - HG13
5936   5947      1 ;     CA - CD1
5936   5954      1 ;     CA - H
5936   5955      1 ;     CA - CA
5937   5939      1 ;     HA - HB
5937   5940      1 ;     HA - CG2
5937   5944      1 ;     HA - CG1
5937   5952      1 ;     HA - O
5937   5953      1 ;     HA - N
5938   5948      1 ;     CB - HD11
5938   5949      1 ;     CB - HD12
5938   5950      1 ;     CB - HD13
5938   5952      1 ;     CB - O
5938   5953      1 ;     CB - N
5939   5941      1 ;     HB - HG21
5939   5942      1 ;     HB - HG22
5939   5943      1 ;     HB - HG23
5939   5945      1 ;     HB - HG12
5939   5946      1 ;     HB - HG13
5939   5947      1 ;     HB - CD1
5939   5951      1 ;     HB - C
5940   5945      1 ;    CG2 - HG12
5940   5946      1 ;    CG2 - HG13
5940   5947      1 ;    CG2 - CD1
5940   5951      1 ;    CG2 - C
5941   5944      1 ;   HG21 - CG1
5942   5944      1 ;   HG22 - CG1
5943   5944      1 ;   HG23 - CG1
5944   5951      1 ;    CG1 - C
5945   5948      1 ;   HG12 - HD11
5945   5949      1 ;   HG12 - HD12
5945   5950      1 ;   HG12 - HD13
5946   5948      1 ;   HG13 - HD11
5946   5949      1 ;   HG13 - HD12
5946   5950      1 ;   HG13 - HD13
5951   5956      1 ;      C - HA
5951   5957      1 ;      C - CB
5951   5969      1 ;      C - C
5952   5954      1 ;      O - H
5952   5955      1 ;      O - CA
```

| | | | | | |
|---|---|---|---|---|---|
| 5953 | 5958 | 1 ; | N | - | HB2 |
| 5953 | 5959 | 1 ; | N | - | HB3 |
| 5953 | 5960 | 1 ; | N | - | CG |
| 5953 | 5970 | 1 ; | N | - | O |
| 5953 | 5971 | 1 ; | N | - | N |
| 5954 | 5956 | 1 ; | H | - | HA |
| 5954 | 5957 | 1 ; | H | - | CB |
| 5954 | 5969 | 1 ; | H | - | C |
| 5955 | 5961 | 1 ; | CA | - | ND1 |
| 5955 | 5967 | 1 ; | CA | - | CD2 |
| 5955 | 5972 | 1 ; | CA | - | H |
| 5955 | 5973 | 1 ; | CA | - | CA |
| 5956 | 5958 | 1 ; | HA | - | HB2 |
| 5956 | 5959 | 1 ; | HA | - | HB3 |
| 5956 | 5960 | 1 ; | HA | - | CG |
| 5956 | 5970 | 1 ; | HA | - | O |
| 5956 | 5971 | 1 ; | HA | - | N |
| 5957 | 5962 | 1 ; | CB | - | HD1 |
| 5957 | 5963 | 1 ; | CB | - | CE1 |
| 5957 | 5965 | 1 ; | CB | - | NE2 |
| 5957 | 5968 | 1 ; | CB | - | HD2 |
| 5957 | 5970 | 1 ; | CB | - | O |
| 5957 | 5971 | 1 ; | CB | - | N |
| 5958 | 5961 | 1 ; | HB2 | - | ND1 |
| 5958 | 5967 | 1 ; | HB2 | - | CD2 |
| 5958 | 5969 | 1 ; | HB2 | - | C |
| 5959 | 5961 | 1 ; | HB3 | - | ND1 |
| 5959 | 5967 | 1 ; | HB3 | - | CD2 |
| 5959 | 5969 | 1 ; | HB3 | - | C |
| 5960 | 5964 | 1 ; | CG | - | HE1 |
| 5960 | 5966 | 1 ; | CG | - | HE2 |
| 5960 | 5969 | 1 ; | CG | - | C |
| 5961 | 5966 | 1 ; | ND1 | - | HE2 |
| 5961 | 5968 | 1 ; | ND1 | - | HD2 |
| 5962 | 5964 | 1 ; | HD1 | - | HE1 |
| 5962 | 5965 | 1 ; | HD1 | - | NE2 |
| 5962 | 5967 | 1 ; | HD1 | - | CD2 |
| 5963 | 5968 | 1 ; | CE1 | - | HD2 |
| 5964 | 5966 | 1 ; | HE1 | - | HE2 |
| 5964 | 5967 | 1 ; | HE1 | - | CD2 |
| 5966 | 5968 | 1 ; | HE2 | - | HD2 |
| 5969 | 5974 | 1 ; | C | - | HA |
| 5969 | 5975 | 1 ; | C | - | CB |
| 5969 | 5990 | 1 ; | C | - | C |
| 5970 | 5972 | 1 ; | O | - | H |
| 5970 | 5973 | 1 ; | O | - | CA |
| 5971 | 5976 | 1 ; | N | - | HB2 |
| 5971 | 5977 | 1 ; | N | - | HB3 |
| 5971 | 5978 | 1 ; | N | - | CG |
| 5971 | 5991 | 1 ; | N | - | O |
| 5971 | 5992 | 1 ; | N | - | N |
| 5972 | 5974 | 1 ; | H | - | HA |

```
5972   5975       1 ;      H - CB
5972   5990       1 ;      H - C
5973   5979       1 ;     CA - CD1
5973   5988       1 ;     CA - CD2
5973   5993       1 ;     CA - H
5973   5994       1 ;     CA - CA
5974   5976       1 ;     HA - HB2
5974   5977       1 ;     HA - HB3
5974   5978       1 ;     HA - CG
5974   5991       1 ;     HA - O
5974   5992       1 ;     HA - N
5975   5980       1 ;     CB - HD1
5975   5981       1 ;     CB - CE1
5975   5986       1 ;     CB - CE2
5975   5989       1 ;     CB - HD2
5975   5991       1 ;     CB - O
5975   5992       1 ;     CB - N
5976   5979       1 ;    HB2 - CD1
5976   5988       1 ;    HB2 - CD2
5976   5990       1 ;    HB2 - C
5977   5979       1 ;    HB3 - CD1
5977   5988       1 ;    HB3 - CD2
5977   5990       1 ;    HB3 - C
5978   5982       1 ;     CG - HE1
5978   5983       1 ;     CG - CZ
5978   5987       1 ;     CG - HE2
5978   5990       1 ;     CG - C
5979   5984       1 ;    CD1 - OH
5979   5986       1 ;    CD1 - CE2
5979   5989       1 ;    CD1 - HD2
5980   5982       1 ;    HD1 - HE1
5980   5983       1 ;    HD1 - CZ
5980   5988       1 ;    HD1 - CD2
5981   5985       1 ;    CE1 - HH
5981   5987       1 ;    CE1 - HE2
5981   5988       1 ;    CE1 - CD2
5982   5984       1 ;    HE1 - OH
5982   5986       1 ;    HE1 - CE2
5983   5989       1 ;     CZ - HD2
5984   5987       1 ;     OH - HE2
5984   5988       1 ;     OH - CD2
5985   5986       1 ;     HH - CE2
5987   5989       1 ;    HE2 - HD2
5990   5995       1 ;      C - HA
5990   5996       1 ;      C - CB
5990   6004       1 ;      C - C
5991   5993       1 ;      O - H
5991   5994       1 ;      O - CA
5992   5997       1 ;      N - HB
5992   5998       1 ;      N - CG2
5992   6002       1 ;      N - OG1
5992   6005       1 ;      N - O
```

```
5992   6006      1 ;      N - N
5993   5995      1 ;      H - HA
5993   5996      1 ;      H - CB
5993   6004      1 ;      H - C
5994   5999      1 ;     CA - HG21
5994   6000      1 ;     CA - HG22
5994   6001      1 ;     CA - HG23
5994   6003      1 ;     CA - HG1
5994   6007      1 ;     CA - H
5994   6008      1 ;     CA - CA
5995   5997      1 ;     HA - HB
5995   5998      1 ;     HA - CG2
5995   6002      1 ;     HA - OG1
5995   6005      1 ;     HA - O
5995   6006      1 ;     HA - N
5996   6005      1 ;     CB - O
5996   6006      1 ;     CB - N
5997   5999      1 ;     HB - HG21
5997   6000      1 ;     HB - HG22
5997   6001      1 ;     HB - HG23
5997   6003      1 ;     HB - HG1
5997   6004      1 ;     HB - C
5998   6003      1 ;    CG2 - HG1
5998   6004      1 ;    CG2 - C
5999   6002      1 ;   HG21 - OG1
6000   6002      1 ;   HG22 - OG1
6001   6002      1 ;   HG23 - OG1
6002   6004      1 ;    OG1 - C
6004   6009      1 ;      C - HA
6004   6010      1 ;      C - CB
6004   6016      1 ;      C - C
6005   6007      1 ;      O - H
6005   6008      1 ;      O - CA
6006   6011      1 ;      N - HB2
6006   6012      1 ;      N - HB3
6006   6013      1 ;      N - CG
6006   6017      1 ;      N - O
6006   6018      1 ;      N - N
6007   6009      1 ;      H - HA
6007   6010      1 ;      H - CB
6007   6016      1 ;      H - C
6008   6014      1 ;     CA - OD1
6008   6015      1 ;     CA - OD2
6008   6019      1 ;     CA - H
6008   6020      1 ;     CA - CA
6009   6011      1 ;     HA - HB2
6009   6012      1 ;     HA - HB3
6009   6013      1 ;     HA - CG
6009   6017      1 ;     HA - O
6009   6018      1 ;     HA - N
6010   6017      1 ;     CB - O
6010   6018      1 ;     CB - N
```

```
6011   6014       1 ;     HB2 - OD1
6011   6015       1 ;     HB2 - OD2
6011   6016       1 ;     HB2 - C
6012   6014       1 ;     HB3 - OD1
6012   6015       1 ;     HB3 - OD2
6012   6016       1 ;     HB3 - C
6013   6016       1 ;      CG - C
6016   6021       1 ;       C - HA
6016   6022       1 ;       C - CB
6016   6036       1 ;       C - C
6017   6019       1 ;       O - H
6017   6020       1 ;       O - CA
6018   6023       1 ;       N - HB2
6018   6024       1 ;       N - HB3
6018   6025       1 ;       N - CG
6018   6037       1 ;       N - O
6018   6038       1 ;       N - N
6019   6021       1 ;       H - HA
6019   6022       1 ;       H - CB
6019   6036       1 ;       H - C
6020   6026       1 ;      CA - CD1
6020   6034       1 ;      CA - CD2
6020   6039       1 ;      CA - H
6020   6040       1 ;      CA - CA
6021   6023       1 ;      HA - HB2
6021   6024       1 ;      HA - HB3
6021   6025       1 ;      HA - CG
6021   6037       1 ;      HA - O
6021   6038       1 ;      HA - N
6022   6027       1 ;      CB - HD1
6022   6028       1 ;      CB - CE1
6022   6032       1 ;      CB - CE2
6022   6035       1 ;      CB - HD2
6022   6037       1 ;      CB - O
6022   6038       1 ;      CB - N
6023   6026       1 ;     HB2 - CD1
6023   6034       1 ;     HB2 - CD2
6023   6036       1 ;     HB2 - C
6024   6026       1 ;     HB3 - CD1
6024   6034       1 ;     HB3 - CD2
6024   6036       1 ;     HB3 - C
6025   6029       1 ;      CG - HE1
6025   6030       1 ;      CG - CZ
6025   6033       1 ;      CG - HE2
6025   6036       1 ;      CG - C
6026   6031       1 ;     CD1 - HZ
6026   6032       1 ;     CD1 - CE2
6026   6035       1 ;     CD1 - HD2
6027   6029       1 ;     HD1 - HE1
6027   6030       1 ;     HD1 - CZ
6027   6034       1 ;     HD1 - CD2
6028   6033       1 ;     CE1 - HE2
```

```
6028   6034      1 ;    CE1 - CD2
6029   6031      1 ;    HE1 - HZ
6029   6032      1 ;    HE1 - CE2
6030   6035      1 ;     CZ - HD2
6031   6033      1 ;     HZ - HE2
6031   6034      1 ;     HZ - CD2
6033   6035      1 ;    HE2 - HD2
6036   6041      1 ;      C - HA
6036   6042      1 ;      C - CB
6036   6055      1 ;      C - C
6037   6039      1 ;      O - H
6037   6040      1 ;      O - CA
6038   6043      1 ;      N - HB2
6038   6044      1 ;      N - HB3
6038   6045      1 ;      N - CG
6038   6056      1 ;      N - O
6038   6057      1 ;      N - N
6039   6041      1 ;      H - HA
6039   6042      1 ;      H - CB
6039   6055      1 ;      H - C
6040   6046      1 ;     CA - HG
6040   6047      1 ;     CA - CD1
6040   6051      1 ;     CA - CD2
6040   6058      1 ;     CA - H
6040   6059      1 ;     CA - CA
6041   6043      1 ;     HA - HB2
6041   6044      1 ;     HA - HB3
6041   6045      1 ;     HA - CG
6041   6056      1 ;     HA - O
6041   6057      1 ;     HA - N
6042   6048      1 ;     CB - HD11
6042   6049      1 ;     CB - HD12
6042   6050      1 ;     CB - HD13
6042   6052      1 ;     CB - HD21
6042   6053      1 ;     CB - HD22
6042   6054      1 ;     CB - HD23
6042   6056      1 ;     CB - O
6042   6057      1 ;     CB - N
6043   6046      1 ;    HB2 - HG
6043   6047      1 ;    HB2 - CD1
6043   6051      1 ;    HB2 - CD2
6043   6055      1 ;    HB2 - C
6044   6046      1 ;    HB3 - HG
6044   6047      1 ;    HB3 - CD1
6044   6051      1 ;    HB3 - CD2
6044   6055      1 ;    HB3 - C
6045   6055      1 ;     CG - C
6046   6048      1 ;     HG - HD11
6046   6049      1 ;     HG - HD12
6046   6050      1 ;     HG - HD13
6046   6052      1 ;     HG - HD21
6046   6053      1 ;     HG - HD22
```

| | | | | | | |
|---|---|---|---|---|---|---|
| 6046 | 6054 | 1 | ; | HG | - | HD23 |
| 6047 | 6052 | 1 | ; | CD1 | - | HD21 |
| 6047 | 6053 | 1 | ; | CD1 | - | HD22 |
| 6047 | 6054 | 1 | ; | CD1 | - | HD23 |
| 6048 | 6051 | 1 | ; | HD11 | - | CD2 |
| 6049 | 6051 | 1 | ; | HD12 | - | CD2 |
| 6050 | 6051 | 1 | ; | HD13 | - | CD2 |
| 6055 | 6060 | 1 | ; | C | - | HA |
| 6055 | 6061 | 1 | ; | C | - | CB |
| 6055 | 6065 | 1 | ; | C | - | C |
| 6056 | 6058 | 1 | ; | O | - | H |
| 6056 | 6059 | 1 | ; | O | - | CA |
| 6057 | 6062 | 1 | ; | N | - | HB1 |
| 6057 | 6063 | 1 | ; | N | - | HB2 |
| 6057 | 6064 | 1 | ; | N | - | HB3 |
| 6057 | 6066 | 1 | ; | N | - | O |
| 6057 | 6067 | 1 | ; | N | - | N |
| 6058 | 6060 | 1 | ; | H | - | HA |
| 6058 | 6061 | 1 | ; | H | - | CB |
| 6058 | 6065 | 1 | ; | H | - | C |
| 6059 | 6068 | 1 | ; | CA | - | H |
| 6059 | 6069 | 1 | ; | CA | - | CA |
| 6060 | 6062 | 1 | ; | HA | - | HB1 |
| 6060 | 6063 | 1 | ; | HA | - | HB2 |
| 6060 | 6064 | 1 | ; | HA | - | HB3 |
| 6060 | 6066 | 1 | ; | HA | - | O |
| 6060 | 6067 | 1 | ; | HA | - | N |
| 6061 | 6066 | 1 | ; | CB | - | O |
| 6061 | 6067 | 1 | ; | CB | - | N |
| 6062 | 6065 | 1 | ; | HB1 | - | C |
| 6063 | 6065 | 1 | ; | HB2 | - | C |
| 6064 | 6065 | 1 | ; | HB3 | - | C |
| 6065 | 6070 | 1 | ; | C | - | HA |
| 6065 | 6071 | 1 | ; | C | - | CB |
| 6065 | 6075 | 1 | ; | C | - | C |
| 6066 | 6068 | 1 | ; | O | - | H |
| 6066 | 6069 | 1 | ; | O | - | CA |
| 6067 | 6072 | 1 | ; | N | - | HB1 |
| 6067 | 6073 | 1 | ; | N | - | HB2 |
| 6067 | 6074 | 1 | ; | N | - | HB3 |
| 6067 | 6076 | 1 | ; | N | - | O |
| 6067 | 6077 | 1 | ; | N | - | N |
| 6068 | 6070 | 1 | ; | H | - | HA |
| 6068 | 6071 | 1 | ; | H | - | CB |
| 6068 | 6075 | 1 | ; | H | - | C |
| 6069 | 6078 | 1 | ; | CA | - | H |
| 6069 | 6079 | 1 | ; | CA | - | CA |
| 6070 | 6072 | 1 | ; | HA | - | HB1 |
| 6070 | 6073 | 1 | ; | HA | - | HB2 |
| 6070 | 6074 | 1 | ; | HA | - | HB3 |
| 6070 | 6076 | 1 | ; | HA | - | O |
| 6070 | 6077 | 1 | ; | HA | - | N |

| | | | | |
|---|---|---|---|---|
| 6071 | 6076 | 1 ; | CB | - O |
| 6071 | 6077 | 1 ; | CB | - N |
| 6072 | 6075 | 1 ; | HB1 | - C |
| 6073 | 6075 | 1 ; | HB2 | - C |
| 6074 | 6075 | 1 ; | HB3 | - C |
| 6075 | 6080 | 1 ; | C | - HA |
| 6075 | 6081 | 1 ; | C | - CB |
| 6075 | 6089 | 1 ; | C | - C |
| 6076 | 6078 | 1 ; | O | - H |
| 6076 | 6079 | 1 ; | O | - CA |
| 6077 | 6082 | 1 ; | N | - HB |
| 6077 | 6083 | 1 ; | N | - CG2 |
| 6077 | 6087 | 1 ; | N | - OG1 |
| 6077 | 6090 | 1 ; | N | - O |
| 6077 | 6091 | 1 ; | N | - N |
| 6078 | 6080 | 1 ; | H | - HA |
| 6078 | 6081 | 1 ; | H | - CB |
| 6078 | 6089 | 1 ; | H | - C |
| 6079 | 6084 | 1 ; | CA | - HG21 |
| 6079 | 6085 | 1 ; | CA | - HG22 |
| 6079 | 6086 | 1 ; | CA | - HG23 |
| 6079 | 6088 | 1 ; | CA | - HG1 |
| 6079 | 6092 | 1 ; | CA | - H |
| 6079 | 6093 | 1 ; | CA | - CA |
| 6080 | 6082 | 1 ; | HA | - HB |
| 6080 | 6083 | 1 ; | HA | - CG2 |
| 6080 | 6087 | 1 ; | HA | - OG1 |
| 6080 | 6090 | 1 ; | HA | - O |
| 6080 | 6091 | 1 ; | HA | - N |
| 6081 | 6090 | 1 ; | CB | - O |
| 6081 | 6091 | 1 ; | CB | - N |
| 6082 | 6084 | 1 ; | HB | - HG21 |
| 6082 | 6085 | 1 ; | HB | - HG22 |
| 6082 | 6086 | 1 ; | HB | - HG23 |
| 6082 | 6088 | 1 ; | HB | - HG1 |
| 6082 | 6089 | 1 ; | HB | - C |
| 6083 | 6088 | 1 ; | CG2 | - HG1 |
| 6083 | 6089 | 1 ; | CG2 | - C |
| 6084 | 6087 | 1 ; | HG21 | - OG1 |
| 6085 | 6087 | 1 ; | HG22 | - OG1 |
| 6086 | 6087 | 1 ; | HG23 | - OG1 |
| 6087 | 6089 | 1 ; | OG1 | - C |
| 6089 | 6094 | 1 ; | C | - HA |
| 6089 | 6095 | 1 ; | C | - CB |
| 6089 | 6108 | 1 ; | C | - C |
| 6090 | 6092 | 1 ; | O | - H |
| 6090 | 6093 | 1 ; | O | - CA |
| 6091 | 6096 | 1 ; | N | - HB |
| 6091 | 6097 | 1 ; | N | - CG2 |
| 6091 | 6101 | 1 ; | N | - CG1 |
| 6091 | 6109 | 1 ; | N | - O |
| 6091 | 6110 | 1 ; | N | - N |

```
6092   6094       1 ;       H - HA
6092   6095       1 ;       H - CB
6092   6108       1 ;       H - C
6093   6098       1 ;      CA - HG21
6093   6099       1 ;      CA - HG22
6093   6100       1 ;      CA - HG23
6093   6102       1 ;      CA - HG12
6093   6103       1 ;      CA - HG13
6093   6104       1 ;      CA - CD1
6093   6111       1 ;      CA - H
6093   6112       1 ;      CA - CA
6094   6096       1 ;      HA - HB
6094   6097       1 ;      HA - CG2
6094   6101       1 ;      HA - CG1
6094   6109       1 ;      HA - O
6094   6110       1 ;      HA - N
6095   6105       1 ;      CB - HD11
6095   6106       1 ;      CB - HD12
6095   6107       1 ;      CB - HD13
6095   6109       1 ;      CB - O
6095   6110       1 ;      CB - N
6096   6098       1 ;      HB - HG21
6096   6099       1 ;      HB - HG22
6096   6100       1 ;      HB - HG23
6096   6102       1 ;      HB - HG12
6096   6103       1 ;      HB - HG13
6096   6104       1 ;      HB - CD1
6096   6108       1 ;      HB - C
6097   6102       1 ;     CG2 - HG12
6097   6103       1 ;     CG2 - HG13
6097   6104       1 ;     CG2 - CD1
6097   6108       1 ;     CG2 - C
6098   6101       1 ;    HG21 - CG1
6099   6101       1 ;    HG22 - CG1
6100   6101       1 ;    HG23 - CG1
6101   6108       1 ;     CG1 - C
6102   6105       1 ;    HG12 - HD11
6102   6106       1 ;    HG12 - HD12
6102   6107       1 ;    HG12 - HD13
6103   6105       1 ;    HG13 - HD11
6103   6106       1 ;    HG13 - HD12
6103   6107       1 ;    HG13 - HD13
6108   6113       1 ;       C - HA
6108   6114       1 ;       C - CB
6108   6120       1 ;       C - C
6109   6111       1 ;       O - H
6109   6112       1 ;       O - CA
6110   6115       1 ;       N - HB2
6110   6116       1 ;       N - HB3
6110   6117       1 ;       N - CG
6110   6121       1 ;       N - O
6110   6122       1 ;       N - N
```

| | | | | | |
|---|---|---|---|---|---|
| 6111 | 6113 | 1 ; | H | - | HA |
| 6111 | 6114 | 1 ; | H | - | CB |
| 6111 | 6120 | 1 ; | H | - | C |
| 6112 | 6118 | 1 ; | CA | - | OD1 |
| 6112 | 6119 | 1 ; | CA | - | OD2 |
| 6112 | 6123 | 1 ; | CA | - | H |
| 6112 | 6124 | 1 ; | CA | - | CA |
| 6113 | 6115 | 1 ; | HA | - | HB2 |
| 6113 | 6116 | 1 ; | HA | - | HB3 |
| 6113 | 6117 | 1 ; | HA | - | CG |
| 6113 | 6121 | 1 ; | HA | - | O |
| 6113 | 6122 | 1 ; | HA | - | N |
| 6114 | 6121 | 1 ; | CB | - | O |
| 6114 | 6122 | 1 ; | CB | - | N |
| 6115 | 6118 | 1 ; | HB2 | - | OD1 |
| 6115 | 6119 | 1 ; | HB2 | - | OD2 |
| 6115 | 6120 | 1 ; | HB2 | - | C |
| 6116 | 6118 | 1 ; | HB3 | - | OD1 |
| 6116 | 6119 | 1 ; | HB3 | - | OD2 |
| 6116 | 6120 | 1 ; | HB3 | - | C |
| 6117 | 6120 | 1 ; | CG | - | C |
| 6120 | 6125 | 1 ; | C | - | HA |
| 6120 | 6126 | 1 ; | C | - | CB |
| 6120 | 6142 | 1 ; | C | - | C |
| 6121 | 6123 | 1 ; | O | - | H |
| 6121 | 6124 | 1 ; | O | - | CA |
| 6122 | 6127 | 1 ; | N | - | HB2 |
| 6122 | 6128 | 1 ; | N | - | HB3 |
| 6122 | 6129 | 1 ; | N | - | CG |
| 6122 | 6143 | 1 ; | N | - | O |
| 6122 | 6144 | 1 ; | N | - | N |
| 6123 | 6125 | 1 ; | H | - | HA |
| 6123 | 6126 | 1 ; | H | - | CB |
| 6123 | 6142 | 1 ; | H | - | C |
| 6124 | 6130 | 1 ; | CA | - | HG2 |
| 6124 | 6131 | 1 ; | CA | - | HG3 |
| 6124 | 6132 | 1 ; | CA | - | CD |
| 6124 | 6145 | 1 ; | CA | - | H |
| 6124 | 6146 | 1 ; | CA | - | CA |
| 6125 | 6127 | 1 ; | HA | - | HB2 |
| 6125 | 6128 | 1 ; | HA | - | HB3 |
| 6125 | 6129 | 1 ; | HA | - | CG |
| 6125 | 6143 | 1 ; | HA | - | O |
| 6125 | 6144 | 1 ; | HA | - | N |
| 6126 | 6133 | 1 ; | CB | - | HD2 |
| 6126 | 6134 | 1 ; | CB | - | HD3 |
| 6126 | 6135 | 1 ; | CB | - | CE |
| 6126 | 6143 | 1 ; | CB | - | O |
| 6126 | 6144 | 1 ; | CB | - | N |
| 6127 | 6130 | 1 ; | HB2 | - | HG2 |
| 6127 | 6131 | 1 ; | HB2 | - | HG3 |
| 6127 | 6132 | 1 ; | HB2 | - | CD |

| | | | | |
|---|---|---|---|---|
| 6127 | 6142 | 1 ; | HB2 - | C |
| 6128 | 6130 | 1 ; | HB3 - | HG2 |
| 6128 | 6131 | 1 ; | HB3 - | HG3 |
| 6128 | 6132 | 1 ; | HB3 - | CD |
| 6128 | 6142 | 1 ; | HB3 - | C |
| 6129 | 6136 | 1 ; | CG - | HE2 |
| 6129 | 6137 | 1 ; | CG - | HE3 |
| 6129 | 6138 | 1 ; | CG - | NZ |
| 6129 | 6142 | 1 ; | CG - | C |
| 6130 | 6133 | 1 ; | HG2 - | HD2 |
| 6130 | 6134 | 1 ; | HG2 - | HD3 |
| 6130 | 6135 | 1 ; | HG2 - | CE |
| 6131 | 6133 | 1 ; | HG3 - | HD2 |
| 6131 | 6134 | 1 ; | HG3 - | HD3 |
| 6131 | 6135 | 1 ; | HG3 - | CE |
| 6132 | 6139 | 1 ; | CD - | HZ1 |
| 6132 | 6140 | 1 ; | CD - | HZ2 |
| 6132 | 6141 | 1 ; | CD - | HZ3 |
| 6133 | 6136 | 1 ; | HD2 - | HE2 |
| 6133 | 6137 | 1 ; | HD2 - | HE3 |
| 6133 | 6138 | 1 ; | HD2 - | NZ |
| 6134 | 6136 | 1 ; | HD3 - | HE2 |
| 6134 | 6137 | 1 ; | HD3 - | HE3 |
| 6134 | 6138 | 1 ; | HD3 - | NZ |
| 6136 | 6139 | 1 ; | HE2 - | HZ1 |
| 6136 | 6140 | 1 ; | HE2 - | HZ2 |
| 6136 | 6141 | 1 ; | HE2 - | HZ3 |
| 6137 | 6139 | 1 ; | HE3 - | HZ1 |
| 6137 | 6140 | 1 ; | HE3 - | HZ2 |
| 6137 | 6141 | 1 ; | HE3 - | HZ3 |
| 6142 | 6147 | 1 ; | C - | HA |
| 6142 | 6148 | 1 ; | C - | CB |
| 6142 | 6159 | 1 ; | C - | C |
| 6143 | 6145 | 1 ; | O - | H |
| 6143 | 6146 | 1 ; | O - | CA |
| 6144 | 6149 | 1 ; | N - | HB2 |
| 6144 | 6150 | 1 ; | N - | HB3 |
| 6144 | 6151 | 1 ; | N - | CG |
| 6144 | 6160 | 1 ; | N - | O |
| 6144 | 6161 | 1 ; | N - | N |
| 6145 | 6147 | 1 ; | H - | HA |
| 6145 | 6148 | 1 ; | H - | CB |
| 6145 | 6159 | 1 ; | H - | C |
| 6146 | 6152 | 1 ; | CA - | HG2 |
| 6146 | 6153 | 1 ; | CA - | HG3 |
| 6146 | 6154 | 1 ; | CA - | CD |
| 6146 | 6162 | 1 ; | CA - | H |
| 6146 | 6163 | 1 ; | CA - | CA |
| 6147 | 6149 | 1 ; | HA - | HB2 |
| 6147 | 6150 | 1 ; | HA - | HB3 |
| 6147 | 6151 | 1 ; | HA - | CG |
| 6147 | 6160 | 1 ; | HA - | O |

```
6147   6161       1 ;      HA - N
6148   6155       1 ;      CB - OE1
6148   6156       1 ;      CB - NE2
6148   6160       1 ;      CB - O
6148   6161       1 ;      CB - N
6149   6152       1 ;     HB2 - HG2
6149   6153       1 ;     HB2 - HG3
6149   6154       1 ;     HB2 - CD
6149   6159       1 ;     HB2 - C
6150   6152       1 ;     HB3 - HG2
6150   6153       1 ;     HB3 - HG3
6150   6154       1 ;     HB3 - CD
6150   6159       1 ;     HB3 - C
6151   6157       1 ;      CG - HE21
6151   6158       1 ;      CG - HE22
6151   6159       1 ;      CG - C
6152   6155       1 ;     HG2 - OE1
6152   6156       1 ;     HG2 - NE2
6153   6155       1 ;     HG3 - OE1
6153   6156       1 ;     HG3 - NE2
6155   6157       1 ;     OE1 - HE21
6155   6158       1 ;     OE1 - HE22
6159   6164       1 ;       C - HA
6159   6165       1 ;       C - CB
6159   6173       1 ;       C - C
6160   6162       1 ;       O - H
6160   6163       1 ;       O - CA
6161   6166       1 ;       N - HB
6161   6167       1 ;       N - CG2
6161   6171       1 ;       N - OG1
6161   6174       1 ;       N - O
6161   6175       1 ;       N - N
6162   6164       1 ;       H - HA
6162   6165       1 ;       H - CB
6162   6173       1 ;       H - C
6163   6168       1 ;      CA - HG21
6163   6169       1 ;      CA - HG22
6163   6170       1 ;      CA - HG23
6163   6172       1 ;      CA - HG1
6163   6176       1 ;      CA - H
6163   6177       1 ;      CA - CA
6164   6166       1 ;      HA - HB
6164   6167       1 ;      HA - CG2
6164   6171       1 ;      HA - OG1
6164   6174       1 ;      HA - O
6164   6175       1 ;      HA - N
6165   6174       1 ;      CB - O
6165   6175       1 ;      CB - N
6166   6168       1 ;      HB - HG21
6166   6169       1 ;      HB - HG22
6166   6170       1 ;      HB - HG23
6166   6172       1 ;      HB - HG1
```

| | | | | | |
|---|---|---|---|---|---|
| 6166 | 6173 | 1 ; | HB | - | C |
| 6167 | 6172 | 1 ; | CG2 | - | HG1 |
| 6167 | 6173 | 1 ; | CG2 | - | C |
| 6168 | 6171 | 1 ; | HG21 | - | OG1 |
| 6169 | 6171 | 1 ; | HG22 | - | OG1 |
| 6170 | 6171 | 1 ; | HG23 | - | OG1 |
| 6171 | 6173 | 1 ; | OG1 | - | C |
| 6173 | 6178 | 1 ; | C | - | HA |
| 6173 | 6179 | 1 ; | C | - | CB |
| 6173 | 6194 | 1 ; | C | - | C |
| 6174 | 6176 | 1 ; | O | - | H |
| 6174 | 6177 | 1 ; | O | - | CA |
| 6175 | 6180 | 1 ; | N | - | HB2 |
| 6175 | 6181 | 1 ; | N | - | HB3 |
| 6175 | 6182 | 1 ; | N | - | CG |
| 6175 | 6195 | 1 ; | N | - | O |
| 6175 | 6196 | 1 ; | N | - | N |
| 6176 | 6178 | 1 ; | H | - | HA |
| 6176 | 6179 | 1 ; | H | - | CB |
| 6176 | 6194 | 1 ; | H | - | C |
| 6177 | 6183 | 1 ; | CA | - | CD1 |
| 6177 | 6192 | 1 ; | CA | - | CD2 |
| 6177 | 6197 | 1 ; | CA | - | H |
| 6177 | 6198 | 1 ; | CA | - | CA |
| 6178 | 6180 | 1 ; | HA | - | HB2 |
| 6178 | 6181 | 1 ; | HA | - | HB3 |
| 6178 | 6182 | 1 ; | HA | - | CG |
| 6178 | 6195 | 1 ; | HA | - | O |
| 6178 | 6196 | 1 ; | HA | - | N |
| 6179 | 6184 | 1 ; | CB | - | HD1 |
| 6179 | 6185 | 1 ; | CB | - | CE1 |
| 6179 | 6190 | 1 ; | CB | - | CE2 |
| 6179 | 6193 | 1 ; | CB | - | HD2 |
| 6179 | 6195 | 1 ; | CB | - | O |
| 6179 | 6196 | 1 ; | CB | - | N |
| 6180 | 6183 | 1 ; | HB2 | - | CD1 |
| 6180 | 6192 | 1 ; | HB2 | - | CD2 |
| 6180 | 6194 | 1 ; | HB2 | - | C |
| 6181 | 6183 | 1 ; | HB3 | - | CD1 |
| 6181 | 6192 | 1 ; | HB3 | - | CD2 |
| 6181 | 6194 | 1 ; | HB3 | - | C |
| 6182 | 6186 | 1 ; | CG | - | HE1 |
| 6182 | 6187 | 1 ; | CG | - | CZ |
| 6182 | 6191 | 1 ; | CG | - | HE2 |
| 6182 | 6194 | 1 ; | CG | - | C |
| 6183 | 6188 | 1 ; | CD1 | - | OH |
| 6183 | 6190 | 1 ; | CD1 | - | CE2 |
| 6183 | 6193 | 1 ; | CD1 | - | HD2 |
| 6184 | 6186 | 1 ; | HD1 | - | HE1 |
| 6184 | 6187 | 1 ; | HD1 | - | CZ |
| 6184 | 6192 | 1 ; | HD1 | - | CD2 |
| 6185 | 6189 | 1 ; | CE1 | - | HH |

| | | | | |
|---|---|---|---|---|
| 6185 | 6191 | 1 | ; | CE1 - HE2 |
| 6185 | 6192 | 1 | ; | CE1 - CD2 |
| 6186 | 6188 | 1 | ; | HE1 - OH |
| 6186 | 6190 | 1 | ; | HE1 - CE2 |
| 6187 | 6193 | 1 | ; | CZ - HD2 |
| 6188 | 6191 | 1 | ; | OH - HE2 |
| 6188 | 6192 | 1 | ; | OH - CD2 |
| 6189 | 6190 | 1 | ; | HH - CE2 |
| 6191 | 6193 | 1 | ; | HE2 - HD2 |
| 6194 | 6199 | 1 | ; | C - HA |
| 6194 | 6200 | 1 | ; | C - CB |
| 6194 | 6213 | 1 | ; | C - C |
| 6195 | 6197 | 1 | ; | O - H |
| 6195 | 6198 | 1 | ; | O - CA |
| 6196 | 6201 | 1 | ; | N - HB2 |
| 6196 | 6202 | 1 | ; | N - HB3 |
| 6196 | 6203 | 1 | ; | N - CG |
| 6196 | 6214 | 1 | ; | N - O |
| 6196 | 6215 | 1 | ; | N - N |
| 6197 | 6199 | 1 | ; | H - HA |
| 6197 | 6200 | 1 | ; | H - CB |
| 6197 | 6213 | 1 | ; | H - C |
| 6198 | 6204 | 1 | ; | CA - HG |
| 6198 | 6205 | 1 | ; | CA - CD1 |
| 6198 | 6209 | 1 | ; | CA - CD2 |
| 6198 | 6216 | 1 | ; | CA - H |
| 6198 | 6217 | 1 | ; | CA - CA |
| 6199 | 6201 | 1 | ; | HA - HB2 |
| 6199 | 6202 | 1 | ; | HA - HB3 |
| 6199 | 6203 | 1 | ; | HA - CG |
| 6199 | 6214 | 1 | ; | HA - O |
| 6199 | 6215 | 1 | ; | HA - N |
| 6200 | 6206 | 1 | ; | CB - HD11 |
| 6200 | 6207 | 1 | ; | CB - HD12 |
| 6200 | 6208 | 1 | ; | CB - HD13 |
| 6200 | 6210 | 1 | ; | CB - HD21 |
| 6200 | 6211 | 1 | ; | CB - HD22 |
| 6200 | 6212 | 1 | ; | CB - HD23 |
| 6200 | 6214 | 1 | ; | CB - O |
| 6200 | 6215 | 1 | ; | CB - N |
| 6201 | 6204 | 1 | ; | HB2 - HG |
| 6201 | 6205 | 1 | ; | HB2 - CD1 |
| 6201 | 6209 | 1 | ; | HB2 - CD2 |
| 6201 | 6213 | 1 | ; | HB2 - C |
| 6202 | 6204 | 1 | ; | HB3 - HG |
| 6202 | 6205 | 1 | ; | HB3 - CD1 |
| 6202 | 6209 | 1 | ; | HB3 - CD2 |
| 6202 | 6213 | 1 | ; | HB3 - C |
| 6203 | 6213 | 1 | ; | CG - C |
| 6204 | 6206 | 1 | ; | HG - HD11 |
| 6204 | 6207 | 1 | ; | HG - HD12 |
| 6204 | 6208 | 1 | ; | HG - HD13 |

| | | | | | | |
|---|---|---|---|---|---|---|
| 6204 | 6210 | 1 | ; | HG | - | HD21 |
| 6204 | 6211 | 1 | ; | HG | - | HD22 |
| 6204 | 6212 | 1 | ; | HG | - | HD23 |
| 6205 | 6210 | 1 | ; | CD1 | - | HD21 |
| 6205 | 6211 | 1 | ; | CD1 | - | HD22 |
| 6205 | 6212 | 1 | ; | CD1 | - | HD23 |
| 6206 | 6209 | 1 | ; | HD11 | - | CD2 |
| 6207 | 6209 | 1 | ; | HD12 | - | CD2 |
| 6208 | 6209 | 1 | ; | HD13 | - | CD2 |
| 6213 | 6218 | 1 | ; | C | - | HA |
| 6213 | 6219 | 1 | ; | C | - | CB |
| 6213 | 6235 | 1 | ; | C | - | C |
| 6214 | 6216 | 1 | ; | O | - | H |
| 6214 | 6217 | 1 | ; | O | - | CA |
| 6215 | 6220 | 1 | ; | N | - | HB2 |
| 6215 | 6221 | 1 | ; | N | - | HB3 |
| 6215 | 6222 | 1 | ; | N | - | CG |
| 6215 | 6236 | 1 | ; | N | - | O |
| 6215 | 6237 | 1 | ; | N | - | N |
| 6216 | 6218 | 1 | ; | H | - | HA |
| 6216 | 6219 | 1 | ; | H | - | CB |
| 6216 | 6235 | 1 | ; | H | - | C |
| 6217 | 6223 | 1 | ; | CA | - | HG2 |
| 6217 | 6224 | 1 | ; | CA | - | HG3 |
| 6217 | 6225 | 1 | ; | CA | - | CD |
| 6217 | 6238 | 1 | ; | CA | - | H |
| 6217 | 6239 | 1 | ; | CA | - | CA |
| 6218 | 6220 | 1 | ; | HA | - | HB2 |
| 6218 | 6221 | 1 | ; | HA | - | HB3 |
| 6218 | 6222 | 1 | ; | HA | - | CG |
| 6218 | 6236 | 1 | ; | HA | - | O |
| 6218 | 6237 | 1 | ; | HA | - | N |
| 6219 | 6226 | 1 | ; | CB | - | HD2 |
| 6219 | 6227 | 1 | ; | CB | - | HD3 |
| 6219 | 6228 | 1 | ; | CB | - | CE |
| 6219 | 6236 | 1 | ; | CB | - | O |
| 6219 | 6237 | 1 | ; | CB | - | N |
| 6220 | 6223 | 1 | ; | HB2 | - | HG2 |
| 6220 | 6224 | 1 | ; | HB2 | - | HG3 |
| 6220 | 6225 | 1 | ; | HB2 | - | CD |
| 6220 | 6235 | 1 | ; | HB2 | - | C |
| 6221 | 6223 | 1 | ; | HB3 | - | HG2 |
| 6221 | 6224 | 1 | ; | HB3 | - | HG3 |
| 6221 | 6225 | 1 | ; | HB3 | - | CD |
| 6221 | 6235 | 1 | ; | HB3 | - | C |
| 6222 | 6229 | 1 | ; | CG | - | HE2 |
| 6222 | 6230 | 1 | ; | CG | - | HE3 |
| 6222 | 6231 | 1 | ; | CG | - | NZ |
| 6222 | 6235 | 1 | ; | CG | - | C |
| 6223 | 6226 | 1 | ; | HG2 | - | HD2 |
| 6223 | 6227 | 1 | ; | HG2 | - | HD3 |
| 6223 | 6228 | 1 | ; | HG2 | - | CE |

```
6224   6226      1 ;    HG3 - HD2
6224   6227      1 ;    HG3 - HD3
6224   6228      1 ;    HG3 - CE
6225   6232      1 ;     CD - HZ1
6225   6233      1 ;     CD - HZ2
6225   6234      1 ;     CD - HZ3
6226   6229      1 ;    HD2 - HE2
6226   6230      1 ;    HD2 - HE3
6226   6231      1 ;    HD2 - NZ
6227   6229      1 ;    HD3 - HE2
6227   6230      1 ;    HD3 - HE3
6227   6231      1 ;    HD3 - NZ
6229   6232      1 ;    HE2 - HZ1
6229   6233      1 ;    HE2 - HZ2
6229   6234      1 ;    HE2 - HZ3
6230   6232      1 ;    HE3 - HZ1
6230   6233      1 ;    HE3 - HZ2
6230   6234      1 ;    HE3 - HZ3
6235   6240      1 ;      C - HA
6235   6241      1 ;      C - CB
6235   6257      1 ;      C - C
6236   6238      1 ;      O - H
6236   6239      1 ;      O - CA
6237   6242      1 ;      N - HB2
6237   6243      1 ;      N - HB3
6237   6244      1 ;      N - CG
6237   6258      1 ;      N - O
6237   6259      1 ;      N - N
6238   6240      1 ;      H - HA
6238   6241      1 ;      H - CB
6238   6257      1 ;      H - C
6239   6245      1 ;     CA - HG2
6239   6246      1 ;     CA - HG3
6239   6247      1 ;     CA - CD
6239   6260      1 ;     CA - H
6239   6261      1 ;     CA - CA
6240   6242      1 ;     HA - HB2
6240   6243      1 ;     HA - HB3
6240   6244      1 ;     HA - CG
6240   6258      1 ;     HA - O
6240   6259      1 ;     HA - N
6241   6248      1 ;     CB - HD2
6241   6249      1 ;     CB - HD3
6241   6250      1 ;     CB - CE
6241   6258      1 ;     CB - O
6241   6259      1 ;     CB - N
6242   6245      1 ;    HB2 - HG2
6242   6246      1 ;    HB2 - HG3
6242   6247      1 ;    HB2 - CD
6242   6257      1 ;    HB2 - C
6243   6245      1 ;    HB3 - HG2
6243   6246      1 ;    HB3 - HG3
```

```
6243   6247       1 ;    HB3 - CD
6243   6257       1 ;    HB3 - C
6244   6251       1 ;     CG - HE2
6244   6252       1 ;     CG - HE3
6244   6253       1 ;     CG - NZ
6244   6257       1 ;     CG - C
6245   6248       1 ;    HG2 - HD2
6245   6249       1 ;    HG2 - HD3
6245   6250       1 ;    HG2 - CE
6246   6248       1 ;    HG3 - HD2
6246   6249       1 ;    HG3 - HD3
6246   6250       1 ;    HG3 - CE
6247   6254       1 ;     CD - HZ1
6247   6255       1 ;     CD - HZ2
6247   6256       1 ;     CD - HZ3
6248   6251       1 ;    HD2 - HE2
6248   6252       1 ;    HD2 - HE3
6248   6253       1 ;    HD2 - NZ
6249   6251       1 ;    HD3 - HE2
6249   6252       1 ;    HD3 - HE3
6249   6253       1 ;    HD3 - NZ
6251   6254       1 ;    HE2 - HZ1
6251   6255       1 ;    HE2 - HZ2
6251   6256       1 ;    HE2 - HZ3
6252   6254       1 ;    HE3 - HZ1
6252   6255       1 ;    HE3 - HZ2
6252   6256       1 ;    HE3 - HZ3
6257   6262       1 ;      C - HA
6257   6263       1 ;      C - CB
6257   6272       1 ;      C - C
6258   6260       1 ;      O - H
6258   6261       1 ;      O - CA
6259   6264       1 ;      N - HB2
6259   6265       1 ;      N - HB3
6259   6266       1 ;      N - CG
6259   6273       1 ;      N - O
6259   6274       1 ;      N - N
6260   6262       1 ;      H - HA
6260   6263       1 ;      H - CB
6260   6272       1 ;      H - C
6261   6267       1 ;     CA - HG2
6261   6268       1 ;     CA - HG3
6261   6269       1 ;     CA - CD
6261   6275       1 ;     CA - H
6261   6276       1 ;     CA - CA
6262   6264       1 ;     HA - HB2
6262   6265       1 ;     HA - HB3
6262   6266       1 ;     HA - CG
6262   6273       1 ;     HA - O
6262   6274       1 ;     HA - N
6263   6270       1 ;     CB - OE1
6263   6271       1 ;     CB - OE2
```

```
6263   6273       1 ;      CB - O
6263   6274       1 ;      CB - N
6264   6267       1 ;     HB2 - HG2
6264   6268       1 ;     HB2 - HG3
6264   6269       1 ;     HB2 - CD
6264   6272       1 ;     HB2 - C
6265   6267       1 ;     HB3 - HG2
6265   6268       1 ;     HB3 - HG3
6265   6269       1 ;     HB3 - CD
6265   6272       1 ;     HB3 - C
6266   6272       1 ;      CG - C
6267   6270       1 ;     HG2 - OE1
6267   6271       1 ;     HG2 - OE2
6268   6270       1 ;     HG3 - OE1
6268   6271       1 ;     HG3 - OE2
6272   6277       1 ;       C - HA
6272   6278       1 ;       C - CB
6272   6288       1 ;       C - C
6273   6275       1 ;       O - H
6273   6276       1 ;       O - CA
6274   6279       1 ;       N - HB
6274   6280       1 ;       N - CG1
6274   6284       1 ;       N - CG2
6274   6289       1 ;       N - O
6274   6290       1 ;       N - N
6275   6277       1 ;       H - HA
6275   6278       1 ;       H - CB
6275   6288       1 ;       H - C
6276   6281       1 ;      CA - HG11
6276   6282       1 ;      CA - HG12
6276   6283       1 ;      CA - HG13
6276   6285       1 ;      CA - HG21
6276   6286       1 ;      CA - HG22
6276   6287       1 ;      CA - HG23
6276   6291       1 ;      CA - H
6276   6292       1 ;      CA - CA
6277   6279       1 ;      HA - HB
6277   6280       1 ;      HA - CG1
6277   6284       1 ;      HA - CG2
6277   6289       1 ;      HA - O
6277   6290       1 ;      HA - N
6278   6289       1 ;      CB - O
6278   6290       1 ;      CB - N
6279   6281       1 ;      HB - HG11
6279   6282       1 ;      HB - HG12
6279   6283       1 ;      HB - HG13
6279   6285       1 ;      HB - HG21
6279   6286       1 ;      HB - HG22
6279   6287       1 ;      HB - HG23
6279   6288       1 ;      HB - C
6280   6285       1 ;     CG1 - HG21
6280   6286       1 ;     CG1 - HG22
```

| | | | | | |
|---|---|---|---|---|---|
| 6280 | 6287 | 1 ; | CG1 | - | HG23 |
| 6280 | 6288 | 1 ; | CG1 | - | C |
| 6281 | 6284 | 1 ; | HG11 | - | CG2 |
| 6282 | 6284 | 1 ; | HG12 | - | CG2 |
| 6283 | 6284 | 1 ; | HG13 | - | CG2 |
| 6284 | 6288 | 1 ; | CG2 | - | C |
| 6288 | 6293 | 1 ; | C | - | HA |
| 6288 | 6294 | 1 ; | C | - | CB |
| 6288 | 6299 | 1 ; | C | - | C |
| 6289 | 6291 | 1 ; | O | - | H |
| 6289 | 6292 | 1 ; | O | - | CA |
| 6290 | 6295 | 1 ; | N | - | HB2 |
| 6290 | 6296 | 1 ; | N | - | HB3 |
| 6290 | 6297 | 1 ; | N | - | SG |
| 6290 | 6300 | 1 ; | N | - | O |
| 6290 | 6301 | 1 ; | N | - | N |
| 6291 | 6293 | 1 ; | H | - | HA |
| 6291 | 6294 | 1 ; | H | - | CB |
| 6291 | 6299 | 1 ; | H | - | C |
| 6292 | 6298 | 1 ; | CA | - | HG |
| 6292 | 6302 | 1 ; | CA | - | H |
| 6292 | 6303 | 1 ; | CA | - | CA |
| 6293 | 6295 | 1 ; | HA | - | HB2 |
| 6293 | 6296 | 1 ; | HA | - | HB3 |
| 6293 | 6297 | 1 ; | HA | - | SG |
| 6293 | 6300 | 1 ; | HA | - | O |
| 6293 | 6301 | 1 ; | HA | - | N |
| 6294 | 6300 | 1 ; | CB | - | O |
| 6294 | 6301 | 1 ; | CB | - | N |
| 6295 | 6298 | 1 ; | HB2 | - | HG |
| 6295 | 6299 | 1 ; | HB2 | - | C |
| 6296 | 6298 | 1 ; | HB3 | - | HG |
| 6296 | 6299 | 1 ; | HB3 | - | C |
| 6297 | 6299 | 1 ; | SG | - | C |
| 6299 | 6304 | 1 ; | C | - | HA |
| 6299 | 6305 | 1 ; | C | - | CB |
| 6299 | 6318 | 1 ; | C | - | C |
| 6300 | 6302 | 1 ; | O | - | H |
| 6300 | 6303 | 1 ; | O | - | CA |
| 6301 | 6306 | 1 ; | N | - | HB2 |
| 6301 | 6307 | 1 ; | N | - | HB3 |
| 6301 | 6308 | 1 ; | N | - | CG |
| 6301 | 6319 | 1 ; | N | - | O |
| 6301 | 6320 | 1 ; | N | - | N |
| 6302 | 6304 | 1 ; | H | - | HA |
| 6302 | 6305 | 1 ; | H | - | CB |
| 6302 | 6318 | 1 ; | H | - | C |
| 6303 | 6309 | 1 ; | CA | - | HG |
| 6303 | 6310 | 1 ; | CA | - | CD1 |
| 6303 | 6314 | 1 ; | CA | - | CD2 |
| 6303 | 6321 | 1 ; | CA | - | H |
| 6303 | 6322 | 1 ; | CA | - | CA |

| | | | | |
|---|---|---|---|---|
| 6304 | 6306 | 1 | ; | HA - HB2 |
| 6304 | 6307 | 1 | ; | HA - HB3 |
| 6304 | 6308 | 1 | ; | HA - CG |
| 6304 | 6319 | 1 | ; | HA - O |
| 6304 | 6320 | 1 | ; | HA - N |
| 6305 | 6311 | 1 | ; | CB - HD11 |
| 6305 | 6312 | 1 | ; | CB - HD12 |
| 6305 | 6313 | 1 | ; | CB - HD13 |
| 6305 | 6315 | 1 | ; | CB - HD21 |
| 6305 | 6316 | 1 | ; | CB - HD22 |
| 6305 | 6317 | 1 | ; | CB - HD23 |
| 6305 | 6319 | 1 | ; | CB - O |
| 6305 | 6320 | 1 | ; | CB - N |
| 6306 | 6309 | 1 | ; | HB2 - HG |
| 6306 | 6310 | 1 | ; | HB2 - CD1 |
| 6306 | 6314 | 1 | ; | HB2 - CD2 |
| 6306 | 6318 | 1 | ; | HB2 - C |
| 6307 | 6309 | 1 | ; | HB3 - HG |
| 6307 | 6310 | 1 | ; | HB3 - CD1 |
| 6307 | 6314 | 1 | ; | HB3 - CD2 |
| 6307 | 6318 | 1 | ; | HB3 - C |
| 6308 | 6318 | 1 | ; | CG - C |
| 6309 | 6311 | 1 | ; | HG - HD11 |
| 6309 | 6312 | 1 | ; | HG - HD12 |
| 6309 | 6313 | 1 | ; | HG - HD13 |
| 6309 | 6315 | 1 | ; | HG - HD21 |
| 6309 | 6316 | 1 | ; | HG - HD22 |
| 6309 | 6317 | 1 | ; | HG - HD23 |
| 6310 | 6315 | 1 | ; | CD1 - HD21 |
| 6310 | 6316 | 1 | ; | CD1 - HD22 |
| 6310 | 6317 | 1 | ; | CD1 - HD23 |
| 6311 | 6314 | 1 | ; | HD11 - CD2 |
| 6312 | 6314 | 1 | ; | HD12 - CD2 |
| 6313 | 6314 | 1 | ; | HD13 - CD2 |
| 6318 | 6323 | 1 | ; | C - HA |
| 6318 | 6324 | 1 | ; | C - CB |
| 6318 | 6337 | 1 | ; | C - C |
| 6319 | 6321 | 1 | ; | O - H |
| 6319 | 6322 | 1 | ; | O - CA |
| 6320 | 6325 | 1 | ; | N - HB |
| 6320 | 6326 | 1 | ; | N - CG2 |
| 6320 | 6330 | 1 | ; | N - CG1 |
| 6320 | 6338 | 1 | ; | N - O |
| 6320 | 6339 | 1 | ; | N - N |
| 6321 | 6323 | 1 | ; | H - HA |
| 6321 | 6324 | 1 | ; | H - CB |
| 6321 | 6337 | 1 | ; | H - C |
| 6322 | 6327 | 1 | ; | CA - HG21 |
| 6322 | 6328 | 1 | ; | CA - HG22 |
| 6322 | 6329 | 1 | ; | CA - HG23 |
| 6322 | 6331 | 1 | ; | CA - HG12 |
| 6322 | 6332 | 1 | ; | CA - HG13 |

```
6322   6333      1 ;     CA - CD1
6322   6340      1 ;     CA - CD
6322   6349      1 ;     CA - CA
6323   6325      1 ;     HA - HB
6323   6326      1 ;     HA - CG2
6323   6330      1 ;     HA - CG1
6323   6338      1 ;     HA - O
6323   6339      1 ;     HA - N
6324   6334      1 ;     CB - HD11
6324   6335      1 ;     CB - HD12
6324   6336      1 ;     CB - HD13
6324   6338      1 ;     CB - O
6324   6339      1 ;     CB - N
6325   6327      1 ;     HB - HG21
6325   6328      1 ;     HB - HG22
6325   6329      1 ;     HB - HG23
6325   6331      1 ;     HB - HG12
6325   6332      1 ;     HB - HG13
6325   6333      1 ;     HB - CD1
6325   6337      1 ;     HB - C
6326   6331      1 ;    CG2 - HG12
6326   6332      1 ;    CG2 - HG13
6326   6333      1 ;    CG2 - CD1
6326   6337      1 ;    CG2 - C
6327   6330      1 ;   HG21 - CG1
6328   6330      1 ;   HG22 - CG1
6329   6330      1 ;   HG23 - CG1
6330   6337      1 ;    CG1 - C
6331   6334      1 ;   HG12 - HD11
6331   6335      1 ;   HG12 - HD12
6331   6336      1 ;   HG12 - HD13
6332   6334      1 ;   HG13 - HD11
6332   6335      1 ;   HG13 - HD12
6332   6336      1 ;   HG13 - HD13
6337   6341      1 ;      C - HD2
6337   6342      1 ;      C - HD3
6337   6343      1 ;      C - CG
6337   6346      1 ;      C - CB
6337   6350      1 ;      C - HA
6337   6351      1 ;      C - C
6338   6340      1 ;      O - CD
6338   6349      1 ;      O - CA
6339   6344      1 ;      N - HG2
6339   6345      1 ;      N - HG3
6339   6347      1 ;      N - HB2
6339   6348      1 ;      N - HB3
6339   6352      1 ;      N - O
6339   6353      1 ;      N - N
6340   6347      1 ;     CD - HB2
6340   6348      1 ;     CD - HB3
6340   6350      1 ;     CD - HA
6340   6351      1 ;     CD - C
```

```
6341   6344      1 ;    HD2 - HG2
6341   6345      1 ;    HD2 - HG3
6341   6346      1 ;    HD2 - CB
6341   6349      1 ;    HD2 - CA
6342   6344      1 ;    HD3 - HG2
6342   6345      1 ;    HD3 - HG3
6342   6346      1 ;    HD3 - CB
6342   6349      1 ;    HD3 - CA
6343   6350      1 ;     CG - HA
6343   6351      1 ;     CG - C
6344   6347      1 ;    HG2 - HB2
6344   6348      1 ;    HG2 - HB3
6344   6349      1 ;    HG2 - CA
6345   6347      1 ;    HG3 - HB2
6345   6348      1 ;    HG3 - HB3
6345   6349      1 ;    HG3 - CA
6346   6352      1 ;     CB - O
6346   6353      1 ;     CB - N
6347   6350      1 ;    HB2 - HA
6347   6351      1 ;    HB2 - C
6348   6350      1 ;    HB3 - HA
6348   6351      1 ;    HB3 - C
6349   6354      1 ;     CA - H
6349   6355      1 ;     CA - CA
6350   6352      1 ;     HA - O
6350   6353      1 ;     HA - N
6351   6356      1 ;      C - HA
6351   6357      1 ;      C - CB
6351   6371      1 ;      C - C
6352   6354      1 ;      O - H
6352   6355      1 ;      O - CA
6353   6358      1 ;      N - HB2
6353   6359      1 ;      N - HB3
6353   6360      1 ;      N - CG
6353   6372      1 ;      N - O
6353   6373      1 ;      N - N
6354   6356      1 ;      H - HA
6354   6357      1 ;      H - CB
6354   6371      1 ;      H - C
6355   6361      1 ;     CA - CD1
6355   6369      1 ;     CA - CD2
6355   6374      1 ;     CA - H
6355   6375      1 ;     CA - CA
6356   6358      1 ;     HA - HB2
6356   6359      1 ;     HA - HB3
6356   6360      1 ;     HA - CG
6356   6372      1 ;     HA - O
6356   6373      1 ;     HA - N
6357   6362      1 ;     CB - HD1
6357   6363      1 ;     CB - CE1
6357   6367      1 ;     CB - CE2
6357   6370      1 ;     CB - HD2
```

| | | | | | | |
|---|---|---|---|---|---|---|
| 6357 | 6372 | 1 | ; | CB | - | O |
| 6357 | 6373 | 1 | ; | CB | - | N |
| 6358 | 6361 | 1 | ; | HB2 | - | CD1 |
| 6358 | 6369 | 1 | ; | HB2 | - | CD2 |
| 6358 | 6371 | 1 | ; | HB2 | - | C |
| 6359 | 6361 | 1 | ; | HB3 | - | CD1 |
| 6359 | 6369 | 1 | ; | HB3 | - | CD2 |
| 6359 | 6371 | 1 | ; | HB3 | - | C |
| 6360 | 6364 | 1 | ; | CG | - | HE1 |
| 6360 | 6365 | 1 | ; | CG | - | CZ |
| 6360 | 6368 | 1 | ; | CG | - | HE2 |
| 6360 | 6371 | 1 | ; | CG | - | C |
| 6361 | 6366 | 1 | ; | CD1 | - | HZ |
| 6361 | 6367 | 1 | ; | CD1 | - | CE2 |
| 6361 | 6370 | 1 | ; | CD1 | - | HD2 |
| 6362 | 6364 | 1 | ; | HD1 | - | HE1 |
| 6362 | 6365 | 1 | ; | HD1 | - | CZ |
| 6362 | 6369 | 1 | ; | HD1 | - | CD2 |
| 6363 | 6368 | 1 | ; | CE1 | - | HE2 |
| 6363 | 6369 | 1 | ; | CE1 | - | CD2 |
| 6364 | 6366 | 1 | ; | HE1 | - | HZ |
| 6364 | 6367 | 1 | ; | HE1 | - | CE2 |
| 6365 | 6370 | 1 | ; | CZ | - | HD2 |
| 6366 | 6368 | 1 | ; | HZ | - | HE2 |
| 6366 | 6369 | 1 | ; | HZ | - | CD2 |
| 6368 | 6370 | 1 | ; | HE2 | - | HD2 |
| 6371 | 6376 | 1 | ; | C | - | HA |
| 6371 | 6377 | 1 | ; | C | - | CB |
| 6371 | 6393 | 1 | ; | C | - | C |
| 6372 | 6374 | 1 | ; | O | - | H |
| 6372 | 6375 | 1 | ; | O | - | CA |
| 6373 | 6378 | 1 | ; | N | - | HB2 |
| 6373 | 6379 | 1 | ; | N | - | HB3 |
| 6373 | 6380 | 1 | ; | N | - | CG |
| 6373 | 6394 | 1 | ; | N | - | O |
| 6373 | 6395 | 1 | ; | N | - | N |
| 6374 | 6376 | 1 | ; | H | - | HA |
| 6374 | 6377 | 1 | ; | H | - | CB |
| 6374 | 6393 | 1 | ; | H | - | C |
| 6375 | 6381 | 1 | ; | CA | - | HG2 |
| 6375 | 6382 | 1 | ; | CA | - | HG3 |
| 6375 | 6383 | 1 | ; | CA | - | CD |
| 6375 | 6396 | 1 | ; | CA | - | H |
| 6375 | 6397 | 1 | ; | CA | - | CA |
| 6376 | 6378 | 1 | ; | HA | - | HB2 |
| 6376 | 6379 | 1 | ; | HA | - | HB3 |
| 6376 | 6380 | 1 | ; | HA | - | CG |
| 6376 | 6394 | 1 | ; | HA | - | O |
| 6376 | 6395 | 1 | ; | HA | - | N |
| 6377 | 6384 | 1 | ; | CB | - | HD2 |
| 6377 | 6385 | 1 | ; | CB | - | HD3 |
| 6377 | 6386 | 1 | ; | CB | - | CE |

| | | | |
|---|---|---|---|
| 6377 | 6394 | 1 ; | CB - O |
| 6377 | 6395 | 1 ; | CB - N |
| 6378 | 6381 | 1 ; | HB2 - HG2 |
| 6378 | 6382 | 1 ; | HB2 - HG3 |
| 6378 | 6383 | 1 ; | HB2 - CD |
| 6378 | 6393 | 1 ; | HB2 - C |
| 6379 | 6381 | 1 ; | HB3 - HG2 |
| 6379 | 6382 | 1 ; | HB3 - HG3 |
| 6379 | 6383 | 1 ; | HB3 - CD |
| 6379 | 6393 | 1 ; | HB3 - C |
| 6380 | 6387 | 1 ; | CG - HE2 |
| 6380 | 6388 | 1 ; | CG - HE3 |
| 6380 | 6389 | 1 ; | CG - NZ |
| 6380 | 6393 | 1 ; | CG - C |
| 6381 | 6384 | 1 ; | HG2 - HD2 |
| 6381 | 6385 | 1 ; | HG2 - HD3 |
| 6381 | 6386 | 1 ; | HG2 - CE |
| 6382 | 6384 | 1 ; | HG3 - HD2 |
| 6382 | 6385 | 1 ; | HG3 - HD3 |
| 6382 | 6386 | 1 ; | HG3 - CE |
| 6383 | 6390 | 1 ; | CD - HZ1 |
| 6383 | 6391 | 1 ; | CD - HZ2 |
| 6383 | 6392 | 1 ; | CD - HZ3 |
| 6384 | 6387 | 1 ; | HD2 - HE2 |
| 6384 | 6388 | 1 ; | HD2 - HE3 |
| 6384 | 6389 | 1 ; | HD2 - NZ |
| 6385 | 6387 | 1 ; | HD3 - HE2 |
| 6385 | 6388 | 1 ; | HD3 - HE3 |
| 6385 | 6389 | 1 ; | HD3 - NZ |
| 6387 | 6390 | 1 ; | HE2 - HZ1 |
| 6387 | 6391 | 1 ; | HE2 - HZ2 |
| 6387 | 6392 | 1 ; | HE2 - HZ3 |
| 6388 | 6390 | 1 ; | HE3 - HZ1 |
| 6388 | 6391 | 1 ; | HE3 - HZ2 |
| 6388 | 6392 | 1 ; | HE3 - HZ3 |
| 6393 | 6398 | 1 ; | C - HA |
| 6393 | 6399 | 1 ; | C - CB |
| 6393 | 6413 | 1 ; | C - C |
| 6394 | 6396 | 1 ; | O - H |
| 6394 | 6397 | 1 ; | O - CA |
| 6395 | 6400 | 1 ; | N - HB2 |
| 6395 | 6401 | 1 ; | N - HB3 |
| 6395 | 6402 | 1 ; | N - CG |
| 6395 | 6414 | 1 ; | N - O |
| 6395 | 6415 | 1 ; | N - N |
| 6396 | 6398 | 1 ; | H - HA |
| 6396 | 6399 | 1 ; | H - CB |
| 6396 | 6413 | 1 ; | H - C |
| 6397 | 6403 | 1 ; | CA - CD1 |
| 6397 | 6411 | 1 ; | CA - CD2 |
| 6397 | 6416 | 1 ; | CA - H |
| 6397 | 6417 | 1 ; | CA - CA |

| | | | | |
|---|---|---|---|---|
| 6398 | 6400 | 1 | ; | HA - HB2 |
| 6398 | 6401 | 1 | ; | HA - HB3 |
| 6398 | 6402 | 1 | ; | HA - CG |
| 6398 | 6414 | 1 | ; | HA - O |
| 6398 | 6415 | 1 | ; | HA - N |
| 6399 | 6404 | 1 | ; | CB - HD1 |
| 6399 | 6405 | 1 | ; | CB - CE1 |
| 6399 | 6409 | 1 | ; | CB - CE2 |
| 6399 | 6412 | 1 | ; | CB - HD2 |
| 6399 | 6414 | 1 | ; | CB - O |
| 6399 | 6415 | 1 | ; | CB - N |
| 6400 | 6403 | 1 | ; | HB2 - CD1 |
| 6400 | 6411 | 1 | ; | HB2 - CD2 |
| 6400 | 6413 | 1 | ; | HB2 - C |
| 6401 | 6403 | 1 | ; | HB3 - CD1 |
| 6401 | 6411 | 1 | ; | HB3 - CD2 |
| 6401 | 6413 | 1 | ; | HB3 - C |
| 6402 | 6406 | 1 | ; | CG - HE1 |
| 6402 | 6407 | 1 | ; | CG - CZ |
| 6402 | 6410 | 1 | ; | CG - HE2 |
| 6402 | 6413 | 1 | ; | CG - C |
| 6403 | 6408 | 1 | ; | CD1 - HZ |
| 6403 | 6409 | 1 | ; | CD1 - CE2 |
| 6403 | 6412 | 1 | ; | CD1 - HD2 |
| 6404 | 6406 | 1 | ; | HD1 - HE1 |
| 6404 | 6407 | 1 | ; | HD1 - CZ |
| 6404 | 6411 | 1 | ; | HD1 - CD2 |
| 6405 | 6410 | 1 | ; | CE1 - HE2 |
| 6405 | 6411 | 1 | ; | CE1 - CD2 |
| 6406 | 6408 | 1 | ; | HE1 - HZ |
| 6406 | 6409 | 1 | ; | HE1 - CE2 |
| 6407 | 6412 | 1 | ; | CZ - HD2 |
| 6408 | 6410 | 1 | ; | HZ - HE2 |
| 6408 | 6411 | 1 | ; | HZ - CD2 |
| 6410 | 6412 | 1 | ; | HE2 - HD2 |
| 6413 | 6418 | 1 | ; | C - HA |
| 6413 | 6419 | 1 | ; | C - CB |
| 6413 | 6433 | 1 | ; | C - C |
| 6414 | 6416 | 1 | ; | O - H |
| 6414 | 6417 | 1 | ; | O - CA |
| 6415 | 6420 | 1 | ; | N - HB2 |
| 6415 | 6421 | 1 | ; | N - HB3 |
| 6415 | 6422 | 1 | ; | N - CG |
| 6415 | 6434 | 1 | ; | N - O |
| 6415 | 6435 | 1 | ; | N - N |
| 6416 | 6418 | 1 | ; | H - HA |
| 6416 | 6419 | 1 | ; | H - CB |
| 6416 | 6433 | 1 | ; | H - C |
| 6417 | 6423 | 1 | ; | CA - CD1 |
| 6417 | 6431 | 1 | ; | CA - CD2 |
| 6417 | 6436 | 1 | ; | CA - H |
| 6417 | 6437 | 1 | ; | CA - CA |

| | | | |
|---|---|---|---|
| 6418 | 6420 | 1 ; | HA - HB2 |
| 6418 | 6421 | 1 ; | HA - HB3 |
| 6418 | 6422 | 1 ; | HA - CG |
| 6418 | 6434 | 1 ; | HA - O |
| 6418 | 6435 | 1 ; | HA - N |
| 6419 | 6424 | 1 ; | CB - HD1 |
| 6419 | 6425 | 1 ; | CB - CE1 |
| 6419 | 6429 | 1 ; | CB - CE2 |
| 6419 | 6432 | 1 ; | CB - HD2 |
| 6419 | 6434 | 1 ; | CB - O |
| 6419 | 6435 | 1 ; | CB - N |
| 6420 | 6423 | 1 ; | HB2 - CD1 |
| 6420 | 6431 | 1 ; | HB2 - CD2 |
| 6420 | 6433 | 1 ; | HB2 - C |
| 6421 | 6423 | 1 ; | HB3 - CD1 |
| 6421 | 6431 | 1 ; | HB3 - CD2 |
| 6421 | 6433 | 1 ; | HB3 - C |
| 6422 | 6426 | 1 ; | CG - HE1 |
| 6422 | 6427 | 1 ; | CG - CZ |
| 6422 | 6430 | 1 ; | CG - HE2 |
| 6422 | 6433 | 1 ; | CG - C |
| 6423 | 6428 | 1 ; | CD1 - HZ |
| 6423 | 6429 | 1 ; | CD1 - CE2 |
| 6423 | 6432 | 1 ; | CD1 - HD2 |
| 6424 | 6426 | 1 ; | HD1 - HE1 |
| 6424 | 6427 | 1 ; | HD1 - CZ |
| 6424 | 6431 | 1 ; | HD1 - CD2 |
| 6425 | 6430 | 1 ; | CE1 - HE2 |
| 6425 | 6431 | 1 ; | CE1 - CD2 |
| 6426 | 6428 | 1 ; | HE1 - HZ |
| 6426 | 6429 | 1 ; | HE1 - CE2 |
| 6427 | 6432 | 1 ; | CZ - HD2 |
| 6428 | 6430 | 1 ; | HZ - HE2 |
| 6428 | 6431 | 1 ; | HZ - CD2 |
| 6430 | 6432 | 1 ; | HE2 - HD2 |
| 6433 | 6438 | 1 ; | C - HA |
| 6433 | 6439 | 1 ; | C - CB |
| 6433 | 6445 | 1 ; | C - C |
| 6434 | 6436 | 1 ; | O - H |
| 6434 | 6437 | 1 ; | O - CA |
| 6435 | 6440 | 1 ; | N - HB2 |
| 6435 | 6441 | 1 ; | N - HB3 |
| 6435 | 6442 | 1 ; | N - CG |
| 6435 | 6446 | 1 ; | N - O |
| 6435 | 6447 | 1 ; | N - N |
| 6436 | 6438 | 1 ; | H - HA |
| 6436 | 6439 | 1 ; | H - CB |
| 6436 | 6445 | 1 ; | H - C |
| 6437 | 6443 | 1 ; | CA - OD1 |
| 6437 | 6444 | 1 ; | CA - OD2 |
| 6437 | 6448 | 1 ; | CA - H |
| 6437 | 6449 | 1 ; | CA - CA |

| | | | |
|---|---|---|---|
| 6438 | 6440 | 1 ; | HA - HB2 |
| 6438 | 6441 | 1 ; | HA - HB3 |
| 6438 | 6442 | 1 ; | HA - CG |
| 6438 | 6446 | 1 ; | HA - O |
| 6438 | 6447 | 1 ; | HA - N |
| 6439 | 6446 | 1 ; | CB - O |
| 6439 | 6447 | 1 ; | CB - N |
| 6440 | 6443 | 1 ; | HB2 - OD1 |
| 6440 | 6444 | 1 ; | HB2 - OD2 |
| 6440 | 6445 | 1 ; | HB2 - C |
| 6441 | 6443 | 1 ; | HB3 - OD1 |
| 6441 | 6444 | 1 ; | HB3 - OD2 |
| 6441 | 6445 | 1 ; | HB3 - C |
| 6442 | 6445 | 1 ; | CG - C |
| 6445 | 6450 | 1 ; | C - HA |
| 6445 | 6451 | 1 ; | C - CB |
| 6445 | 6464 | 1 ; | C - C |
| 6446 | 6448 | 1 ; | O - H |
| 6446 | 6449 | 1 ; | O - CA |
| 6447 | 6452 | 1 ; | N - HB |
| 6447 | 6453 | 1 ; | N - CG2 |
| 6447 | 6457 | 1 ; | N - CG1 |
| 6447 | 6465 | 1 ; | N - O |
| 6447 | 6466 | 1 ; | N - N |
| 6448 | 6450 | 1 ; | H - HA |
| 6448 | 6451 | 1 ; | H - CB |
| 6448 | 6464 | 1 ; | H - C |
| 6449 | 6454 | 1 ; | CA - HG21 |
| 6449 | 6455 | 1 ; | CA - HG22 |
| 6449 | 6456 | 1 ; | CA - HG23 |
| 6449 | 6458 | 1 ; | CA - HG12 |
| 6449 | 6459 | 1 ; | CA - HG13 |
| 6449 | 6460 | 1 ; | CA - CD1 |
| 6449 | 6467 | 1 ; | CA - H |
| 6449 | 6468 | 1 ; | CA - CA |
| 6450 | 6452 | 1 ; | HA - HB |
| 6450 | 6453 | 1 ; | HA - CG2 |
| 6450 | 6457 | 1 ; | HA - CG1 |
| 6450 | 6465 | 1 ; | HA - O |
| 6450 | 6466 | 1 ; | HA - N |
| 6451 | 6461 | 1 ; | CB - HD11 |
| 6451 | 6462 | 1 ; | CB - HD12 |
| 6451 | 6463 | 1 ; | CB - HD13 |
| 6451 | 6465 | 1 ; | CB - O |
| 6451 | 6466 | 1 ; | CB - N |
| 6452 | 6454 | 1 ; | HB - HG21 |
| 6452 | 6455 | 1 ; | HB - HG22 |
| 6452 | 6456 | 1 ; | HB - HG23 |
| 6452 | 6458 | 1 ; | HB - HG12 |
| 6452 | 6459 | 1 ; | HB - HG13 |
| 6452 | 6460 | 1 ; | HB - CD1 |
| 6452 | 6464 | 1 ; | HB - C |

| | | | | | | |
|---|---|---|---|---|---|---|
| 6453 | 6458 | 1 | ; | CG2 | - | HG12 |
| 6453 | 6459 | 1 | ; | CG2 | - | HG13 |
| 6453 | 6460 | 1 | ; | CG2 | - | CD1 |
| 6453 | 6464 | 1 | ; | CG2 | - | C |
| 6454 | 6457 | 1 | ; | HG21 | - | CG1 |
| 6455 | 6457 | 1 | ; | HG22 | - | CG1 |
| 6456 | 6457 | 1 | ; | HG23 | - | CG1 |
| 6457 | 6464 | 1 | ; | CG1 | - | C |
| 6458 | 6461 | 1 | ; | HG12 | - | HD11 |
| 6458 | 6462 | 1 | ; | HG12 | - | HD12 |
| 6458 | 6463 | 1 | ; | HG12 | - | HD13 |
| 6459 | 6461 | 1 | ; | HG13 | - | HD11 |
| 6459 | 6462 | 1 | ; | HG13 | - | HD12 |
| 6459 | 6463 | 1 | ; | HG13 | - | HD13 |
| 6464 | 6469 | 1 | ; | C | - | HA |
| 6464 | 6470 | 1 | ; | C | - | CB |
| 6464 | 6476 | 1 | ; | C | - | C |
| 6465 | 6467 | 1 | ; | O | - | H |
| 6465 | 6468 | 1 | ; | O | - | CA |
| 6466 | 6471 | 1 | ; | N | - | HB2 |
| 6466 | 6472 | 1 | ; | N | - | HB3 |
| 6466 | 6473 | 1 | ; | N | - | CG |
| 6466 | 6477 | 1 | ; | N | - | O |
| 6466 | 6478 | 1 | ; | N | - | N |
| 6467 | 6469 | 1 | ; | H | - | HA |
| 6467 | 6470 | 1 | ; | H | - | CB |
| 6467 | 6476 | 1 | ; | H | - | C |
| 6468 | 6474 | 1 | ; | CA | - | OD1 |
| 6468 | 6475 | 1 | ; | CA | - | OD2 |
| 6468 | 6479 | 1 | ; | CA | - | H |
| 6468 | 6480 | 1 | ; | CA | - | CA |
| 6469 | 6471 | 1 | ; | HA | - | HB2 |
| 6469 | 6472 | 1 | ; | HA | - | HB3 |
| 6469 | 6473 | 1 | ; | HA | - | CG |
| 6469 | 6477 | 1 | ; | HA | - | O |
| 6469 | 6478 | 1 | ; | HA | - | N |
| 6470 | 6477 | 1 | ; | CB | - | O |
| 6470 | 6478 | 1 | ; | CB | - | N |
| 6471 | 6474 | 1 | ; | HB2 | - | OD1 |
| 6471 | 6475 | 1 | ; | HB2 | - | OD2 |
| 6471 | 6476 | 1 | ; | HB2 | - | C |
| 6472 | 6474 | 1 | ; | HB3 | - | OD1 |
| 6472 | 6475 | 1 | ; | HB3 | - | OD2 |
| 6472 | 6476 | 1 | ; | HB3 | - | C |
| 6473 | 6476 | 1 | ; | CG | - | C |
| 6476 | 6481 | 1 | ; | C | - | HA2 |
| 6476 | 6482 | 1 | ; | C | - | HA3 |
| 6476 | 6483 | 1 | ; | C | - | C |
| 6477 | 6479 | 1 | ; | O | - | H |
| 6477 | 6480 | 1 | ; | O | - | CA |
| 6478 | 6484 | 1 | ; | N | - | O |
| 6478 | 6485 | 1 | ; | N | - | N |

```
6479   6481       1 ;       H - HA2
6479   6482       1 ;       H - HA3
6479   6483       1 ;       H - C
6480   6486       1 ;      CA - H
6480   6487       1 ;      CA - CA
6481   6484       1 ;     HA2 - O
6481   6485       1 ;     HA2 - N
6482   6484       1 ;     HA3 - O
6482   6485       1 ;     HA3 - N
6483   6488       1 ;       C - HA
6483   6489       1 ;       C - CB
6483   6497       1 ;       C - C
6484   6486       1 ;       O - H
6484   6487       1 ;       O - CA
6485   6490       1 ;       N - HB2
6485   6491       1 ;       N - HB3
6485   6492       1 ;       N - CG
6485   6498       1 ;       N - O
6485   6499       1 ;       N - N
6486   6488       1 ;       H - HA
6486   6489       1 ;       H - CB
6486   6497       1 ;       H - C
6487   6493       1 ;      CA - OD1
6487   6494       1 ;      CA - ND2
6487   6500       1 ;      CA - H
6487   6501       1 ;      CA - CA
6488   6490       1 ;      HA - HB2
6488   6491       1 ;      HA - HB3
6488   6492       1 ;      HA - CG
6488   6498       1 ;      HA - O
6488   6499       1 ;      HA - N
6489   6495       1 ;      CB - HD21
6489   6496       1 ;      CB - HD22
6489   6498       1 ;      CB - O
6489   6499       1 ;      CB - N
6490   6493       1 ;     HB2 - OD1
6490   6494       1 ;     HB2 - ND2
6490   6497       1 ;     HB2 - C
6491   6493       1 ;     HB3 - OD1
6491   6494       1 ;     HB3 - ND2
6491   6497       1 ;     HB3 - C
6492   6497       1 ;      CG - C
6493   6495       1 ;     OD1 - HD21
6493   6496       1 ;     OD1 - HD22
6497   6502       1 ;       C - HA2
6497   6503       1 ;       C - HA3
6497   6504       1 ;       C - C
6498   6500       1 ;       O - H
6498   6501       1 ;       O - CA
6499   6505       1 ;       N - O
6499   6506       1 ;       N - N
6500   6502       1 ;       H - HA2
```

| | | | | | |
|---|---|---|---|---|---|
| 6500 | 6503 | 1 ; | H | - | HA3 |
| 6500 | 6504 | 1 ; | H | - | C |
| 6501 | 6507 | 1 ; | CA | - | H |
| 6501 | 6508 | 1 ; | CA | - | CA |
| 6502 | 6505 | 1 ; | HA2 | - | O |
| 6502 | 6506 | 1 ; | HA2 | - | N |
| 6503 | 6505 | 1 ; | HA3 | - | O |
| 6503 | 6506 | 1 ; | HA3 | - | N |
| 6504 | 6509 | 1 ; | C | - | HA |
| 6504 | 6510 | 1 ; | C | - | CB |
| 6504 | 6526 | 1 ; | C | - | C |
| 6505 | 6507 | 1 ; | O | - | H |
| 6505 | 6508 | 1 ; | O | - | CA |
| 6506 | 6511 | 1 ; | N | - | HB2 |
| 6506 | 6512 | 1 ; | N | - | HB3 |
| 6506 | 6513 | 1 ; | N | - | CG |
| 6506 | 6527 | 1 ; | N | - | O |
| 6506 | 6528 | 1 ; | N | - | N |
| 6507 | 6509 | 1 ; | H | - | HA |
| 6507 | 6510 | 1 ; | H | - | CB |
| 6507 | 6526 | 1 ; | H | - | C |
| 6508 | 6514 | 1 ; | CA | - | HG2 |
| 6508 | 6515 | 1 ; | CA | - | HG3 |
| 6508 | 6516 | 1 ; | CA | - | CD |
| 6508 | 6529 | 1 ; | CA | - | H |
| 6508 | 6530 | 1 ; | CA | - | CA |
| 6509 | 6511 | 1 ; | HA | - | HB2 |
| 6509 | 6512 | 1 ; | HA | - | HB3 |
| 6509 | 6513 | 1 ; | HA | - | CG |
| 6509 | 6527 | 1 ; | HA | - | O |
| 6509 | 6528 | 1 ; | HA | - | N |
| 6510 | 6517 | 1 ; | CB | - | HD2 |
| 6510 | 6518 | 1 ; | CB | - | HD3 |
| 6510 | 6519 | 1 ; | CB | - | CE |
| 6510 | 6527 | 1 ; | CB | - | O |
| 6510 | 6528 | 1 ; | CB | - | N |
| 6511 | 6514 | 1 ; | HB2 | - | HG2 |
| 6511 | 6515 | 1 ; | HB2 | - | HG3 |
| 6511 | 6516 | 1 ; | HB2 | - | CD |
| 6511 | 6526 | 1 ; | HB2 | - | C |
| 6512 | 6514 | 1 ; | HB3 | - | HG2 |
| 6512 | 6515 | 1 ; | HB3 | - | HG3 |
| 6512 | 6516 | 1 ; | HB3 | - | CD |
| 6512 | 6526 | 1 ; | HB3 | - | C |
| 6513 | 6520 | 1 ; | CG | - | HE2 |
| 6513 | 6521 | 1 ; | CG | - | HE3 |
| 6513 | 6522 | 1 ; | CG | - | NZ |
| 6513 | 6526 | 1 ; | CG | - | C |
| 6514 | 6517 | 1 ; | HG2 | - | HD2 |
| 6514 | 6518 | 1 ; | HG2 | - | HD3 |
| 6514 | 6519 | 1 ; | HG2 | - | CE |
| 6515 | 6517 | 1 ; | HG3 | - | HD2 |

```
6515   6518       1 ;     HG3 - HD3
6515   6519       1 ;     HG3 - CE
6516   6523       1 ;      CD - HZ1
6516   6524       1 ;      CD - HZ2
6516   6525       1 ;      CD - HZ3
6517   6520       1 ;     HD2 - HE2
6517   6521       1 ;     HD2 - HE3
6517   6522       1 ;     HD2 - NZ
6518   6520       1 ;     HD3 - HE2
6518   6521       1 ;     HD3 - HE3
6518   6522       1 ;     HD3 - NZ
6520   6523       1 ;     HE2 - HZ1
6520   6524       1 ;     HE2 - HZ2
6520   6525       1 ;     HE2 - HZ3
6521   6523       1 ;     HE3 - HZ1
6521   6524       1 ;     HE3 - HZ2
6521   6525       1 ;     HE3 - HZ3
6526   6531       1 ;       C - HA
6526   6532       1 ;       C - CB
6526   6545       1 ;       C - C
6527   6529       1 ;       O - H
6527   6530       1 ;       O - CA
6528   6533       1 ;       N - HB
6528   6534       1 ;       N - CG2
6528   6538       1 ;       N - CG1
6528   6546       1 ;       N - O
6528   6547       1 ;       N - N
6529   6531       1 ;       H - HA
6529   6532       1 ;       H - CB
6529   6545       1 ;       H - C
6530   6535       1 ;      CA - HG21
6530   6536       1 ;      CA - HG22
6530   6537       1 ;      CA - HG23
6530   6539       1 ;      CA - HG12
6530   6540       1 ;      CA - HG13
6530   6541       1 ;      CA - CD1
6530   6548       1 ;      CA - H
6530   6549       1 ;      CA - CA
6531   6533       1 ;      HA - HB
6531   6534       1 ;      HA - CG2
6531   6538       1 ;      HA - CG1
6531   6546       1 ;      HA - O
6531   6547       1 ;      HA - N
6532   6542       1 ;      CB - HD11
6532   6543       1 ;      CB - HD12
6532   6544       1 ;      CB - HD13
6532   6546       1 ;      CB - O
6532   6547       1 ;      CB - N
6533   6535       1 ;      HB - HG21
6533   6536       1 ;      HB - HG22
6533   6537       1 ;      HB - HG23
6533   6539       1 ;      HB - HG12
```

| | | | | | |
|---|---|---|---|---|---|
| 6533 | 6540 | 1 ; | HB | - | HG13 |
| 6533 | 6541 | 1 ; | HB | - | CD1 |
| 6533 | 6545 | 1 ; | HB | - | C |
| 6534 | 6539 | 1 ; | CG2 | - | HG12 |
| 6534 | 6540 | 1 ; | CG2 | - | HG13 |
| 6534 | 6541 | 1 ; | CG2 | - | CD1 |
| 6534 | 6545 | 1 ; | CG2 | - | C |
| 6535 | 6538 | 1 ; | HG21 | - | CG1 |
| 6536 | 6538 | 1 ; | HG22 | - | CG1 |
| 6537 | 6538 | 1 ; | HG23 | - | CG1 |
| 6538 | 6545 | 1 ; | CG1 | - | C |
| 6539 | 6542 | 1 ; | HG12 | - | HD11 |
| 6539 | 6543 | 1 ; | HG12 | - | HD12 |
| 6539 | 6544 | 1 ; | HG12 | - | HD13 |
| 6540 | 6542 | 1 ; | HG13 | - | HD11 |
| 6540 | 6543 | 1 ; | HG13 | - | HD12 |
| 6540 | 6544 | 1 ; | HG13 | - | HD13 |
| 6545 | 6550 | 1 ; | C | - | HA |
| 6545 | 6551 | 1 ; | C | - | CB |
| 6545 | 6556 | 1 ; | C | - | C |
| 6546 | 6548 | 1 ; | O | - | H |
| 6546 | 6549 | 1 ; | O | - | CA |
| 6547 | 6552 | 1 ; | N | - | HB2 |
| 6547 | 6553 | 1 ; | N | - | HB3 |
| 6547 | 6554 | 1 ; | N | - | OG |
| 6547 | 6557 | 1 ; | N | - | O |
| 6547 | 6558 | 1 ; | N | - | N |
| 6548 | 6550 | 1 ; | H | - | HA |
| 6548 | 6551 | 1 ; | H | - | CB |
| 6548 | 6556 | 1 ; | H | - | C |
| 6549 | 6555 | 1 ; | CA | - | HG |
| 6549 | 6559 | 1 ; | CA | - | H |
| 6549 | 6560 | 1 ; | CA | - | CA |
| 6550 | 6552 | 1 ; | HA | - | HB2 |
| 6550 | 6553 | 1 ; | HA | - | HB3 |
| 6550 | 6554 | 1 ; | HA | - | OG |
| 6550 | 6557 | 1 ; | HA | - | O |
| 6550 | 6558 | 1 ; | HA | - | N |
| 6551 | 6557 | 1 ; | CB | - | O |
| 6551 | 6558 | 1 ; | CB | - | N |
| 6552 | 6555 | 1 ; | HB2 | - | HG |
| 6552 | 6556 | 1 ; | HB2 | - | C |
| 6553 | 6555 | 1 ; | HB3 | - | HG |
| 6553 | 6556 | 1 ; | HB3 | - | C |
| 6554 | 6556 | 1 ; | OG | - | C |
| 6556 | 6561 | 1 ; | C | - | HA |
| 6556 | 6562 | 1 ; | C | - | CB |
| 6556 | 6572 | 1 ; | C | - | C |
| 6557 | 6559 | 1 ; | O | - | H |
| 6557 | 6560 | 1 ; | O | - | CA |
| 6558 | 6563 | 1 ; | N | - | HB |
| 6558 | 6564 | 1 ; | N | - | CG1 |

| | | | | |
|---|---|---|---|---|
| 6558 | 6568 | 1 ; | N - | CG2 |
| 6558 | 6573 | 1 ; | N - | O |
| 6558 | 6574 | 1 ; | N - | N |
| 6559 | 6561 | 1 ; | H - | HA |
| 6559 | 6562 | 1 ; | H - | CB |
| 6559 | 6572 | 1 ; | H - | C |
| 6560 | 6565 | 1 ; | CA - | HG11 |
| 6560 | 6566 | 1 ; | CA - | HG12 |
| 6560 | 6567 | 1 ; | CA - | HG13 |
| 6560 | 6569 | 1 ; | CA - | HG21 |
| 6560 | 6570 | 1 ; | CA - | HG22 |
| 6560 | 6571 | 1 ; | CA - | HG23 |
| 6560 | 6575 | 1 ; | CA - | H |
| 6560 | 6576 | 1 ; | CA - | CA |
| 6561 | 6563 | 1 ; | HA - | HB |
| 6561 | 6564 | 1 ; | HA - | CG1 |
| 6561 | 6568 | 1 ; | HA - | CG2 |
| 6561 | 6573 | 1 ; | HA - | O |
| 6561 | 6574 | 1 ; | HA - | N |
| 6562 | 6573 | 1 ; | CB - | O |
| 6562 | 6574 | 1 ; | CB - | N |
| 6563 | 6565 | 1 ; | HB - | HG11 |
| 6563 | 6566 | 1 ; | HB - | HG12 |
| 6563 | 6567 | 1 ; | HB - | HG13 |
| 6563 | 6569 | 1 ; | HB - | HG21 |
| 6563 | 6570 | 1 ; | HB - | HG22 |
| 6563 | 6571 | 1 ; | HB - | HG23 |
| 6563 | 6572 | 1 ; | HB - | C |
| 6564 | 6569 | 1 ; | CG1 - | HG21 |
| 6564 | 6570 | 1 ; | CG1 - | HG22 |
| 6564 | 6571 | 1 ; | CG1 - | HG23 |
| 6564 | 6572 | 1 ; | CG1 - | C |
| 6565 | 6568 | 1 ; | HG11 - | CG2 |
| 6566 | 6568 | 1 ; | HG12 - | CG2 |
| 6567 | 6568 | 1 ; | HG13 - | CG2 |
| 6568 | 6572 | 1 ; | CG2 - | C |
| 6572 | 6577 | 1 ; | C - | HA |
| 6572 | 6578 | 1 ; | C - | CB |
| 6572 | 6587 | 1 ; | C - | C |
| 6573 | 6575 | 1 ; | O - | H |
| 6573 | 6576 | 1 ; | O - | CA |
| 6574 | 6579 | 1 ; | N - | HB2 |
| 6574 | 6580 | 1 ; | N - | HB3 |
| 6574 | 6581 | 1 ; | N - | CG |
| 6574 | 6588 | 1 ; | N - | O |
| 6574 | 6589 | 1 ; | N - | N |
| 6575 | 6577 | 1 ; | H - | HA |
| 6575 | 6578 | 1 ; | H - | CB |
| 6575 | 6587 | 1 ; | H - | C |
| 6576 | 6582 | 1 ; | CA - | HG2 |
| 6576 | 6583 | 1 ; | CA - | HG3 |
| 6576 | 6584 | 1 ; | CA - | CD |

| | | | | | |
|---|---|---|---|---|---|
| 6576 | 6590 | 1 ; | CA | - | H |
| 6576 | 6591 | 1 ; | CA | - | CA |
| 6577 | 6579 | 1 ; | HA | - | HB2 |
| 6577 | 6580 | 1 ; | HA | - | HB3 |
| 6577 | 6581 | 1 ; | HA | - | CG |
| 6577 | 6588 | 1 ; | HA | - | O |
| 6577 | 6589 | 1 ; | HA | - | N |
| 6578 | 6585 | 1 ; | CB | - | OE1 |
| 6578 | 6586 | 1 ; | CB | - | OE2 |
| 6578 | 6588 | 1 ; | CB | - | O |
| 6578 | 6589 | 1 ; | CB | - | N |
| 6579 | 6582 | 1 ; | HB2 | - | HG2 |
| 6579 | 6583 | 1 ; | HB2 | - | HG3 |
| 6579 | 6584 | 1 ; | HB2 | - | CD |
| 6579 | 6587 | 1 ; | HB2 | - | C |
| 6580 | 6582 | 1 ; | HB3 | - | HG2 |
| 6580 | 6583 | 1 ; | HB3 | - | HG3 |
| 6580 | 6584 | 1 ; | HB3 | - | CD |
| 6580 | 6587 | 1 ; | HB3 | - | C |
| 6581 | 6587 | 1 ; | CG | - | C |
| 6582 | 6585 | 1 ; | HG2 | - | OE1 |
| 6582 | 6586 | 1 ; | HG2 | - | OE2 |
| 6583 | 6585 | 1 ; | HG3 | - | OE1 |
| 6583 | 6586 | 1 ; | HG3 | - | OE2 |
| 6587 | 6592 | 1 ; | C | - | HA |
| 6587 | 6593 | 1 ; | C | - | CB |
| 6587 | 6603 | 1 ; | C | - | C |
| 6588 | 6590 | 1 ; | O | - | H |
| 6588 | 6591 | 1 ; | O | - | CA |
| 6589 | 6594 | 1 ; | N | - | HB2 |
| 6589 | 6595 | 1 ; | N | - | HB3 |
| 6589 | 6596 | 1 ; | N | - | CG |
| 6589 | 6604 | 1 ; | N | - | O |
| 6589 | 6605 | 1 ; | N | - | N |
| 6590 | 6592 | 1 ; | H | - | HA |
| 6590 | 6593 | 1 ; | H | - | CB |
| 6590 | 6603 | 1 ; | H | - | C |
| 6591 | 6597 | 1 ; | CA | - | HG2 |
| 6591 | 6598 | 1 ; | CA | - | HG3 |
| 6591 | 6599 | 1 ; | CA | - | CD |
| 6591 | 6606 | 1 ; | CA | - | H |
| 6591 | 6607 | 1 ; | CA | - | CA |
| 6592 | 6594 | 1 ; | HA | - | HB2 |
| 6592 | 6595 | 1 ; | HA | - | HB3 |
| 6592 | 6596 | 1 ; | HA | - | CG |
| 6592 | 6604 | 1 ; | HA | - | O |
| 6592 | 6605 | 1 ; | HA | - | N |
| 6593 | 6600 | 1 ; | CB | - | OE1 |
| 6593 | 6601 | 1 ; | CB | - | OE2 |
| 6593 | 6604 | 1 ; | CB | - | O |
| 6593 | 6605 | 1 ; | CB | - | N |
| 6594 | 6597 | 1 ; | HB2 | - | HG2 |

| | | | | |
|---|---|---|---|---|
| 6594 | 6598 | 1 | ; | HB2 - HG3 |
| 6594 | 6599 | 1 | ; | HB2 - CD |
| 6594 | 6603 | 1 | ; | HB2 - C |
| 6595 | 6597 | 1 | ; | HB3 - HG2 |
| 6595 | 6598 | 1 | ; | HB3 - HG3 |
| 6595 | 6599 | 1 | ; | HB3 - CD |
| 6595 | 6603 | 1 | ; | HB3 - C |
| 6596 | 6602 | 1 | ; | CG - HE2 |
| 6596 | 6603 | 1 | ; | CG - C |
| 6597 | 6600 | 1 | ; | HG2 - OE1 |
| 6597 | 6601 | 1 | ; | HG2 - OE2 |
| 6598 | 6600 | 1 | ; | HG3 - OE1 |
| 6598 | 6601 | 1 | ; | HG3 - OE2 |
| 6600 | 6602 | 1 | ; | OE1 - HE2 |
| 6603 | 6608 | 1 | ; | C - HA |
| 6603 | 6609 | 1 | ; | C - CB |
| 6603 | 6622 | 1 | ; | C - C |
| 6604 | 6606 | 1 | ; | O - H |
| 6604 | 6607 | 1 | ; | O - CA |
| 6605 | 6610 | 1 | ; | N - HB2 |
| 6605 | 6611 | 1 | ; | N - HB3 |
| 6605 | 6612 | 1 | ; | N - CG |
| 6605 | 6623 | 1 | ; | N - O |
| 6605 | 6624 | 1 | ; | N - N |
| 6606 | 6608 | 1 | ; | H - HA |
| 6606 | 6609 | 1 | ; | H - CB |
| 6606 | 6622 | 1 | ; | H - C |
| 6607 | 6613 | 1 | ; | CA - HG |
| 6607 | 6614 | 1 | ; | CA - CD1 |
| 6607 | 6618 | 1 | ; | CA - CD2 |
| 6607 | 6625 | 1 | ; | CA - H |
| 6607 | 6626 | 1 | ; | CA - CA |
| 6608 | 6610 | 1 | ; | HA - HB2 |
| 6608 | 6611 | 1 | ; | HA - HB3 |
| 6608 | 6612 | 1 | ; | HA - CG |
| 6608 | 6623 | 1 | ; | HA - O |
| 6608 | 6624 | 1 | ; | HA - N |
| 6609 | 6615 | 1 | ; | CB - HD11 |
| 6609 | 6616 | 1 | ; | CB - HD12 |
| 6609 | 6617 | 1 | ; | CB - HD13 |
| 6609 | 6619 | 1 | ; | CB - HD21 |
| 6609 | 6620 | 1 | ; | CB - HD22 |
| 6609 | 6621 | 1 | ; | CB - HD23 |
| 6609 | 6623 | 1 | ; | CB - O |
| 6609 | 6624 | 1 | ; | CB - N |
| 6610 | 6613 | 1 | ; | HB2 - HG |
| 6610 | 6614 | 1 | ; | HB2 - CD1 |
| 6610 | 6618 | 1 | ; | HB2 - CD2 |
| 6610 | 6622 | 1 | ; | HB2 - C |
| 6611 | 6613 | 1 | ; | HB3 - HG |
| 6611 | 6614 | 1 | ; | HB3 - CD1 |
| 6611 | 6618 | 1 | ; | HB3 - CD2 |

| | | | | | |
|---|---|---|---|---|---|
| 6611 | 6622 | 1 ; | HB3 | - | C |
| 6612 | 6622 | 1 ; | CG | - | C |
| 6613 | 6615 | 1 ; | HG | - | HD11 |
| 6613 | 6616 | 1 ; | HG | - | HD12 |
| 6613 | 6617 | 1 ; | HG | - | HD13 |
| 6613 | 6619 | 1 ; | HG | - | HD21 |
| 6613 | 6620 | 1 ; | HG | - | HD22 |
| 6613 | 6621 | 1 ; | HG | - | HD23 |
| 6614 | 6619 | 1 ; | CD1 | - | HD21 |
| 6614 | 6620 | 1 ; | CD1 | - | HD22 |
| 6614 | 6621 | 1 ; | CD1 | - | HD23 |
| 6615 | 6618 | 1 ; | HD11 | - | CD2 |
| 6616 | 6618 | 1 ; | HD12 | - | CD2 |
| 6617 | 6618 | 1 ; | HD13 | - | CD2 |
| 6622 | 6627 | 1 ; | C | - | HA |
| 6622 | 6628 | 1 ; | C | - | CB |
| 6622 | 6644 | 1 ; | C | - | C |
| 6623 | 6625 | 1 ; | O | - | H |
| 6623 | 6626 | 1 ; | O | - | CA |
| 6624 | 6629 | 1 ; | N | - | HB2 |
| 6624 | 6630 | 1 ; | N | - | HB3 |
| 6624 | 6631 | 1 ; | N | - | CG |
| 6624 | 6645 | 1 ; | N | - | O |
| 6624 | 6646 | 1 ; | N | - | N |
| 6625 | 6627 | 1 ; | H | - | HA |
| 6625 | 6628 | 1 ; | H | - | CB |
| 6625 | 6644 | 1 ; | H | - | C |
| 6626 | 6632 | 1 ; | CA | - | HG2 |
| 6626 | 6633 | 1 ; | CA | - | HG3 |
| 6626 | 6634 | 1 ; | CA | - | CD |
| 6626 | 6647 | 1 ; | CA | - | H |
| 6626 | 6648 | 1 ; | CA | - | CA |
| 6627 | 6629 | 1 ; | HA | - | HB2 |
| 6627 | 6630 | 1 ; | HA | - | HB3 |
| 6627 | 6631 | 1 ; | HA | - | CG |
| 6627 | 6645 | 1 ; | HA | - | O |
| 6627 | 6646 | 1 ; | HA | - | N |
| 6628 | 6635 | 1 ; | CB | - | HD2 |
| 6628 | 6636 | 1 ; | CB | - | HD3 |
| 6628 | 6637 | 1 ; | CB | - | CE |
| 6628 | 6645 | 1 ; | CB | - | O |
| 6628 | 6646 | 1 ; | CB | - | N |
| 6629 | 6632 | 1 ; | HB2 | - | HG2 |
| 6629 | 6633 | 1 ; | HB2 | - | HG3 |
| 6629 | 6634 | 1 ; | HB2 | - | CD |
| 6629 | 6644 | 1 ; | HB2 | - | C |
| 6630 | 6632 | 1 ; | HB3 | - | HG2 |
| 6630 | 6633 | 1 ; | HB3 | - | HG3 |
| 6630 | 6634 | 1 ; | HB3 | - | CD |
| 6630 | 6644 | 1 ; | HB3 | - | C |
| 6631 | 6638 | 1 ; | CG | - | HE2 |
| 6631 | 6639 | 1 ; | CG | - | HE3 |

| | | | | | | |
|---|---|---|---|---|---|---|
| 6631 | 6640 | 1 | ; | CG | - | NZ |
| 6631 | 6644 | 1 | ; | CG | - | C |
| 6632 | 6635 | 1 | ; | HG2 | - | HD2 |
| 6632 | 6636 | 1 | ; | HG2 | - | HD3 |
| 6632 | 6637 | 1 | ; | HG2 | - | CE |
| 6633 | 6635 | 1 | ; | HG3 | - | HD2 |
| 6633 | 6636 | 1 | ; | HG3 | - | HD3 |
| 6633 | 6637 | 1 | ; | HG3 | - | CE |
| 6634 | 6641 | 1 | ; | CD | - | HZ1 |
| 6634 | 6642 | 1 | ; | CD | - | HZ2 |
| 6634 | 6643 | 1 | ; | CD | - | HZ3 |
| 6635 | 6638 | 1 | ; | HD2 | - | HE2 |
| 6635 | 6639 | 1 | ; | HD2 | - | HE3 |
| 6635 | 6640 | 1 | ; | HD2 | - | NZ |
| 6636 | 6638 | 1 | ; | HD3 | - | HE2 |
| 6636 | 6639 | 1 | ; | HD3 | - | HE3 |
| 6636 | 6640 | 1 | ; | HD3 | - | NZ |
| 6638 | 6641 | 1 | ; | HE2 | - | HZ1 |
| 6638 | 6642 | 1 | ; | HE2 | - | HZ2 |
| 6638 | 6643 | 1 | ; | HE2 | - | HZ3 |
| 6639 | 6641 | 1 | ; | HE3 | - | HZ1 |
| 6639 | 6642 | 1 | ; | HE3 | - | HZ2 |
| 6639 | 6643 | 1 | ; | HE3 | - | HZ3 |
| 6644 | 6649 | 1 | ; | C | - | HA |
| 6644 | 6650 | 1 | ; | C | - | CB |
| 6644 | 6668 | 1 | ; | C | - | C |
| 6645 | 6647 | 1 | ; | O | - | H |
| 6645 | 6648 | 1 | ; | O | - | CA |
| 6646 | 6651 | 1 | ; | N | - | HB2 |
| 6646 | 6652 | 1 | ; | N | - | HB3 |
| 6646 | 6653 | 1 | ; | N | - | CG |
| 6646 | 6669 | 1 | ; | N | - | O |
| 6646 | 6670 | 1 | ; | N | - | N |
| 6647 | 6649 | 1 | ; | H | - | HA |
| 6647 | 6650 | 1 | ; | H | - | CB |
| 6647 | 6668 | 1 | ; | H | - | C |
| 6648 | 6654 | 1 | ; | CA | - | HG2 |
| 6648 | 6655 | 1 | ; | CA | - | HG3 |
| 6648 | 6656 | 1 | ; | CA | - | CD |
| 6648 | 6671 | 1 | ; | CA | - | H |
| 6648 | 6672 | 1 | ; | CA | - | CA |
| 6649 | 6651 | 1 | ; | HA | - | HB2 |
| 6649 | 6652 | 1 | ; | HA | - | HB3 |
| 6649 | 6653 | 1 | ; | HA | - | CG |
| 6649 | 6669 | 1 | ; | HA | - | O |
| 6649 | 6670 | 1 | ; | HA | - | N |
| 6650 | 6657 | 1 | ; | CB | - | HD2 |
| 6650 | 6658 | 1 | ; | CB | - | HD3 |
| 6650 | 6659 | 1 | ; | CB | - | NE |
| 6650 | 6669 | 1 | ; | CB | - | O |
| 6650 | 6670 | 1 | ; | CB | - | N |
| 6651 | 6654 | 1 | ; | HB2 | - | HG2 |

| | | | | | |
|---|---|---|---|---|---|
| 6651 | 6655 | 1 ; | HB2 | - | HG3 |
| 6651 | 6656 | 1 ; | HB2 | - | CD |
| 6651 | 6668 | 1 ; | HB2 | - | C |
| 6652 | 6654 | 1 ; | HB3 | - | HG2 |
| 6652 | 6655 | 1 ; | HB3 | - | HG3 |
| 6652 | 6656 | 1 ; | HB3 | - | CD |
| 6652 | 6668 | 1 ; | HB3 | - | C |
| 6653 | 6660 | 1 ; | CG | - | HE |
| 6653 | 6661 | 1 ; | CG | - | CZ |
| 6653 | 6668 | 1 ; | CG | - | C |
| 6654 | 6657 | 1 ; | HG2 | - | HD2 |
| 6654 | 6658 | 1 ; | HG2 | - | HD3 |
| 6654 | 6659 | 1 ; | HG2 | - | NE |
| 6655 | 6657 | 1 ; | HG3 | - | HD2 |
| 6655 | 6658 | 1 ; | HG3 | - | HD3 |
| 6655 | 6659 | 1 ; | HG3 | - | NE |
| 6656 | 6662 | 1 ; | CD | - | NH1 |
| 6656 | 6665 | 1 ; | CD | - | NH2 |
| 6657 | 6660 | 1 ; | HD2 | - | HE |
| 6657 | 6661 | 1 ; | HD2 | - | CZ |
| 6658 | 6660 | 1 ; | HD3 | - | HE |
| 6658 | 6661 | 1 ; | HD3 | - | CZ |
| 6659 | 6663 | 1 ; | NE | - | HH11 |
| 6659 | 6664 | 1 ; | NE | - | HH12 |
| 6659 | 6666 | 1 ; | NE | - | HH21 |
| 6659 | 6667 | 1 ; | NE | - | HH22 |
| 6660 | 6662 | 1 ; | HE | - | NH1 |
| 6660 | 6665 | 1 ; | HE | - | NH2 |
| 6662 | 6666 | 1 ; | NH1 | - | HH21 |
| 6662 | 6667 | 1 ; | NH1 | - | HH22 |
| 6663 | 6665 | 1 ; | HH11 | - | NH2 |
| 6664 | 6665 | 1 ; | HH12 | - | NH2 |
| 6668 | 6673 | 1 ; | C | - | HA |
| 6668 | 6674 | 1 ; | C | - | CB |
| 6668 | 6687 | 1 ; | C | - | C |
| 6669 | 6671 | 1 ; | O | - | H |
| 6669 | 6672 | 1 ; | O | - | CA |
| 6670 | 6675 | 1 ; | N | - | HB |
| 6670 | 6676 | 1 ; | N | - | CG2 |
| 6670 | 6680 | 1 ; | N | - | CG1 |
| 6670 | 6688 | 1 ; | N | - | O |
| 6670 | 6689 | 1 ; | N | - | N |
| 6671 | 6673 | 1 ; | H | - | HA |
| 6671 | 6674 | 1 ; | H | - | CB |
| 6671 | 6687 | 1 ; | H | - | C |
| 6672 | 6677 | 1 ; | CA | - | HG21 |
| 6672 | 6678 | 1 ; | CA | - | HG22 |
| 6672 | 6679 | 1 ; | CA | - | HG23 |
| 6672 | 6681 | 1 ; | CA | - | HG12 |
| 6672 | 6682 | 1 ; | CA | - | HG13 |
| 6672 | 6683 | 1 ; | CA | - | CD1 |
| 6672 | 6690 | 1 ; | CA | - | H |

| | | | |
|---|---|---|---|
| 6672 | 6691 | 1 ; | CA - CA |
| 6673 | 6675 | 1 ; | HA - HB |
| 6673 | 6676 | 1 ; | HA - CG2 |
| 6673 | 6680 | 1 ; | HA - CG1 |
| 6673 | 6688 | 1 ; | HA - O |
| 6673 | 6689 | 1 ; | HA - N |
| 6674 | 6684 | 1 ; | CB - HD11 |
| 6674 | 6685 | 1 ; | CB - HD12 |
| 6674 | 6686 | 1 ; | CB - HD13 |
| 6674 | 6688 | 1 ; | CB - O |
| 6674 | 6689 | 1 ; | CB - N |
| 6675 | 6677 | 1 ; | HB - HG21 |
| 6675 | 6678 | 1 ; | HB - HG22 |
| 6675 | 6679 | 1 ; | HB - HG23 |
| 6675 | 6681 | 1 ; | HB - HG12 |
| 6675 | 6682 | 1 ; | HB - HG13 |
| 6675 | 6683 | 1 ; | HB - CD1 |
| 6675 | 6687 | 1 ; | HB - C |
| 6676 | 6681 | 1 ; | CG2 - HG12 |
| 6676 | 6682 | 1 ; | CG2 - HG13 |
| 6676 | 6683 | 1 ; | CG2 - CD1 |
| 6676 | 6687 | 1 ; | CG2 - C |
| 6677 | 6680 | 1 ; | HG21 - CG1 |
| 6678 | 6680 | 1 ; | HG22 - CG1 |
| 6679 | 6680 | 1 ; | HG23 - CG1 |
| 6680 | 6687 | 1 ; | CG1 - C |
| 6681 | 6684 | 1 ; | HG12 - HD11 |
| 6681 | 6685 | 1 ; | HG12 - HD12 |
| 6681 | 6686 | 1 ; | HG12 - HD13 |
| 6682 | 6684 | 1 ; | HG13 - HD11 |
| 6682 | 6685 | 1 ; | HG13 - HD12 |
| 6682 | 6686 | 1 ; | HG13 - HD13 |
| 6687 | 6692 | 1 ; | C - HA |
| 6687 | 6693 | 1 ; | C - CB |
| 6687 | 6707 | 1 ; | C - C |
| 6688 | 6690 | 1 ; | O - H |
| 6688 | 6691 | 1 ; | O - CA |
| 6689 | 6694 | 1 ; | N - HB2 |
| 6689 | 6695 | 1 ; | N - HB3 |
| 6689 | 6696 | 1 ; | N - CG |
| 6689 | 6708 | 1 ; | N - O |
| 6689 | 6709 | 1 ; | N - N |
| 6690 | 6692 | 1 ; | H - HA |
| 6690 | 6693 | 1 ; | H - CB |
| 6690 | 6707 | 1 ; | H - C |
| 6691 | 6697 | 1 ; | CA - CD1 |
| 6691 | 6705 | 1 ; | CA - CD2 |
| 6691 | 6710 | 1 ; | CA - H |
| 6691 | 6711 | 1 ; | CA - CA |
| 6692 | 6694 | 1 ; | HA - HB2 |
| 6692 | 6695 | 1 ; | HA - HB3 |
| 6692 | 6696 | 1 ; | HA - CG |

```
6692   6708       1 ;      HA - O
6692   6709       1 ;      HA - N
6693   6698       1 ;      CB - HD1
6693   6699       1 ;      CB - CE1
6693   6703       1 ;      CB - CE2
6693   6706       1 ;      CB - HD2
6693   6708       1 ;      CB - O
6693   6709       1 ;      CB - N
6694   6697       1 ;     HB2 - CD1
6694   6705       1 ;     HB2 - CD2
6694   6707       1 ;     HB2 - C
6695   6697       1 ;     HB3 - CD1
6695   6705       1 ;     HB3 - CD2
6695   6707       1 ;     HB3 - C
6696   6700       1 ;      CG - HE1
6696   6701       1 ;      CG - CZ
6696   6704       1 ;      CG - HE2
6696   6707       1 ;      CG - C
6697   6702       1 ;     CD1 - HZ
6697   6703       1 ;     CD1 - CE2
6697   6706       1 ;     CD1 - HD2
6698   6700       1 ;     HD1 - HE1
6698   6701       1 ;     HD1 - CZ
6698   6705       1 ;     HD1 - CD2
6699   6704       1 ;     CE1 - HE2
6699   6705       1 ;     CE1 - CD2
6700   6702       1 ;     HE1 - HZ
6700   6703       1 ;     HE1 - CE2
6701   6706       1 ;      CZ - HD2
6702   6704       1 ;      HZ - HE2
6702   6705       1 ;      HZ - CD2
6704   6706       1 ;     HE2 - HD2
6707   6712       1 ;       C - HA2
6707   6713       1 ;       C - HA3
6707   6714       1 ;       C - C
6708   6710       1 ;       O - H
6708   6711       1 ;       O - CA
6709   6715       1 ;       N - O
6709   6716       1 ;       N - N
6710   6712       1 ;       H - HA2
6710   6713       1 ;       H - HA3
6710   6714       1 ;       H - C
6711   6717       1 ;      CA - H
6711   6718       1 ;      CA - CA
6712   6715       1 ;     HA2 - O
6712   6716       1 ;     HA2 - N
6713   6715       1 ;     HA3 - O
6713   6716       1 ;     HA3 - N
6714   6719       1 ;       C - HA
6714   6720       1 ;       C - CB
6714   6738       1 ;       C - C
6715   6717       1 ;       O - H
```

```
6715   6718       1 ;       O - CA
6716   6721       1 ;       N - HB2
6716   6722       1 ;       N - HB3
6716   6723       1 ;       N - CG
6716   6739       1 ;       N - O
6716   6740       1 ;       N - N
6717   6719       1 ;       H - HA
6717   6720       1 ;       H - CB
6717   6738       1 ;       H - C
6718   6724       1 ;      CA - HG2
6718   6725       1 ;      CA - HG3
6718   6726       1 ;      CA - CD
6718   6741       1 ;      CA - H
6718   6742       1 ;      CA - CA
6719   6721       1 ;      HA - HB2
6719   6722       1 ;      HA - HB3
6719   6723       1 ;      HA - CG
6719   6739       1 ;      HA - O
6719   6740       1 ;      HA - N
6720   6727       1 ;      CB - HD2
6720   6728       1 ;      CB - HD3
6720   6729       1 ;      CB - NE
6720   6739       1 ;      CB - O
6720   6740       1 ;      CB - N
6721   6724       1 ;     HB2 - HG2
6721   6725       1 ;     HB2 - HG3
6721   6726       1 ;     HB2 - CD
6721   6738       1 ;     HB2 - C
6722   6724       1 ;     HB3 - HG2
6722   6725       1 ;     HB3 - HG3
6722   6726       1 ;     HB3 - CD
6722   6738       1 ;     HB3 - C
6723   6730       1 ;      CG - HE
6723   6731       1 ;      CG - CZ
6723   6738       1 ;      CG - C
6724   6727       1 ;     HG2 - HD2
6724   6728       1 ;     HG2 - HD3
6724   6729       1 ;     HG2 - NE
6725   6727       1 ;     HG3 - HD2
6725   6728       1 ;     HG3 - HD3
6725   6729       1 ;     HG3 - NE
6726   6732       1 ;      CD - NH1
6726   6735       1 ;      CD - NH2
6727   6730       1 ;     HD2 - HE
6727   6731       1 ;     HD2 - CZ
6728   6730       1 ;     HD3 - HE
6728   6731       1 ;     HD3 - CZ
6729   6733       1 ;      NE - HH11
6729   6734       1 ;      NE - HH12
6729   6736       1 ;      NE - HH21
6729   6737       1 ;      NE - HH22
6730   6732       1 ;      HE - NH1
```

```
6730   6735       1 ;      HE - NH2
6732   6736       1 ;     NH1 - HH21
6732   6737       1 ;     NH1 - HH22
6733   6735       1 ;    HH11 - NH2
6734   6735       1 ;    HH12 - NH2
6738   6743       1 ;       C - HA
6738   6744       1 ;       C - CB
6738   6750       1 ;       C - C
6739   6741       1 ;       O - H
6739   6742       1 ;       O - CA
6740   6745       1 ;       N - HB2
6740   6746       1 ;       N - HB3
6740   6747       1 ;       N - CG
6740   6751       1 ;       N - O
6740   6752       1 ;       N - N
6741   6743       1 ;       H - HA
6741   6744       1 ;       H - CB
6741   6750       1 ;       H - C
6742   6748       1 ;      CA - OD1
6742   6749       1 ;      CA - OD2
6742   6753       1 ;      CA - H
6742   6754       1 ;      CA - CA
6743   6745       1 ;      HA - HB2
6743   6746       1 ;      HA - HB3
6743   6747       1 ;      HA - CG
6743   6751       1 ;      HA - O
6743   6752       1 ;      HA - N
6744   6751       1 ;      CB - O
6744   6752       1 ;      CB - N
6745   6748       1 ;     HB2 - OD1
6745   6749       1 ;     HB2 - OD2
6745   6750       1 ;     HB2 - C
6746   6748       1 ;     HB3 - OD1
6746   6749       1 ;     HB3 - OD2
6746   6750       1 ;     HB3 - C
6747   6750       1 ;      CG - C
6750   6755       1 ;       C - HA
6750   6756       1 ;       C - CB
6750   6762       1 ;       C - C
6751   6753       1 ;       O - H
6751   6754       1 ;       O - CA
6752   6757       1 ;       N - HB2
6752   6758       1 ;       N - HB3
6752   6759       1 ;       N - CG
6752   6763       1 ;       N - O
6752   6764       1 ;       N - N
6753   6755       1 ;       H - HA
6753   6756       1 ;       H - CB
6753   6762       1 ;       H - C
6754   6760       1 ;      CA - OD1
6754   6761       1 ;      CA - OD2
6754   6765       1 ;      CA - H
```

```
6754   6766      1 ;     CA - CA
6755   6757      1 ;     HA - HB2
6755   6758      1 ;     HA - HB3
6755   6759      1 ;     HA - CG
6755   6763      1 ;     HA - O
6755   6764      1 ;     HA - N
6756   6763      1 ;     CB - O
6756   6764      1 ;     CB - N
6757   6760      1 ;    HB2 - OD1
6757   6761      1 ;    HB2 - OD2
6757   6762      1 ;    HB2 - C
6758   6760      1 ;    HB3 - OD1
6758   6761      1 ;    HB3 - OD2
6758   6762      1 ;    HB3 - C
6759   6762      1 ;     CG - C
6762   6767      1 ;      C - HA
6762   6768      1 ;      C - CB
6762   6781      1 ;      C - C
6763   6765      1 ;      O - H
6763   6766      1 ;      O - CA
6764   6769      1 ;      N - HB
6764   6770      1 ;      N - CG2
6764   6774      1 ;      N - CG1
6764   6782      1 ;      N - O
6764   6783      1 ;      N - N
6765   6767      1 ;      H - HA
6765   6768      1 ;      H - CB
6765   6781      1 ;      H - C
6766   6771      1 ;     CA - HG21
6766   6772      1 ;     CA - HG22
6766   6773      1 ;     CA - HG23
6766   6775      1 ;     CA - HG12
6766   6776      1 ;     CA - HG13
6766   6777      1 ;     CA - CD1
6766   6784      1 ;     CA - H
6766   6785      1 ;     CA - CA
6767   6769      1 ;     HA - HB
6767   6770      1 ;     HA - CG2
6767   6774      1 ;     HA - CG1
6767   6782      1 ;     HA - O
6767   6783      1 ;     HA - N
6768   6778      1 ;     CB - HD11
6768   6779      1 ;     CB - HD12
6768   6780      1 ;     CB - HD13
6768   6782      1 ;     CB - O
6768   6783      1 ;     CB - N
6769   6771      1 ;     HB - HG21
6769   6772      1 ;     HB - HG22
6769   6773      1 ;     HB - HG23
6769   6775      1 ;     HB - HG12
6769   6776      1 ;     HB - HG13
6769   6777      1 ;     HB - CD1
```

```
6769   6781       1 ;      HB - C
6770   6775       1 ;     CG2 - HG12
6770   6776       1 ;     CG2 - HG13
6770   6777       1 ;     CG2 - CD1
6770   6781       1 ;     CG2 - C
6771   6774       1 ;    HG21 - CG1
6772   6774       1 ;    HG22 - CG1
6773   6774       1 ;    HG23 - CG1
6774   6781       1 ;     CG1 - C
6775   6778       1 ;    HG12 - HD11
6775   6779       1 ;    HG12 - HD12
6775   6780       1 ;    HG12 - HD13
6776   6778       1 ;    HG13 - HD11
6776   6779       1 ;    HG13 - HD12
6776   6780       1 ;    HG13 - HD13
6781   6786       1 ;       C - HA
6781   6787       1 ;       C - CB
6781   6796       1 ;       C - C
6782   6784       1 ;       O - H
6782   6785       1 ;       O - CA
6783   6788       1 ;       N - HB2
6783   6789       1 ;       N - HB3
6783   6790       1 ;       N - CG
6783   6797       1 ;       N - O
6783   6798       1 ;       N - N
6784   6786       1 ;       H - HA
6784   6787       1 ;       H - CB
6784   6796       1 ;       H - C
6785   6791       1 ;      CA - HG2
6785   6792       1 ;      CA - HG3
6785   6793       1 ;      CA - CD
6785   6799       1 ;      CA - H
6785   6800       1 ;      CA - CA
6786   6788       1 ;      HA - HB2
6786   6789       1 ;      HA - HB3
6786   6790       1 ;      HA - CG
6786   6797       1 ;      HA - O
6786   6798       1 ;      HA - N
6787   6794       1 ;      CB - OE1
6787   6795       1 ;      CB - OE2
6787   6797       1 ;      CB - O
6787   6798       1 ;      CB - N
6788   6791       1 ;     HB2 - HG2
6788   6792       1 ;     HB2 - HG3
6788   6793       1 ;     HB2 - CD
6788   6796       1 ;     HB2 - C
6789   6791       1 ;     HB3 - HG2
6789   6792       1 ;     HB3 - HG3
6789   6793       1 ;     HB3 - CD
6789   6796       1 ;     HB3 - C
6790   6796       1 ;      CG - C
6791   6794       1 ;     HG2 - OE1
```

| | | | | |
|---|---|---|---|---|
| 6791 | 6795 | 1 | ; | HG2 - OE2 |
| 6792 | 6794 | 1 | ; | HG3 - OE1 |
| 6792 | 6795 | 1 | ; | HG3 - OE2 |
| 6796 | 6801 | 1 | ; | C - HA |
| 6796 | 6802 | 1 | ; | C - CB |
| 6796 | 6810 | 1 | ; | C - C |
| 6797 | 6799 | 1 | ; | O - H |
| 6797 | 6800 | 1 | ; | O - CA |
| 6798 | 6803 | 1 | ; | N - HB2 |
| 6798 | 6804 | 1 | ; | N - HB3 |
| 6798 | 6805 | 1 | ; | N - CG |
| 6798 | 6811 | 1 | ; | N - O |
| 6798 | 6812 | 1 | ; | N - N |
| 6799 | 6801 | 1 | ; | H - HA |
| 6799 | 6802 | 1 | ; | H - CB |
| 6799 | 6810 | 1 | ; | H - C |
| 6800 | 6806 | 1 | ; | CA - OD1 |
| 6800 | 6807 | 1 | ; | CA - ND2 |
| 6800 | 6813 | 1 | ; | CA - CD |
| 6800 | 6822 | 1 | ; | CA - CA |
| 6801 | 6803 | 1 | ; | HA - HB2 |
| 6801 | 6804 | 1 | ; | HA - HB3 |
| 6801 | 6805 | 1 | ; | HA - CG |
| 6801 | 6811 | 1 | ; | HA - O |
| 6801 | 6812 | 1 | ; | HA - N |
| 6802 | 6808 | 1 | ; | CB - HD21 |
| 6802 | 6809 | 1 | ; | CB - HD22 |
| 6802 | 6811 | 1 | ; | CB - O |
| 6802 | 6812 | 1 | ; | CB - N |
| 6803 | 6806 | 1 | ; | HB2 - OD1 |
| 6803 | 6807 | 1 | ; | HB2 - ND2 |
| 6803 | 6810 | 1 | ; | HB2 - C |
| 6804 | 6806 | 1 | ; | HB3 - OD1 |
| 6804 | 6807 | 1 | ; | HB3 - ND2 |
| 6804 | 6810 | 1 | ; | HB3 - C |
| 6805 | 6810 | 1 | ; | CG - C |
| 6806 | 6808 | 1 | ; | OD1 - HD21 |
| 6806 | 6809 | 1 | ; | OD1 - HD22 |
| 6810 | 6814 | 1 | ; | C - HD2 |
| 6810 | 6815 | 1 | ; | C - HD3 |
| 6810 | 6816 | 1 | ; | C - CG |
| 6810 | 6819 | 1 | ; | C - CB |
| 6810 | 6823 | 1 | ; | C - HA |
| 6810 | 6824 | 1 | ; | C - C |
| 6811 | 6813 | 1 | ; | O - CD |
| 6811 | 6822 | 1 | ; | O - CA |
| 6812 | 6817 | 1 | ; | N - HG2 |
| 6812 | 6818 | 1 | ; | N - HG3 |
| 6812 | 6820 | 1 | ; | N - HB2 |
| 6812 | 6821 | 1 | ; | N - HB3 |
| 6812 | 6825 | 1 | ; | N - O |
| 6812 | 6826 | 1 | ; | N - N |

| | | | |
|---|---|---|---|
| 6813 | 6820 | 1 ; | CD - HB2 |
| 6813 | 6821 | 1 ; | CD - HB3 |
| 6813 | 6823 | 1 ; | CD - HA |
| 6813 | 6824 | 1 ; | CD - C |
| 6814 | 6817 | 1 ; | HD2 - HG2 |
| 6814 | 6818 | 1 ; | HD2 - HG3 |
| 6814 | 6819 | 1 ; | HD2 - CB |
| 6814 | 6822 | 1 ; | HD2 - CA |
| 6815 | 6817 | 1 ; | HD3 - HG2 |
| 6815 | 6818 | 1 ; | HD3 - HG3 |
| 6815 | 6819 | 1 ; | HD3 - CB |
| 6815 | 6822 | 1 ; | HD3 - CA |
| 6816 | 6823 | 1 ; | CG - HA |
| 6816 | 6824 | 1 ; | CG - C |
| 6817 | 6820 | 1 ; | HG2 - HB2 |
| 6817 | 6821 | 1 ; | HG2 - HB3 |
| 6817 | 6822 | 1 ; | HG2 - CA |
| 6818 | 6820 | 1 ; | HG3 - HB2 |
| 6818 | 6821 | 1 ; | HG3 - HB3 |
| 6818 | 6822 | 1 ; | HG3 - CA |
| 6819 | 6825 | 1 ; | CB - O |
| 6819 | 6826 | 1 ; | CB - N |
| 6820 | 6823 | 1 ; | HB2 - HA |
| 6820 | 6824 | 1 ; | HB2 - C |
| 6821 | 6823 | 1 ; | HB3 - HA |
| 6821 | 6824 | 1 ; | HB3 - C |
| 6822 | 6827 | 1 ; | CA - H |
| 6822 | 6828 | 1 ; | CA - CA |
| 6823 | 6825 | 1 ; | HA - O |
| 6823 | 6826 | 1 ; | HA - N |
| 6824 | 6829 | 1 ; | C - HA |
| 6824 | 6830 | 1 ; | C - CB |
| 6824 | 6843 | 1 ; | C - C |
| 6825 | 6827 | 1 ; | O - H |
| 6825 | 6828 | 1 ; | O - CA |
| 6826 | 6831 | 1 ; | N - HB2 |
| 6826 | 6832 | 1 ; | N - HB3 |
| 6826 | 6833 | 1 ; | N - CG |
| 6826 | 6844 | 1 ; | N - O |
| 6826 | 6845 | 1 ; | N - N |
| 6827 | 6829 | 1 ; | H - HA |
| 6827 | 6830 | 1 ; | H - CB |
| 6827 | 6843 | 1 ; | H - C |
| 6828 | 6834 | 1 ; | CA - HG |
| 6828 | 6835 | 1 ; | CA - CD1 |
| 6828 | 6839 | 1 ; | CA - CD2 |
| 6828 | 6846 | 1 ; | CA - H |
| 6828 | 6847 | 1 ; | CA - CA |
| 6829 | 6831 | 1 ; | HA - HB2 |
| 6829 | 6832 | 1 ; | HA - HB3 |
| 6829 | 6833 | 1 ; | HA - CG |
| 6829 | 6844 | 1 ; | HA - O |

| | | | | | | |
|---|---|---|---|---|---|---|
| 6829 | 6845 | 1 | ; | HA | - | N |
| 6830 | 6836 | 1 | ; | CB | - | HD11 |
| 6830 | 6837 | 1 | ; | CB | - | HD12 |
| 6830 | 6838 | 1 | ; | CB | - | HD13 |
| 6830 | 6840 | 1 | ; | CB | - | HD21 |
| 6830 | 6841 | 1 | ; | CB | - | HD22 |
| 6830 | 6842 | 1 | ; | CB | - | HD23 |
| 6830 | 6844 | 1 | ; | CB | - | O |
| 6830 | 6845 | 1 | ; | CB | - | N |
| 6831 | 6834 | 1 | ; | HB2 | - | HG |
| 6831 | 6835 | 1 | ; | HB2 | - | CD1 |
| 6831 | 6839 | 1 | ; | HB2 | - | CD2 |
| 6831 | 6843 | 1 | ; | HB2 | - | C |
| 6832 | 6834 | 1 | ; | HB3 | - | HG |
| 6832 | 6835 | 1 | ; | HB3 | - | CD1 |
| 6832 | 6839 | 1 | ; | HB3 | - | CD2 |
| 6832 | 6843 | 1 | ; | HB3 | - | C |
| 6833 | 6843 | 1 | ; | CG | - | C |
| 6834 | 6836 | 1 | ; | HG | - | HD11 |
| 6834 | 6837 | 1 | ; | HG | - | HD12 |
| 6834 | 6838 | 1 | ; | HG | - | HD13 |
| 6834 | 6840 | 1 | ; | HG | - | HD21 |
| 6834 | 6841 | 1 | ; | HG | - | HD22 |
| 6834 | 6842 | 1 | ; | HG | - | HD23 |
| 6835 | 6840 | 1 | ; | CD1 | - | HD21 |
| 6835 | 6841 | 1 | ; | CD1 | - | HD22 |
| 6835 | 6842 | 1 | ; | CD1 | - | HD23 |
| 6836 | 6839 | 1 | ; | HD11 | - | CD2 |
| 6837 | 6839 | 1 | ; | HD12 | - | CD2 |
| 6838 | 6839 | 1 | ; | HD13 | - | CD2 |
| 6843 | 6848 | 1 | ; | C | - | HA |
| 6843 | 6849 | 1 | ; | C | - | CB |
| 6843 | 6862 | 1 | ; | C | - | C |
| 6844 | 6846 | 1 | ; | O | - | H |
| 6844 | 6847 | 1 | ; | O | - | CA |
| 6845 | 6850 | 1 | ; | N | - | HB |
| 6845 | 6851 | 1 | ; | N | - | CG2 |
| 6845 | 6855 | 1 | ; | N | - | CG1 |
| 6845 | 6863 | 1 | ; | N | - | O |
| 6845 | 6864 | 1 | ; | N | - | N |
| 6846 | 6848 | 1 | ; | H | - | HA |
| 6846 | 6849 | 1 | ; | H | - | CB |
| 6846 | 6862 | 1 | ; | H | - | C |
| 6847 | 6852 | 1 | ; | CA | - | HG21 |
| 6847 | 6853 | 1 | ; | CA | - | HG22 |
| 6847 | 6854 | 1 | ; | CA | - | HG23 |
| 6847 | 6856 | 1 | ; | CA | - | HG12 |
| 6847 | 6857 | 1 | ; | CA | - | HG13 |
| 6847 | 6858 | 1 | ; | CA | - | CD1 |
| 6847 | 6865 | 1 | ; | CA | - | H |
| 6847 | 6866 | 1 | ; | CA | - | CA |
| 6848 | 6850 | 1 | ; | HA | - | HB |

```
6848   6851      1 ;     HA - CG2
6848   6855      1 ;     HA - CG1
6848   6863      1 ;     HA - O
6848   6864      1 ;     HA - N
6849   6859      1 ;     CB - HD11
6849   6860      1 ;     CB - HD12
6849   6861      1 ;     CB - HD13
6849   6863      1 ;     CB - O
6849   6864      1 ;     CB - N
6850   6852      1 ;     HB - HG21
6850   6853      1 ;     HB - HG22
6850   6854      1 ;     HB - HG23
6850   6856      1 ;     HB - HG12
6850   6857      1 ;     HB - HG13
6850   6858      1 ;     HB - CD1
6850   6862      1 ;     HB - C
6851   6856      1 ;    CG2 - HG12
6851   6857      1 ;    CG2 - HG13
6851   6858      1 ;    CG2 - CD1
6851   6862      1 ;    CG2 - C
6852   6855      1 ;   HG21 - CG1
6853   6855      1 ;   HG22 - CG1
6854   6855      1 ;   HG23 - CG1
6855   6862      1 ;    CG1 - C
6856   6859      1 ;   HG12 - HD11
6856   6860      1 ;   HG12 - HD12
6856   6861      1 ;   HG12 - HD13
6857   6859      1 ;   HG13 - HD11
6857   6860      1 ;   HG13 - HD12
6857   6861      1 ;   HG13 - HD13
6862   6867      1 ;      C - HA
6862   6868      1 ;      C - CB
6862   6874      1 ;      C - C
6863   6865      1 ;      O - H
6863   6866      1 ;      O - CA
6864   6869      1 ;      N - HB2
6864   6870      1 ;      N - HB3
6864   6871      1 ;      N - CG
6864   6875      1 ;      N - O
6864   6876      1 ;      N - N
6865   6867      1 ;      H - HA
6865   6868      1 ;      H - CB
6865   6874      1 ;      H - C
6866   6872      1 ;     CA - OD1
6866   6873      1 ;     CA - OD2
6866   6877      1 ;     CA - H
6866   6878      1 ;     CA - CA
6867   6869      1 ;     HA - HB2
6867   6870      1 ;     HA - HB3
6867   6871      1 ;     HA - CG
6867   6875      1 ;     HA - O
6867   6876      1 ;     HA - N
```

| | | | |
|---|---|---|---|
| 6868 | 6875 | 1 ; | CB - O |
| 6868 | 6876 | 1 ; | CB - N |
| 6869 | 6872 | 1 ; | HB2 - OD1 |
| 6869 | 6873 | 1 ; | HB2 - OD2 |
| 6869 | 6874 | 1 ; | HB2 - C |
| 6870 | 6872 | 1 ; | HB3 - OD1 |
| 6870 | 6873 | 1 ; | HB3 - OD2 |
| 6870 | 6874 | 1 ; | HB3 - C |
| 6871 | 6874 | 1 ; | CG - C |
| 6874 | 6879 | 1 ; | C - HA |
| 6874 | 6880 | 1 ; | C - CB |
| 6874 | 6896 | 1 ; | C - C |
| 6875 | 6877 | 1 ; | O - H |
| 6875 | 6878 | 1 ; | O - CA |
| 6876 | 6881 | 1 ; | N - HB2 |
| 6876 | 6882 | 1 ; | N - HB3 |
| 6876 | 6883 | 1 ; | N - CG |
| 6876 | 6897 | 1 ; | N - O |
| 6876 | 6898 | 1 ; | N - N |
| 6877 | 6879 | 1 ; | H - HA |
| 6877 | 6880 | 1 ; | H - CB |
| 6877 | 6896 | 1 ; | H - C |
| 6878 | 6884 | 1 ; | CA - HG2 |
| 6878 | 6885 | 1 ; | CA - HG3 |
| 6878 | 6886 | 1 ; | CA - CD |
| 6878 | 6899 | 1 ; | CA - H |
| 6878 | 6900 | 1 ; | CA - CA |
| 6879 | 6881 | 1 ; | HA - HB2 |
| 6879 | 6882 | 1 ; | HA - HB3 |
| 6879 | 6883 | 1 ; | HA - CG |
| 6879 | 6897 | 1 ; | HA - O |
| 6879 | 6898 | 1 ; | HA - N |
| 6880 | 6887 | 1 ; | CB - HD2 |
| 6880 | 6888 | 1 ; | CB - HD3 |
| 6880 | 6889 | 1 ; | CB - CE |
| 6880 | 6897 | 1 ; | CB - O |
| 6880 | 6898 | 1 ; | CB - N |
| 6881 | 6884 | 1 ; | HB2 - HG2 |
| 6881 | 6885 | 1 ; | HB2 - HG3 |
| 6881 | 6886 | 1 ; | HB2 - CD |
| 6881 | 6896 | 1 ; | HB2 - C |
| 6882 | 6884 | 1 ; | HB3 - HG2 |
| 6882 | 6885 | 1 ; | HB3 - HG3 |
| 6882 | 6886 | 1 ; | HB3 - CD |
| 6882 | 6896 | 1 ; | HB3 - C |
| 6883 | 6890 | 1 ; | CG - HE2 |
| 6883 | 6891 | 1 ; | CG - HE3 |
| 6883 | 6892 | 1 ; | CG - NZ |
| 6883 | 6896 | 1 ; | CG - C |
| 6884 | 6887 | 1 ; | HG2 - HD2 |
| 6884 | 6888 | 1 ; | HG2 - HD3 |
| 6884 | 6889 | 1 ; | HG2 - CE |

| | | | |
|---|---|---|---|
| 6885 | 6887 | 1 ; | HG3 - HD2 |
| 6885 | 6888 | 1 ; | HG3 - HD3 |
| 6885 | 6889 | 1 ; | HG3 - CE |
| 6886 | 6893 | 1 ; | CD - HZ1 |
| 6886 | 6894 | 1 ; | CD - HZ2 |
| 6886 | 6895 | 1 ; | CD - HZ3 |
| 6887 | 6890 | 1 ; | HD2 - HE2 |
| 6887 | 6891 | 1 ; | HD2 - HE3 |
| 6887 | 6892 | 1 ; | HD2 - NZ |
| 6888 | 6890 | 1 ; | HD3 - HE2 |
| 6888 | 6891 | 1 ; | HD3 - HE3 |
| 6888 | 6892 | 1 ; | HD3 - NZ |
| 6890 | 6893 | 1 ; | HE2 - HZ1 |
| 6890 | 6894 | 1 ; | HE2 - HZ2 |
| 6890 | 6895 | 1 ; | HE2 - HZ3 |
| 6891 | 6893 | 1 ; | HE3 - HZ1 |
| 6891 | 6894 | 1 ; | HE3 - HZ2 |
| 6891 | 6895 | 1 ; | HE3 - HZ3 |
| 6896 | 6901 | 1 ; | C - HA |
| 6896 | 6902 | 1 ; | C - CB |
| 6896 | 6906 | 1 ; | C - C |
| 6897 | 6899 | 1 ; | O - H |
| 6897 | 6900 | 1 ; | O - CA |
| 6898 | 6903 | 1 ; | N - HB1 |
| 6898 | 6904 | 1 ; | N - HB2 |
| 6898 | 6905 | 1 ; | N - HB3 |
| 6898 | 6907 | 1 ; | N - O |
| 6898 | 6908 | 1 ; | N - N |
| 6899 | 6901 | 1 ; | H - HA |
| 6899 | 6902 | 1 ; | H - CB |
| 6899 | 6906 | 1 ; | H - C |
| 6900 | 6909 | 1 ; | CA - H |
| 6900 | 6910 | 1 ; | CA - CA |
| 6901 | 6903 | 1 ; | HA - HB1 |
| 6901 | 6904 | 1 ; | HA - HB2 |
| 6901 | 6905 | 1 ; | HA - HB3 |
| 6901 | 6907 | 1 ; | HA - O |
| 6901 | 6908 | 1 ; | HA - N |
| 6902 | 6907 | 1 ; | CB - O |
| 6902 | 6908 | 1 ; | CB - N |
| 6903 | 6906 | 1 ; | HB1 - C |
| 6904 | 6906 | 1 ; | HB2 - C |
| 6905 | 6906 | 1 ; | HB3 - C |
| 6906 | 6911 | 1 ; | C - HA |
| 6906 | 6912 | 1 ; | C - CB |
| 6906 | 6925 | 1 ; | C - C |
| 6907 | 6909 | 1 ; | O - H |
| 6907 | 6910 | 1 ; | O - CA |
| 6908 | 6913 | 1 ; | N - HB |
| 6908 | 6914 | 1 ; | N - CG2 |
| 6908 | 6918 | 1 ; | N - CG1 |
| 6908 | 6926 | 1 ; | N - O |

| | | | | | |
|---|---|---|---|---|---|
| 6908 | 6927 | 1 ; | N | - | N |
| 6909 | 6911 | 1 ; | H | - | HA |
| 6909 | 6912 | 1 ; | H | - | CB |
| 6909 | 6925 | 1 ; | H | - | C |
| 6910 | 6915 | 1 ; | CA | - | HG21 |
| 6910 | 6916 | 1 ; | CA | - | HG22 |
| 6910 | 6917 | 1 ; | CA | - | HG23 |
| 6910 | 6919 | 1 ; | CA | - | HG12 |
| 6910 | 6920 | 1 ; | CA | - | HG13 |
| 6910 | 6921 | 1 ; | CA | - | CD1 |
| 6910 | 6928 | 1 ; | CA | - | H |
| 6910 | 6929 | 1 ; | CA | - | CA |
| 6911 | 6913 | 1 ; | HA | - | HB |
| 6911 | 6914 | 1 ; | HA | - | CG2 |
| 6911 | 6918 | 1 ; | HA | - | CG1 |
| 6911 | 6926 | 1 ; | HA | - | O |
| 6911 | 6927 | 1 ; | HA | - | N |
| 6912 | 6922 | 1 ; | CB | - | HD11 |
| 6912 | 6923 | 1 ; | CB | - | HD12 |
| 6912 | 6924 | 1 ; | CB | - | HD13 |
| 6912 | 6926 | 1 ; | CB | - | O |
| 6912 | 6927 | 1 ; | CB | - | N |
| 6913 | 6915 | 1 ; | HB | - | HG21 |
| 6913 | 6916 | 1 ; | HB | - | HG22 |
| 6913 | 6917 | 1 ; | HB | - | HG23 |
| 6913 | 6919 | 1 ; | HB | - | HG12 |
| 6913 | 6920 | 1 ; | HB | - | HG13 |
| 6913 | 6921 | 1 ; | HB | - | CD1 |
| 6913 | 6925 | 1 ; | HB | - | C |
| 6914 | 6919 | 1 ; | CG2 | - | HG12 |
| 6914 | 6920 | 1 ; | CG2 | - | HG13 |
| 6914 | 6921 | 1 ; | CG2 | - | CD1 |
| 6914 | 6925 | 1 ; | CG2 | - | C |
| 6915 | 6918 | 1 ; | HG21 | - | CG1 |
| 6916 | 6918 | 1 ; | HG22 | - | CG1 |
| 6917 | 6918 | 1 ; | HG23 | - | CG1 |
| 6918 | 6925 | 1 ; | CG1 | - | C |
| 6919 | 6922 | 1 ; | HG12 | - | HD11 |
| 6919 | 6923 | 1 ; | HG12 | - | HD12 |
| 6919 | 6924 | 1 ; | HG12 | - | HD13 |
| 6920 | 6922 | 1 ; | HG13 | - | HD11 |
| 6920 | 6923 | 1 ; | HG13 | - | HD12 |
| 6920 | 6924 | 1 ; | HG13 | - | HD13 |
| 6925 | 6930 | 1 ; | C | - | HA |
| 6925 | 6931 | 1 ; | C | - | CB |
| 6925 | 6937 | 1 ; | C | - | C |
| 6926 | 6928 | 1 ; | O | - | H |
| 6926 | 6929 | 1 ; | O | - | CA |
| 6927 | 6932 | 1 ; | N | - | HB2 |
| 6927 | 6933 | 1 ; | N | - | HB3 |
| 6927 | 6934 | 1 ; | N | - | CG |
| 6927 | 6938 | 1 ; | N | - | O |

| | | | |
|---|---|---|---|
| 6927 | 6939 | 1 ; | N - N |
| 6928 | 6930 | 1 ; | H - HA |
| 6928 | 6931 | 1 ; | H - CB |
| 6928 | 6937 | 1 ; | H - C |
| 6929 | 6935 | 1 ; | CA - OD1 |
| 6929 | 6936 | 1 ; | CA - OD2 |
| 6929 | 6940 | 1 ; | CA - H |
| 6929 | 6941 | 1 ; | CA - CA |
| 6930 | 6932 | 1 ; | HA - HB2 |
| 6930 | 6933 | 1 ; | HA - HB3 |
| 6930 | 6934 | 1 ; | HA - CG |
| 6930 | 6938 | 1 ; | HA - O |
| 6930 | 6939 | 1 ; | HA - N |
| 6931 | 6938 | 1 ; | CB - O |
| 6931 | 6939 | 1 ; | CB - N |
| 6932 | 6935 | 1 ; | HB2 - OD1 |
| 6932 | 6936 | 1 ; | HB2 - OD2 |
| 6932 | 6937 | 1 ; | HB2 - C |
| 6933 | 6935 | 1 ; | HB3 - OD1 |
| 6933 | 6936 | 1 ; | HB3 - OD2 |
| 6933 | 6937 | 1 ; | HB3 - C |
| 6934 | 6937 | 1 ; | CG - C |
| 6937 | 6942 | 1 ; | C - HA |
| 6937 | 6943 | 1 ; | C - CB |
| 6937 | 6948 | 1 ; | C - C |
| 6938 | 6940 | 1 ; | O - H |
| 6938 | 6941 | 1 ; | O - CA |
| 6939 | 6944 | 1 ; | N - HB2 |
| 6939 | 6945 | 1 ; | N - HB3 |
| 6939 | 6946 | 1 ; | N - OG |
| 6939 | 6949 | 1 ; | N - O |
| 6939 | 6950 | 1 ; | N - N |
| 6940 | 6942 | 1 ; | H - HA |
| 6940 | 6943 | 1 ; | H - CB |
| 6940 | 6948 | 1 ; | H - C |
| 6941 | 6947 | 1 ; | CA - HG |
| 6941 | 6951 | 1 ; | CA - H |
| 6941 | 6952 | 1 ; | CA - CA |
| 6942 | 6944 | 1 ; | HA - HB2 |
| 6942 | 6945 | 1 ; | HA - HB3 |
| 6942 | 6946 | 1 ; | HA - OG |
| 6942 | 6949 | 1 ; | HA - O |
| 6942 | 6950 | 1 ; | HA - N |
| 6943 | 6949 | 1 ; | CB - O |
| 6943 | 6950 | 1 ; | CB - N |
| 6944 | 6947 | 1 ; | HB2 - HG |
| 6944 | 6948 | 1 ; | HB2 - C |
| 6945 | 6947 | 1 ; | HB3 - HG |
| 6945 | 6948 | 1 ; | HB3 - C |
| 6946 | 6948 | 1 ; | OG - C |
| 6948 | 6953 | 1 ; | C - HA |
| 6948 | 6954 | 1 ; | C - CB |

| | | | | | |
|---|---|---|---|---|---|
| 6948 | 6967 | 1 ; | C | - | C |
| 6949 | 6951 | 1 ; | O | - | H |
| 6949 | 6952 | 1 ; | O | - | CA |
| 6950 | 6955 | 1 ; | N | - | HB2 |
| 6950 | 6956 | 1 ; | N | - | HB3 |
| 6950 | 6957 | 1 ; | N | - | CG |
| 6950 | 6968 | 1 ; | N | - | O |
| 6950 | 6969 | 1 ; | N | - | N |
| 6951 | 6953 | 1 ; | H | - | HA |
| 6951 | 6954 | 1 ; | H | - | CB |
| 6951 | 6967 | 1 ; | H | - | C |
| 6952 | 6958 | 1 ; | CA | - | HG |
| 6952 | 6959 | 1 ; | CA | - | CD1 |
| 6952 | 6963 | 1 ; | CA | - | CD2 |
| 6952 | 6970 | 1 ; | CA | - | H |
| 6952 | 6971 | 1 ; | CA | - | CA |
| 6953 | 6955 | 1 ; | HA | - | HB2 |
| 6953 | 6956 | 1 ; | HA | - | HB3 |
| 6953 | 6957 | 1 ; | HA | - | CG |
| 6953 | 6968 | 1 ; | HA | - | O |
| 6953 | 6969 | 1 ; | HA | - | N |
| 6954 | 6960 | 1 ; | CB | - | HD11 |
| 6954 | 6961 | 1 ; | CB | - | HD12 |
| 6954 | 6962 | 1 ; | CB | - | HD13 |
| 6954 | 6964 | 1 ; | CB | - | HD21 |
| 6954 | 6965 | 1 ; | CB | - | HD22 |
| 6954 | 6966 | 1 ; | CB | - | HD23 |
| 6954 | 6968 | 1 ; | CB | - | O |
| 6954 | 6969 | 1 ; | CB | - | N |
| 6955 | 6958 | 1 ; | HB2 | - | HG |
| 6955 | 6959 | 1 ; | HB2 | - | CD1 |
| 6955 | 6963 | 1 ; | HB2 | - | CD2 |
| 6955 | 6967 | 1 ; | HB2 | - | C |
| 6956 | 6958 | 1 ; | HB3 | - | HG |
| 6956 | 6959 | 1 ; | HB3 | - | CD1 |
| 6956 | 6963 | 1 ; | HB3 | - | CD2 |
| 6956 | 6967 | 1 ; | HB3 | - | C |
| 6957 | 6967 | 1 ; | CG | - | C |
| 6958 | 6960 | 1 ; | HG | - | HD11 |
| 6958 | 6961 | 1 ; | HG | - | HD12 |
| 6958 | 6962 | 1 ; | HG | - | HD13 |
| 6958 | 6964 | 1 ; | HG | - | HD21 |
| 6958 | 6965 | 1 ; | HG | - | HD22 |
| 6958 | 6966 | 1 ; | HG | - | HD23 |
| 6959 | 6964 | 1 ; | CD1 | - | HD21 |
| 6959 | 6965 | 1 ; | CD1 | - | HD22 |
| 6959 | 6966 | 1 ; | CD1 | - | HD23 |
| 6960 | 6963 | 1 ; | HD11 | - | CD2 |
| 6961 | 6963 | 1 ; | HD12 | - | CD2 |
| 6962 | 6963 | 1 ; | HD13 | - | CD2 |
| 6967 | 6972 | 1 ; | C | - | HA |
| 6967 | 6973 | 1 ; | C | - | CB |

| | | | | | |
|---|---|---|---|---|---|
| 6967 | 6986 | 1 ; | C | - | C |
| 6968 | 6970 | 1 ; | O | - | H |
| 6968 | 6971 | 1 ; | O | - | CA |
| 6969 | 6974 | 1 ; | N | - | HB2 |
| 6969 | 6975 | 1 ; | N | - | HB3 |
| 6969 | 6976 | 1 ; | N | - | CG |
| 6969 | 6987 | 1 ; | N | - | O |
| 6969 | 6988 | 1 ; | N | - | N |
| 6970 | 6972 | 1 ; | H | - | HA |
| 6970 | 6973 | 1 ; | H | - | CB |
| 6970 | 6986 | 1 ; | H | - | C |
| 6971 | 6977 | 1 ; | CA | - | HG |
| 6971 | 6978 | 1 ; | CA | - | CD1 |
| 6971 | 6982 | 1 ; | CA | - | CD2 |
| 6971 | 6989 | 1 ; | CA | - | H |
| 6971 | 6990 | 1 ; | CA | - | CA |
| 6972 | 6974 | 1 ; | HA | - | HB2 |
| 6972 | 6975 | 1 ; | HA | - | HB3 |
| 6972 | 6976 | 1 ; | HA | - | CG |
| 6972 | 6987 | 1 ; | HA | - | O |
| 6972 | 6988 | 1 ; | HA | - | N |
| 6973 | 6979 | 1 ; | CB | - | HD11 |
| 6973 | 6980 | 1 ; | CB | - | HD12 |
| 6973 | 6981 | 1 ; | CB | - | HD13 |
| 6973 | 6983 | 1 ; | CB | - | HD21 |
| 6973 | 6984 | 1 ; | CB | - | HD22 |
| 6973 | 6985 | 1 ; | CB | - | HD23 |
| 6973 | 6987 | 1 ; | CB | - | O |
| 6973 | 6988 | 1 ; | CB | - | N |
| 6974 | 6977 | 1 ; | HB2 | - | HG |
| 6974 | 6978 | 1 ; | HB2 | - | CD1 |
| 6974 | 6982 | 1 ; | HB2 | - | CD2 |
| 6974 | 6986 | 1 ; | HB2 | - | C |
| 6975 | 6977 | 1 ; | HB3 | - | HG |
| 6975 | 6978 | 1 ; | HB3 | - | CD1 |
| 6975 | 6982 | 1 ; | HB3 | - | CD2 |
| 6975 | 6986 | 1 ; | HB3 | - | C |
| 6976 | 6986 | 1 ; | CG | - | C |
| 6977 | 6979 | 1 ; | HG | - | HD11 |
| 6977 | 6980 | 1 ; | HG | - | HD12 |
| 6977 | 6981 | 1 ; | HG | - | HD13 |
| 6977 | 6983 | 1 ; | HG | - | HD21 |
| 6977 | 6984 | 1 ; | HG | - | HD22 |
| 6977 | 6985 | 1 ; | HG | - | HD23 |
| 6978 | 6983 | 1 ; | CD1 | - | HD21 |
| 6978 | 6984 | 1 ; | CD1 | - | HD22 |
| 6978 | 6985 | 1 ; | CD1 | - | HD23 |
| 6979 | 6982 | 1 ; | HD11 | - | CD2 |
| 6980 | 6982 | 1 ; | HD12 | - | CD2 |
| 6981 | 6982 | 1 ; | HD13 | - | CD2 |
| 6986 | 6991 | 1 ; | C | - | HA |
| 6986 | 6992 | 1 ; | C | - | CB |

| | | | | | |
|---|---|---|---|---|---|
| 6986 | 7003 | 1 ; | C | - | C |
| 6987 | 6989 | 1 ; | O | - | H |
| 6987 | 6990 | 1 ; | O | - | CA |
| 6988 | 6993 | 1 ; | N | - | HB2 |
| 6988 | 6994 | 1 ; | N | - | HB3 |
| 6988 | 6995 | 1 ; | N | - | CG |
| 6988 | 7004 | 1 ; | N | - | O |
| 6988 | 7005 | 1 ; | N | - | N |
| 6989 | 6991 | 1 ; | H | - | HA |
| 6989 | 6992 | 1 ; | H | - | CB |
| 6989 | 7003 | 1 ; | H | - | C |
| 6990 | 6996 | 1 ; | CA | - | HG2 |
| 6990 | 6997 | 1 ; | CA | - | HG3 |
| 6990 | 6998 | 1 ; | CA | - | CD |
| 6990 | 7006 | 1 ; | CA | - | H |
| 6990 | 7007 | 1 ; | CA | - | CA |
| 6991 | 6993 | 1 ; | HA | - | HB2 |
| 6991 | 6994 | 1 ; | HA | - | HB3 |
| 6991 | 6995 | 1 ; | HA | - | CG |
| 6991 | 7004 | 1 ; | HA | - | O |
| 6991 | 7005 | 1 ; | HA | - | N |
| 6992 | 6999 | 1 ; | CB | - | OE1 |
| 6992 | 7000 | 1 ; | CB | - | NE2 |
| 6992 | 7004 | 1 ; | CB | - | O |
| 6992 | 7005 | 1 ; | CB | - | N |
| 6993 | 6996 | 1 ; | HB2 | - | HG2 |
| 6993 | 6997 | 1 ; | HB2 | - | HG3 |
| 6993 | 6998 | 1 ; | HB2 | - | CD |
| 6993 | 7003 | 1 ; | HB2 | - | C |
| 6994 | 6996 | 1 ; | HB3 | - | HG2 |
| 6994 | 6997 | 1 ; | HB3 | - | HG3 |
| 6994 | 6998 | 1 ; | HB3 | - | CD |
| 6994 | 7003 | 1 ; | HB3 | - | C |
| 6995 | 7001 | 1 ; | CG | - | HE21 |
| 6995 | 7002 | 1 ; | CG | - | HE22 |
| 6995 | 7003 | 1 ; | CG | - | C |
| 6996 | 6999 | 1 ; | HG2 | - | OE1 |
| 6996 | 7000 | 1 ; | HG2 | - | NE2 |
| 6997 | 6999 | 1 ; | HG3 | - | OE1 |
| 6997 | 7000 | 1 ; | HG3 | - | NE2 |
| 6999 | 7001 | 1 ; | OE1 | - | HE21 |
| 6999 | 7002 | 1 ; | OE1 | - | HE22 |
| 7003 | 7008 | 1 ; | C | - | HA |
| 7003 | 7009 | 1 ; | C | - | CB |
| 7003 | 7018 | 1 ; | C | - | C |
| 7004 | 7006 | 1 ; | O | - | H |
| 7004 | 7007 | 1 ; | O | - | CA |
| 7005 | 7010 | 1 ; | N | - | HB2 |
| 7005 | 7011 | 1 ; | N | - | HB3 |
| 7005 | 7012 | 1 ; | N | - | CG |
| 7005 | 7019 | 1 ; | N | - | O |
| 7005 | 7020 | 1 ; | N | - | N |

```
7006   7008       1 ;       H - HA
7006   7009       1 ;       H - CB
7006   7018       1 ;       H - C
7007   7013       1 ;      CA - HG2
7007   7014       1 ;      CA - HG3
7007   7015       1 ;      CA - CD
7007   7021       1 ;      CA - H
7007   7022       1 ;      CA - CA
7008   7010       1 ;      HA - HB2
7008   7011       1 ;      HA - HB3
7008   7012       1 ;      HA - CG
7008   7019       1 ;      HA - O
7008   7020       1 ;      HA - N
7009   7016       1 ;      CB - OE1
7009   7017       1 ;      CB - OE2
7009   7019       1 ;      CB - O
7009   7020       1 ;      CB - N
7010   7013       1 ;     HB2 - HG2
7010   7014       1 ;     HB2 - HG3
7010   7015       1 ;     HB2 - CD
7010   7018       1 ;     HB2 - C
7011   7013       1 ;     HB3 - HG2
7011   7014       1 ;     HB3 - HG3
7011   7015       1 ;     HB3 - CD
7011   7018       1 ;     HB3 - C
7012   7018       1 ;      CG - C
7013   7016       1 ;     HG2 - OE1
7013   7017       1 ;     HG2 - OE2
7014   7016       1 ;     HG3 - OE1
7014   7017       1 ;     HG3 - OE2
7018   7023       1 ;       C - HA
7018   7024       1 ;       C - CB
7018   7034       1 ;       C - C
7019   7021       1 ;       O - H
7019   7022       1 ;       O - CA
7020   7025       1 ;       N - HB
7020   7026       1 ;       N - CG1
7020   7030       1 ;       N - CG2
7020   7035       1 ;       N - O
7020   7036       1 ;       N - N
7021   7023       1 ;       H - HA
7021   7024       1 ;       H - CB
7021   7034       1 ;       H - C
7022   7027       1 ;      CA - HG11
7022   7028       1 ;      CA - HG12
7022   7029       1 ;      CA - HG13
7022   7031       1 ;      CA - HG21
7022   7032       1 ;      CA - HG22
7022   7033       1 ;      CA - HG23
7022   7037       1 ;      CA - H
7022   7038       1 ;      CA - CA
7023   7025       1 ;      HA - HB
```

| | | | | | | |
|---|---|---|---|---|---|---|
| 7023 | 7026 | 1 | ; | HA | - | CG1 |
| 7023 | 7030 | 1 | ; | HA | - | CG2 |
| 7023 | 7035 | 1 | ; | HA | - | O |
| 7023 | 7036 | 1 | ; | HA | - | N |
| 7024 | 7035 | 1 | ; | CB | - | O |
| 7024 | 7036 | 1 | ; | CB | - | N |
| 7025 | 7027 | 1 | ; | HB | - | HG11 |
| 7025 | 7028 | 1 | ; | HB | - | HG12 |
| 7025 | 7029 | 1 | ; | HB | - | HG13 |
| 7025 | 7031 | 1 | ; | HB | - | HG21 |
| 7025 | 7032 | 1 | ; | HB | - | HG22 |
| 7025 | 7033 | 1 | ; | HB | - | HG23 |
| 7025 | 7034 | 1 | ; | HB | - | C |
| 7026 | 7031 | 1 | ; | CG1 | - | HG21 |
| 7026 | 7032 | 1 | ; | CG1 | - | HG22 |
| 7026 | 7033 | 1 | ; | CG1 | - | HG23 |
| 7026 | 7034 | 1 | ; | CG1 | - | C |
| 7027 | 7030 | 1 | ; | HG11 | - | CG2 |
| 7028 | 7030 | 1 | ; | HG12 | - | CG2 |
| 7029 | 7030 | 1 | ; | HG13 | - | CG2 |
| 7030 | 7034 | 1 | ; | CG2 | - | C |
| 7034 | 7039 | 1 | ; | C | - | HA |
| 7034 | 7040 | 1 | ; | C | - | CB |
| 7034 | 7046 | 1 | ; | C | - | C |
| 7035 | 7037 | 1 | ; | O | - | H |
| 7035 | 7038 | 1 | ; | O | - | CA |
| 7036 | 7041 | 1 | ; | N | - | HB2 |
| 7036 | 7042 | 1 | ; | N | - | HB3 |
| 7036 | 7043 | 1 | ; | N | - | CG |
| 7036 | 7047 | 1 | ; | N | - | O |
| 7036 | 7048 | 1 | ; | N | - | N |
| 7037 | 7039 | 1 | ; | H | - | HA |
| 7037 | 7040 | 1 | ; | H | - | CB |
| 7037 | 7046 | 1 | ; | H | - | C |
| 7038 | 7044 | 1 | ; | CA | - | OD1 |
| 7038 | 7045 | 1 | ; | CA | - | OD2 |
| 7038 | 7049 | 1 | ; | CA | - | H |
| 7038 | 7050 | 1 | ; | CA | - | CA |
| 7039 | 7041 | 1 | ; | HA | - | HB2 |
| 7039 | 7042 | 1 | ; | HA | - | HB3 |
| 7039 | 7043 | 1 | ; | HA | - | CG |
| 7039 | 7047 | 1 | ; | HA | - | O |
| 7039 | 7048 | 1 | ; | HA | - | N |
| 7040 | 7047 | 1 | ; | CB | - | O |
| 7040 | 7048 | 1 | ; | CB | - | N |
| 7041 | 7044 | 1 | ; | HB2 | - | OD1 |
| 7041 | 7045 | 1 | ; | HB2 | - | OD2 |
| 7041 | 7046 | 1 | ; | HB2 | - | C |
| 7042 | 7044 | 1 | ; | HB3 | - | OD1 |
| 7042 | 7045 | 1 | ; | HB3 | - | OD2 |
| 7042 | 7046 | 1 | ; | HB3 | - | C |
| 7043 | 7046 | 1 | ; | CG | - | C |

| | | | |
|---|---|---|---|
| 7046 | 7051 | 1 ; | C - HA |
| 7046 | 7052 | 1 ; | C - CB |
| 7046 | 7065 | 1 ; | C - C |
| 7047 | 7049 | 1 ; | O - H |
| 7047 | 7050 | 1 ; | O - CA |
| 7048 | 7053 | 1 ; | N - HB2 |
| 7048 | 7054 | 1 ; | N - HB3 |
| 7048 | 7055 | 1 ; | N - CG |
| 7048 | 7066 | 1 ; | N - O |
| 7048 | 7067 | 1 ; | N - N |
| 7049 | 7051 | 1 ; | H - HA |
| 7049 | 7052 | 1 ; | H - CB |
| 7049 | 7065 | 1 ; | H - C |
| 7050 | 7056 | 1 ; | CA - HG |
| 7050 | 7057 | 1 ; | CA - CD1 |
| 7050 | 7061 | 1 ; | CA - CD2 |
| 7050 | 7068 | 1 ; | CA - H |
| 7050 | 7069 | 1 ; | CA - CA |
| 7051 | 7053 | 1 ; | HA - HB2 |
| 7051 | 7054 | 1 ; | HA - HB3 |
| 7051 | 7055 | 1 ; | HA - CG |
| 7051 | 7066 | 1 ; | HA - O |
| 7051 | 7067 | 1 ; | HA - N |
| 7052 | 7058 | 1 ; | CB - HD11 |
| 7052 | 7059 | 1 ; | CB - HD12 |
| 7052 | 7060 | 1 ; | CB - HD13 |
| 7052 | 7062 | 1 ; | CB - HD21 |
| 7052 | 7063 | 1 ; | CB - HD22 |
| 7052 | 7064 | 1 ; | CB - HD23 |
| 7052 | 7066 | 1 ; | CB - O |
| 7052 | 7067 | 1 ; | CB - N |
| 7053 | 7056 | 1 ; | HB2 - HG |
| 7053 | 7057 | 1 ; | HB2 - CD1 |
| 7053 | 7061 | 1 ; | HB2 - CD2 |
| 7053 | 7065 | 1 ; | HB2 - C |
| 7054 | 7056 | 1 ; | HB3 - HG |
| 7054 | 7057 | 1 ; | HB3 - CD1 |
| 7054 | 7061 | 1 ; | HB3 - CD2 |
| 7054 | 7065 | 1 ; | HB3 - C |
| 7055 | 7065 | 1 ; | CG - C |
| 7056 | 7058 | 1 ; | HG - HD11 |
| 7056 | 7059 | 1 ; | HG - HD12 |
| 7056 | 7060 | 1 ; | HG - HD13 |
| 7056 | 7062 | 1 ; | HG - HD21 |
| 7056 | 7063 | 1 ; | HG - HD22 |
| 7056 | 7064 | 1 ; | HG - HD23 |
| 7057 | 7062 | 1 ; | CD1 - HD21 |
| 7057 | 7063 | 1 ; | CD1 - HD22 |
| 7057 | 7064 | 1 ; | CD1 - HD23 |
| 7058 | 7061 | 1 ; | HD11 - CD2 |
| 7059 | 7061 | 1 ; | HD12 - CD2 |
| 7060 | 7061 | 1 ; | HD13 - CD2 |

```
7065   7070       1 ;      C - HA
7065   7071       1 ;      C - CB
7065   7079       1 ;      C - C
7066   7068       1 ;      O - H
7066   7069       1 ;      O - CA
7067   7072       1 ;      N - HB2
7067   7073       1 ;      N - HB3
7067   7074       1 ;      N - CG
7067   7080       1 ;      N - O
7067   7081       1 ;      N - N
7068   7070       1 ;      H - HA
7068   7071       1 ;      H - CB
7068   7079       1 ;      H - C
7069   7075       1 ;     CA - OD1
7069   7076       1 ;     CA - ND2
7069   7082       1 ;     CA - H
7069   7083       1 ;     CA - CA
7070   7072       1 ;     HA - HB2
7070   7073       1 ;     HA - HB3
7070   7074       1 ;     HA - CG
7070   7080       1 ;     HA - O
7070   7081       1 ;     HA - N
7071   7077       1 ;     CB - HD21
7071   7078       1 ;     CB - HD22
7071   7080       1 ;     CB - O
7071   7081       1 ;     CB - N
7072   7075       1 ;    HB2 - OD1
7072   7076       1 ;    HB2 - ND2
7072   7079       1 ;    HB2 - C
7073   7075       1 ;    HB3 - OD1
7073   7076       1 ;    HB3 - ND2
7073   7079       1 ;    HB3 - C
7074   7079       1 ;     CG - C
7075   7077       1 ;    OD1 - HD21
7075   7078       1 ;    OD1 - HD22
7079   7084       1 ;      C - HA2
7079   7085       1 ;      C - HA3
7079   7086       1 ;      C - C
7080   7082       1 ;      O - H
7080   7083       1 ;      O - CA
7081   7087       1 ;      N - O
7081   7088       1 ;      N - N
7082   7084       1 ;      H - HA2
7082   7085       1 ;      H - HA3
7082   7086       1 ;      H - C
7083   7089       1 ;     CA - H
7083   7090       1 ;     CA - CA
7084   7087       1 ;    HA2 - O
7084   7088       1 ;    HA2 - N
7085   7087       1 ;    HA3 - O
7085   7088       1 ;    HA3 - N
7086   7091       1 ;      C - HA
```

| | | | | | |
|---|---|---|---|---|---|
| 7086 | 7092 | 1 ; | C | - | CB |
| 7086 | 7099 | 1 ; | C | - | C |
| 7087 | 7089 | 1 ; | O | - | H |
| 7087 | 7090 | 1 ; | O | - | CA |
| 7088 | 7093 | 1 ; | N | - | HB2 |
| 7088 | 7094 | 1 ; | N | - | HB3 |
| 7088 | 7095 | 1 ; | N | - | CG |
| 7088 | 7100 | 1 ; | N | - | O |
| 7088 | 7101 | 1 ; | N | - | N |
| 7089 | 7091 | 1 ; | H | - | HA |
| 7089 | 7092 | 1 ; | H | - | CB |
| 7089 | 7099 | 1 ; | H | - | C |
| 7090 | 7096 | 1 ; | CA | - | OD1 |
| 7090 | 7097 | 1 ; | CA | - | OD2 |
| 7090 | 7102 | 1 ; | CA | - | H |
| 7090 | 7103 | 1 ; | CA | - | CA |
| 7091 | 7093 | 1 ; | HA | - | HB2 |
| 7091 | 7094 | 1 ; | HA | - | HB3 |
| 7091 | 7095 | 1 ; | HA | - | CG |
| 7091 | 7100 | 1 ; | HA | - | O |
| 7091 | 7101 | 1 ; | HA | - | N |
| 7092 | 7098 | 1 ; | CB | - | HD2 |
| 7092 | 7100 | 1 ; | CB | - | O |
| 7092 | 7101 | 1 ; | CB | - | N |
| 7093 | 7096 | 1 ; | HB2 | - | OD1 |
| 7093 | 7097 | 1 ; | HB2 | - | OD2 |
| 7093 | 7099 | 1 ; | HB2 | - | C |
| 7094 | 7096 | 1 ; | HB3 | - | OD1 |
| 7094 | 7097 | 1 ; | HB3 | - | OD2 |
| 7094 | 7099 | 1 ; | HB3 | - | C |
| 7095 | 7099 | 1 ; | CG | - | C |
| 7096 | 7098 | 1 ; | OD1 | - | HD2 |
| 7099 | 7104 | 1 ; | C | - | HA2 |
| 7099 | 7105 | 1 ; | C | - | HA3 |
| 7099 | 7106 | 1 ; | C | - | C |
| 7100 | 7102 | 1 ; | O | - | H |
| 7100 | 7103 | 1 ; | O | - | CA |
| 7101 | 7107 | 1 ; | N | - | O |
| 7101 | 7108 | 1 ; | N | - | N |
| 7102 | 7104 | 1 ; | H | - | HA2 |
| 7102 | 7105 | 1 ; | H | - | HA3 |
| 7102 | 7106 | 1 ; | H | - | C |
| 7103 | 7109 | 1 ; | CA | - | H |
| 7103 | 7110 | 1 ; | CA | - | CA |
| 7104 | 7107 | 1 ; | HA2 | - | O |
| 7104 | 7108 | 1 ; | HA2 | - | N |
| 7105 | 7107 | 1 ; | HA3 | - | O |
| 7105 | 7108 | 1 ; | HA3 | - | N |
| 7106 | 7111 | 1 ; | C | - | HA |
| 7106 | 7112 | 1 ; | C | - | CB |
| 7106 | 7121 | 1 ; | C | - | C |
| 7107 | 7109 | 1 ; | O | - | H |

| | | | | |
|---|---|---|---|---|
| 7107 | 7110 | 1 ; | O - | CA |
| 7108 | 7113 | 1 ; | N - | HB2 |
| 7108 | 7114 | 1 ; | N - | HB3 |
| 7108 | 7115 | 1 ; | N - | CG |
| 7108 | 7122 | 1 ; | N - | O |
| 7108 | 7123 | 1 ; | N - | N |
| 7109 | 7111 | 1 ; | H - | HA |
| 7109 | 7112 | 1 ; | H - | CB |
| 7109 | 7121 | 1 ; | H - | C |
| 7110 | 7116 | 1 ; | CA - | HG2 |
| 7110 | 7117 | 1 ; | CA - | HG3 |
| 7110 | 7118 | 1 ; | CA - | CD |
| 7110 | 7124 | 1 ; | CA - | H |
| 7110 | 7125 | 1 ; | CA - | CA |
| 7111 | 7113 | 1 ; | HA - | HB2 |
| 7111 | 7114 | 1 ; | HA - | HB3 |
| 7111 | 7115 | 1 ; | HA - | CG |
| 7111 | 7122 | 1 ; | HA - | O |
| 7111 | 7123 | 1 ; | HA - | N |
| 7112 | 7119 | 1 ; | CB - | OE1 |
| 7112 | 7120 | 1 ; | CB - | OE2 |
| 7112 | 7122 | 1 ; | CB - | O |
| 7112 | 7123 | 1 ; | CB - | N |
| 7113 | 7116 | 1 ; | HB2 - | HG2 |
| 7113 | 7117 | 1 ; | HB2 - | HG3 |
| 7113 | 7118 | 1 ; | HB2 - | CD |
| 7113 | 7121 | 1 ; | HB2 - | C |
| 7114 | 7116 | 1 ; | HB3 - | HG2 |
| 7114 | 7117 | 1 ; | HB3 - | HG3 |
| 7114 | 7118 | 1 ; | HB3 - | CD |
| 7114 | 7121 | 1 ; | HB3 - | C |
| 7115 | 7121 | 1 ; | CG - | C |
| 7116 | 7119 | 1 ; | HG2 - | OE1 |
| 7116 | 7120 | 1 ; | HG2 - | OE2 |
| 7117 | 7119 | 1 ; | HG3 - | OE1 |
| 7117 | 7120 | 1 ; | HG3 - | OE2 |
| 7121 | 7126 | 1 ; | C - | HA |
| 7121 | 7127 | 1 ; | C - | CB |
| 7121 | 7140 | 1 ; | C - | C |
| 7122 | 7124 | 1 ; | O - | H |
| 7122 | 7125 | 1 ; | O - | CA |
| 7123 | 7128 | 1 ; | N - | HB |
| 7123 | 7129 | 1 ; | N - | CG2 |
| 7123 | 7133 | 1 ; | N - | CG1 |
| 7123 | 7141 | 1 ; | N - | O |
| 7123 | 7142 | 1 ; | N - | N |
| 7124 | 7126 | 1 ; | H - | HA |
| 7124 | 7127 | 1 ; | H - | CB |
| 7124 | 7140 | 1 ; | H - | C |
| 7125 | 7130 | 1 ; | CA - | HG21 |
| 7125 | 7131 | 1 ; | CA - | HG22 |
| 7125 | 7132 | 1 ; | CA - | HG23 |

| | | | | | |
|---|---|---|---|---|---|
| 7125 | 7134 | 1 ; | CA | - | HG12 |
| 7125 | 7135 | 1 ; | CA | - | HG13 |
| 7125 | 7136 | 1 ; | CA | - | CD1 |
| 7125 | 7143 | 1 ; | CA | - | H |
| 7125 | 7144 | 1 ; | CA | - | CA |
| 7126 | 7128 | 1 ; | HA | - | HB |
| 7126 | 7129 | 1 ; | HA | - | CG2 |
| 7126 | 7133 | 1 ; | HA | - | CG1 |
| 7126 | 7141 | 1 ; | HA | - | O |
| 7126 | 7142 | 1 ; | HA | - | N |
| 7127 | 7137 | 1 ; | CB | - | HD11 |
| 7127 | 7138 | 1 ; | CB | - | HD12 |
| 7127 | 7139 | 1 ; | CB | - | HD13 |
| 7127 | 7141 | 1 ; | CB | - | O |
| 7127 | 7142 | 1 ; | CB | - | N |
| 7128 | 7130 | 1 ; | HB | - | HG21 |
| 7128 | 7131 | 1 ; | HB | - | HG22 |
| 7128 | 7132 | 1 ; | HB | - | HG23 |
| 7128 | 7134 | 1 ; | HB | - | HG12 |
| 7128 | 7135 | 1 ; | HB | - | HG13 |
| 7128 | 7136 | 1 ; | HB | - | CD1 |
| 7128 | 7140 | 1 ; | HB | - | C |
| 7129 | 7134 | 1 ; | CG2 | - | HG12 |
| 7129 | 7135 | 1 ; | CG2 | - | HG13 |
| 7129 | 7136 | 1 ; | CG2 | - | CD1 |
| 7129 | 7140 | 1 ; | CG2 | - | C |
| 7130 | 7133 | 1 ; | HG21 | - | CG1 |
| 7131 | 7133 | 1 ; | HG22 | - | CG1 |
| 7132 | 7133 | 1 ; | HG23 | - | CG1 |
| 7133 | 7140 | 1 ; | CG1 | - | C |
| 7134 | 7137 | 1 ; | HG12 | - | HD11 |
| 7134 | 7138 | 1 ; | HG12 | - | HD12 |
| 7134 | 7139 | 1 ; | HG12 | - | HD13 |
| 7135 | 7137 | 1 ; | HG13 | - | HD11 |
| 7135 | 7138 | 1 ; | HG13 | - | HD12 |
| 7135 | 7139 | 1 ; | HG13 | - | HD13 |
| 7140 | 7145 | 1 ; | C | - | HA |
| 7140 | 7146 | 1 ; | C | - | CB |
| 7140 | 7152 | 1 ; | C | - | C |
| 7141 | 7143 | 1 ; | O | - | H |
| 7141 | 7144 | 1 ; | O | - | CA |
| 7142 | 7147 | 1 ; | N | - | HB2 |
| 7142 | 7148 | 1 ; | N | - | HB3 |
| 7142 | 7149 | 1 ; | N | - | CG |
| 7142 | 7153 | 1 ; | N | - | O |
| 7142 | 7154 | 1 ; | N | - | N |
| 7143 | 7145 | 1 ; | H | - | HA |
| 7143 | 7146 | 1 ; | H | - | CB |
| 7143 | 7152 | 1 ; | H | - | C |
| 7144 | 7150 | 1 ; | CA | - | OD1 |
| 7144 | 7151 | 1 ; | CA | - | OD2 |
| 7144 | 7155 | 1 ; | CA | - | H |

| | | | | | |
|---|---|---|---|---|---|
| 7144 | 7156 | 1 ; | CA | - | CA |
| 7145 | 7147 | 1 ; | HA | - | HB2 |
| 7145 | 7148 | 1 ; | HA | - | HB3 |
| 7145 | 7149 | 1 ; | HA | - | CG |
| 7145 | 7153 | 1 ; | HA | - | O |
| 7145 | 7154 | 1 ; | HA | - | N |
| 7146 | 7153 | 1 ; | CB | - | O |
| 7146 | 7154 | 1 ; | CB | - | N |
| 7147 | 7150 | 1 ; | HB2 | - | OD1 |
| 7147 | 7151 | 1 ; | HB2 | - | OD2 |
| 7147 | 7152 | 1 ; | HB2 | - | C |
| 7148 | 7150 | 1 ; | HB3 | - | OD1 |
| 7148 | 7151 | 1 ; | HB3 | - | OD2 |
| 7148 | 7152 | 1 ; | HB3 | - | C |
| 7149 | 7152 | 1 ; | CG | - | C |
| 7152 | 7157 | 1 ; | C | - | HA |
| 7152 | 7158 | 1 ; | C | - | CB |
| 7152 | 7172 | 1 ; | C | - | C |
| 7153 | 7155 | 1 ; | O | - | H |
| 7153 | 7156 | 1 ; | O | - | CA |
| 7154 | 7159 | 1 ; | N | - | HB2 |
| 7154 | 7160 | 1 ; | N | - | HB3 |
| 7154 | 7161 | 1 ; | N | - | CG |
| 7154 | 7173 | 1 ; | N | - | O |
| 7154 | 7174 | 1 ; | N | - | N |
| 7155 | 7157 | 1 ; | H | - | HA |
| 7155 | 7158 | 1 ; | H | - | CB |
| 7155 | 7172 | 1 ; | H | - | C |
| 7156 | 7162 | 1 ; | CA | - | CD1 |
| 7156 | 7170 | 1 ; | CA | - | CD2 |
| 7156 | 7175 | 1 ; | CA | - | H |
| 7156 | 7176 | 1 ; | CA | - | CA |
| 7157 | 7159 | 1 ; | HA | - | HB2 |
| 7157 | 7160 | 1 ; | HA | - | HB3 |
| 7157 | 7161 | 1 ; | HA | - | CG |
| 7157 | 7173 | 1 ; | HA | - | O |
| 7157 | 7174 | 1 ; | HA | - | N |
| 7158 | 7163 | 1 ; | CB | - | HD1 |
| 7158 | 7164 | 1 ; | CB | - | CE1 |
| 7158 | 7168 | 1 ; | CB | - | CE2 |
| 7158 | 7171 | 1 ; | CB | - | HD2 |
| 7158 | 7173 | 1 ; | CB | - | O |
| 7158 | 7174 | 1 ; | CB | - | N |
| 7159 | 7162 | 1 ; | HB2 | - | CD1 |
| 7159 | 7170 | 1 ; | HB2 | - | CD2 |
| 7159 | 7172 | 1 ; | HB2 | - | C |
| 7160 | 7162 | 1 ; | HB3 | - | CD1 |
| 7160 | 7170 | 1 ; | HB3 | - | CD2 |
| 7160 | 7172 | 1 ; | HB3 | - | C |
| 7161 | 7165 | 1 ; | CG | - | HE1 |
| 7161 | 7166 | 1 ; | CG | - | CZ |
| 7161 | 7169 | 1 ; | CG | - | HE2 |

| | | | | | |
|---|---|---|---|---|---|
| 7161 | 7172 | 1 ; | CG | - | C |
| 7162 | 7167 | 1 ; | CD1 | - | HZ |
| 7162 | 7168 | 1 ; | CD1 | - | CE2 |
| 7162 | 7171 | 1 ; | CD1 | - | HD2 |
| 7163 | 7165 | 1 ; | HD1 | - | HE1 |
| 7163 | 7166 | 1 ; | HD1 | - | CZ |
| 7163 | 7170 | 1 ; | HD1 | - | CD2 |
| 7164 | 7169 | 1 ; | CE1 | - | HE2 |
| 7164 | 7170 | 1 ; | CE1 | - | CD2 |
| 7165 | 7167 | 1 ; | HE1 | - | HZ |
| 7165 | 7168 | 1 ; | HE1 | - | CE2 |
| 7166 | 7171 | 1 ; | CZ | - | HD2 |
| 7167 | 7169 | 1 ; | HZ | - | HE2 |
| 7167 | 7170 | 1 ; | HZ | - | CD2 |
| 7169 | 7171 | 1 ; | HE2 | - | HD2 |
| 7172 | 7177 | 1 ; | C | - | HA |
| 7172 | 7178 | 1 ; | C | - | CB |
| 7172 | 7189 | 1 ; | C | - | C |
| 7173 | 7175 | 1 ; | O | - | H |
| 7173 | 7176 | 1 ; | O | - | CA |
| 7174 | 7179 | 1 ; | N | - | HB2 |
| 7174 | 7180 | 1 ; | N | - | HB3 |
| 7174 | 7181 | 1 ; | N | - | CG |
| 7174 | 7190 | 1 ; | N | - | O |
| 7174 | 7191 | 1 ; | N | - | N |
| 7175 | 7177 | 1 ; | H | - | HA |
| 7175 | 7178 | 1 ; | H | - | CB |
| 7175 | 7189 | 1 ; | H | - | C |
| 7176 | 7182 | 1 ; | CA | - | ND1 |
| 7176 | 7187 | 1 ; | CA | - | CD2 |
| 7176 | 7192 | 1 ; | CA | - | H |
| 7176 | 7193 | 1 ; | CA | - | CA |
| 7177 | 7179 | 1 ; | HA | - | HB2 |
| 7177 | 7180 | 1 ; | HA | - | HB3 |
| 7177 | 7181 | 1 ; | HA | - | CG |
| 7177 | 7190 | 1 ; | HA | - | O |
| 7177 | 7191 | 1 ; | HA | - | N |
| 7178 | 7183 | 1 ; | CB | - | CE1 |
| 7178 | 7185 | 1 ; | CB | - | NE2 |
| 7178 | 7188 | 1 ; | CB | - | HD2 |
| 7178 | 7190 | 1 ; | CB | - | O |
| 7178 | 7191 | 1 ; | CB | - | N |
| 7179 | 7182 | 1 ; | HB2 | - | ND1 |
| 7179 | 7187 | 1 ; | HB2 | - | CD2 |
| 7179 | 7189 | 1 ; | HB2 | - | C |
| 7180 | 7182 | 1 ; | HB3 | - | ND1 |
| 7180 | 7187 | 1 ; | HB3 | - | CD2 |
| 7180 | 7189 | 1 ; | HB3 | - | C |
| 7181 | 7184 | 1 ; | CG | - | HE1 |
| 7181 | 7186 | 1 ; | CG | - | HE2 |
| 7181 | 7189 | 1 ; | CG | - | C |
| 7182 | 7186 | 1 ; | ND1 | - | HE2 |

| | | | | | | |
|---|---|---|---|---|---|---|
| 7182 | 7188 | 1 | ; | ND1 | - | HD2 |
| 7183 | 7188 | 1 | ; | CE1 | - | HD2 |
| 7184 | 7186 | 1 | ; | HE1 | - | HE2 |
| 7184 | 7187 | 1 | ; | HE1 | - | CD2 |
| 7186 | 7188 | 1 | ; | HE2 | - | HD2 |
| 7189 | 7194 | 1 | ; | C | - | HA |
| 7189 | 7195 | 1 | ; | C | - | CB |
| 7189 | 7205 | 1 | ; | C | - | C |
| 7190 | 7192 | 1 | ; | O | - | H |
| 7190 | 7193 | 1 | ; | O | - | CA |
| 7191 | 7196 | 1 | ; | N | - | HB2 |
| 7191 | 7197 | 1 | ; | N | - | HB3 |
| 7191 | 7198 | 1 | ; | N | - | CG |
| 7191 | 7206 | 1 | ; | N | - | O |
| 7191 | 7207 | 1 | ; | N | - | N |
| 7192 | 7194 | 1 | ; | H | - | HA |
| 7192 | 7195 | 1 | ; | H | - | CB |
| 7192 | 7205 | 1 | ; | H | - | C |
| 7193 | 7199 | 1 | ; | CA | - | HG2 |
| 7193 | 7200 | 1 | ; | CA | - | HG3 |
| 7193 | 7201 | 1 | ; | CA | - | CD |
| 7193 | 7208 | 1 | ; | CA | - | H |
| 7193 | 7209 | 1 | ; | CA | - | CA |
| 7194 | 7196 | 1 | ; | HA | - | HB2 |
| 7194 | 7197 | 1 | ; | HA | - | HB3 |
| 7194 | 7198 | 1 | ; | HA | - | CG |
| 7194 | 7206 | 1 | ; | HA | - | O |
| 7194 | 7207 | 1 | ; | HA | - | N |
| 7195 | 7202 | 1 | ; | CB | - | OE1 |
| 7195 | 7203 | 1 | ; | CB | - | OE2 |
| 7195 | 7206 | 1 | ; | CB | - | O |
| 7195 | 7207 | 1 | ; | CB | - | N |
| 7196 | 7199 | 1 | ; | HB2 | - | HG2 |
| 7196 | 7200 | 1 | ; | HB2 | - | HG3 |
| 7196 | 7201 | 1 | ; | HB2 | - | CD |
| 7196 | 7205 | 1 | ; | HB2 | - | C |
| 7197 | 7199 | 1 | ; | HB3 | - | HG2 |
| 7197 | 7200 | 1 | ; | HB3 | - | HG3 |
| 7197 | 7201 | 1 | ; | HB3 | - | CD |
| 7197 | 7205 | 1 | ; | HB3 | - | C |
| 7198 | 7204 | 1 | ; | CG | - | HE2 |
| 7198 | 7205 | 1 | ; | CG | - | C |
| 7199 | 7202 | 1 | ; | HG2 | - | OE1 |
| 7199 | 7203 | 1 | ; | HG2 | - | OE2 |
| 7200 | 7202 | 1 | ; | HG3 | - | OE1 |
| 7200 | 7203 | 1 | ; | HG3 | - | OE2 |
| 7202 | 7204 | 1 | ; | OE1 | - | HE2 |
| 7205 | 7210 | 1 | ; | C | - | HA |
| 7205 | 7211 | 1 | ; | C | - | CB |
| 7205 | 7225 | 1 | ; | C | - | C |
| 7206 | 7208 | 1 | ; | O | - | H |
| 7206 | 7209 | 1 | ; | O | - | CA |

| | | | |
|---|---|---|---|
| 7207 | 7212 | 1 ; | N - HB2 |
| 7207 | 7213 | 1 ; | N - HB3 |
| 7207 | 7214 | 1 ; | N - CG |
| 7207 | 7226 | 1 ; | N - O |
| 7207 | 7227 | 1 ; | N - N |
| 7208 | 7210 | 1 ; | H - HA |
| 7208 | 7211 | 1 ; | H - CB |
| 7208 | 7225 | 1 ; | H - C |
| 7209 | 7215 | 1 ; | CA - CD1 |
| 7209 | 7223 | 1 ; | CA - CD2 |
| 7209 | 7228 | 1 ; | CA - H |
| 7209 | 7229 | 1 ; | CA - CA |
| 7210 | 7212 | 1 ; | HA - HB2 |
| 7210 | 7213 | 1 ; | HA - HB3 |
| 7210 | 7214 | 1 ; | HA - CG |
| 7210 | 7226 | 1 ; | HA - O |
| 7210 | 7227 | 1 ; | HA - N |
| 7211 | 7216 | 1 ; | CB - HD1 |
| 7211 | 7217 | 1 ; | CB - CE1 |
| 7211 | 7221 | 1 ; | CB - CE2 |
| 7211 | 7224 | 1 ; | CB - HD2 |
| 7211 | 7226 | 1 ; | CB - O |
| 7211 | 7227 | 1 ; | CB - N |
| 7212 | 7215 | 1 ; | HB2 - CD1 |
| 7212 | 7223 | 1 ; | HB2 - CD2 |
| 7212 | 7225 | 1 ; | HB2 - C |
| 7213 | 7215 | 1 ; | HB3 - CD1 |
| 7213 | 7223 | 1 ; | HB3 - CD2 |
| 7213 | 7225 | 1 ; | HB3 - C |
| 7214 | 7218 | 1 ; | CG - HE1 |
| 7214 | 7219 | 1 ; | CG - CZ |
| 7214 | 7222 | 1 ; | CG - HE2 |
| 7214 | 7225 | 1 ; | CG - C |
| 7215 | 7220 | 1 ; | CD1 - HZ |
| 7215 | 7221 | 1 ; | CD1 - CE2 |
| 7215 | 7224 | 1 ; | CD1 - HD2 |
| 7216 | 7218 | 1 ; | HD1 - HE1 |
| 7216 | 7219 | 1 ; | HD1 - CZ |
| 7216 | 7223 | 1 ; | HD1 - CD2 |
| 7217 | 7222 | 1 ; | CE1 - HE2 |
| 7217 | 7223 | 1 ; | CE1 - CD2 |
| 7218 | 7220 | 1 ; | HE1 - HZ |
| 7218 | 7221 | 1 ; | HE1 - CE2 |
| 7219 | 7224 | 1 ; | CZ - HD2 |
| 7220 | 7222 | 1 ; | HZ - HE2 |
| 7220 | 7223 | 1 ; | HZ - CD2 |
| 7222 | 7224 | 1 ; | HE2 - HD2 |
| 7225 | 7230 | 1 ; | C - HA |
| 7225 | 7231 | 1 ; | C - CB |
| 7225 | 7242 | 1 ; | C - C |
| 7226 | 7228 | 1 ; | O - H |
| 7226 | 7229 | 1 ; | O - CA |

```
7227   7232       1 ;       N - HB2
7227   7233       1 ;       N - HB3
7227   7234       1 ;       N - CG
7227   7243       1 ;       N - O
7227   7244       1 ;       N - N
7228   7230       1 ;       H - HA
7228   7231       1 ;       H - CB
7228   7242       1 ;       H - C
7229   7235       1 ;      CA - HG2
7229   7236       1 ;      CA - HG3
7229   7237       1 ;      CA - SD
7229   7245       1 ;      CA - H
7229   7246       1 ;      CA - CA
7230   7232       1 ;      HA - HB2
7230   7233       1 ;      HA - HB3
7230   7234       1 ;      HA - CG
7230   7243       1 ;      HA - O
7230   7244       1 ;      HA - N
7231   7238       1 ;      CB - CE
7231   7243       1 ;      CB - O
7231   7244       1 ;      CB - N
7232   7235       1 ;     HB2 - HG2
7232   7236       1 ;     HB2 - HG3
7232   7237       1 ;     HB2 - SD
7232   7242       1 ;     HB2 - C
7233   7235       1 ;     HB3 - HG2
7233   7236       1 ;     HB3 - HG3
7233   7237       1 ;     HB3 - SD
7233   7242       1 ;     HB3 - C
7234   7239       1 ;      CG - HE1
7234   7240       1 ;      CG - HE2
7234   7241       1 ;      CG - HE3
7234   7242       1 ;      CG - C
7235   7238       1 ;     HG2 - CE
7236   7238       1 ;     HG3 - CE
7242   7247       1 ;       C - HA
7242   7248       1 ;       C - CB
7242   7261       1 ;       C - C
7243   7245       1 ;       O - H
7243   7246       1 ;       O - CA
7244   7249       1 ;       N - HB2
7244   7250       1 ;       N - HB3
7244   7251       1 ;       N - CG
7244   7262       1 ;       N - O
7244   7263       1 ;       N - N
7245   7247       1 ;       H - HA
7245   7248       1 ;       H - CB
7245   7261       1 ;       H - C
7246   7252       1 ;      CA - HG
7246   7253       1 ;      CA - CD1
7246   7257       1 ;      CA - CD2
7246   7264       1 ;      CA - H
```

| | | | | | |
|---|---|---|---|---|---|
| 7246 | 7265 | 1 ; | CA | - | CA |
| 7247 | 7249 | 1 ; | HA | - | HB2 |
| 7247 | 7250 | 1 ; | HA | - | HB3 |
| 7247 | 7251 | 1 ; | HA | - | CG |
| 7247 | 7262 | 1 ; | HA | - | O |
| 7247 | 7263 | 1 ; | HA | - | N |
| 7248 | 7254 | 1 ; | CB | - | HD11 |
| 7248 | 7255 | 1 ; | CB | - | HD12 |
| 7248 | 7256 | 1 ; | CB | - | HD13 |
| 7248 | 7258 | 1 ; | CB | - | HD21 |
| 7248 | 7259 | 1 ; | CB | - | HD22 |
| 7248 | 7260 | 1 ; | CB | - | HD23 |
| 7248 | 7262 | 1 ; | CB | - | O |
| 7248 | 7263 | 1 ; | CB | - | N |
| 7249 | 7252 | 1 ; | HB2 | - | HG |
| 7249 | 7253 | 1 ; | HB2 | - | CD1 |
| 7249 | 7257 | 1 ; | HB2 | - | CD2 |
| 7249 | 7261 | 1 ; | HB2 | - | C |
| 7250 | 7252 | 1 ; | HB3 | - | HG |
| 7250 | 7253 | 1 ; | HB3 | - | CD1 |
| 7250 | 7257 | 1 ; | HB3 | - | CD2 |
| 7250 | 7261 | 1 ; | HB3 | - | C |
| 7251 | 7261 | 1 ; | CG | - | C |
| 7252 | 7254 | 1 ; | HG | - | HD11 |
| 7252 | 7255 | 1 ; | HG | - | HD12 |
| 7252 | 7256 | 1 ; | HG | - | HD13 |
| 7252 | 7258 | 1 ; | HG | - | HD21 |
| 7252 | 7259 | 1 ; | HG | - | HD22 |
| 7252 | 7260 | 1 ; | HG | - | HD23 |
| 7253 | 7258 | 1 ; | CD1 | - | HD21 |
| 7253 | 7259 | 1 ; | CD1 | - | HD22 |
| 7253 | 7260 | 1 ; | CD1 | - | HD23 |
| 7254 | 7257 | 1 ; | HD11 | - | CD2 |
| 7255 | 7257 | 1 ; | HD12 | - | CD2 |
| 7256 | 7257 | 1 ; | HD13 | - | CD2 |
| 7261 | 7266 | 1 ; | C | - | HA |
| 7261 | 7267 | 1 ; | C | - | CB |
| 7261 | 7278 | 1 ; | C | - | C |
| 7262 | 7264 | 1 ; | O | - | H |
| 7262 | 7265 | 1 ; | O | - | CA |
| 7263 | 7268 | 1 ; | N | - | HB2 |
| 7263 | 7269 | 1 ; | N | - | HB3 |
| 7263 | 7270 | 1 ; | N | - | CG |
| 7263 | 7279 | 1 ; | N | - | O |
| 7263 | 7280 | 1 ; | N | - | N |
| 7264 | 7266 | 1 ; | H | - | HA |
| 7264 | 7267 | 1 ; | H | - | CB |
| 7264 | 7278 | 1 ; | H | - | C |
| 7265 | 7271 | 1 ; | CA | - | HG2 |
| 7265 | 7272 | 1 ; | CA | - | HG3 |
| 7265 | 7273 | 1 ; | CA | - | SD |
| 7265 | 7281 | 1 ; | CA | - | H |

| | | | | | |
|---|---|---|---|---|---|
| 7265 | 7282 | 1 ; | CA | - | CA |
| 7266 | 7268 | 1 ; | HA | - | HB2 |
| 7266 | 7269 | 1 ; | HA | - | HB3 |
| 7266 | 7270 | 1 ; | HA | - | CG |
| 7266 | 7279 | 1 ; | HA | - | O |
| 7266 | 7280 | 1 ; | HA | - | N |
| 7267 | 7274 | 1 ; | CB | - | CE |
| 7267 | 7279 | 1 ; | CB | - | O |
| 7267 | 7280 | 1 ; | CB | - | N |
| 7268 | 7271 | 1 ; | HB2 | - | HG2 |
| 7268 | 7272 | 1 ; | HB2 | - | HG3 |
| 7268 | 7273 | 1 ; | HB2 | - | SD |
| 7268 | 7278 | 1 ; | HB2 | - | C |
| 7269 | 7271 | 1 ; | HB3 | - | HG2 |
| 7269 | 7272 | 1 ; | HB3 | - | HG3 |
| 7269 | 7273 | 1 ; | HB3 | - | SD |
| 7269 | 7278 | 1 ; | HB3 | - | C |
| 7270 | 7275 | 1 ; | CG | - | HE1 |
| 7270 | 7276 | 1 ; | CG | - | HE2 |
| 7270 | 7277 | 1 ; | CG | - | HE3 |
| 7270 | 7278 | 1 ; | CG | - | C |
| 7271 | 7274 | 1 ; | HG2 | - | CE |
| 7272 | 7274 | 1 ; | HG3 | - | CE |
| 7278 | 7283 | 1 ; | C | - | HA |
| 7278 | 7284 | 1 ; | C | - | CB |
| 7278 | 7295 | 1 ; | C | - | C |
| 7279 | 7281 | 1 ; | O | - | H |
| 7279 | 7282 | 1 ; | O | - | CA |
| 7280 | 7285 | 1 ; | N | - | HB2 |
| 7280 | 7286 | 1 ; | N | - | HB3 |
| 7280 | 7287 | 1 ; | N | - | CG |
| 7280 | 7296 | 1 ; | N | - | O |
| 7280 | 7297 | 1 ; | N | - | N |
| 7281 | 7283 | 1 ; | H | - | HA |
| 7281 | 7284 | 1 ; | H | - | CB |
| 7281 | 7295 | 1 ; | H | - | C |
| 7282 | 7288 | 1 ; | CA | - | HG2 |
| 7282 | 7289 | 1 ; | CA | - | HG3 |
| 7282 | 7290 | 1 ; | CA | - | SD |
| 7282 | 7298 | 1 ; | CA | - | H |
| 7282 | 7299 | 1 ; | CA | - | CA |
| 7283 | 7285 | 1 ; | HA | - | HB2 |
| 7283 | 7286 | 1 ; | HA | - | HB3 |
| 7283 | 7287 | 1 ; | HA | - | CG |
| 7283 | 7296 | 1 ; | HA | - | O |
| 7283 | 7297 | 1 ; | HA | - | N |
| 7284 | 7291 | 1 ; | CB | - | CE |
| 7284 | 7296 | 1 ; | CB | - | O |
| 7284 | 7297 | 1 ; | CB | - | N |
| 7285 | 7288 | 1 ; | HB2 | - | HG2 |
| 7285 | 7289 | 1 ; | HB2 | - | HG3 |
| 7285 | 7290 | 1 ; | HB2 | - | SD |

| | | | | | |
|---|---|---|---|---|---|
| 7285 | 7295 | 1 ; | HB2 | - | C |
| 7286 | 7288 | 1 ; | HB3 | - | HG2 |
| 7286 | 7289 | 1 ; | HB3 | - | HG3 |
| 7286 | 7290 | 1 ; | HB3 | - | SD |
| 7286 | 7295 | 1 ; | HB3 | - | C |
| 7287 | 7292 | 1 ; | CG | - | HE1 |
| 7287 | 7293 | 1 ; | CG | - | HE2 |
| 7287 | 7294 | 1 ; | CG | - | HE3 |
| 7287 | 7295 | 1 ; | CG | - | C |
| 7288 | 7291 | 1 ; | HG2 | - | CE |
| 7289 | 7291 | 1 ; | HG3 | - | CE |
| 7295 | 7300 | 1 ; | C | - | HA |
| 7295 | 7301 | 1 ; | C | - | CB |
| 7295 | 7306 | 1 ; | C | - | C |
| 7296 | 7298 | 1 ; | O | - | H |
| 7296 | 7299 | 1 ; | O | - | CA |
| 7297 | 7302 | 1 ; | N | - | HB2 |
| 7297 | 7303 | 1 ; | N | - | HB3 |
| 7297 | 7304 | 1 ; | N | - | OG |
| 7297 | 7307 | 1 ; | N | - | O |
| 7297 | 7308 | 1 ; | N | - | N |
| 7298 | 7300 | 1 ; | H | - | HA |
| 7298 | 7301 | 1 ; | H | - | CB |
| 7298 | 7306 | 1 ; | H | - | C |
| 7299 | 7305 | 1 ; | CA | - | HG |
| 7299 | 7309 | 1 ; | CA | - | H |
| 7299 | 7310 | 1 ; | CA | - | CA |
| 7300 | 7302 | 1 ; | HA | - | HB2 |
| 7300 | 7303 | 1 ; | HA | - | HB3 |
| 7300 | 7304 | 1 ; | HA | - | OG |
| 7300 | 7307 | 1 ; | HA | - | O |
| 7300 | 7308 | 1 ; | HA | - | N |
| 7301 | 7307 | 1 ; | CB | - | O |
| 7301 | 7308 | 1 ; | CB | - | N |
| 7302 | 7305 | 1 ; | HB2 | - | HG |
| 7302 | 7306 | 1 ; | HB2 | - | C |
| 7303 | 7305 | 1 ; | HB3 | - | HG |
| 7303 | 7306 | 1 ; | HB3 | - | C |
| 7304 | 7306 | 1 ; | OG | - | C |
| 7306 | 7311 | 1 ; | C | - | HA |
| 7306 | 7312 | 1 ; | C | - | CB |
| 7306 | 7328 | 1 ; | C | - | C |
| 7307 | 7309 | 1 ; | O | - | H |
| 7307 | 7310 | 1 ; | O | - | CA |
| 7308 | 7313 | 1 ; | N | - | HB2 |
| 7308 | 7314 | 1 ; | N | - | HB3 |
| 7308 | 7315 | 1 ; | N | - | CG |
| 7308 | 7329 | 1 ; | N | - | O |
| 7308 | 7330 | 1 ; | N | - | N |
| 7309 | 7311 | 1 ; | H | - | HA |
| 7309 | 7312 | 1 ; | H | - | CB |
| 7309 | 7328 | 1 ; | H | - | C |

```
7310   7316       1 ;      CA - HG2
7310   7317       1 ;      CA - HG3
7310   7318       1 ;      CA - CD
7310   7331       1 ;      CA - H
7310   7332       1 ;      CA - CA
7311   7313       1 ;      HA - HB2
7311   7314       1 ;      HA - HB3
7311   7315       1 ;      HA - CG
7311   7329       1 ;      HA - O
7311   7330       1 ;      HA - N
7312   7319       1 ;      CB - HD2
7312   7320       1 ;      CB - HD3
7312   7321       1 ;      CB - CE
7312   7329       1 ;      CB - O
7312   7330       1 ;      CB - N
7313   7316       1 ;     HB2 - HG2
7313   7317       1 ;     HB2 - HG3
7313   7318       1 ;     HB2 - CD
7313   7328       1 ;     HB2 - C
7314   7316       1 ;     HB3 - HG2
7314   7317       1 ;     HB3 - HG3
7314   7318       1 ;     HB3 - CD
7314   7328       1 ;     HB3 - C
7315   7322       1 ;      CG - HE2
7315   7323       1 ;      CG - HE3
7315   7324       1 ;      CG - NZ
7315   7328       1 ;      CG - C
7316   7319       1 ;     HG2 - HD2
7316   7320       1 ;     HG2 - HD3
7316   7321       1 ;     HG2 - CE
7317   7319       1 ;     HG3 - HD2
7317   7320       1 ;     HG3 - HD3
7317   7321       1 ;     HG3 - CE
7318   7325       1 ;      CD - HZ1
7318   7326       1 ;      CD - HZ2
7318   7327       1 ;      CD - HZ3
7319   7322       1 ;     HD2 - HE2
7319   7323       1 ;     HD2 - HE3
7319   7324       1 ;     HD2 - NZ
7320   7322       1 ;     HD3 - HE2
7320   7323       1 ;     HD3 - HE3
7320   7324       1 ;     HD3 - NZ
7322   7325       1 ;     HE2 - HZ1
7322   7326       1 ;     HE2 - HZ2
7322   7327       1 ;     HE2 - HZ3
7323   7325       1 ;     HE3 - HZ1
7323   7326       1 ;     HE3 - HZ2
7323   7327       1 ;     HE3 - HZ3
7328   7333       1 ;       C - HA
7328   7334       1 ;       C - CB
7328   7350       1 ;       C - C
7329   7331       1 ;       O - H
```

| | | | | |
|---|---|---|---|---|
| 7329 | 7332 | 1 ; | O - | CA |
| 7330 | 7335 | 1 ; | N - | HB2 |
| 7330 | 7336 | 1 ; | N - | HB3 |
| 7330 | 7337 | 1 ; | N - | CG |
| 7330 | 7351 | 1 ; | N - | O |
| 7330 | 7352 | 1 ; | N - | N |
| 7331 | 7333 | 1 ; | H - | HA |
| 7331 | 7334 | 1 ; | H - | CB |
| 7331 | 7350 | 1 ; | H - | C |
| 7332 | 7338 | 1 ; | CA - | HG2 |
| 7332 | 7339 | 1 ; | CA - | HG3 |
| 7332 | 7340 | 1 ; | CA - | CD |
| 7332 | 7353 | 1 ; | CA - | H |
| 7332 | 7354 | 1 ; | CA - | CA |
| 7333 | 7335 | 1 ; | HA - | HB2 |
| 7333 | 7336 | 1 ; | HA - | HB3 |
| 7333 | 7337 | 1 ; | HA - | CG |
| 7333 | 7351 | 1 ; | HA - | O |
| 7333 | 7352 | 1 ; | HA - | N |
| 7334 | 7341 | 1 ; | CB - | HD2 |
| 7334 | 7342 | 1 ; | CB - | HD3 |
| 7334 | 7343 | 1 ; | CB - | CE |
| 7334 | 7351 | 1 ; | CB - | O |
| 7334 | 7352 | 1 ; | CB - | N |
| 7335 | 7338 | 1 ; | HB2 - | HG2 |
| 7335 | 7339 | 1 ; | HB2 - | HG3 |
| 7335 | 7340 | 1 ; | HB2 - | CD |
| 7335 | 7350 | 1 ; | HB2 - | C |
| 7336 | 7338 | 1 ; | HB3 - | HG2 |
| 7336 | 7339 | 1 ; | HB3 - | HG3 |
| 7336 | 7340 | 1 ; | HB3 - | CD |
| 7336 | 7350 | 1 ; | HB3 - | C |
| 7337 | 7344 | 1 ; | CG - | HE2 |
| 7337 | 7345 | 1 ; | CG - | HE3 |
| 7337 | 7346 | 1 ; | CG - | NZ |
| 7337 | 7350 | 1 ; | CG - | C |
| 7338 | 7341 | 1 ; | HG2 - | HD2 |
| 7338 | 7342 | 1 ; | HG2 - | HD3 |
| 7338 | 7343 | 1 ; | HG2 - | CE |
| 7339 | 7341 | 1 ; | HG3 - | HD2 |
| 7339 | 7342 | 1 ; | HG3 - | HD3 |
| 7339 | 7343 | 1 ; | HG3 - | CE |
| 7340 | 7347 | 1 ; | CD - | HZ1 |
| 7340 | 7348 | 1 ; | CD - | HZ2 |
| 7340 | 7349 | 1 ; | CD - | HZ3 |
| 7341 | 7344 | 1 ; | HD2 - | HE2 |
| 7341 | 7345 | 1 ; | HD2 - | HE3 |
| 7341 | 7346 | 1 ; | HD2 - | NZ |
| 7342 | 7344 | 1 ; | HD3 - | HE2 |
| 7342 | 7345 | 1 ; | HD3 - | HE3 |
| 7342 | 7346 | 1 ; | HD3 - | NZ |
| 7344 | 7347 | 1 ; | HE2 - | HZ1 |

```
7344   7348       1 ;     HE2 - HZ2
7344   7349       1 ;     HE2 - HZ3
7345   7347       1 ;     HE3 - HZ1
7345   7348       1 ;     HE3 - HZ2
7345   7349       1 ;     HE3 - HZ3
7350   7355       1 ;       C - HA
7350   7356       1 ;       C - CB
7350   7372       1 ;       C - C
7351   7353       1 ;       O - H
7351   7354       1 ;       O - CA
7352   7357       1 ;       N - HB2
7352   7358       1 ;       N - HB3
7352   7359       1 ;       N - CG
7352   7373       1 ;       N - O
7352   7374       1 ;       N - OXT
7353   7355       1 ;       H - HA
7353   7356       1 ;       H - CB
7353   7372       1 ;       H - C
7354   7360       1 ;      CA - HG2
7354   7361       1 ;      CA - HG3
7354   7362       1 ;      CA - CD
7355   7357       1 ;      HA - HB2
7355   7358       1 ;      HA - HB3
7355   7359       1 ;      HA - CG
7355   7373       1 ;      HA - O
7355   7374       1 ;      HA - OXT
7356   7363       1 ;      CB - HD2
7356   7364       1 ;      CB - HD3
7356   7365       1 ;      CB - CE
7356   7373       1 ;      CB - O
7356   7374       1 ;      CB - OXT
7357   7360       1 ;     HB2 - HG2
7357   7361       1 ;     HB2 - HG3
7357   7362       1 ;     HB2 - CD
7357   7372       1 ;     HB2 - C
7358   7360       1 ;     HB3 - HG2
7358   7361       1 ;     HB3 - HG3
7358   7362       1 ;     HB3 - CD
7358   7372       1 ;     HB3 - C
7359   7366       1 ;      CG - HE2
7359   7367       1 ;      CG - HE3
7359   7368       1 ;      CG - NZ
7359   7372       1 ;      CG - C
7360   7363       1 ;     HG2 - HD2
7360   7364       1 ;     HG2 - HD3
7360   7365       1 ;     HG2 - CE
7361   7363       1 ;     HG3 - HD2
7361   7364       1 ;     HG3 - HD3
7361   7365       1 ;     HG3 - CE
7362   7369       1 ;      CD - HZ1
7362   7370       1 ;      CD - HZ2
7362   7371       1 ;      CD - HZ3
```

| | | | |
|---|---|---|---|
| 7363 | 7366 | 1 ; | HD2 - HE2 |
| 7363 | 7367 | 1 ; | HD2 - HE3 |
| 7363 | 7368 | 1 ; | HD2 - NZ |
| 7364 | 7366 | 1 ; | HD3 - HE2 |
| 7364 | 7367 | 1 ; | HD3 - HE3 |
| 7364 | 7368 | 1 ; | HD3 - NZ |
| 7366 | 7369 | 1 ; | HE2 - HZ1 |
| 7366 | 7370 | 1 ; | HE2 - HZ2 |
| 7366 | 7371 | 1 ; | HE2 - HZ3 |
| 7367 | 7369 | 1 ; | HE3 - HZ1 |
| 7367 | 7370 | 1 ; | HE3 - HZ2 |
| 7367 | 7371 | 1 ; | HE3 - HZ3 |
| 7375 | 7378 | 1 ; | F2 - N23 |
| 7375 | 7384 | 1 ; | F2 - C36 |
| 7375 | 7423 | 1 ; | F2 - H39 |
| 7375 | 7424 | 1 ; | F2 - H37 |
| 7376 | 7385 | 1 ; | C38 - C40 |
| 7376 | 7386 | 1 ; | C38 - C35 |
| 7376 | 7433 | 1 ; | C38 - RU |
| 7377 | 7379 | 1 ; | C39 - N21 |
| 7377 | 7380 | 1 ; | C39 - C13 |
| 7377 | 7382 | 1 ; | C39 - CL17 |
| 7377 | 7384 | 1 ; | C39 - C36 |
| 7377 | 7395 | 1 ; | C39 - C24 |
| 7377 | 7402 | 1 ; | C39 - N15 |
| 7377 | 7424 | 1 ; | C39 - H37 |
| 7378 | 7381 | 1 ; | N23 - O14 |
| 7378 | 7383 | 1 ; | N23 - C37 |
| 7378 | 7386 | 1 ; | N23 - C35 |
| 7378 | 7394 | 1 ; | N23 - C31 |
| 7378 | 7397 | 1 ; | N23 - C25 |
| 7378 | 7403 | 1 ; | N23 - C7 |
| 7378 | 7404 | 1 ; | N23 - C18 |
| 7379 | 7381 | 1 ; | N21 - O14 |
| 7379 | 7384 | 1 ; | N21 - C36 |
| 7379 | 7391 | 1 ; | N21 - C32 |
| 7379 | 7398 | 1 ; | N21 - C29 |
| 7379 | 7400 | 1 ; | N21 - C27 |
| 7379 | 7403 | 1 ; | N21 - C7 |
| 7379 | 7404 | 1 ; | N21 - C18 |
| 7379 | 7428 | 1 ; | N21 - H26 |
| 7380 | 7385 | 1 ; | C13 - C40 |
| 7380 | 7395 | 1 ; | C13 - C24 |
| 7380 | 7397 | 1 ; | C13 - C25 |
| 7380 | 7403 | 1 ; | C13 - C7 |
| 7380 | 7404 | 1 ; | C13 - C18 |
| 7381 | 7382 | 1 ; | O14 - CL17 |
| 7381 | 7402 | 1 ; | O14 - N15 |
| 7382 | 7385 | 1 ; | CL17 - C40 |
| 7382 | 7395 | 1 ; | CL17 - C24 |
| 7382 | 7397 | 1 ; | CL17 - C25 |
| 7382 | 7403 | 1 ; | CL17 - C7 |

| | | | |
|---|---|---|---|
| 7382 | 7404 | 1 ; | CL17 - C18 |
| 7383 | 7387 | 1 ; | C37 - C34 |
| 7383 | 7391 | 1 ; | C37 - C32 |
| 7383 | 7395 | 1 ; | C37 - C24 |
| 7383 | 7423 | 1 ; | C37 - H39 |
| 7384 | 7388 | 1 ; | C36 - N22 |
| 7384 | 7390 | 1 ; | C36 - O42 |
| 7384 | 7392 | 1 ; | C36 - C33 |
| 7384 | 7394 | 1 ; | C36 - C31 |
| 7384 | 7433 | 1 ; | C36 - RU |
| 7385 | 7387 | 1 ; | C40 - C34 |
| 7385 | 7391 | 1 ; | C40 - C32 |
| 7385 | 7396 | 1 ; | C40 - C30 |
| 7385 | 7397 | 1 ; | C40 - C25 |
| 7385 | 7402 | 1 ; | C40 - N15 |
| 7385 | 7423 | 1 ; | C40 - H39 |
| 7385 | 7424 | 1 ; | C40 - H37 |
| 7386 | 7389 | 1 ; | C35 - H22 |
| 7386 | 7393 | 1 ; | C35 - O41 |
| 7386 | 7395 | 1 ; | C35 - C24 |
| 7386 | 7396 | 1 ; | C35 - C30 |
| 7386 | 7424 | 1 ; | C35 - H37 |
| 7387 | 7393 | 1 ; | C34 - O41 |
| 7387 | 7394 | 1 ; | C34 - C31 |
| 7388 | 7394 | 1 ; | N22 - C31 |
| 7389 | 7390 | 1 ; | H22 - O42 |
| 7389 | 7391 | 1 ; | H22 - C32 |
| 7389 | 7393 | 1 ; | H22 - O41 |
| 7390 | 7391 | 1 ; | O42 - C32 |
| 7390 | 7392 | 1 ; | O42 - C33 |
| 7391 | 7397 | 1 ; | C32 - C25 |
| 7391 | 7398 | 1 ; | C32 - C29 |
| 7392 | 7395 | 1 ; | C33 - C24 |
| 7392 | 7396 | 1 ; | C33 - C30 |
| 7393 | 7394 | 1 ; | O41 - C31 |
| 7394 | 7399 | 1 ; | C31 - C28 |
| 7394 | 7401 | 1 ; | C31 - C26 |
| 7394 | 7426 | 1 ; | C31 - H29 |
| 7394 | 7433 | 1 ; | C31 - RU |
| 7395 | 7398 | 1 ; | C24 - C29 |
| 7395 | 7401 | 1 ; | C24 - C26 |
| 7395 | 7402 | 1 ; | C24 - N15 |
| 7396 | 7400 | 1 ; | C30 - C27 |
| 7396 | 7416 | 1 ; | C30 - O1 |
| 7396 | 7428 | 1 ; | C30 - H26 |
| 7396 | 7433 | 1 ; | C30 - RU |
| 7397 | 7399 | 1 ; | C25 - C28 |
| 7397 | 7402 | 1 ; | C25 - N15 |
| 7397 | 7426 | 1 ; | C25 - H29 |
| 7397 | 7427 | 1 ; | C25 - H27 |
| 7398 | 7401 | 1 ; | C29 - C26 |
| 7398 | 7417 | 1 ; | C29 - H1 |

| | | | | |
|---|---|---|---|---|
| 7398 | 7427 | 1 | ; | C29 - H27 |
| 7399 | 7428 | 1 | ; | C28 - H26 |
| 7400 | 7417 | 1 | ; | C27 - H1 |
| 7400 | 7426 | 1 | ; | C27 - H29 |
| 7401 | 7416 | 1 | ; | C26 - O1 |
| 7401 | 7433 | 1 | ; | C26 - RU |
| 7402 | 7406 | 1 | ; | N15 - C6 |
| 7402 | 7408 | 1 | ; | N15 - C3 |
| 7402 | 7411 | 1 | ; | N15 - C9 |
| 7402 | 7415 | 1 | ; | N15 - N16 |
| 7402 | 7421 | 1 | ; | N15 - H5 |
| 7402 | 7429 | 1 | ; | N15 - H19 |
| 7403 | 7405 | 1 | ; | C7 - C5 |
| 7403 | 7409 | 1 | ; | C7 - C19 |
| 7403 | 7412 | 1 | ; | C7 - C10 |
| 7403 | 7414 | 1 | ; | C7 - C11 |
| 7403 | 7418 | 1 | ; | C7 - H9 |
| 7404 | 7407 | 1 | ; | C18 - C4 |
| 7404 | 7410 | 1 | ; | C18 - C8 |
| 7404 | 7419 | 1 | ; | C18 - H7A |
| 7404 | 7420 | 1 | ; | C18 - H6 |
| 7404 | 7425 | 1 | ; | C18 - H3 |
| 7405 | 7408 | 1 | ; | C5 - C3 |
| 7405 | 7422 | 1 | ; | C5 - H4 |
| 7405 | 7429 | 1 | ; | C5 - H19 |
| 7405 | 7433 | 1 | ; | C5 - RU |
| 7406 | 7409 | 1 | ; | C6 - C19 |
| 7406 | 7425 | 1 | ; | C6 - H3 |
| 7407 | 7421 | 1 | ; | C4 - H5 |
| 7407 | 7429 | 1 | ; | C4 - H19 |
| 7408 | 7420 | 1 | ; | C3 - H6 |
| 7409 | 7421 | 1 | ; | C19 - H5 |
| 7409 | 7422 | 1 | ; | C19 - H4 |
| 7409 | 7433 | 1 | ; | C19 - RU |
| 7410 | 7413 | 1 | ; | C8 - C12 |
| 7410 | 7431 | 1 | ; | C8 - H11 |
| 7410 | 7432 | 1 | ; | C8 - H10 |
| 7410 | 7433 | 1 | ; | C8 - RU |
| 7411 | 7414 | 1 | ; | C9 - C11 |
| 7411 | 7419 | 1 | ; | C9 - H7A |
| 7411 | 7430 | 1 | ; | C9 - H12 |
| 7412 | 7415 | 1 | ; | C10 - N16 |
| 7412 | 7431 | 1 | ; | C10 - H11 |
| 7413 | 7418 | 1 | ; | C12 - H9 |
| 7414 | 7432 | 1 | ; | C11 - H10 |
| 7415 | 7418 | 1 | ; | N16 - H9 |
| 7415 | 7419 | 1 | ; | N16 - H7A |
| 7415 | 7430 | 1 | ; | N16 - H12 |
| 7416 | 7426 | 1 | ; | O1 - H29 |
| 7416 | 7427 | 1 | ; | O1 - H27 |
| 7418 | 7432 | 1 | ; | H9 - H10 |
| 7419 | 7433 | 1 | ; | H7A - RU |

```
  7420   7421       1 ;      H6 - H5
  7420   7422       1 ;      H6 - H4
  7422   7425       1 ;      H4 - H3
  7423   7433       1 ;     H39 - RU
  7425   7429       1 ;      H3 - H19
  7427   7428       1 ;     H27 - H26
  7430   7431       1 ;     H12 - H11
  7430   7432       1 ;     H12 - H10

[ angles ]
;   ai     aj     ak    funct   theta         cth
     1      5      6      1    1.0950e+02    4.1840e+02 ;       N - CA
- HA
     1      5      7      1    1.1120e+02    6.6944e+02 ;       N - CA
- CB
     1      5     20      1    1.1120e+02    6.6944e+02 ;       N - CA
- C
     2      1      3      1    1.0950e+02    2.9288e+02 ;      H1 - N
- H2
     2      1      4      1    1.0950e+02    2.9288e+02 ;      H1 - N
- H3
     2      1      5      1    1.0950e+02    4.1840e+02 ;      H1 - N
- CA
     3      1      4      1    1.0950e+02    2.9288e+02 ;      H2 - N
- H3
     3      1      5      1    1.0950e+02    4.1840e+02 ;      H2 - N
- CA
     4      1      5      1    1.0950e+02    4.1840e+02 ;      H3 - N
- CA
     5      7      8      1    1.0950e+02    4.1840e+02 ;      CA - CB
- HB2
     5      7      9      1    1.0950e+02    4.1840e+02 ;      CA - CB
- HB3
     5      7     10      1    1.0950e+02    3.3472e+02 ;      CA - CB
- CG
     5     20     21      1    1.2040e+02    6.6944e+02 ;      CA - C
- O
     5     20     22      1    1.1660e+02    5.8576e+02 ;      CA - C
- N
     6      5      7      1    1.0950e+02    4.1840e+02 ;      HA - CA
- CB
     6      5     20      1    1.0950e+02    4.1840e+02 ;      HA - CA
- C
     7      5     20      1    1.1110e+02    5.2718e+02 ;      CB - CA
- C
     7     10     11      1    1.0950e+02    4.1840e+02 ;      CB - CG
- HG
     7     10     12      1    1.0950e+02    3.3472e+02 ;      CB - CG
- CD1
     7     10     16      1    1.0950e+02    3.3472e+02 ;      CB - CG
- CD2
```

```
     8       7       9       1     1.0950e+02    2.9288e+02 ;    HB2 - CB
- HB3
     8       7      10       1     1.0950e+02    4.1840e+02 ;    HB2 - CB
- CG
     9       7      10       1     1.0950e+02    4.1840e+02 ;    HB3 - CB
- CG
    10      12      13       1     1.0950e+02    4.1840e+02 ;     CG - CD1
- HD11
    10      12      14       1     1.0950e+02    4.1840e+02 ;     CG - CD1
- HD12
    10      12      15       1     1.0950e+02    4.1840e+02 ;     CG - CD1
- HD13
    10      16      17       1     1.0950e+02    4.1840e+02 ;     CG - CD2
- HD21
    10      16      18       1     1.0950e+02    4.1840e+02 ;     CG - CD2
- HD22
    10      16      19       1     1.0950e+02    4.1840e+02 ;     CG - CD2
- HD23
    11      10      12       1     1.0950e+02    4.1840e+02 ;     HG - CG
- CD1
    11      10      16       1     1.0950e+02    4.1840e+02 ;     HG - CG
- CD2
    12      10      16       1     1.0950e+02    3.3472e+02 ;    CD1 - CG
- CD2
    13      12      14       1     1.0950e+02    2.9288e+02 ;   HD11 - CD1
- HD12
    13      12      15       1     1.0950e+02    2.9288e+02 ;   HD11 - CD1
- HD13
    14      12      15       1     1.0950e+02    2.9288e+02 ;   HD12 - CD1
- HD13
    17      16      18       1     1.0950e+02    2.9288e+02 ;   HD21 - CD2
- HD22
    17      16      19       1     1.0950e+02    2.9288e+02 ;   HD21 - CD2
- HD23
    18      16      19       1     1.0950e+02    2.9288e+02 ;   HD22 - CD2
- HD23
    20      22      23       1     1.2000e+02    4.1840e+02 ;      C - N
- H
    20      22      24       1     1.2190e+02    4.1840e+02 ;      C - N
- CA
    21      20      22       1     1.2290e+02    6.6944e+02 ;      O - C
- N
    22      24      25       1     1.0950e+02    4.1840e+02 ;      N - CA
- HA
    22      24      26       1     1.0970e+02    6.6944e+02 ;      N - CA
- CB
    22      24      39       1     1.1010e+02    5.2718e+02 ;      N - CA
- C
    23      22      24       1     1.1804e+02    4.1840e+02 ;      H - N
- CA
    24      26      27       1     1.0950e+02    4.1840e+02 ;     CA - CB
- HB
```

24 26 28 1 1.0950e+02 3.3472e+02 ; CA - CB
- CG2
24 26 32 1 1.0950e+02 3.3472e+02 ; CA - CB
- CG1
24 39 40 1 1.2040e+02 6.6944e+02 ; CA - C
- O
24 39 41 1 1.1660e+02 5.8576e+02 ; CA - C
- N
25 24 26 1 1.0950e+02 4.1840e+02 ; HA - CA
- CB
25 24 39 1 1.0950e+02 4.1840e+02 ; HA - CA
- C
26 24 39 1 1.1110e+02 5.2718e+02 ; CB - CA
- C
26 28 29 1 1.0950e+02 4.1840e+02 ; CB - CG2
- HG21
26 28 30 1 1.0950e+02 4.1840e+02 ; CB - CG2
- HG22
26 28 31 1 1.0950e+02 4.1840e+02 ; CB - CG2
- HG23
26 32 33 1 1.0950e+02 4.1840e+02 ; CB - CG1
- HG12
26 32 34 1 1.0950e+02 4.1840e+02 ; CB - CG1
- HG13
26 32 35 1 1.0950e+02 3.3472e+02 ; CB - CG1
- CD1
27 26 28 1 1.0950e+02 4.1840e+02 ; HB - CB
- CG2
27 26 32 1 1.0950e+02 4.1840e+02 ; HB - CB
- CG1
28 26 32 1 1.0950e+02 3.3472e+02 ; CG2 - CB
- CG1
29 28 30 1 1.0950e+02 2.9288e+02 ; HG21 - CG2
- HG22
29 28 31 1 1.0950e+02 2.9288e+02 ; HG21 - CG2
- HG23
30 28 31 1 1.0950e+02 2.9288e+02 ; HG22 - CG2
- HG23
32 35 36 1 1.0950e+02 4.1840e+02 ; CG1 - CD1
- HD11
32 35 37 1 1.0950e+02 4.1840e+02 ; CG1 - CD1
- HD12
32 35 38 1 1.0950e+02 4.1840e+02 ; CG1 - CD1
- HD13
33 32 34 1 1.0950e+02 2.9288e+02 ; HG12 - CG1
- HG13
33 32 35 1 1.0950e+02 4.1840e+02 ; HG12 - CG1
- CD1
34 32 35 1 1.0950e+02 4.1840e+02 ; HG13 - CG1
- CD1
36 35 37 1 1.0950e+02 2.9288e+02 ; HD11 - CD1
- HD12

```
    36      35      38       1     1.0950e+02    2.9288e+02 ;   HD11 - CD1
 - HD13
    37      35      38       1     1.0950e+02    2.9288e+02 ;   HD12 - CD1
 - HD13
    39      41      42       1     1.2000e+02    4.1840e+02 ;      C - N
 - H
    39      41      43       1     1.2190e+02    4.1840e+02 ;      C - N
 - CA
    40      39      41       1     1.2290e+02    6.6944e+02 ;      O - C
 - N
    41      43      44       1     1.0950e+02    4.1840e+02 ;      N - CA
 - HA
    41      43      45       1     1.0970e+02    6.6944e+02 ;      N - CA
 - CB
    41      43      58       1     1.1010e+02    5.2718e+02 ;      N - CA
 - C
    42      41      43       1     1.1804e+02    4.1840e+02 ;      H - N
 - CA
    43      45      46       1     1.0950e+02    4.1840e+02 ;     CA - CB
 - HB2
    43      45      47       1     1.0950e+02    4.1840e+02 ;     CA - CB
 - HB3
    43      45      48       1     1.0950e+02    3.3472e+02 ;     CA - CB
 - CG
    43      58      59       1     1.2040e+02    6.6944e+02 ;     CA - C
 - O
    43      58      60       1     1.1660e+02    5.8576e+02 ;     CA - C
 - N
    44      43      45       1     1.0950e+02    4.1840e+02 ;     HA - CA
 - CB
    44      43      58       1     1.0950e+02    4.1840e+02 ;     HA - CA
 - C
    45      43      58       1     1.1110e+02    5.2718e+02 ;     CB - CA
 - C
    45      48      49       1     1.0950e+02    4.1840e+02 ;     CB - CG
 - HG
    45      48      50       1     1.0950e+02    3.3472e+02 ;     CB - CG
 - CD1
    45      48      54       1     1.0950e+02    3.3472e+02 ;     CB - CG
 - CD2
    46      45      47       1     1.0950e+02    2.9288e+02 ;    HB2 - CB
 - HB3
    46      45      48       1     1.0950e+02    4.1840e+02 ;    HB2 - CB
 - CG
    47      45      48       1     1.0950e+02    4.1840e+02 ;    HB3 - CB
 - CG
    48      50      51       1     1.0950e+02    4.1840e+02 ;     CG - CD1
 - HD11
    48      50      52       1     1.0950e+02    4.1840e+02 ;     CG - CD1
 - HD12
    48      50      53       1     1.0950e+02    4.1840e+02 ;     CG - CD1
 - HD13
```

```
   48     54     55      1    1.0950e+02    4.1840e+02 ;     CG - CD2
- HD21
   48     54     56      1    1.0950e+02    4.1840e+02 ;     CG - CD2
- HD22
   48     54     57      1    1.0950e+02    4.1840e+02 ;     CG - CD2
- HD23
   49     48     50      1    1.0950e+02    4.1840e+02 ;     HG - CG
- CD1
   49     48     54      1    1.0950e+02    4.1840e+02 ;     HG - CG
- CD2
   50     48     54      1    1.0950e+02    3.3472e+02 ;    CD1 - CG
- CD2
   51     50     52      1    1.0950e+02    2.9288e+02 ;   HD11 - CD1
- HD12
   51     50     53      1    1.0950e+02    2.9288e+02 ;   HD11 - CD1
- HD13
   52     50     53      1    1.0950e+02    2.9288e+02 ;   HD12 - CD1
- HD13
   55     54     56      1    1.0950e+02    2.9288e+02 ;   HD21 - CD2
- HD22
   55     54     57      1    1.0950e+02    2.9288e+02 ;   HD21 - CD2
- HD23
   56     54     57      1    1.0950e+02    2.9288e+02 ;   HD22 - CD2
- HD23
   58     60     61      1    1.2000e+02    4.1840e+02 ;      C - N
- H
   58     60     62      1    1.2190e+02    4.1840e+02 ;      C - N
- CA
   59     58     60      1    1.2290e+02    6.6944e+02 ;      O - C
- N
   60     62     63      1    1.0950e+02    4.1840e+02 ;      N - CA
- HA
   60     62     64      1    1.0970e+02    6.6944e+02 ;      N - CA
- CB
   60     62     69      1    1.1010e+02    5.2718e+02 ;      N - CA
- C
   61     60     62      1    1.1804e+02    4.1840e+02 ;      H - N
- CA
   62     64     65      1    1.0950e+02    4.1840e+02 ;     CA - CB
- HB2
   62     64     66      1    1.0950e+02    4.1840e+02 ;     CA - CB
- HB3
   62     64     67      1    1.0860e+02    4.1840e+02 ;     CA - CB
- SG
   62     69     70      1    1.2040e+02    6.6944e+02 ;     CA - C
- O
   62     69     71      1    1.1660e+02    5.8576e+02 ;     CA - C
- N
   63     62     64      1    1.0950e+02    4.1840e+02 ;     HA - CA
- CB
   63     62     69      1    1.0950e+02    4.1840e+02 ;     HA - CA
- C
```

```
    64    62    69     1    1.1110e+02    5.2718e+02 ;     CB - CA
 - C
    64    67    68     1    9.6000e+01    3.5982e+02 ;     CB - SG
 - HG
    65    64    66     1    1.0950e+02    2.9288e+02 ;    HB2 - CB
 - HB3
    65    64    67     1    1.0950e+02    4.1840e+02 ;    HB2 - CB
 - SG
    66    64    67     1    1.0950e+02    4.1840e+02 ;    HB3 - CB
 - SG
    69    71    72     1    1.2000e+02    4.1840e+02 ;      C - N
 - H
    69    71    73     1    1.2190e+02    4.1840e+02 ;      C - N
 - CA
    70    69    71     1    1.2290e+02    6.6944e+02 ;      O - C
 - N
    71    73    74     1    1.0950e+02    4.1840e+02 ;      N - CA
 - HA
    71    73    75     1    1.0970e+02    6.6944e+02 ;      N - CA
 - CB
    71    73    86     1    1.1010e+02    5.2718e+02 ;      N - CA
 - C
    72    71    73     1    1.1804e+02    4.1840e+02 ;      H - N
 - CA
    73    75    76     1    1.0950e+02    4.1840e+02 ;     CA - CB
 - HB2
    73    75    77     1    1.0950e+02    4.1840e+02 ;     CA - CB
 - HB3
    73    75    78     1    1.1310e+02    5.2718e+02 ;     CA - CB
 - CG
    73    86    87     1    1.2040e+02    6.6944e+02 ;     CA - C
 - O
    73    86    88     1    1.1660e+02    5.8576e+02 ;     CA - C
 - N
    74    73    75     1    1.0950e+02    4.1840e+02 ;     HA - CA
 - CB
    74    73    86     1    1.0950e+02    4.1840e+02 ;     HA - CA
 - C
    75    73    86     1    1.1110e+02    5.2718e+02 ;     CB - CA
 - C
    75    78    79     1    1.2000e+02    5.8576e+02 ;     CB - CG
 - ND1
    75    78    84     1    1.2000e+02    5.8576e+02 ;     CB - CG
 - CD2
    76    75    77     1    1.0950e+02    2.9288e+02 ;    HB2 - CB
 - HB3
    76    75    78     1    1.0950e+02    4.1840e+02 ;    HB2 - CB
 - CG
    77    75    78     1    1.0950e+02    4.1840e+02 ;    HB3 - CB
 - CG
    78    79    80     1    1.1700e+02    5.8576e+02 ;     CG - ND1
 - CE1
```

```
    78      84      82       1    1.2000e+02    5.8576e+02 ;     CG - CD2
- NE2
    78      84      85       1    1.2000e+02    4.1840e+02 ;     CG - CD2
- HD2
    79      78      84       1    1.2000e+02    5.8576e+02 ;    ND1 - CG
- CD2
    79      80      81       1    1.2000e+02    4.1840e+02 ;    ND1 - CE1
- HE1
    79      80      82       1    1.2000e+02    5.8576e+02 ;    ND1 - CE1
- NE2
    80      82      83       1    1.2000e+02    4.1840e+02 ;    CE1 - NE2
- HE2
    80      82      84       1    1.2000e+02    5.8576e+02 ;    CE1 - NE2
- CD2
    81      80      82       1    1.2000e+02    4.1840e+02 ;    HE1 - CE1
- NE2
    82      84      85       1    1.2000e+02    4.1840e+02 ;    NE2 - CD2
- HD2
    83      82      84       1    1.2000e+02    4.1840e+02 ;    HE2 - NE2
- CD2
    86      88      89       1    1.2000e+02    4.1840e+02 ;      C - N
- H
    86      88      90       1    1.2190e+02    4.1840e+02 ;      C - N
- CA
    87      86      88       1    1.2290e+02    6.6944e+02 ;      O - C
- N
    88      90      91       1    1.0950e+02    4.1840e+02 ;      N - CA
- HA
    88      90      92       1    1.0970e+02    6.6944e+02 ;      N - CA
- CB
    88      90      97       1    1.1010e+02    5.2718e+02 ;      N - CA
- C
    89      88      90       1    1.1804e+02    4.1840e+02 ;      H - N
- CA
    90      92      93       1    1.0950e+02    4.1840e+02 ;     CA - CB
- HB2
    90      92      94       1    1.0950e+02    4.1840e+02 ;     CA - CB
- HB3
    90      92      95       1    1.0950e+02    4.1840e+02 ;     CA - CB
- OG
    90      97      98       1    1.2040e+02    6.6944e+02 ;     CA - C
- O
    90      97      99       1    1.1660e+02    5.8576e+02 ;     CA - C
- N
    91      90      92       1    1.0950e+02    4.1840e+02 ;     HA - CA
- CB
    91      90      97       1    1.0950e+02    4.1840e+02 ;     HA - CA
- C
    92      90      97       1    1.1110e+02    5.2718e+02 ;     CB - CA
- C
    92      95      96       1    1.0850e+02    4.6024e+02 ;     CB - OG
- HG
```

```
    93     92     94      1    1.0950e+02    2.9288e+02 ;    HB2 - CB
- HB3
    93     92     95      1    1.0950e+02    4.1840e+02 ;    HB2 - CB
- OG
    94     92     95      1    1.0950e+02    4.1840e+02 ;    HB3 - CB
- OG
    97     99    100      1    1.2000e+02    4.1840e+02 ;      C - N
- H
    97     99    101      1    1.2190e+02    4.1840e+02 ;      C - N
- CA
    98     97     99      1    1.2290e+02    6.6944e+02 ;      O - C
- N
    99    101    102      1    1.0950e+02    4.1840e+02 ;      N - CA
- HA2
    99    101    103      1    1.0950e+02    4.1840e+02 ;      N - CA
- HA3
    99    101    104      1    1.1010e+02    5.2718e+02 ;      N - CA
- C
   100     99    101      1    1.1804e+02    4.1840e+02 ;      H - N
- CA
   101    104    105      1    1.2040e+02    6.6944e+02 ;     CA - C
- O
   101    104    106      1    1.1660e+02    5.8576e+02 ;     CA - C
- N
   102    101    103      1    1.0950e+02    2.9288e+02 ;    HA2 - CA
- HA3
   102    101    104      1    1.0950e+02    4.1840e+02 ;    HA2 - CA
- C
   103    101    104      1    1.0950e+02    4.1840e+02 ;    HA3 - CA
- C
   104    106    107      1    1.2000e+02    4.1840e+02 ;      C - N
- H
   104    106    108      1    1.2190e+02    4.1840e+02 ;      C - N
- CA
   105    104    106      1    1.2290e+02    6.6944e+02 ;      O - C
- N
   106    108    109      1    1.0950e+02    4.1840e+02 ;      N - CA
- HA
   106    108    110      1    1.0970e+02    6.6944e+02 ;      N - CA
- CB
   106    108    126      1    1.1010e+02    5.2718e+02 ;      N - CA
- C
   107    106    108      1    1.1804e+02    4.1840e+02 ;      H - N
- CA
   108    110    111      1    1.0950e+02    4.1840e+02 ;     CA - CB
- HB2
   108    110    112      1    1.0950e+02    4.1840e+02 ;     CA - CB
- HB3
   108    110    113      1    1.0950e+02    3.3472e+02 ;     CA - CB
- CG
   108    126    127      1    1.2040e+02    6.6944e+02 ;     CA - C
- O
```

```
   108    126    128       1    1.1660e+02    5.8576e+02 ;     CA - C
- N
   109    108    110       1    1.0950e+02    4.1840e+02 ;     HA - CA
- CB
   109    108    126       1    1.0950e+02    4.1840e+02 ;     HA - CA
- C
   110    108    126       1    1.1110e+02    5.2718e+02 ;     CB - CA
- C
   110    113    114       1    1.0950e+02    4.1840e+02 ;     CB - CG
- HG2
   110    113    115       1    1.0950e+02    4.1840e+02 ;     CB - CG
- HG3
   110    113    116       1    1.0950e+02    3.3472e+02 ;     CB - CG
- CD
   111    110    112       1    1.0950e+02    2.9288e+02 ;    HB2 - CB
- HB3
   111    110    113       1    1.0950e+02    4.1840e+02 ;    HB2 - CB
- CG
   112    110    113       1    1.0950e+02    4.1840e+02 ;    HB3 - CB
- CG
   113    116    117       1    1.0950e+02    4.1840e+02 ;     CG - CD
- HD2
   113    116    118       1    1.0950e+02    4.1840e+02 ;     CG - CD
- HD3
   113    116    119       1    1.0950e+02    3.3472e+02 ;     CG - CD
- CE
   114    113    115       1    1.0950e+02    2.9288e+02 ;    HG2 - CG
- HG3
   114    113    116       1    1.0950e+02    4.1840e+02 ;    HG2 - CG
- CD
   115    113    116       1    1.0950e+02    4.1840e+02 ;    HG3 - CG
- CD
   116    119    120       1    1.0950e+02    4.1840e+02 ;     CD - CE
- HE2
   116    119    121       1    1.0950e+02    4.1840e+02 ;     CD - CE
- HE3
   116    119    122       1    1.1120e+02    6.6944e+02 ;     CD - CE
- NZ
   117    116    118       1    1.0950e+02    2.9288e+02 ;    HD2 - CD
- HD3
   117    116    119       1    1.0950e+02    4.1840e+02 ;    HD2 - CD
- CE
   118    116    119       1    1.0950e+02    4.1840e+02 ;    HD3 - CD
- CE
   119    122    123       1    1.0950e+02    4.1840e+02 ;     CE - NZ
- HZ1
   119    122    124       1    1.0950e+02    4.1840e+02 ;     CE - NZ
- HZ2
   119    122    125       1    1.0950e+02    4.1840e+02 ;     CE - NZ
- HZ3
   120    119    121       1    1.0950e+02    2.9288e+02 ;    HE2 - CE
- HE3
```

120 119 122 1 1.0950e+02 4.1840e+02 ; HE2 - CE
- NZ
121 119 122 1 1.0950e+02 4.1840e+02 ; HE3 - CE
- NZ
123 122 124 1 1.0950e+02 2.9288e+02 ; HZ1 - NZ
- HZ2
123 122 125 1 1.0950e+02 2.9288e+02 ; HZ1 - NZ
- HZ3
124 122 125 1 1.0950e+02 2.9288e+02 ; HZ2 - NZ
- HZ3
126 128 129 1 1.2000e+02 4.1840e+02 ; C - N
- H
126 128 130 1 1.2190e+02 4.1840e+02 ; C - N
- CA
127 126 128 1 1.2290e+02 6.6944e+02 ; O - C
- N
128 130 131 1 1.0950e+02 4.1840e+02 ; N - CA
- HA
128 130 132 1 1.0970e+02 6.6944e+02 ; N - CA
- CB
128 130 145 1 1.1010e+02 5.2718e+02 ; N - CA
- C
129 128 130 1 1.1804e+02 4.1840e+02 ; H - N
- CA
130 132 133 1 1.0950e+02 4.1840e+02 ; CA - CB
- HB2
130 132 134 1 1.0950e+02 4.1840e+02 ; CA - CB
- HB3
130 132 135 1 1.0950e+02 3.3472e+02 ; CA - CB
- CG
130 145 146 1 1.2040e+02 6.6944e+02 ; CA - C
- O
130 145 147 1 1.1660e+02 5.8576e+02 ; CA - C
- N
131 130 132 1 1.0950e+02 4.1840e+02 ; HA - CA
- CB
131 130 145 1 1.0950e+02 4.1840e+02 ; HA - CA
- C
132 130 145 1 1.1110e+02 5.2718e+02 ; CB - CA
- C
132 135 136 1 1.0950e+02 4.1840e+02 ; CB - CG
- HG
132 135 137 1 1.0950e+02 3.3472e+02 ; CB - CG
- CD1
132 135 141 1 1.0950e+02 3.3472e+02 ; CB - CG
- CD2
133 132 134 1 1.0950e+02 2.9288e+02 ; HB2 - CB
- HB3
133 132 135 1 1.0950e+02 4.1840e+02 ; HB2 - CB
- CG
134 132 135 1 1.0950e+02 4.1840e+02 ; HB3 - CB
- CG

135 137 138 1 1.0950e+02 4.1840e+02 ; CG - CD1
- HD11
135 137 139 1 1.0950e+02 4.1840e+02 ; CG - CD1
- HD12
135 137 140 1 1.0950e+02 4.1840e+02 ; CG - CD1
- HD13
135 141 142 1 1.0950e+02 4.1840e+02 ; CG - CD2
- HD21
135 141 143 1 1.0950e+02 4.1840e+02 ; CG - CD2
- HD22
135 141 144 1 1.0950e+02 4.1840e+02 ; CG - CD2
- HD23
136 135 137 1 1.0950e+02 4.1840e+02 ; HG - CG
- CD1
136 135 141 1 1.0950e+02 4.1840e+02 ; HG - CG
- CD2
137 135 141 1 1.0950e+02 3.3472e+02 ; CD1 - CG
- CD2
138 137 139 1 1.0950e+02 2.9288e+02 ; HD11 - CD1
- HD12
138 137 140 1 1.0950e+02 2.9288e+02 ; HD11 - CD1
- HD13
139 137 140 1 1.0950e+02 2.9288e+02 ; HD12 - CD1
- HD13
142 141 143 1 1.0950e+02 2.9288e+02 ; HD21 - CD2
- HD22
142 141 144 1 1.0950e+02 2.9288e+02 ; HD21 - CD2
- HD23
143 141 144 1 1.0950e+02 2.9288e+02 ; HD22 - CD2
- HD23
145 147 148 1 1.2000e+02 4.1840e+02 ; C - N
- H
145 147 149 1 1.2190e+02 4.1840e+02 ; C - N
- CA
146 145 147 1 1.2290e+02 6.6944e+02 ; O - C
- N
147 149 150 1 1.0950e+02 4.1840e+02 ; N - CA
- HA
147 149 151 1 1.0970e+02 6.6944e+02 ; N - CA
- CB
147 149 160 1 1.1010e+02 5.2718e+02 ; N - CA
- C
148 147 149 1 1.1804e+02 4.1840e+02 ; H - N
- CA
149 151 152 1 1.0950e+02 4.1840e+02 ; CA - CB
- HB2
149 151 153 1 1.0950e+02 4.1840e+02 ; CA - CB
- HB3
149 151 154 1 1.0950e+02 3.3472e+02 ; CA - CB
- CG
149 160 161 1 1.2040e+02 6.6944e+02 ; CA - C
- O

```
   149    160    162      1    1.1660e+02    5.8576e+02 ;     CA - C
- N
   150    149    151      1    1.0950e+02    4.1840e+02 ;     HA - CA
- CB
   150    149    160      1    1.0950e+02    4.1840e+02 ;     HA - CA
- C
   151    149    160      1    1.1110e+02    5.2718e+02 ;     CB - CA
- C
   151    154    155      1    1.0950e+02    4.1840e+02 ;     CB - CG
- HG2
   151    154    156      1    1.0950e+02    4.1840e+02 ;     CB - CG
- HG3
   151    154    157      1    1.1110e+02    5.2718e+02 ;     CB - CG
- CD
   152    151    153      1    1.0950e+02    2.9288e+02 ;    HB2 - CB
- HB3
   152    151    154      1    1.0950e+02    4.1840e+02 ;    HB2 - CB
- CG
   153    151    154      1    1.0950e+02    4.1840e+02 ;    HB3 - CB
- CG
   154    157    158      1    1.1700e+02    5.8576e+02 ;     CG - CD
- OE1
   154    157    159      1    1.1700e+02    5.8576e+02 ;     CG - CD
- OE2
   155    154    156      1    1.0950e+02    2.9288e+02 ;    HG2 - CG
- HG3
   155    154    157      1    1.0950e+02    4.1840e+02 ;    HG2 - CG
- CD
   156    154    157      1    1.0950e+02    4.1840e+02 ;    HG3 - CG
- CD
   158    157    159      1    1.2600e+02    6.6944e+02 ;    OE1 - CD
- OE2
   160    162    163      1    1.2000e+02    4.1840e+02 ;      C - N
- H
   160    162    164      1    1.2190e+02    4.1840e+02 ;      C - N
- CA
   161    160    162      1    1.2290e+02    6.6944e+02 ;      O - C
- N
   162    164    165      1    1.0950e+02    4.1840e+02 ;      N - CA
- HA
   162    164    166      1    1.0970e+02    6.6944e+02 ;      N - CA
- CB
   162    164    172      1    1.1010e+02    5.2718e+02 ;      N - CA
- C
   163    162    164      1    1.1804e+02    4.1840e+02 ;      H - N
- CA
   164    166    167      1    1.0950e+02    4.1840e+02 ;     CA - CB
- HB2
   164    166    168      1    1.0950e+02    4.1840e+02 ;     CA - CB
- HB3
   164    166    169      1    1.1110e+02    5.2718e+02 ;     CA - CB
- CG
```

```
  164    172    173      1    1.2040e+02    6.6944e+02 ;     CA - C
- O
  164    172    174      1    1.1660e+02    5.8576e+02 ;     CA - C
- N
  165    164    166      1    1.0950e+02    4.1840e+02 ;     HA - CA
- CB
  165    164    172      1    1.0950e+02    4.1840e+02 ;     HA - CA
- C
  166    164    172      1    1.1110e+02    5.2718e+02 ;     CB - CA
- C
  166    169    170      1    1.1700e+02    5.8576e+02 ;     CB - CG
- OD1
  166    169    171      1    1.1700e+02    5.8576e+02 ;     CB - CG
- OD2
  167    166    168      1    1.0950e+02    2.9288e+02 ;    HB2 - CB
- HB3
  167    166    169      1    1.0950e+02    4.1840e+02 ;    HB2 - CB
- CG
  168    166    169      1    1.0950e+02    4.1840e+02 ;    HB3 - CB
- CG
  170    169    171      1    1.2600e+02    6.6944e+02 ;    OD1 - CG
- OD2
  172    174    175      1    1.2000e+02    4.1840e+02 ;      C - N
- H
  172    174    176      1    1.2190e+02    4.1840e+02 ;      C - N
- CA
  173    172    174      1    1.2290e+02    6.6944e+02 ;      O - C
- N
  174    176    177      1    1.0950e+02    4.1840e+02 ;      N - CA
- HA
  174    176    178      1    1.0970e+02    6.6944e+02 ;      N - CA
- CB
  174    176    194      1    1.1010e+02    5.2718e+02 ;      N - CA
- C
  175    174    176      1    1.1804e+02    4.1840e+02 ;      H - N
- CA
  176    178    179      1    1.0950e+02    4.1840e+02 ;     CA - CB
- HB2
  176    178    180      1    1.0950e+02    4.1840e+02 ;     CA - CB
- HB3
  176    178    181      1    1.0950e+02    3.3472e+02 ;     CA - CB
- CG
  176    194    195      1    1.2040e+02    6.6944e+02 ;     CA - C
- O
  176    194    196      1    1.1660e+02    5.8576e+02 ;     CA - C
- N
  177    176    178      1    1.0950e+02    4.1840e+02 ;     HA - CA
- CB
  177    176    194      1    1.0950e+02    4.1840e+02 ;     HA - CA
- C
  178    176    194      1    1.1110e+02    5.2718e+02 ;     CB - CA
- C
```

178 181 182 1 1.0950e+02 4.1840e+02 ; CB - CG
- HG2
178 181 183 1 1.0950e+02 4.1840e+02 ; CB - CG
- HG3
178 181 184 1 1.0950e+02 3.3472e+02 ; CB - CG
- CD
179 178 180 1 1.0950e+02 2.9288e+02 ; HB2 - CB
- HB3
179 178 181 1 1.0950e+02 4.1840e+02 ; HB2 - CB
- CG
180 178 181 1 1.0950e+02 4.1840e+02 ; HB3 - CB
- CG
181 184 185 1 1.0950e+02 4.1840e+02 ; CG - CD
- HD2
181 184 186 1 1.0950e+02 4.1840e+02 ; CG - CD
- HD3
181 184 187 1 1.0950e+02 3.3472e+02 ; CG - CD
- CE
182 181 183 1 1.0950e+02 2.9288e+02 ; HG2 - CG
- HG3
182 181 184 1 1.0950e+02 4.1840e+02 ; HG2 - CG
- CD
183 181 184 1 1.0950e+02 4.1840e+02 ; HG3 - CG
- CD
184 187 188 1 1.0950e+02 4.1840e+02 ; CD - CE
- HE2
184 187 189 1 1.0950e+02 4.1840e+02 ; CD - CE
- HE3
184 187 190 1 1.1120e+02 6.6944e+02 ; CD - CE
- NZ
185 184 186 1 1.0950e+02 2.9288e+02 ; HD2 - CD
- HD3
185 184 187 1 1.0950e+02 4.1840e+02 ; HD2 - CD
- CE
186 184 187 1 1.0950e+02 4.1840e+02 ; HD3 - CD
- CE
187 190 191 1 1.0950e+02 4.1840e+02 ; CE - NZ
- HZ1
187 190 192 1 1.0950e+02 4.1840e+02 ; CE - NZ
- HZ2
187 190 193 1 1.0950e+02 4.1840e+02 ; CE - NZ
- HZ3
188 187 189 1 1.0950e+02 2.9288e+02 ; HE2 - CE
- HE3
188 187 190 1 1.0950e+02 4.1840e+02 ; HE2 - CE
- NZ
189 187 190 1 1.0950e+02 4.1840e+02 ; HE3 - CE
- NZ
191 190 192 1 1.0950e+02 2.9288e+02 ; HZ1 - NZ
- HZ2
191 190 193 1 1.0950e+02 2.9288e+02 ; HZ1 - NZ
- HZ3

```
  192    190    193       1    1.0950e+02    2.9288e+02 ;    HZ2 - NZ
- HZ3
  194    196    197       1    1.2000e+02    4.1840e+02 ;      C - N
- H
  194    196    198       1    1.2190e+02    4.1840e+02 ;      C - N
- CA
  195    194    196       1    1.2290e+02    6.6944e+02 ;      O - C
- N
  196    198    199       1    1.0950e+02    4.1840e+02 ;      N - CA
- HA
  196    198    200       1    1.0970e+02    6.6944e+02 ;      N - CA
- CB
  196    198    215       1    1.1010e+02    5.2718e+02 ;      N - CA
- C
  197    196    198       1    1.1804e+02    4.1840e+02 ;      H - N
- CA
  198    200    201       1    1.0950e+02    4.1840e+02 ;     CA - CB
- HB2
  198    200    202       1    1.0950e+02    4.1840e+02 ;     CA - CB
- HB3
  198    200    203       1    1.1400e+02    5.2718e+02 ;     CA - CB
- CG
  198    215    216       1    1.2040e+02    6.6944e+02 ;     CA - C
- O
  198    215    217       1    1.1660e+02    5.8576e+02 ;     CA - C
- N
  199    198    200       1    1.0950e+02    4.1840e+02 ;     HA - CA
- CB
  199    198    215       1    1.0950e+02    4.1840e+02 ;     HA - CA
- C
  200    198    215       1    1.1110e+02    5.2718e+02 ;     CB - CA
- C
  200    203    204       1    1.2000e+02    5.8576e+02 ;     CB - CG
- CD1
  200    203    213       1    1.2000e+02    5.8576e+02 ;     CB - CG
- CD2
  201    200    202       1    1.0950e+02    2.9288e+02 ;    HB2 - CB
- HB3
  201    200    203       1    1.0950e+02    4.1840e+02 ;    HB2 - CB
- CG
  202    200    203       1    1.0950e+02    4.1840e+02 ;    HB3 - CB
- CG
  203    204    205       1    1.2000e+02    4.1840e+02 ;     CG - CD1
- HD1
  203    204    206       1    1.2000e+02    5.2718e+02 ;     CG - CD1
- CE1
  203    213    211       1    1.2000e+02    5.2718e+02 ;     CG - CD2
- CE2
  203    213    214       1    1.2000e+02    4.1840e+02 ;     CG - CD2
- HD2
  204    203    213       1    1.2000e+02    5.2718e+02 ;    CD1 - CG
- CD2
```

```
   204    206    207      1    1.2000e+02    4.1840e+02 ;    CD1 - CE1
- HE1
   204    206    208      1    1.2000e+02    5.2718e+02 ;    CD1 - CE1
- CZ
   205    204    206      1    1.2000e+02    4.1840e+02 ;    HD1 - CD1
- CE1
   206    208    209      1    1.2000e+02    5.8576e+02 ;    CE1 - CZ
- OH
   206    208    211      1    1.2000e+02    5.2718e+02 ;    CE1 - CZ
- CE2
   207    206    208      1    1.2000e+02    4.1840e+02 ;    HE1 - CE1
- CZ
   208    209    210      1    1.1300e+02    4.1840e+02 ;     CZ - OH
- HH
   208    211    212      1    1.2000e+02    4.1840e+02 ;     CZ - CE2
- HE2
   208    211    213      1    1.2000e+02    5.2718e+02 ;     CZ - CE2
- CD2
   209    208    211      1    1.2000e+02    5.8576e+02 ;     OH - CZ
- CE2
   211    213    214      1    1.2000e+02    4.1840e+02 ;    CE2 - CD2
- HD2
   212    211    213      1    1.2000e+02    4.1840e+02 ;    HE2 - CE2
- CD2
   215    217    218      1    1.2000e+02    4.1840e+02 ;      C - N
- H
   215    217    219      1    1.2190e+02    4.1840e+02 ;      C - N
- CA
   216    215    217      1    1.2290e+02    6.6944e+02 ;      O - C
- N
   217    219    220      1    1.0950e+02    4.1840e+02 ;      N - CA
- HA
   217    219    221      1    1.0970e+02    6.6944e+02 ;      N - CA
- CB
   217    219    234      1    1.1010e+02    5.2718e+02 ;      N - CA
- C
   218    217    219      1    1.1804e+02    4.1840e+02 ;      H - N
- CA
   219    221    222      1    1.0950e+02    4.1840e+02 ;     CA - CB
- HB
   219    221    223      1    1.0950e+02    3.3472e+02 ;     CA - CB
- CG2
   219    221    227      1    1.0950e+02    3.3472e+02 ;     CA - CB
- CG1
   219    234    235      1    1.2040e+02    6.6944e+02 ;     CA - C
- O
   219    234    236      1    1.1660e+02    5.8576e+02 ;     CA - C
- N
   220    219    221      1    1.0950e+02    4.1840e+02 ;     HA - CA
- CB
   220    219    234      1    1.0950e+02    4.1840e+02 ;     HA - CA
- C
```

```
   221    219    234      1    1.1110e+02    5.2718e+02 ;     CB - CA
 - C
   221    223    224      1    1.0950e+02    4.1840e+02 ;     CB - CG2
 - HG21
   221    223    225      1    1.0950e+02    4.1840e+02 ;     CB - CG2
 - HG22
   221    223    226      1    1.0950e+02    4.1840e+02 ;     CB - CG2
 - HG23
   221    227    228      1    1.0950e+02    4.1840e+02 ;     CB - CG1
 - HG12
   221    227    229      1    1.0950e+02    4.1840e+02 ;     CB - CG1
 - HG13
   221    227    230      1    1.0950e+02    3.3472e+02 ;     CB - CG1
 - CD1
   222    221    223      1    1.0950e+02    4.1840e+02 ;     HB - CB
 - CG2
   222    221    227      1    1.0950e+02    4.1840e+02 ;     HB - CB
 - CG1
   223    221    227      1    1.0950e+02    3.3472e+02 ;    CG2 - CB
 - CG1
   224    223    225      1    1.0950e+02    2.9288e+02 ;   HG21 - CG2
 - HG22
   224    223    226      1    1.0950e+02    2.9288e+02 ;   HG21 - CG2
 - HG23
   225    223    226      1    1.0950e+02    2.9288e+02 ;   HG22 - CG2
 - HG23
   227    230    231      1    1.0950e+02    4.1840e+02 ;    CG1 - CD1
 - HD11
   227    230    232      1    1.0950e+02    4.1840e+02 ;    CG1 - CD1
 - HD12
   227    230    233      1    1.0950e+02    4.1840e+02 ;    CG1 - CD1
 - HD13
   228    227    229      1    1.0950e+02    2.9288e+02 ;   HG12 - CG1
 - HG13
   228    227    230      1    1.0950e+02    4.1840e+02 ;   HG12 - CG1
 - CD1
   229    227    230      1    1.0950e+02    4.1840e+02 ;   HG13 - CG1
 - CD1
   231    230    232      1    1.0950e+02    2.9288e+02 ;   HD11 - CD1
 - HD12
   231    230    233      1    1.0950e+02    2.9288e+02 ;   HD11 - CD1
 - HD13
   232    230    233      1    1.0950e+02    2.9288e+02 ;   HD12 - CD1
 - HD13
   234    236    237      1    1.2000e+02    4.1840e+02 ;      C - N
 - H
   234    236    238      1    1.2190e+02    4.1840e+02 ;      C - N
 - CA
   235    234    236      1    1.2290e+02    6.6944e+02 ;      O - C
 - N
   236    238    239      1    1.0950e+02    4.1840e+02 ;      N - CA
 - HA
```

```
   236    238    240      1    1.0970e+02    6.6944e+02 ;      N - CA
- CB
   236    238    253      1    1.1010e+02    5.2718e+02 ;      N - CA
- C
   237    236    238      1    1.1804e+02    4.1840e+02 ;      H - N
- CA
   238    240    241      1    1.0950e+02    4.1840e+02 ;     CA - CB
- HB
   238    240    242      1    1.0950e+02    3.3472e+02 ;     CA - CB
- CG2
   238    240    246      1    1.0950e+02    3.3472e+02 ;     CA - CB
- CG1
   238    253    254      1    1.2040e+02    6.6944e+02 ;     CA - C
- O
   238    253    255      1    1.1660e+02    5.8576e+02 ;     CA - C
- N
   239    238    240      1    1.0950e+02    4.1840e+02 ;     HA - CA
- CB
   239    238    253      1    1.0950e+02    4.1840e+02 ;     HA - CA
- C
   240    238    253      1    1.1110e+02    5.2718e+02 ;     CB - CA
- C
   240    242    243      1    1.0950e+02    4.1840e+02 ;     CB - CG2
- HG21
   240    242    244      1    1.0950e+02    4.1840e+02 ;     CB - CG2
- HG22
   240    242    245      1    1.0950e+02    4.1840e+02 ;     CB - CG2
- HG23
   240    246    247      1    1.0950e+02    4.1840e+02 ;     CB - CG1
- HG12
   240    246    248      1    1.0950e+02    4.1840e+02 ;     CB - CG1
- HG13
   240    246    249      1    1.0950e+02    3.3472e+02 ;     CB - CG1
- CD1
   241    240    242      1    1.0950e+02    4.1840e+02 ;     HB - CB
- CG2
   241    240    246      1    1.0950e+02    4.1840e+02 ;     HB - CB
- CG1
   242    240    246      1    1.0950e+02    3.3472e+02 ;    CG2 - CB
- CG1
   243    242    244      1    1.0950e+02    2.9288e+02 ;   HG21 - CG2
- HG22
   243    242    245      1    1.0950e+02    2.9288e+02 ;   HG21 - CG2
- HG23
   244    242    245      1    1.0950e+02    2.9288e+02 ;   HG22 - CG2
- HG23
   246    249    250      1    1.0950e+02    4.1840e+02 ;    CG1 - CD1
- HD11
   246    249    251      1    1.0950e+02    4.1840e+02 ;    CG1 - CD1
- HD12
   246    249    252      1    1.0950e+02    4.1840e+02 ;    CG1 - CD1
- HD13
```

247 246 248 1 1.0950e+02 2.9288e+02 ; HG12 - CG1
- HG13
247 246 249 1 1.0950e+02 4.1840e+02 ; HG12 - CG1
- CD1
248 246 249 1 1.0950e+02 4.1840e+02 ; HG13 - CG1
- CD1
250 249 251 1 1.0950e+02 2.9288e+02 ; HD11 - CD1
- HD12
250 249 252 1 1.0950e+02 2.9288e+02 ; HD11 - CD1
- HD13
251 249 252 1 1.0950e+02 2.9288e+02 ; HD12 - CD1
- HD13
253 255 256 1 1.2000e+02 4.1840e+02 ; C - N
- H
253 255 257 1 1.2190e+02 4.1840e+02 ; C - N
- CA
254 253 255 1 1.2290e+02 6.6944e+02 ; O - C
- N
255 257 258 1 1.0950e+02 4.1840e+02 ; N - CA
- HA
255 257 259 1 1.0970e+02 6.6944e+02 ; N - CA
- CB
255 257 265 1 1.1010e+02 5.2718e+02 ; N - CA
- C
256 255 257 1 1.1804e+02 4.1840e+02 ; H - N
- CA
257 259 260 1 1.0950e+02 4.1840e+02 ; CA - CB
- HB2
257 259 261 1 1.0950e+02 4.1840e+02 ; CA - CB
- HB3
257 259 262 1 1.1110e+02 5.2718e+02 ; CA - CB
- CG
257 265 266 1 1.2040e+02 6.6944e+02 ; CA - C
- O
257 265 267 1 1.1660e+02 5.8576e+02 ; CA - C
- N
258 257 259 1 1.0950e+02 4.1840e+02 ; HA - CA
- CB
258 257 265 1 1.0950e+02 4.1840e+02 ; HA - CA
- C
259 257 265 1 1.1110e+02 5.2718e+02 ; CB - CA
- C
259 262 263 1 1.1700e+02 5.8576e+02 ; CB - CG
- OD1
259 262 264 1 1.1700e+02 5.8576e+02 ; CB - CG
- OD2
260 259 261 1 1.0950e+02 2.9288e+02 ; HB2 - CB
- HB3
260 259 262 1 1.0950e+02 4.1840e+02 ; HB2 - CB
- CG
261 259 262 1 1.0950e+02 4.1840e+02 ; HB3 - CB
- CG

```
   263    262    264      1    1.2600e+02    6.6944e+02 ;    OD1 - CG
- OD2
   265    267    268      1    1.2000e+02    4.1840e+02 ;      C - N
- H
   265    267    269      1    1.2190e+02    4.1840e+02 ;      C - N
- CA
   266    265    267      1    1.2290e+02    6.6944e+02 ;      O - C
- N
   267    269    270      1    1.0950e+02    4.1840e+02 ;      N - CA
- HA
   267    269    271      1    1.0970e+02    6.6944e+02 ;      N - CA
- CB
   267    269    280      1    1.1010e+02    5.2718e+02 ;      N - CA
- C
   268    267    269      1    1.1804e+02    4.1840e+02 ;      H - N
- CA
   269    271    272      1    1.0950e+02    4.1840e+02 ;     CA - CB
- HB2
   269    271    273      1    1.0950e+02    4.1840e+02 ;     CA - CB
- HB3
   269    271    274      1    1.0950e+02    3.3472e+02 ;     CA - CB
- CG
   269    280    281      1    1.2040e+02    6.6944e+02 ;     CA - C
- O
   269    280    282      1    1.1660e+02    5.8576e+02 ;     CA - C
- N
   270    269    271      1    1.0950e+02    4.1840e+02 ;     HA - CA
- CB
   270    269    280      1    1.0950e+02    4.1840e+02 ;     HA - CA
- C
   271    269    280      1    1.1110e+02    5.2718e+02 ;     CB - CA
- C
   271    274    275      1    1.0950e+02    4.1840e+02 ;     CB - CG
- HG2
   271    274    276      1    1.0950e+02    4.1840e+02 ;     CB - CG
- HG3
   271    274    277      1    1.1110e+02    5.2718e+02 ;     CB - CG
- CD
   272    271    273      1    1.0950e+02    2.9288e+02 ;    HB2 - CB
- HB3
   272    271    274      1    1.0950e+02    4.1840e+02 ;    HB2 - CB
- CG
   273    271    274      1    1.0950e+02    4.1840e+02 ;    HB3 - CB
- CG
   274    277    278      1    1.1700e+02    5.8576e+02 ;     CG - CD
- OE1
   274    277    279      1    1.1700e+02    5.8576e+02 ;     CG - CD
- OE2
   275    274    276      1    1.0950e+02    2.9288e+02 ;    HG2 - CG
- HG3
   275    274    277      1    1.0950e+02    4.1840e+02 ;    HG2 - CG
- CD
```

```
  276    274    277      1    1.0950e+02    4.1840e+02 ;    HG3 - CG
- CD
  278    277    279      1    1.2600e+02    6.6944e+02 ;    OE1 - CD
- OE2
  280    282    283      1    1.2000e+02    4.1840e+02 ;      C - N
- H
  280    282    284      1    1.2190e+02    4.1840e+02 ;      C - N
- CA
  281    280    282      1    1.2290e+02    6.6944e+02 ;      O - C
- N
  282    284    285      1    1.0950e+02    4.1840e+02 ;      N - CA
- HA
  282    284    286      1    1.0970e+02    6.6944e+02 ;      N - CA
- CB
  282    284    302      1    1.1010e+02    5.2718e+02 ;      N - CA
- C
  283    282    284      1    1.1804e+02    4.1840e+02 ;      H - N
- CA
  284    286    287      1    1.0950e+02    4.1840e+02 ;     CA - CB
- HB2
  284    286    288      1    1.0950e+02    4.1840e+02 ;     CA - CB
- HB3
  284    286    289      1    1.0950e+02    3.3472e+02 ;     CA - CB
- CG
  284    302    303      1    1.2040e+02    6.6944e+02 ;     CA - C
- O
  284    302    304      1    1.1660e+02    5.8576e+02 ;     CA - C
- N
  285    284    286      1    1.0950e+02    4.1840e+02 ;     HA - CA
- CB
  285    284    302      1    1.0950e+02    4.1840e+02 ;     HA - CA
- C
  286    284    302      1    1.1110e+02    5.2718e+02 ;     CB - CA
- C
  286    289    290      1    1.0950e+02    4.1840e+02 ;     CB - CG
- HG2
  286    289    291      1    1.0950e+02    4.1840e+02 ;     CB - CG
- HG3
  286    289    292      1    1.0950e+02    3.3472e+02 ;     CB - CG
- CD
  287    286    288      1    1.0950e+02    2.9288e+02 ;    HB2 - CB
- HB3
  287    286    289      1    1.0950e+02    4.1840e+02 ;    HB2 - CB
- CG
  288    286    289      1    1.0950e+02    4.1840e+02 ;    HB3 - CB
- CG
  289    292    293      1    1.0950e+02    4.1840e+02 ;     CG - CD
- HD2
  289    292    294      1    1.0950e+02    4.1840e+02 ;     CG - CD
- HD3
  289    292    295      1    1.0950e+02    3.3472e+02 ;     CG - CD
- CE
```

290 289 291 1 1.0950e+02 2.9288e+02 ; HG2 - CG
- HG3
290 289 292 1 1.0950e+02 4.1840e+02 ; HG2 - CG
- CD
291 289 292 1 1.0950e+02 4.1840e+02 ; HG3 - CG
- CD
292 295 296 1 1.0950e+02 4.1840e+02 ; CD - CE
- HE2
292 295 297 1 1.0950e+02 4.1840e+02 ; CD - CE
- HE3
292 295 298 1 1.1120e+02 6.6944e+02 ; CD - CE
- NZ
293 292 294 1 1.0950e+02 2.9288e+02 ; HD2 - CD
- HD3
293 292 295 1 1.0950e+02 4.1840e+02 ; HD2 - CD
- CE
294 292 295 1 1.0950e+02 4.1840e+02 ; HD3 - CD
- CE
295 298 299 1 1.0950e+02 4.1840e+02 ; CE - NZ
- HZ1
295 298 300 1 1.0950e+02 4.1840e+02 ; CE - NZ
- HZ2
295 298 301 1 1.0950e+02 4.1840e+02 ; CE - NZ
- HZ3
296 295 297 1 1.0950e+02 2.9288e+02 ; HE2 - CE
- HE3
296 295 298 1 1.0950e+02 4.1840e+02 ; HE2 - CE
- NZ
297 295 298 1 1.0950e+02 4.1840e+02 ; HE3 - CE
- NZ
299 298 300 1 1.0950e+02 2.9288e+02 ; HZ1 - NZ
- HZ2
299 298 301 1 1.0950e+02 2.9288e+02 ; HZ1 - NZ
- HZ3
300 298 301 1 1.0950e+02 2.9288e+02 ; HZ2 - NZ
- HZ3
302 304 305 1 1.2000e+02 4.1840e+02 ; C - N
- H
302 304 306 1 1.2190e+02 4.1840e+02 ; C - N
- CA
303 302 304 1 1.2290e+02 6.6944e+02 ; O - C
- N
304 306 307 1 1.0950e+02 4.1840e+02 ; N - CA
- HA
304 306 308 1 1.0970e+02 6.6944e+02 ; N - CA
- CB
304 306 321 1 1.1010e+02 5.2718e+02 ; N - CA
- C
305 304 306 1 1.1804e+02 4.1840e+02 ; H - N
- CA
306 308 309 1 1.0950e+02 4.1840e+02 ; CA - CB
- HB2

306 308 310 1 1.0950e+02 4.1840e+02 ; CA - CB
- HB3
306 308 311 1 1.0950e+02 3.3472e+02 ; CA - CB
- CG
306 321 322 1 1.2040e+02 6.6944e+02 ; CA - C
- O
306 321 323 1 1.1660e+02 5.8576e+02 ; CA - C
- N
307 306 308 1 1.0950e+02 4.1840e+02 ; HA - CA
- CB
307 306 321 1 1.0950e+02 4.1840e+02 ; HA - CA
- C
308 306 321 1 1.1110e+02 5.2718e+02 ; CB - CA
- C
308 311 312 1 1.0950e+02 4.1840e+02 ; CB - CG
- HG
308 311 313 1 1.0950e+02 3.3472e+02 ; CB - CG
- CD1
308 311 317 1 1.0950e+02 3.3472e+02 ; CB - CG
- CD2
309 308 310 1 1.0950e+02 2.9288e+02 ; HB2 - CB
- HB3
309 308 311 1 1.0950e+02 4.1840e+02 ; HB2 - CB
- CG
310 308 311 1 1.0950e+02 4.1840e+02 ; HB3 - CB
- CG
311 313 314 1 1.0950e+02 4.1840e+02 ; CG - CD1
- HD11
311 313 315 1 1.0950e+02 4.1840e+02 ; CG - CD1
- HD12
311 313 316 1 1.0950e+02 4.1840e+02 ; CG - CD1
- HD13
311 317 318 1 1.0950e+02 4.1840e+02 ; CG - CD2
- HD21
311 317 319 1 1.0950e+02 4.1840e+02 ; CG - CD2
- HD22
311 317 320 1 1.0950e+02 4.1840e+02 ; CG - CD2
- HD23
312 311 313 1 1.0950e+02 4.1840e+02 ; HG - CG
- CD1
312 311 317 1 1.0950e+02 4.1840e+02 ; HG - CG
- CD2
313 311 317 1 1.0950e+02 3.3472e+02 ; CD1 - CG
- CD2
314 313 315 1 1.0950e+02 2.9288e+02 ; HD11 - CD1
- HD12
314 313 316 1 1.0950e+02 2.9288e+02 ; HD11 - CD1
- HD13
315 313 316 1 1.0950e+02 2.9288e+02 ; HD12 - CD1
- HD13
318 317 319 1 1.0950e+02 2.9288e+02 ; HD21 - CD2
- HD22

```
   318    317    320      1    1.0950e+02    2.9288e+02 ;   HD21 - CD2
 - HD23
   319    317    320      1    1.0950e+02    2.9288e+02 ;   HD22 - CD2
 - HD23
   321    323    324      1    1.2000e+02    4.1840e+02 ;      C - N
 - H
   321    323    325      1    1.2190e+02    4.1840e+02 ;      C - N
 - CA
   322    321    323      1    1.2290e+02    6.6944e+02 ;      O - C
 - N
   323    325    326      1    1.0950e+02    4.1840e+02 ;      N - CA
 - HA2
   323    325    327      1    1.0950e+02    4.1840e+02 ;      N - CA
 - HA3
   323    325    328      1    1.1010e+02    5.2718e+02 ;      N - CA
 - C
   324    323    325      1    1.1804e+02    4.1840e+02 ;      H - N
 - CA
   325    328    329      1    1.2040e+02    6.6944e+02 ;     CA - C
 - O
   325    328    330      1    1.1660e+02    5.8576e+02 ;     CA - C
 - N
   326    325    327      1    1.0950e+02    2.9288e+02 ;    HA2 - CA
 - HA3
   326    325    328      1    1.0950e+02    4.1840e+02 ;    HA2 - CA
 - C
   327    325    328      1    1.0950e+02    4.1840e+02 ;    HA3 - CA
 - C
   328    330    331      1    1.2000e+02    4.1840e+02 ;      C - N
 - H
   328    330    332      1    1.2190e+02    4.1840e+02 ;      C - N
 - CA
   329    328    330      1    1.2290e+02    6.6944e+02 ;      O - C
 - N
   330    332    333      1    1.0950e+02    4.1840e+02 ;      N - CA
 - HA
   330    332    334      1    1.0970e+02    6.6944e+02 ;      N - CA
 - CB
   330    332    345      1    1.1010e+02    5.2718e+02 ;      N - CA
 - C
   331    330    332      1    1.1804e+02    4.1840e+02 ;      H - N
 - CA
   332    334    335      1    1.0950e+02    4.1840e+02 ;     CA - CB
 - HB2
   332    334    336      1    1.0950e+02    4.1840e+02 ;     CA - CB
 - HB3
   332    334    337      1    1.0950e+02    3.3472e+02 ;     CA - CB
 - CG
   332    345    346      1    1.2040e+02    6.6944e+02 ;     CA - C
 - O
   332    345    347      1    1.1660e+02    5.8576e+02 ;     CA - C
 - N
```

```
  333    332    334      1    1.0950e+02    4.1840e+02 ;     HA - CA
- CB
  333    332    345      1    1.0950e+02    4.1840e+02 ;     HA - CA
- C
  334    332    345      1    1.1110e+02    5.2718e+02 ;     CB - CA
- C
  334    337    338      1    1.0950e+02    4.1840e+02 ;     CB - CG
- HG2
  334    337    339      1    1.0950e+02    4.1840e+02 ;     CB - CG
- HG3
  334    337    340      1    1.1110e+02    5.2718e+02 ;     CB - CG
- CD
  335    334    336      1    1.0950e+02    2.9288e+02 ;    HB2 - CB
- HB3
  335    334    337      1    1.0950e+02    4.1840e+02 ;    HB2 - CB
- CG
  336    334    337      1    1.0950e+02    4.1840e+02 ;    HB3 - CB
- CG
  337    340    341      1    1.2040e+02    6.6944e+02 ;     CG - CD
- OE1
  337    340    342      1    1.1660e+02    5.8576e+02 ;     CG - CD
- NE2
  338    337    339      1    1.0950e+02    2.9288e+02 ;    HG2 - CG
- HG3
  338    337    340      1    1.0950e+02    4.1840e+02 ;    HG2 - CG
- CD
  339    337    340      1    1.0950e+02    4.1840e+02 ;    HG3 - CG
- CD
  340    342    343      1    1.2000e+02    4.1840e+02 ;     CD - NE2
- HE21
  340    342    344      1    1.2000e+02    4.1840e+02 ;     CD - NE2
- HE22
  341    340    342      1    1.2290e+02    6.6944e+02 ;    OE1 - CD
- NE2
  343    342    344      1    1.2000e+02    2.9288e+02 ;   HE21 - NE2
- HE22
  345    347    348      1    1.2000e+02    4.1840e+02 ;      C - N
- H
  345    347    349      1    1.2190e+02    4.1840e+02 ;      C - N
- CA
  346    345    347      1    1.2290e+02    6.6944e+02 ;      O - C
- N
  347    349    350      1    1.0950e+02    4.1840e+02 ;      N - CA
- HA2
  347    349    351      1    1.0950e+02    4.1840e+02 ;      N - CA
- HA3
  347    349    352      1    1.1010e+02    5.2718e+02 ;      N - CA
- C
  348    347    349      1    1.1804e+02    4.1840e+02 ;      H - N
- CA
  349    352    353      1    1.2040e+02    6.6944e+02 ;     CA - C
- O
```

349 352 354 1 1.1660e+02 5.8576e+02 ; CA - C
- N
350 349 351 1 1.0950e+02 2.9288e+02 ; HA2 - CA
- HA3
350 349 352 1 1.0950e+02 4.1840e+02 ; HA2 - CA
- C
351 349 352 1 1.0950e+02 4.1840e+02 ; HA3 - CA
- C
352 354 355 1 1.2000e+02 4.1840e+02 ; C - N
- H
352 354 356 1 1.2190e+02 4.1840e+02 ; C - N
- CA
353 352 354 1 1.2290e+02 6.6944e+02 ; O - C
- N
354 356 357 1 1.0950e+02 4.1840e+02 ; N - CA
- HA
354 356 358 1 1.0970e+02 6.6944e+02 ; N - CA
- CB
354 356 366 1 1.1010e+02 5.2718e+02 ; N - CA
- C
355 354 356 1 1.1804e+02 4.1840e+02 ; H - N
- CA
356 358 359 1 1.0950e+02 4.1840e+02 ; CA - CB
- HB
356 358 360 1 1.0950e+02 3.3472e+02 ; CA - CB
- CG2
356 358 364 1 1.0950e+02 4.1840e+02 ; CA - CB
- OG1
356 366 367 1 1.2040e+02 6.6944e+02 ; CA - C
- O
356 366 368 1 1.1660e+02 5.8576e+02 ; CA - C
- N
357 356 358 1 1.0950e+02 4.1840e+02 ; HA - CA
- CB
357 356 366 1 1.0950e+02 4.1840e+02 ; HA - CA
- C
358 356 366 1 1.1110e+02 5.2718e+02 ; CB - CA
- C
358 360 361 1 1.0950e+02 4.1840e+02 ; CB - CG2
- HG21
358 360 362 1 1.0950e+02 4.1840e+02 ; CB - CG2
- HG22
358 360 363 1 1.0950e+02 4.1840e+02 ; CB - CG2
- HG23
358 364 365 1 1.0850e+02 4.6024e+02 ; CB - OG1
- HG1
359 358 360 1 1.0950e+02 4.1840e+02 ; HB - CB
- CG2
359 358 364 1 1.0950e+02 4.1840e+02 ; HB - CB
- OG1
360 358 364 1 1.0950e+02 4.1840e+02 ; CG2 - CB
- OG1

361 360 362 1 1.0950e+02 2.9288e+02 ; HG21 - CG2
- HG22
361 360 363 1 1.0950e+02 2.9288e+02 ; HG21 - CG2
- HG23
362 360 363 1 1.0950e+02 2.9288e+02 ; HG22 - CG2
- HG23
366 368 369 1 1.2000e+02 4.1840e+02 ; C - N
- H
366 368 370 1 1.2190e+02 4.1840e+02 ; C - N
- CA
367 366 368 1 1.2290e+02 6.6944e+02 ; O - C
- N
368 370 371 1 1.0950e+02 4.1840e+02 ; N - CA
- HA
368 370 372 1 1.0970e+02 6.6944e+02 ; N - CA
- CB
368 370 387 1 1.1010e+02 5.2718e+02 ; N - CA
- C
369 368 370 1 1.1804e+02 4.1840e+02 ; H - N
- CA
370 372 373 1 1.0950e+02 4.1840e+02 ; CA - CB
- HB2
370 372 374 1 1.0950e+02 4.1840e+02 ; CA - CB
- HB3
370 372 375 1 1.1400e+02 5.2718e+02 ; CA - CB
- CG
370 387 388 1 1.2040e+02 6.6944e+02 ; CA - C
- O
370 387 389 1 1.1660e+02 5.8576e+02 ; CA - C
- N
371 370 372 1 1.0950e+02 4.1840e+02 ; HA - CA
- CB
371 370 387 1 1.0950e+02 4.1840e+02 ; HA - CA
- C
372 370 387 1 1.1110e+02 5.2718e+02 ; CB - CA
- C
372 375 376 1 1.2000e+02 5.8576e+02 ; CB - CG
- CD1
372 375 385 1 1.2000e+02 5.8576e+02 ; CB - CG
- CD2
373 372 374 1 1.0950e+02 2.9288e+02 ; HB2 - CB
- HB3
373 372 375 1 1.0950e+02 4.1840e+02 ; HB2 - CB
- CG
374 372 375 1 1.0950e+02 4.1840e+02 ; HB3 - CB
- CG
375 376 377 1 1.2000e+02 4.1840e+02 ; CG - CD1
- HD1
375 376 378 1 1.2000e+02 5.2718e+02 ; CG - CD1
- CE1
375 385 383 1 1.2000e+02 5.2718e+02 ; CG - CD2
- CE2

```
  375    385    386       1    1.2000e+02    4.1840e+02 ;     CG - CD2
- HD2
  376    375    385       1    1.2000e+02    5.2718e+02 ;    CD1 - CG
- CD2
  376    378    379       1    1.2000e+02    4.1840e+02 ;    CD1 - CE1
- HE1
  376    378    380       1    1.2000e+02    5.2718e+02 ;    CD1 - CE1
- CZ
  377    376    378       1    1.2000e+02    4.1840e+02 ;    HD1 - CD1
- CE1
  378    380    381       1    1.2000e+02    5.8576e+02 ;    CE1 - CZ
- OH
  378    380    383       1    1.2000e+02    5.2718e+02 ;    CE1 - CZ
- CE2
  379    378    380       1    1.2000e+02    4.1840e+02 ;    HE1 - CE1
- CZ
  380    381    382       1    1.1300e+02    4.1840e+02 ;     CZ - OH
- HH
  380    383    384       1    1.2000e+02    4.1840e+02 ;     CZ - CE2
- HE2
  380    383    385       1    1.2000e+02    5.2718e+02 ;     CZ - CE2
- CD2
  381    380    383       1    1.2000e+02    5.8576e+02 ;     OH - CZ
- CE2
  383    385    386       1    1.2000e+02    4.1840e+02 ;    CE2 - CD2
- HD2
  384    383    385       1    1.2000e+02    4.1840e+02 ;    HE2 - CE2
- CD2
  387    389    390       1    1.2000e+02    4.1840e+02 ;      C - N
- H
  387    389    391       1    1.2190e+02    4.1840e+02 ;      C - N
- CA
  388    387    389       1    1.2290e+02    6.6944e+02 ;      O - C
- N
  389    391    392       1    1.0950e+02    4.1840e+02 ;      N - CA
- HA2
  389    391    393       1    1.0950e+02    4.1840e+02 ;      N - CA
- HA3
  389    391    394       1    1.1010e+02    5.2718e+02 ;      N - CA
- C
  390    389    391       1    1.1804e+02    4.1840e+02 ;      H - N
- CA
  391    394    395       1    1.2040e+02    6.6944e+02 ;     CA - C
- O
  391    394    396       1    1.1660e+02    5.8576e+02 ;     CA - C
- N
  392    391    393       1    1.0950e+02    2.9288e+02 ;    HA2 - CA
- HA3
  392    391    394       1    1.0950e+02    4.1840e+02 ;    HA2 - CA
- C
  393    391    394       1    1.0950e+02    4.1840e+02 ;    HA3 - CA
- C
```

```
  394    396    397      1    1.2000e+02    4.1840e+02 ;      C - N
- H
  394    396    398      1    1.2190e+02    4.1840e+02 ;      C - N
- CA
  395    394    396      1    1.2290e+02    6.6944e+02 ;      O - C
- N
  396    398    399      1    1.0950e+02    4.1840e+02 ;      N - CA
- HA
  396    398    400      1    1.0970e+02    6.6944e+02 ;      N - CA
- CB
  396    398    405      1    1.1010e+02    5.2718e+02 ;      N - CA
- C
  397    396    398      1    1.1804e+02    4.1840e+02 ;      H - N
- CA
  398    400    401      1    1.0950e+02    4.1840e+02 ;     CA - CB
- HB2
  398    400    402      1    1.0950e+02    4.1840e+02 ;     CA - CB
- HB3
  398    400    403      1    1.0860e+02    4.1840e+02 ;     CA - CB
- SG
  398    405    406      1    1.2040e+02    6.6944e+02 ;     CA - C
- O
  398    405    407      1    1.1660e+02    5.8576e+02 ;     CA - C
- N
  399    398    400      1    1.0950e+02    4.1840e+02 ;     HA - CA
- CB
  399    398    405      1    1.0950e+02    4.1840e+02 ;     HA - CA
- C
  400    398    405      1    1.1110e+02    5.2718e+02 ;     CB - CA
- C
  400    403    404      1    9.6000e+01    3.5982e+02 ;     CB - SG
- HG
  401    400    402      1    1.0950e+02    2.9288e+02 ;    HB2 - CB
- HB3
  401    400    403      1    1.0950e+02    4.1840e+02 ;    HB2 - CB
- SG
  402    400    403      1    1.0950e+02    4.1840e+02 ;    HB3 - CB
- SG
  405    407    408      1    1.2000e+02    4.1840e+02 ;      C - N
- H
  405    407    409      1    1.2190e+02    4.1840e+02 ;      C - N
- CA
  406    405    407      1    1.2290e+02    6.6944e+02 ;      O - C
- N
  407    409    410      1    1.0950e+02    4.1840e+02 ;      N - CA
- HA
  407    409    411      1    1.0970e+02    6.6944e+02 ;      N - CA
- CB
  407    409    421      1    1.1010e+02    5.2718e+02 ;      N - CA
- C
  408    407    409      1    1.1804e+02    4.1840e+02 ;      H - N
- CA
```

409 411 412 1 1.0950e+02 4.1840e+02 ; CA - CB
- HB
409 411 413 1 1.0950e+02 3.3472e+02 ; CA - CB
- CG1
409 411 417 1 1.0950e+02 3.3472e+02 ; CA - CB
- CG2
409 421 422 1 1.2040e+02 6.6944e+02 ; CA - C
- O
409 421 423 1 1.1660e+02 5.8576e+02 ; CA - C
- N
410 409 411 1 1.0950e+02 4.1840e+02 ; HA - CA
- CB
410 409 421 1 1.0950e+02 4.1840e+02 ; HA - CA
- C
411 409 421 1 1.1110e+02 5.2718e+02 ; CB - CA
- C
411 413 414 1 1.0950e+02 4.1840e+02 ; CB - CG1
- HG11
411 413 415 1 1.0950e+02 4.1840e+02 ; CB - CG1
- HG12
411 413 416 1 1.0950e+02 4.1840e+02 ; CB - CG1
- HG13
411 417 418 1 1.0950e+02 4.1840e+02 ; CB - CG2
- HG21
411 417 419 1 1.0950e+02 4.1840e+02 ; CB - CG2
- HG22
411 417 420 1 1.0950e+02 4.1840e+02 ; CB - CG2
- HG23
412 411 413 1 1.0950e+02 4.1840e+02 ; HB - CB
- CG1
412 411 417 1 1.0950e+02 4.1840e+02 ; HB - CB
- CG2
413 411 417 1 1.0950e+02 3.3472e+02 ; CG1 - CB
- CG2
414 413 415 1 1.0950e+02 2.9288e+02 ; HG11 - CG1
- HG12
414 413 416 1 1.0950e+02 2.9288e+02 ; HG11 - CG1
- HG13
415 413 416 1 1.0950e+02 2.9288e+02 ; HG12 - CG1
- HG13
418 417 419 1 1.0950e+02 2.9288e+02 ; HG21 - CG2
- HG22
418 417 420 1 1.0950e+02 2.9288e+02 ; HG21 - CG2
- HG23
419 417 420 1 1.0950e+02 2.9288e+02 ; HG22 - CG2
- HG23
421 423 424 1 1.2000e+02 4.1840e+02 ; C - N
- H
421 423 425 1 1.2190e+02 4.1840e+02 ; C - N
- CA
422 421 423 1 1.2290e+02 6.6944e+02 ; O - C
- N

```
   423    425    426       1    1.0950e+02    4.1840e+02 ;      N - CA
- HA
   423    425    427       1    1.0970e+02    6.6944e+02 ;      N - CA
- CB
   423    425    442       1    1.1010e+02    5.2718e+02 ;      N - CA
- C
   424    423    425       1    1.1804e+02    4.1840e+02 ;      H - N
- CA
   425    427    428       1    1.0950e+02    4.1840e+02 ;     CA - CB
- HB2
   425    427    429       1    1.0950e+02    4.1840e+02 ;     CA - CB
- HB3
   425    427    430       1    1.1400e+02    5.2718e+02 ;     CA - CB
- CG
   425    442    443       1    1.2040e+02    6.6944e+02 ;     CA - C
- O
   425    442    444       1    1.1660e+02    5.8576e+02 ;     CA - C
- N
   426    425    427       1    1.0950e+02    4.1840e+02 ;     HA - CA
- CB
   426    425    442       1    1.0950e+02    4.1840e+02 ;     HA - CA
- C
   427    425    442       1    1.1110e+02    5.2718e+02 ;     CB - CA
- C
   427    430    431       1    1.2000e+02    5.8576e+02 ;     CB - CG
- CD1
   427    430    440       1    1.2000e+02    5.8576e+02 ;     CB - CG
- CD2
   428    427    429       1    1.0950e+02    2.9288e+02 ;    HB2 - CB
- HB3
   428    427    430       1    1.0950e+02    4.1840e+02 ;    HB2 - CB
- CG
   429    427    430       1    1.0950e+02    4.1840e+02 ;    HB3 - CB
- CG
   430    431    432       1    1.2000e+02    4.1840e+02 ;     CG - CD1
- HD1
   430    431    433       1    1.2000e+02    5.2718e+02 ;     CG - CD1
- CE1
   430    440    438       1    1.2000e+02    5.2718e+02 ;     CG - CD2
- CE2
   430    440    441       1    1.2000e+02    4.1840e+02 ;     CG - CD2
- HD2
   431    430    440       1    1.2000e+02    5.2718e+02 ;    CD1 - CG
- CD2
   431    433    434       1    1.2000e+02    4.1840e+02 ;    CD1 - CE1
- HE1
   431    433    435       1    1.2000e+02    5.2718e+02 ;    CD1 - CE1
- CZ
   432    431    433       1    1.2000e+02    4.1840e+02 ;    HD1 - CD1
- CE1
   433    435    436       1    1.2000e+02    5.8576e+02 ;    CE1 - CZ
- OH
```

433 435 438 1 1.2000e+02 5.2718e+02 ; CE1 - CZ
- CE2
434 433 435 1 1.2000e+02 4.1840e+02 ; HE1 - CE1
- CZ
435 436 437 1 1.1300e+02 4.1840e+02 ; CZ - OH
- HH
435 438 439 1 1.2000e+02 4.1840e+02 ; CZ - CE2
- HE2
435 438 440 1 1.2000e+02 5.2718e+02 ; CZ - CE2
- CD2
436 435 438 1 1.2000e+02 5.8576e+02 ; OH - CZ
- CE2
438 440 441 1 1.2000e+02 4.1840e+02 ; CE2 - CD2
- HD2
439 438 440 1 1.2000e+02 4.1840e+02 ; HE2 - CE2
- CD2
442 444 445 1 1.2000e+02 4.1840e+02 ; C - N
- H
442 444 446 1 1.2190e+02 4.1840e+02 ; C - N
- CA
443 442 444 1 1.2290e+02 6.6944e+02 ; O - C
- N
444 446 447 1 1.0950e+02 4.1840e+02 ; N - CA
- HA
444 446 448 1 1.0970e+02 6.6944e+02 ; N - CA
- CB
444 446 464 1 1.1010e+02 5.2718e+02 ; N - CA
- C
445 444 446 1 1.1804e+02 4.1840e+02 ; H - N
- CA
446 448 449 1 1.0950e+02 4.1840e+02 ; CA - CB
- HB2
446 448 450 1 1.0950e+02 4.1840e+02 ; CA - CB
- HB3
446 448 451 1 1.0950e+02 3.3472e+02 ; CA - CB
- CG
446 464 465 1 1.2040e+02 6.6944e+02 ; CA - C
- O
446 464 466 1 1.1660e+02 5.8576e+02 ; CA - C
- N
447 446 448 1 1.0950e+02 4.1840e+02 ; HA - CA
- CB
447 446 464 1 1.0950e+02 4.1840e+02 ; HA - CA
- C
448 446 464 1 1.1110e+02 5.2718e+02 ; CB - CA
- C
448 451 452 1 1.0950e+02 4.1840e+02 ; CB - CG
- HG2
448 451 453 1 1.0950e+02 4.1840e+02 ; CB - CG
- HG3
448 451 454 1 1.0950e+02 3.3472e+02 ; CB - CG
- CD

```
  449    448    450      1    1.0950e+02    2.9288e+02 ;    HB2 - CB
- HB3
  449    448    451      1    1.0950e+02    4.1840e+02 ;    HB2 - CB
- CG
  450    448    451      1    1.0950e+02    4.1840e+02 ;    HB3 - CB
- CG
  451    454    455      1    1.0950e+02    4.1840e+02 ;     CG - CD
- HD2
  451    454    456      1    1.0950e+02    4.1840e+02 ;     CG - CD
- HD3
  451    454    457      1    1.0950e+02    3.3472e+02 ;     CG - CD
- CE
  452    451    453      1    1.0950e+02    2.9288e+02 ;    HG2 - CG
- HG3
  452    451    454      1    1.0950e+02    4.1840e+02 ;    HG2 - CG
- CD
  453    451    454      1    1.0950e+02    4.1840e+02 ;    HG3 - CG
- CD
  454    457    458      1    1.0950e+02    4.1840e+02 ;     CD - CE
- HE2
  454    457    459      1    1.0950e+02    4.1840e+02 ;     CD - CE
- HE3
  454    457    460      1    1.1120e+02    6.6944e+02 ;     CD - CE
- NZ
  455    454    456      1    1.0950e+02    2.9288e+02 ;    HD2 - CD
- HD3
  455    454    457      1    1.0950e+02    4.1840e+02 ;    HD2 - CD
- CE
  456    454    457      1    1.0950e+02    4.1840e+02 ;    HD3 - CD
- CE
  457    460    461      1    1.0950e+02    4.1840e+02 ;     CE - NZ
- HZ1
  457    460    462      1    1.0950e+02    4.1840e+02 ;     CE - NZ
- HZ2
  457    460    463      1    1.0950e+02    4.1840e+02 ;     CE - NZ
- HZ3
  458    457    459      1    1.0950e+02    2.9288e+02 ;    HE2 - CE
- HE3
  458    457    460      1    1.0950e+02    4.1840e+02 ;    HE2 - CE
- NZ
  459    457    460      1    1.0950e+02    4.1840e+02 ;    HE3 - CE
- NZ
  461    460    462      1    1.0950e+02    2.9288e+02 ;    HZ1 - NZ
- HZ2
  461    460    463      1    1.0950e+02    2.9288e+02 ;    HZ1 - NZ
- HZ3
  462    460    463      1    1.0950e+02    2.9288e+02 ;    HZ2 - NZ
- HZ3
  464    466    467      1    1.2000e+02    4.1840e+02 ;      C - N
- H
  464    466    468      1    1.2190e+02    4.1840e+02 ;      C - N
- CA
```

465 464 466 1 1.2290e+02 6.6944e+02 ; O - C
- N
466 468 469 1 1.0950e+02 4.1840e+02 ; N - CA
- HA2
466 468 470 1 1.0950e+02 4.1840e+02 ; N - CA
- HA3
466 468 471 1 1.1010e+02 5.2718e+02 ; N - CA
- C
467 466 468 1 1.1804e+02 4.1840e+02 ; H - N
- CA
468 471 472 1 1.2040e+02 6.6944e+02 ; CA - C
- O
468 471 473 1 1.1660e+02 5.8576e+02 ; CA - C
- N
469 468 470 1 1.0950e+02 2.9288e+02 ; HA2 - CA
- HA3
469 468 471 1 1.0950e+02 4.1840e+02 ; HA2 - CA
- C
470 468 471 1 1.0950e+02 4.1840e+02 ; HA3 - CA
- C
471 473 474 1 1.2000e+02 4.1840e+02 ; C - N
- H
471 473 475 1 1.2190e+02 4.1840e+02 ; C - N
- CA
472 471 473 1 1.2290e+02 6.6944e+02 ; O - C
- N
473 475 476 1 1.0950e+02 4.1840e+02 ; N - CA
- HA
473 475 477 1 1.0970e+02 6.6944e+02 ; N - CA
- CB
473 475 490 1 1.1010e+02 5.2718e+02 ; N - CA
- C
474 473 475 1 1.1804e+02 4.1840e+02 ; H - N
- CA
475 477 478 1 1.0950e+02 4.1840e+02 ; CA - CB
- HB
475 477 479 1 1.0950e+02 3.3472e+02 ; CA - CB
- CG2
475 477 483 1 1.0950e+02 3.3472e+02 ; CA - CB
- CG1
475 490 491 1 1.2040e+02 6.6944e+02 ; CA - C
- O
475 490 492 1 1.1660e+02 5.8576e+02 ; CA - C
- N
476 475 477 1 1.0950e+02 4.1840e+02 ; HA - CA
- CB
476 475 490 1 1.0950e+02 4.1840e+02 ; HA - CA
- C
477 475 490 1 1.1110e+02 5.2718e+02 ; CB - CA
- C
477 479 480 1 1.0950e+02 4.1840e+02 ; CB - CG2
- HG21

477 479 481 1 1.0950e+02 4.1840e+02 ; CB - CG2
- HG22
477 479 482 1 1.0950e+02 4.1840e+02 ; CB - CG2
- HG23
477 483 484 1 1.0950e+02 4.1840e+02 ; CB - CG1
- HG12
477 483 485 1 1.0950e+02 4.1840e+02 ; CB - CG1
- HG13
477 483 486 1 1.0950e+02 3.3472e+02 ; CB - CG1
- CD1
478 477 479 1 1.0950e+02 4.1840e+02 ; HB - CB
- CG2
478 477 483 1 1.0950e+02 4.1840e+02 ; HB - CB
- CG1
479 477 483 1 1.0950e+02 3.3472e+02 ; CG2 - CB
- CG1
480 479 481 1 1.0950e+02 2.9288e+02 ; HG21 - CG2
- HG22
480 479 482 1 1.0950e+02 2.9288e+02 ; HG21 - CG2
- HG23
481 479 482 1 1.0950e+02 2.9288e+02 ; HG22 - CG2
- HG23
483 486 487 1 1.0950e+02 4.1840e+02 ; CG1 - CD1
- HD11
483 486 488 1 1.0950e+02 4.1840e+02 ; CG1 - CD1
- HD12
483 486 489 1 1.0950e+02 4.1840e+02 ; CG1 - CD1
- HD13
484 483 485 1 1.0950e+02 2.9288e+02 ; HG12 - CG1
- HG13
484 483 486 1 1.0950e+02 4.1840e+02 ; HG12 - CG1
- CD1
485 483 486 1 1.0950e+02 4.1840e+02 ; HG13 - CG1
- CD1
487 486 488 1 1.0950e+02 2.9288e+02 ; HD11 - CD1
- HD12
487 486 489 1 1.0950e+02 2.9288e+02 ; HD11 - CD1
- HD13
488 486 489 1 1.0950e+02 2.9288e+02 ; HD12 - CD1
- HD13
490 492 493 1 1.2000e+02 4.1840e+02 ; C - N
- H
490 492 494 1 1.2190e+02 4.1840e+02 ; C - N
- CA
491 490 492 1 1.2290e+02 6.6944e+02 ; O - C
- N
492 494 495 1 1.0950e+02 4.1840e+02 ; N - CA
- HA
492 494 496 1 1.0970e+02 6.6944e+02 ; N - CA
- CB
492 494 502 1 1.1010e+02 5.2718e+02 ; N - CA
- C

493 492 494 1 1.1804e+02 4.1840e+02 ; H - N
- CA
494 496 497 1 1.0950e+02 4.1840e+02 ; CA - CB
- HB2
494 496 498 1 1.0950e+02 4.1840e+02 ; CA - CB
- HB3
494 496 499 1 1.1110e+02 5.2718e+02 ; CA - CB
- CG
494 502 503 1 1.2040e+02 6.6944e+02 ; CA - C
- O
494 502 504 1 1.1660e+02 5.8576e+02 ; CA - C
- N
495 494 496 1 1.0950e+02 4.1840e+02 ; HA - CA
- CB
495 494 502 1 1.0950e+02 4.1840e+02 ; HA - CA
- C
496 494 502 1 1.1110e+02 5.2718e+02 ; CB - CA
- C
496 499 500 1 1.1700e+02 5.8576e+02 ; CB - CG
- OD1
496 499 501 1 1.1700e+02 5.8576e+02 ; CB - CG
- OD2
497 496 498 1 1.0950e+02 2.9288e+02 ; HB2 - CB
- HB3
497 496 499 1 1.0950e+02 4.1840e+02 ; HB2 - CB
- CG
498 496 499 1 1.0950e+02 4.1840e+02 ; HB3 - CB
- CG
500 499 501 1 1.2600e+02 6.6944e+02 ; OD1 - CG
- OD2
502 504 505 1 1.2000e+02 4.1840e+02 ; C - N
- H
502 504 506 1 1.2190e+02 4.1840e+02 ; C - N
- CA
503 502 504 1 1.2290e+02 6.6944e+02 ; O - C
- N
504 506 507 1 1.0950e+02 4.1840e+02 ; N - CA
- HA
504 506 508 1 1.0970e+02 6.6944e+02 ; N - CA
- CB
504 506 524 1 1.1010e+02 5.2718e+02 ; N - CA
- C
505 504 506 1 1.1804e+02 4.1840e+02 ; H - N
- CA
506 508 509 1 1.0950e+02 4.1840e+02 ; CA - CB
- HB2
506 508 510 1 1.0950e+02 4.1840e+02 ; CA - CB
- HB3
506 508 511 1 1.0950e+02 3.3472e+02 ; CA - CB
- CG
506 524 525 1 1.2040e+02 6.6944e+02 ; CA - C
- O

```
   506    524    526      1    1.1660e+02    5.8576e+02 ;      CA - C
- N
   507    506    508      1    1.0950e+02    4.1840e+02 ;      HA - CA
- CB
   507    506    524      1    1.0950e+02    4.1840e+02 ;      HA - CA
- C
   508    506    524      1    1.1110e+02    5.2718e+02 ;      CB - CA
- C
   508    511    512      1    1.0950e+02    4.1840e+02 ;      CB - CG
- HG2
   508    511    513      1    1.0950e+02    4.1840e+02 ;      CB - CG
- HG3
   508    511    514      1    1.0950e+02    3.3472e+02 ;      CB - CG
- CD
   509    508    510      1    1.0950e+02    2.9288e+02 ;     HB2 - CB
- HB3
   509    508    511      1    1.0950e+02    4.1840e+02 ;     HB2 - CB
- CG
   510    508    511      1    1.0950e+02    4.1840e+02 ;     HB3 - CB
- CG
   511    514    515      1    1.0950e+02    4.1840e+02 ;      CG - CD
- HD2
   511    514    516      1    1.0950e+02    4.1840e+02 ;      CG - CD
- HD3
   511    514    517      1    1.0950e+02    3.3472e+02 ;      CG - CD
- CE
   512    511    513      1    1.0950e+02    2.9288e+02 ;     HG2 - CG
- HG3
   512    511    514      1    1.0950e+02    4.1840e+02 ;     HG2 - CG
- CD
   513    511    514      1    1.0950e+02    4.1840e+02 ;     HG3 - CG
- CD
   514    517    518      1    1.0950e+02    4.1840e+02 ;      CD - CE
- HE2
   514    517    519      1    1.0950e+02    4.1840e+02 ;      CD - CE
- HE3
   514    517    520      1    1.1120e+02    6.6944e+02 ;      CD - CE
- NZ
   515    514    516      1    1.0950e+02    2.9288e+02 ;     HD2 - CD
- HD3
   515    514    517      1    1.0950e+02    4.1840e+02 ;     HD2 - CD
- CE
   516    514    517      1    1.0950e+02    4.1840e+02 ;     HD3 - CD
- CE
   517    520    521      1    1.0950e+02    4.1840e+02 ;      CE - NZ
- HZ1
   517    520    522      1    1.0950e+02    4.1840e+02 ;      CE - NZ
- HZ2
   517    520    523      1    1.0950e+02    4.1840e+02 ;      CE - NZ
- HZ3
   518    517    519      1    1.0950e+02    2.9288e+02 ;     HE2 - CE
- HE3
```

```
  518    517    520       1    1.0950e+02    4.1840e+02 ;    HE2 - CE
- NZ
  519    517    520       1    1.0950e+02    4.1840e+02 ;    HE3 - CE
- NZ
  521    520    522       1    1.0950e+02    2.9288e+02 ;    HZ1 - NZ
- HZ2
  521    520    523       1    1.0950e+02    2.9288e+02 ;    HZ1 - NZ
- HZ3
  522    520    523       1    1.0950e+02    2.9288e+02 ;    HZ2 - NZ
- HZ3
  524    526    527       1    1.2000e+02    4.1840e+02 ;      C - N
- H
  524    526    528       1    1.2190e+02    4.1840e+02 ;      C - N
- CA
  525    524    526       1    1.2290e+02    6.6944e+02 ;      O - C
- N
  526    528    529       1    1.0950e+02    4.1840e+02 ;      N - CA
- HA
  526    528    530       1    1.0970e+02    6.6944e+02 ;      N - CA
- CB
  526    528    540       1    1.1010e+02    5.2718e+02 ;      N - CA
- C
  527    526    528       1    1.1804e+02    4.1840e+02 ;      H - N
- CA
  528    530    531       1    1.0950e+02    4.1840e+02 ;     CA - CB
- HB
  528    530    532       1    1.0950e+02    3.3472e+02 ;     CA - CB
- CG1
  528    530    536       1    1.0950e+02    3.3472e+02 ;     CA - CB
- CG2
  528    540    541       1    1.2040e+02    6.6944e+02 ;     CA - C
- O
  528    540    542       1    1.1660e+02    5.8576e+02 ;     CA - C
- N
  529    528    530       1    1.0950e+02    4.1840e+02 ;     HA - CA
- CB
  529    528    540       1    1.0950e+02    4.1840e+02 ;     HA - CA
- C
  530    528    540       1    1.1110e+02    5.2718e+02 ;     CB - CA
- C
  530    532    533       1    1.0950e+02    4.1840e+02 ;     CB - CG1
- HG11
  530    532    534       1    1.0950e+02    4.1840e+02 ;     CB - CG1
- HG12
  530    532    535       1    1.0950e+02    4.1840e+02 ;     CB - CG1
- HG13
  530    536    537       1    1.0950e+02    4.1840e+02 ;     CB - CG2
- HG21
  530    536    538       1    1.0950e+02    4.1840e+02 ;     CB - CG2
- HG22
  530    536    539       1    1.0950e+02    4.1840e+02 ;     CB - CG2
- HG23
```

| | | | | | | | |
|---|---|---|---|---|---|---|---|
| 531 | 530 | 532 | 1 | 1.0950e+02 | 4.1840e+02 | ; | HB - CB - CG1 |
| 531 | 530 | 536 | 1 | 1.0950e+02 | 4.1840e+02 | ; | HB - CB - CG2 |
| 532 | 530 | 536 | 1 | 1.0950e+02 | 3.3472e+02 | ; | CG1 - CB - CG2 |
| 533 | 532 | 534 | 1 | 1.0950e+02 | 2.9288e+02 | ; | HG11 - CG1 - HG12 |
| 533 | 532 | 535 | 1 | 1.0950e+02 | 2.9288e+02 | ; | HG11 - CG1 - HG13 |
| 534 | 532 | 535 | 1 | 1.0950e+02 | 2.9288e+02 | ; | HG12 - CG1 - HG13 |
| 537 | 536 | 538 | 1 | 1.0950e+02 | 2.9288e+02 | ; | HG21 - CG2 - HG22 |
| 537 | 536 | 539 | 1 | 1.0950e+02 | 2.9288e+02 | ; | HG21 - CG2 - HG23 |
| 538 | 536 | 539 | 1 | 1.0950e+02 | 2.9288e+02 | ; | HG22 - CG2 - HG23 |
| 540 | 542 | 543 | 1 | 1.2000e+02 | 4.1840e+02 | ; | C - N - H |
| 540 | 542 | 544 | 1 | 1.2190e+02 | 4.1840e+02 | ; | C - N - CA |
| 541 | 540 | 542 | 1 | 1.2290e+02 | 6.6944e+02 | ; | O - C - N |
| 542 | 544 | 545 | 1 | 1.0950e+02 | 4.1840e+02 | ; | N - CA - HA |
| 542 | 544 | 546 | 1 | 1.0970e+02 | 6.6944e+02 | ; | N - CA - CB |
| 542 | 544 | 554 | 1 | 1.1010e+02 | 5.2718e+02 | ; | N - CA - C |
| 543 | 542 | 544 | 1 | 1.1804e+02 | 4.1840e+02 | ; | H - N - CA |
| 544 | 546 | 547 | 1 | 1.0950e+02 | 4.1840e+02 | ; | CA - CB - HB |
| 544 | 546 | 548 | 1 | 1.0950e+02 | 3.3472e+02 | ; | CA - CB - CG2 |
| 544 | 546 | 552 | 1 | 1.0950e+02 | 4.1840e+02 | ; | CA - CB - OG1 |
| 544 | 554 | 555 | 1 | 1.2040e+02 | 6.6944e+02 | ; | CA - C - O |
| 544 | 554 | 556 | 1 | 1.1660e+02 | 5.8576e+02 | ; | CA - C - N |
| 545 | 544 | 546 | 1 | 1.0950e+02 | 4.1840e+02 | ; | HA - CA - CB |
| 545 | 544 | 554 | 1 | 1.0950e+02 | 4.1840e+02 | ; | HA - CA - C |
| 546 | 544 | 554 | 1 | 1.1110e+02 | 5.2718e+02 | ; | CB - CA - C |
| 546 | 548 | 549 | 1 | 1.0950e+02 | 4.1840e+02 | ; | CB - CG2 - HG21 |
| 546 | 548 | 550 | 1 | 1.0950e+02 | 4.1840e+02 | ; | CB - CG2 - HG22 |

```
   546    548    551      1    1.0950e+02    4.1840e+02 ;     CB - CG2
- HG23
   546    552    553      1    1.0850e+02    4.6024e+02 ;     CB - OG1
- HG1
   547    546    548      1    1.0950e+02    4.1840e+02 ;     HB - CB
- CG2
   547    546    552      1    1.0950e+02    4.1840e+02 ;     HB - CB
- OG1
   548    546    552      1    1.0950e+02    4.1840e+02 ;    CG2 - CB
- OG1
   549    548    550      1    1.0950e+02    2.9288e+02 ;   HG21 - CG2
- HG22
   549    548    551      1    1.0950e+02    2.9288e+02 ;   HG21 - CG2
- HG23
   550    548    551      1    1.0950e+02    2.9288e+02 ;   HG22 - CG2
- HG23
   554    556    557      1    1.2000e+02    4.1840e+02 ;      C - N
- H
   554    556    558      1    1.2190e+02    4.1840e+02 ;      C - N
- CA
   555    554    556      1    1.2290e+02    6.6944e+02 ;      O - C
- N
   556    558    559      1    1.0950e+02    4.1840e+02 ;      N - CA
- HA
   556    558    560      1    1.0970e+02    6.6944e+02 ;      N - CA
- CB
   556    558    568      1    1.1010e+02    5.2718e+02 ;      N - CA
- C
   557    556    558      1    1.1804e+02    4.1840e+02 ;      H - N
- CA
   558    560    561      1    1.0950e+02    4.1840e+02 ;     CA - CB
- HB2
   558    560    562      1    1.0950e+02    4.1840e+02 ;     CA - CB
- HB3
   558    560    563      1    1.1110e+02    5.2718e+02 ;     CA - CB
- CG
   558    568    569      1    1.2040e+02    6.6944e+02 ;     CA - C
- O
   558    568    570      1    1.1660e+02    5.8576e+02 ;     CA - C
- N
   559    558    560      1    1.0950e+02    4.1840e+02 ;     HA - CA
- CB
   559    558    568      1    1.0950e+02    4.1840e+02 ;     HA - CA
- C
   560    558    568      1    1.1110e+02    5.2718e+02 ;     CB - CA
- C
   560    563    564      1    1.2040e+02    6.6944e+02 ;     CB - CG
- OD1
   560    563    565      1    1.1660e+02    5.8576e+02 ;     CB - CG
- ND2
   561    560    562      1    1.0950e+02    2.9288e+02 ;    HB2 - CB
- HB3
```

```
   561     560     563       1     1.0950e+02    4.1840e+02 ;    HB2 - CB
 - CG
   562     560     563       1     1.0950e+02    4.1840e+02 ;    HB3 - CB
 - CG
   563     565     566       1     1.2000e+02    4.1840e+02 ;     CG - ND2
 - HD21
   563     565     567       1     1.2000e+02    4.1840e+02 ;     CG - ND2
 - HD22
   564     563     565       1     1.2290e+02    6.6944e+02 ;    OD1 - CG
 - ND2
   566     565     567       1     1.2000e+02    2.9288e+02 ;   HD21 - ND2
 - HD22
   568     570     571       1     1.2000e+02    4.1840e+02 ;      C - N
 - H
   568     570     572       1     1.2190e+02    4.1840e+02 ;      C - N
 - CA
   569     568     570       1     1.2290e+02    6.6944e+02 ;      O - C
 - N
   570     572     573       1     1.0950e+02    4.1840e+02 ;      N - CA
 - HA
   570     572     574       1     1.0970e+02    6.6944e+02 ;      N - CA
 - CB
   570     572     585       1     1.1010e+02    5.2718e+02 ;      N - CA
 - C
   571     570     572       1     1.1804e+02    4.1840e+02 ;      H - N
 - CA
   572     574     575       1     1.0950e+02    4.1840e+02 ;     CA - CB
 - HB2
   572     574     576       1     1.0950e+02    4.1840e+02 ;     CA - CB
 - HB3
   572     574     577       1     1.0950e+02    3.3472e+02 ;     CA - CB
 - CG
   572     585     586       1     1.2040e+02    6.6944e+02 ;     CA - C
 - O
   572     585     587       1     1.1660e+02    5.8576e+02 ;     CA - C
 - N
   573     572     574       1     1.0950e+02    4.1840e+02 ;     HA - CA
 - CB
   573     572     585       1     1.0950e+02    4.1840e+02 ;     HA - CA
 - C
   574     572     585       1     1.1110e+02    5.2718e+02 ;     CB - CA
 - C
   574     577     578       1     1.0950e+02    4.1840e+02 ;     CB - CG
 - HG2
   574     577     579       1     1.0950e+02    4.1840e+02 ;     CB - CG
 - HG3
   574     577     580       1     1.1110e+02    5.2718e+02 ;     CB - CG
 - CD
   575     574     576       1     1.0950e+02    2.9288e+02 ;    HB2 - CB
 - HB3
   575     574     577       1     1.0950e+02    4.1840e+02 ;    HB2 - CB
 - CG
```

```
   576    574    577      1    1.0950e+02    4.1840e+02 ;    HB3 - CB
 - CG
   577    580    581      1    1.2040e+02    6.6944e+02 ;     CG - CD
 - OE1
   577    580    582      1    1.1660e+02    5.8576e+02 ;     CG - CD
 - NE2
   578    577    579      1    1.0950e+02    2.9288e+02 ;    HG2 - CG
 - HG3
   578    577    580      1    1.0950e+02    4.1840e+02 ;    HG2 - CG
 - CD
   579    577    580      1    1.0950e+02    4.1840e+02 ;    HG3 - CG
 - CD
   580    582    583      1    1.2000e+02    4.1840e+02 ;     CD - NE2
 - HE21
   580    582    584      1    1.2000e+02    4.1840e+02 ;     CD - NE2
 - HE22
   581    580    582      1    1.2290e+02    6.6944e+02 ;    OE1 - CD
 - NE2
   583    582    584      1    1.2000e+02    2.9288e+02 ;   HE21 - NE2
 - HE22
   585    587    588      1    1.2000e+02    4.1840e+02 ;      C - N
 - H
   585    587    589      1    1.2190e+02    4.1840e+02 ;      C - N
 - CA
   586    585    587      1    1.2290e+02    6.6944e+02 ;      O - C
 - N
   587    589    590      1    1.0950e+02    4.1840e+02 ;      N - CA
 - HA
   587    589    591      1    1.0970e+02    6.6944e+02 ;      N - CA
 - CB
   587    589    604      1    1.1010e+02    5.2718e+02 ;      N - CA
 - C
   588    587    589      1    1.1804e+02    4.1840e+02 ;      H - N
 - CA
   589    591    592      1    1.0950e+02    4.1840e+02 ;     CA - CB
 - HB2
   589    591    593      1    1.0950e+02    4.1840e+02 ;     CA - CB
 - HB3
   589    591    594      1    1.0950e+02    3.3472e+02 ;     CA - CB
 - CG
   589    604    605      1    1.2040e+02    6.6944e+02 ;     CA - C
 - O
   589    604    606      1    1.1660e+02    5.8576e+02 ;     CA - C
 - N
   590    589    591      1    1.0950e+02    4.1840e+02 ;     HA - CA
 - CB
   590    589    604      1    1.0950e+02    4.1840e+02 ;     HA - CA
 - C
   591    589    604      1    1.1110e+02    5.2718e+02 ;     CB - CA
 - C
   591    594    595      1    1.0950e+02    4.1840e+02 ;     CB - CG
 - HG
```

591 594 596 1 1.0950e+02 3.3472e+02 ; CB - CG
- CD1
591 594 600 1 1.0950e+02 3.3472e+02 ; CB - CG
- CD2
592 591 593 1 1.0950e+02 2.9288e+02 ; HB2 - CB
- HB3
592 591 594 1 1.0950e+02 4.1840e+02 ; HB2 - CB
- CG
593 591 594 1 1.0950e+02 4.1840e+02 ; HB3 - CB
- CG
594 596 597 1 1.0950e+02 4.1840e+02 ; CG - CD1
- HD11
594 596 598 1 1.0950e+02 4.1840e+02 ; CG - CD1
- HD12
594 596 599 1 1.0950e+02 4.1840e+02 ; CG - CD1
- HD13
594 600 601 1 1.0950e+02 4.1840e+02 ; CG - CD2
- HD21
594 600 602 1 1.0950e+02 4.1840e+02 ; CG - CD2
- HD22
594 600 603 1 1.0950e+02 4.1840e+02 ; CG - CD2
- HD23
595 594 596 1 1.0950e+02 4.1840e+02 ; HG - CG
- CD1
595 594 600 1 1.0950e+02 4.1840e+02 ; HG - CG
- CD2
596 594 600 1 1.0950e+02 3.3472e+02 ; CD1 - CG
- CD2
597 596 598 1 1.0950e+02 2.9288e+02 ; HD11 - CD1
- HD12
597 596 599 1 1.0950e+02 2.9288e+02 ; HD11 - CD1
- HD13
598 596 599 1 1.0950e+02 2.9288e+02 ; HD12 - CD1
- HD13
601 600 602 1 1.0950e+02 2.9288e+02 ; HD21 - CD2
- HD22
601 600 603 1 1.0950e+02 2.9288e+02 ; HD21 - CD2
- HD23
602 600 603 1 1.0950e+02 2.9288e+02 ; HD22 - CD2
- HD23
604 606 607 1 1.2000e+02 4.1840e+02 ; C - N
- H
604 606 608 1 1.2190e+02 4.1840e+02 ; C - N
- CA
605 604 606 1 1.2290e+02 6.6944e+02 ; O - C
- N
606 608 609 1 1.0950e+02 4.1840e+02 ; N - CA
- HA
606 608 610 1 1.0970e+02 6.6944e+02 ; N - CA
- CB
606 608 625 1 1.1010e+02 5.2718e+02 ; N - CA
- C

607 606 608 1 1.1804e+02 4.1840e+02 ; H - N
- CA
608 610 611 1 1.0950e+02 4.1840e+02 ; CA - CB
- HB2
608 610 612 1 1.0950e+02 4.1840e+02 ; CA - CB
- HB3
608 610 613 1 1.1400e+02 5.2718e+02 ; CA - CB
- CG
608 625 626 1 1.2040e+02 6.6944e+02 ; CA - C
- O
608 625 627 1 1.1660e+02 5.8576e+02 ; CA - C
- N
609 608 610 1 1.0950e+02 4.1840e+02 ; HA - CA
- CB
609 608 625 1 1.0950e+02 4.1840e+02 ; HA - CA
- C
610 608 625 1 1.1110e+02 5.2718e+02 ; CB - CA
- C
610 613 614 1 1.2000e+02 5.8576e+02 ; CB - CG
- CD1
610 613 623 1 1.2000e+02 5.8576e+02 ; CB - CG
- CD2
611 610 612 1 1.0950e+02 2.9288e+02 ; HB2 - CB
- HB3
611 610 613 1 1.0950e+02 4.1840e+02 ; HB2 - CB
- CG
612 610 613 1 1.0950e+02 4.1840e+02 ; HB3 - CB
- CG
613 614 615 1 1.2000e+02 4.1840e+02 ; CG - CD1
- HD1
613 614 616 1 1.2000e+02 5.2718e+02 ; CG - CD1
- CE1
613 623 621 1 1.2000e+02 5.2718e+02 ; CG - CD2
- CE2
613 623 624 1 1.2000e+02 4.1840e+02 ; CG - CD2
- HD2
614 613 623 1 1.2000e+02 5.2718e+02 ; CD1 - CG
- CD2
614 616 617 1 1.2000e+02 4.1840e+02 ; CD1 - CE1
- HE1
614 616 618 1 1.2000e+02 5.2718e+02 ; CD1 - CE1
- CZ
615 614 616 1 1.2000e+02 4.1840e+02 ; HD1 - CD1
- CE1
616 618 619 1 1.2000e+02 5.8576e+02 ; CE1 - CZ
- OH
616 618 621 1 1.2000e+02 5.2718e+02 ; CE1 - CZ
- CE2
617 616 618 1 1.2000e+02 4.1840e+02 ; HE1 - CE1
- CZ
618 619 620 1 1.1300e+02 4.1840e+02 ; CZ - OH
- HH

```
   618    621    622      1    1.2000e+02    4.1840e+02 ;     CZ - CE2
- HE2
   618    621    623      1    1.2000e+02    5.2718e+02 ;     CZ - CE2
- CD2
   619    618    621      1    1.2000e+02    5.8576e+02 ;     OH - CZ
- CE2
   621    623    624      1    1.2000e+02    4.1840e+02 ;    CE2 - CD2
- HD2
   622    621    623      1    1.2000e+02    4.1840e+02 ;    HE2 - CE2
- CD2
   625    627    628      1    1.2000e+02    4.1840e+02 ;      C - N
- H
   625    627    629      1    1.2190e+02    4.1840e+02 ;      C - N
- CA
   626    625    627      1    1.2290e+02    6.6944e+02 ;      O - C
- N
   627    629    630      1    1.0950e+02    4.1840e+02 ;      N - CA
- HA
   627    629    631      1    1.0970e+02    6.6944e+02 ;      N - CA
- CB
   627    629    635      1    1.1010e+02    5.2718e+02 ;      N - CA
- C
   628    627    629      1    1.1804e+02    4.1840e+02 ;      H - N
- CA
   629    631    632      1    1.0950e+02    4.1840e+02 ;     CA - CB
- HB1
   629    631    633      1    1.0950e+02    4.1840e+02 ;     CA - CB
- HB2
   629    631    634      1    1.0950e+02    4.1840e+02 ;     CA - CB
- HB3
   629    635    636      1    1.2040e+02    6.6944e+02 ;     CA - C
- O
   629    635    637      1    1.1660e+02    5.8576e+02 ;     CA - C
- N
   630    629    631      1    1.0950e+02    4.1840e+02 ;     HA - CA
- CB
   630    629    635      1    1.0950e+02    4.1840e+02 ;     HA - CA
- C
   631    629    635      1    1.1110e+02    5.2718e+02 ;     CB - CA
- C
   632    631    633      1    1.0950e+02    2.9288e+02 ;    HB1 - CB
- HB2
   632    631    634      1    1.0950e+02    2.9288e+02 ;    HB1 - CB
- HB3
   633    631    634      1    1.0950e+02    2.9288e+02 ;    HB2 - CB
- HB3
   635    637    638      1    1.2000e+02    4.1840e+02 ;      C - N
- H
   635    637    639      1    1.2190e+02    4.1840e+02 ;      C - N
- CA
   636    635    637      1    1.2290e+02    6.6944e+02 ;      O - C
- N
```

637 639 640 1 1.0950e+02 4.1840e+02 ; N - CA
- HA
637 639 641 1 1.0970e+02 6.6944e+02 ; N - CA
- CB
637 639 654 1 1.1010e+02 5.2718e+02 ; N - CA
- C
638 637 639 1 1.1804e+02 4.1840e+02 ; H - N
- CA
639 641 642 1 1.0950e+02 4.1840e+02 ; CA - CB
- HB
639 641 643 1 1.0950e+02 3.3472e+02 ; CA - CB
- CG2
639 641 647 1 1.0950e+02 3.3472e+02 ; CA - CB
- CG1
639 654 655 1 1.2040e+02 6.6944e+02 ; CA - C
- O
639 654 656 1 1.1660e+02 5.8576e+02 ; CA - C
- N
640 639 641 1 1.0950e+02 4.1840e+02 ; HA - CA
- CB
640 639 654 1 1.0950e+02 4.1840e+02 ; HA - CA
- C
641 639 654 1 1.1110e+02 5.2718e+02 ; CB - CA
- C
641 643 644 1 1.0950e+02 4.1840e+02 ; CB - CG2
- HG21
641 643 645 1 1.0950e+02 4.1840e+02 ; CB - CG2
- HG22
641 643 646 1 1.0950e+02 4.1840e+02 ; CB - CG2
- HG23
641 647 648 1 1.0950e+02 4.1840e+02 ; CB - CG1
- HG12
641 647 649 1 1.0950e+02 4.1840e+02 ; CB - CG1
- HG13
641 647 650 1 1.0950e+02 3.3472e+02 ; CB - CG1
- CD1
642 641 643 1 1.0950e+02 4.1840e+02 ; HB - CB
- CG2
642 641 647 1 1.0950e+02 4.1840e+02 ; HB - CB
- CG1
643 641 647 1 1.0950e+02 3.3472e+02 ; CG2 - CB
- CG1
644 643 645 1 1.0950e+02 2.9288e+02 ; HG21 - CG2
- HG22
644 643 646 1 1.0950e+02 2.9288e+02 ; HG21 - CG2
- HG23
645 643 646 1 1.0950e+02 2.9288e+02 ; HG22 - CG2
- HG23
647 650 651 1 1.0950e+02 4.1840e+02 ; CG1 - CD1
- HD11
647 650 652 1 1.0950e+02 4.1840e+02 ; CG1 - CD1
- HD12

647 650 653 1 1.0950e+02 4.1840e+02 ; CG1 - CD1
- HD13
648 647 649 1 1.0950e+02 2.9288e+02 ; HG12 - CG1
- HG13
648 647 650 1 1.0950e+02 4.1840e+02 ; HG12 - CG1
- CD1
649 647 650 1 1.0950e+02 4.1840e+02 ; HG13 - CG1
- CD1
651 650 652 1 1.0950e+02 2.9288e+02 ; HD11 - CD1
- HD12
651 650 653 1 1.0950e+02 2.9288e+02 ; HD11 - CD1
- HD13
652 650 653 1 1.0950e+02 2.9288e+02 ; HD12 - CD1
- HD13
654 656 657 1 1.2000e+02 4.1840e+02 ; C - N
- H
654 656 658 1 1.2190e+02 4.1840e+02 ; C - N
- CA
655 654 656 1 1.2290e+02 6.6944e+02 ; O - C
- N
656 658 659 1 1.0950e+02 4.1840e+02 ; N - CA
- HA
656 658 660 1 1.0970e+02 6.6944e+02 ; N - CA
- CB
656 658 676 1 1.1010e+02 5.2718e+02 ; N - CA
- C
657 656 658 1 1.1804e+02 4.1840e+02 ; H - N
- CA
658 660 661 1 1.0950e+02 4.1840e+02 ; CA - CB
- HB2
658 660 662 1 1.0950e+02 4.1840e+02 ; CA - CB
- HB3
658 660 663 1 1.0950e+02 3.3472e+02 ; CA - CB
- CG
658 676 677 1 1.2040e+02 6.6944e+02 ; CA - C
- O
658 676 678 1 1.1660e+02 5.8576e+02 ; CA - C
- N
659 658 660 1 1.0950e+02 4.1840e+02 ; HA - CA
- CB
659 658 676 1 1.0950e+02 4.1840e+02 ; HA - CA
- C
660 658 676 1 1.1110e+02 5.2718e+02 ; CB - CA
- C
660 663 664 1 1.0950e+02 4.1840e+02 ; CB - CG
- HG2
660 663 665 1 1.0950e+02 4.1840e+02 ; CB - CG
- HG3
660 663 666 1 1.0950e+02 3.3472e+02 ; CB - CG
- CD
661 660 662 1 1.0950e+02 2.9288e+02 ; HB2 - CB
- HB3

661 660 663 1 1.0950e+02 4.1840e+02 ; HB2 - CB
- CG
662 660 663 1 1.0950e+02 4.1840e+02 ; HB3 - CB
- CG
663 666 667 1 1.0950e+02 4.1840e+02 ; CG - CD
- HD2
663 666 668 1 1.0950e+02 4.1840e+02 ; CG - CD
- HD3
663 666 669 1 1.0950e+02 3.3472e+02 ; CG - CD
- CE
664 663 665 1 1.0950e+02 2.9288e+02 ; HG2 - CG
- HG3
664 663 666 1 1.0950e+02 4.1840e+02 ; HG2 - CG
- CD
665 663 666 1 1.0950e+02 4.1840e+02 ; HG3 - CG
- CD
666 669 670 1 1.0950e+02 4.1840e+02 ; CD - CE
- HE2
666 669 671 1 1.0950e+02 4.1840e+02 ; CD - CE
- HE3
666 669 672 1 1.1120e+02 6.6944e+02 ; CD - CE
- NZ
667 666 668 1 1.0950e+02 2.9288e+02 ; HD2 - CD
- HD3
667 666 669 1 1.0950e+02 4.1840e+02 ; HD2 - CD
- CE
668 666 669 1 1.0950e+02 4.1840e+02 ; HD3 - CD
- CE
669 672 673 1 1.0950e+02 4.1840e+02 ; CE - NZ
- HZ1
669 672 674 1 1.0950e+02 4.1840e+02 ; CE - NZ
- HZ2
669 672 675 1 1.0950e+02 4.1840e+02 ; CE - NZ
- HZ3
670 669 671 1 1.0950e+02 2.9288e+02 ; HE2 - CE
- HE3
670 669 672 1 1.0950e+02 4.1840e+02 ; HE2 - CE
- NZ
671 669 672 1 1.0950e+02 4.1840e+02 ; HE3 - CE
- NZ
673 672 674 1 1.0950e+02 2.9288e+02 ; HZ1 - NZ
- HZ2
673 672 675 1 1.0950e+02 2.9288e+02 ; HZ1 - NZ
- HZ3
674 672 675 1 1.0950e+02 2.9288e+02 ; HZ2 - NZ
- HZ3
676 678 679 1 1.2000e+02 4.1840e+02 ; C - N
- H
676 678 680 1 1.2190e+02 4.1840e+02 ; C - N
- CA
677 676 678 1 1.2290e+02 6.6944e+02 ; O - C
- N

```
  678    680    681      1    1.0950e+02    4.1840e+02 ;      N - CA
- HA
  678    680    682      1    1.0970e+02    6.6944e+02 ;      N - CA
- CB
  678    680    691      1    1.1010e+02    5.2718e+02 ;      N - CA
- C
  679    678    680      1    1.1804e+02    4.1840e+02 ;      H - N
- CA
  680    682    683      1    1.0950e+02    4.1840e+02 ;     CA - CB
- HB2
  680    682    684      1    1.0950e+02    4.1840e+02 ;     CA - CB
- HB3
  680    682    685      1    1.0950e+02    3.3472e+02 ;     CA - CB
- CG
  680    691    692      1    1.2040e+02    6.6944e+02 ;     CA - C
- O
  680    691    693      1    1.1660e+02    5.8576e+02 ;     CA - C
- N
  681    680    682      1    1.0950e+02    4.1840e+02 ;     HA - CA
- CB
  681    680    691      1    1.0950e+02    4.1840e+02 ;     HA - CA
- C
  682    680    691      1    1.1110e+02    5.2718e+02 ;     CB - CA
- C
  682    685    686      1    1.0950e+02    4.1840e+02 ;     CB - CG
- HG2
  682    685    687      1    1.0950e+02    4.1840e+02 ;     CB - CG
- HG3
  682    685    688      1    1.1110e+02    5.2718e+02 ;     CB - CG
- CD
  683    682    684      1    1.0950e+02    2.9288e+02 ;    HB2 - CB
- HB3
  683    682    685      1    1.0950e+02    4.1840e+02 ;    HB2 - CB
- CG
  684    682    685      1    1.0950e+02    4.1840e+02 ;    HB3 - CB
- CG
  685    688    689      1    1.1700e+02    5.8576e+02 ;     CG - CD
- OE1
  685    688    690      1    1.1700e+02    5.8576e+02 ;     CG - CD
- OE2
  686    685    687      1    1.0950e+02    2.9288e+02 ;    HG2 - CG
- HG3
  686    685    688      1    1.0950e+02    4.1840e+02 ;    HG2 - CG
- CD
  687    685    688      1    1.0950e+02    4.1840e+02 ;    HG3 - CG
- CD
  689    688    690      1    1.2600e+02    6.6944e+02 ;    OE1 - CD
- OE2
  691    693    694      1    1.2000e+02    4.1840e+02 ;      C - N
- H
  691    693    695      1    1.2190e+02    4.1840e+02 ;      C - N
- CA
```

```
   692    691    693      1    1.2290e+02    6.6944e+02 ;      O - C
- N
   693    695    696      1    1.0950e+02    4.1840e+02 ;      N - CA
- HA
   693    695    697      1    1.0970e+02    6.6944e+02 ;      N - CA
- CB
   693    695    707      1    1.1010e+02    5.2718e+02 ;      N - CA
- C
   694    693    695      1    1.1804e+02    4.1840e+02 ;      H - N
- CA
   695    697    698      1    1.0950e+02    4.1840e+02 ;     CA - CB
- HB2
   695    697    699      1    1.0950e+02    4.1840e+02 ;     CA - CB
- HB3
   695    697    700      1    1.0950e+02    3.3472e+02 ;     CA - CB
- CG
   695    707    708      1    1.2040e+02    6.6944e+02 ;     CA - C
- O
   695    707    709      1    1.1660e+02    5.8576e+02 ;     CA - C
- N
   696    695    697      1    1.0950e+02    4.1840e+02 ;     HA - CA
- CB
   696    695    707      1    1.0950e+02    4.1840e+02 ;     HA - CA
- C
   697    695    707      1    1.1110e+02    5.2718e+02 ;     CB - CA
- C
   697    700    701      1    1.0950e+02    4.1840e+02 ;     CB - CG
- HG2
   697    700    702      1    1.0950e+02    4.1840e+02 ;     CB - CG
- HG3
   697    700    703      1    1.1110e+02    5.2718e+02 ;     CB - CG
- CD
   698    697    699      1    1.0950e+02    2.9288e+02 ;    HB2 - CB
- HB3
   698    697    700      1    1.0950e+02    4.1840e+02 ;    HB2 - CB
- CG
   699    697    700      1    1.0950e+02    4.1840e+02 ;    HB3 - CB
- CG
   700    703    704      1    1.2040e+02    6.6944e+02 ;     CG - CD
- OE1
   700    703    705      1    1.1000e+02    6.6944e+02 ;     CG - CD
- OE2
   701    700    702      1    1.0950e+02    2.9288e+02 ;    HG2 - CG
- HG3
   701    700    703      1    1.0950e+02    4.1840e+02 ;    HG2 - CG
- CD
   702    700    703      1    1.0950e+02    4.1840e+02 ;    HG3 - CG
- CD
   703    705    706      1    1.1300e+02    4.1840e+02 ;     CD - OE2
- HE2
   704    703    705      1    1.2000e+02    6.6944e+02 ;    OE1 - CD
- OE2
```

```
   707    709    710       1    1.2000e+02    4.1840e+02 ;      C - N
 - H
   707    709    711       1    1.2190e+02    4.1840e+02 ;      C - N
 - CA
   708    707    709       1    1.2290e+02    6.6944e+02 ;      O - C
 - N
   709    711    712       1    1.0950e+02    4.1840e+02 ;      N - CA
 - HA
   709    711    713       1    1.0970e+02    6.6944e+02 ;      N - CA
 - CB
   709    711    729       1    1.1010e+02    5.2718e+02 ;      N - CA
 - C
   710    709    711       1    1.1804e+02    4.1840e+02 ;      H - N
 - CA
   711    713    714       1    1.0950e+02    4.1840e+02 ;     CA - CB
 - HB2
   711    713    715       1    1.0950e+02    4.1840e+02 ;     CA - CB
 - HB3
   711    713    716       1    1.0950e+02    3.3472e+02 ;     CA - CB
 - CG
   711    729    730       1    1.2040e+02    6.6944e+02 ;     CA - C
 - O
   711    729    731       1    1.1660e+02    5.8576e+02 ;     CA - C
 - N
   712    711    713       1    1.0950e+02    4.1840e+02 ;     HA - CA
 - CB
   712    711    729       1    1.0950e+02    4.1840e+02 ;     HA - CA
 - C
   713    711    729       1    1.1110e+02    5.2718e+02 ;     CB - CA
 - C
   713    716    717       1    1.0950e+02    4.1840e+02 ;     CB - CG
 - HG2
   713    716    718       1    1.0950e+02    4.1840e+02 ;     CB - CG
 - HG3
   713    716    719       1    1.0950e+02    3.3472e+02 ;     CB - CG
 - CD
   714    713    715       1    1.0950e+02    2.9288e+02 ;    HB2 - CB
 - HB3
   714    713    716       1    1.0950e+02    4.1840e+02 ;    HB2 - CB
 - CG
   715    713    716       1    1.0950e+02    4.1840e+02 ;    HB3 - CB
 - CG
   716    719    720       1    1.0950e+02    4.1840e+02 ;     CG - CD
 - HD2
   716    719    721       1    1.0950e+02    4.1840e+02 ;     CG - CD
 - HD3
   716    719    722       1    1.0950e+02    3.3472e+02 ;     CG - CD
 - CE
   717    716    718       1    1.0950e+02    2.9288e+02 ;    HG2 - CG
 - HG3
   717    716    719       1    1.0950e+02    4.1840e+02 ;    HG2 - CG
 - CD
```

```
   718    716    719      1    1.0950e+02    4.1840e+02 ;    HG3 - CG
 - CD
   719    722    723      1    1.0950e+02    4.1840e+02 ;     CD - CE
 - HE2
   719    722    724      1    1.0950e+02    4.1840e+02 ;     CD - CE
 - HE3
   719    722    725      1    1.1120e+02    6.6944e+02 ;     CD - CE
 - NZ
   720    719    721      1    1.0950e+02    2.9288e+02 ;    HD2 - CD
 - HD3
   720    719    722      1    1.0950e+02    4.1840e+02 ;    HD2 - CD
 - CE
   721    719    722      1    1.0950e+02    4.1840e+02 ;    HD3 - CD
 - CE
   722    725    726      1    1.0950e+02    4.1840e+02 ;     CE - NZ
 - HZ1
   722    725    727      1    1.0950e+02    4.1840e+02 ;     CE - NZ
 - HZ2
   722    725    728      1    1.0950e+02    4.1840e+02 ;     CE - NZ
 - HZ3
   723    722    724      1    1.0950e+02    2.9288e+02 ;    HE2 - CE
 - HE3
   723    722    725      1    1.0950e+02    4.1840e+02 ;    HE2 - CE
 - NZ
   724    722    725      1    1.0950e+02    4.1840e+02 ;    HE3 - CE
 - NZ
   726    725    727      1    1.0950e+02    2.9288e+02 ;    HZ1 - NZ
 - HZ2
   726    725    728      1    1.0950e+02    2.9288e+02 ;    HZ1 - NZ
 - HZ3
   727    725    728      1    1.0950e+02    2.9288e+02 ;    HZ2 - NZ
 - HZ3
   729    731    732      1    1.2000e+02    4.1840e+02 ;      C - N
 - H
   729    731    733      1    1.2190e+02    4.1840e+02 ;      C - N
 - CA
   730    729    731      1    1.2290e+02    6.6944e+02 ;      O - C
 - N
   731    733    734      1    1.0950e+02    4.1840e+02 ;      N - CA
 - HA
   731    733    735      1    1.0970e+02    6.6944e+02 ;      N - CA
 - CB
   731    733    751      1    1.1010e+02    5.2718e+02 ;      N - CA
 - C
   732    731    733      1    1.1804e+02    4.1840e+02 ;      H - N
 - CA
   733    735    736      1    1.0950e+02    4.1840e+02 ;     CA - CB
 - HB2
   733    735    737      1    1.0950e+02    4.1840e+02 ;     CA - CB
 - HB3
   733    735    738      1    1.0950e+02    3.3472e+02 ;     CA - CB
 - CG
```

```
   733    751    752       1    1.2040e+02    6.6944e+02 ;      CA - C
- O
   733    751    753       1    1.1660e+02    5.8576e+02 ;      CA - C
- N
   734    733    735       1    1.0950e+02    4.1840e+02 ;      HA - CA
- CB
   734    733    751       1    1.0950e+02    4.1840e+02 ;      HA - CA
- C
   735    733    751       1    1.1110e+02    5.2718e+02 ;      CB - CA
- C
   735    738    739       1    1.0950e+02    4.1840e+02 ;      CB - CG
- HG2
   735    738    740       1    1.0950e+02    4.1840e+02 ;      CB - CG
- HG3
   735    738    741       1    1.0950e+02    3.3472e+02 ;      CB - CG
- CD
   736    735    737       1    1.0950e+02    2.9288e+02 ;     HB2 - CB
- HB3
   736    735    738       1    1.0950e+02    4.1840e+02 ;     HB2 - CB
- CG
   737    735    738       1    1.0950e+02    4.1840e+02 ;     HB3 - CB
- CG
   738    741    742       1    1.0950e+02    4.1840e+02 ;      CG - CD
- HD2
   738    741    743       1    1.0950e+02    4.1840e+02 ;      CG - CD
- HD3
   738    741    744       1    1.0950e+02    3.3472e+02 ;      CG - CD
- CE
   739    738    740       1    1.0950e+02    2.9288e+02 ;     HG2 - CG
- HG3
   739    738    741       1    1.0950e+02    4.1840e+02 ;     HG2 - CG
- CD
   740    738    741       1    1.0950e+02    4.1840e+02 ;     HG3 - CG
- CD
   741    744    745       1    1.0950e+02    4.1840e+02 ;      CD - CE
- HE2
   741    744    746       1    1.0950e+02    4.1840e+02 ;      CD - CE
- HE3
   741    744    747       1    1.1120e+02    6.6944e+02 ;      CD - CE
- NZ
   742    741    743       1    1.0950e+02    2.9288e+02 ;     HD2 - CD
- HD3
   742    741    744       1    1.0950e+02    4.1840e+02 ;     HD2 - CD
- CE
   743    741    744       1    1.0950e+02    4.1840e+02 ;     HD3 - CD
- CE
   744    747    748       1    1.0950e+02    4.1840e+02 ;      CE - NZ
- HZ1
   744    747    749       1    1.0950e+02    4.1840e+02 ;      CE - NZ
- HZ2
   744    747    750       1    1.0950e+02    4.1840e+02 ;      CE - NZ
- HZ3
```

745 744 746 1 1.0950e+02 2.9288e+02 ; HE2 - CE
- HE3
745 744 747 1 1.0950e+02 4.1840e+02 ; HE2 - CE
- NZ
746 744 747 1 1.0950e+02 4.1840e+02 ; HE3 - CE
- NZ
748 747 749 1 1.0950e+02 2.9288e+02 ; HZ1 - NZ
- HZ2
748 747 750 1 1.0950e+02 2.9288e+02 ; HZ1 - NZ
- HZ3
749 747 750 1 1.0950e+02 2.9288e+02 ; HZ2 - NZ
- HZ3
751 753 754 1 1.2000e+02 4.1840e+02 ; C - N
- H
751 753 755 1 1.2190e+02 4.1840e+02 ; C - N
- CA
752 751 753 1 1.2290e+02 6.6944e+02 ; O - C
- N
753 755 756 1 1.0950e+02 4.1840e+02 ; N - CA
- HA
753 755 757 1 1.0970e+02 6.6944e+02 ; N - CA
- CB
753 755 763 1 1.1010e+02 5.2718e+02 ; N - CA
- C
754 753 755 1 1.1804e+02 4.1840e+02 ; H - N
- CA
755 757 758 1 1.0950e+02 4.1840e+02 ; CA - CB
- HB2
755 757 759 1 1.0950e+02 4.1840e+02 ; CA - CB
- HB3
755 757 760 1 1.1110e+02 5.2718e+02 ; CA - CB
- CG
755 763 764 1 1.2040e+02 6.6944e+02 ; CA - C
- O
755 763 765 1 1.1660e+02 5.8576e+02 ; CA - C
- N
756 755 757 1 1.0950e+02 4.1840e+02 ; HA - CA
- CB
756 755 763 1 1.0950e+02 4.1840e+02 ; HA - CA
- C
757 755 763 1 1.1110e+02 5.2718e+02 ; CB - CA
- C
757 760 761 1 1.1700e+02 5.8576e+02 ; CB - CG
- OD1
757 760 762 1 1.1700e+02 5.8576e+02 ; CB - CG
- OD2
758 757 759 1 1.0950e+02 2.9288e+02 ; HB2 - CB
- HB3
758 757 760 1 1.0950e+02 4.1840e+02 ; HB2 - CB
- CG
759 757 760 1 1.0950e+02 4.1840e+02 ; HB3 - CB
- CG

```
   761     760     762       1     1.2600e+02     6.6944e+02 ;    OD1 - CG
- OD2
   763     765     766       1     1.2000e+02     4.1840e+02 ;      C - N
- H
   763     765     767       1     1.2190e+02     4.1840e+02 ;      C - N
- CA
   764     763     765       1     1.2290e+02     6.6944e+02 ;      O - C
- N
   765     767     768       1     1.0950e+02     4.1840e+02 ;      N - CA
- HA
   765     767     769       1     1.0970e+02     6.6944e+02 ;      N - CA
- CB
   765     767     787       1     1.1010e+02     5.2718e+02 ;      N - CA
- C
   766     765     767       1     1.1804e+02     4.1840e+02 ;      H - N
- CA
   767     769     770       1     1.0950e+02     4.1840e+02 ;     CA - CB
- HB2
   767     769     771       1     1.0950e+02     4.1840e+02 ;     CA - CB
- HB3
   767     769     772       1     1.0950e+02     3.3472e+02 ;     CA - CB
- CG
   767     787     788       1     1.2040e+02     6.6944e+02 ;     CA - C
- O
   767     787     789       1     1.1660e+02     5.8576e+02 ;     CA - C
- N
   768     767     769       1     1.0950e+02     4.1840e+02 ;     HA - CA
- CB
   768     767     787       1     1.0950e+02     4.1840e+02 ;     HA - CA
- C
   769     767     787       1     1.1110e+02     5.2718e+02 ;     CB - CA
- C
   769     772     773       1     1.0950e+02     4.1840e+02 ;     CB - CG
- HG2
   769     772     774       1     1.0950e+02     4.1840e+02 ;     CB - CG
- HG3
   769     772     775       1     1.0950e+02     3.3472e+02 ;     CB - CG
- CD
   770     769     771       1     1.0950e+02     2.9288e+02 ;    HB2 - CB
- HB3
   770     769     772       1     1.0950e+02     4.1840e+02 ;    HB2 - CB
- CG
   771     769     772       1     1.0950e+02     4.1840e+02 ;    HB3 - CB
- CG
   772     775     776       1     1.0950e+02     4.1840e+02 ;     CG - CD
- HD2
   772     775     777       1     1.0950e+02     4.1840e+02 ;     CG - CD
- HD3
   772     775     778       1     1.1120e+02     6.6944e+02 ;     CG - CD
- NE
   773     772     774       1     1.0950e+02     2.9288e+02 ;    HG2 - CG
- HG3
```

```
   773    772    775      1    1.0950e+02    4.1840e+02 ;    HG2 - CG
- CD
   774    772    775      1    1.0950e+02    4.1840e+02 ;    HG3 - CG
- CD
   775    778    779      1    1.1840e+02    4.1840e+02 ;     CD - NE
- HE
   775    778    780      1    1.2320e+02    4.1840e+02 ;     CD - NE
- CZ
   776    775    777      1    1.0950e+02    2.9288e+02 ;    HD2 - CD
- HD3
   776    775    778      1    1.0950e+02    4.1840e+02 ;    HD2 - CD
- NE
   777    775    778      1    1.0950e+02    4.1840e+02 ;    HD3 - CD
- NE
   778    780    781      1    1.2000e+02    5.8576e+02 ;     NE - CZ
- NH1
   778    780    784      1    1.2000e+02    5.8576e+02 ;     NE - CZ
- NH2
   779    778    780      1    1.2000e+02    4.1840e+02 ;     HE - NE
- CZ
   780    781    782      1    1.2000e+02    4.1840e+02 ;     CZ - NH1
- HH11
   780    781    783      1    1.2000e+02    4.1840e+02 ;     CZ - NH1
- HH12
   780    784    785      1    1.2000e+02    4.1840e+02 ;     CZ - NH2
- HH21
   780    784    786      1    1.2000e+02    4.1840e+02 ;     CZ - NH2
- HH22
   781    780    784      1    1.2000e+02    5.8576e+02 ;    NH1 - CZ
- NH2
   782    781    783      1    1.2000e+02    2.9288e+02 ;   HH11 - NH1
- HH12
   785    784    786      1    1.2000e+02    2.9288e+02 ;   HH21 - NH2
- HH22
   787    789    790      1    1.2000e+02    4.1840e+02 ;      C - N
- H
   787    789    791      1    1.2190e+02    4.1840e+02 ;      C - N
- CA
   788    787    789      1    1.2290e+02    6.6944e+02 ;      O - C
- N
   789    791    792      1    1.0950e+02    4.1840e+02 ;      N - CA
- HA
   789    791    793      1    1.0970e+02    6.6944e+02 ;      N - CA
- CB
   789    791    806      1    1.1010e+02    5.2718e+02 ;      N - CA
- C
   790    789    791      1    1.1804e+02    4.1840e+02 ;      H - N
- CA
   791    793    794      1    1.0950e+02    4.1840e+02 ;     CA - CB
- HB2
   791    793    795      1    1.0950e+02    4.1840e+02 ;     CA - CB
- HB3
```

791 793 796 1 1.0950e+02 3.3472e+02 ; CA - CB
- CG
791 806 807 1 1.2040e+02 6.6944e+02 ; CA - C
- O
791 806 808 1 1.1660e+02 5.8576e+02 ; CA - C
- N
792 791 793 1 1.0950e+02 4.1840e+02 ; HA - CA
- CB
792 791 806 1 1.0950e+02 4.1840e+02 ; HA - CA
- C
793 791 806 1 1.1110e+02 5.2718e+02 ; CB - CA
- C
793 796 797 1 1.0950e+02 4.1840e+02 ; CB - CG
- HG
793 796 798 1 1.0950e+02 3.3472e+02 ; CB - CG
- CD1
793 796 802 1 1.0950e+02 3.3472e+02 ; CB - CG
- CD2
794 793 795 1 1.0950e+02 2.9288e+02 ; HB2 - CB
- HB3
794 793 796 1 1.0950e+02 4.1840e+02 ; HB2 - CB
- CG
795 793 796 1 1.0950e+02 4.1840e+02 ; HB3 - CB
- CG
796 798 799 1 1.0950e+02 4.1840e+02 ; CG - CD1
- HD11
796 798 800 1 1.0950e+02 4.1840e+02 ; CG - CD1
- HD12
796 798 801 1 1.0950e+02 4.1840e+02 ; CG - CD1
- HD13
796 802 803 1 1.0950e+02 4.1840e+02 ; CG - CD2
- HD21
796 802 804 1 1.0950e+02 4.1840e+02 ; CG - CD2
- HD22
796 802 805 1 1.0950e+02 4.1840e+02 ; CG - CD2
- HD23
797 796 798 1 1.0950e+02 4.1840e+02 ; HG - CG
- CD1
797 796 802 1 1.0950e+02 4.1840e+02 ; HG - CG
- CD2
798 796 802 1 1.0950e+02 3.3472e+02 ; CD1 - CG
- CD2
799 798 800 1 1.0950e+02 2.9288e+02 ; HD11 - CD1
- HD12
799 798 801 1 1.0950e+02 2.9288e+02 ; HD11 - CD1
- HD13
800 798 801 1 1.0950e+02 2.9288e+02 ; HD12 - CD1
- HD13
803 802 804 1 1.0950e+02 2.9288e+02 ; HD21 - CD2
- HD22
803 802 805 1 1.0950e+02 2.9288e+02 ; HD21 - CD2
- HD23

804    802    805    1    1.0950e+02    2.9288e+02 ;   HD22 - CD2
- HD23
806    808    809    1    1.2000e+02    4.1840e+02 ;     C - N
- H
806    808    810    1    1.2190e+02    4.1840e+02 ;     C - N
- CA
807    806    808    1    1.2290e+02    6.6944e+02 ;     O - C
- N
808    810    811    1    1.0950e+02    4.1840e+02 ;     N - CA
- HA
808    810    812    1    1.0970e+02    6.6944e+02 ;     N - CA
- CB
808    810    828    1    1.1010e+02    5.2718e+02 ;     N - CA
- C
809    808    810    1    1.1804e+02    4.1840e+02 ;     H - N
- CA
810    812    813    1    1.0950e+02    4.1840e+02 ;    CA - CB
- HB2
810    812    814    1    1.0950e+02    4.1840e+02 ;    CA - CB
- HB3
810    812    815    1    1.0950e+02    3.3472e+02 ;    CA - CB
- CG
810    828    829    1    1.2040e+02    6.6944e+02 ;    CA - C
- O
810    828    830    1    1.1660e+02    5.8576e+02 ;    CA - C
- N
811    810    812    1    1.0950e+02    4.1840e+02 ;    HA - CA
- CB
811    810    828    1    1.0950e+02    4.1840e+02 ;    HA - CA
- C
812    810    828    1    1.1110e+02    5.2718e+02 ;    CB - CA
- C
812    815    816    1    1.0950e+02    4.1840e+02 ;    CB - CG
- HG2
812    815    817    1    1.0950e+02    4.1840e+02 ;    CB - CG
- HG3
812    815    818    1    1.0950e+02    3.3472e+02 ;    CB - CG
- CD
813    812    814    1    1.0950e+02    2.9288e+02 ;   HB2 - CB
- HB3
813    812    815    1    1.0950e+02    4.1840e+02 ;   HB2 - CB
- CG
814    812    815    1    1.0950e+02    4.1840e+02 ;   HB3 - CB
- CG
815    818    819    1    1.0950e+02    4.1840e+02 ;    CG - CD
- HD2
815    818    820    1    1.0950e+02    4.1840e+02 ;    CG - CD
- HD3
815    818    821    1    1.0950e+02    3.3472e+02 ;    CG - CD
- CE
816    815    817    1    1.0950e+02    2.9288e+02 ;   HG2 - CG
- HG3

816 815 818 1 1.0950e+02 4.1840e+02 ; HG2 - CG
- CD
817 815 818 1 1.0950e+02 4.1840e+02 ; HG3 - CG
- CD
818 821 822 1 1.0950e+02 4.1840e+02 ; CD - CE
- HE2
818 821 823 1 1.0950e+02 4.1840e+02 ; CD - CE
- HE3
818 821 824 1 1.1120e+02 6.6944e+02 ; CD - CE
- NZ
819 818 820 1 1.0950e+02 2.9288e+02 ; HD2 - CD
- HD3
819 818 821 1 1.0950e+02 4.1840e+02 ; HD2 - CD
- CE
820 818 821 1 1.0950e+02 4.1840e+02 ; HD3 - CD
- CE
821 824 825 1 1.0950e+02 4.1840e+02 ; CE - NZ
- HZ1
821 824 826 1 1.0950e+02 4.1840e+02 ; CE - NZ
- HZ2
821 824 827 1 1.0950e+02 4.1840e+02 ; CE - NZ
- HZ3
822 821 823 1 1.0950e+02 2.9288e+02 ; HE2 - CE
- HE3
822 821 824 1 1.0950e+02 4.1840e+02 ; HE2 - CE
- NZ
823 821 824 1 1.0950e+02 4.1840e+02 ; HE3 - CE
- NZ
825 824 826 1 1.0950e+02 2.9288e+02 ; HZ1 - NZ
- HZ2
825 824 827 1 1.0950e+02 2.9288e+02 ; HZ1 - NZ
- HZ3
826 824 827 1 1.0950e+02 2.9288e+02 ; HZ2 - NZ
- HZ3
828 830 831 1 1.2000e+02 4.1840e+02 ; C - N
- H
828 830 832 1 1.2190e+02 4.1840e+02 ; C - N
- CA
829 828 830 1 1.2290e+02 6.6944e+02 ; O - C
- N
830 832 833 1 1.0950e+02 4.1840e+02 ; N - CA
- HA
830 832 834 1 1.0970e+02 6.6944e+02 ; N - CA
- CB
830 832 842 1 1.1010e+02 5.2718e+02 ; N - CA
- C
831 830 832 1 1.1804e+02 4.1840e+02 ; H - N
- CA
832 834 835 1 1.0950e+02 4.1840e+02 ; CA - CB
- HB2
832 834 836 1 1.0950e+02 4.1840e+02 ; CA - CB
- HB3

```
  832    834    837      1    1.1110e+02    5.2718e+02 ;     CA - CB
- CG
  832    842    843      1    1.2040e+02    6.6944e+02 ;     CA - C
- O
  832    842    844      1    1.1660e+02    5.8576e+02 ;     CA - C
- N
  833    832    834      1    1.0950e+02    4.1840e+02 ;     HA - CA
- CB
  833    832    842      1    1.0950e+02    4.1840e+02 ;     HA - CA
- C
  834    832    842      1    1.1110e+02    5.2718e+02 ;     CB - CA
- C
  834    837    838      1    1.2040e+02    6.6944e+02 ;     CB - CG
- OD1
  834    837    839      1    1.1660e+02    5.8576e+02 ;     CB - CG
- ND2
  835    834    836      1    1.0950e+02    2.9288e+02 ;    HB2 - CB
- HB3
  835    834    837      1    1.0950e+02    4.1840e+02 ;    HB2 - CB
- CG
  836    834    837      1    1.0950e+02    4.1840e+02 ;    HB3 - CB
- CG
  837    839    840      1    1.2000e+02    4.1840e+02 ;     CG - ND2
- HD21
  837    839    841      1    1.2000e+02    4.1840e+02 ;     CG - ND2
- HD22
  838    837    839      1    1.2290e+02    6.6944e+02 ;    OD1 - CG
- ND2
  840    839    841      1    1.2000e+02    2.9288e+02 ;   HD21 - ND2
- HD22
  842    844    845      1    1.2000e+02    4.1840e+02 ;      C - N
- H
  842    844    846      1    1.2190e+02    4.1840e+02 ;      C - N
- CA
  843    842    844      1    1.2290e+02    6.6944e+02 ;      O - C
- N
  844    846    847      1    1.0950e+02    4.1840e+02 ;      N - CA
- HA
  844    846    848      1    1.0970e+02    6.6944e+02 ;      N - CA
- CB
  844    846    861      1    1.1010e+02    5.2718e+02 ;      N - CA
- C
  845    844    846      1    1.1804e+02    4.1840e+02 ;      H - N
- CA
  846    848    849      1    1.0950e+02    4.1840e+02 ;     CA - CB
- HB
  846    848    850      1    1.0950e+02    3.3472e+02 ;     CA - CB
- CG2
  846    848    854      1    1.0950e+02    3.3472e+02 ;     CA - CB
- CG1
  846    861    862      1    1.2040e+02    6.6944e+02 ;     CA - C
- O
```

```
   846    861    863      1    1.1660e+02    5.8576e+02 ;     CA - C
 - N
   847    846    848      1    1.0950e+02    4.1840e+02 ;     HA - CA
 - CB
   847    846    861      1    1.0950e+02    4.1840e+02 ;     HA - CA
 - C
   848    846    861      1    1.1110e+02    5.2718e+02 ;     CB - CA
 - C
   848    850    851      1    1.0950e+02    4.1840e+02 ;     CB - CG2
 - HG21
   848    850    852      1    1.0950e+02    4.1840e+02 ;     CB - CG2
 - HG22
   848    850    853      1    1.0950e+02    4.1840e+02 ;     CB - CG2
 - HG23
   848    854    855      1    1.0950e+02    4.1840e+02 ;     CB - CG1
 - HG12
   848    854    856      1    1.0950e+02    4.1840e+02 ;     CB - CG1
 - HG13
   848    854    857      1    1.0950e+02    3.3472e+02 ;     CB - CG1
 - CD1
   849    848    850      1    1.0950e+02    4.1840e+02 ;     HB - CB
 - CG2
   849    848    854      1    1.0950e+02    4.1840e+02 ;     HB - CB
 - CG1
   850    848    854      1    1.0950e+02    3.3472e+02 ;    CG2 - CB
 - CG1
   851    850    852      1    1.0950e+02    2.9288e+02 ;   HG21 - CG2
 - HG22
   851    850    853      1    1.0950e+02    2.9288e+02 ;   HG21 - CG2
 - HG23
   852    850    853      1    1.0950e+02    2.9288e+02 ;   HG22 - CG2
 - HG23
   854    857    858      1    1.0950e+02    4.1840e+02 ;    CG1 - CD1
 - HD11
   854    857    859      1    1.0950e+02    4.1840e+02 ;    CG1 - CD1
 - HD12
   854    857    860      1    1.0950e+02    4.1840e+02 ;    CG1 - CD1
 - HD13
   855    854    856      1    1.0950e+02    2.9288e+02 ;   HG12 - CG1
 - HG13
   855    854    857      1    1.0950e+02    4.1840e+02 ;   HG12 - CG1
 - CD1
   856    854    857      1    1.0950e+02    4.1840e+02 ;   HG13 - CG1
 - CD1
   858    857    859      1    1.0950e+02    2.9288e+02 ;   HD11 - CD1
 - HD12
   858    857    860      1    1.0950e+02    2.9288e+02 ;   HD11 - CD1
 - HD13
   859    857    860      1    1.0950e+02    2.9288e+02 ;   HD12 - CD1
 - HD13
   861    863    864      1    1.2000e+02    4.1840e+02 ;      C - N
 - H
```

```
   861    863    865      1    1.2190e+02    4.1840e+02 ;      C - N
- CA
   862    861    863      1    1.2290e+02    6.6944e+02 ;      O - C
- N
   863    865    866      1    1.0950e+02    4.1840e+02 ;      N - CA
- HA
   863    865    867      1    1.0970e+02    6.6944e+02 ;      N - CA
- CB
   863    865    875      1    1.1010e+02    5.2718e+02 ;      N - CA
- C
   864    863    865      1    1.1804e+02    4.1840e+02 ;      H - N
- CA
   865    867    868      1    1.0950e+02    4.1840e+02 ;     CA - CB
- HB2
   865    867    869      1    1.0950e+02    4.1840e+02 ;     CA - CB
- HB3
   865    867    870      1    1.1110e+02    5.2718e+02 ;     CA - CB
- CG
   865    875    876      1    1.2040e+02    6.6944e+02 ;     CA - C
- O
   865    875    877      1    1.1660e+02    5.8576e+02 ;     CA - C
- N
   866    865    867      1    1.0950e+02    4.1840e+02 ;     HA - CA
- CB
   866    865    875      1    1.0950e+02    4.1840e+02 ;     HA - CA
- C
   867    865    875      1    1.1110e+02    5.2718e+02 ;     CB - CA
- C
   867    870    871      1    1.2040e+02    6.6944e+02 ;     CB - CG
- OD1
   867    870    872      1    1.1660e+02    5.8576e+02 ;     CB - CG
- ND2
   868    867    869      1    1.0950e+02    2.9288e+02 ;    HB2 - CB
- HB3
   868    867    870      1    1.0950e+02    4.1840e+02 ;    HB2 - CB
- CG
   869    867    870      1    1.0950e+02    4.1840e+02 ;    HB3 - CB
- CG
   870    872    873      1    1.2000e+02    4.1840e+02 ;     CG - ND2
- HD21
   870    872    874      1    1.2000e+02    4.1840e+02 ;     CG - ND2
- HD22
   871    870    872      1    1.2290e+02    6.6944e+02 ;    OD1 - CG
- ND2
   873    872    874      1    1.2000e+02    2.9288e+02 ;   HD21 - ND2
- HD22
   875    877    878      1    1.2000e+02    4.1840e+02 ;      C - N
- H
   875    877    879      1    1.2190e+02    4.1840e+02 ;      C - N
- CA
   876    875    877      1    1.2290e+02    6.6944e+02 ;      O - C
- N
```

877 879 880 1 1.0950e+02 4.1840e+02 ; N - CA
- HA
877 879 881 1 1.0970e+02 6.6944e+02 ; N - CA
- CB
877 879 899 1 1.1010e+02 5.2718e+02 ; N - CA
- C
878 877 879 1 1.1804e+02 4.1840e+02 ; H - N
- CA
879 881 882 1 1.0950e+02 4.1840e+02 ; CA - CB
- HB2
879 881 883 1 1.0950e+02 4.1840e+02 ; CA - CB
- HB3
879 881 884 1 1.0950e+02 3.3472e+02 ; CA - CB
- CG
879 899 900 1 1.2040e+02 6.6944e+02 ; CA - C
- O
879 899 901 1 1.1660e+02 5.8576e+02 ; CA - C
- N
880 879 881 1 1.0950e+02 4.1840e+02 ; HA - CA
- CB
880 879 899 1 1.0950e+02 4.1840e+02 ; HA - CA
- C
881 879 899 1 1.1110e+02 5.2718e+02 ; CB - CA
- C
881 884 885 1 1.0950e+02 4.1840e+02 ; CB - CG
- HG2
881 884 886 1 1.0950e+02 4.1840e+02 ; CB - CG
- HG3
881 884 887 1 1.0950e+02 3.3472e+02 ; CB - CG
- CD
882 881 883 1 1.0950e+02 2.9288e+02 ; HB2 - CB
- HB3
882 881 884 1 1.0950e+02 4.1840e+02 ; HB2 - CB
- CG
883 881 884 1 1.0950e+02 4.1840e+02 ; HB3 - CB
- CG
884 887 888 1 1.0950e+02 4.1840e+02 ; CG - CD
- HD2
884 887 889 1 1.0950e+02 4.1840e+02 ; CG - CD
- HD3
884 887 890 1 1.1120e+02 6.6944e+02 ; CG - CD
- NE
885 884 886 1 1.0950e+02 2.9288e+02 ; HG2 - CG
- HG3
885 884 887 1 1.0950e+02 4.1840e+02 ; HG2 - CG
- CD
886 884 887 1 1.0950e+02 4.1840e+02 ; HG3 - CG
- CD
887 890 891 1 1.1840e+02 4.1840e+02 ; CD - NE
- HE
887 890 892 1 1.2320e+02 4.1840e+02 ; CD - NE
- CZ

888 887 889 1 1.0950e+02 2.9288e+02 ; HD2 - CD
- HD3
888 887 890 1 1.0950e+02 4.1840e+02 ; HD2 - CD
- NE
889 887 890 1 1.0950e+02 4.1840e+02 ; HD3 - CD
- NE
890 892 893 1 1.2000e+02 5.8576e+02 ; NE - CZ
- NH1
890 892 896 1 1.2000e+02 5.8576e+02 ; NE - CZ
- NH2
891 890 892 1 1.2000e+02 4.1840e+02 ; HE - NE
- CZ
892 893 894 1 1.2000e+02 4.1840e+02 ; CZ - NH1
- HH11
892 893 895 1 1.2000e+02 4.1840e+02 ; CZ - NH1
- HH12
892 896 897 1 1.2000e+02 4.1840e+02 ; CZ - NH2
- HH21
892 896 898 1 1.2000e+02 4.1840e+02 ; CZ - NH2
- HH22
893 892 896 1 1.2000e+02 5.8576e+02 ; NH1 - CZ
- NH2
894 893 895 1 1.2000e+02 2.9288e+02 ; HH11 - NH1
- HH12
897 896 898 1 1.2000e+02 2.9288e+02 ; HH21 - NH2
- HH22
899 901 902 1 1.2000e+02 4.1840e+02 ; C - N
- H
899 901 903 1 1.2190e+02 4.1840e+02 ; C - N
- CA
900 899 901 1 1.2290e+02 6.6944e+02 ; O - C
- N
901 903 904 1 1.0950e+02 4.1840e+02 ; N - CA
- HA
901 903 905 1 1.0970e+02 6.6944e+02 ; N - CA
- CB
901 903 919 1 1.1010e+02 5.2718e+02 ; N - CA
- C
902 901 903 1 1.1804e+02 4.1840e+02 ; H - N
- CA
903 905 906 1 1.0950e+02 4.1840e+02 ; CA - CB
- HB2
903 905 907 1 1.0950e+02 4.1840e+02 ; CA - CB
- HB3
903 905 908 1 1.1400e+02 5.2718e+02 ; CA - CB
- CG
903 919 920 1 1.2040e+02 6.6944e+02 ; CA - C
- O
903 919 921 1 1.1660e+02 5.8576e+02 ; CA - C
- N
904 903 905 1 1.0950e+02 4.1840e+02 ; HA - CA
- CB

```
   904    903    919      1    1.0950e+02    4.1840e+02 ;     HA - CA
- C
   905    903    919      1    1.1110e+02    5.2718e+02 ;     CB - CA
- C
   905    908    909      1    1.2000e+02    5.8576e+02 ;     CB - CG
- CD1
   905    908    917      1    1.2000e+02    5.8576e+02 ;     CB - CG
- CD2
   906    905    907      1    1.0950e+02    2.9288e+02 ;    HB2 - CB
- HB3
   906    905    908      1    1.0950e+02    4.1840e+02 ;    HB2 - CB
- CG
   907    905    908      1    1.0950e+02    4.1840e+02 ;    HB3 - CB
- CG
   908    909    910      1    1.2000e+02    4.1840e+02 ;     CG - CD1
- HD1
   908    909    911      1    1.2000e+02    5.2718e+02 ;     CG - CD1
- CE1
   908    917    915      1    1.2000e+02    5.2718e+02 ;     CG - CD2
- CE2
   908    917    918      1    1.2000e+02    4.1840e+02 ;     CG - CD2
- HD2
   909    908    917      1    1.2000e+02    5.2718e+02 ;    CD1 - CG
- CD2
   909    911    912      1    1.2000e+02    4.1840e+02 ;    CD1 - CE1
- HE1
   909    911    913      1    1.2000e+02    5.2718e+02 ;    CD1 - CE1
- CZ
   910    909    911      1    1.2000e+02    4.1840e+02 ;    HD1 - CD1
- CE1
   911    913    914      1    1.2000e+02    4.1840e+02 ;    CE1 - CZ
- HZ
   911    913    915      1    1.2000e+02    5.2718e+02 ;    CE1 - CZ
- CE2
   912    911    913      1    1.2000e+02    4.1840e+02 ;    HE1 - CE1
- CZ
   913    915    916      1    1.2000e+02    4.1840e+02 ;     CZ - CE2
- HE2
   913    915    917      1    1.2000e+02    5.2718e+02 ;     CZ - CE2
- CD2
   914    913    915      1    1.2000e+02    4.1840e+02 ;     HZ - CZ
- CE2
   915    917    918      1    1.2000e+02    4.1840e+02 ;    CE2 - CD2
- HD2
   916    915    917      1    1.2000e+02    4.1840e+02 ;    HE2 - CE2
- CD2
   919    921    922      1    1.2000e+02    4.1840e+02 ;      C - N
- H
   919    921    923      1    1.2190e+02    4.1840e+02 ;      C - N
- CA
   920    919    921      1    1.2290e+02    6.6944e+02 ;      O - C
- N
```

```
   921     923     924       1    1.0950e+02    4.1840e+02 ;      N - CA
 - HA
   921     923     925       1    1.0970e+02    6.6944e+02 ;      N - CA
 - CB
   921     923     939       1    1.1010e+02    5.2718e+02 ;      N - CA
 - C
   922     921     923       1    1.1804e+02    4.1840e+02 ;      H - N
 - CA
   923     925     926       1    1.0950e+02    4.1840e+02 ;     CA - CB
 - HB2
   923     925     927       1    1.0950e+02    4.1840e+02 ;     CA - CB
 - HB3
   923     925     928       1    1.1400e+02    5.2718e+02 ;     CA - CB
 - CG
   923     939     940       1    1.2040e+02    6.6944e+02 ;     CA - C
 - O
   923     939     941       1    1.1660e+02    5.8576e+02 ;     CA - C
 - N
   924     923     925       1    1.0950e+02    4.1840e+02 ;     HA - CA
 - CB
   924     923     939       1    1.0950e+02    4.1840e+02 ;     HA - CA
 - C
   925     923     939       1    1.1110e+02    5.2718e+02 ;     CB - CA
 - C
   925     928     929       1    1.2000e+02    5.8576e+02 ;     CB - CG
 - CD1
   925     928     937       1    1.2000e+02    5.8576e+02 ;     CB - CG
 - CD2
   926     925     927       1    1.0950e+02    2.9288e+02 ;    HB2 - CB
 - HB3
   926     925     928       1    1.0950e+02    4.1840e+02 ;    HB2 - CB
 - CG
   927     925     928       1    1.0950e+02    4.1840e+02 ;    HB3 - CB
 - CG
   928     929     930       1    1.2000e+02    4.1840e+02 ;     CG - CD1
 - HD1
   928     929     931       1    1.2000e+02    5.2718e+02 ;     CG - CD1
 - CE1
   928     937     935       1    1.2000e+02    5.2718e+02 ;     CG - CD2
 - CE2
   928     937     938       1    1.2000e+02    4.1840e+02 ;     CG - CD2
 - HD2
   929     928     937       1    1.2000e+02    5.2718e+02 ;    CD1 - CG
 - CD2
   929     931     932       1    1.2000e+02    4.1840e+02 ;    CD1 - CE1
 - HE1
   929     931     933       1    1.2000e+02    5.2718e+02 ;    CD1 - CE1
 - CZ
   930     929     931       1    1.2000e+02    4.1840e+02 ;    HD1 - CD1
 - CE1
   931     933     934       1    1.2000e+02    4.1840e+02 ;    CE1 - CZ
 - HZ
```

```
   931    933    935     1    1.2000e+02    5.2718e+02 ;    CE1 - CZ
- CE2
   932    931    933     1    1.2000e+02    4.1840e+02 ;    HE1 - CE1
- CZ
   933    935    936     1    1.2000e+02    4.1840e+02 ;     CZ - CE2
- HE2
   933    935    937     1    1.2000e+02    5.2718e+02 ;     CZ - CE2
- CD2
   934    933    935     1    1.2000e+02    4.1840e+02 ;     HZ - CZ
- CE2
   935    937    938     1    1.2000e+02    4.1840e+02 ;    CE2 - CD2
- HD2
   936    935    937     1    1.2000e+02    4.1840e+02 ;    HE2 - CE2
- CD2
   939    941    942     1    1.2000e+02    4.1840e+02 ;      C - N
- H
   939    941    943     1    1.2190e+02    4.1840e+02 ;      C - N
- CA
   940    939    941     1    1.2290e+02    6.6944e+02 ;      O - C
- N
   941    943    944     1    1.0950e+02    4.1840e+02 ;      N - CA
- HA
   941    943    945     1    1.0970e+02    6.6944e+02 ;      N - CA
- CB
   941    943    956     1    1.1010e+02    5.2718e+02 ;      N - CA
- C
   942    941    943     1    1.1804e+02    4.1840e+02 ;      H - N
- CA
   943    945    946     1    1.0950e+02    4.1840e+02 ;     CA - CB
- HB2
   943    945    947     1    1.0950e+02    4.1840e+02 ;     CA - CB
- HB3
   943    945    948     1    1.0950e+02    3.3472e+02 ;     CA - CB
- CG
   943    956    957     1    1.2040e+02    6.6944e+02 ;     CA - C
- O
   943    956    958     1    1.1660e+02    5.8576e+02 ;     CA - C
- N
   944    943    945     1    1.0950e+02    4.1840e+02 ;     HA - CA
- CB
   944    943    956     1    1.0950e+02    4.1840e+02 ;     HA - CA
- C
   945    943    956     1    1.1110e+02    5.2718e+02 ;     CB - CA
- C
   945    948    949     1    1.0950e+02    4.1840e+02 ;     CB - CG
- HG2
   945    948    950     1    1.0950e+02    4.1840e+02 ;     CB - CG
- HG3
   945    948    951     1    1.1110e+02    5.2718e+02 ;     CB - CG
- CD
   946    945    947     1    1.0950e+02    2.9288e+02 ;    HB2 - CB
- HB3
```

| | | | | | | | |
|---|---|---|---|---|---|---|---|
| 946 | 945 | 948 | 1 | 1.0950e+02 | 4.1840e+02 | ; | HB2 - CB - CG |
| 947 | 945 | 948 | 1 | 1.0950e+02 | 4.1840e+02 | ; | HB3 - CB - CG |
| 948 | 951 | 952 | 1 | 1.2040e+02 | 6.6944e+02 | ; | CG - CD - OE1 |
| 948 | 951 | 953 | 1 | 1.1660e+02 | 5.8576e+02 | ; | CG - CD - NE2 |
| 949 | 948 | 950 | 1 | 1.0950e+02 | 2.9288e+02 | ; | HG2 - CG - HG3 |
| 949 | 948 | 951 | 1 | 1.0950e+02 | 4.1840e+02 | ; | HG2 - CG - CD |
| 950 | 948 | 951 | 1 | 1.0950e+02 | 4.1840e+02 | ; | HG3 - CG - CD |
| 951 | 953 | 954 | 1 | 1.2000e+02 | 4.1840e+02 | ; | CD - NE2 - HE21 |
| 951 | 953 | 955 | 1 | 1.2000e+02 | 4.1840e+02 | ; | CD - NE2 - HE22 |
| 952 | 951 | 953 | 1 | 1.2290e+02 | 6.6944e+02 | ; | OE1 - CD - NE2 |
| 954 | 953 | 955 | 1 | 1.2000e+02 | 2.9288e+02 | ; | HE21 - NE2 - HE22 |
| 956 | 958 | 959 | 1 | 1.2000e+02 | 4.1840e+02 | ; | C - N - H |
| 956 | 958 | 960 | 1 | 1.2190e+02 | 4.1840e+02 | ; | C - N - CA |
| 957 | 956 | 958 | 1 | 1.2290e+02 | 6.6944e+02 | ; | O - C - N |
| 958 | 960 | 961 | 1 | 1.0950e+02 | 4.1840e+02 | ; | N - CA - HA |
| 958 | 960 | 962 | 1 | 1.0970e+02 | 6.6944e+02 | ; | N - CA - CB |
| 958 | 960 | 971 | 1 | 1.1010e+02 | 5.2718e+02 | ; | N - CA - C |
| 959 | 958 | 960 | 1 | 1.1804e+02 | 4.1840e+02 | ; | H - N - CA |
| 960 | 962 | 963 | 1 | 1.0950e+02 | 4.1840e+02 | ; | CA - CB - HB2 |
| 960 | 962 | 964 | 1 | 1.0950e+02 | 4.1840e+02 | ; | CA - CB - HB3 |
| 960 | 962 | 965 | 1 | 1.0950e+02 | 3.3472e+02 | ; | CA - CB - CG |
| 960 | 971 | 972 | 1 | 1.2040e+02 | 6.6944e+02 | ; | CA - C - O |
| 960 | 971 | 973 | 1 | 1.1660e+02 | 5.8576e+02 | ; | CA - C - N |
| 961 | 960 | 962 | 1 | 1.0950e+02 | 4.1840e+02 | ; | HA - CA - CB |
| 961 | 960 | 971 | 1 | 1.0950e+02 | 4.1840e+02 | ; | HA - CA - C |
| 962 | 960 | 971 | 1 | 1.1110e+02 | 5.2718e+02 | ; | CB - CA - C |

962 965 966 1 1.0950e+02 4.1840e+02 ; CB - CG
- HG2
962 965 967 1 1.0950e+02 4.1840e+02 ; CB - CG
- HG3
962 965 968 1 1.1110e+02 5.2718e+02 ; CB - CG
- CD
963 962 964 1 1.0950e+02 2.9288e+02 ; HB2 - CB
- HB3
963 962 965 1 1.0950e+02 4.1840e+02 ; HB2 - CB
- CG
964 962 965 1 1.0950e+02 4.1840e+02 ; HB3 - CB
- CG
965 968 969 1 1.1700e+02 5.8576e+02 ; CG - CD
- OE1
965 968 970 1 1.1700e+02 5.8576e+02 ; CG - CD
- OE2
966 965 967 1 1.0950e+02 2.9288e+02 ; HG2 - CG
- HG3
966 965 968 1 1.0950e+02 4.1840e+02 ; HG2 - CG
- CD
967 965 968 1 1.0950e+02 4.1840e+02 ; HG3 - CG
- CD
969 968 970 1 1.2600e+02 6.6944e+02 ; OE1 - CD
- OE2
971 973 974 1 1.2000e+02 4.1840e+02 ; C - N
- H
971 973 975 1 1.2190e+02 4.1840e+02 ; C - N
- CA
972 971 973 1 1.2290e+02 6.6944e+02 ; O - C
- N
973 975 976 1 1.0950e+02 4.1840e+02 ; N - CA
- HA
973 975 977 1 1.0970e+02 6.6944e+02 ; N - CA
- CB
973 975 990 1 1.1010e+02 5.2718e+02 ; N - CA
- C
974 973 975 1 1.1804e+02 4.1840e+02 ; H - N
- CA
975 977 978 1 1.0950e+02 4.1840e+02 ; CA - CB
- HB
975 977 979 1 1.0950e+02 3.3472e+02 ; CA - CB
- CG2
975 977 983 1 1.0950e+02 3.3472e+02 ; CA - CB
- CG1
975 990 991 1 1.2040e+02 6.6944e+02 ; CA - C
- O
975 990 992 1 1.1660e+02 5.8576e+02 ; CA - C
- N
976 975 977 1 1.0950e+02 4.1840e+02 ; HA - CA
- CB
976 975 990 1 1.0950e+02 4.1840e+02 ; HA - CA
- C

```
   977    975    990     1    1.1110e+02    5.2718e+02 ;     CB - CA
- C
   977    979    980     1    1.0950e+02    4.1840e+02 ;     CB - CG2
- HG21
   977    979    981     1    1.0950e+02    4.1840e+02 ;     CB - CG2
- HG22
   977    979    982     1    1.0950e+02    4.1840e+02 ;     CB - CG2
- HG23
   977    983    984     1    1.0950e+02    4.1840e+02 ;     CB - CG1
- HG12
   977    983    985     1    1.0950e+02    4.1840e+02 ;     CB - CG1
- HG13
   977    983    986     1    1.0950e+02    3.3472e+02 ;     CB - CG1
- CD1
   978    977    979     1    1.0950e+02    4.1840e+02 ;     HB - CB
- CG2
   978    977    983     1    1.0950e+02    4.1840e+02 ;     HB - CB
- CG1
   979    977    983     1    1.0950e+02    3.3472e+02 ;    CG2 - CB
- CG1
   980    979    981     1    1.0950e+02    2.9288e+02 ;   HG21 - CG2
- HG22
   980    979    982     1    1.0950e+02    2.9288e+02 ;   HG21 - CG2
- HG23
   981    979    982     1    1.0950e+02    2.9288e+02 ;   HG22 - CG2
- HG23
   983    986    987     1    1.0950e+02    4.1840e+02 ;    CG1 - CD1
- HD11
   983    986    988     1    1.0950e+02    4.1840e+02 ;    CG1 - CD1
- HD12
   983    986    989     1    1.0950e+02    4.1840e+02 ;    CG1 - CD1
- HD13
   984    983    985     1    1.0950e+02    2.9288e+02 ;   HG12 - CG1
- HG13
   984    983    986     1    1.0950e+02    4.1840e+02 ;   HG12 - CG1
- CD1
   985    983    986     1    1.0950e+02    4.1840e+02 ;   HG13 - CG1
- CD1
   987    986    988     1    1.0950e+02    2.9288e+02 ;   HD11 - CD1
- HD12
   987    986    989     1    1.0950e+02    2.9288e+02 ;   HD11 - CD1
- HD13
   988    986    989     1    1.0950e+02    2.9288e+02 ;   HD12 - CD1
- HD13
   990    992    993     1    1.2000e+02    4.1840e+02 ;      C - N
- H
   990    992    994     1    1.2190e+02    4.1840e+02 ;      C - N
- CA
   991    990    992     1    1.2290e+02    6.6944e+02 ;      O - C
- N
   992    994    995     1    1.0950e+02    4.1840e+02 ;      N - CA
- HA
```

```
   992    994    996       1    1.0970e+02    6.6944e+02 ;       N - CA
 - CB
   992    994   1005       1    1.1010e+02    5.2718e+02 ;       N - CA
 - C
   993    992    994       1    1.1804e+02    4.1840e+02 ;       H - N
 - CA
   994    996    997       1    1.0950e+02    4.1840e+02 ;      CA - CB
 - HB2
   994    996    998       1    1.0950e+02    4.1840e+02 ;      CA - CB
 - HB3
   994    996    999       1    1.0950e+02    3.3472e+02 ;      CA - CB
 - CG
   994   1005   1006       1    1.2040e+02    6.6944e+02 ;      CA - C
 - O
   994   1005   1007       1    1.1660e+02    5.8576e+02 ;      CA - C
 - N
   995    994    996       1    1.0950e+02    4.1840e+02 ;      HA - CA
 - CB
   995    994   1005       1    1.0950e+02    4.1840e+02 ;      HA - CA
 - C
   996    994   1005       1    1.1110e+02    5.2718e+02 ;      CB - CA
 - C
   996    999   1000       1    1.0950e+02    4.1840e+02 ;      CB - CG
 - HG2
   996    999   1001       1    1.0950e+02    4.1840e+02 ;      CB - CG
 - HG3
   996    999   1002       1    1.1110e+02    5.2718e+02 ;      CB - CG
 - CD
   997    996    998       1    1.0950e+02    2.9288e+02 ;     HB2 - CB
 - HB3
   997    996    999       1    1.0950e+02    4.1840e+02 ;     HB2 - CB
 - CG
   998    996    999       1    1.0950e+02    4.1840e+02 ;     HB3 - CB
 - CG
   999   1002   1003       1    1.1700e+02    5.8576e+02 ;      CG - CD
 - OE1
   999   1002   1004       1    1.1700e+02    5.8576e+02 ;      CG - CD
 - OE2
  1000    999   1001       1    1.0950e+02    2.9288e+02 ;     HG2 - CG
 - HG3
  1000    999   1002       1    1.0950e+02    4.1840e+02 ;     HG2 - CG
 - CD
  1001    999   1002       1    1.0950e+02    4.1840e+02 ;     HG3 - CG
 - CD
  1003   1002   1004       1    1.2600e+02    6.6944e+02 ;     OE1 - CD
 - OE2
  1005   1007   1008       1    1.2000e+02    4.1840e+02 ;       C - N
 - H
  1005   1007   1009       1    1.2190e+02    4.1840e+02 ;       C - N
 - CA
  1006   1005   1007       1    1.2290e+02    6.6944e+02 ;       O - C
 - N
```

1007 1009 1010 1 1.0950e+02 4.1840e+02 ; N - CA
- HA
1007 1009 1011 1 1.0970e+02 6.6944e+02 ; N - CA
- CB
1007 1009 1024 1 1.1010e+02 5.2718e+02 ; N - CA
- C
1008 1007 1009 1 1.1804e+02 4.1840e+02 ; H - N
- CA
1009 1011 1012 1 1.0950e+02 4.1840e+02 ; CA - CB
- HB
1009 1011 1013 1 1.0950e+02 3.3472e+02 ; CA - CB
- CG2
1009 1011 1017 1 1.0950e+02 3.3472e+02 ; CA - CB
- CG1
1009 1024 1025 1 1.2040e+02 6.6944e+02 ; CA - C
- O
1009 1024 1026 1 1.1660e+02 5.8576e+02 ; CA - C
- N
1010 1009 1011 1 1.0950e+02 4.1840e+02 ; HA - CA
- CB
1010 1009 1024 1 1.0950e+02 4.1840e+02 ; HA - CA
- C
1011 1009 1024 1 1.1110e+02 5.2718e+02 ; CB - CA
- C
1011 1013 1014 1 1.0950e+02 4.1840e+02 ; CB - CG2
- HG21
1011 1013 1015 1 1.0950e+02 4.1840e+02 ; CB - CG2
- HG22
1011 1013 1016 1 1.0950e+02 4.1840e+02 ; CB - CG2
- HG23
1011 1017 1018 1 1.0950e+02 4.1840e+02 ; CB - CG1
- HG12
1011 1017 1019 1 1.0950e+02 4.1840e+02 ; CB - CG1
- HG13
1011 1017 1020 1 1.0950e+02 3.3472e+02 ; CB - CG1
- CD1
1012 1011 1013 1 1.0950e+02 4.1840e+02 ; HB - CB
- CG2
1012 1011 1017 1 1.0950e+02 4.1840e+02 ; HB - CB
- CG1
1013 1011 1017 1 1.0950e+02 3.3472e+02 ; CG2 - CB
- CG1
1014 1013 1015 1 1.0950e+02 2.9288e+02 ; HG21 - CG2
- HG22
1014 1013 1016 1 1.0950e+02 2.9288e+02 ; HG21 - CG2
- HG23
1015 1013 1016 1 1.0950e+02 2.9288e+02 ; HG22 - CG2
- HG23
1017 1020 1021 1 1.0950e+02 4.1840e+02 ; CG1 - CD1
- HD11
1017 1020 1022 1 1.0950e+02 4.1840e+02 ; CG1 - CD1
- HD12

```
  1017   1020   1023       1    1.0950e+02    4.1840e+02 ;    CG1 - CD1
- HD13
  1018   1017   1019       1    1.0950e+02    2.9288e+02 ;   HG12 - CG1
- HG13
  1018   1017   1020       1    1.0950e+02    4.1840e+02 ;   HG12 - CG1
- CD1
  1019   1017   1020       1    1.0950e+02    4.1840e+02 ;   HG13 - CG1
- CD1
  1021   1020   1022       1    1.0950e+02    2.9288e+02 ;   HD11 - CD1
- HD12
  1021   1020   1023       1    1.0950e+02    2.9288e+02 ;   HD11 - CD1
- HD13
  1022   1020   1023       1    1.0950e+02    2.9288e+02 ;   HD12 - CD1
- HD13
  1024   1026   1027       1    1.2000e+02    4.1840e+02 ;      C - N
- H
  1024   1026   1028       1    1.2190e+02    4.1840e+02 ;      C - N
- CA
  1025   1024   1026       1    1.2290e+02    6.6944e+02 ;      O - C
- N
  1026   1028   1029       1    1.0950e+02    4.1840e+02 ;      N - CA
- HA
  1026   1028   1030       1    1.0970e+02    6.6944e+02 ;      N - CA
- CB
  1026   1028   1041       1    1.1010e+02    5.2718e+02 ;      N - CA
- C
  1027   1026   1028       1    1.1804e+02    4.1840e+02 ;      H - N
- CA
  1028   1030   1031       1    1.0950e+02    4.1840e+02 ;     CA - CB
- HB2
  1028   1030   1032       1    1.0950e+02    4.1840e+02 ;     CA - CB
- HB3
  1028   1030   1033       1    1.0950e+02    3.3472e+02 ;     CA - CB
- CG
  1028   1041   1042       1    1.2040e+02    6.6944e+02 ;     CA - C
- O
  1028   1041   1043       1    1.1660e+02    5.8576e+02 ;     CA - C
- N
  1029   1028   1030       1    1.0950e+02    4.1840e+02 ;     HA - CA
- CB
  1029   1028   1041       1    1.0950e+02    4.1840e+02 ;     HA - CA
- C
  1030   1028   1041       1    1.1110e+02    5.2718e+02 ;     CB - CA
- C
  1030   1033   1034       1    1.0950e+02    4.1840e+02 ;     CB - CG
- HG2
  1030   1033   1035       1    1.0950e+02    4.1840e+02 ;     CB - CG
- HG3
  1030   1033   1036       1    1.1470e+02    4.1840e+02 ;     CB - CG
- SD
  1031   1030   1032       1    1.0950e+02    2.9288e+02 ;    HB2 - CB
- HB3
```

1031 1030 1033 1 1.0950e+02 4.1840e+02 ; HB2 - CB
- CG
1032 1030 1033 1 1.0950e+02 4.1840e+02 ; HB3 - CB
- CG
1033 1036 1037 1 9.8900e+01 5.1882e+02 ; CG - SD
- CE
1034 1033 1035 1 1.0950e+02 2.9288e+02 ; HG2 - CG
- HG3
1034 1033 1036 1 1.0950e+02 4.1840e+02 ; HG2 - CG
- SD
1035 1033 1036 1 1.0950e+02 4.1840e+02 ; HG3 - CG
- SD
1036 1037 1038 1 1.0950e+02 4.1840e+02 ; SD - CE
- HE1
1036 1037 1039 1 1.0950e+02 4.1840e+02 ; SD - CE
- HE2
1036 1037 1040 1 1.0950e+02 4.1840e+02 ; SD - CE
- HE3
1038 1037 1039 1 1.0950e+02 2.9288e+02 ; HE1 - CE
- HE2
1038 1037 1040 1 1.0950e+02 2.9288e+02 ; HE1 - CE
- HE3
1039 1037 1040 1 1.0950e+02 2.9288e+02 ; HE2 - CE
- HE3
1041 1043 1044 1 1.2000e+02 4.1840e+02 ; C - N
- H
1041 1043 1045 1 1.2190e+02 4.1840e+02 ; C - N
- CA
1042 1041 1043 1 1.2290e+02 6.6944e+02 ; O - C
- N
1043 1045 1046 1 1.0950e+02 4.1840e+02 ; N - CA
- HA
1043 1045 1047 1 1.0970e+02 6.6944e+02 ; N - CA
- CB
1043 1045 1063 1 1.1010e+02 5.2718e+02 ; N - CA
- C
1044 1043 1045 1 1.1804e+02 4.1840e+02 ; H - N
- CA
1045 1047 1048 1 1.0950e+02 4.1840e+02 ; CA - CB
- HB2
1045 1047 1049 1 1.0950e+02 4.1840e+02 ; CA - CB
- HB3
1045 1047 1050 1 1.0950e+02 3.3472e+02 ; CA - CB
- CG
1045 1063 1064 1 1.2040e+02 6.6944e+02 ; CA - C
- O
1045 1063 1065 1 1.1660e+02 5.8576e+02 ; CA - C
- N
1046 1045 1047 1 1.0950e+02 4.1840e+02 ; HA - CA
- CB
1046 1045 1063 1 1.0950e+02 4.1840e+02 ; HA - CA
- C

```
  1047   1045   1063       1    1.1110e+02    5.2718e+02 ;      CB - CA
- C
  1047   1050   1051       1    1.0950e+02    4.1840e+02 ;      CB - CG
- HG2
  1047   1050   1052       1    1.0950e+02    4.1840e+02 ;      CB - CG
- HG3
  1047   1050   1053       1    1.0950e+02    3.3472e+02 ;      CB - CG
- CD
  1048   1047   1049       1    1.0950e+02    2.9288e+02 ;     HB2 - CB
- HB3
  1048   1047   1050       1    1.0950e+02    4.1840e+02 ;     HB2 - CB
- CG
  1049   1047   1050       1    1.0950e+02    4.1840e+02 ;     HB3 - CB
- CG
  1050   1053   1054       1    1.0950e+02    4.1840e+02 ;      CG - CD
- HD2
  1050   1053   1055       1    1.0950e+02    4.1840e+02 ;      CG - CD
- HD3
  1050   1053   1056       1    1.0950e+02    3.3472e+02 ;      CG - CD
- CE
  1051   1050   1052       1    1.0950e+02    2.9288e+02 ;     HG2 - CG
- HG3
  1051   1050   1053       1    1.0950e+02    4.1840e+02 ;     HG2 - CG
- CD
  1052   1050   1053       1    1.0950e+02    4.1840e+02 ;     HG3 - CG
- CD
  1053   1056   1057       1    1.0950e+02    4.1840e+02 ;      CD - CE
- HE2
  1053   1056   1058       1    1.0950e+02    4.1840e+02 ;      CD - CE
- HE3
  1053   1056   1059       1    1.1120e+02    6.6944e+02 ;      CD - CE
- NZ
  1054   1053   1055       1    1.0950e+02    2.9288e+02 ;     HD2 - CD
- HD3
  1054   1053   1056       1    1.0950e+02    4.1840e+02 ;     HD2 - CD
- CE
  1055   1053   1056       1    1.0950e+02    4.1840e+02 ;     HD3 - CD
- CE
  1056   1059   1060       1    1.0950e+02    4.1840e+02 ;      CE - NZ
- HZ1
  1056   1059   1061       1    1.0950e+02    4.1840e+02 ;      CE - NZ
- HZ2
  1056   1059   1062       1    1.0950e+02    4.1840e+02 ;      CE - NZ
- HZ3
  1057   1056   1058       1    1.0950e+02    2.9288e+02 ;     HE2 - CE
- HE3
  1057   1056   1059       1    1.0950e+02    4.1840e+02 ;     HE2 - CE
- NZ
  1058   1056   1059       1    1.0950e+02    4.1840e+02 ;     HE3 - CE
- NZ
  1060   1059   1061       1    1.0950e+02    2.9288e+02 ;     HZ1 - NZ
- HZ2
```

1060 1059 1062 1 1.0950e+02 2.9288e+02 ; HZ1 - NZ
- HZ3
1061 1059 1062 1 1.0950e+02 2.9288e+02 ; HZ2 - NZ
- HZ3
1063 1065 1066 1 1.2000e+02 4.1840e+02 ; C - N
- H
1063 1065 1067 1 1.2190e+02 4.1840e+02 ; C - N
- CA
1064 1063 1065 1 1.2290e+02 6.6944e+02 ; O - C
- N
1065 1067 1068 1 1.0950e+02 4.1840e+02 ; N - CA
- HA
1065 1067 1069 1 1.0970e+02 6.6944e+02 ; N - CA
- CB
1065 1067 1085 1 1.1010e+02 5.2718e+02 ; N - CA
- C
1066 1065 1067 1 1.1804e+02 4.1840e+02 ; H - N
- CA
1067 1069 1070 1 1.0950e+02 4.1840e+02 ; CA - CB
- HB2
1067 1069 1071 1 1.0950e+02 4.1840e+02 ; CA - CB
- HB3
1067 1069 1072 1 1.0950e+02 3.3472e+02 ; CA - CB
- CG
1067 1085 1086 1 1.2040e+02 6.6944e+02 ; CA - C
- O
1067 1085 1087 1 1.1660e+02 5.8576e+02 ; CA - C
- N
1068 1067 1069 1 1.0950e+02 4.1840e+02 ; HA - CA
- CB
1068 1067 1085 1 1.0950e+02 4.1840e+02 ; HA - CA
- C
1069 1067 1085 1 1.1110e+02 5.2718e+02 ; CB - CA
- C
1069 1072 1073 1 1.0950e+02 4.1840e+02 ; CB - CG
- HG2
1069 1072 1074 1 1.0950e+02 4.1840e+02 ; CB - CG
- HG3
1069 1072 1075 1 1.0950e+02 3.3472e+02 ; CB - CG
- CD
1070 1069 1071 1 1.0950e+02 2.9288e+02 ; HB2 - CB
- HB3
1070 1069 1072 1 1.0950e+02 4.1840e+02 ; HB2 - CB
- CG
1071 1069 1072 1 1.0950e+02 4.1840e+02 ; HB3 - CB
- CG
1072 1075 1076 1 1.0950e+02 4.1840e+02 ; CG - CD
- HD2
1072 1075 1077 1 1.0950e+02 4.1840e+02 ; CG - CD
- HD3
1072 1075 1078 1 1.0950e+02 3.3472e+02 ; CG - CD
- CE

```
  1073   1072   1074       1    1.0950e+02    2.9288e+02 ;    HG2 - CG
- HG3
  1073   1072   1075       1    1.0950e+02    4.1840e+02 ;    HG2 - CG
- CD
  1074   1072   1075       1    1.0950e+02    4.1840e+02 ;    HG3 - CG
- CD
  1075   1078   1079       1    1.0950e+02    4.1840e+02 ;     CD - CE
- HE2
  1075   1078   1080       1    1.0950e+02    4.1840e+02 ;     CD - CE
- HE3
  1075   1078   1081       1    1.1120e+02    6.6944e+02 ;     CD - CE
- NZ
  1076   1075   1077       1    1.0950e+02    2.9288e+02 ;    HD2 - CD
- HD3
  1076   1075   1078       1    1.0950e+02    4.1840e+02 ;    HD2 - CD
- CE
  1077   1075   1078       1    1.0950e+02    4.1840e+02 ;    HD3 - CD
- CE
  1078   1081   1082       1    1.0950e+02    4.1840e+02 ;     CE - NZ
- HZ1
  1078   1081   1083       1    1.0950e+02    4.1840e+02 ;     CE - NZ
- HZ2
  1078   1081   1084       1    1.0950e+02    4.1840e+02 ;     CE - NZ
- HZ3
  1079   1078   1080       1    1.0950e+02    2.9288e+02 ;    HE2 - CE
- HE3
  1079   1078   1081       1    1.0950e+02    4.1840e+02 ;    HE2 - CE
- NZ
  1080   1078   1081       1    1.0950e+02    4.1840e+02 ;    HE3 - CE
- NZ
  1082   1081   1083       1    1.0950e+02    2.9288e+02 ;    HZ1 - NZ
- HZ2
  1082   1081   1084       1    1.0950e+02    2.9288e+02 ;    HZ1 - NZ
- HZ3
  1083   1081   1084       1    1.0950e+02    2.9288e+02 ;    HZ2 - NZ
- HZ3
  1085   1087   1088       1    1.2000e+02    4.1840e+02 ;      C - N
- H
  1085   1087   1089       1    1.2190e+02    4.1840e+02 ;      C - N
- CA
  1086   1085   1087       1    1.2290e+02    6.6944e+02 ;      O - C
- N
  1087   1089   1090       1    1.0950e+02    4.1840e+02 ;      N - CA
- HA
  1087   1089   1091       1    1.0970e+02    6.6944e+02 ;      N - CA
- CB
  1087   1089   1104       1    1.1010e+02    5.2718e+02 ;      N - CA
- C
  1088   1087   1089       1    1.1804e+02    4.1840e+02 ;      H - N
- CA
  1089   1091   1092       1    1.0950e+02    4.1840e+02 ;     CA - CB
- HB2
```

```
  1089   1091   1093       1    1.0950e+02    4.1840e+02 ;     CA - CB
- HB3
  1089   1091   1094       1    1.0950e+02    3.3472e+02 ;     CA - CB
- CG
  1089   1104   1105       1    1.2040e+02    6.6944e+02 ;     CA - C
- O
  1089   1104   1106       1    1.1660e+02    5.8576e+02 ;     CA - C
- N
  1090   1089   1091       1    1.0950e+02    4.1840e+02 ;     HA - CA
- CB
  1090   1089   1104       1    1.0950e+02    4.1840e+02 ;     HA - CA
- C
  1091   1089   1104       1    1.1110e+02    5.2718e+02 ;     CB - CA
- C
  1091   1094   1095       1    1.0950e+02    4.1840e+02 ;     CB - CG
- HG
  1091   1094   1096       1    1.0950e+02    3.3472e+02 ;     CB - CG
- CD1
  1091   1094   1100       1    1.0950e+02    3.3472e+02 ;     CB - CG
- CD2
  1092   1091   1093       1    1.0950e+02    2.9288e+02 ;    HB2 - CB
- HB3
  1092   1091   1094       1    1.0950e+02    4.1840e+02 ;    HB2 - CB
- CG
  1093   1091   1094       1    1.0950e+02    4.1840e+02 ;    HB3 - CB
- CG
  1094   1096   1097       1    1.0950e+02    4.1840e+02 ;     CG - CD1
- HD11
  1094   1096   1098       1    1.0950e+02    4.1840e+02 ;     CG - CD1
- HD12
  1094   1096   1099       1    1.0950e+02    4.1840e+02 ;     CG - CD1
- HD13
  1094   1100   1101       1    1.0950e+02    4.1840e+02 ;     CG - CD2
- HD21
  1094   1100   1102       1    1.0950e+02    4.1840e+02 ;     CG - CD2
- HD22
  1094   1100   1103       1    1.0950e+02    4.1840e+02 ;     CG - CD2
- HD23
  1095   1094   1096       1    1.0950e+02    4.1840e+02 ;     HG - CG
- CD1
  1095   1094   1100       1    1.0950e+02    4.1840e+02 ;     HG - CG
- CD2
  1096   1094   1100       1    1.0950e+02    3.3472e+02 ;    CD1 - CG
- CD2
  1097   1096   1098       1    1.0950e+02    2.9288e+02 ;   HD11 - CD1
- HD12
  1097   1096   1099       1    1.0950e+02    2.9288e+02 ;   HD11 - CD1
- HD13
  1098   1096   1099       1    1.0950e+02    2.9288e+02 ;   HD12 - CD1
- HD13
  1101   1100   1102       1    1.0950e+02    2.9288e+02 ;   HD21 - CD2
- HD22
```

1101 1100 1103 1 1.0950e+02 2.9288e+02 ; HD21 - CD2
- HD23
1102 1100 1103 1 1.0950e+02 2.9288e+02 ; HD22 - CD2
- HD23
1104 1106 1107 1 1.2000e+02 4.1840e+02 ; C - N
- H
1104 1106 1108 1 1.2190e+02 4.1840e+02 ; C - N
- CA
1105 1104 1106 1 1.2290e+02 6.6944e+02 ; O - C
- N
1106 1108 1109 1 1.0950e+02 4.1840e+02 ; N - CA
- HA
1106 1108 1110 1 1.0970e+02 6.6944e+02 ; N - CA
- CB
1106 1108 1116 1 1.1010e+02 5.2718e+02 ; N - CA
- C
1107 1106 1108 1 1.1804e+02 4.1840e+02 ; H - N
- CA
1108 1110 1111 1 1.0950e+02 4.1840e+02 ; CA - CB
- HB2
1108 1110 1112 1 1.0950e+02 4.1840e+02 ; CA - CB
- HB3
1108 1110 1113 1 1.1110e+02 5.2718e+02 ; CA - CB
- CG
1108 1116 1117 1 1.2040e+02 6.6944e+02 ; CA - C
- O
1108 1116 1118 1 1.1660e+02 5.8576e+02 ; CA - C
- N
1109 1108 1110 1 1.0950e+02 4.1840e+02 ; HA - CA
- CB
1109 1108 1116 1 1.0950e+02 4.1840e+02 ; HA - CA
- C
1110 1108 1116 1 1.1110e+02 5.2718e+02 ; CB - CA
- C
1110 1113 1114 1 1.1700e+02 5.8576e+02 ; CB - CG
- OD1
1110 1113 1115 1 1.1700e+02 5.8576e+02 ; CB - CG
- OD2
1111 1110 1112 1 1.0950e+02 2.9288e+02 ; HB2 - CB
- HB3
1111 1110 1113 1 1.0950e+02 4.1840e+02 ; HB2 - CB
- CG
1112 1110 1113 1 1.0950e+02 4.1840e+02 ; HB3 - CB
- CG
1114 1113 1115 1 1.2600e+02 6.6944e+02 ; OD1 - CG
- OD2
1116 1118 1119 1 1.2000e+02 4.1840e+02 ; C - N
- H
1116 1118 1120 1 1.2190e+02 4.1840e+02 ; C - N
- CA
1117 1116 1118 1 1.2290e+02 6.6944e+02 ; O - C
- N

```
  1118   1120   1121      1    1.0950e+02    4.1840e+02 ;      N - CA
- HA
  1118   1120   1122      1    1.0970e+02    6.6944e+02 ;      N - CA
- CB
  1118   1120   1133      1    1.1010e+02    5.2718e+02 ;      N - CA
- C
  1119   1118   1120      1    1.1804e+02    4.1840e+02 ;      H - N
- CA
  1120   1122   1123      1    1.0950e+02    4.1840e+02 ;     CA - CB
- HB2
  1120   1122   1124      1    1.0950e+02    4.1840e+02 ;     CA - CB
- HB3
  1120   1122   1125      1    1.1310e+02    5.2718e+02 ;     CA - CB
- CG
  1120   1133   1134      1    1.2040e+02    6.6944e+02 ;     CA - C
- O
  1120   1133   1135      1    1.1660e+02    5.8576e+02 ;     CA - C
- N
  1121   1120   1122      1    1.0950e+02    4.1840e+02 ;     HA - CA
- CB
  1121   1120   1133      1    1.0950e+02    4.1840e+02 ;     HA - CA
- C
  1122   1120   1133      1    1.1110e+02    5.2718e+02 ;     CB - CA
- C
  1122   1125   1126      1    1.2000e+02    5.8576e+02 ;     CB - CG
- ND1
  1122   1125   1131      1    1.2000e+02    5.8576e+02 ;     CB - CG
- CD2
  1123   1122   1124      1    1.0950e+02    2.9288e+02 ;    HB2 - CB
- HB3
  1123   1122   1125      1    1.0950e+02    4.1840e+02 ;    HB2 - CB
- CG
  1124   1122   1125      1    1.0950e+02    4.1840e+02 ;    HB3 - CB
- CG
  1125   1126   1127      1    1.1700e+02    5.8576e+02 ;     CG - ND1
- CE1
  1125   1131   1129      1    1.2000e+02    5.8576e+02 ;     CG - CD2
- NE2
  1125   1131   1132      1    1.2000e+02    4.1840e+02 ;     CG - CD2
- HD2
  1126   1125   1131      1    1.2000e+02    5.8576e+02 ;    ND1 - CG
- CD2
  1126   1127   1128      1    1.2000e+02    4.1840e+02 ;    ND1 - CE1
- HE1
  1126   1127   1129      1    1.2000e+02    5.8576e+02 ;    ND1 - CE1
- NE2
  1127   1129   1130      1    1.2000e+02    4.1840e+02 ;    CE1 - NE2
- HE2
  1127   1129   1131      1    1.2000e+02    5.8576e+02 ;    CE1 - NE2
- CD2
  1128   1127   1129      1    1.2000e+02    4.1840e+02 ;    HE1 - CE1
- NE2
```

1129 1131 1132 1 1.2000e+02 4.1840e+02 ; NE2 - CD2
- HD2
1130 1129 1131 1 1.2000e+02 4.1840e+02 ; HE2 - NE2
- CD2
1133 1135 1136 1 1.2190e+02 4.1840e+02 ; C - N
- CD
1133 1135 1145 1 1.2190e+02 4.1840e+02 ; C - N
- CA
1134 1133 1135 1 1.2290e+02 6.6944e+02 ; O - C
- N
1135 1136 1137 1 1.0950e+02 4.1840e+02 ; N - CD
- HD2
1135 1136 1138 1 1.0950e+02 4.1840e+02 ; N - CD
- HD3
1135 1136 1139 1 1.0970e+02 6.6944e+02 ; N - CD
- CG
1135 1145 1142 1 1.0970e+02 6.6944e+02 ; N - CA
- CB
1135 1145 1146 1 1.0950e+02 4.1840e+02 ; N - CA
- HA
1135 1145 1147 1 1.1010e+02 5.2718e+02 ; N - CA
- C
1136 1135 1145 1 1.1800e+02 4.1840e+02 ; CD - N
- CA
1136 1139 1140 1 1.0950e+02 4.1840e+02 ; CD - CG
- HG2
1136 1139 1141 1 1.0950e+02 4.1840e+02 ; CD - CG
- HG3
1136 1139 1142 1 1.0950e+02 3.3472e+02 ; CD - CG
- CB
1137 1136 1138 1 1.0950e+02 2.9288e+02 ; HD2 - CD
- HD3
1137 1136 1139 1 1.0950e+02 4.1840e+02 ; HD2 - CD
- CG
1138 1136 1139 1 1.0950e+02 4.1840e+02 ; HD3 - CD
- CG
1139 1142 1143 1 1.0950e+02 4.1840e+02 ; CG - CB
- HB2
1139 1142 1144 1 1.0950e+02 4.1840e+02 ; CG - CB
- HB3
1139 1142 1145 1 1.0950e+02 3.3472e+02 ; CG - CB
- CA
1140 1139 1141 1 1.0950e+02 2.9288e+02 ; HG2 - CG
- HG3
1140 1139 1142 1 1.0950e+02 4.1840e+02 ; HG2 - CG
- CB
1141 1139 1142 1 1.0950e+02 4.1840e+02 ; HG3 - CG
- CB
1142 1145 1146 1 1.0950e+02 4.1840e+02 ; CB - CA
- HA
1142 1145 1147 1 1.1110e+02 5.2718e+02 ; CB - CA
- C

1143 1142 1144 1 1.0950e+02 2.9288e+02 ; HB2 - CB
- HB3
1143 1142 1145 1 1.0950e+02 4.1840e+02 ; HB2 - CB
- CA
1144 1142 1145 1 1.0950e+02 4.1840e+02 ; HB3 - CB
- CA
1145 1147 1148 1 1.2040e+02 6.6944e+02 ; CA - C
- O
1145 1147 1149 1 1.1660e+02 5.8576e+02 ; CA - C
- N
1146 1145 1147 1 1.0950e+02 4.1840e+02 ; HA - CA
- C
1147 1149 1150 1 1.2000e+02 4.1840e+02 ; C - N
- H
1147 1149 1151 1 1.2190e+02 4.1840e+02 ; C - N
- CA
1148 1147 1149 1 1.2290e+02 6.6944e+02 ; O - C
- N
1149 1151 1152 1 1.0950e+02 4.1840e+02 ; N - CA
- HA
1149 1151 1153 1 1.0970e+02 6.6944e+02 ; N - CA
- CB
1149 1151 1161 1 1.1010e+02 5.2718e+02 ; N - CA
- C
1150 1149 1151 1 1.1804e+02 4.1840e+02 ; H - N
- CA
1151 1153 1154 1 1.0950e+02 4.1840e+02 ; CA - CB
- HB2
1151 1153 1155 1 1.0950e+02 4.1840e+02 ; CA - CB
- HB3
1151 1153 1156 1 1.1110e+02 5.2718e+02 ; CA - CB
- CG
1151 1161 1162 1 1.2040e+02 6.6944e+02 ; CA - C
- O
1151 1161 1163 1 1.1660e+02 5.8576e+02 ; CA - C
- N
1152 1151 1153 1 1.0950e+02 4.1840e+02 ; HA - CA
- CB
1152 1151 1161 1 1.0950e+02 4.1840e+02 ; HA - CA
- C
1153 1151 1161 1 1.1110e+02 5.2718e+02 ; CB - CA
- C
1153 1156 1157 1 1.2040e+02 6.6944e+02 ; CB - CG
- OD1
1153 1156 1158 1 1.1660e+02 5.8576e+02 ; CB - CG
- ND2
1154 1153 1155 1 1.0950e+02 2.9288e+02 ; HB2 - CB
- HB3
1154 1153 1156 1 1.0950e+02 4.1840e+02 ; HB2 - CB
- CG
1155 1153 1156 1 1.0950e+02 4.1840e+02 ; HB3 - CB
- CG

```
  1156   1158   1159      1    1.2000e+02    4.1840e+02 ;     CG - ND2
- HD21
  1156   1158   1160      1    1.2000e+02    4.1840e+02 ;     CG - ND2
- HD22
  1157   1156   1158      1    1.2290e+02    6.6944e+02 ;    OD1 - CG
- ND2
  1159   1158   1160      1    1.2000e+02    2.9288e+02 ;   HD21 - ND2
- HD22
  1161   1163   1164      1    1.2000e+02    4.1840e+02 ;      C - N
- H
  1161   1163   1165      1    1.2190e+02    4.1840e+02 ;      C - N
- CA
  1162   1161   1163      1    1.2290e+02    6.6944e+02 ;      O - C
- N
  1163   1165   1166      1    1.0950e+02    4.1840e+02 ;      N - CA
- HA
  1163   1165   1167      1    1.0970e+02    6.6944e+02 ;      N - CA
- CB
  1163   1165   1180      1    1.1010e+02    5.2718e+02 ;      N - CA
- C
  1164   1163   1165      1    1.1804e+02    4.1840e+02 ;      H - N
- CA
  1165   1167   1168      1    1.0950e+02    4.1840e+02 ;     CA - CB
- HB
  1165   1167   1169      1    1.0950e+02    3.3472e+02 ;     CA - CB
- CG2
  1165   1167   1173      1    1.0950e+02    3.3472e+02 ;     CA - CB
- CG1
  1165   1180   1181      1    1.2040e+02    6.6944e+02 ;     CA - C
- O
  1165   1180   1182      1    1.1660e+02    5.8576e+02 ;     CA - C
- N
  1166   1165   1167      1    1.0950e+02    4.1840e+02 ;     HA - CA
- CB
  1166   1165   1180      1    1.0950e+02    4.1840e+02 ;     HA - CA
- C
  1167   1165   1180      1    1.1110e+02    5.2718e+02 ;     CB - CA
- C
  1167   1169   1170      1    1.0950e+02    4.1840e+02 ;     CB - CG2
- HG21
  1167   1169   1171      1    1.0950e+02    4.1840e+02 ;     CB - CG2
- HG22
  1167   1169   1172      1    1.0950e+02    4.1840e+02 ;     CB - CG2
- HG23
  1167   1173   1174      1    1.0950e+02    4.1840e+02 ;     CB - CG1
- HG12
  1167   1173   1175      1    1.0950e+02    4.1840e+02 ;     CB - CG1
- HG13
  1167   1173   1176      1    1.0950e+02    3.3472e+02 ;     CB - CG1
- CD1
  1168   1167   1169      1    1.0950e+02    4.1840e+02 ;     HB - CB
- CG2
```

1168 1167 1173 1 1.0950e+02 4.1840e+02 ; HB - CB
- CG1
1169 1167 1173 1 1.0950e+02 3.3472e+02 ; CG2 - CB
- CG1
1170 1169 1171 1 1.0950e+02 2.9288e+02 ; HG21 - CG2
- HG22
1170 1169 1172 1 1.0950e+02 2.9288e+02 ; HG21 - CG2
- HG23
1171 1169 1172 1 1.0950e+02 2.9288e+02 ; HG22 - CG2
- HG23
1173 1176 1177 1 1.0950e+02 4.1840e+02 ; CG1 - CD1
- HD11
1173 1176 1178 1 1.0950e+02 4.1840e+02 ; CG1 - CD1
- HD12
1173 1176 1179 1 1.0950e+02 4.1840e+02 ; CG1 - CD1
- HD13
1174 1173 1175 1 1.0950e+02 2.9288e+02 ; HG12 - CG1
- HG13
1174 1173 1176 1 1.0950e+02 4.1840e+02 ; HG12 - CG1
- CD1
1175 1173 1176 1 1.0950e+02 4.1840e+02 ; HG13 - CG1
- CD1
1177 1176 1178 1 1.0950e+02 2.9288e+02 ; HD11 - CD1
- HD12
1177 1176 1179 1 1.0950e+02 2.9288e+02 ; HD11 - CD1
- HD13
1178 1176 1179 1 1.0950e+02 2.9288e+02 ; HD12 - CD1
- HD13
1180 1182 1183 1 1.2000e+02 4.1840e+02 ; C - N
- H
1180 1182 1184 1 1.2190e+02 4.1840e+02 ; C - N
- CA
1181 1180 1182 1 1.2290e+02 6.6944e+02 ; O - C
- N
1182 1184 1185 1 1.0950e+02 4.1840e+02 ; N - CA
- HA
1182 1184 1186 1 1.0970e+02 6.6944e+02 ; N - CA
- CB
1182 1184 1196 1 1.1010e+02 5.2718e+02 ; N - CA
- C
1183 1182 1184 1 1.1804e+02 4.1840e+02 ; H - N
- CA
1184 1186 1187 1 1.0950e+02 4.1840e+02 ; CA - CB
- HB
1184 1186 1188 1 1.0950e+02 3.3472e+02 ; CA - CB
- CG1
1184 1186 1192 1 1.0950e+02 3.3472e+02 ; CA - CB
- CG2
1184 1196 1197 1 1.2040e+02 6.6944e+02 ; CA - C
- O
1184 1196 1198 1 1.1660e+02 5.8576e+02 ; CA - C
- N

1185 1184 1186 1 1.0950e+02 4.1840e+02 ; HA - CA
- CB
1185 1184 1196 1 1.0950e+02 4.1840e+02 ; HA - CA
- C
1186 1184 1196 1 1.1110e+02 5.2718e+02 ; CB - CA
- C
1186 1188 1189 1 1.0950e+02 4.1840e+02 ; CB - CG1
- HG11
1186 1188 1190 1 1.0950e+02 4.1840e+02 ; CB - CG1
- HG12
1186 1188 1191 1 1.0950e+02 4.1840e+02 ; CB - CG1
- HG13
1186 1192 1193 1 1.0950e+02 4.1840e+02 ; CB - CG2
- HG21
1186 1192 1194 1 1.0950e+02 4.1840e+02 ; CB - CG2
- HG22
1186 1192 1195 1 1.0950e+02 4.1840e+02 ; CB - CG2
- HG23
1187 1186 1188 1 1.0950e+02 4.1840e+02 ; HB - CB
- CG1
1187 1186 1192 1 1.0950e+02 4.1840e+02 ; HB - CB
- CG2
1188 1186 1192 1 1.0950e+02 3.3472e+02 ; CG1 - CB
- CG2
1189 1188 1190 1 1.0950e+02 2.9288e+02 ; HG11 - CG1
- HG12
1189 1188 1191 1 1.0950e+02 2.9288e+02 ; HG11 - CG1
- HG13
1190 1188 1191 1 1.0950e+02 2.9288e+02 ; HG12 - CG1
- HG13
1193 1192 1194 1 1.0950e+02 2.9288e+02 ; HG21 - CG2
- HG22
1193 1192 1195 1 1.0950e+02 2.9288e+02 ; HG21 - CG2
- HG23
1194 1192 1195 1 1.0950e+02 2.9288e+02 ; HG22 - CG2
- HG23
1196 1198 1199 1 1.2000e+02 4.1840e+02 ; C - N
- H
1196 1198 1200 1 1.2190e+02 4.1840e+02 ; C - N
- CA
1197 1196 1198 1 1.2290e+02 6.6944e+02 ; O - C
- N
1198 1200 1201 1 1.0950e+02 4.1840e+02 ; N - CA
- HA
1198 1200 1202 1 1.0970e+02 6.6944e+02 ; N - CA
- CB
1198 1200 1218 1 1.1010e+02 5.2718e+02 ; N - CA
- C
1199 1198 1200 1 1.1804e+02 4.1840e+02 ; H - N
- CA
1200 1202 1203 1 1.0950e+02 4.1840e+02 ; CA - CB
- HB2

1200 1202 1204 1 1.0950e+02 4.1840e+02 ; CA - CB
- HB3
1200 1202 1205 1 1.0950e+02 3.3472e+02 ; CA - CB
- CG
1200 1218 1219 1 1.2040e+02 6.6944e+02 ; CA - C
- O
1200 1218 1220 1 1.1660e+02 5.8576e+02 ; CA - C
- N
1201 1200 1202 1 1.0950e+02 4.1840e+02 ; HA - CA
- CB
1201 1200 1218 1 1.0950e+02 4.1840e+02 ; HA - CA
- C
1202 1200 1218 1 1.1110e+02 5.2718e+02 ; CB - CA
- C
1202 1205 1206 1 1.0950e+02 4.1840e+02 ; CB - CG
- HG2
1202 1205 1207 1 1.0950e+02 4.1840e+02 ; CB - CG
- HG3
1202 1205 1208 1 1.0950e+02 3.3472e+02 ; CB - CG
- CD
1203 1202 1204 1 1.0950e+02 2.9288e+02 ; HB2 - CB
- HB3
1203 1202 1205 1 1.0950e+02 4.1840e+02 ; HB2 - CB
- CG
1204 1202 1205 1 1.0950e+02 4.1840e+02 ; HB3 - CB
- CG
1205 1208 1209 1 1.0950e+02 4.1840e+02 ; CG - CD
- HD2
1205 1208 1210 1 1.0950e+02 4.1840e+02 ; CG - CD
- HD3
1205 1208 1211 1 1.0950e+02 3.3472e+02 ; CG - CD
- CE
1206 1205 1207 1 1.0950e+02 2.9288e+02 ; HG2 - CG
- HG3
1206 1205 1208 1 1.0950e+02 4.1840e+02 ; HG2 - CG
- CD
1207 1205 1208 1 1.0950e+02 4.1840e+02 ; HG3 - CG
- CD
1208 1211 1212 1 1.0950e+02 4.1840e+02 ; CD - CE
- HE2
1208 1211 1213 1 1.0950e+02 4.1840e+02 ; CD - CE
- HE3
1208 1211 1214 1 1.1120e+02 6.6944e+02 ; CD - CE
- NZ
1209 1208 1210 1 1.0950e+02 2.9288e+02 ; HD2 - CD
- HD3
1209 1208 1211 1 1.0950e+02 4.1840e+02 ; HD2 - CD
- CE
1210 1208 1211 1 1.0950e+02 4.1840e+02 ; HD3 - CD
- CE
1211 1214 1215 1 1.0950e+02 4.1840e+02 ; CE - NZ
- HZ1

```
  1211   1214   1216       1    1.0950e+02    4.1840e+02 ;     CE - NZ
- HZ2
  1211   1214   1217       1    1.0950e+02    4.1840e+02 ;     CE - NZ
- HZ3
  1212   1211   1213       1    1.0950e+02    2.9288e+02 ;    HE2 - CE
- HE3
  1212   1211   1214       1    1.0950e+02    4.1840e+02 ;    HE2 - CE
- NZ
  1213   1211   1214       1    1.0950e+02    4.1840e+02 ;    HE3 - CE
- NZ
  1215   1214   1216       1    1.0950e+02    2.9288e+02 ;    HZ1 - NZ
- HZ2
  1215   1214   1217       1    1.0950e+02    2.9288e+02 ;    HZ1 - NZ
- HZ3
  1216   1214   1217       1    1.0950e+02    2.9288e+02 ;    HZ2 - NZ
- HZ3
  1218   1220   1221       1    1.2000e+02    4.1840e+02 ;      C - N
- H
  1218   1220   1222       1    1.2190e+02    4.1840e+02 ;      C - N
- CA
  1219   1218   1220       1    1.2290e+02    6.6944e+02 ;      O - C
- N
  1220   1222   1223       1    1.0950e+02    4.1840e+02 ;      N - CA
- HA
  1220   1222   1224       1    1.0970e+02    6.6944e+02 ;      N - CA
- CB
  1220   1222   1237       1    1.1010e+02    5.2718e+02 ;      N - CA
- C
  1221   1220   1222       1    1.1804e+02    4.1840e+02 ;      H - N
- CA
  1222   1224   1225       1    1.0950e+02    4.1840e+02 ;     CA - CB
- HB2
  1222   1224   1226       1    1.0950e+02    4.1840e+02 ;     CA - CB
- HB3
  1222   1224   1227       1    1.0950e+02    3.3472e+02 ;     CA - CB
- CG
  1222   1237   1238       1    1.2040e+02    6.6944e+02 ;     CA - C
- O
  1222   1237   1239       1    1.1660e+02    5.8576e+02 ;     CA - C
- N
  1223   1222   1224       1    1.0950e+02    4.1840e+02 ;     HA - CA
- CB
  1223   1222   1237       1    1.0950e+02    4.1840e+02 ;     HA - CA
- C
  1224   1222   1237       1    1.1110e+02    5.2718e+02 ;     CB - CA
- C
  1224   1227   1228       1    1.0950e+02    4.1840e+02 ;     CB - CG
- HG
  1224   1227   1229       1    1.0950e+02    3.3472e+02 ;     CB - CG
- CD1
  1224   1227   1233       1    1.0950e+02    3.3472e+02 ;     CB - CG
- CD2
```

1225 1224 1226 1 1.0950e+02 2.9288e+02 ; HB2 - CB
- HB3
1225 1224 1227 1 1.0950e+02 4.1840e+02 ; HB2 - CB
- CG
1226 1224 1227 1 1.0950e+02 4.1840e+02 ; HB3 - CB
- CG
1227 1229 1230 1 1.0950e+02 4.1840e+02 ; CG - CD1
- HD11
1227 1229 1231 1 1.0950e+02 4.1840e+02 ; CG - CD1
- HD12
1227 1229 1232 1 1.0950e+02 4.1840e+02 ; CG - CD1
- HD13
1227 1233 1234 1 1.0950e+02 4.1840e+02 ; CG - CD2
- HD21
1227 1233 1235 1 1.0950e+02 4.1840e+02 ; CG - CD2
- HD22
1227 1233 1236 1 1.0950e+02 4.1840e+02 ; CG - CD2
- HD23
1228 1227 1229 1 1.0950e+02 4.1840e+02 ; HG - CG
- CD1
1228 1227 1233 1 1.0950e+02 4.1840e+02 ; HG - CG
- CD2
1229 1227 1233 1 1.0950e+02 3.3472e+02 ; CD1 - CG
- CD2
1230 1229 1231 1 1.0950e+02 2.9288e+02 ; HD11 - CD1
- HD12
1230 1229 1232 1 1.0950e+02 2.9288e+02 ; HD11 - CD1
- HD13
1231 1229 1232 1 1.0950e+02 2.9288e+02 ; HD12 - CD1
- HD13
1234 1233 1235 1 1.0950e+02 2.9288e+02 ; HD21 - CD2
- HD22
1234 1233 1236 1 1.0950e+02 2.9288e+02 ; HD21 - CD2
- HD23
1235 1233 1236 1 1.0950e+02 2.9288e+02 ; HD22 - CD2
- HD23
1237 1239 1240 1 1.2000e+02 4.1840e+02 ; C - N
- H
1237 1239 1241 1 1.2190e+02 4.1840e+02 ; C - N
- CA
1238 1237 1239 1 1.2290e+02 6.6944e+02 ; O - C
- N
1239 1241 1242 1 1.0950e+02 4.1840e+02 ; N - CA
- HA
1239 1241 1243 1 1.0970e+02 6.6944e+02 ; N - CA
- CB
1239 1241 1258 1 1.1010e+02 5.2718e+02 ; N - CA
- C
1240 1239 1241 1 1.1804e+02 4.1840e+02 ; H - N
- CA
1241 1243 1244 1 1.0950e+02 4.1840e+02 ; CA - CB
- HB2

```
  1241   1243   1245      1    1.0950e+02    4.1840e+02 ;     CA - CB
- HB3
  1241   1243   1246      1    1.1400e+02    5.2718e+02 ;     CA - CB
- CG
  1241   1258   1259      1    1.2040e+02    6.6944e+02 ;     CA - C
- O
  1241   1258   1260      1    1.1660e+02    5.8576e+02 ;     CA - C
- N
  1242   1241   1243      1    1.0950e+02    4.1840e+02 ;     HA - CA
- CB
  1242   1241   1258      1    1.0950e+02    4.1840e+02 ;     HA - CA
- C
  1243   1241   1258      1    1.1110e+02    5.2718e+02 ;     CB - CA
- C
  1243   1246   1247      1    1.2000e+02    5.8576e+02 ;     CB - CG
- CD1
  1243   1246   1256      1    1.2000e+02    5.8576e+02 ;     CB - CG
- CD2
  1244   1243   1245      1    1.0950e+02    2.9288e+02 ;    HB2 - CB
- HB3
  1244   1243   1246      1    1.0950e+02    4.1840e+02 ;    HB2 - CB
- CG
  1245   1243   1246      1    1.0950e+02    4.1840e+02 ;    HB3 - CB
- CG
  1246   1247   1248      1    1.2000e+02    4.1840e+02 ;     CG - CD1
- HD1
  1246   1247   1249      1    1.2000e+02    5.2718e+02 ;     CG - CD1
- CE1
  1246   1256   1254      1    1.2000e+02    5.2718e+02 ;     CG - CD2
- CE2
  1246   1256   1257      1    1.2000e+02    4.1840e+02 ;     CG - CD2
- HD2
  1247   1246   1256      1    1.2000e+02    5.2718e+02 ;    CD1 - CG
- CD2
  1247   1249   1250      1    1.2000e+02    4.1840e+02 ;    CD1 - CE1
- HE1
  1247   1249   1251      1    1.2000e+02    5.2718e+02 ;    CD1 - CE1
- CZ
  1248   1247   1249      1    1.2000e+02    4.1840e+02 ;    HD1 - CD1
- CE1
  1249   1251   1252      1    1.2000e+02    5.8576e+02 ;    CE1 - CZ
- OH
  1249   1251   1254      1    1.2000e+02    5.2718e+02 ;    CE1 - CZ
- CE2
  1250   1249   1251      1    1.2000e+02    4.1840e+02 ;    HE1 - CE1
- CZ
  1251   1252   1253      1    1.1300e+02    4.1840e+02 ;     CZ - OH
- HH
  1251   1254   1255      1    1.2000e+02    4.1840e+02 ;     CZ - CE2
- HE2
  1251   1254   1256      1    1.2000e+02    5.2718e+02 ;     CZ - CE2
- CD2
```

```
  1252   1251   1254       1    1.2000e+02    5.8576e+02 ;     OH - CZ
- CE2
  1254   1256   1257       1    1.2000e+02    4.1840e+02 ;    CE2 - CD2
- HD2
  1255   1254   1256       1    1.2000e+02    4.1840e+02 ;    HE2 - CE2
- CD2
  1258   1260   1261       1    1.2000e+02    4.1840e+02 ;      C - N
- H
  1258   1260   1262       1    1.2190e+02    4.1840e+02 ;      C - N
- CA
  1259   1258   1260       1    1.2290e+02    6.6944e+02 ;      O - C
- N
  1260   1262   1263       1    1.0950e+02    4.1840e+02 ;      N - CA
- HA
  1260   1262   1264       1    1.0970e+02    6.6944e+02 ;      N - CA
- CB
  1260   1262   1273       1    1.1010e+02    5.2718e+02 ;      N - CA
- C
  1261   1260   1262       1    1.1804e+02    4.1840e+02 ;      H - N
- CA
  1262   1264   1265       1    1.0950e+02    4.1840e+02 ;     CA - CB
- HB2
  1262   1264   1266       1    1.0950e+02    4.1840e+02 ;     CA - CB
- HB3
  1262   1264   1267       1    1.0950e+02    3.3472e+02 ;     CA - CB
- CG
  1262   1273   1274       1    1.2040e+02    6.6944e+02 ;     CA - C
- O
  1262   1273   1275       1    1.1660e+02    5.8576e+02 ;     CA - C
- N
  1263   1262   1264       1    1.0950e+02    4.1840e+02 ;     HA - CA
- CB
  1263   1262   1273       1    1.0950e+02    4.1840e+02 ;     HA - CA
- C
  1264   1262   1273       1    1.1110e+02    5.2718e+02 ;     CB - CA
- C
  1264   1267   1268       1    1.0950e+02    4.1840e+02 ;     CB - CG
- HG2
  1264   1267   1269       1    1.0950e+02    4.1840e+02 ;     CB - CG
- HG3
  1264   1267   1270       1    1.1110e+02    5.2718e+02 ;     CB - CG
- CD
  1265   1264   1266       1    1.0950e+02    2.9288e+02 ;    HB2 - CB
- HB3
  1265   1264   1267       1    1.0950e+02    4.1840e+02 ;    HB2 - CB
- CG
  1266   1264   1267       1    1.0950e+02    4.1840e+02 ;    HB3 - CB
- CG
  1267   1270   1271       1    1.1700e+02    5.8576e+02 ;     CG - CD
- OE1
  1267   1270   1272       1    1.1700e+02    5.8576e+02 ;     CG - CD
- OE2
```

```
  1268   1267   1269       1    1.0950e+02    2.9288e+02 ;    HG2 - CG
- HG3
  1268   1267   1270       1    1.0950e+02    4.1840e+02 ;    HG2 - CG
- CD
  1269   1267   1270       1    1.0950e+02    4.1840e+02 ;    HG3 - CG
- CD
  1271   1270   1272       1    1.2600e+02    6.6944e+02 ;    OE1 - CD
- OE2
  1273   1275   1276       1    1.2000e+02    4.1840e+02 ;      C - N
- H
  1273   1275   1277       1    1.2190e+02    4.1840e+02 ;      C - N
- CA
  1274   1273   1275       1    1.2290e+02    6.6944e+02 ;      O - C
- N
  1275   1277   1278       1    1.0950e+02    4.1840e+02 ;      N - CA
- HA
  1275   1277   1279       1    1.0970e+02    6.6944e+02 ;      N - CA
- CB
  1275   1277   1287       1    1.1010e+02    5.2718e+02 ;      N - CA
- C
  1276   1275   1277       1    1.1804e+02    4.1840e+02 ;      H - N
- CA
  1277   1279   1280       1    1.0950e+02    4.1840e+02 ;     CA - CB
- HB
  1277   1279   1281       1    1.0950e+02    3.3472e+02 ;     CA - CB
- CG2
  1277   1279   1285       1    1.0950e+02    4.1840e+02 ;     CA - CB
- OG1
  1277   1287   1288       1    1.2040e+02    6.6944e+02 ;     CA - C
- O
  1277   1287   1289       1    1.1660e+02    5.8576e+02 ;     CA - C
- N
  1278   1277   1279       1    1.0950e+02    4.1840e+02 ;     HA - CA
- CB
  1278   1277   1287       1    1.0950e+02    4.1840e+02 ;     HA - CA
- C
  1279   1277   1287       1    1.1110e+02    5.2718e+02 ;     CB - CA
- C
  1279   1281   1282       1    1.0950e+02    4.1840e+02 ;     CB - CG2
- HG21
  1279   1281   1283       1    1.0950e+02    4.1840e+02 ;     CB - CG2
- HG22
  1279   1281   1284       1    1.0950e+02    4.1840e+02 ;     CB - CG2
- HG23
  1279   1285   1286       1    1.0850e+02    4.6024e+02 ;     CB - OG1
- HG1
  1280   1279   1281       1    1.0950e+02    4.1840e+02 ;     HB - CB
- CG2
  1280   1279   1285       1    1.0950e+02    4.1840e+02 ;     HB - CB
- OG1
  1281   1279   1285       1    1.0950e+02    4.1840e+02 ;    CG2 - CB
- OG1
```

```
  1282   1281   1283       1    1.0950e+02    2.9288e+02 ;   HG21 - CG2
- HG22
  1282   1281   1284       1    1.0950e+02    2.9288e+02 ;   HG21 - CG2
- HG23
  1283   1281   1284       1    1.0950e+02    2.9288e+02 ;   HG22 - CG2
- HG23
  1287   1289   1290       1    1.2000e+02    4.1840e+02 ;      C - N
- H
  1287   1289   1291       1    1.2190e+02    4.1840e+02 ;      C - N
- CA
  1288   1287   1289       1    1.2290e+02    6.6944e+02 ;      O - C
- N
  1289   1291   1292       1    1.0950e+02    4.1840e+02 ;      N - CA
- HA
  1289   1291   1293       1    1.0970e+02    6.6944e+02 ;      N - CA
- CB
  1289   1291   1308       1    1.1010e+02    5.2718e+02 ;      N - CA
- C
  1290   1289   1291       1    1.1804e+02    4.1840e+02 ;      H - N
- CA
  1291   1293   1294       1    1.0950e+02    4.1840e+02 ;     CA - CB
- HB2
  1291   1293   1295       1    1.0950e+02    4.1840e+02 ;     CA - CB
- HB3
  1291   1293   1296       1    1.1400e+02    5.2718e+02 ;     CA - CB
- CG
  1291   1308   1309       1    1.2040e+02    6.6944e+02 ;     CA - C
- O
  1291   1308   1310       1    1.1660e+02    5.8576e+02 ;     CA - C
- N
  1292   1291   1293       1    1.0950e+02    4.1840e+02 ;     HA - CA
- CB
  1292   1291   1308       1    1.0950e+02    4.1840e+02 ;     HA - CA
- C
  1293   1291   1308       1    1.1110e+02    5.2718e+02 ;     CB - CA
- C
  1293   1296   1297       1    1.2000e+02    5.8576e+02 ;     CB - CG
- CD1
  1293   1296   1306       1    1.2000e+02    5.8576e+02 ;     CB - CG
- CD2
  1294   1293   1295       1    1.0950e+02    2.9288e+02 ;    HB2 - CB
- HB3
  1294   1293   1296       1    1.0950e+02    4.1840e+02 ;    HB2 - CB
- CG
  1295   1293   1296       1    1.0950e+02    4.1840e+02 ;    HB3 - CB
- CG
  1296   1297   1298       1    1.2000e+02    4.1840e+02 ;     CG - CD1
- HD1
  1296   1297   1299       1    1.2000e+02    5.2718e+02 ;     CG - CD1
- CE1
  1296   1306   1304       1    1.2000e+02    5.2718e+02 ;     CG - CD2
- CE2
```

```
  1296   1306   1307       1    1.2000e+02    4.1840e+02 ;      CG - CD2
- HD2
  1297   1296   1306       1    1.2000e+02    5.2718e+02 ;     CD1 - CG
- CD2
  1297   1299   1300       1    1.2000e+02    4.1840e+02 ;     CD1 - CE1
- HE1
  1297   1299   1301       1    1.2000e+02    5.2718e+02 ;     CD1 - CE1
- CZ
  1298   1297   1299       1    1.2000e+02    4.1840e+02 ;     HD1 - CD1
- CE1
  1299   1301   1302       1    1.2000e+02    5.8576e+02 ;     CE1 - CZ
- OH
  1299   1301   1304       1    1.2000e+02    5.2718e+02 ;     CE1 - CZ
- CE2
  1300   1299   1301       1    1.2000e+02    4.1840e+02 ;     HE1 - CE1
- CZ
  1301   1302   1303       1    1.1300e+02    4.1840e+02 ;      CZ - OH
- HH
  1301   1304   1305       1    1.2000e+02    4.1840e+02 ;      CZ - CE2
- HE2
  1301   1304   1306       1    1.2000e+02    5.2718e+02 ;      CZ - CE2
- CD2
  1302   1301   1304       1    1.2000e+02    5.8576e+02 ;      OH - CZ
- CE2
  1304   1306   1307       1    1.2000e+02    4.1840e+02 ;     CE2 - CD2
- HD2
  1305   1304   1306       1    1.2000e+02    4.1840e+02 ;     HE2 - CE2
- CD2
  1308   1310   1311       1    1.2000e+02    4.1840e+02 ;       C - N
- H
  1308   1310   1312       1    1.2190e+02    4.1840e+02 ;       C - N
- CA
  1309   1308   1310       1    1.2290e+02    6.6944e+02 ;       O - C
- N
  1310   1312   1313       1    1.0950e+02    4.1840e+02 ;       N - CA
- HA
  1310   1312   1314       1    1.0970e+02    6.6944e+02 ;       N - CA
- CB
  1310   1312   1323       1    1.1010e+02    5.2718e+02 ;       N - CA
- C
  1311   1310   1312       1    1.1804e+02    4.1840e+02 ;       H - N
- CA
  1312   1314   1315       1    1.0950e+02    4.1840e+02 ;      CA - CB
- HB2
  1312   1314   1316       1    1.0950e+02    4.1840e+02 ;      CA - CB
- HB3
  1312   1314   1317       1    1.0950e+02    3.3472e+02 ;      CA - CB
- CG
  1312   1323   1324       1    1.2040e+02    6.6944e+02 ;      CA - C
- O
  1312   1323   1325       1    1.1660e+02    5.8576e+02 ;      CA - C
- N
```

```
  1313   1312   1314       1    1.0950e+02    4.1840e+02 ;     HA - CA
- CB
  1313   1312   1323       1    1.0950e+02    4.1840e+02 ;     HA - CA
- C
  1314   1312   1323       1    1.1110e+02    5.2718e+02 ;     CB - CA
- C
  1314   1317   1318       1    1.0950e+02    4.1840e+02 ;     CB - CG
- HG2
  1314   1317   1319       1    1.0950e+02    4.1840e+02 ;     CB - CG
- HG3
  1314   1317   1320       1    1.1110e+02    5.2718e+02 ;     CB - CG
- CD
  1315   1314   1316       1    1.0950e+02    2.9288e+02 ;    HB2 - CB
- HB3
  1315   1314   1317       1    1.0950e+02    4.1840e+02 ;    HB2 - CB
- CG
  1316   1314   1317       1    1.0950e+02    4.1840e+02 ;    HB3 - CB
- CG
  1317   1320   1321       1    1.1700e+02    5.8576e+02 ;     CG - CD
- OE1
  1317   1320   1322       1    1.1700e+02    5.8576e+02 ;     CG - CD
- OE2
  1318   1317   1319       1    1.0950e+02    2.9288e+02 ;    HG2 - CG
- HG3
  1318   1317   1320       1    1.0950e+02    4.1840e+02 ;    HG2 - CG
- CD
  1319   1317   1320       1    1.0950e+02    4.1840e+02 ;    HG3 - CG
- CD
  1321   1320   1322       1    1.2600e+02    6.6944e+02 ;    OE1 - CD
- OE2
  1323   1325   1326       1    1.2000e+02    4.1840e+02 ;      C - N
- H
  1323   1325   1327       1    1.2190e+02    4.1840e+02 ;      C - N
- CA
  1324   1323   1325       1    1.2290e+02    6.6944e+02 ;      O - C
- N
  1325   1327   1328       1    1.0950e+02    4.1840e+02 ;      N - CA
- HA
  1325   1327   1329       1    1.0970e+02    6.6944e+02 ;      N - CA
- CB
  1325   1327   1337       1    1.1010e+02    5.2718e+02 ;      N - CA
- C
  1326   1325   1327       1    1.1804e+02    4.1840e+02 ;      H - N
- CA
  1327   1329   1330       1    1.0950e+02    4.1840e+02 ;     CA - CB
- HB2
  1327   1329   1331       1    1.0950e+02    4.1840e+02 ;     CA - CB
- HB3
  1327   1329   1332       1    1.1110e+02    5.2718e+02 ;     CA - CB
- CG
  1327   1337   1338       1    1.2040e+02    6.6944e+02 ;     CA - C
- O
```

```
 1327   1337   1339      1    1.1660e+02    5.8576e+02 ;     CA - C
- N
 1328   1327   1329      1    1.0950e+02    4.1840e+02 ;     HA - CA
- CB
 1328   1327   1337      1    1.0950e+02    4.1840e+02 ;     HA - CA
- C
 1329   1327   1337      1    1.1110e+02    5.2718e+02 ;     CB - CA
- C
 1329   1332   1333      1    1.2040e+02    6.6944e+02 ;     CB - CG
- OD1
 1329   1332   1334      1    1.1660e+02    5.8576e+02 ;     CB - CG
- ND2
 1330   1329   1331      1    1.0950e+02    2.9288e+02 ;    HB2 - CB
- HB3
 1330   1329   1332      1    1.0950e+02    4.1840e+02 ;    HB2 - CB
- CG
 1331   1329   1332      1    1.0950e+02    4.1840e+02 ;    HB3 - CB
- CG
 1332   1334   1335      1    1.2000e+02    4.1840e+02 ;     CG - ND2
- HD21
 1332   1334   1336      1    1.2000e+02    4.1840e+02 ;     CG - ND2
- HD22
 1333   1332   1334      1    1.2290e+02    6.6944e+02 ;    OD1 - CG
- ND2
 1335   1334   1336      1    1.2000e+02    2.9288e+02 ;   HD21 - ND2
- HD22
 1337   1339   1340      1    1.2000e+02    4.1840e+02 ;      C - N
- H
 1337   1339   1341      1    1.2190e+02    4.1840e+02 ;      C - N
- CA
 1338   1337   1339      1    1.2290e+02    6.6944e+02 ;      O - C
- N
 1339   1341   1342      1    1.0950e+02    4.1840e+02 ;      N - CA
- HA
 1339   1341   1343      1    1.0970e+02    6.6944e+02 ;      N - CA
- CB
 1339   1341   1349      1    1.1010e+02    5.2718e+02 ;      N - CA
- C
 1340   1339   1341      1    1.1804e+02    4.1840e+02 ;      H - N
- CA
 1341   1343   1344      1    1.0950e+02    4.1840e+02 ;     CA - CB
- HB2
 1341   1343   1345      1    1.0950e+02    4.1840e+02 ;     CA - CB
- HB3
 1341   1343   1346      1    1.1110e+02    5.2718e+02 ;     CA - CB
- CG
 1341   1349   1350      1    1.2040e+02    6.6944e+02 ;     CA - C
- O
 1341   1349   1351      1    1.1660e+02    5.8576e+02 ;     CA - C
- N
 1342   1341   1343      1    1.0950e+02    4.1840e+02 ;     HA - CA
- CB
```

```
  1342   1341   1349      1    1.0950e+02    4.1840e+02 ;     HA - CA
- C
  1343   1341   1349      1    1.1110e+02    5.2718e+02 ;     CB - CA
- C
  1343   1346   1347      1    1.1700e+02    5.8576e+02 ;     CB - CG
- OD1
  1343   1346   1348      1    1.1700e+02    5.8576e+02 ;     CB - CG
- OD2
  1344   1343   1345      1    1.0950e+02    2.9288e+02 ;    HB2 - CB
- HB3
  1344   1343   1346      1    1.0950e+02    4.1840e+02 ;    HB2 - CB
- CG
  1345   1343   1346      1    1.0950e+02    4.1840e+02 ;    HB3 - CB
- CG
  1347   1346   1348      1    1.2600e+02    6.6944e+02 ;    OD1 - CG
- OD2
  1349   1351   1352      1    1.2000e+02    4.1840e+02 ;      C - N
- H
  1349   1351   1353      1    1.2190e+02    4.1840e+02 ;      C - N
- CA
  1350   1349   1351      1    1.2290e+02    6.6944e+02 ;      O - C
- N
  1351   1353   1354      1    1.0950e+02    4.1840e+02 ;      N - CA
- HA
  1351   1353   1355      1    1.0970e+02    6.6944e+02 ;      N - CA
- CB
  1351   1353   1363      1    1.1010e+02    5.2718e+02 ;      N - CA
- C
  1352   1351   1353      1    1.1804e+02    4.1840e+02 ;      H - N
- CA
  1353   1355   1356      1    1.0950e+02    4.1840e+02 ;     CA - CB
- HB2
  1353   1355   1357      1    1.0950e+02    4.1840e+02 ;     CA - CB
- HB3
  1353   1355   1358      1    1.1110e+02    5.2718e+02 ;     CA - CB
- CG
  1353   1363   1364      1    1.2040e+02    6.6944e+02 ;     CA - C
- O
  1353   1363   1365      1    1.1660e+02    5.8576e+02 ;     CA - C
- N
  1354   1353   1355      1    1.0950e+02    4.1840e+02 ;     HA - CA
- CB
  1354   1353   1363      1    1.0950e+02    4.1840e+02 ;     HA - CA
- C
  1355   1353   1363      1    1.1110e+02    5.2718e+02 ;     CB - CA
- C
  1355   1358   1359      1    1.2040e+02    6.6944e+02 ;     CB - CG
- OD1
  1355   1358   1360      1    1.1660e+02    5.8576e+02 ;     CB - CG
- ND2
  1356   1355   1357      1    1.0950e+02    2.9288e+02 ;    HB2 - CB
- HB3
```

1356 1355 1358 1 1.0950e+02 4.1840e+02 ; HB2 - CB
- CG
1357 1355 1358 1 1.0950e+02 4.1840e+02 ; HB3 - CB
- CG
1358 1360 1361 1 1.2000e+02 4.1840e+02 ; CG - ND2
- HD21
1358 1360 1362 1 1.2000e+02 4.1840e+02 ; CG - ND2
- HD22
1359 1358 1360 1 1.2290e+02 6.6944e+02 ; OD1 - CG
- ND2
1361 1360 1362 1 1.2000e+02 2.9288e+02 ; HD21 - ND2
- HD22
1363 1365 1366 1 1.2000e+02 4.1840e+02 ; C - N
- H
1363 1365 1367 1 1.2190e+02 4.1840e+02 ; C - N
- CA
1364 1363 1365 1 1.2290e+02 6.6944e+02 ; O - C
- N
1365 1367 1368 1 1.0950e+02 4.1840e+02 ; N - CA
- HA
1365 1367 1369 1 1.0970e+02 6.6944e+02 ; N - CA
- CB
1365 1367 1384 1 1.1010e+02 5.2718e+02 ; N - CA
- C
1366 1365 1367 1 1.1804e+02 4.1840e+02 ; H - N
- CA
1367 1369 1370 1 1.0950e+02 4.1840e+02 ; CA - CB
- HB2
1367 1369 1371 1 1.0950e+02 4.1840e+02 ; CA - CB
- HB3
1367 1369 1372 1 1.1400e+02 5.2718e+02 ; CA - CB
- CG
1367 1384 1385 1 1.2040e+02 6.6944e+02 ; CA - C
- O
1367 1384 1386 1 1.1660e+02 5.8576e+02 ; CA - C
- N
1368 1367 1369 1 1.0950e+02 4.1840e+02 ; HA - CA
- CB
1368 1367 1384 1 1.0950e+02 4.1840e+02 ; HA - CA
- C
1369 1367 1384 1 1.1110e+02 5.2718e+02 ; CB - CA
- C
1369 1372 1373 1 1.2000e+02 5.8576e+02 ; CB - CG
- CD1
1369 1372 1382 1 1.2000e+02 5.8576e+02 ; CB - CG
- CD2
1370 1369 1371 1 1.0950e+02 2.9288e+02 ; HB2 - CB
- HB3
1370 1369 1372 1 1.0950e+02 4.1840e+02 ; HB2 - CB
- CG
1371 1369 1372 1 1.0950e+02 4.1840e+02 ; HB3 - CB
- CG

1372 1373 1374 1 1.2000e+02 4.1840e+02 ; CG - CD1
- HD1
1372 1373 1375 1 1.2000e+02 5.2718e+02 ; CG - CD1
- CE1
1372 1382 1380 1 1.2000e+02 5.2718e+02 ; CG - CD2
- CE2
1372 1382 1383 1 1.2000e+02 4.1840e+02 ; CG - CD2
- HD2
1373 1372 1382 1 1.2000e+02 5.2718e+02 ; CD1 - CG
- CD2
1373 1375 1376 1 1.2000e+02 4.1840e+02 ; CD1 - CE1
- HE1
1373 1375 1377 1 1.2000e+02 5.2718e+02 ; CD1 - CE1
- CZ
1374 1373 1375 1 1.2000e+02 4.1840e+02 ; HD1 - CD1
- CE1
1375 1377 1378 1 1.2000e+02 5.8576e+02 ; CE1 - CZ
- OH
1375 1377 1380 1 1.2000e+02 5.2718e+02 ; CE1 - CZ
- CE2
1376 1375 1377 1 1.2000e+02 4.1840e+02 ; HE1 - CE1
- CZ
1377 1378 1379 1 1.1300e+02 4.1840e+02 ; CZ - OH
- HH
1377 1380 1381 1 1.2000e+02 4.1840e+02 ; CZ - CE2
- HE2
1377 1380 1382 1 1.2000e+02 5.2718e+02 ; CZ - CE2
- CD2
1378 1377 1380 1 1.2000e+02 5.8576e+02 ; OH - CZ
- CE2
1380 1382 1383 1 1.2000e+02 4.1840e+02 ; CE2 - CD2
- HD2
1381 1380 1382 1 1.2000e+02 4.1840e+02 ; HE2 - CE2
- CD2
1384 1386 1387 1 1.2000e+02 4.1840e+02 ; C - N
- H
1384 1386 1388 1 1.2190e+02 4.1840e+02 ; C - N
- CA
1385 1384 1386 1 1.2290e+02 6.6944e+02 ; O - C
- N
1386 1388 1389 1 1.0950e+02 4.1840e+02 ; N - CA
- HA
1386 1388 1390 1 1.0970e+02 6.6944e+02 ; N - CA
- CB
1386 1388 1403 1 1.1010e+02 5.2718e+02 ; N - CA
- C
1387 1386 1388 1 1.1804e+02 4.1840e+02 ; H - N
- CA
1388 1390 1391 1 1.0950e+02 4.1840e+02 ; CA - CB
- HB
1388 1390 1392 1 1.0950e+02 3.3472e+02 ; CA - CB
- CG2

1388 1390 1396 1 1.0950e+02 3.3472e+02 ; CA - CB
- CG1
1388 1403 1404 1 1.2040e+02 6.6944e+02 ; CA - C
- O
1388 1403 1405 1 1.1660e+02 5.8576e+02 ; CA - C
- N
1389 1388 1390 1 1.0950e+02 4.1840e+02 ; HA - CA
- CB
1389 1388 1403 1 1.0950e+02 4.1840e+02 ; HA - CA
- C
1390 1388 1403 1 1.1110e+02 5.2718e+02 ; CB - CA
- C
1390 1392 1393 1 1.0950e+02 4.1840e+02 ; CB - CG2
- HG21
1390 1392 1394 1 1.0950e+02 4.1840e+02 ; CB - CG2
- HG22
1390 1392 1395 1 1.0950e+02 4.1840e+02 ; CB - CG2
- HG23
1390 1396 1397 1 1.0950e+02 4.1840e+02 ; CB - CG1
- HG12
1390 1396 1398 1 1.0950e+02 4.1840e+02 ; CB - CG1
- HG13
1390 1396 1399 1 1.0950e+02 3.3472e+02 ; CB - CG1
- CD1
1391 1390 1392 1 1.0950e+02 4.1840e+02 ; HB - CB
- CG2
1391 1390 1396 1 1.0950e+02 4.1840e+02 ; HB - CB
- CG1
1392 1390 1396 1 1.0950e+02 3.3472e+02 ; CG2 - CB
- CG1
1393 1392 1394 1 1.0950e+02 2.9288e+02 ; HG21 - CG2
- HG22
1393 1392 1395 1 1.0950e+02 2.9288e+02 ; HG21 - CG2
- HG23
1394 1392 1395 1 1.0950e+02 2.9288e+02 ; HG22 - CG2
- HG23
1396 1399 1400 1 1.0950e+02 4.1840e+02 ; CG1 - CD1
- HD11
1396 1399 1401 1 1.0950e+02 4.1840e+02 ; CG1 - CD1
- HD12
1396 1399 1402 1 1.0950e+02 4.1840e+02 ; CG1 - CD1
- HD13
1397 1396 1398 1 1.0950e+02 2.9288e+02 ; HG12 - CG1
- HG13
1397 1396 1399 1 1.0950e+02 4.1840e+02 ; HG12 - CG1
- CD1
1398 1396 1399 1 1.0950e+02 4.1840e+02 ; HG13 - CG1
- CD1
1400 1399 1401 1 1.0950e+02 2.9288e+02 ; HD11 - CD1
- HD12
1400 1399 1402 1 1.0950e+02 2.9288e+02 ; HD11 - CD1
- HD13

1401 1399 1402 1 1.0950e+02 2.9288e+02 ; HD12 - CD1
- HD13
1403 1405 1406 1 1.2000e+02 4.1840e+02 ; C - N
- H
1403 1405 1407 1 1.2190e+02 4.1840e+02 ; C - N
- CA
1404 1403 1405 1 1.2290e+02 6.6944e+02 ; O - C
- N
1405 1407 1408 1 1.0950e+02 4.1840e+02 ; N - CA
- HA
1405 1407 1409 1 1.0970e+02 6.6944e+02 ; N - CA
- CB
1405 1407 1424 1 1.1010e+02 5.2718e+02 ; N - CA
- C
1406 1405 1407 1 1.1804e+02 4.1840e+02 ; H - N
- CA
1407 1409 1410 1 1.0950e+02 4.1840e+02 ; CA - CB
- HB2
1407 1409 1411 1 1.0950e+02 4.1840e+02 ; CA - CB
- HB3
1407 1409 1412 1 1.1400e+02 5.2718e+02 ; CA - CB
- CG
1407 1424 1425 1 1.2040e+02 6.6944e+02 ; CA - C
- O
1407 1424 1426 1 1.1660e+02 5.8576e+02 ; CA - C
- N
1408 1407 1409 1 1.0950e+02 4.1840e+02 ; HA - CA
- CB
1408 1407 1424 1 1.0950e+02 4.1840e+02 ; HA - CA
- C
1409 1407 1424 1 1.1110e+02 5.2718e+02 ; CB - CA
- C
1409 1412 1413 1 1.2000e+02 5.8576e+02 ; CB - CG
- CD1
1409 1412 1422 1 1.2000e+02 5.8576e+02 ; CB - CG
- CD2
1410 1409 1411 1 1.0950e+02 2.9288e+02 ; HB2 - CB
- HB3
1410 1409 1412 1 1.0950e+02 4.1840e+02 ; HB2 - CB
- CG
1411 1409 1412 1 1.0950e+02 4.1840e+02 ; HB3 - CB
- CG
1412 1413 1414 1 1.2000e+02 4.1840e+02 ; CG - CD1
- HD1
1412 1413 1415 1 1.2000e+02 5.2718e+02 ; CG - CD1
- CE1
1412 1422 1420 1 1.2000e+02 5.2718e+02 ; CG - CD2
- CE2
1412 1422 1423 1 1.2000e+02 4.1840e+02 ; CG - CD2
- HD2
1413 1412 1422 1 1.2000e+02 5.2718e+02 ; CD1 - CG
- CD2

1413 1415 1416 1 1.2000e+02 4.1840e+02 ; CD1 - CE1 - HE1
1413 1415 1417 1 1.2000e+02 5.2718e+02 ; CD1 - CE1 - CZ
1414 1413 1415 1 1.2000e+02 4.1840e+02 ; HD1 - CD1 - CE1
1415 1417 1418 1 1.2000e+02 5.8576e+02 ; CE1 - CZ - OH
1415 1417 1420 1 1.2000e+02 5.2718e+02 ; CE1 - CZ - CE2
1416 1415 1417 1 1.2000e+02 4.1840e+02 ; HE1 - CE1 - CZ
1417 1418 1419 1 1.1300e+02 4.1840e+02 ; CZ - OH - HH
1417 1420 1421 1 1.2000e+02 4.1840e+02 ; CZ - CE2 - HE2
1417 1420 1422 1 1.2000e+02 5.2718e+02 ; CZ - CE2 - CD2
1418 1417 1420 1 1.2000e+02 5.8576e+02 ; OH - CZ - CE2
1420 1422 1423 1 1.2000e+02 4.1840e+02 ; CE2 - CD2 - HD2
1421 1420 1422 1 1.2000e+02 4.1840e+02 ; HE2 - CE2 - CD2
1424 1426 1427 1 1.2000e+02 4.1840e+02 ; C - N - H
1424 1426 1428 1 1.2190e+02 4.1840e+02 ; C - N - CA
1425 1424 1426 1 1.2290e+02 6.6944e+02 ; O - C - N
1426 1428 1429 1 1.0950e+02 4.1840e+02 ; N - CA - HA
1426 1428 1430 1 1.0970e+02 6.6944e+02 ; N - CA - CB
1426 1428 1443 1 1.1010e+02 5.2718e+02 ; N - CA - C
1427 1426 1428 1 1.1804e+02 4.1840e+02 ; H - N - CA
1428 1430 1431 1 1.0950e+02 4.1840e+02 ; CA - CB - HB2
1428 1430 1432 1 1.0950e+02 4.1840e+02 ; CA - CB - HB3
1428 1430 1433 1 1.0950e+02 3.3472e+02 ; CA - CB - CG
1428 1443 1444 1 1.2040e+02 6.6944e+02 ; CA - C - O
1428 1443 1445 1 1.1660e+02 5.8576e+02 ; CA - C - N
1429 1428 1430 1 1.0950e+02 4.1840e+02 ; HA - CA - CB
1429 1428 1443 1 1.0950e+02 4.1840e+02 ; HA - CA - C

```
  1430   1428   1443      1    1.1110e+02    5.2718e+02 ;     CB - CA
- C
  1430   1433   1434      1    1.0950e+02    4.1840e+02 ;     CB - CG
- HG
  1430   1433   1435      1    1.0950e+02    3.3472e+02 ;     CB - CG
- CD1
  1430   1433   1439      1    1.0950e+02    3.3472e+02 ;     CB - CG
- CD2
  1431   1430   1432      1    1.0950e+02    2.9288e+02 ;    HB2 - CB
- HB3
  1431   1430   1433      1    1.0950e+02    4.1840e+02 ;    HB2 - CB
- CG
  1432   1430   1433      1    1.0950e+02    4.1840e+02 ;    HB3 - CB
- CG
  1433   1435   1436      1    1.0950e+02    4.1840e+02 ;     CG - CD1
- HD11
  1433   1435   1437      1    1.0950e+02    4.1840e+02 ;     CG - CD1
- HD12
  1433   1435   1438      1    1.0950e+02    4.1840e+02 ;     CG - CD1
- HD13
  1433   1439   1440      1    1.0950e+02    4.1840e+02 ;     CG - CD2
- HD21
  1433   1439   1441      1    1.0950e+02    4.1840e+02 ;     CG - CD2
- HD22
  1433   1439   1442      1    1.0950e+02    4.1840e+02 ;     CG - CD2
- HD23
  1434   1433   1435      1    1.0950e+02    4.1840e+02 ;     HG - CG
- CD1
  1434   1433   1439      1    1.0950e+02    4.1840e+02 ;     HG - CG
- CD2
  1435   1433   1439      1    1.0950e+02    3.3472e+02 ;    CD1 - CG
- CD2
  1436   1435   1437      1    1.0950e+02    2.9288e+02 ;   HD11 - CD1
- HD12
  1436   1435   1438      1    1.0950e+02    2.9288e+02 ;   HD11 - CD1
- HD13
  1437   1435   1438      1    1.0950e+02    2.9288e+02 ;   HD12 - CD1
- HD13
  1440   1439   1441      1    1.0950e+02    2.9288e+02 ;   HD21 - CD2
- HD22
  1440   1439   1442      1    1.0950e+02    2.9288e+02 ;   HD21 - CD2
- HD23
  1441   1439   1442      1    1.0950e+02    2.9288e+02 ;   HD22 - CD2
- HD23
  1443   1445   1446      1    1.2000e+02    4.1840e+02 ;      C - N
- H
  1443   1445   1447      1    1.2190e+02    4.1840e+02 ;      C - N
- CA
  1444   1443   1445      1    1.2290e+02    6.6944e+02 ;      O - C
- N
  1445   1447   1448      1    1.0950e+02    4.1840e+02 ;      N - CA
- HA
```

1445 1447 1449 1 1.0970e+02 6.6944e+02 ; N - CA
- CB
1445 1447 1462 1 1.1010e+02 5.2718e+02 ; N - CA
- C
1446 1445 1447 1 1.1804e+02 4.1840e+02 ; H - N
- CA
1447 1449 1450 1 1.0950e+02 4.1840e+02 ; CA - CB
- HB
1447 1449 1451 1 1.0950e+02 3.3472e+02 ; CA - CB
- CG2
1447 1449 1455 1 1.0950e+02 3.3472e+02 ; CA - CB
- CG1
1447 1462 1463 1 1.2040e+02 6.6944e+02 ; CA - C
- O
1447 1462 1464 1 1.1660e+02 5.8576e+02 ; CA - C
- N
1448 1447 1449 1 1.0950e+02 4.1840e+02 ; HA - CA
- CB
1448 1447 1462 1 1.0950e+02 4.1840e+02 ; HA - CA
- C
1449 1447 1462 1 1.1110e+02 5.2718e+02 ; CB - CA
- C
1449 1451 1452 1 1.0950e+02 4.1840e+02 ; CB - CG2
- HG21
1449 1451 1453 1 1.0950e+02 4.1840e+02 ; CB - CG2
- HG22
1449 1451 1454 1 1.0950e+02 4.1840e+02 ; CB - CG2
- HG23
1449 1455 1456 1 1.0950e+02 4.1840e+02 ; CB - CG1
- HG12
1449 1455 1457 1 1.0950e+02 4.1840e+02 ; CB - CG1
- HG13
1449 1455 1458 1 1.0950e+02 3.3472e+02 ; CB - CG1
- CD1
1450 1449 1451 1 1.0950e+02 4.1840e+02 ; HB - CB
- CG2
1450 1449 1455 1 1.0950e+02 4.1840e+02 ; HB - CB
- CG1
1451 1449 1455 1 1.0950e+02 3.3472e+02 ; CG2 - CB
- CG1
1452 1451 1453 1 1.0950e+02 2.9288e+02 ; HG21 - CG2
- HG22
1452 1451 1454 1 1.0950e+02 2.9288e+02 ; HG21 - CG2
- HG23
1453 1451 1454 1 1.0950e+02 2.9288e+02 ; HG22 - CG2
- HG23
1455 1458 1459 1 1.0950e+02 4.1840e+02 ; CG1 - CD1
- HD11
1455 1458 1460 1 1.0950e+02 4.1840e+02 ; CG1 - CD1
- HD12
1455 1458 1461 1 1.0950e+02 4.1840e+02 ; CG1 - CD1
- HD13

1456 1455 1457 1 1.0950e+02 2.9288e+02 ; HG12 - CG1
- HG13
1456 1455 1458 1 1.0950e+02 4.1840e+02 ; HG12 - CG1
- CD1
1457 1455 1458 1 1.0950e+02 4.1840e+02 ; HG13 - CG1
- CD1
1459 1458 1460 1 1.0950e+02 2.9288e+02 ; HD11 - CD1
- HD12
1459 1458 1461 1 1.0950e+02 2.9288e+02 ; HD11 - CD1
- HD13
1460 1458 1461 1 1.0950e+02 2.9288e+02 ; HD12 - CD1
- HD13
1462 1464 1465 1 1.2000e+02 4.1840e+02 ; C - N
- H
1462 1464 1466 1 1.2190e+02 4.1840e+02 ; C - N
- CA
1463 1462 1464 1 1.2290e+02 6.6944e+02 ; O - C
- N
1464 1466 1467 1 1.0950e+02 4.1840e+02 ; N - CA
- HA
1464 1466 1468 1 1.0970e+02 6.6944e+02 ; N - CA
- CB
1464 1466 1479 1 1.1010e+02 5.2718e+02 ; N - CA
- C
1465 1464 1466 1 1.1804e+02 4.1840e+02 ; H - N
- CA
1466 1468 1469 1 1.0950e+02 4.1840e+02 ; CA - CB
- HB2
1466 1468 1470 1 1.0950e+02 4.1840e+02 ; CA - CB
- HB3
1466 1468 1471 1 1.0950e+02 3.3472e+02 ; CA - CB
- CG
1466 1479 1480 1 1.2040e+02 6.6944e+02 ; CA - C
- O
1466 1479 1481 1 1.1660e+02 5.8576e+02 ; CA - C
- N
1467 1466 1468 1 1.0950e+02 4.1840e+02 ; HA - CA
- CB
1467 1466 1479 1 1.0950e+02 4.1840e+02 ; HA - CA
- C
1468 1466 1479 1 1.1110e+02 5.2718e+02 ; CB - CA
- C
1468 1471 1472 1 1.0950e+02 4.1840e+02 ; CB - CG
- HG2
1468 1471 1473 1 1.0950e+02 4.1840e+02 ; CB - CG
- HG3
1468 1471 1474 1 1.1470e+02 4.1840e+02 ; CB - CG
- SD
1469 1468 1470 1 1.0950e+02 2.9288e+02 ; HB2 - CB
- HB3
1469 1468 1471 1 1.0950e+02 4.1840e+02 ; HB2 - CB
- CG

1470 1468 1471 1 1.0950e+02 4.1840e+02 ; HB3 - CB
- CG
1471 1474 1475 1 9.8900e+01 5.1882e+02 ; CG - SD
- CE
1472 1471 1473 1 1.0950e+02 2.9288e+02 ; HG2 - CG
- HG3
1472 1471 1474 1 1.0950e+02 4.1840e+02 ; HG2 - CG
- SD
1473 1471 1474 1 1.0950e+02 4.1840e+02 ; HG3 - CG
- SD
1474 1475 1476 1 1.0950e+02 4.1840e+02 ; SD - CE
- HE1
1474 1475 1477 1 1.0950e+02 4.1840e+02 ; SD - CE
- HE2
1474 1475 1478 1 1.0950e+02 4.1840e+02 ; SD - CE
- HE3
1476 1475 1477 1 1.0950e+02 2.9288e+02 ; HE1 - CE
- HE2
1476 1475 1478 1 1.0950e+02 2.9288e+02 ; HE1 - CE
- HE3
1477 1475 1478 1 1.0950e+02 2.9288e+02 ; HE2 - CE
- HE3
1479 1481 1482 1 1.2000e+02 4.1840e+02 ; C - N
- H
1479 1481 1483 1 1.2190e+02 4.1840e+02 ; C - N
- CA
1480 1479 1481 1 1.2290e+02 6.6944e+02 ; O - C
- N
1481 1483 1484 1 1.0950e+02 4.1840e+02 ; N - CA
- HA
1481 1483 1485 1 1.0970e+02 6.6944e+02 ; N - CA
- CB
1481 1483 1494 1 1.1010e+02 5.2718e+02 ; N - CA
- C
1482 1481 1483 1 1.1804e+02 4.1840e+02 ; H - N
- CA
1483 1485 1486 1 1.0950e+02 4.1840e+02 ; CA - CB
- HB2
1483 1485 1487 1 1.0950e+02 4.1840e+02 ; CA - CB
- HB3
1483 1485 1488 1 1.0950e+02 3.3472e+02 ; CA - CB
- CG
1483 1494 1495 1 1.2040e+02 6.6944e+02 ; CA - C
- O
1483 1494 1496 1 1.1660e+02 5.8576e+02 ; CA - C
- N
1484 1483 1485 1 1.0950e+02 4.1840e+02 ; HA - CA
- CB
1484 1483 1494 1 1.0950e+02 4.1840e+02 ; HA - CA
- C
1485 1483 1494 1 1.1110e+02 5.2718e+02 ; CB - CA
- C

|  |  |  |  |  |  |  |  |
|---|---|---|---|---|---|---|---|
| 1485 | 1488 | 1489 | 1 | 1.0950e+02 | 4.1840e+02 | ; | CB - CG - HG2 |
| 1485 | 1488 | 1490 | 1 | 1.0950e+02 | 4.1840e+02 | ; | CB - CG - HG3 |
| 1485 | 1488 | 1491 | 1 | 1.1110e+02 | 5.2718e+02 | ; | CB - CG - CD |
| 1486 | 1485 | 1487 | 1 | 1.0950e+02 | 2.9288e+02 | ; | HB2 - CB - HB3 |
| 1486 | 1485 | 1488 | 1 | 1.0950e+02 | 4.1840e+02 | ; | HB2 - CB - CG |
| 1487 | 1485 | 1488 | 1 | 1.0950e+02 | 4.1840e+02 | ; | HB3 - CB - CG |
| 1488 | 1491 | 1492 | 1 | 1.1700e+02 | 5.8576e+02 | ; | CG - CD - OE1 |
| 1488 | 1491 | 1493 | 1 | 1.1700e+02 | 5.8576e+02 | ; | CG - CD - OE2 |
| 1489 | 1488 | 1490 | 1 | 1.0950e+02 | 2.9288e+02 | ; | HG2 - CG - HG3 |
| 1489 | 1488 | 1491 | 1 | 1.0950e+02 | 4.1840e+02 | ; | HG2 - CG - CD |
| 1490 | 1488 | 1491 | 1 | 1.0950e+02 | 4.1840e+02 | ; | HG3 - CG - CD |
| 1492 | 1491 | 1493 | 1 | 1.2600e+02 | 6.6944e+02 | ; | OE1 - CD - OE2 |
| 1494 | 1496 | 1497 | 1 | 1.2000e+02 | 4.1840e+02 | ; | C - N - H |
| 1494 | 1496 | 1498 | 1 | 1.2190e+02 | 4.1840e+02 | ; | C - N - CA |
| 1495 | 1494 | 1496 | 1 | 1.2290e+02 | 6.6944e+02 | ; | O - C - N |
| 1496 | 1498 | 1499 | 1 | 1.0950e+02 | 4.1840e+02 | ; | N - CA - HA |
| 1496 | 1498 | 1500 | 1 | 1.0970e+02 | 6.6944e+02 | ; | N - CA - CB |
| 1496 | 1498 | 1513 | 1 | 1.1010e+02 | 5.2718e+02 | ; | N - CA - C |
| 1497 | 1496 | 1498 | 1 | 1.1804e+02 | 4.1840e+02 | ; | H - N - CA |
| 1498 | 1500 | 1501 | 1 | 1.0950e+02 | 4.1840e+02 | ; | CA - CB - HB2 |
| 1498 | 1500 | 1502 | 1 | 1.0950e+02 | 4.1840e+02 | ; | CA - CB - HB3 |
| 1498 | 1500 | 1503 | 1 | 1.0950e+02 | 3.3472e+02 | ; | CA - CB - CG |
| 1498 | 1513 | 1514 | 1 | 1.2040e+02 | 6.6944e+02 | ; | CA - C - O |
| 1498 | 1513 | 1515 | 1 | 1.1660e+02 | 5.8576e+02 | ; | CA - C - N |
| 1499 | 1498 | 1500 | 1 | 1.0950e+02 | 4.1840e+02 | ; | HA - CA - CB |
| 1499 | 1498 | 1513 | 1 | 1.0950e+02 | 4.1840e+02 | ; | HA - CA - C |

```
  1500   1498   1513       1    1.1110e+02    5.2718e+02 ;     CB - CA
- C
  1500   1503   1504       1    1.0950e+02    4.1840e+02 ;     CB - CG
- HG
  1500   1503   1505       1    1.0950e+02    3.3472e+02 ;     CB - CG
- CD1
  1500   1503   1509       1    1.0950e+02    3.3472e+02 ;     CB - CG
- CD2
  1501   1500   1502       1    1.0950e+02    2.9288e+02 ;    HB2 - CB
- HB3
  1501   1500   1503       1    1.0950e+02    4.1840e+02 ;    HB2 - CB
- CG
  1502   1500   1503       1    1.0950e+02    4.1840e+02 ;    HB3 - CB
- CG
  1503   1505   1506       1    1.0950e+02    4.1840e+02 ;     CG - CD1
- HD11
  1503   1505   1507       1    1.0950e+02    4.1840e+02 ;     CG - CD1
- HD12
  1503   1505   1508       1    1.0950e+02    4.1840e+02 ;     CG - CD1
- HD13
  1503   1509   1510       1    1.0950e+02    4.1840e+02 ;     CG - CD2
- HD21
  1503   1509   1511       1    1.0950e+02    4.1840e+02 ;     CG - CD2
- HD22
  1503   1509   1512       1    1.0950e+02    4.1840e+02 ;     CG - CD2
- HD23
  1504   1503   1505       1    1.0950e+02    4.1840e+02 ;     HG - CG
- CD1
  1504   1503   1509       1    1.0950e+02    4.1840e+02 ;     HG - CG
- CD2
  1505   1503   1509       1    1.0950e+02    3.3472e+02 ;    CD1 - CG
- CD2
  1506   1505   1507       1    1.0950e+02    2.9288e+02 ;   HD11 - CD1
- HD12
  1506   1505   1508       1    1.0950e+02    2.9288e+02 ;   HD11 - CD1
- HD13
  1507   1505   1508       1    1.0950e+02    2.9288e+02 ;   HD12 - CD1
- HD13
  1510   1509   1511       1    1.0950e+02    2.9288e+02 ;   HD21 - CD2
- HD22
  1510   1509   1512       1    1.0950e+02    2.9288e+02 ;   HD21 - CD2
- HD23
  1511   1509   1512       1    1.0950e+02    2.9288e+02 ;   HD22 - CD2
- HD23
  1513   1515   1516       1    1.2000e+02    4.1840e+02 ;      C - N
- H
  1513   1515   1517       1    1.2190e+02    4.1840e+02 ;      C - N
- CA
  1514   1513   1515       1    1.2290e+02    6.6944e+02 ;      O - C
- N
  1515   1517   1518       1    1.0950e+02    4.1840e+02 ;      N - CA
- HA
```

1515 1517 1519 1 1.0970e+02 6.6944e+02 ; N - CA
- CB
1515 1517 1524 1 1.1010e+02 5.2718e+02 ; N - CA
- C
1516 1515 1517 1 1.1804e+02 4.1840e+02 ; H - N
- CA
1517 1519 1520 1 1.0950e+02 4.1840e+02 ; CA - CB
- HB2
1517 1519 1521 1 1.0950e+02 4.1840e+02 ; CA - CB
- HB3
1517 1519 1522 1 1.0860e+02 4.1840e+02 ; CA - CB
- SG
1517 1524 1525 1 1.2040e+02 6.6944e+02 ; CA - C
- O
1517 1524 1526 1 1.1660e+02 5.8576e+02 ; CA - C
- N
1518 1517 1519 1 1.0950e+02 4.1840e+02 ; HA - CA
- CB
1518 1517 1524 1 1.0950e+02 4.1840e+02 ; HA - CA
- C
1519 1517 1524 1 1.1110e+02 5.2718e+02 ; CB - CA
- C
1519 1522 1523 1 9.6000e+01 3.5982e+02 ; CB - SG
- HG
1520 1519 1521 1 1.0950e+02 2.9288e+02 ; HB2 - CB
- HB3
1520 1519 1522 1 1.0950e+02 4.1840e+02 ; HB2 - CB
- SG
1521 1519 1522 1 1.0950e+02 4.1840e+02 ; HB3 - CB
- SG
1524 1526 1527 1 1.2000e+02 4.1840e+02 ; C - N
- H
1524 1526 1528 1 1.2190e+02 4.1840e+02 ; C - N
- CA
1525 1524 1526 1 1.2290e+02 6.6944e+02 ; O - C
- N
1526 1528 1529 1 1.0950e+02 4.1840e+02 ; N - CA
- HA
1526 1528 1530 1 1.0970e+02 6.6944e+02 ; N - CA
- CB
1526 1528 1535 1 1.1010e+02 5.2718e+02 ; N - CA
- C
1527 1526 1528 1 1.1804e+02 4.1840e+02 ; H - N
- CA
1528 1530 1531 1 1.0950e+02 4.1840e+02 ; CA - CB
- HB2
1528 1530 1532 1 1.0950e+02 4.1840e+02 ; CA - CB
- HB3
1528 1530 1533 1 1.0950e+02 4.1840e+02 ; CA - CB
- OG
1528 1535 1536 1 1.2040e+02 6.6944e+02 ; CA - C
- O

```
  1528   1535   1537       1    1.1660e+02    5.8576e+02 ;     CA - C
- N
  1529   1528   1530       1    1.0950e+02    4.1840e+02 ;     HA - CA
- CB
  1529   1528   1535       1    1.0950e+02    4.1840e+02 ;     HA - CA
- C
  1530   1528   1535       1    1.1110e+02    5.2718e+02 ;     CB - CA
- C
  1530   1533   1534       1    1.0850e+02    4.6024e+02 ;     CB - OG
- HG
  1531   1530   1532       1    1.0950e+02    2.9288e+02 ;    HB2 - CB
- HB3
  1531   1530   1533       1    1.0950e+02    4.1840e+02 ;    HB2 - CB
- OG
  1532   1530   1533       1    1.0950e+02    4.1840e+02 ;    HB3 - CB
- OG
  1535   1537   1538       1    1.2000e+02    4.1840e+02 ;      C - N
- H
  1535   1537   1539       1    1.2190e+02    4.1840e+02 ;      C - N
- CA
  1536   1535   1537       1    1.2290e+02    6.6944e+02 ;      O - C
- N
  1537   1539   1540       1    1.0950e+02    4.1840e+02 ;      N - CA
- HA2
  1537   1539   1541       1    1.0950e+02    4.1840e+02 ;      N - CA
- HA3
  1537   1539   1542       1    1.1010e+02    5.2718e+02 ;      N - CA
- C
  1538   1537   1539       1    1.1804e+02    4.1840e+02 ;      H - N
- CA
  1539   1542   1543       1    1.2040e+02    6.6944e+02 ;     CA - C
- O
  1539   1542   1544       1    1.1660e+02    5.8576e+02 ;     CA - C
- N
  1540   1539   1541       1    1.0950e+02    2.9288e+02 ;    HA2 - CA
- HA3
  1540   1539   1542       1    1.0950e+02    4.1840e+02 ;    HA2 - CA
- C
  1541   1539   1542       1    1.0950e+02    4.1840e+02 ;    HA3 - CA
- C
  1542   1544   1545       1    1.2000e+02    4.1840e+02 ;      C - N
- H
  1542   1544   1546       1    1.2190e+02    4.1840e+02 ;      C - N
- CA
  1543   1542   1544       1    1.2290e+02    6.6944e+02 ;      O - C
- N
  1544   1546   1547       1    1.0950e+02    4.1840e+02 ;      N - CA
- HA
  1544   1546   1548       1    1.0970e+02    6.6944e+02 ;      N - CA
- CB
  1544   1546   1566       1    1.1010e+02    5.2718e+02 ;      N - CA
- C
```

1545 1544 1546 1 1.1804e+02 4.1840e+02 ; H - N
- CA
1546 1548 1549 1 1.0950e+02 4.1840e+02 ; CA - CB
- HB2
1546 1548 1550 1 1.0950e+02 4.1840e+02 ; CA - CB
- HB3
1546 1548 1551 1 1.0950e+02 3.3472e+02 ; CA - CB
- CG
1546 1566 1567 1 1.2040e+02 6.6944e+02 ; CA - C
- O
1546 1566 1568 1 1.1660e+02 5.8576e+02 ; CA - C
- N
1547 1546 1548 1 1.0950e+02 4.1840e+02 ; HA - CA
- CB
1547 1546 1566 1 1.0950e+02 4.1840e+02 ; HA - CA
- C
1548 1546 1566 1 1.1110e+02 5.2718e+02 ; CB - CA
- C
1548 1551 1552 1 1.0950e+02 4.1840e+02 ; CB - CG
- HG2
1548 1551 1553 1 1.0950e+02 4.1840e+02 ; CB - CG
- HG3
1548 1551 1554 1 1.0950e+02 3.3472e+02 ; CB - CG
- CD
1549 1548 1550 1 1.0950e+02 2.9288e+02 ; HB2 - CB
- HB3
1549 1548 1551 1 1.0950e+02 4.1840e+02 ; HB2 - CB
- CG
1550 1548 1551 1 1.0950e+02 4.1840e+02 ; HB3 - CB
- CG
1551 1554 1555 1 1.0950e+02 4.1840e+02 ; CG - CD
- HD2
1551 1554 1556 1 1.0950e+02 4.1840e+02 ; CG - CD
- HD3
1551 1554 1557 1 1.1120e+02 6.6944e+02 ; CG - CD
- NE
1552 1551 1553 1 1.0950e+02 2.9288e+02 ; HG2 - CG
- HG3
1552 1551 1554 1 1.0950e+02 4.1840e+02 ; HG2 - CG
- CD
1553 1551 1554 1 1.0950e+02 4.1840e+02 ; HG3 - CG
- CD
1554 1557 1558 1 1.1840e+02 4.1840e+02 ; CD - NE
- HE
1554 1557 1559 1 1.2320e+02 4.1840e+02 ; CD - NE
- CZ
1555 1554 1556 1 1.0950e+02 2.9288e+02 ; HD2 - CD
- HD3
1555 1554 1557 1 1.0950e+02 4.1840e+02 ; HD2 - CD
- NE
1556 1554 1557 1 1.0950e+02 4.1840e+02 ; HD3 - CD
- NE

```
  1557   1559   1560      1    1.2000e+02    5.8576e+02 ;     NE - CZ
- NH1
  1557   1559   1563      1    1.2000e+02    5.8576e+02 ;     NE - CZ
- NH2
  1558   1557   1559      1    1.2000e+02    4.1840e+02 ;     HE - NE
- CZ
  1559   1560   1561      1    1.2000e+02    4.1840e+02 ;     CZ - NH1
- HH11
  1559   1560   1562      1    1.2000e+02    4.1840e+02 ;     CZ - NH1
- HH12
  1559   1563   1564      1    1.2000e+02    4.1840e+02 ;     CZ - NH2
- HH21
  1559   1563   1565      1    1.2000e+02    4.1840e+02 ;     CZ - NH2
- HH22
  1560   1559   1563      1    1.2000e+02    5.8576e+02 ;    NH1 - CZ
- NH2
  1561   1560   1562      1    1.2000e+02    2.9288e+02 ;   HH11 - NH1
- HH12
  1564   1563   1565      1    1.2000e+02    2.9288e+02 ;   HH21 - NH2
- HH22
  1566   1568   1569      1    1.2000e+02    4.1840e+02 ;      C - N
- H
  1566   1568   1570      1    1.2190e+02    4.1840e+02 ;      C - N
- CA
  1567   1566   1568      1    1.2290e+02    6.6944e+02 ;      O - C
- N
  1568   1570   1571      1    1.0950e+02    4.1840e+02 ;      N - CA
- HA
  1568   1570   1572      1    1.0970e+02    6.6944e+02 ;      N - CA
- CB
  1568   1570   1581      1    1.1010e+02    5.2718e+02 ;      N - CA
- C
  1569   1568   1570      1    1.1804e+02    4.1840e+02 ;      H - N
- CA
  1570   1572   1573      1    1.0950e+02    4.1840e+02 ;     CA - CB
- HB2
  1570   1572   1574      1    1.0950e+02    4.1840e+02 ;     CA - CB
- HB3
  1570   1572   1575      1    1.0950e+02    3.3472e+02 ;     CA - CB
- CG
  1570   1581   1582      1    1.2040e+02    6.6944e+02 ;     CA - C
- O
  1570   1581   1583      1    1.1660e+02    5.8576e+02 ;     CA - C
- N
  1571   1570   1572      1    1.0950e+02    4.1840e+02 ;     HA - CA
- CB
  1571   1570   1581      1    1.0950e+02    4.1840e+02 ;     HA - CA
- C
  1572   1570   1581      1    1.1110e+02    5.2718e+02 ;     CB - CA
- C
  1572   1575   1576      1    1.0950e+02    4.1840e+02 ;     CB - CG
- HG2
```

```
  1572   1575   1577      1    1.0950e+02    4.1840e+02 ;     CB - CG
- HG3
  1572   1575   1578      1    1.1110e+02    5.2718e+02 ;     CB - CG
- CD
  1573   1572   1574      1    1.0950e+02    2.9288e+02 ;    HB2 - CB
- HB3
  1573   1572   1575      1    1.0950e+02    4.1840e+02 ;    HB2 - CB
- CG
  1574   1572   1575      1    1.0950e+02    4.1840e+02 ;    HB3 - CB
- CG
  1575   1578   1579      1    1.1700e+02    5.8576e+02 ;     CG - CD
- OE1
  1575   1578   1580      1    1.1700e+02    5.8576e+02 ;     CG - CD
- OE2
  1576   1575   1577      1    1.0950e+02    2.9288e+02 ;    HG2 - CG
- HG3
  1576   1575   1578      1    1.0950e+02    4.1840e+02 ;    HG2 - CG
- CD
  1577   1575   1578      1    1.0950e+02    4.1840e+02 ;    HG3 - CG
- CD
  1579   1578   1580      1    1.2600e+02    6.6944e+02 ;    OE1 - CD
- OE2
  1581   1583   1584      1    1.2000e+02    4.1840e+02 ;      C - N
- H
  1581   1583   1585      1    1.2190e+02    4.1840e+02 ;      C - N
- CA
  1582   1581   1583      1    1.2290e+02    6.6944e+02 ;      O - C
- N
  1583   1585   1586      1    1.0950e+02    4.1840e+02 ;      N - CA
- HA
  1583   1585   1587      1    1.0970e+02    6.6944e+02 ;      N - CA
- CB
  1583   1585   1600      1    1.1010e+02    5.2718e+02 ;      N - CA
- C
  1584   1583   1585      1    1.1804e+02    4.1840e+02 ;      H - N
- CA
  1585   1587   1588      1    1.0950e+02    4.1840e+02 ;     CA - CB
- HB2
  1585   1587   1589      1    1.0950e+02    4.1840e+02 ;     CA - CB
- HB3
  1585   1587   1590      1    1.0950e+02    3.3472e+02 ;     CA - CB
- CG
  1585   1600   1601      1    1.2040e+02    6.6944e+02 ;     CA - C
- O
  1585   1600   1602      1    1.1660e+02    5.8576e+02 ;     CA - C
- N
  1586   1585   1587      1    1.0950e+02    4.1840e+02 ;     HA - CA
- CB
  1586   1585   1600      1    1.0950e+02    4.1840e+02 ;     HA - CA
- C
  1587   1585   1600      1    1.1110e+02    5.2718e+02 ;     CB - CA
- C
```

1587 1590 1591 1 1.0950e+02 4.1840e+02 ; CB - CG
- HG
1587 1590 1592 1 1.0950e+02 3.3472e+02 ; CB - CG
- CD1
1587 1590 1596 1 1.0950e+02 3.3472e+02 ; CB - CG
- CD2
1588 1587 1589 1 1.0950e+02 2.9288e+02 ; HB2 - CB
- HB3
1588 1587 1590 1 1.0950e+02 4.1840e+02 ; HB2 - CB
- CG
1589 1587 1590 1 1.0950e+02 4.1840e+02 ; HB3 - CB
- CG
1590 1592 1593 1 1.0950e+02 4.1840e+02 ; CG - CD1
- HD11
1590 1592 1594 1 1.0950e+02 4.1840e+02 ; CG - CD1
- HD12
1590 1592 1595 1 1.0950e+02 4.1840e+02 ; CG - CD1
- HD13
1590 1596 1597 1 1.0950e+02 4.1840e+02 ; CG - CD2
- HD21
1590 1596 1598 1 1.0950e+02 4.1840e+02 ; CG - CD2
- HD22
1590 1596 1599 1 1.0950e+02 4.1840e+02 ; CG - CD2
- HD23
1591 1590 1592 1 1.0950e+02 4.1840e+02 ; HG - CG
- CD1
1591 1590 1596 1 1.0950e+02 4.1840e+02 ; HG - CG
- CD2
1592 1590 1596 1 1.0950e+02 3.3472e+02 ; CD1 - CG
- CD2
1593 1592 1594 1 1.0950e+02 2.9288e+02 ; HD11 - CD1
- HD12
1593 1592 1595 1 1.0950e+02 2.9288e+02 ; HD11 - CD1
- HD13
1594 1592 1595 1 1.0950e+02 2.9288e+02 ; HD12 - CD1
- HD13
1597 1596 1598 1 1.0950e+02 2.9288e+02 ; HD21 - CD2
- HD22
1597 1596 1599 1 1.0950e+02 2.9288e+02 ; HD21 - CD2
- HD23
1598 1596 1599 1 1.0950e+02 2.9288e+02 ; HD22 - CD2
- HD23
1600 1602 1603 1 1.2000e+02 4.1840e+02 ; C - N
- H
1600 1602 1604 1 1.2190e+02 4.1840e+02 ; C - N
- CA
1601 1600 1602 1 1.2290e+02 6.6944e+02 ; O - C
- N
1602 1604 1605 1 1.0950e+02 4.1840e+02 ; N - CA
- HA
1602 1604 1606 1 1.0970e+02 6.6944e+02 ; N - CA
- CB

```
  1602   1604   1620      1    1.1010e+02    5.2718e+02 ;      N - CA
- C
  1603   1602   1604      1    1.1804e+02    4.1840e+02 ;      H - N
- CA
  1604   1606   1607      1    1.0950e+02    4.1840e+02 ;     CA - CB
- HB2
  1604   1606   1608      1    1.0950e+02    4.1840e+02 ;     CA - CB
- HB3
  1604   1606   1609      1    1.1400e+02    5.2718e+02 ;     CA - CB
- CG
  1604   1620   1621      1    1.2040e+02    6.6944e+02 ;     CA - C
- O
  1604   1620   1622      1    1.1660e+02    5.8576e+02 ;     CA - C
- N
  1605   1604   1606      1    1.0950e+02    4.1840e+02 ;     HA - CA
- CB
  1605   1604   1620      1    1.0950e+02    4.1840e+02 ;     HA - CA
- C
  1606   1604   1620      1    1.1110e+02    5.2718e+02 ;     CB - CA
- C
  1606   1609   1610      1    1.2000e+02    5.8576e+02 ;     CB - CG
- CD1
  1606   1609   1618      1    1.2000e+02    5.8576e+02 ;     CB - CG
- CD2
  1607   1606   1608      1    1.0950e+02    2.9288e+02 ;    HB2 - CB
- HB3
  1607   1606   1609      1    1.0950e+02    4.1840e+02 ;    HB2 - CB
- CG
  1608   1606   1609      1    1.0950e+02    4.1840e+02 ;    HB3 - CB
- CG
  1609   1610   1611      1    1.2000e+02    4.1840e+02 ;     CG - CD1
- HD1
  1609   1610   1612      1    1.2000e+02    5.2718e+02 ;     CG - CD1
- CE1
  1609   1618   1616      1    1.2000e+02    5.2718e+02 ;     CG - CD2
- CE2
  1609   1618   1619      1    1.2000e+02    4.1840e+02 ;     CG - CD2
- HD2
  1610   1609   1618      1    1.2000e+02    5.2718e+02 ;    CD1 - CG
- CD2
  1610   1612   1613      1    1.2000e+02    4.1840e+02 ;    CD1 - CE1
- HE1
  1610   1612   1614      1    1.2000e+02    5.2718e+02 ;    CD1 - CE1
- CZ
  1611   1610   1612      1    1.2000e+02    4.1840e+02 ;    HD1 - CD1
- CE1
  1612   1614   1615      1    1.2000e+02    4.1840e+02 ;    CE1 - CZ
- HZ
  1612   1614   1616      1    1.2000e+02    5.2718e+02 ;    CE1 - CZ
- CE2
  1613   1612   1614      1    1.2000e+02    4.1840e+02 ;    HE1 - CE1
- CZ
```

```
  1614   1616   1617      1    1.2000e+02    4.1840e+02 ;     CZ - CE2
- HE2
  1614   1616   1618      1    1.2000e+02    5.2718e+02 ;     CZ - CE2
- CD2
  1615   1614   1616      1    1.2000e+02    4.1840e+02 ;     HZ - CZ
- CE2
  1616   1618   1619      1    1.2000e+02    4.1840e+02 ;    CE2 - CD2
- HD2
  1617   1616   1618      1    1.2000e+02    4.1840e+02 ;    HE2 - CE2
- CD2
  1620   1622   1623      1    1.2000e+02    4.1840e+02 ;      C - N
- H
  1620   1622   1624      1    1.2190e+02    4.1840e+02 ;      C - N
- CA
  1621   1620   1622      1    1.2290e+02    6.6944e+02 ;      O - C
- N
  1622   1624   1625      1    1.0950e+02    4.1840e+02 ;      N - CA
- HA
  1622   1624   1626      1    1.0970e+02    6.6944e+02 ;      N - CA
- CB
  1622   1624   1632      1    1.1010e+02    5.2718e+02 ;      N - CA
- C
  1623   1622   1624      1    1.1804e+02    4.1840e+02 ;      H - N
- CA
  1624   1626   1627      1    1.0950e+02    4.1840e+02 ;     CA - CB
- HB2
  1624   1626   1628      1    1.0950e+02    4.1840e+02 ;     CA - CB
- HB3
  1624   1626   1629      1    1.1110e+02    5.2718e+02 ;     CA - CB
- CG
  1624   1632   1633      1    1.2040e+02    6.6944e+02 ;     CA - C
- O
  1624   1632   1634      1    1.1660e+02    5.8576e+02 ;     CA - C
- N
  1625   1624   1626      1    1.0950e+02    4.1840e+02 ;     HA - CA
- CB
  1625   1624   1632      1    1.0950e+02    4.1840e+02 ;     HA - CA
- C
  1626   1624   1632      1    1.1110e+02    5.2718e+02 ;     CB - CA
- C
  1626   1629   1630      1    1.1700e+02    5.8576e+02 ;     CB - CG
- OD1
  1626   1629   1631      1    1.1700e+02    5.8576e+02 ;     CB - CG
- OD2
  1627   1626   1628      1    1.0950e+02    2.9288e+02 ;    HB2 - CB
- HB3
  1627   1626   1629      1    1.0950e+02    4.1840e+02 ;    HB2 - CB
- CG
  1628   1626   1629      1    1.0950e+02    4.1840e+02 ;    HB3 - CB
- CG
  1630   1629   1631      1    1.2600e+02    6.6944e+02 ;    OD1 - CG
- OD2
```

1632 1634 1635 1 1.2000e+02 4.1840e+02 ; C - N
- H
1632 1634 1636 1 1.2190e+02 4.1840e+02 ; C - N
- CA
1633 1632 1634 1 1.2290e+02 6.6944e+02 ; O - C
- N
1634 1636 1637 1 1.0950e+02 4.1840e+02 ; N - CA
- HA
1634 1636 1638 1 1.0970e+02 6.6944e+02 ; N - CA
- CB
1634 1636 1643 1 1.1010e+02 5.2718e+02 ; N - CA
- C
1635 1634 1636 1 1.1804e+02 4.1840e+02 ; H - N
- CA
1636 1638 1639 1 1.0950e+02 4.1840e+02 ; CA - CB
- HB2
1636 1638 1640 1 1.0950e+02 4.1840e+02 ; CA - CB
- HB3
1636 1638 1641 1 1.0950e+02 4.1840e+02 ; CA - CB
- OG
1636 1643 1644 1 1.2040e+02 6.6944e+02 ; CA - C
- O
1636 1643 1645 1 1.1660e+02 5.8576e+02 ; CA - C
- N
1637 1636 1638 1 1.0950e+02 4.1840e+02 ; HA - CA
- CB
1637 1636 1643 1 1.0950e+02 4.1840e+02 ; HA - CA
- C
1638 1636 1643 1 1.1110e+02 5.2718e+02 ; CB - CA
- C
1638 1641 1642 1 1.0850e+02 4.6024e+02 ; CB - OG
- HG
1639 1638 1640 1 1.0950e+02 2.9288e+02 ; HB2 - CB
- HB3
1639 1638 1641 1 1.0950e+02 4.1840e+02 ; HB2 - CB
- OG
1640 1638 1641 1 1.0950e+02 4.1840e+02 ; HB3 - CB
- OG
1643 1645 1646 1 1.2000e+02 4.1840e+02 ; C - N
- H
1643 1645 1647 1 1.2190e+02 4.1840e+02 ; C - N
- CA
1644 1643 1645 1 1.2290e+02 6.6944e+02 ; O - C
- N
1645 1647 1648 1 1.0950e+02 4.1840e+02 ; N - CA
- HA
1645 1647 1649 1 1.0970e+02 6.6944e+02 ; N - CA
- CB
1645 1647 1662 1 1.1010e+02 5.2718e+02 ; N - CA
- C
1646 1645 1647 1 1.1804e+02 4.1840e+02 ; H - N
- CA

1647 1649 1650 1 1.0950e+02 4.1840e+02 ; CA - CB
- HB
1647 1649 1651 1 1.0950e+02 3.3472e+02 ; CA - CB
- CG2
1647 1649 1655 1 1.0950e+02 3.3472e+02 ; CA - CB
- CG1
1647 1662 1663 1 1.2040e+02 6.6944e+02 ; CA - C
- O
1647 1662 1664 1 1.1660e+02 5.8576e+02 ; CA - C
- N
1648 1647 1649 1 1.0950e+02 4.1840e+02 ; HA - CA
- CB
1648 1647 1662 1 1.0950e+02 4.1840e+02 ; HA - CA
- C
1649 1647 1662 1 1.1110e+02 5.2718e+02 ; CB - CA
- C
1649 1651 1652 1 1.0950e+02 4.1840e+02 ; CB - CG2
- HG21
1649 1651 1653 1 1.0950e+02 4.1840e+02 ; CB - CG2
- HG22
1649 1651 1654 1 1.0950e+02 4.1840e+02 ; CB - CG2
- HG23
1649 1655 1656 1 1.0950e+02 4.1840e+02 ; CB - CG1
- HG12
1649 1655 1657 1 1.0950e+02 4.1840e+02 ; CB - CG1
- HG13
1649 1655 1658 1 1.0950e+02 3.3472e+02 ; CB - CG1
- CD1
1650 1649 1651 1 1.0950e+02 4.1840e+02 ; HB - CB
- CG2
1650 1649 1655 1 1.0950e+02 4.1840e+02 ; HB - CB
- CG1
1651 1649 1655 1 1.0950e+02 3.3472e+02 ; CG2 - CB
- CG1
1652 1651 1653 1 1.0950e+02 2.9288e+02 ; HG21 - CG2
- HG22
1652 1651 1654 1 1.0950e+02 2.9288e+02 ; HG21 - CG2
- HG23
1653 1651 1654 1 1.0950e+02 2.9288e+02 ; HG22 - CG2
- HG23
1655 1658 1659 1 1.0950e+02 4.1840e+02 ; CG1 - CD1
- HD11
1655 1658 1660 1 1.0950e+02 4.1840e+02 ; CG1 - CD1
- HD12
1655 1658 1661 1 1.0950e+02 4.1840e+02 ; CG1 - CD1
- HD13
1656 1655 1657 1 1.0950e+02 2.9288e+02 ; HG12 - CG1
- HG13
1656 1655 1658 1 1.0950e+02 4.1840e+02 ; HG12 - CG1
- CD1
1657 1655 1658 1 1.0950e+02 4.1840e+02 ; HG13 - CG1
- CD1

1659 1658 1660 1 1.0950e+02 2.9288e+02 ; HD11 - CD1
- HD12
1659 1658 1661 1 1.0950e+02 2.9288e+02 ; HD11 - CD1
- HD13
1660 1658 1661 1 1.0950e+02 2.9288e+02 ; HD12 - CD1
- HD13
1662 1664 1665 1 1.2000e+02 4.1840e+02 ; C - N
- H
1662 1664 1666 1 1.2190e+02 4.1840e+02 ; C - N
- CA
1663 1662 1664 1 1.2290e+02 6.6944e+02 ; O - C
- N
1664 1666 1667 1 1.0950e+02 4.1840e+02 ; N - CA
- HA
1664 1666 1668 1 1.0970e+02 6.6944e+02 ; N - CA
- CB
1664 1666 1681 1 1.1010e+02 5.2718e+02 ; N - CA
- C
1665 1664 1666 1 1.1804e+02 4.1840e+02 ; H - N
- CA
1666 1668 1669 1 1.0950e+02 4.1840e+02 ; CA - CB
- HB
1666 1668 1670 1 1.0950e+02 3.3472e+02 ; CA - CB
- CG2
1666 1668 1674 1 1.0950e+02 3.3472e+02 ; CA - CB
- CG1
1666 1681 1682 1 1.2040e+02 6.6944e+02 ; CA - C
- O
1666 1681 1683 1 1.1660e+02 5.8576e+02 ; CA - C
- N
1667 1666 1668 1 1.0950e+02 4.1840e+02 ; HA - CA
- CB
1667 1666 1681 1 1.0950e+02 4.1840e+02 ; HA - CA
- C
1668 1666 1681 1 1.1110e+02 5.2718e+02 ; CB - CA
- C
1668 1670 1671 1 1.0950e+02 4.1840e+02 ; CB - CG2
- HG21
1668 1670 1672 1 1.0950e+02 4.1840e+02 ; CB - CG2
- HG22
1668 1670 1673 1 1.0950e+02 4.1840e+02 ; CB - CG2
- HG23
1668 1674 1675 1 1.0950e+02 4.1840e+02 ; CB - CG1
- HG12
1668 1674 1676 1 1.0950e+02 4.1840e+02 ; CB - CG1
- HG13
1668 1674 1677 1 1.0950e+02 3.3472e+02 ; CB - CG1
- CD1
1669 1668 1670 1 1.0950e+02 4.1840e+02 ; HB - CB
- CG2
1669 1668 1674 1 1.0950e+02 4.1840e+02 ; HB - CB
- CG1

```
  1670   1668   1674      1    1.0950e+02    3.3472e+02 ;    CG2 - CB
- CG1
  1671   1670   1672      1    1.0950e+02    2.9288e+02 ;   HG21 - CG2
- HG22
  1671   1670   1673      1    1.0950e+02    2.9288e+02 ;   HG21 - CG2
- HG23
  1672   1670   1673      1    1.0950e+02    2.9288e+02 ;   HG22 - CG2
- HG23
  1674   1677   1678      1    1.0950e+02    4.1840e+02 ;    CG1 - CD1
- HD11
  1674   1677   1679      1    1.0950e+02    4.1840e+02 ;    CG1 - CD1
- HD12
  1674   1677   1680      1    1.0950e+02    4.1840e+02 ;    CG1 - CD1
- HD13
  1675   1674   1676      1    1.0950e+02    2.9288e+02 ;   HG12 - CG1
- HG13
  1675   1674   1677      1    1.0950e+02    4.1840e+02 ;   HG12 - CG1
- CD1
  1676   1674   1677      1    1.0950e+02    4.1840e+02 ;   HG13 - CG1
- CD1
  1678   1677   1679      1    1.0950e+02    2.9288e+02 ;   HD11 - CD1
- HD12
  1678   1677   1680      1    1.0950e+02    2.9288e+02 ;   HD11 - CD1
- HD13
  1679   1677   1680      1    1.0950e+02    2.9288e+02 ;   HD12 - CD1
- HD13
  1681   1683   1684      1    1.2000e+02    4.1840e+02 ;      C - N
- H
  1681   1683   1685      1    1.2190e+02    4.1840e+02 ;      C - N
- CA
  1682   1681   1683      1    1.2290e+02    6.6944e+02 ;      O - C
- N
  1683   1685   1686      1    1.0950e+02    4.1840e+02 ;      N - CA
- HA
  1683   1685   1687      1    1.0970e+02    6.6944e+02 ;      N - CA
- CB
  1683   1685   1696      1    1.1010e+02    5.2718e+02 ;      N - CA
- C
  1684   1683   1685      1    1.1804e+02    4.1840e+02 ;      H - N
- CA
  1685   1687   1688      1    1.0950e+02    4.1840e+02 ;     CA - CB
- HB2
  1685   1687   1689      1    1.0950e+02    4.1840e+02 ;     CA - CB
- HB3
  1685   1687   1690      1    1.0950e+02    3.3472e+02 ;     CA - CB
- CG
  1685   1696   1697      1    1.2040e+02    6.6944e+02 ;     CA - C
- O
  1685   1696   1698      1    1.1660e+02    5.8576e+02 ;     CA - C
- N
  1686   1685   1687      1    1.0950e+02    4.1840e+02 ;     HA - CA
- CB
```

1686 1685 1696 1 1.0950e+02 4.1840e+02 ; HA - CA
- C
1687 1685 1696 1 1.1110e+02 5.2718e+02 ; CB - CA
- C
1687 1690 1691 1 1.0950e+02 4.1840e+02 ; CB - CG
- HG2
1687 1690 1692 1 1.0950e+02 4.1840e+02 ; CB - CG
- HG3
1687 1690 1693 1 1.1110e+02 5.2718e+02 ; CB - CG
- CD
1688 1687 1689 1 1.0950e+02 2.9288e+02 ; HB2 - CB
- HB3
1688 1687 1690 1 1.0950e+02 4.1840e+02 ; HB2 - CB
- CG
1689 1687 1690 1 1.0950e+02 4.1840e+02 ; HB3 - CB
- CG
1690 1693 1694 1 1.1700e+02 5.8576e+02 ; CG - CD
- OE1
1690 1693 1695 1 1.1700e+02 5.8576e+02 ; CG - CD
- OE2
1691 1690 1692 1 1.0950e+02 2.9288e+02 ; HG2 - CG
- HG3
1691 1690 1693 1 1.0950e+02 4.1840e+02 ; HG2 - CG
- CD
1692 1690 1693 1 1.0950e+02 4.1840e+02 ; HG3 - CG
- CD
1694 1693 1695 1 1.2600e+02 6.6944e+02 ; OE1 - CD
- OE2
1696 1698 1699 1 1.2000e+02 4.1840e+02 ; C - N
- H
1696 1698 1700 1 1.2190e+02 4.1840e+02 ; C - N
- CA
1697 1696 1698 1 1.2290e+02 6.6944e+02 ; O - C
- N
1698 1700 1701 1 1.0950e+02 4.1840e+02 ; N - CA
- HA
1698 1700 1702 1 1.0970e+02 6.6944e+02 ; N - CA
- CB
1698 1700 1710 1 1.1010e+02 5.2718e+02 ; N - CA
- C
1699 1698 1700 1 1.1804e+02 4.1840e+02 ; H - N
- CA
1700 1702 1703 1 1.0950e+02 4.1840e+02 ; CA - CB
- HB2
1700 1702 1704 1 1.0950e+02 4.1840e+02 ; CA - CB
- HB3
1700 1702 1705 1 1.1110e+02 5.2718e+02 ; CA - CB
- CG
1700 1710 1711 1 1.2040e+02 6.6944e+02 ; CA - C
- O
1700 1710 1712 1 1.1660e+02 5.8576e+02 ; CA - C
- N

1701 1700 1702 1 1.0950e+02 4.1840e+02 ; HA - CA
- CB
1701 1700 1710 1 1.0950e+02 4.1840e+02 ; HA - CA
- C
1702 1700 1710 1 1.1110e+02 5.2718e+02 ; CB - CA
- C
1702 1705 1706 1 1.2040e+02 6.6944e+02 ; CB - CG
- OD1
1702 1705 1707 1 1.1660e+02 5.8576e+02 ; CB - CG
- ND2
1703 1702 1704 1 1.0950e+02 2.9288e+02 ; HB2 - CB
- HB3
1703 1702 1705 1 1.0950e+02 4.1840e+02 ; HB2 - CB
- CG
1704 1702 1705 1 1.0950e+02 4.1840e+02 ; HB3 - CB
- CG
1705 1707 1708 1 1.2000e+02 4.1840e+02 ; CG - ND2
- HD21
1705 1707 1709 1 1.2000e+02 4.1840e+02 ; CG - ND2
- HD22
1706 1705 1707 1 1.2290e+02 6.6944e+02 ; OD1 - CG
- ND2
1708 1707 1709 1 1.2000e+02 2.9288e+02 ; HD21 - ND2
- HD22
1710 1712 1713 1 1.2000e+02 4.1840e+02 ; C - N
- H
1710 1712 1714 1 1.2190e+02 4.1840e+02 ; C - N
- CA
1711 1710 1712 1 1.2290e+02 6.6944e+02 ; O - C
- N
1712 1714 1715 1 1.0950e+02 4.1840e+02 ; N - CA
- HA2
1712 1714 1716 1 1.0950e+02 4.1840e+02 ; N - CA
- HA3
1712 1714 1717 1 1.1010e+02 5.2718e+02 ; N - CA
- C
1713 1712 1714 1 1.1804e+02 4.1840e+02 ; H - N
- CA
1714 1717 1718 1 1.2040e+02 6.6944e+02 ; CA - C
- O
1714 1717 1719 1 1.1660e+02 5.8576e+02 ; CA - C
- N
1715 1714 1716 1 1.0950e+02 2.9288e+02 ; HA2 - CA
- HA3
1715 1714 1717 1 1.0950e+02 4.1840e+02 ; HA2 - CA
- C
1716 1714 1717 1 1.0950e+02 4.1840e+02 ; HA3 - CA
- C
1717 1719 1720 1 1.2000e+02 4.1840e+02 ; C - N
- H
1717 1719 1721 1 1.2190e+02 4.1840e+02 ; C - N
- CA

1718 1717 1719 1 1.2290e+02 6.6944e+02 ; O - C
- N
1719 1721 1722 1 1.0950e+02 4.1840e+02 ; N - CA
- HA
1719 1721 1723 1 1.0970e+02 6.6944e+02 ; N - CA
- CB
1719 1721 1728 1 1.1010e+02 5.2718e+02 ; N - CA
- C
1720 1719 1721 1 1.1804e+02 4.1840e+02 ; H - N
- CA
1721 1723 1724 1 1.0950e+02 4.1840e+02 ; CA - CB
- HB2
1721 1723 1725 1 1.0950e+02 4.1840e+02 ; CA - CB
- HB3
1721 1723 1726 1 1.0950e+02 4.1840e+02 ; CA - CB
- OG
1721 1728 1729 1 1.2040e+02 6.6944e+02 ; CA - C
- O
1721 1728 1730 1 1.1660e+02 5.8576e+02 ; CA - C
- N
1722 1721 1723 1 1.0950e+02 4.1840e+02 ; HA - CA
- CB
1722 1721 1728 1 1.0950e+02 4.1840e+02 ; HA - CA
- C
1723 1721 1728 1 1.1110e+02 5.2718e+02 ; CB - CA
- C
1723 1726 1727 1 1.0850e+02 4.6024e+02 ; CB - OG
- HG
1724 1723 1725 1 1.0950e+02 2.9288e+02 ; HB2 - CB
- HB3
1724 1723 1726 1 1.0950e+02 4.1840e+02 ; HB2 - CB
- OG
1725 1723 1726 1 1.0950e+02 4.1840e+02 ; HB3 - CB
- OG
1728 1730 1731 1 1.2000e+02 4.1840e+02 ; C - N
- H
1728 1730 1732 1 1.2190e+02 4.1840e+02 ; C - N
- CA
1729 1728 1730 1 1.2290e+02 6.6944e+02 ; O - C
- N
1730 1732 1733 1 1.0950e+02 4.1840e+02 ; N - CA
- HA
1730 1732 1734 1 1.0970e+02 6.6944e+02 ; N - CA
- CB
1730 1732 1748 1 1.1010e+02 5.2718e+02 ; N - CA
- C
1731 1730 1732 1 1.1804e+02 4.1840e+02 ; H - N
- CA
1732 1734 1735 1 1.0950e+02 4.1840e+02 ; CA - CB
- HB2
1732 1734 1736 1 1.0950e+02 4.1840e+02 ; CA - CB
- HB3

1732 1734 1737 1 1.1400e+02 5.2718e+02 ; CA - CB
- CG
1732 1748 1749 1 1.2040e+02 6.6944e+02 ; CA - C
- O
1732 1748 1750 1 1.1660e+02 5.8576e+02 ; CA - C
- N
1733 1732 1734 1 1.0950e+02 4.1840e+02 ; HA - CA
- CB
1733 1732 1748 1 1.0950e+02 4.1840e+02 ; HA - CA
- C
1734 1732 1748 1 1.1110e+02 5.2718e+02 ; CB - CA
- C
1734 1737 1738 1 1.2000e+02 5.8576e+02 ; CB - CG
- CD1
1734 1737 1746 1 1.2000e+02 5.8576e+02 ; CB - CG
- CD2
1735 1734 1736 1 1.0950e+02 2.9288e+02 ; HB2 - CB
- HB3
1735 1734 1737 1 1.0950e+02 4.1840e+02 ; HB2 - CB
- CG
1736 1734 1737 1 1.0950e+02 4.1840e+02 ; HB3 - CB
- CG
1737 1738 1739 1 1.2000e+02 4.1840e+02 ; CG - CD1
- HD1
1737 1738 1740 1 1.2000e+02 5.2718e+02 ; CG - CD1
- CE1
1737 1746 1744 1 1.2000e+02 5.2718e+02 ; CG - CD2
- CE2
1737 1746 1747 1 1.2000e+02 4.1840e+02 ; CG - CD2
- HD2
1738 1737 1746 1 1.2000e+02 5.2718e+02 ; CD1 - CG
- CD2
1738 1740 1741 1 1.2000e+02 4.1840e+02 ; CD1 - CE1
- HE1
1738 1740 1742 1 1.2000e+02 5.2718e+02 ; CD1 - CE1
- CZ
1739 1738 1740 1 1.2000e+02 4.1840e+02 ; HD1 - CD1
- CE1
1740 1742 1743 1 1.2000e+02 4.1840e+02 ; CE1 - CZ
- HZ
1740 1742 1744 1 1.2000e+02 5.2718e+02 ; CE1 - CZ
- CE2
1741 1740 1742 1 1.2000e+02 4.1840e+02 ; HE1 - CE1
- CZ
1742 1744 1745 1 1.2000e+02 4.1840e+02 ; CZ - CE2
- HE2
1742 1744 1746 1 1.2000e+02 5.2718e+02 ; CZ - CE2
- CD2
1743 1742 1744 1 1.2000e+02 4.1840e+02 ; HZ - CZ
- CE2
1744 1746 1747 1 1.2000e+02 4.1840e+02 ; CE2 - CD2
- HD2

```
  1745   1744   1746       1    1.2000e+02    4.1840e+02 ;    HE2 - CE2
- CD2
  1748   1750   1751       1    1.2000e+02    4.1840e+02 ;      C - N
- H
  1748   1750   1752       1    1.2190e+02    4.1840e+02 ;      C - N
- CA
  1749   1748   1750       1    1.2290e+02    6.6944e+02 ;      O - C
- N
  1750   1752   1753       1    1.0950e+02    4.1840e+02 ;      N - CA
- HA
  1750   1752   1754       1    1.0970e+02    6.6944e+02 ;      N - CA
- CB
  1750   1752   1762       1    1.1010e+02    5.2718e+02 ;      N - CA
- C
  1751   1750   1752       1    1.1804e+02    4.1840e+02 ;      H - N
- CA
  1752   1754   1755       1    1.0950e+02    4.1840e+02 ;     CA - CB
- HB
  1752   1754   1756       1    1.0950e+02    3.3472e+02 ;     CA - CB
- CG2
  1752   1754   1760       1    1.0950e+02    4.1840e+02 ;     CA - CB
- OG1
  1752   1762   1763       1    1.2040e+02    6.6944e+02 ;     CA - C
- O
  1752   1762   1764       1    1.1660e+02    5.8576e+02 ;     CA - C
- N
  1753   1752   1754       1    1.0950e+02    4.1840e+02 ;     HA - CA
- CB
  1753   1752   1762       1    1.0950e+02    4.1840e+02 ;     HA - CA
- C
  1754   1752   1762       1    1.1110e+02    5.2718e+02 ;     CB - CA
- C
  1754   1756   1757       1    1.0950e+02    4.1840e+02 ;     CB - CG2
- HG21
  1754   1756   1758       1    1.0950e+02    4.1840e+02 ;     CB - CG2
- HG22
  1754   1756   1759       1    1.0950e+02    4.1840e+02 ;     CB - CG2
- HG23
  1754   1760   1761       1    1.0850e+02    4.6024e+02 ;     CB - OG1
- HG1
  1755   1754   1756       1    1.0950e+02    4.1840e+02 ;     HB - CB
- CG2
  1755   1754   1760       1    1.0950e+02    4.1840e+02 ;     HB - CB
- OG1
  1756   1754   1760       1    1.0950e+02    4.1840e+02 ;    CG2 - CB
- OG1
  1757   1756   1758       1    1.0950e+02    2.9288e+02 ;   HG21 - CG2
- HG22
  1757   1756   1759       1    1.0950e+02    2.9288e+02 ;   HG21 - CG2
- HG23
  1758   1756   1759       1    1.0950e+02    2.9288e+02 ;   HG22 - CG2
- HG23
```

```
  1762   1764   1765      1    1.2000e+02    4.1840e+02 ;      C - N 
- H
  1762   1764   1766      1    1.2190e+02    4.1840e+02 ;      C - N 
- CA
  1763   1762   1764      1    1.2290e+02    6.6944e+02 ;      O - C 
- N
  1764   1766   1767      1    1.0950e+02    4.1840e+02 ;      N - CA
- HA
  1764   1766   1768      1    1.0970e+02    6.6944e+02 ;      N - CA
- CB
  1764   1766   1777      1    1.1010e+02    5.2718e+02 ;      N - CA
- C
  1765   1764   1766      1    1.1804e+02    4.1840e+02 ;      H - N 
- CA
  1766   1768   1769      1    1.0950e+02    4.1840e+02 ;     CA - CB
- HB2
  1766   1768   1770      1    1.0950e+02    4.1840e+02 ;     CA - CB
- HB3
  1766   1768   1771      1    1.0950e+02    3.3472e+02 ;     CA - CB
- CG
  1766   1777   1778      1    1.2040e+02    6.6944e+02 ;     CA - C 
- O
  1766   1777   1779      1    1.1660e+02    5.8576e+02 ;     CA - C 
- N
  1767   1766   1768      1    1.0950e+02    4.1840e+02 ;     HA - CA
- CB
  1767   1766   1777      1    1.0950e+02    4.1840e+02 ;     HA - CA
- C
  1768   1766   1777      1    1.1110e+02    5.2718e+02 ;     CB - CA
- C
  1768   1771   1772      1    1.0950e+02    4.1840e+02 ;     CB - CG
- HG2
  1768   1771   1773      1    1.0950e+02    4.1840e+02 ;     CB - CG
- HG3
  1768   1771   1774      1    1.1110e+02    5.2718e+02 ;     CB - CG
- CD
  1769   1768   1770      1    1.0950e+02    2.9288e+02 ;    HB2 - CB
- HB3
  1769   1768   1771      1    1.0950e+02    4.1840e+02 ;    HB2 - CB
- CG
  1770   1768   1771      1    1.0950e+02    4.1840e+02 ;    HB3 - CB
- CG
  1771   1774   1775      1    1.1700e+02    5.8576e+02 ;     CG - CD
- OE1
  1771   1774   1776      1    1.1700e+02    5.8576e+02 ;     CG - CD
- OE2
  1772   1771   1773      1    1.0950e+02    2.9288e+02 ;    HG2 - CG
- HG3
  1772   1771   1774      1    1.0950e+02    4.1840e+02 ;    HG2 - CG
- CD
  1773   1771   1774      1    1.0950e+02    4.1840e+02 ;    HG3 - CG
- CD
```

1775 1774 1776 1 1.2600e+02 6.6944e+02 ; OE1 - CD
- OE2
1777 1779 1780 1 1.2000e+02 4.1840e+02 ; C - N
- H
1777 1779 1781 1 1.2190e+02 4.1840e+02 ; C - N
- CA
1778 1777 1779 1 1.2290e+02 6.6944e+02 ; O - C
- N
1779 1781 1782 1 1.0950e+02 4.1840e+02 ; N - CA
- HA
1779 1781 1783 1 1.0970e+02 6.6944e+02 ; N - CA
- CB
1779 1781 1799 1 1.1010e+02 5.2718e+02 ; N - CA
- C
1780 1779 1781 1 1.1804e+02 4.1840e+02 ; H - N
- CA
1781 1783 1784 1 1.0950e+02 4.1840e+02 ; CA - CB
- HB2
1781 1783 1785 1 1.0950e+02 4.1840e+02 ; CA - CB
- HB3
1781 1783 1786 1 1.0950e+02 3.3472e+02 ; CA - CB
- CG
1781 1799 1800 1 1.2040e+02 6.6944e+02 ; CA - C
- O
1781 1799 1801 1 1.1660e+02 5.8576e+02 ; CA - C
- N
1782 1781 1783 1 1.0950e+02 4.1840e+02 ; HA - CA
- CB
1782 1781 1799 1 1.0950e+02 4.1840e+02 ; HA - CA
- C
1783 1781 1799 1 1.1110e+02 5.2718e+02 ; CB - CA
- C
1783 1786 1787 1 1.0950e+02 4.1840e+02 ; CB - CG
- HG2
1783 1786 1788 1 1.0950e+02 4.1840e+02 ; CB - CG
- HG3
1783 1786 1789 1 1.0950e+02 3.3472e+02 ; CB - CG
- CD
1784 1783 1785 1 1.0950e+02 2.9288e+02 ; HB2 - CB
- HB3
1784 1783 1786 1 1.0950e+02 4.1840e+02 ; HB2 - CB
- CG
1785 1783 1786 1 1.0950e+02 4.1840e+02 ; HB3 - CB
- CG
1786 1789 1790 1 1.0950e+02 4.1840e+02 ; CG - CD
- HD2
1786 1789 1791 1 1.0950e+02 4.1840e+02 ; CG - CD
- HD3
1786 1789 1792 1 1.0950e+02 3.3472e+02 ; CG - CD
- CE
1787 1786 1788 1 1.0950e+02 2.9288e+02 ; HG2 - CG
- HG3

```
  1787   1786   1789       1    1.0950e+02    4.1840e+02 ;    HG2 - CG
- CD
  1788   1786   1789       1    1.0950e+02    4.1840e+02 ;    HG3 - CG
- CD
  1789   1792   1793       1    1.0950e+02    4.1840e+02 ;     CD - CE
- HE2
  1789   1792   1794       1    1.0950e+02    4.1840e+02 ;     CD - CE
- HE3
  1789   1792   1795       1    1.1120e+02    6.6944e+02 ;     CD - CE
- NZ
  1790   1789   1791       1    1.0950e+02    2.9288e+02 ;    HD2 - CD
- HD3
  1790   1789   1792       1    1.0950e+02    4.1840e+02 ;    HD2 - CD
- CE
  1791   1789   1792       1    1.0950e+02    4.1840e+02 ;    HD3 - CD
- CE
  1792   1795   1796       1    1.0950e+02    4.1840e+02 ;     CE - NZ
- HZ1
  1792   1795   1797       1    1.0950e+02    4.1840e+02 ;     CE - NZ
- HZ2
  1792   1795   1798       1    1.0950e+02    4.1840e+02 ;     CE - NZ
- HZ3
  1793   1792   1794       1    1.0950e+02    2.9288e+02 ;    HE2 - CE
- HE3
  1793   1792   1795       1    1.0950e+02    4.1840e+02 ;    HE2 - CE
- NZ
  1794   1792   1795       1    1.0950e+02    4.1840e+02 ;    HE3 - CE
- NZ
  1796   1795   1797       1    1.0950e+02    2.9288e+02 ;    HZ1 - NZ
- HZ2
  1796   1795   1798       1    1.0950e+02    2.9288e+02 ;    HZ1 - NZ
- HZ3
  1797   1795   1798       1    1.0950e+02    2.9288e+02 ;    HZ2 - NZ
- HZ3
  1799   1801   1802       1    1.2000e+02    4.1840e+02 ;      C - N
- H
  1799   1801   1803       1    1.2190e+02    4.1840e+02 ;      C - N
- CA
  1800   1799   1801       1    1.2290e+02    6.6944e+02 ;      O - C
- N
  1801   1803   1804       1    1.0950e+02    4.1840e+02 ;      N - CA
- HA
  1801   1803   1805       1    1.0970e+02    6.6944e+02 ;      N - CA
- CB
  1801   1803   1813       1    1.1010e+02    5.2718e+02 ;      N - CA
- C
  1802   1801   1803       1    1.1804e+02    4.1840e+02 ;      H - N
- CA
  1803   1805   1806       1    1.0950e+02    4.1840e+02 ;     CA - CB
- HB2
  1803   1805   1807       1    1.0950e+02    4.1840e+02 ;     CA - CB
- HB3
```

1803 1805 1808 1 1.1110e+02 5.2718e+02 ; CA - CB
- CG
1803 1813 1814 1 1.2040e+02 6.6944e+02 ; CA - C
- O
1803 1813 1815 1 1.1660e+02 5.8576e+02 ; CA - C
- N
1804 1803 1805 1 1.0950e+02 4.1840e+02 ; HA - CA
- CB
1804 1803 1813 1 1.0950e+02 4.1840e+02 ; HA - CA
- C
1805 1803 1813 1 1.1110e+02 5.2718e+02 ; CB - CA
- C
1805 1808 1809 1 1.2040e+02 6.6944e+02 ; CB - CG
- OD1
1805 1808 1810 1 1.1660e+02 5.8576e+02 ; CB - CG
- ND2
1806 1805 1807 1 1.0950e+02 2.9288e+02 ; HB2 - CB
- HB3
1806 1805 1808 1 1.0950e+02 4.1840e+02 ; HB2 - CB
- CG
1807 1805 1808 1 1.0950e+02 4.1840e+02 ; HB3 - CB
- CG
1808 1810 1811 1 1.2000e+02 4.1840e+02 ; CG - ND2
- HD21
1808 1810 1812 1 1.2000e+02 4.1840e+02 ; CG - ND2
- HD22
1809 1808 1810 1 1.2290e+02 6.6944e+02 ; OD1 - CG
- ND2
1811 1810 1812 1 1.2000e+02 2.9288e+02 ; HD21 - ND2
- HD22
1813 1815 1816 1 1.2000e+02 4.1840e+02 ; C - N
- H
1813 1815 1817 1 1.2190e+02 4.1840e+02 ; C - N
- CA
1814 1813 1815 1 1.2290e+02 6.6944e+02 ; O - C
- N
1815 1817 1818 1 1.0950e+02 4.1840e+02 ; N - CA
- HA
1815 1817 1819 1 1.0970e+02 6.6944e+02 ; N - CA
- CB
1815 1817 1823 1 1.1010e+02 5.2718e+02 ; N - CA
- C
1816 1815 1817 1 1.1804e+02 4.1840e+02 ; H - N
- CA
1817 1819 1820 1 1.0950e+02 4.1840e+02 ; CA - CB
- HB1
1817 1819 1821 1 1.0950e+02 4.1840e+02 ; CA - CB
- HB2
1817 1819 1822 1 1.0950e+02 4.1840e+02 ; CA - CB
- HB3
1817 1823 1824 1 1.2040e+02 6.6944e+02 ; CA - C
- O

1817 1823 1825 1 1.1660e+02 5.8576e+02 ; CA - C
- N
1818 1817 1819 1 1.0950e+02 4.1840e+02 ; HA - CA
- CB
1818 1817 1823 1 1.0950e+02 4.1840e+02 ; HA - CA
- C
1819 1817 1823 1 1.1110e+02 5.2718e+02 ; CB - CA
- C
1820 1819 1821 1 1.0950e+02 2.9288e+02 ; HB1 - CB
- HB2
1820 1819 1822 1 1.0950e+02 2.9288e+02 ; HB1 - CB
- HB3
1821 1819 1822 1 1.0950e+02 2.9288e+02 ; HB2 - CB
- HB3
1823 1825 1826 1 1.2000e+02 4.1840e+02 ; C - N
- H
1823 1825 1827 1 1.2190e+02 4.1840e+02 ; C - N
- CA
1824 1823 1825 1 1.2290e+02 6.6944e+02 ; O - C
- N
1825 1827 1828 1 1.0950e+02 4.1840e+02 ; N - CA
- HA
1825 1827 1829 1 1.0970e+02 6.6944e+02 ; N - CA
- CB
1825 1827 1833 1 1.1010e+02 5.2718e+02 ; N - CA
- C
1826 1825 1827 1 1.1804e+02 4.1840e+02 ; H - N
- CA
1827 1829 1830 1 1.0950e+02 4.1840e+02 ; CA - CB
- HB1
1827 1829 1831 1 1.0950e+02 4.1840e+02 ; CA - CB
- HB2
1827 1829 1832 1 1.0950e+02 4.1840e+02 ; CA - CB
- HB3
1827 1833 1834 1 1.2040e+02 6.6944e+02 ; CA - C
- O
1827 1833 1835 1 1.1660e+02 5.8576e+02 ; CA - C
- N
1828 1827 1829 1 1.0950e+02 4.1840e+02 ; HA - CA
- CB
1828 1827 1833 1 1.0950e+02 4.1840e+02 ; HA - CA
- C
1829 1827 1833 1 1.1110e+02 5.2718e+02 ; CB - CA
- C
1830 1829 1831 1 1.0950e+02 2.9288e+02 ; HB1 - CB
- HB2
1830 1829 1832 1 1.0950e+02 2.9288e+02 ; HB1 - CB
- HB3
1831 1829 1832 1 1.0950e+02 2.9288e+02 ; HB2 - CB
- HB3
1833 1835 1836 1 1.2000e+02 4.1840e+02 ; C - N
- H

1833 1835 1837 1 1.2190e+02 4.1840e+02 ; C - N
- CA
1834 1833 1835 1 1.2290e+02 6.6944e+02 ; O - C
- N
1835 1837 1838 1 1.0950e+02 4.1840e+02 ; N - CA
- HA
1835 1837 1839 1 1.0970e+02 6.6944e+02 ; N - CA
- CB
1835 1837 1847 1 1.1010e+02 5.2718e+02 ; N - CA
- C
1836 1835 1837 1 1.1804e+02 4.1840e+02 ; H - N
- CA
1837 1839 1840 1 1.0950e+02 4.1840e+02 ; CA - CB
- HB
1837 1839 1841 1 1.0950e+02 3.3472e+02 ; CA - CB
- CG2
1837 1839 1845 1 1.0950e+02 4.1840e+02 ; CA - CB
- OG1
1837 1847 1848 1 1.2040e+02 6.6944e+02 ; CA - C
- O
1837 1847 1849 1 1.1660e+02 5.8576e+02 ; CA - C
- N
1838 1837 1839 1 1.0950e+02 4.1840e+02 ; HA - CA
- CB
1838 1837 1847 1 1.0950e+02 4.1840e+02 ; HA - CA
- C
1839 1837 1847 1 1.1110e+02 5.2718e+02 ; CB - CA
- C
1839 1841 1842 1 1.0950e+02 4.1840e+02 ; CB - CG2
- HG21
1839 1841 1843 1 1.0950e+02 4.1840e+02 ; CB - CG2
- HG22
1839 1841 1844 1 1.0950e+02 4.1840e+02 ; CB - CG2
- HG23
1839 1845 1846 1 1.0850e+02 4.6024e+02 ; CB - OG1
- HG1
1840 1839 1841 1 1.0950e+02 4.1840e+02 ; HB - CB
- CG2
1840 1839 1845 1 1.0950e+02 4.1840e+02 ; HB - CB
- OG1
1841 1839 1845 1 1.0950e+02 4.1840e+02 ; CG2 - CB
- OG1
1842 1841 1843 1 1.0950e+02 2.9288e+02 ; HG21 - CG2
- HG22
1842 1841 1844 1 1.0950e+02 2.9288e+02 ; HG21 - CG2
- HG23
1843 1841 1844 1 1.0950e+02 2.9288e+02 ; HG22 - CG2
- HG23
1847 1849 1850 1 1.2000e+02 4.1840e+02 ; C - N
- H
1847 1849 1851 1 1.2190e+02 4.1840e+02 ; C - N
- CA

| | | | | | | | |
|---|---|---|---|---|---|---|---|
| 1848 | 1847 | 1849 | 1 | 1.2290e+02 | 6.6944e+02 | ; | O - C - N |
| 1849 | 1851 | 1852 | 1 | 1.0950e+02 | 4.1840e+02 | ; | N - CA - HA |
| 1849 | 1851 | 1853 | 1 | 1.0970e+02 | 6.6944e+02 | ; | N - CA - CB |
| 1849 | 1851 | 1866 | 1 | 1.1010e+02 | 5.2718e+02 | ; | N - CA - C |
| 1850 | 1849 | 1851 | 1 | 1.1804e+02 | 4.1840e+02 | ; | H - N - CA |
| 1851 | 1853 | 1854 | 1 | 1.0950e+02 | 4.1840e+02 | ; | CA - CB - HB |
| 1851 | 1853 | 1855 | 1 | 1.0950e+02 | 3.3472e+02 | ; | CA - CB - CG2 |
| 1851 | 1853 | 1859 | 1 | 1.0950e+02 | 3.3472e+02 | ; | CA - CB - CG1 |
| 1851 | 1866 | 1867 | 1 | 1.2040e+02 | 6.6944e+02 | ; | CA - C - O |
| 1851 | 1866 | 1868 | 1 | 1.1660e+02 | 5.8576e+02 | ; | CA - C - N |
| 1852 | 1851 | 1853 | 1 | 1.0950e+02 | 4.1840e+02 | ; | HA - CA - CB |
| 1852 | 1851 | 1866 | 1 | 1.0950e+02 | 4.1840e+02 | ; | HA - CA - C |
| 1853 | 1851 | 1866 | 1 | 1.1110e+02 | 5.2718e+02 | ; | CB - CA - C |
| 1853 | 1855 | 1856 | 1 | 1.0950e+02 | 4.1840e+02 | ; | CB - CG2 - HG21 |
| 1853 | 1855 | 1857 | 1 | 1.0950e+02 | 4.1840e+02 | ; | CB - CG2 - HG22 |
| 1853 | 1855 | 1858 | 1 | 1.0950e+02 | 4.1840e+02 | ; | CB - CG2 - HG23 |
| 1853 | 1859 | 1860 | 1 | 1.0950e+02 | 4.1840e+02 | ; | CB - CG1 - HG12 |
| 1853 | 1859 | 1861 | 1 | 1.0950e+02 | 4.1840e+02 | ; | CB - CG1 - HG13 |
| 1853 | 1859 | 1862 | 1 | 1.0950e+02 | 3.3472e+02 | ; | CB - CG1 - CD1 |
| 1854 | 1853 | 1855 | 1 | 1.0950e+02 | 4.1840e+02 | ; | HB - CB - CG2 |
| 1854 | 1853 | 1859 | 1 | 1.0950e+02 | 4.1840e+02 | ; | HB - CB - CG1 |
| 1855 | 1853 | 1859 | 1 | 1.0950e+02 | 3.3472e+02 | ; | CG2 - CB - CG1 |
| 1856 | 1855 | 1857 | 1 | 1.0950e+02 | 2.9288e+02 | ; | HG21 - CG2 - HG22 |
| 1856 | 1855 | 1858 | 1 | 1.0950e+02 | 2.9288e+02 | ; | HG21 - CG2 - HG23 |
| 1857 | 1855 | 1858 | 1 | 1.0950e+02 | 2.9288e+02 | ; | HG22 - CG2 - HG23 |
| 1859 | 1862 | 1863 | 1 | 1.0950e+02 | 4.1840e+02 | ; | CG1 - CD1 - HD11 |

1859 1862 1864 1 1.0950e+02 4.1840e+02 ; CG1 - CD1
- HD12
1859 1862 1865 1 1.0950e+02 4.1840e+02 ; CG1 - CD1
- HD13
1860 1859 1861 1 1.0950e+02 2.9288e+02 ; HG12 - CG1
- HG13
1860 1859 1862 1 1.0950e+02 4.1840e+02 ; HG12 - CG1
- CD1
1861 1859 1862 1 1.0950e+02 4.1840e+02 ; HG13 - CG1
- CD1
1863 1862 1864 1 1.0950e+02 2.9288e+02 ; HD11 - CD1
- HD12
1863 1862 1865 1 1.0950e+02 2.9288e+02 ; HD11 - CD1
- HD13
1864 1862 1865 1 1.0950e+02 2.9288e+02 ; HD12 - CD1
- HD13
1866 1868 1869 1 1.2000e+02 4.1840e+02 ; C - N
- H
1866 1868 1870 1 1.2190e+02 4.1840e+02 ; C - N
- CA
1867 1866 1868 1 1.2290e+02 6.6944e+02 ; O - C
- N
1868 1870 1871 1 1.0950e+02 4.1840e+02 ; N - CA
- HA
1868 1870 1872 1 1.0970e+02 6.6944e+02 ; N - CA
- CB
1868 1870 1883 1 1.1010e+02 5.2718e+02 ; N - CA
- C
1869 1868 1870 1 1.1804e+02 4.1840e+02 ; H - N
- CA
1870 1872 1873 1 1.0950e+02 4.1840e+02 ; CA - CB
- HB2
1870 1872 1874 1 1.0950e+02 4.1840e+02 ; CA - CB
- HB3
1870 1872 1875 1 1.0950e+02 3.3472e+02 ; CA - CB
- CG
1870 1883 1884 1 1.2040e+02 6.6944e+02 ; CA - C
- O
1870 1883 1885 1 1.1660e+02 5.8576e+02 ; CA - C
- N
1871 1870 1872 1 1.0950e+02 4.1840e+02 ; HA - CA
- CB
1871 1870 1883 1 1.0950e+02 4.1840e+02 ; HA - CA
- C
1872 1870 1883 1 1.1110e+02 5.2718e+02 ; CB - CA
- C
1872 1875 1876 1 1.0950e+02 4.1840e+02 ; CB - CG
- HG2
1872 1875 1877 1 1.0950e+02 4.1840e+02 ; CB - CG
- HG3
1872 1875 1878 1 1.1470e+02 4.1840e+02 ; CB - CG
- SD

1873 1872 1874 1 1.0950e+02 2.9288e+02 ; HB2 - CB
- HB3
1873 1872 1875 1 1.0950e+02 4.1840e+02 ; HB2 - CB
- CG
1874 1872 1875 1 1.0950e+02 4.1840e+02 ; HB3 - CB
- CG
1875 1878 1879 1 9.8900e+01 5.1882e+02 ; CG - SD
- CE
1876 1875 1877 1 1.0950e+02 2.9288e+02 ; HG2 - CG
- HG3
1876 1875 1878 1 1.0950e+02 4.1840e+02 ; HG2 - CG
- SD
1877 1875 1878 1 1.0950e+02 4.1840e+02 ; HG3 - CG
- SD
1878 1879 1880 1 1.0950e+02 4.1840e+02 ; SD - CE
- HE1
1878 1879 1881 1 1.0950e+02 4.1840e+02 ; SD - CE
- HE2
1878 1879 1882 1 1.0950e+02 4.1840e+02 ; SD - CE
- HE3
1880 1879 1881 1 1.0950e+02 2.9288e+02 ; HE1 - CE
- HE2
1880 1879 1882 1 1.0950e+02 2.9288e+02 ; HE1 - CE
- HE3
1881 1879 1882 1 1.0950e+02 2.9288e+02 ; HE2 - CE
- HE3
1883 1885 1886 1 1.2000e+02 4.1840e+02 ; C - N
- H
1883 1885 1887 1 1.2190e+02 4.1840e+02 ; C - N
- CA
1884 1883 1885 1 1.2290e+02 6.6944e+02 ; O - C
- N
1885 1887 1888 1 1.0950e+02 4.1840e+02 ; N - CA
- HA
1885 1887 1889 1 1.0970e+02 6.6944e+02 ; N - CA
- CB
1885 1887 1905 1 1.1010e+02 5.2718e+02 ; N - CA
- C
1886 1885 1887 1 1.1804e+02 4.1840e+02 ; H - N
- CA
1887 1889 1890 1 1.0950e+02 4.1840e+02 ; CA - CB
- HB2
1887 1889 1891 1 1.0950e+02 4.1840e+02 ; CA - CB
- HB3
1887 1889 1892 1 1.0950e+02 3.3472e+02 ; CA - CB
- CG
1887 1905 1906 1 1.2040e+02 6.6944e+02 ; CA - C
- O
1887 1905 1907 1 1.1660e+02 5.8576e+02 ; CA - C
- N
1888 1887 1889 1 1.0950e+02 4.1840e+02 ; HA - CA
- CB

1888 1887 1905 1 1.0950e+02 4.1840e+02 ; HA - CA
- C
1889 1887 1905 1 1.1110e+02 5.2718e+02 ; CB - CA
- C
1889 1892 1893 1 1.0950e+02 4.1840e+02 ; CB - CG
- HG2
1889 1892 1894 1 1.0950e+02 4.1840e+02 ; CB - CG
- HG3
1889 1892 1895 1 1.0950e+02 3.3472e+02 ; CB - CG
- CD
1890 1889 1891 1 1.0950e+02 2.9288e+02 ; HB2 - CB
- HB3
1890 1889 1892 1 1.0950e+02 4.1840e+02 ; HB2 - CB
- CG
1891 1889 1892 1 1.0950e+02 4.1840e+02 ; HB3 - CB
- CG
1892 1895 1896 1 1.0950e+02 4.1840e+02 ; CG - CD
- HD2
1892 1895 1897 1 1.0950e+02 4.1840e+02 ; CG - CD
- HD3
1892 1895 1898 1 1.0950e+02 3.3472e+02 ; CG - CD
- CE
1893 1892 1894 1 1.0950e+02 2.9288e+02 ; HG2 - CG
- HG3
1893 1892 1895 1 1.0950e+02 4.1840e+02 ; HG2 - CG
- CD
1894 1892 1895 1 1.0950e+02 4.1840e+02 ; HG3 - CG
- CD
1895 1898 1899 1 1.0950e+02 4.1840e+02 ; CD - CE
- HE2
1895 1898 1900 1 1.0950e+02 4.1840e+02 ; CD - CE
- HE3
1895 1898 1901 1 1.1120e+02 6.6944e+02 ; CD - CE
- NZ
1896 1895 1897 1 1.0950e+02 2.9288e+02 ; HD2 - CD
- HD3
1896 1895 1898 1 1.0950e+02 4.1840e+02 ; HD2 - CD
- CE
1897 1895 1898 1 1.0950e+02 4.1840e+02 ; HD3 - CD
- CE
1898 1901 1902 1 1.0950e+02 4.1840e+02 ; CE - NZ
- HZ1
1898 1901 1903 1 1.0950e+02 4.1840e+02 ; CE - NZ
- HZ2
1898 1901 1904 1 1.0950e+02 4.1840e+02 ; CE - NZ
- HZ3
1899 1898 1900 1 1.0950e+02 2.9288e+02 ; HE2 - CE
- HE3
1899 1898 1901 1 1.0950e+02 4.1840e+02 ; HE2 - CE
- NZ
1900 1898 1901 1 1.0950e+02 4.1840e+02 ; HE3 - CE
- NZ

1902 1901 1903 1 1.0950e+02 2.9288e+02 ; HZ1 - NZ
- HZ2
1902 1901 1904 1 1.0950e+02 2.9288e+02 ; HZ1 - NZ
- HZ3
1903 1901 1904 1 1.0950e+02 2.9288e+02 ; HZ2 - NZ
- HZ3
1905 1907 1908 1 1.2000e+02 4.1840e+02 ; C - N
- H
1905 1907 1909 1 1.2190e+02 4.1840e+02 ; C - N
- CA
1906 1905 1907 1 1.2290e+02 6.6944e+02 ; O - C
- N
1907 1909 1910 1 1.0950e+02 4.1840e+02 ; N - CA
- HA
1907 1909 1911 1 1.0970e+02 6.6944e+02 ; N - CA
- CB
1907 1909 1922 1 1.1010e+02 5.2718e+02 ; N - CA
- C
1908 1907 1909 1 1.1804e+02 4.1840e+02 ; H - N
- CA
1909 1911 1912 1 1.0950e+02 4.1840e+02 ; CA - CB
- HB2
1909 1911 1913 1 1.0950e+02 4.1840e+02 ; CA - CB
- HB3
1909 1911 1914 1 1.0950e+02 3.3472e+02 ; CA - CB
- CG
1909 1922 1923 1 1.2040e+02 6.6944e+02 ; CA - C
- O
1909 1922 1924 1 1.1660e+02 5.8576e+02 ; CA - C
- N
1910 1909 1911 1 1.0950e+02 4.1840e+02 ; HA - CA
- CB
1910 1909 1922 1 1.0950e+02 4.1840e+02 ; HA - CA
- C
1911 1909 1922 1 1.1110e+02 5.2718e+02 ; CB - CA
- C
1911 1914 1915 1 1.0950e+02 4.1840e+02 ; CB - CG
- HG2
1911 1914 1916 1 1.0950e+02 4.1840e+02 ; CB - CG
- HG3
1911 1914 1917 1 1.1110e+02 5.2718e+02 ; CB - CG
- CD
1912 1911 1913 1 1.0950e+02 2.9288e+02 ; HB2 - CB
- HB3
1912 1911 1914 1 1.0950e+02 4.1840e+02 ; HB2 - CB
- CG
1913 1911 1914 1 1.0950e+02 4.1840e+02 ; HB3 - CB
- CG
1914 1917 1918 1 1.2040e+02 6.6944e+02 ; CG - CD
- OE1
1914 1917 1919 1 1.1660e+02 5.8576e+02 ; CG - CD
- NE2

1915 1914 1916 1 1.0950e+02 2.9288e+02 ; HG2 - CG
- HG3
1915 1914 1917 1 1.0950e+02 4.1840e+02 ; HG2 - CG
- CD
1916 1914 1917 1 1.0950e+02 4.1840e+02 ; HG3 - CG
- CD
1917 1919 1920 1 1.2000e+02 4.1840e+02 ; CD - NE2
- HE21
1917 1919 1921 1 1.2000e+02 4.1840e+02 ; CD - NE2
- HE22
1918 1917 1919 1 1.2290e+02 6.6944e+02 ; OE1 - CD
- NE2
1920 1919 1921 1 1.2000e+02 2.9288e+02 ; HE21 - NE2
- HE22
1922 1924 1925 1 1.2000e+02 4.1840e+02 ; C - N
- H
1922 1924 1926 1 1.2190e+02 4.1840e+02 ; C - N
- CA
1923 1922 1924 1 1.2290e+02 6.6944e+02 ; O - C
- N
1924 1926 1927 1 1.0950e+02 4.1840e+02 ; N - CA
- HA
1924 1926 1928 1 1.0970e+02 6.6944e+02 ; N - CA
- CB
1924 1926 1941 1 1.1010e+02 5.2718e+02 ; N - CA
- C
1925 1924 1926 1 1.1804e+02 4.1840e+02 ; H - N
- CA
1926 1928 1929 1 1.0950e+02 4.1840e+02 ; CA - CB
- HB
1926 1928 1930 1 1.0950e+02 3.3472e+02 ; CA - CB
- CG2
1926 1928 1934 1 1.0950e+02 3.3472e+02 ; CA - CB
- CG1
1926 1941 1942 1 1.2040e+02 6.6944e+02 ; CA - C
- O
1926 1941 1943 1 1.1660e+02 5.8576e+02 ; CA - C
- N
1927 1926 1928 1 1.0950e+02 4.1840e+02 ; HA - CA
- CB
1927 1926 1941 1 1.0950e+02 4.1840e+02 ; HA - CA
- C
1928 1926 1941 1 1.1110e+02 5.2718e+02 ; CB - CA
- C
1928 1930 1931 1 1.0950e+02 4.1840e+02 ; CB - CG2
- HG21
1928 1930 1932 1 1.0950e+02 4.1840e+02 ; CB - CG2
- HG22
1928 1930 1933 1 1.0950e+02 4.1840e+02 ; CB - CG2
- HG23
1928 1934 1935 1 1.0950e+02 4.1840e+02 ; CB - CG1
- HG12

```
  1928   1934   1936      1    1.0950e+02    4.1840e+02 ;     CB - CG1
- HG13
  1928   1934   1937      1    1.0950e+02    3.3472e+02 ;     CB - CG1
- CD1
  1929   1928   1930      1    1.0950e+02    4.1840e+02 ;     HB - CB
- CG2
  1929   1928   1934      1    1.0950e+02    4.1840e+02 ;     HB - CB
- CG1
  1930   1928   1934      1    1.0950e+02    3.3472e+02 ;    CG2 - CB
- CG1
  1931   1930   1932      1    1.0950e+02    2.9288e+02 ;   HG21 - CG2
- HG22
  1931   1930   1933      1    1.0950e+02    2.9288e+02 ;   HG21 - CG2
- HG23
  1932   1930   1933      1    1.0950e+02    2.9288e+02 ;   HG22 - CG2
- HG23
  1934   1937   1938      1    1.0950e+02    4.1840e+02 ;    CG1 - CD1
- HD11
  1934   1937   1939      1    1.0950e+02    4.1840e+02 ;    CG1 - CD1
- HD12
  1934   1937   1940      1    1.0950e+02    4.1840e+02 ;    CG1 - CD1
- HD13
  1935   1934   1936      1    1.0950e+02    2.9288e+02 ;   HG12 - CG1
- HG13
  1935   1934   1937      1    1.0950e+02    4.1840e+02 ;   HG12 - CG1
- CD1
  1936   1934   1937      1    1.0950e+02    4.1840e+02 ;   HG13 - CG1
- CD1
  1938   1937   1939      1    1.0950e+02    2.9288e+02 ;   HD11 - CD1
- HD12
  1938   1937   1940      1    1.0950e+02    2.9288e+02 ;   HD11 - CD1
- HD13
  1939   1937   1940      1    1.0950e+02    2.9288e+02 ;   HD12 - CD1
- HD13
  1941   1943   1944      1    1.2000e+02    4.1840e+02 ;      C - N
- H
  1941   1943   1945      1    1.2190e+02    4.1840e+02 ;      C - N
- CA
  1942   1941   1943      1    1.2290e+02    6.6944e+02 ;      O - C
- N
  1943   1945   1946      1    1.0950e+02    4.1840e+02 ;      N - CA
- HA
  1943   1945   1947      1    1.0970e+02    6.6944e+02 ;      N - CA
- CB
  1943   1945   1961      1    1.1010e+02    5.2718e+02 ;      N - CA
- C
  1944   1943   1945      1    1.1804e+02    4.1840e+02 ;      H - N
- CA
  1945   1947   1948      1    1.0950e+02    4.1840e+02 ;     CA - CB
- HB2
  1945   1947   1949      1    1.0950e+02    4.1840e+02 ;     CA - CB
- HB3
```

1945 1947 1950 1 1.1400e+02 5.2718e+02 ; CA - CB
- CG
1945 1961 1962 1 1.2040e+02 6.6944e+02 ; CA - C
- O
1945 1961 1963 1 1.1660e+02 5.8576e+02 ; CA - C
- N
1946 1945 1947 1 1.0950e+02 4.1840e+02 ; HA - CA
- CB
1946 1945 1961 1 1.0950e+02 4.1840e+02 ; HA - CA
- C
1947 1945 1961 1 1.1110e+02 5.2718e+02 ; CB - CA
- C
1947 1950 1951 1 1.2000e+02 5.8576e+02 ; CB - CG
- CD1
1947 1950 1959 1 1.2000e+02 5.8576e+02 ; CB - CG
- CD2
1948 1947 1949 1 1.0950e+02 2.9288e+02 ; HB2 - CB
- HB3
1948 1947 1950 1 1.0950e+02 4.1840e+02 ; HB2 - CB
- CG
1949 1947 1950 1 1.0950e+02 4.1840e+02 ; HB3 - CB
- CG
1950 1951 1952 1 1.2000e+02 4.1840e+02 ; CG - CD1
- HD1
1950 1951 1953 1 1.2000e+02 5.2718e+02 ; CG - CD1
- CE1
1950 1959 1957 1 1.2000e+02 5.2718e+02 ; CG - CD2
- CE2
1950 1959 1960 1 1.2000e+02 4.1840e+02 ; CG - CD2
- HD2
1951 1950 1959 1 1.2000e+02 5.2718e+02 ; CD1 - CG
- CD2
1951 1953 1954 1 1.2000e+02 4.1840e+02 ; CD1 - CE1
- HE1
1951 1953 1955 1 1.2000e+02 5.2718e+02 ; CD1 - CE1
- CZ
1952 1951 1953 1 1.2000e+02 4.1840e+02 ; HD1 - CD1
- CE1
1953 1955 1956 1 1.2000e+02 4.1840e+02 ; CE1 - CZ
- HZ
1953 1955 1957 1 1.2000e+02 5.2718e+02 ; CE1 - CZ
- CE2
1954 1953 1955 1 1.2000e+02 4.1840e+02 ; HE1 - CE1
- CZ
1955 1957 1958 1 1.2000e+02 4.1840e+02 ; CZ - CE2
- HE2
1955 1957 1959 1 1.2000e+02 5.2718e+02 ; CZ - CE2
- CD2
1956 1955 1957 1 1.2000e+02 4.1840e+02 ; HZ - CZ
- CE2
1957 1959 1960 1 1.2000e+02 4.1840e+02 ; CE2 - CD2
- HD2

1958 1957 1959 1 1.2000e+02 4.1840e+02 ; HE2 - CE2
- CD2
1961 1963 1964 1 1.2000e+02 4.1840e+02 ; C - N
- H
1961 1963 1965 1 1.2190e+02 4.1840e+02 ; C - N
- CA
1962 1961 1963 1 1.2290e+02 6.6944e+02 ; O - C
- N
1963 1965 1966 1 1.0950e+02 4.1840e+02 ; N - CA
- HA
1963 1965 1967 1 1.0970e+02 6.6944e+02 ; N - CA
- CB
1963 1965 1972 1 1.1010e+02 5.2718e+02 ; N - CA
- C
1964 1963 1965 1 1.1804e+02 4.1840e+02 ; H - N
- CA
1965 1967 1968 1 1.0950e+02 4.1840e+02 ; CA - CB
- HB2
1965 1967 1969 1 1.0950e+02 4.1840e+02 ; CA - CB
- HB3
1965 1967 1970 1 1.0950e+02 4.1840e+02 ; CA - CB
- OG
1965 1972 1973 1 1.2040e+02 6.6944e+02 ; CA - C
- O
1965 1972 1974 1 1.1660e+02 5.8576e+02 ; CA - C
- N
1966 1965 1967 1 1.0950e+02 4.1840e+02 ; HA - CA
- CB
1966 1965 1972 1 1.0950e+02 4.1840e+02 ; HA - CA
- C
1967 1965 1972 1 1.1110e+02 5.2718e+02 ; CB - CA
- C
1967 1970 1971 1 1.0850e+02 4.6024e+02 ; CB - OG
- HG
1968 1967 1969 1 1.0950e+02 2.9288e+02 ; HB2 - CB
- HB3
1968 1967 1970 1 1.0950e+02 4.1840e+02 ; HB2 - CB
- OG
1969 1967 1970 1 1.0950e+02 4.1840e+02 ; HB3 - CB
- OG
1972 1974 1975 1 1.2000e+02 4.1840e+02 ; C - N
- H
1972 1974 1976 1 1.2190e+02 4.1840e+02 ; C - N
- CA
1973 1972 1974 1 1.2290e+02 6.6944e+02 ; O - C
- N
1974 1976 1977 1 1.0950e+02 4.1840e+02 ; N - CA
- HA
1974 1976 1978 1 1.0970e+02 6.6944e+02 ; N - CA
- CB
1974 1976 1982 1 1.1010e+02 5.2718e+02 ; N - CA
- C

1975 1974 1976 1 1.1804e+02 4.1840e+02 ; H - N
- CA
1976 1978 1979 1 1.0950e+02 4.1840e+02 ; CA - CB
- HB1
1976 1978 1980 1 1.0950e+02 4.1840e+02 ; CA - CB
- HB2
1976 1978 1981 1 1.0950e+02 4.1840e+02 ; CA - CB
- HB3
1976 1982 1983 1 1.2040e+02 6.6944e+02 ; CA - C
- O
1976 1982 1984 1 1.1660e+02 5.8576e+02 ; CA - C
- N
1977 1976 1978 1 1.0950e+02 4.1840e+02 ; HA - CA
- CB
1977 1976 1982 1 1.0950e+02 4.1840e+02 ; HA - CA
- C
1978 1976 1982 1 1.1110e+02 5.2718e+02 ; CB - CA
- C
1979 1978 1980 1 1.0950e+02 2.9288e+02 ; HB1 - CB
- HB2
1979 1978 1981 1 1.0950e+02 2.9288e+02 ; HB1 - CB
- HB3
1980 1978 1981 1 1.0950e+02 2.9288e+02 ; HB2 - CB
- HB3
1982 1984 1985 1 1.2000e+02 4.1840e+02 ; C - N
- H
1982 1984 1986 1 1.2190e+02 4.1840e+02 ; C - N
- CA
1983 1982 1984 1 1.2290e+02 6.6944e+02 ; O - C
- N
1984 1986 1987 1 1.0950e+02 4.1840e+02 ; N - CA
- HA
1984 1986 1988 1 1.0970e+02 6.6944e+02 ; N - CA
- CB
1984 1986 2001 1 1.1010e+02 5.2718e+02 ; N - CA
- C
1985 1984 1986 1 1.1804e+02 4.1840e+02 ; H - N
- CA
1986 1988 1989 1 1.0950e+02 4.1840e+02 ; CA - CB
- HB
1986 1988 1990 1 1.0950e+02 3.3472e+02 ; CA - CB
- CG2
1986 1988 1994 1 1.0950e+02 3.3472e+02 ; CA - CB
- CG1
1986 2001 2002 1 1.2040e+02 6.6944e+02 ; CA - C
- O
1986 2001 2003 1 1.1660e+02 5.8576e+02 ; CA - C
- N
1987 1986 1988 1 1.0950e+02 4.1840e+02 ; HA - CA
- CB
1987 1986 2001 1 1.0950e+02 4.1840e+02 ; HA - CA
- C

```
  1988   1986   2001      1    1.1110e+02    5.2718e+02 ;     CB - CA
- C
  1988   1990   1991      1    1.0950e+02    4.1840e+02 ;     CB - CG2
- HG21
  1988   1990   1992      1    1.0950e+02    4.1840e+02 ;     CB - CG2
- HG22
  1988   1990   1993      1    1.0950e+02    4.1840e+02 ;     CB - CG2
- HG23
  1988   1994   1995      1    1.0950e+02    4.1840e+02 ;     CB - CG1
- HG12
  1988   1994   1996      1    1.0950e+02    4.1840e+02 ;     CB - CG1
- HG13
  1988   1994   1997      1    1.0950e+02    3.3472e+02 ;     CB - CG1
- CD1
  1989   1988   1990      1    1.0950e+02    4.1840e+02 ;     HB - CB
- CG2
  1989   1988   1994      1    1.0950e+02    4.1840e+02 ;     HB - CB
- CG1
  1990   1988   1994      1    1.0950e+02    3.3472e+02 ;    CG2 - CB
- CG1
  1991   1990   1992      1    1.0950e+02    2.9288e+02 ;   HG21 - CG2
- HG22
  1991   1990   1993      1    1.0950e+02    2.9288e+02 ;   HG21 - CG2
- HG23
  1992   1990   1993      1    1.0950e+02    2.9288e+02 ;   HG22 - CG2
- HG23
  1994   1997   1998      1    1.0950e+02    4.1840e+02 ;    CG1 - CD1
- HD11
  1994   1997   1999      1    1.0950e+02    4.1840e+02 ;    CG1 - CD1
- HD12
  1994   1997   2000      1    1.0950e+02    4.1840e+02 ;    CG1 - CD1
- HD13
  1995   1994   1996      1    1.0950e+02    2.9288e+02 ;   HG12 - CG1
- HG13
  1995   1994   1997      1    1.0950e+02    4.1840e+02 ;   HG12 - CG1
- CD1
  1996   1994   1997      1    1.0950e+02    4.1840e+02 ;   HG13 - CG1
- CD1
  1998   1997   1999      1    1.0950e+02    2.9288e+02 ;   HD11 - CD1
- HD12
  1998   1997   2000      1    1.0950e+02    2.9288e+02 ;   HD11 - CD1
- HD13
  1999   1997   2000      1    1.0950e+02    2.9288e+02 ;   HD12 - CD1
- HD13
  2001   2003   2004      1    1.2000e+02    4.1840e+02 ;      C - N
- H
  2001   2003   2005      1    1.2190e+02    4.1840e+02 ;      C - N
- CA
  2002   2001   2003      1    1.2290e+02    6.6944e+02 ;      O - C
- N
  2003   2005   2006      1    1.0950e+02    4.1840e+02 ;      N - CA
- HA
```

```
  2003   2005   2007       1    1.0970e+02    6.6944e+02 ;      N - CA
- CB
  2003   2005   2021       1    1.1010e+02    5.2718e+02 ;      N - CA
- C
  2004   2003   2005       1    1.1804e+02    4.1840e+02 ;      H - N
- CA
  2005   2007   2008       1    1.0950e+02    4.1840e+02 ;     CA - CB
- HB2
  2005   2007   2009       1    1.0950e+02    4.1840e+02 ;     CA - CB
- HB3
  2005   2007   2010       1    1.1400e+02    5.2718e+02 ;     CA - CB
- CG
  2005   2021   2022       1    1.2040e+02    6.6944e+02 ;     CA - C
- O
  2005   2021   2023       1    1.1660e+02    5.8576e+02 ;     CA - C
- N
  2006   2005   2007       1    1.0950e+02    4.1840e+02 ;     HA - CA
- CB
  2006   2005   2021       1    1.0950e+02    4.1840e+02 ;     HA - CA
- C
  2007   2005   2021       1    1.1110e+02    5.2718e+02 ;     CB - CA
- C
  2007   2010   2011       1    1.2000e+02    5.8576e+02 ;     CB - CG
- CD1
  2007   2010   2019       1    1.2000e+02    5.8576e+02 ;     CB - CG
- CD2
  2008   2007   2009       1    1.0950e+02    2.9288e+02 ;    HB2 - CB
- HB3
  2008   2007   2010       1    1.0950e+02    4.1840e+02 ;    HB2 - CB
- CG
  2009   2007   2010       1    1.0950e+02    4.1840e+02 ;    HB3 - CB
- CG
  2010   2011   2012       1    1.2000e+02    4.1840e+02 ;     CG - CD1
- HD1
  2010   2011   2013       1    1.2000e+02    5.2718e+02 ;     CG - CD1
- CE1
  2010   2019   2017       1    1.2000e+02    5.2718e+02 ;     CG - CD2
- CE2
  2010   2019   2020       1    1.2000e+02    4.1840e+02 ;     CG - CD2
- HD2
  2011   2010   2019       1    1.2000e+02    5.2718e+02 ;    CD1 - CG
- CD2
  2011   2013   2014       1    1.2000e+02    4.1840e+02 ;    CD1 - CE1
- HE1
  2011   2013   2015       1    1.2000e+02    5.2718e+02 ;    CD1 - CE1
- CZ
  2012   2011   2013       1    1.2000e+02    4.1840e+02 ;    HD1 - CD1
- CE1
  2013   2015   2016       1    1.2000e+02    4.1840e+02 ;    CE1 - CZ
- HZ
  2013   2015   2017       1    1.2000e+02    5.2718e+02 ;    CE1 - CZ
- CE2
```

2014 2013 2015 1 1.2000e+02 4.1840e+02 ; HE1 - CE1 - CZ
2015 2017 2018 1 1.2000e+02 4.1840e+02 ; CZ - CE2 - HE2
2015 2017 2019 1 1.2000e+02 5.2718e+02 ; CZ - CE2 - CD2
2016 2015 2017 1 1.2000e+02 4.1840e+02 ; HZ - CZ - CE2
2017 2019 2020 1 1.2000e+02 4.1840e+02 ; CE2 - CD2 - HD2
2018 2017 2019 1 1.2000e+02 4.1840e+02 ; HE2 - CE2 - CD2
2021 2023 2024 1 1.2000e+02 4.1840e+02 ; C - N - H
2021 2023 2025 1 1.2190e+02 4.1840e+02 ; C - N - CA
2022 2021 2023 1 1.2290e+02 6.6944e+02 ; O - C - N
2023 2025 2026 1 1.0950e+02 4.1840e+02 ; N - CA - HA
2023 2025 2027 1 1.0970e+02 6.6944e+02 ; N - CA - CB
2023 2025 2042 1 1.1010e+02 5.2718e+02 ; N - CA - C
2024 2023 2025 1 1.1804e+02 4.1840e+02 ; H - N - CA
2025 2027 2028 1 1.0950e+02 4.1840e+02 ; CA - CB - HB2
2025 2027 2029 1 1.0950e+02 4.1840e+02 ; CA - CB - HB3
2025 2027 2030 1 1.1400e+02 5.2718e+02 ; CA - CB - CG
2025 2042 2043 1 1.2040e+02 6.6944e+02 ; CA - C - O
2025 2042 2044 1 1.1660e+02 5.8576e+02 ; CA - C - N
2026 2025 2027 1 1.0950e+02 4.1840e+02 ; HA - CA - CB
2026 2025 2042 1 1.0950e+02 4.1840e+02 ; HA - CA - C
2027 2025 2042 1 1.1110e+02 5.2718e+02 ; CB - CA - C
2027 2030 2031 1 1.2000e+02 5.8576e+02 ; CB - CG - CD1
2027 2030 2040 1 1.2000e+02 5.8576e+02 ; CB - CG - CD2
2028 2027 2029 1 1.0950e+02 2.9288e+02 ; HB2 - CB - HB3
2028 2027 2030 1 1.0950e+02 4.1840e+02 ; HB2 - CB - CG
2029 2027 2030 1 1.0950e+02 4.1840e+02 ; HB3 - CB - CG

2030 2031 2032 1 1.2000e+02 4.1840e+02 ; CG - CD1
- HD1
2030 2031 2033 1 1.2000e+02 5.2718e+02 ; CG - CD1
- CE1
2030 2040 2038 1 1.2000e+02 5.2718e+02 ; CG - CD2
- CE2
2030 2040 2041 1 1.2000e+02 4.1840e+02 ; CG - CD2
- HD2
2031 2030 2040 1 1.2000e+02 5.2718e+02 ; CD1 - CG
- CD2
2031 2033 2034 1 1.2000e+02 4.1840e+02 ; CD1 - CE1
- HE1
2031 2033 2035 1 1.2000e+02 5.2718e+02 ; CD1 - CE1
- CZ
2032 2031 2033 1 1.2000e+02 4.1840e+02 ; HD1 - CD1
- CE1
2033 2035 2036 1 1.2000e+02 5.8576e+02 ; CE1 - CZ
- OH
2033 2035 2038 1 1.2000e+02 5.2718e+02 ; CE1 - CZ
- CE2
2034 2033 2035 1 1.2000e+02 4.1840e+02 ; HE1 - CE1
- CZ
2035 2036 2037 1 1.1300e+02 4.1840e+02 ; CZ - OH
- HH
2035 2038 2039 1 1.2000e+02 4.1840e+02 ; CZ - CE2
- HE2
2035 2038 2040 1 1.2000e+02 5.2718e+02 ; CZ - CE2
- CD2
2036 2035 2038 1 1.2000e+02 5.8576e+02 ; OH - CZ
- CE2
2038 2040 2041 1 1.2000e+02 4.1840e+02 ; CE2 - CD2
- HD2
2039 2038 2040 1 1.2000e+02 4.1840e+02 ; HE2 - CE2
- CD2
2042 2044 2045 1 1.2000e+02 4.1840e+02 ; C - N
- H
2042 2044 2046 1 1.2190e+02 4.1840e+02 ; C - N
- CA
2043 2042 2044 1 1.2290e+02 6.6944e+02 ; O - C
- N
2044 2046 2047 1 1.0950e+02 4.1840e+02 ; N - CA
- HA
2044 2046 2048 1 1.0970e+02 6.6944e+02 ; N - CA
- CB
2044 2046 2061 1 1.1010e+02 5.2718e+02 ; N - CA
- C
2045 2044 2046 1 1.1804e+02 4.1840e+02 ; H - N
- CA
2046 2048 2049 1 1.0950e+02 4.1840e+02 ; CA - CB
- HB2
2046 2048 2050 1 1.0950e+02 4.1840e+02 ; CA - CB
- HB3

2046 2048 2051 1 1.0950e+02 3.3472e+02 ; CA - CB
- CG
2046 2061 2062 1 1.2040e+02 6.6944e+02 ; CA - C
- O
2046 2061 2063 1 1.1660e+02 5.8576e+02 ; CA - C
- N
2047 2046 2048 1 1.0950e+02 4.1840e+02 ; HA - CA
- CB
2047 2046 2061 1 1.0950e+02 4.1840e+02 ; HA - CA
- C
2048 2046 2061 1 1.1110e+02 5.2718e+02 ; CB - CA
- C
2048 2051 2052 1 1.0950e+02 4.1840e+02 ; CB - CG
- HG
2048 2051 2053 1 1.0950e+02 3.3472e+02 ; CB - CG
- CD1
2048 2051 2057 1 1.0950e+02 3.3472e+02 ; CB - CG
- CD2
2049 2048 2050 1 1.0950e+02 2.9288e+02 ; HB2 - CB
- HB3
2049 2048 2051 1 1.0950e+02 4.1840e+02 ; HB2 - CB
- CG
2050 2048 2051 1 1.0950e+02 4.1840e+02 ; HB3 - CB
- CG
2051 2053 2054 1 1.0950e+02 4.1840e+02 ; CG - CD1
- HD11
2051 2053 2055 1 1.0950e+02 4.1840e+02 ; CG - CD1
- HD12
2051 2053 2056 1 1.0950e+02 4.1840e+02 ; CG - CD1
- HD13
2051 2057 2058 1 1.0950e+02 4.1840e+02 ; CG - CD2
- HD21
2051 2057 2059 1 1.0950e+02 4.1840e+02 ; CG - CD2
- HD22
2051 2057 2060 1 1.0950e+02 4.1840e+02 ; CG - CD2
- HD23
2052 2051 2053 1 1.0950e+02 4.1840e+02 ; HG - CG
- CD1
2052 2051 2057 1 1.0950e+02 4.1840e+02 ; HG - CG
- CD2
2053 2051 2057 1 1.0950e+02 3.3472e+02 ; CD1 - CG
- CD2
2054 2053 2055 1 1.0950e+02 2.9288e+02 ; HD11 - CD1
- HD12
2054 2053 2056 1 1.0950e+02 2.9288e+02 ; HD11 - CD1
- HD13
2055 2053 2056 1 1.0950e+02 2.9288e+02 ; HD12 - CD1
- HD13
2058 2057 2059 1 1.0950e+02 2.9288e+02 ; HD21 - CD2
- HD22
2058 2057 2060 1 1.0950e+02 2.9288e+02 ; HD21 - CD2
- HD23

2059 2057 2060 1 1.0950e+02 2.9288e+02 ; HD22 - CD2
- HD23
2061 2063 2064 1 1.2000e+02 4.1840e+02 ; C - N
- H
2061 2063 2065 1 1.2190e+02 4.1840e+02 ; C - N
- CA
2062 2061 2063 1 1.2290e+02 6.6944e+02 ; O - C
- N
2063 2065 2066 1 1.0950e+02 4.1840e+02 ; N - CA
- HA
2063 2065 2067 1 1.0970e+02 6.6944e+02 ; N - CA
- CB
2063 2065 2079 1 1.1010e+02 5.2718e+02 ; N - CA
- C
2064 2063 2065 1 1.1804e+02 4.1840e+02 ; H - N
- CA
2065 2067 2068 1 1.0950e+02 4.1840e+02 ; CA - CB
- HB2
2065 2067 2069 1 1.0950e+02 4.1840e+02 ; CA - CB
- HB3
2065 2067 2070 1 1.1310e+02 5.2718e+02 ; CA - CB
- CG
2065 2079 2080 1 1.2040e+02 6.6944e+02 ; CA - C
- O
2065 2079 2081 1 1.1660e+02 5.8576e+02 ; CA - C
- N
2066 2065 2067 1 1.0950e+02 4.1840e+02 ; HA - CA
- CB
2066 2065 2079 1 1.0950e+02 4.1840e+02 ; HA - CA
- C
2067 2065 2079 1 1.1110e+02 5.2718e+02 ; CB - CA
- C
2067 2070 2071 1 1.2000e+02 5.8576e+02 ; CB - CG
- ND1
2067 2070 2077 1 1.2000e+02 5.8576e+02 ; CB - CG
- CD2
2068 2067 2069 1 1.0950e+02 2.9288e+02 ; HB2 - CB
- HB3
2068 2067 2070 1 1.0950e+02 4.1840e+02 ; HB2 - CB
- CG
2069 2067 2070 1 1.0950e+02 4.1840e+02 ; HB3 - CB
- CG
2070 2071 2072 1 1.2000e+02 4.1840e+02 ; CG - ND1
- HD1
2070 2071 2073 1 1.2000e+02 5.8576e+02 ; CG - ND1
- CE1
2070 2077 2075 1 1.2000e+02 5.8576e+02 ; CG - CD2
- NE2
2070 2077 2078 1 1.2000e+02 4.1840e+02 ; CG - CD2
- HD2
2071 2070 2077 1 1.2000e+02 5.8576e+02 ; ND1 - CG
- CD2

2071   2073   2074   1   1.2000e+02   4.1840e+02 ;   ND1 - CE1
- HE1
2071   2073   2075   1   1.2000e+02   5.8576e+02 ;   ND1 - CE1
- NE2
2072   2071   2073   1   1.2000e+02   4.1840e+02 ;   HD1 - ND1
- CE1
2073   2075   2076   1   1.2000e+02   4.1840e+02 ;   CE1 - NE2
- HE2
2073   2075   2077   1   1.2000e+02   5.8576e+02 ;   CE1 - NE2
- CD2
2074   2073   2075   1   1.2000e+02   4.1840e+02 ;   HE1 - CE1
- NE2
2075   2077   2078   1   1.2000e+02   4.1840e+02 ;   NE2 - CD2
- HD2
2076   2075   2077   1   1.2000e+02   4.1840e+02 ;   HE2 - NE2
- CD2
2079   2081   2082   1   1.2000e+02   4.1840e+02 ;   C - N
- H
2079   2081   2083   1   1.2190e+02   4.1840e+02 ;   C - N
- CA
2080   2079   2081   1   1.2290e+02   6.6944e+02 ;   O - C
- N
2081   2083   2084   1   1.0950e+02   4.1840e+02 ;   N - CA
- HA
2081   2083   2085   1   1.0970e+02   6.6944e+02 ;   N - CA
- CB
2081   2083   2090   1   1.1010e+02   5.2718e+02 ;   N - CA
- C
2082   2081   2083   1   1.1804e+02   4.1840e+02 ;   H - N
- CA
2083   2085   2086   1   1.0950e+02   4.1840e+02 ;   CA - CB
- HB2
2083   2085   2087   1   1.0950e+02   4.1840e+02 ;   CA - CB
- HB3
2083   2085   2088   1   1.0950e+02   4.1840e+02 ;   CA - CB
- OG
2083   2090   2091   1   1.2040e+02   6.6944e+02 ;   CA - C
- O
2083   2090   2092   1   1.1660e+02   5.8576e+02 ;   CA - C
- N
2084   2083   2085   1   1.0950e+02   4.1840e+02 ;   HA - CA
- CB
2084   2083   2090   1   1.0950e+02   4.1840e+02 ;   HA - CA
- C
2085   2083   2090   1   1.1110e+02   5.2718e+02 ;   CB - CA
- C
2085   2088   2089   1   1.0850e+02   4.6024e+02 ;   CB - OG
- HG
2086   2085   2087   1   1.0950e+02   2.9288e+02 ;   HB2 - CB
- HB3
2086   2085   2088   1   1.0950e+02   4.1840e+02 ;   HB2 - CB
- OG

```
  2087   2085   2088      1    1.0950e+02    4.1840e+02 ;    HB3 - CB
- OG
  2090   2092   2093      1    1.2000e+02    4.1840e+02 ;      C - N
- H
  2090   2092   2094      1    1.2190e+02    4.1840e+02 ;      C - N
- CA
  2091   2090   2092      1    1.2290e+02    6.6944e+02 ;      O - C
- N
  2092   2094   2095      1    1.0950e+02    4.1840e+02 ;      N - CA
- HA
  2092   2094   2096      1    1.0970e+02    6.6944e+02 ;      N - CA
- CB
  2092   2094   2109      1    1.1010e+02    5.2718e+02 ;      N - CA
- C
  2093   2092   2094      1    1.1804e+02    4.1840e+02 ;      H - N
- CA
  2094   2096   2097      1    1.0950e+02    4.1840e+02 ;     CA - CB
- HB2
  2094   2096   2098      1    1.0950e+02    4.1840e+02 ;     CA - CB
- HB3
  2094   2096   2099      1    1.0950e+02    3.3472e+02 ;     CA - CB
- CG
  2094   2109   2110      1    1.2040e+02    6.6944e+02 ;     CA - C
- O
  2094   2109   2111      1    1.1660e+02    5.8576e+02 ;     CA - C
- N
  2095   2094   2096      1    1.0950e+02    4.1840e+02 ;     HA - CA
- CB
  2095   2094   2109      1    1.0950e+02    4.1840e+02 ;     HA - CA
- C
  2096   2094   2109      1    1.1110e+02    5.2718e+02 ;     CB - CA
- C
  2096   2099   2100      1    1.0950e+02    4.1840e+02 ;     CB - CG
- HG
  2096   2099   2101      1    1.0950e+02    3.3472e+02 ;     CB - CG
- CD1
  2096   2099   2105      1    1.0950e+02    3.3472e+02 ;     CB - CG
- CD2
  2097   2096   2098      1    1.0950e+02    2.9288e+02 ;    HB2 - CB
- HB3
  2097   2096   2099      1    1.0950e+02    4.1840e+02 ;    HB2 - CB
- CG
  2098   2096   2099      1    1.0950e+02    4.1840e+02 ;    HB3 - CB
- CG
  2099   2101   2102      1    1.0950e+02    4.1840e+02 ;     CG - CD1
- HD11
  2099   2101   2103      1    1.0950e+02    4.1840e+02 ;     CG - CD1
- HD12
  2099   2101   2104      1    1.0950e+02    4.1840e+02 ;     CG - CD1
- HD13
  2099   2105   2106      1    1.0950e+02    4.1840e+02 ;     CG - CD2
- HD21
```

2099 2105 2107 1 1.0950e+02 4.1840e+02 ; CG - CD2
- HD22
2099 2105 2108 1 1.0950e+02 4.1840e+02 ; CG - CD2
- HD23
2100 2099 2101 1 1.0950e+02 4.1840e+02 ; HG - CG
- CD1
2100 2099 2105 1 1.0950e+02 4.1840e+02 ; HG - CG
- CD2
2101 2099 2105 1 1.0950e+02 3.3472e+02 ; CD1 - CG
- CD2
2102 2101 2103 1 1.0950e+02 2.9288e+02 ; HD11 - CD1
- HD12
2102 2101 2104 1 1.0950e+02 2.9288e+02 ; HD11 - CD1
- HD13
2103 2101 2104 1 1.0950e+02 2.9288e+02 ; HD12 - CD1
- HD13
2106 2105 2107 1 1.0950e+02 2.9288e+02 ; HD21 - CD2
- HD22
2106 2105 2108 1 1.0950e+02 2.9288e+02 ; HD21 - CD2
- HD23
2107 2105 2108 1 1.0950e+02 2.9288e+02 ; HD22 - CD2
- HD23
2109 2111 2112 1 1.2000e+02 4.1840e+02 ; C - N
- H
2109 2111 2113 1 1.2190e+02 4.1840e+02 ; C - N
- CA
2110 2109 2111 1 1.2290e+02 6.6944e+02 ; O - C
- N
2111 2113 2114 1 1.0950e+02 4.1840e+02 ; N - CA
- HA
2111 2113 2115 1 1.0970e+02 6.6944e+02 ; N - CA
- CB
2111 2113 2123 1 1.1010e+02 5.2718e+02 ; N - CA
- C
2112 2111 2113 1 1.1804e+02 4.1840e+02 ; H - N
- CA
2113 2115 2116 1 1.0950e+02 4.1840e+02 ; CA - CB
- HB2
2113 2115 2117 1 1.0950e+02 4.1840e+02 ; CA - CB
- HB3
2113 2115 2118 1 1.1110e+02 5.2718e+02 ; CA - CB
- CG
2113 2123 2124 1 1.2040e+02 6.6944e+02 ; CA - C
- O
2113 2123 2125 1 1.1660e+02 5.8576e+02 ; CA - C
- N
2114 2113 2115 1 1.0950e+02 4.1840e+02 ; HA - CA
- CB
2114 2113 2123 1 1.0950e+02 4.1840e+02 ; HA - CA
- C
2115 2113 2123 1 1.1110e+02 5.2718e+02 ; CB - CA
- C

2115 2118 2119 1 1.2040e+02 6.6944e+02 ; CB - CG
- OD1
2115 2118 2120 1 1.1660e+02 5.8576e+02 ; CB - CG
- ND2
2116 2115 2117 1 1.0950e+02 2.9288e+02 ; HB2 - CB
- HB3
2116 2115 2118 1 1.0950e+02 4.1840e+02 ; HB2 - CB
- CG
2117 2115 2118 1 1.0950e+02 4.1840e+02 ; HB3 - CB
- CG
2118 2120 2121 1 1.2000e+02 4.1840e+02 ; CG - ND2
- HD21
2118 2120 2122 1 1.2000e+02 4.1840e+02 ; CG - ND2
- HD22
2119 2118 2120 1 1.2290e+02 6.6944e+02 ; OD1 - CG
- ND2
2121 2120 2122 1 1.2000e+02 2.9288e+02 ; HD21 - ND2
- HD22
2123 2125 2126 1 1.2000e+02 4.1840e+02 ; C - N
- H
2123 2125 2127 1 1.2190e+02 4.1840e+02 ; C - N
- CA
2124 2123 2125 1 1.2290e+02 6.6944e+02 ; O - C
- N
2125 2127 2128 1 1.0950e+02 4.1840e+02 ; N - CA
- HA
2125 2127 2129 1 1.0970e+02 6.6944e+02 ; N - CA
- CB
2125 2127 2142 1 1.1010e+02 5.2718e+02 ; N - CA
- C
2126 2125 2127 1 1.1804e+02 4.1840e+02 ; H - N
- CA
2127 2129 2130 1 1.0950e+02 4.1840e+02 ; CA - CB
- HB
2127 2129 2131 1 1.0950e+02 3.3472e+02 ; CA - CB
- CG2
2127 2129 2135 1 1.0950e+02 3.3472e+02 ; CA - CB
- CG1
2127 2142 2143 1 1.2040e+02 6.6944e+02 ; CA - C
- O
2127 2142 2144 1 1.1660e+02 5.8576e+02 ; CA - C
- N
2128 2127 2129 1 1.0950e+02 4.1840e+02 ; HA - CA
- CB
2128 2127 2142 1 1.0950e+02 4.1840e+02 ; HA - CA
- C
2129 2127 2142 1 1.1110e+02 5.2718e+02 ; CB - CA
- C
2129 2131 2132 1 1.0950e+02 4.1840e+02 ; CB - CG2
- HG21
2129 2131 2133 1 1.0950e+02 4.1840e+02 ; CB - CG2
- HG22

```
  2129   2131   2134       1    1.0950e+02    4.1840e+02 ;     CB - CG2
- HG23
  2129   2135   2136       1    1.0950e+02    4.1840e+02 ;     CB - CG1
- HG12
  2129   2135   2137       1    1.0950e+02    4.1840e+02 ;     CB - CG1
- HG13
  2129   2135   2138       1    1.0950e+02    3.3472e+02 ;     CB - CG1
- CD1
  2130   2129   2131       1    1.0950e+02    4.1840e+02 ;     HB - CB
- CG2
  2130   2129   2135       1    1.0950e+02    4.1840e+02 ;     HB - CB
- CG1
  2131   2129   2135       1    1.0950e+02    3.3472e+02 ;    CG2 - CB
- CG1
  2132   2131   2133       1    1.0950e+02    2.9288e+02 ;   HG21 - CG2
- HG22
  2132   2131   2134       1    1.0950e+02    2.9288e+02 ;   HG21 - CG2
- HG23
  2133   2131   2134       1    1.0950e+02    2.9288e+02 ;   HG22 - CG2
- HG23
  2135   2138   2139       1    1.0950e+02    4.1840e+02 ;    CG1 - CD1
- HD11
  2135   2138   2140       1    1.0950e+02    4.1840e+02 ;    CG1 - CD1
- HD12
  2135   2138   2141       1    1.0950e+02    4.1840e+02 ;    CG1 - CD1
- HD13
  2136   2135   2137       1    1.0950e+02    2.9288e+02 ;   HG12 - CG1
- HG13
  2136   2135   2138       1    1.0950e+02    4.1840e+02 ;   HG12 - CG1
- CD1
  2137   2135   2138       1    1.0950e+02    4.1840e+02 ;   HG13 - CG1
- CD1
  2139   2138   2140       1    1.0950e+02    2.9288e+02 ;   HD11 - CD1
- HD12
  2139   2138   2141       1    1.0950e+02    2.9288e+02 ;   HD11 - CD1
- HD13
  2140   2138   2141       1    1.0950e+02    2.9288e+02 ;   HD12 - CD1
- HD13
  2142   2144   2145       1    1.2000e+02    4.1840e+02 ;      C - N
- H
  2142   2144   2146       1    1.2190e+02    4.1840e+02 ;      C - N
- CA
  2143   2142   2144       1    1.2290e+02    6.6944e+02 ;      O - C
- N
  2144   2146   2147       1    1.0950e+02    4.1840e+02 ;      N - CA
- HA
  2144   2146   2148       1    1.0970e+02    6.6944e+02 ;      N - CA
- CB
  2144   2146   2158       1    1.1010e+02    5.2718e+02 ;      N - CA
- C
  2145   2144   2146       1    1.1804e+02    4.1840e+02 ;      H - N
- CA
```

2146 2148 2149 1 1.0950e+02 4.1840e+02 ; CA - CB
- HB
2146 2148 2150 1 1.0950e+02 3.3472e+02 ; CA - CB
- CG1
2146 2148 2154 1 1.0950e+02 3.3472e+02 ; CA - CB
- CG2
2146 2158 2159 1 1.2040e+02 6.6944e+02 ; CA - C
- O
2146 2158 2160 1 1.1660e+02 5.8576e+02 ; CA - C
- N
2147 2146 2148 1 1.0950e+02 4.1840e+02 ; HA - CA
- CB
2147 2146 2158 1 1.0950e+02 4.1840e+02 ; HA - CA
- C
2148 2146 2158 1 1.1110e+02 5.2718e+02 ; CB - CA
- C
2148 2150 2151 1 1.0950e+02 4.1840e+02 ; CB - CG1
- HG11
2148 2150 2152 1 1.0950e+02 4.1840e+02 ; CB - CG1
- HG12
2148 2150 2153 1 1.0950e+02 4.1840e+02 ; CB - CG1
- HG13
2148 2154 2155 1 1.0950e+02 4.1840e+02 ; CB - CG2
- HG21
2148 2154 2156 1 1.0950e+02 4.1840e+02 ; CB - CG2
- HG22
2148 2154 2157 1 1.0950e+02 4.1840e+02 ; CB - CG2
- HG23
2149 2148 2150 1 1.0950e+02 4.1840e+02 ; HB - CB
- CG1
2149 2148 2154 1 1.0950e+02 4.1840e+02 ; HB - CB
- CG2
2150 2148 2154 1 1.0950e+02 3.3472e+02 ; CG1 - CB
- CG2
2151 2150 2152 1 1.0950e+02 2.9288e+02 ; HG11 - CG1
- HG12
2151 2150 2153 1 1.0950e+02 2.9288e+02 ; HG11 - CG1
- HG13
2152 2150 2153 1 1.0950e+02 2.9288e+02 ; HG12 - CG1
- HG13
2155 2154 2156 1 1.0950e+02 2.9288e+02 ; HG21 - CG2
- HG22
2155 2154 2157 1 1.0950e+02 2.9288e+02 ; HG21 - CG2
- HG23
2156 2154 2157 1 1.0950e+02 2.9288e+02 ; HG22 - CG2
- HG23
2158 2160 2161 1 1.2000e+02 4.1840e+02 ; C - N
- H
2158 2160 2162 1 1.2190e+02 4.1840e+02 ; C - N
- CA
2159 2158 2160 1 1.2290e+02 6.6944e+02 ; O - C
- N

2160 2162 2163 1 1.0950e+02 4.1840e+02 ; N - CA
- HA
2160 2162 2164 1 1.0970e+02 6.6944e+02 ; N - CA
- CB
2160 2162 2175 1 1.1010e+02 5.2718e+02 ; N - CA
- C
2161 2160 2162 1 1.1804e+02 4.1840e+02 ; H - N
- CA
2162 2164 2165 1 1.0950e+02 4.1840e+02 ; CA - CB
- HB2
2162 2164 2166 1 1.0950e+02 4.1840e+02 ; CA - CB
- HB3
2162 2164 2167 1 1.1310e+02 5.2718e+02 ; CA - CB
- CG
2162 2175 2176 1 1.2040e+02 6.6944e+02 ; CA - C
- O
2162 2175 2177 1 1.1660e+02 5.8576e+02 ; CA - C
- N
2163 2162 2164 1 1.0950e+02 4.1840e+02 ; HA - CA
- CB
2163 2162 2175 1 1.0950e+02 4.1840e+02 ; HA - CA
- C
2164 2162 2175 1 1.1110e+02 5.2718e+02 ; CB - CA
- C
2164 2167 2168 1 1.2000e+02 5.8576e+02 ; CB - CG
- ND1
2164 2167 2173 1 1.2000e+02 5.8576e+02 ; CB - CG
- CD2
2165 2164 2166 1 1.0950e+02 2.9288e+02 ; HB2 - CB
- HB3
2165 2164 2167 1 1.0950e+02 4.1840e+02 ; HB2 - CB
- CG
2166 2164 2167 1 1.0950e+02 4.1840e+02 ; HB3 - CB
- CG
2167 2168 2169 1 1.1700e+02 5.8576e+02 ; CG - ND1
- CE1
2167 2173 2171 1 1.2000e+02 5.8576e+02 ; CG - CD2
- NE2
2167 2173 2174 1 1.2000e+02 4.1840e+02 ; CG - CD2
- HD2
2168 2167 2173 1 1.2000e+02 5.8576e+02 ; ND1 - CG
- CD2
2168 2169 2170 1 1.2000e+02 4.1840e+02 ; ND1 - CE1
- HE1
2168 2169 2171 1 1.2000e+02 5.8576e+02 ; ND1 - CE1
- NE2
2169 2171 2172 1 1.2000e+02 4.1840e+02 ; CE1 - NE2
- HE2
2169 2171 2173 1 1.2000e+02 5.8576e+02 ; CE1 - NE2
- CD2
2170 2169 2171 1 1.2000e+02 4.1840e+02 ; HE1 - CE1
- NE2

2171 2173 2174 1 1.2000e+02 4.1840e+02 ; NE2 - CD2
- HD2
2172 2171 2173 1 1.2000e+02 4.1840e+02 ; HE2 - NE2
- CD2
2175 2177 2178 1 1.2000e+02 4.1840e+02 ; C - N
- H
2175 2177 2179 1 1.2190e+02 4.1840e+02 ; C - N
- CA
2176 2175 2177 1 1.2290e+02 6.6944e+02 ; O - C
- N
2177 2179 2180 1 1.0950e+02 4.1840e+02 ; N - CA
- HA
2177 2179 2181 1 1.0970e+02 6.6944e+02 ; N - CA
- CB
2177 2179 2199 1 1.1010e+02 5.2718e+02 ; N - CA
- C
2178 2177 2179 1 1.1804e+02 4.1840e+02 ; H - N
- CA
2179 2181 2182 1 1.0950e+02 4.1840e+02 ; CA - CB
- HB2
2179 2181 2183 1 1.0950e+02 4.1840e+02 ; CA - CB
- HB3
2179 2181 2184 1 1.0950e+02 3.3472e+02 ; CA - CB
- CG
2179 2199 2200 1 1.2040e+02 6.6944e+02 ; CA - C
- O
2179 2199 2201 1 1.1660e+02 5.8576e+02 ; CA - C
- N
2180 2179 2181 1 1.0950e+02 4.1840e+02 ; HA - CA
- CB
2180 2179 2199 1 1.0950e+02 4.1840e+02 ; HA - CA
- C
2181 2179 2199 1 1.1110e+02 5.2718e+02 ; CB - CA
- C
2181 2184 2185 1 1.0950e+02 4.1840e+02 ; CB - CG
- HG2
2181 2184 2186 1 1.0950e+02 4.1840e+02 ; CB - CG
- HG3
2181 2184 2187 1 1.0950e+02 3.3472e+02 ; CB - CG
- CD
2182 2181 2183 1 1.0950e+02 2.9288e+02 ; HB2 - CB
- HB3
2182 2181 2184 1 1.0950e+02 4.1840e+02 ; HB2 - CB
- CG
2183 2181 2184 1 1.0950e+02 4.1840e+02 ; HB3 - CB
- CG
2184 2187 2188 1 1.0950e+02 4.1840e+02 ; CG - CD
- HD2
2184 2187 2189 1 1.0950e+02 4.1840e+02 ; CG - CD
- HD3
2184 2187 2190 1 1.1120e+02 6.6944e+02 ; CG - CD
- NE

2185 2184 2186 1 1.0950e+02 2.9288e+02 ; HG2 - CG
- HG3
2185 2184 2187 1 1.0950e+02 4.1840e+02 ; HG2 - CG
- CD
2186 2184 2187 1 1.0950e+02 4.1840e+02 ; HG3 - CG
- CD
2187 2190 2191 1 1.1840e+02 4.1840e+02 ; CD - NE
- HE
2187 2190 2192 1 1.2320e+02 4.1840e+02 ; CD - NE
- CZ
2188 2187 2189 1 1.0950e+02 2.9288e+02 ; HD2 - CD
- HD3
2188 2187 2190 1 1.0950e+02 4.1840e+02 ; HD2 - CD
- NE
2189 2187 2190 1 1.0950e+02 4.1840e+02 ; HD3 - CD
- NE
2190 2192 2193 1 1.2000e+02 5.8576e+02 ; NE - CZ
- NH1
2190 2192 2196 1 1.2000e+02 5.8576e+02 ; NE - CZ
- NH2
2191 2190 2192 1 1.2000e+02 4.1840e+02 ; HE - NE
- CZ
2192 2193 2194 1 1.2000e+02 4.1840e+02 ; CZ - NH1
- HH11
2192 2193 2195 1 1.2000e+02 4.1840e+02 ; CZ - NH1
- HH12
2192 2196 2197 1 1.2000e+02 4.1840e+02 ; CZ - NH2
- HH21
2192 2196 2198 1 1.2000e+02 4.1840e+02 ; CZ - NH2
- HH22
2193 2192 2196 1 1.2000e+02 5.8576e+02 ; NH1 - CZ
- NH2
2194 2193 2195 1 1.2000e+02 2.9288e+02 ; HH11 - NH1
- HH12
2197 2196 2198 1 1.2000e+02 2.9288e+02 ; HH21 - NH2
- HH22
2199 2201 2202 1 1.2000e+02 4.1840e+02 ; C - N
- H
2199 2201 2203 1 1.2190e+02 4.1840e+02 ; C - N
- CA
2200 2199 2201 1 1.2290e+02 6.6944e+02 ; O - C
- N
2201 2203 2204 1 1.0950e+02 4.1840e+02 ; N - CA
- HA
2201 2203 2205 1 1.0970e+02 6.6944e+02 ; N - CA
- CB
2201 2203 2211 1 1.1010e+02 5.2718e+02 ; N - CA
- C
2202 2201 2203 1 1.1804e+02 4.1840e+02 ; H - N
- CA
2203 2205 2206 1 1.0950e+02 4.1840e+02 ; CA - CB
- HB2

2203 2205 2207 1 1.0950e+02 4.1840e+02 ; CA - CB
- HB3
2203 2205 2208 1 1.1110e+02 5.2718e+02 ; CA - CB
- CG
2203 2211 2212 1 1.2040e+02 6.6944e+02 ; CA - C
- O
2203 2211 2213 1 1.1660e+02 5.8576e+02 ; CA - C
- N
2204 2203 2205 1 1.0950e+02 4.1840e+02 ; HA - CA
- CB
2204 2203 2211 1 1.0950e+02 4.1840e+02 ; HA - CA
- C
2205 2203 2211 1 1.1110e+02 5.2718e+02 ; CB - CA
- C
2205 2208 2209 1 1.1700e+02 5.8576e+02 ; CB - CG
- OD1
2205 2208 2210 1 1.1700e+02 5.8576e+02 ; CB - CG
- OD2
2206 2205 2207 1 1.0950e+02 2.9288e+02 ; HB2 - CB
- HB3
2206 2205 2208 1 1.0950e+02 4.1840e+02 ; HB2 - CB
- CG
2207 2205 2208 1 1.0950e+02 4.1840e+02 ; HB3 - CB
- CG
2209 2208 2210 1 1.2600e+02 6.6944e+02 ; OD1 - CG
- OD2
2211 2213 2214 1 1.2000e+02 4.1840e+02 ; C - N
- H
2211 2213 2215 1 1.2190e+02 4.1840e+02 ; C - N
- CA
2212 2211 2213 1 1.2290e+02 6.6944e+02 ; O - C
- N
2213 2215 2216 1 1.0950e+02 4.1840e+02 ; N - CA
- HA
2213 2215 2217 1 1.0970e+02 6.6944e+02 ; N - CA
- CB
2213 2215 2230 1 1.1010e+02 5.2718e+02 ; N - CA
- C
2214 2213 2215 1 1.1804e+02 4.1840e+02 ; H - N
- CA
2215 2217 2218 1 1.0950e+02 4.1840e+02 ; CA - CB
- HB2
2215 2217 2219 1 1.0950e+02 4.1840e+02 ; CA - CB
- HB3
2215 2217 2220 1 1.0950e+02 3.3472e+02 ; CA - CB
- CG
2215 2230 2231 1 1.2040e+02 6.6944e+02 ; CA - C
- O
2215 2230 2232 1 1.1660e+02 5.8576e+02 ; CA - C
- N
2216 2215 2217 1 1.0950e+02 4.1840e+02 ; HA - CA
- CB

```
  2216   2215   2230       1    1.0950e+02    4.1840e+02 ;     HA - CA
- C
  2217   2215   2230       1    1.1110e+02    5.2718e+02 ;     CB - CA
- C
  2217   2220   2221       1    1.0950e+02    4.1840e+02 ;     CB - CG
- HG
  2217   2220   2222       1    1.0950e+02    3.3472e+02 ;     CB - CG
- CD1
  2217   2220   2226       1    1.0950e+02    3.3472e+02 ;     CB - CG
- CD2
  2218   2217   2219       1    1.0950e+02    2.9288e+02 ;    HB2 - CB
- HB3
  2218   2217   2220       1    1.0950e+02    4.1840e+02 ;    HB2 - CB
- CG
  2219   2217   2220       1    1.0950e+02    4.1840e+02 ;    HB3 - CB
- CG
  2220   2222   2223       1    1.0950e+02    4.1840e+02 ;     CG - CD1
- HD11
  2220   2222   2224       1    1.0950e+02    4.1840e+02 ;     CG - CD1
- HD12
  2220   2222   2225       1    1.0950e+02    4.1840e+02 ;     CG - CD1
- HD13
  2220   2226   2227       1    1.0950e+02    4.1840e+02 ;     CG - CD2
- HD21
  2220   2226   2228       1    1.0950e+02    4.1840e+02 ;     CG - CD2
- HD22
  2220   2226   2229       1    1.0950e+02    4.1840e+02 ;     CG - CD2
- HD23
  2221   2220   2222       1    1.0950e+02    4.1840e+02 ;     HG - CG
- CD1
  2221   2220   2226       1    1.0950e+02    4.1840e+02 ;     HG - CG
- CD2
  2222   2220   2226       1    1.0950e+02    3.3472e+02 ;    CD1 - CG
- CD2
  2223   2222   2224       1    1.0950e+02    2.9288e+02 ;   HD11 - CD1
- HD12
  2223   2222   2225       1    1.0950e+02    2.9288e+02 ;   HD11 - CD1
- HD13
  2224   2222   2225       1    1.0950e+02    2.9288e+02 ;   HD12 - CD1
- HD13
  2227   2226   2228       1    1.0950e+02    2.9288e+02 ;   HD21 - CD2
- HD22
  2227   2226   2229       1    1.0950e+02    2.9288e+02 ;   HD21 - CD2
- HD23
  2228   2226   2229       1    1.0950e+02    2.9288e+02 ;   HD22 - CD2
- HD23
  2230   2232   2233       1    1.2000e+02    4.1840e+02 ;      C - N
- H
  2230   2232   2234       1    1.2190e+02    4.1840e+02 ;      C - N
- CA
  2231   2230   2232       1    1.2290e+02    6.6944e+02 ;      O - C
- N
```

```
  2232   2234   2235      1    1.0950e+02    4.1840e+02 ;      N - CA
- HA
  2232   2234   2236      1    1.0970e+02    6.6944e+02 ;      N - CA
- CB
  2232   2234   2252      1    1.1010e+02    5.2718e+02 ;      N - CA
- C
  2233   2232   2234      1    1.1804e+02    4.1840e+02 ;      H - N
- CA
  2234   2236   2237      1    1.0950e+02    4.1840e+02 ;     CA - CB
- HB2
  2234   2236   2238      1    1.0950e+02    4.1840e+02 ;     CA - CB
- HB3
  2234   2236   2239      1    1.0950e+02    3.3472e+02 ;     CA - CB
- CG
  2234   2252   2253      1    1.2040e+02    6.6944e+02 ;     CA - C
- O
  2234   2252   2254      1    1.1660e+02    5.8576e+02 ;     CA - C
- N
  2235   2234   2236      1    1.0950e+02    4.1840e+02 ;     HA - CA
- CB
  2235   2234   2252      1    1.0950e+02    4.1840e+02 ;     HA - CA
- C
  2236   2234   2252      1    1.1110e+02    5.2718e+02 ;     CB - CA
- C
  2236   2239   2240      1    1.0950e+02    4.1840e+02 ;     CB - CG
- HG2
  2236   2239   2241      1    1.0950e+02    4.1840e+02 ;     CB - CG
- HG3
  2236   2239   2242      1    1.0950e+02    3.3472e+02 ;     CB - CG
- CD
  2237   2236   2238      1    1.0950e+02    2.9288e+02 ;    HB2 - CB
- HB3
  2237   2236   2239      1    1.0950e+02    4.1840e+02 ;    HB2 - CB
- CG
  2238   2236   2239      1    1.0950e+02    4.1840e+02 ;    HB3 - CB
- CG
  2239   2242   2243      1    1.0950e+02    4.1840e+02 ;     CG - CD
- HD2
  2239   2242   2244      1    1.0950e+02    4.1840e+02 ;     CG - CD
- HD3
  2239   2242   2245      1    1.0950e+02    3.3472e+02 ;     CG - CD
- CE
  2240   2239   2241      1    1.0950e+02    2.9288e+02 ;    HG2 - CG
- HG3
  2240   2239   2242      1    1.0950e+02    4.1840e+02 ;    HG2 - CG
- CD
  2241   2239   2242      1    1.0950e+02    4.1840e+02 ;    HG3 - CG
- CD
  2242   2245   2246      1    1.0950e+02    4.1840e+02 ;     CD - CE
- HE2
  2242   2245   2247      1    1.0950e+02    4.1840e+02 ;     CD - CE
- HE3
```

2242 2245 2248 1 1.1120e+02 6.6944e+02 ; CD - CE
- NZ
2243 2242 2244 1 1.0950e+02 2.9288e+02 ; HD2 - CD
- HD3
2243 2242 2245 1 1.0950e+02 4.1840e+02 ; HD2 - CD
- CE
2244 2242 2245 1 1.0950e+02 4.1840e+02 ; HD3 - CD
- CE
2245 2248 2249 1 1.0950e+02 4.1840e+02 ; CE - NZ
- HZ1
2245 2248 2250 1 1.0950e+02 4.1840e+02 ; CE - NZ
- HZ2
2245 2248 2251 1 1.0950e+02 4.1840e+02 ; CE - NZ
- HZ3
2246 2245 2247 1 1.0950e+02 2.9288e+02 ; HE2 - CE
- HE3
2246 2245 2248 1 1.0950e+02 4.1840e+02 ; HE2 - CE
- NZ
2247 2245 2248 1 1.0950e+02 4.1840e+02 ; HE3 - CE
- NZ
2249 2248 2250 1 1.0950e+02 2.9288e+02 ; HZ1 - NZ
- HZ2
2249 2248 2251 1 1.0950e+02 2.9288e+02 ; HZ1 - NZ
- HZ3
2250 2248 2251 1 1.0950e+02 2.9288e+02 ; HZ2 - NZ
- HZ3
2252 2254 2255 1 1.2190e+02 4.1840e+02 ; C - N
- CD
2252 2254 2264 1 1.2190e+02 4.1840e+02 ; C - N
- CA
2253 2252 2254 1 1.2290e+02 6.6944e+02 ; O - C
- N
2254 2255 2256 1 1.0950e+02 4.1840e+02 ; N - CD
- HD2
2254 2255 2257 1 1.0950e+02 4.1840e+02 ; N - CD
- HD3
2254 2255 2258 1 1.0970e+02 6.6944e+02 ; N - CD
- CG
2254 2264 2261 1 1.0970e+02 6.6944e+02 ; N - CA
- CB
2254 2264 2265 1 1.0950e+02 4.1840e+02 ; N - CA
- HA
2254 2264 2266 1 1.1010e+02 5.2718e+02 ; N - CA
- C
2255 2254 2264 1 1.1800e+02 4.1840e+02 ; CD - N
- CA
2255 2258 2259 1 1.0950e+02 4.1840e+02 ; CD - CG
- HG2
2255 2258 2260 1 1.0950e+02 4.1840e+02 ; CD - CG
- HG3
2255 2258 2261 1 1.0950e+02 3.3472e+02 ; CD - CG
- CB

2256 2255 2257 1 1.0950e+02 2.9288e+02 ; HD2 - CD
- HD3
2256 2255 2258 1 1.0950e+02 4.1840e+02 ; HD2 - CD
- CG
2257 2255 2258 1 1.0950e+02 4.1840e+02 ; HD3 - CD
- CG
2258 2261 2262 1 1.0950e+02 4.1840e+02 ; CG - CB
- HB2
2258 2261 2263 1 1.0950e+02 4.1840e+02 ; CG - CB
- HB3
2258 2261 2264 1 1.0950e+02 3.3472e+02 ; CG - CB
- CA
2259 2258 2260 1 1.0950e+02 2.9288e+02 ; HG2 - CG
- HG3
2259 2258 2261 1 1.0950e+02 4.1840e+02 ; HG2 - CG
- CB
2260 2258 2261 1 1.0950e+02 4.1840e+02 ; HG3 - CG
- CB
2261 2264 2265 1 1.0950e+02 4.1840e+02 ; CB - CA
- HA
2261 2264 2266 1 1.1110e+02 5.2718e+02 ; CB - CA
- C
2262 2261 2263 1 1.0950e+02 2.9288e+02 ; HB2 - CB
- HB3
2262 2261 2264 1 1.0950e+02 4.1840e+02 ; HB2 - CB
- CA
2263 2261 2264 1 1.0950e+02 4.1840e+02 ; HB3 - CB
- CA
2264 2266 2267 1 1.2040e+02 6.6944e+02 ; CA - C
- O
2264 2266 2268 1 1.1660e+02 5.8576e+02 ; CA - C
- N
2265 2264 2266 1 1.0950e+02 4.1840e+02 ; HA - CA
- C
2266 2268 2269 1 1.2000e+02 4.1840e+02 ; C - N
- H
2266 2268 2270 1 1.2190e+02 4.1840e+02 ; C - N
- CA
2267 2266 2268 1 1.2290e+02 6.6944e+02 ; O - C
- N
2268 2270 2271 1 1.0950e+02 4.1840e+02 ; N - CA
- HA
2268 2270 2272 1 1.0970e+02 6.6944e+02 ; N - CA
- CB
2268 2270 2281 1 1.1010e+02 5.2718e+02 ; N - CA
- C
2269 2268 2270 1 1.1804e+02 4.1840e+02 ; H - N
- CA
2270 2272 2273 1 1.0950e+02 4.1840e+02 ; CA - CB
- HB2
2270 2272 2274 1 1.0950e+02 4.1840e+02 ; CA - CB
- HB3

2270 2272 2275 1 1.0950e+02 3.3472e+02 ; CA - CB
- CG
2270 2281 2282 1 1.2040e+02 6.6944e+02 ; CA - C
- O
2270 2281 2283 1 1.1660e+02 5.8576e+02 ; CA - C
- N
2271 2270 2272 1 1.0950e+02 4.1840e+02 ; HA - CA
- CB
2271 2270 2281 1 1.0950e+02 4.1840e+02 ; HA - CA
- C
2272 2270 2281 1 1.1110e+02 5.2718e+02 ; CB - CA
- C
2272 2275 2276 1 1.0950e+02 4.1840e+02 ; CB - CG
- HG2
2272 2275 2277 1 1.0950e+02 4.1840e+02 ; CB - CG
- HG3
2272 2275 2278 1 1.1110e+02 5.2718e+02 ; CB - CG
- CD
2273 2272 2274 1 1.0950e+02 2.9288e+02 ; HB2 - CB
- HB3
2273 2272 2275 1 1.0950e+02 4.1840e+02 ; HB2 - CB
- CG
2274 2272 2275 1 1.0950e+02 4.1840e+02 ; HB3 - CB
- CG
2275 2278 2279 1 1.1700e+02 5.8576e+02 ; CG - CD
- OE1
2275 2278 2280 1 1.1700e+02 5.8576e+02 ; CG - CD
- OE2
2276 2275 2277 1 1.0950e+02 2.9288e+02 ; HG2 - CG
- HG3
2276 2275 2278 1 1.0950e+02 4.1840e+02 ; HG2 - CG
- CD
2277 2275 2278 1 1.0950e+02 4.1840e+02 ; HG3 - CG
- CD
2279 2278 2280 1 1.2600e+02 6.6944e+02 ; OE1 - CD
- OE2
2281 2283 2284 1 1.2000e+02 4.1840e+02 ; C - N
- H
2281 2283 2285 1 1.2190e+02 4.1840e+02 ; C - N
- CA
2282 2281 2283 1 1.2290e+02 6.6944e+02 ; O - C
- N
2283 2285 2286 1 1.0950e+02 4.1840e+02 ; N - CA
- HA
2283 2285 2287 1 1.0970e+02 6.6944e+02 ; N - CA
- CB
2283 2285 2295 1 1.1010e+02 5.2718e+02 ; N - CA
- C
2284 2283 2285 1 1.1804e+02 4.1840e+02 ; H - N
- CA
2285 2287 2288 1 1.0950e+02 4.1840e+02 ; CA - CB
- HB2

```
  2285   2287   2289      1    1.0950e+02    4.1840e+02 ;     CA - CB
- HB3
  2285   2287   2290      1    1.1110e+02    5.2718e+02 ;     CA - CB
- CG
  2285   2295   2296      1    1.2040e+02    6.6944e+02 ;     CA - C
- O
  2285   2295   2297      1    1.1660e+02    5.8576e+02 ;     CA - C
- N
  2286   2285   2287      1    1.0950e+02    4.1840e+02 ;     HA - CA
- CB
  2286   2285   2295      1    1.0950e+02    4.1840e+02 ;     HA - CA
- C
  2287   2285   2295      1    1.1110e+02    5.2718e+02 ;     CB - CA
- C
  2287   2290   2291      1    1.2040e+02    6.6944e+02 ;     CB - CG
- OD1
  2287   2290   2292      1    1.1660e+02    5.8576e+02 ;     CB - CG
- ND2
  2288   2287   2289      1    1.0950e+02    2.9288e+02 ;    HB2 - CB
- HB3
  2288   2287   2290      1    1.0950e+02    4.1840e+02 ;    HB2 - CB
- CG
  2289   2287   2290      1    1.0950e+02    4.1840e+02 ;    HB3 - CB
- CG
  2290   2292   2293      1    1.2000e+02    4.1840e+02 ;     CG - ND2
- HD21
  2290   2292   2294      1    1.2000e+02    4.1840e+02 ;     CG - ND2
- HD22
  2291   2290   2292      1    1.2290e+02    6.6944e+02 ;    OD1 - CG
- ND2
  2293   2292   2294      1    1.2000e+02    2.9288e+02 ;   HD21 - ND2
- HD22
  2295   2297   2298      1    1.2000e+02    4.1840e+02 ;      C - N
- H
  2295   2297   2299      1    1.2190e+02    4.1840e+02 ;      C - N
- CA
  2296   2295   2297      1    1.2290e+02    6.6944e+02 ;      O - C
- N
  2297   2299   2300      1    1.0950e+02    4.1840e+02 ;      N - CA
- HA
  2297   2299   2301      1    1.0970e+02    6.6944e+02 ;      N - CA
- CB
  2297   2299   2315      1    1.1010e+02    5.2718e+02 ;      N - CA
- C
  2298   2297   2299      1    1.1804e+02    4.1840e+02 ;      H - N
- CA
  2299   2301   2302      1    1.0950e+02    4.1840e+02 ;     CA - CB
- HB2
  2299   2301   2303      1    1.0950e+02    4.1840e+02 ;     CA - CB
- HB3
  2299   2301   2304      1    1.1400e+02    5.2718e+02 ;     CA - CB
- CG
```

```
  2299   2315   2316       1    1.2040e+02    6.6944e+02 ;     CA - C
- O
  2299   2315   2317       1    1.1660e+02    5.8576e+02 ;     CA - C
- N
  2300   2299   2301       1    1.0950e+02    4.1840e+02 ;     HA - CA
- CB
  2300   2299   2315       1    1.0950e+02    4.1840e+02 ;     HA - CA
- C
  2301   2299   2315       1    1.1110e+02    5.2718e+02 ;     CB - CA
- C
  2301   2304   2305       1    1.2000e+02    5.8576e+02 ;     CB - CG
- CD1
  2301   2304   2313       1    1.2000e+02    5.8576e+02 ;     CB - CG
- CD2
  2302   2301   2303       1    1.0950e+02    2.9288e+02 ;    HB2 - CB
- HB3
  2302   2301   2304       1    1.0950e+02    4.1840e+02 ;    HB2 - CB
- CG
  2303   2301   2304       1    1.0950e+02    4.1840e+02 ;    HB3 - CB
- CG
  2304   2305   2306       1    1.2000e+02    4.1840e+02 ;     CG - CD1
- HD1
  2304   2305   2307       1    1.2000e+02    5.2718e+02 ;     CG - CD1
- CE1
  2304   2313   2311       1    1.2000e+02    5.2718e+02 ;     CG - CD2
- CE2
  2304   2313   2314       1    1.2000e+02    4.1840e+02 ;     CG - CD2
- HD2
  2305   2304   2313       1    1.2000e+02    5.2718e+02 ;    CD1 - CG
- CD2
  2305   2307   2308       1    1.2000e+02    4.1840e+02 ;    CD1 - CE1
- HE1
  2305   2307   2309       1    1.2000e+02    5.2718e+02 ;    CD1 - CE1
- CZ
  2306   2305   2307       1    1.2000e+02    4.1840e+02 ;    HD1 - CD1
- CE1
  2307   2309   2310       1    1.2000e+02    4.1840e+02 ;    CE1 - CZ
- HZ
  2307   2309   2311       1    1.2000e+02    5.2718e+02 ;    CE1 - CZ
- CE2
  2308   2307   2309       1    1.2000e+02    4.1840e+02 ;    HE1 - CE1
- CZ
  2309   2311   2312       1    1.2000e+02    4.1840e+02 ;     CZ - CE2
- HE2
  2309   2311   2313       1    1.2000e+02    5.2718e+02 ;     CZ - CE2
- CD2
  2310   2309   2311       1    1.2000e+02    4.1840e+02 ;     HZ - CZ
- CE2
  2311   2313   2314       1    1.2000e+02    4.1840e+02 ;    CE2 - CD2
- HD2
  2312   2311   2313       1    1.2000e+02    4.1840e+02 ;    HE2 - CE2
- CD2
```

```
  2315   2317   2318      1    1.2000e+02    4.1840e+02 ;      C - N 
- H
  2315   2317   2319      1    1.2190e+02    4.1840e+02 ;      C - N 
- CA
  2316   2315   2317      1    1.2290e+02    6.6944e+02 ;      O - C 
- N
  2317   2319   2320      1    1.0950e+02    4.1840e+02 ;      N - CA
- HA
  2317   2319   2321      1    1.0970e+02    6.6944e+02 ;      N - CA
- CB
  2317   2319   2334      1    1.1010e+02    5.2718e+02 ;      N - CA
- C
  2318   2317   2319      1    1.1804e+02    4.1840e+02 ;      H - N 
- CA
  2319   2321   2322      1    1.0950e+02    4.1840e+02 ;     CA - CB
- HB2
  2319   2321   2323      1    1.0950e+02    4.1840e+02 ;     CA - CB
- HB3
  2319   2321   2324      1    1.0950e+02    3.3472e+02 ;     CA - CB
- CG
  2319   2334   2335      1    1.2040e+02    6.6944e+02 ;     CA - C 
- O
  2319   2334   2336      1    1.1660e+02    5.8576e+02 ;     CA - C 
- N
  2320   2319   2321      1    1.0950e+02    4.1840e+02 ;     HA - CA
- CB
  2320   2319   2334      1    1.0950e+02    4.1840e+02 ;     HA - CA
- C
  2321   2319   2334      1    1.1110e+02    5.2718e+02 ;     CB - CA
- C
  2321   2324   2325      1    1.0950e+02    4.1840e+02 ;     CB - CG
- HG
  2321   2324   2326      1    1.0950e+02    3.3472e+02 ;     CB - CG
- CD1
  2321   2324   2330      1    1.0950e+02    3.3472e+02 ;     CB - CG
- CD2
  2322   2321   2323      1    1.0950e+02    2.9288e+02 ;    HB2 - CB
- HB3
  2322   2321   2324      1    1.0950e+02    4.1840e+02 ;    HB2 - CB
- CG
  2323   2321   2324      1    1.0950e+02    4.1840e+02 ;    HB3 - CB
- CG
  2324   2326   2327      1    1.0950e+02    4.1840e+02 ;     CG - CD1
- HD11
  2324   2326   2328      1    1.0950e+02    4.1840e+02 ;     CG - CD1
- HD12
  2324   2326   2329      1    1.0950e+02    4.1840e+02 ;     CG - CD1
- HD13
  2324   2330   2331      1    1.0950e+02    4.1840e+02 ;     CG - CD2
- HD21
  2324   2330   2332      1    1.0950e+02    4.1840e+02 ;     CG - CD2
- HD22
```

```
  2324   2330   2333       1    1.0950e+02    4.1840e+02 ;     CG - CD2
- HD23
  2325   2324   2326       1    1.0950e+02    4.1840e+02 ;     HG - CG
- CD1
  2325   2324   2330       1    1.0950e+02    4.1840e+02 ;     HG - CG
- CD2
  2326   2324   2330       1    1.0950e+02    3.3472e+02 ;    CD1 - CG
- CD2
  2327   2326   2328       1    1.0950e+02    2.9288e+02 ;   HD11 - CD1
- HD12
  2327   2326   2329       1    1.0950e+02    2.9288e+02 ;   HD11 - CD1
- HD13
  2328   2326   2329       1    1.0950e+02    2.9288e+02 ;   HD12 - CD1
- HD13
  2331   2330   2332       1    1.0950e+02    2.9288e+02 ;   HD21 - CD2
- HD22
  2331   2330   2333       1    1.0950e+02    2.9288e+02 ;   HD21 - CD2
- HD23
  2332   2330   2333       1    1.0950e+02    2.9288e+02 ;   HD22 - CD2
- HD23
  2334   2336   2337       1    1.2000e+02    4.1840e+02 ;      C - N
- H
  2334   2336   2338       1    1.2190e+02    4.1840e+02 ;      C - N
- CA
  2335   2334   2336       1    1.2290e+02    6.6944e+02 ;      O - C
- N
  2336   2338   2339       1    1.0950e+02    4.1840e+02 ;      N - CA
- HA
  2336   2338   2340       1    1.0970e+02    6.6944e+02 ;      N - CA
- CB
  2336   2338   2354       1    1.1010e+02    5.2718e+02 ;      N - CA
- C
  2337   2336   2338       1    1.1804e+02    4.1840e+02 ;      H - N
- CA
  2338   2340   2341       1    1.0950e+02    4.1840e+02 ;     CA - CB
- HB2
  2338   2340   2342       1    1.0950e+02    4.1840e+02 ;     CA - CB
- HB3
  2338   2340   2343       1    1.1400e+02    5.2718e+02 ;     CA - CB
- CG
  2338   2354   2355       1    1.2040e+02    6.6944e+02 ;     CA - C
- O
  2338   2354   2356       1    1.1660e+02    5.8576e+02 ;     CA - C
- N
  2339   2338   2340       1    1.0950e+02    4.1840e+02 ;     HA - CA
- CB
  2339   2338   2354       1    1.0950e+02    4.1840e+02 ;     HA - CA
- C
  2340   2338   2354       1    1.1110e+02    5.2718e+02 ;     CB - CA
- C
  2340   2343   2344       1    1.2000e+02    5.8576e+02 ;     CB - CG
- CD1
```

2340 2343 2352 1 1.2000e+02 5.8576e+02 ; CB - CG
- CD2
2341 2340 2342 1 1.0950e+02 2.9288e+02 ; HB2 - CB
- HB3
2341 2340 2343 1 1.0950e+02 4.1840e+02 ; HB2 - CB
- CG
2342 2340 2343 1 1.0950e+02 4.1840e+02 ; HB3 - CB
- CG
2343 2344 2345 1 1.2000e+02 4.1840e+02 ; CG - CD1
- HD1
2343 2344 2346 1 1.2000e+02 5.2718e+02 ; CG - CD1
- CE1
2343 2352 2350 1 1.2000e+02 5.2718e+02 ; CG - CD2
- CE2
2343 2352 2353 1 1.2000e+02 4.1840e+02 ; CG - CD2
- HD2
2344 2343 2352 1 1.2000e+02 5.2718e+02 ; CD1 - CG
- CD2
2344 2346 2347 1 1.2000e+02 4.1840e+02 ; CD1 - CE1
- HE1
2344 2346 2348 1 1.2000e+02 5.2718e+02 ; CD1 - CE1
- CZ
2345 2344 2346 1 1.2000e+02 4.1840e+02 ; HD1 - CD1
- CE1
2346 2348 2349 1 1.2000e+02 4.1840e+02 ; CE1 - CZ
- HZ
2346 2348 2350 1 1.2000e+02 5.2718e+02 ; CE1 - CZ
- CE2
2347 2346 2348 1 1.2000e+02 4.1840e+02 ; HE1 - CE1
- CZ
2348 2350 2351 1 1.2000e+02 4.1840e+02 ; CZ - CE2
- HE2
2348 2350 2352 1 1.2000e+02 5.2718e+02 ; CZ - CE2
- CD2
2349 2348 2350 1 1.2000e+02 4.1840e+02 ; HZ - CZ
- CE2
2350 2352 2353 1 1.2000e+02 4.1840e+02 ; CE2 - CD2
- HD2
2351 2350 2352 1 1.2000e+02 4.1840e+02 ; HE2 - CE2
- CD2
2354 2356 2357 1 1.2000e+02 4.1840e+02 ; C - N
- H
2354 2356 2358 1 1.2190e+02 4.1840e+02 ; C - N
- CA
2355 2354 2356 1 1.2290e+02 6.6944e+02 ; O - C
- N
2356 2358 2359 1 1.0950e+02 4.1840e+02 ; N - CA
- HA
2356 2358 2360 1 1.0970e+02 6.6944e+02 ; N - CA
- CB
2356 2358 2371 1 1.1010e+02 5.2718e+02 ; N - CA
- C

```
  2357   2356   2358       1    1.1804e+02    4.1840e+02 ;      H - N
- CA
  2358   2360   2361       1    1.0950e+02    4.1840e+02 ;     CA - CB
- HB2
  2358   2360   2362       1    1.0950e+02    4.1840e+02 ;     CA - CB
- HB3
  2358   2360   2363       1    1.0950e+02    3.3472e+02 ;     CA - CB
- CG
  2358   2371   2372       1    1.2040e+02    6.6944e+02 ;     CA - C
- O
  2358   2371   2373       1    1.1660e+02    5.8576e+02 ;     CA - C
- N
  2359   2358   2360       1    1.0950e+02    4.1840e+02 ;     HA - CA
- CB
  2359   2358   2371       1    1.0950e+02    4.1840e+02 ;     HA - CA
- C
  2360   2358   2371       1    1.1110e+02    5.2718e+02 ;     CB - CA
- C
  2360   2363   2364       1    1.0950e+02    4.1840e+02 ;     CB - CG
- HG2
  2360   2363   2365       1    1.0950e+02    4.1840e+02 ;     CB - CG
- HG3
  2360   2363   2366       1    1.1110e+02    5.2718e+02 ;     CB - CG
- CD
  2361   2360   2362       1    1.0950e+02    2.9288e+02 ;    HB2 - CB
- HB3
  2361   2360   2363       1    1.0950e+02    4.1840e+02 ;    HB2 - CB
- CG
  2362   2360   2363       1    1.0950e+02    4.1840e+02 ;    HB3 - CB
- CG
  2363   2366   2367       1    1.2040e+02    6.6944e+02 ;     CG - CD
- OE1
  2363   2366   2368       1    1.1660e+02    5.8576e+02 ;     CG - CD
- NE2
  2364   2363   2365       1    1.0950e+02    2.9288e+02 ;    HG2 - CG
- HG3
  2364   2363   2366       1    1.0950e+02    4.1840e+02 ;    HG2 - CG
- CD
  2365   2363   2366       1    1.0950e+02    4.1840e+02 ;    HG3 - CG
- CD
  2366   2368   2369       1    1.2000e+02    4.1840e+02 ;     CD - NE2
- HE21
  2366   2368   2370       1    1.2000e+02    4.1840e+02 ;     CD - NE2
- HE22
  2367   2366   2368       1    1.2290e+02    6.6944e+02 ;    OE1 - CD
- NE2
  2369   2368   2370       1    1.2000e+02    2.9288e+02 ;   HE21 - NE2
- HE22
  2371   2373   2374       1    1.2000e+02    4.1840e+02 ;      C - N
- H
  2371   2373   2375       1    1.2190e+02    4.1840e+02 ;      C - N
- CA
```

2372 2371 2373 1 1.2290e+02 6.6944e+02 ; O - C
- N
2373 2375 2376 1 1.0950e+02 4.1840e+02 ; N - CA
- HA
2373 2375 2377 1 1.0970e+02 6.6944e+02 ; N - CA
- CB
2373 2375 2382 1 1.1010e+02 5.2718e+02 ; N - CA
- C
2374 2373 2375 1 1.1804e+02 4.1840e+02 ; H - N
- CA
2375 2377 2378 1 1.0950e+02 4.1840e+02 ; CA - CB
- HB2
2375 2377 2379 1 1.0950e+02 4.1840e+02 ; CA - CB
- HB3
2375 2377 2380 1 1.0950e+02 4.1840e+02 ; CA - CB
- OG
2375 2382 2383 1 1.2040e+02 6.6944e+02 ; CA - C
- O
2375 2382 2384 1 1.1660e+02 5.8576e+02 ; CA - C
- N
2376 2375 2377 1 1.0950e+02 4.1840e+02 ; HA - CA
- CB
2376 2375 2382 1 1.0950e+02 4.1840e+02 ; HA - CA
- C
2377 2375 2382 1 1.1110e+02 5.2718e+02 ; CB - CA
- C
2377 2380 2381 1 1.0850e+02 4.6024e+02 ; CB - OG
- HG
2378 2377 2379 1 1.0950e+02 2.9288e+02 ; HB2 - CB
- HB3
2378 2377 2380 1 1.0950e+02 4.1840e+02 ; HB2 - CB
- OG
2379 2377 2380 1 1.0950e+02 4.1840e+02 ; HB3 - CB
- OG
2382 2384 2385 1 1.2000e+02 4.1840e+02 ; C - N
- H
2382 2384 2386 1 1.2190e+02 4.1840e+02 ; C - N
- CA
2383 2382 2384 1 1.2290e+02 6.6944e+02 ; O - C
- N
2384 2386 2387 1 1.0950e+02 4.1840e+02 ; N - CA
- HA
2384 2386 2388 1 1.0970e+02 6.6944e+02 ; N - CA
- CB
2384 2386 2397 1 1.1010e+02 5.2718e+02 ; N - CA
- C
2385 2384 2386 1 1.1804e+02 4.1840e+02 ; H - N
- CA
2386 2388 2389 1 1.0950e+02 4.1840e+02 ; CA - CB
- HB2
2386 2388 2390 1 1.0950e+02 4.1840e+02 ; CA - CB
- HB3

2386 2388 2391 1 1.0950e+02 3.3472e+02 ; CA - CB
- CG
2386 2397 2398 1 1.2040e+02 6.6944e+02 ; CA - C
- O
2386 2397 2399 1 1.1660e+02 5.8576e+02 ; CA - C
- N
2387 2386 2388 1 1.0950e+02 4.1840e+02 ; HA - CA
- CB
2387 2386 2397 1 1.0950e+02 4.1840e+02 ; HA - CA
- C
2388 2386 2397 1 1.1110e+02 5.2718e+02 ; CB - CA
- C
2388 2391 2392 1 1.0950e+02 4.1840e+02 ; CB - CG
- HG2
2388 2391 2393 1 1.0950e+02 4.1840e+02 ; CB - CG
- HG3
2388 2391 2394 1 1.1110e+02 5.2718e+02 ; CB - CG
- CD
2389 2388 2390 1 1.0950e+02 2.9288e+02 ; HB2 - CB
- HB3
2389 2388 2391 1 1.0950e+02 4.1840e+02 ; HB2 - CB
- CG
2390 2388 2391 1 1.0950e+02 4.1840e+02 ; HB3 - CB
- CG
2391 2394 2395 1 1.1700e+02 5.8576e+02 ; CG - CD
- OE1
2391 2394 2396 1 1.1700e+02 5.8576e+02 ; CG - CD
- OE2
2392 2391 2393 1 1.0950e+02 2.9288e+02 ; HG2 - CG
- HG3
2392 2391 2394 1 1.0950e+02 4.1840e+02 ; HG2 - CG
- CD
2393 2391 2394 1 1.0950e+02 4.1840e+02 ; HG3 - CG
- CD
2395 2394 2396 1 1.2600e+02 6.6944e+02 ; OE1 - CD
- OE2
2397 2399 2400 1 1.2000e+02 4.1840e+02 ; C - N
- H
2397 2399 2401 1 1.2190e+02 4.1840e+02 ; C - N
- CA
2398 2397 2399 1 1.2290e+02 6.6944e+02 ; O - C
- N
2399 2401 2402 1 1.0950e+02 4.1840e+02 ; N - CA
- HA
2399 2401 2403 1 1.0970e+02 6.6944e+02 ; N - CA
- CB
2399 2401 2411 1 1.1010e+02 5.2718e+02 ; N - CA
- C
2400 2399 2401 1 1.1804e+02 4.1840e+02 ; H - N
- CA
2401 2403 2404 1 1.0950e+02 4.1840e+02 ; CA - CB
- HB2

```
  2401   2403   2405       1    1.0950e+02    4.1840e+02 ;      CA - CB
- HB3
  2401   2403   2406       1    1.1110e+02    5.2718e+02 ;      CA - CB
- CG
  2401   2411   2412       1    1.2040e+02    6.6944e+02 ;      CA - C
- O
  2401   2411   2413       1    1.1660e+02    5.8576e+02 ;      CA - C
- N
  2402   2401   2403       1    1.0950e+02    4.1840e+02 ;      HA - CA
- CB
  2402   2401   2411       1    1.0950e+02    4.1840e+02 ;      HA - CA
- C
  2403   2401   2411       1    1.1110e+02    5.2718e+02 ;      CB - CA
- C
  2403   2406   2407       1    1.2040e+02    6.6944e+02 ;      CB - CG
- OD1
  2403   2406   2408       1    1.1660e+02    5.8576e+02 ;      CB - CG
- ND2
  2404   2403   2405       1    1.0950e+02    2.9288e+02 ;     HB2 - CB
- HB3
  2404   2403   2406       1    1.0950e+02    4.1840e+02 ;     HB2 - CB
- CG
  2405   2403   2406       1    1.0950e+02    4.1840e+02 ;     HB3 - CB
- CG
  2406   2408   2409       1    1.2000e+02    4.1840e+02 ;      CG - ND2
- HD21
  2406   2408   2410       1    1.2000e+02    4.1840e+02 ;      CG - ND2
- HD22
  2407   2406   2408       1    1.2290e+02    6.6944e+02 ;     OD1 - CG
- ND2
  2409   2408   2410       1    1.2000e+02    2.9288e+02 ;    HD21 - ND2
- HD22
  2411   2413   2414       1    1.2000e+02    4.1840e+02 ;       C - N
- H
  2411   2413   2415       1    1.2190e+02    4.1840e+02 ;       C - N
- CA
  2412   2411   2413       1    1.2290e+02    6.6944e+02 ;       O - C
- N
  2413   2415   2416       1    1.0950e+02    4.1840e+02 ;       N - CA
- HA
  2413   2415   2417       1    1.0970e+02    6.6944e+02 ;       N - CA
- CB
  2413   2415   2433       1    1.1010e+02    5.2718e+02 ;       N - CA
- C
  2414   2413   2415       1    1.1804e+02    4.1840e+02 ;       H - N
- CA
  2415   2417   2418       1    1.0950e+02    4.1840e+02 ;      CA - CB
- HB2
  2415   2417   2419       1    1.0950e+02    4.1840e+02 ;      CA - CB
- HB3
  2415   2417   2420       1    1.0950e+02    3.3472e+02 ;      CA - CB
- CG
```

2415 2433 2434 1 1.2040e+02 6.6944e+02 ; CA - C
- O
2415 2433 2435 1 1.1660e+02 5.8576e+02 ; CA - C
- N
2416 2415 2417 1 1.0950e+02 4.1840e+02 ; HA - CA
- CB
2416 2415 2433 1 1.0950e+02 4.1840e+02 ; HA - CA
- C
2417 2415 2433 1 1.1110e+02 5.2718e+02 ; CB - CA
- C
2417 2420 2421 1 1.0950e+02 4.1840e+02 ; CB - CG
- HG2
2417 2420 2422 1 1.0950e+02 4.1840e+02 ; CB - CG
- HG3
2417 2420 2423 1 1.0950e+02 3.3472e+02 ; CB - CG
- CD
2418 2417 2419 1 1.0950e+02 2.9288e+02 ; HB2 - CB
- HB3
2418 2417 2420 1 1.0950e+02 4.1840e+02 ; HB2 - CB
- CG
2419 2417 2420 1 1.0950e+02 4.1840e+02 ; HB3 - CB
- CG
2420 2423 2424 1 1.0950e+02 4.1840e+02 ; CG - CD
- HD2
2420 2423 2425 1 1.0950e+02 4.1840e+02 ; CG - CD
- HD3
2420 2423 2426 1 1.0950e+02 3.3472e+02 ; CG - CD
- CE
2421 2420 2422 1 1.0950e+02 2.9288e+02 ; HG2 - CG
- HG3
2421 2420 2423 1 1.0950e+02 4.1840e+02 ; HG2 - CG
- CD
2422 2420 2423 1 1.0950e+02 4.1840e+02 ; HG3 - CG
- CD
2423 2426 2427 1 1.0950e+02 4.1840e+02 ; CD - CE
- HE2
2423 2426 2428 1 1.0950e+02 4.1840e+02 ; CD - CE
- HE3
2423 2426 2429 1 1.1120e+02 6.6944e+02 ; CD - CE
- NZ
2424 2423 2425 1 1.0950e+02 2.9288e+02 ; HD2 - CD
- HD3
2424 2423 2426 1 1.0950e+02 4.1840e+02 ; HD2 - CD
- CE
2425 2423 2426 1 1.0950e+02 4.1840e+02 ; HD3 - CD
- CE
2426 2429 2430 1 1.0950e+02 4.1840e+02 ; CE - NZ
- HZ1
2426 2429 2431 1 1.0950e+02 4.1840e+02 ; CE - NZ
- HZ2
2426 2429 2432 1 1.0950e+02 4.1840e+02 ; CE - NZ
- HZ3

2427   2426   2428      1    1.0950e+02    2.9288e+02 ;    HE2 - CE
- HE3
2427   2426   2429      1    1.0950e+02    4.1840e+02 ;    HE2 - CE
- NZ
2428   2426   2429      1    1.0950e+02    4.1840e+02 ;    HE3 - CE
- NZ
2430   2429   2431      1    1.0950e+02    2.9288e+02 ;    HZ1 - NZ
- HZ2
2430   2429   2432      1    1.0950e+02    2.9288e+02 ;    HZ1 - NZ
- HZ3
2431   2429   2432      1    1.0950e+02    2.9288e+02 ;    HZ2 - NZ
- HZ3
2433   2435   2436      1    1.2000e+02    4.1840e+02 ;      C - N
- H
2433   2435   2437      1    1.2190e+02    4.1840e+02 ;      C - N
- CA
2434   2433   2435      1    1.2290e+02    6.6944e+02 ;      O - C
- N
2435   2437   2438      1    1.0950e+02    4.1840e+02 ;      N - CA
- HA
2435   2437   2439      1    1.0970e+02    6.6944e+02 ;      N - CA
- CB
2435   2437   2445      1    1.1010e+02    5.2718e+02 ;      N - CA
- C
2436   2435   2437      1    1.1804e+02    4.1840e+02 ;      H - N
- CA
2437   2439   2440      1    1.0950e+02    4.1840e+02 ;     CA - CB
- HB2
2437   2439   2441      1    1.0950e+02    4.1840e+02 ;     CA - CB
- HB3
2437   2439   2442      1    1.1110e+02    5.2718e+02 ;     CA - CB
- CG
2437   2445   2446      1    1.2040e+02    6.6944e+02 ;     CA - C
- O
2437   2445   2447      1    1.1660e+02    5.8576e+02 ;     CA - C
- N
2438   2437   2439      1    1.0950e+02    4.1840e+02 ;     HA - CA
- CB
2438   2437   2445      1    1.0950e+02    4.1840e+02 ;     HA - CA
- C
2439   2437   2445      1    1.1110e+02    5.2718e+02 ;     CB - CA
- C
2439   2442   2443      1    1.1700e+02    5.8576e+02 ;     CB - CG
- OD1
2439   2442   2444      1    1.1700e+02    5.8576e+02 ;     CB - CG
- OD2
2440   2439   2441      1    1.0950e+02    2.9288e+02 ;    HB2 - CB
- HB3
2440   2439   2442      1    1.0950e+02    4.1840e+02 ;    HB2 - CB
- CG
2441   2439   2442      1    1.0950e+02    4.1840e+02 ;    HB3 - CB
- CG

2443 2442 2444 1 1.2600e+02 6.6944e+02 ; OD1 - CG
- OD2
2445 2447 2448 1 1.2000e+02 4.1840e+02 ; C - N
- H
2445 2447 2449 1 1.2190e+02 4.1840e+02 ; C - N
- CA
2446 2445 2447 1 1.2290e+02 6.6944e+02 ; O - C
- N
2447 2449 2450 1 1.0950e+02 4.1840e+02 ; N - CA
- HA
2447 2449 2451 1 1.0970e+02 6.6944e+02 ; N - CA
- CB
2447 2449 2456 1 1.1010e+02 5.2718e+02 ; N - CA
- C
2448 2447 2449 1 1.1804e+02 4.1840e+02 ; H - N
- CA
2449 2451 2452 1 1.0950e+02 4.1840e+02 ; CA - CB
- HB2
2449 2451 2453 1 1.0950e+02 4.1840e+02 ; CA - CB
- HB3
2449 2451 2454 1 1.0950e+02 4.1840e+02 ; CA - CB
- OG
2449 2456 2457 1 1.2040e+02 6.6944e+02 ; CA - C
- O
2449 2456 2458 1 1.1660e+02 5.8576e+02 ; CA - C
- N
2450 2449 2451 1 1.0950e+02 4.1840e+02 ; HA - CA
- CB
2450 2449 2456 1 1.0950e+02 4.1840e+02 ; HA - CA
- C
2451 2449 2456 1 1.1110e+02 5.2718e+02 ; CB - CA
- C
2451 2454 2455 1 1.0850e+02 4.6024e+02 ; CB - OG
- HG
2452 2451 2453 1 1.0950e+02 2.9288e+02 ; HB2 - CB
- HB3
2452 2451 2454 1 1.0950e+02 4.1840e+02 ; HB2 - CB
- OG
2453 2451 2454 1 1.0950e+02 4.1840e+02 ; HB3 - CB
- OG
2456 2458 2459 1 1.2000e+02 4.1840e+02 ; C - N
- H
2456 2458 2460 1 1.2190e+02 4.1840e+02 ; C - N
- CA
2457 2456 2458 1 1.2290e+02 6.6944e+02 ; O - C
- N
2458 2460 2461 1 1.0950e+02 4.1840e+02 ; N - CA
- HA
2458 2460 2462 1 1.0970e+02 6.6944e+02 ; N - CA
- CB
2458 2460 2475 1 1.1010e+02 5.2718e+02 ; N - CA
- C

2459 2458 2460 1 1.1804e+02 4.1840e+02 ; H - N
- CA
2460 2462 2463 1 1.0950e+02 4.1840e+02 ; CA - CB
- HB2
2460 2462 2464 1 1.0950e+02 4.1840e+02 ; CA - CB
- HB3
2460 2462 2465 1 1.0950e+02 3.3472e+02 ; CA - CB
- CG
2460 2475 2476 1 1.2040e+02 6.6944e+02 ; CA - C
- O
2460 2475 2477 1 1.1660e+02 5.8576e+02 ; CA - C
- N
2461 2460 2462 1 1.0950e+02 4.1840e+02 ; HA - CA
- CB
2461 2460 2475 1 1.0950e+02 4.1840e+02 ; HA - CA
- C
2462 2460 2475 1 1.1110e+02 5.2718e+02 ; CB - CA
- C
2462 2465 2466 1 1.0950e+02 4.1840e+02 ; CB - CG
- HG
2462 2465 2467 1 1.0950e+02 3.3472e+02 ; CB - CG
- CD1
2462 2465 2471 1 1.0950e+02 3.3472e+02 ; CB - CG
- CD2
2463 2462 2464 1 1.0950e+02 2.9288e+02 ; HB2 - CB
- HB3
2463 2462 2465 1 1.0950e+02 4.1840e+02 ; HB2 - CB
- CG
2464 2462 2465 1 1.0950e+02 4.1840e+02 ; HB3 - CB
- CG
2465 2467 2468 1 1.0950e+02 4.1840e+02 ; CG - CD1
- HD11
2465 2467 2469 1 1.0950e+02 4.1840e+02 ; CG - CD1
- HD12
2465 2467 2470 1 1.0950e+02 4.1840e+02 ; CG - CD1
- HD13
2465 2471 2472 1 1.0950e+02 4.1840e+02 ; CG - CD2
- HD21
2465 2471 2473 1 1.0950e+02 4.1840e+02 ; CG - CD2
- HD22
2465 2471 2474 1 1.0950e+02 4.1840e+02 ; CG - CD2
- HD23
2466 2465 2467 1 1.0950e+02 4.1840e+02 ; HG - CG
- CD1
2466 2465 2471 1 1.0950e+02 4.1840e+02 ; HG - CG
- CD2
2467 2465 2471 1 1.0950e+02 3.3472e+02 ; CD1 - CG
- CD2
2468 2467 2469 1 1.0950e+02 2.9288e+02 ; HD11 - CD1
- HD12
2468 2467 2470 1 1.0950e+02 2.9288e+02 ; HD11 - CD1
- HD13

2469 2467 2470 1 1.0950e+02 2.9288e+02 ; HD12 - CD1
- HD13
2472 2471 2473 1 1.0950e+02 2.9288e+02 ; HD21 - CD2
- HD22
2472 2471 2474 1 1.0950e+02 2.9288e+02 ; HD21 - CD2
- HD23
2473 2471 2474 1 1.0950e+02 2.9288e+02 ; HD22 - CD2
- HD23
2475 2477 2478 1 1.2000e+02 4.1840e+02 ; C - N
- H
2475 2477 2479 1 1.2190e+02 4.1840e+02 ; C - N
- CA
2476 2475 2477 1 1.2290e+02 6.6944e+02 ; O - C
- N
2477 2479 2480 1 1.0950e+02 4.1840e+02 ; N - CA
- HA
2477 2479 2481 1 1.0970e+02 6.6944e+02 ; N - CA
- CB
2477 2479 2494 1 1.1010e+02 5.2718e+02 ; N - CA
- C
2478 2477 2479 1 1.1804e+02 4.1840e+02 ; H - N
- CA
2479 2481 2482 1 1.0950e+02 4.1840e+02 ; CA - CB
- HB2
2479 2481 2483 1 1.0950e+02 4.1840e+02 ; CA - CB
- HB3
2479 2481 2484 1 1.0950e+02 3.3472e+02 ; CA - CB
- CG
2479 2494 2495 1 1.2040e+02 6.6944e+02 ; CA - C
- O
2479 2494 2496 1 1.1660e+02 5.8576e+02 ; CA - C
- N
2480 2479 2481 1 1.0950e+02 4.1840e+02 ; HA - CA
- CB
2480 2479 2494 1 1.0950e+02 4.1840e+02 ; HA - CA
- C
2481 2479 2494 1 1.1110e+02 5.2718e+02 ; CB - CA
- C
2481 2484 2485 1 1.0950e+02 4.1840e+02 ; CB - CG
- HG
2481 2484 2486 1 1.0950e+02 3.3472e+02 ; CB - CG
- CD1
2481 2484 2490 1 1.0950e+02 3.3472e+02 ; CB - CG
- CD2
2482 2481 2483 1 1.0950e+02 2.9288e+02 ; HB2 - CB
- HB3
2482 2481 2484 1 1.0950e+02 4.1840e+02 ; HB2 - CB
- CG
2483 2481 2484 1 1.0950e+02 4.1840e+02 ; HB3 - CB
- CG
2484 2486 2487 1 1.0950e+02 4.1840e+02 ; CG - CD1
- HD11

2484 2486 2488 1 1.0950e+02 4.1840e+02 ; CG - CD1
- HD12
2484 2486 2489 1 1.0950e+02 4.1840e+02 ; CG - CD1
- HD13
2484 2490 2491 1 1.0950e+02 4.1840e+02 ; CG - CD2
- HD21
2484 2490 2492 1 1.0950e+02 4.1840e+02 ; CG - CD2
- HD22
2484 2490 2493 1 1.0950e+02 4.1840e+02 ; CG - CD2
- HD23
2485 2484 2486 1 1.0950e+02 4.1840e+02 ; HG - CG
- CD1
2485 2484 2490 1 1.0950e+02 4.1840e+02 ; HG - CG
- CD2
2486 2484 2490 1 1.0950e+02 3.3472e+02 ; CD1 - CG
- CD2
2487 2486 2488 1 1.0950e+02 2.9288e+02 ; HD11 - CD1
- HD12
2487 2486 2489 1 1.0950e+02 2.9288e+02 ; HD11 - CD1
- HD13
2488 2486 2489 1 1.0950e+02 2.9288e+02 ; HD12 - CD1
- HD13
2491 2490 2492 1 1.0950e+02 2.9288e+02 ; HD21 - CD2
- HD22
2491 2490 2493 1 1.0950e+02 2.9288e+02 ; HD21 - CD2
- HD23
2492 2490 2493 1 1.0950e+02 2.9288e+02 ; HD22 - CD2
- HD23
2494 2496 2497 1 1.2000e+02 4.1840e+02 ; C - N
- H
2494 2496 2498 1 1.2190e+02 4.1840e+02 ; C - N
- CA
2495 2494 2496 1 1.2290e+02 6.6944e+02 ; O - C
- N
2496 2498 2499 1 1.0950e+02 4.1840e+02 ; N - CA
- HA
2496 2498 2500 1 1.0970e+02 6.6944e+02 ; N - CA
- CB
2496 2498 2516 1 1.1010e+02 5.2718e+02 ; N - CA
- C
2497 2496 2498 1 1.1804e+02 4.1840e+02 ; H - N
- CA
2498 2500 2501 1 1.0950e+02 4.1840e+02 ; CA - CB
- HB2
2498 2500 2502 1 1.0950e+02 4.1840e+02 ; CA - CB
- HB3
2498 2500 2503 1 1.0950e+02 3.3472e+02 ; CA - CB
- CG
2498 2516 2517 1 1.2040e+02 6.6944e+02 ; CA - C
- O
2498 2516 2518 1 1.1660e+02 5.8576e+02 ; CA - C
- N

2499 2498 2500 1 1.0950e+02 4.1840e+02 ; HA - CA
- CB
2499 2498 2516 1 1.0950e+02 4.1840e+02 ; HA - CA
- C
2500 2498 2516 1 1.1110e+02 5.2718e+02 ; CB - CA
- C
2500 2503 2504 1 1.0950e+02 4.1840e+02 ; CB - CG
- HG2
2500 2503 2505 1 1.0950e+02 4.1840e+02 ; CB - CG
- HG3
2500 2503 2506 1 1.0950e+02 3.3472e+02 ; CB - CG
- CD
2501 2500 2502 1 1.0950e+02 2.9288e+02 ; HB2 - CB
- HB3
2501 2500 2503 1 1.0950e+02 4.1840e+02 ; HB2 - CB
- CG
2502 2500 2503 1 1.0950e+02 4.1840e+02 ; HB3 - CB
- CG
2503 2506 2507 1 1.0950e+02 4.1840e+02 ; CG - CD
- HD2
2503 2506 2508 1 1.0950e+02 4.1840e+02 ; CG - CD
- HD3
2503 2506 2509 1 1.0950e+02 3.3472e+02 ; CG - CD
- CE
2504 2503 2505 1 1.0950e+02 2.9288e+02 ; HG2 - CG
- HG3
2504 2503 2506 1 1.0950e+02 4.1840e+02 ; HG2 - CG
- CD
2505 2503 2506 1 1.0950e+02 4.1840e+02 ; HG3 - CG
- CD
2506 2509 2510 1 1.0950e+02 4.1840e+02 ; CD - CE
- HE2
2506 2509 2511 1 1.0950e+02 4.1840e+02 ; CD - CE
- HE3
2506 2509 2512 1 1.1120e+02 6.6944e+02 ; CD - CE
- NZ
2507 2506 2508 1 1.0950e+02 2.9288e+02 ; HD2 - CD
- HD3
2507 2506 2509 1 1.0950e+02 4.1840e+02 ; HD2 - CD
- CE
2508 2506 2509 1 1.0950e+02 4.1840e+02 ; HD3 - CD
- CE
2509 2512 2513 1 1.0950e+02 4.1840e+02 ; CE - NZ
- HZ1
2509 2512 2514 1 1.0950e+02 4.1840e+02 ; CE - NZ
- HZ2
2509 2512 2515 1 1.0950e+02 4.1840e+02 ; CE - NZ
- HZ3
2510 2509 2511 1 1.0950e+02 2.9288e+02 ; HE2 - CE
- HE3
2510 2509 2512 1 1.0950e+02 4.1840e+02 ; HE2 - CE
- NZ

2511 2509 2512 1 1.0950e+02 4.1840e+02 ; HE3 - CE - NZ
2513 2512 2514 1 1.0950e+02 2.9288e+02 ; HZ1 - NZ - HZ2
2513 2512 2515 1 1.0950e+02 2.9288e+02 ; HZ1 - NZ - HZ3
2514 2512 2515 1 1.0950e+02 2.9288e+02 ; HZ2 - NZ - HZ3
2516 2518 2519 1 1.2000e+02 4.1840e+02 ; C - N - H
2516 2518 2520 1 1.2190e+02 4.1840e+02 ; C - N - CA
2517 2516 2518 1 1.2290e+02 6.6944e+02 ; O - C - N
2518 2520 2521 1 1.0950e+02 4.1840e+02 ; N - CA - HA
2518 2520 2522 1 1.0970e+02 6.6944e+02 ; N - CA - CB
2518 2520 2535 1 1.1010e+02 5.2718e+02 ; N - CA - C
2519 2518 2520 1 1.1804e+02 4.1840e+02 ; H - N - CA
2520 2522 2523 1 1.0950e+02 4.1840e+02 ; CA - CB - HB
2520 2522 2524 1 1.0950e+02 3.3472e+02 ; CA - CB - CG2
2520 2522 2528 1 1.0950e+02 3.3472e+02 ; CA - CB - CG1
2520 2535 2536 1 1.2040e+02 6.6944e+02 ; CA - C - O
2520 2535 2537 1 1.1660e+02 5.8576e+02 ; CA - C - N
2521 2520 2522 1 1.0950e+02 4.1840e+02 ; HA - CA - CB
2521 2520 2535 1 1.0950e+02 4.1840e+02 ; HA - CA - C
2522 2520 2535 1 1.1110e+02 5.2718e+02 ; CB - CA - C
2522 2524 2525 1 1.0950e+02 4.1840e+02 ; CB - CG2 - HG21
2522 2524 2526 1 1.0950e+02 4.1840e+02 ; CB - CG2 - HG22
2522 2524 2527 1 1.0950e+02 4.1840e+02 ; CB - CG2 - HG23
2522 2528 2529 1 1.0950e+02 4.1840e+02 ; CB - CG1 - HG12
2522 2528 2530 1 1.0950e+02 4.1840e+02 ; CB - CG1 - HG13
2522 2528 2531 1 1.0950e+02 3.3472e+02 ; CB - CG1 - CD1
2523 2522 2524 1 1.0950e+02 4.1840e+02 ; HB - CB - CG2

2523 2522 2528 1 1.0950e+02 4.1840e+02 ; HB - CB
- CG1
2524 2522 2528 1 1.0950e+02 3.3472e+02 ; CG2 - CB
- CG1
2525 2524 2526 1 1.0950e+02 2.9288e+02 ; HG21 - CG2
- HG22
2525 2524 2527 1 1.0950e+02 2.9288e+02 ; HG21 - CG2
- HG23
2526 2524 2527 1 1.0950e+02 2.9288e+02 ; HG22 - CG2
- HG23
2528 2531 2532 1 1.0950e+02 4.1840e+02 ; CG1 - CD1
- HD11
2528 2531 2533 1 1.0950e+02 4.1840e+02 ; CG1 - CD1
- HD12
2528 2531 2534 1 1.0950e+02 4.1840e+02 ; CG1 - CD1
- HD13
2529 2528 2530 1 1.0950e+02 2.9288e+02 ; HG12 - CG1
- HG13
2529 2528 2531 1 1.0950e+02 4.1840e+02 ; HG12 - CG1
- CD1
2530 2528 2531 1 1.0950e+02 4.1840e+02 ; HG13 - CG1
- CD1
2532 2531 2533 1 1.0950e+02 2.9288e+02 ; HD11 - CD1
- HD12
2532 2531 2534 1 1.0950e+02 2.9288e+02 ; HD11 - CD1
- HD13
2533 2531 2534 1 1.0950e+02 2.9288e+02 ; HD12 - CD1
- HD13
2535 2537 2538 1 1.2000e+02 4.1840e+02 ; C - N
- H
2535 2537 2539 1 1.2190e+02 4.1840e+02 ; C - N
- CA
2536 2535 2537 1 1.2290e+02 6.6944e+02 ; O - C
- N
2537 2539 2540 1 1.0950e+02 4.1840e+02 ; N - CA
- HA
2537 2539 2541 1 1.0970e+02 6.6944e+02 ; N - CA
- CB
2537 2539 2554 1 1.1010e+02 5.2718e+02 ; N - CA
- C
2538 2537 2539 1 1.1804e+02 4.1840e+02 ; H - N
- CA
2539 2541 2542 1 1.0950e+02 4.1840e+02 ; CA - CB
- HB
2539 2541 2543 1 1.0950e+02 3.3472e+02 ; CA - CB
- CG2
2539 2541 2547 1 1.0950e+02 3.3472e+02 ; CA - CB
- CG1
2539 2554 2555 1 1.2040e+02 6.6944e+02 ; CA - C
- O
2539 2554 2556 1 1.1660e+02 5.8576e+02 ; CA - C
- N

2540 2539 2541 1 1.0950e+02 4.1840e+02 ; HA - CA
- CB
2540 2539 2554 1 1.0950e+02 4.1840e+02 ; HA - CA
- C
2541 2539 2554 1 1.1110e+02 5.2718e+02 ; CB - CA
- C
2541 2543 2544 1 1.0950e+02 4.1840e+02 ; CB - CG2
- HG21
2541 2543 2545 1 1.0950e+02 4.1840e+02 ; CB - CG2
- HG22
2541 2543 2546 1 1.0950e+02 4.1840e+02 ; CB - CG2
- HG23
2541 2547 2548 1 1.0950e+02 4.1840e+02 ; CB - CG1
- HG12
2541 2547 2549 1 1.0950e+02 4.1840e+02 ; CB - CG1
- HG13
2541 2547 2550 1 1.0950e+02 3.3472e+02 ; CB - CG1
- CD1
2542 2541 2543 1 1.0950e+02 4.1840e+02 ; HB - CB
- CG2
2542 2541 2547 1 1.0950e+02 4.1840e+02 ; HB - CB
- CG1
2543 2541 2547 1 1.0950e+02 3.3472e+02 ; CG2 - CB
- CG1
2544 2543 2545 1 1.0950e+02 2.9288e+02 ; HG21 - CG2
- HG22
2544 2543 2546 1 1.0950e+02 2.9288e+02 ; HG21 - CG2
- HG23
2545 2543 2546 1 1.0950e+02 2.9288e+02 ; HG22 - CG2
- HG23
2547 2550 2551 1 1.0950e+02 4.1840e+02 ; CG1 - CD1
- HD11
2547 2550 2552 1 1.0950e+02 4.1840e+02 ; CG1 - CD1
- HD12
2547 2550 2553 1 1.0950e+02 4.1840e+02 ; CG1 - CD1
- HD13
2548 2547 2549 1 1.0950e+02 2.9288e+02 ; HG12 - CG1
- HG13
2548 2547 2550 1 1.0950e+02 4.1840e+02 ; HG12 - CG1
- CD1
2549 2547 2550 1 1.0950e+02 4.1840e+02 ; HG13 - CG1
- CD1
2551 2550 2552 1 1.0950e+02 2.9288e+02 ; HD11 - CD1
- HD12
2551 2550 2553 1 1.0950e+02 2.9288e+02 ; HD11 - CD1
- HD13
2552 2550 2553 1 1.0950e+02 2.9288e+02 ; HD12 - CD1
- HD13
2554 2556 2557 1 1.2000e+02 4.1840e+02 ; C - N
- H
2554 2556 2558 1 1.2190e+02 4.1840e+02 ; C - N
- CA

2555 2554 2556 1 1.2290e+02 6.6944e+02 ; O - C
- N
2556 2558 2559 1 1.0950e+02 4.1840e+02 ; N - CA
- HA
2556 2558 2560 1 1.0970e+02 6.6944e+02 ; N - CA
- CB
2556 2558 2566 1 1.1010e+02 5.2718e+02 ; N - CA
- C
2557 2556 2558 1 1.1804e+02 4.1840e+02 ; H - N
- CA
2558 2560 2561 1 1.0950e+02 4.1840e+02 ; CA - CB
- HB2
2558 2560 2562 1 1.0950e+02 4.1840e+02 ; CA - CB
- HB3
2558 2560 2563 1 1.1110e+02 5.2718e+02 ; CA - CB
- CG
2558 2566 2567 1 1.2040e+02 6.6944e+02 ; CA - C
- O
2558 2566 2568 1 1.1660e+02 5.8576e+02 ; CA - C
- N
2559 2558 2560 1 1.0950e+02 4.1840e+02 ; HA - CA
- CB
2559 2558 2566 1 1.0950e+02 4.1840e+02 ; HA - CA
- C
2560 2558 2566 1 1.1110e+02 5.2718e+02 ; CB - CA
- C
2560 2563 2564 1 1.1700e+02 5.8576e+02 ; CB - CG
- OD1
2560 2563 2565 1 1.1700e+02 5.8576e+02 ; CB - CG
- OD2
2561 2560 2562 1 1.0950e+02 2.9288e+02 ; HB2 - CB
- HB3
2561 2560 2563 1 1.0950e+02 4.1840e+02 ; HB2 - CB
- CG
2562 2560 2563 1 1.0950e+02 4.1840e+02 ; HB3 - CB
- CG
2564 2563 2565 1 1.2600e+02 6.6944e+02 ; OD1 - CG
- OD2
2566 2568 2569 1 1.2000e+02 4.1840e+02 ; C - N
- H
2566 2568 2570 1 1.2190e+02 4.1840e+02 ; C - N
- CA
2567 2566 2568 1 1.2290e+02 6.6944e+02 ; O - C
- N
2568 2570 2571 1 1.0950e+02 4.1840e+02 ; N - CA
- HA
2568 2570 2572 1 1.0970e+02 6.6944e+02 ; N - CA
- CB
2568 2570 2586 1 1.1010e+02 5.2718e+02 ; N - CA
- C
2569 2568 2570 1 1.1804e+02 4.1840e+02 ; H - N
- CA

```
  2570   2572   2573       1    1.0950e+02    4.1840e+02 ;     CA - CB
- HB2
  2570   2572   2574       1    1.0950e+02    4.1840e+02 ;     CA - CB
- HB3
  2570   2572   2575       1    1.1400e+02    5.2718e+02 ;     CA - CB
- CG
  2570   2586   2587       1    1.2040e+02    6.6944e+02 ;     CA - C
- O
  2570   2586   2588       1    1.1660e+02    5.8576e+02 ;     CA - C
- N
  2571   2570   2572       1    1.0950e+02    4.1840e+02 ;     HA - CA
- CB
  2571   2570   2586       1    1.0950e+02    4.1840e+02 ;     HA - CA
- C
  2572   2570   2586       1    1.1110e+02    5.2718e+02 ;     CB - CA
- C
  2572   2575   2576       1    1.2000e+02    5.8576e+02 ;     CB - CG
- CD1
  2572   2575   2584       1    1.2000e+02    5.8576e+02 ;     CB - CG
- CD2
  2573   2572   2574       1    1.0950e+02    2.9288e+02 ;    HB2 - CB
- HB3
  2573   2572   2575       1    1.0950e+02    4.1840e+02 ;    HB2 - CB
- CG
  2574   2572   2575       1    1.0950e+02    4.1840e+02 ;    HB3 - CB
- CG
  2575   2576   2577       1    1.2000e+02    4.1840e+02 ;     CG - CD1
- HD1
  2575   2576   2578       1    1.2000e+02    5.2718e+02 ;     CG - CD1
- CE1
  2575   2584   2582       1    1.2000e+02    5.2718e+02 ;     CG - CD2
- CE2
  2575   2584   2585       1    1.2000e+02    4.1840e+02 ;     CG - CD2
- HD2
  2576   2575   2584       1    1.2000e+02    5.2718e+02 ;    CD1 - CG
- CD2
  2576   2578   2579       1    1.2000e+02    4.1840e+02 ;    CD1 - CE1
- HE1
  2576   2578   2580       1    1.2000e+02    5.2718e+02 ;    CD1 - CE1
- CZ
  2577   2576   2578       1    1.2000e+02    4.1840e+02 ;    HD1 - CD1
- CE1
  2578   2580   2581       1    1.2000e+02    4.1840e+02 ;    CE1 - CZ
- HZ
  2578   2580   2582       1    1.2000e+02    5.2718e+02 ;    CE1 - CZ
- CE2
  2579   2578   2580       1    1.2000e+02    4.1840e+02 ;    HE1 - CE1
- CZ
  2580   2582   2583       1    1.2000e+02    4.1840e+02 ;     CZ - CE2
- HE2
  2580   2582   2584       1    1.2000e+02    5.2718e+02 ;     CZ - CE2
- CD2
```

2581 2580 2582 1 1.2000e+02 4.1840e+02 ; HZ - CZ
- CE2
2582 2584 2585 1 1.2000e+02 4.1840e+02 ; CE2 - CD2
- HD2
2583 2582 2584 1 1.2000e+02 4.1840e+02 ; HE2 - CE2
- CD2
2586 2588 2589 1 1.2000e+02 4.1840e+02 ; C - N
- H
2586 2588 2590 1 1.2190e+02 4.1840e+02 ; C - N
- CA
2587 2586 2588 1 1.2290e+02 6.6944e+02 ; O - C
- N
2588 2590 2591 1 1.0950e+02 4.1840e+02 ; N - CA
- HA2
2588 2590 2592 1 1.0950e+02 4.1840e+02 ; N - CA
- HA3
2588 2590 2593 1 1.1010e+02 5.2718e+02 ; N - CA
- C
2589 2588 2590 1 1.1804e+02 4.1840e+02 ; H - N
- CA
2590 2593 2594 1 1.2040e+02 6.6944e+02 ; CA - C
- O
2590 2593 2595 1 1.1660e+02 5.8576e+02 ; CA - C
- N
2591 2590 2592 1 1.0950e+02 2.9288e+02 ; HA2 - CA
- HA3
2591 2590 2593 1 1.0950e+02 4.1840e+02 ; HA2 - CA
- C
2592 2590 2593 1 1.0950e+02 4.1840e+02 ; HA3 - CA
- C
2593 2595 2596 1 1.2000e+02 4.1840e+02 ; C - N
- H
2593 2595 2597 1 1.2190e+02 4.1840e+02 ; C - N
- CA
2594 2593 2595 1 1.2290e+02 6.6944e+02 ; O - C
- N
2595 2597 2598 1 1.0950e+02 4.1840e+02 ; N - CA
- HA
2595 2597 2599 1 1.0970e+02 6.6944e+02 ; N - CA
- CB
2595 2597 2612 1 1.1010e+02 5.2718e+02 ; N - CA
- C
2596 2595 2597 1 1.1804e+02 4.1840e+02 ; H - N
- CA
2597 2599 2600 1 1.0950e+02 4.1840e+02 ; CA - CB
- HB2
2597 2599 2601 1 1.0950e+02 4.1840e+02 ; CA - CB
- HB3
2597 2599 2602 1 1.0950e+02 3.3472e+02 ; CA - CB
- CG
2597 2612 2613 1 1.2040e+02 6.6944e+02 ; CA - C
- O

2597 2612 2614 1 1.1660e+02 5.8576e+02 ; CA - C
- N
2598 2597 2599 1 1.0950e+02 4.1840e+02 ; HA - CA
- CB
2598 2597 2612 1 1.0950e+02 4.1840e+02 ; HA - CA
- C
2599 2597 2612 1 1.1110e+02 5.2718e+02 ; CB - CA
- C
2599 2602 2603 1 1.0950e+02 4.1840e+02 ; CB - CG
- HG
2599 2602 2604 1 1.0950e+02 3.3472e+02 ; CB - CG
- CD1
2599 2602 2608 1 1.0950e+02 3.3472e+02 ; CB - CG
- CD2
2600 2599 2601 1 1.0950e+02 2.9288e+02 ; HB2 - CB
- HB3
2600 2599 2602 1 1.0950e+02 4.1840e+02 ; HB2 - CB
- CG
2601 2599 2602 1 1.0950e+02 4.1840e+02 ; HB3 - CB
- CG
2602 2604 2605 1 1.0950e+02 4.1840e+02 ; CG - CD1
- HD11
2602 2604 2606 1 1.0950e+02 4.1840e+02 ; CG - CD1
- HD12
2602 2604 2607 1 1.0950e+02 4.1840e+02 ; CG - CD1
- HD13
2602 2608 2609 1 1.0950e+02 4.1840e+02 ; CG - CD2
- HD21
2602 2608 2610 1 1.0950e+02 4.1840e+02 ; CG - CD2
- HD22
2602 2608 2611 1 1.0950e+02 4.1840e+02 ; CG - CD2
- HD23
2603 2602 2604 1 1.0950e+02 4.1840e+02 ; HG - CG
- CD1
2603 2602 2608 1 1.0950e+02 4.1840e+02 ; HG - CG
- CD2
2604 2602 2608 1 1.0950e+02 3.3472e+02 ; CD1 - CG
- CD2
2605 2604 2606 1 1.0950e+02 2.9288e+02 ; HD11 - CD1
- HD12
2605 2604 2607 1 1.0950e+02 2.9288e+02 ; HD11 - CD1
- HD13
2606 2604 2607 1 1.0950e+02 2.9288e+02 ; HD12 - CD1
- HD13
2609 2608 2610 1 1.0950e+02 2.9288e+02 ; HD21 - CD2
- HD22
2609 2608 2611 1 1.0950e+02 2.9288e+02 ; HD21 - CD2
- HD23
2610 2608 2611 1 1.0950e+02 2.9288e+02 ; HD22 - CD2
- HD23
2612 2614 2615 1 1.2000e+02 4.1840e+02 ; C - N
- H

| | | | | | | |
|---|---|---|---|---|---|---|
| 2612 | 2614 | 2616 | 1 | 1.2190e+02 | 4.1840e+02 ; | C - N - CA |
| 2613 | 2612 | 2614 | 1 | 1.2290e+02 | 6.6944e+02 ; | O - C - N |
| 2614 | 2616 | 2617 | 1 | 1.0950e+02 | 4.1840e+02 ; | N - CA - HA |
| 2614 | 2616 | 2618 | 1 | 1.0970e+02 | 6.6944e+02 ; | N - CA - CB |
| 2614 | 2616 | 2623 | 1 | 1.1010e+02 | 5.2718e+02 ; | N - CA - C |
| 2615 | 2614 | 2616 | 1 | 1.1804e+02 | 4.1840e+02 ; | H - N - CA |
| 2616 | 2618 | 2619 | 1 | 1.0950e+02 | 4.1840e+02 ; | CA - CB - HB2 |
| 2616 | 2618 | 2620 | 1 | 1.0950e+02 | 4.1840e+02 ; | CA - CB - HB3 |
| 2616 | 2618 | 2621 | 1 | 1.0950e+02 | 4.1840e+02 ; | CA - CB - OG |
| 2616 | 2623 | 2624 | 1 | 1.2040e+02 | 6.6944e+02 ; | CA - C - O |
| 2616 | 2623 | 2625 | 1 | 1.1660e+02 | 5.8576e+02 ; | CA - C - N |
| 2617 | 2616 | 2618 | 1 | 1.0950e+02 | 4.1840e+02 ; | HA - CA - CB |
| 2617 | 2616 | 2623 | 1 | 1.0950e+02 | 4.1840e+02 ; | HA - CA - C |
| 2618 | 2616 | 2623 | 1 | 1.1110e+02 | 5.2718e+02 ; | CB - CA - C |
| 2618 | 2621 | 2622 | 1 | 1.0850e+02 | 4.6024e+02 ; | CB - OG - HG |
| 2619 | 2618 | 2620 | 1 | 1.0950e+02 | 2.9288e+02 ; | HB2 - CB - HB3 |
| 2619 | 2618 | 2621 | 1 | 1.0950e+02 | 4.1840e+02 ; | HB2 - CB - OG |
| 2620 | 2618 | 2621 | 1 | 1.0950e+02 | 4.1840e+02 ; | HB3 - CB - OG |
| 2623 | 2625 | 2626 | 1 | 1.2000e+02 | 4.1840e+02 ; | C - N - H |
| 2623 | 2625 | 2627 | 1 | 1.2190e+02 | 4.1840e+02 ; | C - N - CA |
| 2624 | 2623 | 2625 | 1 | 1.2290e+02 | 6.6944e+02 ; | O - C - N |
| 2625 | 2627 | 2628 | 1 | 1.0950e+02 | 4.1840e+02 ; | N - CA - HA |
| 2625 | 2627 | 2629 | 1 | 1.0970e+02 | 6.6944e+02 ; | N - CA - CB |
| 2625 | 2627 | 2645 | 1 | 1.1010e+02 | 5.2718e+02 ; | N - CA - C |
| 2626 | 2625 | 2627 | 1 | 1.1804e+02 | 4.1840e+02 ; | H - N - CA |
| 2627 | 2629 | 2630 | 1 | 1.0950e+02 | 4.1840e+02 ; | CA - CB - HB2 |

2627 2629 2631 1 1.0950e+02 4.1840e+02 ; CA - CB
- HB3
2627 2629 2632 1 1.0950e+02 3.3472e+02 ; CA - CB
- CG
2627 2645 2646 1 1.2040e+02 6.6944e+02 ; CA - C
- O
2627 2645 2647 1 1.1660e+02 5.8576e+02 ; CA - C
- N
2628 2627 2629 1 1.0950e+02 4.1840e+02 ; HA - CA
- CB
2628 2627 2645 1 1.0950e+02 4.1840e+02 ; HA - CA
- C
2629 2627 2645 1 1.1110e+02 5.2718e+02 ; CB - CA
- C
2629 2632 2633 1 1.0950e+02 4.1840e+02 ; CB - CG
- HG2
2629 2632 2634 1 1.0950e+02 4.1840e+02 ; CB - CG
- HG3
2629 2632 2635 1 1.0950e+02 3.3472e+02 ; CB - CG
- CD
2630 2629 2631 1 1.0950e+02 2.9288e+02 ; HB2 - CB
- HB3
2630 2629 2632 1 1.0950e+02 4.1840e+02 ; HB2 - CB
- CG
2631 2629 2632 1 1.0950e+02 4.1840e+02 ; HB3 - CB
- CG
2632 2635 2636 1 1.0950e+02 4.1840e+02 ; CG - CD
- HD2
2632 2635 2637 1 1.0950e+02 4.1840e+02 ; CG - CD
- HD3
2632 2635 2638 1 1.0950e+02 3.3472e+02 ; CG - CD
- CE
2633 2632 2634 1 1.0950e+02 2.9288e+02 ; HG2 - CG
- HG3
2633 2632 2635 1 1.0950e+02 4.1840e+02 ; HG2 - CG
- CD
2634 2632 2635 1 1.0950e+02 4.1840e+02 ; HG3 - CG
- CD
2635 2638 2639 1 1.0950e+02 4.1840e+02 ; CD - CE
- HE2
2635 2638 2640 1 1.0950e+02 4.1840e+02 ; CD - CE
- HE3
2635 2638 2641 1 1.1120e+02 6.6944e+02 ; CD - CE
- NZ
2636 2635 2637 1 1.0950e+02 2.9288e+02 ; HD2 - CD
- HD3
2636 2635 2638 1 1.0950e+02 4.1840e+02 ; HD2 - CD
- CE
2637 2635 2638 1 1.0950e+02 4.1840e+02 ; HD3 - CD
- CE
2638 2641 2642 1 1.0950e+02 4.1840e+02 ; CE - NZ
- HZ1

```
  2638   2641   2643      1    1.0950e+02    4.1840e+02 ;     CE - NZ
- HZ2
  2638   2641   2644      1    1.0950e+02    4.1840e+02 ;     CE - NZ
- HZ3
  2639   2638   2640      1    1.0950e+02    2.9288e+02 ;    HE2 - CE
- HE3
  2639   2638   2641      1    1.0950e+02    4.1840e+02 ;    HE2 - CE
- NZ
  2640   2638   2641      1    1.0950e+02    4.1840e+02 ;    HE3 - CE
- NZ
  2642   2641   2643      1    1.0950e+02    2.9288e+02 ;    HZ1 - NZ
- HZ2
  2642   2641   2644      1    1.0950e+02    2.9288e+02 ;    HZ1 - NZ
- HZ3
  2643   2641   2644      1    1.0950e+02    2.9288e+02 ;    HZ2 - NZ
- HZ3
  2645   2647   2648      1    1.2000e+02    4.1840e+02 ;      C - N
- H
  2645   2647   2649      1    1.2190e+02    4.1840e+02 ;      C - N
- CA
  2646   2645   2647      1    1.2290e+02    6.6944e+02 ;      O - C
- N
  2647   2649   2650      1    1.0950e+02    4.1840e+02 ;      N - CA
- HA
  2647   2649   2651      1    1.0970e+02    6.6944e+02 ;      N - CA
- CB
  2647   2649   2659      1    1.1010e+02    5.2718e+02 ;      N - CA
- C
  2648   2647   2649      1    1.1804e+02    4.1840e+02 ;      H - N
- CA
  2649   2651   2652      1    1.0950e+02    4.1840e+02 ;     CA - CB
- HB2
  2649   2651   2653      1    1.0950e+02    4.1840e+02 ;     CA - CB
- HB3
  2649   2651   2654      1    1.1110e+02    5.2718e+02 ;     CA - CB
- CG
  2649   2659   2660      1    1.2040e+02    6.6944e+02 ;     CA - C
- O
  2649   2659   2661      1    1.1660e+02    5.8576e+02 ;     CA - C
- N
  2650   2649   2651      1    1.0950e+02    4.1840e+02 ;     HA - CA
- CB
  2650   2649   2659      1    1.0950e+02    4.1840e+02 ;     HA - CA
- C
  2651   2649   2659      1    1.1110e+02    5.2718e+02 ;     CB - CA
- C
  2651   2654   2655      1    1.2040e+02    6.6944e+02 ;     CB - CG
- OD1
  2651   2654   2656      1    1.1660e+02    5.8576e+02 ;     CB - CG
- ND2
  2652   2651   2653      1    1.0950e+02    2.9288e+02 ;    HB2 - CB
- HB3
```

2652 2651 2654 1 1.0950e+02 4.1840e+02 ; HB2 - CB
- CG
2653 2651 2654 1 1.0950e+02 4.1840e+02 ; HB3 - CB
- CG
2654 2656 2657 1 1.2000e+02 4.1840e+02 ; CG - ND2
- HD21
2654 2656 2658 1 1.2000e+02 4.1840e+02 ; CG - ND2
- HD22
2655 2654 2656 1 1.2290e+02 6.6944e+02 ; OD1 - CG
- ND2
2657 2656 2658 1 1.2000e+02 2.9288e+02 ; HD21 - ND2
- HD22
2659 2661 2662 1 1.2000e+02 4.1840e+02 ; C - N
- H
2659 2661 2663 1 1.2190e+02 4.1840e+02 ; C - N
- CA
2660 2659 2661 1 1.2290e+02 6.6944e+02 ; O - C
- N
2661 2663 2664 1 1.0950e+02 4.1840e+02 ; N - CA
- HA
2661 2663 2665 1 1.0970e+02 6.6944e+02 ; N - CA
- CB
2661 2663 2678 1 1.1010e+02 5.2718e+02 ; N - CA
- C
2662 2661 2663 1 1.1804e+02 4.1840e+02 ; H - N
- CA
2663 2665 2666 1 1.0950e+02 4.1840e+02 ; CA - CB
- HB2
2663 2665 2667 1 1.0950e+02 4.1840e+02 ; CA - CB
- HB3
2663 2665 2668 1 1.0950e+02 3.3472e+02 ; CA - CB
- CG
2663 2678 2679 1 1.2040e+02 6.6944e+02 ; CA - C
- O
2663 2678 2680 1 1.1660e+02 5.8576e+02 ; CA - C
- N
2664 2663 2665 1 1.0950e+02 4.1840e+02 ; HA - CA
- CB
2664 2663 2678 1 1.0950e+02 4.1840e+02 ; HA - CA
- C
2665 2663 2678 1 1.1110e+02 5.2718e+02 ; CB - CA
- C
2665 2668 2669 1 1.0950e+02 4.1840e+02 ; CB - CG
- HG
2665 2668 2670 1 1.0950e+02 3.3472e+02 ; CB - CG
- CD1
2665 2668 2674 1 1.0950e+02 3.3472e+02 ; CB - CG
- CD2
2666 2665 2667 1 1.0950e+02 2.9288e+02 ; HB2 - CB
- HB3
2666 2665 2668 1 1.0950e+02 4.1840e+02 ; HB2 - CB
- CG

2667 2665 2668 1 1.0950e+02 4.1840e+02 ; HB3 - CB
- CG
2668 2670 2671 1 1.0950e+02 4.1840e+02 ; CG - CD1
- HD11
2668 2670 2672 1 1.0950e+02 4.1840e+02 ; CG - CD1
- HD12
2668 2670 2673 1 1.0950e+02 4.1840e+02 ; CG - CD1
- HD13
2668 2674 2675 1 1.0950e+02 4.1840e+02 ; CG - CD2
- HD21
2668 2674 2676 1 1.0950e+02 4.1840e+02 ; CG - CD2
- HD22
2668 2674 2677 1 1.0950e+02 4.1840e+02 ; CG - CD2
- HD23
2669 2668 2670 1 1.0950e+02 4.1840e+02 ; HG - CG
- CD1
2669 2668 2674 1 1.0950e+02 4.1840e+02 ; HG - CG
- CD2
2670 2668 2674 1 1.0950e+02 3.3472e+02 ; CD1 - CG
- CD2
2671 2670 2672 1 1.0950e+02 2.9288e+02 ; HD11 - CD1
- HD12
2671 2670 2673 1 1.0950e+02 2.9288e+02 ; HD11 - CD1
- HD13
2672 2670 2673 1 1.0950e+02 2.9288e+02 ; HD12 - CD1
- HD13
2675 2674 2676 1 1.0950e+02 2.9288e+02 ; HD21 - CD2
- HD22
2675 2674 2677 1 1.0950e+02 2.9288e+02 ; HD21 - CD2
- HD23
2676 2674 2677 1 1.0950e+02 2.9288e+02 ; HD22 - CD2
- HD23
2678 2680 2681 1 1.2000e+02 4.1840e+02 ; C - N
- H
2678 2680 2682 1 1.2190e+02 4.1840e+02 ; C - N
- CA
2679 2678 2680 1 1.2290e+02 6.6944e+02 ; O - C
- N
2680 2682 2683 1 1.0950e+02 4.1840e+02 ; N - CA
- HA2
2680 2682 2684 1 1.0950e+02 4.1840e+02 ; N - CA
- HA3
2680 2682 2685 1 1.1010e+02 5.2718e+02 ; N - CA
- C
2681 2680 2682 1 1.1804e+02 4.1840e+02 ; H - N
- CA
2682 2685 2686 1 1.2040e+02 6.6944e+02 ; CA - C
- O
2682 2685 2687 1 1.1660e+02 5.8576e+02 ; CA - C
- N
2683 2682 2684 1 1.0950e+02 2.9288e+02 ; HA2 - CA
- HA3

```
  2683   2682   2685       1    1.0950e+02    4.1840e+02 ;    HA2 - CA
- C
  2684   2682   2685       1    1.0950e+02    4.1840e+02 ;    HA3 - CA
- C
  2685   2687   2688       1    1.2000e+02    4.1840e+02 ;      C - N
- H
  2685   2687   2689       1    1.2190e+02    4.1840e+02 ;      C - N
- CA
  2686   2685   2687       1    1.2290e+02    6.6944e+02 ;      O - C
- N
  2687   2689   2690       1    1.0950e+02    4.1840e+02 ;      N - CA
- HA
  2687   2689   2691       1    1.0970e+02    6.6944e+02 ;      N - CA
- CB
  2687   2689   2699       1    1.1010e+02    5.2718e+02 ;      N - CA
- C
  2688   2687   2689       1    1.1804e+02    4.1840e+02 ;      H - N
- CA
  2689   2691   2692       1    1.0950e+02    4.1840e+02 ;     CA - CB
- HB
  2689   2691   2693       1    1.0950e+02    3.3472e+02 ;     CA - CB
- CG2
  2689   2691   2697       1    1.0950e+02    4.1840e+02 ;     CA - CB
- OG1
  2689   2699   2700       1    1.2040e+02    6.6944e+02 ;     CA - C
- O
  2689   2699   2701       1    1.1660e+02    5.8576e+02 ;     CA - C
- N
  2690   2689   2691       1    1.0950e+02    4.1840e+02 ;     HA - CA
- CB
  2690   2689   2699       1    1.0950e+02    4.1840e+02 ;     HA - CA
- C
  2691   2689   2699       1    1.1110e+02    5.2718e+02 ;     CB - CA
- C
  2691   2693   2694       1    1.0950e+02    4.1840e+02 ;     CB - CG2
- HG21
  2691   2693   2695       1    1.0950e+02    4.1840e+02 ;     CB - CG2
- HG22
  2691   2693   2696       1    1.0950e+02    4.1840e+02 ;     CB - CG2
- HG23
  2691   2697   2698       1    1.0850e+02    4.6024e+02 ;     CB - OG1
- HG1
  2692   2691   2693       1    1.0950e+02    4.1840e+02 ;     HB - CB
- CG2
  2692   2691   2697       1    1.0950e+02    4.1840e+02 ;     HB - CB
- OG1
  2693   2691   2697       1    1.0950e+02    4.1840e+02 ;    CG2 - CB
- OG1
  2694   2693   2695       1    1.0950e+02    2.9288e+02 ;   HG21 - CG2
- HG22
  2694   2693   2696       1    1.0950e+02    2.9288e+02 ;   HG21 - CG2
- HG23
```

2695 2693 2696 1 1.0950e+02 2.9288e+02 ; HG22 - CG2
- HG23
2699 2701 2702 1 1.2000e+02 4.1840e+02 ; C - N
- H
2699 2701 2703 1 1.2190e+02 4.1840e+02 ; C - N
- CA
2700 2699 2701 1 1.2290e+02 6.6944e+02 ; O - C
- N
2701 2703 2704 1 1.0950e+02 4.1840e+02 ; N - CA
- HA2
2701 2703 2705 1 1.0950e+02 4.1840e+02 ; N - CA
- HA3
2701 2703 2706 1 1.1010e+02 5.2718e+02 ; N - CA
- C
2702 2701 2703 1 1.1804e+02 4.1840e+02 ; H - N
- CA
2703 2706 2707 1 1.2040e+02 6.6944e+02 ; CA - C
- O
2703 2706 2708 1 1.1660e+02 5.8576e+02 ; CA - C
- N
2704 2703 2705 1 1.0950e+02 2.9288e+02 ; HA2 - CA
- HA3
2704 2703 2706 1 1.0950e+02 4.1840e+02 ; HA2 - CA
- C
2705 2703 2706 1 1.0950e+02 4.1840e+02 ; HA3 - CA
- C
2706 2708 2709 1 1.2000e+02 4.1840e+02 ; C - N
- H
2706 2708 2710 1 1.2190e+02 4.1840e+02 ; C - N
- CA
2707 2706 2708 1 1.2290e+02 6.6944e+02 ; O - C
- N
2708 2710 2711 1 1.0950e+02 4.1840e+02 ; N - CA
- HA
2708 2710 2712 1 1.0970e+02 6.6944e+02 ; N - CA
- CB
2708 2710 2721 1 1.1010e+02 5.2718e+02 ; N - CA
- C
2709 2708 2710 1 1.1804e+02 4.1840e+02 ; H - N
- CA
2710 2712 2713 1 1.0950e+02 4.1840e+02 ; CA - CB
- HB2
2710 2712 2714 1 1.0950e+02 4.1840e+02 ; CA - CB
- HB3
2710 2712 2715 1 1.0950e+02 3.3472e+02 ; CA - CB
- CG
2710 2721 2722 1 1.2040e+02 6.6944e+02 ; CA - C
- O
2710 2721 2723 1 1.1660e+02 5.8576e+02 ; CA - C
- N
2711 2710 2712 1 1.0950e+02 4.1840e+02 ; HA - CA
- CB

2711 2710 2721 1 1.0950e+02 4.1840e+02 ; HA - CA
- C
2712 2710 2721 1 1.1110e+02 5.2718e+02 ; CB - CA
- C
2712 2715 2716 1 1.0950e+02 4.1840e+02 ; CB - CG
- HG2
2712 2715 2717 1 1.0950e+02 4.1840e+02 ; CB - CG
- HG3
2712 2715 2718 1 1.1110e+02 5.2718e+02 ; CB - CG
- CD
2713 2712 2714 1 1.0950e+02 2.9288e+02 ; HB2 - CB
- HB3
2713 2712 2715 1 1.0950e+02 4.1840e+02 ; HB2 - CB
- CG
2714 2712 2715 1 1.0950e+02 4.1840e+02 ; HB3 - CB
- CG
2715 2718 2719 1 1.1700e+02 5.8576e+02 ; CG - CD
- OE1
2715 2718 2720 1 1.1700e+02 5.8576e+02 ; CG - CD
- OE2
2716 2715 2717 1 1.0950e+02 2.9288e+02 ; HG2 - CG
- HG3
2716 2715 2718 1 1.0950e+02 4.1840e+02 ; HG2 - CG
- CD
2717 2715 2718 1 1.0950e+02 4.1840e+02 ; HG3 - CG
- CD
2719 2718 2720 1 1.2600e+02 6.6944e+02 ; OE1 - CD
- OE2
2721 2723 2724 1 1.2000e+02 4.1840e+02 ; C - N
- H
2721 2723 2725 1 1.2190e+02 4.1840e+02 ; C - N
- CA
2722 2721 2723 1 1.2290e+02 6.6944e+02 ; O - C
- N
2723 2725 2726 1 1.0950e+02 4.1840e+02 ; N - CA
- HA
2723 2725 2727 1 1.0970e+02 6.6944e+02 ; N - CA
- CB
2723 2725 2741 1 1.1010e+02 5.2718e+02 ; N - CA
- C
2724 2723 2725 1 1.1804e+02 4.1840e+02 ; H - N
- CA
2725 2727 2728 1 1.0950e+02 4.1840e+02 ; CA - CB
- HB2
2725 2727 2729 1 1.0950e+02 4.1840e+02 ; CA - CB
- HB3
2725 2727 2730 1 1.1400e+02 5.2718e+02 ; CA - CB
- CG
2725 2741 2742 1 1.2040e+02 6.6944e+02 ; CA - C
- O
2725 2741 2743 1 1.1660e+02 5.8576e+02 ; CA - C
- N

2726 2725 2727 1 1.0950e+02 4.1840e+02 ; HA - CA
- CB
2726 2725 2741 1 1.0950e+02 4.1840e+02 ; HA - CA
- C
2727 2725 2741 1 1.1110e+02 5.2718e+02 ; CB - CA
- C
2727 2730 2731 1 1.2000e+02 5.8576e+02 ; CB - CG
- CD1
2727 2730 2739 1 1.2000e+02 5.8576e+02 ; CB - CG
- CD2
2728 2727 2729 1 1.0950e+02 2.9288e+02 ; HB2 - CB
- HB3
2728 2727 2730 1 1.0950e+02 4.1840e+02 ; HB2 - CB
- CG
2729 2727 2730 1 1.0950e+02 4.1840e+02 ; HB3 - CB
- CG
2730 2731 2732 1 1.2000e+02 4.1840e+02 ; CG - CD1
- HD1
2730 2731 2733 1 1.2000e+02 5.2718e+02 ; CG - CD1
- CE1
2730 2739 2737 1 1.2000e+02 5.2718e+02 ; CG - CD2
- CE2
2730 2739 2740 1 1.2000e+02 4.1840e+02 ; CG - CD2
- HD2
2731 2730 2739 1 1.2000e+02 5.2718e+02 ; CD1 - CG
- CD2
2731 2733 2734 1 1.2000e+02 4.1840e+02 ; CD1 - CE1
- HE1
2731 2733 2735 1 1.2000e+02 5.2718e+02 ; CD1 - CE1
- CZ
2732 2731 2733 1 1.2000e+02 4.1840e+02 ; HD1 - CD1
- CE1
2733 2735 2736 1 1.2000e+02 4.1840e+02 ; CE1 - CZ
- HZ
2733 2735 2737 1 1.2000e+02 5.2718e+02 ; CE1 - CZ
- CE2
2734 2733 2735 1 1.2000e+02 4.1840e+02 ; HE1 - CE1
- CZ
2735 2737 2738 1 1.2000e+02 4.1840e+02 ; CZ - CE2
- HE2
2735 2737 2739 1 1.2000e+02 5.2718e+02 ; CZ - CE2
- CD2
2736 2735 2737 1 1.2000e+02 4.1840e+02 ; HZ - CZ
- CE2
2737 2739 2740 1 1.2000e+02 4.1840e+02 ; CE2 - CD2
- HD2
2738 2737 2739 1 1.2000e+02 4.1840e+02 ; HE2 - CE2
- CD2
2741 2743 2744 1 1.2000e+02 4.1840e+02 ; C - N
- H
2741 2743 2745 1 1.2190e+02 4.1840e+02 ; C - N
- CA

2742 2741 2743 1 1.2290e+02 6.6944e+02 ; O - C
- N
2743 2745 2746 1 1.0950e+02 4.1840e+02 ; N - CA
- HA
2743 2745 2747 1 1.0970e+02 6.6944e+02 ; N - CA
- CB
2743 2745 2755 1 1.1010e+02 5.2718e+02 ; N - CA
- C
2744 2743 2745 1 1.1804e+02 4.1840e+02 ; H - N
- CA
2745 2747 2748 1 1.0950e+02 4.1840e+02 ; CA - CB
- HB
2745 2747 2749 1 1.0950e+02 3.3472e+02 ; CA - CB
- CG2
2745 2747 2753 1 1.0950e+02 4.1840e+02 ; CA - CB
- OG1
2745 2755 2756 1 1.2040e+02 6.6944e+02 ; CA - C
- O
2745 2755 2757 1 1.1660e+02 5.8576e+02 ; CA - C
- N
2746 2745 2747 1 1.0950e+02 4.1840e+02 ; HA - CA
- CB
2746 2745 2755 1 1.0950e+02 4.1840e+02 ; HA - CA
- C
2747 2745 2755 1 1.1110e+02 5.2718e+02 ; CB - CA
- C
2747 2749 2750 1 1.0950e+02 4.1840e+02 ; CB - CG2
- HG21
2747 2749 2751 1 1.0950e+02 4.1840e+02 ; CB - CG2
- HG22
2747 2749 2752 1 1.0950e+02 4.1840e+02 ; CB - CG2
- HG23
2747 2753 2754 1 1.0850e+02 4.6024e+02 ; CB - OG1
- HG1
2748 2747 2749 1 1.0950e+02 4.1840e+02 ; HB - CB
- CG2
2748 2747 2753 1 1.0950e+02 4.1840e+02 ; HB - CB
- OG1
2749 2747 2753 1 1.0950e+02 4.1840e+02 ; CG2 - CB
- OG1
2750 2749 2751 1 1.0950e+02 2.9288e+02 ; HG21 - CG2
- HG22
2750 2749 2752 1 1.0950e+02 2.9288e+02 ; HG21 - CG2
- HG23
2751 2749 2752 1 1.0950e+02 2.9288e+02 ; HG22 - CG2
- HG23
2755 2757 2758 1 1.2000e+02 4.1840e+02 ; C - N
- H
2755 2757 2759 1 1.2190e+02 4.1840e+02 ; C - N
- CA
2756 2755 2757 1 1.2290e+02 6.6944e+02 ; O - C
- N

2757 2759 2760 1 1.0950e+02 4.1840e+02 ; N - CA
- HA
2757 2759 2761 1 1.0970e+02 6.6944e+02 ; N - CA
- CB
2757 2759 2769 1 1.1010e+02 5.2718e+02 ; N - CA
- C
2758 2757 2759 1 1.1804e+02 4.1840e+02 ; H - N
- CA
2759 2761 2762 1 1.0950e+02 4.1840e+02 ; CA - CB
- HB
2759 2761 2763 1 1.0950e+02 3.3472e+02 ; CA - CB
- CG2
2759 2761 2767 1 1.0950e+02 4.1840e+02 ; CA - CB
- OG1
2759 2769 2770 1 1.2040e+02 6.6944e+02 ; CA - C
- O
2759 2769 2771 1 1.1660e+02 5.8576e+02 ; CA - C
- N
2760 2759 2761 1 1.0950e+02 4.1840e+02 ; HA - CA
- CB
2760 2759 2769 1 1.0950e+02 4.1840e+02 ; HA - CA
- C
2761 2759 2769 1 1.1110e+02 5.2718e+02 ; CB - CA
- C
2761 2763 2764 1 1.0950e+02 4.1840e+02 ; CB - CG2
- HG21
2761 2763 2765 1 1.0950e+02 4.1840e+02 ; CB - CG2
- HG22
2761 2763 2766 1 1.0950e+02 4.1840e+02 ; CB - CG2
- HG23
2761 2767 2768 1 1.0850e+02 4.6024e+02 ; CB - OG1
- HG1
2762 2761 2763 1 1.0950e+02 4.1840e+02 ; HB - CB
- CG2
2762 2761 2767 1 1.0950e+02 4.1840e+02 ; HB - CB
- OG1
2763 2761 2767 1 1.0950e+02 4.1840e+02 ; CG2 - CB
- OG1
2764 2763 2765 1 1.0950e+02 2.9288e+02 ; HG21 - CG2
- HG22
2764 2763 2766 1 1.0950e+02 2.9288e+02 ; HG21 - CG2
- HG23
2765 2763 2766 1 1.0950e+02 2.9288e+02 ; HG22 - CG2
- HG23
2769 2771 2772 1 1.2000e+02 4.1840e+02 ; C - N
- H
2769 2771 2773 1 1.2190e+02 4.1840e+02 ; C - N
- CA
2770 2769 2771 1 1.2290e+02 6.6944e+02 ; O - C
- N
2771 2773 2774 1 1.0950e+02 4.1840e+02 ; N - CA
- HA

2771 2773 2775 1 1.0970e+02 6.6944e+02 ; N - CA
- CB
2771 2773 2783 1 1.1010e+02 5.2718e+02 ; N - CA
- C
2772 2771 2773 1 1.1804e+02 4.1840e+02 ; H - N
- CA
2773 2775 2776 1 1.0950e+02 4.1840e+02 ; CA - CB
- HB
2773 2775 2777 1 1.0950e+02 3.3472e+02 ; CA - CB
- CG2
2773 2775 2781 1 1.0950e+02 4.1840e+02 ; CA - CB
- OG1
2773 2783 2784 1 1.2040e+02 6.6944e+02 ; CA - C
- O
2773 2783 2785 1 1.1660e+02 5.8576e+02 ; CA - C
- N
2774 2773 2775 1 1.0950e+02 4.1840e+02 ; HA - CA
- CB
2774 2773 2783 1 1.0950e+02 4.1840e+02 ; HA - CA
- C
2775 2773 2783 1 1.1110e+02 5.2718e+02 ; CB - CA
- C
2775 2777 2778 1 1.0950e+02 4.1840e+02 ; CB - CG2
- HG21
2775 2777 2779 1 1.0950e+02 4.1840e+02 ; CB - CG2
- HG22
2775 2777 2780 1 1.0950e+02 4.1840e+02 ; CB - CG2
- HG23
2775 2781 2782 1 1.0850e+02 4.6024e+02 ; CB - OG1
- HG1
2776 2775 2777 1 1.0950e+02 4.1840e+02 ; HB - CB
- CG2
2776 2775 2781 1 1.0950e+02 4.1840e+02 ; HB - CB
- OG1
2777 2775 2781 1 1.0950e+02 4.1840e+02 ; CG2 - CB
- OG1
2778 2777 2779 1 1.0950e+02 2.9288e+02 ; HG21 - CG2
- HG22
2778 2777 2780 1 1.0950e+02 2.9288e+02 ; HG21 - CG2
- HG23
2779 2777 2780 1 1.0950e+02 2.9288e+02 ; HG22 - CG2
- HG23
2783 2785 2786 1 1.2000e+02 4.1840e+02 ; C - N
- H
2783 2785 2787 1 1.2190e+02 4.1840e+02 ; C - N
- CA
2784 2783 2785 1 1.2290e+02 6.6944e+02 ; O - C
- N
2785 2787 2788 1 1.0950e+02 4.1840e+02 ; N - CA
- HA
2785 2787 2789 1 1.0970e+02 6.6944e+02 ; N - CA
- CB

2785 2787 2805 1 1.1010e+02 5.2718e+02 ; N - CA
- C
2786 2785 2787 1 1.1804e+02 4.1840e+02 ; H - N
- CA
2787 2789 2790 1 1.0950e+02 4.1840e+02 ; CA - CB
- HB2
2787 2789 2791 1 1.0950e+02 4.1840e+02 ; CA - CB
- HB3
2787 2789 2792 1 1.0950e+02 3.3472e+02 ; CA - CB
- CG
2787 2805 2806 1 1.2040e+02 6.6944e+02 ; CA - C
- O
2787 2805 2807 1 1.1660e+02 5.8576e+02 ; CA - C
- N
2788 2787 2789 1 1.0950e+02 4.1840e+02 ; HA - CA
- CB
2788 2787 2805 1 1.0950e+02 4.1840e+02 ; HA - CA
- C
2789 2787 2805 1 1.1110e+02 5.2718e+02 ; CB - CA
- C
2789 2792 2793 1 1.0950e+02 4.1840e+02 ; CB - CG
- HG2
2789 2792 2794 1 1.0950e+02 4.1840e+02 ; CB - CG
- HG3
2789 2792 2795 1 1.0950e+02 3.3472e+02 ; CB - CG
- CD
2790 2789 2791 1 1.0950e+02 2.9288e+02 ; HB2 - CB
- HB3
2790 2789 2792 1 1.0950e+02 4.1840e+02 ; HB2 - CB
- CG
2791 2789 2792 1 1.0950e+02 4.1840e+02 ; HB3 - CB
- CG
2792 2795 2796 1 1.0950e+02 4.1840e+02 ; CG - CD
- HD2
2792 2795 2797 1 1.0950e+02 4.1840e+02 ; CG - CD
- HD3
2792 2795 2798 1 1.0950e+02 3.3472e+02 ; CG - CD
- CE
2793 2792 2794 1 1.0950e+02 2.9288e+02 ; HG2 - CG
- HG3
2793 2792 2795 1 1.0950e+02 4.1840e+02 ; HG2 - CG
- CD
2794 2792 2795 1 1.0950e+02 4.1840e+02 ; HG3 - CG
- CD
2795 2798 2799 1 1.0950e+02 4.1840e+02 ; CD - CE
- HE2
2795 2798 2800 1 1.0950e+02 4.1840e+02 ; CD - CE
- HE3
2795 2798 2801 1 1.1120e+02 6.6944e+02 ; CD - CE
- NZ
2796 2795 2797 1 1.0950e+02 2.9288e+02 ; HD2 - CD
- HD3

2796 2795 2798 1 1.0950e+02 4.1840e+02 ; HD2 - CD
- CE
2797 2795 2798 1 1.0950e+02 4.1840e+02 ; HD3 - CD
- CE
2798 2801 2802 1 1.0950e+02 4.1840e+02 ; CE - NZ
- HZ1
2798 2801 2803 1 1.0950e+02 4.1840e+02 ; CE - NZ
- HZ2
2798 2801 2804 1 1.0950e+02 4.1840e+02 ; CE - NZ
- HZ3
2799 2798 2800 1 1.0950e+02 2.9288e+02 ; HE2 - CE
- HE3
2799 2798 2801 1 1.0950e+02 4.1840e+02 ; HE2 - CE
- NZ
2800 2798 2801 1 1.0950e+02 4.1840e+02 ; HE3 - CE
- NZ
2802 2801 2803 1 1.0950e+02 2.9288e+02 ; HZ1 - NZ
- HZ2
2802 2801 2804 1 1.0950e+02 2.9288e+02 ; HZ1 - NZ
- HZ3
2803 2801 2804 1 1.0950e+02 2.9288e+02 ; HZ2 - NZ
- HZ3
2805 2807 2808 1 1.2000e+02 4.1840e+02 ; C - N
- H
2805 2807 2809 1 1.2190e+02 4.1840e+02 ; C - N
- CA
2806 2805 2807 1 1.2290e+02 6.6944e+02 ; O - C
- N
2807 2809 2810 1 1.0950e+02 4.1840e+02 ; N - CA
- HA
2807 2809 2811 1 1.0970e+02 6.6944e+02 ; N - CA
- CB
2807 2809 2815 1 1.1010e+02 5.2718e+02 ; N - CA
- C
2808 2807 2809 1 1.1804e+02 4.1840e+02 ; H - N
- CA
2809 2811 2812 1 1.0950e+02 4.1840e+02 ; CA - CB
- HB1
2809 2811 2813 1 1.0950e+02 4.1840e+02 ; CA - CB
- HB2
2809 2811 2814 1 1.0950e+02 4.1840e+02 ; CA - CB
- HB3
2809 2815 2816 1 1.2040e+02 6.6944e+02 ; CA - C
- O
2809 2815 2817 1 1.1660e+02 5.8576e+02 ; CA - C
- N
2810 2809 2811 1 1.0950e+02 4.1840e+02 ; HA - CA
- CB
2810 2809 2815 1 1.0950e+02 4.1840e+02 ; HA - CA
- C
2811 2809 2815 1 1.1110e+02 5.2718e+02 ; CB - CA
- C

2812 2811 2813 1 1.0950e+02 2.9288e+02 ; HB1 - CB
- HB2
2812 2811 2814 1 1.0950e+02 2.9288e+02 ; HB1 - CB
- HB3
2813 2811 2814 1 1.0950e+02 2.9288e+02 ; HB2 - CB
- HB3
2815 2817 2818 1 1.2000e+02 4.1840e+02 ; C - N
- H
2815 2817 2819 1 1.2190e+02 4.1840e+02 ; C - N
- CA
2816 2815 2817 1 1.2290e+02 6.6944e+02 ; O - C
- N
2817 2819 2820 1 1.0950e+02 4.1840e+02 ; N - CA
- HA2
2817 2819 2821 1 1.0950e+02 4.1840e+02 ; N - CA
- HA3
2817 2819 2822 1 1.1010e+02 5.2718e+02 ; N - CA
- C
2818 2817 2819 1 1.1804e+02 4.1840e+02 ; H - N
- CA
2819 2822 2823 1 1.2040e+02 6.6944e+02 ; CA - C
- O
2819 2822 2824 1 1.1660e+02 5.8576e+02 ; CA - C
- N
2820 2819 2821 1 1.0950e+02 2.9288e+02 ; HA2 - CA
- HA3
2820 2819 2822 1 1.0950e+02 4.1840e+02 ; HA2 - CA
- C
2821 2819 2822 1 1.0950e+02 4.1840e+02 ; HA3 - CA
- C
2822 2824 2825 1 1.2000e+02 4.1840e+02 ; C - N
- H
2822 2824 2826 1 1.2190e+02 4.1840e+02 ; C - N
- CA
2823 2822 2824 1 1.2290e+02 6.6944e+02 ; O - C
- N
2824 2826 2827 1 1.0950e+02 4.1840e+02 ; N - CA
- HA
2824 2826 2828 1 1.0970e+02 6.6944e+02 ; N - CA
- CB
2824 2826 2836 1 1.1010e+02 5.2718e+02 ; N - CA
- C
2825 2824 2826 1 1.1804e+02 4.1840e+02 ; H - N
- CA
2826 2828 2829 1 1.0950e+02 4.1840e+02 ; CA - CB
- HB
2826 2828 2830 1 1.0950e+02 3.3472e+02 ; CA - CB
- CG2
2826 2828 2834 1 1.0950e+02 4.1840e+02 ; CA - CB
- OG1
2826 2836 2837 1 1.2040e+02 6.6944e+02 ; CA - C
- O

2826 2836 2838 1 1.1660e+02 5.8576e+02 ; CA - C
- N
2827 2826 2828 1 1.0950e+02 4.1840e+02 ; HA - CA
- CB
2827 2826 2836 1 1.0950e+02 4.1840e+02 ; HA - CA
- C
2828 2826 2836 1 1.1110e+02 5.2718e+02 ; CB - CA
- C
2828 2830 2831 1 1.0950e+02 4.1840e+02 ; CB - CG2
- HG21
2828 2830 2832 1 1.0950e+02 4.1840e+02 ; CB - CG2
- HG22
2828 2830 2833 1 1.0950e+02 4.1840e+02 ; CB - CG2
- HG23
2828 2834 2835 1 1.0850e+02 4.6024e+02 ; CB - OG1
- HG1
2829 2828 2830 1 1.0950e+02 4.1840e+02 ; HB - CB
- CG2
2829 2828 2834 1 1.0950e+02 4.1840e+02 ; HB - CB
- OG1
2830 2828 2834 1 1.0950e+02 4.1840e+02 ; CG2 - CB
- OG1
2831 2830 2832 1 1.0950e+02 2.9288e+02 ; HG21 - CG2
- HG22
2831 2830 2833 1 1.0950e+02 2.9288e+02 ; HG21 - CG2
- HG23
2832 2830 2833 1 1.0950e+02 2.9288e+02 ; HG22 - CG2
- HG23
2836 2838 2839 1 1.2190e+02 4.1840e+02 ; C - N
- CD
2836 2838 2848 1 1.2190e+02 4.1840e+02 ; C - N
- CA
2837 2836 2838 1 1.2290e+02 6.6944e+02 ; O - C
- N
2838 2839 2840 1 1.0950e+02 4.1840e+02 ; N - CD
- HD2
2838 2839 2841 1 1.0950e+02 4.1840e+02 ; N - CD
- HD3
2838 2839 2842 1 1.0970e+02 6.6944e+02 ; N - CD
- CG
2838 2848 2845 1 1.0970e+02 6.6944e+02 ; N - CA
- CB
2838 2848 2849 1 1.0950e+02 4.1840e+02 ; N - CA
- HA
2838 2848 2850 1 1.1010e+02 5.2718e+02 ; N - CA
- C
2839 2838 2848 1 1.1800e+02 4.1840e+02 ; CD - N
- CA
2839 2842 2843 1 1.0950e+02 4.1840e+02 ; CD - CG
- HG2
2839 2842 2844 1 1.0950e+02 4.1840e+02 ; CD - CG
- HG3

2839 2842 2845 1 1.0950e+02 3.3472e+02 ; CD - CG - CB
2840 2839 2841 1 1.0950e+02 2.9288e+02 ; HD2 - CD - HD3
2840 2839 2842 1 1.0950e+02 4.1840e+02 ; HD2 - CD - CG
2841 2839 2842 1 1.0950e+02 4.1840e+02 ; HD3 - CD - CG
2842 2845 2846 1 1.0950e+02 4.1840e+02 ; CG - CB - HB2
2842 2845 2847 1 1.0950e+02 4.1840e+02 ; CG - CB - HB3
2842 2845 2848 1 1.0950e+02 3.3472e+02 ; CG - CB - CA
2843 2842 2844 1 1.0950e+02 2.9288e+02 ; HG2 - CG - HG3
2843 2842 2845 1 1.0950e+02 4.1840e+02 ; HG2 - CG - CB
2844 2842 2845 1 1.0950e+02 4.1840e+02 ; HG3 - CG - CB
2845 2848 2849 1 1.0950e+02 4.1840e+02 ; CB - CA - HA
2845 2848 2850 1 1.1110e+02 5.2718e+02 ; CB - CA - C
2846 2845 2847 1 1.0950e+02 2.9288e+02 ; HB2 - CB - HB3
2846 2845 2848 1 1.0950e+02 4.1840e+02 ; HB2 - CB - CA
2847 2845 2848 1 1.0950e+02 4.1840e+02 ; HB3 - CB - CA
2848 2850 2851 1 1.2040e+02 6.6944e+02 ; CA - C - O
2848 2850 2852 1 1.1660e+02 5.8576e+02 ; CA - C - N
2849 2848 2850 1 1.0950e+02 4.1840e+02 ; HA - CA - C
2850 2852 2853 1 1.2000e+02 4.1840e+02 ; C - N - H
2850 2852 2854 1 1.2190e+02 4.1840e+02 ; C - N - CA
2851 2850 2852 1 1.2290e+02 6.6944e+02 ; O - C - N
2852 2854 2855 1 1.0950e+02 4.1840e+02 ; N - CA - HA
2852 2854 2856 1 1.0970e+02 6.6944e+02 ; N - CA - CB
2852 2854 2871 1 1.1010e+02 5.2718e+02 ; N - CA - C
2853 2852 2854 1 1.1804e+02 4.1840e+02 ; H - N - CA
2854 2856 2857 1 1.0950e+02 4.1840e+02 ; CA - CB - HB2

2854 2856 2858 1 1.0950e+02 4.1840e+02 ; CA - CB
- HB3
2854 2856 2859 1 1.1400e+02 5.2718e+02 ; CA - CB
- CG
2854 2871 2872 1 1.2040e+02 6.6944e+02 ; CA - C
- O
2854 2871 2873 1 1.1660e+02 5.8576e+02 ; CA - C
- N
2855 2854 2856 1 1.0950e+02 4.1840e+02 ; HA - CA
- CB
2855 2854 2871 1 1.0950e+02 4.1840e+02 ; HA - CA
- C
2856 2854 2871 1 1.1110e+02 5.2718e+02 ; CB - CA
- C
2856 2859 2860 1 1.2000e+02 5.8576e+02 ; CB - CG
- CD1
2856 2859 2869 1 1.2000e+02 5.8576e+02 ; CB - CG
- CD2
2857 2856 2858 1 1.0950e+02 2.9288e+02 ; HB2 - CB
- HB3
2857 2856 2859 1 1.0950e+02 4.1840e+02 ; HB2 - CB
- CG
2858 2856 2859 1 1.0950e+02 4.1840e+02 ; HB3 - CB
- CG
2859 2860 2861 1 1.2000e+02 4.1840e+02 ; CG - CD1
- HD1
2859 2860 2862 1 1.2000e+02 5.2718e+02 ; CG - CD1
- CE1
2859 2869 2867 1 1.2000e+02 5.2718e+02 ; CG - CD2
- CE2
2859 2869 2870 1 1.2000e+02 4.1840e+02 ; CG - CD2
- HD2
2860 2859 2869 1 1.2000e+02 5.2718e+02 ; CD1 - CG
- CD2
2860 2862 2863 1 1.2000e+02 4.1840e+02 ; CD1 - CE1
- HE1
2860 2862 2864 1 1.2000e+02 5.2718e+02 ; CD1 - CE1
- CZ
2861 2860 2862 1 1.2000e+02 4.1840e+02 ; HD1 - CD1
- CE1
2862 2864 2865 1 1.2000e+02 5.8576e+02 ; CE1 - CZ
- OH
2862 2864 2867 1 1.2000e+02 5.2718e+02 ; CE1 - CZ
- CE2
2863 2862 2864 1 1.2000e+02 4.1840e+02 ; HE1 - CE1
- CZ
2864 2865 2866 1 1.1300e+02 4.1840e+02 ; CZ - OH
- HH
2864 2867 2868 1 1.2000e+02 4.1840e+02 ; CZ - CE2
- HE2
2864 2867 2869 1 1.2000e+02 5.2718e+02 ; CZ - CE2
- CD2

2865 2864 2867 1 1.2000e+02 5.8576e+02 ; OH - CZ
- CE2
2867 2869 2870 1 1.2000e+02 4.1840e+02 ; CE2 - CD2
- HD2
2868 2867 2869 1 1.2000e+02 4.1840e+02 ; HE2 - CE2
- CD2
2871 2873 2874 1 1.2000e+02 4.1840e+02 ; C - N
- H
2871 2873 2875 1 1.2190e+02 4.1840e+02 ; C - N
- CA
2872 2871 2873 1 1.2290e+02 6.6944e+02 ; O - C
- N
2873 2875 2876 1 1.0950e+02 4.1840e+02 ; N - CA
- HA
2873 2875 2877 1 1.0970e+02 6.6944e+02 ; N - CA
- CB
2873 2875 2892 1 1.1010e+02 5.2718e+02 ; N - CA
- C
2874 2873 2875 1 1.1804e+02 4.1840e+02 ; H - N
- CA
2875 2877 2878 1 1.0950e+02 4.1840e+02 ; CA - CB
- HB2
2875 2877 2879 1 1.0950e+02 4.1840e+02 ; CA - CB
- HB3
2875 2877 2880 1 1.1400e+02 5.2718e+02 ; CA - CB
- CG
2875 2892 2893 1 1.2040e+02 6.6944e+02 ; CA - C
- O
2875 2892 2894 1 1.1660e+02 5.8576e+02 ; CA - C
- N
2876 2875 2877 1 1.0950e+02 4.1840e+02 ; HA - CA
- CB
2876 2875 2892 1 1.0950e+02 4.1840e+02 ; HA - CA
- C
2877 2875 2892 1 1.1110e+02 5.2718e+02 ; CB - CA
- C
2877 2880 2881 1 1.2000e+02 5.8576e+02 ; CB - CG
- CD1
2877 2880 2890 1 1.2000e+02 5.8576e+02 ; CB - CG
- CD2
2878 2877 2879 1 1.0950e+02 2.9288e+02 ; HB2 - CB
- HB3
2878 2877 2880 1 1.0950e+02 4.1840e+02 ; HB2 - CB
- CG
2879 2877 2880 1 1.0950e+02 4.1840e+02 ; HB3 - CB
- CG
2880 2881 2882 1 1.2000e+02 4.1840e+02 ; CG - CD1
- HD1
2880 2881 2883 1 1.2000e+02 5.2718e+02 ; CG - CD1
- CE1
2880 2890 2888 1 1.2000e+02 5.2718e+02 ; CG - CD2
- CE2

```
  2880   2890   2891      1    1.2000e+02    4.1840e+02 ;     CG - CD2
- HD2
  2881   2880   2890      1    1.2000e+02    5.2718e+02 ;    CD1 - CG
- CD2
  2881   2883   2884      1    1.2000e+02    4.1840e+02 ;    CD1 - CE1
- HE1
  2881   2883   2885      1    1.2000e+02    5.2718e+02 ;    CD1 - CE1
- CZ
  2882   2881   2883      1    1.2000e+02    4.1840e+02 ;    HD1 - CD1
- CE1
  2883   2885   2886      1    1.2000e+02    5.8576e+02 ;    CE1 - CZ
- OH
  2883   2885   2888      1    1.2000e+02    5.2718e+02 ;    CE1 - CZ
- CE2
  2884   2883   2885      1    1.2000e+02    4.1840e+02 ;    HE1 - CE1
- CZ
  2885   2886   2887      1    1.1300e+02    4.1840e+02 ;     CZ - OH
- HH
  2885   2888   2889      1    1.2000e+02    4.1840e+02 ;     CZ - CE2
- HE2
  2885   2888   2890      1    1.2000e+02    5.2718e+02 ;     CZ - CE2
- CD2
  2886   2885   2888      1    1.2000e+02    5.8576e+02 ;     OH - CZ
- CE2
  2888   2890   2891      1    1.2000e+02    4.1840e+02 ;    CE2 - CD2
- HD2
  2889   2888   2890      1    1.2000e+02    4.1840e+02 ;    HE2 - CE2
- CD2
  2892   2894   2895      1    1.2000e+02    4.1840e+02 ;      C - N
- H
  2892   2894   2896      1    1.2190e+02    4.1840e+02 ;      C - N
- CA
  2893   2892   2894      1    1.2290e+02    6.6944e+02 ;      O - C
- N
  2894   2896   2897      1    1.0950e+02    4.1840e+02 ;      N - CA
- HA
  2894   2896   2898      1    1.0970e+02    6.6944e+02 ;      N - CA
- CB
  2894   2896   2908      1    1.1010e+02    5.2718e+02 ;      N - CA
- C
  2895   2894   2896      1    1.1804e+02    4.1840e+02 ;      H - N
- CA
  2896   2898   2899      1    1.0950e+02    4.1840e+02 ;     CA - CB
- HB
  2896   2898   2900      1    1.0950e+02    3.3472e+02 ;     CA - CB
- CG1
  2896   2898   2904      1    1.0950e+02    3.3472e+02 ;     CA - CB
- CG2
  2896   2908   2909      1    1.2040e+02    6.6944e+02 ;     CA - C
- O
  2896   2908   2910      1    1.1660e+02    5.8576e+02 ;     CA - C
- N
```

2897 2896 2898 1 1.0950e+02 4.1840e+02 ; HA - CA
- CB
2897 2896 2908 1 1.0950e+02 4.1840e+02 ; HA - CA
- C
2898 2896 2908 1 1.1110e+02 5.2718e+02 ; CB - CA
- C
2898 2900 2901 1 1.0950e+02 4.1840e+02 ; CB - CG1
- HG11
2898 2900 2902 1 1.0950e+02 4.1840e+02 ; CB - CG1
- HG12
2898 2900 2903 1 1.0950e+02 4.1840e+02 ; CB - CG1
- HG13
2898 2904 2905 1 1.0950e+02 4.1840e+02 ; CB - CG2
- HG21
2898 2904 2906 1 1.0950e+02 4.1840e+02 ; CB - CG2
- HG22
2898 2904 2907 1 1.0950e+02 4.1840e+02 ; CB - CG2
- HG23
2899 2898 2900 1 1.0950e+02 4.1840e+02 ; HB - CB
- CG1
2899 2898 2904 1 1.0950e+02 4.1840e+02 ; HB - CB
- CG2
2900 2898 2904 1 1.0950e+02 3.3472e+02 ; CG1 - CB
- CG2
2901 2900 2902 1 1.0950e+02 2.9288e+02 ; HG11 - CG1
- HG12
2901 2900 2903 1 1.0950e+02 2.9288e+02 ; HG11 - CG1
- HG13
2902 2900 2903 1 1.0950e+02 2.9288e+02 ; HG12 - CG1
- HG13
2905 2904 2906 1 1.0950e+02 2.9288e+02 ; HG21 - CG2
- HG22
2905 2904 2907 1 1.0950e+02 2.9288e+02 ; HG21 - CG2
- HG23
2906 2904 2907 1 1.0950e+02 2.9288e+02 ; HG22 - CG2
- HG23
2908 2910 2911 1 1.2000e+02 4.1840e+02 ; C - N
- H
2908 2910 2912 1 1.2190e+02 4.1840e+02 ; C - N
- CA
2909 2908 2910 1 1.2290e+02 6.6944e+02 ; O - C
- N
2910 2912 2913 1 1.0950e+02 4.1840e+02 ; N - CA
- HA
2910 2912 2914 1 1.0970e+02 6.6944e+02 ; N - CA
- CB
2910 2912 2918 1 1.1010e+02 5.2718e+02 ; N - CA
- C
2911 2910 2912 1 1.1804e+02 4.1840e+02 ; H - N
- CA
2912 2914 2915 1 1.0950e+02 4.1840e+02 ; CA - CB
- HB1

```
  2912   2914   2916      1    1.0950e+02    4.1840e+02 ;     CA - CB
- HB2
  2912   2914   2917      1    1.0950e+02    4.1840e+02 ;     CA - CB
- HB3
  2912   2918   2919      1    1.2040e+02    6.6944e+02 ;     CA - C
- O
  2912   2918   2920      1    1.1660e+02    5.8576e+02 ;     CA - C
- N
  2913   2912   2914      1    1.0950e+02    4.1840e+02 ;     HA - CA
- CB
  2913   2912   2918      1    1.0950e+02    4.1840e+02 ;     HA - CA
- C
  2914   2912   2918      1    1.1110e+02    5.2718e+02 ;     CB - CA
- C
  2915   2914   2916      1    1.0950e+02    2.9288e+02 ;    HB1 - CB
- HB2
  2915   2914   2917      1    1.0950e+02    2.9288e+02 ;    HB1 - CB
- HB3
  2916   2914   2917      1    1.0950e+02    2.9288e+02 ;    HB2 - CB
- HB3
  2918   2920   2921      1    1.2190e+02    4.1840e+02 ;      C - N
- CD
  2918   2920   2930      1    1.2190e+02    4.1840e+02 ;      C - N
- CA
  2919   2918   2920      1    1.2290e+02    6.6944e+02 ;      O - C
- N
  2920   2921   2922      1    1.0950e+02    4.1840e+02 ;      N - CD
- HD2
  2920   2921   2923      1    1.0950e+02    4.1840e+02 ;      N - CD
- HD3
  2920   2921   2924      1    1.0970e+02    6.6944e+02 ;      N - CD
- CG
  2920   2930   2927      1    1.0970e+02    6.6944e+02 ;      N - CA
- CB
  2920   2930   2931      1    1.0950e+02    4.1840e+02 ;      N - CA
- HA
  2920   2930   2932      1    1.1010e+02    5.2718e+02 ;      N - CA
- C
  2921   2920   2930      1    1.1800e+02    4.1840e+02 ;     CD - N
- CA
  2921   2924   2925      1    1.0950e+02    4.1840e+02 ;     CD - CG
- HG2
  2921   2924   2926      1    1.0950e+02    4.1840e+02 ;     CD - CG
- HG3
  2921   2924   2927      1    1.0950e+02    3.3472e+02 ;     CD - CG
- CB
  2922   2921   2923      1    1.0950e+02    2.9288e+02 ;    HD2 - CD
- HD3
  2922   2921   2924      1    1.0950e+02    4.1840e+02 ;    HD2 - CD
- CG
  2923   2921   2924      1    1.0950e+02    4.1840e+02 ;    HD3 - CD
- CG
```

2924 2927 2928 1 1.0950e+02 4.1840e+02 ; CG - CB
- HB2
2924 2927 2929 1 1.0950e+02 4.1840e+02 ; CG - CB
- HB3
2924 2927 2930 1 1.0950e+02 3.3472e+02 ; CG - CB
- CA
2925 2924 2926 1 1.0950e+02 2.9288e+02 ; HG2 - CG
- HG3
2925 2924 2927 1 1.0950e+02 4.1840e+02 ; HG2 - CG
- CB
2926 2924 2927 1 1.0950e+02 4.1840e+02 ; HG3 - CG
- CB
2927 2930 2931 1 1.0950e+02 4.1840e+02 ; CB - CA
- HA
2927 2930 2932 1 1.1110e+02 5.2718e+02 ; CB - CA
- C
2928 2927 2929 1 1.0950e+02 2.9288e+02 ; HB2 - CB
- HB3
2928 2927 2930 1 1.0950e+02 4.1840e+02 ; HB2 - CB
- CA
2929 2927 2930 1 1.0950e+02 4.1840e+02 ; HB3 - CB
- CA
2930 2932 2933 1 1.2040e+02 6.6944e+02 ; CA - C
- O
2930 2932 2934 1 1.1660e+02 5.8576e+02 ; CA - C
- N
2931 2930 2932 1 1.0950e+02 4.1840e+02 ; HA - CA
- C
2932 2934 2935 1 1.2000e+02 4.1840e+02 ; C - N
- H
2932 2934 2936 1 1.2190e+02 4.1840e+02 ; C - N
- CA
2933 2932 2934 1 1.2290e+02 6.6944e+02 ; O - C
- N
2934 2936 2937 1 1.0950e+02 4.1840e+02 ; N - CA
- HA
2934 2936 2938 1 1.0970e+02 6.6944e+02 ; N - CA
- CB
2934 2936 2949 1 1.1010e+02 5.2718e+02 ; N - CA
- C
2935 2934 2936 1 1.1804e+02 4.1840e+02 ; H - N
- CA
2936 2938 2939 1 1.0950e+02 4.1840e+02 ; CA - CB
- HB2
2936 2938 2940 1 1.0950e+02 4.1840e+02 ; CA - CB
- HB3
2936 2938 2941 1 1.0950e+02 3.3472e+02 ; CA - CB
- CG
2936 2949 2950 1 1.2040e+02 6.6944e+02 ; CA - C
- O
2936 2949 2951 1 1.1660e+02 5.8576e+02 ; CA - C
- N

2937 2936 2938 1 1.0950e+02 4.1840e+02 ; HA - CA
- CB
2937 2936 2949 1 1.0950e+02 4.1840e+02 ; HA - CA
- C
2938 2936 2949 1 1.1110e+02 5.2718e+02 ; CB - CA
- C
2938 2941 2942 1 1.0950e+02 4.1840e+02 ; CB - CG
- HG2
2938 2941 2943 1 1.0950e+02 4.1840e+02 ; CB - CG
- HG3
2938 2941 2944 1 1.1110e+02 5.2718e+02 ; CB - CG
- CD
2939 2938 2940 1 1.0950e+02 2.9288e+02 ; HB2 - CB
- HB3
2939 2938 2941 1 1.0950e+02 4.1840e+02 ; HB2 - CB
- CG
2940 2938 2941 1 1.0950e+02 4.1840e+02 ; HB3 - CB
- CG
2941 2944 2945 1 1.2040e+02 6.6944e+02 ; CG - CD
- OE1
2941 2944 2946 1 1.1660e+02 5.8576e+02 ; CG - CD
- NE2
2942 2941 2943 1 1.0950e+02 2.9288e+02 ; HG2 - CG
- HG3
2942 2941 2944 1 1.0950e+02 4.1840e+02 ; HG2 - CG
- CD
2943 2941 2944 1 1.0950e+02 4.1840e+02 ; HG3 - CG
- CD
2944 2946 2947 1 1.2000e+02 4.1840e+02 ; CD - NE2
- HE21
2944 2946 2948 1 1.2000e+02 4.1840e+02 ; CD - NE2
- HE22
2945 2944 2946 1 1.2290e+02 6.6944e+02 ; OE1 - CD
- NE2
2947 2946 2948 1 1.2000e+02 2.9288e+02 ; HE21 - NE2
- HE22
2949 2951 2952 1 1.2000e+02 4.1840e+02 ; C - N
- H
2949 2951 2953 1 1.2190e+02 4.1840e+02 ; C - N
- CA
2950 2949 2951 1 1.2290e+02 6.6944e+02 ; O - C
- N
2951 2953 2954 1 1.0950e+02 4.1840e+02 ; N - CA
- HA
2951 2953 2955 1 1.0970e+02 6.6944e+02 ; N - CA
- CB
2951 2953 2965 1 1.1010e+02 5.2718e+02 ; N - CA
- C
2952 2951 2953 1 1.1804e+02 4.1840e+02 ; H - N
- CA
2953 2955 2956 1 1.0950e+02 4.1840e+02 ; CA - CB
- HB

```
  2953   2955   2957      1    1.0950e+02    3.3472e+02 ;     CA - CB
- CG1
  2953   2955   2961      1    1.0950e+02    3.3472e+02 ;     CA - CB
- CG2
  2953   2965   2966      1    1.2040e+02    6.6944e+02 ;     CA - C
- O
  2953   2965   2967      1    1.1660e+02    5.8576e+02 ;     CA - C
- N
  2954   2953   2955      1    1.0950e+02    4.1840e+02 ;     HA - CA
- CB
  2954   2953   2965      1    1.0950e+02    4.1840e+02 ;     HA - CA
- C
  2955   2953   2965      1    1.1110e+02    5.2718e+02 ;     CB - CA
- C
  2955   2957   2958      1    1.0950e+02    4.1840e+02 ;     CB - CG1
- HG11
  2955   2957   2959      1    1.0950e+02    4.1840e+02 ;     CB - CG1
- HG12
  2955   2957   2960      1    1.0950e+02    4.1840e+02 ;     CB - CG1
- HG13
  2955   2961   2962      1    1.0950e+02    4.1840e+02 ;     CB - CG2
- HG21
  2955   2961   2963      1    1.0950e+02    4.1840e+02 ;     CB - CG2
- HG22
  2955   2961   2964      1    1.0950e+02    4.1840e+02 ;     CB - CG2
- HG23
  2956   2955   2957      1    1.0950e+02    4.1840e+02 ;     HB - CB
- CG1
  2956   2955   2961      1    1.0950e+02    4.1840e+02 ;     HB - CB
- CG2
  2957   2955   2961      1    1.0950e+02    3.3472e+02 ;    CG1 - CB
- CG2
  2958   2957   2959      1    1.0950e+02    2.9288e+02 ;   HG11 - CG1
- HG12
  2958   2957   2960      1    1.0950e+02    2.9288e+02 ;   HG11 - CG1
- HG13
  2959   2957   2960      1    1.0950e+02    2.9288e+02 ;   HG12 - CG1
- HG13
  2962   2961   2963      1    1.0950e+02    2.9288e+02 ;   HG21 - CG2
- HG22
  2962   2961   2964      1    1.0950e+02    2.9288e+02 ;   HG21 - CG2
- HG23
  2963   2961   2964      1    1.0950e+02    2.9288e+02 ;   HG22 - CG2
- HG23
  2965   2967   2968      1    1.2000e+02    4.1840e+02 ;      C - N
- H
  2965   2967   2969      1    1.2190e+02    4.1840e+02 ;      C - N
- CA
  2966   2965   2967      1    1.2290e+02    6.6944e+02 ;      O - C
- N
  2967   2969   2970      1    1.0950e+02    4.1840e+02 ;      N - CA
- HA
```

2967 2969 2971 1 1.0970e+02 6.6944e+02 ; N - CA
- CB
2967 2969 2984 1 1.1010e+02 5.2718e+02 ; N - CA
- C
2968 2967 2969 1 1.1804e+02 4.1840e+02 ; H - N
- CA
2969 2971 2972 1 1.0950e+02 4.1840e+02 ; CA - CB
- HB2
2969 2971 2973 1 1.0950e+02 4.1840e+02 ; CA - CB
- HB3
2969 2971 2974 1 1.0950e+02 3.3472e+02 ; CA - CB
- CG
2969 2984 2985 1 1.2040e+02 6.6944e+02 ; CA - C
- O
2969 2984 2986 1 1.1660e+02 5.8576e+02 ; CA - C
- N
2970 2969 2971 1 1.0950e+02 4.1840e+02 ; HA - CA
- CB
2970 2969 2984 1 1.0950e+02 4.1840e+02 ; HA - CA
- C
2971 2969 2984 1 1.1110e+02 5.2718e+02 ; CB - CA
- C
2971 2974 2975 1 1.0950e+02 4.1840e+02 ; CB - CG
- HG
2971 2974 2976 1 1.0950e+02 3.3472e+02 ; CB - CG
- CD1
2971 2974 2980 1 1.0950e+02 3.3472e+02 ; CB - CG
- CD2
2972 2971 2973 1 1.0950e+02 2.9288e+02 ; HB2 - CB
- HB3
2972 2971 2974 1 1.0950e+02 4.1840e+02 ; HB2 - CB
- CG
2973 2971 2974 1 1.0950e+02 4.1840e+02 ; HB3 - CB
- CG
2974 2976 2977 1 1.0950e+02 4.1840e+02 ; CG - CD1
- HD11
2974 2976 2978 1 1.0950e+02 4.1840e+02 ; CG - CD1
- HD12
2974 2976 2979 1 1.0950e+02 4.1840e+02 ; CG - CD1
- HD13
2974 2980 2981 1 1.0950e+02 4.1840e+02 ; CG - CD2
- HD21
2974 2980 2982 1 1.0950e+02 4.1840e+02 ; CG - CD2
- HD22
2974 2980 2983 1 1.0950e+02 4.1840e+02 ; CG - CD2
- HD23
2975 2974 2976 1 1.0950e+02 4.1840e+02 ; HG - CG
- CD1
2975 2974 2980 1 1.0950e+02 4.1840e+02 ; HG - CG
- CD2
2976 2974 2980 1 1.0950e+02 3.3472e+02 ; CD1 - CG
- CD2

2977 2976 2978 1 1.0950e+02 2.9288e+02 ; HD11 - CD1
- HD12
2977 2976 2979 1 1.0950e+02 2.9288e+02 ; HD11 - CD1
- HD13
2978 2976 2979 1 1.0950e+02 2.9288e+02 ; HD12 - CD1
- HD13
2981 2980 2982 1 1.0950e+02 2.9288e+02 ; HD21 - CD2
- HD22
2981 2980 2983 1 1.0950e+02 2.9288e+02 ; HD21 - CD2
- HD23
2982 2980 2983 1 1.0950e+02 2.9288e+02 ; HD22 - CD2
- HD23
2984 2986 2987 1 1.2000e+02 4.1840e+02 ; C - N
- H
2984 2986 2988 1 1.2190e+02 4.1840e+02 ; C - N
- CA
2985 2984 2986 1 1.2290e+02 6.6944e+02 ; O - C
- N
2986 2988 2989 1 1.0950e+02 4.1840e+02 ; N - CA
- HA
2986 2988 2990 1 1.0970e+02 6.6944e+02 ; N - CA
- CB
2986 2988 2996 1 1.1010e+02 5.2718e+02 ; N - CA
- C
2987 2986 2988 1 1.1804e+02 4.1840e+02 ; H - N
- CA
2988 2990 2991 1 1.0950e+02 4.1840e+02 ; CA - CB
- HB2
2988 2990 2992 1 1.0950e+02 4.1840e+02 ; CA - CB
- HB3
2988 2990 2993 1 1.1110e+02 5.2718e+02 ; CA - CB
- CG
2988 2996 2997 1 1.2040e+02 6.6944e+02 ; CA - C
- O
2988 2996 2998 1 1.1660e+02 5.8576e+02 ; CA - C
- N
2989 2988 2990 1 1.0950e+02 4.1840e+02 ; HA - CA
- CB
2989 2988 2996 1 1.0950e+02 4.1840e+02 ; HA - CA
- C
2990 2988 2996 1 1.1110e+02 5.2718e+02 ; CB - CA
- C
2990 2993 2994 1 1.1700e+02 5.8576e+02 ; CB - CG
- OD1
2990 2993 2995 1 1.1700e+02 5.8576e+02 ; CB - CG
- OD2
2991 2990 2992 1 1.0950e+02 2.9288e+02 ; HB2 - CB
- HB3
2991 2990 2993 1 1.0950e+02 4.1840e+02 ; HB2 - CB
- CG
2992 2990 2993 1 1.0950e+02 4.1840e+02 ; HB3 - CB
- CG

2994 2993 2995 1 1.2600e+02 6.6944e+02 ; OD1 - CG
- OD2
2996 2998 2999 1 1.2000e+02 4.1840e+02 ; C - N
- H
2996 2998 3000 1 1.2190e+02 4.1840e+02 ; C - N
- CA
2997 2996 2998 1 1.2290e+02 6.6944e+02 ; O - C
- N
2998 3000 3001 1 1.0950e+02 4.1840e+02 ; N - CA
- HA2
2998 3000 3002 1 1.0950e+02 4.1840e+02 ; N - CA
- HA3
2998 3000 3003 1 1.1010e+02 5.2718e+02 ; N - CA
- C
2999 2998 3000 1 1.1804e+02 4.1840e+02 ; H - N
- CA
3000 3003 3004 1 1.2040e+02 6.6944e+02 ; CA - C
- O
3000 3003 3005 1 1.1660e+02 5.8576e+02 ; CA - C
- N
3001 3000 3002 1 1.0950e+02 2.9288e+02 ; HA2 - CA
- HA3
3001 3000 3003 1 1.0950e+02 4.1840e+02 ; HA2 - CA
- C
3002 3000 3003 1 1.0950e+02 4.1840e+02 ; HA3 - CA
- C
3003 3005 3006 1 1.2000e+02 4.1840e+02 ; C - N
- H
3003 3005 3007 1 1.2190e+02 4.1840e+02 ; C - N
- CA
3004 3003 3005 1 1.2290e+02 6.6944e+02 ; O - C
- N
3005 3007 3008 1 1.0950e+02 4.1840e+02 ; N - CA
- HA
3005 3007 3009 1 1.0970e+02 6.6944e+02 ; N - CA
- CB
3005 3007 3025 1 1.1010e+02 5.2718e+02 ; N - CA
- C
3006 3005 3007 1 1.1804e+02 4.1840e+02 ; H - N
- CA
3007 3009 3010 1 1.0950e+02 4.1840e+02 ; CA - CB
- HB2
3007 3009 3011 1 1.0950e+02 4.1840e+02 ; CA - CB
- HB3
3007 3009 3012 1 1.0950e+02 3.3472e+02 ; CA - CB
- CG
3007 3025 3026 1 1.2040e+02 6.6944e+02 ; CA - C
- O
3007 3025 3027 1 1.1660e+02 5.8576e+02 ; CA - C
- N
3008 3007 3009 1 1.0950e+02 4.1840e+02 ; HA - CA
- CB

3008 3007 3025 1 1.0950e+02 4.1840e+02 ; HA - CA
- C
3009 3007 3025 1 1.1110e+02 5.2718e+02 ; CB - CA
- C
3009 3012 3013 1 1.0950e+02 4.1840e+02 ; CB - CG
- HG2
3009 3012 3014 1 1.0950e+02 4.1840e+02 ; CB - CG
- HG3
3009 3012 3015 1 1.0950e+02 3.3472e+02 ; CB - CG
- CD
3010 3009 3011 1 1.0950e+02 2.9288e+02 ; HB2 - CB
- HB3
3010 3009 3012 1 1.0950e+02 4.1840e+02 ; HB2 - CB
- CG
3011 3009 3012 1 1.0950e+02 4.1840e+02 ; HB3 - CB
- CG
3012 3015 3016 1 1.0950e+02 4.1840e+02 ; CG - CD
- HD2
3012 3015 3017 1 1.0950e+02 4.1840e+02 ; CG - CD
- HD3
3012 3015 3018 1 1.0950e+02 3.3472e+02 ; CG - CD
- CE
3013 3012 3014 1 1.0950e+02 2.9288e+02 ; HG2 - CG
- HG3
3013 3012 3015 1 1.0950e+02 4.1840e+02 ; HG2 - CG
- CD
3014 3012 3015 1 1.0950e+02 4.1840e+02 ; HG3 - CG
- CD
3015 3018 3019 1 1.0950e+02 4.1840e+02 ; CD - CE
- HE2
3015 3018 3020 1 1.0950e+02 4.1840e+02 ; CD - CE
- HE3
3015 3018 3021 1 1.1120e+02 6.6944e+02 ; CD - CE
- NZ
3016 3015 3017 1 1.0950e+02 2.9288e+02 ; HD2 - CD
- HD3
3016 3015 3018 1 1.0950e+02 4.1840e+02 ; HD2 - CD
- CE
3017 3015 3018 1 1.0950e+02 4.1840e+02 ; HD3 - CD
- CE
3018 3021 3022 1 1.0950e+02 4.1840e+02 ; CE - NZ
- HZ1
3018 3021 3023 1 1.0950e+02 4.1840e+02 ; CE - NZ
- HZ2
3018 3021 3024 1 1.0950e+02 4.1840e+02 ; CE - NZ
- HZ3
3019 3018 3020 1 1.0950e+02 2.9288e+02 ; HE2 - CE
- HE3
3019 3018 3021 1 1.0950e+02 4.1840e+02 ; HE2 - CE
- NZ
3020 3018 3021 1 1.0950e+02 4.1840e+02 ; HE3 - CE
- NZ

3022 3021 3023 1 1.0950e+02 2.9288e+02 ; HZ1 - NZ
- HZ2
3022 3021 3024 1 1.0950e+02 2.9288e+02 ; HZ1 - NZ
- HZ3
3023 3021 3024 1 1.0950e+02 2.9288e+02 ; HZ2 - NZ
- HZ3
3025 3027 3028 1 1.2000e+02 4.1840e+02 ; C - N
- H
3025 3027 3029 1 1.2190e+02 4.1840e+02 ; C - N
- CA
3026 3025 3027 1 1.2290e+02 6.6944e+02 ; O - C
- N
3027 3029 3030 1 1.0950e+02 4.1840e+02 ; N - CA
- HA
3027 3029 3031 1 1.0970e+02 6.6944e+02 ; N - CA
- CB
3027 3029 3046 1 1.1010e+02 5.2718e+02 ; N - CA
- C
3028 3027 3029 1 1.1804e+02 4.1840e+02 ; H - N
- CA
3029 3031 3032 1 1.0950e+02 4.1840e+02 ; CA - CB
- HB2
3029 3031 3033 1 1.0950e+02 4.1840e+02 ; CA - CB
- HB3
3029 3031 3034 1 1.1400e+02 5.2718e+02 ; CA - CB
- CG
3029 3046 3047 1 1.2040e+02 6.6944e+02 ; CA - C
- O
3029 3046 3048 1 1.1660e+02 5.8576e+02 ; CA - C
- N
3030 3029 3031 1 1.0950e+02 4.1840e+02 ; HA - CA
- CB
3030 3029 3046 1 1.0950e+02 4.1840e+02 ; HA - CA
- C
3031 3029 3046 1 1.1110e+02 5.2718e+02 ; CB - CA
- C
3031 3034 3035 1 1.2000e+02 5.8576e+02 ; CB - CG
- CD1
3031 3034 3044 1 1.2000e+02 5.8576e+02 ; CB - CG
- CD2
3032 3031 3033 1 1.0950e+02 2.9288e+02 ; HB2 - CB
- HB3
3032 3031 3034 1 1.0950e+02 4.1840e+02 ; HB2 - CB
- CG
3033 3031 3034 1 1.0950e+02 4.1840e+02 ; HB3 - CB
- CG
3034 3035 3036 1 1.2000e+02 4.1840e+02 ; CG - CD1
- HD1
3034 3035 3037 1 1.2000e+02 5.2718e+02 ; CG - CD1
- CE1
3034 3044 3042 1 1.2000e+02 5.2718e+02 ; CG - CD2
- CE2

3034 3044 3045 1 1.2000e+02 4.1840e+02 ; CG - CD2
- HD2
3035 3034 3044 1 1.2000e+02 5.2718e+02 ; CD1 - CG
- CD2
3035 3037 3038 1 1.2000e+02 4.1840e+02 ; CD1 - CE1
- HE1
3035 3037 3039 1 1.2000e+02 5.2718e+02 ; CD1 - CE1
- CZ
3036 3035 3037 1 1.2000e+02 4.1840e+02 ; HD1 - CD1
- CE1
3037 3039 3040 1 1.2000e+02 5.8576e+02 ; CE1 - CZ
- OH
3037 3039 3042 1 1.2000e+02 5.2718e+02 ; CE1 - CZ
- CE2
3038 3037 3039 1 1.2000e+02 4.1840e+02 ; HE1 - CE1
- CZ
3039 3040 3041 1 1.1300e+02 4.1840e+02 ; CZ - OH
- HH
3039 3042 3043 1 1.2000e+02 4.1840e+02 ; CZ - CE2
- HE2
3039 3042 3044 1 1.2000e+02 5.2718e+02 ; CZ - CE2
- CD2
3040 3039 3042 1 1.2000e+02 5.8576e+02 ; OH - CZ
- CE2
3042 3044 3045 1 1.2000e+02 4.1840e+02 ; CE2 - CD2
- HD2
3043 3042 3044 1 1.2000e+02 4.1840e+02 ; HE2 - CE2
- CD2
3046 3048 3049 1 1.2000e+02 4.1840e+02 ; C - N
- H
3046 3048 3050 1 1.2190e+02 4.1840e+02 ; C - N
- CA
3047 3046 3048 1 1.2290e+02 6.6944e+02 ; O - C
- N
3048 3050 3051 1 1.0950e+02 4.1840e+02 ; N - CA
- HA
3048 3050 3052 1 1.0970e+02 6.6944e+02 ; N - CA
- CB
3048 3050 3058 1 1.1010e+02 5.2718e+02 ; N - CA
- C
3049 3048 3050 1 1.1804e+02 4.1840e+02 ; H - N
- CA
3050 3052 3053 1 1.0950e+02 4.1840e+02 ; CA - CB
- HB2
3050 3052 3054 1 1.0950e+02 4.1840e+02 ; CA - CB
- HB3
3050 3052 3055 1 1.1110e+02 5.2718e+02 ; CA - CB
- CG
3050 3058 3059 1 1.2040e+02 6.6944e+02 ; CA - C
- O
3050 3058 3060 1 1.1660e+02 5.8576e+02 ; CA - C
- N

3051 3050 3052 1 1.0950e+02 4.1840e+02 ; HA - CA
- CB
3051 3050 3058 1 1.0950e+02 4.1840e+02 ; HA - CA
- C
3052 3050 3058 1 1.1110e+02 5.2718e+02 ; CB - CA
- C
3052 3055 3056 1 1.1700e+02 5.8576e+02 ; CB - CG
- OD1
3052 3055 3057 1 1.1700e+02 5.8576e+02 ; CB - CG
- OD2
3053 3052 3054 1 1.0950e+02 2.9288e+02 ; HB2 - CB
- HB3
3053 3052 3055 1 1.0950e+02 4.1840e+02 ; HB2 - CB
- CG
3054 3052 3055 1 1.0950e+02 4.1840e+02 ; HB3 - CB
- CG
3056 3055 3057 1 1.2600e+02 6.6944e+02 ; OD1 - CG
- OD2
3058 3060 3061 1 1.2000e+02 4.1840e+02 ; C - N
- H
3058 3060 3062 1 1.2190e+02 4.1840e+02 ; C - N
- CA
3059 3058 3060 1 1.2290e+02 6.6944e+02 ; O - C
- N
3060 3062 3063 1 1.0950e+02 4.1840e+02 ; N - CA
- HA
3060 3062 3064 1 1.0970e+02 6.6944e+02 ; N - CA
- CB
3060 3062 3080 1 1.1010e+02 5.2718e+02 ; N - CA
- C
3061 3060 3062 1 1.1804e+02 4.1840e+02 ; H - N
- CA
3062 3064 3065 1 1.0950e+02 4.1840e+02 ; CA - CB
- HB2
3062 3064 3066 1 1.0950e+02 4.1840e+02 ; CA - CB
- HB3
3062 3064 3067 1 1.0950e+02 3.3472e+02 ; CA - CB
- CG
3062 3080 3081 1 1.2040e+02 6.6944e+02 ; CA - C
- O
3062 3080 3082 1 1.1660e+02 5.8576e+02 ; CA - C
- N
3063 3062 3064 1 1.0950e+02 4.1840e+02 ; HA - CA
- CB
3063 3062 3080 1 1.0950e+02 4.1840e+02 ; HA - CA
- C
3064 3062 3080 1 1.1110e+02 5.2718e+02 ; CB - CA
- C
3064 3067 3068 1 1.0950e+02 4.1840e+02 ; CB - CG
- HG2
3064 3067 3069 1 1.0950e+02 4.1840e+02 ; CB - CG
- HG3

3064 3067 3070 1 1.0950e+02 3.3472e+02 ; CB - CG
- CD
3065 3064 3066 1 1.0950e+02 2.9288e+02 ; HB2 - CB
- HB3
3065 3064 3067 1 1.0950e+02 4.1840e+02 ; HB2 - CB
- CG
3066 3064 3067 1 1.0950e+02 4.1840e+02 ; HB3 - CB
- CG
3067 3070 3071 1 1.0950e+02 4.1840e+02 ; CG - CD
- HD2
3067 3070 3072 1 1.0950e+02 4.1840e+02 ; CG - CD
- HD3
3067 3070 3073 1 1.0950e+02 3.3472e+02 ; CG - CD
- CE
3068 3067 3069 1 1.0950e+02 2.9288e+02 ; HG2 - CG
- HG3
3068 3067 3070 1 1.0950e+02 4.1840e+02 ; HG2 - CG
- CD
3069 3067 3070 1 1.0950e+02 4.1840e+02 ; HG3 - CG
- CD
3070 3073 3074 1 1.0950e+02 4.1840e+02 ; CD - CE
- HE2
3070 3073 3075 1 1.0950e+02 4.1840e+02 ; CD - CE
- HE3
3070 3073 3076 1 1.1120e+02 6.6944e+02 ; CD - CE
- NZ
3071 3070 3072 1 1.0950e+02 2.9288e+02 ; HD2 - CD
- HD3
3071 3070 3073 1 1.0950e+02 4.1840e+02 ; HD2 - CD
- CE
3072 3070 3073 1 1.0950e+02 4.1840e+02 ; HD3 - CD
- CE
3073 3076 3077 1 1.0950e+02 4.1840e+02 ; CE - NZ
- HZ1
3073 3076 3078 1 1.0950e+02 4.1840e+02 ; CE - NZ
- HZ2
3073 3076 3079 1 1.0950e+02 4.1840e+02 ; CE - NZ
- HZ3
3074 3073 3075 1 1.0950e+02 2.9288e+02 ; HE2 - CE
- HE3
3074 3073 3076 1 1.0950e+02 4.1840e+02 ; HE2 - CE
- NZ
3075 3073 3076 1 1.0950e+02 4.1840e+02 ; HE3 - CE
- NZ
3077 3076 3078 1 1.0950e+02 2.9288e+02 ; HZ1 - NZ
- HZ2
3077 3076 3079 1 1.0950e+02 2.9288e+02 ; HZ1 - NZ
- HZ3
3078 3076 3079 1 1.0950e+02 2.9288e+02 ; HZ2 - NZ
- HZ3
3080 3082 3083 1 1.2000e+02 4.1840e+02 ; C - N
- H

3080 3082 3084 1 1.2190e+02 4.1840e+02 ; C - N
- CA
3081 3080 3082 1 1.2290e+02 6.6944e+02 ; O - C
- N
3082 3084 3085 1 1.0950e+02 4.1840e+02 ; N - CA
- HA
3082 3084 3086 1 1.0970e+02 6.6944e+02 ; N - CA
- CB
3082 3084 3102 1 1.1010e+02 5.2718e+02 ; N - CA
- C
3083 3082 3084 1 1.1804e+02 4.1840e+02 ; H - N
- CA
3084 3086 3087 1 1.0950e+02 4.1840e+02 ; CA - CB
- HB2
3084 3086 3088 1 1.0950e+02 4.1840e+02 ; CA - CB
- HB3
3084 3086 3089 1 1.0950e+02 3.3472e+02 ; CA - CB
- CG
3084 3102 3103 1 1.2040e+02 6.6944e+02 ; CA - C
- O
3084 3102 3104 1 1.1660e+02 5.8576e+02 ; CA - C
- N
3085 3084 3086 1 1.0950e+02 4.1840e+02 ; HA - CA
- CB
3085 3084 3102 1 1.0950e+02 4.1840e+02 ; HA - CA
- C
3086 3084 3102 1 1.1110e+02 5.2718e+02 ; CB - CA
- C
3086 3089 3090 1 1.0950e+02 4.1840e+02 ; CB - CG
- HG2
3086 3089 3091 1 1.0950e+02 4.1840e+02 ; CB - CG
- HG3
3086 3089 3092 1 1.0950e+02 3.3472e+02 ; CB - CG
- CD
3087 3086 3088 1 1.0950e+02 2.9288e+02 ; HB2 - CB
- HB3
3087 3086 3089 1 1.0950e+02 4.1840e+02 ; HB2 - CB
- CG
3088 3086 3089 1 1.0950e+02 4.1840e+02 ; HB3 - CB
- CG
3089 3092 3093 1 1.0950e+02 4.1840e+02 ; CG - CD
- HD2
3089 3092 3094 1 1.0950e+02 4.1840e+02 ; CG - CD
- HD3
3089 3092 3095 1 1.0950e+02 3.3472e+02 ; CG - CD
- CE
3090 3089 3091 1 1.0950e+02 2.9288e+02 ; HG2 - CG
- HG3
3090 3089 3092 1 1.0950e+02 4.1840e+02 ; HG2 - CG
- CD
3091 3089 3092 1 1.0950e+02 4.1840e+02 ; HG3 - CG
- CD

3092 3095 3096 1 1.0950e+02 4.1840e+02 ; CD - CE
- HE2
3092 3095 3097 1 1.0950e+02 4.1840e+02 ; CD - CE
- HE3
3092 3095 3098 1 1.1120e+02 6.6944e+02 ; CD - CE
- NZ
3093 3092 3094 1 1.0950e+02 2.9288e+02 ; HD2 - CD
- HD3
3093 3092 3095 1 1.0950e+02 4.1840e+02 ; HD2 - CD
- CE
3094 3092 3095 1 1.0950e+02 4.1840e+02 ; HD3 - CD
- CE
3095 3098 3099 1 1.0950e+02 4.1840e+02 ; CE - NZ
- HZ1
3095 3098 3100 1 1.0950e+02 4.1840e+02 ; CE - NZ
- HZ2
3095 3098 3101 1 1.0950e+02 4.1840e+02 ; CE - NZ
- HZ3
3096 3095 3097 1 1.0950e+02 2.9288e+02 ; HE2 - CE
- HE3
3096 3095 3098 1 1.0950e+02 4.1840e+02 ; HE2 - CE
- NZ
3097 3095 3098 1 1.0950e+02 4.1840e+02 ; HE3 - CE
- NZ
3099 3098 3100 1 1.0950e+02 2.9288e+02 ; HZ1 - NZ
- HZ2
3099 3098 3101 1 1.0950e+02 2.9288e+02 ; HZ1 - NZ
- HZ3
3100 3098 3101 1 1.0950e+02 2.9288e+02 ; HZ2 - NZ
- HZ3
3102 3104 3105 1 1.2000e+02 4.1840e+02 ; C - N
- H
3102 3104 3106 1 1.2190e+02 4.1840e+02 ; C - N
- CA
3103 3102 3104 1 1.2290e+02 6.6944e+02 ; O - C
- N
3104 3106 3107 1 1.0950e+02 4.1840e+02 ; N - CA
- HA
3104 3106 3108 1 1.0970e+02 6.6944e+02 ; N - CA
- CB
3104 3106 3113 1 1.1010e+02 5.2718e+02 ; N - CA
- C
3105 3104 3106 1 1.1804e+02 4.1840e+02 ; H - N
- CA
3106 3108 3109 1 1.0950e+02 4.1840e+02 ; CA - CB
- HB2
3106 3108 3110 1 1.0950e+02 4.1840e+02 ; CA - CB
- HB3
3106 3108 3111 1 1.0860e+02 4.1840e+02 ; CA - CB
- SG
3106 3113 3114 1 1.2040e+02 6.6944e+02 ; CA - C
- O

3106 3113 3115 1 1.1660e+02 5.8576e+02 ; CA - C
- N
3107 3106 3108 1 1.0950e+02 4.1840e+02 ; HA - CA
- CB
3107 3106 3113 1 1.0950e+02 4.1840e+02 ; HA - CA
- C
3108 3106 3113 1 1.1110e+02 5.2718e+02 ; CB - CA
- C
3108 3111 3112 1 9.6000e+01 3.5982e+02 ; CB - SG
- HG
3109 3108 3110 1 1.0950e+02 2.9288e+02 ; HB2 - CB
- HB3
3109 3108 3111 1 1.0950e+02 4.1840e+02 ; HB2 - CB
- SG
3110 3108 3111 1 1.0950e+02 4.1840e+02 ; HB3 - CB
- SG
3113 3115 3116 1 1.2000e+02 4.1840e+02 ; C - N
- H
3113 3115 3117 1 1.2190e+02 4.1840e+02 ; C - N
- CA
3114 3113 3115 1 1.2290e+02 6.6944e+02 ; O - C
- N
3115 3117 3118 1 1.0950e+02 4.1840e+02 ; N - CA
- HA
3115 3117 3119 1 1.0970e+02 6.6944e+02 ; N - CA
- CB
3115 3117 3125 1 1.1010e+02 5.2718e+02 ; N - CA
- C
3116 3115 3117 1 1.1804e+02 4.1840e+02 ; H - N
- CA
3117 3119 3120 1 1.0950e+02 4.1840e+02 ; CA - CB
- HB2
3117 3119 3121 1 1.0950e+02 4.1840e+02 ; CA - CB
- HB3
3117 3119 3122 1 1.1110e+02 5.2718e+02 ; CA - CB
- CG
3117 3125 3126 1 1.2040e+02 6.6944e+02 ; CA - C
- O
3117 3125 3127 1 1.1660e+02 5.8576e+02 ; CA - C
- N
3118 3117 3119 1 1.0950e+02 4.1840e+02 ; HA - CA
- CB
3118 3117 3125 1 1.0950e+02 4.1840e+02 ; HA - CA
- C
3119 3117 3125 1 1.1110e+02 5.2718e+02 ; CB - CA
- C
3119 3122 3123 1 1.1700e+02 5.8576e+02 ; CB - CG
- OD1
3119 3122 3124 1 1.1700e+02 5.8576e+02 ; CB - CG
- OD2
3120 3119 3121 1 1.0950e+02 2.9288e+02 ; HB2 - CB
- HB3

3120 3119 3122 1 1.0950e+02 4.1840e+02 ; HB2 - CB
- CG
3121 3119 3122 1 1.0950e+02 4.1840e+02 ; HB3 - CB
- CG
3123 3122 3124 1 1.2600e+02 6.6944e+02 ; OD1 - CG
- OD2
3125 3127 3128 1 1.2000e+02 4.1840e+02 ; C - N
- H
3125 3127 3129 1 1.2190e+02 4.1840e+02 ; C - N
- CA
3126 3125 3127 1 1.2290e+02 6.6944e+02 ; O - C
- N
3127 3129 3130 1 1.0950e+02 4.1840e+02 ; N - CA
- HA
3127 3129 3131 1 1.0970e+02 6.6944e+02 ; N - CA
- CB
3127 3129 3144 1 1.1010e+02 5.2718e+02 ; N - CA
- C
3128 3127 3129 1 1.1804e+02 4.1840e+02 ; H - N
- CA
3129 3131 3132 1 1.0950e+02 4.1840e+02 ; CA - CB
- HB
3129 3131 3133 1 1.0950e+02 3.3472e+02 ; CA - CB
- CG2
3129 3131 3137 1 1.0950e+02 3.3472e+02 ; CA - CB
- CG1
3129 3144 3145 1 1.2040e+02 6.6944e+02 ; CA - C
- O
3129 3144 3146 1 1.1660e+02 5.8576e+02 ; CA - C
- N
3130 3129 3131 1 1.0950e+02 4.1840e+02 ; HA - CA
- CB
3130 3129 3144 1 1.0950e+02 4.1840e+02 ; HA - CA
- C
3131 3129 3144 1 1.1110e+02 5.2718e+02 ; CB - CA
- C
3131 3133 3134 1 1.0950e+02 4.1840e+02 ; CB - CG2
- HG21
3131 3133 3135 1 1.0950e+02 4.1840e+02 ; CB - CG2
- HG22
3131 3133 3136 1 1.0950e+02 4.1840e+02 ; CB - CG2
- HG23
3131 3137 3138 1 1.0950e+02 4.1840e+02 ; CB - CG1
- HG12
3131 3137 3139 1 1.0950e+02 4.1840e+02 ; CB - CG1
- HG13
3131 3137 3140 1 1.0950e+02 3.3472e+02 ; CB - CG1
- CD1
3132 3131 3133 1 1.0950e+02 4.1840e+02 ; HB - CB
- CG2
3132 3131 3137 1 1.0950e+02 4.1840e+02 ; HB - CB
- CG1

```
  3133   3131   3137      1    1.0950e+02    3.3472e+02 ;    CG2 - CB
- CG1
  3134   3133   3135      1    1.0950e+02    2.9288e+02 ;   HG21 - CG2
- HG22
  3134   3133   3136      1    1.0950e+02    2.9288e+02 ;   HG21 - CG2
- HG23
  3135   3133   3136      1    1.0950e+02    2.9288e+02 ;   HG22 - CG2
- HG23
  3137   3140   3141      1    1.0950e+02    4.1840e+02 ;    CG1 - CD1
- HD11
  3137   3140   3142      1    1.0950e+02    4.1840e+02 ;    CG1 - CD1
- HD12
  3137   3140   3143      1    1.0950e+02    4.1840e+02 ;    CG1 - CD1
- HD13
  3138   3137   3139      1    1.0950e+02    2.9288e+02 ;   HG12 - CG1
- HG13
  3138   3137   3140      1    1.0950e+02    4.1840e+02 ;   HG12 - CG1
- CD1
  3139   3137   3140      1    1.0950e+02    4.1840e+02 ;   HG13 - CG1
- CD1
  3141   3140   3142      1    1.0950e+02    2.9288e+02 ;   HD11 - CD1
- HD12
  3141   3140   3143      1    1.0950e+02    2.9288e+02 ;   HD11 - CD1
- HD13
  3142   3140   3143      1    1.0950e+02    2.9288e+02 ;   HD12 - CD1
- HD13
  3144   3146   3147      1    1.2000e+02    4.1840e+02 ;      C - N
- H
  3144   3146   3148      1    1.2190e+02    4.1840e+02 ;      C - N
- CA
  3145   3144   3146      1    1.2290e+02    6.6944e+02 ;      O - C
- N
  3146   3148   3149      1    1.0950e+02    4.1840e+02 ;      N - CA
- HA
  3146   3148   3150      1    1.0970e+02    6.6944e+02 ;      N - CA
- CB
  3146   3148   3168      1    1.1010e+02    5.2718e+02 ;      N - CA
- C
  3147   3146   3148      1    1.1804e+02    4.1840e+02 ;      H - N
- CA
  3148   3150   3151      1    1.0950e+02    4.1840e+02 ;     CA - CB
- HB2
  3148   3150   3152      1    1.0950e+02    4.1840e+02 ;     CA - CB
- HB3
  3148   3150   3153      1    1.1560e+02    5.2718e+02 ;     CA - CB
- CG
  3148   3168   3169      1    1.2040e+02    6.6944e+02 ;     CA - C
- O
  3148   3168   3170      1    1.1660e+02    5.8576e+02 ;     CA - C
- N
  3149   3148   3150      1    1.0950e+02    4.1840e+02 ;     HA - CA
- CB
```

```
  3149   3148   3168       1    1.0950e+02    4.1840e+02 ;     HA - CA
- C
  3150   3148   3168       1    1.1110e+02    5.2718e+02 ;     CB - CA
- C
  3150   3153   3154       1    1.2500e+02    5.8576e+02 ;     CB - CG
- CD1
  3150   3153   3167       1    1.2860e+02    5.8576e+02 ;     CB - CG
- CD2
  3151   3150   3152       1    1.0950e+02    2.9288e+02 ;    HB2 - CB
- HB3
  3151   3150   3153       1    1.0950e+02    4.1840e+02 ;    HB2 - CB
- CG
  3152   3150   3153       1    1.0950e+02    4.1840e+02 ;    HB3 - CB
- CG
  3153   3154   3155       1    1.2000e+02    4.1840e+02 ;     CG - CD1
- HD1
  3153   3154   3156       1    1.0870e+02    5.8576e+02 ;     CG - CD1
- NE1
  3153   3167   3158       1    1.0880e+02    5.2718e+02 ;     CG - CD2
- CE2
  3153   3167   3165       1    1.3490e+02    5.2718e+02 ;     CG - CD2
- CE3
  3154   3153   3167       1    1.0640e+02    5.2718e+02 ;    CD1 - CG
- CD2
  3154   3156   3157       1    1.2000e+02    4.1840e+02 ;    CD1 - NE1
- HE1
  3154   3156   3158       1    1.1160e+02    5.8576e+02 ;    CD1 - NE1
- CE2
  3155   3154   3156       1    1.2000e+02    4.1840e+02 ;    HD1 - CD1
- NE1
  3156   3158   3159       1    1.3280e+02    5.8576e+02 ;    NE1 - CE2
- CZ2
  3156   3158   3167       1    1.0440e+02    5.8576e+02 ;    NE1 - CE2
- CD2
  3157   3156   3158       1    1.2310e+02    4.1840e+02 ;    HE1 - NE1
- CE2
  3158   3159   3160       1    1.2000e+02    4.1840e+02 ;    CE2 - CZ2
- HZ2
  3158   3159   3161       1    1.2000e+02    5.2718e+02 ;    CE2 - CZ2
- CH2
  3158   3167   3165       1    1.1620e+02    5.2718e+02 ;    CE2 - CD2
- CE3
  3159   3158   3167       1    1.2270e+02    5.2718e+02 ;    CZ2 - CE2
- CD2
  3159   3161   3162       1    1.2000e+02    4.1840e+02 ;    CZ2 - CH2
- HH2
  3159   3161   3163       1    1.2000e+02    5.2718e+02 ;    CZ2 - CH2
- CZ3
  3160   3159   3161       1    1.2000e+02    4.1840e+02 ;    HZ2 - CZ2
- CH2
  3161   3163   3164       1    1.2000e+02    4.1840e+02 ;    CH2 - CZ3
- HZ3
```

```
  3161   3163   3165       1    1.2000e+02    5.2718e+02 ;    CH2 - CZ3
- CE3
  3162   3161   3163       1    1.2000e+02    4.1840e+02 ;    HH2 - CH2
- CZ3
  3163   3165   3166       1    1.2000e+02    4.1840e+02 ;    CZ3 - CE3
- HE3
  3163   3165   3167       1    1.2000e+02    5.2718e+02 ;    CZ3 - CE3
- CD2
  3164   3163   3165       1    1.2000e+02    4.1840e+02 ;    HZ3 - CZ3
- CE3
  3166   3165   3167       1    1.2000e+02    4.1840e+02 ;    HE3 - CE3
- CD2
  3168   3170   3171       1    1.2000e+02    4.1840e+02 ;      C - N
- H
  3168   3170   3172       1    1.2190e+02    4.1840e+02 ;      C - N
- CA
  3169   3168   3170       1    1.2290e+02    6.6944e+02 ;      O - C
- N
  3170   3172   3173       1    1.0950e+02    4.1840e+02 ;      N - CA
- HA
  3170   3172   3174       1    1.0970e+02    6.6944e+02 ;      N - CA
- CB
  3170   3172   3179       1    1.1010e+02    5.2718e+02 ;      N - CA
- C
  3171   3170   3172       1    1.1804e+02    4.1840e+02 ;      H - N
- CA
  3172   3174   3175       1    1.0950e+02    4.1840e+02 ;     CA - CB
- HB2
  3172   3174   3176       1    1.0950e+02    4.1840e+02 ;     CA - CB
- HB3
  3172   3174   3177       1    1.0950e+02    4.1840e+02 ;     CA - CB
- OG
  3172   3179   3180       1    1.2040e+02    6.6944e+02 ;     CA - C
- O
  3172   3179   3181       1    1.1660e+02    5.8576e+02 ;     CA - C
- N
  3173   3172   3174       1    1.0950e+02    4.1840e+02 ;     HA - CA
- CB
  3173   3172   3179       1    1.0950e+02    4.1840e+02 ;     HA - CA
- C
  3174   3172   3179       1    1.1110e+02    5.2718e+02 ;     CB - CA
- C
  3174   3177   3178       1    1.0850e+02    4.6024e+02 ;     CB - OG
- HG
  3175   3174   3176       1    1.0950e+02    2.9288e+02 ;    HB2 - CB
- HB3
  3175   3174   3177       1    1.0950e+02    4.1840e+02 ;    HB2 - CB
- OG
  3176   3174   3177       1    1.0950e+02    4.1840e+02 ;    HB3 - CB
- OG
  3179   3181   3182       1    1.2000e+02    4.1840e+02 ;      C - N
- H
```

3179 3181 3183 1 1.2190e+02 4.1840e+02 ; C - N
- CA
3180 3179 3181 1 1.2290e+02 6.6944e+02 ; O - C
- N
3181 3183 3184 1 1.0950e+02 4.1840e+02 ; N - CA
- HA
3181 3183 3185 1 1.0970e+02 6.6944e+02 ; N - CA
- CB
3181 3183 3190 1 1.1010e+02 5.2718e+02 ; N - CA
- C
3182 3181 3183 1 1.1804e+02 4.1840e+02 ; H - N
- CA
3183 3185 3186 1 1.0950e+02 4.1840e+02 ; CA - CB
- HB2
3183 3185 3187 1 1.0950e+02 4.1840e+02 ; CA - CB
- HB3
3183 3185 3188 1 1.0950e+02 4.1840e+02 ; CA - CB
- OG
3183 3190 3191 1 1.2040e+02 6.6944e+02 ; CA - C
- O
3183 3190 3192 1 1.1660e+02 5.8576e+02 ; CA - C
- N
3184 3183 3185 1 1.0950e+02 4.1840e+02 ; HA - CA
- CB
3184 3183 3190 1 1.0950e+02 4.1840e+02 ; HA - CA
- C
3185 3183 3190 1 1.1110e+02 5.2718e+02 ; CB - CA
- C
3185 3188 3189 1 1.0850e+02 4.6024e+02 ; CB - OG
- HG
3186 3185 3187 1 1.0950e+02 2.9288e+02 ; HB2 - CB
- HB3
3186 3185 3188 1 1.0950e+02 4.1840e+02 ; HB2 - CB
- OG
3187 3185 3188 1 1.0950e+02 4.1840e+02 ; HB3 - CB
- OG
3190 3192 3193 1 1.2000e+02 4.1840e+02 ; C - N
- H
3190 3192 3194 1 1.2190e+02 4.1840e+02 ; C - N
- CA
3191 3190 3192 1 1.2290e+02 6.6944e+02 ; O - C
- N
3192 3194 3195 1 1.0950e+02 4.1840e+02 ; N - CA
- HA2
3192 3194 3196 1 1.0950e+02 4.1840e+02 ; N - CA
- HA3
3192 3194 3197 1 1.1010e+02 5.2718e+02 ; N - CA
- C
3193 3192 3194 1 1.1804e+02 4.1840e+02 ; H - N
- CA
3194 3197 3198 1 1.2040e+02 6.6944e+02 ; CA - C
- O

3194 3197 3199 1 1.1660e+02 5.8576e+02 ; CA - C
- N
3195 3194 3196 1 1.0950e+02 2.9288e+02 ; HA2 - CA
- HA3
3195 3194 3197 1 1.0950e+02 4.1840e+02 ; HA2 - CA
- C
3196 3194 3197 1 1.0950e+02 4.1840e+02 ; HA3 - CA
- C
3197 3199 3200 1 1.2000e+02 4.1840e+02 ; C - N
- H
3197 3199 3201 1 1.2190e+02 4.1840e+02 ; C - N
- CA
3198 3197 3199 1 1.2290e+02 6.6944e+02 ; O - C
- N
3199 3201 3202 1 1.0950e+02 4.1840e+02 ; N - CA
- HA
3199 3201 3203 1 1.0970e+02 6.6944e+02 ; N - CA
- CB
3199 3201 3213 1 1.1010e+02 5.2718e+02 ; N - CA
- C
3200 3199 3201 1 1.1804e+02 4.1840e+02 ; H - N
- CA
3201 3203 3204 1 1.0950e+02 4.1840e+02 ; CA - CB
- HB
3201 3203 3205 1 1.0950e+02 3.3472e+02 ; CA - CB
- CG1
3201 3203 3209 1 1.0950e+02 3.3472e+02 ; CA - CB
- CG2
3201 3213 3214 1 1.2040e+02 6.6944e+02 ; CA - C
- O
3201 3213 3215 1 1.1660e+02 5.8576e+02 ; CA - C
- N
3202 3201 3203 1 1.0950e+02 4.1840e+02 ; HA - CA
- CB
3202 3201 3213 1 1.0950e+02 4.1840e+02 ; HA - CA
- C
3203 3201 3213 1 1.1110e+02 5.2718e+02 ; CB - CA
- C
3203 3205 3206 1 1.0950e+02 4.1840e+02 ; CB - CG1
- HG11
3203 3205 3207 1 1.0950e+02 4.1840e+02 ; CB - CG1
- HG12
3203 3205 3208 1 1.0950e+02 4.1840e+02 ; CB - CG1
- HG13
3203 3209 3210 1 1.0950e+02 4.1840e+02 ; CB - CG2
- HG21
3203 3209 3211 1 1.0950e+02 4.1840e+02 ; CB - CG2
- HG22
3203 3209 3212 1 1.0950e+02 4.1840e+02 ; CB - CG2
- HG23
3204 3203 3205 1 1.0950e+02 4.1840e+02 ; HB - CB
- CG1

```
  3204   3203   3209       1    1.0950e+02    4.1840e+02 ;     HB - CB
- CG2
  3205   3203   3209       1    1.0950e+02    3.3472e+02 ;    CG1 - CB
- CG2
  3206   3205   3207       1    1.0950e+02    2.9288e+02 ;   HG11 - CG1
- HG12
  3206   3205   3208       1    1.0950e+02    2.9288e+02 ;   HG11 - CG1
- HG13
  3207   3205   3208       1    1.0950e+02    2.9288e+02 ;   HG12 - CG1
- HG13
  3210   3209   3211       1    1.0950e+02    2.9288e+02 ;   HG21 - CG2
- HG22
  3210   3209   3212       1    1.0950e+02    2.9288e+02 ;   HG21 - CG2
- HG23
  3211   3209   3212       1    1.0950e+02    2.9288e+02 ;   HG22 - CG2
- HG23
  3213   3215   3216       1    1.2000e+02    4.1840e+02 ;      C - N
- H
  3213   3215   3217       1    1.2190e+02    4.1840e+02 ;      C - N
- CA
  3214   3213   3215       1    1.2290e+02    6.6944e+02 ;      O - C
- N
  3215   3217   3218       1    1.0950e+02    4.1840e+02 ;      N - CA
- HA
  3215   3217   3219       1    1.0970e+02    6.6944e+02 ;      N - CA
- CB
  3215   3217   3232       1    1.1010e+02    5.2718e+02 ;      N - CA
- C
  3216   3215   3217       1    1.1804e+02    4.1840e+02 ;      H - N
- CA
  3217   3219   3220       1    1.0950e+02    4.1840e+02 ;     CA - CB
- HB
  3217   3219   3221       1    1.0950e+02    3.3472e+02 ;     CA - CB
- CG2
  3217   3219   3225       1    1.0950e+02    3.3472e+02 ;     CA - CB
- CG1
  3217   3232   3233       1    1.2040e+02    6.6944e+02 ;     CA - C
- O
  3217   3232   3234       1    1.1660e+02    5.8576e+02 ;     CA - C
- N
  3218   3217   3219       1    1.0950e+02    4.1840e+02 ;     HA - CA
- CB
  3218   3217   3232       1    1.0950e+02    4.1840e+02 ;     HA - CA
- C
  3219   3217   3232       1    1.1110e+02    5.2718e+02 ;     CB - CA
- C
  3219   3221   3222       1    1.0950e+02    4.1840e+02 ;     CB - CG2
- HG21
  3219   3221   3223       1    1.0950e+02    4.1840e+02 ;     CB - CG2
- HG22
  3219   3221   3224       1    1.0950e+02    4.1840e+02 ;     CB - CG2
- HG23
```

3219 3225 3226 1 1.0950e+02 4.1840e+02 ; CB - CG1
- HG12
3219 3225 3227 1 1.0950e+02 4.1840e+02 ; CB - CG1
- HG13
3219 3225 3228 1 1.0950e+02 3.3472e+02 ; CB - CG1
- CD1
3220 3219 3221 1 1.0950e+02 4.1840e+02 ; HB - CB
- CG2
3220 3219 3225 1 1.0950e+02 4.1840e+02 ; HB - CB
- CG1
3221 3219 3225 1 1.0950e+02 3.3472e+02 ; CG2 - CB
- CG1
3222 3221 3223 1 1.0950e+02 2.9288e+02 ; HG21 - CG2
- HG22
3222 3221 3224 1 1.0950e+02 2.9288e+02 ; HG21 - CG2
- HG23
3223 3221 3224 1 1.0950e+02 2.9288e+02 ; HG22 - CG2
- HG23
3225 3228 3229 1 1.0950e+02 4.1840e+02 ; CG1 - CD1
- HD11
3225 3228 3230 1 1.0950e+02 4.1840e+02 ; CG1 - CD1
- HD12
3225 3228 3231 1 1.0950e+02 4.1840e+02 ; CG1 - CD1
- HD13
3226 3225 3227 1 1.0950e+02 2.9288e+02 ; HG12 - CG1
- HG13
3226 3225 3228 1 1.0950e+02 4.1840e+02 ; HG12 - CG1
- CD1
3227 3225 3228 1 1.0950e+02 4.1840e+02 ; HG13 - CG1
- CD1
3229 3228 3230 1 1.0950e+02 2.9288e+02 ; HD11 - CD1
- HD12
3229 3228 3231 1 1.0950e+02 2.9288e+02 ; HD11 - CD1
- HD13
3230 3228 3231 1 1.0950e+02 2.9288e+02 ; HD12 - CD1
- HD13
3232 3234 3235 1 1.2000e+02 4.1840e+02 ; C - N
- H
3232 3234 3236 1 1.2190e+02 4.1840e+02 ; C - N
- CA
3233 3232 3234 1 1.2290e+02 6.6944e+02 ; O - C
- N
3234 3236 3237 1 1.0950e+02 4.1840e+02 ; N - CA
- HA
3234 3236 3238 1 1.0970e+02 6.6944e+02 ; N - CA
- CB
3234 3236 3249 1 1.1010e+02 5.2718e+02 ; N - CA
- C
3235 3234 3236 1 1.1804e+02 4.1840e+02 ; H - N
- CA
3236 3238 3239 1 1.0950e+02 4.1840e+02 ; CA - CB
- HB2

3236 3238 3240 1 1.0950e+02 4.1840e+02 ; CA - CB
- HB3
3236 3238 3241 1 1.0950e+02 3.3472e+02 ; CA - CB
- CG
3236 3249 3250 1 1.2040e+02 6.6944e+02 ; CA - C
- O
3236 3249 3251 1 1.1660e+02 5.8576e+02 ; CA - C
- N
3237 3236 3238 1 1.0950e+02 4.1840e+02 ; HA - CA
- CB
3237 3236 3249 1 1.0950e+02 4.1840e+02 ; HA - CA
- C
3238 3236 3249 1 1.1110e+02 5.2718e+02 ; CB - CA
- C
3238 3241 3242 1 1.0950e+02 4.1840e+02 ; CB - CG
- HG2
3238 3241 3243 1 1.0950e+02 4.1840e+02 ; CB - CG
- HG3
3238 3241 3244 1 1.1470e+02 4.1840e+02 ; CB - CG
- SD
3239 3238 3240 1 1.0950e+02 2.9288e+02 ; HB2 - CB
- HB3
3239 3238 3241 1 1.0950e+02 4.1840e+02 ; HB2 - CB
- CG
3240 3238 3241 1 1.0950e+02 4.1840e+02 ; HB3 - CB
- CG
3241 3244 3245 1 9.8900e+01 5.1882e+02 ; CG - SD
- CE
3242 3241 3243 1 1.0950e+02 2.9288e+02 ; HG2 - CG
- HG3
3242 3241 3244 1 1.0950e+02 4.1840e+02 ; HG2 - CG
- SD
3243 3241 3244 1 1.0950e+02 4.1840e+02 ; HG3 - CG
- SD
3244 3245 3246 1 1.0950e+02 4.1840e+02 ; SD - CE
- HE1
3244 3245 3247 1 1.0950e+02 4.1840e+02 ; SD - CE
- HE2
3244 3245 3248 1 1.0950e+02 4.1840e+02 ; SD - CE
- HE3
3246 3245 3247 1 1.0950e+02 2.9288e+02 ; HE1 - CE
- HE2
3246 3245 3248 1 1.0950e+02 2.9288e+02 ; HE1 - CE
- HE3
3247 3245 3248 1 1.0950e+02 2.9288e+02 ; HE2 - CE
- HE3
3249 3251 3252 1 1.2000e+02 4.1840e+02 ; C - N
- H
3249 3251 3253 1 1.2190e+02 4.1840e+02 ; C - N
- CA
3250 3249 3251 1 1.2290e+02 6.6944e+02 ; O - C
- N

3251 3253 3254 1 1.0950e+02 4.1840e+02 ; N - CA
- HA
3251 3253 3255 1 1.0970e+02 6.6944e+02 ; N - CA
- CB
3251 3253 3270 1 1.1010e+02 5.2718e+02 ; N - CA
- C
3252 3251 3253 1 1.1804e+02 4.1840e+02 ; H - N
- CA
3253 3255 3256 1 1.0950e+02 4.1840e+02 ; CA - CB
- HB2
3253 3255 3257 1 1.0950e+02 4.1840e+02 ; CA - CB
- HB3
3253 3255 3258 1 1.1400e+02 5.2718e+02 ; CA - CB
- CG
3253 3270 3271 1 1.2040e+02 6.6944e+02 ; CA - C
- O
3253 3270 3272 1 1.1660e+02 5.8576e+02 ; CA - C
- N
3254 3253 3255 1 1.0950e+02 4.1840e+02 ; HA - CA
- CB
3254 3253 3270 1 1.0950e+02 4.1840e+02 ; HA - CA
- C
3255 3253 3270 1 1.1110e+02 5.2718e+02 ; CB - CA
- C
3255 3258 3259 1 1.2000e+02 5.8576e+02 ; CB - CG
- CD1
3255 3258 3268 1 1.2000e+02 5.8576e+02 ; CB - CG
- CD2
3256 3255 3257 1 1.0950e+02 2.9288e+02 ; HB2 - CB
- HB3
3256 3255 3258 1 1.0950e+02 4.1840e+02 ; HB2 - CB
- CG
3257 3255 3258 1 1.0950e+02 4.1840e+02 ; HB3 - CB
- CG
3258 3259 3260 1 1.2000e+02 4.1840e+02 ; CG - CD1
- HD1
3258 3259 3261 1 1.2000e+02 5.2718e+02 ; CG - CD1
- CE1
3258 3268 3266 1 1.2000e+02 5.2718e+02 ; CG - CD2
- CE2
3258 3268 3269 1 1.2000e+02 4.1840e+02 ; CG - CD2
- HD2
3259 3258 3268 1 1.2000e+02 5.2718e+02 ; CD1 - CG
- CD2
3259 3261 3262 1 1.2000e+02 4.1840e+02 ; CD1 - CE1
- HE1
3259 3261 3263 1 1.2000e+02 5.2718e+02 ; CD1 - CE1
- CZ
3260 3259 3261 1 1.2000e+02 4.1840e+02 ; HD1 - CD1
- CE1
3261 3263 3264 1 1.2000e+02 5.8576e+02 ; CE1 - CZ
- OH

3261 3263 3266 1 1.2000e+02 5.2718e+02 ; CE1 - CZ
- CE2
3262 3261 3263 1 1.2000e+02 4.1840e+02 ; HE1 - CE1
- CZ
3263 3264 3265 1 1.1300e+02 4.1840e+02 ; CZ - OH
- HH
3263 3266 3267 1 1.2000e+02 4.1840e+02 ; CZ - CE2
- HE2
3263 3266 3268 1 1.2000e+02 5.2718e+02 ; CZ - CE2
- CD2
3264 3263 3266 1 1.2000e+02 5.8576e+02 ; OH - CZ
- CE2
3266 3268 3269 1 1.2000e+02 4.1840e+02 ; CE2 - CD2
- HD2
3267 3266 3268 1 1.2000e+02 4.1840e+02 ; HE2 - CE2
- CD2
3270 3272 3273 1 1.2000e+02 4.1840e+02 ; C - N
- H
3270 3272 3274 1 1.2190e+02 4.1840e+02 ; C - N
- CA
3271 3270 3272 1 1.2290e+02 6.6944e+02 ; O - C
- N
3272 3274 3275 1 1.0950e+02 4.1840e+02 ; N - CA
- HA
3272 3274 3276 1 1.0970e+02 6.6944e+02 ; N - CA
- CB
3272 3274 3284 1 1.1010e+02 5.2718e+02 ; N - CA
- C
3273 3272 3274 1 1.1804e+02 4.1840e+02 ; H - N
- CA
3274 3276 3277 1 1.0950e+02 4.1840e+02 ; CA - CB
- HB
3274 3276 3278 1 1.0950e+02 3.3472e+02 ; CA - CB
- CG2
3274 3276 3282 1 1.0950e+02 4.1840e+02 ; CA - CB
- OG1
3274 3284 3285 1 1.2040e+02 6.6944e+02 ; CA - C
- O
3274 3284 3286 1 1.1660e+02 5.8576e+02 ; CA - C
- N
3275 3274 3276 1 1.0950e+02 4.1840e+02 ; HA - CA
- CB
3275 3274 3284 1 1.0950e+02 4.1840e+02 ; HA - CA
- C
3276 3274 3284 1 1.1110e+02 5.2718e+02 ; CB - CA
- C
3276 3278 3279 1 1.0950e+02 4.1840e+02 ; CB - CG2
- HG21
3276 3278 3280 1 1.0950e+02 4.1840e+02 ; CB - CG2
- HG22
3276 3278 3281 1 1.0950e+02 4.1840e+02 ; CB - CG2
- HG23

3276 3282 3283 1 1.0850e+02 4.6024e+02 ; CB - OG1
- HG1
3277 3276 3278 1 1.0950e+02 4.1840e+02 ; HB - CB
- CG2
3277 3276 3282 1 1.0950e+02 4.1840e+02 ; HB - CB
- OG1
3278 3276 3282 1 1.0950e+02 4.1840e+02 ; CG2 - CB
- OG1
3279 3278 3280 1 1.0950e+02 2.9288e+02 ; HG21 - CG2
- HG22
3279 3278 3281 1 1.0950e+02 2.9288e+02 ; HG21 - CG2
- HG23
3280 3278 3281 1 1.0950e+02 2.9288e+02 ; HG22 - CG2
- HG23
3284 3286 3287 1 1.2000e+02 4.1840e+02 ; C - N
- H
3284 3286 3288 1 1.2190e+02 4.1840e+02 ; C - N
- CA
3285 3284 3286 1 1.2290e+02 6.6944e+02 ; O - C
- N
3286 3288 3289 1 1.0950e+02 4.1840e+02 ; N - CA
- HA
3286 3288 3290 1 1.0970e+02 6.6944e+02 ; N - CA
- CB
3286 3288 3303 1 1.1010e+02 5.2718e+02 ; N - CA
- C
3287 3286 3288 1 1.1804e+02 4.1840e+02 ; H - N
- CA
3288 3290 3291 1 1.0950e+02 4.1840e+02 ; CA - CB
- HB2
3288 3290 3292 1 1.0950e+02 4.1840e+02 ; CA - CB
- HB3
3288 3290 3293 1 1.0950e+02 3.3472e+02 ; CA - CB
- CG
3288 3303 3304 1 1.2040e+02 6.6944e+02 ; CA - C
- O
3288 3303 3305 1 1.1660e+02 5.8576e+02 ; CA - C
- N
3289 3288 3290 1 1.0950e+02 4.1840e+02 ; HA - CA
- CB
3289 3288 3303 1 1.0950e+02 4.1840e+02 ; HA - CA
- C
3290 3288 3303 1 1.1110e+02 5.2718e+02 ; CB - CA
- C
3290 3293 3294 1 1.0950e+02 4.1840e+02 ; CB - CG
- HG
3290 3293 3295 1 1.0950e+02 3.3472e+02 ; CB - CG
- CD1
3290 3293 3299 1 1.0950e+02 3.3472e+02 ; CB - CG
- CD2
3291 3290 3292 1 1.0950e+02 2.9288e+02 ; HB2 - CB
- HB3

3291 3290 3293 1 1.0950e+02 4.1840e+02 ; HB2 - CB
- CG
3292 3290 3293 1 1.0950e+02 4.1840e+02 ; HB3 - CB
- CG
3293 3295 3296 1 1.0950e+02 4.1840e+02 ; CG - CD1
- HD11
3293 3295 3297 1 1.0950e+02 4.1840e+02 ; CG - CD1
- HD12
3293 3295 3298 1 1.0950e+02 4.1840e+02 ; CG - CD1
- HD13
3293 3299 3300 1 1.0950e+02 4.1840e+02 ; CG - CD2
- HD21
3293 3299 3301 1 1.0950e+02 4.1840e+02 ; CG - CD2
- HD22
3293 3299 3302 1 1.0950e+02 4.1840e+02 ; CG - CD2
- HD23
3294 3293 3295 1 1.0950e+02 4.1840e+02 ; HG - CG
- CD1
3294 3293 3299 1 1.0950e+02 4.1840e+02 ; HG - CG
- CD2
3295 3293 3299 1 1.0950e+02 3.3472e+02 ; CD1 - CG
- CD2
3296 3295 3297 1 1.0950e+02 2.9288e+02 ; HD11 - CD1
- HD12
3296 3295 3298 1 1.0950e+02 2.9288e+02 ; HD11 - CD1
- HD13
3297 3295 3298 1 1.0950e+02 2.9288e+02 ; HD12 - CD1
- HD13
3300 3299 3301 1 1.0950e+02 2.9288e+02 ; HD21 - CD2
- HD22
3300 3299 3302 1 1.0950e+02 2.9288e+02 ; HD21 - CD2
- HD23
3301 3299 3302 1 1.0950e+02 2.9288e+02 ; HD22 - CD2
- HD23
3303 3305 3306 1 1.2000e+02 4.1840e+02 ; C - N
- H
3303 3305 3307 1 1.2190e+02 4.1840e+02 ; C - N
- CA
3304 3303 3305 1 1.2290e+02 6.6944e+02 ; O - C
- N
3305 3307 3308 1 1.0950e+02 4.1840e+02 ; N - CA
- HA
3305 3307 3309 1 1.0970e+02 6.6944e+02 ; N - CA
- CB
3305 3307 3322 1 1.1010e+02 5.2718e+02 ; N - CA
- C
3306 3305 3307 1 1.1804e+02 4.1840e+02 ; H - N
- CA
3307 3309 3310 1 1.0950e+02 4.1840e+02 ; CA - CB
- HB2
3307 3309 3311 1 1.0950e+02 4.1840e+02 ; CA - CB
- HB3

3307 3309 3312 1 1.0950e+02 3.3472e+02 ; CA - CB
- CG
3307 3322 3323 1 1.2040e+02 6.6944e+02 ; CA - C
- O
3307 3322 3324 1 1.1660e+02 5.8576e+02 ; CA - C
- N
3308 3307 3309 1 1.0950e+02 4.1840e+02 ; HA - CA
- CB
3308 3307 3322 1 1.0950e+02 4.1840e+02 ; HA - CA
- C
3309 3307 3322 1 1.1110e+02 5.2718e+02 ; CB - CA
- C
3309 3312 3313 1 1.0950e+02 4.1840e+02 ; CB - CG
- HG
3309 3312 3314 1 1.0950e+02 3.3472e+02 ; CB - CG
- CD1
3309 3312 3318 1 1.0950e+02 3.3472e+02 ; CB - CG
- CD2
3310 3309 3311 1 1.0950e+02 2.9288e+02 ; HB2 - CB
- HB3
3310 3309 3312 1 1.0950e+02 4.1840e+02 ; HB2 - CB
- CG
3311 3309 3312 1 1.0950e+02 4.1840e+02 ; HB3 - CB
- CG
3312 3314 3315 1 1.0950e+02 4.1840e+02 ; CG - CD1
- HD11
3312 3314 3316 1 1.0950e+02 4.1840e+02 ; CG - CD1
- HD12
3312 3314 3317 1 1.0950e+02 4.1840e+02 ; CG - CD1
- HD13
3312 3318 3319 1 1.0950e+02 4.1840e+02 ; CG - CD2
- HD21
3312 3318 3320 1 1.0950e+02 4.1840e+02 ; CG - CD2
- HD22
3312 3318 3321 1 1.0950e+02 4.1840e+02 ; CG - CD2
- HD23
3313 3312 3314 1 1.0950e+02 4.1840e+02 ; HG - CG
- CD1
3313 3312 3318 1 1.0950e+02 4.1840e+02 ; HG - CG
- CD2
3314 3312 3318 1 1.0950e+02 3.3472e+02 ; CD1 - CG
- CD2
3315 3314 3316 1 1.0950e+02 2.9288e+02 ; HD11 - CD1
- HD12
3315 3314 3317 1 1.0950e+02 2.9288e+02 ; HD11 - CD1
- HD13
3316 3314 3317 1 1.0950e+02 2.9288e+02 ; HD12 - CD1
- HD13
3319 3318 3320 1 1.0950e+02 2.9288e+02 ; HD21 - CD2
- HD22
3319 3318 3321 1 1.0950e+02 2.9288e+02 ; HD21 - CD2
- HD23

3320 3318 3321 1 1.0950e+02 2.9288e+02 ; HD22 - CD2
- HD23
3322 3324 3325 1 1.2000e+02 4.1840e+02 ; C - N
- H
3322 3324 3326 1 1.2190e+02 4.1840e+02 ; C - N
- CA
3323 3322 3324 1 1.2290e+02 6.6944e+02 ; O - C
- N
3324 3326 3327 1 1.0950e+02 4.1840e+02 ; N - CA
- HA
3324 3326 3328 1 1.0970e+02 6.6944e+02 ; N - CA
- CB
3324 3326 3333 1 1.1010e+02 5.2718e+02 ; N - CA
- C
3325 3324 3326 1 1.1804e+02 4.1840e+02 ; H - N
- CA
3326 3328 3329 1 1.0950e+02 4.1840e+02 ; CA - CB
- HB2
3326 3328 3330 1 1.0950e+02 4.1840e+02 ; CA - CB
- HB3
3326 3328 3331 1 1.0860e+02 4.1840e+02 ; CA - CB
- SG
3326 3333 3334 1 1.2040e+02 6.6944e+02 ; CA - C
- O
3326 3333 3335 1 1.1660e+02 5.8576e+02 ; CA - C
- N
3327 3326 3328 1 1.0950e+02 4.1840e+02 ; HA - CA
- CB
3327 3326 3333 1 1.0950e+02 4.1840e+02 ; HA - CA
- C
3328 3326 3333 1 1.1110e+02 5.2718e+02 ; CB - CA
- C
3328 3331 3332 1 9.6000e+01 3.5982e+02 ; CB - SG
- HG
3329 3328 3330 1 1.0950e+02 2.9288e+02 ; HB2 - CB
- HB3
3329 3328 3331 1 1.0950e+02 4.1840e+02 ; HB2 - CB
- SG
3330 3328 3331 1 1.0950e+02 4.1840e+02 ; HB3 - CB
- SG
3333 3335 3336 1 1.2000e+02 4.1840e+02 ; C - N
- H
3333 3335 3337 1 1.2190e+02 4.1840e+02 ; C - N
- CA
3334 3333 3335 1 1.2290e+02 6.6944e+02 ; O - C
- N
3335 3337 3338 1 1.0950e+02 4.1840e+02 ; N - CA
- HA2
3335 3337 3339 1 1.0950e+02 4.1840e+02 ; N - CA
- HA3
3335 3337 3340 1 1.1010e+02 5.2718e+02 ; N - CA
- C

3336 3335 3337 1 1.1804e+02 4.1840e+02 ; H - N
- CA
3337 3340 3341 1 1.2040e+02 6.6944e+02 ; CA - C
- O
3337 3340 3342 1 1.1660e+02 5.8576e+02 ; CA - C
- N
3338 3337 3339 1 1.0950e+02 2.9288e+02 ; HA2 - CA
- HA3
3338 3337 3340 1 1.0950e+02 4.1840e+02 ; HA2 - CA
- C
3339 3337 3340 1 1.0950e+02 4.1840e+02 ; HA3 - CA
- C
3340 3342 3343 1 1.2000e+02 4.1840e+02 ; C - N
- H
3340 3342 3344 1 1.2190e+02 4.1840e+02 ; C - N
- CA
3341 3340 3342 1 1.2290e+02 6.6944e+02 ; O - C
- N
3342 3344 3345 1 1.0950e+02 4.1840e+02 ; N - CA
- HA
3342 3344 3346 1 1.0970e+02 6.6944e+02 ; N - CA
- CB
3342 3344 3361 1 1.1010e+02 5.2718e+02 ; N - CA
- C
3343 3342 3344 1 1.1804e+02 4.1840e+02 ; H - N
- CA
3344 3346 3347 1 1.0950e+02 4.1840e+02 ; CA - CB
- HB2
3344 3346 3348 1 1.0950e+02 4.1840e+02 ; CA - CB
- HB3
3344 3346 3349 1 1.1400e+02 5.2718e+02 ; CA - CB
- CG
3344 3361 3362 1 1.2040e+02 6.6944e+02 ; CA - C
- O
3344 3361 3363 1 1.1660e+02 5.8576e+02 ; CA - C
- N
3345 3344 3346 1 1.0950e+02 4.1840e+02 ; HA - CA
- CB
3345 3344 3361 1 1.0950e+02 4.1840e+02 ; HA - CA
- C
3346 3344 3361 1 1.1110e+02 5.2718e+02 ; CB - CA
- C
3346 3349 3350 1 1.2000e+02 5.8576e+02 ; CB - CG
- CD1
3346 3349 3359 1 1.2000e+02 5.8576e+02 ; CB - CG
- CD2
3347 3346 3348 1 1.0950e+02 2.9288e+02 ; HB2 - CB
- HB3
3347 3346 3349 1 1.0950e+02 4.1840e+02 ; HB2 - CB
- CG
3348 3346 3349 1 1.0950e+02 4.1840e+02 ; HB3 - CB
- CG

3349 3350 3351 1 1.2000e+02 4.1840e+02 ; CG - CD1
- HD1
3349 3350 3352 1 1.2000e+02 5.2718e+02 ; CG - CD1
- CE1
3349 3359 3357 1 1.2000e+02 5.2718e+02 ; CG - CD2
- CE2
3349 3359 3360 1 1.2000e+02 4.1840e+02 ; CG - CD2
- HD2
3350 3349 3359 1 1.2000e+02 5.2718e+02 ; CD1 - CG
- CD2
3350 3352 3353 1 1.2000e+02 4.1840e+02 ; CD1 - CE1
- HE1
3350 3352 3354 1 1.2000e+02 5.2718e+02 ; CD1 - CE1
- CZ
3351 3350 3352 1 1.2000e+02 4.1840e+02 ; HD1 - CD1
- CE1
3352 3354 3355 1 1.2000e+02 5.8576e+02 ; CE1 - CZ
- OH
3352 3354 3357 1 1.2000e+02 5.2718e+02 ; CE1 - CZ
- CE2
3353 3352 3354 1 1.2000e+02 4.1840e+02 ; HE1 - CE1
- CZ
3354 3355 3356 1 1.1300e+02 4.1840e+02 ; CZ - OH
- HH
3354 3357 3358 1 1.2000e+02 4.1840e+02 ; CZ - CE2
- HE2
3354 3357 3359 1 1.2000e+02 5.2718e+02 ; CZ - CE2
- CD2
3355 3354 3357 1 1.2000e+02 5.8576e+02 ; OH - CZ
- CE2
3357 3359 3360 1 1.2000e+02 4.1840e+02 ; CE2 - CD2
- HD2
3358 3357 3359 1 1.2000e+02 4.1840e+02 ; HE2 - CE2
- CD2
3361 3363 3364 1 1.2190e+02 4.1840e+02 ; C - N
- CD
3361 3363 3373 1 1.2190e+02 4.1840e+02 ; C - N
- CA
3362 3361 3363 1 1.2290e+02 6.6944e+02 ; O - C
- N
3363 3364 3365 1 1.0950e+02 4.1840e+02 ; N - CD
- HD2
3363 3364 3366 1 1.0950e+02 4.1840e+02 ; N - CD
- HD3
3363 3364 3367 1 1.0970e+02 6.6944e+02 ; N - CD
- CG
3363 3373 3370 1 1.0970e+02 6.6944e+02 ; N - CA
- CB
3363 3373 3374 1 1.0950e+02 4.1840e+02 ; N - CA
- HA
3363 3373 3375 1 1.1010e+02 5.2718e+02 ; N - CA
- C

3364 3363 3373 1 1.1800e+02 4.1840e+02 ; CD - N
- CA
3364 3367 3368 1 1.0950e+02 4.1840e+02 ; CD - CG
- HG2
3364 3367 3369 1 1.0950e+02 4.1840e+02 ; CD - CG
- HG3
3364 3367 3370 1 1.0950e+02 3.3472e+02 ; CD - CG
- CB
3365 3364 3366 1 1.0950e+02 2.9288e+02 ; HD2 - CD
- HD3
3365 3364 3367 1 1.0950e+02 4.1840e+02 ; HD2 - CD
- CG
3366 3364 3367 1 1.0950e+02 4.1840e+02 ; HD3 - CD
- CG
3367 3370 3371 1 1.0950e+02 4.1840e+02 ; CG - CB
- HB2
3367 3370 3372 1 1.0950e+02 4.1840e+02 ; CG - CB
- HB3
3367 3370 3373 1 1.0950e+02 3.3472e+02 ; CG - CB
- CA
3368 3367 3369 1 1.0950e+02 2.9288e+02 ; HG2 - CG
- HG3
3368 3367 3370 1 1.0950e+02 4.1840e+02 ; HG2 - CG
- CB
3369 3367 3370 1 1.0950e+02 4.1840e+02 ; HG3 - CG
- CB
3370 3373 3374 1 1.0950e+02 4.1840e+02 ; CB - CA
- HA
3370 3373 3375 1 1.1110e+02 5.2718e+02 ; CB - CA
- C
3371 3370 3372 1 1.0950e+02 2.9288e+02 ; HB2 - CB
- HB3
3371 3370 3373 1 1.0950e+02 4.1840e+02 ; HB2 - CB
- CA
3372 3370 3373 1 1.0950e+02 4.1840e+02 ; HB3 - CB
- CA
3373 3375 3376 1 1.2040e+02 6.6944e+02 ; CA - C
- O
3373 3375 3377 1 1.1660e+02 5.8576e+02 ; CA - C
- N
3374 3373 3375 1 1.0950e+02 4.1840e+02 ; HA - CA
- C
3375 3377 3378 1 1.2190e+02 4.1840e+02 ; C - N
- CD
3375 3377 3387 1 1.2190e+02 4.1840e+02 ; C - N
- CA
3376 3375 3377 1 1.2290e+02 6.6944e+02 ; O - C
- N
3377 3378 3379 1 1.0950e+02 4.1840e+02 ; N - CD
- HD2
3377 3378 3380 1 1.0950e+02 4.1840e+02 ; N - CD
- HD3

3377 3378 3381 1 1.0970e+02 6.6944e+02 ; N - CD - CG
3377 3387 3384 1 1.0970e+02 6.6944e+02 ; N - CA - CB
3377 3387 3388 1 1.0950e+02 4.1840e+02 ; N - CA - HA
3377 3387 3389 1 1.1010e+02 5.2718e+02 ; N - CA - C
3378 3377 3387 1 1.1800e+02 4.1840e+02 ; CD - N - CA
3378 3381 3382 1 1.0950e+02 4.1840e+02 ; CD - CG - HG2
3378 3381 3383 1 1.0950e+02 4.1840e+02 ; CD - CG - HG3
3378 3381 3384 1 1.0950e+02 3.3472e+02 ; CD - CG - CB
3379 3378 3380 1 1.0950e+02 2.9288e+02 ; HD2 - CD - HD3
3379 3378 3381 1 1.0950e+02 4.1840e+02 ; HD2 - CD - CG
3380 3378 3381 1 1.0950e+02 4.1840e+02 ; HD3 - CD - CG
3381 3384 3385 1 1.0950e+02 4.1840e+02 ; CG - CB - HB2
3381 3384 3386 1 1.0950e+02 4.1840e+02 ; CG - CB - HB3
3381 3384 3387 1 1.0950e+02 3.3472e+02 ; CG - CB - CA
3382 3381 3383 1 1.0950e+02 2.9288e+02 ; HG2 - CG - HG3
3382 3381 3384 1 1.0950e+02 4.1840e+02 ; HG2 - CG - CB
3383 3381 3384 1 1.0950e+02 4.1840e+02 ; HG3 - CG - CB
3384 3387 3388 1 1.0950e+02 4.1840e+02 ; CB - CA - HA
3384 3387 3389 1 1.1110e+02 5.2718e+02 ; CB - CA - C
3385 3384 3386 1 1.0950e+02 2.9288e+02 ; HB2 - CB - HB3
3385 3384 3387 1 1.0950e+02 4.1840e+02 ; HB2 - CB - CA
3386 3384 3387 1 1.0950e+02 4.1840e+02 ; HB3 - CB - CA
3387 3389 3390 1 1.2040e+02 6.6944e+02 ; CA - C - O
3387 3389 3391 1 1.1660e+02 5.8576e+02 ; CA - C - N
3388 3387 3389 1 1.0950e+02 4.1840e+02 ; HA - CA - C
3389 3391 3392 1 1.2000e+02 4.1840e+02 ; C - N - H

3389 3391 3393 1 1.2190e+02 4.1840e+02 ; C - N
- CA
3390 3389 3391 1 1.2290e+02 6.6944e+02 ; O - C
- N
3391 3393 3394 1 1.0950e+02 4.1840e+02 ; N - CA
- HA
3391 3393 3395 1 1.0970e+02 6.6944e+02 ; N - CA
- CB
3391 3393 3409 1 1.1010e+02 5.2718e+02 ; N - CA
- C
3392 3391 3393 1 1.1804e+02 4.1840e+02 ; H - N
- CA
3393 3395 3396 1 1.0950e+02 4.1840e+02 ; CA - CB
- HB2
3393 3395 3397 1 1.0950e+02 4.1840e+02 ; CA - CB
- HB3
3393 3395 3398 1 1.1400e+02 5.2718e+02 ; CA - CB
- CG
3393 3409 3410 1 1.2040e+02 6.6944e+02 ; CA - C
- O
3393 3409 3411 1 1.1660e+02 5.8576e+02 ; CA - C
- N
3394 3393 3395 1 1.0950e+02 4.1840e+02 ; HA - CA
- CB
3394 3393 3409 1 1.0950e+02 4.1840e+02 ; HA - CA
- C
3395 3393 3409 1 1.1110e+02 5.2718e+02 ; CB - CA
- C
3395 3398 3399 1 1.2000e+02 5.8576e+02 ; CB - CG
- CD1
3395 3398 3407 1 1.2000e+02 5.8576e+02 ; CB - CG
- CD2
3396 3395 3397 1 1.0950e+02 2.9288e+02 ; HB2 - CB
- HB3
3396 3395 3398 1 1.0950e+02 4.1840e+02 ; HB2 - CB
- CG
3397 3395 3398 1 1.0950e+02 4.1840e+02 ; HB3 - CB
- CG
3398 3399 3400 1 1.2000e+02 4.1840e+02 ; CG - CD1
- HD1
3398 3399 3401 1 1.2000e+02 5.2718e+02 ; CG - CD1
- CE1
3398 3407 3405 1 1.2000e+02 5.2718e+02 ; CG - CD2
- CE2
3398 3407 3408 1 1.2000e+02 4.1840e+02 ; CG - CD2
- HD2
3399 3398 3407 1 1.2000e+02 5.2718e+02 ; CD1 - CG
- CD2
3399 3401 3402 1 1.2000e+02 4.1840e+02 ; CD1 - CE1
- HE1
3399 3401 3403 1 1.2000e+02 5.2718e+02 ; CD1 - CE1
- CZ

3400 3399 3401 1 1.2000e+02 4.1840e+02 ; HD1 - CD1
- CE1
3401 3403 3404 1 1.2000e+02 4.1840e+02 ; CE1 - CZ
- HZ
3401 3403 3405 1 1.2000e+02 5.2718e+02 ; CE1 - CZ
- CE2
3402 3401 3403 1 1.2000e+02 4.1840e+02 ; HE1 - CE1
- CZ
3403 3405 3406 1 1.2000e+02 4.1840e+02 ; CZ - CE2
- HE2
3403 3405 3407 1 1.2000e+02 5.2718e+02 ; CZ - CE2
- CD2
3404 3403 3405 1 1.2000e+02 4.1840e+02 ; HZ - CZ
- CE2
3405 3407 3408 1 1.2000e+02 4.1840e+02 ; CE2 - CD2
- HD2
3406 3405 3407 1 1.2000e+02 4.1840e+02 ; HE2 - CE2
- CD2
3409 3411 3412 1 1.2000e+02 4.1840e+02 ; C - N
- H
3409 3411 3413 1 1.2190e+02 4.1840e+02 ; C - N
- CA
3410 3409 3411 1 1.2290e+02 6.6944e+02 ; O - C
- N
3411 3413 3414 1 1.0950e+02 4.1840e+02 ; N - CA
- HA
3411 3413 3415 1 1.0970e+02 6.6944e+02 ; N - CA
- CB
3411 3413 3430 1 1.1010e+02 5.2718e+02 ; N - CA
- C
3412 3411 3413 1 1.1804e+02 4.1840e+02 ; H - N
- CA
3413 3415 3416 1 1.0950e+02 4.1840e+02 ; CA - CB
- HB2
3413 3415 3417 1 1.0950e+02 4.1840e+02 ; CA - CB
- HB3
3413 3415 3418 1 1.1400e+02 5.2718e+02 ; CA - CB
- CG
3413 3430 3431 1 1.2040e+02 6.6944e+02 ; CA - C
- O
3413 3430 3432 1 1.1660e+02 5.8576e+02 ; CA - C
- N
3414 3413 3415 1 1.0950e+02 4.1840e+02 ; HA - CA
- CB
3414 3413 3430 1 1.0950e+02 4.1840e+02 ; HA - CA
- C
3415 3413 3430 1 1.1110e+02 5.2718e+02 ; CB - CA
- C
3415 3418 3419 1 1.2000e+02 5.8576e+02 ; CB - CG
- CD1
3415 3418 3428 1 1.2000e+02 5.8576e+02 ; CB - CG
- CD2

3416 3415 3417 1 1.0950e+02 2.9288e+02 ; HB2 - CB
- HB3
3416 3415 3418 1 1.0950e+02 4.1840e+02 ; HB2 - CB
- CG
3417 3415 3418 1 1.0950e+02 4.1840e+02 ; HB3 - CB
- CG
3418 3419 3420 1 1.2000e+02 4.1840e+02 ; CG - CD1
- HD1
3418 3419 3421 1 1.2000e+02 5.2718e+02 ; CG - CD1
- CE1
3418 3428 3426 1 1.2000e+02 5.2718e+02 ; CG - CD2
- CE2
3418 3428 3429 1 1.2000e+02 4.1840e+02 ; CG - CD2
- HD2
3419 3418 3428 1 1.2000e+02 5.2718e+02 ; CD1 - CG
- CD2
3419 3421 3422 1 1.2000e+02 4.1840e+02 ; CD1 - CE1
- HE1
3419 3421 3423 1 1.2000e+02 5.2718e+02 ; CD1 - CE1
- CZ
3420 3419 3421 1 1.2000e+02 4.1840e+02 ; HD1 - CD1
- CE1
3421 3423 3424 1 1.2000e+02 5.8576e+02 ; CE1 - CZ
- OH
3421 3423 3426 1 1.2000e+02 5.2718e+02 ; CE1 - CZ
- CE2
3422 3421 3423 1 1.2000e+02 4.1840e+02 ; HE1 - CE1
- CZ
3423 3424 3425 1 1.1300e+02 4.1840e+02 ; CZ - OH
- HH
3423 3426 3427 1 1.2000e+02 4.1840e+02 ; CZ - CE2
- HE2
3423 3426 3428 1 1.2000e+02 5.2718e+02 ; CZ - CE2
- CD2
3424 3423 3426 1 1.2000e+02 5.8576e+02 ; OH - CZ
- CE2
3426 3428 3429 1 1.2000e+02 4.1840e+02 ; CE2 - CD2
- HD2
3427 3426 3428 1 1.2000e+02 4.1840e+02 ; HE2 - CE2
- CD2
3430 3432 3433 1 1.2000e+02 4.1840e+02 ; C - N
- H
3430 3432 3434 1 1.2190e+02 4.1840e+02 ; C - N
- CA
3431 3430 3432 1 1.2290e+02 6.6944e+02 ; O - C
- N
3432 3434 3435 1 1.0950e+02 4.1840e+02 ; N - CA
- HA2
3432 3434 3436 1 1.0950e+02 4.1840e+02 ; N - CA
- HA3
3432 3434 3437 1 1.1010e+02 5.2718e+02 ; N - CA
- C

3433 3432 3434 1 1.1804e+02 4.1840e+02 ; H - N
- CA
3434 3437 3438 1 1.2040e+02 6.6944e+02 ; CA - C
- O
3434 3437 3439 1 1.1660e+02 5.8576e+02 ; CA - C
- N
3435 3434 3436 1 1.0950e+02 2.9288e+02 ; HA2 - CA
- HA3
3435 3434 3437 1 1.0950e+02 4.1840e+02 ; HA2 - CA
- C
3436 3434 3437 1 1.0950e+02 4.1840e+02 ; HA3 - CA
- C
3437 3439 3440 1 1.2000e+02 4.1840e+02 ; C - N
- H
3437 3439 3441 1 1.2190e+02 4.1840e+02 ; C - N
- CA
3438 3437 3439 1 1.2290e+02 6.6944e+02 ; O - C
- N
3439 3441 3442 1 1.0950e+02 4.1840e+02 ; N - CA
- HA
3439 3441 3443 1 1.0970e+02 6.6944e+02 ; N - CA
- CB
3439 3441 3449 1 1.1010e+02 5.2718e+02 ; N - CA
- C
3440 3439 3441 1 1.1804e+02 4.1840e+02 ; H - N
- CA
3441 3443 3444 1 1.0950e+02 4.1840e+02 ; CA - CB
- HB2
3441 3443 3445 1 1.0950e+02 4.1840e+02 ; CA - CB
- HB3
3441 3443 3446 1 1.1110e+02 5.2718e+02 ; CA - CB
- CG
3441 3449 3450 1 1.2040e+02 6.6944e+02 ; CA - C
- O
3441 3449 3451 1 1.1660e+02 5.8576e+02 ; CA - C
- N
3442 3441 3443 1 1.0950e+02 4.1840e+02 ; HA - CA
- CB
3442 3441 3449 1 1.0950e+02 4.1840e+02 ; HA - CA
- C
3443 3441 3449 1 1.1110e+02 5.2718e+02 ; CB - CA
- C
3443 3446 3447 1 1.1700e+02 5.8576e+02 ; CB - CG
- OD1
3443 3446 3448 1 1.1700e+02 5.8576e+02 ; CB - CG
- OD2
3444 3443 3445 1 1.0950e+02 2.9288e+02 ; HB2 - CB
- HB3
3444 3443 3446 1 1.0950e+02 4.1840e+02 ; HB2 - CB
- CG
3445 3443 3446 1 1.0950e+02 4.1840e+02 ; HB3 - CB
- CG

3447 3446 3448 1 1.2600e+02 6.6944e+02 ; OD1 - CG
- OD2
3449 3451 3452 1 1.2000e+02 4.1840e+02 ; C - N
- H
3449 3451 3453 1 1.2190e+02 4.1840e+02 ; C - N
- CA
3450 3449 3451 1 1.2290e+02 6.6944e+02 ; O - C
- N
3451 3453 3454 1 1.0950e+02 4.1840e+02 ; N - CA
- HA
3451 3453 3455 1 1.0970e+02 6.6944e+02 ; N - CA
- CB
3451 3453 3463 1 1.1010e+02 5.2718e+02 ; N - CA
- C
3452 3451 3453 1 1.1804e+02 4.1840e+02 ; H - N
- CA
3453 3455 3456 1 1.0950e+02 4.1840e+02 ; CA - CB
- HB
3453 3455 3457 1 1.0950e+02 3.3472e+02 ; CA - CB
- CG2
3453 3455 3461 1 1.0950e+02 4.1840e+02 ; CA - CB
- OG1
3453 3463 3464 1 1.2040e+02 6.6944e+02 ; CA - C
- O
3453 3463 3465 1 1.1660e+02 5.8576e+02 ; CA - C
- N
3454 3453 3455 1 1.0950e+02 4.1840e+02 ; HA - CA
- CB
3454 3453 3463 1 1.0950e+02 4.1840e+02 ; HA - CA
- C
3455 3453 3463 1 1.1110e+02 5.2718e+02 ; CB - CA
- C
3455 3457 3458 1 1.0950e+02 4.1840e+02 ; CB - CG2
- HG21
3455 3457 3459 1 1.0950e+02 4.1840e+02 ; CB - CG2
- HG22
3455 3457 3460 1 1.0950e+02 4.1840e+02 ; CB - CG2
- HG23
3455 3461 3462 1 1.0850e+02 4.6024e+02 ; CB - OG1
- HG1
3456 3455 3457 1 1.0950e+02 4.1840e+02 ; HB - CB
- CG2
3456 3455 3461 1 1.0950e+02 4.1840e+02 ; HB - CB
- OG1
3457 3455 3461 1 1.0950e+02 4.1840e+02 ; CG2 - CB
- OG1
3458 3457 3459 1 1.0950e+02 2.9288e+02 ; HG21 - CG2
- HG22
3458 3457 3460 1 1.0950e+02 2.9288e+02 ; HG21 - CG2
- HG23
3459 3457 3460 1 1.0950e+02 2.9288e+02 ; HG22 - CG2
- HG23

3463 3465 3466 1 1.2000e+02 4.1840e+02 ; C - N
- H
3463 3465 3467 1 1.2190e+02 4.1840e+02 ; C - N
- CA
3464 3463 3465 1 1.2290e+02 6.6944e+02 ; O - C
- N
3465 3467 3468 1 1.0950e+02 4.1840e+02 ; N - CA
- HA
3465 3467 3469 1 1.0970e+02 6.6944e+02 ; N - CA
- CB
3465 3467 3475 1 1.1010e+02 5.2718e+02 ; N - CA
- C
3466 3465 3467 1 1.1804e+02 4.1840e+02 ; H - N
- CA
3467 3469 3470 1 1.0950e+02 4.1840e+02 ; CA - CB
- HB2
3467 3469 3471 1 1.0950e+02 4.1840e+02 ; CA - CB
- HB3
3467 3469 3472 1 1.1110e+02 5.2718e+02 ; CA - CB
- CG
3467 3475 3476 1 1.2040e+02 6.6944e+02 ; CA - C
- O
3467 3475 3477 1 1.1660e+02 5.8576e+02 ; CA - C
- N
3468 3467 3469 1 1.0950e+02 4.1840e+02 ; HA - CA
- CB
3468 3467 3475 1 1.0950e+02 4.1840e+02 ; HA - CA
- C
3469 3467 3475 1 1.1110e+02 5.2718e+02 ; CB - CA
- C
3469 3472 3473 1 1.1700e+02 5.8576e+02 ; CB - CG
- OD1
3469 3472 3474 1 1.1700e+02 5.8576e+02 ; CB - CG
- OD2
3470 3469 3471 1 1.0950e+02 2.9288e+02 ; HB2 - CB
- HB3
3470 3469 3472 1 1.0950e+02 4.1840e+02 ; HB2 - CB
- CG
3471 3469 3472 1 1.0950e+02 4.1840e+02 ; HB3 - CB
- CG
3473 3472 3474 1 1.2600e+02 6.6944e+02 ; OD1 - CG
- OD2
3475 3477 3478 1 1.2000e+02 4.1840e+02 ; C - N
- H
3475 3477 3479 1 1.2190e+02 4.1840e+02 ; C - N
- CA
3476 3475 3477 1 1.2290e+02 6.6944e+02 ; O - C
- N
3477 3479 3480 1 1.0950e+02 4.1840e+02 ; N - CA
- HA
3477 3479 3481 1 1.0970e+02 6.6944e+02 ; N - CA
- CB

3477 3479 3489 1 1.1010e+02 5.2718e+02 ; N - CA
- C
3478 3477 3479 1 1.1804e+02 4.1840e+02 ; H - N
- CA
3479 3481 3482 1 1.0950e+02 4.1840e+02 ; CA - CB
- HB2
3479 3481 3483 1 1.0950e+02 4.1840e+02 ; CA - CB
- HB3
3479 3481 3484 1 1.1110e+02 5.2718e+02 ; CA - CB
- CG
3479 3489 3490 1 1.2040e+02 6.6944e+02 ; CA - C
- O
3479 3489 3491 1 1.1660e+02 5.8576e+02 ; CA - C
- N
3480 3479 3481 1 1.0950e+02 4.1840e+02 ; HA - CA
- CB
3480 3479 3489 1 1.0950e+02 4.1840e+02 ; HA - CA
- C
3481 3479 3489 1 1.1110e+02 5.2718e+02 ; CB - CA
- C
3481 3484 3485 1 1.2040e+02 6.6944e+02 ; CB - CG
- OD1
3481 3484 3486 1 1.1660e+02 5.8576e+02 ; CB - CG
- ND2
3482 3481 3483 1 1.0950e+02 2.9288e+02 ; HB2 - CB
- HB3
3482 3481 3484 1 1.0950e+02 4.1840e+02 ; HB2 - CB
- CG
3483 3481 3484 1 1.0950e+02 4.1840e+02 ; HB3 - CB
- CG
3484 3486 3487 1 1.2000e+02 4.1840e+02 ; CG - ND2
- HD21
3484 3486 3488 1 1.2000e+02 4.1840e+02 ; CG - ND2
- HD22
3485 3484 3486 1 1.2290e+02 6.6944e+02 ; OD1 - CG
- ND2
3487 3486 3488 1 1.2000e+02 2.9288e+02 ; HD21 - ND2
- HD22
3489 3491 3492 1 1.2000e+02 4.1840e+02 ; C - N
- H
3489 3491 3493 1 1.2190e+02 4.1840e+02 ; C - N
- CA
3490 3489 3491 1 1.2290e+02 6.6944e+02 ; O - C
- N
3491 3493 3494 1 1.0950e+02 4.1840e+02 ; N - CA
- HA
3491 3493 3495 1 1.0970e+02 6.6944e+02 ; N - CA
- CB
3491 3493 3504 1 1.1010e+02 5.2718e+02 ; N - CA
- C
3492 3491 3493 1 1.1804e+02 4.1840e+02 ; H - N
- CA

3493 3495 3496 1 1.0950e+02 4.1840e+02 ; CA - CB
- HB2
3493 3495 3497 1 1.0950e+02 4.1840e+02 ; CA - CB
- HB3
3493 3495 3498 1 1.0950e+02 3.3472e+02 ; CA - CB
- CG
3493 3504 3505 1 1.2040e+02 6.6944e+02 ; CA - C
- O
3493 3504 3506 1 1.1660e+02 5.8576e+02 ; CA - C
- N
3494 3493 3495 1 1.0950e+02 4.1840e+02 ; HA - CA
- CB
3494 3493 3504 1 1.0950e+02 4.1840e+02 ; HA - CA
- C
3495 3493 3504 1 1.1110e+02 5.2718e+02 ; CB - CA
- C
3495 3498 3499 1 1.0950e+02 4.1840e+02 ; CB - CG
- HG2
3495 3498 3500 1 1.0950e+02 4.1840e+02 ; CB - CG
- HG3
3495 3498 3501 1 1.1110e+02 5.2718e+02 ; CB - CG
- CD
3496 3495 3497 1 1.0950e+02 2.9288e+02 ; HB2 - CB
- HB3
3496 3495 3498 1 1.0950e+02 4.1840e+02 ; HB2 - CB
- CG
3497 3495 3498 1 1.0950e+02 4.1840e+02 ; HB3 - CB
- CG
3498 3501 3502 1 1.1700e+02 5.8576e+02 ; CG - CD
- OE1
3498 3501 3503 1 1.1700e+02 5.8576e+02 ; CG - CD
- OE2
3499 3498 3500 1 1.0950e+02 2.9288e+02 ; HG2 - CG
- HG3
3499 3498 3501 1 1.0950e+02 4.1840e+02 ; HG2 - CG
- CD
3500 3498 3501 1 1.0950e+02 4.1840e+02 ; HG3 - CG
- CD
3502 3501 3503 1 1.2600e+02 6.6944e+02 ; OE1 - CD
- OE2
3504 3506 3507 1 1.2000e+02 4.1840e+02 ; C - N
- H
3504 3506 3508 1 1.2190e+02 4.1840e+02 ; C - N
- CA
3505 3504 3506 1 1.2290e+02 6.6944e+02 ; O - C
- N
3506 3508 3509 1 1.0950e+02 4.1840e+02 ; N - CA
- HA
3506 3508 3510 1 1.0970e+02 6.6944e+02 ; N - CA
- CB
3506 3508 3520 1 1.1010e+02 5.2718e+02 ; N - CA
- C

3507 3506 3508 1 1.1804e+02 4.1840e+02 ; H - N
- CA
3508 3510 3511 1 1.0950e+02 4.1840e+02 ; CA - CB
- HB
3508 3510 3512 1 1.0950e+02 3.3472e+02 ; CA - CB
- CG1
3508 3510 3516 1 1.0950e+02 3.3472e+02 ; CA - CB
- CG2
3508 3520 3521 1 1.2040e+02 6.6944e+02 ; CA - C
- O
3508 3520 3522 1 1.1660e+02 5.8576e+02 ; CA - C
- N
3509 3508 3510 1 1.0950e+02 4.1840e+02 ; HA - CA
- CB
3509 3508 3520 1 1.0950e+02 4.1840e+02 ; HA - CA
- C
3510 3508 3520 1 1.1110e+02 5.2718e+02 ; CB - CA
- C
3510 3512 3513 1 1.0950e+02 4.1840e+02 ; CB - CG1
- HG11
3510 3512 3514 1 1.0950e+02 4.1840e+02 ; CB - CG1
- HG12
3510 3512 3515 1 1.0950e+02 4.1840e+02 ; CB - CG1
- HG13
3510 3516 3517 1 1.0950e+02 4.1840e+02 ; CB - CG2
- HG21
3510 3516 3518 1 1.0950e+02 4.1840e+02 ; CB - CG2
- HG22
3510 3516 3519 1 1.0950e+02 4.1840e+02 ; CB - CG2
- HG23
3511 3510 3512 1 1.0950e+02 4.1840e+02 ; HB - CB
- CG1
3511 3510 3516 1 1.0950e+02 4.1840e+02 ; HB - CB
- CG2
3512 3510 3516 1 1.0950e+02 3.3472e+02 ; CG1 - CB
- CG2
3513 3512 3514 1 1.0950e+02 2.9288e+02 ; HG11 - CG1
- HG12
3513 3512 3515 1 1.0950e+02 2.9288e+02 ; HG11 - CG1
- HG13
3514 3512 3515 1 1.0950e+02 2.9288e+02 ; HG12 - CG1
- HG13
3517 3516 3518 1 1.0950e+02 2.9288e+02 ; HG21 - CG2
- HG22
3517 3516 3519 1 1.0950e+02 2.9288e+02 ; HG21 - CG2
- HG23
3518 3516 3519 1 1.0950e+02 2.9288e+02 ; HG22 - CG2
- HG23
3520 3522 3523 1 1.2000e+02 4.1840e+02 ; C - N
- H
3520 3522 3524 1 1.2190e+02 4.1840e+02 ; C - N
- CA

3521 3520 3522 1 1.2290e+02 6.6944e+02 ; O - C
- N
3522 3524 3525 1 1.0950e+02 4.1840e+02 ; N - CA
- HA
3522 3524 3526 1 1.0970e+02 6.6944e+02 ; N - CA
- CB
3522 3524 3539 1 1.1010e+02 5.2718e+02 ; N - CA
- C
3523 3522 3524 1 1.1804e+02 4.1840e+02 ; H - N
- CA
3524 3526 3527 1 1.0950e+02 4.1840e+02 ; CA - CB
- HB2
3524 3526 3528 1 1.0950e+02 4.1840e+02 ; CA - CB
- HB3
3524 3526 3529 1 1.0950e+02 3.3472e+02 ; CA - CB
- CG
3524 3539 3540 1 1.2040e+02 6.6944e+02 ; CA - C
- O
3524 3539 3541 1 1.1660e+02 5.8576e+02 ; CA - C
- N
3525 3524 3526 1 1.0950e+02 4.1840e+02 ; HA - CA
- CB
3525 3524 3539 1 1.0950e+02 4.1840e+02 ; HA - CA
- C
3526 3524 3539 1 1.1110e+02 5.2718e+02 ; CB - CA
- C
3526 3529 3530 1 1.0950e+02 4.1840e+02 ; CB - CG
- HG
3526 3529 3531 1 1.0950e+02 3.3472e+02 ; CB - CG
- CD1
3526 3529 3535 1 1.0950e+02 3.3472e+02 ; CB - CG
- CD2
3527 3526 3528 1 1.0950e+02 2.9288e+02 ; HB2 - CB
- HB3
3527 3526 3529 1 1.0950e+02 4.1840e+02 ; HB2 - CB
- CG
3528 3526 3529 1 1.0950e+02 4.1840e+02 ; HB3 - CB
- CG
3529 3531 3532 1 1.0950e+02 4.1840e+02 ; CG - CD1
- HD11
3529 3531 3533 1 1.0950e+02 4.1840e+02 ; CG - CD1
- HD12
3529 3531 3534 1 1.0950e+02 4.1840e+02 ; CG - CD1
- HD13
3529 3535 3536 1 1.0950e+02 4.1840e+02 ; CG - CD2
- HD21
3529 3535 3537 1 1.0950e+02 4.1840e+02 ; CG - CD2
- HD22
3529 3535 3538 1 1.0950e+02 4.1840e+02 ; CG - CD2
- HD23
3530 3529 3531 1 1.0950e+02 4.1840e+02 ; HG - CG
- CD1

3530 3529 3535 1 1.0950e+02 4.1840e+02 ; HG - CG
- CD2
3531 3529 3535 1 1.0950e+02 3.3472e+02 ; CD1 - CG
- CD2
3532 3531 3533 1 1.0950e+02 2.9288e+02 ; HD11 - CD1
- HD12
3532 3531 3534 1 1.0950e+02 2.9288e+02 ; HD11 - CD1
- HD13
3533 3531 3534 1 1.0950e+02 2.9288e+02 ; HD12 - CD1
- HD13
3536 3535 3537 1 1.0950e+02 2.9288e+02 ; HD21 - CD2
- HD22
3536 3535 3538 1 1.0950e+02 2.9288e+02 ; HD21 - CD2
- HD23
3537 3535 3538 1 1.0950e+02 2.9288e+02 ; HD22 - CD2
- HD23
3539 3541 3542 1 1.2000e+02 4.1840e+02 ; C - N
- H
3539 3541 3543 1 1.2190e+02 4.1840e+02 ; C - N
- CA
3540 3539 3541 1 1.2290e+02 6.6944e+02 ; O - C
- N
3541 3543 3544 1 1.0950e+02 4.1840e+02 ; N - CA
- HA
3541 3543 3545 1 1.0970e+02 6.6944e+02 ; N - CA
- CB
3541 3543 3561 1 1.1010e+02 5.2718e+02 ; N - CA
- C
3542 3541 3543 1 1.1804e+02 4.1840e+02 ; H - N
- CA
3543 3545 3546 1 1.0950e+02 4.1840e+02 ; CA - CB
- HB2
3543 3545 3547 1 1.0950e+02 4.1840e+02 ; CA - CB
- HB3
3543 3545 3548 1 1.0950e+02 3.3472e+02 ; CA - CB
- CG
3543 3561 3562 1 1.2040e+02 6.6944e+02 ; CA - C
- O
3543 3561 3563 1 1.1660e+02 5.8576e+02 ; CA - C
- N
3544 3543 3545 1 1.0950e+02 4.1840e+02 ; HA - CA
- CB
3544 3543 3561 1 1.0950e+02 4.1840e+02 ; HA - CA
- C
3545 3543 3561 1 1.1110e+02 5.2718e+02 ; CB - CA
- C
3545 3548 3549 1 1.0950e+02 4.1840e+02 ; CB - CG
- HG2
3545 3548 3550 1 1.0950e+02 4.1840e+02 ; CB - CG
- HG3
3545 3548 3551 1 1.0950e+02 3.3472e+02 ; CB - CG
- CD

3546 3545 3547 1 1.0950e+02 2.9288e+02 ; HB2 - CB
- HB3
3546 3545 3548 1 1.0950e+02 4.1840e+02 ; HB2 - CB
- CG
3547 3545 3548 1 1.0950e+02 4.1840e+02 ; HB3 - CB
- CG
3548 3551 3552 1 1.0950e+02 4.1840e+02 ; CG - CD
- HD2
3548 3551 3553 1 1.0950e+02 4.1840e+02 ; CG - CD
- HD3
3548 3551 3554 1 1.0950e+02 3.3472e+02 ; CG - CD
- CE
3549 3548 3550 1 1.0950e+02 2.9288e+02 ; HG2 - CG
- HG3
3549 3548 3551 1 1.0950e+02 4.1840e+02 ; HG2 - CG
- CD
3550 3548 3551 1 1.0950e+02 4.1840e+02 ; HG3 - CG
- CD
3551 3554 3555 1 1.0950e+02 4.1840e+02 ; CD - CE
- HE2
3551 3554 3556 1 1.0950e+02 4.1840e+02 ; CD - CE
- HE3
3551 3554 3557 1 1.1120e+02 6.6944e+02 ; CD - CE
- NZ
3552 3551 3553 1 1.0950e+02 2.9288e+02 ; HD2 - CD
- HD3
3552 3551 3554 1 1.0950e+02 4.1840e+02 ; HD2 - CD
- CE
3553 3551 3554 1 1.0950e+02 4.1840e+02 ; HD3 - CD
- CE
3554 3557 3558 1 1.0950e+02 4.1840e+02 ; CE - NZ
- HZ1
3554 3557 3559 1 1.0950e+02 4.1840e+02 ; CE - NZ
- HZ2
3554 3557 3560 1 1.0950e+02 4.1840e+02 ; CE - NZ
- HZ3
3555 3554 3556 1 1.0950e+02 2.9288e+02 ; HE2 - CE
- HE3
3555 3554 3557 1 1.0950e+02 4.1840e+02 ; HE2 - CE
- NZ
3556 3554 3557 1 1.0950e+02 4.1840e+02 ; HE3 - CE
- NZ
3558 3557 3559 1 1.0950e+02 2.9288e+02 ; HZ1 - NZ
- HZ2
3558 3557 3560 1 1.0950e+02 2.9288e+02 ; HZ1 - NZ
- HZ3
3559 3557 3560 1 1.0950e+02 2.9288e+02 ; HZ2 - NZ
- HZ3
3561 3563 3564 1 1.2000e+02 4.1840e+02 ; C - N
- H
3561 3563 3565 1 1.2190e+02 4.1840e+02 ; C - N
- CA

3562 3561 3563 1 1.2290e+02 6.6944e+02 ; O - C
- N
3563 3565 3566 1 1.0950e+02 4.1840e+02 ; N - CA
- HA
3563 3565 3567 1 1.0970e+02 6.6944e+02 ; N - CA
- CB
3563 3565 3583 1 1.1010e+02 5.2718e+02 ; N - CA
- C
3564 3563 3565 1 1.1804e+02 4.1840e+02 ; H - N
- CA
3565 3567 3568 1 1.0950e+02 4.1840e+02 ; CA - CB
- HB2
3565 3567 3569 1 1.0950e+02 4.1840e+02 ; CA - CB
- HB3
3565 3567 3570 1 1.0950e+02 3.3472e+02 ; CA - CB
- CG
3565 3583 3584 1 1.2040e+02 6.6944e+02 ; CA - C
- O
3565 3583 3585 1 1.1660e+02 5.8576e+02 ; CA - C
- N
3566 3565 3567 1 1.0950e+02 4.1840e+02 ; HA - CA
- CB
3566 3565 3583 1 1.0950e+02 4.1840e+02 ; HA - CA
- C
3567 3565 3583 1 1.1110e+02 5.2718e+02 ; CB - CA
- C
3567 3570 3571 1 1.0950e+02 4.1840e+02 ; CB - CG
- HG2
3567 3570 3572 1 1.0950e+02 4.1840e+02 ; CB - CG
- HG3
3567 3570 3573 1 1.0950e+02 3.3472e+02 ; CB - CG
- CD
3568 3567 3569 1 1.0950e+02 2.9288e+02 ; HB2 - CB
- HB3
3568 3567 3570 1 1.0950e+02 4.1840e+02 ; HB2 - CB
- CG
3569 3567 3570 1 1.0950e+02 4.1840e+02 ; HB3 - CB
- CG
3570 3573 3574 1 1.0950e+02 4.1840e+02 ; CG - CD
- HD2
3570 3573 3575 1 1.0950e+02 4.1840e+02 ; CG - CD
- HD3
3570 3573 3576 1 1.0950e+02 3.3472e+02 ; CG - CD
- CE
3571 3570 3572 1 1.0950e+02 2.9288e+02 ; HG2 - CG
- HG3
3571 3570 3573 1 1.0950e+02 4.1840e+02 ; HG2 - CG
- CD
3572 3570 3573 1 1.0950e+02 4.1840e+02 ; HG3 - CG
- CD
3573 3576 3577 1 1.0950e+02 4.1840e+02 ; CD - CE
- HE2

3573 3576 3578 1 1.0950e+02 4.1840e+02 ; CD - CE
- HE3
3573 3576 3579 1 1.1120e+02 6.6944e+02 ; CD - CE
- NZ
3574 3573 3575 1 1.0950e+02 2.9288e+02 ; HD2 - CD
- HD3
3574 3573 3576 1 1.0950e+02 4.1840e+02 ; HD2 - CD
- CE
3575 3573 3576 1 1.0950e+02 4.1840e+02 ; HD3 - CD
- CE
3576 3579 3580 1 1.0950e+02 4.1840e+02 ; CE - NZ
- HZ1
3576 3579 3581 1 1.0950e+02 4.1840e+02 ; CE - NZ
- HZ2
3576 3579 3582 1 1.0950e+02 4.1840e+02 ; CE - NZ
- HZ3
3577 3576 3578 1 1.0950e+02 2.9288e+02 ; HE2 - CE
- HE3
3577 3576 3579 1 1.0950e+02 4.1840e+02 ; HE2 - CE
- NZ
3578 3576 3579 1 1.0950e+02 4.1840e+02 ; HE3 - CE
- NZ
3580 3579 3581 1 1.0950e+02 2.9288e+02 ; HZ1 - NZ
- HZ2
3580 3579 3582 1 1.0950e+02 2.9288e+02 ; HZ1 - NZ
- HZ3
3581 3579 3582 1 1.0950e+02 2.9288e+02 ; HZ2 - NZ
- HZ3
3583 3585 3586 1 1.2000e+02 4.1840e+02 ; C - N
- H
3583 3585 3587 1 1.2190e+02 4.1840e+02 ; C - N
- CA
3584 3583 3585 1 1.2290e+02 6.6944e+02 ; O - C
- N
3585 3587 3588 1 1.0950e+02 4.1840e+02 ; N - CA
- HA
3585 3587 3589 1 1.0970e+02 6.6944e+02 ; N - CA
- CB
3585 3587 3599 1 1.1010e+02 5.2718e+02 ; N - CA
- C
3586 3585 3587 1 1.1804e+02 4.1840e+02 ; H - N
- CA
3587 3589 3590 1 1.0950e+02 4.1840e+02 ; CA - CB
- HB
3587 3589 3591 1 1.0950e+02 3.3472e+02 ; CA - CB
- CG1
3587 3589 3595 1 1.0950e+02 3.3472e+02 ; CA - CB
- CG2
3587 3599 3600 1 1.2040e+02 6.6944e+02 ; CA - C
- O
3587 3599 3601 1 1.1660e+02 5.8576e+02 ; CA - C
- N

3588 3587 3589 1 1.0950e+02 4.1840e+02 ; HA - CA
- CB
3588 3587 3599 1 1.0950e+02 4.1840e+02 ; HA - CA
- C
3589 3587 3599 1 1.1110e+02 5.2718e+02 ; CB - CA
- C
3589 3591 3592 1 1.0950e+02 4.1840e+02 ; CB - CG1
- HG11
3589 3591 3593 1 1.0950e+02 4.1840e+02 ; CB - CG1
- HG12
3589 3591 3594 1 1.0950e+02 4.1840e+02 ; CB - CG1
- HG13
3589 3595 3596 1 1.0950e+02 4.1840e+02 ; CB - CG2
- HG21
3589 3595 3597 1 1.0950e+02 4.1840e+02 ; CB - CG2
- HG22
3589 3595 3598 1 1.0950e+02 4.1840e+02 ; CB - CG2
- HG23
3590 3589 3591 1 1.0950e+02 4.1840e+02 ; HB - CB
- CG1
3590 3589 3595 1 1.0950e+02 4.1840e+02 ; HB - CB
- CG2
3591 3589 3595 1 1.0950e+02 3.3472e+02 ; CG1 - CB
- CG2
3592 3591 3593 1 1.0950e+02 2.9288e+02 ; HG11 - CG1
- HG12
3592 3591 3594 1 1.0950e+02 2.9288e+02 ; HG11 - CG1
- HG13
3593 3591 3594 1 1.0950e+02 2.9288e+02 ; HG12 - CG1
- HG13
3596 3595 3597 1 1.0950e+02 2.9288e+02 ; HG21 - CG2
- HG22
3596 3595 3598 1 1.0950e+02 2.9288e+02 ; HG21 - CG2
- HG23
3597 3595 3598 1 1.0950e+02 2.9288e+02 ; HG22 - CG2
- HG23
3599 3601 3602 1 1.2000e+02 4.1840e+02 ; C - N
- H
3599 3601 3603 1 1.2190e+02 4.1840e+02 ; C - N
- CA
3600 3599 3601 1 1.2290e+02 6.6944e+02 ; O - C
- N
3601 3603 3604 1 1.0950e+02 4.1840e+02 ; N - CA
- HA
3601 3603 3605 1 1.0970e+02 6.6944e+02 ; N - CA
- CB
3601 3603 3621 1 1.1010e+02 5.2718e+02 ; N - CA
- C
3602 3601 3603 1 1.1804e+02 4.1840e+02 ; H - N
- CA
3603 3605 3606 1 1.0950e+02 4.1840e+02 ; CA - CB
- HB2

3603 3605 3607 1 1.0950e+02 4.1840e+02 ; CA - CB
- HB3
3603 3605 3608 1 1.0950e+02 3.3472e+02 ; CA - CB
- CG
3603 3621 3622 1 1.2040e+02 6.6944e+02 ; CA - C
- O
3603 3621 3623 1 1.1660e+02 5.8576e+02 ; CA - C
- N
3604 3603 3605 1 1.0950e+02 4.1840e+02 ; HA - CA
- CB
3604 3603 3621 1 1.0950e+02 4.1840e+02 ; HA - CA
- C
3605 3603 3621 1 1.1110e+02 5.2718e+02 ; CB - CA
- C
3605 3608 3609 1 1.0950e+02 4.1840e+02 ; CB - CG
- HG2
3605 3608 3610 1 1.0950e+02 4.1840e+02 ; CB - CG
- HG3
3605 3608 3611 1 1.0950e+02 3.3472e+02 ; CB - CG
- CD
3606 3605 3607 1 1.0950e+02 2.9288e+02 ; HB2 - CB
- HB3
3606 3605 3608 1 1.0950e+02 4.1840e+02 ; HB2 - CB
- CG
3607 3605 3608 1 1.0950e+02 4.1840e+02 ; HB3 - CB
- CG
3608 3611 3612 1 1.0950e+02 4.1840e+02 ; CG - CD
- HD2
3608 3611 3613 1 1.0950e+02 4.1840e+02 ; CG - CD
- HD3
3608 3611 3614 1 1.0950e+02 3.3472e+02 ; CG - CD
- CE
3609 3608 3610 1 1.0950e+02 2.9288e+02 ; HG2 - CG
- HG3
3609 3608 3611 1 1.0950e+02 4.1840e+02 ; HG2 - CG
- CD
3610 3608 3611 1 1.0950e+02 4.1840e+02 ; HG3 - CG
- CD
3611 3614 3615 1 1.0950e+02 4.1840e+02 ; CD - CE
- HE2
3611 3614 3616 1 1.0950e+02 4.1840e+02 ; CD - CE
- HE3
3611 3614 3617 1 1.1120e+02 6.6944e+02 ; CD - CE
- NZ
3612 3611 3613 1 1.0950e+02 2.9288e+02 ; HD2 - CD
- HD3
3612 3611 3614 1 1.0950e+02 4.1840e+02 ; HD2 - CD
- CE
3613 3611 3614 1 1.0950e+02 4.1840e+02 ; HD3 - CD
- CE
3614 3617 3618 1 1.0950e+02 4.1840e+02 ; CE - NZ
- HZ1

3614 3617 3619 1 1.0950e+02 4.1840e+02 ; CE - NZ
- HZ2
3614 3617 3620 1 1.0950e+02 4.1840e+02 ; CE - NZ
- HZ3
3615 3614 3616 1 1.0950e+02 2.9288e+02 ; HE2 - CE
- HE3
3615 3614 3617 1 1.0950e+02 4.1840e+02 ; HE2 - CE
- NZ
3616 3614 3617 1 1.0950e+02 4.1840e+02 ; HE3 - CE
- NZ
3618 3617 3619 1 1.0950e+02 2.9288e+02 ; HZ1 - NZ
- HZ2
3618 3617 3620 1 1.0950e+02 2.9288e+02 ; HZ1 - NZ
- HZ3
3619 3617 3620 1 1.0950e+02 2.9288e+02 ; HZ2 - NZ
- HZ3
3621 3623 3624 1 1.2000e+02 4.1840e+02 ; C - N
- H
3621 3623 3625 1 1.2190e+02 4.1840e+02 ; C - N
- CA
3622 3621 3623 1 1.2290e+02 6.6944e+02 ; O - C
- N
3623 3625 3626 1 1.0950e+02 4.1840e+02 ; N - CA
- HA
3623 3625 3627 1 1.0970e+02 6.6944e+02 ; N - CA
- CB
3623 3625 3643 1 1.1010e+02 5.2718e+02 ; N - CA
- C
3624 3623 3625 1 1.1804e+02 4.1840e+02 ; H - N
- CA
3625 3627 3628 1 1.0950e+02 4.1840e+02 ; CA - CB
- HB2
3625 3627 3629 1 1.0950e+02 4.1840e+02 ; CA - CB
- HB3
3625 3627 3630 1 1.0950e+02 3.3472e+02 ; CA - CB
- CG
3625 3643 3644 1 1.2040e+02 6.6944e+02 ; CA - C
- O
3625 3643 3645 1 1.1660e+02 5.8576e+02 ; CA - C
- N
3626 3625 3627 1 1.0950e+02 4.1840e+02 ; HA - CA
- CB
3626 3625 3643 1 1.0950e+02 4.1840e+02 ; HA - CA
- C
3627 3625 3643 1 1.1110e+02 5.2718e+02 ; CB - CA
- C
3627 3630 3631 1 1.0950e+02 4.1840e+02 ; CB - CG
- HG2
3627 3630 3632 1 1.0950e+02 4.1840e+02 ; CB - CG
- HG3
3627 3630 3633 1 1.0950e+02 3.3472e+02 ; CB - CG
- CD

3628 3627 3629 1 1.0950e+02 2.9288e+02 ; HB2 - CB
- HB3
3628 3627 3630 1 1.0950e+02 4.1840e+02 ; HB2 - CB
- CG
3629 3627 3630 1 1.0950e+02 4.1840e+02 ; HB3 - CB
- CG
3630 3633 3634 1 1.0950e+02 4.1840e+02 ; CG - CD
- HD2
3630 3633 3635 1 1.0950e+02 4.1840e+02 ; CG - CD
- HD3
3630 3633 3636 1 1.0950e+02 3.3472e+02 ; CG - CD
- CE
3631 3630 3632 1 1.0950e+02 2.9288e+02 ; HG2 - CG
- HG3
3631 3630 3633 1 1.0950e+02 4.1840e+02 ; HG2 - CG
- CD
3632 3630 3633 1 1.0950e+02 4.1840e+02 ; HG3 - CG
- CD
3633 3636 3637 1 1.0950e+02 4.1840e+02 ; CD - CE
- HE2
3633 3636 3638 1 1.0950e+02 4.1840e+02 ; CD - CE
- HE3
3633 3636 3639 1 1.1120e+02 6.6944e+02 ; CD - CE
- NZ
3634 3633 3635 1 1.0950e+02 2.9288e+02 ; HD2 - CD
- HD3
3634 3633 3636 1 1.0950e+02 4.1840e+02 ; HD2 - CD
- CE
3635 3633 3636 1 1.0950e+02 4.1840e+02 ; HD3 - CD
- CE
3636 3639 3640 1 1.0950e+02 4.1840e+02 ; CE - NZ
- HZ1
3636 3639 3641 1 1.0950e+02 4.1840e+02 ; CE - NZ
- HZ2
3636 3639 3642 1 1.0950e+02 4.1840e+02 ; CE - NZ
- HZ3
3637 3636 3638 1 1.0950e+02 2.9288e+02 ; HE2 - CE
- HE3
3637 3636 3639 1 1.0950e+02 4.1840e+02 ; HE2 - CE
- NZ
3638 3636 3639 1 1.0950e+02 4.1840e+02 ; HE3 - CE
- NZ
3640 3639 3641 1 1.0950e+02 2.9288e+02 ; HZ1 - NZ
- HZ2
3640 3639 3642 1 1.0950e+02 2.9288e+02 ; HZ1 - NZ
- HZ3
3641 3639 3642 1 1.0950e+02 2.9288e+02 ; HZ2 - NZ
- HZ3
3643 3645 3646 1 1.2000e+02 4.1840e+02 ; C - N
- H
3643 3645 3647 1 1.2190e+02 4.1840e+02 ; C - N
- CA

3644 3643 3645 1 1.2290e+02 6.6944e+02 ; O - C
- N
3645 3647 3648 1 1.0950e+02 4.1840e+02 ; N - CA
- HA2
3645 3647 3649 1 1.0950e+02 4.1840e+02 ; N - CA
- HA3
3645 3647 3650 1 1.1010e+02 5.2718e+02 ; N - CA
- C
3646 3645 3647 1 1.1804e+02 4.1840e+02 ; H - N
- CA
3647 3650 3651 1 1.2040e+02 6.6944e+02 ; CA - C
- O
3647 3650 3652 1 1.1660e+02 5.8576e+02 ; CA - C
- N
3648 3647 3649 1 1.0950e+02 2.9288e+02 ; HA2 - CA
- HA3
3648 3647 3650 1 1.0950e+02 4.1840e+02 ; HA2 - CA
- C
3649 3647 3650 1 1.0950e+02 4.1840e+02 ; HA3 - CA
- C
3650 3652 3653 1 1.2000e+02 4.1840e+02 ; C - N
- H
3650 3652 3654 1 1.2190e+02 4.1840e+02 ; C - N
- CA
3651 3650 3652 1 1.2290e+02 6.6944e+02 ; O - C
- N
3652 3654 3655 1 1.0950e+02 4.1840e+02 ; N - CA
- HA
3652 3654 3656 1 1.0970e+02 6.6944e+02 ; N - CA
- CB
3652 3654 3665 1 1.1010e+02 5.2718e+02 ; N - CA
- C
3653 3652 3654 1 1.1804e+02 4.1840e+02 ; H - N
- CA
3654 3656 3657 1 1.0950e+02 4.1840e+02 ; CA - CB
- HB2
3654 3656 3658 1 1.0950e+02 4.1840e+02 ; CA - CB
- HB3
3654 3656 3659 1 1.0950e+02 3.3472e+02 ; CA - CB
- CG
3654 3665 3666 1 1.2040e+02 6.6944e+02 ; CA - C
- O
3654 3665 3667 1 1.1660e+02 5.8576e+02 ; CA - C
- N
3655 3654 3656 1 1.0950e+02 4.1840e+02 ; HA - CA
- CB
3655 3654 3665 1 1.0950e+02 4.1840e+02 ; HA - CA
- C
3656 3654 3665 1 1.1110e+02 5.2718e+02 ; CB - CA
- C
3656 3659 3660 1 1.0950e+02 4.1840e+02 ; CB - CG
- HG2

3656 3659 3661 1 1.0950e+02 4.1840e+02 ; CB - CG
- HG3
3656 3659 3662 1 1.1110e+02 5.2718e+02 ; CB - CG
- CD
3657 3656 3658 1 1.0950e+02 2.9288e+02 ; HB2 - CB
- HB3
3657 3656 3659 1 1.0950e+02 4.1840e+02 ; HB2 - CB
- CG
3658 3656 3659 1 1.0950e+02 4.1840e+02 ; HB3 - CB
- CG
3659 3662 3663 1 1.1700e+02 5.8576e+02 ; CG - CD
- OE1
3659 3662 3664 1 1.1700e+02 5.8576e+02 ; CG - CD
- OE2
3660 3659 3661 1 1.0950e+02 2.9288e+02 ; HG2 - CG
- HG3
3660 3659 3662 1 1.0950e+02 4.1840e+02 ; HG2 - CG
- CD
3661 3659 3662 1 1.0950e+02 4.1840e+02 ; HG3 - CG
- CD
3663 3662 3664 1 1.2600e+02 6.6944e+02 ; OE1 - CD
- OE2
3665 3667 3668 1 1.2000e+02 4.1840e+02 ; C - N
- H
3665 3667 3669 1 1.2190e+02 4.1840e+02 ; C - N
- CA
3666 3665 3667 1 1.2290e+02 6.6944e+02 ; O - C
- N
3667 3669 3670 1 1.0950e+02 4.1840e+02 ; N - CA
- HA
3667 3669 3671 1 1.0970e+02 6.6944e+02 ; N - CA
- CB
3667 3669 3685 1 1.1010e+02 5.2718e+02 ; N - CA
- C
3668 3667 3669 1 1.1804e+02 4.1840e+02 ; H - N
- CA
3669 3671 3672 1 1.0950e+02 4.1840e+02 ; CA - CB
- HB2
3669 3671 3673 1 1.0950e+02 4.1840e+02 ; CA - CB
- HB3
3669 3671 3674 1 1.1400e+02 5.2718e+02 ; CA - CB
- CG
3669 3685 3686 1 1.2040e+02 6.6944e+02 ; CA - C
- O
3669 3685 3687 1 1.1660e+02 5.8576e+02 ; CA - C
- N
3670 3669 3671 1 1.0950e+02 4.1840e+02 ; HA - CA
- CB
3670 3669 3685 1 1.0950e+02 4.1840e+02 ; HA - CA
- C
3671 3669 3685 1 1.1110e+02 5.2718e+02 ; CB - CA
- C

3671 3674 3675 1 1.2000e+02 5.8576e+02 ; CB - CG
- CD1
3671 3674 3683 1 1.2000e+02 5.8576e+02 ; CB - CG
- CD2
3672 3671 3673 1 1.0950e+02 2.9288e+02 ; HB2 - CB
- HB3
3672 3671 3674 1 1.0950e+02 4.1840e+02 ; HB2 - CB
- CG
3673 3671 3674 1 1.0950e+02 4.1840e+02 ; HB3 - CB
- CG
3674 3675 3676 1 1.2000e+02 4.1840e+02 ; CG - CD1
- HD1
3674 3675 3677 1 1.2000e+02 5.2718e+02 ; CG - CD1
- CE1
3674 3683 3681 1 1.2000e+02 5.2718e+02 ; CG - CD2
- CE2
3674 3683 3684 1 1.2000e+02 4.1840e+02 ; CG - CD2
- HD2
3675 3674 3683 1 1.2000e+02 5.2718e+02 ; CD1 - CG
- CD2
3675 3677 3678 1 1.2000e+02 4.1840e+02 ; CD1 - CE1
- HE1
3675 3677 3679 1 1.2000e+02 5.2718e+02 ; CD1 - CE1
- CZ
3676 3675 3677 1 1.2000e+02 4.1840e+02 ; HD1 - CD1
- CE1
3677 3679 3680 1 1.2000e+02 4.1840e+02 ; CE1 - CZ
- HZ
3677 3679 3681 1 1.2000e+02 5.2718e+02 ; CE1 - CZ
- CE2
3678 3677 3679 1 1.2000e+02 4.1840e+02 ; HE1 - CE1
- CZ
3679 3681 3682 1 1.2000e+02 4.1840e+02 ; CZ - CE2
- HE2
3679 3681 3683 1 1.2000e+02 5.2718e+02 ; CZ - CE2
- CD2
3680 3679 3681 1 1.2000e+02 4.1840e+02 ; HZ - CZ
- CE2
3681 3683 3684 1 1.2000e+02 4.1840e+02 ; CE2 - CD2
- HD2
3682 3681 3683 1 1.2000e+02 4.1840e+02 ; HE2 - CE2
- CD2
3685 3687 3688 1 1.2000e+02 4.1840e+02 ; C - N
- H
3685 3687 3689 1 1.2190e+02 4.1840e+02 ; C - N
- CA
3686 3685 3687 1 1.2290e+02 6.6944e+02 ; O - C
- N
3687 3689 3690 1 1.0950e+02 4.1840e+02 ; N - CA
- HA
3687 3689 3691 1 1.0970e+02 6.6944e+02 ; N - CA
- CB

3687 3689 3696 1 1.1010e+02 5.2718e+02 ; N - CA
- C
3688 3687 3689 1 1.1804e+02 4.1840e+02 ; H - N
- CA
3689 3691 3692 1 1.0950e+02 4.1840e+02 ; CA - CB
- HB2
3689 3691 3693 1 1.0950e+02 4.1840e+02 ; CA - CB
- HB3
3689 3691 3694 1 1.0860e+02 4.1840e+02 ; CA - CB
- SG
3689 3696 3697 1 1.2040e+02 6.6944e+02 ; CA - C
- O
3689 3696 3698 1 1.1660e+02 5.8576e+02 ; CA - C
- N
3690 3689 3691 1 1.0950e+02 4.1840e+02 ; HA - CA
- CB
3690 3689 3696 1 1.0950e+02 4.1840e+02 ; HA - CA
- C
3691 3689 3696 1 1.1110e+02 5.2718e+02 ; CB - CA
- C
3691 3694 3695 1 9.6000e+01 3.5982e+02 ; CB - SG
- HG
3692 3691 3693 1 1.0950e+02 2.9288e+02 ; HB2 - CB
- HB3
3692 3691 3694 1 1.0950e+02 4.1840e+02 ; HB2 - CB
- SG
3693 3691 3694 1 1.0950e+02 4.1840e+02 ; HB3 - CB
- SG
3696 3698 3699 1 1.2000e+02 4.1840e+02 ; C - N
- H
3696 3698 3700 1 1.2190e+02 4.1840e+02 ; C - N
- CA
3697 3696 3698 1 1.2290e+02 6.6944e+02 ; O - C
- N
3698 3700 3701 1 1.0950e+02 4.1840e+02 ; N - CA
- HA
3698 3700 3702 1 1.0970e+02 6.6944e+02 ; N - CA
- CB
3698 3700 3716 1 1.1010e+02 5.2718e+02 ; N - CA
- C
3699 3698 3700 1 1.1804e+02 4.1840e+02 ; H - N
- CA
3700 3702 3703 1 1.0950e+02 4.1840e+02 ; CA - CB
- HB2
3700 3702 3704 1 1.0950e+02 4.1840e+02 ; CA - CB
- HB3
3700 3702 3705 1 1.1400e+02 5.2718e+02 ; CA - CB
- CG
3700 3716 3717 1 1.2040e+02 6.6944e+02 ; CA - C
- O
3700 3716 3718 1 1.1660e+02 5.8576e+02 ; CA - C
- N

3701 3700 3702 1 1.0950e+02 4.1840e+02 ; HA - CA
- CB
3701 3700 3716 1 1.0950e+02 4.1840e+02 ; HA - CA
- C
3702 3700 3716 1 1.1110e+02 5.2718e+02 ; CB - CA
- C
3702 3705 3706 1 1.2000e+02 5.8576e+02 ; CB - CG
- CD1
3702 3705 3714 1 1.2000e+02 5.8576e+02 ; CB - CG
- CD2
3703 3702 3704 1 1.0950e+02 2.9288e+02 ; HB2 - CB
- HB3
3703 3702 3705 1 1.0950e+02 4.1840e+02 ; HB2 - CB
- CG
3704 3702 3705 1 1.0950e+02 4.1840e+02 ; HB3 - CB
- CG
3705 3706 3707 1 1.2000e+02 4.1840e+02 ; CG - CD1
- HD1
3705 3706 3708 1 1.2000e+02 5.2718e+02 ; CG - CD1
- CE1
3705 3714 3712 1 1.2000e+02 5.2718e+02 ; CG - CD2
- CE2
3705 3714 3715 1 1.2000e+02 4.1840e+02 ; CG - CD2
- HD2
3706 3705 3714 1 1.2000e+02 5.2718e+02 ; CD1 - CG
- CD2
3706 3708 3709 1 1.2000e+02 4.1840e+02 ; CD1 - CE1
- HE1
3706 3708 3710 1 1.2000e+02 5.2718e+02 ; CD1 - CE1
- CZ
3707 3706 3708 1 1.2000e+02 4.1840e+02 ; HD1 - CD1
- CE1
3708 3710 3711 1 1.2000e+02 4.1840e+02 ; CE1 - CZ
- HZ
3708 3710 3712 1 1.2000e+02 5.2718e+02 ; CE1 - CZ
- CE2
3709 3708 3710 1 1.2000e+02 4.1840e+02 ; HE1 - CE1
- CZ
3710 3712 3713 1 1.2000e+02 4.1840e+02 ; CZ - CE2
- HE2
3710 3712 3714 1 1.2000e+02 5.2718e+02 ; CZ - CE2
- CD2
3711 3710 3712 1 1.2000e+02 4.1840e+02 ; HZ - CZ
- CE2
3712 3714 3715 1 1.2000e+02 4.1840e+02 ; CE2 - CD2
- HD2
3713 3712 3714 1 1.2000e+02 4.1840e+02 ; HE2 - CE2
- CD2
3716 3718 3719 1 1.2000e+02 4.1840e+02 ; C - N
- H
3716 3718 3720 1 1.2190e+02 4.1840e+02 ; C - N
- CA

3717 3716 3718 1 1.2290e+02 6.6944e+02 ; O - C
- N
3718 3720 3721 1 1.0950e+02 4.1840e+02 ; N - CA
- HA
3718 3720 3722 1 1.0970e+02 6.6944e+02 ; N - CA
- CB
3718 3720 3737 1 1.1010e+02 5.2718e+02 ; N - CA
- C
3719 3718 3720 1 1.1804e+02 4.1840e+02 ; H - N
- CA
3720 3722 3723 1 1.0950e+02 4.1840e+02 ; CA - CB
- HB2
3720 3722 3724 1 1.0950e+02 4.1840e+02 ; CA - CB
- HB3
3720 3722 3725 1 1.1400e+02 5.2718e+02 ; CA - CB
- CG
3720 3737 3738 1 1.2040e+02 6.6944e+02 ; CA - C
- O
3720 3737 3739 1 1.1660e+02 5.8576e+02 ; CA - C
- N
3721 3720 3722 1 1.0950e+02 4.1840e+02 ; HA - CA
- CB
3721 3720 3737 1 1.0950e+02 4.1840e+02 ; HA - CA
- C
3722 3720 3737 1 1.1110e+02 5.2718e+02 ; CB - CA
- C
3722 3725 3726 1 1.2000e+02 5.8576e+02 ; CB - CG
- CD1
3722 3725 3735 1 1.2000e+02 5.8576e+02 ; CB - CG
- CD2
3723 3722 3724 1 1.0950e+02 2.9288e+02 ; HB2 - CB
- HB3
3723 3722 3725 1 1.0950e+02 4.1840e+02 ; HB2 - CB
- CG
3724 3722 3725 1 1.0950e+02 4.1840e+02 ; HB3 - CB
- CG
3725 3726 3727 1 1.2000e+02 4.1840e+02 ; CG - CD1
- HD1
3725 3726 3728 1 1.2000e+02 5.2718e+02 ; CG - CD1
- CE1
3725 3735 3733 1 1.2000e+02 5.2718e+02 ; CG - CD2
- CE2
3725 3735 3736 1 1.2000e+02 4.1840e+02 ; CG - CD2
- HD2
3726 3725 3735 1 1.2000e+02 5.2718e+02 ; CD1 - CG
- CD2
3726 3728 3729 1 1.2000e+02 4.1840e+02 ; CD1 - CE1
- HE1
3726 3728 3730 1 1.2000e+02 5.2718e+02 ; CD1 - CE1
- CZ
3727 3726 3728 1 1.2000e+02 4.1840e+02 ; HD1 - CD1
- CE1

3728 3730 3731 1 1.2000e+02 5.8576e+02 ; CE1 - CZ
- OH
3728 3730 3733 1 1.2000e+02 5.2718e+02 ; CE1 - CZ
- CE2
3729 3728 3730 1 1.2000e+02 4.1840e+02 ; HE1 - CE1
- CZ
3730 3731 3732 1 1.1300e+02 4.1840e+02 ; CZ - OH
- HH
3730 3733 3734 1 1.2000e+02 4.1840e+02 ; CZ - CE2
- HE2
3730 3733 3735 1 1.2000e+02 5.2718e+02 ; CZ - CE2
- CD2
3731 3730 3733 1 1.2000e+02 5.8576e+02 ; OH - CZ
- CE2
3733 3735 3736 1 1.2000e+02 4.1840e+02 ; CE2 - CD2
- HD2
3734 3733 3735 1 1.2000e+02 4.1840e+02 ; HE2 - CE2
- CD2
3737 3739 3740 1 1.2000e+02 4.1840e+02 ; C - N
- H
3737 3739 3741 1 1.2190e+02 4.1840e+02 ; C - N
- CA
3738 3737 3739 1 1.2290e+02 6.6944e+02 ; O - C
- N
3739 3741 3742 1 1.0950e+02 4.1840e+02 ; N - CA
- HA
3739 3741 3743 1 1.0970e+02 6.6944e+02 ; N - CA
- CB
3739 3741 3752 1 1.1010e+02 5.2718e+02 ; N - CA
- C
3740 3739 3741 1 1.1804e+02 4.1840e+02 ; H - N
- CA
3741 3743 3744 1 1.0950e+02 4.1840e+02 ; CA - CB
- HB2
3741 3743 3745 1 1.0950e+02 4.1840e+02 ; CA - CB
- HB3
3741 3743 3746 1 1.0950e+02 3.3472e+02 ; CA - CB
- CG
3741 3752 3753 1 1.2040e+02 6.6944e+02 ; CA - C
- O
3741 3752 3754 1 1.1660e+02 5.8576e+02 ; CA - C
- N
3742 3741 3743 1 1.0950e+02 4.1840e+02 ; HA - CA
- CB
3742 3741 3752 1 1.0950e+02 4.1840e+02 ; HA - CA
- C
3743 3741 3752 1 1.1110e+02 5.2718e+02 ; CB - CA
- C
3743 3746 3747 1 1.0950e+02 4.1840e+02 ; CB - CG
- HG2
3743 3746 3748 1 1.0950e+02 4.1840e+02 ; CB - CG
- HG3

3743 3746 3749 1 1.1110e+02 5.2718e+02 ; CB - CG
- CD
3744 3743 3745 1 1.0950e+02 2.9288e+02 ; HB2 - CB
- HB3
3744 3743 3746 1 1.0950e+02 4.1840e+02 ; HB2 - CB
- CG
3745 3743 3746 1 1.0950e+02 4.1840e+02 ; HB3 - CB
- CG
3746 3749 3750 1 1.1700e+02 5.8576e+02 ; CG - CD
- OE1
3746 3749 3751 1 1.1700e+02 5.8576e+02 ; CG - CD
- OE2
3747 3746 3748 1 1.0950e+02 2.9288e+02 ; HG2 - CG
- HG3
3747 3746 3749 1 1.0950e+02 4.1840e+02 ; HG2 - CG
- CD
3748 3746 3749 1 1.0950e+02 4.1840e+02 ; HG3 - CG
- CD
3750 3749 3751 1 1.2600e+02 6.6944e+02 ; OE1 - CD
- OE2
3752 3754 3755 1 1.2000e+02 4.1840e+02 ; C - N
- H
3752 3754 3756 1 1.2190e+02 4.1840e+02 ; C - N
- CA
3753 3752 3754 1 1.2290e+02 6.6944e+02 ; O - C
- N
3754 3756 3757 1 1.0950e+02 4.1840e+02 ; N - CA
- HA
3754 3756 3758 1 1.0970e+02 6.6944e+02 ; N - CA
- CB
3754 3756 3766 1 1.1010e+02 5.2718e+02 ; N - CA
- C
3755 3754 3756 1 1.1804e+02 4.1840e+02 ; H - N
- CA
3756 3758 3759 1 1.0950e+02 4.1840e+02 ; CA - CB
- HB2
3756 3758 3760 1 1.0950e+02 4.1840e+02 ; CA - CB
- HB3
3756 3758 3761 1 1.1110e+02 5.2718e+02 ; CA - CB
- CG
3756 3766 3767 1 1.2040e+02 6.6944e+02 ; CA - C
- O
3756 3766 3768 1 1.1660e+02 5.8576e+02 ; CA - C
- N
3757 3756 3758 1 1.0950e+02 4.1840e+02 ; HA - CA
- CB
3757 3756 3766 1 1.0950e+02 4.1840e+02 ; HA - CA
- C
3758 3756 3766 1 1.1110e+02 5.2718e+02 ; CB - CA
- C
3758 3761 3762 1 1.2040e+02 6.6944e+02 ; CB - CG
- OD1

3758 3761 3763 1 1.1660e+02 5.8576e+02 ; CB - CG
- ND2
3759 3758 3760 1 1.0950e+02 2.9288e+02 ; HB2 - CB
- HB3
3759 3758 3761 1 1.0950e+02 4.1840e+02 ; HB2 - CB
- CG
3760 3758 3761 1 1.0950e+02 4.1840e+02 ; HB3 - CB
- CG
3761 3763 3764 1 1.2000e+02 4.1840e+02 ; CG - ND2
- HD21
3761 3763 3765 1 1.2000e+02 4.1840e+02 ; CG - ND2
- HD22
3762 3761 3763 1 1.2290e+02 6.6944e+02 ; OD1 - CG
- ND2
3764 3763 3765 1 1.2000e+02 2.9288e+02 ; HD21 - ND2
- HD22
3766 3768 3769 1 1.2000e+02 4.1840e+02 ; C - N
- H
3766 3768 3770 1 1.2190e+02 4.1840e+02 ; C - N
- CA
3767 3766 3768 1 1.2290e+02 6.6944e+02 ; O - C
- N
3768 3770 3771 1 1.0950e+02 4.1840e+02 ; N - CA
- HA
3768 3770 3772 1 1.0970e+02 6.6944e+02 ; N - CA
- CB
3768 3770 3778 1 1.1010e+02 5.2718e+02 ; N - CA
- C
3769 3768 3770 1 1.1804e+02 4.1840e+02 ; H - N
- CA
3770 3772 3773 1 1.0950e+02 4.1840e+02 ; CA - CB
- HB2
3770 3772 3774 1 1.0950e+02 4.1840e+02 ; CA - CB
- HB3
3770 3772 3775 1 1.1110e+02 5.2718e+02 ; CA - CB
- CG
3770 3778 3779 1 1.2040e+02 6.6944e+02 ; CA - C
- O
3770 3778 3780 1 1.1660e+02 5.8576e+02 ; CA - C
- N
3771 3770 3772 1 1.0950e+02 4.1840e+02 ; HA - CA
- CB
3771 3770 3778 1 1.0950e+02 4.1840e+02 ; HA - CA
- C
3772 3770 3778 1 1.1110e+02 5.2718e+02 ; CB - CA
- C
3772 3775 3776 1 1.1700e+02 5.8576e+02 ; CB - CG
- OD1
3772 3775 3777 1 1.1700e+02 5.8576e+02 ; CB - CG
- OD2
3773 3772 3774 1 1.0950e+02 2.9288e+02 ; HB2 - CB
- HB3

3773 3772 3775 1 1.0950e+02 4.1840e+02 ; HB2 - CB
- CG
3774 3772 3775 1 1.0950e+02 4.1840e+02 ; HB3 - CB
- CG
3776 3775 3777 1 1.2600e+02 6.6944e+02 ; OD1 - CG
- OD2
3778 3780 3781 1 1.2000e+02 4.1840e+02 ; C - N
- H
3778 3780 3782 1 1.2190e+02 4.1840e+02 ; C - N
- CA
3779 3778 3780 1 1.2290e+02 6.6944e+02 ; O - C
- N
3780 3782 3783 1 1.0950e+02 4.1840e+02 ; N - CA
- HA
3780 3782 3784 1 1.0970e+02 6.6944e+02 ; N - CA
- CB
3780 3782 3802 1 1.1010e+02 5.2718e+02 ; N - CA
- C
3781 3780 3782 1 1.1804e+02 4.1840e+02 ; H - N
- CA
3782 3784 3785 1 1.0950e+02 4.1840e+02 ; CA - CB
- HB2
3782 3784 3786 1 1.0950e+02 4.1840e+02 ; CA - CB
- HB3
3782 3784 3787 1 1.1560e+02 5.2718e+02 ; CA - CB
- CG
3782 3802 3803 1 1.2040e+02 6.6944e+02 ; CA - C
- O
3782 3802 3804 1 1.1660e+02 5.8576e+02 ; CA - C
- N
3783 3782 3784 1 1.0950e+02 4.1840e+02 ; HA - CA
- CB
3783 3782 3802 1 1.0950e+02 4.1840e+02 ; HA - CA
- C
3784 3782 3802 1 1.1110e+02 5.2718e+02 ; CB - CA
- C
3784 3787 3788 1 1.2500e+02 5.8576e+02 ; CB - CG
- CD1
3784 3787 3801 1 1.2860e+02 5.8576e+02 ; CB - CG
- CD2
3785 3784 3786 1 1.0950e+02 2.9288e+02 ; HB2 - CB
- HB3
3785 3784 3787 1 1.0950e+02 4.1840e+02 ; HB2 - CB
- CG
3786 3784 3787 1 1.0950e+02 4.1840e+02 ; HB3 - CB
- CG
3787 3788 3789 1 1.2000e+02 4.1840e+02 ; CG - CD1
- HD1
3787 3788 3790 1 1.0870e+02 5.8576e+02 ; CG - CD1
- NE1
3787 3801 3792 1 1.0880e+02 5.2718e+02 ; CG - CD2
- CE2

3787 3801 3799 1 1.3490e+02 5.2718e+02 ; CG - CD2
- CE3
3788 3787 3801 1 1.0640e+02 5.2718e+02 ; CD1 - CG
- CD2
3788 3790 3791 1 1.2000e+02 4.1840e+02 ; CD1 - NE1
- HE1
3788 3790 3792 1 1.1160e+02 5.8576e+02 ; CD1 - NE1
- CE2
3789 3788 3790 1 1.2000e+02 4.1840e+02 ; HD1 - CD1
- NE1
3790 3792 3793 1 1.3280e+02 5.8576e+02 ; NE1 - CE2
- CZ2
3790 3792 3801 1 1.0440e+02 5.8576e+02 ; NE1 - CE2
- CD2
3791 3790 3792 1 1.2310e+02 4.1840e+02 ; HE1 - NE1
- CE2
3792 3793 3794 1 1.2000e+02 4.1840e+02 ; CE2 - CZ2
- HZ2
3792 3793 3795 1 1.2000e+02 5.2718e+02 ; CE2 - CZ2
- CH2
3792 3801 3799 1 1.1620e+02 5.2718e+02 ; CE2 - CD2
- CE3
3793 3792 3801 1 1.2270e+02 5.2718e+02 ; CZ2 - CE2
- CD2
3793 3795 3796 1 1.2000e+02 4.1840e+02 ; CZ2 - CH2
- HH2
3793 3795 3797 1 1.2000e+02 5.2718e+02 ; CZ2 - CH2
- CZ3
3794 3793 3795 1 1.2000e+02 4.1840e+02 ; HZ2 - CZ2
- CH2
3795 3797 3798 1 1.2000e+02 4.1840e+02 ; CH2 - CZ3
- HZ3
3795 3797 3799 1 1.2000e+02 5.2718e+02 ; CH2 - CZ3
- CE3
3796 3795 3797 1 1.2000e+02 4.1840e+02 ; HH2 - CH2
- CZ3
3797 3799 3800 1 1.2000e+02 4.1840e+02 ; CZ3 - CE3
- HE3
3797 3799 3801 1 1.2000e+02 5.2718e+02 ; CZ3 - CE3
- CD2
3798 3797 3799 1 1.2000e+02 4.1840e+02 ; HZ3 - CZ3
- CE3
3800 3799 3801 1 1.2000e+02 4.1840e+02 ; HE3 - CE3
- CD2
3802 3804 3805 1 1.2000e+02 4.1840e+02 ; C - N
- H
3802 3804 3806 1 1.2190e+02 4.1840e+02 ; C - N
- CA
3803 3802 3804 1 1.2290e+02 6.6944e+02 ; O - C
- N
3804 3806 3807 1 1.0950e+02 4.1840e+02 ; N - CA
- HA2

3804 3806 3808 1 1.0950e+02 4.1840e+02 ; N - CA
- HA3
3804 3806 3809 1 1.1010e+02 5.2718e+02 ; N - CA
- C
3805 3804 3806 1 1.1804e+02 4.1840e+02 ; H - N
- CA
3806 3809 3810 1 1.2040e+02 6.6944e+02 ; CA - C
- O
3806 3809 3811 1 1.1660e+02 5.8576e+02 ; CA - C
- N
3807 3806 3808 1 1.0950e+02 2.9288e+02 ; HA2 - CA
- HA3
3807 3806 3809 1 1.0950e+02 4.1840e+02 ; HA2 - CA
- C
3808 3806 3809 1 1.0950e+02 4.1840e+02 ; HA3 - CA
- C
3809 3811 3812 1 1.2000e+02 4.1840e+02 ; C - N
- H
3809 3811 3813 1 1.2190e+02 4.1840e+02 ; C - N
- CA
3810 3809 3811 1 1.2290e+02 6.6944e+02 ; O - C
- N
3811 3813 3814 1 1.0950e+02 4.1840e+02 ; N - CA
- HA
3811 3813 3815 1 1.0970e+02 6.6944e+02 ; N - CA
- CB
3811 3813 3820 1 1.1010e+02 5.2718e+02 ; N - CA
- C
3812 3811 3813 1 1.1804e+02 4.1840e+02 ; H - N
- CA
3813 3815 3816 1 1.0950e+02 4.1840e+02 ; CA - CB
- HB2
3813 3815 3817 1 1.0950e+02 4.1840e+02 ; CA - CB
- HB3
3813 3815 3818 1 1.0950e+02 4.1840e+02 ; CA - CB
- OG
3813 3820 3821 1 1.2040e+02 6.6944e+02 ; CA - C
- O
3813 3820 3822 1 1.1660e+02 5.8576e+02 ; CA - C
- N
3814 3813 3815 1 1.0950e+02 4.1840e+02 ; HA - CA
- CB
3814 3813 3820 1 1.0950e+02 4.1840e+02 ; HA - CA
- C
3815 3813 3820 1 1.1110e+02 5.2718e+02 ; CB - CA
- C
3815 3818 3819 1 1.0850e+02 4.6024e+02 ; CB - OG
- HG
3816 3815 3817 1 1.0950e+02 2.9288e+02 ; HB2 - CB
- HB3
3816 3815 3818 1 1.0950e+02 4.1840e+02 ; HB2 - CB
- OG

3817 3815 3818 1 1.0950e+02 4.1840e+02 ; HB3 - CB
- OG
3820 3822 3823 1 1.2000e+02 4.1840e+02 ; C - N
- H
3820 3822 3824 1 1.2190e+02 4.1840e+02 ; C - N
- CA
3821 3820 3822 1 1.2290e+02 6.6944e+02 ; O - C
- N
3822 3824 3825 1 1.0950e+02 4.1840e+02 ; N - CA
- HA
3822 3824 3826 1 1.0970e+02 6.6944e+02 ; N - CA
- CB
3822 3824 3839 1 1.1010e+02 5.2718e+02 ; N - CA
- C
3823 3822 3824 1 1.1804e+02 4.1840e+02 ; H - N
- CA
3824 3826 3827 1 1.0950e+02 4.1840e+02 ; CA - CB
- HB
3824 3826 3828 1 1.0950e+02 3.3472e+02 ; CA - CB
- CG2
3824 3826 3832 1 1.0950e+02 3.3472e+02 ; CA - CB
- CG1
3824 3839 3840 1 1.2040e+02 6.6944e+02 ; CA - C
- O
3824 3839 3841 1 1.1660e+02 5.8576e+02 ; CA - C
- N
3825 3824 3826 1 1.0950e+02 4.1840e+02 ; HA - CA
- CB
3825 3824 3839 1 1.0950e+02 4.1840e+02 ; HA - CA
- C
3826 3824 3839 1 1.1110e+02 5.2718e+02 ; CB - CA
- C
3826 3828 3829 1 1.0950e+02 4.1840e+02 ; CB - CG2
- HG21
3826 3828 3830 1 1.0950e+02 4.1840e+02 ; CB - CG2
- HG22
3826 3828 3831 1 1.0950e+02 4.1840e+02 ; CB - CG2
- HG23
3826 3832 3833 1 1.0950e+02 4.1840e+02 ; CB - CG1
- HG12
3826 3832 3834 1 1.0950e+02 4.1840e+02 ; CB - CG1
- HG13
3826 3832 3835 1 1.0950e+02 3.3472e+02 ; CB - CG1
- CD1
3827 3826 3828 1 1.0950e+02 4.1840e+02 ; HB - CB
- CG2
3827 3826 3832 1 1.0950e+02 4.1840e+02 ; HB - CB
- CG1
3828 3826 3832 1 1.0950e+02 3.3472e+02 ; CG2 - CB
- CG1
3829 3828 3830 1 1.0950e+02 2.9288e+02 ; HG21 - CG2
- HG22

3829 3828 3831 1 1.0950e+02 2.9288e+02 ; HG21 - CG2
- HG23
3830 3828 3831 1 1.0950e+02 2.9288e+02 ; HG22 - CG2
- HG23
3832 3835 3836 1 1.0950e+02 4.1840e+02 ; CG1 - CD1
- HD11
3832 3835 3837 1 1.0950e+02 4.1840e+02 ; CG1 - CD1
- HD12
3832 3835 3838 1 1.0950e+02 4.1840e+02 ; CG1 - CD1
- HD13
3833 3832 3834 1 1.0950e+02 2.9288e+02 ; HG12 - CG1
- HG13
3833 3832 3835 1 1.0950e+02 4.1840e+02 ; HG12 - CG1
- CD1
3834 3832 3835 1 1.0950e+02 4.1840e+02 ; HG13 - CG1
- CD1
3836 3835 3837 1 1.0950e+02 2.9288e+02 ; HD11 - CD1
- HD12
3836 3835 3838 1 1.0950e+02 2.9288e+02 ; HD11 - CD1
- HD13
3837 3835 3838 1 1.0950e+02 2.9288e+02 ; HD12 - CD1
- HD13
3839 3841 3842 1 1.2000e+02 4.1840e+02 ; C - N
- H
3839 3841 3843 1 1.2190e+02 4.1840e+02 ; C - N
- CA
3840 3839 3841 1 1.2290e+02 6.6944e+02 ; O - C
- N
3841 3843 3844 1 1.0950e+02 4.1840e+02 ; N - CA
- HA
3841 3843 3845 1 1.0970e+02 6.6944e+02 ; N - CA
- CB
3841 3843 3850 1 1.1010e+02 5.2718e+02 ; N - CA
- C
3842 3841 3843 1 1.1804e+02 4.1840e+02 ; H - N
- CA
3843 3845 3846 1 1.0950e+02 4.1840e+02 ; CA - CB
- HB2
3843 3845 3847 1 1.0950e+02 4.1840e+02 ; CA - CB
- HB3
3843 3845 3848 1 1.0950e+02 4.1840e+02 ; CA - CB
- OG
3843 3850 3851 1 1.2040e+02 6.6944e+02 ; CA - C
- O
3843 3850 3852 1 1.1660e+02 5.8576e+02 ; CA - C
- N
3844 3843 3845 1 1.0950e+02 4.1840e+02 ; HA - CA
- CB
3844 3843 3850 1 1.0950e+02 4.1840e+02 ; HA - CA
- C
3845 3843 3850 1 1.1110e+02 5.2718e+02 ; CB - CA
- C

```
  3845   3848   3849      1    1.0850e+02    4.6024e+02 ;     CB - OG
- HG
  3846   3845   3847      1    1.0950e+02    2.9288e+02 ;    HB2 - CB
- HB3
  3846   3845   3848      1    1.0950e+02    4.1840e+02 ;    HB2 - CB
- OG
  3847   3845   3848      1    1.0950e+02    4.1840e+02 ;    HB3 - CB
- OG
  3850   3852   3853      1    1.2000e+02    4.1840e+02 ;      C - N
- H
  3850   3852   3854      1    1.2190e+02    4.1840e+02 ;      C - N
- CA
  3851   3850   3852      1    1.2290e+02    6.6944e+02 ;      O - C
- N
  3852   3854   3855      1    1.0950e+02    4.1840e+02 ;      N - CA
- HA
  3852   3854   3856      1    1.0970e+02    6.6944e+02 ;      N - CA
- CB
  3852   3854   3861      1    1.1010e+02    5.2718e+02 ;      N - CA
- C
  3853   3852   3854      1    1.1804e+02    4.1840e+02 ;      H - N
- CA
  3854   3856   3857      1    1.0950e+02    4.1840e+02 ;     CA - CB
- HB2
  3854   3856   3858      1    1.0950e+02    4.1840e+02 ;     CA - CB
- HB3
  3854   3856   3859      1    1.0950e+02    4.1840e+02 ;     CA - CB
- OG
  3854   3861   3862      1    1.2040e+02    6.6944e+02 ;     CA - C
- O
  3854   3861   3863      1    1.1660e+02    5.8576e+02 ;     CA - C
- N
  3855   3854   3856      1    1.0950e+02    4.1840e+02 ;     HA - CA
- CB
  3855   3854   3861      1    1.0950e+02    4.1840e+02 ;     HA - CA
- C
  3856   3854   3861      1    1.1110e+02    5.2718e+02 ;     CB - CA
- C
  3856   3859   3860      1    1.0850e+02    4.6024e+02 ;     CB - OG
- HG
  3857   3856   3858      1    1.0950e+02    2.9288e+02 ;    HB2 - CB
- HB3
  3857   3856   3859      1    1.0950e+02    4.1840e+02 ;    HB2 - CB
- OG
  3858   3856   3859      1    1.0950e+02    4.1840e+02 ;    HB3 - CB
- OG
  3861   3863   3864      1    1.2000e+02    4.1840e+02 ;      C - N
- H
  3861   3863   3865      1    1.2190e+02    4.1840e+02 ;      C - N
- CA
  3862   3861   3863      1    1.2290e+02    6.6944e+02 ;      O - C
- N
```

3863 3865 3866 1 1.0950e+02 4.1840e+02 ; N - CA
- HA
3863 3865 3867 1 1.0970e+02 6.6944e+02 ; N - CA
- CB
3863 3865 3873 1 1.1010e+02 5.2718e+02 ; N - CA
- C
3864 3863 3865 1 1.1804e+02 4.1840e+02 ; H - N
- CA
3865 3867 3868 1 1.0950e+02 4.1840e+02 ; CA - CB
- HB2
3865 3867 3869 1 1.0950e+02 4.1840e+02 ; CA - CB
- HB3
3865 3867 3870 1 1.1110e+02 5.2718e+02 ; CA - CB
- CG
3865 3873 3874 1 1.2040e+02 6.6944e+02 ; CA - C
- O
3865 3873 3875 1 1.1660e+02 5.8576e+02 ; CA - C
- N
3866 3865 3867 1 1.0950e+02 4.1840e+02 ; HA - CA
- CB
3866 3865 3873 1 1.0950e+02 4.1840e+02 ; HA - CA
- C
3867 3865 3873 1 1.1110e+02 5.2718e+02 ; CB - CA
- C
3867 3870 3871 1 1.1700e+02 5.8576e+02 ; CB - CG
- OD1
3867 3870 3872 1 1.1700e+02 5.8576e+02 ; CB - CG
- OD2
3868 3867 3869 1 1.0950e+02 2.9288e+02 ; HB2 - CB
- HB3
3868 3867 3870 1 1.0950e+02 4.1840e+02 ; HB2 - CB
- CG
3869 3867 3870 1 1.0950e+02 4.1840e+02 ; HB3 - CB
- CG
3871 3870 3872 1 1.2600e+02 6.6944e+02 ; OD1 - CG
- OD2
3873 3875 3876 1 1.2000e+02 4.1840e+02 ; C - N
- H
3873 3875 3877 1 1.2190e+02 4.1840e+02 ; C - N
- CA
3874 3873 3875 1 1.2290e+02 6.6944e+02 ; O - C
- N
3875 3877 3878 1 1.0950e+02 4.1840e+02 ; N - CA
- HA
3875 3877 3879 1 1.0970e+02 6.6944e+02 ; N - CA
- CB
3875 3877 3883 1 1.1010e+02 5.2718e+02 ; N - CA
- C
3876 3875 3877 1 1.1804e+02 4.1840e+02 ; H - N
- CA
3877 3879 3880 1 1.0950e+02 4.1840e+02 ; CA - CB
- HB1

3877 3879 3881 1 1.0950e+02 4.1840e+02 ; CA - CB
- HB2
3877 3879 3882 1 1.0950e+02 4.1840e+02 ; CA - CB
- HB3
3877 3883 3884 1 1.2040e+02 6.6944e+02 ; CA - C
- O
3877 3883 3885 1 1.1660e+02 5.8576e+02 ; CA - C
- N
3878 3877 3879 1 1.0950e+02 4.1840e+02 ; HA - CA
- CB
3878 3877 3883 1 1.0950e+02 4.1840e+02 ; HA - CA
- C
3879 3877 3883 1 1.1110e+02 5.2718e+02 ; CB - CA
- C
3880 3879 3881 1 1.0950e+02 2.9288e+02 ; HB1 - CB
- HB2
3880 3879 3882 1 1.0950e+02 2.9288e+02 ; HB1 - CB
- HB3
3881 3879 3882 1 1.0950e+02 2.9288e+02 ; HB2 - CB
- HB3
3883 3885 3886 1 1.2000e+02 4.1840e+02 ; C - N
- H
3883 3885 3887 1 1.2190e+02 4.1840e+02 ; C - N
- CA
3884 3883 3885 1 1.2290e+02 6.6944e+02 ; O - C
- N
3885 3887 3888 1 1.0950e+02 4.1840e+02 ; N - CA
- HA
3885 3887 3889 1 1.0970e+02 6.6944e+02 ; N - CA
- CB
3885 3887 3905 1 1.1010e+02 5.2718e+02 ; N - CA
- C
3886 3885 3887 1 1.1804e+02 4.1840e+02 ; H - N
- CA
3887 3889 3890 1 1.0950e+02 4.1840e+02 ; CA - CB
- HB2
3887 3889 3891 1 1.0950e+02 4.1840e+02 ; CA - CB
- HB3
3887 3889 3892 1 1.0950e+02 3.3472e+02 ; CA - CB
- CG
3887 3905 3906 1 1.2040e+02 6.6944e+02 ; CA - C
- O
3887 3905 3907 1 1.1660e+02 5.8576e+02 ; CA - C
- N
3888 3887 3889 1 1.0950e+02 4.1840e+02 ; HA - CA
- CB
3888 3887 3905 1 1.0950e+02 4.1840e+02 ; HA - CA
- C
3889 3887 3905 1 1.1110e+02 5.2718e+02 ; CB - CA
- C
3889 3892 3893 1 1.0950e+02 4.1840e+02 ; CB - CG
- HG2

```
  3889   3892   3894      1    1.0950e+02    4.1840e+02 ;     CB - CG
- HG3
  3889   3892   3895      1    1.0950e+02    3.3472e+02 ;     CB - CG
- CD
  3890   3889   3891      1    1.0950e+02    2.9288e+02 ;    HB2 - CB
- HB3
  3890   3889   3892      1    1.0950e+02    4.1840e+02 ;    HB2 - CB
- CG
  3891   3889   3892      1    1.0950e+02    4.1840e+02 ;    HB3 - CB
- CG
  3892   3895   3896      1    1.0950e+02    4.1840e+02 ;     CG - CD
- HD2
  3892   3895   3897      1    1.0950e+02    4.1840e+02 ;     CG - CD
- HD3
  3892   3895   3898      1    1.0950e+02    3.3472e+02 ;     CG - CD
- CE
  3893   3892   3894      1    1.0950e+02    2.9288e+02 ;    HG2 - CG
- HG3
  3893   3892   3895      1    1.0950e+02    4.1840e+02 ;    HG2 - CG
- CD
  3894   3892   3895      1    1.0950e+02    4.1840e+02 ;    HG3 - CG
- CD
  3895   3898   3899      1    1.0950e+02    4.1840e+02 ;     CD - CE
- HE2
  3895   3898   3900      1    1.0950e+02    4.1840e+02 ;     CD - CE
- HE3
  3895   3898   3901      1    1.1120e+02    6.6944e+02 ;     CD - CE
- NZ
  3896   3895   3897      1    1.0950e+02    2.9288e+02 ;    HD2 - CD
- HD3
  3896   3895   3898      1    1.0950e+02    4.1840e+02 ;    HD2 - CD
- CE
  3897   3895   3898      1    1.0950e+02    4.1840e+02 ;    HD3 - CD
- CE
  3898   3901   3902      1    1.0950e+02    4.1840e+02 ;     CE - NZ
- HZ1
  3898   3901   3903      1    1.0950e+02    4.1840e+02 ;     CE - NZ
- HZ2
  3898   3901   3904      1    1.0950e+02    4.1840e+02 ;     CE - NZ
- HZ3
  3899   3898   3900      1    1.0950e+02    2.9288e+02 ;    HE2 - CE
- HE3
  3899   3898   3901      1    1.0950e+02    4.1840e+02 ;    HE2 - CE
- NZ
  3900   3898   3901      1    1.0950e+02    4.1840e+02 ;    HE3 - CE
- NZ
  3902   3901   3903      1    1.0950e+02    2.9288e+02 ;    HZ1 - NZ
- HZ2
  3902   3901   3904      1    1.0950e+02    2.9288e+02 ;    HZ1 - NZ
- HZ3
  3903   3901   3904      1    1.0950e+02    2.9288e+02 ;    HZ2 - NZ
- HZ3
```

```
  3905   3907   3908      1    1.2000e+02    4.1840e+02 ;      C - N
- H
  3905   3907   3909      1    1.2190e+02    4.1840e+02 ;      C - N
- CA
  3906   3905   3907      1    1.2290e+02    6.6944e+02 ;      O - C
- N
  3907   3909   3910      1    1.0950e+02    4.1840e+02 ;      N - CA
- HA
  3907   3909   3911      1    1.0970e+02    6.6944e+02 ;      N - CA
- CB
  3907   3909   3919      1    1.1010e+02    5.2718e+02 ;      N - CA
- C
  3908   3907   3909      1    1.1804e+02    4.1840e+02 ;      H - N
- CA
  3909   3911   3912      1    1.0950e+02    4.1840e+02 ;     CA - CB
- HB2
  3909   3911   3913      1    1.0950e+02    4.1840e+02 ;     CA - CB
- HB3
  3909   3911   3914      1    1.1110e+02    5.2718e+02 ;     CA - CB
- CG
  3909   3919   3920      1    1.2040e+02    6.6944e+02 ;     CA - C
- O
  3909   3919   3921      1    1.1660e+02    5.8576e+02 ;     CA - C
- N
  3910   3909   3911      1    1.0950e+02    4.1840e+02 ;     HA - CA
- CB
  3910   3909   3919      1    1.0950e+02    4.1840e+02 ;     HA - CA
- C
  3911   3909   3919      1    1.1110e+02    5.2718e+02 ;     CB - CA
- C
  3911   3914   3915      1    1.2040e+02    6.6944e+02 ;     CB - CG
- OD1
  3911   3914   3916      1    1.1660e+02    5.8576e+02 ;     CB - CG
- ND2
  3912   3911   3913      1    1.0950e+02    2.9288e+02 ;    HB2 - CB
- HB3
  3912   3911   3914      1    1.0950e+02    4.1840e+02 ;    HB2 - CB
- CG
  3913   3911   3914      1    1.0950e+02    4.1840e+02 ;    HB3 - CB
- CG
  3914   3916   3917      1    1.2000e+02    4.1840e+02 ;     CG - ND2
- HD21
  3914   3916   3918      1    1.2000e+02    4.1840e+02 ;     CG - ND2
- HD22
  3915   3914   3916      1    1.2290e+02    6.6944e+02 ;    OD1 - CG
- ND2
  3917   3916   3918      1    1.2000e+02    2.9288e+02 ;   HD21 - ND2
- HD22
  3919   3921   3922      1    1.2000e+02    4.1840e+02 ;      C - N
- H
  3919   3921   3923      1    1.2190e+02    4.1840e+02 ;      C - N
- CA
```

3920 3919 3921 1 1.2290e+02 6.6944e+02 ; O - C
- N
3921 3923 3924 1 1.0950e+02 4.1840e+02 ; N - CA
- HA
3921 3923 3925 1 1.0970e+02 6.6944e+02 ; N - CA
- CB
3921 3923 3938 1 1.1010e+02 5.2718e+02 ; N - CA
- C
3922 3921 3923 1 1.1804e+02 4.1840e+02 ; H - N
- CA
3923 3925 3926 1 1.0950e+02 4.1840e+02 ; CA - CB
- HB2
3923 3925 3927 1 1.0950e+02 4.1840e+02 ; CA - CB
- HB3
3923 3925 3928 1 1.0950e+02 3.3472e+02 ; CA - CB
- CG
3923 3938 3939 1 1.2040e+02 6.6944e+02 ; CA - C
- O
3923 3938 3940 1 1.1660e+02 5.8576e+02 ; CA - C
- N
3924 3923 3925 1 1.0950e+02 4.1840e+02 ; HA - CA
- CB
3924 3923 3938 1 1.0950e+02 4.1840e+02 ; HA - CA
- C
3925 3923 3938 1 1.1110e+02 5.2718e+02 ; CB - CA
- C
3925 3928 3929 1 1.0950e+02 4.1840e+02 ; CB - CG
- HG
3925 3928 3930 1 1.0950e+02 3.3472e+02 ; CB - CG
- CD1
3925 3928 3934 1 1.0950e+02 3.3472e+02 ; CB - CG
- CD2
3926 3925 3927 1 1.0950e+02 2.9288e+02 ; HB2 - CB
- HB3
3926 3925 3928 1 1.0950e+02 4.1840e+02 ; HB2 - CB
- CG
3927 3925 3928 1 1.0950e+02 4.1840e+02 ; HB3 - CB
- CG
3928 3930 3931 1 1.0950e+02 4.1840e+02 ; CG - CD1
- HD11
3928 3930 3932 1 1.0950e+02 4.1840e+02 ; CG - CD1
- HD12
3928 3930 3933 1 1.0950e+02 4.1840e+02 ; CG - CD1
- HD13
3928 3934 3935 1 1.0950e+02 4.1840e+02 ; CG - CD2
- HD21
3928 3934 3936 1 1.0950e+02 4.1840e+02 ; CG - CD2
- HD22
3928 3934 3937 1 1.0950e+02 4.1840e+02 ; CG - CD2
- HD23
3929 3928 3930 1 1.0950e+02 4.1840e+02 ; HG - CG
- CD1

```
  3929   3928   3934       1    1.0950e+02    4.1840e+02 ;     HG - CG
- CD2
  3930   3928   3934       1    1.0950e+02    3.3472e+02 ;    CD1 - CG
- CD2
  3931   3930   3932       1    1.0950e+02    2.9288e+02 ;   HD11 - CD1
- HD12
  3931   3930   3933       1    1.0950e+02    2.9288e+02 ;   HD11 - CD1
- HD13
  3932   3930   3933       1    1.0950e+02    2.9288e+02 ;   HD12 - CD1
- HD13
  3935   3934   3936       1    1.0950e+02    2.9288e+02 ;   HD21 - CD2
- HD22
  3935   3934   3937       1    1.0950e+02    2.9288e+02 ;   HD21 - CD2
- HD23
  3936   3934   3937       1    1.0950e+02    2.9288e+02 ;   HD22 - CD2
- HD23
  3938   3940   3941       1    1.2000e+02    4.1840e+02 ;      C - N
- H
  3938   3940   3942       1    1.2190e+02    4.1840e+02 ;      C - N
- CA
  3939   3938   3940       1    1.2290e+02    6.6944e+02 ;      O - C
- N
  3940   3942   3943       1    1.0950e+02    4.1840e+02 ;      N - CA
- HA
  3940   3942   3944       1    1.0970e+02    6.6944e+02 ;      N - CA
- CB
  3940   3942   3957       1    1.1010e+02    5.2718e+02 ;      N - CA
- C
  3941   3940   3942       1    1.1804e+02    4.1840e+02 ;      H - N
- CA
  3942   3944   3945       1    1.0950e+02    4.1840e+02 ;     CA - CB
- HB
  3942   3944   3946       1    1.0950e+02    3.3472e+02 ;     CA - CB
- CG2
  3942   3944   3950       1    1.0950e+02    3.3472e+02 ;     CA - CB
- CG1
  3942   3957   3958       1    1.2040e+02    6.6944e+02 ;     CA - C
- O
  3942   3957   3959       1    1.1660e+02    5.8576e+02 ;     CA - C
- N
  3943   3942   3944       1    1.0950e+02    4.1840e+02 ;     HA - CA
- CB
  3943   3942   3957       1    1.0950e+02    4.1840e+02 ;     HA - CA
- C
  3944   3942   3957       1    1.1110e+02    5.2718e+02 ;     CB - CA
- C
  3944   3946   3947       1    1.0950e+02    4.1840e+02 ;     CB - CG2
- HG21
  3944   3946   3948       1    1.0950e+02    4.1840e+02 ;     CB - CG2
- HG22
  3944   3946   3949       1    1.0950e+02    4.1840e+02 ;     CB - CG2
- HG23
```

3944 3950 3951 1 1.0950e+02 4.1840e+02 ; CB - CG1
- HG12
3944 3950 3952 1 1.0950e+02 4.1840e+02 ; CB - CG1
- HG13
3944 3950 3953 1 1.0950e+02 3.3472e+02 ; CB - CG1
- CD1
3945 3944 3946 1 1.0950e+02 4.1840e+02 ; HB - CB
- CG2
3945 3944 3950 1 1.0950e+02 4.1840e+02 ; HB - CB
- CG1
3946 3944 3950 1 1.0950e+02 3.3472e+02 ; CG2 - CB
- CG1
3947 3946 3948 1 1.0950e+02 2.9288e+02 ; HG21 - CG2
- HG22
3947 3946 3949 1 1.0950e+02 2.9288e+02 ; HG21 - CG2
- HG23
3948 3946 3949 1 1.0950e+02 2.9288e+02 ; HG22 - CG2
- HG23
3950 3953 3954 1 1.0950e+02 4.1840e+02 ; CG1 - CD1
- HD11
3950 3953 3955 1 1.0950e+02 4.1840e+02 ; CG1 - CD1
- HD12
3950 3953 3956 1 1.0950e+02 4.1840e+02 ; CG1 - CD1
- HD13
3951 3950 3952 1 1.0950e+02 2.9288e+02 ; HG12 - CG1
- HG13
3951 3950 3953 1 1.0950e+02 4.1840e+02 ; HG12 - CG1
- CD1
3952 3950 3953 1 1.0950e+02 4.1840e+02 ; HG13 - CG1
- CD1
3954 3953 3955 1 1.0950e+02 2.9288e+02 ; HD11 - CD1
- HD12
3954 3953 3956 1 1.0950e+02 2.9288e+02 ; HD11 - CD1
- HD13
3955 3953 3956 1 1.0950e+02 2.9288e+02 ; HD12 - CD1
- HD13
3957 3959 3960 1 1.2000e+02 4.1840e+02 ; C - N
- H
3957 3959 3961 1 1.2190e+02 4.1840e+02 ; C - N
- CA
3958 3957 3959 1 1.2290e+02 6.6944e+02 ; O - C
- N
3959 3961 3962 1 1.0950e+02 4.1840e+02 ; N - CA
- HA
3959 3961 3963 1 1.0970e+02 6.6944e+02 ; N - CA
- CB
3959 3961 3971 1 1.1010e+02 5.2718e+02 ; N - CA
- C
3960 3959 3961 1 1.1804e+02 4.1840e+02 ; H - N
- CA
3961 3963 3964 1 1.0950e+02 4.1840e+02 ; CA - CB
- HB

3961 3963 3965 1 1.0950e+02 3.3472e+02 ; CA - CB
- CG2
3961 3963 3969 1 1.0950e+02 4.1840e+02 ; CA - CB
- OG1
3961 3971 3972 1 1.2040e+02 6.6944e+02 ; CA - C
- O
3961 3971 3973 1 1.1660e+02 5.8576e+02 ; CA - C
- N
3962 3961 3963 1 1.0950e+02 4.1840e+02 ; HA - CA
- CB
3962 3961 3971 1 1.0950e+02 4.1840e+02 ; HA - CA
- C
3963 3961 3971 1 1.1110e+02 5.2718e+02 ; CB - CA
- C
3963 3965 3966 1 1.0950e+02 4.1840e+02 ; CB - CG2
- HG21
3963 3965 3967 1 1.0950e+02 4.1840e+02 ; CB - CG2
- HG22
3963 3965 3968 1 1.0950e+02 4.1840e+02 ; CB - CG2
- HG23
3963 3969 3970 1 1.0850e+02 4.6024e+02 ; CB - OG1
- HG1
3964 3963 3965 1 1.0950e+02 4.1840e+02 ; HB - CB
- CG2
3964 3963 3969 1 1.0950e+02 4.1840e+02 ; HB - CB
- OG1
3965 3963 3969 1 1.0950e+02 4.1840e+02 ; CG2 - CB
- OG1
3966 3965 3967 1 1.0950e+02 2.9288e+02 ; HG21 - CG2
- HG22
3966 3965 3968 1 1.0950e+02 2.9288e+02 ; HG21 - CG2
- HG23
3967 3965 3968 1 1.0950e+02 2.9288e+02 ; HG22 - CG2
- HG23
3971 3973 3974 1 1.2000e+02 4.1840e+02 ; C - N
- H
3971 3973 3975 1 1.2190e+02 4.1840e+02 ; C - N
- CA
3972 3971 3973 1 1.2290e+02 6.6944e+02 ; O - C
- N
3973 3975 3976 1 1.0950e+02 4.1840e+02 ; N - CA
- HA
3973 3975 3977 1 1.0970e+02 6.6944e+02 ; N - CA
- CB
3973 3975 3993 1 1.1010e+02 5.2718e+02 ; N - CA
- C
3974 3973 3975 1 1.1804e+02 4.1840e+02 ; H - N
- CA
3975 3977 3978 1 1.0950e+02 4.1840e+02 ; CA - CB
- HB2
3975 3977 3979 1 1.0950e+02 4.1840e+02 ; CA - CB
- HB3

```
  3975   3977   3980       1    1.0950e+02    3.3472e+02 ;      CA - CB
- CG
  3975   3993   3994       1    1.2040e+02    6.6944e+02 ;      CA - C
- O
  3975   3993   3995       1    1.1660e+02    5.8576e+02 ;      CA - C
- N
  3976   3975   3977       1    1.0950e+02    4.1840e+02 ;      HA - CA
- CB
  3976   3975   3993       1    1.0950e+02    4.1840e+02 ;      HA - CA
- C
  3977   3975   3993       1    1.1110e+02    5.2718e+02 ;      CB - CA
- C
  3977   3980   3981       1    1.0950e+02    4.1840e+02 ;      CB - CG
- HG2
  3977   3980   3982       1    1.0950e+02    4.1840e+02 ;      CB - CG
- HG3
  3977   3980   3983       1    1.0950e+02    3.3472e+02 ;      CB - CG
- CD
  3978   3977   3979       1    1.0950e+02    2.9288e+02 ;     HB2 - CB
- HB3
  3978   3977   3980       1    1.0950e+02    4.1840e+02 ;     HB2 - CB
- CG
  3979   3977   3980       1    1.0950e+02    4.1840e+02 ;     HB3 - CB
- CG
  3980   3983   3984       1    1.0950e+02    4.1840e+02 ;      CG - CD
- HD2
  3980   3983   3985       1    1.0950e+02    4.1840e+02 ;      CG - CD
- HD3
  3980   3983   3986       1    1.0950e+02    3.3472e+02 ;      CG - CD
- CE
  3981   3980   3982       1    1.0950e+02    2.9288e+02 ;     HG2 - CG
- HG3
  3981   3980   3983       1    1.0950e+02    4.1840e+02 ;     HG2 - CG
- CD
  3982   3980   3983       1    1.0950e+02    4.1840e+02 ;     HG3 - CG
- CD
  3983   3986   3987       1    1.0950e+02    4.1840e+02 ;      CD - CE
- HE2
  3983   3986   3988       1    1.0950e+02    4.1840e+02 ;      CD - CE
- HE3
  3983   3986   3989       1    1.1120e+02    6.6944e+02 ;      CD - CE
- NZ
  3984   3983   3985       1    1.0950e+02    2.9288e+02 ;     HD2 - CD
- HD3
  3984   3983   3986       1    1.0950e+02    4.1840e+02 ;     HD2 - CD
- CE
  3985   3983   3986       1    1.0950e+02    4.1840e+02 ;     HD3 - CD
- CE
  3986   3989   3990       1    1.0950e+02    4.1840e+02 ;      CE - NZ
- HZ1
  3986   3989   3991       1    1.0950e+02    4.1840e+02 ;      CE - NZ
- HZ2
```

```
  3986   3989   3992      1    1.0950e+02    4.1840e+02 ;     CE - NZ
- HZ3
  3987   3986   3988      1    1.0950e+02    2.9288e+02 ;    HE2 - CE
- HE3
  3987   3986   3989      1    1.0950e+02    4.1840e+02 ;    HE2 - CE
- NZ
  3988   3986   3989      1    1.0950e+02    4.1840e+02 ;    HE3 - CE
- NZ
  3990   3989   3991      1    1.0950e+02    2.9288e+02 ;    HZ1 - NZ
- HZ2
  3990   3989   3992      1    1.0950e+02    2.9288e+02 ;    HZ1 - NZ
- HZ3
  3991   3989   3992      1    1.0950e+02    2.9288e+02 ;    HZ2 - NZ
- HZ3
  3993   3995   3996      1    1.2000e+02    4.1840e+02 ;      C - N
- H
  3993   3995   3997      1    1.2190e+02    4.1840e+02 ;      C - N
- CA
  3994   3993   3995      1    1.2290e+02    6.6944e+02 ;      O - C
- N
  3995   3997   3998      1    1.0950e+02    4.1840e+02 ;      N - CA
- HA
  3995   3997   3999      1    1.0970e+02    6.6944e+02 ;      N - CA
- CB
  3995   3997   4012      1    1.1010e+02    5.2718e+02 ;      N - CA
- C
  3996   3995   3997      1    1.1804e+02    4.1840e+02 ;      H - N
- CA
  3997   3999   4000      1    1.0950e+02    4.1840e+02 ;     CA - CB
- HB2
  3997   3999   4001      1    1.0950e+02    4.1840e+02 ;     CA - CB
- HB3
  3997   3999   4002      1    1.0950e+02    3.3472e+02 ;     CA - CB
- CG
  3997   4012   4013      1    1.2040e+02    6.6944e+02 ;     CA - C
- O
  3997   4012   4014      1    1.1660e+02    5.8576e+02 ;     CA - C
- N
  3998   3997   3999      1    1.0950e+02    4.1840e+02 ;     HA - CA
- CB
  3998   3997   4012      1    1.0950e+02    4.1840e+02 ;     HA - CA
- C
  3999   3997   4012      1    1.1110e+02    5.2718e+02 ;     CB - CA
- C
  3999   4002   4003      1    1.0950e+02    4.1840e+02 ;     CB - CG
- HG
  3999   4002   4004      1    1.0950e+02    3.3472e+02 ;     CB - CG
- CD1
  3999   4002   4008      1    1.0950e+02    3.3472e+02 ;     CB - CG
- CD2
  4000   3999   4001      1    1.0950e+02    2.9288e+02 ;    HB2 - CB
- HB3
```

4000 3999 4002 1 1.0950e+02 4.1840e+02 ; HB2 - CB
- CG
4001 3999 4002 1 1.0950e+02 4.1840e+02 ; HB3 - CB
- CG
4002 4004 4005 1 1.0950e+02 4.1840e+02 ; CG - CD1
- HD11
4002 4004 4006 1 1.0950e+02 4.1840e+02 ; CG - CD1
- HD12
4002 4004 4007 1 1.0950e+02 4.1840e+02 ; CG - CD1
- HD13
4002 4008 4009 1 1.0950e+02 4.1840e+02 ; CG - CD2
- HD21
4002 4008 4010 1 1.0950e+02 4.1840e+02 ; CG - CD2
- HD22
4002 4008 4011 1 1.0950e+02 4.1840e+02 ; CG - CD2
- HD23
4003 4002 4004 1 1.0950e+02 4.1840e+02 ; HG - CG
- CD1
4003 4002 4008 1 1.0950e+02 4.1840e+02 ; HG - CG
- CD2
4004 4002 4008 1 1.0950e+02 3.3472e+02 ; CD1 - CG
- CD2
4005 4004 4006 1 1.0950e+02 2.9288e+02 ; HD11 - CD1
- HD12
4005 4004 4007 1 1.0950e+02 2.9288e+02 ; HD11 - CD1
- HD13
4006 4004 4007 1 1.0950e+02 2.9288e+02 ; HD12 - CD1
- HD13
4009 4008 4010 1 1.0950e+02 2.9288e+02 ; HD21 - CD2
- HD22
4009 4008 4011 1 1.0950e+02 2.9288e+02 ; HD21 - CD2
- HD23
4010 4008 4011 1 1.0950e+02 2.9288e+02 ; HD22 - CD2
- HD23
4012 4014 4015 1 1.2000e+02 4.1840e+02 ; C - N
- H
4012 4014 4016 1 1.2190e+02 4.1840e+02 ; C - N
- CA
4013 4012 4014 1 1.2290e+02 6.6944e+02 ; O - C
- N
4014 4016 4017 1 1.0950e+02 4.1840e+02 ; N - CA
- HA
4014 4016 4018 1 1.0970e+02 6.6944e+02 ; N - CA
- CB
4014 4016 4031 1 1.1010e+02 5.2718e+02 ; N - CA
- C
4015 4014 4016 1 1.1804e+02 4.1840e+02 ; H - N
- CA
4016 4018 4019 1 1.0950e+02 4.1840e+02 ; CA - CB
- HB2
4016 4018 4020 1 1.0950e+02 4.1840e+02 ; CA - CB
- HB3

4016 4018 4021 1 1.0950e+02 3.3472e+02 ; CA - CB
- CG
4016 4031 4032 1 1.2040e+02 6.6944e+02 ; CA - C
- O
4016 4031 4033 1 1.1660e+02 5.8576e+02 ; CA - C
- N
4017 4016 4018 1 1.0950e+02 4.1840e+02 ; HA - CA
- CB
4017 4016 4031 1 1.0950e+02 4.1840e+02 ; HA - CA
- C
4018 4016 4031 1 1.1110e+02 5.2718e+02 ; CB - CA
- C
4018 4021 4022 1 1.0950e+02 4.1840e+02 ; CB - CG
- HG
4018 4021 4023 1 1.0950e+02 3.3472e+02 ; CB - CG
- CD1
4018 4021 4027 1 1.0950e+02 3.3472e+02 ; CB - CG
- CD2
4019 4018 4020 1 1.0950e+02 2.9288e+02 ; HB2 - CB
- HB3
4019 4018 4021 1 1.0950e+02 4.1840e+02 ; HB2 - CB
- CG
4020 4018 4021 1 1.0950e+02 4.1840e+02 ; HB3 - CB
- CG
4021 4023 4024 1 1.0950e+02 4.1840e+02 ; CG - CD1
- HD11
4021 4023 4025 1 1.0950e+02 4.1840e+02 ; CG - CD1
- HD12
4021 4023 4026 1 1.0950e+02 4.1840e+02 ; CG - CD1
- HD13
4021 4027 4028 1 1.0950e+02 4.1840e+02 ; CG - CD2
- HD21
4021 4027 4029 1 1.0950e+02 4.1840e+02 ; CG - CD2
- HD22
4021 4027 4030 1 1.0950e+02 4.1840e+02 ; CG - CD2
- HD23
4022 4021 4023 1 1.0950e+02 4.1840e+02 ; HG - CG
- CD1
4022 4021 4027 1 1.0950e+02 4.1840e+02 ; HG - CG
- CD2
4023 4021 4027 1 1.0950e+02 3.3472e+02 ; CD1 - CG
- CD2
4024 4023 4025 1 1.0950e+02 2.9288e+02 ; HD11 - CD1
- HD12
4024 4023 4026 1 1.0950e+02 2.9288e+02 ; HD11 - CD1
- HD13
4025 4023 4026 1 1.0950e+02 2.9288e+02 ; HD12 - CD1
- HD13
4028 4027 4029 1 1.0950e+02 2.9288e+02 ; HD21 - CD2
- HD22
4028 4027 4030 1 1.0950e+02 2.9288e+02 ; HD21 - CD2
- HD23

4029 4027 4030 1 1.0950e+02 2.9288e+02 ; HD22 - CD2 - HD23
4031 4033 4034 1 1.2000e+02 4.1840e+02 ; C - N - H
4031 4033 4035 1 1.2190e+02 4.1840e+02 ; C - N - CA
4032 4031 4033 1 1.2290e+02 6.6944e+02 ; O - C - N
4033 4035 4036 1 1.0950e+02 4.1840e+02 ; N - CA - HA
4033 4035 4037 1 1.0970e+02 6.6944e+02 ; N - CA - CB
4033 4035 4045 1 1.1010e+02 5.2718e+02 ; N - CA - C
4034 4033 4035 1 1.1804e+02 4.1840e+02 ; H - N - CA
4035 4037 4038 1 1.0950e+02 4.1840e+02 ; CA - CB - HB
4035 4037 4039 1 1.0950e+02 3.3472e+02 ; CA - CB - CG2
4035 4037 4043 1 1.0950e+02 4.1840e+02 ; CA - CB - OG1
4035 4045 4046 1 1.2040e+02 6.6944e+02 ; CA - C - O
4035 4045 4047 1 1.1660e+02 5.8576e+02 ; CA - C - N
4036 4035 4037 1 1.0950e+02 4.1840e+02 ; HA - CA - CB
4036 4035 4045 1 1.0950e+02 4.1840e+02 ; HA - CA - C
4037 4035 4045 1 1.1110e+02 5.2718e+02 ; CB - CA - C
4037 4039 4040 1 1.0950e+02 4.1840e+02 ; CB - CG2 - HG21
4037 4039 4041 1 1.0950e+02 4.1840e+02 ; CB - CG2 - HG22
4037 4039 4042 1 1.0950e+02 4.1840e+02 ; CB - CG2 - HG23
4037 4043 4044 1 1.0850e+02 4.6024e+02 ; CB - OG1 - HG1
4038 4037 4039 1 1.0950e+02 4.1840e+02 ; HB - CB - CG2
4038 4037 4043 1 1.0950e+02 4.1840e+02 ; HB - CB - OG1
4039 4037 4043 1 1.0950e+02 4.1840e+02 ; CG2 - CB - OG1
4040 4039 4041 1 1.0950e+02 2.9288e+02 ; HG21 - CG2 - HG22
4040 4039 4042 1 1.0950e+02 2.9288e+02 ; HG21 - CG2 - HG23
4041 4039 4042 1 1.0950e+02 2.9288e+02 ; HG22 - CG2 - HG23

```
  4045   4047   4048      1    1.2000e+02    4.1840e+02 ;      C - N
- H
  4045   4047   4049      1    1.2190e+02    4.1840e+02 ;      C - N
- CA
  4046   4045   4047      1    1.2290e+02    6.6944e+02 ;      O - C
- N
  4047   4049   4050      1    1.0950e+02    4.1840e+02 ;      N - CA
- HA
  4047   4049   4051      1    1.0970e+02    6.6944e+02 ;      N - CA
- CB
  4047   4049   4066      1    1.1010e+02    5.2718e+02 ;      N - CA
- C
  4048   4047   4049      1    1.1804e+02    4.1840e+02 ;      H - N
- CA
  4049   4051   4052      1    1.0950e+02    4.1840e+02 ;     CA - CB
- HB2
  4049   4051   4053      1    1.0950e+02    4.1840e+02 ;     CA - CB
- HB3
  4049   4051   4054      1    1.1400e+02    5.2718e+02 ;     CA - CB
- CG
  4049   4066   4067      1    1.2040e+02    6.6944e+02 ;     CA - C
- O
  4049   4066   4068      1    1.1660e+02    5.8576e+02 ;     CA - C
- N
  4050   4049   4051      1    1.0950e+02    4.1840e+02 ;     HA - CA
- CB
  4050   4049   4066      1    1.0950e+02    4.1840e+02 ;     HA - CA
- C
  4051   4049   4066      1    1.1110e+02    5.2718e+02 ;     CB - CA
- C
  4051   4054   4055      1    1.2000e+02    5.8576e+02 ;     CB - CG
- CD1
  4051   4054   4064      1    1.2000e+02    5.8576e+02 ;     CB - CG
- CD2
  4052   4051   4053      1    1.0950e+02    2.9288e+02 ;    HB2 - CB
- HB3
  4052   4051   4054      1    1.0950e+02    4.1840e+02 ;    HB2 - CB
- CG
  4053   4051   4054      1    1.0950e+02    4.1840e+02 ;    HB3 - CB
- CG
  4054   4055   4056      1    1.2000e+02    4.1840e+02 ;     CG - CD1
- HD1
  4054   4055   4057      1    1.2000e+02    5.2718e+02 ;     CG - CD1
- CE1
  4054   4064   4062      1    1.2000e+02    5.2718e+02 ;     CG - CD2
- CE2
  4054   4064   4065      1    1.2000e+02    4.1840e+02 ;     CG - CD2
- HD2
  4055   4054   4064      1    1.2000e+02    5.2718e+02 ;    CD1 - CG
- CD2
  4055   4057   4058      1    1.2000e+02    4.1840e+02 ;    CD1 - CE1
- HE1
```

4055 4057 4059 1 1.2000e+02 5.2718e+02 ; CD1 - CE1
- CZ
4056 4055 4057 1 1.2000e+02 4.1840e+02 ; HD1 - CD1
- CE1
4057 4059 4060 1 1.2000e+02 5.8576e+02 ; CE1 - CZ
- OH
4057 4059 4062 1 1.2000e+02 5.2718e+02 ; CE1 - CZ
- CE2
4058 4057 4059 1 1.2000e+02 4.1840e+02 ; HE1 - CE1
- CZ
4059 4060 4061 1 1.1300e+02 4.1840e+02 ; CZ - OH
- HH
4059 4062 4063 1 1.2000e+02 4.1840e+02 ; CZ - CE2
- HE2
4059 4062 4064 1 1.2000e+02 5.2718e+02 ; CZ - CE2
- CD2
4060 4059 4062 1 1.2000e+02 5.8576e+02 ; OH - CZ
- CE2
4062 4064 4065 1 1.2000e+02 4.1840e+02 ; CE2 - CD2
- HD2
4063 4062 4064 1 1.2000e+02 4.1840e+02 ; HE2 - CE2
- CD2
4066 4068 4069 1 1.2000e+02 4.1840e+02 ; C - N
- H
4066 4068 4070 1 1.2190e+02 4.1840e+02 ; C - N
- CA
4067 4066 4068 1 1.2290e+02 6.6944e+02 ; O - C
- N
4068 4070 4071 1 1.0950e+02 4.1840e+02 ; N - CA
- HA
4068 4070 4072 1 1.0970e+02 6.6944e+02 ; N - CA
- CB
4068 4070 4080 1 1.1010e+02 5.2718e+02 ; N - CA
- C
4069 4068 4070 1 1.1804e+02 4.1840e+02 ; H - N
- CA
4070 4072 4073 1 1.0950e+02 4.1840e+02 ; CA - CB
- HB2
4070 4072 4074 1 1.0950e+02 4.1840e+02 ; CA - CB
- HB3
4070 4072 4075 1 1.1110e+02 5.2718e+02 ; CA - CB
- CG
4070 4080 4081 1 1.2040e+02 6.6944e+02 ; CA - C
- O
4070 4080 4082 1 1.1660e+02 5.8576e+02 ; CA - C
- N
4071 4070 4072 1 1.0950e+02 4.1840e+02 ; HA - CA
- CB
4071 4070 4080 1 1.0950e+02 4.1840e+02 ; HA - CA
- C
4072 4070 4080 1 1.1110e+02 5.2718e+02 ; CB - CA
- C

4072 4075 4076 1 1.2040e+02 6.6944e+02 ; CB - CG
- OD1
4072 4075 4077 1 1.1660e+02 5.8576e+02 ; CB - CG
- ND2
4073 4072 4074 1 1.0950e+02 2.9288e+02 ; HB2 - CB
- HB3
4073 4072 4075 1 1.0950e+02 4.1840e+02 ; HB2 - CB
- CG
4074 4072 4075 1 1.0950e+02 4.1840e+02 ; HB3 - CB
- CG
4075 4077 4078 1 1.2000e+02 4.1840e+02 ; CG - ND2
- HD21
4075 4077 4079 1 1.2000e+02 4.1840e+02 ; CG - ND2
- HD22
4076 4075 4077 1 1.2290e+02 6.6944e+02 ; OD1 - CG
- ND2
4078 4077 4079 1 1.2000e+02 2.9288e+02 ; HD21 - ND2
- HD22
4080 4082 4083 1 1.2190e+02 4.1840e+02 ; C - N
- CD
4080 4082 4092 1 1.2190e+02 4.1840e+02 ; C - N
- CA
4081 4080 4082 1 1.2290e+02 6.6944e+02 ; O - C
- N
4082 4083 4084 1 1.0950e+02 4.1840e+02 ; N - CD
- HD2
4082 4083 4085 1 1.0950e+02 4.1840e+02 ; N - CD
- HD3
4082 4083 4086 1 1.0970e+02 6.6944e+02 ; N - CD
- CG
4082 4092 4089 1 1.0970e+02 6.6944e+02 ; N - CA
- CB
4082 4092 4093 1 1.0950e+02 4.1840e+02 ; N - CA
- HA
4082 4092 4094 1 1.1010e+02 5.2718e+02 ; N - CA
- C
4083 4082 4092 1 1.1800e+02 4.1840e+02 ; CD - N
- CA
4083 4086 4087 1 1.0950e+02 4.1840e+02 ; CD - CG
- HG2
4083 4086 4088 1 1.0950e+02 4.1840e+02 ; CD - CG
- HG3
4083 4086 4089 1 1.0950e+02 3.3472e+02 ; CD - CG
- CB
4084 4083 4085 1 1.0950e+02 2.9288e+02 ; HD2 - CD
- HD3
4084 4083 4086 1 1.0950e+02 4.1840e+02 ; HD2 - CD
- CG
4085 4083 4086 1 1.0950e+02 4.1840e+02 ; HD3 - CD
- CG
4086 4089 4090 1 1.0950e+02 4.1840e+02 ; CG - CB
- HB2

```
  4086   4089   4091       1    1.0950e+02    4.1840e+02 ;     CG - CB
- HB3
  4086   4089   4092       1    1.0950e+02    3.3472e+02 ;     CG - CB
- CA
  4087   4086   4088       1    1.0950e+02    2.9288e+02 ;    HG2 - CG
- HG3
  4087   4086   4089       1    1.0950e+02    4.1840e+02 ;    HG2 - CG
- CB
  4088   4086   4089       1    1.0950e+02    4.1840e+02 ;    HG3 - CG
- CB
  4089   4092   4093       1    1.0950e+02    4.1840e+02 ;     CB - CA
- HA
  4089   4092   4094       1    1.1110e+02    5.2718e+02 ;     CB - CA
- C
  4090   4089   4091       1    1.0950e+02    2.9288e+02 ;    HB2 - CB
- HB3
  4090   4089   4092       1    1.0950e+02    4.1840e+02 ;    HB2 - CB
- CA
  4091   4089   4092       1    1.0950e+02    4.1840e+02 ;    HB3 - CB
- CA
  4092   4094   4095       1    1.2040e+02    6.6944e+02 ;     CA - C
- O
  4092   4094   4096       1    1.1660e+02    5.8576e+02 ;     CA - C
- N
  4093   4092   4094       1    1.0950e+02    4.1840e+02 ;     HA - CA
- C
  4094   4096   4097       1    1.2000e+02    4.1840e+02 ;      C - N
- H
  4094   4096   4098       1    1.2190e+02    4.1840e+02 ;      C - N
- CA
  4095   4094   4096       1    1.2290e+02    6.6944e+02 ;      O - C
- N
  4096   4098   4099       1    1.0950e+02    4.1840e+02 ;      N - CA
- HA
  4096   4098   4100       1    1.0970e+02    6.6944e+02 ;      N - CA
- CB
  4096   4098   4108       1    1.1010e+02    5.2718e+02 ;      N - CA
- C
  4097   4096   4098       1    1.1804e+02    4.1840e+02 ;      H - N
- CA
  4098   4100   4101       1    1.0950e+02    4.1840e+02 ;     CA - CB
- HB2
  4098   4100   4102       1    1.0950e+02    4.1840e+02 ;     CA - CB
- HB3
  4098   4100   4103       1    1.1110e+02    5.2718e+02 ;     CA - CB
- CG
  4098   4108   4109       1    1.2040e+02    6.6944e+02 ;     CA - C
- O
  4098   4108   4110       1    1.1660e+02    5.8576e+02 ;     CA - C
- N
  4099   4098   4100       1    1.0950e+02    4.1840e+02 ;     HA - CA
- CB
```

4099 4098 4108 1 1.0950e+02 4.1840e+02 ; HA - CA
- C
4100 4098 4108 1 1.1110e+02 5.2718e+02 ; CB - CA
- C
4100 4103 4104 1 1.2040e+02 6.6944e+02 ; CB - CG
- OD1
4100 4103 4105 1 1.1660e+02 5.8576e+02 ; CB - CG
- ND2
4101 4100 4102 1 1.0950e+02 2.9288e+02 ; HB2 - CB
- HB3
4101 4100 4103 1 1.0950e+02 4.1840e+02 ; HB2 - CB
- CG
4102 4100 4103 1 1.0950e+02 4.1840e+02 ; HB3 - CB
- CG
4103 4105 4106 1 1.2000e+02 4.1840e+02 ; CG - ND2
- HD21
4103 4105 4107 1 1.2000e+02 4.1840e+02 ; CG - ND2
- HD22
4104 4103 4105 1 1.2290e+02 6.6944e+02 ; OD1 - CG
- ND2
4106 4105 4107 1 1.2000e+02 2.9288e+02 ; HD21 - ND2
- HD22
4108 4110 4111 1 1.2000e+02 4.1840e+02 ; C - N
- H
4108 4110 4112 1 1.2190e+02 4.1840e+02 ; C - N
- CA
4109 4108 4110 1 1.2290e+02 6.6944e+02 ; O - C
- N
4110 4112 4113 1 1.0950e+02 4.1840e+02 ; N - CA
- HA
4110 4112 4114 1 1.0970e+02 6.6944e+02 ; N - CA
- CB
4110 4112 4123 1 1.1010e+02 5.2718e+02 ; N - CA
- C
4111 4110 4112 1 1.1804e+02 4.1840e+02 ; H - N
- CA
4112 4114 4115 1 1.0950e+02 4.1840e+02 ; CA - CB
- HB2
4112 4114 4116 1 1.0950e+02 4.1840e+02 ; CA - CB
- HB3
4112 4114 4117 1 1.0950e+02 3.3472e+02 ; CA - CB
- CG
4112 4123 4124 1 1.2040e+02 6.6944e+02 ; CA - C
- O
4112 4123 4125 1 1.1660e+02 5.8576e+02 ; CA - C
- N
4113 4112 4114 1 1.0950e+02 4.1840e+02 ; HA - CA
- CB
4113 4112 4123 1 1.0950e+02 4.1840e+02 ; HA - CA
- C
4114 4112 4123 1 1.1110e+02 5.2718e+02 ; CB - CA
- C

4114 4117 4118 1 1.0950e+02 4.1840e+02 ; CB - CG
- HG2
4114 4117 4119 1 1.0950e+02 4.1840e+02 ; CB - CG
- HG3
4114 4117 4120 1 1.1110e+02 5.2718e+02 ; CB - CG
- CD
4115 4114 4116 1 1.0950e+02 2.9288e+02 ; HB2 - CB
- HB3
4115 4114 4117 1 1.0950e+02 4.1840e+02 ; HB2 - CB
- CG
4116 4114 4117 1 1.0950e+02 4.1840e+02 ; HB3 - CB
- CG
4117 4120 4121 1 1.1700e+02 5.8576e+02 ; CG - CD
- OE1
4117 4120 4122 1 1.1700e+02 5.8576e+02 ; CG - CD
- OE2
4118 4117 4119 1 1.0950e+02 2.9288e+02 ; HG2 - CG
- HG3
4118 4117 4120 1 1.0950e+02 4.1840e+02 ; HG2 - CG
- CD
4119 4117 4120 1 1.0950e+02 4.1840e+02 ; HG3 - CG
- CD
4121 4120 4122 1 1.2600e+02 6.6944e+02 ; OE1 - CD
- OE2
4123 4125 4126 1 1.2000e+02 4.1840e+02 ; C - N
- H
4123 4125 4127 1 1.2190e+02 4.1840e+02 ; C - N
- CA
4124 4123 4125 1 1.2290e+02 6.6944e+02 ; O - C
- N
4125 4127 4128 1 1.0950e+02 4.1840e+02 ; N - CA
- HA
4125 4127 4129 1 1.0970e+02 6.6944e+02 ; N - CA
- CB
4125 4127 4147 1 1.1010e+02 5.2718e+02 ; N - CA
- C
4126 4125 4127 1 1.1804e+02 4.1840e+02 ; H - N
- CA
4127 4129 4130 1 1.0950e+02 4.1840e+02 ; CA - CB
- HB2
4127 4129 4131 1 1.0950e+02 4.1840e+02 ; CA - CB
- HB3
4127 4129 4132 1 1.0950e+02 3.3472e+02 ; CA - CB
- CG
4127 4147 4148 1 1.2040e+02 6.6944e+02 ; CA - C
- O
4127 4147 4149 1 1.1660e+02 5.8576e+02 ; CA - C
- N
4128 4127 4129 1 1.0950e+02 4.1840e+02 ; HA - CA
- CB
4128 4127 4147 1 1.0950e+02 4.1840e+02 ; HA - CA
- C

```
  4129   4127   4147       1    1.1110e+02    5.2718e+02 ;     CB - CA
- C
  4129   4132   4133       1    1.0950e+02    4.1840e+02 ;     CB - CG
- HG2
  4129   4132   4134       1    1.0950e+02    4.1840e+02 ;     CB - CG
- HG3
  4129   4132   4135       1    1.0950e+02    3.3472e+02 ;     CB - CG
- CD
  4130   4129   4131       1    1.0950e+02    2.9288e+02 ;    HB2 - CB
- HB3
  4130   4129   4132       1    1.0950e+02    4.1840e+02 ;    HB2 - CB
- CG
  4131   4129   4132       1    1.0950e+02    4.1840e+02 ;    HB3 - CB
- CG
  4132   4135   4136       1    1.0950e+02    4.1840e+02 ;     CG - CD
- HD2
  4132   4135   4137       1    1.0950e+02    4.1840e+02 ;     CG - CD
- HD3
  4132   4135   4138       1    1.1120e+02    6.6944e+02 ;     CG - CD
- NE
  4133   4132   4134       1    1.0950e+02    2.9288e+02 ;    HG2 - CG
- HG3
  4133   4132   4135       1    1.0950e+02    4.1840e+02 ;    HG2 - CG
- CD
  4134   4132   4135       1    1.0950e+02    4.1840e+02 ;    HG3 - CG
- CD
  4135   4138   4139       1    1.1840e+02    4.1840e+02 ;     CD - NE
- HE
  4135   4138   4140       1    1.2320e+02    4.1840e+02 ;     CD - NE
- CZ
  4136   4135   4137       1    1.0950e+02    2.9288e+02 ;    HD2 - CD
- HD3
  4136   4135   4138       1    1.0950e+02    4.1840e+02 ;    HD2 - CD
- NE
  4137   4135   4138       1    1.0950e+02    4.1840e+02 ;    HD3 - CD
- NE
  4138   4140   4141       1    1.2000e+02    5.8576e+02 ;     NE - CZ
- NH1
  4138   4140   4144       1    1.2000e+02    5.8576e+02 ;     NE - CZ
- NH2
  4139   4138   4140       1    1.2000e+02    4.1840e+02 ;     HE - NE
- CZ
  4140   4141   4142       1    1.2000e+02    4.1840e+02 ;     CZ - NH1
- HH11
  4140   4141   4143       1    1.2000e+02    4.1840e+02 ;     CZ - NH1
- HH12
  4140   4144   4145       1    1.2000e+02    4.1840e+02 ;     CZ - NH2
- HH21
  4140   4144   4146       1    1.2000e+02    4.1840e+02 ;     CZ - NH2
- HH22
  4141   4140   4144       1    1.2000e+02    5.8576e+02 ;    NH1 - CZ
- NH2
```

4142 4141 4143 1 1.2000e+02 2.9288e+02 ; HH11 - NH1 - HH12
4145 4144 4146 1 1.2000e+02 2.9288e+02 ; HH21 - NH2 - HH22
4147 4149 4150 1 1.2000e+02 4.1840e+02 ; C - N - H
4147 4149 4151 1 1.2190e+02 4.1840e+02 ; C - N - CA
4148 4147 4149 1 1.2290e+02 6.6944e+02 ; O - C - N
4149 4151 4152 1 1.0950e+02 4.1840e+02 ; N - CA - HA
4149 4151 4153 1 1.0970e+02 6.6944e+02 ; N - CA - CB
4149 4151 4158 1 1.1010e+02 5.2718e+02 ; N - CA - C
4150 4149 4151 1 1.1804e+02 4.1840e+02 ; H - N - CA
4151 4153 4154 1 1.0950e+02 4.1840e+02 ; CA - CB - HB2
4151 4153 4155 1 1.0950e+02 4.1840e+02 ; CA - CB - HB3
4151 4153 4156 1 1.0860e+02 4.1840e+02 ; CA - CB - SG
4151 4158 4159 1 1.2040e+02 6.6944e+02 ; CA - C - O
4151 4158 4160 1 1.1660e+02 5.8576e+02 ; CA - C - N
4152 4151 4153 1 1.0950e+02 4.1840e+02 ; HA - CA - CB
4152 4151 4158 1 1.0950e+02 4.1840e+02 ; HA - CA - C
4153 4151 4158 1 1.1110e+02 5.2718e+02 ; CB - CA - C
4153 4156 4157 1 9.6000e+01 3.5982e+02 ; CB - SG - HG
4154 4153 4155 1 1.0950e+02 2.9288e+02 ; HB2 - CB - HB3
4154 4153 4156 1 1.0950e+02 4.1840e+02 ; HB2 - CB - SG
4155 4153 4156 1 1.0950e+02 4.1840e+02 ; HB3 - CB - SG
4158 4160 4161 1 1.2000e+02 4.1840e+02 ; C - N - H
4158 4160 4162 1 1.2190e+02 4.1840e+02 ; C - N - CA
4159 4158 4160 1 1.2290e+02 6.6944e+02 ; O - C - N
4160 4162 4163 1 1.0950e+02 4.1840e+02 ; N - CA - HA
4160 4162 4164 1 1.0970e+02 6.6944e+02 ; N - CA - CB

```
  4160   4162   4172       1    1.1010e+02    5.2718e+02 ;      N - CA
- C
  4161   4160   4162       1    1.1804e+02    4.1840e+02 ;      H - N
- CA
  4162   4164   4165       1    1.0950e+02    4.1840e+02 ;     CA - CB
- HB
  4162   4164   4166       1    1.0950e+02    3.3472e+02 ;     CA - CB
- CG2
  4162   4164   4170       1    1.0950e+02    4.1840e+02 ;     CA - CB
- OG1
  4162   4172   4173       1    1.2040e+02    6.6944e+02 ;     CA - C
- O
  4162   4172   4174       1    1.1660e+02    5.8576e+02 ;     CA - C
- N
  4163   4162   4164       1    1.0950e+02    4.1840e+02 ;     HA - CA
- CB
  4163   4162   4172       1    1.0950e+02    4.1840e+02 ;     HA - CA
- C
  4164   4162   4172       1    1.1110e+02    5.2718e+02 ;     CB - CA
- C
  4164   4166   4167       1    1.0950e+02    4.1840e+02 ;     CB - CG2
- HG21
  4164   4166   4168       1    1.0950e+02    4.1840e+02 ;     CB - CG2
- HG22
  4164   4166   4169       1    1.0950e+02    4.1840e+02 ;     CB - CG2
- HG23
  4164   4170   4171       1    1.0850e+02    4.6024e+02 ;     CB - OG1
- HG1
  4165   4164   4166       1    1.0950e+02    4.1840e+02 ;     HB - CB
- CG2
  4165   4164   4170       1    1.0950e+02    4.1840e+02 ;     HB - CB
- OG1
  4166   4164   4170       1    1.0950e+02    4.1840e+02 ;    CG2 - CB
- OG1
  4167   4166   4168       1    1.0950e+02    2.9288e+02 ;   HG21 - CG2
- HG22
  4167   4166   4169       1    1.0950e+02    2.9288e+02 ;   HG21 - CG2
- HG23
  4168   4166   4169       1    1.0950e+02    2.9288e+02 ;   HG22 - CG2
- HG23
  4172   4174   4175       1    1.2000e+02    4.1840e+02 ;      C - N
- H
  4172   4174   4176       1    1.2190e+02    4.1840e+02 ;      C - N
- CA
  4173   4172   4174       1    1.2290e+02    6.6944e+02 ;      O - C
- N
  4174   4176   4177       1    1.0950e+02    4.1840e+02 ;      N - CA
- HA
  4174   4176   4178       1    1.0970e+02    6.6944e+02 ;      N - CA
- CB
  4174   4176   4191       1    1.1010e+02    5.2718e+02 ;      N - CA
- C
```

4175 4174 4176 1 1.1804e+02 4.1840e+02 ; H - N
- CA
4176 4178 4179 1 1.0950e+02 4.1840e+02 ; CA - CB
- HB
4176 4178 4180 1 1.0950e+02 3.3472e+02 ; CA - CB
- CG2
4176 4178 4184 1 1.0950e+02 3.3472e+02 ; CA - CB
- CG1
4176 4191 4192 1 1.2040e+02 6.6944e+02 ; CA - C
- O
4176 4191 4193 1 1.1660e+02 5.8576e+02 ; CA - C
- N
4177 4176 4178 1 1.0950e+02 4.1840e+02 ; HA - CA
- CB
4177 4176 4191 1 1.0950e+02 4.1840e+02 ; HA - CA
- C
4178 4176 4191 1 1.1110e+02 5.2718e+02 ; CB - CA
- C
4178 4180 4181 1 1.0950e+02 4.1840e+02 ; CB - CG2
- HG21
4178 4180 4182 1 1.0950e+02 4.1840e+02 ; CB - CG2
- HG22
4178 4180 4183 1 1.0950e+02 4.1840e+02 ; CB - CG2
- HG23
4178 4184 4185 1 1.0950e+02 4.1840e+02 ; CB - CG1
- HG12
4178 4184 4186 1 1.0950e+02 4.1840e+02 ; CB - CG1
- HG13
4178 4184 4187 1 1.0950e+02 3.3472e+02 ; CB - CG1
- CD1
4179 4178 4180 1 1.0950e+02 4.1840e+02 ; HB - CB
- CG2
4179 4178 4184 1 1.0950e+02 4.1840e+02 ; HB - CB
- CG1
4180 4178 4184 1 1.0950e+02 3.3472e+02 ; CG2 - CB
- CG1
4181 4180 4182 1 1.0950e+02 2.9288e+02 ; HG21 - CG2
- HG22
4181 4180 4183 1 1.0950e+02 2.9288e+02 ; HG21 - CG2
- HG23
4182 4180 4183 1 1.0950e+02 2.9288e+02 ; HG22 - CG2
- HG23
4184 4187 4188 1 1.0950e+02 4.1840e+02 ; CG1 - CD1
- HD11
4184 4187 4189 1 1.0950e+02 4.1840e+02 ; CG1 - CD1
- HD12
4184 4187 4190 1 1.0950e+02 4.1840e+02 ; CG1 - CD1
- HD13
4185 4184 4186 1 1.0950e+02 2.9288e+02 ; HG12 - CG1
- HG13
4185 4184 4187 1 1.0950e+02 4.1840e+02 ; HG12 - CG1
- CD1

4186 4184 4187 1 1.0950e+02 4.1840e+02 ; HG13 - CG1 - CD1
4188 4187 4189 1 1.0950e+02 2.9288e+02 ; HD11 - CD1 - HD12
4188 4187 4190 1 1.0950e+02 2.9288e+02 ; HD11 - CD1 - HD13
4189 4187 4190 1 1.0950e+02 2.9288e+02 ; HD12 - CD1 - HD13
4191 4193 4194 1 1.2000e+02 4.1840e+02 ; C - N - H
4191 4193 4195 1 1.2190e+02 4.1840e+02 ; C - N - CA
4192 4191 4193 1 1.2290e+02 6.6944e+02 ; O - C - N
4193 4195 4196 1 1.0950e+02 4.1840e+02 ; N - CA - HA
4193 4195 4197 1 1.0970e+02 6.6944e+02 ; N - CA - CB
4193 4195 4206 1 1.1010e+02 5.2718e+02 ; N - CA - C
4194 4193 4195 1 1.1804e+02 4.1840e+02 ; H - N - CA
4195 4197 4198 1 1.0950e+02 4.1840e+02 ; CA - CB - HB2
4195 4197 4199 1 1.0950e+02 4.1840e+02 ; CA - CB - HB3
4195 4197 4200 1 1.0950e+02 3.3472e+02 ; CA - CB - CG
4195 4206 4207 1 1.2040e+02 6.6944e+02 ; CA - C - O
4195 4206 4208 1 1.1660e+02 5.8576e+02 ; CA - C - N
4196 4195 4197 1 1.0950e+02 4.1840e+02 ; HA - CA - CB
4196 4195 4206 1 1.0950e+02 4.1840e+02 ; HA - CA - C
4197 4195 4206 1 1.1110e+02 5.2718e+02 ; CB - CA - C
4197 4200 4201 1 1.0950e+02 4.1840e+02 ; CB - CG - HG2
4197 4200 4202 1 1.0950e+02 4.1840e+02 ; CB - CG - HG3
4197 4200 4203 1 1.1110e+02 5.2718e+02 ; CB - CG - CD
4198 4197 4199 1 1.0950e+02 2.9288e+02 ; HB2 - CB - HB3
4198 4197 4200 1 1.0950e+02 4.1840e+02 ; HB2 - CB - CG
4199 4197 4200 1 1.0950e+02 4.1840e+02 ; HB3 - CB - CG
4200 4203 4204 1 1.1700e+02 5.8576e+02 ; CG - CD - OE1

4200 4203 4205 1 1.1700e+02 5.8576e+02 ; CG - CD
- OE2
4201 4200 4202 1 1.0950e+02 2.9288e+02 ; HG2 - CG
- HG3
4201 4200 4203 1 1.0950e+02 4.1840e+02 ; HG2 - CG
- CD
4202 4200 4203 1 1.0950e+02 4.1840e+02 ; HG3 - CG
- CD
4204 4203 4205 1 1.2600e+02 6.6944e+02 ; OE1 - CD
- OE2
4206 4208 4209 1 1.2000e+02 4.1840e+02 ; C - N
- H
4206 4208 4210 1 1.2190e+02 4.1840e+02 ; C - N
- CA
4207 4206 4208 1 1.2290e+02 6.6944e+02 ; O - C
- N
4208 4210 4211 1 1.0950e+02 4.1840e+02 ; N - CA
- HA
4208 4210 4212 1 1.0970e+02 6.6944e+02 ; N - CA
- CB
4208 4210 4221 1 1.1010e+02 5.2718e+02 ; N - CA
- C
4209 4208 4210 1 1.1804e+02 4.1840e+02 ; H - N
- CA
4210 4212 4213 1 1.0950e+02 4.1840e+02 ; CA - CB
- HB2
4210 4212 4214 1 1.0950e+02 4.1840e+02 ; CA - CB
- HB3
4210 4212 4215 1 1.0950e+02 3.3472e+02 ; CA - CB
- CG
4210 4221 4222 1 1.2040e+02 6.6944e+02 ; CA - C
- O
4210 4221 4223 1 1.1660e+02 5.8576e+02 ; CA - C
- N
4211 4210 4212 1 1.0950e+02 4.1840e+02 ; HA - CA
- CB
4211 4210 4221 1 1.0950e+02 4.1840e+02 ; HA - CA
- C
4212 4210 4221 1 1.1110e+02 5.2718e+02 ; CB - CA
- C
4212 4215 4216 1 1.0950e+02 4.1840e+02 ; CB - CG
- HG2
4212 4215 4217 1 1.0950e+02 4.1840e+02 ; CB - CG
- HG3
4212 4215 4218 1 1.1110e+02 5.2718e+02 ; CB - CG
- CD
4213 4212 4214 1 1.0950e+02 2.9288e+02 ; HB2 - CB
- HB3
4213 4212 4215 1 1.0950e+02 4.1840e+02 ; HB2 - CB
- CG
4214 4212 4215 1 1.0950e+02 4.1840e+02 ; HB3 - CB
- CG

```
  4215   4218   4219      1    1.1700e+02    5.8576e+02 ;     CG - CD
- OE1
  4215   4218   4220      1    1.1700e+02    5.8576e+02 ;     CG - CD
- OE2
  4216   4215   4217      1    1.0950e+02    2.9288e+02 ;    HG2 - CG
- HG3
  4216   4215   4218      1    1.0950e+02    4.1840e+02 ;    HG2 - CG
- CD
  4217   4215   4218      1    1.0950e+02    4.1840e+02 ;    HG3 - CG
- CD
  4219   4218   4220      1    1.2600e+02    6.6944e+02 ;    OE1 - CD
- OE2
  4221   4223   4224      1    1.2000e+02    4.1840e+02 ;      C - N
- H
  4221   4223   4225      1    1.2190e+02    4.1840e+02 ;      C - N
- CA
  4222   4221   4223      1    1.2290e+02    6.6944e+02 ;      O - C
- N
  4223   4225   4226      1    1.0950e+02    4.1840e+02 ;      N - CA
- HA
  4223   4225   4227      1    1.0970e+02    6.6944e+02 ;      N - CA
- CB
  4223   4225   4231      1    1.1010e+02    5.2718e+02 ;      N - CA
- C
  4224   4223   4225      1    1.1804e+02    4.1840e+02 ;      H - N
- CA
  4225   4227   4228      1    1.0950e+02    4.1840e+02 ;     CA - CB
- HB1
  4225   4227   4229      1    1.0950e+02    4.1840e+02 ;     CA - CB
- HB2
  4225   4227   4230      1    1.0950e+02    4.1840e+02 ;     CA - CB
- HB3
  4225   4231   4232      1    1.2040e+02    6.6944e+02 ;     CA - C
- O
  4225   4231   4233      1    1.1660e+02    5.8576e+02 ;     CA - C
- N
  4226   4225   4227      1    1.0950e+02    4.1840e+02 ;     HA - CA
- CB
  4226   4225   4231      1    1.0950e+02    4.1840e+02 ;     HA - CA
- C
  4227   4225   4231      1    1.1110e+02    5.2718e+02 ;     CB - CA
- C
  4228   4227   4229      1    1.0950e+02    2.9288e+02 ;    HB1 - CB
- HB2
  4228   4227   4230      1    1.0950e+02    2.9288e+02 ;    HB1 - CB
- HB3
  4229   4227   4230      1    1.0950e+02    2.9288e+02 ;    HB2 - CB
- HB3
  4231   4233   4234      1    1.2000e+02    4.1840e+02 ;      C - N
- H
  4231   4233   4235      1    1.2190e+02    4.1840e+02 ;      C - N
- CA
```

```
  4232   4231   4233      1    1.2290e+02    6.6944e+02 ;      O - C
- N
  4233   4235   4236      1    1.0950e+02    4.1840e+02 ;      N - CA
- HA
  4233   4235   4237      1    1.0970e+02    6.6944e+02 ;      N - CA
- CB
  4233   4235   4250      1    1.1010e+02    5.2718e+02 ;      N - CA
- C
  4234   4233   4235      1    1.1804e+02    4.1840e+02 ;      H - N
- CA
  4235   4237   4238      1    1.0950e+02    4.1840e+02 ;     CA - CB
- HB2
  4235   4237   4239      1    1.0950e+02    4.1840e+02 ;     CA - CB
- HB3
  4235   4237   4240      1    1.0950e+02    3.3472e+02 ;     CA - CB
- CG
  4235   4250   4251      1    1.2040e+02    6.6944e+02 ;     CA - C
- O
  4235   4250   4252      1    1.1660e+02    5.8576e+02 ;     CA - C
- N
  4236   4235   4237      1    1.0950e+02    4.1840e+02 ;     HA - CA
- CB
  4236   4235   4250      1    1.0950e+02    4.1840e+02 ;     HA - CA
- C
  4237   4235   4250      1    1.1110e+02    5.2718e+02 ;     CB - CA
- C
  4237   4240   4241      1    1.0950e+02    4.1840e+02 ;     CB - CG
- HG
  4237   4240   4242      1    1.0950e+02    3.3472e+02 ;     CB - CG
- CD1
  4237   4240   4246      1    1.0950e+02    3.3472e+02 ;     CB - CG
- CD2
  4238   4237   4239      1    1.0950e+02    2.9288e+02 ;    HB2 - CB
- HB3
  4238   4237   4240      1    1.0950e+02    4.1840e+02 ;    HB2 - CB
- CG
  4239   4237   4240      1    1.0950e+02    4.1840e+02 ;    HB3 - CB
- CG
  4240   4242   4243      1    1.0950e+02    4.1840e+02 ;     CG - CD1
- HD11
  4240   4242   4244      1    1.0950e+02    4.1840e+02 ;     CG - CD1
- HD12
  4240   4242   4245      1    1.0950e+02    4.1840e+02 ;     CG - CD1
- HD13
  4240   4246   4247      1    1.0950e+02    4.1840e+02 ;     CG - CD2
- HD21
  4240   4246   4248      1    1.0950e+02    4.1840e+02 ;     CG - CD2
- HD22
  4240   4246   4249      1    1.0950e+02    4.1840e+02 ;     CG - CD2
- HD23
  4241   4240   4242      1    1.0950e+02    4.1840e+02 ;     HG - CG
- CD1
```

```
  4241   4240   4246      1    1.0950e+02    4.1840e+02 ;     HG - CG
- CD2
  4242   4240   4246      1    1.0950e+02    3.3472e+02 ;    CD1 - CG
- CD2
  4243   4242   4244      1    1.0950e+02    2.9288e+02 ;   HD11 - CD1
- HD12
  4243   4242   4245      1    1.0950e+02    2.9288e+02 ;   HD11 - CD1
- HD13
  4244   4242   4245      1    1.0950e+02    2.9288e+02 ;   HD12 - CD1
- HD13
  4247   4246   4248      1    1.0950e+02    2.9288e+02 ;   HD21 - CD2
- HD22
  4247   4246   4249      1    1.0950e+02    2.9288e+02 ;   HD21 - CD2
- HD23
  4248   4246   4249      1    1.0950e+02    2.9288e+02 ;   HD22 - CD2
- HD23
  4250   4252   4253      1    1.2000e+02    4.1840e+02 ;      C - N
- H
  4250   4252   4254      1    1.2190e+02    4.1840e+02 ;      C - N
- CA
  4251   4250   4252      1    1.2290e+02    6.6944e+02 ;      O - C
- N
  4252   4254   4255      1    1.0950e+02    4.1840e+02 ;      N - CA
- HA
  4252   4254   4256      1    1.0970e+02    6.6944e+02 ;      N - CA
- CB
  4252   4254   4264      1    1.1010e+02    5.2718e+02 ;      N - CA
- C
  4253   4252   4254      1    1.1804e+02    4.1840e+02 ;      H - N
- CA
  4254   4256   4257      1    1.0950e+02    4.1840e+02 ;     CA - CB
- HB2
  4254   4256   4258      1    1.0950e+02    4.1840e+02 ;     CA - CB
- HB3
  4254   4256   4259      1    1.1110e+02    5.2718e+02 ;     CA - CB
- CG
  4254   4264   4265      1    1.2040e+02    6.6944e+02 ;     CA - C
- O
  4254   4264   4266      1    1.1660e+02    5.8576e+02 ;     CA - C
- N
  4255   4254   4256      1    1.0950e+02    4.1840e+02 ;     HA - CA
- CB
  4255   4254   4264      1    1.0950e+02    4.1840e+02 ;     HA - CA
- C
  4256   4254   4264      1    1.1110e+02    5.2718e+02 ;     CB - CA
- C
  4256   4259   4260      1    1.2040e+02    6.6944e+02 ;     CB - CG
- OD1
  4256   4259   4261      1    1.1660e+02    5.8576e+02 ;     CB - CG
- ND2
  4257   4256   4258      1    1.0950e+02    2.9288e+02 ;    HB2 - CB
- HB3
```

4257 4256 4259 1 1.0950e+02 4.1840e+02 ; HB2 - CB
- CG
4258 4256 4259 1 1.0950e+02 4.1840e+02 ; HB3 - CB
- CG
4259 4261 4262 1 1.2000e+02 4.1840e+02 ; CG - ND2
- HD21
4259 4261 4263 1 1.2000e+02 4.1840e+02 ; CG - ND2
- HD22
4260 4259 4261 1 1.2290e+02 6.6944e+02 ; OD1 - CG
- ND2
4262 4261 4263 1 1.2000e+02 2.9288e+02 ; HD21 - ND2
- HD22
4264 4266 4267 1 1.2000e+02 4.1840e+02 ; C - N
- H
4264 4266 4268 1 1.2190e+02 4.1840e+02 ; C - N
- CA
4265 4264 4266 1 1.2290e+02 6.6944e+02 ; O - C
- N
4266 4268 4269 1 1.0950e+02 4.1840e+02 ; N - CA
- HA
4266 4268 4270 1 1.0970e+02 6.6944e+02 ; N - CA
- CB
4266 4268 4281 1 1.1010e+02 5.2718e+02 ; N - CA
- C
4267 4266 4268 1 1.1804e+02 4.1840e+02 ; H - N
- CA
4268 4270 4271 1 1.0950e+02 4.1840e+02 ; CA - CB
- HB2
4268 4270 4272 1 1.0950e+02 4.1840e+02 ; CA - CB
- HB3
4268 4270 4273 1 1.1310e+02 5.2718e+02 ; CA - CB
- CG
4268 4281 4282 1 1.2040e+02 6.6944e+02 ; CA - C
- O
4268 4281 4283 1 1.1660e+02 5.8576e+02 ; CA - C
- N
4269 4268 4270 1 1.0950e+02 4.1840e+02 ; HA - CA
- CB
4269 4268 4281 1 1.0950e+02 4.1840e+02 ; HA - CA
- C
4270 4268 4281 1 1.1110e+02 5.2718e+02 ; CB - CA
- C
4270 4273 4274 1 1.2000e+02 5.8576e+02 ; CB - CG
- ND1
4270 4273 4279 1 1.2000e+02 5.8576e+02 ; CB - CG
- CD2
4271 4270 4272 1 1.0950e+02 2.9288e+02 ; HB2 - CB
- HB3
4271 4270 4273 1 1.0950e+02 4.1840e+02 ; HB2 - CB
- CG
4272 4270 4273 1 1.0950e+02 4.1840e+02 ; HB3 - CB
- CG

```
  4273   4274   4275      1    1.1700e+02    5.8576e+02 ;      CG - ND1
- CE1
  4273   4279   4277      1    1.2000e+02    5.8576e+02 ;      CG - CD2
- NE2
  4273   4279   4280      1    1.2000e+02    4.1840e+02 ;      CG - CD2
- HD2
  4274   4273   4279      1    1.2000e+02    5.8576e+02 ;     ND1 - CG
- CD2
  4274   4275   4276      1    1.2000e+02    4.1840e+02 ;     ND1 - CE1
- HE1
  4274   4275   4277      1    1.2000e+02    5.8576e+02 ;     ND1 - CE1
- NE2
  4275   4277   4278      1    1.2000e+02    4.1840e+02 ;     CE1 - NE2
- HE2
  4275   4277   4279      1    1.2000e+02    5.8576e+02 ;     CE1 - NE2
- CD2
  4276   4275   4277      1    1.2000e+02    4.1840e+02 ;     HE1 - CE1
- NE2
  4277   4279   4280      1    1.2000e+02    4.1840e+02 ;     NE2 - CD2
- HD2
  4278   4277   4279      1    1.2000e+02    4.1840e+02 ;     HE2 - NE2
- CD2
  4281   4283   4284      1    1.2190e+02    4.1840e+02 ;       C - N
- CD
  4281   4283   4293      1    1.2190e+02    4.1840e+02 ;       C - N
- CA
  4282   4281   4283      1    1.2290e+02    6.6944e+02 ;       O - C
- N
  4283   4284   4285      1    1.0950e+02    4.1840e+02 ;       N - CD
- HD2
  4283   4284   4286      1    1.0950e+02    4.1840e+02 ;       N - CD
- HD3
  4283   4284   4287      1    1.0970e+02    6.6944e+02 ;       N - CD
- CG
  4283   4293   4290      1    1.0970e+02    6.6944e+02 ;       N - CA
- CB
  4283   4293   4294      1    1.0950e+02    4.1840e+02 ;       N - CA
- HA
  4283   4293   4295      1    1.1010e+02    5.2718e+02 ;       N - CA
- C
  4284   4283   4293      1    1.1800e+02    4.1840e+02 ;      CD - N
- CA
  4284   4287   4288      1    1.0950e+02    4.1840e+02 ;      CD - CG
- HG2
  4284   4287   4289      1    1.0950e+02    4.1840e+02 ;      CD - CG
- HG3
  4284   4287   4290      1    1.0950e+02    3.3472e+02 ;      CD - CG
- CB
  4285   4284   4286      1    1.0950e+02    2.9288e+02 ;     HD2 - CD
- HD3
  4285   4284   4287      1    1.0950e+02    4.1840e+02 ;     HD2 - CD
- CG
```

4286 4284 4287 1 1.0950e+02 4.1840e+02 ; HD3 - CD
- CG
4287 4290 4291 1 1.0950e+02 4.1840e+02 ; CG - CB
- HB2
4287 4290 4292 1 1.0950e+02 4.1840e+02 ; CG - CB
- HB3
4287 4290 4293 1 1.0950e+02 3.3472e+02 ; CG - CB
- CA
4288 4287 4289 1 1.0950e+02 2.9288e+02 ; HG2 - CG
- HG3
4288 4287 4290 1 1.0950e+02 4.1840e+02 ; HG2 - CG
- CB
4289 4287 4290 1 1.0950e+02 4.1840e+02 ; HG3 - CG
- CB
4290 4293 4294 1 1.0950e+02 4.1840e+02 ; CB - CA
- HA
4290 4293 4295 1 1.1110e+02 5.2718e+02 ; CB - CA
- C
4291 4290 4292 1 1.0950e+02 2.9288e+02 ; HB2 - CB
- HB3
4291 4290 4293 1 1.0950e+02 4.1840e+02 ; HB2 - CB
- CA
4292 4290 4293 1 1.0950e+02 4.1840e+02 ; HB3 - CB
- CA
4293 4295 4296 1 1.2040e+02 6.6944e+02 ; CA - C
- O
4293 4295 4297 1 1.1660e+02 5.8576e+02 ; CA - C
- N
4294 4293 4295 1 1.0950e+02 4.1840e+02 ; HA - CA
- C
4295 4297 4298 1 1.2000e+02 4.1840e+02 ; C - N
- H
4295 4297 4299 1 1.2190e+02 4.1840e+02 ; C - N
- CA
4296 4295 4297 1 1.2290e+02 6.6944e+02 ; O - C
- N
4297 4299 4300 1 1.0950e+02 4.1840e+02 ; N - CA
- HA
4297 4299 4301 1 1.0970e+02 6.6944e+02 ; N - CA
- CB
4297 4299 4319 1 1.1010e+02 5.2718e+02 ; N - CA
- C
4298 4297 4299 1 1.1804e+02 4.1840e+02 ; H - N
- CA
4299 4301 4302 1 1.0950e+02 4.1840e+02 ; CA - CB
- HB2
4299 4301 4303 1 1.0950e+02 4.1840e+02 ; CA - CB
- HB3
4299 4301 4304 1 1.1560e+02 5.2718e+02 ; CA - CB
- CG
4299 4319 4320 1 1.2040e+02 6.6944e+02 ; CA - C
- O

4299 4319 4321 1 1.1660e+02 5.8576e+02 ; CA - C
- N
4300 4299 4301 1 1.0950e+02 4.1840e+02 ; HA - CA
- CB
4300 4299 4319 1 1.0950e+02 4.1840e+02 ; HA - CA
- C
4301 4299 4319 1 1.1110e+02 5.2718e+02 ; CB - CA
- C
4301 4304 4305 1 1.2500e+02 5.8576e+02 ; CB - CG
- CD1
4301 4304 4318 1 1.2860e+02 5.8576e+02 ; CB - CG
- CD2
4302 4301 4303 1 1.0950e+02 2.9288e+02 ; HB2 - CB
- HB3
4302 4301 4304 1 1.0950e+02 4.1840e+02 ; HB2 - CB
- CG
4303 4301 4304 1 1.0950e+02 4.1840e+02 ; HB3 - CB
- CG
4304 4305 4306 1 1.2000e+02 4.1840e+02 ; CG - CD1
- HD1
4304 4305 4307 1 1.0870e+02 5.8576e+02 ; CG - CD1
- NE1
4304 4318 4309 1 1.0880e+02 5.2718e+02 ; CG - CD2
- CE2
4304 4318 4316 1 1.3490e+02 5.2718e+02 ; CG - CD2
- CE3
4305 4304 4318 1 1.0640e+02 5.2718e+02 ; CD1 - CG
- CD2
4305 4307 4308 1 1.2000e+02 4.1840e+02 ; CD1 - NE1
- HE1
4305 4307 4309 1 1.1160e+02 5.8576e+02 ; CD1 - NE1
- CE2
4306 4305 4307 1 1.2000e+02 4.1840e+02 ; HD1 - CD1
- NE1
4307 4309 4310 1 1.3280e+02 5.8576e+02 ; NE1 - CE2
- CZ2
4307 4309 4318 1 1.0440e+02 5.8576e+02 ; NE1 - CE2
- CD2
4308 4307 4309 1 1.2310e+02 4.1840e+02 ; HE1 - NE1
- CE2
4309 4310 4311 1 1.2000e+02 4.1840e+02 ; CE2 - CZ2
- HZ2
4309 4310 4312 1 1.2000e+02 5.2718e+02 ; CE2 - CZ2
- CH2
4309 4318 4316 1 1.1620e+02 5.2718e+02 ; CE2 - CD2
- CE3
4310 4309 4318 1 1.2270e+02 5.2718e+02 ; CZ2 - CE2
- CD2
4310 4312 4313 1 1.2000e+02 4.1840e+02 ; CZ2 - CH2
- HH2
4310 4312 4314 1 1.2000e+02 5.2718e+02 ; CZ2 - CH2
- CZ3

4311 4310 4312 1 1.2000e+02 4.1840e+02 ; HZ2 - CZ2 - CH2
4312 4314 4315 1 1.2000e+02 4.1840e+02 ; CH2 - CZ3 - HZ3
4312 4314 4316 1 1.2000e+02 5.2718e+02 ; CH2 - CZ3 - CE3
4313 4312 4314 1 1.2000e+02 4.1840e+02 ; HH2 - CH2 - CZ3
4314 4316 4317 1 1.2000e+02 4.1840e+02 ; CZ3 - CE3 - HE3
4314 4316 4318 1 1.2000e+02 5.2718e+02 ; CZ3 - CE3 - CD2
4315 4314 4316 1 1.2000e+02 4.1840e+02 ; HZ3 - CZ3 - CE3
4317 4316 4318 1 1.2000e+02 4.1840e+02 ; HE3 - CE3 - CD2
4319 4321 4322 1 1.2000e+02 4.1840e+02 ; C - N - H
4319 4321 4323 1 1.2190e+02 4.1840e+02 ; C - N - CA
4320 4319 4321 1 1.2290e+02 6.6944e+02 ; O - C - N
4321 4323 4324 1 1.0950e+02 4.1840e+02 ; N - CA - HA
4321 4323 4325 1 1.0970e+02 6.6944e+02 ; N - CA - CB
4321 4323 4338 1 1.1010e+02 5.2718e+02 ; N - CA - C
4322 4321 4323 1 1.1804e+02 4.1840e+02 ; H - N - CA
4323 4325 4326 1 1.0950e+02 4.1840e+02 ; CA - CB - HB
4323 4325 4327 1 1.0950e+02 3.3472e+02 ; CA - CB - CG2
4323 4325 4331 1 1.0950e+02 3.3472e+02 ; CA - CB - CG1
4323 4338 4339 1 1.2040e+02 6.6944e+02 ; CA - C - O
4323 4338 4340 1 1.1660e+02 5.8576e+02 ; CA - C - N
4324 4323 4325 1 1.0950e+02 4.1840e+02 ; HA - CA - CB
4324 4323 4338 1 1.0950e+02 4.1840e+02 ; HA - CA - C
4325 4323 4338 1 1.1110e+02 5.2718e+02 ; CB - CA - C
4325 4327 4328 1 1.0950e+02 4.1840e+02 ; CB - CG2 - HG21
4325 4327 4329 1 1.0950e+02 4.1840e+02 ; CB - CG2 - HG22
4325 4327 4330 1 1.0950e+02 4.1840e+02 ; CB - CG2 - HG23

```
  4325   4331   4332      1    1.0950e+02    4.1840e+02 ;     CB - CG1
- HG12
  4325   4331   4333      1    1.0950e+02    4.1840e+02 ;     CB - CG1
- HG13
  4325   4331   4334      1    1.0950e+02    3.3472e+02 ;     CB - CG1
- CD1
  4326   4325   4327      1    1.0950e+02    4.1840e+02 ;     HB - CB
- CG2
  4326   4325   4331      1    1.0950e+02    4.1840e+02 ;     HB - CB
- CG1
  4327   4325   4331      1    1.0950e+02    3.3472e+02 ;    CG2 - CB
- CG1
  4328   4327   4329      1    1.0950e+02    2.9288e+02 ;   HG21 - CG2
- HG22
  4328   4327   4330      1    1.0950e+02    2.9288e+02 ;   HG21 - CG2
- HG23
  4329   4327   4330      1    1.0950e+02    2.9288e+02 ;   HG22 - CG2
- HG23
  4331   4334   4335      1    1.0950e+02    4.1840e+02 ;    CG1 - CD1
- HD11
  4331   4334   4336      1    1.0950e+02    4.1840e+02 ;    CG1 - CD1
- HD12
  4331   4334   4337      1    1.0950e+02    4.1840e+02 ;    CG1 - CD1
- HD13
  4332   4331   4333      1    1.0950e+02    2.9288e+02 ;   HG12 - CG1
- HG13
  4332   4331   4334      1    1.0950e+02    4.1840e+02 ;   HG12 - CG1
- CD1
  4333   4331   4334      1    1.0950e+02    4.1840e+02 ;   HG13 - CG1
- CD1
  4335   4334   4336      1    1.0950e+02    2.9288e+02 ;   HD11 - CD1
- HD12
  4335   4334   4337      1    1.0950e+02    2.9288e+02 ;   HD11 - CD1
- HD13
  4336   4334   4337      1    1.0950e+02    2.9288e+02 ;   HD12 - CD1
- HD13
  4338   4340   4341      1    1.2000e+02    4.1840e+02 ;      C - N
- H
  4338   4340   4342      1    1.2190e+02    4.1840e+02 ;      C - N
- CA
  4339   4338   4340      1    1.2290e+02    6.6944e+02 ;      O - C
- N
  4340   4342   4343      1    1.0950e+02    4.1840e+02 ;      N - CA
- HA
  4340   4342   4344      1    1.0970e+02    6.6944e+02 ;      N - CA
- CB
  4340   4342   4352      1    1.1010e+02    5.2718e+02 ;      N - CA
- C
  4341   4340   4342      1    1.1804e+02    4.1840e+02 ;      H - N
- CA
  4342   4344   4345      1    1.0950e+02    4.1840e+02 ;     CA - CB
- HB
```

```
  4342   4344   4346      1    1.0950e+02    3.3472e+02 ;     CA - CB
- CG2
  4342   4344   4350      1    1.0950e+02    4.1840e+02 ;     CA - CB
- OG1
  4342   4352   4353      1    1.2040e+02    6.6944e+02 ;     CA - C
- O
  4342   4352   4354      1    1.1660e+02    5.8576e+02 ;     CA - C
- N
  4343   4342   4344      1    1.0950e+02    4.1840e+02 ;     HA - CA
- CB
  4343   4342   4352      1    1.0950e+02    4.1840e+02 ;     HA - CA
- C
  4344   4342   4352      1    1.1110e+02    5.2718e+02 ;     CB - CA
- C
  4344   4346   4347      1    1.0950e+02    4.1840e+02 ;     CB - CG2
- HG21
  4344   4346   4348      1    1.0950e+02    4.1840e+02 ;     CB - CG2
- HG22
  4344   4346   4349      1    1.0950e+02    4.1840e+02 ;     CB - CG2
- HG23
  4344   4350   4351      1    1.0850e+02    4.6024e+02 ;     CB - OG1
- HG1
  4345   4344   4346      1    1.0950e+02    4.1840e+02 ;     HB - CB
- CG2
  4345   4344   4350      1    1.0950e+02    4.1840e+02 ;     HB - CB
- OG1
  4346   4344   4350      1    1.0950e+02    4.1840e+02 ;    CG2 - CB
- OG1
  4347   4346   4348      1    1.0950e+02    2.9288e+02 ;   HG21 - CG2
- HG22
  4347   4346   4349      1    1.0950e+02    2.9288e+02 ;   HG21 - CG2
- HG23
  4348   4346   4349      1    1.0950e+02    2.9288e+02 ;   HG22 - CG2
- HG23
  4352   4354   4355      1    1.2000e+02    4.1840e+02 ;      C - N
- H
  4352   4354   4356      1    1.2190e+02    4.1840e+02 ;      C - N
- CA
  4353   4352   4354      1    1.2290e+02    6.6944e+02 ;      O - C
- N
  4354   4356   4357      1    1.0950e+02    4.1840e+02 ;      N - CA
- HA
  4354   4356   4358      1    1.0970e+02    6.6944e+02 ;      N - CA
- CB
  4354   4356   4369      1    1.1010e+02    5.2718e+02 ;      N - CA
- C
  4355   4354   4356      1    1.1804e+02    4.1840e+02 ;      H - N
- CA
  4356   4358   4359      1    1.0950e+02    4.1840e+02 ;     CA - CB
- HB2
  4356   4358   4360      1    1.0950e+02    4.1840e+02 ;     CA - CB
- HB3
```

4356 4358 4361 1 1.0950e+02 3.3472e+02 ; CA - CB
- CG
4356 4369 4370 1 1.2040e+02 6.6944e+02 ; CA - C
- O
4356 4369 4371 1 1.1660e+02 5.8576e+02 ; CA - C
- N
4357 4356 4358 1 1.0950e+02 4.1840e+02 ; HA - CA
- CB
4357 4356 4369 1 1.0950e+02 4.1840e+02 ; HA - CA
- C
4358 4356 4369 1 1.1110e+02 5.2718e+02 ; CB - CA
- C
4358 4361 4362 1 1.0950e+02 4.1840e+02 ; CB - CG
- HG2
4358 4361 4363 1 1.0950e+02 4.1840e+02 ; CB - CG
- HG3
4358 4361 4364 1 1.1110e+02 5.2718e+02 ; CB - CG
- CD
4359 4358 4360 1 1.0950e+02 2.9288e+02 ; HB2 - CB
- HB3
4359 4358 4361 1 1.0950e+02 4.1840e+02 ; HB2 - CB
- CG
4360 4358 4361 1 1.0950e+02 4.1840e+02 ; HB3 - CB
- CG
4361 4364 4365 1 1.2040e+02 6.6944e+02 ; CG - CD
- OE1
4361 4364 4366 1 1.1660e+02 5.8576e+02 ; CG - CD
- NE2
4362 4361 4363 1 1.0950e+02 2.9288e+02 ; HG2 - CG
- HG3
4362 4361 4364 1 1.0950e+02 4.1840e+02 ; HG2 - CG
- CD
4363 4361 4364 1 1.0950e+02 4.1840e+02 ; HG3 - CG
- CD
4364 4366 4367 1 1.2000e+02 4.1840e+02 ; CD - NE2
- HE21
4364 4366 4368 1 1.2000e+02 4.1840e+02 ; CD - NE2
- HE22
4365 4364 4366 1 1.2290e+02 6.6944e+02 ; OE1 - CD
- NE2
4367 4366 4368 1 1.2000e+02 2.9288e+02 ; HE21 - NE2
- HE22
4369 4371 4372 1 1.2000e+02 4.1840e+02 ; C - N
- H
4369 4371 4373 1 1.2190e+02 4.1840e+02 ; C - N
- CA
4370 4369 4371 1 1.2290e+02 6.6944e+02 ; O - C
- N
4371 4373 4374 1 1.0950e+02 4.1840e+02 ; N - CA
- HA
4371 4373 4375 1 1.0970e+02 6.6944e+02 ; N - CA
- CB

4371 4373 4386 1 1.1010e+02 5.2718e+02 ; N - CA
- C
4372 4371 4373 1 1.1804e+02 4.1840e+02 ; H - N
- CA
4373 4375 4376 1 1.0950e+02 4.1840e+02 ; CA - CB
- HB2
4373 4375 4377 1 1.0950e+02 4.1840e+02 ; CA - CB
- HB3
4373 4375 4378 1 1.0950e+02 3.3472e+02 ; CA - CB
- CG
4373 4386 4387 1 1.2040e+02 6.6944e+02 ; CA - C
- O
4373 4386 4388 1 1.1660e+02 5.8576e+02 ; CA - C
- N
4374 4373 4375 1 1.0950e+02 4.1840e+02 ; HA - CA
- CB
4374 4373 4386 1 1.0950e+02 4.1840e+02 ; HA - CA
- C
4375 4373 4386 1 1.1110e+02 5.2718e+02 ; CB - CA
- C
4375 4378 4379 1 1.0950e+02 4.1840e+02 ; CB - CG
- HG2
4375 4378 4380 1 1.0950e+02 4.1840e+02 ; CB - CG
- HG3
4375 4378 4381 1 1.1470e+02 4.1840e+02 ; CB - CG
- SD
4376 4375 4377 1 1.0950e+02 2.9288e+02 ; HB2 - CB
- HB3
4376 4375 4378 1 1.0950e+02 4.1840e+02 ; HB2 - CB
- CG
4377 4375 4378 1 1.0950e+02 4.1840e+02 ; HB3 - CB
- CG
4378 4381 4382 1 9.8900e+01 5.1882e+02 ; CG - SD
- CE
4379 4378 4380 1 1.0950e+02 2.9288e+02 ; HG2 - CG
- HG3
4379 4378 4381 1 1.0950e+02 4.1840e+02 ; HG2 - CG
- SD
4380 4378 4381 1 1.0950e+02 4.1840e+02 ; HG3 - CG
- SD
4381 4382 4383 1 1.0950e+02 4.1840e+02 ; SD - CE
- HE1
4381 4382 4384 1 1.0950e+02 4.1840e+02 ; SD - CE
- HE2
4381 4382 4385 1 1.0950e+02 4.1840e+02 ; SD - CE
- HE3
4383 4382 4384 1 1.0950e+02 2.9288e+02 ; HE1 - CE
- HE2
4383 4382 4385 1 1.0950e+02 2.9288e+02 ; HE1 - CE
- HE3
4384 4382 4385 1 1.0950e+02 2.9288e+02 ; HE2 - CE
- HE3

4386 4388 4389 1 1.2000e+02 4.1840e+02 ; C - N
- H
4386 4388 4390 1 1.2190e+02 4.1840e+02 ; C - N
- CA
4387 4386 4388 1 1.2290e+02 6.6944e+02 ; O - C
- N
4388 4390 4391 1 1.0950e+02 4.1840e+02 ; N - CA
- HA
4388 4390 4392 1 1.0970e+02 6.6944e+02 ; N - CA
- CB
4388 4390 4400 1 1.1010e+02 5.2718e+02 ; N - CA
- C
4389 4388 4390 1 1.1804e+02 4.1840e+02 ; H - N
- CA
4390 4392 4393 1 1.0950e+02 4.1840e+02 ; CA - CB
- HB
4390 4392 4394 1 1.0950e+02 3.3472e+02 ; CA - CB
- CG2
4390 4392 4398 1 1.0950e+02 4.1840e+02 ; CA - CB
- OG1
4390 4400 4401 1 1.2040e+02 6.6944e+02 ; CA - C
- O
4390 4400 4402 1 1.1660e+02 5.8576e+02 ; CA - C
- N
4391 4390 4392 1 1.0950e+02 4.1840e+02 ; HA - CA
- CB
4391 4390 4400 1 1.0950e+02 4.1840e+02 ; HA - CA
- C
4392 4390 4400 1 1.1110e+02 5.2718e+02 ; CB - CA
- C
4392 4394 4395 1 1.0950e+02 4.1840e+02 ; CB - CG2
- HG21
4392 4394 4396 1 1.0950e+02 4.1840e+02 ; CB - CG2
- HG22
4392 4394 4397 1 1.0950e+02 4.1840e+02 ; CB - CG2
- HG23
4392 4398 4399 1 1.0850e+02 4.6024e+02 ; CB - OG1
- HG1
4393 4392 4394 1 1.0950e+02 4.1840e+02 ; HB - CB
- CG2
4393 4392 4398 1 1.0950e+02 4.1840e+02 ; HB - CB
- OG1
4394 4392 4398 1 1.0950e+02 4.1840e+02 ; CG2 - CB
- OG1
4395 4394 4396 1 1.0950e+02 2.9288e+02 ; HG21 - CG2
- HG22
4395 4394 4397 1 1.0950e+02 2.9288e+02 ; HG21 - CG2
- HG23
4396 4394 4397 1 1.0950e+02 2.9288e+02 ; HG22 - CG2
- HG23
4400 4402 4403 1 1.2000e+02 4.1840e+02 ; C - N
- H

4400 4402 4404 1 1.2190e+02 4.1840e+02 ; C - N
- CA
4401 4400 4402 1 1.2290e+02 6.6944e+02 ; O - C
- N
4402 4404 4405 1 1.0950e+02 4.1840e+02 ; N - CA
- HA
4402 4404 4406 1 1.0970e+02 6.6944e+02 ; N - CA
- CB
4402 4404 4422 1 1.1010e+02 5.2718e+02 ; N - CA
- C
4403 4402 4404 1 1.1804e+02 4.1840e+02 ; H - N
- CA
4404 4406 4407 1 1.0950e+02 4.1840e+02 ; CA - CB
- HB2
4404 4406 4408 1 1.0950e+02 4.1840e+02 ; CA - CB
- HB3
4404 4406 4409 1 1.0950e+02 3.3472e+02 ; CA - CB
- CG
4404 4422 4423 1 1.2040e+02 6.6944e+02 ; CA - C
- O
4404 4422 4424 1 1.1660e+02 5.8576e+02 ; CA - C
- N
4405 4404 4406 1 1.0950e+02 4.1840e+02 ; HA - CA
- CB
4405 4404 4422 1 1.0950e+02 4.1840e+02 ; HA - CA
- C
4406 4404 4422 1 1.1110e+02 5.2718e+02 ; CB - CA
- C
4406 4409 4410 1 1.0950e+02 4.1840e+02 ; CB - CG
- HG2
4406 4409 4411 1 1.0950e+02 4.1840e+02 ; CB - CG
- HG3
4406 4409 4412 1 1.0950e+02 3.3472e+02 ; CB - CG
- CD
4407 4406 4408 1 1.0950e+02 2.9288e+02 ; HB2 - CB
- HB3
4407 4406 4409 1 1.0950e+02 4.1840e+02 ; HB2 - CB
- CG
4408 4406 4409 1 1.0950e+02 4.1840e+02 ; HB3 - CB
- CG
4409 4412 4413 1 1.0950e+02 4.1840e+02 ; CG - CD
- HD2
4409 4412 4414 1 1.0950e+02 4.1840e+02 ; CG - CD
- HD3
4409 4412 4415 1 1.0950e+02 3.3472e+02 ; CG - CD
- CE
4410 4409 4411 1 1.0950e+02 2.9288e+02 ; HG2 - CG
- HG3
4410 4409 4412 1 1.0950e+02 4.1840e+02 ; HG2 - CG
- CD
4411 4409 4412 1 1.0950e+02 4.1840e+02 ; HG3 - CG
- CD

4412 4415 4416 1 1.0950e+02 4.1840e+02 ; CD - CE
- HE2
4412 4415 4417 1 1.0950e+02 4.1840e+02 ; CD - CE
- HE3
4412 4415 4418 1 1.1120e+02 6.6944e+02 ; CD - CE
- NZ
4413 4412 4414 1 1.0950e+02 2.9288e+02 ; HD2 - CD
- HD3
4413 4412 4415 1 1.0950e+02 4.1840e+02 ; HD2 - CD
- CE
4414 4412 4415 1 1.0950e+02 4.1840e+02 ; HD3 - CD
- CE
4415 4418 4419 1 1.0950e+02 4.1840e+02 ; CE - NZ
- HZ1
4415 4418 4420 1 1.0950e+02 4.1840e+02 ; CE - NZ
- HZ2
4415 4418 4421 1 1.0950e+02 4.1840e+02 ; CE - NZ
- HZ3
4416 4415 4417 1 1.0950e+02 2.9288e+02 ; HE2 - CE
- HE3
4416 4415 4418 1 1.0950e+02 4.1840e+02 ; HE2 - CE
- NZ
4417 4415 4418 1 1.0950e+02 4.1840e+02 ; HE3 - CE
- NZ
4419 4418 4420 1 1.0950e+02 2.9288e+02 ; HZ1 - NZ
- HZ2
4419 4418 4421 1 1.0950e+02 2.9288e+02 ; HZ1 - NZ
- HZ3
4420 4418 4421 1 1.0950e+02 2.9288e+02 ; HZ2 - NZ
- HZ3
4422 4424 4425 1 1.2000e+02 4.1840e+02 ; C - N
- H
4422 4424 4426 1 1.2190e+02 4.1840e+02 ; C - N
- CA
4423 4422 4424 1 1.2290e+02 6.6944e+02 ; O - C
- N
4424 4426 4427 1 1.0950e+02 4.1840e+02 ; N - CA
- HA
4424 4426 4428 1 1.0970e+02 6.6944e+02 ; N - CA
- CB
4424 4426 4433 1 1.1010e+02 5.2718e+02 ; N - CA
- C
4425 4424 4426 1 1.1804e+02 4.1840e+02 ; H - N
- CA
4426 4428 4429 1 1.0950e+02 4.1840e+02 ; CA - CB
- HB2
4426 4428 4430 1 1.0950e+02 4.1840e+02 ; CA - CB
- HB3
4426 4428 4431 1 1.0950e+02 4.1840e+02 ; CA - CB
- OG
4426 4433 4434 1 1.2040e+02 6.6944e+02 ; CA - C
- O

4426 4433 4435 1 1.1660e+02 5.8576e+02 ; CA - C
- N
4427 4426 4428 1 1.0950e+02 4.1840e+02 ; HA - CA
- CB
4427 4426 4433 1 1.0950e+02 4.1840e+02 ; HA - CA
- C
4428 4426 4433 1 1.1110e+02 5.2718e+02 ; CB - CA
- C
4428 4431 4432 1 1.0850e+02 4.6024e+02 ; CB - OG
- HG
4429 4428 4430 1 1.0950e+02 2.9288e+02 ; HB2 - CB
- HB3
4429 4428 4431 1 1.0950e+02 4.1840e+02 ; HB2 - CB
- OG
4430 4428 4431 1 1.0950e+02 4.1840e+02 ; HB3 - CB
- OG
4433 4435 4436 1 1.2000e+02 4.1840e+02 ; C - N
- H
4433 4435 4437 1 1.2190e+02 4.1840e+02 ; C - N
- CA
4434 4433 4435 1 1.2290e+02 6.6944e+02 ; O - C
- N
4435 4437 4438 1 1.0950e+02 4.1840e+02 ; N - CA
- HA
4435 4437 4439 1 1.0970e+02 6.6944e+02 ; N - CA
- CB
4435 4437 4450 1 1.1010e+02 5.2718e+02 ; N - CA
- C
4436 4435 4437 1 1.1804e+02 4.1840e+02 ; H - N
- CA
4437 4439 4440 1 1.0950e+02 4.1840e+02 ; CA - CB
- HB2
4437 4439 4441 1 1.0950e+02 4.1840e+02 ; CA - CB
- HB3
4437 4439 4442 1 1.1310e+02 5.2718e+02 ; CA - CB
- CG
4437 4450 4451 1 1.2040e+02 6.6944e+02 ; CA - C
- O
4437 4450 4452 1 1.1660e+02 5.8576e+02 ; CA - C
- N
4438 4437 4439 1 1.0950e+02 4.1840e+02 ; HA - CA
- CB
4438 4437 4450 1 1.0950e+02 4.1840e+02 ; HA - CA
- C
4439 4437 4450 1 1.1110e+02 5.2718e+02 ; CB - CA
- C
4439 4442 4443 1 1.2000e+02 5.8576e+02 ; CB - CG
- ND1
4439 4442 4448 1 1.2000e+02 5.8576e+02 ; CB - CG
- CD2
4440 4439 4441 1 1.0950e+02 2.9288e+02 ; HB2 - CB
- HB3

```
  4440   4439   4442      1    1.0950e+02    4.1840e+02 ;    HB2 - CB
- CG
  4441   4439   4442      1    1.0950e+02    4.1840e+02 ;    HB3 - CB
- CG
  4442   4443   4444      1    1.1700e+02    5.8576e+02 ;     CG - ND1
- CE1
  4442   4448   4446      1    1.2000e+02    5.8576e+02 ;     CG - CD2
- NE2
  4442   4448   4449      1    1.2000e+02    4.1840e+02 ;     CG - CD2
- HD2
  4443   4442   4448      1    1.2000e+02    5.8576e+02 ;    ND1 - CG
- CD2
  4443   4444   4445      1    1.2000e+02    4.1840e+02 ;    ND1 - CE1
- HE1
  4443   4444   4446      1    1.2000e+02    5.8576e+02 ;    ND1 - CE1
- NE2
  4444   4446   4447      1    1.2000e+02    4.1840e+02 ;    CE1 - NE2
- HE2
  4444   4446   4448      1    1.2000e+02    5.8576e+02 ;    CE1 - NE2
- CD2
  4445   4444   4446      1    1.2000e+02    4.1840e+02 ;    HE1 - CE1
- NE2
  4446   4448   4449      1    1.2000e+02    4.1840e+02 ;    NE2 - CD2
- HD2
  4447   4446   4448      1    1.2000e+02    4.1840e+02 ;    HE2 - NE2
- CD2
  4450   4452   4453      1    1.2000e+02    4.1840e+02 ;      C - N
- H
  4450   4452   4454      1    1.2190e+02    4.1840e+02 ;      C - N
- CA
  4451   4450   4452      1    1.2290e+02    6.6944e+02 ;      O - C
- N
  4452   4454   4455      1    1.0950e+02    4.1840e+02 ;      N - CA
- HA
  4452   4454   4456      1    1.0970e+02    6.6944e+02 ;      N - CA
- CB
  4452   4454   4465      1    1.1010e+02    5.2718e+02 ;      N - CA
- C
  4453   4452   4454      1    1.1804e+02    4.1840e+02 ;      H - N
- CA
  4454   4456   4457      1    1.0950e+02    4.1840e+02 ;     CA - CB
- HB2
  4454   4456   4458      1    1.0950e+02    4.1840e+02 ;     CA - CB
- HB3
  4454   4456   4459      1    1.0950e+02    3.3472e+02 ;     CA - CB
- CG
  4454   4465   4466      1    1.2040e+02    6.6944e+02 ;     CA - C
- O
  4454   4465   4467      1    1.1660e+02    5.8576e+02 ;     CA - C
- N
  4455   4454   4456      1    1.0950e+02    4.1840e+02 ;     HA - CA
- CB
```

4455 4454 4465 1 1.0950e+02 4.1840e+02 ; HA - CA
- C
4456 4454 4465 1 1.1110e+02 5.2718e+02 ; CB - CA
- C
4456 4459 4460 1 1.0950e+02 4.1840e+02 ; CB - CG
- HG2
4456 4459 4461 1 1.0950e+02 4.1840e+02 ; CB - CG
- HG3
4456 4459 4462 1 1.1110e+02 5.2718e+02 ; CB - CG
- CD
4457 4456 4458 1 1.0950e+02 2.9288e+02 ; HB2 - CB
- HB3
4457 4456 4459 1 1.0950e+02 4.1840e+02 ; HB2 - CB
- CG
4458 4456 4459 1 1.0950e+02 4.1840e+02 ; HB3 - CB
- CG
4459 4462 4463 1 1.1700e+02 5.8576e+02 ; CG - CD
- OE1
4459 4462 4464 1 1.1700e+02 5.8576e+02 ; CG - CD
- OE2
4460 4459 4461 1 1.0950e+02 2.9288e+02 ; HG2 - CG
- HG3
4460 4459 4462 1 1.0950e+02 4.1840e+02 ; HG2 - CG
- CD
4461 4459 4462 1 1.0950e+02 4.1840e+02 ; HG3 - CG
- CD
4463 4462 4464 1 1.2600e+02 6.6944e+02 ; OE1 - CD
- OE2
4465 4467 4468 1 1.2000e+02 4.1840e+02 ; C - N
- H
4465 4467 4469 1 1.2190e+02 4.1840e+02 ; C - N
- CA
4466 4465 4467 1 1.2290e+02 6.6944e+02 ; O - C
- N
4467 4469 4470 1 1.0950e+02 4.1840e+02 ; N - CA
- HA
4467 4469 4471 1 1.0970e+02 6.6944e+02 ; N - CA
- CB
4467 4469 4482 1 1.1010e+02 5.2718e+02 ; N - CA
- C
4468 4467 4469 1 1.1804e+02 4.1840e+02 ; H - N
- CA
4469 4471 4472 1 1.0950e+02 4.1840e+02 ; CA - CB
- HB2
4469 4471 4473 1 1.0950e+02 4.1840e+02 ; CA - CB
- HB3
4469 4471 4474 1 1.1310e+02 5.2718e+02 ; CA - CB
- CG
4469 4482 4483 1 1.2040e+02 6.6944e+02 ; CA - C
- O
4469 4482 4484 1 1.1660e+02 5.8576e+02 ; CA - C
- N

4470 4469 4471 1 1.0950e+02 4.1840e+02 ; HA - CA
- CB
4470 4469 4482 1 1.0950e+02 4.1840e+02 ; HA - CA
- C
4471 4469 4482 1 1.1110e+02 5.2718e+02 ; CB - CA
- C
4471 4474 4475 1 1.2000e+02 5.8576e+02 ; CB - CG
- ND1
4471 4474 4480 1 1.2000e+02 5.8576e+02 ; CB - CG
- CD2
4472 4471 4473 1 1.0950e+02 2.9288e+02 ; HB2 - CB
- HB3
4472 4471 4474 1 1.0950e+02 4.1840e+02 ; HB2 - CB
- CG
4473 4471 4474 1 1.0950e+02 4.1840e+02 ; HB3 - CB
- CG
4474 4475 4476 1 1.1700e+02 5.8576e+02 ; CG - ND1
- CE1
4474 4480 4478 1 1.2000e+02 5.8576e+02 ; CG - CD2
- NE2
4474 4480 4481 1 1.2000e+02 4.1840e+02 ; CG - CD2
- HD2
4475 4474 4480 1 1.2000e+02 5.8576e+02 ; ND1 - CG
- CD2
4475 4476 4477 1 1.2000e+02 4.1840e+02 ; ND1 - CE1
- HE1
4475 4476 4478 1 1.2000e+02 5.8576e+02 ; ND1 - CE1
- NE2
4476 4478 4479 1 1.2000e+02 4.1840e+02 ; CE1 - NE2
- HE2
4476 4478 4480 1 1.2000e+02 5.8576e+02 ; CE1 - NE2
- CD2
4477 4476 4478 1 1.2000e+02 4.1840e+02 ; HE1 - CE1
- NE2
4478 4480 4481 1 1.2000e+02 4.1840e+02 ; NE2 - CD2
- HD2
4479 4478 4480 1 1.2000e+02 4.1840e+02 ; HE2 - NE2
- CD2
4482 4484 4485 1 1.2000e+02 4.1840e+02 ; C - N
- H
4482 4484 4486 1 1.2190e+02 4.1840e+02 ; C - N
- CA
4483 4482 4484 1 1.2290e+02 6.6944e+02 ; O - C
- N
4484 4486 4487 1 1.0950e+02 4.1840e+02 ; N - CA
- HA
4484 4486 4488 1 1.0970e+02 6.6944e+02 ; N - CA
- CB
4484 4486 4498 1 1.1010e+02 5.2718e+02 ; N - CA
- C
4485 4484 4486 1 1.1804e+02 4.1840e+02 ; H - N
- CA

4486 4488 4489 1 1.0950e+02 4.1840e+02 ; CA - CB
- HB
4486 4488 4490 1 1.0950e+02 3.3472e+02 ; CA - CB
- CG1
4486 4488 4494 1 1.0950e+02 3.3472e+02 ; CA - CB
- CG2
4486 4498 4499 1 1.2040e+02 6.6944e+02 ; CA - C
- O
4486 4498 4500 1 1.1660e+02 5.8576e+02 ; CA - C
- N
4487 4486 4488 1 1.0950e+02 4.1840e+02 ; HA - CA
- CB
4487 4486 4498 1 1.0950e+02 4.1840e+02 ; HA - CA
- C
4488 4486 4498 1 1.1110e+02 5.2718e+02 ; CB - CA
- C
4488 4490 4491 1 1.0950e+02 4.1840e+02 ; CB - CG1
- HG11
4488 4490 4492 1 1.0950e+02 4.1840e+02 ; CB - CG1
- HG12
4488 4490 4493 1 1.0950e+02 4.1840e+02 ; CB - CG1
- HG13
4488 4494 4495 1 1.0950e+02 4.1840e+02 ; CB - CG2
- HG21
4488 4494 4496 1 1.0950e+02 4.1840e+02 ; CB - CG2
- HG22
4488 4494 4497 1 1.0950e+02 4.1840e+02 ; CB - CG2
- HG23
4489 4488 4490 1 1.0950e+02 4.1840e+02 ; HB - CB
- CG1
4489 4488 4494 1 1.0950e+02 4.1840e+02 ; HB - CB
- CG2
4490 4488 4494 1 1.0950e+02 3.3472e+02 ; CG1 - CB
- CG2
4491 4490 4492 1 1.0950e+02 2.9288e+02 ; HG11 - CG1
- HG12
4491 4490 4493 1 1.0950e+02 2.9288e+02 ; HG11 - CG1
- HG13
4492 4490 4493 1 1.0950e+02 2.9288e+02 ; HG12 - CG1
- HG13
4495 4494 4496 1 1.0950e+02 2.9288e+02 ; HG21 - CG2
- HG22
4495 4494 4497 1 1.0950e+02 2.9288e+02 ; HG21 - CG2
- HG23
4496 4494 4497 1 1.0950e+02 2.9288e+02 ; HG22 - CG2
- HG23
4498 4500 4501 1 1.2000e+02 4.1840e+02 ; C - N
- H
4498 4500 4502 1 1.2190e+02 4.1840e+02 ; C - N
- CA
4499 4498 4500 1 1.2290e+02 6.6944e+02 ; O - C
- N

```
  4500   4502   4503       1    1.0950e+02    4.1840e+02 ;       N - CA
- HA
  4500   4502   4504       1    1.0970e+02    6.6944e+02 ;       N - CA
- CB
  4500   4502   4513       1    1.1010e+02    5.2718e+02 ;       N - CA
- C
  4501   4500   4502       1    1.1804e+02    4.1840e+02 ;       H - N
- CA
  4502   4504   4505       1    1.0950e+02    4.1840e+02 ;      CA - CB
- HB2
  4502   4504   4506       1    1.0950e+02    4.1840e+02 ;      CA - CB
- HB3
  4502   4504   4507       1    1.0950e+02    3.3472e+02 ;      CA - CB
- CG
  4502   4513   4514       1    1.2040e+02    6.6944e+02 ;      CA - C
- O
  4502   4513   4515       1    1.1660e+02    5.8576e+02 ;      CA - C
- N
  4503   4502   4504       1    1.0950e+02    4.1840e+02 ;      HA - CA
- CB
  4503   4502   4513       1    1.0950e+02    4.1840e+02 ;      HA - CA
- C
  4504   4502   4513       1    1.1110e+02    5.2718e+02 ;      CB - CA
- C
  4504   4507   4508       1    1.0950e+02    4.1840e+02 ;      CB - CG
- HG2
  4504   4507   4509       1    1.0950e+02    4.1840e+02 ;      CB - CG
- HG3
  4504   4507   4510       1    1.1110e+02    5.2718e+02 ;      CB - CG
- CD
  4505   4504   4506       1    1.0950e+02    2.9288e+02 ;     HB2 - CB
- HB3
  4505   4504   4507       1    1.0950e+02    4.1840e+02 ;     HB2 - CB
- CG
  4506   4504   4507       1    1.0950e+02    4.1840e+02 ;     HB3 - CB
- CG
  4507   4510   4511       1    1.1700e+02    5.8576e+02 ;      CG - CD
- OE1
  4507   4510   4512       1    1.1700e+02    5.8576e+02 ;      CG - CD
- OE2
  4508   4507   4509       1    1.0950e+02    2.9288e+02 ;     HG2 - CG
- HG3
  4508   4507   4510       1    1.0950e+02    4.1840e+02 ;     HG2 - CG
- CD
  4509   4507   4510       1    1.0950e+02    4.1840e+02 ;     HG3 - CG
- CD
  4511   4510   4512       1    1.2600e+02    6.6944e+02 ;     OE1 - CD
- OE2
  4513   4515   4516       1    1.2000e+02    4.1840e+02 ;       C - N
- H
  4513   4515   4517       1    1.2190e+02    4.1840e+02 ;       C - N
- CA
```

4514 4513 4515 1 1.2290e+02 6.6944e+02 ; O - C
- N
4515 4517 4518 1 1.0950e+02 4.1840e+02 ; N - CA
- HA
4515 4517 4519 1 1.0970e+02 6.6944e+02 ; N - CA
- CB
4515 4517 4532 1 1.1010e+02 5.2718e+02 ; N - CA
- C
4516 4515 4517 1 1.1804e+02 4.1840e+02 ; H - N
- CA
4517 4519 4520 1 1.0950e+02 4.1840e+02 ; CA - CB
- HB2
4517 4519 4521 1 1.0950e+02 4.1840e+02 ; CA - CB
- HB3
4517 4519 4522 1 1.0950e+02 3.3472e+02 ; CA - CB
- CG
4517 4532 4533 1 1.2040e+02 6.6944e+02 ; CA - C
- O
4517 4532 4534 1 1.1660e+02 5.8576e+02 ; CA - C
- N
4518 4517 4519 1 1.0950e+02 4.1840e+02 ; HA - CA
- CB
4518 4517 4532 1 1.0950e+02 4.1840e+02 ; HA - CA
- C
4519 4517 4532 1 1.1110e+02 5.2718e+02 ; CB - CA
- C
4519 4522 4523 1 1.0950e+02 4.1840e+02 ; CB - CG
- HG
4519 4522 4524 1 1.0950e+02 3.3472e+02 ; CB - CG
- CD1
4519 4522 4528 1 1.0950e+02 3.3472e+02 ; CB - CG
- CD2
4520 4519 4521 1 1.0950e+02 2.9288e+02 ; HB2 - CB
- HB3
4520 4519 4522 1 1.0950e+02 4.1840e+02 ; HB2 - CB
- CG
4521 4519 4522 1 1.0950e+02 4.1840e+02 ; HB3 - CB
- CG
4522 4524 4525 1 1.0950e+02 4.1840e+02 ; CG - CD1
- HD11
4522 4524 4526 1 1.0950e+02 4.1840e+02 ; CG - CD1
- HD12
4522 4524 4527 1 1.0950e+02 4.1840e+02 ; CG - CD1
- HD13
4522 4528 4529 1 1.0950e+02 4.1840e+02 ; CG - CD2
- HD21
4522 4528 4530 1 1.0950e+02 4.1840e+02 ; CG - CD2
- HD22
4522 4528 4531 1 1.0950e+02 4.1840e+02 ; CG - CD2
- HD23
4523 4522 4524 1 1.0950e+02 4.1840e+02 ; HG - CG
- CD1

4523 4522 4528 1 1.0950e+02 4.1840e+02 ; HG - CG
- CD2
4524 4522 4528 1 1.0950e+02 3.3472e+02 ; CD1 - CG
- CD2
4525 4524 4526 1 1.0950e+02 2.9288e+02 ; HD11 - CD1
- HD12
4525 4524 4527 1 1.0950e+02 2.9288e+02 ; HD11 - CD1
- HD13
4526 4524 4527 1 1.0950e+02 2.9288e+02 ; HD12 - CD1
- HD13
4529 4528 4530 1 1.0950e+02 2.9288e+02 ; HD21 - CD2
- HD22
4529 4528 4531 1 1.0950e+02 2.9288e+02 ; HD21 - CD2
- HD23
4530 4528 4531 1 1.0950e+02 2.9288e+02 ; HD22 - CD2
- HD23
4532 4534 4535 1 1.2000e+02 4.1840e+02 ; C - N
- H
4532 4534 4536 1 1.2190e+02 4.1840e+02 ; C - N
- CA
4533 4532 4534 1 1.2290e+02 6.6944e+02 ; O - C
- N
4534 4536 4537 1 1.0950e+02 4.1840e+02 ; N - CA
- HA
4534 4536 4538 1 1.0970e+02 6.6944e+02 ; N - CA
- CB
4534 4536 4543 1 1.1010e+02 5.2718e+02 ; N - CA
- C
4535 4534 4536 1 1.1804e+02 4.1840e+02 ; H - N
- CA
4536 4538 4539 1 1.0950e+02 4.1840e+02 ; CA - CB
- HB2
4536 4538 4540 1 1.0950e+02 4.1840e+02 ; CA - CB
- HB3
4536 4538 4541 1 1.0950e+02 4.1840e+02 ; CA - CB
- OG
4536 4543 4544 1 1.2040e+02 6.6944e+02 ; CA - C
- O
4536 4543 4545 1 1.1660e+02 5.8576e+02 ; CA - C
- N
4537 4536 4538 1 1.0950e+02 4.1840e+02 ; HA - CA
- CB
4537 4536 4543 1 1.0950e+02 4.1840e+02 ; HA - CA
- C
4538 4536 4543 1 1.1110e+02 5.2718e+02 ; CB - CA
- C
4538 4541 4542 1 1.0850e+02 4.6024e+02 ; CB - OG
- HG
4539 4538 4540 1 1.0950e+02 2.9288e+02 ; HB2 - CB
- HB3
4539 4538 4541 1 1.0950e+02 4.1840e+02 ; HB2 - CB
- OG

4540 4538 4541 1 1.0950e+02 4.1840e+02 ; HB3 - CB
- OG
4543 4545 4546 1 1.2000e+02 4.1840e+02 ; C - N
- H
4543 4545 4547 1 1.2190e+02 4.1840e+02 ; C - N
- CA
4544 4543 4545 1 1.2290e+02 6.6944e+02 ; O - C
- N
4545 4547 4548 1 1.0950e+02 4.1840e+02 ; N - CA
- HA
4545 4547 4549 1 1.0970e+02 6.6944e+02 ; N - CA
- CB
4545 4547 4554 1 1.1010e+02 5.2718e+02 ; N - CA
- C
4546 4545 4547 1 1.1804e+02 4.1840e+02 ; H - N
- CA
4547 4549 4550 1 1.0950e+02 4.1840e+02 ; CA - CB
- HB2
4547 4549 4551 1 1.0950e+02 4.1840e+02 ; CA - CB
- HB3
4547 4549 4552 1 1.0950e+02 4.1840e+02 ; CA - CB
- OG
4547 4554 4555 1 1.2040e+02 6.6944e+02 ; CA - C
- O
4547 4554 4556 1 1.1660e+02 5.8576e+02 ; CA - C
- N
4548 4547 4549 1 1.0950e+02 4.1840e+02 ; HA - CA
- CB
4548 4547 4554 1 1.0950e+02 4.1840e+02 ; HA - CA
- C
4549 4547 4554 1 1.1110e+02 5.2718e+02 ; CB - CA
- C
4549 4552 4553 1 1.0850e+02 4.6024e+02 ; CB - OG
- HG
4550 4549 4551 1 1.0950e+02 2.9288e+02 ; HB2 - CB
- HB3
4550 4549 4552 1 1.0950e+02 4.1840e+02 ; HB2 - CB
- OG
4551 4549 4552 1 1.0950e+02 4.1840e+02 ; HB3 - CB
- OG
4554 4556 4557 1 1.2000e+02 4.1840e+02 ; C - N
- H
4554 4556 4558 1 1.2190e+02 4.1840e+02 ; C - N
- CA
4555 4554 4556 1 1.2290e+02 6.6944e+02 ; O - C
- N
4556 4558 4559 1 1.0950e+02 4.1840e+02 ; N - CA
- HA
4556 4558 4560 1 1.0970e+02 6.6944e+02 ; N - CA
- CB
4556 4558 4568 1 1.1010e+02 5.2718e+02 ; N - CA
- C

```
  4557   4556   4558       1    1.1804e+02    4.1840e+02 ;      H - N
- CA
  4558   4560   4561       1    1.0950e+02    4.1840e+02 ;     CA - CB
- HB
  4558   4560   4562       1    1.0950e+02    3.3472e+02 ;     CA - CB
- CG2
  4558   4560   4566       1    1.0950e+02    4.1840e+02 ;     CA - CB
- OG1
  4558   4568   4569       1    1.2040e+02    6.6944e+02 ;     CA - C
- O
  4558   4568   4570       1    1.1660e+02    5.8576e+02 ;     CA - C
- N
  4559   4558   4560       1    1.0950e+02    4.1840e+02 ;     HA - CA
- CB
  4559   4558   4568       1    1.0950e+02    4.1840e+02 ;     HA - CA
- C
  4560   4558   4568       1    1.1110e+02    5.2718e+02 ;     CB - CA
- C
  4560   4562   4563       1    1.0950e+02    4.1840e+02 ;     CB - CG2
- HG21
  4560   4562   4564       1    1.0950e+02    4.1840e+02 ;     CB - CG2
- HG22
  4560   4562   4565       1    1.0950e+02    4.1840e+02 ;     CB - CG2
- HG23
  4560   4566   4567       1    1.0850e+02    4.6024e+02 ;     CB - OG1
- HG1
  4561   4560   4562       1    1.0950e+02    4.1840e+02 ;     HB - CB
- CG2
  4561   4560   4566       1    1.0950e+02    4.1840e+02 ;     HB - CB
- OG1
  4562   4560   4566       1    1.0950e+02    4.1840e+02 ;    CG2 - CB
- OG1
  4563   4562   4564       1    1.0950e+02    2.9288e+02 ;   HG21 - CG2
- HG22
  4563   4562   4565       1    1.0950e+02    2.9288e+02 ;   HG21 - CG2
- HG23
  4564   4562   4565       1    1.0950e+02    2.9288e+02 ;   HG22 - CG2
- HG23
  4568   4570   4571       1    1.2000e+02    4.1840e+02 ;      C - N
- H
  4568   4570   4572       1    1.2190e+02    4.1840e+02 ;      C - N
- CA
  4569   4568   4570       1    1.2290e+02    6.6944e+02 ;      O - C
- N
  4570   4572   4573       1    1.0950e+02    4.1840e+02 ;      N - CA
- HA
  4570   4572   4574       1    1.0970e+02    6.6944e+02 ;      N - CA
- CB
  4570   4572   4587       1    1.1010e+02    5.2718e+02 ;      N - CA
- C
  4571   4570   4572       1    1.1804e+02    4.1840e+02 ;      H - N
- CA
```

4572 4574 4575 1 1.0950e+02 4.1840e+02 ; CA - CB
- HB2
4572 4574 4576 1 1.0950e+02 4.1840e+02 ; CA - CB
- HB3
4572 4574 4577 1 1.0950e+02 3.3472e+02 ; CA - CB
- CG
4572 4587 4588 1 1.2040e+02 6.6944e+02 ; CA - C
- O
4572 4587 4589 1 1.1660e+02 5.8576e+02 ; CA - C
- N
4573 4572 4574 1 1.0950e+02 4.1840e+02 ; HA - CA
- CB
4573 4572 4587 1 1.0950e+02 4.1840e+02 ; HA - CA
- C
4574 4572 4587 1 1.1110e+02 5.2718e+02 ; CB - CA
- C
4574 4577 4578 1 1.0950e+02 4.1840e+02 ; CB - CG
- HG
4574 4577 4579 1 1.0950e+02 3.3472e+02 ; CB - CG
- CD1
4574 4577 4583 1 1.0950e+02 3.3472e+02 ; CB - CG
- CD2
4575 4574 4576 1 1.0950e+02 2.9288e+02 ; HB2 - CB
- HB3
4575 4574 4577 1 1.0950e+02 4.1840e+02 ; HB2 - CB
- CG
4576 4574 4577 1 1.0950e+02 4.1840e+02 ; HB3 - CB
- CG
4577 4579 4580 1 1.0950e+02 4.1840e+02 ; CG - CD1
- HD11
4577 4579 4581 1 1.0950e+02 4.1840e+02 ; CG - CD1
- HD12
4577 4579 4582 1 1.0950e+02 4.1840e+02 ; CG - CD1
- HD13
4577 4583 4584 1 1.0950e+02 4.1840e+02 ; CG - CD2
- HD21
4577 4583 4585 1 1.0950e+02 4.1840e+02 ; CG - CD2
- HD22
4577 4583 4586 1 1.0950e+02 4.1840e+02 ; CG - CD2
- HD23
4578 4577 4579 1 1.0950e+02 4.1840e+02 ; HG - CG
- CD1
4578 4577 4583 1 1.0950e+02 4.1840e+02 ; HG - CG
- CD2
4579 4577 4583 1 1.0950e+02 3.3472e+02 ; CD1 - CG
- CD2
4580 4579 4581 1 1.0950e+02 2.9288e+02 ; HD11 - CD1
- HD12
4580 4579 4582 1 1.0950e+02 2.9288e+02 ; HD11 - CD1
- HD13
4581 4579 4582 1 1.0950e+02 2.9288e+02 ; HD12 - CD1
- HD13

```
  4584   4583   4585      1    1.0950e+02    2.9288e+02 ;   HD21 - CD2
- HD22
  4584   4583   4586      1    1.0950e+02    2.9288e+02 ;   HD21 - CD2
- HD23
  4585   4583   4586      1    1.0950e+02    2.9288e+02 ;   HD22 - CD2
- HD23
  4587   4589   4590      1    1.2000e+02    4.1840e+02 ;      C - N
- H
  4587   4589   4591      1    1.2190e+02    4.1840e+02 ;      C - N
- CA
  4588   4587   4589      1    1.2290e+02    6.6944e+02 ;      O - C
- N
  4589   4591   4592      1    1.0950e+02    4.1840e+02 ;      N - CA
- HA
  4589   4591   4593      1    1.0970e+02    6.6944e+02 ;      N - CA
- CB
  4589   4591   4606      1    1.1010e+02    5.2718e+02 ;      N - CA
- C
  4590   4589   4591      1    1.1804e+02    4.1840e+02 ;      H - N
- CA
  4591   4593   4594      1    1.0950e+02    4.1840e+02 ;     CA - CB
- HB2
  4591   4593   4595      1    1.0950e+02    4.1840e+02 ;     CA - CB
- HB3
  4591   4593   4596      1    1.0950e+02    3.3472e+02 ;     CA - CB
- CG
  4591   4606   4607      1    1.2040e+02    6.6944e+02 ;     CA - C
- O
  4591   4606   4608      1    1.1660e+02    5.8576e+02 ;     CA - C
- N
  4592   4591   4593      1    1.0950e+02    4.1840e+02 ;     HA - CA
- CB
  4592   4591   4606      1    1.0950e+02    4.1840e+02 ;     HA - CA
- C
  4593   4591   4606      1    1.1110e+02    5.2718e+02 ;     CB - CA
- C
  4593   4596   4597      1    1.0950e+02    4.1840e+02 ;     CB - CG
- HG
  4593   4596   4598      1    1.0950e+02    3.3472e+02 ;     CB - CG
- CD1
  4593   4596   4602      1    1.0950e+02    3.3472e+02 ;     CB - CG
- CD2
  4594   4593   4595      1    1.0950e+02    2.9288e+02 ;    HB2 - CB
- HB3
  4594   4593   4596      1    1.0950e+02    4.1840e+02 ;    HB2 - CB
- CG
  4595   4593   4596      1    1.0950e+02    4.1840e+02 ;    HB3 - CB
- CG
  4596   4598   4599      1    1.0950e+02    4.1840e+02 ;     CG - CD1
- HD11
  4596   4598   4600      1    1.0950e+02    4.1840e+02 ;     CG - CD1
- HD12
```

4596 4598 4601 1 1.0950e+02 4.1840e+02 ; CG - CD1
- HD13
4596 4602 4603 1 1.0950e+02 4.1840e+02 ; CG - CD2
- HD21
4596 4602 4604 1 1.0950e+02 4.1840e+02 ; CG - CD2
- HD22
4596 4602 4605 1 1.0950e+02 4.1840e+02 ; CG - CD2
- HD23
4597 4596 4598 1 1.0950e+02 4.1840e+02 ; HG - CG
- CD1
4597 4596 4602 1 1.0950e+02 4.1840e+02 ; HG - CG
- CD2
4598 4596 4602 1 1.0950e+02 3.3472e+02 ; CD1 - CG
- CD2
4599 4598 4600 1 1.0950e+02 2.9288e+02 ; HD11 - CD1
- HD12
4599 4598 4601 1 1.0950e+02 2.9288e+02 ; HD11 - CD1
- HD13
4600 4598 4601 1 1.0950e+02 2.9288e+02 ; HD12 - CD1
- HD13
4603 4602 4604 1 1.0950e+02 2.9288e+02 ; HD21 - CD2
- HD22
4603 4602 4605 1 1.0950e+02 2.9288e+02 ; HD21 - CD2
- HD23
4604 4602 4605 1 1.0950e+02 2.9288e+02 ; HD22 - CD2
- HD23
4606 4608 4609 1 1.2000e+02 4.1840e+02 ; C - N
- H
4606 4608 4610 1 1.2190e+02 4.1840e+02 ; C - N
- CA
4607 4606 4608 1 1.2290e+02 6.6944e+02 ; O - C
- N
4608 4610 4611 1 1.0950e+02 4.1840e+02 ; N - CA
- HA
4608 4610 4612 1 1.0970e+02 6.6944e+02 ; N - CA
- CB
4608 4610 4628 1 1.1010e+02 5.2718e+02 ; N - CA
- C
4609 4608 4610 1 1.1804e+02 4.1840e+02 ; H - N
- CA
4610 4612 4613 1 1.0950e+02 4.1840e+02 ; CA - CB
- HB2
4610 4612 4614 1 1.0950e+02 4.1840e+02 ; CA - CB
- HB3
4610 4612 4615 1 1.0950e+02 3.3472e+02 ; CA - CB
- CG
4610 4628 4629 1 1.2040e+02 6.6944e+02 ; CA - C
- O
4610 4628 4630 1 1.1660e+02 5.8576e+02 ; CA - C
- N
4611 4610 4612 1 1.0950e+02 4.1840e+02 ; HA - CA
- CB

4611 4610 4628 1 1.0950e+02 4.1840e+02 ; HA - CA - C
4612 4610 4628 1 1.1110e+02 5.2718e+02 ; CB - CA - C
4612 4615 4616 1 1.0950e+02 4.1840e+02 ; CB - CG - HG2
4612 4615 4617 1 1.0950e+02 4.1840e+02 ; CB - CG - HG3
4612 4615 4618 1 1.0950e+02 3.3472e+02 ; CB - CG - CD
4613 4612 4614 1 1.0950e+02 2.9288e+02 ; HB2 - CB - HB3
4613 4612 4615 1 1.0950e+02 4.1840e+02 ; HB2 - CB - CG
4614 4612 4615 1 1.0950e+02 4.1840e+02 ; HB3 - CB - CG
4615 4618 4619 1 1.0950e+02 4.1840e+02 ; CG - CD - HD2
4615 4618 4620 1 1.0950e+02 4.1840e+02 ; CG - CD - HD3
4615 4618 4621 1 1.0950e+02 3.3472e+02 ; CG - CD - CE
4616 4615 4617 1 1.0950e+02 2.9288e+02 ; HG2 - CG - HG3
4616 4615 4618 1 1.0950e+02 4.1840e+02 ; HG2 - CG - CD
4617 4615 4618 1 1.0950e+02 4.1840e+02 ; HG3 - CG - CD
4618 4621 4622 1 1.0950e+02 4.1840e+02 ; CD - CE - HE2
4618 4621 4623 1 1.0950e+02 4.1840e+02 ; CD - CE - HE3
4618 4621 4624 1 1.1120e+02 6.6944e+02 ; CD - CE - NZ
4619 4618 4620 1 1.0950e+02 2.9288e+02 ; HD2 - CD - HD3
4619 4618 4621 1 1.0950e+02 4.1840e+02 ; HD2 - CD - CE
4620 4618 4621 1 1.0950e+02 4.1840e+02 ; HD3 - CD - CE
4621 4624 4625 1 1.0950e+02 4.1840e+02 ; CE - NZ - HZ1
4621 4624 4626 1 1.0950e+02 4.1840e+02 ; CE - NZ - HZ2
4621 4624 4627 1 1.0950e+02 4.1840e+02 ; CE - NZ - HZ3
4622 4621 4623 1 1.0950e+02 2.9288e+02 ; HE2 - CE - HE3
4622 4621 4624 1 1.0950e+02 4.1840e+02 ; HE2 - CE - NZ
4623 4621 4624 1 1.0950e+02 4.1840e+02 ; HE3 - CE - NZ

4625 4624 4626 1 1.0950e+02 2.9288e+02 ; HZ1 - NZ
- HZ2
4625 4624 4627 1 1.0950e+02 2.9288e+02 ; HZ1 - NZ
- HZ3
4626 4624 4627 1 1.0950e+02 2.9288e+02 ; HZ2 - NZ
- HZ3
4628 4630 4631 1 1.2000e+02 4.1840e+02 ; C - N
- H
4628 4630 4632 1 1.2190e+02 4.1840e+02 ; C - N
- CA
4629 4628 4630 1 1.2290e+02 6.6944e+02 ; O - C
- N
4630 4632 4633 1 1.0950e+02 4.1840e+02 ; N - CA
- HA
4630 4632 4634 1 1.0970e+02 6.6944e+02 ; N - CA
- CB
4630 4632 4642 1 1.1010e+02 5.2718e+02 ; N - CA
- C
4631 4630 4632 1 1.1804e+02 4.1840e+02 ; H - N
- CA
4632 4634 4635 1 1.0950e+02 4.1840e+02 ; CA - CB
- HB2
4632 4634 4636 1 1.0950e+02 4.1840e+02 ; CA - CB
- HB3
4632 4634 4637 1 1.1110e+02 5.2718e+02 ; CA - CB
- CG
4632 4642 4643 1 1.2040e+02 6.6944e+02 ; CA - C
- O
4632 4642 4644 1 1.1660e+02 5.8576e+02 ; CA - C
- N
4633 4632 4634 1 1.0950e+02 4.1840e+02 ; HA - CA
- CB
4633 4632 4642 1 1.0950e+02 4.1840e+02 ; HA - CA
- C
4634 4632 4642 1 1.1110e+02 5.2718e+02 ; CB - CA
- C
4634 4637 4638 1 1.2040e+02 6.6944e+02 ; CB - CG
- OD1
4634 4637 4639 1 1.1660e+02 5.8576e+02 ; CB - CG
- ND2
4635 4634 4636 1 1.0950e+02 2.9288e+02 ; HB2 - CB
- HB3
4635 4634 4637 1 1.0950e+02 4.1840e+02 ; HB2 - CB
- CG
4636 4634 4637 1 1.0950e+02 4.1840e+02 ; HB3 - CB
- CG
4637 4639 4640 1 1.2000e+02 4.1840e+02 ; CG - ND2
- HD21
4637 4639 4641 1 1.2000e+02 4.1840e+02 ; CG - ND2
- HD22
4638 4637 4639 1 1.2290e+02 6.6944e+02 ; OD1 - CG
- ND2

4640 4639 4641 1 1.2000e+02 2.9288e+02 ; HD21 - ND2
- HD22
4642 4644 4645 1 1.2000e+02 4.1840e+02 ; C - N
- H
4642 4644 4646 1 1.2190e+02 4.1840e+02 ; C - N
- CA
4643 4642 4644 1 1.2290e+02 6.6944e+02 ; O - C
- N
4644 4646 4647 1 1.0950e+02 4.1840e+02 ; N - CA
- HA
4644 4646 4648 1 1.0970e+02 6.6944e+02 ; N - CA
- CB
4644 4646 4661 1 1.1010e+02 5.2718e+02 ; N - CA
- C
4645 4644 4646 1 1.1804e+02 4.1840e+02 ; H - N
- CA
4646 4648 4649 1 1.0950e+02 4.1840e+02 ; CA - CB
- HB2
4646 4648 4650 1 1.0950e+02 4.1840e+02 ; CA - CB
- HB3
4646 4648 4651 1 1.0950e+02 3.3472e+02 ; CA - CB
- CG
4646 4661 4662 1 1.2040e+02 6.6944e+02 ; CA - C
- O
4646 4661 4663 1 1.1660e+02 5.8576e+02 ; CA - C
- N
4647 4646 4648 1 1.0950e+02 4.1840e+02 ; HA - CA
- CB
4647 4646 4661 1 1.0950e+02 4.1840e+02 ; HA - CA
- C
4648 4646 4661 1 1.1110e+02 5.2718e+02 ; CB - CA
- C
4648 4651 4652 1 1.0950e+02 4.1840e+02 ; CB - CG
- HG
4648 4651 4653 1 1.0950e+02 3.3472e+02 ; CB - CG
- CD1
4648 4651 4657 1 1.0950e+02 3.3472e+02 ; CB - CG
- CD2
4649 4648 4650 1 1.0950e+02 2.9288e+02 ; HB2 - CB
- HB3
4649 4648 4651 1 1.0950e+02 4.1840e+02 ; HB2 - CB
- CG
4650 4648 4651 1 1.0950e+02 4.1840e+02 ; HB3 - CB
- CG
4651 4653 4654 1 1.0950e+02 4.1840e+02 ; CG - CD1
- HD11
4651 4653 4655 1 1.0950e+02 4.1840e+02 ; CG - CD1
- HD12
4651 4653 4656 1 1.0950e+02 4.1840e+02 ; CG - CD1
- HD13
4651 4657 4658 1 1.0950e+02 4.1840e+02 ; CG - CD2
- HD21

4651 4657 4659 1 1.0950e+02 4.1840e+02 ; CG - CD2
- HD22
4651 4657 4660 1 1.0950e+02 4.1840e+02 ; CG - CD2
- HD23
4652 4651 4653 1 1.0950e+02 4.1840e+02 ; HG - CG
- CD1
4652 4651 4657 1 1.0950e+02 4.1840e+02 ; HG - CG
- CD2
4653 4651 4657 1 1.0950e+02 3.3472e+02 ; CD1 - CG
- CD2
4654 4653 4655 1 1.0950e+02 2.9288e+02 ; HD11 - CD1
- HD12
4654 4653 4656 1 1.0950e+02 2.9288e+02 ; HD11 - CD1
- HD13
4655 4653 4656 1 1.0950e+02 2.9288e+02 ; HD12 - CD1
- HD13
4658 4657 4659 1 1.0950e+02 2.9288e+02 ; HD21 - CD2
- HD22
4658 4657 4660 1 1.0950e+02 2.9288e+02 ; HD21 - CD2
- HD23
4659 4657 4660 1 1.0950e+02 2.9288e+02 ; HD22 - CD2
- HD23
4661 4663 4664 1 1.2000e+02 4.1840e+02 ; C - N
- H
4661 4663 4665 1 1.2190e+02 4.1840e+02 ; C - N
- CA
4662 4661 4663 1 1.2290e+02 6.6944e+02 ; O - C
- N
4663 4665 4666 1 1.0950e+02 4.1840e+02 ; N - CA
- HA
4663 4665 4667 1 1.0970e+02 6.6944e+02 ; N - CA
- CB
4663 4665 4683 1 1.1010e+02 5.2718e+02 ; N - CA
- C
4664 4663 4665 1 1.1804e+02 4.1840e+02 ; H - N
- CA
4665 4667 4668 1 1.0950e+02 4.1840e+02 ; CA - CB
- HB2
4665 4667 4669 1 1.0950e+02 4.1840e+02 ; CA - CB
- HB3
4665 4667 4670 1 1.0950e+02 3.3472e+02 ; CA - CB
- CG
4665 4683 4684 1 1.2040e+02 6.6944e+02 ; CA - C
- O
4665 4683 4685 1 1.1660e+02 5.8576e+02 ; CA - C
- N
4666 4665 4667 1 1.0950e+02 4.1840e+02 ; HA - CA
- CB
4666 4665 4683 1 1.0950e+02 4.1840e+02 ; HA - CA
- C
4667 4665 4683 1 1.1110e+02 5.2718e+02 ; CB - CA
- C

4667 4670 4671 1 1.0950e+02 4.1840e+02 ; CB - CG
- HG2
4667 4670 4672 1 1.0950e+02 4.1840e+02 ; CB - CG
- HG3
4667 4670 4673 1 1.0950e+02 3.3472e+02 ; CB - CG
- CD
4668 4667 4669 1 1.0950e+02 2.9288e+02 ; HB2 - CB
- HB3
4668 4667 4670 1 1.0950e+02 4.1840e+02 ; HB2 - CB
- CG
4669 4667 4670 1 1.0950e+02 4.1840e+02 ; HB3 - CB
- CG
4670 4673 4674 1 1.0950e+02 4.1840e+02 ; CG - CD
- HD2
4670 4673 4675 1 1.0950e+02 4.1840e+02 ; CG - CD
- HD3
4670 4673 4676 1 1.0950e+02 3.3472e+02 ; CG - CD
- CE
4671 4670 4672 1 1.0950e+02 2.9288e+02 ; HG2 - CG
- HG3
4671 4670 4673 1 1.0950e+02 4.1840e+02 ; HG2 - CG
- CD
4672 4670 4673 1 1.0950e+02 4.1840e+02 ; HG3 - CG
- CD
4673 4676 4677 1 1.0950e+02 4.1840e+02 ; CD - CE
- HE2
4673 4676 4678 1 1.0950e+02 4.1840e+02 ; CD - CE
- HE3
4673 4676 4679 1 1.1120e+02 6.6944e+02 ; CD - CE
- NZ
4674 4673 4675 1 1.0950e+02 2.9288e+02 ; HD2 - CD
- HD3
4674 4673 4676 1 1.0950e+02 4.1840e+02 ; HD2 - CD
- CE
4675 4673 4676 1 1.0950e+02 4.1840e+02 ; HD3 - CD
- CE
4676 4679 4680 1 1.0950e+02 4.1840e+02 ; CE - NZ
- HZ1
4676 4679 4681 1 1.0950e+02 4.1840e+02 ; CE - NZ
- HZ2
4676 4679 4682 1 1.0950e+02 4.1840e+02 ; CE - NZ
- HZ3
4677 4676 4678 1 1.0950e+02 2.9288e+02 ; HE2 - CE
- HE3
4677 4676 4679 1 1.0950e+02 4.1840e+02 ; HE2 - CE
- NZ
4678 4676 4679 1 1.0950e+02 4.1840e+02 ; HE3 - CE
- NZ
4680 4679 4681 1 1.0950e+02 2.9288e+02 ; HZ1 - NZ
- HZ2
4680 4679 4682 1 1.0950e+02 2.9288e+02 ; HZ1 - NZ
- HZ3

```
  4681   4679   4682      1    1.0950e+02    2.9288e+02 ;    HZ2 - NZ - HZ3
  4683   4685   4686      1    1.2000e+02    4.1840e+02 ;      C - N - H
  4683   4685   4687      1    1.2190e+02    4.1840e+02 ;      C - N - CA
  4684   4683   4685      1    1.2290e+02    6.6944e+02 ;      O - C - N
  4685   4687   4688      1    1.0950e+02    4.1840e+02 ;      N - CA - HA
  4685   4687   4689      1    1.0970e+02    6.6944e+02 ;      N - CA - CB
  4685   4687   4697      1    1.1010e+02    5.2718e+02 ;      N - CA - C
  4686   4685   4687      1    1.1804e+02    4.1840e+02 ;      H - N - CA
  4687   4689   4690      1    1.0950e+02    4.1840e+02 ;     CA - CB - HB2
  4687   4689   4691      1    1.0950e+02    4.1840e+02 ;     CA - CB - HB3
  4687   4689   4692      1    1.1110e+02    5.2718e+02 ;     CA - CB - CG
  4687   4697   4698      1    1.2040e+02    6.6944e+02 ;     CA - C - O
  4687   4697   4699      1    1.1660e+02    5.8576e+02 ;     CA - C - N
  4688   4687   4689      1    1.0950e+02    4.1840e+02 ;     HA - CA - CB
  4688   4687   4697      1    1.0950e+02    4.1840e+02 ;     HA - CA - C
  4689   4687   4697      1    1.1110e+02    5.2718e+02 ;     CB - CA - C
  4689   4692   4693      1    1.2040e+02    6.6944e+02 ;     CB - CG - OD1
  4689   4692   4694      1    1.1660e+02    5.8576e+02 ;     CB - CG - ND2
  4690   4689   4691      1    1.0950e+02    2.9288e+02 ;    HB2 - CB - HB3
  4690   4689   4692      1    1.0950e+02    4.1840e+02 ;    HB2 - CB - CG
  4691   4689   4692      1    1.0950e+02    4.1840e+02 ;    HB3 - CB - CG
  4692   4694   4695      1    1.2000e+02    4.1840e+02 ;     CG - ND2 - HD21
  4692   4694   4696      1    1.2000e+02    4.1840e+02 ;     CG - ND2 - HD22
  4693   4692   4694      1    1.2290e+02    6.6944e+02 ;    OD1 - CG - ND2
  4695   4694   4696      1    1.2000e+02    2.9288e+02 ;   HD21 - ND2 - HD22
  4697   4699   4700      1    1.2000e+02    4.1840e+02 ;      C - N - H
```

4697 4699 4701 1 1.2190e+02 4.1840e+02 ; C - N
- CA
4698 4697 4699 1 1.2290e+02 6.6944e+02 ; O - C
- N
4699 4701 4702 1 1.0950e+02 4.1840e+02 ; N - CA
- HA
4699 4701 4703 1 1.0970e+02 6.6944e+02 ; N - CA
- CB
4699 4701 4717 1 1.1010e+02 5.2718e+02 ; N - CA
- C
4700 4699 4701 1 1.1804e+02 4.1840e+02 ; H - N
- CA
4701 4703 4704 1 1.0950e+02 4.1840e+02 ; CA - CB
- HB2
4701 4703 4705 1 1.0950e+02 4.1840e+02 ; CA - CB
- HB3
4701 4703 4706 1 1.1400e+02 5.2718e+02 ; CA - CB
- CG
4701 4717 4718 1 1.2040e+02 6.6944e+02 ; CA - C
- O
4701 4717 4719 1 1.1660e+02 5.8576e+02 ; CA - C
- N
4702 4701 4703 1 1.0950e+02 4.1840e+02 ; HA - CA
- CB
4702 4701 4717 1 1.0950e+02 4.1840e+02 ; HA - CA
- C
4703 4701 4717 1 1.1110e+02 5.2718e+02 ; CB - CA
- C
4703 4706 4707 1 1.2000e+02 5.8576e+02 ; CB - CG
- CD1
4703 4706 4715 1 1.2000e+02 5.8576e+02 ; CB - CG
- CD2
4704 4703 4705 1 1.0950e+02 2.9288e+02 ; HB2 - CB
- HB3
4704 4703 4706 1 1.0950e+02 4.1840e+02 ; HB2 - CB
- CG
4705 4703 4706 1 1.0950e+02 4.1840e+02 ; HB3 - CB
- CG
4706 4707 4708 1 1.2000e+02 4.1840e+02 ; CG - CD1
- HD1
4706 4707 4709 1 1.2000e+02 5.2718e+02 ; CG - CD1
- CE1
4706 4715 4713 1 1.2000e+02 5.2718e+02 ; CG - CD2
- CE2
4706 4715 4716 1 1.2000e+02 4.1840e+02 ; CG - CD2
- HD2
4707 4706 4715 1 1.2000e+02 5.2718e+02 ; CD1 - CG
- CD2
4707 4709 4710 1 1.2000e+02 4.1840e+02 ; CD1 - CE1
- HE1
4707 4709 4711 1 1.2000e+02 5.2718e+02 ; CD1 - CE1
- CZ

4708 4707 4709 1 1.2000e+02 4.1840e+02 ; HD1 - CD1
- CE1
4709 4711 4712 1 1.2000e+02 4.1840e+02 ; CE1 - CZ
- HZ
4709 4711 4713 1 1.2000e+02 5.2718e+02 ; CE1 - CZ
- CE2
4710 4709 4711 1 1.2000e+02 4.1840e+02 ; HE1 - CE1
- CZ
4711 4713 4714 1 1.2000e+02 4.1840e+02 ; CZ - CE2
- HE2
4711 4713 4715 1 1.2000e+02 5.2718e+02 ; CZ - CE2
- CD2
4712 4711 4713 1 1.2000e+02 4.1840e+02 ; HZ - CZ
- CE2
4713 4715 4716 1 1.2000e+02 4.1840e+02 ; CE2 - CD2
- HD2
4714 4713 4715 1 1.2000e+02 4.1840e+02 ; HE2 - CE2
- CD2
4717 4719 4720 1 1.2000e+02 4.1840e+02 ; C - N
- H
4717 4719 4721 1 1.2190e+02 4.1840e+02 ; C - N
- CA
4718 4717 4719 1 1.2290e+02 6.6944e+02 ; O - C
- N
4719 4721 4722 1 1.0950e+02 4.1840e+02 ; N - CA
- HA
4719 4721 4723 1 1.0970e+02 6.6944e+02 ; N - CA
- CB
4719 4721 4739 1 1.1010e+02 5.2718e+02 ; N - CA
- C
4720 4719 4721 1 1.1804e+02 4.1840e+02 ; H - N
- CA
4721 4723 4724 1 1.0950e+02 4.1840e+02 ; CA - CB
- HB2
4721 4723 4725 1 1.0950e+02 4.1840e+02 ; CA - CB
- HB3
4721 4723 4726 1 1.0950e+02 3.3472e+02 ; CA - CB
- CG
4721 4739 4740 1 1.2040e+02 6.6944e+02 ; CA - C
- O
4721 4739 4741 1 1.1660e+02 5.8576e+02 ; CA - C
- N
4722 4721 4723 1 1.0950e+02 4.1840e+02 ; HA - CA
- CB
4722 4721 4739 1 1.0950e+02 4.1840e+02 ; HA - CA
- C
4723 4721 4739 1 1.1110e+02 5.2718e+02 ; CB - CA
- C
4723 4726 4727 1 1.0950e+02 4.1840e+02 ; CB - CG
- HG2
4723 4726 4728 1 1.0950e+02 4.1840e+02 ; CB - CG
- HG3

4723 4726 4729 1 1.0950e+02 3.3472e+02 ; CB - CG
- CD
4724 4723 4725 1 1.0950e+02 2.9288e+02 ; HB2 - CB
- HB3
4724 4723 4726 1 1.0950e+02 4.1840e+02 ; HB2 - CB
- CG
4725 4723 4726 1 1.0950e+02 4.1840e+02 ; HB3 - CB
- CG
4726 4729 4730 1 1.0950e+02 4.1840e+02 ; CG - CD
- HD2
4726 4729 4731 1 1.0950e+02 4.1840e+02 ; CG - CD
- HD3
4726 4729 4732 1 1.0950e+02 3.3472e+02 ; CG - CD
- CE
4727 4726 4728 1 1.0950e+02 2.9288e+02 ; HG2 - CG
- HG3
4727 4726 4729 1 1.0950e+02 4.1840e+02 ; HG2 - CG
- CD
4728 4726 4729 1 1.0950e+02 4.1840e+02 ; HG3 - CG
- CD
4729 4732 4733 1 1.0950e+02 4.1840e+02 ; CD - CE
- HE2
4729 4732 4734 1 1.0950e+02 4.1840e+02 ; CD - CE
- HE3
4729 4732 4735 1 1.1120e+02 6.6944e+02 ; CD - CE
- NZ
4730 4729 4731 1 1.0950e+02 2.9288e+02 ; HD2 - CD
- HD3
4730 4729 4732 1 1.0950e+02 4.1840e+02 ; HD2 - CD
- CE
4731 4729 4732 1 1.0950e+02 4.1840e+02 ; HD3 - CD
- CE
4732 4735 4736 1 1.0950e+02 4.1840e+02 ; CE - NZ
- HZ1
4732 4735 4737 1 1.0950e+02 4.1840e+02 ; CE - NZ
- HZ2
4732 4735 4738 1 1.0950e+02 4.1840e+02 ; CE - NZ
- HZ3
4733 4732 4734 1 1.0950e+02 2.9288e+02 ; HE2 - CE
- HE3
4733 4732 4735 1 1.0950e+02 4.1840e+02 ; HE2 - CE
- NZ
4734 4732 4735 1 1.0950e+02 4.1840e+02 ; HE3 - CE
- NZ
4736 4735 4737 1 1.0950e+02 2.9288e+02 ; HZ1 - NZ
- HZ2
4736 4735 4738 1 1.0950e+02 2.9288e+02 ; HZ1 - NZ
- HZ3
4737 4735 4738 1 1.0950e+02 2.9288e+02 ; HZ2 - NZ
- HZ3
4739 4741 4742 1 1.2000e+02 4.1840e+02 ; C - N
- H

```
  4739   4741   4743      1    1.2190e+02    4.1840e+02 ;      C - N
- CA
  4740   4739   4741      1    1.2290e+02    6.6944e+02 ;      O - C
- N
  4741   4743   4744      1    1.0950e+02    4.1840e+02 ;      N - CA
- HA
  4741   4743   4745      1    1.0970e+02    6.6944e+02 ;      N - CA
- CB
  4741   4743   4761      1    1.1010e+02    5.2718e+02 ;      N - CA
- C
  4742   4741   4743      1    1.1804e+02    4.1840e+02 ;      H - N
- CA
  4743   4745   4746      1    1.0950e+02    4.1840e+02 ;     CA - CB
- HB2
  4743   4745   4747      1    1.0950e+02    4.1840e+02 ;     CA - CB
- HB3
  4743   4745   4748      1    1.0950e+02    3.3472e+02 ;     CA - CB
- CG
  4743   4761   4762      1    1.2040e+02    6.6944e+02 ;     CA - C
- O
  4743   4761   4763      1    1.1660e+02    5.8576e+02 ;     CA - C
- N
  4744   4743   4745      1    1.0950e+02    4.1840e+02 ;     HA - CA
- CB
  4744   4743   4761      1    1.0950e+02    4.1840e+02 ;     HA - CA
- C
  4745   4743   4761      1    1.1110e+02    5.2718e+02 ;     CB - CA
- C
  4745   4748   4749      1    1.0950e+02    4.1840e+02 ;     CB - CG
- HG2
  4745   4748   4750      1    1.0950e+02    4.1840e+02 ;     CB - CG
- HG3
  4745   4748   4751      1    1.0950e+02    3.3472e+02 ;     CB - CG
- CD
  4746   4745   4747      1    1.0950e+02    2.9288e+02 ;    HB2 - CB
- HB3
  4746   4745   4748      1    1.0950e+02    4.1840e+02 ;    HB2 - CB
- CG
  4747   4745   4748      1    1.0950e+02    4.1840e+02 ;    HB3 - CB
- CG
  4748   4751   4752      1    1.0950e+02    4.1840e+02 ;     CG - CD
- HD2
  4748   4751   4753      1    1.0950e+02    4.1840e+02 ;     CG - CD
- HD3
  4748   4751   4754      1    1.0950e+02    3.3472e+02 ;     CG - CD
- CE
  4749   4748   4750      1    1.0950e+02    2.9288e+02 ;    HG2 - CG
- HG3
  4749   4748   4751      1    1.0950e+02    4.1840e+02 ;    HG2 - CG
- CD
  4750   4748   4751      1    1.0950e+02    4.1840e+02 ;    HG3 - CG
- CD
```

4751 4754 4755 1 1.0950e+02 4.1840e+02 ; CD - CE
- HE2
4751 4754 4756 1 1.0950e+02 4.1840e+02 ; CD - CE
- HE3
4751 4754 4757 1 1.1120e+02 6.6944e+02 ; CD - CE
- NZ
4752 4751 4753 1 1.0950e+02 2.9288e+02 ; HD2 - CD
- HD3
4752 4751 4754 1 1.0950e+02 4.1840e+02 ; HD2 - CD
- CE
4753 4751 4754 1 1.0950e+02 4.1840e+02 ; HD3 - CD
- CE
4754 4757 4758 1 1.0950e+02 4.1840e+02 ; CE - NZ
- HZ1
4754 4757 4759 1 1.0950e+02 4.1840e+02 ; CE - NZ
- HZ2
4754 4757 4760 1 1.0950e+02 4.1840e+02 ; CE - NZ
- HZ3
4755 4754 4756 1 1.0950e+02 2.9288e+02 ; HE2 - CE
- HE3
4755 4754 4757 1 1.0950e+02 4.1840e+02 ; HE2 - CE
- NZ
4756 4754 4757 1 1.0950e+02 4.1840e+02 ; HE3 - CE
- NZ
4758 4757 4759 1 1.0950e+02 2.9288e+02 ; HZ1 - NZ
- HZ2
4758 4757 4760 1 1.0950e+02 2.9288e+02 ; HZ1 - NZ
- HZ3
4759 4757 4760 1 1.0950e+02 2.9288e+02 ; HZ2 - NZ
- HZ3
4761 4763 4764 1 1.2000e+02 4.1840e+02 ; C - N
- H
4761 4763 4765 1 1.2190e+02 4.1840e+02 ; C - N
- CA
4762 4761 4763 1 1.2290e+02 6.6944e+02 ; O - C
- N
4763 4765 4766 1 1.0950e+02 4.1840e+02 ; N - CA
- HA
4763 4765 4767 1 1.0970e+02 6.6944e+02 ; N - CA
- CB
4763 4765 4776 1 1.1010e+02 5.2718e+02 ; N - CA
- C
4764 4763 4765 1 1.1804e+02 4.1840e+02 ; H - N
- CA
4765 4767 4768 1 1.0950e+02 4.1840e+02 ; CA - CB
- HB2
4765 4767 4769 1 1.0950e+02 4.1840e+02 ; CA - CB
- HB3
4765 4767 4770 1 1.0950e+02 3.3472e+02 ; CA - CB
- CG
4765 4776 4777 1 1.2040e+02 6.6944e+02 ; CA - C
- O

4765 4776 4778 1 1.1660e+02 5.8576e+02 ; CA - C
- N
4766 4765 4767 1 1.0950e+02 4.1840e+02 ; HA - CA
- CB
4766 4765 4776 1 1.0950e+02 4.1840e+02 ; HA - CA
- C
4767 4765 4776 1 1.1110e+02 5.2718e+02 ; CB - CA
- C
4767 4770 4771 1 1.0950e+02 4.1840e+02 ; CB - CG
- HG2
4767 4770 4772 1 1.0950e+02 4.1840e+02 ; CB - CG
- HG3
4767 4770 4773 1 1.1110e+02 5.2718e+02 ; CB - CG
- CD
4768 4767 4769 1 1.0950e+02 2.9288e+02 ; HB2 - CB
- HB3
4768 4767 4770 1 1.0950e+02 4.1840e+02 ; HB2 - CB
- CG
4769 4767 4770 1 1.0950e+02 4.1840e+02 ; HB3 - CB
- CG
4770 4773 4774 1 1.1700e+02 5.8576e+02 ; CG - CD
- OE1
4770 4773 4775 1 1.1700e+02 5.8576e+02 ; CG - CD
- OE2
4771 4770 4772 1 1.0950e+02 2.9288e+02 ; HG2 - CG
- HG3
4771 4770 4773 1 1.0950e+02 4.1840e+02 ; HG2 - CG
- CD
4772 4770 4773 1 1.0950e+02 4.1840e+02 ; HG3 - CG
- CD
4774 4773 4775 1 1.2600e+02 6.6944e+02 ; OE1 - CD
- OE2
4776 4778 4779 1 1.2000e+02 4.1840e+02 ; C - N
- H
4776 4778 4780 1 1.2190e+02 4.1840e+02 ; C - N
- CA
4777 4776 4778 1 1.2290e+02 6.6944e+02 ; O - C
- N
4778 4780 4781 1 1.0950e+02 4.1840e+02 ; N - CA
- HA
4778 4780 4782 1 1.0970e+02 6.6944e+02 ; N - CA
- CB
4778 4780 4790 1 1.1010e+02 5.2718e+02 ; N - CA
- C
4779 4778 4780 1 1.1804e+02 4.1840e+02 ; H - N
- CA
4780 4782 4783 1 1.0950e+02 4.1840e+02 ; CA - CB
- HB2
4780 4782 4784 1 1.0950e+02 4.1840e+02 ; CA - CB
- HB3
4780 4782 4785 1 1.1110e+02 5.2718e+02 ; CA - CB
- CG

```
  4780   4790   4791      1    1.2040e+02    6.6944e+02 ;     CA - C
- O
  4780   4790   4792      1    1.1660e+02    5.8576e+02 ;     CA - C
- N
  4781   4780   4782      1    1.0950e+02    4.1840e+02 ;     HA - CA
- CB
  4781   4780   4790      1    1.0950e+02    4.1840e+02 ;     HA - CA
- C
  4782   4780   4790      1    1.1110e+02    5.2718e+02 ;     CB - CA
- C
  4782   4785   4786      1    1.2040e+02    6.6944e+02 ;     CB - CG
- OD1
  4782   4785   4787      1    1.1660e+02    5.8576e+02 ;     CB - CG
- ND2
  4783   4782   4784      1    1.0950e+02    2.9288e+02 ;    HB2 - CB
- HB3
  4783   4782   4785      1    1.0950e+02    4.1840e+02 ;    HB2 - CB
- CG
  4784   4782   4785      1    1.0950e+02    4.1840e+02 ;    HB3 - CB
- CG
  4785   4787   4788      1    1.2000e+02    4.1840e+02 ;     CG - ND2
- HD21
  4785   4787   4789      1    1.2000e+02    4.1840e+02 ;     CG - ND2
- HD22
  4786   4785   4787      1    1.2290e+02    6.6944e+02 ;    OD1 - CG
- ND2
  4788   4787   4789      1    1.2000e+02    2.9288e+02 ;   HD21 - ND2
- HD22
  4790   4792   4793      1    1.2000e+02    4.1840e+02 ;      C - N
- H
  4790   4792   4794      1    1.2190e+02    4.1840e+02 ;      C - N
- CA
  4791   4790   4792      1    1.2290e+02    6.6944e+02 ;      O - C
- N
  4792   4794   4795      1    1.0950e+02    4.1840e+02 ;      N - CA
- HA
  4792   4794   4796      1    1.0970e+02    6.6944e+02 ;      N - CA
- CB
  4792   4794   4805      1    1.1010e+02    5.2718e+02 ;      N - CA
- C
  4793   4792   4794      1    1.1804e+02    4.1840e+02 ;      H - N
- CA
  4794   4796   4797      1    1.0950e+02    4.1840e+02 ;     CA - CB
- HB2
  4794   4796   4798      1    1.0950e+02    4.1840e+02 ;     CA - CB
- HB3
  4794   4796   4799      1    1.0950e+02    3.3472e+02 ;     CA - CB
- CG
  4794   4805   4806      1    1.2040e+02    6.6944e+02 ;     CA - C
- O
  4794   4805   4807      1    1.1660e+02    5.8576e+02 ;     CA - C
- N
```

4795 4794 4796 1 1.0950e+02 4.1840e+02 ; HA - CA
- CB
4795 4794 4805 1 1.0950e+02 4.1840e+02 ; HA - CA
- C
4796 4794 4805 1 1.1110e+02 5.2718e+02 ; CB - CA
- C
4796 4799 4800 1 1.0950e+02 4.1840e+02 ; CB - CG
- HG2
4796 4799 4801 1 1.0950e+02 4.1840e+02 ; CB - CG
- HG3
4796 4799 4802 1 1.1110e+02 5.2718e+02 ; CB - CG
- CD
4797 4796 4798 1 1.0950e+02 2.9288e+02 ; HB2 - CB
- HB3
4797 4796 4799 1 1.0950e+02 4.1840e+02 ; HB2 - CB
- CG
4798 4796 4799 1 1.0950e+02 4.1840e+02 ; HB3 - CB
- CG
4799 4802 4803 1 1.1700e+02 5.8576e+02 ; CG - CD
- OE1
4799 4802 4804 1 1.1700e+02 5.8576e+02 ; CG - CD
- OE2
4800 4799 4801 1 1.0950e+02 2.9288e+02 ; HG2 - CG
- HG3
4800 4799 4802 1 1.0950e+02 4.1840e+02 ; HG2 - CG
- CD
4801 4799 4802 1 1.0950e+02 4.1840e+02 ; HG3 - CG
- CD
4803 4802 4804 1 1.2600e+02 6.6944e+02 ; OE1 - CD
- OE2
4805 4807 4808 1 1.2000e+02 4.1840e+02 ; C - N
- H
4805 4807 4809 1 1.2190e+02 4.1840e+02 ; C - N
- CA
4806 4805 4807 1 1.2290e+02 6.6944e+02 ; O - C
- N
4807 4809 4810 1 1.0950e+02 4.1840e+02 ; N - CA
- HA
4807 4809 4811 1 1.0970e+02 6.6944e+02 ; N - CA
- CB
4807 4809 4824 1 1.1010e+02 5.2718e+02 ; N - CA
- C
4808 4807 4809 1 1.1804e+02 4.1840e+02 ; H - N
- CA
4809 4811 4812 1 1.0950e+02 4.1840e+02 ; CA - CB
- HB2
4809 4811 4813 1 1.0950e+02 4.1840e+02 ; CA - CB
- HB3
4809 4811 4814 1 1.0950e+02 3.3472e+02 ; CA - CB
- CG
4809 4824 4825 1 1.2040e+02 6.6944e+02 ; CA - C
- O

```
  4809   4824   4826      1    1.1660e+02    5.8576e+02 ;     CA - C
- N
  4810   4809   4811      1    1.0950e+02    4.1840e+02 ;     HA - CA
- CB
  4810   4809   4824      1    1.0950e+02    4.1840e+02 ;     HA - CA
- C
  4811   4809   4824      1    1.1110e+02    5.2718e+02 ;     CB - CA
- C
  4811   4814   4815      1    1.0950e+02    4.1840e+02 ;     CB - CG
- HG
  4811   4814   4816      1    1.0950e+02    3.3472e+02 ;     CB - CG
- CD1
  4811   4814   4820      1    1.0950e+02    3.3472e+02 ;     CB - CG
- CD2
  4812   4811   4813      1    1.0950e+02    2.9288e+02 ;    HB2 - CB
- HB3
  4812   4811   4814      1    1.0950e+02    4.1840e+02 ;    HB2 - CB
- CG
  4813   4811   4814      1    1.0950e+02    4.1840e+02 ;    HB3 - CB
- CG
  4814   4816   4817      1    1.0950e+02    4.1840e+02 ;     CG - CD1
- HD11
  4814   4816   4818      1    1.0950e+02    4.1840e+02 ;     CG - CD1
- HD12
  4814   4816   4819      1    1.0950e+02    4.1840e+02 ;     CG - CD1
- HD13
  4814   4820   4821      1    1.0950e+02    4.1840e+02 ;     CG - CD2
- HD21
  4814   4820   4822      1    1.0950e+02    4.1840e+02 ;     CG - CD2
- HD22
  4814   4820   4823      1    1.0950e+02    4.1840e+02 ;     CG - CD2
- HD23
  4815   4814   4816      1    1.0950e+02    4.1840e+02 ;     HG - CG
- CD1
  4815   4814   4820      1    1.0950e+02    4.1840e+02 ;     HG - CG
- CD2
  4816   4814   4820      1    1.0950e+02    3.3472e+02 ;    CD1 - CG
- CD2
  4817   4816   4818      1    1.0950e+02    2.9288e+02 ;   HD11 - CD1
- HD12
  4817   4816   4819      1    1.0950e+02    2.9288e+02 ;   HD11 - CD1
- HD13
  4818   4816   4819      1    1.0950e+02    2.9288e+02 ;   HD12 - CD1
- HD13
  4821   4820   4822      1    1.0950e+02    2.9288e+02 ;   HD21 - CD2
- HD22
  4821   4820   4823      1    1.0950e+02    2.9288e+02 ;   HD21 - CD2
- HD23
  4822   4820   4823      1    1.0950e+02    2.9288e+02 ;   HD22 - CD2
- HD23
  4824   4826   4827      1    1.2000e+02    4.1840e+02 ;      C - N
- H
```

4824 4826 4828 1 1.2190e+02 4.1840e+02 ; C - N
- CA
4825 4824 4826 1 1.2290e+02 6.6944e+02 ; O - C
- N
4826 4828 4829 1 1.0950e+02 4.1840e+02 ; N - CA
- HA
4826 4828 4830 1 1.0970e+02 6.6944e+02 ; N - CA
- CB
4826 4828 4846 1 1.1010e+02 5.2718e+02 ; N - CA
- C
4827 4826 4828 1 1.1804e+02 4.1840e+02 ; H - N
- CA
4828 4830 4831 1 1.0950e+02 4.1840e+02 ; CA - CB
- HB2
4828 4830 4832 1 1.0950e+02 4.1840e+02 ; CA - CB
- HB3
4828 4830 4833 1 1.0950e+02 3.3472e+02 ; CA - CB
- CG
4828 4846 4847 1 1.2040e+02 6.6944e+02 ; CA - C
- O
4828 4846 4848 1 1.1660e+02 5.8576e+02 ; CA - C
- N
4829 4828 4830 1 1.0950e+02 4.1840e+02 ; HA - CA
- CB
4829 4828 4846 1 1.0950e+02 4.1840e+02 ; HA - CA
- C
4830 4828 4846 1 1.1110e+02 5.2718e+02 ; CB - CA
- C
4830 4833 4834 1 1.0950e+02 4.1840e+02 ; CB - CG
- HG2
4830 4833 4835 1 1.0950e+02 4.1840e+02 ; CB - CG
- HG3
4830 4833 4836 1 1.0950e+02 3.3472e+02 ; CB - CG
- CD
4831 4830 4832 1 1.0950e+02 2.9288e+02 ; HB2 - CB
- HB3
4831 4830 4833 1 1.0950e+02 4.1840e+02 ; HB2 - CB
- CG
4832 4830 4833 1 1.0950e+02 4.1840e+02 ; HB3 - CB
- CG
4833 4836 4837 1 1.0950e+02 4.1840e+02 ; CG - CD
- HD2
4833 4836 4838 1 1.0950e+02 4.1840e+02 ; CG - CD
- HD3
4833 4836 4839 1 1.0950e+02 3.3472e+02 ; CG - CD
- CE
4834 4833 4835 1 1.0950e+02 2.9288e+02 ; HG2 - CG
- HG3
4834 4833 4836 1 1.0950e+02 4.1840e+02 ; HG2 - CG
- CD
4835 4833 4836 1 1.0950e+02 4.1840e+02 ; HG3 - CG
- CD

4836 4839 4840 1 1.0950e+02 4.1840e+02 ; CD - CE
- HE2
4836 4839 4841 1 1.0950e+02 4.1840e+02 ; CD - CE
- HE3
4836 4839 4842 1 1.1120e+02 6.6944e+02 ; CD - CE
- NZ
4837 4836 4838 1 1.0950e+02 2.9288e+02 ; HD2 - CD
- HD3
4837 4836 4839 1 1.0950e+02 4.1840e+02 ; HD2 - CD
- CE
4838 4836 4839 1 1.0950e+02 4.1840e+02 ; HD3 - CD
- CE
4839 4842 4843 1 1.0950e+02 4.1840e+02 ; CE - NZ
- HZ1
4839 4842 4844 1 1.0950e+02 4.1840e+02 ; CE - NZ
- HZ2
4839 4842 4845 1 1.0950e+02 4.1840e+02 ; CE - NZ
- HZ3
4840 4839 4841 1 1.0950e+02 2.9288e+02 ; HE2 - CE
- HE3
4840 4839 4842 1 1.0950e+02 4.1840e+02 ; HE2 - CE
- NZ
4841 4839 4842 1 1.0950e+02 4.1840e+02 ; HE3 - CE
- NZ
4843 4842 4844 1 1.0950e+02 2.9288e+02 ; HZ1 - NZ
- HZ2
4843 4842 4845 1 1.0950e+02 2.9288e+02 ; HZ1 - NZ
- HZ3
4844 4842 4845 1 1.0950e+02 2.9288e+02 ; HZ2 - NZ
- HZ3
4846 4848 4849 1 1.2000e+02 4.1840e+02 ; C - N
- H
4846 4848 4850 1 1.2190e+02 4.1840e+02 ; C - N
- CA
4847 4846 4848 1 1.2290e+02 6.6944e+02 ; O - C
- N
4848 4850 4851 1 1.0950e+02 4.1840e+02 ; N - CA
- HA
4848 4850 4852 1 1.0970e+02 6.6944e+02 ; N - CA
- CB
4848 4850 4868 1 1.1010e+02 5.2718e+02 ; N - CA
- C
4849 4848 4850 1 1.1804e+02 4.1840e+02 ; H - N
- CA
4850 4852 4853 1 1.0950e+02 4.1840e+02 ; CA - CB
- HB2
4850 4852 4854 1 1.0950e+02 4.1840e+02 ; CA - CB
- HB3
4850 4852 4855 1 1.0950e+02 3.3472e+02 ; CA - CB
- CG
4850 4868 4869 1 1.2040e+02 6.6944e+02 ; CA - C
- O

4850 4868 4870 1 1.1660e+02 5.8576e+02 ; CA - C
- N
4851 4850 4852 1 1.0950e+02 4.1840e+02 ; HA - CA
- CB
4851 4850 4868 1 1.0950e+02 4.1840e+02 ; HA - CA
- C
4852 4850 4868 1 1.1110e+02 5.2718e+02 ; CB - CA
- C
4852 4855 4856 1 1.0950e+02 4.1840e+02 ; CB - CG
- HG2
4852 4855 4857 1 1.0950e+02 4.1840e+02 ; CB - CG
- HG3
4852 4855 4858 1 1.0950e+02 3.3472e+02 ; CB - CG
- CD
4853 4852 4854 1 1.0950e+02 2.9288e+02 ; HB2 - CB
- HB3
4853 4852 4855 1 1.0950e+02 4.1840e+02 ; HB2 - CB
- CG
4854 4852 4855 1 1.0950e+02 4.1840e+02 ; HB3 - CB
- CG
4855 4858 4859 1 1.0950e+02 4.1840e+02 ; CG - CD
- HD2
4855 4858 4860 1 1.0950e+02 4.1840e+02 ; CG - CD
- HD3
4855 4858 4861 1 1.0950e+02 3.3472e+02 ; CG - CD
- CE
4856 4855 4857 1 1.0950e+02 2.9288e+02 ; HG2 - CG
- HG3
4856 4855 4858 1 1.0950e+02 4.1840e+02 ; HG2 - CG
- CD
4857 4855 4858 1 1.0950e+02 4.1840e+02 ; HG3 - CG
- CD
4858 4861 4862 1 1.0950e+02 4.1840e+02 ; CD - CE
- HE2
4858 4861 4863 1 1.0950e+02 4.1840e+02 ; CD - CE
- HE3
4858 4861 4864 1 1.1120e+02 6.6944e+02 ; CD - CE
- NZ
4859 4858 4860 1 1.0950e+02 2.9288e+02 ; HD2 - CD
- HD3
4859 4858 4861 1 1.0950e+02 4.1840e+02 ; HD2 - CD
- CE
4860 4858 4861 1 1.0950e+02 4.1840e+02 ; HD3 - CD
- CE
4861 4864 4865 1 1.0950e+02 4.1840e+02 ; CE - NZ
- HZ1
4861 4864 4866 1 1.0950e+02 4.1840e+02 ; CE - NZ
- HZ2
4861 4864 4867 1 1.0950e+02 4.1840e+02 ; CE - NZ
- HZ3
4862 4861 4863 1 1.0950e+02 2.9288e+02 ; HE2 - CE
- HE3

4862 4861 4864 1 1.0950e+02 4.1840e+02 ; HE2 - CE
- NZ
4863 4861 4864 1 1.0950e+02 4.1840e+02 ; HE3 - CE
- NZ
4865 4864 4866 1 1.0950e+02 2.9288e+02 ; HZ1 - NZ
- HZ2
4865 4864 4867 1 1.0950e+02 2.9288e+02 ; HZ1 - NZ
- HZ3
4866 4864 4867 1 1.0950e+02 2.9288e+02 ; HZ2 - NZ
- HZ3
4868 4870 4871 1 1.2000e+02 4.1840e+02 ; C - N
- H
4868 4870 4872 1 1.2190e+02 4.1840e+02 ; C - N
- CA
4869 4868 4870 1 1.2290e+02 6.6944e+02 ; O - C
- N
4870 4872 4873 1 1.0950e+02 4.1840e+02 ; N - CA
- HA
4870 4872 4874 1 1.0970e+02 6.6944e+02 ; N - CA
- CB
4870 4872 4887 1 1.1010e+02 5.2718e+02 ; N - CA
- C
4871 4870 4872 1 1.1804e+02 4.1840e+02 ; H - N
- CA
4872 4874 4875 1 1.0950e+02 4.1840e+02 ; CA - CB
- HB
4872 4874 4876 1 1.0950e+02 3.3472e+02 ; CA - CB
- CG2
4872 4874 4880 1 1.0950e+02 3.3472e+02 ; CA - CB
- CG1
4872 4887 4888 1 1.2040e+02 6.6944e+02 ; CA - C
- O
4872 4887 4889 1 1.1660e+02 5.8576e+02 ; CA - C
- N
4873 4872 4874 1 1.0950e+02 4.1840e+02 ; HA - CA
- CB
4873 4872 4887 1 1.0950e+02 4.1840e+02 ; HA - CA
- C
4874 4872 4887 1 1.1110e+02 5.2718e+02 ; CB - CA
- C
4874 4876 4877 1 1.0950e+02 4.1840e+02 ; CB - CG2
- HG21
4874 4876 4878 1 1.0950e+02 4.1840e+02 ; CB - CG2
- HG22
4874 4876 4879 1 1.0950e+02 4.1840e+02 ; CB - CG2
- HG23
4874 4880 4881 1 1.0950e+02 4.1840e+02 ; CB - CG1
- HG12
4874 4880 4882 1 1.0950e+02 4.1840e+02 ; CB - CG1
- HG13
4874 4880 4883 1 1.0950e+02 3.3472e+02 ; CB - CG1
- CD1

4875 4874 4876 1 1.0950e+02 4.1840e+02 ; HB - CB
- CG2
4875 4874 4880 1 1.0950e+02 4.1840e+02 ; HB - CB
- CG1
4876 4874 4880 1 1.0950e+02 3.3472e+02 ; CG2 - CB
- CG1
4877 4876 4878 1 1.0950e+02 2.9288e+02 ; HG21 - CG2
- HG22
4877 4876 4879 1 1.0950e+02 2.9288e+02 ; HG21 - CG2
- HG23
4878 4876 4879 1 1.0950e+02 2.9288e+02 ; HG22 - CG2
- HG23
4880 4883 4884 1 1.0950e+02 4.1840e+02 ; CG1 - CD1
- HD11
4880 4883 4885 1 1.0950e+02 4.1840e+02 ; CG1 - CD1
- HD12
4880 4883 4886 1 1.0950e+02 4.1840e+02 ; CG1 - CD1
- HD13
4881 4880 4882 1 1.0950e+02 2.9288e+02 ; HG12 - CG1
- HG13
4881 4880 4883 1 1.0950e+02 4.1840e+02 ; HG12 - CG1
- CD1
4882 4880 4883 1 1.0950e+02 4.1840e+02 ; HG13 - CG1
- CD1
4884 4883 4885 1 1.0950e+02 2.9288e+02 ; HD11 - CD1
- HD12
4884 4883 4886 1 1.0950e+02 2.9288e+02 ; HD11 - CD1
- HD13
4885 4883 4886 1 1.0950e+02 2.9288e+02 ; HD12 - CD1
- HD13
4887 4889 4890 1 1.2000e+02 4.1840e+02 ; C - N
- H
4887 4889 4891 1 1.2190e+02 4.1840e+02 ; C - N
- CA
4888 4887 4889 1 1.2290e+02 6.6944e+02 ; O - C
- N
4889 4891 4892 1 1.0950e+02 4.1840e+02 ; N - CA
- HA
4889 4891 4893 1 1.0970e+02 6.6944e+02 ; N - CA
- CB
4889 4891 4897 1 1.1010e+02 5.2718e+02 ; N - CA
- C
4890 4889 4891 1 1.1804e+02 4.1840e+02 ; H - N
- CA
4891 4893 4894 1 1.0950e+02 4.1840e+02 ; CA - CB
- HB1
4891 4893 4895 1 1.0950e+02 4.1840e+02 ; CA - CB
- HB2
4891 4893 4896 1 1.0950e+02 4.1840e+02 ; CA - CB
- HB3
4891 4897 4898 1 1.2040e+02 6.6944e+02 ; CA - C
- O

4891 4897 4899 1 1.1660e+02 5.8576e+02 ; CA - C
- N
4892 4891 4893 1 1.0950e+02 4.1840e+02 ; HA - CA
- CB
4892 4891 4897 1 1.0950e+02 4.1840e+02 ; HA - CA
- C
4893 4891 4897 1 1.1110e+02 5.2718e+02 ; CB - CA
- C
4894 4893 4895 1 1.0950e+02 2.9288e+02 ; HB1 - CB
- HB2
4894 4893 4896 1 1.0950e+02 2.9288e+02 ; HB1 - CB
- HB3
4895 4893 4896 1 1.0950e+02 2.9288e+02 ; HB2 - CB
- HB3
4897 4899 4900 1 1.2000e+02 4.1840e+02 ; C - N
- H
4897 4899 4901 1 1.2190e+02 4.1840e+02 ; C - N
- CA
4898 4897 4899 1 1.2290e+02 6.6944e+02 ; O - C
- N
4899 4901 4902 1 1.0950e+02 4.1840e+02 ; N - CA
- HA
4899 4901 4903 1 1.0970e+02 6.6944e+02 ; N - CA
- CB
4899 4901 4916 1 1.1010e+02 5.2718e+02 ; N - CA
- C
4900 4899 4901 1 1.1804e+02 4.1840e+02 ; H - N
- CA
4901 4903 4904 1 1.0950e+02 4.1840e+02 ; CA - CB
- HB2
4901 4903 4905 1 1.0950e+02 4.1840e+02 ; CA - CB
- HB3
4901 4903 4906 1 1.0950e+02 3.3472e+02 ; CA - CB
- CG
4901 4916 4917 1 1.2040e+02 6.6944e+02 ; CA - C
- O
4901 4916 4918 1 1.1660e+02 5.8576e+02 ; CA - C
- N
4902 4901 4903 1 1.0950e+02 4.1840e+02 ; HA - CA
- CB
4902 4901 4916 1 1.0950e+02 4.1840e+02 ; HA - CA
- C
4903 4901 4916 1 1.1110e+02 5.2718e+02 ; CB - CA
- C
4903 4906 4907 1 1.0950e+02 4.1840e+02 ; CB - CG
- HG
4903 4906 4908 1 1.0950e+02 3.3472e+02 ; CB - CG
- CD1
4903 4906 4912 1 1.0950e+02 3.3472e+02 ; CB - CG
- CD2
4904 4903 4905 1 1.0950e+02 2.9288e+02 ; HB2 - CB
- HB3

```
  4904   4903   4906      1    1.0950e+02    4.1840e+02 ;    HB2 - CB
- CG
  4905   4903   4906      1    1.0950e+02    4.1840e+02 ;    HB3 - CB
- CG
  4906   4908   4909      1    1.0950e+02    4.1840e+02 ;     CG - CD1
- HD11
  4906   4908   4910      1    1.0950e+02    4.1840e+02 ;     CG - CD1
- HD12
  4906   4908   4911      1    1.0950e+02    4.1840e+02 ;     CG - CD1
- HD13
  4906   4912   4913      1    1.0950e+02    4.1840e+02 ;     CG - CD2
- HD21
  4906   4912   4914      1    1.0950e+02    4.1840e+02 ;     CG - CD2
- HD22
  4906   4912   4915      1    1.0950e+02    4.1840e+02 ;     CG - CD2
- HD23
  4907   4906   4908      1    1.0950e+02    4.1840e+02 ;     HG - CG
- CD1
  4907   4906   4912      1    1.0950e+02    4.1840e+02 ;     HG - CG
- CD2
  4908   4906   4912      1    1.0950e+02    3.3472e+02 ;    CD1 - CG
- CD2
  4909   4908   4910      1    1.0950e+02    2.9288e+02 ;   HD11 - CD1
- HD12
  4909   4908   4911      1    1.0950e+02    2.9288e+02 ;   HD11 - CD1
- HD13
  4910   4908   4911      1    1.0950e+02    2.9288e+02 ;   HD12 - CD1
- HD13
  4913   4912   4914      1    1.0950e+02    2.9288e+02 ;   HD21 - CD2
- HD22
  4913   4912   4915      1    1.0950e+02    2.9288e+02 ;   HD21 - CD2
- HD23
  4914   4912   4915      1    1.0950e+02    2.9288e+02 ;   HD22 - CD2
- HD23
  4916   4918   4919      1    1.2000e+02    4.1840e+02 ;      C - N
- H
  4916   4918   4920      1    1.2190e+02    4.1840e+02 ;      C - N
- CA
  4917   4916   4918      1    1.2290e+02    6.6944e+02 ;      O - C
- N
  4918   4920   4921      1    1.0950e+02    4.1840e+02 ;      N - CA
- HA
  4918   4920   4922      1    1.0970e+02    6.6944e+02 ;      N - CA
- CB
  4918   4920   4930      1    1.1010e+02    5.2718e+02 ;      N - CA
- C
  4919   4918   4920      1    1.1804e+02    4.1840e+02 ;      H - N
- CA
  4920   4922   4923      1    1.0950e+02    4.1840e+02 ;     CA - CB
- HB
  4920   4922   4924      1    1.0950e+02    3.3472e+02 ;     CA - CB
- CG2
```

4920 4922 4928 1 1.0950e+02 4.1840e+02 ; CA - CB
- OG1
4920 4930 4931 1 1.2040e+02 6.6944e+02 ; CA - C
- O
4920 4930 4932 1 1.1660e+02 5.8576e+02 ; CA - C
- N
4921 4920 4922 1 1.0950e+02 4.1840e+02 ; HA - CA
- CB
4921 4920 4930 1 1.0950e+02 4.1840e+02 ; HA - CA
- C
4922 4920 4930 1 1.1110e+02 5.2718e+02 ; CB - CA
- C
4922 4924 4925 1 1.0950e+02 4.1840e+02 ; CB - CG2
- HG21
4922 4924 4926 1 1.0950e+02 4.1840e+02 ; CB - CG2
- HG22
4922 4924 4927 1 1.0950e+02 4.1840e+02 ; CB - CG2
- HG23
4922 4928 4929 1 1.0850e+02 4.6024e+02 ; CB - OG1
- HG1
4923 4922 4924 1 1.0950e+02 4.1840e+02 ; HB - CB
- CG2
4923 4922 4928 1 1.0950e+02 4.1840e+02 ; HB - CB
- OG1
4924 4922 4928 1 1.0950e+02 4.1840e+02 ; CG2 - CB
- OG1
4925 4924 4926 1 1.0950e+02 2.9288e+02 ; HG21 - CG2
- HG22
4925 4924 4927 1 1.0950e+02 2.9288e+02 ; HG21 - CG2
- HG23
4926 4924 4927 1 1.0950e+02 2.9288e+02 ; HG22 - CG2
- HG23
4930 4932 4933 1 1.2000e+02 4.1840e+02 ; C - N
- H
4930 4932 4934 1 1.2190e+02 4.1840e+02 ; C - N
- CA
4931 4930 4932 1 1.2290e+02 6.6944e+02 ; O - C
- N
4932 4934 4935 1 1.0950e+02 4.1840e+02 ; N - CA
- HA
4932 4934 4936 1 1.0970e+02 6.6944e+02 ; N - CA
- CB
4932 4934 4949 1 1.1010e+02 5.2718e+02 ; N - CA
- C
4933 4932 4934 1 1.1804e+02 4.1840e+02 ; H - N
- CA
4934 4936 4937 1 1.0950e+02 4.1840e+02 ; CA - CB
- HB
4934 4936 4938 1 1.0950e+02 3.3472e+02 ; CA - CB
- CG2
4934 4936 4942 1 1.0950e+02 3.3472e+02 ; CA - CB
- CG1

4934 4949 4950 1 1.2040e+02 6.6944e+02 ; CA - C
- O
4934 4949 4951 1 1.1660e+02 5.8576e+02 ; CA - C
- N
4935 4934 4936 1 1.0950e+02 4.1840e+02 ; HA - CA
- CB
4935 4934 4949 1 1.0950e+02 4.1840e+02 ; HA - CA
- C
4936 4934 4949 1 1.1110e+02 5.2718e+02 ; CB - CA
- C
4936 4938 4939 1 1.0950e+02 4.1840e+02 ; CB - CG2
- HG21
4936 4938 4940 1 1.0950e+02 4.1840e+02 ; CB - CG2
- HG22
4936 4938 4941 1 1.0950e+02 4.1840e+02 ; CB - CG2
- HG23
4936 4942 4943 1 1.0950e+02 4.1840e+02 ; CB - CG1
- HG12
4936 4942 4944 1 1.0950e+02 4.1840e+02 ; CB - CG1
- HG13
4936 4942 4945 1 1.0950e+02 3.3472e+02 ; CB - CG1
- CD1
4937 4936 4938 1 1.0950e+02 4.1840e+02 ; HB - CB
- CG2
4937 4936 4942 1 1.0950e+02 4.1840e+02 ; HB - CB
- CG1
4938 4936 4942 1 1.0950e+02 3.3472e+02 ; CG2 - CB
- CG1
4939 4938 4940 1 1.0950e+02 2.9288e+02 ; HG21 - CG2
- HG22
4939 4938 4941 1 1.0950e+02 2.9288e+02 ; HG21 - CG2
- HG23
4940 4938 4941 1 1.0950e+02 2.9288e+02 ; HG22 - CG2
- HG23
4942 4945 4946 1 1.0950e+02 4.1840e+02 ; CG1 - CD1
- HD11
4942 4945 4947 1 1.0950e+02 4.1840e+02 ; CG1 - CD1
- HD12
4942 4945 4948 1 1.0950e+02 4.1840e+02 ; CG1 - CD1
- HD13
4943 4942 4944 1 1.0950e+02 2.9288e+02 ; HG12 - CG1
- HG13
4943 4942 4945 1 1.0950e+02 4.1840e+02 ; HG12 - CG1
- CD1
4944 4942 4945 1 1.0950e+02 4.1840e+02 ; HG13 - CG1
- CD1
4946 4945 4947 1 1.0950e+02 2.9288e+02 ; HD11 - CD1
- HD12
4946 4945 4948 1 1.0950e+02 2.9288e+02 ; HD11 - CD1
- HD13
4947 4945 4948 1 1.0950e+02 2.9288e+02 ; HD12 - CD1
- HD13

4949 4951 4952 1 1.2000e+02 4.1840e+02 ; C - N
- H
4949 4951 4953 1 1.2190e+02 4.1840e+02 ; C - N
- CA
4950 4949 4951 1 1.2290e+02 6.6944e+02 ; O - C
- N
4951 4953 4954 1 1.0950e+02 4.1840e+02 ; N - CA
- HA
4951 4953 4955 1 1.0970e+02 6.6944e+02 ; N - CA
- CB
4951 4953 4968 1 1.1010e+02 5.2718e+02 ; N - CA
- C
4952 4951 4953 1 1.1804e+02 4.1840e+02 ; H - N
- CA
4953 4955 4956 1 1.0950e+02 4.1840e+02 ; CA - CB
- HB
4953 4955 4957 1 1.0950e+02 3.3472e+02 ; CA - CB
- CG2
4953 4955 4961 1 1.0950e+02 3.3472e+02 ; CA - CB
- CG1
4953 4968 4969 1 1.2040e+02 6.6944e+02 ; CA - C
- O
4953 4968 4970 1 1.1660e+02 5.8576e+02 ; CA - C
- N
4954 4953 4955 1 1.0950e+02 4.1840e+02 ; HA - CA
- CB
4954 4953 4968 1 1.0950e+02 4.1840e+02 ; HA - CA
- C
4955 4953 4968 1 1.1110e+02 5.2718e+02 ; CB - CA
- C
4955 4957 4958 1 1.0950e+02 4.1840e+02 ; CB - CG2
- HG21
4955 4957 4959 1 1.0950e+02 4.1840e+02 ; CB - CG2
- HG22
4955 4957 4960 1 1.0950e+02 4.1840e+02 ; CB - CG2
- HG23
4955 4961 4962 1 1.0950e+02 4.1840e+02 ; CB - CG1
- HG12
4955 4961 4963 1 1.0950e+02 4.1840e+02 ; CB - CG1
- HG13
4955 4961 4964 1 1.0950e+02 3.3472e+02 ; CB - CG1
- CD1
4956 4955 4957 1 1.0950e+02 4.1840e+02 ; HB - CB
- CG2
4956 4955 4961 1 1.0950e+02 4.1840e+02 ; HB - CB
- CG1
4957 4955 4961 1 1.0950e+02 3.3472e+02 ; CG2 - CB
- CG1
4958 4957 4959 1 1.0950e+02 2.9288e+02 ; HG21 - CG2
- HG22
4958 4957 4960 1 1.0950e+02 2.9288e+02 ; HG21 - CG2
- HG23

4959 4957 4960 1 1.0950e+02 2.9288e+02 ; HG22 - CG2 - HG23
4961 4964 4965 1 1.0950e+02 4.1840e+02 ; CG1 - CD1 - HD11
4961 4964 4966 1 1.0950e+02 4.1840e+02 ; CG1 - CD1 - HD12
4961 4964 4967 1 1.0950e+02 4.1840e+02 ; CG1 - CD1 - HD13
4962 4961 4963 1 1.0950e+02 2.9288e+02 ; HG12 - CG1 - HG13
4962 4961 4964 1 1.0950e+02 4.1840e+02 ; HG12 - CG1 - CD1
4963 4961 4964 1 1.0950e+02 4.1840e+02 ; HG13 - CG1 - CD1
4965 4964 4966 1 1.0950e+02 2.9288e+02 ; HD11 - CD1 - HD12
4965 4964 4967 1 1.0950e+02 2.9288e+02 ; HD11 - CD1 - HD13
4966 4964 4967 1 1.0950e+02 2.9288e+02 ; HD12 - CD1 - HD13
4968 4970 4971 1 1.2000e+02 4.1840e+02 ; C - N - H
4968 4970 4972 1 1.2190e+02 4.1840e+02 ; C - N - CA
4969 4968 4970 1 1.2290e+02 6.6944e+02 ; O - C - N
4970 4972 4973 1 1.0950e+02 4.1840e+02 ; N - CA - HA
4970 4972 4974 1 1.0970e+02 6.6944e+02 ; N - CA - CB
4970 4972 4978 1 1.1010e+02 5.2718e+02 ; N - CA - C
4971 4970 4972 1 1.1804e+02 4.1840e+02 ; H - N - CA
4972 4974 4975 1 1.0950e+02 4.1840e+02 ; CA - CB - HB1
4972 4974 4976 1 1.0950e+02 4.1840e+02 ; CA - CB - HB2
4972 4974 4977 1 1.0950e+02 4.1840e+02 ; CA - CB - HB3
4972 4978 4979 1 1.2040e+02 6.6944e+02 ; CA - C - O
4972 4978 4980 1 1.1660e+02 5.8576e+02 ; CA - C - N
4973 4972 4974 1 1.0950e+02 4.1840e+02 ; HA - CA - CB
4973 4972 4978 1 1.0950e+02 4.1840e+02 ; HA - CA - C
4974 4972 4978 1 1.1110e+02 5.2718e+02 ; CB - CA - C
4975 4974 4976 1 1.0950e+02 2.9288e+02 ; HB1 - CB - HB2

4975 4974 4977 1 1.0950e+02 2.9288e+02 ; HB1 - CB
- HB3
4976 4974 4977 1 1.0950e+02 2.9288e+02 ; HB2 - CB
- HB3
4978 4980 4981 1 1.2000e+02 4.1840e+02 ; C - N
- H
4978 4980 4982 1 1.2190e+02 4.1840e+02 ; C - N
- CA
4979 4978 4980 1 1.2290e+02 6.6944e+02 ; O - C
- N
4980 4982 4983 1 1.0950e+02 4.1840e+02 ; N - CA
- HA
4980 4982 4984 1 1.0970e+02 6.6944e+02 ; N - CA
- CB
4980 4982 5000 1 1.1010e+02 5.2718e+02 ; N - CA
- C
4981 4980 4982 1 1.1804e+02 4.1840e+02 ; H - N
- CA
4982 4984 4985 1 1.0950e+02 4.1840e+02 ; CA - CB
- HB2
4982 4984 4986 1 1.0950e+02 4.1840e+02 ; CA - CB
- HB3
4982 4984 4987 1 1.0950e+02 3.3472e+02 ; CA - CB
- CG
4982 5000 5001 1 1.2040e+02 6.6944e+02 ; CA - C
- O
4982 5000 5002 1 1.1660e+02 5.8576e+02 ; CA - C
- N
4983 4982 4984 1 1.0950e+02 4.1840e+02 ; HA - CA
- CB
4983 4982 5000 1 1.0950e+02 4.1840e+02 ; HA - CA
- C
4984 4982 5000 1 1.1110e+02 5.2718e+02 ; CB - CA
- C
4984 4987 4988 1 1.0950e+02 4.1840e+02 ; CB - CG
- HG2
4984 4987 4989 1 1.0950e+02 4.1840e+02 ; CB - CG
- HG3
4984 4987 4990 1 1.0950e+02 3.3472e+02 ; CB - CG
- CD
4985 4984 4986 1 1.0950e+02 2.9288e+02 ; HB2 - CB
- HB3
4985 4984 4987 1 1.0950e+02 4.1840e+02 ; HB2 - CB
- CG
4986 4984 4987 1 1.0950e+02 4.1840e+02 ; HB3 - CB
- CG
4987 4990 4991 1 1.0950e+02 4.1840e+02 ; CG - CD
- HD2
4987 4990 4992 1 1.0950e+02 4.1840e+02 ; CG - CD
- HD3
4987 4990 4993 1 1.0950e+02 3.3472e+02 ; CG - CD
- CE

```
  4988   4987   4989      1    1.0950e+02    2.9288e+02 ;    HG2 - CG
- HG3
  4988   4987   4990      1    1.0950e+02    4.1840e+02 ;    HG2 - CG
- CD
  4989   4987   4990      1    1.0950e+02    4.1840e+02 ;    HG3 - CG
- CD
  4990   4993   4994      1    1.0950e+02    4.1840e+02 ;     CD - CE
- HE2
  4990   4993   4995      1    1.0950e+02    4.1840e+02 ;     CD - CE
- HE3
  4990   4993   4996      1    1.1120e+02    6.6944e+02 ;     CD - CE
- NZ
  4991   4990   4992      1    1.0950e+02    2.9288e+02 ;    HD2 - CD
- HD3
  4991   4990   4993      1    1.0950e+02    4.1840e+02 ;    HD2 - CD
- CE
  4992   4990   4993      1    1.0950e+02    4.1840e+02 ;    HD3 - CD
- CE
  4993   4996   4997      1    1.0950e+02    4.1840e+02 ;     CE - NZ
- HZ1
  4993   4996   4998      1    1.0950e+02    4.1840e+02 ;     CE - NZ
- HZ2
  4993   4996   4999      1    1.0950e+02    4.1840e+02 ;     CE - NZ
- HZ3
  4994   4993   4995      1    1.0950e+02    2.9288e+02 ;    HE2 - CE
- HE3
  4994   4993   4996      1    1.0950e+02    4.1840e+02 ;    HE2 - CE
- NZ
  4995   4993   4996      1    1.0950e+02    4.1840e+02 ;    HE3 - CE
- NZ
  4997   4996   4998      1    1.0950e+02    2.9288e+02 ;    HZ1 - NZ
- HZ2
  4997   4996   4999      1    1.0950e+02    2.9288e+02 ;    HZ1 - NZ
- HZ3
  4998   4996   4999      1    1.0950e+02    2.9288e+02 ;    HZ2 - NZ
- HZ3
  5000   5002   5003      1    1.2000e+02    4.1840e+02 ;      C - N
- H
  5000   5002   5004      1    1.2190e+02    4.1840e+02 ;      C - N
- CA
  5001   5000   5002      1    1.2290e+02    6.6944e+02 ;      O - C
- N
  5002   5004   5005      1    1.0950e+02    4.1840e+02 ;      N - CA
- HA
  5002   5004   5006      1    1.0970e+02    6.6944e+02 ;      N - CA
- CB
  5002   5004   5018      1    1.1010e+02    5.2718e+02 ;      N - CA
- C
  5003   5002   5004      1    1.1804e+02    4.1840e+02 ;      H - N
- CA
  5004   5006   5007      1    1.0950e+02    4.1840e+02 ;     CA - CB
- HB2
```

```
  5004   5006   5008       1    1.0950e+02    4.1840e+02 ;     CA - CB
- HB3
  5004   5006   5009       1    1.1310e+02    5.2718e+02 ;     CA - CB
- CG
  5004   5018   5019       1    1.2040e+02    6.6944e+02 ;     CA - C
- O
  5004   5018   5020       1    1.1660e+02    5.8576e+02 ;     CA - C
- N
  5005   5004   5006       1    1.0950e+02    4.1840e+02 ;     HA - CA
- CB
  5005   5004   5018       1    1.0950e+02    4.1840e+02 ;     HA - CA
- C
  5006   5004   5018       1    1.1110e+02    5.2718e+02 ;     CB - CA
- C
  5006   5009   5010       1    1.2000e+02    5.8576e+02 ;     CB - CG
- ND1
  5006   5009   5016       1    1.2000e+02    5.8576e+02 ;     CB - CG
- CD2
  5007   5006   5008       1    1.0950e+02    2.9288e+02 ;    HB2 - CB
- HB3
  5007   5006   5009       1    1.0950e+02    4.1840e+02 ;    HB2 - CB
- CG
  5008   5006   5009       1    1.0950e+02    4.1840e+02 ;    HB3 - CB
- CG
  5009   5010   5011       1    1.2000e+02    4.1840e+02 ;     CG - ND1
- HD1
  5009   5010   5012       1    1.2000e+02    5.8576e+02 ;     CG - ND1
- CE1
  5009   5016   5014       1    1.2000e+02    5.8576e+02 ;     CG - CD2
- NE2
  5009   5016   5017       1    1.2000e+02    4.1840e+02 ;     CG - CD2
- HD2
  5010   5009   5016       1    1.2000e+02    5.8576e+02 ;    ND1 - CG
- CD2
  5010   5012   5013       1    1.2000e+02    4.1840e+02 ;    ND1 - CE1
- HE1
  5010   5012   5014       1    1.2000e+02    5.8576e+02 ;    ND1 - CE1
- NE2
  5011   5010   5012       1    1.2000e+02    4.1840e+02 ;    HD1 - ND1
- CE1
  5012   5014   5015       1    1.2000e+02    4.1840e+02 ;    CE1 - NE2
- HE2
  5012   5014   5016       1    1.2000e+02    5.8576e+02 ;    CE1 - NE2
- CD2
  5013   5012   5014       1    1.2000e+02    4.1840e+02 ;    HE1 - CE1
- NE2
  5014   5016   5017       1    1.2000e+02    4.1840e+02 ;    NE2 - CD2
- HD2
  5015   5014   5016       1    1.2000e+02    4.1840e+02 ;    HE2 - NE2
- CD2
  5018   5020   5021       1    1.2000e+02    4.1840e+02 ;      C - N
- H
```

5018 5020 5022 1 1.2190e+02 4.1840e+02 ; C - N
- CA
5019 5018 5020 1 1.2290e+02 6.6944e+02 ; O - C
- N
5020 5022 5023 1 1.0950e+02 4.1840e+02 ; N - CA
- HA
5020 5022 5024 1 1.0970e+02 6.6944e+02 ; N - CA
- CB
5020 5022 5037 1 1.1010e+02 5.2718e+02 ; N - CA
- C
5021 5020 5022 1 1.1804e+02 4.1840e+02 ; H - N
- CA
5022 5024 5025 1 1.0950e+02 4.1840e+02 ; CA - CB
- HB2
5022 5024 5026 1 1.0950e+02 4.1840e+02 ; CA - CB
- HB3
5022 5024 5027 1 1.0950e+02 3.3472e+02 ; CA - CB
- CG
5022 5037 5038 1 1.2040e+02 6.6944e+02 ; CA - C
- O
5022 5037 5039 1 1.1660e+02 5.8576e+02 ; CA - C
- N
5023 5022 5024 1 1.0950e+02 4.1840e+02 ; HA - CA
- CB
5023 5022 5037 1 1.0950e+02 4.1840e+02 ; HA - CA
- C
5024 5022 5037 1 1.1110e+02 5.2718e+02 ; CB - CA
- C
5024 5027 5028 1 1.0950e+02 4.1840e+02 ; CB - CG
- HG
5024 5027 5029 1 1.0950e+02 3.3472e+02 ; CB - CG
- CD1
5024 5027 5033 1 1.0950e+02 3.3472e+02 ; CB - CG
- CD2
5025 5024 5026 1 1.0950e+02 2.9288e+02 ; HB2 - CB
- HB3
5025 5024 5027 1 1.0950e+02 4.1840e+02 ; HB2 - CB
- CG
5026 5024 5027 1 1.0950e+02 4.1840e+02 ; HB3 - CB
- CG
5027 5029 5030 1 1.0950e+02 4.1840e+02 ; CG - CD1
- HD11
5027 5029 5031 1 1.0950e+02 4.1840e+02 ; CG - CD1
- HD12
5027 5029 5032 1 1.0950e+02 4.1840e+02 ; CG - CD1
- HD13
5027 5033 5034 1 1.0950e+02 4.1840e+02 ; CG - CD2
- HD21
5027 5033 5035 1 1.0950e+02 4.1840e+02 ; CG - CD2
- HD22
5027 5033 5036 1 1.0950e+02 4.1840e+02 ; CG - CD2
- HD23

5028 5027 5029 1 1.0950e+02 4.1840e+02 ; HG - CG
- CD1
5028 5027 5033 1 1.0950e+02 4.1840e+02 ; HG - CG
- CD2
5029 5027 5033 1 1.0950e+02 3.3472e+02 ; CD1 - CG
- CD2
5030 5029 5031 1 1.0950e+02 2.9288e+02 ; HD11 - CD1
- HD12
5030 5029 5032 1 1.0950e+02 2.9288e+02 ; HD11 - CD1
- HD13
5031 5029 5032 1 1.0950e+02 2.9288e+02 ; HD12 - CD1
- HD13
5034 5033 5035 1 1.0950e+02 2.9288e+02 ; HD21 - CD2
- HD22
5034 5033 5036 1 1.0950e+02 2.9288e+02 ; HD21 - CD2
- HD23
5035 5033 5036 1 1.0950e+02 2.9288e+02 ; HD22 - CD2
- HD23
5037 5039 5040 1 1.2000e+02 4.1840e+02 ; C - N
- H
5037 5039 5041 1 1.2190e+02 4.1840e+02 ; C - N
- CA
5038 5037 5039 1 1.2290e+02 6.6944e+02 ; O - C
- N
5039 5041 5042 1 1.0950e+02 4.1840e+02 ; N - CA
- HA
5039 5041 5043 1 1.0970e+02 6.6944e+02 ; N - CA
- CB
5039 5041 5048 1 1.1010e+02 5.2718e+02 ; N - CA
- C
5040 5039 5041 1 1.1804e+02 4.1840e+02 ; H - N
- CA
5041 5043 5044 1 1.0950e+02 4.1840e+02 ; CA - CB
- HB2
5041 5043 5045 1 1.0950e+02 4.1840e+02 ; CA - CB
- HB3
5041 5043 5046 1 1.0860e+02 4.1840e+02 ; CA - CB
- SG
5041 5048 5049 1 1.2040e+02 6.6944e+02 ; CA - C
- O
5041 5048 5050 1 1.1660e+02 5.8576e+02 ; CA - C
- N
5042 5041 5043 1 1.0950e+02 4.1840e+02 ; HA - CA
- CB
5042 5041 5048 1 1.0950e+02 4.1840e+02 ; HA - CA
- C
5043 5041 5048 1 1.1110e+02 5.2718e+02 ; CB - CA
- C
5043 5046 5047 1 9.6000e+01 3.5982e+02 ; CB - SG
- HG
5044 5043 5045 1 1.0950e+02 2.9288e+02 ; HB2 - CB
- HB3

5044 5043 5046 1 1.0950e+02 4.1840e+02 ; HB2 - CB
- SG
5045 5043 5046 1 1.0950e+02 4.1840e+02 ; HB3 - CB
- SG
5048 5050 5051 1 1.2000e+02 4.1840e+02 ; C - N
- H
5048 5050 5052 1 1.2190e+02 4.1840e+02 ; C - N
- CA
5049 5048 5050 1 1.2290e+02 6.6944e+02 ; O - C
- N
5050 5052 5053 1 1.0950e+02 4.1840e+02 ; N - CA
- HA
5050 5052 5054 1 1.0970e+02 6.6944e+02 ; N - CA
- CB
5050 5052 5060 1 1.1010e+02 5.2718e+02 ; N - CA
- C
5051 5050 5052 1 1.1804e+02 4.1840e+02 ; H - N
- CA
5052 5054 5055 1 1.0950e+02 4.1840e+02 ; CA - CB
- HB2
5052 5054 5056 1 1.0950e+02 4.1840e+02 ; CA - CB
- HB3
5052 5054 5057 1 1.1110e+02 5.2718e+02 ; CA - CB
- CG
5052 5060 5061 1 1.2040e+02 6.6944e+02 ; CA - C
- O
5052 5060 5062 1 1.1660e+02 5.8576e+02 ; CA - C
- N
5053 5052 5054 1 1.0950e+02 4.1840e+02 ; HA - CA
- CB
5053 5052 5060 1 1.0950e+02 4.1840e+02 ; HA - CA
- C
5054 5052 5060 1 1.1110e+02 5.2718e+02 ; CB - CA
- C
5054 5057 5058 1 1.1700e+02 5.8576e+02 ; CB - CG
- OD1
5054 5057 5059 1 1.1700e+02 5.8576e+02 ; CB - CG
- OD2
5055 5054 5056 1 1.0950e+02 2.9288e+02 ; HB2 - CB
- HB3
5055 5054 5057 1 1.0950e+02 4.1840e+02 ; HB2 - CB
- CG
5056 5054 5057 1 1.0950e+02 4.1840e+02 ; HB3 - CB
- CG
5058 5057 5059 1 1.2600e+02 6.6944e+02 ; OD1 - CG
- OD2
5060 5062 5063 1 1.2000e+02 4.1840e+02 ; C - N
- H
5060 5062 5064 1 1.2190e+02 4.1840e+02 ; C - N
- CA
5061 5060 5062 1 1.2290e+02 6.6944e+02 ; O - C
- N

5062 5064 5065 1 1.0950e+02 4.1840e+02 ; N - CA - HA
5062 5064 5066 1 1.0970e+02 6.6944e+02 ; N - CA - CB
5062 5064 5076 1 1.1010e+02 5.2718e+02 ; N - CA - C
5063 5062 5064 1 1.1804e+02 4.1840e+02 ; H - N - CA
5064 5066 5067 1 1.0950e+02 4.1840e+02 ; CA - CB - HB
5064 5066 5068 1 1.0950e+02 3.3472e+02 ; CA - CB - CG1
5064 5066 5072 1 1.0950e+02 3.3472e+02 ; CA - CB - CG2
5064 5076 5077 1 1.2040e+02 6.6944e+02 ; CA - C - O
5064 5076 5078 1 1.1660e+02 5.8576e+02 ; CA - C - N
5065 5064 5066 1 1.0950e+02 4.1840e+02 ; HA - CA - CB
5065 5064 5076 1 1.0950e+02 4.1840e+02 ; HA - CA - C
5066 5064 5076 1 1.1110e+02 5.2718e+02 ; CB - CA - C
5066 5068 5069 1 1.0950e+02 4.1840e+02 ; CB - CG1 - HG11
5066 5068 5070 1 1.0950e+02 4.1840e+02 ; CB - CG1 - HG12
5066 5068 5071 1 1.0950e+02 4.1840e+02 ; CB - CG1 - HG13
5066 5072 5073 1 1.0950e+02 4.1840e+02 ; CB - CG2 - HG21
5066 5072 5074 1 1.0950e+02 4.1840e+02 ; CB - CG2 - HG22
5066 5072 5075 1 1.0950e+02 4.1840e+02 ; CB - CG2 - HG23
5067 5066 5068 1 1.0950e+02 4.1840e+02 ; HB - CB - CG1
5067 5066 5072 1 1.0950e+02 4.1840e+02 ; HB - CB - CG2
5068 5066 5072 1 1.0950e+02 3.3472e+02 ; CG1 - CB - CG2
5069 5068 5070 1 1.0950e+02 2.9288e+02 ; HG11 - CG1 - HG12
5069 5068 5071 1 1.0950e+02 2.9288e+02 ; HG11 - CG1 - HG13
5070 5068 5071 1 1.0950e+02 2.9288e+02 ; HG12 - CG1 - HG13
5073 5072 5074 1 1.0950e+02 2.9288e+02 ; HG21 - CG2 - HG22
5073 5072 5075 1 1.0950e+02 2.9288e+02 ; HG21 - CG2 - HG23

5074 5072 5075 1 1.0950e+02 2.9288e+02 ; HG22 - CG2
- HG23
5076 5078 5079 1 1.2000e+02 4.1840e+02 ; C - N
- H
5076 5078 5080 1 1.2190e+02 4.1840e+02 ; C - N
- CA
5077 5076 5078 1 1.2290e+02 6.6944e+02 ; O - C
- N
5078 5080 5081 1 1.0950e+02 4.1840e+02 ; N - CA
- HA
5078 5080 5082 1 1.0970e+02 6.6944e+02 ; N - CA
- CB
5078 5080 5091 1 1.1010e+02 5.2718e+02 ; N - CA
- C
5079 5078 5080 1 1.1804e+02 4.1840e+02 ; H - N
- CA
5080 5082 5083 1 1.0950e+02 4.1840e+02 ; CA - CB
- HB2
5080 5082 5084 1 1.0950e+02 4.1840e+02 ; CA - CB
- HB3
5080 5082 5085 1 1.0950e+02 3.3472e+02 ; CA - CB
- CG
5080 5091 5092 1 1.2040e+02 6.6944e+02 ; CA - C
- O
5080 5091 5093 1 1.1660e+02 5.8576e+02 ; CA - C
- N
5081 5080 5082 1 1.0950e+02 4.1840e+02 ; HA - CA
- CB
5081 5080 5091 1 1.0950e+02 4.1840e+02 ; HA - CA
- C
5082 5080 5091 1 1.1110e+02 5.2718e+02 ; CB - CA
- C
5082 5085 5086 1 1.0950e+02 4.1840e+02 ; CB - CG
- HG2
5082 5085 5087 1 1.0950e+02 4.1840e+02 ; CB - CG
- HG3
5082 5085 5088 1 1.1110e+02 5.2718e+02 ; CB - CG
- CD
5083 5082 5084 1 1.0950e+02 2.9288e+02 ; HB2 - CB
- HB3
5083 5082 5085 1 1.0950e+02 4.1840e+02 ; HB2 - CB
- CG
5084 5082 5085 1 1.0950e+02 4.1840e+02 ; HB3 - CB
- CG
5085 5088 5089 1 1.1700e+02 5.8576e+02 ; CG - CD
- OE1
5085 5088 5090 1 1.1700e+02 5.8576e+02 ; CG - CD
- OE2
5086 5085 5087 1 1.0950e+02 2.9288e+02 ; HG2 - CG
- HG3
5086 5085 5088 1 1.0950e+02 4.1840e+02 ; HG2 - CG
- CD

5087 5085 5088 1 1.0950e+02 4.1840e+02 ; HG3 - CG
- CD
5089 5088 5090 1 1.2600e+02 6.6944e+02 ; OE1 - CD
- OE2
5091 5093 5094 1 1.2000e+02 4.1840e+02 ; C - N
- H
5091 5093 5095 1 1.2190e+02 4.1840e+02 ; C - N
- CA
5092 5091 5093 1 1.2290e+02 6.6944e+02 ; O - C
- N
5093 5095 5096 1 1.0950e+02 4.1840e+02 ; N - CA
- HA
5093 5095 5097 1 1.0970e+02 6.6944e+02 ; N - CA
- CB
5093 5095 5110 1 1.1010e+02 5.2718e+02 ; N - CA
- C
5094 5093 5095 1 1.1804e+02 4.1840e+02 ; H - N
- CA
5095 5097 5098 1 1.0950e+02 4.1840e+02 ; CA - CB
- HB
5095 5097 5099 1 1.0950e+02 3.3472e+02 ; CA - CB
- CG2
5095 5097 5103 1 1.0950e+02 3.3472e+02 ; CA - CB
- CG1
5095 5110 5111 1 1.2040e+02 6.6944e+02 ; CA - C
- O
5095 5110 5112 1 1.1660e+02 5.8576e+02 ; CA - C
- N
5096 5095 5097 1 1.0950e+02 4.1840e+02 ; HA - CA
- CB
5096 5095 5110 1 1.0950e+02 4.1840e+02 ; HA - CA
- C
5097 5095 5110 1 1.1110e+02 5.2718e+02 ; CB - CA
- C
5097 5099 5100 1 1.0950e+02 4.1840e+02 ; CB - CG2
- HG21
5097 5099 5101 1 1.0950e+02 4.1840e+02 ; CB - CG2
- HG22
5097 5099 5102 1 1.0950e+02 4.1840e+02 ; CB - CG2
- HG23
5097 5103 5104 1 1.0950e+02 4.1840e+02 ; CB - CG1
- HG12
5097 5103 5105 1 1.0950e+02 4.1840e+02 ; CB - CG1
- HG13
5097 5103 5106 1 1.0950e+02 3.3472e+02 ; CB - CG1
- CD1
5098 5097 5099 1 1.0950e+02 4.1840e+02 ; HB - CB
- CG2
5098 5097 5103 1 1.0950e+02 4.1840e+02 ; HB - CB
- CG1
5099 5097 5103 1 1.0950e+02 3.3472e+02 ; CG2 - CB
- CG1

```
  5100   5099   5101      1    1.0950e+02    2.9288e+02 ;   HG21 - CG2
- HG22
  5100   5099   5102      1    1.0950e+02    2.9288e+02 ;   HG21 - CG2
- HG23
  5101   5099   5102      1    1.0950e+02    2.9288e+02 ;   HG22 - CG2
- HG23
  5103   5106   5107      1    1.0950e+02    4.1840e+02 ;    CG1 - CD1
- HD11
  5103   5106   5108      1    1.0950e+02    4.1840e+02 ;    CG1 - CD1
- HD12
  5103   5106   5109      1    1.0950e+02    4.1840e+02 ;    CG1 - CD1
- HD13
  5104   5103   5105      1    1.0950e+02    2.9288e+02 ;   HG12 - CG1
- HG13
  5104   5103   5106      1    1.0950e+02    4.1840e+02 ;   HG12 - CG1
- CD1
  5105   5103   5106      1    1.0950e+02    4.1840e+02 ;   HG13 - CG1
- CD1
  5107   5106   5108      1    1.0950e+02    2.9288e+02 ;   HD11 - CD1
- HD12
  5107   5106   5109      1    1.0950e+02    2.9288e+02 ;   HD11 - CD1
- HD13
  5108   5106   5109      1    1.0950e+02    2.9288e+02 ;   HD12 - CD1
- HD13
  5110   5112   5113      1    1.2000e+02    4.1840e+02 ;      C - N
- H
  5110   5112   5114      1    1.2190e+02    4.1840e+02 ;      C - N
- CA
  5111   5110   5112      1    1.2290e+02    6.6944e+02 ;      O - C
- N
  5112   5114   5115      1    1.0950e+02    4.1840e+02 ;      N - CA
- HA
  5112   5114   5116      1    1.0970e+02    6.6944e+02 ;      N - CA
- CB
  5112   5114   5124      1    1.1010e+02    5.2718e+02 ;      N - CA
- C
  5113   5112   5114      1    1.1804e+02    4.1840e+02 ;      H - N
- CA
  5114   5116   5117      1    1.0950e+02    4.1840e+02 ;     CA - CB
- HB2
  5114   5116   5118      1    1.0950e+02    4.1840e+02 ;     CA - CB
- HB3
  5114   5116   5119      1    1.1110e+02    5.2718e+02 ;     CA - CB
- CG
  5114   5124   5125      1    1.2040e+02    6.6944e+02 ;     CA - C
- O
  5114   5124   5126      1    1.1660e+02    5.8576e+02 ;     CA - C
- N
  5115   5114   5116      1    1.0950e+02    4.1840e+02 ;     HA - CA
- CB
  5115   5114   5124      1    1.0950e+02    4.1840e+02 ;     HA - CA
- C
```

5116 5114 5124 1 1.1110e+02 5.2718e+02 ; CB - CA
- C
5116 5119 5120 1 1.2040e+02 6.6944e+02 ; CB - CG
- OD1
5116 5119 5121 1 1.1660e+02 5.8576e+02 ; CB - CG
- ND2
5117 5116 5118 1 1.0950e+02 2.9288e+02 ; HB2 - CB
- HB3
5117 5116 5119 1 1.0950e+02 4.1840e+02 ; HB2 - CB
- CG
5118 5116 5119 1 1.0950e+02 4.1840e+02 ; HB3 - CB
- CG
5119 5121 5122 1 1.2000e+02 4.1840e+02 ; CG - ND2
- HD21
5119 5121 5123 1 1.2000e+02 4.1840e+02 ; CG - ND2
- HD22
5120 5119 5121 1 1.2290e+02 6.6944e+02 ; OD1 - CG
- ND2
5122 5121 5123 1 1.2000e+02 2.9288e+02 ; HD21 - ND2
- HD22
5124 5126 5127 1 1.2000e+02 4.1840e+02 ; C - N
- H
5124 5126 5128 1 1.2190e+02 4.1840e+02 ; C - N
- CA
5125 5124 5126 1 1.2290e+02 6.6944e+02 ; O - C
- N
5126 5128 5129 1 1.0950e+02 4.1840e+02 ; N - CA
- HA
5126 5128 5130 1 1.0970e+02 6.6944e+02 ; N - CA
- CB
5126 5128 5138 1 1.1010e+02 5.2718e+02 ; N - CA
- C
5127 5126 5128 1 1.1804e+02 4.1840e+02 ; H - N
- CA
5128 5130 5131 1 1.0950e+02 4.1840e+02 ; CA - CB
- HB2
5128 5130 5132 1 1.0950e+02 4.1840e+02 ; CA - CB
- HB3
5128 5130 5133 1 1.1110e+02 5.2718e+02 ; CA - CB
- CG
5128 5138 5139 1 1.2040e+02 6.6944e+02 ; CA - C
- O
5128 5138 5140 1 1.1660e+02 5.8576e+02 ; CA - C
- N
5129 5128 5130 1 1.0950e+02 4.1840e+02 ; HA - CA
- CB
5129 5128 5138 1 1.0950e+02 4.1840e+02 ; HA - CA
- C
5130 5128 5138 1 1.1110e+02 5.2718e+02 ; CB - CA
- C
5130 5133 5134 1 1.2040e+02 6.6944e+02 ; CB - CG
- OD1

5130 5133 5135 1 1.1660e+02 5.8576e+02 ; CB - CG
- ND2
5131 5130 5132 1 1.0950e+02 2.9288e+02 ; HB2 - CB
- HB3
5131 5130 5133 1 1.0950e+02 4.1840e+02 ; HB2 - CB
- CG
5132 5130 5133 1 1.0950e+02 4.1840e+02 ; HB3 - CB
- CG
5133 5135 5136 1 1.2000e+02 4.1840e+02 ; CG - ND2
- HD21
5133 5135 5137 1 1.2000e+02 4.1840e+02 ; CG - ND2
- HD22
5134 5133 5135 1 1.2290e+02 6.6944e+02 ; OD1 - CG
- ND2
5136 5135 5137 1 1.2000e+02 2.9288e+02 ; HD21 - ND2
- HD22
5138 5140 5141 1 1.2000e+02 4.1840e+02 ; C - N
- H
5138 5140 5142 1 1.2190e+02 4.1840e+02 ; C - N
- CA
5139 5138 5140 1 1.2290e+02 6.6944e+02 ; O - C
- N
5140 5142 5143 1 1.0950e+02 4.1840e+02 ; N - CA
- HA
5140 5142 5144 1 1.0970e+02 6.6944e+02 ; N - CA
- CB
5140 5142 5157 1 1.1010e+02 5.2718e+02 ; N - CA
- C
5141 5140 5142 1 1.1804e+02 4.1840e+02 ; H - N
- CA
5142 5144 5145 1 1.0950e+02 4.1840e+02 ; CA - CB
- HB2
5142 5144 5146 1 1.0950e+02 4.1840e+02 ; CA - CB
- HB3
5142 5144 5147 1 1.0950e+02 3.3472e+02 ; CA - CB
- CG
5142 5157 5158 1 1.2040e+02 6.6944e+02 ; CA - C
- O
5142 5157 5159 1 1.1660e+02 5.8576e+02 ; CA - C
- N
5143 5142 5144 1 1.0950e+02 4.1840e+02 ; HA - CA
- CB
5143 5142 5157 1 1.0950e+02 4.1840e+02 ; HA - CA
- C
5144 5142 5157 1 1.1110e+02 5.2718e+02 ; CB - CA
- C
5144 5147 5148 1 1.0950e+02 4.1840e+02 ; CB - CG
- HG
5144 5147 5149 1 1.0950e+02 3.3472e+02 ; CB - CG
- CD1
5144 5147 5153 1 1.0950e+02 3.3472e+02 ; CB - CG
- CD2

5145 5144 5146 1 1.0950e+02 2.9288e+02 ; HB2 - CB
- HB3
5145 5144 5147 1 1.0950e+02 4.1840e+02 ; HB2 - CB
- CG
5146 5144 5147 1 1.0950e+02 4.1840e+02 ; HB3 - CB
- CG
5147 5149 5150 1 1.0950e+02 4.1840e+02 ; CG - CD1
- HD11
5147 5149 5151 1 1.0950e+02 4.1840e+02 ; CG - CD1
- HD12
5147 5149 5152 1 1.0950e+02 4.1840e+02 ; CG - CD1
- HD13
5147 5153 5154 1 1.0950e+02 4.1840e+02 ; CG - CD2
- HD21
5147 5153 5155 1 1.0950e+02 4.1840e+02 ; CG - CD2
- HD22
5147 5153 5156 1 1.0950e+02 4.1840e+02 ; CG - CD2
- HD23
5148 5147 5149 1 1.0950e+02 4.1840e+02 ; HG - CG
- CD1
5148 5147 5153 1 1.0950e+02 4.1840e+02 ; HG - CG
- CD2
5149 5147 5153 1 1.0950e+02 3.3472e+02 ; CD1 - CG
- CD2
5150 5149 5151 1 1.0950e+02 2.9288e+02 ; HD11 - CD1
- HD12
5150 5149 5152 1 1.0950e+02 2.9288e+02 ; HD11 - CD1
- HD13
5151 5149 5152 1 1.0950e+02 2.9288e+02 ; HD12 - CD1
- HD13
5154 5153 5155 1 1.0950e+02 2.9288e+02 ; HD21 - CD2
- HD22
5154 5153 5156 1 1.0950e+02 2.9288e+02 ; HD21 - CD2
- HD23
5155 5153 5156 1 1.0950e+02 2.9288e+02 ; HD22 - CD2
- HD23
5157 5159 5160 1 1.2000e+02 4.1840e+02 ; C - N
- H
5157 5159 5161 1 1.2190e+02 4.1840e+02 ; C - N
- CA
5158 5157 5159 1 1.2290e+02 6.6944e+02 ; O - C
- N
5159 5161 5162 1 1.0950e+02 4.1840e+02 ; N - CA
- HA
5159 5161 5163 1 1.0970e+02 6.6944e+02 ; N - CA
- CB
5159 5161 5181 1 1.1010e+02 5.2718e+02 ; N - CA
- C
5160 5159 5161 1 1.1804e+02 4.1840e+02 ; H - N
- CA
5161 5163 5164 1 1.0950e+02 4.1840e+02 ; CA - CB
- HB2

5161 5163 5165 1 1.0950e+02 4.1840e+02 ; CA - CB
- HB3
5161 5163 5166 1 1.0950e+02 3.3472e+02 ; CA - CB
- CG
5161 5181 5182 1 1.2040e+02 6.6944e+02 ; CA - C
- O
5161 5181 5183 1 1.1660e+02 5.8576e+02 ; CA - C
- N
5162 5161 5163 1 1.0950e+02 4.1840e+02 ; HA - CA
- CB
5162 5161 5181 1 1.0950e+02 4.1840e+02 ; HA - CA
- C
5163 5161 5181 1 1.1110e+02 5.2718e+02 ; CB - CA
- C
5163 5166 5167 1 1.0950e+02 4.1840e+02 ; CB - CG
- HG2
5163 5166 5168 1 1.0950e+02 4.1840e+02 ; CB - CG
- HG3
5163 5166 5169 1 1.0950e+02 3.3472e+02 ; CB - CG
- CD
5164 5163 5165 1 1.0950e+02 2.9288e+02 ; HB2 - CB
- HB3
5164 5163 5166 1 1.0950e+02 4.1840e+02 ; HB2 - CB
- CG
5165 5163 5166 1 1.0950e+02 4.1840e+02 ; HB3 - CB
- CG
5166 5169 5170 1 1.0950e+02 4.1840e+02 ; CG - CD
- HD2
5166 5169 5171 1 1.0950e+02 4.1840e+02 ; CG - CD
- HD3
5166 5169 5172 1 1.1120e+02 6.6944e+02 ; CG - CD
- NE
5167 5166 5168 1 1.0950e+02 2.9288e+02 ; HG2 - CG
- HG3
5167 5166 5169 1 1.0950e+02 4.1840e+02 ; HG2 - CG
- CD
5168 5166 5169 1 1.0950e+02 4.1840e+02 ; HG3 - CG
- CD
5169 5172 5173 1 1.1840e+02 4.1840e+02 ; CD - NE
- HE
5169 5172 5174 1 1.2320e+02 4.1840e+02 ; CD - NE
- CZ
5170 5169 5171 1 1.0950e+02 2.9288e+02 ; HD2 - CD
- HD3
5170 5169 5172 1 1.0950e+02 4.1840e+02 ; HD2 - CD
- NE
5171 5169 5172 1 1.0950e+02 4.1840e+02 ; HD3 - CD
- NE
5172 5174 5175 1 1.2000e+02 5.8576e+02 ; NE - CZ
- NH1
5172 5174 5178 1 1.2000e+02 5.8576e+02 ; NE - CZ
- NH2

5173 5172 5174 1 1.2000e+02 4.1840e+02 ; HE - NE
- CZ
5174 5175 5176 1 1.2000e+02 4.1840e+02 ; CZ - NH1
- HH11
5174 5175 5177 1 1.2000e+02 4.1840e+02 ; CZ - NH1
- HH12
5174 5178 5179 1 1.2000e+02 4.1840e+02 ; CZ - NH2
- HH21
5174 5178 5180 1 1.2000e+02 4.1840e+02 ; CZ - NH2
- HH22
5175 5174 5178 1 1.2000e+02 5.8576e+02 ; NH1 - CZ
- NH2
5176 5175 5177 1 1.2000e+02 2.9288e+02 ; HH11 - NH1
- HH12
5179 5178 5180 1 1.2000e+02 2.9288e+02 ; HH21 - NH2
- HH22
5181 5183 5184 1 1.2000e+02 4.1840e+02 ; C - N
- H
5181 5183 5185 1 1.2190e+02 4.1840e+02 ; C - N
- CA
5182 5181 5183 1 1.2290e+02 6.6944e+02 ; O - C
- N
5183 5185 5186 1 1.0950e+02 4.1840e+02 ; N - CA
- HA
5183 5185 5187 1 1.0970e+02 6.6944e+02 ; N - CA
- CB
5183 5185 5195 1 1.1010e+02 5.2718e+02 ; N - CA
- C
5184 5183 5185 1 1.1804e+02 4.1840e+02 ; H - N
- CA
5185 5187 5188 1 1.0950e+02 4.1840e+02 ; CA - CB
- HB2
5185 5187 5189 1 1.0950e+02 4.1840e+02 ; CA - CB
- HB3
5185 5187 5190 1 1.1110e+02 5.2718e+02 ; CA - CB
- CG
5185 5195 5196 1 1.2040e+02 6.6944e+02 ; CA - C
- O
5185 5195 5197 1 1.1660e+02 5.8576e+02 ; CA - C
- N
5186 5185 5187 1 1.0950e+02 4.1840e+02 ; HA - CA
- CB
5186 5185 5195 1 1.0950e+02 4.1840e+02 ; HA - CA
- C
5187 5185 5195 1 1.1110e+02 5.2718e+02 ; CB - CA
- C
5187 5190 5191 1 1.2040e+02 6.6944e+02 ; CB - CG
- OD1
5187 5190 5192 1 1.1660e+02 5.8576e+02 ; CB - CG
- ND2
5188 5187 5189 1 1.0950e+02 2.9288e+02 ; HB2 - CB
- HB3

5188 5187 5190 1 1.0950e+02 4.1840e+02 ; HB2 - CB
- CG
5189 5187 5190 1 1.0950e+02 4.1840e+02 ; HB3 - CB
- CG
5190 5192 5193 1 1.2000e+02 4.1840e+02 ; CG - ND2
- HD21
5190 5192 5194 1 1.2000e+02 4.1840e+02 ; CG - ND2
- HD22
5191 5190 5192 1 1.2290e+02 6.6944e+02 ; OD1 - CG
- ND2
5193 5192 5194 1 1.2000e+02 2.9288e+02 ; HD21 - ND2
- HD22
5195 5197 5198 1 1.2000e+02 4.1840e+02 ; C - N
- H
5195 5197 5199 1 1.2190e+02 4.1840e+02 ; C - N
- CA
5196 5195 5197 1 1.2290e+02 6.6944e+02 ; O - C
- N
5197 5199 5200 1 1.0950e+02 4.1840e+02 ; N - CA
- HA
5197 5199 5201 1 1.0970e+02 6.6944e+02 ; N - CA
- CB
5197 5199 5214 1 1.1010e+02 5.2718e+02 ; N - CA
- C
5198 5197 5199 1 1.1804e+02 4.1840e+02 ; H - N
- CA
5199 5201 5202 1 1.0950e+02 4.1840e+02 ; CA - CB
- HB
5199 5201 5203 1 1.0950e+02 3.3472e+02 ; CA - CB
- CG2
5199 5201 5207 1 1.0950e+02 3.3472e+02 ; CA - CB
- CG1
5199 5214 5215 1 1.2040e+02 6.6944e+02 ; CA - C
- O
5199 5214 5216 1 1.1660e+02 5.8576e+02 ; CA - C
- N
5200 5199 5201 1 1.0950e+02 4.1840e+02 ; HA - CA
- CB
5200 5199 5214 1 1.0950e+02 4.1840e+02 ; HA - CA
- C
5201 5199 5214 1 1.1110e+02 5.2718e+02 ; CB - CA
- C
5201 5203 5204 1 1.0950e+02 4.1840e+02 ; CB - CG2
- HG21
5201 5203 5205 1 1.0950e+02 4.1840e+02 ; CB - CG2
- HG22
5201 5203 5206 1 1.0950e+02 4.1840e+02 ; CB - CG2
- HG23
5201 5207 5208 1 1.0950e+02 4.1840e+02 ; CB - CG1
- HG12
5201 5207 5209 1 1.0950e+02 4.1840e+02 ; CB - CG1
- HG13

```
  5201   5207   5210      1    1.0950e+02    3.3472e+02 ;     CB - CG1
- CD1
  5202   5201   5203      1    1.0950e+02    4.1840e+02 ;     HB - CB
- CG2
  5202   5201   5207      1    1.0950e+02    4.1840e+02 ;     HB - CB
- CG1
  5203   5201   5207      1    1.0950e+02    3.3472e+02 ;    CG2 - CB
- CG1
  5204   5203   5205      1    1.0950e+02    2.9288e+02 ;   HG21 - CG2
- HG22
  5204   5203   5206      1    1.0950e+02    2.9288e+02 ;   HG21 - CG2
- HG23
  5205   5203   5206      1    1.0950e+02    2.9288e+02 ;   HG22 - CG2
- HG23
  5207   5210   5211      1    1.0950e+02    4.1840e+02 ;    CG1 - CD1
- HD11
  5207   5210   5212      1    1.0950e+02    4.1840e+02 ;    CG1 - CD1
- HD12
  5207   5210   5213      1    1.0950e+02    4.1840e+02 ;    CG1 - CD1
- HD13
  5208   5207   5209      1    1.0950e+02    2.9288e+02 ;   HG12 - CG1
- HG13
  5208   5207   5210      1    1.0950e+02    4.1840e+02 ;   HG12 - CG1
- CD1
  5209   5207   5210      1    1.0950e+02    4.1840e+02 ;   HG13 - CG1
- CD1
  5211   5210   5212      1    1.0950e+02    2.9288e+02 ;   HD11 - CD1
- HD12
  5211   5210   5213      1    1.0950e+02    2.9288e+02 ;   HD11 - CD1
- HD13
  5212   5210   5213      1    1.0950e+02    2.9288e+02 ;   HD12 - CD1
- HD13
  5214   5216   5217      1    1.2000e+02    4.1840e+02 ;      C - N
- H
  5214   5216   5218      1    1.2190e+02    4.1840e+02 ;      C - N
- CA
  5215   5214   5216      1    1.2290e+02    6.6944e+02 ;      O - C
- N
  5216   5218   5219      1    1.0950e+02    4.1840e+02 ;      N - CA
- HA
  5216   5218   5220      1    1.0970e+02    6.6944e+02 ;      N - CA
- CB
  5216   5218   5234      1    1.1010e+02    5.2718e+02 ;      N - CA
- C
  5217   5216   5218      1    1.1804e+02    4.1840e+02 ;      H - N
- CA
  5218   5220   5221      1    1.0950e+02    4.1840e+02 ;     CA - CB
- HB2
  5218   5220   5222      1    1.0950e+02    4.1840e+02 ;     CA - CB
- HB3
  5218   5220   5223      1    1.1400e+02    5.2718e+02 ;     CA - CB
- CG
```

```
  5218   5234   5235       1    1.2040e+02    6.6944e+02 ;      CA - C
- O
  5218   5234   5236       1    1.1660e+02    5.8576e+02 ;      CA - C
- N
  5219   5218   5220       1    1.0950e+02    4.1840e+02 ;      HA - CA
- CB
  5219   5218   5234       1    1.0950e+02    4.1840e+02 ;      HA - CA
- C
  5220   5218   5234       1    1.1110e+02    5.2718e+02 ;      CB - CA
- C
  5220   5223   5224       1    1.2000e+02    5.8576e+02 ;      CB - CG
- CD1
  5220   5223   5232       1    1.2000e+02    5.8576e+02 ;      CB - CG
- CD2
  5221   5220   5222       1    1.0950e+02    2.9288e+02 ;     HB2 - CB
- HB3
  5221   5220   5223       1    1.0950e+02    4.1840e+02 ;     HB2 - CB
- CG
  5222   5220   5223       1    1.0950e+02    4.1840e+02 ;     HB3 - CB
- CG
  5223   5224   5225       1    1.2000e+02    4.1840e+02 ;      CG - CD1
- HD1
  5223   5224   5226       1    1.2000e+02    5.2718e+02 ;      CG - CD1
- CE1
  5223   5232   5230       1    1.2000e+02    5.2718e+02 ;      CG - CD2
- CE2
  5223   5232   5233       1    1.2000e+02    4.1840e+02 ;      CG - CD2
- HD2
  5224   5223   5232       1    1.2000e+02    5.2718e+02 ;     CD1 - CG
- CD2
  5224   5226   5227       1    1.2000e+02    4.1840e+02 ;     CD1 - CE1
- HE1
  5224   5226   5228       1    1.2000e+02    5.2718e+02 ;     CD1 - CE1
- CZ
  5225   5224   5226       1    1.2000e+02    4.1840e+02 ;     HD1 - CD1
- CE1
  5226   5228   5229       1    1.2000e+02    4.1840e+02 ;     CE1 - CZ
- HZ
  5226   5228   5230       1    1.2000e+02    5.2718e+02 ;     CE1 - CZ
- CE2
  5227   5226   5228       1    1.2000e+02    4.1840e+02 ;     HE1 - CE1
- CZ
  5228   5230   5231       1    1.2000e+02    4.1840e+02 ;      CZ - CE2
- HE2
  5228   5230   5232       1    1.2000e+02    5.2718e+02 ;      CZ - CE2
- CD2
  5229   5228   5230       1    1.2000e+02    4.1840e+02 ;      HZ - CZ
- CE2
  5230   5232   5233       1    1.2000e+02    4.1840e+02 ;     CE2 - CD2
- HD2
  5231   5230   5232       1    1.2000e+02    4.1840e+02 ;     HE2 - CE2
- CD2
```

5234 5236 5237 1 1.2000e+02 4.1840e+02 ; C - N
- H
5234 5236 5238 1 1.2190e+02 4.1840e+02 ; C - N
- CA
5235 5234 5236 1 1.2290e+02 6.6944e+02 ; O - C
- N
5236 5238 5239 1 1.0950e+02 4.1840e+02 ; N - CA
- HA
5236 5238 5240 1 1.0970e+02 6.6944e+02 ; N - CA
- CB
5236 5238 5253 1 1.1010e+02 5.2718e+02 ; N - CA
- C
5237 5236 5238 1 1.1804e+02 4.1840e+02 ; H - N
- CA
5238 5240 5241 1 1.0950e+02 4.1840e+02 ; CA - CB
- HB
5238 5240 5242 1 1.0950e+02 3.3472e+02 ; CA - CB
- CG2
5238 5240 5246 1 1.0950e+02 3.3472e+02 ; CA - CB
- CG1
5238 5253 5254 1 1.2040e+02 6.6944e+02 ; CA - C
- O
5238 5253 5255 1 1.1660e+02 5.8576e+02 ; CA - C
- N
5239 5238 5240 1 1.0950e+02 4.1840e+02 ; HA - CA
- CB
5239 5238 5253 1 1.0950e+02 4.1840e+02 ; HA - CA
- C
5240 5238 5253 1 1.1110e+02 5.2718e+02 ; CB - CA
- C
5240 5242 5243 1 1.0950e+02 4.1840e+02 ; CB - CG2
- HG21
5240 5242 5244 1 1.0950e+02 4.1840e+02 ; CB - CG2
- HG22
5240 5242 5245 1 1.0950e+02 4.1840e+02 ; CB - CG2
- HG23
5240 5246 5247 1 1.0950e+02 4.1840e+02 ; CB - CG1
- HG12
5240 5246 5248 1 1.0950e+02 4.1840e+02 ; CB - CG1
- HG13
5240 5246 5249 1 1.0950e+02 3.3472e+02 ; CB - CG1
- CD1
5241 5240 5242 1 1.0950e+02 4.1840e+02 ; HB - CB
- CG2
5241 5240 5246 1 1.0950e+02 4.1840e+02 ; HB - CB
- CG1
5242 5240 5246 1 1.0950e+02 3.3472e+02 ; CG2 - CB
- CG1
5243 5242 5244 1 1.0950e+02 2.9288e+02 ; HG21 - CG2
- HG22
5243 5242 5245 1 1.0950e+02 2.9288e+02 ; HG21 - CG2
- HG23

5244 5242 5245 1 1.0950e+02 2.9288e+02 ; HG22 - CG2
- HG23
5246 5249 5250 1 1.0950e+02 4.1840e+02 ; CG1 - CD1
- HD11
5246 5249 5251 1 1.0950e+02 4.1840e+02 ; CG1 - CD1
- HD12
5246 5249 5252 1 1.0950e+02 4.1840e+02 ; CG1 - CD1
- HD13
5247 5246 5248 1 1.0950e+02 2.9288e+02 ; HG12 - CG1
- HG13
5247 5246 5249 1 1.0950e+02 4.1840e+02 ; HG12 - CG1
- CD1
5248 5246 5249 1 1.0950e+02 4.1840e+02 ; HG13 - CG1
- CD1
5250 5249 5251 1 1.0950e+02 2.9288e+02 ; HD11 - CD1
- HD12
5250 5249 5252 1 1.0950e+02 2.9288e+02 ; HD11 - CD1
- HD13
5251 5249 5252 1 1.0950e+02 2.9288e+02 ; HD12 - CD1
- HD13
5253 5255 5256 1 1.2000e+02 4.1840e+02 ; C - N
- H
5253 5255 5257 1 1.2190e+02 4.1840e+02 ; C - N
- CA
5254 5253 5255 1 1.2290e+02 6.6944e+02 ; O - C
- N
5255 5257 5258 1 1.0950e+02 4.1840e+02 ; N - CA
- HA
5255 5257 5259 1 1.0970e+02 6.6944e+02 ; N - CA
- CB
5255 5257 5263 1 1.1010e+02 5.2718e+02 ; N - CA
- C
5256 5255 5257 1 1.1804e+02 4.1840e+02 ; H - N
- CA
5257 5259 5260 1 1.0950e+02 4.1840e+02 ; CA - CB
- HB1
5257 5259 5261 1 1.0950e+02 4.1840e+02 ; CA - CB
- HB2
5257 5259 5262 1 1.0950e+02 4.1840e+02 ; CA - CB
- HB3
5257 5263 5264 1 1.2040e+02 6.6944e+02 ; CA - C
- O
5257 5263 5265 1 1.1660e+02 5.8576e+02 ; CA - C
- N
5258 5257 5259 1 1.0950e+02 4.1840e+02 ; HA - CA
- CB
5258 5257 5263 1 1.0950e+02 4.1840e+02 ; HA - CA
- C
5259 5257 5263 1 1.1110e+02 5.2718e+02 ; CB - CA
- C
5260 5259 5261 1 1.0950e+02 2.9288e+02 ; HB1 - CB
- HB2

5260 5259 5262 1 1.0950e+02 2.9288e+02 ; HB1 - CB
- HB3
5261 5259 5262 1 1.0950e+02 2.9288e+02 ; HB2 - CB
- HB3
5263 5265 5266 1 1.2000e+02 4.1840e+02 ; C - N
- H
5263 5265 5267 1 1.2190e+02 4.1840e+02 ; C - N
- CA
5264 5263 5265 1 1.2290e+02 6.6944e+02 ; O - C
- N
5265 5267 5268 1 1.0950e+02 4.1840e+02 ; N - CA
- HA
5265 5267 5269 1 1.0970e+02 6.6944e+02 ; N - CA
- CB
5265 5267 5282 1 1.1010e+02 5.2718e+02 ; N - CA
- C
5266 5265 5267 1 1.1804e+02 4.1840e+02 ; H - N
- CA
5267 5269 5270 1 1.0950e+02 4.1840e+02 ; CA - CB
- HB2
5267 5269 5271 1 1.0950e+02 4.1840e+02 ; CA - CB
- HB3
5267 5269 5272 1 1.0950e+02 3.3472e+02 ; CA - CB
- CG
5267 5282 5283 1 1.2040e+02 6.6944e+02 ; CA - C
- O
5267 5282 5284 1 1.1660e+02 5.8576e+02 ; CA - C
- N
5268 5267 5269 1 1.0950e+02 4.1840e+02 ; HA - CA
- CB
5268 5267 5282 1 1.0950e+02 4.1840e+02 ; HA - CA
- C
5269 5267 5282 1 1.1110e+02 5.2718e+02 ; CB - CA
- C
5269 5272 5273 1 1.0950e+02 4.1840e+02 ; CB - CG
- HG
5269 5272 5274 1 1.0950e+02 3.3472e+02 ; CB - CG
- CD1
5269 5272 5278 1 1.0950e+02 3.3472e+02 ; CB - CG
- CD2
5270 5269 5271 1 1.0950e+02 2.9288e+02 ; HB2 - CB
- HB3
5270 5269 5272 1 1.0950e+02 4.1840e+02 ; HB2 - CB
- CG
5271 5269 5272 1 1.0950e+02 4.1840e+02 ; HB3 - CB
- CG
5272 5274 5275 1 1.0950e+02 4.1840e+02 ; CG - CD1
- HD11
5272 5274 5276 1 1.0950e+02 4.1840e+02 ; CG - CD1
- HD12
5272 5274 5277 1 1.0950e+02 4.1840e+02 ; CG - CD1
- HD13

```
  5272   5278   5279      1    1.0950e+02    4.1840e+02 ;     CG - CD2
- HD21
  5272   5278   5280      1    1.0950e+02    4.1840e+02 ;     CG - CD2
- HD22
  5272   5278   5281      1    1.0950e+02    4.1840e+02 ;     CG - CD2
- HD23
  5273   5272   5274      1    1.0950e+02    4.1840e+02 ;     HG - CG
- CD1
  5273   5272   5278      1    1.0950e+02    4.1840e+02 ;     HG - CG
- CD2
  5274   5272   5278      1    1.0950e+02    3.3472e+02 ;    CD1 - CG
- CD2
  5275   5274   5276      1    1.0950e+02    2.9288e+02 ;   HD11 - CD1
- HD12
  5275   5274   5277      1    1.0950e+02    2.9288e+02 ;   HD11 - CD1
- HD13
  5276   5274   5277      1    1.0950e+02    2.9288e+02 ;   HD12 - CD1
- HD13
  5279   5278   5280      1    1.0950e+02    2.9288e+02 ;   HD21 - CD2
- HD22
  5279   5278   5281      1    1.0950e+02    2.9288e+02 ;   HD21 - CD2
- HD23
  5280   5278   5281      1    1.0950e+02    2.9288e+02 ;   HD22 - CD2
- HD23
  5282   5284   5285      1    1.2000e+02    4.1840e+02 ;      C - N
- H
  5282   5284   5286      1    1.2190e+02    4.1840e+02 ;      C - N
- CA
  5283   5282   5284      1    1.2290e+02    6.6944e+02 ;      O - C
- N
  5284   5286   5287      1    1.0950e+02    4.1840e+02 ;      N - CA
- HA
  5284   5286   5288      1    1.0970e+02    6.6944e+02 ;      N - CA
- CB
  5284   5286   5294      1    1.1010e+02    5.2718e+02 ;      N - CA
- C
  5285   5284   5286      1    1.1804e+02    4.1840e+02 ;      H - N
- CA
  5286   5288   5289      1    1.0950e+02    4.1840e+02 ;     CA - CB
- HB2
  5286   5288   5290      1    1.0950e+02    4.1840e+02 ;     CA - CB
- HB3
  5286   5288   5291      1    1.1110e+02    5.2718e+02 ;     CA - CB
- CG
  5286   5294   5295      1    1.2040e+02    6.6944e+02 ;     CA - C
- O
  5286   5294   5296      1    1.1660e+02    5.8576e+02 ;     CA - C
- N
  5287   5286   5288      1    1.0950e+02    4.1840e+02 ;     HA - CA
- CB
  5287   5286   5294      1    1.0950e+02    4.1840e+02 ;     HA - CA
- C
```

5288 5286 5294 1 1.1110e+02 5.2718e+02 ; CB - CA
- C
5288 5291 5292 1 1.1700e+02 5.8576e+02 ; CB - CG
- OD1
5288 5291 5293 1 1.1700e+02 5.8576e+02 ; CB - CG
- OD2
5289 5288 5290 1 1.0950e+02 2.9288e+02 ; HB2 - CB
- HB3
5289 5288 5291 1 1.0950e+02 4.1840e+02 ; HB2 - CB
- CG
5290 5288 5291 1 1.0950e+02 4.1840e+02 ; HB3 - CB
- CG
5292 5291 5293 1 1.2600e+02 6.6944e+02 ; OD1 - CG
- OD2
5294 5296 5297 1 1.2000e+02 4.1840e+02 ; C - N
- H
5294 5296 5298 1 1.2190e+02 4.1840e+02 ; C - N
- CA
5295 5294 5296 1 1.2290e+02 6.6944e+02 ; O - C
- N
5296 5298 5299 1 1.0950e+02 4.1840e+02 ; N - CA
- HA
5296 5298 5300 1 1.0970e+02 6.6944e+02 ; N - CA
- CB
5296 5298 5310 1 1.1010e+02 5.2718e+02 ; N - CA
- C
5297 5296 5298 1 1.1804e+02 4.1840e+02 ; H - N
- CA
5298 5300 5301 1 1.0950e+02 4.1840e+02 ; CA - CB
- HB
5298 5300 5302 1 1.0950e+02 3.3472e+02 ; CA - CB
- CG1
5298 5300 5306 1 1.0950e+02 3.3472e+02 ; CA - CB
- CG2
5298 5310 5311 1 1.2040e+02 6.6944e+02 ; CA - C
- O
5298 5310 5312 1 1.1660e+02 5.8576e+02 ; CA - C
- N
5299 5298 5300 1 1.0950e+02 4.1840e+02 ; HA - CA
- CB
5299 5298 5310 1 1.0950e+02 4.1840e+02 ; HA - CA
- C
5300 5298 5310 1 1.1110e+02 5.2718e+02 ; CB - CA
- C
5300 5302 5303 1 1.0950e+02 4.1840e+02 ; CB - CG1
- HG11
5300 5302 5304 1 1.0950e+02 4.1840e+02 ; CB - CG1
- HG12
5300 5302 5305 1 1.0950e+02 4.1840e+02 ; CB - CG1
- HG13
5300 5306 5307 1 1.0950e+02 4.1840e+02 ; CB - CG2
- HG21

5300 5306 5308 1 1.0950e+02 4.1840e+02 ; CB - CG2
- HG22
5300 5306 5309 1 1.0950e+02 4.1840e+02 ; CB - CG2
- HG23
5301 5300 5302 1 1.0950e+02 4.1840e+02 ; HB - CB
- CG1
5301 5300 5306 1 1.0950e+02 4.1840e+02 ; HB - CB
- CG2
5302 5300 5306 1 1.0950e+02 3.3472e+02 ; CG1 - CB
- CG2
5303 5302 5304 1 1.0950e+02 2.9288e+02 ; HG11 - CG1
- HG12
5303 5302 5305 1 1.0950e+02 2.9288e+02 ; HG11 - CG1
- HG13
5304 5302 5305 1 1.0950e+02 2.9288e+02 ; HG12 - CG1
- HG13
5307 5306 5308 1 1.0950e+02 2.9288e+02 ; HG21 - CG2
- HG22
5307 5306 5309 1 1.0950e+02 2.9288e+02 ; HG21 - CG2
- HG23
5308 5306 5309 1 1.0950e+02 2.9288e+02 ; HG22 - CG2
- HG23
5310 5312 5313 1 1.2000e+02 4.1840e+02 ; C - N
- H
5310 5312 5314 1 1.2190e+02 4.1840e+02 ; C - N
- CA
5311 5310 5312 1 1.2290e+02 6.6944e+02 ; O - C
- N
5312 5314 5315 1 1.0950e+02 4.1840e+02 ; N - CA
- HA
5312 5314 5316 1 1.0970e+02 6.6944e+02 ; N - CA
- CB
5312 5314 5322 1 1.1010e+02 5.2718e+02 ; N - CA
- C
5313 5312 5314 1 1.1804e+02 4.1840e+02 ; H - N
- CA
5314 5316 5317 1 1.0950e+02 4.1840e+02 ; CA - CB
- HB2
5314 5316 5318 1 1.0950e+02 4.1840e+02 ; CA - CB
- HB3
5314 5316 5319 1 1.1110e+02 5.2718e+02 ; CA - CB
- CG
5314 5322 5323 1 1.2040e+02 6.6944e+02 ; CA - C
- O
5314 5322 5324 1 1.1660e+02 5.8576e+02 ; CA - C
- N
5315 5314 5316 1 1.0950e+02 4.1840e+02 ; HA - CA
- CB
5315 5314 5322 1 1.0950e+02 4.1840e+02 ; HA - CA
- C
5316 5314 5322 1 1.1110e+02 5.2718e+02 ; CB - CA
- C

```
  5316   5319   5320       1    1.1700e+02    5.8576e+02 ;     CB - CG 
- OD1
  5316   5319   5321       1    1.1700e+02    5.8576e+02 ;     CB - CG 
- OD2
  5317   5316   5318       1    1.0950e+02    2.9288e+02 ;    HB2 - CB 
- HB3
  5317   5316   5319       1    1.0950e+02    4.1840e+02 ;    HB2 - CB 
- CG
  5318   5316   5319       1    1.0950e+02    4.1840e+02 ;    HB3 - CB 
- CG
  5320   5319   5321       1    1.2600e+02    6.6944e+02 ;    OD1 - CG 
- OD2
  5322   5324   5325       1    1.2000e+02    4.1840e+02 ;      C - N 
- H
  5322   5324   5326       1    1.2190e+02    4.1840e+02 ;      C - N 
- CA
  5323   5322   5324       1    1.2290e+02    6.6944e+02 ;      O - C 
- N
  5324   5326   5327       1    1.0950e+02    4.1840e+02 ;      N - CA 
- HA
  5324   5326   5328       1    1.0970e+02    6.6944e+02 ;      N - CA 
- CB
  5324   5326   5336       1    1.1010e+02    5.2718e+02 ;      N - CA 
- C
  5325   5324   5326       1    1.1804e+02    4.1840e+02 ;      H - N 
- CA
  5326   5328   5329       1    1.0950e+02    4.1840e+02 ;     CA - CB 
- HB2
  5326   5328   5330       1    1.0950e+02    4.1840e+02 ;     CA - CB 
- HB3
  5326   5328   5331       1    1.1110e+02    5.2718e+02 ;     CA - CB 
- CG
  5326   5336   5337       1    1.2040e+02    6.6944e+02 ;     CA - C 
- O
  5326   5336   5338       1    1.1660e+02    5.8576e+02 ;     CA - C 
- N
  5327   5326   5328       1    1.0950e+02    4.1840e+02 ;     HA - CA 
- CB
  5327   5326   5336       1    1.0950e+02    4.1840e+02 ;     HA - CA 
- C
  5328   5326   5336       1    1.1110e+02    5.2718e+02 ;     CB - CA 
- C
  5328   5331   5332       1    1.2040e+02    6.6944e+02 ;     CB - CG 
- OD1
  5328   5331   5333       1    1.1660e+02    5.8576e+02 ;     CB - CG 
- ND2
  5329   5328   5330       1    1.0950e+02    2.9288e+02 ;    HB2 - CB 
- HB3
  5329   5328   5331       1    1.0950e+02    4.1840e+02 ;    HB2 - CB 
- CG
  5330   5328   5331       1    1.0950e+02    4.1840e+02 ;    HB3 - CB 
- CG
```

5331 5333 5334 1 1.2000e+02 4.1840e+02 ; CG - ND2
- HD21
5331 5333 5335 1 1.2000e+02 4.1840e+02 ; CG - ND2
- HD22
5332 5331 5333 1 1.2290e+02 6.6944e+02 ; OD1 - CG
- ND2
5334 5333 5335 1 1.2000e+02 2.9288e+02 ; HD21 - ND2
- HD22
5336 5338 5339 1 1.2000e+02 4.1840e+02 ; C - N
- H
5336 5338 5340 1 1.2190e+02 4.1840e+02 ; C - N
- CA
5337 5336 5338 1 1.2290e+02 6.6944e+02 ; O - C
- N
5338 5340 5341 1 1.0950e+02 4.1840e+02 ; N - CA
- HA
5338 5340 5342 1 1.0970e+02 6.6944e+02 ; N - CA
- CB
5338 5340 5347 1 1.1010e+02 5.2718e+02 ; N - CA
- C
5339 5338 5340 1 1.1804e+02 4.1840e+02 ; H - N
- CA
5340 5342 5343 1 1.0950e+02 4.1840e+02 ; CA - CB
- HB2
5340 5342 5344 1 1.0950e+02 4.1840e+02 ; CA - CB
- HB3
5340 5342 5345 1 1.0950e+02 4.1840e+02 ; CA - CB
- OG
5340 5347 5348 1 1.2040e+02 6.6944e+02 ; CA - C
- O
5340 5347 5349 1 1.1660e+02 5.8576e+02 ; CA - C
- N
5341 5340 5342 1 1.0950e+02 4.1840e+02 ; HA - CA
- CB
5341 5340 5347 1 1.0950e+02 4.1840e+02 ; HA - CA
- C
5342 5340 5347 1 1.1110e+02 5.2718e+02 ; CB - CA
- C
5342 5345 5346 1 1.0850e+02 4.6024e+02 ; CB - OG
- HG
5343 5342 5344 1 1.0950e+02 2.9288e+02 ; HB2 - CB
- HB3
5343 5342 5345 1 1.0950e+02 4.1840e+02 ; HB2 - CB
- OG
5344 5342 5345 1 1.0950e+02 4.1840e+02 ; HB3 - CB
- OG
5347 5349 5350 1 1.2000e+02 4.1840e+02 ; C - N
- H
5347 5349 5351 1 1.2190e+02 4.1840e+02 ; C - N
- CA
5348 5347 5349 1 1.2290e+02 6.6944e+02 ; O - C
- N

5349 5351 5352 1 1.0950e+02 4.1840e+02 ; N - CA
- HA2
5349 5351 5353 1 1.0950e+02 4.1840e+02 ; N - CA
- HA3
5349 5351 5354 1 1.1010e+02 5.2718e+02 ; N - CA
- C
5350 5349 5351 1 1.1804e+02 4.1840e+02 ; H - N
- CA
5351 5354 5355 1 1.2040e+02 6.6944e+02 ; CA - C
- O
5351 5354 5356 1 1.1660e+02 5.8576e+02 ; CA - C
- N
5352 5351 5353 1 1.0950e+02 2.9288e+02 ; HA2 - CA
- HA3
5352 5351 5354 1 1.0950e+02 4.1840e+02 ; HA2 - CA
- C
5353 5351 5354 1 1.0950e+02 4.1840e+02 ; HA3 - CA
- C
5354 5356 5357 1 1.2000e+02 4.1840e+02 ; C - N
- H
5354 5356 5358 1 1.2190e+02 4.1840e+02 ; C - N
- CA
5355 5354 5356 1 1.2290e+02 6.6944e+02 ; O - C
- N
5356 5358 5359 1 1.0950e+02 4.1840e+02 ; N - CA
- HA
5356 5358 5360 1 1.0970e+02 6.6944e+02 ; N - CA
- CB
5356 5358 5368 1 1.1010e+02 5.2718e+02 ; N - CA
- C
5357 5356 5358 1 1.1804e+02 4.1840e+02 ; H - N
- CA
5358 5360 5361 1 1.0950e+02 4.1840e+02 ; CA - CB
- HB
5358 5360 5362 1 1.0950e+02 3.3472e+02 ; CA - CB
- CG2
5358 5360 5366 1 1.0950e+02 4.1840e+02 ; CA - CB
- OG1
5358 5368 5369 1 1.2040e+02 6.6944e+02 ; CA - C
- O
5358 5368 5370 1 1.1660e+02 5.8576e+02 ; CA - C
- N
5359 5358 5360 1 1.0950e+02 4.1840e+02 ; HA - CA
- CB
5359 5358 5368 1 1.0950e+02 4.1840e+02 ; HA - CA
- C
5360 5358 5368 1 1.1110e+02 5.2718e+02 ; CB - CA
- C
5360 5362 5363 1 1.0950e+02 4.1840e+02 ; CB - CG2
- HG21
5360 5362 5364 1 1.0950e+02 4.1840e+02 ; CB - CG2
- HG22

```
  5360   5362   5365      1    1.0950e+02    4.1840e+02 ;     CB - CG2
- HG23
  5360   5366   5367      1    1.0850e+02    4.6024e+02 ;     CB - OG1
- HG1
  5361   5360   5362      1    1.0950e+02    4.1840e+02 ;     HB - CB
- CG2
  5361   5360   5366      1    1.0950e+02    4.1840e+02 ;     HB - CB
- OG1
  5362   5360   5366      1    1.0950e+02    4.1840e+02 ;    CG2 - CB
- OG1
  5363   5362   5364      1    1.0950e+02    2.9288e+02 ;   HG21 - CG2
- HG22
  5363   5362   5365      1    1.0950e+02    2.9288e+02 ;   HG21 - CG2
- HG23
  5364   5362   5365      1    1.0950e+02    2.9288e+02 ;   HG22 - CG2
- HG23
  5368   5370   5371      1    1.2000e+02    4.1840e+02 ;      C - N
- H
  5368   5370   5372      1    1.2190e+02    4.1840e+02 ;      C - N
- CA
  5369   5368   5370      1    1.2290e+02    6.6944e+02 ;      O - C
- N
  5370   5372   5373      1    1.0950e+02    4.1840e+02 ;      N - CA
- HA
  5370   5372   5374      1    1.0970e+02    6.6944e+02 ;      N - CA
- CB
  5370   5372   5387      1    1.1010e+02    5.2718e+02 ;      N - CA
- C
  5371   5370   5372      1    1.1804e+02    4.1840e+02 ;      H - N
- CA
  5372   5374   5375      1    1.0950e+02    4.1840e+02 ;     CA - CB
- HB2
  5372   5374   5376      1    1.0950e+02    4.1840e+02 ;     CA - CB
- HB3
  5372   5374   5377      1    1.0950e+02    3.3472e+02 ;     CA - CB
- CG
  5372   5387   5388      1    1.2040e+02    6.6944e+02 ;     CA - C
- O
  5372   5387   5389      1    1.1660e+02    5.8576e+02 ;     CA - C
- N
  5373   5372   5374      1    1.0950e+02    4.1840e+02 ;     HA - CA
- CB
  5373   5372   5387      1    1.0950e+02    4.1840e+02 ;     HA - CA
- C
  5374   5372   5387      1    1.1110e+02    5.2718e+02 ;     CB - CA
- C
  5374   5377   5378      1    1.0950e+02    4.1840e+02 ;     CB - CG
- HG
  5374   5377   5379      1    1.0950e+02    3.3472e+02 ;     CB - CG
- CD1
  5374   5377   5383      1    1.0950e+02    3.3472e+02 ;     CB - CG
- CD2
```

5375 5374 5376 1 1.0950e+02 2.9288e+02 ; HB2 - CB
- HB3
5375 5374 5377 1 1.0950e+02 4.1840e+02 ; HB2 - CB
- CG
5376 5374 5377 1 1.0950e+02 4.1840e+02 ; HB3 - CB
- CG
5377 5379 5380 1 1.0950e+02 4.1840e+02 ; CG - CD1
- HD11
5377 5379 5381 1 1.0950e+02 4.1840e+02 ; CG - CD1
- HD12
5377 5379 5382 1 1.0950e+02 4.1840e+02 ; CG - CD1
- HD13
5377 5383 5384 1 1.0950e+02 4.1840e+02 ; CG - CD2
- HD21
5377 5383 5385 1 1.0950e+02 4.1840e+02 ; CG - CD2
- HD22
5377 5383 5386 1 1.0950e+02 4.1840e+02 ; CG - CD2
- HD23
5378 5377 5379 1 1.0950e+02 4.1840e+02 ; HG - CG
- CD1
5378 5377 5383 1 1.0950e+02 4.1840e+02 ; HG - CG
- CD2
5379 5377 5383 1 1.0950e+02 3.3472e+02 ; CD1 - CG
- CD2
5380 5379 5381 1 1.0950e+02 2.9288e+02 ; HD11 - CD1
- HD12
5380 5379 5382 1 1.0950e+02 2.9288e+02 ; HD11 - CD1
- HD13
5381 5379 5382 1 1.0950e+02 2.9288e+02 ; HD12 - CD1
- HD13
5384 5383 5385 1 1.0950e+02 2.9288e+02 ; HD21 - CD2
- HD22
5384 5383 5386 1 1.0950e+02 2.9288e+02 ; HD21 - CD2
- HD23
5385 5383 5386 1 1.0950e+02 2.9288e+02 ; HD22 - CD2
- HD23
5387 5389 5390 1 1.2000e+02 4.1840e+02 ; C - N
- H
5387 5389 5391 1 1.2190e+02 4.1840e+02 ; C - N
- CA
5388 5387 5389 1 1.2290e+02 6.6944e+02 ; O - C
- N
5389 5391 5392 1 1.0950e+02 4.1840e+02 ; N - CA
- HA
5389 5391 5393 1 1.0970e+02 6.6944e+02 ; N - CA
- CB
5389 5391 5398 1 1.1010e+02 5.2718e+02 ; N - CA
- C
5390 5389 5391 1 1.1804e+02 4.1840e+02 ; H - N
- CA
5391 5393 5394 1 1.0950e+02 4.1840e+02 ; CA - CB
- HB2

5391 5393 5395 1 1.0950e+02 4.1840e+02 ; CA - CB
- HB3
5391 5393 5396 1 1.0950e+02 4.1840e+02 ; CA - CB
- OG
5391 5398 5399 1 1.2040e+02 6.6944e+02 ; CA - C
- O
5391 5398 5400 1 1.1660e+02 5.8576e+02 ; CA - C
- N
5392 5391 5393 1 1.0950e+02 4.1840e+02 ; HA - CA
- CB
5392 5391 5398 1 1.0950e+02 4.1840e+02 ; HA - CA
- C
5393 5391 5398 1 1.1110e+02 5.2718e+02 ; CB - CA
- C
5393 5396 5397 1 1.0850e+02 4.6024e+02 ; CB - OG
- HG
5394 5393 5395 1 1.0950e+02 2.9288e+02 ; HB2 - CB
- HB3
5394 5393 5396 1 1.0950e+02 4.1840e+02 ; HB2 - CB
- OG
5395 5393 5396 1 1.0950e+02 4.1840e+02 ; HB3 - CB
- OG
5398 5400 5401 1 1.2000e+02 4.1840e+02 ; C - N
- H
5398 5400 5402 1 1.2190e+02 4.1840e+02 ; C - N
- CA
5399 5398 5400 1 1.2290e+02 6.6944e+02 ; O - C
- N
5400 5402 5403 1 1.0950e+02 4.1840e+02 ; N - CA
- HA
5400 5402 5404 1 1.0970e+02 6.6944e+02 ; N - CA
- CB
5400 5402 5409 1 1.1010e+02 5.2718e+02 ; N - CA
- C
5401 5400 5402 1 1.1804e+02 4.1840e+02 ; H - N
- CA
5402 5404 5405 1 1.0950e+02 4.1840e+02 ; CA - CB
- HB2
5402 5404 5406 1 1.0950e+02 4.1840e+02 ; CA - CB
- HB3
5402 5404 5407 1 1.0950e+02 4.1840e+02 ; CA - CB
- OG
5402 5409 5410 1 1.2040e+02 6.6944e+02 ; CA - C
- O
5402 5409 5411 1 1.1660e+02 5.8576e+02 ; CA - C
- N
5403 5402 5404 1 1.0950e+02 4.1840e+02 ; HA - CA
- CB
5403 5402 5409 1 1.0950e+02 4.1840e+02 ; HA - CA
- C
5404 5402 5409 1 1.1110e+02 5.2718e+02 ; CB - CA
- C

5404 5407 5408 1 1.0850e+02 4.6024e+02 ; CB - OG
- HG
5405 5404 5406 1 1.0950e+02 2.9288e+02 ; HB2 - CB
- HB3
5405 5404 5407 1 1.0950e+02 4.1840e+02 ; HB2 - CB
- OG
5406 5404 5407 1 1.0950e+02 4.1840e+02 ; HB3 - CB
- OG
5409 5411 5412 1 1.2000e+02 4.1840e+02 ; C - N
- H
5409 5411 5413 1 1.2190e+02 4.1840e+02 ; C - N
- CA
5410 5409 5411 1 1.2290e+02 6.6944e+02 ; O - C
- N
5411 5413 5414 1 1.0950e+02 4.1840e+02 ; N - CA
- HA
5411 5413 5415 1 1.0970e+02 6.6944e+02 ; N - CA
- CB
5411 5413 5426 1 1.1010e+02 5.2718e+02 ; N - CA
- C
5412 5411 5413 1 1.1804e+02 4.1840e+02 ; H - N
- CA
5413 5415 5416 1 1.0950e+02 4.1840e+02 ; CA - CB
- HB2
5413 5415 5417 1 1.0950e+02 4.1840e+02 ; CA - CB
- HB3
5413 5415 5418 1 1.0950e+02 3.3472e+02 ; CA - CB
- CG
5413 5426 5427 1 1.2040e+02 6.6944e+02 ; CA - C
- O
5413 5426 5428 1 1.1660e+02 5.8576e+02 ; CA - C
- N
5414 5413 5415 1 1.0950e+02 4.1840e+02 ; HA - CA
- CB
5414 5413 5426 1 1.0950e+02 4.1840e+02 ; HA - CA
- C
5415 5413 5426 1 1.1110e+02 5.2718e+02 ; CB - CA
- C
5415 5418 5419 1 1.0950e+02 4.1840e+02 ; CB - CG
- HG2
5415 5418 5420 1 1.0950e+02 4.1840e+02 ; CB - CG
- HG3
5415 5418 5421 1 1.1110e+02 5.2718e+02 ; CB - CG
- CD
5416 5415 5417 1 1.0950e+02 2.9288e+02 ; HB2 - CB
- HB3
5416 5415 5418 1 1.0950e+02 4.1840e+02 ; HB2 - CB
- CG
5417 5415 5418 1 1.0950e+02 4.1840e+02 ; HB3 - CB
- CG
5418 5421 5422 1 1.2040e+02 6.6944e+02 ; CG - CD
- OE1

5418 5421 5423 1 1.1660e+02 5.8576e+02 ; CG - CD
- NE2
5419 5418 5420 1 1.0950e+02 2.9288e+02 ; HG2 - CG
- HG3
5419 5418 5421 1 1.0950e+02 4.1840e+02 ; HG2 - CG
- CD
5420 5418 5421 1 1.0950e+02 4.1840e+02 ; HG3 - CG
- CD
5421 5423 5424 1 1.2000e+02 4.1840e+02 ; CD - NE2
- HE21
5421 5423 5425 1 1.2000e+02 4.1840e+02 ; CD - NE2
- HE22
5422 5421 5423 1 1.2290e+02 6.6944e+02 ; OE1 - CD
- NE2
5424 5423 5425 1 1.2000e+02 2.9288e+02 ; HE21 - NE2
- HE22
5426 5428 5429 1 1.2000e+02 4.1840e+02 ; C - N
- H
5426 5428 5430 1 1.2190e+02 4.1840e+02 ; C - N
- CA
5427 5426 5428 1 1.2290e+02 6.6944e+02 ; O - C
- N
5428 5430 5431 1 1.0950e+02 4.1840e+02 ; N - CA
- HA
5428 5430 5432 1 1.0970e+02 6.6944e+02 ; N - CA
- CB
5428 5430 5442 1 1.1010e+02 5.2718e+02 ; N - CA
- C
5429 5428 5430 1 1.1804e+02 4.1840e+02 ; H - N
- CA
5430 5432 5433 1 1.0950e+02 4.1840e+02 ; CA - CB
- HB2
5430 5432 5434 1 1.0950e+02 4.1840e+02 ; CA - CB
- HB3
5430 5432 5435 1 1.0950e+02 3.3472e+02 ; CA - CB
- CG
5430 5442 5443 1 1.2040e+02 6.6944e+02 ; CA - C
- O
5430 5442 5444 1 1.1660e+02 5.8576e+02 ; CA - C
- N
5431 5430 5432 1 1.0950e+02 4.1840e+02 ; HA - CA
- CB
5431 5430 5442 1 1.0950e+02 4.1840e+02 ; HA - CA
- C
5432 5430 5442 1 1.1110e+02 5.2718e+02 ; CB - CA
- C
5432 5435 5436 1 1.0950e+02 4.1840e+02 ; CB - CG
- HG2
5432 5435 5437 1 1.0950e+02 4.1840e+02 ; CB - CG
- HG3
5432 5435 5438 1 1.1110e+02 5.2718e+02 ; CB - CG
- CD

5433 5432 5434 1 1.0950e+02 2.9288e+02 ; HB2 - CB
- HB3
5433 5432 5435 1 1.0950e+02 4.1840e+02 ; HB2 - CB
- CG
5434 5432 5435 1 1.0950e+02 4.1840e+02 ; HB3 - CB
- CG
5435 5438 5439 1 1.2040e+02 6.6944e+02 ; CG - CD
- OE1
5435 5438 5440 1 1.1000e+02 6.6944e+02 ; CG - CD
- OE2
5436 5435 5437 1 1.0950e+02 2.9288e+02 ; HG2 - CG
- HG3
5436 5435 5438 1 1.0950e+02 4.1840e+02 ; HG2 - CG
- CD
5437 5435 5438 1 1.0950e+02 4.1840e+02 ; HG3 - CG
- CD
5438 5440 5441 1 1.1300e+02 4.1840e+02 ; CD - OE2
- HE2
5439 5438 5440 1 1.2000e+02 6.6944e+02 ; OE1 - CD
- OE2
5442 5444 5445 1 1.2000e+02 4.1840e+02 ; C - N
- H
5442 5444 5446 1 1.2190e+02 4.1840e+02 ; C - N
- CA
5443 5442 5444 1 1.2290e+02 6.6944e+02 ; O - C
- N
5444 5446 5447 1 1.0950e+02 4.1840e+02 ; N - CA
- HA
5444 5446 5448 1 1.0970e+02 6.6944e+02 ; N - CA
- CB
5444 5446 5461 1 1.1010e+02 5.2718e+02 ; N - CA
- C
5445 5444 5446 1 1.1804e+02 4.1840e+02 ; H - N
- CA
5446 5448 5449 1 1.0950e+02 4.1840e+02 ; CA - CB
- HB
5446 5448 5450 1 1.0950e+02 3.3472e+02 ; CA - CB
- CG2
5446 5448 5454 1 1.0950e+02 3.3472e+02 ; CA - CB
- CG1
5446 5461 5462 1 1.2040e+02 6.6944e+02 ; CA - C
- O
5446 5461 5463 1 1.1660e+02 5.8576e+02 ; CA - C
- N
5447 5446 5448 1 1.0950e+02 4.1840e+02 ; HA - CA
- CB
5447 5446 5461 1 1.0950e+02 4.1840e+02 ; HA - CA
- C
5448 5446 5461 1 1.1110e+02 5.2718e+02 ; CB - CA
- C
5448 5450 5451 1 1.0950e+02 4.1840e+02 ; CB - CG2
- HG21

```
  5448   5450   5452       1    1.0950e+02    4.1840e+02 ;     CB - CG2
- HG22
  5448   5450   5453       1    1.0950e+02    4.1840e+02 ;     CB - CG2
- HG23
  5448   5454   5455       1    1.0950e+02    4.1840e+02 ;     CB - CG1
- HG12
  5448   5454   5456       1    1.0950e+02    4.1840e+02 ;     CB - CG1
- HG13
  5448   5454   5457       1    1.0950e+02    3.3472e+02 ;     CB - CG1
- CD1
  5449   5448   5450       1    1.0950e+02    4.1840e+02 ;     HB - CB
- CG2
  5449   5448   5454       1    1.0950e+02    4.1840e+02 ;     HB - CB
- CG1
  5450   5448   5454       1    1.0950e+02    3.3472e+02 ;    CG2 - CB
- CG1
  5451   5450   5452       1    1.0950e+02    2.9288e+02 ;   HG21 - CG2
- HG22
  5451   5450   5453       1    1.0950e+02    2.9288e+02 ;   HG21 - CG2
- HG23
  5452   5450   5453       1    1.0950e+02    2.9288e+02 ;   HG22 - CG2
- HG23
  5454   5457   5458       1    1.0950e+02    4.1840e+02 ;    CG1 - CD1
- HD11
  5454   5457   5459       1    1.0950e+02    4.1840e+02 ;    CG1 - CD1
- HD12
  5454   5457   5460       1    1.0950e+02    4.1840e+02 ;    CG1 - CD1
- HD13
  5455   5454   5456       1    1.0950e+02    2.9288e+02 ;   HG12 - CG1
- HG13
  5455   5454   5457       1    1.0950e+02    4.1840e+02 ;   HG12 - CG1
- CD1
  5456   5454   5457       1    1.0950e+02    4.1840e+02 ;   HG13 - CG1
- CD1
  5458   5457   5459       1    1.0950e+02    2.9288e+02 ;   HD11 - CD1
- HD12
  5458   5457   5460       1    1.0950e+02    2.9288e+02 ;   HD11 - CD1
- HD13
  5459   5457   5460       1    1.0950e+02    2.9288e+02 ;   HD12 - CD1
- HD13
  5461   5463   5464       1    1.2000e+02    4.1840e+02 ;      C - N
- H
  5461   5463   5465       1    1.2190e+02    4.1840e+02 ;      C - N
- CA
  5462   5461   5463       1    1.2290e+02    6.6944e+02 ;      O - C
- N
  5463   5465   5466       1    1.0950e+02    4.1840e+02 ;      N - CA
- HA
  5463   5465   5467       1    1.0970e+02    6.6944e+02 ;      N - CA
- CB
  5463   5465   5480       1    1.1010e+02    5.2718e+02 ;      N - CA
- C
```

5464 5463 5465 1 1.1804e+02 4.1840e+02 ; H - N
- CA
5465 5467 5468 1 1.0950e+02 4.1840e+02 ; CA - CB
- HB2
5465 5467 5469 1 1.0950e+02 4.1840e+02 ; CA - CB
- HB3
5465 5467 5470 1 1.0950e+02 3.3472e+02 ; CA - CB
- CG
5465 5480 5481 1 1.2040e+02 6.6944e+02 ; CA - C
- O
5465 5480 5482 1 1.1660e+02 5.8576e+02 ; CA - C
- N
5466 5465 5467 1 1.0950e+02 4.1840e+02 ; HA - CA
- CB
5466 5465 5480 1 1.0950e+02 4.1840e+02 ; HA - CA
- C
5467 5465 5480 1 1.1110e+02 5.2718e+02 ; CB - CA
- C
5467 5470 5471 1 1.0950e+02 4.1840e+02 ; CB - CG
- HG
5467 5470 5472 1 1.0950e+02 3.3472e+02 ; CB - CG
- CD1
5467 5470 5476 1 1.0950e+02 3.3472e+02 ; CB - CG
- CD2
5468 5467 5469 1 1.0950e+02 2.9288e+02 ; HB2 - CB
- HB3
5468 5467 5470 1 1.0950e+02 4.1840e+02 ; HB2 - CB
- CG
5469 5467 5470 1 1.0950e+02 4.1840e+02 ; HB3 - CB
- CG
5470 5472 5473 1 1.0950e+02 4.1840e+02 ; CG - CD1
- HD11
5470 5472 5474 1 1.0950e+02 4.1840e+02 ; CG - CD1
- HD12
5470 5472 5475 1 1.0950e+02 4.1840e+02 ; CG - CD1
- HD13
5470 5476 5477 1 1.0950e+02 4.1840e+02 ; CG - CD2
- HD21
5470 5476 5478 1 1.0950e+02 4.1840e+02 ; CG - CD2
- HD22
5470 5476 5479 1 1.0950e+02 4.1840e+02 ; CG - CD2
- HD23
5471 5470 5472 1 1.0950e+02 4.1840e+02 ; HG - CG
- CD1
5471 5470 5476 1 1.0950e+02 4.1840e+02 ; HG - CG
- CD2
5472 5470 5476 1 1.0950e+02 3.3472e+02 ; CD1 - CG
- CD2
5473 5472 5474 1 1.0950e+02 2.9288e+02 ; HD11 - CD1
- HD12
5473 5472 5475 1 1.0950e+02 2.9288e+02 ; HD11 - CD1
- HD13

```
  5474   5472   5475       1    1.0950e+02    2.9288e+02 ;   HD12 - CD1
- HD13
  5477   5476   5478       1    1.0950e+02    2.9288e+02 ;   HD21 - CD2
- HD22
  5477   5476   5479       1    1.0950e+02    2.9288e+02 ;   HD21 - CD2
- HD23
  5478   5476   5479       1    1.0950e+02    2.9288e+02 ;   HD22 - CD2
- HD23
  5480   5482   5483       1    1.2000e+02    4.1840e+02 ;      C - N
- H
  5480   5482   5484       1    1.2190e+02    4.1840e+02 ;      C - N
- CA
  5481   5480   5482       1    1.2290e+02    6.6944e+02 ;      O - C
- N
  5482   5484   5485       1    1.0950e+02    4.1840e+02 ;      N - CA
- HA
  5482   5484   5486       1    1.0970e+02    6.6944e+02 ;      N - CA
- CB
  5482   5484   5492       1    1.1010e+02    5.2718e+02 ;      N - CA
- C
  5483   5482   5484       1    1.1804e+02    4.1840e+02 ;      H - N
- CA
  5484   5486   5487       1    1.0950e+02    4.1840e+02 ;     CA - CB
- HB2
  5484   5486   5488       1    1.0950e+02    4.1840e+02 ;     CA - CB
- HB3
  5484   5486   5489       1    1.1110e+02    5.2718e+02 ;     CA - CB
- CG
  5484   5492   5493       1    1.2040e+02    6.6944e+02 ;     CA - C
- O
  5484   5492   5494       1    1.1660e+02    5.8576e+02 ;     CA - C
- N
  5485   5484   5486       1    1.0950e+02    4.1840e+02 ;     HA - CA
- CB
  5485   5484   5492       1    1.0950e+02    4.1840e+02 ;     HA - CA
- C
  5486   5484   5492       1    1.1110e+02    5.2718e+02 ;     CB - CA
- C
  5486   5489   5490       1    1.1700e+02    5.8576e+02 ;     CB - CG
- OD1
  5486   5489   5491       1    1.1700e+02    5.8576e+02 ;     CB - CG
- OD2
  5487   5486   5488       1    1.0950e+02    2.9288e+02 ;    HB2 - CB
- HB3
  5487   5486   5489       1    1.0950e+02    4.1840e+02 ;    HB2 - CB
- CG
  5488   5486   5489       1    1.0950e+02    4.1840e+02 ;    HB3 - CB
- CG
  5490   5489   5491       1    1.2600e+02    6.6944e+02 ;    OD1 - CG
- OD2
  5492   5494   5495       1    1.2000e+02    4.1840e+02 ;      C - N
- H
```

```
  5492   5494   5496       1    1.2190e+02    4.1840e+02 ;       C - N
- CA
  5493   5492   5494       1    1.2290e+02    6.6944e+02 ;       O - C
- N
  5494   5496   5497       1    1.0950e+02    4.1840e+02 ;       N - CA
- HA2
  5494   5496   5498       1    1.0950e+02    4.1840e+02 ;       N - CA
- HA3
  5494   5496   5499       1    1.1010e+02    5.2718e+02 ;       N - CA
- C
  5495   5494   5496       1    1.1804e+02    4.1840e+02 ;       H - N
- CA
  5496   5499   5500       1    1.2040e+02    6.6944e+02 ;      CA - C
- O
  5496   5499   5501       1    1.1660e+02    5.8576e+02 ;      CA - C
- N
  5497   5496   5498       1    1.0950e+02    2.9288e+02 ;     HA2 - CA
- HA3
  5497   5496   5499       1    1.0950e+02    4.1840e+02 ;     HA2 - CA
- C
  5498   5496   5499       1    1.0950e+02    4.1840e+02 ;     HA3 - CA
- C
  5499   5501   5502       1    1.2000e+02    4.1840e+02 ;       C - N
- H
  5499   5501   5503       1    1.2190e+02    4.1840e+02 ;       C - N
- CA
  5500   5499   5501       1    1.2290e+02    6.6944e+02 ;       O - C
- N
  5501   5503   5504       1    1.0950e+02    4.1840e+02 ;       N - CA
- HA
  5501   5503   5505       1    1.0970e+02    6.6944e+02 ;       N - CA
- CB
  5501   5503   5518       1    1.1010e+02    5.2718e+02 ;       N - CA
- C
  5502   5501   5503       1    1.1804e+02    4.1840e+02 ;       H - N
- CA
  5503   5505   5506       1    1.0950e+02    4.1840e+02 ;      CA - CB
- HB2
  5503   5505   5507       1    1.0950e+02    4.1840e+02 ;      CA - CB
- HB3
  5503   5505   5508       1    1.0950e+02    3.3472e+02 ;      CA - CB
- CG
  5503   5518   5519       1    1.2040e+02    6.6944e+02 ;      CA - C
- O
  5503   5518   5520       1    1.1660e+02    5.8576e+02 ;      CA - C
- N
  5504   5503   5505       1    1.0950e+02    4.1840e+02 ;      HA - CA
- CB
  5504   5503   5518       1    1.0950e+02    4.1840e+02 ;      HA - CA
- C
  5505   5503   5518       1    1.1110e+02    5.2718e+02 ;      CB - CA
- C
```

5505 5508 5509 1 1.0950e+02 4.1840e+02 ; CB - CG
- HG
5505 5508 5510 1 1.0950e+02 3.3472e+02 ; CB - CG
- CD1
5505 5508 5514 1 1.0950e+02 3.3472e+02 ; CB - CG
- CD2
5506 5505 5507 1 1.0950e+02 2.9288e+02 ; HB2 - CB
- HB3
5506 5505 5508 1 1.0950e+02 4.1840e+02 ; HB2 - CB
- CG
5507 5505 5508 1 1.0950e+02 4.1840e+02 ; HB3 - CB
- CG
5508 5510 5511 1 1.0950e+02 4.1840e+02 ; CG - CD1
- HD11
5508 5510 5512 1 1.0950e+02 4.1840e+02 ; CG - CD1
- HD12
5508 5510 5513 1 1.0950e+02 4.1840e+02 ; CG - CD1
- HD13
5508 5514 5515 1 1.0950e+02 4.1840e+02 ; CG - CD2
- HD21
5508 5514 5516 1 1.0950e+02 4.1840e+02 ; CG - CD2
- HD22
5508 5514 5517 1 1.0950e+02 4.1840e+02 ; CG - CD2
- HD23
5509 5508 5510 1 1.0950e+02 4.1840e+02 ; HG - CG
- CD1
5509 5508 5514 1 1.0950e+02 4.1840e+02 ; HG - CG
- CD2
5510 5508 5514 1 1.0950e+02 3.3472e+02 ; CD1 - CG
- CD2
5511 5510 5512 1 1.0950e+02 2.9288e+02 ; HD11 - CD1
- HD12
5511 5510 5513 1 1.0950e+02 2.9288e+02 ; HD11 - CD1
- HD13
5512 5510 5513 1 1.0950e+02 2.9288e+02 ; HD12 - CD1
- HD13
5515 5514 5516 1 1.0950e+02 2.9288e+02 ; HD21 - CD2
- HD22
5515 5514 5517 1 1.0950e+02 2.9288e+02 ; HD21 - CD2
- HD23
5516 5514 5517 1 1.0950e+02 2.9288e+02 ; HD22 - CD2
- HD23
5518 5520 5521 1 1.2000e+02 4.1840e+02 ; C - N
- H
5518 5520 5522 1 1.2190e+02 4.1840e+02 ; C - N
- CA
5519 5518 5520 1 1.2290e+02 6.6944e+02 ; O - C
- N
5520 5522 5523 1 1.0950e+02 4.1840e+02 ; N - CA
- HA
5520 5522 5524 1 1.0970e+02 6.6944e+02 ; N - CA
- CB

5520 5522 5540 1 1.1010e+02 5.2718e+02 ; N - CA
- C
5521 5520 5522 1 1.1804e+02 4.1840e+02 ; H - N
- CA
5522 5524 5525 1 1.0950e+02 4.1840e+02 ; CA - CB
- HB2
5522 5524 5526 1 1.0950e+02 4.1840e+02 ; CA - CB
- HB3
5522 5524 5527 1 1.0950e+02 3.3472e+02 ; CA - CB
- CG
5522 5540 5541 1 1.2040e+02 6.6944e+02 ; CA - C
- O
5522 5540 5542 1 1.1660e+02 5.8576e+02 ; CA - C
- N
5523 5522 5524 1 1.0950e+02 4.1840e+02 ; HA - CA
- CB
5523 5522 5540 1 1.0950e+02 4.1840e+02 ; HA - CA
- C
5524 5522 5540 1 1.1110e+02 5.2718e+02 ; CB - CA
- C
5524 5527 5528 1 1.0950e+02 4.1840e+02 ; CB - CG
- HG2
5524 5527 5529 1 1.0950e+02 4.1840e+02 ; CB - CG
- HG3
5524 5527 5530 1 1.0950e+02 3.3472e+02 ; CB - CG
- CD
5525 5524 5526 1 1.0950e+02 2.9288e+02 ; HB2 - CB
- HB3
5525 5524 5527 1 1.0950e+02 4.1840e+02 ; HB2 - CB
- CG
5526 5524 5527 1 1.0950e+02 4.1840e+02 ; HB3 - CB
- CG
5527 5530 5531 1 1.0950e+02 4.1840e+02 ; CG - CD
- HD2
5527 5530 5532 1 1.0950e+02 4.1840e+02 ; CG - CD
- HD3
5527 5530 5533 1 1.0950e+02 3.3472e+02 ; CG - CD
- CE
5528 5527 5529 1 1.0950e+02 2.9288e+02 ; HG2 - CG
- HG3
5528 5527 5530 1 1.0950e+02 4.1840e+02 ; HG2 - CG
- CD
5529 5527 5530 1 1.0950e+02 4.1840e+02 ; HG3 - CG
- CD
5530 5533 5534 1 1.0950e+02 4.1840e+02 ; CD - CE
- HE2
5530 5533 5535 1 1.0950e+02 4.1840e+02 ; CD - CE
- HE3
5530 5533 5536 1 1.1120e+02 6.6944e+02 ; CD - CE
- NZ
5531 5530 5532 1 1.0950e+02 2.9288e+02 ; HD2 - CD
- HD3

```
  5531   5530   5533       1    1.0950e+02    4.1840e+02 ;    HD2 - CD
- CE
  5532   5530   5533       1    1.0950e+02    4.1840e+02 ;    HD3 - CD
- CE
  5533   5536   5537       1    1.0950e+02    4.1840e+02 ;     CE - NZ
- HZ1
  5533   5536   5538       1    1.0950e+02    4.1840e+02 ;     CE - NZ
- HZ2
  5533   5536   5539       1    1.0950e+02    4.1840e+02 ;     CE - NZ
- HZ3
  5534   5533   5535       1    1.0950e+02    2.9288e+02 ;    HE2 - CE
- HE3
  5534   5533   5536       1    1.0950e+02    4.1840e+02 ;    HE2 - CE
- NZ
  5535   5533   5536       1    1.0950e+02    4.1840e+02 ;    HE3 - CE
- NZ
  5537   5536   5538       1    1.0950e+02    2.9288e+02 ;    HZ1 - NZ
- HZ2
  5537   5536   5539       1    1.0950e+02    2.9288e+02 ;    HZ1 - NZ
- HZ3
  5538   5536   5539       1    1.0950e+02    2.9288e+02 ;    HZ2 - NZ
- HZ3
  5540   5542   5543       1    1.2000e+02    4.1840e+02 ;      C - N
- H
  5540   5542   5544       1    1.2190e+02    4.1840e+02 ;      C - N
- CA
  5541   5540   5542       1    1.2290e+02    6.6944e+02 ;      O - C
- N
  5542   5544   5545       1    1.0950e+02    4.1840e+02 ;      N - CA
- HA
  5542   5544   5546       1    1.0970e+02    6.6944e+02 ;      N - CA
- CB
  5542   5544   5562       1    1.1010e+02    5.2718e+02 ;      N - CA
- C
  5543   5542   5544       1    1.1804e+02    4.1840e+02 ;      H - N
- CA
  5544   5546   5547       1    1.0950e+02    4.1840e+02 ;     CA - CB
- HB2
  5544   5546   5548       1    1.0950e+02    4.1840e+02 ;     CA - CB
- HB3
  5544   5546   5549       1    1.0950e+02    3.3472e+02 ;     CA - CB
- CG
  5544   5562   5563       1    1.2040e+02    6.6944e+02 ;     CA - C
- O
  5544   5562   5564       1    1.1660e+02    5.8576e+02 ;     CA - C
- N
  5545   5544   5546       1    1.0950e+02    4.1840e+02 ;     HA - CA
- CB
  5545   5544   5562       1    1.0950e+02    4.1840e+02 ;     HA - CA
- C
  5546   5544   5562       1    1.1110e+02    5.2718e+02 ;     CB - CA
- C
```

5546 5549 5550 1 1.0950e+02 4.1840e+02 ; CB - CG
- HG2
5546 5549 5551 1 1.0950e+02 4.1840e+02 ; CB - CG
- HG3
5546 5549 5552 1 1.0950e+02 3.3472e+02 ; CB - CG
- CD
5547 5546 5548 1 1.0950e+02 2.9288e+02 ; HB2 - CB
- HB3
5547 5546 5549 1 1.0950e+02 4.1840e+02 ; HB2 - CB
- CG
5548 5546 5549 1 1.0950e+02 4.1840e+02 ; HB3 - CB
- CG
5549 5552 5553 1 1.0950e+02 4.1840e+02 ; CG - CD
- HD2
5549 5552 5554 1 1.0950e+02 4.1840e+02 ; CG - CD
- HD3
5549 5552 5555 1 1.0950e+02 3.3472e+02 ; CG - CD
- CE
5550 5549 5551 1 1.0950e+02 2.9288e+02 ; HG2 - CG
- HG3
5550 5549 5552 1 1.0950e+02 4.1840e+02 ; HG2 - CG
- CD
5551 5549 5552 1 1.0950e+02 4.1840e+02 ; HG3 - CG
- CD
5552 5555 5556 1 1.0950e+02 4.1840e+02 ; CD - CE
- HE2
5552 5555 5557 1 1.0950e+02 4.1840e+02 ; CD - CE
- HE3
5552 5555 5558 1 1.1120e+02 6.6944e+02 ; CD - CE
- NZ
5553 5552 5554 1 1.0950e+02 2.9288e+02 ; HD2 - CD
- HD3
5553 5552 5555 1 1.0950e+02 4.1840e+02 ; HD2 - CD
- CE
5554 5552 5555 1 1.0950e+02 4.1840e+02 ; HD3 - CD
- CE
5555 5558 5559 1 1.0950e+02 4.1840e+02 ; CE - NZ
- HZ1
5555 5558 5560 1 1.0950e+02 4.1840e+02 ; CE - NZ
- HZ2
5555 5558 5561 1 1.0950e+02 4.1840e+02 ; CE - NZ
- HZ3
5556 5555 5557 1 1.0950e+02 2.9288e+02 ; HE2 - CE
- HE3
5556 5555 5558 1 1.0950e+02 4.1840e+02 ; HE2 - CE
- NZ
5557 5555 5558 1 1.0950e+02 4.1840e+02 ; HE3 - CE
- NZ
5559 5558 5560 1 1.0950e+02 2.9288e+02 ; HZ1 - NZ
- HZ2
5559 5558 5561 1 1.0950e+02 2.9288e+02 ; HZ1 - NZ
- HZ3

5560 5558 5561 1 1.0950e+02 2.9288e+02 ; HZ2 - NZ
- HZ3
5562 5564 5565 1 1.2000e+02 4.1840e+02 ; C - N
- H
5562 5564 5566 1 1.2190e+02 4.1840e+02 ; C - N
- CA
5563 5562 5564 1 1.2290e+02 6.6944e+02 ; O - C
- N
5564 5566 5567 1 1.0950e+02 4.1840e+02 ; N - CA
- HA
5564 5566 5568 1 1.0970e+02 6.6944e+02 ; N - CA
- CB
5564 5566 5581 1 1.1010e+02 5.2718e+02 ; N - CA
- C
5565 5564 5566 1 1.1804e+02 4.1840e+02 ; H - N
- CA
5566 5568 5569 1 1.0950e+02 4.1840e+02 ; CA - CB
- HB
5566 5568 5570 1 1.0950e+02 3.3472e+02 ; CA - CB
- CG2
5566 5568 5574 1 1.0950e+02 3.3472e+02 ; CA - CB
- CG1
5566 5581 5582 1 1.2040e+02 6.6944e+02 ; CA - C
- O
5566 5581 5583 1 1.1660e+02 5.8576e+02 ; CA - C
- N
5567 5566 5568 1 1.0950e+02 4.1840e+02 ; HA - CA
- CB
5567 5566 5581 1 1.0950e+02 4.1840e+02 ; HA - CA
- C
5568 5566 5581 1 1.1110e+02 5.2718e+02 ; CB - CA
- C
5568 5570 5571 1 1.0950e+02 4.1840e+02 ; CB - CG2
- HG21
5568 5570 5572 1 1.0950e+02 4.1840e+02 ; CB - CG2
- HG22
5568 5570 5573 1 1.0950e+02 4.1840e+02 ; CB - CG2
- HG23
5568 5574 5575 1 1.0950e+02 4.1840e+02 ; CB - CG1
- HG12
5568 5574 5576 1 1.0950e+02 4.1840e+02 ; CB - CG1
- HG13
5568 5574 5577 1 1.0950e+02 3.3472e+02 ; CB - CG1
- CD1
5569 5568 5570 1 1.0950e+02 4.1840e+02 ; HB - CB
- CG2
5569 5568 5574 1 1.0950e+02 4.1840e+02 ; HB - CB
- CG1
5570 5568 5574 1 1.0950e+02 3.3472e+02 ; CG2 - CB
- CG1
5571 5570 5572 1 1.0950e+02 2.9288e+02 ; HG21 - CG2
- HG22

```
  5571   5570   5573      1    1.0950e+02    2.9288e+02 ;   HG21 - CG2
- HG23
  5572   5570   5573      1    1.0950e+02    2.9288e+02 ;   HG22 - CG2
- HG23
  5574   5577   5578      1    1.0950e+02    4.1840e+02 ;    CG1 - CD1
- HD11
  5574   5577   5579      1    1.0950e+02    4.1840e+02 ;    CG1 - CD1
- HD12
  5574   5577   5580      1    1.0950e+02    4.1840e+02 ;    CG1 - CD1
- HD13
  5575   5574   5576      1    1.0950e+02    2.9288e+02 ;   HG12 - CG1
- HG13
  5575   5574   5577      1    1.0950e+02    4.1840e+02 ;   HG12 - CG1
- CD1
  5576   5574   5577      1    1.0950e+02    4.1840e+02 ;   HG13 - CG1
- CD1
  5578   5577   5579      1    1.0950e+02    2.9288e+02 ;   HD11 - CD1
- HD12
  5578   5577   5580      1    1.0950e+02    2.9288e+02 ;   HD11 - CD1
- HD13
  5579   5577   5580      1    1.0950e+02    2.9288e+02 ;   HD12 - CD1
- HD13
  5581   5583   5584      1    1.2000e+02    4.1840e+02 ;      C - N
- H
  5581   5583   5585      1    1.2190e+02    4.1840e+02 ;      C - N
- CA
  5582   5581   5583      1    1.2290e+02    6.6944e+02 ;      O - C
- N
  5583   5585   5586      1    1.0950e+02    4.1840e+02 ;      N - CA
- HA2
  5583   5585   5587      1    1.0950e+02    4.1840e+02 ;      N - CA
- HA3
  5583   5585   5588      1    1.1010e+02    5.2718e+02 ;      N - CA
- C
  5584   5583   5585      1    1.1804e+02    4.1840e+02 ;      H - N
- CA
  5585   5588   5589      1    1.2040e+02    6.6944e+02 ;     CA - C
- O
  5585   5588   5590      1    1.1660e+02    5.8576e+02 ;     CA - C
- N
  5586   5585   5587      1    1.0950e+02    2.9288e+02 ;    HA2 - CA
- HA3
  5586   5585   5588      1    1.0950e+02    4.1840e+02 ;    HA2 - CA
- C
  5587   5585   5588      1    1.0950e+02    4.1840e+02 ;    HA3 - CA
- C
  5588   5590   5591      1    1.2000e+02    4.1840e+02 ;      C - N
- H
  5588   5590   5592      1    1.2190e+02    4.1840e+02 ;      C - N
- CA
  5589   5588   5590      1    1.2290e+02    6.6944e+02 ;      O - C
- N
```

5590 5592 5593 1 1.0950e+02 4.1840e+02 ; N - CA
- HA
5590 5592 5594 1 1.0970e+02 6.6944e+02 ; N - CA
- CB
5590 5592 5609 1 1.1010e+02 5.2718e+02 ; N - CA
- C
5591 5590 5592 1 1.1804e+02 4.1840e+02 ; H - N
- CA
5592 5594 5595 1 1.0950e+02 4.1840e+02 ; CA - CB
- HB2
5592 5594 5596 1 1.0950e+02 4.1840e+02 ; CA - CB
- HB3
5592 5594 5597 1 1.1400e+02 5.2718e+02 ; CA - CB
- CG
5592 5609 5610 1 1.2040e+02 6.6944e+02 ; CA - C
- O
5592 5609 5611 1 1.1660e+02 5.8576e+02 ; CA - C
- N
5593 5592 5594 1 1.0950e+02 4.1840e+02 ; HA - CA
- CB
5593 5592 5609 1 1.0950e+02 4.1840e+02 ; HA - CA
- C
5594 5592 5609 1 1.1110e+02 5.2718e+02 ; CB - CA
- C
5594 5597 5598 1 1.2000e+02 5.8576e+02 ; CB - CG
- CD1
5594 5597 5607 1 1.2000e+02 5.8576e+02 ; CB - CG
- CD2
5595 5594 5596 1 1.0950e+02 2.9288e+02 ; HB2 - CB
- HB3
5595 5594 5597 1 1.0950e+02 4.1840e+02 ; HB2 - CB
- CG
5596 5594 5597 1 1.0950e+02 4.1840e+02 ; HB3 - CB
- CG
5597 5598 5599 1 1.2000e+02 4.1840e+02 ; CG - CD1
- HD1
5597 5598 5600 1 1.2000e+02 5.2718e+02 ; CG - CD1
- CE1
5597 5607 5605 1 1.2000e+02 5.2718e+02 ; CG - CD2
- CE2
5597 5607 5608 1 1.2000e+02 4.1840e+02 ; CG - CD2
- HD2
5598 5597 5607 1 1.2000e+02 5.2718e+02 ; CD1 - CG
- CD2
5598 5600 5601 1 1.2000e+02 4.1840e+02 ; CD1 - CE1
- HE1
5598 5600 5602 1 1.2000e+02 5.2718e+02 ; CD1 - CE1
- CZ
5599 5598 5600 1 1.2000e+02 4.1840e+02 ; HD1 - CD1
- CE1
5600 5602 5603 1 1.2000e+02 5.8576e+02 ; CE1 - CZ
- OH

```
  5600   5602   5605      1    1.2000e+02    5.2718e+02 ;    CE1 - CZ
- CE2
  5601   5600   5602      1    1.2000e+02    4.1840e+02 ;    HE1 - CE1
- CZ
  5602   5603   5604      1    1.1300e+02    4.1840e+02 ;     CZ - OH
- HH
  5602   5605   5606      1    1.2000e+02    4.1840e+02 ;     CZ - CE2
- HE2
  5602   5605   5607      1    1.2000e+02    5.2718e+02 ;     CZ - CE2
- CD2
  5603   5602   5605      1    1.2000e+02    5.8576e+02 ;     OH - CZ
- CE2
  5605   5607   5608      1    1.2000e+02    4.1840e+02 ;    CE2 - CD2
- HD2
  5606   5605   5607      1    1.2000e+02    4.1840e+02 ;    HE2 - CE2
- CD2
  5609   5611   5612      1    1.2000e+02    4.1840e+02 ;      C - N
- H
  5609   5611   5613      1    1.2190e+02    4.1840e+02 ;      C - N
- CA
  5610   5609   5611      1    1.2290e+02    6.6944e+02 ;      O - C
- N
  5611   5613   5614      1    1.0950e+02    4.1840e+02 ;      N - CA
- HA
  5611   5613   5615      1    1.0970e+02    6.6944e+02 ;      N - CA
- CB
  5611   5613   5626      1    1.1010e+02    5.2718e+02 ;      N - CA
- C
  5612   5611   5613      1    1.1804e+02    4.1840e+02 ;      H - N
- CA
  5613   5615   5616      1    1.0950e+02    4.1840e+02 ;     CA - CB
- HB2
  5613   5615   5617      1    1.0950e+02    4.1840e+02 ;     CA - CB
- HB3
  5613   5615   5618      1    1.0950e+02    3.3472e+02 ;     CA - CB
- CG
  5613   5626   5627      1    1.2040e+02    6.6944e+02 ;     CA - C
- O
  5613   5626   5628      1    1.1660e+02    5.8576e+02 ;     CA - C
- N
  5614   5613   5615      1    1.0950e+02    4.1840e+02 ;     HA - CA
- CB
  5614   5613   5626      1    1.0950e+02    4.1840e+02 ;     HA - CA
- C
  5615   5613   5626      1    1.1110e+02    5.2718e+02 ;     CB - CA
- C
  5615   5618   5619      1    1.0950e+02    4.1840e+02 ;     CB - CG
- HG2
  5615   5618   5620      1    1.0950e+02    4.1840e+02 ;     CB - CG
- HG3
  5615   5618   5621      1    1.1110e+02    5.2718e+02 ;     CB - CG
- CD
```

5616 5615 5617 1 1.0950e+02 2.9288e+02 ; HB2 - CB
- HB3
5616 5615 5618 1 1.0950e+02 4.1840e+02 ; HB2 - CB
- CG
5617 5615 5618 1 1.0950e+02 4.1840e+02 ; HB3 - CB
- CG
5618 5621 5622 1 1.2040e+02 6.6944e+02 ; CG - CD
- OE1
5618 5621 5623 1 1.1660e+02 5.8576e+02 ; CG - CD
- NE2
5619 5618 5620 1 1.0950e+02 2.9288e+02 ; HG2 - CG
- HG3
5619 5618 5621 1 1.0950e+02 4.1840e+02 ; HG2 - CG
- CD
5620 5618 5621 1 1.0950e+02 4.1840e+02 ; HG3 - CG
- CD
5621 5623 5624 1 1.2000e+02 4.1840e+02 ; CD - NE2
- HE21
5621 5623 5625 1 1.2000e+02 4.1840e+02 ; CD - NE2
- HE22
5622 5621 5623 1 1.2290e+02 6.6944e+02 ; OE1 - CD
- NE2
5624 5623 5625 1 1.2000e+02 2.9288e+02 ; HE21 - NE2
- HE22
5626 5628 5629 1 1.2000e+02 4.1840e+02 ; C - N
- H
5626 5628 5630 1 1.2190e+02 4.1840e+02 ; C - N
- CA
5627 5626 5628 1 1.2290e+02 6.6944e+02 ; O - C
- N
5628 5630 5631 1 1.0950e+02 4.1840e+02 ; N - CA
- HA
5628 5630 5632 1 1.0970e+02 6.6944e+02 ; N - CA
- CB
5628 5630 5648 1 1.1010e+02 5.2718e+02 ; N - CA
- C
5629 5628 5630 1 1.1804e+02 4.1840e+02 ; H - N
- CA
5630 5632 5633 1 1.0950e+02 4.1840e+02 ; CA - CB
- HB2
5630 5632 5634 1 1.0950e+02 4.1840e+02 ; CA - CB
- HB3
5630 5632 5635 1 1.0950e+02 3.3472e+02 ; CA - CB
- CG
5630 5648 5649 1 1.2040e+02 6.6944e+02 ; CA - C
- O
5630 5648 5650 1 1.1660e+02 5.8576e+02 ; CA - C
- N
5631 5630 5632 1 1.0950e+02 4.1840e+02 ; HA - CA
- CB
5631 5630 5648 1 1.0950e+02 4.1840e+02 ; HA - CA
- C

```
  5632   5630   5648      1    1.1110e+02    5.2718e+02 ;     CB - CA
- C
  5632   5635   5636      1    1.0950e+02    4.1840e+02 ;     CB - CG
- HG2
  5632   5635   5637      1    1.0950e+02    4.1840e+02 ;     CB - CG
- HG3
  5632   5635   5638      1    1.0950e+02    3.3472e+02 ;     CB - CG
- CD
  5633   5632   5634      1    1.0950e+02    2.9288e+02 ;    HB2 - CB
- HB3
  5633   5632   5635      1    1.0950e+02    4.1840e+02 ;    HB2 - CB
- CG
  5634   5632   5635      1    1.0950e+02    4.1840e+02 ;    HB3 - CB
- CG
  5635   5638   5639      1    1.0950e+02    4.1840e+02 ;     CG - CD
- HD2
  5635   5638   5640      1    1.0950e+02    4.1840e+02 ;     CG - CD
- HD3
  5635   5638   5641      1    1.0950e+02    3.3472e+02 ;     CG - CD
- CE
  5636   5635   5637      1    1.0950e+02    2.9288e+02 ;    HG2 - CG
- HG3
  5636   5635   5638      1    1.0950e+02    4.1840e+02 ;    HG2 - CG
- CD
  5637   5635   5638      1    1.0950e+02    4.1840e+02 ;    HG3 - CG
- CD
  5638   5641   5642      1    1.0950e+02    4.1840e+02 ;     CD - CE
- HE2
  5638   5641   5643      1    1.0950e+02    4.1840e+02 ;     CD - CE
- HE3
  5638   5641   5644      1    1.1120e+02    6.6944e+02 ;     CD - CE
- NZ
  5639   5638   5640      1    1.0950e+02    2.9288e+02 ;    HD2 - CD
- HD3
  5639   5638   5641      1    1.0950e+02    4.1840e+02 ;    HD2 - CD
- CE
  5640   5638   5641      1    1.0950e+02    4.1840e+02 ;    HD3 - CD
- CE
  5641   5644   5645      1    1.0950e+02    4.1840e+02 ;     CE - NZ
- HZ1
  5641   5644   5646      1    1.0950e+02    4.1840e+02 ;     CE - NZ
- HZ2
  5641   5644   5647      1    1.0950e+02    4.1840e+02 ;     CE - NZ
- HZ3
  5642   5641   5643      1    1.0950e+02    2.9288e+02 ;    HE2 - CE
- HE3
  5642   5641   5644      1    1.0950e+02    4.1840e+02 ;    HE2 - CE
- NZ
  5643   5641   5644      1    1.0950e+02    4.1840e+02 ;    HE3 - CE
- NZ
  5645   5644   5646      1    1.0950e+02    2.9288e+02 ;    HZ1 - NZ
- HZ2
```

5645 5644 5647 1 1.0950e+02 2.9288e+02 ; HZ1 - NZ
- HZ3
5646 5644 5647 1 1.0950e+02 2.9288e+02 ; HZ2 - NZ
- HZ3
5648 5650 5651 1 1.2000e+02 4.1840e+02 ; C - N
- H
5648 5650 5652 1 1.2190e+02 4.1840e+02 ; C - N
- CA
5649 5648 5650 1 1.2290e+02 6.6944e+02 ; O - C
- N
5650 5652 5653 1 1.0950e+02 4.1840e+02 ; N - CA
- HA
5650 5652 5654 1 1.0970e+02 6.6944e+02 ; N - CA
- CB
5650 5652 5667 1 1.1010e+02 5.2718e+02 ; N - CA
- C
5651 5650 5652 1 1.1804e+02 4.1840e+02 ; H - N
- CA
5652 5654 5655 1 1.0950e+02 4.1840e+02 ; CA - CB
- HB
5652 5654 5656 1 1.0950e+02 3.3472e+02 ; CA - CB
- CG2
5652 5654 5660 1 1.0950e+02 3.3472e+02 ; CA - CB
- CG1
5652 5667 5668 1 1.2040e+02 6.6944e+02 ; CA - C
- O
5652 5667 5669 1 1.1660e+02 5.8576e+02 ; CA - C
- N
5653 5652 5654 1 1.0950e+02 4.1840e+02 ; HA - CA
- CB
5653 5652 5667 1 1.0950e+02 4.1840e+02 ; HA - CA
- C
5654 5652 5667 1 1.1110e+02 5.2718e+02 ; CB - CA
- C
5654 5656 5657 1 1.0950e+02 4.1840e+02 ; CB - CG2
- HG21
5654 5656 5658 1 1.0950e+02 4.1840e+02 ; CB - CG2
- HG22
5654 5656 5659 1 1.0950e+02 4.1840e+02 ; CB - CG2
- HG23
5654 5660 5661 1 1.0950e+02 4.1840e+02 ; CB - CG1
- HG12
5654 5660 5662 1 1.0950e+02 4.1840e+02 ; CB - CG1
- HG13
5654 5660 5663 1 1.0950e+02 3.3472e+02 ; CB - CG1
- CD1
5655 5654 5656 1 1.0950e+02 4.1840e+02 ; HB - CB
- CG2
5655 5654 5660 1 1.0950e+02 4.1840e+02 ; HB - CB
- CG1
5656 5654 5660 1 1.0950e+02 3.3472e+02 ; CG2 - CB
- CG1

5657 5656 5658 1 1.0950e+02 2.9288e+02 ; HG21 - CG2
- HG22
5657 5656 5659 1 1.0950e+02 2.9288e+02 ; HG21 - CG2
- HG23
5658 5656 5659 1 1.0950e+02 2.9288e+02 ; HG22 - CG2
- HG23
5660 5663 5664 1 1.0950e+02 4.1840e+02 ; CG1 - CD1
- HD11
5660 5663 5665 1 1.0950e+02 4.1840e+02 ; CG1 - CD1
- HD12
5660 5663 5666 1 1.0950e+02 4.1840e+02 ; CG1 - CD1
- HD13
5661 5660 5662 1 1.0950e+02 2.9288e+02 ; HG12 - CG1
- HG13
5661 5660 5663 1 1.0950e+02 4.1840e+02 ; HG12 - CG1
- CD1
5662 5660 5663 1 1.0950e+02 4.1840e+02 ; HG13 - CG1
- CD1
5664 5663 5665 1 1.0950e+02 2.9288e+02 ; HD11 - CD1
- HD12
5664 5663 5666 1 1.0950e+02 2.9288e+02 ; HD11 - CD1
- HD13
5665 5663 5666 1 1.0950e+02 2.9288e+02 ; HD12 - CD1
- HD13
5667 5669 5670 1 1.2190e+02 4.1840e+02 ; C - N
- CD
5667 5669 5679 1 1.2190e+02 4.1840e+02 ; C - N
- CA
5668 5667 5669 1 1.2290e+02 6.6944e+02 ; O - C
- N
5669 5670 5671 1 1.0950e+02 4.1840e+02 ; N - CD
- HD2
5669 5670 5672 1 1.0950e+02 4.1840e+02 ; N - CD
- HD3
5669 5670 5673 1 1.0970e+02 6.6944e+02 ; N - CD
- CG
5669 5679 5676 1 1.0970e+02 6.6944e+02 ; N - CA
- CB
5669 5679 5680 1 1.0950e+02 4.1840e+02 ; N - CA
- HA
5669 5679 5681 1 1.1010e+02 5.2718e+02 ; N - CA
- C
5670 5669 5679 1 1.1800e+02 4.1840e+02 ; CD - N
- CA
5670 5673 5674 1 1.0950e+02 4.1840e+02 ; CD - CG
- HG2
5670 5673 5675 1 1.0950e+02 4.1840e+02 ; CD - CG
- HG3
5670 5673 5676 1 1.0950e+02 3.3472e+02 ; CD - CG
- CB
5671 5670 5672 1 1.0950e+02 2.9288e+02 ; HD2 - CD
- HD3

```
  5671   5670   5673       1    1.0950e+02    4.1840e+02 ;    HD2 - CD
- CG
  5672   5670   5673       1    1.0950e+02    4.1840e+02 ;    HD3 - CD
- CG
  5673   5676   5677       1    1.0950e+02    4.1840e+02 ;     CG - CB
- HB2
  5673   5676   5678       1    1.0950e+02    4.1840e+02 ;     CG - CB
- HB3
  5673   5676   5679       1    1.0950e+02    3.3472e+02 ;     CG - CB
- CA
  5674   5673   5675       1    1.0950e+02    2.9288e+02 ;    HG2 - CG
- HG3
  5674   5673   5676       1    1.0950e+02    4.1840e+02 ;    HG2 - CG
- CB
  5675   5673   5676       1    1.0950e+02    4.1840e+02 ;    HG3 - CG
- CB
  5676   5679   5680       1    1.0950e+02    4.1840e+02 ;     CB - CA
- HA
  5676   5679   5681       1    1.1110e+02    5.2718e+02 ;     CB - CA
- C
  5677   5676   5678       1    1.0950e+02    2.9288e+02 ;    HB2 - CB
- HB3
  5677   5676   5679       1    1.0950e+02    4.1840e+02 ;    HB2 - CB
- CA
  5678   5676   5679       1    1.0950e+02    4.1840e+02 ;    HB3 - CB
- CA
  5679   5681   5682       1    1.2040e+02    6.6944e+02 ;     CA - C
- O
  5679   5681   5683       1    1.1660e+02    5.8576e+02 ;     CA - C
- N
  5680   5679   5681       1    1.0950e+02    4.1840e+02 ;     HA - CA
- C
  5681   5683   5684       1    1.2190e+02    4.1840e+02 ;      C - N
- CD
  5681   5683   5693       1    1.2190e+02    4.1840e+02 ;      C - N
- CA
  5682   5681   5683       1    1.2290e+02    6.6944e+02 ;      O - C
- N
  5683   5684   5685       1    1.0950e+02    4.1840e+02 ;      N - CD
- HD2
  5683   5684   5686       1    1.0950e+02    4.1840e+02 ;      N - CD
- HD3
  5683   5684   5687       1    1.0970e+02    6.6944e+02 ;      N - CD
- CG
  5683   5693   5690       1    1.0970e+02    6.6944e+02 ;      N - CA
- CB
  5683   5693   5694       1    1.0950e+02    4.1840e+02 ;      N - CA
- HA
  5683   5693   5695       1    1.1010e+02    5.2718e+02 ;      N - CA
- C
  5684   5683   5693       1    1.1800e+02    4.1840e+02 ;     CD - N
- CA
```

5684 5687 5688 1 1.0950e+02 4.1840e+02 ; CD - CG
- HG2
5684 5687 5689 1 1.0950e+02 4.1840e+02 ; CD - CG
- HG3
5684 5687 5690 1 1.0950e+02 3.3472e+02 ; CD - CG
- CB
5685 5684 5686 1 1.0950e+02 2.9288e+02 ; HD2 - CD
- HD3
5685 5684 5687 1 1.0950e+02 4.1840e+02 ; HD2 - CD
- CG
5686 5684 5687 1 1.0950e+02 4.1840e+02 ; HD3 - CD
- CG
5687 5690 5691 1 1.0950e+02 4.1840e+02 ; CG - CB
- HB2
5687 5690 5692 1 1.0950e+02 4.1840e+02 ; CG - CB
- HB3
5687 5690 5693 1 1.0950e+02 3.3472e+02 ; CG - CB
- CA
5688 5687 5689 1 1.0950e+02 2.9288e+02 ; HG2 - CG
- HG3
5688 5687 5690 1 1.0950e+02 4.1840e+02 ; HG2 - CG
- CB
5689 5687 5690 1 1.0950e+02 4.1840e+02 ; HG3 - CG
- CB
5690 5693 5694 1 1.0950e+02 4.1840e+02 ; CB - CA
- HA
5690 5693 5695 1 1.1110e+02 5.2718e+02 ; CB - CA
- C
5691 5690 5692 1 1.0950e+02 2.9288e+02 ; HB2 - CB
- HB3
5691 5690 5693 1 1.0950e+02 4.1840e+02 ; HB2 - CB
- CA
5692 5690 5693 1 1.0950e+02 4.1840e+02 ; HB3 - CB
- CA
5693 5695 5696 1 1.2040e+02 6.6944e+02 ; CA - C
- O
5693 5695 5697 1 1.1660e+02 5.8576e+02 ; CA - C
- N
5694 5693 5695 1 1.0950e+02 4.1840e+02 ; HA - CA
- C
5695 5697 5698 1 1.2000e+02 4.1840e+02 ; C - N
- H
5695 5697 5699 1 1.2190e+02 4.1840e+02 ; C - N
- CA
5696 5695 5697 1 1.2290e+02 6.6944e+02 ; O - C
- N
5697 5699 5700 1 1.0950e+02 4.1840e+02 ; N - CA
- HA
5697 5699 5701 1 1.0970e+02 6.6944e+02 ; N - CA
- CB
5697 5699 5707 1 1.1010e+02 5.2718e+02 ; N - CA
- C

5698 5697 5699 1 1.1804e+02 4.1840e+02 ; H - N
- CA
5699 5701 5702 1 1.0950e+02 4.1840e+02 ; CA - CB
- HB2
5699 5701 5703 1 1.0950e+02 4.1840e+02 ; CA - CB
- HB3
5699 5701 5704 1 1.1110e+02 5.2718e+02 ; CA - CB
- CG
5699 5707 5708 1 1.2040e+02 6.6944e+02 ; CA - C
- O
5699 5707 5709 1 1.1660e+02 5.8576e+02 ; CA - C
- N
5700 5699 5701 1 1.0950e+02 4.1840e+02 ; HA - CA
- CB
5700 5699 5707 1 1.0950e+02 4.1840e+02 ; HA - CA
- C
5701 5699 5707 1 1.1110e+02 5.2718e+02 ; CB - CA
- C
5701 5704 5705 1 1.1700e+02 5.8576e+02 ; CB - CG
- OD1
5701 5704 5706 1 1.1700e+02 5.8576e+02 ; CB - CG
- OD2
5702 5701 5703 1 1.0950e+02 2.9288e+02 ; HB2 - CB
- HB3
5702 5701 5704 1 1.0950e+02 4.1840e+02 ; HB2 - CB
- CG
5703 5701 5704 1 1.0950e+02 4.1840e+02 ; HB3 - CB
- CG
5705 5704 5706 1 1.2600e+02 6.6944e+02 ; OD1 - CG
- OD2
5707 5709 5710 1 1.2000e+02 4.1840e+02 ; C - N
- H
5707 5709 5711 1 1.2190e+02 4.1840e+02 ; C - N
- CA
5708 5707 5709 1 1.2290e+02 6.6944e+02 ; O - C
- N
5709 5711 5712 1 1.0950e+02 4.1840e+02 ; N - CA
- HA
5709 5711 5713 1 1.0970e+02 6.6944e+02 ; N - CA
- CB
5709 5711 5726 1 1.1010e+02 5.2718e+02 ; N - CA
- C
5710 5709 5711 1 1.1804e+02 4.1840e+02 ; H - N
- CA
5711 5713 5714 1 1.0950e+02 4.1840e+02 ; CA - CB
- HB
5711 5713 5715 1 1.0950e+02 3.3472e+02 ; CA - CB
- CG2
5711 5713 5719 1 1.0950e+02 3.3472e+02 ; CA - CB
- CG1
5711 5726 5727 1 1.2040e+02 6.6944e+02 ; CA - C
- O

5711 5726 5728 1 1.1660e+02 5.8576e+02 ; CA - C
- N
5712 5711 5713 1 1.0950e+02 4.1840e+02 ; HA - CA
- CB
5712 5711 5726 1 1.0950e+02 4.1840e+02 ; HA - CA
- C
5713 5711 5726 1 1.1110e+02 5.2718e+02 ; CB - CA
- C
5713 5715 5716 1 1.0950e+02 4.1840e+02 ; CB - CG2
- HG21
5713 5715 5717 1 1.0950e+02 4.1840e+02 ; CB - CG2
- HG22
5713 5715 5718 1 1.0950e+02 4.1840e+02 ; CB - CG2
- HG23
5713 5719 5720 1 1.0950e+02 4.1840e+02 ; CB - CG1
- HG12
5713 5719 5721 1 1.0950e+02 4.1840e+02 ; CB - CG1
- HG13
5713 5719 5722 1 1.0950e+02 3.3472e+02 ; CB - CG1
- CD1
5714 5713 5715 1 1.0950e+02 4.1840e+02 ; HB - CB
- CG2
5714 5713 5719 1 1.0950e+02 4.1840e+02 ; HB - CB
- CG1
5715 5713 5719 1 1.0950e+02 3.3472e+02 ; CG2 - CB
- CG1
5716 5715 5717 1 1.0950e+02 2.9288e+02 ; HG21 - CG2
- HG22
5716 5715 5718 1 1.0950e+02 2.9288e+02 ; HG21 - CG2
- HG23
5717 5715 5718 1 1.0950e+02 2.9288e+02 ; HG22 - CG2
- HG23
5719 5722 5723 1 1.0950e+02 4.1840e+02 ; CG1 - CD1
- HD11
5719 5722 5724 1 1.0950e+02 4.1840e+02 ; CG1 - CD1
- HD12
5719 5722 5725 1 1.0950e+02 4.1840e+02 ; CG1 - CD1
- HD13
5720 5719 5721 1 1.0950e+02 2.9288e+02 ; HG12 - CG1
- HG13
5720 5719 5722 1 1.0950e+02 4.1840e+02 ; HG12 - CG1
- CD1
5721 5719 5722 1 1.0950e+02 4.1840e+02 ; HG13 - CG1
- CD1
5723 5722 5724 1 1.0950e+02 2.9288e+02 ; HD11 - CD1
- HD12
5723 5722 5725 1 1.0950e+02 2.9288e+02 ; HD11 - CD1
- HD13
5724 5722 5725 1 1.0950e+02 2.9288e+02 ; HD12 - CD1
- HD13
5726 5728 5729 1 1.2000e+02 4.1840e+02 ; C - N
- H

```
  5726   5728   5730      1    1.2190e+02    4.1840e+02 ;       C - N
- CA
  5727   5726   5728      1    1.2290e+02    6.6944e+02 ;       O - C
- N
  5728   5730   5731      1    1.0950e+02    4.1840e+02 ;       N - CA
- HA
  5728   5730   5732      1    1.0970e+02    6.6944e+02 ;       N - CA
- CB
  5728   5730   5743      1    1.1010e+02    5.2718e+02 ;       N - CA
- C
  5729   5728   5730      1    1.1804e+02    4.1840e+02 ;       H - N
- CA
  5730   5732   5733      1    1.0950e+02    4.1840e+02 ;      CA - CB
- HB2
  5730   5732   5734      1    1.0950e+02    4.1840e+02 ;      CA - CB
- HB3
  5730   5732   5735      1    1.1310e+02    5.2718e+02 ;      CA - CB
- CG
  5730   5743   5744      1    1.2040e+02    6.6944e+02 ;      CA - C
- O
  5730   5743   5745      1    1.1660e+02    5.8576e+02 ;      CA - C
- N
  5731   5730   5732      1    1.0950e+02    4.1840e+02 ;      HA - CA
- CB
  5731   5730   5743      1    1.0950e+02    4.1840e+02 ;      HA - CA
- C
  5732   5730   5743      1    1.1110e+02    5.2718e+02 ;      CB - CA
- C
  5732   5735   5736      1    1.2000e+02    5.8576e+02 ;      CB - CG
- ND1
  5732   5735   5741      1    1.2000e+02    5.8576e+02 ;      CB - CG
- CD2
  5733   5732   5734      1    1.0950e+02    2.9288e+02 ;     HB2 - CB
- HB3
  5733   5732   5735      1    1.0950e+02    4.1840e+02 ;     HB2 - CB
- CG
  5734   5732   5735      1    1.0950e+02    4.1840e+02 ;     HB3 - CB
- CG
  5735   5736   5737      1    1.1700e+02    5.8576e+02 ;      CG - ND1
- CE1
  5735   5741   5739      1    1.2000e+02    5.8576e+02 ;      CG - CD2
- NE2
  5735   5741   5742      1    1.2000e+02    4.1840e+02 ;      CG - CD2
- HD2
  5736   5735   5741      1    1.2000e+02    5.8576e+02 ;     ND1 - CG
- CD2
  5736   5737   5738      1    1.2000e+02    4.1840e+02 ;     ND1 - CE1
- HE1
  5736   5737   5739      1    1.2000e+02    5.8576e+02 ;     ND1 - CE1
- NE2
  5737   5739   5740      1    1.2000e+02    4.1840e+02 ;     CE1 - NE2
- HE2
```

5737 5739 5741 1 1.2000e+02 5.8576e+02 ; CE1 - NE2
- CD2
5738 5737 5739 1 1.2000e+02 4.1840e+02 ; HE1 - CE1
- NE2
5739 5741 5742 1 1.2000e+02 4.1840e+02 ; NE2 - CD2
- HD2
5740 5739 5741 1 1.2000e+02 4.1840e+02 ; HE2 - NE2
- CD2
5743 5745 5746 1 1.2000e+02 4.1840e+02 ; C - N
- H
5743 5745 5747 1 1.2190e+02 4.1840e+02 ; C - N
- CA
5744 5743 5745 1 1.2290e+02 6.6944e+02 ; O - C
- N
5745 5747 5748 1 1.0950e+02 4.1840e+02 ; N - CA
- HA
5745 5747 5749 1 1.0970e+02 6.6944e+02 ; N - CA
- CB
5745 5747 5760 1 1.1010e+02 5.2718e+02 ; N - CA
- C
5746 5745 5747 1 1.1804e+02 4.1840e+02 ; H - N
- CA
5747 5749 5750 1 1.0950e+02 4.1840e+02 ; CA - CB
- HB2
5747 5749 5751 1 1.0950e+02 4.1840e+02 ; CA - CB
- HB3
5747 5749 5752 1 1.0950e+02 3.3472e+02 ; CA - CB
- CG
5747 5760 5761 1 1.2040e+02 6.6944e+02 ; CA - C
- O
5747 5760 5762 1 1.1660e+02 5.8576e+02 ; CA - C
- N
5748 5747 5749 1 1.0950e+02 4.1840e+02 ; HA - CA
- CB
5748 5747 5760 1 1.0950e+02 4.1840e+02 ; HA - CA
- C
5749 5747 5760 1 1.1110e+02 5.2718e+02 ; CB - CA
- C
5749 5752 5753 1 1.0950e+02 4.1840e+02 ; CB - CG
- HG2
5749 5752 5754 1 1.0950e+02 4.1840e+02 ; CB - CG
- HG3
5749 5752 5755 1 1.1110e+02 5.2718e+02 ; CB - CG
- CD
5750 5749 5751 1 1.0950e+02 2.9288e+02 ; HB2 - CB
- HB3
5750 5749 5752 1 1.0950e+02 4.1840e+02 ; HB2 - CB
- CG
5751 5749 5752 1 1.0950e+02 4.1840e+02 ; HB3 - CB
- CG
5752 5755 5756 1 1.2040e+02 6.6944e+02 ; CG - CD
- OE1

5752 5755 5757 1 1.1660e+02 5.8576e+02 ; CG - CD
- NE2
5753 5752 5754 1 1.0950e+02 2.9288e+02 ; HG2 - CG
- HG3
5753 5752 5755 1 1.0950e+02 4.1840e+02 ; HG2 - CG
- CD
5754 5752 5755 1 1.0950e+02 4.1840e+02 ; HG3 - CG
- CD
5755 5757 5758 1 1.2000e+02 4.1840e+02 ; CD - NE2
- HE21
5755 5757 5759 1 1.2000e+02 4.1840e+02 ; CD - NE2
- HE22
5756 5755 5757 1 1.2290e+02 6.6944e+02 ; OE1 - CD
- NE2
5758 5757 5759 1 1.2000e+02 2.9288e+02 ; HE21 - NE2
- HE22
5760 5762 5763 1 1.2000e+02 4.1840e+02 ; C - N
- H
5760 5762 5764 1 1.2190e+02 4.1840e+02 ; C - N
- CA
5761 5760 5762 1 1.2290e+02 6.6944e+02 ; O - C
- N
5762 5764 5765 1 1.0950e+02 4.1840e+02 ; N - CA
- HA
5762 5764 5766 1 1.0970e+02 6.6944e+02 ; N - CA
- CB
5762 5764 5776 1 1.1010e+02 5.2718e+02 ; N - CA
- C
5763 5762 5764 1 1.1804e+02 4.1840e+02 ; H - N
- CA
5764 5766 5767 1 1.0950e+02 4.1840e+02 ; CA - CB
- HB
5764 5766 5768 1 1.0950e+02 3.3472e+02 ; CA - CB
- CG1
5764 5766 5772 1 1.0950e+02 3.3472e+02 ; CA - CB
- CG2
5764 5776 5777 1 1.2040e+02 6.6944e+02 ; CA - C
- O
5764 5776 5778 1 1.1660e+02 5.8576e+02 ; CA - C
- N
5765 5764 5766 1 1.0950e+02 4.1840e+02 ; HA - CA
- CB
5765 5764 5776 1 1.0950e+02 4.1840e+02 ; HA - CA
- C
5766 5764 5776 1 1.1110e+02 5.2718e+02 ; CB - CA
- C
5766 5768 5769 1 1.0950e+02 4.1840e+02 ; CB - CG1
- HG11
5766 5768 5770 1 1.0950e+02 4.1840e+02 ; CB - CG1
- HG12
5766 5768 5771 1 1.0950e+02 4.1840e+02 ; CB - CG1
- HG13

5766 5772 5773 1 1.0950e+02 4.1840e+02 ; CB - CG2
- HG21
5766 5772 5774 1 1.0950e+02 4.1840e+02 ; CB - CG2
- HG22
5766 5772 5775 1 1.0950e+02 4.1840e+02 ; CB - CG2
- HG23
5767 5766 5768 1 1.0950e+02 4.1840e+02 ; HB - CB
- CG1
5767 5766 5772 1 1.0950e+02 4.1840e+02 ; HB - CB
- CG2
5768 5766 5772 1 1.0950e+02 3.3472e+02 ; CG1 - CB
- CG2
5769 5768 5770 1 1.0950e+02 2.9288e+02 ; HG11 - CG1
- HG12
5769 5768 5771 1 1.0950e+02 2.9288e+02 ; HG11 - CG1
- HG13
5770 5768 5771 1 1.0950e+02 2.9288e+02 ; HG12 - CG1
- HG13
5773 5772 5774 1 1.0950e+02 2.9288e+02 ; HG21 - CG2
- HG22
5773 5772 5775 1 1.0950e+02 2.9288e+02 ; HG21 - CG2
- HG23
5774 5772 5775 1 1.0950e+02 2.9288e+02 ; HG22 - CG2
- HG23
5776 5778 5779 1 1.2000e+02 4.1840e+02 ; C - N
- H
5776 5778 5780 1 1.2190e+02 4.1840e+02 ; C - N
- CA
5777 5776 5778 1 1.2290e+02 6.6944e+02 ; O - C
- N
5778 5780 5781 1 1.0950e+02 4.1840e+02 ; N - CA
- HA
5778 5780 5782 1 1.0970e+02 6.6944e+02 ; N - CA
- CB
5778 5780 5795 1 1.1010e+02 5.2718e+02 ; N - CA
- C
5779 5778 5780 1 1.1804e+02 4.1840e+02 ; H - N
- CA
5780 5782 5783 1 1.0950e+02 4.1840e+02 ; CA - CB
- HB2
5780 5782 5784 1 1.0950e+02 4.1840e+02 ; CA - CB
- HB3
5780 5782 5785 1 1.0950e+02 3.3472e+02 ; CA - CB
- CG
5780 5795 5796 1 1.2040e+02 6.6944e+02 ; CA - C
- O
5780 5795 5797 1 1.1660e+02 5.8576e+02 ; CA - C
- N
5781 5780 5782 1 1.0950e+02 4.1840e+02 ; HA - CA
- CB
5781 5780 5795 1 1.0950e+02 4.1840e+02 ; HA - CA
- C

5782 5780 5795 1 1.1110e+02 5.2718e+02 ; CB - CA
- C
5782 5785 5786 1 1.0950e+02 4.1840e+02 ; CB - CG
- HG
5782 5785 5787 1 1.0950e+02 3.3472e+02 ; CB - CG
- CD1
5782 5785 5791 1 1.0950e+02 3.3472e+02 ; CB - CG
- CD2
5783 5782 5784 1 1.0950e+02 2.9288e+02 ; HB2 - CB
- HB3
5783 5782 5785 1 1.0950e+02 4.1840e+02 ; HB2 - CB
- CG
5784 5782 5785 1 1.0950e+02 4.1840e+02 ; HB3 - CB
- CG
5785 5787 5788 1 1.0950e+02 4.1840e+02 ; CG - CD1
- HD11
5785 5787 5789 1 1.0950e+02 4.1840e+02 ; CG - CD1
- HD12
5785 5787 5790 1 1.0950e+02 4.1840e+02 ; CG - CD1
- HD13
5785 5791 5792 1 1.0950e+02 4.1840e+02 ; CG - CD2
- HD21
5785 5791 5793 1 1.0950e+02 4.1840e+02 ; CG - CD2
- HD22
5785 5791 5794 1 1.0950e+02 4.1840e+02 ; CG - CD2
- HD23
5786 5785 5787 1 1.0950e+02 4.1840e+02 ; HG - CG
- CD1
5786 5785 5791 1 1.0950e+02 4.1840e+02 ; HG - CG
- CD2
5787 5785 5791 1 1.0950e+02 3.3472e+02 ; CD1 - CG
- CD2
5788 5787 5789 1 1.0950e+02 2.9288e+02 ; HD11 - CD1
- HD12
5788 5787 5790 1 1.0950e+02 2.9288e+02 ; HD11 - CD1
- HD13
5789 5787 5790 1 1.0950e+02 2.9288e+02 ; HD12 - CD1
- HD13
5792 5791 5793 1 1.0950e+02 2.9288e+02 ; HD21 - CD2
- HD22
5792 5791 5794 1 1.0950e+02 2.9288e+02 ; HD21 - CD2
- HD23
5793 5791 5794 1 1.0950e+02 2.9288e+02 ; HD22 - CD2
- HD23
5795 5797 5798 1 1.2000e+02 4.1840e+02 ; C - N
- H
5795 5797 5799 1 1.2190e+02 4.1840e+02 ; C - N
- CA
5796 5795 5797 1 1.2290e+02 6.6944e+02 ; O - C
- N
5797 5799 5800 1 1.0950e+02 4.1840e+02 ; N - CA
- HA

5797 5799 5801 1 1.0970e+02 6.6944e+02 ; N - CA
- CB
5797 5799 5819 1 1.1010e+02 5.2718e+02 ; N - CA
- C
5798 5797 5799 1 1.1804e+02 4.1840e+02 ; H - N
- CA
5799 5801 5802 1 1.0950e+02 4.1840e+02 ; CA - CB
- HB2
5799 5801 5803 1 1.0950e+02 4.1840e+02 ; CA - CB
- HB3
5799 5801 5804 1 1.0950e+02 3.3472e+02 ; CA - CB
- CG
5799 5819 5820 1 1.2040e+02 6.6944e+02 ; CA - C
- O
5799 5819 5821 1 1.1660e+02 5.8576e+02 ; CA - C
- N
5800 5799 5801 1 1.0950e+02 4.1840e+02 ; HA - CA
- CB
5800 5799 5819 1 1.0950e+02 4.1840e+02 ; HA - CA
- C
5801 5799 5819 1 1.1110e+02 5.2718e+02 ; CB - CA
- C
5801 5804 5805 1 1.0950e+02 4.1840e+02 ; CB - CG
- HG2
5801 5804 5806 1 1.0950e+02 4.1840e+02 ; CB - CG
- HG3
5801 5804 5807 1 1.0950e+02 3.3472e+02 ; CB - CG
- CD
5802 5801 5803 1 1.0950e+02 2.9288e+02 ; HB2 - CB
- HB3
5802 5801 5804 1 1.0950e+02 4.1840e+02 ; HB2 - CB
- CG
5803 5801 5804 1 1.0950e+02 4.1840e+02 ; HB3 - CB
- CG
5804 5807 5808 1 1.0950e+02 4.1840e+02 ; CG - CD
- HD2
5804 5807 5809 1 1.0950e+02 4.1840e+02 ; CG - CD
- HD3
5804 5807 5810 1 1.1120e+02 6.6944e+02 ; CG - CD
- NE
5805 5804 5806 1 1.0950e+02 2.9288e+02 ; HG2 - CG
- HG3
5805 5804 5807 1 1.0950e+02 4.1840e+02 ; HG2 - CG
- CD
5806 5804 5807 1 1.0950e+02 4.1840e+02 ; HG3 - CG
- CD
5807 5810 5811 1 1.1840e+02 4.1840e+02 ; CD - NE
- HE
5807 5810 5812 1 1.2320e+02 4.1840e+02 ; CD - NE
- CZ
5808 5807 5809 1 1.0950e+02 2.9288e+02 ; HD2 - CD
- HD3

```
  5808   5807   5810       1    1.0950e+02    4.1840e+02 ;    HD2 - CD
- NE
  5809   5807   5810       1    1.0950e+02    4.1840e+02 ;    HD3 - CD
- NE
  5810   5812   5813       1    1.2000e+02    5.8576e+02 ;     NE - CZ
- NH1
  5810   5812   5816       1    1.2000e+02    5.8576e+02 ;     NE - CZ
- NH2
  5811   5810   5812       1    1.2000e+02    4.1840e+02 ;     HE - NE
- CZ
  5812   5813   5814       1    1.2000e+02    4.1840e+02 ;     CZ - NH1
- HH11
  5812   5813   5815       1    1.2000e+02    4.1840e+02 ;     CZ - NH1
- HH12
  5812   5816   5817       1    1.2000e+02    4.1840e+02 ;     CZ - NH2
- HH21
  5812   5816   5818       1    1.2000e+02    4.1840e+02 ;     CZ - NH2
- HH22
  5813   5812   5816       1    1.2000e+02    5.8576e+02 ;    NH1 - CZ
- NH2
  5814   5813   5815       1    1.2000e+02    2.9288e+02 ;   HH11 - NH1
- HH12
  5817   5816   5818       1    1.2000e+02    2.9288e+02 ;   HH21 - NH2
- HH22
  5819   5821   5822       1    1.2000e+02    4.1840e+02 ;      C - N
- H
  5819   5821   5823       1    1.2190e+02    4.1840e+02 ;      C - N
- CA
  5820   5819   5821       1    1.2290e+02    6.6944e+02 ;      O - C
- N
  5821   5823   5824       1    1.0950e+02    4.1840e+02 ;      N - CA
- HA
  5821   5823   5825       1    1.0970e+02    6.6944e+02 ;      N - CA
- CB
  5821   5823   5831       1    1.1010e+02    5.2718e+02 ;      N - CA
- C
  5822   5821   5823       1    1.1804e+02    4.1840e+02 ;      H - N
- CA
  5823   5825   5826       1    1.0950e+02    4.1840e+02 ;     CA - CB
- HB2
  5823   5825   5827       1    1.0950e+02    4.1840e+02 ;     CA - CB
- HB3
  5823   5825   5828       1    1.1110e+02    5.2718e+02 ;     CA - CB
- CG
  5823   5831   5832       1    1.2040e+02    6.6944e+02 ;     CA - C
- O
  5823   5831   5833       1    1.1660e+02    5.8576e+02 ;     CA - C
- N
  5824   5823   5825       1    1.0950e+02    4.1840e+02 ;     HA - CA
- CB
  5824   5823   5831       1    1.0950e+02    4.1840e+02 ;     HA - CA
- C
```

5825 5823 5831 1 1.1110e+02 5.2718e+02 ; CB - CA
- C
5825 5828 5829 1 1.1700e+02 5.8576e+02 ; CB - CG
- OD1
5825 5828 5830 1 1.1700e+02 5.8576e+02 ; CB - CG
- OD2
5826 5825 5827 1 1.0950e+02 2.9288e+02 ; HB2 - CB
- HB3
5826 5825 5828 1 1.0950e+02 4.1840e+02 ; HB2 - CB
- CG
5827 5825 5828 1 1.0950e+02 4.1840e+02 ; HB3 - CB
- CG
5829 5828 5830 1 1.2600e+02 6.6944e+02 ; OD1 - CG
- OD2
5831 5833 5834 1 1.2000e+02 4.1840e+02 ; C - N
- H
5831 5833 5835 1 1.2190e+02 4.1840e+02 ; C - N
- CA
5832 5831 5833 1 1.2290e+02 6.6944e+02 ; O - C
- N
5833 5835 5836 1 1.0950e+02 4.1840e+02 ; N - CA
- HA
5833 5835 5837 1 1.0970e+02 6.6944e+02 ; N - CA
- CB
5833 5835 5850 1 1.1010e+02 5.2718e+02 ; N - CA
- C
5834 5833 5835 1 1.1804e+02 4.1840e+02 ; H - N
- CA
5835 5837 5838 1 1.0950e+02 4.1840e+02 ; CA - CB
- HB
5835 5837 5839 1 1.0950e+02 3.3472e+02 ; CA - CB
- CG2
5835 5837 5843 1 1.0950e+02 3.3472e+02 ; CA - CB
- CG1
5835 5850 5851 1 1.2040e+02 6.6944e+02 ; CA - C
- O
5835 5850 5852 1 1.1660e+02 5.8576e+02 ; CA - C
- N
5836 5835 5837 1 1.0950e+02 4.1840e+02 ; HA - CA
- CB
5836 5835 5850 1 1.0950e+02 4.1840e+02 ; HA - CA
- C
5837 5835 5850 1 1.1110e+02 5.2718e+02 ; CB - CA
- C
5837 5839 5840 1 1.0950e+02 4.1840e+02 ; CB - CG2
- HG21
5837 5839 5841 1 1.0950e+02 4.1840e+02 ; CB - CG2
- HG22
5837 5839 5842 1 1.0950e+02 4.1840e+02 ; CB - CG2
- HG23
5837 5843 5844 1 1.0950e+02 4.1840e+02 ; CB - CG1
- HG12

5837 5843 5845 1 1.0950e+02 4.1840e+02 ; CB - CG1
- HG13
5837 5843 5846 1 1.0950e+02 3.3472e+02 ; CB - CG1
- CD1
5838 5837 5839 1 1.0950e+02 4.1840e+02 ; HB - CB
- CG2
5838 5837 5843 1 1.0950e+02 4.1840e+02 ; HB - CB
- CG1
5839 5837 5843 1 1.0950e+02 3.3472e+02 ; CG2 - CB
- CG1
5840 5839 5841 1 1.0950e+02 2.9288e+02 ; HG21 - CG2
- HG22
5840 5839 5842 1 1.0950e+02 2.9288e+02 ; HG21 - CG2
- HG23
5841 5839 5842 1 1.0950e+02 2.9288e+02 ; HG22 - CG2
- HG23
5843 5846 5847 1 1.0950e+02 4.1840e+02 ; CG1 - CD1
- HD11
5843 5846 5848 1 1.0950e+02 4.1840e+02 ; CG1 - CD1
- HD12
5843 5846 5849 1 1.0950e+02 4.1840e+02 ; CG1 - CD1
- HD13
5844 5843 5845 1 1.0950e+02 2.9288e+02 ; HG12 - CG1
- HG13
5844 5843 5846 1 1.0950e+02 4.1840e+02 ; HG12 - CG1
- CD1
5845 5843 5846 1 1.0950e+02 4.1840e+02 ; HG13 - CG1
- CD1
5847 5846 5848 1 1.0950e+02 2.9288e+02 ; HD11 - CD1
- HD12
5847 5846 5849 1 1.0950e+02 2.9288e+02 ; HD11 - CD1
- HD13
5848 5846 5849 1 1.0950e+02 2.9288e+02 ; HD12 - CD1
- HD13
5850 5852 5853 1 1.2000e+02 4.1840e+02 ; C - N
- H
5850 5852 5854 1 1.2190e+02 4.1840e+02 ; C - N
- CA
5851 5850 5852 1 1.2290e+02 6.6944e+02 ; O - C
- N
5852 5854 5855 1 1.0950e+02 4.1840e+02 ; N - CA
- HA
5852 5854 5856 1 1.0970e+02 6.6944e+02 ; N - CA
- CB
5852 5854 5862 1 1.1010e+02 5.2718e+02 ; N - CA
- C
5853 5852 5854 1 1.1804e+02 4.1840e+02 ; H - N
- CA
5854 5856 5857 1 1.0950e+02 4.1840e+02 ; CA - CB
- HB2
5854 5856 5858 1 1.0950e+02 4.1840e+02 ; CA - CB
- HB3

5854 5856 5859 1 1.1110e+02 5.2718e+02 ; CA - CB
- CG
5854 5862 5863 1 1.2040e+02 6.6944e+02 ; CA - C
- O
5854 5862 5864 1 1.1660e+02 5.8576e+02 ; CA - C
- N
5855 5854 5856 1 1.0950e+02 4.1840e+02 ; HA - CA
- CB
5855 5854 5862 1 1.0950e+02 4.1840e+02 ; HA - CA
- C
5856 5854 5862 1 1.1110e+02 5.2718e+02 ; CB - CA
- C
5856 5859 5860 1 1.1700e+02 5.8576e+02 ; CB - CG
- OD1
5856 5859 5861 1 1.1700e+02 5.8576e+02 ; CB - CG
- OD2
5857 5856 5858 1 1.0950e+02 2.9288e+02 ; HB2 - CB
- HB3
5857 5856 5859 1 1.0950e+02 4.1840e+02 ; HB2 - CB
- CG
5858 5856 5859 1 1.0950e+02 4.1840e+02 ; HB3 - CB
- CG
5860 5859 5861 1 1.2600e+02 6.6944e+02 ; OD1 - CG
- OD2
5862 5864 5865 1 1.2000e+02 4.1840e+02 ; C - N
- H
5862 5864 5866 1 1.2190e+02 4.1840e+02 ; C - N
- CA
5863 5862 5864 1 1.2290e+02 6.6944e+02 ; O - C
- N
5864 5866 5867 1 1.0950e+02 4.1840e+02 ; N - CA
- HA
5864 5866 5868 1 1.0970e+02 6.6944e+02 ; N - CA
- CB
5864 5866 5873 1 1.1010e+02 5.2718e+02 ; N - CA
- C
5865 5864 5866 1 1.1804e+02 4.1840e+02 ; H - N
- CA
5866 5868 5869 1 1.0950e+02 4.1840e+02 ; CA - CB
- HB2
5866 5868 5870 1 1.0950e+02 4.1840e+02 ; CA - CB
- HB3
5866 5868 5871 1 1.0950e+02 4.1840e+02 ; CA - CB
- OG
5866 5873 5874 1 1.2040e+02 6.6944e+02 ; CA - C
- O
5866 5873 5875 1 1.1660e+02 5.8576e+02 ; CA - C
- N
5867 5866 5868 1 1.0950e+02 4.1840e+02 ; HA - CA
- CB
5867 5866 5873 1 1.0950e+02 4.1840e+02 ; HA - CA
- C

5868 5866 5873 1 1.1110e+02 5.2718e+02 ; CB - CA
- C
5868 5871 5872 1 1.0850e+02 4.6024e+02 ; CB - OG
- HG
5869 5868 5870 1 1.0950e+02 2.9288e+02 ; HB2 - CB
- HB3
5869 5868 5871 1 1.0950e+02 4.1840e+02 ; HB2 - CB
- OG
5870 5868 5871 1 1.0950e+02 4.1840e+02 ; HB3 - CB
- OG
5873 5875 5876 1 1.2000e+02 4.1840e+02 ; C - N
- H
5873 5875 5877 1 1.2190e+02 4.1840e+02 ; C - N
- CA
5874 5873 5875 1 1.2290e+02 6.6944e+02 ; O - C
- N
5875 5877 5878 1 1.0950e+02 4.1840e+02 ; N - CA
- HA
5875 5877 5879 1 1.0970e+02 6.6944e+02 ; N - CA
- CB
5875 5877 5887 1 1.1010e+02 5.2718e+02 ; N - CA
- C
5876 5875 5877 1 1.1804e+02 4.1840e+02 ; H - N
- CA
5877 5879 5880 1 1.0950e+02 4.1840e+02 ; CA - CB
- HB2
5877 5879 5881 1 1.0950e+02 4.1840e+02 ; CA - CB
- HB3
5877 5879 5882 1 1.1110e+02 5.2718e+02 ; CA - CB
- CG
5877 5887 5888 1 1.2040e+02 6.6944e+02 ; CA - C
- O
5877 5887 5889 1 1.1660e+02 5.8576e+02 ; CA - C
- N
5878 5877 5879 1 1.0950e+02 4.1840e+02 ; HA - CA
- CB
5878 5877 5887 1 1.0950e+02 4.1840e+02 ; HA - CA
- C
5879 5877 5887 1 1.1110e+02 5.2718e+02 ; CB - CA
- C
5879 5882 5883 1 1.2040e+02 6.6944e+02 ; CB - CG
- OD1
5879 5882 5884 1 1.1660e+02 5.8576e+02 ; CB - CG
- ND2
5880 5879 5881 1 1.0950e+02 2.9288e+02 ; HB2 - CB
- HB3
5880 5879 5882 1 1.0950e+02 4.1840e+02 ; HB2 - CB
- CG
5881 5879 5882 1 1.0950e+02 4.1840e+02 ; HB3 - CB
- CG
5882 5884 5885 1 1.2000e+02 4.1840e+02 ; CG - ND2
- HD21

5882 5884 5886 1 1.2000e+02 4.1840e+02 ; CG - ND2
- HD22
5883 5882 5884 1 1.2290e+02 6.6944e+02 ; OD1 - CG
- ND2
5885 5884 5886 1 1.2000e+02 2.9288e+02 ; HD21 - ND2
- HD22
5887 5889 5890 1 1.2000e+02 4.1840e+02 ; C - N
- H
5887 5889 5891 1 1.2190e+02 4.1840e+02 ; C - N
- CA
5888 5887 5889 1 1.2290e+02 6.6944e+02 ; O - C
- N
5889 5891 5892 1 1.0950e+02 4.1840e+02 ; N - CA
- HA
5889 5891 5893 1 1.0970e+02 6.6944e+02 ; N - CA
- CB
5889 5891 5897 1 1.1010e+02 5.2718e+02 ; N - CA
- C
5890 5889 5891 1 1.1804e+02 4.1840e+02 ; H - N
- CA
5891 5893 5894 1 1.0950e+02 4.1840e+02 ; CA - CB
- HB1
5891 5893 5895 1 1.0950e+02 4.1840e+02 ; CA - CB
- HB2
5891 5893 5896 1 1.0950e+02 4.1840e+02 ; CA - CB
- HB3
5891 5897 5898 1 1.2040e+02 6.6944e+02 ; CA - C
- O
5891 5897 5899 1 1.1660e+02 5.8576e+02 ; CA - C
- N
5892 5891 5893 1 1.0950e+02 4.1840e+02 ; HA - CA
- CB
5892 5891 5897 1 1.0950e+02 4.1840e+02 ; HA - CA
- C
5893 5891 5897 1 1.1110e+02 5.2718e+02 ; CB - CA
- C
5894 5893 5895 1 1.0950e+02 2.9288e+02 ; HB1 - CB
- HB2
5894 5893 5896 1 1.0950e+02 2.9288e+02 ; HB1 - CB
- HB3
5895 5893 5896 1 1.0950e+02 2.9288e+02 ; HB2 - CB
- HB3
5897 5899 5900 1 1.2000e+02 4.1840e+02 ; C - N
- H
5897 5899 5901 1 1.2190e+02 4.1840e+02 ; C - N
- CA
5898 5897 5899 1 1.2290e+02 6.6944e+02 ; O - C
- N
5899 5901 5902 1 1.0950e+02 4.1840e+02 ; N - CA
- HA
5899 5901 5903 1 1.0970e+02 6.6944e+02 ; N - CA
- CB

```
  5899   5901   5908       1    1.1010e+02    5.2718e+02 ;      N - CA
- C
  5900   5899   5901       1    1.1804e+02    4.1840e+02 ;      H - N
- CA
  5901   5903   5904       1    1.0950e+02    4.1840e+02 ;     CA - CB
- HB2
  5901   5903   5905       1    1.0950e+02    4.1840e+02 ;     CA - CB
- HB3
  5901   5903   5906       1    1.0950e+02    4.1840e+02 ;     CA - CB
- OG
  5901   5908   5909       1    1.2040e+02    6.6944e+02 ;     CA - C
- O
  5901   5908   5910       1    1.1660e+02    5.8576e+02 ;     CA - C
- N
  5902   5901   5903       1    1.0950e+02    4.1840e+02 ;     HA - CA
- CB
  5902   5901   5908       1    1.0950e+02    4.1840e+02 ;     HA - CA
- C
  5903   5901   5908       1    1.1110e+02    5.2718e+02 ;     CB - CA
- C
  5903   5906   5907       1    1.0850e+02    4.6024e+02 ;     CB - OG
- HG
  5904   5903   5905       1    1.0950e+02    2.9288e+02 ;    HB2 - CB
- HB3
  5904   5903   5906       1    1.0950e+02    4.1840e+02 ;    HB2 - CB
- OG
  5905   5903   5906       1    1.0950e+02    4.1840e+02 ;    HB3 - CB
- OG
  5908   5910   5911       1    1.2000e+02    4.1840e+02 ;      C - N
- H
  5908   5910   5912       1    1.2190e+02    4.1840e+02 ;      C - N
- CA
  5909   5908   5910       1    1.2290e+02    6.6944e+02 ;      O - C
- N
  5910   5912   5913       1    1.0950e+02    4.1840e+02 ;      N - CA
- HA2
  5910   5912   5914       1    1.0950e+02    4.1840e+02 ;      N - CA
- HA3
  5910   5912   5915       1    1.1010e+02    5.2718e+02 ;      N - CA
- C
  5911   5910   5912       1    1.1804e+02    4.1840e+02 ;      H - N
- CA
  5912   5915   5916       1    1.2040e+02    6.6944e+02 ;     CA - C
- O
  5912   5915   5917       1    1.1660e+02    5.8576e+02 ;     CA - C
- N
  5913   5912   5914       1    1.0950e+02    2.9288e+02 ;    HA2 - CA
- HA3
  5913   5912   5915       1    1.0950e+02    4.1840e+02 ;    HA2 - CA
- C
  5914   5912   5915       1    1.0950e+02    4.1840e+02 ;    HA3 - CA
- C
```

```
  5915   5917   5918       1    1.2000e+02    4.1840e+02 ;      C - N
- H
  5915   5917   5919       1    1.2190e+02    4.1840e+02 ;      C - N
- CA
  5916   5915   5917       1    1.2290e+02    6.6944e+02 ;      O - C
- N
  5917   5919   5920       1    1.0950e+02    4.1840e+02 ;      N - CA
- HA
  5917   5919   5921       1    1.0970e+02    6.6944e+02 ;      N - CA
- CB
  5917   5919   5932       1    1.1010e+02    5.2718e+02 ;      N - CA
- C
  5918   5917   5919       1    1.1804e+02    4.1840e+02 ;      H - N
- CA
  5919   5921   5922       1    1.0950e+02    4.1840e+02 ;     CA - CB
- HB2
  5919   5921   5923       1    1.0950e+02    4.1840e+02 ;     CA - CB
- HB3
  5919   5921   5924       1    1.0950e+02    3.3472e+02 ;     CA - CB
- CG
  5919   5932   5933       1    1.2040e+02    6.6944e+02 ;     CA - C
- O
  5919   5932   5934       1    1.1660e+02    5.8576e+02 ;     CA - C
- N
  5920   5919   5921       1    1.0950e+02    4.1840e+02 ;     HA - CA
- CB
  5920   5919   5932       1    1.0950e+02    4.1840e+02 ;     HA - CA
- C
  5921   5919   5932       1    1.1110e+02    5.2718e+02 ;     CB - CA
- C
  5921   5924   5925       1    1.0950e+02    4.1840e+02 ;     CB - CG
- HG2
  5921   5924   5926       1    1.0950e+02    4.1840e+02 ;     CB - CG
- HG3
  5921   5924   5927       1    1.1110e+02    5.2718e+02 ;     CB - CG
- CD
  5922   5921   5923       1    1.0950e+02    2.9288e+02 ;    HB2 - CB
- HB3
  5922   5921   5924       1    1.0950e+02    4.1840e+02 ;    HB2 - CB
- CG
  5923   5921   5924       1    1.0950e+02    4.1840e+02 ;    HB3 - CB
- CG
  5924   5927   5928       1    1.2040e+02    6.6944e+02 ;     CG - CD
- OE1
  5924   5927   5929       1    1.1660e+02    5.8576e+02 ;     CG - CD
- NE2
  5925   5924   5926       1    1.0950e+02    2.9288e+02 ;    HG2 - CG
- HG3
  5925   5924   5927       1    1.0950e+02    4.1840e+02 ;    HG2 - CG
- CD
  5926   5924   5927       1    1.0950e+02    4.1840e+02 ;    HG3 - CG
- CD
```

```
  5927   5929   5930      1    1.2000e+02    4.1840e+02 ;     CD - NE2
- HE21
  5927   5929   5931      1    1.2000e+02    4.1840e+02 ;     CD - NE2
- HE22
  5928   5927   5929      1    1.2290e+02    6.6944e+02 ;    OE1 - CD
- NE2
  5930   5929   5931      1    1.2000e+02    2.9288e+02 ;   HE21 - NE2
- HE22
  5932   5934   5935      1    1.2000e+02    4.1840e+02 ;      C - N
- H
  5932   5934   5936      1    1.2190e+02    4.1840e+02 ;      C - N
- CA
  5933   5932   5934      1    1.2290e+02    6.6944e+02 ;      O - C
- N
  5934   5936   5937      1    1.0950e+02    4.1840e+02 ;      N - CA
- HA
  5934   5936   5938      1    1.0970e+02    6.6944e+02 ;      N - CA
- CB
  5934   5936   5951      1    1.1010e+02    5.2718e+02 ;      N - CA
- C
  5935   5934   5936      1    1.1804e+02    4.1840e+02 ;      H - N
- CA
  5936   5938   5939      1    1.0950e+02    4.1840e+02 ;     CA - CB
- HB
  5936   5938   5940      1    1.0950e+02    3.3472e+02 ;     CA - CB
- CG2
  5936   5938   5944      1    1.0950e+02    3.3472e+02 ;     CA - CB
- CG1
  5936   5951   5952      1    1.2040e+02    6.6944e+02 ;     CA - C
- O
  5936   5951   5953      1    1.1660e+02    5.8576e+02 ;     CA - C
- N
  5937   5936   5938      1    1.0950e+02    4.1840e+02 ;     HA - CA
- CB
  5937   5936   5951      1    1.0950e+02    4.1840e+02 ;     HA - CA
- C
  5938   5936   5951      1    1.1110e+02    5.2718e+02 ;     CB - CA
- C
  5938   5940   5941      1    1.0950e+02    4.1840e+02 ;     CB - CG2
- HG21
  5938   5940   5942      1    1.0950e+02    4.1840e+02 ;     CB - CG2
- HG22
  5938   5940   5943      1    1.0950e+02    4.1840e+02 ;     CB - CG2
- HG23
  5938   5944   5945      1    1.0950e+02    4.1840e+02 ;     CB - CG1
- HG12
  5938   5944   5946      1    1.0950e+02    4.1840e+02 ;     CB - CG1
- HG13
  5938   5944   5947      1    1.0950e+02    3.3472e+02 ;     CB - CG1
- CD1
  5939   5938   5940      1    1.0950e+02    4.1840e+02 ;     HB - CB
- CG2
```

5939 5938 5944 1 1.0950e+02 4.1840e+02 ; HB - CB
- CG1
5940 5938 5944 1 1.0950e+02 3.3472e+02 ; CG2 - CB
- CG1
5941 5940 5942 1 1.0950e+02 2.9288e+02 ; HG21 - CG2
- HG22
5941 5940 5943 1 1.0950e+02 2.9288e+02 ; HG21 - CG2
- HG23
5942 5940 5943 1 1.0950e+02 2.9288e+02 ; HG22 - CG2
- HG23
5944 5947 5948 1 1.0950e+02 4.1840e+02 ; CG1 - CD1
- HD11
5944 5947 5949 1 1.0950e+02 4.1840e+02 ; CG1 - CD1
- HD12
5944 5947 5950 1 1.0950e+02 4.1840e+02 ; CG1 - CD1
- HD13
5945 5944 5946 1 1.0950e+02 2.9288e+02 ; HG12 - CG1
- HG13
5945 5944 5947 1 1.0950e+02 4.1840e+02 ; HG12 - CG1
- CD1
5946 5944 5947 1 1.0950e+02 4.1840e+02 ; HG13 - CG1
- CD1
5948 5947 5949 1 1.0950e+02 2.9288e+02 ; HD11 - CD1
- HD12
5948 5947 5950 1 1.0950e+02 2.9288e+02 ; HD11 - CD1
- HD13
5949 5947 5950 1 1.0950e+02 2.9288e+02 ; HD12 - CD1
- HD13
5951 5953 5954 1 1.2000e+02 4.1840e+02 ; C - N
- H
5951 5953 5955 1 1.2190e+02 4.1840e+02 ; C - N
- CA
5952 5951 5953 1 1.2290e+02 6.6944e+02 ; O - C
- N
5953 5955 5956 1 1.0950e+02 4.1840e+02 ; N - CA
- HA
5953 5955 5957 1 1.0970e+02 6.6944e+02 ; N - CA
- CB
5953 5955 5969 1 1.1010e+02 5.2718e+02 ; N - CA
- C
5954 5953 5955 1 1.1804e+02 4.1840e+02 ; H - N
- CA
5955 5957 5958 1 1.0950e+02 4.1840e+02 ; CA - CB
- HB2
5955 5957 5959 1 1.0950e+02 4.1840e+02 ; CA - CB
- HB3
5955 5957 5960 1 1.1310e+02 5.2718e+02 ; CA - CB
- CG
5955 5969 5970 1 1.2040e+02 6.6944e+02 ; CA - C
- O
5955 5969 5971 1 1.1660e+02 5.8576e+02 ; CA - C
- N

5956 5955 5957 1 1.0950e+02 4.1840e+02 ; HA - CA
- CB
5956 5955 5969 1 1.0950e+02 4.1840e+02 ; HA - CA
- C
5957 5955 5969 1 1.1110e+02 5.2718e+02 ; CB - CA
- C
5957 5960 5961 1 1.2000e+02 5.8576e+02 ; CB - CG
- ND1
5957 5960 5967 1 1.2000e+02 5.8576e+02 ; CB - CG
- CD2
5958 5957 5959 1 1.0950e+02 2.9288e+02 ; HB2 - CB
- HB3
5958 5957 5960 1 1.0950e+02 4.1840e+02 ; HB2 - CB
- CG
5959 5957 5960 1 1.0950e+02 4.1840e+02 ; HB3 - CB
- CG
5960 5961 5962 1 1.2000e+02 4.1840e+02 ; CG - ND1
- HD1
5960 5961 5963 1 1.2000e+02 5.8576e+02 ; CG - ND1
- CE1
5960 5967 5965 1 1.2000e+02 5.8576e+02 ; CG - CD2
- NE2
5960 5967 5968 1 1.2000e+02 4.1840e+02 ; CG - CD2
- HD2
5961 5960 5967 1 1.2000e+02 5.8576e+02 ; ND1 - CG
- CD2
5961 5963 5964 1 1.2000e+02 4.1840e+02 ; ND1 - CE1
- HE1
5961 5963 5965 1 1.2000e+02 5.8576e+02 ; ND1 - CE1
- NE2
5962 5961 5963 1 1.2000e+02 4.1840e+02 ; HD1 - ND1
- CE1
5963 5965 5966 1 1.2000e+02 4.1840e+02 ; CE1 - NE2
- HE2
5963 5965 5967 1 1.2000e+02 5.8576e+02 ; CE1 - NE2
- CD2
5964 5963 5965 1 1.2000e+02 4.1840e+02 ; HE1 - CE1
- NE2
5965 5967 5968 1 1.2000e+02 4.1840e+02 ; NE2 - CD2
- HD2
5966 5965 5967 1 1.2000e+02 4.1840e+02 ; HE2 - NE2
- CD2
5969 5971 5972 1 1.2000e+02 4.1840e+02 ; C - N
- H
5969 5971 5973 1 1.2190e+02 4.1840e+02 ; C - N
- CA
5970 5969 5971 1 1.2290e+02 6.6944e+02 ; O - C
- N
5971 5973 5974 1 1.0950e+02 4.1840e+02 ; N - CA
- HA
5971 5973 5975 1 1.0970e+02 6.6944e+02 ; N - CA
- CB

5971 5973 5990 1 1.1010e+02 5.2718e+02 ; N - CA
- C
5972 5971 5973 1 1.1804e+02 4.1840e+02 ; H - N
- CA
5973 5975 5976 1 1.0950e+02 4.1840e+02 ; CA - CB
- HB2
5973 5975 5977 1 1.0950e+02 4.1840e+02 ; CA - CB
- HB3
5973 5975 5978 1 1.1400e+02 5.2718e+02 ; CA - CB
- CG
5973 5990 5991 1 1.2040e+02 6.6944e+02 ; CA - C
- O
5973 5990 5992 1 1.1660e+02 5.8576e+02 ; CA - C
- N
5974 5973 5975 1 1.0950e+02 4.1840e+02 ; HA - CA
- CB
5974 5973 5990 1 1.0950e+02 4.1840e+02 ; HA - CA
- C
5975 5973 5990 1 1.1110e+02 5.2718e+02 ; CB - CA
- C
5975 5978 5979 1 1.2000e+02 5.8576e+02 ; CB - CG
- CD1
5975 5978 5988 1 1.2000e+02 5.8576e+02 ; CB - CG
- CD2
5976 5975 5977 1 1.0950e+02 2.9288e+02 ; HB2 - CB
- HB3
5976 5975 5978 1 1.0950e+02 4.1840e+02 ; HB2 - CB
- CG
5977 5975 5978 1 1.0950e+02 4.1840e+02 ; HB3 - CB
- CG
5978 5979 5980 1 1.2000e+02 4.1840e+02 ; CG - CD1
- HD1
5978 5979 5981 1 1.2000e+02 5.2718e+02 ; CG - CD1
- CE1
5978 5988 5986 1 1.2000e+02 5.2718e+02 ; CG - CD2
- CE2
5978 5988 5989 1 1.2000e+02 4.1840e+02 ; CG - CD2
- HD2
5979 5978 5988 1 1.2000e+02 5.2718e+02 ; CD1 - CG
- CD2
5979 5981 5982 1 1.2000e+02 4.1840e+02 ; CD1 - CE1
- HE1
5979 5981 5983 1 1.2000e+02 5.2718e+02 ; CD1 - CE1
- CZ
5980 5979 5981 1 1.2000e+02 4.1840e+02 ; HD1 - CD1
- CE1
5981 5983 5984 1 1.2000e+02 5.8576e+02 ; CE1 - CZ
- OH
5981 5983 5986 1 1.2000e+02 5.2718e+02 ; CE1 - CZ
- CE2
5982 5981 5983 1 1.2000e+02 4.1840e+02 ; HE1 - CE1
- CZ

5983 5984 5985 1 1.1300e+02 4.1840e+02 ; CZ - OH
- HH
5983 5986 5987 1 1.2000e+02 4.1840e+02 ; CZ - CE2
- HE2
5983 5986 5988 1 1.2000e+02 5.2718e+02 ; CZ - CE2
- CD2
5984 5983 5986 1 1.2000e+02 5.8576e+02 ; OH - CZ
- CE2
5986 5988 5989 1 1.2000e+02 4.1840e+02 ; CE2 - CD2
- HD2
5987 5986 5988 1 1.2000e+02 4.1840e+02 ; HE2 - CE2
- CD2
5990 5992 5993 1 1.2000e+02 4.1840e+02 ; C - N
- H
5990 5992 5994 1 1.2190e+02 4.1840e+02 ; C - N
- CA
5991 5990 5992 1 1.2290e+02 6.6944e+02 ; O - C
- N
5992 5994 5995 1 1.0950e+02 4.1840e+02 ; N - CA
- HA
5992 5994 5996 1 1.0970e+02 6.6944e+02 ; N - CA
- CB
5992 5994 6004 1 1.1010e+02 5.2718e+02 ; N - CA
- C
5993 5992 5994 1 1.1804e+02 4.1840e+02 ; H - N
- CA
5994 5996 5997 1 1.0950e+02 4.1840e+02 ; CA - CB
- HB
5994 5996 5998 1 1.0950e+02 3.3472e+02 ; CA - CB
- CG2
5994 5996 6002 1 1.0950e+02 4.1840e+02 ; CA - CB
- OG1
5994 6004 6005 1 1.2040e+02 6.6944e+02 ; CA - C
- O
5994 6004 6006 1 1.1660e+02 5.8576e+02 ; CA - C
- N
5995 5994 5996 1 1.0950e+02 4.1840e+02 ; HA - CA
- CB
5995 5994 6004 1 1.0950e+02 4.1840e+02 ; HA - CA
- C
5996 5994 6004 1 1.1110e+02 5.2718e+02 ; CB - CA
- C
5996 5998 5999 1 1.0950e+02 4.1840e+02 ; CB - CG2
- HG21
5996 5998 6000 1 1.0950e+02 4.1840e+02 ; CB - CG2
- HG22
5996 5998 6001 1 1.0950e+02 4.1840e+02 ; CB - CG2
- HG23
5996 6002 6003 1 1.0850e+02 4.6024e+02 ; CB - OG1
- HG1
5997 5996 5998 1 1.0950e+02 4.1840e+02 ; HB - CB
- CG2

5997 5996 6002 1 1.0950e+02 4.1840e+02 ; HB - CB
- OG1
5998 5996 6002 1 1.0950e+02 4.1840e+02 ; CG2 - CB
- OG1
5999 5998 6000 1 1.0950e+02 2.9288e+02 ; HG21 - CG2
- HG22
5999 5998 6001 1 1.0950e+02 2.9288e+02 ; HG21 - CG2
- HG23
6000 5998 6001 1 1.0950e+02 2.9288e+02 ; HG22 - CG2
- HG23
6004 6006 6007 1 1.2000e+02 4.1840e+02 ; C - N
- H
6004 6006 6008 1 1.2190e+02 4.1840e+02 ; C - N
- CA
6005 6004 6006 1 1.2290e+02 6.6944e+02 ; O - C
- N
6006 6008 6009 1 1.0950e+02 4.1840e+02 ; N - CA
- HA
6006 6008 6010 1 1.0970e+02 6.6944e+02 ; N - CA
- CB
6006 6008 6016 1 1.1010e+02 5.2718e+02 ; N - CA
- C
6007 6006 6008 1 1.1804e+02 4.1840e+02 ; H - N
- CA
6008 6010 6011 1 1.0950e+02 4.1840e+02 ; CA - CB
- HB2
6008 6010 6012 1 1.0950e+02 4.1840e+02 ; CA - CB
- HB3
6008 6010 6013 1 1.1110e+02 5.2718e+02 ; CA - CB
- CG
6008 6016 6017 1 1.2040e+02 6.6944e+02 ; CA - C
- O
6008 6016 6018 1 1.1660e+02 5.8576e+02 ; CA - C
- N
6009 6008 6010 1 1.0950e+02 4.1840e+02 ; HA - CA
- CB
6009 6008 6016 1 1.0950e+02 4.1840e+02 ; HA - CA
- C
6010 6008 6016 1 1.1110e+02 5.2718e+02 ; CB - CA
- C
6010 6013 6014 1 1.1700e+02 5.8576e+02 ; CB - CG
- OD1
6010 6013 6015 1 1.1700e+02 5.8576e+02 ; CB - CG
- OD2
6011 6010 6012 1 1.0950e+02 2.9288e+02 ; HB2 - CB
- HB3
6011 6010 6013 1 1.0950e+02 4.1840e+02 ; HB2 - CB
- CG
6012 6010 6013 1 1.0950e+02 4.1840e+02 ; HB3 - CB
- CG
6014 6013 6015 1 1.2600e+02 6.6944e+02 ; OD1 - CG
- OD2

6016 6018 6019 1 1.2000e+02 4.1840e+02 ; C - N
- H
6016 6018 6020 1 1.2190e+02 4.1840e+02 ; C - N
- CA
6017 6016 6018 1 1.2290e+02 6.6944e+02 ; O - C
- N
6018 6020 6021 1 1.0950e+02 4.1840e+02 ; N - CA
- HA
6018 6020 6022 1 1.0970e+02 6.6944e+02 ; N - CA
- CB
6018 6020 6036 1 1.1010e+02 5.2718e+02 ; N - CA
- C
6019 6018 6020 1 1.1804e+02 4.1840e+02 ; H - N
- CA
6020 6022 6023 1 1.0950e+02 4.1840e+02 ; CA - CB
- HB2
6020 6022 6024 1 1.0950e+02 4.1840e+02 ; CA - CB
- HB3
6020 6022 6025 1 1.1400e+02 5.2718e+02 ; CA - CB
- CG
6020 6036 6037 1 1.2040e+02 6.6944e+02 ; CA - C
- O
6020 6036 6038 1 1.1660e+02 5.8576e+02 ; CA - C
- N
6021 6020 6022 1 1.0950e+02 4.1840e+02 ; HA - CA
- CB
6021 6020 6036 1 1.0950e+02 4.1840e+02 ; HA - CA
- C
6022 6020 6036 1 1.1110e+02 5.2718e+02 ; CB - CA
- C
6022 6025 6026 1 1.2000e+02 5.8576e+02 ; CB - CG
- CD1
6022 6025 6034 1 1.2000e+02 5.8576e+02 ; CB - CG
- CD2
6023 6022 6024 1 1.0950e+02 2.9288e+02 ; HB2 - CB
- HB3
6023 6022 6025 1 1.0950e+02 4.1840e+02 ; HB2 - CB
- CG
6024 6022 6025 1 1.0950e+02 4.1840e+02 ; HB3 - CB
- CG
6025 6026 6027 1 1.2000e+02 4.1840e+02 ; CG - CD1
- HD1
6025 6026 6028 1 1.2000e+02 5.2718e+02 ; CG - CD1
- CE1
6025 6034 6032 1 1.2000e+02 5.2718e+02 ; CG - CD2
- CE2
6025 6034 6035 1 1.2000e+02 4.1840e+02 ; CG - CD2
- HD2
6026 6025 6034 1 1.2000e+02 5.2718e+02 ; CD1 - CG
- CD2
6026 6028 6029 1 1.2000e+02 4.1840e+02 ; CD1 - CE1
- HE1

```
  6026   6028   6030       1    1.2000e+02    5.2718e+02 ;    CD1 - CE1
- CZ
  6027   6026   6028       1    1.2000e+02    4.1840e+02 ;    HD1 - CD1
- CE1
  6028   6030   6031       1    1.2000e+02    4.1840e+02 ;    CE1 - CZ
- HZ
  6028   6030   6032       1    1.2000e+02    5.2718e+02 ;    CE1 - CZ
- CE2
  6029   6028   6030       1    1.2000e+02    4.1840e+02 ;    HE1 - CE1
- CZ
  6030   6032   6033       1    1.2000e+02    4.1840e+02 ;     CZ - CE2
- HE2
  6030   6032   6034       1    1.2000e+02    5.2718e+02 ;     CZ - CE2
- CD2
  6031   6030   6032       1    1.2000e+02    4.1840e+02 ;     HZ - CZ
- CE2
  6032   6034   6035       1    1.2000e+02    4.1840e+02 ;    CE2 - CD2
- HD2
  6033   6032   6034       1    1.2000e+02    4.1840e+02 ;    HE2 - CE2
- CD2
  6036   6038   6039       1    1.2000e+02    4.1840e+02 ;      C - N
- H
  6036   6038   6040       1    1.2190e+02    4.1840e+02 ;      C - N
- CA
  6037   6036   6038       1    1.2290e+02    6.6944e+02 ;      O - C
- N
  6038   6040   6041       1    1.0950e+02    4.1840e+02 ;      N - CA
- HA
  6038   6040   6042       1    1.0970e+02    6.6944e+02 ;      N - CA
- CB
  6038   6040   6055       1    1.1010e+02    5.2718e+02 ;      N - CA
- C
  6039   6038   6040       1    1.1804e+02    4.1840e+02 ;      H - N
- CA
  6040   6042   6043       1    1.0950e+02    4.1840e+02 ;     CA - CB
- HB2
  6040   6042   6044       1    1.0950e+02    4.1840e+02 ;     CA - CB
- HB3
  6040   6042   6045       1    1.0950e+02    3.3472e+02 ;     CA - CB
- CG
  6040   6055   6056       1    1.2040e+02    6.6944e+02 ;     CA - C
- O
  6040   6055   6057       1    1.1660e+02    5.8576e+02 ;     CA - C
- N
  6041   6040   6042       1    1.0950e+02    4.1840e+02 ;     HA - CA
- CB
  6041   6040   6055       1    1.0950e+02    4.1840e+02 ;     HA - CA
- C
  6042   6040   6055       1    1.1110e+02    5.2718e+02 ;     CB - CA
- C
  6042   6045   6046       1    1.0950e+02    4.1840e+02 ;     CB - CG
- HG
```

```
  6042   6045   6047       1    1.0950e+02    3.3472e+02 ;     CB - CG
- CD1
  6042   6045   6051       1    1.0950e+02    3.3472e+02 ;     CB - CG
- CD2
  6043   6042   6044       1    1.0950e+02    2.9288e+02 ;    HB2 - CB
- HB3
  6043   6042   6045       1    1.0950e+02    4.1840e+02 ;    HB2 - CB
- CG
  6044   6042   6045       1    1.0950e+02    4.1840e+02 ;    HB3 - CB
- CG
  6045   6047   6048       1    1.0950e+02    4.1840e+02 ;     CG - CD1
- HD11
  6045   6047   6049       1    1.0950e+02    4.1840e+02 ;     CG - CD1
- HD12
  6045   6047   6050       1    1.0950e+02    4.1840e+02 ;     CG - CD1
- HD13
  6045   6051   6052       1    1.0950e+02    4.1840e+02 ;     CG - CD2
- HD21
  6045   6051   6053       1    1.0950e+02    4.1840e+02 ;     CG - CD2
- HD22
  6045   6051   6054       1    1.0950e+02    4.1840e+02 ;     CG - CD2
- HD23
  6046   6045   6047       1    1.0950e+02    4.1840e+02 ;     HG - CG
- CD1
  6046   6045   6051       1    1.0950e+02    4.1840e+02 ;     HG - CG
- CD2
  6047   6045   6051       1    1.0950e+02    3.3472e+02 ;    CD1 - CG
- CD2
  6048   6047   6049       1    1.0950e+02    2.9288e+02 ;   HD11 - CD1
- HD12
  6048   6047   6050       1    1.0950e+02    2.9288e+02 ;   HD11 - CD1
- HD13
  6049   6047   6050       1    1.0950e+02    2.9288e+02 ;   HD12 - CD1
- HD13
  6052   6051   6053       1    1.0950e+02    2.9288e+02 ;   HD21 - CD2
- HD22
  6052   6051   6054       1    1.0950e+02    2.9288e+02 ;   HD21 - CD2
- HD23
  6053   6051   6054       1    1.0950e+02    2.9288e+02 ;   HD22 - CD2
- HD23
  6055   6057   6058       1    1.2000e+02    4.1840e+02 ;      C - N
- H
  6055   6057   6059       1    1.2190e+02    4.1840e+02 ;      C - N
- CA
  6056   6055   6057       1    1.2290e+02    6.6944e+02 ;      O - C
- N
  6057   6059   6060       1    1.0950e+02    4.1840e+02 ;      N - CA
- HA
  6057   6059   6061       1    1.0970e+02    6.6944e+02 ;      N - CA
- CB
  6057   6059   6065       1    1.1010e+02    5.2718e+02 ;      N - CA
- C
```

| | | | | | | | |
|---|---|---|---|---|---|---|---|
| 6058 | 6057 | 6059 | 1 | 1.1804e+02 | 4.1840e+02 | ; | H - N - CA |
| 6059 | 6061 | 6062 | 1 | 1.0950e+02 | 4.1840e+02 | ; | CA - CB - HB1 |
| 6059 | 6061 | 6063 | 1 | 1.0950e+02 | 4.1840e+02 | ; | CA - CB - HB2 |
| 6059 | 6061 | 6064 | 1 | 1.0950e+02 | 4.1840e+02 | ; | CA - CB - HB3 |
| 6059 | 6065 | 6066 | 1 | 1.2040e+02 | 6.6944e+02 | ; | CA - C - O |
| 6059 | 6065 | 6067 | 1 | 1.1660e+02 | 5.8576e+02 | ; | CA - C - N |
| 6060 | 6059 | 6061 | 1 | 1.0950e+02 | 4.1840e+02 | ; | HA - CA - CB |
| 6060 | 6059 | 6065 | 1 | 1.0950e+02 | 4.1840e+02 | ; | HA - CA - C |
| 6061 | 6059 | 6065 | 1 | 1.1110e+02 | 5.2718e+02 | ; | CB - CA - C |
| 6062 | 6061 | 6063 | 1 | 1.0950e+02 | 2.9288e+02 | ; | HB1 - CB - HB2 |
| 6062 | 6061 | 6064 | 1 | 1.0950e+02 | 2.9288e+02 | ; | HB1 - CB - HB3 |
| 6063 | 6061 | 6064 | 1 | 1.0950e+02 | 2.9288e+02 | ; | HB2 - CB - HB3 |
| 6065 | 6067 | 6068 | 1 | 1.2000e+02 | 4.1840e+02 | ; | C - N - H |
| 6065 | 6067 | 6069 | 1 | 1.2190e+02 | 4.1840e+02 | ; | C - N - CA |
| 6066 | 6065 | 6067 | 1 | 1.2290e+02 | 6.6944e+02 | ; | O - C - N |
| 6067 | 6069 | 6070 | 1 | 1.0950e+02 | 4.1840e+02 | ; | N - CA - HA |
| 6067 | 6069 | 6071 | 1 | 1.0970e+02 | 6.6944e+02 | ; | N - CA - CB |
| 6067 | 6069 | 6075 | 1 | 1.1010e+02 | 5.2718e+02 | ; | N - CA - C |
| 6068 | 6067 | 6069 | 1 | 1.1804e+02 | 4.1840e+02 | ; | H - N - CA |
| 6069 | 6071 | 6072 | 1 | 1.0950e+02 | 4.1840e+02 | ; | CA - CB - HB1 |
| 6069 | 6071 | 6073 | 1 | 1.0950e+02 | 4.1840e+02 | ; | CA - CB - HB2 |
| 6069 | 6071 | 6074 | 1 | 1.0950e+02 | 4.1840e+02 | ; | CA - CB - HB3 |
| 6069 | 6075 | 6076 | 1 | 1.2040e+02 | 6.6944e+02 | ; | CA - C - O |
| 6069 | 6075 | 6077 | 1 | 1.1660e+02 | 5.8576e+02 | ; | CA - C - N |
| 6070 | 6069 | 6071 | 1 | 1.0950e+02 | 4.1840e+02 | ; | HA - CA - CB |
| 6070 | 6069 | 6075 | 1 | 1.0950e+02 | 4.1840e+02 | ; | HA - CA - C |

6071 6069 6075 1 1.1110e+02 5.2718e+02 ; CB - CA
- C
6072 6071 6073 1 1.0950e+02 2.9288e+02 ; HB1 - CB
- HB2
6072 6071 6074 1 1.0950e+02 2.9288e+02 ; HB1 - CB
- HB3
6073 6071 6074 1 1.0950e+02 2.9288e+02 ; HB2 - CB
- HB3
6075 6077 6078 1 1.2000e+02 4.1840e+02 ; C - N
- H
6075 6077 6079 1 1.2190e+02 4.1840e+02 ; C - N
- CA
6076 6075 6077 1 1.2290e+02 6.6944e+02 ; O - C
- N
6077 6079 6080 1 1.0950e+02 4.1840e+02 ; N - CA
- HA
6077 6079 6081 1 1.0970e+02 6.6944e+02 ; N - CA
- CB
6077 6079 6089 1 1.1010e+02 5.2718e+02 ; N - CA
- C
6078 6077 6079 1 1.1804e+02 4.1840e+02 ; H - N
- CA
6079 6081 6082 1 1.0950e+02 4.1840e+02 ; CA - CB
- HB
6079 6081 6083 1 1.0950e+02 3.3472e+02 ; CA - CB
- CG2
6079 6081 6087 1 1.0950e+02 4.1840e+02 ; CA - CB
- OG1
6079 6089 6090 1 1.2040e+02 6.6944e+02 ; CA - C
- O
6079 6089 6091 1 1.1660e+02 5.8576e+02 ; CA - C
- N
6080 6079 6081 1 1.0950e+02 4.1840e+02 ; HA - CA
- CB
6080 6079 6089 1 1.0950e+02 4.1840e+02 ; HA - CA
- C
6081 6079 6089 1 1.1110e+02 5.2718e+02 ; CB - CA
- C
6081 6083 6084 1 1.0950e+02 4.1840e+02 ; CB - CG2
- HG21
6081 6083 6085 1 1.0950e+02 4.1840e+02 ; CB - CG2
- HG22
6081 6083 6086 1 1.0950e+02 4.1840e+02 ; CB - CG2
- HG23
6081 6087 6088 1 1.0850e+02 4.6024e+02 ; CB - OG1
- HG1
6082 6081 6083 1 1.0950e+02 4.1840e+02 ; HB - CB
- CG2
6082 6081 6087 1 1.0950e+02 4.1840e+02 ; HB - CB
- OG1
6083 6081 6087 1 1.0950e+02 4.1840e+02 ; CG2 - CB
- OG1

6084 6083 6085 1 1.0950e+02 2.9288e+02 ; HG21 - CG2
- HG22
6084 6083 6086 1 1.0950e+02 2.9288e+02 ; HG21 - CG2
- HG23
6085 6083 6086 1 1.0950e+02 2.9288e+02 ; HG22 - CG2
- HG23
6089 6091 6092 1 1.2000e+02 4.1840e+02 ; C - N
- H
6089 6091 6093 1 1.2190e+02 4.1840e+02 ; C - N
- CA
6090 6089 6091 1 1.2290e+02 6.6944e+02 ; O - C
- N
6091 6093 6094 1 1.0950e+02 4.1840e+02 ; N - CA
- HA
6091 6093 6095 1 1.0970e+02 6.6944e+02 ; N - CA
- CB
6091 6093 6108 1 1.1010e+02 5.2718e+02 ; N - CA
- C
6092 6091 6093 1 1.1804e+02 4.1840e+02 ; H - N
- CA
6093 6095 6096 1 1.0950e+02 4.1840e+02 ; CA - CB
- HB
6093 6095 6097 1 1.0950e+02 3.3472e+02 ; CA - CB
- CG2
6093 6095 6101 1 1.0950e+02 3.3472e+02 ; CA - CB
- CG1
6093 6108 6109 1 1.2040e+02 6.6944e+02 ; CA - C
- O
6093 6108 6110 1 1.1660e+02 5.8576e+02 ; CA - C
- N
6094 6093 6095 1 1.0950e+02 4.1840e+02 ; HA - CA
- CB
6094 6093 6108 1 1.0950e+02 4.1840e+02 ; HA - CA
- C
6095 6093 6108 1 1.1110e+02 5.2718e+02 ; CB - CA
- C
6095 6097 6098 1 1.0950e+02 4.1840e+02 ; CB - CG2
- HG21
6095 6097 6099 1 1.0950e+02 4.1840e+02 ; CB - CG2
- HG22
6095 6097 6100 1 1.0950e+02 4.1840e+02 ; CB - CG2
- HG23
6095 6101 6102 1 1.0950e+02 4.1840e+02 ; CB - CG1
- HG12
6095 6101 6103 1 1.0950e+02 4.1840e+02 ; CB - CG1
- HG13
6095 6101 6104 1 1.0950e+02 3.3472e+02 ; CB - CG1
- CD1
6096 6095 6097 1 1.0950e+02 4.1840e+02 ; HB - CB
- CG2
6096 6095 6101 1 1.0950e+02 4.1840e+02 ; HB - CB
- CG1

6097 6095 6101 1 1.0950e+02 3.3472e+02 ; CG2 - CB
- CG1
6098 6097 6099 1 1.0950e+02 2.9288e+02 ; HG21 - CG2
- HG22
6098 6097 6100 1 1.0950e+02 2.9288e+02 ; HG21 - CG2
- HG23
6099 6097 6100 1 1.0950e+02 2.9288e+02 ; HG22 - CG2
- HG23
6101 6104 6105 1 1.0950e+02 4.1840e+02 ; CG1 - CD1
- HD11
6101 6104 6106 1 1.0950e+02 4.1840e+02 ; CG1 - CD1
- HD12
6101 6104 6107 1 1.0950e+02 4.1840e+02 ; CG1 - CD1
- HD13
6102 6101 6103 1 1.0950e+02 2.9288e+02 ; HG12 - CG1
- HG13
6102 6101 6104 1 1.0950e+02 4.1840e+02 ; HG12 - CG1
- CD1
6103 6101 6104 1 1.0950e+02 4.1840e+02 ; HG13 - CG1
- CD1
6105 6104 6106 1 1.0950e+02 2.9288e+02 ; HD11 - CD1
- HD12
6105 6104 6107 1 1.0950e+02 2.9288e+02 ; HD11 - CD1
- HD13
6106 6104 6107 1 1.0950e+02 2.9288e+02 ; HD12 - CD1
- HD13
6108 6110 6111 1 1.2000e+02 4.1840e+02 ; C - N
- H
6108 6110 6112 1 1.2190e+02 4.1840e+02 ; C - N
- CA
6109 6108 6110 1 1.2290e+02 6.6944e+02 ; O - C
- N
6110 6112 6113 1 1.0950e+02 4.1840e+02 ; N - CA
- HA
6110 6112 6114 1 1.0970e+02 6.6944e+02 ; N - CA
- CB
6110 6112 6120 1 1.1010e+02 5.2718e+02 ; N - CA
- C
6111 6110 6112 1 1.1804e+02 4.1840e+02 ; H - N
- CA
6112 6114 6115 1 1.0950e+02 4.1840e+02 ; CA - CB
- HB2
6112 6114 6116 1 1.0950e+02 4.1840e+02 ; CA - CB
- HB3
6112 6114 6117 1 1.1110e+02 5.2718e+02 ; CA - CB
- CG
6112 6120 6121 1 1.2040e+02 6.6944e+02 ; CA - C
- O
6112 6120 6122 1 1.1660e+02 5.8576e+02 ; CA - C
- N
6113 6112 6114 1 1.0950e+02 4.1840e+02 ; HA - CA
- CB

6113 6112 6120 1 1.0950e+02 4.1840e+02 ; HA - CA
- C
6114 6112 6120 1 1.1110e+02 5.2718e+02 ; CB - CA
- C
6114 6117 6118 1 1.1700e+02 5.8576e+02 ; CB - CG
- OD1
6114 6117 6119 1 1.1700e+02 5.8576e+02 ; CB - CG
- OD2
6115 6114 6116 1 1.0950e+02 2.9288e+02 ; HB2 - CB
- HB3
6115 6114 6117 1 1.0950e+02 4.1840e+02 ; HB2 - CB
- CG
6116 6114 6117 1 1.0950e+02 4.1840e+02 ; HB3 - CB
- CG
6118 6117 6119 1 1.2600e+02 6.6944e+02 ; OD1 - CG
- OD2
6120 6122 6123 1 1.2000e+02 4.1840e+02 ; C - N
- H
6120 6122 6124 1 1.2190e+02 4.1840e+02 ; C - N
- CA
6121 6120 6122 1 1.2290e+02 6.6944e+02 ; O - C
- N
6122 6124 6125 1 1.0950e+02 4.1840e+02 ; N - CA
- HA
6122 6124 6126 1 1.0970e+02 6.6944e+02 ; N - CA
- CB
6122 6124 6142 1 1.1010e+02 5.2718e+02 ; N - CA
- C
6123 6122 6124 1 1.1804e+02 4.1840e+02 ; H - N
- CA
6124 6126 6127 1 1.0950e+02 4.1840e+02 ; CA - CB
- HB2
6124 6126 6128 1 1.0950e+02 4.1840e+02 ; CA - CB
- HB3
6124 6126 6129 1 1.0950e+02 3.3472e+02 ; CA - CB
- CG
6124 6142 6143 1 1.2040e+02 6.6944e+02 ; CA - C
- O
6124 6142 6144 1 1.1660e+02 5.8576e+02 ; CA - C
- N
6125 6124 6126 1 1.0950e+02 4.1840e+02 ; HA - CA
- CB
6125 6124 6142 1 1.0950e+02 4.1840e+02 ; HA - CA
- C
6126 6124 6142 1 1.1110e+02 5.2718e+02 ; CB - CA
- C
6126 6129 6130 1 1.0950e+02 4.1840e+02 ; CB - CG
- HG2
6126 6129 6131 1 1.0950e+02 4.1840e+02 ; CB - CG
- HG3
6126 6129 6132 1 1.0950e+02 3.3472e+02 ; CB - CG
- CD

```
  6127   6126   6128       1    1.0950e+02    2.9288e+02 ;    HB2 - CB
- HB3
  6127   6126   6129       1    1.0950e+02    4.1840e+02 ;    HB2 - CB
- CG
  6128   6126   6129       1    1.0950e+02    4.1840e+02 ;    HB3 - CB
- CG
  6129   6132   6133       1    1.0950e+02    4.1840e+02 ;     CG - CD
- HD2
  6129   6132   6134       1    1.0950e+02    4.1840e+02 ;     CG - CD
- HD3
  6129   6132   6135       1    1.0950e+02    3.3472e+02 ;     CG - CD
- CE
  6130   6129   6131       1    1.0950e+02    2.9288e+02 ;    HG2 - CG
- HG3
  6130   6129   6132       1    1.0950e+02    4.1840e+02 ;    HG2 - CG
- CD
  6131   6129   6132       1    1.0950e+02    4.1840e+02 ;    HG3 - CG
- CD
  6132   6135   6136       1    1.0950e+02    4.1840e+02 ;     CD - CE
- HE2
  6132   6135   6137       1    1.0950e+02    4.1840e+02 ;     CD - CE
- HE3
  6132   6135   6138       1    1.1120e+02    6.6944e+02 ;     CD - CE
- NZ
  6133   6132   6134       1    1.0950e+02    2.9288e+02 ;    HD2 - CD
- HD3
  6133   6132   6135       1    1.0950e+02    4.1840e+02 ;    HD2 - CD
- CE
  6134   6132   6135       1    1.0950e+02    4.1840e+02 ;    HD3 - CD
- CE
  6135   6138   6139       1    1.0950e+02    4.1840e+02 ;     CE - NZ
- HZ1
  6135   6138   6140       1    1.0950e+02    4.1840e+02 ;     CE - NZ
- HZ2
  6135   6138   6141       1    1.0950e+02    4.1840e+02 ;     CE - NZ
- HZ3
  6136   6135   6137       1    1.0950e+02    2.9288e+02 ;    HE2 - CE
- HE3
  6136   6135   6138       1    1.0950e+02    4.1840e+02 ;    HE2 - CE
- NZ
  6137   6135   6138       1    1.0950e+02    4.1840e+02 ;    HE3 - CE
- NZ
  6139   6138   6140       1    1.0950e+02    2.9288e+02 ;    HZ1 - NZ
- HZ2
  6139   6138   6141       1    1.0950e+02    2.9288e+02 ;    HZ1 - NZ
- HZ3
  6140   6138   6141       1    1.0950e+02    2.9288e+02 ;    HZ2 - NZ
- HZ3
  6142   6144   6145       1    1.2000e+02    4.1840e+02 ;      C - N
- H
  6142   6144   6146       1    1.2190e+02    4.1840e+02 ;      C - N
- CA
```

6143 6142 6144 1 1.2290e+02 6.6944e+02 ; O - C
- N
6144 6146 6147 1 1.0950e+02 4.1840e+02 ; N - CA
- HA
6144 6146 6148 1 1.0970e+02 6.6944e+02 ; N - CA
- CB
6144 6146 6159 1 1.1010e+02 5.2718e+02 ; N - CA
- C
6145 6144 6146 1 1.1804e+02 4.1840e+02 ; H - N
- CA
6146 6148 6149 1 1.0950e+02 4.1840e+02 ; CA - CB
- HB2
6146 6148 6150 1 1.0950e+02 4.1840e+02 ; CA - CB
- HB3
6146 6148 6151 1 1.0950e+02 3.3472e+02 ; CA - CB
- CG
6146 6159 6160 1 1.2040e+02 6.6944e+02 ; CA - C
- O
6146 6159 6161 1 1.1660e+02 5.8576e+02 ; CA - C
- N
6147 6146 6148 1 1.0950e+02 4.1840e+02 ; HA - CA
- CB
6147 6146 6159 1 1.0950e+02 4.1840e+02 ; HA - CA
- C
6148 6146 6159 1 1.1110e+02 5.2718e+02 ; CB - CA
- C
6148 6151 6152 1 1.0950e+02 4.1840e+02 ; CB - CG
- HG2
6148 6151 6153 1 1.0950e+02 4.1840e+02 ; CB - CG
- HG3
6148 6151 6154 1 1.1110e+02 5.2718e+02 ; CB - CG
- CD
6149 6148 6150 1 1.0950e+02 2.9288e+02 ; HB2 - CB
- HB3
6149 6148 6151 1 1.0950e+02 4.1840e+02 ; HB2 - CB
- CG
6150 6148 6151 1 1.0950e+02 4.1840e+02 ; HB3 - CB
- CG
6151 6154 6155 1 1.2040e+02 6.6944e+02 ; CG - CD
- OE1
6151 6154 6156 1 1.1660e+02 5.8576e+02 ; CG - CD
- NE2
6152 6151 6153 1 1.0950e+02 2.9288e+02 ; HG2 - CG
- HG3
6152 6151 6154 1 1.0950e+02 4.1840e+02 ; HG2 - CG
- CD
6153 6151 6154 1 1.0950e+02 4.1840e+02 ; HG3 - CG
- CD
6154 6156 6157 1 1.2000e+02 4.1840e+02 ; CD - NE2
- HE21
6154 6156 6158 1 1.2000e+02 4.1840e+02 ; CD - NE2
- HE22

6155 6154 6156 1 1.2290e+02 6.6944e+02 ; OE1 - CD
- NE2
6157 6156 6158 1 1.2000e+02 2.9288e+02 ; HE21 - NE2
- HE22
6159 6161 6162 1 1.2000e+02 4.1840e+02 ; C - N
- H
6159 6161 6163 1 1.2190e+02 4.1840e+02 ; C - N
- CA
6160 6159 6161 1 1.2290e+02 6.6944e+02 ; O - C
- N
6161 6163 6164 1 1.0950e+02 4.1840e+02 ; N - CA
- HA
6161 6163 6165 1 1.0970e+02 6.6944e+02 ; N - CA
- CB
6161 6163 6173 1 1.1010e+02 5.2718e+02 ; N - CA
- C
6162 6161 6163 1 1.1804e+02 4.1840e+02 ; H - N
- CA
6163 6165 6166 1 1.0950e+02 4.1840e+02 ; CA - CB
- HB
6163 6165 6167 1 1.0950e+02 3.3472e+02 ; CA - CB
- CG2
6163 6165 6171 1 1.0950e+02 4.1840e+02 ; CA - CB
- OG1
6163 6173 6174 1 1.2040e+02 6.6944e+02 ; CA - C
- O
6163 6173 6175 1 1.1660e+02 5.8576e+02 ; CA - C
- N
6164 6163 6165 1 1.0950e+02 4.1840e+02 ; HA - CA
- CB
6164 6163 6173 1 1.0950e+02 4.1840e+02 ; HA - CA
- C
6165 6163 6173 1 1.1110e+02 5.2718e+02 ; CB - CA
- C
6165 6167 6168 1 1.0950e+02 4.1840e+02 ; CB - CG2
- HG21
6165 6167 6169 1 1.0950e+02 4.1840e+02 ; CB - CG2
- HG22
6165 6167 6170 1 1.0950e+02 4.1840e+02 ; CB - CG2
- HG23
6165 6171 6172 1 1.0850e+02 4.6024e+02 ; CB - OG1
- HG1
6166 6165 6167 1 1.0950e+02 4.1840e+02 ; HB - CB
- CG2
6166 6165 6171 1 1.0950e+02 4.1840e+02 ; HB - CB
- OG1
6167 6165 6171 1 1.0950e+02 4.1840e+02 ; CG2 - CB
- OG1
6168 6167 6169 1 1.0950e+02 2.9288e+02 ; HG21 - CG2
- HG22
6168 6167 6170 1 1.0950e+02 2.9288e+02 ; HG21 - CG2
- HG23

6169 6167 6170 1 1.0950e+02 2.9288e+02 ; HG22 - CG2
- HG23
6173 6175 6176 1 1.2000e+02 4.1840e+02 ; C - N
- H
6173 6175 6177 1 1.2190e+02 4.1840e+02 ; C - N
- CA
6174 6173 6175 1 1.2290e+02 6.6944e+02 ; O - C
- N
6175 6177 6178 1 1.0950e+02 4.1840e+02 ; N - CA
- HA
6175 6177 6179 1 1.0970e+02 6.6944e+02 ; N - CA
- CB
6175 6177 6194 1 1.1010e+02 5.2718e+02 ; N - CA
- C
6176 6175 6177 1 1.1804e+02 4.1840e+02 ; H - N
- CA
6177 6179 6180 1 1.0950e+02 4.1840e+02 ; CA - CB
- HB2
6177 6179 6181 1 1.0950e+02 4.1840e+02 ; CA - CB
- HB3
6177 6179 6182 1 1.1400e+02 5.2718e+02 ; CA - CB
- CG
6177 6194 6195 1 1.2040e+02 6.6944e+02 ; CA - C
- O
6177 6194 6196 1 1.1660e+02 5.8576e+02 ; CA - C
- N
6178 6177 6179 1 1.0950e+02 4.1840e+02 ; HA - CA
- CB
6178 6177 6194 1 1.0950e+02 4.1840e+02 ; HA - CA
- C
6179 6177 6194 1 1.1110e+02 5.2718e+02 ; CB - CA
- C
6179 6182 6183 1 1.2000e+02 5.8576e+02 ; CB - CG
- CD1
6179 6182 6192 1 1.2000e+02 5.8576e+02 ; CB - CG
- CD2
6180 6179 6181 1 1.0950e+02 2.9288e+02 ; HB2 - CB
- HB3
6180 6179 6182 1 1.0950e+02 4.1840e+02 ; HB2 - CB
- CG
6181 6179 6182 1 1.0950e+02 4.1840e+02 ; HB3 - CB
- CG
6182 6183 6184 1 1.2000e+02 4.1840e+02 ; CG - CD1
- HD1
6182 6183 6185 1 1.2000e+02 5.2718e+02 ; CG - CD1
- CE1
6182 6192 6190 1 1.2000e+02 5.2718e+02 ; CG - CD2
- CE2
6182 6192 6193 1 1.2000e+02 4.1840e+02 ; CG - CD2
- HD2
6183 6182 6192 1 1.2000e+02 5.2718e+02 ; CD1 - CG
- CD2

6183 6185 6186 1 1.2000e+02 4.1840e+02 ; CD1 - CE1
- HE1
6183 6185 6187 1 1.2000e+02 5.2718e+02 ; CD1 - CE1
- CZ
6184 6183 6185 1 1.2000e+02 4.1840e+02 ; HD1 - CD1
- CE1
6185 6187 6188 1 1.2000e+02 5.8576e+02 ; CE1 - CZ
- OH
6185 6187 6190 1 1.2000e+02 5.2718e+02 ; CE1 - CZ
- CE2
6186 6185 6187 1 1.2000e+02 4.1840e+02 ; HE1 - CE1
- CZ
6187 6188 6189 1 1.1300e+02 4.1840e+02 ; CZ - OH
- HH
6187 6190 6191 1 1.2000e+02 4.1840e+02 ; CZ - CE2
- HE2
6187 6190 6192 1 1.2000e+02 5.2718e+02 ; CZ - CE2
- CD2
6188 6187 6190 1 1.2000e+02 5.8576e+02 ; OH - CZ
- CE2
6190 6192 6193 1 1.2000e+02 4.1840e+02 ; CE2 - CD2
- HD2
6191 6190 6192 1 1.2000e+02 4.1840e+02 ; HE2 - CE2
- CD2
6194 6196 6197 1 1.2000e+02 4.1840e+02 ; C - N
- H
6194 6196 6198 1 1.2190e+02 4.1840e+02 ; C - N
- CA
6195 6194 6196 1 1.2290e+02 6.6944e+02 ; O - C
- N
6196 6198 6199 1 1.0950e+02 4.1840e+02 ; N - CA
- HA
6196 6198 6200 1 1.0970e+02 6.6944e+02 ; N - CA
- CB
6196 6198 6213 1 1.1010e+02 5.2718e+02 ; N - CA
- C
6197 6196 6198 1 1.1804e+02 4.1840e+02 ; H - N
- CA
6198 6200 6201 1 1.0950e+02 4.1840e+02 ; CA - CB
- HB2
6198 6200 6202 1 1.0950e+02 4.1840e+02 ; CA - CB
- HB3
6198 6200 6203 1 1.0950e+02 3.3472e+02 ; CA - CB
- CG
6198 6213 6214 1 1.2040e+02 6.6944e+02 ; CA - C
- O
6198 6213 6215 1 1.1660e+02 5.8576e+02 ; CA - C
- N
6199 6198 6200 1 1.0950e+02 4.1840e+02 ; HA - CA
- CB
6199 6198 6213 1 1.0950e+02 4.1840e+02 ; HA - CA
- C

```
  6200   6198   6213      1    1.1110e+02    5.2718e+02 ;      CB - CA
- C
  6200   6203   6204      1    1.0950e+02    4.1840e+02 ;      CB - CG
- HG
  6200   6203   6205      1    1.0950e+02    3.3472e+02 ;      CB - CG
- CD1
  6200   6203   6209      1    1.0950e+02    3.3472e+02 ;      CB - CG
- CD2
  6201   6200   6202      1    1.0950e+02    2.9288e+02 ;     HB2 - CB
- HB3
  6201   6200   6203      1    1.0950e+02    4.1840e+02 ;     HB2 - CB
- CG
  6202   6200   6203      1    1.0950e+02    4.1840e+02 ;     HB3 - CB
- CG
  6203   6205   6206      1    1.0950e+02    4.1840e+02 ;      CG - CD1
- HD11
  6203   6205   6207      1    1.0950e+02    4.1840e+02 ;      CG - CD1
- HD12
  6203   6205   6208      1    1.0950e+02    4.1840e+02 ;      CG - CD1
- HD13
  6203   6209   6210      1    1.0950e+02    4.1840e+02 ;      CG - CD2
- HD21
  6203   6209   6211      1    1.0950e+02    4.1840e+02 ;      CG - CD2
- HD22
  6203   6209   6212      1    1.0950e+02    4.1840e+02 ;      CG - CD2
- HD23
  6204   6203   6205      1    1.0950e+02    4.1840e+02 ;      HG - CG
- CD1
  6204   6203   6209      1    1.0950e+02    4.1840e+02 ;      HG - CG
- CD2
  6205   6203   6209      1    1.0950e+02    3.3472e+02 ;     CD1 - CG
- CD2
  6206   6205   6207      1    1.0950e+02    2.9288e+02 ;    HD11 - CD1
- HD12
  6206   6205   6208      1    1.0950e+02    2.9288e+02 ;    HD11 - CD1
- HD13
  6207   6205   6208      1    1.0950e+02    2.9288e+02 ;    HD12 - CD1
- HD13
  6210   6209   6211      1    1.0950e+02    2.9288e+02 ;    HD21 - CD2
- HD22
  6210   6209   6212      1    1.0950e+02    2.9288e+02 ;    HD21 - CD2
- HD23
  6211   6209   6212      1    1.0950e+02    2.9288e+02 ;    HD22 - CD2
- HD23
  6213   6215   6216      1    1.2000e+02    4.1840e+02 ;       C - N
- H
  6213   6215   6217      1    1.2190e+02    4.1840e+02 ;       C - N
- CA
  6214   6213   6215      1    1.2290e+02    6.6944e+02 ;       O - C
- N
  6215   6217   6218      1    1.0950e+02    4.1840e+02 ;       N - CA
- HA
```

6215 6217 6219 1 1.0970e+02 6.6944e+02 ; N - CA
- CB
6215 6217 6235 1 1.1010e+02 5.2718e+02 ; N - CA
- C
6216 6215 6217 1 1.1804e+02 4.1840e+02 ; H - N
- CA
6217 6219 6220 1 1.0950e+02 4.1840e+02 ; CA - CB
- HB2
6217 6219 6221 1 1.0950e+02 4.1840e+02 ; CA - CB
- HB3
6217 6219 6222 1 1.0950e+02 3.3472e+02 ; CA - CB
- CG
6217 6235 6236 1 1.2040e+02 6.6944e+02 ; CA - C
- O
6217 6235 6237 1 1.1660e+02 5.8576e+02 ; CA - C
- N
6218 6217 6219 1 1.0950e+02 4.1840e+02 ; HA - CA
- CB
6218 6217 6235 1 1.0950e+02 4.1840e+02 ; HA - CA
- C
6219 6217 6235 1 1.1110e+02 5.2718e+02 ; CB - CA
- C
6219 6222 6223 1 1.0950e+02 4.1840e+02 ; CB - CG
- HG2
6219 6222 6224 1 1.0950e+02 4.1840e+02 ; CB - CG
- HG3
6219 6222 6225 1 1.0950e+02 3.3472e+02 ; CB - CG
- CD
6220 6219 6221 1 1.0950e+02 2.9288e+02 ; HB2 - CB
- HB3
6220 6219 6222 1 1.0950e+02 4.1840e+02 ; HB2 - CB
- CG
6221 6219 6222 1 1.0950e+02 4.1840e+02 ; HB3 - CB
- CG
6222 6225 6226 1 1.0950e+02 4.1840e+02 ; CG - CD
- HD2
6222 6225 6227 1 1.0950e+02 4.1840e+02 ; CG - CD
- HD3
6222 6225 6228 1 1.0950e+02 3.3472e+02 ; CG - CD
- CE
6223 6222 6224 1 1.0950e+02 2.9288e+02 ; HG2 - CG
- HG3
6223 6222 6225 1 1.0950e+02 4.1840e+02 ; HG2 - CG
- CD
6224 6222 6225 1 1.0950e+02 4.1840e+02 ; HG3 - CG
- CD
6225 6228 6229 1 1.0950e+02 4.1840e+02 ; CD - CE
- HE2
6225 6228 6230 1 1.0950e+02 4.1840e+02 ; CD - CE
- HE3
6225 6228 6231 1 1.1120e+02 6.6944e+02 ; CD - CE
- NZ

6226 6225 6227 1 1.0950e+02 2.9288e+02 ; HD2 - CD
- HD3
6226 6225 6228 1 1.0950e+02 4.1840e+02 ; HD2 - CD
- CE
6227 6225 6228 1 1.0950e+02 4.1840e+02 ; HD3 - CD
- CE
6228 6231 6232 1 1.0950e+02 4.1840e+02 ; CE - NZ
- HZ1
6228 6231 6233 1 1.0950e+02 4.1840e+02 ; CE - NZ
- HZ2
6228 6231 6234 1 1.0950e+02 4.1840e+02 ; CE - NZ
- HZ3
6229 6228 6230 1 1.0950e+02 2.9288e+02 ; HE2 - CE
- HE3
6229 6228 6231 1 1.0950e+02 4.1840e+02 ; HE2 - CE
- NZ
6230 6228 6231 1 1.0950e+02 4.1840e+02 ; HE3 - CE
- NZ
6232 6231 6233 1 1.0950e+02 2.9288e+02 ; HZ1 - NZ
- HZ2
6232 6231 6234 1 1.0950e+02 2.9288e+02 ; HZ1 - NZ
- HZ3
6233 6231 6234 1 1.0950e+02 2.9288e+02 ; HZ2 - NZ
- HZ3
6235 6237 6238 1 1.2000e+02 4.1840e+02 ; C - N
- H
6235 6237 6239 1 1.2190e+02 4.1840e+02 ; C - N
- CA
6236 6235 6237 1 1.2290e+02 6.6944e+02 ; O - C
- N
6237 6239 6240 1 1.0950e+02 4.1840e+02 ; N - CA
- HA
6237 6239 6241 1 1.0970e+02 6.6944e+02 ; N - CA
- CB
6237 6239 6257 1 1.1010e+02 5.2718e+02 ; N - CA
- C
6238 6237 6239 1 1.1804e+02 4.1840e+02 ; H - N
- CA
6239 6241 6242 1 1.0950e+02 4.1840e+02 ; CA - CB
- HB2
6239 6241 6243 1 1.0950e+02 4.1840e+02 ; CA - CB
- HB3
6239 6241 6244 1 1.0950e+02 3.3472e+02 ; CA - CB
- CG
6239 6257 6258 1 1.2040e+02 6.6944e+02 ; CA - C
- O
6239 6257 6259 1 1.1660e+02 5.8576e+02 ; CA - C
- N
6240 6239 6241 1 1.0950e+02 4.1840e+02 ; HA - CA
- CB
6240 6239 6257 1 1.0950e+02 4.1840e+02 ; HA - CA
- C

6241 6239 6257 1 1.1110e+02 5.2718e+02 ; CB - CA
- C
6241 6244 6245 1 1.0950e+02 4.1840e+02 ; CB - CG
- HG2
6241 6244 6246 1 1.0950e+02 4.1840e+02 ; CB - CG
- HG3
6241 6244 6247 1 1.0950e+02 3.3472e+02 ; CB - CG
- CD
6242 6241 6243 1 1.0950e+02 2.9288e+02 ; HB2 - CB
- HB3
6242 6241 6244 1 1.0950e+02 4.1840e+02 ; HB2 - CB
- CG
6243 6241 6244 1 1.0950e+02 4.1840e+02 ; HB3 - CB
- CG
6244 6247 6248 1 1.0950e+02 4.1840e+02 ; CG - CD
- HD2
6244 6247 6249 1 1.0950e+02 4.1840e+02 ; CG - CD
- HD3
6244 6247 6250 1 1.0950e+02 3.3472e+02 ; CG - CD
- CE
6245 6244 6246 1 1.0950e+02 2.9288e+02 ; HG2 - CG
- HG3
6245 6244 6247 1 1.0950e+02 4.1840e+02 ; HG2 - CG
- CD
6246 6244 6247 1 1.0950e+02 4.1840e+02 ; HG3 - CG
- CD
6247 6250 6251 1 1.0950e+02 4.1840e+02 ; CD - CE
- HE2
6247 6250 6252 1 1.0950e+02 4.1840e+02 ; CD - CE
- HE3
6247 6250 6253 1 1.1120e+02 6.6944e+02 ; CD - CE
- NZ
6248 6247 6249 1 1.0950e+02 2.9288e+02 ; HD2 - CD
- HD3
6248 6247 6250 1 1.0950e+02 4.1840e+02 ; HD2 - CD
- CE
6249 6247 6250 1 1.0950e+02 4.1840e+02 ; HD3 - CD
- CE
6250 6253 6254 1 1.0950e+02 4.1840e+02 ; CE - NZ
- HZ1
6250 6253 6255 1 1.0950e+02 4.1840e+02 ; CE - NZ
- HZ2
6250 6253 6256 1 1.0950e+02 4.1840e+02 ; CE - NZ
- HZ3
6251 6250 6252 1 1.0950e+02 2.9288e+02 ; HE2 - CE
- HE3
6251 6250 6253 1 1.0950e+02 4.1840e+02 ; HE2 - CE
- NZ
6252 6250 6253 1 1.0950e+02 4.1840e+02 ; HE3 - CE
- NZ
6254 6253 6255 1 1.0950e+02 2.9288e+02 ; HZ1 - NZ
- HZ2

6254 6253 6256 1 1.0950e+02 2.9288e+02 ; HZ1 - NZ
- HZ3
6255 6253 6256 1 1.0950e+02 2.9288e+02 ; HZ2 - NZ
- HZ3
6257 6259 6260 1 1.2000e+02 4.1840e+02 ; C - N
- H
6257 6259 6261 1 1.2190e+02 4.1840e+02 ; C - N
- CA
6258 6257 6259 1 1.2290e+02 6.6944e+02 ; O - C
- N
6259 6261 6262 1 1.0950e+02 4.1840e+02 ; N - CA
- HA
6259 6261 6263 1 1.0970e+02 6.6944e+02 ; N - CA
- CB
6259 6261 6272 1 1.1010e+02 5.2718e+02 ; N - CA
- C
6260 6259 6261 1 1.1804e+02 4.1840e+02 ; H - N
- CA
6261 6263 6264 1 1.0950e+02 4.1840e+02 ; CA - CB
- HB2
6261 6263 6265 1 1.0950e+02 4.1840e+02 ; CA - CB
- HB3
6261 6263 6266 1 1.0950e+02 3.3472e+02 ; CA - CB
- CG
6261 6272 6273 1 1.2040e+02 6.6944e+02 ; CA - C
- O
6261 6272 6274 1 1.1660e+02 5.8576e+02 ; CA - C
- N
6262 6261 6263 1 1.0950e+02 4.1840e+02 ; HA - CA
- CB
6262 6261 6272 1 1.0950e+02 4.1840e+02 ; HA - CA
- C
6263 6261 6272 1 1.1110e+02 5.2718e+02 ; CB - CA
- C
6263 6266 6267 1 1.0950e+02 4.1840e+02 ; CB - CG
- HG2
6263 6266 6268 1 1.0950e+02 4.1840e+02 ; CB - CG
- HG3
6263 6266 6269 1 1.1110e+02 5.2718e+02 ; CB - CG
- CD
6264 6263 6265 1 1.0950e+02 2.9288e+02 ; HB2 - CB
- HB3
6264 6263 6266 1 1.0950e+02 4.1840e+02 ; HB2 - CB
- CG
6265 6263 6266 1 1.0950e+02 4.1840e+02 ; HB3 - CB
- CG
6266 6269 6270 1 1.1700e+02 5.8576e+02 ; CG - CD
- OE1
6266 6269 6271 1 1.1700e+02 5.8576e+02 ; CG - CD
- OE2
6267 6266 6268 1 1.0950e+02 2.9288e+02 ; HG2 - CG
- HG3

6267 6266 6269 1 1.0950e+02 4.1840e+02 ; HG2 - CG
- CD
6268 6266 6269 1 1.0950e+02 4.1840e+02 ; HG3 - CG
- CD
6270 6269 6271 1 1.2600e+02 6.6944e+02 ; OE1 - CD
- OE2
6272 6274 6275 1 1.2000e+02 4.1840e+02 ; C - N
- H
6272 6274 6276 1 1.2190e+02 4.1840e+02 ; C - N
- CA
6273 6272 6274 1 1.2290e+02 6.6944e+02 ; O - C
- N
6274 6276 6277 1 1.0950e+02 4.1840e+02 ; N - CA
- HA
6274 6276 6278 1 1.0970e+02 6.6944e+02 ; N - CA
- CB
6274 6276 6288 1 1.1010e+02 5.2718e+02 ; N - CA
- C
6275 6274 6276 1 1.1804e+02 4.1840e+02 ; H - N
- CA
6276 6278 6279 1 1.0950e+02 4.1840e+02 ; CA - CB
- HB
6276 6278 6280 1 1.0950e+02 3.3472e+02 ; CA - CB
- CG1
6276 6278 6284 1 1.0950e+02 3.3472e+02 ; CA - CB
- CG2
6276 6288 6289 1 1.2040e+02 6.6944e+02 ; CA - C
- O
6276 6288 6290 1 1.1660e+02 5.8576e+02 ; CA - C
- N
6277 6276 6278 1 1.0950e+02 4.1840e+02 ; HA - CA
- CB
6277 6276 6288 1 1.0950e+02 4.1840e+02 ; HA - CA
- C
6278 6276 6288 1 1.1110e+02 5.2718e+02 ; CB - CA
- C
6278 6280 6281 1 1.0950e+02 4.1840e+02 ; CB - CG1
- HG11
6278 6280 6282 1 1.0950e+02 4.1840e+02 ; CB - CG1
- HG12
6278 6280 6283 1 1.0950e+02 4.1840e+02 ; CB - CG1
- HG13
6278 6284 6285 1 1.0950e+02 4.1840e+02 ; CB - CG2
- HG21
6278 6284 6286 1 1.0950e+02 4.1840e+02 ; CB - CG2
- HG22
6278 6284 6287 1 1.0950e+02 4.1840e+02 ; CB - CG2
- HG23
6279 6278 6280 1 1.0950e+02 4.1840e+02 ; HB - CB
- CG1
6279 6278 6284 1 1.0950e+02 4.1840e+02 ; HB - CB
- CG2

6280 6278 6284 1 1.0950e+02 3.3472e+02 ; CG1 - CB
- CG2
6281 6280 6282 1 1.0950e+02 2.9288e+02 ; HG11 - CG1
- HG12
6281 6280 6283 1 1.0950e+02 2.9288e+02 ; HG11 - CG1
- HG13
6282 6280 6283 1 1.0950e+02 2.9288e+02 ; HG12 - CG1
- HG13
6285 6284 6286 1 1.0950e+02 2.9288e+02 ; HG21 - CG2
- HG22
6285 6284 6287 1 1.0950e+02 2.9288e+02 ; HG21 - CG2
- HG23
6286 6284 6287 1 1.0950e+02 2.9288e+02 ; HG22 - CG2
- HG23
6288 6290 6291 1 1.2000e+02 4.1840e+02 ; C - N
- H
6288 6290 6292 1 1.2190e+02 4.1840e+02 ; C - N
- CA
6289 6288 6290 1 1.2290e+02 6.6944e+02 ; O - C
- N
6290 6292 6293 1 1.0950e+02 4.1840e+02 ; N - CA
- HA
6290 6292 6294 1 1.0970e+02 6.6944e+02 ; N - CA
- CB
6290 6292 6299 1 1.1010e+02 5.2718e+02 ; N - CA
- C
6291 6290 6292 1 1.1804e+02 4.1840e+02 ; H - N
- CA
6292 6294 6295 1 1.0950e+02 4.1840e+02 ; CA - CB
- HB2
6292 6294 6296 1 1.0950e+02 4.1840e+02 ; CA - CB
- HB3
6292 6294 6297 1 1.0860e+02 4.1840e+02 ; CA - CB
- SG
6292 6299 6300 1 1.2040e+02 6.6944e+02 ; CA - C
- O
6292 6299 6301 1 1.1660e+02 5.8576e+02 ; CA - C
- N
6293 6292 6294 1 1.0950e+02 4.1840e+02 ; HA - CA
- CB
6293 6292 6299 1 1.0950e+02 4.1840e+02 ; HA - CA
- C
6294 6292 6299 1 1.1110e+02 5.2718e+02 ; CB - CA
- C
6294 6297 6298 1 9.6000e+01 3.5982e+02 ; CB - SG
- HG
6295 6294 6296 1 1.0950e+02 2.9288e+02 ; HB2 - CB
- HB3
6295 6294 6297 1 1.0950e+02 4.1840e+02 ; HB2 - CB
- SG
6296 6294 6297 1 1.0950e+02 4.1840e+02 ; HB3 - CB
- SG

6299 6301 6302 1 1.2000e+02 4.1840e+02 ; C - N
- H
6299 6301 6303 1 1.2190e+02 4.1840e+02 ; C - N
- CA
6300 6299 6301 1 1.2290e+02 6.6944e+02 ; O - C
- N
6301 6303 6304 1 1.0950e+02 4.1840e+02 ; N - CA
- HA
6301 6303 6305 1 1.0970e+02 6.6944e+02 ; N - CA
- CB
6301 6303 6318 1 1.1010e+02 5.2718e+02 ; N - CA
- C
6302 6301 6303 1 1.1804e+02 4.1840e+02 ; H - N
- CA
6303 6305 6306 1 1.0950e+02 4.1840e+02 ; CA - CB
- HB2
6303 6305 6307 1 1.0950e+02 4.1840e+02 ; CA - CB
- HB3
6303 6305 6308 1 1.0950e+02 3.3472e+02 ; CA - CB
- CG
6303 6318 6319 1 1.2040e+02 6.6944e+02 ; CA - C
- O
6303 6318 6320 1 1.1660e+02 5.8576e+02 ; CA - C
- N
6304 6303 6305 1 1.0950e+02 4.1840e+02 ; HA - CA
- CB
6304 6303 6318 1 1.0950e+02 4.1840e+02 ; HA - CA
- C
6305 6303 6318 1 1.1110e+02 5.2718e+02 ; CB - CA
- C
6305 6308 6309 1 1.0950e+02 4.1840e+02 ; CB - CG
- HG
6305 6308 6310 1 1.0950e+02 3.3472e+02 ; CB - CG
- CD1
6305 6308 6314 1 1.0950e+02 3.3472e+02 ; CB - CG
- CD2
6306 6305 6307 1 1.0950e+02 2.9288e+02 ; HB2 - CB
- HB3
6306 6305 6308 1 1.0950e+02 4.1840e+02 ; HB2 - CB
- CG
6307 6305 6308 1 1.0950e+02 4.1840e+02 ; HB3 - CB
- CG
6308 6310 6311 1 1.0950e+02 4.1840e+02 ; CG - CD1
- HD11
6308 6310 6312 1 1.0950e+02 4.1840e+02 ; CG - CD1
- HD12
6308 6310 6313 1 1.0950e+02 4.1840e+02 ; CG - CD1
- HD13
6308 6314 6315 1 1.0950e+02 4.1840e+02 ; CG - CD2
- HD21
6308 6314 6316 1 1.0950e+02 4.1840e+02 ; CG - CD2
- HD22

```
  6308   6314   6317      1    1.0950e+02    4.1840e+02 ;     CG - CD2
- HD23
  6309   6308   6310      1    1.0950e+02    4.1840e+02 ;     HG - CG
- CD1
  6309   6308   6314      1    1.0950e+02    4.1840e+02 ;     HG - CG
- CD2
  6310   6308   6314      1    1.0950e+02    3.3472e+02 ;    CD1 - CG
- CD2
  6311   6310   6312      1    1.0950e+02    2.9288e+02 ;   HD11 - CD1
- HD12
  6311   6310   6313      1    1.0950e+02    2.9288e+02 ;   HD11 - CD1
- HD13
  6312   6310   6313      1    1.0950e+02    2.9288e+02 ;   HD12 - CD1
- HD13
  6315   6314   6316      1    1.0950e+02    2.9288e+02 ;   HD21 - CD2
- HD22
  6315   6314   6317      1    1.0950e+02    2.9288e+02 ;   HD21 - CD2
- HD23
  6316   6314   6317      1    1.0950e+02    2.9288e+02 ;   HD22 - CD2
- HD23
  6318   6320   6321      1    1.2000e+02    4.1840e+02 ;      C - N
- H
  6318   6320   6322      1    1.2190e+02    4.1840e+02 ;      C - N
- CA
  6319   6318   6320      1    1.2290e+02    6.6944e+02 ;      O - C
- N
  6320   6322   6323      1    1.0950e+02    4.1840e+02 ;      N - CA
- HA
  6320   6322   6324      1    1.0970e+02    6.6944e+02 ;      N - CA
- CB
  6320   6322   6337      1    1.1010e+02    5.2718e+02 ;      N - CA
- C
  6321   6320   6322      1    1.1804e+02    4.1840e+02 ;      H - N
- CA
  6322   6324   6325      1    1.0950e+02    4.1840e+02 ;     CA - CB
- HB
  6322   6324   6326      1    1.0950e+02    3.3472e+02 ;     CA - CB
- CG2
  6322   6324   6330      1    1.0950e+02    3.3472e+02 ;     CA - CB
- CG1
  6322   6337   6338      1    1.2040e+02    6.6944e+02 ;     CA - C
- O
  6322   6337   6339      1    1.1660e+02    5.8576e+02 ;     CA - C
- N
  6323   6322   6324      1    1.0950e+02    4.1840e+02 ;     HA - CA
- CB
  6323   6322   6337      1    1.0950e+02    4.1840e+02 ;     HA - CA
- C
  6324   6322   6337      1    1.1110e+02    5.2718e+02 ;     CB - CA
- C
  6324   6326   6327      1    1.0950e+02    4.1840e+02 ;     CB - CG2
- HG21
```

6324 6326 6328 1 1.0950e+02 4.1840e+02 ; CB - CG2
- HG22
6324 6326 6329 1 1.0950e+02 4.1840e+02 ; CB - CG2
- HG23
6324 6330 6331 1 1.0950e+02 4.1840e+02 ; CB - CG1
- HG12
6324 6330 6332 1 1.0950e+02 4.1840e+02 ; CB - CG1
- HG13
6324 6330 6333 1 1.0950e+02 3.3472e+02 ; CB - CG1
- CD1
6325 6324 6326 1 1.0950e+02 4.1840e+02 ; HB - CB
- CG2
6325 6324 6330 1 1.0950e+02 4.1840e+02 ; HB - CB
- CG1
6326 6324 6330 1 1.0950e+02 3.3472e+02 ; CG2 - CB
- CG1
6327 6326 6328 1 1.0950e+02 2.9288e+02 ; HG21 - CG2
- HG22
6327 6326 6329 1 1.0950e+02 2.9288e+02 ; HG21 - CG2
- HG23
6328 6326 6329 1 1.0950e+02 2.9288e+02 ; HG22 - CG2
- HG23
6330 6333 6334 1 1.0950e+02 4.1840e+02 ; CG1 - CD1
- HD11
6330 6333 6335 1 1.0950e+02 4.1840e+02 ; CG1 - CD1
- HD12
6330 6333 6336 1 1.0950e+02 4.1840e+02 ; CG1 - CD1
- HD13
6331 6330 6332 1 1.0950e+02 2.9288e+02 ; HG12 - CG1
- HG13
6331 6330 6333 1 1.0950e+02 4.1840e+02 ; HG12 - CG1
- CD1
6332 6330 6333 1 1.0950e+02 4.1840e+02 ; HG13 - CG1
- CD1
6334 6333 6335 1 1.0950e+02 2.9288e+02 ; HD11 - CD1
- HD12
6334 6333 6336 1 1.0950e+02 2.9288e+02 ; HD11 - CD1
- HD13
6335 6333 6336 1 1.0950e+02 2.9288e+02 ; HD12 - CD1
- HD13
6337 6339 6340 1 1.2190e+02 4.1840e+02 ; C - N
- CD
6337 6339 6349 1 1.2190e+02 4.1840e+02 ; C - N
- CA
6338 6337 6339 1 1.2290e+02 6.6944e+02 ; O - C
- N
6339 6340 6341 1 1.0950e+02 4.1840e+02 ; N - CD
- HD2
6339 6340 6342 1 1.0950e+02 4.1840e+02 ; N - CD
- HD3
6339 6340 6343 1 1.0970e+02 6.6944e+02 ; N - CD
- CG

| | | | | | | | |
|---|---|---|---|---|---|---|---|
| 6339 | 6349 | 6346 | 1 | 1.0970e+02 | 6.6944e+02 | ; | N - CA - CB |
| 6339 | 6349 | 6350 | 1 | 1.0950e+02 | 4.1840e+02 | ; | N - CA - HA |
| 6339 | 6349 | 6351 | 1 | 1.1010e+02 | 5.2718e+02 | ; | N - CA - C |
| 6340 | 6339 | 6349 | 1 | 1.1800e+02 | 4.1840e+02 | ; | CD - N - CA |
| 6340 | 6343 | 6344 | 1 | 1.0950e+02 | 4.1840e+02 | ; | CD - CG - HG2 |
| 6340 | 6343 | 6345 | 1 | 1.0950e+02 | 4.1840e+02 | ; | CD - CG - HG3 |
| 6340 | 6343 | 6346 | 1 | 1.0950e+02 | 3.3472e+02 | ; | CD - CG - CB |
| 6341 | 6340 | 6342 | 1 | 1.0950e+02 | 2.9288e+02 | ; | HD2 - CD - HD3 |
| 6341 | 6340 | 6343 | 1 | 1.0950e+02 | 4.1840e+02 | ; | HD2 - CD - CG |
| 6342 | 6340 | 6343 | 1 | 1.0950e+02 | 4.1840e+02 | ; | HD3 - CD - CG |
| 6343 | 6346 | 6347 | 1 | 1.0950e+02 | 4.1840e+02 | ; | CG - CB - HB2 |
| 6343 | 6346 | 6348 | 1 | 1.0950e+02 | 4.1840e+02 | ; | CG - CB - HB3 |
| 6343 | 6346 | 6349 | 1 | 1.0950e+02 | 3.3472e+02 | ; | CG - CB - CA |
| 6344 | 6343 | 6345 | 1 | 1.0950e+02 | 2.9288e+02 | ; | HG2 - CG - HG3 |
| 6344 | 6343 | 6346 | 1 | 1.0950e+02 | 4.1840e+02 | ; | HG2 - CG - CB |
| 6345 | 6343 | 6346 | 1 | 1.0950e+02 | 4.1840e+02 | ; | HG3 - CG - CB |
| 6346 | 6349 | 6350 | 1 | 1.0950e+02 | 4.1840e+02 | ; | CB - CA - HA |
| 6346 | 6349 | 6351 | 1 | 1.1110e+02 | 5.2718e+02 | ; | CB - CA - C |
| 6347 | 6346 | 6348 | 1 | 1.0950e+02 | 2.9288e+02 | ; | HB2 - CB - HB3 |
| 6347 | 6346 | 6349 | 1 | 1.0950e+02 | 4.1840e+02 | ; | HB2 - CB - CA |
| 6348 | 6346 | 6349 | 1 | 1.0950e+02 | 4.1840e+02 | ; | HB3 - CB - CA |
| 6349 | 6351 | 6352 | 1 | 1.2040e+02 | 6.6944e+02 | ; | CA - C - O |
| 6349 | 6351 | 6353 | 1 | 1.1660e+02 | 5.8576e+02 | ; | CA - C - N |
| 6350 | 6349 | 6351 | 1 | 1.0950e+02 | 4.1840e+02 | ; | HA - CA - C |
| 6351 | 6353 | 6354 | 1 | 1.2000e+02 | 4.1840e+02 | ; | C - N - H |
| 6351 | 6353 | 6355 | 1 | 1.2190e+02 | 4.1840e+02 | ; | C - N - CA |

```
  6352   6351   6353       1    1.2290e+02    6.6944e+02 ;       O - C
- N
  6353   6355   6356       1    1.0950e+02    4.1840e+02 ;       N - CA
- HA
  6353   6355   6357       1    1.0970e+02    6.6944e+02 ;       N - CA
- CB
  6353   6355   6371       1    1.1010e+02    5.2718e+02 ;       N - CA
- C
  6354   6353   6355       1    1.1804e+02    4.1840e+02 ;       H - N
- CA
  6355   6357   6358       1    1.0950e+02    4.1840e+02 ;      CA - CB
- HB2
  6355   6357   6359       1    1.0950e+02    4.1840e+02 ;      CA - CB
- HB3
  6355   6357   6360       1    1.1400e+02    5.2718e+02 ;      CA - CB
- CG
  6355   6371   6372       1    1.2040e+02    6.6944e+02 ;      CA - C
- O
  6355   6371   6373       1    1.1660e+02    5.8576e+02 ;      CA - C
- N
  6356   6355   6357       1    1.0950e+02    4.1840e+02 ;      HA - CA
- CB
  6356   6355   6371       1    1.0950e+02    4.1840e+02 ;      HA - CA
- C
  6357   6355   6371       1    1.1110e+02    5.2718e+02 ;      CB - CA
- C
  6357   6360   6361       1    1.2000e+02    5.8576e+02 ;      CB - CG
- CD1
  6357   6360   6369       1    1.2000e+02    5.8576e+02 ;      CB - CG
- CD2
  6358   6357   6359       1    1.0950e+02    2.9288e+02 ;     HB2 - CB
- HB3
  6358   6357   6360       1    1.0950e+02    4.1840e+02 ;     HB2 - CB
- CG
  6359   6357   6360       1    1.0950e+02    4.1840e+02 ;     HB3 - CB
- CG
  6360   6361   6362       1    1.2000e+02    4.1840e+02 ;      CG - CD1
- HD1
  6360   6361   6363       1    1.2000e+02    5.2718e+02 ;      CG - CD1
- CE1
  6360   6369   6367       1    1.2000e+02    5.2718e+02 ;      CG - CD2
- CE2
  6360   6369   6370       1    1.2000e+02    4.1840e+02 ;      CG - CD2
- HD2
  6361   6360   6369       1    1.2000e+02    5.2718e+02 ;     CD1 - CG
- CD2
  6361   6363   6364       1    1.2000e+02    4.1840e+02 ;     CD1 - CE1
- HE1
  6361   6363   6365       1    1.2000e+02    5.2718e+02 ;     CD1 - CE1
- CZ
  6362   6361   6363       1    1.2000e+02    4.1840e+02 ;     HD1 - CD1
- CE1
```

```
  6363   6365   6366      1    1.2000e+02    4.1840e+02 ;    CE1 - CZ
- HZ
  6363   6365   6367      1    1.2000e+02    5.2718e+02 ;    CE1 - CZ
- CE2
  6364   6363   6365      1    1.2000e+02    4.1840e+02 ;    HE1 - CE1
- CZ
  6365   6367   6368      1    1.2000e+02    4.1840e+02 ;     CZ - CE2
- HE2
  6365   6367   6369      1    1.2000e+02    5.2718e+02 ;     CZ - CE2
- CD2
  6366   6365   6367      1    1.2000e+02    4.1840e+02 ;     HZ - CZ
- CE2
  6367   6369   6370      1    1.2000e+02    4.1840e+02 ;    CE2 - CD2
- HD2
  6368   6367   6369      1    1.2000e+02    4.1840e+02 ;    HE2 - CE2
- CD2
  6371   6373   6374      1    1.2000e+02    4.1840e+02 ;      C - N
- H
  6371   6373   6375      1    1.2190e+02    4.1840e+02 ;      C - N
- CA
  6372   6371   6373      1    1.2290e+02    6.6944e+02 ;      O - C
- N
  6373   6375   6376      1    1.0950e+02    4.1840e+02 ;      N - CA
- HA
  6373   6375   6377      1    1.0970e+02    6.6944e+02 ;      N - CA
- CB
  6373   6375   6393      1    1.1010e+02    5.2718e+02 ;      N - CA
- C
  6374   6373   6375      1    1.1804e+02    4.1840e+02 ;      H - N
- CA
  6375   6377   6378      1    1.0950e+02    4.1840e+02 ;     CA - CB
- HB2
  6375   6377   6379      1    1.0950e+02    4.1840e+02 ;     CA - CB
- HB3
  6375   6377   6380      1    1.0950e+02    3.3472e+02 ;     CA - CB
- CG
  6375   6393   6394      1    1.2040e+02    6.6944e+02 ;     CA - C
- O
  6375   6393   6395      1    1.1660e+02    5.8576e+02 ;     CA - C
- N
  6376   6375   6377      1    1.0950e+02    4.1840e+02 ;     HA - CA
- CB
  6376   6375   6393      1    1.0950e+02    4.1840e+02 ;     HA - CA
- C
  6377   6375   6393      1    1.1110e+02    5.2718e+02 ;     CB - CA
- C
  6377   6380   6381      1    1.0950e+02    4.1840e+02 ;     CB - CG
- HG2
  6377   6380   6382      1    1.0950e+02    4.1840e+02 ;     CB - CG
- HG3
  6377   6380   6383      1    1.0950e+02    3.3472e+02 ;     CB - CG
- CD
```

6378 6377 6379 1 1.0950e+02 2.9288e+02 ; HB2 - CB
- HB3
6378 6377 6380 1 1.0950e+02 4.1840e+02 ; HB2 - CB
- CG
6379 6377 6380 1 1.0950e+02 4.1840e+02 ; HB3 - CB
- CG
6380 6383 6384 1 1.0950e+02 4.1840e+02 ; CG - CD
- HD2
6380 6383 6385 1 1.0950e+02 4.1840e+02 ; CG - CD
- HD3
6380 6383 6386 1 1.0950e+02 3.3472e+02 ; CG - CD
- CE
6381 6380 6382 1 1.0950e+02 2.9288e+02 ; HG2 - CG
- HG3
6381 6380 6383 1 1.0950e+02 4.1840e+02 ; HG2 - CG
- CD
6382 6380 6383 1 1.0950e+02 4.1840e+02 ; HG3 - CG
- CD
6383 6386 6387 1 1.0950e+02 4.1840e+02 ; CD - CE
- HE2
6383 6386 6388 1 1.0950e+02 4.1840e+02 ; CD - CE
- HE3
6383 6386 6389 1 1.1120e+02 6.6944e+02 ; CD - CE
- NZ
6384 6383 6385 1 1.0950e+02 2.9288e+02 ; HD2 - CD
- HD3
6384 6383 6386 1 1.0950e+02 4.1840e+02 ; HD2 - CD
- CE
6385 6383 6386 1 1.0950e+02 4.1840e+02 ; HD3 - CD
- CE
6386 6389 6390 1 1.0950e+02 4.1840e+02 ; CE - NZ
- HZ1
6386 6389 6391 1 1.0950e+02 4.1840e+02 ; CE - NZ
- HZ2
6386 6389 6392 1 1.0950e+02 4.1840e+02 ; CE - NZ
- HZ3
6387 6386 6388 1 1.0950e+02 2.9288e+02 ; HE2 - CE
- HE3
6387 6386 6389 1 1.0950e+02 4.1840e+02 ; HE2 - CE
- NZ
6388 6386 6389 1 1.0950e+02 4.1840e+02 ; HE3 - CE
- NZ
6390 6389 6391 1 1.0950e+02 2.9288e+02 ; HZ1 - NZ
- HZ2
6390 6389 6392 1 1.0950e+02 2.9288e+02 ; HZ1 - NZ
- HZ3
6391 6389 6392 1 1.0950e+02 2.9288e+02 ; HZ2 - NZ
- HZ3
6393 6395 6396 1 1.2000e+02 4.1840e+02 ; C - N
- H
6393 6395 6397 1 1.2190e+02 4.1840e+02 ; C - N
- CA

6394 6393 6395 1 1.2290e+02 6.6944e+02 ; O - C
- N
6395 6397 6398 1 1.0950e+02 4.1840e+02 ; N - CA
- HA
6395 6397 6399 1 1.0970e+02 6.6944e+02 ; N - CA
- CB
6395 6397 6413 1 1.1010e+02 5.2718e+02 ; N - CA
- C
6396 6395 6397 1 1.1804e+02 4.1840e+02 ; H - N
- CA
6397 6399 6400 1 1.0950e+02 4.1840e+02 ; CA - CB
- HB2
6397 6399 6401 1 1.0950e+02 4.1840e+02 ; CA - CB
- HB3
6397 6399 6402 1 1.1400e+02 5.2718e+02 ; CA - CB
- CG
6397 6413 6414 1 1.2040e+02 6.6944e+02 ; CA - C
- O
6397 6413 6415 1 1.1660e+02 5.8576e+02 ; CA - C
- N
6398 6397 6399 1 1.0950e+02 4.1840e+02 ; HA - CA
- CB
6398 6397 6413 1 1.0950e+02 4.1840e+02 ; HA - CA
- C
6399 6397 6413 1 1.1110e+02 5.2718e+02 ; CB - CA
- C
6399 6402 6403 1 1.2000e+02 5.8576e+02 ; CB - CG
- CD1
6399 6402 6411 1 1.2000e+02 5.8576e+02 ; CB - CG
- CD2
6400 6399 6401 1 1.0950e+02 2.9288e+02 ; HB2 - CB
- HB3
6400 6399 6402 1 1.0950e+02 4.1840e+02 ; HB2 - CB
- CG
6401 6399 6402 1 1.0950e+02 4.1840e+02 ; HB3 - CB
- CG
6402 6403 6404 1 1.2000e+02 4.1840e+02 ; CG - CD1
- HD1
6402 6403 6405 1 1.2000e+02 5.2718e+02 ; CG - CD1
- CE1
6402 6411 6409 1 1.2000e+02 5.2718e+02 ; CG - CD2
- CE2
6402 6411 6412 1 1.2000e+02 4.1840e+02 ; CG - CD2
- HD2
6403 6402 6411 1 1.2000e+02 5.2718e+02 ; CD1 - CG
- CD2
6403 6405 6406 1 1.2000e+02 4.1840e+02 ; CD1 - CE1
- HE1
6403 6405 6407 1 1.2000e+02 5.2718e+02 ; CD1 - CE1
- CZ
6404 6403 6405 1 1.2000e+02 4.1840e+02 ; HD1 - CD1
- CE1

6405 6407 6408 1 1.2000e+02 4.1840e+02 ; CE1 - CZ
- HZ
6405 6407 6409 1 1.2000e+02 5.2718e+02 ; CE1 - CZ
- CE2
6406 6405 6407 1 1.2000e+02 4.1840e+02 ; HE1 - CE1
- CZ
6407 6409 6410 1 1.2000e+02 4.1840e+02 ; CZ - CE2
- HE2
6407 6409 6411 1 1.2000e+02 5.2718e+02 ; CZ - CE2
- CD2
6408 6407 6409 1 1.2000e+02 4.1840e+02 ; HZ - CZ
- CE2
6409 6411 6412 1 1.2000e+02 4.1840e+02 ; CE2 - CD2
- HD2
6410 6409 6411 1 1.2000e+02 4.1840e+02 ; HE2 - CE2
- CD2
6413 6415 6416 1 1.2000e+02 4.1840e+02 ; C - N
- H
6413 6415 6417 1 1.2190e+02 4.1840e+02 ; C - N
- CA
6414 6413 6415 1 1.2290e+02 6.6944e+02 ; O - C
- N
6415 6417 6418 1 1.0950e+02 4.1840e+02 ; N - CA
- HA
6415 6417 6419 1 1.0970e+02 6.6944e+02 ; N - CA
- CB
6415 6417 6433 1 1.1010e+02 5.2718e+02 ; N - CA
- C
6416 6415 6417 1 1.1804e+02 4.1840e+02 ; H - N
- CA
6417 6419 6420 1 1.0950e+02 4.1840e+02 ; CA - CB
- HB2
6417 6419 6421 1 1.0950e+02 4.1840e+02 ; CA - CB
- HB3
6417 6419 6422 1 1.1400e+02 5.2718e+02 ; CA - CB
- CG
6417 6433 6434 1 1.2040e+02 6.6944e+02 ; CA - C
- O
6417 6433 6435 1 1.1660e+02 5.8576e+02 ; CA - C
- N
6418 6417 6419 1 1.0950e+02 4.1840e+02 ; HA - CA
- CB
6418 6417 6433 1 1.0950e+02 4.1840e+02 ; HA - CA
- C
6419 6417 6433 1 1.1110e+02 5.2718e+02 ; CB - CA
- C
6419 6422 6423 1 1.2000e+02 5.8576e+02 ; CB - CG
- CD1
6419 6422 6431 1 1.2000e+02 5.8576e+02 ; CB - CG
- CD2
6420 6419 6421 1 1.0950e+02 2.9288e+02 ; HB2 - CB
- HB3

```
  6420   6419   6422      1    1.0950e+02    4.1840e+02 ;    HB2 - CB
- CG
  6421   6419   6422      1    1.0950e+02    4.1840e+02 ;    HB3 - CB
- CG
  6422   6423   6424      1    1.2000e+02    4.1840e+02 ;     CG - CD1
- HD1
  6422   6423   6425      1    1.2000e+02    5.2718e+02 ;     CG - CD1
- CE1
  6422   6431   6429      1    1.2000e+02    5.2718e+02 ;     CG - CD2
- CE2
  6422   6431   6432      1    1.2000e+02    4.1840e+02 ;     CG - CD2
- HD2
  6423   6422   6431      1    1.2000e+02    5.2718e+02 ;    CD1 - CG
- CD2
  6423   6425   6426      1    1.2000e+02    4.1840e+02 ;    CD1 - CE1
- HE1
  6423   6425   6427      1    1.2000e+02    5.2718e+02 ;    CD1 - CE1
- CZ
  6424   6423   6425      1    1.2000e+02    4.1840e+02 ;    HD1 - CD1
- CE1
  6425   6427   6428      1    1.2000e+02    4.1840e+02 ;    CE1 - CZ
- HZ
  6425   6427   6429      1    1.2000e+02    5.2718e+02 ;    CE1 - CZ
- CE2
  6426   6425   6427      1    1.2000e+02    4.1840e+02 ;    HE1 - CE1
- CZ
  6427   6429   6430      1    1.2000e+02    4.1840e+02 ;     CZ - CE2
- HE2
  6427   6429   6431      1    1.2000e+02    5.2718e+02 ;     CZ - CE2
- CD2
  6428   6427   6429      1    1.2000e+02    4.1840e+02 ;     HZ - CZ
- CE2
  6429   6431   6432      1    1.2000e+02    4.1840e+02 ;    CE2 - CD2
- HD2
  6430   6429   6431      1    1.2000e+02    4.1840e+02 ;    HE2 - CE2
- CD2
  6433   6435   6436      1    1.2000e+02    4.1840e+02 ;      C - N
- H
  6433   6435   6437      1    1.2190e+02    4.1840e+02 ;      C - N
- CA
  6434   6433   6435      1    1.2290e+02    6.6944e+02 ;      O - C
- N
  6435   6437   6438      1    1.0950e+02    4.1840e+02 ;      N - CA
- HA
  6435   6437   6439      1    1.0970e+02    6.6944e+02 ;      N - CA
- CB
  6435   6437   6445      1    1.1010e+02    5.2718e+02 ;      N - CA
- C
  6436   6435   6437      1    1.1804e+02    4.1840e+02 ;      H - N
- CA
  6437   6439   6440      1    1.0950e+02    4.1840e+02 ;     CA - CB
- HB2
```

6437 6439 6441 1 1.0950e+02 4.1840e+02 ; CA - CB
- HB3
6437 6439 6442 1 1.1110e+02 5.2718e+02 ; CA - CB
- CG
6437 6445 6446 1 1.2040e+02 6.6944e+02 ; CA - C
- O
6437 6445 6447 1 1.1660e+02 5.8576e+02 ; CA - C
- N
6438 6437 6439 1 1.0950e+02 4.1840e+02 ; HA - CA
- CB
6438 6437 6445 1 1.0950e+02 4.1840e+02 ; HA - CA
- C
6439 6437 6445 1 1.1110e+02 5.2718e+02 ; CB - CA
- C
6439 6442 6443 1 1.1700e+02 5.8576e+02 ; CB - CG
- OD1
6439 6442 6444 1 1.1700e+02 5.8576e+02 ; CB - CG
- OD2
6440 6439 6441 1 1.0950e+02 2.9288e+02 ; HB2 - CB
- HB3
6440 6439 6442 1 1.0950e+02 4.1840e+02 ; HB2 - CB
- CG
6441 6439 6442 1 1.0950e+02 4.1840e+02 ; HB3 - CB
- CG
6443 6442 6444 1 1.2600e+02 6.6944e+02 ; OD1 - CG
- OD2
6445 6447 6448 1 1.2000e+02 4.1840e+02 ; C - N
- H
6445 6447 6449 1 1.2190e+02 4.1840e+02 ; C - N
- CA
6446 6445 6447 1 1.2290e+02 6.6944e+02 ; O - C
- N
6447 6449 6450 1 1.0950e+02 4.1840e+02 ; N - CA
- HA
6447 6449 6451 1 1.0970e+02 6.6944e+02 ; N - CA
- CB
6447 6449 6464 1 1.1010e+02 5.2718e+02 ; N - CA
- C
6448 6447 6449 1 1.1804e+02 4.1840e+02 ; H - N
- CA
6449 6451 6452 1 1.0950e+02 4.1840e+02 ; CA - CB
- HB
6449 6451 6453 1 1.0950e+02 3.3472e+02 ; CA - CB
- CG2
6449 6451 6457 1 1.0950e+02 3.3472e+02 ; CA - CB
- CG1
6449 6464 6465 1 1.2040e+02 6.6944e+02 ; CA - C
- O
6449 6464 6466 1 1.1660e+02 5.8576e+02 ; CA - C
- N
6450 6449 6451 1 1.0950e+02 4.1840e+02 ; HA - CA
- CB

```
  6450   6449   6464       1    1.0950e+02    4.1840e+02 ;     HA - CA
- C
  6451   6449   6464       1    1.1110e+02    5.2718e+02 ;     CB - CA
- C
  6451   6453   6454       1    1.0950e+02    4.1840e+02 ;     CB - CG2
- HG21
  6451   6453   6455       1    1.0950e+02    4.1840e+02 ;     CB - CG2
- HG22
  6451   6453   6456       1    1.0950e+02    4.1840e+02 ;     CB - CG2
- HG23
  6451   6457   6458       1    1.0950e+02    4.1840e+02 ;     CB - CG1
- HG12
  6451   6457   6459       1    1.0950e+02    4.1840e+02 ;     CB - CG1
- HG13
  6451   6457   6460       1    1.0950e+02    3.3472e+02 ;     CB - CG1
- CD1
  6452   6451   6453       1    1.0950e+02    4.1840e+02 ;     HB - CB
- CG2
  6452   6451   6457       1    1.0950e+02    4.1840e+02 ;     HB - CB
- CG1
  6453   6451   6457       1    1.0950e+02    3.3472e+02 ;    CG2 - CB
- CG1
  6454   6453   6455       1    1.0950e+02    2.9288e+02 ;   HG21 - CG2
- HG22
  6454   6453   6456       1    1.0950e+02    2.9288e+02 ;   HG21 - CG2
- HG23
  6455   6453   6456       1    1.0950e+02    2.9288e+02 ;   HG22 - CG2
- HG23
  6457   6460   6461       1    1.0950e+02    4.1840e+02 ;    CG1 - CD1
- HD11
  6457   6460   6462       1    1.0950e+02    4.1840e+02 ;    CG1 - CD1
- HD12
  6457   6460   6463       1    1.0950e+02    4.1840e+02 ;    CG1 - CD1
- HD13
  6458   6457   6459       1    1.0950e+02    2.9288e+02 ;   HG12 - CG1
- HG13
  6458   6457   6460       1    1.0950e+02    4.1840e+02 ;   HG12 - CG1
- CD1
  6459   6457   6460       1    1.0950e+02    4.1840e+02 ;   HG13 - CG1
- CD1
  6461   6460   6462       1    1.0950e+02    2.9288e+02 ;   HD11 - CD1
- HD12
  6461   6460   6463       1    1.0950e+02    2.9288e+02 ;   HD11 - CD1
- HD13
  6462   6460   6463       1    1.0950e+02    2.9288e+02 ;   HD12 - CD1
- HD13
  6464   6466   6467       1    1.2000e+02    4.1840e+02 ;      C - N
- H
  6464   6466   6468       1    1.2190e+02    4.1840e+02 ;      C - N
- CA
  6465   6464   6466       1    1.2290e+02    6.6944e+02 ;      O - C
- N
```

```
  6466   6468   6469      1    1.0950e+02    4.1840e+02 ;      N - CA
- HA
  6466   6468   6470      1    1.0970e+02    6.6944e+02 ;      N - CA
- CB
  6466   6468   6476      1    1.1010e+02    5.2718e+02 ;      N - CA
- C
  6467   6466   6468      1    1.1804e+02    4.1840e+02 ;      H - N
- CA
  6468   6470   6471      1    1.0950e+02    4.1840e+02 ;     CA - CB
- HB2
  6468   6470   6472      1    1.0950e+02    4.1840e+02 ;     CA - CB
- HB3
  6468   6470   6473      1    1.1110e+02    5.2718e+02 ;     CA - CB
- CG
  6468   6476   6477      1    1.2040e+02    6.6944e+02 ;     CA - C
- O
  6468   6476   6478      1    1.1660e+02    5.8576e+02 ;     CA - C
- N
  6469   6468   6470      1    1.0950e+02    4.1840e+02 ;     HA - CA
- CB
  6469   6468   6476      1    1.0950e+02    4.1840e+02 ;     HA - CA
- C
  6470   6468   6476      1    1.1110e+02    5.2718e+02 ;     CB - CA
- C
  6470   6473   6474      1    1.1700e+02    5.8576e+02 ;     CB - CG
- OD1
  6470   6473   6475      1    1.1700e+02    5.8576e+02 ;     CB - CG
- OD2
  6471   6470   6472      1    1.0950e+02    2.9288e+02 ;    HB2 - CB
- HB3
  6471   6470   6473      1    1.0950e+02    4.1840e+02 ;    HB2 - CB
- CG
  6472   6470   6473      1    1.0950e+02    4.1840e+02 ;    HB3 - CB
- CG
  6474   6473   6475      1    1.2600e+02    6.6944e+02 ;    OD1 - CG
- OD2
  6476   6478   6479      1    1.2000e+02    4.1840e+02 ;      C - N
- H
  6476   6478   6480      1    1.2190e+02    4.1840e+02 ;      C - N
- CA
  6477   6476   6478      1    1.2290e+02    6.6944e+02 ;      O - C
- N
  6478   6480   6481      1    1.0950e+02    4.1840e+02 ;      N - CA
- HA2
  6478   6480   6482      1    1.0950e+02    4.1840e+02 ;      N - CA
- HA3
  6478   6480   6483      1    1.1010e+02    5.2718e+02 ;      N - CA
- C
  6479   6478   6480      1    1.1804e+02    4.1840e+02 ;      H - N
- CA
  6480   6483   6484      1    1.2040e+02    6.6944e+02 ;     CA - C
- O
```

6480 6483 6485 1 1.1660e+02 5.8576e+02 ; CA - C
- N
6481 6480 6482 1 1.0950e+02 2.9288e+02 ; HA2 - CA
- HA3
6481 6480 6483 1 1.0950e+02 4.1840e+02 ; HA2 - CA
- C
6482 6480 6483 1 1.0950e+02 4.1840e+02 ; HA3 - CA
- C
6483 6485 6486 1 1.2000e+02 4.1840e+02 ; C - N
- H
6483 6485 6487 1 1.2190e+02 4.1840e+02 ; C - N
- CA
6484 6483 6485 1 1.2290e+02 6.6944e+02 ; O - C
- N
6485 6487 6488 1 1.0950e+02 4.1840e+02 ; N - CA
- HA
6485 6487 6489 1 1.0970e+02 6.6944e+02 ; N - CA
- CB
6485 6487 6497 1 1.1010e+02 5.2718e+02 ; N - CA
- C
6486 6485 6487 1 1.1804e+02 4.1840e+02 ; H - N
- CA
6487 6489 6490 1 1.0950e+02 4.1840e+02 ; CA - CB
- HB2
6487 6489 6491 1 1.0950e+02 4.1840e+02 ; CA - CB
- HB3
6487 6489 6492 1 1.1110e+02 5.2718e+02 ; CA - CB
- CG
6487 6497 6498 1 1.2040e+02 6.6944e+02 ; CA - C
- O
6487 6497 6499 1 1.1660e+02 5.8576e+02 ; CA - C
- N
6488 6487 6489 1 1.0950e+02 4.1840e+02 ; HA - CA
- CB
6488 6487 6497 1 1.0950e+02 4.1840e+02 ; HA - CA
- C
6489 6487 6497 1 1.1110e+02 5.2718e+02 ; CB - CA
- C
6489 6492 6493 1 1.2040e+02 6.6944e+02 ; CB - CG
- OD1
6489 6492 6494 1 1.1660e+02 5.8576e+02 ; CB - CG
- ND2
6490 6489 6491 1 1.0950e+02 2.9288e+02 ; HB2 - CB
- HB3
6490 6489 6492 1 1.0950e+02 4.1840e+02 ; HB2 - CB
- CG
6491 6489 6492 1 1.0950e+02 4.1840e+02 ; HB3 - CB
- CG
6492 6494 6495 1 1.2000e+02 4.1840e+02 ; CG - ND2
- HD21
6492 6494 6496 1 1.2000e+02 4.1840e+02 ; CG - ND2
- HD22

```
  6493   6492   6494      1    1.2290e+02    6.6944e+02 ;    OD1 - CG
- ND2
  6495   6494   6496      1    1.2000e+02    2.9288e+02 ;   HD21 - ND2
- HD22
  6497   6499   6500      1    1.2000e+02    4.1840e+02 ;      C - N
- H
  6497   6499   6501      1    1.2190e+02    4.1840e+02 ;      C - N
- CA
  6498   6497   6499      1    1.2290e+02    6.6944e+02 ;      O - C
- N
  6499   6501   6502      1    1.0950e+02    4.1840e+02 ;      N - CA
- HA2
  6499   6501   6503      1    1.0950e+02    4.1840e+02 ;      N - CA
- HA3
  6499   6501   6504      1    1.1010e+02    5.2718e+02 ;      N - CA
- C
  6500   6499   6501      1    1.1804e+02    4.1840e+02 ;      H - N
- CA
  6501   6504   6505      1    1.2040e+02    6.6944e+02 ;     CA - C
- O
  6501   6504   6506      1    1.1660e+02    5.8576e+02 ;     CA - C
- N
  6502   6501   6503      1    1.0950e+02    2.9288e+02 ;    HA2 - CA
- HA3
  6502   6501   6504      1    1.0950e+02    4.1840e+02 ;    HA2 - CA
- C
  6503   6501   6504      1    1.0950e+02    4.1840e+02 ;    HA3 - CA
- C
  6504   6506   6507      1    1.2000e+02    4.1840e+02 ;      C - N
- H
  6504   6506   6508      1    1.2190e+02    4.1840e+02 ;      C - N
- CA
  6505   6504   6506      1    1.2290e+02    6.6944e+02 ;      O - C
- N
  6506   6508   6509      1    1.0950e+02    4.1840e+02 ;      N - CA
- HA
  6506   6508   6510      1    1.0970e+02    6.6944e+02 ;      N - CA
- CB
  6506   6508   6526      1    1.1010e+02    5.2718e+02 ;      N - CA
- C
  6507   6506   6508      1    1.1804e+02    4.1840e+02 ;      H - N
- CA
  6508   6510   6511      1    1.0950e+02    4.1840e+02 ;     CA - CB
- HB2
  6508   6510   6512      1    1.0950e+02    4.1840e+02 ;     CA - CB
- HB3
  6508   6510   6513      1    1.0950e+02    3.3472e+02 ;     CA - CB
- CG
  6508   6526   6527      1    1.2040e+02    6.6944e+02 ;     CA - C
- O
  6508   6526   6528      1    1.1660e+02    5.8576e+02 ;     CA - C
- N
```

6509 6508 6510 1 1.0950e+02 4.1840e+02 ; HA - CA
- CB
6509 6508 6526 1 1.0950e+02 4.1840e+02 ; HA - CA
- C
6510 6508 6526 1 1.1110e+02 5.2718e+02 ; CB - CA
- C
6510 6513 6514 1 1.0950e+02 4.1840e+02 ; CB - CG
- HG2
6510 6513 6515 1 1.0950e+02 4.1840e+02 ; CB - CG
- HG3
6510 6513 6516 1 1.0950e+02 3.3472e+02 ; CB - CG
- CD
6511 6510 6512 1 1.0950e+02 2.9288e+02 ; HB2 - CB
- HB3
6511 6510 6513 1 1.0950e+02 4.1840e+02 ; HB2 - CB
- CG
6512 6510 6513 1 1.0950e+02 4.1840e+02 ; HB3 - CB
- CG
6513 6516 6517 1 1.0950e+02 4.1840e+02 ; CG - CD
- HD2
6513 6516 6518 1 1.0950e+02 4.1840e+02 ; CG - CD
- HD3
6513 6516 6519 1 1.0950e+02 3.3472e+02 ; CG - CD
- CE
6514 6513 6515 1 1.0950e+02 2.9288e+02 ; HG2 - CG
- HG3
6514 6513 6516 1 1.0950e+02 4.1840e+02 ; HG2 - CG
- CD
6515 6513 6516 1 1.0950e+02 4.1840e+02 ; HG3 - CG
- CD
6516 6519 6520 1 1.0950e+02 4.1840e+02 ; CD - CE
- HE2
6516 6519 6521 1 1.0950e+02 4.1840e+02 ; CD - CE
- HE3
6516 6519 6522 1 1.1120e+02 6.6944e+02 ; CD - CE
- NZ
6517 6516 6518 1 1.0950e+02 2.9288e+02 ; HD2 - CD
- HD3
6517 6516 6519 1 1.0950e+02 4.1840e+02 ; HD2 - CD
- CE
6518 6516 6519 1 1.0950e+02 4.1840e+02 ; HD3 - CD
- CE
6519 6522 6523 1 1.0950e+02 4.1840e+02 ; CE - NZ
- HZ1
6519 6522 6524 1 1.0950e+02 4.1840e+02 ; CE - NZ
- HZ2
6519 6522 6525 1 1.0950e+02 4.1840e+02 ; CE - NZ
- HZ3
6520 6519 6521 1 1.0950e+02 2.9288e+02 ; HE2 - CE
- HE3
6520 6519 6522 1 1.0950e+02 4.1840e+02 ; HE2 - CE
- NZ

6521 6519 6522 1 1.0950e+02 4.1840e+02 ; HE3 - CE
- NZ
6523 6522 6524 1 1.0950e+02 2.9288e+02 ; HZ1 - NZ
- HZ2
6523 6522 6525 1 1.0950e+02 2.9288e+02 ; HZ1 - NZ
- HZ3
6524 6522 6525 1 1.0950e+02 2.9288e+02 ; HZ2 - NZ
- HZ3
6526 6528 6529 1 1.2000e+02 4.1840e+02 ; C - N
- H
6526 6528 6530 1 1.2190e+02 4.1840e+02 ; C - N
- CA
6527 6526 6528 1 1.2290e+02 6.6944e+02 ; O - C
- N
6528 6530 6531 1 1.0950e+02 4.1840e+02 ; N - CA
- HA
6528 6530 6532 1 1.0970e+02 6.6944e+02 ; N - CA
- CB
6528 6530 6545 1 1.1010e+02 5.2718e+02 ; N - CA
- C
6529 6528 6530 1 1.1804e+02 4.1840e+02 ; H - N
- CA
6530 6532 6533 1 1.0950e+02 4.1840e+02 ; CA - CB
- HB
6530 6532 6534 1 1.0950e+02 3.3472e+02 ; CA - CB
- CG2
6530 6532 6538 1 1.0950e+02 3.3472e+02 ; CA - CB
- CG1
6530 6545 6546 1 1.2040e+02 6.6944e+02 ; CA - C
- O
6530 6545 6547 1 1.1660e+02 5.8576e+02 ; CA - C
- N
6531 6530 6532 1 1.0950e+02 4.1840e+02 ; HA - CA
- CB
6531 6530 6545 1 1.0950e+02 4.1840e+02 ; HA - CA
- C
6532 6530 6545 1 1.1110e+02 5.2718e+02 ; CB - CA
- C
6532 6534 6535 1 1.0950e+02 4.1840e+02 ; CB - CG2
- HG21
6532 6534 6536 1 1.0950e+02 4.1840e+02 ; CB - CG2
- HG22
6532 6534 6537 1 1.0950e+02 4.1840e+02 ; CB - CG2
- HG23
6532 6538 6539 1 1.0950e+02 4.1840e+02 ; CB - CG1
- HG12
6532 6538 6540 1 1.0950e+02 4.1840e+02 ; CB - CG1
- HG13
6532 6538 6541 1 1.0950e+02 3.3472e+02 ; CB - CG1
- CD1
6533 6532 6534 1 1.0950e+02 4.1840e+02 ; HB - CB
- CG2

6533 6532 6538 1 1.0950e+02 4.1840e+02 ; HB - CB
- CG1
6534 6532 6538 1 1.0950e+02 3.3472e+02 ; CG2 - CB
- CG1
6535 6534 6536 1 1.0950e+02 2.9288e+02 ; HG21 - CG2
- HG22
6535 6534 6537 1 1.0950e+02 2.9288e+02 ; HG21 - CG2
- HG23
6536 6534 6537 1 1.0950e+02 2.9288e+02 ; HG22 - CG2
- HG23
6538 6541 6542 1 1.0950e+02 4.1840e+02 ; CG1 - CD1
- HD11
6538 6541 6543 1 1.0950e+02 4.1840e+02 ; CG1 - CD1
- HD12
6538 6541 6544 1 1.0950e+02 4.1840e+02 ; CG1 - CD1
- HD13
6539 6538 6540 1 1.0950e+02 2.9288e+02 ; HG12 - CG1
- HG13
6539 6538 6541 1 1.0950e+02 4.1840e+02 ; HG12 - CG1
- CD1
6540 6538 6541 1 1.0950e+02 4.1840e+02 ; HG13 - CG1
- CD1
6542 6541 6543 1 1.0950e+02 2.9288e+02 ; HD11 - CD1
- HD12
6542 6541 6544 1 1.0950e+02 2.9288e+02 ; HD11 - CD1
- HD13
6543 6541 6544 1 1.0950e+02 2.9288e+02 ; HD12 - CD1
- HD13
6545 6547 6548 1 1.2000e+02 4.1840e+02 ; C - N
- H
6545 6547 6549 1 1.2190e+02 4.1840e+02 ; C - N
- CA
6546 6545 6547 1 1.2290e+02 6.6944e+02 ; O - C
- N
6547 6549 6550 1 1.0950e+02 4.1840e+02 ; N - CA
- HA
6547 6549 6551 1 1.0970e+02 6.6944e+02 ; N - CA
- CB
6547 6549 6556 1 1.1010e+02 5.2718e+02 ; N - CA
- C
6548 6547 6549 1 1.1804e+02 4.1840e+02 ; H - N
- CA
6549 6551 6552 1 1.0950e+02 4.1840e+02 ; CA - CB
- HB2
6549 6551 6553 1 1.0950e+02 4.1840e+02 ; CA - CB
- HB3
6549 6551 6554 1 1.0950e+02 4.1840e+02 ; CA - CB
- OG
6549 6556 6557 1 1.2040e+02 6.6944e+02 ; CA - C
- O
6549 6556 6558 1 1.1660e+02 5.8576e+02 ; CA - C
- N

6550 6549 6551 1 1.0950e+02 4.1840e+02 ; HA - CA
- CB
6550 6549 6556 1 1.0950e+02 4.1840e+02 ; HA - CA
- C
6551 6549 6556 1 1.1110e+02 5.2718e+02 ; CB - CA
- C
6551 6554 6555 1 1.0850e+02 4.6024e+02 ; CB - OG
- HG
6552 6551 6553 1 1.0950e+02 2.9288e+02 ; HB2 - CB
- HB3
6552 6551 6554 1 1.0950e+02 4.1840e+02 ; HB2 - CB
- OG
6553 6551 6554 1 1.0950e+02 4.1840e+02 ; HB3 - CB
- OG
6556 6558 6559 1 1.2000e+02 4.1840e+02 ; C - N
- H
6556 6558 6560 1 1.2190e+02 4.1840e+02 ; C - N
- CA
6557 6556 6558 1 1.2290e+02 6.6944e+02 ; O - C
- N
6558 6560 6561 1 1.0950e+02 4.1840e+02 ; N - CA
- HA
6558 6560 6562 1 1.0970e+02 6.6944e+02 ; N - CA
- CB
6558 6560 6572 1 1.1010e+02 5.2718e+02 ; N - CA
- C
6559 6558 6560 1 1.1804e+02 4.1840e+02 ; H - N
- CA
6560 6562 6563 1 1.0950e+02 4.1840e+02 ; CA - CB
- HB
6560 6562 6564 1 1.0950e+02 3.3472e+02 ; CA - CB
- CG1
6560 6562 6568 1 1.0950e+02 3.3472e+02 ; CA - CB
- CG2
6560 6572 6573 1 1.2040e+02 6.6944e+02 ; CA - C
- O
6560 6572 6574 1 1.1660e+02 5.8576e+02 ; CA - C
- N
6561 6560 6562 1 1.0950e+02 4.1840e+02 ; HA - CA
- CB
6561 6560 6572 1 1.0950e+02 4.1840e+02 ; HA - CA
- C
6562 6560 6572 1 1.1110e+02 5.2718e+02 ; CB - CA
- C
6562 6564 6565 1 1.0950e+02 4.1840e+02 ; CB - CG1
- HG11
6562 6564 6566 1 1.0950e+02 4.1840e+02 ; CB - CG1
- HG12
6562 6564 6567 1 1.0950e+02 4.1840e+02 ; CB - CG1
- HG13
6562 6568 6569 1 1.0950e+02 4.1840e+02 ; CB - CG2
- HG21

6562 6568 6570 1 1.0950e+02 4.1840e+02 ; CB - CG2
- HG22
6562 6568 6571 1 1.0950e+02 4.1840e+02 ; CB - CG2
- HG23
6563 6562 6564 1 1.0950e+02 4.1840e+02 ; HB - CB
- CG1
6563 6562 6568 1 1.0950e+02 4.1840e+02 ; HB - CB
- CG2
6564 6562 6568 1 1.0950e+02 3.3472e+02 ; CG1 - CB
- CG2
6565 6564 6566 1 1.0950e+02 2.9288e+02 ; HG11 - CG1
- HG12
6565 6564 6567 1 1.0950e+02 2.9288e+02 ; HG11 - CG1
- HG13
6566 6564 6567 1 1.0950e+02 2.9288e+02 ; HG12 - CG1
- HG13
6569 6568 6570 1 1.0950e+02 2.9288e+02 ; HG21 - CG2
- HG22
6569 6568 6571 1 1.0950e+02 2.9288e+02 ; HG21 - CG2
- HG23
6570 6568 6571 1 1.0950e+02 2.9288e+02 ; HG22 - CG2
- HG23
6572 6574 6575 1 1.2000e+02 4.1840e+02 ; C - N
- H
6572 6574 6576 1 1.2190e+02 4.1840e+02 ; C - N
- CA
6573 6572 6574 1 1.2290e+02 6.6944e+02 ; O - C
- N
6574 6576 6577 1 1.0950e+02 4.1840e+02 ; N - CA
- HA
6574 6576 6578 1 1.0970e+02 6.6944e+02 ; N - CA
- CB
6574 6576 6587 1 1.1010e+02 5.2718e+02 ; N - CA
- C
6575 6574 6576 1 1.1804e+02 4.1840e+02 ; H - N
- CA
6576 6578 6579 1 1.0950e+02 4.1840e+02 ; CA - CB
- HB2
6576 6578 6580 1 1.0950e+02 4.1840e+02 ; CA - CB
- HB3
6576 6578 6581 1 1.0950e+02 3.3472e+02 ; CA - CB
- CG
6576 6587 6588 1 1.2040e+02 6.6944e+02 ; CA - C
- O
6576 6587 6589 1 1.1660e+02 5.8576e+02 ; CA - C
- N
6577 6576 6578 1 1.0950e+02 4.1840e+02 ; HA - CA
- CB
6577 6576 6587 1 1.0950e+02 4.1840e+02 ; HA - CA
- C
6578 6576 6587 1 1.1110e+02 5.2718e+02 ; CB - CA
- C

6578 6581 6582 1 1.0950e+02 4.1840e+02 ; CB - CG
- HG2
6578 6581 6583 1 1.0950e+02 4.1840e+02 ; CB - CG
- HG3
6578 6581 6584 1 1.1110e+02 5.2718e+02 ; CB - CG
- CD
6579 6578 6580 1 1.0950e+02 2.9288e+02 ; HB2 - CB
- HB3
6579 6578 6581 1 1.0950e+02 4.1840e+02 ; HB2 - CB
- CG
6580 6578 6581 1 1.0950e+02 4.1840e+02 ; HB3 - CB
- CG
6581 6584 6585 1 1.1700e+02 5.8576e+02 ; CG - CD
- OE1
6581 6584 6586 1 1.1700e+02 5.8576e+02 ; CG - CD
- OE2
6582 6581 6583 1 1.0950e+02 2.9288e+02 ; HG2 - CG
- HG3
6582 6581 6584 1 1.0950e+02 4.1840e+02 ; HG2 - CG
- CD
6583 6581 6584 1 1.0950e+02 4.1840e+02 ; HG3 - CG
- CD
6585 6584 6586 1 1.2600e+02 6.6944e+02 ; OE1 - CD
- OE2
6587 6589 6590 1 1.2000e+02 4.1840e+02 ; C - N
- H
6587 6589 6591 1 1.2190e+02 4.1840e+02 ; C - N
- CA
6588 6587 6589 1 1.2290e+02 6.6944e+02 ; O - C
- N
6589 6591 6592 1 1.0950e+02 4.1840e+02 ; N - CA
- HA
6589 6591 6593 1 1.0970e+02 6.6944e+02 ; N - CA
- CB
6589 6591 6603 1 1.1010e+02 5.2718e+02 ; N - CA
- C
6590 6589 6591 1 1.1804e+02 4.1840e+02 ; H - N
- CA
6591 6593 6594 1 1.0950e+02 4.1840e+02 ; CA - CB
- HB2
6591 6593 6595 1 1.0950e+02 4.1840e+02 ; CA - CB
- HB3
6591 6593 6596 1 1.0950e+02 3.3472e+02 ; CA - CB
- CG
6591 6603 6604 1 1.2040e+02 6.6944e+02 ; CA - C
- O
6591 6603 6605 1 1.1660e+02 5.8576e+02 ; CA - C
- N
6592 6591 6593 1 1.0950e+02 4.1840e+02 ; HA - CA
- CB
6592 6591 6603 1 1.0950e+02 4.1840e+02 ; HA - CA
- C

```
  6593   6591   6603      1    1.1110e+02    5.2718e+02 ;     CB - CA
- C
  6593   6596   6597      1    1.0950e+02    4.1840e+02 ;     CB - CG
- HG2
  6593   6596   6598      1    1.0950e+02    4.1840e+02 ;     CB - CG
- HG3
  6593   6596   6599      1    1.1110e+02    5.2718e+02 ;     CB - CG
- CD
  6594   6593   6595      1    1.0950e+02    2.9288e+02 ;    HB2 - CB
- HB3
  6594   6593   6596      1    1.0950e+02    4.1840e+02 ;    HB2 - CB
- CG
  6595   6593   6596      1    1.0950e+02    4.1840e+02 ;    HB3 - CB
- CG
  6596   6599   6600      1    1.2040e+02    6.6944e+02 ;     CG - CD
- OE1
  6596   6599   6601      1    1.1000e+02    6.6944e+02 ;     CG - CD
- OE2
  6597   6596   6598      1    1.0950e+02    2.9288e+02 ;    HG2 - CG
- HG3
  6597   6596   6599      1    1.0950e+02    4.1840e+02 ;    HG2 - CG
- CD
  6598   6596   6599      1    1.0950e+02    4.1840e+02 ;    HG3 - CG
- CD
  6599   6601   6602      1    1.1300e+02    4.1840e+02 ;     CD - OE2
- HE2
  6600   6599   6601      1    1.2000e+02    6.6944e+02 ;    OE1 - CD
- OE2
  6603   6605   6606      1    1.2000e+02    4.1840e+02 ;      C - N
- H
  6603   6605   6607      1    1.2190e+02    4.1840e+02 ;      C - N
- CA
  6604   6603   6605      1    1.2290e+02    6.6944e+02 ;      O - C
- N
  6605   6607   6608      1    1.0950e+02    4.1840e+02 ;      N - CA
- HA
  6605   6607   6609      1    1.0970e+02    6.6944e+02 ;      N - CA
- CB
  6605   6607   6622      1    1.1010e+02    5.2718e+02 ;      N - CA
- C
  6606   6605   6607      1    1.1804e+02    4.1840e+02 ;      H - N
- CA
  6607   6609   6610      1    1.0950e+02    4.1840e+02 ;     CA - CB
- HB2
  6607   6609   6611      1    1.0950e+02    4.1840e+02 ;     CA - CB
- HB3
  6607   6609   6612      1    1.0950e+02    3.3472e+02 ;     CA - CB
- CG
  6607   6622   6623      1    1.2040e+02    6.6944e+02 ;     CA - C
- O
  6607   6622   6624      1    1.1660e+02    5.8576e+02 ;     CA - C
- N
```

```
  6608   6607   6609      1    1.0950e+02    4.1840e+02 ;     HA - CA
- CB
  6608   6607   6622      1    1.0950e+02    4.1840e+02 ;     HA - CA
- C
  6609   6607   6622      1    1.1110e+02    5.2718e+02 ;     CB - CA
- C
  6609   6612   6613      1    1.0950e+02    4.1840e+02 ;     CB - CG
- HG
  6609   6612   6614      1    1.0950e+02    3.3472e+02 ;     CB - CG
- CD1
  6609   6612   6618      1    1.0950e+02    3.3472e+02 ;     CB - CG
- CD2
  6610   6609   6611      1    1.0950e+02    2.9288e+02 ;    HB2 - CB
- HB3
  6610   6609   6612      1    1.0950e+02    4.1840e+02 ;    HB2 - CB
- CG
  6611   6609   6612      1    1.0950e+02    4.1840e+02 ;    HB3 - CB
- CG
  6612   6614   6615      1    1.0950e+02    4.1840e+02 ;     CG - CD1
- HD11
  6612   6614   6616      1    1.0950e+02    4.1840e+02 ;     CG - CD1
- HD12
  6612   6614   6617      1    1.0950e+02    4.1840e+02 ;     CG - CD1
- HD13
  6612   6618   6619      1    1.0950e+02    4.1840e+02 ;     CG - CD2
- HD21
  6612   6618   6620      1    1.0950e+02    4.1840e+02 ;     CG - CD2
- HD22
  6612   6618   6621      1    1.0950e+02    4.1840e+02 ;     CG - CD2
- HD23
  6613   6612   6614      1    1.0950e+02    4.1840e+02 ;     HG - CG
- CD1
  6613   6612   6618      1    1.0950e+02    4.1840e+02 ;     HG - CG
- CD2
  6614   6612   6618      1    1.0950e+02    3.3472e+02 ;    CD1 - CG
- CD2
  6615   6614   6616      1    1.0950e+02    2.9288e+02 ;   HD11 - CD1
- HD12
  6615   6614   6617      1    1.0950e+02    2.9288e+02 ;   HD11 - CD1
- HD13
  6616   6614   6617      1    1.0950e+02    2.9288e+02 ;   HD12 - CD1
- HD13
  6619   6618   6620      1    1.0950e+02    2.9288e+02 ;   HD21 - CD2
- HD22
  6619   6618   6621      1    1.0950e+02    2.9288e+02 ;   HD21 - CD2
- HD23
  6620   6618   6621      1    1.0950e+02    2.9288e+02 ;   HD22 - CD2
- HD23
  6622   6624   6625      1    1.2000e+02    4.1840e+02 ;      C - N
- H
  6622   6624   6626      1    1.2190e+02    4.1840e+02 ;      C - N
- CA
```

6623 6622 6624 1 1.2290e+02 6.6944e+02 ; O - C
- N
6624 6626 6627 1 1.0950e+02 4.1840e+02 ; N - CA
- HA
6624 6626 6628 1 1.0970e+02 6.6944e+02 ; N - CA
- CB
6624 6626 6644 1 1.1010e+02 5.2718e+02 ; N - CA
- C
6625 6624 6626 1 1.1804e+02 4.1840e+02 ; H - N
- CA
6626 6628 6629 1 1.0950e+02 4.1840e+02 ; CA - CB
- HB2
6626 6628 6630 1 1.0950e+02 4.1840e+02 ; CA - CB
- HB3
6626 6628 6631 1 1.0950e+02 3.3472e+02 ; CA - CB
- CG
6626 6644 6645 1 1.2040e+02 6.6944e+02 ; CA - C
- O
6626 6644 6646 1 1.1660e+02 5.8576e+02 ; CA - C
- N
6627 6626 6628 1 1.0950e+02 4.1840e+02 ; HA - CA
- CB
6627 6626 6644 1 1.0950e+02 4.1840e+02 ; HA - CA
- C
6628 6626 6644 1 1.1110e+02 5.2718e+02 ; CB - CA
- C
6628 6631 6632 1 1.0950e+02 4.1840e+02 ; CB - CG
- HG2
6628 6631 6633 1 1.0950e+02 4.1840e+02 ; CB - CG
- HG3
6628 6631 6634 1 1.0950e+02 3.3472e+02 ; CB - CG
- CD
6629 6628 6630 1 1.0950e+02 2.9288e+02 ; HB2 - CB
- HB3
6629 6628 6631 1 1.0950e+02 4.1840e+02 ; HB2 - CB
- CG
6630 6628 6631 1 1.0950e+02 4.1840e+02 ; HB3 - CB
- CG
6631 6634 6635 1 1.0950e+02 4.1840e+02 ; CG - CD
- HD2
6631 6634 6636 1 1.0950e+02 4.1840e+02 ; CG - CD
- HD3
6631 6634 6637 1 1.0950e+02 3.3472e+02 ; CG - CD
- CE
6632 6631 6633 1 1.0950e+02 2.9288e+02 ; HG2 - CG
- HG3
6632 6631 6634 1 1.0950e+02 4.1840e+02 ; HG2 - CG
- CD
6633 6631 6634 1 1.0950e+02 4.1840e+02 ; HG3 - CG
- CD
6634 6637 6638 1 1.0950e+02 4.1840e+02 ; CD - CE
- HE2

```
  6634   6637   6639       1    1.0950e+02    4.1840e+02 ;     CD - CE
- HE3
  6634   6637   6640       1    1.1120e+02    6.6944e+02 ;     CD - CE
- NZ
  6635   6634   6636       1    1.0950e+02    2.9288e+02 ;    HD2 - CD
- HD3
  6635   6634   6637       1    1.0950e+02    4.1840e+02 ;    HD2 - CD
- CE
  6636   6634   6637       1    1.0950e+02    4.1840e+02 ;    HD3 - CD
- CE
  6637   6640   6641       1    1.0950e+02    4.1840e+02 ;     CE - NZ
- HZ1
  6637   6640   6642       1    1.0950e+02    4.1840e+02 ;     CE - NZ
- HZ2
  6637   6640   6643       1    1.0950e+02    4.1840e+02 ;     CE - NZ
- HZ3
  6638   6637   6639       1    1.0950e+02    2.9288e+02 ;    HE2 - CE
- HE3
  6638   6637   6640       1    1.0950e+02    4.1840e+02 ;    HE2 - CE
- NZ
  6639   6637   6640       1    1.0950e+02    4.1840e+02 ;    HE3 - CE
- NZ
  6641   6640   6642       1    1.0950e+02    2.9288e+02 ;    HZ1 - NZ
- HZ2
  6641   6640   6643       1    1.0950e+02    2.9288e+02 ;    HZ1 - NZ
- HZ3
  6642   6640   6643       1    1.0950e+02    2.9288e+02 ;    HZ2 - NZ
- HZ3
  6644   6646   6647       1    1.2000e+02    4.1840e+02 ;      C - N
- H
  6644   6646   6648       1    1.2190e+02    4.1840e+02 ;      C - N
- CA
  6645   6644   6646       1    1.2290e+02    6.6944e+02 ;      O - C
- N
  6646   6648   6649       1    1.0950e+02    4.1840e+02 ;      N - CA
- HA
  6646   6648   6650       1    1.0970e+02    6.6944e+02 ;      N - CA
- CB
  6646   6648   6668       1    1.1010e+02    5.2718e+02 ;      N - CA
- C
  6647   6646   6648       1    1.1804e+02    4.1840e+02 ;      H - N
- CA
  6648   6650   6651       1    1.0950e+02    4.1840e+02 ;     CA - CB
- HB2
  6648   6650   6652       1    1.0950e+02    4.1840e+02 ;     CA - CB
- HB3
  6648   6650   6653       1    1.0950e+02    3.3472e+02 ;     CA - CB
- CG
  6648   6668   6669       1    1.2040e+02    6.6944e+02 ;     CA - C
- O
  6648   6668   6670       1    1.1660e+02    5.8576e+02 ;     CA - C
- N
```

```
  6649   6648   6650      1    1.0950e+02    4.1840e+02 ;     HA - CA
- CB
  6649   6648   6668      1    1.0950e+02    4.1840e+02 ;     HA - CA
- C
  6650   6648   6668      1    1.1110e+02    5.2718e+02 ;     CB - CA
- C
  6650   6653   6654      1    1.0950e+02    4.1840e+02 ;     CB - CG
- HG2
  6650   6653   6655      1    1.0950e+02    4.1840e+02 ;     CB - CG
- HG3
  6650   6653   6656      1    1.0950e+02    3.3472e+02 ;     CB - CG
- CD
  6651   6650   6652      1    1.0950e+02    2.9288e+02 ;    HB2 - CB
- HB3
  6651   6650   6653      1    1.0950e+02    4.1840e+02 ;    HB2 - CB
- CG
  6652   6650   6653      1    1.0950e+02    4.1840e+02 ;    HB3 - CB
- CG
  6653   6656   6657      1    1.0950e+02    4.1840e+02 ;     CG - CD
- HD2
  6653   6656   6658      1    1.0950e+02    4.1840e+02 ;     CG - CD
- HD3
  6653   6656   6659      1    1.1120e+02    6.6944e+02 ;     CG - CD
- NE
  6654   6653   6655      1    1.0950e+02    2.9288e+02 ;    HG2 - CG
- HG3
  6654   6653   6656      1    1.0950e+02    4.1840e+02 ;    HG2 - CG
- CD
  6655   6653   6656      1    1.0950e+02    4.1840e+02 ;    HG3 - CG
- CD
  6656   6659   6660      1    1.1840e+02    4.1840e+02 ;     CD - NE
- HE
  6656   6659   6661      1    1.2320e+02    4.1840e+02 ;     CD - NE
- CZ
  6657   6656   6658      1    1.0950e+02    2.9288e+02 ;    HD2 - CD
- HD3
  6657   6656   6659      1    1.0950e+02    4.1840e+02 ;    HD2 - CD
- NE
  6658   6656   6659      1    1.0950e+02    4.1840e+02 ;    HD3 - CD
- NE
  6659   6661   6662      1    1.2000e+02    5.8576e+02 ;     NE - CZ
- NH1
  6659   6661   6665      1    1.2000e+02    5.8576e+02 ;     NE - CZ
- NH2
  6660   6659   6661      1    1.2000e+02    4.1840e+02 ;     HE - NE
- CZ
  6661   6662   6663      1    1.2000e+02    4.1840e+02 ;     CZ - NH1
- HH11
  6661   6662   6664      1    1.2000e+02    4.1840e+02 ;     CZ - NH1
- HH12
  6661   6665   6666      1    1.2000e+02    4.1840e+02 ;     CZ - NH2
- HH21
```

6661 6665 6667 1 1.2000e+02 4.1840e+02 ; CZ - NH2
- HH22
6662 6661 6665 1 1.2000e+02 5.8576e+02 ; NH1 - CZ
- NH2
6663 6662 6664 1 1.2000e+02 2.9288e+02 ; HH11 - NH1
- HH12
6666 6665 6667 1 1.2000e+02 2.9288e+02 ; HH21 - NH2
- HH22
6668 6670 6671 1 1.2000e+02 4.1840e+02 ; C - N
- H
6668 6670 6672 1 1.2190e+02 4.1840e+02 ; C - N
- CA
6669 6668 6670 1 1.2290e+02 6.6944e+02 ; O - C
- N
6670 6672 6673 1 1.0950e+02 4.1840e+02 ; N - CA
- HA
6670 6672 6674 1 1.0970e+02 6.6944e+02 ; N - CA
- CB
6670 6672 6687 1 1.1010e+02 5.2718e+02 ; N - CA
- C
6671 6670 6672 1 1.1804e+02 4.1840e+02 ; H - N
- CA
6672 6674 6675 1 1.0950e+02 4.1840e+02 ; CA - CB
- HB
6672 6674 6676 1 1.0950e+02 3.3472e+02 ; CA - CB
- CG2
6672 6674 6680 1 1.0950e+02 3.3472e+02 ; CA - CB
- CG1
6672 6687 6688 1 1.2040e+02 6.6944e+02 ; CA - C
- O
6672 6687 6689 1 1.1660e+02 5.8576e+02 ; CA - C
- N
6673 6672 6674 1 1.0950e+02 4.1840e+02 ; HA - CA
- CB
6673 6672 6687 1 1.0950e+02 4.1840e+02 ; HA - CA
- C
6674 6672 6687 1 1.1110e+02 5.2718e+02 ; CB - CA
- C
6674 6676 6677 1 1.0950e+02 4.1840e+02 ; CB - CG2
- HG21
6674 6676 6678 1 1.0950e+02 4.1840e+02 ; CB - CG2
- HG22
6674 6676 6679 1 1.0950e+02 4.1840e+02 ; CB - CG2
- HG23
6674 6680 6681 1 1.0950e+02 4.1840e+02 ; CB - CG1
- HG12
6674 6680 6682 1 1.0950e+02 4.1840e+02 ; CB - CG1
- HG13
6674 6680 6683 1 1.0950e+02 3.3472e+02 ; CB - CG1
- CD1
6675 6674 6676 1 1.0950e+02 4.1840e+02 ; HB - CB
- CG2

```
  6675   6674   6680      1    1.0950e+02    4.1840e+02 ;     HB - CB
- CG1
  6676   6674   6680      1    1.0950e+02    3.3472e+02 ;    CG2 - CB
- CG1
  6677   6676   6678      1    1.0950e+02    2.9288e+02 ;   HG21 - CG2
- HG22
  6677   6676   6679      1    1.0950e+02    2.9288e+02 ;   HG21 - CG2
- HG23
  6678   6676   6679      1    1.0950e+02    2.9288e+02 ;   HG22 - CG2
- HG23
  6680   6683   6684      1    1.0950e+02    4.1840e+02 ;    CG1 - CD1
- HD11
  6680   6683   6685      1    1.0950e+02    4.1840e+02 ;    CG1 - CD1
- HD12
  6680   6683   6686      1    1.0950e+02    4.1840e+02 ;    CG1 - CD1
- HD13
  6681   6680   6682      1    1.0950e+02    2.9288e+02 ;   HG12 - CG1
- HG13
  6681   6680   6683      1    1.0950e+02    4.1840e+02 ;   HG12 - CG1
- CD1
  6682   6680   6683      1    1.0950e+02    4.1840e+02 ;   HG13 - CG1
- CD1
  6684   6683   6685      1    1.0950e+02    2.9288e+02 ;   HD11 - CD1
- HD12
  6684   6683   6686      1    1.0950e+02    2.9288e+02 ;   HD11 - CD1
- HD13
  6685   6683   6686      1    1.0950e+02    2.9288e+02 ;   HD12 - CD1
- HD13
  6687   6689   6690      1    1.2000e+02    4.1840e+02 ;      C - N
- H
  6687   6689   6691      1    1.2190e+02    4.1840e+02 ;      C - N
- CA
  6688   6687   6689      1    1.2290e+02    6.6944e+02 ;      O - C
- N
  6689   6691   6692      1    1.0950e+02    4.1840e+02 ;      N - CA
- HA
  6689   6691   6693      1    1.0970e+02    6.6944e+02 ;      N - CA
- CB
  6689   6691   6707      1    1.1010e+02    5.2718e+02 ;      N - CA
- C
  6690   6689   6691      1    1.1804e+02    4.1840e+02 ;      H - N
- CA
  6691   6693   6694      1    1.0950e+02    4.1840e+02 ;     CA - CB
- HB2
  6691   6693   6695      1    1.0950e+02    4.1840e+02 ;     CA - CB
- HB3
  6691   6693   6696      1    1.1400e+02    5.2718e+02 ;     CA - CB
- CG
  6691   6707   6708      1    1.2040e+02    6.6944e+02 ;     CA - C
- O
  6691   6707   6709      1    1.1660e+02    5.8576e+02 ;     CA - C
- N
```

6692 6691 6693 1 1.0950e+02 4.1840e+02 ; HA - CA
- CB
6692 6691 6707 1 1.0950e+02 4.1840e+02 ; HA - CA
- C
6693 6691 6707 1 1.1110e+02 5.2718e+02 ; CB - CA
- C
6693 6696 6697 1 1.2000e+02 5.8576e+02 ; CB - CG
- CD1
6693 6696 6705 1 1.2000e+02 5.8576e+02 ; CB - CG
- CD2
6694 6693 6695 1 1.0950e+02 2.9288e+02 ; HB2 - CB
- HB3
6694 6693 6696 1 1.0950e+02 4.1840e+02 ; HB2 - CB
- CG
6695 6693 6696 1 1.0950e+02 4.1840e+02 ; HB3 - CB
- CG
6696 6697 6698 1 1.2000e+02 4.1840e+02 ; CG - CD1
- HD1
6696 6697 6699 1 1.2000e+02 5.2718e+02 ; CG - CD1
- CE1
6696 6705 6703 1 1.2000e+02 5.2718e+02 ; CG - CD2
- CE2
6696 6705 6706 1 1.2000e+02 4.1840e+02 ; CG - CD2
- HD2
6697 6696 6705 1 1.2000e+02 5.2718e+02 ; CD1 - CG
- CD2
6697 6699 6700 1 1.2000e+02 4.1840e+02 ; CD1 - CE1
- HE1
6697 6699 6701 1 1.2000e+02 5.2718e+02 ; CD1 - CE1
- CZ
6698 6697 6699 1 1.2000e+02 4.1840e+02 ; HD1 - CD1
- CE1
6699 6701 6702 1 1.2000e+02 4.1840e+02 ; CE1 - CZ
- HZ
6699 6701 6703 1 1.2000e+02 5.2718e+02 ; CE1 - CZ
- CE2
6700 6699 6701 1 1.2000e+02 4.1840e+02 ; HE1 - CE1
- CZ
6701 6703 6704 1 1.2000e+02 4.1840e+02 ; CZ - CE2
- HE2
6701 6703 6705 1 1.2000e+02 5.2718e+02 ; CZ - CE2
- CD2
6702 6701 6703 1 1.2000e+02 4.1840e+02 ; HZ - CZ
- CE2
6703 6705 6706 1 1.2000e+02 4.1840e+02 ; CE2 - CD2
- HD2
6704 6703 6705 1 1.2000e+02 4.1840e+02 ; HE2 - CE2
- CD2
6707 6709 6710 1 1.2000e+02 4.1840e+02 ; C - N
- H
6707 6709 6711 1 1.2190e+02 4.1840e+02 ; C - N
- CA

6708 6707 6709 1 1.2290e+02 6.6944e+02 ; O - C
- N
6709 6711 6712 1 1.0950e+02 4.1840e+02 ; N - CA
- HA2
6709 6711 6713 1 1.0950e+02 4.1840e+02 ; N - CA
- HA3
6709 6711 6714 1 1.1010e+02 5.2718e+02 ; N - CA
- C
6710 6709 6711 1 1.1804e+02 4.1840e+02 ; H - N
- CA
6711 6714 6715 1 1.2040e+02 6.6944e+02 ; CA - C
- O
6711 6714 6716 1 1.1660e+02 5.8576e+02 ; CA - C
- N
6712 6711 6713 1 1.0950e+02 2.9288e+02 ; HA2 - CA
- HA3
6712 6711 6714 1 1.0950e+02 4.1840e+02 ; HA2 - CA
- C
6713 6711 6714 1 1.0950e+02 4.1840e+02 ; HA3 - CA
- C
6714 6716 6717 1 1.2000e+02 4.1840e+02 ; C - N
- H
6714 6716 6718 1 1.2190e+02 4.1840e+02 ; C - N
- CA
6715 6714 6716 1 1.2290e+02 6.6944e+02 ; O - C
- N
6716 6718 6719 1 1.0950e+02 4.1840e+02 ; N - CA
- HA
6716 6718 6720 1 1.0970e+02 6.6944e+02 ; N - CA
- CB
6716 6718 6738 1 1.1010e+02 5.2718e+02 ; N - CA
- C
6717 6716 6718 1 1.1804e+02 4.1840e+02 ; H - N
- CA
6718 6720 6721 1 1.0950e+02 4.1840e+02 ; CA - CB
- HB2
6718 6720 6722 1 1.0950e+02 4.1840e+02 ; CA - CB
- HB3
6718 6720 6723 1 1.0950e+02 3.3472e+02 ; CA - CB
- CG
6718 6738 6739 1 1.2040e+02 6.6944e+02 ; CA - C
- O
6718 6738 6740 1 1.1660e+02 5.8576e+02 ; CA - C
- N
6719 6718 6720 1 1.0950e+02 4.1840e+02 ; HA - CA
- CB
6719 6718 6738 1 1.0950e+02 4.1840e+02 ; HA - CA
- C
6720 6718 6738 1 1.1110e+02 5.2718e+02 ; CB - CA
- C
6720 6723 6724 1 1.0950e+02 4.1840e+02 ; CB - CG
- HG2

6720 6723 6725 1 1.0950e+02 4.1840e+02 ; CB - CG
- HG3
6720 6723 6726 1 1.0950e+02 3.3472e+02 ; CB - CG
- CD
6721 6720 6722 1 1.0950e+02 2.9288e+02 ; HB2 - CB
- HB3
6721 6720 6723 1 1.0950e+02 4.1840e+02 ; HB2 - CB
- CG
6722 6720 6723 1 1.0950e+02 4.1840e+02 ; HB3 - CB
- CG
6723 6726 6727 1 1.0950e+02 4.1840e+02 ; CG - CD
- HD2
6723 6726 6728 1 1.0950e+02 4.1840e+02 ; CG - CD
- HD3
6723 6726 6729 1 1.1120e+02 6.6944e+02 ; CG - CD
- NE
6724 6723 6725 1 1.0950e+02 2.9288e+02 ; HG2 - CG
- HG3
6724 6723 6726 1 1.0950e+02 4.1840e+02 ; HG2 - CG
- CD
6725 6723 6726 1 1.0950e+02 4.1840e+02 ; HG3 - CG
- CD
6726 6729 6730 1 1.1840e+02 4.1840e+02 ; CD - NE
- HE
6726 6729 6731 1 1.2320e+02 4.1840e+02 ; CD - NE
- CZ
6727 6726 6728 1 1.0950e+02 2.9288e+02 ; HD2 - CD
- HD3
6727 6726 6729 1 1.0950e+02 4.1840e+02 ; HD2 - CD
- NE
6728 6726 6729 1 1.0950e+02 4.1840e+02 ; HD3 - CD
- NE
6729 6731 6732 1 1.2000e+02 5.8576e+02 ; NE - CZ
- NH1
6729 6731 6735 1 1.2000e+02 5.8576e+02 ; NE - CZ
- NH2
6730 6729 6731 1 1.2000e+02 4.1840e+02 ; HE - NE
- CZ
6731 6732 6733 1 1.2000e+02 4.1840e+02 ; CZ - NH1
- HH11
6731 6732 6734 1 1.2000e+02 4.1840e+02 ; CZ - NH1
- HH12
6731 6735 6736 1 1.2000e+02 4.1840e+02 ; CZ - NH2
- HH21
6731 6735 6737 1 1.2000e+02 4.1840e+02 ; CZ - NH2
- HH22
6732 6731 6735 1 1.2000e+02 5.8576e+02 ; NH1 - CZ
- NH2
6733 6732 6734 1 1.2000e+02 2.9288e+02 ; HH11 - NH1
- HH12
6736 6735 6737 1 1.2000e+02 2.9288e+02 ; HH21 - NH2
- HH22

6738 6740 6741 1 1.2000e+02 4.1840e+02 ; C - N
- H
6738 6740 6742 1 1.2190e+02 4.1840e+02 ; C - N
- CA
6739 6738 6740 1 1.2290e+02 6.6944e+02 ; O - C
- N
6740 6742 6743 1 1.0950e+02 4.1840e+02 ; N - CA
- HA
6740 6742 6744 1 1.0970e+02 6.6944e+02 ; N - CA
- CB
6740 6742 6750 1 1.1010e+02 5.2718e+02 ; N - CA
- C
6741 6740 6742 1 1.1804e+02 4.1840e+02 ; H - N
- CA
6742 6744 6745 1 1.0950e+02 4.1840e+02 ; CA - CB
- HB2
6742 6744 6746 1 1.0950e+02 4.1840e+02 ; CA - CB
- HB3
6742 6744 6747 1 1.1110e+02 5.2718e+02 ; CA - CB
- CG
6742 6750 6751 1 1.2040e+02 6.6944e+02 ; CA - C
- O
6742 6750 6752 1 1.1660e+02 5.8576e+02 ; CA - C
- N
6743 6742 6744 1 1.0950e+02 4.1840e+02 ; HA - CA
- CB
6743 6742 6750 1 1.0950e+02 4.1840e+02 ; HA - CA
- C
6744 6742 6750 1 1.1110e+02 5.2718e+02 ; CB - CA
- C
6744 6747 6748 1 1.1700e+02 5.8576e+02 ; CB - CG
- OD1
6744 6747 6749 1 1.1700e+02 5.8576e+02 ; CB - CG
- OD2
6745 6744 6746 1 1.0950e+02 2.9288e+02 ; HB2 - CB
- HB3
6745 6744 6747 1 1.0950e+02 4.1840e+02 ; HB2 - CB
- CG
6746 6744 6747 1 1.0950e+02 4.1840e+02 ; HB3 - CB
- CG
6748 6747 6749 1 1.2600e+02 6.6944e+02 ; OD1 - CG
- OD2
6750 6752 6753 1 1.2000e+02 4.1840e+02 ; C - N
- H
6750 6752 6754 1 1.2190e+02 4.1840e+02 ; C - N
- CA
6751 6750 6752 1 1.2290e+02 6.6944e+02 ; O - C
- N
6752 6754 6755 1 1.0950e+02 4.1840e+02 ; N - CA
- HA
6752 6754 6756 1 1.0970e+02 6.6944e+02 ; N - CA
- CB

6752 6754 6762 1 1.1010e+02 5.2718e+02 ; N - CA
- C
6753 6752 6754 1 1.1804e+02 4.1840e+02 ; H - N
- CA
6754 6756 6757 1 1.0950e+02 4.1840e+02 ; CA - CB
- HB2
6754 6756 6758 1 1.0950e+02 4.1840e+02 ; CA - CB
- HB3
6754 6756 6759 1 1.1110e+02 5.2718e+02 ; CA - CB
- CG
6754 6762 6763 1 1.2040e+02 6.6944e+02 ; CA - C
- O
6754 6762 6764 1 1.1660e+02 5.8576e+02 ; CA - C
- N
6755 6754 6756 1 1.0950e+02 4.1840e+02 ; HA - CA
- CB
6755 6754 6762 1 1.0950e+02 4.1840e+02 ; HA - CA
- C
6756 6754 6762 1 1.1110e+02 5.2718e+02 ; CB - CA
- C
6756 6759 6760 1 1.1700e+02 5.8576e+02 ; CB - CG
- OD1
6756 6759 6761 1 1.1700e+02 5.8576e+02 ; CB - CG
- OD2
6757 6756 6758 1 1.0950e+02 2.9288e+02 ; HB2 - CB
- HB3
6757 6756 6759 1 1.0950e+02 4.1840e+02 ; HB2 - CB
- CG
6758 6756 6759 1 1.0950e+02 4.1840e+02 ; HB3 - CB
- CG
6760 6759 6761 1 1.2600e+02 6.6944e+02 ; OD1 - CG
- OD2
6762 6764 6765 1 1.2000e+02 4.1840e+02 ; C - N
- H
6762 6764 6766 1 1.2190e+02 4.1840e+02 ; C - N
- CA
6763 6762 6764 1 1.2290e+02 6.6944e+02 ; O - C
- N
6764 6766 6767 1 1.0950e+02 4.1840e+02 ; N - CA
- HA
6764 6766 6768 1 1.0970e+02 6.6944e+02 ; N - CA
- CB
6764 6766 6781 1 1.1010e+02 5.2718e+02 ; N - CA
- C
6765 6764 6766 1 1.1804e+02 4.1840e+02 ; H - N
- CA
6766 6768 6769 1 1.0950e+02 4.1840e+02 ; CA - CB
- HB
6766 6768 6770 1 1.0950e+02 3.3472e+02 ; CA - CB
- CG2
6766 6768 6774 1 1.0950e+02 3.3472e+02 ; CA - CB
- CG1

6766 6781 6782 1 1.2040e+02 6.6944e+02 ; CA - C
- O
6766 6781 6783 1 1.1660e+02 5.8576e+02 ; CA - C
- N
6767 6766 6768 1 1.0950e+02 4.1840e+02 ; HA - CA
- CB
6767 6766 6781 1 1.0950e+02 4.1840e+02 ; HA - CA
- C
6768 6766 6781 1 1.1110e+02 5.2718e+02 ; CB - CA
- C
6768 6770 6771 1 1.0950e+02 4.1840e+02 ; CB - CG2
- HG21
6768 6770 6772 1 1.0950e+02 4.1840e+02 ; CB - CG2
- HG22
6768 6770 6773 1 1.0950e+02 4.1840e+02 ; CB - CG2
- HG23
6768 6774 6775 1 1.0950e+02 4.1840e+02 ; CB - CG1
- HG12
6768 6774 6776 1 1.0950e+02 4.1840e+02 ; CB - CG1
- HG13
6768 6774 6777 1 1.0950e+02 3.3472e+02 ; CB - CG1
- CD1
6769 6768 6770 1 1.0950e+02 4.1840e+02 ; HB - CB
- CG2
6769 6768 6774 1 1.0950e+02 4.1840e+02 ; HB - CB
- CG1
6770 6768 6774 1 1.0950e+02 3.3472e+02 ; CG2 - CB
- CG1
6771 6770 6772 1 1.0950e+02 2.9288e+02 ; HG21 - CG2
- HG22
6771 6770 6773 1 1.0950e+02 2.9288e+02 ; HG21 - CG2
- HG23
6772 6770 6773 1 1.0950e+02 2.9288e+02 ; HG22 - CG2
- HG23
6774 6777 6778 1 1.0950e+02 4.1840e+02 ; CG1 - CD1
- HD11
6774 6777 6779 1 1.0950e+02 4.1840e+02 ; CG1 - CD1
- HD12
6774 6777 6780 1 1.0950e+02 4.1840e+02 ; CG1 - CD1
- HD13
6775 6774 6776 1 1.0950e+02 2.9288e+02 ; HG12 - CG1
- HG13
6775 6774 6777 1 1.0950e+02 4.1840e+02 ; HG12 - CG1
- CD1
6776 6774 6777 1 1.0950e+02 4.1840e+02 ; HG13 - CG1
- CD1
6778 6777 6779 1 1.0950e+02 2.9288e+02 ; HD11 - CD1
- HD12
6778 6777 6780 1 1.0950e+02 2.9288e+02 ; HD11 - CD1
- HD13
6779 6777 6780 1 1.0950e+02 2.9288e+02 ; HD12 - CD1
- HD13

```
  6781   6783   6784       1    1.2000e+02    4.1840e+02 ;      C - N 
- H
  6781   6783   6785       1    1.2190e+02    4.1840e+02 ;      C - N 
- CA
  6782   6781   6783       1    1.2290e+02    6.6944e+02 ;      O - C 
- N
  6783   6785   6786       1    1.0950e+02    4.1840e+02 ;      N - CA
- HA
  6783   6785   6787       1    1.0970e+02    6.6944e+02 ;      N - CA
- CB
  6783   6785   6796       1    1.1010e+02    5.2718e+02 ;      N - CA
- C
  6784   6783   6785       1    1.1804e+02    4.1840e+02 ;      H - N 
- CA
  6785   6787   6788       1    1.0950e+02    4.1840e+02 ;     CA - CB
- HB2
  6785   6787   6789       1    1.0950e+02    4.1840e+02 ;     CA - CB
- HB3
  6785   6787   6790       1    1.0950e+02    3.3472e+02 ;     CA - CB
- CG
  6785   6796   6797       1    1.2040e+02    6.6944e+02 ;     CA - C 
- O
  6785   6796   6798       1    1.1660e+02    5.8576e+02 ;     CA - C 
- N
  6786   6785   6787       1    1.0950e+02    4.1840e+02 ;     HA - CA
- CB
  6786   6785   6796       1    1.0950e+02    4.1840e+02 ;     HA - CA
- C
  6787   6785   6796       1    1.1110e+02    5.2718e+02 ;     CB - CA
- C
  6787   6790   6791       1    1.0950e+02    4.1840e+02 ;     CB - CG
- HG2
  6787   6790   6792       1    1.0950e+02    4.1840e+02 ;     CB - CG
- HG3
  6787   6790   6793       1    1.1110e+02    5.2718e+02 ;     CB - CG
- CD
  6788   6787   6789       1    1.0950e+02    2.9288e+02 ;    HB2 - CB
- HB3
  6788   6787   6790       1    1.0950e+02    4.1840e+02 ;    HB2 - CB
- CG
  6789   6787   6790       1    1.0950e+02    4.1840e+02 ;    HB3 - CB
- CG
  6790   6793   6794       1    1.1700e+02    5.8576e+02 ;     CG - CD
- OE1
  6790   6793   6795       1    1.1700e+02    5.8576e+02 ;     CG - CD
- OE2
  6791   6790   6792       1    1.0950e+02    2.9288e+02 ;    HG2 - CG
- HG3
  6791   6790   6793       1    1.0950e+02    4.1840e+02 ;    HG2 - CG
- CD
  6792   6790   6793       1    1.0950e+02    4.1840e+02 ;    HG3 - CG
- CD
```

6794 6793 6795 1 1.2600e+02 6.6944e+02 ; OE1 - CD
- OE2
6796 6798 6799 1 1.2000e+02 4.1840e+02 ; C - N
- H
6796 6798 6800 1 1.2190e+02 4.1840e+02 ; C - N
- CA
6797 6796 6798 1 1.2290e+02 6.6944e+02 ; O - C
- N
6798 6800 6801 1 1.0950e+02 4.1840e+02 ; N - CA
- HA
6798 6800 6802 1 1.0970e+02 6.6944e+02 ; N - CA
- CB
6798 6800 6810 1 1.1010e+02 5.2718e+02 ; N - CA
- C
6799 6798 6800 1 1.1804e+02 4.1840e+02 ; H - N
- CA
6800 6802 6803 1 1.0950e+02 4.1840e+02 ; CA - CB
- HB2
6800 6802 6804 1 1.0950e+02 4.1840e+02 ; CA - CB
- HB3
6800 6802 6805 1 1.1110e+02 5.2718e+02 ; CA - CB
- CG
6800 6810 6811 1 1.2040e+02 6.6944e+02 ; CA - C
- O
6800 6810 6812 1 1.1660e+02 5.8576e+02 ; CA - C
- N
6801 6800 6802 1 1.0950e+02 4.1840e+02 ; HA - CA
- CB
6801 6800 6810 1 1.0950e+02 4.1840e+02 ; HA - CA
- C
6802 6800 6810 1 1.1110e+02 5.2718e+02 ; CB - CA
- C
6802 6805 6806 1 1.2040e+02 6.6944e+02 ; CB - CG
- OD1
6802 6805 6807 1 1.1660e+02 5.8576e+02 ; CB - CG
- ND2
6803 6802 6804 1 1.0950e+02 2.9288e+02 ; HB2 - CB
- HB3
6803 6802 6805 1 1.0950e+02 4.1840e+02 ; HB2 - CB
- CG
6804 6802 6805 1 1.0950e+02 4.1840e+02 ; HB3 - CB
- CG
6805 6807 6808 1 1.2000e+02 4.1840e+02 ; CG - ND2
- HD21
6805 6807 6809 1 1.2000e+02 4.1840e+02 ; CG - ND2
- HD22
6806 6805 6807 1 1.2290e+02 6.6944e+02 ; OD1 - CG
- ND2
6808 6807 6809 1 1.2000e+02 2.9288e+02 ; HD21 - ND2
- HD22
6810 6812 6813 1 1.2190e+02 4.1840e+02 ; C - N
- CD

6810 6812 6822 1 1.2190e+02 4.1840e+02 ; C - N
- CA
6811 6810 6812 1 1.2290e+02 6.6944e+02 ; O - C
- N
6812 6813 6814 1 1.0950e+02 4.1840e+02 ; N - CD
- HD2
6812 6813 6815 1 1.0950e+02 4.1840e+02 ; N - CD
- HD3
6812 6813 6816 1 1.0970e+02 6.6944e+02 ; N - CD
- CG
6812 6822 6819 1 1.0970e+02 6.6944e+02 ; N - CA
- CB
6812 6822 6823 1 1.0950e+02 4.1840e+02 ; N - CA
- HA
6812 6822 6824 1 1.1010e+02 5.2718e+02 ; N - CA
- C
6813 6812 6822 1 1.1800e+02 4.1840e+02 ; CD - N
- CA
6813 6816 6817 1 1.0950e+02 4.1840e+02 ; CD - CG
- HG2
6813 6816 6818 1 1.0950e+02 4.1840e+02 ; CD - CG
- HG3
6813 6816 6819 1 1.0950e+02 3.3472e+02 ; CD - CG
- CB
6814 6813 6815 1 1.0950e+02 2.9288e+02 ; HD2 - CD
- HD3
6814 6813 6816 1 1.0950e+02 4.1840e+02 ; HD2 - CD
- CG
6815 6813 6816 1 1.0950e+02 4.1840e+02 ; HD3 - CD
- CG
6816 6819 6820 1 1.0950e+02 4.1840e+02 ; CG - CB
- HB2
6816 6819 6821 1 1.0950e+02 4.1840e+02 ; CG - CB
- HB3
6816 6819 6822 1 1.0950e+02 3.3472e+02 ; CG - CB
- CA
6817 6816 6818 1 1.0950e+02 2.9288e+02 ; HG2 - CG
- HG3
6817 6816 6819 1 1.0950e+02 4.1840e+02 ; HG2 - CG
- CB
6818 6816 6819 1 1.0950e+02 4.1840e+02 ; HG3 - CG
- CB
6819 6822 6823 1 1.0950e+02 4.1840e+02 ; CB - CA
- HA
6819 6822 6824 1 1.1110e+02 5.2718e+02 ; CB - CA
- C
6820 6819 6821 1 1.0950e+02 2.9288e+02 ; HB2 - CB
- HB3
6820 6819 6822 1 1.0950e+02 4.1840e+02 ; HB2 - CB
- CA
6821 6819 6822 1 1.0950e+02 4.1840e+02 ; HB3 - CB
- CA

6822 6824 6825 1 1.2040e+02 6.6944e+02 ; CA - C
- O
6822 6824 6826 1 1.1660e+02 5.8576e+02 ; CA - C
- N
6823 6822 6824 1 1.0950e+02 4.1840e+02 ; HA - CA
- C
6824 6826 6827 1 1.2000e+02 4.1840e+02 ; C - N
- H
6824 6826 6828 1 1.2190e+02 4.1840e+02 ; C - N
- CA
6825 6824 6826 1 1.2290e+02 6.6944e+02 ; O - C
- N
6826 6828 6829 1 1.0950e+02 4.1840e+02 ; N - CA
- HA
6826 6828 6830 1 1.0970e+02 6.6944e+02 ; N - CA
- CB
6826 6828 6843 1 1.1010e+02 5.2718e+02 ; N - CA
- C
6827 6826 6828 1 1.1804e+02 4.1840e+02 ; H - N
- CA
6828 6830 6831 1 1.0950e+02 4.1840e+02 ; CA - CB
- HB2
6828 6830 6832 1 1.0950e+02 4.1840e+02 ; CA - CB
- HB3
6828 6830 6833 1 1.0950e+02 3.3472e+02 ; CA - CB
- CG
6828 6843 6844 1 1.2040e+02 6.6944e+02 ; CA - C
- O
6828 6843 6845 1 1.1660e+02 5.8576e+02 ; CA - C
- N
6829 6828 6830 1 1.0950e+02 4.1840e+02 ; HA - CA
- CB
6829 6828 6843 1 1.0950e+02 4.1840e+02 ; HA - CA
- C
6830 6828 6843 1 1.1110e+02 5.2718e+02 ; CB - CA
- C
6830 6833 6834 1 1.0950e+02 4.1840e+02 ; CB - CG
- HG
6830 6833 6835 1 1.0950e+02 3.3472e+02 ; CB - CG
- CD1
6830 6833 6839 1 1.0950e+02 3.3472e+02 ; CB - CG
- CD2
6831 6830 6832 1 1.0950e+02 2.9288e+02 ; HB2 - CB
- HB3
6831 6830 6833 1 1.0950e+02 4.1840e+02 ; HB2 - CB
- CG
6832 6830 6833 1 1.0950e+02 4.1840e+02 ; HB3 - CB
- CG
6833 6835 6836 1 1.0950e+02 4.1840e+02 ; CG - CD1
- HD11
6833 6835 6837 1 1.0950e+02 4.1840e+02 ; CG - CD1
- HD12

```
  6833   6835   6838      1    1.0950e+02    4.1840e+02 ;     CG - CD1
- HD13
  6833   6839   6840      1    1.0950e+02    4.1840e+02 ;     CG - CD2
- HD21
  6833   6839   6841      1    1.0950e+02    4.1840e+02 ;     CG - CD2
- HD22
  6833   6839   6842      1    1.0950e+02    4.1840e+02 ;     CG - CD2
- HD23
  6834   6833   6835      1    1.0950e+02    4.1840e+02 ;     HG - CG
- CD1
  6834   6833   6839      1    1.0950e+02    4.1840e+02 ;     HG - CG
- CD2
  6835   6833   6839      1    1.0950e+02    3.3472e+02 ;    CD1 - CG
- CD2
  6836   6835   6837      1    1.0950e+02    2.9288e+02 ;   HD11 - CD1
- HD12
  6836   6835   6838      1    1.0950e+02    2.9288e+02 ;   HD11 - CD1
- HD13
  6837   6835   6838      1    1.0950e+02    2.9288e+02 ;   HD12 - CD1
- HD13
  6840   6839   6841      1    1.0950e+02    2.9288e+02 ;   HD21 - CD2
- HD22
  6840   6839   6842      1    1.0950e+02    2.9288e+02 ;   HD21 - CD2
- HD23
  6841   6839   6842      1    1.0950e+02    2.9288e+02 ;   HD22 - CD2
- HD23
  6843   6845   6846      1    1.2000e+02    4.1840e+02 ;      C - N
- H
  6843   6845   6847      1    1.2190e+02    4.1840e+02 ;      C - N
- CA
  6844   6843   6845      1    1.2290e+02    6.6944e+02 ;      O - C
- N
  6845   6847   6848      1    1.0950e+02    4.1840e+02 ;      N - CA
- HA
  6845   6847   6849      1    1.0970e+02    6.6944e+02 ;      N - CA
- CB
  6845   6847   6862      1    1.1010e+02    5.2718e+02 ;      N - CA
- C
  6846   6845   6847      1    1.1804e+02    4.1840e+02 ;      H - N
- CA
  6847   6849   6850      1    1.0950e+02    4.1840e+02 ;     CA - CB
- HB
  6847   6849   6851      1    1.0950e+02    3.3472e+02 ;     CA - CB
- CG2
  6847   6849   6855      1    1.0950e+02    3.3472e+02 ;     CA - CB
- CG1
  6847   6862   6863      1    1.2040e+02    6.6944e+02 ;     CA - C
- O
  6847   6862   6864      1    1.1660e+02    5.8576e+02 ;     CA - C
- N
  6848   6847   6849      1    1.0950e+02    4.1840e+02 ;     HA - CA
- CB
```

```
  6848   6847   6862      1    1.0950e+02    4.1840e+02 ;     HA - CA
- C
  6849   6847   6862      1    1.1110e+02    5.2718e+02 ;     CB - CA
- C
  6849   6851   6852      1    1.0950e+02    4.1840e+02 ;     CB - CG2
- HG21
  6849   6851   6853      1    1.0950e+02    4.1840e+02 ;     CB - CG2
- HG22
  6849   6851   6854      1    1.0950e+02    4.1840e+02 ;     CB - CG2
- HG23
  6849   6855   6856      1    1.0950e+02    4.1840e+02 ;     CB - CG1
- HG12
  6849   6855   6857      1    1.0950e+02    4.1840e+02 ;     CB - CG1
- HG13
  6849   6855   6858      1    1.0950e+02    3.3472e+02 ;     CB - CG1
- CD1
  6850   6849   6851      1    1.0950e+02    4.1840e+02 ;     HB - CB
- CG2
  6850   6849   6855      1    1.0950e+02    4.1840e+02 ;     HB - CB
- CG1
  6851   6849   6855      1    1.0950e+02    3.3472e+02 ;    CG2 - CB
- CG1
  6852   6851   6853      1    1.0950e+02    2.9288e+02 ;   HG21 - CG2
- HG22
  6852   6851   6854      1    1.0950e+02    2.9288e+02 ;   HG21 - CG2
- HG23
  6853   6851   6854      1    1.0950e+02    2.9288e+02 ;   HG22 - CG2
- HG23
  6855   6858   6859      1    1.0950e+02    4.1840e+02 ;    CG1 - CD1
- HD11
  6855   6858   6860      1    1.0950e+02    4.1840e+02 ;    CG1 - CD1
- HD12
  6855   6858   6861      1    1.0950e+02    4.1840e+02 ;    CG1 - CD1
- HD13
  6856   6855   6857      1    1.0950e+02    2.9288e+02 ;   HG12 - CG1
- HG13
  6856   6855   6858      1    1.0950e+02    4.1840e+02 ;   HG12 - CG1
- CD1
  6857   6855   6858      1    1.0950e+02    4.1840e+02 ;   HG13 - CG1
- CD1
  6859   6858   6860      1    1.0950e+02    2.9288e+02 ;   HD11 - CD1
- HD12
  6859   6858   6861      1    1.0950e+02    2.9288e+02 ;   HD11 - CD1
- HD13
  6860   6858   6861      1    1.0950e+02    2.9288e+02 ;   HD12 - CD1
- HD13
  6862   6864   6865      1    1.2000e+02    4.1840e+02 ;      C - N
- H
  6862   6864   6866      1    1.2190e+02    4.1840e+02 ;      C - N
- CA
  6863   6862   6864      1    1.2290e+02    6.6944e+02 ;      O - C
- N
```

```
  6864   6866   6867      1    1.0950e+02    4.1840e+02 ;      N - CA
- HA
  6864   6866   6868      1    1.0970e+02    6.6944e+02 ;      N - CA
- CB
  6864   6866   6874      1    1.1010e+02    5.2718e+02 ;      N - CA
- C
  6865   6864   6866      1    1.1804e+02    4.1840e+02 ;      H - N
- CA
  6866   6868   6869      1    1.0950e+02    4.1840e+02 ;     CA - CB
- HB2
  6866   6868   6870      1    1.0950e+02    4.1840e+02 ;     CA - CB
- HB3
  6866   6868   6871      1    1.1110e+02    5.2718e+02 ;     CA - CB
- CG
  6866   6874   6875      1    1.2040e+02    6.6944e+02 ;     CA - C
- O
  6866   6874   6876      1    1.1660e+02    5.8576e+02 ;     CA - C
- N
  6867   6866   6868      1    1.0950e+02    4.1840e+02 ;     HA - CA
- CB
  6867   6866   6874      1    1.0950e+02    4.1840e+02 ;     HA - CA
- C
  6868   6866   6874      1    1.1110e+02    5.2718e+02 ;     CB - CA
- C
  6868   6871   6872      1    1.1700e+02    5.8576e+02 ;     CB - CG
- OD1
  6868   6871   6873      1    1.1700e+02    5.8576e+02 ;     CB - CG
- OD2
  6869   6868   6870      1    1.0950e+02    2.9288e+02 ;    HB2 - CB
- HB3
  6869   6868   6871      1    1.0950e+02    4.1840e+02 ;    HB2 - CB
- CG
  6870   6868   6871      1    1.0950e+02    4.1840e+02 ;    HB3 - CB
- CG
  6872   6871   6873      1    1.2600e+02    6.6944e+02 ;    OD1 - CG
- OD2
  6874   6876   6877      1    1.2000e+02    4.1840e+02 ;      C - N
- H
  6874   6876   6878      1    1.2190e+02    4.1840e+02 ;      C - N
- CA
  6875   6874   6876      1    1.2290e+02    6.6944e+02 ;      O - C
- N
  6876   6878   6879      1    1.0950e+02    4.1840e+02 ;      N - CA
- HA
  6876   6878   6880      1    1.0970e+02    6.6944e+02 ;      N - CA
- CB
  6876   6878   6896      1    1.1010e+02    5.2718e+02 ;      N - CA
- C
  6877   6876   6878      1    1.1804e+02    4.1840e+02 ;      H - N
- CA
  6878   6880   6881      1    1.0950e+02    4.1840e+02 ;     CA - CB
- HB2
```

6878 6880 6882 1 1.0950e+02 4.1840e+02 ; CA - CB
- HB3
6878 6880 6883 1 1.0950e+02 3.3472e+02 ; CA - CB
- CG
6878 6896 6897 1 1.2040e+02 6.6944e+02 ; CA - C
- O
6878 6896 6898 1 1.1660e+02 5.8576e+02 ; CA - C
- N
6879 6878 6880 1 1.0950e+02 4.1840e+02 ; HA - CA
- CB
6879 6878 6896 1 1.0950e+02 4.1840e+02 ; HA - CA
- C
6880 6878 6896 1 1.1110e+02 5.2718e+02 ; CB - CA
- C
6880 6883 6884 1 1.0950e+02 4.1840e+02 ; CB - CG
- HG2
6880 6883 6885 1 1.0950e+02 4.1840e+02 ; CB - CG
- HG3
6880 6883 6886 1 1.0950e+02 3.3472e+02 ; CB - CG
- CD
6881 6880 6882 1 1.0950e+02 2.9288e+02 ; HB2 - CB
- HB3
6881 6880 6883 1 1.0950e+02 4.1840e+02 ; HB2 - CB
- CG
6882 6880 6883 1 1.0950e+02 4.1840e+02 ; HB3 - CB
- CG
6883 6886 6887 1 1.0950e+02 4.1840e+02 ; CG - CD
- HD2
6883 6886 6888 1 1.0950e+02 4.1840e+02 ; CG - CD
- HD3
6883 6886 6889 1 1.0950e+02 3.3472e+02 ; CG - CD
- CE
6884 6883 6885 1 1.0950e+02 2.9288e+02 ; HG2 - CG
- HG3
6884 6883 6886 1 1.0950e+02 4.1840e+02 ; HG2 - CG
- CD
6885 6883 6886 1 1.0950e+02 4.1840e+02 ; HG3 - CG
- CD
6886 6889 6890 1 1.0950e+02 4.1840e+02 ; CD - CE
- HE2
6886 6889 6891 1 1.0950e+02 4.1840e+02 ; CD - CE
- HE3
6886 6889 6892 1 1.1120e+02 6.6944e+02 ; CD - CE
- NZ
6887 6886 6888 1 1.0950e+02 2.9288e+02 ; HD2 - CD
- HD3
6887 6886 6889 1 1.0950e+02 4.1840e+02 ; HD2 - CD
- CE
6888 6886 6889 1 1.0950e+02 4.1840e+02 ; HD3 - CD
- CE
6889 6892 6893 1 1.0950e+02 4.1840e+02 ; CE - NZ
- HZ1

6889 6892 6894 1 1.0950e+02 4.1840e+02 ; CE - NZ
- HZ2
6889 6892 6895 1 1.0950e+02 4.1840e+02 ; CE - NZ
- HZ3
6890 6889 6891 1 1.0950e+02 2.9288e+02 ; HE2 - CE
- HE3
6890 6889 6892 1 1.0950e+02 4.1840e+02 ; HE2 - CE
- NZ
6891 6889 6892 1 1.0950e+02 4.1840e+02 ; HE3 - CE
- NZ
6893 6892 6894 1 1.0950e+02 2.9288e+02 ; HZ1 - NZ
- HZ2
6893 6892 6895 1 1.0950e+02 2.9288e+02 ; HZ1 - NZ
- HZ3
6894 6892 6895 1 1.0950e+02 2.9288e+02 ; HZ2 - NZ
- HZ3
6896 6898 6899 1 1.2000e+02 4.1840e+02 ; C - N
- H
6896 6898 6900 1 1.2190e+02 4.1840e+02 ; C - N
- CA
6897 6896 6898 1 1.2290e+02 6.6944e+02 ; O - C
- N
6898 6900 6901 1 1.0950e+02 4.1840e+02 ; N - CA
- HA
6898 6900 6902 1 1.0970e+02 6.6944e+02 ; N - CA
- CB
6898 6900 6906 1 1.1010e+02 5.2718e+02 ; N - CA
- C
6899 6898 6900 1 1.1804e+02 4.1840e+02 ; H - N
- CA
6900 6902 6903 1 1.0950e+02 4.1840e+02 ; CA - CB
- HB1
6900 6902 6904 1 1.0950e+02 4.1840e+02 ; CA - CB
- HB2
6900 6902 6905 1 1.0950e+02 4.1840e+02 ; CA - CB
- HB3
6900 6906 6907 1 1.2040e+02 6.6944e+02 ; CA - C
- O
6900 6906 6908 1 1.1660e+02 5.8576e+02 ; CA - C
- N
6901 6900 6902 1 1.0950e+02 4.1840e+02 ; HA - CA
- CB
6901 6900 6906 1 1.0950e+02 4.1840e+02 ; HA - CA
- C
6902 6900 6906 1 1.1110e+02 5.2718e+02 ; CB - CA
- C
6903 6902 6904 1 1.0950e+02 2.9288e+02 ; HB1 - CB
- HB2
6903 6902 6905 1 1.0950e+02 2.9288e+02 ; HB1 - CB
- HB3
6904 6902 6905 1 1.0950e+02 2.9288e+02 ; HB2 - CB
- HB3

6906 6908 6909 1 1.2000e+02 4.1840e+02 ; C - N
- H
6906 6908 6910 1 1.2190e+02 4.1840e+02 ; C - N
- CA
6907 6906 6908 1 1.2290e+02 6.6944e+02 ; O - C
- N
6908 6910 6911 1 1.0950e+02 4.1840e+02 ; N - CA
- HA
6908 6910 6912 1 1.0970e+02 6.6944e+02 ; N - CA
- CB
6908 6910 6925 1 1.1010e+02 5.2718e+02 ; N - CA
- C
6909 6908 6910 1 1.1804e+02 4.1840e+02 ; H - N
- CA
6910 6912 6913 1 1.0950e+02 4.1840e+02 ; CA - CB
- HB
6910 6912 6914 1 1.0950e+02 3.3472e+02 ; CA - CB
- CG2
6910 6912 6918 1 1.0950e+02 3.3472e+02 ; CA - CB
- CG1
6910 6925 6926 1 1.2040e+02 6.6944e+02 ; CA - C
- O
6910 6925 6927 1 1.1660e+02 5.8576e+02 ; CA - C
- N
6911 6910 6912 1 1.0950e+02 4.1840e+02 ; HA - CA
- CB
6911 6910 6925 1 1.0950e+02 4.1840e+02 ; HA - CA
- C
6912 6910 6925 1 1.1110e+02 5.2718e+02 ; CB - CA
- C
6912 6914 6915 1 1.0950e+02 4.1840e+02 ; CB - CG2
- HG21
6912 6914 6916 1 1.0950e+02 4.1840e+02 ; CB - CG2
- HG22
6912 6914 6917 1 1.0950e+02 4.1840e+02 ; CB - CG2
- HG23
6912 6918 6919 1 1.0950e+02 4.1840e+02 ; CB - CG1
- HG12
6912 6918 6920 1 1.0950e+02 4.1840e+02 ; CB - CG1
- HG13
6912 6918 6921 1 1.0950e+02 3.3472e+02 ; CB - CG1
- CD1
6913 6912 6914 1 1.0950e+02 4.1840e+02 ; HB - CB
- CG2
6913 6912 6918 1 1.0950e+02 4.1840e+02 ; HB - CB
- CG1
6914 6912 6918 1 1.0950e+02 3.3472e+02 ; CG2 - CB
- CG1
6915 6914 6916 1 1.0950e+02 2.9288e+02 ; HG21 - CG2
- HG22
6915 6914 6917 1 1.0950e+02 2.9288e+02 ; HG21 - CG2
- HG23

6916 6914 6917 1 1.0950e+02 2.9288e+02 ; HG22 - CG2
- HG23
6918 6921 6922 1 1.0950e+02 4.1840e+02 ; CG1 - CD1
- HD11
6918 6921 6923 1 1.0950e+02 4.1840e+02 ; CG1 - CD1
- HD12
6918 6921 6924 1 1.0950e+02 4.1840e+02 ; CG1 - CD1
- HD13
6919 6918 6920 1 1.0950e+02 2.9288e+02 ; HG12 - CG1
- HG13
6919 6918 6921 1 1.0950e+02 4.1840e+02 ; HG12 - CG1
- CD1
6920 6918 6921 1 1.0950e+02 4.1840e+02 ; HG13 - CG1
- CD1
6922 6921 6923 1 1.0950e+02 2.9288e+02 ; HD11 - CD1
- HD12
6922 6921 6924 1 1.0950e+02 2.9288e+02 ; HD11 - CD1
- HD13
6923 6921 6924 1 1.0950e+02 2.9288e+02 ; HD12 - CD1
- HD13
6925 6927 6928 1 1.2000e+02 4.1840e+02 ; C - N
- H
6925 6927 6929 1 1.2190e+02 4.1840e+02 ; C - N
- CA
6926 6925 6927 1 1.2290e+02 6.6944e+02 ; O - C
- N
6927 6929 6930 1 1.0950e+02 4.1840e+02 ; N - CA
- HA
6927 6929 6931 1 1.0970e+02 6.6944e+02 ; N - CA
- CB
6927 6929 6937 1 1.1010e+02 5.2718e+02 ; N - CA
- C
6928 6927 6929 1 1.1804e+02 4.1840e+02 ; H - N
- CA
6929 6931 6932 1 1.0950e+02 4.1840e+02 ; CA - CB
- HB2
6929 6931 6933 1 1.0950e+02 4.1840e+02 ; CA - CB
- HB3
6929 6931 6934 1 1.1110e+02 5.2718e+02 ; CA - CB
- CG
6929 6937 6938 1 1.2040e+02 6.6944e+02 ; CA - C
- O
6929 6937 6939 1 1.1660e+02 5.8576e+02 ; CA - C
- N
6930 6929 6931 1 1.0950e+02 4.1840e+02 ; HA - CA
- CB
6930 6929 6937 1 1.0950e+02 4.1840e+02 ; HA - CA
- C
6931 6929 6937 1 1.1110e+02 5.2718e+02 ; CB - CA
- C
6931 6934 6935 1 1.1700e+02 5.8576e+02 ; CB - CG
- OD1

6931 6934 6936 1 1.1700e+02 5.8576e+02 ; CB - CG
- OD2
6932 6931 6933 1 1.0950e+02 2.9288e+02 ; HB2 - CB
- HB3
6932 6931 6934 1 1.0950e+02 4.1840e+02 ; HB2 - CB
- CG
6933 6931 6934 1 1.0950e+02 4.1840e+02 ; HB3 - CB
- CG
6935 6934 6936 1 1.2600e+02 6.6944e+02 ; OD1 - CG
- OD2
6937 6939 6940 1 1.2000e+02 4.1840e+02 ; C - N
- H
6937 6939 6941 1 1.2190e+02 4.1840e+02 ; C - N
- CA
6938 6937 6939 1 1.2290e+02 6.6944e+02 ; O - C
- N
6939 6941 6942 1 1.0950e+02 4.1840e+02 ; N - CA
- HA
6939 6941 6943 1 1.0970e+02 6.6944e+02 ; N - CA
- CB
6939 6941 6948 1 1.1010e+02 5.2718e+02 ; N - CA
- C
6940 6939 6941 1 1.1804e+02 4.1840e+02 ; H - N
- CA
6941 6943 6944 1 1.0950e+02 4.1840e+02 ; CA - CB
- HB2
6941 6943 6945 1 1.0950e+02 4.1840e+02 ; CA - CB
- HB3
6941 6943 6946 1 1.0950e+02 4.1840e+02 ; CA - CB
- OG
6941 6948 6949 1 1.2040e+02 6.6944e+02 ; CA - C
- O
6941 6948 6950 1 1.1660e+02 5.8576e+02 ; CA - C
- N
6942 6941 6943 1 1.0950e+02 4.1840e+02 ; HA - CA
- CB
6942 6941 6948 1 1.0950e+02 4.1840e+02 ; HA - CA
- C
6943 6941 6948 1 1.1110e+02 5.2718e+02 ; CB - CA
- C
6943 6946 6947 1 1.0850e+02 4.6024e+02 ; CB - OG
- HG
6944 6943 6945 1 1.0950e+02 2.9288e+02 ; HB2 - CB
- HB3
6944 6943 6946 1 1.0950e+02 4.1840e+02 ; HB2 - CB
- OG
6945 6943 6946 1 1.0950e+02 4.1840e+02 ; HB3 - CB
- OG
6948 6950 6951 1 1.2000e+02 4.1840e+02 ; C - N
- H
6948 6950 6952 1 1.2190e+02 4.1840e+02 ; C - N
- CA

6949 6948 6950 1 1.2290e+02 6.6944e+02 ; O - C
- N
6950 6952 6953 1 1.0950e+02 4.1840e+02 ; N - CA
- HA
6950 6952 6954 1 1.0970e+02 6.6944e+02 ; N - CA
- CB
6950 6952 6967 1 1.1010e+02 5.2718e+02 ; N - CA
- C
6951 6950 6952 1 1.1804e+02 4.1840e+02 ; H - N
- CA
6952 6954 6955 1 1.0950e+02 4.1840e+02 ; CA - CB
- HB2
6952 6954 6956 1 1.0950e+02 4.1840e+02 ; CA - CB
- HB3
6952 6954 6957 1 1.0950e+02 3.3472e+02 ; CA - CB
- CG
6952 6967 6968 1 1.2040e+02 6.6944e+02 ; CA - C
- O
6952 6967 6969 1 1.1660e+02 5.8576e+02 ; CA - C
- N
6953 6952 6954 1 1.0950e+02 4.1840e+02 ; HA - CA
- CB
6953 6952 6967 1 1.0950e+02 4.1840e+02 ; HA - CA
- C
6954 6952 6967 1 1.1110e+02 5.2718e+02 ; CB - CA
- C
6954 6957 6958 1 1.0950e+02 4.1840e+02 ; CB - CG
- HG
6954 6957 6959 1 1.0950e+02 3.3472e+02 ; CB - CG
- CD1
6954 6957 6963 1 1.0950e+02 3.3472e+02 ; CB - CG
- CD2
6955 6954 6956 1 1.0950e+02 2.9288e+02 ; HB2 - CB
- HB3
6955 6954 6957 1 1.0950e+02 4.1840e+02 ; HB2 - CB
- CG
6956 6954 6957 1 1.0950e+02 4.1840e+02 ; HB3 - CB
- CG
6957 6959 6960 1 1.0950e+02 4.1840e+02 ; CG - CD1
- HD11
6957 6959 6961 1 1.0950e+02 4.1840e+02 ; CG - CD1
- HD12
6957 6959 6962 1 1.0950e+02 4.1840e+02 ; CG - CD1
- HD13
6957 6963 6964 1 1.0950e+02 4.1840e+02 ; CG - CD2
- HD21
6957 6963 6965 1 1.0950e+02 4.1840e+02 ; CG - CD2
- HD22
6957 6963 6966 1 1.0950e+02 4.1840e+02 ; CG - CD2
- HD23
6958 6957 6959 1 1.0950e+02 4.1840e+02 ; HG - CG
- CD1

6958 6957 6963 1 1.0950e+02 4.1840e+02 ; HG - CG
- CD2
6959 6957 6963 1 1.0950e+02 3.3472e+02 ; CD1 - CG
- CD2
6960 6959 6961 1 1.0950e+02 2.9288e+02 ; HD11 - CD1
- HD12
6960 6959 6962 1 1.0950e+02 2.9288e+02 ; HD11 - CD1
- HD13
6961 6959 6962 1 1.0950e+02 2.9288e+02 ; HD12 - CD1
- HD13
6964 6963 6965 1 1.0950e+02 2.9288e+02 ; HD21 - CD2
- HD22
6964 6963 6966 1 1.0950e+02 2.9288e+02 ; HD21 - CD2
- HD23
6965 6963 6966 1 1.0950e+02 2.9288e+02 ; HD22 - CD2
- HD23
6967 6969 6970 1 1.2000e+02 4.1840e+02 ; C - N
- H
6967 6969 6971 1 1.2190e+02 4.1840e+02 ; C - N
- CA
6968 6967 6969 1 1.2290e+02 6.6944e+02 ; O - C
- N
6969 6971 6972 1 1.0950e+02 4.1840e+02 ; N - CA
- HA
6969 6971 6973 1 1.0970e+02 6.6944e+02 ; N - CA
- CB
6969 6971 6986 1 1.1010e+02 5.2718e+02 ; N - CA
- C
6970 6969 6971 1 1.1804e+02 4.1840e+02 ; H - N
- CA
6971 6973 6974 1 1.0950e+02 4.1840e+02 ; CA - CB
- HB2
6971 6973 6975 1 1.0950e+02 4.1840e+02 ; CA - CB
- HB3
6971 6973 6976 1 1.0950e+02 3.3472e+02 ; CA - CB
- CG
6971 6986 6987 1 1.2040e+02 6.6944e+02 ; CA - C
- O
6971 6986 6988 1 1.1660e+02 5.8576e+02 ; CA - C
- N
6972 6971 6973 1 1.0950e+02 4.1840e+02 ; HA - CA
- CB
6972 6971 6986 1 1.0950e+02 4.1840e+02 ; HA - CA
- C
6973 6971 6986 1 1.1110e+02 5.2718e+02 ; CB - CA
- C
6973 6976 6977 1 1.0950e+02 4.1840e+02 ; CB - CG
- HG
6973 6976 6978 1 1.0950e+02 3.3472e+02 ; CB - CG
- CD1
6973 6976 6982 1 1.0950e+02 3.3472e+02 ; CB - CG
- CD2

6974 6973 6975 1 1.0950e+02 2.9288e+02 ; HB2 - CB
- HB3
6974 6973 6976 1 1.0950e+02 4.1840e+02 ; HB2 - CB
- CG
6975 6973 6976 1 1.0950e+02 4.1840e+02 ; HB3 - CB
- CG
6976 6978 6979 1 1.0950e+02 4.1840e+02 ; CG - CD1
- HD11
6976 6978 6980 1 1.0950e+02 4.1840e+02 ; CG - CD1
- HD12
6976 6978 6981 1 1.0950e+02 4.1840e+02 ; CG - CD1
- HD13
6976 6982 6983 1 1.0950e+02 4.1840e+02 ; CG - CD2
- HD21
6976 6982 6984 1 1.0950e+02 4.1840e+02 ; CG - CD2
- HD22
6976 6982 6985 1 1.0950e+02 4.1840e+02 ; CG - CD2
- HD23
6977 6976 6978 1 1.0950e+02 4.1840e+02 ; HG - CG
- CD1
6977 6976 6982 1 1.0950e+02 4.1840e+02 ; HG - CG
- CD2
6978 6976 6982 1 1.0950e+02 3.3472e+02 ; CD1 - CG
- CD2
6979 6978 6980 1 1.0950e+02 2.9288e+02 ; HD11 - CD1
- HD12
6979 6978 6981 1 1.0950e+02 2.9288e+02 ; HD11 - CD1
- HD13
6980 6978 6981 1 1.0950e+02 2.9288e+02 ; HD12 - CD1
- HD13
6983 6982 6984 1 1.0950e+02 2.9288e+02 ; HD21 - CD2
- HD22
6983 6982 6985 1 1.0950e+02 2.9288e+02 ; HD21 - CD2
- HD23
6984 6982 6985 1 1.0950e+02 2.9288e+02 ; HD22 - CD2
- HD23
6986 6988 6989 1 1.2000e+02 4.1840e+02 ; C - N
- H
6986 6988 6990 1 1.2190e+02 4.1840e+02 ; C - N
- CA
6987 6986 6988 1 1.2290e+02 6.6944e+02 ; O - C
- N
6988 6990 6991 1 1.0950e+02 4.1840e+02 ; N - CA
- HA
6988 6990 6992 1 1.0970e+02 6.6944e+02 ; N - CA
- CB
6988 6990 7003 1 1.1010e+02 5.2718e+02 ; N - CA
- C
6989 6988 6990 1 1.1804e+02 4.1840e+02 ; H - N
- CA
6990 6992 6993 1 1.0950e+02 4.1840e+02 ; CA - CB
- HB2

6990 6992 6994 1 1.0950e+02 4.1840e+02 ; CA - CB
- HB3
6990 6992 6995 1 1.0950e+02 3.3472e+02 ; CA - CB
- CG
6990 7003 7004 1 1.2040e+02 6.6944e+02 ; CA - C
- O
6990 7003 7005 1 1.1660e+02 5.8576e+02 ; CA - C
- N
6991 6990 6992 1 1.0950e+02 4.1840e+02 ; HA - CA
- CB
6991 6990 7003 1 1.0950e+02 4.1840e+02 ; HA - CA
- C
6992 6990 7003 1 1.1110e+02 5.2718e+02 ; CB - CA
- C
6992 6995 6996 1 1.0950e+02 4.1840e+02 ; CB - CG
- HG2
6992 6995 6997 1 1.0950e+02 4.1840e+02 ; CB - CG
- HG3
6992 6995 6998 1 1.1110e+02 5.2718e+02 ; CB - CG
- CD
6993 6992 6994 1 1.0950e+02 2.9288e+02 ; HB2 - CB
- HB3
6993 6992 6995 1 1.0950e+02 4.1840e+02 ; HB2 - CB
- CG
6994 6992 6995 1 1.0950e+02 4.1840e+02 ; HB3 - CB
- CG
6995 6998 6999 1 1.2040e+02 6.6944e+02 ; CG - CD
- OE1
6995 6998 7000 1 1.1660e+02 5.8576e+02 ; CG - CD
- NE2
6996 6995 6997 1 1.0950e+02 2.9288e+02 ; HG2 - CG
- HG3
6996 6995 6998 1 1.0950e+02 4.1840e+02 ; HG2 - CG
- CD
6997 6995 6998 1 1.0950e+02 4.1840e+02 ; HG3 - CG
- CD
6998 7000 7001 1 1.2000e+02 4.1840e+02 ; CD - NE2
- HE21
6998 7000 7002 1 1.2000e+02 4.1840e+02 ; CD - NE2
- HE22
6999 6998 7000 1 1.2290e+02 6.6944e+02 ; OE1 - CD
- NE2
7001 7000 7002 1 1.2000e+02 2.9288e+02 ; HE21 - NE2
- HE22
7003 7005 7006 1 1.2000e+02 4.1840e+02 ; C - N
- H
7003 7005 7007 1 1.2190e+02 4.1840e+02 ; C - N
- CA
7004 7003 7005 1 1.2290e+02 6.6944e+02 ; O - C
- N
7005 7007 7008 1 1.0950e+02 4.1840e+02 ; N - CA
- HA

7005 7007 7009 1 1.0970e+02 6.6944e+02 ; N - CA
- CB
7005 7007 7018 1 1.1010e+02 5.2718e+02 ; N - CA
- C
7006 7005 7007 1 1.1804e+02 4.1840e+02 ; H - N
- CA
7007 7009 7010 1 1.0950e+02 4.1840e+02 ; CA - CB
- HB2
7007 7009 7011 1 1.0950e+02 4.1840e+02 ; CA - CB
- HB3
7007 7009 7012 1 1.0950e+02 3.3472e+02 ; CA - CB
- CG
7007 7018 7019 1 1.2040e+02 6.6944e+02 ; CA - C
- O
7007 7018 7020 1 1.1660e+02 5.8576e+02 ; CA - C
- N
7008 7007 7009 1 1.0950e+02 4.1840e+02 ; HA - CA
- CB
7008 7007 7018 1 1.0950e+02 4.1840e+02 ; HA - CA
- C
7009 7007 7018 1 1.1110e+02 5.2718e+02 ; CB - CA
- C
7009 7012 7013 1 1.0950e+02 4.1840e+02 ; CB - CG
- HG2
7009 7012 7014 1 1.0950e+02 4.1840e+02 ; CB - CG
- HG3
7009 7012 7015 1 1.1110e+02 5.2718e+02 ; CB - CG
- CD
7010 7009 7011 1 1.0950e+02 2.9288e+02 ; HB2 - CB
- HB3
7010 7009 7012 1 1.0950e+02 4.1840e+02 ; HB2 - CB
- CG
7011 7009 7012 1 1.0950e+02 4.1840e+02 ; HB3 - CB
- CG
7012 7015 7016 1 1.1700e+02 5.8576e+02 ; CG - CD
- OE1
7012 7015 7017 1 1.1700e+02 5.8576e+02 ; CG - CD
- OE2
7013 7012 7014 1 1.0950e+02 2.9288e+02 ; HG2 - CG
- HG3
7013 7012 7015 1 1.0950e+02 4.1840e+02 ; HG2 - CG
- CD
7014 7012 7015 1 1.0950e+02 4.1840e+02 ; HG3 - CG
- CD
7016 7015 7017 1 1.2600e+02 6.6944e+02 ; OE1 - CD
- OE2
7018 7020 7021 1 1.2000e+02 4.1840e+02 ; C - N
- H
7018 7020 7022 1 1.2190e+02 4.1840e+02 ; C - N
- CA
7019 7018 7020 1 1.2290e+02 6.6944e+02 ; O - C
- N

7020 7022 7023 1 1.0950e+02 4.1840e+02 ; N - CA
- HA
7020 7022 7024 1 1.0970e+02 6.6944e+02 ; N - CA
- CB
7020 7022 7034 1 1.1010e+02 5.2718e+02 ; N - CA
- C
7021 7020 7022 1 1.1804e+02 4.1840e+02 ; H - N
- CA
7022 7024 7025 1 1.0950e+02 4.1840e+02 ; CA - CB
- HB
7022 7024 7026 1 1.0950e+02 3.3472e+02 ; CA - CB
- CG1
7022 7024 7030 1 1.0950e+02 3.3472e+02 ; CA - CB
- CG2
7022 7034 7035 1 1.2040e+02 6.6944e+02 ; CA - C
- O
7022 7034 7036 1 1.1660e+02 5.8576e+02 ; CA - C
- N
7023 7022 7024 1 1.0950e+02 4.1840e+02 ; HA - CA
- CB
7023 7022 7034 1 1.0950e+02 4.1840e+02 ; HA - CA
- C
7024 7022 7034 1 1.1110e+02 5.2718e+02 ; CB - CA
- C
7024 7026 7027 1 1.0950e+02 4.1840e+02 ; CB - CG1
- HG11
7024 7026 7028 1 1.0950e+02 4.1840e+02 ; CB - CG1
- HG12
7024 7026 7029 1 1.0950e+02 4.1840e+02 ; CB - CG1
- HG13
7024 7030 7031 1 1.0950e+02 4.1840e+02 ; CB - CG2
- HG21
7024 7030 7032 1 1.0950e+02 4.1840e+02 ; CB - CG2
- HG22
7024 7030 7033 1 1.0950e+02 4.1840e+02 ; CB - CG2
- HG23
7025 7024 7026 1 1.0950e+02 4.1840e+02 ; HB - CB
- CG1
7025 7024 7030 1 1.0950e+02 4.1840e+02 ; HB - CB
- CG2
7026 7024 7030 1 1.0950e+02 3.3472e+02 ; CG1 - CB
- CG2
7027 7026 7028 1 1.0950e+02 2.9288e+02 ; HG11 - CG1
- HG12
7027 7026 7029 1 1.0950e+02 2.9288e+02 ; HG11 - CG1
- HG13
7028 7026 7029 1 1.0950e+02 2.9288e+02 ; HG12 - CG1
- HG13
7031 7030 7032 1 1.0950e+02 2.9288e+02 ; HG21 - CG2
- HG22
7031 7030 7033 1 1.0950e+02 2.9288e+02 ; HG21 - CG2
- HG23

```
  7032   7030   7033       1    1.0950e+02    2.9288e+02 ;   HG22 - CG2
- HG23
  7034   7036   7037       1    1.2000e+02    4.1840e+02 ;      C - N
- H
  7034   7036   7038       1    1.2190e+02    4.1840e+02 ;      C - N
- CA
  7035   7034   7036       1    1.2290e+02    6.6944e+02 ;      O - C
- N
  7036   7038   7039       1    1.0950e+02    4.1840e+02 ;      N - CA
- HA
  7036   7038   7040       1    1.0970e+02    6.6944e+02 ;      N - CA
- CB
  7036   7038   7046       1    1.1010e+02    5.2718e+02 ;      N - CA
- C
  7037   7036   7038       1    1.1804e+02    4.1840e+02 ;      H - N
- CA
  7038   7040   7041       1    1.0950e+02    4.1840e+02 ;     CA - CB
- HB2
  7038   7040   7042       1    1.0950e+02    4.1840e+02 ;     CA - CB
- HB3
  7038   7040   7043       1    1.1110e+02    5.2718e+02 ;     CA - CB
- CG
  7038   7046   7047       1    1.2040e+02    6.6944e+02 ;     CA - C
- O
  7038   7046   7048       1    1.1660e+02    5.8576e+02 ;     CA - C
- N
  7039   7038   7040       1    1.0950e+02    4.1840e+02 ;     HA - CA
- CB
  7039   7038   7046       1    1.0950e+02    4.1840e+02 ;     HA - CA
- C
  7040   7038   7046       1    1.1110e+02    5.2718e+02 ;     CB - CA
- C
  7040   7043   7044       1    1.1700e+02    5.8576e+02 ;     CB - CG
- OD1
  7040   7043   7045       1    1.1700e+02    5.8576e+02 ;     CB - CG
- OD2
  7041   7040   7042       1    1.0950e+02    2.9288e+02 ;    HB2 - CB
- HB3
  7041   7040   7043       1    1.0950e+02    4.1840e+02 ;    HB2 - CB
- CG
  7042   7040   7043       1    1.0950e+02    4.1840e+02 ;    HB3 - CB
- CG
  7044   7043   7045       1    1.2600e+02    6.6944e+02 ;    OD1 - CG
- OD2
  7046   7048   7049       1    1.2000e+02    4.1840e+02 ;      C - N
- H
  7046   7048   7050       1    1.2190e+02    4.1840e+02 ;      C - N
- CA
  7047   7046   7048       1    1.2290e+02    6.6944e+02 ;      O - C
- N
  7048   7050   7051       1    1.0950e+02    4.1840e+02 ;      N - CA
- HA
```

7048 7050 7052 1 1.0970e+02 6.6944e+02 ; N - CA
- CB
7048 7050 7065 1 1.1010e+02 5.2718e+02 ; N - CA
- C
7049 7048 7050 1 1.1804e+02 4.1840e+02 ; H - N
- CA
7050 7052 7053 1 1.0950e+02 4.1840e+02 ; CA - CB
- HB2
7050 7052 7054 1 1.0950e+02 4.1840e+02 ; CA - CB
- HB3
7050 7052 7055 1 1.0950e+02 3.3472e+02 ; CA - CB
- CG
7050 7065 7066 1 1.2040e+02 6.6944e+02 ; CA - C
- O
7050 7065 7067 1 1.1660e+02 5.8576e+02 ; CA - C
- N
7051 7050 7052 1 1.0950e+02 4.1840e+02 ; HA - CA
- CB
7051 7050 7065 1 1.0950e+02 4.1840e+02 ; HA - CA
- C
7052 7050 7065 1 1.1110e+02 5.2718e+02 ; CB - CA
- C
7052 7055 7056 1 1.0950e+02 4.1840e+02 ; CB - CG
- HG
7052 7055 7057 1 1.0950e+02 3.3472e+02 ; CB - CG
- CD1
7052 7055 7061 1 1.0950e+02 3.3472e+02 ; CB - CG
- CD2
7053 7052 7054 1 1.0950e+02 2.9288e+02 ; HB2 - CB
- HB3
7053 7052 7055 1 1.0950e+02 4.1840e+02 ; HB2 - CB
- CG
7054 7052 7055 1 1.0950e+02 4.1840e+02 ; HB3 - CB
- CG
7055 7057 7058 1 1.0950e+02 4.1840e+02 ; CG - CD1
- HD11
7055 7057 7059 1 1.0950e+02 4.1840e+02 ; CG - CD1
- HD12
7055 7057 7060 1 1.0950e+02 4.1840e+02 ; CG - CD1
- HD13
7055 7061 7062 1 1.0950e+02 4.1840e+02 ; CG - CD2
- HD21
7055 7061 7063 1 1.0950e+02 4.1840e+02 ; CG - CD2
- HD22
7055 7061 7064 1 1.0950e+02 4.1840e+02 ; CG - CD2
- HD23
7056 7055 7057 1 1.0950e+02 4.1840e+02 ; HG - CG
- CD1
7056 7055 7061 1 1.0950e+02 4.1840e+02 ; HG - CG
- CD2
7057 7055 7061 1 1.0950e+02 3.3472e+02 ; CD1 - CG
- CD2

7058 7057 7059 1 1.0950e+02 2.9288e+02 ; HD11 - CD1 - HD12
7058 7057 7060 1 1.0950e+02 2.9288e+02 ; HD11 - CD1 - HD13
7059 7057 7060 1 1.0950e+02 2.9288e+02 ; HD12 - CD1 - HD13
7062 7061 7063 1 1.0950e+02 2.9288e+02 ; HD21 - CD2 - HD22
7062 7061 7064 1 1.0950e+02 2.9288e+02 ; HD21 - CD2 - HD23
7063 7061 7064 1 1.0950e+02 2.9288e+02 ; HD22 - CD2 - HD23
7065 7067 7068 1 1.2000e+02 4.1840e+02 ; C - N - H
7065 7067 7069 1 1.2190e+02 4.1840e+02 ; C - N - CA
7066 7065 7067 1 1.2290e+02 6.6944e+02 ; O - C - N
7067 7069 7070 1 1.0950e+02 4.1840e+02 ; N - CA - HA
7067 7069 7071 1 1.0970e+02 6.6944e+02 ; N - CA - CB
7067 7069 7079 1 1.1010e+02 5.2718e+02 ; N - CA - C
7068 7067 7069 1 1.1804e+02 4.1840e+02 ; H - N - CA
7069 7071 7072 1 1.0950e+02 4.1840e+02 ; CA - CB - HB2
7069 7071 7073 1 1.0950e+02 4.1840e+02 ; CA - CB - HB3
7069 7071 7074 1 1.1110e+02 5.2718e+02 ; CA - CB - CG
7069 7079 7080 1 1.2040e+02 6.6944e+02 ; CA - C - O
7069 7079 7081 1 1.1660e+02 5.8576e+02 ; CA - C - N
7070 7069 7071 1 1.0950e+02 4.1840e+02 ; HA - CA - CB
7070 7069 7079 1 1.0950e+02 4.1840e+02 ; HA - CA - C
7071 7069 7079 1 1.1110e+02 5.2718e+02 ; CB - CA - C
7071 7074 7075 1 1.2040e+02 6.6944e+02 ; CB - CG - OD1
7071 7074 7076 1 1.1660e+02 5.8576e+02 ; CB - CG - ND2
7072 7071 7073 1 1.0950e+02 2.9288e+02 ; HB2 - CB - HB3
7072 7071 7074 1 1.0950e+02 4.1840e+02 ; HB2 - CB - CG
7073 7071 7074 1 1.0950e+02 4.1840e+02 ; HB3 - CB - CG

7074 7076 7077 1 1.2000e+02 4.1840e+02 ; CG - ND2
- HD21
7074 7076 7078 1 1.2000e+02 4.1840e+02 ; CG - ND2
- HD22
7075 7074 7076 1 1.2290e+02 6.6944e+02 ; OD1 - CG
- ND2
7077 7076 7078 1 1.2000e+02 2.9288e+02 ; HD21 - ND2
- HD22
7079 7081 7082 1 1.2000e+02 4.1840e+02 ; C - N
- H
7079 7081 7083 1 1.2190e+02 4.1840e+02 ; C - N
- CA
7080 7079 7081 1 1.2290e+02 6.6944e+02 ; O - C
- N
7081 7083 7084 1 1.0950e+02 4.1840e+02 ; N - CA
- HA2
7081 7083 7085 1 1.0950e+02 4.1840e+02 ; N - CA
- HA3
7081 7083 7086 1 1.1010e+02 5.2718e+02 ; N - CA
- C
7082 7081 7083 1 1.1804e+02 4.1840e+02 ; H - N
- CA
7083 7086 7087 1 1.2040e+02 6.6944e+02 ; CA - C
- O
7083 7086 7088 1 1.1660e+02 5.8576e+02 ; CA - C
- N
7084 7083 7085 1 1.0950e+02 2.9288e+02 ; HA2 - CA
- HA3
7084 7083 7086 1 1.0950e+02 4.1840e+02 ; HA2 - CA
- C
7085 7083 7086 1 1.0950e+02 4.1840e+02 ; HA3 - CA
- C
7086 7088 7089 1 1.2000e+02 4.1840e+02 ; C - N
- H
7086 7088 7090 1 1.2190e+02 4.1840e+02 ; C - N
- CA
7087 7086 7088 1 1.2290e+02 6.6944e+02 ; O - C
- N
7088 7090 7091 1 1.0950e+02 4.1840e+02 ; N - CA
- HA
7088 7090 7092 1 1.0970e+02 6.6944e+02 ; N - CA
- CB
7088 7090 7099 1 1.1010e+02 5.2718e+02 ; N - CA
- C
7089 7088 7090 1 1.1804e+02 4.1840e+02 ; H - N
- CA
7090 7092 7093 1 1.0950e+02 4.1840e+02 ; CA - CB
- HB2
7090 7092 7094 1 1.0950e+02 4.1840e+02 ; CA - CB
- HB3
7090 7092 7095 1 1.1110e+02 5.2718e+02 ; CA - CB
- CG

```
  7090   7099   7100       1    1.2040e+02    6.6944e+02 ;     CA - C
- O
  7090   7099   7101       1    1.1660e+02    5.8576e+02 ;     CA - C
- N
  7091   7090   7092       1    1.0950e+02    4.1840e+02 ;     HA - CA
- CB
  7091   7090   7099       1    1.0950e+02    4.1840e+02 ;     HA - CA
- C
  7092   7090   7099       1    1.1110e+02    5.2718e+02 ;     CB - CA
- C
  7092   7095   7096       1    1.2040e+02    6.6944e+02 ;     CB - CG
- OD1
  7092   7095   7097       1    1.1000e+02    6.6944e+02 ;     CB - CG
- OD2
  7093   7092   7094       1    1.0950e+02    2.9288e+02 ;    HB2 - CB
- HB3
  7093   7092   7095       1    1.0950e+02    4.1840e+02 ;    HB2 - CB
- CG
  7094   7092   7095       1    1.0950e+02    4.1840e+02 ;    HB3 - CB
- CG
  7095   7097   7098       1    1.1300e+02    4.1840e+02 ;     CG - OD2
- HD2
  7096   7095   7097       1    1.2000e+02    6.6944e+02 ;    OD1 - CG
- OD2
  7099   7101   7102       1    1.2000e+02    4.1840e+02 ;      C - N
- H
  7099   7101   7103       1    1.2190e+02    4.1840e+02 ;      C - N
- CA
  7100   7099   7101       1    1.2290e+02    6.6944e+02 ;      O - C
- N
  7101   7103   7104       1    1.0950e+02    4.1840e+02 ;      N - CA
- HA2
  7101   7103   7105       1    1.0950e+02    4.1840e+02 ;      N - CA
- HA3
  7101   7103   7106       1    1.1010e+02    5.2718e+02 ;      N - CA
- C
  7102   7101   7103       1    1.1804e+02    4.1840e+02 ;      H - N
- CA
  7103   7106   7107       1    1.2040e+02    6.6944e+02 ;     CA - C
- O
  7103   7106   7108       1    1.1660e+02    5.8576e+02 ;     CA - C
- N
  7104   7103   7105       1    1.0950e+02    2.9288e+02 ;    HA2 - CA
- HA3
  7104   7103   7106       1    1.0950e+02    4.1840e+02 ;    HA2 - CA
- C
  7105   7103   7106       1    1.0950e+02    4.1840e+02 ;    HA3 - CA
- C
  7106   7108   7109       1    1.2000e+02    4.1840e+02 ;      C - N
- H
  7106   7108   7110       1    1.2190e+02    4.1840e+02 ;      C - N
- CA
```

7107 7106 7108 1 1.2290e+02 6.6944e+02 ; O - C
- N
7108 7110 7111 1 1.0950e+02 4.1840e+02 ; N - CA
- HA
7108 7110 7112 1 1.0970e+02 6.6944e+02 ; N - CA
- CB
7108 7110 7121 1 1.1010e+02 5.2718e+02 ; N - CA
- C
7109 7108 7110 1 1.1804e+02 4.1840e+02 ; H - N
- CA
7110 7112 7113 1 1.0950e+02 4.1840e+02 ; CA - CB
- HB2
7110 7112 7114 1 1.0950e+02 4.1840e+02 ; CA - CB
- HB3
7110 7112 7115 1 1.0950e+02 3.3472e+02 ; CA - CB
- CG
7110 7121 7122 1 1.2040e+02 6.6944e+02 ; CA - C
- O
7110 7121 7123 1 1.1660e+02 5.8576e+02 ; CA - C
- N
7111 7110 7112 1 1.0950e+02 4.1840e+02 ; HA - CA
- CB
7111 7110 7121 1 1.0950e+02 4.1840e+02 ; HA - CA
- C
7112 7110 7121 1 1.1110e+02 5.2718e+02 ; CB - CA
- C
7112 7115 7116 1 1.0950e+02 4.1840e+02 ; CB - CG
- HG2
7112 7115 7117 1 1.0950e+02 4.1840e+02 ; CB - CG
- HG3
7112 7115 7118 1 1.1110e+02 5.2718e+02 ; CB - CG
- CD
7113 7112 7114 1 1.0950e+02 2.9288e+02 ; HB2 - CB
- HB3
7113 7112 7115 1 1.0950e+02 4.1840e+02 ; HB2 - CB
- CG
7114 7112 7115 1 1.0950e+02 4.1840e+02 ; HB3 - CB
- CG
7115 7118 7119 1 1.1700e+02 5.8576e+02 ; CG - CD
- OE1
7115 7118 7120 1 1.1700e+02 5.8576e+02 ; CG - CD
- OE2
7116 7115 7117 1 1.0950e+02 2.9288e+02 ; HG2 - CG
- HG3
7116 7115 7118 1 1.0950e+02 4.1840e+02 ; HG2 - CG
- CD
7117 7115 7118 1 1.0950e+02 4.1840e+02 ; HG3 - CG
- CD
7119 7118 7120 1 1.2600e+02 6.6944e+02 ; OE1 - CD
- OE2
7121 7123 7124 1 1.2000e+02 4.1840e+02 ; C - N
- H

7121 7123 7125 1 1.2190e+02 4.1840e+02 ; C - N
- CA
7122 7121 7123 1 1.2290e+02 6.6944e+02 ; O - C
- N
7123 7125 7126 1 1.0950e+02 4.1840e+02 ; N - CA
- HA
7123 7125 7127 1 1.0970e+02 6.6944e+02 ; N - CA
- CB
7123 7125 7140 1 1.1010e+02 5.2718e+02 ; N - CA
- C
7124 7123 7125 1 1.1804e+02 4.1840e+02 ; H - N
- CA
7125 7127 7128 1 1.0950e+02 4.1840e+02 ; CA - CB
- HB
7125 7127 7129 1 1.0950e+02 3.3472e+02 ; CA - CB
- CG2
7125 7127 7133 1 1.0950e+02 3.3472e+02 ; CA - CB
- CG1
7125 7140 7141 1 1.2040e+02 6.6944e+02 ; CA - C
- O
7125 7140 7142 1 1.1660e+02 5.8576e+02 ; CA - C
- N
7126 7125 7127 1 1.0950e+02 4.1840e+02 ; HA - CA
- CB
7126 7125 7140 1 1.0950e+02 4.1840e+02 ; HA - CA
- C
7127 7125 7140 1 1.1110e+02 5.2718e+02 ; CB - CA
- C
7127 7129 7130 1 1.0950e+02 4.1840e+02 ; CB - CG2
- HG21
7127 7129 7131 1 1.0950e+02 4.1840e+02 ; CB - CG2
- HG22
7127 7129 7132 1 1.0950e+02 4.1840e+02 ; CB - CG2
- HG23
7127 7133 7134 1 1.0950e+02 4.1840e+02 ; CB - CG1
- HG12
7127 7133 7135 1 1.0950e+02 4.1840e+02 ; CB - CG1
- HG13
7127 7133 7136 1 1.0950e+02 3.3472e+02 ; CB - CG1
- CD1
7128 7127 7129 1 1.0950e+02 4.1840e+02 ; HB - CB
- CG2
7128 7127 7133 1 1.0950e+02 4.1840e+02 ; HB - CB
- CG1
7129 7127 7133 1 1.0950e+02 3.3472e+02 ; CG2 - CB
- CG1
7130 7129 7131 1 1.0950e+02 2.9288e+02 ; HG21 - CG2
- HG22
7130 7129 7132 1 1.0950e+02 2.9288e+02 ; HG21 - CG2
- HG23
7131 7129 7132 1 1.0950e+02 2.9288e+02 ; HG22 - CG2
- HG23

```
  7133   7136   7137       1    1.0950e+02    4.1840e+02 ;    CG1 - CD1
- HD11
  7133   7136   7138       1    1.0950e+02    4.1840e+02 ;    CG1 - CD1
- HD12
  7133   7136   7139       1    1.0950e+02    4.1840e+02 ;    CG1 - CD1
- HD13
  7134   7133   7135       1    1.0950e+02    2.9288e+02 ;   HG12 - CG1
- HG13
  7134   7133   7136       1    1.0950e+02    4.1840e+02 ;   HG12 - CG1
- CD1
  7135   7133   7136       1    1.0950e+02    4.1840e+02 ;   HG13 - CG1
- CD1
  7137   7136   7138       1    1.0950e+02    2.9288e+02 ;   HD11 - CD1
- HD12
  7137   7136   7139       1    1.0950e+02    2.9288e+02 ;   HD11 - CD1
- HD13
  7138   7136   7139       1    1.0950e+02    2.9288e+02 ;   HD12 - CD1
- HD13
  7140   7142   7143       1    1.2000e+02    4.1840e+02 ;      C - N
- H
  7140   7142   7144       1    1.2190e+02    4.1840e+02 ;      C - N
- CA
  7141   7140   7142       1    1.2290e+02    6.6944e+02 ;      O - C
- N
  7142   7144   7145       1    1.0950e+02    4.1840e+02 ;      N - CA
- HA
  7142   7144   7146       1    1.0970e+02    6.6944e+02 ;      N - CA
- CB
  7142   7144   7152       1    1.1010e+02    5.2718e+02 ;      N - CA
- C
  7143   7142   7144       1    1.1804e+02    4.1840e+02 ;      H - N
- CA
  7144   7146   7147       1    1.0950e+02    4.1840e+02 ;     CA - CB
- HB2
  7144   7146   7148       1    1.0950e+02    4.1840e+02 ;     CA - CB
- HB3
  7144   7146   7149       1    1.1110e+02    5.2718e+02 ;     CA - CB
- CG
  7144   7152   7153       1    1.2040e+02    6.6944e+02 ;     CA - C
- O
  7144   7152   7154       1    1.1660e+02    5.8576e+02 ;     CA - C
- N
  7145   7144   7146       1    1.0950e+02    4.1840e+02 ;     HA - CA
- CB
  7145   7144   7152       1    1.0950e+02    4.1840e+02 ;     HA - CA
- C
  7146   7144   7152       1    1.1110e+02    5.2718e+02 ;     CB - CA
- C
  7146   7149   7150       1    1.1700e+02    5.8576e+02 ;     CB - CG
- OD1
  7146   7149   7151       1    1.1700e+02    5.8576e+02 ;     CB - CG
- OD2
```

7147 7146 7148 1 1.0950e+02 2.9288e+02 ; HB2 - CB
- HB3
7147 7146 7149 1 1.0950e+02 4.1840e+02 ; HB2 - CB
- CG
7148 7146 7149 1 1.0950e+02 4.1840e+02 ; HB3 - CB
- CG
7150 7149 7151 1 1.2600e+02 6.6944e+02 ; OD1 - CG
- OD2
7152 7154 7155 1 1.2000e+02 4.1840e+02 ; C - N
- H
7152 7154 7156 1 1.2190e+02 4.1840e+02 ; C - N
- CA
7153 7152 7154 1 1.2290e+02 6.6944e+02 ; O - C
- N
7154 7156 7157 1 1.0950e+02 4.1840e+02 ; N - CA
- HA
7154 7156 7158 1 1.0970e+02 6.6944e+02 ; N - CA
- CB
7154 7156 7172 1 1.1010e+02 5.2718e+02 ; N - CA
- C
7155 7154 7156 1 1.1804e+02 4.1840e+02 ; H - N
- CA
7156 7158 7159 1 1.0950e+02 4.1840e+02 ; CA - CB
- HB2
7156 7158 7160 1 1.0950e+02 4.1840e+02 ; CA - CB
- HB3
7156 7158 7161 1 1.1400e+02 5.2718e+02 ; CA - CB
- CG
7156 7172 7173 1 1.2040e+02 6.6944e+02 ; CA - C
- O
7156 7172 7174 1 1.1660e+02 5.8576e+02 ; CA - C
- N
7157 7156 7158 1 1.0950e+02 4.1840e+02 ; HA - CA
- CB
7157 7156 7172 1 1.0950e+02 4.1840e+02 ; HA - CA
- C
7158 7156 7172 1 1.1110e+02 5.2718e+02 ; CB - CA
- C
7158 7161 7162 1 1.2000e+02 5.8576e+02 ; CB - CG
- CD1
7158 7161 7170 1 1.2000e+02 5.8576e+02 ; CB - CG
- CD2
7159 7158 7160 1 1.0950e+02 2.9288e+02 ; HB2 - CB
- HB3
7159 7158 7161 1 1.0950e+02 4.1840e+02 ; HB2 - CB
- CG
7160 7158 7161 1 1.0950e+02 4.1840e+02 ; HB3 - CB
- CG
7161 7162 7163 1 1.2000e+02 4.1840e+02 ; CG - CD1
- HD1
7161 7162 7164 1 1.2000e+02 5.2718e+02 ; CG - CD1
- CE1

7161 7170 7168 1 1.2000e+02 5.2718e+02 ; CG - CD2
- CE2
7161 7170 7171 1 1.2000e+02 4.1840e+02 ; CG - CD2
- HD2
7162 7161 7170 1 1.2000e+02 5.2718e+02 ; CD1 - CG
- CD2
7162 7164 7165 1 1.2000e+02 4.1840e+02 ; CD1 - CE1
- HE1
7162 7164 7166 1 1.2000e+02 5.2718e+02 ; CD1 - CE1
- CZ
7163 7162 7164 1 1.2000e+02 4.1840e+02 ; HD1 - CD1
- CE1
7164 7166 7167 1 1.2000e+02 4.1840e+02 ; CE1 - CZ
- HZ
7164 7166 7168 1 1.2000e+02 5.2718e+02 ; CE1 - CZ
- CE2
7165 7164 7166 1 1.2000e+02 4.1840e+02 ; HE1 - CE1
- CZ
7166 7168 7169 1 1.2000e+02 4.1840e+02 ; CZ - CE2
- HE2
7166 7168 7170 1 1.2000e+02 5.2718e+02 ; CZ - CE2
- CD2
7167 7166 7168 1 1.2000e+02 4.1840e+02 ; HZ - CZ
- CE2
7168 7170 7171 1 1.2000e+02 4.1840e+02 ; CE2 - CD2
- HD2
7169 7168 7170 1 1.2000e+02 4.1840e+02 ; HE2 - CE2
- CD2
7172 7174 7175 1 1.2000e+02 4.1840e+02 ; C - N
- H
7172 7174 7176 1 1.2190e+02 4.1840e+02 ; C - N
- CA
7173 7172 7174 1 1.2290e+02 6.6944e+02 ; O - C
- N
7174 7176 7177 1 1.0950e+02 4.1840e+02 ; N - CA
- HA
7174 7176 7178 1 1.0970e+02 6.6944e+02 ; N - CA
- CB
7174 7176 7189 1 1.1010e+02 5.2718e+02 ; N - CA
- C
7175 7174 7176 1 1.1804e+02 4.1840e+02 ; H - N
- CA
7176 7178 7179 1 1.0950e+02 4.1840e+02 ; CA - CB
- HB2
7176 7178 7180 1 1.0950e+02 4.1840e+02 ; CA - CB
- HB3
7176 7178 7181 1 1.1310e+02 5.2718e+02 ; CA - CB
- CG
7176 7189 7190 1 1.2040e+02 6.6944e+02 ; CA - C
- O
7176 7189 7191 1 1.1660e+02 5.8576e+02 ; CA - C
- N

7177 7176 7178 1 1.0950e+02 4.1840e+02 ; HA - CA
- CB
7177 7176 7189 1 1.0950e+02 4.1840e+02 ; HA - CA
- C
7178 7176 7189 1 1.1110e+02 5.2718e+02 ; CB - CA
- C
7178 7181 7182 1 1.2000e+02 5.8576e+02 ; CB - CG
- ND1
7178 7181 7187 1 1.2000e+02 5.8576e+02 ; CB - CG
- CD2
7179 7178 7180 1 1.0950e+02 2.9288e+02 ; HB2 - CB
- HB3
7179 7178 7181 1 1.0950e+02 4.1840e+02 ; HB2 - CB
- CG
7180 7178 7181 1 1.0950e+02 4.1840e+02 ; HB3 - CB
- CG
7181 7182 7183 1 1.1700e+02 5.8576e+02 ; CG - ND1
- CE1
7181 7187 7185 1 1.2000e+02 5.8576e+02 ; CG - CD2
- NE2
7181 7187 7188 1 1.2000e+02 4.1840e+02 ; CG - CD2
- HD2
7182 7181 7187 1 1.2000e+02 5.8576e+02 ; ND1 - CG
- CD2
7182 7183 7184 1 1.2000e+02 4.1840e+02 ; ND1 - CE1
- HE1
7182 7183 7185 1 1.2000e+02 5.8576e+02 ; ND1 - CE1
- NE2
7183 7185 7186 1 1.2000e+02 4.1840e+02 ; CE1 - NE2
- HE2
7183 7185 7187 1 1.2000e+02 5.8576e+02 ; CE1 - NE2
- CD2
7184 7183 7185 1 1.2000e+02 4.1840e+02 ; HE1 - CE1
- NE2
7185 7187 7188 1 1.2000e+02 4.1840e+02 ; NE2 - CD2
- HD2
7186 7185 7187 1 1.2000e+02 4.1840e+02 ; HE2 - NE2
- CD2
7189 7191 7192 1 1.2000e+02 4.1840e+02 ; C - N
- H
7189 7191 7193 1 1.2190e+02 4.1840e+02 ; C - N
- CA
7190 7189 7191 1 1.2290e+02 6.6944e+02 ; O - C
- N
7191 7193 7194 1 1.0950e+02 4.1840e+02 ; N - CA
- HA
7191 7193 7195 1 1.0970e+02 6.6944e+02 ; N - CA
- CB
7191 7193 7205 1 1.1010e+02 5.2718e+02 ; N - CA
- C
7192 7191 7193 1 1.1804e+02 4.1840e+02 ; H - N
- CA

7193 7195 7196 1 1.0950e+02 4.1840e+02 ; CA - CB - HB2
7193 7195 7197 1 1.0950e+02 4.1840e+02 ; CA - CB - HB3
7193 7195 7198 1 1.0950e+02 3.3472e+02 ; CA - CB - CG
7193 7205 7206 1 1.2040e+02 6.6944e+02 ; CA - C - O
7193 7205 7207 1 1.1660e+02 5.8576e+02 ; CA - C - N
7194 7193 7195 1 1.0950e+02 4.1840e+02 ; HA - CA - CB
7194 7193 7205 1 1.0950e+02 4.1840e+02 ; HA - CA - C
7195 7193 7205 1 1.1110e+02 5.2718e+02 ; CB - CA - C
7195 7198 7199 1 1.0950e+02 4.1840e+02 ; CB - CG - HG2
7195 7198 7200 1 1.0950e+02 4.1840e+02 ; CB - CG - HG3
7195 7198 7201 1 1.1110e+02 5.2718e+02 ; CB - CG - CD
7196 7195 7197 1 1.0950e+02 2.9288e+02 ; HB2 - CB - HB3
7196 7195 7198 1 1.0950e+02 4.1840e+02 ; HB2 - CB - CG
7197 7195 7198 1 1.0950e+02 4.1840e+02 ; HB3 - CB - CG
7198 7201 7202 1 1.2040e+02 6.6944e+02 ; CG - CD - OE1
7198 7201 7203 1 1.1000e+02 6.6944e+02 ; CG - CD - OE2
7199 7198 7200 1 1.0950e+02 2.9288e+02 ; HG2 - CG - HG3
7199 7198 7201 1 1.0950e+02 4.1840e+02 ; HG2 - CG - CD
7200 7198 7201 1 1.0950e+02 4.1840e+02 ; HG3 - CG - CD
7201 7203 7204 1 1.1300e+02 4.1840e+02 ; CD - OE2 - HE2
7202 7201 7203 1 1.2000e+02 6.6944e+02 ; OE1 - CD - OE2
7205 7207 7208 1 1.2000e+02 4.1840e+02 ; C - N - H
7205 7207 7209 1 1.2190e+02 4.1840e+02 ; C - N - CA
7206 7205 7207 1 1.2290e+02 6.6944e+02 ; O - C - N
7207 7209 7210 1 1.0950e+02 4.1840e+02 ; N - CA - HA
7207 7209 7211 1 1.0970e+02 6.6944e+02 ; N - CA - CB

7207 7209 7225 1 1.1010e+02 5.2718e+02 ; N - CA
- C
7208 7207 7209 1 1.1804e+02 4.1840e+02 ; H - N
- CA
7209 7211 7212 1 1.0950e+02 4.1840e+02 ; CA - CB
- HB2
7209 7211 7213 1 1.0950e+02 4.1840e+02 ; CA - CB
- HB3
7209 7211 7214 1 1.1400e+02 5.2718e+02 ; CA - CB
- CG
7209 7225 7226 1 1.2040e+02 6.6944e+02 ; CA - C
- O
7209 7225 7227 1 1.1660e+02 5.8576e+02 ; CA - C
- N
7210 7209 7211 1 1.0950e+02 4.1840e+02 ; HA - CA
- CB
7210 7209 7225 1 1.0950e+02 4.1840e+02 ; HA - CA
- C
7211 7209 7225 1 1.1110e+02 5.2718e+02 ; CB - CA
- C
7211 7214 7215 1 1.2000e+02 5.8576e+02 ; CB - CG
- CD1
7211 7214 7223 1 1.2000e+02 5.8576e+02 ; CB - CG
- CD2
7212 7211 7213 1 1.0950e+02 2.9288e+02 ; HB2 - CB
- HB3
7212 7211 7214 1 1.0950e+02 4.1840e+02 ; HB2 - CB
- CG
7213 7211 7214 1 1.0950e+02 4.1840e+02 ; HB3 - CB
- CG
7214 7215 7216 1 1.2000e+02 4.1840e+02 ; CG - CD1
- HD1
7214 7215 7217 1 1.2000e+02 5.2718e+02 ; CG - CD1
- CE1
7214 7223 7221 1 1.2000e+02 5.2718e+02 ; CG - CD2
- CE2
7214 7223 7224 1 1.2000e+02 4.1840e+02 ; CG - CD2
- HD2
7215 7214 7223 1 1.2000e+02 5.2718e+02 ; CD1 - CG
- CD2
7215 7217 7218 1 1.2000e+02 4.1840e+02 ; CD1 - CE1
- HE1
7215 7217 7219 1 1.2000e+02 5.2718e+02 ; CD1 - CE1
- CZ
7216 7215 7217 1 1.2000e+02 4.1840e+02 ; HD1 - CD1
- CE1
7217 7219 7220 1 1.2000e+02 4.1840e+02 ; CE1 - CZ
- HZ
7217 7219 7221 1 1.2000e+02 5.2718e+02 ; CE1 - CZ
- CE2
7218 7217 7219 1 1.2000e+02 4.1840e+02 ; HE1 - CE1
- CZ

7219 7221 7222 1 1.2000e+02 4.1840e+02 ; CZ - CE2
- HE2
7219 7221 7223 1 1.2000e+02 5.2718e+02 ; CZ - CE2
- CD2
7220 7219 7221 1 1.2000e+02 4.1840e+02 ; HZ - CZ
- CE2
7221 7223 7224 1 1.2000e+02 4.1840e+02 ; CE2 - CD2
- HD2
7222 7221 7223 1 1.2000e+02 4.1840e+02 ; HE2 - CE2
- CD2
7225 7227 7228 1 1.2000e+02 4.1840e+02 ; C - N
- H
7225 7227 7229 1 1.2190e+02 4.1840e+02 ; C - N
- CA
7226 7225 7227 1 1.2290e+02 6.6944e+02 ; O - C
- N
7227 7229 7230 1 1.0950e+02 4.1840e+02 ; N - CA
- HA
7227 7229 7231 1 1.0970e+02 6.6944e+02 ; N - CA
- CB
7227 7229 7242 1 1.1010e+02 5.2718e+02 ; N - CA
- C
7228 7227 7229 1 1.1804e+02 4.1840e+02 ; H - N
- CA
7229 7231 7232 1 1.0950e+02 4.1840e+02 ; CA - CB
- HB2
7229 7231 7233 1 1.0950e+02 4.1840e+02 ; CA - CB
- HB3
7229 7231 7234 1 1.0950e+02 3.3472e+02 ; CA - CB
- CG
7229 7242 7243 1 1.2040e+02 6.6944e+02 ; CA - C
- O
7229 7242 7244 1 1.1660e+02 5.8576e+02 ; CA - C
- N
7230 7229 7231 1 1.0950e+02 4.1840e+02 ; HA - CA
- CB
7230 7229 7242 1 1.0950e+02 4.1840e+02 ; HA - CA
- C
7231 7229 7242 1 1.1110e+02 5.2718e+02 ; CB - CA
- C
7231 7234 7235 1 1.0950e+02 4.1840e+02 ; CB - CG
- HG2
7231 7234 7236 1 1.0950e+02 4.1840e+02 ; CB - CG
- HG3
7231 7234 7237 1 1.1470e+02 4.1840e+02 ; CB - CG
- SD
7232 7231 7233 1 1.0950e+02 2.9288e+02 ; HB2 - CB
- HB3
7232 7231 7234 1 1.0950e+02 4.1840e+02 ; HB2 - CB
- CG
7233 7231 7234 1 1.0950e+02 4.1840e+02 ; HB3 - CB
- CG

7234 7237 7238 1 9.8900e+01 5.1882e+02 ; CG - SD
- CE
7235 7234 7236 1 1.0950e+02 2.9288e+02 ; HG2 - CG
- HG3
7235 7234 7237 1 1.0950e+02 4.1840e+02 ; HG2 - CG
- SD
7236 7234 7237 1 1.0950e+02 4.1840e+02 ; HG3 - CG
- SD
7237 7238 7239 1 1.0950e+02 4.1840e+02 ; SD - CE
- HE1
7237 7238 7240 1 1.0950e+02 4.1840e+02 ; SD - CE
- HE2
7237 7238 7241 1 1.0950e+02 4.1840e+02 ; SD - CE
- HE3
7239 7238 7240 1 1.0950e+02 2.9288e+02 ; HE1 - CE
- HE2
7239 7238 7241 1 1.0950e+02 2.9288e+02 ; HE1 - CE
- HE3
7240 7238 7241 1 1.0950e+02 2.9288e+02 ; HE2 - CE
- HE3
7242 7244 7245 1 1.2000e+02 4.1840e+02 ; C - N
- H
7242 7244 7246 1 1.2190e+02 4.1840e+02 ; C - N
- CA
7243 7242 7244 1 1.2290e+02 6.6944e+02 ; O - C
- N
7244 7246 7247 1 1.0950e+02 4.1840e+02 ; N - CA
- HA
7244 7246 7248 1 1.0970e+02 6.6944e+02 ; N - CA
- CB
7244 7246 7261 1 1.1010e+02 5.2718e+02 ; N - CA
- C
7245 7244 7246 1 1.1804e+02 4.1840e+02 ; H - N
- CA
7246 7248 7249 1 1.0950e+02 4.1840e+02 ; CA - CB
- HB2
7246 7248 7250 1 1.0950e+02 4.1840e+02 ; CA - CB
- HB3
7246 7248 7251 1 1.0950e+02 3.3472e+02 ; CA - CB
- CG
7246 7261 7262 1 1.2040e+02 6.6944e+02 ; CA - C
- O
7246 7261 7263 1 1.1660e+02 5.8576e+02 ; CA - C
- N
7247 7246 7248 1 1.0950e+02 4.1840e+02 ; HA - CA
- CB
7247 7246 7261 1 1.0950e+02 4.1840e+02 ; HA - CA
- C
7248 7246 7261 1 1.1110e+02 5.2718e+02 ; CB - CA
- C
7248 7251 7252 1 1.0950e+02 4.1840e+02 ; CB - CG
- HG

```
  7248   7251   7253      1    1.0950e+02    3.3472e+02 ;     CB - CG
- CD1
  7248   7251   7257      1    1.0950e+02    3.3472e+02 ;     CB - CG
- CD2
  7249   7248   7250      1    1.0950e+02    2.9288e+02 ;    HB2 - CB
- HB3
  7249   7248   7251      1    1.0950e+02    4.1840e+02 ;    HB2 - CB
- CG
  7250   7248   7251      1    1.0950e+02    4.1840e+02 ;    HB3 - CB
- CG
  7251   7253   7254      1    1.0950e+02    4.1840e+02 ;     CG - CD1
- HD11
  7251   7253   7255      1    1.0950e+02    4.1840e+02 ;     CG - CD1
- HD12
  7251   7253   7256      1    1.0950e+02    4.1840e+02 ;     CG - CD1
- HD13
  7251   7257   7258      1    1.0950e+02    4.1840e+02 ;     CG - CD2
- HD21
  7251   7257   7259      1    1.0950e+02    4.1840e+02 ;     CG - CD2
- HD22
  7251   7257   7260      1    1.0950e+02    4.1840e+02 ;     CG - CD2
- HD23
  7252   7251   7253      1    1.0950e+02    4.1840e+02 ;     HG - CG
- CD1
  7252   7251   7257      1    1.0950e+02    4.1840e+02 ;     HG - CG
- CD2
  7253   7251   7257      1    1.0950e+02    3.3472e+02 ;    CD1 - CG
- CD2
  7254   7253   7255      1    1.0950e+02    2.9288e+02 ;   HD11 - CD1
- HD12
  7254   7253   7256      1    1.0950e+02    2.9288e+02 ;   HD11 - CD1
- HD13
  7255   7253   7256      1    1.0950e+02    2.9288e+02 ;   HD12 - CD1
- HD13
  7258   7257   7259      1    1.0950e+02    2.9288e+02 ;   HD21 - CD2
- HD22
  7258   7257   7260      1    1.0950e+02    2.9288e+02 ;   HD21 - CD2
- HD23
  7259   7257   7260      1    1.0950e+02    2.9288e+02 ;   HD22 - CD2
- HD23
  7261   7263   7264      1    1.2000e+02    4.1840e+02 ;      C - N
- H
  7261   7263   7265      1    1.2190e+02    4.1840e+02 ;      C - N
- CA
  7262   7261   7263      1    1.2290e+02    6.6944e+02 ;      O - C
- N
  7263   7265   7266      1    1.0950e+02    4.1840e+02 ;      N - CA
- HA
  7263   7265   7267      1    1.0970e+02    6.6944e+02 ;      N - CA
- CB
  7263   7265   7278      1    1.1010e+02    5.2718e+02 ;      N - CA
- C
```

7264 7263 7265 1 1.1804e+02 4.1840e+02 ; H - N
- CA
7265 7267 7268 1 1.0950e+02 4.1840e+02 ; CA - CB
- HB2
7265 7267 7269 1 1.0950e+02 4.1840e+02 ; CA - CB
- HB3
7265 7267 7270 1 1.0950e+02 3.3472e+02 ; CA - CB
- CG
7265 7278 7279 1 1.2040e+02 6.6944e+02 ; CA - C
- O
7265 7278 7280 1 1.1660e+02 5.8576e+02 ; CA - C
- N
7266 7265 7267 1 1.0950e+02 4.1840e+02 ; HA - CA
- CB
7266 7265 7278 1 1.0950e+02 4.1840e+02 ; HA - CA
- C
7267 7265 7278 1 1.1110e+02 5.2718e+02 ; CB - CA
- C
7267 7270 7271 1 1.0950e+02 4.1840e+02 ; CB - CG
- HG2
7267 7270 7272 1 1.0950e+02 4.1840e+02 ; CB - CG
- HG3
7267 7270 7273 1 1.1470e+02 4.1840e+02 ; CB - CG
- SD
7268 7267 7269 1 1.0950e+02 2.9288e+02 ; HB2 - CB
- HB3
7268 7267 7270 1 1.0950e+02 4.1840e+02 ; HB2 - CB
- CG
7269 7267 7270 1 1.0950e+02 4.1840e+02 ; HB3 - CB
- CG
7270 7273 7274 1 9.8900e+01 5.1882e+02 ; CG - SD
- CE
7271 7270 7272 1 1.0950e+02 2.9288e+02 ; HG2 - CG
- HG3
7271 7270 7273 1 1.0950e+02 4.1840e+02 ; HG2 - CG
- SD
7272 7270 7273 1 1.0950e+02 4.1840e+02 ; HG3 - CG
- SD
7273 7274 7275 1 1.0950e+02 4.1840e+02 ; SD - CE
- HE1
7273 7274 7276 1 1.0950e+02 4.1840e+02 ; SD - CE
- HE2
7273 7274 7277 1 1.0950e+02 4.1840e+02 ; SD - CE
- HE3
7275 7274 7276 1 1.0950e+02 2.9288e+02 ; HE1 - CE
- HE2
7275 7274 7277 1 1.0950e+02 2.9288e+02 ; HE1 - CE
- HE3
7276 7274 7277 1 1.0950e+02 2.9288e+02 ; HE2 - CE
- HE3
7278 7280 7281 1 1.2000e+02 4.1840e+02 ; C - N
- H

7278 7280 7282 1 1.2190e+02 4.1840e+02 ; C - N
- CA
7279 7278 7280 1 1.2290e+02 6.6944e+02 ; O - C
- N
7280 7282 7283 1 1.0950e+02 4.1840e+02 ; N - CA
- HA
7280 7282 7284 1 1.0970e+02 6.6944e+02 ; N - CA
- CB
7280 7282 7295 1 1.1010e+02 5.2718e+02 ; N - CA
- C
7281 7280 7282 1 1.1804e+02 4.1840e+02 ; H - N
- CA
7282 7284 7285 1 1.0950e+02 4.1840e+02 ; CA - CB
- HB2
7282 7284 7286 1 1.0950e+02 4.1840e+02 ; CA - CB
- HB3
7282 7284 7287 1 1.0950e+02 3.3472e+02 ; CA - CB
- CG
7282 7295 7296 1 1.2040e+02 6.6944e+02 ; CA - C
- O
7282 7295 7297 1 1.1660e+02 5.8576e+02 ; CA - C
- N
7283 7282 7284 1 1.0950e+02 4.1840e+02 ; HA - CA
- CB
7283 7282 7295 1 1.0950e+02 4.1840e+02 ; HA - CA
- C
7284 7282 7295 1 1.1110e+02 5.2718e+02 ; CB - CA
- C
7284 7287 7288 1 1.0950e+02 4.1840e+02 ; CB - CG
- HG2
7284 7287 7289 1 1.0950e+02 4.1840e+02 ; CB - CG
- HG3
7284 7287 7290 1 1.1470e+02 4.1840e+02 ; CB - CG
- SD
7285 7284 7286 1 1.0950e+02 2.9288e+02 ; HB2 - CB
- HB3
7285 7284 7287 1 1.0950e+02 4.1840e+02 ; HB2 - CB
- CG
7286 7284 7287 1 1.0950e+02 4.1840e+02 ; HB3 - CB
- CG
7287 7290 7291 1 9.8900e+01 5.1882e+02 ; CG - SD
- CE
7288 7287 7289 1 1.0950e+02 2.9288e+02 ; HG2 - CG
- HG3
7288 7287 7290 1 1.0950e+02 4.1840e+02 ; HG2 - CG
- SD
7289 7287 7290 1 1.0950e+02 4.1840e+02 ; HG3 - CG
- SD
7290 7291 7292 1 1.0950e+02 4.1840e+02 ; SD - CE
- HE1
7290 7291 7293 1 1.0950e+02 4.1840e+02 ; SD - CE
- HE2

7290 7291 7294 1 1.0950e+02 4.1840e+02 ; SD - CE
- HE3
7292 7291 7293 1 1.0950e+02 2.9288e+02 ; HE1 - CE
- HE2
7292 7291 7294 1 1.0950e+02 2.9288e+02 ; HE1 - CE
- HE3
7293 7291 7294 1 1.0950e+02 2.9288e+02 ; HE2 - CE
- HE3
7295 7297 7298 1 1.2000e+02 4.1840e+02 ; C - N
- H
7295 7297 7299 1 1.2190e+02 4.1840e+02 ; C - N
- CA
7296 7295 7297 1 1.2290e+02 6.6944e+02 ; O - C
- N
7297 7299 7300 1 1.0950e+02 4.1840e+02 ; N - CA
- HA
7297 7299 7301 1 1.0970e+02 6.6944e+02 ; N - CA
- CB
7297 7299 7306 1 1.1010e+02 5.2718e+02 ; N - CA
- C
7298 7297 7299 1 1.1804e+02 4.1840e+02 ; H - N
- CA
7299 7301 7302 1 1.0950e+02 4.1840e+02 ; CA - CB
- HB2
7299 7301 7303 1 1.0950e+02 4.1840e+02 ; CA - CB
- HB3
7299 7301 7304 1 1.0950e+02 4.1840e+02 ; CA - CB
- OG
7299 7306 7307 1 1.2040e+02 6.6944e+02 ; CA - C
- O
7299 7306 7308 1 1.1660e+02 5.8576e+02 ; CA - C
- N
7300 7299 7301 1 1.0950e+02 4.1840e+02 ; HA - CA
- CB
7300 7299 7306 1 1.0950e+02 4.1840e+02 ; HA - CA
- C
7301 7299 7306 1 1.1110e+02 5.2718e+02 ; CB - CA
- C
7301 7304 7305 1 1.0850e+02 4.6024e+02 ; CB - OG
- HG
7302 7301 7303 1 1.0950e+02 2.9288e+02 ; HB2 - CB
- HB3
7302 7301 7304 1 1.0950e+02 4.1840e+02 ; HB2 - CB
- OG
7303 7301 7304 1 1.0950e+02 4.1840e+02 ; HB3 - CB
- OG
7306 7308 7309 1 1.2000e+02 4.1840e+02 ; C - N
- H
7306 7308 7310 1 1.2190e+02 4.1840e+02 ; C - N
- CA
7307 7306 7308 1 1.2290e+02 6.6944e+02 ; O - C
- N

7308 7310 7311 1 1.0950e+02 4.1840e+02 ; N - CA
- HA
7308 7310 7312 1 1.0970e+02 6.6944e+02 ; N - CA
- CB
7308 7310 7328 1 1.1010e+02 5.2718e+02 ; N - CA
- C
7309 7308 7310 1 1.1804e+02 4.1840e+02 ; H - N
- CA
7310 7312 7313 1 1.0950e+02 4.1840e+02 ; CA - CB
- HB2
7310 7312 7314 1 1.0950e+02 4.1840e+02 ; CA - CB
- HB3
7310 7312 7315 1 1.0950e+02 3.3472e+02 ; CA - CB
- CG
7310 7328 7329 1 1.2040e+02 6.6944e+02 ; CA - C
- O
7310 7328 7330 1 1.1660e+02 5.8576e+02 ; CA - C
- N
7311 7310 7312 1 1.0950e+02 4.1840e+02 ; HA - CA
- CB
7311 7310 7328 1 1.0950e+02 4.1840e+02 ; HA - CA
- C
7312 7310 7328 1 1.1110e+02 5.2718e+02 ; CB - CA
- C
7312 7315 7316 1 1.0950e+02 4.1840e+02 ; CB - CG
- HG2
7312 7315 7317 1 1.0950e+02 4.1840e+02 ; CB - CG
- HG3
7312 7315 7318 1 1.0950e+02 3.3472e+02 ; CB - CG
- CD
7313 7312 7314 1 1.0950e+02 2.9288e+02 ; HB2 - CB
- HB3
7313 7312 7315 1 1.0950e+02 4.1840e+02 ; HB2 - CB
- CG
7314 7312 7315 1 1.0950e+02 4.1840e+02 ; HB3 - CB
- CG
7315 7318 7319 1 1.0950e+02 4.1840e+02 ; CG - CD
- HD2
7315 7318 7320 1 1.0950e+02 4.1840e+02 ; CG - CD
- HD3
7315 7318 7321 1 1.0950e+02 3.3472e+02 ; CG - CD
- CE
7316 7315 7317 1 1.0950e+02 2.9288e+02 ; HG2 - CG
- HG3
7316 7315 7318 1 1.0950e+02 4.1840e+02 ; HG2 - CG
- CD
7317 7315 7318 1 1.0950e+02 4.1840e+02 ; HG3 - CG
- CD
7318 7321 7322 1 1.0950e+02 4.1840e+02 ; CD - CE
- HE2
7318 7321 7323 1 1.0950e+02 4.1840e+02 ; CD - CE
- HE3

7318 7321 7324 1 1.1120e+02 6.6944e+02 ; CD - CE
- NZ
7319 7318 7320 1 1.0950e+02 2.9288e+02 ; HD2 - CD
- HD3
7319 7318 7321 1 1.0950e+02 4.1840e+02 ; HD2 - CD
- CE
7320 7318 7321 1 1.0950e+02 4.1840e+02 ; HD3 - CD
- CE
7321 7324 7325 1 1.0950e+02 4.1840e+02 ; CE - NZ
- HZ1
7321 7324 7326 1 1.0950e+02 4.1840e+02 ; CE - NZ
- HZ2
7321 7324 7327 1 1.0950e+02 4.1840e+02 ; CE - NZ
- HZ3
7322 7321 7323 1 1.0950e+02 2.9288e+02 ; HE2 - CE
- HE3
7322 7321 7324 1 1.0950e+02 4.1840e+02 ; HE2 - CE
- NZ
7323 7321 7324 1 1.0950e+02 4.1840e+02 ; HE3 - CE
- NZ
7325 7324 7326 1 1.0950e+02 2.9288e+02 ; HZ1 - NZ
- HZ2
7325 7324 7327 1 1.0950e+02 2.9288e+02 ; HZ1 - NZ
- HZ3
7326 7324 7327 1 1.0950e+02 2.9288e+02 ; HZ2 - NZ
- HZ3
7328 7330 7331 1 1.2000e+02 4.1840e+02 ; C - N
- H
7328 7330 7332 1 1.2190e+02 4.1840e+02 ; C - N
- CA
7329 7328 7330 1 1.2290e+02 6.6944e+02 ; O - C
- N
7330 7332 7333 1 1.0950e+02 4.1840e+02 ; N - CA
- HA
7330 7332 7334 1 1.0970e+02 6.6944e+02 ; N - CA
- CB
7330 7332 7350 1 1.1010e+02 5.2718e+02 ; N - CA
- C
7331 7330 7332 1 1.1804e+02 4.1840e+02 ; H - N
- CA
7332 7334 7335 1 1.0950e+02 4.1840e+02 ; CA - CB
- HB2
7332 7334 7336 1 1.0950e+02 4.1840e+02 ; CA - CB
- HB3
7332 7334 7337 1 1.0950e+02 3.3472e+02 ; CA - CB
- CG
7332 7350 7351 1 1.2040e+02 6.6944e+02 ; CA - C
- O
7332 7350 7352 1 1.1660e+02 5.8576e+02 ; CA - C
- N
7333 7332 7334 1 1.0950e+02 4.1840e+02 ; HA - CA
- CB

7333 7332 7350 1 1.0950e+02 4.1840e+02 ; HA - CA
- C
7334 7332 7350 1 1.1110e+02 5.2718e+02 ; CB - CA
- C
7334 7337 7338 1 1.0950e+02 4.1840e+02 ; CB - CG
- HG2
7334 7337 7339 1 1.0950e+02 4.1840e+02 ; CB - CG
- HG3
7334 7337 7340 1 1.0950e+02 3.3472e+02 ; CB - CG
- CD
7335 7334 7336 1 1.0950e+02 2.9288e+02 ; HB2 - CB
- HB3
7335 7334 7337 1 1.0950e+02 4.1840e+02 ; HB2 - CB
- CG
7336 7334 7337 1 1.0950e+02 4.1840e+02 ; HB3 - CB
- CG
7337 7340 7341 1 1.0950e+02 4.1840e+02 ; CG - CD
- HD2
7337 7340 7342 1 1.0950e+02 4.1840e+02 ; CG - CD
- HD3
7337 7340 7343 1 1.0950e+02 3.3472e+02 ; CG - CD
- CE
7338 7337 7339 1 1.0950e+02 2.9288e+02 ; HG2 - CG
- HG3
7338 7337 7340 1 1.0950e+02 4.1840e+02 ; HG2 - CG
- CD
7339 7337 7340 1 1.0950e+02 4.1840e+02 ; HG3 - CG
- CD
7340 7343 7344 1 1.0950e+02 4.1840e+02 ; CD - CE
- HE2
7340 7343 7345 1 1.0950e+02 4.1840e+02 ; CD - CE
- HE3
7340 7343 7346 1 1.1120e+02 6.6944e+02 ; CD - CE
- NZ
7341 7340 7342 1 1.0950e+02 2.9288e+02 ; HD2 - CD
- HD3
7341 7340 7343 1 1.0950e+02 4.1840e+02 ; HD2 - CD
- CE
7342 7340 7343 1 1.0950e+02 4.1840e+02 ; HD3 - CD
- CE
7343 7346 7347 1 1.0950e+02 4.1840e+02 ; CE - NZ
- HZ1
7343 7346 7348 1 1.0950e+02 4.1840e+02 ; CE - NZ
- HZ2
7343 7346 7349 1 1.0950e+02 4.1840e+02 ; CE - NZ
- HZ3
7344 7343 7345 1 1.0950e+02 2.9288e+02 ; HE2 - CE
- HE3
7344 7343 7346 1 1.0950e+02 4.1840e+02 ; HE2 - CE
- NZ
7345 7343 7346 1 1.0950e+02 4.1840e+02 ; HE3 - CE
- NZ

```
  7347   7346   7348      1    1.0950e+02    2.9288e+02 ;    HZ1 - NZ
- HZ2
  7347   7346   7349      1    1.0950e+02    2.9288e+02 ;    HZ1 - NZ
- HZ3
  7348   7346   7349      1    1.0950e+02    2.9288e+02 ;    HZ2 - NZ
- HZ3
  7350   7352   7353      1    1.2000e+02    4.1840e+02 ;      C - N
- H
  7350   7352   7354      1    1.2190e+02    4.1840e+02 ;      C - N
- CA
  7351   7350   7352      1    1.2290e+02    6.6944e+02 ;      O - C
- N
  7352   7354   7355      1    1.0950e+02    4.1840e+02 ;      N - CA
- HA
  7352   7354   7356      1    1.0970e+02    6.6944e+02 ;      N - CA
- CB
  7352   7354   7372      1    1.1010e+02    5.2718e+02 ;      N - CA
- C
  7353   7352   7354      1    1.1804e+02    4.1840e+02 ;      H - N
- CA
  7354   7356   7357      1    1.0950e+02    4.1840e+02 ;     CA - CB
- HB2
  7354   7356   7358      1    1.0950e+02    4.1840e+02 ;     CA - CB
- HB3
  7354   7356   7359      1    1.0950e+02    3.3472e+02 ;     CA - CB
- CG
  7354   7372   7373      1    1.1700e+02    5.8576e+02 ;     CA - C
- O
  7354   7372   7374      1    1.1700e+02    5.8576e+02 ;     CA - C
- OXT
  7355   7354   7356      1    1.0950e+02    4.1840e+02 ;     HA - CA
- CB
  7355   7354   7372      1    1.0950e+02    4.1840e+02 ;     HA - CA
- C
  7356   7354   7372      1    1.1110e+02    5.2718e+02 ;     CB - CA
- C
  7356   7359   7360      1    1.0950e+02    4.1840e+02 ;     CB - CG
- HG2
  7356   7359   7361      1    1.0950e+02    4.1840e+02 ;     CB - CG
- HG3
  7356   7359   7362      1    1.0950e+02    3.3472e+02 ;     CB - CG
- CD
  7357   7356   7358      1    1.0950e+02    2.9288e+02 ;    HB2 - CB
- HB3
  7357   7356   7359      1    1.0950e+02    4.1840e+02 ;    HB2 - CB
- CG
  7358   7356   7359      1    1.0950e+02    4.1840e+02 ;    HB3 - CB
- CG
  7359   7362   7363      1    1.0950e+02    4.1840e+02 ;     CG - CD
- HD2
  7359   7362   7364      1    1.0950e+02    4.1840e+02 ;     CG - CD
- HD3
```

7359 7362 7365 1 1.0950e+02 3.3472e+02 ; CG - CD
- CE
7360 7359 7361 1 1.0950e+02 2.9288e+02 ; HG2 - CG
- HG3
7360 7359 7362 1 1.0950e+02 4.1840e+02 ; HG2 - CG
- CD
7361 7359 7362 1 1.0950e+02 4.1840e+02 ; HG3 - CG
- CD
7362 7365 7366 1 1.0950e+02 4.1840e+02 ; CD - CE
- HE2
7362 7365 7367 1 1.0950e+02 4.1840e+02 ; CD - CE
- HE3
7362 7365 7368 1 1.1120e+02 6.6944e+02 ; CD - CE
- NZ
7363 7362 7364 1 1.0950e+02 2.9288e+02 ; HD2 - CD
- HD3
7363 7362 7365 1 1.0950e+02 4.1840e+02 ; HD2 - CD
- CE
7364 7362 7365 1 1.0950e+02 4.1840e+02 ; HD3 - CD
- CE
7365 7368 7369 1 1.0950e+02 4.1840e+02 ; CE - NZ
- HZ1
7365 7368 7370 1 1.0950e+02 4.1840e+02 ; CE - NZ
- HZ2
7365 7368 7371 1 1.0950e+02 4.1840e+02 ; CE - NZ
- HZ3
7366 7365 7367 1 1.0950e+02 2.9288e+02 ; HE2 - CE
- HE3
7366 7365 7368 1 1.0950e+02 4.1840e+02 ; HE2 - CE
- NZ
7367 7365 7368 1 1.0950e+02 4.1840e+02 ; HE3 - CE
- NZ
7369 7368 7370 1 1.0950e+02 2.9288e+02 ; HZ1 - NZ
- HZ2
7369 7368 7371 1 1.0950e+02 2.9288e+02 ; HZ1 - NZ
- HZ3
7370 7368 7371 1 1.0950e+02 2.9288e+02 ; HZ2 - NZ
- HZ3
7373 7372 7374 1 1.2600e+02 6.6944e+02 ; O - C
- OXT
7375 7376 7377 1 1.2287e+02 5.6819e+02 ; F2 - C38
- C39
7375 7376 7383 1 1.2287e+02 5.6819e+02 ; F2 - C38
- C37
7376 7377 7378 1 1.2601e+02 5.9580e+02 ; C38 - C39
- N23
7376 7377 7423 1 1.2267e+02 4.1338e+02 ; C38 - C39
- H39
7376 7383 7384 1 1.1700e+02 5.8074e+02 ; C38 - C37
- C36
7376 7383 7424 1 1.2043e+02 4.1756e+02 ; C38 - C37
- H37

| | | | | | | | |
|---|---|---|---|---|---|---|---|
| 7377 | 7376 | 7383 | 1 | 1.2181e+02 | 5.7990e+02 | ; | C39 - C38 - C37 |
| 7377 | 7378 | 7385 | 1 | 1.1994e+02 | 5.8576e+02 | ; | C39 - N23 - C40 |
| 7377 | 7378 | 7433 | 1 | 1.3089e+02 | 7.6467e+02 | ; | C39 - N23 - RU |
| 7378 | 7377 | 7423 | 1 | 1.2099e+02 | 4.3932e+02 | ; | N23 - C39 - H39 |
| 7378 | 7385 | 7384 | 1 | 1.1957e+02 | 5.9078e+02 | ; | N23 - C40 - C36 |
| 7378 | 7385 | 7395 | 1 | 1.1957e+02 | 5.9078e+02 | ; | N23 - C40 - C24 |
| 7378 | 7433 | 7379 | 1 | 8.0080e+01 | 1.5944e+03 | ; | N23 - RU - N21 |
| 7378 | 7433 | 7380 | 1 | 9.4800e+01 | 1.1312e+03 | ; | N23 - RU - C13 |
| 7378 | 7433 | 7382 | 1 | 1.7362e+02 | 6.5655e+02 | ; | N23 - RU - CL17 |
| 7378 | 7433 | 7402 | 1 | 9.7200e+01 | 1.5943e+03 | ; | N23 - RU - N15 |
| 7379 | 7395 | 7385 | 1 | 1.1957e+02 | 5.9078e+02 | ; | N21 - C24 - C40 |
| 7379 | 7395 | 7394 | 1 | 1.1957e+02 | 5.9078e+02 | ; | N21 - C24 - C31 |
| 7379 | 7397 | 7396 | 1 | 1.1957e+02 | 5.9078e+02 | ; | N21 - C25 - C30 |
| 7379 | 7397 | 7401 | 1 | 1.1957e+02 | 5.9078e+02 | ; | N21 - C25 - C26 |
| 7379 | 7433 | 7380 | 1 | 9.0220e+01 | 1.1077e+03 | ; | N21 - RU - C13 |
| 7379 | 7433 | 7382 | 1 | 9.4710e+01 | 7.7822e+02 | ; | N21 - RU - CL17 |
| 7379 | 7433 | 7402 | 1 | 1.7121e+02 | 1.5575e+03 | ; | N21 - RU - N15 |
| 7380 | 7433 | 7382 | 1 | 8.8880e+01 | 1.1400e+03 | ; | C13 - RU - CL17 |
| 7380 | 7433 | 7402 | 1 | 9.8350e+01 | 1.7278e+03 | ; | C13 - RU - N15 |
| 7381 | 7380 | 7433 | 1 | 1.7733e+02 | 7.1546e+02 | ; | O14 - C13 - RU |
| 7382 | 7433 | 7402 | 1 | 8.7400e+01 | 8.3513e+02 | ; | CL17 - RU - N15 |
| 7383 | 7384 | 7385 | 1 | 1.2060e+02 | 5.5898e+02 | ; | C37 - C36 - C40 |
| 7383 | 7384 | 7386 | 1 | 1.2060e+02 | 5.5898e+02 | ; | C37 - C36 - C35 |
| 7384 | 7383 | 7424 | 1 | 1.2330e+02 | 4.0083e+02 | ; | C36 - C37 - H37 |
| 7384 | 7385 | 7395 | 1 | 1.2002e+02 | 5.5731e+02 | ; | C36 - C40 - C24 |
| 7384 | 7386 | 7387 | 1 | 1.2060e+02 | 5.5898e+02 | ; | C36 - C35 - C34 |

7384 7386 7391 1 1.2002e+02 5.5731e+02 ; C36 - C35 - C32
7385 7378 7433 1 1.1072e+02 8.2492e+02 ; C40 - N23 - RU
7385 7384 7386 1 1.2002e+02 5.5731e+02 ; C40 - C36 - C35
7385 7395 7394 1 1.2069e+02 5.5480e+02 ; C40 - C24 - C31
7386 7387 7388 1 1.2367e+02 5.7488e+02 ; C35 - C34 - N22
7386 7387 7390 1 1.2359e+02 6.0919e+02 ; C35 - C34 - O42
7386 7391 7392 1 1.2060e+02 5.5898e+02 ; C35 - C32 - C33
7386 7391 7394 1 1.2069e+02 5.5480e+02 ; C35 - C32 - C31
7387 7386 7391 1 1.2060e+02 5.5898e+02 ; C34 - C35 - C32
7387 7388 7389 1 1.1790e+02 3.9832e+02 ; C34 - N22 - H22
7387 7388 7392 1 1.1675e+02 5.4810e+02 ; C34 - N22 - C33
7388 7387 7390 1 1.1746e+02 6.2843e+02 ; N22 - C34 - O42
7388 7392 7391 1 1.2367e+02 5.7488e+02 ; N22 - C33 - C32
7388 7392 7393 1 1.1746e+02 6.2843e+02 ; N22 - C33 - O41
7389 7388 7392 1 1.1790e+02 3.9832e+02 ; H22 - N22 - C33
7391 7392 7393 1 1.2359e+02 6.0919e+02 ; C32 - C33 - O41
7391 7394 7395 1 1.1838e+02 5.5815e+02 ; C32 - C31 - C24
7391 7394 7396 1 1.2111e+02 5.3555e+02 ; C32 - C31 - C30
7392 7391 7394 1 1.2000e+02 5.6317e+02 ; C33 - C32 - C31
7394 7396 7397 1 1.2111e+02 5.3555e+02 ; C31 - C30 - C25
7394 7396 7398 1 1.2111e+02 5.3555e+02 ; C31 - C30 - C29
7395 7379 7397 1 1.1220e+02 6.0082e+02 ; C24 - N21 - C25
7395 7379 7433 1 1.7519e+02 8.8952e+01 ; C24 - N21 - RU
7395 7394 7396 1 1.2111e+02 5.3555e+02 ; C24 - C31 - C30
7396 7397 7401 1 1.2069e+02 5.5480e+02 ; C30 - C25 - C26
7396 7398 7399 1 1.2069e+02 5.5480e+02 ; C30 - C29 - C28

7396 7398 7426 1 1.1986e+02 4.0166e+02 ; C30 - C29
- H29
7397 7379 7433 1 1.7519e+02 8.8952e+01 ; C25 - N21
- RU
7397 7396 7398 1 1.1838e+02 5.5815e+02 ; C25 - C30
- C29
7397 7401 7400 1 1.2002e+02 5.5731e+02 ; C25 - C26
- C27
7397 7401 7428 1 1.1988e+02 4.0334e+02 ; C25 - C26
- H26
7398 7399 7400 1 1.2002e+02 5.5731e+02 ; C29 - C28
- C27
7398 7399 7416 1 1.1990e+02 5.8158e+02 ; C29 - C28
- O1
7399 7398 7426 1 1.1988e+02 4.0334e+02 ; C28 - C29
- H29
7399 7400 7401 1 1.2002e+02 5.5731e+02 ; C28 - C27
- C26
7399 7400 7427 1 1.1988e+02 4.0334e+02 ; C28 - C27
- H27
7399 7416 7417 1 1.0858e+02 4.1003e+02 ; C28 - O1
- H1
7400 7399 7416 1 1.1990e+02 5.8158e+02 ; C27 - C28
- O1
7400 7401 7428 1 1.1988e+02 4.0334e+02 ; C27 - C26
- H26
7401 7400 7427 1 1.1988e+02 4.0334e+02 ; C26 - C27
- H27
7402 7403 7410 1 1.2601e+02 5.9580e+02 ; N15 - C7
- C8
7402 7403 7419 1 1.2099e+02 4.3932e+02 ; N15 - C7
- H7A
7402 7404 7405 1 1.1957e+02 5.9078e+02 ; N15 - C18
- C5
7402 7404 7409 1 1.1957e+02 5.9078e+02 ; N15 - C18
- C19
7403 7402 7404 1 1.1994e+02 5.8576e+02 ; C7 - N15
- C18
7403 7402 7433 1 1.1544e+02 1.4209e+03 ; C7 - N15
- RU
7403 7410 7411 1 1.2181e+02 5.7990e+02 ; C7 - C8
- C9
7403 7410 7415 1 1.2601e+02 5.9580e+02 ; C7 - C8
- N16
7404 7402 7433 1 1.2478e+02 1.3001e+03 ; C18 - N15
- RU
7404 7405 7406 1 1.2002e+02 5.5731e+02 ; C18 - C5
- C6
7404 7405 7421 1 1.1988e+02 4.0334e+02 ; C18 - C5
- H5
7404 7409 7408 1 1.2002e+02 5.5731e+02 ; C18 - C19
- C3

7404 7409 7429 1 1.1988e+02 4.0334e+02 ; C18 - C19
- H19
7405 7404 7409 1 1.2002e+02 5.5731e+02 ; C5 - C18
- C19
7405 7406 7407 1 1.2002e+02 5.5731e+02 ; C5 - C6
- C4
7405 7406 7420 1 1.1988e+02 4.0334e+02 ; C5 - C6
- H6
7406 7405 7421 1 1.1988e+02 4.0334e+02 ; C6 - C5
- H5
7406 7407 7408 1 1.2002e+02 5.5731e+02 ; C6 - C4
- C3
7406 7407 7422 1 1.1988e+02 4.0334e+02 ; C6 - C4
- H4
7407 7406 7420 1 1.1988e+02 4.0334e+02 ; C4 - C6
- H6
7407 7408 7409 1 1.2002e+02 5.5731e+02 ; C4 - C3
- C19
7407 7408 7425 1 1.1988e+02 4.0334e+02 ; C4 - C3
- H3
7408 7407 7422 1 1.1988e+02 4.0334e+02 ; C3 - C4
- H4
7408 7409 7429 1 1.1988e+02 4.0334e+02 ; C3 - C19
- H19
7409 7408 7425 1 1.1988e+02 4.0334e+02 ; C19 - C3
- H3
7410 7403 7419 1 1.2267e+02 4.1338e+02 ; C8 - C7
- H7A
7410 7411 7412 1 1.2181e+02 5.7990e+02 ; C8 - C9
- C10
7410 7411 7418 1 1.2043e+02 4.1756e+02 ; C8 - C9
- H9
7410 7415 7414 1 1.1818e+02 5.9496e+02 ; C8 - N16
- C11
7411 7410 7415 1 1.2601e+02 5.9580e+02 ; C9 - C8
- N16
7411 7412 7413 1 1.2181e+02 5.7990e+02 ; C9 - C10
- C12
7411 7412 7432 1 1.2043e+02 4.1756e+02 ; C9 - C10
- H10
7412 7411 7418 1 1.2043e+02 4.1756e+02 ; C10 - C9
- H9
7412 7413 7414 1 1.2181e+02 5.7990e+02 ; C10 - C12
- C11
7412 7413 7430 1 1.2043e+02 4.1756e+02 ; C10 - C12
- H12
7413 7412 7432 1 1.2043e+02 4.1756e+02 ; C12 - C10
- H10
7413 7414 7415 1 1.2601e+02 5.9580e+02 ; C12 - C11
- N16
7413 7414 7431 1 1.2267e+02 4.1338e+02 ; C12 - C11
- H11

```
  7414   7413   7430      1    1.2043e+02    4.1756e+02 ;    C11 - C12
- H12
  7415   7414   7431      1    1.2099e+02    4.3932e+02 ;    N16 - C11
- H11

[ dihedrals ] ; propers
; treated as RBs in GROMACS to use combine multiple AMBER torsions per
quartet
;    i      j      k      l   func     C0          C1          C2
C3          C4          C5
     1      5      7      8      3    0.65084    1.95253    0.00000
-2.60338    0.00000    0.00000 ;      N-    CA-    CB-   HB2
     1      5      7      9      3    0.65084    1.95253    0.00000
-2.60338    0.00000    0.00000 ;      N-    CA-    CB-   HB3
     1      5      7     10      3    3.79070    1.39746   -4.61077
-2.40998    2.44346    0.00000 ;      N-    CA-    CB-    CG
     1      5     20     21      3    0.00000    0.00000    0.00000
0.00000    0.00000    0.00000 ;      N-    CA-     C-     O
     1      5     20     22      3    0.00000    0.00000    0.00000
0.00000    0.00000    0.00000 ;      N-    CA-     C-     N
     2      1      5      6      3    0.65084    1.95253    0.00000
-2.60338    0.00000    0.00000 ;     H1-     N-    CA-    HA
     2      1      5      7      3    0.65084    1.95253    0.00000
-2.60338    0.00000    0.00000 ;     H1-     N-    CA-    CB
     2      1      5     20      3    0.65084    1.95253    0.00000
-2.60338    0.00000    0.00000 ;     H1-     N-    CA-     C
     3      1      5      6      3    0.65084    1.95253    0.00000
-2.60338    0.00000    0.00000 ;     H2-     N-    CA-    HA
     3      1      5      7      3    0.65084    1.95253    0.00000
-2.60338    0.00000    0.00000 ;     H2-     N-    CA-    CB
     3      1      5     20      3    0.65084    1.95253    0.00000
-2.60338    0.00000    0.00000 ;     H2-     N-    CA-     C
     4      1      5      6      3    0.65084    1.95253    0.00000
-2.60338    0.00000    0.00000 ;     H3-     N-    CA-    HA
     4      1      5      7      3    0.65084    1.95253    0.00000
-2.60338    0.00000    0.00000 ;     H3-     N-    CA-    CB
     4      1      5     20      3    0.65084    1.95253    0.00000
-2.60338    0.00000    0.00000 ;     H3-     N-    CA-     C
     5      7     10     11      3    0.66944    2.00832    0.00000
-2.67776    0.00000    0.00000 ;     CA-    CB-    CG-    HG
     5      7     10     12      3    3.90367    0.19665   -6.21742
-2.37651    5.99149    0.00000 ;     CA-    CB-    CG-   CD1
     5      7     10     16      3    3.90367    0.19665   -6.21742
-2.37651    5.99149    0.00000 ;     CA-    CB-    CG-   CD2
     5     20     22     23      3   20.92000    0.00000  -20.92000
0.00000    0.00000    0.00000 ;     CA-     C-     N-     H
     5     20     22     24      3   20.92000    0.00000  -20.92000
0.00000    0.00000    0.00000 ;     CA-     C-     N-    CA
     6      5      7      8      3    0.65084    1.95253    0.00000
-2.60338    0.00000    0.00000 ;     HA-    CA-    CB-   HB2
     6      5      7      9      3    0.65084    1.95253    0.00000
-2.60338    0.00000    0.00000 ;     HA-    CA-    CB-   HB3
```

```
    6     5     7    10     3   0.65084   1.95253   0.00000
-2.60338   0.00000   0.00000 ;    HA-   CA-   CB-   CG
    6     5    20    21     3   0.00000   0.00000   0.00000
0.00000   0.00000   0.00000 ;    HA-   CA-    C-     O
    6     5    20    22     3   0.00000   0.00000   0.00000
0.00000   0.00000   0.00000 ;    HA-   CA-    C-     N
    7     5    20    21     3   0.00000   0.00000   0.00000
0.00000   0.00000   0.00000 ;    CB-   CA-    C-     O
    7     5    20    22     3   2.51040   4.18400   1.67360
-6.69440   0.00000   0.00000 ;    CB-   CA-    C-     N
    7    10    12    13     3   0.66944   2.00832   0.00000
-2.67776   0.00000   0.00000 ;    CB-   CG-  CD1-  HD11
    7    10    12    14     3   0.66944   2.00832   0.00000
-2.67776   0.00000   0.00000 ;    CB-   CG-  CD1-  HD12
    7    10    12    15     3   0.66944   2.00832   0.00000
-2.67776   0.00000   0.00000 ;    CB-   CG-  CD1-  HD13
    7    10    16    17     3   0.66944   2.00832   0.00000
-2.67776   0.00000   0.00000 ;    CB-   CG-  CD2-  HD21
    7    10    16    18     3   0.66944   2.00832   0.00000
-2.67776   0.00000   0.00000 ;    CB-   CG-  CD2-  HD22
    7    10    16    19     3   0.66944   2.00832   0.00000
-2.67776   0.00000   0.00000 ;    CB-   CG-  CD2-  HD23
    8     7     5    20     3   0.65084   1.95253   0.00000
-2.60338   0.00000   0.00000 ;   HB2-   CB-   CA-     C
    8     7    10    11     3   0.62760   1.88280   0.00000
-2.51040   0.00000   0.00000 ;   HB2-   CB-   CG-    HG
    8     7    10    12     3   0.66944   2.00832   0.00000
-2.67776   0.00000   0.00000 ;   HB2-   CB-   CG-   CD1
    8     7    10    16     3   0.66944   2.00832   0.00000
-2.67776   0.00000   0.00000 ;   HB2-   CB-   CG-   CD2
    9     7     5    20     3   0.65084   1.95253   0.00000
-2.60338   0.00000   0.00000 ;   HB3-   CB-   CA-     C
    9     7    10    11     3   0.62760   1.88280   0.00000
-2.51040   0.00000   0.00000 ;   HB3-   CB-   CG-    HG
    9     7    10    12     3   0.66944   2.00832   0.00000
-2.67776   0.00000   0.00000 ;   HB3-   CB-   CG-   CD1
    9     7    10    16     3   0.66944   2.00832   0.00000
-2.67776   0.00000   0.00000 ;   HB3-   CB-   CG-   CD2
   10     7     5    20     3  10.33448  -1.14642  -11.54784
-2.40998   6.35968   0.00000 ;    CG-   CB-   CA-     C
   11    10    12    13     3   0.62760   1.88280   0.00000
-2.51040   0.00000   0.00000 ;    HG-   CG-  CD1-  HD11
   11    10    12    14     3   0.62760   1.88280   0.00000
-2.51040   0.00000   0.00000 ;    HG-   CG-  CD1-  HD12
   11    10    12    15     3   0.62760   1.88280   0.00000
-2.51040   0.00000   0.00000 ;    HG-   CG-  CD1-  HD13
   11    10    16    17     3   0.62760   1.88280   0.00000
-2.51040   0.00000   0.00000 ;    HG-   CG-  CD2-  HD21
   11    10    16    18     3   0.62760   1.88280   0.00000
-2.51040   0.00000   0.00000 ;    HG-   CG-  CD2-  HD22
   11    10    16    19     3   0.62760   1.88280   0.00000
-2.51040   0.00000   0.00000 ;    HG-   CG-  CD2-  HD23
```

12 10 16 17 3 0.66944 2.00832 0.00000
-2.67776 0.00000 0.00000 ; CD1- CG- CD2- HD21
12 10 16 18 3 0.66944 2.00832 0.00000
-2.67776 0.00000 0.00000 ; CD1- CG- CD2- HD22
12 10 16 19 3 0.66944 2.00832 0.00000
-2.67776 0.00000 0.00000 ; CD1- CG- CD2- HD23
13 12 10 16 3 0.66944 2.00832 0.00000
-2.67776 0.00000 0.00000 ; HD11- CD1- CG- CD2
14 12 10 16 3 0.66944 2.00832 0.00000
-2.67776 0.00000 0.00000 ; HD12- CD1- CG- CD2
15 12 10 16 3 0.66944 2.00832 0.00000
-2.67776 0.00000 0.00000 ; HD13- CD1- CG- CD2
20 22 24 25 3 0.00000 0.00000 0.00000
0.00000 0.00000 0.00000 ; C- N- CA- HA
20 22 24 26 3 11.71520 1.67360 15.06240 -
13.38880 0.00000 0.00000 ; C- N- CA- CB
20 22 24 39 3 1.75728 5.27184 2.25936
-7.02912 0.00000 0.00000 ; C- N- CA- C
21 20 22 23 3 29.28800 -8.36800 -20.92000
0.00000 0.00000 0.00000 ; O- C- N- H
21 20 22 24 3 20.92000 0.00000 -20.92000
0.00000 0.00000 0.00000 ; O- C- N- CA
22 24 26 27 3 0.65084 1.95253 0.00000
-2.60338 0.00000 0.00000 ; N- CA- CB- HB
22 24 26 28 3 3.83673 0.44769 -1.77402
-2.47693 -0.03347 0.00000 ; N- CA- CB- CG2
22 24 26 32 3 2.97482 0.12134 2.04179
-1.89117 -3.24678 0.00000 ; N- CA- CB- CG1
22 24 39 40 3 0.00000 0.00000 0.00000
0.00000 0.00000 0.00000 ; N- CA- C- O
22 24 39 41 3 17.40544 -5.02080 -13.22144
9.20480 0.00000 0.00000 ; N- CA- C- N
23 22 24 25 3 0.00000 0.00000 0.00000
0.00000 0.00000 0.00000 ; H- N- CA- HA
23 22 24 26 3 0.00000 0.00000 0.00000
0.00000 0.00000 0.00000 ; H- N- CA- CB
23 22 24 39 3 0.00000 0.00000 0.00000
0.00000 0.00000 0.00000 ; H- N- CA- C
24 26 28 29 3 0.66944 2.00832 0.00000
-2.67776 0.00000 0.00000 ; CA- CB- CG2- HG21
24 26 28 30 3 0.66944 2.00832 0.00000
-2.67776 0.00000 0.00000 ; CA- CB- CG2- HG22
24 26 28 31 3 0.66944 2.00832 0.00000
-2.67776 0.00000 0.00000 ; CA- CB- CG2- HG23
24 26 32 33 3 0.66944 2.00832 0.00000
-2.67776 0.00000 0.00000 ; CA- CB- CG1- HG12
24 26 32 34 3 0.66944 2.00832 0.00000
-2.67776 0.00000 0.00000 ; CA- CB- CG1- HG13
24 26 32 35 3 4.24258 -0.52718 -7.25506
-1.79075 7.69856 0.00000 ; CA- CB- CG1- CD1
24 39 41 42 3 20.92000 0.00000 -20.92000
0.00000 0.00000 0.00000 ; CA- C- N- H

24 39 41 43 3 20.92000 0.00000 -20.92000
0.00000 0.00000 0.00000 ; CA- C- N- CA
25 24 26 27 3 0.65084 1.95253 0.00000
-2.60338 0.00000 0.00000 ; HA- CA- CB- HB
25 24 26 28 3 0.65084 1.95253 0.00000
-2.60338 0.00000 0.00000 ; HA- CA- CB- CG2
25 24 26 32 3 0.65084 1.95253 0.00000
-2.60338 0.00000 0.00000 ; HA- CA- CB- CG1
25 24 39 40 3 3.68192 -4.35136 0.00000
1.33888 0.00000 0.00000 ; HA- CA- C- O
25 24 39 41 3 0.00000 0.00000 0.00000
0.00000 0.00000 0.00000 ; HA- CA- C- N
26 24 39 40 3 0.00000 0.00000 0.00000
0.00000 0.00000 0.00000 ; CB- CA- C- O
26 24 39 41 3 2.51040 4.18400 1.67360
-6.69440 0.00000 0.00000 ; CB- CA- C- N
26 32 35 36 3 0.66944 2.00832 0.00000
-2.67776 0.00000 0.00000 ; CB- CG1- CD1- HD11
26 32 35 37 3 0.66944 2.00832 0.00000
-2.67776 0.00000 0.00000 ; CB- CG1- CD1- HD12
26 32 35 38 3 0.66944 2.00832 0.00000
-2.67776 0.00000 0.00000 ; CB- CG1- CD1- HD13
27 26 24 39 3 0.65084 1.95253 0.00000
-2.60338 0.00000 0.00000 ; HB- CB- CA- C
27 26 28 29 3 0.62760 1.88280 0.00000
-2.51040 0.00000 0.00000 ; HB- CB- CG2- HG21
27 26 28 30 3 0.62760 1.88280 0.00000
-2.51040 0.00000 0.00000 ; HB- CB- CG2- HG22
27 26 28 31 3 0.62760 1.88280 0.00000
-2.51040 0.00000 0.00000 ; HB- CB- CG2- HG23
27 26 32 33 3 0.62760 1.88280 0.00000
-2.51040 0.00000 0.00000 ; HB- CB- CG1- HG12
27 26 32 34 3 0.62760 1.88280 0.00000
-2.51040 0.00000 0.00000 ; HB- CB- CG1- HG13
27 26 32 35 3 0.66944 2.00832 0.00000
-2.67776 0.00000 0.00000 ; HB- CB- CG1- CD1
28 26 24 39 3 5.67350 3.55640 -6.16722
-2.47693 3.74886 0.00000 ; CG2- CB- CA- C
28 26 32 33 3 0.66944 2.00832 0.00000
-2.67776 0.00000 0.00000 ; CG2- CB- CG1- HG12
28 26 32 34 3 0.66944 2.00832 0.00000
-2.67776 0.00000 0.00000 ; CG2- CB- CG1- HG13
28 26 32 35 3 3.81162 0.49790 -8.14206
-1.79075 7.49773 0.00000 ; CG2- CB- CG1- CD1
29 28 26 32 3 0.66944 2.00832 0.00000
-2.67776 0.00000 0.00000 ; HG21- CG2- CB- CG1
30 28 26 32 3 0.66944 2.00832 0.00000
-2.67776 0.00000 0.00000 ; HG22- CG2- CB- CG1
31 28 26 32 3 0.66944 2.00832 0.00000
-2.67776 0.00000 0.00000 ; HG23- CG2- CB- CG1
32 26 24 39 3 8.26340 0.74057 -9.99976
-1.89117 3.84928 0.00000 ; CG1- CB- CA- C

33 32 35 36 3 0.62760 1.88280 0.00000
-2.51040 0.00000 0.00000 ; HG12- CG1- CD1- HD11
33 32 35 37 3 0.62760 1.88280 0.00000
-2.51040 0.00000 0.00000 ; HG12- CG1- CD1- HD12
33 32 35 38 3 0.62760 1.88280 0.00000
-2.51040 0.00000 0.00000 ; HG12- CG1- CD1- HD13
34 32 35 36 3 0.62760 1.88280 0.00000
-2.51040 0.00000 0.00000 ; HG13- CG1- CD1- HD11
34 32 35 37 3 0.62760 1.88280 0.00000
-2.51040 0.00000 0.00000 ; HG13- CG1- CD1- HD12
34 32 35 38 3 0.62760 1.88280 0.00000
-2.51040 0.00000 0.00000 ; HG13- CG1- CD1- HD13
39 41 43 44 3 0.00000 0.00000 0.00000
0.00000 0.00000 0.00000 ; C- N- CA- HA
39 41 43 45 3 11.71520 1.67360 15.06240 -
13.38880 0.00000 0.00000 ; C- N- CA- CB
39 41 43 58 3 1.75728 5.27184 2.25936
-7.02912 0.00000 0.00000 ; C- N- CA- C
40 39 41 42 3 29.28800 -8.36800 -20.92000
0.00000 0.00000 0.00000 ; O- C- N- H
40 39 41 43 3 20.92000 0.00000 -20.92000
0.00000 0.00000 0.00000 ; O- C- N- CA
41 43 45 46 3 0.65084 1.95253 0.00000
-2.60338 0.00000 0.00000 ; N- CA- CB- HB2
41 43 45 47 3 0.65084 1.95253 0.00000
-2.60338 0.00000 0.00000 ; N- CA- CB- HB3
41 43 45 48 3 3.79070 1.39746 -4.61077
-2.40998 2.44346 0.00000 ; N- CA- CB- CG
41 43 58 59 3 0.00000 0.00000 0.00000
0.00000 0.00000 0.00000 ; N- CA- C- O
41 43 58 60 3 17.40544 -5.02080 -13.22144
9.20480 0.00000 0.00000 ; N- CA- C- N
42 41 43 44 3 0.00000 0.00000 0.00000
0.00000 0.00000 0.00000 ; H- N- CA- HA
42 41 43 45 3 0.00000 0.00000 0.00000
0.00000 0.00000 0.00000 ; H- N- CA- CB
42 41 43 58 3 0.00000 0.00000 0.00000
0.00000 0.00000 0.00000 ; H- N- CA- C
43 45 48 49 3 0.66944 2.00832 0.00000
-2.67776 0.00000 0.00000 ; CA- CB- CG- HG
43 45 48 50 3 3.90367 0.19665 -6.21742
-2.37651 5.99149 0.00000 ; CA- CB- CG- CD1
43 45 48 54 3 3.90367 0.19665 -6.21742
-2.37651 5.99149 0.00000 ; CA- CB- CG- CD2
43 58 60 61 3 20.92000 0.00000 -20.92000
0.00000 0.00000 0.00000 ; CA- C- N- H
43 58 60 62 3 20.92000 0.00000 -20.92000
0.00000 0.00000 0.00000 ; CA- C- N- CA
44 43 45 46 3 0.65084 1.95253 0.00000
-2.60338 0.00000 0.00000 ; HA- CA- CB- HB2
44 43 45 47 3 0.65084 1.95253 0.00000
-2.60338 0.00000 0.00000 ; HA- CA- CB- HB3

44 43 45 48 3 0.65084 1.95253 0.00000
-2.60338 0.00000 0.00000 ; HA- CA- CB- CG
44 43 58 59 3 3.68192 -4.35136 0.00000
1.33888 0.00000 0.00000 ; HA- CA- C- O
44 43 58 60 3 0.00000 0.00000 0.00000
0.00000 0.00000 0.00000 ; HA- CA- C- N
45 43 58 59 3 0.00000 0.00000 0.00000
0.00000 0.00000 0.00000 ; CB- CA- C- O
45 43 58 60 3 2.51040 4.18400 1.67360
-6.69440 0.00000 0.00000 ; CB- CA- C- N
45 48 50 51 3 0.66944 2.00832 0.00000
-2.67776 0.00000 0.00000 ; CB- CG- CD1- HD11
45 48 50 52 3 0.66944 2.00832 0.00000
-2.67776 0.00000 0.00000 ; CB- CG- CD1- HD12
45 48 50 53 3 0.66944 2.00832 0.00000
-2.67776 0.00000 0.00000 ; CB- CG- CD1- HD13
45 48 54 55 3 0.66944 2.00832 0.00000
-2.67776 0.00000 0.00000 ; CB- CG- CD2- HD21
45 48 54 56 3 0.66944 2.00832 0.00000
-2.67776 0.00000 0.00000 ; CB- CG- CD2- HD22
45 48 54 57 3 0.66944 2.00832 0.00000
-2.67776 0.00000 0.00000 ; CB- CG- CD2- HD23
46 45 43 58 3 0.65084 1.95253 0.00000
-2.60338 0.00000 0.00000 ; HB2- CB- CA- C
46 45 48 49 3 0.62760 1.88280 0.00000
-2.51040 0.00000 0.00000 ; HB2- CB- CG- HG
46 45 48 50 3 0.66944 2.00832 0.00000
-2.67776 0.00000 0.00000 ; HB2- CB- CG- CD1
46 45 48 54 3 0.66944 2.00832 0.00000
-2.67776 0.00000 0.00000 ; HB2- CB- CG- CD2
47 45 43 58 3 0.65084 1.95253 0.00000
-2.60338 0.00000 0.00000 ; HB3- CB- CA- C
47 45 48 49 3 0.62760 1.88280 0.00000
-2.51040 0.00000 0.00000 ; HB3- CB- CG- HG
47 45 48 50 3 0.66944 2.00832 0.00000
-2.67776 0.00000 0.00000 ; HB3- CB- CG- CD1
47 45 48 54 3 0.66944 2.00832 0.00000
-2.67776 0.00000 0.00000 ; HB3- CB- CG- CD2
48 45 43 58 3 10.33448 -1.14642 -11.54784
-2.40998 6.35968 0.00000 ; CG- CB- CA- C
49 48 50 51 3 0.62760 1.88280 0.00000
-2.51040 0.00000 0.00000 ; HG- CG- CD1- HD11
49 48 50 52 3 0.62760 1.88280 0.00000
-2.51040 0.00000 0.00000 ; HG- CG- CD1- HD12
49 48 50 53 3 0.62760 1.88280 0.00000
-2.51040 0.00000 0.00000 ; HG- CG- CD1- HD13
49 48 54 55 3 0.62760 1.88280 0.00000
-2.51040 0.00000 0.00000 ; HG- CG- CD2- HD21
49 48 54 56 3 0.62760 1.88280 0.00000
-2.51040 0.00000 0.00000 ; HG- CG- CD2- HD22
49 48 54 57 3 0.62760 1.88280 0.00000
-2.51040 0.00000 0.00000 ; HG- CG- CD2- HD23

50 48 54 55 3 0.66944 2.00832 0.00000
-2.67776 0.00000 0.00000 ; CD1- CG- CD2- HD21
50 48 54 56 3 0.66944 2.00832 0.00000
-2.67776 0.00000 0.00000 ; CD1- CG- CD2- HD22
50 48 54 57 3 0.66944 2.00832 0.00000
-2.67776 0.00000 0.00000 ; CD1- CG- CD2- HD23
51 50 48 54 3 0.66944 2.00832 0.00000
-2.67776 0.00000 0.00000 ; HD11- CD1- CG- CD2
52 50 48 54 3 0.66944 2.00832 0.00000
-2.67776 0.00000 0.00000 ; HD12- CD1- CG- CD2
53 50 48 54 3 0.66944 2.00832 0.00000
-2.67776 0.00000 0.00000 ; HD13- CD1- CG- CD2
58 60 62 63 3 0.00000 0.00000 0.00000
0.00000 0.00000 0.00000 ; C- N- CA- HA
58 60 62 64 3 11.71520 1.67360 15.06240 -
13.38880 0.00000 0.00000 ; C- N- CA- CB
58 60 62 69 3 1.75728 5.27184 2.25936
-7.02912 0.00000 0.00000 ; C- N- CA- C
59 58 60 61 3 29.28800 -8.36800 -20.92000
0.00000 0.00000 0.00000 ; O- C- N- H
59 58 60 62 3 20.92000 0.00000 -20.92000
0.00000 0.00000 0.00000 ; O- C- N- CA
60 62 64 65 3 0.65084 1.95253 0.00000
-2.60338 0.00000 0.00000 ; N- CA- CB- HB2
60 62 64 66 3 0.65084 1.95253 0.00000
-2.60338 0.00000 0.00000 ; N- CA- CB- HB3
60 62 64 67 3 6.03751 2.50622 -5.17142
-4.20074 1.10458 0.00000 ; N- CA- CB- SG
60 62 69 70 3 0.00000 0.00000 0.00000
0.00000 0.00000 0.00000 ; N- CA- C- O
60 62 69 71 3 17.40544 -5.02080 -13.22144
9.20480 0.00000 0.00000 ; N- CA- C- N
61 60 62 63 3 0.00000 0.00000 0.00000
0.00000 0.00000 0.00000 ; H- N- CA- HA
61 60 62 64 3 0.00000 0.00000 0.00000
0.00000 0.00000 0.00000 ; H- N- CA- CB
61 60 62 69 3 0.00000 0.00000 0.00000
0.00000 0.00000 0.00000 ; H- N- CA- C
62 64 67 68 3 1.69034 2.77818 4.11706
-4.21747 1.00416 0.00000 ; CA- CB- SG- HG
62 69 71 72 3 20.92000 0.00000 -20.92000
0.00000 0.00000 0.00000 ; CA- C- N- H
62 69 71 73 3 20.92000 0.00000 -20.92000
0.00000 0.00000 0.00000 ; CA- C- N- CA
63 62 64 65 3 0.65084 1.95253 0.00000
-2.60338 0.00000 0.00000 ; HA- CA- CB- HB2
63 62 64 66 3 0.65084 1.95253 0.00000
-2.60338 0.00000 0.00000 ; HA- CA- CB- HB3
63 62 64 67 3 0.65084 1.95253 0.00000
-2.60338 0.00000 0.00000 ; HA- CA- CB- SG
63 62 69 70 3 3.68192 -4.35136 0.00000
1.33888 0.00000 0.00000 ; HA- CA- C- O

63 62 69 71 3 0.00000 0.00000 0.00000
0.00000 0.00000 0.00000 ; HA- CA- C- N
64 62 69 70 3 0.00000 0.00000 0.00000
0.00000 0.00000 0.00000 ; CB- CA- C- O
64 62 69 71 3 2.51040 4.18400 1.67360
-6.69440 0.00000 0.00000 ; CB- CA- C- N
65 64 62 69 3 0.65084 1.95253 0.00000
-2.60338 0.00000 0.00000 ; HB2- CB- CA- C
65 64 67 68 3 1.04600 3.13800 0.00000
-4.18400 0.00000 0.00000 ; HB2- CB- SG- HG
66 64 62 69 3 0.65084 1.95253 0.00000
-2.60338 0.00000 0.00000 ; HB3- CB- CA- C
66 64 67 68 3 1.04600 3.13800 0.00000
-4.18400 0.00000 0.00000 ; HB3- CB- SG- HG
67 64 62 69 3 5.62330 4.27605 -5.33042
-4.20074 2.51040 0.00000 ; SG- CB- CA- C
69 71 73 74 3 0.00000 0.00000 0.00000
0.00000 0.00000 0.00000 ; C- N- CA- HA
69 71 73 75 3 11.71520 1.67360 15.06240 -
13.38880 0.00000 0.00000 ; C- N- CA- CB
69 71 73 86 3 1.75728 5.27184 2.25936
-7.02912 0.00000 0.00000 ; C- N- CA- C
70 69 71 72 3 29.28800 -8.36800 -20.92000
0.00000 0.00000 0.00000 ; O- C- N- H
70 69 71 73 3 20.92000 0.00000 -20.92000
0.00000 0.00000 0.00000 ; O- C- N- CA
71 73 75 76 3 0.65084 1.95253 0.00000
-2.60338 0.00000 0.00000 ; N- CA- CB- HB2
71 73 75 77 3 0.65084 1.95253 0.00000
-2.60338 0.00000 0.00000 ; N- CA- CB- HB3
71 73 75 78 3 4.79068 4.02919 -4.82834
-3.66518 2.97901 0.00000 ; N- CA- CB- CG
71 73 86 87 3 0.00000 0.00000 0.00000
0.00000 0.00000 0.00000 ; N- CA- C- O
71 73 86 88 3 17.40544 -5.02080 -13.22144
9.20480 0.00000 0.00000 ; N- CA- C- N
72 71 73 74 3 0.00000 0.00000 0.00000
0.00000 0.00000 0.00000 ; H- N- CA- HA
72 71 73 75 3 0.00000 0.00000 0.00000
0.00000 0.00000 0.00000 ; H- N- CA- CB
72 71 73 86 3 0.00000 0.00000 0.00000
0.00000 0.00000 0.00000 ; H- N- CA- C
73 75 78 79 3 5.98312 6.40152 3.28026 -
12.38464 -1.57318 0.00000 ; CA- CB- CG- ND1
73 75 78 84 3 0.00000 0.00000 0.00000
0.00000 0.00000 0.00000 ; CA- CB- CG- CD2
73 86 88 89 3 20.92000 0.00000 -20.92000
0.00000 0.00000 0.00000 ; CA- C- N- H
73 86 88 90 3 20.92000 0.00000 -20.92000
0.00000 0.00000 0.00000 ; CA- C- N- CA
74 73 75 76 3 0.65084 1.95253 0.00000
-2.60338 0.00000 0.00000 ; HA- CA- CB- HB2

74 73 75 77 3 0.65084 1.95253 0.00000
-2.60338 0.00000 0.00000 ; HA- CA- CB- HB3
74 73 75 78 3 0.65084 1.95253 0.00000
-2.60338 0.00000 0.00000 ; HA- CA- CB- CG
74 73 86 87 3 3.68192 -4.35136 0.00000
1.33888 0.00000 0.00000 ; HA- CA- C- O
74 73 86 88 3 0.00000 0.00000 0.00000
0.00000 0.00000 0.00000 ; HA- CA- C- N
75 73 86 87 3 0.00000 0.00000 0.00000
0.00000 0.00000 0.00000 ; CB- CA- C- O
75 73 86 88 3 2.51040 4.18400 1.67360
-6.69440 0.00000 0.00000 ; CB- CA- C- N
75 78 79 80 3 20.08320 0.00000 -20.08320
0.00000 0.00000 0.00000 ; CB- CG- ND1- CE1
75 78 84 82 3 44.97800 0.00000 -44.97800
0.00000 0.00000 0.00000 ; CB- CG- CD2- NE2
75 78 84 85 3 44.97800 0.00000 -44.97800
0.00000 0.00000 0.00000 ; CB- CG- CD2- HD2
76 75 73 86 3 0.65084 1.95253 0.00000
-2.60338 0.00000 0.00000 ; HB2- CB- CA- C
76 75 78 79 3 0.00000 0.00000 0.00000
0.00000 0.00000 0.00000 ; HB2- CB- CG- ND1
76 75 78 84 3 0.00000 0.00000 0.00000
0.00000 0.00000 0.00000 ; HB2- CB- CG- CD2
77 75 73 86 3 0.65084 1.95253 0.00000
-2.60338 0.00000 0.00000 ; HB3- CB- CA- C
77 75 78 79 3 0.00000 0.00000 0.00000
0.00000 0.00000 0.00000 ; HB3- CB- CG- ND1
77 75 78 84 3 0.00000 0.00000 0.00000
0.00000 0.00000 0.00000 ; HB3- CB- CG- CD2
78 75 73 86 3 3.76560 3.34720 -2.87859
-3.66518 0.83680 0.00000 ; CG- CB- CA- C
78 79 80 81 3 41.84000 0.00000 -41.84000
0.00000 0.00000 0.00000 ; CG- ND1- CE1- HE1
78 79 80 82 3 41.84000 0.00000 -41.84000
0.00000 0.00000 0.00000 ; CG- ND1- CE1- NE2
78 84 82 80 3 12.55200 0.00000 -12.55200
0.00000 0.00000 0.00000 ; CG- CD2- NE2- CE1
78 84 82 83 3 12.55200 0.00000 -12.55200
0.00000 0.00000 0.00000 ; CG- CD2- NE2- HE2
79 78 84 82 3 44.97800 0.00000 -44.97800
0.00000 0.00000 0.00000 ; ND1- CG- CD2- NE2
79 78 84 85 3 44.97800 0.00000 -44.97800
0.00000 0.00000 0.00000 ; ND1- CG- CD2- HD2
79 80 82 83 3 19.45560 0.00000 -19.45560
0.00000 0.00000 0.00000 ; ND1- CE1- NE2- HE2
79 80 82 84 3 19.45560 0.00000 -19.45560
0.00000 0.00000 0.00000 ; ND1- CE1- NE2- CD2
80 79 78 84 3 20.08320 0.00000 -20.08320
0.00000 0.00000 0.00000 ; CE1- ND1- CG- CD2
80 82 84 85 3 12.55200 0.00000 -12.55200
0.00000 0.00000 0.00000 ; CE1- NE2- CD2- HD2

```
    81      80      82      83       3   19.45560    0.00000  -19.45560
0.00000    0.00000    0.00000 ;    HE1-   CE1-   NE2-   HE2
    81      80      82      84       3   19.45560    0.00000  -19.45560
0.00000    0.00000    0.00000 ;    HE1-   CE1-   NE2-   CD2
    83      82      84      85       3   12.55200    0.00000  -12.55200
0.00000    0.00000    0.00000 ;    HE2-   NE2-   CD2-   HD2
    86      88      90      91       3    0.00000    0.00000    0.00000
0.00000    0.00000    0.00000 ;      C-     N-    CA-    HA
    86      88      90      92       3   11.71520    1.67360   15.06240  -
13.38880    0.00000    0.00000 ;      C-     N-    CA-    CB
    86      88      90      97       3    1.75728    5.27184    2.25936
-7.02912    0.00000    0.00000 ;      C-     N-    CA-     C
    87      86      88      89       3   29.28800   -8.36800  -20.92000
0.00000    0.00000    0.00000 ;      O-     C-     N-     H
    87      86      88      90       3   20.92000    0.00000  -20.92000
0.00000    0.00000    0.00000 ;      O-     C-     N-    CA
    88      90      92      93       3    0.65084    1.95253    0.00000
-2.60338    0.00000    0.00000 ;      N-    CA-    CB-   HB2
    88      90      92      94       3    0.65084    1.95253    0.00000
-2.60338    0.00000    0.00000 ;      N-    CA-    CB-   HB3
    88      90      92      95       3    7.86174    2.24681   -7.41405
-6.71114    5.35552    0.00000 ;      N-    CA-    CB-    OG
    88      90      97      98       3    0.00000    0.00000    0.00000
0.00000    0.00000    0.00000 ;      N-    CA-     C-     O
    88      90      97      99       3   17.40544   -5.02080  -13.22144
9.20480    0.00000    0.00000 ;      N-    CA-     C-     N
    89      88      90      91       3    0.00000    0.00000    0.00000
0.00000    0.00000    0.00000 ;      H-     N-    CA-    HA
    89      88      90      92       3    0.00000    0.00000    0.00000
0.00000    0.00000    0.00000 ;      H-     N-    CA-    CB
    89      88      90      97       3    0.00000    0.00000    0.00000
0.00000    0.00000    0.00000 ;      H-     N-    CA-     C
    90      92      95      96       3    2.05853    2.46856    3.48109
-4.46851    0.23430    0.00000 ;     CA-    CB-    OG-    HG
    90      97      99     100       3   20.92000    0.00000  -20.92000
0.00000    0.00000    0.00000 ;     CA-     C-     N-     H
    90      97      99     101       3   20.92000    0.00000  -20.92000
0.00000    0.00000    0.00000 ;     CA-     C-     N-    CA
    91      90      92      93       3    0.65084    1.95253    0.00000
-2.60338    0.00000    0.00000 ;     HA-    CA-    CB-   HB2
    91      90      92      94       3    0.65084    1.95253    0.00000
-2.60338    0.00000    0.00000 ;     HA-    CA-    CB-   HB3
    91      90      92      95       3    1.04600   -1.04600    0.00000
0.00000    0.00000    0.00000 ;     HA-    CA-    CB-    OG
    91      90      97      98       3    3.68192   -4.35136    0.00000
1.33888    0.00000    0.00000 ;     HA-    CA-     C-     O
    91      90      97      99       3    0.00000    0.00000    0.00000
0.00000    0.00000    0.00000 ;     HA-    CA-     C-     N
    92      90      97      98       3    0.00000    0.00000    0.00000
0.00000    0.00000    0.00000 ;     CB-    CA-     C-     O
    92      90      97      99       3    2.51040    4.18400    1.67360
-6.69440    0.00000    0.00000 ;     CB-    CA-     C-     N
```

93 92 90 97 3 0.65084 1.95253 0.00000
-2.60338 0.00000 0.00000 ; HB2- CB- CA- C
93 92 95 96 3 0.69733 2.09200 0.00000
-2.78933 0.00000 0.00000 ; HB2- CB- OG- HG
94 92 90 97 3 0.65084 1.95253 0.00000
-2.60338 0.00000 0.00000 ; HB3- CB- CA- C
94 92 95 96 3 0.69733 2.09200 0.00000
-2.78933 0.00000 0.00000 ; HB3- CB- OG- HG
95 92 90 97 3 7.34710 7.79898 -6.14211
-6.71114 4.31789 0.00000 ; OG- CB- CA- C
97 99 101 102 3 0.00000 0.00000 0.00000
0.00000 0.00000 0.00000 ; C- N- CA- HA2
97 99 101 103 3 0.00000 0.00000 0.00000
0.00000 0.00000 0.00000 ; C- N- CA- HA3
97 99 101 104 3 1.75728 5.27184 2.25936
-7.02912 0.00000 0.00000 ; C- N- CA- C
98 97 99 100 3 29.28800 -8.36800 -20.92000
0.00000 0.00000 0.00000 ; O- C- N- H
98 97 99 101 3 20.92000 0.00000 -20.92000
0.00000 0.00000 0.00000 ; O- C- N- CA
99 101 104 105 3 0.00000 0.00000 0.00000
0.00000 0.00000 0.00000 ; N- CA- C- O
99 101 104 106 3 17.40544 -5.02080 -13.22144
9.20480 0.00000 0.00000 ; N- CA- C- N
100 99 101 102 3 0.00000 0.00000 0.00000
0.00000 0.00000 0.00000 ; H- N- CA- HA2
100 99 101 103 3 0.00000 0.00000 0.00000
0.00000 0.00000 0.00000 ; H- N- CA- HA3
100 99 101 104 3 0.00000 0.00000 0.00000
0.00000 0.00000 0.00000 ; H- N- CA- C
101 104 106 107 3 20.92000 0.00000 -20.92000
0.00000 0.00000 0.00000 ; CA- C- N- H
101 104 106 108 3 20.92000 0.00000 -20.92000
0.00000 0.00000 0.00000 ; CA- C- N- CA
102 101 104 105 3 3.68192 -4.35136 0.00000
1.33888 0.00000 0.00000 ; HA2- CA- C- O
102 101 104 106 3 0.00000 0.00000 0.00000
0.00000 0.00000 0.00000 ; HA2- CA- C- N
103 101 104 105 3 3.68192 -4.35136 0.00000
1.33888 0.00000 0.00000 ; HA3- CA- C- O
103 101 104 106 3 0.00000 0.00000 0.00000
0.00000 0.00000 0.00000 ; HA3- CA- C- N
104 106 108 109 3 0.00000 0.00000 0.00000
0.00000 0.00000 0.00000 ; C- N- CA- HA
104 106 108 110 3 11.71520 1.67360 15.06240 -
13.38880 0.00000 0.00000 ; C- N- CA- CB
104 106 108 126 3 1.75728 5.27184 2.25936
-7.02912 0.00000 0.00000 ; C- N- CA- C
105 104 106 107 3 29.28800 -8.36800 -20.92000
0.00000 0.00000 0.00000 ; O- C- N- H
105 104 106 108 3 20.92000 0.00000 -20.92000
0.00000 0.00000 0.00000 ; O- C- N- CA

```
   106    108    110    111      3    0.65084    1.95253    0.00000
-2.60338    0.00000    0.00000 ;      N-    CA-    CB-   HB2
   106    108    110    112      3    0.65084    1.95253    0.00000
-2.60338    0.00000    0.00000 ;      N-    CA-    CB-   HB3
   106    108    110    113      3    0.65084    1.95253    0.00000
-2.60338    0.00000    0.00000 ;      N-    CA-    CB-    CG
   106    108    126    127      3    0.00000    0.00000    0.00000
0.00000    0.00000    0.00000 ;      N-    CA-     C-     O
   106    108    126    128      3   17.40544   -5.02080  -13.22144
9.20480    0.00000    0.00000 ;      N-    CA-     C-     N
   107    106    108    109      3    0.00000    0.00000    0.00000
0.00000    0.00000    0.00000 ;      H-     N-    CA-    HA
   107    106    108    110      3    0.00000    0.00000    0.00000
0.00000    0.00000    0.00000 ;      H-     N-    CA-    CB
   107    106    108    126      3    0.00000    0.00000    0.00000
0.00000    0.00000    0.00000 ;      H-     N-    CA-     C
   108    110    113    114      3    0.66944    2.00832    0.00000
-2.67776    0.00000    0.00000 ;     CA-    CB-    CG-   HG2
   108    110    113    115      3    0.66944    2.00832    0.00000
-2.67776    0.00000    0.00000 ;     CA-    CB-    CG-   HG3
   108    110    113    116      3    3.68192    3.09616   -2.09200
-3.01248    0.00000    0.00000 ;     CA-    CB-    CG-    CD
   108    126    128    129      3   20.92000    0.00000  -20.92000
0.00000    0.00000    0.00000 ;     CA-     C-     N-     H
   108    126    128    130      3   20.92000    0.00000  -20.92000
0.00000    0.00000    0.00000 ;     CA-     C-     N-    CA
   109    108    110    111      3    0.65084    1.95253    0.00000
-2.60338    0.00000    0.00000 ;     HA-    CA-    CB-   HB2
   109    108    110    112      3    0.65084    1.95253    0.00000
-2.60338    0.00000    0.00000 ;     HA-    CA-    CB-   HB3
   109    108    110    113      3    0.65084    1.95253    0.00000
-2.60338    0.00000    0.00000 ;     HA-    CA-    CB-    CG
   109    108    126    127      3    3.68192   -4.35136    0.00000
1.33888    0.00000    0.00000 ;     HA-    CA-     C-     O
   109    108    126    128      3    0.00000    0.00000    0.00000
0.00000    0.00000    0.00000 ;     HA-    CA-     C-     N
   110    108    126    127      3    0.00000    0.00000    0.00000
0.00000    0.00000    0.00000 ;     CB-    CA-     C-     O
   110    108    126    128      3    2.51040    4.18400    1.67360
-6.69440    0.00000    0.00000 ;     CB-    CA-     C-     N
   110    113    116    117      3    0.66944    2.00832    0.00000
-2.67776    0.00000    0.00000 ;     CB-    CG-    CD-   HD2
   110    113    116    118      3    0.66944    2.00832    0.00000
-2.67776    0.00000    0.00000 ;     CB-    CG-    CD-   HD3
   110    113    116    119      3    3.68192    3.09616   -2.09200
-3.01248    0.00000    0.00000 ;     CB-    CG-    CD-    CE
   111    110    108    126      3    0.65084    1.95253    0.00000
-2.60338    0.00000    0.00000 ;    HB2-    CB-    CA-     C
   111    110    113    114      3    0.62760    1.88280    0.00000
-2.51040    0.00000    0.00000 ;    HB2-    CB-    CG-   HG2
   111    110    113    115      3    0.62760    1.88280    0.00000
-2.51040    0.00000    0.00000 ;    HB2-    CB-    CG-   HG3
```

```
   111     110     113     116       3    0.66944    2.00832    0.00000
-2.67776    0.00000    0.00000 ;    HB2-    CB-    CG-    CD
   112     110     108     126       3    0.65084    1.95253    0.00000
-2.60338    0.00000    0.00000 ;    HB3-    CB-    CA-     C
   112     110     113     114       3    0.62760    1.88280    0.00000
-2.51040    0.00000    0.00000 ;    HB3-    CB-    CG-   HG2
   112     110     113     115       3    0.62760    1.88280    0.00000
-2.51040    0.00000    0.00000 ;    HB3-    CB-    CG-   HG3
   112     110     113     116       3    0.66944    2.00832    0.00000
-2.67776    0.00000    0.00000 ;    HB3-    CB-    CG-    CD
   113     110     108     126       3    0.65084    1.95253    0.00000
-2.60338    0.00000    0.00000 ;     CG-    CB-    CA-     C
   113     116     119     120       3    0.65084    1.95253    0.00000
-2.60338    0.00000    0.00000 ;     CG-    CD-    CE-   HE2
   113     116     119     121       3    0.65084    1.95253    0.00000
-2.60338    0.00000    0.00000 ;     CG-    CD-    CE-   HE3
   113     116     119     122       3    0.65084    1.95253    0.00000
-2.60338    0.00000    0.00000 ;     CG-    CD-    CE-    NZ
   114     113     116     117       3    0.62760    1.88280    0.00000
-2.51040    0.00000    0.00000 ;    HG2-    CG-    CD-   HD2
   114     113     116     118       3    0.62760    1.88280    0.00000
-2.51040    0.00000    0.00000 ;    HG2-    CG-    CD-   HD3
   114     113     116     119       3    0.66944    2.00832    0.00000
-2.67776    0.00000    0.00000 ;    HG2-    CG-    CD-    CE
   115     113     116     117       3    0.62760    1.88280    0.00000
-2.51040    0.00000    0.00000 ;    HG3-    CG-    CD-   HD2
   115     113     116     118       3    0.62760    1.88280    0.00000
-2.51040    0.00000    0.00000 ;    HG3-    CG-    CD-   HD3
   115     113     116     119       3    0.66944    2.00832    0.00000
-2.67776    0.00000    0.00000 ;    HG3-    CG-    CD-    CE
   116     119     122     123       3    0.65084    1.95253    0.00000
-2.60338    0.00000    0.00000 ;     CD-    CE-    NZ-   HZ1
   116     119     122     124       3    0.65084    1.95253    0.00000
-2.60338    0.00000    0.00000 ;     CD-    CE-    NZ-   HZ2
   116     119     122     125       3    0.65084    1.95253    0.00000
-2.60338    0.00000    0.00000 ;     CD-    CE-    NZ-   HZ3
   117     116     119     120       3    0.65084    1.95253    0.00000
-2.60338    0.00000    0.00000 ;    HD2-    CD-    CE-   HE2
   117     116     119     121       3    0.65084    1.95253    0.00000
-2.60338    0.00000    0.00000 ;    HD2-    CD-    CE-   HE3
   117     116     119     122       3    0.65084    1.95253    0.00000
-2.60338    0.00000    0.00000 ;    HD2-    CD-    CE-    NZ
   118     116     119     120       3    0.65084    1.95253    0.00000
-2.60338    0.00000    0.00000 ;    HD3-    CD-    CE-   HE2
   118     116     119     121       3    0.65084    1.95253    0.00000
-2.60338    0.00000    0.00000 ;    HD3-    CD-    CE-   HE3
   118     116     119     122       3    0.65084    1.95253    0.00000
-2.60338    0.00000    0.00000 ;    HD3-    CD-    CE-    NZ
   120     119     122     123       3    0.65084    1.95253    0.00000
-2.60338    0.00000    0.00000 ;    HE2-    CE-    NZ-   HZ1
   120     119     122     124       3    0.65084    1.95253    0.00000
-2.60338    0.00000    0.00000 ;    HE2-    CE-    NZ-   HZ2
```

120 119 122 125 3 0.65084 1.95253 0.00000
-2.60338 0.00000 0.00000 ; HE2- CE- NZ- HZ3
121 119 122 123 3 0.65084 1.95253 0.00000
-2.60338 0.00000 0.00000 ; HE3- CE- NZ- HZ1
121 119 122 124 3 0.65084 1.95253 0.00000
-2.60338 0.00000 0.00000 ; HE3- CE- NZ- HZ2
121 119 122 125 3 0.65084 1.95253 0.00000
-2.60338 0.00000 0.00000 ; HE3- CE- NZ- HZ3
126 128 130 131 3 0.00000 0.00000 0.00000
0.00000 0.00000 0.00000 ; C- N- CA- HA
126 128 130 132 3 11.71520 1.67360 15.06240 -
13.38880 0.00000 0.00000 ; C- N- CA- CB
126 128 130 145 3 1.75728 5.27184 2.25936
-7.02912 0.00000 0.00000 ; C- N- CA- C
127 126 128 129 3 29.28800 -8.36800 -20.92000
0.00000 0.00000 0.00000 ; O- C- N- H
127 126 128 130 3 20.92000 0.00000 -20.92000
0.00000 0.00000 0.00000 ; O- C- N- CA
128 130 132 133 3 0.65084 1.95253 0.00000
-2.60338 0.00000 0.00000 ; N- CA- CB- HB2
128 130 132 134 3 0.65084 1.95253 0.00000
-2.60338 0.00000 0.00000 ; N- CA- CB- HB3
128 130 132 135 3 3.79070 1.39746 -4.61077
-2.40998 2.44346 0.00000 ; N- CA- CB- CG
128 130 145 146 3 0.00000 0.00000 0.00000
0.00000 0.00000 0.00000 ; N- CA- C- O
128 130 145 147 3 17.40544 -5.02080 -13.22144
9.20480 0.00000 0.00000 ; N- CA- C- N
129 128 130 131 3 0.00000 0.00000 0.00000
0.00000 0.00000 0.00000 ; H- N- CA- HA
129 128 130 132 3 0.00000 0.00000 0.00000
0.00000 0.00000 0.00000 ; H- N- CA- CB
129 128 130 145 3 0.00000 0.00000 0.00000
0.00000 0.00000 0.00000 ; H- N- CA- C
130 132 135 136 3 0.66944 2.00832 0.00000
-2.67776 0.00000 0.00000 ; CA- CB- CG- HG
130 132 135 137 3 3.90367 0.19665 -6.21742
-2.37651 5.99149 0.00000 ; CA- CB- CG- CD1
130 132 135 141 3 3.90367 0.19665 -6.21742
-2.37651 5.99149 0.00000 ; CA- CB- CG- CD2
130 145 147 148 3 20.92000 0.00000 -20.92000
0.00000 0.00000 0.00000 ; CA- C- N- H
130 145 147 149 3 20.92000 0.00000 -20.92000
0.00000 0.00000 0.00000 ; CA- C- N- CA
131 130 132 133 3 0.65084 1.95253 0.00000
-2.60338 0.00000 0.00000 ; HA- CA- CB- HB2
131 130 132 134 3 0.65084 1.95253 0.00000
-2.60338 0.00000 0.00000 ; HA- CA- CB- HB3
131 130 132 135 3 0.65084 1.95253 0.00000
-2.60338 0.00000 0.00000 ; HA- CA- CB- CG
131 130 145 146 3 3.68192 -4.35136 0.00000
1.33888 0.00000 0.00000 ; HA- CA- C- O

131 130 145 147 3 0.00000 0.00000 0.00000
0.00000 0.00000 0.00000 ; HA- CA- C- N
132 130 145 146 3 0.00000 0.00000 0.00000
0.00000 0.00000 0.00000 ; CB- CA- C- O
132 130 145 147 3 2.51040 4.18400 1.67360
-6.69440 0.00000 0.00000 ; CB- CA- C- N
132 135 137 138 3 0.66944 2.00832 0.00000
-2.67776 0.00000 0.00000 ; CB- CG- CD1- HD11
132 135 137 139 3 0.66944 2.00832 0.00000
-2.67776 0.00000 0.00000 ; CB- CG- CD1- HD12
132 135 137 140 3 0.66944 2.00832 0.00000
-2.67776 0.00000 0.00000 ; CB- CG- CD1- HD13
132 135 141 142 3 0.66944 2.00832 0.00000
-2.67776 0.00000 0.00000 ; CB- CG- CD2- HD21
132 135 141 143 3 0.66944 2.00832 0.00000
-2.67776 0.00000 0.00000 ; CB- CG- CD2- HD22
132 135 141 144 3 0.66944 2.00832 0.00000
-2.67776 0.00000 0.00000 ; CB- CG- CD2- HD23
133 132 130 145 3 0.65084 1.95253 0.00000
-2.60338 0.00000 0.00000 ; HB2- CB- CA- C
133 132 135 136 3 0.62760 1.88280 0.00000
-2.51040 0.00000 0.00000 ; HB2- CB- CG- HG
133 132 135 137 3 0.66944 2.00832 0.00000
-2.67776 0.00000 0.00000 ; HB2- CB- CG- CD1
133 132 135 141 3 0.66944 2.00832 0.00000
-2.67776 0.00000 0.00000 ; HB2- CB- CG- CD2
134 132 130 145 3 0.65084 1.95253 0.00000
-2.60338 0.00000 0.00000 ; HB3- CB- CA- C
134 132 135 136 3 0.62760 1.88280 0.00000
-2.51040 0.00000 0.00000 ; HB3- CB- CG- HG
134 132 135 137 3 0.66944 2.00832 0.00000
-2.67776 0.00000 0.00000 ; HB3- CB- CG- CD1
134 132 135 141 3 0.66944 2.00832 0.00000
-2.67776 0.00000 0.00000 ; HB3- CB- CG- CD2
135 132 130 145 3 10.33448 -1.14642 -11.54784
-2.40998 6.35968 0.00000 ; CG- CB- CA- C
136 135 137 138 3 0.62760 1.88280 0.00000
-2.51040 0.00000 0.00000 ; HG- CG- CD1- HD11
136 135 137 139 3 0.62760 1.88280 0.00000
-2.51040 0.00000 0.00000 ; HG- CG- CD1- HD12
136 135 137 140 3 0.62760 1.88280 0.00000
-2.51040 0.00000 0.00000 ; HG- CG- CD1- HD13
136 135 141 142 3 0.62760 1.88280 0.00000
-2.51040 0.00000 0.00000 ; HG- CG- CD2- HD21
136 135 141 143 3 0.62760 1.88280 0.00000
-2.51040 0.00000 0.00000 ; HG- CG- CD2- HD22
136 135 141 144 3 0.62760 1.88280 0.00000
-2.51040 0.00000 0.00000 ; HG- CG- CD2- HD23
137 135 141 142 3 0.66944 2.00832 0.00000
-2.67776 0.00000 0.00000 ; CD1- CG- CD2- HD21
137 135 141 143 3 0.66944 2.00832 0.00000
-2.67776 0.00000 0.00000 ; CD1- CG- CD2- HD22

```
   137    135    141    144      3    0.66944    2.00832    0.00000
-2.67776    0.00000    0.00000 ;   CD1-   CG-   CD2-  HD23
   138    137    135    141      3    0.66944    2.00832    0.00000
-2.67776    0.00000    0.00000 ;   HD11-   CD1-    CG-   CD2
   139    137    135    141      3    0.66944    2.00832    0.00000
-2.67776    0.00000    0.00000 ;   HD12-   CD1-    CG-   CD2
   140    137    135    141      3    0.66944    2.00832    0.00000
-2.67776    0.00000    0.00000 ;   HD13-   CD1-    CG-   CD2
   145    147    149    150      3    0.00000    0.00000    0.00000
0.00000    0.00000    0.00000 ;     C-     N-    CA-    HA
   145    147    149    151      3   11.71520    1.67360   15.06240  -
13.38880    0.00000    0.00000 ;     C-     N-    CA-    CB
   145    147    149    160      3    1.75728    5.27184    2.25936
-7.02912    0.00000    0.00000 ;     C-     N-    CA-     C
   146    145    147    148      3   29.28800   -8.36800  -20.92000
0.00000    0.00000    0.00000 ;     O-     C-     N-     H
   146    145    147    149      3   20.92000    0.00000  -20.92000
0.00000    0.00000    0.00000 ;     O-     C-     N-    CA
   147    149    151    152      3    0.65084    1.95253    0.00000
-2.60338    0.00000    0.00000 ;     N-    CA-    CB-   HB2
   147    149    151    153      3    0.65084    1.95253    0.00000
-2.60338    0.00000    0.00000 ;     N-    CA-    CB-   HB3
   147    149    151    154      3    3.21331    2.22589   -4.15053
-2.40998    2.61082    0.00000 ;     N-    CA-    CB-    CG
   147    149    160    161      3    0.00000    0.00000    0.00000
0.00000    0.00000    0.00000 ;     N-    CA-     C-     O
   147    149    160    162      3   17.40544   -5.02080  -13.22144
9.20480    0.00000    0.00000 ;     N-    CA-     C-     N
   148    147    149    150      3    0.00000    0.00000    0.00000
0.00000    0.00000    0.00000 ;     H-     N-    CA-    HA
   148    147    149    151      3    0.00000    0.00000    0.00000
0.00000    0.00000    0.00000 ;     H-     N-    CA-    CB
   148    147    149    160      3    0.00000    0.00000    0.00000
0.00000    0.00000    0.00000 ;     H-     N-    CA-     C
   149    151    154    155      3    0.66944    2.00832    0.00000
-2.67776    0.00000    0.00000 ;    CA-    CB-    CG-   HG2
   149    151    154    156      3    0.66944    2.00832    0.00000
-2.67776    0.00000    0.00000 ;    CA-    CB-    CG-   HG3
   149    151    154    157      3   10.12110   -1.91209    0.01674
10.17549   -1.87443    0.00000 ;    CA-    CB-    CG-    CD
   149    160    162    163      3   20.92000    0.00000  -20.92000
0.00000    0.00000    0.00000 ;    CA-     C-     N-     H
   149    160    162    164      3   20.92000    0.00000  -20.92000
0.00000    0.00000    0.00000 ;    CA-     C-     N-    CA
   150    149    151    152      3    0.65084    1.95253    0.00000
-2.60338    0.00000    0.00000 ;    HA-    CA-    CB-   HB2
   150    149    151    153      3    0.65084    1.95253    0.00000
-2.60338    0.00000    0.00000 ;    HA-    CA-    CB-   HB3
   150    149    151    154      3    0.65084    1.95253    0.00000
-2.60338    0.00000    0.00000 ;    HA-    CA-    CB-    CG
   150    149    160    161      3    3.68192   -4.35136    0.00000
1.33888    0.00000    0.00000 ;    HA-    CA-     C-     O
```

150 149 160 162 3 0.00000 0.00000 0.00000
0.00000 0.00000 0.00000 ; HA- CA- C- N
151 149 160 161 3 0.00000 0.00000 0.00000
0.00000 0.00000 0.00000 ; CB- CA- C- O
151 149 160 162 3 2.51040 4.18400 1.67360
-6.69440 0.00000 0.00000 ; CB- CA- C- N
151 154 157 158 3 3.79907 0.00000 -5.40573
0.00000 2.14221 0.00000 ; CB- CG- CD- OE1
151 154 157 159 3 3.79907 0.00000 -5.40573
0.00000 2.14221 0.00000 ; CB- CG- CD- OE2
152 151 149 160 3 0.65084 1.95253 0.00000
-2.60338 0.00000 0.00000 ; HB2- CB- CA- C
152 151 154 155 3 0.62760 1.88280 0.00000
-2.51040 0.00000 0.00000 ; HB2- CB- CG- HG2
152 151 154 156 3 0.62760 1.88280 0.00000
-2.51040 0.00000 0.00000 ; HB2- CB- CG- HG3
152 151 154 157 3 0.65084 1.95253 0.00000
-2.60338 0.00000 0.00000 ; HB2- CB- CG- CD
153 151 149 160 3 0.65084 1.95253 0.00000
-2.60338 0.00000 0.00000 ; HB3- CB- CA- C
153 151 154 155 3 0.62760 1.88280 0.00000
-2.51040 0.00000 0.00000 ; HB3- CB- CG- HG2
153 151 154 156 3 0.62760 1.88280 0.00000
-2.51040 0.00000 0.00000 ; HB3- CB- CG- HG3
153 151 154 157 3 0.65084 1.95253 0.00000
-2.60338 0.00000 0.00000 ; HB3- CB- CG- CD
154 151 149 160 3 6.86594 3.56895 -8.14206
-2.40998 4.85344 0.00000 ; CG- CB- CA- C
155 154 157 158 3 0.00000 0.00000 0.00000
0.00000 0.00000 0.00000 ; HG2- CG- CD- OE1
155 154 157 159 3 0.00000 0.00000 0.00000
0.00000 0.00000 0.00000 ; HG2- CG- CD- OE2
156 154 157 158 3 0.00000 0.00000 0.00000
0.00000 0.00000 0.00000 ; HG3- CG- CD- OE1
156 154 157 159 3 0.00000 0.00000 0.00000
0.00000 0.00000 0.00000 ; HG3- CG- CD- OE2
160 162 164 165 3 0.00000 0.00000 0.00000
0.00000 0.00000 0.00000 ; C- N- CA- HA
160 162 164 166 3 11.71520 1.67360 15.06240 -
13.38880 0.00000 0.00000 ; C- N- CA- CB
160 162 164 172 3 1.75728 5.27184 2.25936
-7.02912 0.00000 0.00000 ; C- N- CA- C
161 160 162 163 3 29.28800 -8.36800 -20.92000
0.00000 0.00000 0.00000 ; O- C- N- H
161 160 162 164 3 20.92000 0.00000 -20.92000
0.00000 0.00000 0.00000 ; O- C- N- CA
162 164 166 167 3 0.65084 1.95253 0.00000
-2.60338 0.00000 0.00000 ; N- CA- CB- HB2
162 164 166 168 3 0.65084 1.95253 0.00000
-2.60338 0.00000 0.00000 ; N- CA- CB- HB3
162 164 166 169 3 15.41386 9.74035 -8.39310
-0.97069 2.97901 0.00000 ; N- CA- CB- CG

162 164 172 173 3 0.00000 0.00000 0.00000
0.00000 0.00000 0.00000 ; N- CA- C- O
162 164 172 174 3 17.40544 -5.02080 -13.22144
9.20480 0.00000 0.00000 ; N- CA- C- N
163 162 164 165 3 0.00000 0.00000 0.00000
0.00000 0.00000 0.00000 ; H- N- CA- HA
163 162 164 166 3 0.00000 0.00000 0.00000
0.00000 0.00000 0.00000 ; H- N- CA- CB
163 162 164 172 3 0.00000 0.00000 0.00000
0.00000 0.00000 0.00000 ; H- N- CA- C
164 166 169 170 3 6.43499 0.00000 -5.39736
0.00000 -1.03763 0.00000 ; CA- CB- CG- OD1
164 166 169 171 3 6.43499 0.00000 -5.39736
0.00000 -1.03763 0.00000 ; CA- CB- CG- OD2
164 172 174 175 3 20.92000 0.00000 -20.92000
0.00000 0.00000 0.00000 ; CA- C- N- H
164 172 174 176 3 20.92000 0.00000 -20.92000
0.00000 0.00000 0.00000 ; CA- C- N- CA
165 164 166 167 3 0.65084 1.95253 0.00000
-2.60338 0.00000 0.00000 ; HA- CA- CB- HB2
165 164 166 168 3 0.65084 1.95253 0.00000
-2.60338 0.00000 0.00000 ; HA- CA- CB- HB3
165 164 166 169 3 0.65084 1.95253 0.00000
-2.60338 0.00000 0.00000 ; HA- CA- CB- CG
165 164 172 173 3 3.68192 -4.35136 0.00000
1.33888 0.00000 0.00000 ; HA- CA- C- O
165 164 172 174 3 0.00000 0.00000 0.00000
0.00000 0.00000 0.00000 ; HA- CA- C- N
166 164 172 173 3 0.00000 0.00000 0.00000
0.00000 0.00000 0.00000 ; CB- CA- C- O
166 164 172 174 3 2.51040 4.18400 1.67360
-6.69440 0.00000 0.00000 ; CB- CA- C- N
167 166 164 172 3 0.65084 1.95253 0.00000
-2.60338 0.00000 0.00000 ; HB2- CB- CA- C
167 166 169 170 3 0.00000 0.00000 0.00000
0.00000 0.00000 0.00000 ; HB2- CB- CG- OD1
167 166 169 171 3 0.00000 0.00000 0.00000
0.00000 0.00000 0.00000 ; HB2- CB- CG- OD2
168 166 164 172 3 0.65084 1.95253 0.00000
-2.60338 0.00000 0.00000 ; HB3- CB- CA- C
168 166 169 170 3 0.00000 0.00000 0.00000
0.00000 0.00000 0.00000 ; HB3- CB- CG- OD1
168 166 169 171 3 0.00000 0.00000 0.00000
0.00000 0.00000 0.00000 ; HB3- CB- CG- OD2
169 166 164 172 3 7.14627 -1.04600 -8.99560
-0.97069 5.15469 0.00000 ; CG- CB- CA- C
172 174 176 177 3 0.00000 0.00000 0.00000
0.00000 0.00000 0.00000 ; C- N- CA- HA
172 174 176 178 3 11.71520 1.67360 15.06240 -
13.38880 0.00000 0.00000 ; C- N- CA- CB
172 174 176 194 3 1.75728 5.27184 2.25936
-7.02912 0.00000 0.00000 ; C- N- CA- C

173 172 174 175 3 29.28800 -8.36800 -20.92000
0.00000 0.00000 0.00000 ; O- C- N- H
173 172 174 176 3 20.92000 0.00000 -20.92000
0.00000 0.00000 0.00000 ; O- C- N- CA
174 176 178 179 3 0.65084 1.95253 0.00000
-2.60338 0.00000 0.00000 ; N- CA- CB- HB2
174 176 178 180 3 0.65084 1.95253 0.00000
-2.60338 0.00000 0.00000 ; N- CA- CB- HB3
174 176 178 181 3 0.65084 1.95253 0.00000
-2.60338 0.00000 0.00000 ; N- CA- CB- CG
174 176 194 195 3 0.00000 0.00000 0.00000
0.00000 0.00000 0.00000 ; N- CA- C- O
174 176 194 196 3 17.40544 -5.02080 -13.22144
9.20480 0.00000 0.00000 ; N- CA- C- N
175 174 176 177 3 0.00000 0.00000 0.00000
0.00000 0.00000 0.00000 ; H- N- CA- HA
175 174 176 178 3 0.00000 0.00000 0.00000
0.00000 0.00000 0.00000 ; H- N- CA- CB
175 174 176 194 3 0.00000 0.00000 0.00000
0.00000 0.00000 0.00000 ; H- N- CA- C
176 178 181 182 3 0.66944 2.00832 0.00000
-2.67776 0.00000 0.00000 ; CA- CB- CG- HG2
176 178 181 183 3 0.66944 2.00832 0.00000
-2.67776 0.00000 0.00000 ; CA- CB- CG- HG3
176 178 181 184 3 3.68192 3.09616 -2.09200
-3.01248 0.00000 0.00000 ; CA- CB- CG- CD
176 194 196 197 3 20.92000 0.00000 -20.92000
0.00000 0.00000 0.00000 ; CA- C- N- H
176 194 196 198 3 20.92000 0.00000 -20.92000
0.00000 0.00000 0.00000 ; CA- C- N- CA
177 176 178 179 3 0.65084 1.95253 0.00000
-2.60338 0.00000 0.00000 ; HA- CA- CB- HB2
177 176 178 180 3 0.65084 1.95253 0.00000
-2.60338 0.00000 0.00000 ; HA- CA- CB- HB3
177 176 178 181 3 0.65084 1.95253 0.00000
-2.60338 0.00000 0.00000 ; HA- CA- CB- CG
177 176 194 195 3 3.68192 -4.35136 0.00000
1.33888 0.00000 0.00000 ; HA- CA- C- O
177 176 194 196 3 0.00000 0.00000 0.00000
0.00000 0.00000 0.00000 ; HA- CA- C- N
178 176 194 195 3 0.00000 0.00000 0.00000
0.00000 0.00000 0.00000 ; CB- CA- C- O
178 176 194 196 3 2.51040 4.18400 1.67360
-6.69440 0.00000 0.00000 ; CB- CA- C- N
178 181 184 185 3 0.66944 2.00832 0.00000
-2.67776 0.00000 0.00000 ; CB- CG- CD- HD2
178 181 184 186 3 0.66944 2.00832 0.00000
-2.67776 0.00000 0.00000 ; CB- CG- CD- HD3
178 181 184 187 3 3.68192 3.09616 -2.09200
-3.01248 0.00000 0.00000 ; CB- CG- CD- CE
179 178 176 194 3 0.65084 1.95253 0.00000
-2.60338 0.00000 0.00000 ; HB2- CB- CA- C

```
   179    178    181    182      3    0.62760    1.88280    0.00000
-2.51040    0.00000    0.00000 ;    HB2-    CB-    CG-   HG2
   179    178    181    183      3    0.62760    1.88280    0.00000
-2.51040    0.00000    0.00000 ;    HB2-    CB-    CG-   HG3
   179    178    181    184      3    0.66944    2.00832    0.00000
-2.67776    0.00000    0.00000 ;    HB2-    CB-    CG-    CD
   180    178    176    194      3    0.65084    1.95253    0.00000
-2.60338    0.00000    0.00000 ;    HB3-    CB-    CA-     C
   180    178    181    182      3    0.62760    1.88280    0.00000
-2.51040    0.00000    0.00000 ;    HB3-    CB-    CG-   HG2
   180    178    181    183      3    0.62760    1.88280    0.00000
-2.51040    0.00000    0.00000 ;    HB3-    CB-    CG-   HG3
   180    178    181    184      3    0.66944    2.00832    0.00000
-2.67776    0.00000    0.00000 ;    HB3-    CB-    CG-    CD
   181    178    176    194      3    0.65084    1.95253    0.00000
-2.60338    0.00000    0.00000 ;     CG-    CB-    CA-     C
   181    184    187    188      3    0.65084    1.95253    0.00000
-2.60338    0.00000    0.00000 ;     CG-    CD-    CE-   HE2
   181    184    187    189      3    0.65084    1.95253    0.00000
-2.60338    0.00000    0.00000 ;     CG-    CD-    CE-   HE3
   181    184    187    190      3    0.65084    1.95253    0.00000
-2.60338    0.00000    0.00000 ;     CG-    CD-    CE-    NZ
   182    181    184    185      3    0.62760    1.88280    0.00000
-2.51040    0.00000    0.00000 ;    HG2-    CG-    CD-   HD2
   182    181    184    186      3    0.62760    1.88280    0.00000
-2.51040    0.00000    0.00000 ;    HG2-    CG-    CD-   HD3
   182    181    184    187      3    0.66944    2.00832    0.00000
-2.67776    0.00000    0.00000 ;    HG2-    CG-    CD-    CE
   183    181    184    185      3    0.62760    1.88280    0.00000
-2.51040    0.00000    0.00000 ;    HG3-    CG-    CD-   HD2
   183    181    184    186      3    0.62760    1.88280    0.00000
-2.51040    0.00000    0.00000 ;    HG3-    CG-    CD-   HD3
   183    181    184    187      3    0.66944    2.00832    0.00000
-2.67776    0.00000    0.00000 ;    HG3-    CG-    CD-    CE
   184    187    190    191      3    0.65084    1.95253    0.00000
-2.60338    0.00000    0.00000 ;     CD-    CE-    NZ-   HZ1
   184    187    190    192      3    0.65084    1.95253    0.00000
-2.60338    0.00000    0.00000 ;     CD-    CE-    NZ-   HZ2
   184    187    190    193      3    0.65084    1.95253    0.00000
-2.60338    0.00000    0.00000 ;     CD-    CE-    NZ-   HZ3
   185    184    187    188      3    0.65084    1.95253    0.00000
-2.60338    0.00000    0.00000 ;    HD2-    CD-    CE-   HE2
   185    184    187    189      3    0.65084    1.95253    0.00000
-2.60338    0.00000    0.00000 ;    HD2-    CD-    CE-   HE3
   185    184    187    190      3    0.65084    1.95253    0.00000
-2.60338    0.00000    0.00000 ;    HD2-    CD-    CE-    NZ
   186    184    187    188      3    0.65084    1.95253    0.00000
-2.60338    0.00000    0.00000 ;    HD3-    CD-    CE-   HE2
   186    184    187    189      3    0.65084    1.95253    0.00000
-2.60338    0.00000    0.00000 ;    HD3-    CD-    CE-   HE3
   186    184    187    190      3    0.65084    1.95253    0.00000
-2.60338    0.00000    0.00000 ;    HD3-    CD-    CE-    NZ
```

188 187 190 191 3 0.65084 1.95253 0.00000
-2.60338 0.00000 0.00000 ; HE2- CE- NZ- HZ1
188 187 190 192 3 0.65084 1.95253 0.00000
-2.60338 0.00000 0.00000 ; HE2- CE- NZ- HZ2
188 187 190 193 3 0.65084 1.95253 0.00000
-2.60338 0.00000 0.00000 ; HE2- CE- NZ- HZ3
189 187 190 191 3 0.65084 1.95253 0.00000
-2.60338 0.00000 0.00000 ; HE3- CE- NZ- HZ1
189 187 190 192 3 0.65084 1.95253 0.00000
-2.60338 0.00000 0.00000 ; HE3- CE- NZ- HZ2
189 187 190 193 3 0.65084 1.95253 0.00000
-2.60338 0.00000 0.00000 ; HE3- CE- NZ- HZ3
194 196 198 199 3 0.00000 0.00000 0.00000
0.00000 0.00000 0.00000 ; C- N- CA- HA
194 196 198 200 3 11.71520 1.67360 15.06240 -
13.38880 0.00000 0.00000 ; C- N- CA- CB
194 196 198 215 3 1.75728 5.27184 2.25936
-7.02912 0.00000 0.00000 ; C- N- CA- C
195 194 196 197 3 29.28800 -8.36800 -20.92000
0.00000 0.00000 0.00000 ; O- C- N- H
195 194 196 198 3 20.92000 0.00000 -20.92000
0.00000 0.00000 0.00000 ; O- C- N- CA
196 198 200 201 3 0.65084 1.95253 0.00000
-2.60338 0.00000 0.00000 ; N- CA- CB- HB2
196 198 200 202 3 0.65084 1.95253 0.00000
-2.60338 0.00000 0.00000 ; N- CA- CB- HB3
196 198 200 203 3 3.28026 2.46019 -2.19242
-3.21331 -0.23430 0.00000 ; N- CA- CB- CG
196 198 215 216 3 0.00000 0.00000 0.00000
0.00000 0.00000 0.00000 ; N- CA- C- O
196 198 215 217 3 17.40544 -5.02080 -13.22144
9.20480 0.00000 0.00000 ; N- CA- C- N
197 196 198 199 3 0.00000 0.00000 0.00000
0.00000 0.00000 0.00000 ; H- N- CA- HA
197 196 198 200 3 0.00000 0.00000 0.00000
0.00000 0.00000 0.00000 ; H- N- CA- CB
197 196 198 215 3 0.00000 0.00000 0.00000
0.00000 0.00000 0.00000 ; H- N- CA- C
198 200 203 204 3 0.57739 0.00000 1.02926
0.00000 -1.60666 0.00000 ; CA- CB- CG- CD1
198 200 203 213 3 0.57739 0.00000 1.02926
0.00000 -1.60666 0.00000 ; CA- CB- CG- CD2
198 215 217 218 3 20.92000 0.00000 -20.92000
0.00000 0.00000 0.00000 ; CA- C- N- H
198 215 217 219 3 20.92000 0.00000 -20.92000
0.00000 0.00000 0.00000 ; CA- C- N- CA
199 198 200 201 3 0.65084 1.95253 0.00000
-2.60338 0.00000 0.00000 ; HA- CA- CB- HB2
199 198 200 202 3 0.65084 1.95253 0.00000
-2.60338 0.00000 0.00000 ; HA- CA- CB- HB3
199 198 200 203 3 0.65084 1.95253 0.00000
-2.60338 0.00000 0.00000 ; HA- CA- CB- CG

199 198 215 216 3 3.68192 -4.35136 0.00000
1.33888 0.00000 0.00000 ; HA- CA- C- O
199 198 215 217 3 0.00000 0.00000 0.00000
0.00000 0.00000 0.00000 ; HA- CA- C- N
200 198 215 216 3 0.00000 0.00000 0.00000
0.00000 0.00000 0.00000 ; CB- CA- C- O
200 198 215 217 3 2.51040 4.18400 1.67360
-6.69440 0.00000 0.00000 ; CB- CA- C- N
200 203 204 205 3 30.33400 0.00000 -30.33400
0.00000 0.00000 0.00000 ; CB- CG- CD1- HD1
200 203 204 206 3 30.33400 0.00000 -30.33400
0.00000 0.00000 0.00000 ; CB- CG- CD1- CE1
200 203 213 211 3 30.33400 0.00000 -30.33400
0.00000 0.00000 0.00000 ; CB- CG- CD2- CE2
200 203 213 214 3 30.33400 0.00000 -30.33400
0.00000 0.00000 0.00000 ; CB- CG- CD2- HD2
201 200 198 215 3 0.65084 1.95253 0.00000
-2.60338 0.00000 0.00000 ; HB2- CB- CA- C
201 200 203 204 3 0.00000 0.00000 0.00000
0.00000 0.00000 0.00000 ; HB2- CB- CG- CD1
201 200 203 213 3 0.00000 0.00000 0.00000
0.00000 0.00000 0.00000 ; HB2- CB- CG- CD2
202 200 198 215 3 0.65084 1.95253 0.00000
-2.60338 0.00000 0.00000 ; HB3- CB- CA- C
202 200 203 204 3 0.00000 0.00000 0.00000
0.00000 0.00000 0.00000 ; HB3- CB- CG- CD1
202 200 203 213 3 0.00000 0.00000 0.00000
0.00000 0.00000 0.00000 ; HB3- CB- CG- CD2
203 200 198 215 3 4.95804 2.17986 -3.52293
-3.21331 -0.40166 0.00000 ; CG- CB- CA- C
203 204 206 207 3 30.33400 0.00000 -30.33400
0.00000 0.00000 0.00000 ; CG- CD1- CE1- HE1
203 204 206 208 3 30.33400 0.00000 -30.33400
0.00000 0.00000 0.00000 ; CG- CD1- CE1- CZ
203 213 211 208 3 30.33400 0.00000 -30.33400
0.00000 0.00000 0.00000 ; CG- CD2- CE2- CZ
203 213 211 212 3 30.33400 0.00000 -30.33400
0.00000 0.00000 0.00000 ; CG- CD2- CE2- HE2
204 203 213 211 3 30.33400 0.00000 -30.33400
0.00000 0.00000 0.00000 ; CD1- CG- CD2- CE2
204 203 213 214 3 30.33400 0.00000 -30.33400
0.00000 0.00000 0.00000 ; CD1- CG- CD2- HD2
204 206 208 209 3 30.33400 0.00000 -30.33400
0.00000 0.00000 0.00000 ; CD1- CE1- CZ- OH
204 206 208 211 3 30.33400 0.00000 -30.33400
0.00000 0.00000 0.00000 ; CD1- CE1- CZ- CE2
205 204 203 213 3 30.33400 0.00000 -30.33400
0.00000 0.00000 0.00000 ; HD1- CD1- CG- CD2
205 204 206 207 3 30.33400 0.00000 -30.33400
0.00000 0.00000 0.00000 ; HD1- CD1- CE1- HE1
205 204 206 208 3 30.33400 0.00000 -30.33400
0.00000 0.00000 0.00000 ; HD1- CD1- CE1- CZ

206 204 203 213 3 30.33400 0.00000 -30.33400
0.00000 0.00000 0.00000 ; CE1- CD1- CG- CD2
206 208 209 210 3 7.93286 0.00000 -9.56462
0.00000 2.17568 0.00000 ; CE1- CZ- OH- HH
206 208 211 212 3 30.33400 0.00000 -30.33400
0.00000 0.00000 0.00000 ; CE1- CZ- CE2- HE2
206 208 211 213 3 30.33400 0.00000 -30.33400
0.00000 0.00000 0.00000 ; CE1- CZ- CE2- CD2
207 206 208 209 3 30.33400 0.00000 -30.33400
0.00000 0.00000 0.00000 ; HE1- CE1- CZ- OH
207 206 208 211 3 30.33400 0.00000 -30.33400
0.00000 0.00000 0.00000 ; HE1- CE1- CZ- CE2
208 211 213 214 3 30.33400 0.00000 -30.33400
0.00000 0.00000 0.00000 ; CZ- CE2- CD2- HD2
209 208 211 212 3 30.33400 0.00000 -30.33400
0.00000 0.00000 0.00000 ; OH- CZ- CE2- HE2
209 208 211 213 3 30.33400 0.00000 -30.33400
0.00000 0.00000 0.00000 ; OH- CZ- CE2- CD2
210 209 208 211 3 7.93286 0.00000 -9.56462
0.00000 2.17568 0.00000 ; HH- OH- CZ- CE2
212 211 213 214 3 30.33400 0.00000 -30.33400
0.00000 0.00000 0.00000 ; HE2- CE2- CD2- HD2
215 217 219 220 3 0.00000 0.00000 0.00000
0.00000 0.00000 0.00000 ; C- N- CA- HA
215 217 219 221 3 11.71520 1.67360 15.06240 -
13.38880 0.00000 0.00000 ; C- N- CA- CB
215 217 219 234 3 1.75728 5.27184 2.25936
-7.02912 0.00000 0.00000 ; C- N- CA- C
216 215 217 218 3 29.28800 -8.36800 -20.92000
0.00000 0.00000 0.00000 ; O- C- N- H
216 215 217 219 3 20.92000 0.00000 -20.92000
0.00000 0.00000 0.00000 ; O- C- N- CA
217 219 221 222 3 0.65084 1.95253 0.00000
-2.60338 0.00000 0.00000 ; N- CA- CB- HB
217 219 221 223 3 3.83673 0.44769 -1.77402
-2.47693 -0.03347 0.00000 ; N- CA- CB- CG2
217 219 221 227 3 2.97482 0.12134 2.04179
-1.89117 -3.24678 0.00000 ; N- CA- CB- CG1
217 219 234 235 3 0.00000 0.00000 0.00000
0.00000 0.00000 0.00000 ; N- CA- C- O
217 219 234 236 3 17.40544 -5.02080 -13.22144
9.20480 0.00000 0.00000 ; N- CA- C- N
218 217 219 220 3 0.00000 0.00000 0.00000
0.00000 0.00000 0.00000 ; H- N- CA- HA
218 217 219 221 3 0.00000 0.00000 0.00000
0.00000 0.00000 0.00000 ; H- N- CA- CB
218 217 219 234 3 0.00000 0.00000 0.00000
0.00000 0.00000 0.00000 ; H- N- CA- C
219 221 223 224 3 0.66944 2.00832 0.00000
-2.67776 0.00000 0.00000 ; CA- CB- CG2- HG21
219 221 223 225 3 0.66944 2.00832 0.00000
-2.67776 0.00000 0.00000 ; CA- CB- CG2- HG22

```
   219    221    223    226      3    0.66944    2.00832    0.00000
-2.67776    0.00000    0.00000 ;    CA-   CB-   CG2-  HG23
   219    221    227    228      3    0.66944    2.00832    0.00000
-2.67776    0.00000    0.00000 ;    CA-   CB-   CG1-  HG12
   219    221    227    229      3    0.66944    2.00832    0.00000
-2.67776    0.00000    0.00000 ;    CA-   CB-   CG1-  HG13
   219    221    227    230      3    4.24258   -0.52718   -7.25506
-1.79075    7.69856    0.00000 ;    CA-   CB-   CG1-   CD1
   219    234    236    237      3   20.92000    0.00000  -20.92000
0.00000    0.00000    0.00000 ;    CA-    C-    N-    H
   219    234    236    238      3   20.92000    0.00000  -20.92000
0.00000    0.00000    0.00000 ;    CA-    C-    N-   CA
   220    219    221    222      3    0.65084    1.95253    0.00000
-2.60338    0.00000    0.00000 ;    HA-   CA-   CB-    HB
   220    219    221    223      3    0.65084    1.95253    0.00000
-2.60338    0.00000    0.00000 ;    HA-   CA-   CB-   CG2
   220    219    221    227      3    0.65084    1.95253    0.00000
-2.60338    0.00000    0.00000 ;    HA-   CA-   CB-   CG1
   220    219    234    235      3    3.68192   -4.35136    0.00000
1.33888    0.00000    0.00000 ;    HA-   CA-    C-    O
   220    219    234    236      3    0.00000    0.00000    0.00000
0.00000    0.00000    0.00000 ;    HA-   CA-    C-    N
   221    219    234    235      3    0.00000    0.00000    0.00000
0.00000    0.00000    0.00000 ;    CB-   CA-    C-    O
   221    219    234    236      3    2.51040    4.18400    1.67360
-6.69440    0.00000    0.00000 ;    CB-   CA-    C-    N
   221    227    230    231      3    0.66944    2.00832    0.00000
-2.67776    0.00000    0.00000 ;    CB-   CG1-   CD1-  HD11
   221    227    230    232      3    0.66944    2.00832    0.00000
-2.67776    0.00000    0.00000 ;    CB-   CG1-   CD1-  HD12
   221    227    230    233      3    0.66944    2.00832    0.00000
-2.67776    0.00000    0.00000 ;    CB-   CG1-   CD1-  HD13
   222    221    219    234      3    0.65084    1.95253    0.00000
-2.60338    0.00000    0.00000 ;    HB-   CB-   CA-    C
   222    221    223    224      3    0.62760    1.88280    0.00000
-2.51040    0.00000    0.00000 ;    HB-   CB-   CG2-  HG21
   222    221    223    225      3    0.62760    1.88280    0.00000
-2.51040    0.00000    0.00000 ;    HB-   CB-   CG2-  HG22
   222    221    223    226      3    0.62760    1.88280    0.00000
-2.51040    0.00000    0.00000 ;    HB-   CB-   CG2-  HG23
   222    221    227    228      3    0.62760    1.88280    0.00000
-2.51040    0.00000    0.00000 ;    HB-   CB-   CG1-  HG12
   222    221    227    229      3    0.62760    1.88280    0.00000
-2.51040    0.00000    0.00000 ;    HB-   CB-   CG1-  HG13
   222    221    227    230      3    0.66944    2.00832    0.00000
-2.67776    0.00000    0.00000 ;    HB-   CB-   CG1-   CD1
   223    221    219    234      3    5.67350    3.55640   -6.16722
-2.47693    3.74886    0.00000 ;   CG2-   CB-   CA-    C
   223    221    227    228      3    0.66944    2.00832    0.00000
-2.67776    0.00000    0.00000 ;   CG2-   CB-   CG1-  HG12
   223    221    227    229      3    0.66944    2.00832    0.00000
-2.67776    0.00000    0.00000 ;   CG2-   CB-   CG1-  HG13
```

223 221 227 230 3 3.81162 0.49790 -8.14206
-1.79075 7.49773 0.00000 ; CG2- CB- CG1- CD1
224 223 221 227 3 0.66944 2.00832 0.00000
-2.67776 0.00000 0.00000 ; HG21- CG2- CB- CG1
225 223 221 227 3 0.66944 2.00832 0.00000
-2.67776 0.00000 0.00000 ; HG22- CG2- CB- CG1
226 223 221 227 3 0.66944 2.00832 0.00000
-2.67776 0.00000 0.00000 ; HG23- CG2- CB- CG1
227 221 219 234 3 8.26340 0.74057 -9.99976
-1.89117 3.84928 0.00000 ; CG1- CB- CA- C
228 227 230 231 3 0.62760 1.88280 0.00000
-2.51040 0.00000 0.00000 ; HG12- CG1- CD1- HD11
228 227 230 232 3 0.62760 1.88280 0.00000
-2.51040 0.00000 0.00000 ; HG12- CG1- CD1- HD12
228 227 230 233 3 0.62760 1.88280 0.00000
-2.51040 0.00000 0.00000 ; HG12- CG1- CD1- HD13
229 227 230 231 3 0.62760 1.88280 0.00000
-2.51040 0.00000 0.00000 ; HG13- CG1- CD1- HD11
229 227 230 232 3 0.62760 1.88280 0.00000
-2.51040 0.00000 0.00000 ; HG13- CG1- CD1- HD12
229 227 230 233 3 0.62760 1.88280 0.00000
-2.51040 0.00000 0.00000 ; HG13- CG1- CD1- HD13
234 236 238 239 3 0.00000 0.00000 0.00000
0.00000 0.00000 0.00000 ; C- N- CA- HA
234 236 238 240 3 11.71520 1.67360 15.06240 -
13.38880 0.00000 0.00000 ; C- N- CA- CB
234 236 238 253 3 1.75728 5.27184 2.25936
-7.02912 0.00000 0.00000 ; C- N- CA- C
235 234 236 237 3 29.28800 -8.36800 -20.92000
0.00000 0.00000 0.00000 ; O- C- N- H
235 234 236 238 3 20.92000 0.00000 -20.92000
0.00000 0.00000 0.00000 ; O- C- N- CA
236 238 240 241 3 0.65084 1.95253 0.00000
-2.60338 0.00000 0.00000 ; N- CA- CB- HB
236 238 240 242 3 3.83673 0.44769 -1.77402
-2.47693 -0.03347 0.00000 ; N- CA- CB- CG2
236 238 240 246 3 2.97482 0.12134 2.04179
-1.89117 -3.24678 0.00000 ; N- CA- CB- CG1
236 238 253 254 3 0.00000 0.00000 0.00000
0.00000 0.00000 0.00000 ; N- CA- C- O
236 238 253 255 3 17.40544 -5.02080 -13.22144
9.20480 0.00000 0.00000 ; N- CA- C- N
237 236 238 239 3 0.00000 0.00000 0.00000
0.00000 0.00000 0.00000 ; H- N- CA- HA
237 236 238 240 3 0.00000 0.00000 0.00000
0.00000 0.00000 0.00000 ; H- N- CA- CB
237 236 238 253 3 0.00000 0.00000 0.00000
0.00000 0.00000 0.00000 ; H- N- CA- C
238 240 242 243 3 0.66944 2.00832 0.00000
-2.67776 0.00000 0.00000 ; CA- CB- CG2- HG21
238 240 242 244 3 0.66944 2.00832 0.00000
-2.67776 0.00000 0.00000 ; CA- CB- CG2- HG22

238 240 242 245 3 0.66944 2.00832 0.00000
-2.67776 0.00000 0.00000 ; CA- CB- CG2- HG23
238 240 246 247 3 0.66944 2.00832 0.00000
-2.67776 0.00000 0.00000 ; CA- CB- CG1- HG12
238 240 246 248 3 0.66944 2.00832 0.00000
-2.67776 0.00000 0.00000 ; CA- CB- CG1- HG13
238 240 246 249 3 4.24258 -0.52718 -7.25506
-1.79075 7.69856 0.00000 ; CA- CB- CG1- CD1
238 253 255 256 3 20.92000 0.00000 -20.92000
0.00000 0.00000 0.00000 ; CA- C- N- H
238 253 255 257 3 20.92000 0.00000 -20.92000
0.00000 0.00000 0.00000 ; CA- C- N- CA
239 238 240 241 3 0.65084 1.95253 0.00000
-2.60338 0.00000 0.00000 ; HA- CA- CB- HB
239 238 240 242 3 0.65084 1.95253 0.00000
-2.60338 0.00000 0.00000 ; HA- CA- CB- CG2
239 238 240 246 3 0.65084 1.95253 0.00000
-2.60338 0.00000 0.00000 ; HA- CA- CB- CG1
239 238 253 254 3 3.68192 -4.35136 0.00000
1.33888 0.00000 0.00000 ; HA- CA- C- O
239 238 253 255 3 0.00000 0.00000 0.00000
0.00000 0.00000 0.00000 ; HA- CA- C- N
240 238 253 254 3 0.00000 0.00000 0.00000
0.00000 0.00000 0.00000 ; CB- CA- C- O
240 238 253 255 3 2.51040 4.18400 1.67360
-6.69440 0.00000 0.00000 ; CB- CA- C- N
240 246 249 250 3 0.66944 2.00832 0.00000
-2.67776 0.00000 0.00000 ; CB- CG1- CD1- HD11
240 246 249 251 3 0.66944 2.00832 0.00000
-2.67776 0.00000 0.00000 ; CB- CG1- CD1- HD12
240 246 249 252 3 0.66944 2.00832 0.00000
-2.67776 0.00000 0.00000 ; CB- CG1- CD1- HD13
241 240 238 253 3 0.65084 1.95253 0.00000
-2.60338 0.00000 0.00000 ; HB- CB- CA- C
241 240 242 243 3 0.62760 1.88280 0.00000
-2.51040 0.00000 0.00000 ; HB- CB- CG2- HG21
241 240 242 244 3 0.62760 1.88280 0.00000
-2.51040 0.00000 0.00000 ; HB- CB- CG2- HG22
241 240 242 245 3 0.62760 1.88280 0.00000
-2.51040 0.00000 0.00000 ; HB- CB- CG2- HG23
241 240 246 247 3 0.62760 1.88280 0.00000
-2.51040 0.00000 0.00000 ; HB- CB- CG1- HG12
241 240 246 248 3 0.62760 1.88280 0.00000
-2.51040 0.00000 0.00000 ; HB- CB- CG1- HG13
241 240 246 249 3 0.66944 2.00832 0.00000
-2.67776 0.00000 0.00000 ; HB- CB- CG1- CD1
242 240 238 253 3 5.67350 3.55640 -6.16722
-2.47693 3.74886 0.00000 ; CG2- CB- CA- C
242 240 246 247 3 0.66944 2.00832 0.00000
-2.67776 0.00000 0.00000 ; CG2- CB- CG1- HG12
242 240 246 248 3 0.66944 2.00832 0.00000
-2.67776 0.00000 0.00000 ; CG2- CB- CG1- HG13

```
   242    240    246    249      3    3.81162    0.49790   -8.14206
-1.79075    7.49773    0.00000 ;    CG2-    CB-   CG1-   CD1
   243    242    240    246      3    0.66944    2.00832    0.00000
-2.67776    0.00000    0.00000 ;   HG21-   CG2-    CB-   CG1
   244    242    240    246      3    0.66944    2.00832    0.00000
-2.67776    0.00000    0.00000 ;   HG22-   CG2-    CB-   CG1
   245    242    240    246      3    0.66944    2.00832    0.00000
-2.67776    0.00000    0.00000 ;   HG23-   CG2-    CB-   CG1
   246    240    238    253      3    8.26340    0.74057   -9.99976
-1.89117    3.84928    0.00000 ;    CG1-    CB-    CA-     C
   247    246    249    250      3    0.62760    1.88280    0.00000
-2.51040    0.00000    0.00000 ;   HG12-   CG1-   CD1-  HD11
   247    246    249    251      3    0.62760    1.88280    0.00000
-2.51040    0.00000    0.00000 ;   HG12-   CG1-   CD1-  HD12
   247    246    249    252      3    0.62760    1.88280    0.00000
-2.51040    0.00000    0.00000 ;   HG12-   CG1-   CD1-  HD13
   248    246    249    250      3    0.62760    1.88280    0.00000
-2.51040    0.00000    0.00000 ;   HG13-   CG1-   CD1-  HD11
   248    246    249    251      3    0.62760    1.88280    0.00000
-2.51040    0.00000    0.00000 ;   HG13-   CG1-   CD1-  HD12
   248    246    249    252      3    0.62760    1.88280    0.00000
-2.51040    0.00000    0.00000 ;   HG13-   CG1-   CD1-  HD13
   253    255    257    258      3    0.00000    0.00000    0.00000
0.00000    0.00000    0.00000 ;      C-     N-    CA-    HA
   253    255    257    259      3   11.71520    1.67360   15.06240  -
13.38880    0.00000    0.00000 ;      C-     N-    CA-    CB
   253    255    257    265      3    1.75728    5.27184    2.25936
-7.02912    0.00000    0.00000 ;      C-     N-    CA-     C
   254    253    255    256      3   29.28800   -8.36800  -20.92000
0.00000    0.00000    0.00000 ;      O-     C-     N-     H
   254    253    255    257      3   20.92000    0.00000  -20.92000
0.00000    0.00000    0.00000 ;      O-     C-     N-    CA
   255    257    259    260      3    0.65084    1.95253    0.00000
-2.60338    0.00000    0.00000 ;      N-    CA-    CB-   HB2
   255    257    259    261      3    0.65084    1.95253    0.00000
-2.60338    0.00000    0.00000 ;      N-    CA-    CB-   HB3
   255    257    259    262      3   15.41386    9.74035   -8.39310
-0.97069    2.97901    0.00000 ;      N-    CA-    CB-    CG
   255    257    265    266      3    0.00000    0.00000    0.00000
0.00000    0.00000    0.00000 ;      N-    CA-     C-     O
   255    257    265    267      3   17.40544   -5.02080  -13.22144
9.20480    0.00000    0.00000 ;      N-    CA-     C-     N
   256    255    257    258      3    0.00000    0.00000    0.00000
0.00000    0.00000    0.00000 ;      H-     N-    CA-    HA
   256    255    257    259      3    0.00000    0.00000    0.00000
0.00000    0.00000    0.00000 ;      H-     N-    CA-    CB
   256    255    257    265      3    0.00000    0.00000    0.00000
0.00000    0.00000    0.00000 ;      H-     N-    CA-     C
   257    259    262    263      3    6.43499    0.00000   -5.39736
0.00000   -1.03763    0.00000 ;     CA-    CB-    CG-   OD1
   257    259    262    264      3    6.43499    0.00000   -5.39736
0.00000   -1.03763    0.00000 ;     CA-    CB-    CG-   OD2
```

257 265 267 268 3 20.92000 0.00000 -20.92000
0.00000 0.00000 0.00000 ; CA- C- N- H
257 265 267 269 3 20.92000 0.00000 -20.92000
0.00000 0.00000 0.00000 ; CA- C- N- CA
258 257 259 260 3 0.65084 1.95253 0.00000
-2.60338 0.00000 0.00000 ; HA- CA- CB- HB2
258 257 259 261 3 0.65084 1.95253 0.00000
-2.60338 0.00000 0.00000 ; HA- CA- CB- HB3
258 257 259 262 3 0.65084 1.95253 0.00000
-2.60338 0.00000 0.00000 ; HA- CA- CB- CG
258 257 265 266 3 3.68192 -4.35136 0.00000
1.33888 0.00000 0.00000 ; HA- CA- C- O
258 257 265 267 3 0.00000 0.00000 0.00000
0.00000 0.00000 0.00000 ; HA- CA- C- N
259 257 265 266 3 0.00000 0.00000 0.00000
0.00000 0.00000 0.00000 ; CB- CA- C- O
259 257 265 267 3 2.51040 4.18400 1.67360
-6.69440 0.00000 0.00000 ; CB- CA- C- N
260 259 257 265 3 0.65084 1.95253 0.00000
-2.60338 0.00000 0.00000 ; HB2- CB- CA- C
260 259 262 263 3 0.00000 0.00000 0.00000
0.00000 0.00000 0.00000 ; HB2- CB- CG- OD1
260 259 262 264 3 0.00000 0.00000 0.00000
0.00000 0.00000 0.00000 ; HB2- CB- CG- OD2
261 259 257 265 3 0.65084 1.95253 0.00000
-2.60338 0.00000 0.00000 ; HB3- CB- CA- C
261 259 262 263 3 0.00000 0.00000 0.00000
0.00000 0.00000 0.00000 ; HB3- CB- CG- OD1
261 259 262 264 3 0.00000 0.00000 0.00000
0.00000 0.00000 0.00000 ; HB3- CB- CG- OD2
262 259 257 265 3 7.14627 -1.04600 -8.99560
-0.97069 5.15469 0.00000 ; CG- CB- CA- C
265 267 269 270 3 0.00000 0.00000 0.00000
0.00000 0.00000 0.00000 ; C- N- CA- HA
265 267 269 271 3 11.71520 1.67360 15.06240 -
13.38880 0.00000 0.00000 ; C- N- CA- CB
265 267 269 280 3 1.75728 5.27184 2.25936
-7.02912 0.00000 0.00000 ; C- N- CA- C
266 265 267 268 3 29.28800 -8.36800 -20.92000
0.00000 0.00000 0.00000 ; O- C- N- H
266 265 267 269 3 20.92000 0.00000 -20.92000
0.00000 0.00000 0.00000 ; O- C- N- CA
267 269 271 272 3 0.65084 1.95253 0.00000
-2.60338 0.00000 0.00000 ; N- CA- CB- HB2
267 269 271 273 3 0.65084 1.95253 0.00000
-2.60338 0.00000 0.00000 ; N- CA- CB- HB3
267 269 271 274 3 3.21331 2.22589 -4.15053
-2.40998 2.61082 0.00000 ; N- CA- CB- CG
267 269 280 281 3 0.00000 0.00000 0.00000
0.00000 0.00000 0.00000 ; N- CA- C- O
267 269 280 282 3 17.40544 -5.02080 -13.22144
9.20480 0.00000 0.00000 ; N- CA- C- N

268 267 269 270 3 0.00000 0.00000 0.00000
0.00000 0.00000 0.00000 ; H- N- CA- HA
268 267 269 271 3 0.00000 0.00000 0.00000
0.00000 0.00000 0.00000 ; H- N- CA- CB
268 267 269 280 3 0.00000 0.00000 0.00000
0.00000 0.00000 0.00000 ; H- N- CA- C
269 271 274 275 3 0.66944 2.00832 0.00000
-2.67776 0.00000 0.00000 ; CA- CB- CG- HG2
269 271 274 276 3 0.66944 2.00832 0.00000
-2.67776 0.00000 0.00000 ; CA- CB- CG- HG3
269 271 274 277 3 10.12110 -1.91209 0.01674
10.17549 -1.87443 0.00000 ; CA- CB- CG- CD
269 280 282 283 3 20.92000 0.00000 -20.92000
0.00000 0.00000 0.00000 ; CA- C- N- H
269 280 282 284 3 20.92000 0.00000 -20.92000
0.00000 0.00000 0.00000 ; CA- C- N- CA
270 269 271 272 3 0.65084 1.95253 0.00000
-2.60338 0.00000 0.00000 ; HA- CA- CB- HB2
270 269 271 273 3 0.65084 1.95253 0.00000
-2.60338 0.00000 0.00000 ; HA- CA- CB- HB3
270 269 271 274 3 0.65084 1.95253 0.00000
-2.60338 0.00000 0.00000 ; HA- CA- CB- CG
270 269 280 281 3 3.68192 -4.35136 0.00000
1.33888 0.00000 0.00000 ; HA- CA- C- O
270 269 280 282 3 0.00000 0.00000 0.00000
0.00000 0.00000 0.00000 ; HA- CA- C- N
271 269 280 281 3 0.00000 0.00000 0.00000
0.00000 0.00000 0.00000 ; CB- CA- C- O
271 269 280 282 3 2.51040 4.18400 1.67360
-6.69440 0.00000 0.00000 ; CB- CA- C- N
271 274 277 278 3 3.79907 0.00000 -5.40573
0.00000 2.14221 0.00000 ; CB- CG- CD- OE1
271 274 277 279 3 3.79907 0.00000 -5.40573
0.00000 2.14221 0.00000 ; CB- CG- CD- OE2
272 271 269 280 3 0.65084 1.95253 0.00000
-2.60338 0.00000 0.00000 ; HB2- CB- CA- C
272 271 274 275 3 0.62760 1.88280 0.00000
-2.51040 0.00000 0.00000 ; HB2- CB- CG- HG2
272 271 274 276 3 0.62760 1.88280 0.00000
-2.51040 0.00000 0.00000 ; HB2- CB- CG- HG3
272 271 274 277 3 0.65084 1.95253 0.00000
-2.60338 0.00000 0.00000 ; HB2- CB- CG- CD
273 271 269 280 3 0.65084 1.95253 0.00000
-2.60338 0.00000 0.00000 ; HB3- CB- CA- C
273 271 274 275 3 0.62760 1.88280 0.00000
-2.51040 0.00000 0.00000 ; HB3- CB- CG- HG2
273 271 274 276 3 0.62760 1.88280 0.00000
-2.51040 0.00000 0.00000 ; HB3- CB- CG- HG3
273 271 274 277 3 0.65084 1.95253 0.00000
-2.60338 0.00000 0.00000 ; HB3- CB- CG- CD
274 271 269 280 3 6.86594 3.56895 -8.14206
-2.40998 4.85344 0.00000 ; CG- CB- CA- C

```
   275    274    277    278      3    0.00000    0.00000    0.00000
0.00000   0.00000   0.00000 ;    HG2-    CG-    CD-   OE1
   275    274    277    279      3    0.00000    0.00000    0.00000
0.00000   0.00000   0.00000 ;    HG2-    CG-    CD-   OE2
   276    274    277    278      3    0.00000    0.00000    0.00000
0.00000   0.00000   0.00000 ;    HG3-    CG-    CD-   OE1
   276    274    277    279      3    0.00000    0.00000    0.00000
0.00000   0.00000   0.00000 ;    HG3-    CG-    CD-   OE2
   280    282    284    285      3    0.00000    0.00000    0.00000
0.00000   0.00000   0.00000 ;     C-     N-    CA-    HA
   280    282    284    286      3   11.71520    1.67360   15.06240  -
13.38880   0.00000   0.00000 ;     C-     N-    CA-    CB
   280    282    284    302      3    1.75728    5.27184    2.25936
-7.02912   0.00000   0.00000 ;     C-     N-    CA-     C
   281    280    282    283      3   29.28800   -8.36800  -20.92000
0.00000   0.00000   0.00000 ;     O-     C-     N-     H
   281    280    282    284      3   20.92000    0.00000  -20.92000
0.00000   0.00000   0.00000 ;     O-     C-     N-    CA
   282    284    286    287      3    0.65084    1.95253    0.00000
-2.60338   0.00000   0.00000 ;     N-    CA-    CB-   HB2
   282    284    286    288      3    0.65084    1.95253    0.00000
-2.60338   0.00000   0.00000 ;     N-    CA-    CB-   HB3
   282    284    286    289      3    0.65084    1.95253    0.00000
-2.60338   0.00000   0.00000 ;     N-    CA-    CB-    CG
   282    284    302    303      3    0.00000    0.00000    0.00000
0.00000   0.00000   0.00000 ;     N-    CA-     C-     O
   282    284    302    304      3   17.40544   -5.02080  -13.22144
9.20480   0.00000   0.00000 ;     N-    CA-     C-     N
   283    282    284    285      3    0.00000    0.00000    0.00000
0.00000   0.00000   0.00000 ;     H-     N-    CA-    HA
   283    282    284    286      3    0.00000    0.00000    0.00000
0.00000   0.00000   0.00000 ;     H-     N-    CA-    CB
   283    282    284    302      3    0.00000    0.00000    0.00000
0.00000   0.00000   0.00000 ;     H-     N-    CA-     C
   284    286    289    290      3    0.66944    2.00832    0.00000
-2.67776   0.00000   0.00000 ;    CA-    CB-    CG-   HG2
   284    286    289    291      3    0.66944    2.00832    0.00000
-2.67776   0.00000   0.00000 ;    CA-    CB-    CG-   HG3
   284    286    289    292      3    3.68192    3.09616   -2.09200
-3.01248   0.00000   0.00000 ;    CA-    CB-    CG-    CD
   284    302    304    305      3   20.92000    0.00000  -20.92000
0.00000   0.00000   0.00000 ;    CA-     C-     N-     H
   284    302    304    306      3   20.92000    0.00000  -20.92000
0.00000   0.00000   0.00000 ;    CA-     C-     N-    CA
   285    284    286    287      3    0.65084    1.95253    0.00000
-2.60338   0.00000   0.00000 ;    HA-    CA-    CB-   HB2
   285    284    286    288      3    0.65084    1.95253    0.00000
-2.60338   0.00000   0.00000 ;    HA-    CA-    CB-   HB3
   285    284    286    289      3    0.65084    1.95253    0.00000
-2.60338   0.00000   0.00000 ;    HA-    CA-    CB-    CG
   285    284    302    303      3    3.68192   -4.35136    0.00000
1.33888   0.00000   0.00000 ;    HA-    CA-     C-     O
```

285 284 302 304 3 0.00000 0.00000 0.00000
0.00000 0.00000 0.00000 ; HA- CA- C- N
286 284 302 303 3 0.00000 0.00000 0.00000
0.00000 0.00000 0.00000 ; CB- CA- C- O
286 284 302 304 3 2.51040 4.18400 1.67360
-6.69440 0.00000 0.00000 ; CB- CA- C- N
286 289 292 293 3 0.66944 2.00832 0.00000
-2.67776 0.00000 0.00000 ; CB- CG- CD- HD2
286 289 292 294 3 0.66944 2.00832 0.00000
-2.67776 0.00000 0.00000 ; CB- CG- CD- HD3
286 289 292 295 3 3.68192 3.09616 -2.09200
-3.01248 0.00000 0.00000 ; CB- CG- CD- CE
287 286 284 302 3 0.65084 1.95253 0.00000
-2.60338 0.00000 0.00000 ; HB2- CB- CA- C
287 286 289 290 3 0.62760 1.88280 0.00000
-2.51040 0.00000 0.00000 ; HB2- CB- CG- HG2
287 286 289 291 3 0.62760 1.88280 0.00000
-2.51040 0.00000 0.00000 ; HB2- CB- CG- HG3
287 286 289 292 3 0.66944 2.00832 0.00000
-2.67776 0.00000 0.00000 ; HB2- CB- CG- CD
288 286 284 302 3 0.65084 1.95253 0.00000
-2.60338 0.00000 0.00000 ; HB3- CB- CA- C
288 286 289 290 3 0.62760 1.88280 0.00000
-2.51040 0.00000 0.00000 ; HB3- CB- CG- HG2
288 286 289 291 3 0.62760 1.88280 0.00000
-2.51040 0.00000 0.00000 ; HB3- CB- CG- HG3
288 286 289 292 3 0.66944 2.00832 0.00000
-2.67776 0.00000 0.00000 ; HB3- CB- CG- CD
289 286 284 302 3 0.65084 1.95253 0.00000
-2.60338 0.00000 0.00000 ; CG- CB- CA- C
289 292 295 296 3 0.65084 1.95253 0.00000
-2.60338 0.00000 0.00000 ; CG- CD- CE- HE2
289 292 295 297 3 0.65084 1.95253 0.00000
-2.60338 0.00000 0.00000 ; CG- CD- CE- HE3
289 292 295 298 3 0.65084 1.95253 0.00000
-2.60338 0.00000 0.00000 ; CG- CD- CE- NZ
290 289 292 293 3 0.62760 1.88280 0.00000
-2.51040 0.00000 0.00000 ; HG2- CG- CD- HD2
290 289 292 294 3 0.62760 1.88280 0.00000
-2.51040 0.00000 0.00000 ; HG2- CG- CD- HD3
290 289 292 295 3 0.66944 2.00832 0.00000
-2.67776 0.00000 0.00000 ; HG2- CG- CD- CE
291 289 292 293 3 0.62760 1.88280 0.00000
-2.51040 0.00000 0.00000 ; HG3- CG- CD- HD2
291 289 292 294 3 0.62760 1.88280 0.00000
-2.51040 0.00000 0.00000 ; HG3- CG- CD- HD3
291 289 292 295 3 0.66944 2.00832 0.00000
-2.67776 0.00000 0.00000 ; HG3- CG- CD- CE
292 295 298 299 3 0.65084 1.95253 0.00000
-2.60338 0.00000 0.00000 ; CD- CE- NZ- HZ1
292 295 298 300 3 0.65084 1.95253 0.00000
-2.60338 0.00000 0.00000 ; CD- CE- NZ- HZ2

292 295 298 301 3 0.65084 1.95253 0.00000
-2.60338 0.00000 0.00000 ; CD- CE- NZ- HZ3
293 292 295 296 3 0.65084 1.95253 0.00000
-2.60338 0.00000 0.00000 ; HD2- CD- CE- HE2
293 292 295 297 3 0.65084 1.95253 0.00000
-2.60338 0.00000 0.00000 ; HD2- CD- CE- HE3
293 292 295 298 3 0.65084 1.95253 0.00000
-2.60338 0.00000 0.00000 ; HD2- CD- CE- NZ
294 292 295 296 3 0.65084 1.95253 0.00000
-2.60338 0.00000 0.00000 ; HD3- CD- CE- HE2
294 292 295 297 3 0.65084 1.95253 0.00000
-2.60338 0.00000 0.00000 ; HD3- CD- CE- HE3
294 292 295 298 3 0.65084 1.95253 0.00000
-2.60338 0.00000 0.00000 ; HD3- CD- CE- NZ
296 295 298 299 3 0.65084 1.95253 0.00000
-2.60338 0.00000 0.00000 ; HE2- CE- NZ- HZ1
296 295 298 300 3 0.65084 1.95253 0.00000
-2.60338 0.00000 0.00000 ; HE2- CE- NZ- HZ2
296 295 298 301 3 0.65084 1.95253 0.00000
-2.60338 0.00000 0.00000 ; HE2- CE- NZ- HZ3
297 295 298 299 3 0.65084 1.95253 0.00000
-2.60338 0.00000 0.00000 ; HE3- CE- NZ- HZ1
297 295 298 300 3 0.65084 1.95253 0.00000
-2.60338 0.00000 0.00000 ; HE3- CE- NZ- HZ2
297 295 298 301 3 0.65084 1.95253 0.00000
-2.60338 0.00000 0.00000 ; HE3- CE- NZ- HZ3
302 304 306 307 3 0.00000 0.00000 0.00000
0.00000 0.00000 0.00000 ; C- N- CA- HA
302 304 306 308 3 11.71520 1.67360 15.06240 -
13.38880 0.00000 0.00000 ; C- N- CA- CB
302 304 306 321 3 1.75728 5.27184 2.25936
-7.02912 0.00000 0.00000 ; C- N- CA- C
303 302 304 305 3 29.28800 -8.36800 -20.92000
0.00000 0.00000 0.00000 ; O- C- N- H
303 302 304 306 3 20.92000 0.00000 -20.92000
0.00000 0.00000 0.00000 ; O- C- N- CA
304 306 308 309 3 0.65084 1.95253 0.00000
-2.60338 0.00000 0.00000 ; N- CA- CB- HB2
304 306 308 310 3 0.65084 1.95253 0.00000
-2.60338 0.00000 0.00000 ; N- CA- CB- HB3
304 306 308 311 3 3.79070 1.39746 -4.61077
-2.40998 2.44346 0.00000 ; N- CA- CB- CG
304 306 321 322 3 0.00000 0.00000 0.00000
0.00000 0.00000 0.00000 ; N- CA- C- O
304 306 321 323 3 17.40544 -5.02080 -13.22144
9.20480 0.00000 0.00000 ; N- CA- C- N
305 304 306 307 3 0.00000 0.00000 0.00000
0.00000 0.00000 0.00000 ; H- N- CA- HA
305 304 306 308 3 0.00000 0.00000 0.00000
0.00000 0.00000 0.00000 ; H- N- CA- CB
305 304 306 321 3 0.00000 0.00000 0.00000
0.00000 0.00000 0.00000 ; H- N- CA- C

```
   306    308    311    312      3    0.66944    2.00832    0.00000
-2.67776    0.00000    0.00000 ;     CA-    CB-    CG-    HG
   306    308    311    313      3    3.90367    0.19665   -6.21742
-2.37651    5.99149    0.00000 ;     CA-    CB-    CG-   CD1
   306    308    311    317      3    3.90367    0.19665   -6.21742
-2.37651    5.99149    0.00000 ;     CA-    CB-    CG-   CD2
   306    321    323    324      3   20.92000    0.00000  -20.92000
0.00000    0.00000    0.00000 ;     CA-     C-     N-     H
   306    321    323    325      3   20.92000    0.00000  -20.92000
0.00000    0.00000    0.00000 ;     CA-     C-     N-    CA
   307    306    308    309      3    0.65084    1.95253    0.00000
-2.60338    0.00000    0.00000 ;     HA-    CA-    CB-   HB2
   307    306    308    310      3    0.65084    1.95253    0.00000
-2.60338    0.00000    0.00000 ;     HA-    CA-    CB-   HB3
   307    306    308    311      3    0.65084    1.95253    0.00000
-2.60338    0.00000    0.00000 ;     HA-    CA-    CB-    CG
   307    306    321    322      3    3.68192   -4.35136    0.00000
1.33888    0.00000    0.00000 ;     HA-    CA-     C-     O
   307    306    321    323      3    0.00000    0.00000    0.00000
0.00000    0.00000    0.00000 ;     HA-    CA-     C-     N
   308    306    321    322      3    0.00000    0.00000    0.00000
0.00000    0.00000    0.00000 ;     CB-    CA-     C-     O
   308    306    321    323      3    2.51040    4.18400    1.67360
-6.69440    0.00000    0.00000 ;     CB-    CA-     C-     N
   308    311    313    314      3    0.66944    2.00832    0.00000
-2.67776    0.00000    0.00000 ;     CB-    CG-   CD1-  HD11
   308    311    313    315      3    0.66944    2.00832    0.00000
-2.67776    0.00000    0.00000 ;     CB-    CG-   CD1-  HD12
   308    311    313    316      3    0.66944    2.00832    0.00000
-2.67776    0.00000    0.00000 ;     CB-    CG-   CD1-  HD13
   308    311    317    318      3    0.66944    2.00832    0.00000
-2.67776    0.00000    0.00000 ;     CB-    CG-   CD2-  HD21
   308    311    317    319      3    0.66944    2.00832    0.00000
-2.67776    0.00000    0.00000 ;     CB-    CG-   CD2-  HD22
   308    311    317    320      3    0.66944    2.00832    0.00000
-2.67776    0.00000    0.00000 ;     CB-    CG-   CD2-  HD23
   309    308    306    321      3    0.65084    1.95253    0.00000
-2.60338    0.00000    0.00000 ;    HB2-    CB-    CA-     C
   309    308    311    312      3    0.62760    1.88280    0.00000
-2.51040    0.00000    0.00000 ;    HB2-    CB-    CG-    HG
   309    308    311    313      3    0.66944    2.00832    0.00000
-2.67776    0.00000    0.00000 ;    HB2-    CB-    CG-   CD1
   309    308    311    317      3    0.66944    2.00832    0.00000
-2.67776    0.00000    0.00000 ;    HB2-    CB-    CG-   CD2
   310    308    306    321      3    0.65084    1.95253    0.00000
-2.60338    0.00000    0.00000 ;    HB3-    CB-    CA-     C
   310    308    311    312      3    0.62760    1.88280    0.00000
-2.51040    0.00000    0.00000 ;    HB3-    CB-    CG-    HG
   310    308    311    313      3    0.66944    2.00832    0.00000
-2.67776    0.00000    0.00000 ;    HB3-    CB-    CG-   CD1
   310    308    311    317      3    0.66944    2.00832    0.00000
-2.67776    0.00000    0.00000 ;    HB3-    CB-    CG-   CD2
```

311 308 306 321 3 10.33448 -1.14642 -11.54784
-2.40998 6.35968 0.00000 ; CG- CB- CA- C
312 311 313 314 3 0.62760 1.88280 0.00000
-2.51040 0.00000 0.00000 ; HG- CG- CD1- HD11
312 311 313 315 3 0.62760 1.88280 0.00000
-2.51040 0.00000 0.00000 ; HG- CG- CD1- HD12
312 311 313 316 3 0.62760 1.88280 0.00000
-2.51040 0.00000 0.00000 ; HG- CG- CD1- HD13
312 311 317 318 3 0.62760 1.88280 0.00000
-2.51040 0.00000 0.00000 ; HG- CG- CD2- HD21
312 311 317 319 3 0.62760 1.88280 0.00000
-2.51040 0.00000 0.00000 ; HG- CG- CD2- HD22
312 311 317 320 3 0.62760 1.88280 0.00000
-2.51040 0.00000 0.00000 ; HG- CG- CD2- HD23
313 311 317 318 3 0.66944 2.00832 0.00000
-2.67776 0.00000 0.00000 ; CD1- CG- CD2- HD21
313 311 317 319 3 0.66944 2.00832 0.00000
-2.67776 0.00000 0.00000 ; CD1- CG- CD2- HD22
313 311 317 320 3 0.66944 2.00832 0.00000
-2.67776 0.00000 0.00000 ; CD1- CG- CD2- HD23
314 313 311 317 3 0.66944 2.00832 0.00000
-2.67776 0.00000 0.00000 ; HD11- CD1- CG- CD2
315 313 311 317 3 0.66944 2.00832 0.00000
-2.67776 0.00000 0.00000 ; HD12- CD1- CG- CD2
316 313 311 317 3 0.66944 2.00832 0.00000
-2.67776 0.00000 0.00000 ; HD13- CD1- CG- CD2
321 323 325 326 3 0.00000 0.00000 0.00000
0.00000 0.00000 0.00000 ; C- N- CA- HA2
321 323 325 327 3 0.00000 0.00000 0.00000
0.00000 0.00000 0.00000 ; C- N- CA- HA3
321 323 325 328 3 1.75728 5.27184 2.25936
-7.02912 0.00000 0.00000 ; C- N- CA- C
322 321 323 324 3 29.28800 -8.36800 -20.92000
0.00000 0.00000 0.00000 ; O- C- N- H
322 321 323 325 3 20.92000 0.00000 -20.92000
0.00000 0.00000 0.00000 ; O- C- N- CA
323 325 328 329 3 0.00000 0.00000 0.00000
0.00000 0.00000 0.00000 ; N- CA- C- O
323 325 328 330 3 17.40544 -5.02080 -13.22144
9.20480 0.00000 0.00000 ; N- CA- C- N
324 323 325 326 3 0.00000 0.00000 0.00000
0.00000 0.00000 0.00000 ; H- N- CA- HA2
324 323 325 327 3 0.00000 0.00000 0.00000
0.00000 0.00000 0.00000 ; H- N- CA- HA3
324 323 325 328 3 0.00000 0.00000 0.00000
0.00000 0.00000 0.00000 ; H- N- CA- C
325 328 330 331 3 20.92000 0.00000 -20.92000
0.00000 0.00000 0.00000 ; CA- C- N- H
325 328 330 332 3 20.92000 0.00000 -20.92000
0.00000 0.00000 0.00000 ; CA- C- N- CA
326 325 328 329 3 3.68192 -4.35136 0.00000
1.33888 0.00000 0.00000 ; HA2- CA- C- O

326 325 328 330 3 0.00000 0.00000 0.00000
0.00000 0.00000 0.00000 ; HA2- CA- C- N
327 325 328 329 3 3.68192 -4.35136 0.00000
1.33888 0.00000 0.00000 ; HA3- CA- C- O
327 325 328 330 3 0.00000 0.00000 0.00000
0.00000 0.00000 0.00000 ; HA3- CA- C- N
328 330 332 333 3 0.00000 0.00000 0.00000
0.00000 0.00000 0.00000 ; C- N- CA- HA
328 330 332 334 3 11.71520 1.67360 15.06240 -
13.38880 0.00000 0.00000 ; C- N- CA- CB
328 330 332 345 3 1.75728 5.27184 2.25936
-7.02912 0.00000 0.00000 ; C- N- CA- C
329 328 330 331 3 29.28800 -8.36800 -20.92000
0.00000 0.00000 0.00000 ; O- C- N- H
329 328 330 332 3 20.92000 0.00000 -20.92000
0.00000 0.00000 0.00000 ; O- C- N- CA
330 332 334 335 3 0.65084 1.95253 0.00000
-2.60338 0.00000 0.00000 ; N- CA- CB- HB2
330 332 334 336 3 0.65084 1.95253 0.00000
-2.60338 0.00000 0.00000 ; N- CA- CB- HB3
330 332 334 337 3 3.21331 2.22589 -4.15053
-2.40998 2.61082 0.00000 ; N- CA- CB- CG
330 332 345 346 3 0.00000 0.00000 0.00000
0.00000 0.00000 0.00000 ; N- CA- C- O
330 332 345 347 3 17.40544 -5.02080 -13.22144
9.20480 0.00000 0.00000 ; N- CA- C- N
331 330 332 333 3 0.00000 0.00000 0.00000
0.00000 0.00000 0.00000 ; H- N- CA- HA
331 330 332 334 3 0.00000 0.00000 0.00000
0.00000 0.00000 0.00000 ; H- N- CA- CB
331 330 332 345 3 0.00000 0.00000 0.00000
0.00000 0.00000 0.00000 ; H- N- CA- C
332 334 337 338 3 0.66944 2.00832 0.00000
-2.67776 0.00000 0.00000 ; CA- CB- CG- HG2
332 334 337 339 3 0.66944 2.00832 0.00000
-2.67776 0.00000 0.00000 ; CA- CB- CG- HG3
332 334 337 340 3 3.69866 -4.35136 -3.92459
6.89523 4.61914 0.00000 ; CA- CB- CG- CD
332 345 347 348 3 20.92000 0.00000 -20.92000
0.00000 0.00000 0.00000 ; CA- C- N- H
332 345 347 349 3 20.92000 0.00000 -20.92000
0.00000 0.00000 0.00000 ; CA- C- N- CA
333 332 334 335 3 0.65084 1.95253 0.00000
-2.60338 0.00000 0.00000 ; HA- CA- CB- HB2
333 332 334 336 3 0.65084 1.95253 0.00000
-2.60338 0.00000 0.00000 ; HA- CA- CB- HB3
333 332 334 337 3 0.65084 1.95253 0.00000
-2.60338 0.00000 0.00000 ; HA- CA- CB- CG
333 332 345 346 3 3.68192 -4.35136 0.00000
1.33888 0.00000 0.00000 ; HA- CA- C- O
333 332 345 347 3 0.00000 0.00000 0.00000
0.00000 0.00000 0.00000 ; HA- CA- C- N

334 332 345 346 3 0.00000 0.00000 0.00000
0.00000 0.00000 0.00000 ; CB- CA- C- O
334 332 345 347 3 2.51040 4.18400 1.67360
-6.69440 0.00000 0.00000 ; CB- CA- C- N
334 337 340 341 3 0.00000 0.00000 0.00000
0.00000 0.00000 0.00000 ; CB- CG- CD- OE1
334 337 340 342 3 10.32611 1.48114 -8.47678
1.42256 1.40582 0.00000 ; CB- CG- CD- NE2
335 334 332 345 3 0.65084 1.95253 0.00000
-2.60338 0.00000 0.00000 ; HB2- CB- CA- C
335 334 337 338 3 0.62760 1.88280 0.00000
-2.51040 0.00000 0.00000 ; HB2- CB- CG- HG2
335 334 337 339 3 0.62760 1.88280 0.00000
-2.51040 0.00000 0.00000 ; HB2- CB- CG- HG3
335 334 337 340 3 0.65084 1.95253 0.00000
-2.60338 0.00000 0.00000 ; HB2- CB- CG- CD
336 334 332 345 3 0.65084 1.95253 0.00000
-2.60338 0.00000 0.00000 ; HB3- CB- CA- C
336 334 337 338 3 0.62760 1.88280 0.00000
-2.51040 0.00000 0.00000 ; HB3- CB- CG- HG2
336 334 337 339 3 0.62760 1.88280 0.00000
-2.51040 0.00000 0.00000 ; HB3- CB- CG- HG3
336 334 337 340 3 0.65084 1.95253 0.00000
-2.60338 0.00000 0.00000 ; HB3- CB- CG- CD
337 334 332 345 3 6.86594 3.56895 -8.14206
-2.40998 4.85344 0.00000 ; CG- CB- CA- C
337 340 342 343 3 20.92000 0.00000 -20.92000
0.00000 0.00000 0.00000 ; CG- CD- NE2- HE21
337 340 342 344 3 20.92000 0.00000 -20.92000
0.00000 0.00000 0.00000 ; CG- CD- NE2- HE22
338 337 340 341 3 3.68192 -4.35136 0.00000
1.33888 0.00000 0.00000 ; HG2- CG- CD- OE1
338 337 340 342 3 0.00000 0.00000 0.00000
0.00000 0.00000 0.00000 ; HG2- CG- CD- NE2
339 337 340 341 3 3.68192 -4.35136 0.00000
1.33888 0.00000 0.00000 ; HG3- CG- CD- OE1
339 337 340 342 3 0.00000 0.00000 0.00000
0.00000 0.00000 0.00000 ; HG3- CG- CD- NE2
341 340 342 343 3 29.28800 -8.36800 -20.92000
0.00000 0.00000 0.00000 ; OE1- CD- NE2- HE21
341 340 342 344 3 29.28800 -8.36800 -20.92000
0.00000 0.00000 0.00000 ; OE1- CD- NE2- HE22
345 347 349 350 3 0.00000 0.00000 0.00000
0.00000 0.00000 0.00000 ; C- N- CA- HA2
345 347 349 351 3 0.00000 0.00000 0.00000
0.00000 0.00000 0.00000 ; C- N- CA- HA3
345 347 349 352 3 1.75728 5.27184 2.25936
-7.02912 0.00000 0.00000 ; C- N- CA- C
346 345 347 348 3 29.28800 -8.36800 -20.92000
0.00000 0.00000 0.00000 ; O- C- N- H
346 345 347 349 3 20.92000 0.00000 -20.92000
0.00000 0.00000 0.00000 ; O- C- N- CA

347 349 352 353 3 0.00000 0.00000 0.00000
0.00000 0.00000 0.00000 ; N- CA- C- O
347 349 352 354 3 17.40544 -5.02080 -13.22144
9.20480 0.00000 0.00000 ; N- CA- C- N
348 347 349 350 3 0.00000 0.00000 0.00000
0.00000 0.00000 0.00000 ; H- N- CA- HA2
348 347 349 351 3 0.00000 0.00000 0.00000
0.00000 0.00000 0.00000 ; H- N- CA- HA3
348 347 349 352 3 0.00000 0.00000 0.00000
0.00000 0.00000 0.00000 ; H- N- CA- C
349 352 354 355 3 20.92000 0.00000 -20.92000
0.00000 0.00000 0.00000 ; CA- C- N- H
349 352 354 356 3 20.92000 0.00000 -20.92000
0.00000 0.00000 0.00000 ; CA- C- N- CA
350 349 352 353 3 3.68192 -4.35136 0.00000
1.33888 0.00000 0.00000 ; HA2- CA- C- O
350 349 352 354 3 0.00000 0.00000 0.00000
0.00000 0.00000 0.00000 ; HA2- CA- C- N
351 349 352 353 3 3.68192 -4.35136 0.00000
1.33888 0.00000 0.00000 ; HA3- CA- C- O
351 349 352 354 3 0.00000 0.00000 0.00000
0.00000 0.00000 0.00000 ; HA3- CA- C- N
352 354 356 357 3 0.00000 0.00000 0.00000
0.00000 0.00000 0.00000 ; C- N- CA- HA
352 354 356 358 3 11.71520 1.67360 15.06240 -
13.38880 0.00000 0.00000 ; C- N- CA- CB
352 354 356 366 3 1.75728 5.27184 2.25936
-7.02912 0.00000 0.00000 ; C- N- CA- C
353 352 354 355 3 29.28800 -8.36800 -20.92000
0.00000 0.00000 0.00000 ; O- C- N- H
353 352 354 356 3 20.92000 0.00000 -20.92000
0.00000 0.00000 0.00000 ; O- C- N- CA
354 356 358 359 3 0.65084 1.95253 0.00000
-2.60338 0.00000 0.00000 ; N- CA- CB- HB
354 356 358 360 3 3.83673 0.44769 -1.77402
-2.47693 -0.03347 0.00000 ; N- CA- CB- CG2
354 356 358 364 3 4.93294 1.13386 -3.12963
-5.27184 3.17984 0.00000 ; N- CA- CB- OG1
354 356 366 367 3 0.00000 0.00000 0.00000
0.00000 0.00000 0.00000 ; N- CA- C- O
354 356 366 368 3 17.40544 -5.02080 -13.22144
9.20480 0.00000 0.00000 ; N- CA- C- N
355 354 356 357 3 0.00000 0.00000 0.00000
0.00000 0.00000 0.00000 ; H- N- CA- HA
355 354 356 358 3 0.00000 0.00000 0.00000
0.00000 0.00000 0.00000 ; H- N- CA- CB
355 354 356 366 3 0.00000 0.00000 0.00000
0.00000 0.00000 0.00000 ; H- N- CA- C
356 358 360 361 3 0.66944 2.00832 0.00000
-2.67776 0.00000 0.00000 ; CA- CB- CG2- HG21
356 358 360 362 3 0.66944 2.00832 0.00000
-2.67776 0.00000 0.00000 ; CA- CB- CG2- HG22

356 358 360 363 3 0.66944 2.00832 0.00000
-2.67776 0.00000 0.00000 ; CA- CB- CG2- HG23
356 358 364 365 3 1.12131 2.98738 1.66523
-3.94970 0.43514 0.00000 ; CA- CB- OG1- HG1
356 366 368 369 3 20.92000 0.00000 -20.92000
0.00000 0.00000 0.00000 ; CA- C- N- H
356 366 368 370 3 20.92000 0.00000 -20.92000
0.00000 0.00000 0.00000 ; CA- C- N- CA
357 356 358 359 3 0.65084 1.95253 0.00000
-2.60338 0.00000 0.00000 ; HA- CA- CB- HB
357 356 358 360 3 0.65084 1.95253 0.00000
-2.60338 0.00000 0.00000 ; HA- CA- CB- CG2
357 356 358 364 3 1.04600 -1.04600 0.00000
0.00000 0.00000 0.00000 ; HA- CA- CB- OG1
357 356 366 367 3 3.68192 -4.35136 0.00000
1.33888 0.00000 0.00000 ; HA- CA- C- O
357 356 366 368 3 0.00000 0.00000 0.00000
0.00000 0.00000 0.00000 ; HA- CA- C- N
358 356 366 367 3 0.00000 0.00000 0.00000
0.00000 0.00000 0.00000 ; CB- CA- C- O
358 356 366 368 3 2.51040 4.18400 1.67360
-6.69440 0.00000 0.00000 ; CB- CA- C- N
359 358 356 366 3 0.65084 1.95253 0.00000
-2.60338 0.00000 0.00000 ; HB- CB- CA- C
359 358 360 361 3 0.65084 1.95253 0.00000
-2.60338 0.00000 0.00000 ; HB- CB- CG2- HG21
359 358 360 362 3 0.65084 1.95253 0.00000
-2.60338 0.00000 0.00000 ; HB- CB- CG2- HG22
359 358 360 363 3 0.65084 1.95253 0.00000
-2.60338 0.00000 0.00000 ; HB- CB- CG2- HG23
359 358 364 365 3 0.69733 2.09200 0.00000
-2.78933 0.00000 0.00000 ; HB- CB- OG1- HG1
360 358 356 366 3 5.67350 3.55640 -6.16722
-2.47693 3.74886 0.00000 ; CG2- CB- CA- C
360 358 364 365 3 4.74047 0.27196 -2.26773
-3.94970 1.60666 0.00000 ; CG2- CB- OG1- HG1
361 360 358 364 3 1.04600 -1.04600 0.00000
0.00000 0.00000 0.00000 ; HG21- CG2- CB- OG1
362 360 358 364 3 1.04600 -1.04600 0.00000
0.00000 0.00000 0.00000 ; HG22- CG2- CB- OG1
363 360 358 364 3 1.04600 -1.04600 0.00000
0.00000 0.00000 0.00000 ; HG23- CG2- CB- OG1
364 358 356 366 3 6.53541 6.87013 -6.21742
-5.27184 5.22163 0.00000 ; OG1- CB- CA- C
366 368 370 371 3 0.00000 0.00000 0.00000
0.00000 0.00000 0.00000 ; C- N- CA- HA
366 368 370 372 3 11.71520 1.67360 15.06240 -
13.38880 0.00000 0.00000 ; C- N- CA- CB
366 368 370 387 3 1.75728 5.27184 2.25936
-7.02912 0.00000 0.00000 ; C- N- CA- C
367 366 368 369 3 29.28800 -8.36800 -20.92000
0.00000 0.00000 0.00000 ; O- C- N- H

367 366 368 370 3 20.92000 0.00000 -20.92000
0.00000 0.00000 0.00000 ; O- C- N- CA
368 370 372 373 3 0.65084 1.95253 0.00000
-2.60338 0.00000 0.00000 ; N- CA- CB- HB2
368 370 372 374 3 0.65084 1.95253 0.00000
-2.60338 0.00000 0.00000 ; N- CA- CB- HB3
368 370 372 375 3 3.28026 2.46019 -2.19242
-3.21331 -0.23430 0.00000 ; N- CA- CB- CG
368 370 387 388 3 0.00000 0.00000 0.00000
0.00000 0.00000 0.00000 ; N- CA- C- O
368 370 387 389 3 17.40544 -5.02080 -13.22144
9.20480 0.00000 0.00000 ; N- CA- C- N
369 368 370 371 3 0.00000 0.00000 0.00000
0.00000 0.00000 0.00000 ; H- N- CA- HA
369 368 370 372 3 0.00000 0.00000 0.00000
0.00000 0.00000 0.00000 ; H- N- CA- CB
369 368 370 387 3 0.00000 0.00000 0.00000
0.00000 0.00000 0.00000 ; H- N- CA- C
370 372 375 376 3 0.57739 0.00000 1.02926
0.00000 -1.60666 0.00000 ; CA- CB- CG- CD1
370 372 375 385 3 0.57739 0.00000 1.02926
0.00000 -1.60666 0.00000 ; CA- CB- CG- CD2
370 387 389 390 3 20.92000 0.00000 -20.92000
0.00000 0.00000 0.00000 ; CA- C- N- H
370 387 389 391 3 20.92000 0.00000 -20.92000
0.00000 0.00000 0.00000 ; CA- C- N- CA
371 370 372 373 3 0.65084 1.95253 0.00000
-2.60338 0.00000 0.00000 ; HA- CA- CB- HB2
371 370 372 374 3 0.65084 1.95253 0.00000
-2.60338 0.00000 0.00000 ; HA- CA- CB- HB3
371 370 372 375 3 0.65084 1.95253 0.00000
-2.60338 0.00000 0.00000 ; HA- CA- CB- CG
371 370 387 388 3 3.68192 -4.35136 0.00000
1.33888 0.00000 0.00000 ; HA- CA- C- O
371 370 387 389 3 0.00000 0.00000 0.00000
0.00000 0.00000 0.00000 ; HA- CA- C- N
372 370 387 388 3 0.00000 0.00000 0.00000
0.00000 0.00000 0.00000 ; CB- CA- C- O
372 370 387 389 3 2.51040 4.18400 1.67360
-6.69440 0.00000 0.00000 ; CB- CA- C- N
372 375 376 377 3 30.33400 0.00000 -30.33400
0.00000 0.00000 0.00000 ; CB- CG- CD1- HD1
372 375 376 378 3 30.33400 0.00000 -30.33400
0.00000 0.00000 0.00000 ; CB- CG- CD1- CE1
372 375 385 383 3 30.33400 0.00000 -30.33400
0.00000 0.00000 0.00000 ; CB- CG- CD2- CE2
372 375 385 386 3 30.33400 0.00000 -30.33400
0.00000 0.00000 0.00000 ; CB- CG- CD2- HD2
373 372 370 387 3 0.65084 1.95253 0.00000
-2.60338 0.00000 0.00000 ; HB2- CB- CA- C
373 372 375 376 3 0.00000 0.00000 0.00000
0.00000 0.00000 0.00000 ; HB2- CB- CG- CD1

373 372 375 385 3 0.00000 0.00000 0.00000
0.00000 0.00000 0.00000 ; HB2- CB- CG- CD2
374 372 370 387 3 0.65084 1.95253 0.00000
-2.60338 0.00000 0.00000 ; HB3- CB- CA- C
374 372 375 376 3 0.00000 0.00000 0.00000
0.00000 0.00000 0.00000 ; HB3- CB- CG- CD1
374 372 375 385 3 0.00000 0.00000 0.00000
0.00000 0.00000 0.00000 ; HB3- CB- CG- CD2
375 372 370 387 3 4.95804 2.17986 -3.52293
-3.21331 -0.40166 0.00000 ; CG- CB- CA- C
375 376 378 379 3 30.33400 0.00000 -30.33400
0.00000 0.00000 0.00000 ; CG- CD1- CE1- HE1
375 376 378 380 3 30.33400 0.00000 -30.33400
0.00000 0.00000 0.00000 ; CG- CD1- CE1- CZ
375 385 383 380 3 30.33400 0.00000 -30.33400
0.00000 0.00000 0.00000 ; CG- CD2- CE2- CZ
375 385 383 384 3 30.33400 0.00000 -30.33400
0.00000 0.00000 0.00000 ; CG- CD2- CE2- HE2
376 375 385 383 3 30.33400 0.00000 -30.33400
0.00000 0.00000 0.00000 ; CD1- CG- CD2- CE2
376 375 385 386 3 30.33400 0.00000 -30.33400
0.00000 0.00000 0.00000 ; CD1- CG- CD2- HD2
376 378 380 381 3 30.33400 0.00000 -30.33400
0.00000 0.00000 0.00000 ; CD1- CE1- CZ- OH
376 378 380 383 3 30.33400 0.00000 -30.33400
0.00000 0.00000 0.00000 ; CD1- CE1- CZ- CE2
377 376 375 385 3 30.33400 0.00000 -30.33400
0.00000 0.00000 0.00000 ; HD1- CD1- CG- CD2
377 376 378 379 3 30.33400 0.00000 -30.33400
0.00000 0.00000 0.00000 ; HD1- CD1- CE1- HE1
377 376 378 380 3 30.33400 0.00000 -30.33400
0.00000 0.00000 0.00000 ; HD1- CD1- CE1- CZ
378 376 375 385 3 30.33400 0.00000 -30.33400
0.00000 0.00000 0.00000 ; CE1- CD1- CG- CD2
378 380 381 382 3 7.93286 0.00000 -9.56462
0.00000 2.17568 0.00000 ; CE1- CZ- OH- HH
378 380 383 384 3 30.33400 0.00000 -30.33400
0.00000 0.00000 0.00000 ; CE1- CZ- CE2- HE2
378 380 383 385 3 30.33400 0.00000 -30.33400
0.00000 0.00000 0.00000 ; CE1- CZ- CE2- CD2
379 378 380 381 3 30.33400 0.00000 -30.33400
0.00000 0.00000 0.00000 ; HE1- CE1- CZ- OH
379 378 380 383 3 30.33400 0.00000 -30.33400
0.00000 0.00000 0.00000 ; HE1- CE1- CZ- CE2
380 383 385 386 3 30.33400 0.00000 -30.33400
0.00000 0.00000 0.00000 ; CZ- CE2- CD2- HD2
381 380 383 384 3 30.33400 0.00000 -30.33400
0.00000 0.00000 0.00000 ; OH- CZ- CE2- HE2
381 380 383 385 3 30.33400 0.00000 -30.33400
0.00000 0.00000 0.00000 ; OH- CZ- CE2- CD2
382 381 380 383 3 7.93286 0.00000 -9.56462
0.00000 2.17568 0.00000 ; HH- OH- CZ- CE2

384 383 385 386 3 30.33400 0.00000 -30.33400
0.00000 0.00000 0.00000 ; HE2- CE2- CD2- HD2
387 389 391 392 3 0.00000 0.00000 0.00000
0.00000 0.00000 0.00000 ; C- N- CA- HA2
387 389 391 393 3 0.00000 0.00000 0.00000
0.00000 0.00000 0.00000 ; C- N- CA- HA3
387 389 391 394 3 1.75728 5.27184 2.25936
-7.02912 0.00000 0.00000 ; C- N- CA- C
388 387 389 390 3 29.28800 -8.36800 -20.92000
0.00000 0.00000 0.00000 ; O- C- N- H
388 387 389 391 3 20.92000 0.00000 -20.92000
0.00000 0.00000 0.00000 ; O- C- N- CA
389 391 394 395 3 0.00000 0.00000 0.00000
0.00000 0.00000 0.00000 ; N- CA- C- O
389 391 394 396 3 17.40544 -5.02080 -13.22144
9.20480 0.00000 0.00000 ; N- CA- C- N
390 389 391 392 3 0.00000 0.00000 0.00000
0.00000 0.00000 0.00000 ; H- N- CA- HA2
390 389 391 393 3 0.00000 0.00000 0.00000
0.00000 0.00000 0.00000 ; H- N- CA- HA3
390 389 391 394 3 0.00000 0.00000 0.00000
0.00000 0.00000 0.00000 ; H- N- CA- C
391 394 396 397 3 20.92000 0.00000 -20.92000
0.00000 0.00000 0.00000 ; CA- C- N- H
391 394 396 398 3 20.92000 0.00000 -20.92000
0.00000 0.00000 0.00000 ; CA- C- N- CA
392 391 394 395 3 3.68192 -4.35136 0.00000
1.33888 0.00000 0.00000 ; HA2- CA- C- O
392 391 394 396 3 0.00000 0.00000 0.00000
0.00000 0.00000 0.00000 ; HA2- CA- C- N
393 391 394 395 3 3.68192 -4.35136 0.00000
1.33888 0.00000 0.00000 ; HA3- CA- C- O
393 391 394 396 3 0.00000 0.00000 0.00000
0.00000 0.00000 0.00000 ; HA3- CA- C- N
394 396 398 399 3 0.00000 0.00000 0.00000
0.00000 0.00000 0.00000 ; C- N- CA- HA
394 396 398 400 3 11.71520 1.67360 15.06240 -
13.38880 0.00000 0.00000 ; C- N- CA- CB
394 396 398 405 3 1.75728 5.27184 2.25936
-7.02912 0.00000 0.00000 ; C- N- CA- C
395 394 396 397 3 29.28800 -8.36800 -20.92000
0.00000 0.00000 0.00000 ; O- C- N- H
395 394 396 398 3 20.92000 0.00000 -20.92000
0.00000 0.00000 0.00000 ; O- C- N- CA
396 398 400 401 3 0.65084 1.95253 0.00000
-2.60338 0.00000 0.00000 ; N- CA- CB- HB2
396 398 400 402 3 0.65084 1.95253 0.00000
-2.60338 0.00000 0.00000 ; N- CA- CB- HB3
396 398 400 403 3 6.03751 2.50622 -5.17142
-4.20074 1.10458 0.00000 ; N- CA- CB- SG
396 398 405 406 3 0.00000 0.00000 0.00000
0.00000 0.00000 0.00000 ; N- CA- C- O

```
   396    398    405    407      3   17.40544   -5.02080  -13.22144
9.20480    0.00000    0.00000 ;      N-    CA-     C-     N
   397    396    398    399      3    0.00000    0.00000    0.00000
0.00000    0.00000    0.00000 ;      H-     N-    CA-    HA
   397    396    398    400      3    0.00000    0.00000    0.00000
0.00000    0.00000    0.00000 ;      H-     N-    CA-    CB
   397    396    398    405      3    0.00000    0.00000    0.00000
0.00000    0.00000    0.00000 ;      H-     N-    CA-     C
   398    400    403    404      3    1.69034    2.77818    4.11706
-4.21747    1.00416    0.00000 ;     CA-    CB-    SG-    HG
   398    405    407    408      3   20.92000    0.00000  -20.92000
0.00000    0.00000    0.00000 ;     CA-     C-     N-     H
   398    405    407    409      3   20.92000    0.00000  -20.92000
0.00000    0.00000    0.00000 ;     CA-     C-     N-    CA
   399    398    400    401      3    0.65084    1.95253    0.00000
-2.60338    0.00000    0.00000 ;     HA-    CA-    CB-   HB2
   399    398    400    402      3    0.65084    1.95253    0.00000
-2.60338    0.00000    0.00000 ;     HA-    CA-    CB-   HB3
   399    398    400    403      3    0.65084    1.95253    0.00000
-2.60338    0.00000    0.00000 ;     HA-    CA-    CB-    SG
   399    398    405    406      3    3.68192   -4.35136    0.00000
1.33888    0.00000    0.00000 ;     HA-    CA-     C-     O
   399    398    405    407      3    0.00000    0.00000    0.00000
0.00000    0.00000    0.00000 ;     HA-    CA-     C-     N
   400    398    405    406      3    0.00000    0.00000    0.00000
0.00000    0.00000    0.00000 ;     CB-    CA-     C-     O
   400    398    405    407      3    2.51040    4.18400    1.67360
-6.69440    0.00000    0.00000 ;     CB-    CA-     C-     N
   401    400    398    405      3    0.65084    1.95253    0.00000
-2.60338    0.00000    0.00000 ;    HB2-    CB-    CA-     C
   401    400    403    404      3    1.04600    3.13800    0.00000
-4.18400    0.00000    0.00000 ;    HB2-    CB-    SG-    HG
   402    400    398    405      3    0.65084    1.95253    0.00000
-2.60338    0.00000    0.00000 ;    HB3-    CB-    CA-     C
   402    400    403    404      3    1.04600    3.13800    0.00000
-4.18400    0.00000    0.00000 ;    HB3-    CB-    SG-    HG
   403    400    398    405      3    5.62330    4.27605   -5.33042
-4.20074    2.51040    0.00000 ;     SG-    CB-    CA-     C
   405    407    409    410      3    0.00000    0.00000    0.00000
0.00000    0.00000    0.00000 ;      C-     N-    CA-    HA
   405    407    409    411      3   11.71520    1.67360   15.06240  -
13.38880    0.00000    0.00000 ;      C-     N-    CA-    CB
   405    407    409    421      3    1.75728    5.27184    2.25936
-7.02912    0.00000    0.00000 ;      C-     N-    CA-     C
   406    405    407    408      3   29.28800   -8.36800  -20.92000
0.00000    0.00000    0.00000 ;      O-     C-     N-     H
   406    405    407    409      3   20.92000    0.00000  -20.92000
0.00000    0.00000    0.00000 ;      O-     C-     N-    CA
   407    409    411    412      3    0.65084    1.95253    0.00000
-2.60338    0.00000    0.00000 ;      N-    CA-    CB-    HB
   407    409    411    413      3    3.83673    0.44769   -1.77402
-2.47693   -0.03347    0.00000 ;      N-    CA-    CB-   CG1
```

407 409 411 417 3 3.83673 0.44769 -1.77402
-2.47693 -0.03347 0.00000 ; N- CA- CB- CG2
407 409 421 422 3 0.00000 0.00000 0.00000
0.00000 0.00000 0.00000 ; N- CA- C- O
407 409 421 423 3 17.40544 -5.02080 -13.22144
9.20480 0.00000 0.00000 ; N- CA- C- N
408 407 409 410 3 0.00000 0.00000 0.00000
0.00000 0.00000 0.00000 ; H- N- CA- HA
408 407 409 411 3 0.00000 0.00000 0.00000
0.00000 0.00000 0.00000 ; H- N- CA- CB
408 407 409 421 3 0.00000 0.00000 0.00000
0.00000 0.00000 0.00000 ; H- N- CA- C
409 411 413 414 3 0.66944 2.00832 0.00000
-2.67776 0.00000 0.00000 ; CA- CB- CG1- HG11
409 411 413 415 3 0.66944 2.00832 0.00000
-2.67776 0.00000 0.00000 ; CA- CB- CG1- HG12
409 411 413 416 3 0.66944 2.00832 0.00000
-2.67776 0.00000 0.00000 ; CA- CB- CG1- HG13
409 411 417 418 3 0.66944 2.00832 0.00000
-2.67776 0.00000 0.00000 ; CA- CB- CG2- HG21
409 411 417 419 3 0.66944 2.00832 0.00000
-2.67776 0.00000 0.00000 ; CA- CB- CG2- HG22
409 411 417 420 3 0.66944 2.00832 0.00000
-2.67776 0.00000 0.00000 ; CA- CB- CG2- HG23
409 421 423 424 3 20.92000 0.00000 -20.92000
0.00000 0.00000 0.00000 ; CA- C- N- H
409 421 423 425 3 20.92000 0.00000 -20.92000
0.00000 0.00000 0.00000 ; CA- C- N- CA
410 409 411 412 3 0.65084 1.95253 0.00000
-2.60338 0.00000 0.00000 ; HA- CA- CB- HB
410 409 411 413 3 0.65084 1.95253 0.00000
-2.60338 0.00000 0.00000 ; HA- CA- CB- CG1
410 409 411 417 3 0.65084 1.95253 0.00000
-2.60338 0.00000 0.00000 ; HA- CA- CB- CG2
410 409 421 422 3 3.68192 -4.35136 0.00000
1.33888 0.00000 0.00000 ; HA- CA- C- O
410 409 421 423 3 0.00000 0.00000 0.00000
0.00000 0.00000 0.00000 ; HA- CA- C- N
411 409 421 422 3 0.00000 0.00000 0.00000
0.00000 0.00000 0.00000 ; CB- CA- C- O
411 409 421 423 3 2.51040 4.18400 1.67360
-6.69440 0.00000 0.00000 ; CB- CA- C- N
412 411 409 421 3 0.65084 1.95253 0.00000
-2.60338 0.00000 0.00000 ; HB- CB- CA- C
412 411 413 414 3 0.62760 1.88280 0.00000
-2.51040 0.00000 0.00000 ; HB- CB- CG1- HG11
412 411 413 415 3 0.62760 1.88280 0.00000
-2.51040 0.00000 0.00000 ; HB- CB- CG1- HG12
412 411 413 416 3 0.62760 1.88280 0.00000
-2.51040 0.00000 0.00000 ; HB- CB- CG1- HG13
412 411 417 418 3 0.62760 1.88280 0.00000
-2.51040 0.00000 0.00000 ; HB- CB- CG2- HG21

```
   412    411    417    419      3    0.62760    1.88280    0.00000
-2.51040    0.00000    0.00000 ;     HB-    CB-   CG2-  HG22
   412    411    417    420      3    0.62760    1.88280    0.00000
-2.51040    0.00000    0.00000 ;     HB-    CB-   CG2-  HG23
   413    411    409    421      3    5.67350    3.55640   -6.16722
-2.47693    3.74886    0.00000 ;    CG1-    CB-    CA-     C
   413    411    417    418      3    0.66944    2.00832    0.00000
-2.67776    0.00000    0.00000 ;    CG1-    CB-   CG2-  HG21
   413    411    417    419      3    0.66944    2.00832    0.00000
-2.67776    0.00000    0.00000 ;    CG1-    CB-   CG2-  HG22
   413    411    417    420      3    0.66944    2.00832    0.00000
-2.67776    0.00000    0.00000 ;    CG1-    CB-   CG2-  HG23
   414    413    411    417      3    0.66944    2.00832    0.00000
-2.67776    0.00000    0.00000 ;   HG11-   CG1-    CB-   CG2
   415    413    411    417      3    0.66944    2.00832    0.00000
-2.67776    0.00000    0.00000 ;   HG12-   CG1-    CB-   CG2
   416    413    411    417      3    0.66944    2.00832    0.00000
-2.67776    0.00000    0.00000 ;   HG13-   CG1-    CB-   CG2
   417    411    409    421      3    5.67350    3.55640   -6.16722
-2.47693    3.74886    0.00000 ;    CG2-    CB-    CA-     C
   421    423    425    426      3    0.00000    0.00000    0.00000
0.00000    0.00000    0.00000 ;      C-     N-    CA-    HA
   421    423    425    427      3   11.71520    1.67360   15.06240  -
13.38880    0.00000    0.00000 ;      C-     N-    CA-    CB
   421    423    425    442      3    1.75728    5.27184    2.25936
-7.02912    0.00000    0.00000 ;      C-     N-    CA-     C
   422    421    423    424      3   29.28800   -8.36800  -20.92000
0.00000    0.00000    0.00000 ;      O-     C-     N-     H
   422    421    423    425      3   20.92000    0.00000  -20.92000
0.00000    0.00000    0.00000 ;      O-     C-     N-    CA
   423    425    427    428      3    0.65084    1.95253    0.00000
-2.60338    0.00000    0.00000 ;      N-    CA-    CB-   HB2
   423    425    427    429      3    0.65084    1.95253    0.00000
-2.60338    0.00000    0.00000 ;      N-    CA-    CB-   HB3
   423    425    427    430      3    3.28026    2.46019   -2.19242
-3.21331   -0.23430    0.00000 ;      N-    CA-    CB-    CG
   423    425    442    443      3    0.00000    0.00000    0.00000
0.00000    0.00000    0.00000 ;      N-    CA-     C-     O
   423    425    442    444      3   17.40544   -5.02080  -13.22144
9.20480    0.00000    0.00000 ;      N-    CA-     C-     N
   424    423    425    426      3    0.00000    0.00000    0.00000
0.00000    0.00000    0.00000 ;      H-     N-    CA-    HA
   424    423    425    427      3    0.00000    0.00000    0.00000
0.00000    0.00000    0.00000 ;      H-     N-    CA-    CB
   424    423    425    442      3    0.00000    0.00000    0.00000
0.00000    0.00000    0.00000 ;      H-     N-    CA-     C
   425    427    430    431      3    0.57739    0.00000    1.02926
0.00000   -1.60666    0.00000 ;     CA-    CB-    CG-   CD1
   425    427    430    440      3    0.57739    0.00000    1.02926
0.00000   -1.60666    0.00000 ;     CA-    CB-    CG-   CD2
   425    442    444    445      3   20.92000    0.00000  -20.92000
0.00000    0.00000    0.00000 ;     CA-     C-     N-     H
```

425 442 444 446 3 20.92000 0.00000 -20.92000
0.00000 0.00000 0.00000 ; CA- C- N- CA
426 425 427 428 3 0.65084 1.95253 0.00000
-2.60338 0.00000 0.00000 ; HA- CA- CB- HB2
426 425 427 429 3 0.65084 1.95253 0.00000
-2.60338 0.00000 0.00000 ; HA- CA- CB- HB3
426 425 427 430 3 0.65084 1.95253 0.00000
-2.60338 0.00000 0.00000 ; HA- CA- CB- CG
426 425 442 443 3 3.68192 -4.35136 0.00000
1.33888 0.00000 0.00000 ; HA- CA- C- O
426 425 442 444 3 0.00000 0.00000 0.00000
0.00000 0.00000 0.00000 ; HA- CA- C- N
427 425 442 443 3 0.00000 0.00000 0.00000
0.00000 0.00000 0.00000 ; CB- CA- C- O
427 425 442 444 3 2.51040 4.18400 1.67360
-6.69440 0.00000 0.00000 ; CB- CA- C- N
427 430 431 432 3 30.33400 0.00000 -30.33400
0.00000 0.00000 0.00000 ; CB- CG- CD1- HD1
427 430 431 433 3 30.33400 0.00000 -30.33400
0.00000 0.00000 0.00000 ; CB- CG- CD1- CE1
427 430 440 438 3 30.33400 0.00000 -30.33400
0.00000 0.00000 0.00000 ; CB- CG- CD2- CE2
427 430 440 441 3 30.33400 0.00000 -30.33400
0.00000 0.00000 0.00000 ; CB- CG- CD2- HD2
428 427 425 442 3 0.65084 1.95253 0.00000
-2.60338 0.00000 0.00000 ; HB2- CB- CA- C
428 427 430 431 3 0.00000 0.00000 0.00000
0.00000 0.00000 0.00000 ; HB2- CB- CG- CD1
428 427 430 440 3 0.00000 0.00000 0.00000
0.00000 0.00000 0.00000 ; HB2- CB- CG- CD2
429 427 425 442 3 0.65084 1.95253 0.00000
-2.60338 0.00000 0.00000 ; HB3- CB- CA- C
429 427 430 431 3 0.00000 0.00000 0.00000
0.00000 0.00000 0.00000 ; HB3- CB- CG- CD1
429 427 430 440 3 0.00000 0.00000 0.00000
0.00000 0.00000 0.00000 ; HB3- CB- CG- CD2
430 427 425 442 3 4.95804 2.17986 -3.52293
-3.21331 -0.40166 0.00000 ; CG- CB- CA- C
430 431 433 434 3 30.33400 0.00000 -30.33400
0.00000 0.00000 0.00000 ; CG- CD1- CE1- HE1
430 431 433 435 3 30.33400 0.00000 -30.33400
0.00000 0.00000 0.00000 ; CG- CD1- CE1- CZ
430 440 438 435 3 30.33400 0.00000 -30.33400
0.00000 0.00000 0.00000 ; CG- CD2- CE2- CZ
430 440 438 439 3 30.33400 0.00000 -30.33400
0.00000 0.00000 0.00000 ; CG- CD2- CE2- HE2
431 430 440 438 3 30.33400 0.00000 -30.33400
0.00000 0.00000 0.00000 ; CD1- CG- CD2- CE2
431 430 440 441 3 30.33400 0.00000 -30.33400
0.00000 0.00000 0.00000 ; CD1- CG- CD2- HD2
431 433 435 436 3 30.33400 0.00000 -30.33400
0.00000 0.00000 0.00000 ; CD1- CE1- CZ- OH

431 433 435 438 3 30.33400 0.00000 -30.33400
0.00000 0.00000 0.00000 ; CD1- CE1- CZ- CE2
432 431 430 440 3 30.33400 0.00000 -30.33400
0.00000 0.00000 0.00000 ; HD1- CD1- CG- CD2
432 431 433 434 3 30.33400 0.00000 -30.33400
0.00000 0.00000 0.00000 ; HD1- CD1- CE1- HE1
432 431 433 435 3 30.33400 0.00000 -30.33400
0.00000 0.00000 0.00000 ; HD1- CD1- CE1- CZ
433 431 430 440 3 30.33400 0.00000 -30.33400
0.00000 0.00000 0.00000 ; CE1- CD1- CG- CD2
433 435 436 437 3 7.93286 0.00000 -9.56462
0.00000 2.17568 0.00000 ; CE1- CZ- OH- HH
433 435 438 439 3 30.33400 0.00000 -30.33400
0.00000 0.00000 0.00000 ; CE1- CZ- CE2- HE2
433 435 438 440 3 30.33400 0.00000 -30.33400
0.00000 0.00000 0.00000 ; CE1- CZ- CE2- CD2
434 433 435 436 3 30.33400 0.00000 -30.33400
0.00000 0.00000 0.00000 ; HE1- CE1- CZ- OH
434 433 435 438 3 30.33400 0.00000 -30.33400
0.00000 0.00000 0.00000 ; HE1- CE1- CZ- CE2
435 438 440 441 3 30.33400 0.00000 -30.33400
0.00000 0.00000 0.00000 ; CZ- CE2- CD2- HD2
436 435 438 439 3 30.33400 0.00000 -30.33400
0.00000 0.00000 0.00000 ; OH- CZ- CE2- HE2
436 435 438 440 3 30.33400 0.00000 -30.33400
0.00000 0.00000 0.00000 ; OH- CZ- CE2- CD2
437 436 435 438 3 7.93286 0.00000 -9.56462
0.00000 2.17568 0.00000 ; HH- OH- CZ- CE2
439 438 440 441 3 30.33400 0.00000 -30.33400
0.00000 0.00000 0.00000 ; HE2- CE2- CD2- HD2
442 444 446 447 3 0.00000 0.00000 0.00000
0.00000 0.00000 0.00000 ; C- N- CA- HA
442 444 446 448 3 11.71520 1.67360 15.06240 -
13.38880 0.00000 0.00000 ; C- N- CA- CB
442 444 446 464 3 1.75728 5.27184 2.25936
-7.02912 0.00000 0.00000 ; C- N- CA- C
443 442 444 445 3 29.28800 -8.36800 -20.92000
0.00000 0.00000 0.00000 ; O- C- N- H
443 442 444 446 3 20.92000 0.00000 -20.92000
0.00000 0.00000 0.00000 ; O- C- N- CA
444 446 448 449 3 0.65084 1.95253 0.00000
-2.60338 0.00000 0.00000 ; N- CA- CB- HB2
444 446 448 450 3 0.65084 1.95253 0.00000
-2.60338 0.00000 0.00000 ; N- CA- CB- HB3
444 446 448 451 3 0.65084 1.95253 0.00000
-2.60338 0.00000 0.00000 ; N- CA- CB- CG
444 446 464 465 3 0.00000 0.00000 0.00000
0.00000 0.00000 0.00000 ; N- CA- C- O
444 446 464 466 3 17.40544 -5.02080 -13.22144
9.20480 0.00000 0.00000 ; N- CA- C- N
445 444 446 447 3 0.00000 0.00000 0.00000
0.00000 0.00000 0.00000 ; H- N- CA- HA

445 444 446 448 3 0.00000 0.00000 0.00000
0.00000 0.00000 0.00000 ; H- N- CA- CB
445 444 446 464 3 0.00000 0.00000 0.00000
0.00000 0.00000 0.00000 ; H- N- CA- C
446 448 451 452 3 0.66944 2.00832 0.00000
-2.67776 0.00000 0.00000 ; CA- CB- CG- HG2
446 448 451 453 3 0.66944 2.00832 0.00000
-2.67776 0.00000 0.00000 ; CA- CB- CG- HG3
446 448 451 454 3 3.68192 3.09616 -2.09200
-3.01248 0.00000 0.00000 ; CA- CB- CG- CD
446 464 466 467 3 20.92000 0.00000 -20.92000
0.00000 0.00000 0.00000 ; CA- C- N- H
446 464 466 468 3 20.92000 0.00000 -20.92000
0.00000 0.00000 0.00000 ; CA- C- N- CA
447 446 448 449 3 0.65084 1.95253 0.00000
-2.60338 0.00000 0.00000 ; HA- CA- CB- HB2
447 446 448 450 3 0.65084 1.95253 0.00000
-2.60338 0.00000 0.00000 ; HA- CA- CB- HB3
447 446 448 451 3 0.65084 1.95253 0.00000
-2.60338 0.00000 0.00000 ; HA- CA- CB- CG
447 446 464 465 3 3.68192 -4.35136 0.00000
1.33888 0.00000 0.00000 ; HA- CA- C- O
447 446 464 466 3 0.00000 0.00000 0.00000
0.00000 0.00000 0.00000 ; HA- CA- C- N
448 446 464 465 3 0.00000 0.00000 0.00000
0.00000 0.00000 0.00000 ; CB- CA- C- O
448 446 464 466 3 2.51040 4.18400 1.67360
-6.69440 0.00000 0.00000 ; CB- CA- C- N
448 451 454 455 3 0.66944 2.00832 0.00000
-2.67776 0.00000 0.00000 ; CB- CG- CD- HD2
448 451 454 456 3 0.66944 2.00832 0.00000
-2.67776 0.00000 0.00000 ; CB- CG- CD- HD3
448 451 454 457 3 3.68192 3.09616 -2.09200
-3.01248 0.00000 0.00000 ; CB- CG- CD- CE
449 448 446 464 3 0.65084 1.95253 0.00000
-2.60338 0.00000 0.00000 ; HB2- CB- CA- C
449 448 451 452 3 0.62760 1.88280 0.00000
-2.51040 0.00000 0.00000 ; HB2- CB- CG- HG2
449 448 451 453 3 0.62760 1.88280 0.00000
-2.51040 0.00000 0.00000 ; HB2- CB- CG- HG3
449 448 451 454 3 0.66944 2.00832 0.00000
-2.67776 0.00000 0.00000 ; HB2- CB- CG- CD
450 448 446 464 3 0.65084 1.95253 0.00000
-2.60338 0.00000 0.00000 ; HB3- CB- CA- C
450 448 451 452 3 0.62760 1.88280 0.00000
-2.51040 0.00000 0.00000 ; HB3- CB- CG- HG2
450 448 451 453 3 0.62760 1.88280 0.00000
-2.51040 0.00000 0.00000 ; HB3- CB- CG- HG3
450 448 451 454 3 0.66944 2.00832 0.00000
-2.67776 0.00000 0.00000 ; HB3- CB- CG- CD
451 448 446 464 3 0.65084 1.95253 0.00000
-2.60338 0.00000 0.00000 ; CG- CB- CA- C

451 454 457 458 3 0.65084 1.95253 0.00000
-2.60338 0.00000 0.00000 ; CG- CD- CE- HE2
451 454 457 459 3 0.65084 1.95253 0.00000
-2.60338 0.00000 0.00000 ; CG- CD- CE- HE3
451 454 457 460 3 0.65084 1.95253 0.00000
-2.60338 0.00000 0.00000 ; CG- CD- CE- NZ
452 451 454 455 3 0.62760 1.88280 0.00000
-2.51040 0.00000 0.00000 ; HG2- CG- CD- HD2
452 451 454 456 3 0.62760 1.88280 0.00000
-2.51040 0.00000 0.00000 ; HG2- CG- CD- HD3
452 451 454 457 3 0.66944 2.00832 0.00000
-2.67776 0.00000 0.00000 ; HG2- CG- CD- CE
453 451 454 455 3 0.62760 1.88280 0.00000
-2.51040 0.00000 0.00000 ; HG3- CG- CD- HD2
453 451 454 456 3 0.62760 1.88280 0.00000
-2.51040 0.00000 0.00000 ; HG3- CG- CD- HD3
453 451 454 457 3 0.66944 2.00832 0.00000
-2.67776 0.00000 0.00000 ; HG3- CG- CD- CE
454 457 460 461 3 0.65084 1.95253 0.00000
-2.60338 0.00000 0.00000 ; CD- CE- NZ- HZ1
454 457 460 462 3 0.65084 1.95253 0.00000
-2.60338 0.00000 0.00000 ; CD- CE- NZ- HZ2
454 457 460 463 3 0.65084 1.95253 0.00000
-2.60338 0.00000 0.00000 ; CD- CE- NZ- HZ3
455 454 457 458 3 0.65084 1.95253 0.00000
-2.60338 0.00000 0.00000 ; HD2- CD- CE- HE2
455 454 457 459 3 0.65084 1.95253 0.00000
-2.60338 0.00000 0.00000 ; HD2- CD- CE- HE3
455 454 457 460 3 0.65084 1.95253 0.00000
-2.60338 0.00000 0.00000 ; HD2- CD- CE- NZ
456 454 457 458 3 0.65084 1.95253 0.00000
-2.60338 0.00000 0.00000 ; HD3- CD- CE- HE2
456 454 457 459 3 0.65084 1.95253 0.00000
-2.60338 0.00000 0.00000 ; HD3- CD- CE- HE3
456 454 457 460 3 0.65084 1.95253 0.00000
-2.60338 0.00000 0.00000 ; HD3- CD- CE- NZ
458 457 460 461 3 0.65084 1.95253 0.00000
-2.60338 0.00000 0.00000 ; HE2- CE- NZ- HZ1
458 457 460 462 3 0.65084 1.95253 0.00000
-2.60338 0.00000 0.00000 ; HE2- CE- NZ- HZ2
458 457 460 463 3 0.65084 1.95253 0.00000
-2.60338 0.00000 0.00000 ; HE2- CE- NZ- HZ3
459 457 460 461 3 0.65084 1.95253 0.00000
-2.60338 0.00000 0.00000 ; HE3- CE- NZ- HZ1
459 457 460 462 3 0.65084 1.95253 0.00000
-2.60338 0.00000 0.00000 ; HE3- CE- NZ- HZ2
459 457 460 463 3 0.65084 1.95253 0.00000
-2.60338 0.00000 0.00000 ; HE3- CE- NZ- HZ3
464 466 468 469 3 0.00000 0.00000 0.00000
0.00000 0.00000 0.00000 ; C- N- CA- HA2
464 466 468 470 3 0.00000 0.00000 0.00000
0.00000 0.00000 0.00000 ; C- N- CA- HA3

464 466 468 471 3 1.75728 5.27184 2.25936
-7.02912 0.00000 0.00000 ; C- N- CA- C
465 464 466 467 3 29.28800 -8.36800 -20.92000
0.00000 0.00000 0.00000 ; O- C- N- H
465 464 466 468 3 20.92000 0.00000 -20.92000
0.00000 0.00000 0.00000 ; O- C- N- CA
466 468 471 472 3 0.00000 0.00000 0.00000
0.00000 0.00000 0.00000 ; N- CA- C- O
466 468 471 473 3 17.40544 -5.02080 -13.22144
9.20480 0.00000 0.00000 ; N- CA- C- N
467 466 468 469 3 0.00000 0.00000 0.00000
0.00000 0.00000 0.00000 ; H- N- CA- HA2
467 466 468 470 3 0.00000 0.00000 0.00000
0.00000 0.00000 0.00000 ; H- N- CA- HA3
467 466 468 471 3 0.00000 0.00000 0.00000
0.00000 0.00000 0.00000 ; H- N- CA- C
468 471 473 474 3 20.92000 0.00000 -20.92000
0.00000 0.00000 0.00000 ; CA- C- N- H
468 471 473 475 3 20.92000 0.00000 -20.92000
0.00000 0.00000 0.00000 ; CA- C- N- CA
469 468 471 472 3 3.68192 -4.35136 0.00000
1.33888 0.00000 0.00000 ; HA2- CA- C- O
469 468 471 473 3 0.00000 0.00000 0.00000
0.00000 0.00000 0.00000 ; HA2- CA- C- N
470 468 471 472 3 3.68192 -4.35136 0.00000
1.33888 0.00000 0.00000 ; HA3- CA- C- O
470 468 471 473 3 0.00000 0.00000 0.00000
0.00000 0.00000 0.00000 ; HA3- CA- C- N
471 473 475 476 3 0.00000 0.00000 0.00000
0.00000 0.00000 0.00000 ; C- N- CA- HA
471 473 475 477 3 11.71520 1.67360 15.06240 -
13.38880 0.00000 0.00000 ; C- N- CA- CB
471 473 475 490 3 1.75728 5.27184 2.25936
-7.02912 0.00000 0.00000 ; C- N- CA- C
472 471 473 474 3 29.28800 -8.36800 -20.92000
0.00000 0.00000 0.00000 ; O- C- N- H
472 471 473 475 3 20.92000 0.00000 -20.92000
0.00000 0.00000 0.00000 ; O- C- N- CA
473 475 477 478 3 0.65084 1.95253 0.00000
-2.60338 0.00000 0.00000 ; N- CA- CB- HB
473 475 477 479 3 3.83673 0.44769 -1.77402
-2.47693 -0.03347 0.00000 ; N- CA- CB- CG2
473 475 477 483 3 2.97482 0.12134 2.04179
-1.89117 -3.24678 0.00000 ; N- CA- CB- CG1
473 475 490 491 3 0.00000 0.00000 0.00000
0.00000 0.00000 0.00000 ; N- CA- C- O
473 475 490 492 3 17.40544 -5.02080 -13.22144
9.20480 0.00000 0.00000 ; N- CA- C- N
474 473 475 476 3 0.00000 0.00000 0.00000
0.00000 0.00000 0.00000 ; H- N- CA- HA
474 473 475 477 3 0.00000 0.00000 0.00000
0.00000 0.00000 0.00000 ; H- N- CA- CB

474 473 475 490 3 0.00000 0.00000 0.00000
0.00000 0.00000 0.00000 ; H- N- CA- C
475 477 479 480 3 0.66944 2.00832 0.00000
-2.67776 0.00000 0.00000 ; CA- CB- CG2- HG21
475 477 479 481 3 0.66944 2.00832 0.00000
-2.67776 0.00000 0.00000 ; CA- CB- CG2- HG22
475 477 479 482 3 0.66944 2.00832 0.00000
-2.67776 0.00000 0.00000 ; CA- CB- CG2- HG23
475 477 483 484 3 0.66944 2.00832 0.00000
-2.67776 0.00000 0.00000 ; CA- CB- CG1- HG12
475 477 483 485 3 0.66944 2.00832 0.00000
-2.67776 0.00000 0.00000 ; CA- CB- CG1- HG13
475 477 483 486 3 4.24258 -0.52718 -7.25506
-1.79075 7.69856 0.00000 ; CA- CB- CG1- CD1
475 490 492 493 3 20.92000 0.00000 -20.92000
0.00000 0.00000 0.00000 ; CA- C- N- H
475 490 492 494 3 20.92000 0.00000 -20.92000
0.00000 0.00000 0.00000 ; CA- C- N- CA
476 475 477 478 3 0.65084 1.95253 0.00000
-2.60338 0.00000 0.00000 ; HA- CA- CB- HB
476 475 477 479 3 0.65084 1.95253 0.00000
-2.60338 0.00000 0.00000 ; HA- CA- CB- CG2
476 475 477 483 3 0.65084 1.95253 0.00000
-2.60338 0.00000 0.00000 ; HA- CA- CB- CG1
476 475 490 491 3 3.68192 -4.35136 0.00000
1.33888 0.00000 0.00000 ; HA- CA- C- O
476 475 490 492 3 0.00000 0.00000 0.00000
0.00000 0.00000 0.00000 ; HA- CA- C- N
477 475 490 491 3 0.00000 0.00000 0.00000
0.00000 0.00000 0.00000 ; CB- CA- C- O
477 475 490 492 3 2.51040 4.18400 1.67360
-6.69440 0.00000 0.00000 ; CB- CA- C- N
477 483 486 487 3 0.66944 2.00832 0.00000
-2.67776 0.00000 0.00000 ; CB- CG1- CD1- HD11
477 483 486 488 3 0.66944 2.00832 0.00000
-2.67776 0.00000 0.00000 ; CB- CG1- CD1- HD12
477 483 486 489 3 0.66944 2.00832 0.00000
-2.67776 0.00000 0.00000 ; CB- CG1- CD1- HD13
478 477 475 490 3 0.65084 1.95253 0.00000
-2.60338 0.00000 0.00000 ; HB- CB- CA- C
478 477 479 480 3 0.62760 1.88280 0.00000
-2.51040 0.00000 0.00000 ; HB- CB- CG2- HG21
478 477 479 481 3 0.62760 1.88280 0.00000
-2.51040 0.00000 0.00000 ; HB- CB- CG2- HG22
478 477 479 482 3 0.62760 1.88280 0.00000
-2.51040 0.00000 0.00000 ; HB- CB- CG2- HG23
478 477 483 484 3 0.62760 1.88280 0.00000
-2.51040 0.00000 0.00000 ; HB- CB- CG1- HG12
478 477 483 485 3 0.62760 1.88280 0.00000
-2.51040 0.00000 0.00000 ; HB- CB- CG1- HG13
478 477 483 486 3 0.66944 2.00832 0.00000
-2.67776 0.00000 0.00000 ; HB- CB- CG1- CD1

479 477 475 490 3 5.67350 3.55640 -6.16722
-2.47693 3.74886 0.00000 ; CG2- CB- CA- C
479 477 483 484 3 0.66944 2.00832 0.00000
-2.67776 0.00000 0.00000 ; CG2- CB- CG1- HG12
479 477 483 485 3 0.66944 2.00832 0.00000
-2.67776 0.00000 0.00000 ; CG2- CB- CG1- HG13
479 477 483 486 3 3.81162 0.49790 -8.14206
-1.79075 7.49773 0.00000 ; CG2- CB- CG1- CD1
480 479 477 483 3 0.66944 2.00832 0.00000
-2.67776 0.00000 0.00000 ; HG21- CG2- CB- CG1
481 479 477 483 3 0.66944 2.00832 0.00000
-2.67776 0.00000 0.00000 ; HG22- CG2- CB- CG1
482 479 477 483 3 0.66944 2.00832 0.00000
-2.67776 0.00000 0.00000 ; HG23- CG2- CB- CG1
483 477 475 490 3 8.26340 0.74057 -9.99976
-1.89117 3.84928 0.00000 ; CG1- CB- CA- C
484 483 486 487 3 0.62760 1.88280 0.00000
-2.51040 0.00000 0.00000 ; HG12- CG1- CD1- HD11
484 483 486 488 3 0.62760 1.88280 0.00000
-2.51040 0.00000 0.00000 ; HG12- CG1- CD1- HD12
484 483 486 489 3 0.62760 1.88280 0.00000
-2.51040 0.00000 0.00000 ; HG12- CG1- CD1- HD13
485 483 486 487 3 0.62760 1.88280 0.00000
-2.51040 0.00000 0.00000 ; HG13- CG1- CD1- HD11
485 483 486 488 3 0.62760 1.88280 0.00000
-2.51040 0.00000 0.00000 ; HG13- CG1- CD1- HD12
485 483 486 489 3 0.62760 1.88280 0.00000
-2.51040 0.00000 0.00000 ; HG13- CG1- CD1- HD13
490 492 494 495 3 0.00000 0.00000 0.00000
0.00000 0.00000 0.00000 ; C- N- CA- HA
490 492 494 496 3 11.71520 1.67360 15.06240 -
13.38880 0.00000 0.00000 ; C- N- CA- CB
490 492 494 502 3 1.75728 5.27184 2.25936
-7.02912 0.00000 0.00000 ; C- N- CA- C
491 490 492 493 3 29.28800 -8.36800 -20.92000
0.00000 0.00000 0.00000 ; O- C- N- H
491 490 492 494 3 20.92000 0.00000 -20.92000
0.00000 0.00000 0.00000 ; O- C- N- CA
492 494 496 497 3 0.65084 1.95253 0.00000
-2.60338 0.00000 0.00000 ; N- CA- CB- HB2
492 494 496 498 3 0.65084 1.95253 0.00000
-2.60338 0.00000 0.00000 ; N- CA- CB- HB3
492 494 496 499 3 15.41386 9.74035 -8.39310
-0.97069 2.97901 0.00000 ; N- CA- CB- CG
492 494 502 503 3 0.00000 0.00000 0.00000
0.00000 0.00000 0.00000 ; N- CA- C- O
492 494 502 504 3 17.40544 -5.02080 -13.22144
9.20480 0.00000 0.00000 ; N- CA- C- N
493 492 494 495 3 0.00000 0.00000 0.00000
0.00000 0.00000 0.00000 ; H- N- CA- HA
493 492 494 496 3 0.00000 0.00000 0.00000
0.00000 0.00000 0.00000 ; H- N- CA- CB

493 492 494 502 3 0.00000 0.00000 0.00000
0.00000 0.00000 0.00000 ; H- N- CA- C
494 496 499 500 3 6.43499 0.00000 -5.39736
0.00000 -1.03763 0.00000 ; CA- CB- CG- OD1
494 496 499 501 3 6.43499 0.00000 -5.39736
0.00000 -1.03763 0.00000 ; CA- CB- CG- OD2
494 502 504 505 3 20.92000 0.00000 -20.92000
0.00000 0.00000 0.00000 ; CA- C- N- H
494 502 504 506 3 20.92000 0.00000 -20.92000
0.00000 0.00000 0.00000 ; CA- C- N- CA
495 494 496 497 3 0.65084 1.95253 0.00000
-2.60338 0.00000 0.00000 ; HA- CA- CB- HB2
495 494 496 498 3 0.65084 1.95253 0.00000
-2.60338 0.00000 0.00000 ; HA- CA- CB- HB3
495 494 496 499 3 0.65084 1.95253 0.00000
-2.60338 0.00000 0.00000 ; HA- CA- CB- CG
495 494 502 503 3 3.68192 -4.35136 0.00000
1.33888 0.00000 0.00000 ; HA- CA- C- O
495 494 502 504 3 0.00000 0.00000 0.00000
0.00000 0.00000 0.00000 ; HA- CA- C- N
496 494 502 503 3 0.00000 0.00000 0.00000
0.00000 0.00000 0.00000 ; CB- CA- C- O
496 494 502 504 3 2.51040 4.18400 1.67360
-6.69440 0.00000 0.00000 ; CB- CA- C- N
497 496 494 502 3 0.65084 1.95253 0.00000
-2.60338 0.00000 0.00000 ; HB2- CB- CA- C
497 496 499 500 3 0.00000 0.00000 0.00000
0.00000 0.00000 0.00000 ; HB2- CB- CG- OD1
497 496 499 501 3 0.00000 0.00000 0.00000
0.00000 0.00000 0.00000 ; HB2- CB- CG- OD2
498 496 494 502 3 0.65084 1.95253 0.00000
-2.60338 0.00000 0.00000 ; HB3- CB- CA- C
498 496 499 500 3 0.00000 0.00000 0.00000
0.00000 0.00000 0.00000 ; HB3- CB- CG- OD1
498 496 499 501 3 0.00000 0.00000 0.00000
0.00000 0.00000 0.00000 ; HB3- CB- CG- OD2
499 496 494 502 3 7.14627 -1.04600 -8.99560
-0.97069 5.15469 0.00000 ; CG- CB- CA- C
502 504 506 507 3 0.00000 0.00000 0.00000
0.00000 0.00000 0.00000 ; C- N- CA- HA
502 504 506 508 3 11.71520 1.67360 15.06240 -
13.38880 0.00000 0.00000 ; C- N- CA- CB
502 504 506 524 3 1.75728 5.27184 2.25936
-7.02912 0.00000 0.00000 ; C- N- CA- C
503 502 504 505 3 29.28800 -8.36800 -20.92000
0.00000 0.00000 0.00000 ; O- C- N- H
503 502 504 506 3 20.92000 0.00000 -20.92000
0.00000 0.00000 0.00000 ; O- C- N- CA
504 506 508 509 3 0.65084 1.95253 0.00000
-2.60338 0.00000 0.00000 ; N- CA- CB- HB2
504 506 508 510 3 0.65084 1.95253 0.00000
-2.60338 0.00000 0.00000 ; N- CA- CB- HB3

```
   504    506    508    511      3    0.65084    1.95253    0.00000
-2.60338    0.00000    0.00000 ;     N-    CA-    CB-    CG
   504    506    524    525      3    0.00000    0.00000    0.00000
0.00000    0.00000    0.00000 ;     N-    CA-    C-    O
   504    506    524    526      3   17.40544   -5.02080  -13.22144
9.20480    0.00000    0.00000 ;     N-    CA-    C-    N
   505    504    506    507      3    0.00000    0.00000    0.00000
0.00000    0.00000    0.00000 ;     H-    N-    CA-    HA
   505    504    506    508      3    0.00000    0.00000    0.00000
0.00000    0.00000    0.00000 ;     H-    N-    CA-    CB
   505    504    506    524      3    0.00000    0.00000    0.00000
0.00000    0.00000    0.00000 ;     H-    N-    CA-    C
   506    508    511    512      3    0.66944    2.00832    0.00000
-2.67776    0.00000    0.00000 ;    CA-    CB-    CG-   HG2
   506    508    511    513      3    0.66944    2.00832    0.00000
-2.67776    0.00000    0.00000 ;    CA-    CB-    CG-   HG3
   506    508    511    514      3    3.68192    3.09616   -2.09200
-3.01248    0.00000    0.00000 ;    CA-    CB-    CG-    CD
   506    524    526    527      3   20.92000    0.00000  -20.92000
0.00000    0.00000    0.00000 ;    CA-    C-    N-    H
   506    524    526    528      3   20.92000    0.00000  -20.92000
0.00000    0.00000    0.00000 ;    CA-    C-    N-    CA
   507    506    508    509      3    0.65084    1.95253    0.00000
-2.60338    0.00000    0.00000 ;    HA-    CA-    CB-   HB2
   507    506    508    510      3    0.65084    1.95253    0.00000
-2.60338    0.00000    0.00000 ;    HA-    CA-    CB-   HB3
   507    506    508    511      3    0.65084    1.95253    0.00000
-2.60338    0.00000    0.00000 ;    HA-    CA-    CB-    CG
   507    506    524    525      3    3.68192   -4.35136    0.00000
1.33888    0.00000    0.00000 ;    HA-    CA-    C-    O
   507    506    524    526      3    0.00000    0.00000    0.00000
0.00000    0.00000    0.00000 ;    HA-    CA-    C-    N
   508    506    524    525      3    0.00000    0.00000    0.00000
0.00000    0.00000    0.00000 ;    CB-    CA-    C-    O
   508    506    524    526      3    2.51040    4.18400    1.67360
-6.69440    0.00000    0.00000 ;    CB-    CA-    C-    N
   508    511    514    515      3    0.66944    2.00832    0.00000
-2.67776    0.00000    0.00000 ;    CB-    CG-    CD-   HD2
   508    511    514    516      3    0.66944    2.00832    0.00000
-2.67776    0.00000    0.00000 ;    CB-    CG-    CD-   HD3
   508    511    514    517      3    3.68192    3.09616   -2.09200
-3.01248    0.00000    0.00000 ;    CB-    CG-    CD-    CE
   509    508    506    524      3    0.65084    1.95253    0.00000
-2.60338    0.00000    0.00000 ;   HB2-    CB-    CA-    C
   509    508    511    512      3    0.62760    1.88280    0.00000
-2.51040    0.00000    0.00000 ;   HB2-    CB-    CG-   HG2
   509    508    511    513      3    0.62760    1.88280    0.00000
-2.51040    0.00000    0.00000 ;   HB2-    CB-    CG-   HG3
   509    508    511    514      3    0.66944    2.00832    0.00000
-2.67776    0.00000    0.00000 ;   HB2-    CB-    CG-    CD
   510    508    506    524      3    0.65084    1.95253    0.00000
-2.60338    0.00000    0.00000 ;   HB3-    CB-    CA-    C
```

```
  510    508    511    512      3    0.62760    1.88280    0.00000
-2.51040    0.00000    0.00000 ;    HB3-   CB-    CG-   HG2
  510    508    511    513      3    0.62760    1.88280    0.00000
-2.51040    0.00000    0.00000 ;    HB3-   CB-    CG-   HG3
  510    508    511    514      3    0.66944    2.00832    0.00000
-2.67776    0.00000    0.00000 ;    HB3-   CB-    CG-    CD
  511    508    506    524      3    0.65084    1.95253    0.00000
-2.60338    0.00000    0.00000 ;     CG-   CB-    CA-     C
  511    514    517    518      3    0.65084    1.95253    0.00000
-2.60338    0.00000    0.00000 ;     CG-   CD-    CE-   HE2
  511    514    517    519      3    0.65084    1.95253    0.00000
-2.60338    0.00000    0.00000 ;     CG-   CD-    CE-   HE3
  511    514    517    520      3    0.65084    1.95253    0.00000
-2.60338    0.00000    0.00000 ;     CG-   CD-    CE-    NZ
  512    511    514    515      3    0.62760    1.88280    0.00000
-2.51040    0.00000    0.00000 ;    HG2-   CG-    CD-   HD2
  512    511    514    516      3    0.62760    1.88280    0.00000
-2.51040    0.00000    0.00000 ;    HG2-   CG-    CD-   HD3
  512    511    514    517      3    0.66944    2.00832    0.00000
-2.67776    0.00000    0.00000 ;    HG2-   CG-    CD-    CE
  513    511    514    515      3    0.62760    1.88280    0.00000
-2.51040    0.00000    0.00000 ;    HG3-   CG-    CD-   HD2
  513    511    514    516      3    0.62760    1.88280    0.00000
-2.51040    0.00000    0.00000 ;    HG3-   CG-    CD-   HD3
  513    511    514    517      3    0.66944    2.00832    0.00000
-2.67776    0.00000    0.00000 ;    HG3-   CG-    CD-    CE
  514    517    520    521      3    0.65084    1.95253    0.00000
-2.60338    0.00000    0.00000 ;     CD-   CE-    NZ-   HZ1
  514    517    520    522      3    0.65084    1.95253    0.00000
-2.60338    0.00000    0.00000 ;     CD-   CE-    NZ-   HZ2
  514    517    520    523      3    0.65084    1.95253    0.00000
-2.60338    0.00000    0.00000 ;     CD-   CE-    NZ-   HZ3
  515    514    517    518      3    0.65084    1.95253    0.00000
-2.60338    0.00000    0.00000 ;    HD2-   CD-    CE-   HE2
  515    514    517    519      3    0.65084    1.95253    0.00000
-2.60338    0.00000    0.00000 ;    HD2-   CD-    CE-   HE3
  515    514    517    520      3    0.65084    1.95253    0.00000
-2.60338    0.00000    0.00000 ;    HD2-   CD-    CE-    NZ
  516    514    517    518      3    0.65084    1.95253    0.00000
-2.60338    0.00000    0.00000 ;    HD3-   CD-    CE-   HE2
  516    514    517    519      3    0.65084    1.95253    0.00000
-2.60338    0.00000    0.00000 ;    HD3-   CD-    CE-   HE3
  516    514    517    520      3    0.65084    1.95253    0.00000
-2.60338    0.00000    0.00000 ;    HD3-   CD-    CE-    NZ
  518    517    520    521      3    0.65084    1.95253    0.00000
-2.60338    0.00000    0.00000 ;    HE2-   CE-    NZ-   HZ1
  518    517    520    522      3    0.65084    1.95253    0.00000
-2.60338    0.00000    0.00000 ;    HE2-   CE-    NZ-   HZ2
  518    517    520    523      3    0.65084    1.95253    0.00000
-2.60338    0.00000    0.00000 ;    HE2-   CE-    NZ-   HZ3
  519    517    520    521      3    0.65084    1.95253    0.00000
-2.60338    0.00000    0.00000 ;    HE3-   CE-    NZ-   HZ1
```

519 517 520 522 3 0.65084 1.95253 0.00000
-2.60338 0.00000 0.00000 ; HE3- CE- NZ- HZ2
519 517 520 523 3 0.65084 1.95253 0.00000
-2.60338 0.00000 0.00000 ; HE3- CE- NZ- HZ3
524 526 528 529 3 0.00000 0.00000 0.00000
0.00000 0.00000 0.00000 ; C- N- CA- HA
524 526 528 530 3 11.71520 1.67360 15.06240 -
13.38880 0.00000 0.00000 ; C- N- CA- CB
524 526 528 540 3 1.75728 5.27184 2.25936
-7.02912 0.00000 0.00000 ; C- N- CA- C
525 524 526 527 3 29.28800 -8.36800 -20.92000
0.00000 0.00000 0.00000 ; O- C- N- H
525 524 526 528 3 20.92000 0.00000 -20.92000
0.00000 0.00000 0.00000 ; O- C- N- CA
526 528 530 531 3 0.65084 1.95253 0.00000
-2.60338 0.00000 0.00000 ; N- CA- CB- HB
526 528 530 532 3 3.83673 0.44769 -1.77402
-2.47693 -0.03347 0.00000 ; N- CA- CB- CG1
526 528 530 536 3 3.83673 0.44769 -1.77402
-2.47693 -0.03347 0.00000 ; N- CA- CB- CG2
526 528 540 541 3 0.00000 0.00000 0.00000
0.00000 0.00000 0.00000 ; N- CA- C- O
526 528 540 542 3 17.40544 -5.02080 -13.22144
9.20480 0.00000 0.00000 ; N- CA- C- N
527 526 528 529 3 0.00000 0.00000 0.00000
0.00000 0.00000 0.00000 ; H- N- CA- HA
527 526 528 530 3 0.00000 0.00000 0.00000
0.00000 0.00000 0.00000 ; H- N- CA- CB
527 526 528 540 3 0.00000 0.00000 0.00000
0.00000 0.00000 0.00000 ; H- N- CA- C
528 530 532 533 3 0.66944 2.00832 0.00000
-2.67776 0.00000 0.00000 ; CA- CB- CG1- HG11
528 530 532 534 3 0.66944 2.00832 0.00000
-2.67776 0.00000 0.00000 ; CA- CB- CG1- HG12
528 530 532 535 3 0.66944 2.00832 0.00000
-2.67776 0.00000 0.00000 ; CA- CB- CG1- HG13
528 530 536 537 3 0.66944 2.00832 0.00000
-2.67776 0.00000 0.00000 ; CA- CB- CG2- HG21
528 530 536 538 3 0.66944 2.00832 0.00000
-2.67776 0.00000 0.00000 ; CA- CB- CG2- HG22
528 530 536 539 3 0.66944 2.00832 0.00000
-2.67776 0.00000 0.00000 ; CA- CB- CG2- HG23
528 540 542 543 3 20.92000 0.00000 -20.92000
0.00000 0.00000 0.00000 ; CA- C- N- H
528 540 542 544 3 20.92000 0.00000 -20.92000
0.00000 0.00000 0.00000 ; CA- C- N- CA
529 528 530 531 3 0.65084 1.95253 0.00000
-2.60338 0.00000 0.00000 ; HA- CA- CB- HB
529 528 530 532 3 0.65084 1.95253 0.00000
-2.60338 0.00000 0.00000 ; HA- CA- CB- CG1
529 528 530 536 3 0.65084 1.95253 0.00000
-2.60338 0.00000 0.00000 ; HA- CA- CB- CG2

529 528 540 541 3 3.68192 -4.35136 0.00000
1.33888 0.00000 0.00000 ; HA- CA- C- O
529 528 540 542 3 0.00000 0.00000 0.00000
0.00000 0.00000 0.00000 ; HA- CA- C- N
530 528 540 541 3 0.00000 0.00000 0.00000
0.00000 0.00000 0.00000 ; CB- CA- C- O
530 528 540 542 3 2.51040 4.18400 1.67360
-6.69440 0.00000 0.00000 ; CB- CA- C- N
531 530 528 540 3 0.65084 1.95253 0.00000
-2.60338 0.00000 0.00000 ; HB- CB- CA- C
531 530 532 533 3 0.62760 1.88280 0.00000
-2.51040 0.00000 0.00000 ; HB- CB- CG1- HG11
531 530 532 534 3 0.62760 1.88280 0.00000
-2.51040 0.00000 0.00000 ; HB- CB- CG1- HG12
531 530 532 535 3 0.62760 1.88280 0.00000
-2.51040 0.00000 0.00000 ; HB- CB- CG1- HG13
531 530 536 537 3 0.62760 1.88280 0.00000
-2.51040 0.00000 0.00000 ; HB- CB- CG2- HG21
531 530 536 538 3 0.62760 1.88280 0.00000
-2.51040 0.00000 0.00000 ; HB- CB- CG2- HG22
531 530 536 539 3 0.62760 1.88280 0.00000
-2.51040 0.00000 0.00000 ; HB- CB- CG2- HG23
532 530 528 540 3 5.67350 3.55640 -6.16722
-2.47693 3.74886 0.00000 ; CG1- CB- CA- C
532 530 536 537 3 0.66944 2.00832 0.00000
-2.67776 0.00000 0.00000 ; CG1- CB- CG2- HG21
532 530 536 538 3 0.66944 2.00832 0.00000
-2.67776 0.00000 0.00000 ; CG1- CB- CG2- HG22
532 530 536 539 3 0.66944 2.00832 0.00000
-2.67776 0.00000 0.00000 ; CG1- CB- CG2- HG23
533 532 530 536 3 0.66944 2.00832 0.00000
-2.67776 0.00000 0.00000 ; HG11- CG1- CB- CG2
534 532 530 536 3 0.66944 2.00832 0.00000
-2.67776 0.00000 0.00000 ; HG12- CG1- CB- CG2
535 532 530 536 3 0.66944 2.00832 0.00000
-2.67776 0.00000 0.00000 ; HG13- CG1- CB- CG2
536 530 528 540 3 5.67350 3.55640 -6.16722
-2.47693 3.74886 0.00000 ; CG2- CB- CA- C
540 542 544 545 3 0.00000 0.00000 0.00000
0.00000 0.00000 0.00000 ; C- N- CA- HA
540 542 544 546 3 11.71520 1.67360 15.06240 -
13.38880 0.00000 0.00000 ; C- N- CA- CB
540 542 544 554 3 1.75728 5.27184 2.25936
-7.02912 0.00000 0.00000 ; C- N- CA- C
541 540 542 543 3 29.28800 -8.36800 -20.92000
0.00000 0.00000 0.00000 ; O- C- N- H
541 540 542 544 3 20.92000 0.00000 -20.92000
0.00000 0.00000 0.00000 ; O- C- N- CA
542 544 546 547 3 0.65084 1.95253 0.00000
-2.60338 0.00000 0.00000 ; N- CA- CB- HB
542 544 546 548 3 3.83673 0.44769 -1.77402
-2.47693 -0.03347 0.00000 ; N- CA- CB- CG2

542 544 546 552 3 4.93294 1.13386 -3.12963
-5.27184 3.17984 0.00000 ; N- CA- CB- OG1
542 544 554 555 3 0.00000 0.00000 0.00000
0.00000 0.00000 0.00000 ; N- CA- C- O
542 544 554 556 3 17.40544 -5.02080 -13.22144
9.20480 0.00000 0.00000 ; N- CA- C- N
543 542 544 545 3 0.00000 0.00000 0.00000
0.00000 0.00000 0.00000 ; H- N- CA- HA
543 542 544 546 3 0.00000 0.00000 0.00000
0.00000 0.00000 0.00000 ; H- N- CA- CB
543 542 544 554 3 0.00000 0.00000 0.00000
0.00000 0.00000 0.00000 ; H- N- CA- C
544 546 548 549 3 0.66944 2.00832 0.00000
-2.67776 0.00000 0.00000 ; CA- CB- CG2- HG21
544 546 548 550 3 0.66944 2.00832 0.00000
-2.67776 0.00000 0.00000 ; CA- CB- CG2- HG22
544 546 548 551 3 0.66944 2.00832 0.00000
-2.67776 0.00000 0.00000 ; CA- CB- CG2- HG23
544 546 552 553 3 1.12131 2.98738 1.66523
-3.94970 0.43514 0.00000 ; CA- CB- OG1- HG1
544 554 556 557 3 20.92000 0.00000 -20.92000
0.00000 0.00000 0.00000 ; CA- C- N- H
544 554 556 558 3 20.92000 0.00000 -20.92000
0.00000 0.00000 0.00000 ; CA- C- N- CA
545 544 546 547 3 0.65084 1.95253 0.00000
-2.60338 0.00000 0.00000 ; HA- CA- CB- HB
545 544 546 548 3 0.65084 1.95253 0.00000
-2.60338 0.00000 0.00000 ; HA- CA- CB- CG2
545 544 546 552 3 1.04600 -1.04600 0.00000
0.00000 0.00000 0.00000 ; HA- CA- CB- OG1
545 544 554 555 3 3.68192 -4.35136 0.00000
1.33888 0.00000 0.00000 ; HA- CA- C- O
545 544 554 556 3 0.00000 0.00000 0.00000
0.00000 0.00000 0.00000 ; HA- CA- C- N
546 544 554 555 3 0.00000 0.00000 0.00000
0.00000 0.00000 0.00000 ; CB- CA- C- O
546 544 554 556 3 2.51040 4.18400 1.67360
-6.69440 0.00000 0.00000 ; CB- CA- C- N
547 546 544 554 3 0.65084 1.95253 0.00000
-2.60338 0.00000 0.00000 ; HB- CB- CA- C
547 546 548 549 3 0.65084 1.95253 0.00000
-2.60338 0.00000 0.00000 ; HB- CB- CG2- HG21
547 546 548 550 3 0.65084 1.95253 0.00000
-2.60338 0.00000 0.00000 ; HB- CB- CG2- HG22
547 546 548 551 3 0.65084 1.95253 0.00000
-2.60338 0.00000 0.00000 ; HB- CB- CG2- HG23
547 546 552 553 3 0.69733 2.09200 0.00000
-2.78933 0.00000 0.00000 ; HB- CB- OG1- HG1
548 546 544 554 3 5.67350 3.55640 -6.16722
-2.47693 3.74886 0.00000 ; CG2- CB- CA- C
548 546 552 553 3 4.74047 0.27196 -2.26773
-3.94970 1.60666 0.00000 ; CG2- CB- OG1- HG1

549 548 546 552 3 1.04600 -1.04600 0.00000
0.00000 0.00000 0.00000 ; HG21- CG2- CB- OG1
550 548 546 552 3 1.04600 -1.04600 0.00000
0.00000 0.00000 0.00000 ; HG22- CG2- CB- OG1
551 548 546 552 3 1.04600 -1.04600 0.00000
0.00000 0.00000 0.00000 ; HG23- CG2- CB- OG1
552 546 544 554 3 6.53541 6.87013 -6.21742
-5.27184 5.22163 0.00000 ; OG1- CB- CA- C
554 556 558 559 3 0.00000 0.00000 0.00000
0.00000 0.00000 0.00000 ; C- N- CA- HA
554 556 558 560 3 11.71520 1.67360 15.06240 -
13.38880 0.00000 0.00000 ; C- N- CA- CB
554 556 558 568 3 1.75728 5.27184 2.25936
-7.02912 0.00000 0.00000 ; C- N- CA- C
555 554 556 557 3 29.28800 -8.36800 -20.92000
0.00000 0.00000 0.00000 ; O- C- N- H
555 554 556 558 3 20.92000 0.00000 -20.92000
0.00000 0.00000 0.00000 ; O- C- N- CA
556 558 560 561 3 0.65084 1.95253 0.00000
-2.60338 0.00000 0.00000 ; N- CA- CB- HB2
556 558 560 562 3 0.65084 1.95253 0.00000
-2.60338 0.00000 0.00000 ; N- CA- CB- HB3
556 558 560 563 3 5.99567 3.29281 -4.46014
-0.55229 1.97485 0.00000 ; N- CA- CB- CG
556 558 568 569 3 0.00000 0.00000 0.00000
0.00000 0.00000 0.00000 ; N- CA- C- O
556 558 568 570 3 17.40544 -5.02080 -13.22144
9.20480 0.00000 0.00000 ; N- CA- C- N
557 556 558 559 3 0.00000 0.00000 0.00000
0.00000 0.00000 0.00000 ; H- N- CA- HA
557 556 558 560 3 0.00000 0.00000 0.00000
0.00000 0.00000 0.00000 ; H- N- CA- CB
557 556 558 568 3 0.00000 0.00000 0.00000
0.00000 0.00000 0.00000 ; H- N- CA- C
558 560 563 564 3 0.00000 0.00000 0.00000
0.00000 0.00000 0.00000 ; CA- CB- CG- OD1
558 560 563 565 3 8.84916 -0.31380 -4.32626
5.03754 0.26778 0.00000 ; CA- CB- CG- ND2
558 568 570 571 3 20.92000 0.00000 -20.92000
0.00000 0.00000 0.00000 ; CA- C- N- H
558 568 570 572 3 20.92000 0.00000 -20.92000
0.00000 0.00000 0.00000 ; CA- C- N- CA
559 558 560 561 3 0.65084 1.95253 0.00000
-2.60338 0.00000 0.00000 ; HA- CA- CB- HB2
559 558 560 562 3 0.65084 1.95253 0.00000
-2.60338 0.00000 0.00000 ; HA- CA- CB- HB3
559 558 560 563 3 0.65084 1.95253 0.00000
-2.60338 0.00000 0.00000 ; HA- CA- CB- CG
559 558 568 569 3 3.68192 -4.35136 0.00000
1.33888 0.00000 0.00000 ; HA- CA- C- O
559 558 568 570 3 0.00000 0.00000 0.00000
0.00000 0.00000 0.00000 ; HA- CA- C- N

```
   560    558    568    569      3    0.00000    0.00000    0.00000
0.00000    0.00000    0.00000 ;     CB-    CA-     C-     O
   560    558    568    570      3    2.51040    4.18400    1.67360
-6.69440    0.00000    0.00000 ;     CB-    CA-     C-     N
   560    563    565    566      3   20.92000    0.00000  -20.92000
0.00000    0.00000    0.00000 ;     CB-    CG-   ND2-  HD21
   560    563    565    567      3   20.92000    0.00000  -20.92000
0.00000    0.00000    0.00000 ;     CB-    CG-   ND2-  HD22
   561    560    558    568      3    0.65084    1.95253    0.00000
-2.60338    0.00000    0.00000 ;    HB2-    CB-    CA-     C
   561    560    563    564      3    3.68192   -4.35136    0.00000
1.33888    0.00000    0.00000 ;    HB2-    CB-    CG-   OD1
   561    560    563    565      3    0.00000    0.00000    0.00000
0.00000    0.00000    0.00000 ;    HB2-    CB-    CG-   ND2
   562    560    558    568      3    0.65084    1.95253    0.00000
-2.60338    0.00000    0.00000 ;    HB3-    CB-    CA-     C
   562    560    563    564      3    3.68192   -4.35136    0.00000
1.33888    0.00000    0.00000 ;    HB3-    CB-    CG-   OD1
   562    560    563    565      3    0.00000    0.00000    0.00000
0.00000    0.00000    0.00000 ;    HB3-    CB-    CG-   ND2
   563    560    558    568      3    7.94542   -3.96225   -6.11701
-0.55229    3.58150    0.00000 ;     CG-    CB-    CA-     C
   564    563    565    566      3   29.28800   -8.36800  -20.92000
0.00000    0.00000    0.00000 ;    OD1-    CG-   ND2-  HD21
   564    563    565    567      3   29.28800   -8.36800  -20.92000
0.00000    0.00000    0.00000 ;    OD1-    CG-   ND2-  HD22
   568    570    572    573      3    0.00000    0.00000    0.00000
0.00000    0.00000    0.00000 ;      C-     N-    CA-    HA
   568    570    572    574      3   11.71520    1.67360   15.06240  -
13.38880    0.00000    0.00000 ;      C-     N-    CA-    CB
   568    570    572    585      3    1.75728    5.27184    2.25936
-7.02912    0.00000    0.00000 ;      C-     N-    CA-     C
   569    568    570    571      3   29.28800   -8.36800  -20.92000
0.00000    0.00000    0.00000 ;      O-     C-     N-     H
   569    568    570    572      3   20.92000    0.00000  -20.92000
0.00000    0.00000    0.00000 ;      O-     C-     N-    CA
   570    572    574    575      3    0.65084    1.95253    0.00000
-2.60338    0.00000    0.00000 ;      N-    CA-    CB-   HB2
   570    572    574    576      3    0.65084    1.95253    0.00000
-2.60338    0.00000    0.00000 ;      N-    CA-    CB-   HB3
   570    572    574    577      3    3.21331    2.22589   -4.15053
-2.40998    2.61082    0.00000 ;      N-    CA-    CB-    CG
   570    572    585    586      3    0.00000    0.00000    0.00000
0.00000    0.00000    0.00000 ;      N-    CA-     C-     O
   570    572    585    587      3   17.40544   -5.02080  -13.22144
9.20480    0.00000    0.00000 ;      N-    CA-     C-     N
   571    570    572    573      3    0.00000    0.00000    0.00000
0.00000    0.00000    0.00000 ;      H-     N-    CA-    HA
   571    570    572    574      3    0.00000    0.00000    0.00000
0.00000    0.00000    0.00000 ;      H-     N-    CA-    CB
   571    570    572    585      3    0.00000    0.00000    0.00000
0.00000    0.00000    0.00000 ;      H-     N-    CA-     C
```

572 574 577 578 3 0.66944 2.00832 0.00000
-2.67776 0.00000 0.00000 ; CA- CB- CG- HG2
572 574 577 579 3 0.66944 2.00832 0.00000
-2.67776 0.00000 0.00000 ; CA- CB- CG- HG3
572 574 577 580 3 3.69866 -4.35136 -3.92459
6.89523 4.61914 0.00000 ; CA- CB- CG- CD
572 585 587 588 3 20.92000 0.00000 -20.92000
0.00000 0.00000 0.00000 ; CA- C- N- H
572 585 587 589 3 20.92000 0.00000 -20.92000
0.00000 0.00000 0.00000 ; CA- C- N- CA
573 572 574 575 3 0.65084 1.95253 0.00000
-2.60338 0.00000 0.00000 ; HA- CA- CB- HB2
573 572 574 576 3 0.65084 1.95253 0.00000
-2.60338 0.00000 0.00000 ; HA- CA- CB- HB3
573 572 574 577 3 0.65084 1.95253 0.00000
-2.60338 0.00000 0.00000 ; HA- CA- CB- CG
573 572 585 586 3 3.68192 -4.35136 0.00000
1.33888 0.00000 0.00000 ; HA- CA- C- O
573 572 585 587 3 0.00000 0.00000 0.00000
0.00000 0.00000 0.00000 ; HA- CA- C- N
574 572 585 586 3 0.00000 0.00000 0.00000
0.00000 0.00000 0.00000 ; CB- CA- C- O
574 572 585 587 3 2.51040 4.18400 1.67360
-6.69440 0.00000 0.00000 ; CB- CA- C- N
574 577 580 581 3 0.00000 0.00000 0.00000
0.00000 0.00000 0.00000 ; CB- CG- CD- OE1
574 577 580 582 3 10.32611 1.48114 -8.47678
1.42256 1.40582 0.00000 ; CB- CG- CD- NE2
575 574 572 585 3 0.65084 1.95253 0.00000
-2.60338 0.00000 0.00000 ; HB2- CB- CA- C
575 574 577 578 3 0.62760 1.88280 0.00000
-2.51040 0.00000 0.00000 ; HB2- CB- CG- HG2
575 574 577 579 3 0.62760 1.88280 0.00000
-2.51040 0.00000 0.00000 ; HB2- CB- CG- HG3
575 574 577 580 3 0.65084 1.95253 0.00000
-2.60338 0.00000 0.00000 ; HB2- CB- CG- CD
576 574 572 585 3 0.65084 1.95253 0.00000
-2.60338 0.00000 0.00000 ; HB3- CB- CA- C
576 574 577 578 3 0.62760 1.88280 0.00000
-2.51040 0.00000 0.00000 ; HB3- CB- CG- HG2
576 574 577 579 3 0.62760 1.88280 0.00000
-2.51040 0.00000 0.00000 ; HB3- CB- CG- HG3
576 574 577 580 3 0.65084 1.95253 0.00000
-2.60338 0.00000 0.00000 ; HB3- CB- CG- CD
577 574 572 585 3 6.86594 3.56895 -8.14206
-2.40998 4.85344 0.00000 ; CG- CB- CA- C
577 580 582 583 3 20.92000 0.00000 -20.92000
0.00000 0.00000 0.00000 ; CG- CD- NE2- HE21
577 580 582 584 3 20.92000 0.00000 -20.92000
0.00000 0.00000 0.00000 ; CG- CD- NE2- HE22
578 577 580 581 3 3.68192 -4.35136 0.00000
1.33888 0.00000 0.00000 ; HG2- CG- CD- OE1

578 577 580 582 3 0.00000 0.00000 0.00000
0.00000 0.00000 0.00000 ; HG2- CG- CD- NE2
579 577 580 581 3 3.68192 -4.35136 0.00000
1.33888 0.00000 0.00000 ; HG3- CG- CD- OE1
579 577 580 582 3 0.00000 0.00000 0.00000
0.00000 0.00000 0.00000 ; HG3- CG- CD- NE2
581 580 582 583 3 29.28800 -8.36800 -20.92000
0.00000 0.00000 0.00000 ; OE1- CD- NE2- HE21
581 580 582 584 3 29.28800 -8.36800 -20.92000
0.00000 0.00000 0.00000 ; OE1- CD- NE2- HE22
585 587 589 590 3 0.00000 0.00000 0.00000
0.00000 0.00000 0.00000 ; C- N- CA- HA
585 587 589 591 3 11.71520 1.67360 15.06240 -
13.38880 0.00000 0.00000 ; C- N- CA- CB
585 587 589 604 3 1.75728 5.27184 2.25936
-7.02912 0.00000 0.00000 ; C- N- CA- C
586 585 587 588 3 29.28800 -8.36800 -20.92000
0.00000 0.00000 0.00000 ; O- C- N- H
586 585 587 589 3 20.92000 0.00000 -20.92000
0.00000 0.00000 0.00000 ; O- C- N- CA
587 589 591 592 3 0.65084 1.95253 0.00000
-2.60338 0.00000 0.00000 ; N- CA- CB- HB2
587 589 591 593 3 0.65084 1.95253 0.00000
-2.60338 0.00000 0.00000 ; N- CA- CB- HB3
587 589 591 594 3 3.79070 1.39746 -4.61077
-2.40998 2.44346 0.00000 ; N- CA- CB- CG
587 589 604 605 3 0.00000 0.00000 0.00000
0.00000 0.00000 0.00000 ; N- CA- C- O
587 589 604 606 3 17.40544 -5.02080 -13.22144
9.20480 0.00000 0.00000 ; N- CA- C- N
588 587 589 590 3 0.00000 0.00000 0.00000
0.00000 0.00000 0.00000 ; H- N- CA- HA
588 587 589 591 3 0.00000 0.00000 0.00000
0.00000 0.00000 0.00000 ; H- N- CA- CB
588 587 589 604 3 0.00000 0.00000 0.00000
0.00000 0.00000 0.00000 ; H- N- CA- C
589 591 594 595 3 0.66944 2.00832 0.00000
-2.67776 0.00000 0.00000 ; CA- CB- CG- HG
589 591 594 596 3 3.90367 0.19665 -6.21742
-2.37651 5.99149 0.00000 ; CA- CB- CG- CD1
589 591 594 600 3 3.90367 0.19665 -6.21742
-2.37651 5.99149 0.00000 ; CA- CB- CG- CD2
589 604 606 607 3 20.92000 0.00000 -20.92000
0.00000 0.00000 0.00000 ; CA- C- N- H
589 604 606 608 3 20.92000 0.00000 -20.92000
0.00000 0.00000 0.00000 ; CA- C- N- CA
590 589 591 592 3 0.65084 1.95253 0.00000
-2.60338 0.00000 0.00000 ; HA- CA- CB- HB2
590 589 591 593 3 0.65084 1.95253 0.00000
-2.60338 0.00000 0.00000 ; HA- CA- CB- HB3
590 589 591 594 3 0.65084 1.95253 0.00000
-2.60338 0.00000 0.00000 ; HA- CA- CB- CG

590 589 604 605 3 3.68192 -4.35136 0.00000
1.33888 0.00000 0.00000 ; HA- CA- C- O
590 589 604 606 3 0.00000 0.00000 0.00000
0.00000 0.00000 0.00000 ; HA- CA- C- N
591 589 604 605 3 0.00000 0.00000 0.00000
0.00000 0.00000 0.00000 ; CB- CA- C- O
591 589 604 606 3 2.51040 4.18400 1.67360
-6.69440 0.00000 0.00000 ; CB- CA- C- N
591 594 596 597 3 0.66944 2.00832 0.00000
-2.67776 0.00000 0.00000 ; CB- CG- CD1- HD11
591 594 596 598 3 0.66944 2.00832 0.00000
-2.67776 0.00000 0.00000 ; CB- CG- CD1- HD12
591 594 596 599 3 0.66944 2.00832 0.00000
-2.67776 0.00000 0.00000 ; CB- CG- CD1- HD13
591 594 600 601 3 0.66944 2.00832 0.00000
-2.67776 0.00000 0.00000 ; CB- CG- CD2- HD21
591 594 600 602 3 0.66944 2.00832 0.00000
-2.67776 0.00000 0.00000 ; CB- CG- CD2- HD22
591 594 600 603 3 0.66944 2.00832 0.00000
-2.67776 0.00000 0.00000 ; CB- CG- CD2- HD23
592 591 589 604 3 0.65084 1.95253 0.00000
-2.60338 0.00000 0.00000 ; HB2- CB- CA- C
592 591 594 595 3 0.62760 1.88280 0.00000
-2.51040 0.00000 0.00000 ; HB2- CB- CG- HG
592 591 594 596 3 0.66944 2.00832 0.00000
-2.67776 0.00000 0.00000 ; HB2- CB- CG- CD1
592 591 594 600 3 0.66944 2.00832 0.00000
-2.67776 0.00000 0.00000 ; HB2- CB- CG- CD2
593 591 589 604 3 0.65084 1.95253 0.00000
-2.60338 0.00000 0.00000 ; HB3- CB- CA- C
593 591 594 595 3 0.62760 1.88280 0.00000
-2.51040 0.00000 0.00000 ; HB3- CB- CG- HG
593 591 594 596 3 0.66944 2.00832 0.00000
-2.67776 0.00000 0.00000 ; HB3- CB- CG- CD1
593 591 594 600 3 0.66944 2.00832 0.00000
-2.67776 0.00000 0.00000 ; HB3- CB- CG- CD2
594 591 589 604 3 10.33448 -1.14642 -11.54784
-2.40998 6.35968 0.00000 ; CG- CB- CA- C
595 594 596 597 3 0.62760 1.88280 0.00000
-2.51040 0.00000 0.00000 ; HG- CG- CD1- HD11
595 594 596 598 3 0.62760 1.88280 0.00000
-2.51040 0.00000 0.00000 ; HG- CG- CD1- HD12
595 594 596 599 3 0.62760 1.88280 0.00000
-2.51040 0.00000 0.00000 ; HG- CG- CD1- HD13
595 594 600 601 3 0.62760 1.88280 0.00000
-2.51040 0.00000 0.00000 ; HG- CG- CD2- HD21
595 594 600 602 3 0.62760 1.88280 0.00000
-2.51040 0.00000 0.00000 ; HG- CG- CD2- HD22
595 594 600 603 3 0.62760 1.88280 0.00000
-2.51040 0.00000 0.00000 ; HG- CG- CD2- HD23
596 594 600 601 3 0.66944 2.00832 0.00000
-2.67776 0.00000 0.00000 ; CD1- CG- CD2- HD21

596 594 600 602 3 0.66944 2.00832 0.00000
-2.67776 0.00000 0.00000 ; CD1- CG- CD2- HD22
596 594 600 603 3 0.66944 2.00832 0.00000
-2.67776 0.00000 0.00000 ; CD1- CG- CD2- HD23
597 596 594 600 3 0.66944 2.00832 0.00000
-2.67776 0.00000 0.00000 ; HD11- CD1- CG- CD2
598 596 594 600 3 0.66944 2.00832 0.00000
-2.67776 0.00000 0.00000 ; HD12- CD1- CG- CD2
599 596 594 600 3 0.66944 2.00832 0.00000
-2.67776 0.00000 0.00000 ; HD13- CD1- CG- CD2
604 606 608 609 3 0.00000 0.00000 0.00000
0.00000 0.00000 0.00000 ; C- N- CA- HA
604 606 608 610 3 11.71520 1.67360 15.06240 -
13.38880 0.00000 0.00000 ; C- N- CA- CB
604 606 608 625 3 1.75728 5.27184 2.25936
-7.02912 0.00000 0.00000 ; C- N- CA- C
605 604 606 607 3 29.28800 -8.36800 -20.92000
0.00000 0.00000 0.00000 ; O- C- N- H
605 604 606 608 3 20.92000 0.00000 -20.92000
0.00000 0.00000 0.00000 ; O- C- N- CA
606 608 610 611 3 0.65084 1.95253 0.00000
-2.60338 0.00000 0.00000 ; N- CA- CB- HB2
606 608 610 612 3 0.65084 1.95253 0.00000
-2.60338 0.00000 0.00000 ; N- CA- CB- HB3
606 608 610 613 3 3.28026 2.46019 -2.19242
-3.21331 -0.23430 0.00000 ; N- CA- CB- CG
606 608 625 626 3 0.00000 0.00000 0.00000
0.00000 0.00000 0.00000 ; N- CA- C- O
606 608 625 627 3 17.40544 -5.02080 -13.22144
9.20480 0.00000 0.00000 ; N- CA- C- N
607 606 608 609 3 0.00000 0.00000 0.00000
0.00000 0.00000 0.00000 ; H- N- CA- HA
607 606 608 610 3 0.00000 0.00000 0.00000
0.00000 0.00000 0.00000 ; H- N- CA- CB
607 606 608 625 3 0.00000 0.00000 0.00000
0.00000 0.00000 0.00000 ; H- N- CA- C
608 610 613 614 3 0.57739 0.00000 1.02926
0.00000 -1.60666 0.00000 ; CA- CB- CG- CD1
608 610 613 623 3 0.57739 0.00000 1.02926
0.00000 -1.60666 0.00000 ; CA- CB- CG- CD2
608 625 627 628 3 20.92000 0.00000 -20.92000
0.00000 0.00000 0.00000 ; CA- C- N- H
608 625 627 629 3 20.92000 0.00000 -20.92000
0.00000 0.00000 0.00000 ; CA- C- N- CA
609 608 610 611 3 0.65084 1.95253 0.00000
-2.60338 0.00000 0.00000 ; HA- CA- CB- HB2
609 608 610 612 3 0.65084 1.95253 0.00000
-2.60338 0.00000 0.00000 ; HA- CA- CB- HB3
609 608 610 613 3 0.65084 1.95253 0.00000
-2.60338 0.00000 0.00000 ; HA- CA- CB- CG
609 608 625 626 3 3.68192 -4.35136 0.00000
1.33888 0.00000 0.00000 ; HA- CA- C- O

609 608 625 627 3 0.00000 0.00000 0.00000
0.00000 0.00000 0.00000 ; HA- CA- C- N
610 608 625 626 3 0.00000 0.00000 0.00000
0.00000 0.00000 0.00000 ; CB- CA- C- O
610 608 625 627 3 2.51040 4.18400 1.67360
-6.69440 0.00000 0.00000 ; CB- CA- C- N
610 613 614 615 3 30.33400 0.00000 -30.33400
0.00000 0.00000 0.00000 ; CB- CG- CD1- HD1
610 613 614 616 3 30.33400 0.00000 -30.33400
0.00000 0.00000 0.00000 ; CB- CG- CD1- CE1
610 613 623 621 3 30.33400 0.00000 -30.33400
0.00000 0.00000 0.00000 ; CB- CG- CD2- CE2
610 613 623 624 3 30.33400 0.00000 -30.33400
0.00000 0.00000 0.00000 ; CB- CG- CD2- HD2
611 610 608 625 3 0.65084 1.95253 0.00000
-2.60338 0.00000 0.00000 ; HB2- CB- CA- C
611 610 613 614 3 0.00000 0.00000 0.00000
0.00000 0.00000 0.00000 ; HB2- CB- CG- CD1
611 610 613 623 3 0.00000 0.00000 0.00000
0.00000 0.00000 0.00000 ; HB2- CB- CG- CD2
612 610 608 625 3 0.65084 1.95253 0.00000
-2.60338 0.00000 0.00000 ; HB3- CB- CA- C
612 610 613 614 3 0.00000 0.00000 0.00000
0.00000 0.00000 0.00000 ; HB3- CB- CG- CD1
612 610 613 623 3 0.00000 0.00000 0.00000
0.00000 0.00000 0.00000 ; HB3- CB- CG- CD2
613 610 608 625 3 4.95804 2.17986 -3.52293
-3.21331 -0.40166 0.00000 ; CG- CB- CA- C
613 614 616 617 3 30.33400 0.00000 -30.33400
0.00000 0.00000 0.00000 ; CG- CD1- CE1- HE1
613 614 616 618 3 30.33400 0.00000 -30.33400
0.00000 0.00000 0.00000 ; CG- CD1- CE1- CZ
613 623 621 618 3 30.33400 0.00000 -30.33400
0.00000 0.00000 0.00000 ; CG- CD2- CE2- CZ
613 623 621 622 3 30.33400 0.00000 -30.33400
0.00000 0.00000 0.00000 ; CG- CD2- CE2- HE2
614 613 623 621 3 30.33400 0.00000 -30.33400
0.00000 0.00000 0.00000 ; CD1- CG- CD2- CE2
614 613 623 624 3 30.33400 0.00000 -30.33400
0.00000 0.00000 0.00000 ; CD1- CG- CD2- HD2
614 616 618 619 3 30.33400 0.00000 -30.33400
0.00000 0.00000 0.00000 ; CD1- CE1- CZ- OH
614 616 618 621 3 30.33400 0.00000 -30.33400
0.00000 0.00000 0.00000 ; CD1- CE1- CZ- CE2
615 614 613 623 3 30.33400 0.00000 -30.33400
0.00000 0.00000 0.00000 ; HD1- CD1- CG- CD2
615 614 616 617 3 30.33400 0.00000 -30.33400
0.00000 0.00000 0.00000 ; HD1- CD1- CE1- HE1
615 614 616 618 3 30.33400 0.00000 -30.33400
0.00000 0.00000 0.00000 ; HD1- CD1- CE1- CZ
616 614 613 623 3 30.33400 0.00000 -30.33400
0.00000 0.00000 0.00000 ; CE1- CD1- CG- CD2

616 618 619 620 3 7.93286 0.00000 -9.56462
0.00000 2.17568 0.00000 ; CE1- CZ- OH- HH
616 618 621 622 3 30.33400 0.00000 -30.33400
0.00000 0.00000 0.00000 ; CE1- CZ- CE2- HE2
616 618 621 623 3 30.33400 0.00000 -30.33400
0.00000 0.00000 0.00000 ; CE1- CZ- CE2- CD2
617 616 618 619 3 30.33400 0.00000 -30.33400
0.00000 0.00000 0.00000 ; HE1- CE1- CZ- OH
617 616 618 621 3 30.33400 0.00000 -30.33400
0.00000 0.00000 0.00000 ; HE1- CE1- CZ- CE2
618 621 623 624 3 30.33400 0.00000 -30.33400
0.00000 0.00000 0.00000 ; CZ- CE2- CD2- HD2
619 618 621 622 3 30.33400 0.00000 -30.33400
0.00000 0.00000 0.00000 ; OH- CZ- CE2- HE2
619 618 621 623 3 30.33400 0.00000 -30.33400
0.00000 0.00000 0.00000 ; OH- CZ- CE2- CD2
620 619 618 621 3 7.93286 0.00000 -9.56462
0.00000 2.17568 0.00000 ; HH- OH- CZ- CE2
622 621 623 624 3 30.33400 0.00000 -30.33400
0.00000 0.00000 0.00000 ; HE2- CE2- CD2- HD2
625 627 629 630 3 0.00000 0.00000 0.00000
0.00000 0.00000 0.00000 ; C- N- CA- HA
625 627 629 631 3 11.71520 1.67360 15.06240 -
13.38880 0.00000 0.00000 ; C- N- CA- CB
625 627 629 635 3 1.75728 5.27184 2.25936
-7.02912 0.00000 0.00000 ; C- N- CA- C
626 625 627 628 3 29.28800 -8.36800 -20.92000
0.00000 0.00000 0.00000 ; O- C- N- H
626 625 627 629 3 20.92000 0.00000 -20.92000
0.00000 0.00000 0.00000 ; O- C- N- CA
627 629 631 632 3 0.65084 1.95253 0.00000
-2.60338 0.00000 0.00000 ; N- CA- CB- HB1
627 629 631 633 3 0.65084 1.95253 0.00000
-2.60338 0.00000 0.00000 ; N- CA- CB- HB2
627 629 631 634 3 0.65084 1.95253 0.00000
-2.60338 0.00000 0.00000 ; N- CA- CB- HB3
627 629 635 636 3 0.00000 0.00000 0.00000
0.00000 0.00000 0.00000 ; N- CA- C- O
627 629 635 637 3 17.40544 -5.02080 -13.22144
9.20480 0.00000 0.00000 ; N- CA- C- N
628 627 629 630 3 0.00000 0.00000 0.00000
0.00000 0.00000 0.00000 ; H- N- CA- HA
628 627 629 631 3 0.00000 0.00000 0.00000
0.00000 0.00000 0.00000 ; H- N- CA- CB
628 627 629 635 3 0.00000 0.00000 0.00000
0.00000 0.00000 0.00000 ; H- N- CA- C
629 635 637 638 3 20.92000 0.00000 -20.92000
0.00000 0.00000 0.00000 ; CA- C- N- H
629 635 637 639 3 20.92000 0.00000 -20.92000
0.00000 0.00000 0.00000 ; CA- C- N- CA
630 629 631 632 3 0.65084 1.95253 0.00000
-2.60338 0.00000 0.00000 ; HA- CA- CB- HB1

630 629 631 633 3 0.65084 1.95253 0.00000
-2.60338 0.00000 0.00000 ; HA- CA- CB- HB2
630 629 631 634 3 0.65084 1.95253 0.00000
-2.60338 0.00000 0.00000 ; HA- CA- CB- HB3
630 629 635 636 3 3.68192 -4.35136 0.00000
1.33888 0.00000 0.00000 ; HA- CA- C- O
630 629 635 637 3 0.00000 0.00000 0.00000
0.00000 0.00000 0.00000 ; HA- CA- C- N
631 629 635 636 3 0.00000 0.00000 0.00000
0.00000 0.00000 0.00000 ; CB- CA- C- O
631 629 635 637 3 2.51040 4.18400 1.67360
-6.69440 0.00000 0.00000 ; CB- CA- C- N
632 631 629 635 3 0.65084 1.95253 0.00000
-2.60338 0.00000 0.00000 ; HB1- CB- CA- C
633 631 629 635 3 0.65084 1.95253 0.00000
-2.60338 0.00000 0.00000 ; HB2- CB- CA- C
634 631 629 635 3 0.65084 1.95253 0.00000
-2.60338 0.00000 0.00000 ; HB3- CB- CA- C
635 637 639 640 3 0.00000 0.00000 0.00000
0.00000 0.00000 0.00000 ; C- N- CA- HA
635 637 639 641 3 11.71520 1.67360 15.06240 -
13.38880 0.00000 0.00000 ; C- N- CA- CB
635 637 639 654 3 1.75728 5.27184 2.25936
-7.02912 0.00000 0.00000 ; C- N- CA- C
636 635 637 638 3 29.28800 -8.36800 -20.92000
0.00000 0.00000 0.00000 ; O- C- N- H
636 635 637 639 3 20.92000 0.00000 -20.92000
0.00000 0.00000 0.00000 ; O- C- N- CA
637 639 641 642 3 0.65084 1.95253 0.00000
-2.60338 0.00000 0.00000 ; N- CA- CB- HB
637 639 641 643 3 3.83673 0.44769 -1.77402
-2.47693 -0.03347 0.00000 ; N- CA- CB- CG2
637 639 641 647 3 2.97482 0.12134 2.04179
-1.89117 -3.24678 0.00000 ; N- CA- CB- CG1
637 639 654 655 3 0.00000 0.00000 0.00000
0.00000 0.00000 0.00000 ; N- CA- C- O
637 639 654 656 3 17.40544 -5.02080 -13.22144
9.20480 0.00000 0.00000 ; N- CA- C- N
638 637 639 640 3 0.00000 0.00000 0.00000
0.00000 0.00000 0.00000 ; H- N- CA- HA
638 637 639 641 3 0.00000 0.00000 0.00000
0.00000 0.00000 0.00000 ; H- N- CA- CB
638 637 639 654 3 0.00000 0.00000 0.00000
0.00000 0.00000 0.00000 ; H- N- CA- C
639 641 643 644 3 0.66944 2.00832 0.00000
-2.67776 0.00000 0.00000 ; CA- CB- CG2- HG21
639 641 643 645 3 0.66944 2.00832 0.00000
-2.67776 0.00000 0.00000 ; CA- CB- CG2- HG22
639 641 643 646 3 0.66944 2.00832 0.00000
-2.67776 0.00000 0.00000 ; CA- CB- CG2- HG23
639 641 647 648 3 0.66944 2.00832 0.00000
-2.67776 0.00000 0.00000 ; CA- CB- CG1- HG12

```
   639    641    647    649      3    0.66944    2.00832    0.00000
-2.67776    0.00000    0.00000 ;     CA-    CB-   CG1-  HG13
   639    641    647    650      3    4.24258   -0.52718   -7.25506
-1.79075    7.69856    0.00000 ;     CA-    CB-   CG1-   CD1
   639    654    656    657      3   20.92000    0.00000  -20.92000
0.00000    0.00000    0.00000 ;     CA-     C-     N-     H
   639    654    656    658      3   20.92000    0.00000  -20.92000
0.00000    0.00000    0.00000 ;     CA-     C-     N-    CA
   640    639    641    642      3    0.65084    1.95253    0.00000
-2.60338    0.00000    0.00000 ;     HA-    CA-    CB-    HB
   640    639    641    643      3    0.65084    1.95253    0.00000
-2.60338    0.00000    0.00000 ;     HA-    CA-    CB-   CG2
   640    639    641    647      3    0.65084    1.95253    0.00000
-2.60338    0.00000    0.00000 ;     HA-    CA-    CB-   CG1
   640    639    654    655      3    3.68192   -4.35136    0.00000
1.33888    0.00000    0.00000 ;     HA-    CA-     C-     O
   640    639    654    656      3    0.00000    0.00000    0.00000
0.00000    0.00000    0.00000 ;     HA-    CA-     C-     N
   641    639    654    655      3    0.00000    0.00000    0.00000
0.00000    0.00000    0.00000 ;     CB-    CA-     C-     O
   641    639    654    656      3    2.51040    4.18400    1.67360
-6.69440    0.00000    0.00000 ;     CB-    CA-     C-     N
   641    647    650    651      3    0.66944    2.00832    0.00000
-2.67776    0.00000    0.00000 ;     CB-   CG1-   CD1-  HD11
   641    647    650    652      3    0.66944    2.00832    0.00000
-2.67776    0.00000    0.00000 ;     CB-   CG1-   CD1-  HD12
   641    647    650    653      3    0.66944    2.00832    0.00000
-2.67776    0.00000    0.00000 ;     CB-   CG1-   CD1-  HD13
   642    641    639    654      3    0.65084    1.95253    0.00000
-2.60338    0.00000    0.00000 ;     HB-    CB-    CA-     C
   642    641    643    644      3    0.62760    1.88280    0.00000
-2.51040    0.00000    0.00000 ;     HB-    CB-   CG2-  HG21
   642    641    643    645      3    0.62760    1.88280    0.00000
-2.51040    0.00000    0.00000 ;     HB-    CB-   CG2-  HG22
   642    641    643    646      3    0.62760    1.88280    0.00000
-2.51040    0.00000    0.00000 ;     HB-    CB-   CG2-  HG23
   642    641    647    648      3    0.62760    1.88280    0.00000
-2.51040    0.00000    0.00000 ;     HB-    CB-   CG1-  HG12
   642    641    647    649      3    0.62760    1.88280    0.00000
-2.51040    0.00000    0.00000 ;     HB-    CB-   CG1-  HG13
   642    641    647    650      3    0.66944    2.00832    0.00000
-2.67776    0.00000    0.00000 ;     HB-    CB-   CG1-   CD1
   643    641    639    654      3    5.67350    3.55640   -6.16722
-2.47693    3.74886    0.00000 ;    CG2-    CB-    CA-     C
   643    641    647    648      3    0.66944    2.00832    0.00000
-2.67776    0.00000    0.00000 ;    CG2-    CB-   CG1-  HG12
   643    641    647    649      3    0.66944    2.00832    0.00000
-2.67776    0.00000    0.00000 ;    CG2-    CB-   CG1-  HG13
   643    641    647    650      3    3.81162    0.49790   -8.14206
-1.79075    7.49773    0.00000 ;    CG2-    CB-   CG1-   CD1
   644    643    641    647      3    0.66944    2.00832    0.00000
-2.67776    0.00000    0.00000 ;   HG21-   CG2-    CB-   CG1
```

645 643 641 647 3 0.66944 2.00832 0.00000
-2.67776 0.00000 0.00000 ; HG22- CG2- CB- CG1
646 643 641 647 3 0.66944 2.00832 0.00000
-2.67776 0.00000 0.00000 ; HG23- CG2- CB- CG1
647 641 639 654 3 8.26340 0.74057 -9.99976
-1.89117 3.84928 0.00000 ; CG1- CB- CA- C
648 647 650 651 3 0.62760 1.88280 0.00000
-2.51040 0.00000 0.00000 ; HG12- CG1- CD1- HD11
648 647 650 652 3 0.62760 1.88280 0.00000
-2.51040 0.00000 0.00000 ; HG12- CG1- CD1- HD12
648 647 650 653 3 0.62760 1.88280 0.00000
-2.51040 0.00000 0.00000 ; HG12- CG1- CD1- HD13
649 647 650 651 3 0.62760 1.88280 0.00000
-2.51040 0.00000 0.00000 ; HG13- CG1- CD1- HD11
649 647 650 652 3 0.62760 1.88280 0.00000
-2.51040 0.00000 0.00000 ; HG13- CG1- CD1- HD12
649 647 650 653 3 0.62760 1.88280 0.00000
-2.51040 0.00000 0.00000 ; HG13- CG1- CD1- HD13
654 656 658 659 3 0.00000 0.00000 0.00000
0.00000 0.00000 0.00000 ; C- N- CA- HA
654 656 658 660 3 11.71520 1.67360 15.06240 -
13.38880 0.00000 0.00000 ; C- N- CA- CB
654 656 658 676 3 1.75728 5.27184 2.25936
-7.02912 0.00000 0.00000 ; C- N- CA- C
655 654 656 657 3 29.28800 -8.36800 -20.92000
0.00000 0.00000 0.00000 ; O- C- N- H
655 654 656 658 3 20.92000 0.00000 -20.92000
0.00000 0.00000 0.00000 ; O- C- N- CA
656 658 660 661 3 0.65084 1.95253 0.00000
-2.60338 0.00000 0.00000 ; N- CA- CB- HB2
656 658 660 662 3 0.65084 1.95253 0.00000
-2.60338 0.00000 0.00000 ; N- CA- CB- HB3
656 658 660 663 3 0.65084 1.95253 0.00000
-2.60338 0.00000 0.00000 ; N- CA- CB- CG
656 658 676 677 3 0.00000 0.00000 0.00000
0.00000 0.00000 0.00000 ; N- CA- C- O
656 658 676 678 3 17.40544 -5.02080 -13.22144
9.20480 0.00000 0.00000 ; N- CA- C- N
657 656 658 659 3 0.00000 0.00000 0.00000
0.00000 0.00000 0.00000 ; H- N- CA- HA
657 656 658 660 3 0.00000 0.00000 0.00000
0.00000 0.00000 0.00000 ; H- N- CA- CB
657 656 658 676 3 0.00000 0.00000 0.00000
0.00000 0.00000 0.00000 ; H- N- CA- C
658 660 663 664 3 0.66944 2.00832 0.00000
-2.67776 0.00000 0.00000 ; CA- CB- CG- HG2
658 660 663 665 3 0.66944 2.00832 0.00000
-2.67776 0.00000 0.00000 ; CA- CB- CG- HG3
658 660 663 666 3 3.68192 3.09616 -2.09200
-3.01248 0.00000 0.00000 ; CA- CB- CG- CD
658 676 678 679 3 20.92000 0.00000 -20.92000
0.00000 0.00000 0.00000 ; CA- C- N- H

658 676 678 680 3 20.92000 0.00000 -20.92000
0.00000 0.00000 0.00000 ; CA- C- N- CA
659 658 660 661 3 0.65084 1.95253 0.00000
-2.60338 0.00000 0.00000 ; HA- CA- CB- HB2
659 658 660 662 3 0.65084 1.95253 0.00000
-2.60338 0.00000 0.00000 ; HA- CA- CB- HB3
659 658 660 663 3 0.65084 1.95253 0.00000
-2.60338 0.00000 0.00000 ; HA- CA- CB- CG
659 658 676 677 3 3.68192 -4.35136 0.00000
1.33888 0.00000 0.00000 ; HA- CA- C- O
659 658 676 678 3 0.00000 0.00000 0.00000
0.00000 0.00000 0.00000 ; HA- CA- C- N
660 658 676 677 3 0.00000 0.00000 0.00000
0.00000 0.00000 0.00000 ; CB- CA- C- O
660 658 676 678 3 2.51040 4.18400 1.67360
-6.69440 0.00000 0.00000 ; CB- CA- C- N
660 663 666 667 3 0.66944 2.00832 0.00000
-2.67776 0.00000 0.00000 ; CB- CG- CD- HD2
660 663 666 668 3 0.66944 2.00832 0.00000
-2.67776 0.00000 0.00000 ; CB- CG- CD- HD3
660 663 666 669 3 3.68192 3.09616 -2.09200
-3.01248 0.00000 0.00000 ; CB- CG- CD- CE
661 660 658 676 3 0.65084 1.95253 0.00000
-2.60338 0.00000 0.00000 ; HB2- CB- CA- C
661 660 663 664 3 0.62760 1.88280 0.00000
-2.51040 0.00000 0.00000 ; HB2- CB- CG- HG2
661 660 663 665 3 0.62760 1.88280 0.00000
-2.51040 0.00000 0.00000 ; HB2- CB- CG- HG3
661 660 663 666 3 0.66944 2.00832 0.00000
-2.67776 0.00000 0.00000 ; HB2- CB- CG- CD
662 660 658 676 3 0.65084 1.95253 0.00000
-2.60338 0.00000 0.00000 ; HB3- CB- CA- C
662 660 663 664 3 0.62760 1.88280 0.00000
-2.51040 0.00000 0.00000 ; HB3- CB- CG- HG2
662 660 663 665 3 0.62760 1.88280 0.00000
-2.51040 0.00000 0.00000 ; HB3- CB- CG- HG3
662 660 663 666 3 0.66944 2.00832 0.00000
-2.67776 0.00000 0.00000 ; HB3- CB- CG- CD
663 660 658 676 3 0.65084 1.95253 0.00000
-2.60338 0.00000 0.00000 ; CG- CB- CA- C
663 666 669 670 3 0.65084 1.95253 0.00000
-2.60338 0.00000 0.00000 ; CG- CD- CE- HE2
663 666 669 671 3 0.65084 1.95253 0.00000
-2.60338 0.00000 0.00000 ; CG- CD- CE- HE3
663 666 669 672 3 0.65084 1.95253 0.00000
-2.60338 0.00000 0.00000 ; CG- CD- CE- NZ
664 663 666 667 3 0.62760 1.88280 0.00000
-2.51040 0.00000 0.00000 ; HG2- CG- CD- HD2
664 663 666 668 3 0.62760 1.88280 0.00000
-2.51040 0.00000 0.00000 ; HG2- CG- CD- HD3
664 663 666 669 3 0.66944 2.00832 0.00000
-2.67776 0.00000 0.00000 ; HG2- CG- CD- CE

665 663 666 667 3 0.62760 1.88280 0.00000
-2.51040 0.00000 0.00000 ; HG3- CG- CD- HD2
665 663 666 668 3 0.62760 1.88280 0.00000
-2.51040 0.00000 0.00000 ; HG3- CG- CD- HD3
665 663 666 669 3 0.66944 2.00832 0.00000
-2.67776 0.00000 0.00000 ; HG3- CG- CD- CE
666 669 672 673 3 0.65084 1.95253 0.00000
-2.60338 0.00000 0.00000 ; CD- CE- NZ- HZ1
666 669 672 674 3 0.65084 1.95253 0.00000
-2.60338 0.00000 0.00000 ; CD- CE- NZ- HZ2
666 669 672 675 3 0.65084 1.95253 0.00000
-2.60338 0.00000 0.00000 ; CD- CE- NZ- HZ3
667 666 669 670 3 0.65084 1.95253 0.00000
-2.60338 0.00000 0.00000 ; HD2- CD- CE- HE2
667 666 669 671 3 0.65084 1.95253 0.00000
-2.60338 0.00000 0.00000 ; HD2- CD- CE- HE3
667 666 669 672 3 0.65084 1.95253 0.00000
-2.60338 0.00000 0.00000 ; HD2- CD- CE- NZ
668 666 669 670 3 0.65084 1.95253 0.00000
-2.60338 0.00000 0.00000 ; HD3- CD- CE- HE2
668 666 669 671 3 0.65084 1.95253 0.00000
-2.60338 0.00000 0.00000 ; HD3- CD- CE- HE3
668 666 669 672 3 0.65084 1.95253 0.00000
-2.60338 0.00000 0.00000 ; HD3- CD- CE- NZ
670 669 672 673 3 0.65084 1.95253 0.00000
-2.60338 0.00000 0.00000 ; HE2- CE- NZ- HZ1
670 669 672 674 3 0.65084 1.95253 0.00000
-2.60338 0.00000 0.00000 ; HE2- CE- NZ- HZ2
670 669 672 675 3 0.65084 1.95253 0.00000
-2.60338 0.00000 0.00000 ; HE2- CE- NZ- HZ3
671 669 672 673 3 0.65084 1.95253 0.00000
-2.60338 0.00000 0.00000 ; HE3- CE- NZ- HZ1
671 669 672 674 3 0.65084 1.95253 0.00000
-2.60338 0.00000 0.00000 ; HE3- CE- NZ- HZ2
671 669 672 675 3 0.65084 1.95253 0.00000
-2.60338 0.00000 0.00000 ; HE3- CE- NZ- HZ3
676 678 680 681 3 0.00000 0.00000 0.00000
0.00000 0.00000 0.00000 ; C- N- CA- HA
676 678 680 682 3 11.71520 1.67360 15.06240 -
13.38880 0.00000 0.00000 ; C- N- CA- CB
676 678 680 691 3 1.75728 5.27184 2.25936
-7.02912 0.00000 0.00000 ; C- N- CA- C
677 676 678 679 3 29.28800 -8.36800 -20.92000
0.00000 0.00000 0.00000 ; O- C- N- H
677 676 678 680 3 20.92000 0.00000 -20.92000
0.00000 0.00000 0.00000 ; O- C- N- CA
678 680 682 683 3 0.65084 1.95253 0.00000
-2.60338 0.00000 0.00000 ; N- CA- CB- HB2
678 680 682 684 3 0.65084 1.95253 0.00000
-2.60338 0.00000 0.00000 ; N- CA- CB- HB3
678 680 682 685 3 3.21331 2.22589 -4.15053
-2.40998 2.61082 0.00000 ; N- CA- CB- CG

678 680 691 692 3 0.00000 0.00000 0.00000
0.00000 0.00000 0.00000 ; N- CA- C- O
678 680 691 693 3 17.40544 -5.02080 -13.22144
9.20480 0.00000 0.00000 ; N- CA- C- N
679 678 680 681 3 0.00000 0.00000 0.00000
0.00000 0.00000 0.00000 ; H- N- CA- HA
679 678 680 682 3 0.00000 0.00000 0.00000
0.00000 0.00000 0.00000 ; H- N- CA- CB
679 678 680 691 3 0.00000 0.00000 0.00000
0.00000 0.00000 0.00000 ; H- N- CA- C
680 682 685 686 3 0.66944 2.00832 0.00000
-2.67776 0.00000 0.00000 ; CA- CB- CG- HG2
680 682 685 687 3 0.66944 2.00832 0.00000
-2.67776 0.00000 0.00000 ; CA- CB- CG- HG3
680 682 685 688 3 10.12110 -1.91209 0.01674
10.17549 -1.87443 0.00000 ; CA- CB- CG- CD
680 691 693 694 3 20.92000 0.00000 -20.92000
0.00000 0.00000 0.00000 ; CA- C- N- H
680 691 693 695 3 20.92000 0.00000 -20.92000
0.00000 0.00000 0.00000 ; CA- C- N- CA
681 680 682 683 3 0.65084 1.95253 0.00000
-2.60338 0.00000 0.00000 ; HA- CA- CB- HB2
681 680 682 684 3 0.65084 1.95253 0.00000
-2.60338 0.00000 0.00000 ; HA- CA- CB- HB3
681 680 682 685 3 0.65084 1.95253 0.00000
-2.60338 0.00000 0.00000 ; HA- CA- CB- CG
681 680 691 692 3 3.68192 -4.35136 0.00000
1.33888 0.00000 0.00000 ; HA- CA- C- O
681 680 691 693 3 0.00000 0.00000 0.00000
0.00000 0.00000 0.00000 ; HA- CA- C- N
682 680 691 692 3 0.00000 0.00000 0.00000
0.00000 0.00000 0.00000 ; CB- CA- C- O
682 680 691 693 3 2.51040 4.18400 1.67360
-6.69440 0.00000 0.00000 ; CB- CA- C- N
682 685 688 689 3 3.79907 0.00000 -5.40573
0.00000 2.14221 0.00000 ; CB- CG- CD- OE1
682 685 688 690 3 3.79907 0.00000 -5.40573
0.00000 2.14221 0.00000 ; CB- CG- CD- OE2
683 682 680 691 3 0.65084 1.95253 0.00000
-2.60338 0.00000 0.00000 ; HB2- CB- CA- C
683 682 685 686 3 0.62760 1.88280 0.00000
-2.51040 0.00000 0.00000 ; HB2- CB- CG- HG2
683 682 685 687 3 0.62760 1.88280 0.00000
-2.51040 0.00000 0.00000 ; HB2- CB- CG- HG3
683 682 685 688 3 0.65084 1.95253 0.00000
-2.60338 0.00000 0.00000 ; HB2- CB- CG- CD
684 682 680 691 3 0.65084 1.95253 0.00000
-2.60338 0.00000 0.00000 ; HB3- CB- CA- C
684 682 685 686 3 0.62760 1.88280 0.00000
-2.51040 0.00000 0.00000 ; HB3- CB- CG- HG2
684 682 685 687 3 0.62760 1.88280 0.00000
-2.51040 0.00000 0.00000 ; HB3- CB- CG- HG3

684 682 685 688 3 0.65084 1.95253 0.00000
-2.60338 0.00000 0.00000 ; HB3- CB- CG- CD
685 682 680 691 3 6.86594 3.56895 -8.14206
-2.40998 4.85344 0.00000 ; CG- CB- CA- C
686 685 688 689 3 0.00000 0.00000 0.00000
0.00000 0.00000 0.00000 ; HG2- CG- CD- OE1
686 685 688 690 3 0.00000 0.00000 0.00000
0.00000 0.00000 0.00000 ; HG2- CG- CD- OE2
687 685 688 689 3 0.00000 0.00000 0.00000
0.00000 0.00000 0.00000 ; HG3- CG- CD- OE1
687 685 688 690 3 0.00000 0.00000 0.00000
0.00000 0.00000 0.00000 ; HG3- CG- CD- OE2
691 693 695 696 3 0.00000 0.00000 0.00000
0.00000 0.00000 0.00000 ; C- N- CA- HA
691 693 695 697 3 11.71520 1.67360 15.06240 -
13.38880 0.00000 0.00000 ; C- N- CA- CB
691 693 695 707 3 1.75728 5.27184 2.25936
-7.02912 0.00000 0.00000 ; C- N- CA- C
692 691 693 694 3 29.28800 -8.36800 -20.92000
0.00000 0.00000 0.00000 ; O- C- N- H
692 691 693 695 3 20.92000 0.00000 -20.92000
0.00000 0.00000 0.00000 ; O- C- N- CA
693 695 697 698 3 0.65084 1.95253 0.00000
-2.60338 0.00000 0.00000 ; N- CA- CB- HB2
693 695 697 699 3 0.65084 1.95253 0.00000
-2.60338 0.00000 0.00000 ; N- CA- CB- HB3
693 695 697 700 3 3.21331 2.22589 -4.15053
-2.40998 2.61082 0.00000 ; N- CA- CB- CG
693 695 707 708 3 0.00000 0.00000 0.00000
0.00000 0.00000 0.00000 ; N- CA- C- O
693 695 707 709 3 17.40544 -5.02080 -13.22144
9.20480 0.00000 0.00000 ; N- CA- C- N
694 693 695 696 3 0.00000 0.00000 0.00000
0.00000 0.00000 0.00000 ; H- N- CA- HA
694 693 695 697 3 0.00000 0.00000 0.00000
0.00000 0.00000 0.00000 ; H- N- CA- CB
694 693 695 707 3 0.00000 0.00000 0.00000
0.00000 0.00000 0.00000 ; H- N- CA- C
695 697 700 701 3 0.66944 2.00832 0.00000
-2.67776 0.00000 0.00000 ; CA- CB- CG- HG2
695 697 700 702 3 0.66944 2.00832 0.00000
-2.67776 0.00000 0.00000 ; CA- CB- CG- HG3
695 697 700 703 3 3.69866 -4.35136 -3.92459
6.89523 4.61914 0.00000 ; CA- CB- CG- CD
695 707 709 710 3 20.92000 0.00000 -20.92000
0.00000 0.00000 0.00000 ; CA- C- N- H
695 707 709 711 3 20.92000 0.00000 -20.92000
0.00000 0.00000 0.00000 ; CA- C- N- CA
696 695 697 698 3 0.65084 1.95253 0.00000
-2.60338 0.00000 0.00000 ; HA- CA- CB- HB2
696 695 697 699 3 0.65084 1.95253 0.00000
-2.60338 0.00000 0.00000 ; HA- CA- CB- HB3

696 695 697 700 3 0.65084 1.95253 0.00000
-2.60338 0.00000 0.00000 ; HA- CA- CB- CG
696 695 707 708 3 3.68192 -4.35136 0.00000
1.33888 0.00000 0.00000 ; HA- CA- C- O
696 695 707 709 3 0.00000 0.00000 0.00000
0.00000 0.00000 0.00000 ; HA- CA- C- N
697 695 707 708 3 0.00000 0.00000 0.00000
0.00000 0.00000 0.00000 ; CB- CA- C- O
697 695 707 709 3 2.51040 4.18400 1.67360
-6.69440 0.00000 0.00000 ; CB- CA- C- N
697 700 703 704 3 0.00000 0.00000 0.00000
0.00000 0.00000 0.00000 ; CB- CG- CD- OE1
697 700 703 705 3 12.79049 3.13382 -7.02912
0.41840 -2.20915 0.00000 ; CB- CG- CD- OE2
698 697 695 707 3 0.65084 1.95253 0.00000
-2.60338 0.00000 0.00000 ; HB2- CB- CA- C
698 697 700 701 3 0.62760 1.88280 0.00000
-2.51040 0.00000 0.00000 ; HB2- CB- CG- HG2
698 697 700 702 3 0.62760 1.88280 0.00000
-2.51040 0.00000 0.00000 ; HB2- CB- CG- HG3
698 697 700 703 3 0.65084 1.95253 0.00000
-2.60338 0.00000 0.00000 ; HB2- CB- CG- CD
699 697 695 707 3 0.65084 1.95253 0.00000
-2.60338 0.00000 0.00000 ; HB3- CB- CA- C
699 697 700 701 3 0.62760 1.88280 0.00000
-2.51040 0.00000 0.00000 ; HB3- CB- CG- HG2
699 697 700 702 3 0.62760 1.88280 0.00000
-2.51040 0.00000 0.00000 ; HB3- CB- CG- HG3
699 697 700 703 3 0.65084 1.95253 0.00000
-2.60338 0.00000 0.00000 ; HB3- CB- CG- CD
700 697 695 707 3 6.86594 3.56895 -8.14206
-2.40998 4.85344 0.00000 ; CG- CB- CA- C
700 703 705 706 3 27.46796 7.88684 -26.42614
-8.01654 3.78234 0.00000 ; CG- CD- OE2- HE2
701 700 703 704 3 3.68192 -4.35136 0.00000
1.33888 0.00000 0.00000 ; HG2- CG- CD- OE1
701 700 703 705 3 0.00000 0.00000 0.00000
0.00000 0.00000 0.00000 ; HG2- CG- CD- OE2
702 700 703 704 3 3.68192 -4.35136 0.00000
1.33888 0.00000 0.00000 ; HG3- CG- CD- OE1
702 700 703 705 3 0.00000 0.00000 0.00000
0.00000 0.00000 0.00000 ; HG3- CG- CD- OE2
704 703 705 706 3 27.19600 -7.94960 -19.24640
0.00000 0.00000 0.00000 ; OE1- CD- OE2- HE2
707 709 711 712 3 0.00000 0.00000 0.00000
0.00000 0.00000 0.00000 ; C- N- CA- HA
707 709 711 713 3 11.71520 1.67360 15.06240 -
13.38880 0.00000 0.00000 ; C- N- CA- CB
707 709 711 729 3 1.75728 5.27184 2.25936
-7.02912 0.00000 0.00000 ; C- N- CA- C
708 707 709 710 3 29.28800 -8.36800 -20.92000
0.00000 0.00000 0.00000 ; O- C- N- H

708 707 709 711 3 20.92000 0.00000 -20.92000
0.00000 0.00000 0.00000 ; O- C- N- CA
709 711 713 714 3 0.65084 1.95253 0.00000
-2.60338 0.00000 0.00000 ; N- CA- CB- HB2
709 711 713 715 3 0.65084 1.95253 0.00000
-2.60338 0.00000 0.00000 ; N- CA- CB- HB3
709 711 713 716 3 0.65084 1.95253 0.00000
-2.60338 0.00000 0.00000 ; N- CA- CB- CG
709 711 729 730 3 0.00000 0.00000 0.00000
0.00000 0.00000 0.00000 ; N- CA- C- O
709 711 729 731 3 17.40544 -5.02080 -13.22144
9.20480 0.00000 0.00000 ; N- CA- C- N
710 709 711 712 3 0.00000 0.00000 0.00000
0.00000 0.00000 0.00000 ; H- N- CA- HA
710 709 711 713 3 0.00000 0.00000 0.00000
0.00000 0.00000 0.00000 ; H- N- CA- CB
710 709 711 729 3 0.00000 0.00000 0.00000
0.00000 0.00000 0.00000 ; H- N- CA- C
711 713 716 717 3 0.66944 2.00832 0.00000
-2.67776 0.00000 0.00000 ; CA- CB- CG- HG2
711 713 716 718 3 0.66944 2.00832 0.00000
-2.67776 0.00000 0.00000 ; CA- CB- CG- HG3
711 713 716 719 3 3.68192 3.09616 -2.09200
-3.01248 0.00000 0.00000 ; CA- CB- CG- CD
711 729 731 732 3 20.92000 0.00000 -20.92000
0.00000 0.00000 0.00000 ; CA- C- N- H
711 729 731 733 3 20.92000 0.00000 -20.92000
0.00000 0.00000 0.00000 ; CA- C- N- CA
712 711 713 714 3 0.65084 1.95253 0.00000
-2.60338 0.00000 0.00000 ; HA- CA- CB- HB2
712 711 713 715 3 0.65084 1.95253 0.00000
-2.60338 0.00000 0.00000 ; HA- CA- CB- HB3
712 711 713 716 3 0.65084 1.95253 0.00000
-2.60338 0.00000 0.00000 ; HA- CA- CB- CG
712 711 729 730 3 3.68192 -4.35136 0.00000
1.33888 0.00000 0.00000 ; HA- CA- C- O
712 711 729 731 3 0.00000 0.00000 0.00000
0.00000 0.00000 0.00000 ; HA- CA- C- N
713 711 729 730 3 0.00000 0.00000 0.00000
0.00000 0.00000 0.00000 ; CB- CA- C- O
713 711 729 731 3 2.51040 4.18400 1.67360
-6.69440 0.00000 0.00000 ; CB- CA- C- N
713 716 719 720 3 0.66944 2.00832 0.00000
-2.67776 0.00000 0.00000 ; CB- CG- CD- HD2
713 716 719 721 3 0.66944 2.00832 0.00000
-2.67776 0.00000 0.00000 ; CB- CG- CD- HD3
713 716 719 722 3 3.68192 3.09616 -2.09200
-3.01248 0.00000 0.00000 ; CB- CG- CD- CE
714 713 711 729 3 0.65084 1.95253 0.00000
-2.60338 0.00000 0.00000 ; HB2- CB- CA- C
714 713 716 717 3 0.62760 1.88280 0.00000
-2.51040 0.00000 0.00000 ; HB2- CB- CG- HG2

```
   714    713    716    718      3    0.62760    1.88280    0.00000
-2.51040    0.00000    0.00000 ;    HB2-    CB-    CG-   HG3
   714    713    716    719      3    0.66944    2.00832    0.00000
-2.67776    0.00000    0.00000 ;    HB2-    CB-    CG-    CD
   715    713    711    729      3    0.65084    1.95253    0.00000
-2.60338    0.00000    0.00000 ;    HB3-    CB-    CA-     C
   715    713    716    717      3    0.62760    1.88280    0.00000
-2.51040    0.00000    0.00000 ;    HB3-    CB-    CG-   HG2
   715    713    716    718      3    0.62760    1.88280    0.00000
-2.51040    0.00000    0.00000 ;    HB3-    CB-    CG-   HG3
   715    713    716    719      3    0.66944    2.00832    0.00000
-2.67776    0.00000    0.00000 ;    HB3-    CB-    CG-    CD
   716    713    711    729      3    0.65084    1.95253    0.00000
-2.60338    0.00000    0.00000 ;     CG-    CB-    CA-     C
   716    719    722    723      3    0.65084    1.95253    0.00000
-2.60338    0.00000    0.00000 ;     CG-    CD-    CE-   HE2
   716    719    722    724      3    0.65084    1.95253    0.00000
-2.60338    0.00000    0.00000 ;     CG-    CD-    CE-   HE3
   716    719    722    725      3    0.65084    1.95253    0.00000
-2.60338    0.00000    0.00000 ;     CG-    CD-    CE-    NZ
   717    716    719    720      3    0.62760    1.88280    0.00000
-2.51040    0.00000    0.00000 ;    HG2-    CG-    CD-   HD2
   717    716    719    721      3    0.62760    1.88280    0.00000
-2.51040    0.00000    0.00000 ;    HG2-    CG-    CD-   HD3
   717    716    719    722      3    0.66944    2.00832    0.00000
-2.67776    0.00000    0.00000 ;    HG2-    CG-    CD-    CE
   718    716    719    720      3    0.62760    1.88280    0.00000
-2.51040    0.00000    0.00000 ;    HG3-    CG-    CD-   HD2
   718    716    719    721      3    0.62760    1.88280    0.00000
-2.51040    0.00000    0.00000 ;    HG3-    CG-    CD-   HD3
   718    716    719    722      3    0.66944    2.00832    0.00000
-2.67776    0.00000    0.00000 ;    HG3-    CG-    CD-    CE
   719    722    725    726      3    0.65084    1.95253    0.00000
-2.60338    0.00000    0.00000 ;     CD-    CE-    NZ-   HZ1
   719    722    725    727      3    0.65084    1.95253    0.00000
-2.60338    0.00000    0.00000 ;     CD-    CE-    NZ-   HZ2
   719    722    725    728      3    0.65084    1.95253    0.00000
-2.60338    0.00000    0.00000 ;     CD-    CE-    NZ-   HZ3
   720    719    722    723      3    0.65084    1.95253    0.00000
-2.60338    0.00000    0.00000 ;    HD2-    CD-    CE-   HE2
   720    719    722    724      3    0.65084    1.95253    0.00000
-2.60338    0.00000    0.00000 ;    HD2-    CD-    CE-   HE3
   720    719    722    725      3    0.65084    1.95253    0.00000
-2.60338    0.00000    0.00000 ;    HD2-    CD-    CE-    NZ
   721    719    722    723      3    0.65084    1.95253    0.00000
-2.60338    0.00000    0.00000 ;    HD3-    CD-    CE-   HE2
   721    719    722    724      3    0.65084    1.95253    0.00000
-2.60338    0.00000    0.00000 ;    HD3-    CD-    CE-   HE3
   721    719    722    725      3    0.65084    1.95253    0.00000
-2.60338    0.00000    0.00000 ;    HD3-    CD-    CE-    NZ
   723    722    725    726      3    0.65084    1.95253    0.00000
-2.60338    0.00000    0.00000 ;    HE2-    CE-    NZ-   HZ1
```

723 722 725 727 3 0.65084 1.95253 0.00000
-2.60338 0.00000 0.00000 ; HE2- CE- NZ- HZ2
723 722 725 728 3 0.65084 1.95253 0.00000
-2.60338 0.00000 0.00000 ; HE2- CE- NZ- HZ3
724 722 725 726 3 0.65084 1.95253 0.00000
-2.60338 0.00000 0.00000 ; HE3- CE- NZ- HZ1
724 722 725 727 3 0.65084 1.95253 0.00000
-2.60338 0.00000 0.00000 ; HE3- CE- NZ- HZ2
724 722 725 728 3 0.65084 1.95253 0.00000
-2.60338 0.00000 0.00000 ; HE3- CE- NZ- HZ3
729 731 733 734 3 0.00000 0.00000 0.00000
0.00000 0.00000 0.00000 ; C- N- CA- HA
729 731 733 735 3 11.71520 1.67360 15.06240 -
13.38880 0.00000 0.00000 ; C- N- CA- CB
729 731 733 751 3 1.75728 5.27184 2.25936
-7.02912 0.00000 0.00000 ; C- N- CA- C
730 729 731 732 3 29.28800 -8.36800 -20.92000
0.00000 0.00000 0.00000 ; O- C- N- H
730 729 731 733 3 20.92000 0.00000 -20.92000
0.00000 0.00000 0.00000 ; O- C- N- CA
731 733 735 736 3 0.65084 1.95253 0.00000
-2.60338 0.00000 0.00000 ; N- CA- CB- HB2
731 733 735 737 3 0.65084 1.95253 0.00000
-2.60338 0.00000 0.00000 ; N- CA- CB- HB3
731 733 735 738 3 0.65084 1.95253 0.00000
-2.60338 0.00000 0.00000 ; N- CA- CB- CG
731 733 751 752 3 0.00000 0.00000 0.00000
0.00000 0.00000 0.00000 ; N- CA- C- O
731 733 751 753 3 17.40544 -5.02080 -13.22144
9.20480 0.00000 0.00000 ; N- CA- C- N
732 731 733 734 3 0.00000 0.00000 0.00000
0.00000 0.00000 0.00000 ; H- N- CA- HA
732 731 733 735 3 0.00000 0.00000 0.00000
0.00000 0.00000 0.00000 ; H- N- CA- CB
732 731 733 751 3 0.00000 0.00000 0.00000
0.00000 0.00000 0.00000 ; H- N- CA- C
733 735 738 739 3 0.66944 2.00832 0.00000
-2.67776 0.00000 0.00000 ; CA- CB- CG- HG2
733 735 738 740 3 0.66944 2.00832 0.00000
-2.67776 0.00000 0.00000 ; CA- CB- CG- HG3
733 735 738 741 3 3.68192 3.09616 -2.09200
-3.01248 0.00000 0.00000 ; CA- CB- CG- CD
733 751 753 754 3 20.92000 0.00000 -20.92000
0.00000 0.00000 0.00000 ; CA- C- N- H
733 751 753 755 3 20.92000 0.00000 -20.92000
0.00000 0.00000 0.00000 ; CA- C- N- CA
734 733 735 736 3 0.65084 1.95253 0.00000
-2.60338 0.00000 0.00000 ; HA- CA- CB- HB2
734 733 735 737 3 0.65084 1.95253 0.00000
-2.60338 0.00000 0.00000 ; HA- CA- CB- HB3
734 733 735 738 3 0.65084 1.95253 0.00000
-2.60338 0.00000 0.00000 ; HA- CA- CB- CG

734 733 751 752 3 3.68192 -4.35136 0.00000
1.33888 0.00000 0.00000 ; HA- CA- C- O
734 733 751 753 3 0.00000 0.00000 0.00000
0.00000 0.00000 0.00000 ; HA- CA- C- N
735 733 751 752 3 0.00000 0.00000 0.00000
0.00000 0.00000 0.00000 ; CB- CA- C- O
735 733 751 753 3 2.51040 4.18400 1.67360
-6.69440 0.00000 0.00000 ; CB- CA- C- N
735 738 741 742 3 0.66944 2.00832 0.00000
-2.67776 0.00000 0.00000 ; CB- CG- CD- HD2
735 738 741 743 3 0.66944 2.00832 0.00000
-2.67776 0.00000 0.00000 ; CB- CG- CD- HD3
735 738 741 744 3 3.68192 3.09616 -2.09200
-3.01248 0.00000 0.00000 ; CB- CG- CD- CE
736 735 733 751 3 0.65084 1.95253 0.00000
-2.60338 0.00000 0.00000 ; HB2- CB- CA- C
736 735 738 739 3 0.62760 1.88280 0.00000
-2.51040 0.00000 0.00000 ; HB2- CB- CG- HG2
736 735 738 740 3 0.62760 1.88280 0.00000
-2.51040 0.00000 0.00000 ; HB2- CB- CG- HG3
736 735 738 741 3 0.66944 2.00832 0.00000
-2.67776 0.00000 0.00000 ; HB2- CB- CG- CD
737 735 733 751 3 0.65084 1.95253 0.00000
-2.60338 0.00000 0.00000 ; HB3- CB- CA- C
737 735 738 739 3 0.62760 1.88280 0.00000
-2.51040 0.00000 0.00000 ; HB3- CB- CG- HG2
737 735 738 740 3 0.62760 1.88280 0.00000
-2.51040 0.00000 0.00000 ; HB3- CB- CG- HG3
737 735 738 741 3 0.66944 2.00832 0.00000
-2.67776 0.00000 0.00000 ; HB3- CB- CG- CD
738 735 733 751 3 0.65084 1.95253 0.00000
-2.60338 0.00000 0.00000 ; CG- CB- CA- C
738 741 744 745 3 0.65084 1.95253 0.00000
-2.60338 0.00000 0.00000 ; CG- CD- CE- HE2
738 741 744 746 3 0.65084 1.95253 0.00000
-2.60338 0.00000 0.00000 ; CG- CD- CE- HE3
738 741 744 747 3 0.65084 1.95253 0.00000
-2.60338 0.00000 0.00000 ; CG- CD- CE- NZ
739 738 741 742 3 0.62760 1.88280 0.00000
-2.51040 0.00000 0.00000 ; HG2- CG- CD- HD2
739 738 741 743 3 0.62760 1.88280 0.00000
-2.51040 0.00000 0.00000 ; HG2- CG- CD- HD3
739 738 741 744 3 0.66944 2.00832 0.00000
-2.67776 0.00000 0.00000 ; HG2- CG- CD- CE
740 738 741 742 3 0.62760 1.88280 0.00000
-2.51040 0.00000 0.00000 ; HG3- CG- CD- HD2
740 738 741 743 3 0.62760 1.88280 0.00000
-2.51040 0.00000 0.00000 ; HG3- CG- CD- HD3
740 738 741 744 3 0.66944 2.00832 0.00000
-2.67776 0.00000 0.00000 ; HG3- CG- CD- CE
741 744 747 748 3 0.65084 1.95253 0.00000
-2.60338 0.00000 0.00000 ; CD- CE- NZ- HZ1

741 744 747 749 3 0.65084 1.95253 0.00000
-2.60338 0.00000 0.00000 ; CD- CE- NZ- HZ2
741 744 747 750 3 0.65084 1.95253 0.00000
-2.60338 0.00000 0.00000 ; CD- CE- NZ- HZ3
742 741 744 745 3 0.65084 1.95253 0.00000
-2.60338 0.00000 0.00000 ; HD2- CD- CE- HE2
742 741 744 746 3 0.65084 1.95253 0.00000
-2.60338 0.00000 0.00000 ; HD2- CD- CE- HE3
742 741 744 747 3 0.65084 1.95253 0.00000
-2.60338 0.00000 0.00000 ; HD2- CD- CE- NZ
743 741 744 745 3 0.65084 1.95253 0.00000
-2.60338 0.00000 0.00000 ; HD3- CD- CE- HE2
743 741 744 746 3 0.65084 1.95253 0.00000
-2.60338 0.00000 0.00000 ; HD3- CD- CE- HE3
743 741 744 747 3 0.65084 1.95253 0.00000
-2.60338 0.00000 0.00000 ; HD3- CD- CE- NZ
745 744 747 748 3 0.65084 1.95253 0.00000
-2.60338 0.00000 0.00000 ; HE2- CE- NZ- HZ1
745 744 747 749 3 0.65084 1.95253 0.00000
-2.60338 0.00000 0.00000 ; HE2- CE- NZ- HZ2
745 744 747 750 3 0.65084 1.95253 0.00000
-2.60338 0.00000 0.00000 ; HE2- CE- NZ- HZ3
746 744 747 748 3 0.65084 1.95253 0.00000
-2.60338 0.00000 0.00000 ; HE3- CE- NZ- HZ1
746 744 747 749 3 0.65084 1.95253 0.00000
-2.60338 0.00000 0.00000 ; HE3- CE- NZ- HZ2
746 744 747 750 3 0.65084 1.95253 0.00000
-2.60338 0.00000 0.00000 ; HE3- CE- NZ- HZ3
751 753 755 756 3 0.00000 0.00000 0.00000
0.00000 0.00000 0.00000 ; C- N- CA- HA
751 753 755 757 3 11.71520 1.67360 15.06240 -
13.38880 0.00000 0.00000 ; C- N- CA- CB
751 753 755 763 3 1.75728 5.27184 2.25936
-7.02912 0.00000 0.00000 ; C- N- CA- C
752 751 753 754 3 29.28800 -8.36800 -20.92000
0.00000 0.00000 0.00000 ; O- C- N- H
752 751 753 755 3 20.92000 0.00000 -20.92000
0.00000 0.00000 0.00000 ; O- C- N- CA
753 755 757 758 3 0.65084 1.95253 0.00000
-2.60338 0.00000 0.00000 ; N- CA- CB- HB2
753 755 757 759 3 0.65084 1.95253 0.00000
-2.60338 0.00000 0.00000 ; N- CA- CB- HB3
753 755 757 760 3 15.41386 9.74035 -8.39310
-0.97069 2.97901 0.00000 ; N- CA- CB- CG
753 755 763 764 3 0.00000 0.00000 0.00000
0.00000 0.00000 0.00000 ; N- CA- C- O
753 755 763 765 3 17.40544 -5.02080 -13.22144
9.20480 0.00000 0.00000 ; N- CA- C- N
754 753 755 756 3 0.00000 0.00000 0.00000
0.00000 0.00000 0.00000 ; H- N- CA- HA
754 753 755 757 3 0.00000 0.00000 0.00000
0.00000 0.00000 0.00000 ; H- N- CA- CB

754 753 755 763 3 0.00000 0.00000 0.00000
0.00000 0.00000 0.00000 ; H- N- CA- C
755 757 760 761 3 6.43499 0.00000 -5.39736
0.00000 -1.03763 0.00000 ; CA- CB- CG- OD1
755 757 760 762 3 6.43499 0.00000 -5.39736
0.00000 -1.03763 0.00000 ; CA- CB- CG- OD2
755 763 765 766 3 20.92000 0.00000 -20.92000
0.00000 0.00000 0.00000 ; CA- C- N- H
755 763 765 767 3 20.92000 0.00000 -20.92000
0.00000 0.00000 0.00000 ; CA- C- N- CA
756 755 757 758 3 0.65084 1.95253 0.00000
-2.60338 0.00000 0.00000 ; HA- CA- CB- HB2
756 755 757 759 3 0.65084 1.95253 0.00000
-2.60338 0.00000 0.00000 ; HA- CA- CB- HB3
756 755 757 760 3 0.65084 1.95253 0.00000
-2.60338 0.00000 0.00000 ; HA- CA- CB- CG
756 755 763 764 3 3.68192 -4.35136 0.00000
1.33888 0.00000 0.00000 ; HA- CA- C- O
756 755 763 765 3 0.00000 0.00000 0.00000
0.00000 0.00000 0.00000 ; HA- CA- C- N
757 755 763 764 3 0.00000 0.00000 0.00000
0.00000 0.00000 0.00000 ; CB- CA- C- O
757 755 763 765 3 2.51040 4.18400 1.67360
-6.69440 0.00000 0.00000 ; CB- CA- C- N
758 757 755 763 3 0.65084 1.95253 0.00000
-2.60338 0.00000 0.00000 ; HB2- CB- CA- C
758 757 760 761 3 0.00000 0.00000 0.00000
0.00000 0.00000 0.00000 ; HB2- CB- CG- OD1
758 757 760 762 3 0.00000 0.00000 0.00000
0.00000 0.00000 0.00000 ; HB2- CB- CG- OD2
759 757 755 763 3 0.65084 1.95253 0.00000
-2.60338 0.00000 0.00000 ; HB3- CB- CA- C
759 757 760 761 3 0.00000 0.00000 0.00000
0.00000 0.00000 0.00000 ; HB3- CB- CG- OD1
759 757 760 762 3 0.00000 0.00000 0.00000
0.00000 0.00000 0.00000 ; HB3- CB- CG- OD2
760 757 755 763 3 7.14627 -1.04600 -8.99560
-0.97069 5.15469 0.00000 ; CG- CB- CA- C
763 765 767 768 3 0.00000 0.00000 0.00000
0.00000 0.00000 0.00000 ; C- N- CA- HA
763 765 767 769 3 11.71520 1.67360 15.06240 -
13.38880 0.00000 0.00000 ; C- N- CA- CB
763 765 767 787 3 1.75728 5.27184 2.25936
-7.02912 0.00000 0.00000 ; C- N- CA- C
764 763 765 766 3 29.28800 -8.36800 -20.92000
0.00000 0.00000 0.00000 ; O- C- N- H
764 763 765 767 3 20.92000 0.00000 -20.92000
0.00000 0.00000 0.00000 ; O- C- N- CA
765 767 769 770 3 0.65084 1.95253 0.00000
-2.60338 0.00000 0.00000 ; N- CA- CB- HB2
765 767 769 771 3 0.65084 1.95253 0.00000
-2.60338 0.00000 0.00000 ; N- CA- CB- HB3

765 767 769 772 3 0.65084 1.95253 0.00000
-2.60338 0.00000 0.00000 ; N- CA- CB- CG
765 767 787 788 3 0.00000 0.00000 0.00000
0.00000 0.00000 0.00000 ; N- CA- C- O
765 767 787 789 3 17.40544 -5.02080 -13.22144
9.20480 0.00000 0.00000 ; N- CA- C- N
766 765 767 768 3 0.00000 0.00000 0.00000
0.00000 0.00000 0.00000 ; H- N- CA- HA
766 765 767 769 3 0.00000 0.00000 0.00000
0.00000 0.00000 0.00000 ; H- N- CA- CB
766 765 767 787 3 0.00000 0.00000 0.00000
0.00000 0.00000 0.00000 ; H- N- CA- C
767 769 772 773 3 0.66944 2.00832 0.00000
-2.67776 0.00000 0.00000 ; CA- CB- CG- HG2
767 769 772 774 3 0.66944 2.00832 0.00000
-2.67776 0.00000 0.00000 ; CA- CB- CG- HG3
767 769 772 775 3 3.68192 3.09616 -2.09200
-3.01248 0.00000 0.00000 ; CA- CB- CG- CD
767 787 789 790 3 20.92000 0.00000 -20.92000
0.00000 0.00000 0.00000 ; CA- C- N- H
767 787 789 791 3 20.92000 0.00000 -20.92000
0.00000 0.00000 0.00000 ; CA- C- N- CA
768 767 769 770 3 0.65084 1.95253 0.00000
-2.60338 0.00000 0.00000 ; HA- CA- CB- HB2
768 767 769 771 3 0.65084 1.95253 0.00000
-2.60338 0.00000 0.00000 ; HA- CA- CB- HB3
768 767 769 772 3 0.65084 1.95253 0.00000
-2.60338 0.00000 0.00000 ; HA- CA- CB- CG
768 767 787 788 3 3.68192 -4.35136 0.00000
1.33888 0.00000 0.00000 ; HA- CA- C- O
768 767 787 789 3 0.00000 0.00000 0.00000
0.00000 0.00000 0.00000 ; HA- CA- C- N
769 767 787 788 3 0.00000 0.00000 0.00000
0.00000 0.00000 0.00000 ; CB- CA- C- O
769 767 787 789 3 2.51040 4.18400 1.67360
-6.69440 0.00000 0.00000 ; CB- CA- C- N
769 772 775 776 3 0.65084 1.95253 0.00000
-2.60338 0.00000 0.00000 ; CB- CG- CD- HD2
769 772 775 777 3 0.65084 1.95253 0.00000
-2.60338 0.00000 0.00000 ; CB- CG- CD- HD3
769 772 775 778 3 0.65084 1.95253 0.00000
-2.60338 0.00000 0.00000 ; CB- CG- CD- NE
770 769 767 787 3 0.65084 1.95253 0.00000
-2.60338 0.00000 0.00000 ; HB2- CB- CA- C
770 769 772 773 3 0.62760 1.88280 0.00000
-2.51040 0.00000 0.00000 ; HB2- CB- CG- HG2
770 769 772 774 3 0.62760 1.88280 0.00000
-2.51040 0.00000 0.00000 ; HB2- CB- CG- HG3
770 769 772 775 3 0.66944 2.00832 0.00000
-2.67776 0.00000 0.00000 ; HB2- CB- CG- CD
771 769 767 787 3 0.65084 1.95253 0.00000
-2.60338 0.00000 0.00000 ; HB3- CB- CA- C

```
   771     769     772     773       3    0.62760    1.88280    0.00000
-2.51040    0.00000    0.00000 ;    HB3-    CB-    CG-   HG2
   771     769     772     774       3    0.62760    1.88280    0.00000
-2.51040    0.00000    0.00000 ;    HB3-    CB-    CG-   HG3
   771     769     772     775       3    0.66944    2.00832    0.00000
-2.67776    0.00000    0.00000 ;    HB3-    CB-    CG-    CD
   772     769     767     787       3    0.65084    1.95253    0.00000
-2.60338    0.00000    0.00000 ;     CG-    CB-    CA-     C
   772     775     778     779       3    0.00000    0.00000    0.00000
0.00000    0.00000    0.00000 ;     CG-    CD-    NE-    HE
   772     775     778     780       3    0.00000    0.00000    0.00000
0.00000    0.00000    0.00000 ;     CG-    CD-    NE-    CZ
   773     772     775     776       3    0.65084    1.95253    0.00000
-2.60338    0.00000    0.00000 ;    HG2-    CG-    CD-   HD2
   773     772     775     777       3    0.65084    1.95253    0.00000
-2.60338    0.00000    0.00000 ;    HG2-    CG-    CD-   HD3
   773     772     775     778       3    0.65084    1.95253    0.00000
-2.60338    0.00000    0.00000 ;    HG2-    CG-    CD-    NE
   774     772     775     776       3    0.65084    1.95253    0.00000
-2.60338    0.00000    0.00000 ;    HG3-    CG-    CD-   HD2
   774     772     775     777       3    0.65084    1.95253    0.00000
-2.60338    0.00000    0.00000 ;    HG3-    CG-    CD-   HD3
   774     772     775     778       3    0.65084    1.95253    0.00000
-2.60338    0.00000    0.00000 ;    HG3-    CG-    CD-    NE
   775     778     780     781       3   20.08320    0.00000  -20.08320
0.00000    0.00000    0.00000 ;     CD-    NE-    CZ-   NH1
   775     778     780     784       3   20.08320    0.00000  -20.08320
0.00000    0.00000    0.00000 ;     CD-    NE-    CZ-   NH2
   776     775     778     779       3    0.00000    0.00000    0.00000
0.00000    0.00000    0.00000 ;    HD2-    CD-    NE-    HE
   776     775     778     780       3    0.00000    0.00000    0.00000
0.00000    0.00000    0.00000 ;    HD2-    CD-    NE-    CZ
   777     775     778     779       3    0.00000    0.00000    0.00000
0.00000    0.00000    0.00000 ;    HD3-    CD-    NE-    HE
   777     775     778     780       3    0.00000    0.00000    0.00000
0.00000    0.00000    0.00000 ;    HD3-    CD-    NE-    CZ
   778     780     781     782       3   20.08320    0.00000  -20.08320
0.00000    0.00000    0.00000 ;     NE-    CZ-   NH1-  HH11
   778     780     781     783       3   20.08320    0.00000  -20.08320
0.00000    0.00000    0.00000 ;     NE-    CZ-   NH1-  HH12
   778     780     784     785       3   20.08320    0.00000  -20.08320
0.00000    0.00000    0.00000 ;     NE-    CZ-   NH2-  HH21
   778     780     784     786       3   20.08320    0.00000  -20.08320
0.00000    0.00000    0.00000 ;     NE-    CZ-   NH2-  HH22
   779     778     780     781       3   20.08320    0.00000  -20.08320
0.00000    0.00000    0.00000 ;     HE-    NE-    CZ-   NH1
   779     778     780     784       3   20.08320    0.00000  -20.08320
0.00000    0.00000    0.00000 ;     HE-    NE-    CZ-   NH2
   781     780     784     785       3   20.08320    0.00000  -20.08320
0.00000    0.00000    0.00000 ;    NH1-    CZ-   NH2-  HH21
   781     780     784     786       3   20.08320    0.00000  -20.08320
0.00000    0.00000    0.00000 ;    NH1-    CZ-   NH2-  HH22
```

782 781 780 784 3 20.08320 0.00000 -20.08320
0.00000 0.00000 0.00000 ; HH11- NH1- CZ- NH2
783 781 780 784 3 20.08320 0.00000 -20.08320
0.00000 0.00000 0.00000 ; HH12- NH1- CZ- NH2
787 789 791 792 3 0.00000 0.00000 0.00000
0.00000 0.00000 0.00000 ; C- N- CA- HA
787 789 791 793 3 11.71520 1.67360 15.06240 -
13.38880 0.00000 0.00000 ; C- N- CA- CB
787 789 791 806 3 1.75728 5.27184 2.25936
-7.02912 0.00000 0.00000 ; C- N- CA- C
788 787 789 790 3 29.28800 -8.36800 -20.92000
0.00000 0.00000 0.00000 ; O- C- N- H
788 787 789 791 3 20.92000 0.00000 -20.92000
0.00000 0.00000 0.00000 ; O- C- N- CA
789 791 793 794 3 0.65084 1.95253 0.00000
-2.60338 0.00000 0.00000 ; N- CA- CB- HB2
789 791 793 795 3 0.65084 1.95253 0.00000
-2.60338 0.00000 0.00000 ; N- CA- CB- HB3
789 791 793 796 3 3.79070 1.39746 -4.61077
-2.40998 2.44346 0.00000 ; N- CA- CB- CG
789 791 806 807 3 0.00000 0.00000 0.00000
0.00000 0.00000 0.00000 ; N- CA- C- O
789 791 806 808 3 17.40544 -5.02080 -13.22144
9.20480 0.00000 0.00000 ; N- CA- C- N
790 789 791 792 3 0.00000 0.00000 0.00000
0.00000 0.00000 0.00000 ; H- N- CA- HA
790 789 791 793 3 0.00000 0.00000 0.00000
0.00000 0.00000 0.00000 ; H- N- CA- CB
790 789 791 806 3 0.00000 0.00000 0.00000
0.00000 0.00000 0.00000 ; H- N- CA- C
791 793 796 797 3 0.66944 2.00832 0.00000
-2.67776 0.00000 0.00000 ; CA- CB- CG- HG
791 793 796 798 3 3.90367 0.19665 -6.21742
-2.37651 5.99149 0.00000 ; CA- CB- CG- CD1
791 793 796 802 3 3.90367 0.19665 -6.21742
-2.37651 5.99149 0.00000 ; CA- CB- CG- CD2
791 806 808 809 3 20.92000 0.00000 -20.92000
0.00000 0.00000 0.00000 ; CA- C- N- H
791 806 808 810 3 20.92000 0.00000 -20.92000
0.00000 0.00000 0.00000 ; CA- C- N- CA
792 791 793 794 3 0.65084 1.95253 0.00000
-2.60338 0.00000 0.00000 ; HA- CA- CB- HB2
792 791 793 795 3 0.65084 1.95253 0.00000
-2.60338 0.00000 0.00000 ; HA- CA- CB- HB3
792 791 793 796 3 0.65084 1.95253 0.00000
-2.60338 0.00000 0.00000 ; HA- CA- CB- CG
792 791 806 807 3 3.68192 -4.35136 0.00000
1.33888 0.00000 0.00000 ; HA- CA- C- O
792 791 806 808 3 0.00000 0.00000 0.00000
0.00000 0.00000 0.00000 ; HA- CA- C- N
793 791 806 807 3 0.00000 0.00000 0.00000
0.00000 0.00000 0.00000 ; CB- CA- C- O

```
   793    791    806    808      3    2.51040    4.18400    1.67360
 -6.69440    0.00000    0.00000 ;     CB-    CA-     C-     N
   793    796    798    799      3    0.66944    2.00832    0.00000
 -2.67776    0.00000    0.00000 ;     CB-    CG-   CD1-  HD11
   793    796    798    800      3    0.66944    2.00832    0.00000
 -2.67776    0.00000    0.00000 ;     CB-    CG-   CD1-  HD12
   793    796    798    801      3    0.66944    2.00832    0.00000
 -2.67776    0.00000    0.00000 ;     CB-    CG-   CD1-  HD13
   793    796    802    803      3    0.66944    2.00832    0.00000
 -2.67776    0.00000    0.00000 ;     CB-    CG-   CD2-  HD21
   793    796    802    804      3    0.66944    2.00832    0.00000
 -2.67776    0.00000    0.00000 ;     CB-    CG-   CD2-  HD22
   793    796    802    805      3    0.66944    2.00832    0.00000
 -2.67776    0.00000    0.00000 ;     CB-    CG-   CD2-  HD23
   794    793    791    806      3    0.65084    1.95253    0.00000
 -2.60338    0.00000    0.00000 ;    HB2-    CB-    CA-     C
   794    793    796    797      3    0.62760    1.88280    0.00000
 -2.51040    0.00000    0.00000 ;    HB2-    CB-    CG-    HG
   794    793    796    798      3    0.66944    2.00832    0.00000
 -2.67776    0.00000    0.00000 ;    HB2-    CB-    CG-   CD1
   794    793    796    802      3    0.66944    2.00832    0.00000
 -2.67776    0.00000    0.00000 ;    HB2-    CB-    CG-   CD2
   795    793    791    806      3    0.65084    1.95253    0.00000
 -2.60338    0.00000    0.00000 ;    HB3-    CB-    CA-     C
   795    793    796    797      3    0.62760    1.88280    0.00000
 -2.51040    0.00000    0.00000 ;    HB3-    CB-    CG-    HG
   795    793    796    798      3    0.66944    2.00832    0.00000
 -2.67776    0.00000    0.00000 ;    HB3-    CB-    CG-   CD1
   795    793    796    802      3    0.66944    2.00832    0.00000
 -2.67776    0.00000    0.00000 ;    HB3-    CB-    CG-   CD2
   796    793    791    806      3   10.33448   -1.14642  -11.54784
 -2.40998    6.35968    0.00000 ;     CG-    CB-    CA-     C
   797    796    798    799      3    0.62760    1.88280    0.00000
 -2.51040    0.00000    0.00000 ;     HG-    CG-   CD1-  HD11
   797    796    798    800      3    0.62760    1.88280    0.00000
 -2.51040    0.00000    0.00000 ;     HG-    CG-   CD1-  HD12
   797    796    798    801      3    0.62760    1.88280    0.00000
 -2.51040    0.00000    0.00000 ;     HG-    CG-   CD1-  HD13
   797    796    802    803      3    0.62760    1.88280    0.00000
 -2.51040    0.00000    0.00000 ;     HG-    CG-   CD2-  HD21
   797    796    802    804      3    0.62760    1.88280    0.00000
 -2.51040    0.00000    0.00000 ;     HG-    CG-   CD2-  HD22
   797    796    802    805      3    0.62760    1.88280    0.00000
 -2.51040    0.00000    0.00000 ;     HG-    CG-   CD2-  HD23
   798    796    802    803      3    0.66944    2.00832    0.00000
 -2.67776    0.00000    0.00000 ;    CD1-    CG-   CD2-  HD21
   798    796    802    804      3    0.66944    2.00832    0.00000
 -2.67776    0.00000    0.00000 ;    CD1-    CG-   CD2-  HD22
   798    796    802    805      3    0.66944    2.00832    0.00000
 -2.67776    0.00000    0.00000 ;    CD1-    CG-   CD2-  HD23
   799    798    796    802      3    0.66944    2.00832    0.00000
 -2.67776    0.00000    0.00000 ;   HD11-   CD1-    CG-   CD2
```

```
   800     798     796     802       3    0.66944    2.00832    0.00000
-2.67776    0.00000    0.00000 ;   HD12-   CD1-    CG-   CD2
   801     798     796     802       3    0.66944    2.00832    0.00000
-2.67776    0.00000    0.00000 ;   HD13-   CD1-    CG-   CD2
   806     808     810     811       3    0.00000    0.00000    0.00000
0.00000    0.00000    0.00000 ;      C-     N-    CA-    HA
   806     808     810     812       3   11.71520    1.67360   15.06240  -
13.38880    0.00000    0.00000 ;      C-     N-    CA-    CB
   806     808     810     828       3    1.75728    5.27184    2.25936
-7.02912    0.00000    0.00000 ;      C-     N-    CA-     C
   807     806     808     809       3   29.28800   -8.36800  -20.92000
0.00000    0.00000    0.00000 ;      O-     C-     N-     H
   807     806     808     810       3   20.92000    0.00000  -20.92000
0.00000    0.00000    0.00000 ;      O-     C-     N-    CA
   808     810     812     813       3    0.65084    1.95253    0.00000
-2.60338    0.00000    0.00000 ;      N-    CA-    CB-   HB2
   808     810     812     814       3    0.65084    1.95253    0.00000
-2.60338    0.00000    0.00000 ;      N-    CA-    CB-   HB3
   808     810     812     815       3    0.65084    1.95253    0.00000
-2.60338    0.00000    0.00000 ;      N-    CA-    CB-    CG
   808     810     828     829       3    0.00000    0.00000    0.00000
0.00000    0.00000    0.00000 ;      N-    CA-     C-     O
   808     810     828     830       3   17.40544   -5.02080  -13.22144
9.20480    0.00000    0.00000 ;      N-    CA-     C-     N
   809     808     810     811       3    0.00000    0.00000    0.00000
0.00000    0.00000    0.00000 ;      H-     N-    CA-    HA
   809     808     810     812       3    0.00000    0.00000    0.00000
0.00000    0.00000    0.00000 ;      H-     N-    CA-    CB
   809     808     810     828       3    0.00000    0.00000    0.00000
0.00000    0.00000    0.00000 ;      H-     N-    CA-     C
   810     812     815     816       3    0.66944    2.00832    0.00000
-2.67776    0.00000    0.00000 ;     CA-    CB-    CG-   HG2
   810     812     815     817       3    0.66944    2.00832    0.00000
-2.67776    0.00000    0.00000 ;     CA-    CB-    CG-   HG3
   810     812     815     818       3    3.68192    3.09616   -2.09200
-3.01248    0.00000    0.00000 ;     CA-    CB-    CG-    CD
   810     828     830     831       3   20.92000    0.00000  -20.92000
0.00000    0.00000    0.00000 ;     CA-     C-     N-     H
   810     828     830     832       3   20.92000    0.00000  -20.92000
0.00000    0.00000    0.00000 ;     CA-     C-     N-    CA
   811     810     812     813       3    0.65084    1.95253    0.00000
-2.60338    0.00000    0.00000 ;     HA-    CA-    CB-   HB2
   811     810     812     814       3    0.65084    1.95253    0.00000
-2.60338    0.00000    0.00000 ;     HA-    CA-    CB-   HB3
   811     810     812     815       3    0.65084    1.95253    0.00000
-2.60338    0.00000    0.00000 ;     HA-    CA-    CB-    CG
   811     810     828     829       3    3.68192   -4.35136    0.00000
1.33888    0.00000    0.00000 ;     HA-    CA-     C-     O
   811     810     828     830       3    0.00000    0.00000    0.00000
0.00000    0.00000    0.00000 ;     HA-    CA-     C-     N
   812     810     828     829       3    0.00000    0.00000    0.00000
0.00000    0.00000    0.00000 ;     CB-    CA-     C-     O
```

```
   812    810    828    830      3    2.51040    4.18400    1.67360
-6.69440    0.00000    0.00000 ;     CB-    CA-     C-     N
   812    815    818    819      3    0.66944    2.00832    0.00000
-2.67776    0.00000    0.00000 ;     CB-    CG-    CD-   HD2
   812    815    818    820      3    0.66944    2.00832    0.00000
-2.67776    0.00000    0.00000 ;     CB-    CG-    CD-   HD3
   812    815    818    821      3    3.68192    3.09616   -2.09200
-3.01248    0.00000    0.00000 ;     CB-    CG-    CD-    CE
   813    812    810    828      3    0.65084    1.95253    0.00000
-2.60338    0.00000    0.00000 ;    HB2-    CB-    CA-     C
   813    812    815    816      3    0.62760    1.88280    0.00000
-2.51040    0.00000    0.00000 ;    HB2-    CB-    CG-   HG2
   813    812    815    817      3    0.62760    1.88280    0.00000
-2.51040    0.00000    0.00000 ;    HB2-    CB-    CG-   HG3
   813    812    815    818      3    0.66944    2.00832    0.00000
-2.67776    0.00000    0.00000 ;    HB2-    CB-    CG-    CD
   814    812    810    828      3    0.65084    1.95253    0.00000
-2.60338    0.00000    0.00000 ;    HB3-    CB-    CA-     C
   814    812    815    816      3    0.62760    1.88280    0.00000
-2.51040    0.00000    0.00000 ;    HB3-    CB-    CG-   HG2
   814    812    815    817      3    0.62760    1.88280    0.00000
-2.51040    0.00000    0.00000 ;    HB3-    CB-    CG-   HG3
   814    812    815    818      3    0.66944    2.00832    0.00000
-2.67776    0.00000    0.00000 ;    HB3-    CB-    CG-    CD
   815    812    810    828      3    0.65084    1.95253    0.00000
-2.60338    0.00000    0.00000 ;     CG-    CB-    CA-     C
   815    818    821    822      3    0.65084    1.95253    0.00000
-2.60338    0.00000    0.00000 ;     CG-    CD-    CE-   HE2
   815    818    821    823      3    0.65084    1.95253    0.00000
-2.60338    0.00000    0.00000 ;     CG-    CD-    CE-   HE3
   815    818    821    824      3    0.65084    1.95253    0.00000
-2.60338    0.00000    0.00000 ;     CG-    CD-    CE-    NZ
   816    815    818    819      3    0.62760    1.88280    0.00000
-2.51040    0.00000    0.00000 ;    HG2-    CG-    CD-   HD2
   816    815    818    820      3    0.62760    1.88280    0.00000
-2.51040    0.00000    0.00000 ;    HG2-    CG-    CD-   HD3
   816    815    818    821      3    0.66944    2.00832    0.00000
-2.67776    0.00000    0.00000 ;    HG2-    CG-    CD-    CE
   817    815    818    819      3    0.62760    1.88280    0.00000
-2.51040    0.00000    0.00000 ;    HG3-    CG-    CD-   HD2
   817    815    818    820      3    0.62760    1.88280    0.00000
-2.51040    0.00000    0.00000 ;    HG3-    CG-    CD-   HD3
   817    815    818    821      3    0.66944    2.00832    0.00000
-2.67776    0.00000    0.00000 ;    HG3-    CG-    CD-    CE
   818    821    824    825      3    0.65084    1.95253    0.00000
-2.60338    0.00000    0.00000 ;     CD-    CE-    NZ-   HZ1
   818    821    824    826      3    0.65084    1.95253    0.00000
-2.60338    0.00000    0.00000 ;     CD-    CE-    NZ-   HZ2
   818    821    824    827      3    0.65084    1.95253    0.00000
-2.60338    0.00000    0.00000 ;     CD-    CE-    NZ-   HZ3
   819    818    821    822      3    0.65084    1.95253    0.00000
-2.60338    0.00000    0.00000 ;    HD2-    CD-    CE-   HE2
```

```
   819    818    821    823      3    0.65084    1.95253    0.00000
-2.60338    0.00000    0.00000 ;    HD2-    CD-    CE-   HE3
   819    818    821    824      3    0.65084    1.95253    0.00000
-2.60338    0.00000    0.00000 ;    HD2-    CD-    CE-    NZ
   820    818    821    822      3    0.65084    1.95253    0.00000
-2.60338    0.00000    0.00000 ;    HD3-    CD-    CE-   HE2
   820    818    821    823      3    0.65084    1.95253    0.00000
-2.60338    0.00000    0.00000 ;    HD3-    CD-    CE-   HE3
   820    818    821    824      3    0.65084    1.95253    0.00000
-2.60338    0.00000    0.00000 ;    HD3-    CD-    CE-    NZ
   822    821    824    825      3    0.65084    1.95253    0.00000
-2.60338    0.00000    0.00000 ;    HE2-    CE-    NZ-   HZ1
   822    821    824    826      3    0.65084    1.95253    0.00000
-2.60338    0.00000    0.00000 ;    HE2-    CE-    NZ-   HZ2
   822    821    824    827      3    0.65084    1.95253    0.00000
-2.60338    0.00000    0.00000 ;    HE2-    CE-    NZ-   HZ3
   823    821    824    825      3    0.65084    1.95253    0.00000
-2.60338    0.00000    0.00000 ;    HE3-    CE-    NZ-   HZ1
   823    821    824    826      3    0.65084    1.95253    0.00000
-2.60338    0.00000    0.00000 ;    HE3-    CE-    NZ-   HZ2
   823    821    824    827      3    0.65084    1.95253    0.00000
-2.60338    0.00000    0.00000 ;    HE3-    CE-    NZ-   HZ3
   828    830    832    833      3    0.00000    0.00000    0.00000
0.00000    0.00000    0.00000 ;      C-     N-    CA-    HA
   828    830    832    834      3   11.71520    1.67360   15.06240  -
13.38880    0.00000    0.00000 ;      C-     N-    CA-    CB
   828    830    832    842      3    1.75728    5.27184    2.25936
-7.02912    0.00000    0.00000 ;      C-     N-    CA-     C
   829    828    830    831      3   29.28800   -8.36800  -20.92000
0.00000    0.00000    0.00000 ;      O-     C-     N-     H
   829    828    830    832      3   20.92000    0.00000  -20.92000
0.00000    0.00000    0.00000 ;      O-     C-     N-    CA
   830    832    834    835      3    0.65084    1.95253    0.00000
-2.60338    0.00000    0.00000 ;      N-    CA-    CB-   HB2
   830    832    834    836      3    0.65084    1.95253    0.00000
-2.60338    0.00000    0.00000 ;      N-    CA-    CB-   HB3
   830    832    834    837      3    5.99567    3.29281   -4.46014
-0.55229    1.97485    0.00000 ;      N-    CA-    CB-    CG
   830    832    842    843      3    0.00000    0.00000    0.00000
0.00000    0.00000    0.00000 ;      N-    CA-     C-     O
   830    832    842    844      3   17.40544   -5.02080  -13.22144
9.20480    0.00000    0.00000 ;      N-    CA-     C-     N
   831    830    832    833      3    0.00000    0.00000    0.00000
0.00000    0.00000    0.00000 ;      H-     N-    CA-    HA
   831    830    832    834      3    0.00000    0.00000    0.00000
0.00000    0.00000    0.00000 ;      H-     N-    CA-    CB
   831    830    832    842      3    0.00000    0.00000    0.00000
0.00000    0.00000    0.00000 ;      H-     N-    CA-     C
   832    834    837    838      3    0.00000    0.00000    0.00000
0.00000    0.00000    0.00000 ;    CA-    CB-    CG-   OD1
   832    834    837    839      3    8.84916   -0.31380   -4.32626
5.03754    0.26778    0.00000 ;    CA-    CB-    CG-   ND2
```

832 842 844 845 3 20.92000 0.00000 -20.92000
0.00000 0.00000 0.00000 ; CA- C- N- H
832 842 844 846 3 20.92000 0.00000 -20.92000
0.00000 0.00000 0.00000 ; CA- C- N- CA
833 832 834 835 3 0.65084 1.95253 0.00000
-2.60338 0.00000 0.00000 ; HA- CA- CB- HB2
833 832 834 836 3 0.65084 1.95253 0.00000
-2.60338 0.00000 0.00000 ; HA- CA- CB- HB3
833 832 834 837 3 0.65084 1.95253 0.00000
-2.60338 0.00000 0.00000 ; HA- CA- CB- CG
833 832 842 843 3 3.68192 -4.35136 0.00000
1.33888 0.00000 0.00000 ; HA- CA- C- O
833 832 842 844 3 0.00000 0.00000 0.00000
0.00000 0.00000 0.00000 ; HA- CA- C- N
834 832 842 843 3 0.00000 0.00000 0.00000
0.00000 0.00000 0.00000 ; CB- CA- C- O
834 832 842 844 3 2.51040 4.18400 1.67360
-6.69440 0.00000 0.00000 ; CB- CA- C- N
834 837 839 840 3 20.92000 0.00000 -20.92000
0.00000 0.00000 0.00000 ; CB- CG- ND2- HD21
834 837 839 841 3 20.92000 0.00000 -20.92000
0.00000 0.00000 0.00000 ; CB- CG- ND2- HD22
835 834 832 842 3 0.65084 1.95253 0.00000
-2.60338 0.00000 0.00000 ; HB2- CB- CA- C
835 834 837 838 3 3.68192 -4.35136 0.00000
1.33888 0.00000 0.00000 ; HB2- CB- CG- OD1
835 834 837 839 3 0.00000 0.00000 0.00000
0.00000 0.00000 0.00000 ; HB2- CB- CG- ND2
836 834 832 842 3 0.65084 1.95253 0.00000
-2.60338 0.00000 0.00000 ; HB3- CB- CA- C
836 834 837 838 3 3.68192 -4.35136 0.00000
1.33888 0.00000 0.00000 ; HB3- CB- CG- OD1
836 834 837 839 3 0.00000 0.00000 0.00000
0.00000 0.00000 0.00000 ; HB3- CB- CG- ND2
837 834 832 842 3 7.94542 -3.96225 -6.11701
-0.55229 3.58150 0.00000 ; CG- CB- CA- C
838 837 839 840 3 29.28800 -8.36800 -20.92000
0.00000 0.00000 0.00000 ; OD1- CG- ND2- HD21
838 837 839 841 3 29.28800 -8.36800 -20.92000
0.00000 0.00000 0.00000 ; OD1- CG- ND2- HD22
842 844 846 847 3 0.00000 0.00000 0.00000
0.00000 0.00000 0.00000 ; C- N- CA- HA
842 844 846 848 3 11.71520 1.67360 15.06240 -
13.38880 0.00000 0.00000 ; C- N- CA- CB
842 844 846 861 3 1.75728 5.27184 2.25936
-7.02912 0.00000 0.00000 ; C- N- CA- C
843 842 844 845 3 29.28800 -8.36800 -20.92000
0.00000 0.00000 0.00000 ; O- C- N- H
843 842 844 846 3 20.92000 0.00000 -20.92000
0.00000 0.00000 0.00000 ; O- C- N- CA
844 846 848 849 3 0.65084 1.95253 0.00000
-2.60338 0.00000 0.00000 ; N- CA- CB- HB

844 846 848 850 3 3.83673 0.44769 -1.77402
-2.47693 -0.03347 0.00000 ; N- CA- CB- CG2
844 846 848 854 3 2.97482 0.12134 2.04179
-1.89117 -3.24678 0.00000 ; N- CA- CB- CG1
844 846 861 862 3 0.00000 0.00000 0.00000
0.00000 0.00000 0.00000 ; N- CA- C- O
844 846 861 863 3 17.40544 -5.02080 -13.22144
9.20480 0.00000 0.00000 ; N- CA- C- N
845 844 846 847 3 0.00000 0.00000 0.00000
0.00000 0.00000 0.00000 ; H- N- CA- HA
845 844 846 848 3 0.00000 0.00000 0.00000
0.00000 0.00000 0.00000 ; H- N- CA- CB
845 844 846 861 3 0.00000 0.00000 0.00000
0.00000 0.00000 0.00000 ; H- N- CA- C
846 848 850 851 3 0.66944 2.00832 0.00000
-2.67776 0.00000 0.00000 ; CA- CB- CG2- HG21
846 848 850 852 3 0.66944 2.00832 0.00000
-2.67776 0.00000 0.00000 ; CA- CB- CG2- HG22
846 848 850 853 3 0.66944 2.00832 0.00000
-2.67776 0.00000 0.00000 ; CA- CB- CG2- HG23
846 848 854 855 3 0.66944 2.00832 0.00000
-2.67776 0.00000 0.00000 ; CA- CB- CG1- HG12
846 848 854 856 3 0.66944 2.00832 0.00000
-2.67776 0.00000 0.00000 ; CA- CB- CG1- HG13
846 848 854 857 3 4.24258 -0.52718 -7.25506
-1.79075 7.69856 0.00000 ; CA- CB- CG1- CD1
846 861 863 864 3 20.92000 0.00000 -20.92000
0.00000 0.00000 0.00000 ; CA- C- N- H
846 861 863 865 3 20.92000 0.00000 -20.92000
0.00000 0.00000 0.00000 ; CA- C- N- CA
847 846 848 849 3 0.65084 1.95253 0.00000
-2.60338 0.00000 0.00000 ; HA- CA- CB- HB
847 846 848 850 3 0.65084 1.95253 0.00000
-2.60338 0.00000 0.00000 ; HA- CA- CB- CG2
847 846 848 854 3 0.65084 1.95253 0.00000
-2.60338 0.00000 0.00000 ; HA- CA- CB- CG1
847 846 861 862 3 3.68192 -4.35136 0.00000
1.33888 0.00000 0.00000 ; HA- CA- C- O
847 846 861 863 3 0.00000 0.00000 0.00000
0.00000 0.00000 0.00000 ; HA- CA- C- N
848 846 861 862 3 0.00000 0.00000 0.00000
0.00000 0.00000 0.00000 ; CB- CA- C- O
848 846 861 863 3 2.51040 4.18400 1.67360
-6.69440 0.00000 0.00000 ; CB- CA- C- N
848 854 857 858 3 0.66944 2.00832 0.00000
-2.67776 0.00000 0.00000 ; CB- CG1- CD1- HD11
848 854 857 859 3 0.66944 2.00832 0.00000
-2.67776 0.00000 0.00000 ; CB- CG1- CD1- HD12
848 854 857 860 3 0.66944 2.00832 0.00000
-2.67776 0.00000 0.00000 ; CB- CG1- CD1- HD13
849 848 846 861 3 0.65084 1.95253 0.00000
-2.60338 0.00000 0.00000 ; HB- CB- CA- C

849 848 850 851 3 0.62760 1.88280 0.00000
-2.51040 0.00000 0.00000 ; HB- CB- CG2- HG21
849 848 850 852 3 0.62760 1.88280 0.00000
-2.51040 0.00000 0.00000 ; HB- CB- CG2- HG22
849 848 850 853 3 0.62760 1.88280 0.00000
-2.51040 0.00000 0.00000 ; HB- CB- CG2- HG23
849 848 854 855 3 0.62760 1.88280 0.00000
-2.51040 0.00000 0.00000 ; HB- CB- CG1- HG12
849 848 854 856 3 0.62760 1.88280 0.00000
-2.51040 0.00000 0.00000 ; HB- CB- CG1- HG13
849 848 854 857 3 0.66944 2.00832 0.00000
-2.67776 0.00000 0.00000 ; HB- CB- CG1- CD1
850 848 846 861 3 5.67350 3.55640 -6.16722
-2.47693 3.74886 0.00000 ; CG2- CB- CA- C
850 848 854 855 3 0.66944 2.00832 0.00000
-2.67776 0.00000 0.00000 ; CG2- CB- CG1- HG12
850 848 854 856 3 0.66944 2.00832 0.00000
-2.67776 0.00000 0.00000 ; CG2- CB- CG1- HG13
850 848 854 857 3 3.81162 0.49790 -8.14206
-1.79075 7.49773 0.00000 ; CG2- CB- CG1- CD1
851 850 848 854 3 0.66944 2.00832 0.00000
-2.67776 0.00000 0.00000 ; HG21- CG2- CB- CG1
852 850 848 854 3 0.66944 2.00832 0.00000
-2.67776 0.00000 0.00000 ; HG22- CG2- CB- CG1
853 850 848 854 3 0.66944 2.00832 0.00000
-2.67776 0.00000 0.00000 ; HG23- CG2- CB- CG1
854 848 846 861 3 8.26340 0.74057 -9.99976
-1.89117 3.84928 0.00000 ; CG1- CB- CA- C
855 854 857 858 3 0.62760 1.88280 0.00000
-2.51040 0.00000 0.00000 ; HG12- CG1- CD1- HD11
855 854 857 859 3 0.62760 1.88280 0.00000
-2.51040 0.00000 0.00000 ; HG12- CG1- CD1- HD12
855 854 857 860 3 0.62760 1.88280 0.00000
-2.51040 0.00000 0.00000 ; HG12- CG1- CD1- HD13
856 854 857 858 3 0.62760 1.88280 0.00000
-2.51040 0.00000 0.00000 ; HG13- CG1- CD1- HD11
856 854 857 859 3 0.62760 1.88280 0.00000
-2.51040 0.00000 0.00000 ; HG13- CG1- CD1- HD12
856 854 857 860 3 0.62760 1.88280 0.00000
-2.51040 0.00000 0.00000 ; HG13- CG1- CD1- HD13
861 863 865 866 3 0.00000 0.00000 0.00000
0.00000 0.00000 0.00000 ; C- N- CA- HA
861 863 865 867 3 11.71520 1.67360 15.06240 -
13.38880 0.00000 0.00000 ; C- N- CA- CB
861 863 865 875 3 1.75728 5.27184 2.25936
-7.02912 0.00000 0.00000 ; C- N- CA- C
862 861 863 864 3 29.28800 -8.36800 -20.92000
0.00000 0.00000 0.00000 ; O- C- N- H
862 861 863 865 3 20.92000 0.00000 -20.92000
0.00000 0.00000 0.00000 ; O- C- N- CA
863 865 867 868 3 0.65084 1.95253 0.00000
-2.60338 0.00000 0.00000 ; N- CA- CB- HB2

863 865 867 869 3 0.65084 1.95253 0.00000
-2.60338 0.00000 0.00000 ; N- CA- CB- HB3
863 865 867 870 3 5.99567 3.29281 -4.46014
-0.55229 1.97485 0.00000 ; N- CA- CB- CG
863 865 875 876 3 0.00000 0.00000 0.00000
0.00000 0.00000 0.00000 ; N- CA- C- O
863 865 875 877 3 17.40544 -5.02080 -13.22144
9.20480 0.00000 0.00000 ; N- CA- C- N
864 863 865 866 3 0.00000 0.00000 0.00000
0.00000 0.00000 0.00000 ; H- N- CA- HA
864 863 865 867 3 0.00000 0.00000 0.00000
0.00000 0.00000 0.00000 ; H- N- CA- CB
864 863 865 875 3 0.00000 0.00000 0.00000
0.00000 0.00000 0.00000 ; H- N- CA- C
865 867 870 871 3 0.00000 0.00000 0.00000
0.00000 0.00000 0.00000 ; CA- CB- CG- OD1
865 867 870 872 3 8.84916 -0.31380 -4.32626
5.03754 0.26778 0.00000 ; CA- CB- CG- ND2
865 875 877 878 3 20.92000 0.00000 -20.92000
0.00000 0.00000 0.00000 ; CA- C- N- H
865 875 877 879 3 20.92000 0.00000 -20.92000
0.00000 0.00000 0.00000 ; CA- C- N- CA
866 865 867 868 3 0.65084 1.95253 0.00000
-2.60338 0.00000 0.00000 ; HA- CA- CB- HB2
866 865 867 869 3 0.65084 1.95253 0.00000
-2.60338 0.00000 0.00000 ; HA- CA- CB- HB3
866 865 867 870 3 0.65084 1.95253 0.00000
-2.60338 0.00000 0.00000 ; HA- CA- CB- CG
866 865 875 876 3 3.68192 -4.35136 0.00000
1.33888 0.00000 0.00000 ; HA- CA- C- O
866 865 875 877 3 0.00000 0.00000 0.00000
0.00000 0.00000 0.00000 ; HA- CA- C- N
867 865 875 876 3 0.00000 0.00000 0.00000
0.00000 0.00000 0.00000 ; CB- CA- C- O
867 865 875 877 3 2.51040 4.18400 1.67360
-6.69440 0.00000 0.00000 ; CB- CA- C- N
867 870 872 873 3 20.92000 0.00000 -20.92000
0.00000 0.00000 0.00000 ; CB- CG- ND2- HD21
867 870 872 874 3 20.92000 0.00000 -20.92000
0.00000 0.00000 0.00000 ; CB- CG- ND2- HD22
868 867 865 875 3 0.65084 1.95253 0.00000
-2.60338 0.00000 0.00000 ; HB2- CB- CA- C
868 867 870 871 3 3.68192 -4.35136 0.00000
1.33888 0.00000 0.00000 ; HB2- CB- CG- OD1
868 867 870 872 3 0.00000 0.00000 0.00000
0.00000 0.00000 0.00000 ; HB2- CB- CG- ND2
869 867 865 875 3 0.65084 1.95253 0.00000
-2.60338 0.00000 0.00000 ; HB3- CB- CA- C
869 867 870 871 3 3.68192 -4.35136 0.00000
1.33888 0.00000 0.00000 ; HB3- CB- CG- OD1
869 867 870 872 3 0.00000 0.00000 0.00000
0.00000 0.00000 0.00000 ; HB3- CB- CG- ND2

870 867 865 875 3 7.94542 -3.96225 -6.11701
-0.55229 3.58150 0.00000 ; CG- CB- CA- C
871 870 872 873 3 29.28800 -8.36800 -20.92000
0.00000 0.00000 0.00000 ; OD1- CG- ND2- HD21
871 870 872 874 3 29.28800 -8.36800 -20.92000
0.00000 0.00000 0.00000 ; OD1- CG- ND2- HD22
875 877 879 880 3 0.00000 0.00000 0.00000
0.00000 0.00000 0.00000 ; C- N- CA- HA
875 877 879 881 3 11.71520 1.67360 15.06240 -
13.38880 0.00000 0.00000 ; C- N- CA- CB
875 877 879 899 3 1.75728 5.27184 2.25936
-7.02912 0.00000 0.00000 ; C- N- CA- C
876 875 877 878 3 29.28800 -8.36800 -20.92000
0.00000 0.00000 0.00000 ; O- C- N- H
876 875 877 879 3 20.92000 0.00000 -20.92000
0.00000 0.00000 0.00000 ; O- C- N- CA
877 879 881 882 3 0.65084 1.95253 0.00000
-2.60338 0.00000 0.00000 ; N- CA- CB- HB2
877 879 881 883 3 0.65084 1.95253 0.00000
-2.60338 0.00000 0.00000 ; N- CA- CB- HB3
877 879 881 884 3 0.65084 1.95253 0.00000
-2.60338 0.00000 0.00000 ; N- CA- CB- CG
877 879 899 900 3 0.00000 0.00000 0.00000
0.00000 0.00000 0.00000 ; N- CA- C- O
877 879 899 901 3 17.40544 -5.02080 -13.22144
9.20480 0.00000 0.00000 ; N- CA- C- N
878 877 879 880 3 0.00000 0.00000 0.00000
0.00000 0.00000 0.00000 ; H- N- CA- HA
878 877 879 881 3 0.00000 0.00000 0.00000
0.00000 0.00000 0.00000 ; H- N- CA- CB
878 877 879 899 3 0.00000 0.00000 0.00000
0.00000 0.00000 0.00000 ; H- N- CA- C
879 881 884 885 3 0.66944 2.00832 0.00000
-2.67776 0.00000 0.00000 ; CA- CB- CG- HG2
879 881 884 886 3 0.66944 2.00832 0.00000
-2.67776 0.00000 0.00000 ; CA- CB- CG- HG3
879 881 884 887 3 3.68192 3.09616 -2.09200
-3.01248 0.00000 0.00000 ; CA- CB- CG- CD
879 899 901 902 3 20.92000 0.00000 -20.92000
0.00000 0.00000 0.00000 ; CA- C- N- H
879 899 901 903 3 20.92000 0.00000 -20.92000
0.00000 0.00000 0.00000 ; CA- C- N- CA
880 879 881 882 3 0.65084 1.95253 0.00000
-2.60338 0.00000 0.00000 ; HA- CA- CB- HB2
880 879 881 883 3 0.65084 1.95253 0.00000
-2.60338 0.00000 0.00000 ; HA- CA- CB- HB3
880 879 881 884 3 0.65084 1.95253 0.00000
-2.60338 0.00000 0.00000 ; HA- CA- CB- CG
880 879 899 900 3 3.68192 -4.35136 0.00000
1.33888 0.00000 0.00000 ; HA- CA- C- O
880 879 899 901 3 0.00000 0.00000 0.00000
0.00000 0.00000 0.00000 ; HA- CA- C- N

881 879 899 900 3 0.00000 0.00000 0.00000
0.00000 0.00000 0.00000 ; CB- CA- C- O
881 879 899 901 3 2.51040 4.18400 1.67360
-6.69440 0.00000 0.00000 ; CB- CA- C- N
881 884 887 888 3 0.65084 1.95253 0.00000
-2.60338 0.00000 0.00000 ; CB- CG- CD- HD2
881 884 887 889 3 0.65084 1.95253 0.00000
-2.60338 0.00000 0.00000 ; CB- CG- CD- HD3
881 884 887 890 3 0.65084 1.95253 0.00000
-2.60338 0.00000 0.00000 ; CB- CG- CD- NE
882 881 879 899 3 0.65084 1.95253 0.00000
-2.60338 0.00000 0.00000 ; HB2- CB- CA- C
882 881 884 885 3 0.62760 1.88280 0.00000
-2.51040 0.00000 0.00000 ; HB2- CB- CG- HG2
882 881 884 886 3 0.62760 1.88280 0.00000
-2.51040 0.00000 0.00000 ; HB2- CB- CG- HG3
882 881 884 887 3 0.66944 2.00832 0.00000
-2.67776 0.00000 0.00000 ; HB2- CB- CG- CD
883 881 879 899 3 0.65084 1.95253 0.00000
-2.60338 0.00000 0.00000 ; HB3- CB- CA- C
883 881 884 885 3 0.62760 1.88280 0.00000
-2.51040 0.00000 0.00000 ; HB3- CB- CG- HG2
883 881 884 886 3 0.62760 1.88280 0.00000
-2.51040 0.00000 0.00000 ; HB3- CB- CG- HG3
883 881 884 887 3 0.66944 2.00832 0.00000
-2.67776 0.00000 0.00000 ; HB3- CB- CG- CD
884 881 879 899 3 0.65084 1.95253 0.00000
-2.60338 0.00000 0.00000 ; CG- CB- CA- C
884 887 890 891 3 0.00000 0.00000 0.00000
0.00000 0.00000 0.00000 ; CG- CD- NE- HE
884 887 890 892 3 0.00000 0.00000 0.00000
0.00000 0.00000 0.00000 ; CG- CD- NE- CZ
885 884 887 888 3 0.65084 1.95253 0.00000
-2.60338 0.00000 0.00000 ; HG2- CG- CD- HD2
885 884 887 889 3 0.65084 1.95253 0.00000
-2.60338 0.00000 0.00000 ; HG2- CG- CD- HD3
885 884 887 890 3 0.65084 1.95253 0.00000
-2.60338 0.00000 0.00000 ; HG2- CG- CD- NE
886 884 887 888 3 0.65084 1.95253 0.00000
-2.60338 0.00000 0.00000 ; HG3- CG- CD- HD2
886 884 887 889 3 0.65084 1.95253 0.00000
-2.60338 0.00000 0.00000 ; HG3- CG- CD- HD3
886 884 887 890 3 0.65084 1.95253 0.00000
-2.60338 0.00000 0.00000 ; HG3- CG- CD- NE
887 890 892 893 3 20.08320 0.00000 -20.08320
0.00000 0.00000 0.00000 ; CD- NE- CZ- NH1
887 890 892 896 3 20.08320 0.00000 -20.08320
0.00000 0.00000 0.00000 ; CD- NE- CZ- NH2
888 887 890 891 3 0.00000 0.00000 0.00000
0.00000 0.00000 0.00000 ; HD2- CD- NE- HE
888 887 890 892 3 0.00000 0.00000 0.00000
0.00000 0.00000 0.00000 ; HD2- CD- NE- CZ

889 887 890 891 3 0.00000 0.00000 0.00000
0.00000 0.00000 0.00000 ; HD3- CD- NE- HE
889 887 890 892 3 0.00000 0.00000 0.00000
0.00000 0.00000 0.00000 ; HD3- CD- NE- CZ
890 892 893 894 3 20.08320 0.00000 -20.08320
0.00000 0.00000 0.00000 ; NE- CZ- NH1- HH11
890 892 893 895 3 20.08320 0.00000 -20.08320
0.00000 0.00000 0.00000 ; NE- CZ- NH1- HH12
890 892 896 897 3 20.08320 0.00000 -20.08320
0.00000 0.00000 0.00000 ; NE- CZ- NH2- HH21
890 892 896 898 3 20.08320 0.00000 -20.08320
0.00000 0.00000 0.00000 ; NE- CZ- NH2- HH22
891 890 892 893 3 20.08320 0.00000 -20.08320
0.00000 0.00000 0.00000 ; HE- NE- CZ- NH1
891 890 892 896 3 20.08320 0.00000 -20.08320
0.00000 0.00000 0.00000 ; HE- NE- CZ- NH2
893 892 896 897 3 20.08320 0.00000 -20.08320
0.00000 0.00000 0.00000 ; NH1- CZ- NH2- HH21
893 892 896 898 3 20.08320 0.00000 -20.08320
0.00000 0.00000 0.00000 ; NH1- CZ- NH2- HH22
894 893 892 896 3 20.08320 0.00000 -20.08320
0.00000 0.00000 0.00000 ; HH11- NH1- CZ- NH2
895 893 892 896 3 20.08320 0.00000 -20.08320
0.00000 0.00000 0.00000 ; HH12- NH1- CZ- NH2
899 901 903 904 3 0.00000 0.00000 0.00000
0.00000 0.00000 0.00000 ; C- N- CA- HA
899 901 903 905 3 11.71520 1.67360 15.06240 -
13.38880 0.00000 0.00000 ; C- N- CA- CB
899 901 903 919 3 1.75728 5.27184 2.25936
-7.02912 0.00000 0.00000 ; C- N- CA- C
900 899 901 902 3 29.28800 -8.36800 -20.92000
0.00000 0.00000 0.00000 ; O- C- N- H
900 899 901 903 3 20.92000 0.00000 -20.92000
0.00000 0.00000 0.00000 ; O- C- N- CA
901 903 905 906 3 0.65084 1.95253 0.00000
-2.60338 0.00000 0.00000 ; N- CA- CB- HB2
901 903 905 907 3 0.65084 1.95253 0.00000
-2.60338 0.00000 0.00000 ; N- CA- CB- HB3
901 903 905 908 3 3.28026 2.46019 -2.19242
-3.21331 -0.23430 0.00000 ; N- CA- CB- CG
901 903 919 920 3 0.00000 0.00000 0.00000
0.00000 0.00000 0.00000 ; N- CA- C- O
901 903 919 921 3 17.40544 -5.02080 -13.22144
9.20480 0.00000 0.00000 ; N- CA- C- N
902 901 903 904 3 0.00000 0.00000 0.00000
0.00000 0.00000 0.00000 ; H- N- CA- HA
902 901 903 905 3 0.00000 0.00000 0.00000
0.00000 0.00000 0.00000 ; H- N- CA- CB
902 901 903 919 3 0.00000 0.00000 0.00000
0.00000 0.00000 0.00000 ; H- N- CA- C
903 905 908 909 3 0.57739 0.00000 1.02926
0.00000 -1.60666 0.00000 ; CA- CB- CG- CD1

903 905 908 917 3 0.57739 0.00000 1.02926
0.00000 -1.60666 0.00000 ; CA- CB- CG- CD2
903 919 921 922 3 20.92000 0.00000 -20.92000
0.00000 0.00000 0.00000 ; CA- C- N- H
903 919 921 923 3 20.92000 0.00000 -20.92000
0.00000 0.00000 0.00000 ; CA- C- N- CA
904 903 905 906 3 0.65084 1.95253 0.00000
-2.60338 0.00000 0.00000 ; HA- CA- CB- HB2
904 903 905 907 3 0.65084 1.95253 0.00000
-2.60338 0.00000 0.00000 ; HA- CA- CB- HB3
904 903 905 908 3 0.65084 1.95253 0.00000
-2.60338 0.00000 0.00000 ; HA- CA- CB- CG
904 903 919 920 3 3.68192 -4.35136 0.00000
1.33888 0.00000 0.00000 ; HA- CA- C- O
904 903 919 921 3 0.00000 0.00000 0.00000
0.00000 0.00000 0.00000 ; HA- CA- C- N
905 903 919 920 3 0.00000 0.00000 0.00000
0.00000 0.00000 0.00000 ; CB- CA- C- O
905 903 919 921 3 2.51040 4.18400 1.67360
-6.69440 0.00000 0.00000 ; CB- CA- C- N
905 908 909 910 3 30.33400 0.00000 -30.33400
0.00000 0.00000 0.00000 ; CB- CG- CD1- HD1
905 908 909 911 3 30.33400 0.00000 -30.33400
0.00000 0.00000 0.00000 ; CB- CG- CD1- CE1
905 908 917 915 3 30.33400 0.00000 -30.33400
0.00000 0.00000 0.00000 ; CB- CG- CD2- CE2
905 908 917 918 3 30.33400 0.00000 -30.33400
0.00000 0.00000 0.00000 ; CB- CG- CD2- HD2
906 905 903 919 3 0.65084 1.95253 0.00000
-2.60338 0.00000 0.00000 ; HB2- CB- CA- C
906 905 908 909 3 0.00000 0.00000 0.00000
0.00000 0.00000 0.00000 ; HB2- CB- CG- CD1
906 905 908 917 3 0.00000 0.00000 0.00000
0.00000 0.00000 0.00000 ; HB2- CB- CG- CD2
907 905 903 919 3 0.65084 1.95253 0.00000
-2.60338 0.00000 0.00000 ; HB3- CB- CA- C
907 905 908 909 3 0.00000 0.00000 0.00000
0.00000 0.00000 0.00000 ; HB3- CB- CG- CD1
907 905 908 917 3 0.00000 0.00000 0.00000
0.00000 0.00000 0.00000 ; HB3- CB- CG- CD2
908 905 903 919 3 4.95804 2.17986 -3.52293
-3.21331 -0.40166 0.00000 ; CG- CB- CA- C
908 909 911 912 3 30.33400 0.00000 -30.33400
0.00000 0.00000 0.00000 ; CG- CD1- CE1- HE1
908 909 911 913 3 30.33400 0.00000 -30.33400
0.00000 0.00000 0.00000 ; CG- CD1- CE1- CZ
908 917 915 913 3 30.33400 0.00000 -30.33400
0.00000 0.00000 0.00000 ; CG- CD2- CE2- CZ
908 917 915 916 3 30.33400 0.00000 -30.33400
0.00000 0.00000 0.00000 ; CG- CD2- CE2- HE2
909 908 917 915 3 30.33400 0.00000 -30.33400
0.00000 0.00000 0.00000 ; CD1- CG- CD2- CE2

909 908 917 918 3 30.33400 0.00000 -30.33400
0.00000 0.00000 0.00000 ; CD1- CG- CD2- HD2
909 911 913 914 3 30.33400 0.00000 -30.33400
0.00000 0.00000 0.00000 ; CD1- CE1- CZ- HZ
909 911 913 915 3 30.33400 0.00000 -30.33400
0.00000 0.00000 0.00000 ; CD1- CE1- CZ- CE2
910 909 908 917 3 30.33400 0.00000 -30.33400
0.00000 0.00000 0.00000 ; HD1- CD1- CG- CD2
910 909 911 912 3 30.33400 0.00000 -30.33400
0.00000 0.00000 0.00000 ; HD1- CD1- CE1- HE1
910 909 911 913 3 30.33400 0.00000 -30.33400
0.00000 0.00000 0.00000 ; HD1- CD1- CE1- CZ
911 909 908 917 3 30.33400 0.00000 -30.33400
0.00000 0.00000 0.00000 ; CE1- CD1- CG- CD2
911 913 915 916 3 30.33400 0.00000 -30.33400
0.00000 0.00000 0.00000 ; CE1- CZ- CE2- HE2
911 913 915 917 3 30.33400 0.00000 -30.33400
0.00000 0.00000 0.00000 ; CE1- CZ- CE2- CD2
912 911 913 914 3 30.33400 0.00000 -30.33400
0.00000 0.00000 0.00000 ; HE1- CE1- CZ- HZ
912 911 913 915 3 30.33400 0.00000 -30.33400
0.00000 0.00000 0.00000 ; HE1- CE1- CZ- CE2
913 915 917 918 3 30.33400 0.00000 -30.33400
0.00000 0.00000 0.00000 ; CZ- CE2- CD2- HD2
914 913 915 916 3 30.33400 0.00000 -30.33400
0.00000 0.00000 0.00000 ; HZ- CZ- CE2- HE2
914 913 915 917 3 30.33400 0.00000 -30.33400
0.00000 0.00000 0.00000 ; HZ- CZ- CE2- CD2
916 915 917 918 3 30.33400 0.00000 -30.33400
0.00000 0.00000 0.00000 ; HE2- CE2- CD2- HD2
919 921 923 924 3 0.00000 0.00000 0.00000
0.00000 0.00000 0.00000 ; C- N- CA- HA
919 921 923 925 3 11.71520 1.67360 15.06240 -
13.38880 0.00000 0.00000 ; C- N- CA- CB
919 921 923 939 3 1.75728 5.27184 2.25936
-7.02912 0.00000 0.00000 ; C- N- CA- C
920 919 921 922 3 29.28800 -8.36800 -20.92000
0.00000 0.00000 0.00000 ; O- C- N- H
920 919 921 923 3 20.92000 0.00000 -20.92000
0.00000 0.00000 0.00000 ; O- C- N- CA
921 923 925 926 3 0.65084 1.95253 0.00000
-2.60338 0.00000 0.00000 ; N- CA- CB- HB2
921 923 925 927 3 0.65084 1.95253 0.00000
-2.60338 0.00000 0.00000 ; N- CA- CB- HB3
921 923 925 928 3 3.28026 2.46019 -2.19242
-3.21331 -0.23430 0.00000 ; N- CA- CB- CG
921 923 939 940 3 0.00000 0.00000 0.00000
0.00000 0.00000 0.00000 ; N- CA- C- O
921 923 939 941 3 17.40544 -5.02080 -13.22144
9.20480 0.00000 0.00000 ; N- CA- C- N
922 921 923 924 3 0.00000 0.00000 0.00000
0.00000 0.00000 0.00000 ; H- N- CA- HA

922 921 923 925 3 0.00000 0.00000 0.00000
0.00000 0.00000 0.00000 ; H- N- CA- CB
922 921 923 939 3 0.00000 0.00000 0.00000
0.00000 0.00000 0.00000 ; H- N- CA- C
923 925 928 929 3 0.57739 0.00000 1.02926
0.00000 -1.60666 0.00000 ; CA- CB- CG- CD1
923 925 928 937 3 0.57739 0.00000 1.02926
0.00000 -1.60666 0.00000 ; CA- CB- CG- CD2
923 939 941 942 3 20.92000 0.00000 -20.92000
0.00000 0.00000 0.00000 ; CA- C- N- H
923 939 941 943 3 20.92000 0.00000 -20.92000
0.00000 0.00000 0.00000 ; CA- C- N- CA
924 923 925 926 3 0.65084 1.95253 0.00000
-2.60338 0.00000 0.00000 ; HA- CA- CB- HB2
924 923 925 927 3 0.65084 1.95253 0.00000
-2.60338 0.00000 0.00000 ; HA- CA- CB- HB3
924 923 925 928 3 0.65084 1.95253 0.00000
-2.60338 0.00000 0.00000 ; HA- CA- CB- CG
924 923 939 940 3 3.68192 -4.35136 0.00000
1.33888 0.00000 0.00000 ; HA- CA- C- O
924 923 939 941 3 0.00000 0.00000 0.00000
0.00000 0.00000 0.00000 ; HA- CA- C- N
925 923 939 940 3 0.00000 0.00000 0.00000
0.00000 0.00000 0.00000 ; CB- CA- C- O
925 923 939 941 3 2.51040 4.18400 1.67360
-6.69440 0.00000 0.00000 ; CB- CA- C- N
925 928 929 930 3 30.33400 0.00000 -30.33400
0.00000 0.00000 0.00000 ; CB- CG- CD1- HD1
925 928 929 931 3 30.33400 0.00000 -30.33400
0.00000 0.00000 0.00000 ; CB- CG- CD1- CE1
925 928 937 935 3 30.33400 0.00000 -30.33400
0.00000 0.00000 0.00000 ; CB- CG- CD2- CE2
925 928 937 938 3 30.33400 0.00000 -30.33400
0.00000 0.00000 0.00000 ; CB- CG- CD2- HD2
926 925 923 939 3 0.65084 1.95253 0.00000
-2.60338 0.00000 0.00000 ; HB2- CB- CA- C
926 925 928 929 3 0.00000 0.00000 0.00000
0.00000 0.00000 0.00000 ; HB2- CB- CG- CD1
926 925 928 937 3 0.00000 0.00000 0.00000
0.00000 0.00000 0.00000 ; HB2- CB- CG- CD2
927 925 923 939 3 0.65084 1.95253 0.00000
-2.60338 0.00000 0.00000 ; HB3- CB- CA- C
927 925 928 929 3 0.00000 0.00000 0.00000
0.00000 0.00000 0.00000 ; HB3- CB- CG- CD1
927 925 928 937 3 0.00000 0.00000 0.00000
0.00000 0.00000 0.00000 ; HB3- CB- CG- CD2
928 925 923 939 3 4.95804 2.17986 -3.52293
-3.21331 -0.40166 0.00000 ; CG- CB- CA- C
928 929 931 932 3 30.33400 0.00000 -30.33400
0.00000 0.00000 0.00000 ; CG- CD1- CE1- HE1
928 929 931 933 3 30.33400 0.00000 -30.33400
0.00000 0.00000 0.00000 ; CG- CD1- CE1- CZ

928 937 935 933 3 30.33400 0.00000 -30.33400
0.00000 0.00000 0.00000 ; CG- CD2- CE2- CZ
928 937 935 936 3 30.33400 0.00000 -30.33400
0.00000 0.00000 0.00000 ; CG- CD2- CE2- HE2
929 928 937 935 3 30.33400 0.00000 -30.33400
0.00000 0.00000 0.00000 ; CD1- CG- CD2- CE2
929 928 937 938 3 30.33400 0.00000 -30.33400
0.00000 0.00000 0.00000 ; CD1- CG- CD2- HD2
929 931 933 934 3 30.33400 0.00000 -30.33400
0.00000 0.00000 0.00000 ; CD1- CE1- CZ- HZ
929 931 933 935 3 30.33400 0.00000 -30.33400
0.00000 0.00000 0.00000 ; CD1- CE1- CZ- CE2
930 929 928 937 3 30.33400 0.00000 -30.33400
0.00000 0.00000 0.00000 ; HD1- CD1- CG- CD2
930 929 931 932 3 30.33400 0.00000 -30.33400
0.00000 0.00000 0.00000 ; HD1- CD1- CE1- HE1
930 929 931 933 3 30.33400 0.00000 -30.33400
0.00000 0.00000 0.00000 ; HD1- CD1- CE1- CZ
931 929 928 937 3 30.33400 0.00000 -30.33400
0.00000 0.00000 0.00000 ; CE1- CD1- CG- CD2
931 933 935 936 3 30.33400 0.00000 -30.33400
0.00000 0.00000 0.00000 ; CE1- CZ- CE2- HE2
931 933 935 937 3 30.33400 0.00000 -30.33400
0.00000 0.00000 0.00000 ; CE1- CZ- CE2- CD2
932 931 933 934 3 30.33400 0.00000 -30.33400
0.00000 0.00000 0.00000 ; HE1- CE1- CZ- HZ
932 931 933 935 3 30.33400 0.00000 -30.33400
0.00000 0.00000 0.00000 ; HE1- CE1- CZ- CE2
933 935 937 938 3 30.33400 0.00000 -30.33400
0.00000 0.00000 0.00000 ; CZ- CE2- CD2- HD2
934 933 935 936 3 30.33400 0.00000 -30.33400
0.00000 0.00000 0.00000 ; HZ- CZ- CE2- HE2
934 933 935 937 3 30.33400 0.00000 -30.33400
0.00000 0.00000 0.00000 ; HZ- CZ- CE2- CD2
936 935 937 938 3 30.33400 0.00000 -30.33400
0.00000 0.00000 0.00000 ; HE2- CE2- CD2- HD2
939 941 943 944 3 0.00000 0.00000 0.00000
0.00000 0.00000 0.00000 ; C- N- CA- HA
939 941 943 945 3 11.71520 1.67360 15.06240 -
13.38880 0.00000 0.00000 ; C- N- CA- CB
939 941 943 956 3 1.75728 5.27184 2.25936
-7.02912 0.00000 0.00000 ; C- N- CA- C
940 939 941 942 3 29.28800 -8.36800 -20.92000
0.00000 0.00000 0.00000 ; O- C- N- H
940 939 941 943 3 20.92000 0.00000 -20.92000
0.00000 0.00000 0.00000 ; O- C- N- CA
941 943 945 946 3 0.65084 1.95253 0.00000
-2.60338 0.00000 0.00000 ; N- CA- CB- HB2
941 943 945 947 3 0.65084 1.95253 0.00000
-2.60338 0.00000 0.00000 ; N- CA- CB- HB3
941 943 945 948 3 3.21331 2.22589 -4.15053
-2.40998 2.61082 0.00000 ; N- CA- CB- CG

941 943 956 957 3 0.00000 0.00000 0.00000
0.00000 0.00000 0.00000 ; N- CA- C- O
941 943 956 958 3 17.40544 -5.02080 -13.22144
9.20480 0.00000 0.00000 ; N- CA- C- N
942 941 943 944 3 0.00000 0.00000 0.00000
0.00000 0.00000 0.00000 ; H- N- CA- HA
942 941 943 945 3 0.00000 0.00000 0.00000
0.00000 0.00000 0.00000 ; H- N- CA- CB
942 941 943 956 3 0.00000 0.00000 0.00000
0.00000 0.00000 0.00000 ; H- N- CA- C
943 945 948 949 3 0.66944 2.00832 0.00000
-2.67776 0.00000 0.00000 ; CA- CB- CG- HG2
943 945 948 950 3 0.66944 2.00832 0.00000
-2.67776 0.00000 0.00000 ; CA- CB- CG- HG3
943 945 948 951 3 3.69866 -4.35136 -3.92459
6.89523 4.61914 0.00000 ; CA- CB- CG- CD
943 956 958 959 3 20.92000 0.00000 -20.92000
0.00000 0.00000 0.00000 ; CA- C- N- H
943 956 958 960 3 20.92000 0.00000 -20.92000
0.00000 0.00000 0.00000 ; CA- C- N- CA
944 943 945 946 3 0.65084 1.95253 0.00000
-2.60338 0.00000 0.00000 ; HA- CA- CB- HB2
944 943 945 947 3 0.65084 1.95253 0.00000
-2.60338 0.00000 0.00000 ; HA- CA- CB- HB3
944 943 945 948 3 0.65084 1.95253 0.00000
-2.60338 0.00000 0.00000 ; HA- CA- CB- CG
944 943 956 957 3 3.68192 -4.35136 0.00000
1.33888 0.00000 0.00000 ; HA- CA- C- O
944 943 956 958 3 0.00000 0.00000 0.00000
0.00000 0.00000 0.00000 ; HA- CA- C- N
945 943 956 957 3 0.00000 0.00000 0.00000
0.00000 0.00000 0.00000 ; CB- CA- C- O
945 943 956 958 3 2.51040 4.18400 1.67360
-6.69440 0.00000 0.00000 ; CB- CA- C- N
945 948 951 952 3 0.00000 0.00000 0.00000
0.00000 0.00000 0.00000 ; CB- CG- CD- OE1
945 948 951 953 3 10.32611 1.48114 -8.47678
1.42256 1.40582 0.00000 ; CB- CG- CD- NE2
946 945 943 956 3 0.65084 1.95253 0.00000
-2.60338 0.00000 0.00000 ; HB2- CB- CA- C
946 945 948 949 3 0.62760 1.88280 0.00000
-2.51040 0.00000 0.00000 ; HB2- CB- CG- HG2
946 945 948 950 3 0.62760 1.88280 0.00000
-2.51040 0.00000 0.00000 ; HB2- CB- CG- HG3
946 945 948 951 3 0.65084 1.95253 0.00000
-2.60338 0.00000 0.00000 ; HB2- CB- CG- CD
947 945 943 956 3 0.65084 1.95253 0.00000
-2.60338 0.00000 0.00000 ; HB3- CB- CA- C
947 945 948 949 3 0.62760 1.88280 0.00000
-2.51040 0.00000 0.00000 ; HB3- CB- CG- HG2
947 945 948 950 3 0.62760 1.88280 0.00000
-2.51040 0.00000 0.00000 ; HB3- CB- CG- HG3

947 945 948 951 3 0.65084 1.95253 0.00000
-2.60338 0.00000 0.00000 ; HB3- CB- CG- CD
948 945 943 956 3 6.86594 3.56895 -8.14206
-2.40998 4.85344 0.00000 ; CG- CB- CA- C
948 951 953 954 3 20.92000 0.00000 -20.92000
0.00000 0.00000 0.00000 ; CG- CD- NE2- HE21
948 951 953 955 3 20.92000 0.00000 -20.92000
0.00000 0.00000 0.00000 ; CG- CD- NE2- HE22
949 948 951 952 3 3.68192 -4.35136 0.00000
1.33888 0.00000 0.00000 ; HG2- CG- CD- OE1
949 948 951 953 3 0.00000 0.00000 0.00000
0.00000 0.00000 0.00000 ; HG2- CG- CD- NE2
950 948 951 952 3 3.68192 -4.35136 0.00000
1.33888 0.00000 0.00000 ; HG3- CG- CD- OE1
950 948 951 953 3 0.00000 0.00000 0.00000
0.00000 0.00000 0.00000 ; HG3- CG- CD- NE2
952 951 953 954 3 29.28800 -8.36800 -20.92000
0.00000 0.00000 0.00000 ; OE1- CD- NE2- HE21
952 951 953 955 3 29.28800 -8.36800 -20.92000
0.00000 0.00000 0.00000 ; OE1- CD- NE2- HE22
956 958 960 961 3 0.00000 0.00000 0.00000
0.00000 0.00000 0.00000 ; C- N- CA- HA
956 958 960 962 3 11.71520 1.67360 15.06240 -
13.38880 0.00000 0.00000 ; C- N- CA- CB
956 958 960 971 3 1.75728 5.27184 2.25936
-7.02912 0.00000 0.00000 ; C- N- CA- C
957 956 958 959 3 29.28800 -8.36800 -20.92000
0.00000 0.00000 0.00000 ; O- C- N- H
957 956 958 960 3 20.92000 0.00000 -20.92000
0.00000 0.00000 0.00000 ; O- C- N- CA
958 960 962 963 3 0.65084 1.95253 0.00000
-2.60338 0.00000 0.00000 ; N- CA- CB- HB2
958 960 962 964 3 0.65084 1.95253 0.00000
-2.60338 0.00000 0.00000 ; N- CA- CB- HB3
958 960 962 965 3 3.21331 2.22589 -4.15053
-2.40998 2.61082 0.00000 ; N- CA- CB- CG
958 960 971 972 3 0.00000 0.00000 0.00000
0.00000 0.00000 0.00000 ; N- CA- C- O
958 960 971 973 3 17.40544 -5.02080 -13.22144
9.20480 0.00000 0.00000 ; N- CA- C- N
959 958 960 961 3 0.00000 0.00000 0.00000
0.00000 0.00000 0.00000 ; H- N- CA- HA
959 958 960 962 3 0.00000 0.00000 0.00000
0.00000 0.00000 0.00000 ; H- N- CA- CB
959 958 960 971 3 0.00000 0.00000 0.00000
0.00000 0.00000 0.00000 ; H- N- CA- C
960 962 965 966 3 0.66944 2.00832 0.00000
-2.67776 0.00000 0.00000 ; CA- CB- CG- HG2
960 962 965 967 3 0.66944 2.00832 0.00000
-2.67776 0.00000 0.00000 ; CA- CB- CG- HG3
960 962 965 968 3 10.12110 -1.91209 0.01674
10.17549 -1.87443 0.00000 ; CA- CB- CG- CD

960 971 973 974 3 20.92000 0.00000 -20.92000
0.00000 0.00000 0.00000 ; CA- C- N- H
960 971 973 975 3 20.92000 0.00000 -20.92000
0.00000 0.00000 0.00000 ; CA- C- N- CA
961 960 962 963 3 0.65084 1.95253 0.00000
-2.60338 0.00000 0.00000 ; HA- CA- CB- HB2
961 960 962 964 3 0.65084 1.95253 0.00000
-2.60338 0.00000 0.00000 ; HA- CA- CB- HB3
961 960 962 965 3 0.65084 1.95253 0.00000
-2.60338 0.00000 0.00000 ; HA- CA- CB- CG
961 960 971 972 3 3.68192 -4.35136 0.00000
1.33888 0.00000 0.00000 ; HA- CA- C- O
961 960 971 973 3 0.00000 0.00000 0.00000
0.00000 0.00000 0.00000 ; HA- CA- C- N
962 960 971 972 3 0.00000 0.00000 0.00000
0.00000 0.00000 0.00000 ; CB- CA- C- O
962 960 971 973 3 2.51040 4.18400 1.67360
-6.69440 0.00000 0.00000 ; CB- CA- C- N
962 965 968 969 3 3.79907 0.00000 -5.40573
0.00000 2.14221 0.00000 ; CB- CG- CD- OE1
962 965 968 970 3 3.79907 0.00000 -5.40573
0.00000 2.14221 0.00000 ; CB- CG- CD- OE2
963 962 960 971 3 0.65084 1.95253 0.00000
-2.60338 0.00000 0.00000 ; HB2- CB- CA- C
963 962 965 966 3 0.62760 1.88280 0.00000
-2.51040 0.00000 0.00000 ; HB2- CB- CG- HG2
963 962 965 967 3 0.62760 1.88280 0.00000
-2.51040 0.00000 0.00000 ; HB2- CB- CG- HG3
963 962 965 968 3 0.65084 1.95253 0.00000
-2.60338 0.00000 0.00000 ; HB2- CB- CG- CD
964 962 960 971 3 0.65084 1.95253 0.00000
-2.60338 0.00000 0.00000 ; HB3- CB- CA- C
964 962 965 966 3 0.62760 1.88280 0.00000
-2.51040 0.00000 0.00000 ; HB3- CB- CG- HG2
964 962 965 967 3 0.62760 1.88280 0.00000
-2.51040 0.00000 0.00000 ; HB3- CB- CG- HG3
964 962 965 968 3 0.65084 1.95253 0.00000
-2.60338 0.00000 0.00000 ; HB3- CB- CG- CD
965 962 960 971 3 6.86594 3.56895 -8.14206
-2.40998 4.85344 0.00000 ; CG- CB- CA- C
966 965 968 969 3 0.00000 0.00000 0.00000
0.00000 0.00000 0.00000 ; HG2- CG- CD- OE1
966 965 968 970 3 0.00000 0.00000 0.00000
0.00000 0.00000 0.00000 ; HG2- CG- CD- OE2
967 965 968 969 3 0.00000 0.00000 0.00000
0.00000 0.00000 0.00000 ; HG3- CG- CD- OE1
967 965 968 970 3 0.00000 0.00000 0.00000
0.00000 0.00000 0.00000 ; HG3- CG- CD- OE2
971 973 975 976 3 0.00000 0.00000 0.00000
0.00000 0.00000 0.00000 ; C- N- CA- HA
971 973 975 977 3 11.71520 1.67360 15.06240 -
13.38880 0.00000 0.00000 ; C- N- CA- CB

```
   971    973    975    990      3    1.75728    5.27184    2.25936
-7.02912    0.00000    0.00000 ;      C-     N-    CA-     C
   972    971    973    974      3   29.28800   -8.36800  -20.92000
0.00000    0.00000    0.00000 ;      O-     C-     N-     H
   972    971    973    975      3   20.92000    0.00000  -20.92000
0.00000    0.00000    0.00000 ;      O-     C-     N-    CA
   973    975    977    978      3    0.65084    1.95253    0.00000
-2.60338    0.00000    0.00000 ;      N-    CA-    CB-    HB
   973    975    977    979      3    3.83673    0.44769   -1.77402
-2.47693   -0.03347    0.00000 ;      N-    CA-    CB-   CG2
   973    975    977    983      3    2.97482    0.12134    2.04179
-1.89117   -3.24678    0.00000 ;      N-    CA-    CB-   CG1
   973    975    990    991      3    0.00000    0.00000    0.00000
0.00000    0.00000    0.00000 ;      N-    CA-     C-     O
   973    975    990    992      3   17.40544   -5.02080  -13.22144
9.20480    0.00000    0.00000 ;      N-    CA-     C-     N
   974    973    975    976      3    0.00000    0.00000    0.00000
0.00000    0.00000    0.00000 ;      H-     N-    CA-    HA
   974    973    975    977      3    0.00000    0.00000    0.00000
0.00000    0.00000    0.00000 ;      H-     N-    CA-    CB
   974    973    975    990      3    0.00000    0.00000    0.00000
0.00000    0.00000    0.00000 ;      H-     N-    CA-     C
   975    977    979    980      3    0.66944    2.00832    0.00000
-2.67776    0.00000    0.00000 ;     CA-    CB-   CG2-  HG21
   975    977    979    981      3    0.66944    2.00832    0.00000
-2.67776    0.00000    0.00000 ;     CA-    CB-   CG2-  HG22
   975    977    979    982      3    0.66944    2.00832    0.00000
-2.67776    0.00000    0.00000 ;     CA-    CB-   CG2-  HG23
   975    977    983    984      3    0.66944    2.00832    0.00000
-2.67776    0.00000    0.00000 ;     CA-    CB-   CG1-  HG12
   975    977    983    985      3    0.66944    2.00832    0.00000
-2.67776    0.00000    0.00000 ;     CA-    CB-   CG1-  HG13
   975    977    983    986      3    4.24258   -0.52718   -7.25506
-1.79075    7.69856    0.00000 ;     CA-    CB-   CG1-   CD1
   975    990    992    993      3   20.92000    0.00000  -20.92000
0.00000    0.00000    0.00000 ;     CA-     C-     N-     H
   975    990    992    994      3   20.92000    0.00000  -20.92000
0.00000    0.00000    0.00000 ;     CA-     C-     N-    CA
   976    975    977    978      3    0.65084    1.95253    0.00000
-2.60338    0.00000    0.00000 ;     HA-    CA-    CB-    HB
   976    975    977    979      3    0.65084    1.95253    0.00000
-2.60338    0.00000    0.00000 ;     HA-    CA-    CB-   CG2
   976    975    977    983      3    0.65084    1.95253    0.00000
-2.60338    0.00000    0.00000 ;     HA-    CA-    CB-   CG1
   976    975    990    991      3    3.68192   -4.35136    0.00000
1.33888    0.00000    0.00000 ;     HA-    CA-     C-     O
   976    975    990    992      3    0.00000    0.00000    0.00000
0.00000    0.00000    0.00000 ;     HA-    CA-     C-     N
   977    975    990    991      3    0.00000    0.00000    0.00000
0.00000    0.00000    0.00000 ;     CB-    CA-     C-     O
   977    975    990    992      3    2.51040    4.18400    1.67360
-6.69440    0.00000    0.00000 ;     CB-    CA-     C-     N
```

977 983 986 987 3 0.66944 2.00832 0.00000
-2.67776 0.00000 0.00000 ; CB- CG1- CD1- HD11
977 983 986 988 3 0.66944 2.00832 0.00000
-2.67776 0.00000 0.00000 ; CB- CG1- CD1- HD12
977 983 986 989 3 0.66944 2.00832 0.00000
-2.67776 0.00000 0.00000 ; CB- CG1- CD1- HD13
978 977 975 990 3 0.65084 1.95253 0.00000
-2.60338 0.00000 0.00000 ; HB- CB- CA- C
978 977 979 980 3 0.62760 1.88280 0.00000
-2.51040 0.00000 0.00000 ; HB- CB- CG2- HG21
978 977 979 981 3 0.62760 1.88280 0.00000
-2.51040 0.00000 0.00000 ; HB- CB- CG2- HG22
978 977 979 982 3 0.62760 1.88280 0.00000
-2.51040 0.00000 0.00000 ; HB- CB- CG2- HG23
978 977 983 984 3 0.62760 1.88280 0.00000
-2.51040 0.00000 0.00000 ; HB- CB- CG1- HG12
978 977 983 985 3 0.62760 1.88280 0.00000
-2.51040 0.00000 0.00000 ; HB- CB- CG1- HG13
978 977 983 986 3 0.66944 2.00832 0.00000
-2.67776 0.00000 0.00000 ; HB- CB- CG1- CD1
979 977 975 990 3 5.67350 3.55640 -6.16722
-2.47693 3.74886 0.00000 ; CG2- CB- CA- C
979 977 983 984 3 0.66944 2.00832 0.00000
-2.67776 0.00000 0.00000 ; CG2- CB- CG1- HG12
979 977 983 985 3 0.66944 2.00832 0.00000
-2.67776 0.00000 0.00000 ; CG2- CB- CG1- HG13
979 977 983 986 3 3.81162 0.49790 -8.14206
-1.79075 7.49773 0.00000 ; CG2- CB- CG1- CD1
980 979 977 983 3 0.66944 2.00832 0.00000
-2.67776 0.00000 0.00000 ; HG21- CG2- CB- CG1
981 979 977 983 3 0.66944 2.00832 0.00000
-2.67776 0.00000 0.00000 ; HG22- CG2- CB- CG1
982 979 977 983 3 0.66944 2.00832 0.00000
-2.67776 0.00000 0.00000 ; HG23- CG2- CB- CG1
983 977 975 990 3 8.26340 0.74057 -9.99976
-1.89117 3.84928 0.00000 ; CG1- CB- CA- C
984 983 986 987 3 0.62760 1.88280 0.00000
-2.51040 0.00000 0.00000 ; HG12- CG1- CD1- HD11
984 983 986 988 3 0.62760 1.88280 0.00000
-2.51040 0.00000 0.00000 ; HG12- CG1- CD1- HD12
984 983 986 989 3 0.62760 1.88280 0.00000
-2.51040 0.00000 0.00000 ; HG12- CG1- CD1- HD13
985 983 986 987 3 0.62760 1.88280 0.00000
-2.51040 0.00000 0.00000 ; HG13- CG1- CD1- HD11
985 983 986 988 3 0.62760 1.88280 0.00000
-2.51040 0.00000 0.00000 ; HG13- CG1- CD1- HD12
985 983 986 989 3 0.62760 1.88280 0.00000
-2.51040 0.00000 0.00000 ; HG13- CG1- CD1- HD13
990 992 994 995 3 0.00000 0.00000 0.00000
0.00000 0.00000 0.00000 ; C- N- CA- HA
990 992 994 996 3 11.71520 1.67360 15.06240 -
13.38880 0.00000 0.00000 ; C- N- CA- CB

```
   990    992    994   1005       3    1.75728    5.27184    2.25936
-7.02912    0.00000    0.00000 ;      C-     N-    CA-     C
   991    990    992    993       3   29.28800   -8.36800  -20.92000
0.00000    0.00000    0.00000 ;      O-     C-     N-     H
   991    990    992    994       3   20.92000    0.00000  -20.92000
0.00000    0.00000    0.00000 ;      O-     C-     N-    CA
   992    994    996    997       3    0.65084    1.95253    0.00000
-2.60338    0.00000    0.00000 ;      N-    CA-    CB-   HB2
   992    994    996    998       3    0.65084    1.95253    0.00000
-2.60338    0.00000    0.00000 ;      N-    CA-    CB-   HB3
   992    994    996    999       3    3.21331    2.22589   -4.15053
-2.40998    2.61082    0.00000 ;      N-    CA-    CB-    CG
   992    994   1005   1006       3    0.00000    0.00000    0.00000
0.00000    0.00000    0.00000 ;      N-    CA-     C-     O
   992    994   1005   1007       3   17.40544   -5.02080  -13.22144
9.20480    0.00000    0.00000 ;      N-    CA-     C-     N
   993    992    994    995       3    0.00000    0.00000    0.00000
0.00000    0.00000    0.00000 ;      H-     N-    CA-    HA
   993    992    994    996       3    0.00000    0.00000    0.00000
0.00000    0.00000    0.00000 ;      H-     N-    CA-    CB
   993    992    994   1005       3    0.00000    0.00000    0.00000
0.00000    0.00000    0.00000 ;      H-     N-    CA-     C
   994    996    999   1000       3    0.66944    2.00832    0.00000
-2.67776    0.00000    0.00000 ;     CA-    CB-    CG-   HG2
   994    996    999   1001       3    0.66944    2.00832    0.00000
-2.67776    0.00000    0.00000 ;     CA-    CB-    CG-   HG3
   994    996    999   1002       3   10.12110   -1.91209    0.01674
10.17549   -1.87443    0.00000 ;     CA-    CB-    CG-    CD
   994   1005   1007   1008       3   20.92000    0.00000  -20.92000
0.00000    0.00000    0.00000 ;     CA-     C-     N-     H
   994   1005   1007   1009       3   20.92000    0.00000  -20.92000
0.00000    0.00000    0.00000 ;     CA-     C-     N-    CA
   995    994    996    997       3    0.65084    1.95253    0.00000
-2.60338    0.00000    0.00000 ;     HA-    CA-    CB-   HB2
   995    994    996    998       3    0.65084    1.95253    0.00000
-2.60338    0.00000    0.00000 ;     HA-    CA-    CB-   HB3
   995    994    996    999       3    0.65084    1.95253    0.00000
-2.60338    0.00000    0.00000 ;     HA-    CA-    CB-    CG
   995    994   1005   1006       3    3.68192   -4.35136    0.00000
1.33888    0.00000    0.00000 ;     HA-    CA-     C-     O
   995    994   1005   1007       3    0.00000    0.00000    0.00000
0.00000    0.00000    0.00000 ;     HA-    CA-     C-     N
   996    994   1005   1006       3    0.00000    0.00000    0.00000
0.00000    0.00000    0.00000 ;     CB-    CA-     C-     O
   996    994   1005   1007       3    2.51040    4.18400    1.67360
-6.69440    0.00000    0.00000 ;     CB-    CA-     C-     N
   996    999   1002   1003       3    3.79907    0.00000   -5.40573
0.00000    2.14221    0.00000 ;     CB-    CG-    CD-   OE1
   996    999   1002   1004       3    3.79907    0.00000   -5.40573
0.00000    2.14221    0.00000 ;     CB-    CG-    CD-   OE2
   997    996    994   1005       3    0.65084    1.95253    0.00000
-2.60338    0.00000    0.00000 ;    HB2-    CB-    CA-     C
```

```
   997    996    999   1000      3    0.62760    1.88280    0.00000
-2.51040    0.00000    0.00000 ;    HB2-    CB-    CG-   HG2
   997    996    999   1001      3    0.62760    1.88280    0.00000
-2.51040    0.00000    0.00000 ;    HB2-    CB-    CG-   HG3
   997    996    999   1002      3    0.65084    1.95253    0.00000
-2.60338    0.00000    0.00000 ;    HB2-    CB-    CG-    CD
   998    996    994   1005      3    0.65084    1.95253    0.00000
-2.60338    0.00000    0.00000 ;    HB3-    CB-    CA-     C
   998    996    999   1000      3    0.62760    1.88280    0.00000
-2.51040    0.00000    0.00000 ;    HB3-    CB-    CG-   HG2
   998    996    999   1001      3    0.62760    1.88280    0.00000
-2.51040    0.00000    0.00000 ;    HB3-    CB-    CG-   HG3
   998    996    999   1002      3    0.65084    1.95253    0.00000
-2.60338    0.00000    0.00000 ;    HB3-    CB-    CG-    CD
   999    996    994   1005      3    6.86594    3.56895   -8.14206
-2.40998    4.85344    0.00000 ;     CG-    CB-    CA-     C
  1000    999   1002   1003      3    0.00000    0.00000    0.00000
0.00000    0.00000    0.00000 ;    HG2-    CG-    CD-   OE1
  1000    999   1002   1004      3    0.00000    0.00000    0.00000
0.00000    0.00000    0.00000 ;    HG2-    CG-    CD-   OE2
  1001    999   1002   1003      3    0.00000    0.00000    0.00000
0.00000    0.00000    0.00000 ;    HG3-    CG-    CD-   OE1
  1001    999   1002   1004      3    0.00000    0.00000    0.00000
0.00000    0.00000    0.00000 ;    HG3-    CG-    CD-   OE2
  1005   1007   1009   1010      3    0.00000    0.00000    0.00000
0.00000    0.00000    0.00000 ;     C-     N-    CA-    HA
  1005   1007   1009   1011      3   11.71520    1.67360   15.06240  -
13.38880    0.00000    0.00000 ;     C-     N-    CA-    CB
  1005   1007   1009   1024      3    1.75728    5.27184    2.25936
-7.02912    0.00000    0.00000 ;     C-     N-    CA-     C
  1006   1005   1007   1008      3   29.28800   -8.36800  -20.92000
0.00000    0.00000    0.00000 ;     O-     C-     N-     H
  1006   1005   1007   1009      3   20.92000    0.00000  -20.92000
0.00000    0.00000    0.00000 ;     O-     C-     N-    CA
  1007   1009   1011   1012      3    0.65084    1.95253    0.00000
-2.60338    0.00000    0.00000 ;     N-    CA-    CB-    HB
  1007   1009   1011   1013      3    3.83673    0.44769   -1.77402
-2.47693   -0.03347    0.00000 ;     N-    CA-    CB-   CG2
  1007   1009   1011   1017      3    2.97482    0.12134    2.04179
-1.89117   -3.24678    0.00000 ;     N-    CA-    CB-   CG1
  1007   1009   1024   1025      3    0.00000    0.00000    0.00000
0.00000    0.00000    0.00000 ;     N-    CA-     C-     O
  1007   1009   1024   1026      3   17.40544   -5.02080  -13.22144
9.20480    0.00000    0.00000 ;     N-    CA-     C-     N
  1008   1007   1009   1010      3    0.00000    0.00000    0.00000
0.00000    0.00000    0.00000 ;     H-     N-    CA-    HA
  1008   1007   1009   1011      3    0.00000    0.00000    0.00000
0.00000    0.00000    0.00000 ;     H-     N-    CA-    CB
  1008   1007   1009   1024      3    0.00000    0.00000    0.00000
0.00000    0.00000    0.00000 ;     H-     N-    CA-     C
  1009   1011   1013   1014      3    0.66944    2.00832    0.00000
-2.67776    0.00000    0.00000 ;    CA-    CB-   CG2-  HG21
```

```
  1009   1011   1013   1015      3    0.66944    2.00832    0.00000
-2.67776    0.00000    0.00000 ;     CA-    CB-   CG2-  HG22
  1009   1011   1013   1016      3    0.66944    2.00832    0.00000
-2.67776    0.00000    0.00000 ;     CA-    CB-   CG2-  HG23
  1009   1011   1017   1018      3    0.66944    2.00832    0.00000
-2.67776    0.00000    0.00000 ;     CA-    CB-   CG1-  HG12
  1009   1011   1017   1019      3    0.66944    2.00832    0.00000
-2.67776    0.00000    0.00000 ;     CA-    CB-   CG1-  HG13
  1009   1011   1017   1020      3    4.24258   -0.52718   -7.25506
-1.79075    7.69856    0.00000 ;     CA-    CB-   CG1-   CD1
  1009   1024   1026   1027      3   20.92000    0.00000  -20.92000
0.00000    0.00000    0.00000 ;     CA-     C-     N-     H
  1009   1024   1026   1028      3   20.92000    0.00000  -20.92000
0.00000    0.00000    0.00000 ;     CA-     C-     N-    CA
  1010   1009   1011   1012      3    0.65084    1.95253    0.00000
-2.60338    0.00000    0.00000 ;     HA-    CA-    CB-    HB
  1010   1009   1011   1013      3    0.65084    1.95253    0.00000
-2.60338    0.00000    0.00000 ;     HA-    CA-    CB-   CG2
  1010   1009   1011   1017      3    0.65084    1.95253    0.00000
-2.60338    0.00000    0.00000 ;     HA-    CA-    CB-   CG1
  1010   1009   1024   1025      3    3.68192   -4.35136    0.00000
1.33888    0.00000    0.00000 ;     HA-    CA-     C-     O
  1010   1009   1024   1026      3    0.00000    0.00000    0.00000
0.00000    0.00000    0.00000 ;     HA-    CA-     C-     N
  1011   1009   1024   1025      3    0.00000    0.00000    0.00000
0.00000    0.00000    0.00000 ;     CB-    CA-     C-     O
  1011   1009   1024   1026      3    2.51040    4.18400    1.67360
-6.69440    0.00000    0.00000 ;     CB-    CA-     C-     N
  1011   1017   1020   1021      3    0.66944    2.00832    0.00000
-2.67776    0.00000    0.00000 ;     CB-   CG1-   CD1-  HD11
  1011   1017   1020   1022      3    0.66944    2.00832    0.00000
-2.67776    0.00000    0.00000 ;     CB-   CG1-   CD1-  HD12
  1011   1017   1020   1023      3    0.66944    2.00832    0.00000
-2.67776    0.00000    0.00000 ;     CB-   CG1-   CD1-  HD13
  1012   1011   1009   1024      3    0.65084    1.95253    0.00000
-2.60338    0.00000    0.00000 ;     HB-    CB-    CA-     C
  1012   1011   1013   1014      3    0.62760    1.88280    0.00000
-2.51040    0.00000    0.00000 ;     HB-    CB-   CG2-  HG21
  1012   1011   1013   1015      3    0.62760    1.88280    0.00000
-2.51040    0.00000    0.00000 ;     HB-    CB-   CG2-  HG22
  1012   1011   1013   1016      3    0.62760    1.88280    0.00000
-2.51040    0.00000    0.00000 ;     HB-    CB-   CG2-  HG23
  1012   1011   1017   1018      3    0.62760    1.88280    0.00000
-2.51040    0.00000    0.00000 ;     HB-    CB-   CG1-  HG12
  1012   1011   1017   1019      3    0.62760    1.88280    0.00000
-2.51040    0.00000    0.00000 ;     HB-    CB-   CG1-  HG13
  1012   1011   1017   1020      3    0.66944    2.00832    0.00000
-2.67776    0.00000    0.00000 ;     HB-    CB-   CG1-   CD1
  1013   1011   1009   1024      3    5.67350    3.55640   -6.16722
-2.47693    3.74886    0.00000 ;    CG2-    CB-    CA-     C
  1013   1011   1017   1018      3    0.66944    2.00832    0.00000
-2.67776    0.00000    0.00000 ;    CG2-    CB-   CG1-  HG12
```

```
  1013   1011   1017   1019      3    0.66944    2.00832    0.00000
-2.67776    0.00000    0.00000 ;    CG2-    CB-   CG1-  HG13
  1013   1011   1017   1020      3    3.81162    0.49790   -8.14206
-1.79075    7.49773    0.00000 ;    CG2-    CB-   CG1-   CD1
  1014   1013   1011   1017      3    0.66944    2.00832    0.00000
-2.67776    0.00000    0.00000 ;   HG21-   CG2-    CB-   CG1
  1015   1013   1011   1017      3    0.66944    2.00832    0.00000
-2.67776    0.00000    0.00000 ;   HG22-   CG2-    CB-   CG1
  1016   1013   1011   1017      3    0.66944    2.00832    0.00000
-2.67776    0.00000    0.00000 ;   HG23-   CG2-    CB-   CG1
  1017   1011   1009   1024      3    8.26340    0.74057   -9.99976
-1.89117    3.84928    0.00000 ;    CG1-    CB-    CA-     C
  1018   1017   1020   1021      3    0.62760    1.88280    0.00000
-2.51040    0.00000    0.00000 ;   HG12-   CG1-   CD1-  HD11
  1018   1017   1020   1022      3    0.62760    1.88280    0.00000
-2.51040    0.00000    0.00000 ;   HG12-   CG1-   CD1-  HD12
  1018   1017   1020   1023      3    0.62760    1.88280    0.00000
-2.51040    0.00000    0.00000 ;   HG12-   CG1-   CD1-  HD13
  1019   1017   1020   1021      3    0.62760    1.88280    0.00000
-2.51040    0.00000    0.00000 ;   HG13-   CG1-   CD1-  HD11
  1019   1017   1020   1022      3    0.62760    1.88280    0.00000
-2.51040    0.00000    0.00000 ;   HG13-   CG1-   CD1-  HD12
  1019   1017   1020   1023      3    0.62760    1.88280    0.00000
-2.51040    0.00000    0.00000 ;   HG13-   CG1-   CD1-  HD13
  1024   1026   1028   1029      3    0.00000    0.00000    0.00000
0.00000    0.00000    0.00000 ;      C-     N-    CA-    HA
  1024   1026   1028   1030      3   11.71520    1.67360   15.06240  -
13.38880    0.00000    0.00000 ;      C-     N-    CA-    CB
  1024   1026   1028   1041      3    1.75728    5.27184    2.25936
-7.02912    0.00000    0.00000 ;      C-     N-    CA-     C
  1025   1024   1026   1027      3   29.28800   -8.36800  -20.92000
0.00000    0.00000    0.00000 ;      O-     C-     N-     H
  1025   1024   1026   1028      3   20.92000    0.00000  -20.92000
0.00000    0.00000    0.00000 ;      O-     C-     N-    CA
  1026   1028   1030   1031      3    0.65084    1.95253    0.00000
-2.60338    0.00000    0.00000 ;      N-    CA-    CB-   HB2
  1026   1028   1030   1032      3    0.65084    1.95253    0.00000
-2.60338    0.00000    0.00000 ;      N-    CA-    CB-   HB3
  1026   1028   1030   1033      3    3.21331    2.22589   -4.15053
-2.40998    2.61082    0.00000 ;      N-    CA-    CB-    CG
  1026   1028   1041   1042      3    0.00000    0.00000    0.00000
0.00000    0.00000    0.00000 ;      N-    CA-     C-     O
  1026   1028   1041   1043      3   17.40544   -5.02080  -13.22144
9.20480    0.00000    0.00000 ;      N-    CA-     C-     N
  1027   1026   1028   1029      3    0.00000    0.00000    0.00000
0.00000    0.00000    0.00000 ;      H-     N-    CA-    HA
  1027   1026   1028   1030      3    0.00000    0.00000    0.00000
0.00000    0.00000    0.00000 ;      H-     N-    CA-    CB
  1027   1026   1028   1041      3    0.00000    0.00000    0.00000
0.00000    0.00000    0.00000 ;      H-     N-    CA-     C
  1028   1030   1033   1034      3    0.65084    1.95253    0.00000
-2.60338    0.00000    0.00000 ;     CA-    CB-    CG-   HG2
```

```
  1028   1030   1033   1035      3    0.65084    1.95253    0.00000
-2.60338    0.00000    0.00000 ;    CA-    CB-    CG-   HG3
  1028   1030   1033   1036      3    2.04598   -1.54390    1.11294
-0.26778    0.93722    0.00000 ;    CA-    CB-    CG-    SD
  1028   1041   1043   1044      3   20.92000    0.00000  -20.92000
0.00000    0.00000    0.00000 ;    CA-     C-     N-     H
  1028   1041   1043   1045      3   20.92000    0.00000  -20.92000
0.00000    0.00000    0.00000 ;    CA-     C-     N-    CA
  1029   1028   1030   1031      3    0.65084    1.95253    0.00000
-2.60338    0.00000    0.00000 ;    HA-    CA-    CB-   HB2
  1029   1028   1030   1032      3    0.65084    1.95253    0.00000
-2.60338    0.00000    0.00000 ;    HA-    CA-    CB-   HB3
  1029   1028   1030   1033      3    0.65084    1.95253    0.00000
-2.60338    0.00000    0.00000 ;    HA-    CA-    CB-    CG
  1029   1028   1041   1042      3    3.68192   -4.35136    0.00000
1.33888    0.00000    0.00000 ;    HA-    CA-     C-     O
  1029   1028   1041   1043      3    0.00000    0.00000    0.00000
0.00000    0.00000    0.00000 ;    HA-    CA-     C-     N
  1030   1028   1041   1042      3    0.00000    0.00000    0.00000
0.00000    0.00000    0.00000 ;    CB-    CA-     C-     O
  1030   1028   1041   1043      3    2.51040    4.18400    1.67360
-6.69440    0.00000    0.00000 ;    CB-    CA-     C-     N
  1030   1033   1036   1037      3    3.24260    6.22998    1.79075
-6.92870    1.90790    0.00000 ;    CB-    CG-    SD-    CE
  1031   1030   1028   1041      3    0.65084    1.95253    0.00000
-2.60338    0.00000    0.00000 ;   HB2-    CB-    CA-     C
  1031   1030   1033   1034      3    0.65084    1.95253    0.00000
-2.60338    0.00000    0.00000 ;   HB2-    CB-    CG-   HG2
  1031   1030   1033   1035      3    0.65084    1.95253    0.00000
-2.60338    0.00000    0.00000 ;   HB2-    CB-    CG-   HG3
  1031   1030   1033   1036      3    0.65084    1.95253    0.00000
-2.60338    0.00000    0.00000 ;   HB2-    CB-    CG-    SD
  1032   1030   1028   1041      3    0.65084    1.95253    0.00000
-2.60338    0.00000    0.00000 ;   HB3-    CB-    CA-     C
  1032   1030   1033   1034      3    0.65084    1.95253    0.00000
-2.60338    0.00000    0.00000 ;   HB3-    CB-    CG-   HG2
  1032   1030   1033   1035      3    0.65084    1.95253    0.00000
-2.60338    0.00000    0.00000 ;   HB3-    CB-    CG-   HG3
  1032   1030   1033   1036      3    0.65084    1.95253    0.00000
-2.60338    0.00000    0.00000 ;   HB3-    CB-    CG-    SD
  1033   1030   1028   1041      3    6.86594    3.56895   -8.14206
-2.40998    4.85344    0.00000 ;    CG-    CB-    CA-     C
  1033   1036   1037   1038      3    1.39467    4.18400    0.00000
-5.57867    0.00000    0.00000 ;    CG-    SD-    CE-   HE1
  1033   1036   1037   1039      3    1.39467    4.18400    0.00000
-5.57867    0.00000    0.00000 ;    CG-    SD-    CE-   HE2
  1033   1036   1037   1040      3    1.39467    4.18400    0.00000
-5.57867    0.00000    0.00000 ;    CG-    SD-    CE-   HE3
  1034   1033   1036   1037      3    1.39467    4.18400    0.00000
-5.57867    0.00000    0.00000 ;   HG2-    CG-    SD-    CE
  1035   1033   1036   1037      3    1.39467    4.18400    0.00000
-5.57867    0.00000    0.00000 ;   HG3-    CG-    SD-    CE
```

```
  1041   1043   1045   1046      3    0.00000    0.00000    0.00000
0.00000    0.00000    0.00000 ;     C-     N-    CA-    HA
  1041   1043   1045   1047      3   11.71520    1.67360   15.06240  -
13.38880    0.00000    0.00000 ;     C-     N-    CA-    CB
  1041   1043   1045   1063      3    1.75728    5.27184    2.25936
-7.02912    0.00000    0.00000 ;     C-     N-    CA-     C
  1042   1041   1043   1044      3   29.28800   -8.36800  -20.92000
0.00000    0.00000    0.00000 ;     O-     C-     N-     H
  1042   1041   1043   1045      3   20.92000    0.00000  -20.92000
0.00000    0.00000    0.00000 ;     O-     C-     N-    CA
  1043   1045   1047   1048      3    0.65084    1.95253    0.00000
-2.60338    0.00000    0.00000 ;     N-    CA-    CB-   HB2
  1043   1045   1047   1049      3    0.65084    1.95253    0.00000
-2.60338    0.00000    0.00000 ;     N-    CA-    CB-   HB3
  1043   1045   1047   1050      3    0.65084    1.95253    0.00000
-2.60338    0.00000    0.00000 ;     N-    CA-    CB-    CG
  1043   1045   1063   1064      3    0.00000    0.00000    0.00000
0.00000    0.00000    0.00000 ;     N-    CA-     C-     O
  1043   1045   1063   1065      3   17.40544   -5.02080  -13.22144
9.20480    0.00000    0.00000 ;     N-    CA-     C-     N
  1044   1043   1045   1046      3    0.00000    0.00000    0.00000
0.00000    0.00000    0.00000 ;     H-     N-    CA-    HA
  1044   1043   1045   1047      3    0.00000    0.00000    0.00000
0.00000    0.00000    0.00000 ;     H-     N-    CA-    CB
  1044   1043   1045   1063      3    0.00000    0.00000    0.00000
0.00000    0.00000    0.00000 ;     H-     N-    CA-     C
  1045   1047   1050   1051      3    0.66944    2.00832    0.00000
-2.67776    0.00000    0.00000 ;    CA-    CB-    CG-   HG2
  1045   1047   1050   1052      3    0.66944    2.00832    0.00000
-2.67776    0.00000    0.00000 ;    CA-    CB-    CG-   HG3
  1045   1047   1050   1053      3    3.68192    3.09616   -2.09200
-3.01248    0.00000    0.00000 ;    CA-    CB-    CG-    CD
  1045   1063   1065   1066      3   20.92000    0.00000  -20.92000
0.00000    0.00000    0.00000 ;    CA-     C-     N-     H
  1045   1063   1065   1067      3   20.92000    0.00000  -20.92000
0.00000    0.00000    0.00000 ;    CA-     C-     N-    CA
  1046   1045   1047   1048      3    0.65084    1.95253    0.00000
-2.60338    0.00000    0.00000 ;    HA-    CA-    CB-   HB2
  1046   1045   1047   1049      3    0.65084    1.95253    0.00000
-2.60338    0.00000    0.00000 ;    HA-    CA-    CB-   HB3
  1046   1045   1047   1050      3    0.65084    1.95253    0.00000
-2.60338    0.00000    0.00000 ;    HA-    CA-    CB-    CG
  1046   1045   1063   1064      3    3.68192   -4.35136    0.00000
1.33888    0.00000    0.00000 ;    HA-    CA-     C-     O
  1046   1045   1063   1065      3    0.00000    0.00000    0.00000
0.00000    0.00000    0.00000 ;    HA-    CA-     C-     N
  1047   1045   1063   1064      3    0.00000    0.00000    0.00000
0.00000    0.00000    0.00000 ;    CB-    CA-     C-     O
  1047   1045   1063   1065      3    2.51040    4.18400    1.67360
-6.69440    0.00000    0.00000 ;    CB-    CA-     C-     N
  1047   1050   1053   1054      3    0.66944    2.00832    0.00000
-2.67776    0.00000    0.00000 ;    CB-    CG-    CD-   HD2
```

```
  1047   1050   1053   1055       3    0.66944    2.00832    0.00000
-2.67776    0.00000    0.00000 ;     CB-    CG-    CD-   HD3
  1047   1050   1053   1056       3    3.68192    3.09616   -2.09200
-3.01248    0.00000    0.00000 ;     CB-    CG-    CD-    CE
  1048   1047   1045   1063       3    0.65084    1.95253    0.00000
-2.60338    0.00000    0.00000 ;    HB2-    CB-    CA-     C
  1048   1047   1050   1051       3    0.62760    1.88280    0.00000
-2.51040    0.00000    0.00000 ;    HB2-    CB-    CG-   HG2
  1048   1047   1050   1052       3    0.62760    1.88280    0.00000
-2.51040    0.00000    0.00000 ;    HB2-    CB-    CG-   HG3
  1048   1047   1050   1053       3    0.66944    2.00832    0.00000
-2.67776    0.00000    0.00000 ;    HB2-    CB-    CG-    CD
  1049   1047   1045   1063       3    0.65084    1.95253    0.00000
-2.60338    0.00000    0.00000 ;    HB3-    CB-    CA-     C
  1049   1047   1050   1051       3    0.62760    1.88280    0.00000
-2.51040    0.00000    0.00000 ;    HB3-    CB-    CG-   HG2
  1049   1047   1050   1052       3    0.62760    1.88280    0.00000
-2.51040    0.00000    0.00000 ;    HB3-    CB-    CG-   HG3
  1049   1047   1050   1053       3    0.66944    2.00832    0.00000
-2.67776    0.00000    0.00000 ;    HB3-    CB-    CG-    CD
  1050   1047   1045   1063       3    0.65084    1.95253    0.00000
-2.60338    0.00000    0.00000 ;     CG-    CB-    CA-     C
  1050   1053   1056   1057       3    0.65084    1.95253    0.00000
-2.60338    0.00000    0.00000 ;     CG-    CD-    CE-   HE2
  1050   1053   1056   1058       3    0.65084    1.95253    0.00000
-2.60338    0.00000    0.00000 ;     CG-    CD-    CE-   HE3
  1050   1053   1056   1059       3    0.65084    1.95253    0.00000
-2.60338    0.00000    0.00000 ;     CG-    CD-    CE-    NZ
  1051   1050   1053   1054       3    0.62760    1.88280    0.00000
-2.51040    0.00000    0.00000 ;    HG2-    CG-    CD-   HD2
  1051   1050   1053   1055       3    0.62760    1.88280    0.00000
-2.51040    0.00000    0.00000 ;    HG2-    CG-    CD-   HD3
  1051   1050   1053   1056       3    0.66944    2.00832    0.00000
-2.67776    0.00000    0.00000 ;    HG2-    CG-    CD-    CE
  1052   1050   1053   1054       3    0.62760    1.88280    0.00000
-2.51040    0.00000    0.00000 ;    HG3-    CG-    CD-   HD2
  1052   1050   1053   1055       3    0.62760    1.88280    0.00000
-2.51040    0.00000    0.00000 ;    HG3-    CG-    CD-   HD3
  1052   1050   1053   1056       3    0.66944    2.00832    0.00000
-2.67776    0.00000    0.00000 ;    HG3-    CG-    CD-    CE
  1053   1056   1059   1060       3    0.65084    1.95253    0.00000
-2.60338    0.00000    0.00000 ;     CD-    CE-    NZ-   HZ1
  1053   1056   1059   1061       3    0.65084    1.95253    0.00000
-2.60338    0.00000    0.00000 ;     CD-    CE-    NZ-   HZ2
  1053   1056   1059   1062       3    0.65084    1.95253    0.00000
-2.60338    0.00000    0.00000 ;     CD-    CE-    NZ-   HZ3
  1054   1053   1056   1057       3    0.65084    1.95253    0.00000
-2.60338    0.00000    0.00000 ;    HD2-    CD-    CE-   HE2
  1054   1053   1056   1058       3    0.65084    1.95253    0.00000
-2.60338    0.00000    0.00000 ;    HD2-    CD-    CE-   HE3
  1054   1053   1056   1059       3    0.65084    1.95253    0.00000
-2.60338    0.00000    0.00000 ;    HD2-    CD-    CE-    NZ
```

1055 1053 1056 1057 3 0.65084 1.95253 0.00000
-2.60338 0.00000 0.00000 ; HD3- CD- CE- HE2
1055 1053 1056 1058 3 0.65084 1.95253 0.00000
-2.60338 0.00000 0.00000 ; HD3- CD- CE- HE3
1055 1053 1056 1059 3 0.65084 1.95253 0.00000
-2.60338 0.00000 0.00000 ; HD3- CD- CE- NZ
1057 1056 1059 1060 3 0.65084 1.95253 0.00000
-2.60338 0.00000 0.00000 ; HE2- CE- NZ- HZ1
1057 1056 1059 1061 3 0.65084 1.95253 0.00000
-2.60338 0.00000 0.00000 ; HE2- CE- NZ- HZ2
1057 1056 1059 1062 3 0.65084 1.95253 0.00000
-2.60338 0.00000 0.00000 ; HE2- CE- NZ- HZ3
1058 1056 1059 1060 3 0.65084 1.95253 0.00000
-2.60338 0.00000 0.00000 ; HE3- CE- NZ- HZ1
1058 1056 1059 1061 3 0.65084 1.95253 0.00000
-2.60338 0.00000 0.00000 ; HE3- CE- NZ- HZ2
1058 1056 1059 1062 3 0.65084 1.95253 0.00000
-2.60338 0.00000 0.00000 ; HE3- CE- NZ- HZ3
1063 1065 1067 1068 3 0.00000 0.00000 0.00000
0.00000 0.00000 0.00000 ; C- N- CA- HA
1063 1065 1067 1069 3 11.71520 1.67360 15.06240 -
13.38880 0.00000 0.00000 ; C- N- CA- CB
1063 1065 1067 1085 3 1.75728 5.27184 2.25936
-7.02912 0.00000 0.00000 ; C- N- CA- C
1064 1063 1065 1066 3 29.28800 -8.36800 -20.92000
0.00000 0.00000 0.00000 ; O- C- N- H
1064 1063 1065 1067 3 20.92000 0.00000 -20.92000
0.00000 0.00000 0.00000 ; O- C- N- CA
1065 1067 1069 1070 3 0.65084 1.95253 0.00000
-2.60338 0.00000 0.00000 ; N- CA- CB- HB2
1065 1067 1069 1071 3 0.65084 1.95253 0.00000
-2.60338 0.00000 0.00000 ; N- CA- CB- HB3
1065 1067 1069 1072 3 0.65084 1.95253 0.00000
-2.60338 0.00000 0.00000 ; N- CA- CB- CG
1065 1067 1085 1086 3 0.00000 0.00000 0.00000
0.00000 0.00000 0.00000 ; N- CA- C- O
1065 1067 1085 1087 3 17.40544 -5.02080 -13.22144
9.20480 0.00000 0.00000 ; N- CA- C- N
1066 1065 1067 1068 3 0.00000 0.00000 0.00000
0.00000 0.00000 0.00000 ; H- N- CA- HA
1066 1065 1067 1069 3 0.00000 0.00000 0.00000
0.00000 0.00000 0.00000 ; H- N- CA- CB
1066 1065 1067 1085 3 0.00000 0.00000 0.00000
0.00000 0.00000 0.00000 ; H- N- CA- C
1067 1069 1072 1073 3 0.66944 2.00832 0.00000
-2.67776 0.00000 0.00000 ; CA- CB- CG- HG2
1067 1069 1072 1074 3 0.66944 2.00832 0.00000
-2.67776 0.00000 0.00000 ; CA- CB- CG- HG3
1067 1069 1072 1075 3 3.68192 3.09616 -2.09200
-3.01248 0.00000 0.00000 ; CA- CB- CG- CD
1067 1085 1087 1088 3 20.92000 0.00000 -20.92000
0.00000 0.00000 0.00000 ; CA- C- N- H

1067 1085 1087 1089 3 20.92000 0.00000 -20.92000
0.00000 0.00000 0.00000 ; CA- C- N- CA
1068 1067 1069 1070 3 0.65084 1.95253 0.00000
-2.60338 0.00000 0.00000 ; HA- CA- CB- HB2
1068 1067 1069 1071 3 0.65084 1.95253 0.00000
-2.60338 0.00000 0.00000 ; HA- CA- CB- HB3
1068 1067 1069 1072 3 0.65084 1.95253 0.00000
-2.60338 0.00000 0.00000 ; HA- CA- CB- CG
1068 1067 1085 1086 3 3.68192 -4.35136 0.00000
1.33888 0.00000 0.00000 ; HA- CA- C- O
1068 1067 1085 1087 3 0.00000 0.00000 0.00000
0.00000 0.00000 0.00000 ; HA- CA- C- N
1069 1067 1085 1086 3 0.00000 0.00000 0.00000
0.00000 0.00000 0.00000 ; CB- CA- C- O
1069 1067 1085 1087 3 2.51040 4.18400 1.67360
-6.69440 0.00000 0.00000 ; CB- CA- C- N
1069 1072 1075 1076 3 0.66944 2.00832 0.00000
-2.67776 0.00000 0.00000 ; CB- CG- CD- HD2
1069 1072 1075 1077 3 0.66944 2.00832 0.00000
-2.67776 0.00000 0.00000 ; CB- CG- CD- HD3
1069 1072 1075 1078 3 3.68192 3.09616 -2.09200
-3.01248 0.00000 0.00000 ; CB- CG- CD- CE
1070 1069 1067 1085 3 0.65084 1.95253 0.00000
-2.60338 0.00000 0.00000 ; HB2- CB- CA- C
1070 1069 1072 1073 3 0.62760 1.88280 0.00000
-2.51040 0.00000 0.00000 ; HB2- CB- CG- HG2
1070 1069 1072 1074 3 0.62760 1.88280 0.00000
-2.51040 0.00000 0.00000 ; HB2- CB- CG- HG3
1070 1069 1072 1075 3 0.66944 2.00832 0.00000
-2.67776 0.00000 0.00000 ; HB2- CB- CG- CD
1071 1069 1067 1085 3 0.65084 1.95253 0.00000
-2.60338 0.00000 0.00000 ; HB3- CB- CA- C
1071 1069 1072 1073 3 0.62760 1.88280 0.00000
-2.51040 0.00000 0.00000 ; HB3- CB- CG- HG2
1071 1069 1072 1074 3 0.62760 1.88280 0.00000
-2.51040 0.00000 0.00000 ; HB3- CB- CG- HG3
1071 1069 1072 1075 3 0.66944 2.00832 0.00000
-2.67776 0.00000 0.00000 ; HB3- CB- CG- CD
1072 1069 1067 1085 3 0.65084 1.95253 0.00000
-2.60338 0.00000 0.00000 ; CG- CB- CA- C
1072 1075 1078 1079 3 0.65084 1.95253 0.00000
-2.60338 0.00000 0.00000 ; CG- CD- CE- HE2
1072 1075 1078 1080 3 0.65084 1.95253 0.00000
-2.60338 0.00000 0.00000 ; CG- CD- CE- HE3
1072 1075 1078 1081 3 0.65084 1.95253 0.00000
-2.60338 0.00000 0.00000 ; CG- CD- CE- NZ
1073 1072 1075 1076 3 0.62760 1.88280 0.00000
-2.51040 0.00000 0.00000 ; HG2- CG- CD- HD2
1073 1072 1075 1077 3 0.62760 1.88280 0.00000
-2.51040 0.00000 0.00000 ; HG2- CG- CD- HD3
1073 1072 1075 1078 3 0.66944 2.00832 0.00000
-2.67776 0.00000 0.00000 ; HG2- CG- CD- CE

1074 1072 1075 1076 3 0.62760 1.88280 0.00000
-2.51040 0.00000 0.00000 ; HG3- CG- CD- HD2
1074 1072 1075 1077 3 0.62760 1.88280 0.00000
-2.51040 0.00000 0.00000 ; HG3- CG- CD- HD3
1074 1072 1075 1078 3 0.66944 2.00832 0.00000
-2.67776 0.00000 0.00000 ; HG3- CG- CD- CE
1075 1078 1081 1082 3 0.65084 1.95253 0.00000
-2.60338 0.00000 0.00000 ; CD- CE- NZ- HZ1
1075 1078 1081 1083 3 0.65084 1.95253 0.00000
-2.60338 0.00000 0.00000 ; CD- CE- NZ- HZ2
1075 1078 1081 1084 3 0.65084 1.95253 0.00000
-2.60338 0.00000 0.00000 ; CD- CE- NZ- HZ3
1076 1075 1078 1079 3 0.65084 1.95253 0.00000
-2.60338 0.00000 0.00000 ; HD2- CD- CE- HE2
1076 1075 1078 1080 3 0.65084 1.95253 0.00000
-2.60338 0.00000 0.00000 ; HD2- CD- CE- HE3
1076 1075 1078 1081 3 0.65084 1.95253 0.00000
-2.60338 0.00000 0.00000 ; HD2- CD- CE- NZ
1077 1075 1078 1079 3 0.65084 1.95253 0.00000
-2.60338 0.00000 0.00000 ; HD3- CD- CE- HE2
1077 1075 1078 1080 3 0.65084 1.95253 0.00000
-2.60338 0.00000 0.00000 ; HD3- CD- CE- HE3
1077 1075 1078 1081 3 0.65084 1.95253 0.00000
-2.60338 0.00000 0.00000 ; HD3- CD- CE- NZ
1079 1078 1081 1082 3 0.65084 1.95253 0.00000
-2.60338 0.00000 0.00000 ; HE2- CE- NZ- HZ1
1079 1078 1081 1083 3 0.65084 1.95253 0.00000
-2.60338 0.00000 0.00000 ; HE2- CE- NZ- HZ2
1079 1078 1081 1084 3 0.65084 1.95253 0.00000
-2.60338 0.00000 0.00000 ; HE2- CE- NZ- HZ3
1080 1078 1081 1082 3 0.65084 1.95253 0.00000
-2.60338 0.00000 0.00000 ; HE3- CE- NZ- HZ1
1080 1078 1081 1083 3 0.65084 1.95253 0.00000
-2.60338 0.00000 0.00000 ; HE3- CE- NZ- HZ2
1080 1078 1081 1084 3 0.65084 1.95253 0.00000
-2.60338 0.00000 0.00000 ; HE3- CE- NZ- HZ3
1085 1087 1089 1090 3 0.00000 0.00000 0.00000
0.00000 0.00000 0.00000 ; C- N- CA- HA
1085 1087 1089 1091 3 11.71520 1.67360 15.06240 -
13.38880 0.00000 0.00000 ; C- N- CA- CB
1085 1087 1089 1104 3 1.75728 5.27184 2.25936
-7.02912 0.00000 0.00000 ; C- N- CA- C
1086 1085 1087 1088 3 29.28800 -8.36800 -20.92000
0.00000 0.00000 0.00000 ; O- C- N- H
1086 1085 1087 1089 3 20.92000 0.00000 -20.92000
0.00000 0.00000 0.00000 ; O- C- N- CA
1087 1089 1091 1092 3 0.65084 1.95253 0.00000
-2.60338 0.00000 0.00000 ; N- CA- CB- HB2
1087 1089 1091 1093 3 0.65084 1.95253 0.00000
-2.60338 0.00000 0.00000 ; N- CA- CB- HB3
1087 1089 1091 1094 3 3.79070 1.39746 -4.61077
-2.40998 2.44346 0.00000 ; N- CA- CB- CG

```
  1087   1089   1104   1105      3    0.00000    0.00000    0.00000
0.00000    0.00000    0.00000 ;      N-    CA-     C-     O
  1087   1089   1104   1106      3   17.40544   -5.02080  -13.22144
9.20480    0.00000    0.00000 ;      N-    CA-     C-     N
  1088   1087   1089   1090      3    0.00000    0.00000    0.00000
0.00000    0.00000    0.00000 ;      H-     N-    CA-    HA
  1088   1087   1089   1091      3    0.00000    0.00000    0.00000
0.00000    0.00000    0.00000 ;      H-     N-    CA-    CB
  1088   1087   1089   1104      3    0.00000    0.00000    0.00000
0.00000    0.00000    0.00000 ;      H-     N-    CA-     C
  1089   1091   1094   1095      3    0.66944    2.00832    0.00000
-2.67776    0.00000    0.00000 ;     CA-    CB-    CG-    HG
  1089   1091   1094   1096      3    3.90367    0.19665   -6.21742
-2.37651    5.99149    0.00000 ;     CA-    CB-    CG-   CD1
  1089   1091   1094   1100      3    3.90367    0.19665   -6.21742
-2.37651    5.99149    0.00000 ;     CA-    CB-    CG-   CD2
  1089   1104   1106   1107      3   20.92000    0.00000  -20.92000
0.00000    0.00000    0.00000 ;     CA-     C-     N-     H
  1089   1104   1106   1108      3   20.92000    0.00000  -20.92000
0.00000    0.00000    0.00000 ;     CA-     C-     N-    CA
  1090   1089   1091   1092      3    0.65084    1.95253    0.00000
-2.60338    0.00000    0.00000 ;     HA-    CA-    CB-   HB2
  1090   1089   1091   1093      3    0.65084    1.95253    0.00000
-2.60338    0.00000    0.00000 ;     HA-    CA-    CB-   HB3
  1090   1089   1091   1094      3    0.65084    1.95253    0.00000
-2.60338    0.00000    0.00000 ;     HA-    CA-    CB-    CG
  1090   1089   1104   1105      3    3.68192   -4.35136    0.00000
1.33888    0.00000    0.00000 ;     HA-    CA-     C-     O
  1090   1089   1104   1106      3    0.00000    0.00000    0.00000
0.00000    0.00000    0.00000 ;     HA-    CA-     C-     N
  1091   1089   1104   1105      3    0.00000    0.00000    0.00000
0.00000    0.00000    0.00000 ;     CB-    CA-     C-     O
  1091   1089   1104   1106      3    2.51040    4.18400    1.67360
-6.69440    0.00000    0.00000 ;     CB-    CA-     C-     N
  1091   1094   1096   1097      3    0.66944    2.00832    0.00000
-2.67776    0.00000    0.00000 ;     CB-    CG-   CD1-  HD11
  1091   1094   1096   1098      3    0.66944    2.00832    0.00000
-2.67776    0.00000    0.00000 ;     CB-    CG-   CD1-  HD12
  1091   1094   1096   1099      3    0.66944    2.00832    0.00000
-2.67776    0.00000    0.00000 ;     CB-    CG-   CD1-  HD13
  1091   1094   1100   1101      3    0.66944    2.00832    0.00000
-2.67776    0.00000    0.00000 ;     CB-    CG-   CD2-  HD21
  1091   1094   1100   1102      3    0.66944    2.00832    0.00000
-2.67776    0.00000    0.00000 ;     CB-    CG-   CD2-  HD22
  1091   1094   1100   1103      3    0.66944    2.00832    0.00000
-2.67776    0.00000    0.00000 ;     CB-    CG-   CD2-  HD23
  1092   1091   1089   1104      3    0.65084    1.95253    0.00000
-2.60338    0.00000    0.00000 ;    HB2-    CB-    CA-     C
  1092   1091   1094   1095      3    0.62760    1.88280    0.00000
-2.51040    0.00000    0.00000 ;    HB2-    CB-    CG-    HG
  1092   1091   1094   1096      3    0.66944    2.00832    0.00000
-2.67776    0.00000    0.00000 ;    HB2-    CB-    CG-   CD1
```

1092 1091 1094 1100 3 0.66944 2.00832 0.00000
-2.67776 0.00000 0.00000 ; HB2- CB- CG- CD2
1093 1091 1089 1104 3 0.65084 1.95253 0.00000
-2.60338 0.00000 0.00000 ; HB3- CB- CA- C
1093 1091 1094 1095 3 0.62760 1.88280 0.00000
-2.51040 0.00000 0.00000 ; HB3- CB- CG- HG
1093 1091 1094 1096 3 0.66944 2.00832 0.00000
-2.67776 0.00000 0.00000 ; HB3- CB- CG- CD1
1093 1091 1094 1100 3 0.66944 2.00832 0.00000
-2.67776 0.00000 0.00000 ; HB3- CB- CG- CD2
1094 1091 1089 1104 3 10.33448 -1.14642 -11.54784
-2.40998 6.35968 0.00000 ; CG- CB- CA- C
1095 1094 1096 1097 3 0.62760 1.88280 0.00000
-2.51040 0.00000 0.00000 ; HG- CG- CD1- HD11
1095 1094 1096 1098 3 0.62760 1.88280 0.00000
-2.51040 0.00000 0.00000 ; HG- CG- CD1- HD12
1095 1094 1096 1099 3 0.62760 1.88280 0.00000
-2.51040 0.00000 0.00000 ; HG- CG- CD1- HD13
1095 1094 1100 1101 3 0.62760 1.88280 0.00000
-2.51040 0.00000 0.00000 ; HG- CG- CD2- HD21
1095 1094 1100 1102 3 0.62760 1.88280 0.00000
-2.51040 0.00000 0.00000 ; HG- CG- CD2- HD22
1095 1094 1100 1103 3 0.62760 1.88280 0.00000
-2.51040 0.00000 0.00000 ; HG- CG- CD2- HD23
1096 1094 1100 1101 3 0.66944 2.00832 0.00000
-2.67776 0.00000 0.00000 ; CD1- CG- CD2- HD21
1096 1094 1100 1102 3 0.66944 2.00832 0.00000
-2.67776 0.00000 0.00000 ; CD1- CG- CD2- HD22
1096 1094 1100 1103 3 0.66944 2.00832 0.00000
-2.67776 0.00000 0.00000 ; CD1- CG- CD2- HD23
1097 1096 1094 1100 3 0.66944 2.00832 0.00000
-2.67776 0.00000 0.00000 ; HD11- CD1- CG- CD2
1098 1096 1094 1100 3 0.66944 2.00832 0.00000
-2.67776 0.00000 0.00000 ; HD12- CD1- CG- CD2
1099 1096 1094 1100 3 0.66944 2.00832 0.00000
-2.67776 0.00000 0.00000 ; HD13- CD1- CG- CD2
1104 1106 1108 1109 3 0.00000 0.00000 0.00000
0.00000 0.00000 0.00000 ; C- N- CA- HA
1104 1106 1108 1110 3 11.71520 1.67360 15.06240 -
13.38880 0.00000 0.00000 ; C- N- CA- CB
1104 1106 1108 1116 3 1.75728 5.27184 2.25936
-7.02912 0.00000 0.00000 ; C- N- CA- C
1105 1104 1106 1107 3 29.28800 -8.36800 -20.92000
0.00000 0.00000 0.00000 ; O- C- N- H
1105 1104 1106 1108 3 20.92000 0.00000 -20.92000
0.00000 0.00000 0.00000 ; O- C- N- CA
1106 1108 1110 1111 3 0.65084 1.95253 0.00000
-2.60338 0.00000 0.00000 ; N- CA- CB- HB2
1106 1108 1110 1112 3 0.65084 1.95253 0.00000
-2.60338 0.00000 0.00000 ; N- CA- CB- HB3
1106 1108 1110 1113 3 15.41386 9.74035 -8.39310
-0.97069 2.97901 0.00000 ; N- CA- CB- CG

1106 1108 1116 1117 3 0.00000 0.00000 0.00000
0.00000 0.00000 0.00000 ; N- CA- C- O
1106 1108 1116 1118 3 17.40544 -5.02080 -13.22144
9.20480 0.00000 0.00000 ; N- CA- C- N
1107 1106 1108 1109 3 0.00000 0.00000 0.00000
0.00000 0.00000 0.00000 ; H- N- CA- HA
1107 1106 1108 1110 3 0.00000 0.00000 0.00000
0.00000 0.00000 0.00000 ; H- N- CA- CB
1107 1106 1108 1116 3 0.00000 0.00000 0.00000
0.00000 0.00000 0.00000 ; H- N- CA- C
1108 1110 1113 1114 3 6.43499 0.00000 -5.39736
0.00000 -1.03763 0.00000 ; CA- CB- CG- OD1
1108 1110 1113 1115 3 6.43499 0.00000 -5.39736
0.00000 -1.03763 0.00000 ; CA- CB- CG- OD2
1108 1116 1118 1119 3 20.92000 0.00000 -20.92000
0.00000 0.00000 0.00000 ; CA- C- N- H
1108 1116 1118 1120 3 20.92000 0.00000 -20.92000
0.00000 0.00000 0.00000 ; CA- C- N- CA
1109 1108 1110 1111 3 0.65084 1.95253 0.00000
-2.60338 0.00000 0.00000 ; HA- CA- CB- HB2
1109 1108 1110 1112 3 0.65084 1.95253 0.00000
-2.60338 0.00000 0.00000 ; HA- CA- CB- HB3
1109 1108 1110 1113 3 0.65084 1.95253 0.00000
-2.60338 0.00000 0.00000 ; HA- CA- CB- CG
1109 1108 1116 1117 3 3.68192 -4.35136 0.00000
1.33888 0.00000 0.00000 ; HA- CA- C- O
1109 1108 1116 1118 3 0.00000 0.00000 0.00000
0.00000 0.00000 0.00000 ; HA- CA- C- N
1110 1108 1116 1117 3 0.00000 0.00000 0.00000
0.00000 0.00000 0.00000 ; CB- CA- C- O
1110 1108 1116 1118 3 2.51040 4.18400 1.67360
-6.69440 0.00000 0.00000 ; CB- CA- C- N
1111 1110 1108 1116 3 0.65084 1.95253 0.00000
-2.60338 0.00000 0.00000 ; HB2- CB- CA- C
1111 1110 1113 1114 3 0.00000 0.00000 0.00000
0.00000 0.00000 0.00000 ; HB2- CB- CG- OD1
1111 1110 1113 1115 3 0.00000 0.00000 0.00000
0.00000 0.00000 0.00000 ; HB2- CB- CG- OD2
1112 1110 1108 1116 3 0.65084 1.95253 0.00000
-2.60338 0.00000 0.00000 ; HB3- CB- CA- C
1112 1110 1113 1114 3 0.00000 0.00000 0.00000
0.00000 0.00000 0.00000 ; HB3- CB- CG- OD1
1112 1110 1113 1115 3 0.00000 0.00000 0.00000
0.00000 0.00000 0.00000 ; HB3- CB- CG- OD2
1113 1110 1108 1116 3 7.14627 -1.04600 -8.99560
-0.97069 5.15469 0.00000 ; CG- CB- CA- C
1116 1118 1120 1121 3 0.00000 0.00000 0.00000
0.00000 0.00000 0.00000 ; C- N- CA- HA
1116 1118 1120 1122 3 11.71520 1.67360 15.06240 -
13.38880 0.00000 0.00000 ; C- N- CA- CB
1116 1118 1120 1133 3 1.75728 5.27184 2.25936
-7.02912 0.00000 0.00000 ; C- N- CA- C

1117 1116 1118 1119 3 29.28800 -8.36800 -20.92000
0.00000 0.00000 0.00000 ; O- C- N- H
1117 1116 1118 1120 3 20.92000 0.00000 -20.92000
0.00000 0.00000 0.00000 ; O- C- N- CA
1118 1120 1122 1123 3 0.65084 1.95253 0.00000
-2.60338 0.00000 0.00000 ; N- CA- CB- HB2
1118 1120 1122 1124 3 0.65084 1.95253 0.00000
-2.60338 0.00000 0.00000 ; N- CA- CB- HB3
1118 1120 1122 1125 3 4.79068 4.02919 -4.82834
-3.66518 2.97901 0.00000 ; N- CA- CB- CG
1118 1120 1133 1134 3 0.00000 0.00000 0.00000
0.00000 0.00000 0.00000 ; N- CA- C- O
1118 1120 1133 1135 3 17.40544 -5.02080 -13.22144
9.20480 0.00000 0.00000 ; N- CA- C- N
1119 1118 1120 1121 3 0.00000 0.00000 0.00000
0.00000 0.00000 0.00000 ; H- N- CA- HA
1119 1118 1120 1122 3 0.00000 0.00000 0.00000
0.00000 0.00000 0.00000 ; H- N- CA- CB
1119 1118 1120 1133 3 0.00000 0.00000 0.00000
0.00000 0.00000 0.00000 ; H- N- CA- C
1120 1122 1125 1126 3 5.98312 6.40152 3.28026 -
12.38464 -1.57318 0.00000 ; CA- CB- CG- ND1
1120 1122 1125 1131 3 0.00000 0.00000 0.00000
0.00000 0.00000 0.00000 ; CA- CB- CG- CD2
1120 1133 1135 1136 3 20.92000 0.00000 -20.92000
0.00000 0.00000 0.00000 ; CA- C- N- CD
1120 1133 1135 1145 3 20.92000 0.00000 -20.92000
0.00000 0.00000 0.00000 ; CA- C- N- CA
1121 1120 1122 1123 3 0.65084 1.95253 0.00000
-2.60338 0.00000 0.00000 ; HA- CA- CB- HB2
1121 1120 1122 1124 3 0.65084 1.95253 0.00000
-2.60338 0.00000 0.00000 ; HA- CA- CB- HB3
1121 1120 1122 1125 3 0.65084 1.95253 0.00000
-2.60338 0.00000 0.00000 ; HA- CA- CB- CG
1121 1120 1133 1134 3 3.68192 -4.35136 0.00000
1.33888 0.00000 0.00000 ; HA- CA- C- O
1121 1120 1133 1135 3 0.00000 0.00000 0.00000
0.00000 0.00000 0.00000 ; HA- CA- C- N
1122 1120 1133 1134 3 0.00000 0.00000 0.00000
0.00000 0.00000 0.00000 ; CB- CA- C- O
1122 1120 1133 1135 3 2.51040 4.18400 1.67360
-6.69440 0.00000 0.00000 ; CB- CA- C- N
1122 1125 1126 1127 3 20.08320 0.00000 -20.08320
0.00000 0.00000 0.00000 ; CB- CG- ND1- CE1
1122 1125 1131 1129 3 44.97800 0.00000 -44.97800
0.00000 0.00000 0.00000 ; CB- CG- CD2- NE2
1122 1125 1131 1132 3 44.97800 0.00000 -44.97800
0.00000 0.00000 0.00000 ; CB- CG- CD2- HD2
1123 1122 1120 1133 3 0.65084 1.95253 0.00000
-2.60338 0.00000 0.00000 ; HB2- CB- CA- C
1123 1122 1125 1126 3 0.00000 0.00000 0.00000
0.00000 0.00000 0.00000 ; HB2- CB- CG- ND1

1123 1122 1125 1131 3 0.00000 0.00000 0.00000
0.00000 0.00000 0.00000 ; HB2- CB- CG- CD2
1124 1122 1120 1133 3 0.65084 1.95253 0.00000
-2.60338 0.00000 0.00000 ; HB3- CB- CA- C
1124 1122 1125 1126 3 0.00000 0.00000 0.00000
0.00000 0.00000 0.00000 ; HB3- CB- CG- ND1
1124 1122 1125 1131 3 0.00000 0.00000 0.00000
0.00000 0.00000 0.00000 ; HB3- CB- CG- CD2
1125 1122 1120 1133 3 3.76560 3.34720 -2.87859
-3.66518 0.83680 0.00000 ; CG- CB- CA- C
1125 1126 1127 1128 3 41.84000 0.00000 -41.84000
0.00000 0.00000 0.00000 ; CG- ND1- CE1- HE1
1125 1126 1127 1129 3 41.84000 0.00000 -41.84000
0.00000 0.00000 0.00000 ; CG- ND1- CE1- NE2
1125 1131 1129 1127 3 12.55200 0.00000 -12.55200
0.00000 0.00000 0.00000 ; CG- CD2- NE2- CE1
1125 1131 1129 1130 3 12.55200 0.00000 -12.55200
0.00000 0.00000 0.00000 ; CG- CD2- NE2- HE2
1126 1125 1131 1129 3 44.97800 0.00000 -44.97800
0.00000 0.00000 0.00000 ; ND1- CG- CD2- NE2
1126 1125 1131 1132 3 44.97800 0.00000 -44.97800
0.00000 0.00000 0.00000 ; ND1- CG- CD2- HD2
1126 1127 1129 1130 3 19.45560 0.00000 -19.45560
0.00000 0.00000 0.00000 ; ND1- CE1- NE2- HE2
1126 1127 1129 1131 3 19.45560 0.00000 -19.45560
0.00000 0.00000 0.00000 ; ND1- CE1- NE2- CD2
1127 1126 1125 1131 3 20.08320 0.00000 -20.08320
0.00000 0.00000 0.00000 ; CE1- ND1- CG- CD2
1127 1129 1131 1132 3 12.55200 0.00000 -12.55200
0.00000 0.00000 0.00000 ; CE1- NE2- CD2- HD2
1128 1127 1129 1130 3 19.45560 0.00000 -19.45560
0.00000 0.00000 0.00000 ; HE1- CE1- NE2- HE2
1128 1127 1129 1131 3 19.45560 0.00000 -19.45560
0.00000 0.00000 0.00000 ; HE1- CE1- NE2- CD2
1130 1129 1131 1132 3 12.55200 0.00000 -12.55200
0.00000 0.00000 0.00000 ; HE2- NE2- CD2- HD2
1133 1135 1136 1137 3 0.00000 0.00000 0.00000
0.00000 0.00000 0.00000 ; C- N- CD- HD2
1133 1135 1136 1138 3 0.00000 0.00000 0.00000
0.00000 0.00000 0.00000 ; C- N- CD- HD3
1133 1135 1136 1139 3 10.04160 -3.34720 16.73600
-6.69440 0.00000 0.00000 ; C- N- CD- CG
1133 1135 1145 1142 3 11.71520 1.67360 15.06240 -
13.38880 0.00000 0.00000 ; C- N- CA- CB
1133 1135 1145 1146 3 0.00000 0.00000 0.00000
0.00000 0.00000 0.00000 ; C- N- CA- HA
1133 1135 1145 1147 3 1.75728 5.27184 2.25936
-7.02912 0.00000 0.00000 ; C- N- CA- C
1134 1133 1135 1136 3 20.92000 0.00000 -20.92000
0.00000 0.00000 0.00000 ; O- C- N- CD
1134 1133 1135 1145 3 20.92000 0.00000 -20.92000
0.00000 0.00000 0.00000 ; O- C- N- CA

```
  1135   1136   1139   1140      3    0.65084    1.95253    0.00000
-2.60338    0.00000    0.00000 ;      N-    CD-    CG-   HG2
  1135   1136   1139   1141      3    0.65084    1.95253    0.00000
-2.60338    0.00000    0.00000 ;      N-    CD-    CG-   HG3
  1135   1136   1139   1142      3    0.65084    1.95253    0.00000
-2.60338    0.00000    0.00000 ;      N-    CD-    CG-    CB
  1135   1145   1142   1139      3    0.65084    1.95253    0.00000
-2.60338    0.00000    0.00000 ;      N-    CA-    CB-    CG
  1135   1145   1142   1143      3    0.65084    1.95253    0.00000
-2.60338    0.00000    0.00000 ;      N-    CA-    CB-   HB2
  1135   1145   1142   1144      3    0.65084    1.95253    0.00000
-2.60338    0.00000    0.00000 ;      N-    CA-    CB-   HB3
  1135   1145   1147   1148      3    0.00000    0.00000    0.00000
0.00000    0.00000    0.00000 ;      N-    CA-     C-     O
  1135   1145   1147   1149      3   17.40544   -5.02080  -13.22144
9.20480    0.00000    0.00000 ;      N-    CA-     C-     N
  1136   1135   1145   1142      3    0.00000    0.00000    0.00000
0.00000    0.00000    0.00000 ;     CD-     N-    CA-    CB
  1136   1135   1145   1146      3    0.00000    0.00000    0.00000
0.00000    0.00000    0.00000 ;     CD-     N-    CA-    HA
  1136   1135   1145   1147      3    0.00000    0.00000    0.00000
0.00000    0.00000    0.00000 ;     CD-     N-    CA-     C
  1136   1139   1142   1143      3    0.66944    2.00832    0.00000
-2.67776    0.00000    0.00000 ;     CD-    CG-    CB-   HB2
  1136   1139   1142   1144      3    0.66944    2.00832    0.00000
-2.67776    0.00000    0.00000 ;     CD-    CG-    CB-   HB3
  1136   1139   1142   1145      3    3.68192    3.09616   -2.09200
-3.01248    0.00000    0.00000 ;     CD-    CG-    CB-    CA
  1137   1136   1135   1145      3    0.00000    0.00000    0.00000
0.00000    0.00000    0.00000 ;    HD2-    CD-     N-    CA
  1137   1136   1139   1140      3    0.65084    1.95253    0.00000
-2.60338    0.00000    0.00000 ;    HD2-    CD-    CG-   HG2
  1137   1136   1139   1141      3    0.65084    1.95253    0.00000
-2.60338    0.00000    0.00000 ;    HD2-    CD-    CG-   HG3
  1137   1136   1139   1142      3    0.65084    1.95253    0.00000
-2.60338    0.00000    0.00000 ;    HD2-    CD-    CG-    CB
  1138   1136   1135   1145      3    0.00000    0.00000    0.00000
0.00000    0.00000    0.00000 ;    HD3-    CD-     N-    CA
  1138   1136   1139   1140      3    0.65084    1.95253    0.00000
-2.60338    0.00000    0.00000 ;    HD3-    CD-    CG-   HG2
  1138   1136   1139   1141      3    0.65084    1.95253    0.00000
-2.60338    0.00000    0.00000 ;    HD3-    CD-    CG-   HG3
  1138   1136   1139   1142      3    0.65084    1.95253    0.00000
-2.60338    0.00000    0.00000 ;    HD3-    CD-    CG-    CB
  1139   1136   1135   1145      3    0.00000    0.00000    0.00000
0.00000    0.00000    0.00000 ;     CG-    CD-     N-    CA
  1139   1142   1145   1146      3    0.65084    1.95253    0.00000
-2.60338    0.00000    0.00000 ;     CG-    CB-    CA-    HA
  1139   1142   1145   1147      3    0.65084    1.95253    0.00000
-2.60338    0.00000    0.00000 ;     CG-    CB-    CA-     C
  1140   1139   1142   1143      3    0.62760    1.88280    0.00000
-2.51040    0.00000    0.00000 ;    HG2-    CG-    CB-   HB2
```

1140 1139 1142 1144 3 0.62760 1.88280 0.00000
-2.51040 0.00000 0.00000 ; HG2- CG- CB- HB3
1140 1139 1142 1145 3 0.66944 2.00832 0.00000
-2.67776 0.00000 0.00000 ; HG2- CG- CB- CA
1141 1139 1142 1143 3 0.62760 1.88280 0.00000
-2.51040 0.00000 0.00000 ; HG3- CG- CB- HB2
1141 1139 1142 1144 3 0.62760 1.88280 0.00000
-2.51040 0.00000 0.00000 ; HG3- CG- CB- HB3
1141 1139 1142 1145 3 0.66944 2.00832 0.00000
-2.67776 0.00000 0.00000 ; HG3- CG- CB- CA
1142 1145 1147 1148 3 0.00000 0.00000 0.00000
0.00000 0.00000 0.00000 ; CB- CA- C- O
1142 1145 1147 1149 3 2.51040 4.18400 1.67360
-6.69440 0.00000 0.00000 ; CB- CA- C- N
1143 1142 1145 1146 3 0.65084 1.95253 0.00000
-2.60338 0.00000 0.00000 ; HB2- CB- CA- HA
1143 1142 1145 1147 3 0.65084 1.95253 0.00000
-2.60338 0.00000 0.00000 ; HB2- CB- CA- C
1144 1142 1145 1146 3 0.65084 1.95253 0.00000
-2.60338 0.00000 0.00000 ; HB3- CB- CA- HA
1144 1142 1145 1147 3 0.65084 1.95253 0.00000
-2.60338 0.00000 0.00000 ; HB3- CB- CA- C
1145 1147 1149 1150 3 20.92000 0.00000 -20.92000
0.00000 0.00000 0.00000 ; CA- C- N- H
1145 1147 1149 1151 3 20.92000 0.00000 -20.92000
0.00000 0.00000 0.00000 ; CA- C- N- CA
1146 1145 1147 1148 3 3.68192 -4.35136 0.00000
1.33888 0.00000 0.00000 ; HA- CA- C- O
1146 1145 1147 1149 3 0.00000 0.00000 0.00000
0.00000 0.00000 0.00000 ; HA- CA- C- N
1147 1149 1151 1152 3 0.00000 0.00000 0.00000
0.00000 0.00000 0.00000 ; C- N- CA- HA
1147 1149 1151 1153 3 11.71520 1.67360 15.06240 -
13.38880 0.00000 0.00000 ; C- N- CA- CB
1147 1149 1151 1161 3 1.75728 5.27184 2.25936
-7.02912 0.00000 0.00000 ; C- N- CA- C
1148 1147 1149 1150 3 29.28800 -8.36800 -20.92000
0.00000 0.00000 0.00000 ; O- C- N- H
1148 1147 1149 1151 3 20.92000 0.00000 -20.92000
0.00000 0.00000 0.00000 ; O- C- N- CA
1149 1151 1153 1154 3 0.65084 1.95253 0.00000
-2.60338 0.00000 0.00000 ; N- CA- CB- HB2
1149 1151 1153 1155 3 0.65084 1.95253 0.00000
-2.60338 0.00000 0.00000 ; N- CA- CB- HB3
1149 1151 1153 1156 3 5.99567 3.29281 -4.46014
-0.55229 1.97485 0.00000 ; N- CA- CB- CG
1149 1151 1161 1162 3 0.00000 0.00000 0.00000
0.00000 0.00000 0.00000 ; N- CA- C- O
1149 1151 1161 1163 3 17.40544 -5.02080 -13.22144
9.20480 0.00000 0.00000 ; N- CA- C- N
1150 1149 1151 1152 3 0.00000 0.00000 0.00000
0.00000 0.00000 0.00000 ; H- N- CA- HA

1150 1149 1151 1153 3 0.00000 0.00000 0.00000
0.00000 0.00000 0.00000 ; H- N- CA- CB
1150 1149 1151 1161 3 0.00000 0.00000 0.00000
0.00000 0.00000 0.00000 ; H- N- CA- C
1151 1153 1156 1157 3 0.00000 0.00000 0.00000
0.00000 0.00000 0.00000 ; CA- CB- CG- OD1
1151 1153 1156 1158 3 8.84916 -0.31380 -4.32626
5.03754 0.26778 0.00000 ; CA- CB- CG- ND2
1151 1161 1163 1164 3 20.92000 0.00000 -20.92000
0.00000 0.00000 0.00000 ; CA- C- N- H
1151 1161 1163 1165 3 20.92000 0.00000 -20.92000
0.00000 0.00000 0.00000 ; CA- C- N- CA
1152 1151 1153 1154 3 0.65084 1.95253 0.00000
-2.60338 0.00000 0.00000 ; HA- CA- CB- HB2
1152 1151 1153 1155 3 0.65084 1.95253 0.00000
-2.60338 0.00000 0.00000 ; HA- CA- CB- HB3
1152 1151 1153 1156 3 0.65084 1.95253 0.00000
-2.60338 0.00000 0.00000 ; HA- CA- CB- CG
1152 1151 1161 1162 3 3.68192 -4.35136 0.00000
1.33888 0.00000 0.00000 ; HA- CA- C- O
1152 1151 1161 1163 3 0.00000 0.00000 0.00000
0.00000 0.00000 0.00000 ; HA- CA- C- N
1153 1151 1161 1162 3 0.00000 0.00000 0.00000
0.00000 0.00000 0.00000 ; CB- CA- C- O
1153 1151 1161 1163 3 2.51040 4.18400 1.67360
-6.69440 0.00000 0.00000 ; CB- CA- C- N
1153 1156 1158 1159 3 20.92000 0.00000 -20.92000
0.00000 0.00000 0.00000 ; CB- CG- ND2- HD21
1153 1156 1158 1160 3 20.92000 0.00000 -20.92000
0.00000 0.00000 0.00000 ; CB- CG- ND2- HD22
1154 1153 1151 1161 3 0.65084 1.95253 0.00000
-2.60338 0.00000 0.00000 ; HB2- CB- CA- C
1154 1153 1156 1157 3 3.68192 -4.35136 0.00000
1.33888 0.00000 0.00000 ; HB2- CB- CG- OD1
1154 1153 1156 1158 3 0.00000 0.00000 0.00000
0.00000 0.00000 0.00000 ; HB2- CB- CG- ND2
1155 1153 1151 1161 3 0.65084 1.95253 0.00000
-2.60338 0.00000 0.00000 ; HB3- CB- CA- C
1155 1153 1156 1157 3 3.68192 -4.35136 0.00000
1.33888 0.00000 0.00000 ; HB3- CB- CG- OD1
1155 1153 1156 1158 3 0.00000 0.00000 0.00000
0.00000 0.00000 0.00000 ; HB3- CB- CG- ND2
1156 1153 1151 1161 3 7.94542 -3.96225 -6.11701
-0.55229 3.58150 0.00000 ; CG- CB- CA- C
1157 1156 1158 1159 3 29.28800 -8.36800 -20.92000
0.00000 0.00000 0.00000 ; OD1- CG- ND2- HD21
1157 1156 1158 1160 3 29.28800 -8.36800 -20.92000
0.00000 0.00000 0.00000 ; OD1- CG- ND2- HD22
1161 1163 1165 1166 3 0.00000 0.00000 0.00000
0.00000 0.00000 0.00000 ; C- N- CA- HA
1161 1163 1165 1167 3 11.71520 1.67360 15.06240 -
13.38880 0.00000 0.00000 ; C- N- CA- CB

1161 1163 1165 1180 3 1.75728 5.27184 2.25936
-7.02912 0.00000 0.00000 ; C- N- CA- C
1162 1161 1163 1164 3 29.28800 -8.36800 -20.92000
0.00000 0.00000 0.00000 ; O- C- N- H
1162 1161 1163 1165 3 20.92000 0.00000 -20.92000
0.00000 0.00000 0.00000 ; O- C- N- CA
1163 1165 1167 1168 3 0.65084 1.95253 0.00000
-2.60338 0.00000 0.00000 ; N- CA- CB- HB
1163 1165 1167 1169 3 3.83673 0.44769 -1.77402
-2.47693 -0.03347 0.00000 ; N- CA- CB- CG2
1163 1165 1167 1173 3 2.97482 0.12134 2.04179
-1.89117 -3.24678 0.00000 ; N- CA- CB- CG1
1163 1165 1180 1181 3 0.00000 0.00000 0.00000
0.00000 0.00000 0.00000 ; N- CA- C- O
1163 1165 1180 1182 3 17.40544 -5.02080 -13.22144
9.20480 0.00000 0.00000 ; N- CA- C- N
1164 1163 1165 1166 3 0.00000 0.00000 0.00000
0.00000 0.00000 0.00000 ; H- N- CA- HA
1164 1163 1165 1167 3 0.00000 0.00000 0.00000
0.00000 0.00000 0.00000 ; H- N- CA- CB
1164 1163 1165 1180 3 0.00000 0.00000 0.00000
0.00000 0.00000 0.00000 ; H- N- CA- C
1165 1167 1169 1170 3 0.66944 2.00832 0.00000
-2.67776 0.00000 0.00000 ; CA- CB- CG2- HG21
1165 1167 1169 1171 3 0.66944 2.00832 0.00000
-2.67776 0.00000 0.00000 ; CA- CB- CG2- HG22
1165 1167 1169 1172 3 0.66944 2.00832 0.00000
-2.67776 0.00000 0.00000 ; CA- CB- CG2- HG23
1165 1167 1173 1174 3 0.66944 2.00832 0.00000
-2.67776 0.00000 0.00000 ; CA- CB- CG1- HG12
1165 1167 1173 1175 3 0.66944 2.00832 0.00000
-2.67776 0.00000 0.00000 ; CA- CB- CG1- HG13
1165 1167 1173 1176 3 4.24258 -0.52718 -7.25506
-1.79075 7.69856 0.00000 ; CA- CB- CG1- CD1
1165 1180 1182 1183 3 20.92000 0.00000 -20.92000
0.00000 0.00000 0.00000 ; CA- C- N- H
1165 1180 1182 1184 3 20.92000 0.00000 -20.92000
0.00000 0.00000 0.00000 ; CA- C- N- CA
1166 1165 1167 1168 3 0.65084 1.95253 0.00000
-2.60338 0.00000 0.00000 ; HA- CA- CB- HB
1166 1165 1167 1169 3 0.65084 1.95253 0.00000
-2.60338 0.00000 0.00000 ; HA- CA- CB- CG2
1166 1165 1167 1173 3 0.65084 1.95253 0.00000
-2.60338 0.00000 0.00000 ; HA- CA- CB- CG1
1166 1165 1180 1181 3 3.68192 -4.35136 0.00000
1.33888 0.00000 0.00000 ; HA- CA- C- O
1166 1165 1180 1182 3 0.00000 0.00000 0.00000
0.00000 0.00000 0.00000 ; HA- CA- C- N
1167 1165 1180 1181 3 0.00000 0.00000 0.00000
0.00000 0.00000 0.00000 ; CB- CA- C- O
1167 1165 1180 1182 3 2.51040 4.18400 1.67360
-6.69440 0.00000 0.00000 ; CB- CA- C- N

1167 1173 1176 1177 3 0.66944 2.00832 0.00000
-2.67776 0.00000 0.00000 ; CB- CG1- CD1- HD11
1167 1173 1176 1178 3 0.66944 2.00832 0.00000
-2.67776 0.00000 0.00000 ; CB- CG1- CD1- HD12
1167 1173 1176 1179 3 0.66944 2.00832 0.00000
-2.67776 0.00000 0.00000 ; CB- CG1- CD1- HD13
1168 1167 1165 1180 3 0.65084 1.95253 0.00000
-2.60338 0.00000 0.00000 ; HB- CB- CA- C
1168 1167 1169 1170 3 0.62760 1.88280 0.00000
-2.51040 0.00000 0.00000 ; HB- CB- CG2- HG21
1168 1167 1169 1171 3 0.62760 1.88280 0.00000
-2.51040 0.00000 0.00000 ; HB- CB- CG2- HG22
1168 1167 1169 1172 3 0.62760 1.88280 0.00000
-2.51040 0.00000 0.00000 ; HB- CB- CG2- HG23
1168 1167 1173 1174 3 0.62760 1.88280 0.00000
-2.51040 0.00000 0.00000 ; HB- CB- CG1- HG12
1168 1167 1173 1175 3 0.62760 1.88280 0.00000
-2.51040 0.00000 0.00000 ; HB- CB- CG1- HG13
1168 1167 1173 1176 3 0.66944 2.00832 0.00000
-2.67776 0.00000 0.00000 ; HB- CB- CG1- CD1
1169 1167 1165 1180 3 5.67350 3.55640 -6.16722
-2.47693 3.74886 0.00000 ; CG2- CB- CA- C
1169 1167 1173 1174 3 0.66944 2.00832 0.00000
-2.67776 0.00000 0.00000 ; CG2- CB- CG1- HG12
1169 1167 1173 1175 3 0.66944 2.00832 0.00000
-2.67776 0.00000 0.00000 ; CG2- CB- CG1- HG13
1169 1167 1173 1176 3 3.81162 0.49790 -8.14206
-1.79075 7.49773 0.00000 ; CG2- CB- CG1- CD1
1170 1169 1167 1173 3 0.66944 2.00832 0.00000
-2.67776 0.00000 0.00000 ; HG21- CG2- CB- CG1
1171 1169 1167 1173 3 0.66944 2.00832 0.00000
-2.67776 0.00000 0.00000 ; HG22- CG2- CB- CG1
1172 1169 1167 1173 3 0.66944 2.00832 0.00000
-2.67776 0.00000 0.00000 ; HG23- CG2- CB- CG1
1173 1167 1165 1180 3 8.26340 0.74057 -9.99976
-1.89117 3.84928 0.00000 ; CG1- CB- CA- C
1174 1173 1176 1177 3 0.62760 1.88280 0.00000
-2.51040 0.00000 0.00000 ; HG12- CG1- CD1- HD11
1174 1173 1176 1178 3 0.62760 1.88280 0.00000
-2.51040 0.00000 0.00000 ; HG12- CG1- CD1- HD12
1174 1173 1176 1179 3 0.62760 1.88280 0.00000
-2.51040 0.00000 0.00000 ; HG12- CG1- CD1- HD13
1175 1173 1176 1177 3 0.62760 1.88280 0.00000
-2.51040 0.00000 0.00000 ; HG13- CG1- CD1- HD11
1175 1173 1176 1178 3 0.62760 1.88280 0.00000
-2.51040 0.00000 0.00000 ; HG13- CG1- CD1- HD12
1175 1173 1176 1179 3 0.62760 1.88280 0.00000
-2.51040 0.00000 0.00000 ; HG13- CG1- CD1- HD13
1180 1182 1184 1185 3 0.00000 0.00000 0.00000
0.00000 0.00000 0.00000 ; C- N- CA- HA
1180 1182 1184 1186 3 11.71520 1.67360 15.06240 -
13.38880 0.00000 0.00000 ; C- N- CA- CB

1180 1182 1184 1196 3 1.75728 5.27184 2.25936
-7.02912 0.00000 0.00000 ; C- N- CA- C
1181 1180 1182 1183 3 29.28800 -8.36800 -20.92000
0.00000 0.00000 0.00000 ; O- C- N- H
1181 1180 1182 1184 3 20.92000 0.00000 -20.92000
0.00000 0.00000 0.00000 ; O- C- N- CA
1182 1184 1186 1187 3 0.65084 1.95253 0.00000
-2.60338 0.00000 0.00000 ; N- CA- CB- HB
1182 1184 1186 1188 3 3.83673 0.44769 -1.77402
-2.47693 -0.03347 0.00000 ; N- CA- CB- CG1
1182 1184 1186 1192 3 3.83673 0.44769 -1.77402
-2.47693 -0.03347 0.00000 ; N- CA- CB- CG2
1182 1184 1196 1197 3 0.00000 0.00000 0.00000
0.00000 0.00000 0.00000 ; N- CA- C- O
1182 1184 1196 1198 3 17.40544 -5.02080 -13.22144
9.20480 0.00000 0.00000 ; N- CA- C- N
1183 1182 1184 1185 3 0.00000 0.00000 0.00000
0.00000 0.00000 0.00000 ; H- N- CA- HA
1183 1182 1184 1186 3 0.00000 0.00000 0.00000
0.00000 0.00000 0.00000 ; H- N- CA- CB
1183 1182 1184 1196 3 0.00000 0.00000 0.00000
0.00000 0.00000 0.00000 ; H- N- CA- C
1184 1186 1188 1189 3 0.66944 2.00832 0.00000
-2.67776 0.00000 0.00000 ; CA- CB- CG1- HG11
1184 1186 1188 1190 3 0.66944 2.00832 0.00000
-2.67776 0.00000 0.00000 ; CA- CB- CG1- HG12
1184 1186 1188 1191 3 0.66944 2.00832 0.00000
-2.67776 0.00000 0.00000 ; CA- CB- CG1- HG13
1184 1186 1192 1193 3 0.66944 2.00832 0.00000
-2.67776 0.00000 0.00000 ; CA- CB- CG2- HG21
1184 1186 1192 1194 3 0.66944 2.00832 0.00000
-2.67776 0.00000 0.00000 ; CA- CB- CG2- HG22
1184 1186 1192 1195 3 0.66944 2.00832 0.00000
-2.67776 0.00000 0.00000 ; CA- CB- CG2- HG23
1184 1196 1198 1199 3 20.92000 0.00000 -20.92000
0.00000 0.00000 0.00000 ; CA- C- N- H
1184 1196 1198 1200 3 20.92000 0.00000 -20.92000
0.00000 0.00000 0.00000 ; CA- C- N- CA
1185 1184 1186 1187 3 0.65084 1.95253 0.00000
-2.60338 0.00000 0.00000 ; HA- CA- CB- HB
1185 1184 1186 1188 3 0.65084 1.95253 0.00000
-2.60338 0.00000 0.00000 ; HA- CA- CB- CG1
1185 1184 1186 1192 3 0.65084 1.95253 0.00000
-2.60338 0.00000 0.00000 ; HA- CA- CB- CG2
1185 1184 1196 1197 3 3.68192 -4.35136 0.00000
1.33888 0.00000 0.00000 ; HA- CA- C- O
1185 1184 1196 1198 3 0.00000 0.00000 0.00000
0.00000 0.00000 0.00000 ; HA- CA- C- N
1186 1184 1196 1197 3 0.00000 0.00000 0.00000
0.00000 0.00000 0.00000 ; CB- CA- C- O
1186 1184 1196 1198 3 2.51040 4.18400 1.67360
-6.69440 0.00000 0.00000 ; CB- CA- C- N

1187 1186 1184 1196 3 0.65084 1.95253 0.00000
-2.60338 0.00000 0.00000 ; HB- CB- CA- C
1187 1186 1188 1189 3 0.62760 1.88280 0.00000
-2.51040 0.00000 0.00000 ; HB- CB- CG1- HG11
1187 1186 1188 1190 3 0.62760 1.88280 0.00000
-2.51040 0.00000 0.00000 ; HB- CB- CG1- HG12
1187 1186 1188 1191 3 0.62760 1.88280 0.00000
-2.51040 0.00000 0.00000 ; HB- CB- CG1- HG13
1187 1186 1192 1193 3 0.62760 1.88280 0.00000
-2.51040 0.00000 0.00000 ; HB- CB- CG2- HG21
1187 1186 1192 1194 3 0.62760 1.88280 0.00000
-2.51040 0.00000 0.00000 ; HB- CB- CG2- HG22
1187 1186 1192 1195 3 0.62760 1.88280 0.00000
-2.51040 0.00000 0.00000 ; HB- CB- CG2- HG23
1188 1186 1184 1196 3 5.67350 3.55640 -6.16722
-2.47693 3.74886 0.00000 ; CG1- CB- CA- C
1188 1186 1192 1193 3 0.66944 2.00832 0.00000
-2.67776 0.00000 0.00000 ; CG1- CB- CG2- HG21
1188 1186 1192 1194 3 0.66944 2.00832 0.00000
-2.67776 0.00000 0.00000 ; CG1- CB- CG2- HG22
1188 1186 1192 1195 3 0.66944 2.00832 0.00000
-2.67776 0.00000 0.00000 ; CG1- CB- CG2- HG23
1189 1188 1186 1192 3 0.66944 2.00832 0.00000
-2.67776 0.00000 0.00000 ; HG11- CG1- CB- CG2
1190 1188 1186 1192 3 0.66944 2.00832 0.00000
-2.67776 0.00000 0.00000 ; HG12- CG1- CB- CG2
1191 1188 1186 1192 3 0.66944 2.00832 0.00000
-2.67776 0.00000 0.00000 ; HG13- CG1- CB- CG2
1192 1186 1184 1196 3 5.67350 3.55640 -6.16722
-2.47693 3.74886 0.00000 ; CG2- CB- CA- C
1196 1198 1200 1201 3 0.00000 0.00000 0.00000
0.00000 0.00000 0.00000 ; C- N- CA- HA
1196 1198 1200 1202 3 11.71520 1.67360 15.06240 -
13.38880 0.00000 0.00000 ; C- N- CA- CB
1196 1198 1200 1218 3 1.75728 5.27184 2.25936
-7.02912 0.00000 0.00000 ; C- N- CA- C
1197 1196 1198 1199 3 29.28800 -8.36800 -20.92000
0.00000 0.00000 0.00000 ; O- C- N- H
1197 1196 1198 1200 3 20.92000 0.00000 -20.92000
0.00000 0.00000 0.00000 ; O- C- N- CA
1198 1200 1202 1203 3 0.65084 1.95253 0.00000
-2.60338 0.00000 0.00000 ; N- CA- CB- HB2
1198 1200 1202 1204 3 0.65084 1.95253 0.00000
-2.60338 0.00000 0.00000 ; N- CA- CB- HB3
1198 1200 1202 1205 3 0.65084 1.95253 0.00000
-2.60338 0.00000 0.00000 ; N- CA- CB- CG
1198 1200 1218 1219 3 0.00000 0.00000 0.00000
0.00000 0.00000 0.00000 ; N- CA- C- O
1198 1200 1218 1220 3 17.40544 -5.02080 -13.22144
9.20480 0.00000 0.00000 ; N- CA- C- N
1199 1198 1200 1201 3 0.00000 0.00000 0.00000
0.00000 0.00000 0.00000 ; H- N- CA- HA

1199 1198 1200 1202 3 0.00000 0.00000 0.00000
0.00000 0.00000 0.00000 ; H- N- CA- CB
1199 1198 1200 1218 3 0.00000 0.00000 0.00000
0.00000 0.00000 0.00000 ; H- N- CA- C
1200 1202 1205 1206 3 0.66944 2.00832 0.00000
-2.67776 0.00000 0.00000 ; CA- CB- CG- HG2
1200 1202 1205 1207 3 0.66944 2.00832 0.00000
-2.67776 0.00000 0.00000 ; CA- CB- CG- HG3
1200 1202 1205 1208 3 3.68192 3.09616 -2.09200
-3.01248 0.00000 0.00000 ; CA- CB- CG- CD
1200 1218 1220 1221 3 20.92000 0.00000 -20.92000
0.00000 0.00000 0.00000 ; CA- C- N- H
1200 1218 1220 1222 3 20.92000 0.00000 -20.92000
0.00000 0.00000 0.00000 ; CA- C- N- CA
1201 1200 1202 1203 3 0.65084 1.95253 0.00000
-2.60338 0.00000 0.00000 ; HA- CA- CB- HB2
1201 1200 1202 1204 3 0.65084 1.95253 0.00000
-2.60338 0.00000 0.00000 ; HA- CA- CB- HB3
1201 1200 1202 1205 3 0.65084 1.95253 0.00000
-2.60338 0.00000 0.00000 ; HA- CA- CB- CG
1201 1200 1218 1219 3 3.68192 -4.35136 0.00000
1.33888 0.00000 0.00000 ; HA- CA- C- O
1201 1200 1218 1220 3 0.00000 0.00000 0.00000
0.00000 0.00000 0.00000 ; HA- CA- C- N
1202 1200 1218 1219 3 0.00000 0.00000 0.00000
0.00000 0.00000 0.00000 ; CB- CA- C- O
1202 1200 1218 1220 3 2.51040 4.18400 1.67360
-6.69440 0.00000 0.00000 ; CB- CA- C- N
1202 1205 1208 1209 3 0.66944 2.00832 0.00000
-2.67776 0.00000 0.00000 ; CB- CG- CD- HD2
1202 1205 1208 1210 3 0.66944 2.00832 0.00000
-2.67776 0.00000 0.00000 ; CB- CG- CD- HD3
1202 1205 1208 1211 3 3.68192 3.09616 -2.09200
-3.01248 0.00000 0.00000 ; CB- CG- CD- CE
1203 1202 1200 1218 3 0.65084 1.95253 0.00000
-2.60338 0.00000 0.00000 ; HB2- CB- CA- C
1203 1202 1205 1206 3 0.62760 1.88280 0.00000
-2.51040 0.00000 0.00000 ; HB2- CB- CG- HG2
1203 1202 1205 1207 3 0.62760 1.88280 0.00000
-2.51040 0.00000 0.00000 ; HB2- CB- CG- HG3
1203 1202 1205 1208 3 0.66944 2.00832 0.00000
-2.67776 0.00000 0.00000 ; HB2- CB- CG- CD
1204 1202 1200 1218 3 0.65084 1.95253 0.00000
-2.60338 0.00000 0.00000 ; HB3- CB- CA- C
1204 1202 1205 1206 3 0.62760 1.88280 0.00000
-2.51040 0.00000 0.00000 ; HB3- CB- CG- HG2
1204 1202 1205 1207 3 0.62760 1.88280 0.00000
-2.51040 0.00000 0.00000 ; HB3- CB- CG- HG3
1204 1202 1205 1208 3 0.66944 2.00832 0.00000
-2.67776 0.00000 0.00000 ; HB3- CB- CG- CD
1205 1202 1200 1218 3 0.65084 1.95253 0.00000
-2.60338 0.00000 0.00000 ; CG- CB- CA- C

1205 1208 1211 1212 3 0.65084 1.95253 0.00000
-2.60338 0.00000 0.00000 ; CG- CD- CE- HE2
1205 1208 1211 1213 3 0.65084 1.95253 0.00000
-2.60338 0.00000 0.00000 ; CG- CD- CE- HE3
1205 1208 1211 1214 3 0.65084 1.95253 0.00000
-2.60338 0.00000 0.00000 ; CG- CD- CE- NZ
1206 1205 1208 1209 3 0.62760 1.88280 0.00000
-2.51040 0.00000 0.00000 ; HG2- CG- CD- HD2
1206 1205 1208 1210 3 0.62760 1.88280 0.00000
-2.51040 0.00000 0.00000 ; HG2- CG- CD- HD3
1206 1205 1208 1211 3 0.66944 2.00832 0.00000
-2.67776 0.00000 0.00000 ; HG2- CG- CD- CE
1207 1205 1208 1209 3 0.62760 1.88280 0.00000
-2.51040 0.00000 0.00000 ; HG3- CG- CD- HD2
1207 1205 1208 1210 3 0.62760 1.88280 0.00000
-2.51040 0.00000 0.00000 ; HG3- CG- CD- HD3
1207 1205 1208 1211 3 0.66944 2.00832 0.00000
-2.67776 0.00000 0.00000 ; HG3- CG- CD- CE
1208 1211 1214 1215 3 0.65084 1.95253 0.00000
-2.60338 0.00000 0.00000 ; CD- CE- NZ- HZ1
1208 1211 1214 1216 3 0.65084 1.95253 0.00000
-2.60338 0.00000 0.00000 ; CD- CE- NZ- HZ2
1208 1211 1214 1217 3 0.65084 1.95253 0.00000
-2.60338 0.00000 0.00000 ; CD- CE- NZ- HZ3
1209 1208 1211 1212 3 0.65084 1.95253 0.00000
-2.60338 0.00000 0.00000 ; HD2- CD- CE- HE2
1209 1208 1211 1213 3 0.65084 1.95253 0.00000
-2.60338 0.00000 0.00000 ; HD2- CD- CE- HE3
1209 1208 1211 1214 3 0.65084 1.95253 0.00000
-2.60338 0.00000 0.00000 ; HD2- CD- CE- NZ
1210 1208 1211 1212 3 0.65084 1.95253 0.00000
-2.60338 0.00000 0.00000 ; HD3- CD- CE- HE2
1210 1208 1211 1213 3 0.65084 1.95253 0.00000
-2.60338 0.00000 0.00000 ; HD3- CD- CE- HE3
1210 1208 1211 1214 3 0.65084 1.95253 0.00000
-2.60338 0.00000 0.00000 ; HD3- CD- CE- NZ
1212 1211 1214 1215 3 0.65084 1.95253 0.00000
-2.60338 0.00000 0.00000 ; HE2- CE- NZ- HZ1
1212 1211 1214 1216 3 0.65084 1.95253 0.00000
-2.60338 0.00000 0.00000 ; HE2- CE- NZ- HZ2
1212 1211 1214 1217 3 0.65084 1.95253 0.00000
-2.60338 0.00000 0.00000 ; HE2- CE- NZ- HZ3
1213 1211 1214 1215 3 0.65084 1.95253 0.00000
-2.60338 0.00000 0.00000 ; HE3- CE- NZ- HZ1
1213 1211 1214 1216 3 0.65084 1.95253 0.00000
-2.60338 0.00000 0.00000 ; HE3- CE- NZ- HZ2
1213 1211 1214 1217 3 0.65084 1.95253 0.00000
-2.60338 0.00000 0.00000 ; HE3- CE- NZ- HZ3
1218 1220 1222 1223 3 0.00000 0.00000 0.00000
0.00000 0.00000 0.00000 ; C- N- CA- HA
1218 1220 1222 1224 3 11.71520 1.67360 15.06240 -
13.38880 0.00000 0.00000 ; C- N- CA- CB

1218 1220 1222 1237 3 1.75728 5.27184 2.25936
-7.02912 0.00000 0.00000 ; C- N- CA- C
1219 1218 1220 1221 3 29.28800 -8.36800 -20.92000
0.00000 0.00000 0.00000 ; O- C- N- H
1219 1218 1220 1222 3 20.92000 0.00000 -20.92000
0.00000 0.00000 0.00000 ; O- C- N- CA
1220 1222 1224 1225 3 0.65084 1.95253 0.00000
-2.60338 0.00000 0.00000 ; N- CA- CB- HB2
1220 1222 1224 1226 3 0.65084 1.95253 0.00000
-2.60338 0.00000 0.00000 ; N- CA- CB- HB3
1220 1222 1224 1227 3 3.79070 1.39746 -4.61077
-2.40998 2.44346 0.00000 ; N- CA- CB- CG
1220 1222 1237 1238 3 0.00000 0.00000 0.00000
0.00000 0.00000 0.00000 ; N- CA- C- O
1220 1222 1237 1239 3 17.40544 -5.02080 -13.22144
9.20480 0.00000 0.00000 ; N- CA- C- N
1221 1220 1222 1223 3 0.00000 0.00000 0.00000
0.00000 0.00000 0.00000 ; H- N- CA- HA
1221 1220 1222 1224 3 0.00000 0.00000 0.00000
0.00000 0.00000 0.00000 ; H- N- CA- CB
1221 1220 1222 1237 3 0.00000 0.00000 0.00000
0.00000 0.00000 0.00000 ; H- N- CA- C
1222 1224 1227 1228 3 0.66944 2.00832 0.00000
-2.67776 0.00000 0.00000 ; CA- CB- CG- HG
1222 1224 1227 1229 3 3.90367 0.19665 -6.21742
-2.37651 5.99149 0.00000 ; CA- CB- CG- CD1
1222 1224 1227 1233 3 3.90367 0.19665 -6.21742
-2.37651 5.99149 0.00000 ; CA- CB- CG- CD2
1222 1237 1239 1240 3 20.92000 0.00000 -20.92000
0.00000 0.00000 0.00000 ; CA- C- N- H
1222 1237 1239 1241 3 20.92000 0.00000 -20.92000
0.00000 0.00000 0.00000 ; CA- C- N- CA
1223 1222 1224 1225 3 0.65084 1.95253 0.00000
-2.60338 0.00000 0.00000 ; HA- CA- CB- HB2
1223 1222 1224 1226 3 0.65084 1.95253 0.00000
-2.60338 0.00000 0.00000 ; HA- CA- CB- HB3
1223 1222 1224 1227 3 0.65084 1.95253 0.00000
-2.60338 0.00000 0.00000 ; HA- CA- CB- CG
1223 1222 1237 1238 3 3.68192 -4.35136 0.00000
1.33888 0.00000 0.00000 ; HA- CA- C- O
1223 1222 1237 1239 3 0.00000 0.00000 0.00000
0.00000 0.00000 0.00000 ; HA- CA- C- N
1224 1222 1237 1238 3 0.00000 0.00000 0.00000
0.00000 0.00000 0.00000 ; CB- CA- C- O
1224 1222 1237 1239 3 2.51040 4.18400 1.67360
-6.69440 0.00000 0.00000 ; CB- CA- C- N
1224 1227 1229 1230 3 0.66944 2.00832 0.00000
-2.67776 0.00000 0.00000 ; CB- CG- CD1- HD11
1224 1227 1229 1231 3 0.66944 2.00832 0.00000
-2.67776 0.00000 0.00000 ; CB- CG- CD1- HD12
1224 1227 1229 1232 3 0.66944 2.00832 0.00000
-2.67776 0.00000 0.00000 ; CB- CG- CD1- HD13

1224 1227 1233 1234 3 0.66944 2.00832 0.00000
-2.67776 0.00000 0.00000 ; CB- CG- CD2- HD21
1224 1227 1233 1235 3 0.66944 2.00832 0.00000
-2.67776 0.00000 0.00000 ; CB- CG- CD2- HD22
1224 1227 1233 1236 3 0.66944 2.00832 0.00000
-2.67776 0.00000 0.00000 ; CB- CG- CD2- HD23
1225 1224 1222 1237 3 0.65084 1.95253 0.00000
-2.60338 0.00000 0.00000 ; HB2- CB- CA- C
1225 1224 1227 1228 3 0.62760 1.88280 0.00000
-2.51040 0.00000 0.00000 ; HB2- CB- CG- HG
1225 1224 1227 1229 3 0.66944 2.00832 0.00000
-2.67776 0.00000 0.00000 ; HB2- CB- CG- CD1
1225 1224 1227 1233 3 0.66944 2.00832 0.00000
-2.67776 0.00000 0.00000 ; HB2- CB- CG- CD2
1226 1224 1222 1237 3 0.65084 1.95253 0.00000
-2.60338 0.00000 0.00000 ; HB3- CB- CA- C
1226 1224 1227 1228 3 0.62760 1.88280 0.00000
-2.51040 0.00000 0.00000 ; HB3- CB- CG- HG
1226 1224 1227 1229 3 0.66944 2.00832 0.00000
-2.67776 0.00000 0.00000 ; HB3- CB- CG- CD1
1226 1224 1227 1233 3 0.66944 2.00832 0.00000
-2.67776 0.00000 0.00000 ; HB3- CB- CG- CD2
1227 1224 1222 1237 3 10.33448 -1.14642 -11.54784
-2.40998 6.35968 0.00000 ; CG- CB- CA- C
1228 1227 1229 1230 3 0.62760 1.88280 0.00000
-2.51040 0.00000 0.00000 ; HG- CG- CD1- HD11
1228 1227 1229 1231 3 0.62760 1.88280 0.00000
-2.51040 0.00000 0.00000 ; HG- CG- CD1- HD12
1228 1227 1229 1232 3 0.62760 1.88280 0.00000
-2.51040 0.00000 0.00000 ; HG- CG- CD1- HD13
1228 1227 1233 1234 3 0.62760 1.88280 0.00000
-2.51040 0.00000 0.00000 ; HG- CG- CD2- HD21
1228 1227 1233 1235 3 0.62760 1.88280 0.00000
-2.51040 0.00000 0.00000 ; HG- CG- CD2- HD22
1228 1227 1233 1236 3 0.62760 1.88280 0.00000
-2.51040 0.00000 0.00000 ; HG- CG- CD2- HD23
1229 1227 1233 1234 3 0.66944 2.00832 0.00000
-2.67776 0.00000 0.00000 ; CD1- CG- CD2- HD21
1229 1227 1233 1235 3 0.66944 2.00832 0.00000
-2.67776 0.00000 0.00000 ; CD1- CG- CD2- HD22
1229 1227 1233 1236 3 0.66944 2.00832 0.00000
-2.67776 0.00000 0.00000 ; CD1- CG- CD2- HD23
1230 1229 1227 1233 3 0.66944 2.00832 0.00000
-2.67776 0.00000 0.00000 ; HD11- CD1- CG- CD2
1231 1229 1227 1233 3 0.66944 2.00832 0.00000
-2.67776 0.00000 0.00000 ; HD12- CD1- CG- CD2
1232 1229 1227 1233 3 0.66944 2.00832 0.00000
-2.67776 0.00000 0.00000 ; HD13- CD1- CG- CD2
1237 1239 1241 1242 3 0.00000 0.00000 0.00000
0.00000 0.00000 0.00000 ; C- N- CA- HA
1237 1239 1241 1243 3 11.71520 1.67360 15.06240 -
13.38880 0.00000 0.00000 ; C- N- CA- CB

1237 1239 1241 1258 3 1.75728 5.27184 2.25936
-7.02912 0.00000 0.00000 ; C- N- CA- C
1238 1237 1239 1240 3 29.28800 -8.36800 -20.92000
0.00000 0.00000 0.00000 ; O- C- N- H
1238 1237 1239 1241 3 20.92000 0.00000 -20.92000
0.00000 0.00000 0.00000 ; O- C- N- CA
1239 1241 1243 1244 3 0.65084 1.95253 0.00000
-2.60338 0.00000 0.00000 ; N- CA- CB- HB2
1239 1241 1243 1245 3 0.65084 1.95253 0.00000
-2.60338 0.00000 0.00000 ; N- CA- CB- HB3
1239 1241 1243 1246 3 3.28026 2.46019 -2.19242
-3.21331 -0.23430 0.00000 ; N- CA- CB- CG
1239 1241 1258 1259 3 0.00000 0.00000 0.00000
0.00000 0.00000 0.00000 ; N- CA- C- O
1239 1241 1258 1260 3 17.40544 -5.02080 -13.22144
9.20480 0.00000 0.00000 ; N- CA- C- N
1240 1239 1241 1242 3 0.00000 0.00000 0.00000
0.00000 0.00000 0.00000 ; H- N- CA- HA
1240 1239 1241 1243 3 0.00000 0.00000 0.00000
0.00000 0.00000 0.00000 ; H- N- CA- CB
1240 1239 1241 1258 3 0.00000 0.00000 0.00000
0.00000 0.00000 0.00000 ; H- N- CA- C
1241 1243 1246 1247 3 0.57739 0.00000 1.02926
0.00000 -1.60666 0.00000 ; CA- CB- CG- CD1
1241 1243 1246 1256 3 0.57739 0.00000 1.02926
0.00000 -1.60666 0.00000 ; CA- CB- CG- CD2
1241 1258 1260 1261 3 20.92000 0.00000 -20.92000
0.00000 0.00000 0.00000 ; CA- C- N- H
1241 1258 1260 1262 3 20.92000 0.00000 -20.92000
0.00000 0.00000 0.00000 ; CA- C- N- CA
1242 1241 1243 1244 3 0.65084 1.95253 0.00000
-2.60338 0.00000 0.00000 ; HA- CA- CB- HB2
1242 1241 1243 1245 3 0.65084 1.95253 0.00000
-2.60338 0.00000 0.00000 ; HA- CA- CB- HB3
1242 1241 1243 1246 3 0.65084 1.95253 0.00000
-2.60338 0.00000 0.00000 ; HA- CA- CB- CG
1242 1241 1258 1259 3 3.68192 -4.35136 0.00000
1.33888 0.00000 0.00000 ; HA- CA- C- O
1242 1241 1258 1260 3 0.00000 0.00000 0.00000
0.00000 0.00000 0.00000 ; HA- CA- C- N
1243 1241 1258 1259 3 0.00000 0.00000 0.00000
0.00000 0.00000 0.00000 ; CB- CA- C- O
1243 1241 1258 1260 3 2.51040 4.18400 1.67360
-6.69440 0.00000 0.00000 ; CB- CA- C- N
1243 1246 1247 1248 3 30.33400 0.00000 -30.33400
0.00000 0.00000 0.00000 ; CB- CG- CD1- HD1
1243 1246 1247 1249 3 30.33400 0.00000 -30.33400
0.00000 0.00000 0.00000 ; CB- CG- CD1- CE1
1243 1246 1256 1254 3 30.33400 0.00000 -30.33400
0.00000 0.00000 0.00000 ; CB- CG- CD2- CE2
1243 1246 1256 1257 3 30.33400 0.00000 -30.33400
0.00000 0.00000 0.00000 ; CB- CG- CD2- HD2

1244 1243 1241 1258 3 0.65084 1.95253 0.00000
-2.60338 0.00000 0.00000 ; HB2- CB- CA- C
1244 1243 1246 1247 3 0.00000 0.00000 0.00000
0.00000 0.00000 0.00000 ; HB2- CB- CG- CD1
1244 1243 1246 1256 3 0.00000 0.00000 0.00000
0.00000 0.00000 0.00000 ; HB2- CB- CG- CD2
1245 1243 1241 1258 3 0.65084 1.95253 0.00000
-2.60338 0.00000 0.00000 ; HB3- CB- CA- C
1245 1243 1246 1247 3 0.00000 0.00000 0.00000
0.00000 0.00000 0.00000 ; HB3- CB- CG- CD1
1245 1243 1246 1256 3 0.00000 0.00000 0.00000
0.00000 0.00000 0.00000 ; HB3- CB- CG- CD2
1246 1243 1241 1258 3 4.95804 2.17986 -3.52293
-3.21331 -0.40166 0.00000 ; CG- CB- CA- C
1246 1247 1249 1250 3 30.33400 0.00000 -30.33400
0.00000 0.00000 0.00000 ; CG- CD1- CE1- HE1
1246 1247 1249 1251 3 30.33400 0.00000 -30.33400
0.00000 0.00000 0.00000 ; CG- CD1- CE1- CZ
1246 1256 1254 1251 3 30.33400 0.00000 -30.33400
0.00000 0.00000 0.00000 ; CG- CD2- CE2- CZ
1246 1256 1254 1255 3 30.33400 0.00000 -30.33400
0.00000 0.00000 0.00000 ; CG- CD2- CE2- HE2
1247 1246 1256 1254 3 30.33400 0.00000 -30.33400
0.00000 0.00000 0.00000 ; CD1- CG- CD2- CE2
1247 1246 1256 1257 3 30.33400 0.00000 -30.33400
0.00000 0.00000 0.00000 ; CD1- CG- CD2- HD2
1247 1249 1251 1252 3 30.33400 0.00000 -30.33400
0.00000 0.00000 0.00000 ; CD1- CE1- CZ- OH
1247 1249 1251 1254 3 30.33400 0.00000 -30.33400
0.00000 0.00000 0.00000 ; CD1- CE1- CZ- CE2
1248 1247 1246 1256 3 30.33400 0.00000 -30.33400
0.00000 0.00000 0.00000 ; HD1- CD1- CG- CD2
1248 1247 1249 1250 3 30.33400 0.00000 -30.33400
0.00000 0.00000 0.00000 ; HD1- CD1- CE1- HE1
1248 1247 1249 1251 3 30.33400 0.00000 -30.33400
0.00000 0.00000 0.00000 ; HD1- CD1- CE1- CZ
1249 1247 1246 1256 3 30.33400 0.00000 -30.33400
0.00000 0.00000 0.00000 ; CE1- CD1- CG- CD2
1249 1251 1252 1253 3 7.93286 0.00000 -9.56462
0.00000 2.17568 0.00000 ; CE1- CZ- OH- HH
1249 1251 1254 1255 3 30.33400 0.00000 -30.33400
0.00000 0.00000 0.00000 ; CE1- CZ- CE2- HE2
1249 1251 1254 1256 3 30.33400 0.00000 -30.33400
0.00000 0.00000 0.00000 ; CE1- CZ- CE2- CD2
1250 1249 1251 1252 3 30.33400 0.00000 -30.33400
0.00000 0.00000 0.00000 ; HE1- CE1- CZ- OH
1250 1249 1251 1254 3 30.33400 0.00000 -30.33400
0.00000 0.00000 0.00000 ; HE1- CE1- CZ- CE2
1251 1254 1256 1257 3 30.33400 0.00000 -30.33400
0.00000 0.00000 0.00000 ; CZ- CE2- CD2- HD2
1252 1251 1254 1255 3 30.33400 0.00000 -30.33400
0.00000 0.00000 0.00000 ; OH- CZ- CE2- HE2

1252 1251 1254 1256 3 30.33400 0.00000 -30.33400
0.00000 0.00000 0.00000 ; OH- CZ- CE2- CD2
1253 1252 1251 1254 3 7.93286 0.00000 -9.56462
0.00000 2.17568 0.00000 ; HH- OH- CZ- CE2
1255 1254 1256 1257 3 30.33400 0.00000 -30.33400
0.00000 0.00000 0.00000 ; HE2- CE2- CD2- HD2
1258 1260 1262 1263 3 0.00000 0.00000 0.00000
0.00000 0.00000 0.00000 ; C- N- CA- HA
1258 1260 1262 1264 3 11.71520 1.67360 15.06240 -
13.38880 0.00000 0.00000 ; C- N- CA- CB
1258 1260 1262 1273 3 1.75728 5.27184 2.25936
-7.02912 0.00000 0.00000 ; C- N- CA- C
1259 1258 1260 1261 3 29.28800 -8.36800 -20.92000
0.00000 0.00000 0.00000 ; O- C- N- H
1259 1258 1260 1262 3 20.92000 0.00000 -20.92000
0.00000 0.00000 0.00000 ; O- C- N- CA
1260 1262 1264 1265 3 0.65084 1.95253 0.00000
-2.60338 0.00000 0.00000 ; N- CA- CB- HB2
1260 1262 1264 1266 3 0.65084 1.95253 0.00000
-2.60338 0.00000 0.00000 ; N- CA- CB- HB3
1260 1262 1264 1267 3 3.21331 2.22589 -4.15053
-2.40998 2.61082 0.00000 ; N- CA- CB- CG
1260 1262 1273 1274 3 0.00000 0.00000 0.00000
0.00000 0.00000 0.00000 ; N- CA- C- O
1260 1262 1273 1275 3 17.40544 -5.02080 -13.22144
9.20480 0.00000 0.00000 ; N- CA- C- N
1261 1260 1262 1263 3 0.00000 0.00000 0.00000
0.00000 0.00000 0.00000 ; H- N- CA- HA
1261 1260 1262 1264 3 0.00000 0.00000 0.00000
0.00000 0.00000 0.00000 ; H- N- CA- CB
1261 1260 1262 1273 3 0.00000 0.00000 0.00000
0.00000 0.00000 0.00000 ; H- N- CA- C
1262 1264 1267 1268 3 0.66944 2.00832 0.00000
-2.67776 0.00000 0.00000 ; CA- CB- CG- HG2
1262 1264 1267 1269 3 0.66944 2.00832 0.00000
-2.67776 0.00000 0.00000 ; CA- CB- CG- HG3
1262 1264 1267 1270 3 10.12110 -1.91209 0.01674
10.17549 -1.87443 0.00000 ; CA- CB- CG- CD
1262 1273 1275 1276 3 20.92000 0.00000 -20.92000
0.00000 0.00000 0.00000 ; CA- C- N- H
1262 1273 1275 1277 3 20.92000 0.00000 -20.92000
0.00000 0.00000 0.00000 ; CA- C- N- CA
1263 1262 1264 1265 3 0.65084 1.95253 0.00000
-2.60338 0.00000 0.00000 ; HA- CA- CB- HB2
1263 1262 1264 1266 3 0.65084 1.95253 0.00000
-2.60338 0.00000 0.00000 ; HA- CA- CB- HB3
1263 1262 1264 1267 3 0.65084 1.95253 0.00000
-2.60338 0.00000 0.00000 ; HA- CA- CB- CG
1263 1262 1273 1274 3 3.68192 -4.35136 0.00000
1.33888 0.00000 0.00000 ; HA- CA- C- O
1263 1262 1273 1275 3 0.00000 0.00000 0.00000
0.00000 0.00000 0.00000 ; HA- CA- C- N

1264 1262 1273 1274 3 0.00000 0.00000 0.00000
0.00000 0.00000 0.00000 ; CB- CA- C- O
1264 1262 1273 1275 3 2.51040 4.18400 1.67360
-6.69440 0.00000 0.00000 ; CB- CA- C- N
1264 1267 1270 1271 3 3.79907 0.00000 -5.40573
0.00000 2.14221 0.00000 ; CB- CG- CD- OE1
1264 1267 1270 1272 3 3.79907 0.00000 -5.40573
0.00000 2.14221 0.00000 ; CB- CG- CD- OE2
1265 1264 1262 1273 3 0.65084 1.95253 0.00000
-2.60338 0.00000 0.00000 ; HB2- CB- CA- C
1265 1264 1267 1268 3 0.62760 1.88280 0.00000
-2.51040 0.00000 0.00000 ; HB2- CB- CG- HG2
1265 1264 1267 1269 3 0.62760 1.88280 0.00000
-2.51040 0.00000 0.00000 ; HB2- CB- CG- HG3
1265 1264 1267 1270 3 0.65084 1.95253 0.00000
-2.60338 0.00000 0.00000 ; HB2- CB- CG- CD
1266 1264 1262 1273 3 0.65084 1.95253 0.00000
-2.60338 0.00000 0.00000 ; HB3- CB- CA- C
1266 1264 1267 1268 3 0.62760 1.88280 0.00000
-2.51040 0.00000 0.00000 ; HB3- CB- CG- HG2
1266 1264 1267 1269 3 0.62760 1.88280 0.00000
-2.51040 0.00000 0.00000 ; HB3- CB- CG- HG3
1266 1264 1267 1270 3 0.65084 1.95253 0.00000
-2.60338 0.00000 0.00000 ; HB3- CB- CG- CD
1267 1264 1262 1273 3 6.86594 3.56895 -8.14206
-2.40998 4.85344 0.00000 ; CG- CB- CA- C
1268 1267 1270 1271 3 0.00000 0.00000 0.00000
0.00000 0.00000 0.00000 ; HG2- CG- CD- OE1
1268 1267 1270 1272 3 0.00000 0.00000 0.00000
0.00000 0.00000 0.00000 ; HG2- CG- CD- OE2
1269 1267 1270 1271 3 0.00000 0.00000 0.00000
0.00000 0.00000 0.00000 ; HG3- CG- CD- OE1
1269 1267 1270 1272 3 0.00000 0.00000 0.00000
0.00000 0.00000 0.00000 ; HG3- CG- CD- OE2
1273 1275 1277 1278 3 0.00000 0.00000 0.00000
0.00000 0.00000 0.00000 ; C- N- CA- HA
1273 1275 1277 1279 3 11.71520 1.67360 15.06240 -
13.38880 0.00000 0.00000 ; C- N- CA- CB
1273 1275 1277 1287 3 1.75728 5.27184 2.25936
-7.02912 0.00000 0.00000 ; C- N- CA- C
1274 1273 1275 1276 3 29.28800 -8.36800 -20.92000
0.00000 0.00000 0.00000 ; O- C- N- H
1274 1273 1275 1277 3 20.92000 0.00000 -20.92000
0.00000 0.00000 0.00000 ; O- C- N- CA
1275 1277 1279 1280 3 0.65084 1.95253 0.00000
-2.60338 0.00000 0.00000 ; N- CA- CB- HB
1275 1277 1279 1281 3 3.83673 0.44769 -1.77402
-2.47693 -0.03347 0.00000 ; N- CA- CB- CG2
1275 1277 1279 1285 3 4.93294 1.13386 -3.12963
-5.27184 3.17984 0.00000 ; N- CA- CB- OG1
1275 1277 1287 1288 3 0.00000 0.00000 0.00000
0.00000 0.00000 0.00000 ; N- CA- C- O

1275 1277 1287 1289 3 17.40544 -5.02080 -13.22144
9.20480 0.00000 0.00000 ; N- CA- C- N
1276 1275 1277 1278 3 0.00000 0.00000 0.00000
0.00000 0.00000 0.00000 ; H- N- CA- HA
1276 1275 1277 1279 3 0.00000 0.00000 0.00000
0.00000 0.00000 0.00000 ; H- N- CA- CB
1276 1275 1277 1287 3 0.00000 0.00000 0.00000
0.00000 0.00000 0.00000 ; H- N- CA- C
1277 1279 1281 1282 3 0.66944 2.00832 0.00000
-2.67776 0.00000 0.00000 ; CA- CB- CG2- HG21
1277 1279 1281 1283 3 0.66944 2.00832 0.00000
-2.67776 0.00000 0.00000 ; CA- CB- CG2- HG22
1277 1279 1281 1284 3 0.66944 2.00832 0.00000
-2.67776 0.00000 0.00000 ; CA- CB- CG2- HG23
1277 1279 1285 1286 3 1.12131 2.98738 1.66523
-3.94970 0.43514 0.00000 ; CA- CB- OG1- HG1
1277 1287 1289 1290 3 20.92000 0.00000 -20.92000
0.00000 0.00000 0.00000 ; CA- C- N- H
1277 1287 1289 1291 3 20.92000 0.00000 -20.92000
0.00000 0.00000 0.00000 ; CA- C- N- CA
1278 1277 1279 1280 3 0.65084 1.95253 0.00000
-2.60338 0.00000 0.00000 ; HA- CA- CB- HB
1278 1277 1279 1281 3 0.65084 1.95253 0.00000
-2.60338 0.00000 0.00000 ; HA- CA- CB- CG2
1278 1277 1279 1285 3 1.04600 -1.04600 0.00000
0.00000 0.00000 0.00000 ; HA- CA- CB- OG1
1278 1277 1287 1288 3 3.68192 -4.35136 0.00000
1.33888 0.00000 0.00000 ; HA- CA- C- O
1278 1277 1287 1289 3 0.00000 0.00000 0.00000
0.00000 0.00000 0.00000 ; HA- CA- C- N
1279 1277 1287 1288 3 0.00000 0.00000 0.00000
0.00000 0.00000 0.00000 ; CB- CA- C- O
1279 1277 1287 1289 3 2.51040 4.18400 1.67360
-6.69440 0.00000 0.00000 ; CB- CA- C- N
1280 1279 1277 1287 3 0.65084 1.95253 0.00000
-2.60338 0.00000 0.00000 ; HB- CB- CA- C
1280 1279 1281 1282 3 0.65084 1.95253 0.00000
-2.60338 0.00000 0.00000 ; HB- CB- CG2- HG21
1280 1279 1281 1283 3 0.65084 1.95253 0.00000
-2.60338 0.00000 0.00000 ; HB- CB- CG2- HG22
1280 1279 1281 1284 3 0.65084 1.95253 0.00000
-2.60338 0.00000 0.00000 ; HB- CB- CG2- HG23
1280 1279 1285 1286 3 0.69733 2.09200 0.00000
-2.78933 0.00000 0.00000 ; HB- CB- OG1- HG1
1281 1279 1277 1287 3 5.67350 3.55640 -6.16722
-2.47693 3.74886 0.00000 ; CG2- CB- CA- C
1281 1279 1285 1286 3 4.74047 0.27196 -2.26773
-3.94970 1.60666 0.00000 ; CG2- CB- OG1- HG1
1282 1281 1279 1285 3 1.04600 -1.04600 0.00000
0.00000 0.00000 0.00000 ; HG21- CG2- CB- OG1
1283 1281 1279 1285 3 1.04600 -1.04600 0.00000
0.00000 0.00000 0.00000 ; HG22- CG2- CB- OG1

1284 1281 1279 1285 3 1.04600 -1.04600 0.00000
0.00000 0.00000 0.00000 ; HG23- CG2- CB- OG1
1285 1279 1277 1287 3 6.53541 6.87013 -6.21742
-5.27184 5.22163 0.00000 ; OG1- CB- CA- C
1287 1289 1291 1292 3 0.00000 0.00000 0.00000
0.00000 0.00000 0.00000 ; C- N- CA- HA
1287 1289 1291 1293 3 11.71520 1.67360 15.06240 -
13.38880 0.00000 0.00000 ; C- N- CA- CB
1287 1289 1291 1308 3 1.75728 5.27184 2.25936
-7.02912 0.00000 0.00000 ; C- N- CA- C
1288 1287 1289 1290 3 29.28800 -8.36800 -20.92000
0.00000 0.00000 0.00000 ; O- C- N- H
1288 1287 1289 1291 3 20.92000 0.00000 -20.92000
0.00000 0.00000 0.00000 ; O- C- N- CA
1289 1291 1293 1294 3 0.65084 1.95253 0.00000
-2.60338 0.00000 0.00000 ; N- CA- CB- HB2
1289 1291 1293 1295 3 0.65084 1.95253 0.00000
-2.60338 0.00000 0.00000 ; N- CA- CB- HB3
1289 1291 1293 1296 3 3.28026 2.46019 -2.19242
-3.21331 -0.23430 0.00000 ; N- CA- CB- CG
1289 1291 1308 1309 3 0.00000 0.00000 0.00000
0.00000 0.00000 0.00000 ; N- CA- C- O
1289 1291 1308 1310 3 17.40544 -5.02080 -13.22144
9.20480 0.00000 0.00000 ; N- CA- C- N
1290 1289 1291 1292 3 0.00000 0.00000 0.00000
0.00000 0.00000 0.00000 ; H- N- CA- HA
1290 1289 1291 1293 3 0.00000 0.00000 0.00000
0.00000 0.00000 0.00000 ; H- N- CA- CB
1290 1289 1291 1308 3 0.00000 0.00000 0.00000
0.00000 0.00000 0.00000 ; H- N- CA- C
1291 1293 1296 1297 3 0.57739 0.00000 1.02926
0.00000 -1.60666 0.00000 ; CA- CB- CG- CD1
1291 1293 1296 1306 3 0.57739 0.00000 1.02926
0.00000 -1.60666 0.00000 ; CA- CB- CG- CD2
1291 1308 1310 1311 3 20.92000 0.00000 -20.92000
0.00000 0.00000 0.00000 ; CA- C- N- H
1291 1308 1310 1312 3 20.92000 0.00000 -20.92000
0.00000 0.00000 0.00000 ; CA- C- N- CA
1292 1291 1293 1294 3 0.65084 1.95253 0.00000
-2.60338 0.00000 0.00000 ; HA- CA- CB- HB2
1292 1291 1293 1295 3 0.65084 1.95253 0.00000
-2.60338 0.00000 0.00000 ; HA- CA- CB- HB3
1292 1291 1293 1296 3 0.65084 1.95253 0.00000
-2.60338 0.00000 0.00000 ; HA- CA- CB- CG
1292 1291 1308 1309 3 3.68192 -4.35136 0.00000
1.33888 0.00000 0.00000 ; HA- CA- C- O
1292 1291 1308 1310 3 0.00000 0.00000 0.00000
0.00000 0.00000 0.00000 ; HA- CA- C- N
1293 1291 1308 1309 3 0.00000 0.00000 0.00000
0.00000 0.00000 0.00000 ; CB- CA- C- O
1293 1291 1308 1310 3 2.51040 4.18400 1.67360
-6.69440 0.00000 0.00000 ; CB- CA- C- N

1293 1296 1297 1298 3 30.33400 0.00000 -30.33400
0.00000 0.00000 0.00000 ; CB- CG- CD1- HD1
1293 1296 1297 1299 3 30.33400 0.00000 -30.33400
0.00000 0.00000 0.00000 ; CB- CG- CD1- CE1
1293 1296 1306 1304 3 30.33400 0.00000 -30.33400
0.00000 0.00000 0.00000 ; CB- CG- CD2- CE2
1293 1296 1306 1307 3 30.33400 0.00000 -30.33400
0.00000 0.00000 0.00000 ; CB- CG- CD2- HD2
1294 1293 1291 1308 3 0.65084 1.95253 0.00000
-2.60338 0.00000 0.00000 ; HB2- CB- CA- C
1294 1293 1296 1297 3 0.00000 0.00000 0.00000
0.00000 0.00000 0.00000 ; HB2- CB- CG- CD1
1294 1293 1296 1306 3 0.00000 0.00000 0.00000
0.00000 0.00000 0.00000 ; HB2- CB- CG- CD2
1295 1293 1291 1308 3 0.65084 1.95253 0.00000
-2.60338 0.00000 0.00000 ; HB3- CB- CA- C
1295 1293 1296 1297 3 0.00000 0.00000 0.00000
0.00000 0.00000 0.00000 ; HB3- CB- CG- CD1
1295 1293 1296 1306 3 0.00000 0.00000 0.00000
0.00000 0.00000 0.00000 ; HB3- CB- CG- CD2
1296 1293 1291 1308 3 4.95804 2.17986 -3.52293
-3.21331 -0.40166 0.00000 ; CG- CB- CA- C
1296 1297 1299 1300 3 30.33400 0.00000 -30.33400
0.00000 0.00000 0.00000 ; CG- CD1- CE1- HE1
1296 1297 1299 1301 3 30.33400 0.00000 -30.33400
0.00000 0.00000 0.00000 ; CG- CD1- CE1- CZ
1296 1306 1304 1301 3 30.33400 0.00000 -30.33400
0.00000 0.00000 0.00000 ; CG- CD2- CE2- CZ
1296 1306 1304 1305 3 30.33400 0.00000 -30.33400
0.00000 0.00000 0.00000 ; CG- CD2- CE2- HE2
1297 1296 1306 1304 3 30.33400 0.00000 -30.33400
0.00000 0.00000 0.00000 ; CD1- CG- CD2- CE2
1297 1296 1306 1307 3 30.33400 0.00000 -30.33400
0.00000 0.00000 0.00000 ; CD1- CG- CD2- HD2
1297 1299 1301 1302 3 30.33400 0.00000 -30.33400
0.00000 0.00000 0.00000 ; CD1- CE1- CZ- OH
1297 1299 1301 1304 3 30.33400 0.00000 -30.33400
0.00000 0.00000 0.00000 ; CD1- CE1- CZ- CE2
1298 1297 1296 1306 3 30.33400 0.00000 -30.33400
0.00000 0.00000 0.00000 ; HD1- CD1- CG- CD2
1298 1297 1299 1300 3 30.33400 0.00000 -30.33400
0.00000 0.00000 0.00000 ; HD1- CD1- CE1- HE1
1298 1297 1299 1301 3 30.33400 0.00000 -30.33400
0.00000 0.00000 0.00000 ; HD1- CD1- CE1- CZ
1299 1297 1296 1306 3 30.33400 0.00000 -30.33400
0.00000 0.00000 0.00000 ; CE1- CD1- CG- CD2
1299 1301 1302 1303 3 7.93286 0.00000 -9.56462
0.00000 2.17568 0.00000 ; CE1- CZ- OH- HH
1299 1301 1304 1305 3 30.33400 0.00000 -30.33400
0.00000 0.00000 0.00000 ; CE1- CZ- CE2- HE2
1299 1301 1304 1306 3 30.33400 0.00000 -30.33400
0.00000 0.00000 0.00000 ; CE1- CZ- CE2- CD2

```
  1300   1299   1301   1302      3   30.33400    0.00000  -30.33400
0.00000    0.00000    0.00000 ;    HE1-   CE1-    CZ-    OH
  1300   1299   1301   1304      3   30.33400    0.00000  -30.33400
0.00000    0.00000    0.00000 ;    HE1-   CE1-    CZ-   CE2
  1301   1304   1306   1307      3   30.33400    0.00000  -30.33400
0.00000    0.00000    0.00000 ;     CZ-   CE2-   CD2-   HD2
  1302   1301   1304   1305      3   30.33400    0.00000  -30.33400
0.00000    0.00000    0.00000 ;     OH-    CZ-   CE2-   HE2
  1302   1301   1304   1306      3   30.33400    0.00000  -30.33400
0.00000    0.00000    0.00000 ;     OH-    CZ-   CE2-   CD2
  1303   1302   1301   1304      3    7.93286    0.00000   -9.56462
0.00000    2.17568    0.00000 ;     HH-    OH-    CZ-   CE2
  1305   1304   1306   1307      3   30.33400    0.00000  -30.33400
0.00000    0.00000    0.00000 ;    HE2-   CE2-   CD2-   HD2
  1308   1310   1312   1313      3    0.00000    0.00000    0.00000
0.00000    0.00000    0.00000 ;      C-     N-    CA-    HA
  1308   1310   1312   1314      3   11.71520    1.67360   15.06240  -
13.38880    0.00000    0.00000 ;      C-     N-    CA-    CB
  1308   1310   1312   1323      3    1.75728    5.27184    2.25936
-7.02912    0.00000    0.00000 ;      C-     N-    CA-     C
  1309   1308   1310   1311      3   29.28800   -8.36800  -20.92000
0.00000    0.00000    0.00000 ;      O-     C-     N-     H
  1309   1308   1310   1312      3   20.92000    0.00000  -20.92000
0.00000    0.00000    0.00000 ;      O-     C-     N-    CA
  1310   1312   1314   1315      3    0.65084    1.95253    0.00000
-2.60338    0.00000    0.00000 ;      N-    CA-    CB-   HB2
  1310   1312   1314   1316      3    0.65084    1.95253    0.00000
-2.60338    0.00000    0.00000 ;      N-    CA-    CB-   HB3
  1310   1312   1314   1317      3    3.21331    2.22589   -4.15053
-2.40998    2.61082    0.00000 ;      N-    CA-    CB-    CG
  1310   1312   1323   1324      3    0.00000    0.00000    0.00000
0.00000    0.00000    0.00000 ;      N-    CA-     C-     O
  1310   1312   1323   1325      3   17.40544   -5.02080  -13.22144
9.20480    0.00000    0.00000 ;      N-    CA-     C-     N
  1311   1310   1312   1313      3    0.00000    0.00000    0.00000
0.00000    0.00000    0.00000 ;      H-     N-    CA-    HA
  1311   1310   1312   1314      3    0.00000    0.00000    0.00000
0.00000    0.00000    0.00000 ;      H-     N-    CA-    CB
  1311   1310   1312   1323      3    0.00000    0.00000    0.00000
0.00000    0.00000    0.00000 ;      H-     N-    CA-     C
  1312   1314   1317   1318      3    0.66944    2.00832    0.00000
-2.67776    0.00000    0.00000 ;     CA-    CB-    CG-   HG2
  1312   1314   1317   1319      3    0.66944    2.00832    0.00000
-2.67776    0.00000    0.00000 ;     CA-    CB-    CG-   HG3
  1312   1314   1317   1320      3   10.12110   -1.91209    0.01674
10.17549   -1.87443    0.00000 ;     CA-    CB-    CG-    CD
  1312   1323   1325   1326      3   20.92000    0.00000  -20.92000
0.00000    0.00000    0.00000 ;     CA-     C-     N-     H
  1312   1323   1325   1327      3   20.92000    0.00000  -20.92000
0.00000    0.00000    0.00000 ;     CA-     C-     N-    CA
  1313   1312   1314   1315      3    0.65084    1.95253    0.00000
-2.60338    0.00000    0.00000 ;     HA-    CA-    CB-   HB2
```

1313 1312 1314 1316 3 0.65084 1.95253 0.00000
-2.60338 0.00000 0.00000 ; HA- CA- CB- HB3
1313 1312 1314 1317 3 0.65084 1.95253 0.00000
-2.60338 0.00000 0.00000 ; HA- CA- CB- CG
1313 1312 1323 1324 3 3.68192 -4.35136 0.00000
1.33888 0.00000 0.00000 ; HA- CA- C- O
1313 1312 1323 1325 3 0.00000 0.00000 0.00000
0.00000 0.00000 0.00000 ; HA- CA- C- N
1314 1312 1323 1324 3 0.00000 0.00000 0.00000
0.00000 0.00000 0.00000 ; CB- CA- C- O
1314 1312 1323 1325 3 2.51040 4.18400 1.67360
-6.69440 0.00000 0.00000 ; CB- CA- C- N
1314 1317 1320 1321 3 3.79907 0.00000 -5.40573
0.00000 2.14221 0.00000 ; CB- CG- CD- OE1
1314 1317 1320 1322 3 3.79907 0.00000 -5.40573
0.00000 2.14221 0.00000 ; CB- CG- CD- OE2
1315 1314 1312 1323 3 0.65084 1.95253 0.00000
-2.60338 0.00000 0.00000 ; HB2- CB- CA- C
1315 1314 1317 1318 3 0.62760 1.88280 0.00000
-2.51040 0.00000 0.00000 ; HB2- CB- CG- HG2
1315 1314 1317 1319 3 0.62760 1.88280 0.00000
-2.51040 0.00000 0.00000 ; HB2- CB- CG- HG3
1315 1314 1317 1320 3 0.65084 1.95253 0.00000
-2.60338 0.00000 0.00000 ; HB2- CB- CG- CD
1316 1314 1312 1323 3 0.65084 1.95253 0.00000
-2.60338 0.00000 0.00000 ; HB3- CB- CA- C
1316 1314 1317 1318 3 0.62760 1.88280 0.00000
-2.51040 0.00000 0.00000 ; HB3- CB- CG- HG2
1316 1314 1317 1319 3 0.62760 1.88280 0.00000
-2.51040 0.00000 0.00000 ; HB3- CB- CG- HG3
1316 1314 1317 1320 3 0.65084 1.95253 0.00000
-2.60338 0.00000 0.00000 ; HB3- CB- CG- CD
1317 1314 1312 1323 3 6.86594 3.56895 -8.14206
-2.40998 4.85344 0.00000 ; CG- CB- CA- C
1318 1317 1320 1321 3 0.00000 0.00000 0.00000
0.00000 0.00000 0.00000 ; HG2- CG- CD- OE1
1318 1317 1320 1322 3 0.00000 0.00000 0.00000
0.00000 0.00000 0.00000 ; HG2- CG- CD- OE2
1319 1317 1320 1321 3 0.00000 0.00000 0.00000
0.00000 0.00000 0.00000 ; HG3- CG- CD- OE1
1319 1317 1320 1322 3 0.00000 0.00000 0.00000
0.00000 0.00000 0.00000 ; HG3- CG- CD- OE2
1323 1325 1327 1328 3 0.00000 0.00000 0.00000
0.00000 0.00000 0.00000 ; C- N- CA- HA
1323 1325 1327 1329 3 11.71520 1.67360 15.06240 -
13.38880 0.00000 0.00000 ; C- N- CA- CB
1323 1325 1327 1337 3 1.75728 5.27184 2.25936
-7.02912 0.00000 0.00000 ; C- N- CA- C
1324 1323 1325 1326 3 29.28800 -8.36800 -20.92000
0.00000 0.00000 0.00000 ; O- C- N- H
1324 1323 1325 1327 3 20.92000 0.00000 -20.92000
0.00000 0.00000 0.00000 ; O- C- N- CA

1325 1327 1329 1330 3 0.65084 1.95253 0.00000
-2.60338 0.00000 0.00000 ; N- CA- CB- HB2
1325 1327 1329 1331 3 0.65084 1.95253 0.00000
-2.60338 0.00000 0.00000 ; N- CA- CB- HB3
1325 1327 1329 1332 3 5.99567 3.29281 -4.46014
-0.55229 1.97485 0.00000 ; N- CA- CB- CG
1325 1327 1337 1338 3 0.00000 0.00000 0.00000
0.00000 0.00000 0.00000 ; N- CA- C- O
1325 1327 1337 1339 3 17.40544 -5.02080 -13.22144
9.20480 0.00000 0.00000 ; N- CA- C- N
1326 1325 1327 1328 3 0.00000 0.00000 0.00000
0.00000 0.00000 0.00000 ; H- N- CA- HA
1326 1325 1327 1329 3 0.00000 0.00000 0.00000
0.00000 0.00000 0.00000 ; H- N- CA- CB
1326 1325 1327 1337 3 0.00000 0.00000 0.00000
0.00000 0.00000 0.00000 ; H- N- CA- C
1327 1329 1332 1333 3 0.00000 0.00000 0.00000
0.00000 0.00000 0.00000 ; CA- CB- CG- OD1
1327 1329 1332 1334 3 8.84916 -0.31380 -4.32626
5.03754 0.26778 0.00000 ; CA- CB- CG- ND2
1327 1337 1339 1340 3 20.92000 0.00000 -20.92000
0.00000 0.00000 0.00000 ; CA- C- N- H
1327 1337 1339 1341 3 20.92000 0.00000 -20.92000
0.00000 0.00000 0.00000 ; CA- C- N- CA
1328 1327 1329 1330 3 0.65084 1.95253 0.00000
-2.60338 0.00000 0.00000 ; HA- CA- CB- HB2
1328 1327 1329 1331 3 0.65084 1.95253 0.00000
-2.60338 0.00000 0.00000 ; HA- CA- CB- HB3
1328 1327 1329 1332 3 0.65084 1.95253 0.00000
-2.60338 0.00000 0.00000 ; HA- CA- CB- CG
1328 1327 1337 1338 3 3.68192 -4.35136 0.00000
1.33888 0.00000 0.00000 ; HA- CA- C- O
1328 1327 1337 1339 3 0.00000 0.00000 0.00000
0.00000 0.00000 0.00000 ; HA- CA- C- N
1329 1327 1337 1338 3 0.00000 0.00000 0.00000
0.00000 0.00000 0.00000 ; CB- CA- C- O
1329 1327 1337 1339 3 2.51040 4.18400 1.67360
-6.69440 0.00000 0.00000 ; CB- CA- C- N
1329 1332 1334 1335 3 20.92000 0.00000 -20.92000
0.00000 0.00000 0.00000 ; CB- CG- ND2- HD21
1329 1332 1334 1336 3 20.92000 0.00000 -20.92000
0.00000 0.00000 0.00000 ; CB- CG- ND2- HD22
1330 1329 1327 1337 3 0.65084 1.95253 0.00000
-2.60338 0.00000 0.00000 ; HB2- CB- CA- C
1330 1329 1332 1333 3 3.68192 -4.35136 0.00000
1.33888 0.00000 0.00000 ; HB2- CB- CG- OD1
1330 1329 1332 1334 3 0.00000 0.00000 0.00000
0.00000 0.00000 0.00000 ; HB2- CB- CG- ND2
1331 1329 1327 1337 3 0.65084 1.95253 0.00000
-2.60338 0.00000 0.00000 ; HB3- CB- CA- C
1331 1329 1332 1333 3 3.68192 -4.35136 0.00000
1.33888 0.00000 0.00000 ; HB3- CB- CG- OD1

1331 1329 1332 1334 3 0.00000 0.00000 0.00000
0.00000 0.00000 0.00000 ; HB3- CB- CG- ND2
1332 1329 1327 1337 3 7.94542 -3.96225 -6.11701
-0.55229 3.58150 0.00000 ; CG- CB- CA- C
1333 1332 1334 1335 3 29.28800 -8.36800 -20.92000
0.00000 0.00000 0.00000 ; OD1- CG- ND2- HD21
1333 1332 1334 1336 3 29.28800 -8.36800 -20.92000
0.00000 0.00000 0.00000 ; OD1- CG- ND2- HD22
1337 1339 1341 1342 3 0.00000 0.00000 0.00000
0.00000 0.00000 0.00000 ; C- N- CA- HA
1337 1339 1341 1343 3 11.71520 1.67360 15.06240 -
13.38880 0.00000 0.00000 ; C- N- CA- CB
1337 1339 1341 1349 3 1.75728 5.27184 2.25936
-7.02912 0.00000 0.00000 ; C- N- CA- C
1338 1337 1339 1340 3 29.28800 -8.36800 -20.92000
0.00000 0.00000 0.00000 ; O- C- N- H
1338 1337 1339 1341 3 20.92000 0.00000 -20.92000
0.00000 0.00000 0.00000 ; O- C- N- CA
1339 1341 1343 1344 3 0.65084 1.95253 0.00000
-2.60338 0.00000 0.00000 ; N- CA- CB- HB2
1339 1341 1343 1345 3 0.65084 1.95253 0.00000
-2.60338 0.00000 0.00000 ; N- CA- CB- HB3
1339 1341 1343 1346 3 15.41386 9.74035 -8.39310
-0.97069 2.97901 0.00000 ; N- CA- CB- CG
1339 1341 1349 1350 3 0.00000 0.00000 0.00000
0.00000 0.00000 0.00000 ; N- CA- C- O
1339 1341 1349 1351 3 17.40544 -5.02080 -13.22144
9.20480 0.00000 0.00000 ; N- CA- C- N
1340 1339 1341 1342 3 0.00000 0.00000 0.00000
0.00000 0.00000 0.00000 ; H- N- CA- HA
1340 1339 1341 1343 3 0.00000 0.00000 0.00000
0.00000 0.00000 0.00000 ; H- N- CA- CB
1340 1339 1341 1349 3 0.00000 0.00000 0.00000
0.00000 0.00000 0.00000 ; H- N- CA- C
1341 1343 1346 1347 3 6.43499 0.00000 -5.39736
0.00000 -1.03763 0.00000 ; CA- CB- CG- OD1
1341 1343 1346 1348 3 6.43499 0.00000 -5.39736
0.00000 -1.03763 0.00000 ; CA- CB- CG- OD2
1341 1349 1351 1352 3 20.92000 0.00000 -20.92000
0.00000 0.00000 0.00000 ; CA- C- N- H
1341 1349 1351 1353 3 20.92000 0.00000 -20.92000
0.00000 0.00000 0.00000 ; CA- C- N- CA
1342 1341 1343 1344 3 0.65084 1.95253 0.00000
-2.60338 0.00000 0.00000 ; HA- CA- CB- HB2
1342 1341 1343 1345 3 0.65084 1.95253 0.00000
-2.60338 0.00000 0.00000 ; HA- CA- CB- HB3
1342 1341 1343 1346 3 0.65084 1.95253 0.00000
-2.60338 0.00000 0.00000 ; HA- CA- CB- CG
1342 1341 1349 1350 3 3.68192 -4.35136 0.00000
1.33888 0.00000 0.00000 ; HA- CA- C- O
1342 1341 1349 1351 3 0.00000 0.00000 0.00000
0.00000 0.00000 0.00000 ; HA- CA- C- N

1343 1341 1349 1350 3 0.00000 0.00000 0.00000
0.00000 0.00000 0.00000 ; CB- CA- C- O
1343 1341 1349 1351 3 2.51040 4.18400 1.67360
-6.69440 0.00000 0.00000 ; CB- CA- C- N
1344 1343 1341 1349 3 0.65084 1.95253 0.00000
-2.60338 0.00000 0.00000 ; HB2- CB- CA- C
1344 1343 1346 1347 3 0.00000 0.00000 0.00000
0.00000 0.00000 0.00000 ; HB2- CB- CG- OD1
1344 1343 1346 1348 3 0.00000 0.00000 0.00000
0.00000 0.00000 0.00000 ; HB2- CB- CG- OD2
1345 1343 1341 1349 3 0.65084 1.95253 0.00000
-2.60338 0.00000 0.00000 ; HB3- CB- CA- C
1345 1343 1346 1347 3 0.00000 0.00000 0.00000
0.00000 0.00000 0.00000 ; HB3- CB- CG- OD1
1345 1343 1346 1348 3 0.00000 0.00000 0.00000
0.00000 0.00000 0.00000 ; HB3- CB- CG- OD2
1346 1343 1341 1349 3 7.14627 -1.04600 -8.99560
-0.97069 5.15469 0.00000 ; CG- CB- CA- C
1349 1351 1353 1354 3 0.00000 0.00000 0.00000
0.00000 0.00000 0.00000 ; C- N- CA- HA
1349 1351 1353 1355 3 11.71520 1.67360 15.06240 -
13.38880 0.00000 0.00000 ; C- N- CA- CB
1349 1351 1353 1363 3 1.75728 5.27184 2.25936
-7.02912 0.00000 0.00000 ; C- N- CA- C
1350 1349 1351 1352 3 29.28800 -8.36800 -20.92000
0.00000 0.00000 0.00000 ; O- C- N- H
1350 1349 1351 1353 3 20.92000 0.00000 -20.92000
0.00000 0.00000 0.00000 ; O- C- N- CA
1351 1353 1355 1356 3 0.65084 1.95253 0.00000
-2.60338 0.00000 0.00000 ; N- CA- CB- HB2
1351 1353 1355 1357 3 0.65084 1.95253 0.00000
-2.60338 0.00000 0.00000 ; N- CA- CB- HB3
1351 1353 1355 1358 3 5.99567 3.29281 -4.46014
-0.55229 1.97485 0.00000 ; N- CA- CB- CG
1351 1353 1363 1364 3 0.00000 0.00000 0.00000
0.00000 0.00000 0.00000 ; N- CA- C- O
1351 1353 1363 1365 3 17.40544 -5.02080 -13.22144
9.20480 0.00000 0.00000 ; N- CA- C- N
1352 1351 1353 1354 3 0.00000 0.00000 0.00000
0.00000 0.00000 0.00000 ; H- N- CA- HA
1352 1351 1353 1355 3 0.00000 0.00000 0.00000
0.00000 0.00000 0.00000 ; H- N- CA- CB
1352 1351 1353 1363 3 0.00000 0.00000 0.00000
0.00000 0.00000 0.00000 ; H- N- CA- C
1353 1355 1358 1359 3 0.00000 0.00000 0.00000
0.00000 0.00000 0.00000 ; CA- CB- CG- OD1
1353 1355 1358 1360 3 8.84916 -0.31380 -4.32626
5.03754 0.26778 0.00000 ; CA- CB- CG- ND2
1353 1363 1365 1366 3 20.92000 0.00000 -20.92000
0.00000 0.00000 0.00000 ; CA- C- N- H
1353 1363 1365 1367 3 20.92000 0.00000 -20.92000
0.00000 0.00000 0.00000 ; CA- C- N- CA

1354 1353 1355 1356 3 0.65084 1.95253 0.00000
-2.60338 0.00000 0.00000 ; HA- CA- CB- HB2
1354 1353 1355 1357 3 0.65084 1.95253 0.00000
-2.60338 0.00000 0.00000 ; HA- CA- CB- HB3
1354 1353 1355 1358 3 0.65084 1.95253 0.00000
-2.60338 0.00000 0.00000 ; HA- CA- CB- CG
1354 1353 1363 1364 3 3.68192 -4.35136 0.00000
1.33888 0.00000 0.00000 ; HA- CA- C- O
1354 1353 1363 1365 3 0.00000 0.00000 0.00000
0.00000 0.00000 0.00000 ; HA- CA- C- N
1355 1353 1363 1364 3 0.00000 0.00000 0.00000
0.00000 0.00000 0.00000 ; CB- CA- C- O
1355 1353 1363 1365 3 2.51040 4.18400 1.67360
-6.69440 0.00000 0.00000 ; CB- CA- C- N
1355 1358 1360 1361 3 20.92000 0.00000 -20.92000
0.00000 0.00000 0.00000 ; CB- CG- ND2- HD21
1355 1358 1360 1362 3 20.92000 0.00000 -20.92000
0.00000 0.00000 0.00000 ; CB- CG- ND2- HD22
1356 1355 1353 1363 3 0.65084 1.95253 0.00000
-2.60338 0.00000 0.00000 ; HB2- CB- CA- C
1356 1355 1358 1359 3 3.68192 -4.35136 0.00000
1.33888 0.00000 0.00000 ; HB2- CB- CG- OD1
1356 1355 1358 1360 3 0.00000 0.00000 0.00000
0.00000 0.00000 0.00000 ; HB2- CB- CG- ND2
1357 1355 1353 1363 3 0.65084 1.95253 0.00000
-2.60338 0.00000 0.00000 ; HB3- CB- CA- C
1357 1355 1358 1359 3 3.68192 -4.35136 0.00000
1.33888 0.00000 0.00000 ; HB3- CB- CG- OD1
1357 1355 1358 1360 3 0.00000 0.00000 0.00000
0.00000 0.00000 0.00000 ; HB3- CB- CG- ND2
1358 1355 1353 1363 3 7.94542 -3.96225 -6.11701
-0.55229 3.58150 0.00000 ; CG- CB- CA- C
1359 1358 1360 1361 3 29.28800 -8.36800 -20.92000
0.00000 0.00000 0.00000 ; OD1- CG- ND2- HD21
1359 1358 1360 1362 3 29.28800 -8.36800 -20.92000
0.00000 0.00000 0.00000 ; OD1- CG- ND2- HD22
1363 1365 1367 1368 3 0.00000 0.00000 0.00000
0.00000 0.00000 0.00000 ; C- N- CA- HA
1363 1365 1367 1369 3 11.71520 1.67360 15.06240 -
13.38880 0.00000 0.00000 ; C- N- CA- CB
1363 1365 1367 1384 3 1.75728 5.27184 2.25936
-7.02912 0.00000 0.00000 ; C- N- CA- C
1364 1363 1365 1366 3 29.28800 -8.36800 -20.92000
0.00000 0.00000 0.00000 ; O- C- N- H
1364 1363 1365 1367 3 20.92000 0.00000 -20.92000
0.00000 0.00000 0.00000 ; O- C- N- CA
1365 1367 1369 1370 3 0.65084 1.95253 0.00000
-2.60338 0.00000 0.00000 ; N- CA- CB- HB2
1365 1367 1369 1371 3 0.65084 1.95253 0.00000
-2.60338 0.00000 0.00000 ; N- CA- CB- HB3
1365 1367 1369 1372 3 3.28026 2.46019 -2.19242
-3.21331 -0.23430 0.00000 ; N- CA- CB- CG

```
  1365   1367   1384   1385      3    0.00000    0.00000    0.00000
0.00000    0.00000    0.00000 ;      N-    CA-     C-     O
  1365   1367   1384   1386      3   17.40544   -5.02080  -13.22144
9.20480    0.00000    0.00000 ;      N-    CA-     C-     N
  1366   1365   1367   1368      3    0.00000    0.00000    0.00000
0.00000    0.00000    0.00000 ;      H-     N-    CA-    HA
  1366   1365   1367   1369      3    0.00000    0.00000    0.00000
0.00000    0.00000    0.00000 ;      H-     N-    CA-    CB
  1366   1365   1367   1384      3    0.00000    0.00000    0.00000
0.00000    0.00000    0.00000 ;      H-     N-    CA-     C
  1367   1369   1372   1373      3    0.57739    0.00000    1.02926
0.00000   -1.60666    0.00000 ;     CA-    CB-    CG-   CD1
  1367   1369   1372   1382      3    0.57739    0.00000    1.02926
0.00000   -1.60666    0.00000 ;     CA-    CB-    CG-   CD2
  1367   1384   1386   1387      3   20.92000    0.00000  -20.92000
0.00000    0.00000    0.00000 ;     CA-     C-     N-     H
  1367   1384   1386   1388      3   20.92000    0.00000  -20.92000
0.00000    0.00000    0.00000 ;     CA-     C-     N-    CA
  1368   1367   1369   1370      3    0.65084    1.95253    0.00000
-2.60338    0.00000    0.00000 ;     HA-    CA-    CB-   HB2
  1368   1367   1369   1371      3    0.65084    1.95253    0.00000
-2.60338    0.00000    0.00000 ;     HA-    CA-    CB-   HB3
  1368   1367   1369   1372      3    0.65084    1.95253    0.00000
-2.60338    0.00000    0.00000 ;     HA-    CA-    CB-    CG
  1368   1367   1384   1385      3    3.68192   -4.35136    0.00000
1.33888    0.00000    0.00000 ;     HA-    CA-     C-     O
  1368   1367   1384   1386      3    0.00000    0.00000    0.00000
0.00000    0.00000    0.00000 ;     HA-    CA-     C-     N
  1369   1367   1384   1385      3    0.00000    0.00000    0.00000
0.00000    0.00000    0.00000 ;     CB-    CA-     C-     O
  1369   1367   1384   1386      3    2.51040    4.18400    1.67360
-6.69440    0.00000    0.00000 ;     CB-    CA-     C-     N
  1369   1372   1373   1374      3   30.33400    0.00000  -30.33400
0.00000    0.00000    0.00000 ;     CB-    CG-   CD1-   HD1
  1369   1372   1373   1375      3   30.33400    0.00000  -30.33400
0.00000    0.00000    0.00000 ;     CB-    CG-   CD1-   CE1
  1369   1372   1382   1380      3   30.33400    0.00000  -30.33400
0.00000    0.00000    0.00000 ;     CB-    CG-   CD2-   CE2
  1369   1372   1382   1383      3   30.33400    0.00000  -30.33400
0.00000    0.00000    0.00000 ;     CB-    CG-   CD2-   HD2
  1370   1369   1367   1384      3    0.65084    1.95253    0.00000
-2.60338    0.00000    0.00000 ;    HB2-    CB-    CA-     C
  1370   1369   1372   1373      3    0.00000    0.00000    0.00000
0.00000    0.00000    0.00000 ;    HB2-    CB-    CG-   CD1
  1370   1369   1372   1382      3    0.00000    0.00000    0.00000
0.00000    0.00000    0.00000 ;    HB2-    CB-    CG-   CD2
  1371   1369   1367   1384      3    0.65084    1.95253    0.00000
-2.60338    0.00000    0.00000 ;    HB3-    CB-    CA-     C
  1371   1369   1372   1373      3    0.00000    0.00000    0.00000
0.00000    0.00000    0.00000 ;    HB3-    CB-    CG-   CD1
  1371   1369   1372   1382      3    0.00000    0.00000    0.00000
0.00000    0.00000    0.00000 ;    HB3-    CB-    CG-   CD2
```

1372 1369 1367 1384 3 4.95804 2.17986 -3.52293
-3.21331 -0.40166 0.00000 ; CG- CB- CA- C
1372 1373 1375 1376 3 30.33400 0.00000 -30.33400
0.00000 0.00000 0.00000 ; CG- CD1- CE1- HE1
1372 1373 1375 1377 3 30.33400 0.00000 -30.33400
0.00000 0.00000 0.00000 ; CG- CD1- CE1- CZ
1372 1382 1380 1377 3 30.33400 0.00000 -30.33400
0.00000 0.00000 0.00000 ; CG- CD2- CE2- CZ
1372 1382 1380 1381 3 30.33400 0.00000 -30.33400
0.00000 0.00000 0.00000 ; CG- CD2- CE2- HE2
1373 1372 1382 1380 3 30.33400 0.00000 -30.33400
0.00000 0.00000 0.00000 ; CD1- CG- CD2- CE2
1373 1372 1382 1383 3 30.33400 0.00000 -30.33400
0.00000 0.00000 0.00000 ; CD1- CG- CD2- HD2
1373 1375 1377 1378 3 30.33400 0.00000 -30.33400
0.00000 0.00000 0.00000 ; CD1- CE1- CZ- OH
1373 1375 1377 1380 3 30.33400 0.00000 -30.33400
0.00000 0.00000 0.00000 ; CD1- CE1- CZ- CE2
1374 1373 1372 1382 3 30.33400 0.00000 -30.33400
0.00000 0.00000 0.00000 ; HD1- CD1- CG- CD2
1374 1373 1375 1376 3 30.33400 0.00000 -30.33400
0.00000 0.00000 0.00000 ; HD1- CD1- CE1- HE1
1374 1373 1375 1377 3 30.33400 0.00000 -30.33400
0.00000 0.00000 0.00000 ; HD1- CD1- CE1- CZ
1375 1373 1372 1382 3 30.33400 0.00000 -30.33400
0.00000 0.00000 0.00000 ; CE1- CD1- CG- CD2
1375 1377 1378 1379 3 7.93286 0.00000 -9.56462
0.00000 2.17568 0.00000 ; CE1- CZ- OH- HH
1375 1377 1380 1381 3 30.33400 0.00000 -30.33400
0.00000 0.00000 0.00000 ; CE1- CZ- CE2- HE2
1375 1377 1380 1382 3 30.33400 0.00000 -30.33400
0.00000 0.00000 0.00000 ; CE1- CZ- CE2- CD2
1376 1375 1377 1378 3 30.33400 0.00000 -30.33400
0.00000 0.00000 0.00000 ; HE1- CE1- CZ- OH
1376 1375 1377 1380 3 30.33400 0.00000 -30.33400
0.00000 0.00000 0.00000 ; HE1- CE1- CZ- CE2
1377 1380 1382 1383 3 30.33400 0.00000 -30.33400
0.00000 0.00000 0.00000 ; CZ- CE2- CD2- HD2
1378 1377 1380 1381 3 30.33400 0.00000 -30.33400
0.00000 0.00000 0.00000 ; OH- CZ- CE2- HE2
1378 1377 1380 1382 3 30.33400 0.00000 -30.33400
0.00000 0.00000 0.00000 ; OH- CZ- CE2- CD2
1379 1378 1377 1380 3 7.93286 0.00000 -9.56462
0.00000 2.17568 0.00000 ; HH- OH- CZ- CE2
1381 1380 1382 1383 3 30.33400 0.00000 -30.33400
0.00000 0.00000 0.00000 ; HE2- CE2- CD2- HD2
1384 1386 1388 1389 3 0.00000 0.00000 0.00000
0.00000 0.00000 0.00000 ; C- N- CA- HA
1384 1386 1388 1390 3 11.71520 1.67360 15.06240 -
13.38880 0.00000 0.00000 ; C- N- CA- CB
1384 1386 1388 1403 3 1.75728 5.27184 2.25936
-7.02912 0.00000 0.00000 ; C- N- CA- C

1385 1384 1386 1387 3 29.28800 -8.36800 -20.92000
0.00000 0.00000 0.00000 ; O- C- N- H
1385 1384 1386 1388 3 20.92000 0.00000 -20.92000
0.00000 0.00000 0.00000 ; O- C- N- CA
1386 1388 1390 1391 3 0.65084 1.95253 0.00000
-2.60338 0.00000 0.00000 ; N- CA- CB- HB
1386 1388 1390 1392 3 3.83673 0.44769 -1.77402
-2.47693 -0.03347 0.00000 ; N- CA- CB- CG2
1386 1388 1390 1396 3 2.97482 0.12134 2.04179
-1.89117 -3.24678 0.00000 ; N- CA- CB- CG1
1386 1388 1403 1404 3 0.00000 0.00000 0.00000
0.00000 0.00000 0.00000 ; N- CA- C- O
1386 1388 1403 1405 3 17.40544 -5.02080 -13.22144
9.20480 0.00000 0.00000 ; N- CA- C- N
1387 1386 1388 1389 3 0.00000 0.00000 0.00000
0.00000 0.00000 0.00000 ; H- N- CA- HA
1387 1386 1388 1390 3 0.00000 0.00000 0.00000
0.00000 0.00000 0.00000 ; H- N- CA- CB
1387 1386 1388 1403 3 0.00000 0.00000 0.00000
0.00000 0.00000 0.00000 ; H- N- CA- C
1388 1390 1392 1393 3 0.66944 2.00832 0.00000
-2.67776 0.00000 0.00000 ; CA- CB- CG2- HG21
1388 1390 1392 1394 3 0.66944 2.00832 0.00000
-2.67776 0.00000 0.00000 ; CA- CB- CG2- HG22
1388 1390 1392 1395 3 0.66944 2.00832 0.00000
-2.67776 0.00000 0.00000 ; CA- CB- CG2- HG23
1388 1390 1396 1397 3 0.66944 2.00832 0.00000
-2.67776 0.00000 0.00000 ; CA- CB- CG1- HG12
1388 1390 1396 1398 3 0.66944 2.00832 0.00000
-2.67776 0.00000 0.00000 ; CA- CB- CG1- HG13
1388 1390 1396 1399 3 4.24258 -0.52718 -7.25506
-1.79075 7.69856 0.00000 ; CA- CB- CG1- CD1
1388 1403 1405 1406 3 20.92000 0.00000 -20.92000
0.00000 0.00000 0.00000 ; CA- C- N- H
1388 1403 1405 1407 3 20.92000 0.00000 -20.92000
0.00000 0.00000 0.00000 ; CA- C- N- CA
1389 1388 1390 1391 3 0.65084 1.95253 0.00000
-2.60338 0.00000 0.00000 ; HA- CA- CB- HB
1389 1388 1390 1392 3 0.65084 1.95253 0.00000
-2.60338 0.00000 0.00000 ; HA- CA- CB- CG2
1389 1388 1390 1396 3 0.65084 1.95253 0.00000
-2.60338 0.00000 0.00000 ; HA- CA- CB- CG1
1389 1388 1403 1404 3 3.68192 -4.35136 0.00000
1.33888 0.00000 0.00000 ; HA- CA- C- O
1389 1388 1403 1405 3 0.00000 0.00000 0.00000
0.00000 0.00000 0.00000 ; HA- CA- C- N
1390 1388 1403 1404 3 0.00000 0.00000 0.00000
0.00000 0.00000 0.00000 ; CB- CA- C- O
1390 1388 1403 1405 3 2.51040 4.18400 1.67360
-6.69440 0.00000 0.00000 ; CB- CA- C- N
1390 1396 1399 1400 3 0.66944 2.00832 0.00000
-2.67776 0.00000 0.00000 ; CB- CG1- CD1- HD11

1390 1396 1399 1401 3 0.66944 2.00832 0.00000
-2.67776 0.00000 0.00000 ; CB- CG1- CD1- HD12
1390 1396 1399 1402 3 0.66944 2.00832 0.00000
-2.67776 0.00000 0.00000 ; CB- CG1- CD1- HD13
1391 1390 1388 1403 3 0.65084 1.95253 0.00000
-2.60338 0.00000 0.00000 ; HB- CB- CA- C
1391 1390 1392 1393 3 0.62760 1.88280 0.00000
-2.51040 0.00000 0.00000 ; HB- CB- CG2- HG21
1391 1390 1392 1394 3 0.62760 1.88280 0.00000
-2.51040 0.00000 0.00000 ; HB- CB- CG2- HG22
1391 1390 1392 1395 3 0.62760 1.88280 0.00000
-2.51040 0.00000 0.00000 ; HB- CB- CG2- HG23
1391 1390 1396 1397 3 0.62760 1.88280 0.00000
-2.51040 0.00000 0.00000 ; HB- CB- CG1- HG12
1391 1390 1396 1398 3 0.62760 1.88280 0.00000
-2.51040 0.00000 0.00000 ; HB- CB- CG1- HG13
1391 1390 1396 1399 3 0.66944 2.00832 0.00000
-2.67776 0.00000 0.00000 ; HB- CB- CG1- CD1
1392 1390 1388 1403 3 5.67350 3.55640 -6.16722
-2.47693 3.74886 0.00000 ; CG2- CB- CA- C
1392 1390 1396 1397 3 0.66944 2.00832 0.00000
-2.67776 0.00000 0.00000 ; CG2- CB- CG1- HG12
1392 1390 1396 1398 3 0.66944 2.00832 0.00000
-2.67776 0.00000 0.00000 ; CG2- CB- CG1- HG13
1392 1390 1396 1399 3 3.81162 0.49790 -8.14206
-1.79075 7.49773 0.00000 ; CG2- CB- CG1- CD1
1393 1392 1390 1396 3 0.66944 2.00832 0.00000
-2.67776 0.00000 0.00000 ; HG21- CG2- CB- CG1
1394 1392 1390 1396 3 0.66944 2.00832 0.00000
-2.67776 0.00000 0.00000 ; HG22- CG2- CB- CG1
1395 1392 1390 1396 3 0.66944 2.00832 0.00000
-2.67776 0.00000 0.00000 ; HG23- CG2- CB- CG1
1396 1390 1388 1403 3 8.26340 0.74057 -9.99976
-1.89117 3.84928 0.00000 ; CG1- CB- CA- C
1397 1396 1399 1400 3 0.62760 1.88280 0.00000
-2.51040 0.00000 0.00000 ; HG12- CG1- CD1- HD11
1397 1396 1399 1401 3 0.62760 1.88280 0.00000
-2.51040 0.00000 0.00000 ; HG12- CG1- CD1- HD12
1397 1396 1399 1402 3 0.62760 1.88280 0.00000
-2.51040 0.00000 0.00000 ; HG12- CG1- CD1- HD13
1398 1396 1399 1400 3 0.62760 1.88280 0.00000
-2.51040 0.00000 0.00000 ; HG13- CG1- CD1- HD11
1398 1396 1399 1401 3 0.62760 1.88280 0.00000
-2.51040 0.00000 0.00000 ; HG13- CG1- CD1- HD12
1398 1396 1399 1402 3 0.62760 1.88280 0.00000
-2.51040 0.00000 0.00000 ; HG13- CG1- CD1- HD13
1403 1405 1407 1408 3 0.00000 0.00000 0.00000
0.00000 0.00000 0.00000 ; C- N- CA- HA
1403 1405 1407 1409 3 11.71520 1.67360 15.06240 -
13.38880 0.00000 0.00000 ; C- N- CA- CB
1403 1405 1407 1424 3 1.75728 5.27184 2.25936
-7.02912 0.00000 0.00000 ; C- N- CA- C

1404 1403 1405 1406 3 29.28800 -8.36800 -20.92000
0.00000 0.00000 0.00000 ; O- C- N- H
1404 1403 1405 1407 3 20.92000 0.00000 -20.92000
0.00000 0.00000 0.00000 ; O- C- N- CA
1405 1407 1409 1410 3 0.65084 1.95253 0.00000
-2.60338 0.00000 0.00000 ; N- CA- CB- HB2
1405 1407 1409 1411 3 0.65084 1.95253 0.00000
-2.60338 0.00000 0.00000 ; N- CA- CB- HB3
1405 1407 1409 1412 3 3.28026 2.46019 -2.19242
-3.21331 -0.23430 0.00000 ; N- CA- CB- CG
1405 1407 1424 1425 3 0.00000 0.00000 0.00000
0.00000 0.00000 0.00000 ; N- CA- C- O
1405 1407 1424 1426 3 17.40544 -5.02080 -13.22144
9.20480 0.00000 0.00000 ; N- CA- C- N
1406 1405 1407 1408 3 0.00000 0.00000 0.00000
0.00000 0.00000 0.00000 ; H- N- CA- HA
1406 1405 1407 1409 3 0.00000 0.00000 0.00000
0.00000 0.00000 0.00000 ; H- N- CA- CB
1406 1405 1407 1424 3 0.00000 0.00000 0.00000
0.00000 0.00000 0.00000 ; H- N- CA- C
1407 1409 1412 1413 3 0.57739 0.00000 1.02926
0.00000 -1.60666 0.00000 ; CA- CB- CG- CD1
1407 1409 1412 1422 3 0.57739 0.00000 1.02926
0.00000 -1.60666 0.00000 ; CA- CB- CG- CD2
1407 1424 1426 1427 3 20.92000 0.00000 -20.92000
0.00000 0.00000 0.00000 ; CA- C- N- H
1407 1424 1426 1428 3 20.92000 0.00000 -20.92000
0.00000 0.00000 0.00000 ; CA- C- N- CA
1408 1407 1409 1410 3 0.65084 1.95253 0.00000
-2.60338 0.00000 0.00000 ; HA- CA- CB- HB2
1408 1407 1409 1411 3 0.65084 1.95253 0.00000
-2.60338 0.00000 0.00000 ; HA- CA- CB- HB3
1408 1407 1409 1412 3 0.65084 1.95253 0.00000
-2.60338 0.00000 0.00000 ; HA- CA- CB- CG
1408 1407 1424 1425 3 3.68192 -4.35136 0.00000
1.33888 0.00000 0.00000 ; HA- CA- C- O
1408 1407 1424 1426 3 0.00000 0.00000 0.00000
0.00000 0.00000 0.00000 ; HA- CA- C- N
1409 1407 1424 1425 3 0.00000 0.00000 0.00000
0.00000 0.00000 0.00000 ; CB- CA- C- O
1409 1407 1424 1426 3 2.51040 4.18400 1.67360
-6.69440 0.00000 0.00000 ; CB- CA- C- N
1409 1412 1413 1414 3 30.33400 0.00000 -30.33400
0.00000 0.00000 0.00000 ; CB- CG- CD1- HD1
1409 1412 1413 1415 3 30.33400 0.00000 -30.33400
0.00000 0.00000 0.00000 ; CB- CG- CD1- CE1
1409 1412 1422 1420 3 30.33400 0.00000 -30.33400
0.00000 0.00000 0.00000 ; CB- CG- CD2- CE2
1409 1412 1422 1423 3 30.33400 0.00000 -30.33400
0.00000 0.00000 0.00000 ; CB- CG- CD2- HD2
1410 1409 1407 1424 3 0.65084 1.95253 0.00000
-2.60338 0.00000 0.00000 ; HB2- CB- CA- C

1410 1409 1412 1413 3 0.00000 0.00000 0.00000
0.00000 0.00000 0.00000 ; HB2- CB- CG- CD1
1410 1409 1412 1422 3 0.00000 0.00000 0.00000
0.00000 0.00000 0.00000 ; HB2- CB- CG- CD2
1411 1409 1407 1424 3 0.65084 1.95253 0.00000
-2.60338 0.00000 0.00000 ; HB3- CB- CA- C
1411 1409 1412 1413 3 0.00000 0.00000 0.00000
0.00000 0.00000 0.00000 ; HB3- CB- CG- CD1
1411 1409 1412 1422 3 0.00000 0.00000 0.00000
0.00000 0.00000 0.00000 ; HB3- CB- CG- CD2
1412 1409 1407 1424 3 4.95804 2.17986 -3.52293
-3.21331 -0.40166 0.00000 ; CG- CB- CA- C
1412 1413 1415 1416 3 30.33400 0.00000 -30.33400
0.00000 0.00000 0.00000 ; CG- CD1- CE1- HE1
1412 1413 1415 1417 3 30.33400 0.00000 -30.33400
0.00000 0.00000 0.00000 ; CG- CD1- CE1- CZ
1412 1422 1420 1417 3 30.33400 0.00000 -30.33400
0.00000 0.00000 0.00000 ; CG- CD2- CE2- CZ
1412 1422 1420 1421 3 30.33400 0.00000 -30.33400
0.00000 0.00000 0.00000 ; CG- CD2- CE2- HE2
1413 1412 1422 1420 3 30.33400 0.00000 -30.33400
0.00000 0.00000 0.00000 ; CD1- CG- CD2- CE2
1413 1412 1422 1423 3 30.33400 0.00000 -30.33400
0.00000 0.00000 0.00000 ; CD1- CG- CD2- HD2
1413 1415 1417 1418 3 30.33400 0.00000 -30.33400
0.00000 0.00000 0.00000 ; CD1- CE1- CZ- OH
1413 1415 1417 1420 3 30.33400 0.00000 -30.33400
0.00000 0.00000 0.00000 ; CD1- CE1- CZ- CE2
1414 1413 1412 1422 3 30.33400 0.00000 -30.33400
0.00000 0.00000 0.00000 ; HD1- CD1- CG- CD2
1414 1413 1415 1416 3 30.33400 0.00000 -30.33400
0.00000 0.00000 0.00000 ; HD1- CD1- CE1- HE1
1414 1413 1415 1417 3 30.33400 0.00000 -30.33400
0.00000 0.00000 0.00000 ; HD1- CD1- CE1- CZ
1415 1413 1412 1422 3 30.33400 0.00000 -30.33400
0.00000 0.00000 0.00000 ; CE1- CD1- CG- CD2
1415 1417 1418 1419 3 7.93286 0.00000 -9.56462
0.00000 2.17568 0.00000 ; CE1- CZ- OH- HH
1415 1417 1420 1421 3 30.33400 0.00000 -30.33400
0.00000 0.00000 0.00000 ; CE1- CZ- CE2- HE2
1415 1417 1420 1422 3 30.33400 0.00000 -30.33400
0.00000 0.00000 0.00000 ; CE1- CZ- CE2- CD2
1416 1415 1417 1418 3 30.33400 0.00000 -30.33400
0.00000 0.00000 0.00000 ; HE1- CE1- CZ- OH
1416 1415 1417 1420 3 30.33400 0.00000 -30.33400
0.00000 0.00000 0.00000 ; HE1- CE1- CZ- CE2
1417 1420 1422 1423 3 30.33400 0.00000 -30.33400
0.00000 0.00000 0.00000 ; CZ- CE2- CD2- HD2
1418 1417 1420 1421 3 30.33400 0.00000 -30.33400
0.00000 0.00000 0.00000 ; OH- CZ- CE2- HE2
1418 1417 1420 1422 3 30.33400 0.00000 -30.33400
0.00000 0.00000 0.00000 ; OH- CZ- CE2- CD2

1419 1418 1417 1420 3 7.93286 0.00000 -9.56462
0.00000 2.17568 0.00000 ; HH- OH- CZ- CE2
1421 1420 1422 1423 3 30.33400 0.00000 -30.33400
0.00000 0.00000 0.00000 ; HE2- CE2- CD2- HD2
1424 1426 1428 1429 3 0.00000 0.00000 0.00000
0.00000 0.00000 0.00000 ; C- N- CA- HA
1424 1426 1428 1430 3 11.71520 1.67360 15.06240 -
13.38880 0.00000 0.00000 ; C- N- CA- CB
1424 1426 1428 1443 3 1.75728 5.27184 2.25936
-7.02912 0.00000 0.00000 ; C- N- CA- C
1425 1424 1426 1427 3 29.28800 -8.36800 -20.92000
0.00000 0.00000 0.00000 ; O- C- N- H
1425 1424 1426 1428 3 20.92000 0.00000 -20.92000
0.00000 0.00000 0.00000 ; O- C- N- CA
1426 1428 1430 1431 3 0.65084 1.95253 0.00000
-2.60338 0.00000 0.00000 ; N- CA- CB- HB2
1426 1428 1430 1432 3 0.65084 1.95253 0.00000
-2.60338 0.00000 0.00000 ; N- CA- CB- HB3
1426 1428 1430 1433 3 3.79070 1.39746 -4.61077
-2.40998 2.44346 0.00000 ; N- CA- CB- CG
1426 1428 1443 1444 3 0.00000 0.00000 0.00000
0.00000 0.00000 0.00000 ; N- CA- C- O
1426 1428 1443 1445 3 17.40544 -5.02080 -13.22144
9.20480 0.00000 0.00000 ; N- CA- C- N
1427 1426 1428 1429 3 0.00000 0.00000 0.00000
0.00000 0.00000 0.00000 ; H- N- CA- HA
1427 1426 1428 1430 3 0.00000 0.00000 0.00000
0.00000 0.00000 0.00000 ; H- N- CA- CB
1427 1426 1428 1443 3 0.00000 0.00000 0.00000
0.00000 0.00000 0.00000 ; H- N- CA- C
1428 1430 1433 1434 3 0.66944 2.00832 0.00000
-2.67776 0.00000 0.00000 ; CA- CB- CG- HG
1428 1430 1433 1435 3 3.90367 0.19665 -6.21742
-2.37651 5.99149 0.00000 ; CA- CB- CG- CD1
1428 1430 1433 1439 3 3.90367 0.19665 -6.21742
-2.37651 5.99149 0.00000 ; CA- CB- CG- CD2
1428 1443 1445 1446 3 20.92000 0.00000 -20.92000
0.00000 0.00000 0.00000 ; CA- C- N- H
1428 1443 1445 1447 3 20.92000 0.00000 -20.92000
0.00000 0.00000 0.00000 ; CA- C- N- CA
1429 1428 1430 1431 3 0.65084 1.95253 0.00000
-2.60338 0.00000 0.00000 ; HA- CA- CB- HB2
1429 1428 1430 1432 3 0.65084 1.95253 0.00000
-2.60338 0.00000 0.00000 ; HA- CA- CB- HB3
1429 1428 1430 1433 3 0.65084 1.95253 0.00000
-2.60338 0.00000 0.00000 ; HA- CA- CB- CG
1429 1428 1443 1444 3 3.68192 -4.35136 0.00000
1.33888 0.00000 0.00000 ; HA- CA- C- O
1429 1428 1443 1445 3 0.00000 0.00000 0.00000
0.00000 0.00000 0.00000 ; HA- CA- C- N
1430 1428 1443 1444 3 0.00000 0.00000 0.00000
0.00000 0.00000 0.00000 ; CB- CA- C- O

```
  1430   1428   1443   1445      3    2.51040    4.18400    1.67360
-6.69440    0.00000    0.00000 ;     CB-    CA-     C-     N
  1430   1433   1435   1436      3    0.66944    2.00832    0.00000
-2.67776    0.00000    0.00000 ;     CB-    CG-   CD1-  HD11
  1430   1433   1435   1437      3    0.66944    2.00832    0.00000
-2.67776    0.00000    0.00000 ;     CB-    CG-   CD1-  HD12
  1430   1433   1435   1438      3    0.66944    2.00832    0.00000
-2.67776    0.00000    0.00000 ;     CB-    CG-   CD1-  HD13
  1430   1433   1439   1440      3    0.66944    2.00832    0.00000
-2.67776    0.00000    0.00000 ;     CB-    CG-   CD2-  HD21
  1430   1433   1439   1441      3    0.66944    2.00832    0.00000
-2.67776    0.00000    0.00000 ;     CB-    CG-   CD2-  HD22
  1430   1433   1439   1442      3    0.66944    2.00832    0.00000
-2.67776    0.00000    0.00000 ;     CB-    CG-   CD2-  HD23
  1431   1430   1428   1443      3    0.65084    1.95253    0.00000
-2.60338    0.00000    0.00000 ;    HB2-    CB-    CA-     C
  1431   1430   1433   1434      3    0.62760    1.88280    0.00000
-2.51040    0.00000    0.00000 ;    HB2-    CB-    CG-    HG
  1431   1430   1433   1435      3    0.66944    2.00832    0.00000
-2.67776    0.00000    0.00000 ;    HB2-    CB-    CG-   CD1
  1431   1430   1433   1439      3    0.66944    2.00832    0.00000
-2.67776    0.00000    0.00000 ;    HB2-    CB-    CG-   CD2
  1432   1430   1428   1443      3    0.65084    1.95253    0.00000
-2.60338    0.00000    0.00000 ;    HB3-    CB-    CA-     C
  1432   1430   1433   1434      3    0.62760    1.88280    0.00000
-2.51040    0.00000    0.00000 ;    HB3-    CB-    CG-    HG
  1432   1430   1433   1435      3    0.66944    2.00832    0.00000
-2.67776    0.00000    0.00000 ;    HB3-    CB-    CG-   CD1
  1432   1430   1433   1439      3    0.66944    2.00832    0.00000
-2.67776    0.00000    0.00000 ;    HB3-    CB-    CG-   CD2
  1433   1430   1428   1443      3   10.33448   -1.14642  -11.54784
-2.40998    6.35968    0.00000 ;     CG-    CB-    CA-     C
  1434   1433   1435   1436      3    0.62760    1.88280    0.00000
-2.51040    0.00000    0.00000 ;     HG-    CG-   CD1-  HD11
  1434   1433   1435   1437      3    0.62760    1.88280    0.00000
-2.51040    0.00000    0.00000 ;     HG-    CG-   CD1-  HD12
  1434   1433   1435   1438      3    0.62760    1.88280    0.00000
-2.51040    0.00000    0.00000 ;     HG-    CG-   CD1-  HD13
  1434   1433   1439   1440      3    0.62760    1.88280    0.00000
-2.51040    0.00000    0.00000 ;     HG-    CG-   CD2-  HD21
  1434   1433   1439   1441      3    0.62760    1.88280    0.00000
-2.51040    0.00000    0.00000 ;     HG-    CG-   CD2-  HD22
  1434   1433   1439   1442      3    0.62760    1.88280    0.00000
-2.51040    0.00000    0.00000 ;     HG-    CG-   CD2-  HD23
  1435   1433   1439   1440      3    0.66944    2.00832    0.00000
-2.67776    0.00000    0.00000 ;    CD1-    CG-   CD2-  HD21
  1435   1433   1439   1441      3    0.66944    2.00832    0.00000
-2.67776    0.00000    0.00000 ;    CD1-    CG-   CD2-  HD22
  1435   1433   1439   1442      3    0.66944    2.00832    0.00000
-2.67776    0.00000    0.00000 ;    CD1-    CG-   CD2-  HD23
  1436   1435   1433   1439      3    0.66944    2.00832    0.00000
-2.67776    0.00000    0.00000 ;   HD11-   CD1-    CG-   CD2
```

1437 1435 1433 1439 3 0.66944 2.00832 0.00000
-2.67776 0.00000 0.00000 ; HD12- CD1- CG- CD2
1438 1435 1433 1439 3 0.66944 2.00832 0.00000
-2.67776 0.00000 0.00000 ; HD13- CD1- CG- CD2
1443 1445 1447 1448 3 0.00000 0.00000 0.00000
0.00000 0.00000 0.00000 ; C- N- CA- HA
1443 1445 1447 1449 3 11.71520 1.67360 15.06240 -
13.38880 0.00000 0.00000 ; C- N- CA- CB
1443 1445 1447 1462 3 1.75728 5.27184 2.25936
-7.02912 0.00000 0.00000 ; C- N- CA- C
1444 1443 1445 1446 3 29.28800 -8.36800 -20.92000
0.00000 0.00000 0.00000 ; O- C- N- H
1444 1443 1445 1447 3 20.92000 0.00000 -20.92000
0.00000 0.00000 0.00000 ; O- C- N- CA
1445 1447 1449 1450 3 0.65084 1.95253 0.00000
-2.60338 0.00000 0.00000 ; N- CA- CB- HB
1445 1447 1449 1451 3 3.83673 0.44769 -1.77402
-2.47693 -0.03347 0.00000 ; N- CA- CB- CG2
1445 1447 1449 1455 3 2.97482 0.12134 2.04179
-1.89117 -3.24678 0.00000 ; N- CA- CB- CG1
1445 1447 1462 1463 3 0.00000 0.00000 0.00000
0.00000 0.00000 0.00000 ; N- CA- C- O
1445 1447 1462 1464 3 17.40544 -5.02080 -13.22144
9.20480 0.00000 0.00000 ; N- CA- C- N
1446 1445 1447 1448 3 0.00000 0.00000 0.00000
0.00000 0.00000 0.00000 ; H- N- CA- HA
1446 1445 1447 1449 3 0.00000 0.00000 0.00000
0.00000 0.00000 0.00000 ; H- N- CA- CB
1446 1445 1447 1462 3 0.00000 0.00000 0.00000
0.00000 0.00000 0.00000 ; H- N- CA- C
1447 1449 1451 1452 3 0.66944 2.00832 0.00000
-2.67776 0.00000 0.00000 ; CA- CB- CG2- HG21
1447 1449 1451 1453 3 0.66944 2.00832 0.00000
-2.67776 0.00000 0.00000 ; CA- CB- CG2- HG22
1447 1449 1451 1454 3 0.66944 2.00832 0.00000
-2.67776 0.00000 0.00000 ; CA- CB- CG2- HG23
1447 1449 1455 1456 3 0.66944 2.00832 0.00000
-2.67776 0.00000 0.00000 ; CA- CB- CG1- HG12
1447 1449 1455 1457 3 0.66944 2.00832 0.00000
-2.67776 0.00000 0.00000 ; CA- CB- CG1- HG13
1447 1449 1455 1458 3 4.24258 -0.52718 -7.25506
-1.79075 7.69856 0.00000 ; CA- CB- CG1- CD1
1447 1462 1464 1465 3 20.92000 0.00000 -20.92000
0.00000 0.00000 0.00000 ; CA- C- N- H
1447 1462 1464 1466 3 20.92000 0.00000 -20.92000
0.00000 0.00000 0.00000 ; CA- C- N- CA
1448 1447 1449 1450 3 0.65084 1.95253 0.00000
-2.60338 0.00000 0.00000 ; HA- CA- CB- HB
1448 1447 1449 1451 3 0.65084 1.95253 0.00000
-2.60338 0.00000 0.00000 ; HA- CA- CB- CG2
1448 1447 1449 1455 3 0.65084 1.95253 0.00000
-2.60338 0.00000 0.00000 ; HA- CA- CB- CG1

1448 1447 1462 1463 3 3.68192 -4.35136 0.00000
1.33888 0.00000 0.00000 ; HA- CA- C- O
1448 1447 1462 1464 3 0.00000 0.00000 0.00000
0.00000 0.00000 0.00000 ; HA- CA- C- N
1449 1447 1462 1463 3 0.00000 0.00000 0.00000
0.00000 0.00000 0.00000 ; CB- CA- C- O
1449 1447 1462 1464 3 2.51040 4.18400 1.67360
-6.69440 0.00000 0.00000 ; CB- CA- C- N
1449 1455 1458 1459 3 0.66944 2.00832 0.00000
-2.67776 0.00000 0.00000 ; CB- CG1- CD1- HD11
1449 1455 1458 1460 3 0.66944 2.00832 0.00000
-2.67776 0.00000 0.00000 ; CB- CG1- CD1- HD12
1449 1455 1458 1461 3 0.66944 2.00832 0.00000
-2.67776 0.00000 0.00000 ; CB- CG1- CD1- HD13
1450 1449 1447 1462 3 0.65084 1.95253 0.00000
-2.60338 0.00000 0.00000 ; HB- CB- CA- C
1450 1449 1451 1452 3 0.62760 1.88280 0.00000
-2.51040 0.00000 0.00000 ; HB- CB- CG2- HG21
1450 1449 1451 1453 3 0.62760 1.88280 0.00000
-2.51040 0.00000 0.00000 ; HB- CB- CG2- HG22
1450 1449 1451 1454 3 0.62760 1.88280 0.00000
-2.51040 0.00000 0.00000 ; HB- CB- CG2- HG23
1450 1449 1455 1456 3 0.62760 1.88280 0.00000
-2.51040 0.00000 0.00000 ; HB- CB- CG1- HG12
1450 1449 1455 1457 3 0.62760 1.88280 0.00000
-2.51040 0.00000 0.00000 ; HB- CB- CG1- HG13
1450 1449 1455 1458 3 0.66944 2.00832 0.00000
-2.67776 0.00000 0.00000 ; HB- CB- CG1- CD1
1451 1449 1447 1462 3 5.67350 3.55640 -6.16722
-2.47693 3.74886 0.00000 ; CG2- CB- CA- C
1451 1449 1455 1456 3 0.66944 2.00832 0.00000
-2.67776 0.00000 0.00000 ; CG2- CB- CG1- HG12
1451 1449 1455 1457 3 0.66944 2.00832 0.00000
-2.67776 0.00000 0.00000 ; CG2- CB- CG1- HG13
1451 1449 1455 1458 3 3.81162 0.49790 -8.14206
-1.79075 7.49773 0.00000 ; CG2- CB- CG1- CD1
1452 1451 1449 1455 3 0.66944 2.00832 0.00000
-2.67776 0.00000 0.00000 ; HG21- CG2- CB- CG1
1453 1451 1449 1455 3 0.66944 2.00832 0.00000
-2.67776 0.00000 0.00000 ; HG22- CG2- CB- CG1
1454 1451 1449 1455 3 0.66944 2.00832 0.00000
-2.67776 0.00000 0.00000 ; HG23- CG2- CB- CG1
1455 1449 1447 1462 3 8.26340 0.74057 -9.99976
-1.89117 3.84928 0.00000 ; CG1- CB- CA- C
1456 1455 1458 1459 3 0.62760 1.88280 0.00000
-2.51040 0.00000 0.00000 ; HG12- CG1- CD1- HD11
1456 1455 1458 1460 3 0.62760 1.88280 0.00000
-2.51040 0.00000 0.00000 ; HG12- CG1- CD1- HD12
1456 1455 1458 1461 3 0.62760 1.88280 0.00000
-2.51040 0.00000 0.00000 ; HG12- CG1- CD1- HD13
1457 1455 1458 1459 3 0.62760 1.88280 0.00000
-2.51040 0.00000 0.00000 ; HG13- CG1- CD1- HD11

1457 1455 1458 1460 3 0.62760 1.88280 0.00000
-2.51040 0.00000 0.00000 ; HG13- CG1- CD1- HD12
1457 1455 1458 1461 3 0.62760 1.88280 0.00000
-2.51040 0.00000 0.00000 ; HG13- CG1- CD1- HD13
1462 1464 1466 1467 3 0.00000 0.00000 0.00000
0.00000 0.00000 0.00000 ; C- N- CA- HA
1462 1464 1466 1468 3 11.71520 1.67360 15.06240 -
13.38880 0.00000 0.00000 ; C- N- CA- CB
1462 1464 1466 1479 3 1.75728 5.27184 2.25936
-7.02912 0.00000 0.00000 ; C- N- CA- C
1463 1462 1464 1465 3 29.28800 -8.36800 -20.92000
0.00000 0.00000 0.00000 ; O- C- N- H
1463 1462 1464 1466 3 20.92000 0.00000 -20.92000
0.00000 0.00000 0.00000 ; O- C- N- CA
1464 1466 1468 1469 3 0.65084 1.95253 0.00000
-2.60338 0.00000 0.00000 ; N- CA- CB- HB2
1464 1466 1468 1470 3 0.65084 1.95253 0.00000
-2.60338 0.00000 0.00000 ; N- CA- CB- HB3
1464 1466 1468 1471 3 3.21331 2.22589 -4.15053
-2.40998 2.61082 0.00000 ; N- CA- CB- CG
1464 1466 1479 1480 3 0.00000 0.00000 0.00000
0.00000 0.00000 0.00000 ; N- CA- C- O
1464 1466 1479 1481 3 17.40544 -5.02080 -13.22144
9.20480 0.00000 0.00000 ; N- CA- C- N
1465 1464 1466 1467 3 0.00000 0.00000 0.00000
0.00000 0.00000 0.00000 ; H- N- CA- HA
1465 1464 1466 1468 3 0.00000 0.00000 0.00000
0.00000 0.00000 0.00000 ; H- N- CA- CB
1465 1464 1466 1479 3 0.00000 0.00000 0.00000
0.00000 0.00000 0.00000 ; H- N- CA- C
1466 1468 1471 1472 3 0.65084 1.95253 0.00000
-2.60338 0.00000 0.00000 ; CA- CB- CG- HG2
1466 1468 1471 1473 3 0.65084 1.95253 0.00000
-2.60338 0.00000 0.00000 ; CA- CB- CG- HG3
1466 1468 1471 1474 3 2.04598 -1.54390 1.11294
-0.26778 0.93722 0.00000 ; CA- CB- CG- SD
1466 1479 1481 1482 3 20.92000 0.00000 -20.92000
0.00000 0.00000 0.00000 ; CA- C- N- H
1466 1479 1481 1483 3 20.92000 0.00000 -20.92000
0.00000 0.00000 0.00000 ; CA- C- N- CA
1467 1466 1468 1469 3 0.65084 1.95253 0.00000
-2.60338 0.00000 0.00000 ; HA- CA- CB- HB2
1467 1466 1468 1470 3 0.65084 1.95253 0.00000
-2.60338 0.00000 0.00000 ; HA- CA- CB- HB3
1467 1466 1468 1471 3 0.65084 1.95253 0.00000
-2.60338 0.00000 0.00000 ; HA- CA- CB- CG
1467 1466 1479 1480 3 3.68192 -4.35136 0.00000
1.33888 0.00000 0.00000 ; HA- CA- C- O
1467 1466 1479 1481 3 0.00000 0.00000 0.00000
0.00000 0.00000 0.00000 ; HA- CA- C- N
1468 1466 1479 1480 3 0.00000 0.00000 0.00000
0.00000 0.00000 0.00000 ; CB- CA- C- O

1468 1466 1479 1481 3 2.51040 4.18400 1.67360
-6.69440 0.00000 0.00000 ; CB- CA- C- N
1468 1471 1474 1475 3 3.24260 6.22998 1.79075
-6.92870 1.90790 0.00000 ; CB- CG- SD- CE
1469 1468 1466 1479 3 0.65084 1.95253 0.00000
-2.60338 0.00000 0.00000 ; HB2- CB- CA- C
1469 1468 1471 1472 3 0.65084 1.95253 0.00000
-2.60338 0.00000 0.00000 ; HB2- CB- CG- HG2
1469 1468 1471 1473 3 0.65084 1.95253 0.00000
-2.60338 0.00000 0.00000 ; HB2- CB- CG- HG3
1469 1468 1471 1474 3 0.65084 1.95253 0.00000
-2.60338 0.00000 0.00000 ; HB2- CB- CG- SD
1470 1468 1466 1479 3 0.65084 1.95253 0.00000
-2.60338 0.00000 0.00000 ; HB3- CB- CA- C
1470 1468 1471 1472 3 0.65084 1.95253 0.00000
-2.60338 0.00000 0.00000 ; HB3- CB- CG- HG2
1470 1468 1471 1473 3 0.65084 1.95253 0.00000
-2.60338 0.00000 0.00000 ; HB3- CB- CG- HG3
1470 1468 1471 1474 3 0.65084 1.95253 0.00000
-2.60338 0.00000 0.00000 ; HB3- CB- CG- SD
1471 1468 1466 1479 3 6.86594 3.56895 -8.14206
-2.40998 4.85344 0.00000 ; CG- CB- CA- C
1471 1474 1475 1476 3 1.39467 4.18400 0.00000
-5.57867 0.00000 0.00000 ; CG- SD- CE- HE1
1471 1474 1475 1477 3 1.39467 4.18400 0.00000
-5.57867 0.00000 0.00000 ; CG- SD- CE- HE2
1471 1474 1475 1478 3 1.39467 4.18400 0.00000
-5.57867 0.00000 0.00000 ; CG- SD- CE- HE3
1472 1471 1474 1475 3 1.39467 4.18400 0.00000
-5.57867 0.00000 0.00000 ; HG2- CG- SD- CE
1473 1471 1474 1475 3 1.39467 4.18400 0.00000
-5.57867 0.00000 0.00000 ; HG3- CG- SD- CE
1479 1481 1483 1484 3 0.00000 0.00000 0.00000
0.00000 0.00000 0.00000 ; C- N- CA- HA
1479 1481 1483 1485 3 11.71520 1.67360 15.06240 -
13.38880 0.00000 0.00000 ; C- N- CA- CB
1479 1481 1483 1494 3 1.75728 5.27184 2.25936
-7.02912 0.00000 0.00000 ; C- N- CA- C
1480 1479 1481 1482 3 29.28800 -8.36800 -20.92000
0.00000 0.00000 0.00000 ; O- C- N- H
1480 1479 1481 1483 3 20.92000 0.00000 -20.92000
0.00000 0.00000 0.00000 ; O- C- N- CA
1481 1483 1485 1486 3 0.65084 1.95253 0.00000
-2.60338 0.00000 0.00000 ; N- CA- CB- HB2
1481 1483 1485 1487 3 0.65084 1.95253 0.00000
-2.60338 0.00000 0.00000 ; N- CA- CB- HB3
1481 1483 1485 1488 3 3.21331 2.22589 -4.15053
-2.40998 2.61082 0.00000 ; N- CA- CB- CG
1481 1483 1494 1495 3 0.00000 0.00000 0.00000
0.00000 0.00000 0.00000 ; N- CA- C- O
1481 1483 1494 1496 3 17.40544 -5.02080 -13.22144
9.20480 0.00000 0.00000 ; N- CA- C- N

1482 1481 1483 1484 3 0.00000 0.00000 0.00000
0.00000 0.00000 0.00000 ; H- N- CA- HA
1482 1481 1483 1485 3 0.00000 0.00000 0.00000
0.00000 0.00000 0.00000 ; H- N- CA- CB
1482 1481 1483 1494 3 0.00000 0.00000 0.00000
0.00000 0.00000 0.00000 ; H- N- CA- C
1483 1485 1488 1489 3 0.66944 2.00832 0.00000
-2.67776 0.00000 0.00000 ; CA- CB- CG- HG2
1483 1485 1488 1490 3 0.66944 2.00832 0.00000
-2.67776 0.00000 0.00000 ; CA- CB- CG- HG3
1483 1485 1488 1491 3 10.12110 -1.91209 0.01674
10.17549 -1.87443 0.00000 ; CA- CB- CG- CD
1483 1494 1496 1497 3 20.92000 0.00000 -20.92000
0.00000 0.00000 0.00000 ; CA- C- N- H
1483 1494 1496 1498 3 20.92000 0.00000 -20.92000
0.00000 0.00000 0.00000 ; CA- C- N- CA
1484 1483 1485 1486 3 0.65084 1.95253 0.00000
-2.60338 0.00000 0.00000 ; HA- CA- CB- HB2
1484 1483 1485 1487 3 0.65084 1.95253 0.00000
-2.60338 0.00000 0.00000 ; HA- CA- CB- HB3
1484 1483 1485 1488 3 0.65084 1.95253 0.00000
-2.60338 0.00000 0.00000 ; HA- CA- CB- CG
1484 1483 1494 1495 3 3.68192 -4.35136 0.00000
1.33888 0.00000 0.00000 ; HA- CA- C- O
1484 1483 1494 1496 3 0.00000 0.00000 0.00000
0.00000 0.00000 0.00000 ; HA- CA- C- N
1485 1483 1494 1495 3 0.00000 0.00000 0.00000
0.00000 0.00000 0.00000 ; CB- CA- C- O
1485 1483 1494 1496 3 2.51040 4.18400 1.67360
-6.69440 0.00000 0.00000 ; CB- CA- C- N
1485 1488 1491 1492 3 3.79907 0.00000 -5.40573
0.00000 2.14221 0.00000 ; CB- CG- CD- OE1
1485 1488 1491 1493 3 3.79907 0.00000 -5.40573
0.00000 2.14221 0.00000 ; CB- CG- CD- OE2
1486 1485 1483 1494 3 0.65084 1.95253 0.00000
-2.60338 0.00000 0.00000 ; HB2- CB- CA- C
1486 1485 1488 1489 3 0.62760 1.88280 0.00000
-2.51040 0.00000 0.00000 ; HB2- CB- CG- HG2
1486 1485 1488 1490 3 0.62760 1.88280 0.00000
-2.51040 0.00000 0.00000 ; HB2- CB- CG- HG3
1486 1485 1488 1491 3 0.65084 1.95253 0.00000
-2.60338 0.00000 0.00000 ; HB2- CB- CG- CD
1487 1485 1483 1494 3 0.65084 1.95253 0.00000
-2.60338 0.00000 0.00000 ; HB3- CB- CA- C
1487 1485 1488 1489 3 0.62760 1.88280 0.00000
-2.51040 0.00000 0.00000 ; HB3- CB- CG- HG2
1487 1485 1488 1490 3 0.62760 1.88280 0.00000
-2.51040 0.00000 0.00000 ; HB3- CB- CG- HG3
1487 1485 1488 1491 3 0.65084 1.95253 0.00000
-2.60338 0.00000 0.00000 ; HB3- CB- CG- CD
1488 1485 1483 1494 3 6.86594 3.56895 -8.14206
-2.40998 4.85344 0.00000 ; CG- CB- CA- C

1489 1488 1491 1492 3 0.00000 0.00000 0.00000
0.00000 0.00000 0.00000 ; HG2- CG- CD- OE1
1489 1488 1491 1493 3 0.00000 0.00000 0.00000
0.00000 0.00000 0.00000 ; HG2- CG- CD- OE2
1490 1488 1491 1492 3 0.00000 0.00000 0.00000
0.00000 0.00000 0.00000 ; HG3- CG- CD- OE1
1490 1488 1491 1493 3 0.00000 0.00000 0.00000
0.00000 0.00000 0.00000 ; HG3- CG- CD- OE2
1494 1496 1498 1499 3 0.00000 0.00000 0.00000
0.00000 0.00000 0.00000 ; C- N- CA- HA
1494 1496 1498 1500 3 11.71520 1.67360 15.06240 -
13.38880 0.00000 0.00000 ; C- N- CA- CB
1494 1496 1498 1513 3 1.75728 5.27184 2.25936
-7.02912 0.00000 0.00000 ; C- N- CA- C
1495 1494 1496 1497 3 29.28800 -8.36800 -20.92000
0.00000 0.00000 0.00000 ; O- C- N- H
1495 1494 1496 1498 3 20.92000 0.00000 -20.92000
0.00000 0.00000 0.00000 ; O- C- N- CA
1496 1498 1500 1501 3 0.65084 1.95253 0.00000
-2.60338 0.00000 0.00000 ; N- CA- CB- HB2
1496 1498 1500 1502 3 0.65084 1.95253 0.00000
-2.60338 0.00000 0.00000 ; N- CA- CB- HB3
1496 1498 1500 1503 3 3.79070 1.39746 -4.61077
-2.40998 2.44346 0.00000 ; N- CA- CB- CG
1496 1498 1513 1514 3 0.00000 0.00000 0.00000
0.00000 0.00000 0.00000 ; N- CA- C- O
1496 1498 1513 1515 3 17.40544 -5.02080 -13.22144
9.20480 0.00000 0.00000 ; N- CA- C- N
1497 1496 1498 1499 3 0.00000 0.00000 0.00000
0.00000 0.00000 0.00000 ; H- N- CA- HA
1497 1496 1498 1500 3 0.00000 0.00000 0.00000
0.00000 0.00000 0.00000 ; H- N- CA- CB
1497 1496 1498 1513 3 0.00000 0.00000 0.00000
0.00000 0.00000 0.00000 ; H- N- CA- C
1498 1500 1503 1504 3 0.66944 2.00832 0.00000
-2.67776 0.00000 0.00000 ; CA- CB- CG- HG
1498 1500 1503 1505 3 3.90367 0.19665 -6.21742
-2.37651 5.99149 0.00000 ; CA- CB- CG- CD1
1498 1500 1503 1509 3 3.90367 0.19665 -6.21742
-2.37651 5.99149 0.00000 ; CA- CB- CG- CD2
1498 1513 1515 1516 3 20.92000 0.00000 -20.92000
0.00000 0.00000 0.00000 ; CA- C- N- H
1498 1513 1515 1517 3 20.92000 0.00000 -20.92000
0.00000 0.00000 0.00000 ; CA- C- N- CA
1499 1498 1500 1501 3 0.65084 1.95253 0.00000
-2.60338 0.00000 0.00000 ; HA- CA- CB- HB2
1499 1498 1500 1502 3 0.65084 1.95253 0.00000
-2.60338 0.00000 0.00000 ; HA- CA- CB- HB3
1499 1498 1500 1503 3 0.65084 1.95253 0.00000
-2.60338 0.00000 0.00000 ; HA- CA- CB- CG
1499 1498 1513 1514 3 3.68192 -4.35136 0.00000
1.33888 0.00000 0.00000 ; HA- CA- C- O

1499 1498 1513 1515 3 0.00000 0.00000 0.00000
0.00000 0.00000 0.00000 ; HA- CA- C- N
1500 1498 1513 1514 3 0.00000 0.00000 0.00000
0.00000 0.00000 0.00000 ; CB- CA- C- O
1500 1498 1513 1515 3 2.51040 4.18400 1.67360
-6.69440 0.00000 0.00000 ; CB- CA- C- N
1500 1503 1505 1506 3 0.66944 2.00832 0.00000
-2.67776 0.00000 0.00000 ; CB- CG- CD1- HD11
1500 1503 1505 1507 3 0.66944 2.00832 0.00000
-2.67776 0.00000 0.00000 ; CB- CG- CD1- HD12
1500 1503 1505 1508 3 0.66944 2.00832 0.00000
-2.67776 0.00000 0.00000 ; CB- CG- CD1- HD13
1500 1503 1509 1510 3 0.66944 2.00832 0.00000
-2.67776 0.00000 0.00000 ; CB- CG- CD2- HD21
1500 1503 1509 1511 3 0.66944 2.00832 0.00000
-2.67776 0.00000 0.00000 ; CB- CG- CD2- HD22
1500 1503 1509 1512 3 0.66944 2.00832 0.00000
-2.67776 0.00000 0.00000 ; CB- CG- CD2- HD23
1501 1500 1498 1513 3 0.65084 1.95253 0.00000
-2.60338 0.00000 0.00000 ; HB2- CB- CA- C
1501 1500 1503 1504 3 0.62760 1.88280 0.00000
-2.51040 0.00000 0.00000 ; HB2- CB- CG- HG
1501 1500 1503 1505 3 0.66944 2.00832 0.00000
-2.67776 0.00000 0.00000 ; HB2- CB- CG- CD1
1501 1500 1503 1509 3 0.66944 2.00832 0.00000
-2.67776 0.00000 0.00000 ; HB2- CB- CG- CD2
1502 1500 1498 1513 3 0.65084 1.95253 0.00000
-2.60338 0.00000 0.00000 ; HB3- CB- CA- C
1502 1500 1503 1504 3 0.62760 1.88280 0.00000
-2.51040 0.00000 0.00000 ; HB3- CB- CG- HG
1502 1500 1503 1505 3 0.66944 2.00832 0.00000
-2.67776 0.00000 0.00000 ; HB3- CB- CG- CD1
1502 1500 1503 1509 3 0.66944 2.00832 0.00000
-2.67776 0.00000 0.00000 ; HB3- CB- CG- CD2
1503 1500 1498 1513 3 10.33448 -1.14642 -11.54784
-2.40998 6.35968 0.00000 ; CG- CB- CA- C
1504 1503 1505 1506 3 0.62760 1.88280 0.00000
-2.51040 0.00000 0.00000 ; HG- CG- CD1- HD11
1504 1503 1505 1507 3 0.62760 1.88280 0.00000
-2.51040 0.00000 0.00000 ; HG- CG- CD1- HD12
1504 1503 1505 1508 3 0.62760 1.88280 0.00000
-2.51040 0.00000 0.00000 ; HG- CG- CD1- HD13
1504 1503 1509 1510 3 0.62760 1.88280 0.00000
-2.51040 0.00000 0.00000 ; HG- CG- CD2- HD21
1504 1503 1509 1511 3 0.62760 1.88280 0.00000
-2.51040 0.00000 0.00000 ; HG- CG- CD2- HD22
1504 1503 1509 1512 3 0.62760 1.88280 0.00000
-2.51040 0.00000 0.00000 ; HG- CG- CD2- HD23
1505 1503 1509 1510 3 0.66944 2.00832 0.00000
-2.67776 0.00000 0.00000 ; CD1- CG- CD2- HD21
1505 1503 1509 1511 3 0.66944 2.00832 0.00000
-2.67776 0.00000 0.00000 ; CD1- CG- CD2- HD22

1505 1503 1509 1512 3 0.66944 2.00832 0.00000
-2.67776 0.00000 0.00000 ; CD1- CG- CD2- HD23
1506 1505 1503 1509 3 0.66944 2.00832 0.00000
-2.67776 0.00000 0.00000 ; HD11- CD1- CG- CD2
1507 1505 1503 1509 3 0.66944 2.00832 0.00000
-2.67776 0.00000 0.00000 ; HD12- CD1- CG- CD2
1508 1505 1503 1509 3 0.66944 2.00832 0.00000
-2.67776 0.00000 0.00000 ; HD13- CD1- CG- CD2
1513 1515 1517 1518 3 0.00000 0.00000 0.00000
0.00000 0.00000 0.00000 ; C- N- CA- HA
1513 1515 1517 1519 3 11.71520 1.67360 15.06240 -
13.38880 0.00000 0.00000 ; C- N- CA- CB
1513 1515 1517 1524 3 1.75728 5.27184 2.25936
-7.02912 0.00000 0.00000 ; C- N- CA- C
1514 1513 1515 1516 3 29.28800 -8.36800 -20.92000
0.00000 0.00000 0.00000 ; O- C- N- H
1514 1513 1515 1517 3 20.92000 0.00000 -20.92000
0.00000 0.00000 0.00000 ; O- C- N- CA
1515 1517 1519 1520 3 0.65084 1.95253 0.00000
-2.60338 0.00000 0.00000 ; N- CA- CB- HB2
1515 1517 1519 1521 3 0.65084 1.95253 0.00000
-2.60338 0.00000 0.00000 ; N- CA- CB- HB3
1515 1517 1519 1522 3 6.03751 2.50622 -5.17142
-4.20074 1.10458 0.00000 ; N- CA- CB- SG
1515 1517 1524 1525 3 0.00000 0.00000 0.00000
0.00000 0.00000 0.00000 ; N- CA- C- O
1515 1517 1524 1526 3 17.40544 -5.02080 -13.22144
9.20480 0.00000 0.00000 ; N- CA- C- N
1516 1515 1517 1518 3 0.00000 0.00000 0.00000
0.00000 0.00000 0.00000 ; H- N- CA- HA
1516 1515 1517 1519 3 0.00000 0.00000 0.00000
0.00000 0.00000 0.00000 ; H- N- CA- CB
1516 1515 1517 1524 3 0.00000 0.00000 0.00000
0.00000 0.00000 0.00000 ; H- N- CA- C
1517 1519 1522 1523 3 1.69034 2.77818 4.11706
-4.21747 1.00416 0.00000 ; CA- CB- SG- HG
1517 1524 1526 1527 3 20.92000 0.00000 -20.92000
0.00000 0.00000 0.00000 ; CA- C- N- H
1517 1524 1526 1528 3 20.92000 0.00000 -20.92000
0.00000 0.00000 0.00000 ; CA- C- N- CA
1518 1517 1519 1520 3 0.65084 1.95253 0.00000
-2.60338 0.00000 0.00000 ; HA- CA- CB- HB2
1518 1517 1519 1521 3 0.65084 1.95253 0.00000
-2.60338 0.00000 0.00000 ; HA- CA- CB- HB3
1518 1517 1519 1522 3 0.65084 1.95253 0.00000
-2.60338 0.00000 0.00000 ; HA- CA- CB- SG
1518 1517 1524 1525 3 3.68192 -4.35136 0.00000
1.33888 0.00000 0.00000 ; HA- CA- C- O
1518 1517 1524 1526 3 0.00000 0.00000 0.00000
0.00000 0.00000 0.00000 ; HA- CA- C- N
1519 1517 1524 1525 3 0.00000 0.00000 0.00000
0.00000 0.00000 0.00000 ; CB- CA- C- O

1519 1517 1524 1526 3 2.51040 4.18400 1.67360
-6.69440 0.00000 0.00000 ; CB- CA- C- N
1520 1519 1517 1524 3 0.65084 1.95253 0.00000
-2.60338 0.00000 0.00000 ; HB2- CB- CA- C
1520 1519 1522 1523 3 1.04600 3.13800 0.00000
-4.18400 0.00000 0.00000 ; HB2- CB- SG- HG
1521 1519 1517 1524 3 0.65084 1.95253 0.00000
-2.60338 0.00000 0.00000 ; HB3- CB- CA- C
1521 1519 1522 1523 3 1.04600 3.13800 0.00000
-4.18400 0.00000 0.00000 ; HB3- CB- SG- HG
1522 1519 1517 1524 3 5.62330 4.27605 -5.33042
-4.20074 2.51040 0.00000 ; SG- CB- CA- C
1524 1526 1528 1529 3 0.00000 0.00000 0.00000
0.00000 0.00000 0.00000 ; C- N- CA- HA
1524 1526 1528 1530 3 11.71520 1.67360 15.06240 -
13.38880 0.00000 0.00000 ; C- N- CA- CB
1524 1526 1528 1535 3 1.75728 5.27184 2.25936
-7.02912 0.00000 0.00000 ; C- N- CA- C
1525 1524 1526 1527 3 29.28800 -8.36800 -20.92000
0.00000 0.00000 0.00000 ; O- C- N- H
1525 1524 1526 1528 3 20.92000 0.00000 -20.92000
0.00000 0.00000 0.00000 ; O- C- N- CA
1526 1528 1530 1531 3 0.65084 1.95253 0.00000
-2.60338 0.00000 0.00000 ; N- CA- CB- HB2
1526 1528 1530 1532 3 0.65084 1.95253 0.00000
-2.60338 0.00000 0.00000 ; N- CA- CB- HB3
1526 1528 1530 1533 3 7.86174 2.24681 -7.41405
-6.71114 5.35552 0.00000 ; N- CA- CB- OG
1526 1528 1535 1536 3 0.00000 0.00000 0.00000
0.00000 0.00000 0.00000 ; N- CA- C- O
1526 1528 1535 1537 3 17.40544 -5.02080 -13.22144
9.20480 0.00000 0.00000 ; N- CA- C- N
1527 1526 1528 1529 3 0.00000 0.00000 0.00000
0.00000 0.00000 0.00000 ; H- N- CA- HA
1527 1526 1528 1530 3 0.00000 0.00000 0.00000
0.00000 0.00000 0.00000 ; H- N- CA- CB
1527 1526 1528 1535 3 0.00000 0.00000 0.00000
0.00000 0.00000 0.00000 ; H- N- CA- C
1528 1530 1533 1534 3 2.05853 2.46856 3.48109
-4.46851 0.23430 0.00000 ; CA- CB- OG- HG
1528 1535 1537 1538 3 20.92000 0.00000 -20.92000
0.00000 0.00000 0.00000 ; CA- C- N- H
1528 1535 1537 1539 3 20.92000 0.00000 -20.92000
0.00000 0.00000 0.00000 ; CA- C- N- CA
1529 1528 1530 1531 3 0.65084 1.95253 0.00000
-2.60338 0.00000 0.00000 ; HA- CA- CB- HB2
1529 1528 1530 1532 3 0.65084 1.95253 0.00000
-2.60338 0.00000 0.00000 ; HA- CA- CB- HB3
1529 1528 1530 1533 3 1.04600 -1.04600 0.00000
0.00000 0.00000 0.00000 ; HA- CA- CB- OG
1529 1528 1535 1536 3 3.68192 -4.35136 0.00000
1.33888 0.00000 0.00000 ; HA- CA- C- O

1529 1528 1535 1537 3 0.00000 0.00000 0.00000
0.00000 0.00000 0.00000 ; HA- CA- C- N
1530 1528 1535 1536 3 0.00000 0.00000 0.00000
0.00000 0.00000 0.00000 ; CB- CA- C- O
1530 1528 1535 1537 3 2.51040 4.18400 1.67360
-6.69440 0.00000 0.00000 ; CB- CA- C- N
1531 1530 1528 1535 3 0.65084 1.95253 0.00000
-2.60338 0.00000 0.00000 ; HB2- CB- CA- C
1531 1530 1533 1534 3 0.69733 2.09200 0.00000
-2.78933 0.00000 0.00000 ; HB2- CB- OG- HG
1532 1530 1528 1535 3 0.65084 1.95253 0.00000
-2.60338 0.00000 0.00000 ; HB3- CB- CA- C
1532 1530 1533 1534 3 0.69733 2.09200 0.00000
-2.78933 0.00000 0.00000 ; HB3- CB- OG- HG
1533 1530 1528 1535 3 7.34710 7.79898 -6.14211
-6.71114 4.31789 0.00000 ; OG- CB- CA- C
1535 1537 1539 1540 3 0.00000 0.00000 0.00000
0.00000 0.00000 0.00000 ; C- N- CA- HA2
1535 1537 1539 1541 3 0.00000 0.00000 0.00000
0.00000 0.00000 0.00000 ; C- N- CA- HA3
1535 1537 1539 1542 3 1.75728 5.27184 2.25936
-7.02912 0.00000 0.00000 ; C- N- CA- C
1536 1535 1537 1538 3 29.28800 -8.36800 -20.92000
0.00000 0.00000 0.00000 ; O- C- N- H
1536 1535 1537 1539 3 20.92000 0.00000 -20.92000
0.00000 0.00000 0.00000 ; O- C- N- CA
1537 1539 1542 1543 3 0.00000 0.00000 0.00000
0.00000 0.00000 0.00000 ; N- CA- C- O
1537 1539 1542 1544 3 17.40544 -5.02080 -13.22144
9.20480 0.00000 0.00000 ; N- CA- C- N
1538 1537 1539 1540 3 0.00000 0.00000 0.00000
0.00000 0.00000 0.00000 ; H- N- CA- HA2
1538 1537 1539 1541 3 0.00000 0.00000 0.00000
0.00000 0.00000 0.00000 ; H- N- CA- HA3
1538 1537 1539 1542 3 0.00000 0.00000 0.00000
0.00000 0.00000 0.00000 ; H- N- CA- C
1539 1542 1544 1545 3 20.92000 0.00000 -20.92000
0.00000 0.00000 0.00000 ; CA- C- N- H
1539 1542 1544 1546 3 20.92000 0.00000 -20.92000
0.00000 0.00000 0.00000 ; CA- C- N- CA
1540 1539 1542 1543 3 3.68192 -4.35136 0.00000
1.33888 0.00000 0.00000 ; HA2- CA- C- O
1540 1539 1542 1544 3 0.00000 0.00000 0.00000
0.00000 0.00000 0.00000 ; HA2- CA- C- N
1541 1539 1542 1543 3 3.68192 -4.35136 0.00000
1.33888 0.00000 0.00000 ; HA3- CA- C- O
1541 1539 1542 1544 3 0.00000 0.00000 0.00000
0.00000 0.00000 0.00000 ; HA3- CA- C- N
1542 1544 1546 1547 3 0.00000 0.00000 0.00000
0.00000 0.00000 0.00000 ; C- N- CA- HA
1542 1544 1546 1548 3 11.71520 1.67360 15.06240 -
13.38880 0.00000 0.00000 ; C- N- CA- CB

1542 1544 1546 1566 3 1.75728 5.27184 2.25936
-7.02912 0.00000 0.00000 ; C- N- CA- C
1543 1542 1544 1545 3 29.28800 -8.36800 -20.92000
0.00000 0.00000 0.00000 ; O- C- N- H
1543 1542 1544 1546 3 20.92000 0.00000 -20.92000
0.00000 0.00000 0.00000 ; O- C- N- CA
1544 1546 1548 1549 3 0.65084 1.95253 0.00000
-2.60338 0.00000 0.00000 ; N- CA- CB- HB2
1544 1546 1548 1550 3 0.65084 1.95253 0.00000
-2.60338 0.00000 0.00000 ; N- CA- CB- HB3
1544 1546 1548 1551 3 0.65084 1.95253 0.00000
-2.60338 0.00000 0.00000 ; N- CA- CB- CG
1544 1546 1566 1567 3 0.00000 0.00000 0.00000
0.00000 0.00000 0.00000 ; N- CA- C- O
1544 1546 1566 1568 3 17.40544 -5.02080 -13.22144
9.20480 0.00000 0.00000 ; N- CA- C- N
1545 1544 1546 1547 3 0.00000 0.00000 0.00000
0.00000 0.00000 0.00000 ; H- N- CA- HA
1545 1544 1546 1548 3 0.00000 0.00000 0.00000
0.00000 0.00000 0.00000 ; H- N- CA- CB
1545 1544 1546 1566 3 0.00000 0.00000 0.00000
0.00000 0.00000 0.00000 ; H- N- CA- C
1546 1548 1551 1552 3 0.66944 2.00832 0.00000
-2.67776 0.00000 0.00000 ; CA- CB- CG- HG2
1546 1548 1551 1553 3 0.66944 2.00832 0.00000
-2.67776 0.00000 0.00000 ; CA- CB- CG- HG3
1546 1548 1551 1554 3 3.68192 3.09616 -2.09200
-3.01248 0.00000 0.00000 ; CA- CB- CG- CD
1546 1566 1568 1569 3 20.92000 0.00000 -20.92000
0.00000 0.00000 0.00000 ; CA- C- N- H
1546 1566 1568 1570 3 20.92000 0.00000 -20.92000
0.00000 0.00000 0.00000 ; CA- C- N- CA
1547 1546 1548 1549 3 0.65084 1.95253 0.00000
-2.60338 0.00000 0.00000 ; HA- CA- CB- HB2
1547 1546 1548 1550 3 0.65084 1.95253 0.00000
-2.60338 0.00000 0.00000 ; HA- CA- CB- HB3
1547 1546 1548 1551 3 0.65084 1.95253 0.00000
-2.60338 0.00000 0.00000 ; HA- CA- CB- CG
1547 1546 1566 1567 3 3.68192 -4.35136 0.00000
1.33888 0.00000 0.00000 ; HA- CA- C- O
1547 1546 1566 1568 3 0.00000 0.00000 0.00000
0.00000 0.00000 0.00000 ; HA- CA- C- N
1548 1546 1566 1567 3 0.00000 0.00000 0.00000
0.00000 0.00000 0.00000 ; CB- CA- C- O
1548 1546 1566 1568 3 2.51040 4.18400 1.67360
-6.69440 0.00000 0.00000 ; CB- CA- C- N
1548 1551 1554 1555 3 0.65084 1.95253 0.00000
-2.60338 0.00000 0.00000 ; CB- CG- CD- HD2
1548 1551 1554 1556 3 0.65084 1.95253 0.00000
-2.60338 0.00000 0.00000 ; CB- CG- CD- HD3
1548 1551 1554 1557 3 0.65084 1.95253 0.00000
-2.60338 0.00000 0.00000 ; CB- CG- CD- NE

```
  1549   1548   1546   1566      3    0.65084    1.95253    0.00000
-2.60338    0.00000    0.00000 ;    HB2-    CB-    CA-     C
  1549   1548   1551   1552      3    0.62760    1.88280    0.00000
-2.51040    0.00000    0.00000 ;    HB2-    CB-    CG-   HG2
  1549   1548   1551   1553      3    0.62760    1.88280    0.00000
-2.51040    0.00000    0.00000 ;    HB2-    CB-    CG-   HG3
  1549   1548   1551   1554      3    0.66944    2.00832    0.00000
-2.67776    0.00000    0.00000 ;    HB2-    CB-    CG-    CD
  1550   1548   1546   1566      3    0.65084    1.95253    0.00000
-2.60338    0.00000    0.00000 ;    HB3-    CB-    CA-     C
  1550   1548   1551   1552      3    0.62760    1.88280    0.00000
-2.51040    0.00000    0.00000 ;    HB3-    CB-    CG-   HG2
  1550   1548   1551   1553      3    0.62760    1.88280    0.00000
-2.51040    0.00000    0.00000 ;    HB3-    CB-    CG-   HG3
  1550   1548   1551   1554      3    0.66944    2.00832    0.00000
-2.67776    0.00000    0.00000 ;    HB3-    CB-    CG-    CD
  1551   1548   1546   1566      3    0.65084    1.95253    0.00000
-2.60338    0.00000    0.00000 ;     CG-    CB-    CA-     C
  1551   1554   1557   1558      3    0.00000    0.00000    0.00000
0.00000    0.00000    0.00000 ;     CG-    CD-    NE-    HE
  1551   1554   1557   1559      3    0.00000    0.00000    0.00000
0.00000    0.00000    0.00000 ;     CG-    CD-    NE-    CZ
  1552   1551   1554   1555      3    0.65084    1.95253    0.00000
-2.60338    0.00000    0.00000 ;    HG2-    CG-    CD-   HD2
  1552   1551   1554   1556      3    0.65084    1.95253    0.00000
-2.60338    0.00000    0.00000 ;    HG2-    CG-    CD-   HD3
  1552   1551   1554   1557      3    0.65084    1.95253    0.00000
-2.60338    0.00000    0.00000 ;    HG2-    CG-    CD-    NE
  1553   1551   1554   1555      3    0.65084    1.95253    0.00000
-2.60338    0.00000    0.00000 ;    HG3-    CG-    CD-   HD2
  1553   1551   1554   1556      3    0.65084    1.95253    0.00000
-2.60338    0.00000    0.00000 ;    HG3-    CG-    CD-   HD3
  1553   1551   1554   1557      3    0.65084    1.95253    0.00000
-2.60338    0.00000    0.00000 ;    HG3-    CG-    CD-    NE
  1554   1557   1559   1560      3   20.08320    0.00000  -20.08320
0.00000    0.00000    0.00000 ;     CD-    NE-    CZ-   NH1
  1554   1557   1559   1563      3   20.08320    0.00000  -20.08320
0.00000    0.00000    0.00000 ;     CD-    NE-    CZ-   NH2
  1555   1554   1557   1558      3    0.00000    0.00000    0.00000
0.00000    0.00000    0.00000 ;    HD2-    CD-    NE-    HE
  1555   1554   1557   1559      3    0.00000    0.00000    0.00000
0.00000    0.00000    0.00000 ;    HD2-    CD-    NE-    CZ
  1556   1554   1557   1558      3    0.00000    0.00000    0.00000
0.00000    0.00000    0.00000 ;    HD3-    CD-    NE-    HE
  1556   1554   1557   1559      3    0.00000    0.00000    0.00000
0.00000    0.00000    0.00000 ;    HD3-    CD-    NE-    CZ
  1557   1559   1560   1561      3   20.08320    0.00000  -20.08320
0.00000    0.00000    0.00000 ;     NE-    CZ-   NH1-  HH11
  1557   1559   1560   1562      3   20.08320    0.00000  -20.08320
0.00000    0.00000    0.00000 ;     NE-    CZ-   NH1-  HH12
  1557   1559   1563   1564      3   20.08320    0.00000  -20.08320
0.00000    0.00000    0.00000 ;     NE-    CZ-   NH2-  HH21
```

1557 1559 1563 1565 3 20.08320 0.00000 -20.08320
0.00000 0.00000 0.00000 ; NE- CZ- NH2- HH22
1558 1557 1559 1560 3 20.08320 0.00000 -20.08320
0.00000 0.00000 0.00000 ; HE- NE- CZ- NH1
1558 1557 1559 1563 3 20.08320 0.00000 -20.08320
0.00000 0.00000 0.00000 ; HE- NE- CZ- NH2
1560 1559 1563 1564 3 20.08320 0.00000 -20.08320
0.00000 0.00000 0.00000 ; NH1- CZ- NH2- HH21
1560 1559 1563 1565 3 20.08320 0.00000 -20.08320
0.00000 0.00000 0.00000 ; NH1- CZ- NH2- HH22
1561 1560 1559 1563 3 20.08320 0.00000 -20.08320
0.00000 0.00000 0.00000 ; HH11- NH1- CZ- NH2
1562 1560 1559 1563 3 20.08320 0.00000 -20.08320
0.00000 0.00000 0.00000 ; HH12- NH1- CZ- NH2
1566 1568 1570 1571 3 0.00000 0.00000 0.00000
0.00000 0.00000 0.00000 ; C- N- CA- HA
1566 1568 1570 1572 3 11.71520 1.67360 15.06240 -
13.38880 0.00000 0.00000 ; C- N- CA- CB
1566 1568 1570 1581 3 1.75728 5.27184 2.25936
-7.02912 0.00000 0.00000 ; C- N- CA- C
1567 1566 1568 1569 3 29.28800 -8.36800 -20.92000
0.00000 0.00000 0.00000 ; O- C- N- H
1567 1566 1568 1570 3 20.92000 0.00000 -20.92000
0.00000 0.00000 0.00000 ; O- C- N- CA
1568 1570 1572 1573 3 0.65084 1.95253 0.00000
-2.60338 0.00000 0.00000 ; N- CA- CB- HB2
1568 1570 1572 1574 3 0.65084 1.95253 0.00000
-2.60338 0.00000 0.00000 ; N- CA- CB- HB3
1568 1570 1572 1575 3 3.21331 2.22589 -4.15053
-2.40998 2.61082 0.00000 ; N- CA- CB- CG
1568 1570 1581 1582 3 0.00000 0.00000 0.00000
0.00000 0.00000 0.00000 ; N- CA- C- O
1568 1570 1581 1583 3 17.40544 -5.02080 -13.22144
9.20480 0.00000 0.00000 ; N- CA- C- N
1569 1568 1570 1571 3 0.00000 0.00000 0.00000
0.00000 0.00000 0.00000 ; H- N- CA- HA
1569 1568 1570 1572 3 0.00000 0.00000 0.00000
0.00000 0.00000 0.00000 ; H- N- CA- CB
1569 1568 1570 1581 3 0.00000 0.00000 0.00000
0.00000 0.00000 0.00000 ; H- N- CA- C
1570 1572 1575 1576 3 0.66944 2.00832 0.00000
-2.67776 0.00000 0.00000 ; CA- CB- CG- HG2
1570 1572 1575 1577 3 0.66944 2.00832 0.00000
-2.67776 0.00000 0.00000 ; CA- CB- CG- HG3
1570 1572 1575 1578 3 10.12110 -1.91209 0.01674
10.17549 -1.87443 0.00000 ; CA- CB- CG- CD
1570 1581 1583 1584 3 20.92000 0.00000 -20.92000
0.00000 0.00000 0.00000 ; CA- C- N- H
1570 1581 1583 1585 3 20.92000 0.00000 -20.92000
0.00000 0.00000 0.00000 ; CA- C- N- CA
1571 1570 1572 1573 3 0.65084 1.95253 0.00000
-2.60338 0.00000 0.00000 ; HA- CA- CB- HB2

1571 1570 1572 1574 3 0.65084 1.95253 0.00000
-2.60338 0.00000 0.00000 ; HA- CA- CB- HB3
1571 1570 1572 1575 3 0.65084 1.95253 0.00000
-2.60338 0.00000 0.00000 ; HA- CA- CB- CG
1571 1570 1581 1582 3 3.68192 -4.35136 0.00000
1.33888 0.00000 0.00000 ; HA- CA- C- O
1571 1570 1581 1583 3 0.00000 0.00000 0.00000
0.00000 0.00000 0.00000 ; HA- CA- C- N
1572 1570 1581 1582 3 0.00000 0.00000 0.00000
0.00000 0.00000 0.00000 ; CB- CA- C- O
1572 1570 1581 1583 3 2.51040 4.18400 1.67360
-6.69440 0.00000 0.00000 ; CB- CA- C- N
1572 1575 1578 1579 3 3.79907 0.00000 -5.40573
0.00000 2.14221 0.00000 ; CB- CG- CD- OE1
1572 1575 1578 1580 3 3.79907 0.00000 -5.40573
0.00000 2.14221 0.00000 ; CB- CG- CD- OE2
1573 1572 1570 1581 3 0.65084 1.95253 0.00000
-2.60338 0.00000 0.00000 ; HB2- CB- CA- C
1573 1572 1575 1576 3 0.62760 1.88280 0.00000
-2.51040 0.00000 0.00000 ; HB2- CB- CG- HG2
1573 1572 1575 1577 3 0.62760 1.88280 0.00000
-2.51040 0.00000 0.00000 ; HB2- CB- CG- HG3
1573 1572 1575 1578 3 0.65084 1.95253 0.00000
-2.60338 0.00000 0.00000 ; HB2- CB- CG- CD
1574 1572 1570 1581 3 0.65084 1.95253 0.00000
-2.60338 0.00000 0.00000 ; HB3- CB- CA- C
1574 1572 1575 1576 3 0.62760 1.88280 0.00000
-2.51040 0.00000 0.00000 ; HB3- CB- CG- HG2
1574 1572 1575 1577 3 0.62760 1.88280 0.00000
-2.51040 0.00000 0.00000 ; HB3- CB- CG- HG3
1574 1572 1575 1578 3 0.65084 1.95253 0.00000
-2.60338 0.00000 0.00000 ; HB3- CB- CG- CD
1575 1572 1570 1581 3 6.86594 3.56895 -8.14206
-2.40998 4.85344 0.00000 ; CG- CB- CA- C
1576 1575 1578 1579 3 0.00000 0.00000 0.00000
0.00000 0.00000 0.00000 ; HG2- CG- CD- OE1
1576 1575 1578 1580 3 0.00000 0.00000 0.00000
0.00000 0.00000 0.00000 ; HG2- CG- CD- OE2
1577 1575 1578 1579 3 0.00000 0.00000 0.00000
0.00000 0.00000 0.00000 ; HG3- CG- CD- OE1
1577 1575 1578 1580 3 0.00000 0.00000 0.00000
0.00000 0.00000 0.00000 ; HG3- CG- CD- OE2
1581 1583 1585 1586 3 0.00000 0.00000 0.00000
0.00000 0.00000 0.00000 ; C- N- CA- HA
1581 1583 1585 1587 3 11.71520 1.67360 15.06240 -
13.38880 0.00000 0.00000 ; C- N- CA- CB
1581 1583 1585 1600 3 1.75728 5.27184 2.25936
-7.02912 0.00000 0.00000 ; C- N- CA- C
1582 1581 1583 1584 3 29.28800 -8.36800 -20.92000
0.00000 0.00000 0.00000 ; O- C- N- H
1582 1581 1583 1585 3 20.92000 0.00000 -20.92000
0.00000 0.00000 0.00000 ; O- C- N- CA

```
  1583   1585   1587   1588      3    0.65084    1.95253    0.00000
-2.60338    0.00000    0.00000 ;      N-    CA-    CB-   HB2
  1583   1585   1587   1589      3    0.65084    1.95253    0.00000
-2.60338    0.00000    0.00000 ;      N-    CA-    CB-   HB3
  1583   1585   1587   1590      3    3.79070    1.39746   -4.61077
-2.40998    2.44346    0.00000 ;      N-    CA-    CB-    CG
  1583   1585   1600   1601      3    0.00000    0.00000    0.00000
0.00000    0.00000    0.00000 ;      N-    CA-     C-     O
  1583   1585   1600   1602      3   17.40544   -5.02080  -13.22144
9.20480    0.00000    0.00000 ;      N-    CA-     C-     N
  1584   1583   1585   1586      3    0.00000    0.00000    0.00000
0.00000    0.00000    0.00000 ;      H-     N-    CA-    HA
  1584   1583   1585   1587      3    0.00000    0.00000    0.00000
0.00000    0.00000    0.00000 ;      H-     N-    CA-    CB
  1584   1583   1585   1600      3    0.00000    0.00000    0.00000
0.00000    0.00000    0.00000 ;      H-     N-    CA-     C
  1585   1587   1590   1591      3    0.66944    2.00832    0.00000
-2.67776    0.00000    0.00000 ;     CA-    CB-    CG-    HG
  1585   1587   1590   1592      3    3.90367    0.19665   -6.21742
-2.37651    5.99149    0.00000 ;     CA-    CB-    CG-   CD1
  1585   1587   1590   1596      3    3.90367    0.19665   -6.21742
-2.37651    5.99149    0.00000 ;     CA-    CB-    CG-   CD2
  1585   1600   1602   1603      3   20.92000    0.00000  -20.92000
0.00000    0.00000    0.00000 ;     CA-     C-     N-     H
  1585   1600   1602   1604      3   20.92000    0.00000  -20.92000
0.00000    0.00000    0.00000 ;     CA-     C-     N-    CA
  1586   1585   1587   1588      3    0.65084    1.95253    0.00000
-2.60338    0.00000    0.00000 ;     HA-    CA-    CB-   HB2
  1586   1585   1587   1589      3    0.65084    1.95253    0.00000
-2.60338    0.00000    0.00000 ;     HA-    CA-    CB-   HB3
  1586   1585   1587   1590      3    0.65084    1.95253    0.00000
-2.60338    0.00000    0.00000 ;     HA-    CA-    CB-    CG
  1586   1585   1600   1601      3    3.68192   -4.35136    0.00000
1.33888    0.00000    0.00000 ;     HA-    CA-     C-     O
  1586   1585   1600   1602      3    0.00000    0.00000    0.00000
0.00000    0.00000    0.00000 ;     HA-    CA-     C-     N
  1587   1585   1600   1601      3    0.00000    0.00000    0.00000
0.00000    0.00000    0.00000 ;     CB-    CA-     C-     O
  1587   1585   1600   1602      3    2.51040    4.18400    1.67360
-6.69440    0.00000    0.00000 ;     CB-    CA-     C-     N
  1587   1590   1592   1593      3    0.66944    2.00832    0.00000
-2.67776    0.00000    0.00000 ;     CB-    CG-   CD1-  HD11
  1587   1590   1592   1594      3    0.66944    2.00832    0.00000
-2.67776    0.00000    0.00000 ;     CB-    CG-   CD1-  HD12
  1587   1590   1592   1595      3    0.66944    2.00832    0.00000
-2.67776    0.00000    0.00000 ;     CB-    CG-   CD1-  HD13
  1587   1590   1596   1597      3    0.66944    2.00832    0.00000
-2.67776    0.00000    0.00000 ;     CB-    CG-   CD2-  HD21
  1587   1590   1596   1598      3    0.66944    2.00832    0.00000
-2.67776    0.00000    0.00000 ;     CB-    CG-   CD2-  HD22
  1587   1590   1596   1599      3    0.66944    2.00832    0.00000
-2.67776    0.00000    0.00000 ;     CB-    CG-   CD2-  HD23
```

1588 1587 1585 1600 3 0.65084 1.95253 0.00000
-2.60338 0.00000 0.00000 ; HB2- CB- CA- C
1588 1587 1590 1591 3 0.62760 1.88280 0.00000
-2.51040 0.00000 0.00000 ; HB2- CB- CG- HG
1588 1587 1590 1592 3 0.66944 2.00832 0.00000
-2.67776 0.00000 0.00000 ; HB2- CB- CG- CD1
1588 1587 1590 1596 3 0.66944 2.00832 0.00000
-2.67776 0.00000 0.00000 ; HB2- CB- CG- CD2
1589 1587 1585 1600 3 0.65084 1.95253 0.00000
-2.60338 0.00000 0.00000 ; HB3- CB- CA- C
1589 1587 1590 1591 3 0.62760 1.88280 0.00000
-2.51040 0.00000 0.00000 ; HB3- CB- CG- HG
1589 1587 1590 1592 3 0.66944 2.00832 0.00000
-2.67776 0.00000 0.00000 ; HB3- CB- CG- CD1
1589 1587 1590 1596 3 0.66944 2.00832 0.00000
-2.67776 0.00000 0.00000 ; HB3- CB- CG- CD2
1590 1587 1585 1600 3 10.33448 -1.14642 -11.54784
-2.40998 6.35968 0.00000 ; CG- CB- CA- C
1591 1590 1592 1593 3 0.62760 1.88280 0.00000
-2.51040 0.00000 0.00000 ; HG- CG- CD1- HD11
1591 1590 1592 1594 3 0.62760 1.88280 0.00000
-2.51040 0.00000 0.00000 ; HG- CG- CD1- HD12
1591 1590 1592 1595 3 0.62760 1.88280 0.00000
-2.51040 0.00000 0.00000 ; HG- CG- CD1- HD13
1591 1590 1596 1597 3 0.62760 1.88280 0.00000
-2.51040 0.00000 0.00000 ; HG- CG- CD2- HD21
1591 1590 1596 1598 3 0.62760 1.88280 0.00000
-2.51040 0.00000 0.00000 ; HG- CG- CD2- HD22
1591 1590 1596 1599 3 0.62760 1.88280 0.00000
-2.51040 0.00000 0.00000 ; HG- CG- CD2- HD23
1592 1590 1596 1597 3 0.66944 2.00832 0.00000
-2.67776 0.00000 0.00000 ; CD1- CG- CD2- HD21
1592 1590 1596 1598 3 0.66944 2.00832 0.00000
-2.67776 0.00000 0.00000 ; CD1- CG- CD2- HD22
1592 1590 1596 1599 3 0.66944 2.00832 0.00000
-2.67776 0.00000 0.00000 ; CD1- CG- CD2- HD23
1593 1592 1590 1596 3 0.66944 2.00832 0.00000
-2.67776 0.00000 0.00000 ; HD11- CD1- CG- CD2
1594 1592 1590 1596 3 0.66944 2.00832 0.00000
-2.67776 0.00000 0.00000 ; HD12- CD1- CG- CD2
1595 1592 1590 1596 3 0.66944 2.00832 0.00000
-2.67776 0.00000 0.00000 ; HD13- CD1- CG- CD2
1600 1602 1604 1605 3 0.00000 0.00000 0.00000
0.00000 0.00000 0.00000 ; C- N- CA- HA
1600 1602 1604 1606 3 11.71520 1.67360 15.06240 -
13.38880 0.00000 0.00000 ; C- N- CA- CB
1600 1602 1604 1620 3 1.75728 5.27184 2.25936
-7.02912 0.00000 0.00000 ; C- N- CA- C
1601 1600 1602 1603 3 29.28800 -8.36800 -20.92000
0.00000 0.00000 0.00000 ; O- C- N- H
1601 1600 1602 1604 3 20.92000 0.00000 -20.92000
0.00000 0.00000 0.00000 ; O- C- N- CA

1602 1604 1606 1607 3 0.65084 1.95253 0.00000
-2.60338 0.00000 0.00000 ; N- CA- CB- HB2
1602 1604 1606 1608 3 0.65084 1.95253 0.00000
-2.60338 0.00000 0.00000 ; N- CA- CB- HB3
1602 1604 1606 1609 3 3.28026 2.46019 -2.19242
-3.21331 -0.23430 0.00000 ; N- CA- CB- CG
1602 1604 1620 1621 3 0.00000 0.00000 0.00000
0.00000 0.00000 0.00000 ; N- CA- C- O
1602 1604 1620 1622 3 17.40544 -5.02080 -13.22144
9.20480 0.00000 0.00000 ; N- CA- C- N
1603 1602 1604 1605 3 0.00000 0.00000 0.00000
0.00000 0.00000 0.00000 ; H- N- CA- HA
1603 1602 1604 1606 3 0.00000 0.00000 0.00000
0.00000 0.00000 0.00000 ; H- N- CA- CB
1603 1602 1604 1620 3 0.00000 0.00000 0.00000
0.00000 0.00000 0.00000 ; H- N- CA- C
1604 1606 1609 1610 3 0.57739 0.00000 1.02926
0.00000 -1.60666 0.00000 ; CA- CB- CG- CD1
1604 1606 1609 1618 3 0.57739 0.00000 1.02926
0.00000 -1.60666 0.00000 ; CA- CB- CG- CD2
1604 1620 1622 1623 3 20.92000 0.00000 -20.92000
0.00000 0.00000 0.00000 ; CA- C- N- H
1604 1620 1622 1624 3 20.92000 0.00000 -20.92000
0.00000 0.00000 0.00000 ; CA- C- N- CA
1605 1604 1606 1607 3 0.65084 1.95253 0.00000
-2.60338 0.00000 0.00000 ; HA- CA- CB- HB2
1605 1604 1606 1608 3 0.65084 1.95253 0.00000
-2.60338 0.00000 0.00000 ; HA- CA- CB- HB3
1605 1604 1606 1609 3 0.65084 1.95253 0.00000
-2.60338 0.00000 0.00000 ; HA- CA- CB- CG
1605 1604 1620 1621 3 3.68192 -4.35136 0.00000
1.33888 0.00000 0.00000 ; HA- CA- C- O
1605 1604 1620 1622 3 0.00000 0.00000 0.00000
0.00000 0.00000 0.00000 ; HA- CA- C- N
1606 1604 1620 1621 3 0.00000 0.00000 0.00000
0.00000 0.00000 0.00000 ; CB- CA- C- O
1606 1604 1620 1622 3 2.51040 4.18400 1.67360
-6.69440 0.00000 0.00000 ; CB- CA- C- N
1606 1609 1610 1611 3 30.33400 0.00000 -30.33400
0.00000 0.00000 0.00000 ; CB- CG- CD1- HD1
1606 1609 1610 1612 3 30.33400 0.00000 -30.33400
0.00000 0.00000 0.00000 ; CB- CG- CD1- CE1
1606 1609 1618 1616 3 30.33400 0.00000 -30.33400
0.00000 0.00000 0.00000 ; CB- CG- CD2- CE2
1606 1609 1618 1619 3 30.33400 0.00000 -30.33400
0.00000 0.00000 0.00000 ; CB- CG- CD2- HD2
1607 1606 1604 1620 3 0.65084 1.95253 0.00000
-2.60338 0.00000 0.00000 ; HB2- CB- CA- C
1607 1606 1609 1610 3 0.00000 0.00000 0.00000
0.00000 0.00000 0.00000 ; HB2- CB- CG- CD1
1607 1606 1609 1618 3 0.00000 0.00000 0.00000
0.00000 0.00000 0.00000 ; HB2- CB- CG- CD2

1608 1606 1604 1620 3 0.65084 1.95253 0.00000
-2.60338 0.00000 0.00000 ; HB3- CB- CA- C
1608 1606 1609 1610 3 0.00000 0.00000 0.00000
0.00000 0.00000 0.00000 ; HB3- CB- CG- CD1
1608 1606 1609 1618 3 0.00000 0.00000 0.00000
0.00000 0.00000 0.00000 ; HB3- CB- CG- CD2
1609 1606 1604 1620 3 4.95804 2.17986 -3.52293
-3.21331 -0.40166 0.00000 ; CG- CB- CA- C
1609 1610 1612 1613 3 30.33400 0.00000 -30.33400
0.00000 0.00000 0.00000 ; CG- CD1- CE1- HE1
1609 1610 1612 1614 3 30.33400 0.00000 -30.33400
0.00000 0.00000 0.00000 ; CG- CD1- CE1- CZ
1609 1618 1616 1614 3 30.33400 0.00000 -30.33400
0.00000 0.00000 0.00000 ; CG- CD2- CE2- CZ
1609 1618 1616 1617 3 30.33400 0.00000 -30.33400
0.00000 0.00000 0.00000 ; CG- CD2- CE2- HE2
1610 1609 1618 1616 3 30.33400 0.00000 -30.33400
0.00000 0.00000 0.00000 ; CD1- CG- CD2- CE2
1610 1609 1618 1619 3 30.33400 0.00000 -30.33400
0.00000 0.00000 0.00000 ; CD1- CG- CD2- HD2
1610 1612 1614 1615 3 30.33400 0.00000 -30.33400
0.00000 0.00000 0.00000 ; CD1- CE1- CZ- HZ
1610 1612 1614 1616 3 30.33400 0.00000 -30.33400
0.00000 0.00000 0.00000 ; CD1- CE1- CZ- CE2
1611 1610 1609 1618 3 30.33400 0.00000 -30.33400
0.00000 0.00000 0.00000 ; HD1- CD1- CG- CD2
1611 1610 1612 1613 3 30.33400 0.00000 -30.33400
0.00000 0.00000 0.00000 ; HD1- CD1- CE1- HE1
1611 1610 1612 1614 3 30.33400 0.00000 -30.33400
0.00000 0.00000 0.00000 ; HD1- CD1- CE1- CZ
1612 1610 1609 1618 3 30.33400 0.00000 -30.33400
0.00000 0.00000 0.00000 ; CE1- CD1- CG- CD2
1612 1614 1616 1617 3 30.33400 0.00000 -30.33400
0.00000 0.00000 0.00000 ; CE1- CZ- CE2- HE2
1612 1614 1616 1618 3 30.33400 0.00000 -30.33400
0.00000 0.00000 0.00000 ; CE1- CZ- CE2- CD2
1613 1612 1614 1615 3 30.33400 0.00000 -30.33400
0.00000 0.00000 0.00000 ; HE1- CE1- CZ- HZ
1613 1612 1614 1616 3 30.33400 0.00000 -30.33400
0.00000 0.00000 0.00000 ; HE1- CE1- CZ- CE2
1614 1616 1618 1619 3 30.33400 0.00000 -30.33400
0.00000 0.00000 0.00000 ; CZ- CE2- CD2- HD2
1615 1614 1616 1617 3 30.33400 0.00000 -30.33400
0.00000 0.00000 0.00000 ; HZ- CZ- CE2- HE2
1615 1614 1616 1618 3 30.33400 0.00000 -30.33400
0.00000 0.00000 0.00000 ; HZ- CZ- CE2- CD2
1617 1616 1618 1619 3 30.33400 0.00000 -30.33400
0.00000 0.00000 0.00000 ; HE2- CE2- CD2- HD2
1620 1622 1624 1625 3 0.00000 0.00000 0.00000
0.00000 0.00000 0.00000 ; C- N- CA- HA
1620 1622 1624 1626 3 11.71520 1.67360 15.06240 -
13.38880 0.00000 0.00000 ; C- N- CA- CB

```
  1620   1622   1624   1632       3    1.75728    5.27184    2.25936
-7.02912    0.00000    0.00000 ;       C-     N-    CA-     C
  1621   1620   1622   1623       3   29.28800   -8.36800  -20.92000
0.00000    0.00000    0.00000 ;       O-     C-     N-     H
  1621   1620   1622   1624       3   20.92000    0.00000  -20.92000
0.00000    0.00000    0.00000 ;       O-     C-     N-    CA
  1622   1624   1626   1627       3    0.65084    1.95253    0.00000
-2.60338    0.00000    0.00000 ;       N-    CA-    CB-   HB2
  1622   1624   1626   1628       3    0.65084    1.95253    0.00000
-2.60338    0.00000    0.00000 ;       N-    CA-    CB-   HB3
  1622   1624   1626   1629       3   15.41386    9.74035   -8.39310
-0.97069    2.97901    0.00000 ;       N-    CA-    CB-    CG
  1622   1624   1632   1633       3    0.00000    0.00000    0.00000
0.00000    0.00000    0.00000 ;      N-    CA-     C-     O
  1622   1624   1632   1634       3   17.40544   -5.02080  -13.22144
9.20480    0.00000    0.00000 ;      N-    CA-     C-     N
  1623   1622   1624   1625       3    0.00000    0.00000    0.00000
0.00000    0.00000    0.00000 ;      H-     N-    CA-    HA
  1623   1622   1624   1626       3    0.00000    0.00000    0.00000
0.00000    0.00000    0.00000 ;      H-     N-    CA-    CB
  1623   1622   1624   1632       3    0.00000    0.00000    0.00000
0.00000    0.00000    0.00000 ;      H-     N-    CA-     C
  1624   1626   1629   1630       3    6.43499    0.00000   -5.39736
0.00000   -1.03763    0.00000 ;     CA-    CB-    CG-   OD1
  1624   1626   1629   1631       3    6.43499    0.00000   -5.39736
0.00000   -1.03763    0.00000 ;     CA-    CB-    CG-   OD2
  1624   1632   1634   1635       3   20.92000    0.00000  -20.92000
0.00000    0.00000    0.00000 ;     CA-     C-     N-     H
  1624   1632   1634   1636       3   20.92000    0.00000  -20.92000
0.00000    0.00000    0.00000 ;     CA-     C-     N-    CA
  1625   1624   1626   1627       3    0.65084    1.95253    0.00000
-2.60338    0.00000    0.00000 ;     HA-    CA-    CB-   HB2
  1625   1624   1626   1628       3    0.65084    1.95253    0.00000
-2.60338    0.00000    0.00000 ;     HA-    CA-    CB-   HB3
  1625   1624   1626   1629       3    0.65084    1.95253    0.00000
-2.60338    0.00000    0.00000 ;     HA-    CA-    CB-    CG
  1625   1624   1632   1633       3    3.68192   -4.35136    0.00000
1.33888    0.00000    0.00000 ;     HA-    CA-     C-     O
  1625   1624   1632   1634       3    0.00000    0.00000    0.00000
0.00000    0.00000    0.00000 ;     HA-    CA-     C-     N
  1626   1624   1632   1633       3    0.00000    0.00000    0.00000
0.00000    0.00000    0.00000 ;     CB-    CA-     C-     O
  1626   1624   1632   1634       3    2.51040    4.18400    1.67360
-6.69440    0.00000    0.00000 ;     CB-    CA-     C-     N
  1627   1626   1624   1632       3    0.65084    1.95253    0.00000
-2.60338    0.00000    0.00000 ;    HB2-    CB-    CA-     C
  1627   1626   1629   1630       3    0.00000    0.00000    0.00000
0.00000    0.00000    0.00000 ;    HB2-    CB-    CG-   OD1
  1627   1626   1629   1631       3    0.00000    0.00000    0.00000
0.00000    0.00000    0.00000 ;    HB2-    CB-    CG-   OD2
  1628   1626   1624   1632       3    0.65084    1.95253    0.00000
-2.60338    0.00000    0.00000 ;    HB3-    CB-    CA-     C
```

```
  1628   1626   1629   1630      3    0.00000    0.00000    0.00000
0.00000    0.00000    0.00000 ;    HB3-    CB-    CG-   OD1
  1628   1626   1629   1631      3    0.00000    0.00000    0.00000
0.00000    0.00000    0.00000 ;    HB3-    CB-    CG-   OD2
  1629   1626   1624   1632      3    7.14627   -1.04600   -8.99560
-0.97069    5.15469    0.00000 ;     CG-    CB-    CA-     C
  1632   1634   1636   1637      3    0.00000    0.00000    0.00000
0.00000    0.00000    0.00000 ;     C-     N-    CA-    HA
  1632   1634   1636   1638      3   11.71520    1.67360   15.06240  -
13.38880    0.00000    0.00000 ;     C-     N-    CA-    CB
  1632   1634   1636   1643      3    1.75728    5.27184    2.25936
-7.02912    0.00000    0.00000 ;     C-     N-    CA-     C
  1633   1632   1634   1635      3   29.28800   -8.36800  -20.92000
0.00000    0.00000    0.00000 ;     O-     C-     N-     H
  1633   1632   1634   1636      3   20.92000    0.00000  -20.92000
0.00000    0.00000    0.00000 ;     O-     C-     N-    CA
  1634   1636   1638   1639      3    0.65084    1.95253    0.00000
-2.60338    0.00000    0.00000 ;     N-    CA-    CB-   HB2
  1634   1636   1638   1640      3    0.65084    1.95253    0.00000
-2.60338    0.00000    0.00000 ;     N-    CA-    CB-   HB3
  1634   1636   1638   1641      3    7.86174    2.24681   -7.41405
-6.71114    5.35552    0.00000 ;     N-    CA-    CB-    OG
  1634   1636   1643   1644      3    0.00000    0.00000    0.00000
0.00000    0.00000    0.00000 ;     N-    CA-     C-     O
  1634   1636   1643   1645      3   17.40544   -5.02080  -13.22144
9.20480    0.00000    0.00000 ;     N-    CA-     C-     N
  1635   1634   1636   1637      3    0.00000    0.00000    0.00000
0.00000    0.00000    0.00000 ;     H-     N-    CA-    HA
  1635   1634   1636   1638      3    0.00000    0.00000    0.00000
0.00000    0.00000    0.00000 ;     H-     N-    CA-    CB
  1635   1634   1636   1643      3    0.00000    0.00000    0.00000
0.00000    0.00000    0.00000 ;     H-     N-    CA-     C
  1636   1638   1641   1642      3    2.05853    2.46856    3.48109
-4.46851    0.23430    0.00000 ;    CA-    CB-    OG-    HG
  1636   1643   1645   1646      3   20.92000    0.00000  -20.92000
0.00000    0.00000    0.00000 ;    CA-     C-     N-     H
  1636   1643   1645   1647      3   20.92000    0.00000  -20.92000
0.00000    0.00000    0.00000 ;    CA-     C-     N-    CA
  1637   1636   1638   1639      3    0.65084    1.95253    0.00000
-2.60338    0.00000    0.00000 ;    HA-    CA-    CB-   HB2
  1637   1636   1638   1640      3    0.65084    1.95253    0.00000
-2.60338    0.00000    0.00000 ;    HA-    CA-    CB-   HB3
  1637   1636   1638   1641      3    1.04600   -1.04600    0.00000
0.00000    0.00000    0.00000 ;    HA-    CA-    CB-    OG
  1637   1636   1643   1644      3    3.68192   -4.35136    0.00000
1.33888    0.00000    0.00000 ;    HA-    CA-     C-     O
  1637   1636   1643   1645      3    0.00000    0.00000    0.00000
0.00000    0.00000    0.00000 ;    HA-    CA-     C-     N
  1638   1636   1643   1644      3    0.00000    0.00000    0.00000
0.00000    0.00000    0.00000 ;    CB-    CA-     C-     O
  1638   1636   1643   1645      3    2.51040    4.18400    1.67360
-6.69440    0.00000    0.00000 ;    CB-    CA-     C-     N
```

1639 1638 1636 1643 3 0.65084 1.95253 0.00000
-2.60338 0.00000 0.00000 ; HB2- CB- CA- C
1639 1638 1641 1642 3 0.69733 2.09200 0.00000
-2.78933 0.00000 0.00000 ; HB2- CB- OG- HG
1640 1638 1636 1643 3 0.65084 1.95253 0.00000
-2.60338 0.00000 0.00000 ; HB3- CB- CA- C
1640 1638 1641 1642 3 0.69733 2.09200 0.00000
-2.78933 0.00000 0.00000 ; HB3- CB- OG- HG
1641 1638 1636 1643 3 7.34710 7.79898 -6.14211
-6.71114 4.31789 0.00000 ; OG- CB- CA- C
1643 1645 1647 1648 3 0.00000 0.00000 0.00000
0.00000 0.00000 0.00000 ; C- N- CA- HA
1643 1645 1647 1649 3 11.71520 1.67360 15.06240 -
13.38880 0.00000 0.00000 ; C- N- CA- CB
1643 1645 1647 1662 3 1.75728 5.27184 2.25936
-7.02912 0.00000 0.00000 ; C- N- CA- C
1644 1643 1645 1646 3 29.28800 -8.36800 -20.92000
0.00000 0.00000 0.00000 ; O- C- N- H
1644 1643 1645 1647 3 20.92000 0.00000 -20.92000
0.00000 0.00000 0.00000 ; O- C- N- CA
1645 1647 1649 1650 3 0.65084 1.95253 0.00000
-2.60338 0.00000 0.00000 ; N- CA- CB- HB
1645 1647 1649 1651 3 3.83673 0.44769 -1.77402
-2.47693 -0.03347 0.00000 ; N- CA- CB- CG2
1645 1647 1649 1655 3 2.97482 0.12134 2.04179
-1.89117 -3.24678 0.00000 ; N- CA- CB- CG1
1645 1647 1662 1663 3 0.00000 0.00000 0.00000
0.00000 0.00000 0.00000 ; N- CA- C- O
1645 1647 1662 1664 3 17.40544 -5.02080 -13.22144
9.20480 0.00000 0.00000 ; N- CA- C- N
1646 1645 1647 1648 3 0.00000 0.00000 0.00000
0.00000 0.00000 0.00000 ; H- N- CA- HA
1646 1645 1647 1649 3 0.00000 0.00000 0.00000
0.00000 0.00000 0.00000 ; H- N- CA- CB
1646 1645 1647 1662 3 0.00000 0.00000 0.00000
0.00000 0.00000 0.00000 ; H- N- CA- C
1647 1649 1651 1652 3 0.66944 2.00832 0.00000
-2.67776 0.00000 0.00000 ; CA- CB- CG2- HG21
1647 1649 1651 1653 3 0.66944 2.00832 0.00000
-2.67776 0.00000 0.00000 ; CA- CB- CG2- HG22
1647 1649 1651 1654 3 0.66944 2.00832 0.00000
-2.67776 0.00000 0.00000 ; CA- CB- CG2- HG23
1647 1649 1655 1656 3 0.66944 2.00832 0.00000
-2.67776 0.00000 0.00000 ; CA- CB- CG1- HG12
1647 1649 1655 1657 3 0.66944 2.00832 0.00000
-2.67776 0.00000 0.00000 ; CA- CB- CG1- HG13
1647 1649 1655 1658 3 4.24258 -0.52718 -7.25506
-1.79075 7.69856 0.00000 ; CA- CB- CG1- CD1
1647 1662 1664 1665 3 20.92000 0.00000 -20.92000
0.00000 0.00000 0.00000 ; CA- C- N- H
1647 1662 1664 1666 3 20.92000 0.00000 -20.92000
0.00000 0.00000 0.00000 ; CA- C- N- CA

1648 1647 1649 1650 3 0.65084 1.95253 0.00000
-2.60338 0.00000 0.00000 ; HA- CA- CB- HB
1648 1647 1649 1651 3 0.65084 1.95253 0.00000
-2.60338 0.00000 0.00000 ; HA- CA- CB- CG2
1648 1647 1649 1655 3 0.65084 1.95253 0.00000
-2.60338 0.00000 0.00000 ; HA- CA- CB- CG1
1648 1647 1662 1663 3 3.68192 -4.35136 0.00000
1.33888 0.00000 0.00000 ; HA- CA- C- O
1648 1647 1662 1664 3 0.00000 0.00000 0.00000
0.00000 0.00000 0.00000 ; HA- CA- C- N
1649 1647 1662 1663 3 0.00000 0.00000 0.00000
0.00000 0.00000 0.00000 ; CB- CA- C- O
1649 1647 1662 1664 3 2.51040 4.18400 1.67360
-6.69440 0.00000 0.00000 ; CB- CA- C- N
1649 1655 1658 1659 3 0.66944 2.00832 0.00000
-2.67776 0.00000 0.00000 ; CB- CG1- CD1- HD11
1649 1655 1658 1660 3 0.66944 2.00832 0.00000
-2.67776 0.00000 0.00000 ; CB- CG1- CD1- HD12
1649 1655 1658 1661 3 0.66944 2.00832 0.00000
-2.67776 0.00000 0.00000 ; CB- CG1- CD1- HD13
1650 1649 1647 1662 3 0.65084 1.95253 0.00000
-2.60338 0.00000 0.00000 ; HB- CB- CA- C
1650 1649 1651 1652 3 0.62760 1.88280 0.00000
-2.51040 0.00000 0.00000 ; HB- CB- CG2- HG21
1650 1649 1651 1653 3 0.62760 1.88280 0.00000
-2.51040 0.00000 0.00000 ; HB- CB- CG2- HG22
1650 1649 1651 1654 3 0.62760 1.88280 0.00000
-2.51040 0.00000 0.00000 ; HB- CB- CG2- HG23
1650 1649 1655 1656 3 0.62760 1.88280 0.00000
-2.51040 0.00000 0.00000 ; HB- CB- CG1- HG12
1650 1649 1655 1657 3 0.62760 1.88280 0.00000
-2.51040 0.00000 0.00000 ; HB- CB- CG1- HG13
1650 1649 1655 1658 3 0.66944 2.00832 0.00000
-2.67776 0.00000 0.00000 ; HB- CB- CG1- CD1
1651 1649 1647 1662 3 5.67350 3.55640 -6.16722
-2.47693 3.74886 0.00000 ; CG2- CB- CA- C
1651 1649 1655 1656 3 0.66944 2.00832 0.00000
-2.67776 0.00000 0.00000 ; CG2- CB- CG1- HG12
1651 1649 1655 1657 3 0.66944 2.00832 0.00000
-2.67776 0.00000 0.00000 ; CG2- CB- CG1- HG13
1651 1649 1655 1658 3 3.81162 0.49790 -8.14206
-1.79075 7.49773 0.00000 ; CG2- CB- CG1- CD1
1652 1651 1649 1655 3 0.66944 2.00832 0.00000
-2.67776 0.00000 0.00000 ; HG21- CG2- CB- CG1
1653 1651 1649 1655 3 0.66944 2.00832 0.00000
-2.67776 0.00000 0.00000 ; HG22- CG2- CB- CG1
1654 1651 1649 1655 3 0.66944 2.00832 0.00000
-2.67776 0.00000 0.00000 ; HG23- CG2- CB- CG1
1655 1649 1647 1662 3 8.26340 0.74057 -9.99976
-1.89117 3.84928 0.00000 ; CG1- CB- CA- C
1656 1655 1658 1659 3 0.62760 1.88280 0.00000
-2.51040 0.00000 0.00000 ; HG12- CG1- CD1- HD11

```
  1656   1655   1658   1660       3    0.62760    1.88280    0.00000
-2.51040    0.00000    0.00000 ;   HG12-   CG1-   CD1-  HD12
  1656   1655   1658   1661       3    0.62760    1.88280    0.00000
-2.51040    0.00000    0.00000 ;   HG12-   CG1-   CD1-  HD13
  1657   1655   1658   1659       3    0.62760    1.88280    0.00000
-2.51040    0.00000    0.00000 ;   HG13-   CG1-   CD1-  HD11
  1657   1655   1658   1660       3    0.62760    1.88280    0.00000
-2.51040    0.00000    0.00000 ;   HG13-   CG1-   CD1-  HD12
  1657   1655   1658   1661       3    0.62760    1.88280    0.00000
-2.51040    0.00000    0.00000 ;   HG13-   CG1-   CD1-  HD13
  1662   1664   1666   1667       3    0.00000    0.00000    0.00000
0.00000    0.00000    0.00000 ;      C-     N-    CA-    HA
  1662   1664   1666   1668       3   11.71520    1.67360   15.06240  -
13.38880    0.00000    0.00000 ;      C-     N-    CA-    CB
  1662   1664   1666   1681       3    1.75728    5.27184    2.25936
-7.02912    0.00000    0.00000 ;      C-     N-    CA-     C
  1663   1662   1664   1665       3   29.28800   -8.36800  -20.92000
0.00000    0.00000    0.00000 ;      O-     C-     N-     H
  1663   1662   1664   1666       3   20.92000    0.00000  -20.92000
0.00000    0.00000    0.00000 ;      O-     C-     N-    CA
  1664   1666   1668   1669       3    0.65084    1.95253    0.00000
-2.60338    0.00000    0.00000 ;      N-    CA-    CB-    HB
  1664   1666   1668   1670       3    3.83673    0.44769   -1.77402
-2.47693   -0.03347    0.00000 ;      N-    CA-    CB-   CG2
  1664   1666   1668   1674       3    2.97482    0.12134    2.04179
-1.89117   -3.24678    0.00000 ;      N-    CA-    CB-   CG1
  1664   1666   1681   1682       3    0.00000    0.00000    0.00000
0.00000    0.00000    0.00000 ;      N-    CA-     C-     O
  1664   1666   1681   1683       3   17.40544   -5.02080  -13.22144
9.20480    0.00000    0.00000 ;      N-    CA-     C-     N
  1665   1664   1666   1667       3    0.00000    0.00000    0.00000
0.00000    0.00000    0.00000 ;      H-     N-    CA-    HA
  1665   1664   1666   1668       3    0.00000    0.00000    0.00000
0.00000    0.00000    0.00000 ;      H-     N-    CA-    CB
  1665   1664   1666   1681       3    0.00000    0.00000    0.00000
0.00000    0.00000    0.00000 ;      H-     N-    CA-     C
  1666   1668   1670   1671       3    0.66944    2.00832    0.00000
-2.67776    0.00000    0.00000 ;     CA-    CB-   CG2-  HG21
  1666   1668   1670   1672       3    0.66944    2.00832    0.00000
-2.67776    0.00000    0.00000 ;     CA-    CB-   CG2-  HG22
  1666   1668   1670   1673       3    0.66944    2.00832    0.00000
-2.67776    0.00000    0.00000 ;     CA-    CB-   CG2-  HG23
  1666   1668   1674   1675       3    0.66944    2.00832    0.00000
-2.67776    0.00000    0.00000 ;     CA-    CB-   CG1-  HG12
  1666   1668   1674   1676       3    0.66944    2.00832    0.00000
-2.67776    0.00000    0.00000 ;     CA-    CB-   CG1-  HG13
  1666   1668   1674   1677       3    4.24258   -0.52718   -7.25506
-1.79075    7.69856    0.00000 ;     CA-    CB-   CG1-   CD1
  1666   1681   1683   1684       3   20.92000    0.00000  -20.92000
0.00000    0.00000    0.00000 ;     CA-     C-     N-     H
  1666   1681   1683   1685       3   20.92000    0.00000  -20.92000
0.00000    0.00000    0.00000 ;     CA-     C-     N-    CA
```

```
  1667   1666   1668   1669      3    0.65084    1.95253    0.00000
-2.60338    0.00000    0.00000 ;     HA-    CA-    CB-    HB
  1667   1666   1668   1670      3    0.65084    1.95253    0.00000
-2.60338    0.00000    0.00000 ;     HA-    CA-    CB-   CG2
  1667   1666   1668   1674      3    0.65084    1.95253    0.00000
-2.60338    0.00000    0.00000 ;     HA-    CA-    CB-   CG1
  1667   1666   1681   1682      3    3.68192   -4.35136    0.00000
1.33888    0.00000    0.00000 ;     HA-    CA-     C-     O
  1667   1666   1681   1683      3    0.00000    0.00000    0.00000
0.00000    0.00000    0.00000 ;     HA-    CA-     C-     N
  1668   1666   1681   1682      3    0.00000    0.00000    0.00000
0.00000    0.00000    0.00000 ;     CB-    CA-     C-     O
  1668   1666   1681   1683      3    2.51040    4.18400    1.67360
-6.69440    0.00000    0.00000 ;     CB-    CA-     C-     N
  1668   1674   1677   1678      3    0.66944    2.00832    0.00000
-2.67776    0.00000    0.00000 ;     CB-   CG1-   CD1-  HD11
  1668   1674   1677   1679      3    0.66944    2.00832    0.00000
-2.67776    0.00000    0.00000 ;     CB-   CG1-   CD1-  HD12
  1668   1674   1677   1680      3    0.66944    2.00832    0.00000
-2.67776    0.00000    0.00000 ;     CB-   CG1-   CD1-  HD13
  1669   1668   1666   1681      3    0.65084    1.95253    0.00000
-2.60338    0.00000    0.00000 ;     HB-    CB-    CA-     C
  1669   1668   1670   1671      3    0.62760    1.88280    0.00000
-2.51040    0.00000    0.00000 ;     HB-    CB-   CG2-  HG21
  1669   1668   1670   1672      3    0.62760    1.88280    0.00000
-2.51040    0.00000    0.00000 ;     HB-    CB-   CG2-  HG22
  1669   1668   1670   1673      3    0.62760    1.88280    0.00000
-2.51040    0.00000    0.00000 ;     HB-    CB-   CG2-  HG23
  1669   1668   1674   1675      3    0.62760    1.88280    0.00000
-2.51040    0.00000    0.00000 ;     HB-    CB-   CG1-  HG12
  1669   1668   1674   1676      3    0.62760    1.88280    0.00000
-2.51040    0.00000    0.00000 ;     HB-    CB-   CG1-  HG13
  1669   1668   1674   1677      3    0.66944    2.00832    0.00000
-2.67776    0.00000    0.00000 ;     HB-    CB-   CG1-   CD1
  1670   1668   1666   1681      3    5.67350    3.55640   -6.16722
-2.47693    3.74886    0.00000 ;    CG2-    CB-    CA-     C
  1670   1668   1674   1675      3    0.66944    2.00832    0.00000
-2.67776    0.00000    0.00000 ;    CG2-    CB-   CG1-  HG12
  1670   1668   1674   1676      3    0.66944    2.00832    0.00000
-2.67776    0.00000    0.00000 ;    CG2-    CB-   CG1-  HG13
  1670   1668   1674   1677      3    3.81162    0.49790   -8.14206
-1.79075    7.49773    0.00000 ;    CG2-    CB-   CG1-   CD1
  1671   1670   1668   1674      3    0.66944    2.00832    0.00000
-2.67776    0.00000    0.00000 ;   HG21-   CG2-    CB-   CG1
  1672   1670   1668   1674      3    0.66944    2.00832    0.00000
-2.67776    0.00000    0.00000 ;   HG22-   CG2-    CB-   CG1
  1673   1670   1668   1674      3    0.66944    2.00832    0.00000
-2.67776    0.00000    0.00000 ;   HG23-   CG2-    CB-   CG1
  1674   1668   1666   1681      3    8.26340    0.74057   -9.99976
-1.89117    3.84928    0.00000 ;    CG1-    CB-    CA-     C
  1675   1674   1677   1678      3    0.62760    1.88280    0.00000
-2.51040    0.00000    0.00000 ;   HG12-   CG1-   CD1-  HD11
```

1675 1674 1677 1679 3 0.62760 1.88280 0.00000
-2.51040 0.00000 0.00000 ; HG12- CG1- CD1- HD12
1675 1674 1677 1680 3 0.62760 1.88280 0.00000
-2.51040 0.00000 0.00000 ; HG12- CG1- CD1- HD13
1676 1674 1677 1678 3 0.62760 1.88280 0.00000
-2.51040 0.00000 0.00000 ; HG13- CG1- CD1- HD11
1676 1674 1677 1679 3 0.62760 1.88280 0.00000
-2.51040 0.00000 0.00000 ; HG13- CG1- CD1- HD12
1676 1674 1677 1680 3 0.62760 1.88280 0.00000
-2.51040 0.00000 0.00000 ; HG13- CG1- CD1- HD13
1681 1683 1685 1686 3 0.00000 0.00000 0.00000
0.00000 0.00000 0.00000 ; C- N- CA- HA
1681 1683 1685 1687 3 11.71520 1.67360 15.06240 -
13.38880 0.00000 0.00000 ; C- N- CA- CB
1681 1683 1685 1696 3 1.75728 5.27184 2.25936
-7.02912 0.00000 0.00000 ; C- N- CA- C
1682 1681 1683 1684 3 29.28800 -8.36800 -20.92000
0.00000 0.00000 0.00000 ; O- C- N- H
1682 1681 1683 1685 3 20.92000 0.00000 -20.92000
0.00000 0.00000 0.00000 ; O- C- N- CA
1683 1685 1687 1688 3 0.65084 1.95253 0.00000
-2.60338 0.00000 0.00000 ; N- CA- CB- HB2
1683 1685 1687 1689 3 0.65084 1.95253 0.00000
-2.60338 0.00000 0.00000 ; N- CA- CB- HB3
1683 1685 1687 1690 3 3.21331 2.22589 -4.15053
-2.40998 2.61082 0.00000 ; N- CA- CB- CG
1683 1685 1696 1697 3 0.00000 0.00000 0.00000
0.00000 0.00000 0.00000 ; N- CA- C- O
1683 1685 1696 1698 3 17.40544 -5.02080 -13.22144
9.20480 0.00000 0.00000 ; N- CA- C- N
1684 1683 1685 1686 3 0.00000 0.00000 0.00000
0.00000 0.00000 0.00000 ; H- N- CA- HA
1684 1683 1685 1687 3 0.00000 0.00000 0.00000
0.00000 0.00000 0.00000 ; H- N- CA- CB
1684 1683 1685 1696 3 0.00000 0.00000 0.00000
0.00000 0.00000 0.00000 ; H- N- CA- C
1685 1687 1690 1691 3 0.66944 2.00832 0.00000
-2.67776 0.00000 0.00000 ; CA- CB- CG- HG2
1685 1687 1690 1692 3 0.66944 2.00832 0.00000
-2.67776 0.00000 0.00000 ; CA- CB- CG- HG3
1685 1687 1690 1693 3 10.12110 -1.91209 0.01674
10.17549 -1.87443 0.00000 ; CA- CB- CG- CD
1685 1696 1698 1699 3 20.92000 0.00000 -20.92000
0.00000 0.00000 0.00000 ; CA- C- N- H
1685 1696 1698 1700 3 20.92000 0.00000 -20.92000
0.00000 0.00000 0.00000 ; CA- C- N- CA
1686 1685 1687 1688 3 0.65084 1.95253 0.00000
-2.60338 0.00000 0.00000 ; HA- CA- CB- HB2
1686 1685 1687 1689 3 0.65084 1.95253 0.00000
-2.60338 0.00000 0.00000 ; HA- CA- CB- HB3
1686 1685 1687 1690 3 0.65084 1.95253 0.00000
-2.60338 0.00000 0.00000 ; HA- CA- CB- CG

1686 1685 1696 1697 3 3.68192 -4.35136 0.00000
1.33888 0.00000 0.00000 ; HA- CA- C- O
1686 1685 1696 1698 3 0.00000 0.00000 0.00000
0.00000 0.00000 0.00000 ; HA- CA- C- N
1687 1685 1696 1697 3 0.00000 0.00000 0.00000
0.00000 0.00000 0.00000 ; CB- CA- C- O
1687 1685 1696 1698 3 2.51040 4.18400 1.67360
-6.69440 0.00000 0.00000 ; CB- CA- C- N
1687 1690 1693 1694 3 3.79907 0.00000 -5.40573
0.00000 2.14221 0.00000 ; CB- CG- CD- OE1
1687 1690 1693 1695 3 3.79907 0.00000 -5.40573
0.00000 2.14221 0.00000 ; CB- CG- CD- OE2
1688 1687 1685 1696 3 0.65084 1.95253 0.00000
-2.60338 0.00000 0.00000 ; HB2- CB- CA- C
1688 1687 1690 1691 3 0.62760 1.88280 0.00000
-2.51040 0.00000 0.00000 ; HB2- CB- CG- HG2
1688 1687 1690 1692 3 0.62760 1.88280 0.00000
-2.51040 0.00000 0.00000 ; HB2- CB- CG- HG3
1688 1687 1690 1693 3 0.65084 1.95253 0.00000
-2.60338 0.00000 0.00000 ; HB2- CB- CG- CD
1689 1687 1685 1696 3 0.65084 1.95253 0.00000
-2.60338 0.00000 0.00000 ; HB3- CB- CA- C
1689 1687 1690 1691 3 0.62760 1.88280 0.00000
-2.51040 0.00000 0.00000 ; HB3- CB- CG- HG2
1689 1687 1690 1692 3 0.62760 1.88280 0.00000
-2.51040 0.00000 0.00000 ; HB3- CB- CG- HG3
1689 1687 1690 1693 3 0.65084 1.95253 0.00000
-2.60338 0.00000 0.00000 ; HB3- CB- CG- CD
1690 1687 1685 1696 3 6.86594 3.56895 -8.14206
-2.40998 4.85344 0.00000 ; CG- CB- CA- C
1691 1690 1693 1694 3 0.00000 0.00000 0.00000
0.00000 0.00000 0.00000 ; HG2- CG- CD- OE1
1691 1690 1693 1695 3 0.00000 0.00000 0.00000
0.00000 0.00000 0.00000 ; HG2- CG- CD- OE2
1692 1690 1693 1694 3 0.00000 0.00000 0.00000
0.00000 0.00000 0.00000 ; HG3- CG- CD- OE1
1692 1690 1693 1695 3 0.00000 0.00000 0.00000
0.00000 0.00000 0.00000 ; HG3- CG- CD- OE2
1696 1698 1700 1701 3 0.00000 0.00000 0.00000
0.00000 0.00000 0.00000 ; C- N- CA- HA
1696 1698 1700 1702 3 11.71520 1.67360 15.06240 -
13.38880 0.00000 0.00000 ; C- N- CA- CB
1696 1698 1700 1710 3 1.75728 5.27184 2.25936
-7.02912 0.00000 0.00000 ; C- N- CA- C
1697 1696 1698 1699 3 29.28800 -8.36800 -20.92000
0.00000 0.00000 0.00000 ; O- C- N- H
1697 1696 1698 1700 3 20.92000 0.00000 -20.92000
0.00000 0.00000 0.00000 ; O- C- N- CA
1698 1700 1702 1703 3 0.65084 1.95253 0.00000
-2.60338 0.00000 0.00000 ; N- CA- CB- HB2
1698 1700 1702 1704 3 0.65084 1.95253 0.00000
-2.60338 0.00000 0.00000 ; N- CA- CB- HB3

```
  1698   1700   1702   1705      3    5.99567    3.29281   -4.46014
-0.55229    1.97485    0.00000 ;       N-    CA-    CB-    CG
  1698   1700   1710   1711      3    0.00000    0.00000    0.00000
0.00000    0.00000    0.00000 ;       N-    CA-     C-     O
  1698   1700   1710   1712      3   17.40544   -5.02080  -13.22144
9.20480    0.00000    0.00000 ;       N-    CA-     C-     N
  1699   1698   1700   1701      3    0.00000    0.00000    0.00000
0.00000    0.00000    0.00000 ;       H-     N-    CA-    HA
  1699   1698   1700   1702      3    0.00000    0.00000    0.00000
0.00000    0.00000    0.00000 ;       H-     N-    CA-    CB
  1699   1698   1700   1710      3    0.00000    0.00000    0.00000
0.00000    0.00000    0.00000 ;       H-     N-    CA-     C
  1700   1702   1705   1706      3    0.00000    0.00000    0.00000
0.00000    0.00000    0.00000 ;      CA-    CB-    CG-   OD1
  1700   1702   1705   1707      3    8.84916   -0.31380   -4.32626
5.03754    0.26778    0.00000 ;      CA-    CB-    CG-   ND2
  1700   1710   1712   1713      3   20.92000    0.00000  -20.92000
0.00000    0.00000    0.00000 ;      CA-     C-     N-     H
  1700   1710   1712   1714      3   20.92000    0.00000  -20.92000
0.00000    0.00000    0.00000 ;      CA-     C-     N-    CA
  1701   1700   1702   1703      3    0.65084    1.95253    0.00000
-2.60338    0.00000    0.00000 ;      HA-    CA-    CB-   HB2
  1701   1700   1702   1704      3    0.65084    1.95253    0.00000
-2.60338    0.00000    0.00000 ;      HA-    CA-    CB-   HB3
  1701   1700   1702   1705      3    0.65084    1.95253    0.00000
-2.60338    0.00000    0.00000 ;      HA-    CA-    CB-    CG
  1701   1700   1710   1711      3    3.68192   -4.35136    0.00000
1.33888    0.00000    0.00000 ;      HA-    CA-     C-     O
  1701   1700   1710   1712      3    0.00000    0.00000    0.00000
0.00000    0.00000    0.00000 ;      HA-    CA-     C-     N
  1702   1700   1710   1711      3    0.00000    0.00000    0.00000
0.00000    0.00000    0.00000 ;      CB-    CA-     C-     O
  1702   1700   1710   1712      3    2.51040    4.18400    1.67360
-6.69440    0.00000    0.00000 ;      CB-    CA-     C-     N
  1702   1705   1707   1708      3   20.92000    0.00000  -20.92000
0.00000    0.00000    0.00000 ;      CB-    CG-   ND2-  HD21
  1702   1705   1707   1709      3   20.92000    0.00000  -20.92000
0.00000    0.00000    0.00000 ;      CB-    CG-   ND2-  HD22
  1703   1702   1700   1710      3    0.65084    1.95253    0.00000
-2.60338    0.00000    0.00000 ;     HB2-    CB-    CA-     C
  1703   1702   1705   1706      3    3.68192   -4.35136    0.00000
1.33888    0.00000    0.00000 ;     HB2-    CB-    CG-   OD1
  1703   1702   1705   1707      3    0.00000    0.00000    0.00000
0.00000    0.00000    0.00000 ;     HB2-    CB-    CG-   ND2
  1704   1702   1700   1710      3    0.65084    1.95253    0.00000
-2.60338    0.00000    0.00000 ;     HB3-    CB-    CA-     C
  1704   1702   1705   1706      3    3.68192   -4.35136    0.00000
1.33888    0.00000    0.00000 ;     HB3-    CB-    CG-   OD1
  1704   1702   1705   1707      3    0.00000    0.00000    0.00000
0.00000    0.00000    0.00000 ;     HB3-    CB-    CG-   ND2
  1705   1702   1700   1710      3    7.94542   -3.96225   -6.11701
-0.55229    3.58150    0.00000 ;      CG-    CB-    CA-     C
```

1706 1705 1707 1708 3 29.28800 -8.36800 -20.92000
0.00000 0.00000 0.00000 ; OD1- CG- ND2- HD21
1706 1705 1707 1709 3 29.28800 -8.36800 -20.92000
0.00000 0.00000 0.00000 ; OD1- CG- ND2- HD22
1710 1712 1714 1715 3 0.00000 0.00000 0.00000
0.00000 0.00000 0.00000 ; C- N- CA- HA2
1710 1712 1714 1716 3 0.00000 0.00000 0.00000
0.00000 0.00000 0.00000 ; C- N- CA- HA3
1710 1712 1714 1717 3 1.75728 5.27184 2.25936
-7.02912 0.00000 0.00000 ; C- N- CA- C
1711 1710 1712 1713 3 29.28800 -8.36800 -20.92000
0.00000 0.00000 0.00000 ; O- C- N- H
1711 1710 1712 1714 3 20.92000 0.00000 -20.92000
0.00000 0.00000 0.00000 ; O- C- N- CA
1712 1714 1717 1718 3 0.00000 0.00000 0.00000
0.00000 0.00000 0.00000 ; N- CA- C- O
1712 1714 1717 1719 3 17.40544 -5.02080 -13.22144
9.20480 0.00000 0.00000 ; N- CA- C- N
1713 1712 1714 1715 3 0.00000 0.00000 0.00000
0.00000 0.00000 0.00000 ; H- N- CA- HA2
1713 1712 1714 1716 3 0.00000 0.00000 0.00000
0.00000 0.00000 0.00000 ; H- N- CA- HA3
1713 1712 1714 1717 3 0.00000 0.00000 0.00000
0.00000 0.00000 0.00000 ; H- N- CA- C
1714 1717 1719 1720 3 20.92000 0.00000 -20.92000
0.00000 0.00000 0.00000 ; CA- C- N- H
1714 1717 1719 1721 3 20.92000 0.00000 -20.92000
0.00000 0.00000 0.00000 ; CA- C- N- CA
1715 1714 1717 1718 3 3.68192 -4.35136 0.00000
1.33888 0.00000 0.00000 ; HA2- CA- C- O
1715 1714 1717 1719 3 0.00000 0.00000 0.00000
0.00000 0.00000 0.00000 ; HA2- CA- C- N
1716 1714 1717 1718 3 3.68192 -4.35136 0.00000
1.33888 0.00000 0.00000 ; HA3- CA- C- O
1716 1714 1717 1719 3 0.00000 0.00000 0.00000
0.00000 0.00000 0.00000 ; HA3- CA- C- N
1717 1719 1721 1722 3 0.00000 0.00000 0.00000
0.00000 0.00000 0.00000 ; C- N- CA- HA
1717 1719 1721 1723 3 11.71520 1.67360 15.06240 -
13.38880 0.00000 0.00000 ; C- N- CA- CB
1717 1719 1721 1728 3 1.75728 5.27184 2.25936
-7.02912 0.00000 0.00000 ; C- N- CA- C
1718 1717 1719 1720 3 29.28800 -8.36800 -20.92000
0.00000 0.00000 0.00000 ; O- C- N- H
1718 1717 1719 1721 3 20.92000 0.00000 -20.92000
0.00000 0.00000 0.00000 ; O- C- N- CA
1719 1721 1723 1724 3 0.65084 1.95253 0.00000
-2.60338 0.00000 0.00000 ; N- CA- CB- HB2
1719 1721 1723 1725 3 0.65084 1.95253 0.00000
-2.60338 0.00000 0.00000 ; N- CA- CB- HB3
1719 1721 1723 1726 3 7.86174 2.24681 -7.41405
-6.71114 5.35552 0.00000 ; N- CA- CB- OG

1719 1721 1728 1729 3 0.00000 0.00000 0.00000
0.00000 0.00000 0.00000 ; N- CA- C- O
1719 1721 1728 1730 3 17.40544 -5.02080 -13.22144
9.20480 0.00000 0.00000 ; N- CA- C- N
1720 1719 1721 1722 3 0.00000 0.00000 0.00000
0.00000 0.00000 0.00000 ; H- N- CA- HA
1720 1719 1721 1723 3 0.00000 0.00000 0.00000
0.00000 0.00000 0.00000 ; H- N- CA- CB
1720 1719 1721 1728 3 0.00000 0.00000 0.00000
0.00000 0.00000 0.00000 ; H- N- CA- C
1721 1723 1726 1727 3 2.05853 2.46856 3.48109
-4.46851 0.23430 0.00000 ; CA- CB- OG- HG
1721 1728 1730 1731 3 20.92000 0.00000 -20.92000
0.00000 0.00000 0.00000 ; CA- C- N- H
1721 1728 1730 1732 3 20.92000 0.00000 -20.92000
0.00000 0.00000 0.00000 ; CA- C- N- CA
1722 1721 1723 1724 3 0.65084 1.95253 0.00000
-2.60338 0.00000 0.00000 ; HA- CA- CB- HB2
1722 1721 1723 1725 3 0.65084 1.95253 0.00000
-2.60338 0.00000 0.00000 ; HA- CA- CB- HB3
1722 1721 1723 1726 3 1.04600 -1.04600 0.00000
0.00000 0.00000 0.00000 ; HA- CA- CB- OG
1722 1721 1728 1729 3 3.68192 -4.35136 0.00000
1.33888 0.00000 0.00000 ; HA- CA- C- O
1722 1721 1728 1730 3 0.00000 0.00000 0.00000
0.00000 0.00000 0.00000 ; HA- CA- C- N
1723 1721 1728 1729 3 0.00000 0.00000 0.00000
0.00000 0.00000 0.00000 ; CB- CA- C- O
1723 1721 1728 1730 3 2.51040 4.18400 1.67360
-6.69440 0.00000 0.00000 ; CB- CA- C- N
1724 1723 1721 1728 3 0.65084 1.95253 0.00000
-2.60338 0.00000 0.00000 ; HB2- CB- CA- C
1724 1723 1726 1727 3 0.69733 2.09200 0.00000
-2.78933 0.00000 0.00000 ; HB2- CB- OG- HG
1725 1723 1721 1728 3 0.65084 1.95253 0.00000
-2.60338 0.00000 0.00000 ; HB3- CB- CA- C
1725 1723 1726 1727 3 0.69733 2.09200 0.00000
-2.78933 0.00000 0.00000 ; HB3- CB- OG- HG
1726 1723 1721 1728 3 7.34710 7.79898 -6.14211
-6.71114 4.31789 0.00000 ; OG- CB- CA- C
1728 1730 1732 1733 3 0.00000 0.00000 0.00000
0.00000 0.00000 0.00000 ; C- N- CA- HA
1728 1730 1732 1734 3 11.71520 1.67360 15.06240 -
13.38880 0.00000 0.00000 ; C- N- CA- CB
1728 1730 1732 1748 3 1.75728 5.27184 2.25936
-7.02912 0.00000 0.00000 ; C- N- CA- C
1729 1728 1730 1731 3 29.28800 -8.36800 -20.92000
0.00000 0.00000 0.00000 ; O- C- N- H
1729 1728 1730 1732 3 20.92000 0.00000 -20.92000
0.00000 0.00000 0.00000 ; O- C- N- CA
1730 1732 1734 1735 3 0.65084 1.95253 0.00000
-2.60338 0.00000 0.00000 ; N- CA- CB- HB2

1730 1732 1734 1736 3 0.65084 1.95253 0.00000
-2.60338 0.00000 0.00000 ; N- CA- CB- HB3
1730 1732 1734 1737 3 3.28026 2.46019 -2.19242
-3.21331 -0.23430 0.00000 ; N- CA- CB- CG
1730 1732 1748 1749 3 0.00000 0.00000 0.00000
0.00000 0.00000 0.00000 ; N- CA- C- O
1730 1732 1748 1750 3 17.40544 -5.02080 -13.22144
9.20480 0.00000 0.00000 ; N- CA- C- N
1731 1730 1732 1733 3 0.00000 0.00000 0.00000
0.00000 0.00000 0.00000 ; H- N- CA- HA
1731 1730 1732 1734 3 0.00000 0.00000 0.00000
0.00000 0.00000 0.00000 ; H- N- CA- CB
1731 1730 1732 1748 3 0.00000 0.00000 0.00000
0.00000 0.00000 0.00000 ; H- N- CA- C
1732 1734 1737 1738 3 0.57739 0.00000 1.02926
0.00000 -1.60666 0.00000 ; CA- CB- CG- CD1
1732 1734 1737 1746 3 0.57739 0.00000 1.02926
0.00000 -1.60666 0.00000 ; CA- CB- CG- CD2
1732 1748 1750 1751 3 20.92000 0.00000 -20.92000
0.00000 0.00000 0.00000 ; CA- C- N- H
1732 1748 1750 1752 3 20.92000 0.00000 -20.92000
0.00000 0.00000 0.00000 ; CA- C- N- CA
1733 1732 1734 1735 3 0.65084 1.95253 0.00000
-2.60338 0.00000 0.00000 ; HA- CA- CB- HB2
1733 1732 1734 1736 3 0.65084 1.95253 0.00000
-2.60338 0.00000 0.00000 ; HA- CA- CB- HB3
1733 1732 1734 1737 3 0.65084 1.95253 0.00000
-2.60338 0.00000 0.00000 ; HA- CA- CB- CG
1733 1732 1748 1749 3 3.68192 -4.35136 0.00000
1.33888 0.00000 0.00000 ; HA- CA- C- O
1733 1732 1748 1750 3 0.00000 0.00000 0.00000
0.00000 0.00000 0.00000 ; HA- CA- C- N
1734 1732 1748 1749 3 0.00000 0.00000 0.00000
0.00000 0.00000 0.00000 ; CB- CA- C- O
1734 1732 1748 1750 3 2.51040 4.18400 1.67360
-6.69440 0.00000 0.00000 ; CB- CA- C- N
1734 1737 1738 1739 3 30.33400 0.00000 -30.33400
0.00000 0.00000 0.00000 ; CB- CG- CD1- HD1
1734 1737 1738 1740 3 30.33400 0.00000 -30.33400
0.00000 0.00000 0.00000 ; CB- CG- CD1- CE1
1734 1737 1746 1744 3 30.33400 0.00000 -30.33400
0.00000 0.00000 0.00000 ; CB- CG- CD2- CE2
1734 1737 1746 1747 3 30.33400 0.00000 -30.33400
0.00000 0.00000 0.00000 ; CB- CG- CD2- HD2
1735 1734 1732 1748 3 0.65084 1.95253 0.00000
-2.60338 0.00000 0.00000 ; HB2- CB- CA- C
1735 1734 1737 1738 3 0.00000 0.00000 0.00000
0.00000 0.00000 0.00000 ; HB2- CB- CG- CD1
1735 1734 1737 1746 3 0.00000 0.00000 0.00000
0.00000 0.00000 0.00000 ; HB2- CB- CG- CD2
1736 1734 1732 1748 3 0.65084 1.95253 0.00000
-2.60338 0.00000 0.00000 ; HB3- CB- CA- C

1736 1734 1737 1738 3 0.00000 0.00000 0.00000
0.00000 0.00000 0.00000 ; HB3- CB- CG- CD1
1736 1734 1737 1746 3 0.00000 0.00000 0.00000
0.00000 0.00000 0.00000 ; HB3- CB- CG- CD2
1737 1734 1732 1748 3 4.95804 2.17986 -3.52293
-3.21331 -0.40166 0.00000 ; CG- CB- CA- C
1737 1738 1740 1741 3 30.33400 0.00000 -30.33400
0.00000 0.00000 0.00000 ; CG- CD1- CE1- HE1
1737 1738 1740 1742 3 30.33400 0.00000 -30.33400
0.00000 0.00000 0.00000 ; CG- CD1- CE1- CZ
1737 1746 1744 1742 3 30.33400 0.00000 -30.33400
0.00000 0.00000 0.00000 ; CG- CD2- CE2- CZ
1737 1746 1744 1745 3 30.33400 0.00000 -30.33400
0.00000 0.00000 0.00000 ; CG- CD2- CE2- HE2
1738 1737 1746 1744 3 30.33400 0.00000 -30.33400
0.00000 0.00000 0.00000 ; CD1- CG- CD2- CE2
1738 1737 1746 1747 3 30.33400 0.00000 -30.33400
0.00000 0.00000 0.00000 ; CD1- CG- CD2- HD2
1738 1740 1742 1743 3 30.33400 0.00000 -30.33400
0.00000 0.00000 0.00000 ; CD1- CE1- CZ- HZ
1738 1740 1742 1744 3 30.33400 0.00000 -30.33400
0.00000 0.00000 0.00000 ; CD1- CE1- CZ- CE2
1739 1738 1737 1746 3 30.33400 0.00000 -30.33400
0.00000 0.00000 0.00000 ; HD1- CD1- CG- CD2
1739 1738 1740 1741 3 30.33400 0.00000 -30.33400
0.00000 0.00000 0.00000 ; HD1- CD1- CE1- HE1
1739 1738 1740 1742 3 30.33400 0.00000 -30.33400
0.00000 0.00000 0.00000 ; HD1- CD1- CE1- CZ
1740 1738 1737 1746 3 30.33400 0.00000 -30.33400
0.00000 0.00000 0.00000 ; CE1- CD1- CG- CD2
1740 1742 1744 1745 3 30.33400 0.00000 -30.33400
0.00000 0.00000 0.00000 ; CE1- CZ- CE2- HE2
1740 1742 1744 1746 3 30.33400 0.00000 -30.33400
0.00000 0.00000 0.00000 ; CE1- CZ- CE2- CD2
1741 1740 1742 1743 3 30.33400 0.00000 -30.33400
0.00000 0.00000 0.00000 ; HE1- CE1- CZ- HZ
1741 1740 1742 1744 3 30.33400 0.00000 -30.33400
0.00000 0.00000 0.00000 ; HE1- CE1- CZ- CE2
1742 1744 1746 1747 3 30.33400 0.00000 -30.33400
0.00000 0.00000 0.00000 ; CZ- CE2- CD2- HD2
1743 1742 1744 1745 3 30.33400 0.00000 -30.33400
0.00000 0.00000 0.00000 ; HZ- CZ- CE2- HE2
1743 1742 1744 1746 3 30.33400 0.00000 -30.33400
0.00000 0.00000 0.00000 ; HZ- CZ- CE2- CD2
1745 1744 1746 1747 3 30.33400 0.00000 -30.33400
0.00000 0.00000 0.00000 ; HE2- CE2- CD2- HD2
1748 1750 1752 1753 3 0.00000 0.00000 0.00000
0.00000 0.00000 0.00000 ; C- N- CA- HA
1748 1750 1752 1754 3 11.71520 1.67360 15.06240 -
13.38880 0.00000 0.00000 ; C- N- CA- CB
1748 1750 1752 1762 3 1.75728 5.27184 2.25936
-7.02912 0.00000 0.00000 ; C- N- CA- C

1749 1748 1750 1751 3 29.28800 -8.36800 -20.92000
0.00000 0.00000 0.00000 ; O- C- N- H
1749 1748 1750 1752 3 20.92000 0.00000 -20.92000
0.00000 0.00000 0.00000 ; O- C- N- CA
1750 1752 1754 1755 3 0.65084 1.95253 0.00000
-2.60338 0.00000 0.00000 ; N- CA- CB- HB
1750 1752 1754 1756 3 3.83673 0.44769 -1.77402
-2.47693 -0.03347 0.00000 ; N- CA- CB- CG2
1750 1752 1754 1760 3 4.93294 1.13386 -3.12963
-5.27184 3.17984 0.00000 ; N- CA- CB- OG1
1750 1752 1762 1763 3 0.00000 0.00000 0.00000
0.00000 0.00000 0.00000 ; N- CA- C- O
1750 1752 1762 1764 3 17.40544 -5.02080 -13.22144
9.20480 0.00000 0.00000 ; N- CA- C- N
1751 1750 1752 1753 3 0.00000 0.00000 0.00000
0.00000 0.00000 0.00000 ; H- N- CA- HA
1751 1750 1752 1754 3 0.00000 0.00000 0.00000
0.00000 0.00000 0.00000 ; H- N- CA- CB
1751 1750 1752 1762 3 0.00000 0.00000 0.00000
0.00000 0.00000 0.00000 ; H- N- CA- C
1752 1754 1756 1757 3 0.66944 2.00832 0.00000
-2.67776 0.00000 0.00000 ; CA- CB- CG2- HG21
1752 1754 1756 1758 3 0.66944 2.00832 0.00000
-2.67776 0.00000 0.00000 ; CA- CB- CG2- HG22
1752 1754 1756 1759 3 0.66944 2.00832 0.00000
-2.67776 0.00000 0.00000 ; CA- CB- CG2- HG23
1752 1754 1760 1761 3 1.12131 2.98738 1.66523
-3.94970 0.43514 0.00000 ; CA- CB- OG1- HG1
1752 1762 1764 1765 3 20.92000 0.00000 -20.92000
0.00000 0.00000 0.00000 ; CA- C- N- H
1752 1762 1764 1766 3 20.92000 0.00000 -20.92000
0.00000 0.00000 0.00000 ; CA- C- N- CA
1753 1752 1754 1755 3 0.65084 1.95253 0.00000
-2.60338 0.00000 0.00000 ; HA- CA- CB- HB
1753 1752 1754 1756 3 0.65084 1.95253 0.00000
-2.60338 0.00000 0.00000 ; HA- CA- CB- CG2
1753 1752 1754 1760 3 1.04600 -1.04600 0.00000
0.00000 0.00000 0.00000 ; HA- CA- CB- OG1
1753 1752 1762 1763 3 3.68192 -4.35136 0.00000
1.33888 0.00000 0.00000 ; HA- CA- C- O
1753 1752 1762 1764 3 0.00000 0.00000 0.00000
0.00000 0.00000 0.00000 ; HA- CA- C- N
1754 1752 1762 1763 3 0.00000 0.00000 0.00000
0.00000 0.00000 0.00000 ; CB- CA- C- O
1754 1752 1762 1764 3 2.51040 4.18400 1.67360
-6.69440 0.00000 0.00000 ; CB- CA- C- N
1755 1754 1752 1762 3 0.65084 1.95253 0.00000
-2.60338 0.00000 0.00000 ; HB- CB- CA- C
1755 1754 1756 1757 3 0.65084 1.95253 0.00000
-2.60338 0.00000 0.00000 ; HB- CB- CG2- HG21
1755 1754 1756 1758 3 0.65084 1.95253 0.00000
-2.60338 0.00000 0.00000 ; HB- CB- CG2- HG22

1755 1754 1756 1759 3 0.65084 1.95253 0.00000
-2.60338 0.00000 0.00000 ; HB- CB- CG2- HG23
1755 1754 1760 1761 3 0.69733 2.09200 0.00000
-2.78933 0.00000 0.00000 ; HB- CB- OG1- HG1
1756 1754 1752 1762 3 5.67350 3.55640 -6.16722
-2.47693 3.74886 0.00000 ; CG2- CB- CA- C
1756 1754 1760 1761 3 4.74047 0.27196 -2.26773
-3.94970 1.60666 0.00000 ; CG2- CB- OG1- HG1
1757 1756 1754 1760 3 1.04600 -1.04600 0.00000
0.00000 0.00000 0.00000 ; HG21- CG2- CB- OG1
1758 1756 1754 1760 3 1.04600 -1.04600 0.00000
0.00000 0.00000 0.00000 ; HG22- CG2- CB- OG1
1759 1756 1754 1760 3 1.04600 -1.04600 0.00000
0.00000 0.00000 0.00000 ; HG23- CG2- CB- OG1
1760 1754 1752 1762 3 6.53541 6.87013 -6.21742
-5.27184 5.22163 0.00000 ; OG1- CB- CA- C
1762 1764 1766 1767 3 0.00000 0.00000 0.00000
0.00000 0.00000 0.00000 ; C- N- CA- HA
1762 1764 1766 1768 3 11.71520 1.67360 15.06240 -
13.38880 0.00000 0.00000 ; C- N- CA- CB
1762 1764 1766 1777 3 1.75728 5.27184 2.25936
-7.02912 0.00000 0.00000 ; C- N- CA- C
1763 1762 1764 1765 3 29.28800 -8.36800 -20.92000
0.00000 0.00000 0.00000 ; O- C- N- H
1763 1762 1764 1766 3 20.92000 0.00000 -20.92000
0.00000 0.00000 0.00000 ; O- C- N- CA
1764 1766 1768 1769 3 0.65084 1.95253 0.00000
-2.60338 0.00000 0.00000 ; N- CA- CB- HB2
1764 1766 1768 1770 3 0.65084 1.95253 0.00000
-2.60338 0.00000 0.00000 ; N- CA- CB- HB3
1764 1766 1768 1771 3 3.21331 2.22589 -4.15053
-2.40998 2.61082 0.00000 ; N- CA- CB- CG
1764 1766 1777 1778 3 0.00000 0.00000 0.00000
0.00000 0.00000 0.00000 ; N- CA- C- O
1764 1766 1777 1779 3 17.40544 -5.02080 -13.22144
9.20480 0.00000 0.00000 ; N- CA- C- N
1765 1764 1766 1767 3 0.00000 0.00000 0.00000
0.00000 0.00000 0.00000 ; H- N- CA- HA
1765 1764 1766 1768 3 0.00000 0.00000 0.00000
0.00000 0.00000 0.00000 ; H- N- CA- CB
1765 1764 1766 1777 3 0.00000 0.00000 0.00000
0.00000 0.00000 0.00000 ; H- N- CA- C
1766 1768 1771 1772 3 0.66944 2.00832 0.00000
-2.67776 0.00000 0.00000 ; CA- CB- CG- HG2
1766 1768 1771 1773 3 0.66944 2.00832 0.00000
-2.67776 0.00000 0.00000 ; CA- CB- CG- HG3
1766 1768 1771 1774 3 10.12110 -1.91209 0.01674
10.17549 -1.87443 0.00000 ; CA- CB- CG- CD
1766 1777 1779 1780 3 20.92000 0.00000 -20.92000
0.00000 0.00000 0.00000 ; CA- C- N- H
1766 1777 1779 1781 3 20.92000 0.00000 -20.92000
0.00000 0.00000 0.00000 ; CA- C- N- CA

1767 1766 1768 1769 3 0.65084 1.95253 0.00000
-2.60338 0.00000 0.00000 ; HA- CA- CB- HB2
1767 1766 1768 1770 3 0.65084 1.95253 0.00000
-2.60338 0.00000 0.00000 ; HA- CA- CB- HB3
1767 1766 1768 1771 3 0.65084 1.95253 0.00000
-2.60338 0.00000 0.00000 ; HA- CA- CB- CG
1767 1766 1777 1778 3 3.68192 -4.35136 0.00000
1.33888 0.00000 0.00000 ; HA- CA- C- O
1767 1766 1777 1779 3 0.00000 0.00000 0.00000
0.00000 0.00000 0.00000 ; HA- CA- C- N
1768 1766 1777 1778 3 0.00000 0.00000 0.00000
0.00000 0.00000 0.00000 ; CB- CA- C- O
1768 1766 1777 1779 3 2.51040 4.18400 1.67360
-6.69440 0.00000 0.00000 ; CB- CA- C- N
1768 1771 1774 1775 3 3.79907 0.00000 -5.40573
0.00000 2.14221 0.00000 ; CB- CG- CD- OE1
1768 1771 1774 1776 3 3.79907 0.00000 -5.40573
0.00000 2.14221 0.00000 ; CB- CG- CD- OE2
1769 1768 1766 1777 3 0.65084 1.95253 0.00000
-2.60338 0.00000 0.00000 ; HB2- CB- CA- C
1769 1768 1771 1772 3 0.62760 1.88280 0.00000
-2.51040 0.00000 0.00000 ; HB2- CB- CG- HG2
1769 1768 1771 1773 3 0.62760 1.88280 0.00000
-2.51040 0.00000 0.00000 ; HB2- CB- CG- HG3
1769 1768 1771 1774 3 0.65084 1.95253 0.00000
-2.60338 0.00000 0.00000 ; HB2- CB- CG- CD
1770 1768 1766 1777 3 0.65084 1.95253 0.00000
-2.60338 0.00000 0.00000 ; HB3- CB- CA- C
1770 1768 1771 1772 3 0.62760 1.88280 0.00000
-2.51040 0.00000 0.00000 ; HB3- CB- CG- HG2
1770 1768 1771 1773 3 0.62760 1.88280 0.00000
-2.51040 0.00000 0.00000 ; HB3- CB- CG- HG3
1770 1768 1771 1774 3 0.65084 1.95253 0.00000
-2.60338 0.00000 0.00000 ; HB3- CB- CG- CD
1771 1768 1766 1777 3 6.86594 3.56895 -8.14206
-2.40998 4.85344 0.00000 ; CG- CB- CA- C
1772 1771 1774 1775 3 0.00000 0.00000 0.00000
0.00000 0.00000 0.00000 ; HG2- CG- CD- OE1
1772 1771 1774 1776 3 0.00000 0.00000 0.00000
0.00000 0.00000 0.00000 ; HG2- CG- CD- OE2
1773 1771 1774 1775 3 0.00000 0.00000 0.00000
0.00000 0.00000 0.00000 ; HG3- CG- CD- OE1
1773 1771 1774 1776 3 0.00000 0.00000 0.00000
0.00000 0.00000 0.00000 ; HG3- CG- CD- OE2
1777 1779 1781 1782 3 0.00000 0.00000 0.00000
0.00000 0.00000 0.00000 ; C- N- CA- HA
1777 1779 1781 1783 3 11.71520 1.67360 15.06240 -
13.38880 0.00000 0.00000 ; C- N- CA- CB
1777 1779 1781 1799 3 1.75728 5.27184 2.25936
-7.02912 0.00000 0.00000 ; C- N- CA- C
1778 1777 1779 1780 3 29.28800 -8.36800 -20.92000
0.00000 0.00000 0.00000 ; O- C- N- H

1778 1777 1779 1781 3 20.92000 0.00000 -20.92000
0.00000 0.00000 0.00000 ; O- C- N- CA
1779 1781 1783 1784 3 0.65084 1.95253 0.00000
-2.60338 0.00000 0.00000 ; N- CA- CB- HB2
1779 1781 1783 1785 3 0.65084 1.95253 0.00000
-2.60338 0.00000 0.00000 ; N- CA- CB- HB3
1779 1781 1783 1786 3 0.65084 1.95253 0.00000
-2.60338 0.00000 0.00000 ; N- CA- CB- CG
1779 1781 1799 1800 3 0.00000 0.00000 0.00000
0.00000 0.00000 0.00000 ; N- CA- C- O
1779 1781 1799 1801 3 17.40544 -5.02080 -13.22144
9.20480 0.00000 0.00000 ; N- CA- C- N
1780 1779 1781 1782 3 0.00000 0.00000 0.00000
0.00000 0.00000 0.00000 ; H- N- CA- HA
1780 1779 1781 1783 3 0.00000 0.00000 0.00000
0.00000 0.00000 0.00000 ; H- N- CA- CB
1780 1779 1781 1799 3 0.00000 0.00000 0.00000
0.00000 0.00000 0.00000 ; H- N- CA- C
1781 1783 1786 1787 3 0.66944 2.00832 0.00000
-2.67776 0.00000 0.00000 ; CA- CB- CG- HG2
1781 1783 1786 1788 3 0.66944 2.00832 0.00000
-2.67776 0.00000 0.00000 ; CA- CB- CG- HG3
1781 1783 1786 1789 3 3.68192 3.09616 -2.09200
-3.01248 0.00000 0.00000 ; CA- CB- CG- CD
1781 1799 1801 1802 3 20.92000 0.00000 -20.92000
0.00000 0.00000 0.00000 ; CA- C- N- H
1781 1799 1801 1803 3 20.92000 0.00000 -20.92000
0.00000 0.00000 0.00000 ; CA- C- N- CA
1782 1781 1783 1784 3 0.65084 1.95253 0.00000
-2.60338 0.00000 0.00000 ; HA- CA- CB- HB2
1782 1781 1783 1785 3 0.65084 1.95253 0.00000
-2.60338 0.00000 0.00000 ; HA- CA- CB- HB3
1782 1781 1783 1786 3 0.65084 1.95253 0.00000
-2.60338 0.00000 0.00000 ; HA- CA- CB- CG
1782 1781 1799 1800 3 3.68192 -4.35136 0.00000
1.33888 0.00000 0.00000 ; HA- CA- C- O
1782 1781 1799 1801 3 0.00000 0.00000 0.00000
0.00000 0.00000 0.00000 ; HA- CA- C- N
1783 1781 1799 1800 3 0.00000 0.00000 0.00000
0.00000 0.00000 0.00000 ; CB- CA- C- O
1783 1781 1799 1801 3 2.51040 4.18400 1.67360
-6.69440 0.00000 0.00000 ; CB- CA- C- N
1783 1786 1789 1790 3 0.66944 2.00832 0.00000
-2.67776 0.00000 0.00000 ; CB- CG- CD- HD2
1783 1786 1789 1791 3 0.66944 2.00832 0.00000
-2.67776 0.00000 0.00000 ; CB- CG- CD- HD3
1783 1786 1789 1792 3 3.68192 3.09616 -2.09200
-3.01248 0.00000 0.00000 ; CB- CG- CD- CE
1784 1783 1781 1799 3 0.65084 1.95253 0.00000
-2.60338 0.00000 0.00000 ; HB2- CB- CA- C
1784 1783 1786 1787 3 0.62760 1.88280 0.00000
-2.51040 0.00000 0.00000 ; HB2- CB- CG- HG2

```
  1784   1783   1786   1788       3    0.62760    1.88280    0.00000
 -2.51040    0.00000    0.00000 ;    HB2-    CB-    CG-   HG3
  1784   1783   1786   1789       3    0.66944    2.00832    0.00000
 -2.67776    0.00000    0.00000 ;    HB2-    CB-    CG-    CD
  1785   1783   1781   1799       3    0.65084    1.95253    0.00000
 -2.60338    0.00000    0.00000 ;    HB3-    CB-    CA-     C
  1785   1783   1786   1787       3    0.62760    1.88280    0.00000
 -2.51040    0.00000    0.00000 ;    HB3-    CB-    CG-   HG2
  1785   1783   1786   1788       3    0.62760    1.88280    0.00000
 -2.51040    0.00000    0.00000 ;    HB3-    CB-    CG-   HG3
  1785   1783   1786   1789       3    0.66944    2.00832    0.00000
 -2.67776    0.00000    0.00000 ;    HB3-    CB-    CG-    CD
  1786   1783   1781   1799       3    0.65084    1.95253    0.00000
 -2.60338    0.00000    0.00000 ;     CG-    CB-    CA-     C
  1786   1789   1792   1793       3    0.65084    1.95253    0.00000
 -2.60338    0.00000    0.00000 ;     CG-    CD-    CE-   HE2
  1786   1789   1792   1794       3    0.65084    1.95253    0.00000
 -2.60338    0.00000    0.00000 ;     CG-    CD-    CE-   HE3
  1786   1789   1792   1795       3    0.65084    1.95253    0.00000
 -2.60338    0.00000    0.00000 ;     CG-    CD-    CE-    NZ
  1787   1786   1789   1790       3    0.62760    1.88280    0.00000
 -2.51040    0.00000    0.00000 ;    HG2-    CG-    CD-   HD2
  1787   1786   1789   1791       3    0.62760    1.88280    0.00000
 -2.51040    0.00000    0.00000 ;    HG2-    CG-    CD-   HD3
  1787   1786   1789   1792       3    0.66944    2.00832    0.00000
 -2.67776    0.00000    0.00000 ;    HG2-    CG-    CD-    CE
  1788   1786   1789   1790       3    0.62760    1.88280    0.00000
 -2.51040    0.00000    0.00000 ;    HG3-    CG-    CD-   HD2
  1788   1786   1789   1791       3    0.62760    1.88280    0.00000
 -2.51040    0.00000    0.00000 ;    HG3-    CG-    CD-   HD3
  1788   1786   1789   1792       3    0.66944    2.00832    0.00000
 -2.67776    0.00000    0.00000 ;    HG3-    CG-    CD-    CE
  1789   1792   1795   1796       3    0.65084    1.95253    0.00000
 -2.60338    0.00000    0.00000 ;     CD-    CE-    NZ-   HZ1
  1789   1792   1795   1797       3    0.65084    1.95253    0.00000
 -2.60338    0.00000    0.00000 ;     CD-    CE-    NZ-   HZ2
  1789   1792   1795   1798       3    0.65084    1.95253    0.00000
 -2.60338    0.00000    0.00000 ;     CD-    CE-    NZ-   HZ3
  1790   1789   1792   1793       3    0.65084    1.95253    0.00000
 -2.60338    0.00000    0.00000 ;    HD2-    CD-    CE-   HE2
  1790   1789   1792   1794       3    0.65084    1.95253    0.00000
 -2.60338    0.00000    0.00000 ;    HD2-    CD-    CE-   HE3
  1790   1789   1792   1795       3    0.65084    1.95253    0.00000
 -2.60338    0.00000    0.00000 ;    HD2-    CD-    CE-    NZ
  1791   1789   1792   1793       3    0.65084    1.95253    0.00000
 -2.60338    0.00000    0.00000 ;    HD3-    CD-    CE-   HE2
  1791   1789   1792   1794       3    0.65084    1.95253    0.00000
 -2.60338    0.00000    0.00000 ;    HD3-    CD-    CE-   HE3
  1791   1789   1792   1795       3    0.65084    1.95253    0.00000
 -2.60338    0.00000    0.00000 ;    HD3-    CD-    CE-    NZ
  1793   1792   1795   1796       3    0.65084    1.95253    0.00000
 -2.60338    0.00000    0.00000 ;    HE2-    CE-    NZ-   HZ1
```

1793 1792 1795 1797 3 0.65084 1.95253 0.00000
-2.60338 0.00000 0.00000 ; HE2- CE- NZ- HZ2
1793 1792 1795 1798 3 0.65084 1.95253 0.00000
-2.60338 0.00000 0.00000 ; HE2- CE- NZ- HZ3
1794 1792 1795 1796 3 0.65084 1.95253 0.00000
-2.60338 0.00000 0.00000 ; HE3- CE- NZ- HZ1
1794 1792 1795 1797 3 0.65084 1.95253 0.00000
-2.60338 0.00000 0.00000 ; HE3- CE- NZ- HZ2
1794 1792 1795 1798 3 0.65084 1.95253 0.00000
-2.60338 0.00000 0.00000 ; HE3- CE- NZ- HZ3
1799 1801 1803 1804 3 0.00000 0.00000 0.00000
0.00000 0.00000 0.00000 ; C- N- CA- HA
1799 1801 1803 1805 3 11.71520 1.67360 15.06240 -
13.38880 0.00000 0.00000 ; C- N- CA- CB
1799 1801 1803 1813 3 1.75728 5.27184 2.25936
-7.02912 0.00000 0.00000 ; C- N- CA- C
1800 1799 1801 1802 3 29.28800 -8.36800 -20.92000
0.00000 0.00000 0.00000 ; O- C- N- H
1800 1799 1801 1803 3 20.92000 0.00000 -20.92000
0.00000 0.00000 0.00000 ; O- C- N- CA
1801 1803 1805 1806 3 0.65084 1.95253 0.00000
-2.60338 0.00000 0.00000 ; N- CA- CB- HB2
1801 1803 1805 1807 3 0.65084 1.95253 0.00000
-2.60338 0.00000 0.00000 ; N- CA- CB- HB3
1801 1803 1805 1808 3 5.99567 3.29281 -4.46014
-0.55229 1.97485 0.00000 ; N- CA- CB- CG
1801 1803 1813 1814 3 0.00000 0.00000 0.00000
0.00000 0.00000 0.00000 ; N- CA- C- O
1801 1803 1813 1815 3 17.40544 -5.02080 -13.22144
9.20480 0.00000 0.00000 ; N- CA- C- N
1802 1801 1803 1804 3 0.00000 0.00000 0.00000
0.00000 0.00000 0.00000 ; H- N- CA- HA
1802 1801 1803 1805 3 0.00000 0.00000 0.00000
0.00000 0.00000 0.00000 ; H- N- CA- CB
1802 1801 1803 1813 3 0.00000 0.00000 0.00000
0.00000 0.00000 0.00000 ; H- N- CA- C
1803 1805 1808 1809 3 0.00000 0.00000 0.00000
0.00000 0.00000 0.00000 ; CA- CB- CG- OD1
1803 1805 1808 1810 3 8.84916 -0.31380 -4.32626
5.03754 0.26778 0.00000 ; CA- CB- CG- ND2
1803 1813 1815 1816 3 20.92000 0.00000 -20.92000
0.00000 0.00000 0.00000 ; CA- C- N- H
1803 1813 1815 1817 3 20.92000 0.00000 -20.92000
0.00000 0.00000 0.00000 ; CA- C- N- CA
1804 1803 1805 1806 3 0.65084 1.95253 0.00000
-2.60338 0.00000 0.00000 ; HA- CA- CB- HB2
1804 1803 1805 1807 3 0.65084 1.95253 0.00000
-2.60338 0.00000 0.00000 ; HA- CA- CB- HB3
1804 1803 1805 1808 3 0.65084 1.95253 0.00000
-2.60338 0.00000 0.00000 ; HA- CA- CB- CG
1804 1803 1813 1814 3 3.68192 -4.35136 0.00000
1.33888 0.00000 0.00000 ; HA- CA- C- O

1804 1803 1813 1815 3 0.00000 0.00000 0.00000
0.00000 0.00000 0.00000 ; HA- CA- C- N
1805 1803 1813 1814 3 0.00000 0.00000 0.00000
0.00000 0.00000 0.00000 ; CB- CA- C- O
1805 1803 1813 1815 3 2.51040 4.18400 1.67360
-6.69440 0.00000 0.00000 ; CB- CA- C- N
1805 1808 1810 1811 3 20.92000 0.00000 -20.92000
0.00000 0.00000 0.00000 ; CB- CG- ND2- HD21
1805 1808 1810 1812 3 20.92000 0.00000 -20.92000
0.00000 0.00000 0.00000 ; CB- CG- ND2- HD22
1806 1805 1803 1813 3 0.65084 1.95253 0.00000
-2.60338 0.00000 0.00000 ; HB2- CB- CA- C
1806 1805 1808 1809 3 3.68192 -4.35136 0.00000
1.33888 0.00000 0.00000 ; HB2- CB- CG- OD1
1806 1805 1808 1810 3 0.00000 0.00000 0.00000
0.00000 0.00000 0.00000 ; HB2- CB- CG- ND2
1807 1805 1803 1813 3 0.65084 1.95253 0.00000
-2.60338 0.00000 0.00000 ; HB3- CB- CA- C
1807 1805 1808 1809 3 3.68192 -4.35136 0.00000
1.33888 0.00000 0.00000 ; HB3- CB- CG- OD1
1807 1805 1808 1810 3 0.00000 0.00000 0.00000
0.00000 0.00000 0.00000 ; HB3- CB- CG- ND2
1808 1805 1803 1813 3 7.94542 -3.96225 -6.11701
-0.55229 3.58150 0.00000 ; CG- CB- CA- C
1809 1808 1810 1811 3 29.28800 -8.36800 -20.92000
0.00000 0.00000 0.00000 ; OD1- CG- ND2- HD21
1809 1808 1810 1812 3 29.28800 -8.36800 -20.92000
0.00000 0.00000 0.00000 ; OD1- CG- ND2- HD22
1813 1815 1817 1818 3 0.00000 0.00000 0.00000
0.00000 0.00000 0.00000 ; C- N- CA- HA
1813 1815 1817 1819 3 11.71520 1.67360 15.06240 -
13.38880 0.00000 0.00000 ; C- N- CA- CB
1813 1815 1817 1823 3 1.75728 5.27184 2.25936
-7.02912 0.00000 0.00000 ; C- N- CA- C
1814 1813 1815 1816 3 29.28800 -8.36800 -20.92000
0.00000 0.00000 0.00000 ; O- C- N- H
1814 1813 1815 1817 3 20.92000 0.00000 -20.92000
0.00000 0.00000 0.00000 ; O- C- N- CA
1815 1817 1819 1820 3 0.65084 1.95253 0.00000
-2.60338 0.00000 0.00000 ; N- CA- CB- HB1
1815 1817 1819 1821 3 0.65084 1.95253 0.00000
-2.60338 0.00000 0.00000 ; N- CA- CB- HB2
1815 1817 1819 1822 3 0.65084 1.95253 0.00000
-2.60338 0.00000 0.00000 ; N- CA- CB- HB3
1815 1817 1823 1824 3 0.00000 0.00000 0.00000
0.00000 0.00000 0.00000 ; N- CA- C- O
1815 1817 1823 1825 3 17.40544 -5.02080 -13.22144
9.20480 0.00000 0.00000 ; N- CA- C- N
1816 1815 1817 1818 3 0.00000 0.00000 0.00000
0.00000 0.00000 0.00000 ; H- N- CA- HA
1816 1815 1817 1819 3 0.00000 0.00000 0.00000
0.00000 0.00000 0.00000 ; H- N- CA- CB

1816 1815 1817 1823 3 0.00000 0.00000 0.00000
0.00000 0.00000 0.00000 ; H- N- CA- C
1817 1823 1825 1826 3 20.92000 0.00000 -20.92000
0.00000 0.00000 0.00000 ; CA- C- N- H
1817 1823 1825 1827 3 20.92000 0.00000 -20.92000
0.00000 0.00000 0.00000 ; CA- C- N- CA
1818 1817 1819 1820 3 0.65084 1.95253 0.00000
-2.60338 0.00000 0.00000 ; HA- CA- CB- HB1
1818 1817 1819 1821 3 0.65084 1.95253 0.00000
-2.60338 0.00000 0.00000 ; HA- CA- CB- HB2
1818 1817 1819 1822 3 0.65084 1.95253 0.00000
-2.60338 0.00000 0.00000 ; HA- CA- CB- HB3
1818 1817 1823 1824 3 3.68192 -4.35136 0.00000
1.33888 0.00000 0.00000 ; HA- CA- C- O
1818 1817 1823 1825 3 0.00000 0.00000 0.00000
0.00000 0.00000 0.00000 ; HA- CA- C- N
1819 1817 1823 1824 3 0.00000 0.00000 0.00000
0.00000 0.00000 0.00000 ; CB- CA- C- O
1819 1817 1823 1825 3 2.51040 4.18400 1.67360
-6.69440 0.00000 0.00000 ; CB- CA- C- N
1820 1819 1817 1823 3 0.65084 1.95253 0.00000
-2.60338 0.00000 0.00000 ; HB1- CB- CA- C
1821 1819 1817 1823 3 0.65084 1.95253 0.00000
-2.60338 0.00000 0.00000 ; HB2- CB- CA- C
1822 1819 1817 1823 3 0.65084 1.95253 0.00000
-2.60338 0.00000 0.00000 ; HB3- CB- CA- C
1823 1825 1827 1828 3 0.00000 0.00000 0.00000
0.00000 0.00000 0.00000 ; C- N- CA- HA
1823 1825 1827 1829 3 11.71520 1.67360 15.06240 -
13.38880 0.00000 0.00000 ; C- N- CA- CB
1823 1825 1827 1833 3 1.75728 5.27184 2.25936
-7.02912 0.00000 0.00000 ; C- N- CA- C
1824 1823 1825 1826 3 29.28800 -8.36800 -20.92000
0.00000 0.00000 0.00000 ; O- C- N- H
1824 1823 1825 1827 3 20.92000 0.00000 -20.92000
0.00000 0.00000 0.00000 ; O- C- N- CA
1825 1827 1829 1830 3 0.65084 1.95253 0.00000
-2.60338 0.00000 0.00000 ; N- CA- CB- HB1
1825 1827 1829 1831 3 0.65084 1.95253 0.00000
-2.60338 0.00000 0.00000 ; N- CA- CB- HB2
1825 1827 1829 1832 3 0.65084 1.95253 0.00000
-2.60338 0.00000 0.00000 ; N- CA- CB- HB3
1825 1827 1833 1834 3 0.00000 0.00000 0.00000
0.00000 0.00000 0.00000 ; N- CA- C- O
1825 1827 1833 1835 3 17.40544 -5.02080 -13.22144
9.20480 0.00000 0.00000 ; N- CA- C- N
1826 1825 1827 1828 3 0.00000 0.00000 0.00000
0.00000 0.00000 0.00000 ; H- N- CA- HA
1826 1825 1827 1829 3 0.00000 0.00000 0.00000
0.00000 0.00000 0.00000 ; H- N- CA- CB
1826 1825 1827 1833 3 0.00000 0.00000 0.00000
0.00000 0.00000 0.00000 ; H- N- CA- C

1827 1833 1835 1836 3 20.92000 0.00000 -20.92000
0.00000 0.00000 0.00000 ; CA- C- N- H
1827 1833 1835 1837 3 20.92000 0.00000 -20.92000
0.00000 0.00000 0.00000 ; CA- C- N- CA
1828 1827 1829 1830 3 0.65084 1.95253 0.00000
-2.60338 0.00000 0.00000 ; HA- CA- CB- HB1
1828 1827 1829 1831 3 0.65084 1.95253 0.00000
-2.60338 0.00000 0.00000 ; HA- CA- CB- HB2
1828 1827 1829 1832 3 0.65084 1.95253 0.00000
-2.60338 0.00000 0.00000 ; HA- CA- CB- HB3
1828 1827 1833 1834 3 3.68192 -4.35136 0.00000
1.33888 0.00000 0.00000 ; HA- CA- C- O
1828 1827 1833 1835 3 0.00000 0.00000 0.00000
0.00000 0.00000 0.00000 ; HA- CA- C- N
1829 1827 1833 1834 3 0.00000 0.00000 0.00000
0.00000 0.00000 0.00000 ; CB- CA- C- O
1829 1827 1833 1835 3 2.51040 4.18400 1.67360
-6.69440 0.00000 0.00000 ; CB- CA- C- N
1830 1829 1827 1833 3 0.65084 1.95253 0.00000
-2.60338 0.00000 0.00000 ; HB1- CB- CA- C
1831 1829 1827 1833 3 0.65084 1.95253 0.00000
-2.60338 0.00000 0.00000 ; HB2- CB- CA- C
1832 1829 1827 1833 3 0.65084 1.95253 0.00000
-2.60338 0.00000 0.00000 ; HB3- CB- CA- C
1833 1835 1837 1838 3 0.00000 0.00000 0.00000
0.00000 0.00000 0.00000 ; C- N- CA- HA
1833 1835 1837 1839 3 11.71520 1.67360 15.06240 -
13.38880 0.00000 0.00000 ; C- N- CA- CB
1833 1835 1837 1847 3 1.75728 5.27184 2.25936
-7.02912 0.00000 0.00000 ; C- N- CA- C
1834 1833 1835 1836 3 29.28800 -8.36800 -20.92000
0.00000 0.00000 0.00000 ; O- C- N- H
1834 1833 1835 1837 3 20.92000 0.00000 -20.92000
0.00000 0.00000 0.00000 ; O- C- N- CA
1835 1837 1839 1840 3 0.65084 1.95253 0.00000
-2.60338 0.00000 0.00000 ; N- CA- CB- HB
1835 1837 1839 1841 3 3.83673 0.44769 -1.77402
-2.47693 -0.03347 0.00000 ; N- CA- CB- CG2
1835 1837 1839 1845 3 4.93294 1.13386 -3.12963
-5.27184 3.17984 0.00000 ; N- CA- CB- OG1
1835 1837 1847 1848 3 0.00000 0.00000 0.00000
0.00000 0.00000 0.00000 ; N- CA- C- O
1835 1837 1847 1849 3 17.40544 -5.02080 -13.22144
9.20480 0.00000 0.00000 ; N- CA- C- N
1836 1835 1837 1838 3 0.00000 0.00000 0.00000
0.00000 0.00000 0.00000 ; H- N- CA- HA
1836 1835 1837 1839 3 0.00000 0.00000 0.00000
0.00000 0.00000 0.00000 ; H- N- CA- CB
1836 1835 1837 1847 3 0.00000 0.00000 0.00000
0.00000 0.00000 0.00000 ; H- N- CA- C
1837 1839 1841 1842 3 0.66944 2.00832 0.00000
-2.67776 0.00000 0.00000 ; CA- CB- CG2- HG21

1837 1839 1841 1843 3 0.66944 2.00832 0.00000
-2.67776 0.00000 0.00000 ; CA- CB- CG2- HG22
1837 1839 1841 1844 3 0.66944 2.00832 0.00000
-2.67776 0.00000 0.00000 ; CA- CB- CG2- HG23
1837 1839 1845 1846 3 1.12131 2.98738 1.66523
-3.94970 0.43514 0.00000 ; CA- CB- OG1- HG1
1837 1847 1849 1850 3 20.92000 0.00000 -20.92000
0.00000 0.00000 0.00000 ; CA- C- N- H
1837 1847 1849 1851 3 20.92000 0.00000 -20.92000
0.00000 0.00000 0.00000 ; CA- C- N- CA
1838 1837 1839 1840 3 0.65084 1.95253 0.00000
-2.60338 0.00000 0.00000 ; HA- CA- CB- HB
1838 1837 1839 1841 3 0.65084 1.95253 0.00000
-2.60338 0.00000 0.00000 ; HA- CA- CB- CG2
1838 1837 1839 1845 3 1.04600 -1.04600 0.00000
0.00000 0.00000 0.00000 ; HA- CA- CB- OG1
1838 1837 1847 1848 3 3.68192 -4.35136 0.00000
1.33888 0.00000 0.00000 ; HA- CA- C- O
1838 1837 1847 1849 3 0.00000 0.00000 0.00000
0.00000 0.00000 0.00000 ; HA- CA- C- N
1839 1837 1847 1848 3 0.00000 0.00000 0.00000
0.00000 0.00000 0.00000 ; CB- CA- C- O
1839 1837 1847 1849 3 2.51040 4.18400 1.67360
-6.69440 0.00000 0.00000 ; CB- CA- C- N
1840 1839 1837 1847 3 0.65084 1.95253 0.00000
-2.60338 0.00000 0.00000 ; HB- CB- CA- C
1840 1839 1841 1842 3 0.65084 1.95253 0.00000
-2.60338 0.00000 0.00000 ; HB- CB- CG2- HG21
1840 1839 1841 1843 3 0.65084 1.95253 0.00000
-2.60338 0.00000 0.00000 ; HB- CB- CG2- HG22
1840 1839 1841 1844 3 0.65084 1.95253 0.00000
-2.60338 0.00000 0.00000 ; HB- CB- CG2- HG23
1840 1839 1845 1846 3 0.69733 2.09200 0.00000
-2.78933 0.00000 0.00000 ; HB- CB- OG1- HG1
1841 1839 1837 1847 3 5.67350 3.55640 -6.16722
-2.47693 3.74886 0.00000 ; CG2- CB- CA- C
1841 1839 1845 1846 3 4.74047 0.27196 -2.26773
-3.94970 1.60666 0.00000 ; CG2- CB- OG1- HG1
1842 1841 1839 1845 3 1.04600 -1.04600 0.00000
0.00000 0.00000 0.00000 ; HG21- CG2- CB- OG1
1843 1841 1839 1845 3 1.04600 -1.04600 0.00000
0.00000 0.00000 0.00000 ; HG22- CG2- CB- OG1
1844 1841 1839 1845 3 1.04600 -1.04600 0.00000
0.00000 0.00000 0.00000 ; HG23- CG2- CB- OG1
1845 1839 1837 1847 3 6.53541 6.87013 -6.21742
-5.27184 5.22163 0.00000 ; OG1- CB- CA- C
1847 1849 1851 1852 3 0.00000 0.00000 0.00000
0.00000 0.00000 0.00000 ; C- N- CA- HA
1847 1849 1851 1853 3 11.71520 1.67360 15.06240 -
13.38880 0.00000 0.00000 ; C- N- CA- CB
1847 1849 1851 1866 3 1.75728 5.27184 2.25936
-7.02912 0.00000 0.00000 ; C- N- CA- C

1848 1847 1849 1850 3 29.28800 -8.36800 -20.92000
0.00000 0.00000 0.00000 ; O- C- N- H
1848 1847 1849 1851 3 20.92000 0.00000 -20.92000
0.00000 0.00000 0.00000 ; O- C- N- CA
1849 1851 1853 1854 3 0.65084 1.95253 0.00000
-2.60338 0.00000 0.00000 ; N- CA- CB- HB
1849 1851 1853 1855 3 3.83673 0.44769 -1.77402
-2.47693 -0.03347 0.00000 ; N- CA- CB- CG2
1849 1851 1853 1859 3 2.97482 0.12134 2.04179
-1.89117 -3.24678 0.00000 ; N- CA- CB- CG1
1849 1851 1866 1867 3 0.00000 0.00000 0.00000
0.00000 0.00000 0.00000 ; N- CA- C- O
1849 1851 1866 1868 3 17.40544 -5.02080 -13.22144
9.20480 0.00000 0.00000 ; N- CA- C- N
1850 1849 1851 1852 3 0.00000 0.00000 0.00000
0.00000 0.00000 0.00000 ; H- N- CA- HA
1850 1849 1851 1853 3 0.00000 0.00000 0.00000
0.00000 0.00000 0.00000 ; H- N- CA- CB
1850 1849 1851 1866 3 0.00000 0.00000 0.00000
0.00000 0.00000 0.00000 ; H- N- CA- C
1851 1853 1855 1856 3 0.66944 2.00832 0.00000
-2.67776 0.00000 0.00000 ; CA- CB- CG2- HG21
1851 1853 1855 1857 3 0.66944 2.00832 0.00000
-2.67776 0.00000 0.00000 ; CA- CB- CG2- HG22
1851 1853 1855 1858 3 0.66944 2.00832 0.00000
-2.67776 0.00000 0.00000 ; CA- CB- CG2- HG23
1851 1853 1859 1860 3 0.66944 2.00832 0.00000
-2.67776 0.00000 0.00000 ; CA- CB- CG1- HG12
1851 1853 1859 1861 3 0.66944 2.00832 0.00000
-2.67776 0.00000 0.00000 ; CA- CB- CG1- HG13
1851 1853 1859 1862 3 4.24258 -0.52718 -7.25506
-1.79075 7.69856 0.00000 ; CA- CB- CG1- CD1
1851 1866 1868 1869 3 20.92000 0.00000 -20.92000
0.00000 0.00000 0.00000 ; CA- C- N- H
1851 1866 1868 1870 3 20.92000 0.00000 -20.92000
0.00000 0.00000 0.00000 ; CA- C- N- CA
1852 1851 1853 1854 3 0.65084 1.95253 0.00000
-2.60338 0.00000 0.00000 ; HA- CA- CB- HB
1852 1851 1853 1855 3 0.65084 1.95253 0.00000
-2.60338 0.00000 0.00000 ; HA- CA- CB- CG2
1852 1851 1853 1859 3 0.65084 1.95253 0.00000
-2.60338 0.00000 0.00000 ; HA- CA- CB- CG1
1852 1851 1866 1867 3 3.68192 -4.35136 0.00000
1.33888 0.00000 0.00000 ; HA- CA- C- O
1852 1851 1866 1868 3 0.00000 0.00000 0.00000
0.00000 0.00000 0.00000 ; HA- CA- C- N
1853 1851 1866 1867 3 0.00000 0.00000 0.00000
0.00000 0.00000 0.00000 ; CB- CA- C- O
1853 1851 1866 1868 3 2.51040 4.18400 1.67360
-6.69440 0.00000 0.00000 ; CB- CA- C- N
1853 1859 1862 1863 3 0.66944 2.00832 0.00000
-2.67776 0.00000 0.00000 ; CB- CG1- CD1- HD11

1853 1859 1862 1864 3 0.66944 2.00832 0.00000
-2.67776 0.00000 0.00000 ; CB- CG1- CD1- HD12
1853 1859 1862 1865 3 0.66944 2.00832 0.00000
-2.67776 0.00000 0.00000 ; CB- CG1- CD1- HD13
1854 1853 1851 1866 3 0.65084 1.95253 0.00000
-2.60338 0.00000 0.00000 ; HB- CB- CA- C
1854 1853 1855 1856 3 0.62760 1.88280 0.00000
-2.51040 0.00000 0.00000 ; HB- CB- CG2- HG21
1854 1853 1855 1857 3 0.62760 1.88280 0.00000
-2.51040 0.00000 0.00000 ; HB- CB- CG2- HG22
1854 1853 1855 1858 3 0.62760 1.88280 0.00000
-2.51040 0.00000 0.00000 ; HB- CB- CG2- HG23
1854 1853 1859 1860 3 0.62760 1.88280 0.00000
-2.51040 0.00000 0.00000 ; HB- CB- CG1- HG12
1854 1853 1859 1861 3 0.62760 1.88280 0.00000
-2.51040 0.00000 0.00000 ; HB- CB- CG1- HG13
1854 1853 1859 1862 3 0.66944 2.00832 0.00000
-2.67776 0.00000 0.00000 ; HB- CB- CG1- CD1
1855 1853 1851 1866 3 5.67350 3.55640 -6.16722
-2.47693 3.74886 0.00000 ; CG2- CB- CA- C
1855 1853 1859 1860 3 0.66944 2.00832 0.00000
-2.67776 0.00000 0.00000 ; CG2- CB- CG1- HG12
1855 1853 1859 1861 3 0.66944 2.00832 0.00000
-2.67776 0.00000 0.00000 ; CG2- CB- CG1- HG13
1855 1853 1859 1862 3 3.81162 0.49790 -8.14206
-1.79075 7.49773 0.00000 ; CG2- CB- CG1- CD1
1856 1855 1853 1859 3 0.66944 2.00832 0.00000
-2.67776 0.00000 0.00000 ; HG21- CG2- CB- CG1
1857 1855 1853 1859 3 0.66944 2.00832 0.00000
-2.67776 0.00000 0.00000 ; HG22- CG2- CB- CG1
1858 1855 1853 1859 3 0.66944 2.00832 0.00000
-2.67776 0.00000 0.00000 ; HG23- CG2- CB- CG1
1859 1853 1851 1866 3 8.26340 0.74057 -9.99976
-1.89117 3.84928 0.00000 ; CG1- CB- CA- C
1860 1859 1862 1863 3 0.62760 1.88280 0.00000
-2.51040 0.00000 0.00000 ; HG12- CG1- CD1- HD11
1860 1859 1862 1864 3 0.62760 1.88280 0.00000
-2.51040 0.00000 0.00000 ; HG12- CG1- CD1- HD12
1860 1859 1862 1865 3 0.62760 1.88280 0.00000
-2.51040 0.00000 0.00000 ; HG12- CG1- CD1- HD13
1861 1859 1862 1863 3 0.62760 1.88280 0.00000
-2.51040 0.00000 0.00000 ; HG13- CG1- CD1- HD11
1861 1859 1862 1864 3 0.62760 1.88280 0.00000
-2.51040 0.00000 0.00000 ; HG13- CG1- CD1- HD12
1861 1859 1862 1865 3 0.62760 1.88280 0.00000
-2.51040 0.00000 0.00000 ; HG13- CG1- CD1- HD13
1866 1868 1870 1871 3 0.00000 0.00000 0.00000
0.00000 0.00000 0.00000 ; C- N- CA- HA
1866 1868 1870 1872 3 11.71520 1.67360 15.06240 -
13.38880 0.00000 0.00000 ; C- N- CA- CB
1866 1868 1870 1883 3 1.75728 5.27184 2.25936
-7.02912 0.00000 0.00000 ; C- N- CA- C

1867 1866 1868 1869 3 29.28800 -8.36800 -20.92000
0.00000 0.00000 0.00000 ; O- C- N- H
1867 1866 1868 1870 3 20.92000 0.00000 -20.92000
0.00000 0.00000 0.00000 ; O- C- N- CA
1868 1870 1872 1873 3 0.65084 1.95253 0.00000
-2.60338 0.00000 0.00000 ; N- CA- CB- HB2
1868 1870 1872 1874 3 0.65084 1.95253 0.00000
-2.60338 0.00000 0.00000 ; N- CA- CB- HB3
1868 1870 1872 1875 3 3.21331 2.22589 -4.15053
-2.40998 2.61082 0.00000 ; N- CA- CB- CG
1868 1870 1883 1884 3 0.00000 0.00000 0.00000
0.00000 0.00000 0.00000 ; N- CA- C- O
1868 1870 1883 1885 3 17.40544 -5.02080 -13.22144
9.20480 0.00000 0.00000 ; N- CA- C- N
1869 1868 1870 1871 3 0.00000 0.00000 0.00000
0.00000 0.00000 0.00000 ; H- N- CA- HA
1869 1868 1870 1872 3 0.00000 0.00000 0.00000
0.00000 0.00000 0.00000 ; H- N- CA- CB
1869 1868 1870 1883 3 0.00000 0.00000 0.00000
0.00000 0.00000 0.00000 ; H- N- CA- C
1870 1872 1875 1876 3 0.65084 1.95253 0.00000
-2.60338 0.00000 0.00000 ; CA- CB- CG- HG2
1870 1872 1875 1877 3 0.65084 1.95253 0.00000
-2.60338 0.00000 0.00000 ; CA- CB- CG- HG3
1870 1872 1875 1878 3 2.04598 -1.54390 1.11294
-0.26778 0.93722 0.00000 ; CA- CB- CG- SD
1870 1883 1885 1886 3 20.92000 0.00000 -20.92000
0.00000 0.00000 0.00000 ; CA- C- N- H
1870 1883 1885 1887 3 20.92000 0.00000 -20.92000
0.00000 0.00000 0.00000 ; CA- C- N- CA
1871 1870 1872 1873 3 0.65084 1.95253 0.00000
-2.60338 0.00000 0.00000 ; HA- CA- CB- HB2
1871 1870 1872 1874 3 0.65084 1.95253 0.00000
-2.60338 0.00000 0.00000 ; HA- CA- CB- HB3
1871 1870 1872 1875 3 0.65084 1.95253 0.00000
-2.60338 0.00000 0.00000 ; HA- CA- CB- CG
1871 1870 1883 1884 3 3.68192 -4.35136 0.00000
1.33888 0.00000 0.00000 ; HA- CA- C- O
1871 1870 1883 1885 3 0.00000 0.00000 0.00000
0.00000 0.00000 0.00000 ; HA- CA- C- N
1872 1870 1883 1884 3 0.00000 0.00000 0.00000
0.00000 0.00000 0.00000 ; CB- CA- C- O
1872 1870 1883 1885 3 2.51040 4.18400 1.67360
-6.69440 0.00000 0.00000 ; CB- CA- C- N
1872 1875 1878 1879 3 3.24260 6.22998 1.79075
-6.92870 1.90790 0.00000 ; CB- CG- SD- CE
1873 1872 1870 1883 3 0.65084 1.95253 0.00000
-2.60338 0.00000 0.00000 ; HB2- CB- CA- C
1873 1872 1875 1876 3 0.65084 1.95253 0.00000
-2.60338 0.00000 0.00000 ; HB2- CB- CG- HG2
1873 1872 1875 1877 3 0.65084 1.95253 0.00000
-2.60338 0.00000 0.00000 ; HB2- CB- CG- HG3

1873 1872 1875 1878 3 0.65084 1.95253 0.00000
-2.60338 0.00000 0.00000 ; HB2- CB- CG- SD
1874 1872 1870 1883 3 0.65084 1.95253 0.00000
-2.60338 0.00000 0.00000 ; HB3- CB- CA- C
1874 1872 1875 1876 3 0.65084 1.95253 0.00000
-2.60338 0.00000 0.00000 ; HB3- CB- CG- HG2
1874 1872 1875 1877 3 0.65084 1.95253 0.00000
-2.60338 0.00000 0.00000 ; HB3- CB- CG- HG3
1874 1872 1875 1878 3 0.65084 1.95253 0.00000
-2.60338 0.00000 0.00000 ; HB3- CB- CG- SD
1875 1872 1870 1883 3 6.86594 3.56895 -8.14206
-2.40998 4.85344 0.00000 ; CG- CB- CA- C
1875 1878 1879 1880 3 1.39467 4.18400 0.00000
-5.57867 0.00000 0.00000 ; CG- SD- CE- HE1
1875 1878 1879 1881 3 1.39467 4.18400 0.00000
-5.57867 0.00000 0.00000 ; CG- SD- CE- HE2
1875 1878 1879 1882 3 1.39467 4.18400 0.00000
-5.57867 0.00000 0.00000 ; CG- SD- CE- HE3
1876 1875 1878 1879 3 1.39467 4.18400 0.00000
-5.57867 0.00000 0.00000 ; HG2- CG- SD- CE
1877 1875 1878 1879 3 1.39467 4.18400 0.00000
-5.57867 0.00000 0.00000 ; HG3- CG- SD- CE
1883 1885 1887 1888 3 0.00000 0.00000 0.00000
0.00000 0.00000 0.00000 ; C- N- CA- HA
1883 1885 1887 1889 3 11.71520 1.67360 15.06240 -
13.38880 0.00000 0.00000 ; C- N- CA- CB
1883 1885 1887 1905 3 1.75728 5.27184 2.25936
-7.02912 0.00000 0.00000 ; C- N- CA- C
1884 1883 1885 1886 3 29.28800 -8.36800 -20.92000
0.00000 0.00000 0.00000 ; O- C- N- H
1884 1883 1885 1887 3 20.92000 0.00000 -20.92000
0.00000 0.00000 0.00000 ; O- C- N- CA
1885 1887 1889 1890 3 0.65084 1.95253 0.00000
-2.60338 0.00000 0.00000 ; N- CA- CB- HB2
1885 1887 1889 1891 3 0.65084 1.95253 0.00000
-2.60338 0.00000 0.00000 ; N- CA- CB- HB3
1885 1887 1889 1892 3 0.65084 1.95253 0.00000
-2.60338 0.00000 0.00000 ; N- CA- CB- CG
1885 1887 1905 1906 3 0.00000 0.00000 0.00000
0.00000 0.00000 0.00000 ; N- CA- C- O
1885 1887 1905 1907 3 17.40544 -5.02080 -13.22144
9.20480 0.00000 0.00000 ; N- CA- C- N
1886 1885 1887 1888 3 0.00000 0.00000 0.00000
0.00000 0.00000 0.00000 ; H- N- CA- HA
1886 1885 1887 1889 3 0.00000 0.00000 0.00000
0.00000 0.00000 0.00000 ; H- N- CA- CB
1886 1885 1887 1905 3 0.00000 0.00000 0.00000
0.00000 0.00000 0.00000 ; H- N- CA- C
1887 1889 1892 1893 3 0.66944 2.00832 0.00000
-2.67776 0.00000 0.00000 ; CA- CB- CG- HG2
1887 1889 1892 1894 3 0.66944 2.00832 0.00000
-2.67776 0.00000 0.00000 ; CA- CB- CG- HG3

```
  1887   1889   1892   1895      3    3.68192    3.09616   -2.09200
-3.01248    0.00000    0.00000 ;     CA-    CB-    CG-    CD
  1887   1905   1907   1908      3   20.92000    0.00000  -20.92000
0.00000    0.00000    0.00000 ;     CA-     C-     N-     H
  1887   1905   1907   1909      3   20.92000    0.00000  -20.92000
0.00000    0.00000    0.00000 ;     CA-     C-     N-    CA
  1888   1887   1889   1890      3    0.65084    1.95253    0.00000
-2.60338    0.00000    0.00000 ;     HA-    CA-    CB-   HB2
  1888   1887   1889   1891      3    0.65084    1.95253    0.00000
-2.60338    0.00000    0.00000 ;     HA-    CA-    CB-   HB3
  1888   1887   1889   1892      3    0.65084    1.95253    0.00000
-2.60338    0.00000    0.00000 ;     HA-    CA-    CB-    CG
  1888   1887   1905   1906      3    3.68192   -4.35136    0.00000
1.33888    0.00000    0.00000 ;     HA-    CA-     C-     O
  1888   1887   1905   1907      3    0.00000    0.00000    0.00000
0.00000    0.00000    0.00000 ;     HA-    CA-     C-     N
  1889   1887   1905   1906      3    0.00000    0.00000    0.00000
0.00000    0.00000    0.00000 ;     CB-    CA-     C-     O
  1889   1887   1905   1907      3    2.51040    4.18400    1.67360
-6.69440    0.00000    0.00000 ;     CB-    CA-     C-     N
  1889   1892   1895   1896      3    0.66944    2.00832    0.00000
-2.67776    0.00000    0.00000 ;     CB-    CG-    CD-   HD2
  1889   1892   1895   1897      3    0.66944    2.00832    0.00000
-2.67776    0.00000    0.00000 ;     CB-    CG-    CD-   HD3
  1889   1892   1895   1898      3    3.68192    3.09616   -2.09200
-3.01248    0.00000    0.00000 ;     CB-    CG-    CD-    CE
  1890   1889   1887   1905      3    0.65084    1.95253    0.00000
-2.60338    0.00000    0.00000 ;    HB2-    CB-    CA-     C
  1890   1889   1892   1893      3    0.62760    1.88280    0.00000
-2.51040    0.00000    0.00000 ;    HB2-    CB-    CG-   HG2
  1890   1889   1892   1894      3    0.62760    1.88280    0.00000
-2.51040    0.00000    0.00000 ;    HB2-    CB-    CG-   HG3
  1890   1889   1892   1895      3    0.66944    2.00832    0.00000
-2.67776    0.00000    0.00000 ;    HB2-    CB-    CG-    CD
  1891   1889   1887   1905      3    0.65084    1.95253    0.00000
-2.60338    0.00000    0.00000 ;    HB3-    CB-    CA-     C
  1891   1889   1892   1893      3    0.62760    1.88280    0.00000
-2.51040    0.00000    0.00000 ;    HB3-    CB-    CG-   HG2
  1891   1889   1892   1894      3    0.62760    1.88280    0.00000
-2.51040    0.00000    0.00000 ;    HB3-    CB-    CG-   HG3
  1891   1889   1892   1895      3    0.66944    2.00832    0.00000
-2.67776    0.00000    0.00000 ;    HB3-    CB-    CG-    CD
  1892   1889   1887   1905      3    0.65084    1.95253    0.00000
-2.60338    0.00000    0.00000 ;     CG-    CB-    CA-     C
  1892   1895   1898   1899      3    0.65084    1.95253    0.00000
-2.60338    0.00000    0.00000 ;     CG-    CD-    CE-   HE2
  1892   1895   1898   1900      3    0.65084    1.95253    0.00000
-2.60338    0.00000    0.00000 ;     CG-    CD-    CE-   HE3
  1892   1895   1898   1901      3    0.65084    1.95253    0.00000
-2.60338    0.00000    0.00000 ;     CG-    CD-    CE-    NZ
  1893   1892   1895   1896      3    0.62760    1.88280    0.00000
-2.51040    0.00000    0.00000 ;    HG2-    CG-    CD-   HD2
```

```
  1893   1892   1895   1897      3    0.62760    1.88280    0.00000
-2.51040    0.00000    0.00000 ;    HG2-   CG-    CD-   HD3
  1893   1892   1895   1898      3    0.66944    2.00832    0.00000
-2.67776    0.00000    0.00000 ;    HG2-   CG-    CD-    CE
  1894   1892   1895   1896      3    0.62760    1.88280    0.00000
-2.51040    0.00000    0.00000 ;    HG3-   CG-    CD-   HD2
  1894   1892   1895   1897      3    0.62760    1.88280    0.00000
-2.51040    0.00000    0.00000 ;    HG3-   CG-    CD-   HD3
  1894   1892   1895   1898      3    0.66944    2.00832    0.00000
-2.67776    0.00000    0.00000 ;    HG3-   CG-    CD-    CE
  1895   1898   1901   1902      3    0.65084    1.95253    0.00000
-2.60338    0.00000    0.00000 ;     CD-   CE-    NZ-   HZ1
  1895   1898   1901   1903      3    0.65084    1.95253    0.00000
-2.60338    0.00000    0.00000 ;     CD-   CE-    NZ-   HZ2
  1895   1898   1901   1904      3    0.65084    1.95253    0.00000
-2.60338    0.00000    0.00000 ;     CD-   CE-    NZ-   HZ3
  1896   1895   1898   1899      3    0.65084    1.95253    0.00000
-2.60338    0.00000    0.00000 ;    HD2-   CD-    CE-   HE2
  1896   1895   1898   1900      3    0.65084    1.95253    0.00000
-2.60338    0.00000    0.00000 ;    HD2-   CD-    CE-   HE3
  1896   1895   1898   1901      3    0.65084    1.95253    0.00000
-2.60338    0.00000    0.00000 ;    HD2-   CD-    CE-    NZ
  1897   1895   1898   1899      3    0.65084    1.95253    0.00000
-2.60338    0.00000    0.00000 ;    HD3-   CD-    CE-   HE2
  1897   1895   1898   1900      3    0.65084    1.95253    0.00000
-2.60338    0.00000    0.00000 ;    HD3-   CD-    CE-   HE3
  1897   1895   1898   1901      3    0.65084    1.95253    0.00000
-2.60338    0.00000    0.00000 ;    HD3-   CD-    CE-    NZ
  1899   1898   1901   1902      3    0.65084    1.95253    0.00000
-2.60338    0.00000    0.00000 ;    HE2-   CE-    NZ-   HZ1
  1899   1898   1901   1903      3    0.65084    1.95253    0.00000
-2.60338    0.00000    0.00000 ;    HE2-   CE-    NZ-   HZ2
  1899   1898   1901   1904      3    0.65084    1.95253    0.00000
-2.60338    0.00000    0.00000 ;    HE2-   CE-    NZ-   HZ3
  1900   1898   1901   1902      3    0.65084    1.95253    0.00000
-2.60338    0.00000    0.00000 ;    HE3-   CE-    NZ-   HZ1
  1900   1898   1901   1903      3    0.65084    1.95253    0.00000
-2.60338    0.00000    0.00000 ;    HE3-   CE-    NZ-   HZ2
  1900   1898   1901   1904      3    0.65084    1.95253    0.00000
-2.60338    0.00000    0.00000 ;    HE3-   CE-    NZ-   HZ3
  1905   1907   1909   1910      3    0.00000    0.00000    0.00000
0.00000    0.00000    0.00000 ;     C-     N-    CA-    HA
  1905   1907   1909   1911      3   11.71520    1.67360   15.06240  -
13.38880    0.00000    0.00000 ;     C-     N-    CA-    CB
  1905   1907   1909   1922      3    1.75728    5.27184    2.25936
-7.02912    0.00000    0.00000 ;     C-     N-    CA-     C
  1906   1905   1907   1908      3   29.28800   -8.36800  -20.92000
0.00000    0.00000    0.00000 ;     O-     C-     N-     H
  1906   1905   1907   1909      3   20.92000    0.00000  -20.92000
0.00000    0.00000    0.00000 ;     O-     C-     N-    CA
  1907   1909   1911   1912      3    0.65084    1.95253    0.00000
-2.60338    0.00000    0.00000 ;      N-   CA-    CB-   HB2
```

```
  1907   1909   1911   1913      3    0.65084    1.95253    0.00000
-2.60338    0.00000    0.00000 ;     N-    CA-    CB-   HB3
  1907   1909   1911   1914      3    3.21331    2.22589   -4.15053
-2.40998    2.61082    0.00000 ;     N-    CA-    CB-    CG
  1907   1909   1922   1923      3    0.00000    0.00000    0.00000
0.00000    0.00000    0.00000 ;     N-    CA-     C-     O
  1907   1909   1922   1924      3   17.40544   -5.02080  -13.22144
9.20480    0.00000    0.00000 ;     N-    CA-     C-     N
  1908   1907   1909   1910      3    0.00000    0.00000    0.00000
0.00000    0.00000    0.00000 ;     H-     N-    CA-    HA
  1908   1907   1909   1911      3    0.00000    0.00000    0.00000
0.00000    0.00000    0.00000 ;     H-     N-    CA-    CB
  1908   1907   1909   1922      3    0.00000    0.00000    0.00000
0.00000    0.00000    0.00000 ;     H-     N-    CA-     C
  1909   1911   1914   1915      3    0.66944    2.00832    0.00000
-2.67776    0.00000    0.00000 ;    CA-    CB-    CG-   HG2
  1909   1911   1914   1916      3    0.66944    2.00832    0.00000
-2.67776    0.00000    0.00000 ;    CA-    CB-    CG-   HG3
  1909   1911   1914   1917      3    3.69866   -4.35136   -3.92459
6.89523    4.61914    0.00000 ;    CA-    CB-    CG-    CD
  1909   1922   1924   1925      3   20.92000    0.00000  -20.92000
0.00000    0.00000    0.00000 ;    CA-     C-     N-     H
  1909   1922   1924   1926      3   20.92000    0.00000  -20.92000
0.00000    0.00000    0.00000 ;    CA-     C-     N-    CA
  1910   1909   1911   1912      3    0.65084    1.95253    0.00000
-2.60338    0.00000    0.00000 ;    HA-    CA-    CB-   HB2
  1910   1909   1911   1913      3    0.65084    1.95253    0.00000
-2.60338    0.00000    0.00000 ;    HA-    CA-    CB-   HB3
  1910   1909   1911   1914      3    0.65084    1.95253    0.00000
-2.60338    0.00000    0.00000 ;    HA-    CA-    CB-    CG
  1910   1909   1922   1923      3    3.68192   -4.35136    0.00000
1.33888    0.00000    0.00000 ;    HA-    CA-     C-     O
  1910   1909   1922   1924      3    0.00000    0.00000    0.00000
0.00000    0.00000    0.00000 ;    HA-    CA-     C-     N
  1911   1909   1922   1923      3    0.00000    0.00000    0.00000
0.00000    0.00000    0.00000 ;    CB-    CA-     C-     O
  1911   1909   1922   1924      3    2.51040    4.18400    1.67360
-6.69440    0.00000    0.00000 ;    CB-    CA-     C-     N
  1911   1914   1917   1918      3    0.00000    0.00000    0.00000
0.00000    0.00000    0.00000 ;    CB-    CG-    CD-   OE1
  1911   1914   1917   1919      3   10.32611    1.48114   -8.47678
1.42256    1.40582    0.00000 ;    CB-    CG-    CD-   NE2
  1912   1911   1909   1922      3    0.65084    1.95253    0.00000
-2.60338    0.00000    0.00000 ;   HB2-    CB-    CA-     C
  1912   1911   1914   1915      3    0.62760    1.88280    0.00000
-2.51040    0.00000    0.00000 ;   HB2-    CB-    CG-   HG2
  1912   1911   1914   1916      3    0.62760    1.88280    0.00000
-2.51040    0.00000    0.00000 ;   HB2-    CB-    CG-   HG3
  1912   1911   1914   1917      3    0.65084    1.95253    0.00000
-2.60338    0.00000    0.00000 ;   HB2-    CB-    CG-    CD
  1913   1911   1909   1922      3    0.65084    1.95253    0.00000
-2.60338    0.00000    0.00000 ;   HB3-    CB-    CA-     C
```

1913 1911 1914 1915 3 0.62760 1.88280 0.00000
-2.51040 0.00000 0.00000 ; HB3- CB- CG- HG2
1913 1911 1914 1916 3 0.62760 1.88280 0.00000
-2.51040 0.00000 0.00000 ; HB3- CB- CG- HG3
1913 1911 1914 1917 3 0.65084 1.95253 0.00000
-2.60338 0.00000 0.00000 ; HB3- CB- CG- CD
1914 1911 1909 1922 3 6.86594 3.56895 -8.14206
-2.40998 4.85344 0.00000 ; CG- CB- CA- C
1914 1917 1919 1920 3 20.92000 0.00000 -20.92000
0.00000 0.00000 0.00000 ; CG- CD- NE2- HE21
1914 1917 1919 1921 3 20.92000 0.00000 -20.92000
0.00000 0.00000 0.00000 ; CG- CD- NE2- HE22
1915 1914 1917 1918 3 3.68192 -4.35136 0.00000
1.33888 0.00000 0.00000 ; HG2- CG- CD- OE1
1915 1914 1917 1919 3 0.00000 0.00000 0.00000
0.00000 0.00000 0.00000 ; HG2- CG- CD- NE2
1916 1914 1917 1918 3 3.68192 -4.35136 0.00000
1.33888 0.00000 0.00000 ; HG3- CG- CD- OE1
1916 1914 1917 1919 3 0.00000 0.00000 0.00000
0.00000 0.00000 0.00000 ; HG3- CG- CD- NE2
1918 1917 1919 1920 3 29.28800 -8.36800 -20.92000
0.00000 0.00000 0.00000 ; OE1- CD- NE2- HE21
1918 1917 1919 1921 3 29.28800 -8.36800 -20.92000
0.00000 0.00000 0.00000 ; OE1- CD- NE2- HE22
1922 1924 1926 1927 3 0.00000 0.00000 0.00000
0.00000 0.00000 0.00000 ; C- N- CA- HA
1922 1924 1926 1928 3 11.71520 1.67360 15.06240 -
13.38880 0.00000 0.00000 ; C- N- CA- CB
1922 1924 1926 1941 3 1.75728 5.27184 2.25936
-7.02912 0.00000 0.00000 ; C- N- CA- C
1923 1922 1924 1925 3 29.28800 -8.36800 -20.92000
0.00000 0.00000 0.00000 ; O- C- N- H
1923 1922 1924 1926 3 20.92000 0.00000 -20.92000
0.00000 0.00000 0.00000 ; O- C- N- CA
1924 1926 1928 1929 3 0.65084 1.95253 0.00000
-2.60338 0.00000 0.00000 ; N- CA- CB- HB
1924 1926 1928 1930 3 3.83673 0.44769 -1.77402
-2.47693 -0.03347 0.00000 ; N- CA- CB- CG2
1924 1926 1928 1934 3 2.97482 0.12134 2.04179
-1.89117 -3.24678 0.00000 ; N- CA- CB- CG1
1924 1926 1941 1942 3 0.00000 0.00000 0.00000
0.00000 0.00000 0.00000 ; N- CA- C- O
1924 1926 1941 1943 3 17.40544 -5.02080 -13.22144
9.20480 0.00000 0.00000 ; N- CA- C- N
1925 1924 1926 1927 3 0.00000 0.00000 0.00000
0.00000 0.00000 0.00000 ; H- N- CA- HA
1925 1924 1926 1928 3 0.00000 0.00000 0.00000
0.00000 0.00000 0.00000 ; H- N- CA- CB
1925 1924 1926 1941 3 0.00000 0.00000 0.00000
0.00000 0.00000 0.00000 ; H- N- CA- C
1926 1928 1930 1931 3 0.66944 2.00832 0.00000
-2.67776 0.00000 0.00000 ; CA- CB- CG2- HG21

1926 1928 1930 1932 3 0.66944 2.00832 0.00000
-2.67776 0.00000 0.00000 ; CA- CB- CG2- HG22
1926 1928 1930 1933 3 0.66944 2.00832 0.00000
-2.67776 0.00000 0.00000 ; CA- CB- CG2- HG23
1926 1928 1934 1935 3 0.66944 2.00832 0.00000
-2.67776 0.00000 0.00000 ; CA- CB- CG1- HG12
1926 1928 1934 1936 3 0.66944 2.00832 0.00000
-2.67776 0.00000 0.00000 ; CA- CB- CG1- HG13
1926 1928 1934 1937 3 4.24258 -0.52718 -7.25506
-1.79075 7.69856 0.00000 ; CA- CB- CG1- CD1
1926 1941 1943 1944 3 20.92000 0.00000 -20.92000
0.00000 0.00000 0.00000 ; CA- C- N- H
1926 1941 1943 1945 3 20.92000 0.00000 -20.92000
0.00000 0.00000 0.00000 ; CA- C- N- CA
1927 1926 1928 1929 3 0.65084 1.95253 0.00000
-2.60338 0.00000 0.00000 ; HA- CA- CB- HB
1927 1926 1928 1930 3 0.65084 1.95253 0.00000
-2.60338 0.00000 0.00000 ; HA- CA- CB- CG2
1927 1926 1928 1934 3 0.65084 1.95253 0.00000
-2.60338 0.00000 0.00000 ; HA- CA- CB- CG1
1927 1926 1941 1942 3 3.68192 -4.35136 0.00000
1.33888 0.00000 0.00000 ; HA- CA- C- O
1927 1926 1941 1943 3 0.00000 0.00000 0.00000
0.00000 0.00000 0.00000 ; HA- CA- C- N
1928 1926 1941 1942 3 0.00000 0.00000 0.00000
0.00000 0.00000 0.00000 ; CB- CA- C- O
1928 1926 1941 1943 3 2.51040 4.18400 1.67360
-6.69440 0.00000 0.00000 ; CB- CA- C- N
1928 1934 1937 1938 3 0.66944 2.00832 0.00000
-2.67776 0.00000 0.00000 ; CB- CG1- CD1- HD11
1928 1934 1937 1939 3 0.66944 2.00832 0.00000
-2.67776 0.00000 0.00000 ; CB- CG1- CD1- HD12
1928 1934 1937 1940 3 0.66944 2.00832 0.00000
-2.67776 0.00000 0.00000 ; CB- CG1- CD1- HD13
1929 1928 1926 1941 3 0.65084 1.95253 0.00000
-2.60338 0.00000 0.00000 ; HB- CB- CA- C
1929 1928 1930 1931 3 0.62760 1.88280 0.00000
-2.51040 0.00000 0.00000 ; HB- CB- CG2- HG21
1929 1928 1930 1932 3 0.62760 1.88280 0.00000
-2.51040 0.00000 0.00000 ; HB- CB- CG2- HG22
1929 1928 1930 1933 3 0.62760 1.88280 0.00000
-2.51040 0.00000 0.00000 ; HB- CB- CG2- HG23
1929 1928 1934 1935 3 0.62760 1.88280 0.00000
-2.51040 0.00000 0.00000 ; HB- CB- CG1- HG12
1929 1928 1934 1936 3 0.62760 1.88280 0.00000
-2.51040 0.00000 0.00000 ; HB- CB- CG1- HG13
1929 1928 1934 1937 3 0.66944 2.00832 0.00000
-2.67776 0.00000 0.00000 ; HB- CB- CG1- CD1
1930 1928 1926 1941 3 5.67350 3.55640 -6.16722
-2.47693 3.74886 0.00000 ; CG2- CB- CA- C
1930 1928 1934 1935 3 0.66944 2.00832 0.00000
-2.67776 0.00000 0.00000 ; CG2- CB- CG1- HG12

1930 1928 1934 1936 3 0.66944 2.00832 0.00000
-2.67776 0.00000 0.00000 ; CG2- CB- CG1- HG13
1930 1928 1934 1937 3 3.81162 0.49790 -8.14206
-1.79075 7.49773 0.00000 ; CG2- CB- CG1- CD1
1931 1930 1928 1934 3 0.66944 2.00832 0.00000
-2.67776 0.00000 0.00000 ; HG21- CG2- CB- CG1
1932 1930 1928 1934 3 0.66944 2.00832 0.00000
-2.67776 0.00000 0.00000 ; HG22- CG2- CB- CG1
1933 1930 1928 1934 3 0.66944 2.00832 0.00000
-2.67776 0.00000 0.00000 ; HG23- CG2- CB- CG1
1934 1928 1926 1941 3 8.26340 0.74057 -9.99976
-1.89117 3.84928 0.00000 ; CG1- CB- CA- C
1935 1934 1937 1938 3 0.62760 1.88280 0.00000
-2.51040 0.00000 0.00000 ; HG12- CG1- CD1- HD11
1935 1934 1937 1939 3 0.62760 1.88280 0.00000
-2.51040 0.00000 0.00000 ; HG12- CG1- CD1- HD12
1935 1934 1937 1940 3 0.62760 1.88280 0.00000
-2.51040 0.00000 0.00000 ; HG12- CG1- CD1- HD13
1936 1934 1937 1938 3 0.62760 1.88280 0.00000
-2.51040 0.00000 0.00000 ; HG13- CG1- CD1- HD11
1936 1934 1937 1939 3 0.62760 1.88280 0.00000
-2.51040 0.00000 0.00000 ; HG13- CG1- CD1- HD12
1936 1934 1937 1940 3 0.62760 1.88280 0.00000
-2.51040 0.00000 0.00000 ; HG13- CG1- CD1- HD13
1941 1943 1945 1946 3 0.00000 0.00000 0.00000
0.00000 0.00000 0.00000 ; C- N- CA- HA
1941 1943 1945 1947 3 11.71520 1.67360 15.06240 -
13.38880 0.00000 0.00000 ; C- N- CA- CB
1941 1943 1945 1961 3 1.75728 5.27184 2.25936
-7.02912 0.00000 0.00000 ; C- N- CA- C
1942 1941 1943 1944 3 29.28800 -8.36800 -20.92000
0.00000 0.00000 0.00000 ; O- C- N- H
1942 1941 1943 1945 3 20.92000 0.00000 -20.92000
0.00000 0.00000 0.00000 ; O- C- N- CA
1943 1945 1947 1948 3 0.65084 1.95253 0.00000
-2.60338 0.00000 0.00000 ; N- CA- CB- HB2
1943 1945 1947 1949 3 0.65084 1.95253 0.00000
-2.60338 0.00000 0.00000 ; N- CA- CB- HB3
1943 1945 1947 1950 3 3.28026 2.46019 -2.19242
-3.21331 -0.23430 0.00000 ; N- CA- CB- CG
1943 1945 1961 1962 3 0.00000 0.00000 0.00000
0.00000 0.00000 0.00000 ; N- CA- C- O
1943 1945 1961 1963 3 17.40544 -5.02080 -13.22144
9.20480 0.00000 0.00000 ; N- CA- C- N
1944 1943 1945 1946 3 0.00000 0.00000 0.00000
0.00000 0.00000 0.00000 ; H- N- CA- HA
1944 1943 1945 1947 3 0.00000 0.00000 0.00000
0.00000 0.00000 0.00000 ; H- N- CA- CB
1944 1943 1945 1961 3 0.00000 0.00000 0.00000
0.00000 0.00000 0.00000 ; H- N- CA- C
1945 1947 1950 1951 3 0.57739 0.00000 1.02926
0.00000 -1.60666 0.00000 ; CA- CB- CG- CD1

```
  1945   1947   1950   1959      3    0.57739    0.00000    1.02926
0.00000   -1.60666   0.00000 ;     CA-    CB-    CG-   CD2
  1945   1961   1963   1964      3   20.92000    0.00000  -20.92000
0.00000    0.00000   0.00000 ;     CA-     C-     N-     H
  1945   1961   1963   1965      3   20.92000    0.00000  -20.92000
0.00000    0.00000   0.00000 ;     CA-     C-     N-    CA
  1946   1945   1947   1948      3    0.65084    1.95253    0.00000
-2.60338    0.00000    0.00000 ;     HA-    CA-    CB-   HB2
  1946   1945   1947   1949      3    0.65084    1.95253    0.00000
-2.60338    0.00000    0.00000 ;     HA-    CA-    CB-   HB3
  1946   1945   1947   1950      3    0.65084    1.95253    0.00000
-2.60338    0.00000    0.00000 ;     HA-    CA-    CB-    CG
  1946   1945   1961   1962      3    3.68192   -4.35136    0.00000
1.33888    0.00000   0.00000 ;     HA-    CA-     C-     O
  1946   1945   1961   1963      3    0.00000    0.00000    0.00000
0.00000    0.00000   0.00000 ;     HA-    CA-     C-     N
  1947   1945   1961   1962      3    0.00000    0.00000    0.00000
0.00000    0.00000   0.00000 ;     CB-    CA-     C-     O
  1947   1945   1961   1963      3    2.51040    4.18400    1.67360
-6.69440    0.00000    0.00000 ;     CB-    CA-     C-     N
  1947   1950   1951   1952      3   30.33400    0.00000  -30.33400
0.00000    0.00000   0.00000 ;     CB-    CG-   CD1-   HD1
  1947   1950   1951   1953      3   30.33400    0.00000  -30.33400
0.00000    0.00000   0.00000 ;     CB-    CG-   CD1-   CE1
  1947   1950   1959   1957      3   30.33400    0.00000  -30.33400
0.00000    0.00000   0.00000 ;     CB-    CG-   CD2-   CE2
  1947   1950   1959   1960      3   30.33400    0.00000  -30.33400
0.00000    0.00000   0.00000 ;     CB-    CG-   CD2-   HD2
  1948   1947   1945   1961      3    0.65084    1.95253    0.00000
-2.60338    0.00000    0.00000 ;    HB2-    CB-    CA-     C
  1948   1947   1950   1951      3    0.00000    0.00000    0.00000
0.00000    0.00000   0.00000 ;    HB2-    CB-    CG-   CD1
  1948   1947   1950   1959      3    0.00000    0.00000    0.00000
0.00000    0.00000   0.00000 ;    HB2-    CB-    CG-   CD2
  1949   1947   1945   1961      3    0.65084    1.95253    0.00000
-2.60338    0.00000    0.00000 ;    HB3-    CB-    CA-     C
  1949   1947   1950   1951      3    0.00000    0.00000    0.00000
0.00000    0.00000   0.00000 ;    HB3-    CB-    CG-   CD1
  1949   1947   1950   1959      3    0.00000    0.00000    0.00000
0.00000    0.00000   0.00000 ;    HB3-    CB-    CG-   CD2
  1950   1947   1945   1961      3    4.95804    2.17986   -3.52293
-3.21331   -0.40166    0.00000 ;     CG-    CB-    CA-     C
  1950   1951   1953   1954      3   30.33400    0.00000  -30.33400
0.00000    0.00000   0.00000 ;     CG-   CD1-   CE1-   HE1
  1950   1951   1953   1955      3   30.33400    0.00000  -30.33400
0.00000    0.00000   0.00000 ;     CG-   CD1-   CE1-    CZ
  1950   1959   1957   1955      3   30.33400    0.00000  -30.33400
0.00000    0.00000   0.00000 ;     CG-   CD2-   CE2-    CZ
  1950   1959   1957   1958      3   30.33400    0.00000  -30.33400
0.00000    0.00000   0.00000 ;     CG-   CD2-   CE2-   HE2
  1951   1950   1959   1957      3   30.33400    0.00000  -30.33400
0.00000    0.00000   0.00000 ;    CD1-    CG-   CD2-   CE2
```

```
  1951   1950   1959   1960      3   30.33400    0.00000  -30.33400
0.00000    0.00000    0.00000 ;    CD1-   CG-   CD2-   HD2
  1951   1953   1955   1956      3   30.33400    0.00000  -30.33400
0.00000    0.00000    0.00000 ;    CD1-   CE1-    CZ-    HZ
  1951   1953   1955   1957      3   30.33400    0.00000  -30.33400
0.00000    0.00000    0.00000 ;    CD1-   CE1-    CZ-   CE2
  1952   1951   1950   1959      3   30.33400    0.00000  -30.33400
0.00000    0.00000    0.00000 ;    HD1-   CD1-    CG-   CD2
  1952   1951   1953   1954      3   30.33400    0.00000  -30.33400
0.00000    0.00000    0.00000 ;    HD1-   CD1-   CE1-   HE1
  1952   1951   1953   1955      3   30.33400    0.00000  -30.33400
0.00000    0.00000    0.00000 ;    HD1-   CD1-   CE1-    CZ
  1953   1951   1950   1959      3   30.33400    0.00000  -30.33400
0.00000    0.00000    0.00000 ;    CE1-   CD1-    CG-   CD2
  1953   1955   1957   1958      3   30.33400    0.00000  -30.33400
0.00000    0.00000    0.00000 ;    CE1-    CZ-   CE2-   HE2
  1953   1955   1957   1959      3   30.33400    0.00000  -30.33400
0.00000    0.00000    0.00000 ;    CE1-    CZ-   CE2-   CD2
  1954   1953   1955   1956      3   30.33400    0.00000  -30.33400
0.00000    0.00000    0.00000 ;    HE1-   CE1-    CZ-    HZ
  1954   1953   1955   1957      3   30.33400    0.00000  -30.33400
0.00000    0.00000    0.00000 ;    HE1-   CE1-    CZ-   CE2
  1955   1957   1959   1960      3   30.33400    0.00000  -30.33400
0.00000    0.00000    0.00000 ;     CZ-   CE2-   CD2-   HD2
  1956   1955   1957   1958      3   30.33400    0.00000  -30.33400
0.00000    0.00000    0.00000 ;     HZ-    CZ-   CE2-   HE2
  1956   1955   1957   1959      3   30.33400    0.00000  -30.33400
0.00000    0.00000    0.00000 ;     HZ-    CZ-   CE2-   CD2
  1958   1957   1959   1960      3   30.33400    0.00000  -30.33400
0.00000    0.00000    0.00000 ;    HE2-   CE2-   CD2-   HD2
  1961   1963   1965   1966      3    0.00000    0.00000    0.00000
0.00000    0.00000    0.00000 ;      C-     N-    CA-    HA
  1961   1963   1965   1967      3   11.71520    1.67360   15.06240  -
13.38880    0.00000    0.00000 ;      C-     N-    CA-    CB
  1961   1963   1965   1972      3    1.75728    5.27184    2.25936
-7.02912    0.00000    0.00000 ;      C-     N-    CA-     C
  1962   1961   1963   1964      3   29.28800   -8.36800  -20.92000
0.00000    0.00000    0.00000 ;      O-     C-     N-     H
  1962   1961   1963   1965      3   20.92000    0.00000  -20.92000
0.00000    0.00000    0.00000 ;      O-     C-     N-    CA
  1963   1965   1967   1968      3    0.65084    1.95253    0.00000
-2.60338    0.00000    0.00000 ;      N-    CA-    CB-   HB2
  1963   1965   1967   1969      3    0.65084    1.95253    0.00000
-2.60338    0.00000    0.00000 ;      N-    CA-    CB-   HB3
  1963   1965   1967   1970      3    7.86174    2.24681   -7.41405
-6.71114    5.35552    0.00000 ;      N-    CA-    CB-    OG
  1963   1965   1972   1973      3    0.00000    0.00000    0.00000
0.00000    0.00000    0.00000 ;      N-    CA-     C-     O
  1963   1965   1972   1974      3   17.40544   -5.02080  -13.22144
9.20480    0.00000    0.00000 ;      N-    CA-     C-     N
  1964   1963   1965   1966      3    0.00000    0.00000    0.00000
0.00000    0.00000    0.00000 ;      H-     N-    CA-    HA
```

1964 1963 1965 1967 3 0.00000 0.00000 0.00000
0.00000 0.00000 0.00000 ; H- N- CA- CB
1964 1963 1965 1972 3 0.00000 0.00000 0.00000
0.00000 0.00000 0.00000 ; H- N- CA- C
1965 1967 1970 1971 3 2.05853 2.46856 3.48109
-4.46851 0.23430 0.00000 ; CA- CB- OG- HG
1965 1972 1974 1975 3 20.92000 0.00000 -20.92000
0.00000 0.00000 0.00000 ; CA- C- N- H
1965 1972 1974 1976 3 20.92000 0.00000 -20.92000
0.00000 0.00000 0.00000 ; CA- C- N- CA
1966 1965 1967 1968 3 0.65084 1.95253 0.00000
-2.60338 0.00000 0.00000 ; HA- CA- CB- HB2
1966 1965 1967 1969 3 0.65084 1.95253 0.00000
-2.60338 0.00000 0.00000 ; HA- CA- CB- HB3
1966 1965 1967 1970 3 1.04600 -1.04600 0.00000
0.00000 0.00000 0.00000 ; HA- CA- CB- OG
1966 1965 1972 1973 3 3.68192 -4.35136 0.00000
1.33888 0.00000 0.00000 ; HA- CA- C- O
1966 1965 1972 1974 3 0.00000 0.00000 0.00000
0.00000 0.00000 0.00000 ; HA- CA- C- N
1967 1965 1972 1973 3 0.00000 0.00000 0.00000
0.00000 0.00000 0.00000 ; CB- CA- C- O
1967 1965 1972 1974 3 2.51040 4.18400 1.67360
-6.69440 0.00000 0.00000 ; CB- CA- C- N
1968 1967 1965 1972 3 0.65084 1.95253 0.00000
-2.60338 0.00000 0.00000 ; HB2- CB- CA- C
1968 1967 1970 1971 3 0.69733 2.09200 0.00000
-2.78933 0.00000 0.00000 ; HB2- CB- OG- HG
1969 1967 1965 1972 3 0.65084 1.95253 0.00000
-2.60338 0.00000 0.00000 ; HB3- CB- CA- C
1969 1967 1970 1971 3 0.69733 2.09200 0.00000
-2.78933 0.00000 0.00000 ; HB3- CB- OG- HG
1970 1967 1965 1972 3 7.34710 7.79898 -6.14211
-6.71114 4.31789 0.00000 ; OG- CB- CA- C
1972 1974 1976 1977 3 0.00000 0.00000 0.00000
0.00000 0.00000 0.00000 ; C- N- CA- HA
1972 1974 1976 1978 3 11.71520 1.67360 15.06240 -
13.38880 0.00000 0.00000 ; C- N- CA- CB
1972 1974 1976 1982 3 1.75728 5.27184 2.25936
-7.02912 0.00000 0.00000 ; C- N- CA- C
1973 1972 1974 1975 3 29.28800 -8.36800 -20.92000
0.00000 0.00000 0.00000 ; O- C- N- H
1973 1972 1974 1976 3 20.92000 0.00000 -20.92000
0.00000 0.00000 0.00000 ; O- C- N- CA
1974 1976 1978 1979 3 0.65084 1.95253 0.00000
-2.60338 0.00000 0.00000 ; N- CA- CB- HB1
1974 1976 1978 1980 3 0.65084 1.95253 0.00000
-2.60338 0.00000 0.00000 ; N- CA- CB- HB2
1974 1976 1978 1981 3 0.65084 1.95253 0.00000
-2.60338 0.00000 0.00000 ; N- CA- CB- HB3
1974 1976 1982 1983 3 0.00000 0.00000 0.00000
0.00000 0.00000 0.00000 ; N- CA- C- O

1974 1976 1982 1984 3 17.40544 -5.02080 -13.22144
9.20480 0.00000 0.00000 ; N- CA- C- N
1975 1974 1976 1977 3 0.00000 0.00000 0.00000
0.00000 0.00000 0.00000 ; H- N- CA- HA
1975 1974 1976 1978 3 0.00000 0.00000 0.00000
0.00000 0.00000 0.00000 ; H- N- CA- CB
1975 1974 1976 1982 3 0.00000 0.00000 0.00000
0.00000 0.00000 0.00000 ; H- N- CA- C
1976 1982 1984 1985 3 20.92000 0.00000 -20.92000
0.00000 0.00000 0.00000 ; CA- C- N- H
1976 1982 1984 1986 3 20.92000 0.00000 -20.92000
0.00000 0.00000 0.00000 ; CA- C- N- CA
1977 1976 1978 1979 3 0.65084 1.95253 0.00000
-2.60338 0.00000 0.00000 ; HA- CA- CB- HB1
1977 1976 1978 1980 3 0.65084 1.95253 0.00000
-2.60338 0.00000 0.00000 ; HA- CA- CB- HB2
1977 1976 1978 1981 3 0.65084 1.95253 0.00000
-2.60338 0.00000 0.00000 ; HA- CA- CB- HB3
1977 1976 1982 1983 3 3.68192 -4.35136 0.00000
1.33888 0.00000 0.00000 ; HA- CA- C- O
1977 1976 1982 1984 3 0.00000 0.00000 0.00000
0.00000 0.00000 0.00000 ; HA- CA- C- N
1978 1976 1982 1983 3 0.00000 0.00000 0.00000
0.00000 0.00000 0.00000 ; CB- CA- C- O
1978 1976 1982 1984 3 2.51040 4.18400 1.67360
-6.69440 0.00000 0.00000 ; CB- CA- C- N
1979 1978 1976 1982 3 0.65084 1.95253 0.00000
-2.60338 0.00000 0.00000 ; HB1- CB- CA- C
1980 1978 1976 1982 3 0.65084 1.95253 0.00000
-2.60338 0.00000 0.00000 ; HB2- CB- CA- C
1981 1978 1976 1982 3 0.65084 1.95253 0.00000
-2.60338 0.00000 0.00000 ; HB3- CB- CA- C
1982 1984 1986 1987 3 0.00000 0.00000 0.00000
0.00000 0.00000 0.00000 ; C- N- CA- HA
1982 1984 1986 1988 3 11.71520 1.67360 15.06240 -
13.38880 0.00000 0.00000 ; C- N- CA- CB
1982 1984 1986 2001 3 1.75728 5.27184 2.25936
-7.02912 0.00000 0.00000 ; C- N- CA- C
1983 1982 1984 1985 3 29.28800 -8.36800 -20.92000
0.00000 0.00000 0.00000 ; O- C- N- H
1983 1982 1984 1986 3 20.92000 0.00000 -20.92000
0.00000 0.00000 0.00000 ; O- C- N- CA
1984 1986 1988 1989 3 0.65084 1.95253 0.00000
-2.60338 0.00000 0.00000 ; N- CA- CB- HB
1984 1986 1988 1990 3 3.83673 0.44769 -1.77402
-2.47693 -0.03347 0.00000 ; N- CA- CB- CG2
1984 1986 1988 1994 3 2.97482 0.12134 2.04179
-1.89117 -3.24678 0.00000 ; N- CA- CB- CG1
1984 1986 2001 2002 3 0.00000 0.00000 0.00000
0.00000 0.00000 0.00000 ; N- CA- C- O
1984 1986 2001 2003 3 17.40544 -5.02080 -13.22144
9.20480 0.00000 0.00000 ; N- CA- C- N

1985 1984 1986 1987 3 0.00000 0.00000 0.00000
0.00000 0.00000 0.00000 ; H- N- CA- HA
1985 1984 1986 1988 3 0.00000 0.00000 0.00000
0.00000 0.00000 0.00000 ; H- N- CA- CB
1985 1984 1986 2001 3 0.00000 0.00000 0.00000
0.00000 0.00000 0.00000 ; H- N- CA- C
1986 1988 1990 1991 3 0.66944 2.00832 0.00000
-2.67776 0.00000 0.00000 ; CA- CB- CG2- HG21
1986 1988 1990 1992 3 0.66944 2.00832 0.00000
-2.67776 0.00000 0.00000 ; CA- CB- CG2- HG22
1986 1988 1990 1993 3 0.66944 2.00832 0.00000
-2.67776 0.00000 0.00000 ; CA- CB- CG2- HG23
1986 1988 1994 1995 3 0.66944 2.00832 0.00000
-2.67776 0.00000 0.00000 ; CA- CB- CG1- HG12
1986 1988 1994 1996 3 0.66944 2.00832 0.00000
-2.67776 0.00000 0.00000 ; CA- CB- CG1- HG13
1986 1988 1994 1997 3 4.24258 -0.52718 -7.25506
-1.79075 7.69856 0.00000 ; CA- CB- CG1- CD1
1986 2001 2003 2004 3 20.92000 0.00000 -20.92000
0.00000 0.00000 0.00000 ; CA- C- N- H
1986 2001 2003 2005 3 20.92000 0.00000 -20.92000
0.00000 0.00000 0.00000 ; CA- C- N- CA
1987 1986 1988 1989 3 0.65084 1.95253 0.00000
-2.60338 0.00000 0.00000 ; HA- CA- CB- HB
1987 1986 1988 1990 3 0.65084 1.95253 0.00000
-2.60338 0.00000 0.00000 ; HA- CA- CB- CG2
1987 1986 1988 1994 3 0.65084 1.95253 0.00000
-2.60338 0.00000 0.00000 ; HA- CA- CB- CG1
1987 1986 2001 2002 3 3.68192 -4.35136 0.00000
1.33888 0.00000 0.00000 ; HA- CA- C- O
1987 1986 2001 2003 3 0.00000 0.00000 0.00000
0.00000 0.00000 0.00000 ; HA- CA- C- N
1988 1986 2001 2002 3 0.00000 0.00000 0.00000
0.00000 0.00000 0.00000 ; CB- CA- C- O
1988 1986 2001 2003 3 2.51040 4.18400 1.67360
-6.69440 0.00000 0.00000 ; CB- CA- C- N
1988 1994 1997 1998 3 0.66944 2.00832 0.00000
-2.67776 0.00000 0.00000 ; CB- CG1- CD1- HD11
1988 1994 1997 1999 3 0.66944 2.00832 0.00000
-2.67776 0.00000 0.00000 ; CB- CG1- CD1- HD12
1988 1994 1997 2000 3 0.66944 2.00832 0.00000
-2.67776 0.00000 0.00000 ; CB- CG1- CD1- HD13
1989 1988 1986 2001 3 0.65084 1.95253 0.00000
-2.60338 0.00000 0.00000 ; HB- CB- CA- C
1989 1988 1990 1991 3 0.62760 1.88280 0.00000
-2.51040 0.00000 0.00000 ; HB- CB- CG2- HG21
1989 1988 1990 1992 3 0.62760 1.88280 0.00000
-2.51040 0.00000 0.00000 ; HB- CB- CG2- HG22
1989 1988 1990 1993 3 0.62760 1.88280 0.00000
-2.51040 0.00000 0.00000 ; HB- CB- CG2- HG23
1989 1988 1994 1995 3 0.62760 1.88280 0.00000
-2.51040 0.00000 0.00000 ; HB- CB- CG1- HG12

```
  1989   1988   1994   1996      3    0.62760    1.88280    0.00000
-2.51040    0.00000    0.00000 ;     HB-    CB-   CG1-  HG13
  1989   1988   1994   1997      3    0.66944    2.00832    0.00000
-2.67776    0.00000    0.00000 ;     HB-    CB-   CG1-   CD1
  1990   1988   1986   2001      3    5.67350    3.55640   -6.16722
-2.47693    3.74886    0.00000 ;    CG2-    CB-    CA-     C
  1990   1988   1994   1995      3    0.66944    2.00832    0.00000
-2.67776    0.00000    0.00000 ;    CG2-    CB-   CG1-  HG12
  1990   1988   1994   1996      3    0.66944    2.00832    0.00000
-2.67776    0.00000    0.00000 ;    CG2-    CB-   CG1-  HG13
  1990   1988   1994   1997      3    3.81162    0.49790   -8.14206
-1.79075    7.49773    0.00000 ;    CG2-    CB-   CG1-   CD1
  1991   1990   1988   1994      3    0.66944    2.00832    0.00000
-2.67776    0.00000    0.00000 ;   HG21-   CG2-    CB-   CG1
  1992   1990   1988   1994      3    0.66944    2.00832    0.00000
-2.67776    0.00000    0.00000 ;   HG22-   CG2-    CB-   CG1
  1993   1990   1988   1994      3    0.66944    2.00832    0.00000
-2.67776    0.00000    0.00000 ;   HG23-   CG2-    CB-   CG1
  1994   1988   1986   2001      3    8.26340    0.74057   -9.99976
-1.89117    3.84928    0.00000 ;    CG1-    CB-    CA-     C
  1995   1994   1997   1998      3    0.62760    1.88280    0.00000
-2.51040    0.00000    0.00000 ;   HG12-   CG1-   CD1-  HD11
  1995   1994   1997   1999      3    0.62760    1.88280    0.00000
-2.51040    0.00000    0.00000 ;   HG12-   CG1-   CD1-  HD12
  1995   1994   1997   2000      3    0.62760    1.88280    0.00000
-2.51040    0.00000    0.00000 ;   HG12-   CG1-   CD1-  HD13
  1996   1994   1997   1998      3    0.62760    1.88280    0.00000
-2.51040    0.00000    0.00000 ;   HG13-   CG1-   CD1-  HD11
  1996   1994   1997   1999      3    0.62760    1.88280    0.00000
-2.51040    0.00000    0.00000 ;   HG13-   CG1-   CD1-  HD12
  1996   1994   1997   2000      3    0.62760    1.88280    0.00000
-2.51040    0.00000    0.00000 ;   HG13-   CG1-   CD1-  HD13
  2001   2003   2005   2006      3    0.00000    0.00000    0.00000
0.00000    0.00000    0.00000 ;      C-     N-    CA-    HA
  2001   2003   2005   2007      3   11.71520    1.67360   15.06240  -
13.38880    0.00000    0.00000 ;      C-     N-    CA-    CB
  2001   2003   2005   2021      3    1.75728    5.27184    2.25936
-7.02912    0.00000    0.00000 ;      C-     N-    CA-     C
  2002   2001   2003   2004      3   29.28800   -8.36800  -20.92000
0.00000    0.00000    0.00000 ;      O-     C-     N-     H
  2002   2001   2003   2005      3   20.92000    0.00000  -20.92000
0.00000    0.00000    0.00000 ;      O-     C-     N-    CA
  2003   2005   2007   2008      3    0.65084    1.95253    0.00000
-2.60338    0.00000    0.00000 ;      N-    CA-    CB-   HB2
  2003   2005   2007   2009      3    0.65084    1.95253    0.00000
-2.60338    0.00000    0.00000 ;      N-    CA-    CB-   HB3
  2003   2005   2007   2010      3    3.28026    2.46019   -2.19242
-3.21331   -0.23430    0.00000 ;      N-    CA-    CB-    CG
  2003   2005   2021   2022      3    0.00000    0.00000    0.00000
0.00000    0.00000    0.00000 ;      N-    CA-     C-     O
  2003   2005   2021   2023      3   17.40544   -5.02080  -13.22144
9.20480    0.00000    0.00000 ;      N-    CA-     C-     N
```

2004 2003 2005 2006 3 0.00000 0.00000 0.00000
0.00000 0.00000 0.00000 ; H- N- CA- HA
2004 2003 2005 2007 3 0.00000 0.00000 0.00000
0.00000 0.00000 0.00000 ; H- N- CA- CB
2004 2003 2005 2021 3 0.00000 0.00000 0.00000
0.00000 0.00000 0.00000 ; H- N- CA- C
2005 2007 2010 2011 3 0.57739 0.00000 1.02926
0.00000 -1.60666 0.00000 ; CA- CB- CG- CD1
2005 2007 2010 2019 3 0.57739 0.00000 1.02926
0.00000 -1.60666 0.00000 ; CA- CB- CG- CD2
2005 2021 2023 2024 3 20.92000 0.00000 -20.92000
0.00000 0.00000 0.00000 ; CA- C- N- H
2005 2021 2023 2025 3 20.92000 0.00000 -20.92000
0.00000 0.00000 0.00000 ; CA- C- N- CA
2006 2005 2007 2008 3 0.65084 1.95253 0.00000
-2.60338 0.00000 0.00000 ; HA- CA- CB- HB2
2006 2005 2007 2009 3 0.65084 1.95253 0.00000
-2.60338 0.00000 0.00000 ; HA- CA- CB- HB3
2006 2005 2007 2010 3 0.65084 1.95253 0.00000
-2.60338 0.00000 0.00000 ; HA- CA- CB- CG
2006 2005 2021 2022 3 3.68192 -4.35136 0.00000
1.33888 0.00000 0.00000 ; HA- CA- C- O
2006 2005 2021 2023 3 0.00000 0.00000 0.00000
0.00000 0.00000 0.00000 ; HA- CA- C- N
2007 2005 2021 2022 3 0.00000 0.00000 0.00000
0.00000 0.00000 0.00000 ; CB- CA- C- O
2007 2005 2021 2023 3 2.51040 4.18400 1.67360
-6.69440 0.00000 0.00000 ; CB- CA- C- N
2007 2010 2011 2012 3 30.33400 0.00000 -30.33400
0.00000 0.00000 0.00000 ; CB- CG- CD1- HD1
2007 2010 2011 2013 3 30.33400 0.00000 -30.33400
0.00000 0.00000 0.00000 ; CB- CG- CD1- CE1
2007 2010 2019 2017 3 30.33400 0.00000 -30.33400
0.00000 0.00000 0.00000 ; CB- CG- CD2- CE2
2007 2010 2019 2020 3 30.33400 0.00000 -30.33400
0.00000 0.00000 0.00000 ; CB- CG- CD2- HD2
2008 2007 2005 2021 3 0.65084 1.95253 0.00000
-2.60338 0.00000 0.00000 ; HB2- CB- CA- C
2008 2007 2010 2011 3 0.00000 0.00000 0.00000
0.00000 0.00000 0.00000 ; HB2- CB- CG- CD1
2008 2007 2010 2019 3 0.00000 0.00000 0.00000
0.00000 0.00000 0.00000 ; HB2- CB- CG- CD2
2009 2007 2005 2021 3 0.65084 1.95253 0.00000
-2.60338 0.00000 0.00000 ; HB3- CB- CA- C
2009 2007 2010 2011 3 0.00000 0.00000 0.00000
0.00000 0.00000 0.00000 ; HB3- CB- CG- CD1
2009 2007 2010 2019 3 0.00000 0.00000 0.00000
0.00000 0.00000 0.00000 ; HB3- CB- CG- CD2
2010 2007 2005 2021 3 4.95804 2.17986 -3.52293
-3.21331 -0.40166 0.00000 ; CG- CB- CA- C
2010 2011 2013 2014 3 30.33400 0.00000 -30.33400
0.00000 0.00000 0.00000 ; CG- CD1- CE1- HE1

2010 2011 2013 2015 3 30.33400 0.00000 -30.33400
0.00000 0.00000 0.00000 ; CG- CD1- CE1- CZ
2010 2019 2017 2015 3 30.33400 0.00000 -30.33400
0.00000 0.00000 0.00000 ; CG- CD2- CE2- CZ
2010 2019 2017 2018 3 30.33400 0.00000 -30.33400
0.00000 0.00000 0.00000 ; CG- CD2- CE2- HE2
2011 2010 2019 2017 3 30.33400 0.00000 -30.33400
0.00000 0.00000 0.00000 ; CD1- CG- CD2- CE2
2011 2010 2019 2020 3 30.33400 0.00000 -30.33400
0.00000 0.00000 0.00000 ; CD1- CG- CD2- HD2
2011 2013 2015 2016 3 30.33400 0.00000 -30.33400
0.00000 0.00000 0.00000 ; CD1- CE1- CZ- HZ
2011 2013 2015 2017 3 30.33400 0.00000 -30.33400
0.00000 0.00000 0.00000 ; CD1- CE1- CZ- CE2
2012 2011 2010 2019 3 30.33400 0.00000 -30.33400
0.00000 0.00000 0.00000 ; HD1- CD1- CG- CD2
2012 2011 2013 2014 3 30.33400 0.00000 -30.33400
0.00000 0.00000 0.00000 ; HD1- CD1- CE1- HE1
2012 2011 2013 2015 3 30.33400 0.00000 -30.33400
0.00000 0.00000 0.00000 ; HD1- CD1- CE1- CZ
2013 2011 2010 2019 3 30.33400 0.00000 -30.33400
0.00000 0.00000 0.00000 ; CE1- CD1- CG- CD2
2013 2015 2017 2018 3 30.33400 0.00000 -30.33400
0.00000 0.00000 0.00000 ; CE1- CZ- CE2- HE2
2013 2015 2017 2019 3 30.33400 0.00000 -30.33400
0.00000 0.00000 0.00000 ; CE1- CZ- CE2- CD2
2014 2013 2015 2016 3 30.33400 0.00000 -30.33400
0.00000 0.00000 0.00000 ; HE1- CE1- CZ- HZ
2014 2013 2015 2017 3 30.33400 0.00000 -30.33400
0.00000 0.00000 0.00000 ; HE1- CE1- CZ- CE2
2015 2017 2019 2020 3 30.33400 0.00000 -30.33400
0.00000 0.00000 0.00000 ; CZ- CE2- CD2- HD2
2016 2015 2017 2018 3 30.33400 0.00000 -30.33400
0.00000 0.00000 0.00000 ; HZ- CZ- CE2- HE2
2016 2015 2017 2019 3 30.33400 0.00000 -30.33400
0.00000 0.00000 0.00000 ; HZ- CZ- CE2- CD2
2018 2017 2019 2020 3 30.33400 0.00000 -30.33400
0.00000 0.00000 0.00000 ; HE2- CE2- CD2- HD2
2021 2023 2025 2026 3 0.00000 0.00000 0.00000
0.00000 0.00000 0.00000 ; C- N- CA- HA
2021 2023 2025 2027 3 11.71520 1.67360 15.06240 -
13.38880 0.00000 0.00000 ; C- N- CA- CB
2021 2023 2025 2042 3 1.75728 5.27184 2.25936
-7.02912 0.00000 0.00000 ; C- N- CA- C
2022 2021 2023 2024 3 29.28800 -8.36800 -20.92000
0.00000 0.00000 0.00000 ; O- C- N- H
2022 2021 2023 2025 3 20.92000 0.00000 -20.92000
0.00000 0.00000 0.00000 ; O- C- N- CA
2023 2025 2027 2028 3 0.65084 1.95253 0.00000
-2.60338 0.00000 0.00000 ; N- CA- CB- HB2
2023 2025 2027 2029 3 0.65084 1.95253 0.00000
-2.60338 0.00000 0.00000 ; N- CA- CB- HB3

2023 2025 2027 2030 3 3.28026 2.46019 -2.19242
-3.21331 -0.23430 0.00000 ; N- CA- CB- CG
2023 2025 2042 2043 3 0.00000 0.00000 0.00000
0.00000 0.00000 0.00000 ; N- CA- C- O
2023 2025 2042 2044 3 17.40544 -5.02080 -13.22144
9.20480 0.00000 0.00000 ; N- CA- C- N
2024 2023 2025 2026 3 0.00000 0.00000 0.00000
0.00000 0.00000 0.00000 ; H- N- CA- HA
2024 2023 2025 2027 3 0.00000 0.00000 0.00000
0.00000 0.00000 0.00000 ; H- N- CA- CB
2024 2023 2025 2042 3 0.00000 0.00000 0.00000
0.00000 0.00000 0.00000 ; H- N- CA- C
2025 2027 2030 2031 3 0.57739 0.00000 1.02926
0.00000 -1.60666 0.00000 ; CA- CB- CG- CD1
2025 2027 2030 2040 3 0.57739 0.00000 1.02926
0.00000 -1.60666 0.00000 ; CA- CB- CG- CD2
2025 2042 2044 2045 3 20.92000 0.00000 -20.92000
0.00000 0.00000 0.00000 ; CA- C- N- H
2025 2042 2044 2046 3 20.92000 0.00000 -20.92000
0.00000 0.00000 0.00000 ; CA- C- N- CA
2026 2025 2027 2028 3 0.65084 1.95253 0.00000
-2.60338 0.00000 0.00000 ; HA- CA- CB- HB2
2026 2025 2027 2029 3 0.65084 1.95253 0.00000
-2.60338 0.00000 0.00000 ; HA- CA- CB- HB3
2026 2025 2027 2030 3 0.65084 1.95253 0.00000
-2.60338 0.00000 0.00000 ; HA- CA- CB- CG
2026 2025 2042 2043 3 3.68192 -4.35136 0.00000
1.33888 0.00000 0.00000 ; HA- CA- C- O
2026 2025 2042 2044 3 0.00000 0.00000 0.00000
0.00000 0.00000 0.00000 ; HA- CA- C- N
2027 2025 2042 2043 3 0.00000 0.00000 0.00000
0.00000 0.00000 0.00000 ; CB- CA- C- O
2027 2025 2042 2044 3 2.51040 4.18400 1.67360
-6.69440 0.00000 0.00000 ; CB- CA- C- N
2027 2030 2031 2032 3 30.33400 0.00000 -30.33400
0.00000 0.00000 0.00000 ; CB- CG- CD1- HD1
2027 2030 2031 2033 3 30.33400 0.00000 -30.33400
0.00000 0.00000 0.00000 ; CB- CG- CD1- CE1
2027 2030 2040 2038 3 30.33400 0.00000 -30.33400
0.00000 0.00000 0.00000 ; CB- CG- CD2- CE2
2027 2030 2040 2041 3 30.33400 0.00000 -30.33400
0.00000 0.00000 0.00000 ; CB- CG- CD2- HD2
2028 2027 2025 2042 3 0.65084 1.95253 0.00000
-2.60338 0.00000 0.00000 ; HB2- CB- CA- C
2028 2027 2030 2031 3 0.00000 0.00000 0.00000
0.00000 0.00000 0.00000 ; HB2- CB- CG- CD1
2028 2027 2030 2040 3 0.00000 0.00000 0.00000
0.00000 0.00000 0.00000 ; HB2- CB- CG- CD2
2029 2027 2025 2042 3 0.65084 1.95253 0.00000
-2.60338 0.00000 0.00000 ; HB3- CB- CA- C
2029 2027 2030 2031 3 0.00000 0.00000 0.00000
0.00000 0.00000 0.00000 ; HB3- CB- CG- CD1

2029 2027 2030 2040 3 0.00000 0.00000 0.00000
0.00000 0.00000 0.00000 ; HB3- CB- CG- CD2
2030 2027 2025 2042 3 4.95804 2.17986 -3.52293
-3.21331 -0.40166 0.00000 ; CG- CB- CA- C
2030 2031 2033 2034 3 30.33400 0.00000 -30.33400
0.00000 0.00000 0.00000 ; CG- CD1- CE1- HE1
2030 2031 2033 2035 3 30.33400 0.00000 -30.33400
0.00000 0.00000 0.00000 ; CG- CD1- CE1- CZ
2030 2040 2038 2035 3 30.33400 0.00000 -30.33400
0.00000 0.00000 0.00000 ; CG- CD2- CE2- CZ
2030 2040 2038 2039 3 30.33400 0.00000 -30.33400
0.00000 0.00000 0.00000 ; CG- CD2- CE2- HE2
2031 2030 2040 2038 3 30.33400 0.00000 -30.33400
0.00000 0.00000 0.00000 ; CD1- CG- CD2- CE2
2031 2030 2040 2041 3 30.33400 0.00000 -30.33400
0.00000 0.00000 0.00000 ; CD1- CG- CD2- HD2
2031 2033 2035 2036 3 30.33400 0.00000 -30.33400
0.00000 0.00000 0.00000 ; CD1- CE1- CZ- OH
2031 2033 2035 2038 3 30.33400 0.00000 -30.33400
0.00000 0.00000 0.00000 ; CD1- CE1- CZ- CE2
2032 2031 2030 2040 3 30.33400 0.00000 -30.33400
0.00000 0.00000 0.00000 ; HD1- CD1- CG- CD2
2032 2031 2033 2034 3 30.33400 0.00000 -30.33400
0.00000 0.00000 0.00000 ; HD1- CD1- CE1- HE1
2032 2031 2033 2035 3 30.33400 0.00000 -30.33400
0.00000 0.00000 0.00000 ; HD1- CD1- CE1- CZ
2033 2031 2030 2040 3 30.33400 0.00000 -30.33400
0.00000 0.00000 0.00000 ; CE1- CD1- CG- CD2
2033 2035 2036 2037 3 7.93286 0.00000 -9.56462
0.00000 2.17568 0.00000 ; CE1- CZ- OH- HH
2033 2035 2038 2039 3 30.33400 0.00000 -30.33400
0.00000 0.00000 0.00000 ; CE1- CZ- CE2- HE2
2033 2035 2038 2040 3 30.33400 0.00000 -30.33400
0.00000 0.00000 0.00000 ; CE1- CZ- CE2- CD2
2034 2033 2035 2036 3 30.33400 0.00000 -30.33400
0.00000 0.00000 0.00000 ; HE1- CE1- CZ- OH
2034 2033 2035 2038 3 30.33400 0.00000 -30.33400
0.00000 0.00000 0.00000 ; HE1- CE1- CZ- CE2
2035 2038 2040 2041 3 30.33400 0.00000 -30.33400
0.00000 0.00000 0.00000 ; CZ- CE2- CD2- HD2
2036 2035 2038 2039 3 30.33400 0.00000 -30.33400
0.00000 0.00000 0.00000 ; OH- CZ- CE2- HE2
2036 2035 2038 2040 3 30.33400 0.00000 -30.33400
0.00000 0.00000 0.00000 ; OH- CZ- CE2- CD2
2037 2036 2035 2038 3 7.93286 0.00000 -9.56462
0.00000 2.17568 0.00000 ; HH- OH- CZ- CE2
2039 2038 2040 2041 3 30.33400 0.00000 -30.33400
0.00000 0.00000 0.00000 ; HE2- CE2- CD2- HD2
2042 2044 2046 2047 3 0.00000 0.00000 0.00000
0.00000 0.00000 0.00000 ; C- N- CA- HA
2042 2044 2046 2048 3 11.71520 1.67360 15.06240 -
13.38880 0.00000 0.00000 ; C- N- CA- CB

2042 2044 2046 2061 3 1.75728 5.27184 2.25936
-7.02912 0.00000 0.00000 ; C- N- CA- C
2043 2042 2044 2045 3 29.28800 -8.36800 -20.92000
0.00000 0.00000 0.00000 ; O- C- N- H
2043 2042 2044 2046 3 20.92000 0.00000 -20.92000
0.00000 0.00000 0.00000 ; O- C- N- CA
2044 2046 2048 2049 3 0.65084 1.95253 0.00000
-2.60338 0.00000 0.00000 ; N- CA- CB- HB2
2044 2046 2048 2050 3 0.65084 1.95253 0.00000
-2.60338 0.00000 0.00000 ; N- CA- CB- HB3
2044 2046 2048 2051 3 3.79070 1.39746 -4.61077
-2.40998 2.44346 0.00000 ; N- CA- CB- CG
2044 2046 2061 2062 3 0.00000 0.00000 0.00000
0.00000 0.00000 0.00000 ; N- CA- C- O
2044 2046 2061 2063 3 17.40544 -5.02080 -13.22144
9.20480 0.00000 0.00000 ; N- CA- C- N
2045 2044 2046 2047 3 0.00000 0.00000 0.00000
0.00000 0.00000 0.00000 ; H- N- CA- HA
2045 2044 2046 2048 3 0.00000 0.00000 0.00000
0.00000 0.00000 0.00000 ; H- N- CA- CB
2045 2044 2046 2061 3 0.00000 0.00000 0.00000
0.00000 0.00000 0.00000 ; H- N- CA- C
2046 2048 2051 2052 3 0.66944 2.00832 0.00000
-2.67776 0.00000 0.00000 ; CA- CB- CG- HG
2046 2048 2051 2053 3 3.90367 0.19665 -6.21742
-2.37651 5.99149 0.00000 ; CA- CB- CG- CD1
2046 2048 2051 2057 3 3.90367 0.19665 -6.21742
-2.37651 5.99149 0.00000 ; CA- CB- CG- CD2
2046 2061 2063 2064 3 20.92000 0.00000 -20.92000
0.00000 0.00000 0.00000 ; CA- C- N- H
2046 2061 2063 2065 3 20.92000 0.00000 -20.92000
0.00000 0.00000 0.00000 ; CA- C- N- CA
2047 2046 2048 2049 3 0.65084 1.95253 0.00000
-2.60338 0.00000 0.00000 ; HA- CA- CB- HB2
2047 2046 2048 2050 3 0.65084 1.95253 0.00000
-2.60338 0.00000 0.00000 ; HA- CA- CB- HB3
2047 2046 2048 2051 3 0.65084 1.95253 0.00000
-2.60338 0.00000 0.00000 ; HA- CA- CB- CG
2047 2046 2061 2062 3 3.68192 -4.35136 0.00000
1.33888 0.00000 0.00000 ; HA- CA- C- O
2047 2046 2061 2063 3 0.00000 0.00000 0.00000
0.00000 0.00000 0.00000 ; HA- CA- C- N
2048 2046 2061 2062 3 0.00000 0.00000 0.00000
0.00000 0.00000 0.00000 ; CB- CA- C- O
2048 2046 2061 2063 3 2.51040 4.18400 1.67360
-6.69440 0.00000 0.00000 ; CB- CA- C- N
2048 2051 2053 2054 3 0.66944 2.00832 0.00000
-2.67776 0.00000 0.00000 ; CB- CG- CD1- HD11
2048 2051 2053 2055 3 0.66944 2.00832 0.00000
-2.67776 0.00000 0.00000 ; CB- CG- CD1- HD12
2048 2051 2053 2056 3 0.66944 2.00832 0.00000
-2.67776 0.00000 0.00000 ; CB- CG- CD1- HD13

2048 2051 2057 2058 3 0.66944 2.00832 0.00000
-2.67776 0.00000 0.00000 ; CB- CG- CD2- HD21
2048 2051 2057 2059 3 0.66944 2.00832 0.00000
-2.67776 0.00000 0.00000 ; CB- CG- CD2- HD22
2048 2051 2057 2060 3 0.66944 2.00832 0.00000
-2.67776 0.00000 0.00000 ; CB- CG- CD2- HD23
2049 2048 2046 2061 3 0.65084 1.95253 0.00000
-2.60338 0.00000 0.00000 ; HB2- CB- CA- C
2049 2048 2051 2052 3 0.62760 1.88280 0.00000
-2.51040 0.00000 0.00000 ; HB2- CB- CG- HG
2049 2048 2051 2053 3 0.66944 2.00832 0.00000
-2.67776 0.00000 0.00000 ; HB2- CB- CG- CD1
2049 2048 2051 2057 3 0.66944 2.00832 0.00000
-2.67776 0.00000 0.00000 ; HB2- CB- CG- CD2
2050 2048 2046 2061 3 0.65084 1.95253 0.00000
-2.60338 0.00000 0.00000 ; HB3- CB- CA- C
2050 2048 2051 2052 3 0.62760 1.88280 0.00000
-2.51040 0.00000 0.00000 ; HB3- CB- CG- HG
2050 2048 2051 2053 3 0.66944 2.00832 0.00000
-2.67776 0.00000 0.00000 ; HB3- CB- CG- CD1
2050 2048 2051 2057 3 0.66944 2.00832 0.00000
-2.67776 0.00000 0.00000 ; HB3- CB- CG- CD2
2051 2048 2046 2061 3 10.33448 -1.14642 -11.54784
-2.40998 6.35968 0.00000 ; CG- CB- CA- C
2052 2051 2053 2054 3 0.62760 1.88280 0.00000
-2.51040 0.00000 0.00000 ; HG- CG- CD1- HD11
2052 2051 2053 2055 3 0.62760 1.88280 0.00000
-2.51040 0.00000 0.00000 ; HG- CG- CD1- HD12
2052 2051 2053 2056 3 0.62760 1.88280 0.00000
-2.51040 0.00000 0.00000 ; HG- CG- CD1- HD13
2052 2051 2057 2058 3 0.62760 1.88280 0.00000
-2.51040 0.00000 0.00000 ; HG- CG- CD2- HD21
2052 2051 2057 2059 3 0.62760 1.88280 0.00000
-2.51040 0.00000 0.00000 ; HG- CG- CD2- HD22
2052 2051 2057 2060 3 0.62760 1.88280 0.00000
-2.51040 0.00000 0.00000 ; HG- CG- CD2- HD23
2053 2051 2057 2058 3 0.66944 2.00832 0.00000
-2.67776 0.00000 0.00000 ; CD1- CG- CD2- HD21
2053 2051 2057 2059 3 0.66944 2.00832 0.00000
-2.67776 0.00000 0.00000 ; CD1- CG- CD2- HD22
2053 2051 2057 2060 3 0.66944 2.00832 0.00000
-2.67776 0.00000 0.00000 ; CD1- CG- CD2- HD23
2054 2053 2051 2057 3 0.66944 2.00832 0.00000
-2.67776 0.00000 0.00000 ; HD11- CD1- CG- CD2
2055 2053 2051 2057 3 0.66944 2.00832 0.00000
-2.67776 0.00000 0.00000 ; HD12- CD1- CG- CD2
2056 2053 2051 2057 3 0.66944 2.00832 0.00000
-2.67776 0.00000 0.00000 ; HD13- CD1- CG- CD2
2061 2063 2065 2066 3 0.00000 0.00000 0.00000
0.00000 0.00000 0.00000 ; C- N- CA- HA
2061 2063 2065 2067 3 11.71520 1.67360 15.06240 -
13.38880 0.00000 0.00000 ; C- N- CA- CB

2061 2063 2065 2079 3 1.75728 5.27184 2.25936
-7.02912 0.00000 0.00000 ; C- N- CA- C
2062 2061 2063 2064 3 29.28800 -8.36800 -20.92000
0.00000 0.00000 0.00000 ; O- C- N- H
2062 2061 2063 2065 3 20.92000 0.00000 -20.92000
0.00000 0.00000 0.00000 ; O- C- N- CA
2063 2065 2067 2068 3 0.65084 1.95253 0.00000
-2.60338 0.00000 0.00000 ; N- CA- CB- HB2
2063 2065 2067 2069 3 0.65084 1.95253 0.00000
-2.60338 0.00000 0.00000 ; N- CA- CB- HB3
2063 2065 2067 2070 3 4.79068 4.02919 -4.82834
-3.66518 2.97901 0.00000 ; N- CA- CB- CG
2063 2065 2079 2080 3 0.00000 0.00000 0.00000
0.00000 0.00000 0.00000 ; N- CA- C- O
2063 2065 2079 2081 3 17.40544 -5.02080 -13.22144
9.20480 0.00000 0.00000 ; N- CA- C- N
2064 2063 2065 2066 3 0.00000 0.00000 0.00000
0.00000 0.00000 0.00000 ; H- N- CA- HA
2064 2063 2065 2067 3 0.00000 0.00000 0.00000
0.00000 0.00000 0.00000 ; H- N- CA- CB
2064 2063 2065 2079 3 0.00000 0.00000 0.00000
0.00000 0.00000 0.00000 ; H- N- CA- C
2065 2067 2070 2071 3 6.81992 9.28011 -2.04179 -
11.48090 -1.23846 0.00000 ; CA- CB- CG- ND1
2065 2067 2070 2077 3 0.00000 0.00000 0.00000
0.00000 0.00000 0.00000 ; CA- CB- CG- CD2
2065 2079 2081 2082 3 20.92000 0.00000 -20.92000
0.00000 0.00000 0.00000 ; CA- C- N- H
2065 2079 2081 2083 3 20.92000 0.00000 -20.92000
0.00000 0.00000 0.00000 ; CA- C- N- CA
2066 2065 2067 2068 3 0.65084 1.95253 0.00000
-2.60338 0.00000 0.00000 ; HA- CA- CB- HB2
2066 2065 2067 2069 3 0.65084 1.95253 0.00000
-2.60338 0.00000 0.00000 ; HA- CA- CB- HB3
2066 2065 2067 2070 3 0.65084 1.95253 0.00000
-2.60338 0.00000 0.00000 ; HA- CA- CB- CG
2066 2065 2079 2080 3 3.68192 -4.35136 0.00000
1.33888 0.00000 0.00000 ; HA- CA- C- O
2066 2065 2079 2081 3 0.00000 0.00000 0.00000
0.00000 0.00000 0.00000 ; HA- CA- C- N
2067 2065 2079 2080 3 0.00000 0.00000 0.00000
0.00000 0.00000 0.00000 ; CB- CA- C- O
2067 2065 2079 2081 3 2.51040 4.18400 1.67360
-6.69440 0.00000 0.00000 ; CB- CA- C- N
2067 2070 2071 2072 3 11.71520 0.00000 -11.71520
0.00000 0.00000 0.00000 ; CB- CG- ND1- HD1
2067 2070 2071 2073 3 11.71520 0.00000 -11.71520
0.00000 0.00000 0.00000 ; CB- CG- ND1- CE1
2067 2070 2077 2075 3 44.97800 0.00000 -44.97800
0.00000 0.00000 0.00000 ; CB- CG- CD2- NE2
2067 2070 2077 2078 3 44.97800 0.00000 -44.97800
0.00000 0.00000 0.00000 ; CB- CG- CD2- HD2

2068 2067 2065 2079 3 0.65084 1.95253 0.00000
-2.60338 0.00000 0.00000 ; HB2- CB- CA- C
2068 2067 2070 2071 3 0.00000 0.00000 0.00000
0.00000 0.00000 0.00000 ; HB2- CB- CG- ND1
2068 2067 2070 2077 3 0.00000 0.00000 0.00000
0.00000 0.00000 0.00000 ; HB2- CB- CG- CD2
2069 2067 2065 2079 3 0.65084 1.95253 0.00000
-2.60338 0.00000 0.00000 ; HB3- CB- CA- C
2069 2067 2070 2071 3 0.00000 0.00000 0.00000
0.00000 0.00000 0.00000 ; HB3- CB- CG- ND1
2069 2067 2070 2077 3 0.00000 0.00000 0.00000
0.00000 0.00000 0.00000 ; HB3- CB- CG- CD2
2070 2067 2065 2079 3 3.76560 3.34720 -2.87859
-3.66518 0.83680 0.00000 ; CG- CB- CA- C
2070 2071 2073 2074 3 19.45560 0.00000 -19.45560
0.00000 0.00000 0.00000 ; CG- ND1- CE1- HE1
2070 2071 2073 2075 3 19.45560 0.00000 -19.45560
0.00000 0.00000 0.00000 ; CG- ND1- CE1- NE2
2070 2077 2075 2073 3 12.55200 0.00000 -12.55200
0.00000 0.00000 0.00000 ; CG- CD2- NE2- CE1
2070 2077 2075 2076 3 12.55200 0.00000 -12.55200
0.00000 0.00000 0.00000 ; CG- CD2- NE2- HE2
2071 2070 2077 2075 3 44.97800 0.00000 -44.97800
0.00000 0.00000 0.00000 ; ND1- CG- CD2- NE2
2071 2070 2077 2078 3 44.97800 0.00000 -44.97800
0.00000 0.00000 0.00000 ; ND1- CG- CD2- HD2
2071 2073 2075 2076 3 19.45560 0.00000 -19.45560
0.00000 0.00000 0.00000 ; ND1- CE1- NE2- HE2
2071 2073 2075 2077 3 19.45560 0.00000 -19.45560
0.00000 0.00000 0.00000 ; ND1- CE1- NE2- CD2
2072 2071 2070 2077 3 11.71520 0.00000 -11.71520
0.00000 0.00000 0.00000 ; HD1- ND1- CG- CD2
2072 2071 2073 2074 3 19.45560 0.00000 -19.45560
0.00000 0.00000 0.00000 ; HD1- ND1- CE1- HE1
2072 2071 2073 2075 3 19.45560 0.00000 -19.45560
0.00000 0.00000 0.00000 ; HD1- ND1- CE1- NE2
2073 2071 2070 2077 3 11.71520 0.00000 -11.71520
0.00000 0.00000 0.00000 ; CE1- ND1- CG- CD2
2073 2075 2077 2078 3 12.55200 0.00000 -12.55200
0.00000 0.00000 0.00000 ; CE1- NE2- CD2- HD2
2074 2073 2075 2076 3 19.45560 0.00000 -19.45560
0.00000 0.00000 0.00000 ; HE1- CE1- NE2- HE2
2074 2073 2075 2077 3 19.45560 0.00000 -19.45560
0.00000 0.00000 0.00000 ; HE1- CE1- NE2- CD2
2076 2075 2077 2078 3 12.55200 0.00000 -12.55200
0.00000 0.00000 0.00000 ; HE2- NE2- CD2- HD2
2079 2081 2083 2084 3 0.00000 0.00000 0.00000
0.00000 0.00000 0.00000 ; C- N- CA- HA
2079 2081 2083 2085 3 11.71520 1.67360 15.06240 -
13.38880 0.00000 0.00000 ; C- N- CA- CB
2079 2081 2083 2090 3 1.75728 5.27184 2.25936
-7.02912 0.00000 0.00000 ; C- N- CA- C

2080 2079 2081 2082 3 29.28800 -8.36800 -20.92000
0.00000 0.00000 0.00000 ; O- C- N- H
2080 2079 2081 2083 3 20.92000 0.00000 -20.92000
0.00000 0.00000 0.00000 ; O- C- N- CA
2081 2083 2085 2086 3 0.65084 1.95253 0.00000
-2.60338 0.00000 0.00000 ; N- CA- CB- HB2
2081 2083 2085 2087 3 0.65084 1.95253 0.00000
-2.60338 0.00000 0.00000 ; N- CA- CB- HB3
2081 2083 2085 2088 3 7.86174 2.24681 -7.41405
-6.71114 5.35552 0.00000 ; N- CA- CB- OG
2081 2083 2090 2091 3 0.00000 0.00000 0.00000
0.00000 0.00000 0.00000 ; N- CA- C- O
2081 2083 2090 2092 3 17.40544 -5.02080 -13.22144
9.20480 0.00000 0.00000 ; N- CA- C- N
2082 2081 2083 2084 3 0.00000 0.00000 0.00000
0.00000 0.00000 0.00000 ; H- N- CA- HA
2082 2081 2083 2085 3 0.00000 0.00000 0.00000
0.00000 0.00000 0.00000 ; H- N- CA- CB
2082 2081 2083 2090 3 0.00000 0.00000 0.00000
0.00000 0.00000 0.00000 ; H- N- CA- C
2083 2085 2088 2089 3 2.05853 2.46856 3.48109
-4.46851 0.23430 0.00000 ; CA- CB- OG- HG
2083 2090 2092 2093 3 20.92000 0.00000 -20.92000
0.00000 0.00000 0.00000 ; CA- C- N- H
2083 2090 2092 2094 3 20.92000 0.00000 -20.92000
0.00000 0.00000 0.00000 ; CA- C- N- CA
2084 2083 2085 2086 3 0.65084 1.95253 0.00000
-2.60338 0.00000 0.00000 ; HA- CA- CB- HB2
2084 2083 2085 2087 3 0.65084 1.95253 0.00000
-2.60338 0.00000 0.00000 ; HA- CA- CB- HB3
2084 2083 2085 2088 3 1.04600 -1.04600 0.00000
0.00000 0.00000 0.00000 ; HA- CA- CB- OG
2084 2083 2090 2091 3 3.68192 -4.35136 0.00000
1.33888 0.00000 0.00000 ; HA- CA- C- O
2084 2083 2090 2092 3 0.00000 0.00000 0.00000
0.00000 0.00000 0.00000 ; HA- CA- C- N
2085 2083 2090 2091 3 0.00000 0.00000 0.00000
0.00000 0.00000 0.00000 ; CB- CA- C- O
2085 2083 2090 2092 3 2.51040 4.18400 1.67360
-6.69440 0.00000 0.00000 ; CB- CA- C- N
2086 2085 2083 2090 3 0.65084 1.95253 0.00000
-2.60338 0.00000 0.00000 ; HB2- CB- CA- C
2086 2085 2088 2089 3 0.69733 2.09200 0.00000
-2.78933 0.00000 0.00000 ; HB2- CB- OG- HG
2087 2085 2083 2090 3 0.65084 1.95253 0.00000
-2.60338 0.00000 0.00000 ; HB3- CB- CA- C
2087 2085 2088 2089 3 0.69733 2.09200 0.00000
-2.78933 0.00000 0.00000 ; HB3- CB- OG- HG
2088 2085 2083 2090 3 7.34710 7.79898 -6.14211
-6.71114 4.31789 0.00000 ; OG- CB- CA- C
2090 2092 2094 2095 3 0.00000 0.00000 0.00000
0.00000 0.00000 0.00000 ; C- N- CA- HA

2090 2092 2094 2096 3 11.71520 1.67360 15.06240 -
13.38880 0.00000 0.00000 ; C- N- CA- CB
2090 2092 2094 2109 3 1.75728 5.27184 2.25936
-7.02912 0.00000 0.00000 ; C- N- CA- C
2091 2090 2092 2093 3 29.28800 -8.36800 -20.92000
0.00000 0.00000 0.00000 ; O- C- N- H
2091 2090 2092 2094 3 20.92000 0.00000 -20.92000
0.00000 0.00000 0.00000 ; O- C- N- CA
2092 2094 2096 2097 3 0.65084 1.95253 0.00000
-2.60338 0.00000 0.00000 ; N- CA- CB- HB2
2092 2094 2096 2098 3 0.65084 1.95253 0.00000
-2.60338 0.00000 0.00000 ; N- CA- CB- HB3
2092 2094 2096 2099 3 3.79070 1.39746 -4.61077
-2.40998 2.44346 0.00000 ; N- CA- CB- CG
2092 2094 2109 2110 3 0.00000 0.00000 0.00000
0.00000 0.00000 0.00000 ; N- CA- C- O
2092 2094 2109 2111 3 17.40544 -5.02080 -13.22144
9.20480 0.00000 0.00000 ; N- CA- C- N
2093 2092 2094 2095 3 0.00000 0.00000 0.00000
0.00000 0.00000 0.00000 ; H- N- CA- HA
2093 2092 2094 2096 3 0.00000 0.00000 0.00000
0.00000 0.00000 0.00000 ; H- N- CA- CB
2093 2092 2094 2109 3 0.00000 0.00000 0.00000
0.00000 0.00000 0.00000 ; H- N- CA- C
2094 2096 2099 2100 3 0.66944 2.00832 0.00000
-2.67776 0.00000 0.00000 ; CA- CB- CG- HG
2094 2096 2099 2101 3 3.90367 0.19665 -6.21742
-2.37651 5.99149 0.00000 ; CA- CB- CG- CD1
2094 2096 2099 2105 3 3.90367 0.19665 -6.21742
-2.37651 5.99149 0.00000 ; CA- CB- CG- CD2
2094 2109 2111 2112 3 20.92000 0.00000 -20.92000
0.00000 0.00000 0.00000 ; CA- C- N- H
2094 2109 2111 2113 3 20.92000 0.00000 -20.92000
0.00000 0.00000 0.00000 ; CA- C- N- CA
2095 2094 2096 2097 3 0.65084 1.95253 0.00000
-2.60338 0.00000 0.00000 ; HA- CA- CB- HB2
2095 2094 2096 2098 3 0.65084 1.95253 0.00000
-2.60338 0.00000 0.00000 ; HA- CA- CB- HB3
2095 2094 2096 2099 3 0.65084 1.95253 0.00000
-2.60338 0.00000 0.00000 ; HA- CA- CB- CG
2095 2094 2109 2110 3 3.68192 -4.35136 0.00000
1.33888 0.00000 0.00000 ; HA- CA- C- O
2095 2094 2109 2111 3 0.00000 0.00000 0.00000
0.00000 0.00000 0.00000 ; HA- CA- C- N
2096 2094 2109 2110 3 0.00000 0.00000 0.00000
0.00000 0.00000 0.00000 ; CB- CA- C- O
2096 2094 2109 2111 3 2.51040 4.18400 1.67360
-6.69440 0.00000 0.00000 ; CB- CA- C- N
2096 2099 2101 2102 3 0.66944 2.00832 0.00000
-2.67776 0.00000 0.00000 ; CB- CG- CD1- HD11
2096 2099 2101 2103 3 0.66944 2.00832 0.00000
-2.67776 0.00000 0.00000 ; CB- CG- CD1- HD12

2096 2099 2101 2104 3 0.66944 2.00832 0.00000
-2.67776 0.00000 0.00000 ; CB- CG- CD1- HD13
2096 2099 2105 2106 3 0.66944 2.00832 0.00000
-2.67776 0.00000 0.00000 ; CB- CG- CD2- HD21
2096 2099 2105 2107 3 0.66944 2.00832 0.00000
-2.67776 0.00000 0.00000 ; CB- CG- CD2- HD22
2096 2099 2105 2108 3 0.66944 2.00832 0.00000
-2.67776 0.00000 0.00000 ; CB- CG- CD2- HD23
2097 2096 2094 2109 3 0.65084 1.95253 0.00000
-2.60338 0.00000 0.00000 ; HB2- CB- CA- C
2097 2096 2099 2100 3 0.62760 1.88280 0.00000
-2.51040 0.00000 0.00000 ; HB2- CB- CG- HG
2097 2096 2099 2101 3 0.66944 2.00832 0.00000
-2.67776 0.00000 0.00000 ; HB2- CB- CG- CD1
2097 2096 2099 2105 3 0.66944 2.00832 0.00000
-2.67776 0.00000 0.00000 ; HB2- CB- CG- CD2
2098 2096 2094 2109 3 0.65084 1.95253 0.00000
-2.60338 0.00000 0.00000 ; HB3- CB- CA- C
2098 2096 2099 2100 3 0.62760 1.88280 0.00000
-2.51040 0.00000 0.00000 ; HB3- CB- CG- HG
2098 2096 2099 2101 3 0.66944 2.00832 0.00000
-2.67776 0.00000 0.00000 ; HB3- CB- CG- CD1
2098 2096 2099 2105 3 0.66944 2.00832 0.00000
-2.67776 0.00000 0.00000 ; HB3- CB- CG- CD2
2099 2096 2094 2109 3 10.33448 -1.14642 -11.54784
-2.40998 6.35968 0.00000 ; CG- CB- CA- C
2100 2099 2101 2102 3 0.62760 1.88280 0.00000
-2.51040 0.00000 0.00000 ; HG- CG- CD1- HD11
2100 2099 2101 2103 3 0.62760 1.88280 0.00000
-2.51040 0.00000 0.00000 ; HG- CG- CD1- HD12
2100 2099 2101 2104 3 0.62760 1.88280 0.00000
-2.51040 0.00000 0.00000 ; HG- CG- CD1- HD13
2100 2099 2105 2106 3 0.62760 1.88280 0.00000
-2.51040 0.00000 0.00000 ; HG- CG- CD2- HD21
2100 2099 2105 2107 3 0.62760 1.88280 0.00000
-2.51040 0.00000 0.00000 ; HG- CG- CD2- HD22
2100 2099 2105 2108 3 0.62760 1.88280 0.00000
-2.51040 0.00000 0.00000 ; HG- CG- CD2- HD23
2101 2099 2105 2106 3 0.66944 2.00832 0.00000
-2.67776 0.00000 0.00000 ; CD1- CG- CD2- HD21
2101 2099 2105 2107 3 0.66944 2.00832 0.00000
-2.67776 0.00000 0.00000 ; CD1- CG- CD2- HD22
2101 2099 2105 2108 3 0.66944 2.00832 0.00000
-2.67776 0.00000 0.00000 ; CD1- CG- CD2- HD23
2102 2101 2099 2105 3 0.66944 2.00832 0.00000
-2.67776 0.00000 0.00000 ; HD11- CD1- CG- CD2
2103 2101 2099 2105 3 0.66944 2.00832 0.00000
-2.67776 0.00000 0.00000 ; HD12- CD1- CG- CD2
2104 2101 2099 2105 3 0.66944 2.00832 0.00000
-2.67776 0.00000 0.00000 ; HD13- CD1- CG- CD2
2109 2111 2113 2114 3 0.00000 0.00000 0.00000
0.00000 0.00000 0.00000 ; C- N- CA- HA

2109 2111 2113 2115 3 11.71520 1.67360 15.06240 -
13.38880 0.00000 0.00000 ; C- N- CA- CB
2109 2111 2113 2123 3 1.75728 5.27184 2.25936
-7.02912 0.00000 0.00000 ; C- N- CA- C
2110 2109 2111 2112 3 29.28800 -8.36800 -20.92000
0.00000 0.00000 0.00000 ; O- C- N- H
2110 2109 2111 2113 3 20.92000 0.00000 -20.92000
0.00000 0.00000 0.00000 ; O- C- N- CA
2111 2113 2115 2116 3 0.65084 1.95253 0.00000
-2.60338 0.00000 0.00000 ; N- CA- CB- HB2
2111 2113 2115 2117 3 0.65084 1.95253 0.00000
-2.60338 0.00000 0.00000 ; N- CA- CB- HB3
2111 2113 2115 2118 3 5.99567 3.29281 -4.46014
-0.55229 1.97485 0.00000 ; N- CA- CB- CG
2111 2113 2123 2124 3 0.00000 0.00000 0.00000
0.00000 0.00000 0.00000 ; N- CA- C- O
2111 2113 2123 2125 3 17.40544 -5.02080 -13.22144
9.20480 0.00000 0.00000 ; N- CA- C- N
2112 2111 2113 2114 3 0.00000 0.00000 0.00000
0.00000 0.00000 0.00000 ; H- N- CA- HA
2112 2111 2113 2115 3 0.00000 0.00000 0.00000
0.00000 0.00000 0.00000 ; H- N- CA- CB
2112 2111 2113 2123 3 0.00000 0.00000 0.00000
0.00000 0.00000 0.00000 ; H- N- CA- C
2113 2115 2118 2119 3 0.00000 0.00000 0.00000
0.00000 0.00000 0.00000 ; CA- CB- CG- OD1
2113 2115 2118 2120 3 8.84916 -0.31380 -4.32626
5.03754 0.26778 0.00000 ; CA- CB- CG- ND2
2113 2123 2125 2126 3 20.92000 0.00000 -20.92000
0.00000 0.00000 0.00000 ; CA- C- N- H
2113 2123 2125 2127 3 20.92000 0.00000 -20.92000
0.00000 0.00000 0.00000 ; CA- C- N- CA
2114 2113 2115 2116 3 0.65084 1.95253 0.00000
-2.60338 0.00000 0.00000 ; HA- CA- CB- HB2
2114 2113 2115 2117 3 0.65084 1.95253 0.00000
-2.60338 0.00000 0.00000 ; HA- CA- CB- HB3
2114 2113 2115 2118 3 0.65084 1.95253 0.00000
-2.60338 0.00000 0.00000 ; HA- CA- CB- CG
2114 2113 2123 2124 3 3.68192 -4.35136 0.00000
1.33888 0.00000 0.00000 ; HA- CA- C- O
2114 2113 2123 2125 3 0.00000 0.00000 0.00000
0.00000 0.00000 0.00000 ; HA- CA- C- N
2115 2113 2123 2124 3 0.00000 0.00000 0.00000
0.00000 0.00000 0.00000 ; CB- CA- C- O
2115 2113 2123 2125 3 2.51040 4.18400 1.67360
-6.69440 0.00000 0.00000 ; CB- CA- C- N
2115 2118 2120 2121 3 20.92000 0.00000 -20.92000
0.00000 0.00000 0.00000 ; CB- CG- ND2- HD21
2115 2118 2120 2122 3 20.92000 0.00000 -20.92000
0.00000 0.00000 0.00000 ; CB- CG- ND2- HD22
2116 2115 2113 2123 3 0.65084 1.95253 0.00000
-2.60338 0.00000 0.00000 ; HB2- CB- CA- C

2116 2115 2118 2119 3 3.68192 -4.35136 0.00000
1.33888 0.00000 0.00000 ; HB2- CB- CG- OD1
2116 2115 2118 2120 3 0.00000 0.00000 0.00000
0.00000 0.00000 0.00000 ; HB2- CB- CG- ND2
2117 2115 2113 2123 3 0.65084 1.95253 0.00000
-2.60338 0.00000 0.00000 ; HB3- CB- CA- C
2117 2115 2118 2119 3 3.68192 -4.35136 0.00000
1.33888 0.00000 0.00000 ; HB3- CB- CG- OD1
2117 2115 2118 2120 3 0.00000 0.00000 0.00000
0.00000 0.00000 0.00000 ; HB3- CB- CG- ND2
2118 2115 2113 2123 3 7.94542 -3.96225 -6.11701
-0.55229 3.58150 0.00000 ; CG- CB- CA- C
2119 2118 2120 2121 3 29.28800 -8.36800 -20.92000
0.00000 0.00000 0.00000 ; OD1- CG- ND2- HD21
2119 2118 2120 2122 3 29.28800 -8.36800 -20.92000
0.00000 0.00000 0.00000 ; OD1- CG- ND2- HD22
2123 2125 2127 2128 3 0.00000 0.00000 0.00000
0.00000 0.00000 0.00000 ; C- N- CA- HA
2123 2125 2127 2129 3 11.71520 1.67360 15.06240 -
13.38880 0.00000 0.00000 ; C- N- CA- CB
2123 2125 2127 2142 3 1.75728 5.27184 2.25936
-7.02912 0.00000 0.00000 ; C- N- CA- C
2124 2123 2125 2126 3 29.28800 -8.36800 -20.92000
0.00000 0.00000 0.00000 ; O- C- N- H
2124 2123 2125 2127 3 20.92000 0.00000 -20.92000
0.00000 0.00000 0.00000 ; O- C- N- CA
2125 2127 2129 2130 3 0.65084 1.95253 0.00000
-2.60338 0.00000 0.00000 ; N- CA- CB- HB
2125 2127 2129 2131 3 3.83673 0.44769 -1.77402
-2.47693 -0.03347 0.00000 ; N- CA- CB- CG2
2125 2127 2129 2135 3 2.97482 0.12134 2.04179
-1.89117 -3.24678 0.00000 ; N- CA- CB- CG1
2125 2127 2142 2143 3 0.00000 0.00000 0.00000
0.00000 0.00000 0.00000 ; N- CA- C- O
2125 2127 2142 2144 3 17.40544 -5.02080 -13.22144
9.20480 0.00000 0.00000 ; N- CA- C- N
2126 2125 2127 2128 3 0.00000 0.00000 0.00000
0.00000 0.00000 0.00000 ; H- N- CA- HA
2126 2125 2127 2129 3 0.00000 0.00000 0.00000
0.00000 0.00000 0.00000 ; H- N- CA- CB
2126 2125 2127 2142 3 0.00000 0.00000 0.00000
0.00000 0.00000 0.00000 ; H- N- CA- C
2127 2129 2131 2132 3 0.66944 2.00832 0.00000
-2.67776 0.00000 0.00000 ; CA- CB- CG2- HG21
2127 2129 2131 2133 3 0.66944 2.00832 0.00000
-2.67776 0.00000 0.00000 ; CA- CB- CG2- HG22
2127 2129 2131 2134 3 0.66944 2.00832 0.00000
-2.67776 0.00000 0.00000 ; CA- CB- CG2- HG23
2127 2129 2135 2136 3 0.66944 2.00832 0.00000
-2.67776 0.00000 0.00000 ; CA- CB- CG1- HG12
2127 2129 2135 2137 3 0.66944 2.00832 0.00000
-2.67776 0.00000 0.00000 ; CA- CB- CG1- HG13

```
  2127   2129   2135   2138       3    4.24258   -0.52718   -7.25506
-1.79075    7.69856    0.00000 ;     CA-    CB-   CG1-   CD1
  2127   2142   2144   2145       3   20.92000    0.00000  -20.92000
0.00000    0.00000    0.00000 ;     CA-     C-     N-     H
  2127   2142   2144   2146       3   20.92000    0.00000  -20.92000
0.00000    0.00000    0.00000 ;     CA-     C-     N-    CA
  2128   2127   2129   2130       3    0.65084    1.95253    0.00000
-2.60338    0.00000    0.00000 ;     HA-    CA-    CB-    HB
  2128   2127   2129   2131       3    0.65084    1.95253    0.00000
-2.60338    0.00000    0.00000 ;     HA-    CA-    CB-   CG2
  2128   2127   2129   2135       3    0.65084    1.95253    0.00000
-2.60338    0.00000    0.00000 ;     HA-    CA-    CB-   CG1
  2128   2127   2142   2143       3    3.68192   -4.35136    0.00000
1.33888    0.00000    0.00000 ;     HA-    CA-     C-     O
  2128   2127   2142   2144       3    0.00000    0.00000    0.00000
0.00000    0.00000    0.00000 ;     HA-    CA-     C-     N
  2129   2127   2142   2143       3    0.00000    0.00000    0.00000
0.00000    0.00000    0.00000 ;     CB-    CA-     C-     O
  2129   2127   2142   2144       3    2.51040    4.18400    1.67360
-6.69440    0.00000    0.00000 ;     CB-    CA-     C-     N
  2129   2135   2138   2139       3    0.66944    2.00832    0.00000
-2.67776    0.00000    0.00000 ;     CB-   CG1-   CD1-  HD11
  2129   2135   2138   2140       3    0.66944    2.00832    0.00000
-2.67776    0.00000    0.00000 ;     CB-   CG1-   CD1-  HD12
  2129   2135   2138   2141       3    0.66944    2.00832    0.00000
-2.67776    0.00000    0.00000 ;     CB-   CG1-   CD1-  HD13
  2130   2129   2127   2142       3    0.65084    1.95253    0.00000
-2.60338    0.00000    0.00000 ;     HB-    CB-    CA-     C
  2130   2129   2131   2132       3    0.62760    1.88280    0.00000
-2.51040    0.00000    0.00000 ;     HB-    CB-   CG2-  HG21
  2130   2129   2131   2133       3    0.62760    1.88280    0.00000
-2.51040    0.00000    0.00000 ;     HB-    CB-   CG2-  HG22
  2130   2129   2131   2134       3    0.62760    1.88280    0.00000
-2.51040    0.00000    0.00000 ;     HB-    CB-   CG2-  HG23
  2130   2129   2135   2136       3    0.62760    1.88280    0.00000
-2.51040    0.00000    0.00000 ;     HB-    CB-   CG1-  HG12
  2130   2129   2135   2137       3    0.62760    1.88280    0.00000
-2.51040    0.00000    0.00000 ;     HB-    CB-   CG1-  HG13
  2130   2129   2135   2138       3    0.66944    2.00832    0.00000
-2.67776    0.00000    0.00000 ;     HB-    CB-   CG1-   CD1
  2131   2129   2127   2142       3    5.67350    3.55640   -6.16722
-2.47693    3.74886    0.00000 ;    CG2-    CB-    CA-     C
  2131   2129   2135   2136       3    0.66944    2.00832    0.00000
-2.67776    0.00000    0.00000 ;    CG2-    CB-   CG1-  HG12
  2131   2129   2135   2137       3    0.66944    2.00832    0.00000
-2.67776    0.00000    0.00000 ;    CG2-    CB-   CG1-  HG13
  2131   2129   2135   2138       3    3.81162    0.49790   -8.14206
-1.79075    7.49773    0.00000 ;    CG2-    CB-   CG1-   CD1
  2132   2131   2129   2135       3    0.66944    2.00832    0.00000
-2.67776    0.00000    0.00000 ;   HG21-   CG2-    CB-   CG1
  2133   2131   2129   2135       3    0.66944    2.00832    0.00000
-2.67776    0.00000    0.00000 ;   HG22-   CG2-    CB-   CG1
```

2134 2131 2129 2135 3 0.66944 2.00832 0.00000
-2.67776 0.00000 0.00000 ; HG23- CG2- CB- CG1
2135 2129 2127 2142 3 8.26340 0.74057 -9.99976
-1.89117 3.84928 0.00000 ; CG1- CB- CA- C
2136 2135 2138 2139 3 0.62760 1.88280 0.00000
-2.51040 0.00000 0.00000 ; HG12- CG1- CD1- HD11
2136 2135 2138 2140 3 0.62760 1.88280 0.00000
-2.51040 0.00000 0.00000 ; HG12- CG1- CD1- HD12
2136 2135 2138 2141 3 0.62760 1.88280 0.00000
-2.51040 0.00000 0.00000 ; HG12- CG1- CD1- HD13
2137 2135 2138 2139 3 0.62760 1.88280 0.00000
-2.51040 0.00000 0.00000 ; HG13- CG1- CD1- HD11
2137 2135 2138 2140 3 0.62760 1.88280 0.00000
-2.51040 0.00000 0.00000 ; HG13- CG1- CD1- HD12
2137 2135 2138 2141 3 0.62760 1.88280 0.00000
-2.51040 0.00000 0.00000 ; HG13- CG1- CD1- HD13
2142 2144 2146 2147 3 0.00000 0.00000 0.00000
0.00000 0.00000 0.00000 ; C- N- CA- HA
2142 2144 2146 2148 3 11.71520 1.67360 15.06240 -
13.38880 0.00000 0.00000 ; C- N- CA- CB
2142 2144 2146 2158 3 1.75728 5.27184 2.25936
-7.02912 0.00000 0.00000 ; C- N- CA- C
2143 2142 2144 2145 3 29.28800 -8.36800 -20.92000
0.00000 0.00000 0.00000 ; O- C- N- H
2143 2142 2144 2146 3 20.92000 0.00000 -20.92000
0.00000 0.00000 0.00000 ; O- C- N- CA
2144 2146 2148 2149 3 0.65084 1.95253 0.00000
-2.60338 0.00000 0.00000 ; N- CA- CB- HB
2144 2146 2148 2150 3 3.83673 0.44769 -1.77402
-2.47693 -0.03347 0.00000 ; N- CA- CB- CG1
2144 2146 2148 2154 3 3.83673 0.44769 -1.77402
-2.47693 -0.03347 0.00000 ; N- CA- CB- CG2
2144 2146 2158 2159 3 0.00000 0.00000 0.00000
0.00000 0.00000 0.00000 ; N- CA- C- O
2144 2146 2158 2160 3 17.40544 -5.02080 -13.22144
9.20480 0.00000 0.00000 ; N- CA- C- N
2145 2144 2146 2147 3 0.00000 0.00000 0.00000
0.00000 0.00000 0.00000 ; H- N- CA- HA
2145 2144 2146 2148 3 0.00000 0.00000 0.00000
0.00000 0.00000 0.00000 ; H- N- CA- CB
2145 2144 2146 2158 3 0.00000 0.00000 0.00000
0.00000 0.00000 0.00000 ; H- N- CA- C
2146 2148 2150 2151 3 0.66944 2.00832 0.00000
-2.67776 0.00000 0.00000 ; CA- CB- CG1- HG11
2146 2148 2150 2152 3 0.66944 2.00832 0.00000
-2.67776 0.00000 0.00000 ; CA- CB- CG1- HG12
2146 2148 2150 2153 3 0.66944 2.00832 0.00000
-2.67776 0.00000 0.00000 ; CA- CB- CG1- HG13
2146 2148 2154 2155 3 0.66944 2.00832 0.00000
-2.67776 0.00000 0.00000 ; CA- CB- CG2- HG21
2146 2148 2154 2156 3 0.66944 2.00832 0.00000
-2.67776 0.00000 0.00000 ; CA- CB- CG2- HG22

2146 2148 2154 2157 3 0.66944 2.00832 0.00000
-2.67776 0.00000 0.00000 ; CA- CB- CG2- HG23
2146 2158 2160 2161 3 20.92000 0.00000 -20.92000
0.00000 0.00000 0.00000 ; CA- C- N- H
2146 2158 2160 2162 3 20.92000 0.00000 -20.92000
0.00000 0.00000 0.00000 ; CA- C- N- CA
2147 2146 2148 2149 3 0.65084 1.95253 0.00000
-2.60338 0.00000 0.00000 ; HA- CA- CB- HB
2147 2146 2148 2150 3 0.65084 1.95253 0.00000
-2.60338 0.00000 0.00000 ; HA- CA- CB- CG1
2147 2146 2148 2154 3 0.65084 1.95253 0.00000
-2.60338 0.00000 0.00000 ; HA- CA- CB- CG2
2147 2146 2158 2159 3 3.68192 -4.35136 0.00000
1.33888 0.00000 0.00000 ; HA- CA- C- O
2147 2146 2158 2160 3 0.00000 0.00000 0.00000
0.00000 0.00000 0.00000 ; HA- CA- C- N
2148 2146 2158 2159 3 0.00000 0.00000 0.00000
0.00000 0.00000 0.00000 ; CB- CA- C- O
2148 2146 2158 2160 3 2.51040 4.18400 1.67360
-6.69440 0.00000 0.00000 ; CB- CA- C- N
2149 2148 2146 2158 3 0.65084 1.95253 0.00000
-2.60338 0.00000 0.00000 ; HB- CB- CA- C
2149 2148 2150 2151 3 0.62760 1.88280 0.00000
-2.51040 0.00000 0.00000 ; HB- CB- CG1- HG11
2149 2148 2150 2152 3 0.62760 1.88280 0.00000
-2.51040 0.00000 0.00000 ; HB- CB- CG1- HG12
2149 2148 2150 2153 3 0.62760 1.88280 0.00000
-2.51040 0.00000 0.00000 ; HB- CB- CG1- HG13
2149 2148 2154 2155 3 0.62760 1.88280 0.00000
-2.51040 0.00000 0.00000 ; HB- CB- CG2- HG21
2149 2148 2154 2156 3 0.62760 1.88280 0.00000
-2.51040 0.00000 0.00000 ; HB- CB- CG2- HG22
2149 2148 2154 2157 3 0.62760 1.88280 0.00000
-2.51040 0.00000 0.00000 ; HB- CB- CG2- HG23
2150 2148 2146 2158 3 5.67350 3.55640 -6.16722
-2.47693 3.74886 0.00000 ; CG1- CB- CA- C
2150 2148 2154 2155 3 0.66944 2.00832 0.00000
-2.67776 0.00000 0.00000 ; CG1- CB- CG2- HG21
2150 2148 2154 2156 3 0.66944 2.00832 0.00000
-2.67776 0.00000 0.00000 ; CG1- CB- CG2- HG22
2150 2148 2154 2157 3 0.66944 2.00832 0.00000
-2.67776 0.00000 0.00000 ; CG1- CB- CG2- HG23
2151 2150 2148 2154 3 0.66944 2.00832 0.00000
-2.67776 0.00000 0.00000 ; HG11- CG1- CB- CG2
2152 2150 2148 2154 3 0.66944 2.00832 0.00000
-2.67776 0.00000 0.00000 ; HG12- CG1- CB- CG2
2153 2150 2148 2154 3 0.66944 2.00832 0.00000
-2.67776 0.00000 0.00000 ; HG13- CG1- CB- CG2
2154 2148 2146 2158 3 5.67350 3.55640 -6.16722
-2.47693 3.74886 0.00000 ; CG2- CB- CA- C
2158 2160 2162 2163 3 0.00000 0.00000 0.00000
0.00000 0.00000 0.00000 ; C- N- CA- HA

2158 2160 2162 2164 3 11.71520 1.67360 15.06240 -
13.38880 0.00000 0.00000 ; C- N- CA- CB
2158 2160 2162 2175 3 1.75728 5.27184 2.25936
-7.02912 0.00000 0.00000 ; C- N- CA- C
2159 2158 2160 2161 3 29.28800 -8.36800 -20.92000
0.00000 0.00000 0.00000 ; O- C- N- H
2159 2158 2160 2162 3 20.92000 0.00000 -20.92000
0.00000 0.00000 0.00000 ; O- C- N- CA
2160 2162 2164 2165 3 0.65084 1.95253 0.00000
-2.60338 0.00000 0.00000 ; N- CA- CB- HB2
2160 2162 2164 2166 3 0.65084 1.95253 0.00000
-2.60338 0.00000 0.00000 ; N- CA- CB- HB3
2160 2162 2164 2167 3 4.79068 4.02919 -4.82834
-3.66518 2.97901 0.00000 ; N- CA- CB- CG
2160 2162 2175 2176 3 0.00000 0.00000 0.00000
0.00000 0.00000 0.00000 ; N- CA- C- O
2160 2162 2175 2177 3 17.40544 -5.02080 -13.22144
9.20480 0.00000 0.00000 ; N- CA- C- N
2161 2160 2162 2163 3 0.00000 0.00000 0.00000
0.00000 0.00000 0.00000 ; H- N- CA- HA
2161 2160 2162 2164 3 0.00000 0.00000 0.00000
0.00000 0.00000 0.00000 ; H- N- CA- CB
2161 2160 2162 2175 3 0.00000 0.00000 0.00000
0.00000 0.00000 0.00000 ; H- N- CA- C
2162 2164 2167 2168 3 5.98312 6.40152 3.28026 -
12.38464 -1.57318 0.00000 ; CA- CB- CG- ND1
2162 2164 2167 2173 3 0.00000 0.00000 0.00000
0.00000 0.00000 0.00000 ; CA- CB- CG- CD2
2162 2175 2177 2178 3 20.92000 0.00000 -20.92000
0.00000 0.00000 0.00000 ; CA- C- N- H
2162 2175 2177 2179 3 20.92000 0.00000 -20.92000
0.00000 0.00000 0.00000 ; CA- C- N- CA
2163 2162 2164 2165 3 0.65084 1.95253 0.00000
-2.60338 0.00000 0.00000 ; HA- CA- CB- HB2
2163 2162 2164 2166 3 0.65084 1.95253 0.00000
-2.60338 0.00000 0.00000 ; HA- CA- CB- HB3
2163 2162 2164 2167 3 0.65084 1.95253 0.00000
-2.60338 0.00000 0.00000 ; HA- CA- CB- CG
2163 2162 2175 2176 3 3.68192 -4.35136 0.00000
1.33888 0.00000 0.00000 ; HA- CA- C- O
2163 2162 2175 2177 3 0.00000 0.00000 0.00000
0.00000 0.00000 0.00000 ; HA- CA- C- N
2164 2162 2175 2176 3 0.00000 0.00000 0.00000
0.00000 0.00000 0.00000 ; CB- CA- C- O
2164 2162 2175 2177 3 2.51040 4.18400 1.67360
-6.69440 0.00000 0.00000 ; CB- CA- C- N
2164 2167 2168 2169 3 20.08320 0.00000 -20.08320
0.00000 0.00000 0.00000 ; CB- CG- ND1- CE1
2164 2167 2173 2171 3 44.97800 0.00000 -44.97800
0.00000 0.00000 0.00000 ; CB- CG- CD2- NE2
2164 2167 2173 2174 3 44.97800 0.00000 -44.97800
0.00000 0.00000 0.00000 ; CB- CG- CD2- HD2

2165 2164 2162 2175 3 0.65084 1.95253 0.00000
-2.60338 0.00000 0.00000 ; HB2- CB- CA- C
2165 2164 2167 2168 3 0.00000 0.00000 0.00000
0.00000 0.00000 0.00000 ; HB2- CB- CG- ND1
2165 2164 2167 2173 3 0.00000 0.00000 0.00000
0.00000 0.00000 0.00000 ; HB2- CB- CG- CD2
2166 2164 2162 2175 3 0.65084 1.95253 0.00000
-2.60338 0.00000 0.00000 ; HB3- CB- CA- C
2166 2164 2167 2168 3 0.00000 0.00000 0.00000
0.00000 0.00000 0.00000 ; HB3- CB- CG- ND1
2166 2164 2167 2173 3 0.00000 0.00000 0.00000
0.00000 0.00000 0.00000 ; HB3- CB- CG- CD2
2167 2164 2162 2175 3 3.76560 3.34720 -2.87859
-3.66518 0.83680 0.00000 ; CG- CB- CA- C
2167 2168 2169 2170 3 41.84000 0.00000 -41.84000
0.00000 0.00000 0.00000 ; CG- ND1- CE1- HE1
2167 2168 2169 2171 3 41.84000 0.00000 -41.84000
0.00000 0.00000 0.00000 ; CG- ND1- CE1- NE2
2167 2173 2171 2169 3 12.55200 0.00000 -12.55200
0.00000 0.00000 0.00000 ; CG- CD2- NE2- CE1
2167 2173 2171 2172 3 12.55200 0.00000 -12.55200
0.00000 0.00000 0.00000 ; CG- CD2- NE2- HE2
2168 2167 2173 2171 3 44.97800 0.00000 -44.97800
0.00000 0.00000 0.00000 ; ND1- CG- CD2- NE2
2168 2167 2173 2174 3 44.97800 0.00000 -44.97800
0.00000 0.00000 0.00000 ; ND1- CG- CD2- HD2
2168 2169 2171 2172 3 19.45560 0.00000 -19.45560
0.00000 0.00000 0.00000 ; ND1- CE1- NE2- HE2
2168 2169 2171 2173 3 19.45560 0.00000 -19.45560
0.00000 0.00000 0.00000 ; ND1- CE1- NE2- CD2
2169 2168 2167 2173 3 20.08320 0.00000 -20.08320
0.00000 0.00000 0.00000 ; CE1- ND1- CG- CD2
2169 2171 2173 2174 3 12.55200 0.00000 -12.55200
0.00000 0.00000 0.00000 ; CE1- NE2- CD2- HD2
2170 2169 2171 2172 3 19.45560 0.00000 -19.45560
0.00000 0.00000 0.00000 ; HE1- CE1- NE2- HE2
2170 2169 2171 2173 3 19.45560 0.00000 -19.45560
0.00000 0.00000 0.00000 ; HE1- CE1- NE2- CD2
2172 2171 2173 2174 3 12.55200 0.00000 -12.55200
0.00000 0.00000 0.00000 ; HE2- NE2- CD2- HD2
2175 2177 2179 2180 3 0.00000 0.00000 0.00000
0.00000 0.00000 0.00000 ; C- N- CA- HA
2175 2177 2179 2181 3 11.71520 1.67360 15.06240 -
13.38880 0.00000 0.00000 ; C- N- CA- CB
2175 2177 2179 2199 3 1.75728 5.27184 2.25936
-7.02912 0.00000 0.00000 ; C- N- CA- C
2176 2175 2177 2178 3 29.28800 -8.36800 -20.92000
0.00000 0.00000 0.00000 ; O- C- N- H
2176 2175 2177 2179 3 20.92000 0.00000 -20.92000
0.00000 0.00000 0.00000 ; O- C- N- CA
2177 2179 2181 2182 3 0.65084 1.95253 0.00000
-2.60338 0.00000 0.00000 ; N- CA- CB- HB2

```
  2177   2179   2181   2183      3    0.65084    1.95253    0.00000
-2.60338    0.00000    0.00000 ;     N-    CA-    CB-   HB3
  2177   2179   2181   2184      3    0.65084    1.95253    0.00000
-2.60338    0.00000    0.00000 ;     N-    CA-    CB-    CG
  2177   2179   2199   2200      3    0.00000    0.00000    0.00000
0.00000    0.00000    0.00000 ;     N-    CA-     C-     O
  2177   2179   2199   2201      3   17.40544   -5.02080  -13.22144
9.20480    0.00000    0.00000 ;     N-    CA-     C-     N
  2178   2177   2179   2180      3    0.00000    0.00000    0.00000
0.00000    0.00000    0.00000 ;     H-     N-    CA-    HA
  2178   2177   2179   2181      3    0.00000    0.00000    0.00000
0.00000    0.00000    0.00000 ;     H-     N-    CA-    CB
  2178   2177   2179   2199      3    0.00000    0.00000    0.00000
0.00000    0.00000    0.00000 ;     H-     N-    CA-     C
  2179   2181   2184   2185      3    0.66944    2.00832    0.00000
-2.67776    0.00000    0.00000 ;    CA-    CB-    CG-   HG2
  2179   2181   2184   2186      3    0.66944    2.00832    0.00000
-2.67776    0.00000    0.00000 ;    CA-    CB-    CG-   HG3
  2179   2181   2184   2187      3    3.68192    3.09616   -2.09200
-3.01248    0.00000    0.00000 ;    CA-    CB-    CG-    CD
  2179   2199   2201   2202      3   20.92000    0.00000  -20.92000
0.00000    0.00000    0.00000 ;    CA-     C-     N-     H
  2179   2199   2201   2203      3   20.92000    0.00000  -20.92000
0.00000    0.00000    0.00000 ;    CA-     C-     N-    CA
  2180   2179   2181   2182      3    0.65084    1.95253    0.00000
-2.60338    0.00000    0.00000 ;    HA-    CA-    CB-   HB2
  2180   2179   2181   2183      3    0.65084    1.95253    0.00000
-2.60338    0.00000    0.00000 ;    HA-    CA-    CB-   HB3
  2180   2179   2181   2184      3    0.65084    1.95253    0.00000
-2.60338    0.00000    0.00000 ;    HA-    CA-    CB-    CG
  2180   2179   2199   2200      3    3.68192   -4.35136    0.00000
1.33888    0.00000    0.00000 ;    HA-    CA-     C-     O
  2180   2179   2199   2201      3    0.00000    0.00000    0.00000
0.00000    0.00000    0.00000 ;    HA-    CA-     C-     N
  2181   2179   2199   2200      3    0.00000    0.00000    0.00000
0.00000    0.00000    0.00000 ;    CB-    CA-     C-     O
  2181   2179   2199   2201      3    2.51040    4.18400    1.67360
-6.69440    0.00000    0.00000 ;    CB-    CA-     C-     N
  2181   2184   2187   2188      3    0.65084    1.95253    0.00000
-2.60338    0.00000    0.00000 ;    CB-    CG-    CD-   HD2
  2181   2184   2187   2189      3    0.65084    1.95253    0.00000
-2.60338    0.00000    0.00000 ;    CB-    CG-    CD-   HD3
  2181   2184   2187   2190      3    0.65084    1.95253    0.00000
-2.60338    0.00000    0.00000 ;    CB-    CG-    CD-    NE
  2182   2181   2179   2199      3    0.65084    1.95253    0.00000
-2.60338    0.00000    0.00000 ;   HB2-    CB-    CA-     C
  2182   2181   2184   2185      3    0.62760    1.88280    0.00000
-2.51040    0.00000    0.00000 ;   HB2-    CB-    CG-   HG2
  2182   2181   2184   2186      3    0.62760    1.88280    0.00000
-2.51040    0.00000    0.00000 ;   HB2-    CB-    CG-   HG3
  2182   2181   2184   2187      3    0.66944    2.00832    0.00000
-2.67776    0.00000    0.00000 ;   HB2-    CB-    CG-    CD
```

2183 2181 2179 2199 3 0.65084 1.95253 0.00000
-2.60338 0.00000 0.00000 ; HB3- CB- CA- C
2183 2181 2184 2185 3 0.62760 1.88280 0.00000
-2.51040 0.00000 0.00000 ; HB3- CB- CG- HG2
2183 2181 2184 2186 3 0.62760 1.88280 0.00000
-2.51040 0.00000 0.00000 ; HB3- CB- CG- HG3
2183 2181 2184 2187 3 0.66944 2.00832 0.00000
-2.67776 0.00000 0.00000 ; HB3- CB- CG- CD
2184 2181 2179 2199 3 0.65084 1.95253 0.00000
-2.60338 0.00000 0.00000 ; CG- CB- CA- C
2184 2187 2190 2191 3 0.00000 0.00000 0.00000
0.00000 0.00000 0.00000 ; CG- CD- NE- HE
2184 2187 2190 2192 3 0.00000 0.00000 0.00000
0.00000 0.00000 0.00000 ; CG- CD- NE- CZ
2185 2184 2187 2188 3 0.65084 1.95253 0.00000
-2.60338 0.00000 0.00000 ; HG2- CG- CD- HD2
2185 2184 2187 2189 3 0.65084 1.95253 0.00000
-2.60338 0.00000 0.00000 ; HG2- CG- CD- HD3
2185 2184 2187 2190 3 0.65084 1.95253 0.00000
-2.60338 0.00000 0.00000 ; HG2- CG- CD- NE
2186 2184 2187 2188 3 0.65084 1.95253 0.00000
-2.60338 0.00000 0.00000 ; HG3- CG- CD- HD2
2186 2184 2187 2189 3 0.65084 1.95253 0.00000
-2.60338 0.00000 0.00000 ; HG3- CG- CD- HD3
2186 2184 2187 2190 3 0.65084 1.95253 0.00000
-2.60338 0.00000 0.00000 ; HG3- CG- CD- NE
2187 2190 2192 2193 3 20.08320 0.00000 -20.08320
0.00000 0.00000 0.00000 ; CD- NE- CZ- NH1
2187 2190 2192 2196 3 20.08320 0.00000 -20.08320
0.00000 0.00000 0.00000 ; CD- NE- CZ- NH2
2188 2187 2190 2191 3 0.00000 0.00000 0.00000
0.00000 0.00000 0.00000 ; HD2- CD- NE- HE
2188 2187 2190 2192 3 0.00000 0.00000 0.00000
0.00000 0.00000 0.00000 ; HD2- CD- NE- CZ
2189 2187 2190 2191 3 0.00000 0.00000 0.00000
0.00000 0.00000 0.00000 ; HD3- CD- NE- HE
2189 2187 2190 2192 3 0.00000 0.00000 0.00000
0.00000 0.00000 0.00000 ; HD3- CD- NE- CZ
2190 2192 2193 2194 3 20.08320 0.00000 -20.08320
0.00000 0.00000 0.00000 ; NE- CZ- NH1- HH11
2190 2192 2193 2195 3 20.08320 0.00000 -20.08320
0.00000 0.00000 0.00000 ; NE- CZ- NH1- HH12
2190 2192 2196 2197 3 20.08320 0.00000 -20.08320
0.00000 0.00000 0.00000 ; NE- CZ- NH2- HH21
2190 2192 2196 2198 3 20.08320 0.00000 -20.08320
0.00000 0.00000 0.00000 ; NE- CZ- NH2- HH22
2191 2190 2192 2193 3 20.08320 0.00000 -20.08320
0.00000 0.00000 0.00000 ; HE- NE- CZ- NH1
2191 2190 2192 2196 3 20.08320 0.00000 -20.08320
0.00000 0.00000 0.00000 ; HE- NE- CZ- NH2
2193 2192 2196 2197 3 20.08320 0.00000 -20.08320
0.00000 0.00000 0.00000 ; NH1- CZ- NH2- HH21

2193 2192 2196 2198 3 20.08320 0.00000 -20.08320
0.00000 0.00000 0.00000 ; NH1- CZ- NH2- HH22
2194 2193 2192 2196 3 20.08320 0.00000 -20.08320
0.00000 0.00000 0.00000 ; HH11- NH1- CZ- NH2
2195 2193 2192 2196 3 20.08320 0.00000 -20.08320
0.00000 0.00000 0.00000 ; HH12- NH1- CZ- NH2
2199 2201 2203 2204 3 0.00000 0.00000 0.00000
0.00000 0.00000 0.00000 ; C- N- CA- HA
2199 2201 2203 2205 3 11.71520 1.67360 15.06240 -
13.38880 0.00000 0.00000 ; C- N- CA- CB
2199 2201 2203 2211 3 1.75728 5.27184 2.25936
-7.02912 0.00000 0.00000 ; C- N- CA- C
2200 2199 2201 2202 3 29.28800 -8.36800 -20.92000
0.00000 0.00000 0.00000 ; O- C- N- H
2200 2199 2201 2203 3 20.92000 0.00000 -20.92000
0.00000 0.00000 0.00000 ; O- C- N- CA
2201 2203 2205 2206 3 0.65084 1.95253 0.00000
-2.60338 0.00000 0.00000 ; N- CA- CB- HB2
2201 2203 2205 2207 3 0.65084 1.95253 0.00000
-2.60338 0.00000 0.00000 ; N- CA- CB- HB3
2201 2203 2205 2208 3 15.41386 9.74035 -8.39310
-0.97069 2.97901 0.00000 ; N- CA- CB- CG
2201 2203 2211 2212 3 0.00000 0.00000 0.00000
0.00000 0.00000 0.00000 ; N- CA- C- O
2201 2203 2211 2213 3 17.40544 -5.02080 -13.22144
9.20480 0.00000 0.00000 ; N- CA- C- N
2202 2201 2203 2204 3 0.00000 0.00000 0.00000
0.00000 0.00000 0.00000 ; H- N- CA- HA
2202 2201 2203 2205 3 0.00000 0.00000 0.00000
0.00000 0.00000 0.00000 ; H- N- CA- CB
2202 2201 2203 2211 3 0.00000 0.00000 0.00000
0.00000 0.00000 0.00000 ; H- N- CA- C
2203 2205 2208 2209 3 6.43499 0.00000 -5.39736
0.00000 -1.03763 0.00000 ; CA- CB- CG- OD1
2203 2205 2208 2210 3 6.43499 0.00000 -5.39736
0.00000 -1.03763 0.00000 ; CA- CB- CG- OD2
2203 2211 2213 2214 3 20.92000 0.00000 -20.92000
0.00000 0.00000 0.00000 ; CA- C- N- H
2203 2211 2213 2215 3 20.92000 0.00000 -20.92000
0.00000 0.00000 0.00000 ; CA- C- N- CA
2204 2203 2205 2206 3 0.65084 1.95253 0.00000
-2.60338 0.00000 0.00000 ; HA- CA- CB- HB2
2204 2203 2205 2207 3 0.65084 1.95253 0.00000
-2.60338 0.00000 0.00000 ; HA- CA- CB- HB3
2204 2203 2205 2208 3 0.65084 1.95253 0.00000
-2.60338 0.00000 0.00000 ; HA- CA- CB- CG
2204 2203 2211 2212 3 3.68192 -4.35136 0.00000
1.33888 0.00000 0.00000 ; HA- CA- C- O
2204 2203 2211 2213 3 0.00000 0.00000 0.00000
0.00000 0.00000 0.00000 ; HA- CA- C- N
2205 2203 2211 2212 3 0.00000 0.00000 0.00000
0.00000 0.00000 0.00000 ; CB- CA- C- O

2205 2203 2211 2213 3 2.51040 4.18400 1.67360
-6.69440 0.00000 0.00000 ; CB- CA- C- N
2206 2205 2203 2211 3 0.65084 1.95253 0.00000
-2.60338 0.00000 0.00000 ; HB2- CB- CA- C
2206 2205 2208 2209 3 0.00000 0.00000 0.00000
0.00000 0.00000 0.00000 ; HB2- CB- CG- OD1
2206 2205 2208 2210 3 0.00000 0.00000 0.00000
0.00000 0.00000 0.00000 ; HB2- CB- CG- OD2
2207 2205 2203 2211 3 0.65084 1.95253 0.00000
-2.60338 0.00000 0.00000 ; HB3- CB- CA- C
2207 2205 2208 2209 3 0.00000 0.00000 0.00000
0.00000 0.00000 0.00000 ; HB3- CB- CG- OD1
2207 2205 2208 2210 3 0.00000 0.00000 0.00000
0.00000 0.00000 0.00000 ; HB3- CB- CG- OD2
2208 2205 2203 2211 3 7.14627 -1.04600 -8.99560
-0.97069 5.15469 0.00000 ; CG- CB- CA- C
2211 2213 2215 2216 3 0.00000 0.00000 0.00000
0.00000 0.00000 0.00000 ; C- N- CA- HA
2211 2213 2215 2217 3 11.71520 1.67360 15.06240 -
13.38880 0.00000 0.00000 ; C- N- CA- CB
2211 2213 2215 2230 3 1.75728 5.27184 2.25936
-7.02912 0.00000 0.00000 ; C- N- CA- C
2212 2211 2213 2214 3 29.28800 -8.36800 -20.92000
0.00000 0.00000 0.00000 ; O- C- N- H
2212 2211 2213 2215 3 20.92000 0.00000 -20.92000
0.00000 0.00000 0.00000 ; O- C- N- CA
2213 2215 2217 2218 3 0.65084 1.95253 0.00000
-2.60338 0.00000 0.00000 ; N- CA- CB- HB2
2213 2215 2217 2219 3 0.65084 1.95253 0.00000
-2.60338 0.00000 0.00000 ; N- CA- CB- HB3
2213 2215 2217 2220 3 3.79070 1.39746 -4.61077
-2.40998 2.44346 0.00000 ; N- CA- CB- CG
2213 2215 2230 2231 3 0.00000 0.00000 0.00000
0.00000 0.00000 0.00000 ; N- CA- C- O
2213 2215 2230 2232 3 17.40544 -5.02080 -13.22144
9.20480 0.00000 0.00000 ; N- CA- C- N
2214 2213 2215 2216 3 0.00000 0.00000 0.00000
0.00000 0.00000 0.00000 ; H- N- CA- HA
2214 2213 2215 2217 3 0.00000 0.00000 0.00000
0.00000 0.00000 0.00000 ; H- N- CA- CB
2214 2213 2215 2230 3 0.00000 0.00000 0.00000
0.00000 0.00000 0.00000 ; H- N- CA- C
2215 2217 2220 2221 3 0.66944 2.00832 0.00000
-2.67776 0.00000 0.00000 ; CA- CB- CG- HG
2215 2217 2220 2222 3 3.90367 0.19665 -6.21742
-2.37651 5.99149 0.00000 ; CA- CB- CG- CD1
2215 2217 2220 2226 3 3.90367 0.19665 -6.21742
-2.37651 5.99149 0.00000 ; CA- CB- CG- CD2
2215 2230 2232 2233 3 20.92000 0.00000 -20.92000
0.00000 0.00000 0.00000 ; CA- C- N- H
2215 2230 2232 2234 3 20.92000 0.00000 -20.92000
0.00000 0.00000 0.00000 ; CA- C- N- CA

2216 2215 2217 2218 3 0.65084 1.95253 0.00000
-2.60338 0.00000 0.00000 ; HA- CA- CB- HB2
2216 2215 2217 2219 3 0.65084 1.95253 0.00000
-2.60338 0.00000 0.00000 ; HA- CA- CB- HB3
2216 2215 2217 2220 3 0.65084 1.95253 0.00000
-2.60338 0.00000 0.00000 ; HA- CA- CB- CG
2216 2215 2230 2231 3 3.68192 -4.35136 0.00000
1.33888 0.00000 0.00000 ; HA- CA- C- O
2216 2215 2230 2232 3 0.00000 0.00000 0.00000
0.00000 0.00000 0.00000 ; HA- CA- C- N
2217 2215 2230 2231 3 0.00000 0.00000 0.00000
0.00000 0.00000 0.00000 ; CB- CA- C- O
2217 2215 2230 2232 3 2.51040 4.18400 1.67360
-6.69440 0.00000 0.00000 ; CB- CA- C- N
2217 2220 2222 2223 3 0.66944 2.00832 0.00000
-2.67776 0.00000 0.00000 ; CB- CG- CD1- HD11
2217 2220 2222 2224 3 0.66944 2.00832 0.00000
-2.67776 0.00000 0.00000 ; CB- CG- CD1- HD12
2217 2220 2222 2225 3 0.66944 2.00832 0.00000
-2.67776 0.00000 0.00000 ; CB- CG- CD1- HD13
2217 2220 2226 2227 3 0.66944 2.00832 0.00000
-2.67776 0.00000 0.00000 ; CB- CG- CD2- HD21
2217 2220 2226 2228 3 0.66944 2.00832 0.00000
-2.67776 0.00000 0.00000 ; CB- CG- CD2- HD22
2217 2220 2226 2229 3 0.66944 2.00832 0.00000
-2.67776 0.00000 0.00000 ; CB- CG- CD2- HD23
2218 2217 2215 2230 3 0.65084 1.95253 0.00000
-2.60338 0.00000 0.00000 ; HB2- CB- CA- C
2218 2217 2220 2221 3 0.62760 1.88280 0.00000
-2.51040 0.00000 0.00000 ; HB2- CB- CG- HG
2218 2217 2220 2222 3 0.66944 2.00832 0.00000
-2.67776 0.00000 0.00000 ; HB2- CB- CG- CD1
2218 2217 2220 2226 3 0.66944 2.00832 0.00000
-2.67776 0.00000 0.00000 ; HB2- CB- CG- CD2
2219 2217 2215 2230 3 0.65084 1.95253 0.00000
-2.60338 0.00000 0.00000 ; HB3- CB- CA- C
2219 2217 2220 2221 3 0.62760 1.88280 0.00000
-2.51040 0.00000 0.00000 ; HB3- CB- CG- HG
2219 2217 2220 2222 3 0.66944 2.00832 0.00000
-2.67776 0.00000 0.00000 ; HB3- CB- CG- CD1
2219 2217 2220 2226 3 0.66944 2.00832 0.00000
-2.67776 0.00000 0.00000 ; HB3- CB- CG- CD2
2220 2217 2215 2230 3 10.33448 -1.14642 -11.54784
-2.40998 6.35968 0.00000 ; CG- CB- CA- C
2221 2220 2222 2223 3 0.62760 1.88280 0.00000
-2.51040 0.00000 0.00000 ; HG- CG- CD1- HD11
2221 2220 2222 2224 3 0.62760 1.88280 0.00000
-2.51040 0.00000 0.00000 ; HG- CG- CD1- HD12
2221 2220 2222 2225 3 0.62760 1.88280 0.00000
-2.51040 0.00000 0.00000 ; HG- CG- CD1- HD13
2221 2220 2226 2227 3 0.62760 1.88280 0.00000
-2.51040 0.00000 0.00000 ; HG- CG- CD2- HD21

2221 2220 2226 2228 3 0.62760 1.88280 0.00000
-2.51040 0.00000 0.00000 ; HG- CG- CD2- HD22
2221 2220 2226 2229 3 0.62760 1.88280 0.00000
-2.51040 0.00000 0.00000 ; HG- CG- CD2- HD23
2222 2220 2226 2227 3 0.66944 2.00832 0.00000
-2.67776 0.00000 0.00000 ; CD1- CG- CD2- HD21
2222 2220 2226 2228 3 0.66944 2.00832 0.00000
-2.67776 0.00000 0.00000 ; CD1- CG- CD2- HD22
2222 2220 2226 2229 3 0.66944 2.00832 0.00000
-2.67776 0.00000 0.00000 ; CD1- CG- CD2- HD23
2223 2222 2220 2226 3 0.66944 2.00832 0.00000
-2.67776 0.00000 0.00000 ; HD11- CD1- CG- CD2
2224 2222 2220 2226 3 0.66944 2.00832 0.00000
-2.67776 0.00000 0.00000 ; HD12- CD1- CG- CD2
2225 2222 2220 2226 3 0.66944 2.00832 0.00000
-2.67776 0.00000 0.00000 ; HD13- CD1- CG- CD2
2230 2232 2234 2235 3 0.00000 0.00000 0.00000
0.00000 0.00000 0.00000 ; C- N- CA- HA
2230 2232 2234 2236 3 11.71520 1.67360 15.06240 -
13.38880 0.00000 0.00000 ; C- N- CA- CB
2230 2232 2234 2252 3 1.75728 5.27184 2.25936
-7.02912 0.00000 0.00000 ; C- N- CA- C
2231 2230 2232 2233 3 29.28800 -8.36800 -20.92000
0.00000 0.00000 0.00000 ; O- C- N- H
2231 2230 2232 2234 3 20.92000 0.00000 -20.92000
0.00000 0.00000 0.00000 ; O- C- N- CA
2232 2234 2236 2237 3 0.65084 1.95253 0.00000
-2.60338 0.00000 0.00000 ; N- CA- CB- HB2
2232 2234 2236 2238 3 0.65084 1.95253 0.00000
-2.60338 0.00000 0.00000 ; N- CA- CB- HB3
2232 2234 2236 2239 3 0.65084 1.95253 0.00000
-2.60338 0.00000 0.00000 ; N- CA- CB- CG
2232 2234 2252 2253 3 0.00000 0.00000 0.00000
0.00000 0.00000 0.00000 ; N- CA- C- O
2232 2234 2252 2254 3 17.40544 -5.02080 -13.22144
9.20480 0.00000 0.00000 ; N- CA- C- N
2233 2232 2234 2235 3 0.00000 0.00000 0.00000
0.00000 0.00000 0.00000 ; H- N- CA- HA
2233 2232 2234 2236 3 0.00000 0.00000 0.00000
0.00000 0.00000 0.00000 ; H- N- CA- CB
2233 2232 2234 2252 3 0.00000 0.00000 0.00000
0.00000 0.00000 0.00000 ; H- N- CA- C
2234 2236 2239 2240 3 0.66944 2.00832 0.00000
-2.67776 0.00000 0.00000 ; CA- CB- CG- HG2
2234 2236 2239 2241 3 0.66944 2.00832 0.00000
-2.67776 0.00000 0.00000 ; CA- CB- CG- HG3
2234 2236 2239 2242 3 3.68192 3.09616 -2.09200
-3.01248 0.00000 0.00000 ; CA- CB- CG- CD
2234 2252 2254 2255 3 20.92000 0.00000 -20.92000
0.00000 0.00000 0.00000 ; CA- C- N- CD
2234 2252 2254 2264 3 20.92000 0.00000 -20.92000
0.00000 0.00000 0.00000 ; CA- C- N- CA

2235 2234 2236 2237 3 0.65084 1.95253 0.00000
-2.60338 0.00000 0.00000 ; HA- CA- CB- HB2
2235 2234 2236 2238 3 0.65084 1.95253 0.00000
-2.60338 0.00000 0.00000 ; HA- CA- CB- HB3
2235 2234 2236 2239 3 0.65084 1.95253 0.00000
-2.60338 0.00000 0.00000 ; HA- CA- CB- CG
2235 2234 2252 2253 3 3.68192 -4.35136 0.00000
1.33888 0.00000 0.00000 ; HA- CA- C- O
2235 2234 2252 2254 3 0.00000 0.00000 0.00000
0.00000 0.00000 0.00000 ; HA- CA- C- N
2236 2234 2252 2253 3 0.00000 0.00000 0.00000
0.00000 0.00000 0.00000 ; CB- CA- C- O
2236 2234 2252 2254 3 2.51040 4.18400 1.67360
-6.69440 0.00000 0.00000 ; CB- CA- C- N
2236 2239 2242 2243 3 0.66944 2.00832 0.00000
-2.67776 0.00000 0.00000 ; CB- CG- CD- HD2
2236 2239 2242 2244 3 0.66944 2.00832 0.00000
-2.67776 0.00000 0.00000 ; CB- CG- CD- HD3
2236 2239 2242 2245 3 3.68192 3.09616 -2.09200
-3.01248 0.00000 0.00000 ; CB- CG- CD- CE
2237 2236 2234 2252 3 0.65084 1.95253 0.00000
-2.60338 0.00000 0.00000 ; HB2- CB- CA- C
2237 2236 2239 2240 3 0.62760 1.88280 0.00000
-2.51040 0.00000 0.00000 ; HB2- CB- CG- HG2
2237 2236 2239 2241 3 0.62760 1.88280 0.00000
-2.51040 0.00000 0.00000 ; HB2- CB- CG- HG3
2237 2236 2239 2242 3 0.66944 2.00832 0.00000
-2.67776 0.00000 0.00000 ; HB2- CB- CG- CD
2238 2236 2234 2252 3 0.65084 1.95253 0.00000
-2.60338 0.00000 0.00000 ; HB3- CB- CA- C
2238 2236 2239 2240 3 0.62760 1.88280 0.00000
-2.51040 0.00000 0.00000 ; HB3- CB- CG- HG2
2238 2236 2239 2241 3 0.62760 1.88280 0.00000
-2.51040 0.00000 0.00000 ; HB3- CB- CG- HG3
2238 2236 2239 2242 3 0.66944 2.00832 0.00000
-2.67776 0.00000 0.00000 ; HB3- CB- CG- CD
2239 2236 2234 2252 3 0.65084 1.95253 0.00000
-2.60338 0.00000 0.00000 ; CG- CB- CA- C
2239 2242 2245 2246 3 0.65084 1.95253 0.00000
-2.60338 0.00000 0.00000 ; CG- CD- CE- HE2
2239 2242 2245 2247 3 0.65084 1.95253 0.00000
-2.60338 0.00000 0.00000 ; CG- CD- CE- HE3
2239 2242 2245 2248 3 0.65084 1.95253 0.00000
-2.60338 0.00000 0.00000 ; CG- CD- CE- NZ
2240 2239 2242 2243 3 0.62760 1.88280 0.00000
-2.51040 0.00000 0.00000 ; HG2- CG- CD- HD2
2240 2239 2242 2244 3 0.62760 1.88280 0.00000
-2.51040 0.00000 0.00000 ; HG2- CG- CD- HD3
2240 2239 2242 2245 3 0.66944 2.00832 0.00000
-2.67776 0.00000 0.00000 ; HG2- CG- CD- CE
2241 2239 2242 2243 3 0.62760 1.88280 0.00000
-2.51040 0.00000 0.00000 ; HG3- CG- CD- HD2

2241 2239 2242 2244 3 0.62760 1.88280 0.00000
-2.51040 0.00000 0.00000 ; HG3- CG- CD- HD3
2241 2239 2242 2245 3 0.66944 2.00832 0.00000
-2.67776 0.00000 0.00000 ; HG3- CG- CD- CE
2242 2245 2248 2249 3 0.65084 1.95253 0.00000
-2.60338 0.00000 0.00000 ; CD- CE- NZ- HZ1
2242 2245 2248 2250 3 0.65084 1.95253 0.00000
-2.60338 0.00000 0.00000 ; CD- CE- NZ- HZ2
2242 2245 2248 2251 3 0.65084 1.95253 0.00000
-2.60338 0.00000 0.00000 ; CD- CE- NZ- HZ3
2243 2242 2245 2246 3 0.65084 1.95253 0.00000
-2.60338 0.00000 0.00000 ; HD2- CD- CE- HE2
2243 2242 2245 2247 3 0.65084 1.95253 0.00000
-2.60338 0.00000 0.00000 ; HD2- CD- CE- HE3
2243 2242 2245 2248 3 0.65084 1.95253 0.00000
-2.60338 0.00000 0.00000 ; HD2- CD- CE- NZ
2244 2242 2245 2246 3 0.65084 1.95253 0.00000
-2.60338 0.00000 0.00000 ; HD3- CD- CE- HE2
2244 2242 2245 2247 3 0.65084 1.95253 0.00000
-2.60338 0.00000 0.00000 ; HD3- CD- CE- HE3
2244 2242 2245 2248 3 0.65084 1.95253 0.00000
-2.60338 0.00000 0.00000 ; HD3- CD- CE- NZ
2246 2245 2248 2249 3 0.65084 1.95253 0.00000
-2.60338 0.00000 0.00000 ; HE2- CE- NZ- HZ1
2246 2245 2248 2250 3 0.65084 1.95253 0.00000
-2.60338 0.00000 0.00000 ; HE2- CE- NZ- HZ2
2246 2245 2248 2251 3 0.65084 1.95253 0.00000
-2.60338 0.00000 0.00000 ; HE2- CE- NZ- HZ3
2247 2245 2248 2249 3 0.65084 1.95253 0.00000
-2.60338 0.00000 0.00000 ; HE3- CE- NZ- HZ1
2247 2245 2248 2250 3 0.65084 1.95253 0.00000
-2.60338 0.00000 0.00000 ; HE3- CE- NZ- HZ2
2247 2245 2248 2251 3 0.65084 1.95253 0.00000
-2.60338 0.00000 0.00000 ; HE3- CE- NZ- HZ3
2252 2254 2255 2256 3 0.00000 0.00000 0.00000
0.00000 0.00000 0.00000 ; C- N- CD- HD2
2252 2254 2255 2257 3 0.00000 0.00000 0.00000
0.00000 0.00000 0.00000 ; C- N- CD- HD3
2252 2254 2255 2258 3 10.04160 -3.34720 16.73600
-6.69440 0.00000 0.00000 ; C- N- CD- CG
2252 2254 2264 2261 3 11.71520 1.67360 15.06240 -
13.38880 0.00000 0.00000 ; C- N- CA- CB
2252 2254 2264 2265 3 0.00000 0.00000 0.00000
0.00000 0.00000 0.00000 ; C- N- CA- HA
2252 2254 2264 2266 3 1.75728 5.27184 2.25936
-7.02912 0.00000 0.00000 ; C- N- CA- C
2253 2252 2254 2255 3 20.92000 0.00000 -20.92000
0.00000 0.00000 0.00000 ; O- C- N- CD
2253 2252 2254 2264 3 20.92000 0.00000 -20.92000
0.00000 0.00000 0.00000 ; O- C- N- CA
2254 2255 2258 2259 3 0.65084 1.95253 0.00000
-2.60338 0.00000 0.00000 ; N- CD- CG- HG2

2254 2255 2258 2260 3 0.65084 1.95253 0.00000
-2.60338 0.00000 0.00000 ; N- CD- CG- HG3
2254 2255 2258 2261 3 0.65084 1.95253 0.00000
-2.60338 0.00000 0.00000 ; N- CD- CG- CB
2254 2264 2261 2258 3 0.65084 1.95253 0.00000
-2.60338 0.00000 0.00000 ; N- CA- CB- CG
2254 2264 2261 2262 3 0.65084 1.95253 0.00000
-2.60338 0.00000 0.00000 ; N- CA- CB- HB2
2254 2264 2261 2263 3 0.65084 1.95253 0.00000
-2.60338 0.00000 0.00000 ; N- CA- CB- HB3
2254 2264 2266 2267 3 0.00000 0.00000 0.00000
0.00000 0.00000 0.00000 ; N- CA- C- O
2254 2264 2266 2268 3 17.40544 -5.02080 -13.22144
9.20480 0.00000 0.00000 ; N- CA- C- N
2255 2254 2264 2261 3 0.00000 0.00000 0.00000
0.00000 0.00000 0.00000 ; CD- N- CA- CB
2255 2254 2264 2265 3 0.00000 0.00000 0.00000
0.00000 0.00000 0.00000 ; CD- N- CA- HA
2255 2254 2264 2266 3 0.00000 0.00000 0.00000
0.00000 0.00000 0.00000 ; CD- N- CA- C
2255 2258 2261 2262 3 0.66944 2.00832 0.00000
-2.67776 0.00000 0.00000 ; CD- CG- CB- HB2
2255 2258 2261 2263 3 0.66944 2.00832 0.00000
-2.67776 0.00000 0.00000 ; CD- CG- CB- HB3
2255 2258 2261 2264 3 3.68192 3.09616 -2.09200
-3.01248 0.00000 0.00000 ; CD- CG- CB- CA
2256 2255 2254 2264 3 0.00000 0.00000 0.00000
0.00000 0.00000 0.00000 ; HD2- CD- N- CA
2256 2255 2258 2259 3 0.65084 1.95253 0.00000
-2.60338 0.00000 0.00000 ; HD2- CD- CG- HG2
2256 2255 2258 2260 3 0.65084 1.95253 0.00000
-2.60338 0.00000 0.00000 ; HD2- CD- CG- HG3
2256 2255 2258 2261 3 0.65084 1.95253 0.00000
-2.60338 0.00000 0.00000 ; HD2- CD- CG- CB
2257 2255 2254 2264 3 0.00000 0.00000 0.00000
0.00000 0.00000 0.00000 ; HD3- CD- N- CA
2257 2255 2258 2259 3 0.65084 1.95253 0.00000
-2.60338 0.00000 0.00000 ; HD3- CD- CG- HG2
2257 2255 2258 2260 3 0.65084 1.95253 0.00000
-2.60338 0.00000 0.00000 ; HD3- CD- CG- HG3
2257 2255 2258 2261 3 0.65084 1.95253 0.00000
-2.60338 0.00000 0.00000 ; HD3- CD- CG- CB
2258 2255 2254 2264 3 0.00000 0.00000 0.00000
0.00000 0.00000 0.00000 ; CG- CD- N- CA
2258 2261 2264 2265 3 0.65084 1.95253 0.00000
-2.60338 0.00000 0.00000 ; CG- CB- CA- HA
2258 2261 2264 2266 3 0.65084 1.95253 0.00000
-2.60338 0.00000 0.00000 ; CG- CB- CA- C
2259 2258 2261 2262 3 0.62760 1.88280 0.00000
-2.51040 0.00000 0.00000 ; HG2- CG- CB- HB2
2259 2258 2261 2263 3 0.62760 1.88280 0.00000
-2.51040 0.00000 0.00000 ; HG2- CG- CB- HB3

2259 2258 2261 2264 3 0.66944 2.00832 0.00000
-2.67776 0.00000 0.00000 ; HG2- CG- CB- CA
2260 2258 2261 2262 3 0.62760 1.88280 0.00000
-2.51040 0.00000 0.00000 ; HG3- CG- CB- HB2
2260 2258 2261 2263 3 0.62760 1.88280 0.00000
-2.51040 0.00000 0.00000 ; HG3- CG- CB- HB3
2260 2258 2261 2264 3 0.66944 2.00832 0.00000
-2.67776 0.00000 0.00000 ; HG3- CG- CB- CA
2261 2264 2266 2267 3 0.00000 0.00000 0.00000
0.00000 0.00000 0.00000 ; CB- CA- C- O
2261 2264 2266 2268 3 2.51040 4.18400 1.67360
-6.69440 0.00000 0.00000 ; CB- CA- C- N
2262 2261 2264 2265 3 0.65084 1.95253 0.00000
-2.60338 0.00000 0.00000 ; HB2- CB- CA- HA
2262 2261 2264 2266 3 0.65084 1.95253 0.00000
-2.60338 0.00000 0.00000 ; HB2- CB- CA- C
2263 2261 2264 2265 3 0.65084 1.95253 0.00000
-2.60338 0.00000 0.00000 ; HB3- CB- CA- HA
2263 2261 2264 2266 3 0.65084 1.95253 0.00000
-2.60338 0.00000 0.00000 ; HB3- CB- CA- C
2264 2266 2268 2269 3 20.92000 0.00000 -20.92000
0.00000 0.00000 0.00000 ; CA- C- N- H
2264 2266 2268 2270 3 20.92000 0.00000 -20.92000
0.00000 0.00000 0.00000 ; CA- C- N- CA
2265 2264 2266 2267 3 3.68192 -4.35136 0.00000
1.33888 0.00000 0.00000 ; HA- CA- C- O
2265 2264 2266 2268 3 0.00000 0.00000 0.00000
0.00000 0.00000 0.00000 ; HA- CA- C- N
2266 2268 2270 2271 3 0.00000 0.00000 0.00000
0.00000 0.00000 0.00000 ; C- N- CA- HA
2266 2268 2270 2272 3 11.71520 1.67360 15.06240 -
13.38880 0.00000 0.00000 ; C- N- CA- CB
2266 2268 2270 2281 3 1.75728 5.27184 2.25936
-7.02912 0.00000 0.00000 ; C- N- CA- C
2267 2266 2268 2269 3 29.28800 -8.36800 -20.92000
0.00000 0.00000 0.00000 ; O- C- N- H
2267 2266 2268 2270 3 20.92000 0.00000 -20.92000
0.00000 0.00000 0.00000 ; O- C- N- CA
2268 2270 2272 2273 3 0.65084 1.95253 0.00000
-2.60338 0.00000 0.00000 ; N- CA- CB- HB2
2268 2270 2272 2274 3 0.65084 1.95253 0.00000
-2.60338 0.00000 0.00000 ; N- CA- CB- HB3
2268 2270 2272 2275 3 3.21331 2.22589 -4.15053
-2.40998 2.61082 0.00000 ; N- CA- CB- CG
2268 2270 2281 2282 3 0.00000 0.00000 0.00000
0.00000 0.00000 0.00000 ; N- CA- C- O
2268 2270 2281 2283 3 17.40544 -5.02080 -13.22144
9.20480 0.00000 0.00000 ; N- CA- C- N
2269 2268 2270 2271 3 0.00000 0.00000 0.00000
0.00000 0.00000 0.00000 ; H- N- CA- HA
2269 2268 2270 2272 3 0.00000 0.00000 0.00000
0.00000 0.00000 0.00000 ; H- N- CA- CB

2269 2268 2270 2281 3 0.00000 0.00000 0.00000
0.00000 0.00000 0.00000 ; H- N- CA- C
2270 2272 2275 2276 3 0.66944 2.00832 0.00000
-2.67776 0.00000 0.00000 ; CA- CB- CG- HG2
2270 2272 2275 2277 3 0.66944 2.00832 0.00000
-2.67776 0.00000 0.00000 ; CA- CB- CG- HG3
2270 2272 2275 2278 3 10.12110 -1.91209 0.01674
10.17549 -1.87443 0.00000 ; CA- CB- CG- CD
2270 2281 2283 2284 3 20.92000 0.00000 -20.92000
0.00000 0.00000 0.00000 ; CA- C- N- H
2270 2281 2283 2285 3 20.92000 0.00000 -20.92000
0.00000 0.00000 0.00000 ; CA- C- N- CA
2271 2270 2272 2273 3 0.65084 1.95253 0.00000
-2.60338 0.00000 0.00000 ; HA- CA- CB- HB2
2271 2270 2272 2274 3 0.65084 1.95253 0.00000
-2.60338 0.00000 0.00000 ; HA- CA- CB- HB3
2271 2270 2272 2275 3 0.65084 1.95253 0.00000
-2.60338 0.00000 0.00000 ; HA- CA- CB- CG
2271 2270 2281 2282 3 3.68192 -4.35136 0.00000
1.33888 0.00000 0.00000 ; HA- CA- C- O
2271 2270 2281 2283 3 0.00000 0.00000 0.00000
0.00000 0.00000 0.00000 ; HA- CA- C- N
2272 2270 2281 2282 3 0.00000 0.00000 0.00000
0.00000 0.00000 0.00000 ; CB- CA- C- O
2272 2270 2281 2283 3 2.51040 4.18400 1.67360
-6.69440 0.00000 0.00000 ; CB- CA- C- N
2272 2275 2278 2279 3 3.79907 0.00000 -5.40573
0.00000 2.14221 0.00000 ; CB- CG- CD- OE1
2272 2275 2278 2280 3 3.79907 0.00000 -5.40573
0.00000 2.14221 0.00000 ; CB- CG- CD- OE2
2273 2272 2270 2281 3 0.65084 1.95253 0.00000
-2.60338 0.00000 0.00000 ; HB2- CB- CA- C
2273 2272 2275 2276 3 0.62760 1.88280 0.00000
-2.51040 0.00000 0.00000 ; HB2- CB- CG- HG2
2273 2272 2275 2277 3 0.62760 1.88280 0.00000
-2.51040 0.00000 0.00000 ; HB2- CB- CG- HG3
2273 2272 2275 2278 3 0.65084 1.95253 0.00000
-2.60338 0.00000 0.00000 ; HB2- CB- CG- CD
2274 2272 2270 2281 3 0.65084 1.95253 0.00000
-2.60338 0.00000 0.00000 ; HB3- CB- CA- C
2274 2272 2275 2276 3 0.62760 1.88280 0.00000
-2.51040 0.00000 0.00000 ; HB3- CB- CG- HG2
2274 2272 2275 2277 3 0.62760 1.88280 0.00000
-2.51040 0.00000 0.00000 ; HB3- CB- CG- HG3
2274 2272 2275 2278 3 0.65084 1.95253 0.00000
-2.60338 0.00000 0.00000 ; HB3- CB- CG- CD
2275 2272 2270 2281 3 6.86594 3.56895 -8.14206
-2.40998 4.85344 0.00000 ; CG- CB- CA- C
2276 2275 2278 2279 3 0.00000 0.00000 0.00000
0.00000 0.00000 0.00000 ; HG2- CG- CD- OE1
2276 2275 2278 2280 3 0.00000 0.00000 0.00000
0.00000 0.00000 0.00000 ; HG2- CG- CD- OE2

2277 2275 2278 2279 3 0.00000 0.00000 0.00000
0.00000 0.00000 0.00000 ; HG3- CG- CD- OE1
2277 2275 2278 2280 3 0.00000 0.00000 0.00000
0.00000 0.00000 0.00000 ; HG3- CG- CD- OE2
2281 2283 2285 2286 3 0.00000 0.00000 0.00000
0.00000 0.00000 0.00000 ; C- N- CA- HA
2281 2283 2285 2287 3 11.71520 1.67360 15.06240 -
13.38880 0.00000 0.00000 ; C- N- CA- CB
2281 2283 2285 2295 3 1.75728 5.27184 2.25936
-7.02912 0.00000 0.00000 ; C- N- CA- C
2282 2281 2283 2284 3 29.28800 -8.36800 -20.92000
0.00000 0.00000 0.00000 ; O- C- N- H
2282 2281 2283 2285 3 20.92000 0.00000 -20.92000
0.00000 0.00000 0.00000 ; O- C- N- CA
2283 2285 2287 2288 3 0.65084 1.95253 0.00000
-2.60338 0.00000 0.00000 ; N- CA- CB- HB2
2283 2285 2287 2289 3 0.65084 1.95253 0.00000
-2.60338 0.00000 0.00000 ; N- CA- CB- HB3
2283 2285 2287 2290 3 5.99567 3.29281 -4.46014
-0.55229 1.97485 0.00000 ; N- CA- CB- CG
2283 2285 2295 2296 3 0.00000 0.00000 0.00000
0.00000 0.00000 0.00000 ; N- CA- C- O
2283 2285 2295 2297 3 17.40544 -5.02080 -13.22144
9.20480 0.00000 0.00000 ; N- CA- C- N
2284 2283 2285 2286 3 0.00000 0.00000 0.00000
0.00000 0.00000 0.00000 ; H- N- CA- HA
2284 2283 2285 2287 3 0.00000 0.00000 0.00000
0.00000 0.00000 0.00000 ; H- N- CA- CB
2284 2283 2285 2295 3 0.00000 0.00000 0.00000
0.00000 0.00000 0.00000 ; H- N- CA- C
2285 2287 2290 2291 3 0.00000 0.00000 0.00000
0.00000 0.00000 0.00000 ; CA- CB- CG- OD1
2285 2287 2290 2292 3 8.84916 -0.31380 -4.32626
5.03754 0.26778 0.00000 ; CA- CB- CG- ND2
2285 2295 2297 2298 3 20.92000 0.00000 -20.92000
0.00000 0.00000 0.00000 ; CA- C- N- H
2285 2295 2297 2299 3 20.92000 0.00000 -20.92000
0.00000 0.00000 0.00000 ; CA- C- N- CA
2286 2285 2287 2288 3 0.65084 1.95253 0.00000
-2.60338 0.00000 0.00000 ; HA- CA- CB- HB2
2286 2285 2287 2289 3 0.65084 1.95253 0.00000
-2.60338 0.00000 0.00000 ; HA- CA- CB- HB3
2286 2285 2287 2290 3 0.65084 1.95253 0.00000
-2.60338 0.00000 0.00000 ; HA- CA- CB- CG
2286 2285 2295 2296 3 3.68192 -4.35136 0.00000
1.33888 0.00000 0.00000 ; HA- CA- C- O
2286 2285 2295 2297 3 0.00000 0.00000 0.00000
0.00000 0.00000 0.00000 ; HA- CA- C- N
2287 2285 2295 2296 3 0.00000 0.00000 0.00000
0.00000 0.00000 0.00000 ; CB- CA- C- O
2287 2285 2295 2297 3 2.51040 4.18400 1.67360
-6.69440 0.00000 0.00000 ; CB- CA- C- N

2287 2290 2292 2293 3 20.92000 0.00000 -20.92000
0.00000 0.00000 0.00000 ; CB- CG- ND2- HD21
2287 2290 2292 2294 3 20.92000 0.00000 -20.92000
0.00000 0.00000 0.00000 ; CB- CG- ND2- HD22
2288 2287 2285 2295 3 0.65084 1.95253 0.00000
-2.60338 0.00000 0.00000 ; HB2- CB- CA- C
2288 2287 2290 2291 3 3.68192 -4.35136 0.00000
1.33888 0.00000 0.00000 ; HB2- CB- CG- OD1
2288 2287 2290 2292 3 0.00000 0.00000 0.00000
0.00000 0.00000 0.00000 ; HB2- CB- CG- ND2
2289 2287 2285 2295 3 0.65084 1.95253 0.00000
-2.60338 0.00000 0.00000 ; HB3- CB- CA- C
2289 2287 2290 2291 3 3.68192 -4.35136 0.00000
1.33888 0.00000 0.00000 ; HB3- CB- CG- OD1
2289 2287 2290 2292 3 0.00000 0.00000 0.00000
0.00000 0.00000 0.00000 ; HB3- CB- CG- ND2
2290 2287 2285 2295 3 7.94542 -3.96225 -6.11701
-0.55229 3.58150 0.00000 ; CG- CB- CA- C
2291 2290 2292 2293 3 29.28800 -8.36800 -20.92000
0.00000 0.00000 0.00000 ; OD1- CG- ND2- HD21
2291 2290 2292 2294 3 29.28800 -8.36800 -20.92000
0.00000 0.00000 0.00000 ; OD1- CG- ND2- HD22
2295 2297 2299 2300 3 0.00000 0.00000 0.00000
0.00000 0.00000 0.00000 ; C- N- CA- HA
2295 2297 2299 2301 3 11.71520 1.67360 15.06240 -
13.38880 0.00000 0.00000 ; C- N- CA- CB
2295 2297 2299 2315 3 1.75728 5.27184 2.25936
-7.02912 0.00000 0.00000 ; C- N- CA- C
2296 2295 2297 2298 3 29.28800 -8.36800 -20.92000
0.00000 0.00000 0.00000 ; O- C- N- H
2296 2295 2297 2299 3 20.92000 0.00000 -20.92000
0.00000 0.00000 0.00000 ; O- C- N- CA
2297 2299 2301 2302 3 0.65084 1.95253 0.00000
-2.60338 0.00000 0.00000 ; N- CA- CB- HB2
2297 2299 2301 2303 3 0.65084 1.95253 0.00000
-2.60338 0.00000 0.00000 ; N- CA- CB- HB3
2297 2299 2301 2304 3 3.28026 2.46019 -2.19242
-3.21331 -0.23430 0.00000 ; N- CA- CB- CG
2297 2299 2315 2316 3 0.00000 0.00000 0.00000
0.00000 0.00000 0.00000 ; N- CA- C- O
2297 2299 2315 2317 3 17.40544 -5.02080 -13.22144
9.20480 0.00000 0.00000 ; N- CA- C- N
2298 2297 2299 2300 3 0.00000 0.00000 0.00000
0.00000 0.00000 0.00000 ; H- N- CA- HA
2298 2297 2299 2301 3 0.00000 0.00000 0.00000
0.00000 0.00000 0.00000 ; H- N- CA- CB
2298 2297 2299 2315 3 0.00000 0.00000 0.00000
0.00000 0.00000 0.00000 ; H- N- CA- C
2299 2301 2304 2305 3 0.57739 0.00000 1.02926
0.00000 -1.60666 0.00000 ; CA- CB- CG- CD1
2299 2301 2304 2313 3 0.57739 0.00000 1.02926
0.00000 -1.60666 0.00000 ; CA- CB- CG- CD2

2299 2315 2317 2318 3 20.92000 0.00000 -20.92000
0.00000 0.00000 0.00000 ; CA- C- N- H
2299 2315 2317 2319 3 20.92000 0.00000 -20.92000
0.00000 0.00000 0.00000 ; CA- C- N- CA
2300 2299 2301 2302 3 0.65084 1.95253 0.00000
-2.60338 0.00000 0.00000 ; HA- CA- CB- HB2
2300 2299 2301 2303 3 0.65084 1.95253 0.00000
-2.60338 0.00000 0.00000 ; HA- CA- CB- HB3
2300 2299 2301 2304 3 0.65084 1.95253 0.00000
-2.60338 0.00000 0.00000 ; HA- CA- CB- CG
2300 2299 2315 2316 3 3.68192 -4.35136 0.00000
1.33888 0.00000 0.00000 ; HA- CA- C- O
2300 2299 2315 2317 3 0.00000 0.00000 0.00000
0.00000 0.00000 0.00000 ; HA- CA- C- N
2301 2299 2315 2316 3 0.00000 0.00000 0.00000
0.00000 0.00000 0.00000 ; CB- CA- C- O
2301 2299 2315 2317 3 2.51040 4.18400 1.67360
-6.69440 0.00000 0.00000 ; CB- CA- C- N
2301 2304 2305 2306 3 30.33400 0.00000 -30.33400
0.00000 0.00000 0.00000 ; CB- CG- CD1- HD1
2301 2304 2305 2307 3 30.33400 0.00000 -30.33400
0.00000 0.00000 0.00000 ; CB- CG- CD1- CE1
2301 2304 2313 2311 3 30.33400 0.00000 -30.33400
0.00000 0.00000 0.00000 ; CB- CG- CD2- CE2
2301 2304 2313 2314 3 30.33400 0.00000 -30.33400
0.00000 0.00000 0.00000 ; CB- CG- CD2- HD2
2302 2301 2299 2315 3 0.65084 1.95253 0.00000
-2.60338 0.00000 0.00000 ; HB2- CB- CA- C
2302 2301 2304 2305 3 0.00000 0.00000 0.00000
0.00000 0.00000 0.00000 ; HB2- CB- CG- CD1
2302 2301 2304 2313 3 0.00000 0.00000 0.00000
0.00000 0.00000 0.00000 ; HB2- CB- CG- CD2
2303 2301 2299 2315 3 0.65084 1.95253 0.00000
-2.60338 0.00000 0.00000 ; HB3- CB- CA- C
2303 2301 2304 2305 3 0.00000 0.00000 0.00000
0.00000 0.00000 0.00000 ; HB3- CB- CG- CD1
2303 2301 2304 2313 3 0.00000 0.00000 0.00000
0.00000 0.00000 0.00000 ; HB3- CB- CG- CD2
2304 2301 2299 2315 3 4.95804 2.17986 -3.52293
-3.21331 -0.40166 0.00000 ; CG- CB- CA- C
2304 2305 2307 2308 3 30.33400 0.00000 -30.33400
0.00000 0.00000 0.00000 ; CG- CD1- CE1- HE1
2304 2305 2307 2309 3 30.33400 0.00000 -30.33400
0.00000 0.00000 0.00000 ; CG- CD1- CE1- CZ
2304 2313 2311 2309 3 30.33400 0.00000 -30.33400
0.00000 0.00000 0.00000 ; CG- CD2- CE2- CZ
2304 2313 2311 2312 3 30.33400 0.00000 -30.33400
0.00000 0.00000 0.00000 ; CG- CD2- CE2- HE2
2305 2304 2313 2311 3 30.33400 0.00000 -30.33400
0.00000 0.00000 0.00000 ; CD1- CG- CD2- CE2
2305 2304 2313 2314 3 30.33400 0.00000 -30.33400
0.00000 0.00000 0.00000 ; CD1- CG- CD2- HD2

2305 2307 2309 2310 3 30.33400 0.00000 -30.33400
0.00000 0.00000 0.00000 ; CD1- CE1- CZ- HZ
2305 2307 2309 2311 3 30.33400 0.00000 -30.33400
0.00000 0.00000 0.00000 ; CD1- CE1- CZ- CE2
2306 2305 2304 2313 3 30.33400 0.00000 -30.33400
0.00000 0.00000 0.00000 ; HD1- CD1- CG- CD2
2306 2305 2307 2308 3 30.33400 0.00000 -30.33400
0.00000 0.00000 0.00000 ; HD1- CD1- CE1- HE1
2306 2305 2307 2309 3 30.33400 0.00000 -30.33400
0.00000 0.00000 0.00000 ; HD1- CD1- CE1- CZ
2307 2305 2304 2313 3 30.33400 0.00000 -30.33400
0.00000 0.00000 0.00000 ; CE1- CD1- CG- CD2
2307 2309 2311 2312 3 30.33400 0.00000 -30.33400
0.00000 0.00000 0.00000 ; CE1- CZ- CE2- HE2
2307 2309 2311 2313 3 30.33400 0.00000 -30.33400
0.00000 0.00000 0.00000 ; CE1- CZ- CE2- CD2
2308 2307 2309 2310 3 30.33400 0.00000 -30.33400
0.00000 0.00000 0.00000 ; HE1- CE1- CZ- HZ
2308 2307 2309 2311 3 30.33400 0.00000 -30.33400
0.00000 0.00000 0.00000 ; HE1- CE1- CZ- CE2
2309 2311 2313 2314 3 30.33400 0.00000 -30.33400
0.00000 0.00000 0.00000 ; CZ- CE2- CD2- HD2
2310 2309 2311 2312 3 30.33400 0.00000 -30.33400
0.00000 0.00000 0.00000 ; HZ- CZ- CE2- HE2
2310 2309 2311 2313 3 30.33400 0.00000 -30.33400
0.00000 0.00000 0.00000 ; HZ- CZ- CE2- CD2
2312 2311 2313 2314 3 30.33400 0.00000 -30.33400
0.00000 0.00000 0.00000 ; HE2- CE2- CD2- HD2
2315 2317 2319 2320 3 0.00000 0.00000 0.00000
0.00000 0.00000 0.00000 ; C- N- CA- HA
2315 2317 2319 2321 3 11.71520 1.67360 15.06240 -
13.38880 0.00000 0.00000 ; C- N- CA- CB
2315 2317 2319 2334 3 1.75728 5.27184 2.25936
-7.02912 0.00000 0.00000 ; C- N- CA- C
2316 2315 2317 2318 3 29.28800 -8.36800 -20.92000
0.00000 0.00000 0.00000 ; O- C- N- H
2316 2315 2317 2319 3 20.92000 0.00000 -20.92000
0.00000 0.00000 0.00000 ; O- C- N- CA
2317 2319 2321 2322 3 0.65084 1.95253 0.00000
-2.60338 0.00000 0.00000 ; N- CA- CB- HB2
2317 2319 2321 2323 3 0.65084 1.95253 0.00000
-2.60338 0.00000 0.00000 ; N- CA- CB- HB3
2317 2319 2321 2324 3 3.79070 1.39746 -4.61077
-2.40998 2.44346 0.00000 ; N- CA- CB- CG
2317 2319 2334 2335 3 0.00000 0.00000 0.00000
0.00000 0.00000 0.00000 ; N- CA- C- O
2317 2319 2334 2336 3 17.40544 -5.02080 -13.22144
9.20480 0.00000 0.00000 ; N- CA- C- N
2318 2317 2319 2320 3 0.00000 0.00000 0.00000
0.00000 0.00000 0.00000 ; H- N- CA- HA
2318 2317 2319 2321 3 0.00000 0.00000 0.00000
0.00000 0.00000 0.00000 ; H- N- CA- CB

2318 2317 2319 2334 3 0.00000 0.00000 0.00000
0.00000 0.00000 0.00000 ; H- N- CA- C
2319 2321 2324 2325 3 0.66944 2.00832 0.00000
-2.67776 0.00000 0.00000 ; CA- CB- CG- HG
2319 2321 2324 2326 3 3.90367 0.19665 -6.21742
-2.37651 5.99149 0.00000 ; CA- CB- CG- CD1
2319 2321 2324 2330 3 3.90367 0.19665 -6.21742
-2.37651 5.99149 0.00000 ; CA- CB- CG- CD2
2319 2334 2336 2337 3 20.92000 0.00000 -20.92000
0.00000 0.00000 0.00000 ; CA- C- N- H
2319 2334 2336 2338 3 20.92000 0.00000 -20.92000
0.00000 0.00000 0.00000 ; CA- C- N- CA
2320 2319 2321 2322 3 0.65084 1.95253 0.00000
-2.60338 0.00000 0.00000 ; HA- CA- CB- HB2
2320 2319 2321 2323 3 0.65084 1.95253 0.00000
-2.60338 0.00000 0.00000 ; HA- CA- CB- HB3
2320 2319 2321 2324 3 0.65084 1.95253 0.00000
-2.60338 0.00000 0.00000 ; HA- CA- CB- CG
2320 2319 2334 2335 3 3.68192 -4.35136 0.00000
1.33888 0.00000 0.00000 ; HA- CA- C- O
2320 2319 2334 2336 3 0.00000 0.00000 0.00000
0.00000 0.00000 0.00000 ; HA- CA- C- N
2321 2319 2334 2335 3 0.00000 0.00000 0.00000
0.00000 0.00000 0.00000 ; CB- CA- C- O
2321 2319 2334 2336 3 2.51040 4.18400 1.67360
-6.69440 0.00000 0.00000 ; CB- CA- C- N
2321 2324 2326 2327 3 0.66944 2.00832 0.00000
-2.67776 0.00000 0.00000 ; CB- CG- CD1- HD11
2321 2324 2326 2328 3 0.66944 2.00832 0.00000
-2.67776 0.00000 0.00000 ; CB- CG- CD1- HD12
2321 2324 2326 2329 3 0.66944 2.00832 0.00000
-2.67776 0.00000 0.00000 ; CB- CG- CD1- HD13
2321 2324 2330 2331 3 0.66944 2.00832 0.00000
-2.67776 0.00000 0.00000 ; CB- CG- CD2- HD21
2321 2324 2330 2332 3 0.66944 2.00832 0.00000
-2.67776 0.00000 0.00000 ; CB- CG- CD2- HD22
2321 2324 2330 2333 3 0.66944 2.00832 0.00000
-2.67776 0.00000 0.00000 ; CB- CG- CD2- HD23
2322 2321 2319 2334 3 0.65084 1.95253 0.00000
-2.60338 0.00000 0.00000 ; HB2- CB- CA- C
2322 2321 2324 2325 3 0.62760 1.88280 0.00000
-2.51040 0.00000 0.00000 ; HB2- CB- CG- HG
2322 2321 2324 2326 3 0.66944 2.00832 0.00000
-2.67776 0.00000 0.00000 ; HB2- CB- CG- CD1
2322 2321 2324 2330 3 0.66944 2.00832 0.00000
-2.67776 0.00000 0.00000 ; HB2- CB- CG- CD2
2323 2321 2319 2334 3 0.65084 1.95253 0.00000
-2.60338 0.00000 0.00000 ; HB3- CB- CA- C
2323 2321 2324 2325 3 0.62760 1.88280 0.00000
-2.51040 0.00000 0.00000 ; HB3- CB- CG- HG
2323 2321 2324 2326 3 0.66944 2.00832 0.00000
-2.67776 0.00000 0.00000 ; HB3- CB- CG- CD1

2323 2321 2324 2330 3 0.66944 2.00832 0.00000
-2.67776 0.00000 0.00000 ; HB3- CB- CG- CD2
2324 2321 2319 2334 3 10.33448 -1.14642 -11.54784
-2.40998 6.35968 0.00000 ; CG- CB- CA- C
2325 2324 2326 2327 3 0.62760 1.88280 0.00000
-2.51040 0.00000 0.00000 ; HG- CG- CD1- HD11
2325 2324 2326 2328 3 0.62760 1.88280 0.00000
-2.51040 0.00000 0.00000 ; HG- CG- CD1- HD12
2325 2324 2326 2329 3 0.62760 1.88280 0.00000
-2.51040 0.00000 0.00000 ; HG- CG- CD1- HD13
2325 2324 2330 2331 3 0.62760 1.88280 0.00000
-2.51040 0.00000 0.00000 ; HG- CG- CD2- HD21
2325 2324 2330 2332 3 0.62760 1.88280 0.00000
-2.51040 0.00000 0.00000 ; HG- CG- CD2- HD22
2325 2324 2330 2333 3 0.62760 1.88280 0.00000
-2.51040 0.00000 0.00000 ; HG- CG- CD2- HD23
2326 2324 2330 2331 3 0.66944 2.00832 0.00000
-2.67776 0.00000 0.00000 ; CD1- CG- CD2- HD21
2326 2324 2330 2332 3 0.66944 2.00832 0.00000
-2.67776 0.00000 0.00000 ; CD1- CG- CD2- HD22
2326 2324 2330 2333 3 0.66944 2.00832 0.00000
-2.67776 0.00000 0.00000 ; CD1- CG- CD2- HD23
2327 2326 2324 2330 3 0.66944 2.00832 0.00000
-2.67776 0.00000 0.00000 ; HD11- CD1- CG- CD2
2328 2326 2324 2330 3 0.66944 2.00832 0.00000
-2.67776 0.00000 0.00000 ; HD12- CD1- CG- CD2
2329 2326 2324 2330 3 0.66944 2.00832 0.00000
-2.67776 0.00000 0.00000 ; HD13- CD1- CG- CD2
2334 2336 2338 2339 3 0.00000 0.00000 0.00000
0.00000 0.00000 0.00000 ; C- N- CA- HA
2334 2336 2338 2340 3 11.71520 1.67360 15.06240 -
13.38880 0.00000 0.00000 ; C- N- CA- CB
2334 2336 2338 2354 3 1.75728 5.27184 2.25936
-7.02912 0.00000 0.00000 ; C- N- CA- C
2335 2334 2336 2337 3 29.28800 -8.36800 -20.92000
0.00000 0.00000 0.00000 ; O- C- N- H
2335 2334 2336 2338 3 20.92000 0.00000 -20.92000
0.00000 0.00000 0.00000 ; O- C- N- CA
2336 2338 2340 2341 3 0.65084 1.95253 0.00000
-2.60338 0.00000 0.00000 ; N- CA- CB- HB2
2336 2338 2340 2342 3 0.65084 1.95253 0.00000
-2.60338 0.00000 0.00000 ; N- CA- CB- HB3
2336 2338 2340 2343 3 3.28026 2.46019 -2.19242
-3.21331 -0.23430 0.00000 ; N- CA- CB- CG
2336 2338 2354 2355 3 0.00000 0.00000 0.00000
0.00000 0.00000 0.00000 ; N- CA- C- O
2336 2338 2354 2356 3 17.40544 -5.02080 -13.22144
9.20480 0.00000 0.00000 ; N- CA- C- N
2337 2336 2338 2339 3 0.00000 0.00000 0.00000
0.00000 0.00000 0.00000 ; H- N- CA- HA
2337 2336 2338 2340 3 0.00000 0.00000 0.00000
0.00000 0.00000 0.00000 ; H- N- CA- CB

2337 2336 2338 2354 3 0.00000 0.00000 0.00000
0.00000 0.00000 0.00000 ; H- N- CA- C
2338 2340 2343 2344 3 0.57739 0.00000 1.02926
0.00000 -1.60666 0.00000 ; CA- CB- CG- CD1
2338 2340 2343 2352 3 0.57739 0.00000 1.02926
0.00000 -1.60666 0.00000 ; CA- CB- CG- CD2
2338 2354 2356 2357 3 20.92000 0.00000 -20.92000
0.00000 0.00000 0.00000 ; CA- C- N- H
2338 2354 2356 2358 3 20.92000 0.00000 -20.92000
0.00000 0.00000 0.00000 ; CA- C- N- CA
2339 2338 2340 2341 3 0.65084 1.95253 0.00000
-2.60338 0.00000 0.00000 ; HA- CA- CB- HB2
2339 2338 2340 2342 3 0.65084 1.95253 0.00000
-2.60338 0.00000 0.00000 ; HA- CA- CB- HB3
2339 2338 2340 2343 3 0.65084 1.95253 0.00000
-2.60338 0.00000 0.00000 ; HA- CA- CB- CG
2339 2338 2354 2355 3 3.68192 -4.35136 0.00000
1.33888 0.00000 0.00000 ; HA- CA- C- O
2339 2338 2354 2356 3 0.00000 0.00000 0.00000
0.00000 0.00000 0.00000 ; HA- CA- C- N
2340 2338 2354 2355 3 0.00000 0.00000 0.00000
0.00000 0.00000 0.00000 ; CB- CA- C- O
2340 2338 2354 2356 3 2.51040 4.18400 1.67360
-6.69440 0.00000 0.00000 ; CB- CA- C- N
2340 2343 2344 2345 3 30.33400 0.00000 -30.33400
0.00000 0.00000 0.00000 ; CB- CG- CD1- HD1
2340 2343 2344 2346 3 30.33400 0.00000 -30.33400
0.00000 0.00000 0.00000 ; CB- CG- CD1- CE1
2340 2343 2352 2350 3 30.33400 0.00000 -30.33400
0.00000 0.00000 0.00000 ; CB- CG- CD2- CE2
2340 2343 2352 2353 3 30.33400 0.00000 -30.33400
0.00000 0.00000 0.00000 ; CB- CG- CD2- HD2
2341 2340 2338 2354 3 0.65084 1.95253 0.00000
-2.60338 0.00000 0.00000 ; HB2- CB- CA- C
2341 2340 2343 2344 3 0.00000 0.00000 0.00000
0.00000 0.00000 0.00000 ; HB2- CB- CG- CD1
2341 2340 2343 2352 3 0.00000 0.00000 0.00000
0.00000 0.00000 0.00000 ; HB2- CB- CG- CD2
2342 2340 2338 2354 3 0.65084 1.95253 0.00000
-2.60338 0.00000 0.00000 ; HB3- CB- CA- C
2342 2340 2343 2344 3 0.00000 0.00000 0.00000
0.00000 0.00000 0.00000 ; HB3- CB- CG- CD1
2342 2340 2343 2352 3 0.00000 0.00000 0.00000
0.00000 0.00000 0.00000 ; HB3- CB- CG- CD2
2343 2340 2338 2354 3 4.95804 2.17986 -3.52293
-3.21331 -0.40166 0.00000 ; CG- CB- CA- C
2343 2344 2346 2347 3 30.33400 0.00000 -30.33400
0.00000 0.00000 0.00000 ; CG- CD1- CE1- HE1
2343 2344 2346 2348 3 30.33400 0.00000 -30.33400
0.00000 0.00000 0.00000 ; CG- CD1- CE1- CZ
2343 2352 2350 2348 3 30.33400 0.00000 -30.33400
0.00000 0.00000 0.00000 ; CG- CD2- CE2- CZ

2343 2352 2350 2351 3 30.33400 0.00000 -30.33400
0.00000 0.00000 0.00000 ; CG- CD2- CE2- HE2
2344 2343 2352 2350 3 30.33400 0.00000 -30.33400
0.00000 0.00000 0.00000 ; CD1- CG- CD2- CE2
2344 2343 2352 2353 3 30.33400 0.00000 -30.33400
0.00000 0.00000 0.00000 ; CD1- CG- CD2- HD2
2344 2346 2348 2349 3 30.33400 0.00000 -30.33400
0.00000 0.00000 0.00000 ; CD1- CE1- CZ- HZ
2344 2346 2348 2350 3 30.33400 0.00000 -30.33400
0.00000 0.00000 0.00000 ; CD1- CE1- CZ- CE2
2345 2344 2343 2352 3 30.33400 0.00000 -30.33400
0.00000 0.00000 0.00000 ; HD1- CD1- CG- CD2
2345 2344 2346 2347 3 30.33400 0.00000 -30.33400
0.00000 0.00000 0.00000 ; HD1- CD1- CE1- HE1
2345 2344 2346 2348 3 30.33400 0.00000 -30.33400
0.00000 0.00000 0.00000 ; HD1- CD1- CE1- CZ
2346 2344 2343 2352 3 30.33400 0.00000 -30.33400
0.00000 0.00000 0.00000 ; CE1- CD1- CG- CD2
2346 2348 2350 2351 3 30.33400 0.00000 -30.33400
0.00000 0.00000 0.00000 ; CE1- CZ- CE2- HE2
2346 2348 2350 2352 3 30.33400 0.00000 -30.33400
0.00000 0.00000 0.00000 ; CE1- CZ- CE2- CD2
2347 2346 2348 2349 3 30.33400 0.00000 -30.33400
0.00000 0.00000 0.00000 ; HE1- CE1- CZ- HZ
2347 2346 2348 2350 3 30.33400 0.00000 -30.33400
0.00000 0.00000 0.00000 ; HE1- CE1- CZ- CE2
2348 2350 2352 2353 3 30.33400 0.00000 -30.33400
0.00000 0.00000 0.00000 ; CZ- CE2- CD2- HD2
2349 2348 2350 2351 3 30.33400 0.00000 -30.33400
0.00000 0.00000 0.00000 ; HZ- CZ- CE2- HE2
2349 2348 2350 2352 3 30.33400 0.00000 -30.33400
0.00000 0.00000 0.00000 ; HZ- CZ- CE2- CD2
2351 2350 2352 2353 3 30.33400 0.00000 -30.33400
0.00000 0.00000 0.00000 ; HE2- CE2- CD2- HD2
2354 2356 2358 2359 3 0.00000 0.00000 0.00000
0.00000 0.00000 0.00000 ; C- N- CA- HA
2354 2356 2358 2360 3 11.71520 1.67360 15.06240 -
13.38880 0.00000 0.00000 ; C- N- CA- CB
2354 2356 2358 2371 3 1.75728 5.27184 2.25936
-7.02912 0.00000 0.00000 ; C- N- CA- C
2355 2354 2356 2357 3 29.28800 -8.36800 -20.92000
0.00000 0.00000 0.00000 ; O- C- N- H
2355 2354 2356 2358 3 20.92000 0.00000 -20.92000
0.00000 0.00000 0.00000 ; O- C- N- CA
2356 2358 2360 2361 3 0.65084 1.95253 0.00000
-2.60338 0.00000 0.00000 ; N- CA- CB- HB2
2356 2358 2360 2362 3 0.65084 1.95253 0.00000
-2.60338 0.00000 0.00000 ; N- CA- CB- HB3
2356 2358 2360 2363 3 3.21331 2.22589 -4.15053
-2.40998 2.61082 0.00000 ; N- CA- CB- CG
2356 2358 2371 2372 3 0.00000 0.00000 0.00000
0.00000 0.00000 0.00000 ; N- CA- C- O

2356 2358 2371 2373 3 17.40544 -5.02080 -13.22144
9.20480 0.00000 0.00000 ; N- CA- C- N
2357 2356 2358 2359 3 0.00000 0.00000 0.00000
0.00000 0.00000 0.00000 ; H- N- CA- HA
2357 2356 2358 2360 3 0.00000 0.00000 0.00000
0.00000 0.00000 0.00000 ; H- N- CA- CB
2357 2356 2358 2371 3 0.00000 0.00000 0.00000
0.00000 0.00000 0.00000 ; H- N- CA- C
2358 2360 2363 2364 3 0.66944 2.00832 0.00000
-2.67776 0.00000 0.00000 ; CA- CB- CG- HG2
2358 2360 2363 2365 3 0.66944 2.00832 0.00000
-2.67776 0.00000 0.00000 ; CA- CB- CG- HG3
2358 2360 2363 2366 3 3.69866 -4.35136 -3.92459
6.89523 4.61914 0.00000 ; CA- CB- CG- CD
2358 2371 2373 2374 3 20.92000 0.00000 -20.92000
0.00000 0.00000 0.00000 ; CA- C- N- H
2358 2371 2373 2375 3 20.92000 0.00000 -20.92000
0.00000 0.00000 0.00000 ; CA- C- N- CA
2359 2358 2360 2361 3 0.65084 1.95253 0.00000
-2.60338 0.00000 0.00000 ; HA- CA- CB- HB2
2359 2358 2360 2362 3 0.65084 1.95253 0.00000
-2.60338 0.00000 0.00000 ; HA- CA- CB- HB3
2359 2358 2360 2363 3 0.65084 1.95253 0.00000
-2.60338 0.00000 0.00000 ; HA- CA- CB- CG
2359 2358 2371 2372 3 3.68192 -4.35136 0.00000
1.33888 0.00000 0.00000 ; HA- CA- C- O
2359 2358 2371 2373 3 0.00000 0.00000 0.00000
0.00000 0.00000 0.00000 ; HA- CA- C- N
2360 2358 2371 2372 3 0.00000 0.00000 0.00000
0.00000 0.00000 0.00000 ; CB- CA- C- O
2360 2358 2371 2373 3 2.51040 4.18400 1.67360
-6.69440 0.00000 0.00000 ; CB- CA- C- N
2360 2363 2366 2367 3 0.00000 0.00000 0.00000
0.00000 0.00000 0.00000 ; CB- CG- CD- OE1
2360 2363 2366 2368 3 10.32611 1.48114 -8.47678
1.42256 1.40582 0.00000 ; CB- CG- CD- NE2
2361 2360 2358 2371 3 0.65084 1.95253 0.00000
-2.60338 0.00000 0.00000 ; HB2- CB- CA- C
2361 2360 2363 2364 3 0.62760 1.88280 0.00000
-2.51040 0.00000 0.00000 ; HB2- CB- CG- HG2
2361 2360 2363 2365 3 0.62760 1.88280 0.00000
-2.51040 0.00000 0.00000 ; HB2- CB- CG- HG3
2361 2360 2363 2366 3 0.65084 1.95253 0.00000
-2.60338 0.00000 0.00000 ; HB2- CB- CG- CD
2362 2360 2358 2371 3 0.65084 1.95253 0.00000
-2.60338 0.00000 0.00000 ; HB3- CB- CA- C
2362 2360 2363 2364 3 0.62760 1.88280 0.00000
-2.51040 0.00000 0.00000 ; HB3- CB- CG- HG2
2362 2360 2363 2365 3 0.62760 1.88280 0.00000
-2.51040 0.00000 0.00000 ; HB3- CB- CG- HG3
2362 2360 2363 2366 3 0.65084 1.95253 0.00000
-2.60338 0.00000 0.00000 ; HB3- CB- CG- CD

2363 2360 2358 2371 3 6.86594 3.56895 -8.14206
-2.40998 4.85344 0.00000 ; CG- CB- CA- C
2363 2366 2368 2369 3 20.92000 0.00000 -20.92000
0.00000 0.00000 0.00000 ; CG- CD- NE2- HE21
2363 2366 2368 2370 3 20.92000 0.00000 -20.92000
0.00000 0.00000 0.00000 ; CG- CD- NE2- HE22
2364 2363 2366 2367 3 3.68192 -4.35136 0.00000
1.33888 0.00000 0.00000 ; HG2- CG- CD- OE1
2364 2363 2366 2368 3 0.00000 0.00000 0.00000
0.00000 0.00000 0.00000 ; HG2- CG- CD- NE2
2365 2363 2366 2367 3 3.68192 -4.35136 0.00000
1.33888 0.00000 0.00000 ; HG3- CG- CD- OE1
2365 2363 2366 2368 3 0.00000 0.00000 0.00000
0.00000 0.00000 0.00000 ; HG3- CG- CD- NE2
2367 2366 2368 2369 3 29.28800 -8.36800 -20.92000
0.00000 0.00000 0.00000 ; OE1- CD- NE2- HE21
2367 2366 2368 2370 3 29.28800 -8.36800 -20.92000
0.00000 0.00000 0.00000 ; OE1- CD- NE2- HE22
2371 2373 2375 2376 3 0.00000 0.00000 0.00000
0.00000 0.00000 0.00000 ; C- N- CA- HA
2371 2373 2375 2377 3 11.71520 1.67360 15.06240 -
13.38880 0.00000 0.00000 ; C- N- CA- CB
2371 2373 2375 2382 3 1.75728 5.27184 2.25936
-7.02912 0.00000 0.00000 ; C- N- CA- C
2372 2371 2373 2374 3 29.28800 -8.36800 -20.92000
0.00000 0.00000 0.00000 ; O- C- N- H
2372 2371 2373 2375 3 20.92000 0.00000 -20.92000
0.00000 0.00000 0.00000 ; O- C- N- CA
2373 2375 2377 2378 3 0.65084 1.95253 0.00000
-2.60338 0.00000 0.00000 ; N- CA- CB- HB2
2373 2375 2377 2379 3 0.65084 1.95253 0.00000
-2.60338 0.00000 0.00000 ; N- CA- CB- HB3
2373 2375 2377 2380 3 7.86174 2.24681 -7.41405
-6.71114 5.35552 0.00000 ; N- CA- CB- OG
2373 2375 2382 2383 3 0.00000 0.00000 0.00000
0.00000 0.00000 0.00000 ; N- CA- C- O
2373 2375 2382 2384 3 17.40544 -5.02080 -13.22144
9.20480 0.00000 0.00000 ; N- CA- C- N
2374 2373 2375 2376 3 0.00000 0.00000 0.00000
0.00000 0.00000 0.00000 ; H- N- CA- HA
2374 2373 2375 2377 3 0.00000 0.00000 0.00000
0.00000 0.00000 0.00000 ; H- N- CA- CB
2374 2373 2375 2382 3 0.00000 0.00000 0.00000
0.00000 0.00000 0.00000 ; H- N- CA- C
2375 2377 2380 2381 3 2.05853 2.46856 3.48109
-4.46851 0.23430 0.00000 ; CA- CB- OG- HG
2375 2382 2384 2385 3 20.92000 0.00000 -20.92000
0.00000 0.00000 0.00000 ; CA- C- N- H
2375 2382 2384 2386 3 20.92000 0.00000 -20.92000
0.00000 0.00000 0.00000 ; CA- C- N- CA
2376 2375 2377 2378 3 0.65084 1.95253 0.00000
-2.60338 0.00000 0.00000 ; HA- CA- CB- HB2

2376 2375 2377 2379 3 0.65084 1.95253 0.00000
-2.60338 0.00000 0.00000 ; HA- CA- CB- HB3
2376 2375 2377 2380 3 1.04600 -1.04600 0.00000
0.00000 0.00000 0.00000 ; HA- CA- CB- OG
2376 2375 2382 2383 3 3.68192 -4.35136 0.00000
1.33888 0.00000 0.00000 ; HA- CA- C- O
2376 2375 2382 2384 3 0.00000 0.00000 0.00000
0.00000 0.00000 0.00000 ; HA- CA- C- N
2377 2375 2382 2383 3 0.00000 0.00000 0.00000
0.00000 0.00000 0.00000 ; CB- CA- C- O
2377 2375 2382 2384 3 2.51040 4.18400 1.67360
-6.69440 0.00000 0.00000 ; CB- CA- C- N
2378 2377 2375 2382 3 0.65084 1.95253 0.00000
-2.60338 0.00000 0.00000 ; HB2- CB- CA- C
2378 2377 2380 2381 3 0.69733 2.09200 0.00000
-2.78933 0.00000 0.00000 ; HB2- CB- OG- HG
2379 2377 2375 2382 3 0.65084 1.95253 0.00000
-2.60338 0.00000 0.00000 ; HB3- CB- CA- C
2379 2377 2380 2381 3 0.69733 2.09200 0.00000
-2.78933 0.00000 0.00000 ; HB3- CB- OG- HG
2380 2377 2375 2382 3 7.34710 7.79898 -6.14211
-6.71114 4.31789 0.00000 ; OG- CB- CA- C
2382 2384 2386 2387 3 0.00000 0.00000 0.00000
0.00000 0.00000 0.00000 ; C- N- CA- HA
2382 2384 2386 2388 3 11.71520 1.67360 15.06240 -
13.38880 0.00000 0.00000 ; C- N- CA- CB
2382 2384 2386 2397 3 1.75728 5.27184 2.25936
-7.02912 0.00000 0.00000 ; C- N- CA- C
2383 2382 2384 2385 3 29.28800 -8.36800 -20.92000
0.00000 0.00000 0.00000 ; O- C- N- H
2383 2382 2384 2386 3 20.92000 0.00000 -20.92000
0.00000 0.00000 0.00000 ; O- C- N- CA
2384 2386 2388 2389 3 0.65084 1.95253 0.00000
-2.60338 0.00000 0.00000 ; N- CA- CB- HB2
2384 2386 2388 2390 3 0.65084 1.95253 0.00000
-2.60338 0.00000 0.00000 ; N- CA- CB- HB3
2384 2386 2388 2391 3 3.21331 2.22589 -4.15053
-2.40998 2.61082 0.00000 ; N- CA- CB- CG
2384 2386 2397 2398 3 0.00000 0.00000 0.00000
0.00000 0.00000 0.00000 ; N- CA- C- O
2384 2386 2397 2399 3 17.40544 -5.02080 -13.22144
9.20480 0.00000 0.00000 ; N- CA- C- N
2385 2384 2386 2387 3 0.00000 0.00000 0.00000
0.00000 0.00000 0.00000 ; H- N- CA- HA
2385 2384 2386 2388 3 0.00000 0.00000 0.00000
0.00000 0.00000 0.00000 ; H- N- CA- CB
2385 2384 2386 2397 3 0.00000 0.00000 0.00000
0.00000 0.00000 0.00000 ; H- N- CA- C
2386 2388 2391 2392 3 0.66944 2.00832 0.00000
-2.67776 0.00000 0.00000 ; CA- CB- CG- HG2
2386 2388 2391 2393 3 0.66944 2.00832 0.00000
-2.67776 0.00000 0.00000 ; CA- CB- CG- HG3

2386 2388 2391 2394 3 10.12110 -1.91209 0.01674
10.17549 -1.87443 0.00000 ; CA- CB- CG- CD
2386 2397 2399 2400 3 20.92000 0.00000 -20.92000
0.00000 0.00000 0.00000 ; CA- C- N- H
2386 2397 2399 2401 3 20.92000 0.00000 -20.92000
0.00000 0.00000 0.00000 ; CA- C- N- CA
2387 2386 2388 2389 3 0.65084 1.95253 0.00000
-2.60338 0.00000 0.00000 ; HA- CA- CB- HB2
2387 2386 2388 2390 3 0.65084 1.95253 0.00000
-2.60338 0.00000 0.00000 ; HA- CA- CB- HB3
2387 2386 2388 2391 3 0.65084 1.95253 0.00000
-2.60338 0.00000 0.00000 ; HA- CA- CB- CG
2387 2386 2397 2398 3 3.68192 -4.35136 0.00000
1.33888 0.00000 0.00000 ; HA- CA- C- O
2387 2386 2397 2399 3 0.00000 0.00000 0.00000
0.00000 0.00000 0.00000 ; HA- CA- C- N
2388 2386 2397 2398 3 0.00000 0.00000 0.00000
0.00000 0.00000 0.00000 ; CB- CA- C- O
2388 2386 2397 2399 3 2.51040 4.18400 1.67360
-6.69440 0.00000 0.00000 ; CB- CA- C- N
2388 2391 2394 2395 3 3.79907 0.00000 -5.40573
0.00000 2.14221 0.00000 ; CB- CG- CD- OE1
2388 2391 2394 2396 3 3.79907 0.00000 -5.40573
0.00000 2.14221 0.00000 ; CB- CG- CD- OE2
2389 2388 2386 2397 3 0.65084 1.95253 0.00000
-2.60338 0.00000 0.00000 ; HB2- CB- CA- C
2389 2388 2391 2392 3 0.62760 1.88280 0.00000
-2.51040 0.00000 0.00000 ; HB2- CB- CG- HG2
2389 2388 2391 2393 3 0.62760 1.88280 0.00000
-2.51040 0.00000 0.00000 ; HB2- CB- CG- HG3
2389 2388 2391 2394 3 0.65084 1.95253 0.00000
-2.60338 0.00000 0.00000 ; HB2- CB- CG- CD
2390 2388 2386 2397 3 0.65084 1.95253 0.00000
-2.60338 0.00000 0.00000 ; HB3- CB- CA- C
2390 2388 2391 2392 3 0.62760 1.88280 0.00000
-2.51040 0.00000 0.00000 ; HB3- CB- CG- HG2
2390 2388 2391 2393 3 0.62760 1.88280 0.00000
-2.51040 0.00000 0.00000 ; HB3- CB- CG- HG3
2390 2388 2391 2394 3 0.65084 1.95253 0.00000
-2.60338 0.00000 0.00000 ; HB3- CB- CG- CD
2391 2388 2386 2397 3 6.86594 3.56895 -8.14206
-2.40998 4.85344 0.00000 ; CG- CB- CA- C
2392 2391 2394 2395 3 0.00000 0.00000 0.00000
0.00000 0.00000 0.00000 ; HG2- CG- CD- OE1
2392 2391 2394 2396 3 0.00000 0.00000 0.00000
0.00000 0.00000 0.00000 ; HG2- CG- CD- OE2
2393 2391 2394 2395 3 0.00000 0.00000 0.00000
0.00000 0.00000 0.00000 ; HG3- CG- CD- OE1
2393 2391 2394 2396 3 0.00000 0.00000 0.00000
0.00000 0.00000 0.00000 ; HG3- CG- CD- OE2
2397 2399 2401 2402 3 0.00000 0.00000 0.00000
0.00000 0.00000 0.00000 ; C- N- CA- HA

2397 2399 2401 2403 3 11.71520 1.67360 15.06240 -
13.38880 0.00000 0.00000 ; C- N- CA- CB
2397 2399 2401 2411 3 1.75728 5.27184 2.25936
-7.02912 0.00000 0.00000 ; C- N- CA- C
2398 2397 2399 2400 3 29.28800 -8.36800 -20.92000
0.00000 0.00000 0.00000 ; O- C- N- H
2398 2397 2399 2401 3 20.92000 0.00000 -20.92000
0.00000 0.00000 0.00000 ; O- C- N- CA
2399 2401 2403 2404 3 0.65084 1.95253 0.00000
-2.60338 0.00000 0.00000 ; N- CA- CB- HB2
2399 2401 2403 2405 3 0.65084 1.95253 0.00000
-2.60338 0.00000 0.00000 ; N- CA- CB- HB3
2399 2401 2403 2406 3 5.99567 3.29281 -4.46014
-0.55229 1.97485 0.00000 ; N- CA- CB- CG
2399 2401 2411 2412 3 0.00000 0.00000 0.00000
0.00000 0.00000 0.00000 ; N- CA- C- O
2399 2401 2411 2413 3 17.40544 -5.02080 -13.22144
9.20480 0.00000 0.00000 ; N- CA- C- N
2400 2399 2401 2402 3 0.00000 0.00000 0.00000
0.00000 0.00000 0.00000 ; H- N- CA- HA
2400 2399 2401 2403 3 0.00000 0.00000 0.00000
0.00000 0.00000 0.00000 ; H- N- CA- CB
2400 2399 2401 2411 3 0.00000 0.00000 0.00000
0.00000 0.00000 0.00000 ; H- N- CA- C
2401 2403 2406 2407 3 0.00000 0.00000 0.00000
0.00000 0.00000 0.00000 ; CA- CB- CG- OD1
2401 2403 2406 2408 3 8.84916 -0.31380 -4.32626
5.03754 0.26778 0.00000 ; CA- CB- CG- ND2
2401 2411 2413 2414 3 20.92000 0.00000 -20.92000
0.00000 0.00000 0.00000 ; CA- C- N- H
2401 2411 2413 2415 3 20.92000 0.00000 -20.92000
0.00000 0.00000 0.00000 ; CA- C- N- CA
2402 2401 2403 2404 3 0.65084 1.95253 0.00000
-2.60338 0.00000 0.00000 ; HA- CA- CB- HB2
2402 2401 2403 2405 3 0.65084 1.95253 0.00000
-2.60338 0.00000 0.00000 ; HA- CA- CB- HB3
2402 2401 2403 2406 3 0.65084 1.95253 0.00000
-2.60338 0.00000 0.00000 ; HA- CA- CB- CG
2402 2401 2411 2412 3 3.68192 -4.35136 0.00000
1.33888 0.00000 0.00000 ; HA- CA- C- O
2402 2401 2411 2413 3 0.00000 0.00000 0.00000
0.00000 0.00000 0.00000 ; HA- CA- C- N
2403 2401 2411 2412 3 0.00000 0.00000 0.00000
0.00000 0.00000 0.00000 ; CB- CA- C- O
2403 2401 2411 2413 3 2.51040 4.18400 1.67360
-6.69440 0.00000 0.00000 ; CB- CA- C- N
2403 2406 2408 2409 3 20.92000 0.00000 -20.92000
0.00000 0.00000 0.00000 ; CB- CG- ND2- HD21
2403 2406 2408 2410 3 20.92000 0.00000 -20.92000
0.00000 0.00000 0.00000 ; CB- CG- ND2- HD22
2404 2403 2401 2411 3 0.65084 1.95253 0.00000
-2.60338 0.00000 0.00000 ; HB2- CB- CA- C

2404 2403 2406 2407 3 3.68192 -4.35136 0.00000
1.33888 0.00000 0.00000 ; HB2- CB- CG- OD1
2404 2403 2406 2408 3 0.00000 0.00000 0.00000
0.00000 0.00000 0.00000 ; HB2- CB- CG- ND2
2405 2403 2401 2411 3 0.65084 1.95253 0.00000
-2.60338 0.00000 0.00000 ; HB3- CB- CA- C
2405 2403 2406 2407 3 3.68192 -4.35136 0.00000
1.33888 0.00000 0.00000 ; HB3- CB- CG- OD1
2405 2403 2406 2408 3 0.00000 0.00000 0.00000
0.00000 0.00000 0.00000 ; HB3- CB- CG- ND2
2406 2403 2401 2411 3 7.94542 -3.96225 -6.11701
-0.55229 3.58150 0.00000 ; CG- CB- CA- C
2407 2406 2408 2409 3 29.28800 -8.36800 -20.92000
0.00000 0.00000 0.00000 ; OD1- CG- ND2- HD21
2407 2406 2408 2410 3 29.28800 -8.36800 -20.92000
0.00000 0.00000 0.00000 ; OD1- CG- ND2- HD22
2411 2413 2415 2416 3 0.00000 0.00000 0.00000
0.00000 0.00000 0.00000 ; C- N- CA- HA
2411 2413 2415 2417 3 11.71520 1.67360 15.06240 -
13.38880 0.00000 0.00000 ; C- N- CA- CB
2411 2413 2415 2433 3 1.75728 5.27184 2.25936
-7.02912 0.00000 0.00000 ; C- N- CA- C
2412 2411 2413 2414 3 29.28800 -8.36800 -20.92000
0.00000 0.00000 0.00000 ; O- C- N- H
2412 2411 2413 2415 3 20.92000 0.00000 -20.92000
0.00000 0.00000 0.00000 ; O- C- N- CA
2413 2415 2417 2418 3 0.65084 1.95253 0.00000
-2.60338 0.00000 0.00000 ; N- CA- CB- HB2
2413 2415 2417 2419 3 0.65084 1.95253 0.00000
-2.60338 0.00000 0.00000 ; N- CA- CB- HB3
2413 2415 2417 2420 3 0.65084 1.95253 0.00000
-2.60338 0.00000 0.00000 ; N- CA- CB- CG
2413 2415 2433 2434 3 0.00000 0.00000 0.00000
0.00000 0.00000 0.00000 ; N- CA- C- O
2413 2415 2433 2435 3 17.40544 -5.02080 -13.22144
9.20480 0.00000 0.00000 ; N- CA- C- N
2414 2413 2415 2416 3 0.00000 0.00000 0.00000
0.00000 0.00000 0.00000 ; H- N- CA- HA
2414 2413 2415 2417 3 0.00000 0.00000 0.00000
0.00000 0.00000 0.00000 ; H- N- CA- CB
2414 2413 2415 2433 3 0.00000 0.00000 0.00000
0.00000 0.00000 0.00000 ; H- N- CA- C
2415 2417 2420 2421 3 0.66944 2.00832 0.00000
-2.67776 0.00000 0.00000 ; CA- CB- CG- HG2
2415 2417 2420 2422 3 0.66944 2.00832 0.00000
-2.67776 0.00000 0.00000 ; CA- CB- CG- HG3
2415 2417 2420 2423 3 3.68192 3.09616 -2.09200
-3.01248 0.00000 0.00000 ; CA- CB- CG- CD
2415 2433 2435 2436 3 20.92000 0.00000 -20.92000
0.00000 0.00000 0.00000 ; CA- C- N- H
2415 2433 2435 2437 3 20.92000 0.00000 -20.92000
0.00000 0.00000 0.00000 ; CA- C- N- CA

2416 2415 2417 2418 3 0.65084 1.95253 0.00000
-2.60338 0.00000 0.00000 ; HA- CA- CB- HB2
2416 2415 2417 2419 3 0.65084 1.95253 0.00000
-2.60338 0.00000 0.00000 ; HA- CA- CB- HB3
2416 2415 2417 2420 3 0.65084 1.95253 0.00000
-2.60338 0.00000 0.00000 ; HA- CA- CB- CG
2416 2415 2433 2434 3 3.68192 -4.35136 0.00000
1.33888 0.00000 0.00000 ; HA- CA- C- O
2416 2415 2433 2435 3 0.00000 0.00000 0.00000
0.00000 0.00000 0.00000 ; HA- CA- C- N
2417 2415 2433 2434 3 0.00000 0.00000 0.00000
0.00000 0.00000 0.00000 ; CB- CA- C- O
2417 2415 2433 2435 3 2.51040 4.18400 1.67360
-6.69440 0.00000 0.00000 ; CB- CA- C- N
2417 2420 2423 2424 3 0.66944 2.00832 0.00000
-2.67776 0.00000 0.00000 ; CB- CG- CD- HD2
2417 2420 2423 2425 3 0.66944 2.00832 0.00000
-2.67776 0.00000 0.00000 ; CB- CG- CD- HD3
2417 2420 2423 2426 3 3.68192 3.09616 -2.09200
-3.01248 0.00000 0.00000 ; CB- CG- CD- CE
2418 2417 2415 2433 3 0.65084 1.95253 0.00000
-2.60338 0.00000 0.00000 ; HB2- CB- CA- C
2418 2417 2420 2421 3 0.62760 1.88280 0.00000
-2.51040 0.00000 0.00000 ; HB2- CB- CG- HG2
2418 2417 2420 2422 3 0.62760 1.88280 0.00000
-2.51040 0.00000 0.00000 ; HB2- CB- CG- HG3
2418 2417 2420 2423 3 0.66944 2.00832 0.00000
-2.67776 0.00000 0.00000 ; HB2- CB- CG- CD
2419 2417 2415 2433 3 0.65084 1.95253 0.00000
-2.60338 0.00000 0.00000 ; HB3- CB- CA- C
2419 2417 2420 2421 3 0.62760 1.88280 0.00000
-2.51040 0.00000 0.00000 ; HB3- CB- CG- HG2
2419 2417 2420 2422 3 0.62760 1.88280 0.00000
-2.51040 0.00000 0.00000 ; HB3- CB- CG- HG3
2419 2417 2420 2423 3 0.66944 2.00832 0.00000
-2.67776 0.00000 0.00000 ; HB3- CB- CG- CD
2420 2417 2415 2433 3 0.65084 1.95253 0.00000
-2.60338 0.00000 0.00000 ; CG- CB- CA- C
2420 2423 2426 2427 3 0.65084 1.95253 0.00000
-2.60338 0.00000 0.00000 ; CG- CD- CE- HE2
2420 2423 2426 2428 3 0.65084 1.95253 0.00000
-2.60338 0.00000 0.00000 ; CG- CD- CE- HE3
2420 2423 2426 2429 3 0.65084 1.95253 0.00000
-2.60338 0.00000 0.00000 ; CG- CD- CE- NZ
2421 2420 2423 2424 3 0.62760 1.88280 0.00000
-2.51040 0.00000 0.00000 ; HG2- CG- CD- HD2
2421 2420 2423 2425 3 0.62760 1.88280 0.00000
-2.51040 0.00000 0.00000 ; HG2- CG- CD- HD3
2421 2420 2423 2426 3 0.66944 2.00832 0.00000
-2.67776 0.00000 0.00000 ; HG2- CG- CD- CE
2422 2420 2423 2424 3 0.62760 1.88280 0.00000
-2.51040 0.00000 0.00000 ; HG3- CG- CD- HD2

2422 2420 2423 2425 3 0.62760 1.88280 0.00000
-2.51040 0.00000 0.00000 ; HG3- CG- CD- HD3
2422 2420 2423 2426 3 0.66944 2.00832 0.00000
-2.67776 0.00000 0.00000 ; HG3- CG- CD- CE
2423 2426 2429 2430 3 0.65084 1.95253 0.00000
-2.60338 0.00000 0.00000 ; CD- CE- NZ- HZ1
2423 2426 2429 2431 3 0.65084 1.95253 0.00000
-2.60338 0.00000 0.00000 ; CD- CE- NZ- HZ2
2423 2426 2429 2432 3 0.65084 1.95253 0.00000
-2.60338 0.00000 0.00000 ; CD- CE- NZ- HZ3
2424 2423 2426 2427 3 0.65084 1.95253 0.00000
-2.60338 0.00000 0.00000 ; HD2- CD- CE- HE2
2424 2423 2426 2428 3 0.65084 1.95253 0.00000
-2.60338 0.00000 0.00000 ; HD2- CD- CE- HE3
2424 2423 2426 2429 3 0.65084 1.95253 0.00000
-2.60338 0.00000 0.00000 ; HD2- CD- CE- NZ
2425 2423 2426 2427 3 0.65084 1.95253 0.00000
-2.60338 0.00000 0.00000 ; HD3- CD- CE- HE2
2425 2423 2426 2428 3 0.65084 1.95253 0.00000
-2.60338 0.00000 0.00000 ; HD3- CD- CE- HE3
2425 2423 2426 2429 3 0.65084 1.95253 0.00000
-2.60338 0.00000 0.00000 ; HD3- CD- CE- NZ
2427 2426 2429 2430 3 0.65084 1.95253 0.00000
-2.60338 0.00000 0.00000 ; HE2- CE- NZ- HZ1
2427 2426 2429 2431 3 0.65084 1.95253 0.00000
-2.60338 0.00000 0.00000 ; HE2- CE- NZ- HZ2
2427 2426 2429 2432 3 0.65084 1.95253 0.00000
-2.60338 0.00000 0.00000 ; HE2- CE- NZ- HZ3
2428 2426 2429 2430 3 0.65084 1.95253 0.00000
-2.60338 0.00000 0.00000 ; HE3- CE- NZ- HZ1
2428 2426 2429 2431 3 0.65084 1.95253 0.00000
-2.60338 0.00000 0.00000 ; HE3- CE- NZ- HZ2
2428 2426 2429 2432 3 0.65084 1.95253 0.00000
-2.60338 0.00000 0.00000 ; HE3- CE- NZ- HZ3
2433 2435 2437 2438 3 0.00000 0.00000 0.00000
0.00000 0.00000 0.00000 ; C- N- CA- HA
2433 2435 2437 2439 3 11.71520 1.67360 15.06240 -
13.38880 0.00000 0.00000 ; C- N- CA- CB
2433 2435 2437 2445 3 1.75728 5.27184 2.25936
-7.02912 0.00000 0.00000 ; C- N- CA- C
2434 2433 2435 2436 3 29.28800 -8.36800 -20.92000
0.00000 0.00000 0.00000 ; O- C- N- H
2434 2433 2435 2437 3 20.92000 0.00000 -20.92000
0.00000 0.00000 0.00000 ; O- C- N- CA
2435 2437 2439 2440 3 0.65084 1.95253 0.00000
-2.60338 0.00000 0.00000 ; N- CA- CB- HB2
2435 2437 2439 2441 3 0.65084 1.95253 0.00000
-2.60338 0.00000 0.00000 ; N- CA- CB- HB3
2435 2437 2439 2442 3 15.41386 9.74035 -8.39310
-0.97069 2.97901 0.00000 ; N- CA- CB- CG
2435 2437 2445 2446 3 0.00000 0.00000 0.00000
0.00000 0.00000 0.00000 ; N- CA- C- O

2435 2437 2445 2447 3 17.40544 -5.02080 -13.22144
9.20480 0.00000 0.00000 ; N- CA- C- N
2436 2435 2437 2438 3 0.00000 0.00000 0.00000
0.00000 0.00000 0.00000 ; H- N- CA- HA
2436 2435 2437 2439 3 0.00000 0.00000 0.00000
0.00000 0.00000 0.00000 ; H- N- CA- CB
2436 2435 2437 2445 3 0.00000 0.00000 0.00000
0.00000 0.00000 0.00000 ; H- N- CA- C
2437 2439 2442 2443 3 6.43499 0.00000 -5.39736
0.00000 -1.03763 0.00000 ; CA- CB- CG- OD1
2437 2439 2442 2444 3 6.43499 0.00000 -5.39736
0.00000 -1.03763 0.00000 ; CA- CB- CG- OD2
2437 2445 2447 2448 3 20.92000 0.00000 -20.92000
0.00000 0.00000 0.00000 ; CA- C- N- H
2437 2445 2447 2449 3 20.92000 0.00000 -20.92000
0.00000 0.00000 0.00000 ; CA- C- N- CA
2438 2437 2439 2440 3 0.65084 1.95253 0.00000
-2.60338 0.00000 0.00000 ; HA- CA- CB- HB2
2438 2437 2439 2441 3 0.65084 1.95253 0.00000
-2.60338 0.00000 0.00000 ; HA- CA- CB- HB3
2438 2437 2439 2442 3 0.65084 1.95253 0.00000
-2.60338 0.00000 0.00000 ; HA- CA- CB- CG
2438 2437 2445 2446 3 3.68192 -4.35136 0.00000
1.33888 0.00000 0.00000 ; HA- CA- C- O
2438 2437 2445 2447 3 0.00000 0.00000 0.00000
0.00000 0.00000 0.00000 ; HA- CA- C- N
2439 2437 2445 2446 3 0.00000 0.00000 0.00000
0.00000 0.00000 0.00000 ; CB- CA- C- O
2439 2437 2445 2447 3 2.51040 4.18400 1.67360
-6.69440 0.00000 0.00000 ; CB- CA- C- N
2440 2439 2437 2445 3 0.65084 1.95253 0.00000
-2.60338 0.00000 0.00000 ; HB2- CB- CA- C
2440 2439 2442 2443 3 0.00000 0.00000 0.00000
0.00000 0.00000 0.00000 ; HB2- CB- CG- OD1
2440 2439 2442 2444 3 0.00000 0.00000 0.00000
0.00000 0.00000 0.00000 ; HB2- CB- CG- OD2
2441 2439 2437 2445 3 0.65084 1.95253 0.00000
-2.60338 0.00000 0.00000 ; HB3- CB- CA- C
2441 2439 2442 2443 3 0.00000 0.00000 0.00000
0.00000 0.00000 0.00000 ; HB3- CB- CG- OD1
2441 2439 2442 2444 3 0.00000 0.00000 0.00000
0.00000 0.00000 0.00000 ; HB3- CB- CG- OD2
2442 2439 2437 2445 3 7.14627 -1.04600 -8.99560
-0.97069 5.15469 0.00000 ; CG- CB- CA- C
2445 2447 2449 2450 3 0.00000 0.00000 0.00000
0.00000 0.00000 0.00000 ; C- N- CA- HA
2445 2447 2449 2451 3 11.71520 1.67360 15.06240 -
13.38880 0.00000 0.00000 ; C- N- CA- CB
2445 2447 2449 2456 3 1.75728 5.27184 2.25936
-7.02912 0.00000 0.00000 ; C- N- CA- C
2446 2445 2447 2448 3 29.28800 -8.36800 -20.92000
0.00000 0.00000 0.00000 ; O- C- N- H

2446 2445 2447 2449 3 20.92000 0.00000 -20.92000
0.00000 0.00000 0.00000 ; O- C- N- CA
2447 2449 2451 2452 3 0.65084 1.95253 0.00000
-2.60338 0.00000 0.00000 ; N- CA- CB- HB2
2447 2449 2451 2453 3 0.65084 1.95253 0.00000
-2.60338 0.00000 0.00000 ; N- CA- CB- HB3
2447 2449 2451 2454 3 7.86174 2.24681 -7.41405
-6.71114 5.35552 0.00000 ; N- CA- CB- OG
2447 2449 2456 2457 3 0.00000 0.00000 0.00000
0.00000 0.00000 0.00000 ; N- CA- C- O
2447 2449 2456 2458 3 17.40544 -5.02080 -13.22144
9.20480 0.00000 0.00000 ; N- CA- C- N
2448 2447 2449 2450 3 0.00000 0.00000 0.00000
0.00000 0.00000 0.00000 ; H- N- CA- HA
2448 2447 2449 2451 3 0.00000 0.00000 0.00000
0.00000 0.00000 0.00000 ; H- N- CA- CB
2448 2447 2449 2456 3 0.00000 0.00000 0.00000
0.00000 0.00000 0.00000 ; H- N- CA- C
2449 2451 2454 2455 3 2.05853 2.46856 3.48109
-4.46851 0.23430 0.00000 ; CA- CB- OG- HG
2449 2456 2458 2459 3 20.92000 0.00000 -20.92000
0.00000 0.00000 0.00000 ; CA- C- N- H
2449 2456 2458 2460 3 20.92000 0.00000 -20.92000
0.00000 0.00000 0.00000 ; CA- C- N- CA
2450 2449 2451 2452 3 0.65084 1.95253 0.00000
-2.60338 0.00000 0.00000 ; HA- CA- CB- HB2
2450 2449 2451 2453 3 0.65084 1.95253 0.00000
-2.60338 0.00000 0.00000 ; HA- CA- CB- HB3
2450 2449 2451 2454 3 1.04600 -1.04600 0.00000
0.00000 0.00000 0.00000 ; HA- CA- CB- OG
2450 2449 2456 2457 3 3.68192 -4.35136 0.00000
1.33888 0.00000 0.00000 ; HA- CA- C- O
2450 2449 2456 2458 3 0.00000 0.00000 0.00000
0.00000 0.00000 0.00000 ; HA- CA- C- N
2451 2449 2456 2457 3 0.00000 0.00000 0.00000
0.00000 0.00000 0.00000 ; CB- CA- C- O
2451 2449 2456 2458 3 2.51040 4.18400 1.67360
-6.69440 0.00000 0.00000 ; CB- CA- C- N
2452 2451 2449 2456 3 0.65084 1.95253 0.00000
-2.60338 0.00000 0.00000 ; HB2- CB- CA- C
2452 2451 2454 2455 3 0.69733 2.09200 0.00000
-2.78933 0.00000 0.00000 ; HB2- CB- OG- HG
2453 2451 2449 2456 3 0.65084 1.95253 0.00000
-2.60338 0.00000 0.00000 ; HB3- CB- CA- C
2453 2451 2454 2455 3 0.69733 2.09200 0.00000
-2.78933 0.00000 0.00000 ; HB3- CB- OG- HG
2454 2451 2449 2456 3 7.34710 7.79898 -6.14211
-6.71114 4.31789 0.00000 ; OG- CB- CA- C
2456 2458 2460 2461 3 0.00000 0.00000 0.00000
0.00000 0.00000 0.00000 ; C- N- CA- HA
2456 2458 2460 2462 3 11.71520 1.67360 15.06240 -
13.38880 0.00000 0.00000 ; C- N- CA- CB

2456 2458 2460 2475 3 1.75728 5.27184 2.25936
-7.02912 0.00000 0.00000 ; C- N- CA- C
2457 2456 2458 2459 3 29.28800 -8.36800 -20.92000
0.00000 0.00000 0.00000 ; O- C- N- H
2457 2456 2458 2460 3 20.92000 0.00000 -20.92000
0.00000 0.00000 0.00000 ; O- C- N- CA
2458 2460 2462 2463 3 0.65084 1.95253 0.00000
-2.60338 0.00000 0.00000 ; N- CA- CB- HB2
2458 2460 2462 2464 3 0.65084 1.95253 0.00000
-2.60338 0.00000 0.00000 ; N- CA- CB- HB3
2458 2460 2462 2465 3 3.79070 1.39746 -4.61077
-2.40998 2.44346 0.00000 ; N- CA- CB- CG
2458 2460 2475 2476 3 0.00000 0.00000 0.00000
0.00000 0.00000 0.00000 ; N- CA- C- O
2458 2460 2475 2477 3 17.40544 -5.02080 -13.22144
9.20480 0.00000 0.00000 ; N- CA- C- N
2459 2458 2460 2461 3 0.00000 0.00000 0.00000
0.00000 0.00000 0.00000 ; H- N- CA- HA
2459 2458 2460 2462 3 0.00000 0.00000 0.00000
0.00000 0.00000 0.00000 ; H- N- CA- CB
2459 2458 2460 2475 3 0.00000 0.00000 0.00000
0.00000 0.00000 0.00000 ; H- N- CA- C
2460 2462 2465 2466 3 0.66944 2.00832 0.00000
-2.67776 0.00000 0.00000 ; CA- CB- CG- HG
2460 2462 2465 2467 3 3.90367 0.19665 -6.21742
-2.37651 5.99149 0.00000 ; CA- CB- CG- CD1
2460 2462 2465 2471 3 3.90367 0.19665 -6.21742
-2.37651 5.99149 0.00000 ; CA- CB- CG- CD2
2460 2475 2477 2478 3 20.92000 0.00000 -20.92000
0.00000 0.00000 0.00000 ; CA- C- N- H
2460 2475 2477 2479 3 20.92000 0.00000 -20.92000
0.00000 0.00000 0.00000 ; CA- C- N- CA
2461 2460 2462 2463 3 0.65084 1.95253 0.00000
-2.60338 0.00000 0.00000 ; HA- CA- CB- HB2
2461 2460 2462 2464 3 0.65084 1.95253 0.00000
-2.60338 0.00000 0.00000 ; HA- CA- CB- HB3
2461 2460 2462 2465 3 0.65084 1.95253 0.00000
-2.60338 0.00000 0.00000 ; HA- CA- CB- CG
2461 2460 2475 2476 3 3.68192 -4.35136 0.00000
1.33888 0.00000 0.00000 ; HA- CA- C- O
2461 2460 2475 2477 3 0.00000 0.00000 0.00000
0.00000 0.00000 0.00000 ; HA- CA- C- N
2462 2460 2475 2476 3 0.00000 0.00000 0.00000
0.00000 0.00000 0.00000 ; CB- CA- C- O
2462 2460 2475 2477 3 2.51040 4.18400 1.67360
-6.69440 0.00000 0.00000 ; CB- CA- C- N
2462 2465 2467 2468 3 0.66944 2.00832 0.00000
-2.67776 0.00000 0.00000 ; CB- CG- CD1- HD11
2462 2465 2467 2469 3 0.66944 2.00832 0.00000
-2.67776 0.00000 0.00000 ; CB- CG- CD1- HD12
2462 2465 2467 2470 3 0.66944 2.00832 0.00000
-2.67776 0.00000 0.00000 ; CB- CG- CD1- HD13

2462 2465 2471 2472 3 0.66944 2.00832 0.00000
-2.67776 0.00000 0.00000 ; CB- CG- CD2- HD21
2462 2465 2471 2473 3 0.66944 2.00832 0.00000
-2.67776 0.00000 0.00000 ; CB- CG- CD2- HD22
2462 2465 2471 2474 3 0.66944 2.00832 0.00000
-2.67776 0.00000 0.00000 ; CB- CG- CD2- HD23
2463 2462 2460 2475 3 0.65084 1.95253 0.00000
-2.60338 0.00000 0.00000 ; HB2- CB- CA- C
2463 2462 2465 2466 3 0.62760 1.88280 0.00000
-2.51040 0.00000 0.00000 ; HB2- CB- CG- HG
2463 2462 2465 2467 3 0.66944 2.00832 0.00000
-2.67776 0.00000 0.00000 ; HB2- CB- CG- CD1
2463 2462 2465 2471 3 0.66944 2.00832 0.00000
-2.67776 0.00000 0.00000 ; HB2- CB- CG- CD2
2464 2462 2460 2475 3 0.65084 1.95253 0.00000
-2.60338 0.00000 0.00000 ; HB3- CB- CA- C
2464 2462 2465 2466 3 0.62760 1.88280 0.00000
-2.51040 0.00000 0.00000 ; HB3- CB- CG- HG
2464 2462 2465 2467 3 0.66944 2.00832 0.00000
-2.67776 0.00000 0.00000 ; HB3- CB- CG- CD1
2464 2462 2465 2471 3 0.66944 2.00832 0.00000
-2.67776 0.00000 0.00000 ; HB3- CB- CG- CD2
2465 2462 2460 2475 3 10.33448 -1.14642 -11.54784
-2.40998 6.35968 0.00000 ; CG- CB- CA- C
2466 2465 2467 2468 3 0.62760 1.88280 0.00000
-2.51040 0.00000 0.00000 ; HG- CG- CD1- HD11
2466 2465 2467 2469 3 0.62760 1.88280 0.00000
-2.51040 0.00000 0.00000 ; HG- CG- CD1- HD12
2466 2465 2467 2470 3 0.62760 1.88280 0.00000
-2.51040 0.00000 0.00000 ; HG- CG- CD1- HD13
2466 2465 2471 2472 3 0.62760 1.88280 0.00000
-2.51040 0.00000 0.00000 ; HG- CG- CD2- HD21
2466 2465 2471 2473 3 0.62760 1.88280 0.00000
-2.51040 0.00000 0.00000 ; HG- CG- CD2- HD22
2466 2465 2471 2474 3 0.62760 1.88280 0.00000
-2.51040 0.00000 0.00000 ; HG- CG- CD2- HD23
2467 2465 2471 2472 3 0.66944 2.00832 0.00000
-2.67776 0.00000 0.00000 ; CD1- CG- CD2- HD21
2467 2465 2471 2473 3 0.66944 2.00832 0.00000
-2.67776 0.00000 0.00000 ; CD1- CG- CD2- HD22
2467 2465 2471 2474 3 0.66944 2.00832 0.00000
-2.67776 0.00000 0.00000 ; CD1- CG- CD2- HD23
2468 2467 2465 2471 3 0.66944 2.00832 0.00000
-2.67776 0.00000 0.00000 ; HD11- CD1- CG- CD2
2469 2467 2465 2471 3 0.66944 2.00832 0.00000
-2.67776 0.00000 0.00000 ; HD12- CD1- CG- CD2
2470 2467 2465 2471 3 0.66944 2.00832 0.00000
-2.67776 0.00000 0.00000 ; HD13- CD1- CG- CD2
2475 2477 2479 2480 3 0.00000 0.00000 0.00000
0.00000 0.00000 0.00000 ; C- N- CA- HA
2475 2477 2479 2481 3 11.71520 1.67360 15.06240 -
13.38880 0.00000 0.00000 ; C- N- CA- CB

2475 2477 2479 2494 3 1.75728 5.27184 2.25936
-7.02912 0.00000 0.00000 ; C- N- CA- C
2476 2475 2477 2478 3 29.28800 -8.36800 -20.92000
0.00000 0.00000 0.00000 ; O- C- N- H
2476 2475 2477 2479 3 20.92000 0.00000 -20.92000
0.00000 0.00000 0.00000 ; O- C- N- CA
2477 2479 2481 2482 3 0.65084 1.95253 0.00000
-2.60338 0.00000 0.00000 ; N- CA- CB- HB2
2477 2479 2481 2483 3 0.65084 1.95253 0.00000
-2.60338 0.00000 0.00000 ; N- CA- CB- HB3
2477 2479 2481 2484 3 3.79070 1.39746 -4.61077
-2.40998 2.44346 0.00000 ; N- CA- CB- CG
2477 2479 2494 2495 3 0.00000 0.00000 0.00000
0.00000 0.00000 0.00000 ; N- CA- C- O
2477 2479 2494 2496 3 17.40544 -5.02080 -13.22144
9.20480 0.00000 0.00000 ; N- CA- C- N
2478 2477 2479 2480 3 0.00000 0.00000 0.00000
0.00000 0.00000 0.00000 ; H- N- CA- HA
2478 2477 2479 2481 3 0.00000 0.00000 0.00000
0.00000 0.00000 0.00000 ; H- N- CA- CB
2478 2477 2479 2494 3 0.00000 0.00000 0.00000
0.00000 0.00000 0.00000 ; H- N- CA- C
2479 2481 2484 2485 3 0.66944 2.00832 0.00000
-2.67776 0.00000 0.00000 ; CA- CB- CG- HG
2479 2481 2484 2486 3 3.90367 0.19665 -6.21742
-2.37651 5.99149 0.00000 ; CA- CB- CG- CD1
2479 2481 2484 2490 3 3.90367 0.19665 -6.21742
-2.37651 5.99149 0.00000 ; CA- CB- CG- CD2
2479 2494 2496 2497 3 20.92000 0.00000 -20.92000
0.00000 0.00000 0.00000 ; CA- C- N- H
2479 2494 2496 2498 3 20.92000 0.00000 -20.92000
0.00000 0.00000 0.00000 ; CA- C- N- CA
2480 2479 2481 2482 3 0.65084 1.95253 0.00000
-2.60338 0.00000 0.00000 ; HA- CA- CB- HB2
2480 2479 2481 2483 3 0.65084 1.95253 0.00000
-2.60338 0.00000 0.00000 ; HA- CA- CB- HB3
2480 2479 2481 2484 3 0.65084 1.95253 0.00000
-2.60338 0.00000 0.00000 ; HA- CA- CB- CG
2480 2479 2494 2495 3 3.68192 -4.35136 0.00000
1.33888 0.00000 0.00000 ; HA- CA- C- O
2480 2479 2494 2496 3 0.00000 0.00000 0.00000
0.00000 0.00000 0.00000 ; HA- CA- C- N
2481 2479 2494 2495 3 0.00000 0.00000 0.00000
0.00000 0.00000 0.00000 ; CB- CA- C- O
2481 2479 2494 2496 3 2.51040 4.18400 1.67360
-6.69440 0.00000 0.00000 ; CB- CA- C- N
2481 2484 2486 2487 3 0.66944 2.00832 0.00000
-2.67776 0.00000 0.00000 ; CB- CG- CD1- HD11
2481 2484 2486 2488 3 0.66944 2.00832 0.00000
-2.67776 0.00000 0.00000 ; CB- CG- CD1- HD12
2481 2484 2486 2489 3 0.66944 2.00832 0.00000
-2.67776 0.00000 0.00000 ; CB- CG- CD1- HD13

2481 2484 2490 2491 3 0.66944 2.00832 0.00000
-2.67776 0.00000 0.00000 ; CB- CG- CD2- HD21
2481 2484 2490 2492 3 0.66944 2.00832 0.00000
-2.67776 0.00000 0.00000 ; CB- CG- CD2- HD22
2481 2484 2490 2493 3 0.66944 2.00832 0.00000
-2.67776 0.00000 0.00000 ; CB- CG- CD2- HD23
2482 2481 2479 2494 3 0.65084 1.95253 0.00000
-2.60338 0.00000 0.00000 ; HB2- CB- CA- C
2482 2481 2484 2485 3 0.62760 1.88280 0.00000
-2.51040 0.00000 0.00000 ; HB2- CB- CG- HG
2482 2481 2484 2486 3 0.66944 2.00832 0.00000
-2.67776 0.00000 0.00000 ; HB2- CB- CG- CD1
2482 2481 2484 2490 3 0.66944 2.00832 0.00000
-2.67776 0.00000 0.00000 ; HB2- CB- CG- CD2
2483 2481 2479 2494 3 0.65084 1.95253 0.00000
-2.60338 0.00000 0.00000 ; HB3- CB- CA- C
2483 2481 2484 2485 3 0.62760 1.88280 0.00000
-2.51040 0.00000 0.00000 ; HB3- CB- CG- HG
2483 2481 2484 2486 3 0.66944 2.00832 0.00000
-2.67776 0.00000 0.00000 ; HB3- CB- CG- CD1
2483 2481 2484 2490 3 0.66944 2.00832 0.00000
-2.67776 0.00000 0.00000 ; HB3- CB- CG- CD2
2484 2481 2479 2494 3 10.33448 -1.14642 -11.54784
-2.40998 6.35968 0.00000 ; CG- CB- CA- C
2485 2484 2486 2487 3 0.62760 1.88280 0.00000
-2.51040 0.00000 0.00000 ; HG- CG- CD1- HD11
2485 2484 2486 2488 3 0.62760 1.88280 0.00000
-2.51040 0.00000 0.00000 ; HG- CG- CD1- HD12
2485 2484 2486 2489 3 0.62760 1.88280 0.00000
-2.51040 0.00000 0.00000 ; HG- CG- CD1- HD13
2485 2484 2490 2491 3 0.62760 1.88280 0.00000
-2.51040 0.00000 0.00000 ; HG- CG- CD2- HD21
2485 2484 2490 2492 3 0.62760 1.88280 0.00000
-2.51040 0.00000 0.00000 ; HG- CG- CD2- HD22
2485 2484 2490 2493 3 0.62760 1.88280 0.00000
-2.51040 0.00000 0.00000 ; HG- CG- CD2- HD23
2486 2484 2490 2491 3 0.66944 2.00832 0.00000
-2.67776 0.00000 0.00000 ; CD1- CG- CD2- HD21
2486 2484 2490 2492 3 0.66944 2.00832 0.00000
-2.67776 0.00000 0.00000 ; CD1- CG- CD2- HD22
2486 2484 2490 2493 3 0.66944 2.00832 0.00000
-2.67776 0.00000 0.00000 ; CD1- CG- CD2- HD23
2487 2486 2484 2490 3 0.66944 2.00832 0.00000
-2.67776 0.00000 0.00000 ; HD11- CD1- CG- CD2
2488 2486 2484 2490 3 0.66944 2.00832 0.00000
-2.67776 0.00000 0.00000 ; HD12- CD1- CG- CD2
2489 2486 2484 2490 3 0.66944 2.00832 0.00000
-2.67776 0.00000 0.00000 ; HD13- CD1- CG- CD2
2494 2496 2498 2499 3 0.00000 0.00000 0.00000
0.00000 0.00000 0.00000 ; C- N- CA- HA
2494 2496 2498 2500 3 11.71520 1.67360 15.06240 -
13.38880 0.00000 0.00000 ; C- N- CA- CB

2494 2496 2498 2516 3 1.75728 5.27184 2.25936
-7.02912 0.00000 0.00000 ; C- N- CA- C
2495 2494 2496 2497 3 29.28800 -8.36800 -20.92000
0.00000 0.00000 0.00000 ; O- C- N- H
2495 2494 2496 2498 3 20.92000 0.00000 -20.92000
0.00000 0.00000 0.00000 ; O- C- N- CA
2496 2498 2500 2501 3 0.65084 1.95253 0.00000
-2.60338 0.00000 0.00000 ; N- CA- CB- HB2
2496 2498 2500 2502 3 0.65084 1.95253 0.00000
-2.60338 0.00000 0.00000 ; N- CA- CB- HB3
2496 2498 2500 2503 3 0.65084 1.95253 0.00000
-2.60338 0.00000 0.00000 ; N- CA- CB- CG
2496 2498 2516 2517 3 0.00000 0.00000 0.00000
0.00000 0.00000 0.00000 ; N- CA- C- O
2496 2498 2516 2518 3 17.40544 -5.02080 -13.22144
9.20480 0.00000 0.00000 ; N- CA- C- N
2497 2496 2498 2499 3 0.00000 0.00000 0.00000
0.00000 0.00000 0.00000 ; H- N- CA- HA
2497 2496 2498 2500 3 0.00000 0.00000 0.00000
0.00000 0.00000 0.00000 ; H- N- CA- CB
2497 2496 2498 2516 3 0.00000 0.00000 0.00000
0.00000 0.00000 0.00000 ; H- N- CA- C
2498 2500 2503 2504 3 0.66944 2.00832 0.00000
-2.67776 0.00000 0.00000 ; CA- CB- CG- HG2
2498 2500 2503 2505 3 0.66944 2.00832 0.00000
-2.67776 0.00000 0.00000 ; CA- CB- CG- HG3
2498 2500 2503 2506 3 3.68192 3.09616 -2.09200
-3.01248 0.00000 0.00000 ; CA- CB- CG- CD
2498 2516 2518 2519 3 20.92000 0.00000 -20.92000
0.00000 0.00000 0.00000 ; CA- C- N- H
2498 2516 2518 2520 3 20.92000 0.00000 -20.92000
0.00000 0.00000 0.00000 ; CA- C- N- CA
2499 2498 2500 2501 3 0.65084 1.95253 0.00000
-2.60338 0.00000 0.00000 ; HA- CA- CB- HB2
2499 2498 2500 2502 3 0.65084 1.95253 0.00000
-2.60338 0.00000 0.00000 ; HA- CA- CB- HB3
2499 2498 2500 2503 3 0.65084 1.95253 0.00000
-2.60338 0.00000 0.00000 ; HA- CA- CB- CG
2499 2498 2516 2517 3 3.68192 -4.35136 0.00000
1.33888 0.00000 0.00000 ; HA- CA- C- O
2499 2498 2516 2518 3 0.00000 0.00000 0.00000
0.00000 0.00000 0.00000 ; HA- CA- C- N
2500 2498 2516 2517 3 0.00000 0.00000 0.00000
0.00000 0.00000 0.00000 ; CB- CA- C- O
2500 2498 2516 2518 3 2.51040 4.18400 1.67360
-6.69440 0.00000 0.00000 ; CB- CA- C- N
2500 2503 2506 2507 3 0.66944 2.00832 0.00000
-2.67776 0.00000 0.00000 ; CB- CG- CD- HD2
2500 2503 2506 2508 3 0.66944 2.00832 0.00000
-2.67776 0.00000 0.00000 ; CB- CG- CD- HD3
2500 2503 2506 2509 3 3.68192 3.09616 -2.09200
-3.01248 0.00000 0.00000 ; CB- CG- CD- CE

```
  2501   2500   2498   2516      3    0.65084    1.95253    0.00000
-2.60338    0.00000    0.00000 ;    HB2-    CB-    CA-     C
  2501   2500   2503   2504      3    0.62760    1.88280    0.00000
-2.51040    0.00000    0.00000 ;    HB2-    CB-    CG-   HG2
  2501   2500   2503   2505      3    0.62760    1.88280    0.00000
-2.51040    0.00000    0.00000 ;    HB2-    CB-    CG-   HG3
  2501   2500   2503   2506      3    0.66944    2.00832    0.00000
-2.67776    0.00000    0.00000 ;    HB2-    CB-    CG-    CD
  2502   2500   2498   2516      3    0.65084    1.95253    0.00000
-2.60338    0.00000    0.00000 ;    HB3-    CB-    CA-     C
  2502   2500   2503   2504      3    0.62760    1.88280    0.00000
-2.51040    0.00000    0.00000 ;    HB3-    CB-    CG-   HG2
  2502   2500   2503   2505      3    0.62760    1.88280    0.00000
-2.51040    0.00000    0.00000 ;    HB3-    CB-    CG-   HG3
  2502   2500   2503   2506      3    0.66944    2.00832    0.00000
-2.67776    0.00000    0.00000 ;    HB3-    CB-    CG-    CD
  2503   2500   2498   2516      3    0.65084    1.95253    0.00000
-2.60338    0.00000    0.00000 ;     CG-    CB-    CA-     C
  2503   2506   2509   2510      3    0.65084    1.95253    0.00000
-2.60338    0.00000    0.00000 ;     CG-    CD-    CE-   HE2
  2503   2506   2509   2511      3    0.65084    1.95253    0.00000
-2.60338    0.00000    0.00000 ;     CG-    CD-    CE-   HE3
  2503   2506   2509   2512      3    0.65084    1.95253    0.00000
-2.60338    0.00000    0.00000 ;     CG-    CD-    CE-    NZ
  2504   2503   2506   2507      3    0.62760    1.88280    0.00000
-2.51040    0.00000    0.00000 ;    HG2-    CG-    CD-   HD2
  2504   2503   2506   2508      3    0.62760    1.88280    0.00000
-2.51040    0.00000    0.00000 ;    HG2-    CG-    CD-   HD3
  2504   2503   2506   2509      3    0.66944    2.00832    0.00000
-2.67776    0.00000    0.00000 ;    HG2-    CG-    CD-    CE
  2505   2503   2506   2507      3    0.62760    1.88280    0.00000
-2.51040    0.00000    0.00000 ;    HG3-    CG-    CD-   HD2
  2505   2503   2506   2508      3    0.62760    1.88280    0.00000
-2.51040    0.00000    0.00000 ;    HG3-    CG-    CD-   HD3
  2505   2503   2506   2509      3    0.66944    2.00832    0.00000
-2.67776    0.00000    0.00000 ;    HG3-    CG-    CD-    CE
  2506   2509   2512   2513      3    0.65084    1.95253    0.00000
-2.60338    0.00000    0.00000 ;     CD-    CE-    NZ-   HZ1
  2506   2509   2512   2514      3    0.65084    1.95253    0.00000
-2.60338    0.00000    0.00000 ;     CD-    CE-    NZ-   HZ2
  2506   2509   2512   2515      3    0.65084    1.95253    0.00000
-2.60338    0.00000    0.00000 ;     CD-    CE-    NZ-   HZ3
  2507   2506   2509   2510      3    0.65084    1.95253    0.00000
-2.60338    0.00000    0.00000 ;    HD2-    CD-    CE-   HE2
  2507   2506   2509   2511      3    0.65084    1.95253    0.00000
-2.60338    0.00000    0.00000 ;    HD2-    CD-    CE-   HE3
  2507   2506   2509   2512      3    0.65084    1.95253    0.00000
-2.60338    0.00000    0.00000 ;    HD2-    CD-    CE-    NZ
  2508   2506   2509   2510      3    0.65084    1.95253    0.00000
-2.60338    0.00000    0.00000 ;    HD3-    CD-    CE-   HE2
  2508   2506   2509   2511      3    0.65084    1.95253    0.00000
-2.60338    0.00000    0.00000 ;    HD3-    CD-    CE-   HE3
```

2508 2506 2509 2512 3 0.65084 1.95253 0.00000
-2.60338 0.00000 0.00000 ; HD3- CD- CE- NZ
2510 2509 2512 2513 3 0.65084 1.95253 0.00000
-2.60338 0.00000 0.00000 ; HE2- CE- NZ- HZ1
2510 2509 2512 2514 3 0.65084 1.95253 0.00000
-2.60338 0.00000 0.00000 ; HE2- CE- NZ- HZ2
2510 2509 2512 2515 3 0.65084 1.95253 0.00000
-2.60338 0.00000 0.00000 ; HE2- CE- NZ- HZ3
2511 2509 2512 2513 3 0.65084 1.95253 0.00000
-2.60338 0.00000 0.00000 ; HE3- CE- NZ- HZ1
2511 2509 2512 2514 3 0.65084 1.95253 0.00000
-2.60338 0.00000 0.00000 ; HE3- CE- NZ- HZ2
2511 2509 2512 2515 3 0.65084 1.95253 0.00000
-2.60338 0.00000 0.00000 ; HE3- CE- NZ- HZ3
2516 2518 2520 2521 3 0.00000 0.00000 0.00000
0.00000 0.00000 0.00000 ; C- N- CA- HA
2516 2518 2520 2522 3 11.71520 1.67360 15.06240 -
13.38880 0.00000 0.00000 ; C- N- CA- CB
2516 2518 2520 2535 3 1.75728 5.27184 2.25936
-7.02912 0.00000 0.00000 ; C- N- CA- C
2517 2516 2518 2519 3 29.28800 -8.36800 -20.92000
0.00000 0.00000 0.00000 ; O- C- N- H
2517 2516 2518 2520 3 20.92000 0.00000 -20.92000
0.00000 0.00000 0.00000 ; O- C- N- CA
2518 2520 2522 2523 3 0.65084 1.95253 0.00000
-2.60338 0.00000 0.00000 ; N- CA- CB- HB
2518 2520 2522 2524 3 3.83673 0.44769 -1.77402
-2.47693 -0.03347 0.00000 ; N- CA- CB- CG2
2518 2520 2522 2528 3 2.97482 0.12134 2.04179
-1.89117 -3.24678 0.00000 ; N- CA- CB- CG1
2518 2520 2535 2536 3 0.00000 0.00000 0.00000
0.00000 0.00000 0.00000 ; N- CA- C- O
2518 2520 2535 2537 3 17.40544 -5.02080 -13.22144
9.20480 0.00000 0.00000 ; N- CA- C- N
2519 2518 2520 2521 3 0.00000 0.00000 0.00000
0.00000 0.00000 0.00000 ; H- N- CA- HA
2519 2518 2520 2522 3 0.00000 0.00000 0.00000
0.00000 0.00000 0.00000 ; H- N- CA- CB
2519 2518 2520 2535 3 0.00000 0.00000 0.00000
0.00000 0.00000 0.00000 ; H- N- CA- C
2520 2522 2524 2525 3 0.66944 2.00832 0.00000
-2.67776 0.00000 0.00000 ; CA- CB- CG2- HG21
2520 2522 2524 2526 3 0.66944 2.00832 0.00000
-2.67776 0.00000 0.00000 ; CA- CB- CG2- HG22
2520 2522 2524 2527 3 0.66944 2.00832 0.00000
-2.67776 0.00000 0.00000 ; CA- CB- CG2- HG23
2520 2522 2528 2529 3 0.66944 2.00832 0.00000
-2.67776 0.00000 0.00000 ; CA- CB- CG1- HG12
2520 2522 2528 2530 3 0.66944 2.00832 0.00000
-2.67776 0.00000 0.00000 ; CA- CB- CG1- HG13
2520 2522 2528 2531 3 4.24258 -0.52718 -7.25506
-1.79075 7.69856 0.00000 ; CA- CB- CG1- CD1

```
  2520   2535   2537   2538      3   20.92000    0.00000  -20.92000
0.00000    0.00000    0.00000 ;     CA-     C-     N-     H
  2520   2535   2537   2539      3   20.92000    0.00000  -20.92000
0.00000    0.00000    0.00000 ;     CA-     C-     N-    CA
  2521   2520   2522   2523      3    0.65084    1.95253    0.00000
-2.60338    0.00000    0.00000 ;     HA-    CA-    CB-    HB
  2521   2520   2522   2524      3    0.65084    1.95253    0.00000
-2.60338    0.00000    0.00000 ;     HA-    CA-    CB-   CG2
  2521   2520   2522   2528      3    0.65084    1.95253    0.00000
-2.60338    0.00000    0.00000 ;     HA-    CA-    CB-   CG1
  2521   2520   2535   2536      3    3.68192   -4.35136    0.00000
1.33888    0.00000    0.00000 ;     HA-    CA-     C-     O
  2521   2520   2535   2537      3    0.00000    0.00000    0.00000
0.00000    0.00000    0.00000 ;     HA-    CA-     C-     N
  2522   2520   2535   2536      3    0.00000    0.00000    0.00000
0.00000    0.00000    0.00000 ;     CB-    CA-     C-     O
  2522   2520   2535   2537      3    2.51040    4.18400    1.67360
-6.69440    0.00000    0.00000 ;     CB-    CA-     C-     N
  2522   2528   2531   2532      3    0.66944    2.00832    0.00000
-2.67776    0.00000    0.00000 ;     CB-   CG1-   CD1-  HD11
  2522   2528   2531   2533      3    0.66944    2.00832    0.00000
-2.67776    0.00000    0.00000 ;     CB-   CG1-   CD1-  HD12
  2522   2528   2531   2534      3    0.66944    2.00832    0.00000
-2.67776    0.00000    0.00000 ;     CB-   CG1-   CD1-  HD13
  2523   2522   2520   2535      3    0.65084    1.95253    0.00000
-2.60338    0.00000    0.00000 ;     HB-    CB-    CA-     C
  2523   2522   2524   2525      3    0.62760    1.88280    0.00000
-2.51040    0.00000    0.00000 ;     HB-    CB-   CG2-  HG21
  2523   2522   2524   2526      3    0.62760    1.88280    0.00000
-2.51040    0.00000    0.00000 ;     HB-    CB-   CG2-  HG22
  2523   2522   2524   2527      3    0.62760    1.88280    0.00000
-2.51040    0.00000    0.00000 ;     HB-    CB-   CG2-  HG23
  2523   2522   2528   2529      3    0.62760    1.88280    0.00000
-2.51040    0.00000    0.00000 ;     HB-    CB-   CG1-  HG12
  2523   2522   2528   2530      3    0.62760    1.88280    0.00000
-2.51040    0.00000    0.00000 ;     HB-    CB-   CG1-  HG13
  2523   2522   2528   2531      3    0.66944    2.00832    0.00000
-2.67776    0.00000    0.00000 ;     HB-    CB-   CG1-   CD1
  2524   2522   2520   2535      3    5.67350    3.55640   -6.16722
-2.47693    3.74886    0.00000 ;    CG2-    CB-    CA-     C
  2524   2522   2528   2529      3    0.66944    2.00832    0.00000
-2.67776    0.00000    0.00000 ;    CG2-    CB-   CG1-  HG12
  2524   2522   2528   2530      3    0.66944    2.00832    0.00000
-2.67776    0.00000    0.00000 ;    CG2-    CB-   CG1-  HG13
  2524   2522   2528   2531      3    3.81162    0.49790   -8.14206
-1.79075    7.49773    0.00000 ;    CG2-    CB-   CG1-   CD1
  2525   2524   2522   2528      3    0.66944    2.00832    0.00000
-2.67776    0.00000    0.00000 ;   HG21-   CG2-    CB-   CG1
  2526   2524   2522   2528      3    0.66944    2.00832    0.00000
-2.67776    0.00000    0.00000 ;   HG22-   CG2-    CB-   CG1
  2527   2524   2522   2528      3    0.66944    2.00832    0.00000
-2.67776    0.00000    0.00000 ;   HG23-   CG2-    CB-   CG1
```

2528 2522 2520 2535 3 8.26340 0.74057 -9.99976
-1.89117 3.84928 0.00000 ; CG1- CB- CA- C
2529 2528 2531 2532 3 0.62760 1.88280 0.00000
-2.51040 0.00000 0.00000 ; HG12- CG1- CD1- HD11
2529 2528 2531 2533 3 0.62760 1.88280 0.00000
-2.51040 0.00000 0.00000 ; HG12- CG1- CD1- HD12
2529 2528 2531 2534 3 0.62760 1.88280 0.00000
-2.51040 0.00000 0.00000 ; HG12- CG1- CD1- HD13
2530 2528 2531 2532 3 0.62760 1.88280 0.00000
-2.51040 0.00000 0.00000 ; HG13- CG1- CD1- HD11
2530 2528 2531 2533 3 0.62760 1.88280 0.00000
-2.51040 0.00000 0.00000 ; HG13- CG1- CD1- HD12
2530 2528 2531 2534 3 0.62760 1.88280 0.00000
-2.51040 0.00000 0.00000 ; HG13- CG1- CD1- HD13
2535 2537 2539 2540 3 0.00000 0.00000 0.00000
0.00000 0.00000 0.00000 ; C- N- CA- HA
2535 2537 2539 2541 3 11.71520 1.67360 15.06240 -
13.38880 0.00000 0.00000 ; C- N- CA- CB
2535 2537 2539 2554 3 1.75728 5.27184 2.25936
-7.02912 0.00000 0.00000 ; C- N- CA- C
2536 2535 2537 2538 3 29.28800 -8.36800 -20.92000
0.00000 0.00000 0.00000 ; O- C- N- H
2536 2535 2537 2539 3 20.92000 0.00000 -20.92000
0.00000 0.00000 0.00000 ; O- C- N- CA
2537 2539 2541 2542 3 0.65084 1.95253 0.00000
-2.60338 0.00000 0.00000 ; N- CA- CB- HB
2537 2539 2541 2543 3 3.83673 0.44769 -1.77402
-2.47693 -0.03347 0.00000 ; N- CA- CB- CG2
2537 2539 2541 2547 3 2.97482 0.12134 2.04179
-1.89117 -3.24678 0.00000 ; N- CA- CB- CG1
2537 2539 2554 2555 3 0.00000 0.00000 0.00000
0.00000 0.00000 0.00000 ; N- CA- C- O
2537 2539 2554 2556 3 17.40544 -5.02080 -13.22144
9.20480 0.00000 0.00000 ; N- CA- C- N
2538 2537 2539 2540 3 0.00000 0.00000 0.00000
0.00000 0.00000 0.00000 ; H- N- CA- HA
2538 2537 2539 2541 3 0.00000 0.00000 0.00000
0.00000 0.00000 0.00000 ; H- N- CA- CB
2538 2537 2539 2554 3 0.00000 0.00000 0.00000
0.00000 0.00000 0.00000 ; H- N- CA- C
2539 2541 2543 2544 3 0.66944 2.00832 0.00000
-2.67776 0.00000 0.00000 ; CA- CB- CG2- HG21
2539 2541 2543 2545 3 0.66944 2.00832 0.00000
-2.67776 0.00000 0.00000 ; CA- CB- CG2- HG22
2539 2541 2543 2546 3 0.66944 2.00832 0.00000
-2.67776 0.00000 0.00000 ; CA- CB- CG2- HG23
2539 2541 2547 2548 3 0.66944 2.00832 0.00000
-2.67776 0.00000 0.00000 ; CA- CB- CG1- HG12
2539 2541 2547 2549 3 0.66944 2.00832 0.00000
-2.67776 0.00000 0.00000 ; CA- CB- CG1- HG13
2539 2541 2547 2550 3 4.24258 -0.52718 -7.25506
-1.79075 7.69856 0.00000 ; CA- CB- CG1- CD1

```
  2539   2554   2556   2557      3   20.92000    0.00000  -20.92000
0.00000    0.00000    0.00000 ;     CA-     C-     N-     H
  2539   2554   2556   2558      3   20.92000    0.00000  -20.92000
0.00000    0.00000    0.00000 ;     CA-     C-     N-    CA
  2540   2539   2541   2542      3    0.65084    1.95253    0.00000
-2.60338    0.00000    0.00000 ;     HA-    CA-    CB-    HB
  2540   2539   2541   2543      3    0.65084    1.95253    0.00000
-2.60338    0.00000    0.00000 ;     HA-    CA-    CB-   CG2
  2540   2539   2541   2547      3    0.65084    1.95253    0.00000
-2.60338    0.00000    0.00000 ;     HA-    CA-    CB-   CG1
  2540   2539   2554   2555      3    3.68192   -4.35136    0.00000
1.33888    0.00000    0.00000 ;     HA-    CA-     C-     O
  2540   2539   2554   2556      3    0.00000    0.00000    0.00000
0.00000    0.00000    0.00000 ;     HA-    CA-     C-     N
  2541   2539   2554   2555      3    0.00000    0.00000    0.00000
0.00000    0.00000    0.00000 ;     CB-    CA-     C-     O
  2541   2539   2554   2556      3    2.51040    4.18400    1.67360
-6.69440    0.00000    0.00000 ;     CB-    CA-     C-     N
  2541   2547   2550   2551      3    0.66944    2.00832    0.00000
-2.67776    0.00000    0.00000 ;     CB-   CG1-   CD1-  HD11
  2541   2547   2550   2552      3    0.66944    2.00832    0.00000
-2.67776    0.00000    0.00000 ;     CB-   CG1-   CD1-  HD12
  2541   2547   2550   2553      3    0.66944    2.00832    0.00000
-2.67776    0.00000    0.00000 ;     CB-   CG1-   CD1-  HD13
  2542   2541   2539   2554      3    0.65084    1.95253    0.00000
-2.60338    0.00000    0.00000 ;     HB-    CB-    CA-     C
  2542   2541   2543   2544      3    0.62760    1.88280    0.00000
-2.51040    0.00000    0.00000 ;     HB-    CB-   CG2-  HG21
  2542   2541   2543   2545      3    0.62760    1.88280    0.00000
-2.51040    0.00000    0.00000 ;     HB-    CB-   CG2-  HG22
  2542   2541   2543   2546      3    0.62760    1.88280    0.00000
-2.51040    0.00000    0.00000 ;     HB-    CB-   CG2-  HG23
  2542   2541   2547   2548      3    0.62760    1.88280    0.00000
-2.51040    0.00000    0.00000 ;     HB-    CB-   CG1-  HG12
  2542   2541   2547   2549      3    0.62760    1.88280    0.00000
-2.51040    0.00000    0.00000 ;     HB-    CB-   CG1-  HG13
  2542   2541   2547   2550      3    0.66944    2.00832    0.00000
-2.67776    0.00000    0.00000 ;     HB-    CB-   CG1-   CD1
  2543   2541   2539   2554      3    5.67350    3.55640   -6.16722
-2.47693    3.74886    0.00000 ;    CG2-    CB-    CA-     C
  2543   2541   2547   2548      3    0.66944    2.00832    0.00000
-2.67776    0.00000    0.00000 ;    CG2-    CB-   CG1-  HG12
  2543   2541   2547   2549      3    0.66944    2.00832    0.00000
-2.67776    0.00000    0.00000 ;    CG2-    CB-   CG1-  HG13
  2543   2541   2547   2550      3    3.81162    0.49790   -8.14206
-1.79075    7.49773    0.00000 ;    CG2-    CB-   CG1-   CD1
  2544   2543   2541   2547      3    0.66944    2.00832    0.00000
-2.67776    0.00000    0.00000 ;   HG21-   CG2-    CB-   CG1
  2545   2543   2541   2547      3    0.66944    2.00832    0.00000
-2.67776    0.00000    0.00000 ;   HG22-   CG2-    CB-   CG1
  2546   2543   2541   2547      3    0.66944    2.00832    0.00000
-2.67776    0.00000    0.00000 ;   HG23-   CG2-    CB-   CG1
```

```
  2547   2541   2539   2554       3    8.26340    0.74057   -9.99976
-1.89117    3.84928    0.00000 ;    CG1-    CB-    CA-     C
  2548   2547   2550   2551       3    0.62760    1.88280    0.00000
-2.51040    0.00000    0.00000 ;   HG12-   CG1-   CD1-  HD11
  2548   2547   2550   2552       3    0.62760    1.88280    0.00000
-2.51040    0.00000    0.00000 ;   HG12-   CG1-   CD1-  HD12
  2548   2547   2550   2553       3    0.62760    1.88280    0.00000
-2.51040    0.00000    0.00000 ;   HG12-   CG1-   CD1-  HD13
  2549   2547   2550   2551       3    0.62760    1.88280    0.00000
-2.51040    0.00000    0.00000 ;   HG13-   CG1-   CD1-  HD11
  2549   2547   2550   2552       3    0.62760    1.88280    0.00000
-2.51040    0.00000    0.00000 ;   HG13-   CG1-   CD1-  HD12
  2549   2547   2550   2553       3    0.62760    1.88280    0.00000
-2.51040    0.00000    0.00000 ;   HG13-   CG1-   CD1-  HD13
  2554   2556   2558   2559       3    0.00000    0.00000    0.00000
0.00000    0.00000    0.00000 ;      C-     N-    CA-    HA
  2554   2556   2558   2560       3   11.71520    1.67360   15.06240  -
13.38880    0.00000    0.00000 ;      C-      N-    CA-    CB
  2554   2556   2558   2566       3    1.75728    5.27184    2.25936
-7.02912    0.00000    0.00000 ;      C-      N-    CA-     C
  2555   2554   2556   2557       3   29.28800   -8.36800  -20.92000
0.00000    0.00000    0.00000 ;      O-     C-     N-     H
  2555   2554   2556   2558       3   20.92000    0.00000  -20.92000
0.00000    0.00000    0.00000 ;      O-     C-     N-    CA
  2556   2558   2560   2561       3    0.65084    1.95253    0.00000
-2.60338    0.00000    0.00000 ;      N-    CA-    CB-   HB2
  2556   2558   2560   2562       3    0.65084    1.95253    0.00000
-2.60338    0.00000    0.00000 ;      N-    CA-    CB-   HB3
  2556   2558   2560   2563       3   15.41386    9.74035   -8.39310
-0.97069    2.97901    0.00000 ;      N-    CA-    CB-    CG
  2556   2558   2566   2567       3    0.00000    0.00000    0.00000
0.00000    0.00000    0.00000 ;      N-    CA-     C-     O
  2556   2558   2566   2568       3   17.40544   -5.02080  -13.22144
9.20480    0.00000    0.00000 ;      N-    CA-     C-     N
  2557   2556   2558   2559       3    0.00000    0.00000    0.00000
0.00000    0.00000    0.00000 ;      H-     N-    CA-    HA
  2557   2556   2558   2560       3    0.00000    0.00000    0.00000
0.00000    0.00000    0.00000 ;      H-     N-    CA-    CB
  2557   2556   2558   2566       3    0.00000    0.00000    0.00000
0.00000    0.00000    0.00000 ;      H-     N-    CA-     C
  2558   2560   2563   2564       3    6.43499    0.00000   -5.39736
0.00000   -1.03763    0.00000 ;    CA-    CB-    CG-   OD1
  2558   2560   2563   2565       3    6.43499    0.00000   -5.39736
0.00000   -1.03763    0.00000 ;    CA-    CB-    CG-   OD2
  2558   2566   2568   2569       3   20.92000    0.00000  -20.92000
0.00000    0.00000    0.00000 ;    CA-     C-     N-     H
  2558   2566   2568   2570       3   20.92000    0.00000  -20.92000
0.00000    0.00000    0.00000 ;    CA-     C-     N-    CA
  2559   2558   2560   2561       3    0.65084    1.95253    0.00000
-2.60338    0.00000    0.00000 ;     HA-    CA-    CB-   HB2
  2559   2558   2560   2562       3    0.65084    1.95253    0.00000
-2.60338    0.00000    0.00000 ;     HA-    CA-    CB-   HB3
```

2559 2558 2560 2563 3 0.65084 1.95253 0.00000
-2.60338 0.00000 0.00000 ; HA- CA- CB- CG
2559 2558 2566 2567 3 3.68192 -4.35136 0.00000
1.33888 0.00000 0.00000 ; HA- CA- C- O
2559 2558 2566 2568 3 0.00000 0.00000 0.00000
0.00000 0.00000 0.00000 ; HA- CA- C- N
2560 2558 2566 2567 3 0.00000 0.00000 0.00000
0.00000 0.00000 0.00000 ; CB- CA- C- O
2560 2558 2566 2568 3 2.51040 4.18400 1.67360
-6.69440 0.00000 0.00000 ; CB- CA- C- N
2561 2560 2558 2566 3 0.65084 1.95253 0.00000
-2.60338 0.00000 0.00000 ; HB2- CB- CA- C
2561 2560 2563 2564 3 0.00000 0.00000 0.00000
0.00000 0.00000 0.00000 ; HB2- CB- CG- OD1
2561 2560 2563 2565 3 0.00000 0.00000 0.00000
0.00000 0.00000 0.00000 ; HB2- CB- CG- OD2
2562 2560 2558 2566 3 0.65084 1.95253 0.00000
-2.60338 0.00000 0.00000 ; HB3- CB- CA- C
2562 2560 2563 2564 3 0.00000 0.00000 0.00000
0.00000 0.00000 0.00000 ; HB3- CB- CG- OD1
2562 2560 2563 2565 3 0.00000 0.00000 0.00000
0.00000 0.00000 0.00000 ; HB3- CB- CG- OD2
2563 2560 2558 2566 3 7.14627 -1.04600 -8.99560
-0.97069 5.15469 0.00000 ; CG- CB- CA- C
2566 2568 2570 2571 3 0.00000 0.00000 0.00000
0.00000 0.00000 0.00000 ; C- N- CA- HA
2566 2568 2570 2572 3 11.71520 1.67360 15.06240 -
13.38880 0.00000 0.00000 ; C- N- CA- CB
2566 2568 2570 2586 3 1.75728 5.27184 2.25936
-7.02912 0.00000 0.00000 ; C- N- CA- C
2567 2566 2568 2569 3 29.28800 -8.36800 -20.92000
0.00000 0.00000 0.00000 ; O- C- N- H
2567 2566 2568 2570 3 20.92000 0.00000 -20.92000
0.00000 0.00000 0.00000 ; O- C- N- CA
2568 2570 2572 2573 3 0.65084 1.95253 0.00000
-2.60338 0.00000 0.00000 ; N- CA- CB- HB2
2568 2570 2572 2574 3 0.65084 1.95253 0.00000
-2.60338 0.00000 0.00000 ; N- CA- CB- HB3
2568 2570 2572 2575 3 3.28026 2.46019 -2.19242
-3.21331 -0.23430 0.00000 ; N- CA- CB- CG
2568 2570 2586 2587 3 0.00000 0.00000 0.00000
0.00000 0.00000 0.00000 ; N- CA- C- O
2568 2570 2586 2588 3 17.40544 -5.02080 -13.22144
9.20480 0.00000 0.00000 ; N- CA- C- N
2569 2568 2570 2571 3 0.00000 0.00000 0.00000
0.00000 0.00000 0.00000 ; H- N- CA- HA
2569 2568 2570 2572 3 0.00000 0.00000 0.00000
0.00000 0.00000 0.00000 ; H- N- CA- CB
2569 2568 2570 2586 3 0.00000 0.00000 0.00000
0.00000 0.00000 0.00000 ; H- N- CA- C
2570 2572 2575 2576 3 0.57739 0.00000 1.02926
0.00000 -1.60666 0.00000 ; CA- CB- CG- CD1

2570 2572 2575 2584 3 0.57739 0.00000 1.02926
0.00000 -1.60666 0.00000 ; CA- CB- CG- CD2
2570 2586 2588 2589 3 20.92000 0.00000 -20.92000
0.00000 0.00000 0.00000 ; CA- C- N- H
2570 2586 2588 2590 3 20.92000 0.00000 -20.92000
0.00000 0.00000 0.00000 ; CA- C- N- CA
2571 2570 2572 2573 3 0.65084 1.95253 0.00000
-2.60338 0.00000 0.00000 ; HA- CA- CB- HB2
2571 2570 2572 2574 3 0.65084 1.95253 0.00000
-2.60338 0.00000 0.00000 ; HA- CA- CB- HB3
2571 2570 2572 2575 3 0.65084 1.95253 0.00000
-2.60338 0.00000 0.00000 ; HA- CA- CB- CG
2571 2570 2586 2587 3 3.68192 -4.35136 0.00000
1.33888 0.00000 0.00000 ; HA- CA- C- O
2571 2570 2586 2588 3 0.00000 0.00000 0.00000
0.00000 0.00000 0.00000 ; HA- CA- C- N
2572 2570 2586 2587 3 0.00000 0.00000 0.00000
0.00000 0.00000 0.00000 ; CB- CA- C- O
2572 2570 2586 2588 3 2.51040 4.18400 1.67360
-6.69440 0.00000 0.00000 ; CB- CA- C- N
2572 2575 2576 2577 3 30.33400 0.00000 -30.33400
0.00000 0.00000 0.00000 ; CB- CG- CD1- HD1
2572 2575 2576 2578 3 30.33400 0.00000 -30.33400
0.00000 0.00000 0.00000 ; CB- CG- CD1- CE1
2572 2575 2584 2582 3 30.33400 0.00000 -30.33400
0.00000 0.00000 0.00000 ; CB- CG- CD2- CE2
2572 2575 2584 2585 3 30.33400 0.00000 -30.33400
0.00000 0.00000 0.00000 ; CB- CG- CD2- HD2
2573 2572 2570 2586 3 0.65084 1.95253 0.00000
-2.60338 0.00000 0.00000 ; HB2- CB- CA- C
2573 2572 2575 2576 3 0.00000 0.00000 0.00000
0.00000 0.00000 0.00000 ; HB2- CB- CG- CD1
2573 2572 2575 2584 3 0.00000 0.00000 0.00000
0.00000 0.00000 0.00000 ; HB2- CB- CG- CD2
2574 2572 2570 2586 3 0.65084 1.95253 0.00000
-2.60338 0.00000 0.00000 ; HB3- CB- CA- C
2574 2572 2575 2576 3 0.00000 0.00000 0.00000
0.00000 0.00000 0.00000 ; HB3- CB- CG- CD1
2574 2572 2575 2584 3 0.00000 0.00000 0.00000
0.00000 0.00000 0.00000 ; HB3- CB- CG- CD2
2575 2572 2570 2586 3 4.95804 2.17986 -3.52293
-3.21331 -0.40166 0.00000 ; CG- CB- CA- C
2575 2576 2578 2579 3 30.33400 0.00000 -30.33400
0.00000 0.00000 0.00000 ; CG- CD1- CE1- HE1
2575 2576 2578 2580 3 30.33400 0.00000 -30.33400
0.00000 0.00000 0.00000 ; CG- CD1- CE1- CZ
2575 2584 2582 2580 3 30.33400 0.00000 -30.33400
0.00000 0.00000 0.00000 ; CG- CD2- CE2- CZ
2575 2584 2582 2583 3 30.33400 0.00000 -30.33400
0.00000 0.00000 0.00000 ; CG- CD2- CE2- HE2
2576 2575 2584 2582 3 30.33400 0.00000 -30.33400
0.00000 0.00000 0.00000 ; CD1- CG- CD2- CE2

```
  2576   2575   2584   2585       3   30.33400    0.00000  -30.33400
0.00000    0.00000    0.00000 ;    CD1-    CG-   CD2-   HD2
  2576   2578   2580   2581       3   30.33400    0.00000  -30.33400
0.00000    0.00000    0.00000 ;    CD1-   CE1-    CZ-    HZ
  2576   2578   2580   2582       3   30.33400    0.00000  -30.33400
0.00000    0.00000    0.00000 ;    CD1-   CE1-    CZ-   CE2
  2577   2576   2575   2584       3   30.33400    0.00000  -30.33400
0.00000    0.00000    0.00000 ;    HD1-   CD1-    CG-   CD2
  2577   2576   2578   2579       3   30.33400    0.00000  -30.33400
0.00000    0.00000    0.00000 ;    HD1-   CD1-   CE1-   HE1
  2577   2576   2578   2580       3   30.33400    0.00000  -30.33400
0.00000    0.00000    0.00000 ;    HD1-   CD1-   CE1-    CZ
  2578   2576   2575   2584       3   30.33400    0.00000  -30.33400
0.00000    0.00000    0.00000 ;    CE1-   CD1-    CG-   CD2
  2578   2580   2582   2583       3   30.33400    0.00000  -30.33400
0.00000    0.00000    0.00000 ;    CE1-    CZ-   CE2-   HE2
  2578   2580   2582   2584       3   30.33400    0.00000  -30.33400
0.00000    0.00000    0.00000 ;    CE1-    CZ-   CE2-   CD2
  2579   2578   2580   2581       3   30.33400    0.00000  -30.33400
0.00000    0.00000    0.00000 ;    HE1-   CE1-    CZ-    HZ
  2579   2578   2580   2582       3   30.33400    0.00000  -30.33400
0.00000    0.00000    0.00000 ;    HE1-   CE1-    CZ-   CE2
  2580   2582   2584   2585       3   30.33400    0.00000  -30.33400
0.00000    0.00000    0.00000 ;     CZ-   CE2-   CD2-   HD2
  2581   2580   2582   2583       3   30.33400    0.00000  -30.33400
0.00000    0.00000    0.00000 ;     HZ-    CZ-   CE2-   HE2
  2581   2580   2582   2584       3   30.33400    0.00000  -30.33400
0.00000    0.00000    0.00000 ;     HZ-    CZ-   CE2-   CD2
  2583   2582   2584   2585       3   30.33400    0.00000  -30.33400
0.00000    0.00000    0.00000 ;    HE2-   CE2-   CD2-   HD2
  2586   2588   2590   2591       3    0.00000    0.00000    0.00000
0.00000    0.00000    0.00000 ;      C-     N-    CA-   HA2
  2586   2588   2590   2592       3    0.00000    0.00000    0.00000
0.00000    0.00000    0.00000 ;      C-     N-    CA-   HA3
  2586   2588   2590   2593       3    1.75728    5.27184    2.25936
-7.02912    0.00000    0.00000 ;      C-     N-    CA-     C
  2587   2586   2588   2589       3   29.28800   -8.36800  -20.92000
0.00000    0.00000    0.00000 ;      O-     C-     N-     H
  2587   2586   2588   2590       3   20.92000    0.00000  -20.92000
0.00000    0.00000    0.00000 ;      O-     C-     N-    CA
  2588   2590   2593   2594       3    0.00000    0.00000    0.00000
0.00000    0.00000    0.00000 ;      N-    CA-     C-     O
  2588   2590   2593   2595       3   17.40544   -5.02080  -13.22144
9.20480    0.00000    0.00000 ;      N-    CA-     C-     N
  2589   2588   2590   2591       3    0.00000    0.00000    0.00000
0.00000    0.00000    0.00000 ;      H-     N-    CA-   HA2
  2589   2588   2590   2592       3    0.00000    0.00000    0.00000
0.00000    0.00000    0.00000 ;      H-     N-    CA-   HA3
  2589   2588   2590   2593       3    0.00000    0.00000    0.00000
0.00000    0.00000    0.00000 ;      H-     N-    CA-     C
  2590   2593   2595   2596       3   20.92000    0.00000  -20.92000
0.00000    0.00000    0.00000 ;     CA-     C-     N-     H
```

2590 2593 2595 2597 3 20.92000 0.00000 -20.92000
0.00000 0.00000 0.00000 ; CA- C- N- CA
2591 2590 2593 2594 3 3.68192 -4.35136 0.00000
1.33888 0.00000 0.00000 ; HA2- CA- C- O
2591 2590 2593 2595 3 0.00000 0.00000 0.00000
0.00000 0.00000 0.00000 ; HA2- CA- C- N
2592 2590 2593 2594 3 3.68192 -4.35136 0.00000
1.33888 0.00000 0.00000 ; HA3- CA- C- O
2592 2590 2593 2595 3 0.00000 0.00000 0.00000
0.00000 0.00000 0.00000 ; HA3- CA- C- N
2593 2595 2597 2598 3 0.00000 0.00000 0.00000
0.00000 0.00000 0.00000 ; C- N- CA- HA
2593 2595 2597 2599 3 11.71520 1.67360 15.06240 -
13.38880 0.00000 0.00000 ; C- N- CA- CB
2593 2595 2597 2612 3 1.75728 5.27184 2.25936
-7.02912 0.00000 0.00000 ; C- N- CA- C
2594 2593 2595 2596 3 29.28800 -8.36800 -20.92000
0.00000 0.00000 0.00000 ; O- C- N- H
2594 2593 2595 2597 3 20.92000 0.00000 -20.92000
0.00000 0.00000 0.00000 ; O- C- N- CA
2595 2597 2599 2600 3 0.65084 1.95253 0.00000
-2.60338 0.00000 0.00000 ; N- CA- CB- HB2
2595 2597 2599 2601 3 0.65084 1.95253 0.00000
-2.60338 0.00000 0.00000 ; N- CA- CB- HB3
2595 2597 2599 2602 3 3.79070 1.39746 -4.61077
-2.40998 2.44346 0.00000 ; N- CA- CB- CG
2595 2597 2612 2613 3 0.00000 0.00000 0.00000
0.00000 0.00000 0.00000 ; N- CA- C- O
2595 2597 2612 2614 3 17.40544 -5.02080 -13.22144
9.20480 0.00000 0.00000 ; N- CA- C- N
2596 2595 2597 2598 3 0.00000 0.00000 0.00000
0.00000 0.00000 0.00000 ; H- N- CA- HA
2596 2595 2597 2599 3 0.00000 0.00000 0.00000
0.00000 0.00000 0.00000 ; H- N- CA- CB
2596 2595 2597 2612 3 0.00000 0.00000 0.00000
0.00000 0.00000 0.00000 ; H- N- CA- C
2597 2599 2602 2603 3 0.66944 2.00832 0.00000
-2.67776 0.00000 0.00000 ; CA- CB- CG- HG
2597 2599 2602 2604 3 3.90367 0.19665 -6.21742
-2.37651 5.99149 0.00000 ; CA- CB- CG- CD1
2597 2599 2602 2608 3 3.90367 0.19665 -6.21742
-2.37651 5.99149 0.00000 ; CA- CB- CG- CD2
2597 2612 2614 2615 3 20.92000 0.00000 -20.92000
0.00000 0.00000 0.00000 ; CA- C- N- H
2597 2612 2614 2616 3 20.92000 0.00000 -20.92000
0.00000 0.00000 0.00000 ; CA- C- N- CA
2598 2597 2599 2600 3 0.65084 1.95253 0.00000
-2.60338 0.00000 0.00000 ; HA- CA- CB- HB2
2598 2597 2599 2601 3 0.65084 1.95253 0.00000
-2.60338 0.00000 0.00000 ; HA- CA- CB- HB3
2598 2597 2599 2602 3 0.65084 1.95253 0.00000
-2.60338 0.00000 0.00000 ; HA- CA- CB- CG

2598 2597 2612 2613 3 3.68192 -4.35136 0.00000
1.33888 0.00000 0.00000 ; HA- CA- C- O
2598 2597 2612 2614 3 0.00000 0.00000 0.00000
0.00000 0.00000 0.00000 ; HA- CA- C- N
2599 2597 2612 2613 3 0.00000 0.00000 0.00000
0.00000 0.00000 0.00000 ; CB- CA- C- O
2599 2597 2612 2614 3 2.51040 4.18400 1.67360
-6.69440 0.00000 0.00000 ; CB- CA- C- N
2599 2602 2604 2605 3 0.66944 2.00832 0.00000
-2.67776 0.00000 0.00000 ; CB- CG- CD1- HD11
2599 2602 2604 2606 3 0.66944 2.00832 0.00000
-2.67776 0.00000 0.00000 ; CB- CG- CD1- HD12
2599 2602 2604 2607 3 0.66944 2.00832 0.00000
-2.67776 0.00000 0.00000 ; CB- CG- CD1- HD13
2599 2602 2608 2609 3 0.66944 2.00832 0.00000
-2.67776 0.00000 0.00000 ; CB- CG- CD2- HD21
2599 2602 2608 2610 3 0.66944 2.00832 0.00000
-2.67776 0.00000 0.00000 ; CB- CG- CD2- HD22
2599 2602 2608 2611 3 0.66944 2.00832 0.00000
-2.67776 0.00000 0.00000 ; CB- CG- CD2- HD23
2600 2599 2597 2612 3 0.65084 1.95253 0.00000
-2.60338 0.00000 0.00000 ; HB2- CB- CA- C
2600 2599 2602 2603 3 0.62760 1.88280 0.00000
-2.51040 0.00000 0.00000 ; HB2- CB- CG- HG
2600 2599 2602 2604 3 0.66944 2.00832 0.00000
-2.67776 0.00000 0.00000 ; HB2- CB- CG- CD1
2600 2599 2602 2608 3 0.66944 2.00832 0.00000
-2.67776 0.00000 0.00000 ; HB2- CB- CG- CD2
2601 2599 2597 2612 3 0.65084 1.95253 0.00000
-2.60338 0.00000 0.00000 ; HB3- CB- CA- C
2601 2599 2602 2603 3 0.62760 1.88280 0.00000
-2.51040 0.00000 0.00000 ; HB3- CB- CG- HG
2601 2599 2602 2604 3 0.66944 2.00832 0.00000
-2.67776 0.00000 0.00000 ; HB3- CB- CG- CD1
2601 2599 2602 2608 3 0.66944 2.00832 0.00000
-2.67776 0.00000 0.00000 ; HB3- CB- CG- CD2
2602 2599 2597 2612 3 10.33448 -1.14642 -11.54784
-2.40998 6.35968 0.00000 ; CG- CB- CA- C
2603 2602 2604 2605 3 0.62760 1.88280 0.00000
-2.51040 0.00000 0.00000 ; HG- CG- CD1- HD11
2603 2602 2604 2606 3 0.62760 1.88280 0.00000
-2.51040 0.00000 0.00000 ; HG- CG- CD1- HD12
2603 2602 2604 2607 3 0.62760 1.88280 0.00000
-2.51040 0.00000 0.00000 ; HG- CG- CD1- HD13
2603 2602 2608 2609 3 0.62760 1.88280 0.00000
-2.51040 0.00000 0.00000 ; HG- CG- CD2- HD21
2603 2602 2608 2610 3 0.62760 1.88280 0.00000
-2.51040 0.00000 0.00000 ; HG- CG- CD2- HD22
2603 2602 2608 2611 3 0.62760 1.88280 0.00000
-2.51040 0.00000 0.00000 ; HG- CG- CD2- HD23
2604 2602 2608 2609 3 0.66944 2.00832 0.00000
-2.67776 0.00000 0.00000 ; CD1- CG- CD2- HD21

2604 2602 2608 2610 3 0.66944 2.00832 0.00000
-2.67776 0.00000 0.00000 ; CD1- CG- CD2- HD22
2604 2602 2608 2611 3 0.66944 2.00832 0.00000
-2.67776 0.00000 0.00000 ; CD1- CG- CD2- HD23
2605 2604 2602 2608 3 0.66944 2.00832 0.00000
-2.67776 0.00000 0.00000 ; HD11- CD1- CG- CD2
2606 2604 2602 2608 3 0.66944 2.00832 0.00000
-2.67776 0.00000 0.00000 ; HD12- CD1- CG- CD2
2607 2604 2602 2608 3 0.66944 2.00832 0.00000
-2.67776 0.00000 0.00000 ; HD13- CD1- CG- CD2
2612 2614 2616 2617 3 0.00000 0.00000 0.00000
0.00000 0.00000 0.00000 ; C- N- CA- HA
2612 2614 2616 2618 3 11.71520 1.67360 15.06240 -
13.38880 0.00000 0.00000 ; C- N- CA- CB
2612 2614 2616 2623 3 1.75728 5.27184 2.25936
-7.02912 0.00000 0.00000 ; C- N- CA- C
2613 2612 2614 2615 3 29.28800 -8.36800 -20.92000
0.00000 0.00000 0.00000 ; O- C- N- H
2613 2612 2614 2616 3 20.92000 0.00000 -20.92000
0.00000 0.00000 0.00000 ; O- C- N- CA
2614 2616 2618 2619 3 0.65084 1.95253 0.00000
-2.60338 0.00000 0.00000 ; N- CA- CB- HB2
2614 2616 2618 2620 3 0.65084 1.95253 0.00000
-2.60338 0.00000 0.00000 ; N- CA- CB- HB3
2614 2616 2618 2621 3 7.86174 2.24681 -7.41405
-6.71114 5.35552 0.00000 ; N- CA- CB- OG
2614 2616 2623 2624 3 0.00000 0.00000 0.00000
0.00000 0.00000 0.00000 ; N- CA- C- O
2614 2616 2623 2625 3 17.40544 -5.02080 -13.22144
9.20480 0.00000 0.00000 ; N- CA- C- N
2615 2614 2616 2617 3 0.00000 0.00000 0.00000
0.00000 0.00000 0.00000 ; H- N- CA- HA
2615 2614 2616 2618 3 0.00000 0.00000 0.00000
0.00000 0.00000 0.00000 ; H- N- CA- CB
2615 2614 2616 2623 3 0.00000 0.00000 0.00000
0.00000 0.00000 0.00000 ; H- N- CA- C
2616 2618 2621 2622 3 2.05853 2.46856 3.48109
-4.46851 0.23430 0.00000 ; CA- CB- OG- HG
2616 2623 2625 2626 3 20.92000 0.00000 -20.92000
0.00000 0.00000 0.00000 ; CA- C- N- H
2616 2623 2625 2627 3 20.92000 0.00000 -20.92000
0.00000 0.00000 0.00000 ; CA- C- N- CA
2617 2616 2618 2619 3 0.65084 1.95253 0.00000
-2.60338 0.00000 0.00000 ; HA- CA- CB- HB2
2617 2616 2618 2620 3 0.65084 1.95253 0.00000
-2.60338 0.00000 0.00000 ; HA- CA- CB- HB3
2617 2616 2618 2621 3 1.04600 -1.04600 0.00000
0.00000 0.00000 0.00000 ; HA- CA- CB- OG
2617 2616 2623 2624 3 3.68192 -4.35136 0.00000
1.33888 0.00000 0.00000 ; HA- CA- C- O
2617 2616 2623 2625 3 0.00000 0.00000 0.00000
0.00000 0.00000 0.00000 ; HA- CA- C- N

2618 2616 2623 2624 3 0.00000 0.00000 0.00000
0.00000 0.00000 0.00000 ; CB- CA- C- O
2618 2616 2623 2625 3 2.51040 4.18400 1.67360
-6.69440 0.00000 0.00000 ; CB- CA- C- N
2619 2618 2616 2623 3 0.65084 1.95253 0.00000
-2.60338 0.00000 0.00000 ; HB2- CB- CA- C
2619 2618 2621 2622 3 0.69733 2.09200 0.00000
-2.78933 0.00000 0.00000 ; HB2- CB- OG- HG
2620 2618 2616 2623 3 0.65084 1.95253 0.00000
-2.60338 0.00000 0.00000 ; HB3- CB- CA- C
2620 2618 2621 2622 3 0.69733 2.09200 0.00000
-2.78933 0.00000 0.00000 ; HB3- CB- OG- HG
2621 2618 2616 2623 3 7.34710 7.79898 -6.14211
-6.71114 4.31789 0.00000 ; OG- CB- CA- C
2623 2625 2627 2628 3 0.00000 0.00000 0.00000
0.00000 0.00000 0.00000 ; C- N- CA- HA
2623 2625 2627 2629 3 11.71520 1.67360 15.06240 -
13.38880 0.00000 0.00000 ; C- N- CA- CB
2623 2625 2627 2645 3 1.75728 5.27184 2.25936
-7.02912 0.00000 0.00000 ; C- N- CA- C
2624 2623 2625 2626 3 29.28800 -8.36800 -20.92000
0.00000 0.00000 0.00000 ; O- C- N- H
2624 2623 2625 2627 3 20.92000 0.00000 -20.92000
0.00000 0.00000 0.00000 ; O- C- N- CA
2625 2627 2629 2630 3 0.65084 1.95253 0.00000
-2.60338 0.00000 0.00000 ; N- CA- CB- HB2
2625 2627 2629 2631 3 0.65084 1.95253 0.00000
-2.60338 0.00000 0.00000 ; N- CA- CB- HB3
2625 2627 2629 2632 3 0.65084 1.95253 0.00000
-2.60338 0.00000 0.00000 ; N- CA- CB- CG
2625 2627 2645 2646 3 0.00000 0.00000 0.00000
0.00000 0.00000 0.00000 ; N- CA- C- O
2625 2627 2645 2647 3 17.40544 -5.02080 -13.22144
9.20480 0.00000 0.00000 ; N- CA- C- N
2626 2625 2627 2628 3 0.00000 0.00000 0.00000
0.00000 0.00000 0.00000 ; H- N- CA- HA
2626 2625 2627 2629 3 0.00000 0.00000 0.00000
0.00000 0.00000 0.00000 ; H- N- CA- CB
2626 2625 2627 2645 3 0.00000 0.00000 0.00000
0.00000 0.00000 0.00000 ; H- N- CA- C
2627 2629 2632 2633 3 0.66944 2.00832 0.00000
-2.67776 0.00000 0.00000 ; CA- CB- CG- HG2
2627 2629 2632 2634 3 0.66944 2.00832 0.00000
-2.67776 0.00000 0.00000 ; CA- CB- CG- HG3
2627 2629 2632 2635 3 3.68192 3.09616 -2.09200
-3.01248 0.00000 0.00000 ; CA- CB- CG- CD
2627 2645 2647 2648 3 20.92000 0.00000 -20.92000
0.00000 0.00000 0.00000 ; CA- C- N- H
2627 2645 2647 2649 3 20.92000 0.00000 -20.92000
0.00000 0.00000 0.00000 ; CA- C- N- CA
2628 2627 2629 2630 3 0.65084 1.95253 0.00000
-2.60338 0.00000 0.00000 ; HA- CA- CB- HB2

2628 2627 2629 2631 3 0.65084 1.95253 0.00000
-2.60338 0.00000 0.00000 ; HA- CA- CB- HB3
2628 2627 2629 2632 3 0.65084 1.95253 0.00000
-2.60338 0.00000 0.00000 ; HA- CA- CB- CG
2628 2627 2645 2646 3 3.68192 -4.35136 0.00000
1.33888 0.00000 0.00000 ; HA- CA- C- O
2628 2627 2645 2647 3 0.00000 0.00000 0.00000
0.00000 0.00000 0.00000 ; HA- CA- C- N
2629 2627 2645 2646 3 0.00000 0.00000 0.00000
0.00000 0.00000 0.00000 ; CB- CA- C- O
2629 2627 2645 2647 3 2.51040 4.18400 1.67360
-6.69440 0.00000 0.00000 ; CB- CA- C- N
2629 2632 2635 2636 3 0.66944 2.00832 0.00000
-2.67776 0.00000 0.00000 ; CB- CG- CD- HD2
2629 2632 2635 2637 3 0.66944 2.00832 0.00000
-2.67776 0.00000 0.00000 ; CB- CG- CD- HD3
2629 2632 2635 2638 3 3.68192 3.09616 -2.09200
-3.01248 0.00000 0.00000 ; CB- CG- CD- CE
2630 2629 2627 2645 3 0.65084 1.95253 0.00000
-2.60338 0.00000 0.00000 ; HB2- CB- CA- C
2630 2629 2632 2633 3 0.62760 1.88280 0.00000
-2.51040 0.00000 0.00000 ; HB2- CB- CG- HG2
2630 2629 2632 2634 3 0.62760 1.88280 0.00000
-2.51040 0.00000 0.00000 ; HB2- CB- CG- HG3
2630 2629 2632 2635 3 0.66944 2.00832 0.00000
-2.67776 0.00000 0.00000 ; HB2- CB- CG- CD
2631 2629 2627 2645 3 0.65084 1.95253 0.00000
-2.60338 0.00000 0.00000 ; HB3- CB- CA- C
2631 2629 2632 2633 3 0.62760 1.88280 0.00000
-2.51040 0.00000 0.00000 ; HB3- CB- CG- HG2
2631 2629 2632 2634 3 0.62760 1.88280 0.00000
-2.51040 0.00000 0.00000 ; HB3- CB- CG- HG3
2631 2629 2632 2635 3 0.66944 2.00832 0.00000
-2.67776 0.00000 0.00000 ; HB3- CB- CG- CD
2632 2629 2627 2645 3 0.65084 1.95253 0.00000
-2.60338 0.00000 0.00000 ; CG- CB- CA- C
2632 2635 2638 2639 3 0.65084 1.95253 0.00000
-2.60338 0.00000 0.00000 ; CG- CD- CE- HE2
2632 2635 2638 2640 3 0.65084 1.95253 0.00000
-2.60338 0.00000 0.00000 ; CG- CD- CE- HE3
2632 2635 2638 2641 3 0.65084 1.95253 0.00000
-2.60338 0.00000 0.00000 ; CG- CD- CE- NZ
2633 2632 2635 2636 3 0.62760 1.88280 0.00000
-2.51040 0.00000 0.00000 ; HG2- CG- CD- HD2
2633 2632 2635 2637 3 0.62760 1.88280 0.00000
-2.51040 0.00000 0.00000 ; HG2- CG- CD- HD3
2633 2632 2635 2638 3 0.66944 2.00832 0.00000
-2.67776 0.00000 0.00000 ; HG2- CG- CD- CE
2634 2632 2635 2636 3 0.62760 1.88280 0.00000
-2.51040 0.00000 0.00000 ; HG3- CG- CD- HD2
2634 2632 2635 2637 3 0.62760 1.88280 0.00000
-2.51040 0.00000 0.00000 ; HG3- CG- CD- HD3

2634 2632 2635 2638 3 0.66944 2.00832 0.00000
-2.67776 0.00000 0.00000 ; HG3- CG- CD- CE
2635 2638 2641 2642 3 0.65084 1.95253 0.00000
-2.60338 0.00000 0.00000 ; CD- CE- NZ- HZ1
2635 2638 2641 2643 3 0.65084 1.95253 0.00000
-2.60338 0.00000 0.00000 ; CD- CE- NZ- HZ2
2635 2638 2641 2644 3 0.65084 1.95253 0.00000
-2.60338 0.00000 0.00000 ; CD- CE- NZ- HZ3
2636 2635 2638 2639 3 0.65084 1.95253 0.00000
-2.60338 0.00000 0.00000 ; HD2- CD- CE- HE2
2636 2635 2638 2640 3 0.65084 1.95253 0.00000
-2.60338 0.00000 0.00000 ; HD2- CD- CE- HE3
2636 2635 2638 2641 3 0.65084 1.95253 0.00000
-2.60338 0.00000 0.00000 ; HD2- CD- CE- NZ
2637 2635 2638 2639 3 0.65084 1.95253 0.00000
-2.60338 0.00000 0.00000 ; HD3- CD- CE- HE2
2637 2635 2638 2640 3 0.65084 1.95253 0.00000
-2.60338 0.00000 0.00000 ; HD3- CD- CE- HE3
2637 2635 2638 2641 3 0.65084 1.95253 0.00000
-2.60338 0.00000 0.00000 ; HD3- CD- CE- NZ
2639 2638 2641 2642 3 0.65084 1.95253 0.00000
-2.60338 0.00000 0.00000 ; HE2- CE- NZ- HZ1
2639 2638 2641 2643 3 0.65084 1.95253 0.00000
-2.60338 0.00000 0.00000 ; HE2- CE- NZ- HZ2
2639 2638 2641 2644 3 0.65084 1.95253 0.00000
-2.60338 0.00000 0.00000 ; HE2- CE- NZ- HZ3
2640 2638 2641 2642 3 0.65084 1.95253 0.00000
-2.60338 0.00000 0.00000 ; HE3- CE- NZ- HZ1
2640 2638 2641 2643 3 0.65084 1.95253 0.00000
-2.60338 0.00000 0.00000 ; HE3- CE- NZ- HZ2
2640 2638 2641 2644 3 0.65084 1.95253 0.00000
-2.60338 0.00000 0.00000 ; HE3- CE- NZ- HZ3
2645 2647 2649 2650 3 0.00000 0.00000 0.00000
0.00000 0.00000 0.00000 ; C- N- CA- HA
2645 2647 2649 2651 3 11.71520 1.67360 15.06240 -
13.38880 0.00000 0.00000 ; C- N- CA- CB
2645 2647 2649 2659 3 1.75728 5.27184 2.25936
-7.02912 0.00000 0.00000 ; C- N- CA- C
2646 2645 2647 2648 3 29.28800 -8.36800 -20.92000
0.00000 0.00000 0.00000 ; O- C- N- H
2646 2645 2647 2649 3 20.92000 0.00000 -20.92000
0.00000 0.00000 0.00000 ; O- C- N- CA
2647 2649 2651 2652 3 0.65084 1.95253 0.00000
-2.60338 0.00000 0.00000 ; N- CA- CB- HB2
2647 2649 2651 2653 3 0.65084 1.95253 0.00000
-2.60338 0.00000 0.00000 ; N- CA- CB- HB3
2647 2649 2651 2654 3 5.99567 3.29281 -4.46014
-0.55229 1.97485 0.00000 ; N- CA- CB- CG
2647 2649 2659 2660 3 0.00000 0.00000 0.00000
0.00000 0.00000 0.00000 ; N- CA- C- O
2647 2649 2659 2661 3 17.40544 -5.02080 -13.22144
9.20480 0.00000 0.00000 ; N- CA- C- N

2648 2647 2649 2650 3 0.00000 0.00000 0.00000
0.00000 0.00000 0.00000 ; H- N- CA- HA
2648 2647 2649 2651 3 0.00000 0.00000 0.00000
0.00000 0.00000 0.00000 ; H- N- CA- CB
2648 2647 2649 2659 3 0.00000 0.00000 0.00000
0.00000 0.00000 0.00000 ; H- N- CA- C
2649 2651 2654 2655 3 0.00000 0.00000 0.00000
0.00000 0.00000 0.00000 ; CA- CB- CG- OD1
2649 2651 2654 2656 3 8.84916 -0.31380 -4.32626
5.03754 0.26778 0.00000 ; CA- CB- CG- ND2
2649 2659 2661 2662 3 20.92000 0.00000 -20.92000
0.00000 0.00000 0.00000 ; CA- C- N- H
2649 2659 2661 2663 3 20.92000 0.00000 -20.92000
0.00000 0.00000 0.00000 ; CA- C- N- CA
2650 2649 2651 2652 3 0.65084 1.95253 0.00000
-2.60338 0.00000 0.00000 ; HA- CA- CB- HB2
2650 2649 2651 2653 3 0.65084 1.95253 0.00000
-2.60338 0.00000 0.00000 ; HA- CA- CB- HB3
2650 2649 2651 2654 3 0.65084 1.95253 0.00000
-2.60338 0.00000 0.00000 ; HA- CA- CB- CG
2650 2649 2659 2660 3 3.68192 -4.35136 0.00000
1.33888 0.00000 0.00000 ; HA- CA- C- O
2650 2649 2659 2661 3 0.00000 0.00000 0.00000
0.00000 0.00000 0.00000 ; HA- CA- C- N
2651 2649 2659 2660 3 0.00000 0.00000 0.00000
0.00000 0.00000 0.00000 ; CB- CA- C- O
2651 2649 2659 2661 3 2.51040 4.18400 1.67360
-6.69440 0.00000 0.00000 ; CB- CA- C- N
2651 2654 2656 2657 3 20.92000 0.00000 -20.92000
0.00000 0.00000 0.00000 ; CB- CG- ND2- HD21
2651 2654 2656 2658 3 20.92000 0.00000 -20.92000
0.00000 0.00000 0.00000 ; CB- CG- ND2- HD22
2652 2651 2649 2659 3 0.65084 1.95253 0.00000
-2.60338 0.00000 0.00000 ; HB2- CB- CA- C
2652 2651 2654 2655 3 3.68192 -4.35136 0.00000
1.33888 0.00000 0.00000 ; HB2- CB- CG- OD1
2652 2651 2654 2656 3 0.00000 0.00000 0.00000
0.00000 0.00000 0.00000 ; HB2- CB- CG- ND2
2653 2651 2649 2659 3 0.65084 1.95253 0.00000
-2.60338 0.00000 0.00000 ; HB3- CB- CA- C
2653 2651 2654 2655 3 3.68192 -4.35136 0.00000
1.33888 0.00000 0.00000 ; HB3- CB- CG- OD1
2653 2651 2654 2656 3 0.00000 0.00000 0.00000
0.00000 0.00000 0.00000 ; HB3- CB- CG- ND2
2654 2651 2649 2659 3 7.94542 -3.96225 -6.11701
-0.55229 3.58150 0.00000 ; CG- CB- CA- C
2655 2654 2656 2657 3 29.28800 -8.36800 -20.92000
0.00000 0.00000 0.00000 ; OD1- CG- ND2- HD21
2655 2654 2656 2658 3 29.28800 -8.36800 -20.92000
0.00000 0.00000 0.00000 ; OD1- CG- ND2- HD22
2659 2661 2663 2664 3 0.00000 0.00000 0.00000
0.00000 0.00000 0.00000 ; C- N- CA- HA

2659 2661 2663 2665 3 11.71520 1.67360 15.06240 -
13.38880 0.00000 0.00000 ; C- N- CA- CB
2659 2661 2663 2678 3 1.75728 5.27184 2.25936
-7.02912 0.00000 0.00000 ; C- N- CA- C
2660 2659 2661 2662 3 29.28800 -8.36800 -20.92000
0.00000 0.00000 0.00000 ; O- C- N- H
2660 2659 2661 2663 3 20.92000 0.00000 -20.92000
0.00000 0.00000 0.00000 ; O- C- N- CA
2661 2663 2665 2666 3 0.65084 1.95253 0.00000
-2.60338 0.00000 0.00000 ; N- CA- CB- HB2
2661 2663 2665 2667 3 0.65084 1.95253 0.00000
-2.60338 0.00000 0.00000 ; N- CA- CB- HB3
2661 2663 2665 2668 3 3.79070 1.39746 -4.61077
-2.40998 2.44346 0.00000 ; N- CA- CB- CG
2661 2663 2678 2679 3 0.00000 0.00000 0.00000
0.00000 0.00000 0.00000 ; N- CA- C- O
2661 2663 2678 2680 3 17.40544 -5.02080 -13.22144
9.20480 0.00000 0.00000 ; N- CA- C- N
2662 2661 2663 2664 3 0.00000 0.00000 0.00000
0.00000 0.00000 0.00000 ; H- N- CA- HA
2662 2661 2663 2665 3 0.00000 0.00000 0.00000
0.00000 0.00000 0.00000 ; H- N- CA- CB
2662 2661 2663 2678 3 0.00000 0.00000 0.00000
0.00000 0.00000 0.00000 ; H- N- CA- C
2663 2665 2668 2669 3 0.66944 2.00832 0.00000
-2.67776 0.00000 0.00000 ; CA- CB- CG- HG
2663 2665 2668 2670 3 3.90367 0.19665 -6.21742
-2.37651 5.99149 0.00000 ; CA- CB- CG- CD1
2663 2665 2668 2674 3 3.90367 0.19665 -6.21742
-2.37651 5.99149 0.00000 ; CA- CB- CG- CD2
2663 2678 2680 2681 3 20.92000 0.00000 -20.92000
0.00000 0.00000 0.00000 ; CA- C- N- H
2663 2678 2680 2682 3 20.92000 0.00000 -20.92000
0.00000 0.00000 0.00000 ; CA- C- N- CA
2664 2663 2665 2666 3 0.65084 1.95253 0.00000
-2.60338 0.00000 0.00000 ; HA- CA- CB- HB2
2664 2663 2665 2667 3 0.65084 1.95253 0.00000
-2.60338 0.00000 0.00000 ; HA- CA- CB- HB3
2664 2663 2665 2668 3 0.65084 1.95253 0.00000
-2.60338 0.00000 0.00000 ; HA- CA- CB- CG
2664 2663 2678 2679 3 3.68192 -4.35136 0.00000
1.33888 0.00000 0.00000 ; HA- CA- C- O
2664 2663 2678 2680 3 0.00000 0.00000 0.00000
0.00000 0.00000 0.00000 ; HA- CA- C- N
2665 2663 2678 2679 3 0.00000 0.00000 0.00000
0.00000 0.00000 0.00000 ; CB- CA- C- O
2665 2663 2678 2680 3 2.51040 4.18400 1.67360
-6.69440 0.00000 0.00000 ; CB- CA- C- N
2665 2668 2670 2671 3 0.66944 2.00832 0.00000
-2.67776 0.00000 0.00000 ; CB- CG- CD1- HD11
2665 2668 2670 2672 3 0.66944 2.00832 0.00000
-2.67776 0.00000 0.00000 ; CB- CG- CD1- HD12

2665 2668 2670 2673 3 0.66944 2.00832 0.00000
-2.67776 0.00000 0.00000 ; CB- CG- CD1- HD13
2665 2668 2674 2675 3 0.66944 2.00832 0.00000
-2.67776 0.00000 0.00000 ; CB- CG- CD2- HD21
2665 2668 2674 2676 3 0.66944 2.00832 0.00000
-2.67776 0.00000 0.00000 ; CB- CG- CD2- HD22
2665 2668 2674 2677 3 0.66944 2.00832 0.00000
-2.67776 0.00000 0.00000 ; CB- CG- CD2- HD23
2666 2665 2663 2678 3 0.65084 1.95253 0.00000
-2.60338 0.00000 0.00000 ; HB2- CB- CA- C
2666 2665 2668 2669 3 0.62760 1.88280 0.00000
-2.51040 0.00000 0.00000 ; HB2- CB- CG- HG
2666 2665 2668 2670 3 0.66944 2.00832 0.00000
-2.67776 0.00000 0.00000 ; HB2- CB- CG- CD1
2666 2665 2668 2674 3 0.66944 2.00832 0.00000
-2.67776 0.00000 0.00000 ; HB2- CB- CG- CD2
2667 2665 2663 2678 3 0.65084 1.95253 0.00000
-2.60338 0.00000 0.00000 ; HB3- CB- CA- C
2667 2665 2668 2669 3 0.62760 1.88280 0.00000
-2.51040 0.00000 0.00000 ; HB3- CB- CG- HG
2667 2665 2668 2670 3 0.66944 2.00832 0.00000
-2.67776 0.00000 0.00000 ; HB3- CB- CG- CD1
2667 2665 2668 2674 3 0.66944 2.00832 0.00000
-2.67776 0.00000 0.00000 ; HB3- CB- CG- CD2
2668 2665 2663 2678 3 10.33448 -1.14642 -11.54784
-2.40998 6.35968 0.00000 ; CG- CB- CA- C
2669 2668 2670 2671 3 0.62760 1.88280 0.00000
-2.51040 0.00000 0.00000 ; HG- CG- CD1- HD11
2669 2668 2670 2672 3 0.62760 1.88280 0.00000
-2.51040 0.00000 0.00000 ; HG- CG- CD1- HD12
2669 2668 2670 2673 3 0.62760 1.88280 0.00000
-2.51040 0.00000 0.00000 ; HG- CG- CD1- HD13
2669 2668 2674 2675 3 0.62760 1.88280 0.00000
-2.51040 0.00000 0.00000 ; HG- CG- CD2- HD21
2669 2668 2674 2676 3 0.62760 1.88280 0.00000
-2.51040 0.00000 0.00000 ; HG- CG- CD2- HD22
2669 2668 2674 2677 3 0.62760 1.88280 0.00000
-2.51040 0.00000 0.00000 ; HG- CG- CD2- HD23
2670 2668 2674 2675 3 0.66944 2.00832 0.00000
-2.67776 0.00000 0.00000 ; CD1- CG- CD2- HD21
2670 2668 2674 2676 3 0.66944 2.00832 0.00000
-2.67776 0.00000 0.00000 ; CD1- CG- CD2- HD22
2670 2668 2674 2677 3 0.66944 2.00832 0.00000
-2.67776 0.00000 0.00000 ; CD1- CG- CD2- HD23
2671 2670 2668 2674 3 0.66944 2.00832 0.00000
-2.67776 0.00000 0.00000 ; HD11- CD1- CG- CD2
2672 2670 2668 2674 3 0.66944 2.00832 0.00000
-2.67776 0.00000 0.00000 ; HD12- CD1- CG- CD2
2673 2670 2668 2674 3 0.66944 2.00832 0.00000
-2.67776 0.00000 0.00000 ; HD13- CD1- CG- CD2
2678 2680 2682 2683 3 0.00000 0.00000 0.00000
0.00000 0.00000 0.00000 ; C- N- CA- HA2

```
  2678   2680   2682   2684      3    0.00000    0.00000    0.00000
0.00000    0.00000    0.00000 ;      C-     N-    CA-   HA3
  2678   2680   2682   2685      3    1.75728    5.27184    2.25936
-7.02912    0.00000    0.00000 ;      C-     N-    CA-    C
  2679   2678   2680   2681      3   29.28800   -8.36800  -20.92000
0.00000    0.00000    0.00000 ;      O-     C-     N-    H
  2679   2678   2680   2682      3   20.92000    0.00000  -20.92000
0.00000    0.00000    0.00000 ;      O-     C-     N-   CA
  2680   2682   2685   2686      3    0.00000    0.00000    0.00000
0.00000    0.00000    0.00000 ;      N-    CA-     C-    O
  2680   2682   2685   2687      3   17.40544   -5.02080  -13.22144
9.20480    0.00000    0.00000 ;      N-    CA-     C-    N
  2681   2680   2682   2683      3    0.00000    0.00000    0.00000
0.00000    0.00000    0.00000 ;      H-     N-    CA-   HA2
  2681   2680   2682   2684      3    0.00000    0.00000    0.00000
0.00000    0.00000    0.00000 ;      H-     N-    CA-   HA3
  2681   2680   2682   2685      3    0.00000    0.00000    0.00000
0.00000    0.00000    0.00000 ;      H-     N-    CA-    C
  2682   2685   2687   2688      3   20.92000    0.00000  -20.92000
0.00000    0.00000    0.00000 ;     CA-     C-     N-    H
  2682   2685   2687   2689      3   20.92000    0.00000  -20.92000
0.00000    0.00000    0.00000 ;     CA-     C-     N-   CA
  2683   2682   2685   2686      3    3.68192   -4.35136    0.00000
1.33888    0.00000    0.00000 ;    HA2-    CA-     C-    O
  2683   2682   2685   2687      3    0.00000    0.00000    0.00000
0.00000    0.00000    0.00000 ;    HA2-    CA-     C-    N
  2684   2682   2685   2686      3    3.68192   -4.35136    0.00000
1.33888    0.00000    0.00000 ;    HA3-    CA-     C-    O
  2684   2682   2685   2687      3    0.00000    0.00000    0.00000
0.00000    0.00000    0.00000 ;    HA3-    CA-     C-    N
  2685   2687   2689   2690      3    0.00000    0.00000    0.00000
0.00000    0.00000    0.00000 ;      C-     N-    CA-    HA
  2685   2687   2689   2691      3   11.71520    1.67360   15.06240  -
13.38880    0.00000    0.00000 ;      C-     N-    CA-    CB
  2685   2687   2689   2699      3    1.75728    5.27184    2.25936
-7.02912    0.00000    0.00000 ;      C-     N-    CA-    C
  2686   2685   2687   2688      3   29.28800   -8.36800  -20.92000
0.00000    0.00000    0.00000 ;      O-     C-     N-    H
  2686   2685   2687   2689      3   20.92000    0.00000  -20.92000
0.00000    0.00000    0.00000 ;      O-     C-     N-   CA
  2687   2689   2691   2692      3    0.65084    1.95253    0.00000
-2.60338    0.00000    0.00000 ;      N-    CA-    CB-    HB
  2687   2689   2691   2693      3    3.83673    0.44769   -1.77402
-2.47693   -0.03347    0.00000 ;      N-    CA-    CB-   CG2
  2687   2689   2691   2697      3    4.93294    1.13386   -3.12963
-5.27184    3.17984    0.00000 ;      N-    CA-    CB-   OG1
  2687   2689   2699   2700      3    0.00000    0.00000    0.00000
0.00000    0.00000    0.00000 ;      N-    CA-     C-    O
  2687   2689   2699   2701      3   17.40544   -5.02080  -13.22144
9.20480    0.00000    0.00000 ;      N-    CA-     C-    N
  2688   2687   2689   2690      3    0.00000    0.00000    0.00000
0.00000    0.00000    0.00000 ;      H-     N-    CA-    HA
```

2688 2687 2689 2691 3 0.00000 0.00000 0.00000
0.00000 0.00000 0.00000 ; H- N- CA- CB
2688 2687 2689 2699 3 0.00000 0.00000 0.00000
0.00000 0.00000 0.00000 ; H- N- CA- C
2689 2691 2693 2694 3 0.66944 2.00832 0.00000
-2.67776 0.00000 0.00000 ; CA- CB- CG2- HG21
2689 2691 2693 2695 3 0.66944 2.00832 0.00000
-2.67776 0.00000 0.00000 ; CA- CB- CG2- HG22
2689 2691 2693 2696 3 0.66944 2.00832 0.00000
-2.67776 0.00000 0.00000 ; CA- CB- CG2- HG23
2689 2691 2697 2698 3 1.12131 2.98738 1.66523
-3.94970 0.43514 0.00000 ; CA- CB- OG1- HG1
2689 2699 2701 2702 3 20.92000 0.00000 -20.92000
0.00000 0.00000 0.00000 ; CA- C- N- H
2689 2699 2701 2703 3 20.92000 0.00000 -20.92000
0.00000 0.00000 0.00000 ; CA- C- N- CA
2690 2689 2691 2692 3 0.65084 1.95253 0.00000
-2.60338 0.00000 0.00000 ; HA- CA- CB- HB
2690 2689 2691 2693 3 0.65084 1.95253 0.00000
-2.60338 0.00000 0.00000 ; HA- CA- CB- CG2
2690 2689 2691 2697 3 1.04600 -1.04600 0.00000
0.00000 0.00000 0.00000 ; HA- CA- CB- OG1
2690 2689 2699 2700 3 3.68192 -4.35136 0.00000
1.33888 0.00000 0.00000 ; HA- CA- C- O
2690 2689 2699 2701 3 0.00000 0.00000 0.00000
0.00000 0.00000 0.00000 ; HA- CA- C- N
2691 2689 2699 2700 3 0.00000 0.00000 0.00000
0.00000 0.00000 0.00000 ; CB- CA- C- O
2691 2689 2699 2701 3 2.51040 4.18400 1.67360
-6.69440 0.00000 0.00000 ; CB- CA- C- N
2692 2691 2689 2699 3 0.65084 1.95253 0.00000
-2.60338 0.00000 0.00000 ; HB- CB- CA- C
2692 2691 2693 2694 3 0.65084 1.95253 0.00000
-2.60338 0.00000 0.00000 ; HB- CB- CG2- HG21
2692 2691 2693 2695 3 0.65084 1.95253 0.00000
-2.60338 0.00000 0.00000 ; HB- CB- CG2- HG22
2692 2691 2693 2696 3 0.65084 1.95253 0.00000
-2.60338 0.00000 0.00000 ; HB- CB- CG2- HG23
2692 2691 2697 2698 3 0.69733 2.09200 0.00000
-2.78933 0.00000 0.00000 ; HB- CB- OG1- HG1
2693 2691 2689 2699 3 5.67350 3.55640 -6.16722
-2.47693 3.74886 0.00000 ; CG2- CB- CA- C
2693 2691 2697 2698 3 4.74047 0.27196 -2.26773
-3.94970 1.60666 0.00000 ; CG2- CB- OG1- HG1
2694 2693 2691 2697 3 1.04600 -1.04600 0.00000
0.00000 0.00000 0.00000 ; HG21- CG2- CB- OG1
2695 2693 2691 2697 3 1.04600 -1.04600 0.00000
0.00000 0.00000 0.00000 ; HG22- CG2- CB- OG1
2696 2693 2691 2697 3 1.04600 -1.04600 0.00000
0.00000 0.00000 0.00000 ; HG23- CG2- CB- OG1
2697 2691 2689 2699 3 6.53541 6.87013 -6.21742
-5.27184 5.22163 0.00000 ; OG1- CB- CA- C

2699 2701 2703 2704 3 0.00000 0.00000 0.00000
0.00000 0.00000 0.00000 ; C- N- CA- HA2
2699 2701 2703 2705 3 0.00000 0.00000 0.00000
0.00000 0.00000 0.00000 ; C- N- CA- HA3
2699 2701 2703 2706 3 1.75728 5.27184 2.25936
-7.02912 0.00000 0.00000 ; C- N- CA- C
2700 2699 2701 2702 3 29.28800 -8.36800 -20.92000
0.00000 0.00000 0.00000 ; O- C- N- H
2700 2699 2701 2703 3 20.92000 0.00000 -20.92000
0.00000 0.00000 0.00000 ; O- C- N- CA
2701 2703 2706 2707 3 0.00000 0.00000 0.00000
0.00000 0.00000 0.00000 ; N- CA- C- O
2701 2703 2706 2708 3 17.40544 -5.02080 -13.22144
9.20480 0.00000 0.00000 ; N- CA- C- N
2702 2701 2703 2704 3 0.00000 0.00000 0.00000
0.00000 0.00000 0.00000 ; H- N- CA- HA2
2702 2701 2703 2705 3 0.00000 0.00000 0.00000
0.00000 0.00000 0.00000 ; H- N- CA- HA3
2702 2701 2703 2706 3 0.00000 0.00000 0.00000
0.00000 0.00000 0.00000 ; H- N- CA- C
2703 2706 2708 2709 3 20.92000 0.00000 -20.92000
0.00000 0.00000 0.00000 ; CA- C- N- H
2703 2706 2708 2710 3 20.92000 0.00000 -20.92000
0.00000 0.00000 0.00000 ; CA- C- N- CA
2704 2703 2706 2707 3 3.68192 -4.35136 0.00000
1.33888 0.00000 0.00000 ; HA2- CA- C- O
2704 2703 2706 2708 3 0.00000 0.00000 0.00000
0.00000 0.00000 0.00000 ; HA2- CA- C- N
2705 2703 2706 2707 3 3.68192 -4.35136 0.00000
1.33888 0.00000 0.00000 ; HA3- CA- C- O
2705 2703 2706 2708 3 0.00000 0.00000 0.00000
0.00000 0.00000 0.00000 ; HA3- CA- C- N
2706 2708 2710 2711 3 0.00000 0.00000 0.00000
0.00000 0.00000 0.00000 ; C- N- CA- HA
2706 2708 2710 2712 3 11.71520 1.67360 15.06240 -
13.38880 0.00000 0.00000 ; C- N- CA- CB
2706 2708 2710 2721 3 1.75728 5.27184 2.25936
-7.02912 0.00000 0.00000 ; C- N- CA- C
2707 2706 2708 2709 3 29.28800 -8.36800 -20.92000
0.00000 0.00000 0.00000 ; O- C- N- H
2707 2706 2708 2710 3 20.92000 0.00000 -20.92000
0.00000 0.00000 0.00000 ; O- C- N- CA
2708 2710 2712 2713 3 0.65084 1.95253 0.00000
-2.60338 0.00000 0.00000 ; N- CA- CB- HB2
2708 2710 2712 2714 3 0.65084 1.95253 0.00000
-2.60338 0.00000 0.00000 ; N- CA- CB- HB3
2708 2710 2712 2715 3 3.21331 2.22589 -4.15053
-2.40998 2.61082 0.00000 ; N- CA- CB- CG
2708 2710 2721 2722 3 0.00000 0.00000 0.00000
0.00000 0.00000 0.00000 ; N- CA- C- O
2708 2710 2721 2723 3 17.40544 -5.02080 -13.22144
9.20480 0.00000 0.00000 ; N- CA- C- N

```
  2709   2708   2710   2711      3    0.00000    0.00000    0.00000
0.00000    0.00000    0.00000 ;      H-     N-    CA-    HA
  2709   2708   2710   2712      3    0.00000    0.00000    0.00000
0.00000    0.00000    0.00000 ;      H-     N-    CA-    CB
  2709   2708   2710   2721      3    0.00000    0.00000    0.00000
0.00000    0.00000    0.00000 ;      H-     N-    CA-     C
  2710   2712   2715   2716      3    0.66944    2.00832    0.00000
-2.67776    0.00000    0.00000 ;     CA-    CB-    CG-   HG2
  2710   2712   2715   2717      3    0.66944    2.00832    0.00000
-2.67776    0.00000    0.00000 ;     CA-    CB-    CG-   HG3
  2710   2712   2715   2718      3   10.12110   -1.91209    0.01674
10.17549   -1.87443    0.00000 ;     CA-    CB-    CG-    CD
  2710   2721   2723   2724      3   20.92000    0.00000  -20.92000
0.00000    0.00000    0.00000 ;     CA-     C-     N-     H
  2710   2721   2723   2725      3   20.92000    0.00000  -20.92000
0.00000    0.00000    0.00000 ;     CA-     C-     N-    CA
  2711   2710   2712   2713      3    0.65084    1.95253    0.00000
-2.60338    0.00000    0.00000 ;     HA-    CA-    CB-   HB2
  2711   2710   2712   2714      3    0.65084    1.95253    0.00000
-2.60338    0.00000    0.00000 ;     HA-    CA-    CB-   HB3
  2711   2710   2712   2715      3    0.65084    1.95253    0.00000
-2.60338    0.00000    0.00000 ;     HA-    CA-    CB-    CG
  2711   2710   2721   2722      3    3.68192   -4.35136    0.00000
1.33888    0.00000    0.00000 ;     HA-    CA-     C-     O
  2711   2710   2721   2723      3    0.00000    0.00000    0.00000
0.00000    0.00000    0.00000 ;     HA-    CA-     C-     N
  2712   2710   2721   2722      3    0.00000    0.00000    0.00000
0.00000    0.00000    0.00000 ;     CB-    CA-     C-     O
  2712   2710   2721   2723      3    2.51040    4.18400    1.67360
-6.69440    0.00000    0.00000 ;     CB-    CA-     C-     N
  2712   2715   2718   2719      3    3.79907    0.00000   -5.40573
0.00000    2.14221    0.00000 ;     CB-    CG-    CD-   OE1
  2712   2715   2718   2720      3    3.79907    0.00000   -5.40573
0.00000    2.14221    0.00000 ;     CB-    CG-    CD-   OE2
  2713   2712   2710   2721      3    0.65084    1.95253    0.00000
-2.60338    0.00000    0.00000 ;    HB2-    CB-    CA-     C
  2713   2712   2715   2716      3    0.62760    1.88280    0.00000
-2.51040    0.00000    0.00000 ;    HB2-    CB-    CG-   HG2
  2713   2712   2715   2717      3    0.62760    1.88280    0.00000
-2.51040    0.00000    0.00000 ;    HB2-    CB-    CG-   HG3
  2713   2712   2715   2718      3    0.65084    1.95253    0.00000
-2.60338    0.00000    0.00000 ;    HB2-    CB-    CG-    CD
  2714   2712   2710   2721      3    0.65084    1.95253    0.00000
-2.60338    0.00000    0.00000 ;    HB3-    CB-    CA-     C
  2714   2712   2715   2716      3    0.62760    1.88280    0.00000
-2.51040    0.00000    0.00000 ;    HB3-    CB-    CG-   HG2
  2714   2712   2715   2717      3    0.62760    1.88280    0.00000
-2.51040    0.00000    0.00000 ;    HB3-    CB-    CG-   HG3
  2714   2712   2715   2718      3    0.65084    1.95253    0.00000
-2.60338    0.00000    0.00000 ;    HB3-    CB-    CG-    CD
  2715   2712   2710   2721      3    6.86594    3.56895   -8.14206
-2.40998    4.85344    0.00000 ;     CG-    CB-    CA-     C
```

2716 2715 2718 2719 3 0.00000 0.00000 0.00000
0.00000 0.00000 0.00000 ; HG2- CG- CD- OE1
2716 2715 2718 2720 3 0.00000 0.00000 0.00000
0.00000 0.00000 0.00000 ; HG2- CG- CD- OE2
2717 2715 2718 2719 3 0.00000 0.00000 0.00000
0.00000 0.00000 0.00000 ; HG3- CG- CD- OE1
2717 2715 2718 2720 3 0.00000 0.00000 0.00000
0.00000 0.00000 0.00000 ; HG3- CG- CD- OE2
2721 2723 2725 2726 3 0.00000 0.00000 0.00000
0.00000 0.00000 0.00000 ; C- N- CA- HA
2721 2723 2725 2727 3 11.71520 1.67360 15.06240 -
13.38880 0.00000 0.00000 ; C- N- CA- CB
2721 2723 2725 2741 3 1.75728 5.27184 2.25936
-7.02912 0.00000 0.00000 ; C- N- CA- C
2722 2721 2723 2724 3 29.28800 -8.36800 -20.92000
0.00000 0.00000 0.00000 ; O- C- N- H
2722 2721 2723 2725 3 20.92000 0.00000 -20.92000
0.00000 0.00000 0.00000 ; O- C- N- CA
2723 2725 2727 2728 3 0.65084 1.95253 0.00000
-2.60338 0.00000 0.00000 ; N- CA- CB- HB2
2723 2725 2727 2729 3 0.65084 1.95253 0.00000
-2.60338 0.00000 0.00000 ; N- CA- CB- HB3
2723 2725 2727 2730 3 3.28026 2.46019 -2.19242
-3.21331 -0.23430 0.00000 ; N- CA- CB- CG
2723 2725 2741 2742 3 0.00000 0.00000 0.00000
0.00000 0.00000 0.00000 ; N- CA- C- O
2723 2725 2741 2743 3 17.40544 -5.02080 -13.22144
9.20480 0.00000 0.00000 ; N- CA- C- N
2724 2723 2725 2726 3 0.00000 0.00000 0.00000
0.00000 0.00000 0.00000 ; H- N- CA- HA
2724 2723 2725 2727 3 0.00000 0.00000 0.00000
0.00000 0.00000 0.00000 ; H- N- CA- CB
2724 2723 2725 2741 3 0.00000 0.00000 0.00000
0.00000 0.00000 0.00000 ; H- N- CA- C
2725 2727 2730 2731 3 0.57739 0.00000 1.02926
0.00000 -1.60666 0.00000 ; CA- CB- CG- CD1
2725 2727 2730 2739 3 0.57739 0.00000 1.02926
0.00000 -1.60666 0.00000 ; CA- CB- CG- CD2
2725 2741 2743 2744 3 20.92000 0.00000 -20.92000
0.00000 0.00000 0.00000 ; CA- C- N- H
2725 2741 2743 2745 3 20.92000 0.00000 -20.92000
0.00000 0.00000 0.00000 ; CA- C- N- CA
2726 2725 2727 2728 3 0.65084 1.95253 0.00000
-2.60338 0.00000 0.00000 ; HA- CA- CB- HB2
2726 2725 2727 2729 3 0.65084 1.95253 0.00000
-2.60338 0.00000 0.00000 ; HA- CA- CB- HB3
2726 2725 2727 2730 3 0.65084 1.95253 0.00000
-2.60338 0.00000 0.00000 ; HA- CA- CB- CG
2726 2725 2741 2742 3 3.68192 -4.35136 0.00000
1.33888 0.00000 0.00000 ; HA- CA- C- O
2726 2725 2741 2743 3 0.00000 0.00000 0.00000
0.00000 0.00000 0.00000 ; HA- CA- C- N

2727 2725 2741 2742 3 0.00000 0.00000 0.00000
0.00000 0.00000 0.00000 ; CB- CA- C- O
2727 2725 2741 2743 3 2.51040 4.18400 1.67360
-6.69440 0.00000 0.00000 ; CB- CA- C- N
2727 2730 2731 2732 3 30.33400 0.00000 -30.33400
0.00000 0.00000 0.00000 ; CB- CG- CD1- HD1
2727 2730 2731 2733 3 30.33400 0.00000 -30.33400
0.00000 0.00000 0.00000 ; CB- CG- CD1- CE1
2727 2730 2739 2737 3 30.33400 0.00000 -30.33400
0.00000 0.00000 0.00000 ; CB- CG- CD2- CE2
2727 2730 2739 2740 3 30.33400 0.00000 -30.33400
0.00000 0.00000 0.00000 ; CB- CG- CD2- HD2
2728 2727 2725 2741 3 0.65084 1.95253 0.00000
-2.60338 0.00000 0.00000 ; HB2- CB- CA- C
2728 2727 2730 2731 3 0.00000 0.00000 0.00000
0.00000 0.00000 0.00000 ; HB2- CB- CG- CD1
2728 2727 2730 2739 3 0.00000 0.00000 0.00000
0.00000 0.00000 0.00000 ; HB2- CB- CG- CD2
2729 2727 2725 2741 3 0.65084 1.95253 0.00000
-2.60338 0.00000 0.00000 ; HB3- CB- CA- C
2729 2727 2730 2731 3 0.00000 0.00000 0.00000
0.00000 0.00000 0.00000 ; HB3- CB- CG- CD1
2729 2727 2730 2739 3 0.00000 0.00000 0.00000
0.00000 0.00000 0.00000 ; HB3- CB- CG- CD2
2730 2727 2725 2741 3 4.95804 2.17986 -3.52293
-3.21331 -0.40166 0.00000 ; CG- CB- CA- C
2730 2731 2733 2734 3 30.33400 0.00000 -30.33400
0.00000 0.00000 0.00000 ; CG- CD1- CE1- HE1
2730 2731 2733 2735 3 30.33400 0.00000 -30.33400
0.00000 0.00000 0.00000 ; CG- CD1- CE1- CZ
2730 2739 2737 2735 3 30.33400 0.00000 -30.33400
0.00000 0.00000 0.00000 ; CG- CD2- CE2- CZ
2730 2739 2737 2738 3 30.33400 0.00000 -30.33400
0.00000 0.00000 0.00000 ; CG- CD2- CE2- HE2
2731 2730 2739 2737 3 30.33400 0.00000 -30.33400
0.00000 0.00000 0.00000 ; CD1- CG- CD2- CE2
2731 2730 2739 2740 3 30.33400 0.00000 -30.33400
0.00000 0.00000 0.00000 ; CD1- CG- CD2- HD2
2731 2733 2735 2736 3 30.33400 0.00000 -30.33400
0.00000 0.00000 0.00000 ; CD1- CE1- CZ- HZ
2731 2733 2735 2737 3 30.33400 0.00000 -30.33400
0.00000 0.00000 0.00000 ; CD1- CE1- CZ- CE2
2732 2731 2730 2739 3 30.33400 0.00000 -30.33400
0.00000 0.00000 0.00000 ; HD1- CD1- CG- CD2
2732 2731 2733 2734 3 30.33400 0.00000 -30.33400
0.00000 0.00000 0.00000 ; HD1- CD1- CE1- HE1
2732 2731 2733 2735 3 30.33400 0.00000 -30.33400
0.00000 0.00000 0.00000 ; HD1- CD1- CE1- CZ
2733 2731 2730 2739 3 30.33400 0.00000 -30.33400
0.00000 0.00000 0.00000 ; CE1- CD1- CG- CD2
2733 2735 2737 2738 3 30.33400 0.00000 -30.33400
0.00000 0.00000 0.00000 ; CE1- CZ- CE2- HE2

2733 2735 2737 2739 3 30.33400 0.00000 -30.33400
0.00000 0.00000 0.00000 ; CE1- CZ- CE2- CD2
2734 2733 2735 2736 3 30.33400 0.00000 -30.33400
0.00000 0.00000 0.00000 ; HE1- CE1- CZ- HZ
2734 2733 2735 2737 3 30.33400 0.00000 -30.33400
0.00000 0.00000 0.00000 ; HE1- CE1- CZ- CE2
2735 2737 2739 2740 3 30.33400 0.00000 -30.33400
0.00000 0.00000 0.00000 ; CZ- CE2- CD2- HD2
2736 2735 2737 2738 3 30.33400 0.00000 -30.33400
0.00000 0.00000 0.00000 ; HZ- CZ- CE2- HE2
2736 2735 2737 2739 3 30.33400 0.00000 -30.33400
0.00000 0.00000 0.00000 ; HZ- CZ- CE2- CD2
2738 2737 2739 2740 3 30.33400 0.00000 -30.33400
0.00000 0.00000 0.00000 ; HE2- CE2- CD2- HD2
2741 2743 2745 2746 3 0.00000 0.00000 0.00000
0.00000 0.00000 0.00000 ; C- N- CA- HA
2741 2743 2745 2747 3 11.71520 1.67360 15.06240 -
13.38880 0.00000 0.00000 ; C- N- CA- CB
2741 2743 2745 2755 3 1.75728 5.27184 2.25936
-7.02912 0.00000 0.00000 ; C- N- CA- C
2742 2741 2743 2744 3 29.28800 -8.36800 -20.92000
0.00000 0.00000 0.00000 ; O- C- N- H
2742 2741 2743 2745 3 20.92000 0.00000 -20.92000
0.00000 0.00000 0.00000 ; O- C- N- CA
2743 2745 2747 2748 3 0.65084 1.95253 0.00000
-2.60338 0.00000 0.00000 ; N- CA- CB- HB
2743 2745 2747 2749 3 3.83673 0.44769 -1.77402
-2.47693 -0.03347 0.00000 ; N- CA- CB- CG2
2743 2745 2747 2753 3 4.93294 1.13386 -3.12963
-5.27184 3.17984 0.00000 ; N- CA- CB- OG1
2743 2745 2755 2756 3 0.00000 0.00000 0.00000
0.00000 0.00000 0.00000 ; N- CA- C- O
2743 2745 2755 2757 3 17.40544 -5.02080 -13.22144
9.20480 0.00000 0.00000 ; N- CA- C- N
2744 2743 2745 2746 3 0.00000 0.00000 0.00000
0.00000 0.00000 0.00000 ; H- N- CA- HA
2744 2743 2745 2747 3 0.00000 0.00000 0.00000
0.00000 0.00000 0.00000 ; H- N- CA- CB
2744 2743 2745 2755 3 0.00000 0.00000 0.00000
0.00000 0.00000 0.00000 ; H- N- CA- C
2745 2747 2749 2750 3 0.66944 2.00832 0.00000
-2.67776 0.00000 0.00000 ; CA- CB- CG2- HG21
2745 2747 2749 2751 3 0.66944 2.00832 0.00000
-2.67776 0.00000 0.00000 ; CA- CB- CG2- HG22
2745 2747 2749 2752 3 0.66944 2.00832 0.00000
-2.67776 0.00000 0.00000 ; CA- CB- CG2- HG23
2745 2747 2753 2754 3 1.12131 2.98738 1.66523
-3.94970 0.43514 0.00000 ; CA- CB- OG1- HG1
2745 2755 2757 2758 3 20.92000 0.00000 -20.92000
0.00000 0.00000 0.00000 ; CA- C- N- H
2745 2755 2757 2759 3 20.92000 0.00000 -20.92000
0.00000 0.00000 0.00000 ; CA- C- N- CA

2746 2745 2747 2748 3 0.65084 1.95253 0.00000
-2.60338 0.00000 0.00000 ; HA- CA- CB- HB
2746 2745 2747 2749 3 0.65084 1.95253 0.00000
-2.60338 0.00000 0.00000 ; HA- CA- CB- CG2
2746 2745 2747 2753 3 1.04600 -1.04600 0.00000
0.00000 0.00000 0.00000 ; HA- CA- CB- OG1
2746 2745 2755 2756 3 3.68192 -4.35136 0.00000
1.33888 0.00000 0.00000 ; HA- CA- C- O
2746 2745 2755 2757 3 0.00000 0.00000 0.00000
0.00000 0.00000 0.00000 ; HA- CA- C- N
2747 2745 2755 2756 3 0.00000 0.00000 0.00000
0.00000 0.00000 0.00000 ; CB- CA- C- O
2747 2745 2755 2757 3 2.51040 4.18400 1.67360
-6.69440 0.00000 0.00000 ; CB- CA- C- N
2748 2747 2745 2755 3 0.65084 1.95253 0.00000
-2.60338 0.00000 0.00000 ; HB- CB- CA- C
2748 2747 2749 2750 3 0.65084 1.95253 0.00000
-2.60338 0.00000 0.00000 ; HB- CB- CG2- HG21
2748 2747 2749 2751 3 0.65084 1.95253 0.00000
-2.60338 0.00000 0.00000 ; HB- CB- CG2- HG22
2748 2747 2749 2752 3 0.65084 1.95253 0.00000
-2.60338 0.00000 0.00000 ; HB- CB- CG2- HG23
2748 2747 2753 2754 3 0.69733 2.09200 0.00000
-2.78933 0.00000 0.00000 ; HB- CB- OG1- HG1
2749 2747 2745 2755 3 5.67350 3.55640 -6.16722
-2.47693 3.74886 0.00000 ; CG2- CB- CA- C
2749 2747 2753 2754 3 4.74047 0.27196 -2.26773
-3.94970 1.60666 0.00000 ; CG2- CB- OG1- HG1
2750 2749 2747 2753 3 1.04600 -1.04600 0.00000
0.00000 0.00000 0.00000 ; HG21- CG2- CB- OG1
2751 2749 2747 2753 3 1.04600 -1.04600 0.00000
0.00000 0.00000 0.00000 ; HG22- CG2- CB- OG1
2752 2749 2747 2753 3 1.04600 -1.04600 0.00000
0.00000 0.00000 0.00000 ; HG23- CG2- CB- OG1
2753 2747 2745 2755 3 6.53541 6.87013 -6.21742
-5.27184 5.22163 0.00000 ; OG1- CB- CA- C
2755 2757 2759 2760 3 0.00000 0.00000 0.00000
0.00000 0.00000 0.00000 ; C- N- CA- HA
2755 2757 2759 2761 3 11.71520 1.67360 15.06240 -
13.38880 0.00000 0.00000 ; C- N- CA- CB
2755 2757 2759 2769 3 1.75728 5.27184 2.25936
-7.02912 0.00000 0.00000 ; C- N- CA- C
2756 2755 2757 2758 3 29.28800 -8.36800 -20.92000
0.00000 0.00000 0.00000 ; O- C- N- H
2756 2755 2757 2759 3 20.92000 0.00000 -20.92000
0.00000 0.00000 0.00000 ; O- C- N- CA
2757 2759 2761 2762 3 0.65084 1.95253 0.00000
-2.60338 0.00000 0.00000 ; N- CA- CB- HB
2757 2759 2761 2763 3 3.83673 0.44769 -1.77402
-2.47693 -0.03347 0.00000 ; N- CA- CB- CG2
2757 2759 2761 2767 3 4.93294 1.13386 -3.12963
-5.27184 3.17984 0.00000 ; N- CA- CB- OG1

2757 2759 2769 2770 3 0.00000 0.00000 0.00000
0.00000 0.00000 0.00000 ; N- CA- C- O
2757 2759 2769 2771 3 17.40544 -5.02080 -13.22144
9.20480 0.00000 0.00000 ; N- CA- C- N
2758 2757 2759 2760 3 0.00000 0.00000 0.00000
0.00000 0.00000 0.00000 ; H- N- CA- HA
2758 2757 2759 2761 3 0.00000 0.00000 0.00000
0.00000 0.00000 0.00000 ; H- N- CA- CB
2758 2757 2759 2769 3 0.00000 0.00000 0.00000
0.00000 0.00000 0.00000 ; H- N- CA- C
2759 2761 2763 2764 3 0.66944 2.00832 0.00000
-2.67776 0.00000 0.00000 ; CA- CB- CG2- HG21
2759 2761 2763 2765 3 0.66944 2.00832 0.00000
-2.67776 0.00000 0.00000 ; CA- CB- CG2- HG22
2759 2761 2763 2766 3 0.66944 2.00832 0.00000
-2.67776 0.00000 0.00000 ; CA- CB- CG2- HG23
2759 2761 2767 2768 3 1.12131 2.98738 1.66523
-3.94970 0.43514 0.00000 ; CA- CB- OG1- HG1
2759 2769 2771 2772 3 20.92000 0.00000 -20.92000
0.00000 0.00000 0.00000 ; CA- C- N- H
2759 2769 2771 2773 3 20.92000 0.00000 -20.92000
0.00000 0.00000 0.00000 ; CA- C- N- CA
2760 2759 2761 2762 3 0.65084 1.95253 0.00000
-2.60338 0.00000 0.00000 ; HA- CA- CB- HB
2760 2759 2761 2763 3 0.65084 1.95253 0.00000
-2.60338 0.00000 0.00000 ; HA- CA- CB- CG2
2760 2759 2761 2767 3 1.04600 -1.04600 0.00000
0.00000 0.00000 0.00000 ; HA- CA- CB- OG1
2760 2759 2769 2770 3 3.68192 -4.35136 0.00000
1.33888 0.00000 0.00000 ; HA- CA- C- O
2760 2759 2769 2771 3 0.00000 0.00000 0.00000
0.00000 0.00000 0.00000 ; HA- CA- C- N
2761 2759 2769 2770 3 0.00000 0.00000 0.00000
0.00000 0.00000 0.00000 ; CB- CA- C- O
2761 2759 2769 2771 3 2.51040 4.18400 1.67360
-6.69440 0.00000 0.00000 ; CB- CA- C- N
2762 2761 2759 2769 3 0.65084 1.95253 0.00000
-2.60338 0.00000 0.00000 ; HB- CB- CA- C
2762 2761 2763 2764 3 0.65084 1.95253 0.00000
-2.60338 0.00000 0.00000 ; HB- CB- CG2- HG21
2762 2761 2763 2765 3 0.65084 1.95253 0.00000
-2.60338 0.00000 0.00000 ; HB- CB- CG2- HG22
2762 2761 2763 2766 3 0.65084 1.95253 0.00000
-2.60338 0.00000 0.00000 ; HB- CB- CG2- HG23
2762 2761 2767 2768 3 0.69733 2.09200 0.00000
-2.78933 0.00000 0.00000 ; HB- CB- OG1- HG1
2763 2761 2759 2769 3 5.67350 3.55640 -6.16722
-2.47693 3.74886 0.00000 ; CG2- CB- CA- C
2763 2761 2767 2768 3 4.74047 0.27196 -2.26773
-3.94970 1.60666 0.00000 ; CG2- CB- OG1- HG1
2764 2763 2761 2767 3 1.04600 -1.04600 0.00000
0.00000 0.00000 0.00000 ; HG21- CG2- CB- OG1

2765 2763 2761 2767 3 1.04600 -1.04600 0.00000
0.00000 0.00000 0.00000 ; HG22- CG2- CB- OG1
2766 2763 2761 2767 3 1.04600 -1.04600 0.00000
0.00000 0.00000 0.00000 ; HG23- CG2- CB- OG1
2767 2761 2759 2769 3 6.53541 6.87013 -6.21742
-5.27184 5.22163 0.00000 ; OG1- CB- CA- C
2769 2771 2773 2774 3 0.00000 0.00000 0.00000
0.00000 0.00000 0.00000 ; C- N- CA- HA
2769 2771 2773 2775 3 11.71520 1.67360 15.06240 -
13.38880 0.00000 0.00000 ; C- N- CA- CB
2769 2771 2773 2783 3 1.75728 5.27184 2.25936
-7.02912 0.00000 0.00000 ; C- N- CA- C
2770 2769 2771 2772 3 29.28800 -8.36800 -20.92000
0.00000 0.00000 0.00000 ; O- C- N- H
2770 2769 2771 2773 3 20.92000 0.00000 -20.92000
0.00000 0.00000 0.00000 ; O- C- N- CA
2771 2773 2775 2776 3 0.65084 1.95253 0.00000
-2.60338 0.00000 0.00000 ; N- CA- CB- HB
2771 2773 2775 2777 3 3.83673 0.44769 -1.77402
-2.47693 -0.03347 0.00000 ; N- CA- CB- CG2
2771 2773 2775 2781 3 4.93294 1.13386 -3.12963
-5.27184 3.17984 0.00000 ; N- CA- CB- OG1
2771 2773 2783 2784 3 0.00000 0.00000 0.00000
0.00000 0.00000 0.00000 ; N- CA- C- O
2771 2773 2783 2785 3 17.40544 -5.02080 -13.22144
9.20480 0.00000 0.00000 ; N- CA- C- N
2772 2771 2773 2774 3 0.00000 0.00000 0.00000
0.00000 0.00000 0.00000 ; H- N- CA- HA
2772 2771 2773 2775 3 0.00000 0.00000 0.00000
0.00000 0.00000 0.00000 ; H- N- CA- CB
2772 2771 2773 2783 3 0.00000 0.00000 0.00000
0.00000 0.00000 0.00000 ; H- N- CA- C
2773 2775 2777 2778 3 0.66944 2.00832 0.00000
-2.67776 0.00000 0.00000 ; CA- CB- CG2- HG21
2773 2775 2777 2779 3 0.66944 2.00832 0.00000
-2.67776 0.00000 0.00000 ; CA- CB- CG2- HG22
2773 2775 2777 2780 3 0.66944 2.00832 0.00000
-2.67776 0.00000 0.00000 ; CA- CB- CG2- HG23
2773 2775 2781 2782 3 1.12131 2.98738 1.66523
-3.94970 0.43514 0.00000 ; CA- CB- OG1- HG1
2773 2783 2785 2786 3 20.92000 0.00000 -20.92000
0.00000 0.00000 0.00000 ; CA- C- N- H
2773 2783 2785 2787 3 20.92000 0.00000 -20.92000
0.00000 0.00000 0.00000 ; CA- C- N- CA
2774 2773 2775 2776 3 0.65084 1.95253 0.00000
-2.60338 0.00000 0.00000 ; HA- CA- CB- HB
2774 2773 2775 2777 3 0.65084 1.95253 0.00000
-2.60338 0.00000 0.00000 ; HA- CA- CB- CG2
2774 2773 2775 2781 3 1.04600 -1.04600 0.00000
0.00000 0.00000 0.00000 ; HA- CA- CB- OG1
2774 2773 2783 2784 3 3.68192 -4.35136 0.00000
1.33888 0.00000 0.00000 ; HA- CA- C- O

```
  2774   2773   2783   2785       3    0.00000    0.00000    0.00000
0.00000    0.00000    0.00000 ;      HA-    CA-     C-     N
  2775   2773   2783   2784       3    0.00000    0.00000    0.00000
0.00000    0.00000    0.00000 ;      CB-    CA-     C-     O
  2775   2773   2783   2785       3    2.51040    4.18400    1.67360
-6.69440    0.00000    0.00000 ;      CB-    CA-     C-     N
  2776   2775   2773   2783       3    0.65084    1.95253    0.00000
-2.60338    0.00000    0.00000 ;      HB-    CB-    CA-     C
  2776   2775   2777   2778       3    0.65084    1.95253    0.00000
-2.60338    0.00000    0.00000 ;      HB-    CB-   CG2-  HG21
  2776   2775   2777   2779       3    0.65084    1.95253    0.00000
-2.60338    0.00000    0.00000 ;      HB-    CB-   CG2-  HG22
  2776   2775   2777   2780       3    0.65084    1.95253    0.00000
-2.60338    0.00000    0.00000 ;      HB-    CB-   CG2-  HG23
  2776   2775   2781   2782       3    0.69733    2.09200    0.00000
-2.78933    0.00000    0.00000 ;      HB-    CB-   OG1-   HG1
  2777   2775   2773   2783       3    5.67350    3.55640   -6.16722
-2.47693    3.74886    0.00000 ;    CG2-    CB-    CA-     C
  2777   2775   2781   2782       3    4.74047    0.27196   -2.26773
-3.94970    1.60666    0.00000 ;    CG2-    CB-   OG1-   HG1
  2778   2777   2775   2781       3    1.04600   -1.04600    0.00000
0.00000    0.00000    0.00000 ;   HG21-   CG2-    CB-   OG1
  2779   2777   2775   2781       3    1.04600   -1.04600    0.00000
0.00000    0.00000    0.00000 ;   HG22-   CG2-    CB-   OG1
  2780   2777   2775   2781       3    1.04600   -1.04600    0.00000
0.00000    0.00000    0.00000 ;   HG23-   CG2-    CB-   OG1
  2781   2775   2773   2783       3    6.53541    6.87013   -6.21742
-5.27184    5.22163    0.00000 ;    OG1-    CB-    CA-     C
  2783   2785   2787   2788       3    0.00000    0.00000    0.00000
0.00000    0.00000    0.00000 ;      C-     N-    CA-    HA
  2783   2785   2787   2789       3   11.71520    1.67360   15.06240  -
13.38880    0.00000    0.00000 ;      C-     N-    CA-    CB
  2783   2785   2787   2805       3    1.75728    5.27184    2.25936
-7.02912    0.00000    0.00000 ;      C-     N-    CA-     C
  2784   2783   2785   2786       3   29.28800   -8.36800  -20.92000
0.00000    0.00000    0.00000 ;      O-     C-     N-     H
  2784   2783   2785   2787       3   20.92000    0.00000  -20.92000
0.00000    0.00000    0.00000 ;      O-     C-     N-    CA
  2785   2787   2789   2790       3    0.65084    1.95253    0.00000
-2.60338    0.00000    0.00000 ;      N-    CA-    CB-   HB2
  2785   2787   2789   2791       3    0.65084    1.95253    0.00000
-2.60338    0.00000    0.00000 ;      N-    CA-    CB-   HB3
  2785   2787   2789   2792       3    0.65084    1.95253    0.00000
-2.60338    0.00000    0.00000 ;      N-    CA-    CB-    CG
  2785   2787   2805   2806       3    0.00000    0.00000    0.00000
0.00000    0.00000    0.00000 ;      N-    CA-     C-     O
  2785   2787   2805   2807       3   17.40544   -5.02080  -13.22144
9.20480    0.00000    0.00000 ;      N-    CA-     C-     N
  2786   2785   2787   2788       3    0.00000    0.00000    0.00000
0.00000    0.00000    0.00000 ;      H-     N-    CA-    HA
  2786   2785   2787   2789       3    0.00000    0.00000    0.00000
0.00000    0.00000    0.00000 ;      H-     N-    CA-    CB
```

2786 2785 2787 2805 3 0.00000 0.00000 0.00000
0.00000 0.00000 0.00000 ; H- N- CA- C
2787 2789 2792 2793 3 0.66944 2.00832 0.00000
-2.67776 0.00000 0.00000 ; CA- CB- CG- HG2
2787 2789 2792 2794 3 0.66944 2.00832 0.00000
-2.67776 0.00000 0.00000 ; CA- CB- CG- HG3
2787 2789 2792 2795 3 3.68192 3.09616 -2.09200
-3.01248 0.00000 0.00000 ; CA- CB- CG- CD
2787 2805 2807 2808 3 20.92000 0.00000 -20.92000
0.00000 0.00000 0.00000 ; CA- C- N- H
2787 2805 2807 2809 3 20.92000 0.00000 -20.92000
0.00000 0.00000 0.00000 ; CA- C- N- CA
2788 2787 2789 2790 3 0.65084 1.95253 0.00000
-2.60338 0.00000 0.00000 ; HA- CA- CB- HB2
2788 2787 2789 2791 3 0.65084 1.95253 0.00000
-2.60338 0.00000 0.00000 ; HA- CA- CB- HB3
2788 2787 2789 2792 3 0.65084 1.95253 0.00000
-2.60338 0.00000 0.00000 ; HA- CA- CB- CG
2788 2787 2805 2806 3 3.68192 -4.35136 0.00000
1.33888 0.00000 0.00000 ; HA- CA- C- O
2788 2787 2805 2807 3 0.00000 0.00000 0.00000
0.00000 0.00000 0.00000 ; HA- CA- C- N
2789 2787 2805 2806 3 0.00000 0.00000 0.00000
0.00000 0.00000 0.00000 ; CB- CA- C- O
2789 2787 2805 2807 3 2.51040 4.18400 1.67360
-6.69440 0.00000 0.00000 ; CB- CA- C- N
2789 2792 2795 2796 3 0.66944 2.00832 0.00000
-2.67776 0.00000 0.00000 ; CB- CG- CD- HD2
2789 2792 2795 2797 3 0.66944 2.00832 0.00000
-2.67776 0.00000 0.00000 ; CB- CG- CD- HD3
2789 2792 2795 2798 3 3.68192 3.09616 -2.09200
-3.01248 0.00000 0.00000 ; CB- CG- CD- CE
2790 2789 2787 2805 3 0.65084 1.95253 0.00000
-2.60338 0.00000 0.00000 ; HB2- CB- CA- C
2790 2789 2792 2793 3 0.62760 1.88280 0.00000
-2.51040 0.00000 0.00000 ; HB2- CB- CG- HG2
2790 2789 2792 2794 3 0.62760 1.88280 0.00000
-2.51040 0.00000 0.00000 ; HB2- CB- CG- HG3
2790 2789 2792 2795 3 0.66944 2.00832 0.00000
-2.67776 0.00000 0.00000 ; HB2- CB- CG- CD
2791 2789 2787 2805 3 0.65084 1.95253 0.00000
-2.60338 0.00000 0.00000 ; HB3- CB- CA- C
2791 2789 2792 2793 3 0.62760 1.88280 0.00000
-2.51040 0.00000 0.00000 ; HB3- CB- CG- HG2
2791 2789 2792 2794 3 0.62760 1.88280 0.00000
-2.51040 0.00000 0.00000 ; HB3- CB- CG- HG3
2791 2789 2792 2795 3 0.66944 2.00832 0.00000
-2.67776 0.00000 0.00000 ; HB3- CB- CG- CD
2792 2789 2787 2805 3 0.65084 1.95253 0.00000
-2.60338 0.00000 0.00000 ; CG- CB- CA- C
2792 2795 2798 2799 3 0.65084 1.95253 0.00000
-2.60338 0.00000 0.00000 ; CG- CD- CE- HE2

2792 2795 2798 2800 3 0.65084 1.95253 0.00000
-2.60338 0.00000 0.00000 ; CG- CD- CE- HE3
2792 2795 2798 2801 3 0.65084 1.95253 0.00000
-2.60338 0.00000 0.00000 ; CG- CD- CE- NZ
2793 2792 2795 2796 3 0.62760 1.88280 0.00000
-2.51040 0.00000 0.00000 ; HG2- CG- CD- HD2
2793 2792 2795 2797 3 0.62760 1.88280 0.00000
-2.51040 0.00000 0.00000 ; HG2- CG- CD- HD3
2793 2792 2795 2798 3 0.66944 2.00832 0.00000
-2.67776 0.00000 0.00000 ; HG2- CG- CD- CE
2794 2792 2795 2796 3 0.62760 1.88280 0.00000
-2.51040 0.00000 0.00000 ; HG3- CG- CD- HD2
2794 2792 2795 2797 3 0.62760 1.88280 0.00000
-2.51040 0.00000 0.00000 ; HG3- CG- CD- HD3
2794 2792 2795 2798 3 0.66944 2.00832 0.00000
-2.67776 0.00000 0.00000 ; HG3- CG- CD- CE
2795 2798 2801 2802 3 0.65084 1.95253 0.00000
-2.60338 0.00000 0.00000 ; CD- CE- NZ- HZ1
2795 2798 2801 2803 3 0.65084 1.95253 0.00000
-2.60338 0.00000 0.00000 ; CD- CE- NZ- HZ2
2795 2798 2801 2804 3 0.65084 1.95253 0.00000
-2.60338 0.00000 0.00000 ; CD- CE- NZ- HZ3
2796 2795 2798 2799 3 0.65084 1.95253 0.00000
-2.60338 0.00000 0.00000 ; HD2- CD- CE- HE2
2796 2795 2798 2800 3 0.65084 1.95253 0.00000
-2.60338 0.00000 0.00000 ; HD2- CD- CE- HE3
2796 2795 2798 2801 3 0.65084 1.95253 0.00000
-2.60338 0.00000 0.00000 ; HD2- CD- CE- NZ
2797 2795 2798 2799 3 0.65084 1.95253 0.00000
-2.60338 0.00000 0.00000 ; HD3- CD- CE- HE2
2797 2795 2798 2800 3 0.65084 1.95253 0.00000
-2.60338 0.00000 0.00000 ; HD3- CD- CE- HE3
2797 2795 2798 2801 3 0.65084 1.95253 0.00000
-2.60338 0.00000 0.00000 ; HD3- CD- CE- NZ
2799 2798 2801 2802 3 0.65084 1.95253 0.00000
-2.60338 0.00000 0.00000 ; HE2- CE- NZ- HZ1
2799 2798 2801 2803 3 0.65084 1.95253 0.00000
-2.60338 0.00000 0.00000 ; HE2- CE- NZ- HZ2
2799 2798 2801 2804 3 0.65084 1.95253 0.00000
-2.60338 0.00000 0.00000 ; HE2- CE- NZ- HZ3
2800 2798 2801 2802 3 0.65084 1.95253 0.00000
-2.60338 0.00000 0.00000 ; HE3- CE- NZ- HZ1
2800 2798 2801 2803 3 0.65084 1.95253 0.00000
-2.60338 0.00000 0.00000 ; HE3- CE- NZ- HZ2
2800 2798 2801 2804 3 0.65084 1.95253 0.00000
-2.60338 0.00000 0.00000 ; HE3- CE- NZ- HZ3
2805 2807 2809 2810 3 0.00000 0.00000 0.00000
0.00000 0.00000 0.00000 ; C- N- CA- HA
2805 2807 2809 2811 3 11.71520 1.67360 15.06240 -
13.38880 0.00000 0.00000 ; C- N- CA- CB
2805 2807 2809 2815 3 1.75728 5.27184 2.25936
-7.02912 0.00000 0.00000 ; C- N- CA- C

2806 2805 2807 2808 3 29.28800 -8.36800 -20.92000
0.00000 0.00000 0.00000 ; O- C- N- H
2806 2805 2807 2809 3 20.92000 0.00000 -20.92000
0.00000 0.00000 0.00000 ; O- C- N- CA
2807 2809 2811 2812 3 0.65084 1.95253 0.00000
-2.60338 0.00000 0.00000 ; N- CA- CB- HB1
2807 2809 2811 2813 3 0.65084 1.95253 0.00000
-2.60338 0.00000 0.00000 ; N- CA- CB- HB2
2807 2809 2811 2814 3 0.65084 1.95253 0.00000
-2.60338 0.00000 0.00000 ; N- CA- CB- HB3
2807 2809 2815 2816 3 0.00000 0.00000 0.00000
0.00000 0.00000 0.00000 ; N- CA- C- O
2807 2809 2815 2817 3 17.40544 -5.02080 -13.22144
9.20480 0.00000 0.00000 ; N- CA- C- N
2808 2807 2809 2810 3 0.00000 0.00000 0.00000
0.00000 0.00000 0.00000 ; H- N- CA- HA
2808 2807 2809 2811 3 0.00000 0.00000 0.00000
0.00000 0.00000 0.00000 ; H- N- CA- CB
2808 2807 2809 2815 3 0.00000 0.00000 0.00000
0.00000 0.00000 0.00000 ; H- N- CA- C
2809 2815 2817 2818 3 20.92000 0.00000 -20.92000
0.00000 0.00000 0.00000 ; CA- C- N- H
2809 2815 2817 2819 3 20.92000 0.00000 -20.92000
0.00000 0.00000 0.00000 ; CA- C- N- CA
2810 2809 2811 2812 3 0.65084 1.95253 0.00000
-2.60338 0.00000 0.00000 ; HA- CA- CB- HB1
2810 2809 2811 2813 3 0.65084 1.95253 0.00000
-2.60338 0.00000 0.00000 ; HA- CA- CB- HB2
2810 2809 2811 2814 3 0.65084 1.95253 0.00000
-2.60338 0.00000 0.00000 ; HA- CA- CB- HB3
2810 2809 2815 2816 3 3.68192 -4.35136 0.00000
1.33888 0.00000 0.00000 ; HA- CA- C- O
2810 2809 2815 2817 3 0.00000 0.00000 0.00000
0.00000 0.00000 0.00000 ; HA- CA- C- N
2811 2809 2815 2816 3 0.00000 0.00000 0.00000
0.00000 0.00000 0.00000 ; CB- CA- C- O
2811 2809 2815 2817 3 2.51040 4.18400 1.67360
-6.69440 0.00000 0.00000 ; CB- CA- C- N
2812 2811 2809 2815 3 0.65084 1.95253 0.00000
-2.60338 0.00000 0.00000 ; HB1- CB- CA- C
2813 2811 2809 2815 3 0.65084 1.95253 0.00000
-2.60338 0.00000 0.00000 ; HB2- CB- CA- C
2814 2811 2809 2815 3 0.65084 1.95253 0.00000
-2.60338 0.00000 0.00000 ; HB3- CB- CA- C
2815 2817 2819 2820 3 0.00000 0.00000 0.00000
0.00000 0.00000 0.00000 ; C- N- CA- HA2
2815 2817 2819 2821 3 0.00000 0.00000 0.00000
0.00000 0.00000 0.00000 ; C- N- CA- HA3
2815 2817 2819 2822 3 1.75728 5.27184 2.25936
-7.02912 0.00000 0.00000 ; C- N- CA- C
2816 2815 2817 2818 3 29.28800 -8.36800 -20.92000
0.00000 0.00000 0.00000 ; O- C- N- H

```
  2816   2815   2817   2819      3   20.92000    0.00000  -20.92000
0.00000    0.00000    0.00000 ;      O-     C-     N-    CA
  2817   2819   2822   2823      3    0.00000    0.00000    0.00000
0.00000    0.00000    0.00000 ;      N-    CA-     C-     O
  2817   2819   2822   2824      3   17.40544   -5.02080  -13.22144
9.20480    0.00000    0.00000 ;      N-    CA-     C-     N
  2818   2817   2819   2820      3    0.00000    0.00000    0.00000
0.00000    0.00000    0.00000 ;      H-     N-    CA-   HA2
  2818   2817   2819   2821      3    0.00000    0.00000    0.00000
0.00000    0.00000    0.00000 ;      H-     N-    CA-   HA3
  2818   2817   2819   2822      3    0.00000    0.00000    0.00000
0.00000    0.00000    0.00000 ;      H-     N-    CA-     C
  2819   2822   2824   2825      3   20.92000    0.00000  -20.92000
0.00000    0.00000    0.00000 ;     CA-     C-     N-     H
  2819   2822   2824   2826      3   20.92000    0.00000  -20.92000
0.00000    0.00000    0.00000 ;     CA-     C-     N-    CA
  2820   2819   2822   2823      3    3.68192   -4.35136    0.00000
1.33888    0.00000    0.00000 ;    HA2-    CA-     C-     O
  2820   2819   2822   2824      3    0.00000    0.00000    0.00000
0.00000    0.00000    0.00000 ;    HA2-    CA-     C-     N
  2821   2819   2822   2823      3    3.68192   -4.35136    0.00000
1.33888    0.00000    0.00000 ;    HA3-    CA-     C-     O
  2821   2819   2822   2824      3    0.00000    0.00000    0.00000
0.00000    0.00000    0.00000 ;    HA3-    CA-     C-     N
  2822   2824   2826   2827      3    0.00000    0.00000    0.00000
0.00000    0.00000    0.00000 ;      C-     N-    CA-    HA
  2822   2824   2826   2828      3   11.71520    1.67360   15.06240  -
13.38880    0.00000    0.00000 ;      C-     N-    CA-    CB
  2822   2824   2826   2836      3    1.75728    5.27184    2.25936
-7.02912    0.00000    0.00000 ;      C-     N-    CA-     C
  2823   2822   2824   2825      3   29.28800   -8.36800  -20.92000
0.00000    0.00000    0.00000 ;      O-     C-     N-     H
  2823   2822   2824   2826      3   20.92000    0.00000  -20.92000
0.00000    0.00000    0.00000 ;      O-     C-     N-    CA
  2824   2826   2828   2829      3    0.65084    1.95253    0.00000
-2.60338    0.00000    0.00000 ;      N-    CA-    CB-    HB
  2824   2826   2828   2830      3    3.83673    0.44769   -1.77402
-2.47693   -0.03347    0.00000 ;      N-    CA-    CB-   CG2
  2824   2826   2828   2834      3    4.93294    1.13386   -3.12963
-5.27184    3.17984    0.00000 ;      N-    CA-    CB-   OG1
  2824   2826   2836   2837      3    0.00000    0.00000    0.00000
0.00000    0.00000    0.00000 ;      N-    CA-     C-     O
  2824   2826   2836   2838      3   17.40544   -5.02080  -13.22144
9.20480    0.00000    0.00000 ;      N-    CA-     C-     N
  2825   2824   2826   2827      3    0.00000    0.00000    0.00000
0.00000    0.00000    0.00000 ;      H-     N-    CA-    HA
  2825   2824   2826   2828      3    0.00000    0.00000    0.00000
0.00000    0.00000    0.00000 ;      H-     N-    CA-    CB
  2825   2824   2826   2836      3    0.00000    0.00000    0.00000
0.00000    0.00000    0.00000 ;      H-     N-    CA-     C
  2826   2828   2830   2831      3    0.66944    2.00832    0.00000
-2.67776    0.00000    0.00000 ;     CA-    CB-   CG2-  HG21
```

2826 2828 2830 2832 3 0.66944 2.00832 0.00000
-2.67776 0.00000 0.00000 ; CA- CB- CG2- HG22
2826 2828 2830 2833 3 0.66944 2.00832 0.00000
-2.67776 0.00000 0.00000 ; CA- CB- CG2- HG23
2826 2828 2834 2835 3 1.12131 2.98738 1.66523
-3.94970 0.43514 0.00000 ; CA- CB- OG1- HG1
2826 2836 2838 2839 3 20.92000 0.00000 -20.92000
0.00000 0.00000 0.00000 ; CA- C- N- CD
2826 2836 2838 2848 3 20.92000 0.00000 -20.92000
0.00000 0.00000 0.00000 ; CA- C- N- CA
2827 2826 2828 2829 3 0.65084 1.95253 0.00000
-2.60338 0.00000 0.00000 ; HA- CA- CB- HB
2827 2826 2828 2830 3 0.65084 1.95253 0.00000
-2.60338 0.00000 0.00000 ; HA- CA- CB- CG2
2827 2826 2828 2834 3 1.04600 -1.04600 0.00000
0.00000 0.00000 0.00000 ; HA- CA- CB- OG1
2827 2826 2836 2837 3 3.68192 -4.35136 0.00000
1.33888 0.00000 0.00000 ; HA- CA- C- O
2827 2826 2836 2838 3 0.00000 0.00000 0.00000
0.00000 0.00000 0.00000 ; HA- CA- C- N
2828 2826 2836 2837 3 0.00000 0.00000 0.00000
0.00000 0.00000 0.00000 ; CB- CA- C- O
2828 2826 2836 2838 3 2.51040 4.18400 1.67360
-6.69440 0.00000 0.00000 ; CB- CA- C- N
2829 2828 2826 2836 3 0.65084 1.95253 0.00000
-2.60338 0.00000 0.00000 ; HB- CB- CA- C
2829 2828 2830 2831 3 0.65084 1.95253 0.00000
-2.60338 0.00000 0.00000 ; HB- CB- CG2- HG21
2829 2828 2830 2832 3 0.65084 1.95253 0.00000
-2.60338 0.00000 0.00000 ; HB- CB- CG2- HG22
2829 2828 2830 2833 3 0.65084 1.95253 0.00000
-2.60338 0.00000 0.00000 ; HB- CB- CG2- HG23
2829 2828 2834 2835 3 0.69733 2.09200 0.00000
-2.78933 0.00000 0.00000 ; HB- CB- OG1- HG1
2830 2828 2826 2836 3 5.67350 3.55640 -6.16722
-2.47693 3.74886 0.00000 ; CG2- CB- CA- C
2830 2828 2834 2835 3 4.74047 0.27196 -2.26773
-3.94970 1.60666 0.00000 ; CG2- CB- OG1- HG1
2831 2830 2828 2834 3 1.04600 -1.04600 0.00000
0.00000 0.00000 0.00000 ; HG21- CG2- CB- OG1
2832 2830 2828 2834 3 1.04600 -1.04600 0.00000
0.00000 0.00000 0.00000 ; HG22- CG2- CB- OG1
2833 2830 2828 2834 3 1.04600 -1.04600 0.00000
0.00000 0.00000 0.00000 ; HG23- CG2- CB- OG1
2834 2828 2826 2836 3 6.53541 6.87013 -6.21742
-5.27184 5.22163 0.00000 ; OG1- CB- CA- C
2836 2838 2839 2840 3 0.00000 0.00000 0.00000
0.00000 0.00000 0.00000 ; C- N- CD- HD2
2836 2838 2839 2841 3 0.00000 0.00000 0.00000
0.00000 0.00000 0.00000 ; C- N- CD- HD3
2836 2838 2839 2842 3 10.04160 -3.34720 16.73600
-6.69440 0.00000 0.00000 ; C- N- CD- CG

2836 2838 2848 2845 3 11.71520 1.67360 15.06240 -
13.38880 0.00000 0.00000 ; C- N- CA- CB
2836 2838 2848 2849 3 0.00000 0.00000 0.00000
0.00000 0.00000 0.00000 ; C- N- CA- HA
2836 2838 2848 2850 3 1.75728 5.27184 2.25936
-7.02912 0.00000 0.00000 ; C- N- CA- C
2837 2836 2838 2839 3 20.92000 0.00000 -20.92000
0.00000 0.00000 0.00000 ; O- C- N- CD
2837 2836 2838 2848 3 20.92000 0.00000 -20.92000
0.00000 0.00000 0.00000 ; O- C- N- CA
2838 2839 2842 2843 3 0.65084 1.95253 0.00000
-2.60338 0.00000 0.00000 ; N- CD- CG- HG2
2838 2839 2842 2844 3 0.65084 1.95253 0.00000
-2.60338 0.00000 0.00000 ; N- CD- CG- HG3
2838 2839 2842 2845 3 0.65084 1.95253 0.00000
-2.60338 0.00000 0.00000 ; N- CD- CG- CB
2838 2848 2845 2842 3 0.65084 1.95253 0.00000
-2.60338 0.00000 0.00000 ; N- CA- CB- CG
2838 2848 2845 2846 3 0.65084 1.95253 0.00000
-2.60338 0.00000 0.00000 ; N- CA- CB- HB2
2838 2848 2845 2847 3 0.65084 1.95253 0.00000
-2.60338 0.00000 0.00000 ; N- CA- CB- HB3
2838 2848 2850 2851 3 0.00000 0.00000 0.00000
0.00000 0.00000 0.00000 ; N- CA- C- O
2838 2848 2850 2852 3 17.40544 -5.02080 -13.22144
9.20480 0.00000 0.00000 ; N- CA- C- N
2839 2838 2848 2845 3 0.00000 0.00000 0.00000
0.00000 0.00000 0.00000 ; CD- N- CA- CB
2839 2838 2848 2849 3 0.00000 0.00000 0.00000
0.00000 0.00000 0.00000 ; CD- N- CA- HA
2839 2838 2848 2850 3 0.00000 0.00000 0.00000
0.00000 0.00000 0.00000 ; CD- N- CA- C
2839 2842 2845 2846 3 0.66944 2.00832 0.00000
-2.67776 0.00000 0.00000 ; CD- CG- CB- HB2
2839 2842 2845 2847 3 0.66944 2.00832 0.00000
-2.67776 0.00000 0.00000 ; CD- CG- CB- HB3
2839 2842 2845 2848 3 3.68192 3.09616 -2.09200
-3.01248 0.00000 0.00000 ; CD- CG- CB- CA
2840 2839 2838 2848 3 0.00000 0.00000 0.00000
0.00000 0.00000 0.00000 ; HD2- CD- N- CA
2840 2839 2842 2843 3 0.65084 1.95253 0.00000
-2.60338 0.00000 0.00000 ; HD2- CD- CG- HG2
2840 2839 2842 2844 3 0.65084 1.95253 0.00000
-2.60338 0.00000 0.00000 ; HD2- CD- CG- HG3
2840 2839 2842 2845 3 0.65084 1.95253 0.00000
-2.60338 0.00000 0.00000 ; HD2- CD- CG- CB
2841 2839 2838 2848 3 0.00000 0.00000 0.00000
0.00000 0.00000 0.00000 ; HD3- CD- N- CA
2841 2839 2842 2843 3 0.65084 1.95253 0.00000
-2.60338 0.00000 0.00000 ; HD3- CD- CG- HG2
2841 2839 2842 2844 3 0.65084 1.95253 0.00000
-2.60338 0.00000 0.00000 ; HD3- CD- CG- HG3

```
  2841   2839   2842   2845      3    0.65084    1.95253    0.00000
-2.60338    0.00000    0.00000 ;    HD3-    CD-    CG-    CB
  2842   2839   2838   2848      3    0.00000    0.00000    0.00000
0.00000    0.00000    0.00000 ;    CG-    CD-     N-    CA
  2842   2845   2848   2849      3    0.65084    1.95253    0.00000
-2.60338    0.00000    0.00000 ;    CG-    CB-    CA-    HA
  2842   2845   2848   2850      3    0.65084    1.95253    0.00000
-2.60338    0.00000    0.00000 ;    CG-    CB-    CA-     C
  2843   2842   2845   2846      3    0.62760    1.88280    0.00000
-2.51040    0.00000    0.00000 ;    HG2-    CG-    CB-   HB2
  2843   2842   2845   2847      3    0.62760    1.88280    0.00000
-2.51040    0.00000    0.00000 ;    HG2-    CG-    CB-   HB3
  2843   2842   2845   2848      3    0.66944    2.00832    0.00000
-2.67776    0.00000    0.00000 ;    HG2-    CG-    CB-    CA
  2844   2842   2845   2846      3    0.62760    1.88280    0.00000
-2.51040    0.00000    0.00000 ;    HG3-    CG-    CB-   HB2
  2844   2842   2845   2847      3    0.62760    1.88280    0.00000
-2.51040    0.00000    0.00000 ;    HG3-    CG-    CB-   HB3
  2844   2842   2845   2848      3    0.66944    2.00832    0.00000
-2.67776    0.00000    0.00000 ;    HG3-    CG-    CB-    CA
  2845   2848   2850   2851      3    0.00000    0.00000    0.00000
0.00000    0.00000    0.00000 ;    CB-    CA-     C-     O
  2845   2848   2850   2852      3    2.51040    4.18400    1.67360
-6.69440    0.00000    0.00000 ;    CB-    CA-     C-     N
  2846   2845   2848   2849      3    0.65084    1.95253    0.00000
-2.60338    0.00000    0.00000 ;    HB2-    CB-    CA-    HA
  2846   2845   2848   2850      3    0.65084    1.95253    0.00000
-2.60338    0.00000    0.00000 ;    HB2-    CB-    CA-     C
  2847   2845   2848   2849      3    0.65084    1.95253    0.00000
-2.60338    0.00000    0.00000 ;    HB3-    CB-    CA-    HA
  2847   2845   2848   2850      3    0.65084    1.95253    0.00000
-2.60338    0.00000    0.00000 ;    HB3-    CB-    CA-     C
  2848   2850   2852   2853      3   20.92000    0.00000  -20.92000
0.00000    0.00000    0.00000 ;    CA-     C-     N-     H
  2848   2850   2852   2854      3   20.92000    0.00000  -20.92000
0.00000    0.00000    0.00000 ;    CA-     C-     N-    CA
  2849   2848   2850   2851      3    3.68192   -4.35136    0.00000
1.33888    0.00000    0.00000 ;    HA-    CA-     C-     O
  2849   2848   2850   2852      3    0.00000    0.00000    0.00000
0.00000    0.00000    0.00000 ;    HA-    CA-     C-     N
  2850   2852   2854   2855      3    0.00000    0.00000    0.00000
0.00000    0.00000    0.00000 ;     C-     N-    CA-    HA
  2850   2852   2854   2856      3   11.71520    1.67360   15.06240  -
13.38880    0.00000    0.00000 ;     C-     N-    CA-    CB
  2850   2852   2854   2871      3    1.75728    5.27184    2.25936
-7.02912    0.00000    0.00000 ;     C-     N-    CA-     C
  2851   2850   2852   2853      3   29.28800   -8.36800  -20.92000
0.00000    0.00000    0.00000 ;     O-     C-     N-     H
  2851   2850   2852   2854      3   20.92000    0.00000  -20.92000
0.00000    0.00000    0.00000 ;     O-     C-     N-    CA
  2852   2854   2856   2857      3    0.65084    1.95253    0.00000
-2.60338    0.00000    0.00000 ;     N-    CA-    CB-   HB2
```

2852 2854 2856 2858 3 0.65084 1.95253 0.00000
-2.60338 0.00000 0.00000 ; N- CA- CB- HB3
2852 2854 2856 2859 3 3.28026 2.46019 -2.19242
-3.21331 -0.23430 0.00000 ; N- CA- CB- CG
2852 2854 2871 2872 3 0.00000 0.00000 0.00000
0.00000 0.00000 0.00000 ; N- CA- C- O
2852 2854 2871 2873 3 17.40544 -5.02080 -13.22144
9.20480 0.00000 0.00000 ; N- CA- C- N
2853 2852 2854 2855 3 0.00000 0.00000 0.00000
0.00000 0.00000 0.00000 ; H- N- CA- HA
2853 2852 2854 2856 3 0.00000 0.00000 0.00000
0.00000 0.00000 0.00000 ; H- N- CA- CB
2853 2852 2854 2871 3 0.00000 0.00000 0.00000
0.00000 0.00000 0.00000 ; H- N- CA- C
2854 2856 2859 2860 3 0.57739 0.00000 1.02926
0.00000 -1.60666 0.00000 ; CA- CB- CG- CD1
2854 2856 2859 2869 3 0.57739 0.00000 1.02926
0.00000 -1.60666 0.00000 ; CA- CB- CG- CD2
2854 2871 2873 2874 3 20.92000 0.00000 -20.92000
0.00000 0.00000 0.00000 ; CA- C- N- H
2854 2871 2873 2875 3 20.92000 0.00000 -20.92000
0.00000 0.00000 0.00000 ; CA- C- N- CA
2855 2854 2856 2857 3 0.65084 1.95253 0.00000
-2.60338 0.00000 0.00000 ; HA- CA- CB- HB2
2855 2854 2856 2858 3 0.65084 1.95253 0.00000
-2.60338 0.00000 0.00000 ; HA- CA- CB- HB3
2855 2854 2856 2859 3 0.65084 1.95253 0.00000
-2.60338 0.00000 0.00000 ; HA- CA- CB- CG
2855 2854 2871 2872 3 3.68192 -4.35136 0.00000
1.33888 0.00000 0.00000 ; HA- CA- C- O
2855 2854 2871 2873 3 0.00000 0.00000 0.00000
0.00000 0.00000 0.00000 ; HA- CA- C- N
2856 2854 2871 2872 3 0.00000 0.00000 0.00000
0.00000 0.00000 0.00000 ; CB- CA- C- O
2856 2854 2871 2873 3 2.51040 4.18400 1.67360
-6.69440 0.00000 0.00000 ; CB- CA- C- N
2856 2859 2860 2861 3 30.33400 0.00000 -30.33400
0.00000 0.00000 0.00000 ; CB- CG- CD1- HD1
2856 2859 2860 2862 3 30.33400 0.00000 -30.33400
0.00000 0.00000 0.00000 ; CB- CG- CD1- CE1
2856 2859 2869 2867 3 30.33400 0.00000 -30.33400
0.00000 0.00000 0.00000 ; CB- CG- CD2- CE2
2856 2859 2869 2870 3 30.33400 0.00000 -30.33400
0.00000 0.00000 0.00000 ; CB- CG- CD2- HD2
2857 2856 2854 2871 3 0.65084 1.95253 0.00000
-2.60338 0.00000 0.00000 ; HB2- CB- CA- C
2857 2856 2859 2860 3 0.00000 0.00000 0.00000
0.00000 0.00000 0.00000 ; HB2- CB- CG- CD1
2857 2856 2859 2869 3 0.00000 0.00000 0.00000
0.00000 0.00000 0.00000 ; HB2- CB- CG- CD2
2858 2856 2854 2871 3 0.65084 1.95253 0.00000
-2.60338 0.00000 0.00000 ; HB3- CB- CA- C

2858 2856 2859 2860 3 0.00000 0.00000 0.00000
0.00000 0.00000 0.00000 ; HB3- CB- CG- CD1
2858 2856 2859 2869 3 0.00000 0.00000 0.00000
0.00000 0.00000 0.00000 ; HB3- CB- CG- CD2
2859 2856 2854 2871 3 4.95804 2.17986 -3.52293
-3.21331 -0.40166 0.00000 ; CG- CB- CA- C
2859 2860 2862 2863 3 30.33400 0.00000 -30.33400
0.00000 0.00000 0.00000 ; CG- CD1- CE1- HE1
2859 2860 2862 2864 3 30.33400 0.00000 -30.33400
0.00000 0.00000 0.00000 ; CG- CD1- CE1- CZ
2859 2869 2867 2864 3 30.33400 0.00000 -30.33400
0.00000 0.00000 0.00000 ; CG- CD2- CE2- CZ
2859 2869 2867 2868 3 30.33400 0.00000 -30.33400
0.00000 0.00000 0.00000 ; CG- CD2- CE2- HE2
2860 2859 2869 2867 3 30.33400 0.00000 -30.33400
0.00000 0.00000 0.00000 ; CD1- CG- CD2- CE2
2860 2859 2869 2870 3 30.33400 0.00000 -30.33400
0.00000 0.00000 0.00000 ; CD1- CG- CD2- HD2
2860 2862 2864 2865 3 30.33400 0.00000 -30.33400
0.00000 0.00000 0.00000 ; CD1- CE1- CZ- OH
2860 2862 2864 2867 3 30.33400 0.00000 -30.33400
0.00000 0.00000 0.00000 ; CD1- CE1- CZ- CE2
2861 2860 2859 2869 3 30.33400 0.00000 -30.33400
0.00000 0.00000 0.00000 ; HD1- CD1- CG- CD2
2861 2860 2862 2863 3 30.33400 0.00000 -30.33400
0.00000 0.00000 0.00000 ; HD1- CD1- CE1- HE1
2861 2860 2862 2864 3 30.33400 0.00000 -30.33400
0.00000 0.00000 0.00000 ; HD1- CD1- CE1- CZ
2862 2860 2859 2869 3 30.33400 0.00000 -30.33400
0.00000 0.00000 0.00000 ; CE1- CD1- CG- CD2
2862 2864 2865 2866 3 7.93286 0.00000 -9.56462
0.00000 2.17568 0.00000 ; CE1- CZ- OH- HH
2862 2864 2867 2868 3 30.33400 0.00000 -30.33400
0.00000 0.00000 0.00000 ; CE1- CZ- CE2- HE2
2862 2864 2867 2869 3 30.33400 0.00000 -30.33400
0.00000 0.00000 0.00000 ; CE1- CZ- CE2- CD2
2863 2862 2864 2865 3 30.33400 0.00000 -30.33400
0.00000 0.00000 0.00000 ; HE1- CE1- CZ- OH
2863 2862 2864 2867 3 30.33400 0.00000 -30.33400
0.00000 0.00000 0.00000 ; HE1- CE1- CZ- CE2
2864 2867 2869 2870 3 30.33400 0.00000 -30.33400
0.00000 0.00000 0.00000 ; CZ- CE2- CD2- HD2
2865 2864 2867 2868 3 30.33400 0.00000 -30.33400
0.00000 0.00000 0.00000 ; OH- CZ- CE2- HE2
2865 2864 2867 2869 3 30.33400 0.00000 -30.33400
0.00000 0.00000 0.00000 ; OH- CZ- CE2- CD2
2866 2865 2864 2867 3 7.93286 0.00000 -9.56462
0.00000 2.17568 0.00000 ; HH- OH- CZ- CE2
2868 2867 2869 2870 3 30.33400 0.00000 -30.33400
0.00000 0.00000 0.00000 ; HE2- CE2- CD2- HD2
2871 2873 2875 2876 3 0.00000 0.00000 0.00000
0.00000 0.00000 0.00000 ; C- N- CA- HA

2871 2873 2875 2877 3 11.71520 1.67360 15.06240 -
13.38880 0.00000 0.00000 ; C- N- CA- CB
2871 2873 2875 2892 3 1.75728 5.27184 2.25936
-7.02912 0.00000 0.00000 ; C- N- CA- C
2872 2871 2873 2874 3 29.28800 -8.36800 -20.92000
0.00000 0.00000 0.00000 ; O- C- N- H
2872 2871 2873 2875 3 20.92000 0.00000 -20.92000
0.00000 0.00000 0.00000 ; O- C- N- CA
2873 2875 2877 2878 3 0.65084 1.95253 0.00000
-2.60338 0.00000 0.00000 ; N- CA- CB- HB2
2873 2875 2877 2879 3 0.65084 1.95253 0.00000
-2.60338 0.00000 0.00000 ; N- CA- CB- HB3
2873 2875 2877 2880 3 3.28026 2.46019 -2.19242
-3.21331 -0.23430 0.00000 ; N- CA- CB- CG
2873 2875 2892 2893 3 0.00000 0.00000 0.00000
0.00000 0.00000 0.00000 ; N- CA- C- O
2873 2875 2892 2894 3 17.40544 -5.02080 -13.22144
9.20480 0.00000 0.00000 ; N- CA- C- N
2874 2873 2875 2876 3 0.00000 0.00000 0.00000
0.00000 0.00000 0.00000 ; H- N- CA- HA
2874 2873 2875 2877 3 0.00000 0.00000 0.00000
0.00000 0.00000 0.00000 ; H- N- CA- CB
2874 2873 2875 2892 3 0.00000 0.00000 0.00000
0.00000 0.00000 0.00000 ; H- N- CA- C
2875 2877 2880 2881 3 0.57739 0.00000 1.02926
0.00000 -1.60666 0.00000 ; CA- CB- CG- CD1
2875 2877 2880 2890 3 0.57739 0.00000 1.02926
0.00000 -1.60666 0.00000 ; CA- CB- CG- CD2
2875 2892 2894 2895 3 20.92000 0.00000 -20.92000
0.00000 0.00000 0.00000 ; CA- C- N- H
2875 2892 2894 2896 3 20.92000 0.00000 -20.92000
0.00000 0.00000 0.00000 ; CA- C- N- CA
2876 2875 2877 2878 3 0.65084 1.95253 0.00000
-2.60338 0.00000 0.00000 ; HA- CA- CB- HB2
2876 2875 2877 2879 3 0.65084 1.95253 0.00000
-2.60338 0.00000 0.00000 ; HA- CA- CB- HB3
2876 2875 2877 2880 3 0.65084 1.95253 0.00000
-2.60338 0.00000 0.00000 ; HA- CA- CB- CG
2876 2875 2892 2893 3 3.68192 -4.35136 0.00000
1.33888 0.00000 0.00000 ; HA- CA- C- O
2876 2875 2892 2894 3 0.00000 0.00000 0.00000
0.00000 0.00000 0.00000 ; HA- CA- C- N
2877 2875 2892 2893 3 0.00000 0.00000 0.00000
0.00000 0.00000 0.00000 ; CB- CA- C- O
2877 2875 2892 2894 3 2.51040 4.18400 1.67360
-6.69440 0.00000 0.00000 ; CB- CA- C- N
2877 2880 2881 2882 3 30.33400 0.00000 -30.33400
0.00000 0.00000 0.00000 ; CB- CG- CD1- HD1
2877 2880 2881 2883 3 30.33400 0.00000 -30.33400
0.00000 0.00000 0.00000 ; CB- CG- CD1- CE1
2877 2880 2890 2888 3 30.33400 0.00000 -30.33400
0.00000 0.00000 0.00000 ; CB- CG- CD2- CE2

2877 2880 2890 2891 3 30.33400 0.00000 -30.33400
0.00000 0.00000 0.00000 ; CB- CG- CD2- HD2
2878 2877 2875 2892 3 0.65084 1.95253 0.00000
-2.60338 0.00000 0.00000 ; HB2- CB- CA- C
2878 2877 2880 2881 3 0.00000 0.00000 0.00000
0.00000 0.00000 0.00000 ; HB2- CB- CG- CD1
2878 2877 2880 2890 3 0.00000 0.00000 0.00000
0.00000 0.00000 0.00000 ; HB2- CB- CG- CD2
2879 2877 2875 2892 3 0.65084 1.95253 0.00000
-2.60338 0.00000 0.00000 ; HB3- CB- CA- C
2879 2877 2880 2881 3 0.00000 0.00000 0.00000
0.00000 0.00000 0.00000 ; HB3- CB- CG- CD1
2879 2877 2880 2890 3 0.00000 0.00000 0.00000
0.00000 0.00000 0.00000 ; HB3- CB- CG- CD2
2880 2877 2875 2892 3 4.95804 2.17986 -3.52293
-3.21331 -0.40166 0.00000 ; CG- CB- CA- C
2880 2881 2883 2884 3 30.33400 0.00000 -30.33400
0.00000 0.00000 0.00000 ; CG- CD1- CE1- HE1
2880 2881 2883 2885 3 30.33400 0.00000 -30.33400
0.00000 0.00000 0.00000 ; CG- CD1- CE1- CZ
2880 2890 2888 2885 3 30.33400 0.00000 -30.33400
0.00000 0.00000 0.00000 ; CG- CD2- CE2- CZ
2880 2890 2888 2889 3 30.33400 0.00000 -30.33400
0.00000 0.00000 0.00000 ; CG- CD2- CE2- HE2
2881 2880 2890 2888 3 30.33400 0.00000 -30.33400
0.00000 0.00000 0.00000 ; CD1- CG- CD2- CE2
2881 2880 2890 2891 3 30.33400 0.00000 -30.33400
0.00000 0.00000 0.00000 ; CD1- CG- CD2- HD2
2881 2883 2885 2886 3 30.33400 0.00000 -30.33400
0.00000 0.00000 0.00000 ; CD1- CE1- CZ- OH
2881 2883 2885 2888 3 30.33400 0.00000 -30.33400
0.00000 0.00000 0.00000 ; CD1- CE1- CZ- CE2
2882 2881 2880 2890 3 30.33400 0.00000 -30.33400
0.00000 0.00000 0.00000 ; HD1- CD1- CG- CD2
2882 2881 2883 2884 3 30.33400 0.00000 -30.33400
0.00000 0.00000 0.00000 ; HD1- CD1- CE1- HE1
2882 2881 2883 2885 3 30.33400 0.00000 -30.33400
0.00000 0.00000 0.00000 ; HD1- CD1- CE1- CZ
2883 2881 2880 2890 3 30.33400 0.00000 -30.33400
0.00000 0.00000 0.00000 ; CE1- CD1- CG- CD2
2883 2885 2886 2887 3 7.93286 0.00000 -9.56462
0.00000 2.17568 0.00000 ; CE1- CZ- OH- HH
2883 2885 2888 2889 3 30.33400 0.00000 -30.33400
0.00000 0.00000 0.00000 ; CE1- CZ- CE2- HE2
2883 2885 2888 2890 3 30.33400 0.00000 -30.33400
0.00000 0.00000 0.00000 ; CE1- CZ- CE2- CD2
2884 2883 2885 2886 3 30.33400 0.00000 -30.33400
0.00000 0.00000 0.00000 ; HE1- CE1- CZ- OH
2884 2883 2885 2888 3 30.33400 0.00000 -30.33400
0.00000 0.00000 0.00000 ; HE1- CE1- CZ- CE2
2885 2888 2890 2891 3 30.33400 0.00000 -30.33400
0.00000 0.00000 0.00000 ; CZ- CE2- CD2- HD2

2886 2885 2888 2889 3 30.33400 0.00000 -30.33400
0.00000 0.00000 0.00000 ; OH- CZ- CE2- HE2
2886 2885 2888 2890 3 30.33400 0.00000 -30.33400
0.00000 0.00000 0.00000 ; OH- CZ- CE2- CD2
2887 2886 2885 2888 3 7.93286 0.00000 -9.56462
0.00000 2.17568 0.00000 ; HH- OH- CZ- CE2
2889 2888 2890 2891 3 30.33400 0.00000 -30.33400
0.00000 0.00000 0.00000 ; HE2- CE2- CD2- HD2
2892 2894 2896 2897 3 0.00000 0.00000 0.00000
0.00000 0.00000 0.00000 ; C- N- CA- HA
2892 2894 2896 2898 3 11.71520 1.67360 15.06240 -
13.38880 0.00000 0.00000 ; C- N- CA- CB
2892 2894 2896 2908 3 1.75728 5.27184 2.25936
-7.02912 0.00000 0.00000 ; C- N- CA- C
2893 2892 2894 2895 3 29.28800 -8.36800 -20.92000
0.00000 0.00000 0.00000 ; O- C- N- H
2893 2892 2894 2896 3 20.92000 0.00000 -20.92000
0.00000 0.00000 0.00000 ; O- C- N- CA
2894 2896 2898 2899 3 0.65084 1.95253 0.00000
-2.60338 0.00000 0.00000 ; N- CA- CB- HB
2894 2896 2898 2900 3 3.83673 0.44769 -1.77402
-2.47693 -0.03347 0.00000 ; N- CA- CB- CG1
2894 2896 2898 2904 3 3.83673 0.44769 -1.77402
-2.47693 -0.03347 0.00000 ; N- CA- CB- CG2
2894 2896 2908 2909 3 0.00000 0.00000 0.00000
0.00000 0.00000 0.00000 ; N- CA- C- O
2894 2896 2908 2910 3 17.40544 -5.02080 -13.22144
9.20480 0.00000 0.00000 ; N- CA- C- N
2895 2894 2896 2897 3 0.00000 0.00000 0.00000
0.00000 0.00000 0.00000 ; H- N- CA- HA
2895 2894 2896 2898 3 0.00000 0.00000 0.00000
0.00000 0.00000 0.00000 ; H- N- CA- CB
2895 2894 2896 2908 3 0.00000 0.00000 0.00000
0.00000 0.00000 0.00000 ; H- N- CA- C
2896 2898 2900 2901 3 0.66944 2.00832 0.00000
-2.67776 0.00000 0.00000 ; CA- CB- CG1- HG11
2896 2898 2900 2902 3 0.66944 2.00832 0.00000
-2.67776 0.00000 0.00000 ; CA- CB- CG1- HG12
2896 2898 2900 2903 3 0.66944 2.00832 0.00000
-2.67776 0.00000 0.00000 ; CA- CB- CG1- HG13
2896 2898 2904 2905 3 0.66944 2.00832 0.00000
-2.67776 0.00000 0.00000 ; CA- CB- CG2- HG21
2896 2898 2904 2906 3 0.66944 2.00832 0.00000
-2.67776 0.00000 0.00000 ; CA- CB- CG2- HG22
2896 2898 2904 2907 3 0.66944 2.00832 0.00000
-2.67776 0.00000 0.00000 ; CA- CB- CG2- HG23
2896 2908 2910 2911 3 20.92000 0.00000 -20.92000
0.00000 0.00000 0.00000 ; CA- C- N- H
2896 2908 2910 2912 3 20.92000 0.00000 -20.92000
0.00000 0.00000 0.00000 ; CA- C- N- CA
2897 2896 2898 2899 3 0.65084 1.95253 0.00000
-2.60338 0.00000 0.00000 ; HA- CA- CB- HB

2897 2896 2898 2900 3 0.65084 1.95253 0.00000
-2.60338 0.00000 0.00000 ; HA- CA- CB- CG1
2897 2896 2898 2904 3 0.65084 1.95253 0.00000
-2.60338 0.00000 0.00000 ; HA- CA- CB- CG2
2897 2896 2908 2909 3 3.68192 -4.35136 0.00000
1.33888 0.00000 0.00000 ; HA- CA- C- O
2897 2896 2908 2910 3 0.00000 0.00000 0.00000
0.00000 0.00000 0.00000 ; HA- CA- C- N
2898 2896 2908 2909 3 0.00000 0.00000 0.00000
0.00000 0.00000 0.00000 ; CB- CA- C- O
2898 2896 2908 2910 3 2.51040 4.18400 1.67360
-6.69440 0.00000 0.00000 ; CB- CA- C- N
2899 2898 2896 2908 3 0.65084 1.95253 0.00000
-2.60338 0.00000 0.00000 ; HB- CB- CA- C
2899 2898 2900 2901 3 0.62760 1.88280 0.00000
-2.51040 0.00000 0.00000 ; HB- CB- CG1- HG11
2899 2898 2900 2902 3 0.62760 1.88280 0.00000
-2.51040 0.00000 0.00000 ; HB- CB- CG1- HG12
2899 2898 2900 2903 3 0.62760 1.88280 0.00000
-2.51040 0.00000 0.00000 ; HB- CB- CG1- HG13
2899 2898 2904 2905 3 0.62760 1.88280 0.00000
-2.51040 0.00000 0.00000 ; HB- CB- CG2- HG21
2899 2898 2904 2906 3 0.62760 1.88280 0.00000
-2.51040 0.00000 0.00000 ; HB- CB- CG2- HG22
2899 2898 2904 2907 3 0.62760 1.88280 0.00000
-2.51040 0.00000 0.00000 ; HB- CB- CG2- HG23
2900 2898 2896 2908 3 5.67350 3.55640 -6.16722
-2.47693 3.74886 0.00000 ; CG1- CB- CA- C
2900 2898 2904 2905 3 0.66944 2.00832 0.00000
-2.67776 0.00000 0.00000 ; CG1- CB- CG2- HG21
2900 2898 2904 2906 3 0.66944 2.00832 0.00000
-2.67776 0.00000 0.00000 ; CG1- CB- CG2- HG22
2900 2898 2904 2907 3 0.66944 2.00832 0.00000
-2.67776 0.00000 0.00000 ; CG1- CB- CG2- HG23
2901 2900 2898 2904 3 0.66944 2.00832 0.00000
-2.67776 0.00000 0.00000 ; HG11- CG1- CB- CG2
2902 2900 2898 2904 3 0.66944 2.00832 0.00000
-2.67776 0.00000 0.00000 ; HG12- CG1- CB- CG2
2903 2900 2898 2904 3 0.66944 2.00832 0.00000
-2.67776 0.00000 0.00000 ; HG13- CG1- CB- CG2
2904 2898 2896 2908 3 5.67350 3.55640 -6.16722
-2.47693 3.74886 0.00000 ; CG2- CB- CA- C
2908 2910 2912 2913 3 0.00000 0.00000 0.00000
0.00000 0.00000 0.00000 ; C- N- CA- HA
2908 2910 2912 2914 3 11.71520 1.67360 15.06240 -
13.38880 0.00000 0.00000 ; C- N- CA- CB
2908 2910 2912 2918 3 1.75728 5.27184 2.25936
-7.02912 0.00000 0.00000 ; C- N- CA- C
2909 2908 2910 2911 3 29.28800 -8.36800 -20.92000
0.00000 0.00000 0.00000 ; O- C- N- H
2909 2908 2910 2912 3 20.92000 0.00000 -20.92000
0.00000 0.00000 0.00000 ; O- C- N- CA

2910 2912 2914 2915 3 0.65084 1.95253 0.00000
-2.60338 0.00000 0.00000 ; N- CA- CB- HB1
2910 2912 2914 2916 3 0.65084 1.95253 0.00000
-2.60338 0.00000 0.00000 ; N- CA- CB- HB2
2910 2912 2914 2917 3 0.65084 1.95253 0.00000
-2.60338 0.00000 0.00000 ; N- CA- CB- HB3
2910 2912 2918 2919 3 0.00000 0.00000 0.00000
0.00000 0.00000 0.00000 ; N- CA- C- O
2910 2912 2918 2920 3 17.40544 -5.02080 -13.22144
9.20480 0.00000 0.00000 ; N- CA- C- N
2911 2910 2912 2913 3 0.00000 0.00000 0.00000
0.00000 0.00000 0.00000 ; H- N- CA- HA
2911 2910 2912 2914 3 0.00000 0.00000 0.00000
0.00000 0.00000 0.00000 ; H- N- CA- CB
2911 2910 2912 2918 3 0.00000 0.00000 0.00000
0.00000 0.00000 0.00000 ; H- N- CA- C
2912 2918 2920 2921 3 20.92000 0.00000 -20.92000
0.00000 0.00000 0.00000 ; CA- C- N- CD
2912 2918 2920 2930 3 20.92000 0.00000 -20.92000
0.00000 0.00000 0.00000 ; CA- C- N- CA
2913 2912 2914 2915 3 0.65084 1.95253 0.00000
-2.60338 0.00000 0.00000 ; HA- CA- CB- HB1
2913 2912 2914 2916 3 0.65084 1.95253 0.00000
-2.60338 0.00000 0.00000 ; HA- CA- CB- HB2
2913 2912 2914 2917 3 0.65084 1.95253 0.00000
-2.60338 0.00000 0.00000 ; HA- CA- CB- HB3
2913 2912 2918 2919 3 3.68192 -4.35136 0.00000
1.33888 0.00000 0.00000 ; HA- CA- C- O
2913 2912 2918 2920 3 0.00000 0.00000 0.00000
0.00000 0.00000 0.00000 ; HA- CA- C- N
2914 2912 2918 2919 3 0.00000 0.00000 0.00000
0.00000 0.00000 0.00000 ; CB- CA- C- O
2914 2912 2918 2920 3 2.51040 4.18400 1.67360
-6.69440 0.00000 0.00000 ; CB- CA- C- N
2915 2914 2912 2918 3 0.65084 1.95253 0.00000
-2.60338 0.00000 0.00000 ; HB1- CB- CA- C
2916 2914 2912 2918 3 0.65084 1.95253 0.00000
-2.60338 0.00000 0.00000 ; HB2- CB- CA- C
2917 2914 2912 2918 3 0.65084 1.95253 0.00000
-2.60338 0.00000 0.00000 ; HB3- CB- CA- C
2918 2920 2921 2922 3 0.00000 0.00000 0.00000
0.00000 0.00000 0.00000 ; C- N- CD- HD2
2918 2920 2921 2923 3 0.00000 0.00000 0.00000
0.00000 0.00000 0.00000 ; C- N- CD- HD3
2918 2920 2921 2924 3 10.04160 -3.34720 16.73600
-6.69440 0.00000 0.00000 ; C- N- CD- CG
2918 2920 2930 2927 3 11.71520 1.67360 15.06240 -
13.38880 0.00000 0.00000 ; C- N- CA- CB
2918 2920 2930 2931 3 0.00000 0.00000 0.00000
0.00000 0.00000 0.00000 ; C- N- CA- HA
2918 2920 2930 2932 3 1.75728 5.27184 2.25936
-7.02912 0.00000 0.00000 ; C- N- CA- C

2919 2918 2920 2921 3 20.92000 0.00000 -20.92000
0.00000 0.00000 0.00000 ; O- C- N- CD
2919 2918 2920 2930 3 20.92000 0.00000 -20.92000
0.00000 0.00000 0.00000 ; O- C- N- CA
2920 2921 2924 2925 3 0.65084 1.95253 0.00000
-2.60338 0.00000 0.00000 ; N- CD- CG- HG2
2920 2921 2924 2926 3 0.65084 1.95253 0.00000
-2.60338 0.00000 0.00000 ; N- CD- CG- HG3
2920 2921 2924 2927 3 0.65084 1.95253 0.00000
-2.60338 0.00000 0.00000 ; N- CD- CG- CB
2920 2930 2927 2924 3 0.65084 1.95253 0.00000
-2.60338 0.00000 0.00000 ; N- CA- CB- CG
2920 2930 2927 2928 3 0.65084 1.95253 0.00000
-2.60338 0.00000 0.00000 ; N- CA- CB- HB2
2920 2930 2927 2929 3 0.65084 1.95253 0.00000
-2.60338 0.00000 0.00000 ; N- CA- CB- HB3
2920 2930 2932 2933 3 0.00000 0.00000 0.00000
0.00000 0.00000 0.00000 ; N- CA- C- O
2920 2930 2932 2934 3 17.40544 -5.02080 -13.22144
9.20480 0.00000 0.00000 ; N- CA- C- N
2921 2920 2930 2927 3 0.00000 0.00000 0.00000
0.00000 0.00000 0.00000 ; CD- N- CA- CB
2921 2920 2930 2931 3 0.00000 0.00000 0.00000
0.00000 0.00000 0.00000 ; CD- N- CA- HA
2921 2920 2930 2932 3 0.00000 0.00000 0.00000
0.00000 0.00000 0.00000 ; CD- N- CA- C
2921 2924 2927 2928 3 0.66944 2.00832 0.00000
-2.67776 0.00000 0.00000 ; CD- CG- CB- HB2
2921 2924 2927 2929 3 0.66944 2.00832 0.00000
-2.67776 0.00000 0.00000 ; CD- CG- CB- HB3
2921 2924 2927 2930 3 3.68192 3.09616 -2.09200
-3.01248 0.00000 0.00000 ; CD- CG- CB- CA
2922 2921 2920 2930 3 0.00000 0.00000 0.00000
0.00000 0.00000 0.00000 ; HD2- CD- N- CA
2922 2921 2924 2925 3 0.65084 1.95253 0.00000
-2.60338 0.00000 0.00000 ; HD2- CD- CG- HG2
2922 2921 2924 2926 3 0.65084 1.95253 0.00000
-2.60338 0.00000 0.00000 ; HD2- CD- CG- HG3
2922 2921 2924 2927 3 0.65084 1.95253 0.00000
-2.60338 0.00000 0.00000 ; HD2- CD- CG- CB
2923 2921 2920 2930 3 0.00000 0.00000 0.00000
0.00000 0.00000 0.00000 ; HD3- CD- N- CA
2923 2921 2924 2925 3 0.65084 1.95253 0.00000
-2.60338 0.00000 0.00000 ; HD3- CD- CG- HG2
2923 2921 2924 2926 3 0.65084 1.95253 0.00000
-2.60338 0.00000 0.00000 ; HD3- CD- CG- HG3
2923 2921 2924 2927 3 0.65084 1.95253 0.00000
-2.60338 0.00000 0.00000 ; HD3- CD- CG- CB
2924 2921 2920 2930 3 0.00000 0.00000 0.00000
0.00000 0.00000 0.00000 ; CG- CD- N- CA
2924 2927 2930 2931 3 0.65084 1.95253 0.00000
-2.60338 0.00000 0.00000 ; CG- CB- CA- HA

```
  2924   2927   2930   2932       3    0.65084    1.95253    0.00000
-2.60338    0.00000    0.00000 ;     CG-    CB-    CA-     C
  2925   2924   2927   2928       3    0.62760    1.88280    0.00000
-2.51040    0.00000    0.00000 ;    HG2-    CG-    CB-   HB2
  2925   2924   2927   2929       3    0.62760    1.88280    0.00000
-2.51040    0.00000    0.00000 ;    HG2-    CG-    CB-   HB3
  2925   2924   2927   2930       3    0.66944    2.00832    0.00000
-2.67776    0.00000    0.00000 ;    HG2-    CG-    CB-    CA
  2926   2924   2927   2928       3    0.62760    1.88280    0.00000
-2.51040    0.00000    0.00000 ;    HG3-    CG-    CB-   HB2
  2926   2924   2927   2929       3    0.62760    1.88280    0.00000
-2.51040    0.00000    0.00000 ;    HG3-    CG-    CB-   HB3
  2926   2924   2927   2930       3    0.66944    2.00832    0.00000
-2.67776    0.00000    0.00000 ;    HG3-    CG-    CB-    CA
  2927   2930   2932   2933       3    0.00000    0.00000    0.00000
0.00000    0.00000    0.00000 ;     CB-    CA-     C-     O
  2927   2930   2932   2934       3    2.51040    4.18400    1.67360
-6.69440    0.00000    0.00000 ;     CB-    CA-     C-     N
  2928   2927   2930   2931       3    0.65084    1.95253    0.00000
-2.60338    0.00000    0.00000 ;    HB2-    CB-    CA-    HA
  2928   2927   2930   2932       3    0.65084    1.95253    0.00000
-2.60338    0.00000    0.00000 ;    HB2-    CB-    CA-     C
  2929   2927   2930   2931       3    0.65084    1.95253    0.00000
-2.60338    0.00000    0.00000 ;    HB3-    CB-    CA-    HA
  2929   2927   2930   2932       3    0.65084    1.95253    0.00000
-2.60338    0.00000    0.00000 ;    HB3-    CB-    CA-     C
  2930   2932   2934   2935       3   20.92000    0.00000  -20.92000
0.00000    0.00000    0.00000 ;     CA-     C-     N-     H
  2930   2932   2934   2936       3   20.92000    0.00000  -20.92000
0.00000    0.00000    0.00000 ;     CA-     C-     N-    CA
  2931   2930   2932   2933       3    3.68192   -4.35136    0.00000
1.33888    0.00000    0.00000 ;     HA-    CA-     C-     O
  2931   2930   2932   2934       3    0.00000    0.00000    0.00000
0.00000    0.00000    0.00000 ;     HA-    CA-     C-     N
  2932   2934   2936   2937       3    0.00000    0.00000    0.00000
0.00000    0.00000    0.00000 ;      C-     N-    CA-    HA
  2932   2934   2936   2938       3   11.71520    1.67360   15.06240  -
13.38880    0.00000    0.00000 ;      C-     N-    CA-    CB
  2932   2934   2936   2949       3    1.75728    5.27184    2.25936
-7.02912    0.00000    0.00000 ;      C-     N-    CA-     C
  2933   2932   2934   2935       3   29.28800   -8.36800  -20.92000
0.00000    0.00000    0.00000 ;      O-     C-     N-     H
  2933   2932   2934   2936       3   20.92000    0.00000  -20.92000
0.00000    0.00000    0.00000 ;      O-     C-     N-    CA
  2934   2936   2938   2939       3    0.65084    1.95253    0.00000
-2.60338    0.00000    0.00000 ;      N-    CA-    CB-   HB2
  2934   2936   2938   2940       3    0.65084    1.95253    0.00000
-2.60338    0.00000    0.00000 ;      N-    CA-    CB-   HB3
  2934   2936   2938   2941       3    3.21331    2.22589   -4.15053
-2.40998    2.61082    0.00000 ;      N-    CA-    CB-    CG
  2934   2936   2949   2950       3    0.00000    0.00000    0.00000
0.00000    0.00000    0.00000 ;      N-    CA-     C-     O
```

2934 2936 2949 2951 3 17.40544 -5.02080 -13.22144
9.20480 0.00000 0.00000 ; N- CA- C- N
2935 2934 2936 2937 3 0.00000 0.00000 0.00000
0.00000 0.00000 0.00000 ; H- N- CA- HA
2935 2934 2936 2938 3 0.00000 0.00000 0.00000
0.00000 0.00000 0.00000 ; H- N- CA- CB
2935 2934 2936 2949 3 0.00000 0.00000 0.00000
0.00000 0.00000 0.00000 ; H- N- CA- C
2936 2938 2941 2942 3 0.66944 2.00832 0.00000
-2.67776 0.00000 0.00000 ; CA- CB- CG- HG2
2936 2938 2941 2943 3 0.66944 2.00832 0.00000
-2.67776 0.00000 0.00000 ; CA- CB- CG- HG3
2936 2938 2941 2944 3 3.69866 -4.35136 -3.92459
6.89523 4.61914 0.00000 ; CA- CB- CG- CD
2936 2949 2951 2952 3 20.92000 0.00000 -20.92000
0.00000 0.00000 0.00000 ; CA- C- N- H
2936 2949 2951 2953 3 20.92000 0.00000 -20.92000
0.00000 0.00000 0.00000 ; CA- C- N- CA
2937 2936 2938 2939 3 0.65084 1.95253 0.00000
-2.60338 0.00000 0.00000 ; HA- CA- CB- HB2
2937 2936 2938 2940 3 0.65084 1.95253 0.00000
-2.60338 0.00000 0.00000 ; HA- CA- CB- HB3
2937 2936 2938 2941 3 0.65084 1.95253 0.00000
-2.60338 0.00000 0.00000 ; HA- CA- CB- CG
2937 2936 2949 2950 3 3.68192 -4.35136 0.00000
1.33888 0.00000 0.00000 ; HA- CA- C- O
2937 2936 2949 2951 3 0.00000 0.00000 0.00000
0.00000 0.00000 0.00000 ; HA- CA- C- N
2938 2936 2949 2950 3 0.00000 0.00000 0.00000
0.00000 0.00000 0.00000 ; CB- CA- C- O
2938 2936 2949 2951 3 2.51040 4.18400 1.67360
-6.69440 0.00000 0.00000 ; CB- CA- C- N
2938 2941 2944 2945 3 0.00000 0.00000 0.00000
0.00000 0.00000 0.00000 ; CB- CG- CD- OE1
2938 2941 2944 2946 3 10.32611 1.48114 -8.47678
1.42256 1.40582 0.00000 ; CB- CG- CD- NE2
2939 2938 2936 2949 3 0.65084 1.95253 0.00000
-2.60338 0.00000 0.00000 ; HB2- CB- CA- C
2939 2938 2941 2942 3 0.62760 1.88280 0.00000
-2.51040 0.00000 0.00000 ; HB2- CB- CG- HG2
2939 2938 2941 2943 3 0.62760 1.88280 0.00000
-2.51040 0.00000 0.00000 ; HB2- CB- CG- HG3
2939 2938 2941 2944 3 0.65084 1.95253 0.00000
-2.60338 0.00000 0.00000 ; HB2- CB- CG- CD
2940 2938 2936 2949 3 0.65084 1.95253 0.00000
-2.60338 0.00000 0.00000 ; HB3- CB- CA- C
2940 2938 2941 2942 3 0.62760 1.88280 0.00000
-2.51040 0.00000 0.00000 ; HB3- CB- CG- HG2
2940 2938 2941 2943 3 0.62760 1.88280 0.00000
-2.51040 0.00000 0.00000 ; HB3- CB- CG- HG3
2940 2938 2941 2944 3 0.65084 1.95253 0.00000
-2.60338 0.00000 0.00000 ; HB3- CB- CG- CD

2941 2938 2936 2949 3 6.86594 3.56895 -8.14206
-2.40998 4.85344 0.00000 ; CG- CB- CA- C
2941 2944 2946 2947 3 20.92000 0.00000 -20.92000
0.00000 0.00000 0.00000 ; CG- CD- NE2- HE21
2941 2944 2946 2948 3 20.92000 0.00000 -20.92000
0.00000 0.00000 0.00000 ; CG- CD- NE2- HE22
2942 2941 2944 2945 3 3.68192 -4.35136 0.00000
1.33888 0.00000 0.00000 ; HG2- CG- CD- OE1
2942 2941 2944 2946 3 0.00000 0.00000 0.00000
0.00000 0.00000 0.00000 ; HG2- CG- CD- NE2
2943 2941 2944 2945 3 3.68192 -4.35136 0.00000
1.33888 0.00000 0.00000 ; HG3- CG- CD- OE1
2943 2941 2944 2946 3 0.00000 0.00000 0.00000
0.00000 0.00000 0.00000 ; HG3- CG- CD- NE2
2945 2944 2946 2947 3 29.28800 -8.36800 -20.92000
0.00000 0.00000 0.00000 ; OE1- CD- NE2- HE21
2945 2944 2946 2948 3 29.28800 -8.36800 -20.92000
0.00000 0.00000 0.00000 ; OE1- CD- NE2- HE22
2949 2951 2953 2954 3 0.00000 0.00000 0.00000
0.00000 0.00000 0.00000 ; C- N- CA- HA
2949 2951 2953 2955 3 11.71520 1.67360 15.06240 -
13.38880 0.00000 0.00000 ; C- N- CA- CB
2949 2951 2953 2965 3 1.75728 5.27184 2.25936
-7.02912 0.00000 0.00000 ; C- N- CA- C
2950 2949 2951 2952 3 29.28800 -8.36800 -20.92000
0.00000 0.00000 0.00000 ; O- C- N- H
2950 2949 2951 2953 3 20.92000 0.00000 -20.92000
0.00000 0.00000 0.00000 ; O- C- N- CA
2951 2953 2955 2956 3 0.65084 1.95253 0.00000
-2.60338 0.00000 0.00000 ; N- CA- CB- HB
2951 2953 2955 2957 3 3.83673 0.44769 -1.77402
-2.47693 -0.03347 0.00000 ; N- CA- CB- CG1
2951 2953 2955 2961 3 3.83673 0.44769 -1.77402
-2.47693 -0.03347 0.00000 ; N- CA- CB- CG2
2951 2953 2965 2966 3 0.00000 0.00000 0.00000
0.00000 0.00000 0.00000 ; N- CA- C- O
2951 2953 2965 2967 3 17.40544 -5.02080 -13.22144
9.20480 0.00000 0.00000 ; N- CA- C- N
2952 2951 2953 2954 3 0.00000 0.00000 0.00000
0.00000 0.00000 0.00000 ; H- N- CA- HA
2952 2951 2953 2955 3 0.00000 0.00000 0.00000
0.00000 0.00000 0.00000 ; H- N- CA- CB
2952 2951 2953 2965 3 0.00000 0.00000 0.00000
0.00000 0.00000 0.00000 ; H- N- CA- C
2953 2955 2957 2958 3 0.66944 2.00832 0.00000
-2.67776 0.00000 0.00000 ; CA- CB- CG1- HG11
2953 2955 2957 2959 3 0.66944 2.00832 0.00000
-2.67776 0.00000 0.00000 ; CA- CB- CG1- HG12
2953 2955 2957 2960 3 0.66944 2.00832 0.00000
-2.67776 0.00000 0.00000 ; CA- CB- CG1- HG13
2953 2955 2961 2962 3 0.66944 2.00832 0.00000
-2.67776 0.00000 0.00000 ; CA- CB- CG2- HG21

```
  2953   2955   2961   2963       3    0.66944    2.00832    0.00000
-2.67776    0.00000    0.00000 ;     CA-    CB-   CG2-  HG22
  2953   2955   2961   2964       3    0.66944    2.00832    0.00000
-2.67776    0.00000    0.00000 ;     CA-    CB-   CG2-  HG23
  2953   2965   2967   2968       3   20.92000    0.00000  -20.92000
0.00000    0.00000    0.00000 ;     CA-     C-     N-     H
  2953   2965   2967   2969       3   20.92000    0.00000  -20.92000
0.00000    0.00000    0.00000 ;     CA-     C-     N-    CA
  2954   2953   2955   2956       3    0.65084    1.95253    0.00000
-2.60338    0.00000    0.00000 ;     HA-    CA-    CB-    HB
  2954   2953   2955   2957       3    0.65084    1.95253    0.00000
-2.60338    0.00000    0.00000 ;     HA-    CA-    CB-   CG1
  2954   2953   2955   2961       3    0.65084    1.95253    0.00000
-2.60338    0.00000    0.00000 ;     HA-    CA-    CB-   CG2
  2954   2953   2965   2966       3    3.68192   -4.35136    0.00000
1.33888    0.00000    0.00000 ;     HA-    CA-     C-     O
  2954   2953   2965   2967       3    0.00000    0.00000    0.00000
0.00000    0.00000    0.00000 ;     HA-    CA-     C-     N
  2955   2953   2965   2966       3    0.00000    0.00000    0.00000
0.00000    0.00000    0.00000 ;     CB-    CA-     C-     O
  2955   2953   2965   2967       3    2.51040    4.18400    1.67360
-6.69440    0.00000    0.00000 ;     CB-    CA-     C-     N
  2956   2955   2953   2965       3    0.65084    1.95253    0.00000
-2.60338    0.00000    0.00000 ;     HB-    CB-    CA-     C
  2956   2955   2957   2958       3    0.62760    1.88280    0.00000
-2.51040    0.00000    0.00000 ;     HB-    CB-   CG1-  HG11
  2956   2955   2957   2959       3    0.62760    1.88280    0.00000
-2.51040    0.00000    0.00000 ;     HB-    CB-   CG1-  HG12
  2956   2955   2957   2960       3    0.62760    1.88280    0.00000
-2.51040    0.00000    0.00000 ;     HB-    CB-   CG1-  HG13
  2956   2955   2961   2962       3    0.62760    1.88280    0.00000
-2.51040    0.00000    0.00000 ;     HB-    CB-   CG2-  HG21
  2956   2955   2961   2963       3    0.62760    1.88280    0.00000
-2.51040    0.00000    0.00000 ;     HB-    CB-   CG2-  HG22
  2956   2955   2961   2964       3    0.62760    1.88280    0.00000
-2.51040    0.00000    0.00000 ;     HB-    CB-   CG2-  HG23
  2957   2955   2953   2965       3    5.67350    3.55640   -6.16722
-2.47693    3.74886    0.00000 ;    CG1-    CB-    CA-     C
  2957   2955   2961   2962       3    0.66944    2.00832    0.00000
-2.67776    0.00000    0.00000 ;    CG1-    CB-   CG2-  HG21
  2957   2955   2961   2963       3    0.66944    2.00832    0.00000
-2.67776    0.00000    0.00000 ;    CG1-    CB-   CG2-  HG22
  2957   2955   2961   2964       3    0.66944    2.00832    0.00000
-2.67776    0.00000    0.00000 ;    CG1-    CB-   CG2-  HG23
  2958   2957   2955   2961       3    0.66944    2.00832    0.00000
-2.67776    0.00000    0.00000 ;   HG11-   CG1-    CB-   CG2
  2959   2957   2955   2961       3    0.66944    2.00832    0.00000
-2.67776    0.00000    0.00000 ;   HG12-   CG1-    CB-   CG2
  2960   2957   2955   2961       3    0.66944    2.00832    0.00000
-2.67776    0.00000    0.00000 ;   HG13-   CG1-    CB-   CG2
  2961   2955   2953   2965       3    5.67350    3.55640   -6.16722
-2.47693    3.74886    0.00000 ;    CG2-    CB-    CA-     C
```

2965 2967 2969 2970 3 0.00000 0.00000 0.00000
0.00000 0.00000 0.00000 ; C- N- CA- HA
2965 2967 2969 2971 3 11.71520 1.67360 15.06240 -
13.38880 0.00000 0.00000 ; C- N- CA- CB
2965 2967 2969 2984 3 1.75728 5.27184 2.25936
-7.02912 0.00000 0.00000 ; C- N- CA- C
2966 2965 2967 2968 3 29.28800 -8.36800 -20.92000
0.00000 0.00000 0.00000 ; O- C- N- H
2966 2965 2967 2969 3 20.92000 0.00000 -20.92000
0.00000 0.00000 0.00000 ; O- C- N- CA
2967 2969 2971 2972 3 0.65084 1.95253 0.00000
-2.60338 0.00000 0.00000 ; N- CA- CB- HB2
2967 2969 2971 2973 3 0.65084 1.95253 0.00000
-2.60338 0.00000 0.00000 ; N- CA- CB- HB3
2967 2969 2971 2974 3 3.79070 1.39746 -4.61077
-2.40998 2.44346 0.00000 ; N- CA- CB- CG
2967 2969 2984 2985 3 0.00000 0.00000 0.00000
0.00000 0.00000 0.00000 ; N- CA- C- O
2967 2969 2984 2986 3 17.40544 -5.02080 -13.22144
9.20480 0.00000 0.00000 ; N- CA- C- N
2968 2967 2969 2970 3 0.00000 0.00000 0.00000
0.00000 0.00000 0.00000 ; H- N- CA- HA
2968 2967 2969 2971 3 0.00000 0.00000 0.00000
0.00000 0.00000 0.00000 ; H- N- CA- CB
2968 2967 2969 2984 3 0.00000 0.00000 0.00000
0.00000 0.00000 0.00000 ; H- N- CA- C
2969 2971 2974 2975 3 0.66944 2.00832 0.00000
-2.67776 0.00000 0.00000 ; CA- CB- CG- HG
2969 2971 2974 2976 3 3.90367 0.19665 -6.21742
-2.37651 5.99149 0.00000 ; CA- CB- CG- CD1
2969 2971 2974 2980 3 3.90367 0.19665 -6.21742
-2.37651 5.99149 0.00000 ; CA- CB- CG- CD2
2969 2984 2986 2987 3 20.92000 0.00000 -20.92000
0.00000 0.00000 0.00000 ; CA- C- N- H
2969 2984 2986 2988 3 20.92000 0.00000 -20.92000
0.00000 0.00000 0.00000 ; CA- C- N- CA
2970 2969 2971 2972 3 0.65084 1.95253 0.00000
-2.60338 0.00000 0.00000 ; HA- CA- CB- HB2
2970 2969 2971 2973 3 0.65084 1.95253 0.00000
-2.60338 0.00000 0.00000 ; HA- CA- CB- HB3
2970 2969 2971 2974 3 0.65084 1.95253 0.00000
-2.60338 0.00000 0.00000 ; HA- CA- CB- CG
2970 2969 2984 2985 3 3.68192 -4.35136 0.00000
1.33888 0.00000 0.00000 ; HA- CA- C- O
2970 2969 2984 2986 3 0.00000 0.00000 0.00000
0.00000 0.00000 0.00000 ; HA- CA- C- N
2971 2969 2984 2985 3 0.00000 0.00000 0.00000
0.00000 0.00000 0.00000 ; CB- CA- C- O
2971 2969 2984 2986 3 2.51040 4.18400 1.67360
-6.69440 0.00000 0.00000 ; CB- CA- C- N
2971 2974 2976 2977 3 0.66944 2.00832 0.00000
-2.67776 0.00000 0.00000 ; CB- CG- CD1- HD11

2971 2974 2976 2978 3 0.66944 2.00832 0.00000
-2.67776 0.00000 0.00000 ; CB- CG- CD1- HD12
2971 2974 2976 2979 3 0.66944 2.00832 0.00000
-2.67776 0.00000 0.00000 ; CB- CG- CD1- HD13
2971 2974 2980 2981 3 0.66944 2.00832 0.00000
-2.67776 0.00000 0.00000 ; CB- CG- CD2- HD21
2971 2974 2980 2982 3 0.66944 2.00832 0.00000
-2.67776 0.00000 0.00000 ; CB- CG- CD2- HD22
2971 2974 2980 2983 3 0.66944 2.00832 0.00000
-2.67776 0.00000 0.00000 ; CB- CG- CD2- HD23
2972 2971 2969 2984 3 0.65084 1.95253 0.00000
-2.60338 0.00000 0.00000 ; HB2- CB- CA- C
2972 2971 2974 2975 3 0.62760 1.88280 0.00000
-2.51040 0.00000 0.00000 ; HB2- CB- CG- HG
2972 2971 2974 2976 3 0.66944 2.00832 0.00000
-2.67776 0.00000 0.00000 ; HB2- CB- CG- CD1
2972 2971 2974 2980 3 0.66944 2.00832 0.00000
-2.67776 0.00000 0.00000 ; HB2- CB- CG- CD2
2973 2971 2969 2984 3 0.65084 1.95253 0.00000
-2.60338 0.00000 0.00000 ; HB3- CB- CA- C
2973 2971 2974 2975 3 0.62760 1.88280 0.00000
-2.51040 0.00000 0.00000 ; HB3- CB- CG- HG
2973 2971 2974 2976 3 0.66944 2.00832 0.00000
-2.67776 0.00000 0.00000 ; HB3- CB- CG- CD1
2973 2971 2974 2980 3 0.66944 2.00832 0.00000
-2.67776 0.00000 0.00000 ; HB3- CB- CG- CD2
2974 2971 2969 2984 3 10.33448 -1.14642 -11.54784
-2.40998 6.35968 0.00000 ; CG- CB- CA- C
2975 2974 2976 2977 3 0.62760 1.88280 0.00000
-2.51040 0.00000 0.00000 ; HG- CG- CD1- HD11
2975 2974 2976 2978 3 0.62760 1.88280 0.00000
-2.51040 0.00000 0.00000 ; HG- CG- CD1- HD12
2975 2974 2976 2979 3 0.62760 1.88280 0.00000
-2.51040 0.00000 0.00000 ; HG- CG- CD1- HD13
2975 2974 2980 2981 3 0.62760 1.88280 0.00000
-2.51040 0.00000 0.00000 ; HG- CG- CD2- HD21
2975 2974 2980 2982 3 0.62760 1.88280 0.00000
-2.51040 0.00000 0.00000 ; HG- CG- CD2- HD22
2975 2974 2980 2983 3 0.62760 1.88280 0.00000
-2.51040 0.00000 0.00000 ; HG- CG- CD2- HD23
2976 2974 2980 2981 3 0.66944 2.00832 0.00000
-2.67776 0.00000 0.00000 ; CD1- CG- CD2- HD21
2976 2974 2980 2982 3 0.66944 2.00832 0.00000
-2.67776 0.00000 0.00000 ; CD1- CG- CD2- HD22
2976 2974 2980 2983 3 0.66944 2.00832 0.00000
-2.67776 0.00000 0.00000 ; CD1- CG- CD2- HD23
2977 2976 2974 2980 3 0.66944 2.00832 0.00000
-2.67776 0.00000 0.00000 ; HD11- CD1- CG- CD2
2978 2976 2974 2980 3 0.66944 2.00832 0.00000
-2.67776 0.00000 0.00000 ; HD12- CD1- CG- CD2
2979 2976 2974 2980 3 0.66944 2.00832 0.00000
-2.67776 0.00000 0.00000 ; HD13- CD1- CG- CD2

2984 2986 2988 2989 3 0.00000 0.00000 0.00000
0.00000 0.00000 0.00000 ; C- N- CA- HA
2984 2986 2988 2990 3 11.71520 1.67360 15.06240 -
13.38880 0.00000 0.00000 ; C- N- CA- CB
2984 2986 2988 2996 3 1.75728 5.27184 2.25936
-7.02912 0.00000 0.00000 ; C- N- CA- C
2985 2984 2986 2987 3 29.28800 -8.36800 -20.92000
0.00000 0.00000 0.00000 ; O- C- N- H
2985 2984 2986 2988 3 20.92000 0.00000 -20.92000
0.00000 0.00000 0.00000 ; O- C- N- CA
2986 2988 2990 2991 3 0.65084 1.95253 0.00000
-2.60338 0.00000 0.00000 ; N- CA- CB- HB2
2986 2988 2990 2992 3 0.65084 1.95253 0.00000
-2.60338 0.00000 0.00000 ; N- CA- CB- HB3
2986 2988 2990 2993 3 15.41386 9.74035 -8.39310
-0.97069 2.97901 0.00000 ; N- CA- CB- CG
2986 2988 2996 2997 3 0.00000 0.00000 0.00000
0.00000 0.00000 0.00000 ; N- CA- C- O
2986 2988 2996 2998 3 17.40544 -5.02080 -13.22144
9.20480 0.00000 0.00000 ; N- CA- C- N
2987 2986 2988 2989 3 0.00000 0.00000 0.00000
0.00000 0.00000 0.00000 ; H- N- CA- HA
2987 2986 2988 2990 3 0.00000 0.00000 0.00000
0.00000 0.00000 0.00000 ; H- N- CA- CB
2987 2986 2988 2996 3 0.00000 0.00000 0.00000
0.00000 0.00000 0.00000 ; H- N- CA- C
2988 2990 2993 2994 3 6.43499 0.00000 -5.39736
0.00000 -1.03763 0.00000 ; CA- CB- CG- OD1
2988 2990 2993 2995 3 6.43499 0.00000 -5.39736
0.00000 -1.03763 0.00000 ; CA- CB- CG- OD2
2988 2996 2998 2999 3 20.92000 0.00000 -20.92000
0.00000 0.00000 0.00000 ; CA- C- N- H
2988 2996 2998 3000 3 20.92000 0.00000 -20.92000
0.00000 0.00000 0.00000 ; CA- C- N- CA
2989 2988 2990 2991 3 0.65084 1.95253 0.00000
-2.60338 0.00000 0.00000 ; HA- CA- CB- HB2
2989 2988 2990 2992 3 0.65084 1.95253 0.00000
-2.60338 0.00000 0.00000 ; HA- CA- CB- HB3
2989 2988 2990 2993 3 0.65084 1.95253 0.00000
-2.60338 0.00000 0.00000 ; HA- CA- CB- CG
2989 2988 2996 2997 3 3.68192 -4.35136 0.00000
1.33888 0.00000 0.00000 ; HA- CA- C- O
2989 2988 2996 2998 3 0.00000 0.00000 0.00000
0.00000 0.00000 0.00000 ; HA- CA- C- N
2990 2988 2996 2997 3 0.00000 0.00000 0.00000
0.00000 0.00000 0.00000 ; CB- CA- C- O
2990 2988 2996 2998 3 2.51040 4.18400 1.67360
-6.69440 0.00000 0.00000 ; CB- CA- C- N
2991 2990 2988 2996 3 0.65084 1.95253 0.00000
-2.60338 0.00000 0.00000 ; HB2- CB- CA- C
2991 2990 2993 2994 3 0.00000 0.00000 0.00000
0.00000 0.00000 0.00000 ; HB2- CB- CG- OD1

2991 2990 2993 2995 3 0.00000 0.00000 0.00000
0.00000 0.00000 0.00000 ; HB2- CB- CG- OD2
2992 2990 2988 2996 3 0.65084 1.95253 0.00000
-2.60338 0.00000 0.00000 ; HB3- CB- CA- C
2992 2990 2993 2994 3 0.00000 0.00000 0.00000
0.00000 0.00000 0.00000 ; HB3- CB- CG- OD1
2992 2990 2993 2995 3 0.00000 0.00000 0.00000
0.00000 0.00000 0.00000 ; HB3- CB- CG- OD2
2993 2990 2988 2996 3 7.14627 -1.04600 -8.99560
-0.97069 5.15469 0.00000 ; CG- CB- CA- C
2996 2998 3000 3001 3 0.00000 0.00000 0.00000
0.00000 0.00000 0.00000 ; C- N- CA- HA2
2996 2998 3000 3002 3 0.00000 0.00000 0.00000
0.00000 0.00000 0.00000 ; C- N- CA- HA3
2996 2998 3000 3003 3 1.75728 5.27184 2.25936
-7.02912 0.00000 0.00000 ; C- N- CA- C
2997 2996 2998 2999 3 29.28800 -8.36800 -20.92000
0.00000 0.00000 0.00000 ; O- C- N- H
2997 2996 2998 3000 3 20.92000 0.00000 -20.92000
0.00000 0.00000 0.00000 ; O- C- N- CA
2998 3000 3003 3004 3 0.00000 0.00000 0.00000
0.00000 0.00000 0.00000 ; N- CA- C- O
2998 3000 3003 3005 3 17.40544 -5.02080 -13.22144
9.20480 0.00000 0.00000 ; N- CA- C- N
2999 2998 3000 3001 3 0.00000 0.00000 0.00000
0.00000 0.00000 0.00000 ; H- N- CA- HA2
2999 2998 3000 3002 3 0.00000 0.00000 0.00000
0.00000 0.00000 0.00000 ; H- N- CA- HA3
2999 2998 3000 3003 3 0.00000 0.00000 0.00000
0.00000 0.00000 0.00000 ; H- N- CA- C
3000 3003 3005 3006 3 20.92000 0.00000 -20.92000
0.00000 0.00000 0.00000 ; CA- C- N- H
3000 3003 3005 3007 3 20.92000 0.00000 -20.92000
0.00000 0.00000 0.00000 ; CA- C- N- CA
3001 3000 3003 3004 3 3.68192 -4.35136 0.00000
1.33888 0.00000 0.00000 ; HA2- CA- C- O
3001 3000 3003 3005 3 0.00000 0.00000 0.00000
0.00000 0.00000 0.00000 ; HA2- CA- C- N
3002 3000 3003 3004 3 3.68192 -4.35136 0.00000
1.33888 0.00000 0.00000 ; HA3- CA- C- O
3002 3000 3003 3005 3 0.00000 0.00000 0.00000
0.00000 0.00000 0.00000 ; HA3- CA- C- N
3003 3005 3007 3008 3 0.00000 0.00000 0.00000
0.00000 0.00000 0.00000 ; C- N- CA- HA
3003 3005 3007 3009 3 11.71520 1.67360 15.06240 -
13.38880 0.00000 0.00000 ; C- N- CA- CB
3003 3005 3007 3025 3 1.75728 5.27184 2.25936
-7.02912 0.00000 0.00000 ; C- N- CA- C
3004 3003 3005 3006 3 29.28800 -8.36800 -20.92000
0.00000 0.00000 0.00000 ; O- C- N- H
3004 3003 3005 3007 3 20.92000 0.00000 -20.92000
0.00000 0.00000 0.00000 ; O- C- N- CA

3005 3007 3009 3010 3 0.65084 1.95253 0.00000
-2.60338 0.00000 0.00000 ; N- CA- CB- HB2
3005 3007 3009 3011 3 0.65084 1.95253 0.00000
-2.60338 0.00000 0.00000 ; N- CA- CB- HB3
3005 3007 3009 3012 3 0.65084 1.95253 0.00000
-2.60338 0.00000 0.00000 ; N- CA- CB- CG
3005 3007 3025 3026 3 0.00000 0.00000 0.00000
0.00000 0.00000 0.00000 ; N- CA- C- O
3005 3007 3025 3027 3 17.40544 -5.02080 -13.22144
9.20480 0.00000 0.00000 ; N- CA- C- N
3006 3005 3007 3008 3 0.00000 0.00000 0.00000
0.00000 0.00000 0.00000 ; H- N- CA- HA
3006 3005 3007 3009 3 0.00000 0.00000 0.00000
0.00000 0.00000 0.00000 ; H- N- CA- CB
3006 3005 3007 3025 3 0.00000 0.00000 0.00000
0.00000 0.00000 0.00000 ; H- N- CA- C
3007 3009 3012 3013 3 0.66944 2.00832 0.00000
-2.67776 0.00000 0.00000 ; CA- CB- CG- HG2
3007 3009 3012 3014 3 0.66944 2.00832 0.00000
-2.67776 0.00000 0.00000 ; CA- CB- CG- HG3
3007 3009 3012 3015 3 3.68192 3.09616 -2.09200
-3.01248 0.00000 0.00000 ; CA- CB- CG- CD
3007 3025 3027 3028 3 20.92000 0.00000 -20.92000
0.00000 0.00000 0.00000 ; CA- C- N- H
3007 3025 3027 3029 3 20.92000 0.00000 -20.92000
0.00000 0.00000 0.00000 ; CA- C- N- CA
3008 3007 3009 3010 3 0.65084 1.95253 0.00000
-2.60338 0.00000 0.00000 ; HA- CA- CB- HB2
3008 3007 3009 3011 3 0.65084 1.95253 0.00000
-2.60338 0.00000 0.00000 ; HA- CA- CB- HB3
3008 3007 3009 3012 3 0.65084 1.95253 0.00000
-2.60338 0.00000 0.00000 ; HA- CA- CB- CG
3008 3007 3025 3026 3 3.68192 -4.35136 0.00000
1.33888 0.00000 0.00000 ; HA- CA- C- O
3008 3007 3025 3027 3 0.00000 0.00000 0.00000
0.00000 0.00000 0.00000 ; HA- CA- C- N
3009 3007 3025 3026 3 0.00000 0.00000 0.00000
0.00000 0.00000 0.00000 ; CB- CA- C- O
3009 3007 3025 3027 3 2.51040 4.18400 1.67360
-6.69440 0.00000 0.00000 ; CB- CA- C- N
3009 3012 3015 3016 3 0.66944 2.00832 0.00000
-2.67776 0.00000 0.00000 ; CB- CG- CD- HD2
3009 3012 3015 3017 3 0.66944 2.00832 0.00000
-2.67776 0.00000 0.00000 ; CB- CG- CD- HD3
3009 3012 3015 3018 3 3.68192 3.09616 -2.09200
-3.01248 0.00000 0.00000 ; CB- CG- CD- CE
3010 3009 3007 3025 3 0.65084 1.95253 0.00000
-2.60338 0.00000 0.00000 ; HB2- CB- CA- C
3010 3009 3012 3013 3 0.62760 1.88280 0.00000
-2.51040 0.00000 0.00000 ; HB2- CB- CG- HG2
3010 3009 3012 3014 3 0.62760 1.88280 0.00000
-2.51040 0.00000 0.00000 ; HB2- CB- CG- HG3

```
  3010   3009   3012   3015      3    0.66944    2.00832    0.00000
-2.67776    0.00000    0.00000 ;    HB2-    CB-    CG-    CD
  3011   3009   3007   3025      3    0.65084    1.95253    0.00000
-2.60338    0.00000    0.00000 ;    HB3-    CB-    CA-     C
  3011   3009   3012   3013      3    0.62760    1.88280    0.00000
-2.51040    0.00000    0.00000 ;    HB3-    CB-    CG-   HG2
  3011   3009   3012   3014      3    0.62760    1.88280    0.00000
-2.51040    0.00000    0.00000 ;    HB3-    CB-    CG-   HG3
  3011   3009   3012   3015      3    0.66944    2.00832    0.00000
-2.67776    0.00000    0.00000 ;    HB3-    CB-    CG-    CD
  3012   3009   3007   3025      3    0.65084    1.95253    0.00000
-2.60338    0.00000    0.00000 ;     CG-    CB-    CA-     C
  3012   3015   3018   3019      3    0.65084    1.95253    0.00000
-2.60338    0.00000    0.00000 ;     CG-    CD-    CE-   HE2
  3012   3015   3018   3020      3    0.65084    1.95253    0.00000
-2.60338    0.00000    0.00000 ;     CG-    CD-    CE-   HE3
  3012   3015   3018   3021      3    0.65084    1.95253    0.00000
-2.60338    0.00000    0.00000 ;     CG-    CD-    CE-    NZ
  3013   3012   3015   3016      3    0.62760    1.88280    0.00000
-2.51040    0.00000    0.00000 ;    HG2-    CG-    CD-   HD2
  3013   3012   3015   3017      3    0.62760    1.88280    0.00000
-2.51040    0.00000    0.00000 ;    HG2-    CG-    CD-   HD3
  3013   3012   3015   3018      3    0.66944    2.00832    0.00000
-2.67776    0.00000    0.00000 ;    HG2-    CG-    CD-    CE
  3014   3012   3015   3016      3    0.62760    1.88280    0.00000
-2.51040    0.00000    0.00000 ;    HG3-    CG-    CD-   HD2
  3014   3012   3015   3017      3    0.62760    1.88280    0.00000
-2.51040    0.00000    0.00000 ;    HG3-    CG-    CD-   HD3
  3014   3012   3015   3018      3    0.66944    2.00832    0.00000
-2.67776    0.00000    0.00000 ;    HG3-    CG-    CD-    CE
  3015   3018   3021   3022      3    0.65084    1.95253    0.00000
-2.60338    0.00000    0.00000 ;     CD-    CE-    NZ-   HZ1
  3015   3018   3021   3023      3    0.65084    1.95253    0.00000
-2.60338    0.00000    0.00000 ;     CD-    CE-    NZ-   HZ2
  3015   3018   3021   3024      3    0.65084    1.95253    0.00000
-2.60338    0.00000    0.00000 ;     CD-    CE-    NZ-   HZ3
  3016   3015   3018   3019      3    0.65084    1.95253    0.00000
-2.60338    0.00000    0.00000 ;    HD2-    CD-    CE-   HE2
  3016   3015   3018   3020      3    0.65084    1.95253    0.00000
-2.60338    0.00000    0.00000 ;    HD2-    CD-    CE-   HE3
  3016   3015   3018   3021      3    0.65084    1.95253    0.00000
-2.60338    0.00000    0.00000 ;    HD2-    CD-    CE-    NZ
  3017   3015   3018   3019      3    0.65084    1.95253    0.00000
-2.60338    0.00000    0.00000 ;    HD3-    CD-    CE-   HE2
  3017   3015   3018   3020      3    0.65084    1.95253    0.00000
-2.60338    0.00000    0.00000 ;    HD3-    CD-    CE-   HE3
  3017   3015   3018   3021      3    0.65084    1.95253    0.00000
-2.60338    0.00000    0.00000 ;    HD3-    CD-    CE-    NZ
  3019   3018   3021   3022      3    0.65084    1.95253    0.00000
-2.60338    0.00000    0.00000 ;    HE2-    CE-    NZ-   HZ1
  3019   3018   3021   3023      3    0.65084    1.95253    0.00000
-2.60338    0.00000    0.00000 ;    HE2-    CE-    NZ-   HZ2
```

3019 3018 3021 3024 3 0.65084 1.95253 0.00000
-2.60338 0.00000 0.00000 ; HE2- CE- NZ- HZ3
3020 3018 3021 3022 3 0.65084 1.95253 0.00000
-2.60338 0.00000 0.00000 ; HE3- CE- NZ- HZ1
3020 3018 3021 3023 3 0.65084 1.95253 0.00000
-2.60338 0.00000 0.00000 ; HE3- CE- NZ- HZ2
3020 3018 3021 3024 3 0.65084 1.95253 0.00000
-2.60338 0.00000 0.00000 ; HE3- CE- NZ- HZ3
3025 3027 3029 3030 3 0.00000 0.00000 0.00000
0.00000 0.00000 0.00000 ; C- N- CA- HA
3025 3027 3029 3031 3 11.71520 1.67360 15.06240 -
13.38880 0.00000 0.00000 ; C- N- CA- CB
3025 3027 3029 3046 3 1.75728 5.27184 2.25936
-7.02912 0.00000 0.00000 ; C- N- CA- C
3026 3025 3027 3028 3 29.28800 -8.36800 -20.92000
0.00000 0.00000 0.00000 ; O- C- N- H
3026 3025 3027 3029 3 20.92000 0.00000 -20.92000
0.00000 0.00000 0.00000 ; O- C- N- CA
3027 3029 3031 3032 3 0.65084 1.95253 0.00000
-2.60338 0.00000 0.00000 ; N- CA- CB- HB2
3027 3029 3031 3033 3 0.65084 1.95253 0.00000
-2.60338 0.00000 0.00000 ; N- CA- CB- HB3
3027 3029 3031 3034 3 3.28026 2.46019 -2.19242
-3.21331 -0.23430 0.00000 ; N- CA- CB- CG
3027 3029 3046 3047 3 0.00000 0.00000 0.00000
0.00000 0.00000 0.00000 ; N- CA- C- O
3027 3029 3046 3048 3 17.40544 -5.02080 -13.22144
9.20480 0.00000 0.00000 ; N- CA- C- N
3028 3027 3029 3030 3 0.00000 0.00000 0.00000
0.00000 0.00000 0.00000 ; H- N- CA- HA
3028 3027 3029 3031 3 0.00000 0.00000 0.00000
0.00000 0.00000 0.00000 ; H- N- CA- CB
3028 3027 3029 3046 3 0.00000 0.00000 0.00000
0.00000 0.00000 0.00000 ; H- N- CA- C
3029 3031 3034 3035 3 0.57739 0.00000 1.02926
0.00000 -1.60666 0.00000 ; CA- CB- CG- CD1
3029 3031 3034 3044 3 0.57739 0.00000 1.02926
0.00000 -1.60666 0.00000 ; CA- CB- CG- CD2
3029 3046 3048 3049 3 20.92000 0.00000 -20.92000
0.00000 0.00000 0.00000 ; CA- C- N- H
3029 3046 3048 3050 3 20.92000 0.00000 -20.92000
0.00000 0.00000 0.00000 ; CA- C- N- CA
3030 3029 3031 3032 3 0.65084 1.95253 0.00000
-2.60338 0.00000 0.00000 ; HA- CA- CB- HB2
3030 3029 3031 3033 3 0.65084 1.95253 0.00000
-2.60338 0.00000 0.00000 ; HA- CA- CB- HB3
3030 3029 3031 3034 3 0.65084 1.95253 0.00000
-2.60338 0.00000 0.00000 ; HA- CA- CB- CG
3030 3029 3046 3047 3 3.68192 -4.35136 0.00000
1.33888 0.00000 0.00000 ; HA- CA- C- O
3030 3029 3046 3048 3 0.00000 0.00000 0.00000
0.00000 0.00000 0.00000 ; HA- CA- C- N

3031 3029 3046 3047 3 0.00000 0.00000 0.00000
0.00000 0.00000 0.00000 ; CB- CA- C- O
3031 3029 3046 3048 3 2.51040 4.18400 1.67360
-6.69440 0.00000 0.00000 ; CB- CA- C- N
3031 3034 3035 3036 3 30.33400 0.00000 -30.33400
0.00000 0.00000 0.00000 ; CB- CG- CD1- HD1
3031 3034 3035 3037 3 30.33400 0.00000 -30.33400
0.00000 0.00000 0.00000 ; CB- CG- CD1- CE1
3031 3034 3044 3042 3 30.33400 0.00000 -30.33400
0.00000 0.00000 0.00000 ; CB- CG- CD2- CE2
3031 3034 3044 3045 3 30.33400 0.00000 -30.33400
0.00000 0.00000 0.00000 ; CB- CG- CD2- HD2
3032 3031 3029 3046 3 0.65084 1.95253 0.00000
-2.60338 0.00000 0.00000 ; HB2- CB- CA- C
3032 3031 3034 3035 3 0.00000 0.00000 0.00000
0.00000 0.00000 0.00000 ; HB2- CB- CG- CD1
3032 3031 3034 3044 3 0.00000 0.00000 0.00000
0.00000 0.00000 0.00000 ; HB2- CB- CG- CD2
3033 3031 3029 3046 3 0.65084 1.95253 0.00000
-2.60338 0.00000 0.00000 ; HB3- CB- CA- C
3033 3031 3034 3035 3 0.00000 0.00000 0.00000
0.00000 0.00000 0.00000 ; HB3- CB- CG- CD1
3033 3031 3034 3044 3 0.00000 0.00000 0.00000
0.00000 0.00000 0.00000 ; HB3- CB- CG- CD2
3034 3031 3029 3046 3 4.95804 2.17986 -3.52293
-3.21331 -0.40166 0.00000 ; CG- CB- CA- C
3034 3035 3037 3038 3 30.33400 0.00000 -30.33400
0.00000 0.00000 0.00000 ; CG- CD1- CE1- HE1
3034 3035 3037 3039 3 30.33400 0.00000 -30.33400
0.00000 0.00000 0.00000 ; CG- CD1- CE1- CZ
3034 3044 3042 3039 3 30.33400 0.00000 -30.33400
0.00000 0.00000 0.00000 ; CG- CD2- CE2- CZ
3034 3044 3042 3043 3 30.33400 0.00000 -30.33400
0.00000 0.00000 0.00000 ; CG- CD2- CE2- HE2
3035 3034 3044 3042 3 30.33400 0.00000 -30.33400
0.00000 0.00000 0.00000 ; CD1- CG- CD2- CE2
3035 3034 3044 3045 3 30.33400 0.00000 -30.33400
0.00000 0.00000 0.00000 ; CD1- CG- CD2- HD2
3035 3037 3039 3040 3 30.33400 0.00000 -30.33400
0.00000 0.00000 0.00000 ; CD1- CE1- CZ- OH
3035 3037 3039 3042 3 30.33400 0.00000 -30.33400
0.00000 0.00000 0.00000 ; CD1- CE1- CZ- CE2
3036 3035 3034 3044 3 30.33400 0.00000 -30.33400
0.00000 0.00000 0.00000 ; HD1- CD1- CG- CD2
3036 3035 3037 3038 3 30.33400 0.00000 -30.33400
0.00000 0.00000 0.00000 ; HD1- CD1- CE1- HE1
3036 3035 3037 3039 3 30.33400 0.00000 -30.33400
0.00000 0.00000 0.00000 ; HD1- CD1- CE1- CZ
3037 3035 3034 3044 3 30.33400 0.00000 -30.33400
0.00000 0.00000 0.00000 ; CE1- CD1- CG- CD2
3037 3039 3040 3041 3 7.93286 0.00000 -9.56462
0.00000 2.17568 0.00000 ; CE1- CZ- OH- HH

3037 3039 3042 3043 3 30.33400 0.00000 -30.33400
0.00000 0.00000 0.00000 ; CE1- CZ- CE2- HE2
3037 3039 3042 3044 3 30.33400 0.00000 -30.33400
0.00000 0.00000 0.00000 ; CE1- CZ- CE2- CD2
3038 3037 3039 3040 3 30.33400 0.00000 -30.33400
0.00000 0.00000 0.00000 ; HE1- CE1- CZ- OH
3038 3037 3039 3042 3 30.33400 0.00000 -30.33400
0.00000 0.00000 0.00000 ; HE1- CE1- CZ- CE2
3039 3042 3044 3045 3 30.33400 0.00000 -30.33400
0.00000 0.00000 0.00000 ; CZ- CE2- CD2- HD2
3040 3039 3042 3043 3 30.33400 0.00000 -30.33400
0.00000 0.00000 0.00000 ; OH- CZ- CE2- HE2
3040 3039 3042 3044 3 30.33400 0.00000 -30.33400
0.00000 0.00000 0.00000 ; OH- CZ- CE2- CD2
3041 3040 3039 3042 3 7.93286 0.00000 -9.56462
0.00000 2.17568 0.00000 ; HH- OH- CZ- CE2
3043 3042 3044 3045 3 30.33400 0.00000 -30.33400
0.00000 0.00000 0.00000 ; HE2- CE2- CD2- HD2
3046 3048 3050 3051 3 0.00000 0.00000 0.00000
0.00000 0.00000 0.00000 ; C- N- CA- HA
3046 3048 3050 3052 3 11.71520 1.67360 15.06240 -
13.38880 0.00000 0.00000 ; C- N- CA- CB
3046 3048 3050 3058 3 1.75728 5.27184 2.25936
-7.02912 0.00000 0.00000 ; C- N- CA- C
3047 3046 3048 3049 3 29.28800 -8.36800 -20.92000
0.00000 0.00000 0.00000 ; O- C- N- H
3047 3046 3048 3050 3 20.92000 0.00000 -20.92000
0.00000 0.00000 0.00000 ; O- C- N- CA
3048 3050 3052 3053 3 0.65084 1.95253 0.00000
-2.60338 0.00000 0.00000 ; N- CA- CB- HB2
3048 3050 3052 3054 3 0.65084 1.95253 0.00000
-2.60338 0.00000 0.00000 ; N- CA- CB- HB3
3048 3050 3052 3055 3 15.41386 9.74035 -8.39310
-0.97069 2.97901 0.00000 ; N- CA- CB- CG
3048 3050 3058 3059 3 0.00000 0.00000 0.00000
0.00000 0.00000 0.00000 ; N- CA- C- O
3048 3050 3058 3060 3 17.40544 -5.02080 -13.22144
9.20480 0.00000 0.00000 ; N- CA- C- N
3049 3048 3050 3051 3 0.00000 0.00000 0.00000
0.00000 0.00000 0.00000 ; H- N- CA- HA
3049 3048 3050 3052 3 0.00000 0.00000 0.00000
0.00000 0.00000 0.00000 ; H- N- CA- CB
3049 3048 3050 3058 3 0.00000 0.00000 0.00000
0.00000 0.00000 0.00000 ; H- N- CA- C
3050 3052 3055 3056 3 6.43499 0.00000 -5.39736
0.00000 -1.03763 0.00000 ; CA- CB- CG- OD1
3050 3052 3055 3057 3 6.43499 0.00000 -5.39736
0.00000 -1.03763 0.00000 ; CA- CB- CG- OD2
3050 3058 3060 3061 3 20.92000 0.00000 -20.92000
0.00000 0.00000 0.00000 ; CA- C- N- H
3050 3058 3060 3062 3 20.92000 0.00000 -20.92000
0.00000 0.00000 0.00000 ; CA- C- N- CA

3051 3050 3052 3053 3 0.65084 1.95253 0.00000
-2.60338 0.00000 0.00000 ; HA- CA- CB- HB2
3051 3050 3052 3054 3 0.65084 1.95253 0.00000
-2.60338 0.00000 0.00000 ; HA- CA- CB- HB3
3051 3050 3052 3055 3 0.65084 1.95253 0.00000
-2.60338 0.00000 0.00000 ; HA- CA- CB- CG
3051 3050 3058 3059 3 3.68192 -4.35136 0.00000
1.33888 0.00000 0.00000 ; HA- CA- C- O
3051 3050 3058 3060 3 0.00000 0.00000 0.00000
0.00000 0.00000 0.00000 ; HA- CA- C- N
3052 3050 3058 3059 3 0.00000 0.00000 0.00000
0.00000 0.00000 0.00000 ; CB- CA- C- O
3052 3050 3058 3060 3 2.51040 4.18400 1.67360
-6.69440 0.00000 0.00000 ; CB- CA- C- N
3053 3052 3050 3058 3 0.65084 1.95253 0.00000
-2.60338 0.00000 0.00000 ; HB2- CB- CA- C
3053 3052 3055 3056 3 0.00000 0.00000 0.00000
0.00000 0.00000 0.00000 ; HB2- CB- CG- OD1
3053 3052 3055 3057 3 0.00000 0.00000 0.00000
0.00000 0.00000 0.00000 ; HB2- CB- CG- OD2
3054 3052 3050 3058 3 0.65084 1.95253 0.00000
-2.60338 0.00000 0.00000 ; HB3- CB- CA- C
3054 3052 3055 3056 3 0.00000 0.00000 0.00000
0.00000 0.00000 0.00000 ; HB3- CB- CG- OD1
3054 3052 3055 3057 3 0.00000 0.00000 0.00000
0.00000 0.00000 0.00000 ; HB3- CB- CG- OD2
3055 3052 3050 3058 3 7.14627 -1.04600 -8.99560
-0.97069 5.15469 0.00000 ; CG- CB- CA- C
3058 3060 3062 3063 3 0.00000 0.00000 0.00000
0.00000 0.00000 0.00000 ; C- N- CA- HA
3058 3060 3062 3064 3 11.71520 1.67360 15.06240 -
13.38880 0.00000 0.00000 ; C- N- CA- CB
3058 3060 3062 3080 3 1.75728 5.27184 2.25936
-7.02912 0.00000 0.00000 ; C- N- CA- C
3059 3058 3060 3061 3 29.28800 -8.36800 -20.92000
0.00000 0.00000 0.00000 ; O- C- N- H
3059 3058 3060 3062 3 20.92000 0.00000 -20.92000
0.00000 0.00000 0.00000 ; O- C- N- CA
3060 3062 3064 3065 3 0.65084 1.95253 0.00000
-2.60338 0.00000 0.00000 ; N- CA- CB- HB2
3060 3062 3064 3066 3 0.65084 1.95253 0.00000
-2.60338 0.00000 0.00000 ; N- CA- CB- HB3
3060 3062 3064 3067 3 0.65084 1.95253 0.00000
-2.60338 0.00000 0.00000 ; N- CA- CB- CG
3060 3062 3080 3081 3 0.00000 0.00000 0.00000
0.00000 0.00000 0.00000 ; N- CA- C- O
3060 3062 3080 3082 3 17.40544 -5.02080 -13.22144
9.20480 0.00000 0.00000 ; N- CA- C- N
3061 3060 3062 3063 3 0.00000 0.00000 0.00000
0.00000 0.00000 0.00000 ; H- N- CA- HA
3061 3060 3062 3064 3 0.00000 0.00000 0.00000
0.00000 0.00000 0.00000 ; H- N- CA- CB

3061 3060 3062 3080 3 0.00000 0.00000 0.00000
0.00000 0.00000 0.00000 ; H- N- CA- C
3062 3064 3067 3068 3 0.66944 2.00832 0.00000
-2.67776 0.00000 0.00000 ; CA- CB- CG- HG2
3062 3064 3067 3069 3 0.66944 2.00832 0.00000
-2.67776 0.00000 0.00000 ; CA- CB- CG- HG3
3062 3064 3067 3070 3 3.68192 3.09616 -2.09200
-3.01248 0.00000 0.00000 ; CA- CB- CG- CD
3062 3080 3082 3083 3 20.92000 0.00000 -20.92000
0.00000 0.00000 0.00000 ; CA- C- N- H
3062 3080 3082 3084 3 20.92000 0.00000 -20.92000
0.00000 0.00000 0.00000 ; CA- C- N- CA
3063 3062 3064 3065 3 0.65084 1.95253 0.00000
-2.60338 0.00000 0.00000 ; HA- CA- CB- HB2
3063 3062 3064 3066 3 0.65084 1.95253 0.00000
-2.60338 0.00000 0.00000 ; HA- CA- CB- HB3
3063 3062 3064 3067 3 0.65084 1.95253 0.00000
-2.60338 0.00000 0.00000 ; HA- CA- CB- CG
3063 3062 3080 3081 3 3.68192 -4.35136 0.00000
1.33888 0.00000 0.00000 ; HA- CA- C- O
3063 3062 3080 3082 3 0.00000 0.00000 0.00000
0.00000 0.00000 0.00000 ; HA- CA- C- N
3064 3062 3080 3081 3 0.00000 0.00000 0.00000
0.00000 0.00000 0.00000 ; CB- CA- C- O
3064 3062 3080 3082 3 2.51040 4.18400 1.67360
-6.69440 0.00000 0.00000 ; CB- CA- C- N
3064 3067 3070 3071 3 0.66944 2.00832 0.00000
-2.67776 0.00000 0.00000 ; CB- CG- CD- HD2
3064 3067 3070 3072 3 0.66944 2.00832 0.00000
-2.67776 0.00000 0.00000 ; CB- CG- CD- HD3
3064 3067 3070 3073 3 3.68192 3.09616 -2.09200
-3.01248 0.00000 0.00000 ; CB- CG- CD- CE
3065 3064 3062 3080 3 0.65084 1.95253 0.00000
-2.60338 0.00000 0.00000 ; HB2- CB- CA- C
3065 3064 3067 3068 3 0.62760 1.88280 0.00000
-2.51040 0.00000 0.00000 ; HB2- CB- CG- HG2
3065 3064 3067 3069 3 0.62760 1.88280 0.00000
-2.51040 0.00000 0.00000 ; HB2- CB- CG- HG3
3065 3064 3067 3070 3 0.66944 2.00832 0.00000
-2.67776 0.00000 0.00000 ; HB2- CB- CG- CD
3066 3064 3062 3080 3 0.65084 1.95253 0.00000
-2.60338 0.00000 0.00000 ; HB3- CB- CA- C
3066 3064 3067 3068 3 0.62760 1.88280 0.00000
-2.51040 0.00000 0.00000 ; HB3- CB- CG- HG2
3066 3064 3067 3069 3 0.62760 1.88280 0.00000
-2.51040 0.00000 0.00000 ; HB3- CB- CG- HG3
3066 3064 3067 3070 3 0.66944 2.00832 0.00000
-2.67776 0.00000 0.00000 ; HB3- CB- CG- CD
3067 3064 3062 3080 3 0.65084 1.95253 0.00000
-2.60338 0.00000 0.00000 ; CG- CB- CA- C
3067 3070 3073 3074 3 0.65084 1.95253 0.00000
-2.60338 0.00000 0.00000 ; CG- CD- CE- HE2

3067 3070 3073 3075 3 0.65084 1.95253 0.00000
-2.60338 0.00000 0.00000 ; CG- CD- CE- HE3
3067 3070 3073 3076 3 0.65084 1.95253 0.00000
-2.60338 0.00000 0.00000 ; CG- CD- CE- NZ
3068 3067 3070 3071 3 0.62760 1.88280 0.00000
-2.51040 0.00000 0.00000 ; HG2- CG- CD- HD2
3068 3067 3070 3072 3 0.62760 1.88280 0.00000
-2.51040 0.00000 0.00000 ; HG2- CG- CD- HD3
3068 3067 3070 3073 3 0.66944 2.00832 0.00000
-2.67776 0.00000 0.00000 ; HG2- CG- CD- CE
3069 3067 3070 3071 3 0.62760 1.88280 0.00000
-2.51040 0.00000 0.00000 ; HG3- CG- CD- HD2
3069 3067 3070 3072 3 0.62760 1.88280 0.00000
-2.51040 0.00000 0.00000 ; HG3- CG- CD- HD3
3069 3067 3070 3073 3 0.66944 2.00832 0.00000
-2.67776 0.00000 0.00000 ; HG3- CG- CD- CE
3070 3073 3076 3077 3 0.65084 1.95253 0.00000
-2.60338 0.00000 0.00000 ; CD- CE- NZ- HZ1
3070 3073 3076 3078 3 0.65084 1.95253 0.00000
-2.60338 0.00000 0.00000 ; CD- CE- NZ- HZ2
3070 3073 3076 3079 3 0.65084 1.95253 0.00000
-2.60338 0.00000 0.00000 ; CD- CE- NZ- HZ3
3071 3070 3073 3074 3 0.65084 1.95253 0.00000
-2.60338 0.00000 0.00000 ; HD2- CD- CE- HE2
3071 3070 3073 3075 3 0.65084 1.95253 0.00000
-2.60338 0.00000 0.00000 ; HD2- CD- CE- HE3
3071 3070 3073 3076 3 0.65084 1.95253 0.00000
-2.60338 0.00000 0.00000 ; HD2- CD- CE- NZ
3072 3070 3073 3074 3 0.65084 1.95253 0.00000
-2.60338 0.00000 0.00000 ; HD3- CD- CE- HE2
3072 3070 3073 3075 3 0.65084 1.95253 0.00000
-2.60338 0.00000 0.00000 ; HD3- CD- CE- HE3
3072 3070 3073 3076 3 0.65084 1.95253 0.00000
-2.60338 0.00000 0.00000 ; HD3- CD- CE- NZ
3074 3073 3076 3077 3 0.65084 1.95253 0.00000
-2.60338 0.00000 0.00000 ; HE2- CE- NZ- HZ1
3074 3073 3076 3078 3 0.65084 1.95253 0.00000
-2.60338 0.00000 0.00000 ; HE2- CE- NZ- HZ2
3074 3073 3076 3079 3 0.65084 1.95253 0.00000
-2.60338 0.00000 0.00000 ; HE2- CE- NZ- HZ3
3075 3073 3076 3077 3 0.65084 1.95253 0.00000
-2.60338 0.00000 0.00000 ; HE3- CE- NZ- HZ1
3075 3073 3076 3078 3 0.65084 1.95253 0.00000
-2.60338 0.00000 0.00000 ; HE3- CE- NZ- HZ2
3075 3073 3076 3079 3 0.65084 1.95253 0.00000
-2.60338 0.00000 0.00000 ; HE3- CE- NZ- HZ3
3080 3082 3084 3085 3 0.00000 0.00000 0.00000
0.00000 0.00000 0.00000 ; C- N- CA- HA
3080 3082 3084 3086 3 11.71520 1.67360 15.06240 -
13.38880 0.00000 0.00000 ; C- N- CA- CB
3080 3082 3084 3102 3 1.75728 5.27184 2.25936
-7.02912 0.00000 0.00000 ; C- N- CA- C

```
  3081   3080   3082   3083      3   29.28800   -8.36800  -20.92000
0.00000   0.00000   0.00000 ;      O-    C-     N-     H
  3081   3080   3082   3084      3   20.92000    0.00000  -20.92000
0.00000   0.00000   0.00000 ;      O-    C-     N-    CA
  3082   3084   3086   3087      3    0.65084    1.95253    0.00000
-2.60338    0.00000    0.00000 ;     N-    CA-    CB-   HB2
  3082   3084   3086   3088      3    0.65084    1.95253    0.00000
-2.60338    0.00000    0.00000 ;     N-    CA-    CB-   HB3
  3082   3084   3086   3089      3    0.65084    1.95253    0.00000
-2.60338    0.00000    0.00000 ;     N-    CA-    CB-    CG
  3082   3084   3102   3103      3    0.00000    0.00000    0.00000
0.00000   0.00000   0.00000 ;      N-   CA-     C-     O
  3082   3084   3102   3104      3   17.40544   -5.02080  -13.22144
9.20480   0.00000   0.00000 ;      N-    CA-     C-     N
  3083   3082   3084   3085      3    0.00000    0.00000    0.00000
0.00000   0.00000   0.00000 ;      H-     N-    CA-    HA
  3083   3082   3084   3086      3    0.00000    0.00000    0.00000
0.00000   0.00000   0.00000 ;      H-     N-    CA-    CB
  3083   3082   3084   3102      3    0.00000    0.00000    0.00000
0.00000   0.00000   0.00000 ;      H-     N-    CA-     C
  3084   3086   3089   3090      3    0.66944    2.00832    0.00000
-2.67776    0.00000    0.00000 ;     CA-    CB-    CG-   HG2
  3084   3086   3089   3091      3    0.66944    2.00832    0.00000
-2.67776    0.00000    0.00000 ;     CA-    CB-    CG-   HG3
  3084   3086   3089   3092      3    3.68192    3.09616   -2.09200
-3.01248    0.00000    0.00000 ;     CA-    CB-    CG-    CD
  3084   3102   3104   3105      3   20.92000    0.00000  -20.92000
0.00000   0.00000   0.00000 ;     CA-     C-     N-     H
  3084   3102   3104   3106      3   20.92000    0.00000  -20.92000
0.00000   0.00000   0.00000 ;     CA-     C-     N-    CA
  3085   3084   3086   3087      3    0.65084    1.95253    0.00000
-2.60338    0.00000    0.00000 ;     HA-    CA-    CB-   HB2
  3085   3084   3086   3088      3    0.65084    1.95253    0.00000
-2.60338    0.00000    0.00000 ;     HA-    CA-    CB-   HB3
  3085   3084   3086   3089      3    0.65084    1.95253    0.00000
-2.60338    0.00000    0.00000 ;     HA-    CA-    CB-    CG
  3085   3084   3102   3103      3    3.68192   -4.35136    0.00000
1.33888   0.00000   0.00000 ;     HA-    CA-     C-     O
  3085   3084   3102   3104      3    0.00000    0.00000    0.00000
0.00000   0.00000   0.00000 ;     HA-    CA-     C-     N
  3086   3084   3102   3103      3    0.00000    0.00000    0.00000
0.00000   0.00000   0.00000 ;     CB-    CA-     C-     O
  3086   3084   3102   3104      3    2.51040    4.18400    1.67360
-6.69440    0.00000    0.00000 ;     CB-    CA-     C-     N
  3086   3089   3092   3093      3    0.66944    2.00832    0.00000
-2.67776    0.00000    0.00000 ;     CB-    CG-    CD-   HD2
  3086   3089   3092   3094      3    0.66944    2.00832    0.00000
-2.67776    0.00000    0.00000 ;     CB-    CG-    CD-   HD3
  3086   3089   3092   3095      3    3.68192    3.09616   -2.09200
-3.01248    0.00000    0.00000 ;     CB-    CG-    CD-    CE
  3087   3086   3084   3102      3    0.65084    1.95253    0.00000
-2.60338    0.00000    0.00000 ;    HB2-    CB-    CA-     C
```

3087 3086 3089 3090 3 0.62760 1.88280 0.00000
-2.51040 0.00000 0.00000 ; HB2- CB- CG- HG2
3087 3086 3089 3091 3 0.62760 1.88280 0.00000
-2.51040 0.00000 0.00000 ; HB2- CB- CG- HG3
3087 3086 3089 3092 3 0.66944 2.00832 0.00000
-2.67776 0.00000 0.00000 ; HB2- CB- CG- CD
3088 3086 3084 3102 3 0.65084 1.95253 0.00000
-2.60338 0.00000 0.00000 ; HB3- CB- CA- C
3088 3086 3089 3090 3 0.62760 1.88280 0.00000
-2.51040 0.00000 0.00000 ; HB3- CB- CG- HG2
3088 3086 3089 3091 3 0.62760 1.88280 0.00000
-2.51040 0.00000 0.00000 ; HB3- CB- CG- HG3
3088 3086 3089 3092 3 0.66944 2.00832 0.00000
-2.67776 0.00000 0.00000 ; HB3- CB- CG- CD
3089 3086 3084 3102 3 0.65084 1.95253 0.00000
-2.60338 0.00000 0.00000 ; CG- CB- CA- C
3089 3092 3095 3096 3 0.65084 1.95253 0.00000
-2.60338 0.00000 0.00000 ; CG- CD- CE- HE2
3089 3092 3095 3097 3 0.65084 1.95253 0.00000
-2.60338 0.00000 0.00000 ; CG- CD- CE- HE3
3089 3092 3095 3098 3 0.65084 1.95253 0.00000
-2.60338 0.00000 0.00000 ; CG- CD- CE- NZ
3090 3089 3092 3093 3 0.62760 1.88280 0.00000
-2.51040 0.00000 0.00000 ; HG2- CG- CD- HD2
3090 3089 3092 3094 3 0.62760 1.88280 0.00000
-2.51040 0.00000 0.00000 ; HG2- CG- CD- HD3
3090 3089 3092 3095 3 0.66944 2.00832 0.00000
-2.67776 0.00000 0.00000 ; HG2- CG- CD- CE
3091 3089 3092 3093 3 0.62760 1.88280 0.00000
-2.51040 0.00000 0.00000 ; HG3- CG- CD- HD2
3091 3089 3092 3094 3 0.62760 1.88280 0.00000
-2.51040 0.00000 0.00000 ; HG3- CG- CD- HD3
3091 3089 3092 3095 3 0.66944 2.00832 0.00000
-2.67776 0.00000 0.00000 ; HG3- CG- CD- CE
3092 3095 3098 3099 3 0.65084 1.95253 0.00000
-2.60338 0.00000 0.00000 ; CD- CE- NZ- HZ1
3092 3095 3098 3100 3 0.65084 1.95253 0.00000
-2.60338 0.00000 0.00000 ; CD- CE- NZ- HZ2
3092 3095 3098 3101 3 0.65084 1.95253 0.00000
-2.60338 0.00000 0.00000 ; CD- CE- NZ- HZ3
3093 3092 3095 3096 3 0.65084 1.95253 0.00000
-2.60338 0.00000 0.00000 ; HD2- CD- CE- HE2
3093 3092 3095 3097 3 0.65084 1.95253 0.00000
-2.60338 0.00000 0.00000 ; HD2- CD- CE- HE3
3093 3092 3095 3098 3 0.65084 1.95253 0.00000
-2.60338 0.00000 0.00000 ; HD2- CD- CE- NZ
3094 3092 3095 3096 3 0.65084 1.95253 0.00000
-2.60338 0.00000 0.00000 ; HD3- CD- CE- HE2
3094 3092 3095 3097 3 0.65084 1.95253 0.00000
-2.60338 0.00000 0.00000 ; HD3- CD- CE- HE3
3094 3092 3095 3098 3 0.65084 1.95253 0.00000
-2.60338 0.00000 0.00000 ; HD3- CD- CE- NZ

3096 3095 3098 3099 3 0.65084 1.95253 0.00000
-2.60338 0.00000 0.00000 ; HE2- CE- NZ- HZ1
3096 3095 3098 3100 3 0.65084 1.95253 0.00000
-2.60338 0.00000 0.00000 ; HE2- CE- NZ- HZ2
3096 3095 3098 3101 3 0.65084 1.95253 0.00000
-2.60338 0.00000 0.00000 ; HE2- CE- NZ- HZ3
3097 3095 3098 3099 3 0.65084 1.95253 0.00000
-2.60338 0.00000 0.00000 ; HE3- CE- NZ- HZ1
3097 3095 3098 3100 3 0.65084 1.95253 0.00000
-2.60338 0.00000 0.00000 ; HE3- CE- NZ- HZ2
3097 3095 3098 3101 3 0.65084 1.95253 0.00000
-2.60338 0.00000 0.00000 ; HE3- CE- NZ- HZ3
3102 3104 3106 3107 3 0.00000 0.00000 0.00000
0.00000 0.00000 0.00000 ; C- N- CA- HA
3102 3104 3106 3108 3 11.71520 1.67360 15.06240 -
13.38880 0.00000 0.00000 ; C- N- CA- CB
3102 3104 3106 3113 3 1.75728 5.27184 2.25936
-7.02912 0.00000 0.00000 ; C- N- CA- C
3103 3102 3104 3105 3 29.28800 -8.36800 -20.92000
0.00000 0.00000 0.00000 ; O- C- N- H
3103 3102 3104 3106 3 20.92000 0.00000 -20.92000
0.00000 0.00000 0.00000 ; O- C- N- CA
3104 3106 3108 3109 3 0.65084 1.95253 0.00000
-2.60338 0.00000 0.00000 ; N- CA- CB- HB2
3104 3106 3108 3110 3 0.65084 1.95253 0.00000
-2.60338 0.00000 0.00000 ; N- CA- CB- HB3
3104 3106 3108 3111 3 6.03751 2.50622 -5.17142
-4.20074 1.10458 0.00000 ; N- CA- CB- SG
3104 3106 3113 3114 3 0.00000 0.00000 0.00000
0.00000 0.00000 0.00000 ; N- CA- C- O
3104 3106 3113 3115 3 17.40544 -5.02080 -13.22144
9.20480 0.00000 0.00000 ; N- CA- C- N
3105 3104 3106 3107 3 0.00000 0.00000 0.00000
0.00000 0.00000 0.00000 ; H- N- CA- HA
3105 3104 3106 3108 3 0.00000 0.00000 0.00000
0.00000 0.00000 0.00000 ; H- N- CA- CB
3105 3104 3106 3113 3 0.00000 0.00000 0.00000
0.00000 0.00000 0.00000 ; H- N- CA- C
3106 3108 3111 3112 3 1.69034 2.77818 4.11706
-4.21747 1.00416 0.00000 ; CA- CB- SG- HG
3106 3113 3115 3116 3 20.92000 0.00000 -20.92000
0.00000 0.00000 0.00000 ; CA- C- N- H
3106 3113 3115 3117 3 20.92000 0.00000 -20.92000
0.00000 0.00000 0.00000 ; CA- C- N- CA
3107 3106 3108 3109 3 0.65084 1.95253 0.00000
-2.60338 0.00000 0.00000 ; HA- CA- CB- HB2
3107 3106 3108 3110 3 0.65084 1.95253 0.00000
-2.60338 0.00000 0.00000 ; HA- CA- CB- HB3
3107 3106 3108 3111 3 0.65084 1.95253 0.00000
-2.60338 0.00000 0.00000 ; HA- CA- CB- SG
3107 3106 3113 3114 3 3.68192 -4.35136 0.00000
1.33888 0.00000 0.00000 ; HA- CA- C- O

3107 3106 3113 3115 3 0.00000 0.00000 0.00000
0.00000 0.00000 0.00000 ; HA- CA- C- N
3108 3106 3113 3114 3 0.00000 0.00000 0.00000
0.00000 0.00000 0.00000 ; CB- CA- C- O
3108 3106 3113 3115 3 2.51040 4.18400 1.67360
-6.69440 0.00000 0.00000 ; CB- CA- C- N
3109 3108 3106 3113 3 0.65084 1.95253 0.00000
-2.60338 0.00000 0.00000 ; HB2- CB- CA- C
3109 3108 3111 3112 3 1.04600 3.13800 0.00000
-4.18400 0.00000 0.00000 ; HB2- CB- SG- HG
3110 3108 3106 3113 3 0.65084 1.95253 0.00000
-2.60338 0.00000 0.00000 ; HB3- CB- CA- C
3110 3108 3111 3112 3 1.04600 3.13800 0.00000
-4.18400 0.00000 0.00000 ; HB3- CB- SG- HG
3111 3108 3106 3113 3 5.62330 4.27605 -5.33042
-4.20074 2.51040 0.00000 ; SG- CB- CA- C
3113 3115 3117 3118 3 0.00000 0.00000 0.00000
0.00000 0.00000 0.00000 ; C- N- CA- HA
3113 3115 3117 3119 3 11.71520 1.67360 15.06240 -
13.38880 0.00000 0.00000 ; C- N- CA- CB
3113 3115 3117 3125 3 1.75728 5.27184 2.25936
-7.02912 0.00000 0.00000 ; C- N- CA- C
3114 3113 3115 3116 3 29.28800 -8.36800 -20.92000
0.00000 0.00000 0.00000 ; O- C- N- H
3114 3113 3115 3117 3 20.92000 0.00000 -20.92000
0.00000 0.00000 0.00000 ; O- C- N- CA
3115 3117 3119 3120 3 0.65084 1.95253 0.00000
-2.60338 0.00000 0.00000 ; N- CA- CB- HB2
3115 3117 3119 3121 3 0.65084 1.95253 0.00000
-2.60338 0.00000 0.00000 ; N- CA- CB- HB3
3115 3117 3119 3122 3 15.41386 9.74035 -8.39310
-0.97069 2.97901 0.00000 ; N- CA- CB- CG
3115 3117 3125 3126 3 0.00000 0.00000 0.00000
0.00000 0.00000 0.00000 ; N- CA- C- O
3115 3117 3125 3127 3 17.40544 -5.02080 -13.22144
9.20480 0.00000 0.00000 ; N- CA- C- N
3116 3115 3117 3118 3 0.00000 0.00000 0.00000
0.00000 0.00000 0.00000 ; H- N- CA- HA
3116 3115 3117 3119 3 0.00000 0.00000 0.00000
0.00000 0.00000 0.00000 ; H- N- CA- CB
3116 3115 3117 3125 3 0.00000 0.00000 0.00000
0.00000 0.00000 0.00000 ; H- N- CA- C
3117 3119 3122 3123 3 6.43499 0.00000 -5.39736
0.00000 -1.03763 0.00000 ; CA- CB- CG- OD1
3117 3119 3122 3124 3 6.43499 0.00000 -5.39736
0.00000 -1.03763 0.00000 ; CA- CB- CG- OD2
3117 3125 3127 3128 3 20.92000 0.00000 -20.92000
0.00000 0.00000 0.00000 ; CA- C- N- H
3117 3125 3127 3129 3 20.92000 0.00000 -20.92000
0.00000 0.00000 0.00000 ; CA- C- N- CA
3118 3117 3119 3120 3 0.65084 1.95253 0.00000
-2.60338 0.00000 0.00000 ; HA- CA- CB- HB2

3118 3117 3119 3121 3 0.65084 1.95253 0.00000
-2.60338 0.00000 0.00000 ; HA- CA- CB- HB3
3118 3117 3119 3122 3 0.65084 1.95253 0.00000
-2.60338 0.00000 0.00000 ; HA- CA- CB- CG
3118 3117 3125 3126 3 3.68192 -4.35136 0.00000
1.33888 0.00000 0.00000 ; HA- CA- C- O
3118 3117 3125 3127 3 0.00000 0.00000 0.00000
0.00000 0.00000 0.00000 ; HA- CA- C- N
3119 3117 3125 3126 3 0.00000 0.00000 0.00000
0.00000 0.00000 0.00000 ; CB- CA- C- O
3119 3117 3125 3127 3 2.51040 4.18400 1.67360
-6.69440 0.00000 0.00000 ; CB- CA- C- N
3120 3119 3117 3125 3 0.65084 1.95253 0.00000
-2.60338 0.00000 0.00000 ; HB2- CB- CA- C
3120 3119 3122 3123 3 0.00000 0.00000 0.00000
0.00000 0.00000 0.00000 ; HB2- CB- CG- OD1
3120 3119 3122 3124 3 0.00000 0.00000 0.00000
0.00000 0.00000 0.00000 ; HB2- CB- CG- OD2
3121 3119 3117 3125 3 0.65084 1.95253 0.00000
-2.60338 0.00000 0.00000 ; HB3- CB- CA- C
3121 3119 3122 3123 3 0.00000 0.00000 0.00000
0.00000 0.00000 0.00000 ; HB3- CB- CG- OD1
3121 3119 3122 3124 3 0.00000 0.00000 0.00000
0.00000 0.00000 0.00000 ; HB3- CB- CG- OD2
3122 3119 3117 3125 3 7.14627 -1.04600 -8.99560
-0.97069 5.15469 0.00000 ; CG- CB- CA- C
3125 3127 3129 3130 3 0.00000 0.00000 0.00000
0.00000 0.00000 0.00000 ; C- N- CA- HA
3125 3127 3129 3131 3 11.71520 1.67360 15.06240 -
13.38880 0.00000 0.00000 ; C- N- CA- CB
3125 3127 3129 3144 3 1.75728 5.27184 2.25936
-7.02912 0.00000 0.00000 ; C- N- CA- C
3126 3125 3127 3128 3 29.28800 -8.36800 -20.92000
0.00000 0.00000 0.00000 ; O- C- N- H
3126 3125 3127 3129 3 20.92000 0.00000 -20.92000
0.00000 0.00000 0.00000 ; O- C- N- CA
3127 3129 3131 3132 3 0.65084 1.95253 0.00000
-2.60338 0.00000 0.00000 ; N- CA- CB- HB
3127 3129 3131 3133 3 3.83673 0.44769 -1.77402
-2.47693 -0.03347 0.00000 ; N- CA- CB- CG2
3127 3129 3131 3137 3 2.97482 0.12134 2.04179
-1.89117 -3.24678 0.00000 ; N- CA- CB- CG1
3127 3129 3144 3145 3 0.00000 0.00000 0.00000
0.00000 0.00000 0.00000 ; N- CA- C- O
3127 3129 3144 3146 3 17.40544 -5.02080 -13.22144
9.20480 0.00000 0.00000 ; N- CA- C- N
3128 3127 3129 3130 3 0.00000 0.00000 0.00000
0.00000 0.00000 0.00000 ; H- N- CA- HA
3128 3127 3129 3131 3 0.00000 0.00000 0.00000
0.00000 0.00000 0.00000 ; H- N- CA- CB
3128 3127 3129 3144 3 0.00000 0.00000 0.00000
0.00000 0.00000 0.00000 ; H- N- CA- C

```
  3129   3131   3133   3134       3    0.66944    2.00832    0.00000
-2.67776    0.00000    0.00000 ;     CA-    CB-   CG2-  HG21
  3129   3131   3133   3135       3    0.66944    2.00832    0.00000
-2.67776    0.00000    0.00000 ;     CA-    CB-   CG2-  HG22
  3129   3131   3133   3136       3    0.66944    2.00832    0.00000
-2.67776    0.00000    0.00000 ;     CA-    CB-   CG2-  HG23
  3129   3131   3137   3138       3    0.66944    2.00832    0.00000
-2.67776    0.00000    0.00000 ;     CA-    CB-   CG1-  HG12
  3129   3131   3137   3139       3    0.66944    2.00832    0.00000
-2.67776    0.00000    0.00000 ;     CA-    CB-   CG1-  HG13
  3129   3131   3137   3140       3    4.24258   -0.52718   -7.25506
-1.79075    7.69856    0.00000 ;     CA-    CB-   CG1-   CD1
  3129   3144   3146   3147       3   20.92000    0.00000  -20.92000
0.00000    0.00000    0.00000 ;     CA-     C-     N-     H
  3129   3144   3146   3148       3   20.92000    0.00000  -20.92000
0.00000    0.00000    0.00000 ;     CA-     C-     N-    CA
  3130   3129   3131   3132       3    0.65084    1.95253    0.00000
-2.60338    0.00000    0.00000 ;     HA-    CA-    CB-    HB
  3130   3129   3131   3133       3    0.65084    1.95253    0.00000
-2.60338    0.00000    0.00000 ;     HA-    CA-    CB-   CG2
  3130   3129   3131   3137       3    0.65084    1.95253    0.00000
-2.60338    0.00000    0.00000 ;     HA-    CA-    CB-   CG1
  3130   3129   3144   3145       3    3.68192   -4.35136    0.00000
1.33888    0.00000    0.00000 ;     HA-    CA-     C-     O
  3130   3129   3144   3146       3    0.00000    0.00000    0.00000
0.00000    0.00000    0.00000 ;     HA-    CA-     C-     N
  3131   3129   3144   3145       3    0.00000    0.00000    0.00000
0.00000    0.00000    0.00000 ;     CB-    CA-     C-     O
  3131   3129   3144   3146       3    2.51040    4.18400    1.67360
-6.69440    0.00000    0.00000 ;     CB-    CA-     C-     N
  3131   3137   3140   3141       3    0.66944    2.00832    0.00000
-2.67776    0.00000    0.00000 ;     CB-   CG1-   CD1-  HD11
  3131   3137   3140   3142       3    0.66944    2.00832    0.00000
-2.67776    0.00000    0.00000 ;     CB-   CG1-   CD1-  HD12
  3131   3137   3140   3143       3    0.66944    2.00832    0.00000
-2.67776    0.00000    0.00000 ;     CB-   CG1-   CD1-  HD13
  3132   3131   3129   3144       3    0.65084    1.95253    0.00000
-2.60338    0.00000    0.00000 ;     HB-    CB-    CA-     C
  3132   3131   3133   3134       3    0.62760    1.88280    0.00000
-2.51040    0.00000    0.00000 ;     HB-    CB-   CG2-  HG21
  3132   3131   3133   3135       3    0.62760    1.88280    0.00000
-2.51040    0.00000    0.00000 ;     HB-    CB-   CG2-  HG22
  3132   3131   3133   3136       3    0.62760    1.88280    0.00000
-2.51040    0.00000    0.00000 ;     HB-    CB-   CG2-  HG23
  3132   3131   3137   3138       3    0.62760    1.88280    0.00000
-2.51040    0.00000    0.00000 ;     HB-    CB-   CG1-  HG12
  3132   3131   3137   3139       3    0.62760    1.88280    0.00000
-2.51040    0.00000    0.00000 ;     HB-    CB-   CG1-  HG13
  3132   3131   3137   3140       3    0.66944    2.00832    0.00000
-2.67776    0.00000    0.00000 ;     HB-    CB-   CG1-   CD1
  3133   3131   3129   3144       3    5.67350    3.55640   -6.16722
-2.47693    3.74886    0.00000 ;    CG2-    CB-    CA-     C
```

3133 3131 3137 3138 3 0.66944 2.00832 0.00000
-2.67776 0.00000 0.00000 ; CG2- CB- CG1- HG12
3133 3131 3137 3139 3 0.66944 2.00832 0.00000
-2.67776 0.00000 0.00000 ; CG2- CB- CG1- HG13
3133 3131 3137 3140 3 3.81162 0.49790 -8.14206
-1.79075 7.49773 0.00000 ; CG2- CB- CG1- CD1
3134 3133 3131 3137 3 0.66944 2.00832 0.00000
-2.67776 0.00000 0.00000 ; HG21- CG2- CB- CG1
3135 3133 3131 3137 3 0.66944 2.00832 0.00000
-2.67776 0.00000 0.00000 ; HG22- CG2- CB- CG1
3136 3133 3131 3137 3 0.66944 2.00832 0.00000
-2.67776 0.00000 0.00000 ; HG23- CG2- CB- CG1
3137 3131 3129 3144 3 8.26340 0.74057 -9.99976
-1.89117 3.84928 0.00000 ; CG1- CB- CA- C
3138 3137 3140 3141 3 0.62760 1.88280 0.00000
-2.51040 0.00000 0.00000 ; HG12- CG1- CD1- HD11
3138 3137 3140 3142 3 0.62760 1.88280 0.00000
-2.51040 0.00000 0.00000 ; HG12- CG1- CD1- HD12
3138 3137 3140 3143 3 0.62760 1.88280 0.00000
-2.51040 0.00000 0.00000 ; HG12- CG1- CD1- HD13
3139 3137 3140 3141 3 0.62760 1.88280 0.00000
-2.51040 0.00000 0.00000 ; HG13- CG1- CD1- HD11
3139 3137 3140 3142 3 0.62760 1.88280 0.00000
-2.51040 0.00000 0.00000 ; HG13- CG1- CD1- HD12
3139 3137 3140 3143 3 0.62760 1.88280 0.00000
-2.51040 0.00000 0.00000 ; HG13- CG1- CD1- HD13
3144 3146 3148 3149 3 0.00000 0.00000 0.00000
0.00000 0.00000 0.00000 ; C- N- CA- HA
3144 3146 3148 3150 3 11.71520 1.67360 15.06240 -
13.38880 0.00000 0.00000 ; C- N- CA- CB
3144 3146 3148 3168 3 1.75728 5.27184 2.25936
-7.02912 0.00000 0.00000 ; C- N- CA- C
3145 3144 3146 3147 3 29.28800 -8.36800 -20.92000
0.00000 0.00000 0.00000 ; O- C- N- H
3145 3144 3146 3148 3 20.92000 0.00000 -20.92000
0.00000 0.00000 0.00000 ; O- C- N- CA
3146 3148 3150 3151 3 0.65084 1.95253 0.00000
-2.60338 0.00000 0.00000 ; N- CA- CB- HB2
3146 3148 3150 3152 3 0.65084 1.95253 0.00000
-2.60338 0.00000 0.00000 ; N- CA- CB- HB3
3146 3148 3150 3153 3 4.18818 2.60663 -3.65682
-3.91622 1.03763 0.00000 ; N- CA- CB- CG
3146 3148 3168 3169 3 0.00000 0.00000 0.00000
0.00000 0.00000 0.00000 ; N- CA- C- O
3146 3148 3168 3170 3 17.40544 -5.02080 -13.22144
9.20480 0.00000 0.00000 ; N- CA- C- N
3147 3146 3148 3149 3 0.00000 0.00000 0.00000
0.00000 0.00000 0.00000 ; H- N- CA- HA
3147 3146 3148 3150 3 0.00000 0.00000 0.00000
0.00000 0.00000 0.00000 ; H- N- CA- CB
3147 3146 3148 3168 3 0.00000 0.00000 0.00000
0.00000 0.00000 0.00000 ; H- N- CA- C

3148 3150 3153 3154 3 0.00000 0.00000 0.00000
0.00000 0.00000 0.00000 ; CA- CB- CG- CD1
3148 3150 3153 3167 3 4.95386 8.75293 6.59398 -
13.70678 -3.17984 0.00000 ; CA- CB- CG- CD2
3148 3168 3170 3171 3 20.92000 0.00000 -20.92000
0.00000 0.00000 0.00000 ; CA- C- N- H
3148 3168 3170 3172 3 20.92000 0.00000 -20.92000
0.00000 0.00000 0.00000 ; CA- C- N- CA
3149 3148 3150 3151 3 0.65084 1.95253 0.00000
-2.60338 0.00000 0.00000 ; HA- CA- CB- HB2
3149 3148 3150 3152 3 0.65084 1.95253 0.00000
-2.60338 0.00000 0.00000 ; HA- CA- CB- HB3
3149 3148 3150 3153 3 0.65084 1.95253 0.00000
-2.60338 0.00000 0.00000 ; HA- CA- CB- CG
3149 3148 3168 3169 3 3.68192 -4.35136 0.00000
1.33888 0.00000 0.00000 ; HA- CA- C- O
3149 3148 3168 3170 3 0.00000 0.00000 0.00000
0.00000 0.00000 0.00000 ; HA- CA- C- N
3150 3148 3168 3169 3 0.00000 0.00000 0.00000
0.00000 0.00000 0.00000 ; CB- CA- C- O
3150 3148 3168 3170 3 2.51040 4.18400 1.67360
-6.69440 0.00000 0.00000 ; CB- CA- C- N
3150 3153 3154 3155 3 54.60120 0.00000 -54.60120
0.00000 0.00000 0.00000 ; CB- CG- CD1- HD1
3150 3153 3154 3156 3 54.60120 0.00000 -54.60120
0.00000 0.00000 0.00000 ; CB- CG- CD1- NE1
3150 3153 3167 3158 3 14.01640 0.00000 -14.01640
0.00000 0.00000 0.00000 ; CB- CG- CD2- CE2
3150 3153 3167 3165 3 14.01640 0.00000 -14.01640
0.00000 0.00000 0.00000 ; CB- CG- CD2- CE3
3151 3150 3148 3168 3 0.65084 1.95253 0.00000
-2.60338 0.00000 0.00000 ; HB2- CB- CA- C
3151 3150 3153 3154 3 0.00000 0.00000 0.00000
0.00000 0.00000 0.00000 ; HB2- CB- CG- CD1
3151 3150 3153 3167 3 0.00000 0.00000 0.00000
0.00000 0.00000 0.00000 ; HB2- CB- CG- CD2
3152 3150 3148 3168 3 0.65084 1.95253 0.00000
-2.60338 0.00000 0.00000 ; HB3- CB- CA- C
3152 3150 3153 3154 3 0.00000 0.00000 0.00000
0.00000 0.00000 0.00000 ; HB3- CB- CG- CD1
3152 3150 3153 3167 3 0.00000 0.00000 0.00000
0.00000 0.00000 0.00000 ; HB3- CB- CG- CD2
3153 3150 3148 3168 3 4.62332 3.00830 -5.43083
-3.91622 2.47693 0.00000 ; CG- CB- CA- C
3153 3154 3156 3157 3 12.55200 0.00000 -12.55200
0.00000 0.00000 0.00000 ; CG- CD1- NE1- HE1
3153 3154 3156 3158 3 12.55200 0.00000 -12.55200
0.00000 0.00000 0.00000 ; CG- CD1- NE1- CE2
3153 3167 3158 3156 3 25.10400 0.00000 -25.10400
0.00000 0.00000 0.00000 ; CG- CD2- CE2- NE1
3153 3167 3158 3159 3 25.10400 0.00000 -25.10400
0.00000 0.00000 0.00000 ; CG- CD2- CE2- CZ2

```
  3153   3167   3165   3163       3   29.28800    0.00000  -29.28800
0.00000    0.00000    0.00000 ;     CG-   CD2-   CE3-   CZ3
  3153   3167   3165   3166       3   29.28800    0.00000  -29.28800
0.00000    0.00000    0.00000 ;     CG-   CD2-   CE3-   HE3
  3154   3153   3167   3158       3   14.01640    0.00000  -14.01640
0.00000    0.00000    0.00000 ;    CD1-    CG-   CD2-   CE2
  3154   3153   3167   3165       3   14.01640    0.00000  -14.01640
0.00000    0.00000    0.00000 ;    CD1-    CG-   CD2-   CE3
  3154   3156   3158   3159       3   12.76120    0.00000  -12.76120
0.00000    0.00000    0.00000 ;    CD1-   NE1-   CE2-   CZ2
  3154   3156   3158   3167       3   12.76120    0.00000  -12.76120
0.00000    0.00000    0.00000 ;    CD1-   NE1-   CE2-   CD2
  3155   3154   3153   3167       3   54.60120    0.00000  -54.60120
0.00000    0.00000    0.00000 ;    HD1-   CD1-    CG-   CD2
  3155   3154   3156   3157       3   12.55200    0.00000  -12.55200
0.00000    0.00000    0.00000 ;    HD1-   CD1-   NE1-   HE1
  3155   3154   3156   3158       3   12.55200    0.00000  -12.55200
0.00000    0.00000    0.00000 ;    HD1-   CD1-   NE1-   CE2
  3156   3154   3153   3167       3   54.60120    0.00000  -54.60120
0.00000    0.00000    0.00000 ;    NE1-   CD1-    CG-   CD2
  3156   3158   3159   3160       3   30.33400    0.00000  -30.33400
0.00000    0.00000    0.00000 ;    NE1-   CE2-   CZ2-   HZ2
  3156   3158   3159   3161       3   30.33400    0.00000  -30.33400
0.00000    0.00000    0.00000 ;    NE1-   CE2-   CZ2-   CH2
  3156   3158   3167   3165       3   25.10400    0.00000  -25.10400
0.00000    0.00000    0.00000 ;    NE1-   CE2-   CD2-   CE3
  3157   3156   3158   3159       3   12.76120    0.00000  -12.76120
0.00000    0.00000    0.00000 ;    HE1-   NE1-   CE2-   CZ2
  3157   3156   3158   3167       3   12.76120    0.00000  -12.76120
0.00000    0.00000    0.00000 ;    HE1-   NE1-   CE2-   CD2
  3158   3159   3161   3162       3   30.33400    0.00000  -30.33400
0.00000    0.00000    0.00000 ;    CE2-   CZ2-   CH2-   HH2
  3158   3159   3161   3163       3   30.33400    0.00000  -30.33400
0.00000    0.00000    0.00000 ;    CE2-   CZ2-   CH2-   CZ3
  3158   3167   3165   3163       3   29.28800    0.00000  -29.28800
0.00000    0.00000    0.00000 ;    CE2-   CD2-   CE3-   CZ3
  3158   3167   3165   3166       3   29.28800    0.00000  -29.28800
0.00000    0.00000    0.00000 ;    CE2-   CD2-   CE3-   HE3
  3159   3158   3167   3165       3   25.10400    0.00000  -25.10400
0.00000    0.00000    0.00000 ;    CZ2-   CE2-   CD2-   CE3
  3159   3161   3163   3164       3   30.33400    0.00000  -30.33400
0.00000    0.00000    0.00000 ;    CZ2-   CH2-   CZ3-   HZ3
  3159   3161   3163   3165       3   30.33400    0.00000  -30.33400
0.00000    0.00000    0.00000 ;    CZ2-   CH2-   CZ3-   CE3
  3160   3159   3158   3167       3   30.33400    0.00000  -30.33400
0.00000    0.00000    0.00000 ;    HZ2-   CZ2-   CE2-   CD2
  3160   3159   3161   3162       3   30.33400    0.00000  -30.33400
0.00000    0.00000    0.00000 ;    HZ2-   CZ2-   CH2-   HH2
  3160   3159   3161   3163       3   30.33400    0.00000  -30.33400
0.00000    0.00000    0.00000 ;    HZ2-   CZ2-   CH2-   CZ3
  3161   3159   3158   3167       3   30.33400    0.00000  -30.33400
0.00000    0.00000    0.00000 ;    CH2-   CZ2-   CE2-   CD2
```

3161 3163 3165 3166 3 30.33400 0.00000 -30.33400
0.00000 0.00000 0.00000 ; CH2- CZ3- CE3- HE3
3161 3163 3165 3167 3 30.33400 0.00000 -30.33400
0.00000 0.00000 0.00000 ; CH2- CZ3- CE3- CD2
3162 3161 3163 3164 3 30.33400 0.00000 -30.33400
0.00000 0.00000 0.00000 ; HH2- CH2- CZ3- HZ3
3162 3161 3163 3165 3 30.33400 0.00000 -30.33400
0.00000 0.00000 0.00000 ; HH2- CH2- CZ3- CE3
3164 3163 3165 3166 3 30.33400 0.00000 -30.33400
0.00000 0.00000 0.00000 ; HZ3- CZ3- CE3- HE3
3164 3163 3165 3167 3 30.33400 0.00000 -30.33400
0.00000 0.00000 0.00000 ; HZ3- CZ3- CE3- CD2
3168 3170 3172 3173 3 0.00000 0.00000 0.00000
0.00000 0.00000 0.00000 ; C- N- CA- HA
3168 3170 3172 3174 3 11.71520 1.67360 15.06240 -
13.38880 0.00000 0.00000 ; C- N- CA- CB
3168 3170 3172 3179 3 1.75728 5.27184 2.25936
-7.02912 0.00000 0.00000 ; C- N- CA- C
3169 3168 3170 3171 3 29.28800 -8.36800 -20.92000
0.00000 0.00000 0.00000 ; O- C- N- H
3169 3168 3170 3172 3 20.92000 0.00000 -20.92000
0.00000 0.00000 0.00000 ; O- C- N- CA
3170 3172 3174 3175 3 0.65084 1.95253 0.00000
-2.60338 0.00000 0.00000 ; N- CA- CB- HB2
3170 3172 3174 3176 3 0.65084 1.95253 0.00000
-2.60338 0.00000 0.00000 ; N- CA- CB- HB3
3170 3172 3174 3177 3 7.86174 2.24681 -7.41405
-6.71114 5.35552 0.00000 ; N- CA- CB- OG
3170 3172 3179 3180 3 0.00000 0.00000 0.00000
0.00000 0.00000 0.00000 ; N- CA- C- O
3170 3172 3179 3181 3 17.40544 -5.02080 -13.22144
9.20480 0.00000 0.00000 ; N- CA- C- N
3171 3170 3172 3173 3 0.00000 0.00000 0.00000
0.00000 0.00000 0.00000 ; H- N- CA- HA
3171 3170 3172 3174 3 0.00000 0.00000 0.00000
0.00000 0.00000 0.00000 ; H- N- CA- CB
3171 3170 3172 3179 3 0.00000 0.00000 0.00000
0.00000 0.00000 0.00000 ; H- N- CA- C
3172 3174 3177 3178 3 2.05853 2.46856 3.48109
-4.46851 0.23430 0.00000 ; CA- CB- OG- HG
3172 3179 3181 3182 3 20.92000 0.00000 -20.92000
0.00000 0.00000 0.00000 ; CA- C- N- H
3172 3179 3181 3183 3 20.92000 0.00000 -20.92000
0.00000 0.00000 0.00000 ; CA- C- N- CA
3173 3172 3174 3175 3 0.65084 1.95253 0.00000
-2.60338 0.00000 0.00000 ; HA- CA- CB- HB2
3173 3172 3174 3176 3 0.65084 1.95253 0.00000
-2.60338 0.00000 0.00000 ; HA- CA- CB- HB3
3173 3172 3174 3177 3 1.04600 -1.04600 0.00000
0.00000 0.00000 0.00000 ; HA- CA- CB- OG
3173 3172 3179 3180 3 3.68192 -4.35136 0.00000
1.33888 0.00000 0.00000 ; HA- CA- C- O

3173 3172 3179 3181 3 0.00000 0.00000 0.00000
0.00000 0.00000 0.00000 ; HA- CA- C- N
3174 3172 3179 3180 3 0.00000 0.00000 0.00000
0.00000 0.00000 0.00000 ; CB- CA- C- O
3174 3172 3179 3181 3 2.51040 4.18400 1.67360
-6.69440 0.00000 0.00000 ; CB- CA- C- N
3175 3174 3172 3179 3 0.65084 1.95253 0.00000
-2.60338 0.00000 0.00000 ; HB2- CB- CA- C
3175 3174 3177 3178 3 0.69733 2.09200 0.00000
-2.78933 0.00000 0.00000 ; HB2- CB- OG- HG
3176 3174 3172 3179 3 0.65084 1.95253 0.00000
-2.60338 0.00000 0.00000 ; HB3- CB- CA- C
3176 3174 3177 3178 3 0.69733 2.09200 0.00000
-2.78933 0.00000 0.00000 ; HB3- CB- OG- HG
3177 3174 3172 3179 3 7.34710 7.79898 -6.14211
-6.71114 4.31789 0.00000 ; OG- CB- CA- C
3179 3181 3183 3184 3 0.00000 0.00000 0.00000
0.00000 0.00000 0.00000 ; C- N- CA- HA
3179 3181 3183 3185 3 11.71520 1.67360 15.06240 -
13.38880 0.00000 0.00000 ; C- N- CA- CB
3179 3181 3183 3190 3 1.75728 5.27184 2.25936
-7.02912 0.00000 0.00000 ; C- N- CA- C
3180 3179 3181 3182 3 29.28800 -8.36800 -20.92000
0.00000 0.00000 0.00000 ; O- C- N- H
3180 3179 3181 3183 3 20.92000 0.00000 -20.92000
0.00000 0.00000 0.00000 ; O- C- N- CA
3181 3183 3185 3186 3 0.65084 1.95253 0.00000
-2.60338 0.00000 0.00000 ; N- CA- CB- HB2
3181 3183 3185 3187 3 0.65084 1.95253 0.00000
-2.60338 0.00000 0.00000 ; N- CA- CB- HB3
3181 3183 3185 3188 3 7.86174 2.24681 -7.41405
-6.71114 5.35552 0.00000 ; N- CA- CB- OG
3181 3183 3190 3191 3 0.00000 0.00000 0.00000
0.00000 0.00000 0.00000 ; N- CA- C- O
3181 3183 3190 3192 3 17.40544 -5.02080 -13.22144
9.20480 0.00000 0.00000 ; N- CA- C- N
3182 3181 3183 3184 3 0.00000 0.00000 0.00000
0.00000 0.00000 0.00000 ; H- N- CA- HA
3182 3181 3183 3185 3 0.00000 0.00000 0.00000
0.00000 0.00000 0.00000 ; H- N- CA- CB
3182 3181 3183 3190 3 0.00000 0.00000 0.00000
0.00000 0.00000 0.00000 ; H- N- CA- C
3183 3185 3188 3189 3 2.05853 2.46856 3.48109
-4.46851 0.23430 0.00000 ; CA- CB- OG- HG
3183 3190 3192 3193 3 20.92000 0.00000 -20.92000
0.00000 0.00000 0.00000 ; CA- C- N- H
3183 3190 3192 3194 3 20.92000 0.00000 -20.92000
0.00000 0.00000 0.00000 ; CA- C- N- CA
3184 3183 3185 3186 3 0.65084 1.95253 0.00000
-2.60338 0.00000 0.00000 ; HA- CA- CB- HB2
3184 3183 3185 3187 3 0.65084 1.95253 0.00000
-2.60338 0.00000 0.00000 ; HA- CA- CB- HB3

3184 3183 3185 3188 3 1.04600 -1.04600 0.00000
0.00000 0.00000 0.00000 ; HA- CA- CB- OG
3184 3183 3190 3191 3 3.68192 -4.35136 0.00000
1.33888 0.00000 0.00000 ; HA- CA- C- O
3184 3183 3190 3192 3 0.00000 0.00000 0.00000
0.00000 0.00000 0.00000 ; HA- CA- C- N
3185 3183 3190 3191 3 0.00000 0.00000 0.00000
0.00000 0.00000 0.00000 ; CB- CA- C- O
3185 3183 3190 3192 3 2.51040 4.18400 1.67360
-6.69440 0.00000 0.00000 ; CB- CA- C- N
3186 3185 3183 3190 3 0.65084 1.95253 0.00000
-2.60338 0.00000 0.00000 ; HB2- CB- CA- C
3186 3185 3188 3189 3 0.69733 2.09200 0.00000
-2.78933 0.00000 0.00000 ; HB2- CB- OG- HG
3187 3185 3183 3190 3 0.65084 1.95253 0.00000
-2.60338 0.00000 0.00000 ; HB3- CB- CA- C
3187 3185 3188 3189 3 0.69733 2.09200 0.00000
-2.78933 0.00000 0.00000 ; HB3- CB- OG- HG
3188 3185 3183 3190 3 7.34710 7.79898 -6.14211
-6.71114 4.31789 0.00000 ; OG- CB- CA- C
3190 3192 3194 3195 3 0.00000 0.00000 0.00000
0.00000 0.00000 0.00000 ; C- N- CA- HA2
3190 3192 3194 3196 3 0.00000 0.00000 0.00000
0.00000 0.00000 0.00000 ; C- N- CA- HA3
3190 3192 3194 3197 3 1.75728 5.27184 2.25936
-7.02912 0.00000 0.00000 ; C- N- CA- C
3191 3190 3192 3193 3 29.28800 -8.36800 -20.92000
0.00000 0.00000 0.00000 ; O- C- N- H
3191 3190 3192 3194 3 20.92000 0.00000 -20.92000
0.00000 0.00000 0.00000 ; O- C- N- CA
3192 3194 3197 3198 3 0.00000 0.00000 0.00000
0.00000 0.00000 0.00000 ; N- CA- C- O
3192 3194 3197 3199 3 17.40544 -5.02080 -13.22144
9.20480 0.00000 0.00000 ; N- CA- C- N
3193 3192 3194 3195 3 0.00000 0.00000 0.00000
0.00000 0.00000 0.00000 ; H- N- CA- HA2
3193 3192 3194 3196 3 0.00000 0.00000 0.00000
0.00000 0.00000 0.00000 ; H- N- CA- HA3
3193 3192 3194 3197 3 0.00000 0.00000 0.00000
0.00000 0.00000 0.00000 ; H- N- CA- C
3194 3197 3199 3200 3 20.92000 0.00000 -20.92000
0.00000 0.00000 0.00000 ; CA- C- N- H
3194 3197 3199 3201 3 20.92000 0.00000 -20.92000
0.00000 0.00000 0.00000 ; CA- C- N- CA
3195 3194 3197 3198 3 3.68192 -4.35136 0.00000
1.33888 0.00000 0.00000 ; HA2- CA- C- O
3195 3194 3197 3199 3 0.00000 0.00000 0.00000
0.00000 0.00000 0.00000 ; HA2- CA- C- N
3196 3194 3197 3198 3 3.68192 -4.35136 0.00000
1.33888 0.00000 0.00000 ; HA3- CA- C- O
3196 3194 3197 3199 3 0.00000 0.00000 0.00000
0.00000 0.00000 0.00000 ; HA3- CA- C- N

3197 3199 3201 3202 3 0.00000 0.00000 0.00000
0.00000 0.00000 0.00000 ; C- N- CA- HA
3197 3199 3201 3203 3 11.71520 1.67360 15.06240 -
13.38880 0.00000 0.00000 ; C- N- CA- CB
3197 3199 3201 3213 3 1.75728 5.27184 2.25936
-7.02912 0.00000 0.00000 ; C- N- CA- C
3198 3197 3199 3200 3 29.28800 -8.36800 -20.92000
0.00000 0.00000 0.00000 ; O- C- N- H
3198 3197 3199 3201 3 20.92000 0.00000 -20.92000
0.00000 0.00000 0.00000 ; O- C- N- CA
3199 3201 3203 3204 3 0.65084 1.95253 0.00000
-2.60338 0.00000 0.00000 ; N- CA- CB- HB
3199 3201 3203 3205 3 3.83673 0.44769 -1.77402
-2.47693 -0.03347 0.00000 ; N- CA- CB- CG1
3199 3201 3203 3209 3 3.83673 0.44769 -1.77402
-2.47693 -0.03347 0.00000 ; N- CA- CB- CG2
3199 3201 3213 3214 3 0.00000 0.00000 0.00000
0.00000 0.00000 0.00000 ; N- CA- C- O
3199 3201 3213 3215 3 17.40544 -5.02080 -13.22144
9.20480 0.00000 0.00000 ; N- CA- C- N
3200 3199 3201 3202 3 0.00000 0.00000 0.00000
0.00000 0.00000 0.00000 ; H- N- CA- HA
3200 3199 3201 3203 3 0.00000 0.00000 0.00000
0.00000 0.00000 0.00000 ; H- N- CA- CB
3200 3199 3201 3213 3 0.00000 0.00000 0.00000
0.00000 0.00000 0.00000 ; H- N- CA- C
3201 3203 3205 3206 3 0.66944 2.00832 0.00000
-2.67776 0.00000 0.00000 ; CA- CB- CG1- HG11
3201 3203 3205 3207 3 0.66944 2.00832 0.00000
-2.67776 0.00000 0.00000 ; CA- CB- CG1- HG12
3201 3203 3205 3208 3 0.66944 2.00832 0.00000
-2.67776 0.00000 0.00000 ; CA- CB- CG1- HG13
3201 3203 3209 3210 3 0.66944 2.00832 0.00000
-2.67776 0.00000 0.00000 ; CA- CB- CG2- HG21
3201 3203 3209 3211 3 0.66944 2.00832 0.00000
-2.67776 0.00000 0.00000 ; CA- CB- CG2- HG22
3201 3203 3209 3212 3 0.66944 2.00832 0.00000
-2.67776 0.00000 0.00000 ; CA- CB- CG2- HG23
3201 3213 3215 3216 3 20.92000 0.00000 -20.92000
0.00000 0.00000 0.00000 ; CA- C- N- H
3201 3213 3215 3217 3 20.92000 0.00000 -20.92000
0.00000 0.00000 0.00000 ; CA- C- N- CA
3202 3201 3203 3204 3 0.65084 1.95253 0.00000
-2.60338 0.00000 0.00000 ; HA- CA- CB- HB
3202 3201 3203 3205 3 0.65084 1.95253 0.00000
-2.60338 0.00000 0.00000 ; HA- CA- CB- CG1
3202 3201 3203 3209 3 0.65084 1.95253 0.00000
-2.60338 0.00000 0.00000 ; HA- CA- CB- CG2
3202 3201 3213 3214 3 3.68192 -4.35136 0.00000
1.33888 0.00000 0.00000 ; HA- CA- C- O
3202 3201 3213 3215 3 0.00000 0.00000 0.00000
0.00000 0.00000 0.00000 ; HA- CA- C- N

3203 3201 3213 3214 3 0.00000 0.00000 0.00000
0.00000 0.00000 0.00000 ; CB- CA- C- O
3203 3201 3213 3215 3 2.51040 4.18400 1.67360
-6.69440 0.00000 0.00000 ; CB- CA- C- N
3204 3203 3201 3213 3 0.65084 1.95253 0.00000
-2.60338 0.00000 0.00000 ; HB- CB- CA- C
3204 3203 3205 3206 3 0.62760 1.88280 0.00000
-2.51040 0.00000 0.00000 ; HB- CB- CG1- HG11
3204 3203 3205 3207 3 0.62760 1.88280 0.00000
-2.51040 0.00000 0.00000 ; HB- CB- CG1- HG12
3204 3203 3205 3208 3 0.62760 1.88280 0.00000
-2.51040 0.00000 0.00000 ; HB- CB- CG1- HG13
3204 3203 3209 3210 3 0.62760 1.88280 0.00000
-2.51040 0.00000 0.00000 ; HB- CB- CG2- HG21
3204 3203 3209 3211 3 0.62760 1.88280 0.00000
-2.51040 0.00000 0.00000 ; HB- CB- CG2- HG22
3204 3203 3209 3212 3 0.62760 1.88280 0.00000
-2.51040 0.00000 0.00000 ; HB- CB- CG2- HG23
3205 3203 3201 3213 3 5.67350 3.55640 -6.16722
-2.47693 3.74886 0.00000 ; CG1- CB- CA- C
3205 3203 3209 3210 3 0.66944 2.00832 0.00000
-2.67776 0.00000 0.00000 ; CG1- CB- CG2- HG21
3205 3203 3209 3211 3 0.66944 2.00832 0.00000
-2.67776 0.00000 0.00000 ; CG1- CB- CG2- HG22
3205 3203 3209 3212 3 0.66944 2.00832 0.00000
-2.67776 0.00000 0.00000 ; CG1- CB- CG2- HG23
3206 3205 3203 3209 3 0.66944 2.00832 0.00000
-2.67776 0.00000 0.00000 ; HG11- CG1- CB- CG2
3207 3205 3203 3209 3 0.66944 2.00832 0.00000
-2.67776 0.00000 0.00000 ; HG12- CG1- CB- CG2
3208 3205 3203 3209 3 0.66944 2.00832 0.00000
-2.67776 0.00000 0.00000 ; HG13- CG1- CB- CG2
3209 3203 3201 3213 3 5.67350 3.55640 -6.16722
-2.47693 3.74886 0.00000 ; CG2- CB- CA- C
3213 3215 3217 3218 3 0.00000 0.00000 0.00000
0.00000 0.00000 0.00000 ; C- N- CA- HA
3213 3215 3217 3219 3 11.71520 1.67360 15.06240 -
13.38880 0.00000 0.00000 ; C- N- CA- CB
3213 3215 3217 3232 3 1.75728 5.27184 2.25936
-7.02912 0.00000 0.00000 ; C- N- CA- C
3214 3213 3215 3216 3 29.28800 -8.36800 -20.92000
0.00000 0.00000 0.00000 ; O- C- N- H
3214 3213 3215 3217 3 20.92000 0.00000 -20.92000
0.00000 0.00000 0.00000 ; O- C- N- CA
3215 3217 3219 3220 3 0.65084 1.95253 0.00000
-2.60338 0.00000 0.00000 ; N- CA- CB- HB
3215 3217 3219 3221 3 3.83673 0.44769 -1.77402
-2.47693 -0.03347 0.00000 ; N- CA- CB- CG2
3215 3217 3219 3225 3 2.97482 0.12134 2.04179
-1.89117 -3.24678 0.00000 ; N- CA- CB- CG1
3215 3217 3232 3233 3 0.00000 0.00000 0.00000
0.00000 0.00000 0.00000 ; N- CA- C- O

3215 3217 3232 3234 3 17.40544 -5.02080 -13.22144
9.20480 0.00000 0.00000 ; N- CA- C- N
3216 3215 3217 3218 3 0.00000 0.00000 0.00000
0.00000 0.00000 0.00000 ; H- N- CA- HA
3216 3215 3217 3219 3 0.00000 0.00000 0.00000
0.00000 0.00000 0.00000 ; H- N- CA- CB
3216 3215 3217 3232 3 0.00000 0.00000 0.00000
0.00000 0.00000 0.00000 ; H- N- CA- C
3217 3219 3221 3222 3 0.66944 2.00832 0.00000
-2.67776 0.00000 0.00000 ; CA- CB- CG2- HG21
3217 3219 3221 3223 3 0.66944 2.00832 0.00000
-2.67776 0.00000 0.00000 ; CA- CB- CG2- HG22
3217 3219 3221 3224 3 0.66944 2.00832 0.00000
-2.67776 0.00000 0.00000 ; CA- CB- CG2- HG23
3217 3219 3225 3226 3 0.66944 2.00832 0.00000
-2.67776 0.00000 0.00000 ; CA- CB- CG1- HG12
3217 3219 3225 3227 3 0.66944 2.00832 0.00000
-2.67776 0.00000 0.00000 ; CA- CB- CG1- HG13
3217 3219 3225 3228 3 4.24258 -0.52718 -7.25506
-1.79075 7.69856 0.00000 ; CA- CB- CG1- CD1
3217 3232 3234 3235 3 20.92000 0.00000 -20.92000
0.00000 0.00000 0.00000 ; CA- C- N- H
3217 3232 3234 3236 3 20.92000 0.00000 -20.92000
0.00000 0.00000 0.00000 ; CA- C- N- CA
3218 3217 3219 3220 3 0.65084 1.95253 0.00000
-2.60338 0.00000 0.00000 ; HA- CA- CB- HB
3218 3217 3219 3221 3 0.65084 1.95253 0.00000
-2.60338 0.00000 0.00000 ; HA- CA- CB- CG2
3218 3217 3219 3225 3 0.65084 1.95253 0.00000
-2.60338 0.00000 0.00000 ; HA- CA- CB- CG1
3218 3217 3232 3233 3 3.68192 -4.35136 0.00000
1.33888 0.00000 0.00000 ; HA- CA- C- O
3218 3217 3232 3234 3 0.00000 0.00000 0.00000
0.00000 0.00000 0.00000 ; HA- CA- C- N
3219 3217 3232 3233 3 0.00000 0.00000 0.00000
0.00000 0.00000 0.00000 ; CB- CA- C- O
3219 3217 3232 3234 3 2.51040 4.18400 1.67360
-6.69440 0.00000 0.00000 ; CB- CA- C- N
3219 3225 3228 3229 3 0.66944 2.00832 0.00000
-2.67776 0.00000 0.00000 ; CB- CG1- CD1- HD11
3219 3225 3228 3230 3 0.66944 2.00832 0.00000
-2.67776 0.00000 0.00000 ; CB- CG1- CD1- HD12
3219 3225 3228 3231 3 0.66944 2.00832 0.00000
-2.67776 0.00000 0.00000 ; CB- CG1- CD1- HD13
3220 3219 3217 3232 3 0.65084 1.95253 0.00000
-2.60338 0.00000 0.00000 ; HB- CB- CA- C
3220 3219 3221 3222 3 0.62760 1.88280 0.00000
-2.51040 0.00000 0.00000 ; HB- CB- CG2- HG21
3220 3219 3221 3223 3 0.62760 1.88280 0.00000
-2.51040 0.00000 0.00000 ; HB- CB- CG2- HG22
3220 3219 3221 3224 3 0.62760 1.88280 0.00000
-2.51040 0.00000 0.00000 ; HB- CB- CG2- HG23

3220 3219 3225 3226 3 0.62760 1.88280 0.00000
-2.51040 0.00000 0.00000 ; HB- CB- CG1- HG12
3220 3219 3225 3227 3 0.62760 1.88280 0.00000
-2.51040 0.00000 0.00000 ; HB- CB- CG1- HG13
3220 3219 3225 3228 3 0.66944 2.00832 0.00000
-2.67776 0.00000 0.00000 ; HB- CB- CG1- CD1
3221 3219 3217 3232 3 5.67350 3.55640 -6.16722
-2.47693 3.74886 0.00000 ; CG2- CB- CA- C
3221 3219 3225 3226 3 0.66944 2.00832 0.00000
-2.67776 0.00000 0.00000 ; CG2- CB- CG1- HG12
3221 3219 3225 3227 3 0.66944 2.00832 0.00000
-2.67776 0.00000 0.00000 ; CG2- CB- CG1- HG13
3221 3219 3225 3228 3 3.81162 0.49790 -8.14206
-1.79075 7.49773 0.00000 ; CG2- CB- CG1- CD1
3222 3221 3219 3225 3 0.66944 2.00832 0.00000
-2.67776 0.00000 0.00000 ; HG21- CG2- CB- CG1
3223 3221 3219 3225 3 0.66944 2.00832 0.00000
-2.67776 0.00000 0.00000 ; HG22- CG2- CB- CG1
3224 3221 3219 3225 3 0.66944 2.00832 0.00000
-2.67776 0.00000 0.00000 ; HG23- CG2- CB- CG1
3225 3219 3217 3232 3 8.26340 0.74057 -9.99976
-1.89117 3.84928 0.00000 ; CG1- CB- CA- C
3226 3225 3228 3229 3 0.62760 1.88280 0.00000
-2.51040 0.00000 0.00000 ; HG12- CG1- CD1- HD11
3226 3225 3228 3230 3 0.62760 1.88280 0.00000
-2.51040 0.00000 0.00000 ; HG12- CG1- CD1- HD12
3226 3225 3228 3231 3 0.62760 1.88280 0.00000
-2.51040 0.00000 0.00000 ; HG12- CG1- CD1- HD13
3227 3225 3228 3229 3 0.62760 1.88280 0.00000
-2.51040 0.00000 0.00000 ; HG13- CG1- CD1- HD11
3227 3225 3228 3230 3 0.62760 1.88280 0.00000
-2.51040 0.00000 0.00000 ; HG13- CG1- CD1- HD12
3227 3225 3228 3231 3 0.62760 1.88280 0.00000
-2.51040 0.00000 0.00000 ; HG13- CG1- CD1- HD13
3232 3234 3236 3237 3 0.00000 0.00000 0.00000
0.00000 0.00000 0.00000 ; C- N- CA- HA
3232 3234 3236 3238 3 11.71520 1.67360 15.06240 -
13.38880 0.00000 0.00000 ; C- N- CA- CB
3232 3234 3236 3249 3 1.75728 5.27184 2.25936
-7.02912 0.00000 0.00000 ; C- N- CA- C
3233 3232 3234 3235 3 29.28800 -8.36800 -20.92000
0.00000 0.00000 0.00000 ; O- C- N- H
3233 3232 3234 3236 3 20.92000 0.00000 -20.92000
0.00000 0.00000 0.00000 ; O- C- N- CA
3234 3236 3238 3239 3 0.65084 1.95253 0.00000
-2.60338 0.00000 0.00000 ; N- CA- CB- HB2
3234 3236 3238 3240 3 0.65084 1.95253 0.00000
-2.60338 0.00000 0.00000 ; N- CA- CB- HB3
3234 3236 3238 3241 3 3.21331 2.22589 -4.15053
-2.40998 2.61082 0.00000 ; N- CA- CB- CG
3234 3236 3249 3250 3 0.00000 0.00000 0.00000
0.00000 0.00000 0.00000 ; N- CA- C- O

3234 3236 3249 3251 3 17.40544 -5.02080 -13.22144
9.20480 0.00000 0.00000 ; N- CA- C- N
3235 3234 3236 3237 3 0.00000 0.00000 0.00000
0.00000 0.00000 0.00000 ; H- N- CA- HA
3235 3234 3236 3238 3 0.00000 0.00000 0.00000
0.00000 0.00000 0.00000 ; H- N- CA- CB
3235 3234 3236 3249 3 0.00000 0.00000 0.00000
0.00000 0.00000 0.00000 ; H- N- CA- C
3236 3238 3241 3242 3 0.65084 1.95253 0.00000
-2.60338 0.00000 0.00000 ; CA- CB- CG- HG2
3236 3238 3241 3243 3 0.65084 1.95253 0.00000
-2.60338 0.00000 0.00000 ; CA- CB- CG- HG3
3236 3238 3241 3244 3 2.04598 -1.54390 1.11294
-0.26778 0.93722 0.00000 ; CA- CB- CG- SD
3236 3249 3251 3252 3 20.92000 0.00000 -20.92000
0.00000 0.00000 0.00000 ; CA- C- N- H
3236 3249 3251 3253 3 20.92000 0.00000 -20.92000
0.00000 0.00000 0.00000 ; CA- C- N- CA
3237 3236 3238 3239 3 0.65084 1.95253 0.00000
-2.60338 0.00000 0.00000 ; HA- CA- CB- HB2
3237 3236 3238 3240 3 0.65084 1.95253 0.00000
-2.60338 0.00000 0.00000 ; HA- CA- CB- HB3
3237 3236 3238 3241 3 0.65084 1.95253 0.00000
-2.60338 0.00000 0.00000 ; HA- CA- CB- CG
3237 3236 3249 3250 3 3.68192 -4.35136 0.00000
1.33888 0.00000 0.00000 ; HA- CA- C- O
3237 3236 3249 3251 3 0.00000 0.00000 0.00000
0.00000 0.00000 0.00000 ; HA- CA- C- N
3238 3236 3249 3250 3 0.00000 0.00000 0.00000
0.00000 0.00000 0.00000 ; CB- CA- C- O
3238 3236 3249 3251 3 2.51040 4.18400 1.67360
-6.69440 0.00000 0.00000 ; CB- CA- C- N
3238 3241 3244 3245 3 3.24260 6.22998 1.79075
-6.92870 1.90790 0.00000 ; CB- CG- SD- CE
3239 3238 3236 3249 3 0.65084 1.95253 0.00000
-2.60338 0.00000 0.00000 ; HB2- CB- CA- C
3239 3238 3241 3242 3 0.65084 1.95253 0.00000
-2.60338 0.00000 0.00000 ; HB2- CB- CG- HG2
3239 3238 3241 3243 3 0.65084 1.95253 0.00000
-2.60338 0.00000 0.00000 ; HB2- CB- CG- HG3
3239 3238 3241 3244 3 0.65084 1.95253 0.00000
-2.60338 0.00000 0.00000 ; HB2- CB- CG- SD
3240 3238 3236 3249 3 0.65084 1.95253 0.00000
-2.60338 0.00000 0.00000 ; HB3- CB- CA- C
3240 3238 3241 3242 3 0.65084 1.95253 0.00000
-2.60338 0.00000 0.00000 ; HB3- CB- CG- HG2
3240 3238 3241 3243 3 0.65084 1.95253 0.00000
-2.60338 0.00000 0.00000 ; HB3- CB- CG- HG3
3240 3238 3241 3244 3 0.65084 1.95253 0.00000
-2.60338 0.00000 0.00000 ; HB3- CB- CG- SD
3241 3238 3236 3249 3 6.86594 3.56895 -8.14206
-2.40998 4.85344 0.00000 ; CG- CB- CA- C

3241 3244 3245 3246 3 1.39467 4.18400 0.00000
-5.57867 0.00000 0.00000 ; CG- SD- CE- HE1
3241 3244 3245 3247 3 1.39467 4.18400 0.00000
-5.57867 0.00000 0.00000 ; CG- SD- CE- HE2
3241 3244 3245 3248 3 1.39467 4.18400 0.00000
-5.57867 0.00000 0.00000 ; CG- SD- CE- HE3
3242 3241 3244 3245 3 1.39467 4.18400 0.00000
-5.57867 0.00000 0.00000 ; HG2- CG- SD- CE
3243 3241 3244 3245 3 1.39467 4.18400 0.00000
-5.57867 0.00000 0.00000 ; HG3- CG- SD- CE
3249 3251 3253 3254 3 0.00000 0.00000 0.00000
0.00000 0.00000 0.00000 ; C- N- CA- HA
3249 3251 3253 3255 3 11.71520 1.67360 15.06240 -
13.38880 0.00000 0.00000 ; C- N- CA- CB
3249 3251 3253 3270 3 1.75728 5.27184 2.25936
-7.02912 0.00000 0.00000 ; C- N- CA- C
3250 3249 3251 3252 3 29.28800 -8.36800 -20.92000
0.00000 0.00000 0.00000 ; O- C- N- H
3250 3249 3251 3253 3 20.92000 0.00000 -20.92000
0.00000 0.00000 0.00000 ; O- C- N- CA
3251 3253 3255 3256 3 0.65084 1.95253 0.00000
-2.60338 0.00000 0.00000 ; N- CA- CB- HB2
3251 3253 3255 3257 3 0.65084 1.95253 0.00000
-2.60338 0.00000 0.00000 ; N- CA- CB- HB3
3251 3253 3255 3258 3 3.28026 2.46019 -2.19242
-3.21331 -0.23430 0.00000 ; N- CA- CB- CG
3251 3253 3270 3271 3 0.00000 0.00000 0.00000
0.00000 0.00000 0.00000 ; N- CA- C- O
3251 3253 3270 3272 3 17.40544 -5.02080 -13.22144
9.20480 0.00000 0.00000 ; N- CA- C- N
3252 3251 3253 3254 3 0.00000 0.00000 0.00000
0.00000 0.00000 0.00000 ; H- N- CA- HA
3252 3251 3253 3255 3 0.00000 0.00000 0.00000
0.00000 0.00000 0.00000 ; H- N- CA- CB
3252 3251 3253 3270 3 0.00000 0.00000 0.00000
0.00000 0.00000 0.00000 ; H- N- CA- C
3253 3255 3258 3259 3 0.57739 0.00000 1.02926
0.00000 -1.60666 0.00000 ; CA- CB- CG- CD1
3253 3255 3258 3268 3 0.57739 0.00000 1.02926
0.00000 -1.60666 0.00000 ; CA- CB- CG- CD2
3253 3270 3272 3273 3 20.92000 0.00000 -20.92000
0.00000 0.00000 0.00000 ; CA- C- N- H
3253 3270 3272 3274 3 20.92000 0.00000 -20.92000
0.00000 0.00000 0.00000 ; CA- C- N- CA
3254 3253 3255 3256 3 0.65084 1.95253 0.00000
-2.60338 0.00000 0.00000 ; HA- CA- CB- HB2
3254 3253 3255 3257 3 0.65084 1.95253 0.00000
-2.60338 0.00000 0.00000 ; HA- CA- CB- HB3
3254 3253 3255 3258 3 0.65084 1.95253 0.00000
-2.60338 0.00000 0.00000 ; HA- CA- CB- CG
3254 3253 3270 3271 3 3.68192 -4.35136 0.00000
1.33888 0.00000 0.00000 ; HA- CA- C- O

3254 3253 3270 3272 3 0.00000 0.00000 0.00000
0.00000 0.00000 0.00000 ; HA- CA- C- N
3255 3253 3270 3271 3 0.00000 0.00000 0.00000
0.00000 0.00000 0.00000 ; CB- CA- C- O
3255 3253 3270 3272 3 2.51040 4.18400 1.67360
-6.69440 0.00000 0.00000 ; CB- CA- C- N
3255 3258 3259 3260 3 30.33400 0.00000 -30.33400
0.00000 0.00000 0.00000 ; CB- CG- CD1- HD1
3255 3258 3259 3261 3 30.33400 0.00000 -30.33400
0.00000 0.00000 0.00000 ; CB- CG- CD1- CE1
3255 3258 3268 3266 3 30.33400 0.00000 -30.33400
0.00000 0.00000 0.00000 ; CB- CG- CD2- CE2
3255 3258 3268 3269 3 30.33400 0.00000 -30.33400
0.00000 0.00000 0.00000 ; CB- CG- CD2- HD2
3256 3255 3253 3270 3 0.65084 1.95253 0.00000
-2.60338 0.00000 0.00000 ; HB2- CB- CA- C
3256 3255 3258 3259 3 0.00000 0.00000 0.00000
0.00000 0.00000 0.00000 ; HB2- CB- CG- CD1
3256 3255 3258 3268 3 0.00000 0.00000 0.00000
0.00000 0.00000 0.00000 ; HB2- CB- CG- CD2
3257 3255 3253 3270 3 0.65084 1.95253 0.00000
-2.60338 0.00000 0.00000 ; HB3- CB- CA- C
3257 3255 3258 3259 3 0.00000 0.00000 0.00000
0.00000 0.00000 0.00000 ; HB3- CB- CG- CD1
3257 3255 3258 3268 3 0.00000 0.00000 0.00000
0.00000 0.00000 0.00000 ; HB3- CB- CG- CD2
3258 3255 3253 3270 3 4.95804 2.17986 -3.52293
-3.21331 -0.40166 0.00000 ; CG- CB- CA- C
3258 3259 3261 3262 3 30.33400 0.00000 -30.33400
0.00000 0.00000 0.00000 ; CG- CD1- CE1- HE1
3258 3259 3261 3263 3 30.33400 0.00000 -30.33400
0.00000 0.00000 0.00000 ; CG- CD1- CE1- CZ
3258 3268 3266 3263 3 30.33400 0.00000 -30.33400
0.00000 0.00000 0.00000 ; CG- CD2- CE2- CZ
3258 3268 3266 3267 3 30.33400 0.00000 -30.33400
0.00000 0.00000 0.00000 ; CG- CD2- CE2- HE2
3259 3258 3268 3266 3 30.33400 0.00000 -30.33400
0.00000 0.00000 0.00000 ; CD1- CG- CD2- CE2
3259 3258 3268 3269 3 30.33400 0.00000 -30.33400
0.00000 0.00000 0.00000 ; CD1- CG- CD2- HD2
3259 3261 3263 3264 3 30.33400 0.00000 -30.33400
0.00000 0.00000 0.00000 ; CD1- CE1- CZ- OH
3259 3261 3263 3266 3 30.33400 0.00000 -30.33400
0.00000 0.00000 0.00000 ; CD1- CE1- CZ- CE2
3260 3259 3258 3268 3 30.33400 0.00000 -30.33400
0.00000 0.00000 0.00000 ; HD1- CD1- CG- CD2
3260 3259 3261 3262 3 30.33400 0.00000 -30.33400
0.00000 0.00000 0.00000 ; HD1- CD1- CE1- HE1
3260 3259 3261 3263 3 30.33400 0.00000 -30.33400
0.00000 0.00000 0.00000 ; HD1- CD1- CE1- CZ
3261 3259 3258 3268 3 30.33400 0.00000 -30.33400
0.00000 0.00000 0.00000 ; CE1- CD1- CG- CD2

3261 3263 3264 3265 3 7.93286 0.00000 -9.56462
0.00000 2.17568 0.00000 ; CE1- CZ- OH- HH
3261 3263 3266 3267 3 30.33400 0.00000 -30.33400
0.00000 0.00000 0.00000 ; CE1- CZ- CE2- HE2
3261 3263 3266 3268 3 30.33400 0.00000 -30.33400
0.00000 0.00000 0.00000 ; CE1- CZ- CE2- CD2
3262 3261 3263 3264 3 30.33400 0.00000 -30.33400
0.00000 0.00000 0.00000 ; HE1- CE1- CZ- OH
3262 3261 3263 3266 3 30.33400 0.00000 -30.33400
0.00000 0.00000 0.00000 ; HE1- CE1- CZ- CE2
3263 3266 3268 3269 3 30.33400 0.00000 -30.33400
0.00000 0.00000 0.00000 ; CZ- CE2- CD2- HD2
3264 3263 3266 3267 3 30.33400 0.00000 -30.33400
0.00000 0.00000 0.00000 ; OH- CZ- CE2- HE2
3264 3263 3266 3268 3 30.33400 0.00000 -30.33400
0.00000 0.00000 0.00000 ; OH- CZ- CE2- CD2
3265 3264 3263 3266 3 7.93286 0.00000 -9.56462
0.00000 2.17568 0.00000 ; HH- OH- CZ- CE2
3267 3266 3268 3269 3 30.33400 0.00000 -30.33400
0.00000 0.00000 0.00000 ; HE2- CE2- CD2- HD2
3270 3272 3274 3275 3 0.00000 0.00000 0.00000
0.00000 0.00000 0.00000 ; C- N- CA- HA
3270 3272 3274 3276 3 11.71520 1.67360 15.06240 -
13.38880 0.00000 0.00000 ; C- N- CA- CB
3270 3272 3274 3284 3 1.75728 5.27184 2.25936
-7.02912 0.00000 0.00000 ; C- N- CA- C
3271 3270 3272 3273 3 29.28800 -8.36800 -20.92000
0.00000 0.00000 0.00000 ; O- C- N- H
3271 3270 3272 3274 3 20.92000 0.00000 -20.92000
0.00000 0.00000 0.00000 ; O- C- N- CA
3272 3274 3276 3277 3 0.65084 1.95253 0.00000
-2.60338 0.00000 0.00000 ; N- CA- CB- HB
3272 3274 3276 3278 3 3.83673 0.44769 -1.77402
-2.47693 -0.03347 0.00000 ; N- CA- CB- CG2
3272 3274 3276 3282 3 4.93294 1.13386 -3.12963
-5.27184 3.17984 0.00000 ; N- CA- CB- OG1
3272 3274 3284 3285 3 0.00000 0.00000 0.00000
0.00000 0.00000 0.00000 ; N- CA- C- O
3272 3274 3284 3286 3 17.40544 -5.02080 -13.22144
9.20480 0.00000 0.00000 ; N- CA- C- N
3273 3272 3274 3275 3 0.00000 0.00000 0.00000
0.00000 0.00000 0.00000 ; H- N- CA- HA
3273 3272 3274 3276 3 0.00000 0.00000 0.00000
0.00000 0.00000 0.00000 ; H- N- CA- CB
3273 3272 3274 3284 3 0.00000 0.00000 0.00000
0.00000 0.00000 0.00000 ; H- N- CA- C
3274 3276 3278 3279 3 0.66944 2.00832 0.00000
-2.67776 0.00000 0.00000 ; CA- CB- CG2- HG21
3274 3276 3278 3280 3 0.66944 2.00832 0.00000
-2.67776 0.00000 0.00000 ; CA- CB- CG2- HG22
3274 3276 3278 3281 3 0.66944 2.00832 0.00000
-2.67776 0.00000 0.00000 ; CA- CB- CG2- HG23

3274 3276 3282 3283 3 1.12131 2.98738 1.66523
-3.94970 0.43514 0.00000 ; CA- CB- OG1- HG1
3274 3284 3286 3287 3 20.92000 0.00000 -20.92000
0.00000 0.00000 0.00000 ; CA- C- N- H
3274 3284 3286 3288 3 20.92000 0.00000 -20.92000
0.00000 0.00000 0.00000 ; CA- C- N- CA
3275 3274 3276 3277 3 0.65084 1.95253 0.00000
-2.60338 0.00000 0.00000 ; HA- CA- CB- HB
3275 3274 3276 3278 3 0.65084 1.95253 0.00000
-2.60338 0.00000 0.00000 ; HA- CA- CB- CG2
3275 3274 3276 3282 3 1.04600 -1.04600 0.00000
0.00000 0.00000 0.00000 ; HA- CA- CB- OG1
3275 3274 3284 3285 3 3.68192 -4.35136 0.00000
1.33888 0.00000 0.00000 ; HA- CA- C- O
3275 3274 3284 3286 3 0.00000 0.00000 0.00000
0.00000 0.00000 0.00000 ; HA- CA- C- N
3276 3274 3284 3285 3 0.00000 0.00000 0.00000
0.00000 0.00000 0.00000 ; CB- CA- C- O
3276 3274 3284 3286 3 2.51040 4.18400 1.67360
-6.69440 0.00000 0.00000 ; CB- CA- C- N
3277 3276 3274 3284 3 0.65084 1.95253 0.00000
-2.60338 0.00000 0.00000 ; HB- CB- CA- C
3277 3276 3278 3279 3 0.65084 1.95253 0.00000
-2.60338 0.00000 0.00000 ; HB- CB- CG2- HG21
3277 3276 3278 3280 3 0.65084 1.95253 0.00000
-2.60338 0.00000 0.00000 ; HB- CB- CG2- HG22
3277 3276 3278 3281 3 0.65084 1.95253 0.00000
-2.60338 0.00000 0.00000 ; HB- CB- CG2- HG23
3277 3276 3282 3283 3 0.69733 2.09200 0.00000
-2.78933 0.00000 0.00000 ; HB- CB- OG1- HG1
3278 3276 3274 3284 3 5.67350 3.55640 -6.16722
-2.47693 3.74886 0.00000 ; CG2- CB- CA- C
3278 3276 3282 3283 3 4.74047 0.27196 -2.26773
-3.94970 1.60666 0.00000 ; CG2- CB- OG1- HG1
3279 3278 3276 3282 3 1.04600 -1.04600 0.00000
0.00000 0.00000 0.00000 ; HG21- CG2- CB- OG1
3280 3278 3276 3282 3 1.04600 -1.04600 0.00000
0.00000 0.00000 0.00000 ; HG22- CG2- CB- OG1
3281 3278 3276 3282 3 1.04600 -1.04600 0.00000
0.00000 0.00000 0.00000 ; HG23- CG2- CB- OG1
3282 3276 3274 3284 3 6.53541 6.87013 -6.21742
-5.27184 5.22163 0.00000 ; OG1- CB- CA- C
3284 3286 3288 3289 3 0.00000 0.00000 0.00000
0.00000 0.00000 0.00000 ; C- N- CA- HA
3284 3286 3288 3290 3 11.71520 1.67360 15.06240 -
13.38880 0.00000 0.00000 ; C- N- CA- CB
3284 3286 3288 3303 3 1.75728 5.27184 2.25936
-7.02912 0.00000 0.00000 ; C- N- CA- C
3285 3284 3286 3287 3 29.28800 -8.36800 -20.92000
0.00000 0.00000 0.00000 ; O- C- N- H
3285 3284 3286 3288 3 20.92000 0.00000 -20.92000
0.00000 0.00000 0.00000 ; O- C- N- CA

3286 3288 3290 3291 3 0.65084 1.95253 0.00000
-2.60338 0.00000 0.00000 ; N- CA- CB- HB2
3286 3288 3290 3292 3 0.65084 1.95253 0.00000
-2.60338 0.00000 0.00000 ; N- CA- CB- HB3
3286 3288 3290 3293 3 3.79070 1.39746 -4.61077
-2.40998 2.44346 0.00000 ; N- CA- CB- CG
3286 3288 3303 3304 3 0.00000 0.00000 0.00000
0.00000 0.00000 0.00000 ; N- CA- C- O
3286 3288 3303 3305 3 17.40544 -5.02080 -13.22144
9.20480 0.00000 0.00000 ; N- CA- C- N
3287 3286 3288 3289 3 0.00000 0.00000 0.00000
0.00000 0.00000 0.00000 ; H- N- CA- HA
3287 3286 3288 3290 3 0.00000 0.00000 0.00000
0.00000 0.00000 0.00000 ; H- N- CA- CB
3287 3286 3288 3303 3 0.00000 0.00000 0.00000
0.00000 0.00000 0.00000 ; H- N- CA- C
3288 3290 3293 3294 3 0.66944 2.00832 0.00000
-2.67776 0.00000 0.00000 ; CA- CB- CG- HG
3288 3290 3293 3295 3 3.90367 0.19665 -6.21742
-2.37651 5.99149 0.00000 ; CA- CB- CG- CD1
3288 3290 3293 3299 3 3.90367 0.19665 -6.21742
-2.37651 5.99149 0.00000 ; CA- CB- CG- CD2
3288 3303 3305 3306 3 20.92000 0.00000 -20.92000
0.00000 0.00000 0.00000 ; CA- C- N- H
3288 3303 3305 3307 3 20.92000 0.00000 -20.92000
0.00000 0.00000 0.00000 ; CA- C- N- CA
3289 3288 3290 3291 3 0.65084 1.95253 0.00000
-2.60338 0.00000 0.00000 ; HA- CA- CB- HB2
3289 3288 3290 3292 3 0.65084 1.95253 0.00000
-2.60338 0.00000 0.00000 ; HA- CA- CB- HB3
3289 3288 3290 3293 3 0.65084 1.95253 0.00000
-2.60338 0.00000 0.00000 ; HA- CA- CB- CG
3289 3288 3303 3304 3 3.68192 -4.35136 0.00000
1.33888 0.00000 0.00000 ; HA- CA- C- O
3289 3288 3303 3305 3 0.00000 0.00000 0.00000
0.00000 0.00000 0.00000 ; HA- CA- C- N
3290 3288 3303 3304 3 0.00000 0.00000 0.00000
0.00000 0.00000 0.00000 ; CB- CA- C- O
3290 3288 3303 3305 3 2.51040 4.18400 1.67360
-6.69440 0.00000 0.00000 ; CB- CA- C- N
3290 3293 3295 3296 3 0.66944 2.00832 0.00000
-2.67776 0.00000 0.00000 ; CB- CG- CD1- HD11
3290 3293 3295 3297 3 0.66944 2.00832 0.00000
-2.67776 0.00000 0.00000 ; CB- CG- CD1- HD12
3290 3293 3295 3298 3 0.66944 2.00832 0.00000
-2.67776 0.00000 0.00000 ; CB- CG- CD1- HD13
3290 3293 3299 3300 3 0.66944 2.00832 0.00000
-2.67776 0.00000 0.00000 ; CB- CG- CD2- HD21
3290 3293 3299 3301 3 0.66944 2.00832 0.00000
-2.67776 0.00000 0.00000 ; CB- CG- CD2- HD22
3290 3293 3299 3302 3 0.66944 2.00832 0.00000
-2.67776 0.00000 0.00000 ; CB- CG- CD2- HD23

3291 3290 3288 3303 3 0.65084 1.95253 0.00000
-2.60338 0.00000 0.00000 ; HB2- CB- CA- C
3291 3290 3293 3294 3 0.62760 1.88280 0.00000
-2.51040 0.00000 0.00000 ; HB2- CB- CG- HG
3291 3290 3293 3295 3 0.66944 2.00832 0.00000
-2.67776 0.00000 0.00000 ; HB2- CB- CG- CD1
3291 3290 3293 3299 3 0.66944 2.00832 0.00000
-2.67776 0.00000 0.00000 ; HB2- CB- CG- CD2
3292 3290 3288 3303 3 0.65084 1.95253 0.00000
-2.60338 0.00000 0.00000 ; HB3- CB- CA- C
3292 3290 3293 3294 3 0.62760 1.88280 0.00000
-2.51040 0.00000 0.00000 ; HB3- CB- CG- HG
3292 3290 3293 3295 3 0.66944 2.00832 0.00000
-2.67776 0.00000 0.00000 ; HB3- CB- CG- CD1
3292 3290 3293 3299 3 0.66944 2.00832 0.00000
-2.67776 0.00000 0.00000 ; HB3- CB- CG- CD2
3293 3290 3288 3303 3 10.33448 -1.14642 -11.54784
-2.40998 6.35968 0.00000 ; CG- CB- CA- C
3294 3293 3295 3296 3 0.62760 1.88280 0.00000
-2.51040 0.00000 0.00000 ; HG- CG- CD1- HD11
3294 3293 3295 3297 3 0.62760 1.88280 0.00000
-2.51040 0.00000 0.00000 ; HG- CG- CD1- HD12
3294 3293 3295 3298 3 0.62760 1.88280 0.00000
-2.51040 0.00000 0.00000 ; HG- CG- CD1- HD13
3294 3293 3299 3300 3 0.62760 1.88280 0.00000
-2.51040 0.00000 0.00000 ; HG- CG- CD2- HD21
3294 3293 3299 3301 3 0.62760 1.88280 0.00000
-2.51040 0.00000 0.00000 ; HG- CG- CD2- HD22
3294 3293 3299 3302 3 0.62760 1.88280 0.00000
-2.51040 0.00000 0.00000 ; HG- CG- CD2- HD23
3295 3293 3299 3300 3 0.66944 2.00832 0.00000
-2.67776 0.00000 0.00000 ; CD1- CG- CD2- HD21
3295 3293 3299 3301 3 0.66944 2.00832 0.00000
-2.67776 0.00000 0.00000 ; CD1- CG- CD2- HD22
3295 3293 3299 3302 3 0.66944 2.00832 0.00000
-2.67776 0.00000 0.00000 ; CD1- CG- CD2- HD23
3296 3295 3293 3299 3 0.66944 2.00832 0.00000
-2.67776 0.00000 0.00000 ; HD11- CD1- CG- CD2
3297 3295 3293 3299 3 0.66944 2.00832 0.00000
-2.67776 0.00000 0.00000 ; HD12- CD1- CG- CD2
3298 3295 3293 3299 3 0.66944 2.00832 0.00000
-2.67776 0.00000 0.00000 ; HD13- CD1- CG- CD2
3303 3305 3307 3308 3 0.00000 0.00000 0.00000
0.00000 0.00000 0.00000 ; C- N- CA- HA
3303 3305 3307 3309 3 11.71520 1.67360 15.06240 -
13.38880 0.00000 0.00000 ; C- N- CA- CB
3303 3305 3307 3322 3 1.75728 5.27184 2.25936
-7.02912 0.00000 0.00000 ; C- N- CA- C
3304 3303 3305 3306 3 29.28800 -8.36800 -20.92000
0.00000 0.00000 0.00000 ; O- C- N- H
3304 3303 3305 3307 3 20.92000 0.00000 -20.92000
0.00000 0.00000 0.00000 ; O- C- N- CA

3305 3307 3309 3310 3 0.65084 1.95253 0.00000
-2.60338 0.00000 0.00000 ; N- CA- CB- HB2
3305 3307 3309 3311 3 0.65084 1.95253 0.00000
-2.60338 0.00000 0.00000 ; N- CA- CB- HB3
3305 3307 3309 3312 3 3.79070 1.39746 -4.61077
-2.40998 2.44346 0.00000 ; N- CA- CB- CG
3305 3307 3322 3323 3 0.00000 0.00000 0.00000
0.00000 0.00000 0.00000 ; N- CA- C- O
3305 3307 3322 3324 3 17.40544 -5.02080 -13.22144
9.20480 0.00000 0.00000 ; N- CA- C- N
3306 3305 3307 3308 3 0.00000 0.00000 0.00000
0.00000 0.00000 0.00000 ; H- N- CA- HA
3306 3305 3307 3309 3 0.00000 0.00000 0.00000
0.00000 0.00000 0.00000 ; H- N- CA- CB
3306 3305 3307 3322 3 0.00000 0.00000 0.00000
0.00000 0.00000 0.00000 ; H- N- CA- C
3307 3309 3312 3313 3 0.66944 2.00832 0.00000
-2.67776 0.00000 0.00000 ; CA- CB- CG- HG
3307 3309 3312 3314 3 3.90367 0.19665 -6.21742
-2.37651 5.99149 0.00000 ; CA- CB- CG- CD1
3307 3309 3312 3318 3 3.90367 0.19665 -6.21742
-2.37651 5.99149 0.00000 ; CA- CB- CG- CD2
3307 3322 3324 3325 3 20.92000 0.00000 -20.92000
0.00000 0.00000 0.00000 ; CA- C- N- H
3307 3322 3324 3326 3 20.92000 0.00000 -20.92000
0.00000 0.00000 0.00000 ; CA- C- N- CA
3308 3307 3309 3310 3 0.65084 1.95253 0.00000
-2.60338 0.00000 0.00000 ; HA- CA- CB- HB2
3308 3307 3309 3311 3 0.65084 1.95253 0.00000
-2.60338 0.00000 0.00000 ; HA- CA- CB- HB3
3308 3307 3309 3312 3 0.65084 1.95253 0.00000
-2.60338 0.00000 0.00000 ; HA- CA- CB- CG
3308 3307 3322 3323 3 3.68192 -4.35136 0.00000
1.33888 0.00000 0.00000 ; HA- CA- C- O
3308 3307 3322 3324 3 0.00000 0.00000 0.00000
0.00000 0.00000 0.00000 ; HA- CA- C- N
3309 3307 3322 3323 3 0.00000 0.00000 0.00000
0.00000 0.00000 0.00000 ; CB- CA- C- O
3309 3307 3322 3324 3 2.51040 4.18400 1.67360
-6.69440 0.00000 0.00000 ; CB- CA- C- N
3309 3312 3314 3315 3 0.66944 2.00832 0.00000
-2.67776 0.00000 0.00000 ; CB- CG- CD1- HD11
3309 3312 3314 3316 3 0.66944 2.00832 0.00000
-2.67776 0.00000 0.00000 ; CB- CG- CD1- HD12
3309 3312 3314 3317 3 0.66944 2.00832 0.00000
-2.67776 0.00000 0.00000 ; CB- CG- CD1- HD13
3309 3312 3318 3319 3 0.66944 2.00832 0.00000
-2.67776 0.00000 0.00000 ; CB- CG- CD2- HD21
3309 3312 3318 3320 3 0.66944 2.00832 0.00000
-2.67776 0.00000 0.00000 ; CB- CG- CD2- HD22
3309 3312 3318 3321 3 0.66944 2.00832 0.00000
-2.67776 0.00000 0.00000 ; CB- CG- CD2- HD23

3310 3309 3307 3322 3 0.65084 1.95253 0.00000
-2.60338 0.00000 0.00000 ; HB2- CB- CA- C
3310 3309 3312 3313 3 0.62760 1.88280 0.00000
-2.51040 0.00000 0.00000 ; HB2- CB- CG- HG
3310 3309 3312 3314 3 0.66944 2.00832 0.00000
-2.67776 0.00000 0.00000 ; HB2- CB- CG- CD1
3310 3309 3312 3318 3 0.66944 2.00832 0.00000
-2.67776 0.00000 0.00000 ; HB2- CB- CG- CD2
3311 3309 3307 3322 3 0.65084 1.95253 0.00000
-2.60338 0.00000 0.00000 ; HB3- CB- CA- C
3311 3309 3312 3313 3 0.62760 1.88280 0.00000
-2.51040 0.00000 0.00000 ; HB3- CB- CG- HG
3311 3309 3312 3314 3 0.66944 2.00832 0.00000
-2.67776 0.00000 0.00000 ; HB3- CB- CG- CD1
3311 3309 3312 3318 3 0.66944 2.00832 0.00000
-2.67776 0.00000 0.00000 ; HB3- CB- CG- CD2
3312 3309 3307 3322 3 10.33448 -1.14642 -11.54784
-2.40998 6.35968 0.00000 ; CG- CB- CA- C
3313 3312 3314 3315 3 0.62760 1.88280 0.00000
-2.51040 0.00000 0.00000 ; HG- CG- CD1- HD11
3313 3312 3314 3316 3 0.62760 1.88280 0.00000
-2.51040 0.00000 0.00000 ; HG- CG- CD1- HD12
3313 3312 3314 3317 3 0.62760 1.88280 0.00000
-2.51040 0.00000 0.00000 ; HG- CG- CD1- HD13
3313 3312 3318 3319 3 0.62760 1.88280 0.00000
-2.51040 0.00000 0.00000 ; HG- CG- CD2- HD21
3313 3312 3318 3320 3 0.62760 1.88280 0.00000
-2.51040 0.00000 0.00000 ; HG- CG- CD2- HD22
3313 3312 3318 3321 3 0.62760 1.88280 0.00000
-2.51040 0.00000 0.00000 ; HG- CG- CD2- HD23
3314 3312 3318 3319 3 0.66944 2.00832 0.00000
-2.67776 0.00000 0.00000 ; CD1- CG- CD2- HD21
3314 3312 3318 3320 3 0.66944 2.00832 0.00000
-2.67776 0.00000 0.00000 ; CD1- CG- CD2- HD22
3314 3312 3318 3321 3 0.66944 2.00832 0.00000
-2.67776 0.00000 0.00000 ; CD1- CG- CD2- HD23
3315 3314 3312 3318 3 0.66944 2.00832 0.00000
-2.67776 0.00000 0.00000 ; HD11- CD1- CG- CD2
3316 3314 3312 3318 3 0.66944 2.00832 0.00000
-2.67776 0.00000 0.00000 ; HD12- CD1- CG- CD2
3317 3314 3312 3318 3 0.66944 2.00832 0.00000
-2.67776 0.00000 0.00000 ; HD13- CD1- CG- CD2
3322 3324 3326 3327 3 0.00000 0.00000 0.00000
0.00000 0.00000 0.00000 ; C- N- CA- HA
3322 3324 3326 3328 3 11.71520 1.67360 15.06240 -
13.38880 0.00000 0.00000 ; C- N- CA- CB
3322 3324 3326 3333 3 1.75728 5.27184 2.25936
-7.02912 0.00000 0.00000 ; C- N- CA- C
3323 3322 3324 3325 3 29.28800 -8.36800 -20.92000
0.00000 0.00000 0.00000 ; O- C- N- H
3323 3322 3324 3326 3 20.92000 0.00000 -20.92000
0.00000 0.00000 0.00000 ; O- C- N- CA

3324 3326 3328 3329 3 0.65084 1.95253 0.00000
-2.60338 0.00000 0.00000 ; N- CA- CB- HB2
3324 3326 3328 3330 3 0.65084 1.95253 0.00000
-2.60338 0.00000 0.00000 ; N- CA- CB- HB3
3324 3326 3328 3331 3 6.03751 2.50622 -5.17142
-4.20074 1.10458 0.00000 ; N- CA- CB- SG
3324 3326 3333 3334 3 0.00000 0.00000 0.00000
0.00000 0.00000 0.00000 ; N- CA- C- O
3324 3326 3333 3335 3 17.40544 -5.02080 -13.22144
9.20480 0.00000 0.00000 ; N- CA- C- N
3325 3324 3326 3327 3 0.00000 0.00000 0.00000
0.00000 0.00000 0.00000 ; H- N- CA- HA
3325 3324 3326 3328 3 0.00000 0.00000 0.00000
0.00000 0.00000 0.00000 ; H- N- CA- CB
3325 3324 3326 3333 3 0.00000 0.00000 0.00000
0.00000 0.00000 0.00000 ; H- N- CA- C
3326 3328 3331 3332 3 1.69034 2.77818 4.11706
-4.21747 1.00416 0.00000 ; CA- CB- SG- HG
3326 3333 3335 3336 3 20.92000 0.00000 -20.92000
0.00000 0.00000 0.00000 ; CA- C- N- H
3326 3333 3335 3337 3 20.92000 0.00000 -20.92000
0.00000 0.00000 0.00000 ; CA- C- N- CA
3327 3326 3328 3329 3 0.65084 1.95253 0.00000
-2.60338 0.00000 0.00000 ; HA- CA- CB- HB2
3327 3326 3328 3330 3 0.65084 1.95253 0.00000
-2.60338 0.00000 0.00000 ; HA- CA- CB- HB3
3327 3326 3328 3331 3 0.65084 1.95253 0.00000
-2.60338 0.00000 0.00000 ; HA- CA- CB- SG
3327 3326 3333 3334 3 3.68192 -4.35136 0.00000
1.33888 0.00000 0.00000 ; HA- CA- C- O
3327 3326 3333 3335 3 0.00000 0.00000 0.00000
0.00000 0.00000 0.00000 ; HA- CA- C- N
3328 3326 3333 3334 3 0.00000 0.00000 0.00000
0.00000 0.00000 0.00000 ; CB- CA- C- O
3328 3326 3333 3335 3 2.51040 4.18400 1.67360
-6.69440 0.00000 0.00000 ; CB- CA- C- N
3329 3328 3326 3333 3 0.65084 1.95253 0.00000
-2.60338 0.00000 0.00000 ; HB2- CB- CA- C
3329 3328 3331 3332 3 1.04600 3.13800 0.00000
-4.18400 0.00000 0.00000 ; HB2- CB- SG- HG
3330 3328 3326 3333 3 0.65084 1.95253 0.00000
-2.60338 0.00000 0.00000 ; HB3- CB- CA- C
3330 3328 3331 3332 3 1.04600 3.13800 0.00000
-4.18400 0.00000 0.00000 ; HB3- CB- SG- HG
3331 3328 3326 3333 3 5.62330 4.27605 -5.33042
-4.20074 2.51040 0.00000 ; SG- CB- CA- C
3333 3335 3337 3338 3 0.00000 0.00000 0.00000
0.00000 0.00000 0.00000 ; C- N- CA- HA2
3333 3335 3337 3339 3 0.00000 0.00000 0.00000
0.00000 0.00000 0.00000 ; C- N- CA- HA3
3333 3335 3337 3340 3 1.75728 5.27184 2.25936
-7.02912 0.00000 0.00000 ; C- N- CA- C

3334 3333 3335 3336 3 29.28800 -8.36800 -20.92000
0.00000 0.00000 0.00000 ; O- C- N- H
3334 3333 3335 3337 3 20.92000 0.00000 -20.92000
0.00000 0.00000 0.00000 ; O- C- N- CA
3335 3337 3340 3341 3 0.00000 0.00000 0.00000
0.00000 0.00000 0.00000 ; N- CA- C- O
3335 3337 3340 3342 3 17.40544 -5.02080 -13.22144
9.20480 0.00000 0.00000 ; N- CA- C- N
3336 3335 3337 3338 3 0.00000 0.00000 0.00000
0.00000 0.00000 0.00000 ; H- N- CA- HA2
3336 3335 3337 3339 3 0.00000 0.00000 0.00000
0.00000 0.00000 0.00000 ; H- N- CA- HA3
3336 3335 3337 3340 3 0.00000 0.00000 0.00000
0.00000 0.00000 0.00000 ; H- N- CA- C
3337 3340 3342 3343 3 20.92000 0.00000 -20.92000
0.00000 0.00000 0.00000 ; CA- C- N- H
3337 3340 3342 3344 3 20.92000 0.00000 -20.92000
0.00000 0.00000 0.00000 ; CA- C- N- CA
3338 3337 3340 3341 3 3.68192 -4.35136 0.00000
1.33888 0.00000 0.00000 ; HA2- CA- C- O
3338 3337 3340 3342 3 0.00000 0.00000 0.00000
0.00000 0.00000 0.00000 ; HA2- CA- C- N
3339 3337 3340 3341 3 3.68192 -4.35136 0.00000
1.33888 0.00000 0.00000 ; HA3- CA- C- O
3339 3337 3340 3342 3 0.00000 0.00000 0.00000
0.00000 0.00000 0.00000 ; HA3- CA- C- N
3340 3342 3344 3345 3 0.00000 0.00000 0.00000
0.00000 0.00000 0.00000 ; C- N- CA- HA
3340 3342 3344 3346 3 11.71520 1.67360 15.06240 -
13.38880 0.00000 0.00000 ; C- N- CA- CB
3340 3342 3344 3361 3 1.75728 5.27184 2.25936
-7.02912 0.00000 0.00000 ; C- N- CA- C
3341 3340 3342 3343 3 29.28800 -8.36800 -20.92000
0.00000 0.00000 0.00000 ; O- C- N- H
3341 3340 3342 3344 3 20.92000 0.00000 -20.92000
0.00000 0.00000 0.00000 ; O- C- N- CA
3342 3344 3346 3347 3 0.65084 1.95253 0.00000
-2.60338 0.00000 0.00000 ; N- CA- CB- HB2
3342 3344 3346 3348 3 0.65084 1.95253 0.00000
-2.60338 0.00000 0.00000 ; N- CA- CB- HB3
3342 3344 3346 3349 3 3.28026 2.46019 -2.19242
-3.21331 -0.23430 0.00000 ; N- CA- CB- CG
3342 3344 3361 3362 3 0.00000 0.00000 0.00000
0.00000 0.00000 0.00000 ; N- CA- C- O
3342 3344 3361 3363 3 17.40544 -5.02080 -13.22144
9.20480 0.00000 0.00000 ; N- CA- C- N
3343 3342 3344 3345 3 0.00000 0.00000 0.00000
0.00000 0.00000 0.00000 ; H- N- CA- HA
3343 3342 3344 3346 3 0.00000 0.00000 0.00000
0.00000 0.00000 0.00000 ; H- N- CA- CB
3343 3342 3344 3361 3 0.00000 0.00000 0.00000
0.00000 0.00000 0.00000 ; H- N- CA- C

3344 3346 3349 3350 3 0.57739 0.00000 1.02926
0.00000 -1.60666 0.00000 ; CA- CB- CG- CD1
3344 3346 3349 3359 3 0.57739 0.00000 1.02926
0.00000 -1.60666 0.00000 ; CA- CB- CG- CD2
3344 3361 3363 3364 3 20.92000 0.00000 -20.92000
0.00000 0.00000 0.00000 ; CA- C- N- CD
3344 3361 3363 3373 3 20.92000 0.00000 -20.92000
0.00000 0.00000 0.00000 ; CA- C- N- CA
3345 3344 3346 3347 3 0.65084 1.95253 0.00000
-2.60338 0.00000 0.00000 ; HA- CA- CB- HB2
3345 3344 3346 3348 3 0.65084 1.95253 0.00000
-2.60338 0.00000 0.00000 ; HA- CA- CB- HB3
3345 3344 3346 3349 3 0.65084 1.95253 0.00000
-2.60338 0.00000 0.00000 ; HA- CA- CB- CG
3345 3344 3361 3362 3 3.68192 -4.35136 0.00000
1.33888 0.00000 0.00000 ; HA- CA- C- O
3345 3344 3361 3363 3 0.00000 0.00000 0.00000
0.00000 0.00000 0.00000 ; HA- CA- C- N
3346 3344 3361 3362 3 0.00000 0.00000 0.00000
0.00000 0.00000 0.00000 ; CB- CA- C- O
3346 3344 3361 3363 3 2.51040 4.18400 1.67360
-6.69440 0.00000 0.00000 ; CB- CA- C- N
3346 3349 3350 3351 3 30.33400 0.00000 -30.33400
0.00000 0.00000 0.00000 ; CB- CG- CD1- HD1
3346 3349 3350 3352 3 30.33400 0.00000 -30.33400
0.00000 0.00000 0.00000 ; CB- CG- CD1- CE1
3346 3349 3359 3357 3 30.33400 0.00000 -30.33400
0.00000 0.00000 0.00000 ; CB- CG- CD2- CE2
3346 3349 3359 3360 3 30.33400 0.00000 -30.33400
0.00000 0.00000 0.00000 ; CB- CG- CD2- HD2
3347 3346 3344 3361 3 0.65084 1.95253 0.00000
-2.60338 0.00000 0.00000 ; HB2- CB- CA- C
3347 3346 3349 3350 3 0.00000 0.00000 0.00000
0.00000 0.00000 0.00000 ; HB2- CB- CG- CD1
3347 3346 3349 3359 3 0.00000 0.00000 0.00000
0.00000 0.00000 0.00000 ; HB2- CB- CG- CD2
3348 3346 3344 3361 3 0.65084 1.95253 0.00000
-2.60338 0.00000 0.00000 ; HB3- CB- CA- C
3348 3346 3349 3350 3 0.00000 0.00000 0.00000
0.00000 0.00000 0.00000 ; HB3- CB- CG- CD1
3348 3346 3349 3359 3 0.00000 0.00000 0.00000
0.00000 0.00000 0.00000 ; HB3- CB- CG- CD2
3349 3346 3344 3361 3 4.95804 2.17986 -3.52293
-3.21331 -0.40166 0.00000 ; CG- CB- CA- C
3349 3350 3352 3353 3 30.33400 0.00000 -30.33400
0.00000 0.00000 0.00000 ; CG- CD1- CE1- HE1
3349 3350 3352 3354 3 30.33400 0.00000 -30.33400
0.00000 0.00000 0.00000 ; CG- CD1- CE1- CZ
3349 3359 3357 3354 3 30.33400 0.00000 -30.33400
0.00000 0.00000 0.00000 ; CG- CD2- CE2- CZ
3349 3359 3357 3358 3 30.33400 0.00000 -30.33400
0.00000 0.00000 0.00000 ; CG- CD2- CE2- HE2

3350 3349 3359 3357 3 30.33400 0.00000 -30.33400
0.00000 0.00000 0.00000 ; CD1- CG- CD2- CE2
3350 3349 3359 3360 3 30.33400 0.00000 -30.33400
0.00000 0.00000 0.00000 ; CD1- CG- CD2- HD2
3350 3352 3354 3355 3 30.33400 0.00000 -30.33400
0.00000 0.00000 0.00000 ; CD1- CE1- CZ- OH
3350 3352 3354 3357 3 30.33400 0.00000 -30.33400
0.00000 0.00000 0.00000 ; CD1- CE1- CZ- CE2
3351 3350 3349 3359 3 30.33400 0.00000 -30.33400
0.00000 0.00000 0.00000 ; HD1- CD1- CG- CD2
3351 3350 3352 3353 3 30.33400 0.00000 -30.33400
0.00000 0.00000 0.00000 ; HD1- CD1- CE1- HE1
3351 3350 3352 3354 3 30.33400 0.00000 -30.33400
0.00000 0.00000 0.00000 ; HD1- CD1- CE1- CZ
3352 3350 3349 3359 3 30.33400 0.00000 -30.33400
0.00000 0.00000 0.00000 ; CE1- CD1- CG- CD2
3352 3354 3355 3356 3 7.93286 0.00000 -9.56462
0.00000 2.17568 0.00000 ; CE1- CZ- OH- HH
3352 3354 3357 3358 3 30.33400 0.00000 -30.33400
0.00000 0.00000 0.00000 ; CE1- CZ- CE2- HE2
3352 3354 3357 3359 3 30.33400 0.00000 -30.33400
0.00000 0.00000 0.00000 ; CE1- CZ- CE2- CD2
3353 3352 3354 3355 3 30.33400 0.00000 -30.33400
0.00000 0.00000 0.00000 ; HE1- CE1- CZ- OH
3353 3352 3354 3357 3 30.33400 0.00000 -30.33400
0.00000 0.00000 0.00000 ; HE1- CE1- CZ- CE2
3354 3357 3359 3360 3 30.33400 0.00000 -30.33400
0.00000 0.00000 0.00000 ; CZ- CE2- CD2- HD2
3355 3354 3357 3358 3 30.33400 0.00000 -30.33400
0.00000 0.00000 0.00000 ; OH- CZ- CE2- HE2
3355 3354 3357 3359 3 30.33400 0.00000 -30.33400
0.00000 0.00000 0.00000 ; OH- CZ- CE2- CD2
3356 3355 3354 3357 3 7.93286 0.00000 -9.56462
0.00000 2.17568 0.00000 ; HH- OH- CZ- CE2
3358 3357 3359 3360 3 30.33400 0.00000 -30.33400
0.00000 0.00000 0.00000 ; HE2- CE2- CD2- HD2
3361 3363 3364 3365 3 0.00000 0.00000 0.00000
0.00000 0.00000 0.00000 ; C- N- CD- HD2
3361 3363 3364 3366 3 0.00000 0.00000 0.00000
0.00000 0.00000 0.00000 ; C- N- CD- HD3
3361 3363 3364 3367 3 10.04160 -3.34720 16.73600
-6.69440 0.00000 0.00000 ; C- N- CD- CG
3361 3363 3373 3370 3 11.71520 1.67360 15.06240 -
13.38880 0.00000 0.00000 ; C- N- CA- CB
3361 3363 3373 3374 3 0.00000 0.00000 0.00000
0.00000 0.00000 0.00000 ; C- N- CA- HA
3361 3363 3373 3375 3 1.75728 5.27184 2.25936
-7.02912 0.00000 0.00000 ; C- N- CA- C
3362 3361 3363 3364 3 20.92000 0.00000 -20.92000
0.00000 0.00000 0.00000 ; O- C- N- CD
3362 3361 3363 3373 3 20.92000 0.00000 -20.92000
0.00000 0.00000 0.00000 ; O- C- N- CA

3363 3364 3367 3368 3 0.65084 1.95253 0.00000
-2.60338 0.00000 0.00000 ; N- CD- CG- HG2
3363 3364 3367 3369 3 0.65084 1.95253 0.00000
-2.60338 0.00000 0.00000 ; N- CD- CG- HG3
3363 3364 3367 3370 3 0.65084 1.95253 0.00000
-2.60338 0.00000 0.00000 ; N- CD- CG- CB
3363 3373 3370 3367 3 0.65084 1.95253 0.00000
-2.60338 0.00000 0.00000 ; N- CA- CB- CG
3363 3373 3370 3371 3 0.65084 1.95253 0.00000
-2.60338 0.00000 0.00000 ; N- CA- CB- HB2
3363 3373 3370 3372 3 0.65084 1.95253 0.00000
-2.60338 0.00000 0.00000 ; N- CA- CB- HB3
3363 3373 3375 3376 3 0.00000 0.00000 0.00000
0.00000 0.00000 0.00000 ; N- CA- C- O
3363 3373 3375 3377 3 17.40544 -5.02080 -13.22144
9.20480 0.00000 0.00000 ; N- CA- C- N
3364 3363 3373 3370 3 0.00000 0.00000 0.00000
0.00000 0.00000 0.00000 ; CD- N- CA- CB
3364 3363 3373 3374 3 0.00000 0.00000 0.00000
0.00000 0.00000 0.00000 ; CD- N- CA- HA
3364 3363 3373 3375 3 0.00000 0.00000 0.00000
0.00000 0.00000 0.00000 ; CD- N- CA- C
3364 3367 3370 3371 3 0.66944 2.00832 0.00000
-2.67776 0.00000 0.00000 ; CD- CG- CB- HB2
3364 3367 3370 3372 3 0.66944 2.00832 0.00000
-2.67776 0.00000 0.00000 ; CD- CG- CB- HB3
3364 3367 3370 3373 3 3.68192 3.09616 -2.09200
-3.01248 0.00000 0.00000 ; CD- CG- CB- CA
3365 3364 3363 3373 3 0.00000 0.00000 0.00000
0.00000 0.00000 0.00000 ; HD2- CD- N- CA
3365 3364 3367 3368 3 0.65084 1.95253 0.00000
-2.60338 0.00000 0.00000 ; HD2- CD- CG- HG2
3365 3364 3367 3369 3 0.65084 1.95253 0.00000
-2.60338 0.00000 0.00000 ; HD2- CD- CG- HG3
3365 3364 3367 3370 3 0.65084 1.95253 0.00000
-2.60338 0.00000 0.00000 ; HD2- CD- CG- CB
3366 3364 3363 3373 3 0.00000 0.00000 0.00000
0.00000 0.00000 0.00000 ; HD3- CD- N- CA
3366 3364 3367 3368 3 0.65084 1.95253 0.00000
-2.60338 0.00000 0.00000 ; HD3- CD- CG- HG2
3366 3364 3367 3369 3 0.65084 1.95253 0.00000
-2.60338 0.00000 0.00000 ; HD3- CD- CG- HG3
3366 3364 3367 3370 3 0.65084 1.95253 0.00000
-2.60338 0.00000 0.00000 ; HD3- CD- CG- CB
3367 3364 3363 3373 3 0.00000 0.00000 0.00000
0.00000 0.00000 0.00000 ; CG- CD- N- CA
3367 3370 3373 3374 3 0.65084 1.95253 0.00000
-2.60338 0.00000 0.00000 ; CG- CB- CA- HA
3367 3370 3373 3375 3 0.65084 1.95253 0.00000
-2.60338 0.00000 0.00000 ; CG- CB- CA- C
3368 3367 3370 3371 3 0.62760 1.88280 0.00000
-2.51040 0.00000 0.00000 ; HG2- CG- CB- HB2

3368 3367 3370 3372 3 0.62760 1.88280 0.00000
-2.51040 0.00000 0.00000 ; HG2- CG- CB- HB3
3368 3367 3370 3373 3 0.66944 2.00832 0.00000
-2.67776 0.00000 0.00000 ; HG2- CG- CB- CA
3369 3367 3370 3371 3 0.62760 1.88280 0.00000
-2.51040 0.00000 0.00000 ; HG3- CG- CB- HB2
3369 3367 3370 3372 3 0.62760 1.88280 0.00000
-2.51040 0.00000 0.00000 ; HG3- CG- CB- HB3
3369 3367 3370 3373 3 0.66944 2.00832 0.00000
-2.67776 0.00000 0.00000 ; HG3- CG- CB- CA
3370 3373 3375 3376 3 0.00000 0.00000 0.00000
0.00000 0.00000 0.00000 ; CB- CA- C- O
3370 3373 3375 3377 3 2.51040 4.18400 1.67360
-6.69440 0.00000 0.00000 ; CB- CA- C- N
3371 3370 3373 3374 3 0.65084 1.95253 0.00000
-2.60338 0.00000 0.00000 ; HB2- CB- CA- HA
3371 3370 3373 3375 3 0.65084 1.95253 0.00000
-2.60338 0.00000 0.00000 ; HB2- CB- CA- C
3372 3370 3373 3374 3 0.65084 1.95253 0.00000
-2.60338 0.00000 0.00000 ; HB3- CB- CA- HA
3372 3370 3373 3375 3 0.65084 1.95253 0.00000
-2.60338 0.00000 0.00000 ; HB3- CB- CA- C
3373 3375 3377 3378 3 20.92000 0.00000 -20.92000
0.00000 0.00000 0.00000 ; CA- C- N- CD
3373 3375 3377 3387 3 20.92000 0.00000 -20.92000
0.00000 0.00000 0.00000 ; CA- C- N- CA
3374 3373 3375 3376 3 3.68192 -4.35136 0.00000
1.33888 0.00000 0.00000 ; HA- CA- C- O
3374 3373 3375 3377 3 0.00000 0.00000 0.00000
0.00000 0.00000 0.00000 ; HA- CA- C- N
3375 3377 3378 3379 3 0.00000 0.00000 0.00000
0.00000 0.00000 0.00000 ; C- N- CD- HD2
3375 3377 3378 3380 3 0.00000 0.00000 0.00000
0.00000 0.00000 0.00000 ; C- N- CD- HD3
3375 3377 3378 3381 3 10.04160 -3.34720 16.73600
-6.69440 0.00000 0.00000 ; C- N- CD- CG
3375 3377 3387 3384 3 11.71520 1.67360 15.06240 -
13.38880 0.00000 0.00000 ; C- N- CA- CB
3375 3377 3387 3388 3 0.00000 0.00000 0.00000
0.00000 0.00000 0.00000 ; C- N- CA- HA
3375 3377 3387 3389 3 1.75728 5.27184 2.25936
-7.02912 0.00000 0.00000 ; C- N- CA- C
3376 3375 3377 3378 3 20.92000 0.00000 -20.92000
0.00000 0.00000 0.00000 ; O- C- N- CD
3376 3375 3377 3387 3 20.92000 0.00000 -20.92000
0.00000 0.00000 0.00000 ; O- C- N- CA
3377 3378 3381 3382 3 0.65084 1.95253 0.00000
-2.60338 0.00000 0.00000 ; N- CD- CG- HG2
3377 3378 3381 3383 3 0.65084 1.95253 0.00000
-2.60338 0.00000 0.00000 ; N- CD- CG- HG3
3377 3378 3381 3384 3 0.65084 1.95253 0.00000
-2.60338 0.00000 0.00000 ; N- CD- CG- CB

3377 3387 3384 3381 3 0.65084 1.95253 0.00000
-2.60338 0.00000 0.00000 ; N- CA- CB- CG
3377 3387 3384 3385 3 0.65084 1.95253 0.00000
-2.60338 0.00000 0.00000 ; N- CA- CB- HB2
3377 3387 3384 3386 3 0.65084 1.95253 0.00000
-2.60338 0.00000 0.00000 ; N- CA- CB- HB3
3377 3387 3389 3390 3 0.00000 0.00000 0.00000
0.00000 0.00000 0.00000 ; N- CA- C- O
3377 3387 3389 3391 3 17.40544 -5.02080 -13.22144
9.20480 0.00000 0.00000 ; N- CA- C- N
3378 3377 3387 3384 3 0.00000 0.00000 0.00000
0.00000 0.00000 0.00000 ; CD- N- CA- CB
3378 3377 3387 3388 3 0.00000 0.00000 0.00000
0.00000 0.00000 0.00000 ; CD- N- CA- HA
3378 3377 3387 3389 3 0.00000 0.00000 0.00000
0.00000 0.00000 0.00000 ; CD- N- CA- C
3378 3381 3384 3385 3 0.66944 2.00832 0.00000
-2.67776 0.00000 0.00000 ; CD- CG- CB- HB2
3378 3381 3384 3386 3 0.66944 2.00832 0.00000
-2.67776 0.00000 0.00000 ; CD- CG- CB- HB3
3378 3381 3384 3387 3 3.68192 3.09616 -2.09200
-3.01248 0.00000 0.00000 ; CD- CG- CB- CA
3379 3378 3377 3387 3 0.00000 0.00000 0.00000
0.00000 0.00000 0.00000 ; HD2- CD- N- CA
3379 3378 3381 3382 3 0.65084 1.95253 0.00000
-2.60338 0.00000 0.00000 ; HD2- CD- CG- HG2
3379 3378 3381 3383 3 0.65084 1.95253 0.00000
-2.60338 0.00000 0.00000 ; HD2- CD- CG- HG3
3379 3378 3381 3384 3 0.65084 1.95253 0.00000
-2.60338 0.00000 0.00000 ; HD2- CD- CG- CB
3380 3378 3377 3387 3 0.00000 0.00000 0.00000
0.00000 0.00000 0.00000 ; HD3- CD- N- CA
3380 3378 3381 3382 3 0.65084 1.95253 0.00000
-2.60338 0.00000 0.00000 ; HD3- CD- CG- HG2
3380 3378 3381 3383 3 0.65084 1.95253 0.00000
-2.60338 0.00000 0.00000 ; HD3- CD- CG- HG3
3380 3378 3381 3384 3 0.65084 1.95253 0.00000
-2.60338 0.00000 0.00000 ; HD3- CD- CG- CB
3381 3378 3377 3387 3 0.00000 0.00000 0.00000
0.00000 0.00000 0.00000 ; CG- CD- N- CA
3381 3384 3387 3388 3 0.65084 1.95253 0.00000
-2.60338 0.00000 0.00000 ; CG- CB- CA- HA
3381 3384 3387 3389 3 0.65084 1.95253 0.00000
-2.60338 0.00000 0.00000 ; CG- CB- CA- C
3382 3381 3384 3385 3 0.62760 1.88280 0.00000
-2.51040 0.00000 0.00000 ; HG2- CG- CB- HB2
3382 3381 3384 3386 3 0.62760 1.88280 0.00000
-2.51040 0.00000 0.00000 ; HG2- CG- CB- HB3
3382 3381 3384 3387 3 0.66944 2.00832 0.00000
-2.67776 0.00000 0.00000 ; HG2- CG- CB- CA
3383 3381 3384 3385 3 0.62760 1.88280 0.00000
-2.51040 0.00000 0.00000 ; HG3- CG- CB- HB2

3383 3381 3384 3386 3 0.62760 1.88280 0.00000
-2.51040 0.00000 0.00000 ; HG3- CG- CB- HB3
3383 3381 3384 3387 3 0.66944 2.00832 0.00000
-2.67776 0.00000 0.00000 ; HG3- CG- CB- CA
3384 3387 3389 3390 3 0.00000 0.00000 0.00000
0.00000 0.00000 0.00000 ; CB- CA- C- O
3384 3387 3389 3391 3 2.51040 4.18400 1.67360
-6.69440 0.00000 0.00000 ; CB- CA- C- N
3385 3384 3387 3388 3 0.65084 1.95253 0.00000
-2.60338 0.00000 0.00000 ; HB2- CB- CA- HA
3385 3384 3387 3389 3 0.65084 1.95253 0.00000
-2.60338 0.00000 0.00000 ; HB2- CB- CA- C
3386 3384 3387 3388 3 0.65084 1.95253 0.00000
-2.60338 0.00000 0.00000 ; HB3- CB- CA- HA
3386 3384 3387 3389 3 0.65084 1.95253 0.00000
-2.60338 0.00000 0.00000 ; HB3- CB- CA- C
3387 3389 3391 3392 3 20.92000 0.00000 -20.92000
0.00000 0.00000 0.00000 ; CA- C- N- H
3387 3389 3391 3393 3 20.92000 0.00000 -20.92000
0.00000 0.00000 0.00000 ; CA- C- N- CA
3388 3387 3389 3390 3 3.68192 -4.35136 0.00000
1.33888 0.00000 0.00000 ; HA- CA- C- O
3388 3387 3389 3391 3 0.00000 0.00000 0.00000
0.00000 0.00000 0.00000 ; HA- CA- C- N
3389 3391 3393 3394 3 0.00000 0.00000 0.00000
0.00000 0.00000 0.00000 ; C- N- CA- HA
3389 3391 3393 3395 3 11.71520 1.67360 15.06240 -
13.38880 0.00000 0.00000 ; C- N- CA- CB
3389 3391 3393 3409 3 1.75728 5.27184 2.25936
-7.02912 0.00000 0.00000 ; C- N- CA- C
3390 3389 3391 3392 3 29.28800 -8.36800 -20.92000
0.00000 0.00000 0.00000 ; O- C- N- H
3390 3389 3391 3393 3 20.92000 0.00000 -20.92000
0.00000 0.00000 0.00000 ; O- C- N- CA
3391 3393 3395 3396 3 0.65084 1.95253 0.00000
-2.60338 0.00000 0.00000 ; N- CA- CB- HB2
3391 3393 3395 3397 3 0.65084 1.95253 0.00000
-2.60338 0.00000 0.00000 ; N- CA- CB- HB3
3391 3393 3395 3398 3 3.28026 2.46019 -2.19242
-3.21331 -0.23430 0.00000 ; N- CA- CB- CG
3391 3393 3409 3410 3 0.00000 0.00000 0.00000
0.00000 0.00000 0.00000 ; N- CA- C- O
3391 3393 3409 3411 3 17.40544 -5.02080 -13.22144
9.20480 0.00000 0.00000 ; N- CA- C- N
3392 3391 3393 3394 3 0.00000 0.00000 0.00000
0.00000 0.00000 0.00000 ; H- N- CA- HA
3392 3391 3393 3395 3 0.00000 0.00000 0.00000
0.00000 0.00000 0.00000 ; H- N- CA- CB
3392 3391 3393 3409 3 0.00000 0.00000 0.00000
0.00000 0.00000 0.00000 ; H- N- CA- C
3393 3395 3398 3399 3 0.57739 0.00000 1.02926
0.00000 -1.60666 0.00000 ; CA- CB- CG- CD1

3393 3395 3398 3407 3 0.57739 0.00000 1.02926
0.00000 -1.60666 0.00000 ; CA- CB- CG- CD2
3393 3409 3411 3412 3 20.92000 0.00000 -20.92000
0.00000 0.00000 0.00000 ; CA- C- N- H
3393 3409 3411 3413 3 20.92000 0.00000 -20.92000
0.00000 0.00000 0.00000 ; CA- C- N- CA
3394 3393 3395 3396 3 0.65084 1.95253 0.00000
-2.60338 0.00000 0.00000 ; HA- CA- CB- HB2
3394 3393 3395 3397 3 0.65084 1.95253 0.00000
-2.60338 0.00000 0.00000 ; HA- CA- CB- HB3
3394 3393 3395 3398 3 0.65084 1.95253 0.00000
-2.60338 0.00000 0.00000 ; HA- CA- CB- CG
3394 3393 3409 3410 3 3.68192 -4.35136 0.00000
1.33888 0.00000 0.00000 ; HA- CA- C- O
3394 3393 3409 3411 3 0.00000 0.00000 0.00000
0.00000 0.00000 0.00000 ; HA- CA- C- N
3395 3393 3409 3410 3 0.00000 0.00000 0.00000
0.00000 0.00000 0.00000 ; CB- CA- C- O
3395 3393 3409 3411 3 2.51040 4.18400 1.67360
-6.69440 0.00000 0.00000 ; CB- CA- C- N
3395 3398 3399 3400 3 30.33400 0.00000 -30.33400
0.00000 0.00000 0.00000 ; CB- CG- CD1- HD1
3395 3398 3399 3401 3 30.33400 0.00000 -30.33400
0.00000 0.00000 0.00000 ; CB- CG- CD1- CE1
3395 3398 3407 3405 3 30.33400 0.00000 -30.33400
0.00000 0.00000 0.00000 ; CB- CG- CD2- CE2
3395 3398 3407 3408 3 30.33400 0.00000 -30.33400
0.00000 0.00000 0.00000 ; CB- CG- CD2- HD2
3396 3395 3393 3409 3 0.65084 1.95253 0.00000
-2.60338 0.00000 0.00000 ; HB2- CB- CA- C
3396 3395 3398 3399 3 0.00000 0.00000 0.00000
0.00000 0.00000 0.00000 ; HB2- CB- CG- CD1
3396 3395 3398 3407 3 0.00000 0.00000 0.00000
0.00000 0.00000 0.00000 ; HB2- CB- CG- CD2
3397 3395 3393 3409 3 0.65084 1.95253 0.00000
-2.60338 0.00000 0.00000 ; HB3- CB- CA- C
3397 3395 3398 3399 3 0.00000 0.00000 0.00000
0.00000 0.00000 0.00000 ; HB3- CB- CG- CD1
3397 3395 3398 3407 3 0.00000 0.00000 0.00000
0.00000 0.00000 0.00000 ; HB3- CB- CG- CD2
3398 3395 3393 3409 3 4.95804 2.17986 -3.52293
-3.21331 -0.40166 0.00000 ; CG- CB- CA- C
3398 3399 3401 3402 3 30.33400 0.00000 -30.33400
0.00000 0.00000 0.00000 ; CG- CD1- CE1- HE1
3398 3399 3401 3403 3 30.33400 0.00000 -30.33400
0.00000 0.00000 0.00000 ; CG- CD1- CE1- CZ
3398 3407 3405 3403 3 30.33400 0.00000 -30.33400
0.00000 0.00000 0.00000 ; CG- CD2- CE2- CZ
3398 3407 3405 3406 3 30.33400 0.00000 -30.33400
0.00000 0.00000 0.00000 ; CG- CD2- CE2- HE2
3399 3398 3407 3405 3 30.33400 0.00000 -30.33400
0.00000 0.00000 0.00000 ; CD1- CG- CD2- CE2

3399 3398 3407 3408 3 30.33400 0.00000 -30.33400
0.00000 0.00000 0.00000 ; CD1- CG- CD2- HD2
3399 3401 3403 3404 3 30.33400 0.00000 -30.33400
0.00000 0.00000 0.00000 ; CD1- CE1- CZ- HZ
3399 3401 3403 3405 3 30.33400 0.00000 -30.33400
0.00000 0.00000 0.00000 ; CD1- CE1- CZ- CE2
3400 3399 3398 3407 3 30.33400 0.00000 -30.33400
0.00000 0.00000 0.00000 ; HD1- CD1- CG- CD2
3400 3399 3401 3402 3 30.33400 0.00000 -30.33400
0.00000 0.00000 0.00000 ; HD1- CD1- CE1- HE1
3400 3399 3401 3403 3 30.33400 0.00000 -30.33400
0.00000 0.00000 0.00000 ; HD1- CD1- CE1- CZ
3401 3399 3398 3407 3 30.33400 0.00000 -30.33400
0.00000 0.00000 0.00000 ; CE1- CD1- CG- CD2
3401 3403 3405 3406 3 30.33400 0.00000 -30.33400
0.00000 0.00000 0.00000 ; CE1- CZ- CE2- HE2
3401 3403 3405 3407 3 30.33400 0.00000 -30.33400
0.00000 0.00000 0.00000 ; CE1- CZ- CE2- CD2
3402 3401 3403 3404 3 30.33400 0.00000 -30.33400
0.00000 0.00000 0.00000 ; HE1- CE1- CZ- HZ
3402 3401 3403 3405 3 30.33400 0.00000 -30.33400
0.00000 0.00000 0.00000 ; HE1- CE1- CZ- CE2
3403 3405 3407 3408 3 30.33400 0.00000 -30.33400
0.00000 0.00000 0.00000 ; CZ- CE2- CD2- HD2
3404 3403 3405 3406 3 30.33400 0.00000 -30.33400
0.00000 0.00000 0.00000 ; HZ- CZ- CE2- HE2
3404 3403 3405 3407 3 30.33400 0.00000 -30.33400
0.00000 0.00000 0.00000 ; HZ- CZ- CE2- CD2
3406 3405 3407 3408 3 30.33400 0.00000 -30.33400
0.00000 0.00000 0.00000 ; HE2- CE2- CD2- HD2
3409 3411 3413 3414 3 0.00000 0.00000 0.00000
0.00000 0.00000 0.00000 ; C- N- CA- HA
3409 3411 3413 3415 3 11.71520 1.67360 15.06240 -
13.38880 0.00000 0.00000 ; C- N- CA- CB
3409 3411 3413 3430 3 1.75728 5.27184 2.25936
-7.02912 0.00000 0.00000 ; C- N- CA- C
3410 3409 3411 3412 3 29.28800 -8.36800 -20.92000
0.00000 0.00000 0.00000 ; O- C- N- H
3410 3409 3411 3413 3 20.92000 0.00000 -20.92000
0.00000 0.00000 0.00000 ; O- C- N- CA
3411 3413 3415 3416 3 0.65084 1.95253 0.00000
-2.60338 0.00000 0.00000 ; N- CA- CB- HB2
3411 3413 3415 3417 3 0.65084 1.95253 0.00000
-2.60338 0.00000 0.00000 ; N- CA- CB- HB3
3411 3413 3415 3418 3 3.28026 2.46019 -2.19242
-3.21331 -0.23430 0.00000 ; N- CA- CB- CG
3411 3413 3430 3431 3 0.00000 0.00000 0.00000
0.00000 0.00000 0.00000 ; N- CA- C- O
3411 3413 3430 3432 3 17.40544 -5.02080 -13.22144
9.20480 0.00000 0.00000 ; N- CA- C- N
3412 3411 3413 3414 3 0.00000 0.00000 0.00000
0.00000 0.00000 0.00000 ; H- N- CA- HA

3412 3411 3413 3415 3 0.00000 0.00000 0.00000
0.00000 0.00000 0.00000 ; H- N- CA- CB
3412 3411 3413 3430 3 0.00000 0.00000 0.00000
0.00000 0.00000 0.00000 ; H- N- CA- C
3413 3415 3418 3419 3 0.57739 0.00000 1.02926
0.00000 -1.60666 0.00000 ; CA- CB- CG- CD1
3413 3415 3418 3428 3 0.57739 0.00000 1.02926
0.00000 -1.60666 0.00000 ; CA- CB- CG- CD2
3413 3430 3432 3433 3 20.92000 0.00000 -20.92000
0.00000 0.00000 0.00000 ; CA- C- N- H
3413 3430 3432 3434 3 20.92000 0.00000 -20.92000
0.00000 0.00000 0.00000 ; CA- C- N- CA
3414 3413 3415 3416 3 0.65084 1.95253 0.00000
-2.60338 0.00000 0.00000 ; HA- CA- CB- HB2
3414 3413 3415 3417 3 0.65084 1.95253 0.00000
-2.60338 0.00000 0.00000 ; HA- CA- CB- HB3
3414 3413 3415 3418 3 0.65084 1.95253 0.00000
-2.60338 0.00000 0.00000 ; HA- CA- CB- CG
3414 3413 3430 3431 3 3.68192 -4.35136 0.00000
1.33888 0.00000 0.00000 ; HA- CA- C- O
3414 3413 3430 3432 3 0.00000 0.00000 0.00000
0.00000 0.00000 0.00000 ; HA- CA- C- N
3415 3413 3430 3431 3 0.00000 0.00000 0.00000
0.00000 0.00000 0.00000 ; CB- CA- C- O
3415 3413 3430 3432 3 2.51040 4.18400 1.67360
-6.69440 0.00000 0.00000 ; CB- CA- C- N
3415 3418 3419 3420 3 30.33400 0.00000 -30.33400
0.00000 0.00000 0.00000 ; CB- CG- CD1- HD1
3415 3418 3419 3421 3 30.33400 0.00000 -30.33400
0.00000 0.00000 0.00000 ; CB- CG- CD1- CE1
3415 3418 3428 3426 3 30.33400 0.00000 -30.33400
0.00000 0.00000 0.00000 ; CB- CG- CD2- CE2
3415 3418 3428 3429 3 30.33400 0.00000 -30.33400
0.00000 0.00000 0.00000 ; CB- CG- CD2- HD2
3416 3415 3413 3430 3 0.65084 1.95253 0.00000
-2.60338 0.00000 0.00000 ; HB2- CB- CA- C
3416 3415 3418 3419 3 0.00000 0.00000 0.00000
0.00000 0.00000 0.00000 ; HB2- CB- CG- CD1
3416 3415 3418 3428 3 0.00000 0.00000 0.00000
0.00000 0.00000 0.00000 ; HB2- CB- CG- CD2
3417 3415 3413 3430 3 0.65084 1.95253 0.00000
-2.60338 0.00000 0.00000 ; HB3- CB- CA- C
3417 3415 3418 3419 3 0.00000 0.00000 0.00000
0.00000 0.00000 0.00000 ; HB3- CB- CG- CD1
3417 3415 3418 3428 3 0.00000 0.00000 0.00000
0.00000 0.00000 0.00000 ; HB3- CB- CG- CD2
3418 3415 3413 3430 3 4.95804 2.17986 -3.52293
-3.21331 -0.40166 0.00000 ; CG- CB- CA- C
3418 3419 3421 3422 3 30.33400 0.00000 -30.33400
0.00000 0.00000 0.00000 ; CG- CD1- CE1- HE1
3418 3419 3421 3423 3 30.33400 0.00000 -30.33400
0.00000 0.00000 0.00000 ; CG- CD1- CE1- CZ

3418 3428 3426 3423 3 30.33400 0.00000 -30.33400
0.00000 0.00000 0.00000 ; CG- CD2- CE2- CZ
3418 3428 3426 3427 3 30.33400 0.00000 -30.33400
0.00000 0.00000 0.00000 ; CG- CD2- CE2- HE2
3419 3418 3428 3426 3 30.33400 0.00000 -30.33400
0.00000 0.00000 0.00000 ; CD1- CG- CD2- CE2
3419 3418 3428 3429 3 30.33400 0.00000 -30.33400
0.00000 0.00000 0.00000 ; CD1- CG- CD2- HD2
3419 3421 3423 3424 3 30.33400 0.00000 -30.33400
0.00000 0.00000 0.00000 ; CD1- CE1- CZ- OH
3419 3421 3423 3426 3 30.33400 0.00000 -30.33400
0.00000 0.00000 0.00000 ; CD1- CE1- CZ- CE2
3420 3419 3418 3428 3 30.33400 0.00000 -30.33400
0.00000 0.00000 0.00000 ; HD1- CD1- CG- CD2
3420 3419 3421 3422 3 30.33400 0.00000 -30.33400
0.00000 0.00000 0.00000 ; HD1- CD1- CE1- HE1
3420 3419 3421 3423 3 30.33400 0.00000 -30.33400
0.00000 0.00000 0.00000 ; HD1- CD1- CE1- CZ
3421 3419 3418 3428 3 30.33400 0.00000 -30.33400
0.00000 0.00000 0.00000 ; CE1- CD1- CG- CD2
3421 3423 3424 3425 3 7.93286 0.00000 -9.56462
0.00000 2.17568 0.00000 ; CE1- CZ- OH- HH
3421 3423 3426 3427 3 30.33400 0.00000 -30.33400
0.00000 0.00000 0.00000 ; CE1- CZ- CE2- HE2
3421 3423 3426 3428 3 30.33400 0.00000 -30.33400
0.00000 0.00000 0.00000 ; CE1- CZ- CE2- CD2
3422 3421 3423 3424 3 30.33400 0.00000 -30.33400
0.00000 0.00000 0.00000 ; HE1- CE1- CZ- OH
3422 3421 3423 3426 3 30.33400 0.00000 -30.33400
0.00000 0.00000 0.00000 ; HE1- CE1- CZ- CE2
3423 3426 3428 3429 3 30.33400 0.00000 -30.33400
0.00000 0.00000 0.00000 ; CZ- CE2- CD2- HD2
3424 3423 3426 3427 3 30.33400 0.00000 -30.33400
0.00000 0.00000 0.00000 ; OH- CZ- CE2- HE2
3424 3423 3426 3428 3 30.33400 0.00000 -30.33400
0.00000 0.00000 0.00000 ; OH- CZ- CE2- CD2
3425 3424 3423 3426 3 7.93286 0.00000 -9.56462
0.00000 2.17568 0.00000 ; HH- OH- CZ- CE2
3427 3426 3428 3429 3 30.33400 0.00000 -30.33400
0.00000 0.00000 0.00000 ; HE2- CE2- CD2- HD2
3430 3432 3434 3435 3 0.00000 0.00000 0.00000
0.00000 0.00000 0.00000 ; C- N- CA- HA2
3430 3432 3434 3436 3 0.00000 0.00000 0.00000
0.00000 0.00000 0.00000 ; C- N- CA- HA3
3430 3432 3434 3437 3 1.75728 5.27184 2.25936
-7.02912 0.00000 0.00000 ; C- N- CA- C
3431 3430 3432 3433 3 29.28800 -8.36800 -20.92000
0.00000 0.00000 0.00000 ; O- C- N- H
3431 3430 3432 3434 3 20.92000 0.00000 -20.92000
0.00000 0.00000 0.00000 ; O- C- N- CA
3432 3434 3437 3438 3 0.00000 0.00000 0.00000
0.00000 0.00000 0.00000 ; N- CA- C- O

3432 3434 3437 3439 3 17.40544 -5.02080 -13.22144
9.20480 0.00000 0.00000 ; N- CA- C- N
3433 3432 3434 3435 3 0.00000 0.00000 0.00000
0.00000 0.00000 0.00000 ; H- N- CA- HA2
3433 3432 3434 3436 3 0.00000 0.00000 0.00000
0.00000 0.00000 0.00000 ; H- N- CA- HA3
3433 3432 3434 3437 3 0.00000 0.00000 0.00000
0.00000 0.00000 0.00000 ; H- N- CA- C
3434 3437 3439 3440 3 20.92000 0.00000 -20.92000
0.00000 0.00000 0.00000 ; CA- C- N- H
3434 3437 3439 3441 3 20.92000 0.00000 -20.92000
0.00000 0.00000 0.00000 ; CA- C- N- CA
3435 3434 3437 3438 3 3.68192 -4.35136 0.00000
1.33888 0.00000 0.00000 ; HA2- CA- C- O
3435 3434 3437 3439 3 0.00000 0.00000 0.00000
0.00000 0.00000 0.00000 ; HA2- CA- C- N
3436 3434 3437 3438 3 3.68192 -4.35136 0.00000
1.33888 0.00000 0.00000 ; HA3- CA- C- O
3436 3434 3437 3439 3 0.00000 0.00000 0.00000
0.00000 0.00000 0.00000 ; HA3- CA- C- N
3437 3439 3441 3442 3 0.00000 0.00000 0.00000
0.00000 0.00000 0.00000 ; C- N- CA- HA
3437 3439 3441 3443 3 11.71520 1.67360 15.06240 -
13.38880 0.00000 0.00000 ; C- N- CA- CB
3437 3439 3441 3449 3 1.75728 5.27184 2.25936
-7.02912 0.00000 0.00000 ; C- N- CA- C
3438 3437 3439 3440 3 29.28800 -8.36800 -20.92000
0.00000 0.00000 0.00000 ; O- C- N- H
3438 3437 3439 3441 3 20.92000 0.00000 -20.92000
0.00000 0.00000 0.00000 ; O- C- N- CA
3439 3441 3443 3444 3 0.65084 1.95253 0.00000
-2.60338 0.00000 0.00000 ; N- CA- CB- HB2
3439 3441 3443 3445 3 0.65084 1.95253 0.00000
-2.60338 0.00000 0.00000 ; N- CA- CB- HB3
3439 3441 3443 3446 3 15.41386 9.74035 -8.39310
-0.97069 2.97901 0.00000 ; N- CA- CB- CG
3439 3441 3449 3450 3 0.00000 0.00000 0.00000
0.00000 0.00000 0.00000 ; N- CA- C- O
3439 3441 3449 3451 3 17.40544 -5.02080 -13.22144
9.20480 0.00000 0.00000 ; N- CA- C- N
3440 3439 3441 3442 3 0.00000 0.00000 0.00000
0.00000 0.00000 0.00000 ; H- N- CA- HA
3440 3439 3441 3443 3 0.00000 0.00000 0.00000
0.00000 0.00000 0.00000 ; H- N- CA- CB
3440 3439 3441 3449 3 0.00000 0.00000 0.00000
0.00000 0.00000 0.00000 ; H- N- CA- C
3441 3443 3446 3447 3 6.43499 0.00000 -5.39736
0.00000 -1.03763 0.00000 ; CA- CB- CG- OD1
3441 3443 3446 3448 3 6.43499 0.00000 -5.39736
0.00000 -1.03763 0.00000 ; CA- CB- CG- OD2
3441 3449 3451 3452 3 20.92000 0.00000 -20.92000
0.00000 0.00000 0.00000 ; CA- C- N- H

3441 3449 3451 3453 3 20.92000 0.00000 -20.92000
0.00000 0.00000 0.00000 ; CA- C- N- CA
3442 3441 3443 3444 3 0.65084 1.95253 0.00000
-2.60338 0.00000 0.00000 ; HA- CA- CB- HB2
3442 3441 3443 3445 3 0.65084 1.95253 0.00000
-2.60338 0.00000 0.00000 ; HA- CA- CB- HB3
3442 3441 3443 3446 3 0.65084 1.95253 0.00000
-2.60338 0.00000 0.00000 ; HA- CA- CB- CG
3442 3441 3449 3450 3 3.68192 -4.35136 0.00000
1.33888 0.00000 0.00000 ; HA- CA- C- O
3442 3441 3449 3451 3 0.00000 0.00000 0.00000
0.00000 0.00000 0.00000 ; HA- CA- C- N
3443 3441 3449 3450 3 0.00000 0.00000 0.00000
0.00000 0.00000 0.00000 ; CB- CA- C- O
3443 3441 3449 3451 3 2.51040 4.18400 1.67360
-6.69440 0.00000 0.00000 ; CB- CA- C- N
3444 3443 3441 3449 3 0.65084 1.95253 0.00000
-2.60338 0.00000 0.00000 ; HB2- CB- CA- C
3444 3443 3446 3447 3 0.00000 0.00000 0.00000
0.00000 0.00000 0.00000 ; HB2- CB- CG- OD1
3444 3443 3446 3448 3 0.00000 0.00000 0.00000
0.00000 0.00000 0.00000 ; HB2- CB- CG- OD2
3445 3443 3441 3449 3 0.65084 1.95253 0.00000
-2.60338 0.00000 0.00000 ; HB3- CB- CA- C
3445 3443 3446 3447 3 0.00000 0.00000 0.00000
0.00000 0.00000 0.00000 ; HB3- CB- CG- OD1
3445 3443 3446 3448 3 0.00000 0.00000 0.00000
0.00000 0.00000 0.00000 ; HB3- CB- CG- OD2
3446 3443 3441 3449 3 7.14627 -1.04600 -8.99560
-0.97069 5.15469 0.00000 ; CG- CB- CA- C
3449 3451 3453 3454 3 0.00000 0.00000 0.00000
0.00000 0.00000 0.00000 ; C- N- CA- HA
3449 3451 3453 3455 3 11.71520 1.67360 15.06240 -
13.38880 0.00000 0.00000 ; C- N- CA- CB
3449 3451 3453 3463 3 1.75728 5.27184 2.25936
-7.02912 0.00000 0.00000 ; C- N- CA- C
3450 3449 3451 3452 3 29.28800 -8.36800 -20.92000
0.00000 0.00000 0.00000 ; O- C- N- H
3450 3449 3451 3453 3 20.92000 0.00000 -20.92000
0.00000 0.00000 0.00000 ; O- C- N- CA
3451 3453 3455 3456 3 0.65084 1.95253 0.00000
-2.60338 0.00000 0.00000 ; N- CA- CB- HB
3451 3453 3455 3457 3 3.83673 0.44769 -1.77402
-2.47693 -0.03347 0.00000 ; N- CA- CB- CG2
3451 3453 3455 3461 3 4.93294 1.13386 -3.12963
-5.27184 3.17984 0.00000 ; N- CA- CB- OG1
3451 3453 3463 3464 3 0.00000 0.00000 0.00000
0.00000 0.00000 0.00000 ; N- CA- C- O
3451 3453 3463 3465 3 17.40544 -5.02080 -13.22144
9.20480 0.00000 0.00000 ; N- CA- C- N
3452 3451 3453 3454 3 0.00000 0.00000 0.00000
0.00000 0.00000 0.00000 ; H- N- CA- HA

```
  3452   3451   3453   3455      3    0.00000    0.00000    0.00000
0.00000    0.00000    0.00000 ;       H-     N-    CA-    CB
  3452   3451   3453   3463      3    0.00000    0.00000    0.00000
0.00000    0.00000    0.00000 ;       H-     N-    CA-     C
  3453   3455   3457   3458      3    0.66944    2.00832    0.00000
-2.67776    0.00000    0.00000 ;      CA-    CB-   CG2-  HG21
  3453   3455   3457   3459      3    0.66944    2.00832    0.00000
-2.67776    0.00000    0.00000 ;      CA-    CB-   CG2-  HG22
  3453   3455   3457   3460      3    0.66944    2.00832    0.00000
-2.67776    0.00000    0.00000 ;      CA-    CB-   CG2-  HG23
  3453   3455   3461   3462      3    1.12131    2.98738    1.66523
-3.94970    0.43514    0.00000 ;      CA-    CB-   OG1-   HG1
  3453   3463   3465   3466      3   20.92000    0.00000  -20.92000
0.00000    0.00000    0.00000 ;      CA-     C-     N-     H
  3453   3463   3465   3467      3   20.92000    0.00000  -20.92000
0.00000    0.00000    0.00000 ;      CA-     C-     N-    CA
  3454   3453   3455   3456      3    0.65084    1.95253    0.00000
-2.60338    0.00000    0.00000 ;      HA-    CA-    CB-    HB
  3454   3453   3455   3457      3    0.65084    1.95253    0.00000
-2.60338    0.00000    0.00000 ;      HA-    CA-    CB-   CG2
  3454   3453   3455   3461      3    1.04600   -1.04600    0.00000
0.00000    0.00000    0.00000 ;      HA-    CA-    CB-   OG1
  3454   3453   3463   3464      3    3.68192   -4.35136    0.00000
1.33888    0.00000    0.00000 ;      HA-    CA-     C-     O
  3454   3453   3463   3465      3    0.00000    0.00000    0.00000
0.00000    0.00000    0.00000 ;      HA-    CA-     C-     N
  3455   3453   3463   3464      3    0.00000    0.00000    0.00000
0.00000    0.00000    0.00000 ;      CB-    CA-     C-     O
  3455   3453   3463   3465      3    2.51040    4.18400    1.67360
-6.69440    0.00000    0.00000 ;      CB-    CA-     C-     N
  3456   3455   3453   3463      3    0.65084    1.95253    0.00000
-2.60338    0.00000    0.00000 ;      HB-    CB-    CA-     C
  3456   3455   3457   3458      3    0.65084    1.95253    0.00000
-2.60338    0.00000    0.00000 ;      HB-    CB-   CG2-  HG21
  3456   3455   3457   3459      3    0.65084    1.95253    0.00000
-2.60338    0.00000    0.00000 ;      HB-    CB-   CG2-  HG22
  3456   3455   3457   3460      3    0.65084    1.95253    0.00000
-2.60338    0.00000    0.00000 ;      HB-    CB-   CG2-  HG23
  3456   3455   3461   3462      3    0.69733    2.09200    0.00000
-2.78933    0.00000    0.00000 ;      HB-    CB-   OG1-   HG1
  3457   3455   3453   3463      3    5.67350    3.55640   -6.16722
-2.47693    3.74886    0.00000 ;     CG2-    CB-    CA-     C
  3457   3455   3461   3462      3    4.74047    0.27196   -2.26773
-3.94970    1.60666    0.00000 ;     CG2-    CB-   OG1-   HG1
  3458   3457   3455   3461      3    1.04600   -1.04600    0.00000
0.00000    0.00000    0.00000 ;    HG21-   CG2-    CB-   OG1
  3459   3457   3455   3461      3    1.04600   -1.04600    0.00000
0.00000    0.00000    0.00000 ;    HG22-   CG2-    CB-   OG1
  3460   3457   3455   3461      3    1.04600   -1.04600    0.00000
0.00000    0.00000    0.00000 ;    HG23-   CG2-    CB-   OG1
  3461   3455   3453   3463      3    6.53541    6.87013   -6.21742
-5.27184    5.22163    0.00000 ;     OG1-    CB-    CA-     C
```

3463 3465 3467 3468 3 0.00000 0.00000 0.00000
0.00000 0.00000 0.00000 ; C- N- CA- HA
3463 3465 3467 3469 3 11.71520 1.67360 15.06240 -
13.38880 0.00000 0.00000 ; C- N- CA- CB
3463 3465 3467 3475 3 1.75728 5.27184 2.25936
-7.02912 0.00000 0.00000 ; C- N- CA- C
3464 3463 3465 3466 3 29.28800 -8.36800 -20.92000
0.00000 0.00000 0.00000 ; O- C- N- H
3464 3463 3465 3467 3 20.92000 0.00000 -20.92000
0.00000 0.00000 0.00000 ; O- C- N- CA
3465 3467 3469 3470 3 0.65084 1.95253 0.00000
-2.60338 0.00000 0.00000 ; N- CA- CB- HB2
3465 3467 3469 3471 3 0.65084 1.95253 0.00000
-2.60338 0.00000 0.00000 ; N- CA- CB- HB3
3465 3467 3469 3472 3 15.41386 9.74035 -8.39310
-0.97069 2.97901 0.00000 ; N- CA- CB- CG
3465 3467 3475 3476 3 0.00000 0.00000 0.00000
0.00000 0.00000 0.00000 ; N- CA- C- O
3465 3467 3475 3477 3 17.40544 -5.02080 -13.22144
9.20480 0.00000 0.00000 ; N- CA- C- N
3466 3465 3467 3468 3 0.00000 0.00000 0.00000
0.00000 0.00000 0.00000 ; H- N- CA- HA
3466 3465 3467 3469 3 0.00000 0.00000 0.00000
0.00000 0.00000 0.00000 ; H- N- CA- CB
3466 3465 3467 3475 3 0.00000 0.00000 0.00000
0.00000 0.00000 0.00000 ; H- N- CA- C
3467 3469 3472 3473 3 6.43499 0.00000 -5.39736
0.00000 -1.03763 0.00000 ; CA- CB- CG- OD1
3467 3469 3472 3474 3 6.43499 0.00000 -5.39736
0.00000 -1.03763 0.00000 ; CA- CB- CG- OD2
3467 3475 3477 3478 3 20.92000 0.00000 -20.92000
0.00000 0.00000 0.00000 ; CA- C- N- H
3467 3475 3477 3479 3 20.92000 0.00000 -20.92000
0.00000 0.00000 0.00000 ; CA- C- N- CA
3468 3467 3469 3470 3 0.65084 1.95253 0.00000
-2.60338 0.00000 0.00000 ; HA- CA- CB- HB2
3468 3467 3469 3471 3 0.65084 1.95253 0.00000
-2.60338 0.00000 0.00000 ; HA- CA- CB- HB3
3468 3467 3469 3472 3 0.65084 1.95253 0.00000
-2.60338 0.00000 0.00000 ; HA- CA- CB- CG
3468 3467 3475 3476 3 3.68192 -4.35136 0.00000
1.33888 0.00000 0.00000 ; HA- CA- C- O
3468 3467 3475 3477 3 0.00000 0.00000 0.00000
0.00000 0.00000 0.00000 ; HA- CA- C- N
3469 3467 3475 3476 3 0.00000 0.00000 0.00000
0.00000 0.00000 0.00000 ; CB- CA- C- O
3469 3467 3475 3477 3 2.51040 4.18400 1.67360
-6.69440 0.00000 0.00000 ; CB- CA- C- N
3470 3469 3467 3475 3 0.65084 1.95253 0.00000
-2.60338 0.00000 0.00000 ; HB2- CB- CA- C
3470 3469 3472 3473 3 0.00000 0.00000 0.00000
0.00000 0.00000 0.00000 ; HB2- CB- CG- OD1

3470 3469 3472 3474 3 0.00000 0.00000 0.00000
0.00000 0.00000 0.00000 ; HB2- CB- CG- OD2
3471 3469 3467 3475 3 0.65084 1.95253 0.00000
-2.60338 0.00000 0.00000 ; HB3- CB- CA- C
3471 3469 3472 3473 3 0.00000 0.00000 0.00000
0.00000 0.00000 0.00000 ; HB3- CB- CG- OD1
3471 3469 3472 3474 3 0.00000 0.00000 0.00000
0.00000 0.00000 0.00000 ; HB3- CB- CG- OD2
3472 3469 3467 3475 3 7.14627 -1.04600 -8.99560
-0.97069 5.15469 0.00000 ; CG- CB- CA- C
3475 3477 3479 3480 3 0.00000 0.00000 0.00000
0.00000 0.00000 0.00000 ; C- N- CA- HA
3475 3477 3479 3481 3 11.71520 1.67360 15.06240 -
13.38880 0.00000 0.00000 ; C- N- CA- CB
3475 3477 3479 3489 3 1.75728 5.27184 2.25936
-7.02912 0.00000 0.00000 ; C- N- CA- C
3476 3475 3477 3478 3 29.28800 -8.36800 -20.92000
0.00000 0.00000 0.00000 ; O- C- N- H
3476 3475 3477 3479 3 20.92000 0.00000 -20.92000
0.00000 0.00000 0.00000 ; O- C- N- CA
3477 3479 3481 3482 3 0.65084 1.95253 0.00000
-2.60338 0.00000 0.00000 ; N- CA- CB- HB2
3477 3479 3481 3483 3 0.65084 1.95253 0.00000
-2.60338 0.00000 0.00000 ; N- CA- CB- HB3
3477 3479 3481 3484 3 5.99567 3.29281 -4.46014
-0.55229 1.97485 0.00000 ; N- CA- CB- CG
3477 3479 3489 3490 3 0.00000 0.00000 0.00000
0.00000 0.00000 0.00000 ; N- CA- C- O
3477 3479 3489 3491 3 17.40544 -5.02080 -13.22144
9.20480 0.00000 0.00000 ; N- CA- C- N
3478 3477 3479 3480 3 0.00000 0.00000 0.00000
0.00000 0.00000 0.00000 ; H- N- CA- HA
3478 3477 3479 3481 3 0.00000 0.00000 0.00000
0.00000 0.00000 0.00000 ; H- N- CA- CB
3478 3477 3479 3489 3 0.00000 0.00000 0.00000
0.00000 0.00000 0.00000 ; H- N- CA- C
3479 3481 3484 3485 3 0.00000 0.00000 0.00000
0.00000 0.00000 0.00000 ; CA- CB- CG- OD1
3479 3481 3484 3486 3 8.84916 -0.31380 -4.32626
5.03754 0.26778 0.00000 ; CA- CB- CG- ND2
3479 3489 3491 3492 3 20.92000 0.00000 -20.92000
0.00000 0.00000 0.00000 ; CA- C- N- H
3479 3489 3491 3493 3 20.92000 0.00000 -20.92000
0.00000 0.00000 0.00000 ; CA- C- N- CA
3480 3479 3481 3482 3 0.65084 1.95253 0.00000
-2.60338 0.00000 0.00000 ; HA- CA- CB- HB2
3480 3479 3481 3483 3 0.65084 1.95253 0.00000
-2.60338 0.00000 0.00000 ; HA- CA- CB- HB3
3480 3479 3481 3484 3 0.65084 1.95253 0.00000
-2.60338 0.00000 0.00000 ; HA- CA- CB- CG
3480 3479 3489 3490 3 3.68192 -4.35136 0.00000
1.33888 0.00000 0.00000 ; HA- CA- C- O

3480 3479 3489 3491 3 0.00000 0.00000 0.00000
0.00000 0.00000 0.00000 ; HA- CA- C- N
3481 3479 3489 3490 3 0.00000 0.00000 0.00000
0.00000 0.00000 0.00000 ; CB- CA- C- O
3481 3479 3489 3491 3 2.51040 4.18400 1.67360
-6.69440 0.00000 0.00000 ; CB- CA- C- N
3481 3484 3486 3487 3 20.92000 0.00000 -20.92000
0.00000 0.00000 0.00000 ; CB- CG- ND2- HD21
3481 3484 3486 3488 3 20.92000 0.00000 -20.92000
0.00000 0.00000 0.00000 ; CB- CG- ND2- HD22
3482 3481 3479 3489 3 0.65084 1.95253 0.00000
-2.60338 0.00000 0.00000 ; HB2- CB- CA- C
3482 3481 3484 3485 3 3.68192 -4.35136 0.00000
1.33888 0.00000 0.00000 ; HB2- CB- CG- OD1
3482 3481 3484 3486 3 0.00000 0.00000 0.00000
0.00000 0.00000 0.00000 ; HB2- CB- CG- ND2
3483 3481 3479 3489 3 0.65084 1.95253 0.00000
-2.60338 0.00000 0.00000 ; HB3- CB- CA- C
3483 3481 3484 3485 3 3.68192 -4.35136 0.00000
1.33888 0.00000 0.00000 ; HB3- CB- CG- OD1
3483 3481 3484 3486 3 0.00000 0.00000 0.00000
0.00000 0.00000 0.00000 ; HB3- CB- CG- ND2
3484 3481 3479 3489 3 7.94542 -3.96225 -6.11701
-0.55229 3.58150 0.00000 ; CG- CB- CA- C
3485 3484 3486 3487 3 29.28800 -8.36800 -20.92000
0.00000 0.00000 0.00000 ; OD1- CG- ND2- HD21
3485 3484 3486 3488 3 29.28800 -8.36800 -20.92000
0.00000 0.00000 0.00000 ; OD1- CG- ND2- HD22
3489 3491 3493 3494 3 0.00000 0.00000 0.00000
0.00000 0.00000 0.00000 ; C- N- CA- HA
3489 3491 3493 3495 3 11.71520 1.67360 15.06240 -
13.38880 0.00000 0.00000 ; C- N- CA- CB
3489 3491 3493 3504 3 1.75728 5.27184 2.25936
-7.02912 0.00000 0.00000 ; C- N- CA- C
3490 3489 3491 3492 3 29.28800 -8.36800 -20.92000
0.00000 0.00000 0.00000 ; O- C- N- H
3490 3489 3491 3493 3 20.92000 0.00000 -20.92000
0.00000 0.00000 0.00000 ; O- C- N- CA
3491 3493 3495 3496 3 0.65084 1.95253 0.00000
-2.60338 0.00000 0.00000 ; N- CA- CB- HB2
3491 3493 3495 3497 3 0.65084 1.95253 0.00000
-2.60338 0.00000 0.00000 ; N- CA- CB- HB3
3491 3493 3495 3498 3 3.21331 2.22589 -4.15053
-2.40998 2.61082 0.00000 ; N- CA- CB- CG
3491 3493 3504 3505 3 0.00000 0.00000 0.00000
0.00000 0.00000 0.00000 ; N- CA- C- O
3491 3493 3504 3506 3 17.40544 -5.02080 -13.22144
9.20480 0.00000 0.00000 ; N- CA- C- N
3492 3491 3493 3494 3 0.00000 0.00000 0.00000
0.00000 0.00000 0.00000 ; H- N- CA- HA
3492 3491 3493 3495 3 0.00000 0.00000 0.00000
0.00000 0.00000 0.00000 ; H- N- CA- CB

3492 3491 3493 3504 3 0.00000 0.00000 0.00000
0.00000 0.00000 0.00000 ; H- N- CA- C
3493 3495 3498 3499 3 0.66944 2.00832 0.00000
-2.67776 0.00000 0.00000 ; CA- CB- CG- HG2
3493 3495 3498 3500 3 0.66944 2.00832 0.00000
-2.67776 0.00000 0.00000 ; CA- CB- CG- HG3
3493 3495 3498 3501 3 10.12110 -1.91209 0.01674
10.17549 -1.87443 0.00000 ; CA- CB- CG- CD
3493 3504 3506 3507 3 20.92000 0.00000 -20.92000
0.00000 0.00000 0.00000 ; CA- C- N- H
3493 3504 3506 3508 3 20.92000 0.00000 -20.92000
0.00000 0.00000 0.00000 ; CA- C- N- CA
3494 3493 3495 3496 3 0.65084 1.95253 0.00000
-2.60338 0.00000 0.00000 ; HA- CA- CB- HB2
3494 3493 3495 3497 3 0.65084 1.95253 0.00000
-2.60338 0.00000 0.00000 ; HA- CA- CB- HB3
3494 3493 3495 3498 3 0.65084 1.95253 0.00000
-2.60338 0.00000 0.00000 ; HA- CA- CB- CG
3494 3493 3504 3505 3 3.68192 -4.35136 0.00000
1.33888 0.00000 0.00000 ; HA- CA- C- O
3494 3493 3504 3506 3 0.00000 0.00000 0.00000
0.00000 0.00000 0.00000 ; HA- CA- C- N
3495 3493 3504 3505 3 0.00000 0.00000 0.00000
0.00000 0.00000 0.00000 ; CB- CA- C- O
3495 3493 3504 3506 3 2.51040 4.18400 1.67360
-6.69440 0.00000 0.00000 ; CB- CA- C- N
3495 3498 3501 3502 3 3.79907 0.00000 -5.40573
0.00000 2.14221 0.00000 ; CB- CG- CD- OE1
3495 3498 3501 3503 3 3.79907 0.00000 -5.40573
0.00000 2.14221 0.00000 ; CB- CG- CD- OE2
3496 3495 3493 3504 3 0.65084 1.95253 0.00000
-2.60338 0.00000 0.00000 ; HB2- CB- CA- C
3496 3495 3498 3499 3 0.62760 1.88280 0.00000
-2.51040 0.00000 0.00000 ; HB2- CB- CG- HG2
3496 3495 3498 3500 3 0.62760 1.88280 0.00000
-2.51040 0.00000 0.00000 ; HB2- CB- CG- HG3
3496 3495 3498 3501 3 0.65084 1.95253 0.00000
-2.60338 0.00000 0.00000 ; HB2- CB- CG- CD
3497 3495 3493 3504 3 0.65084 1.95253 0.00000
-2.60338 0.00000 0.00000 ; HB3- CB- CA- C
3497 3495 3498 3499 3 0.62760 1.88280 0.00000
-2.51040 0.00000 0.00000 ; HB3- CB- CG- HG2
3497 3495 3498 3500 3 0.62760 1.88280 0.00000
-2.51040 0.00000 0.00000 ; HB3- CB- CG- HG3
3497 3495 3498 3501 3 0.65084 1.95253 0.00000
-2.60338 0.00000 0.00000 ; HB3- CB- CG- CD
3498 3495 3493 3504 3 6.86594 3.56895 -8.14206
-2.40998 4.85344 0.00000 ; CG- CB- CA- C
3499 3498 3501 3502 3 0.00000 0.00000 0.00000
0.00000 0.00000 0.00000 ; HG2- CG- CD- OE1
3499 3498 3501 3503 3 0.00000 0.00000 0.00000
0.00000 0.00000 0.00000 ; HG2- CG- CD- OE2

```
  3500   3498   3501   3502      3    0.00000    0.00000    0.00000
0.00000    0.00000    0.00000 ;    HG3-    CG-    CD-   OE1
  3500   3498   3501   3503      3    0.00000    0.00000    0.00000
0.00000    0.00000    0.00000 ;    HG3-    CG-    CD-   OE2
  3504   3506   3508   3509      3    0.00000    0.00000    0.00000
0.00000    0.00000    0.00000 ;      C-     N-    CA-    HA
  3504   3506   3508   3510      3   11.71520    1.67360   15.06240  -
13.38880    0.00000    0.00000 ;      C-     N-    CA-    CB
  3504   3506   3508   3520      3    1.75728    5.27184    2.25936
-7.02912    0.00000    0.00000 ;      C-     N-    CA-     C
  3505   3504   3506   3507      3   29.28800   -8.36800  -20.92000
0.00000    0.00000    0.00000 ;      O-     C-     N-     H
  3505   3504   3506   3508      3   20.92000    0.00000  -20.92000
0.00000    0.00000    0.00000 ;      O-     C-     N-    CA
  3506   3508   3510   3511      3    0.65084    1.95253    0.00000
-2.60338    0.00000    0.00000 ;      N-    CA-    CB-    HB
  3506   3508   3510   3512      3    3.83673    0.44769   -1.77402
-2.47693   -0.03347    0.00000 ;      N-    CA-    CB-   CG1
  3506   3508   3510   3516      3    3.83673    0.44769   -1.77402
-2.47693   -0.03347    0.00000 ;      N-    CA-    CB-   CG2
  3506   3508   3520   3521      3    0.00000    0.00000    0.00000
0.00000    0.00000    0.00000 ;      N-    CA-     C-     O
  3506   3508   3520   3522      3   17.40544   -5.02080  -13.22144
9.20480    0.00000    0.00000 ;      N-    CA-     C-     N
  3507   3506   3508   3509      3    0.00000    0.00000    0.00000
0.00000    0.00000    0.00000 ;      H-     N-    CA-    HA
  3507   3506   3508   3510      3    0.00000    0.00000    0.00000
0.00000    0.00000    0.00000 ;      H-     N-    CA-    CB
  3507   3506   3508   3520      3    0.00000    0.00000    0.00000
0.00000    0.00000    0.00000 ;      H-     N-    CA-     C
  3508   3510   3512   3513      3    0.66944    2.00832    0.00000
-2.67776    0.00000    0.00000 ;     CA-    CB-   CG1-  HG11
  3508   3510   3512   3514      3    0.66944    2.00832    0.00000
-2.67776    0.00000    0.00000 ;     CA-    CB-   CG1-  HG12
  3508   3510   3512   3515      3    0.66944    2.00832    0.00000
-2.67776    0.00000    0.00000 ;     CA-    CB-   CG1-  HG13
  3508   3510   3516   3517      3    0.66944    2.00832    0.00000
-2.67776    0.00000    0.00000 ;     CA-    CB-   CG2-  HG21
  3508   3510   3516   3518      3    0.66944    2.00832    0.00000
-2.67776    0.00000    0.00000 ;     CA-    CB-   CG2-  HG22
  3508   3510   3516   3519      3    0.66944    2.00832    0.00000
-2.67776    0.00000    0.00000 ;     CA-    CB-   CG2-  HG23
  3508   3520   3522   3523      3   20.92000    0.00000  -20.92000
0.00000    0.00000    0.00000 ;     CA-     C-     N-     H
  3508   3520   3522   3524      3   20.92000    0.00000  -20.92000
0.00000    0.00000    0.00000 ;     CA-     C-     N-    CA
  3509   3508   3510   3511      3    0.65084    1.95253    0.00000
-2.60338    0.00000    0.00000 ;     HA-    CA-    CB-    HB
  3509   3508   3510   3512      3    0.65084    1.95253    0.00000
-2.60338    0.00000    0.00000 ;     HA-    CA-    CB-   CG1
  3509   3508   3510   3516      3    0.65084    1.95253    0.00000
-2.60338    0.00000    0.00000 ;     HA-    CA-    CB-   CG2
```

3509 3508 3520 3521 3 3.68192 -4.35136 0.00000
1.33888 0.00000 0.00000 ; HA- CA- C- O
3509 3508 3520 3522 3 0.00000 0.00000 0.00000
0.00000 0.00000 0.00000 ; HA- CA- C- N
3510 3508 3520 3521 3 0.00000 0.00000 0.00000
0.00000 0.00000 0.00000 ; CB- CA- C- O
3510 3508 3520 3522 3 2.51040 4.18400 1.67360
-6.69440 0.00000 0.00000 ; CB- CA- C- N
3511 3510 3508 3520 3 0.65084 1.95253 0.00000
-2.60338 0.00000 0.00000 ; HB- CB- CA- C
3511 3510 3512 3513 3 0.62760 1.88280 0.00000
-2.51040 0.00000 0.00000 ; HB- CB- CG1- HG11
3511 3510 3512 3514 3 0.62760 1.88280 0.00000
-2.51040 0.00000 0.00000 ; HB- CB- CG1- HG12
3511 3510 3512 3515 3 0.62760 1.88280 0.00000
-2.51040 0.00000 0.00000 ; HB- CB- CG1- HG13
3511 3510 3516 3517 3 0.62760 1.88280 0.00000
-2.51040 0.00000 0.00000 ; HB- CB- CG2- HG21
3511 3510 3516 3518 3 0.62760 1.88280 0.00000
-2.51040 0.00000 0.00000 ; HB- CB- CG2- HG22
3511 3510 3516 3519 3 0.62760 1.88280 0.00000
-2.51040 0.00000 0.00000 ; HB- CB- CG2- HG23
3512 3510 3508 3520 3 5.67350 3.55640 -6.16722
-2.47693 3.74886 0.00000 ; CG1- CB- CA- C
3512 3510 3516 3517 3 0.66944 2.00832 0.00000
-2.67776 0.00000 0.00000 ; CG1- CB- CG2- HG21
3512 3510 3516 3518 3 0.66944 2.00832 0.00000
-2.67776 0.00000 0.00000 ; CG1- CB- CG2- HG22
3512 3510 3516 3519 3 0.66944 2.00832 0.00000
-2.67776 0.00000 0.00000 ; CG1- CB- CG2- HG23
3513 3512 3510 3516 3 0.66944 2.00832 0.00000
-2.67776 0.00000 0.00000 ; HG11- CG1- CB- CG2
3514 3512 3510 3516 3 0.66944 2.00832 0.00000
-2.67776 0.00000 0.00000 ; HG12- CG1- CB- CG2
3515 3512 3510 3516 3 0.66944 2.00832 0.00000
-2.67776 0.00000 0.00000 ; HG13- CG1- CB- CG2
3516 3510 3508 3520 3 5.67350 3.55640 -6.16722
-2.47693 3.74886 0.00000 ; CG2- CB- CA- C
3520 3522 3524 3525 3 0.00000 0.00000 0.00000
0.00000 0.00000 0.00000 ; C- N- CA- HA
3520 3522 3524 3526 3 11.71520 1.67360 15.06240 -
13.38880 0.00000 0.00000 ; C- N- CA- CB
3520 3522 3524 3539 3 1.75728 5.27184 2.25936
-7.02912 0.00000 0.00000 ; C- N- CA- C
3521 3520 3522 3523 3 29.28800 -8.36800 -20.92000
0.00000 0.00000 0.00000 ; O- C- N- H
3521 3520 3522 3524 3 20.92000 0.00000 -20.92000
0.00000 0.00000 0.00000 ; O- C- N- CA
3522 3524 3526 3527 3 0.65084 1.95253 0.00000
-2.60338 0.00000 0.00000 ; N- CA- CB- HB2
3522 3524 3526 3528 3 0.65084 1.95253 0.00000
-2.60338 0.00000 0.00000 ; N- CA- CB- HB3

```
  3522   3524   3526   3529      3    3.79070    1.39746   -4.61077
-2.40998    2.44346    0.00000 ;      N-    CA-    CB-    CG
  3522   3524   3539   3540      3    0.00000    0.00000    0.00000
0.00000    0.00000    0.00000 ;      N-    CA-     C-     O
  3522   3524   3539   3541      3   17.40544   -5.02080  -13.22144
9.20480    0.00000    0.00000 ;      N-    CA-     C-     N
  3523   3522   3524   3525      3    0.00000    0.00000    0.00000
0.00000    0.00000    0.00000 ;      H-     N-    CA-    HA
  3523   3522   3524   3526      3    0.00000    0.00000    0.00000
0.00000    0.00000    0.00000 ;      H-     N-    CA-    CB
  3523   3522   3524   3539      3    0.00000    0.00000    0.00000
0.00000    0.00000    0.00000 ;      H-     N-    CA-     C
  3524   3526   3529   3530      3    0.66944    2.00832    0.00000
-2.67776    0.00000    0.00000 ;     CA-    CB-    CG-    HG
  3524   3526   3529   3531      3    3.90367    0.19665   -6.21742
-2.37651    5.99149    0.00000 ;     CA-    CB-    CG-   CD1
  3524   3526   3529   3535      3    3.90367    0.19665   -6.21742
-2.37651    5.99149    0.00000 ;     CA-    CB-    CG-   CD2
  3524   3539   3541   3542      3   20.92000    0.00000  -20.92000
0.00000    0.00000    0.00000 ;     CA-     C-     N-     H
  3524   3539   3541   3543      3   20.92000    0.00000  -20.92000
0.00000    0.00000    0.00000 ;     CA-     C-     N-    CA
  3525   3524   3526   3527      3    0.65084    1.95253    0.00000
-2.60338    0.00000    0.00000 ;     HA-    CA-    CB-   HB2
  3525   3524   3526   3528      3    0.65084    1.95253    0.00000
-2.60338    0.00000    0.00000 ;     HA-    CA-    CB-   HB3
  3525   3524   3526   3529      3    0.65084    1.95253    0.00000
-2.60338    0.00000    0.00000 ;     HA-    CA-    CB-    CG
  3525   3524   3539   3540      3    3.68192   -4.35136    0.00000
1.33888    0.00000    0.00000 ;     HA-    CA-     C-     O
  3525   3524   3539   3541      3    0.00000    0.00000    0.00000
0.00000    0.00000    0.00000 ;     HA-    CA-     C-     N
  3526   3524   3539   3540      3    0.00000    0.00000    0.00000
0.00000    0.00000    0.00000 ;     CB-    CA-     C-     O
  3526   3524   3539   3541      3    2.51040    4.18400    1.67360
-6.69440    0.00000    0.00000 ;     CB-    CA-     C-     N
  3526   3529   3531   3532      3    0.66944    2.00832    0.00000
-2.67776    0.00000    0.00000 ;     CB-    CG-   CD1-  HD11
  3526   3529   3531   3533      3    0.66944    2.00832    0.00000
-2.67776    0.00000    0.00000 ;     CB-    CG-   CD1-  HD12
  3526   3529   3531   3534      3    0.66944    2.00832    0.00000
-2.67776    0.00000    0.00000 ;     CB-    CG-   CD1-  HD13
  3526   3529   3535   3536      3    0.66944    2.00832    0.00000
-2.67776    0.00000    0.00000 ;     CB-    CG-   CD2-  HD21
  3526   3529   3535   3537      3    0.66944    2.00832    0.00000
-2.67776    0.00000    0.00000 ;     CB-    CG-   CD2-  HD22
  3526   3529   3535   3538      3    0.66944    2.00832    0.00000
-2.67776    0.00000    0.00000 ;     CB-    CG-   CD2-  HD23
  3527   3526   3524   3539      3    0.65084    1.95253    0.00000
-2.60338    0.00000    0.00000 ;    HB2-    CB-    CA-     C
  3527   3526   3529   3530      3    0.62760    1.88280    0.00000
-2.51040    0.00000    0.00000 ;    HB2-    CB-    CG-    HG
```

3527 3526 3529 3531 3 0.66944 2.00832 0.00000
-2.67776 0.00000 0.00000 ; HB2- CB- CG- CD1
3527 3526 3529 3535 3 0.66944 2.00832 0.00000
-2.67776 0.00000 0.00000 ; HB2- CB- CG- CD2
3528 3526 3524 3539 3 0.65084 1.95253 0.00000
-2.60338 0.00000 0.00000 ; HB3- CB- CA- C
3528 3526 3529 3530 3 0.62760 1.88280 0.00000
-2.51040 0.00000 0.00000 ; HB3- CB- CG- HG
3528 3526 3529 3531 3 0.66944 2.00832 0.00000
-2.67776 0.00000 0.00000 ; HB3- CB- CG- CD1
3528 3526 3529 3535 3 0.66944 2.00832 0.00000
-2.67776 0.00000 0.00000 ; HB3- CB- CG- CD2
3529 3526 3524 3539 3 10.33448 -1.14642 -11.54784
-2.40998 6.35968 0.00000 ; CG- CB- CA- C
3530 3529 3531 3532 3 0.62760 1.88280 0.00000
-2.51040 0.00000 0.00000 ; HG- CG- CD1- HD11
3530 3529 3531 3533 3 0.62760 1.88280 0.00000
-2.51040 0.00000 0.00000 ; HG- CG- CD1- HD12
3530 3529 3531 3534 3 0.62760 1.88280 0.00000
-2.51040 0.00000 0.00000 ; HG- CG- CD1- HD13
3530 3529 3535 3536 3 0.62760 1.88280 0.00000
-2.51040 0.00000 0.00000 ; HG- CG- CD2- HD21
3530 3529 3535 3537 3 0.62760 1.88280 0.00000
-2.51040 0.00000 0.00000 ; HG- CG- CD2- HD22
3530 3529 3535 3538 3 0.62760 1.88280 0.00000
-2.51040 0.00000 0.00000 ; HG- CG- CD2- HD23
3531 3529 3535 3536 3 0.66944 2.00832 0.00000
-2.67776 0.00000 0.00000 ; CD1- CG- CD2- HD21
3531 3529 3535 3537 3 0.66944 2.00832 0.00000
-2.67776 0.00000 0.00000 ; CD1- CG- CD2- HD22
3531 3529 3535 3538 3 0.66944 2.00832 0.00000
-2.67776 0.00000 0.00000 ; CD1- CG- CD2- HD23
3532 3531 3529 3535 3 0.66944 2.00832 0.00000
-2.67776 0.00000 0.00000 ; HD11- CD1- CG- CD2
3533 3531 3529 3535 3 0.66944 2.00832 0.00000
-2.67776 0.00000 0.00000 ; HD12- CD1- CG- CD2
3534 3531 3529 3535 3 0.66944 2.00832 0.00000
-2.67776 0.00000 0.00000 ; HD13- CD1- CG- CD2
3539 3541 3543 3544 3 0.00000 0.00000 0.00000
0.00000 0.00000 0.00000 ; C- N- CA- HA
3539 3541 3543 3545 3 11.71520 1.67360 15.06240 -
13.38880 0.00000 0.00000 ; C- N- CA- CB
3539 3541 3543 3561 3 1.75728 5.27184 2.25936
-7.02912 0.00000 0.00000 ; C- N- CA- C
3540 3539 3541 3542 3 29.28800 -8.36800 -20.92000
0.00000 0.00000 0.00000 ; O- C- N- H
3540 3539 3541 3543 3 20.92000 0.00000 -20.92000
0.00000 0.00000 0.00000 ; O- C- N- CA
3541 3543 3545 3546 3 0.65084 1.95253 0.00000
-2.60338 0.00000 0.00000 ; N- CA- CB- HB2
3541 3543 3545 3547 3 0.65084 1.95253 0.00000
-2.60338 0.00000 0.00000 ; N- CA- CB- HB3

3541 3543 3545 3548 3 0.65084 1.95253 0.00000
-2.60338 0.00000 0.00000 ; N- CA- CB- CG
3541 3543 3561 3562 3 0.00000 0.00000 0.00000
0.00000 0.00000 0.00000 ; N- CA- C- O
3541 3543 3561 3563 3 17.40544 -5.02080 -13.22144
9.20480 0.00000 0.00000 ; N- CA- C- N
3542 3541 3543 3544 3 0.00000 0.00000 0.00000
0.00000 0.00000 0.00000 ; H- N- CA- HA
3542 3541 3543 3545 3 0.00000 0.00000 0.00000
0.00000 0.00000 0.00000 ; H- N- CA- CB
3542 3541 3543 3561 3 0.00000 0.00000 0.00000
0.00000 0.00000 0.00000 ; H- N- CA- C
3543 3545 3548 3549 3 0.66944 2.00832 0.00000
-2.67776 0.00000 0.00000 ; CA- CB- CG- HG2
3543 3545 3548 3550 3 0.66944 2.00832 0.00000
-2.67776 0.00000 0.00000 ; CA- CB- CG- HG3
3543 3545 3548 3551 3 3.68192 3.09616 -2.09200
-3.01248 0.00000 0.00000 ; CA- CB- CG- CD
3543 3561 3563 3564 3 20.92000 0.00000 -20.92000
0.00000 0.00000 0.00000 ; CA- C- N- H
3543 3561 3563 3565 3 20.92000 0.00000 -20.92000
0.00000 0.00000 0.00000 ; CA- C- N- CA
3544 3543 3545 3546 3 0.65084 1.95253 0.00000
-2.60338 0.00000 0.00000 ; HA- CA- CB- HB2
3544 3543 3545 3547 3 0.65084 1.95253 0.00000
-2.60338 0.00000 0.00000 ; HA- CA- CB- HB3
3544 3543 3545 3548 3 0.65084 1.95253 0.00000
-2.60338 0.00000 0.00000 ; HA- CA- CB- CG
3544 3543 3561 3562 3 3.68192 -4.35136 0.00000
1.33888 0.00000 0.00000 ; HA- CA- C- O
3544 3543 3561 3563 3 0.00000 0.00000 0.00000
0.00000 0.00000 0.00000 ; HA- CA- C- N
3545 3543 3561 3562 3 0.00000 0.00000 0.00000
0.00000 0.00000 0.00000 ; CB- CA- C- O
3545 3543 3561 3563 3 2.51040 4.18400 1.67360
-6.69440 0.00000 0.00000 ; CB- CA- C- N
3545 3548 3551 3552 3 0.66944 2.00832 0.00000
-2.67776 0.00000 0.00000 ; CB- CG- CD- HD2
3545 3548 3551 3553 3 0.66944 2.00832 0.00000
-2.67776 0.00000 0.00000 ; CB- CG- CD- HD3
3545 3548 3551 3554 3 3.68192 3.09616 -2.09200
-3.01248 0.00000 0.00000 ; CB- CG- CD- CE
3546 3545 3543 3561 3 0.65084 1.95253 0.00000
-2.60338 0.00000 0.00000 ; HB2- CB- CA- C
3546 3545 3548 3549 3 0.62760 1.88280 0.00000
-2.51040 0.00000 0.00000 ; HB2- CB- CG- HG2
3546 3545 3548 3550 3 0.62760 1.88280 0.00000
-2.51040 0.00000 0.00000 ; HB2- CB- CG- HG3
3546 3545 3548 3551 3 0.66944 2.00832 0.00000
-2.67776 0.00000 0.00000 ; HB2- CB- CG- CD
3547 3545 3543 3561 3 0.65084 1.95253 0.00000
-2.60338 0.00000 0.00000 ; HB3- CB- CA- C

3547 3545 3548 3549 3 0.62760 1.88280 0.00000
-2.51040 0.00000 0.00000 ; HB3- CB- CG- HG2
3547 3545 3548 3550 3 0.62760 1.88280 0.00000
-2.51040 0.00000 0.00000 ; HB3- CB- CG- HG3
3547 3545 3548 3551 3 0.66944 2.00832 0.00000
-2.67776 0.00000 0.00000 ; HB3- CB- CG- CD
3548 3545 3543 3561 3 0.65084 1.95253 0.00000
-2.60338 0.00000 0.00000 ; CG- CB- CA- C
3548 3551 3554 3555 3 0.65084 1.95253 0.00000
-2.60338 0.00000 0.00000 ; CG- CD- CE- HE2
3548 3551 3554 3556 3 0.65084 1.95253 0.00000
-2.60338 0.00000 0.00000 ; CG- CD- CE- HE3
3548 3551 3554 3557 3 0.65084 1.95253 0.00000
-2.60338 0.00000 0.00000 ; CG- CD- CE- NZ
3549 3548 3551 3552 3 0.62760 1.88280 0.00000
-2.51040 0.00000 0.00000 ; HG2- CG- CD- HD2
3549 3548 3551 3553 3 0.62760 1.88280 0.00000
-2.51040 0.00000 0.00000 ; HG2- CG- CD- HD3
3549 3548 3551 3554 3 0.66944 2.00832 0.00000
-2.67776 0.00000 0.00000 ; HG2- CG- CD- CE
3550 3548 3551 3552 3 0.62760 1.88280 0.00000
-2.51040 0.00000 0.00000 ; HG3- CG- CD- HD2
3550 3548 3551 3553 3 0.62760 1.88280 0.00000
-2.51040 0.00000 0.00000 ; HG3- CG- CD- HD3
3550 3548 3551 3554 3 0.66944 2.00832 0.00000
-2.67776 0.00000 0.00000 ; HG3- CG- CD- CE
3551 3554 3557 3558 3 0.65084 1.95253 0.00000
-2.60338 0.00000 0.00000 ; CD- CE- NZ- HZ1
3551 3554 3557 3559 3 0.65084 1.95253 0.00000
-2.60338 0.00000 0.00000 ; CD- CE- NZ- HZ2
3551 3554 3557 3560 3 0.65084 1.95253 0.00000
-2.60338 0.00000 0.00000 ; CD- CE- NZ- HZ3
3552 3551 3554 3555 3 0.65084 1.95253 0.00000
-2.60338 0.00000 0.00000 ; HD2- CD- CE- HE2
3552 3551 3554 3556 3 0.65084 1.95253 0.00000
-2.60338 0.00000 0.00000 ; HD2- CD- CE- HE3
3552 3551 3554 3557 3 0.65084 1.95253 0.00000
-2.60338 0.00000 0.00000 ; HD2- CD- CE- NZ
3553 3551 3554 3555 3 0.65084 1.95253 0.00000
-2.60338 0.00000 0.00000 ; HD3- CD- CE- HE2
3553 3551 3554 3556 3 0.65084 1.95253 0.00000
-2.60338 0.00000 0.00000 ; HD3- CD- CE- HE3
3553 3551 3554 3557 3 0.65084 1.95253 0.00000
-2.60338 0.00000 0.00000 ; HD3- CD- CE- NZ
3555 3554 3557 3558 3 0.65084 1.95253 0.00000
-2.60338 0.00000 0.00000 ; HE2- CE- NZ- HZ1
3555 3554 3557 3559 3 0.65084 1.95253 0.00000
-2.60338 0.00000 0.00000 ; HE2- CE- NZ- HZ2
3555 3554 3557 3560 3 0.65084 1.95253 0.00000
-2.60338 0.00000 0.00000 ; HE2- CE- NZ- HZ3
3556 3554 3557 3558 3 0.65084 1.95253 0.00000
-2.60338 0.00000 0.00000 ; HE3- CE- NZ- HZ1

3556 3554 3557 3559 3 0.65084 1.95253 0.00000
-2.60338 0.00000 0.00000 ; HE3- CE- NZ- HZ2
3556 3554 3557 3560 3 0.65084 1.95253 0.00000
-2.60338 0.00000 0.00000 ; HE3- CE- NZ- HZ3
3561 3563 3565 3566 3 0.00000 0.00000 0.00000
0.00000 0.00000 0.00000 ; C- N- CA- HA
3561 3563 3565 3567 3 11.71520 1.67360 15.06240 -
13.38880 0.00000 0.00000 ; C- N- CA- CB
3561 3563 3565 3583 3 1.75728 5.27184 2.25936
-7.02912 0.00000 0.00000 ; C- N- CA- C
3562 3561 3563 3564 3 29.28800 -8.36800 -20.92000
0.00000 0.00000 0.00000 ; O- C- N- H
3562 3561 3563 3565 3 20.92000 0.00000 -20.92000
0.00000 0.00000 0.00000 ; O- C- N- CA
3563 3565 3567 3568 3 0.65084 1.95253 0.00000
-2.60338 0.00000 0.00000 ; N- CA- CB- HB2
3563 3565 3567 3569 3 0.65084 1.95253 0.00000
-2.60338 0.00000 0.00000 ; N- CA- CB- HB3
3563 3565 3567 3570 3 0.65084 1.95253 0.00000
-2.60338 0.00000 0.00000 ; N- CA- CB- CG
3563 3565 3583 3584 3 0.00000 0.00000 0.00000
0.00000 0.00000 0.00000 ; N- CA- C- O
3563 3565 3583 3585 3 17.40544 -5.02080 -13.22144
9.20480 0.00000 0.00000 ; N- CA- C- N
3564 3563 3565 3566 3 0.00000 0.00000 0.00000
0.00000 0.00000 0.00000 ; H- N- CA- HA
3564 3563 3565 3567 3 0.00000 0.00000 0.00000
0.00000 0.00000 0.00000 ; H- N- CA- CB
3564 3563 3565 3583 3 0.00000 0.00000 0.00000
0.00000 0.00000 0.00000 ; H- N- CA- C
3565 3567 3570 3571 3 0.66944 2.00832 0.00000
-2.67776 0.00000 0.00000 ; CA- CB- CG- HG2
3565 3567 3570 3572 3 0.66944 2.00832 0.00000
-2.67776 0.00000 0.00000 ; CA- CB- CG- HG3
3565 3567 3570 3573 3 3.68192 3.09616 -2.09200
-3.01248 0.00000 0.00000 ; CA- CB- CG- CD
3565 3583 3585 3586 3 20.92000 0.00000 -20.92000
0.00000 0.00000 0.00000 ; CA- C- N- H
3565 3583 3585 3587 3 20.92000 0.00000 -20.92000
0.00000 0.00000 0.00000 ; CA- C- N- CA
3566 3565 3567 3568 3 0.65084 1.95253 0.00000
-2.60338 0.00000 0.00000 ; HA- CA- CB- HB2
3566 3565 3567 3569 3 0.65084 1.95253 0.00000
-2.60338 0.00000 0.00000 ; HA- CA- CB- HB3
3566 3565 3567 3570 3 0.65084 1.95253 0.00000
-2.60338 0.00000 0.00000 ; HA- CA- CB- CG
3566 3565 3583 3584 3 3.68192 -4.35136 0.00000
1.33888 0.00000 0.00000 ; HA- CA- C- O
3566 3565 3583 3585 3 0.00000 0.00000 0.00000
0.00000 0.00000 0.00000 ; HA- CA- C- N
3567 3565 3583 3584 3 0.00000 0.00000 0.00000
0.00000 0.00000 0.00000 ; CB- CA- C- O

3567 3565 3583 3585 3 2.51040 4.18400 1.67360
-6.69440 0.00000 0.00000 ; CB- CA- C- N
3567 3570 3573 3574 3 0.66944 2.00832 0.00000
-2.67776 0.00000 0.00000 ; CB- CG- CD- HD2
3567 3570 3573 3575 3 0.66944 2.00832 0.00000
-2.67776 0.00000 0.00000 ; CB- CG- CD- HD3
3567 3570 3573 3576 3 3.68192 3.09616 -2.09200
-3.01248 0.00000 0.00000 ; CB- CG- CD- CE
3568 3567 3565 3583 3 0.65084 1.95253 0.00000
-2.60338 0.00000 0.00000 ; HB2- CB- CA- C
3568 3567 3570 3571 3 0.62760 1.88280 0.00000
-2.51040 0.00000 0.00000 ; HB2- CB- CG- HG2
3568 3567 3570 3572 3 0.62760 1.88280 0.00000
-2.51040 0.00000 0.00000 ; HB2- CB- CG- HG3
3568 3567 3570 3573 3 0.66944 2.00832 0.00000
-2.67776 0.00000 0.00000 ; HB2- CB- CG- CD
3569 3567 3565 3583 3 0.65084 1.95253 0.00000
-2.60338 0.00000 0.00000 ; HB3- CB- CA- C
3569 3567 3570 3571 3 0.62760 1.88280 0.00000
-2.51040 0.00000 0.00000 ; HB3- CB- CG- HG2
3569 3567 3570 3572 3 0.62760 1.88280 0.00000
-2.51040 0.00000 0.00000 ; HB3- CB- CG- HG3
3569 3567 3570 3573 3 0.66944 2.00832 0.00000
-2.67776 0.00000 0.00000 ; HB3- CB- CG- CD
3570 3567 3565 3583 3 0.65084 1.95253 0.00000
-2.60338 0.00000 0.00000 ; CG- CB- CA- C
3570 3573 3576 3577 3 0.65084 1.95253 0.00000
-2.60338 0.00000 0.00000 ; CG- CD- CE- HE2
3570 3573 3576 3578 3 0.65084 1.95253 0.00000
-2.60338 0.00000 0.00000 ; CG- CD- CE- HE3
3570 3573 3576 3579 3 0.65084 1.95253 0.00000
-2.60338 0.00000 0.00000 ; CG- CD- CE- NZ
3571 3570 3573 3574 3 0.62760 1.88280 0.00000
-2.51040 0.00000 0.00000 ; HG2- CG- CD- HD2
3571 3570 3573 3575 3 0.62760 1.88280 0.00000
-2.51040 0.00000 0.00000 ; HG2- CG- CD- HD3
3571 3570 3573 3576 3 0.66944 2.00832 0.00000
-2.67776 0.00000 0.00000 ; HG2- CG- CD- CE
3572 3570 3573 3574 3 0.62760 1.88280 0.00000
-2.51040 0.00000 0.00000 ; HG3- CG- CD- HD2
3572 3570 3573 3575 3 0.62760 1.88280 0.00000
-2.51040 0.00000 0.00000 ; HG3- CG- CD- HD3
3572 3570 3573 3576 3 0.66944 2.00832 0.00000
-2.67776 0.00000 0.00000 ; HG3- CG- CD- CE
3573 3576 3579 3580 3 0.65084 1.95253 0.00000
-2.60338 0.00000 0.00000 ; CD- CE- NZ- HZ1
3573 3576 3579 3581 3 0.65084 1.95253 0.00000
-2.60338 0.00000 0.00000 ; CD- CE- NZ- HZ2
3573 3576 3579 3582 3 0.65084 1.95253 0.00000
-2.60338 0.00000 0.00000 ; CD- CE- NZ- HZ3
3574 3573 3576 3577 3 0.65084 1.95253 0.00000
-2.60338 0.00000 0.00000 ; HD2- CD- CE- HE2

3574 3573 3576 3578 3 0.65084 1.95253 0.00000
-2.60338 0.00000 0.00000 ; HD2- CD- CE- HE3
3574 3573 3576 3579 3 0.65084 1.95253 0.00000
-2.60338 0.00000 0.00000 ; HD2- CD- CE- NZ
3575 3573 3576 3577 3 0.65084 1.95253 0.00000
-2.60338 0.00000 0.00000 ; HD3- CD- CE- HE2
3575 3573 3576 3578 3 0.65084 1.95253 0.00000
-2.60338 0.00000 0.00000 ; HD3- CD- CE- HE3
3575 3573 3576 3579 3 0.65084 1.95253 0.00000
-2.60338 0.00000 0.00000 ; HD3- CD- CE- NZ
3577 3576 3579 3580 3 0.65084 1.95253 0.00000
-2.60338 0.00000 0.00000 ; HE2- CE- NZ- HZ1
3577 3576 3579 3581 3 0.65084 1.95253 0.00000
-2.60338 0.00000 0.00000 ; HE2- CE- NZ- HZ2
3577 3576 3579 3582 3 0.65084 1.95253 0.00000
-2.60338 0.00000 0.00000 ; HE2- CE- NZ- HZ3
3578 3576 3579 3580 3 0.65084 1.95253 0.00000
-2.60338 0.00000 0.00000 ; HE3- CE- NZ- HZ1
3578 3576 3579 3581 3 0.65084 1.95253 0.00000
-2.60338 0.00000 0.00000 ; HE3- CE- NZ- HZ2
3578 3576 3579 3582 3 0.65084 1.95253 0.00000
-2.60338 0.00000 0.00000 ; HE3- CE- NZ- HZ3
3583 3585 3587 3588 3 0.00000 0.00000 0.00000
0.00000 0.00000 0.00000 ; C- N- CA- HA
3583 3585 3587 3589 3 11.71520 1.67360 15.06240 -
13.38880 0.00000 0.00000 ; C- N- CA- CB
3583 3585 3587 3599 3 1.75728 5.27184 2.25936
-7.02912 0.00000 0.00000 ; C- N- CA- C
3584 3583 3585 3586 3 29.28800 -8.36800 -20.92000
0.00000 0.00000 0.00000 ; O- C- N- H
3584 3583 3585 3587 3 20.92000 0.00000 -20.92000
0.00000 0.00000 0.00000 ; O- C- N- CA
3585 3587 3589 3590 3 0.65084 1.95253 0.00000
-2.60338 0.00000 0.00000 ; N- CA- CB- HB
3585 3587 3589 3591 3 3.83673 0.44769 -1.77402
-2.47693 -0.03347 0.00000 ; N- CA- CB- CG1
3585 3587 3589 3595 3 3.83673 0.44769 -1.77402
-2.47693 -0.03347 0.00000 ; N- CA- CB- CG2
3585 3587 3599 3600 3 0.00000 0.00000 0.00000
0.00000 0.00000 0.00000 ; N- CA- C- O
3585 3587 3599 3601 3 17.40544 -5.02080 -13.22144
9.20480 0.00000 0.00000 ; N- CA- C- N
3586 3585 3587 3588 3 0.00000 0.00000 0.00000
0.00000 0.00000 0.00000 ; H- N- CA- HA
3586 3585 3587 3589 3 0.00000 0.00000 0.00000
0.00000 0.00000 0.00000 ; H- N- CA- CB
3586 3585 3587 3599 3 0.00000 0.00000 0.00000
0.00000 0.00000 0.00000 ; H- N- CA- C
3587 3589 3591 3592 3 0.66944 2.00832 0.00000
-2.67776 0.00000 0.00000 ; CA- CB- CG1- HG11
3587 3589 3591 3593 3 0.66944 2.00832 0.00000
-2.67776 0.00000 0.00000 ; CA- CB- CG1- HG12

```
  3587   3589   3591   3594       3    0.66944    2.00832    0.00000
-2.67776    0.00000    0.00000 ;     CA-    CB-   CG1-  HG13
  3587   3589   3595   3596       3    0.66944    2.00832    0.00000
-2.67776    0.00000    0.00000 ;     CA-    CB-   CG2-  HG21
  3587   3589   3595   3597       3    0.66944    2.00832    0.00000
-2.67776    0.00000    0.00000 ;     CA-    CB-   CG2-  HG22
  3587   3589   3595   3598       3    0.66944    2.00832    0.00000
-2.67776    0.00000    0.00000 ;     CA-    CB-   CG2-  HG23
  3587   3599   3601   3602       3   20.92000    0.00000  -20.92000
0.00000    0.00000    0.00000 ;     CA-     C-     N-     H
  3587   3599   3601   3603       3   20.92000    0.00000  -20.92000
0.00000    0.00000    0.00000 ;     CA-     C-     N-    CA
  3588   3587   3589   3590       3    0.65084    1.95253    0.00000
-2.60338    0.00000    0.00000 ;     HA-    CA-    CB-    HB
  3588   3587   3589   3591       3    0.65084    1.95253    0.00000
-2.60338    0.00000    0.00000 ;     HA-    CA-    CB-   CG1
  3588   3587   3589   3595       3    0.65084    1.95253    0.00000
-2.60338    0.00000    0.00000 ;     HA-    CA-    CB-   CG2
  3588   3587   3599   3600       3    3.68192   -4.35136    0.00000
1.33888    0.00000    0.00000 ;     HA-    CA-     C-     O
  3588   3587   3599   3601       3    0.00000    0.00000    0.00000
0.00000    0.00000    0.00000 ;     HA-    CA-     C-     N
  3589   3587   3599   3600       3    0.00000    0.00000    0.00000
0.00000    0.00000    0.00000 ;     CB-    CA-     C-     O
  3589   3587   3599   3601       3    2.51040    4.18400    1.67360
-6.69440    0.00000    0.00000 ;     CB-    CA-     C-     N
  3590   3589   3587   3599       3    0.65084    1.95253    0.00000
-2.60338    0.00000    0.00000 ;     HB-    CB-    CA-     C
  3590   3589   3591   3592       3    0.62760    1.88280    0.00000
-2.51040    0.00000    0.00000 ;     HB-    CB-   CG1-  HG11
  3590   3589   3591   3593       3    0.62760    1.88280    0.00000
-2.51040    0.00000    0.00000 ;     HB-    CB-   CG1-  HG12
  3590   3589   3591   3594       3    0.62760    1.88280    0.00000
-2.51040    0.00000    0.00000 ;     HB-    CB-   CG1-  HG13
  3590   3589   3595   3596       3    0.62760    1.88280    0.00000
-2.51040    0.00000    0.00000 ;     HB-    CB-   CG2-  HG21
  3590   3589   3595   3597       3    0.62760    1.88280    0.00000
-2.51040    0.00000    0.00000 ;     HB-    CB-   CG2-  HG22
  3590   3589   3595   3598       3    0.62760    1.88280    0.00000
-2.51040    0.00000    0.00000 ;     HB-    CB-   CG2-  HG23
  3591   3589   3587   3599       3    5.67350    3.55640   -6.16722
-2.47693    3.74886    0.00000 ;    CG1-    CB-    CA-     C
  3591   3589   3595   3596       3    0.66944    2.00832    0.00000
-2.67776    0.00000    0.00000 ;    CG1-    CB-   CG2-  HG21
  3591   3589   3595   3597       3    0.66944    2.00832    0.00000
-2.67776    0.00000    0.00000 ;    CG1-    CB-   CG2-  HG22
  3591   3589   3595   3598       3    0.66944    2.00832    0.00000
-2.67776    0.00000    0.00000 ;    CG1-    CB-   CG2-  HG23
  3592   3591   3589   3595       3    0.66944    2.00832    0.00000
-2.67776    0.00000    0.00000 ;   HG11-   CG1-    CB-   CG2
  3593   3591   3589   3595       3    0.66944    2.00832    0.00000
-2.67776    0.00000    0.00000 ;   HG12-   CG1-    CB-   CG2
```

3594 3591 3589 3595 3 0.66944 2.00832 0.00000
-2.67776 0.00000 0.00000 ; HG13- CG1- CB- CG2
3595 3589 3587 3599 3 5.67350 3.55640 -6.16722
-2.47693 3.74886 0.00000 ; CG2- CB- CA- C
3599 3601 3603 3604 3 0.00000 0.00000 0.00000
0.00000 0.00000 0.00000 ; C- N- CA- HA
3599 3601 3603 3605 3 11.71520 1.67360 15.06240 -
13.38880 0.00000 0.00000 ; C- N- CA- CB
3599 3601 3603 3621 3 1.75728 5.27184 2.25936
-7.02912 0.00000 0.00000 ; C- N- CA- C
3600 3599 3601 3602 3 29.28800 -8.36800 -20.92000
0.00000 0.00000 0.00000 ; O- C- N- H
3600 3599 3601 3603 3 20.92000 0.00000 -20.92000
0.00000 0.00000 0.00000 ; O- C- N- CA
3601 3603 3605 3606 3 0.65084 1.95253 0.00000
-2.60338 0.00000 0.00000 ; N- CA- CB- HB2
3601 3603 3605 3607 3 0.65084 1.95253 0.00000
-2.60338 0.00000 0.00000 ; N- CA- CB- HB3
3601 3603 3605 3608 3 0.65084 1.95253 0.00000
-2.60338 0.00000 0.00000 ; N- CA- CB- CG
3601 3603 3621 3622 3 0.00000 0.00000 0.00000
0.00000 0.00000 0.00000 ; N- CA- C- O
3601 3603 3621 3623 3 17.40544 -5.02080 -13.22144
9.20480 0.00000 0.00000 ; N- CA- C- N
3602 3601 3603 3604 3 0.00000 0.00000 0.00000
0.00000 0.00000 0.00000 ; H- N- CA- HA
3602 3601 3603 3605 3 0.00000 0.00000 0.00000
0.00000 0.00000 0.00000 ; H- N- CA- CB
3602 3601 3603 3621 3 0.00000 0.00000 0.00000
0.00000 0.00000 0.00000 ; H- N- CA- C
3603 3605 3608 3609 3 0.66944 2.00832 0.00000
-2.67776 0.00000 0.00000 ; CA- CB- CG- HG2
3603 3605 3608 3610 3 0.66944 2.00832 0.00000
-2.67776 0.00000 0.00000 ; CA- CB- CG- HG3
3603 3605 3608 3611 3 3.68192 3.09616 -2.09200
-3.01248 0.00000 0.00000 ; CA- CB- CG- CD
3603 3621 3623 3624 3 20.92000 0.00000 -20.92000
0.00000 0.00000 0.00000 ; CA- C- N- H
3603 3621 3623 3625 3 20.92000 0.00000 -20.92000
0.00000 0.00000 0.00000 ; CA- C- N- CA
3604 3603 3605 3606 3 0.65084 1.95253 0.00000
-2.60338 0.00000 0.00000 ; HA- CA- CB- HB2
3604 3603 3605 3607 3 0.65084 1.95253 0.00000
-2.60338 0.00000 0.00000 ; HA- CA- CB- HB3
3604 3603 3605 3608 3 0.65084 1.95253 0.00000
-2.60338 0.00000 0.00000 ; HA- CA- CB- CG
3604 3603 3621 3622 3 3.68192 -4.35136 0.00000
1.33888 0.00000 0.00000 ; HA- CA- C- O
3604 3603 3621 3623 3 0.00000 0.00000 0.00000
0.00000 0.00000 0.00000 ; HA- CA- C- N
3605 3603 3621 3622 3 0.00000 0.00000 0.00000
0.00000 0.00000 0.00000 ; CB- CA- C- O

```
  3605   3603   3621   3623       3    2.51040    4.18400    1.67360
-6.69440    0.00000    0.00000 ;     CB-    CA-     C-     N
  3605   3608   3611   3612       3    0.66944    2.00832    0.00000
-2.67776    0.00000    0.00000 ;     CB-    CG-    CD-   HD2
  3605   3608   3611   3613       3    0.66944    2.00832    0.00000
-2.67776    0.00000    0.00000 ;     CB-    CG-    CD-   HD3
  3605   3608   3611   3614       3    3.68192    3.09616   -2.09200
-3.01248    0.00000    0.00000 ;     CB-    CG-    CD-    CE
  3606   3605   3603   3621       3    0.65084    1.95253    0.00000
-2.60338    0.00000    0.00000 ;    HB2-    CB-    CA-     C
  3606   3605   3608   3609       3    0.62760    1.88280    0.00000
-2.51040    0.00000    0.00000 ;    HB2-    CB-    CG-   HG2
  3606   3605   3608   3610       3    0.62760    1.88280    0.00000
-2.51040    0.00000    0.00000 ;    HB2-    CB-    CG-   HG3
  3606   3605   3608   3611       3    0.66944    2.00832    0.00000
-2.67776    0.00000    0.00000 ;    HB2-    CB-    CG-    CD
  3607   3605   3603   3621       3    0.65084    1.95253    0.00000
-2.60338    0.00000    0.00000 ;    HB3-    CB-    CA-     C
  3607   3605   3608   3609       3    0.62760    1.88280    0.00000
-2.51040    0.00000    0.00000 ;    HB3-    CB-    CG-   HG2
  3607   3605   3608   3610       3    0.62760    1.88280    0.00000
-2.51040    0.00000    0.00000 ;    HB3-    CB-    CG-   HG3
  3607   3605   3608   3611       3    0.66944    2.00832    0.00000
-2.67776    0.00000    0.00000 ;    HB3-    CB-    CG-    CD
  3608   3605   3603   3621       3    0.65084    1.95253    0.00000
-2.60338    0.00000    0.00000 ;     CG-    CB-    CA-     C
  3608   3611   3614   3615       3    0.65084    1.95253    0.00000
-2.60338    0.00000    0.00000 ;     CG-    CD-    CE-   HE2
  3608   3611   3614   3616       3    0.65084    1.95253    0.00000
-2.60338    0.00000    0.00000 ;     CG-    CD-    CE-   HE3
  3608   3611   3614   3617       3    0.65084    1.95253    0.00000
-2.60338    0.00000    0.00000 ;     CG-    CD-    CE-    NZ
  3609   3608   3611   3612       3    0.62760    1.88280    0.00000
-2.51040    0.00000    0.00000 ;    HG2-    CG-    CD-   HD2
  3609   3608   3611   3613       3    0.62760    1.88280    0.00000
-2.51040    0.00000    0.00000 ;    HG2-    CG-    CD-   HD3
  3609   3608   3611   3614       3    0.66944    2.00832    0.00000
-2.67776    0.00000    0.00000 ;    HG2-    CG-    CD-    CE
  3610   3608   3611   3612       3    0.62760    1.88280    0.00000
-2.51040    0.00000    0.00000 ;    HG3-    CG-    CD-   HD2
  3610   3608   3611   3613       3    0.62760    1.88280    0.00000
-2.51040    0.00000    0.00000 ;    HG3-    CG-    CD-   HD3
  3610   3608   3611   3614       3    0.66944    2.00832    0.00000
-2.67776    0.00000    0.00000 ;    HG3-    CG-    CD-    CE
  3611   3614   3617   3618       3    0.65084    1.95253    0.00000
-2.60338    0.00000    0.00000 ;     CD-    CE-    NZ-   HZ1
  3611   3614   3617   3619       3    0.65084    1.95253    0.00000
-2.60338    0.00000    0.00000 ;     CD-    CE-    NZ-   HZ2
  3611   3614   3617   3620       3    0.65084    1.95253    0.00000
-2.60338    0.00000    0.00000 ;     CD-    CE-    NZ-   HZ3
  3612   3611   3614   3615       3    0.65084    1.95253    0.00000
-2.60338    0.00000    0.00000 ;    HD2-    CD-    CE-   HE2
```

3612 3611 3614 3616 3 0.65084 1.95253 0.00000
-2.60338 0.00000 0.00000 ; HD2- CD- CE- HE3
3612 3611 3614 3617 3 0.65084 1.95253 0.00000
-2.60338 0.00000 0.00000 ; HD2- CD- CE- NZ
3613 3611 3614 3615 3 0.65084 1.95253 0.00000
-2.60338 0.00000 0.00000 ; HD3- CD- CE- HE2
3613 3611 3614 3616 3 0.65084 1.95253 0.00000
-2.60338 0.00000 0.00000 ; HD3- CD- CE- HE3
3613 3611 3614 3617 3 0.65084 1.95253 0.00000
-2.60338 0.00000 0.00000 ; HD3- CD- CE- NZ
3615 3614 3617 3618 3 0.65084 1.95253 0.00000
-2.60338 0.00000 0.00000 ; HE2- CE- NZ- HZ1
3615 3614 3617 3619 3 0.65084 1.95253 0.00000
-2.60338 0.00000 0.00000 ; HE2- CE- NZ- HZ2
3615 3614 3617 3620 3 0.65084 1.95253 0.00000
-2.60338 0.00000 0.00000 ; HE2- CE- NZ- HZ3
3616 3614 3617 3618 3 0.65084 1.95253 0.00000
-2.60338 0.00000 0.00000 ; HE3- CE- NZ- HZ1
3616 3614 3617 3619 3 0.65084 1.95253 0.00000
-2.60338 0.00000 0.00000 ; HE3- CE- NZ- HZ2
3616 3614 3617 3620 3 0.65084 1.95253 0.00000
-2.60338 0.00000 0.00000 ; HE3- CE- NZ- HZ3
3621 3623 3625 3626 3 0.00000 0.00000 0.00000
0.00000 0.00000 0.00000 ; C- N- CA- HA
3621 3623 3625 3627 3 11.71520 1.67360 15.06240 -
13.38880 0.00000 0.00000 ; C- N- CA- CB
3621 3623 3625 3643 3 1.75728 5.27184 2.25936
-7.02912 0.00000 0.00000 ; C- N- CA- C
3622 3621 3623 3624 3 29.28800 -8.36800 -20.92000
0.00000 0.00000 0.00000 ; O- C- N- H
3622 3621 3623 3625 3 20.92000 0.00000 -20.92000
0.00000 0.00000 0.00000 ; O- C- N- CA
3623 3625 3627 3628 3 0.65084 1.95253 0.00000
-2.60338 0.00000 0.00000 ; N- CA- CB- HB2
3623 3625 3627 3629 3 0.65084 1.95253 0.00000
-2.60338 0.00000 0.00000 ; N- CA- CB- HB3
3623 3625 3627 3630 3 0.65084 1.95253 0.00000
-2.60338 0.00000 0.00000 ; N- CA- CB- CG
3623 3625 3643 3644 3 0.00000 0.00000 0.00000
0.00000 0.00000 0.00000 ; N- CA- C- O
3623 3625 3643 3645 3 17.40544 -5.02080 -13.22144
9.20480 0.00000 0.00000 ; N- CA- C- N
3624 3623 3625 3626 3 0.00000 0.00000 0.00000
0.00000 0.00000 0.00000 ; H- N- CA- HA
3624 3623 3625 3627 3 0.00000 0.00000 0.00000
0.00000 0.00000 0.00000 ; H- N- CA- CB
3624 3623 3625 3643 3 0.00000 0.00000 0.00000
0.00000 0.00000 0.00000 ; H- N- CA- C
3625 3627 3630 3631 3 0.66944 2.00832 0.00000
-2.67776 0.00000 0.00000 ; CA- CB- CG- HG2
3625 3627 3630 3632 3 0.66944 2.00832 0.00000
-2.67776 0.00000 0.00000 ; CA- CB- CG- HG3

3625 3627 3630 3633 3 3.68192 3.09616 -2.09200
-3.01248 0.00000 0.00000 ; CA- CB- CG- CD
3625 3643 3645 3646 3 20.92000 0.00000 -20.92000
0.00000 0.00000 0.00000 ; CA- C- N- H
3625 3643 3645 3647 3 20.92000 0.00000 -20.92000
0.00000 0.00000 0.00000 ; CA- C- N- CA
3626 3625 3627 3628 3 0.65084 1.95253 0.00000
-2.60338 0.00000 0.00000 ; HA- CA- CB- HB2
3626 3625 3627 3629 3 0.65084 1.95253 0.00000
-2.60338 0.00000 0.00000 ; HA- CA- CB- HB3
3626 3625 3627 3630 3 0.65084 1.95253 0.00000
-2.60338 0.00000 0.00000 ; HA- CA- CB- CG
3626 3625 3643 3644 3 3.68192 -4.35136 0.00000
1.33888 0.00000 0.00000 ; HA- CA- C- O
3626 3625 3643 3645 3 0.00000 0.00000 0.00000
0.00000 0.00000 0.00000 ; HA- CA- C- N
3627 3625 3643 3644 3 0.00000 0.00000 0.00000
0.00000 0.00000 0.00000 ; CB- CA- C- O
3627 3625 3643 3645 3 2.51040 4.18400 1.67360
-6.69440 0.00000 0.00000 ; CB- CA- C- N
3627 3630 3633 3634 3 0.66944 2.00832 0.00000
-2.67776 0.00000 0.00000 ; CB- CG- CD- HD2
3627 3630 3633 3635 3 0.66944 2.00832 0.00000
-2.67776 0.00000 0.00000 ; CB- CG- CD- HD3
3627 3630 3633 3636 3 3.68192 3.09616 -2.09200
-3.01248 0.00000 0.00000 ; CB- CG- CD- CE
3628 3627 3625 3643 3 0.65084 1.95253 0.00000
-2.60338 0.00000 0.00000 ; HB2- CB- CA- C
3628 3627 3630 3631 3 0.62760 1.88280 0.00000
-2.51040 0.00000 0.00000 ; HB2- CB- CG- HG2
3628 3627 3630 3632 3 0.62760 1.88280 0.00000
-2.51040 0.00000 0.00000 ; HB2- CB- CG- HG3
3628 3627 3630 3633 3 0.66944 2.00832 0.00000
-2.67776 0.00000 0.00000 ; HB2- CB- CG- CD
3629 3627 3625 3643 3 0.65084 1.95253 0.00000
-2.60338 0.00000 0.00000 ; HB3- CB- CA- C
3629 3627 3630 3631 3 0.62760 1.88280 0.00000
-2.51040 0.00000 0.00000 ; HB3- CB- CG- HG2
3629 3627 3630 3632 3 0.62760 1.88280 0.00000
-2.51040 0.00000 0.00000 ; HB3- CB- CG- HG3
3629 3627 3630 3633 3 0.66944 2.00832 0.00000
-2.67776 0.00000 0.00000 ; HB3- CB- CG- CD
3630 3627 3625 3643 3 0.65084 1.95253 0.00000
-2.60338 0.00000 0.00000 ; CG- CB- CA- C
3630 3633 3636 3637 3 0.65084 1.95253 0.00000
-2.60338 0.00000 0.00000 ; CG- CD- CE- HE2
3630 3633 3636 3638 3 0.65084 1.95253 0.00000
-2.60338 0.00000 0.00000 ; CG- CD- CE- HE3
3630 3633 3636 3639 3 0.65084 1.95253 0.00000
-2.60338 0.00000 0.00000 ; CG- CD- CE- NZ
3631 3630 3633 3634 3 0.62760 1.88280 0.00000
-2.51040 0.00000 0.00000 ; HG2- CG- CD- HD2

3631 3630 3633 3635 3 0.62760 1.88280 0.00000
-2.51040 0.00000 0.00000 ; HG2- CG- CD- HD3
3631 3630 3633 3636 3 0.66944 2.00832 0.00000
-2.67776 0.00000 0.00000 ; HG2- CG- CD- CE
3632 3630 3633 3634 3 0.62760 1.88280 0.00000
-2.51040 0.00000 0.00000 ; HG3- CG- CD- HD2
3632 3630 3633 3635 3 0.62760 1.88280 0.00000
-2.51040 0.00000 0.00000 ; HG3- CG- CD- HD3
3632 3630 3633 3636 3 0.66944 2.00832 0.00000
-2.67776 0.00000 0.00000 ; HG3- CG- CD- CE
3633 3636 3639 3640 3 0.65084 1.95253 0.00000
-2.60338 0.00000 0.00000 ; CD- CE- NZ- HZ1
3633 3636 3639 3641 3 0.65084 1.95253 0.00000
-2.60338 0.00000 0.00000 ; CD- CE- NZ- HZ2
3633 3636 3639 3642 3 0.65084 1.95253 0.00000
-2.60338 0.00000 0.00000 ; CD- CE- NZ- HZ3
3634 3633 3636 3637 3 0.65084 1.95253 0.00000
-2.60338 0.00000 0.00000 ; HD2- CD- CE- HE2
3634 3633 3636 3638 3 0.65084 1.95253 0.00000
-2.60338 0.00000 0.00000 ; HD2- CD- CE- HE3
3634 3633 3636 3639 3 0.65084 1.95253 0.00000
-2.60338 0.00000 0.00000 ; HD2- CD- CE- NZ
3635 3633 3636 3637 3 0.65084 1.95253 0.00000
-2.60338 0.00000 0.00000 ; HD3- CD- CE- HE2
3635 3633 3636 3638 3 0.65084 1.95253 0.00000
-2.60338 0.00000 0.00000 ; HD3- CD- CE- HE3
3635 3633 3636 3639 3 0.65084 1.95253 0.00000
-2.60338 0.00000 0.00000 ; HD3- CD- CE- NZ
3637 3636 3639 3640 3 0.65084 1.95253 0.00000
-2.60338 0.00000 0.00000 ; HE2- CE- NZ- HZ1
3637 3636 3639 3641 3 0.65084 1.95253 0.00000
-2.60338 0.00000 0.00000 ; HE2- CE- NZ- HZ2
3637 3636 3639 3642 3 0.65084 1.95253 0.00000
-2.60338 0.00000 0.00000 ; HE2- CE- NZ- HZ3
3638 3636 3639 3640 3 0.65084 1.95253 0.00000
-2.60338 0.00000 0.00000 ; HE3- CE- NZ- HZ1
3638 3636 3639 3641 3 0.65084 1.95253 0.00000
-2.60338 0.00000 0.00000 ; HE3- CE- NZ- HZ2
3638 3636 3639 3642 3 0.65084 1.95253 0.00000
-2.60338 0.00000 0.00000 ; HE3- CE- NZ- HZ3
3643 3645 3647 3648 3 0.00000 0.00000 0.00000
0.00000 0.00000 0.00000 ; C- N- CA- HA2
3643 3645 3647 3649 3 0.00000 0.00000 0.00000
0.00000 0.00000 0.00000 ; C- N- CA- HA3
3643 3645 3647 3650 3 1.75728 5.27184 2.25936
-7.02912 0.00000 0.00000 ; C- N- CA- C
3644 3643 3645 3646 3 29.28800 -8.36800 -20.92000
0.00000 0.00000 0.00000 ; O- C- N- H
3644 3643 3645 3647 3 20.92000 0.00000 -20.92000
0.00000 0.00000 0.00000 ; O- C- N- CA
3645 3647 3650 3651 3 0.00000 0.00000 0.00000
0.00000 0.00000 0.00000 ; N- CA- C- O

3645 3647 3650 3652 3 17.40544 -5.02080 -13.22144
9.20480 0.00000 0.00000 ; N- CA- C- N
3646 3645 3647 3648 3 0.00000 0.00000 0.00000
0.00000 0.00000 0.00000 ; H- N- CA- HA2
3646 3645 3647 3649 3 0.00000 0.00000 0.00000
0.00000 0.00000 0.00000 ; H- N- CA- HA3
3646 3645 3647 3650 3 0.00000 0.00000 0.00000
0.00000 0.00000 0.00000 ; H- N- CA- C
3647 3650 3652 3653 3 20.92000 0.00000 -20.92000
0.00000 0.00000 0.00000 ; CA- C- N- H
3647 3650 3652 3654 3 20.92000 0.00000 -20.92000
0.00000 0.00000 0.00000 ; CA- C- N- CA
3648 3647 3650 3651 3 3.68192 -4.35136 0.00000
1.33888 0.00000 0.00000 ; HA2- CA- C- O
3648 3647 3650 3652 3 0.00000 0.00000 0.00000
0.00000 0.00000 0.00000 ; HA2- CA- C- N
3649 3647 3650 3651 3 3.68192 -4.35136 0.00000
1.33888 0.00000 0.00000 ; HA3- CA- C- O
3649 3647 3650 3652 3 0.00000 0.00000 0.00000
0.00000 0.00000 0.00000 ; HA3- CA- C- N
3650 3652 3654 3655 3 0.00000 0.00000 0.00000
0.00000 0.00000 0.00000 ; C- N- CA- HA
3650 3652 3654 3656 3 11.71520 1.67360 15.06240 -
13.38880 0.00000 0.00000 ; C- N- CA- CB
3650 3652 3654 3665 3 1.75728 5.27184 2.25936
-7.02912 0.00000 0.00000 ; C- N- CA- C
3651 3650 3652 3653 3 29.28800 -8.36800 -20.92000
0.00000 0.00000 0.00000 ; O- C- N- H
3651 3650 3652 3654 3 20.92000 0.00000 -20.92000
0.00000 0.00000 0.00000 ; O- C- N- CA
3652 3654 3656 3657 3 0.65084 1.95253 0.00000
-2.60338 0.00000 0.00000 ; N- CA- CB- HB2
3652 3654 3656 3658 3 0.65084 1.95253 0.00000
-2.60338 0.00000 0.00000 ; N- CA- CB- HB3
3652 3654 3656 3659 3 3.21331 2.22589 -4.15053
-2.40998 2.61082 0.00000 ; N- CA- CB- CG
3652 3654 3665 3666 3 0.00000 0.00000 0.00000
0.00000 0.00000 0.00000 ; N- CA- C- O
3652 3654 3665 3667 3 17.40544 -5.02080 -13.22144
9.20480 0.00000 0.00000 ; N- CA- C- N
3653 3652 3654 3655 3 0.00000 0.00000 0.00000
0.00000 0.00000 0.00000 ; H- N- CA- HA
3653 3652 3654 3656 3 0.00000 0.00000 0.00000
0.00000 0.00000 0.00000 ; H- N- CA- CB
3653 3652 3654 3665 3 0.00000 0.00000 0.00000
0.00000 0.00000 0.00000 ; H- N- CA- C
3654 3656 3659 3660 3 0.66944 2.00832 0.00000
-2.67776 0.00000 0.00000 ; CA- CB- CG- HG2
3654 3656 3659 3661 3 0.66944 2.00832 0.00000
-2.67776 0.00000 0.00000 ; CA- CB- CG- HG3
3654 3656 3659 3662 3 10.12110 -1.91209 0.01674
10.17549 -1.87443 0.00000 ; CA- CB- CG- CD

3654 3665 3667 3668 3 20.92000 0.00000 -20.92000
0.00000 0.00000 0.00000 ; CA- C- N- H
3654 3665 3667 3669 3 20.92000 0.00000 -20.92000
0.00000 0.00000 0.00000 ; CA- C- N- CA
3655 3654 3656 3657 3 0.65084 1.95253 0.00000
-2.60338 0.00000 0.00000 ; HA- CA- CB- HB2
3655 3654 3656 3658 3 0.65084 1.95253 0.00000
-2.60338 0.00000 0.00000 ; HA- CA- CB- HB3
3655 3654 3656 3659 3 0.65084 1.95253 0.00000
-2.60338 0.00000 0.00000 ; HA- CA- CB- CG
3655 3654 3665 3666 3 3.68192 -4.35136 0.00000
1.33888 0.00000 0.00000 ; HA- CA- C- O
3655 3654 3665 3667 3 0.00000 0.00000 0.00000
0.00000 0.00000 0.00000 ; HA- CA- C- N
3656 3654 3665 3666 3 0.00000 0.00000 0.00000
0.00000 0.00000 0.00000 ; CB- CA- C- O
3656 3654 3665 3667 3 2.51040 4.18400 1.67360
-6.69440 0.00000 0.00000 ; CB- CA- C- N
3656 3659 3662 3663 3 3.79907 0.00000 -5.40573
0.00000 2.14221 0.00000 ; CB- CG- CD- OE1
3656 3659 3662 3664 3 3.79907 0.00000 -5.40573
0.00000 2.14221 0.00000 ; CB- CG- CD- OE2
3657 3656 3654 3665 3 0.65084 1.95253 0.00000
-2.60338 0.00000 0.00000 ; HB2- CB- CA- C
3657 3656 3659 3660 3 0.62760 1.88280 0.00000
-2.51040 0.00000 0.00000 ; HB2- CB- CG- HG2
3657 3656 3659 3661 3 0.62760 1.88280 0.00000
-2.51040 0.00000 0.00000 ; HB2- CB- CG- HG3
3657 3656 3659 3662 3 0.65084 1.95253 0.00000
-2.60338 0.00000 0.00000 ; HB2- CB- CG- CD
3658 3656 3654 3665 3 0.65084 1.95253 0.00000
-2.60338 0.00000 0.00000 ; HB3- CB- CA- C
3658 3656 3659 3660 3 0.62760 1.88280 0.00000
-2.51040 0.00000 0.00000 ; HB3- CB- CG- HG2
3658 3656 3659 3661 3 0.62760 1.88280 0.00000
-2.51040 0.00000 0.00000 ; HB3- CB- CG- HG3
3658 3656 3659 3662 3 0.65084 1.95253 0.00000
-2.60338 0.00000 0.00000 ; HB3- CB- CG- CD
3659 3656 3654 3665 3 6.86594 3.56895 -8.14206
-2.40998 4.85344 0.00000 ; CG- CB- CA- C
3660 3659 3662 3663 3 0.00000 0.00000 0.00000
0.00000 0.00000 0.00000 ; HG2- CG- CD- OE1
3660 3659 3662 3664 3 0.00000 0.00000 0.00000
0.00000 0.00000 0.00000 ; HG2- CG- CD- OE2
3661 3659 3662 3663 3 0.00000 0.00000 0.00000
0.00000 0.00000 0.00000 ; HG3- CG- CD- OE1
3661 3659 3662 3664 3 0.00000 0.00000 0.00000
0.00000 0.00000 0.00000 ; HG3- CG- CD- OE2
3665 3667 3669 3670 3 0.00000 0.00000 0.00000
0.00000 0.00000 0.00000 ; C- N- CA- HA
3665 3667 3669 3671 3 11.71520 1.67360 15.06240 -
13.38880 0.00000 0.00000 ; C- N- CA- CB

```
  3665   3667   3669   3685      3    1.75728    5.27184    2.25936
-7.02912    0.00000    0.00000 ;      C-     N-    CA-     C
  3666   3665   3667   3668      3   29.28800   -8.36800  -20.92000
0.00000    0.00000    0.00000 ;      O-     C-     N-     H
  3666   3665   3667   3669      3   20.92000    0.00000  -20.92000
0.00000    0.00000    0.00000 ;      O-     C-     N-    CA
  3667   3669   3671   3672      3    0.65084    1.95253    0.00000
-2.60338    0.00000    0.00000 ;      N-    CA-    CB-   HB2
  3667   3669   3671   3673      3    0.65084    1.95253    0.00000
-2.60338    0.00000    0.00000 ;      N-    CA-    CB-   HB3
  3667   3669   3671   3674      3    3.28026    2.46019   -2.19242
-3.21331   -0.23430    0.00000 ;      N-    CA-    CB-    CG
  3667   3669   3685   3686      3    0.00000    0.00000    0.00000
0.00000    0.00000    0.00000 ;      N-    CA-     C-     O
  3667   3669   3685   3687      3   17.40544   -5.02080  -13.22144
9.20480    0.00000    0.00000 ;      N-    CA-     C-     N
  3668   3667   3669   3670      3    0.00000    0.00000    0.00000
0.00000    0.00000    0.00000 ;      H-     N-    CA-    HA
  3668   3667   3669   3671      3    0.00000    0.00000    0.00000
0.00000    0.00000    0.00000 ;      H-     N-    CA-    CB
  3668   3667   3669   3685      3    0.00000    0.00000    0.00000
0.00000    0.00000    0.00000 ;      H-     N-    CA-     C
  3669   3671   3674   3675      3    0.57739    0.00000    1.02926
0.00000   -1.60666    0.00000 ;     CA-    CB-    CG-   CD1
  3669   3671   3674   3683      3    0.57739    0.00000    1.02926
0.00000   -1.60666    0.00000 ;     CA-    CB-    CG-   CD2
  3669   3685   3687   3688      3   20.92000    0.00000  -20.92000
0.00000    0.00000    0.00000 ;     CA-     C-     N-     H
  3669   3685   3687   3689      3   20.92000    0.00000  -20.92000
0.00000    0.00000    0.00000 ;     CA-     C-     N-    CA
  3670   3669   3671   3672      3    0.65084    1.95253    0.00000
-2.60338    0.00000    0.00000 ;     HA-    CA-    CB-   HB2
  3670   3669   3671   3673      3    0.65084    1.95253    0.00000
-2.60338    0.00000    0.00000 ;     HA-    CA-    CB-   HB3
  3670   3669   3671   3674      3    0.65084    1.95253    0.00000
-2.60338    0.00000    0.00000 ;     HA-    CA-    CB-    CG
  3670   3669   3685   3686      3    3.68192   -4.35136    0.00000
1.33888    0.00000    0.00000 ;     HA-    CA-     C-     O
  3670   3669   3685   3687      3    0.00000    0.00000    0.00000
0.00000    0.00000    0.00000 ;     HA-    CA-     C-     N
  3671   3669   3685   3686      3    0.00000    0.00000    0.00000
0.00000    0.00000    0.00000 ;     CB-    CA-     C-     O
  3671   3669   3685   3687      3    2.51040    4.18400    1.67360
-6.69440    0.00000    0.00000 ;     CB-    CA-     C-     N
  3671   3674   3675   3676      3   30.33400    0.00000  -30.33400
0.00000    0.00000    0.00000 ;     CB-    CG-   CD1-   HD1
  3671   3674   3675   3677      3   30.33400    0.00000  -30.33400
0.00000    0.00000    0.00000 ;     CB-    CG-   CD1-   CE1
  3671   3674   3683   3681      3   30.33400    0.00000  -30.33400
0.00000    0.00000    0.00000 ;     CB-    CG-   CD2-   CE2
  3671   3674   3683   3684      3   30.33400    0.00000  -30.33400
0.00000    0.00000    0.00000 ;     CB-    CG-   CD2-   HD2
```

3672 3671 3669 3685 3 0.65084 1.95253 0.00000
-2.60338 0.00000 0.00000 ; HB2- CB- CA- C
3672 3671 3674 3675 3 0.00000 0.00000 0.00000
0.00000 0.00000 0.00000 ; HB2- CB- CG- CD1
3672 3671 3674 3683 3 0.00000 0.00000 0.00000
0.00000 0.00000 0.00000 ; HB2- CB- CG- CD2
3673 3671 3669 3685 3 0.65084 1.95253 0.00000
-2.60338 0.00000 0.00000 ; HB3- CB- CA- C
3673 3671 3674 3675 3 0.00000 0.00000 0.00000
0.00000 0.00000 0.00000 ; HB3- CB- CG- CD1
3673 3671 3674 3683 3 0.00000 0.00000 0.00000
0.00000 0.00000 0.00000 ; HB3- CB- CG- CD2
3674 3671 3669 3685 3 4.95804 2.17986 -3.52293
-3.21331 -0.40166 0.00000 ; CG- CB- CA- C
3674 3675 3677 3678 3 30.33400 0.00000 -30.33400
0.00000 0.00000 0.00000 ; CG- CD1- CE1- HE1
3674 3675 3677 3679 3 30.33400 0.00000 -30.33400
0.00000 0.00000 0.00000 ; CG- CD1- CE1- CZ
3674 3683 3681 3679 3 30.33400 0.00000 -30.33400
0.00000 0.00000 0.00000 ; CG- CD2- CE2- CZ
3674 3683 3681 3682 3 30.33400 0.00000 -30.33400
0.00000 0.00000 0.00000 ; CG- CD2- CE2- HE2
3675 3674 3683 3681 3 30.33400 0.00000 -30.33400
0.00000 0.00000 0.00000 ; CD1- CG- CD2- CE2
3675 3674 3683 3684 3 30.33400 0.00000 -30.33400
0.00000 0.00000 0.00000 ; CD1- CG- CD2- HD2
3675 3677 3679 3680 3 30.33400 0.00000 -30.33400
0.00000 0.00000 0.00000 ; CD1- CE1- CZ- HZ
3675 3677 3679 3681 3 30.33400 0.00000 -30.33400
0.00000 0.00000 0.00000 ; CD1- CE1- CZ- CE2
3676 3675 3674 3683 3 30.33400 0.00000 -30.33400
0.00000 0.00000 0.00000 ; HD1- CD1- CG- CD2
3676 3675 3677 3678 3 30.33400 0.00000 -30.33400
0.00000 0.00000 0.00000 ; HD1- CD1- CE1- HE1
3676 3675 3677 3679 3 30.33400 0.00000 -30.33400
0.00000 0.00000 0.00000 ; HD1- CD1- CE1- CZ
3677 3675 3674 3683 3 30.33400 0.00000 -30.33400
0.00000 0.00000 0.00000 ; CE1- CD1- CG- CD2
3677 3679 3681 3682 3 30.33400 0.00000 -30.33400
0.00000 0.00000 0.00000 ; CE1- CZ- CE2- HE2
3677 3679 3681 3683 3 30.33400 0.00000 -30.33400
0.00000 0.00000 0.00000 ; CE1- CZ- CE2- CD2
3678 3677 3679 3680 3 30.33400 0.00000 -30.33400
0.00000 0.00000 0.00000 ; HE1- CE1- CZ- HZ
3678 3677 3679 3681 3 30.33400 0.00000 -30.33400
0.00000 0.00000 0.00000 ; HE1- CE1- CZ- CE2
3679 3681 3683 3684 3 30.33400 0.00000 -30.33400
0.00000 0.00000 0.00000 ; CZ- CE2- CD2- HD2
3680 3679 3681 3682 3 30.33400 0.00000 -30.33400
0.00000 0.00000 0.00000 ; HZ- CZ- CE2- HE2
3680 3679 3681 3683 3 30.33400 0.00000 -30.33400
0.00000 0.00000 0.00000 ; HZ- CZ- CE2- CD2

3682 3681 3683 3684 3 30.33400 0.00000 -30.33400
0.00000 0.00000 0.00000 ; HE2- CE2- CD2- HD2
3685 3687 3689 3690 3 0.00000 0.00000 0.00000
0.00000 0.00000 0.00000 ; C- N- CA- HA
3685 3687 3689 3691 3 11.71520 1.67360 15.06240 -
13.38880 0.00000 0.00000 ; C- N- CA- CB
3685 3687 3689 3696 3 1.75728 5.27184 2.25936
-7.02912 0.00000 0.00000 ; C- N- CA- C
3686 3685 3687 3688 3 29.28800 -8.36800 -20.92000
0.00000 0.00000 0.00000 ; O- C- N- H
3686 3685 3687 3689 3 20.92000 0.00000 -20.92000
0.00000 0.00000 0.00000 ; O- C- N- CA
3687 3689 3691 3692 3 0.65084 1.95253 0.00000
-2.60338 0.00000 0.00000 ; N- CA- CB- HB2
3687 3689 3691 3693 3 0.65084 1.95253 0.00000
-2.60338 0.00000 0.00000 ; N- CA- CB- HB3
3687 3689 3691 3694 3 6.03751 2.50622 -5.17142
-4.20074 1.10458 0.00000 ; N- CA- CB- SG
3687 3689 3696 3697 3 0.00000 0.00000 0.00000
0.00000 0.00000 0.00000 ; N- CA- C- O
3687 3689 3696 3698 3 17.40544 -5.02080 -13.22144
9.20480 0.00000 0.00000 ; N- CA- C- N
3688 3687 3689 3690 3 0.00000 0.00000 0.00000
0.00000 0.00000 0.00000 ; H- N- CA- HA
3688 3687 3689 3691 3 0.00000 0.00000 0.00000
0.00000 0.00000 0.00000 ; H- N- CA- CB
3688 3687 3689 3696 3 0.00000 0.00000 0.00000
0.00000 0.00000 0.00000 ; H- N- CA- C
3689 3691 3694 3695 3 1.69034 2.77818 4.11706
-4.21747 1.00416 0.00000 ; CA- CB- SG- HG
3689 3696 3698 3699 3 20.92000 0.00000 -20.92000
0.00000 0.00000 0.00000 ; CA- C- N- H
3689 3696 3698 3700 3 20.92000 0.00000 -20.92000
0.00000 0.00000 0.00000 ; CA- C- N- CA
3690 3689 3691 3692 3 0.65084 1.95253 0.00000
-2.60338 0.00000 0.00000 ; HA- CA- CB- HB2
3690 3689 3691 3693 3 0.65084 1.95253 0.00000
-2.60338 0.00000 0.00000 ; HA- CA- CB- HB3
3690 3689 3691 3694 3 0.65084 1.95253 0.00000
-2.60338 0.00000 0.00000 ; HA- CA- CB- SG
3690 3689 3696 3697 3 3.68192 -4.35136 0.00000
1.33888 0.00000 0.00000 ; HA- CA- C- O
3690 3689 3696 3698 3 0.00000 0.00000 0.00000
0.00000 0.00000 0.00000 ; HA- CA- C- N
3691 3689 3696 3697 3 0.00000 0.00000 0.00000
0.00000 0.00000 0.00000 ; CB- CA- C- O
3691 3689 3696 3698 3 2.51040 4.18400 1.67360
-6.69440 0.00000 0.00000 ; CB- CA- C- N
3692 3691 3689 3696 3 0.65084 1.95253 0.00000
-2.60338 0.00000 0.00000 ; HB2- CB- CA- C
3692 3691 3694 3695 3 1.04600 3.13800 0.00000
-4.18400 0.00000 0.00000 ; HB2- CB- SG- HG

3693 3691 3689 3696 3 0.65084 1.95253 0.00000
-2.60338 0.00000 0.00000 ; HB3- CB- CA- C
3693 3691 3694 3695 3 1.04600 3.13800 0.00000
-4.18400 0.00000 0.00000 ; HB3- CB- SG- HG
3694 3691 3689 3696 3 5.62330 4.27605 -5.33042
-4.20074 2.51040 0.00000 ; SG- CB- CA- C
3696 3698 3700 3701 3 0.00000 0.00000 0.00000
0.00000 0.00000 0.00000 ; C- N- CA- HA
3696 3698 3700 3702 3 11.71520 1.67360 15.06240 -
13.38880 0.00000 0.00000 ; C- N- CA- CB
3696 3698 3700 3716 3 1.75728 5.27184 2.25936
-7.02912 0.00000 0.00000 ; C- N- CA- C
3697 3696 3698 3699 3 29.28800 -8.36800 -20.92000
0.00000 0.00000 0.00000 ; O- C- N- H
3697 3696 3698 3700 3 20.92000 0.00000 -20.92000
0.00000 0.00000 0.00000 ; O- C- N- CA
3698 3700 3702 3703 3 0.65084 1.95253 0.00000
-2.60338 0.00000 0.00000 ; N- CA- CB- HB2
3698 3700 3702 3704 3 0.65084 1.95253 0.00000
-2.60338 0.00000 0.00000 ; N- CA- CB- HB3
3698 3700 3702 3705 3 3.28026 2.46019 -2.19242
-3.21331 -0.23430 0.00000 ; N- CA- CB- CG
3698 3700 3716 3717 3 0.00000 0.00000 0.00000
0.00000 0.00000 0.00000 ; N- CA- C- O
3698 3700 3716 3718 3 17.40544 -5.02080 -13.22144
9.20480 0.00000 0.00000 ; N- CA- C- N
3699 3698 3700 3701 3 0.00000 0.00000 0.00000
0.00000 0.00000 0.00000 ; H- N- CA- HA
3699 3698 3700 3702 3 0.00000 0.00000 0.00000
0.00000 0.00000 0.00000 ; H- N- CA- CB
3699 3698 3700 3716 3 0.00000 0.00000 0.00000
0.00000 0.00000 0.00000 ; H- N- CA- C
3700 3702 3705 3706 3 0.57739 0.00000 1.02926
0.00000 -1.60666 0.00000 ; CA- CB- CG- CD1
3700 3702 3705 3714 3 0.57739 0.00000 1.02926
0.00000 -1.60666 0.00000 ; CA- CB- CG- CD2
3700 3716 3718 3719 3 20.92000 0.00000 -20.92000
0.00000 0.00000 0.00000 ; CA- C- N- H
3700 3716 3718 3720 3 20.92000 0.00000 -20.92000
0.00000 0.00000 0.00000 ; CA- C- N- CA
3701 3700 3702 3703 3 0.65084 1.95253 0.00000
-2.60338 0.00000 0.00000 ; HA- CA- CB- HB2
3701 3700 3702 3704 3 0.65084 1.95253 0.00000
-2.60338 0.00000 0.00000 ; HA- CA- CB- HB3
3701 3700 3702 3705 3 0.65084 1.95253 0.00000
-2.60338 0.00000 0.00000 ; HA- CA- CB- CG
3701 3700 3716 3717 3 3.68192 -4.35136 0.00000
1.33888 0.00000 0.00000 ; HA- CA- C- O
3701 3700 3716 3718 3 0.00000 0.00000 0.00000
0.00000 0.00000 0.00000 ; HA- CA- C- N
3702 3700 3716 3717 3 0.00000 0.00000 0.00000
0.00000 0.00000 0.00000 ; CB- CA- C- O

3702 3700 3716 3718 3 2.51040 4.18400 1.67360
-6.69440 0.00000 0.00000 ; CB- CA- C- N
3702 3705 3706 3707 3 30.33400 0.00000 -30.33400
0.00000 0.00000 0.00000 ; CB- CG- CD1- HD1
3702 3705 3706 3708 3 30.33400 0.00000 -30.33400
0.00000 0.00000 0.00000 ; CB- CG- CD1- CE1
3702 3705 3714 3712 3 30.33400 0.00000 -30.33400
0.00000 0.00000 0.00000 ; CB- CG- CD2- CE2
3702 3705 3714 3715 3 30.33400 0.00000 -30.33400
0.00000 0.00000 0.00000 ; CB- CG- CD2- HD2
3703 3702 3700 3716 3 0.65084 1.95253 0.00000
-2.60338 0.00000 0.00000 ; HB2- CB- CA- C
3703 3702 3705 3706 3 0.00000 0.00000 0.00000
0.00000 0.00000 0.00000 ; HB2- CB- CG- CD1
3703 3702 3705 3714 3 0.00000 0.00000 0.00000
0.00000 0.00000 0.00000 ; HB2- CB- CG- CD2
3704 3702 3700 3716 3 0.65084 1.95253 0.00000
-2.60338 0.00000 0.00000 ; HB3- CB- CA- C
3704 3702 3705 3706 3 0.00000 0.00000 0.00000
0.00000 0.00000 0.00000 ; HB3- CB- CG- CD1
3704 3702 3705 3714 3 0.00000 0.00000 0.00000
0.00000 0.00000 0.00000 ; HB3- CB- CG- CD2
3705 3702 3700 3716 3 4.95804 2.17986 -3.52293
-3.21331 -0.40166 0.00000 ; CG- CB- CA- C
3705 3706 3708 3709 3 30.33400 0.00000 -30.33400
0.00000 0.00000 0.00000 ; CG- CD1- CE1- HE1
3705 3706 3708 3710 3 30.33400 0.00000 -30.33400
0.00000 0.00000 0.00000 ; CG- CD1- CE1- CZ
3705 3714 3712 3710 3 30.33400 0.00000 -30.33400
0.00000 0.00000 0.00000 ; CG- CD2- CE2- CZ
3705 3714 3712 3713 3 30.33400 0.00000 -30.33400
0.00000 0.00000 0.00000 ; CG- CD2- CE2- HE2
3706 3705 3714 3712 3 30.33400 0.00000 -30.33400
0.00000 0.00000 0.00000 ; CD1- CG- CD2- CE2
3706 3705 3714 3715 3 30.33400 0.00000 -30.33400
0.00000 0.00000 0.00000 ; CD1- CG- CD2- HD2
3706 3708 3710 3711 3 30.33400 0.00000 -30.33400
0.00000 0.00000 0.00000 ; CD1- CE1- CZ- HZ
3706 3708 3710 3712 3 30.33400 0.00000 -30.33400
0.00000 0.00000 0.00000 ; CD1- CE1- CZ- CE2
3707 3706 3705 3714 3 30.33400 0.00000 -30.33400
0.00000 0.00000 0.00000 ; HD1- CD1- CG- CD2
3707 3706 3708 3709 3 30.33400 0.00000 -30.33400
0.00000 0.00000 0.00000 ; HD1- CD1- CE1- HE1
3707 3706 3708 3710 3 30.33400 0.00000 -30.33400
0.00000 0.00000 0.00000 ; HD1- CD1- CE1- CZ
3708 3706 3705 3714 3 30.33400 0.00000 -30.33400
0.00000 0.00000 0.00000 ; CE1- CD1- CG- CD2
3708 3710 3712 3713 3 30.33400 0.00000 -30.33400
0.00000 0.00000 0.00000 ; CE1- CZ- CE2- HE2
3708 3710 3712 3714 3 30.33400 0.00000 -30.33400
0.00000 0.00000 0.00000 ; CE1- CZ- CE2- CD2

3709 3708 3710 3711 3 30.33400 0.00000 -30.33400
0.00000 0.00000 0.00000 ; HE1- CE1- CZ- HZ
3709 3708 3710 3712 3 30.33400 0.00000 -30.33400
0.00000 0.00000 0.00000 ; HE1- CE1- CZ- CE2
3710 3712 3714 3715 3 30.33400 0.00000 -30.33400
0.00000 0.00000 0.00000 ; CZ- CE2- CD2- HD2
3711 3710 3712 3713 3 30.33400 0.00000 -30.33400
0.00000 0.00000 0.00000 ; HZ- CZ- CE2- HE2
3711 3710 3712 3714 3 30.33400 0.00000 -30.33400
0.00000 0.00000 0.00000 ; HZ- CZ- CE2- CD2
3713 3712 3714 3715 3 30.33400 0.00000 -30.33400
0.00000 0.00000 0.00000 ; HE2- CE2- CD2- HD2
3716 3718 3720 3721 3 0.00000 0.00000 0.00000
0.00000 0.00000 0.00000 ; C- N- CA- HA
3716 3718 3720 3722 3 11.71520 1.67360 15.06240 -
13.38880 0.00000 0.00000 ; C- N- CA- CB
3716 3718 3720 3737 3 1.75728 5.27184 2.25936
-7.02912 0.00000 0.00000 ; C- N- CA- C
3717 3716 3718 3719 3 29.28800 -8.36800 -20.92000
0.00000 0.00000 0.00000 ; O- C- N- H
3717 3716 3718 3720 3 20.92000 0.00000 -20.92000
0.00000 0.00000 0.00000 ; O- C- N- CA
3718 3720 3722 3723 3 0.65084 1.95253 0.00000
-2.60338 0.00000 0.00000 ; N- CA- CB- HB2
3718 3720 3722 3724 3 0.65084 1.95253 0.00000
-2.60338 0.00000 0.00000 ; N- CA- CB- HB3
3718 3720 3722 3725 3 3.28026 2.46019 -2.19242
-3.21331 -0.23430 0.00000 ; N- CA- CB- CG
3718 3720 3737 3738 3 0.00000 0.00000 0.00000
0.00000 0.00000 0.00000 ; N- CA- C- O
3718 3720 3737 3739 3 17.40544 -5.02080 -13.22144
9.20480 0.00000 0.00000 ; N- CA- C- N
3719 3718 3720 3721 3 0.00000 0.00000 0.00000
0.00000 0.00000 0.00000 ; H- N- CA- HA
3719 3718 3720 3722 3 0.00000 0.00000 0.00000
0.00000 0.00000 0.00000 ; H- N- CA- CB
3719 3718 3720 3737 3 0.00000 0.00000 0.00000
0.00000 0.00000 0.00000 ; H- N- CA- C
3720 3722 3725 3726 3 0.57739 0.00000 1.02926
0.00000 -1.60666 0.00000 ; CA- CB- CG- CD1
3720 3722 3725 3735 3 0.57739 0.00000 1.02926
0.00000 -1.60666 0.00000 ; CA- CB- CG- CD2
3720 3737 3739 3740 3 20.92000 0.00000 -20.92000
0.00000 0.00000 0.00000 ; CA- C- N- H
3720 3737 3739 3741 3 20.92000 0.00000 -20.92000
0.00000 0.00000 0.00000 ; CA- C- N- CA
3721 3720 3722 3723 3 0.65084 1.95253 0.00000
-2.60338 0.00000 0.00000 ; HA- CA- CB- HB2
3721 3720 3722 3724 3 0.65084 1.95253 0.00000
-2.60338 0.00000 0.00000 ; HA- CA- CB- HB3
3721 3720 3722 3725 3 0.65084 1.95253 0.00000
-2.60338 0.00000 0.00000 ; HA- CA- CB- CG

3721 3720 3737 3738 3 3.68192 -4.35136 0.00000
1.33888 0.00000 0.00000 ; HA- CA- C- O
3721 3720 3737 3739 3 0.00000 0.00000 0.00000
0.00000 0.00000 0.00000 ; HA- CA- C- N
3722 3720 3737 3738 3 0.00000 0.00000 0.00000
0.00000 0.00000 0.00000 ; CB- CA- C- O
3722 3720 3737 3739 3 2.51040 4.18400 1.67360
-6.69440 0.00000 0.00000 ; CB- CA- C- N
3722 3725 3726 3727 3 30.33400 0.00000 -30.33400
0.00000 0.00000 0.00000 ; CB- CG- CD1- HD1
3722 3725 3726 3728 3 30.33400 0.00000 -30.33400
0.00000 0.00000 0.00000 ; CB- CG- CD1- CE1
3722 3725 3735 3733 3 30.33400 0.00000 -30.33400
0.00000 0.00000 0.00000 ; CB- CG- CD2- CE2
3722 3725 3735 3736 3 30.33400 0.00000 -30.33400
0.00000 0.00000 0.00000 ; CB- CG- CD2- HD2
3723 3722 3720 3737 3 0.65084 1.95253 0.00000
-2.60338 0.00000 0.00000 ; HB2- CB- CA- C
3723 3722 3725 3726 3 0.00000 0.00000 0.00000
0.00000 0.00000 0.00000 ; HB2- CB- CG- CD1
3723 3722 3725 3735 3 0.00000 0.00000 0.00000
0.00000 0.00000 0.00000 ; HB2- CB- CG- CD2
3724 3722 3720 3737 3 0.65084 1.95253 0.00000
-2.60338 0.00000 0.00000 ; HB3- CB- CA- C
3724 3722 3725 3726 3 0.00000 0.00000 0.00000
0.00000 0.00000 0.00000 ; HB3- CB- CG- CD1
3724 3722 3725 3735 3 0.00000 0.00000 0.00000
0.00000 0.00000 0.00000 ; HB3- CB- CG- CD2
3725 3722 3720 3737 3 4.95804 2.17986 -3.52293
-3.21331 -0.40166 0.00000 ; CG- CB- CA- C
3725 3726 3728 3729 3 30.33400 0.00000 -30.33400
0.00000 0.00000 0.00000 ; CG- CD1- CE1- HE1
3725 3726 3728 3730 3 30.33400 0.00000 -30.33400
0.00000 0.00000 0.00000 ; CG- CD1- CE1- CZ
3725 3735 3733 3730 3 30.33400 0.00000 -30.33400
0.00000 0.00000 0.00000 ; CG- CD2- CE2- CZ
3725 3735 3733 3734 3 30.33400 0.00000 -30.33400
0.00000 0.00000 0.00000 ; CG- CD2- CE2- HE2
3726 3725 3735 3733 3 30.33400 0.00000 -30.33400
0.00000 0.00000 0.00000 ; CD1- CG- CD2- CE2
3726 3725 3735 3736 3 30.33400 0.00000 -30.33400
0.00000 0.00000 0.00000 ; CD1- CG- CD2- HD2
3726 3728 3730 3731 3 30.33400 0.00000 -30.33400
0.00000 0.00000 0.00000 ; CD1- CE1- CZ- OH
3726 3728 3730 3733 3 30.33400 0.00000 -30.33400
0.00000 0.00000 0.00000 ; CD1- CE1- CZ- CE2
3727 3726 3725 3735 3 30.33400 0.00000 -30.33400
0.00000 0.00000 0.00000 ; HD1- CD1- CG- CD2
3727 3726 3728 3729 3 30.33400 0.00000 -30.33400
0.00000 0.00000 0.00000 ; HD1- CD1- CE1- HE1
3727 3726 3728 3730 3 30.33400 0.00000 -30.33400
0.00000 0.00000 0.00000 ; HD1- CD1- CE1- CZ

3728 3726 3725 3735 3 30.33400 0.00000 -30.33400
0.00000 0.00000 0.00000 ; CE1- CD1- CG- CD2
3728 3730 3731 3732 3 7.93286 0.00000 -9.56462
0.00000 2.17568 0.00000 ; CE1- CZ- OH- HH
3728 3730 3733 3734 3 30.33400 0.00000 -30.33400
0.00000 0.00000 0.00000 ; CE1- CZ- CE2- HE2
3728 3730 3733 3735 3 30.33400 0.00000 -30.33400
0.00000 0.00000 0.00000 ; CE1- CZ- CE2- CD2
3729 3728 3730 3731 3 30.33400 0.00000 -30.33400
0.00000 0.00000 0.00000 ; HE1- CE1- CZ- OH
3729 3728 3730 3733 3 30.33400 0.00000 -30.33400
0.00000 0.00000 0.00000 ; HE1- CE1- CZ- CE2
3730 3733 3735 3736 3 30.33400 0.00000 -30.33400
0.00000 0.00000 0.00000 ; CZ- CE2- CD2- HD2
3731 3730 3733 3734 3 30.33400 0.00000 -30.33400
0.00000 0.00000 0.00000 ; OH- CZ- CE2- HE2
3731 3730 3733 3735 3 30.33400 0.00000 -30.33400
0.00000 0.00000 0.00000 ; OH- CZ- CE2- CD2
3732 3731 3730 3733 3 7.93286 0.00000 -9.56462
0.00000 2.17568 0.00000 ; HH- OH- CZ- CE2
3734 3733 3735 3736 3 30.33400 0.00000 -30.33400
0.00000 0.00000 0.00000 ; HE2- CE2- CD2- HD2
3737 3739 3741 3742 3 0.00000 0.00000 0.00000
0.00000 0.00000 0.00000 ; C- N- CA- HA
3737 3739 3741 3743 3 11.71520 1.67360 15.06240 -
13.38880 0.00000 0.00000 ; C- N- CA- CB
3737 3739 3741 3752 3 1.75728 5.27184 2.25936
-7.02912 0.00000 0.00000 ; C- N- CA- C
3738 3737 3739 3740 3 29.28800 -8.36800 -20.92000
0.00000 0.00000 0.00000 ; O- C- N- H
3738 3737 3739 3741 3 20.92000 0.00000 -20.92000
0.00000 0.00000 0.00000 ; O- C- N- CA
3739 3741 3743 3744 3 0.65084 1.95253 0.00000
-2.60338 0.00000 0.00000 ; N- CA- CB- HB2
3739 3741 3743 3745 3 0.65084 1.95253 0.00000
-2.60338 0.00000 0.00000 ; N- CA- CB- HB3
3739 3741 3743 3746 3 3.21331 2.22589 -4.15053
-2.40998 2.61082 0.00000 ; N- CA- CB- CG
3739 3741 3752 3753 3 0.00000 0.00000 0.00000
0.00000 0.00000 0.00000 ; N- CA- C- O
3739 3741 3752 3754 3 17.40544 -5.02080 -13.22144
9.20480 0.00000 0.00000 ; N- CA- C- N
3740 3739 3741 3742 3 0.00000 0.00000 0.00000
0.00000 0.00000 0.00000 ; H- N- CA- HA
3740 3739 3741 3743 3 0.00000 0.00000 0.00000
0.00000 0.00000 0.00000 ; H- N- CA- CB
3740 3739 3741 3752 3 0.00000 0.00000 0.00000
0.00000 0.00000 0.00000 ; H- N- CA- C
3741 3743 3746 3747 3 0.66944 2.00832 0.00000
-2.67776 0.00000 0.00000 ; CA- CB- CG- HG2
3741 3743 3746 3748 3 0.66944 2.00832 0.00000
-2.67776 0.00000 0.00000 ; CA- CB- CG- HG3

3741 3743 3746 3749 3 10.12110 -1.91209 0.01674
10.17549 -1.87443 0.00000 ; CA- CB- CG- CD
3741 3752 3754 3755 3 20.92000 0.00000 -20.92000
0.00000 0.00000 0.00000 ; CA- C- N- H
3741 3752 3754 3756 3 20.92000 0.00000 -20.92000
0.00000 0.00000 0.00000 ; CA- C- N- CA
3742 3741 3743 3744 3 0.65084 1.95253 0.00000
-2.60338 0.00000 0.00000 ; HA- CA- CB- HB2
3742 3741 3743 3745 3 0.65084 1.95253 0.00000
-2.60338 0.00000 0.00000 ; HA- CA- CB- HB3
3742 3741 3743 3746 3 0.65084 1.95253 0.00000
-2.60338 0.00000 0.00000 ; HA- CA- CB- CG
3742 3741 3752 3753 3 3.68192 -4.35136 0.00000
1.33888 0.00000 0.00000 ; HA- CA- C- O
3742 3741 3752 3754 3 0.00000 0.00000 0.00000
0.00000 0.00000 0.00000 ; HA- CA- C- N
3743 3741 3752 3753 3 0.00000 0.00000 0.00000
0.00000 0.00000 0.00000 ; CB- CA- C- O
3743 3741 3752 3754 3 2.51040 4.18400 1.67360
-6.69440 0.00000 0.00000 ; CB- CA- C- N
3743 3746 3749 3750 3 3.79907 0.00000 -5.40573
0.00000 2.14221 0.00000 ; CB- CG- CD- OE1
3743 3746 3749 3751 3 3.79907 0.00000 -5.40573
0.00000 2.14221 0.00000 ; CB- CG- CD- OE2
3744 3743 3741 3752 3 0.65084 1.95253 0.00000
-2.60338 0.00000 0.00000 ; HB2- CB- CA- C
3744 3743 3746 3747 3 0.62760 1.88280 0.00000
-2.51040 0.00000 0.00000 ; HB2- CB- CG- HG2
3744 3743 3746 3748 3 0.62760 1.88280 0.00000
-2.51040 0.00000 0.00000 ; HB2- CB- CG- HG3
3744 3743 3746 3749 3 0.65084 1.95253 0.00000
-2.60338 0.00000 0.00000 ; HB2- CB- CG- CD
3745 3743 3741 3752 3 0.65084 1.95253 0.00000
-2.60338 0.00000 0.00000 ; HB3- CB- CA- C
3745 3743 3746 3747 3 0.62760 1.88280 0.00000
-2.51040 0.00000 0.00000 ; HB3- CB- CG- HG2
3745 3743 3746 3748 3 0.62760 1.88280 0.00000
-2.51040 0.00000 0.00000 ; HB3- CB- CG- HG3
3745 3743 3746 3749 3 0.65084 1.95253 0.00000
-2.60338 0.00000 0.00000 ; HB3- CB- CG- CD
3746 3743 3741 3752 3 6.86594 3.56895 -8.14206
-2.40998 4.85344 0.00000 ; CG- CB- CA- C
3747 3746 3749 3750 3 0.00000 0.00000 0.00000
0.00000 0.00000 0.00000 ; HG2- CG- CD- OE1
3747 3746 3749 3751 3 0.00000 0.00000 0.00000
0.00000 0.00000 0.00000 ; HG2- CG- CD- OE2
3748 3746 3749 3750 3 0.00000 0.00000 0.00000
0.00000 0.00000 0.00000 ; HG3- CG- CD- OE1
3748 3746 3749 3751 3 0.00000 0.00000 0.00000
0.00000 0.00000 0.00000 ; HG3- CG- CD- OE2
3752 3754 3756 3757 3 0.00000 0.00000 0.00000
0.00000 0.00000 0.00000 ; C- N- CA- HA

3752 3754 3756 3758 3 11.71520 1.67360 15.06240 -
13.38880 0.00000 0.00000 ; C- N- CA- CB
3752 3754 3756 3766 3 1.75728 5.27184 2.25936
-7.02912 0.00000 0.00000 ; C- N- CA- C
3753 3752 3754 3755 3 29.28800 -8.36800 -20.92000
0.00000 0.00000 0.00000 ; O- C- N- H
3753 3752 3754 3756 3 20.92000 0.00000 -20.92000
0.00000 0.00000 0.00000 ; O- C- N- CA
3754 3756 3758 3759 3 0.65084 1.95253 0.00000
-2.60338 0.00000 0.00000 ; N- CA- CB- HB2
3754 3756 3758 3760 3 0.65084 1.95253 0.00000
-2.60338 0.00000 0.00000 ; N- CA- CB- HB3
3754 3756 3758 3761 3 5.99567 3.29281 -4.46014
-0.55229 1.97485 0.00000 ; N- CA- CB- CG
3754 3756 3766 3767 3 0.00000 0.00000 0.00000
0.00000 0.00000 0.00000 ; N- CA- C- O
3754 3756 3766 3768 3 17.40544 -5.02080 -13.22144
9.20480 0.00000 0.00000 ; N- CA- C- N
3755 3754 3756 3757 3 0.00000 0.00000 0.00000
0.00000 0.00000 0.00000 ; H- N- CA- HA
3755 3754 3756 3758 3 0.00000 0.00000 0.00000
0.00000 0.00000 0.00000 ; H- N- CA- CB
3755 3754 3756 3766 3 0.00000 0.00000 0.00000
0.00000 0.00000 0.00000 ; H- N- CA- C
3756 3758 3761 3762 3 0.00000 0.00000 0.00000
0.00000 0.00000 0.00000 ; CA- CB- CG- OD1
3756 3758 3761 3763 3 8.84916 -0.31380 -4.32626
5.03754 0.26778 0.00000 ; CA- CB- CG- ND2
3756 3766 3768 3769 3 20.92000 0.00000 -20.92000
0.00000 0.00000 0.00000 ; CA- C- N- H
3756 3766 3768 3770 3 20.92000 0.00000 -20.92000
0.00000 0.00000 0.00000 ; CA- C- N- CA
3757 3756 3758 3759 3 0.65084 1.95253 0.00000
-2.60338 0.00000 0.00000 ; HA- CA- CB- HB2
3757 3756 3758 3760 3 0.65084 1.95253 0.00000
-2.60338 0.00000 0.00000 ; HA- CA- CB- HB3
3757 3756 3758 3761 3 0.65084 1.95253 0.00000
-2.60338 0.00000 0.00000 ; HA- CA- CB- CG
3757 3756 3766 3767 3 3.68192 -4.35136 0.00000
1.33888 0.00000 0.00000 ; HA- CA- C- O
3757 3756 3766 3768 3 0.00000 0.00000 0.00000
0.00000 0.00000 0.00000 ; HA- CA- C- N
3758 3756 3766 3767 3 0.00000 0.00000 0.00000
0.00000 0.00000 0.00000 ; CB- CA- C- O
3758 3756 3766 3768 3 2.51040 4.18400 1.67360
-6.69440 0.00000 0.00000 ; CB- CA- C- N
3758 3761 3763 3764 3 20.92000 0.00000 -20.92000
0.00000 0.00000 0.00000 ; CB- CG- ND2- HD21
3758 3761 3763 3765 3 20.92000 0.00000 -20.92000
0.00000 0.00000 0.00000 ; CB- CG- ND2- HD22
3759 3758 3756 3766 3 0.65084 1.95253 0.00000
-2.60338 0.00000 0.00000 ; HB2- CB- CA- C

3759 3758 3761 3762 3 3.68192 -4.35136 0.00000
1.33888 0.00000 0.00000 ; HB2- CB- CG- OD1
3759 3758 3761 3763 3 0.00000 0.00000 0.00000
0.00000 0.00000 0.00000 ; HB2- CB- CG- ND2
3760 3758 3756 3766 3 0.65084 1.95253 0.00000
-2.60338 0.00000 0.00000 ; HB3- CB- CA- C
3760 3758 3761 3762 3 3.68192 -4.35136 0.00000
1.33888 0.00000 0.00000 ; HB3- CB- CG- OD1
3760 3758 3761 3763 3 0.00000 0.00000 0.00000
0.00000 0.00000 0.00000 ; HB3- CB- CG- ND2
3761 3758 3756 3766 3 7.94542 -3.96225 -6.11701
-0.55229 3.58150 0.00000 ; CG- CB- CA- C
3762 3761 3763 3764 3 29.28800 -8.36800 -20.92000
0.00000 0.00000 0.00000 ; OD1- CG- ND2- HD21
3762 3761 3763 3765 3 29.28800 -8.36800 -20.92000
0.00000 0.00000 0.00000 ; OD1- CG- ND2- HD22
3766 3768 3770 3771 3 0.00000 0.00000 0.00000
0.00000 0.00000 0.00000 ; C- N- CA- HA
3766 3768 3770 3772 3 11.71520 1.67360 15.06240 -
13.38880 0.00000 0.00000 ; C- N- CA- CB
3766 3768 3770 3778 3 1.75728 5.27184 2.25936
-7.02912 0.00000 0.00000 ; C- N- CA- C
3767 3766 3768 3769 3 29.28800 -8.36800 -20.92000
0.00000 0.00000 0.00000 ; O- C- N- H
3767 3766 3768 3770 3 20.92000 0.00000 -20.92000
0.00000 0.00000 0.00000 ; O- C- N- CA
3768 3770 3772 3773 3 0.65084 1.95253 0.00000
-2.60338 0.00000 0.00000 ; N- CA- CB- HB2
3768 3770 3772 3774 3 0.65084 1.95253 0.00000
-2.60338 0.00000 0.00000 ; N- CA- CB- HB3
3768 3770 3772 3775 3 15.41386 9.74035 -8.39310
-0.97069 2.97901 0.00000 ; N- CA- CB- CG
3768 3770 3778 3779 3 0.00000 0.00000 0.00000
0.00000 0.00000 0.00000 ; N- CA- C- O
3768 3770 3778 3780 3 17.40544 -5.02080 -13.22144
9.20480 0.00000 0.00000 ; N- CA- C- N
3769 3768 3770 3771 3 0.00000 0.00000 0.00000
0.00000 0.00000 0.00000 ; H- N- CA- HA
3769 3768 3770 3772 3 0.00000 0.00000 0.00000
0.00000 0.00000 0.00000 ; H- N- CA- CB
3769 3768 3770 3778 3 0.00000 0.00000 0.00000
0.00000 0.00000 0.00000 ; H- N- CA- C
3770 3772 3775 3776 3 6.43499 0.00000 -5.39736
0.00000 -1.03763 0.00000 ; CA- CB- CG- OD1
3770 3772 3775 3777 3 6.43499 0.00000 -5.39736
0.00000 -1.03763 0.00000 ; CA- CB- CG- OD2
3770 3778 3780 3781 3 20.92000 0.00000 -20.92000
0.00000 0.00000 0.00000 ; CA- C- N- H
3770 3778 3780 3782 3 20.92000 0.00000 -20.92000
0.00000 0.00000 0.00000 ; CA- C- N- CA
3771 3770 3772 3773 3 0.65084 1.95253 0.00000
-2.60338 0.00000 0.00000 ; HA- CA- CB- HB2

3771 3770 3772 3774 3 0.65084 1.95253 0.00000
-2.60338 0.00000 0.00000 ; HA- CA- CB- HB3
3771 3770 3772 3775 3 0.65084 1.95253 0.00000
-2.60338 0.00000 0.00000 ; HA- CA- CB- CG
3771 3770 3778 3779 3 3.68192 -4.35136 0.00000
1.33888 0.00000 0.00000 ; HA- CA- C- O
3771 3770 3778 3780 3 0.00000 0.00000 0.00000
0.00000 0.00000 0.00000 ; HA- CA- C- N
3772 3770 3778 3779 3 0.00000 0.00000 0.00000
0.00000 0.00000 0.00000 ; CB- CA- C- O
3772 3770 3778 3780 3 2.51040 4.18400 1.67360
-6.69440 0.00000 0.00000 ; CB- CA- C- N
3773 3772 3770 3778 3 0.65084 1.95253 0.00000
-2.60338 0.00000 0.00000 ; HB2- CB- CA- C
3773 3772 3775 3776 3 0.00000 0.00000 0.00000
0.00000 0.00000 0.00000 ; HB2- CB- CG- OD1
3773 3772 3775 3777 3 0.00000 0.00000 0.00000
0.00000 0.00000 0.00000 ; HB2- CB- CG- OD2
3774 3772 3770 3778 3 0.65084 1.95253 0.00000
-2.60338 0.00000 0.00000 ; HB3- CB- CA- C
3774 3772 3775 3776 3 0.00000 0.00000 0.00000
0.00000 0.00000 0.00000 ; HB3- CB- CG- OD1
3774 3772 3775 3777 3 0.00000 0.00000 0.00000
0.00000 0.00000 0.00000 ; HB3- CB- CG- OD2
3775 3772 3770 3778 3 7.14627 -1.04600 -8.99560
-0.97069 5.15469 0.00000 ; CG- CB- CA- C
3778 3780 3782 3783 3 0.00000 0.00000 0.00000
0.00000 0.00000 0.00000 ; C- N- CA- HA
3778 3780 3782 3784 3 11.71520 1.67360 15.06240 -
13.38880 0.00000 0.00000 ; C- N- CA- CB
3778 3780 3782 3802 3 1.75728 5.27184 2.25936
-7.02912 0.00000 0.00000 ; C- N- CA- C
3779 3778 3780 3781 3 29.28800 -8.36800 -20.92000
0.00000 0.00000 0.00000 ; O- C- N- H
3779 3778 3780 3782 3 20.92000 0.00000 -20.92000
0.00000 0.00000 0.00000 ; O- C- N- CA
3780 3782 3784 3785 3 0.65084 1.95253 0.00000
-2.60338 0.00000 0.00000 ; N- CA- CB- HB2
3780 3782 3784 3786 3 0.65084 1.95253 0.00000
-2.60338 0.00000 0.00000 ; N- CA- CB- HB3
3780 3782 3784 3787 3 4.18818 2.60663 -3.65682
-3.91622 1.03763 0.00000 ; N- CA- CB- CG
3780 3782 3802 3803 3 0.00000 0.00000 0.00000
0.00000 0.00000 0.00000 ; N- CA- C- O
3780 3782 3802 3804 3 17.40544 -5.02080 -13.22144
9.20480 0.00000 0.00000 ; N- CA- C- N
3781 3780 3782 3783 3 0.00000 0.00000 0.00000
0.00000 0.00000 0.00000 ; H- N- CA- HA
3781 3780 3782 3784 3 0.00000 0.00000 0.00000
0.00000 0.00000 0.00000 ; H- N- CA- CB
3781 3780 3782 3802 3 0.00000 0.00000 0.00000
0.00000 0.00000 0.00000 ; H- N- CA- C

3782 3784 3787 3788 3 0.00000 0.00000 0.00000
0.00000 0.00000 0.00000 ; CA- CB- CG- CD1
3782 3784 3787 3801 3 4.95386 8.75293 6.59398 -
13.70678 -3.17984 0.00000 ; CA- CB- CG- CD2
3782 3802 3804 3805 3 20.92000 0.00000 -20.92000
0.00000 0.00000 0.00000 ; CA- C- N- H
3782 3802 3804 3806 3 20.92000 0.00000 -20.92000
0.00000 0.00000 0.00000 ; CA- C- N- CA
3783 3782 3784 3785 3 0.65084 1.95253 0.00000
-2.60338 0.00000 0.00000 ; HA- CA- CB- HB2
3783 3782 3784 3786 3 0.65084 1.95253 0.00000
-2.60338 0.00000 0.00000 ; HA- CA- CB- HB3
3783 3782 3784 3787 3 0.65084 1.95253 0.00000
-2.60338 0.00000 0.00000 ; HA- CA- CB- CG
3783 3782 3802 3803 3 3.68192 -4.35136 0.00000
1.33888 0.00000 0.00000 ; HA- CA- C- O
3783 3782 3802 3804 3 0.00000 0.00000 0.00000
0.00000 0.00000 0.00000 ; HA- CA- C- N
3784 3782 3802 3803 3 0.00000 0.00000 0.00000
0.00000 0.00000 0.00000 ; CB- CA- C- O
3784 3782 3802 3804 3 2.51040 4.18400 1.67360
-6.69440 0.00000 0.00000 ; CB- CA- C- N
3784 3787 3788 3789 3 54.60120 0.00000 -54.60120
0.00000 0.00000 0.00000 ; CB- CG- CD1- HD1
3784 3787 3788 3790 3 54.60120 0.00000 -54.60120
0.00000 0.00000 0.00000 ; CB- CG- CD1- NE1
3784 3787 3801 3792 3 14.01640 0.00000 -14.01640
0.00000 0.00000 0.00000 ; CB- CG- CD2- CE2
3784 3787 3801 3799 3 14.01640 0.00000 -14.01640
0.00000 0.00000 0.00000 ; CB- CG- CD2- CE3
3785 3784 3782 3802 3 0.65084 1.95253 0.00000
-2.60338 0.00000 0.00000 ; HB2- CB- CA- C
3785 3784 3787 3788 3 0.00000 0.00000 0.00000
0.00000 0.00000 0.00000 ; HB2- CB- CG- CD1
3785 3784 3787 3801 3 0.00000 0.00000 0.00000
0.00000 0.00000 0.00000 ; HB2- CB- CG- CD2
3786 3784 3782 3802 3 0.65084 1.95253 0.00000
-2.60338 0.00000 0.00000 ; HB3- CB- CA- C
3786 3784 3787 3788 3 0.00000 0.00000 0.00000
0.00000 0.00000 0.00000 ; HB3- CB- CG- CD1
3786 3784 3787 3801 3 0.00000 0.00000 0.00000
0.00000 0.00000 0.00000 ; HB3- CB- CG- CD2
3787 3784 3782 3802 3 4.62332 3.00830 -5.43083
-3.91622 2.47693 0.00000 ; CG- CB- CA- C
3787 3788 3790 3791 3 12.55200 0.00000 -12.55200
0.00000 0.00000 0.00000 ; CG- CD1- NE1- HE1
3787 3788 3790 3792 3 12.55200 0.00000 -12.55200
0.00000 0.00000 0.00000 ; CG- CD1- NE1- CE2
3787 3801 3792 3790 3 25.10400 0.00000 -25.10400
0.00000 0.00000 0.00000 ; CG- CD2- CE2- NE1
3787 3801 3792 3793 3 25.10400 0.00000 -25.10400
0.00000 0.00000 0.00000 ; CG- CD2- CE2- CZ2

```
  3787   3801   3799   3797      3   29.28800     0.00000  -29.28800
0.00000    0.00000    0.00000 ;     CG-   CD2-    CE3-   CZ3
  3787   3801   3799   3800      3   29.28800     0.00000  -29.28800
0.00000    0.00000    0.00000 ;     CG-   CD2-    CE3-   HE3
  3788   3787   3801   3792      3   14.01640     0.00000  -14.01640
0.00000    0.00000    0.00000 ;    CD1-    CG-    CD2-   CE2
  3788   3787   3801   3799      3   14.01640     0.00000  -14.01640
0.00000    0.00000    0.00000 ;    CD1-    CG-    CD2-   CE3
  3788   3790   3792   3793      3   12.76120     0.00000  -12.76120
0.00000    0.00000    0.00000 ;    CD1-   NE1-    CE2-   CZ2
  3788   3790   3792   3801      3   12.76120     0.00000  -12.76120
0.00000    0.00000    0.00000 ;    CD1-   NE1-    CE2-   CD2
  3789   3788   3787   3801      3   54.60120     0.00000  -54.60120
0.00000    0.00000    0.00000 ;    HD1-   CD1-     CG-   CD2
  3789   3788   3790   3791      3   12.55200     0.00000  -12.55200
0.00000    0.00000    0.00000 ;    HD1-   CD1-    NE1-   HE1
  3789   3788   3790   3792      3   12.55200     0.00000  -12.55200
0.00000    0.00000    0.00000 ;    HD1-   CD1-    NE1-   CE2
  3790   3788   3787   3801      3   54.60120     0.00000  -54.60120
0.00000    0.00000    0.00000 ;    NE1-   CD1-     CG-   CD2
  3790   3792   3793   3794      3   30.33400     0.00000  -30.33400
0.00000    0.00000    0.00000 ;    NE1-   CE2-    CZ2-   HZ2
  3790   3792   3793   3795      3   30.33400     0.00000  -30.33400
0.00000    0.00000    0.00000 ;    NE1-   CE2-    CZ2-   CH2
  3790   3792   3801   3799      3   25.10400     0.00000  -25.10400
0.00000    0.00000    0.00000 ;    NE1-   CE2-    CD2-   CE3
  3791   3790   3792   3793      3   12.76120     0.00000  -12.76120
0.00000    0.00000    0.00000 ;    HE1-   NE1-    CE2-   CZ2
  3791   3790   3792   3801      3   12.76120     0.00000  -12.76120
0.00000    0.00000    0.00000 ;    HE1-   NE1-    CE2-   CD2
  3792   3793   3795   3796      3   30.33400     0.00000  -30.33400
0.00000    0.00000    0.00000 ;    CE2-   CZ2-    CH2-   HH2
  3792   3793   3795   3797      3   30.33400     0.00000  -30.33400
0.00000    0.00000    0.00000 ;    CE2-   CZ2-    CH2-   CZ3
  3792   3801   3799   3797      3   29.28800     0.00000  -29.28800
0.00000    0.00000    0.00000 ;    CE2-   CD2-    CE3-   CZ3
  3792   3801   3799   3800      3   29.28800     0.00000  -29.28800
0.00000    0.00000    0.00000 ;    CE2-   CD2-    CE3-   HE3
  3793   3792   3801   3799      3   25.10400     0.00000  -25.10400
0.00000    0.00000    0.00000 ;    CZ2-   CE2-    CD2-   CE3
  3793   3795   3797   3798      3   30.33400     0.00000  -30.33400
0.00000    0.00000    0.00000 ;    CZ2-   CH2-    CZ3-   HZ3
  3793   3795   3797   3799      3   30.33400     0.00000  -30.33400
0.00000    0.00000    0.00000 ;    CZ2-   CH2-    CZ3-   CE3
  3794   3793   3792   3801      3   30.33400     0.00000  -30.33400
0.00000    0.00000    0.00000 ;    HZ2-   CZ2-    CE2-   CD2
  3794   3793   3795   3796      3   30.33400     0.00000  -30.33400
0.00000    0.00000    0.00000 ;    HZ2-   CZ2-    CH2-   HH2
  3794   3793   3795   3797      3   30.33400     0.00000  -30.33400
0.00000    0.00000    0.00000 ;    HZ2-   CZ2-    CH2-   CZ3
  3795   3793   3792   3801      3   30.33400     0.00000  -30.33400
0.00000    0.00000    0.00000 ;    CH2-   CZ2-    CE2-   CD2
```

3795 3797 3799 3800 3 30.33400 0.00000 -30.33400
0.00000 0.00000 0.00000 ; CH2- CZ3- CE3- HE3
3795 3797 3799 3801 3 30.33400 0.00000 -30.33400
0.00000 0.00000 0.00000 ; CH2- CZ3- CE3- CD2
3796 3795 3797 3798 3 30.33400 0.00000 -30.33400
0.00000 0.00000 0.00000 ; HH2- CH2- CZ3- HZ3
3796 3795 3797 3799 3 30.33400 0.00000 -30.33400
0.00000 0.00000 0.00000 ; HH2- CH2- CZ3- CE3
3798 3797 3799 3800 3 30.33400 0.00000 -30.33400
0.00000 0.00000 0.00000 ; HZ3- CZ3- CE3- HE3
3798 3797 3799 3801 3 30.33400 0.00000 -30.33400
0.00000 0.00000 0.00000 ; HZ3- CZ3- CE3- CD2
3802 3804 3806 3807 3 0.00000 0.00000 0.00000
0.00000 0.00000 0.00000 ; C- N- CA- HA2
3802 3804 3806 3808 3 0.00000 0.00000 0.00000
0.00000 0.00000 0.00000 ; C- N- CA- HA3
3802 3804 3806 3809 3 1.75728 5.27184 2.25936
-7.02912 0.00000 0.00000 ; C- N- CA- C
3803 3802 3804 3805 3 29.28800 -8.36800 -20.92000
0.00000 0.00000 0.00000 ; O- C- N- H
3803 3802 3804 3806 3 20.92000 0.00000 -20.92000
0.00000 0.00000 0.00000 ; O- C- N- CA
3804 3806 3809 3810 3 0.00000 0.00000 0.00000
0.00000 0.00000 0.00000 ; N- CA- C- O
3804 3806 3809 3811 3 17.40544 -5.02080 -13.22144
9.20480 0.00000 0.00000 ; N- CA- C- N
3805 3804 3806 3807 3 0.00000 0.00000 0.00000
0.00000 0.00000 0.00000 ; H- N- CA- HA2
3805 3804 3806 3808 3 0.00000 0.00000 0.00000
0.00000 0.00000 0.00000 ; H- N- CA- HA3
3805 3804 3806 3809 3 0.00000 0.00000 0.00000
0.00000 0.00000 0.00000 ; H- N- CA- C
3806 3809 3811 3812 3 20.92000 0.00000 -20.92000
0.00000 0.00000 0.00000 ; CA- C- N- H
3806 3809 3811 3813 3 20.92000 0.00000 -20.92000
0.00000 0.00000 0.00000 ; CA- C- N- CA
3807 3806 3809 3810 3 3.68192 -4.35136 0.00000
1.33888 0.00000 0.00000 ; HA2- CA- C- O
3807 3806 3809 3811 3 0.00000 0.00000 0.00000
0.00000 0.00000 0.00000 ; HA2- CA- C- N
3808 3806 3809 3810 3 3.68192 -4.35136 0.00000
1.33888 0.00000 0.00000 ; HA3- CA- C- O
3808 3806 3809 3811 3 0.00000 0.00000 0.00000
0.00000 0.00000 0.00000 ; HA3- CA- C- N
3809 3811 3813 3814 3 0.00000 0.00000 0.00000
0.00000 0.00000 0.00000 ; C- N- CA- HA
3809 3811 3813 3815 3 11.71520 1.67360 15.06240 -
13.38880 0.00000 0.00000 ; C- N- CA- CB
3809 3811 3813 3820 3 1.75728 5.27184 2.25936
-7.02912 0.00000 0.00000 ; C- N- CA- C
3810 3809 3811 3812 3 29.28800 -8.36800 -20.92000
0.00000 0.00000 0.00000 ; O- C- N- H

3810 3809 3811 3813 3 20.92000 0.00000 -20.92000
0.00000 0.00000 0.00000 ; O- C- N- CA
3811 3813 3815 3816 3 0.65084 1.95253 0.00000
-2.60338 0.00000 0.00000 ; N- CA- CB- HB2
3811 3813 3815 3817 3 0.65084 1.95253 0.00000
-2.60338 0.00000 0.00000 ; N- CA- CB- HB3
3811 3813 3815 3818 3 7.86174 2.24681 -7.41405
-6.71114 5.35552 0.00000 ; N- CA- CB- OG
3811 3813 3820 3821 3 0.00000 0.00000 0.00000
0.00000 0.00000 0.00000 ; N- CA- C- O
3811 3813 3820 3822 3 17.40544 -5.02080 -13.22144
9.20480 0.00000 0.00000 ; N- CA- C- N
3812 3811 3813 3814 3 0.00000 0.00000 0.00000
0.00000 0.00000 0.00000 ; H- N- CA- HA
3812 3811 3813 3815 3 0.00000 0.00000 0.00000
0.00000 0.00000 0.00000 ; H- N- CA- CB
3812 3811 3813 3820 3 0.00000 0.00000 0.00000
0.00000 0.00000 0.00000 ; H- N- CA- C
3813 3815 3818 3819 3 2.05853 2.46856 3.48109
-4.46851 0.23430 0.00000 ; CA- CB- OG- HG
3813 3820 3822 3823 3 20.92000 0.00000 -20.92000
0.00000 0.00000 0.00000 ; CA- C- N- H
3813 3820 3822 3824 3 20.92000 0.00000 -20.92000
0.00000 0.00000 0.00000 ; CA- C- N- CA
3814 3813 3815 3816 3 0.65084 1.95253 0.00000
-2.60338 0.00000 0.00000 ; HA- CA- CB- HB2
3814 3813 3815 3817 3 0.65084 1.95253 0.00000
-2.60338 0.00000 0.00000 ; HA- CA- CB- HB3
3814 3813 3815 3818 3 1.04600 -1.04600 0.00000
0.00000 0.00000 0.00000 ; HA- CA- CB- OG
3814 3813 3820 3821 3 3.68192 -4.35136 0.00000
1.33888 0.00000 0.00000 ; HA- CA- C- O
3814 3813 3820 3822 3 0.00000 0.00000 0.00000
0.00000 0.00000 0.00000 ; HA- CA- C- N
3815 3813 3820 3821 3 0.00000 0.00000 0.00000
0.00000 0.00000 0.00000 ; CB- CA- C- O
3815 3813 3820 3822 3 2.51040 4.18400 1.67360
-6.69440 0.00000 0.00000 ; CB- CA- C- N
3816 3815 3813 3820 3 0.65084 1.95253 0.00000
-2.60338 0.00000 0.00000 ; HB2- CB- CA- C
3816 3815 3818 3819 3 0.69733 2.09200 0.00000
-2.78933 0.00000 0.00000 ; HB2- CB- OG- HG
3817 3815 3813 3820 3 0.65084 1.95253 0.00000
-2.60338 0.00000 0.00000 ; HB3- CB- CA- C
3817 3815 3818 3819 3 0.69733 2.09200 0.00000
-2.78933 0.00000 0.00000 ; HB3- CB- OG- HG
3818 3815 3813 3820 3 7.34710 7.79898 -6.14211
-6.71114 4.31789 0.00000 ; OG- CB- CA- C
3820 3822 3824 3825 3 0.00000 0.00000 0.00000
0.00000 0.00000 0.00000 ; C- N- CA- HA
3820 3822 3824 3826 3 11.71520 1.67360 15.06240 -
13.38880 0.00000 0.00000 ; C- N- CA- CB

3820 3822 3824 3839 3 1.75728 5.27184 2.25936
-7.02912 0.00000 0.00000 ; C- N- CA- C
3821 3820 3822 3823 3 29.28800 -8.36800 -20.92000
0.00000 0.00000 0.00000 ; O- C- N- H
3821 3820 3822 3824 3 20.92000 0.00000 -20.92000
0.00000 0.00000 0.00000 ; O- C- N- CA
3822 3824 3826 3827 3 0.65084 1.95253 0.00000
-2.60338 0.00000 0.00000 ; N- CA- CB- HB
3822 3824 3826 3828 3 3.83673 0.44769 -1.77402
-2.47693 -0.03347 0.00000 ; N- CA- CB- CG2
3822 3824 3826 3832 3 2.97482 0.12134 2.04179
-1.89117 -3.24678 0.00000 ; N- CA- CB- CG1
3822 3824 3839 3840 3 0.00000 0.00000 0.00000
0.00000 0.00000 0.00000 ; N- CA- C- O
3822 3824 3839 3841 3 17.40544 -5.02080 -13.22144
9.20480 0.00000 0.00000 ; N- CA- C- N
3823 3822 3824 3825 3 0.00000 0.00000 0.00000
0.00000 0.00000 0.00000 ; H- N- CA- HA
3823 3822 3824 3826 3 0.00000 0.00000 0.00000
0.00000 0.00000 0.00000 ; H- N- CA- CB
3823 3822 3824 3839 3 0.00000 0.00000 0.00000
0.00000 0.00000 0.00000 ; H- N- CA- C
3824 3826 3828 3829 3 0.66944 2.00832 0.00000
-2.67776 0.00000 0.00000 ; CA- CB- CG2- HG21
3824 3826 3828 3830 3 0.66944 2.00832 0.00000
-2.67776 0.00000 0.00000 ; CA- CB- CG2- HG22
3824 3826 3828 3831 3 0.66944 2.00832 0.00000
-2.67776 0.00000 0.00000 ; CA- CB- CG2- HG23
3824 3826 3832 3833 3 0.66944 2.00832 0.00000
-2.67776 0.00000 0.00000 ; CA- CB- CG1- HG12
3824 3826 3832 3834 3 0.66944 2.00832 0.00000
-2.67776 0.00000 0.00000 ; CA- CB- CG1- HG13
3824 3826 3832 3835 3 4.24258 -0.52718 -7.25506
-1.79075 7.69856 0.00000 ; CA- CB- CG1- CD1
3824 3839 3841 3842 3 20.92000 0.00000 -20.92000
0.00000 0.00000 0.00000 ; CA- C- N- H
3824 3839 3841 3843 3 20.92000 0.00000 -20.92000
0.00000 0.00000 0.00000 ; CA- C- N- CA
3825 3824 3826 3827 3 0.65084 1.95253 0.00000
-2.60338 0.00000 0.00000 ; HA- CA- CB- HB
3825 3824 3826 3828 3 0.65084 1.95253 0.00000
-2.60338 0.00000 0.00000 ; HA- CA- CB- CG2
3825 3824 3826 3832 3 0.65084 1.95253 0.00000
-2.60338 0.00000 0.00000 ; HA- CA- CB- CG1
3825 3824 3839 3840 3 3.68192 -4.35136 0.00000
1.33888 0.00000 0.00000 ; HA- CA- C- O
3825 3824 3839 3841 3 0.00000 0.00000 0.00000
0.00000 0.00000 0.00000 ; HA- CA- C- N
3826 3824 3839 3840 3 0.00000 0.00000 0.00000
0.00000 0.00000 0.00000 ; CB- CA- C- O
3826 3824 3839 3841 3 2.51040 4.18400 1.67360
-6.69440 0.00000 0.00000 ; CB- CA- C- N

3826 3832 3835 3836 3 0.66944 2.00832 0.00000
-2.67776 0.00000 0.00000 ; CB- CG1- CD1- HD11
3826 3832 3835 3837 3 0.66944 2.00832 0.00000
-2.67776 0.00000 0.00000 ; CB- CG1- CD1- HD12
3826 3832 3835 3838 3 0.66944 2.00832 0.00000
-2.67776 0.00000 0.00000 ; CB- CG1- CD1- HD13
3827 3826 3824 3839 3 0.65084 1.95253 0.00000
-2.60338 0.00000 0.00000 ; HB- CB- CA- C
3827 3826 3828 3829 3 0.62760 1.88280 0.00000
-2.51040 0.00000 0.00000 ; HB- CB- CG2- HG21
3827 3826 3828 3830 3 0.62760 1.88280 0.00000
-2.51040 0.00000 0.00000 ; HB- CB- CG2- HG22
3827 3826 3828 3831 3 0.62760 1.88280 0.00000
-2.51040 0.00000 0.00000 ; HB- CB- CG2- HG23
3827 3826 3832 3833 3 0.62760 1.88280 0.00000
-2.51040 0.00000 0.00000 ; HB- CB- CG1- HG12
3827 3826 3832 3834 3 0.62760 1.88280 0.00000
-2.51040 0.00000 0.00000 ; HB- CB- CG1- HG13
3827 3826 3832 3835 3 0.66944 2.00832 0.00000
-2.67776 0.00000 0.00000 ; HB- CB- CG1- CD1
3828 3826 3824 3839 3 5.67350 3.55640 -6.16722
-2.47693 3.74886 0.00000 ; CG2- CB- CA- C
3828 3826 3832 3833 3 0.66944 2.00832 0.00000
-2.67776 0.00000 0.00000 ; CG2- CB- CG1- HG12
3828 3826 3832 3834 3 0.66944 2.00832 0.00000
-2.67776 0.00000 0.00000 ; CG2- CB- CG1- HG13
3828 3826 3832 3835 3 3.81162 0.49790 -8.14206
-1.79075 7.49773 0.00000 ; CG2- CB- CG1- CD1
3829 3828 3826 3832 3 0.66944 2.00832 0.00000
-2.67776 0.00000 0.00000 ; HG21- CG2- CB- CG1
3830 3828 3826 3832 3 0.66944 2.00832 0.00000
-2.67776 0.00000 0.00000 ; HG22- CG2- CB- CG1
3831 3828 3826 3832 3 0.66944 2.00832 0.00000
-2.67776 0.00000 0.00000 ; HG23- CG2- CB- CG1
3832 3826 3824 3839 3 8.26340 0.74057 -9.99976
-1.89117 3.84928 0.00000 ; CG1- CB- CA- C
3833 3832 3835 3836 3 0.62760 1.88280 0.00000
-2.51040 0.00000 0.00000 ; HG12- CG1- CD1- HD11
3833 3832 3835 3837 3 0.62760 1.88280 0.00000
-2.51040 0.00000 0.00000 ; HG12- CG1- CD1- HD12
3833 3832 3835 3838 3 0.62760 1.88280 0.00000
-2.51040 0.00000 0.00000 ; HG12- CG1- CD1- HD13
3834 3832 3835 3836 3 0.62760 1.88280 0.00000
-2.51040 0.00000 0.00000 ; HG13- CG1- CD1- HD11
3834 3832 3835 3837 3 0.62760 1.88280 0.00000
-2.51040 0.00000 0.00000 ; HG13- CG1- CD1- HD12
3834 3832 3835 3838 3 0.62760 1.88280 0.00000
-2.51040 0.00000 0.00000 ; HG13- CG1- CD1- HD13
3839 3841 3843 3844 3 0.00000 0.00000 0.00000
0.00000 0.00000 0.00000 ; C- N- CA- HA
3839 3841 3843 3845 3 11.71520 1.67360 15.06240 -
13.38880 0.00000 0.00000 ; C- N- CA- CB

3839 3841 3843 3850 3 1.75728 5.27184 2.25936
-7.02912 0.00000 0.00000 ; C- N- CA- C
3840 3839 3841 3842 3 29.28800 -8.36800 -20.92000
0.00000 0.00000 0.00000 ; O- C- N- H
3840 3839 3841 3843 3 20.92000 0.00000 -20.92000
0.00000 0.00000 0.00000 ; O- C- N- CA
3841 3843 3845 3846 3 0.65084 1.95253 0.00000
-2.60338 0.00000 0.00000 ; N- CA- CB- HB2
3841 3843 3845 3847 3 0.65084 1.95253 0.00000
-2.60338 0.00000 0.00000 ; N- CA- CB- HB3
3841 3843 3845 3848 3 7.86174 2.24681 -7.41405
-6.71114 5.35552 0.00000 ; N- CA- CB- OG
3841 3843 3850 3851 3 0.00000 0.00000 0.00000
0.00000 0.00000 0.00000 ; N- CA- C- O
3841 3843 3850 3852 3 17.40544 -5.02080 -13.22144
9.20480 0.00000 0.00000 ; N- CA- C- N
3842 3841 3843 3844 3 0.00000 0.00000 0.00000
0.00000 0.00000 0.00000 ; H- N- CA- HA
3842 3841 3843 3845 3 0.00000 0.00000 0.00000
0.00000 0.00000 0.00000 ; H- N- CA- CB
3842 3841 3843 3850 3 0.00000 0.00000 0.00000
0.00000 0.00000 0.00000 ; H- N- CA- C
3843 3845 3848 3849 3 2.05853 2.46856 3.48109
-4.46851 0.23430 0.00000 ; CA- CB- OG- HG
3843 3850 3852 3853 3 20.92000 0.00000 -20.92000
0.00000 0.00000 0.00000 ; CA- C- N- H
3843 3850 3852 3854 3 20.92000 0.00000 -20.92000
0.00000 0.00000 0.00000 ; CA- C- N- CA
3844 3843 3845 3846 3 0.65084 1.95253 0.00000
-2.60338 0.00000 0.00000 ; HA- CA- CB- HB2
3844 3843 3845 3847 3 0.65084 1.95253 0.00000
-2.60338 0.00000 0.00000 ; HA- CA- CB- HB3
3844 3843 3845 3848 3 1.04600 -1.04600 0.00000
0.00000 0.00000 0.00000 ; HA- CA- CB- OG
3844 3843 3850 3851 3 3.68192 -4.35136 0.00000
1.33888 0.00000 0.00000 ; HA- CA- C- O
3844 3843 3850 3852 3 0.00000 0.00000 0.00000
0.00000 0.00000 0.00000 ; HA- CA- C- N
3845 3843 3850 3851 3 0.00000 0.00000 0.00000
0.00000 0.00000 0.00000 ; CB- CA- C- O
3845 3843 3850 3852 3 2.51040 4.18400 1.67360
-6.69440 0.00000 0.00000 ; CB- CA- C- N
3846 3845 3843 3850 3 0.65084 1.95253 0.00000
-2.60338 0.00000 0.00000 ; HB2- CB- CA- C
3846 3845 3848 3849 3 0.69733 2.09200 0.00000
-2.78933 0.00000 0.00000 ; HB2- CB- OG- HG
3847 3845 3843 3850 3 0.65084 1.95253 0.00000
-2.60338 0.00000 0.00000 ; HB3- CB- CA- C
3847 3845 3848 3849 3 0.69733 2.09200 0.00000
-2.78933 0.00000 0.00000 ; HB3- CB- OG- HG
3848 3845 3843 3850 3 7.34710 7.79898 -6.14211
-6.71114 4.31789 0.00000 ; OG- CB- CA- C

3850 3852 3854 3855 3 0.00000 0.00000 0.00000
0.00000 0.00000 0.00000 ; C- N- CA- HA
3850 3852 3854 3856 3 11.71520 1.67360 15.06240 -
13.38880 0.00000 0.00000 ; C- N- CA- CB
3850 3852 3854 3861 3 1.75728 5.27184 2.25936
-7.02912 0.00000 0.00000 ; C- N- CA- C
3851 3850 3852 3853 3 29.28800 -8.36800 -20.92000
0.00000 0.00000 0.00000 ; O- C- N- H
3851 3850 3852 3854 3 20.92000 0.00000 -20.92000
0.00000 0.00000 0.00000 ; O- C- N- CA
3852 3854 3856 3857 3 0.65084 1.95253 0.00000
-2.60338 0.00000 0.00000 ; N- CA- CB- HB2
3852 3854 3856 3858 3 0.65084 1.95253 0.00000
-2.60338 0.00000 0.00000 ; N- CA- CB- HB3
3852 3854 3856 3859 3 7.86174 2.24681 -7.41405
-6.71114 5.35552 0.00000 ; N- CA- CB- OG
3852 3854 3861 3862 3 0.00000 0.00000 0.00000
0.00000 0.00000 0.00000 ; N- CA- C- O
3852 3854 3861 3863 3 17.40544 -5.02080 -13.22144
9.20480 0.00000 0.00000 ; N- CA- C- N
3853 3852 3854 3855 3 0.00000 0.00000 0.00000
0.00000 0.00000 0.00000 ; H- N- CA- HA
3853 3852 3854 3856 3 0.00000 0.00000 0.00000
0.00000 0.00000 0.00000 ; H- N- CA- CB
3853 3852 3854 3861 3 0.00000 0.00000 0.00000
0.00000 0.00000 0.00000 ; H- N- CA- C
3854 3856 3859 3860 3 2.05853 2.46856 3.48109
-4.46851 0.23430 0.00000 ; CA- CB- OG- HG
3854 3861 3863 3864 3 20.92000 0.00000 -20.92000
0.00000 0.00000 0.00000 ; CA- C- N- H
3854 3861 3863 3865 3 20.92000 0.00000 -20.92000
0.00000 0.00000 0.00000 ; CA- C- N- CA
3855 3854 3856 3857 3 0.65084 1.95253 0.00000
-2.60338 0.00000 0.00000 ; HA- CA- CB- HB2
3855 3854 3856 3858 3 0.65084 1.95253 0.00000
-2.60338 0.00000 0.00000 ; HA- CA- CB- HB3
3855 3854 3856 3859 3 1.04600 -1.04600 0.00000
0.00000 0.00000 0.00000 ; HA- CA- CB- OG
3855 3854 3861 3862 3 3.68192 -4.35136 0.00000
1.33888 0.00000 0.00000 ; HA- CA- C- O
3855 3854 3861 3863 3 0.00000 0.00000 0.00000
0.00000 0.00000 0.00000 ; HA- CA- C- N
3856 3854 3861 3862 3 0.00000 0.00000 0.00000
0.00000 0.00000 0.00000 ; CB- CA- C- O
3856 3854 3861 3863 3 2.51040 4.18400 1.67360
-6.69440 0.00000 0.00000 ; CB- CA- C- N
3857 3856 3854 3861 3 0.65084 1.95253 0.00000
-2.60338 0.00000 0.00000 ; HB2- CB- CA- C
3857 3856 3859 3860 3 0.69733 2.09200 0.00000
-2.78933 0.00000 0.00000 ; HB2- CB- OG- HG
3858 3856 3854 3861 3 0.65084 1.95253 0.00000
-2.60338 0.00000 0.00000 ; HB3- CB- CA- C

3858 3856 3859 3860 3 0.69733 2.09200 0.00000
-2.78933 0.00000 0.00000 ; HB3- CB- OG- HG
3859 3856 3854 3861 3 7.34710 7.79898 -6.14211
-6.71114 4.31789 0.00000 ; OG- CB- CA- C
3861 3863 3865 3866 3 0.00000 0.00000 0.00000
0.00000 0.00000 0.00000 ; C- N- CA- HA
3861 3863 3865 3867 3 11.71520 1.67360 15.06240 -
13.38880 0.00000 0.00000 ; C- N- CA- CB
3861 3863 3865 3873 3 1.75728 5.27184 2.25936
-7.02912 0.00000 0.00000 ; C- N- CA- C
3862 3861 3863 3864 3 29.28800 -8.36800 -20.92000
0.00000 0.00000 0.00000 ; O- C- N- H
3862 3861 3863 3865 3 20.92000 0.00000 -20.92000
0.00000 0.00000 0.00000 ; O- C- N- CA
3863 3865 3867 3868 3 0.65084 1.95253 0.00000
-2.60338 0.00000 0.00000 ; N- CA- CB- HB2
3863 3865 3867 3869 3 0.65084 1.95253 0.00000
-2.60338 0.00000 0.00000 ; N- CA- CB- HB3
3863 3865 3867 3870 3 15.41386 9.74035 -8.39310
-0.97069 2.97901 0.00000 ; N- CA- CB- CG
3863 3865 3873 3874 3 0.00000 0.00000 0.00000
0.00000 0.00000 0.00000 ; N- CA- C- O
3863 3865 3873 3875 3 17.40544 -5.02080 -13.22144
9.20480 0.00000 0.00000 ; N- CA- C- N
3864 3863 3865 3866 3 0.00000 0.00000 0.00000
0.00000 0.00000 0.00000 ; H- N- CA- HA
3864 3863 3865 3867 3 0.00000 0.00000 0.00000
0.00000 0.00000 0.00000 ; H- N- CA- CB
3864 3863 3865 3873 3 0.00000 0.00000 0.00000
0.00000 0.00000 0.00000 ; H- N- CA- C
3865 3867 3870 3871 3 6.43499 0.00000 -5.39736
0.00000 -1.03763 0.00000 ; CA- CB- CG- OD1
3865 3867 3870 3872 3 6.43499 0.00000 -5.39736
0.00000 -1.03763 0.00000 ; CA- CB- CG- OD2
3865 3873 3875 3876 3 20.92000 0.00000 -20.92000
0.00000 0.00000 0.00000 ; CA- C- N- H
3865 3873 3875 3877 3 20.92000 0.00000 -20.92000
0.00000 0.00000 0.00000 ; CA- C- N- CA
3866 3865 3867 3868 3 0.65084 1.95253 0.00000
-2.60338 0.00000 0.00000 ; HA- CA- CB- HB2
3866 3865 3867 3869 3 0.65084 1.95253 0.00000
-2.60338 0.00000 0.00000 ; HA- CA- CB- HB3
3866 3865 3867 3870 3 0.65084 1.95253 0.00000
-2.60338 0.00000 0.00000 ; HA- CA- CB- CG
3866 3865 3873 3874 3 3.68192 -4.35136 0.00000
1.33888 0.00000 0.00000 ; HA- CA- C- O
3866 3865 3873 3875 3 0.00000 0.00000 0.00000
0.00000 0.00000 0.00000 ; HA- CA- C- N
3867 3865 3873 3874 3 0.00000 0.00000 0.00000
0.00000 0.00000 0.00000 ; CB- CA- C- O
3867 3865 3873 3875 3 2.51040 4.18400 1.67360
-6.69440 0.00000 0.00000 ; CB- CA- C- N

3868 3867 3865 3873 3 0.65084 1.95253 0.00000
-2.60338 0.00000 0.00000 ; HB2- CB- CA- C
3868 3867 3870 3871 3 0.00000 0.00000 0.00000
0.00000 0.00000 0.00000 ; HB2- CB- CG- OD1
3868 3867 3870 3872 3 0.00000 0.00000 0.00000
0.00000 0.00000 0.00000 ; HB2- CB- CG- OD2
3869 3867 3865 3873 3 0.65084 1.95253 0.00000
-2.60338 0.00000 0.00000 ; HB3- CB- CA- C
3869 3867 3870 3871 3 0.00000 0.00000 0.00000
0.00000 0.00000 0.00000 ; HB3- CB- CG- OD1
3869 3867 3870 3872 3 0.00000 0.00000 0.00000
0.00000 0.00000 0.00000 ; HB3- CB- CG- OD2
3870 3867 3865 3873 3 7.14627 -1.04600 -8.99560
-0.97069 5.15469 0.00000 ; CG- CB- CA- C
3873 3875 3877 3878 3 0.00000 0.00000 0.00000
0.00000 0.00000 0.00000 ; C- N- CA- HA
3873 3875 3877 3879 3 11.71520 1.67360 15.06240 -
13.38880 0.00000 0.00000 ; C- N- CA- CB
3873 3875 3877 3883 3 1.75728 5.27184 2.25936
-7.02912 0.00000 0.00000 ; C- N- CA- C
3874 3873 3875 3876 3 29.28800 -8.36800 -20.92000
0.00000 0.00000 0.00000 ; O- C- N- H
3874 3873 3875 3877 3 20.92000 0.00000 -20.92000
0.00000 0.00000 0.00000 ; O- C- N- CA
3875 3877 3879 3880 3 0.65084 1.95253 0.00000
-2.60338 0.00000 0.00000 ; N- CA- CB- HB1
3875 3877 3879 3881 3 0.65084 1.95253 0.00000
-2.60338 0.00000 0.00000 ; N- CA- CB- HB2
3875 3877 3879 3882 3 0.65084 1.95253 0.00000
-2.60338 0.00000 0.00000 ; N- CA- CB- HB3
3875 3877 3883 3884 3 0.00000 0.00000 0.00000
0.00000 0.00000 0.00000 ; N- CA- C- O
3875 3877 3883 3885 3 17.40544 -5.02080 -13.22144
9.20480 0.00000 0.00000 ; N- CA- C- N
3876 3875 3877 3878 3 0.00000 0.00000 0.00000
0.00000 0.00000 0.00000 ; H- N- CA- HA
3876 3875 3877 3879 3 0.00000 0.00000 0.00000
0.00000 0.00000 0.00000 ; H- N- CA- CB
3876 3875 3877 3883 3 0.00000 0.00000 0.00000
0.00000 0.00000 0.00000 ; H- N- CA- C
3877 3883 3885 3886 3 20.92000 0.00000 -20.92000
0.00000 0.00000 0.00000 ; CA- C- N- H
3877 3883 3885 3887 3 20.92000 0.00000 -20.92000
0.00000 0.00000 0.00000 ; CA- C- N- CA
3878 3877 3879 3880 3 0.65084 1.95253 0.00000
-2.60338 0.00000 0.00000 ; HA- CA- CB- HB1
3878 3877 3879 3881 3 0.65084 1.95253 0.00000
-2.60338 0.00000 0.00000 ; HA- CA- CB- HB2
3878 3877 3879 3882 3 0.65084 1.95253 0.00000
-2.60338 0.00000 0.00000 ; HA- CA- CB- HB3
3878 3877 3883 3884 3 3.68192 -4.35136 0.00000
1.33888 0.00000 0.00000 ; HA- CA- C- O

3878 3877 3883 3885 3 0.00000 0.00000 0.00000
0.00000 0.00000 0.00000 ; HA- CA- C- N
3879 3877 3883 3884 3 0.00000 0.00000 0.00000
0.00000 0.00000 0.00000 ; CB- CA- C- O
3879 3877 3883 3885 3 2.51040 4.18400 1.67360
-6.69440 0.00000 0.00000 ; CB- CA- C- N
3880 3879 3877 3883 3 0.65084 1.95253 0.00000
-2.60338 0.00000 0.00000 ; HB1- CB- CA- C
3881 3879 3877 3883 3 0.65084 1.95253 0.00000
-2.60338 0.00000 0.00000 ; HB2- CB- CA- C
3882 3879 3877 3883 3 0.65084 1.95253 0.00000
-2.60338 0.00000 0.00000 ; HB3- CB- CA- C
3883 3885 3887 3888 3 0.00000 0.00000 0.00000
0.00000 0.00000 0.00000 ; C- N- CA- HA
3883 3885 3887 3889 3 11.71520 1.67360 15.06240 -
13.38880 0.00000 0.00000 ; C- N- CA- CB
3883 3885 3887 3905 3 1.75728 5.27184 2.25936
-7.02912 0.00000 0.00000 ; C- N- CA- C
3884 3883 3885 3886 3 29.28800 -8.36800 -20.92000
0.00000 0.00000 0.00000 ; O- C- N- H
3884 3883 3885 3887 3 20.92000 0.00000 -20.92000
0.00000 0.00000 0.00000 ; O- C- N- CA
3885 3887 3889 3890 3 0.65084 1.95253 0.00000
-2.60338 0.00000 0.00000 ; N- CA- CB- HB2
3885 3887 3889 3891 3 0.65084 1.95253 0.00000
-2.60338 0.00000 0.00000 ; N- CA- CB- HB3
3885 3887 3889 3892 3 0.65084 1.95253 0.00000
-2.60338 0.00000 0.00000 ; N- CA- CB- CG
3885 3887 3905 3906 3 0.00000 0.00000 0.00000
0.00000 0.00000 0.00000 ; N- CA- C- O
3885 3887 3905 3907 3 17.40544 -5.02080 -13.22144
9.20480 0.00000 0.00000 ; N- CA- C- N
3886 3885 3887 3888 3 0.00000 0.00000 0.00000
0.00000 0.00000 0.00000 ; H- N- CA- HA
3886 3885 3887 3889 3 0.00000 0.00000 0.00000
0.00000 0.00000 0.00000 ; H- N- CA- CB
3886 3885 3887 3905 3 0.00000 0.00000 0.00000
0.00000 0.00000 0.00000 ; H- N- CA- C
3887 3889 3892 3893 3 0.66944 2.00832 0.00000
-2.67776 0.00000 0.00000 ; CA- CB- CG- HG2
3887 3889 3892 3894 3 0.66944 2.00832 0.00000
-2.67776 0.00000 0.00000 ; CA- CB- CG- HG3
3887 3889 3892 3895 3 3.68192 3.09616 -2.09200
-3.01248 0.00000 0.00000 ; CA- CB- CG- CD
3887 3905 3907 3908 3 20.92000 0.00000 -20.92000
0.00000 0.00000 0.00000 ; CA- C- N- H
3887 3905 3907 3909 3 20.92000 0.00000 -20.92000
0.00000 0.00000 0.00000 ; CA- C- N- CA
3888 3887 3889 3890 3 0.65084 1.95253 0.00000
-2.60338 0.00000 0.00000 ; HA- CA- CB- HB2
3888 3887 3889 3891 3 0.65084 1.95253 0.00000
-2.60338 0.00000 0.00000 ; HA- CA- CB- HB3

3888 3887 3889 3892 3 0.65084 1.95253 0.00000
-2.60338 0.00000 0.00000 ; HA- CA- CB- CG
3888 3887 3905 3906 3 3.68192 -4.35136 0.00000
1.33888 0.00000 0.00000 ; HA- CA- C- O
3888 3887 3905 3907 3 0.00000 0.00000 0.00000
0.00000 0.00000 0.00000 ; HA- CA- C- N
3889 3887 3905 3906 3 0.00000 0.00000 0.00000
0.00000 0.00000 0.00000 ; CB- CA- C- O
3889 3887 3905 3907 3 2.51040 4.18400 1.67360
-6.69440 0.00000 0.00000 ; CB- CA- C- N
3889 3892 3895 3896 3 0.66944 2.00832 0.00000
-2.67776 0.00000 0.00000 ; CB- CG- CD- HD2
3889 3892 3895 3897 3 0.66944 2.00832 0.00000
-2.67776 0.00000 0.00000 ; CB- CG- CD- HD3
3889 3892 3895 3898 3 3.68192 3.09616 -2.09200
-3.01248 0.00000 0.00000 ; CB- CG- CD- CE
3890 3889 3887 3905 3 0.65084 1.95253 0.00000
-2.60338 0.00000 0.00000 ; HB2- CB- CA- C
3890 3889 3892 3893 3 0.62760 1.88280 0.00000
-2.51040 0.00000 0.00000 ; HB2- CB- CG- HG2
3890 3889 3892 3894 3 0.62760 1.88280 0.00000
-2.51040 0.00000 0.00000 ; HB2- CB- CG- HG3
3890 3889 3892 3895 3 0.66944 2.00832 0.00000
-2.67776 0.00000 0.00000 ; HB2- CB- CG- CD
3891 3889 3887 3905 3 0.65084 1.95253 0.00000
-2.60338 0.00000 0.00000 ; HB3- CB- CA- C
3891 3889 3892 3893 3 0.62760 1.88280 0.00000
-2.51040 0.00000 0.00000 ; HB3- CB- CG- HG2
3891 3889 3892 3894 3 0.62760 1.88280 0.00000
-2.51040 0.00000 0.00000 ; HB3- CB- CG- HG3
3891 3889 3892 3895 3 0.66944 2.00832 0.00000
-2.67776 0.00000 0.00000 ; HB3- CB- CG- CD
3892 3889 3887 3905 3 0.65084 1.95253 0.00000
-2.60338 0.00000 0.00000 ; CG- CB- CA- C
3892 3895 3898 3899 3 0.65084 1.95253 0.00000
-2.60338 0.00000 0.00000 ; CG- CD- CE- HE2
3892 3895 3898 3900 3 0.65084 1.95253 0.00000
-2.60338 0.00000 0.00000 ; CG- CD- CE- HE3
3892 3895 3898 3901 3 0.65084 1.95253 0.00000
-2.60338 0.00000 0.00000 ; CG- CD- CE- NZ
3893 3892 3895 3896 3 0.62760 1.88280 0.00000
-2.51040 0.00000 0.00000 ; HG2- CG- CD- HD2
3893 3892 3895 3897 3 0.62760 1.88280 0.00000
-2.51040 0.00000 0.00000 ; HG2- CG- CD- HD3
3893 3892 3895 3898 3 0.66944 2.00832 0.00000
-2.67776 0.00000 0.00000 ; HG2- CG- CD- CE
3894 3892 3895 3896 3 0.62760 1.88280 0.00000
-2.51040 0.00000 0.00000 ; HG3- CG- CD- HD2
3894 3892 3895 3897 3 0.62760 1.88280 0.00000
-2.51040 0.00000 0.00000 ; HG3- CG- CD- HD3
3894 3892 3895 3898 3 0.66944 2.00832 0.00000
-2.67776 0.00000 0.00000 ; HG3- CG- CD- CE

3895 3898 3901 3902 3 0.65084 1.95253 0.00000
-2.60338 0.00000 0.00000 ; CD- CE- NZ- HZ1
3895 3898 3901 3903 3 0.65084 1.95253 0.00000
-2.60338 0.00000 0.00000 ; CD- CE- NZ- HZ2
3895 3898 3901 3904 3 0.65084 1.95253 0.00000
-2.60338 0.00000 0.00000 ; CD- CE- NZ- HZ3
3896 3895 3898 3899 3 0.65084 1.95253 0.00000
-2.60338 0.00000 0.00000 ; HD2- CD- CE- HE2
3896 3895 3898 3900 3 0.65084 1.95253 0.00000
-2.60338 0.00000 0.00000 ; HD2- CD- CE- HE3
3896 3895 3898 3901 3 0.65084 1.95253 0.00000
-2.60338 0.00000 0.00000 ; HD2- CD- CE- NZ
3897 3895 3898 3899 3 0.65084 1.95253 0.00000
-2.60338 0.00000 0.00000 ; HD3- CD- CE- HE2
3897 3895 3898 3900 3 0.65084 1.95253 0.00000
-2.60338 0.00000 0.00000 ; HD3- CD- CE- HE3
3897 3895 3898 3901 3 0.65084 1.95253 0.00000
-2.60338 0.00000 0.00000 ; HD3- CD- CE- NZ
3899 3898 3901 3902 3 0.65084 1.95253 0.00000
-2.60338 0.00000 0.00000 ; HE2- CE- NZ- HZ1
3899 3898 3901 3903 3 0.65084 1.95253 0.00000
-2.60338 0.00000 0.00000 ; HE2- CE- NZ- HZ2
3899 3898 3901 3904 3 0.65084 1.95253 0.00000
-2.60338 0.00000 0.00000 ; HE2- CE- NZ- HZ3
3900 3898 3901 3902 3 0.65084 1.95253 0.00000
-2.60338 0.00000 0.00000 ; HE3- CE- NZ- HZ1
3900 3898 3901 3903 3 0.65084 1.95253 0.00000
-2.60338 0.00000 0.00000 ; HE3- CE- NZ- HZ2
3900 3898 3901 3904 3 0.65084 1.95253 0.00000
-2.60338 0.00000 0.00000 ; HE3- CE- NZ- HZ3
3905 3907 3909 3910 3 0.00000 0.00000 0.00000
0.00000 0.00000 0.00000 ; C- N- CA- HA
3905 3907 3909 3911 3 11.71520 1.67360 15.06240 -
13.38880 0.00000 0.00000 ; C- N- CA- CB
3905 3907 3909 3919 3 1.75728 5.27184 2.25936
-7.02912 0.00000 0.00000 ; C- N- CA- C
3906 3905 3907 3908 3 29.28800 -8.36800 -20.92000
0.00000 0.00000 0.00000 ; O- C- N- H
3906 3905 3907 3909 3 20.92000 0.00000 -20.92000
0.00000 0.00000 0.00000 ; O- C- N- CA
3907 3909 3911 3912 3 0.65084 1.95253 0.00000
-2.60338 0.00000 0.00000 ; N- CA- CB- HB2
3907 3909 3911 3913 3 0.65084 1.95253 0.00000
-2.60338 0.00000 0.00000 ; N- CA- CB- HB3
3907 3909 3911 3914 3 5.99567 3.29281 -4.46014
-0.55229 1.97485 0.00000 ; N- CA- CB- CG
3907 3909 3919 3920 3 0.00000 0.00000 0.00000
0.00000 0.00000 0.00000 ; N- CA- C- O
3907 3909 3919 3921 3 17.40544 -5.02080 -13.22144
9.20480 0.00000 0.00000 ; N- CA- C- N
3908 3907 3909 3910 3 0.00000 0.00000 0.00000
0.00000 0.00000 0.00000 ; H- N- CA- HA

```
  3908   3907   3909   3911      3    0.00000    0.00000    0.00000
0.00000    0.00000    0.00000 ;       H-     N-    CA-    CB
  3908   3907   3909   3919      3    0.00000    0.00000    0.00000
0.00000    0.00000    0.00000 ;       H-     N-    CA-     C
  3909   3911   3914   3915      3    0.00000    0.00000    0.00000
0.00000    0.00000    0.00000 ;      CA-    CB-    CG-   OD1
  3909   3911   3914   3916      3    8.84916   -0.31380   -4.32626
5.03754    0.26778    0.00000 ;      CA-    CB-    CG-   ND2
  3909   3919   3921   3922      3   20.92000    0.00000  -20.92000
0.00000    0.00000    0.00000 ;      CA-     C-     N-     H
  3909   3919   3921   3923      3   20.92000    0.00000  -20.92000
0.00000    0.00000    0.00000 ;      CA-     C-     N-    CA
  3910   3909   3911   3912      3    0.65084    1.95253    0.00000
-2.60338    0.00000    0.00000 ;      HA-    CA-    CB-   HB2
  3910   3909   3911   3913      3    0.65084    1.95253    0.00000
-2.60338    0.00000    0.00000 ;      HA-    CA-    CB-   HB3
  3910   3909   3911   3914      3    0.65084    1.95253    0.00000
-2.60338    0.00000    0.00000 ;      HA-    CA-    CB-    CG
  3910   3909   3919   3920      3    3.68192   -4.35136    0.00000
1.33888    0.00000    0.00000 ;      HA-    CA-     C-     O
  3910   3909   3919   3921      3    0.00000    0.00000    0.00000
0.00000    0.00000    0.00000 ;      HA-    CA-     C-     N
  3911   3909   3919   3920      3    0.00000    0.00000    0.00000
0.00000    0.00000    0.00000 ;      CB-    CA-     C-     O
  3911   3909   3919   3921      3    2.51040    4.18400    1.67360
-6.69440    0.00000    0.00000 ;      CB-    CA-     C-     N
  3911   3914   3916   3917      3   20.92000    0.00000  -20.92000
0.00000    0.00000    0.00000 ;      CB-    CG-   ND2-  HD21
  3911   3914   3916   3918      3   20.92000    0.00000  -20.92000
0.00000    0.00000    0.00000 ;      CB-    CG-   ND2-  HD22
  3912   3911   3909   3919      3    0.65084    1.95253    0.00000
-2.60338    0.00000    0.00000 ;     HB2-    CB-    CA-     C
  3912   3911   3914   3915      3    3.68192   -4.35136    0.00000
1.33888    0.00000    0.00000 ;     HB2-    CB-    CG-   OD1
  3912   3911   3914   3916      3    0.00000    0.00000    0.00000
0.00000    0.00000    0.00000 ;     HB2-    CB-    CG-   ND2
  3913   3911   3909   3919      3    0.65084    1.95253    0.00000
-2.60338    0.00000    0.00000 ;     HB3-    CB-    CA-     C
  3913   3911   3914   3915      3    3.68192   -4.35136    0.00000
1.33888    0.00000    0.00000 ;     HB3-    CB-    CG-   OD1
  3913   3911   3914   3916      3    0.00000    0.00000    0.00000
0.00000    0.00000    0.00000 ;     HB3-    CB-    CG-   ND2
  3914   3911   3909   3919      3    7.94542   -3.96225   -6.11701
-0.55229    3.58150    0.00000 ;      CG-    CB-    CA-     C
  3915   3914   3916   3917      3   29.28800   -8.36800  -20.92000
0.00000    0.00000    0.00000 ;     OD1-    CG-   ND2-  HD21
  3915   3914   3916   3918      3   29.28800   -8.36800  -20.92000
0.00000    0.00000    0.00000 ;     OD1-    CG-   ND2-  HD22
  3919   3921   3923   3924      3    0.00000    0.00000    0.00000
0.00000    0.00000    0.00000 ;       C-     N-    CA-    HA
  3919   3921   3923   3925      3   11.71520    1.67360   15.06240  -
13.38880    0.00000    0.00000 ;       C-     N-    CA-    CB
```

3919 3921 3923 3938 3 1.75728 5.27184 2.25936
-7.02912 0.00000 0.00000 ; C- N- CA- C
3920 3919 3921 3922 3 29.28800 -8.36800 -20.92000
0.00000 0.00000 0.00000 ; O- C- N- H
3920 3919 3921 3923 3 20.92000 0.00000 -20.92000
0.00000 0.00000 0.00000 ; O- C- N- CA
3921 3923 3925 3926 3 0.65084 1.95253 0.00000
-2.60338 0.00000 0.00000 ; N- CA- CB- HB2
3921 3923 3925 3927 3 0.65084 1.95253 0.00000
-2.60338 0.00000 0.00000 ; N- CA- CB- HB3
3921 3923 3925 3928 3 3.79070 1.39746 -4.61077
-2.40998 2.44346 0.00000 ; N- CA- CB- CG
3921 3923 3938 3939 3 0.00000 0.00000 0.00000
0.00000 0.00000 0.00000 ; N- CA- C- O
3921 3923 3938 3940 3 17.40544 -5.02080 -13.22144
9.20480 0.00000 0.00000 ; N- CA- C- N
3922 3921 3923 3924 3 0.00000 0.00000 0.00000
0.00000 0.00000 0.00000 ; H- N- CA- HA
3922 3921 3923 3925 3 0.00000 0.00000 0.00000
0.00000 0.00000 0.00000 ; H- N- CA- CB
3922 3921 3923 3938 3 0.00000 0.00000 0.00000
0.00000 0.00000 0.00000 ; H- N- CA- C
3923 3925 3928 3929 3 0.66944 2.00832 0.00000
-2.67776 0.00000 0.00000 ; CA- CB- CG- HG
3923 3925 3928 3930 3 3.90367 0.19665 -6.21742
-2.37651 5.99149 0.00000 ; CA- CB- CG- CD1
3923 3925 3928 3934 3 3.90367 0.19665 -6.21742
-2.37651 5.99149 0.00000 ; CA- CB- CG- CD2
3923 3938 3940 3941 3 20.92000 0.00000 -20.92000
0.00000 0.00000 0.00000 ; CA- C- N- H
3923 3938 3940 3942 3 20.92000 0.00000 -20.92000
0.00000 0.00000 0.00000 ; CA- C- N- CA
3924 3923 3925 3926 3 0.65084 1.95253 0.00000
-2.60338 0.00000 0.00000 ; HA- CA- CB- HB2
3924 3923 3925 3927 3 0.65084 1.95253 0.00000
-2.60338 0.00000 0.00000 ; HA- CA- CB- HB3
3924 3923 3925 3928 3 0.65084 1.95253 0.00000
-2.60338 0.00000 0.00000 ; HA- CA- CB- CG
3924 3923 3938 3939 3 3.68192 -4.35136 0.00000
1.33888 0.00000 0.00000 ; HA- CA- C- O
3924 3923 3938 3940 3 0.00000 0.00000 0.00000
0.00000 0.00000 0.00000 ; HA- CA- C- N
3925 3923 3938 3939 3 0.00000 0.00000 0.00000
0.00000 0.00000 0.00000 ; CB- CA- C- O
3925 3923 3938 3940 3 2.51040 4.18400 1.67360
-6.69440 0.00000 0.00000 ; CB- CA- C- N
3925 3928 3930 3931 3 0.66944 2.00832 0.00000
-2.67776 0.00000 0.00000 ; CB- CG- CD1- HD11
3925 3928 3930 3932 3 0.66944 2.00832 0.00000
-2.67776 0.00000 0.00000 ; CB- CG- CD1- HD12
3925 3928 3930 3933 3 0.66944 2.00832 0.00000
-2.67776 0.00000 0.00000 ; CB- CG- CD1- HD13

3925 3928 3934 3935 3 0.66944 2.00832 0.00000
-2.67776 0.00000 0.00000 ; CB- CG- CD2- HD21
3925 3928 3934 3936 3 0.66944 2.00832 0.00000
-2.67776 0.00000 0.00000 ; CB- CG- CD2- HD22
3925 3928 3934 3937 3 0.66944 2.00832 0.00000
-2.67776 0.00000 0.00000 ; CB- CG- CD2- HD23
3926 3925 3923 3938 3 0.65084 1.95253 0.00000
-2.60338 0.00000 0.00000 ; HB2- CB- CA- C
3926 3925 3928 3929 3 0.62760 1.88280 0.00000
-2.51040 0.00000 0.00000 ; HB2- CB- CG- HG
3926 3925 3928 3930 3 0.66944 2.00832 0.00000
-2.67776 0.00000 0.00000 ; HB2- CB- CG- CD1
3926 3925 3928 3934 3 0.66944 2.00832 0.00000
-2.67776 0.00000 0.00000 ; HB2- CB- CG- CD2
3927 3925 3923 3938 3 0.65084 1.95253 0.00000
-2.60338 0.00000 0.00000 ; HB3- CB- CA- C
3927 3925 3928 3929 3 0.62760 1.88280 0.00000
-2.51040 0.00000 0.00000 ; HB3- CB- CG- HG
3927 3925 3928 3930 3 0.66944 2.00832 0.00000
-2.67776 0.00000 0.00000 ; HB3- CB- CG- CD1
3927 3925 3928 3934 3 0.66944 2.00832 0.00000
-2.67776 0.00000 0.00000 ; HB3- CB- CG- CD2
3928 3925 3923 3938 3 10.33448 -1.14642 -11.54784
-2.40998 6.35968 0.00000 ; CG- CB- CA- C
3929 3928 3930 3931 3 0.62760 1.88280 0.00000
-2.51040 0.00000 0.00000 ; HG- CG- CD1- HD11
3929 3928 3930 3932 3 0.62760 1.88280 0.00000
-2.51040 0.00000 0.00000 ; HG- CG- CD1- HD12
3929 3928 3930 3933 3 0.62760 1.88280 0.00000
-2.51040 0.00000 0.00000 ; HG- CG- CD1- HD13
3929 3928 3934 3935 3 0.62760 1.88280 0.00000
-2.51040 0.00000 0.00000 ; HG- CG- CD2- HD21
3929 3928 3934 3936 3 0.62760 1.88280 0.00000
-2.51040 0.00000 0.00000 ; HG- CG- CD2- HD22
3929 3928 3934 3937 3 0.62760 1.88280 0.00000
-2.51040 0.00000 0.00000 ; HG- CG- CD2- HD23
3930 3928 3934 3935 3 0.66944 2.00832 0.00000
-2.67776 0.00000 0.00000 ; CD1- CG- CD2- HD21
3930 3928 3934 3936 3 0.66944 2.00832 0.00000
-2.67776 0.00000 0.00000 ; CD1- CG- CD2- HD22
3930 3928 3934 3937 3 0.66944 2.00832 0.00000
-2.67776 0.00000 0.00000 ; CD1- CG- CD2- HD23
3931 3930 3928 3934 3 0.66944 2.00832 0.00000
-2.67776 0.00000 0.00000 ; HD11- CD1- CG- CD2
3932 3930 3928 3934 3 0.66944 2.00832 0.00000
-2.67776 0.00000 0.00000 ; HD12- CD1- CG- CD2
3933 3930 3928 3934 3 0.66944 2.00832 0.00000
-2.67776 0.00000 0.00000 ; HD13- CD1- CG- CD2
3938 3940 3942 3943 3 0.00000 0.00000 0.00000
0.00000 0.00000 0.00000 ; C- N- CA- HA
3938 3940 3942 3944 3 11.71520 1.67360 15.06240 -
13.38880 0.00000 0.00000 ; C- N- CA- CB

3938 3940 3942 3957 3 1.75728 5.27184 2.25936
-7.02912 0.00000 0.00000 ; C- N- CA- C
3939 3938 3940 3941 3 29.28800 -8.36800 -20.92000
0.00000 0.00000 0.00000 ; O- C- N- H
3939 3938 3940 3942 3 20.92000 0.00000 -20.92000
0.00000 0.00000 0.00000 ; O- C- N- CA
3940 3942 3944 3945 3 0.65084 1.95253 0.00000
-2.60338 0.00000 0.00000 ; N- CA- CB- HB
3940 3942 3944 3946 3 3.83673 0.44769 -1.77402
-2.47693 -0.03347 0.00000 ; N- CA- CB- CG2
3940 3942 3944 3950 3 2.97482 0.12134 2.04179
-1.89117 -3.24678 0.00000 ; N- CA- CB- CG1
3940 3942 3957 3958 3 0.00000 0.00000 0.00000
0.00000 0.00000 0.00000 ; N- CA- C- O
3940 3942 3957 3959 3 17.40544 -5.02080 -13.22144
9.20480 0.00000 0.00000 ; N- CA- C- N
3941 3940 3942 3943 3 0.00000 0.00000 0.00000
0.00000 0.00000 0.00000 ; H- N- CA- HA
3941 3940 3942 3944 3 0.00000 0.00000 0.00000
0.00000 0.00000 0.00000 ; H- N- CA- CB
3941 3940 3942 3957 3 0.00000 0.00000 0.00000
0.00000 0.00000 0.00000 ; H- N- CA- C
3942 3944 3946 3947 3 0.66944 2.00832 0.00000
-2.67776 0.00000 0.00000 ; CA- CB- CG2- HG21
3942 3944 3946 3948 3 0.66944 2.00832 0.00000
-2.67776 0.00000 0.00000 ; CA- CB- CG2- HG22
3942 3944 3946 3949 3 0.66944 2.00832 0.00000
-2.67776 0.00000 0.00000 ; CA- CB- CG2- HG23
3942 3944 3950 3951 3 0.66944 2.00832 0.00000
-2.67776 0.00000 0.00000 ; CA- CB- CG1- HG12
3942 3944 3950 3952 3 0.66944 2.00832 0.00000
-2.67776 0.00000 0.00000 ; CA- CB- CG1- HG13
3942 3944 3950 3953 3 4.24258 -0.52718 -7.25506
-1.79075 7.69856 0.00000 ; CA- CB- CG1- CD1
3942 3957 3959 3960 3 20.92000 0.00000 -20.92000
0.00000 0.00000 0.00000 ; CA- C- N- H
3942 3957 3959 3961 3 20.92000 0.00000 -20.92000
0.00000 0.00000 0.00000 ; CA- C- N- CA
3943 3942 3944 3945 3 0.65084 1.95253 0.00000
-2.60338 0.00000 0.00000 ; HA- CA- CB- HB
3943 3942 3944 3946 3 0.65084 1.95253 0.00000
-2.60338 0.00000 0.00000 ; HA- CA- CB- CG2
3943 3942 3944 3950 3 0.65084 1.95253 0.00000
-2.60338 0.00000 0.00000 ; HA- CA- CB- CG1
3943 3942 3957 3958 3 3.68192 -4.35136 0.00000
1.33888 0.00000 0.00000 ; HA- CA- C- O
3943 3942 3957 3959 3 0.00000 0.00000 0.00000
0.00000 0.00000 0.00000 ; HA- CA- C- N
3944 3942 3957 3958 3 0.00000 0.00000 0.00000
0.00000 0.00000 0.00000 ; CB- CA- C- O
3944 3942 3957 3959 3 2.51040 4.18400 1.67360
-6.69440 0.00000 0.00000 ; CB- CA- C- N

3944 3950 3953 3954 3 0.66944 2.00832 0.00000
-2.67776 0.00000 0.00000 ; CB- CG1- CD1- HD11
3944 3950 3953 3955 3 0.66944 2.00832 0.00000
-2.67776 0.00000 0.00000 ; CB- CG1- CD1- HD12
3944 3950 3953 3956 3 0.66944 2.00832 0.00000
-2.67776 0.00000 0.00000 ; CB- CG1- CD1- HD13
3945 3944 3942 3957 3 0.65084 1.95253 0.00000
-2.60338 0.00000 0.00000 ; HB- CB- CA- C
3945 3944 3946 3947 3 0.62760 1.88280 0.00000
-2.51040 0.00000 0.00000 ; HB- CB- CG2- HG21
3945 3944 3946 3948 3 0.62760 1.88280 0.00000
-2.51040 0.00000 0.00000 ; HB- CB- CG2- HG22
3945 3944 3946 3949 3 0.62760 1.88280 0.00000
-2.51040 0.00000 0.00000 ; HB- CB- CG2- HG23
3945 3944 3950 3951 3 0.62760 1.88280 0.00000
-2.51040 0.00000 0.00000 ; HB- CB- CG1- HG12
3945 3944 3950 3952 3 0.62760 1.88280 0.00000
-2.51040 0.00000 0.00000 ; HB- CB- CG1- HG13
3945 3944 3950 3953 3 0.66944 2.00832 0.00000
-2.67776 0.00000 0.00000 ; HB- CB- CG1- CD1
3946 3944 3942 3957 3 5.67350 3.55640 -6.16722
-2.47693 3.74886 0.00000 ; CG2- CB- CA- C
3946 3944 3950 3951 3 0.66944 2.00832 0.00000
-2.67776 0.00000 0.00000 ; CG2- CB- CG1- HG12
3946 3944 3950 3952 3 0.66944 2.00832 0.00000
-2.67776 0.00000 0.00000 ; CG2- CB- CG1- HG13
3946 3944 3950 3953 3 3.81162 0.49790 -8.14206
-1.79075 7.49773 0.00000 ; CG2- CB- CG1- CD1
3947 3946 3944 3950 3 0.66944 2.00832 0.00000
-2.67776 0.00000 0.00000 ; HG21- CG2- CB- CG1
3948 3946 3944 3950 3 0.66944 2.00832 0.00000
-2.67776 0.00000 0.00000 ; HG22- CG2- CB- CG1
3949 3946 3944 3950 3 0.66944 2.00832 0.00000
-2.67776 0.00000 0.00000 ; HG23- CG2- CB- CG1
3950 3944 3942 3957 3 8.26340 0.74057 -9.99976
-1.89117 3.84928 0.00000 ; CG1- CB- CA- C
3951 3950 3953 3954 3 0.62760 1.88280 0.00000
-2.51040 0.00000 0.00000 ; HG12- CG1- CD1- HD11
3951 3950 3953 3955 3 0.62760 1.88280 0.00000
-2.51040 0.00000 0.00000 ; HG12- CG1- CD1- HD12
3951 3950 3953 3956 3 0.62760 1.88280 0.00000
-2.51040 0.00000 0.00000 ; HG12- CG1- CD1- HD13
3952 3950 3953 3954 3 0.62760 1.88280 0.00000
-2.51040 0.00000 0.00000 ; HG13- CG1- CD1- HD11
3952 3950 3953 3955 3 0.62760 1.88280 0.00000
-2.51040 0.00000 0.00000 ; HG13- CG1- CD1- HD12
3952 3950 3953 3956 3 0.62760 1.88280 0.00000
-2.51040 0.00000 0.00000 ; HG13- CG1- CD1- HD13
3957 3959 3961 3962 3 0.00000 0.00000 0.00000
0.00000 0.00000 0.00000 ; C- N- CA- HA
3957 3959 3961 3963 3 11.71520 1.67360 15.06240 -
13.38880 0.00000 0.00000 ; C- N- CA- CB

```
  3957   3959   3961   3971       3    1.75728    5.27184    2.25936
-7.02912    0.00000    0.00000 ;      C-     N-    CA-     C
  3958   3957   3959   3960       3   29.28800   -8.36800  -20.92000
0.00000    0.00000    0.00000 ;      O-     C-     N-     H
  3958   3957   3959   3961       3   20.92000    0.00000  -20.92000
0.00000    0.00000    0.00000 ;      O-     C-     N-    CA
  3959   3961   3963   3964       3    0.65084    1.95253    0.00000
-2.60338    0.00000    0.00000 ;      N-    CA-    CB-    HB
  3959   3961   3963   3965       3    3.83673    0.44769   -1.77402
-2.47693   -0.03347    0.00000 ;      N-    CA-    CB-   CG2
  3959   3961   3963   3969       3    4.93294    1.13386   -3.12963
-5.27184    3.17984    0.00000 ;      N-    CA-    CB-   OG1
  3959   3961   3971   3972       3    0.00000    0.00000    0.00000
0.00000    0.00000    0.00000 ;      N-    CA-     C-     O
  3959   3961   3971   3973       3   17.40544   -5.02080  -13.22144
9.20480    0.00000    0.00000 ;      N-    CA-     C-     N
  3960   3959   3961   3962       3    0.00000    0.00000    0.00000
0.00000    0.00000    0.00000 ;      H-     N-    CA-    HA
  3960   3959   3961   3963       3    0.00000    0.00000    0.00000
0.00000    0.00000    0.00000 ;      H-     N-    CA-    CB
  3960   3959   3961   3971       3    0.00000    0.00000    0.00000
0.00000    0.00000    0.00000 ;      H-     N-    CA-     C
  3961   3963   3965   3966       3    0.66944    2.00832    0.00000
-2.67776    0.00000    0.00000 ;     CA-    CB-   CG2-  HG21
  3961   3963   3965   3967       3    0.66944    2.00832    0.00000
-2.67776    0.00000    0.00000 ;     CA-    CB-   CG2-  HG22
  3961   3963   3965   3968       3    0.66944    2.00832    0.00000
-2.67776    0.00000    0.00000 ;     CA-    CB-   CG2-  HG23
  3961   3963   3969   3970       3    1.12131    2.98738    1.66523
-3.94970    0.43514    0.00000 ;     CA-    CB-   OG1-   HG1
  3961   3971   3973   3974       3   20.92000    0.00000  -20.92000
0.00000    0.00000    0.00000 ;     CA-     C-     N-     H
  3961   3971   3973   3975       3   20.92000    0.00000  -20.92000
0.00000    0.00000    0.00000 ;     CA-     C-     N-    CA
  3962   3961   3963   3964       3    0.65084    1.95253    0.00000
-2.60338    0.00000    0.00000 ;     HA-    CA-    CB-    HB
  3962   3961   3963   3965       3    0.65084    1.95253    0.00000
-2.60338    0.00000    0.00000 ;     HA-    CA-    CB-   CG2
  3962   3961   3963   3969       3    1.04600   -1.04600    0.00000
0.00000    0.00000    0.00000 ;     HA-    CA-    CB-   OG1
  3962   3961   3971   3972       3    3.68192   -4.35136    0.00000
1.33888    0.00000    0.00000 ;     HA-    CA-     C-     O
  3962   3961   3971   3973       3    0.00000    0.00000    0.00000
0.00000    0.00000    0.00000 ;     HA-    CA-     C-     N
  3963   3961   3971   3972       3    0.00000    0.00000    0.00000
0.00000    0.00000    0.00000 ;     CB-    CA-     C-     O
  3963   3961   3971   3973       3    2.51040    4.18400    1.67360
-6.69440    0.00000    0.00000 ;     CB-    CA-     C-     N
  3964   3963   3961   3971       3    0.65084    1.95253    0.00000
-2.60338    0.00000    0.00000 ;     HB-    CB-    CA-     C
  3964   3963   3965   3966       3    0.65084    1.95253    0.00000
-2.60338    0.00000    0.00000 ;     HB-    CB-   CG2-  HG21
```

3964 3963 3965 3967 3 0.65084 1.95253 0.00000
-2.60338 0.00000 0.00000 ; HB- CB- CG2- HG22
3964 3963 3965 3968 3 0.65084 1.95253 0.00000
-2.60338 0.00000 0.00000 ; HB- CB- CG2- HG23
3964 3963 3969 3970 3 0.69733 2.09200 0.00000
-2.78933 0.00000 0.00000 ; HB- CB- OG1- HG1
3965 3963 3961 3971 3 5.67350 3.55640 -6.16722
-2.47693 3.74886 0.00000 ; CG2- CB- CA- C
3965 3963 3969 3970 3 4.74047 0.27196 -2.26773
-3.94970 1.60666 0.00000 ; CG2- CB- OG1- HG1
3966 3965 3963 3969 3 1.04600 -1.04600 0.00000
0.00000 0.00000 0.00000 ; HG21- CG2- CB- OG1
3967 3965 3963 3969 3 1.04600 -1.04600 0.00000
0.00000 0.00000 0.00000 ; HG22- CG2- CB- OG1
3968 3965 3963 3969 3 1.04600 -1.04600 0.00000
0.00000 0.00000 0.00000 ; HG23- CG2- CB- OG1
3969 3963 3961 3971 3 6.53541 6.87013 -6.21742
-5.27184 5.22163 0.00000 ; OG1- CB- CA- C
3971 3973 3975 3976 3 0.00000 0.00000 0.00000
0.00000 0.00000 0.00000 ; C- N- CA- HA
3971 3973 3975 3977 3 11.71520 1.67360 15.06240 -
13.38880 0.00000 0.00000 ; C- N- CA- CB
3971 3973 3975 3993 3 1.75728 5.27184 2.25936
-7.02912 0.00000 0.00000 ; C- N- CA- C
3972 3971 3973 3974 3 29.28800 -8.36800 -20.92000
0.00000 0.00000 0.00000 ; O- C- N- H
3972 3971 3973 3975 3 20.92000 0.00000 -20.92000
0.00000 0.00000 0.00000 ; O- C- N- CA
3973 3975 3977 3978 3 0.65084 1.95253 0.00000
-2.60338 0.00000 0.00000 ; N- CA- CB- HB2
3973 3975 3977 3979 3 0.65084 1.95253 0.00000
-2.60338 0.00000 0.00000 ; N- CA- CB- HB3
3973 3975 3977 3980 3 0.65084 1.95253 0.00000
-2.60338 0.00000 0.00000 ; N- CA- CB- CG
3973 3975 3993 3994 3 0.00000 0.00000 0.00000
0.00000 0.00000 0.00000 ; N- CA- C- O
3973 3975 3993 3995 3 17.40544 -5.02080 -13.22144
9.20480 0.00000 0.00000 ; N- CA- C- N
3974 3973 3975 3976 3 0.00000 0.00000 0.00000
0.00000 0.00000 0.00000 ; H- N- CA- HA
3974 3973 3975 3977 3 0.00000 0.00000 0.00000
0.00000 0.00000 0.00000 ; H- N- CA- CB
3974 3973 3975 3993 3 0.00000 0.00000 0.00000
0.00000 0.00000 0.00000 ; H- N- CA- C
3975 3977 3980 3981 3 0.66944 2.00832 0.00000
-2.67776 0.00000 0.00000 ; CA- CB- CG- HG2
3975 3977 3980 3982 3 0.66944 2.00832 0.00000
-2.67776 0.00000 0.00000 ; CA- CB- CG- HG3
3975 3977 3980 3983 3 3.68192 3.09616 -2.09200
-3.01248 0.00000 0.00000 ; CA- CB- CG- CD
3975 3993 3995 3996 3 20.92000 0.00000 -20.92000
0.00000 0.00000 0.00000 ; CA- C- N- H

3975 3993 3995 3997 3 20.92000 0.00000 -20.92000
0.00000 0.00000 0.00000 ; CA- C- N- CA
3976 3975 3977 3978 3 0.65084 1.95253 0.00000
-2.60338 0.00000 0.00000 ; HA- CA- CB- HB2
3976 3975 3977 3979 3 0.65084 1.95253 0.00000
-2.60338 0.00000 0.00000 ; HA- CA- CB- HB3
3976 3975 3977 3980 3 0.65084 1.95253 0.00000
-2.60338 0.00000 0.00000 ; HA- CA- CB- CG
3976 3975 3993 3994 3 3.68192 -4.35136 0.00000
1.33888 0.00000 0.00000 ; HA- CA- C- O
3976 3975 3993 3995 3 0.00000 0.00000 0.00000
0.00000 0.00000 0.00000 ; HA- CA- C- N
3977 3975 3993 3994 3 0.00000 0.00000 0.00000
0.00000 0.00000 0.00000 ; CB- CA- C- O
3977 3975 3993 3995 3 2.51040 4.18400 1.67360
-6.69440 0.00000 0.00000 ; CB- CA- C- N
3977 3980 3983 3984 3 0.66944 2.00832 0.00000
-2.67776 0.00000 0.00000 ; CB- CG- CD- HD2
3977 3980 3983 3985 3 0.66944 2.00832 0.00000
-2.67776 0.00000 0.00000 ; CB- CG- CD- HD3
3977 3980 3983 3986 3 3.68192 3.09616 -2.09200
-3.01248 0.00000 0.00000 ; CB- CG- CD- CE
3978 3977 3975 3993 3 0.65084 1.95253 0.00000
-2.60338 0.00000 0.00000 ; HB2- CB- CA- C
3978 3977 3980 3981 3 0.62760 1.88280 0.00000
-2.51040 0.00000 0.00000 ; HB2- CB- CG- HG2
3978 3977 3980 3982 3 0.62760 1.88280 0.00000
-2.51040 0.00000 0.00000 ; HB2- CB- CG- HG3
3978 3977 3980 3983 3 0.66944 2.00832 0.00000
-2.67776 0.00000 0.00000 ; HB2- CB- CG- CD
3979 3977 3975 3993 3 0.65084 1.95253 0.00000
-2.60338 0.00000 0.00000 ; HB3- CB- CA- C
3979 3977 3980 3981 3 0.62760 1.88280 0.00000
-2.51040 0.00000 0.00000 ; HB3- CB- CG- HG2
3979 3977 3980 3982 3 0.62760 1.88280 0.00000
-2.51040 0.00000 0.00000 ; HB3- CB- CG- HG3
3979 3977 3980 3983 3 0.66944 2.00832 0.00000
-2.67776 0.00000 0.00000 ; HB3- CB- CG- CD
3980 3977 3975 3993 3 0.65084 1.95253 0.00000
-2.60338 0.00000 0.00000 ; CG- CB- CA- C
3980 3983 3986 3987 3 0.65084 1.95253 0.00000
-2.60338 0.00000 0.00000 ; CG- CD- CE- HE2
3980 3983 3986 3988 3 0.65084 1.95253 0.00000
-2.60338 0.00000 0.00000 ; CG- CD- CE- HE3
3980 3983 3986 3989 3 0.65084 1.95253 0.00000
-2.60338 0.00000 0.00000 ; CG- CD- CE- NZ
3981 3980 3983 3984 3 0.62760 1.88280 0.00000
-2.51040 0.00000 0.00000 ; HG2- CG- CD- HD2
3981 3980 3983 3985 3 0.62760 1.88280 0.00000
-2.51040 0.00000 0.00000 ; HG2- CG- CD- HD3
3981 3980 3983 3986 3 0.66944 2.00832 0.00000
-2.67776 0.00000 0.00000 ; HG2- CG- CD- CE

3982 3980 3983 3984 3 0.62760 1.88280 0.00000
-2.51040 0.00000 0.00000 ; HG3- CG- CD- HD2
3982 3980 3983 3985 3 0.62760 1.88280 0.00000
-2.51040 0.00000 0.00000 ; HG3- CG- CD- HD3
3982 3980 3983 3986 3 0.66944 2.00832 0.00000
-2.67776 0.00000 0.00000 ; HG3- CG- CD- CE
3983 3986 3989 3990 3 0.65084 1.95253 0.00000
-2.60338 0.00000 0.00000 ; CD- CE- NZ- HZ1
3983 3986 3989 3991 3 0.65084 1.95253 0.00000
-2.60338 0.00000 0.00000 ; CD- CE- NZ- HZ2
3983 3986 3989 3992 3 0.65084 1.95253 0.00000
-2.60338 0.00000 0.00000 ; CD- CE- NZ- HZ3
3984 3983 3986 3987 3 0.65084 1.95253 0.00000
-2.60338 0.00000 0.00000 ; HD2- CD- CE- HE2
3984 3983 3986 3988 3 0.65084 1.95253 0.00000
-2.60338 0.00000 0.00000 ; HD2- CD- CE- HE3
3984 3983 3986 3989 3 0.65084 1.95253 0.00000
-2.60338 0.00000 0.00000 ; HD2- CD- CE- NZ
3985 3983 3986 3987 3 0.65084 1.95253 0.00000
-2.60338 0.00000 0.00000 ; HD3- CD- CE- HE2
3985 3983 3986 3988 3 0.65084 1.95253 0.00000
-2.60338 0.00000 0.00000 ; HD3- CD- CE- HE3
3985 3983 3986 3989 3 0.65084 1.95253 0.00000
-2.60338 0.00000 0.00000 ; HD3- CD- CE- NZ
3987 3986 3989 3990 3 0.65084 1.95253 0.00000
-2.60338 0.00000 0.00000 ; HE2- CE- NZ- HZ1
3987 3986 3989 3991 3 0.65084 1.95253 0.00000
-2.60338 0.00000 0.00000 ; HE2- CE- NZ- HZ2
3987 3986 3989 3992 3 0.65084 1.95253 0.00000
-2.60338 0.00000 0.00000 ; HE2- CE- NZ- HZ3
3988 3986 3989 3990 3 0.65084 1.95253 0.00000
-2.60338 0.00000 0.00000 ; HE3- CE- NZ- HZ1
3988 3986 3989 3991 3 0.65084 1.95253 0.00000
-2.60338 0.00000 0.00000 ; HE3- CE- NZ- HZ2
3988 3986 3989 3992 3 0.65084 1.95253 0.00000
-2.60338 0.00000 0.00000 ; HE3- CE- NZ- HZ3
3993 3995 3997 3998 3 0.00000 0.00000 0.00000
0.00000 0.00000 0.00000 ; C- N- CA- HA
3993 3995 3997 3999 3 11.71520 1.67360 15.06240 -
13.38880 0.00000 0.00000 ; C- N- CA- CB
3993 3995 3997 4012 3 1.75728 5.27184 2.25936
-7.02912 0.00000 0.00000 ; C- N- CA- C
3994 3993 3995 3996 3 29.28800 -8.36800 -20.92000
0.00000 0.00000 0.00000 ; O- C- N- H
3994 3993 3995 3997 3 20.92000 0.00000 -20.92000
0.00000 0.00000 0.00000 ; O- C- N- CA
3995 3997 3999 4000 3 0.65084 1.95253 0.00000
-2.60338 0.00000 0.00000 ; N- CA- CB- HB2
3995 3997 3999 4001 3 0.65084 1.95253 0.00000
-2.60338 0.00000 0.00000 ; N- CA- CB- HB3
3995 3997 3999 4002 3 3.79070 1.39746 -4.61077
-2.40998 2.44346 0.00000 ; N- CA- CB- CG

3995 3997 4012 4013 3 0.00000 0.00000 0.00000
0.00000 0.00000 0.00000 ; N- CA- C- O
3995 3997 4012 4014 3 17.40544 -5.02080 -13.22144
9.20480 0.00000 0.00000 ; N- CA- C- N
3996 3995 3997 3998 3 0.00000 0.00000 0.00000
0.00000 0.00000 0.00000 ; H- N- CA- HA
3996 3995 3997 3999 3 0.00000 0.00000 0.00000
0.00000 0.00000 0.00000 ; H- N- CA- CB
3996 3995 3997 4012 3 0.00000 0.00000 0.00000
0.00000 0.00000 0.00000 ; H- N- CA- C
3997 3999 4002 4003 3 0.66944 2.00832 0.00000
-2.67776 0.00000 0.00000 ; CA- CB- CG- HG
3997 3999 4002 4004 3 3.90367 0.19665 -6.21742
-2.37651 5.99149 0.00000 ; CA- CB- CG- CD1
3997 3999 4002 4008 3 3.90367 0.19665 -6.21742
-2.37651 5.99149 0.00000 ; CA- CB- CG- CD2
3997 4012 4014 4015 3 20.92000 0.00000 -20.92000
0.00000 0.00000 0.00000 ; CA- C- N- H
3997 4012 4014 4016 3 20.92000 0.00000 -20.92000
0.00000 0.00000 0.00000 ; CA- C- N- CA
3998 3997 3999 4000 3 0.65084 1.95253 0.00000
-2.60338 0.00000 0.00000 ; HA- CA- CB- HB2
3998 3997 3999 4001 3 0.65084 1.95253 0.00000
-2.60338 0.00000 0.00000 ; HA- CA- CB- HB3
3998 3997 3999 4002 3 0.65084 1.95253 0.00000
-2.60338 0.00000 0.00000 ; HA- CA- CB- CG
3998 3997 4012 4013 3 3.68192 -4.35136 0.00000
1.33888 0.00000 0.00000 ; HA- CA- C- O
3998 3997 4012 4014 3 0.00000 0.00000 0.00000
0.00000 0.00000 0.00000 ; HA- CA- C- N
3999 3997 4012 4013 3 0.00000 0.00000 0.00000
0.00000 0.00000 0.00000 ; CB- CA- C- O
3999 3997 4012 4014 3 2.51040 4.18400 1.67360
-6.69440 0.00000 0.00000 ; CB- CA- C- N
3999 4002 4004 4005 3 0.66944 2.00832 0.00000
-2.67776 0.00000 0.00000 ; CB- CG- CD1- HD11
3999 4002 4004 4006 3 0.66944 2.00832 0.00000
-2.67776 0.00000 0.00000 ; CB- CG- CD1- HD12
3999 4002 4004 4007 3 0.66944 2.00832 0.00000
-2.67776 0.00000 0.00000 ; CB- CG- CD1- HD13
3999 4002 4008 4009 3 0.66944 2.00832 0.00000
-2.67776 0.00000 0.00000 ; CB- CG- CD2- HD21
3999 4002 4008 4010 3 0.66944 2.00832 0.00000
-2.67776 0.00000 0.00000 ; CB- CG- CD2- HD22
3999 4002 4008 4011 3 0.66944 2.00832 0.00000
-2.67776 0.00000 0.00000 ; CB- CG- CD2- HD23
4000 3999 3997 4012 3 0.65084 1.95253 0.00000
-2.60338 0.00000 0.00000 ; HB2- CB- CA- C
4000 3999 4002 4003 3 0.62760 1.88280 0.00000
-2.51040 0.00000 0.00000 ; HB2- CB- CG- HG
4000 3999 4002 4004 3 0.66944 2.00832 0.00000
-2.67776 0.00000 0.00000 ; HB2- CB- CG- CD1

```
  4000   3999   4002   4008      3    0.66944    2.00832    0.00000
-2.67776    0.00000    0.00000 ;    HB2-    CB-    CG-   CD2
  4001   3999   3997   4012      3    0.65084    1.95253    0.00000
-2.60338    0.00000    0.00000 ;    HB3-    CB-    CA-     C
  4001   3999   4002   4003      3    0.62760    1.88280    0.00000
-2.51040    0.00000    0.00000 ;    HB3-    CB-    CG-    HG
  4001   3999   4002   4004      3    0.66944    2.00832    0.00000
-2.67776    0.00000    0.00000 ;    HB3-    CB-    CG-   CD1
  4001   3999   4002   4008      3    0.66944    2.00832    0.00000
-2.67776    0.00000    0.00000 ;    HB3-    CB-    CG-   CD2
  4002   3999   3997   4012      3   10.33448   -1.14642  -11.54784
-2.40998    6.35968    0.00000 ;     CG-    CB-    CA-     C
  4003   4002   4004   4005      3    0.62760    1.88280    0.00000
-2.51040    0.00000    0.00000 ;     HG-    CG-   CD1-  HD11
  4003   4002   4004   4006      3    0.62760    1.88280    0.00000
-2.51040    0.00000    0.00000 ;     HG-    CG-   CD1-  HD12
  4003   4002   4004   4007      3    0.62760    1.88280    0.00000
-2.51040    0.00000    0.00000 ;     HG-    CG-   CD1-  HD13
  4003   4002   4008   4009      3    0.62760    1.88280    0.00000
-2.51040    0.00000    0.00000 ;     HG-    CG-   CD2-  HD21
  4003   4002   4008   4010      3    0.62760    1.88280    0.00000
-2.51040    0.00000    0.00000 ;     HG-    CG-   CD2-  HD22
  4003   4002   4008   4011      3    0.62760    1.88280    0.00000
-2.51040    0.00000    0.00000 ;     HG-    CG-   CD2-  HD23
  4004   4002   4008   4009      3    0.66944    2.00832    0.00000
-2.67776    0.00000    0.00000 ;    CD1-    CG-   CD2-  HD21
  4004   4002   4008   4010      3    0.66944    2.00832    0.00000
-2.67776    0.00000    0.00000 ;    CD1-    CG-   CD2-  HD22
  4004   4002   4008   4011      3    0.66944    2.00832    0.00000
-2.67776    0.00000    0.00000 ;    CD1-    CG-   CD2-  HD23
  4005   4004   4002   4008      3    0.66944    2.00832    0.00000
-2.67776    0.00000    0.00000 ;   HD11-   CD1-    CG-   CD2
  4006   4004   4002   4008      3    0.66944    2.00832    0.00000
-2.67776    0.00000    0.00000 ;   HD12-   CD1-    CG-   CD2
  4007   4004   4002   4008      3    0.66944    2.00832    0.00000
-2.67776    0.00000    0.00000 ;   HD13-   CD1-    CG-   CD2
  4012   4014   4016   4017      3    0.00000    0.00000    0.00000
0.00000    0.00000    0.00000 ;     C-     N-    CA-    HA
  4012   4014   4016   4018      3   11.71520    1.67360   15.06240  -
13.38880    0.00000    0.00000 ;      C-     N-    CA-    CB
  4012   4014   4016   4031      3    1.75728    5.27184    2.25936
-7.02912    0.00000    0.00000 ;      C-     N-    CA-     C
  4013   4012   4014   4015      3   29.28800   -8.36800  -20.92000
0.00000    0.00000    0.00000 ;     O-     C-     N-     H
  4013   4012   4014   4016      3   20.92000    0.00000  -20.92000
0.00000    0.00000    0.00000 ;     O-     C-     N-    CA
  4014   4016   4018   4019      3    0.65084    1.95253    0.00000
-2.60338    0.00000    0.00000 ;      N-    CA-    CB-   HB2
  4014   4016   4018   4020      3    0.65084    1.95253    0.00000
-2.60338    0.00000    0.00000 ;      N-    CA-    CB-   HB3
  4014   4016   4018   4021      3    3.79070    1.39746   -4.61077
-2.40998    2.44346    0.00000 ;      N-    CA-    CB-    CG
```

4014 4016 4031 4032 3 0.00000 0.00000 0.00000
0.00000 0.00000 0.00000 ; N- CA- C- O
4014 4016 4031 4033 3 17.40544 -5.02080 -13.22144
9.20480 0.00000 0.00000 ; N- CA- C- N
4015 4014 4016 4017 3 0.00000 0.00000 0.00000
0.00000 0.00000 0.00000 ; H- N- CA- HA
4015 4014 4016 4018 3 0.00000 0.00000 0.00000
0.00000 0.00000 0.00000 ; H- N- CA- CB
4015 4014 4016 4031 3 0.00000 0.00000 0.00000
0.00000 0.00000 0.00000 ; H- N- CA- C
4016 4018 4021 4022 3 0.66944 2.00832 0.00000
-2.67776 0.00000 0.00000 ; CA- CB- CG- HG
4016 4018 4021 4023 3 3.90367 0.19665 -6.21742
-2.37651 5.99149 0.00000 ; CA- CB- CG- CD1
4016 4018 4021 4027 3 3.90367 0.19665 -6.21742
-2.37651 5.99149 0.00000 ; CA- CB- CG- CD2
4016 4031 4033 4034 3 20.92000 0.00000 -20.92000
0.00000 0.00000 0.00000 ; CA- C- N- H
4016 4031 4033 4035 3 20.92000 0.00000 -20.92000
0.00000 0.00000 0.00000 ; CA- C- N- CA
4017 4016 4018 4019 3 0.65084 1.95253 0.00000
-2.60338 0.00000 0.00000 ; HA- CA- CB- HB2
4017 4016 4018 4020 3 0.65084 1.95253 0.00000
-2.60338 0.00000 0.00000 ; HA- CA- CB- HB3
4017 4016 4018 4021 3 0.65084 1.95253 0.00000
-2.60338 0.00000 0.00000 ; HA- CA- CB- CG
4017 4016 4031 4032 3 3.68192 -4.35136 0.00000
1.33888 0.00000 0.00000 ; HA- CA- C- O
4017 4016 4031 4033 3 0.00000 0.00000 0.00000
0.00000 0.00000 0.00000 ; HA- CA- C- N
4018 4016 4031 4032 3 0.00000 0.00000 0.00000
0.00000 0.00000 0.00000 ; CB- CA- C- O
4018 4016 4031 4033 3 2.51040 4.18400 1.67360
-6.69440 0.00000 0.00000 ; CB- CA- C- N
4018 4021 4023 4024 3 0.66944 2.00832 0.00000
-2.67776 0.00000 0.00000 ; CB- CG- CD1- HD11
4018 4021 4023 4025 3 0.66944 2.00832 0.00000
-2.67776 0.00000 0.00000 ; CB- CG- CD1- HD12
4018 4021 4023 4026 3 0.66944 2.00832 0.00000
-2.67776 0.00000 0.00000 ; CB- CG- CD1- HD13
4018 4021 4027 4028 3 0.66944 2.00832 0.00000
-2.67776 0.00000 0.00000 ; CB- CG- CD2- HD21
4018 4021 4027 4029 3 0.66944 2.00832 0.00000
-2.67776 0.00000 0.00000 ; CB- CG- CD2- HD22
4018 4021 4027 4030 3 0.66944 2.00832 0.00000
-2.67776 0.00000 0.00000 ; CB- CG- CD2- HD23
4019 4018 4016 4031 3 0.65084 1.95253 0.00000
-2.60338 0.00000 0.00000 ; HB2- CB- CA- C
4019 4018 4021 4022 3 0.62760 1.88280 0.00000
-2.51040 0.00000 0.00000 ; HB2- CB- CG- HG
4019 4018 4021 4023 3 0.66944 2.00832 0.00000
-2.67776 0.00000 0.00000 ; HB2- CB- CG- CD1

4019 4018 4021 4027 3 0.66944 2.00832 0.00000
-2.67776 0.00000 0.00000 ; HB2- CB- CG- CD2
4020 4018 4016 4031 3 0.65084 1.95253 0.00000
-2.60338 0.00000 0.00000 ; HB3- CB- CA- C
4020 4018 4021 4022 3 0.62760 1.88280 0.00000
-2.51040 0.00000 0.00000 ; HB3- CB- CG- HG
4020 4018 4021 4023 3 0.66944 2.00832 0.00000
-2.67776 0.00000 0.00000 ; HB3- CB- CG- CD1
4020 4018 4021 4027 3 0.66944 2.00832 0.00000
-2.67776 0.00000 0.00000 ; HB3- CB- CG- CD2
4021 4018 4016 4031 3 10.33448 -1.14642 -11.54784
-2.40998 6.35968 0.00000 ; CG- CB- CA- C
4022 4021 4023 4024 3 0.62760 1.88280 0.00000
-2.51040 0.00000 0.00000 ; HG- CG- CD1- HD11
4022 4021 4023 4025 3 0.62760 1.88280 0.00000
-2.51040 0.00000 0.00000 ; HG- CG- CD1- HD12
4022 4021 4023 4026 3 0.62760 1.88280 0.00000
-2.51040 0.00000 0.00000 ; HG- CG- CD1- HD13
4022 4021 4027 4028 3 0.62760 1.88280 0.00000
-2.51040 0.00000 0.00000 ; HG- CG- CD2- HD21
4022 4021 4027 4029 3 0.62760 1.88280 0.00000
-2.51040 0.00000 0.00000 ; HG- CG- CD2- HD22
4022 4021 4027 4030 3 0.62760 1.88280 0.00000
-2.51040 0.00000 0.00000 ; HG- CG- CD2- HD23
4023 4021 4027 4028 3 0.66944 2.00832 0.00000
-2.67776 0.00000 0.00000 ; CD1- CG- CD2- HD21
4023 4021 4027 4029 3 0.66944 2.00832 0.00000
-2.67776 0.00000 0.00000 ; CD1- CG- CD2- HD22
4023 4021 4027 4030 3 0.66944 2.00832 0.00000
-2.67776 0.00000 0.00000 ; CD1- CG- CD2- HD23
4024 4023 4021 4027 3 0.66944 2.00832 0.00000
-2.67776 0.00000 0.00000 ; HD11- CD1- CG- CD2
4025 4023 4021 4027 3 0.66944 2.00832 0.00000
-2.67776 0.00000 0.00000 ; HD12- CD1- CG- CD2
4026 4023 4021 4027 3 0.66944 2.00832 0.00000
-2.67776 0.00000 0.00000 ; HD13- CD1- CG- CD2
4031 4033 4035 4036 3 0.00000 0.00000 0.00000
0.00000 0.00000 0.00000 ; C- N- CA- HA
4031 4033 4035 4037 3 11.71520 1.67360 15.06240 -
13.38880 0.00000 0.00000 ; C- N- CA- CB
4031 4033 4035 4045 3 1.75728 5.27184 2.25936
-7.02912 0.00000 0.00000 ; C- N- CA- C
4032 4031 4033 4034 3 29.28800 -8.36800 -20.92000
0.00000 0.00000 0.00000 ; O- C- N- H
4032 4031 4033 4035 3 20.92000 0.00000 -20.92000
0.00000 0.00000 0.00000 ; O- C- N- CA
4033 4035 4037 4038 3 0.65084 1.95253 0.00000
-2.60338 0.00000 0.00000 ; N- CA- CB- HB
4033 4035 4037 4039 3 3.83673 0.44769 -1.77402
-2.47693 -0.03347 0.00000 ; N- CA- CB- CG2
4033 4035 4037 4043 3 4.93294 1.13386 -3.12963
-5.27184 3.17984 0.00000 ; N- CA- CB- OG1

4033 4035 4045 4046 3 0.00000 0.00000 0.00000
0.00000 0.00000 0.00000 ; N- CA- C- O
4033 4035 4045 4047 3 17.40544 -5.02080 -13.22144
9.20480 0.00000 0.00000 ; N- CA- C- N
4034 4033 4035 4036 3 0.00000 0.00000 0.00000
0.00000 0.00000 0.00000 ; H- N- CA- HA
4034 4033 4035 4037 3 0.00000 0.00000 0.00000
0.00000 0.00000 0.00000 ; H- N- CA- CB
4034 4033 4035 4045 3 0.00000 0.00000 0.00000
0.00000 0.00000 0.00000 ; H- N- CA- C
4035 4037 4039 4040 3 0.66944 2.00832 0.00000
-2.67776 0.00000 0.00000 ; CA- CB- CG2- HG21
4035 4037 4039 4041 3 0.66944 2.00832 0.00000
-2.67776 0.00000 0.00000 ; CA- CB- CG2- HG22
4035 4037 4039 4042 3 0.66944 2.00832 0.00000
-2.67776 0.00000 0.00000 ; CA- CB- CG2- HG23
4035 4037 4043 4044 3 1.12131 2.98738 1.66523
-3.94970 0.43514 0.00000 ; CA- CB- OG1- HG1
4035 4045 4047 4048 3 20.92000 0.00000 -20.92000
0.00000 0.00000 0.00000 ; CA- C- N- H
4035 4045 4047 4049 3 20.92000 0.00000 -20.92000
0.00000 0.00000 0.00000 ; CA- C- N- CA
4036 4035 4037 4038 3 0.65084 1.95253 0.00000
-2.60338 0.00000 0.00000 ; HA- CA- CB- HB
4036 4035 4037 4039 3 0.65084 1.95253 0.00000
-2.60338 0.00000 0.00000 ; HA- CA- CB- CG2
4036 4035 4037 4043 3 1.04600 -1.04600 0.00000
0.00000 0.00000 0.00000 ; HA- CA- CB- OG1
4036 4035 4045 4046 3 3.68192 -4.35136 0.00000
1.33888 0.00000 0.00000 ; HA- CA- C- O
4036 4035 4045 4047 3 0.00000 0.00000 0.00000
0.00000 0.00000 0.00000 ; HA- CA- C- N
4037 4035 4045 4046 3 0.00000 0.00000 0.00000
0.00000 0.00000 0.00000 ; CB- CA- C- O
4037 4035 4045 4047 3 2.51040 4.18400 1.67360
-6.69440 0.00000 0.00000 ; CB- CA- C- N
4038 4037 4035 4045 3 0.65084 1.95253 0.00000
-2.60338 0.00000 0.00000 ; HB- CB- CA- C
4038 4037 4039 4040 3 0.65084 1.95253 0.00000
-2.60338 0.00000 0.00000 ; HB- CB- CG2- HG21
4038 4037 4039 4041 3 0.65084 1.95253 0.00000
-2.60338 0.00000 0.00000 ; HB- CB- CG2- HG22
4038 4037 4039 4042 3 0.65084 1.95253 0.00000
-2.60338 0.00000 0.00000 ; HB- CB- CG2- HG23
4038 4037 4043 4044 3 0.69733 2.09200 0.00000
-2.78933 0.00000 0.00000 ; HB- CB- OG1- HG1
4039 4037 4035 4045 3 5.67350 3.55640 -6.16722
-2.47693 3.74886 0.00000 ; CG2- CB- CA- C
4039 4037 4043 4044 3 4.74047 0.27196 -2.26773
-3.94970 1.60666 0.00000 ; CG2- CB- OG1- HG1
4040 4039 4037 4043 3 1.04600 -1.04600 0.00000
0.00000 0.00000 0.00000 ; HG21- CG2- CB- OG1

4041 4039 4037 4043 3 1.04600 -1.04600 0.00000
0.00000 0.00000 0.00000 ; HG22- CG2- CB- OG1
4042 4039 4037 4043 3 1.04600 -1.04600 0.00000
0.00000 0.00000 0.00000 ; HG23- CG2- CB- OG1
4043 4037 4035 4045 3 6.53541 6.87013 -6.21742
-5.27184 5.22163 0.00000 ; OG1- CB- CA- C
4045 4047 4049 4050 3 0.00000 0.00000 0.00000
0.00000 0.00000 0.00000 ; C- N- CA- HA
4045 4047 4049 4051 3 11.71520 1.67360 15.06240 -
13.38880 0.00000 0.00000 ; C- N- CA- CB
4045 4047 4049 4066 3 1.75728 5.27184 2.25936
-7.02912 0.00000 0.00000 ; C- N- CA- C
4046 4045 4047 4048 3 29.28800 -8.36800 -20.92000
0.00000 0.00000 0.00000 ; O- C- N- H
4046 4045 4047 4049 3 20.92000 0.00000 -20.92000
0.00000 0.00000 0.00000 ; O- C- N- CA
4047 4049 4051 4052 3 0.65084 1.95253 0.00000
-2.60338 0.00000 0.00000 ; N- CA- CB- HB2
4047 4049 4051 4053 3 0.65084 1.95253 0.00000
-2.60338 0.00000 0.00000 ; N- CA- CB- HB3
4047 4049 4051 4054 3 3.28026 2.46019 -2.19242
-3.21331 -0.23430 0.00000 ; N- CA- CB- CG
4047 4049 4066 4067 3 0.00000 0.00000 0.00000
0.00000 0.00000 0.00000 ; N- CA- C- O
4047 4049 4066 4068 3 17.40544 -5.02080 -13.22144
9.20480 0.00000 0.00000 ; N- CA- C- N
4048 4047 4049 4050 3 0.00000 0.00000 0.00000
0.00000 0.00000 0.00000 ; H- N- CA- HA
4048 4047 4049 4051 3 0.00000 0.00000 0.00000
0.00000 0.00000 0.00000 ; H- N- CA- CB
4048 4047 4049 4066 3 0.00000 0.00000 0.00000
0.00000 0.00000 0.00000 ; H- N- CA- C
4049 4051 4054 4055 3 0.57739 0.00000 1.02926
0.00000 -1.60666 0.00000 ; CA- CB- CG- CD1
4049 4051 4054 4064 3 0.57739 0.00000 1.02926
0.00000 -1.60666 0.00000 ; CA- CB- CG- CD2
4049 4066 4068 4069 3 20.92000 0.00000 -20.92000
0.00000 0.00000 0.00000 ; CA- C- N- H
4049 4066 4068 4070 3 20.92000 0.00000 -20.92000
0.00000 0.00000 0.00000 ; CA- C- N- CA
4050 4049 4051 4052 3 0.65084 1.95253 0.00000
-2.60338 0.00000 0.00000 ; HA- CA- CB- HB2
4050 4049 4051 4053 3 0.65084 1.95253 0.00000
-2.60338 0.00000 0.00000 ; HA- CA- CB- HB3
4050 4049 4051 4054 3 0.65084 1.95253 0.00000
-2.60338 0.00000 0.00000 ; HA- CA- CB- CG
4050 4049 4066 4067 3 3.68192 -4.35136 0.00000
1.33888 0.00000 0.00000 ; HA- CA- C- O
4050 4049 4066 4068 3 0.00000 0.00000 0.00000
0.00000 0.00000 0.00000 ; HA- CA- C- N
4051 4049 4066 4067 3 0.00000 0.00000 0.00000
0.00000 0.00000 0.00000 ; CB- CA- C- O

4051 4049 4066 4068 3 2.51040 4.18400 1.67360
-6.69440 0.00000 0.00000 ; CB- CA- C- N
4051 4054 4055 4056 3 30.33400 0.00000 -30.33400
0.00000 0.00000 0.00000 ; CB- CG- CD1- HD1
4051 4054 4055 4057 3 30.33400 0.00000 -30.33400
0.00000 0.00000 0.00000 ; CB- CG- CD1- CE1
4051 4054 4064 4062 3 30.33400 0.00000 -30.33400
0.00000 0.00000 0.00000 ; CB- CG- CD2- CE2
4051 4054 4064 4065 3 30.33400 0.00000 -30.33400
0.00000 0.00000 0.00000 ; CB- CG- CD2- HD2
4052 4051 4049 4066 3 0.65084 1.95253 0.00000
-2.60338 0.00000 0.00000 ; HB2- CB- CA- C
4052 4051 4054 4055 3 0.00000 0.00000 0.00000
0.00000 0.00000 0.00000 ; HB2- CB- CG- CD1
4052 4051 4054 4064 3 0.00000 0.00000 0.00000
0.00000 0.00000 0.00000 ; HB2- CB- CG- CD2
4053 4051 4049 4066 3 0.65084 1.95253 0.00000
-2.60338 0.00000 0.00000 ; HB3- CB- CA- C
4053 4051 4054 4055 3 0.00000 0.00000 0.00000
0.00000 0.00000 0.00000 ; HB3- CB- CG- CD1
4053 4051 4054 4064 3 0.00000 0.00000 0.00000
0.00000 0.00000 0.00000 ; HB3- CB- CG- CD2
4054 4051 4049 4066 3 4.95804 2.17986 -3.52293
-3.21331 -0.40166 0.00000 ; CG- CB- CA- C
4054 4055 4057 4058 3 30.33400 0.00000 -30.33400
0.00000 0.00000 0.00000 ; CG- CD1- CE1- HE1
4054 4055 4057 4059 3 30.33400 0.00000 -30.33400
0.00000 0.00000 0.00000 ; CG- CD1- CE1- CZ
4054 4064 4062 4059 3 30.33400 0.00000 -30.33400
0.00000 0.00000 0.00000 ; CG- CD2- CE2- CZ
4054 4064 4062 4063 3 30.33400 0.00000 -30.33400
0.00000 0.00000 0.00000 ; CG- CD2- CE2- HE2
4055 4054 4064 4062 3 30.33400 0.00000 -30.33400
0.00000 0.00000 0.00000 ; CD1- CG- CD2- CE2
4055 4054 4064 4065 3 30.33400 0.00000 -30.33400
0.00000 0.00000 0.00000 ; CD1- CG- CD2- HD2
4055 4057 4059 4060 3 30.33400 0.00000 -30.33400
0.00000 0.00000 0.00000 ; CD1- CE1- CZ- OH
4055 4057 4059 4062 3 30.33400 0.00000 -30.33400
0.00000 0.00000 0.00000 ; CD1- CE1- CZ- CE2
4056 4055 4054 4064 3 30.33400 0.00000 -30.33400
0.00000 0.00000 0.00000 ; HD1- CD1- CG- CD2
4056 4055 4057 4058 3 30.33400 0.00000 -30.33400
0.00000 0.00000 0.00000 ; HD1- CD1- CE1- HE1
4056 4055 4057 4059 3 30.33400 0.00000 -30.33400
0.00000 0.00000 0.00000 ; HD1- CD1- CE1- CZ
4057 4055 4054 4064 3 30.33400 0.00000 -30.33400
0.00000 0.00000 0.00000 ; CE1- CD1- CG- CD2
4057 4059 4060 4061 3 7.93286 0.00000 -9.56462
0.00000 2.17568 0.00000 ; CE1- CZ- OH- HH
4057 4059 4062 4063 3 30.33400 0.00000 -30.33400
0.00000 0.00000 0.00000 ; CE1- CZ- CE2- HE2

4057 4059 4062 4064 3 30.33400 0.00000 -30.33400
0.00000 0.00000 0.00000 ; CE1- CZ- CE2- CD2
4058 4057 4059 4060 3 30.33400 0.00000 -30.33400
0.00000 0.00000 0.00000 ; HE1- CE1- CZ- OH
4058 4057 4059 4062 3 30.33400 0.00000 -30.33400
0.00000 0.00000 0.00000 ; HE1- CE1- CZ- CE2
4059 4062 4064 4065 3 30.33400 0.00000 -30.33400
0.00000 0.00000 0.00000 ; CZ- CE2- CD2- HD2
4060 4059 4062 4063 3 30.33400 0.00000 -30.33400
0.00000 0.00000 0.00000 ; OH- CZ- CE2- HE2
4060 4059 4062 4064 3 30.33400 0.00000 -30.33400
0.00000 0.00000 0.00000 ; OH- CZ- CE2- CD2
4061 4060 4059 4062 3 7.93286 0.00000 -9.56462
0.00000 2.17568 0.00000 ; HH- OH- CZ- CE2
4063 4062 4064 4065 3 30.33400 0.00000 -30.33400
0.00000 0.00000 0.00000 ; HE2- CE2- CD2- HD2
4066 4068 4070 4071 3 0.00000 0.00000 0.00000
0.00000 0.00000 0.00000 ; C- N- CA- HA
4066 4068 4070 4072 3 11.71520 1.67360 15.06240 -
13.38880 0.00000 0.00000 ; C- N- CA- CB
4066 4068 4070 4080 3 1.75728 5.27184 2.25936
-7.02912 0.00000 0.00000 ; C- N- CA- C
4067 4066 4068 4069 3 29.28800 -8.36800 -20.92000
0.00000 0.00000 0.00000 ; O- C- N- H
4067 4066 4068 4070 3 20.92000 0.00000 -20.92000
0.00000 0.00000 0.00000 ; O- C- N- CA
4068 4070 4072 4073 3 0.65084 1.95253 0.00000
-2.60338 0.00000 0.00000 ; N- CA- CB- HB2
4068 4070 4072 4074 3 0.65084 1.95253 0.00000
-2.60338 0.00000 0.00000 ; N- CA- CB- HB3
4068 4070 4072 4075 3 5.99567 3.29281 -4.46014
-0.55229 1.97485 0.00000 ; N- CA- CB- CG
4068 4070 4080 4081 3 0.00000 0.00000 0.00000
0.00000 0.00000 0.00000 ; N- CA- C- O
4068 4070 4080 4082 3 17.40544 -5.02080 -13.22144
9.20480 0.00000 0.00000 ; N- CA- C- N
4069 4068 4070 4071 3 0.00000 0.00000 0.00000
0.00000 0.00000 0.00000 ; H- N- CA- HA
4069 4068 4070 4072 3 0.00000 0.00000 0.00000
0.00000 0.00000 0.00000 ; H- N- CA- CB
4069 4068 4070 4080 3 0.00000 0.00000 0.00000
0.00000 0.00000 0.00000 ; H- N- CA- C
4070 4072 4075 4076 3 0.00000 0.00000 0.00000
0.00000 0.00000 0.00000 ; CA- CB- CG- OD1
4070 4072 4075 4077 3 8.84916 -0.31380 -4.32626
5.03754 0.26778 0.00000 ; CA- CB- CG- ND2
4070 4080 4082 4083 3 20.92000 0.00000 -20.92000
0.00000 0.00000 0.00000 ; CA- C- N- CD
4070 4080 4082 4092 3 20.92000 0.00000 -20.92000
0.00000 0.00000 0.00000 ; CA- C- N- CA
4071 4070 4072 4073 3 0.65084 1.95253 0.00000
-2.60338 0.00000 0.00000 ; HA- CA- CB- HB2

4071 4070 4072 4074 3 0.65084 1.95253 0.00000
-2.60338 0.00000 0.00000 ; HA- CA- CB- HB3
4071 4070 4072 4075 3 0.65084 1.95253 0.00000
-2.60338 0.00000 0.00000 ; HA- CA- CB- CG
4071 4070 4080 4081 3 3.68192 -4.35136 0.00000
1.33888 0.00000 0.00000 ; HA- CA- C- O
4071 4070 4080 4082 3 0.00000 0.00000 0.00000
0.00000 0.00000 0.00000 ; HA- CA- C- N
4072 4070 4080 4081 3 0.00000 0.00000 0.00000
0.00000 0.00000 0.00000 ; CB- CA- C- O
4072 4070 4080 4082 3 2.51040 4.18400 1.67360
-6.69440 0.00000 0.00000 ; CB- CA- C- N
4072 4075 4077 4078 3 20.92000 0.00000 -20.92000
0.00000 0.00000 0.00000 ; CB- CG- ND2- HD21
4072 4075 4077 4079 3 20.92000 0.00000 -20.92000
0.00000 0.00000 0.00000 ; CB- CG- ND2- HD22
4073 4072 4070 4080 3 0.65084 1.95253 0.00000
-2.60338 0.00000 0.00000 ; HB2- CB- CA- C
4073 4072 4075 4076 3 3.68192 -4.35136 0.00000
1.33888 0.00000 0.00000 ; HB2- CB- CG- OD1
4073 4072 4075 4077 3 0.00000 0.00000 0.00000
0.00000 0.00000 0.00000 ; HB2- CB- CG- ND2
4074 4072 4070 4080 3 0.65084 1.95253 0.00000
-2.60338 0.00000 0.00000 ; HB3- CB- CA- C
4074 4072 4075 4076 3 3.68192 -4.35136 0.00000
1.33888 0.00000 0.00000 ; HB3- CB- CG- OD1
4074 4072 4075 4077 3 0.00000 0.00000 0.00000
0.00000 0.00000 0.00000 ; HB3- CB- CG- ND2
4075 4072 4070 4080 3 7.94542 -3.96225 -6.11701
-0.55229 3.58150 0.00000 ; CG- CB- CA- C
4076 4075 4077 4078 3 29.28800 -8.36800 -20.92000
0.00000 0.00000 0.00000 ; OD1- CG- ND2- HD21
4076 4075 4077 4079 3 29.28800 -8.36800 -20.92000
0.00000 0.00000 0.00000 ; OD1- CG- ND2- HD22
4080 4082 4083 4084 3 0.00000 0.00000 0.00000
0.00000 0.00000 0.00000 ; C- N- CD- HD2
4080 4082 4083 4085 3 0.00000 0.00000 0.00000
0.00000 0.00000 0.00000 ; C- N- CD- HD3
4080 4082 4083 4086 3 10.04160 -3.34720 16.73600
-6.69440 0.00000 0.00000 ; C- N- CD- CG
4080 4082 4092 4089 3 11.71520 1.67360 15.06240 -
13.38880 0.00000 0.00000 ; C- N- CA- CB
4080 4082 4092 4093 3 0.00000 0.00000 0.00000
0.00000 0.00000 0.00000 ; C- N- CA- HA
4080 4082 4092 4094 3 1.75728 5.27184 2.25936
-7.02912 0.00000 0.00000 ; C- N- CA- C
4081 4080 4082 4083 3 20.92000 0.00000 -20.92000
0.00000 0.00000 0.00000 ; O- C- N- CD
4081 4080 4082 4092 3 20.92000 0.00000 -20.92000
0.00000 0.00000 0.00000 ; O- C- N- CA
4082 4083 4086 4087 3 0.65084 1.95253 0.00000
-2.60338 0.00000 0.00000 ; N- CD- CG- HG2

```
  4082   4083   4086   4088       3    0.65084    1.95253    0.00000
-2.60338    0.00000    0.00000 ;       N-    CD-    CG-   HG3
  4082   4083   4086   4089       3    0.65084    1.95253    0.00000
-2.60338    0.00000    0.00000 ;       N-    CD-    CG-    CB
  4082   4092   4089   4086       3    0.65084    1.95253    0.00000
-2.60338    0.00000    0.00000 ;       N-    CA-    CB-    CG
  4082   4092   4089   4090       3    0.65084    1.95253    0.00000
-2.60338    0.00000    0.00000 ;       N-    CA-    CB-   HB2
  4082   4092   4089   4091       3    0.65084    1.95253    0.00000
-2.60338    0.00000    0.00000 ;       N-    CA-    CB-   HB3
  4082   4092   4094   4095       3    0.00000    0.00000    0.00000
0.00000    0.00000    0.00000 ;       N-    CA-     C-     O
  4082   4092   4094   4096       3   17.40544   -5.02080  -13.22144
9.20480    0.00000    0.00000 ;       N-    CA-     C-     N
  4083   4082   4092   4089       3    0.00000    0.00000    0.00000
0.00000    0.00000    0.00000 ;      CD-     N-    CA-    CB
  4083   4082   4092   4093       3    0.00000    0.00000    0.00000
0.00000    0.00000    0.00000 ;      CD-     N-    CA-    HA
  4083   4082   4092   4094       3    0.00000    0.00000    0.00000
0.00000    0.00000    0.00000 ;      CD-     N-    CA-     C
  4083   4086   4089   4090       3    0.66944    2.00832    0.00000
-2.67776    0.00000    0.00000 ;      CD-    CG-    CB-   HB2
  4083   4086   4089   4091       3    0.66944    2.00832    0.00000
-2.67776    0.00000    0.00000 ;      CD-    CG-    CB-   HB3
  4083   4086   4089   4092       3    3.68192    3.09616   -2.09200
-3.01248    0.00000    0.00000 ;      CD-    CG-    CB-    CA
  4084   4083   4082   4092       3    0.00000    0.00000    0.00000
0.00000    0.00000    0.00000 ;     HD2-    CD-     N-    CA
  4084   4083   4086   4087       3    0.65084    1.95253    0.00000
-2.60338    0.00000    0.00000 ;     HD2-    CD-    CG-   HG2
  4084   4083   4086   4088       3    0.65084    1.95253    0.00000
-2.60338    0.00000    0.00000 ;     HD2-    CD-    CG-   HG3
  4084   4083   4086   4089       3    0.65084    1.95253    0.00000
-2.60338    0.00000    0.00000 ;     HD2-    CD-    CG-    CB
  4085   4083   4082   4092       3    0.00000    0.00000    0.00000
0.00000    0.00000    0.00000 ;     HD3-    CD-     N-    CA
  4085   4083   4086   4087       3    0.65084    1.95253    0.00000
-2.60338    0.00000    0.00000 ;     HD3-    CD-    CG-   HG2
  4085   4083   4086   4088       3    0.65084    1.95253    0.00000
-2.60338    0.00000    0.00000 ;     HD3-    CD-    CG-   HG3
  4085   4083   4086   4089       3    0.65084    1.95253    0.00000
-2.60338    0.00000    0.00000 ;     HD3-    CD-    CG-    CB
  4086   4083   4082   4092       3    0.00000    0.00000    0.00000
0.00000    0.00000    0.00000 ;      CG-    CD-     N-    CA
  4086   4089   4092   4093       3    0.65084    1.95253    0.00000
-2.60338    0.00000    0.00000 ;      CG-    CB-    CA-    HA
  4086   4089   4092   4094       3    0.65084    1.95253    0.00000
-2.60338    0.00000    0.00000 ;      CG-    CB-    CA-     C
  4087   4086   4089   4090       3    0.62760    1.88280    0.00000
-2.51040    0.00000    0.00000 ;     HG2-    CG-    CB-   HB2
  4087   4086   4089   4091       3    0.62760    1.88280    0.00000
-2.51040    0.00000    0.00000 ;     HG2-    CG-    CB-   HB3
```

4087 4086 4089 4092 3 0.66944 2.00832 0.00000
-2.67776 0.00000 0.00000 ; HG2- CG- CB- CA
4088 4086 4089 4090 3 0.62760 1.88280 0.00000
-2.51040 0.00000 0.00000 ; HG3- CG- CB- HB2
4088 4086 4089 4091 3 0.62760 1.88280 0.00000
-2.51040 0.00000 0.00000 ; HG3- CG- CB- HB3
4088 4086 4089 4092 3 0.66944 2.00832 0.00000
-2.67776 0.00000 0.00000 ; HG3- CG- CB- CA
4089 4092 4094 4095 3 0.00000 0.00000 0.00000
0.00000 0.00000 0.00000 ; CB- CA- C- O
4089 4092 4094 4096 3 2.51040 4.18400 1.67360
-6.69440 0.00000 0.00000 ; CB- CA- C- N
4090 4089 4092 4093 3 0.65084 1.95253 0.00000
-2.60338 0.00000 0.00000 ; HB2- CB- CA- HA
4090 4089 4092 4094 3 0.65084 1.95253 0.00000
-2.60338 0.00000 0.00000 ; HB2- CB- CA- C
4091 4089 4092 4093 3 0.65084 1.95253 0.00000
-2.60338 0.00000 0.00000 ; HB3- CB- CA- HA
4091 4089 4092 4094 3 0.65084 1.95253 0.00000
-2.60338 0.00000 0.00000 ; HB3- CB- CA- C
4092 4094 4096 4097 3 20.92000 0.00000 -20.92000
0.00000 0.00000 0.00000 ; CA- C- N- H
4092 4094 4096 4098 3 20.92000 0.00000 -20.92000
0.00000 0.00000 0.00000 ; CA- C- N- CA
4093 4092 4094 4095 3 3.68192 -4.35136 0.00000
1.33888 0.00000 0.00000 ; HA- CA- C- O
4093 4092 4094 4096 3 0.00000 0.00000 0.00000
0.00000 0.00000 0.00000 ; HA- CA- C- N
4094 4096 4098 4099 3 0.00000 0.00000 0.00000
0.00000 0.00000 0.00000 ; C- N- CA- HA
4094 4096 4098 4100 3 11.71520 1.67360 15.06240 -
13.38880 0.00000 0.00000 ; C- N- CA- CB
4094 4096 4098 4108 3 1.75728 5.27184 2.25936
-7.02912 0.00000 0.00000 ; C- N- CA- C
4095 4094 4096 4097 3 29.28800 -8.36800 -20.92000
0.00000 0.00000 0.00000 ; O- C- N- H
4095 4094 4096 4098 3 20.92000 0.00000 -20.92000
0.00000 0.00000 0.00000 ; O- C- N- CA
4096 4098 4100 4101 3 0.65084 1.95253 0.00000
-2.60338 0.00000 0.00000 ; N- CA- CB- HB2
4096 4098 4100 4102 3 0.65084 1.95253 0.00000
-2.60338 0.00000 0.00000 ; N- CA- CB- HB3
4096 4098 4100 4103 3 5.99567 3.29281 -4.46014
-0.55229 1.97485 0.00000 ; N- CA- CB- CG
4096 4098 4108 4109 3 0.00000 0.00000 0.00000
0.00000 0.00000 0.00000 ; N- CA- C- O
4096 4098 4108 4110 3 17.40544 -5.02080 -13.22144
9.20480 0.00000 0.00000 ; N- CA- C- N
4097 4096 4098 4099 3 0.00000 0.00000 0.00000
0.00000 0.00000 0.00000 ; H- N- CA- HA
4097 4096 4098 4100 3 0.00000 0.00000 0.00000
0.00000 0.00000 0.00000 ; H- N- CA- CB

4097 4096 4098 4108 3 0.00000 0.00000 0.00000
0.00000 0.00000 0.00000 ; H- N- CA- C
4098 4100 4103 4104 3 0.00000 0.00000 0.00000
0.00000 0.00000 0.00000 ; CA- CB- CG- OD1
4098 4100 4103 4105 3 8.84916 -0.31380 -4.32626
5.03754 0.26778 0.00000 ; CA- CB- CG- ND2
4098 4108 4110 4111 3 20.92000 0.00000 -20.92000
0.00000 0.00000 0.00000 ; CA- C- N- H
4098 4108 4110 4112 3 20.92000 0.00000 -20.92000
0.00000 0.00000 0.00000 ; CA- C- N- CA
4099 4098 4100 4101 3 0.65084 1.95253 0.00000
-2.60338 0.00000 0.00000 ; HA- CA- CB- HB2
4099 4098 4100 4102 3 0.65084 1.95253 0.00000
-2.60338 0.00000 0.00000 ; HA- CA- CB- HB3
4099 4098 4100 4103 3 0.65084 1.95253 0.00000
-2.60338 0.00000 0.00000 ; HA- CA- CB- CG
4099 4098 4108 4109 3 3.68192 -4.35136 0.00000
1.33888 0.00000 0.00000 ; HA- CA- C- O
4099 4098 4108 4110 3 0.00000 0.00000 0.00000
0.00000 0.00000 0.00000 ; HA- CA- C- N
4100 4098 4108 4109 3 0.00000 0.00000 0.00000
0.00000 0.00000 0.00000 ; CB- CA- C- O
4100 4098 4108 4110 3 2.51040 4.18400 1.67360
-6.69440 0.00000 0.00000 ; CB- CA- C- N
4100 4103 4105 4106 3 20.92000 0.00000 -20.92000
0.00000 0.00000 0.00000 ; CB- CG- ND2- HD21
4100 4103 4105 4107 3 20.92000 0.00000 -20.92000
0.00000 0.00000 0.00000 ; CB- CG- ND2- HD22
4101 4100 4098 4108 3 0.65084 1.95253 0.00000
-2.60338 0.00000 0.00000 ; HB2- CB- CA- C
4101 4100 4103 4104 3 3.68192 -4.35136 0.00000
1.33888 0.00000 0.00000 ; HB2- CB- CG- OD1
4101 4100 4103 4105 3 0.00000 0.00000 0.00000
0.00000 0.00000 0.00000 ; HB2- CB- CG- ND2
4102 4100 4098 4108 3 0.65084 1.95253 0.00000
-2.60338 0.00000 0.00000 ; HB3- CB- CA- C
4102 4100 4103 4104 3 3.68192 -4.35136 0.00000
1.33888 0.00000 0.00000 ; HB3- CB- CG- OD1
4102 4100 4103 4105 3 0.00000 0.00000 0.00000
0.00000 0.00000 0.00000 ; HB3- CB- CG- ND2
4103 4100 4098 4108 3 7.94542 -3.96225 -6.11701
-0.55229 3.58150 0.00000 ; CG- CB- CA- C
4104 4103 4105 4106 3 29.28800 -8.36800 -20.92000
0.00000 0.00000 0.00000 ; OD1- CG- ND2- HD21
4104 4103 4105 4107 3 29.28800 -8.36800 -20.92000
0.00000 0.00000 0.00000 ; OD1- CG- ND2- HD22
4108 4110 4112 4113 3 0.00000 0.00000 0.00000
0.00000 0.00000 0.00000 ; C- N- CA- HA
4108 4110 4112 4114 3 11.71520 1.67360 15.06240 -
13.38880 0.00000 0.00000 ; C- N- CA- CB
4108 4110 4112 4123 3 1.75728 5.27184 2.25936
-7.02912 0.00000 0.00000 ; C- N- CA- C

4109 4108 4110 4111 3 29.28800 -8.36800 -20.92000
0.00000 0.00000 0.00000 ; O- C- N- H
4109 4108 4110 4112 3 20.92000 0.00000 -20.92000
0.00000 0.00000 0.00000 ; O- C- N- CA
4110 4112 4114 4115 3 0.65084 1.95253 0.00000
-2.60338 0.00000 0.00000 ; N- CA- CB- HB2
4110 4112 4114 4116 3 0.65084 1.95253 0.00000
-2.60338 0.00000 0.00000 ; N- CA- CB- HB3
4110 4112 4114 4117 3 3.21331 2.22589 -4.15053
-2.40998 2.61082 0.00000 ; N- CA- CB- CG
4110 4112 4123 4124 3 0.00000 0.00000 0.00000
0.00000 0.00000 0.00000 ; N- CA- C- O
4110 4112 4123 4125 3 17.40544 -5.02080 -13.22144
9.20480 0.00000 0.00000 ; N- CA- C- N
4111 4110 4112 4113 3 0.00000 0.00000 0.00000
0.00000 0.00000 0.00000 ; H- N- CA- HA
4111 4110 4112 4114 3 0.00000 0.00000 0.00000
0.00000 0.00000 0.00000 ; H- N- CA- CB
4111 4110 4112 4123 3 0.00000 0.00000 0.00000
0.00000 0.00000 0.00000 ; H- N- CA- C
4112 4114 4117 4118 3 0.66944 2.00832 0.00000
-2.67776 0.00000 0.00000 ; CA- CB- CG- HG2
4112 4114 4117 4119 3 0.66944 2.00832 0.00000
-2.67776 0.00000 0.00000 ; CA- CB- CG- HG3
4112 4114 4117 4120 3 10.12110 -1.91209 0.01674
10.17549 -1.87443 0.00000 ; CA- CB- CG- CD
4112 4123 4125 4126 3 20.92000 0.00000 -20.92000
0.00000 0.00000 0.00000 ; CA- C- N- H
4112 4123 4125 4127 3 20.92000 0.00000 -20.92000
0.00000 0.00000 0.00000 ; CA- C- N- CA
4113 4112 4114 4115 3 0.65084 1.95253 0.00000
-2.60338 0.00000 0.00000 ; HA- CA- CB- HB2
4113 4112 4114 4116 3 0.65084 1.95253 0.00000
-2.60338 0.00000 0.00000 ; HA- CA- CB- HB3
4113 4112 4114 4117 3 0.65084 1.95253 0.00000
-2.60338 0.00000 0.00000 ; HA- CA- CB- CG
4113 4112 4123 4124 3 3.68192 -4.35136 0.00000
1.33888 0.00000 0.00000 ; HA- CA- C- O
4113 4112 4123 4125 3 0.00000 0.00000 0.00000
0.00000 0.00000 0.00000 ; HA- CA- C- N
4114 4112 4123 4124 3 0.00000 0.00000 0.00000
0.00000 0.00000 0.00000 ; CB- CA- C- O
4114 4112 4123 4125 3 2.51040 4.18400 1.67360
-6.69440 0.00000 0.00000 ; CB- CA- C- N
4114 4117 4120 4121 3 3.79907 0.00000 -5.40573
0.00000 2.14221 0.00000 ; CB- CG- CD- OE1
4114 4117 4120 4122 3 3.79907 0.00000 -5.40573
0.00000 2.14221 0.00000 ; CB- CG- CD- OE2
4115 4114 4112 4123 3 0.65084 1.95253 0.00000
-2.60338 0.00000 0.00000 ; HB2- CB- CA- C
4115 4114 4117 4118 3 0.62760 1.88280 0.00000
-2.51040 0.00000 0.00000 ; HB2- CB- CG- HG2

4115 4114 4117 4119 3 0.62760 1.88280 0.00000
-2.51040 0.00000 0.00000 ; HB2- CB- CG- HG3
4115 4114 4117 4120 3 0.65084 1.95253 0.00000
-2.60338 0.00000 0.00000 ; HB2- CB- CG- CD
4116 4114 4112 4123 3 0.65084 1.95253 0.00000
-2.60338 0.00000 0.00000 ; HB3- CB- CA- C
4116 4114 4117 4118 3 0.62760 1.88280 0.00000
-2.51040 0.00000 0.00000 ; HB3- CB- CG- HG2
4116 4114 4117 4119 3 0.62760 1.88280 0.00000
-2.51040 0.00000 0.00000 ; HB3- CB- CG- HG3
4116 4114 4117 4120 3 0.65084 1.95253 0.00000
-2.60338 0.00000 0.00000 ; HB3- CB- CG- CD
4117 4114 4112 4123 3 6.86594 3.56895 -8.14206
-2.40998 4.85344 0.00000 ; CG- CB- CA- C
4118 4117 4120 4121 3 0.00000 0.00000 0.00000
0.00000 0.00000 0.00000 ; HG2- CG- CD- OE1
4118 4117 4120 4122 3 0.00000 0.00000 0.00000
0.00000 0.00000 0.00000 ; HG2- CG- CD- OE2
4119 4117 4120 4121 3 0.00000 0.00000 0.00000
0.00000 0.00000 0.00000 ; HG3- CG- CD- OE1
4119 4117 4120 4122 3 0.00000 0.00000 0.00000
0.00000 0.00000 0.00000 ; HG3- CG- CD- OE2
4123 4125 4127 4128 3 0.00000 0.00000 0.00000
0.00000 0.00000 0.00000 ; C- N- CA- HA
4123 4125 4127 4129 3 11.71520 1.67360 15.06240 -
13.38880 0.00000 0.00000 ; C- N- CA- CB
4123 4125 4127 4147 3 1.75728 5.27184 2.25936
-7.02912 0.00000 0.00000 ; C- N- CA- C
4124 4123 4125 4126 3 29.28800 -8.36800 -20.92000
0.00000 0.00000 0.00000 ; O- C- N- H
4124 4123 4125 4127 3 20.92000 0.00000 -20.92000
0.00000 0.00000 0.00000 ; O- C- N- CA
4125 4127 4129 4130 3 0.65084 1.95253 0.00000
-2.60338 0.00000 0.00000 ; N- CA- CB- HB2
4125 4127 4129 4131 3 0.65084 1.95253 0.00000
-2.60338 0.00000 0.00000 ; N- CA- CB- HB3
4125 4127 4129 4132 3 0.65084 1.95253 0.00000
-2.60338 0.00000 0.00000 ; N- CA- CB- CG
4125 4127 4147 4148 3 0.00000 0.00000 0.00000
0.00000 0.00000 0.00000 ; N- CA- C- O
4125 4127 4147 4149 3 17.40544 -5.02080 -13.22144
9.20480 0.00000 0.00000 ; N- CA- C- N
4126 4125 4127 4128 3 0.00000 0.00000 0.00000
0.00000 0.00000 0.00000 ; H- N- CA- HA
4126 4125 4127 4129 3 0.00000 0.00000 0.00000
0.00000 0.00000 0.00000 ; H- N- CA- CB
4126 4125 4127 4147 3 0.00000 0.00000 0.00000
0.00000 0.00000 0.00000 ; H- N- CA- C
4127 4129 4132 4133 3 0.66944 2.00832 0.00000
-2.67776 0.00000 0.00000 ; CA- CB- CG- HG2
4127 4129 4132 4134 3 0.66944 2.00832 0.00000
-2.67776 0.00000 0.00000 ; CA- CB- CG- HG3

4127 4129 4132 4135 3 3.68192 3.09616 -2.09200
-3.01248 0.00000 0.00000 ; CA- CB- CG- CD
4127 4147 4149 4150 3 20.92000 0.00000 -20.92000
0.00000 0.00000 0.00000 ; CA- C- N- H
4127 4147 4149 4151 3 20.92000 0.00000 -20.92000
0.00000 0.00000 0.00000 ; CA- C- N- CA
4128 4127 4129 4130 3 0.65084 1.95253 0.00000
-2.60338 0.00000 0.00000 ; HA- CA- CB- HB2
4128 4127 4129 4131 3 0.65084 1.95253 0.00000
-2.60338 0.00000 0.00000 ; HA- CA- CB- HB3
4128 4127 4129 4132 3 0.65084 1.95253 0.00000
-2.60338 0.00000 0.00000 ; HA- CA- CB- CG
4128 4127 4147 4148 3 3.68192 -4.35136 0.00000
1.33888 0.00000 0.00000 ; HA- CA- C- O
4128 4127 4147 4149 3 0.00000 0.00000 0.00000
0.00000 0.00000 0.00000 ; HA- CA- C- N
4129 4127 4147 4148 3 0.00000 0.00000 0.00000
0.00000 0.00000 0.00000 ; CB- CA- C- O
4129 4127 4147 4149 3 2.51040 4.18400 1.67360
-6.69440 0.00000 0.00000 ; CB- CA- C- N
4129 4132 4135 4136 3 0.65084 1.95253 0.00000
-2.60338 0.00000 0.00000 ; CB- CG- CD- HD2
4129 4132 4135 4137 3 0.65084 1.95253 0.00000
-2.60338 0.00000 0.00000 ; CB- CG- CD- HD3
4129 4132 4135 4138 3 0.65084 1.95253 0.00000
-2.60338 0.00000 0.00000 ; CB- CG- CD- NE
4130 4129 4127 4147 3 0.65084 1.95253 0.00000
-2.60338 0.00000 0.00000 ; HB2- CB- CA- C
4130 4129 4132 4133 3 0.62760 1.88280 0.00000
-2.51040 0.00000 0.00000 ; HB2- CB- CG- HG2
4130 4129 4132 4134 3 0.62760 1.88280 0.00000
-2.51040 0.00000 0.00000 ; HB2- CB- CG- HG3
4130 4129 4132 4135 3 0.66944 2.00832 0.00000
-2.67776 0.00000 0.00000 ; HB2- CB- CG- CD
4131 4129 4127 4147 3 0.65084 1.95253 0.00000
-2.60338 0.00000 0.00000 ; HB3- CB- CA- C
4131 4129 4132 4133 3 0.62760 1.88280 0.00000
-2.51040 0.00000 0.00000 ; HB3- CB- CG- HG2
4131 4129 4132 4134 3 0.62760 1.88280 0.00000
-2.51040 0.00000 0.00000 ; HB3- CB- CG- HG3
4131 4129 4132 4135 3 0.66944 2.00832 0.00000
-2.67776 0.00000 0.00000 ; HB3- CB- CG- CD
4132 4129 4127 4147 3 0.65084 1.95253 0.00000
-2.60338 0.00000 0.00000 ; CG- CB- CA- C
4132 4135 4138 4139 3 0.00000 0.00000 0.00000
0.00000 0.00000 0.00000 ; CG- CD- NE- HE
4132 4135 4138 4140 3 0.00000 0.00000 0.00000
0.00000 0.00000 0.00000 ; CG- CD- NE- CZ
4133 4132 4135 4136 3 0.65084 1.95253 0.00000
-2.60338 0.00000 0.00000 ; HG2- CG- CD- HD2
4133 4132 4135 4137 3 0.65084 1.95253 0.00000
-2.60338 0.00000 0.00000 ; HG2- CG- CD- HD3

```
  4133   4132   4135   4138       3    0.65084    1.95253    0.00000
-2.60338    0.00000    0.00000 ;    HG2-    CG-    CD-    NE
  4134   4132   4135   4136       3    0.65084    1.95253    0.00000
-2.60338    0.00000    0.00000 ;    HG3-    CG-    CD-   HD2
  4134   4132   4135   4137       3    0.65084    1.95253    0.00000
-2.60338    0.00000    0.00000 ;    HG3-    CG-    CD-   HD3
  4134   4132   4135   4138       3    0.65084    1.95253    0.00000
-2.60338    0.00000    0.00000 ;    HG3-    CG-    CD-    NE
  4135   4138   4140   4141       3   20.08320    0.00000  -20.08320
0.00000    0.00000    0.00000 ;     CD-    NE-    CZ-   NH1
  4135   4138   4140   4144       3   20.08320    0.00000  -20.08320
0.00000    0.00000    0.00000 ;     CD-    NE-    CZ-   NH2
  4136   4135   4138   4139       3    0.00000    0.00000    0.00000
0.00000    0.00000    0.00000 ;    HD2-    CD-    NE-    HE
  4136   4135   4138   4140       3    0.00000    0.00000    0.00000
0.00000    0.00000    0.00000 ;    HD2-    CD-    NE-    CZ
  4137   4135   4138   4139       3    0.00000    0.00000    0.00000
0.00000    0.00000    0.00000 ;    HD3-    CD-    NE-    HE
  4137   4135   4138   4140       3    0.00000    0.00000    0.00000
0.00000    0.00000    0.00000 ;    HD3-    CD-    NE-    CZ
  4138   4140   4141   4142       3   20.08320    0.00000  -20.08320
0.00000    0.00000    0.00000 ;     NE-    CZ-   NH1-  HH11
  4138   4140   4141   4143       3   20.08320    0.00000  -20.08320
0.00000    0.00000    0.00000 ;     NE-    CZ-   NH1-  HH12
  4138   4140   4144   4145       3   20.08320    0.00000  -20.08320
0.00000    0.00000    0.00000 ;     NE-    CZ-   NH2-  HH21
  4138   4140   4144   4146       3   20.08320    0.00000  -20.08320
0.00000    0.00000    0.00000 ;     NE-    CZ-   NH2-  HH22
  4139   4138   4140   4141       3   20.08320    0.00000  -20.08320
0.00000    0.00000    0.00000 ;     HE-    NE-    CZ-   NH1
  4139   4138   4140   4144       3   20.08320    0.00000  -20.08320
0.00000    0.00000    0.00000 ;     HE-    NE-    CZ-   NH2
  4141   4140   4144   4145       3   20.08320    0.00000  -20.08320
0.00000    0.00000    0.00000 ;    NH1-    CZ-   NH2-  HH21
  4141   4140   4144   4146       3   20.08320    0.00000  -20.08320
0.00000    0.00000    0.00000 ;    NH1-    CZ-   NH2-  HH22
  4142   4141   4140   4144       3   20.08320    0.00000  -20.08320
0.00000    0.00000    0.00000 ;   HH11-   NH1-    CZ-   NH2
  4143   4141   4140   4144       3   20.08320    0.00000  -20.08320
0.00000    0.00000    0.00000 ;   HH12-   NH1-    CZ-   NH2
  4147   4149   4151   4152       3    0.00000    0.00000    0.00000
0.00000    0.00000    0.00000 ;      C-     N-    CA-    HA
  4147   4149   4151   4153       3   11.71520    1.67360   15.06240  -
13.38880    0.00000    0.00000 ;      C-     N-    CA-    CB
  4147   4149   4151   4158       3    1.75728    5.27184    2.25936
-7.02912    0.00000    0.00000 ;      C-     N-    CA-     C
  4148   4147   4149   4150       3   29.28800   -8.36800  -20.92000
0.00000    0.00000    0.00000 ;      O-     C-     N-     H
  4148   4147   4149   4151       3   20.92000    0.00000  -20.92000
0.00000    0.00000    0.00000 ;      O-     C-     N-    CA
  4149   4151   4153   4154       3    0.65084    1.95253    0.00000
-2.60338    0.00000    0.00000 ;      N-    CA-    CB-   HB2
```

4149 4151 4153 4155 3 0.65084 1.95253 0.00000
-2.60338 0.00000 0.00000 ; N- CA- CB- HB3
4149 4151 4153 4156 3 6.03751 2.50622 -5.17142
-4.20074 1.10458 0.00000 ; N- CA- CB- SG
4149 4151 4158 4159 3 0.00000 0.00000 0.00000
0.00000 0.00000 0.00000 ; N- CA- C- O
4149 4151 4158 4160 3 17.40544 -5.02080 -13.22144
9.20480 0.00000 0.00000 ; N- CA- C- N
4150 4149 4151 4152 3 0.00000 0.00000 0.00000
0.00000 0.00000 0.00000 ; H- N- CA- HA
4150 4149 4151 4153 3 0.00000 0.00000 0.00000
0.00000 0.00000 0.00000 ; H- N- CA- CB
4150 4149 4151 4158 3 0.00000 0.00000 0.00000
0.00000 0.00000 0.00000 ; H- N- CA- C
4151 4153 4156 4157 3 1.69034 2.77818 4.11706
-4.21747 1.00416 0.00000 ; CA- CB- SG- HG
4151 4158 4160 4161 3 20.92000 0.00000 -20.92000
0.00000 0.00000 0.00000 ; CA- C- N- H
4151 4158 4160 4162 3 20.92000 0.00000 -20.92000
0.00000 0.00000 0.00000 ; CA- C- N- CA
4152 4151 4153 4154 3 0.65084 1.95253 0.00000
-2.60338 0.00000 0.00000 ; HA- CA- CB- HB2
4152 4151 4153 4155 3 0.65084 1.95253 0.00000
-2.60338 0.00000 0.00000 ; HA- CA- CB- HB3
4152 4151 4153 4156 3 0.65084 1.95253 0.00000
-2.60338 0.00000 0.00000 ; HA- CA- CB- SG
4152 4151 4158 4159 3 3.68192 -4.35136 0.00000
1.33888 0.00000 0.00000 ; HA- CA- C- O
4152 4151 4158 4160 3 0.00000 0.00000 0.00000
0.00000 0.00000 0.00000 ; HA- CA- C- N
4153 4151 4158 4159 3 0.00000 0.00000 0.00000
0.00000 0.00000 0.00000 ; CB- CA- C- O
4153 4151 4158 4160 3 2.51040 4.18400 1.67360
-6.69440 0.00000 0.00000 ; CB- CA- C- N
4154 4153 4151 4158 3 0.65084 1.95253 0.00000
-2.60338 0.00000 0.00000 ; HB2- CB- CA- C
4154 4153 4156 4157 3 1.04600 3.13800 0.00000
-4.18400 0.00000 0.00000 ; HB2- CB- SG- HG
4155 4153 4151 4158 3 0.65084 1.95253 0.00000
-2.60338 0.00000 0.00000 ; HB3- CB- CA- C
4155 4153 4156 4157 3 1.04600 3.13800 0.00000
-4.18400 0.00000 0.00000 ; HB3- CB- SG- HG
4156 4153 4151 4158 3 5.62330 4.27605 -5.33042
-4.20074 2.51040 0.00000 ; SG- CB- CA- C
4158 4160 4162 4163 3 0.00000 0.00000 0.00000
0.00000 0.00000 0.00000 ; C- N- CA- HA
4158 4160 4162 4164 3 11.71520 1.67360 15.06240 -
13.38880 0.00000 0.00000 ; C- N- CA- CB
4158 4160 4162 4172 3 1.75728 5.27184 2.25936
-7.02912 0.00000 0.00000 ; C- N- CA- C
4159 4158 4160 4161 3 29.28800 -8.36800 -20.92000
0.00000 0.00000 0.00000 ; O- C- N- H

4159 4158 4160 4162 3 20.92000 0.00000 -20.92000
0.00000 0.00000 0.00000 ; O- C- N- CA
4160 4162 4164 4165 3 0.65084 1.95253 0.00000
-2.60338 0.00000 0.00000 ; N- CA- CB- HB
4160 4162 4164 4166 3 3.83673 0.44769 -1.77402
-2.47693 -0.03347 0.00000 ; N- CA- CB- CG2
4160 4162 4164 4170 3 4.93294 1.13386 -3.12963
-5.27184 3.17984 0.00000 ; N- CA- CB- OG1
4160 4162 4172 4173 3 0.00000 0.00000 0.00000
0.00000 0.00000 0.00000 ; N- CA- C- O
4160 4162 4172 4174 3 17.40544 -5.02080 -13.22144
9.20480 0.00000 0.00000 ; N- CA- C- N
4161 4160 4162 4163 3 0.00000 0.00000 0.00000
0.00000 0.00000 0.00000 ; H- N- CA- HA
4161 4160 4162 4164 3 0.00000 0.00000 0.00000
0.00000 0.00000 0.00000 ; H- N- CA- CB
4161 4160 4162 4172 3 0.00000 0.00000 0.00000
0.00000 0.00000 0.00000 ; H- N- CA- C
4162 4164 4166 4167 3 0.66944 2.00832 0.00000
-2.67776 0.00000 0.00000 ; CA- CB- CG2- HG21
4162 4164 4166 4168 3 0.66944 2.00832 0.00000
-2.67776 0.00000 0.00000 ; CA- CB- CG2- HG22
4162 4164 4166 4169 3 0.66944 2.00832 0.00000
-2.67776 0.00000 0.00000 ; CA- CB- CG2- HG23
4162 4164 4170 4171 3 1.12131 2.98738 1.66523
-3.94970 0.43514 0.00000 ; CA- CB- OG1- HG1
4162 4172 4174 4175 3 20.92000 0.00000 -20.92000
0.00000 0.00000 0.00000 ; CA- C- N- H
4162 4172 4174 4176 3 20.92000 0.00000 -20.92000
0.00000 0.00000 0.00000 ; CA- C- N- CA
4163 4162 4164 4165 3 0.65084 1.95253 0.00000
-2.60338 0.00000 0.00000 ; HA- CA- CB- HB
4163 4162 4164 4166 3 0.65084 1.95253 0.00000
-2.60338 0.00000 0.00000 ; HA- CA- CB- CG2
4163 4162 4164 4170 3 1.04600 -1.04600 0.00000
0.00000 0.00000 0.00000 ; HA- CA- CB- OG1
4163 4162 4172 4173 3 3.68192 -4.35136 0.00000
1.33888 0.00000 0.00000 ; HA- CA- C- O
4163 4162 4172 4174 3 0.00000 0.00000 0.00000
0.00000 0.00000 0.00000 ; HA- CA- C- N
4164 4162 4172 4173 3 0.00000 0.00000 0.00000
0.00000 0.00000 0.00000 ; CB- CA- C- O
4164 4162 4172 4174 3 2.51040 4.18400 1.67360
-6.69440 0.00000 0.00000 ; CB- CA- C- N
4165 4164 4162 4172 3 0.65084 1.95253 0.00000
-2.60338 0.00000 0.00000 ; HB- CB- CA- C
4165 4164 4166 4167 3 0.65084 1.95253 0.00000
-2.60338 0.00000 0.00000 ; HB- CB- CG2- HG21
4165 4164 4166 4168 3 0.65084 1.95253 0.00000
-2.60338 0.00000 0.00000 ; HB- CB- CG2- HG22
4165 4164 4166 4169 3 0.65084 1.95253 0.00000
-2.60338 0.00000 0.00000 ; HB- CB- CG2- HG23

4165 4164 4170 4171 3 0.69733 2.09200 0.00000
-2.78933 0.00000 0.00000 ; HB- CB- OG1- HG1
4166 4164 4162 4172 3 5.67350 3.55640 -6.16722
-2.47693 3.74886 0.00000 ; CG2- CB- CA- C
4166 4164 4170 4171 3 4.74047 0.27196 -2.26773
-3.94970 1.60666 0.00000 ; CG2- CB- OG1- HG1
4167 4166 4164 4170 3 1.04600 -1.04600 0.00000
0.00000 0.00000 0.00000 ; HG21- CG2- CB- OG1
4168 4166 4164 4170 3 1.04600 -1.04600 0.00000
0.00000 0.00000 0.00000 ; HG22- CG2- CB- OG1
4169 4166 4164 4170 3 1.04600 -1.04600 0.00000
0.00000 0.00000 0.00000 ; HG23- CG2- CB- OG1
4170 4164 4162 4172 3 6.53541 6.87013 -6.21742
-5.27184 5.22163 0.00000 ; OG1- CB- CA- C
4172 4174 4176 4177 3 0.00000 0.00000 0.00000
0.00000 0.00000 0.00000 ; C- N- CA- HA
4172 4174 4176 4178 3 11.71520 1.67360 15.06240 -
13.38880 0.00000 0.00000 ; C- N- CA- CB
4172 4174 4176 4191 3 1.75728 5.27184 2.25936
-7.02912 0.00000 0.00000 ; C- N- CA- C
4173 4172 4174 4175 3 29.28800 -8.36800 -20.92000
0.00000 0.00000 0.00000 ; O- C- N- H
4173 4172 4174 4176 3 20.92000 0.00000 -20.92000
0.00000 0.00000 0.00000 ; O- C- N- CA
4174 4176 4178 4179 3 0.65084 1.95253 0.00000
-2.60338 0.00000 0.00000 ; N- CA- CB- HB
4174 4176 4178 4180 3 3.83673 0.44769 -1.77402
-2.47693 -0.03347 0.00000 ; N- CA- CB- CG2
4174 4176 4178 4184 3 2.97482 0.12134 2.04179
-1.89117 -3.24678 0.00000 ; N- CA- CB- CG1
4174 4176 4191 4192 3 0.00000 0.00000 0.00000
0.00000 0.00000 0.00000 ; N- CA- C- O
4174 4176 4191 4193 3 17.40544 -5.02080 -13.22144
9.20480 0.00000 0.00000 ; N- CA- C- N
4175 4174 4176 4177 3 0.00000 0.00000 0.00000
0.00000 0.00000 0.00000 ; H- N- CA- HA
4175 4174 4176 4178 3 0.00000 0.00000 0.00000
0.00000 0.00000 0.00000 ; H- N- CA- CB
4175 4174 4176 4191 3 0.00000 0.00000 0.00000
0.00000 0.00000 0.00000 ; H- N- CA- C
4176 4178 4180 4181 3 0.66944 2.00832 0.00000
-2.67776 0.00000 0.00000 ; CA- CB- CG2- HG21
4176 4178 4180 4182 3 0.66944 2.00832 0.00000
-2.67776 0.00000 0.00000 ; CA- CB- CG2- HG22
4176 4178 4180 4183 3 0.66944 2.00832 0.00000
-2.67776 0.00000 0.00000 ; CA- CB- CG2- HG23
4176 4178 4184 4185 3 0.66944 2.00832 0.00000
-2.67776 0.00000 0.00000 ; CA- CB- CG1- HG12
4176 4178 4184 4186 3 0.66944 2.00832 0.00000
-2.67776 0.00000 0.00000 ; CA- CB- CG1- HG13
4176 4178 4184 4187 3 4.24258 -0.52718 -7.25506
-1.79075 7.69856 0.00000 ; CA- CB- CG1- CD1

4176 4191 4193 4194 3 20.92000 0.00000 -20.92000
0.00000 0.00000 0.00000 ; CA- C- N- H
4176 4191 4193 4195 3 20.92000 0.00000 -20.92000
0.00000 0.00000 0.00000 ; CA- C- N- CA
4177 4176 4178 4179 3 0.65084 1.95253 0.00000
-2.60338 0.00000 0.00000 ; HA- CA- CB- HB
4177 4176 4178 4180 3 0.65084 1.95253 0.00000
-2.60338 0.00000 0.00000 ; HA- CA- CB- CG2
4177 4176 4178 4184 3 0.65084 1.95253 0.00000
-2.60338 0.00000 0.00000 ; HA- CA- CB- CG1
4177 4176 4191 4192 3 3.68192 -4.35136 0.00000
1.33888 0.00000 0.00000 ; HA- CA- C- O
4177 4176 4191 4193 3 0.00000 0.00000 0.00000
0.00000 0.00000 0.00000 ; HA- CA- C- N
4178 4176 4191 4192 3 0.00000 0.00000 0.00000
0.00000 0.00000 0.00000 ; CB- CA- C- O
4178 4176 4191 4193 3 2.51040 4.18400 1.67360
-6.69440 0.00000 0.00000 ; CB- CA- C- N
4178 4184 4187 4188 3 0.66944 2.00832 0.00000
-2.67776 0.00000 0.00000 ; CB- CG1- CD1- HD11
4178 4184 4187 4189 3 0.66944 2.00832 0.00000
-2.67776 0.00000 0.00000 ; CB- CG1- CD1- HD12
4178 4184 4187 4190 3 0.66944 2.00832 0.00000
-2.67776 0.00000 0.00000 ; CB- CG1- CD1- HD13
4179 4178 4176 4191 3 0.65084 1.95253 0.00000
-2.60338 0.00000 0.00000 ; HB- CB- CA- C
4179 4178 4180 4181 3 0.62760 1.88280 0.00000
-2.51040 0.00000 0.00000 ; HB- CB- CG2- HG21
4179 4178 4180 4182 3 0.62760 1.88280 0.00000
-2.51040 0.00000 0.00000 ; HB- CB- CG2- HG22
4179 4178 4180 4183 3 0.62760 1.88280 0.00000
-2.51040 0.00000 0.00000 ; HB- CB- CG2- HG23
4179 4178 4184 4185 3 0.62760 1.88280 0.00000
-2.51040 0.00000 0.00000 ; HB- CB- CG1- HG12
4179 4178 4184 4186 3 0.62760 1.88280 0.00000
-2.51040 0.00000 0.00000 ; HB- CB- CG1- HG13
4179 4178 4184 4187 3 0.66944 2.00832 0.00000
-2.67776 0.00000 0.00000 ; HB- CB- CG1- CD1
4180 4178 4176 4191 3 5.67350 3.55640 -6.16722
-2.47693 3.74886 0.00000 ; CG2- CB- CA- C
4180 4178 4184 4185 3 0.66944 2.00832 0.00000
-2.67776 0.00000 0.00000 ; CG2- CB- CG1- HG12
4180 4178 4184 4186 3 0.66944 2.00832 0.00000
-2.67776 0.00000 0.00000 ; CG2- CB- CG1- HG13
4180 4178 4184 4187 3 3.81162 0.49790 -8.14206
-1.79075 7.49773 0.00000 ; CG2- CB- CG1- CD1
4181 4180 4178 4184 3 0.66944 2.00832 0.00000
-2.67776 0.00000 0.00000 ; HG21- CG2- CB- CG1
4182 4180 4178 4184 3 0.66944 2.00832 0.00000
-2.67776 0.00000 0.00000 ; HG22- CG2- CB- CG1
4183 4180 4178 4184 3 0.66944 2.00832 0.00000
-2.67776 0.00000 0.00000 ; HG23- CG2- CB- CG1

4184 4178 4176 4191 3 8.26340 0.74057 -9.99976
-1.89117 3.84928 0.00000 ; CG1- CB- CA- C
4185 4184 4187 4188 3 0.62760 1.88280 0.00000
-2.51040 0.00000 0.00000 ; HG12- CG1- CD1- HD11
4185 4184 4187 4189 3 0.62760 1.88280 0.00000
-2.51040 0.00000 0.00000 ; HG12- CG1- CD1- HD12
4185 4184 4187 4190 3 0.62760 1.88280 0.00000
-2.51040 0.00000 0.00000 ; HG12- CG1- CD1- HD13
4186 4184 4187 4188 3 0.62760 1.88280 0.00000
-2.51040 0.00000 0.00000 ; HG13- CG1- CD1- HD11
4186 4184 4187 4189 3 0.62760 1.88280 0.00000
-2.51040 0.00000 0.00000 ; HG13- CG1- CD1- HD12
4186 4184 4187 4190 3 0.62760 1.88280 0.00000
-2.51040 0.00000 0.00000 ; HG13- CG1- CD1- HD13
4191 4193 4195 4196 3 0.00000 0.00000 0.00000
0.00000 0.00000 0.00000 ; C- N- CA- HA
4191 4193 4195 4197 3 11.71520 1.67360 15.06240 -
13.38880 0.00000 0.00000 ; C- N- CA- CB
4191 4193 4195 4206 3 1.75728 5.27184 2.25936
-7.02912 0.00000 0.00000 ; C- N- CA- C
4192 4191 4193 4194 3 29.28800 -8.36800 -20.92000
0.00000 0.00000 0.00000 ; O- C- N- H
4192 4191 4193 4195 3 20.92000 0.00000 -20.92000
0.00000 0.00000 0.00000 ; O- C- N- CA
4193 4195 4197 4198 3 0.65084 1.95253 0.00000
-2.60338 0.00000 0.00000 ; N- CA- CB- HB2
4193 4195 4197 4199 3 0.65084 1.95253 0.00000
-2.60338 0.00000 0.00000 ; N- CA- CB- HB3
4193 4195 4197 4200 3 3.21331 2.22589 -4.15053
-2.40998 2.61082 0.00000 ; N- CA- CB- CG
4193 4195 4206 4207 3 0.00000 0.00000 0.00000
0.00000 0.00000 0.00000 ; N- CA- C- O
4193 4195 4206 4208 3 17.40544 -5.02080 -13.22144
9.20480 0.00000 0.00000 ; N- CA- C- N
4194 4193 4195 4196 3 0.00000 0.00000 0.00000
0.00000 0.00000 0.00000 ; H- N- CA- HA
4194 4193 4195 4197 3 0.00000 0.00000 0.00000
0.00000 0.00000 0.00000 ; H- N- CA- CB
4194 4193 4195 4206 3 0.00000 0.00000 0.00000
0.00000 0.00000 0.00000 ; H- N- CA- C
4195 4197 4200 4201 3 0.66944 2.00832 0.00000
-2.67776 0.00000 0.00000 ; CA- CB- CG- HG2
4195 4197 4200 4202 3 0.66944 2.00832 0.00000
-2.67776 0.00000 0.00000 ; CA- CB- CG- HG3
4195 4197 4200 4203 3 10.12110 -1.91209 0.01674
10.17549 -1.87443 0.00000 ; CA- CB- CG- CD
4195 4206 4208 4209 3 20.92000 0.00000 -20.92000
0.00000 0.00000 0.00000 ; CA- C- N- H
4195 4206 4208 4210 3 20.92000 0.00000 -20.92000
0.00000 0.00000 0.00000 ; CA- C- N- CA
4196 4195 4197 4198 3 0.65084 1.95253 0.00000
-2.60338 0.00000 0.00000 ; HA- CA- CB- HB2

4196 4195 4197 4199 3 0.65084 1.95253 0.00000
-2.60338 0.00000 0.00000 ; HA- CA- CB- HB3
4196 4195 4197 4200 3 0.65084 1.95253 0.00000
-2.60338 0.00000 0.00000 ; HA- CA- CB- CG
4196 4195 4206 4207 3 3.68192 -4.35136 0.00000
1.33888 0.00000 0.00000 ; HA- CA- C- O
4196 4195 4206 4208 3 0.00000 0.00000 0.00000
0.00000 0.00000 0.00000 ; HA- CA- C- N
4197 4195 4206 4207 3 0.00000 0.00000 0.00000
0.00000 0.00000 0.00000 ; CB- CA- C- O
4197 4195 4206 4208 3 2.51040 4.18400 1.67360
-6.69440 0.00000 0.00000 ; CB- CA- C- N
4197 4200 4203 4204 3 3.79907 0.00000 -5.40573
0.00000 2.14221 0.00000 ; CB- CG- CD- OE1
4197 4200 4203 4205 3 3.79907 0.00000 -5.40573
0.00000 2.14221 0.00000 ; CB- CG- CD- OE2
4198 4197 4195 4206 3 0.65084 1.95253 0.00000
-2.60338 0.00000 0.00000 ; HB2- CB- CA- C
4198 4197 4200 4201 3 0.62760 1.88280 0.00000
-2.51040 0.00000 0.00000 ; HB2- CB- CG- HG2
4198 4197 4200 4202 3 0.62760 1.88280 0.00000
-2.51040 0.00000 0.00000 ; HB2- CB- CG- HG3
4198 4197 4200 4203 3 0.65084 1.95253 0.00000
-2.60338 0.00000 0.00000 ; HB2- CB- CG- CD
4199 4197 4195 4206 3 0.65084 1.95253 0.00000
-2.60338 0.00000 0.00000 ; HB3- CB- CA- C
4199 4197 4200 4201 3 0.62760 1.88280 0.00000
-2.51040 0.00000 0.00000 ; HB3- CB- CG- HG2
4199 4197 4200 4202 3 0.62760 1.88280 0.00000
-2.51040 0.00000 0.00000 ; HB3- CB- CG- HG3
4199 4197 4200 4203 3 0.65084 1.95253 0.00000
-2.60338 0.00000 0.00000 ; HB3- CB- CG- CD
4200 4197 4195 4206 3 6.86594 3.56895 -8.14206
-2.40998 4.85344 0.00000 ; CG- CB- CA- C
4201 4200 4203 4204 3 0.00000 0.00000 0.00000
0.00000 0.00000 0.00000 ; HG2- CG- CD- OE1
4201 4200 4203 4205 3 0.00000 0.00000 0.00000
0.00000 0.00000 0.00000 ; HG2- CG- CD- OE2
4202 4200 4203 4204 3 0.00000 0.00000 0.00000
0.00000 0.00000 0.00000 ; HG3- CG- CD- OE1
4202 4200 4203 4205 3 0.00000 0.00000 0.00000
0.00000 0.00000 0.00000 ; HG3- CG- CD- OE2
4206 4208 4210 4211 3 0.00000 0.00000 0.00000
0.00000 0.00000 0.00000 ; C- N- CA- HA
4206 4208 4210 4212 3 11.71520 1.67360 15.06240 -
13.38880 0.00000 0.00000 ; C- N- CA- CB
4206 4208 4210 4221 3 1.75728 5.27184 2.25936
-7.02912 0.00000 0.00000 ; C- N- CA- C
4207 4206 4208 4209 3 29.28800 -8.36800 -20.92000
0.00000 0.00000 0.00000 ; O- C- N- H
4207 4206 4208 4210 3 20.92000 0.00000 -20.92000
0.00000 0.00000 0.00000 ; O- C- N- CA

```
  4208   4210   4212   4213       3    0.65084    1.95253    0.00000
-2.60338    0.00000    0.00000 ;       N-    CA-    CB-   HB2
  4208   4210   4212   4214       3    0.65084    1.95253    0.00000
-2.60338    0.00000    0.00000 ;       N-    CA-    CB-   HB3
  4208   4210   4212   4215       3    3.21331    2.22589   -4.15053
-2.40998    2.61082    0.00000 ;       N-    CA-    CB-    CG
  4208   4210   4221   4222       3    0.00000    0.00000    0.00000
0.00000    0.00000    0.00000 ;       N-    CA-     C-     O
  4208   4210   4221   4223       3   17.40544   -5.02080  -13.22144
9.20480    0.00000    0.00000 ;       N-    CA-     C-     N
  4209   4208   4210   4211       3    0.00000    0.00000    0.00000
0.00000    0.00000    0.00000 ;       H-     N-    CA-    HA
  4209   4208   4210   4212       3    0.00000    0.00000    0.00000
0.00000    0.00000    0.00000 ;       H-     N-    CA-    CB
  4209   4208   4210   4221       3    0.00000    0.00000    0.00000
0.00000    0.00000    0.00000 ;       H-     N-    CA-     C
  4210   4212   4215   4216       3    0.66944    2.00832    0.00000
-2.67776    0.00000    0.00000 ;      CA-    CB-    CG-   HG2
  4210   4212   4215   4217       3    0.66944    2.00832    0.00000
-2.67776    0.00000    0.00000 ;      CA-    CB-    CG-   HG3
  4210   4212   4215   4218       3   10.12110   -1.91209    0.01674
10.17549   -1.87443    0.00000 ;      CA-    CB-    CG-    CD
  4210   4221   4223   4224       3   20.92000    0.00000  -20.92000
0.00000    0.00000    0.00000 ;      CA-     C-     N-     H
  4210   4221   4223   4225       3   20.92000    0.00000  -20.92000
0.00000    0.00000    0.00000 ;      CA-     C-     N-    CA
  4211   4210   4212   4213       3    0.65084    1.95253    0.00000
-2.60338    0.00000    0.00000 ;      HA-    CA-    CB-   HB2
  4211   4210   4212   4214       3    0.65084    1.95253    0.00000
-2.60338    0.00000    0.00000 ;      HA-    CA-    CB-   HB3
  4211   4210   4212   4215       3    0.65084    1.95253    0.00000
-2.60338    0.00000    0.00000 ;      HA-    CA-    CB-    CG
  4211   4210   4221   4222       3    3.68192   -4.35136    0.00000
1.33888    0.00000    0.00000 ;      HA-    CA-     C-     O
  4211   4210   4221   4223       3    0.00000    0.00000    0.00000
0.00000    0.00000    0.00000 ;      HA-    CA-     C-     N
  4212   4210   4221   4222       3    0.00000    0.00000    0.00000
0.00000    0.00000    0.00000 ;      CB-    CA-     C-     O
  4212   4210   4221   4223       3    2.51040    4.18400    1.67360
-6.69440    0.00000    0.00000 ;      CB-    CA-     C-     N
  4212   4215   4218   4219       3    3.79907    0.00000   -5.40573
0.00000    2.14221    0.00000 ;      CB-    CG-    CD-   OE1
  4212   4215   4218   4220       3    3.79907    0.00000   -5.40573
0.00000    2.14221    0.00000 ;      CB-    CG-    CD-   OE2
  4213   4212   4210   4221       3    0.65084    1.95253    0.00000
-2.60338    0.00000    0.00000 ;     HB2-    CB-    CA-     C
  4213   4212   4215   4216       3    0.62760    1.88280    0.00000
-2.51040    0.00000    0.00000 ;     HB2-    CB-    CG-   HG2
  4213   4212   4215   4217       3    0.62760    1.88280    0.00000
-2.51040    0.00000    0.00000 ;     HB2-    CB-    CG-   HG3
  4213   4212   4215   4218       3    0.65084    1.95253    0.00000
-2.60338    0.00000    0.00000 ;     HB2-    CB-    CG-    CD
```

4214 4212 4210 4221 3 0.65084 1.95253 0.00000
-2.60338 0.00000 0.00000 ; HB3- CB- CA- C
4214 4212 4215 4216 3 0.62760 1.88280 0.00000
-2.51040 0.00000 0.00000 ; HB3- CB- CG- HG2
4214 4212 4215 4217 3 0.62760 1.88280 0.00000
-2.51040 0.00000 0.00000 ; HB3- CB- CG- HG3
4214 4212 4215 4218 3 0.65084 1.95253 0.00000
-2.60338 0.00000 0.00000 ; HB3- CB- CG- CD
4215 4212 4210 4221 3 6.86594 3.56895 -8.14206
-2.40998 4.85344 0.00000 ; CG- CB- CA- C
4216 4215 4218 4219 3 0.00000 0.00000 0.00000
0.00000 0.00000 0.00000 ; HG2- CG- CD- OE1
4216 4215 4218 4220 3 0.00000 0.00000 0.00000
0.00000 0.00000 0.00000 ; HG2- CG- CD- OE2
4217 4215 4218 4219 3 0.00000 0.00000 0.00000
0.00000 0.00000 0.00000 ; HG3- CG- CD- OE1
4217 4215 4218 4220 3 0.00000 0.00000 0.00000
0.00000 0.00000 0.00000 ; HG3- CG- CD- OE2
4221 4223 4225 4226 3 0.00000 0.00000 0.00000
0.00000 0.00000 0.00000 ; C- N- CA- HA
4221 4223 4225 4227 3 11.71520 1.67360 15.06240 -
13.38880 0.00000 0.00000 ; C- N- CA- CB
4221 4223 4225 4231 3 1.75728 5.27184 2.25936
-7.02912 0.00000 0.00000 ; C- N- CA- C
4222 4221 4223 4224 3 29.28800 -8.36800 -20.92000
0.00000 0.00000 0.00000 ; O- C- N- H
4222 4221 4223 4225 3 20.92000 0.00000 -20.92000
0.00000 0.00000 0.00000 ; O- C- N- CA
4223 4225 4227 4228 3 0.65084 1.95253 0.00000
-2.60338 0.00000 0.00000 ; N- CA- CB- HB1
4223 4225 4227 4229 3 0.65084 1.95253 0.00000
-2.60338 0.00000 0.00000 ; N- CA- CB- HB2
4223 4225 4227 4230 3 0.65084 1.95253 0.00000
-2.60338 0.00000 0.00000 ; N- CA- CB- HB3
4223 4225 4231 4232 3 0.00000 0.00000 0.00000
0.00000 0.00000 0.00000 ; N- CA- C- O
4223 4225 4231 4233 3 17.40544 -5.02080 -13.22144
9.20480 0.00000 0.00000 ; N- CA- C- N
4224 4223 4225 4226 3 0.00000 0.00000 0.00000
0.00000 0.00000 0.00000 ; H- N- CA- HA
4224 4223 4225 4227 3 0.00000 0.00000 0.00000
0.00000 0.00000 0.00000 ; H- N- CA- CB
4224 4223 4225 4231 3 0.00000 0.00000 0.00000
0.00000 0.00000 0.00000 ; H- N- CA- C
4225 4231 4233 4234 3 20.92000 0.00000 -20.92000
0.00000 0.00000 0.00000 ; CA- C- N- H
4225 4231 4233 4235 3 20.92000 0.00000 -20.92000
0.00000 0.00000 0.00000 ; CA- C- N- CA
4226 4225 4227 4228 3 0.65084 1.95253 0.00000
-2.60338 0.00000 0.00000 ; HA- CA- CB- HB1
4226 4225 4227 4229 3 0.65084 1.95253 0.00000
-2.60338 0.00000 0.00000 ; HA- CA- CB- HB2

4226 4225 4227 4230 3 0.65084 1.95253 0.00000
-2.60338 0.00000 0.00000 ; HA- CA- CB- HB3
4226 4225 4231 4232 3 3.68192 -4.35136 0.00000
1.33888 0.00000 0.00000 ; HA- CA- C- O
4226 4225 4231 4233 3 0.00000 0.00000 0.00000
0.00000 0.00000 0.00000 ; HA- CA- C- N
4227 4225 4231 4232 3 0.00000 0.00000 0.00000
0.00000 0.00000 0.00000 ; CB- CA- C- O
4227 4225 4231 4233 3 2.51040 4.18400 1.67360
-6.69440 0.00000 0.00000 ; CB- CA- C- N
4228 4227 4225 4231 3 0.65084 1.95253 0.00000
-2.60338 0.00000 0.00000 ; HB1- CB- CA- C
4229 4227 4225 4231 3 0.65084 1.95253 0.00000
-2.60338 0.00000 0.00000 ; HB2- CB- CA- C
4230 4227 4225 4231 3 0.65084 1.95253 0.00000
-2.60338 0.00000 0.00000 ; HB3- CB- CA- C
4231 4233 4235 4236 3 0.00000 0.00000 0.00000
0.00000 0.00000 0.00000 ; C- N- CA- HA
4231 4233 4235 4237 3 11.71520 1.67360 15.06240 -
13.38880 0.00000 0.00000 ; C- N- CA- CB
4231 4233 4235 4250 3 1.75728 5.27184 2.25936
-7.02912 0.00000 0.00000 ; C- N- CA- C
4232 4231 4233 4234 3 29.28800 -8.36800 -20.92000
0.00000 0.00000 0.00000 ; O- C- N- H
4232 4231 4233 4235 3 20.92000 0.00000 -20.92000
0.00000 0.00000 0.00000 ; O- C- N- CA
4233 4235 4237 4238 3 0.65084 1.95253 0.00000
-2.60338 0.00000 0.00000 ; N- CA- CB- HB2
4233 4235 4237 4239 3 0.65084 1.95253 0.00000
-2.60338 0.00000 0.00000 ; N- CA- CB- HB3
4233 4235 4237 4240 3 3.79070 1.39746 -4.61077
-2.40998 2.44346 0.00000 ; N- CA- CB- CG
4233 4235 4250 4251 3 0.00000 0.00000 0.00000
0.00000 0.00000 0.00000 ; N- CA- C- O
4233 4235 4250 4252 3 17.40544 -5.02080 -13.22144
9.20480 0.00000 0.00000 ; N- CA- C- N
4234 4233 4235 4236 3 0.00000 0.00000 0.00000
0.00000 0.00000 0.00000 ; H- N- CA- HA
4234 4233 4235 4237 3 0.00000 0.00000 0.00000
0.00000 0.00000 0.00000 ; H- N- CA- CB
4234 4233 4235 4250 3 0.00000 0.00000 0.00000
0.00000 0.00000 0.00000 ; H- N- CA- C
4235 4237 4240 4241 3 0.66944 2.00832 0.00000
-2.67776 0.00000 0.00000 ; CA- CB- CG- HG
4235 4237 4240 4242 3 3.90367 0.19665 -6.21742
-2.37651 5.99149 0.00000 ; CA- CB- CG- CD1
4235 4237 4240 4246 3 3.90367 0.19665 -6.21742
-2.37651 5.99149 0.00000 ; CA- CB- CG- CD2
4235 4250 4252 4253 3 20.92000 0.00000 -20.92000
0.00000 0.00000 0.00000 ; CA- C- N- H
4235 4250 4252 4254 3 20.92000 0.00000 -20.92000
0.00000 0.00000 0.00000 ; CA- C- N- CA

```
  4236   4235   4237   4238       3    0.65084    1.95253    0.00000
-2.60338    0.00000    0.00000 ;     HA-    CA-    CB-   HB2
  4236   4235   4237   4239       3    0.65084    1.95253    0.00000
-2.60338    0.00000    0.00000 ;     HA-    CA-    CB-   HB3
  4236   4235   4237   4240       3    0.65084    1.95253    0.00000
-2.60338    0.00000    0.00000 ;     HA-    CA-    CB-    CG
  4236   4235   4250   4251       3    3.68192   -4.35136    0.00000
1.33888    0.00000    0.00000 ;     HA-    CA-     C-     O
  4236   4235   4250   4252       3    0.00000    0.00000    0.00000
0.00000    0.00000    0.00000 ;     HA-    CA-     C-     N
  4237   4235   4250   4251       3    0.00000    0.00000    0.00000
0.00000    0.00000    0.00000 ;     CB-    CA-     C-     O
  4237   4235   4250   4252       3    2.51040    4.18400    1.67360
-6.69440    0.00000    0.00000 ;     CB-    CA-     C-     N
  4237   4240   4242   4243       3    0.66944    2.00832    0.00000
-2.67776    0.00000    0.00000 ;     CB-    CG-   CD1-  HD11
  4237   4240   4242   4244       3    0.66944    2.00832    0.00000
-2.67776    0.00000    0.00000 ;     CB-    CG-   CD1-  HD12
  4237   4240   4242   4245       3    0.66944    2.00832    0.00000
-2.67776    0.00000    0.00000 ;     CB-    CG-   CD1-  HD13
  4237   4240   4246   4247       3    0.66944    2.00832    0.00000
-2.67776    0.00000    0.00000 ;     CB-    CG-   CD2-  HD21
  4237   4240   4246   4248       3    0.66944    2.00832    0.00000
-2.67776    0.00000    0.00000 ;     CB-    CG-   CD2-  HD22
  4237   4240   4246   4249       3    0.66944    2.00832    0.00000
-2.67776    0.00000    0.00000 ;     CB-    CG-   CD2-  HD23
  4238   4237   4235   4250       3    0.65084    1.95253    0.00000
-2.60338    0.00000    0.00000 ;    HB2-    CB-    CA-     C
  4238   4237   4240   4241       3    0.62760    1.88280    0.00000
-2.51040    0.00000    0.00000 ;    HB2-    CB-    CG-    HG
  4238   4237   4240   4242       3    0.66944    2.00832    0.00000
-2.67776    0.00000    0.00000 ;    HB2-    CB-    CG-   CD1
  4238   4237   4240   4246       3    0.66944    2.00832    0.00000
-2.67776    0.00000    0.00000 ;    HB2-    CB-    CG-   CD2
  4239   4237   4235   4250       3    0.65084    1.95253    0.00000
-2.60338    0.00000    0.00000 ;    HB3-    CB-    CA-     C
  4239   4237   4240   4241       3    0.62760    1.88280    0.00000
-2.51040    0.00000    0.00000 ;    HB3-    CB-    CG-    HG
  4239   4237   4240   4242       3    0.66944    2.00832    0.00000
-2.67776    0.00000    0.00000 ;    HB3-    CB-    CG-   CD1
  4239   4237   4240   4246       3    0.66944    2.00832    0.00000
-2.67776    0.00000    0.00000 ;    HB3-    CB-    CG-   CD2
  4240   4237   4235   4250       3   10.33448   -1.14642  -11.54784
-2.40998    6.35968    0.00000 ;     CG-    CB-    CA-     C
  4241   4240   4242   4243       3    0.62760    1.88280    0.00000
-2.51040    0.00000    0.00000 ;     HG-    CG-   CD1-  HD11
  4241   4240   4242   4244       3    0.62760    1.88280    0.00000
-2.51040    0.00000    0.00000 ;     HG-    CG-   CD1-  HD12
  4241   4240   4242   4245       3    0.62760    1.88280    0.00000
-2.51040    0.00000    0.00000 ;     HG-    CG-   CD1-  HD13
  4241   4240   4246   4247       3    0.62760    1.88280    0.00000
-2.51040    0.00000    0.00000 ;     HG-    CG-   CD2-  HD21
```

4241 4240 4246 4248 3 0.62760 1.88280 0.00000
-2.51040 0.00000 0.00000 ; HG- CG- CD2- HD22
4241 4240 4246 4249 3 0.62760 1.88280 0.00000
-2.51040 0.00000 0.00000 ; HG- CG- CD2- HD23
4242 4240 4246 4247 3 0.66944 2.00832 0.00000
-2.67776 0.00000 0.00000 ; CD1- CG- CD2- HD21
4242 4240 4246 4248 3 0.66944 2.00832 0.00000
-2.67776 0.00000 0.00000 ; CD1- CG- CD2- HD22
4242 4240 4246 4249 3 0.66944 2.00832 0.00000
-2.67776 0.00000 0.00000 ; CD1- CG- CD2- HD23
4243 4242 4240 4246 3 0.66944 2.00832 0.00000
-2.67776 0.00000 0.00000 ; HD11- CD1- CG- CD2
4244 4242 4240 4246 3 0.66944 2.00832 0.00000
-2.67776 0.00000 0.00000 ; HD12- CD1- CG- CD2
4245 4242 4240 4246 3 0.66944 2.00832 0.00000
-2.67776 0.00000 0.00000 ; HD13- CD1- CG- CD2
4250 4252 4254 4255 3 0.00000 0.00000 0.00000
0.00000 0.00000 0.00000 ; C- N- CA- HA
4250 4252 4254 4256 3 11.71520 1.67360 15.06240 -
13.38880 0.00000 0.00000 ; C- N- CA- CB
4250 4252 4254 4264 3 1.75728 5.27184 2.25936
-7.02912 0.00000 0.00000 ; C- N- CA- C
4251 4250 4252 4253 3 29.28800 -8.36800 -20.92000
0.00000 0.00000 0.00000 ; O- C- N- H
4251 4250 4252 4254 3 20.92000 0.00000 -20.92000
0.00000 0.00000 0.00000 ; O- C- N- CA
4252 4254 4256 4257 3 0.65084 1.95253 0.00000
-2.60338 0.00000 0.00000 ; N- CA- CB- HB2
4252 4254 4256 4258 3 0.65084 1.95253 0.00000
-2.60338 0.00000 0.00000 ; N- CA- CB- HB3
4252 4254 4256 4259 3 5.99567 3.29281 -4.46014
-0.55229 1.97485 0.00000 ; N- CA- CB- CG
4252 4254 4264 4265 3 0.00000 0.00000 0.00000
0.00000 0.00000 0.00000 ; N- CA- C- O
4252 4254 4264 4266 3 17.40544 -5.02080 -13.22144
9.20480 0.00000 0.00000 ; N- CA- C- N
4253 4252 4254 4255 3 0.00000 0.00000 0.00000
0.00000 0.00000 0.00000 ; H- N- CA- HA
4253 4252 4254 4256 3 0.00000 0.00000 0.00000
0.00000 0.00000 0.00000 ; H- N- CA- CB
4253 4252 4254 4264 3 0.00000 0.00000 0.00000
0.00000 0.00000 0.00000 ; H- N- CA- C
4254 4256 4259 4260 3 0.00000 0.00000 0.00000
0.00000 0.00000 0.00000 ; CA- CB- CG- OD1
4254 4256 4259 4261 3 8.84916 -0.31380 -4.32626
5.03754 0.26778 0.00000 ; CA- CB- CG- ND2
4254 4264 4266 4267 3 20.92000 0.00000 -20.92000
0.00000 0.00000 0.00000 ; CA- C- N- H
4254 4264 4266 4268 3 20.92000 0.00000 -20.92000
0.00000 0.00000 0.00000 ; CA- C- N- CA
4255 4254 4256 4257 3 0.65084 1.95253 0.00000
-2.60338 0.00000 0.00000 ; HA- CA- CB- HB2

4255 4254 4256 4258 3 0.65084 1.95253 0.00000
-2.60338 0.00000 0.00000 ; HA- CA- CB- HB3
4255 4254 4256 4259 3 0.65084 1.95253 0.00000
-2.60338 0.00000 0.00000 ; HA- CA- CB- CG
4255 4254 4264 4265 3 3.68192 -4.35136 0.00000
1.33888 0.00000 0.00000 ; HA- CA- C- O
4255 4254 4264 4266 3 0.00000 0.00000 0.00000
0.00000 0.00000 0.00000 ; HA- CA- C- N
4256 4254 4264 4265 3 0.00000 0.00000 0.00000
0.00000 0.00000 0.00000 ; CB- CA- C- O
4256 4254 4264 4266 3 2.51040 4.18400 1.67360
-6.69440 0.00000 0.00000 ; CB- CA- C- N
4256 4259 4261 4262 3 20.92000 0.00000 -20.92000
0.00000 0.00000 0.00000 ; CB- CG- ND2- HD21
4256 4259 4261 4263 3 20.92000 0.00000 -20.92000
0.00000 0.00000 0.00000 ; CB- CG- ND2- HD22
4257 4256 4254 4264 3 0.65084 1.95253 0.00000
-2.60338 0.00000 0.00000 ; HB2- CB- CA- C
4257 4256 4259 4260 3 3.68192 -4.35136 0.00000
1.33888 0.00000 0.00000 ; HB2- CB- CG- OD1
4257 4256 4259 4261 3 0.00000 0.00000 0.00000
0.00000 0.00000 0.00000 ; HB2- CB- CG- ND2
4258 4256 4254 4264 3 0.65084 1.95253 0.00000
-2.60338 0.00000 0.00000 ; HB3- CB- CA- C
4258 4256 4259 4260 3 3.68192 -4.35136 0.00000
1.33888 0.00000 0.00000 ; HB3- CB- CG- OD1
4258 4256 4259 4261 3 0.00000 0.00000 0.00000
0.00000 0.00000 0.00000 ; HB3- CB- CG- ND2
4259 4256 4254 4264 3 7.94542 -3.96225 -6.11701
-0.55229 3.58150 0.00000 ; CG- CB- CA- C
4260 4259 4261 4262 3 29.28800 -8.36800 -20.92000
0.00000 0.00000 0.00000 ; OD1- CG- ND2- HD21
4260 4259 4261 4263 3 29.28800 -8.36800 -20.92000
0.00000 0.00000 0.00000 ; OD1- CG- ND2- HD22
4264 4266 4268 4269 3 0.00000 0.00000 0.00000
0.00000 0.00000 0.00000 ; C- N- CA- HA
4264 4266 4268 4270 3 11.71520 1.67360 15.06240 -
13.38880 0.00000 0.00000 ; C- N- CA- CB
4264 4266 4268 4281 3 1.75728 5.27184 2.25936
-7.02912 0.00000 0.00000 ; C- N- CA- C
4265 4264 4266 4267 3 29.28800 -8.36800 -20.92000
0.00000 0.00000 0.00000 ; O- C- N- H
4265 4264 4266 4268 3 20.92000 0.00000 -20.92000
0.00000 0.00000 0.00000 ; O- C- N- CA
4266 4268 4270 4271 3 0.65084 1.95253 0.00000
-2.60338 0.00000 0.00000 ; N- CA- CB- HB2
4266 4268 4270 4272 3 0.65084 1.95253 0.00000
-2.60338 0.00000 0.00000 ; N- CA- CB- HB3
4266 4268 4270 4273 3 4.79068 4.02919 -4.82834
-3.66518 2.97901 0.00000 ; N- CA- CB- CG
4266 4268 4281 4282 3 0.00000 0.00000 0.00000
0.00000 0.00000 0.00000 ; N- CA- C- O

4266 4268 4281 4283 3 17.40544 -5.02080 -13.22144
9.20480 0.00000 0.00000 ; N- CA- C- N
4267 4266 4268 4269 3 0.00000 0.00000 0.00000
0.00000 0.00000 0.00000 ; H- N- CA- HA
4267 4266 4268 4270 3 0.00000 0.00000 0.00000
0.00000 0.00000 0.00000 ; H- N- CA- CB
4267 4266 4268 4281 3 0.00000 0.00000 0.00000
0.00000 0.00000 0.00000 ; H- N- CA- C
4268 4270 4273 4274 3 5.98312 6.40152 3.28026 -
12.38464 -1.57318 0.00000 ; CA- CB- CG- ND1
4268 4270 4273 4279 3 0.00000 0.00000 0.00000
0.00000 0.00000 0.00000 ; CA- CB- CG- CD2
4268 4281 4283 4284 3 20.92000 0.00000 -20.92000
0.00000 0.00000 0.00000 ; CA- C- N- CD
4268 4281 4283 4293 3 20.92000 0.00000 -20.92000
0.00000 0.00000 0.00000 ; CA- C- N- CA
4269 4268 4270 4271 3 0.65084 1.95253 0.00000
-2.60338 0.00000 0.00000 ; HA- CA- CB- HB2
4269 4268 4270 4272 3 0.65084 1.95253 0.00000
-2.60338 0.00000 0.00000 ; HA- CA- CB- HB3
4269 4268 4270 4273 3 0.65084 1.95253 0.00000
-2.60338 0.00000 0.00000 ; HA- CA- CB- CG
4269 4268 4281 4282 3 3.68192 -4.35136 0.00000
1.33888 0.00000 0.00000 ; HA- CA- C- O
4269 4268 4281 4283 3 0.00000 0.00000 0.00000
0.00000 0.00000 0.00000 ; HA- CA- C- N
4270 4268 4281 4282 3 0.00000 0.00000 0.00000
0.00000 0.00000 0.00000 ; CB- CA- C- O
4270 4268 4281 4283 3 2.51040 4.18400 1.67360
-6.69440 0.00000 0.00000 ; CB- CA- C- N
4270 4273 4274 4275 3 20.08320 0.00000 -20.08320
0.00000 0.00000 0.00000 ; CB- CG- ND1- CE1
4270 4273 4279 4277 3 44.97800 0.00000 -44.97800
0.00000 0.00000 0.00000 ; CB- CG- CD2- NE2
4270 4273 4279 4280 3 44.97800 0.00000 -44.97800
0.00000 0.00000 0.00000 ; CB- CG- CD2- HD2
4271 4270 4268 4281 3 0.65084 1.95253 0.00000
-2.60338 0.00000 0.00000 ; HB2- CB- CA- C
4271 4270 4273 4274 3 0.00000 0.00000 0.00000
0.00000 0.00000 0.00000 ; HB2- CB- CG- ND1
4271 4270 4273 4279 3 0.00000 0.00000 0.00000
0.00000 0.00000 0.00000 ; HB2- CB- CG- CD2
4272 4270 4268 4281 3 0.65084 1.95253 0.00000
-2.60338 0.00000 0.00000 ; HB3- CB- CA- C
4272 4270 4273 4274 3 0.00000 0.00000 0.00000
0.00000 0.00000 0.00000 ; HB3- CB- CG- ND1
4272 4270 4273 4279 3 0.00000 0.00000 0.00000
0.00000 0.00000 0.00000 ; HB3- CB- CG- CD2
4273 4270 4268 4281 3 3.76560 3.34720 -2.87859
-3.66518 0.83680 0.00000 ; CG- CB- CA- C
4273 4274 4275 4276 3 41.84000 0.00000 -41.84000
0.00000 0.00000 0.00000 ; CG- ND1- CE1- HE1

4273 4274 4275 4277 3 41.84000 0.00000 -41.84000
0.00000 0.00000 0.00000 ; CG- ND1- CE1- NE2
4273 4279 4277 4275 3 12.55200 0.00000 -12.55200
0.00000 0.00000 0.00000 ; CG- CD2- NE2- CE1
4273 4279 4277 4278 3 12.55200 0.00000 -12.55200
0.00000 0.00000 0.00000 ; CG- CD2- NE2- HE2
4274 4273 4279 4277 3 44.97800 0.00000 -44.97800
0.00000 0.00000 0.00000 ; ND1- CG- CD2- NE2
4274 4273 4279 4280 3 44.97800 0.00000 -44.97800
0.00000 0.00000 0.00000 ; ND1- CG- CD2- HD2
4274 4275 4277 4278 3 19.45560 0.00000 -19.45560
0.00000 0.00000 0.00000 ; ND1- CE1- NE2- HE2
4274 4275 4277 4279 3 19.45560 0.00000 -19.45560
0.00000 0.00000 0.00000 ; ND1- CE1- NE2- CD2
4275 4274 4273 4279 3 20.08320 0.00000 -20.08320
0.00000 0.00000 0.00000 ; CE1- ND1- CG- CD2
4275 4277 4279 4280 3 12.55200 0.00000 -12.55200
0.00000 0.00000 0.00000 ; CE1- NE2- CD2- HD2
4276 4275 4277 4278 3 19.45560 0.00000 -19.45560
0.00000 0.00000 0.00000 ; HE1- CE1- NE2- HE2
4276 4275 4277 4279 3 19.45560 0.00000 -19.45560
0.00000 0.00000 0.00000 ; HE1- CE1- NE2- CD2
4278 4277 4279 4280 3 12.55200 0.00000 -12.55200
0.00000 0.00000 0.00000 ; HE2- NE2- CD2- HD2
4281 4283 4284 4285 3 0.00000 0.00000 0.00000
0.00000 0.00000 0.00000 ; C- N- CD- HD2
4281 4283 4284 4286 3 0.00000 0.00000 0.00000
0.00000 0.00000 0.00000 ; C- N- CD- HD3
4281 4283 4284 4287 3 10.04160 -3.34720 16.73600
-6.69440 0.00000 0.00000 ; C- N- CD- CG
4281 4283 4293 4290 3 11.71520 1.67360 15.06240 -
13.38880 0.00000 0.00000 ; C- N- CA- CB
4281 4283 4293 4294 3 0.00000 0.00000 0.00000
0.00000 0.00000 0.00000 ; C- N- CA- HA
4281 4283 4293 4295 3 1.75728 5.27184 2.25936
-7.02912 0.00000 0.00000 ; C- N- CA- C
4282 4281 4283 4284 3 20.92000 0.00000 -20.92000
0.00000 0.00000 0.00000 ; O- C- N- CD
4282 4281 4283 4293 3 20.92000 0.00000 -20.92000
0.00000 0.00000 0.00000 ; O- C- N- CA
4283 4284 4287 4288 3 0.65084 1.95253 0.00000
-2.60338 0.00000 0.00000 ; N- CD- CG- HG2
4283 4284 4287 4289 3 0.65084 1.95253 0.00000
-2.60338 0.00000 0.00000 ; N- CD- CG- HG3
4283 4284 4287 4290 3 0.65084 1.95253 0.00000
-2.60338 0.00000 0.00000 ; N- CD- CG- CB
4283 4293 4290 4287 3 0.65084 1.95253 0.00000
-2.60338 0.00000 0.00000 ; N- CA- CB- CG
4283 4293 4290 4291 3 0.65084 1.95253 0.00000
-2.60338 0.00000 0.00000 ; N- CA- CB- HB2
4283 4293 4290 4292 3 0.65084 1.95253 0.00000
-2.60338 0.00000 0.00000 ; N- CA- CB- HB3

4283 4293 4295 4296 3 0.00000 0.00000 0.00000
0.00000 0.00000 0.00000 ; N- CA- C- O
4283 4293 4295 4297 3 17.40544 -5.02080 -13.22144
9.20480 0.00000 0.00000 ; N- CA- C- N
4284 4283 4293 4290 3 0.00000 0.00000 0.00000
0.00000 0.00000 0.00000 ; CD- N- CA- CB
4284 4283 4293 4294 3 0.00000 0.00000 0.00000
0.00000 0.00000 0.00000 ; CD- N- CA- HA
4284 4283 4293 4295 3 0.00000 0.00000 0.00000
0.00000 0.00000 0.00000 ; CD- N- CA- C
4284 4287 4290 4291 3 0.66944 2.00832 0.00000
-2.67776 0.00000 0.00000 ; CD- CG- CB- HB2
4284 4287 4290 4292 3 0.66944 2.00832 0.00000
-2.67776 0.00000 0.00000 ; CD- CG- CB- HB3
4284 4287 4290 4293 3 3.68192 3.09616 -2.09200
-3.01248 0.00000 0.00000 ; CD- CG- CB- CA
4285 4284 4283 4293 3 0.00000 0.00000 0.00000
0.00000 0.00000 0.00000 ; HD2- CD- N- CA
4285 4284 4287 4288 3 0.65084 1.95253 0.00000
-2.60338 0.00000 0.00000 ; HD2- CD- CG- HG2
4285 4284 4287 4289 3 0.65084 1.95253 0.00000
-2.60338 0.00000 0.00000 ; HD2- CD- CG- HG3
4285 4284 4287 4290 3 0.65084 1.95253 0.00000
-2.60338 0.00000 0.00000 ; HD2- CD- CG- CB
4286 4284 4283 4293 3 0.00000 0.00000 0.00000
0.00000 0.00000 0.00000 ; HD3- CD- N- CA
4286 4284 4287 4288 3 0.65084 1.95253 0.00000
-2.60338 0.00000 0.00000 ; HD3- CD- CG- HG2
4286 4284 4287 4289 3 0.65084 1.95253 0.00000
-2.60338 0.00000 0.00000 ; HD3- CD- CG- HG3
4286 4284 4287 4290 3 0.65084 1.95253 0.00000
-2.60338 0.00000 0.00000 ; HD3- CD- CG- CB
4287 4284 4283 4293 3 0.00000 0.00000 0.00000
0.00000 0.00000 0.00000 ; CG- CD- N- CA
4287 4290 4293 4294 3 0.65084 1.95253 0.00000
-2.60338 0.00000 0.00000 ; CG- CB- CA- HA
4287 4290 4293 4295 3 0.65084 1.95253 0.00000
-2.60338 0.00000 0.00000 ; CG- CB- CA- C
4288 4287 4290 4291 3 0.62760 1.88280 0.00000
-2.51040 0.00000 0.00000 ; HG2- CG- CB- HB2
4288 4287 4290 4292 3 0.62760 1.88280 0.00000
-2.51040 0.00000 0.00000 ; HG2- CG- CB- HB3
4288 4287 4290 4293 3 0.66944 2.00832 0.00000
-2.67776 0.00000 0.00000 ; HG2- CG- CB- CA
4289 4287 4290 4291 3 0.62760 1.88280 0.00000
-2.51040 0.00000 0.00000 ; HG3- CG- CB- HB2
4289 4287 4290 4292 3 0.62760 1.88280 0.00000
-2.51040 0.00000 0.00000 ; HG3- CG- CB- HB3
4289 4287 4290 4293 3 0.66944 2.00832 0.00000
-2.67776 0.00000 0.00000 ; HG3- CG- CB- CA
4290 4293 4295 4296 3 0.00000 0.00000 0.00000
0.00000 0.00000 0.00000 ; CB- CA- C- O

4290 4293 4295 4297 3 2.51040 4.18400 1.67360
-6.69440 0.00000 0.00000 ; CB- CA- C- N
4291 4290 4293 4294 3 0.65084 1.95253 0.00000
-2.60338 0.00000 0.00000 ; HB2- CB- CA- HA
4291 4290 4293 4295 3 0.65084 1.95253 0.00000
-2.60338 0.00000 0.00000 ; HB2- CB- CA- C
4292 4290 4293 4294 3 0.65084 1.95253 0.00000
-2.60338 0.00000 0.00000 ; HB3- CB- CA- HA
4292 4290 4293 4295 3 0.65084 1.95253 0.00000
-2.60338 0.00000 0.00000 ; HB3- CB- CA- C
4293 4295 4297 4298 3 20.92000 0.00000 -20.92000
0.00000 0.00000 0.00000 ; CA- C- N- H
4293 4295 4297 4299 3 20.92000 0.00000 -20.92000
0.00000 0.00000 0.00000 ; CA- C- N- CA
4294 4293 4295 4296 3 3.68192 -4.35136 0.00000
1.33888 0.00000 0.00000 ; HA- CA- C- O
4294 4293 4295 4297 3 0.00000 0.00000 0.00000
0.00000 0.00000 0.00000 ; HA- CA- C- N
4295 4297 4299 4300 3 0.00000 0.00000 0.00000
0.00000 0.00000 0.00000 ; C- N- CA- HA
4295 4297 4299 4301 3 11.71520 1.67360 15.06240 -
13.38880 0.00000 0.00000 ; C- N- CA- CB
4295 4297 4299 4319 3 1.75728 5.27184 2.25936
-7.02912 0.00000 0.00000 ; C- N- CA- C
4296 4295 4297 4298 3 29.28800 -8.36800 -20.92000
0.00000 0.00000 0.00000 ; O- C- N- H
4296 4295 4297 4299 3 20.92000 0.00000 -20.92000
0.00000 0.00000 0.00000 ; O- C- N- CA
4297 4299 4301 4302 3 0.65084 1.95253 0.00000
-2.60338 0.00000 0.00000 ; N- CA- CB- HB2
4297 4299 4301 4303 3 0.65084 1.95253 0.00000
-2.60338 0.00000 0.00000 ; N- CA- CB- HB3
4297 4299 4301 4304 3 4.18818 2.60663 -3.65682
-3.91622 1.03763 0.00000 ; N- CA- CB- CG
4297 4299 4319 4320 3 0.00000 0.00000 0.00000
0.00000 0.00000 0.00000 ; N- CA- C- O
4297 4299 4319 4321 3 17.40544 -5.02080 -13.22144
9.20480 0.00000 0.00000 ; N- CA- C- N
4298 4297 4299 4300 3 0.00000 0.00000 0.00000
0.00000 0.00000 0.00000 ; H- N- CA- HA
4298 4297 4299 4301 3 0.00000 0.00000 0.00000
0.00000 0.00000 0.00000 ; H- N- CA- CB
4298 4297 4299 4319 3 0.00000 0.00000 0.00000
0.00000 0.00000 0.00000 ; H- N- CA- C
4299 4301 4304 4305 3 0.00000 0.00000 0.00000
0.00000 0.00000 0.00000 ; CA- CB- CG- CD1
4299 4301 4304 4318 3 4.95386 8.75293 6.59398 -
13.70678 -3.17984 0.00000 ; CA- CB- CG- CD2
4299 4319 4321 4322 3 20.92000 0.00000 -20.92000
0.00000 0.00000 0.00000 ; CA- C- N- H
4299 4319 4321 4323 3 20.92000 0.00000 -20.92000
0.00000 0.00000 0.00000 ; CA- C- N- CA

4300 4299 4301 4302 3 0.65084 1.95253 0.00000
-2.60338 0.00000 0.00000 ; HA- CA- CB- HB2
4300 4299 4301 4303 3 0.65084 1.95253 0.00000
-2.60338 0.00000 0.00000 ; HA- CA- CB- HB3
4300 4299 4301 4304 3 0.65084 1.95253 0.00000
-2.60338 0.00000 0.00000 ; HA- CA- CB- CG
4300 4299 4319 4320 3 3.68192 -4.35136 0.00000
1.33888 0.00000 0.00000 ; HA- CA- C- O
4300 4299 4319 4321 3 0.00000 0.00000 0.00000
0.00000 0.00000 0.00000 ; HA- CA- C- N
4301 4299 4319 4320 3 0.00000 0.00000 0.00000
0.00000 0.00000 0.00000 ; CB- CA- C- O
4301 4299 4319 4321 3 2.51040 4.18400 1.67360
-6.69440 0.00000 0.00000 ; CB- CA- C- N
4301 4304 4305 4306 3 54.60120 0.00000 -54.60120
0.00000 0.00000 0.00000 ; CB- CG- CD1- HD1
4301 4304 4305 4307 3 54.60120 0.00000 -54.60120
0.00000 0.00000 0.00000 ; CB- CG- CD1- NE1
4301 4304 4318 4309 3 14.01640 0.00000 -14.01640
0.00000 0.00000 0.00000 ; CB- CG- CD2- CE2
4301 4304 4318 4316 3 14.01640 0.00000 -14.01640
0.00000 0.00000 0.00000 ; CB- CG- CD2- CE3
4302 4301 4299 4319 3 0.65084 1.95253 0.00000
-2.60338 0.00000 0.00000 ; HB2- CB- CA- C
4302 4301 4304 4305 3 0.00000 0.00000 0.00000
0.00000 0.00000 0.00000 ; HB2- CB- CG- CD1
4302 4301 4304 4318 3 0.00000 0.00000 0.00000
0.00000 0.00000 0.00000 ; HB2- CB- CG- CD2
4303 4301 4299 4319 3 0.65084 1.95253 0.00000
-2.60338 0.00000 0.00000 ; HB3- CB- CA- C
4303 4301 4304 4305 3 0.00000 0.00000 0.00000
0.00000 0.00000 0.00000 ; HB3- CB- CG- CD1
4303 4301 4304 4318 3 0.00000 0.00000 0.00000
0.00000 0.00000 0.00000 ; HB3- CB- CG- CD2
4304 4301 4299 4319 3 4.62332 3.00830 -5.43083
-3.91622 2.47693 0.00000 ; CG- CB- CA- C
4304 4305 4307 4308 3 12.55200 0.00000 -12.55200
0.00000 0.00000 0.00000 ; CG- CD1- NE1- HE1
4304 4305 4307 4309 3 12.55200 0.00000 -12.55200
0.00000 0.00000 0.00000 ; CG- CD1- NE1- CE2
4304 4318 4309 4307 3 25.10400 0.00000 -25.10400
0.00000 0.00000 0.00000 ; CG- CD2- CE2- NE1
4304 4318 4309 4310 3 25.10400 0.00000 -25.10400
0.00000 0.00000 0.00000 ; CG- CD2- CE2- CZ2
4304 4318 4316 4314 3 29.28800 0.00000 -29.28800
0.00000 0.00000 0.00000 ; CG- CD2- CE3- CZ3
4304 4318 4316 4317 3 29.28800 0.00000 -29.28800
0.00000 0.00000 0.00000 ; CG- CD2- CE3- HE3
4305 4304 4318 4309 3 14.01640 0.00000 -14.01640
0.00000 0.00000 0.00000 ; CD1- CG- CD2- CE2
4305 4304 4318 4316 3 14.01640 0.00000 -14.01640
0.00000 0.00000 0.00000 ; CD1- CG- CD2- CE3

4305 4307 4309 4310 3 12.76120 0.00000 -12.76120
0.00000 0.00000 0.00000 ; CD1- NE1- CE2- CZ2
4305 4307 4309 4318 3 12.76120 0.00000 -12.76120
0.00000 0.00000 0.00000 ; CD1- NE1- CE2- CD2
4306 4305 4304 4318 3 54.60120 0.00000 -54.60120
0.00000 0.00000 0.00000 ; HD1- CD1- CG- CD2
4306 4305 4307 4308 3 12.55200 0.00000 -12.55200
0.00000 0.00000 0.00000 ; HD1- CD1- NE1- HE1
4306 4305 4307 4309 3 12.55200 0.00000 -12.55200
0.00000 0.00000 0.00000 ; HD1- CD1- NE1- CE2
4307 4305 4304 4318 3 54.60120 0.00000 -54.60120
0.00000 0.00000 0.00000 ; NE1- CD1- CG- CD2
4307 4309 4310 4311 3 30.33400 0.00000 -30.33400
0.00000 0.00000 0.00000 ; NE1- CE2- CZ2- HZ2
4307 4309 4310 4312 3 30.33400 0.00000 -30.33400
0.00000 0.00000 0.00000 ; NE1- CE2- CZ2- CH2
4307 4309 4318 4316 3 25.10400 0.00000 -25.10400
0.00000 0.00000 0.00000 ; NE1- CE2- CD2- CE3
4308 4307 4309 4310 3 12.76120 0.00000 -12.76120
0.00000 0.00000 0.00000 ; HE1- NE1- CE2- CZ2
4308 4307 4309 4318 3 12.76120 0.00000 -12.76120
0.00000 0.00000 0.00000 ; HE1- NE1- CE2- CD2
4309 4310 4312 4313 3 30.33400 0.00000 -30.33400
0.00000 0.00000 0.00000 ; CE2- CZ2- CH2- HH2
4309 4310 4312 4314 3 30.33400 0.00000 -30.33400
0.00000 0.00000 0.00000 ; CE2- CZ2- CH2- CZ3
4309 4318 4316 4314 3 29.28800 0.00000 -29.28800
0.00000 0.00000 0.00000 ; CE2- CD2- CE3- CZ3
4309 4318 4316 4317 3 29.28800 0.00000 -29.28800
0.00000 0.00000 0.00000 ; CE2- CD2- CE3- HE3
4310 4309 4318 4316 3 25.10400 0.00000 -25.10400
0.00000 0.00000 0.00000 ; CZ2- CE2- CD2- CE3
4310 4312 4314 4315 3 30.33400 0.00000 -30.33400
0.00000 0.00000 0.00000 ; CZ2- CH2- CZ3- HZ3
4310 4312 4314 4316 3 30.33400 0.00000 -30.33400
0.00000 0.00000 0.00000 ; CZ2- CH2- CZ3- CE3
4311 4310 4309 4318 3 30.33400 0.00000 -30.33400
0.00000 0.00000 0.00000 ; HZ2- CZ2- CE2- CD2
4311 4310 4312 4313 3 30.33400 0.00000 -30.33400
0.00000 0.00000 0.00000 ; HZ2- CZ2- CH2- HH2
4311 4310 4312 4314 3 30.33400 0.00000 -30.33400
0.00000 0.00000 0.00000 ; HZ2- CZ2- CH2- CZ3
4312 4310 4309 4318 3 30.33400 0.00000 -30.33400
0.00000 0.00000 0.00000 ; CH2- CZ2- CE2- CD2
4312 4314 4316 4317 3 30.33400 0.00000 -30.33400
0.00000 0.00000 0.00000 ; CH2- CZ3- CE3- HE3
4312 4314 4316 4318 3 30.33400 0.00000 -30.33400
0.00000 0.00000 0.00000 ; CH2- CZ3- CE3- CD2
4313 4312 4314 4315 3 30.33400 0.00000 -30.33400
0.00000 0.00000 0.00000 ; HH2- CH2- CZ3- HZ3
4313 4312 4314 4316 3 30.33400 0.00000 -30.33400
0.00000 0.00000 0.00000 ; HH2- CH2- CZ3- CE3

4315 4314 4316 4317 3 30.33400 0.00000 -30.33400
0.00000 0.00000 0.00000 ; HZ3- CZ3- CE3- HE3
4315 4314 4316 4318 3 30.33400 0.00000 -30.33400
0.00000 0.00000 0.00000 ; HZ3- CZ3- CE3- CD2
4319 4321 4323 4324 3 0.00000 0.00000 0.00000
0.00000 0.00000 0.00000 ; C- N- CA- HA
4319 4321 4323 4325 3 11.71520 1.67360 15.06240 -
13.38880 0.00000 0.00000 ; C- N- CA- CB
4319 4321 4323 4338 3 1.75728 5.27184 2.25936
-7.02912 0.00000 0.00000 ; C- N- CA- C
4320 4319 4321 4322 3 29.28800 -8.36800 -20.92000
0.00000 0.00000 0.00000 ; O- C- N- H
4320 4319 4321 4323 3 20.92000 0.00000 -20.92000
0.00000 0.00000 0.00000 ; O- C- N- CA
4321 4323 4325 4326 3 0.65084 1.95253 0.00000
-2.60338 0.00000 0.00000 ; N- CA- CB- HB
4321 4323 4325 4327 3 3.83673 0.44769 -1.77402
-2.47693 -0.03347 0.00000 ; N- CA- CB- CG2
4321 4323 4325 4331 3 2.97482 0.12134 2.04179
-1.89117 -3.24678 0.00000 ; N- CA- CB- CG1
4321 4323 4338 4339 3 0.00000 0.00000 0.00000
0.00000 0.00000 0.00000 ; N- CA- C- O
4321 4323 4338 4340 3 17.40544 -5.02080 -13.22144
9.20480 0.00000 0.00000 ; N- CA- C- N
4322 4321 4323 4324 3 0.00000 0.00000 0.00000
0.00000 0.00000 0.00000 ; H- N- CA- HA
4322 4321 4323 4325 3 0.00000 0.00000 0.00000
0.00000 0.00000 0.00000 ; H- N- CA- CB
4322 4321 4323 4338 3 0.00000 0.00000 0.00000
0.00000 0.00000 0.00000 ; H- N- CA- C
4323 4325 4327 4328 3 0.66944 2.00832 0.00000
-2.67776 0.00000 0.00000 ; CA- CB- CG2- HG21
4323 4325 4327 4329 3 0.66944 2.00832 0.00000
-2.67776 0.00000 0.00000 ; CA- CB- CG2- HG22
4323 4325 4327 4330 3 0.66944 2.00832 0.00000
-2.67776 0.00000 0.00000 ; CA- CB- CG2- HG23
4323 4325 4331 4332 3 0.66944 2.00832 0.00000
-2.67776 0.00000 0.00000 ; CA- CB- CG1- HG12
4323 4325 4331 4333 3 0.66944 2.00832 0.00000
-2.67776 0.00000 0.00000 ; CA- CB- CG1- HG13
4323 4325 4331 4334 3 4.24258 -0.52718 -7.25506
-1.79075 7.69856 0.00000 ; CA- CB- CG1- CD1
4323 4338 4340 4341 3 20.92000 0.00000 -20.92000
0.00000 0.00000 0.00000 ; CA- C- N- H
4323 4338 4340 4342 3 20.92000 0.00000 -20.92000
0.00000 0.00000 0.00000 ; CA- C- N- CA
4324 4323 4325 4326 3 0.65084 1.95253 0.00000
-2.60338 0.00000 0.00000 ; HA- CA- CB- HB
4324 4323 4325 4327 3 0.65084 1.95253 0.00000
-2.60338 0.00000 0.00000 ; HA- CA- CB- CG2
4324 4323 4325 4331 3 0.65084 1.95253 0.00000
-2.60338 0.00000 0.00000 ; HA- CA- CB- CG1

4324 4323 4338 4339 3 3.68192 -4.35136 0.00000
1.33888 0.00000 0.00000 ; HA- CA- C- O
4324 4323 4338 4340 3 0.00000 0.00000 0.00000
0.00000 0.00000 0.00000 ; HA- CA- C- N
4325 4323 4338 4339 3 0.00000 0.00000 0.00000
0.00000 0.00000 0.00000 ; CB- CA- C- O
4325 4323 4338 4340 3 2.51040 4.18400 1.67360
-6.69440 0.00000 0.00000 ; CB- CA- C- N
4325 4331 4334 4335 3 0.66944 2.00832 0.00000
-2.67776 0.00000 0.00000 ; CB- CG1- CD1- HD11
4325 4331 4334 4336 3 0.66944 2.00832 0.00000
-2.67776 0.00000 0.00000 ; CB- CG1- CD1- HD12
4325 4331 4334 4337 3 0.66944 2.00832 0.00000
-2.67776 0.00000 0.00000 ; CB- CG1- CD1- HD13
4326 4325 4323 4338 3 0.65084 1.95253 0.00000
-2.60338 0.00000 0.00000 ; HB- CB- CA- C
4326 4325 4327 4328 3 0.62760 1.88280 0.00000
-2.51040 0.00000 0.00000 ; HB- CB- CG2- HG21
4326 4325 4327 4329 3 0.62760 1.88280 0.00000
-2.51040 0.00000 0.00000 ; HB- CB- CG2- HG22
4326 4325 4327 4330 3 0.62760 1.88280 0.00000
-2.51040 0.00000 0.00000 ; HB- CB- CG2- HG23
4326 4325 4331 4332 3 0.62760 1.88280 0.00000
-2.51040 0.00000 0.00000 ; HB- CB- CG1- HG12
4326 4325 4331 4333 3 0.62760 1.88280 0.00000
-2.51040 0.00000 0.00000 ; HB- CB- CG1- HG13
4326 4325 4331 4334 3 0.66944 2.00832 0.00000
-2.67776 0.00000 0.00000 ; HB- CB- CG1- CD1
4327 4325 4323 4338 3 5.67350 3.55640 -6.16722
-2.47693 3.74886 0.00000 ; CG2- CB- CA- C
4327 4325 4331 4332 3 0.66944 2.00832 0.00000
-2.67776 0.00000 0.00000 ; CG2- CB- CG1- HG12
4327 4325 4331 4333 3 0.66944 2.00832 0.00000
-2.67776 0.00000 0.00000 ; CG2- CB- CG1- HG13
4327 4325 4331 4334 3 3.81162 0.49790 -8.14206
-1.79075 7.49773 0.00000 ; CG2- CB- CG1- CD1
4328 4327 4325 4331 3 0.66944 2.00832 0.00000
-2.67776 0.00000 0.00000 ; HG21- CG2- CB- CG1
4329 4327 4325 4331 3 0.66944 2.00832 0.00000
-2.67776 0.00000 0.00000 ; HG22- CG2- CB- CG1
4330 4327 4325 4331 3 0.66944 2.00832 0.00000
-2.67776 0.00000 0.00000 ; HG23- CG2- CB- CG1
4331 4325 4323 4338 3 8.26340 0.74057 -9.99976
-1.89117 3.84928 0.00000 ; CG1- CB- CA- C
4332 4331 4334 4335 3 0.62760 1.88280 0.00000
-2.51040 0.00000 0.00000 ; HG12- CG1- CD1- HD11
4332 4331 4334 4336 3 0.62760 1.88280 0.00000
-2.51040 0.00000 0.00000 ; HG12- CG1- CD1- HD12
4332 4331 4334 4337 3 0.62760 1.88280 0.00000
-2.51040 0.00000 0.00000 ; HG12- CG1- CD1- HD13
4333 4331 4334 4335 3 0.62760 1.88280 0.00000
-2.51040 0.00000 0.00000 ; HG13- CG1- CD1- HD11

4333 4331 4334 4336 3 0.62760 1.88280 0.00000
-2.51040 0.00000 0.00000 ; HG13- CG1- CD1- HD12
4333 4331 4334 4337 3 0.62760 1.88280 0.00000
-2.51040 0.00000 0.00000 ; HG13- CG1- CD1- HD13
4338 4340 4342 4343 3 0.00000 0.00000 0.00000
0.00000 0.00000 0.00000 ; C- N- CA- HA
4338 4340 4342 4344 3 11.71520 1.67360 15.06240 -
13.38880 0.00000 0.00000 ; C- N- CA- CB
4338 4340 4342 4352 3 1.75728 5.27184 2.25936
-7.02912 0.00000 0.00000 ; C- N- CA- C
4339 4338 4340 4341 3 29.28800 -8.36800 -20.92000
0.00000 0.00000 0.00000 ; O- C- N- H
4339 4338 4340 4342 3 20.92000 0.00000 -20.92000
0.00000 0.00000 0.00000 ; O- C- N- CA
4340 4342 4344 4345 3 0.65084 1.95253 0.00000
-2.60338 0.00000 0.00000 ; N- CA- CB- HB
4340 4342 4344 4346 3 3.83673 0.44769 -1.77402
-2.47693 -0.03347 0.00000 ; N- CA- CB- CG2
4340 4342 4344 4350 3 4.93294 1.13386 -3.12963
-5.27184 3.17984 0.00000 ; N- CA- CB- OG1
4340 4342 4352 4353 3 0.00000 0.00000 0.00000
0.00000 0.00000 0.00000 ; N- CA- C- O
4340 4342 4352 4354 3 17.40544 -5.02080 -13.22144
9.20480 0.00000 0.00000 ; N- CA- C- N
4341 4340 4342 4343 3 0.00000 0.00000 0.00000
0.00000 0.00000 0.00000 ; H- N- CA- HA
4341 4340 4342 4344 3 0.00000 0.00000 0.00000
0.00000 0.00000 0.00000 ; H- N- CA- CB
4341 4340 4342 4352 3 0.00000 0.00000 0.00000
0.00000 0.00000 0.00000 ; H- N- CA- C
4342 4344 4346 4347 3 0.66944 2.00832 0.00000
-2.67776 0.00000 0.00000 ; CA- CB- CG2- HG21
4342 4344 4346 4348 3 0.66944 2.00832 0.00000
-2.67776 0.00000 0.00000 ; CA- CB- CG2- HG22
4342 4344 4346 4349 3 0.66944 2.00832 0.00000
-2.67776 0.00000 0.00000 ; CA- CB- CG2- HG23
4342 4344 4350 4351 3 1.12131 2.98738 1.66523
-3.94970 0.43514 0.00000 ; CA- CB- OG1- HG1
4342 4352 4354 4355 3 20.92000 0.00000 -20.92000
0.00000 0.00000 0.00000 ; CA- C- N- H
4342 4352 4354 4356 3 20.92000 0.00000 -20.92000
0.00000 0.00000 0.00000 ; CA- C- N- CA
4343 4342 4344 4345 3 0.65084 1.95253 0.00000
-2.60338 0.00000 0.00000 ; HA- CA- CB- HB
4343 4342 4344 4346 3 0.65084 1.95253 0.00000
-2.60338 0.00000 0.00000 ; HA- CA- CB- CG2
4343 4342 4344 4350 3 1.04600 -1.04600 0.00000
0.00000 0.00000 0.00000 ; HA- CA- CB- OG1
4343 4342 4352 4353 3 3.68192 -4.35136 0.00000
1.33888 0.00000 0.00000 ; HA- CA- C- O
4343 4342 4352 4354 3 0.00000 0.00000 0.00000
0.00000 0.00000 0.00000 ; HA- CA- C- N

4344 4342 4352 4353 3 0.00000 0.00000 0.00000
0.00000 0.00000 0.00000 ; CB- CA- C- O
4344 4342 4352 4354 3 2.51040 4.18400 1.67360
-6.69440 0.00000 0.00000 ; CB- CA- C- N
4345 4344 4342 4352 3 0.65084 1.95253 0.00000
-2.60338 0.00000 0.00000 ; HB- CB- CA- C
4345 4344 4346 4347 3 0.65084 1.95253 0.00000
-2.60338 0.00000 0.00000 ; HB- CB- CG2- HG21
4345 4344 4346 4348 3 0.65084 1.95253 0.00000
-2.60338 0.00000 0.00000 ; HB- CB- CG2- HG22
4345 4344 4346 4349 3 0.65084 1.95253 0.00000
-2.60338 0.00000 0.00000 ; HB- CB- CG2- HG23
4345 4344 4350 4351 3 0.69733 2.09200 0.00000
-2.78933 0.00000 0.00000 ; HB- CB- OG1- HG1
4346 4344 4342 4352 3 5.67350 3.55640 -6.16722
-2.47693 3.74886 0.00000 ; CG2- CB- CA- C
4346 4344 4350 4351 3 4.74047 0.27196 -2.26773
-3.94970 1.60666 0.00000 ; CG2- CB- OG1- HG1
4347 4346 4344 4350 3 1.04600 -1.04600 0.00000
0.00000 0.00000 0.00000 ; HG21- CG2- CB- OG1
4348 4346 4344 4350 3 1.04600 -1.04600 0.00000
0.00000 0.00000 0.00000 ; HG22- CG2- CB- OG1
4349 4346 4344 4350 3 1.04600 -1.04600 0.00000
0.00000 0.00000 0.00000 ; HG23- CG2- CB- OG1
4350 4344 4342 4352 3 6.53541 6.87013 -6.21742
-5.27184 5.22163 0.00000 ; OG1- CB- CA- C
4352 4354 4356 4357 3 0.00000 0.00000 0.00000
0.00000 0.00000 0.00000 ; C- N- CA- HA
4352 4354 4356 4358 3 11.71520 1.67360 15.06240 -
13.38880 0.00000 0.00000 ; C- N- CA- CB
4352 4354 4356 4369 3 1.75728 5.27184 2.25936
-7.02912 0.00000 0.00000 ; C- N- CA- C
4353 4352 4354 4355 3 29.28800 -8.36800 -20.92000
0.00000 0.00000 0.00000 ; O- C- N- H
4353 4352 4354 4356 3 20.92000 0.00000 -20.92000
0.00000 0.00000 0.00000 ; O- C- N- CA
4354 4356 4358 4359 3 0.65084 1.95253 0.00000
-2.60338 0.00000 0.00000 ; N- CA- CB- HB2
4354 4356 4358 4360 3 0.65084 1.95253 0.00000
-2.60338 0.00000 0.00000 ; N- CA- CB- HB3
4354 4356 4358 4361 3 3.21331 2.22589 -4.15053
-2.40998 2.61082 0.00000 ; N- CA- CB- CG
4354 4356 4369 4370 3 0.00000 0.00000 0.00000
0.00000 0.00000 0.00000 ; N- CA- C- O
4354 4356 4369 4371 3 17.40544 -5.02080 -13.22144
9.20480 0.00000 0.00000 ; N- CA- C- N
4355 4354 4356 4357 3 0.00000 0.00000 0.00000
0.00000 0.00000 0.00000 ; H- N- CA- HA
4355 4354 4356 4358 3 0.00000 0.00000 0.00000
0.00000 0.00000 0.00000 ; H- N- CA- CB
4355 4354 4356 4369 3 0.00000 0.00000 0.00000
0.00000 0.00000 0.00000 ; H- N- CA- C

4356 4358 4361 4362 3 0.66944 2.00832 0.00000
-2.67776 0.00000 0.00000 ; CA- CB- CG- HG2
4356 4358 4361 4363 3 0.66944 2.00832 0.00000
-2.67776 0.00000 0.00000 ; CA- CB- CG- HG3
4356 4358 4361 4364 3 3.69866 -4.35136 -3.92459
6.89523 4.61914 0.00000 ; CA- CB- CG- CD
4356 4369 4371 4372 3 20.92000 0.00000 -20.92000
0.00000 0.00000 0.00000 ; CA- C- N- H
4356 4369 4371 4373 3 20.92000 0.00000 -20.92000
0.00000 0.00000 0.00000 ; CA- C- N- CA
4357 4356 4358 4359 3 0.65084 1.95253 0.00000
-2.60338 0.00000 0.00000 ; HA- CA- CB- HB2
4357 4356 4358 4360 3 0.65084 1.95253 0.00000
-2.60338 0.00000 0.00000 ; HA- CA- CB- HB3
4357 4356 4358 4361 3 0.65084 1.95253 0.00000
-2.60338 0.00000 0.00000 ; HA- CA- CB- CG
4357 4356 4369 4370 3 3.68192 -4.35136 0.00000
1.33888 0.00000 0.00000 ; HA- CA- C- O
4357 4356 4369 4371 3 0.00000 0.00000 0.00000
0.00000 0.00000 0.00000 ; HA- CA- C- N
4358 4356 4369 4370 3 0.00000 0.00000 0.00000
0.00000 0.00000 0.00000 ; CB- CA- C- O
4358 4356 4369 4371 3 2.51040 4.18400 1.67360
-6.69440 0.00000 0.00000 ; CB- CA- C- N
4358 4361 4364 4365 3 0.00000 0.00000 0.00000
0.00000 0.00000 0.00000 ; CB- CG- CD- OE1
4358 4361 4364 4366 3 10.32611 1.48114 -8.47678
1.42256 1.40582 0.00000 ; CB- CG- CD- NE2
4359 4358 4356 4369 3 0.65084 1.95253 0.00000
-2.60338 0.00000 0.00000 ; HB2- CB- CA- C
4359 4358 4361 4362 3 0.62760 1.88280 0.00000
-2.51040 0.00000 0.00000 ; HB2- CB- CG- HG2
4359 4358 4361 4363 3 0.62760 1.88280 0.00000
-2.51040 0.00000 0.00000 ; HB2- CB- CG- HG3
4359 4358 4361 4364 3 0.65084 1.95253 0.00000
-2.60338 0.00000 0.00000 ; HB2- CB- CG- CD
4360 4358 4356 4369 3 0.65084 1.95253 0.00000
-2.60338 0.00000 0.00000 ; HB3- CB- CA- C
4360 4358 4361 4362 3 0.62760 1.88280 0.00000
-2.51040 0.00000 0.00000 ; HB3- CB- CG- HG2
4360 4358 4361 4363 3 0.62760 1.88280 0.00000
-2.51040 0.00000 0.00000 ; HB3- CB- CG- HG3
4360 4358 4361 4364 3 0.65084 1.95253 0.00000
-2.60338 0.00000 0.00000 ; HB3- CB- CG- CD
4361 4358 4356 4369 3 6.86594 3.56895 -8.14206
-2.40998 4.85344 0.00000 ; CG- CB- CA- C
4361 4364 4366 4367 3 20.92000 0.00000 -20.92000
0.00000 0.00000 0.00000 ; CG- CD- NE2- HE21
4361 4364 4366 4368 3 20.92000 0.00000 -20.92000
0.00000 0.00000 0.00000 ; CG- CD- NE2- HE22
4362 4361 4364 4365 3 3.68192 -4.35136 0.00000
1.33888 0.00000 0.00000 ; HG2- CG- CD- OE1

4362 4361 4364 4366 3 0.00000 0.00000 0.00000
0.00000 0.00000 0.00000 ; HG2- CG- CD- NE2
4363 4361 4364 4365 3 3.68192 -4.35136 0.00000
1.33888 0.00000 0.00000 ; HG3- CG- CD- OE1
4363 4361 4364 4366 3 0.00000 0.00000 0.00000
0.00000 0.00000 0.00000 ; HG3- CG- CD- NE2
4365 4364 4366 4367 3 29.28800 -8.36800 -20.92000
0.00000 0.00000 0.00000 ; OE1- CD- NE2- HE21
4365 4364 4366 4368 3 29.28800 -8.36800 -20.92000
0.00000 0.00000 0.00000 ; OE1- CD- NE2- HE22
4369 4371 4373 4374 3 0.00000 0.00000 0.00000
0.00000 0.00000 0.00000 ; C- N- CA- HA
4369 4371 4373 4375 3 11.71520 1.67360 15.06240 -
13.38880 0.00000 0.00000 ; C- N- CA- CB
4369 4371 4373 4386 3 1.75728 5.27184 2.25936
-7.02912 0.00000 0.00000 ; C- N- CA- C
4370 4369 4371 4372 3 29.28800 -8.36800 -20.92000
0.00000 0.00000 0.00000 ; O- C- N- H
4370 4369 4371 4373 3 20.92000 0.00000 -20.92000
0.00000 0.00000 0.00000 ; O- C- N- CA
4371 4373 4375 4376 3 0.65084 1.95253 0.00000
-2.60338 0.00000 0.00000 ; N- CA- CB- HB2
4371 4373 4375 4377 3 0.65084 1.95253 0.00000
-2.60338 0.00000 0.00000 ; N- CA- CB- HB3
4371 4373 4375 4378 3 3.21331 2.22589 -4.15053
-2.40998 2.61082 0.00000 ; N- CA- CB- CG
4371 4373 4386 4387 3 0.00000 0.00000 0.00000
0.00000 0.00000 0.00000 ; N- CA- C- O
4371 4373 4386 4388 3 17.40544 -5.02080 -13.22144
9.20480 0.00000 0.00000 ; N- CA- C- N
4372 4371 4373 4374 3 0.00000 0.00000 0.00000
0.00000 0.00000 0.00000 ; H- N- CA- HA
4372 4371 4373 4375 3 0.00000 0.00000 0.00000
0.00000 0.00000 0.00000 ; H- N- CA- CB
4372 4371 4373 4386 3 0.00000 0.00000 0.00000
0.00000 0.00000 0.00000 ; H- N- CA- C
4373 4375 4378 4379 3 0.65084 1.95253 0.00000
-2.60338 0.00000 0.00000 ; CA- CB- CG- HG2
4373 4375 4378 4380 3 0.65084 1.95253 0.00000
-2.60338 0.00000 0.00000 ; CA- CB- CG- HG3
4373 4375 4378 4381 3 2.04598 -1.54390 1.11294
-0.26778 0.93722 0.00000 ; CA- CB- CG- SD
4373 4386 4388 4389 3 20.92000 0.00000 -20.92000
0.00000 0.00000 0.00000 ; CA- C- N- H
4373 4386 4388 4390 3 20.92000 0.00000 -20.92000
0.00000 0.00000 0.00000 ; CA- C- N- CA
4374 4373 4375 4376 3 0.65084 1.95253 0.00000
-2.60338 0.00000 0.00000 ; HA- CA- CB- HB2
4374 4373 4375 4377 3 0.65084 1.95253 0.00000
-2.60338 0.00000 0.00000 ; HA- CA- CB- HB3
4374 4373 4375 4378 3 0.65084 1.95253 0.00000
-2.60338 0.00000 0.00000 ; HA- CA- CB- CG

4374 4373 4386 4387 3 3.68192 -4.35136 0.00000
1.33888 0.00000 0.00000 ; HA- CA- C- O
4374 4373 4386 4388 3 0.00000 0.00000 0.00000
0.00000 0.00000 0.00000 ; HA- CA- C- N
4375 4373 4386 4387 3 0.00000 0.00000 0.00000
0.00000 0.00000 0.00000 ; CB- CA- C- O
4375 4373 4386 4388 3 2.51040 4.18400 1.67360
-6.69440 0.00000 0.00000 ; CB- CA- C- N
4375 4378 4381 4382 3 3.24260 6.22998 1.79075
-6.92870 1.90790 0.00000 ; CB- CG- SD- CE
4376 4375 4373 4386 3 0.65084 1.95253 0.00000
-2.60338 0.00000 0.00000 ; HB2- CB- CA- C
4376 4375 4378 4379 3 0.65084 1.95253 0.00000
-2.60338 0.00000 0.00000 ; HB2- CB- CG- HG2
4376 4375 4378 4380 3 0.65084 1.95253 0.00000
-2.60338 0.00000 0.00000 ; HB2- CB- CG- HG3
4376 4375 4378 4381 3 0.65084 1.95253 0.00000
-2.60338 0.00000 0.00000 ; HB2- CB- CG- SD
4377 4375 4373 4386 3 0.65084 1.95253 0.00000
-2.60338 0.00000 0.00000 ; HB3- CB- CA- C
4377 4375 4378 4379 3 0.65084 1.95253 0.00000
-2.60338 0.00000 0.00000 ; HB3- CB- CG- HG2
4377 4375 4378 4380 3 0.65084 1.95253 0.00000
-2.60338 0.00000 0.00000 ; HB3- CB- CG- HG3
4377 4375 4378 4381 3 0.65084 1.95253 0.00000
-2.60338 0.00000 0.00000 ; HB3- CB- CG- SD
4378 4375 4373 4386 3 6.86594 3.56895 -8.14206
-2.40998 4.85344 0.00000 ; CG- CB- CA- C
4378 4381 4382 4383 3 1.39467 4.18400 0.00000
-5.57867 0.00000 0.00000 ; CG- SD- CE- HE1
4378 4381 4382 4384 3 1.39467 4.18400 0.00000
-5.57867 0.00000 0.00000 ; CG- SD- CE- HE2
4378 4381 4382 4385 3 1.39467 4.18400 0.00000
-5.57867 0.00000 0.00000 ; CG- SD- CE- HE3
4379 4378 4381 4382 3 1.39467 4.18400 0.00000
-5.57867 0.00000 0.00000 ; HG2- CG- SD- CE
4380 4378 4381 4382 3 1.39467 4.18400 0.00000
-5.57867 0.00000 0.00000 ; HG3- CG- SD- CE
4386 4388 4390 4391 3 0.00000 0.00000 0.00000
0.00000 0.00000 0.00000 ; C- N- CA- HA
4386 4388 4390 4392 3 11.71520 1.67360 15.06240 -
13.38880 0.00000 0.00000 ; C- N- CA- CB
4386 4388 4390 4400 3 1.75728 5.27184 2.25936
-7.02912 0.00000 0.00000 ; C- N- CA- C
4387 4386 4388 4389 3 29.28800 -8.36800 -20.92000
0.00000 0.00000 0.00000 ; O- C- N- H
4387 4386 4388 4390 3 20.92000 0.00000 -20.92000
0.00000 0.00000 0.00000 ; O- C- N- CA
4388 4390 4392 4393 3 0.65084 1.95253 0.00000
-2.60338 0.00000 0.00000 ; N- CA- CB- HB
4388 4390 4392 4394 3 3.83673 0.44769 -1.77402
-2.47693 -0.03347 0.00000 ; N- CA- CB- CG2

4388 4390 4392 4398 3 4.93294 1.13386 -3.12963
-5.27184 3.17984 0.00000 ; N- CA- CB- OG1
4388 4390 4400 4401 3 0.00000 0.00000 0.00000
0.00000 0.00000 0.00000 ; N- CA- C- O
4388 4390 4400 4402 3 17.40544 -5.02080 -13.22144
9.20480 0.00000 0.00000 ; N- CA- C- N
4389 4388 4390 4391 3 0.00000 0.00000 0.00000
0.00000 0.00000 0.00000 ; H- N- CA- HA
4389 4388 4390 4392 3 0.00000 0.00000 0.00000
0.00000 0.00000 0.00000 ; H- N- CA- CB
4389 4388 4390 4400 3 0.00000 0.00000 0.00000
0.00000 0.00000 0.00000 ; H- N- CA- C
4390 4392 4394 4395 3 0.66944 2.00832 0.00000
-2.67776 0.00000 0.00000 ; CA- CB- CG2- HG21
4390 4392 4394 4396 3 0.66944 2.00832 0.00000
-2.67776 0.00000 0.00000 ; CA- CB- CG2- HG22
4390 4392 4394 4397 3 0.66944 2.00832 0.00000
-2.67776 0.00000 0.00000 ; CA- CB- CG2- HG23
4390 4392 4398 4399 3 1.12131 2.98738 1.66523
-3.94970 0.43514 0.00000 ; CA- CB- OG1- HG1
4390 4400 4402 4403 3 20.92000 0.00000 -20.92000
0.00000 0.00000 0.00000 ; CA- C- N- H
4390 4400 4402 4404 3 20.92000 0.00000 -20.92000
0.00000 0.00000 0.00000 ; CA- C- N- CA
4391 4390 4392 4393 3 0.65084 1.95253 0.00000
-2.60338 0.00000 0.00000 ; HA- CA- CB- HB
4391 4390 4392 4394 3 0.65084 1.95253 0.00000
-2.60338 0.00000 0.00000 ; HA- CA- CB- CG2
4391 4390 4392 4398 3 1.04600 -1.04600 0.00000
0.00000 0.00000 0.00000 ; HA- CA- CB- OG1
4391 4390 4400 4401 3 3.68192 -4.35136 0.00000
1.33888 0.00000 0.00000 ; HA- CA- C- O
4391 4390 4400 4402 3 0.00000 0.00000 0.00000
0.00000 0.00000 0.00000 ; HA- CA- C- N
4392 4390 4400 4401 3 0.00000 0.00000 0.00000
0.00000 0.00000 0.00000 ; CB- CA- C- O
4392 4390 4400 4402 3 2.51040 4.18400 1.67360
-6.69440 0.00000 0.00000 ; CB- CA- C- N
4393 4392 4390 4400 3 0.65084 1.95253 0.00000
-2.60338 0.00000 0.00000 ; HB- CB- CA- C
4393 4392 4394 4395 3 0.65084 1.95253 0.00000
-2.60338 0.00000 0.00000 ; HB- CB- CG2- HG21
4393 4392 4394 4396 3 0.65084 1.95253 0.00000
-2.60338 0.00000 0.00000 ; HB- CB- CG2- HG22
4393 4392 4394 4397 3 0.65084 1.95253 0.00000
-2.60338 0.00000 0.00000 ; HB- CB- CG2- HG23
4393 4392 4398 4399 3 0.69733 2.09200 0.00000
-2.78933 0.00000 0.00000 ; HB- CB- OG1- HG1
4394 4392 4390 4400 3 5.67350 3.55640 -6.16722
-2.47693 3.74886 0.00000 ; CG2- CB- CA- C
4394 4392 4398 4399 3 4.74047 0.27196 -2.26773
-3.94970 1.60666 0.00000 ; CG2- CB- OG1- HG1

4395 4394 4392 4398 3 1.04600 -1.04600 0.00000
0.00000 0.00000 0.00000 ; HG21- CG2- CB- OG1
4396 4394 4392 4398 3 1.04600 -1.04600 0.00000
0.00000 0.00000 0.00000 ; HG22- CG2- CB- OG1
4397 4394 4392 4398 3 1.04600 -1.04600 0.00000
0.00000 0.00000 0.00000 ; HG23- CG2- CB- OG1
4398 4392 4390 4400 3 6.53541 6.87013 -6.21742
-5.27184 5.22163 0.00000 ; OG1- CB- CA- C
4400 4402 4404 4405 3 0.00000 0.00000 0.00000
0.00000 0.00000 0.00000 ; C- N- CA- HA
4400 4402 4404 4406 3 11.71520 1.67360 15.06240 -
13.38880 0.00000 0.00000 ; C- N- CA- CB
4400 4402 4404 4422 3 1.75728 5.27184 2.25936
-7.02912 0.00000 0.00000 ; C- N- CA- C
4401 4400 4402 4403 3 29.28800 -8.36800 -20.92000
0.00000 0.00000 0.00000 ; O- C- N- H
4401 4400 4402 4404 3 20.92000 0.00000 -20.92000
0.00000 0.00000 0.00000 ; O- C- N- CA
4402 4404 4406 4407 3 0.65084 1.95253 0.00000
-2.60338 0.00000 0.00000 ; N- CA- CB- HB2
4402 4404 4406 4408 3 0.65084 1.95253 0.00000
-2.60338 0.00000 0.00000 ; N- CA- CB- HB3
4402 4404 4406 4409 3 0.65084 1.95253 0.00000
-2.60338 0.00000 0.00000 ; N- CA- CB- CG
4402 4404 4422 4423 3 0.00000 0.00000 0.00000
0.00000 0.00000 0.00000 ; N- CA- C- O
4402 4404 4422 4424 3 17.40544 -5.02080 -13.22144
9.20480 0.00000 0.00000 ; N- CA- C- N
4403 4402 4404 4405 3 0.00000 0.00000 0.00000
0.00000 0.00000 0.00000 ; H- N- CA- HA
4403 4402 4404 4406 3 0.00000 0.00000 0.00000
0.00000 0.00000 0.00000 ; H- N- CA- CB
4403 4402 4404 4422 3 0.00000 0.00000 0.00000
0.00000 0.00000 0.00000 ; H- N- CA- C
4404 4406 4409 4410 3 0.66944 2.00832 0.00000
-2.67776 0.00000 0.00000 ; CA- CB- CG- HG2
4404 4406 4409 4411 3 0.66944 2.00832 0.00000
-2.67776 0.00000 0.00000 ; CA- CB- CG- HG3
4404 4406 4409 4412 3 3.68192 3.09616 -2.09200
-3.01248 0.00000 0.00000 ; CA- CB- CG- CD
4404 4422 4424 4425 3 20.92000 0.00000 -20.92000
0.00000 0.00000 0.00000 ; CA- C- N- H
4404 4422 4424 4426 3 20.92000 0.00000 -20.92000
0.00000 0.00000 0.00000 ; CA- C- N- CA
4405 4404 4406 4407 3 0.65084 1.95253 0.00000
-2.60338 0.00000 0.00000 ; HA- CA- CB- HB2
4405 4404 4406 4408 3 0.65084 1.95253 0.00000
-2.60338 0.00000 0.00000 ; HA- CA- CB- HB3
4405 4404 4406 4409 3 0.65084 1.95253 0.00000
-2.60338 0.00000 0.00000 ; HA- CA- CB- CG
4405 4404 4422 4423 3 3.68192 -4.35136 0.00000
1.33888 0.00000 0.00000 ; HA- CA- C- O

4405 4404 4422 4424 3 0.00000 0.00000 0.00000
0.00000 0.00000 0.00000 ; HA- CA- C- N
4406 4404 4422 4423 3 0.00000 0.00000 0.00000
0.00000 0.00000 0.00000 ; CB- CA- C- O
4406 4404 4422 4424 3 2.51040 4.18400 1.67360
-6.69440 0.00000 0.00000 ; CB- CA- C- N
4406 4409 4412 4413 3 0.66944 2.00832 0.00000
-2.67776 0.00000 0.00000 ; CB- CG- CD- HD2
4406 4409 4412 4414 3 0.66944 2.00832 0.00000
-2.67776 0.00000 0.00000 ; CB- CG- CD- HD3
4406 4409 4412 4415 3 3.68192 3.09616 -2.09200
-3.01248 0.00000 0.00000 ; CB- CG- CD- CE
4407 4406 4404 4422 3 0.65084 1.95253 0.00000
-2.60338 0.00000 0.00000 ; HB2- CB- CA- C
4407 4406 4409 4410 3 0.62760 1.88280 0.00000
-2.51040 0.00000 0.00000 ; HB2- CB- CG- HG2
4407 4406 4409 4411 3 0.62760 1.88280 0.00000
-2.51040 0.00000 0.00000 ; HB2- CB- CG- HG3
4407 4406 4409 4412 3 0.66944 2.00832 0.00000
-2.67776 0.00000 0.00000 ; HB2- CB- CG- CD
4408 4406 4404 4422 3 0.65084 1.95253 0.00000
-2.60338 0.00000 0.00000 ; HB3- CB- CA- C
4408 4406 4409 4410 3 0.62760 1.88280 0.00000
-2.51040 0.00000 0.00000 ; HB3- CB- CG- HG2
4408 4406 4409 4411 3 0.62760 1.88280 0.00000
-2.51040 0.00000 0.00000 ; HB3- CB- CG- HG3
4408 4406 4409 4412 3 0.66944 2.00832 0.00000
-2.67776 0.00000 0.00000 ; HB3- CB- CG- CD
4409 4406 4404 4422 3 0.65084 1.95253 0.00000
-2.60338 0.00000 0.00000 ; CG- CB- CA- C
4409 4412 4415 4416 3 0.65084 1.95253 0.00000
-2.60338 0.00000 0.00000 ; CG- CD- CE- HE2
4409 4412 4415 4417 3 0.65084 1.95253 0.00000
-2.60338 0.00000 0.00000 ; CG- CD- CE- HE3
4409 4412 4415 4418 3 0.65084 1.95253 0.00000
-2.60338 0.00000 0.00000 ; CG- CD- CE- NZ
4410 4409 4412 4413 3 0.62760 1.88280 0.00000
-2.51040 0.00000 0.00000 ; HG2- CG- CD- HD2
4410 4409 4412 4414 3 0.62760 1.88280 0.00000
-2.51040 0.00000 0.00000 ; HG2- CG- CD- HD3
4410 4409 4412 4415 3 0.66944 2.00832 0.00000
-2.67776 0.00000 0.00000 ; HG2- CG- CD- CE
4411 4409 4412 4413 3 0.62760 1.88280 0.00000
-2.51040 0.00000 0.00000 ; HG3- CG- CD- HD2
4411 4409 4412 4414 3 0.62760 1.88280 0.00000
-2.51040 0.00000 0.00000 ; HG3- CG- CD- HD3
4411 4409 4412 4415 3 0.66944 2.00832 0.00000
-2.67776 0.00000 0.00000 ; HG3- CG- CD- CE
4412 4415 4418 4419 3 0.65084 1.95253 0.00000
-2.60338 0.00000 0.00000 ; CD- CE- NZ- HZ1
4412 4415 4418 4420 3 0.65084 1.95253 0.00000
-2.60338 0.00000 0.00000 ; CD- CE- NZ- HZ2

4412 4415 4418 4421 3 0.65084 1.95253 0.00000
-2.60338 0.00000 0.00000 ; CD- CE- NZ- HZ3
4413 4412 4415 4416 3 0.65084 1.95253 0.00000
-2.60338 0.00000 0.00000 ; HD2- CD- CE- HE2
4413 4412 4415 4417 3 0.65084 1.95253 0.00000
-2.60338 0.00000 0.00000 ; HD2- CD- CE- HE3
4413 4412 4415 4418 3 0.65084 1.95253 0.00000
-2.60338 0.00000 0.00000 ; HD2- CD- CE- NZ
4414 4412 4415 4416 3 0.65084 1.95253 0.00000
-2.60338 0.00000 0.00000 ; HD3- CD- CE- HE2
4414 4412 4415 4417 3 0.65084 1.95253 0.00000
-2.60338 0.00000 0.00000 ; HD3- CD- CE- HE3
4414 4412 4415 4418 3 0.65084 1.95253 0.00000
-2.60338 0.00000 0.00000 ; HD3- CD- CE- NZ
4416 4415 4418 4419 3 0.65084 1.95253 0.00000
-2.60338 0.00000 0.00000 ; HE2- CE- NZ- HZ1
4416 4415 4418 4420 3 0.65084 1.95253 0.00000
-2.60338 0.00000 0.00000 ; HE2- CE- NZ- HZ2
4416 4415 4418 4421 3 0.65084 1.95253 0.00000
-2.60338 0.00000 0.00000 ; HE2- CE- NZ- HZ3
4417 4415 4418 4419 3 0.65084 1.95253 0.00000
-2.60338 0.00000 0.00000 ; HE3- CE- NZ- HZ1
4417 4415 4418 4420 3 0.65084 1.95253 0.00000
-2.60338 0.00000 0.00000 ; HE3- CE- NZ- HZ2
4417 4415 4418 4421 3 0.65084 1.95253 0.00000
-2.60338 0.00000 0.00000 ; HE3- CE- NZ- HZ3
4422 4424 4426 4427 3 0.00000 0.00000 0.00000
0.00000 0.00000 0.00000 ; C- N- CA- HA
4422 4424 4426 4428 3 11.71520 1.67360 15.06240 -
13.38880 0.00000 0.00000 ; C- N- CA- CB
4422 4424 4426 4433 3 1.75728 5.27184 2.25936
-7.02912 0.00000 0.00000 ; C- N- CA- C
4423 4422 4424 4425 3 29.28800 -8.36800 -20.92000
0.00000 0.00000 0.00000 ; O- C- N- H
4423 4422 4424 4426 3 20.92000 0.00000 -20.92000
0.00000 0.00000 0.00000 ; O- C- N- CA
4424 4426 4428 4429 3 0.65084 1.95253 0.00000
-2.60338 0.00000 0.00000 ; N- CA- CB- HB2
4424 4426 4428 4430 3 0.65084 1.95253 0.00000
-2.60338 0.00000 0.00000 ; N- CA- CB- HB3
4424 4426 4428 4431 3 7.86174 2.24681 -7.41405
-6.71114 5.35552 0.00000 ; N- CA- CB- OG
4424 4426 4433 4434 3 0.00000 0.00000 0.00000
0.00000 0.00000 0.00000 ; N- CA- C- O
4424 4426 4433 4435 3 17.40544 -5.02080 -13.22144
9.20480 0.00000 0.00000 ; N- CA- C- N
4425 4424 4426 4427 3 0.00000 0.00000 0.00000
0.00000 0.00000 0.00000 ; H- N- CA- HA
4425 4424 4426 4428 3 0.00000 0.00000 0.00000
0.00000 0.00000 0.00000 ; H- N- CA- CB
4425 4424 4426 4433 3 0.00000 0.00000 0.00000
0.00000 0.00000 0.00000 ; H- N- CA- C

4426 4428 4431 4432 3 2.05853 2.46856 3.48109
-4.46851 0.23430 0.00000 ; CA- CB- OG- HG
4426 4433 4435 4436 3 20.92000 0.00000 -20.92000
0.00000 0.00000 0.00000 ; CA- C- N- H
4426 4433 4435 4437 3 20.92000 0.00000 -20.92000
0.00000 0.00000 0.00000 ; CA- C- N- CA
4427 4426 4428 4429 3 0.65084 1.95253 0.00000
-2.60338 0.00000 0.00000 ; HA- CA- CB- HB2
4427 4426 4428 4430 3 0.65084 1.95253 0.00000
-2.60338 0.00000 0.00000 ; HA- CA- CB- HB3
4427 4426 4428 4431 3 1.04600 -1.04600 0.00000
0.00000 0.00000 0.00000 ; HA- CA- CB- OG
4427 4426 4433 4434 3 3.68192 -4.35136 0.00000
1.33888 0.00000 0.00000 ; HA- CA- C- O
4427 4426 4433 4435 3 0.00000 0.00000 0.00000
0.00000 0.00000 0.00000 ; HA- CA- C- N
4428 4426 4433 4434 3 0.00000 0.00000 0.00000
0.00000 0.00000 0.00000 ; CB- CA- C- O
4428 4426 4433 4435 3 2.51040 4.18400 1.67360
-6.69440 0.00000 0.00000 ; CB- CA- C- N
4429 4428 4426 4433 3 0.65084 1.95253 0.00000
-2.60338 0.00000 0.00000 ; HB2- CB- CA- C
4429 4428 4431 4432 3 0.69733 2.09200 0.00000
-2.78933 0.00000 0.00000 ; HB2- CB- OG- HG
4430 4428 4426 4433 3 0.65084 1.95253 0.00000
-2.60338 0.00000 0.00000 ; HB3- CB- CA- C
4430 4428 4431 4432 3 0.69733 2.09200 0.00000
-2.78933 0.00000 0.00000 ; HB3- CB- OG- HG
4431 4428 4426 4433 3 7.34710 7.79898 -6.14211
-6.71114 4.31789 0.00000 ; OG- CB- CA- C
4433 4435 4437 4438 3 0.00000 0.00000 0.00000
0.00000 0.00000 0.00000 ; C- N- CA- HA
4433 4435 4437 4439 3 11.71520 1.67360 15.06240 -
13.38880 0.00000 0.00000 ; C- N- CA- CB
4433 4435 4437 4450 3 1.75728 5.27184 2.25936
-7.02912 0.00000 0.00000 ; C- N- CA- C
4434 4433 4435 4436 3 29.28800 -8.36800 -20.92000
0.00000 0.00000 0.00000 ; O- C- N- H
4434 4433 4435 4437 3 20.92000 0.00000 -20.92000
0.00000 0.00000 0.00000 ; O- C- N- CA
4435 4437 4439 4440 3 0.65084 1.95253 0.00000
-2.60338 0.00000 0.00000 ; N- CA- CB- HB2
4435 4437 4439 4441 3 0.65084 1.95253 0.00000
-2.60338 0.00000 0.00000 ; N- CA- CB- HB3
4435 4437 4439 4442 3 4.79068 4.02919 -4.82834
-3.66518 2.97901 0.00000 ; N- CA- CB- CG
4435 4437 4450 4451 3 0.00000 0.00000 0.00000
0.00000 0.00000 0.00000 ; N- CA- C- O
4435 4437 4450 4452 3 17.40544 -5.02080 -13.22144
9.20480 0.00000 0.00000 ; N- CA- C- N
4436 4435 4437 4438 3 0.00000 0.00000 0.00000
0.00000 0.00000 0.00000 ; H- N- CA- HA

4436 4435 4437 4439 3 0.00000 0.00000 0.00000
0.00000 0.00000 0.00000 ; H- N- CA- CB
4436 4435 4437 4450 3 0.00000 0.00000 0.00000
0.00000 0.00000 0.00000 ; H- N- CA- C
4437 4439 4442 4443 3 5.98312 6.40152 3.28026 -
12.38464 -1.57318 0.00000 ; CA- CB- CG- ND1
4437 4439 4442 4448 3 0.00000 0.00000 0.00000
0.00000 0.00000 0.00000 ; CA- CB- CG- CD2
4437 4450 4452 4453 3 20.92000 0.00000 -20.92000
0.00000 0.00000 0.00000 ; CA- C- N- H
4437 4450 4452 4454 3 20.92000 0.00000 -20.92000
0.00000 0.00000 0.00000 ; CA- C- N- CA
4438 4437 4439 4440 3 0.65084 1.95253 0.00000
-2.60338 0.00000 0.00000 ; HA- CA- CB- HB2
4438 4437 4439 4441 3 0.65084 1.95253 0.00000
-2.60338 0.00000 0.00000 ; HA- CA- CB- HB3
4438 4437 4439 4442 3 0.65084 1.95253 0.00000
-2.60338 0.00000 0.00000 ; HA- CA- CB- CG
4438 4437 4450 4451 3 3.68192 -4.35136 0.00000
1.33888 0.00000 0.00000 ; HA- CA- C- O
4438 4437 4450 4452 3 0.00000 0.00000 0.00000
0.00000 0.00000 0.00000 ; HA- CA- C- N
4439 4437 4450 4451 3 0.00000 0.00000 0.00000
0.00000 0.00000 0.00000 ; CB- CA- C- O
4439 4437 4450 4452 3 2.51040 4.18400 1.67360
-6.69440 0.00000 0.00000 ; CB- CA- C- N
4439 4442 4443 4444 3 20.08320 0.00000 -20.08320
0.00000 0.00000 0.00000 ; CB- CG- ND1- CE1
4439 4442 4448 4446 3 44.97800 0.00000 -44.97800
0.00000 0.00000 0.00000 ; CB- CG- CD2- NE2
4439 4442 4448 4449 3 44.97800 0.00000 -44.97800
0.00000 0.00000 0.00000 ; CB- CG- CD2- HD2
4440 4439 4437 4450 3 0.65084 1.95253 0.00000
-2.60338 0.00000 0.00000 ; HB2- CB- CA- C
4440 4439 4442 4443 3 0.00000 0.00000 0.00000
0.00000 0.00000 0.00000 ; HB2- CB- CG- ND1
4440 4439 4442 4448 3 0.00000 0.00000 0.00000
0.00000 0.00000 0.00000 ; HB2- CB- CG- CD2
4441 4439 4437 4450 3 0.65084 1.95253 0.00000
-2.60338 0.00000 0.00000 ; HB3- CB- CA- C
4441 4439 4442 4443 3 0.00000 0.00000 0.00000
0.00000 0.00000 0.00000 ; HB3- CB- CG- ND1
4441 4439 4442 4448 3 0.00000 0.00000 0.00000
0.00000 0.00000 0.00000 ; HB3- CB- CG- CD2
4442 4439 4437 4450 3 3.76560 3.34720 -2.87859
-3.66518 0.83680 0.00000 ; CG- CB- CA- C
4442 4443 4444 4445 3 41.84000 0.00000 -41.84000
0.00000 0.00000 0.00000 ; CG- ND1- CE1- HE1
4442 4443 4444 4446 3 41.84000 0.00000 -41.84000
0.00000 0.00000 0.00000 ; CG- ND1- CE1- NE2
4442 4448 4446 4444 3 12.55200 0.00000 -12.55200
0.00000 0.00000 0.00000 ; CG- CD2- NE2- CE1

4442 4448 4446 4447 3 12.55200 0.00000 -12.55200
0.00000 0.00000 0.00000 ; CG- CD2- NE2- HE2
4443 4442 4448 4446 3 44.97800 0.00000 -44.97800
0.00000 0.00000 0.00000 ; ND1- CG- CD2- NE2
4443 4442 4448 4449 3 44.97800 0.00000 -44.97800
0.00000 0.00000 0.00000 ; ND1- CG- CD2- HD2
4443 4444 4446 4447 3 19.45560 0.00000 -19.45560
0.00000 0.00000 0.00000 ; ND1- CE1- NE2- HE2
4443 4444 4446 4448 3 19.45560 0.00000 -19.45560
0.00000 0.00000 0.00000 ; ND1- CE1- NE2- CD2
4444 4443 4442 4448 3 20.08320 0.00000 -20.08320
0.00000 0.00000 0.00000 ; CE1- ND1- CG- CD2
4444 4446 4448 4449 3 12.55200 0.00000 -12.55200
0.00000 0.00000 0.00000 ; CE1- NE2- CD2- HD2
4445 4444 4446 4447 3 19.45560 0.00000 -19.45560
0.00000 0.00000 0.00000 ; HE1- CE1- NE2- HE2
4445 4444 4446 4448 3 19.45560 0.00000 -19.45560
0.00000 0.00000 0.00000 ; HE1- CE1- NE2- CD2
4447 4446 4448 4449 3 12.55200 0.00000 -12.55200
0.00000 0.00000 0.00000 ; HE2- NE2- CD2- HD2
4450 4452 4454 4455 3 0.00000 0.00000 0.00000
0.00000 0.00000 0.00000 ; C- N- CA- HA
4450 4452 4454 4456 3 11.71520 1.67360 15.06240 -
13.38880 0.00000 0.00000 ; C- N- CA- CB
4450 4452 4454 4465 3 1.75728 5.27184 2.25936
-7.02912 0.00000 0.00000 ; C- N- CA- C
4451 4450 4452 4453 3 29.28800 -8.36800 -20.92000
0.00000 0.00000 0.00000 ; O- C- N- H
4451 4450 4452 4454 3 20.92000 0.00000 -20.92000
0.00000 0.00000 0.00000 ; O- C- N- CA
4452 4454 4456 4457 3 0.65084 1.95253 0.00000
-2.60338 0.00000 0.00000 ; N- CA- CB- HB2
4452 4454 4456 4458 3 0.65084 1.95253 0.00000
-2.60338 0.00000 0.00000 ; N- CA- CB- HB3
4452 4454 4456 4459 3 3.21331 2.22589 -4.15053
-2.40998 2.61082 0.00000 ; N- CA- CB- CG
4452 4454 4465 4466 3 0.00000 0.00000 0.00000
0.00000 0.00000 0.00000 ; N- CA- C- O
4452 4454 4465 4467 3 17.40544 -5.02080 -13.22144
9.20480 0.00000 0.00000 ; N- CA- C- N
4453 4452 4454 4455 3 0.00000 0.00000 0.00000
0.00000 0.00000 0.00000 ; H- N- CA- HA
4453 4452 4454 4456 3 0.00000 0.00000 0.00000
0.00000 0.00000 0.00000 ; H- N- CA- CB
4453 4452 4454 4465 3 0.00000 0.00000 0.00000
0.00000 0.00000 0.00000 ; H- N- CA- C
4454 4456 4459 4460 3 0.66944 2.00832 0.00000
-2.67776 0.00000 0.00000 ; CA- CB- CG- HG2
4454 4456 4459 4461 3 0.66944 2.00832 0.00000
-2.67776 0.00000 0.00000 ; CA- CB- CG- HG3
4454 4456 4459 4462 3 10.12110 -1.91209 0.01674
10.17549 -1.87443 0.00000 ; CA- CB- CG- CD

4454 4465 4467 4468 3 20.92000 0.00000 -20.92000
0.00000 0.00000 0.00000 ; CA- C- N- H
4454 4465 4467 4469 3 20.92000 0.00000 -20.92000
0.00000 0.00000 0.00000 ; CA- C- N- CA
4455 4454 4456 4457 3 0.65084 1.95253 0.00000
-2.60338 0.00000 0.00000 ; HA- CA- CB- HB2
4455 4454 4456 4458 3 0.65084 1.95253 0.00000
-2.60338 0.00000 0.00000 ; HA- CA- CB- HB3
4455 4454 4456 4459 3 0.65084 1.95253 0.00000
-2.60338 0.00000 0.00000 ; HA- CA- CB- CG
4455 4454 4465 4466 3 3.68192 -4.35136 0.00000
1.33888 0.00000 0.00000 ; HA- CA- C- O
4455 4454 4465 4467 3 0.00000 0.00000 0.00000
0.00000 0.00000 0.00000 ; HA- CA- C- N
4456 4454 4465 4466 3 0.00000 0.00000 0.00000
0.00000 0.00000 0.00000 ; CB- CA- C- O
4456 4454 4465 4467 3 2.51040 4.18400 1.67360
-6.69440 0.00000 0.00000 ; CB- CA- C- N
4456 4459 4462 4463 3 3.79907 0.00000 -5.40573
0.00000 2.14221 0.00000 ; CB- CG- CD- OE1
4456 4459 4462 4464 3 3.79907 0.00000 -5.40573
0.00000 2.14221 0.00000 ; CB- CG- CD- OE2
4457 4456 4454 4465 3 0.65084 1.95253 0.00000
-2.60338 0.00000 0.00000 ; HB2- CB- CA- C
4457 4456 4459 4460 3 0.62760 1.88280 0.00000
-2.51040 0.00000 0.00000 ; HB2- CB- CG- HG2
4457 4456 4459 4461 3 0.62760 1.88280 0.00000
-2.51040 0.00000 0.00000 ; HB2- CB- CG- HG3
4457 4456 4459 4462 3 0.65084 1.95253 0.00000
-2.60338 0.00000 0.00000 ; HB2- CB- CG- CD
4458 4456 4454 4465 3 0.65084 1.95253 0.00000
-2.60338 0.00000 0.00000 ; HB3- CB- CA- C
4458 4456 4459 4460 3 0.62760 1.88280 0.00000
-2.51040 0.00000 0.00000 ; HB3- CB- CG- HG2
4458 4456 4459 4461 3 0.62760 1.88280 0.00000
-2.51040 0.00000 0.00000 ; HB3- CB- CG- HG3
4458 4456 4459 4462 3 0.65084 1.95253 0.00000
-2.60338 0.00000 0.00000 ; HB3- CB- CG- CD
4459 4456 4454 4465 3 6.86594 3.56895 -8.14206
-2.40998 4.85344 0.00000 ; CG- CB- CA- C
4460 4459 4462 4463 3 0.00000 0.00000 0.00000
0.00000 0.00000 0.00000 ; HG2- CG- CD- OE1
4460 4459 4462 4464 3 0.00000 0.00000 0.00000
0.00000 0.00000 0.00000 ; HG2- CG- CD- OE2
4461 4459 4462 4463 3 0.00000 0.00000 0.00000
0.00000 0.00000 0.00000 ; HG3- CG- CD- OE1
4461 4459 4462 4464 3 0.00000 0.00000 0.00000
0.00000 0.00000 0.00000 ; HG3- CG- CD- OE2
4465 4467 4469 4470 3 0.00000 0.00000 0.00000
0.00000 0.00000 0.00000 ; C- N- CA- HA
4465 4467 4469 4471 3 11.71520 1.67360 15.06240 -
13.38880 0.00000 0.00000 ; C- N- CA- CB

4465 4467 4469 4482 3 1.75728 5.27184 2.25936
-7.02912 0.00000 0.00000 ; C- N- CA- C
4466 4465 4467 4468 3 29.28800 -8.36800 -20.92000
0.00000 0.00000 0.00000 ; O- C- N- H
4466 4465 4467 4469 3 20.92000 0.00000 -20.92000
0.00000 0.00000 0.00000 ; O- C- N- CA
4467 4469 4471 4472 3 0.65084 1.95253 0.00000
-2.60338 0.00000 0.00000 ; N- CA- CB- HB2
4467 4469 4471 4473 3 0.65084 1.95253 0.00000
-2.60338 0.00000 0.00000 ; N- CA- CB- HB3
4467 4469 4471 4474 3 4.79068 4.02919 -4.82834
-3.66518 2.97901 0.00000 ; N- CA- CB- CG
4467 4469 4482 4483 3 0.00000 0.00000 0.00000
0.00000 0.00000 0.00000 ; N- CA- C- O
4467 4469 4482 4484 3 17.40544 -5.02080 -13.22144
9.20480 0.00000 0.00000 ; N- CA- C- N
4468 4467 4469 4470 3 0.00000 0.00000 0.00000
0.00000 0.00000 0.00000 ; H- N- CA- HA
4468 4467 4469 4471 3 0.00000 0.00000 0.00000
0.00000 0.00000 0.00000 ; H- N- CA- CB
4468 4467 4469 4482 3 0.00000 0.00000 0.00000
0.00000 0.00000 0.00000 ; H- N- CA- C
4469 4471 4474 4475 3 5.98312 6.40152 3.28026 -
12.38464 -1.57318 0.00000 ; CA- CB- CG- ND1
4469 4471 4474 4480 3 0.00000 0.00000 0.00000
0.00000 0.00000 0.00000 ; CA- CB- CG- CD2
4469 4482 4484 4485 3 20.92000 0.00000 -20.92000
0.00000 0.00000 0.00000 ; CA- C- N- H
4469 4482 4484 4486 3 20.92000 0.00000 -20.92000
0.00000 0.00000 0.00000 ; CA- C- N- CA
4470 4469 4471 4472 3 0.65084 1.95253 0.00000
-2.60338 0.00000 0.00000 ; HA- CA- CB- HB2
4470 4469 4471 4473 3 0.65084 1.95253 0.00000
-2.60338 0.00000 0.00000 ; HA- CA- CB- HB3
4470 4469 4471 4474 3 0.65084 1.95253 0.00000
-2.60338 0.00000 0.00000 ; HA- CA- CB- CG
4470 4469 4482 4483 3 3.68192 -4.35136 0.00000
1.33888 0.00000 0.00000 ; HA- CA- C- O
4470 4469 4482 4484 3 0.00000 0.00000 0.00000
0.00000 0.00000 0.00000 ; HA- CA- C- N
4471 4469 4482 4483 3 0.00000 0.00000 0.00000
0.00000 0.00000 0.00000 ; CB- CA- C- O
4471 4469 4482 4484 3 2.51040 4.18400 1.67360
-6.69440 0.00000 0.00000 ; CB- CA- C- N
4471 4474 4475 4476 3 20.08320 0.00000 -20.08320
0.00000 0.00000 0.00000 ; CB- CG- ND1- CE1
4471 4474 4480 4478 3 44.97800 0.00000 -44.97800
0.00000 0.00000 0.00000 ; CB- CG- CD2- NE2
4471 4474 4480 4481 3 44.97800 0.00000 -44.97800
0.00000 0.00000 0.00000 ; CB- CG- CD2- HD2
4472 4471 4469 4482 3 0.65084 1.95253 0.00000
-2.60338 0.00000 0.00000 ; HB2- CB- CA- C

4472 4471 4474 4475 3 0.00000 0.00000 0.00000
0.00000 0.00000 0.00000 ; HB2- CB- CG- ND1
4472 4471 4474 4480 3 0.00000 0.00000 0.00000
0.00000 0.00000 0.00000 ; HB2- CB- CG- CD2
4473 4471 4469 4482 3 0.65084 1.95253 0.00000
-2.60338 0.00000 0.00000 ; HB3- CB- CA- C
4473 4471 4474 4475 3 0.00000 0.00000 0.00000
0.00000 0.00000 0.00000 ; HB3- CB- CG- ND1
4473 4471 4474 4480 3 0.00000 0.00000 0.00000
0.00000 0.00000 0.00000 ; HB3- CB- CG- CD2
4474 4471 4469 4482 3 3.76560 3.34720 -2.87859
-3.66518 0.83680 0.00000 ; CG- CB- CA- C
4474 4475 4476 4477 3 41.84000 0.00000 -41.84000
0.00000 0.00000 0.00000 ; CG- ND1- CE1- HE1
4474 4475 4476 4478 3 41.84000 0.00000 -41.84000
0.00000 0.00000 0.00000 ; CG- ND1- CE1- NE2
4474 4480 4478 4476 3 12.55200 0.00000 -12.55200
0.00000 0.00000 0.00000 ; CG- CD2- NE2- CE1
4474 4480 4478 4479 3 12.55200 0.00000 -12.55200
0.00000 0.00000 0.00000 ; CG- CD2- NE2- HE2
4475 4474 4480 4478 3 44.97800 0.00000 -44.97800
0.00000 0.00000 0.00000 ; ND1- CG- CD2- NE2
4475 4474 4480 4481 3 44.97800 0.00000 -44.97800
0.00000 0.00000 0.00000 ; ND1- CG- CD2- HD2
4475 4476 4478 4479 3 19.45560 0.00000 -19.45560
0.00000 0.00000 0.00000 ; ND1- CE1- NE2- HE2
4475 4476 4478 4480 3 19.45560 0.00000 -19.45560
0.00000 0.00000 0.00000 ; ND1- CE1- NE2- CD2
4476 4475 4474 4480 3 20.08320 0.00000 -20.08320
0.00000 0.00000 0.00000 ; CE1- ND1- CG- CD2
4476 4478 4480 4481 3 12.55200 0.00000 -12.55200
0.00000 0.00000 0.00000 ; CE1- NE2- CD2- HD2
4477 4476 4478 4479 3 19.45560 0.00000 -19.45560
0.00000 0.00000 0.00000 ; HE1- CE1- NE2- HE2
4477 4476 4478 4480 3 19.45560 0.00000 -19.45560
0.00000 0.00000 0.00000 ; HE1- CE1- NE2- CD2
4479 4478 4480 4481 3 12.55200 0.00000 -12.55200
0.00000 0.00000 0.00000 ; HE2- NE2- CD2- HD2
4482 4484 4486 4487 3 0.00000 0.00000 0.00000
0.00000 0.00000 0.00000 ; C- N- CA- HA
4482 4484 4486 4488 3 11.71520 1.67360 15.06240 -
13.38880 0.00000 0.00000 ; C- N- CA- CB
4482 4484 4486 4498 3 1.75728 5.27184 2.25936
-7.02912 0.00000 0.00000 ; C- N- CA- C
4483 4482 4484 4485 3 29.28800 -8.36800 -20.92000
0.00000 0.00000 0.00000 ; O- C- N- H
4483 4482 4484 4486 3 20.92000 0.00000 -20.92000
0.00000 0.00000 0.00000 ; O- C- N- CA
4484 4486 4488 4489 3 0.65084 1.95253 0.00000
-2.60338 0.00000 0.00000 ; N- CA- CB- HB
4484 4486 4488 4490 3 3.83673 0.44769 -1.77402
-2.47693 -0.03347 0.00000 ; N- CA- CB- CG1

4484 4486 4488 4494 3 3.83673 0.44769 -1.77402
-2.47693 -0.03347 0.00000 ; N- CA- CB- CG2
4484 4486 4498 4499 3 0.00000 0.00000 0.00000
0.00000 0.00000 0.00000 ; N- CA- C- O
4484 4486 4498 4500 3 17.40544 -5.02080 -13.22144
9.20480 0.00000 0.00000 ; N- CA- C- N
4485 4484 4486 4487 3 0.00000 0.00000 0.00000
0.00000 0.00000 0.00000 ; H- N- CA- HA
4485 4484 4486 4488 3 0.00000 0.00000 0.00000
0.00000 0.00000 0.00000 ; H- N- CA- CB
4485 4484 4486 4498 3 0.00000 0.00000 0.00000
0.00000 0.00000 0.00000 ; H- N- CA- C
4486 4488 4490 4491 3 0.66944 2.00832 0.00000
-2.67776 0.00000 0.00000 ; CA- CB- CG1- HG11
4486 4488 4490 4492 3 0.66944 2.00832 0.00000
-2.67776 0.00000 0.00000 ; CA- CB- CG1- HG12
4486 4488 4490 4493 3 0.66944 2.00832 0.00000
-2.67776 0.00000 0.00000 ; CA- CB- CG1- HG13
4486 4488 4494 4495 3 0.66944 2.00832 0.00000
-2.67776 0.00000 0.00000 ; CA- CB- CG2- HG21
4486 4488 4494 4496 3 0.66944 2.00832 0.00000
-2.67776 0.00000 0.00000 ; CA- CB- CG2- HG22
4486 4488 4494 4497 3 0.66944 2.00832 0.00000
-2.67776 0.00000 0.00000 ; CA- CB- CG2- HG23
4486 4498 4500 4501 3 20.92000 0.00000 -20.92000
0.00000 0.00000 0.00000 ; CA- C- N- H
4486 4498 4500 4502 3 20.92000 0.00000 -20.92000
0.00000 0.00000 0.00000 ; CA- C- N- CA
4487 4486 4488 4489 3 0.65084 1.95253 0.00000
-2.60338 0.00000 0.00000 ; HA- CA- CB- HB
4487 4486 4488 4490 3 0.65084 1.95253 0.00000
-2.60338 0.00000 0.00000 ; HA- CA- CB- CG1
4487 4486 4488 4494 3 0.65084 1.95253 0.00000
-2.60338 0.00000 0.00000 ; HA- CA- CB- CG2
4487 4486 4498 4499 3 3.68192 -4.35136 0.00000
1.33888 0.00000 0.00000 ; HA- CA- C- O
4487 4486 4498 4500 3 0.00000 0.00000 0.00000
0.00000 0.00000 0.00000 ; HA- CA- C- N
4488 4486 4498 4499 3 0.00000 0.00000 0.00000
0.00000 0.00000 0.00000 ; CB- CA- C- O
4488 4486 4498 4500 3 2.51040 4.18400 1.67360
-6.69440 0.00000 0.00000 ; CB- CA- C- N
4489 4488 4486 4498 3 0.65084 1.95253 0.00000
-2.60338 0.00000 0.00000 ; HB- CB- CA- C
4489 4488 4490 4491 3 0.62760 1.88280 0.00000
-2.51040 0.00000 0.00000 ; HB- CB- CG1- HG11
4489 4488 4490 4492 3 0.62760 1.88280 0.00000
-2.51040 0.00000 0.00000 ; HB- CB- CG1- HG12
4489 4488 4490 4493 3 0.62760 1.88280 0.00000
-2.51040 0.00000 0.00000 ; HB- CB- CG1- HG13
4489 4488 4494 4495 3 0.62760 1.88280 0.00000
-2.51040 0.00000 0.00000 ; HB- CB- CG2- HG21

4489 4488 4494 4496 3 0.62760 1.88280 0.00000
-2.51040 0.00000 0.00000 ; HB- CB- CG2- HG22
4489 4488 4494 4497 3 0.62760 1.88280 0.00000
-2.51040 0.00000 0.00000 ; HB- CB- CG2- HG23
4490 4488 4486 4498 3 5.67350 3.55640 -6.16722
-2.47693 3.74886 0.00000 ; CG1- CB- CA- C
4490 4488 4494 4495 3 0.66944 2.00832 0.00000
-2.67776 0.00000 0.00000 ; CG1- CB- CG2- HG21
4490 4488 4494 4496 3 0.66944 2.00832 0.00000
-2.67776 0.00000 0.00000 ; CG1- CB- CG2- HG22
4490 4488 4494 4497 3 0.66944 2.00832 0.00000
-2.67776 0.00000 0.00000 ; CG1- CB- CG2- HG23
4491 4490 4488 4494 3 0.66944 2.00832 0.00000
-2.67776 0.00000 0.00000 ; HG11- CG1- CB- CG2
4492 4490 4488 4494 3 0.66944 2.00832 0.00000
-2.67776 0.00000 0.00000 ; HG12- CG1- CB- CG2
4493 4490 4488 4494 3 0.66944 2.00832 0.00000
-2.67776 0.00000 0.00000 ; HG13- CG1- CB- CG2
4494 4488 4486 4498 3 5.67350 3.55640 -6.16722
-2.47693 3.74886 0.00000 ; CG2- CB- CA- C
4498 4500 4502 4503 3 0.00000 0.00000 0.00000
0.00000 0.00000 0.00000 ; C- N- CA- HA
4498 4500 4502 4504 3 11.71520 1.67360 15.06240 -
13.38880 0.00000 0.00000 ; C- N- CA- CB
4498 4500 4502 4513 3 1.75728 5.27184 2.25936
-7.02912 0.00000 0.00000 ; C- N- CA- C
4499 4498 4500 4501 3 29.28800 -8.36800 -20.92000
0.00000 0.00000 0.00000 ; O- C- N- H
4499 4498 4500 4502 3 20.92000 0.00000 -20.92000
0.00000 0.00000 0.00000 ; O- C- N- CA
4500 4502 4504 4505 3 0.65084 1.95253 0.00000
-2.60338 0.00000 0.00000 ; N- CA- CB- HB2
4500 4502 4504 4506 3 0.65084 1.95253 0.00000
-2.60338 0.00000 0.00000 ; N- CA- CB- HB3
4500 4502 4504 4507 3 3.21331 2.22589 -4.15053
-2.40998 2.61082 0.00000 ; N- CA- CB- CG
4500 4502 4513 4514 3 0.00000 0.00000 0.00000
0.00000 0.00000 0.00000 ; N- CA- C- O
4500 4502 4513 4515 3 17.40544 -5.02080 -13.22144
9.20480 0.00000 0.00000 ; N- CA- C- N
4501 4500 4502 4503 3 0.00000 0.00000 0.00000
0.00000 0.00000 0.00000 ; H- N- CA- HA
4501 4500 4502 4504 3 0.00000 0.00000 0.00000
0.00000 0.00000 0.00000 ; H- N- CA- CB
4501 4500 4502 4513 3 0.00000 0.00000 0.00000
0.00000 0.00000 0.00000 ; H- N- CA- C
4502 4504 4507 4508 3 0.66944 2.00832 0.00000
-2.67776 0.00000 0.00000 ; CA- CB- CG- HG2
4502 4504 4507 4509 3 0.66944 2.00832 0.00000
-2.67776 0.00000 0.00000 ; CA- CB- CG- HG3
4502 4504 4507 4510 3 10.12110 -1.91209 0.01674
10.17549 -1.87443 0.00000 ; CA- CB- CG- CD

4502 4513 4515 4516 3 20.92000 0.00000 -20.92000
0.00000 0.00000 0.00000 ; CA- C- N- H
4502 4513 4515 4517 3 20.92000 0.00000 -20.92000
0.00000 0.00000 0.00000 ; CA- C- N- CA
4503 4502 4504 4505 3 0.65084 1.95253 0.00000
-2.60338 0.00000 0.00000 ; HA- CA- CB- HB2
4503 4502 4504 4506 3 0.65084 1.95253 0.00000
-2.60338 0.00000 0.00000 ; HA- CA- CB- HB3
4503 4502 4504 4507 3 0.65084 1.95253 0.00000
-2.60338 0.00000 0.00000 ; HA- CA- CB- CG
4503 4502 4513 4514 3 3.68192 -4.35136 0.00000
1.33888 0.00000 0.00000 ; HA- CA- C- O
4503 4502 4513 4515 3 0.00000 0.00000 0.00000
0.00000 0.00000 0.00000 ; HA- CA- C- N
4504 4502 4513 4514 3 0.00000 0.00000 0.00000
0.00000 0.00000 0.00000 ; CB- CA- C- O
4504 4502 4513 4515 3 2.51040 4.18400 1.67360
-6.69440 0.00000 0.00000 ; CB- CA- C- N
4504 4507 4510 4511 3 3.79907 0.00000 -5.40573
0.00000 2.14221 0.00000 ; CB- CG- CD- OE1
4504 4507 4510 4512 3 3.79907 0.00000 -5.40573
0.00000 2.14221 0.00000 ; CB- CG- CD- OE2
4505 4504 4502 4513 3 0.65084 1.95253 0.00000
-2.60338 0.00000 0.00000 ; HB2- CB- CA- C
4505 4504 4507 4508 3 0.62760 1.88280 0.00000
-2.51040 0.00000 0.00000 ; HB2- CB- CG- HG2
4505 4504 4507 4509 3 0.62760 1.88280 0.00000
-2.51040 0.00000 0.00000 ; HB2- CB- CG- HG3
4505 4504 4507 4510 3 0.65084 1.95253 0.00000
-2.60338 0.00000 0.00000 ; HB2- CB- CG- CD
4506 4504 4502 4513 3 0.65084 1.95253 0.00000
-2.60338 0.00000 0.00000 ; HB3- CB- CA- C
4506 4504 4507 4508 3 0.62760 1.88280 0.00000
-2.51040 0.00000 0.00000 ; HB3- CB- CG- HG2
4506 4504 4507 4509 3 0.62760 1.88280 0.00000
-2.51040 0.00000 0.00000 ; HB3- CB- CG- HG3
4506 4504 4507 4510 3 0.65084 1.95253 0.00000
-2.60338 0.00000 0.00000 ; HB3- CB- CG- CD
4507 4504 4502 4513 3 6.86594 3.56895 -8.14206
-2.40998 4.85344 0.00000 ; CG- CB- CA- C
4508 4507 4510 4511 3 0.00000 0.00000 0.00000
0.00000 0.00000 0.00000 ; HG2- CG- CD- OE1
4508 4507 4510 4512 3 0.00000 0.00000 0.00000
0.00000 0.00000 0.00000 ; HG2- CG- CD- OE2
4509 4507 4510 4511 3 0.00000 0.00000 0.00000
0.00000 0.00000 0.00000 ; HG3- CG- CD- OE1
4509 4507 4510 4512 3 0.00000 0.00000 0.00000
0.00000 0.00000 0.00000 ; HG3- CG- CD- OE2
4513 4515 4517 4518 3 0.00000 0.00000 0.00000
0.00000 0.00000 0.00000 ; C- N- CA- HA
4513 4515 4517 4519 3 11.71520 1.67360 15.06240 -
13.38880 0.00000 0.00000 ; C- N- CA- CB

4513 4515 4517 4532 3 1.75728 5.27184 2.25936
-7.02912 0.00000 0.00000 ; C- N- CA- C
4514 4513 4515 4516 3 29.28800 -8.36800 -20.92000
0.00000 0.00000 0.00000 ; O- C- N- H
4514 4513 4515 4517 3 20.92000 0.00000 -20.92000
0.00000 0.00000 0.00000 ; O- C- N- CA
4515 4517 4519 4520 3 0.65084 1.95253 0.00000
-2.60338 0.00000 0.00000 ; N- CA- CB- HB2
4515 4517 4519 4521 3 0.65084 1.95253 0.00000
-2.60338 0.00000 0.00000 ; N- CA- CB- HB3
4515 4517 4519 4522 3 3.79070 1.39746 -4.61077
-2.40998 2.44346 0.00000 ; N- CA- CB- CG
4515 4517 4532 4533 3 0.00000 0.00000 0.00000
0.00000 0.00000 0.00000 ; N- CA- C- O
4515 4517 4532 4534 3 17.40544 -5.02080 -13.22144
9.20480 0.00000 0.00000 ; N- CA- C- N
4516 4515 4517 4518 3 0.00000 0.00000 0.00000
0.00000 0.00000 0.00000 ; H- N- CA- HA
4516 4515 4517 4519 3 0.00000 0.00000 0.00000
0.00000 0.00000 0.00000 ; H- N- CA- CB
4516 4515 4517 4532 3 0.00000 0.00000 0.00000
0.00000 0.00000 0.00000 ; H- N- CA- C
4517 4519 4522 4523 3 0.66944 2.00832 0.00000
-2.67776 0.00000 0.00000 ; CA- CB- CG- HG
4517 4519 4522 4524 3 3.90367 0.19665 -6.21742
-2.37651 5.99149 0.00000 ; CA- CB- CG- CD1
4517 4519 4522 4528 3 3.90367 0.19665 -6.21742
-2.37651 5.99149 0.00000 ; CA- CB- CG- CD2
4517 4532 4534 4535 3 20.92000 0.00000 -20.92000
0.00000 0.00000 0.00000 ; CA- C- N- H
4517 4532 4534 4536 3 20.92000 0.00000 -20.92000
0.00000 0.00000 0.00000 ; CA- C- N- CA
4518 4517 4519 4520 3 0.65084 1.95253 0.00000
-2.60338 0.00000 0.00000 ; HA- CA- CB- HB2
4518 4517 4519 4521 3 0.65084 1.95253 0.00000
-2.60338 0.00000 0.00000 ; HA- CA- CB- HB3
4518 4517 4519 4522 3 0.65084 1.95253 0.00000
-2.60338 0.00000 0.00000 ; HA- CA- CB- CG
4518 4517 4532 4533 3 3.68192 -4.35136 0.00000
1.33888 0.00000 0.00000 ; HA- CA- C- O
4518 4517 4532 4534 3 0.00000 0.00000 0.00000
0.00000 0.00000 0.00000 ; HA- CA- C- N
4519 4517 4532 4533 3 0.00000 0.00000 0.00000
0.00000 0.00000 0.00000 ; CB- CA- C- O
4519 4517 4532 4534 3 2.51040 4.18400 1.67360
-6.69440 0.00000 0.00000 ; CB- CA- C- N
4519 4522 4524 4525 3 0.66944 2.00832 0.00000
-2.67776 0.00000 0.00000 ; CB- CG- CD1- HD11
4519 4522 4524 4526 3 0.66944 2.00832 0.00000
-2.67776 0.00000 0.00000 ; CB- CG- CD1- HD12
4519 4522 4524 4527 3 0.66944 2.00832 0.00000
-2.67776 0.00000 0.00000 ; CB- CG- CD1- HD13

4519 4522 4528 4529 3 0.66944 2.00832 0.00000
-2.67776 0.00000 0.00000 ; CB- CG- CD2- HD21
4519 4522 4528 4530 3 0.66944 2.00832 0.00000
-2.67776 0.00000 0.00000 ; CB- CG- CD2- HD22
4519 4522 4528 4531 3 0.66944 2.00832 0.00000
-2.67776 0.00000 0.00000 ; CB- CG- CD2- HD23
4520 4519 4517 4532 3 0.65084 1.95253 0.00000
-2.60338 0.00000 0.00000 ; HB2- CB- CA- C
4520 4519 4522 4523 3 0.62760 1.88280 0.00000
-2.51040 0.00000 0.00000 ; HB2- CB- CG- HG
4520 4519 4522 4524 3 0.66944 2.00832 0.00000
-2.67776 0.00000 0.00000 ; HB2- CB- CG- CD1
4520 4519 4522 4528 3 0.66944 2.00832 0.00000
-2.67776 0.00000 0.00000 ; HB2- CB- CG- CD2
4521 4519 4517 4532 3 0.65084 1.95253 0.00000
-2.60338 0.00000 0.00000 ; HB3- CB- CA- C
4521 4519 4522 4523 3 0.62760 1.88280 0.00000
-2.51040 0.00000 0.00000 ; HB3- CB- CG- HG
4521 4519 4522 4524 3 0.66944 2.00832 0.00000
-2.67776 0.00000 0.00000 ; HB3- CB- CG- CD1
4521 4519 4522 4528 3 0.66944 2.00832 0.00000
-2.67776 0.00000 0.00000 ; HB3- CB- CG- CD2
4522 4519 4517 4532 3 10.33448 -1.14642 -11.54784
-2.40998 6.35968 0.00000 ; CG- CB- CA- C
4523 4522 4524 4525 3 0.62760 1.88280 0.00000
-2.51040 0.00000 0.00000 ; HG- CG- CD1- HD11
4523 4522 4524 4526 3 0.62760 1.88280 0.00000
-2.51040 0.00000 0.00000 ; HG- CG- CD1- HD12
4523 4522 4524 4527 3 0.62760 1.88280 0.00000
-2.51040 0.00000 0.00000 ; HG- CG- CD1- HD13
4523 4522 4528 4529 3 0.62760 1.88280 0.00000
-2.51040 0.00000 0.00000 ; HG- CG- CD2- HD21
4523 4522 4528 4530 3 0.62760 1.88280 0.00000
-2.51040 0.00000 0.00000 ; HG- CG- CD2- HD22
4523 4522 4528 4531 3 0.62760 1.88280 0.00000
-2.51040 0.00000 0.00000 ; HG- CG- CD2- HD23
4524 4522 4528 4529 3 0.66944 2.00832 0.00000
-2.67776 0.00000 0.00000 ; CD1- CG- CD2- HD21
4524 4522 4528 4530 3 0.66944 2.00832 0.00000
-2.67776 0.00000 0.00000 ; CD1- CG- CD2- HD22
4524 4522 4528 4531 3 0.66944 2.00832 0.00000
-2.67776 0.00000 0.00000 ; CD1- CG- CD2- HD23
4525 4524 4522 4528 3 0.66944 2.00832 0.00000
-2.67776 0.00000 0.00000 ; HD11- CD1- CG- CD2
4526 4524 4522 4528 3 0.66944 2.00832 0.00000
-2.67776 0.00000 0.00000 ; HD12- CD1- CG- CD2
4527 4524 4522 4528 3 0.66944 2.00832 0.00000
-2.67776 0.00000 0.00000 ; HD13- CD1- CG- CD2
4532 4534 4536 4537 3 0.00000 0.00000 0.00000
0.00000 0.00000 0.00000 ; C- N- CA- HA
4532 4534 4536 4538 3 11.71520 1.67360 15.06240 -
13.38880 0.00000 0.00000 ; C- N- CA- CB

4532 4534 4536 4543 3 1.75728 5.27184 2.25936
-7.02912 0.00000 0.00000 ; C- N- CA- C
4533 4532 4534 4535 3 29.28800 -8.36800 -20.92000
0.00000 0.00000 0.00000 ; O- C- N- H
4533 4532 4534 4536 3 20.92000 0.00000 -20.92000
0.00000 0.00000 0.00000 ; O- C- N- CA
4534 4536 4538 4539 3 0.65084 1.95253 0.00000
-2.60338 0.00000 0.00000 ; N- CA- CB- HB2
4534 4536 4538 4540 3 0.65084 1.95253 0.00000
-2.60338 0.00000 0.00000 ; N- CA- CB- HB3
4534 4536 4538 4541 3 7.86174 2.24681 -7.41405
-6.71114 5.35552 0.00000 ; N- CA- CB- OG
4534 4536 4543 4544 3 0.00000 0.00000 0.00000
0.00000 0.00000 0.00000 ; N- CA- C- O
4534 4536 4543 4545 3 17.40544 -5.02080 -13.22144
9.20480 0.00000 0.00000 ; N- CA- C- N
4535 4534 4536 4537 3 0.00000 0.00000 0.00000
0.00000 0.00000 0.00000 ; H- N- CA- HA
4535 4534 4536 4538 3 0.00000 0.00000 0.00000
0.00000 0.00000 0.00000 ; H- N- CA- CB
4535 4534 4536 4543 3 0.00000 0.00000 0.00000
0.00000 0.00000 0.00000 ; H- N- CA- C
4536 4538 4541 4542 3 2.05853 2.46856 3.48109
-4.46851 0.23430 0.00000 ; CA- CB- OG- HG
4536 4543 4545 4546 3 20.92000 0.00000 -20.92000
0.00000 0.00000 0.00000 ; CA- C- N- H
4536 4543 4545 4547 3 20.92000 0.00000 -20.92000
0.00000 0.00000 0.00000 ; CA- C- N- CA
4537 4536 4538 4539 3 0.65084 1.95253 0.00000
-2.60338 0.00000 0.00000 ; HA- CA- CB- HB2
4537 4536 4538 4540 3 0.65084 1.95253 0.00000
-2.60338 0.00000 0.00000 ; HA- CA- CB- HB3
4537 4536 4538 4541 3 1.04600 -1.04600 0.00000
0.00000 0.00000 0.00000 ; HA- CA- CB- OG
4537 4536 4543 4544 3 3.68192 -4.35136 0.00000
1.33888 0.00000 0.00000 ; HA- CA- C- O
4537 4536 4543 4545 3 0.00000 0.00000 0.00000
0.00000 0.00000 0.00000 ; HA- CA- C- N
4538 4536 4543 4544 3 0.00000 0.00000 0.00000
0.00000 0.00000 0.00000 ; CB- CA- C- O
4538 4536 4543 4545 3 2.51040 4.18400 1.67360
-6.69440 0.00000 0.00000 ; CB- CA- C- N
4539 4538 4536 4543 3 0.65084 1.95253 0.00000
-2.60338 0.00000 0.00000 ; HB2- CB- CA- C
4539 4538 4541 4542 3 0.69733 2.09200 0.00000
-2.78933 0.00000 0.00000 ; HB2- CB- OG- HG
4540 4538 4536 4543 3 0.65084 1.95253 0.00000
-2.60338 0.00000 0.00000 ; HB3- CB- CA- C
4540 4538 4541 4542 3 0.69733 2.09200 0.00000
-2.78933 0.00000 0.00000 ; HB3- CB- OG- HG
4541 4538 4536 4543 3 7.34710 7.79898 -6.14211
-6.71114 4.31789 0.00000 ; OG- CB- CA- C

4543 4545 4547 4548 3 0.00000 0.00000 0.00000
0.00000 0.00000 0.00000 ; C- N- CA- HA
4543 4545 4547 4549 3 11.71520 1.67360 15.06240 -
13.38880 0.00000 0.00000 ; C- N- CA- CB
4543 4545 4547 4554 3 1.75728 5.27184 2.25936
-7.02912 0.00000 0.00000 ; C- N- CA- C
4544 4543 4545 4546 3 29.28800 -8.36800 -20.92000
0.00000 0.00000 0.00000 ; O- C- N- H
4544 4543 4545 4547 3 20.92000 0.00000 -20.92000
0.00000 0.00000 0.00000 ; O- C- N- CA
4545 4547 4549 4550 3 0.65084 1.95253 0.00000
-2.60338 0.00000 0.00000 ; N- CA- CB- HB2
4545 4547 4549 4551 3 0.65084 1.95253 0.00000
-2.60338 0.00000 0.00000 ; N- CA- CB- HB3
4545 4547 4549 4552 3 7.86174 2.24681 -7.41405
-6.71114 5.35552 0.00000 ; N- CA- CB- OG
4545 4547 4554 4555 3 0.00000 0.00000 0.00000
0.00000 0.00000 0.00000 ; N- CA- C- O
4545 4547 4554 4556 3 17.40544 -5.02080 -13.22144
9.20480 0.00000 0.00000 ; N- CA- C- N
4546 4545 4547 4548 3 0.00000 0.00000 0.00000
0.00000 0.00000 0.00000 ; H- N- CA- HA
4546 4545 4547 4549 3 0.00000 0.00000 0.00000
0.00000 0.00000 0.00000 ; H- N- CA- CB
4546 4545 4547 4554 3 0.00000 0.00000 0.00000
0.00000 0.00000 0.00000 ; H- N- CA- C
4547 4549 4552 4553 3 2.05853 2.46856 3.48109
-4.46851 0.23430 0.00000 ; CA- CB- OG- HG
4547 4554 4556 4557 3 20.92000 0.00000 -20.92000
0.00000 0.00000 0.00000 ; CA- C- N- H
4547 4554 4556 4558 3 20.92000 0.00000 -20.92000
0.00000 0.00000 0.00000 ; CA- C- N- CA
4548 4547 4549 4550 3 0.65084 1.95253 0.00000
-2.60338 0.00000 0.00000 ; HA- CA- CB- HB2
4548 4547 4549 4551 3 0.65084 1.95253 0.00000
-2.60338 0.00000 0.00000 ; HA- CA- CB- HB3
4548 4547 4549 4552 3 1.04600 -1.04600 0.00000
0.00000 0.00000 0.00000 ; HA- CA- CB- OG
4548 4547 4554 4555 3 3.68192 -4.35136 0.00000
1.33888 0.00000 0.00000 ; HA- CA- C- O
4548 4547 4554 4556 3 0.00000 0.00000 0.00000
0.00000 0.00000 0.00000 ; HA- CA- C- N
4549 4547 4554 4555 3 0.00000 0.00000 0.00000
0.00000 0.00000 0.00000 ; CB- CA- C- O
4549 4547 4554 4556 3 2.51040 4.18400 1.67360
-6.69440 0.00000 0.00000 ; CB- CA- C- N
4550 4549 4547 4554 3 0.65084 1.95253 0.00000
-2.60338 0.00000 0.00000 ; HB2- CB- CA- C
4550 4549 4552 4553 3 0.69733 2.09200 0.00000
-2.78933 0.00000 0.00000 ; HB2- CB- OG- HG
4551 4549 4547 4554 3 0.65084 1.95253 0.00000
-2.60338 0.00000 0.00000 ; HB3- CB- CA- C

4551 4549 4552 4553 3 0.69733 2.09200 0.00000
-2.78933 0.00000 0.00000 ; HB3- CB- OG- HG
4552 4549 4547 4554 3 7.34710 7.79898 -6.14211
-6.71114 4.31789 0.00000 ; OG- CB- CA- C
4554 4556 4558 4559 3 0.00000 0.00000 0.00000
0.00000 0.00000 0.00000 ; C- N- CA- HA
4554 4556 4558 4560 3 11.71520 1.67360 15.06240 -
13.38880 0.00000 0.00000 ; C- N- CA- CB
4554 4556 4558 4568 3 1.75728 5.27184 2.25936
-7.02912 0.00000 0.00000 ; C- N- CA- C
4555 4554 4556 4557 3 29.28800 -8.36800 -20.92000
0.00000 0.00000 0.00000 ; O- C- N- H
4555 4554 4556 4558 3 20.92000 0.00000 -20.92000
0.00000 0.00000 0.00000 ; O- C- N- CA
4556 4558 4560 4561 3 0.65084 1.95253 0.00000
-2.60338 0.00000 0.00000 ; N- CA- CB- HB
4556 4558 4560 4562 3 3.83673 0.44769 -1.77402
-2.47693 -0.03347 0.00000 ; N- CA- CB- CG2
4556 4558 4560 4566 3 4.93294 1.13386 -3.12963
-5.27184 3.17984 0.00000 ; N- CA- CB- OG1
4556 4558 4568 4569 3 0.00000 0.00000 0.00000
0.00000 0.00000 0.00000 ; N- CA- C- O
4556 4558 4568 4570 3 17.40544 -5.02080 -13.22144
9.20480 0.00000 0.00000 ; N- CA- C- N
4557 4556 4558 4559 3 0.00000 0.00000 0.00000
0.00000 0.00000 0.00000 ; H- N- CA- HA
4557 4556 4558 4560 3 0.00000 0.00000 0.00000
0.00000 0.00000 0.00000 ; H- N- CA- CB
4557 4556 4558 4568 3 0.00000 0.00000 0.00000
0.00000 0.00000 0.00000 ; H- N- CA- C
4558 4560 4562 4563 3 0.66944 2.00832 0.00000
-2.67776 0.00000 0.00000 ; CA- CB- CG2- HG21
4558 4560 4562 4564 3 0.66944 2.00832 0.00000
-2.67776 0.00000 0.00000 ; CA- CB- CG2- HG22
4558 4560 4562 4565 3 0.66944 2.00832 0.00000
-2.67776 0.00000 0.00000 ; CA- CB- CG2- HG23
4558 4560 4566 4567 3 1.12131 2.98738 1.66523
-3.94970 0.43514 0.00000 ; CA- CB- OG1- HG1
4558 4568 4570 4571 3 20.92000 0.00000 -20.92000
0.00000 0.00000 0.00000 ; CA- C- N- H
4558 4568 4570 4572 3 20.92000 0.00000 -20.92000
0.00000 0.00000 0.00000 ; CA- C- N- CA
4559 4558 4560 4561 3 0.65084 1.95253 0.00000
-2.60338 0.00000 0.00000 ; HA- CA- CB- HB
4559 4558 4560 4562 3 0.65084 1.95253 0.00000
-2.60338 0.00000 0.00000 ; HA- CA- CB- CG2
4559 4558 4560 4566 3 1.04600 -1.04600 0.00000
0.00000 0.00000 0.00000 ; HA- CA- CB- OG1
4559 4558 4568 4569 3 3.68192 -4.35136 0.00000
1.33888 0.00000 0.00000 ; HA- CA- C- O
4559 4558 4568 4570 3 0.00000 0.00000 0.00000
0.00000 0.00000 0.00000 ; HA- CA- C- N

4560 4558 4568 4569 3 0.00000 0.00000 0.00000
0.00000 0.00000 0.00000 ; CB- CA- C- O
4560 4558 4568 4570 3 2.51040 4.18400 1.67360
-6.69440 0.00000 0.00000 ; CB- CA- C- N
4561 4560 4558 4568 3 0.65084 1.95253 0.00000
-2.60338 0.00000 0.00000 ; HB- CB- CA- C
4561 4560 4562 4563 3 0.65084 1.95253 0.00000
-2.60338 0.00000 0.00000 ; HB- CB- CG2- HG21
4561 4560 4562 4564 3 0.65084 1.95253 0.00000
-2.60338 0.00000 0.00000 ; HB- CB- CG2- HG22
4561 4560 4562 4565 3 0.65084 1.95253 0.00000
-2.60338 0.00000 0.00000 ; HB- CB- CG2- HG23
4561 4560 4566 4567 3 0.69733 2.09200 0.00000
-2.78933 0.00000 0.00000 ; HB- CB- OG1- HG1
4562 4560 4558 4568 3 5.67350 3.55640 -6.16722
-2.47693 3.74886 0.00000 ; CG2- CB- CA- C
4562 4560 4566 4567 3 4.74047 0.27196 -2.26773
-3.94970 1.60666 0.00000 ; CG2- CB- OG1- HG1
4563 4562 4560 4566 3 1.04600 -1.04600 0.00000
0.00000 0.00000 0.00000 ; HG21- CG2- CB- OG1
4564 4562 4560 4566 3 1.04600 -1.04600 0.00000
0.00000 0.00000 0.00000 ; HG22- CG2- CB- OG1
4565 4562 4560 4566 3 1.04600 -1.04600 0.00000
0.00000 0.00000 0.00000 ; HG23- CG2- CB- OG1
4566 4560 4558 4568 3 6.53541 6.87013 -6.21742
-5.27184 5.22163 0.00000 ; OG1- CB- CA- C
4568 4570 4572 4573 3 0.00000 0.00000 0.00000
0.00000 0.00000 0.00000 ; C- N- CA- HA
4568 4570 4572 4574 3 11.71520 1.67360 15.06240 -
13.38880 0.00000 0.00000 ; C- N- CA- CB
4568 4570 4572 4587 3 1.75728 5.27184 2.25936
-7.02912 0.00000 0.00000 ; C- N- CA- C
4569 4568 4570 4571 3 29.28800 -8.36800 -20.92000
0.00000 0.00000 0.00000 ; O- C- N- H
4569 4568 4570 4572 3 20.92000 0.00000 -20.92000
0.00000 0.00000 0.00000 ; O- C- N- CA
4570 4572 4574 4575 3 0.65084 1.95253 0.00000
-2.60338 0.00000 0.00000 ; N- CA- CB- HB2
4570 4572 4574 4576 3 0.65084 1.95253 0.00000
-2.60338 0.00000 0.00000 ; N- CA- CB- HB3
4570 4572 4574 4577 3 3.79070 1.39746 -4.61077
-2.40998 2.44346 0.00000 ; N- CA- CB- CG
4570 4572 4587 4588 3 0.00000 0.00000 0.00000
0.00000 0.00000 0.00000 ; N- CA- C- O
4570 4572 4587 4589 3 17.40544 -5.02080 -13.22144
9.20480 0.00000 0.00000 ; N- CA- C- N
4571 4570 4572 4573 3 0.00000 0.00000 0.00000
0.00000 0.00000 0.00000 ; H- N- CA- HA
4571 4570 4572 4574 3 0.00000 0.00000 0.00000
0.00000 0.00000 0.00000 ; H- N- CA- CB
4571 4570 4572 4587 3 0.00000 0.00000 0.00000
0.00000 0.00000 0.00000 ; H- N- CA- C

```
  4572   4574   4577   4578       3    0.66944    2.00832    0.00000
-2.67776    0.00000    0.00000 ;     CA-    CB-    CG-    HG
  4572   4574   4577   4579       3    3.90367    0.19665   -6.21742
-2.37651    5.99149    0.00000 ;     CA-    CB-    CG-   CD1
  4572   4574   4577   4583       3    3.90367    0.19665   -6.21742
-2.37651    5.99149    0.00000 ;     CA-    CB-    CG-   CD2
  4572   4587   4589   4590       3   20.92000    0.00000  -20.92000
0.00000    0.00000    0.00000 ;     CA-     C-     N-     H
  4572   4587   4589   4591       3   20.92000    0.00000  -20.92000
0.00000    0.00000    0.00000 ;     CA-     C-     N-    CA
  4573   4572   4574   4575       3    0.65084    1.95253    0.00000
-2.60338    0.00000    0.00000 ;     HA-    CA-    CB-   HB2
  4573   4572   4574   4576       3    0.65084    1.95253    0.00000
-2.60338    0.00000    0.00000 ;     HA-    CA-    CB-   HB3
  4573   4572   4574   4577       3    0.65084    1.95253    0.00000
-2.60338    0.00000    0.00000 ;     HA-    CA-    CB-    CG
  4573   4572   4587   4588       3    3.68192   -4.35136    0.00000
1.33888    0.00000    0.00000 ;     HA-    CA-     C-     O
  4573   4572   4587   4589       3    0.00000    0.00000    0.00000
0.00000    0.00000    0.00000 ;     HA-    CA-     C-     N
  4574   4572   4587   4588       3    0.00000    0.00000    0.00000
0.00000    0.00000    0.00000 ;     CB-    CA-     C-     O
  4574   4572   4587   4589       3    2.51040    4.18400    1.67360
-6.69440    0.00000    0.00000 ;     CB-    CA-     C-     N
  4574   4577   4579   4580       3    0.66944    2.00832    0.00000
-2.67776    0.00000    0.00000 ;     CB-    CG-   CD1-  HD11
  4574   4577   4579   4581       3    0.66944    2.00832    0.00000
-2.67776    0.00000    0.00000 ;     CB-    CG-   CD1-  HD12
  4574   4577   4579   4582       3    0.66944    2.00832    0.00000
-2.67776    0.00000    0.00000 ;     CB-    CG-   CD1-  HD13
  4574   4577   4583   4584       3    0.66944    2.00832    0.00000
-2.67776    0.00000    0.00000 ;     CB-    CG-   CD2-  HD21
  4574   4577   4583   4585       3    0.66944    2.00832    0.00000
-2.67776    0.00000    0.00000 ;     CB-    CG-   CD2-  HD22
  4574   4577   4583   4586       3    0.66944    2.00832    0.00000
-2.67776    0.00000    0.00000 ;     CB-    CG-   CD2-  HD23
  4575   4574   4572   4587       3    0.65084    1.95253    0.00000
-2.60338    0.00000    0.00000 ;    HB2-    CB-    CA-     C
  4575   4574   4577   4578       3    0.62760    1.88280    0.00000
-2.51040    0.00000    0.00000 ;    HB2-    CB-    CG-    HG
  4575   4574   4577   4579       3    0.66944    2.00832    0.00000
-2.67776    0.00000    0.00000 ;    HB2-    CB-    CG-   CD1
  4575   4574   4577   4583       3    0.66944    2.00832    0.00000
-2.67776    0.00000    0.00000 ;    HB2-    CB-    CG-   CD2
  4576   4574   4572   4587       3    0.65084    1.95253    0.00000
-2.60338    0.00000    0.00000 ;    HB3-    CB-    CA-     C
  4576   4574   4577   4578       3    0.62760    1.88280    0.00000
-2.51040    0.00000    0.00000 ;    HB3-    CB-    CG-    HG
  4576   4574   4577   4579       3    0.66944    2.00832    0.00000
-2.67776    0.00000    0.00000 ;    HB3-    CB-    CG-   CD1
  4576   4574   4577   4583       3    0.66944    2.00832    0.00000
-2.67776    0.00000    0.00000 ;    HB3-    CB-    CG-   CD2
```

4577 4574 4572 4587 3 10.33448 -1.14642 -11.54784
-2.40998 6.35968 0.00000 ; CG- CB- CA- C
4578 4577 4579 4580 3 0.62760 1.88280 0.00000
-2.51040 0.00000 0.00000 ; HG- CG- CD1- HD11
4578 4577 4579 4581 3 0.62760 1.88280 0.00000
-2.51040 0.00000 0.00000 ; HG- CG- CD1- HD12
4578 4577 4579 4582 3 0.62760 1.88280 0.00000
-2.51040 0.00000 0.00000 ; HG- CG- CD1- HD13
4578 4577 4583 4584 3 0.62760 1.88280 0.00000
-2.51040 0.00000 0.00000 ; HG- CG- CD2- HD21
4578 4577 4583 4585 3 0.62760 1.88280 0.00000
-2.51040 0.00000 0.00000 ; HG- CG- CD2- HD22
4578 4577 4583 4586 3 0.62760 1.88280 0.00000
-2.51040 0.00000 0.00000 ; HG- CG- CD2- HD23
4579 4577 4583 4584 3 0.66944 2.00832 0.00000
-2.67776 0.00000 0.00000 ; CD1- CG- CD2- HD21
4579 4577 4583 4585 3 0.66944 2.00832 0.00000
-2.67776 0.00000 0.00000 ; CD1- CG- CD2- HD22
4579 4577 4583 4586 3 0.66944 2.00832 0.00000
-2.67776 0.00000 0.00000 ; CD1- CG- CD2- HD23
4580 4579 4577 4583 3 0.66944 2.00832 0.00000
-2.67776 0.00000 0.00000 ; HD11- CD1- CG- CD2
4581 4579 4577 4583 3 0.66944 2.00832 0.00000
-2.67776 0.00000 0.00000 ; HD12- CD1- CG- CD2
4582 4579 4577 4583 3 0.66944 2.00832 0.00000
-2.67776 0.00000 0.00000 ; HD13- CD1- CG- CD2
4587 4589 4591 4592 3 0.00000 0.00000 0.00000
0.00000 0.00000 0.00000 ; C- N- CA- HA
4587 4589 4591 4593 3 11.71520 1.67360 15.06240 -
13.38880 0.00000 0.00000 ; C- N- CA- CB
4587 4589 4591 4606 3 1.75728 5.27184 2.25936
-7.02912 0.00000 0.00000 ; C- N- CA- C
4588 4587 4589 4590 3 29.28800 -8.36800 -20.92000
0.00000 0.00000 0.00000 ; O- C- N- H
4588 4587 4589 4591 3 20.92000 0.00000 -20.92000
0.00000 0.00000 0.00000 ; O- C- N- CA
4589 4591 4593 4594 3 0.65084 1.95253 0.00000
-2.60338 0.00000 0.00000 ; N- CA- CB- HB2
4589 4591 4593 4595 3 0.65084 1.95253 0.00000
-2.60338 0.00000 0.00000 ; N- CA- CB- HB3
4589 4591 4593 4596 3 3.79070 1.39746 -4.61077
-2.40998 2.44346 0.00000 ; N- CA- CB- CG
4589 4591 4606 4607 3 0.00000 0.00000 0.00000
0.00000 0.00000 0.00000 ; N- CA- C- O
4589 4591 4606 4608 3 17.40544 -5.02080 -13.22144
9.20480 0.00000 0.00000 ; N- CA- C- N
4590 4589 4591 4592 3 0.00000 0.00000 0.00000
0.00000 0.00000 0.00000 ; H- N- CA- HA
4590 4589 4591 4593 3 0.00000 0.00000 0.00000
0.00000 0.00000 0.00000 ; H- N- CA- CB
4590 4589 4591 4606 3 0.00000 0.00000 0.00000
0.00000 0.00000 0.00000 ; H- N- CA- C

4591 4593 4596 4597 3 0.66944 2.00832 0.00000
-2.67776 0.00000 0.00000 ; CA- CB- CG- HG
4591 4593 4596 4598 3 3.90367 0.19665 -6.21742
-2.37651 5.99149 0.00000 ; CA- CB- CG- CD1
4591 4593 4596 4602 3 3.90367 0.19665 -6.21742
-2.37651 5.99149 0.00000 ; CA- CB- CG- CD2
4591 4606 4608 4609 3 20.92000 0.00000 -20.92000
0.00000 0.00000 0.00000 ; CA- C- N- H
4591 4606 4608 4610 3 20.92000 0.00000 -20.92000
0.00000 0.00000 0.00000 ; CA- C- N- CA
4592 4591 4593 4594 3 0.65084 1.95253 0.00000
-2.60338 0.00000 0.00000 ; HA- CA- CB- HB2
4592 4591 4593 4595 3 0.65084 1.95253 0.00000
-2.60338 0.00000 0.00000 ; HA- CA- CB- HB3
4592 4591 4593 4596 3 0.65084 1.95253 0.00000
-2.60338 0.00000 0.00000 ; HA- CA- CB- CG
4592 4591 4606 4607 3 3.68192 -4.35136 0.00000
1.33888 0.00000 0.00000 ; HA- CA- C- O
4592 4591 4606 4608 3 0.00000 0.00000 0.00000
0.00000 0.00000 0.00000 ; HA- CA- C- N
4593 4591 4606 4607 3 0.00000 0.00000 0.00000
0.00000 0.00000 0.00000 ; CB- CA- C- O
4593 4591 4606 4608 3 2.51040 4.18400 1.67360
-6.69440 0.00000 0.00000 ; CB- CA- C- N
4593 4596 4598 4599 3 0.66944 2.00832 0.00000
-2.67776 0.00000 0.00000 ; CB- CG- CD1- HD11
4593 4596 4598 4600 3 0.66944 2.00832 0.00000
-2.67776 0.00000 0.00000 ; CB- CG- CD1- HD12
4593 4596 4598 4601 3 0.66944 2.00832 0.00000
-2.67776 0.00000 0.00000 ; CB- CG- CD1- HD13
4593 4596 4602 4603 3 0.66944 2.00832 0.00000
-2.67776 0.00000 0.00000 ; CB- CG- CD2- HD21
4593 4596 4602 4604 3 0.66944 2.00832 0.00000
-2.67776 0.00000 0.00000 ; CB- CG- CD2- HD22
4593 4596 4602 4605 3 0.66944 2.00832 0.00000
-2.67776 0.00000 0.00000 ; CB- CG- CD2- HD23
4594 4593 4591 4606 3 0.65084 1.95253 0.00000
-2.60338 0.00000 0.00000 ; HB2- CB- CA- C
4594 4593 4596 4597 3 0.62760 1.88280 0.00000
-2.51040 0.00000 0.00000 ; HB2- CB- CG- HG
4594 4593 4596 4598 3 0.66944 2.00832 0.00000
-2.67776 0.00000 0.00000 ; HB2- CB- CG- CD1
4594 4593 4596 4602 3 0.66944 2.00832 0.00000
-2.67776 0.00000 0.00000 ; HB2- CB- CG- CD2
4595 4593 4591 4606 3 0.65084 1.95253 0.00000
-2.60338 0.00000 0.00000 ; HB3- CB- CA- C
4595 4593 4596 4597 3 0.62760 1.88280 0.00000
-2.51040 0.00000 0.00000 ; HB3- CB- CG- HG
4595 4593 4596 4598 3 0.66944 2.00832 0.00000
-2.67776 0.00000 0.00000 ; HB3- CB- CG- CD1
4595 4593 4596 4602 3 0.66944 2.00832 0.00000
-2.67776 0.00000 0.00000 ; HB3- CB- CG- CD2

4596 4593 4591 4606 3 10.33448 -1.14642 -11.54784
-2.40998 6.35968 0.00000 ; CG- CB- CA- C
4597 4596 4598 4599 3 0.62760 1.88280 0.00000
-2.51040 0.00000 0.00000 ; HG- CG- CD1- HD11
4597 4596 4598 4600 3 0.62760 1.88280 0.00000
-2.51040 0.00000 0.00000 ; HG- CG- CD1- HD12
4597 4596 4598 4601 3 0.62760 1.88280 0.00000
-2.51040 0.00000 0.00000 ; HG- CG- CD1- HD13
4597 4596 4602 4603 3 0.62760 1.88280 0.00000
-2.51040 0.00000 0.00000 ; HG- CG- CD2- HD21
4597 4596 4602 4604 3 0.62760 1.88280 0.00000
-2.51040 0.00000 0.00000 ; HG- CG- CD2- HD22
4597 4596 4602 4605 3 0.62760 1.88280 0.00000
-2.51040 0.00000 0.00000 ; HG- CG- CD2- HD23
4598 4596 4602 4603 3 0.66944 2.00832 0.00000
-2.67776 0.00000 0.00000 ; CD1- CG- CD2- HD21
4598 4596 4602 4604 3 0.66944 2.00832 0.00000
-2.67776 0.00000 0.00000 ; CD1- CG- CD2- HD22
4598 4596 4602 4605 3 0.66944 2.00832 0.00000
-2.67776 0.00000 0.00000 ; CD1- CG- CD2- HD23
4599 4598 4596 4602 3 0.66944 2.00832 0.00000
-2.67776 0.00000 0.00000 ; HD11- CD1- CG- CD2
4600 4598 4596 4602 3 0.66944 2.00832 0.00000
-2.67776 0.00000 0.00000 ; HD12- CD1- CG- CD2
4601 4598 4596 4602 3 0.66944 2.00832 0.00000
-2.67776 0.00000 0.00000 ; HD13- CD1- CG- CD2
4606 4608 4610 4611 3 0.00000 0.00000 0.00000
0.00000 0.00000 0.00000 ; C- N- CA- HA
4606 4608 4610 4612 3 11.71520 1.67360 15.06240 -
13.38880 0.00000 0.00000 ; C- N- CA- CB
4606 4608 4610 4628 3 1.75728 5.27184 2.25936
-7.02912 0.00000 0.00000 ; C- N- CA- C
4607 4606 4608 4609 3 29.28800 -8.36800 -20.92000
0.00000 0.00000 0.00000 ; O- C- N- H
4607 4606 4608 4610 3 20.92000 0.00000 -20.92000
0.00000 0.00000 0.00000 ; O- C- N- CA
4608 4610 4612 4613 3 0.65084 1.95253 0.00000
-2.60338 0.00000 0.00000 ; N- CA- CB- HB2
4608 4610 4612 4614 3 0.65084 1.95253 0.00000
-2.60338 0.00000 0.00000 ; N- CA- CB- HB3
4608 4610 4612 4615 3 0.65084 1.95253 0.00000
-2.60338 0.00000 0.00000 ; N- CA- CB- CG
4608 4610 4628 4629 3 0.00000 0.00000 0.00000
0.00000 0.00000 0.00000 ; N- CA- C- O
4608 4610 4628 4630 3 17.40544 -5.02080 -13.22144
9.20480 0.00000 0.00000 ; N- CA- C- N
4609 4608 4610 4611 3 0.00000 0.00000 0.00000
0.00000 0.00000 0.00000 ; H- N- CA- HA
4609 4608 4610 4612 3 0.00000 0.00000 0.00000
0.00000 0.00000 0.00000 ; H- N- CA- CB
4609 4608 4610 4628 3 0.00000 0.00000 0.00000
0.00000 0.00000 0.00000 ; H- N- CA- C

```
  4610   4612   4615   4616      3    0.66944    2.00832    0.00000
-2.67776    0.00000    0.00000 ;     CA-    CB-    CG-   HG2
  4610   4612   4615   4617      3    0.66944    2.00832    0.00000
-2.67776    0.00000    0.00000 ;     CA-    CB-    CG-   HG3
  4610   4612   4615   4618      3    3.68192    3.09616   -2.09200
-3.01248    0.00000    0.00000 ;     CA-    CB-    CG-    CD
  4610   4628   4630   4631      3   20.92000    0.00000  -20.92000
0.00000    0.00000    0.00000 ;     CA-     C-     N-     H
  4610   4628   4630   4632      3   20.92000    0.00000  -20.92000
0.00000    0.00000    0.00000 ;     CA-     C-     N-    CA
  4611   4610   4612   4613      3    0.65084    1.95253    0.00000
-2.60338    0.00000    0.00000 ;     HA-    CA-    CB-   HB2
  4611   4610   4612   4614      3    0.65084    1.95253    0.00000
-2.60338    0.00000    0.00000 ;     HA-    CA-    CB-   HB3
  4611   4610   4612   4615      3    0.65084    1.95253    0.00000
-2.60338    0.00000    0.00000 ;     HA-    CA-    CB-    CG
  4611   4610   4628   4629      3    3.68192   -4.35136    0.00000
1.33888    0.00000    0.00000 ;     HA-    CA-     C-     O
  4611   4610   4628   4630      3    0.00000    0.00000    0.00000
0.00000    0.00000    0.00000 ;     HA-    CA-     C-     N
  4612   4610   4628   4629      3    0.00000    0.00000    0.00000
0.00000    0.00000    0.00000 ;     CB-    CA-     C-     O
  4612   4610   4628   4630      3    2.51040    4.18400    1.67360
-6.69440    0.00000    0.00000 ;     CB-    CA-     C-     N
  4612   4615   4618   4619      3    0.66944    2.00832    0.00000
-2.67776    0.00000    0.00000 ;     CB-    CG-    CD-   HD2
  4612   4615   4618   4620      3    0.66944    2.00832    0.00000
-2.67776    0.00000    0.00000 ;     CB-    CG-    CD-   HD3
  4612   4615   4618   4621      3    3.68192    3.09616   -2.09200
-3.01248    0.00000    0.00000 ;     CB-    CG-    CD-    CE
  4613   4612   4610   4628      3    0.65084    1.95253    0.00000
-2.60338    0.00000    0.00000 ;    HB2-    CB-    CA-     C
  4613   4612   4615   4616      3    0.62760    1.88280    0.00000
-2.51040    0.00000    0.00000 ;    HB2-    CB-    CG-   HG2
  4613   4612   4615   4617      3    0.62760    1.88280    0.00000
-2.51040    0.00000    0.00000 ;    HB2-    CB-    CG-   HG3
  4613   4612   4615   4618      3    0.66944    2.00832    0.00000
-2.67776    0.00000    0.00000 ;    HB2-    CB-    CG-    CD
  4614   4612   4610   4628      3    0.65084    1.95253    0.00000
-2.60338    0.00000    0.00000 ;    HB3-    CB-    CA-     C
  4614   4612   4615   4616      3    0.62760    1.88280    0.00000
-2.51040    0.00000    0.00000 ;    HB3-    CB-    CG-   HG2
  4614   4612   4615   4617      3    0.62760    1.88280    0.00000
-2.51040    0.00000    0.00000 ;    HB3-    CB-    CG-   HG3
  4614   4612   4615   4618      3    0.66944    2.00832    0.00000
-2.67776    0.00000    0.00000 ;    HB3-    CB-    CG-    CD
  4615   4612   4610   4628      3    0.65084    1.95253    0.00000
-2.60338    0.00000    0.00000 ;     CG-    CB-    CA-     C
  4615   4618   4621   4622      3    0.65084    1.95253    0.00000
-2.60338    0.00000    0.00000 ;     CG-    CD-    CE-   HE2
  4615   4618   4621   4623      3    0.65084    1.95253    0.00000
-2.60338    0.00000    0.00000 ;     CG-    CD-    CE-   HE3
```

4615 4618 4621 4624 3 0.65084 1.95253 0.00000
-2.60338 0.00000 0.00000 ; CG- CD- CE- NZ
4616 4615 4618 4619 3 0.62760 1.88280 0.00000
-2.51040 0.00000 0.00000 ; HG2- CG- CD- HD2
4616 4615 4618 4620 3 0.62760 1.88280 0.00000
-2.51040 0.00000 0.00000 ; HG2- CG- CD- HD3
4616 4615 4618 4621 3 0.66944 2.00832 0.00000
-2.67776 0.00000 0.00000 ; HG2- CG- CD- CE
4617 4615 4618 4619 3 0.62760 1.88280 0.00000
-2.51040 0.00000 0.00000 ; HG3- CG- CD- HD2
4617 4615 4618 4620 3 0.62760 1.88280 0.00000
-2.51040 0.00000 0.00000 ; HG3- CG- CD- HD3
4617 4615 4618 4621 3 0.66944 2.00832 0.00000
-2.67776 0.00000 0.00000 ; HG3- CG- CD- CE
4618 4621 4624 4625 3 0.65084 1.95253 0.00000
-2.60338 0.00000 0.00000 ; CD- CE- NZ- HZ1
4618 4621 4624 4626 3 0.65084 1.95253 0.00000
-2.60338 0.00000 0.00000 ; CD- CE- NZ- HZ2
4618 4621 4624 4627 3 0.65084 1.95253 0.00000
-2.60338 0.00000 0.00000 ; CD- CE- NZ- HZ3
4619 4618 4621 4622 3 0.65084 1.95253 0.00000
-2.60338 0.00000 0.00000 ; HD2- CD- CE- HE2
4619 4618 4621 4623 3 0.65084 1.95253 0.00000
-2.60338 0.00000 0.00000 ; HD2- CD- CE- HE3
4619 4618 4621 4624 3 0.65084 1.95253 0.00000
-2.60338 0.00000 0.00000 ; HD2- CD- CE- NZ
4620 4618 4621 4622 3 0.65084 1.95253 0.00000
-2.60338 0.00000 0.00000 ; HD3- CD- CE- HE2
4620 4618 4621 4623 3 0.65084 1.95253 0.00000
-2.60338 0.00000 0.00000 ; HD3- CD- CE- HE3
4620 4618 4621 4624 3 0.65084 1.95253 0.00000
-2.60338 0.00000 0.00000 ; HD3- CD- CE- NZ
4622 4621 4624 4625 3 0.65084 1.95253 0.00000
-2.60338 0.00000 0.00000 ; HE2- CE- NZ- HZ1
4622 4621 4624 4626 3 0.65084 1.95253 0.00000
-2.60338 0.00000 0.00000 ; HE2- CE- NZ- HZ2
4622 4621 4624 4627 3 0.65084 1.95253 0.00000
-2.60338 0.00000 0.00000 ; HE2- CE- NZ- HZ3
4623 4621 4624 4625 3 0.65084 1.95253 0.00000
-2.60338 0.00000 0.00000 ; HE3- CE- NZ- HZ1
4623 4621 4624 4626 3 0.65084 1.95253 0.00000
-2.60338 0.00000 0.00000 ; HE3- CE- NZ- HZ2
4623 4621 4624 4627 3 0.65084 1.95253 0.00000
-2.60338 0.00000 0.00000 ; HE3- CE- NZ- HZ3
4628 4630 4632 4633 3 0.00000 0.00000 0.00000
0.00000 0.00000 0.00000 ; C- N- CA- HA
4628 4630 4632 4634 3 11.71520 1.67360 15.06240 -
13.38880 0.00000 0.00000 ; C- N- CA- CB
4628 4630 4632 4642 3 1.75728 5.27184 2.25936
-7.02912 0.00000 0.00000 ; C- N- CA- C
4629 4628 4630 4631 3 29.28800 -8.36800 -20.92000
0.00000 0.00000 0.00000 ; O- C- N- H

4629 4628 4630 4632 3 20.92000 0.00000 -20.92000
0.00000 0.00000 0.00000 ; O- C- N- CA
4630 4632 4634 4635 3 0.65084 1.95253 0.00000
-2.60338 0.00000 0.00000 ; N- CA- CB- HB2
4630 4632 4634 4636 3 0.65084 1.95253 0.00000
-2.60338 0.00000 0.00000 ; N- CA- CB- HB3
4630 4632 4634 4637 3 5.99567 3.29281 -4.46014
-0.55229 1.97485 0.00000 ; N- CA- CB- CG
4630 4632 4642 4643 3 0.00000 0.00000 0.00000
0.00000 0.00000 0.00000 ; N- CA- C- O
4630 4632 4642 4644 3 17.40544 -5.02080 -13.22144
9.20480 0.00000 0.00000 ; N- CA- C- N
4631 4630 4632 4633 3 0.00000 0.00000 0.00000
0.00000 0.00000 0.00000 ; H- N- CA- HA
4631 4630 4632 4634 3 0.00000 0.00000 0.00000
0.00000 0.00000 0.00000 ; H- N- CA- CB
4631 4630 4632 4642 3 0.00000 0.00000 0.00000
0.00000 0.00000 0.00000 ; H- N- CA- C
4632 4634 4637 4638 3 0.00000 0.00000 0.00000
0.00000 0.00000 0.00000 ; CA- CB- CG- OD1
4632 4634 4637 4639 3 8.84916 -0.31380 -4.32626
5.03754 0.26778 0.00000 ; CA- CB- CG- ND2
4632 4642 4644 4645 3 20.92000 0.00000 -20.92000
0.00000 0.00000 0.00000 ; CA- C- N- H
4632 4642 4644 4646 3 20.92000 0.00000 -20.92000
0.00000 0.00000 0.00000 ; CA- C- N- CA
4633 4632 4634 4635 3 0.65084 1.95253 0.00000
-2.60338 0.00000 0.00000 ; HA- CA- CB- HB2
4633 4632 4634 4636 3 0.65084 1.95253 0.00000
-2.60338 0.00000 0.00000 ; HA- CA- CB- HB3
4633 4632 4634 4637 3 0.65084 1.95253 0.00000
-2.60338 0.00000 0.00000 ; HA- CA- CB- CG
4633 4632 4642 4643 3 3.68192 -4.35136 0.00000
1.33888 0.00000 0.00000 ; HA- CA- C- O
4633 4632 4642 4644 3 0.00000 0.00000 0.00000
0.00000 0.00000 0.00000 ; HA- CA- C- N
4634 4632 4642 4643 3 0.00000 0.00000 0.00000
0.00000 0.00000 0.00000 ; CB- CA- C- O
4634 4632 4642 4644 3 2.51040 4.18400 1.67360
-6.69440 0.00000 0.00000 ; CB- CA- C- N
4634 4637 4639 4640 3 20.92000 0.00000 -20.92000
0.00000 0.00000 0.00000 ; CB- CG- ND2- HD21
4634 4637 4639 4641 3 20.92000 0.00000 -20.92000
0.00000 0.00000 0.00000 ; CB- CG- ND2- HD22
4635 4634 4632 4642 3 0.65084 1.95253 0.00000
-2.60338 0.00000 0.00000 ; HB2- CB- CA- C
4635 4634 4637 4638 3 3.68192 -4.35136 0.00000
1.33888 0.00000 0.00000 ; HB2- CB- CG- OD1
4635 4634 4637 4639 3 0.00000 0.00000 0.00000
0.00000 0.00000 0.00000 ; HB2- CB- CG- ND2
4636 4634 4632 4642 3 0.65084 1.95253 0.00000
-2.60338 0.00000 0.00000 ; HB3- CB- CA- C

4636 4634 4637 4638 3 3.68192 -4.35136 0.00000
1.33888 0.00000 0.00000 ; HB3- CB- CG- OD1
4636 4634 4637 4639 3 0.00000 0.00000 0.00000
0.00000 0.00000 0.00000 ; HB3- CB- CG- ND2
4637 4634 4632 4642 3 7.94542 -3.96225 -6.11701
-0.55229 3.58150 0.00000 ; CG- CB- CA- C
4638 4637 4639 4640 3 29.28800 -8.36800 -20.92000
0.00000 0.00000 0.00000 ; OD1- CG- ND2- HD21
4638 4637 4639 4641 3 29.28800 -8.36800 -20.92000
0.00000 0.00000 0.00000 ; OD1- CG- ND2- HD22
4642 4644 4646 4647 3 0.00000 0.00000 0.00000
0.00000 0.00000 0.00000 ; C- N- CA- HA
4642 4644 4646 4648 3 11.71520 1.67360 15.06240 -
13.38880 0.00000 0.00000 ; C- N- CA- CB
4642 4644 4646 4661 3 1.75728 5.27184 2.25936
-7.02912 0.00000 0.00000 ; C- N- CA- C
4643 4642 4644 4645 3 29.28800 -8.36800 -20.92000
0.00000 0.00000 0.00000 ; O- C- N- H
4643 4642 4644 4646 3 20.92000 0.00000 -20.92000
0.00000 0.00000 0.00000 ; O- C- N- CA
4644 4646 4648 4649 3 0.65084 1.95253 0.00000
-2.60338 0.00000 0.00000 ; N- CA- CB- HB2
4644 4646 4648 4650 3 0.65084 1.95253 0.00000
-2.60338 0.00000 0.00000 ; N- CA- CB- HB3
4644 4646 4648 4651 3 3.79070 1.39746 -4.61077
-2.40998 2.44346 0.00000 ; N- CA- CB- CG
4644 4646 4661 4662 3 0.00000 0.00000 0.00000
0.00000 0.00000 0.00000 ; N- CA- C- O
4644 4646 4661 4663 3 17.40544 -5.02080 -13.22144
9.20480 0.00000 0.00000 ; N- CA- C- N
4645 4644 4646 4647 3 0.00000 0.00000 0.00000
0.00000 0.00000 0.00000 ; H- N- CA- HA
4645 4644 4646 4648 3 0.00000 0.00000 0.00000
0.00000 0.00000 0.00000 ; H- N- CA- CB
4645 4644 4646 4661 3 0.00000 0.00000 0.00000
0.00000 0.00000 0.00000 ; H- N- CA- C
4646 4648 4651 4652 3 0.66944 2.00832 0.00000
-2.67776 0.00000 0.00000 ; CA- CB- CG- HG
4646 4648 4651 4653 3 3.90367 0.19665 -6.21742
-2.37651 5.99149 0.00000 ; CA- CB- CG- CD1
4646 4648 4651 4657 3 3.90367 0.19665 -6.21742
-2.37651 5.99149 0.00000 ; CA- CB- CG- CD2
4646 4661 4663 4664 3 20.92000 0.00000 -20.92000
0.00000 0.00000 0.00000 ; CA- C- N- H
4646 4661 4663 4665 3 20.92000 0.00000 -20.92000
0.00000 0.00000 0.00000 ; CA- C- N- CA
4647 4646 4648 4649 3 0.65084 1.95253 0.00000
-2.60338 0.00000 0.00000 ; HA- CA- CB- HB2
4647 4646 4648 4650 3 0.65084 1.95253 0.00000
-2.60338 0.00000 0.00000 ; HA- CA- CB- HB3
4647 4646 4648 4651 3 0.65084 1.95253 0.00000
-2.60338 0.00000 0.00000 ; HA- CA- CB- CG

4647 4646 4661 4662 3 3.68192 -4.35136 0.00000
1.33888 0.00000 0.00000 ; HA- CA- C- O
4647 4646 4661 4663 3 0.00000 0.00000 0.00000
0.00000 0.00000 0.00000 ; HA- CA- C- N
4648 4646 4661 4662 3 0.00000 0.00000 0.00000
0.00000 0.00000 0.00000 ; CB- CA- C- O
4648 4646 4661 4663 3 2.51040 4.18400 1.67360
-6.69440 0.00000 0.00000 ; CB- CA- C- N
4648 4651 4653 4654 3 0.66944 2.00832 0.00000
-2.67776 0.00000 0.00000 ; CB- CG- CD1- HD11
4648 4651 4653 4655 3 0.66944 2.00832 0.00000
-2.67776 0.00000 0.00000 ; CB- CG- CD1- HD12
4648 4651 4653 4656 3 0.66944 2.00832 0.00000
-2.67776 0.00000 0.00000 ; CB- CG- CD1- HD13
4648 4651 4657 4658 3 0.66944 2.00832 0.00000
-2.67776 0.00000 0.00000 ; CB- CG- CD2- HD21
4648 4651 4657 4659 3 0.66944 2.00832 0.00000
-2.67776 0.00000 0.00000 ; CB- CG- CD2- HD22
4648 4651 4657 4660 3 0.66944 2.00832 0.00000
-2.67776 0.00000 0.00000 ; CB- CG- CD2- HD23
4649 4648 4646 4661 3 0.65084 1.95253 0.00000
-2.60338 0.00000 0.00000 ; HB2- CB- CA- C
4649 4648 4651 4652 3 0.62760 1.88280 0.00000
-2.51040 0.00000 0.00000 ; HB2- CB- CG- HG
4649 4648 4651 4653 3 0.66944 2.00832 0.00000
-2.67776 0.00000 0.00000 ; HB2- CB- CG- CD1
4649 4648 4651 4657 3 0.66944 2.00832 0.00000
-2.67776 0.00000 0.00000 ; HB2- CB- CG- CD2
4650 4648 4646 4661 3 0.65084 1.95253 0.00000
-2.60338 0.00000 0.00000 ; HB3- CB- CA- C
4650 4648 4651 4652 3 0.62760 1.88280 0.00000
-2.51040 0.00000 0.00000 ; HB3- CB- CG- HG
4650 4648 4651 4653 3 0.66944 2.00832 0.00000
-2.67776 0.00000 0.00000 ; HB3- CB- CG- CD1
4650 4648 4651 4657 3 0.66944 2.00832 0.00000
-2.67776 0.00000 0.00000 ; HB3- CB- CG- CD2
4651 4648 4646 4661 3 10.33448 -1.14642 -11.54784
-2.40998 6.35968 0.00000 ; CG- CB- CA- C
4652 4651 4653 4654 3 0.62760 1.88280 0.00000
-2.51040 0.00000 0.00000 ; HG- CG- CD1- HD11
4652 4651 4653 4655 3 0.62760 1.88280 0.00000
-2.51040 0.00000 0.00000 ; HG- CG- CD1- HD12
4652 4651 4653 4656 3 0.62760 1.88280 0.00000
-2.51040 0.00000 0.00000 ; HG- CG- CD1- HD13
4652 4651 4657 4658 3 0.62760 1.88280 0.00000
-2.51040 0.00000 0.00000 ; HG- CG- CD2- HD21
4652 4651 4657 4659 3 0.62760 1.88280 0.00000
-2.51040 0.00000 0.00000 ; HG- CG- CD2- HD22
4652 4651 4657 4660 3 0.62760 1.88280 0.00000
-2.51040 0.00000 0.00000 ; HG- CG- CD2- HD23
4653 4651 4657 4658 3 0.66944 2.00832 0.00000
-2.67776 0.00000 0.00000 ; CD1- CG- CD2- HD21

4653 4651 4657 4659 3 0.66944 2.00832 0.00000
-2.67776 0.00000 0.00000 ; CD1- CG- CD2- HD22
4653 4651 4657 4660 3 0.66944 2.00832 0.00000
-2.67776 0.00000 0.00000 ; CD1- CG- CD2- HD23
4654 4653 4651 4657 3 0.66944 2.00832 0.00000
-2.67776 0.00000 0.00000 ; HD11- CD1- CG- CD2
4655 4653 4651 4657 3 0.66944 2.00832 0.00000
-2.67776 0.00000 0.00000 ; HD12- CD1- CG- CD2
4656 4653 4651 4657 3 0.66944 2.00832 0.00000
-2.67776 0.00000 0.00000 ; HD13- CD1- CG- CD2
4661 4663 4665 4666 3 0.00000 0.00000 0.00000
0.00000 0.00000 0.00000 ; C- N- CA- HA
4661 4663 4665 4667 3 11.71520 1.67360 15.06240 -
13.38880 0.00000 0.00000 ; C- N- CA- CB
4661 4663 4665 4683 3 1.75728 5.27184 2.25936
-7.02912 0.00000 0.00000 ; C- N- CA- C
4662 4661 4663 4664 3 29.28800 -8.36800 -20.92000
0.00000 0.00000 0.00000 ; O- C- N- H
4662 4661 4663 4665 3 20.92000 0.00000 -20.92000
0.00000 0.00000 0.00000 ; O- C- N- CA
4663 4665 4667 4668 3 0.65084 1.95253 0.00000
-2.60338 0.00000 0.00000 ; N- CA- CB- HB2
4663 4665 4667 4669 3 0.65084 1.95253 0.00000
-2.60338 0.00000 0.00000 ; N- CA- CB- HB3
4663 4665 4667 4670 3 0.65084 1.95253 0.00000
-2.60338 0.00000 0.00000 ; N- CA- CB- CG
4663 4665 4683 4684 3 0.00000 0.00000 0.00000
0.00000 0.00000 0.00000 ; N- CA- C- O
4663 4665 4683 4685 3 17.40544 -5.02080 -13.22144
9.20480 0.00000 0.00000 ; N- CA- C- N
4664 4663 4665 4666 3 0.00000 0.00000 0.00000
0.00000 0.00000 0.00000 ; H- N- CA- HA
4664 4663 4665 4667 3 0.00000 0.00000 0.00000
0.00000 0.00000 0.00000 ; H- N- CA- CB
4664 4663 4665 4683 3 0.00000 0.00000 0.00000
0.00000 0.00000 0.00000 ; H- N- CA- C
4665 4667 4670 4671 3 0.66944 2.00832 0.00000
-2.67776 0.00000 0.00000 ; CA- CB- CG- HG2
4665 4667 4670 4672 3 0.66944 2.00832 0.00000
-2.67776 0.00000 0.00000 ; CA- CB- CG- HG3
4665 4667 4670 4673 3 3.68192 3.09616 -2.09200
-3.01248 0.00000 0.00000 ; CA- CB- CG- CD
4665 4683 4685 4686 3 20.92000 0.00000 -20.92000
0.00000 0.00000 0.00000 ; CA- C- N- H
4665 4683 4685 4687 3 20.92000 0.00000 -20.92000
0.00000 0.00000 0.00000 ; CA- C- N- CA
4666 4665 4667 4668 3 0.65084 1.95253 0.00000
-2.60338 0.00000 0.00000 ; HA- CA- CB- HB2
4666 4665 4667 4669 3 0.65084 1.95253 0.00000
-2.60338 0.00000 0.00000 ; HA- CA- CB- HB3
4666 4665 4667 4670 3 0.65084 1.95253 0.00000
-2.60338 0.00000 0.00000 ; HA- CA- CB- CG

4666 4665 4683 4684 3 3.68192 -4.35136 0.00000
1.33888 0.00000 0.00000 ; HA- CA- C- O
4666 4665 4683 4685 3 0.00000 0.00000 0.00000
0.00000 0.00000 0.00000 ; HA- CA- C- N
4667 4665 4683 4684 3 0.00000 0.00000 0.00000
0.00000 0.00000 0.00000 ; CB- CA- C- O
4667 4665 4683 4685 3 2.51040 4.18400 1.67360
-6.69440 0.00000 0.00000 ; CB- CA- C- N
4667 4670 4673 4674 3 0.66944 2.00832 0.00000
-2.67776 0.00000 0.00000 ; CB- CG- CD- HD2
4667 4670 4673 4675 3 0.66944 2.00832 0.00000
-2.67776 0.00000 0.00000 ; CB- CG- CD- HD3
4667 4670 4673 4676 3 3.68192 3.09616 -2.09200
-3.01248 0.00000 0.00000 ; CB- CG- CD- CE
4668 4667 4665 4683 3 0.65084 1.95253 0.00000
-2.60338 0.00000 0.00000 ; HB2- CB- CA- C
4668 4667 4670 4671 3 0.62760 1.88280 0.00000
-2.51040 0.00000 0.00000 ; HB2- CB- CG- HG2
4668 4667 4670 4672 3 0.62760 1.88280 0.00000
-2.51040 0.00000 0.00000 ; HB2- CB- CG- HG3
4668 4667 4670 4673 3 0.66944 2.00832 0.00000
-2.67776 0.00000 0.00000 ; HB2- CB- CG- CD
4669 4667 4665 4683 3 0.65084 1.95253 0.00000
-2.60338 0.00000 0.00000 ; HB3- CB- CA- C
4669 4667 4670 4671 3 0.62760 1.88280 0.00000
-2.51040 0.00000 0.00000 ; HB3- CB- CG- HG2
4669 4667 4670 4672 3 0.62760 1.88280 0.00000
-2.51040 0.00000 0.00000 ; HB3- CB- CG- HG3
4669 4667 4670 4673 3 0.66944 2.00832 0.00000
-2.67776 0.00000 0.00000 ; HB3- CB- CG- CD
4670 4667 4665 4683 3 0.65084 1.95253 0.00000
-2.60338 0.00000 0.00000 ; CG- CB- CA- C
4670 4673 4676 4677 3 0.65084 1.95253 0.00000
-2.60338 0.00000 0.00000 ; CG- CD- CE- HE2
4670 4673 4676 4678 3 0.65084 1.95253 0.00000
-2.60338 0.00000 0.00000 ; CG- CD- CE- HE3
4670 4673 4676 4679 3 0.65084 1.95253 0.00000
-2.60338 0.00000 0.00000 ; CG- CD- CE- NZ
4671 4670 4673 4674 3 0.62760 1.88280 0.00000
-2.51040 0.00000 0.00000 ; HG2- CG- CD- HD2
4671 4670 4673 4675 3 0.62760 1.88280 0.00000
-2.51040 0.00000 0.00000 ; HG2- CG- CD- HD3
4671 4670 4673 4676 3 0.66944 2.00832 0.00000
-2.67776 0.00000 0.00000 ; HG2- CG- CD- CE
4672 4670 4673 4674 3 0.62760 1.88280 0.00000
-2.51040 0.00000 0.00000 ; HG3- CG- CD- HD2
4672 4670 4673 4675 3 0.62760 1.88280 0.00000
-2.51040 0.00000 0.00000 ; HG3- CG- CD- HD3
4672 4670 4673 4676 3 0.66944 2.00832 0.00000
-2.67776 0.00000 0.00000 ; HG3- CG- CD- CE
4673 4676 4679 4680 3 0.65084 1.95253 0.00000
-2.60338 0.00000 0.00000 ; CD- CE- NZ- HZ1

4673 4676 4679 4681 3 0.65084 1.95253 0.00000
-2.60338 0.00000 0.00000 ; CD- CE- NZ- HZ2
4673 4676 4679 4682 3 0.65084 1.95253 0.00000
-2.60338 0.00000 0.00000 ; CD- CE- NZ- HZ3
4674 4673 4676 4677 3 0.65084 1.95253 0.00000
-2.60338 0.00000 0.00000 ; HD2- CD- CE- HE2
4674 4673 4676 4678 3 0.65084 1.95253 0.00000
-2.60338 0.00000 0.00000 ; HD2- CD- CE- HE3
4674 4673 4676 4679 3 0.65084 1.95253 0.00000
-2.60338 0.00000 0.00000 ; HD2- CD- CE- NZ
4675 4673 4676 4677 3 0.65084 1.95253 0.00000
-2.60338 0.00000 0.00000 ; HD3- CD- CE- HE2
4675 4673 4676 4678 3 0.65084 1.95253 0.00000
-2.60338 0.00000 0.00000 ; HD3- CD- CE- HE3
4675 4673 4676 4679 3 0.65084 1.95253 0.00000
-2.60338 0.00000 0.00000 ; HD3- CD- CE- NZ
4677 4676 4679 4680 3 0.65084 1.95253 0.00000
-2.60338 0.00000 0.00000 ; HE2- CE- NZ- HZ1
4677 4676 4679 4681 3 0.65084 1.95253 0.00000
-2.60338 0.00000 0.00000 ; HE2- CE- NZ- HZ2
4677 4676 4679 4682 3 0.65084 1.95253 0.00000
-2.60338 0.00000 0.00000 ; HE2- CE- NZ- HZ3
4678 4676 4679 4680 3 0.65084 1.95253 0.00000
-2.60338 0.00000 0.00000 ; HE3- CE- NZ- HZ1
4678 4676 4679 4681 3 0.65084 1.95253 0.00000
-2.60338 0.00000 0.00000 ; HE3- CE- NZ- HZ2
4678 4676 4679 4682 3 0.65084 1.95253 0.00000
-2.60338 0.00000 0.00000 ; HE3- CE- NZ- HZ3
4683 4685 4687 4688 3 0.00000 0.00000 0.00000
0.00000 0.00000 0.00000 ; C- N- CA- HA
4683 4685 4687 4689 3 11.71520 1.67360 15.06240 -
13.38880 0.00000 0.00000 ; C- N- CA- CB
4683 4685 4687 4697 3 1.75728 5.27184 2.25936
-7.02912 0.00000 0.00000 ; C- N- CA- C
4684 4683 4685 4686 3 29.28800 -8.36800 -20.92000
0.00000 0.00000 0.00000 ; O- C- N- H
4684 4683 4685 4687 3 20.92000 0.00000 -20.92000
0.00000 0.00000 0.00000 ; O- C- N- CA
4685 4687 4689 4690 3 0.65084 1.95253 0.00000
-2.60338 0.00000 0.00000 ; N- CA- CB- HB2
4685 4687 4689 4691 3 0.65084 1.95253 0.00000
-2.60338 0.00000 0.00000 ; N- CA- CB- HB3
4685 4687 4689 4692 3 5.99567 3.29281 -4.46014
-0.55229 1.97485 0.00000 ; N- CA- CB- CG
4685 4687 4697 4698 3 0.00000 0.00000 0.00000
0.00000 0.00000 0.00000 ; N- CA- C- O
4685 4687 4697 4699 3 17.40544 -5.02080 -13.22144
9.20480 0.00000 0.00000 ; N- CA- C- N
4686 4685 4687 4688 3 0.00000 0.00000 0.00000
0.00000 0.00000 0.00000 ; H- N- CA- HA
4686 4685 4687 4689 3 0.00000 0.00000 0.00000
0.00000 0.00000 0.00000 ; H- N- CA- CB

4686 4685 4687 4697 3 0.00000 0.00000 0.00000
0.00000 0.00000 0.00000 ; H- N- CA- C
4687 4689 4692 4693 3 0.00000 0.00000 0.00000
0.00000 0.00000 0.00000 ; CA- CB- CG- OD1
4687 4689 4692 4694 3 8.84916 -0.31380 -4.32626
5.03754 0.26778 0.00000 ; CA- CB- CG- ND2
4687 4697 4699 4700 3 20.92000 0.00000 -20.92000
0.00000 0.00000 0.00000 ; CA- C- N- H
4687 4697 4699 4701 3 20.92000 0.00000 -20.92000
0.00000 0.00000 0.00000 ; CA- C- N- CA
4688 4687 4689 4690 3 0.65084 1.95253 0.00000
-2.60338 0.00000 0.00000 ; HA- CA- CB- HB2
4688 4687 4689 4691 3 0.65084 1.95253 0.00000
-2.60338 0.00000 0.00000 ; HA- CA- CB- HB3
4688 4687 4689 4692 3 0.65084 1.95253 0.00000
-2.60338 0.00000 0.00000 ; HA- CA- CB- CG
4688 4687 4697 4698 3 3.68192 -4.35136 0.00000
1.33888 0.00000 0.00000 ; HA- CA- C- O
4688 4687 4697 4699 3 0.00000 0.00000 0.00000
0.00000 0.00000 0.00000 ; HA- CA- C- N
4689 4687 4697 4698 3 0.00000 0.00000 0.00000
0.00000 0.00000 0.00000 ; CB- CA- C- O
4689 4687 4697 4699 3 2.51040 4.18400 1.67360
-6.69440 0.00000 0.00000 ; CB- CA- C- N
4689 4692 4694 4695 3 20.92000 0.00000 -20.92000
0.00000 0.00000 0.00000 ; CB- CG- ND2- HD21
4689 4692 4694 4696 3 20.92000 0.00000 -20.92000
0.00000 0.00000 0.00000 ; CB- CG- ND2- HD22
4690 4689 4687 4697 3 0.65084 1.95253 0.00000
-2.60338 0.00000 0.00000 ; HB2- CB- CA- C
4690 4689 4692 4693 3 3.68192 -4.35136 0.00000
1.33888 0.00000 0.00000 ; HB2- CB- CG- OD1
4690 4689 4692 4694 3 0.00000 0.00000 0.00000
0.00000 0.00000 0.00000 ; HB2- CB- CG- ND2
4691 4689 4687 4697 3 0.65084 1.95253 0.00000
-2.60338 0.00000 0.00000 ; HB3- CB- CA- C
4691 4689 4692 4693 3 3.68192 -4.35136 0.00000
1.33888 0.00000 0.00000 ; HB3- CB- CG- OD1
4691 4689 4692 4694 3 0.00000 0.00000 0.00000
0.00000 0.00000 0.00000 ; HB3- CB- CG- ND2
4692 4689 4687 4697 3 7.94542 -3.96225 -6.11701
-0.55229 3.58150 0.00000 ; CG- CB- CA- C
4693 4692 4694 4695 3 29.28800 -8.36800 -20.92000
0.00000 0.00000 0.00000 ; OD1- CG- ND2- HD21
4693 4692 4694 4696 3 29.28800 -8.36800 -20.92000
0.00000 0.00000 0.00000 ; OD1- CG- ND2- HD22
4697 4699 4701 4702 3 0.00000 0.00000 0.00000
0.00000 0.00000 0.00000 ; C- N- CA- HA
4697 4699 4701 4703 3 11.71520 1.67360 15.06240 -
13.38880 0.00000 0.00000 ; C- N- CA- CB
4697 4699 4701 4717 3 1.75728 5.27184 2.25936
-7.02912 0.00000 0.00000 ; C- N- CA- C

4698 4697 4699 4700 3 29.28800 -8.36800 -20.92000
0.00000 0.00000 0.00000 ; O- C- N- H
4698 4697 4699 4701 3 20.92000 0.00000 -20.92000
0.00000 0.00000 0.00000 ; O- C- N- CA
4699 4701 4703 4704 3 0.65084 1.95253 0.00000
-2.60338 0.00000 0.00000 ; N- CA- CB- HB2
4699 4701 4703 4705 3 0.65084 1.95253 0.00000
-2.60338 0.00000 0.00000 ; N- CA- CB- HB3
4699 4701 4703 4706 3 3.28026 2.46019 -2.19242
-3.21331 -0.23430 0.00000 ; N- CA- CB- CG
4699 4701 4717 4718 3 0.00000 0.00000 0.00000
0.00000 0.00000 0.00000 ; N- CA- C- O
4699 4701 4717 4719 3 17.40544 -5.02080 -13.22144
9.20480 0.00000 0.00000 ; N- CA- C- N
4700 4699 4701 4702 3 0.00000 0.00000 0.00000
0.00000 0.00000 0.00000 ; H- N- CA- HA
4700 4699 4701 4703 3 0.00000 0.00000 0.00000
0.00000 0.00000 0.00000 ; H- N- CA- CB
4700 4699 4701 4717 3 0.00000 0.00000 0.00000
0.00000 0.00000 0.00000 ; H- N- CA- C
4701 4703 4706 4707 3 0.57739 0.00000 1.02926
0.00000 -1.60666 0.00000 ; CA- CB- CG- CD1
4701 4703 4706 4715 3 0.57739 0.00000 1.02926
0.00000 -1.60666 0.00000 ; CA- CB- CG- CD2
4701 4717 4719 4720 3 20.92000 0.00000 -20.92000
0.00000 0.00000 0.00000 ; CA- C- N- H
4701 4717 4719 4721 3 20.92000 0.00000 -20.92000
0.00000 0.00000 0.00000 ; CA- C- N- CA
4702 4701 4703 4704 3 0.65084 1.95253 0.00000
-2.60338 0.00000 0.00000 ; HA- CA- CB- HB2
4702 4701 4703 4705 3 0.65084 1.95253 0.00000
-2.60338 0.00000 0.00000 ; HA- CA- CB- HB3
4702 4701 4703 4706 3 0.65084 1.95253 0.00000
-2.60338 0.00000 0.00000 ; HA- CA- CB- CG
4702 4701 4717 4718 3 3.68192 -4.35136 0.00000
1.33888 0.00000 0.00000 ; HA- CA- C- O
4702 4701 4717 4719 3 0.00000 0.00000 0.00000
0.00000 0.00000 0.00000 ; HA- CA- C- N
4703 4701 4717 4718 3 0.00000 0.00000 0.00000
0.00000 0.00000 0.00000 ; CB- CA- C- O
4703 4701 4717 4719 3 2.51040 4.18400 1.67360
-6.69440 0.00000 0.00000 ; CB- CA- C- N
4703 4706 4707 4708 3 30.33400 0.00000 -30.33400
0.00000 0.00000 0.00000 ; CB- CG- CD1- HD1
4703 4706 4707 4709 3 30.33400 0.00000 -30.33400
0.00000 0.00000 0.00000 ; CB- CG- CD1- CE1
4703 4706 4715 4713 3 30.33400 0.00000 -30.33400
0.00000 0.00000 0.00000 ; CB- CG- CD2- CE2
4703 4706 4715 4716 3 30.33400 0.00000 -30.33400
0.00000 0.00000 0.00000 ; CB- CG- CD2- HD2
4704 4703 4701 4717 3 0.65084 1.95253 0.00000
-2.60338 0.00000 0.00000 ; HB2- CB- CA- C

4704 4703 4706 4707 3 0.00000 0.00000 0.00000
0.00000 0.00000 0.00000 ; HB2- CB- CG- CD1
4704 4703 4706 4715 3 0.00000 0.00000 0.00000
0.00000 0.00000 0.00000 ; HB2- CB- CG- CD2
4705 4703 4701 4717 3 0.65084 1.95253 0.00000
-2.60338 0.00000 0.00000 ; HB3- CB- CA- C
4705 4703 4706 4707 3 0.00000 0.00000 0.00000
0.00000 0.00000 0.00000 ; HB3- CB- CG- CD1
4705 4703 4706 4715 3 0.00000 0.00000 0.00000
0.00000 0.00000 0.00000 ; HB3- CB- CG- CD2
4706 4703 4701 4717 3 4.95804 2.17986 -3.52293
-3.21331 -0.40166 0.00000 ; CG- CB- CA- C
4706 4707 4709 4710 3 30.33400 0.00000 -30.33400
0.00000 0.00000 0.00000 ; CG- CD1- CE1- HE1
4706 4707 4709 4711 3 30.33400 0.00000 -30.33400
0.00000 0.00000 0.00000 ; CG- CD1- CE1- CZ
4706 4715 4713 4711 3 30.33400 0.00000 -30.33400
0.00000 0.00000 0.00000 ; CG- CD2- CE2- CZ
4706 4715 4713 4714 3 30.33400 0.00000 -30.33400
0.00000 0.00000 0.00000 ; CG- CD2- CE2- HE2
4707 4706 4715 4713 3 30.33400 0.00000 -30.33400
0.00000 0.00000 0.00000 ; CD1- CG- CD2- CE2
4707 4706 4715 4716 3 30.33400 0.00000 -30.33400
0.00000 0.00000 0.00000 ; CD1- CG- CD2- HD2
4707 4709 4711 4712 3 30.33400 0.00000 -30.33400
0.00000 0.00000 0.00000 ; CD1- CE1- CZ- HZ
4707 4709 4711 4713 3 30.33400 0.00000 -30.33400
0.00000 0.00000 0.00000 ; CD1- CE1- CZ- CE2
4708 4707 4706 4715 3 30.33400 0.00000 -30.33400
0.00000 0.00000 0.00000 ; HD1- CD1- CG- CD2
4708 4707 4709 4710 3 30.33400 0.00000 -30.33400
0.00000 0.00000 0.00000 ; HD1- CD1- CE1- HE1
4708 4707 4709 4711 3 30.33400 0.00000 -30.33400
0.00000 0.00000 0.00000 ; HD1- CD1- CE1- CZ
4709 4707 4706 4715 3 30.33400 0.00000 -30.33400
0.00000 0.00000 0.00000 ; CE1- CD1- CG- CD2
4709 4711 4713 4714 3 30.33400 0.00000 -30.33400
0.00000 0.00000 0.00000 ; CE1- CZ- CE2- HE2
4709 4711 4713 4715 3 30.33400 0.00000 -30.33400
0.00000 0.00000 0.00000 ; CE1- CZ- CE2- CD2
4710 4709 4711 4712 3 30.33400 0.00000 -30.33400
0.00000 0.00000 0.00000 ; HE1- CE1- CZ- HZ
4710 4709 4711 4713 3 30.33400 0.00000 -30.33400
0.00000 0.00000 0.00000 ; HE1- CE1- CZ- CE2
4711 4713 4715 4716 3 30.33400 0.00000 -30.33400
0.00000 0.00000 0.00000 ; CZ- CE2- CD2- HD2
4712 4711 4713 4714 3 30.33400 0.00000 -30.33400
0.00000 0.00000 0.00000 ; HZ- CZ- CE2- HE2
4712 4711 4713 4715 3 30.33400 0.00000 -30.33400
0.00000 0.00000 0.00000 ; HZ- CZ- CE2- CD2
4714 4713 4715 4716 3 30.33400 0.00000 -30.33400
0.00000 0.00000 0.00000 ; HE2- CE2- CD2- HD2

4717 4719 4721 4722 3 0.00000 0.00000 0.00000
0.00000 0.00000 0.00000 ; C- N- CA- HA
4717 4719 4721 4723 3 11.71520 1.67360 15.06240 -
13.38880 0.00000 0.00000 ; C- N- CA- CB
4717 4719 4721 4739 3 1.75728 5.27184 2.25936
-7.02912 0.00000 0.00000 ; C- N- CA- C
4718 4717 4719 4720 3 29.28800 -8.36800 -20.92000
0.00000 0.00000 0.00000 ; O- C- N- H
4718 4717 4719 4721 3 20.92000 0.00000 -20.92000
0.00000 0.00000 0.00000 ; O- C- N- CA
4719 4721 4723 4724 3 0.65084 1.95253 0.00000
-2.60338 0.00000 0.00000 ; N- CA- CB- HB2
4719 4721 4723 4725 3 0.65084 1.95253 0.00000
-2.60338 0.00000 0.00000 ; N- CA- CB- HB3
4719 4721 4723 4726 3 0.65084 1.95253 0.00000
-2.60338 0.00000 0.00000 ; N- CA- CB- CG
4719 4721 4739 4740 3 0.00000 0.00000 0.00000
0.00000 0.00000 0.00000 ; N- CA- C- O
4719 4721 4739 4741 3 17.40544 -5.02080 -13.22144
9.20480 0.00000 0.00000 ; N- CA- C- N
4720 4719 4721 4722 3 0.00000 0.00000 0.00000
0.00000 0.00000 0.00000 ; H- N- CA- HA
4720 4719 4721 4723 3 0.00000 0.00000 0.00000
0.00000 0.00000 0.00000 ; H- N- CA- CB
4720 4719 4721 4739 3 0.00000 0.00000 0.00000
0.00000 0.00000 0.00000 ; H- N- CA- C
4721 4723 4726 4727 3 0.66944 2.00832 0.00000
-2.67776 0.00000 0.00000 ; CA- CB- CG- HG2
4721 4723 4726 4728 3 0.66944 2.00832 0.00000
-2.67776 0.00000 0.00000 ; CA- CB- CG- HG3
4721 4723 4726 4729 3 3.68192 3.09616 -2.09200
-3.01248 0.00000 0.00000 ; CA- CB- CG- CD
4721 4739 4741 4742 3 20.92000 0.00000 -20.92000
0.00000 0.00000 0.00000 ; CA- C- N- H
4721 4739 4741 4743 3 20.92000 0.00000 -20.92000
0.00000 0.00000 0.00000 ; CA- C- N- CA
4722 4721 4723 4724 3 0.65084 1.95253 0.00000
-2.60338 0.00000 0.00000 ; HA- CA- CB- HB2
4722 4721 4723 4725 3 0.65084 1.95253 0.00000
-2.60338 0.00000 0.00000 ; HA- CA- CB- HB3
4722 4721 4723 4726 3 0.65084 1.95253 0.00000
-2.60338 0.00000 0.00000 ; HA- CA- CB- CG
4722 4721 4739 4740 3 3.68192 -4.35136 0.00000
1.33888 0.00000 0.00000 ; HA- CA- C- O
4722 4721 4739 4741 3 0.00000 0.00000 0.00000
0.00000 0.00000 0.00000 ; HA- CA- C- N
4723 4721 4739 4740 3 0.00000 0.00000 0.00000
0.00000 0.00000 0.00000 ; CB- CA- C- O
4723 4721 4739 4741 3 2.51040 4.18400 1.67360
-6.69440 0.00000 0.00000 ; CB- CA- C- N
4723 4726 4729 4730 3 0.66944 2.00832 0.00000
-2.67776 0.00000 0.00000 ; CB- CG- CD- HD2

```
  4723   4726   4729   4731      3    0.66944    2.00832    0.00000
-2.67776    0.00000    0.00000 ;     CB-    CG-    CD-   HD3
  4723   4726   4729   4732      3    3.68192    3.09616   -2.09200
-3.01248    0.00000    0.00000 ;     CB-    CG-    CD-    CE
  4724   4723   4721   4739      3    0.65084    1.95253    0.00000
-2.60338    0.00000    0.00000 ;    HB2-    CB-    CA-     C
  4724   4723   4726   4727      3    0.62760    1.88280    0.00000
-2.51040    0.00000    0.00000 ;    HB2-    CB-    CG-   HG2
  4724   4723   4726   4728      3    0.62760    1.88280    0.00000
-2.51040    0.00000    0.00000 ;    HB2-    CB-    CG-   HG3
  4724   4723   4726   4729      3    0.66944    2.00832    0.00000
-2.67776    0.00000    0.00000 ;    HB2-    CB-    CG-    CD
  4725   4723   4721   4739      3    0.65084    1.95253    0.00000
-2.60338    0.00000    0.00000 ;    HB3-    CB-    CA-     C
  4725   4723   4726   4727      3    0.62760    1.88280    0.00000
-2.51040    0.00000    0.00000 ;    HB3-    CB-    CG-   HG2
  4725   4723   4726   4728      3    0.62760    1.88280    0.00000
-2.51040    0.00000    0.00000 ;    HB3-    CB-    CG-   HG3
  4725   4723   4726   4729      3    0.66944    2.00832    0.00000
-2.67776    0.00000    0.00000 ;    HB3-    CB-    CG-    CD
  4726   4723   4721   4739      3    0.65084    1.95253    0.00000
-2.60338    0.00000    0.00000 ;     CG-    CB-    CA-     C
  4726   4729   4732   4733      3    0.65084    1.95253    0.00000
-2.60338    0.00000    0.00000 ;     CG-    CD-    CE-   HE2
  4726   4729   4732   4734      3    0.65084    1.95253    0.00000
-2.60338    0.00000    0.00000 ;     CG-    CD-    CE-   HE3
  4726   4729   4732   4735      3    0.65084    1.95253    0.00000
-2.60338    0.00000    0.00000 ;     CG-    CD-    CE-    NZ
  4727   4726   4729   4730      3    0.62760    1.88280    0.00000
-2.51040    0.00000    0.00000 ;    HG2-    CG-    CD-   HD2
  4727   4726   4729   4731      3    0.62760    1.88280    0.00000
-2.51040    0.00000    0.00000 ;    HG2-    CG-    CD-   HD3
  4727   4726   4729   4732      3    0.66944    2.00832    0.00000
-2.67776    0.00000    0.00000 ;    HG2-    CG-    CD-    CE
  4728   4726   4729   4730      3    0.62760    1.88280    0.00000
-2.51040    0.00000    0.00000 ;    HG3-    CG-    CD-   HD2
  4728   4726   4729   4731      3    0.62760    1.88280    0.00000
-2.51040    0.00000    0.00000 ;    HG3-    CG-    CD-   HD3
  4728   4726   4729   4732      3    0.66944    2.00832    0.00000
-2.67776    0.00000    0.00000 ;    HG3-    CG-    CD-    CE
  4729   4732   4735   4736      3    0.65084    1.95253    0.00000
-2.60338    0.00000    0.00000 ;     CD-    CE-    NZ-   HZ1
  4729   4732   4735   4737      3    0.65084    1.95253    0.00000
-2.60338    0.00000    0.00000 ;     CD-    CE-    NZ-   HZ2
  4729   4732   4735   4738      3    0.65084    1.95253    0.00000
-2.60338    0.00000    0.00000 ;     CD-    CE-    NZ-   HZ3
  4730   4729   4732   4733      3    0.65084    1.95253    0.00000
-2.60338    0.00000    0.00000 ;    HD2-    CD-    CE-   HE2
  4730   4729   4732   4734      3    0.65084    1.95253    0.00000
-2.60338    0.00000    0.00000 ;    HD2-    CD-    CE-   HE3
  4730   4729   4732   4735      3    0.65084    1.95253    0.00000
-2.60338    0.00000    0.00000 ;    HD2-    CD-    CE-    NZ
```

4731 4729 4732 4733 3 0.65084 1.95253 0.00000
-2.60338 0.00000 0.00000 ; HD3- CD- CE- HE2
4731 4729 4732 4734 3 0.65084 1.95253 0.00000
-2.60338 0.00000 0.00000 ; HD3- CD- CE- HE3
4731 4729 4732 4735 3 0.65084 1.95253 0.00000
-2.60338 0.00000 0.00000 ; HD3- CD- CE- NZ
4733 4732 4735 4736 3 0.65084 1.95253 0.00000
-2.60338 0.00000 0.00000 ; HE2- CE- NZ- HZ1
4733 4732 4735 4737 3 0.65084 1.95253 0.00000
-2.60338 0.00000 0.00000 ; HE2- CE- NZ- HZ2
4733 4732 4735 4738 3 0.65084 1.95253 0.00000
-2.60338 0.00000 0.00000 ; HE2- CE- NZ- HZ3
4734 4732 4735 4736 3 0.65084 1.95253 0.00000
-2.60338 0.00000 0.00000 ; HE3- CE- NZ- HZ1
4734 4732 4735 4737 3 0.65084 1.95253 0.00000
-2.60338 0.00000 0.00000 ; HE3- CE- NZ- HZ2
4734 4732 4735 4738 3 0.65084 1.95253 0.00000
-2.60338 0.00000 0.00000 ; HE3- CE- NZ- HZ3
4739 4741 4743 4744 3 0.00000 0.00000 0.00000
0.00000 0.00000 0.00000 ; C- N- CA- HA
4739 4741 4743 4745 3 11.71520 1.67360 15.06240 -
13.38880 0.00000 0.00000 ; C- N- CA- CB
4739 4741 4743 4761 3 1.75728 5.27184 2.25936
-7.02912 0.00000 0.00000 ; C- N- CA- C
4740 4739 4741 4742 3 29.28800 -8.36800 -20.92000
0.00000 0.00000 0.00000 ; O- C- N- H
4740 4739 4741 4743 3 20.92000 0.00000 -20.92000
0.00000 0.00000 0.00000 ; O- C- N- CA
4741 4743 4745 4746 3 0.65084 1.95253 0.00000
-2.60338 0.00000 0.00000 ; N- CA- CB- HB2
4741 4743 4745 4747 3 0.65084 1.95253 0.00000
-2.60338 0.00000 0.00000 ; N- CA- CB- HB3
4741 4743 4745 4748 3 0.65084 1.95253 0.00000
-2.60338 0.00000 0.00000 ; N- CA- CB- CG
4741 4743 4761 4762 3 0.00000 0.00000 0.00000
0.00000 0.00000 0.00000 ; N- CA- C- O
4741 4743 4761 4763 3 17.40544 -5.02080 -13.22144
9.20480 0.00000 0.00000 ; N- CA- C- N
4742 4741 4743 4744 3 0.00000 0.00000 0.00000
0.00000 0.00000 0.00000 ; H- N- CA- HA
4742 4741 4743 4745 3 0.00000 0.00000 0.00000
0.00000 0.00000 0.00000 ; H- N- CA- CB
4742 4741 4743 4761 3 0.00000 0.00000 0.00000
0.00000 0.00000 0.00000 ; H- N- CA- C
4743 4745 4748 4749 3 0.66944 2.00832 0.00000
-2.67776 0.00000 0.00000 ; CA- CB- CG- HG2
4743 4745 4748 4750 3 0.66944 2.00832 0.00000
-2.67776 0.00000 0.00000 ; CA- CB- CG- HG3
4743 4745 4748 4751 3 3.68192 3.09616 -2.09200
-3.01248 0.00000 0.00000 ; CA- CB- CG- CD
4743 4761 4763 4764 3 20.92000 0.00000 -20.92000
0.00000 0.00000 0.00000 ; CA- C- N- H

4743 4761 4763 4765 3 20.92000 0.00000 -20.92000
0.00000 0.00000 0.00000 ; CA- C- N- CA
4744 4743 4745 4746 3 0.65084 1.95253 0.00000
-2.60338 0.00000 0.00000 ; HA- CA- CB- HB2
4744 4743 4745 4747 3 0.65084 1.95253 0.00000
-2.60338 0.00000 0.00000 ; HA- CA- CB- HB3
4744 4743 4745 4748 3 0.65084 1.95253 0.00000
-2.60338 0.00000 0.00000 ; HA- CA- CB- CG
4744 4743 4761 4762 3 3.68192 -4.35136 0.00000
1.33888 0.00000 0.00000 ; HA- CA- C- O
4744 4743 4761 4763 3 0.00000 0.00000 0.00000
0.00000 0.00000 0.00000 ; HA- CA- C- N
4745 4743 4761 4762 3 0.00000 0.00000 0.00000
0.00000 0.00000 0.00000 ; CB- CA- C- O
4745 4743 4761 4763 3 2.51040 4.18400 1.67360
-6.69440 0.00000 0.00000 ; CB- CA- C- N
4745 4748 4751 4752 3 0.66944 2.00832 0.00000
-2.67776 0.00000 0.00000 ; CB- CG- CD- HD2
4745 4748 4751 4753 3 0.66944 2.00832 0.00000
-2.67776 0.00000 0.00000 ; CB- CG- CD- HD3
4745 4748 4751 4754 3 3.68192 3.09616 -2.09200
-3.01248 0.00000 0.00000 ; CB- CG- CD- CE
4746 4745 4743 4761 3 0.65084 1.95253 0.00000
-2.60338 0.00000 0.00000 ; HB2- CB- CA- C
4746 4745 4748 4749 3 0.62760 1.88280 0.00000
-2.51040 0.00000 0.00000 ; HB2- CB- CG- HG2
4746 4745 4748 4750 3 0.62760 1.88280 0.00000
-2.51040 0.00000 0.00000 ; HB2- CB- CG- HG3
4746 4745 4748 4751 3 0.66944 2.00832 0.00000
-2.67776 0.00000 0.00000 ; HB2- CB- CG- CD
4747 4745 4743 4761 3 0.65084 1.95253 0.00000
-2.60338 0.00000 0.00000 ; HB3- CB- CA- C
4747 4745 4748 4749 3 0.62760 1.88280 0.00000
-2.51040 0.00000 0.00000 ; HB3- CB- CG- HG2
4747 4745 4748 4750 3 0.62760 1.88280 0.00000
-2.51040 0.00000 0.00000 ; HB3- CB- CG- HG3
4747 4745 4748 4751 3 0.66944 2.00832 0.00000
-2.67776 0.00000 0.00000 ; HB3- CB- CG- CD
4748 4745 4743 4761 3 0.65084 1.95253 0.00000
-2.60338 0.00000 0.00000 ; CG- CB- CA- C
4748 4751 4754 4755 3 0.65084 1.95253 0.00000
-2.60338 0.00000 0.00000 ; CG- CD- CE- HE2
4748 4751 4754 4756 3 0.65084 1.95253 0.00000
-2.60338 0.00000 0.00000 ; CG- CD- CE- HE3
4748 4751 4754 4757 3 0.65084 1.95253 0.00000
-2.60338 0.00000 0.00000 ; CG- CD- CE- NZ
4749 4748 4751 4752 3 0.62760 1.88280 0.00000
-2.51040 0.00000 0.00000 ; HG2- CG- CD- HD2
4749 4748 4751 4753 3 0.62760 1.88280 0.00000
-2.51040 0.00000 0.00000 ; HG2- CG- CD- HD3
4749 4748 4751 4754 3 0.66944 2.00832 0.00000
-2.67776 0.00000 0.00000 ; HG2- CG- CD- CE

4750 4748 4751 4752 3 0.62760 1.88280 0.00000
-2.51040 0.00000 0.00000 ; HG3- CG- CD- HD2
4750 4748 4751 4753 3 0.62760 1.88280 0.00000
-2.51040 0.00000 0.00000 ; HG3- CG- CD- HD3
4750 4748 4751 4754 3 0.66944 2.00832 0.00000
-2.67776 0.00000 0.00000 ; HG3- CG- CD- CE
4751 4754 4757 4758 3 0.65084 1.95253 0.00000
-2.60338 0.00000 0.00000 ; CD- CE- NZ- HZ1
4751 4754 4757 4759 3 0.65084 1.95253 0.00000
-2.60338 0.00000 0.00000 ; CD- CE- NZ- HZ2
4751 4754 4757 4760 3 0.65084 1.95253 0.00000
-2.60338 0.00000 0.00000 ; CD- CE- NZ- HZ3
4752 4751 4754 4755 3 0.65084 1.95253 0.00000
-2.60338 0.00000 0.00000 ; HD2- CD- CE- HE2
4752 4751 4754 4756 3 0.65084 1.95253 0.00000
-2.60338 0.00000 0.00000 ; HD2- CD- CE- HE3
4752 4751 4754 4757 3 0.65084 1.95253 0.00000
-2.60338 0.00000 0.00000 ; HD2- CD- CE- NZ
4753 4751 4754 4755 3 0.65084 1.95253 0.00000
-2.60338 0.00000 0.00000 ; HD3- CD- CE- HE2
4753 4751 4754 4756 3 0.65084 1.95253 0.00000
-2.60338 0.00000 0.00000 ; HD3- CD- CE- HE3
4753 4751 4754 4757 3 0.65084 1.95253 0.00000
-2.60338 0.00000 0.00000 ; HD3- CD- CE- NZ
4755 4754 4757 4758 3 0.65084 1.95253 0.00000
-2.60338 0.00000 0.00000 ; HE2- CE- NZ- HZ1
4755 4754 4757 4759 3 0.65084 1.95253 0.00000
-2.60338 0.00000 0.00000 ; HE2- CE- NZ- HZ2
4755 4754 4757 4760 3 0.65084 1.95253 0.00000
-2.60338 0.00000 0.00000 ; HE2- CE- NZ- HZ3
4756 4754 4757 4758 3 0.65084 1.95253 0.00000
-2.60338 0.00000 0.00000 ; HE3- CE- NZ- HZ1
4756 4754 4757 4759 3 0.65084 1.95253 0.00000
-2.60338 0.00000 0.00000 ; HE3- CE- NZ- HZ2
4756 4754 4757 4760 3 0.65084 1.95253 0.00000
-2.60338 0.00000 0.00000 ; HE3- CE- NZ- HZ3
4761 4763 4765 4766 3 0.00000 0.00000 0.00000
0.00000 0.00000 0.00000 ; C- N- CA- HA
4761 4763 4765 4767 3 11.71520 1.67360 15.06240 -
13.38880 0.00000 0.00000 ; C- N- CA- CB
4761 4763 4765 4776 3 1.75728 5.27184 2.25936
-7.02912 0.00000 0.00000 ; C- N- CA- C
4762 4761 4763 4764 3 29.28800 -8.36800 -20.92000
0.00000 0.00000 0.00000 ; O- C- N- H
4762 4761 4763 4765 3 20.92000 0.00000 -20.92000
0.00000 0.00000 0.00000 ; O- C- N- CA
4763 4765 4767 4768 3 0.65084 1.95253 0.00000
-2.60338 0.00000 0.00000 ; N- CA- CB- HB2
4763 4765 4767 4769 3 0.65084 1.95253 0.00000
-2.60338 0.00000 0.00000 ; N- CA- CB- HB3
4763 4765 4767 4770 3 3.21331 2.22589 -4.15053
-2.40998 2.61082 0.00000 ; N- CA- CB- CG

```
  4763   4765   4776   4777       3    0.00000    0.00000    0.00000
0.00000    0.00000    0.00000 ;      N-    CA-     C-     O
  4763   4765   4776   4778       3   17.40544   -5.02080  -13.22144
9.20480    0.00000    0.00000 ;      N-    CA-     C-     N
  4764   4763   4765   4766       3    0.00000    0.00000    0.00000
0.00000    0.00000    0.00000 ;      H-     N-    CA-    HA
  4764   4763   4765   4767       3    0.00000    0.00000    0.00000
0.00000    0.00000    0.00000 ;      H-     N-    CA-    CB
  4764   4763   4765   4776       3    0.00000    0.00000    0.00000
0.00000    0.00000    0.00000 ;      H-     N-    CA-     C
  4765   4767   4770   4771       3    0.66944    2.00832    0.00000
-2.67776    0.00000    0.00000 ;     CA-    CB-    CG-   HG2
  4765   4767   4770   4772       3    0.66944    2.00832    0.00000
-2.67776    0.00000    0.00000 ;     CA-    CB-    CG-   HG3
  4765   4767   4770   4773       3   10.12110   -1.91209    0.01674
10.17549   -1.87443    0.00000 ;     CA-    CB-    CG-    CD
  4765   4776   4778   4779       3   20.92000    0.00000  -20.92000
0.00000    0.00000    0.00000 ;     CA-     C-     N-     H
  4765   4776   4778   4780       3   20.92000    0.00000  -20.92000
0.00000    0.00000    0.00000 ;     CA-     C-     N-    CA
  4766   4765   4767   4768       3    0.65084    1.95253    0.00000
-2.60338    0.00000    0.00000 ;     HA-    CA-    CB-   HB2
  4766   4765   4767   4769       3    0.65084    1.95253    0.00000
-2.60338    0.00000    0.00000 ;     HA-    CA-    CB-   HB3
  4766   4765   4767   4770       3    0.65084    1.95253    0.00000
-2.60338    0.00000    0.00000 ;     HA-    CA-    CB-    CG
  4766   4765   4776   4777       3    3.68192   -4.35136    0.00000
1.33888    0.00000    0.00000 ;     HA-    CA-     C-     O
  4766   4765   4776   4778       3    0.00000    0.00000    0.00000
0.00000    0.00000    0.00000 ;     HA-    CA-     C-     N
  4767   4765   4776   4777       3    0.00000    0.00000    0.00000
0.00000    0.00000    0.00000 ;     CB-    CA-     C-     O
  4767   4765   4776   4778       3    2.51040    4.18400    1.67360
-6.69440    0.00000    0.00000 ;     CB-    CA-     C-     N
  4767   4770   4773   4774       3    3.79907    0.00000   -5.40573
0.00000    2.14221    0.00000 ;     CB-    CG-    CD-   OE1
  4767   4770   4773   4775       3    3.79907    0.00000   -5.40573
0.00000    2.14221    0.00000 ;     CB-    CG-    CD-   OE2
  4768   4767   4765   4776       3    0.65084    1.95253    0.00000
-2.60338    0.00000    0.00000 ;    HB2-    CB-    CA-     C
  4768   4767   4770   4771       3    0.62760    1.88280    0.00000
-2.51040    0.00000    0.00000 ;    HB2-    CB-    CG-   HG2
  4768   4767   4770   4772       3    0.62760    1.88280    0.00000
-2.51040    0.00000    0.00000 ;    HB2-    CB-    CG-   HG3
  4768   4767   4770   4773       3    0.65084    1.95253    0.00000
-2.60338    0.00000    0.00000 ;    HB2-    CB-    CG-    CD
  4769   4767   4765   4776       3    0.65084    1.95253    0.00000
-2.60338    0.00000    0.00000 ;    HB3-    CB-    CA-     C
  4769   4767   4770   4771       3    0.62760    1.88280    0.00000
-2.51040    0.00000    0.00000 ;    HB3-    CB-    CG-   HG2
  4769   4767   4770   4772       3    0.62760    1.88280    0.00000
-2.51040    0.00000    0.00000 ;    HB3-    CB-    CG-   HG3
```

4769 4767 4770 4773 3 0.65084 1.95253 0.00000
-2.60338 0.00000 0.00000 ; HB3- CB- CG- CD
4770 4767 4765 4776 3 6.86594 3.56895 -8.14206
-2.40998 4.85344 0.00000 ; CG- CB- CA- C
4771 4770 4773 4774 3 0.00000 0.00000 0.00000
0.00000 0.00000 0.00000 ; HG2- CG- CD- OE1
4771 4770 4773 4775 3 0.00000 0.00000 0.00000
0.00000 0.00000 0.00000 ; HG2- CG- CD- OE2
4772 4770 4773 4774 3 0.00000 0.00000 0.00000
0.00000 0.00000 0.00000 ; HG3- CG- CD- OE1
4772 4770 4773 4775 3 0.00000 0.00000 0.00000
0.00000 0.00000 0.00000 ; HG3- CG- CD- OE2
4776 4778 4780 4781 3 0.00000 0.00000 0.00000
0.00000 0.00000 0.00000 ; C- N- CA- HA
4776 4778 4780 4782 3 11.71520 1.67360 15.06240 -
13.38880 0.00000 0.00000 ; C- N- CA- CB
4776 4778 4780 4790 3 1.75728 5.27184 2.25936
-7.02912 0.00000 0.00000 ; C- N- CA- C
4777 4776 4778 4779 3 29.28800 -8.36800 -20.92000
0.00000 0.00000 0.00000 ; O- C- N- H
4777 4776 4778 4780 3 20.92000 0.00000 -20.92000
0.00000 0.00000 0.00000 ; O- C- N- CA
4778 4780 4782 4783 3 0.65084 1.95253 0.00000
-2.60338 0.00000 0.00000 ; N- CA- CB- HB2
4778 4780 4782 4784 3 0.65084 1.95253 0.00000
-2.60338 0.00000 0.00000 ; N- CA- CB- HB3
4778 4780 4782 4785 3 5.99567 3.29281 -4.46014
-0.55229 1.97485 0.00000 ; N- CA- CB- CG
4778 4780 4790 4791 3 0.00000 0.00000 0.00000
0.00000 0.00000 0.00000 ; N- CA- C- O
4778 4780 4790 4792 3 17.40544 -5.02080 -13.22144
9.20480 0.00000 0.00000 ; N- CA- C- N
4779 4778 4780 4781 3 0.00000 0.00000 0.00000
0.00000 0.00000 0.00000 ; H- N- CA- HA
4779 4778 4780 4782 3 0.00000 0.00000 0.00000
0.00000 0.00000 0.00000 ; H- N- CA- CB
4779 4778 4780 4790 3 0.00000 0.00000 0.00000
0.00000 0.00000 0.00000 ; H- N- CA- C
4780 4782 4785 4786 3 0.00000 0.00000 0.00000
0.00000 0.00000 0.00000 ; CA- CB- CG- OD1
4780 4782 4785 4787 3 8.84916 -0.31380 -4.32626
5.03754 0.26778 0.00000 ; CA- CB- CG- ND2
4780 4790 4792 4793 3 20.92000 0.00000 -20.92000
0.00000 0.00000 0.00000 ; CA- C- N- H
4780 4790 4792 4794 3 20.92000 0.00000 -20.92000
0.00000 0.00000 0.00000 ; CA- C- N- CA
4781 4780 4782 4783 3 0.65084 1.95253 0.00000
-2.60338 0.00000 0.00000 ; HA- CA- CB- HB2
4781 4780 4782 4784 3 0.65084 1.95253 0.00000
-2.60338 0.00000 0.00000 ; HA- CA- CB- HB3
4781 4780 4782 4785 3 0.65084 1.95253 0.00000
-2.60338 0.00000 0.00000 ; HA- CA- CB- CG

4781 4780 4790 4791 3 3.68192 -4.35136 0.00000
1.33888 0.00000 0.00000 ; HA- CA- C- O
4781 4780 4790 4792 3 0.00000 0.00000 0.00000
0.00000 0.00000 0.00000 ; HA- CA- C- N
4782 4780 4790 4791 3 0.00000 0.00000 0.00000
0.00000 0.00000 0.00000 ; CB- CA- C- O
4782 4780 4790 4792 3 2.51040 4.18400 1.67360
-6.69440 0.00000 0.00000 ; CB- CA- C- N
4782 4785 4787 4788 3 20.92000 0.00000 -20.92000
0.00000 0.00000 0.00000 ; CB- CG- ND2- HD21
4782 4785 4787 4789 3 20.92000 0.00000 -20.92000
0.00000 0.00000 0.00000 ; CB- CG- ND2- HD22
4783 4782 4780 4790 3 0.65084 1.95253 0.00000
-2.60338 0.00000 0.00000 ; HB2- CB- CA- C
4783 4782 4785 4786 3 3.68192 -4.35136 0.00000
1.33888 0.00000 0.00000 ; HB2- CB- CG- OD1
4783 4782 4785 4787 3 0.00000 0.00000 0.00000
0.00000 0.00000 0.00000 ; HB2- CB- CG- ND2
4784 4782 4780 4790 3 0.65084 1.95253 0.00000
-2.60338 0.00000 0.00000 ; HB3- CB- CA- C
4784 4782 4785 4786 3 3.68192 -4.35136 0.00000
1.33888 0.00000 0.00000 ; HB3- CB- CG- OD1
4784 4782 4785 4787 3 0.00000 0.00000 0.00000
0.00000 0.00000 0.00000 ; HB3- CB- CG- ND2
4785 4782 4780 4790 3 7.94542 -3.96225 -6.11701
-0.55229 3.58150 0.00000 ; CG- CB- CA- C
4786 4785 4787 4788 3 29.28800 -8.36800 -20.92000
0.00000 0.00000 0.00000 ; OD1- CG- ND2- HD21
4786 4785 4787 4789 3 29.28800 -8.36800 -20.92000
0.00000 0.00000 0.00000 ; OD1- CG- ND2- HD22
4790 4792 4794 4795 3 0.00000 0.00000 0.00000
0.00000 0.00000 0.00000 ; C- N- CA- HA
4790 4792 4794 4796 3 11.71520 1.67360 15.06240 -
13.38880 0.00000 0.00000 ; C- N- CA- CB
4790 4792 4794 4805 3 1.75728 5.27184 2.25936
-7.02912 0.00000 0.00000 ; C- N- CA- C
4791 4790 4792 4793 3 29.28800 -8.36800 -20.92000
0.00000 0.00000 0.00000 ; O- C- N- H
4791 4790 4792 4794 3 20.92000 0.00000 -20.92000
0.00000 0.00000 0.00000 ; O- C- N- CA
4792 4794 4796 4797 3 0.65084 1.95253 0.00000
-2.60338 0.00000 0.00000 ; N- CA- CB- HB2
4792 4794 4796 4798 3 0.65084 1.95253 0.00000
-2.60338 0.00000 0.00000 ; N- CA- CB- HB3
4792 4794 4796 4799 3 3.21331 2.22589 -4.15053
-2.40998 2.61082 0.00000 ; N- CA- CB- CG
4792 4794 4805 4806 3 0.00000 0.00000 0.00000
0.00000 0.00000 0.00000 ; N- CA- C- O
4792 4794 4805 4807 3 17.40544 -5.02080 -13.22144
9.20480 0.00000 0.00000 ; N- CA- C- N
4793 4792 4794 4795 3 0.00000 0.00000 0.00000
0.00000 0.00000 0.00000 ; H- N- CA- HA

4793 4792 4794 4796 3 0.00000 0.00000 0.00000
0.00000 0.00000 0.00000 ; H- N- CA- CB
4793 4792 4794 4805 3 0.00000 0.00000 0.00000
0.00000 0.00000 0.00000 ; H- N- CA- C
4794 4796 4799 4800 3 0.66944 2.00832 0.00000
-2.67776 0.00000 0.00000 ; CA- CB- CG- HG2
4794 4796 4799 4801 3 0.66944 2.00832 0.00000
-2.67776 0.00000 0.00000 ; CA- CB- CG- HG3
4794 4796 4799 4802 3 10.12110 -1.91209 0.01674
10.17549 -1.87443 0.00000 ; CA- CB- CG- CD
4794 4805 4807 4808 3 20.92000 0.00000 -20.92000
0.00000 0.00000 0.00000 ; CA- C- N- H
4794 4805 4807 4809 3 20.92000 0.00000 -20.92000
0.00000 0.00000 0.00000 ; CA- C- N- CA
4795 4794 4796 4797 3 0.65084 1.95253 0.00000
-2.60338 0.00000 0.00000 ; HA- CA- CB- HB2
4795 4794 4796 4798 3 0.65084 1.95253 0.00000
-2.60338 0.00000 0.00000 ; HA- CA- CB- HB3
4795 4794 4796 4799 3 0.65084 1.95253 0.00000
-2.60338 0.00000 0.00000 ; HA- CA- CB- CG
4795 4794 4805 4806 3 3.68192 -4.35136 0.00000
1.33888 0.00000 0.00000 ; HA- CA- C- O
4795 4794 4805 4807 3 0.00000 0.00000 0.00000
0.00000 0.00000 0.00000 ; HA- CA- C- N
4796 4794 4805 4806 3 0.00000 0.00000 0.00000
0.00000 0.00000 0.00000 ; CB- CA- C- O
4796 4794 4805 4807 3 2.51040 4.18400 1.67360
-6.69440 0.00000 0.00000 ; CB- CA- C- N
4796 4799 4802 4803 3 3.79907 0.00000 -5.40573
0.00000 2.14221 0.00000 ; CB- CG- CD- OE1
4796 4799 4802 4804 3 3.79907 0.00000 -5.40573
0.00000 2.14221 0.00000 ; CB- CG- CD- OE2
4797 4796 4794 4805 3 0.65084 1.95253 0.00000
-2.60338 0.00000 0.00000 ; HB2- CB- CA- C
4797 4796 4799 4800 3 0.62760 1.88280 0.00000
-2.51040 0.00000 0.00000 ; HB2- CB- CG- HG2
4797 4796 4799 4801 3 0.62760 1.88280 0.00000
-2.51040 0.00000 0.00000 ; HB2- CB- CG- HG3
4797 4796 4799 4802 3 0.65084 1.95253 0.00000
-2.60338 0.00000 0.00000 ; HB2- CB- CG- CD
4798 4796 4794 4805 3 0.65084 1.95253 0.00000
-2.60338 0.00000 0.00000 ; HB3- CB- CA- C
4798 4796 4799 4800 3 0.62760 1.88280 0.00000
-2.51040 0.00000 0.00000 ; HB3- CB- CG- HG2
4798 4796 4799 4801 3 0.62760 1.88280 0.00000
-2.51040 0.00000 0.00000 ; HB3- CB- CG- HG3
4798 4796 4799 4802 3 0.65084 1.95253 0.00000
-2.60338 0.00000 0.00000 ; HB3- CB- CG- CD
4799 4796 4794 4805 3 6.86594 3.56895 -8.14206
-2.40998 4.85344 0.00000 ; CG- CB- CA- C
4800 4799 4802 4803 3 0.00000 0.00000 0.00000
0.00000 0.00000 0.00000 ; HG2- CG- CD- OE1

4800 4799 4802 4804 3 0.00000 0.00000 0.00000
0.00000 0.00000 0.00000 ; HG2- CG- CD- OE2
4801 4799 4802 4803 3 0.00000 0.00000 0.00000
0.00000 0.00000 0.00000 ; HG3- CG- CD- OE1
4801 4799 4802 4804 3 0.00000 0.00000 0.00000
0.00000 0.00000 0.00000 ; HG3- CG- CD- OE2
4805 4807 4809 4810 3 0.00000 0.00000 0.00000
0.00000 0.00000 0.00000 ; C- N- CA- HA
4805 4807 4809 4811 3 11.71520 1.67360 15.06240 -
13.38880 0.00000 0.00000 ; C- N- CA- CB
4805 4807 4809 4824 3 1.75728 5.27184 2.25936
-7.02912 0.00000 0.00000 ; C- N- CA- C
4806 4805 4807 4808 3 29.28800 -8.36800 -20.92000
0.00000 0.00000 0.00000 ; O- C- N- H
4806 4805 4807 4809 3 20.92000 0.00000 -20.92000
0.00000 0.00000 0.00000 ; O- C- N- CA
4807 4809 4811 4812 3 0.65084 1.95253 0.00000
-2.60338 0.00000 0.00000 ; N- CA- CB- HB2
4807 4809 4811 4813 3 0.65084 1.95253 0.00000
-2.60338 0.00000 0.00000 ; N- CA- CB- HB3
4807 4809 4811 4814 3 3.79070 1.39746 -4.61077
-2.40998 2.44346 0.00000 ; N- CA- CB- CG
4807 4809 4824 4825 3 0.00000 0.00000 0.00000
0.00000 0.00000 0.00000 ; N- CA- C- O
4807 4809 4824 4826 3 17.40544 -5.02080 -13.22144
9.20480 0.00000 0.00000 ; N- CA- C- N
4808 4807 4809 4810 3 0.00000 0.00000 0.00000
0.00000 0.00000 0.00000 ; H- N- CA- HA
4808 4807 4809 4811 3 0.00000 0.00000 0.00000
0.00000 0.00000 0.00000 ; H- N- CA- CB
4808 4807 4809 4824 3 0.00000 0.00000 0.00000
0.00000 0.00000 0.00000 ; H- N- CA- C
4809 4811 4814 4815 3 0.66944 2.00832 0.00000
-2.67776 0.00000 0.00000 ; CA- CB- CG- HG
4809 4811 4814 4816 3 3.90367 0.19665 -6.21742
-2.37651 5.99149 0.00000 ; CA- CB- CG- CD1
4809 4811 4814 4820 3 3.90367 0.19665 -6.21742
-2.37651 5.99149 0.00000 ; CA- CB- CG- CD2
4809 4824 4826 4827 3 20.92000 0.00000 -20.92000
0.00000 0.00000 0.00000 ; CA- C- N- H
4809 4824 4826 4828 3 20.92000 0.00000 -20.92000
0.00000 0.00000 0.00000 ; CA- C- N- CA
4810 4809 4811 4812 3 0.65084 1.95253 0.00000
-2.60338 0.00000 0.00000 ; HA- CA- CB- HB2
4810 4809 4811 4813 3 0.65084 1.95253 0.00000
-2.60338 0.00000 0.00000 ; HA- CA- CB- HB3
4810 4809 4811 4814 3 0.65084 1.95253 0.00000
-2.60338 0.00000 0.00000 ; HA- CA- CB- CG
4810 4809 4824 4825 3 3.68192 -4.35136 0.00000
1.33888 0.00000 0.00000 ; HA- CA- C- O
4810 4809 4824 4826 3 0.00000 0.00000 0.00000
0.00000 0.00000 0.00000 ; HA- CA- C- N

```
  4811   4809   4824   4825      3    0.00000    0.00000    0.00000
0.00000    0.00000    0.00000 ;      CB-    CA-     C-     O
  4811   4809   4824   4826      3    2.51040    4.18400    1.67360
-6.69440    0.00000    0.00000 ;      CB-    CA-     C-     N
  4811   4814   4816   4817      3    0.66944    2.00832    0.00000
-2.67776    0.00000    0.00000 ;      CB-    CG-   CD1-  HD11
  4811   4814   4816   4818      3    0.66944    2.00832    0.00000
-2.67776    0.00000    0.00000 ;      CB-    CG-   CD1-  HD12
  4811   4814   4816   4819      3    0.66944    2.00832    0.00000
-2.67776    0.00000    0.00000 ;      CB-    CG-   CD1-  HD13
  4811   4814   4820   4821      3    0.66944    2.00832    0.00000
-2.67776    0.00000    0.00000 ;      CB-    CG-   CD2-  HD21
  4811   4814   4820   4822      3    0.66944    2.00832    0.00000
-2.67776    0.00000    0.00000 ;      CB-    CG-   CD2-  HD22
  4811   4814   4820   4823      3    0.66944    2.00832    0.00000
-2.67776    0.00000    0.00000 ;      CB-    CG-   CD2-  HD23
  4812   4811   4809   4824      3    0.65084    1.95253    0.00000
-2.60338    0.00000    0.00000 ;     HB2-    CB-    CA-     C
  4812   4811   4814   4815      3    0.62760    1.88280    0.00000
-2.51040    0.00000    0.00000 ;     HB2-    CB-    CG-    HG
  4812   4811   4814   4816      3    0.66944    2.00832    0.00000
-2.67776    0.00000    0.00000 ;     HB2-    CB-    CG-   CD1
  4812   4811   4814   4820      3    0.66944    2.00832    0.00000
-2.67776    0.00000    0.00000 ;     HB2-    CB-    CG-   CD2
  4813   4811   4809   4824      3    0.65084    1.95253    0.00000
-2.60338    0.00000    0.00000 ;     HB3-    CB-    CA-     C
  4813   4811   4814   4815      3    0.62760    1.88280    0.00000
-2.51040    0.00000    0.00000 ;     HB3-    CB-    CG-    HG
  4813   4811   4814   4816      3    0.66944    2.00832    0.00000
-2.67776    0.00000    0.00000 ;     HB3-    CB-    CG-   CD1
  4813   4811   4814   4820      3    0.66944    2.00832    0.00000
-2.67776    0.00000    0.00000 ;     HB3-    CB-    CG-   CD2
  4814   4811   4809   4824      3   10.33448   -1.14642  -11.54784
-2.40998    6.35968    0.00000 ;      CG-    CB-    CA-     C
  4815   4814   4816   4817      3    0.62760    1.88280    0.00000
-2.51040    0.00000    0.00000 ;      HG-    CG-   CD1-  HD11
  4815   4814   4816   4818      3    0.62760    1.88280    0.00000
-2.51040    0.00000    0.00000 ;      HG-    CG-   CD1-  HD12
  4815   4814   4816   4819      3    0.62760    1.88280    0.00000
-2.51040    0.00000    0.00000 ;      HG-    CG-   CD1-  HD13
  4815   4814   4820   4821      3    0.62760    1.88280    0.00000
-2.51040    0.00000    0.00000 ;      HG-    CG-   CD2-  HD21
  4815   4814   4820   4822      3    0.62760    1.88280    0.00000
-2.51040    0.00000    0.00000 ;      HG-    CG-   CD2-  HD22
  4815   4814   4820   4823      3    0.62760    1.88280    0.00000
-2.51040    0.00000    0.00000 ;      HG-    CG-   CD2-  HD23
  4816   4814   4820   4821      3    0.66944    2.00832    0.00000
-2.67776    0.00000    0.00000 ;     CD1-    CG-   CD2-  HD21
  4816   4814   4820   4822      3    0.66944    2.00832    0.00000
-2.67776    0.00000    0.00000 ;     CD1-    CG-   CD2-  HD22
  4816   4814   4820   4823      3    0.66944    2.00832    0.00000
-2.67776    0.00000    0.00000 ;     CD1-    CG-   CD2-  HD23
```

4817 4816 4814 4820 3 0.66944 2.00832 0.00000
-2.67776 0.00000 0.00000 ; HD11- CD1- CG- CD2
4818 4816 4814 4820 3 0.66944 2.00832 0.00000
-2.67776 0.00000 0.00000 ; HD12- CD1- CG- CD2
4819 4816 4814 4820 3 0.66944 2.00832 0.00000
-2.67776 0.00000 0.00000 ; HD13- CD1- CG- CD2
4824 4826 4828 4829 3 0.00000 0.00000 0.00000
0.00000 0.00000 0.00000 ; C- N- CA- HA
4824 4826 4828 4830 3 11.71520 1.67360 15.06240 -
13.38880 0.00000 0.00000 ; C- N- CA- CB
4824 4826 4828 4846 3 1.75728 5.27184 2.25936
-7.02912 0.00000 0.00000 ; C- N- CA- C
4825 4824 4826 4827 3 29.28800 -8.36800 -20.92000
0.00000 0.00000 0.00000 ; O- C- N- H
4825 4824 4826 4828 3 20.92000 0.00000 -20.92000
0.00000 0.00000 0.00000 ; O- C- N- CA
4826 4828 4830 4831 3 0.65084 1.95253 0.00000
-2.60338 0.00000 0.00000 ; N- CA- CB- HB2
4826 4828 4830 4832 3 0.65084 1.95253 0.00000
-2.60338 0.00000 0.00000 ; N- CA- CB- HB3
4826 4828 4830 4833 3 0.65084 1.95253 0.00000
-2.60338 0.00000 0.00000 ; N- CA- CB- CG
4826 4828 4846 4847 3 0.00000 0.00000 0.00000
0.00000 0.00000 0.00000 ; N- CA- C- O
4826 4828 4846 4848 3 17.40544 -5.02080 -13.22144
9.20480 0.00000 0.00000 ; N- CA- C- N
4827 4826 4828 4829 3 0.00000 0.00000 0.00000
0.00000 0.00000 0.00000 ; H- N- CA- HA
4827 4826 4828 4830 3 0.00000 0.00000 0.00000
0.00000 0.00000 0.00000 ; H- N- CA- CB
4827 4826 4828 4846 3 0.00000 0.00000 0.00000
0.00000 0.00000 0.00000 ; H- N- CA- C
4828 4830 4833 4834 3 0.66944 2.00832 0.00000
-2.67776 0.00000 0.00000 ; CA- CB- CG- HG2
4828 4830 4833 4835 3 0.66944 2.00832 0.00000
-2.67776 0.00000 0.00000 ; CA- CB- CG- HG3
4828 4830 4833 4836 3 3.68192 3.09616 -2.09200
-3.01248 0.00000 0.00000 ; CA- CB- CG- CD
4828 4846 4848 4849 3 20.92000 0.00000 -20.92000
0.00000 0.00000 0.00000 ; CA- C- N- H
4828 4846 4848 4850 3 20.92000 0.00000 -20.92000
0.00000 0.00000 0.00000 ; CA- C- N- CA
4829 4828 4830 4831 3 0.65084 1.95253 0.00000
-2.60338 0.00000 0.00000 ; HA- CA- CB- HB2
4829 4828 4830 4832 3 0.65084 1.95253 0.00000
-2.60338 0.00000 0.00000 ; HA- CA- CB- HB3
4829 4828 4830 4833 3 0.65084 1.95253 0.00000
-2.60338 0.00000 0.00000 ; HA- CA- CB- CG
4829 4828 4846 4847 3 3.68192 -4.35136 0.00000
1.33888 0.00000 0.00000 ; HA- CA- C- O
4829 4828 4846 4848 3 0.00000 0.00000 0.00000
0.00000 0.00000 0.00000 ; HA- CA- C- N

4830 4828 4846 4847 3 0.00000 0.00000 0.00000
0.00000 0.00000 0.00000 ; CB- CA- C- O
4830 4828 4846 4848 3 2.51040 4.18400 1.67360
-6.69440 0.00000 0.00000 ; CB- CA- C- N
4830 4833 4836 4837 3 0.66944 2.00832 0.00000
-2.67776 0.00000 0.00000 ; CB- CG- CD- HD2
4830 4833 4836 4838 3 0.66944 2.00832 0.00000
-2.67776 0.00000 0.00000 ; CB- CG- CD- HD3
4830 4833 4836 4839 3 3.68192 3.09616 -2.09200
-3.01248 0.00000 0.00000 ; CB- CG- CD- CE
4831 4830 4828 4846 3 0.65084 1.95253 0.00000
-2.60338 0.00000 0.00000 ; HB2- CB- CA- C
4831 4830 4833 4834 3 0.62760 1.88280 0.00000
-2.51040 0.00000 0.00000 ; HB2- CB- CG- HG2
4831 4830 4833 4835 3 0.62760 1.88280 0.00000
-2.51040 0.00000 0.00000 ; HB2- CB- CG- HG3
4831 4830 4833 4836 3 0.66944 2.00832 0.00000
-2.67776 0.00000 0.00000 ; HB2- CB- CG- CD
4832 4830 4828 4846 3 0.65084 1.95253 0.00000
-2.60338 0.00000 0.00000 ; HB3- CB- CA- C
4832 4830 4833 4834 3 0.62760 1.88280 0.00000
-2.51040 0.00000 0.00000 ; HB3- CB- CG- HG2
4832 4830 4833 4835 3 0.62760 1.88280 0.00000
-2.51040 0.00000 0.00000 ; HB3- CB- CG- HG3
4832 4830 4833 4836 3 0.66944 2.00832 0.00000
-2.67776 0.00000 0.00000 ; HB3- CB- CG- CD
4833 4830 4828 4846 3 0.65084 1.95253 0.00000
-2.60338 0.00000 0.00000 ; CG- CB- CA- C
4833 4836 4839 4840 3 0.65084 1.95253 0.00000
-2.60338 0.00000 0.00000 ; CG- CD- CE- HE2
4833 4836 4839 4841 3 0.65084 1.95253 0.00000
-2.60338 0.00000 0.00000 ; CG- CD- CE- HE3
4833 4836 4839 4842 3 0.65084 1.95253 0.00000
-2.60338 0.00000 0.00000 ; CG- CD- CE- NZ
4834 4833 4836 4837 3 0.62760 1.88280 0.00000
-2.51040 0.00000 0.00000 ; HG2- CG- CD- HD2
4834 4833 4836 4838 3 0.62760 1.88280 0.00000
-2.51040 0.00000 0.00000 ; HG2- CG- CD- HD3
4834 4833 4836 4839 3 0.66944 2.00832 0.00000
-2.67776 0.00000 0.00000 ; HG2- CG- CD- CE
4835 4833 4836 4837 3 0.62760 1.88280 0.00000
-2.51040 0.00000 0.00000 ; HG3- CG- CD- HD2
4835 4833 4836 4838 3 0.62760 1.88280 0.00000
-2.51040 0.00000 0.00000 ; HG3- CG- CD- HD3
4835 4833 4836 4839 3 0.66944 2.00832 0.00000
-2.67776 0.00000 0.00000 ; HG3- CG- CD- CE
4836 4839 4842 4843 3 0.65084 1.95253 0.00000
-2.60338 0.00000 0.00000 ; CD- CE- NZ- HZ1
4836 4839 4842 4844 3 0.65084 1.95253 0.00000
-2.60338 0.00000 0.00000 ; CD- CE- NZ- HZ2
4836 4839 4842 4845 3 0.65084 1.95253 0.00000
-2.60338 0.00000 0.00000 ; CD- CE- NZ- HZ3

4837 4836 4839 4840 3 0.65084 1.95253 0.00000
-2.60338 0.00000 0.00000 ; HD2- CD- CE- HE2
4837 4836 4839 4841 3 0.65084 1.95253 0.00000
-2.60338 0.00000 0.00000 ; HD2- CD- CE- HE3
4837 4836 4839 4842 3 0.65084 1.95253 0.00000
-2.60338 0.00000 0.00000 ; HD2- CD- CE- NZ
4838 4836 4839 4840 3 0.65084 1.95253 0.00000
-2.60338 0.00000 0.00000 ; HD3- CD- CE- HE2
4838 4836 4839 4841 3 0.65084 1.95253 0.00000
-2.60338 0.00000 0.00000 ; HD3- CD- CE- HE3
4838 4836 4839 4842 3 0.65084 1.95253 0.00000
-2.60338 0.00000 0.00000 ; HD3- CD- CE- NZ
4840 4839 4842 4843 3 0.65084 1.95253 0.00000
-2.60338 0.00000 0.00000 ; HE2- CE- NZ- HZ1
4840 4839 4842 4844 3 0.65084 1.95253 0.00000
-2.60338 0.00000 0.00000 ; HE2- CE- NZ- HZ2
4840 4839 4842 4845 3 0.65084 1.95253 0.00000
-2.60338 0.00000 0.00000 ; HE2- CE- NZ- HZ3
4841 4839 4842 4843 3 0.65084 1.95253 0.00000
-2.60338 0.00000 0.00000 ; HE3- CE- NZ- HZ1
4841 4839 4842 4844 3 0.65084 1.95253 0.00000
-2.60338 0.00000 0.00000 ; HE3- CE- NZ- HZ2
4841 4839 4842 4845 3 0.65084 1.95253 0.00000
-2.60338 0.00000 0.00000 ; HE3- CE- NZ- HZ3
4846 4848 4850 4851 3 0.00000 0.00000 0.00000
0.00000 0.00000 0.00000 ; C- N- CA- HA
4846 4848 4850 4852 3 11.71520 1.67360 15.06240 -
13.38880 0.00000 0.00000 ; C- N- CA- CB
4846 4848 4850 4868 3 1.75728 5.27184 2.25936
-7.02912 0.00000 0.00000 ; C- N- CA- C
4847 4846 4848 4849 3 29.28800 -8.36800 -20.92000
0.00000 0.00000 0.00000 ; O- C- N- H
4847 4846 4848 4850 3 20.92000 0.00000 -20.92000
0.00000 0.00000 0.00000 ; O- C- N- CA
4848 4850 4852 4853 3 0.65084 1.95253 0.00000
-2.60338 0.00000 0.00000 ; N- CA- CB- HB2
4848 4850 4852 4854 3 0.65084 1.95253 0.00000
-2.60338 0.00000 0.00000 ; N- CA- CB- HB3
4848 4850 4852 4855 3 0.65084 1.95253 0.00000
-2.60338 0.00000 0.00000 ; N- CA- CB- CG
4848 4850 4868 4869 3 0.00000 0.00000 0.00000
0.00000 0.00000 0.00000 ; N- CA- C- O
4848 4850 4868 4870 3 17.40544 -5.02080 -13.22144
9.20480 0.00000 0.00000 ; N- CA- C- N
4849 4848 4850 4851 3 0.00000 0.00000 0.00000
0.00000 0.00000 0.00000 ; H- N- CA- HA
4849 4848 4850 4852 3 0.00000 0.00000 0.00000
0.00000 0.00000 0.00000 ; H- N- CA- CB
4849 4848 4850 4868 3 0.00000 0.00000 0.00000
0.00000 0.00000 0.00000 ; H- N- CA- C
4850 4852 4855 4856 3 0.66944 2.00832 0.00000
-2.67776 0.00000 0.00000 ; CA- CB- CG- HG2

4850 4852 4855 4857 3 0.66944 2.00832 0.00000
-2.67776 0.00000 0.00000 ; CA- CB- CG- HG3
4850 4852 4855 4858 3 3.68192 3.09616 -2.09200
-3.01248 0.00000 0.00000 ; CA- CB- CG- CD
4850 4868 4870 4871 3 20.92000 0.00000 -20.92000
0.00000 0.00000 0.00000 ; CA- C- N- H
4850 4868 4870 4872 3 20.92000 0.00000 -20.92000
0.00000 0.00000 0.00000 ; CA- C- N- CA
4851 4850 4852 4853 3 0.65084 1.95253 0.00000
-2.60338 0.00000 0.00000 ; HA- CA- CB- HB2
4851 4850 4852 4854 3 0.65084 1.95253 0.00000
-2.60338 0.00000 0.00000 ; HA- CA- CB- HB3
4851 4850 4852 4855 3 0.65084 1.95253 0.00000
-2.60338 0.00000 0.00000 ; HA- CA- CB- CG
4851 4850 4868 4869 3 3.68192 -4.35136 0.00000
1.33888 0.00000 0.00000 ; HA- CA- C- O
4851 4850 4868 4870 3 0.00000 0.00000 0.00000
0.00000 0.00000 0.00000 ; HA- CA- C- N
4852 4850 4868 4869 3 0.00000 0.00000 0.00000
0.00000 0.00000 0.00000 ; CB- CA- C- O
4852 4850 4868 4870 3 2.51040 4.18400 1.67360
-6.69440 0.00000 0.00000 ; CB- CA- C- N
4852 4855 4858 4859 3 0.66944 2.00832 0.00000
-2.67776 0.00000 0.00000 ; CB- CG- CD- HD2
4852 4855 4858 4860 3 0.66944 2.00832 0.00000
-2.67776 0.00000 0.00000 ; CB- CG- CD- HD3
4852 4855 4858 4861 3 3.68192 3.09616 -2.09200
-3.01248 0.00000 0.00000 ; CB- CG- CD- CE
4853 4852 4850 4868 3 0.65084 1.95253 0.00000
-2.60338 0.00000 0.00000 ; HB2- CB- CA- C
4853 4852 4855 4856 3 0.62760 1.88280 0.00000
-2.51040 0.00000 0.00000 ; HB2- CB- CG- HG2
4853 4852 4855 4857 3 0.62760 1.88280 0.00000
-2.51040 0.00000 0.00000 ; HB2- CB- CG- HG3
4853 4852 4855 4858 3 0.66944 2.00832 0.00000
-2.67776 0.00000 0.00000 ; HB2- CB- CG- CD
4854 4852 4850 4868 3 0.65084 1.95253 0.00000
-2.60338 0.00000 0.00000 ; HB3- CB- CA- C
4854 4852 4855 4856 3 0.62760 1.88280 0.00000
-2.51040 0.00000 0.00000 ; HB3- CB- CG- HG2
4854 4852 4855 4857 3 0.62760 1.88280 0.00000
-2.51040 0.00000 0.00000 ; HB3- CB- CG- HG3
4854 4852 4855 4858 3 0.66944 2.00832 0.00000
-2.67776 0.00000 0.00000 ; HB3- CB- CG- CD
4855 4852 4850 4868 3 0.65084 1.95253 0.00000
-2.60338 0.00000 0.00000 ; CG- CB- CA- C
4855 4858 4861 4862 3 0.65084 1.95253 0.00000
-2.60338 0.00000 0.00000 ; CG- CD- CE- HE2
4855 4858 4861 4863 3 0.65084 1.95253 0.00000
-2.60338 0.00000 0.00000 ; CG- CD- CE- HE3
4855 4858 4861 4864 3 0.65084 1.95253 0.00000
-2.60338 0.00000 0.00000 ; CG- CD- CE- NZ

```
  4856   4855   4858   4859       3    0.62760    1.88280    0.00000
-2.51040    0.00000    0.00000 ;    HG2-    CG-    CD-   HD2
  4856   4855   4858   4860       3    0.62760    1.88280    0.00000
-2.51040    0.00000    0.00000 ;    HG2-    CG-    CD-   HD3
  4856   4855   4858   4861       3    0.66944    2.00832    0.00000
-2.67776    0.00000    0.00000 ;    HG2-    CG-    CD-    CE
  4857   4855   4858   4859       3    0.62760    1.88280    0.00000
-2.51040    0.00000    0.00000 ;    HG3-    CG-    CD-   HD2
  4857   4855   4858   4860       3    0.62760    1.88280    0.00000
-2.51040    0.00000    0.00000 ;    HG3-    CG-    CD-   HD3
  4857   4855   4858   4861       3    0.66944    2.00832    0.00000
-2.67776    0.00000    0.00000 ;    HG3-    CG-    CD-    CE
  4858   4861   4864   4865       3    0.65084    1.95253    0.00000
-2.60338    0.00000    0.00000 ;     CD-    CE-    NZ-   HZ1
  4858   4861   4864   4866       3    0.65084    1.95253    0.00000
-2.60338    0.00000    0.00000 ;     CD-    CE-    NZ-   HZ2
  4858   4861   4864   4867       3    0.65084    1.95253    0.00000
-2.60338    0.00000    0.00000 ;     CD-    CE-    NZ-   HZ3
  4859   4858   4861   4862       3    0.65084    1.95253    0.00000
-2.60338    0.00000    0.00000 ;    HD2-    CD-    CE-   HE2
  4859   4858   4861   4863       3    0.65084    1.95253    0.00000
-2.60338    0.00000    0.00000 ;    HD2-    CD-    CE-   HE3
  4859   4858   4861   4864       3    0.65084    1.95253    0.00000
-2.60338    0.00000    0.00000 ;    HD2-    CD-    CE-    NZ
  4860   4858   4861   4862       3    0.65084    1.95253    0.00000
-2.60338    0.00000    0.00000 ;    HD3-    CD-    CE-   HE2
  4860   4858   4861   4863       3    0.65084    1.95253    0.00000
-2.60338    0.00000    0.00000 ;    HD3-    CD-    CE-   HE3
  4860   4858   4861   4864       3    0.65084    1.95253    0.00000
-2.60338    0.00000    0.00000 ;    HD3-    CD-    CE-    NZ
  4862   4861   4864   4865       3    0.65084    1.95253    0.00000
-2.60338    0.00000    0.00000 ;    HE2-    CE-    NZ-   HZ1
  4862   4861   4864   4866       3    0.65084    1.95253    0.00000
-2.60338    0.00000    0.00000 ;    HE2-    CE-    NZ-   HZ2
  4862   4861   4864   4867       3    0.65084    1.95253    0.00000
-2.60338    0.00000    0.00000 ;    HE2-    CE-    NZ-   HZ3
  4863   4861   4864   4865       3    0.65084    1.95253    0.00000
-2.60338    0.00000    0.00000 ;    HE3-    CE-    NZ-   HZ1
  4863   4861   4864   4866       3    0.65084    1.95253    0.00000
-2.60338    0.00000    0.00000 ;    HE3-    CE-    NZ-   HZ2
  4863   4861   4864   4867       3    0.65084    1.95253    0.00000
-2.60338    0.00000    0.00000 ;    HE3-    CE-    NZ-   HZ3
  4868   4870   4872   4873       3    0.00000    0.00000    0.00000
0.00000    0.00000    0.00000 ;      C-     N-    CA-    HA
  4868   4870   4872   4874       3   11.71520    1.67360   15.06240  -
13.38880    0.00000    0.00000 ;      C-     N-    CA-    CB
  4868   4870   4872   4887       3    1.75728    5.27184    2.25936
-7.02912    0.00000    0.00000 ;      C-     N-    CA-     C
  4869   4868   4870   4871       3   29.28800   -8.36800  -20.92000
0.00000    0.00000    0.00000 ;      O-     C-     N-     H
  4869   4868   4870   4872       3   20.92000    0.00000  -20.92000
0.00000    0.00000    0.00000 ;      O-     C-     N-    CA
```

4870 4872 4874 4875 3 0.65084 1.95253 0.00000
-2.60338 0.00000 0.00000 ; N- CA- CB- HB
4870 4872 4874 4876 3 3.83673 0.44769 -1.77402
-2.47693 -0.03347 0.00000 ; N- CA- CB- CG2
4870 4872 4874 4880 3 2.97482 0.12134 2.04179
-1.89117 -3.24678 0.00000 ; N- CA- CB- CG1
4870 4872 4887 4888 3 0.00000 0.00000 0.00000
0.00000 0.00000 0.00000 ; N- CA- C- O
4870 4872 4887 4889 3 17.40544 -5.02080 -13.22144
9.20480 0.00000 0.00000 ; N- CA- C- N
4871 4870 4872 4873 3 0.00000 0.00000 0.00000
0.00000 0.00000 0.00000 ; H- N- CA- HA
4871 4870 4872 4874 3 0.00000 0.00000 0.00000
0.00000 0.00000 0.00000 ; H- N- CA- CB
4871 4870 4872 4887 3 0.00000 0.00000 0.00000
0.00000 0.00000 0.00000 ; H- N- CA- C
4872 4874 4876 4877 3 0.66944 2.00832 0.00000
-2.67776 0.00000 0.00000 ; CA- CB- CG2- HG21
4872 4874 4876 4878 3 0.66944 2.00832 0.00000
-2.67776 0.00000 0.00000 ; CA- CB- CG2- HG22
4872 4874 4876 4879 3 0.66944 2.00832 0.00000
-2.67776 0.00000 0.00000 ; CA- CB- CG2- HG23
4872 4874 4880 4881 3 0.66944 2.00832 0.00000
-2.67776 0.00000 0.00000 ; CA- CB- CG1- HG12
4872 4874 4880 4882 3 0.66944 2.00832 0.00000
-2.67776 0.00000 0.00000 ; CA- CB- CG1- HG13
4872 4874 4880 4883 3 4.24258 -0.52718 -7.25506
-1.79075 7.69856 0.00000 ; CA- CB- CG1- CD1
4872 4887 4889 4890 3 20.92000 0.00000 -20.92000
0.00000 0.00000 0.00000 ; CA- C- N- H
4872 4887 4889 4891 3 20.92000 0.00000 -20.92000
0.00000 0.00000 0.00000 ; CA- C- N- CA
4873 4872 4874 4875 3 0.65084 1.95253 0.00000
-2.60338 0.00000 0.00000 ; HA- CA- CB- HB
4873 4872 4874 4876 3 0.65084 1.95253 0.00000
-2.60338 0.00000 0.00000 ; HA- CA- CB- CG2
4873 4872 4874 4880 3 0.65084 1.95253 0.00000
-2.60338 0.00000 0.00000 ; HA- CA- CB- CG1
4873 4872 4887 4888 3 3.68192 -4.35136 0.00000
1.33888 0.00000 0.00000 ; HA- CA- C- O
4873 4872 4887 4889 3 0.00000 0.00000 0.00000
0.00000 0.00000 0.00000 ; HA- CA- C- N
4874 4872 4887 4888 3 0.00000 0.00000 0.00000
0.00000 0.00000 0.00000 ; CB- CA- C- O
4874 4872 4887 4889 3 2.51040 4.18400 1.67360
-6.69440 0.00000 0.00000 ; CB- CA- C- N
4874 4880 4883 4884 3 0.66944 2.00832 0.00000
-2.67776 0.00000 0.00000 ; CB- CG1- CD1- HD11
4874 4880 4883 4885 3 0.66944 2.00832 0.00000
-2.67776 0.00000 0.00000 ; CB- CG1- CD1- HD12
4874 4880 4883 4886 3 0.66944 2.00832 0.00000
-2.67776 0.00000 0.00000 ; CB- CG1- CD1- HD13

4875 4874 4872 4887 3 0.65084 1.95253 0.00000
-2.60338 0.00000 0.00000 ; HB- CB- CA- C
4875 4874 4876 4877 3 0.62760 1.88280 0.00000
-2.51040 0.00000 0.00000 ; HB- CB- CG2- HG21
4875 4874 4876 4878 3 0.62760 1.88280 0.00000
-2.51040 0.00000 0.00000 ; HB- CB- CG2- HG22
4875 4874 4876 4879 3 0.62760 1.88280 0.00000
-2.51040 0.00000 0.00000 ; HB- CB- CG2- HG23
4875 4874 4880 4881 3 0.62760 1.88280 0.00000
-2.51040 0.00000 0.00000 ; HB- CB- CG1- HG12
4875 4874 4880 4882 3 0.62760 1.88280 0.00000
-2.51040 0.00000 0.00000 ; HB- CB- CG1- HG13
4875 4874 4880 4883 3 0.66944 2.00832 0.00000
-2.67776 0.00000 0.00000 ; HB- CB- CG1- CD1
4876 4874 4872 4887 3 5.67350 3.55640 -6.16722
-2.47693 3.74886 0.00000 ; CG2- CB- CA- C
4876 4874 4880 4881 3 0.66944 2.00832 0.00000
-2.67776 0.00000 0.00000 ; CG2- CB- CG1- HG12
4876 4874 4880 4882 3 0.66944 2.00832 0.00000
-2.67776 0.00000 0.00000 ; CG2- CB- CG1- HG13
4876 4874 4880 4883 3 3.81162 0.49790 -8.14206
-1.79075 7.49773 0.00000 ; CG2- CB- CG1- CD1
4877 4876 4874 4880 3 0.66944 2.00832 0.00000
-2.67776 0.00000 0.00000 ; HG21- CG2- CB- CG1
4878 4876 4874 4880 3 0.66944 2.00832 0.00000
-2.67776 0.00000 0.00000 ; HG22- CG2- CB- CG1
4879 4876 4874 4880 3 0.66944 2.00832 0.00000
-2.67776 0.00000 0.00000 ; HG23- CG2- CB- CG1
4880 4874 4872 4887 3 8.26340 0.74057 -9.99976
-1.89117 3.84928 0.00000 ; CG1- CB- CA- C
4881 4880 4883 4884 3 0.62760 1.88280 0.00000
-2.51040 0.00000 0.00000 ; HG12- CG1- CD1- HD11
4881 4880 4883 4885 3 0.62760 1.88280 0.00000
-2.51040 0.00000 0.00000 ; HG12- CG1- CD1- HD12
4881 4880 4883 4886 3 0.62760 1.88280 0.00000
-2.51040 0.00000 0.00000 ; HG12- CG1- CD1- HD13
4882 4880 4883 4884 3 0.62760 1.88280 0.00000
-2.51040 0.00000 0.00000 ; HG13- CG1- CD1- HD11
4882 4880 4883 4885 3 0.62760 1.88280 0.00000
-2.51040 0.00000 0.00000 ; HG13- CG1- CD1- HD12
4882 4880 4883 4886 3 0.62760 1.88280 0.00000
-2.51040 0.00000 0.00000 ; HG13- CG1- CD1- HD13
4887 4889 4891 4892 3 0.00000 0.00000 0.00000
0.00000 0.00000 0.00000 ; C- N- CA- HA
4887 4889 4891 4893 3 11.71520 1.67360 15.06240 -
13.38880 0.00000 0.00000 ; C- N- CA- CB
4887 4889 4891 4897 3 1.75728 5.27184 2.25936
-7.02912 0.00000 0.00000 ; C- N- CA- C
4888 4887 4889 4890 3 29.28800 -8.36800 -20.92000
0.00000 0.00000 0.00000 ; O- C- N- H
4888 4887 4889 4891 3 20.92000 0.00000 -20.92000
0.00000 0.00000 0.00000 ; O- C- N- CA

4889 4891 4893 4894 3 0.65084 1.95253 0.00000
-2.60338 0.00000 0.00000 ; N- CA- CB- HB1
4889 4891 4893 4895 3 0.65084 1.95253 0.00000
-2.60338 0.00000 0.00000 ; N- CA- CB- HB2
4889 4891 4893 4896 3 0.65084 1.95253 0.00000
-2.60338 0.00000 0.00000 ; N- CA- CB- HB3
4889 4891 4897 4898 3 0.00000 0.00000 0.00000
0.00000 0.00000 0.00000 ; N- CA- C- O
4889 4891 4897 4899 3 17.40544 -5.02080 -13.22144
9.20480 0.00000 0.00000 ; N- CA- C- N
4890 4889 4891 4892 3 0.00000 0.00000 0.00000
0.00000 0.00000 0.00000 ; H- N- CA- HA
4890 4889 4891 4893 3 0.00000 0.00000 0.00000
0.00000 0.00000 0.00000 ; H- N- CA- CB
4890 4889 4891 4897 3 0.00000 0.00000 0.00000
0.00000 0.00000 0.00000 ; H- N- CA- C
4891 4897 4899 4900 3 20.92000 0.00000 -20.92000
0.00000 0.00000 0.00000 ; CA- C- N- H
4891 4897 4899 4901 3 20.92000 0.00000 -20.92000
0.00000 0.00000 0.00000 ; CA- C- N- CA
4892 4891 4893 4894 3 0.65084 1.95253 0.00000
-2.60338 0.00000 0.00000 ; HA- CA- CB- HB1
4892 4891 4893 4895 3 0.65084 1.95253 0.00000
-2.60338 0.00000 0.00000 ; HA- CA- CB- HB2
4892 4891 4893 4896 3 0.65084 1.95253 0.00000
-2.60338 0.00000 0.00000 ; HA- CA- CB- HB3
4892 4891 4897 4898 3 3.68192 -4.35136 0.00000
1.33888 0.00000 0.00000 ; HA- CA- C- O
4892 4891 4897 4899 3 0.00000 0.00000 0.00000
0.00000 0.00000 0.00000 ; HA- CA- C- N
4893 4891 4897 4898 3 0.00000 0.00000 0.00000
0.00000 0.00000 0.00000 ; CB- CA- C- O
4893 4891 4897 4899 3 2.51040 4.18400 1.67360
-6.69440 0.00000 0.00000 ; CB- CA- C- N
4894 4893 4891 4897 3 0.65084 1.95253 0.00000
-2.60338 0.00000 0.00000 ; HB1- CB- CA- C
4895 4893 4891 4897 3 0.65084 1.95253 0.00000
-2.60338 0.00000 0.00000 ; HB2- CB- CA- C
4896 4893 4891 4897 3 0.65084 1.95253 0.00000
-2.60338 0.00000 0.00000 ; HB3- CB- CA- C
4897 4899 4901 4902 3 0.00000 0.00000 0.00000
0.00000 0.00000 0.00000 ; C- N- CA- HA
4897 4899 4901 4903 3 11.71520 1.67360 15.06240 -
13.38880 0.00000 0.00000 ; C- N- CA- CB
4897 4899 4901 4916 3 1.75728 5.27184 2.25936
-7.02912 0.00000 0.00000 ; C- N- CA- C
4898 4897 4899 4900 3 29.28800 -8.36800 -20.92000
0.00000 0.00000 0.00000 ; O- C- N- H
4898 4897 4899 4901 3 20.92000 0.00000 -20.92000
0.00000 0.00000 0.00000 ; O- C- N- CA
4899 4901 4903 4904 3 0.65084 1.95253 0.00000
-2.60338 0.00000 0.00000 ; N- CA- CB- HB2

4899 4901 4903 4905 3 0.65084 1.95253 0.00000
-2.60338 0.00000 0.00000 ; N- CA- CB- HB3
4899 4901 4903 4906 3 3.79070 1.39746 -4.61077
-2.40998 2.44346 0.00000 ; N- CA- CB- CG
4899 4901 4916 4917 3 0.00000 0.00000 0.00000
0.00000 0.00000 0.00000 ; N- CA- C- O
4899 4901 4916 4918 3 17.40544 -5.02080 -13.22144
9.20480 0.00000 0.00000 ; N- CA- C- N
4900 4899 4901 4902 3 0.00000 0.00000 0.00000
0.00000 0.00000 0.00000 ; H- N- CA- HA
4900 4899 4901 4903 3 0.00000 0.00000 0.00000
0.00000 0.00000 0.00000 ; H- N- CA- CB
4900 4899 4901 4916 3 0.00000 0.00000 0.00000
0.00000 0.00000 0.00000 ; H- N- CA- C
4901 4903 4906 4907 3 0.66944 2.00832 0.00000
-2.67776 0.00000 0.00000 ; CA- CB- CG- HG
4901 4903 4906 4908 3 3.90367 0.19665 -6.21742
-2.37651 5.99149 0.00000 ; CA- CB- CG- CD1
4901 4903 4906 4912 3 3.90367 0.19665 -6.21742
-2.37651 5.99149 0.00000 ; CA- CB- CG- CD2
4901 4916 4918 4919 3 20.92000 0.00000 -20.92000
0.00000 0.00000 0.00000 ; CA- C- N- H
4901 4916 4918 4920 3 20.92000 0.00000 -20.92000
0.00000 0.00000 0.00000 ; CA- C- N- CA
4902 4901 4903 4904 3 0.65084 1.95253 0.00000
-2.60338 0.00000 0.00000 ; HA- CA- CB- HB2
4902 4901 4903 4905 3 0.65084 1.95253 0.00000
-2.60338 0.00000 0.00000 ; HA- CA- CB- HB3
4902 4901 4903 4906 3 0.65084 1.95253 0.00000
-2.60338 0.00000 0.00000 ; HA- CA- CB- CG
4902 4901 4916 4917 3 3.68192 -4.35136 0.00000
1.33888 0.00000 0.00000 ; HA- CA- C- O
4902 4901 4916 4918 3 0.00000 0.00000 0.00000
0.00000 0.00000 0.00000 ; HA- CA- C- N
4903 4901 4916 4917 3 0.00000 0.00000 0.00000
0.00000 0.00000 0.00000 ; CB- CA- C- O
4903 4901 4916 4918 3 2.51040 4.18400 1.67360
-6.69440 0.00000 0.00000 ; CB- CA- C- N
4903 4906 4908 4909 3 0.66944 2.00832 0.00000
-2.67776 0.00000 0.00000 ; CB- CG- CD1- HD11
4903 4906 4908 4910 3 0.66944 2.00832 0.00000
-2.67776 0.00000 0.00000 ; CB- CG- CD1- HD12
4903 4906 4908 4911 3 0.66944 2.00832 0.00000
-2.67776 0.00000 0.00000 ; CB- CG- CD1- HD13
4903 4906 4912 4913 3 0.66944 2.00832 0.00000
-2.67776 0.00000 0.00000 ; CB- CG- CD2- HD21
4903 4906 4912 4914 3 0.66944 2.00832 0.00000
-2.67776 0.00000 0.00000 ; CB- CG- CD2- HD22
4903 4906 4912 4915 3 0.66944 2.00832 0.00000
-2.67776 0.00000 0.00000 ; CB- CG- CD2- HD23
4904 4903 4901 4916 3 0.65084 1.95253 0.00000
-2.60338 0.00000 0.00000 ; HB2- CB- CA- C

4904 4903 4906 4907 3 0.62760 1.88280 0.00000
-2.51040 0.00000 0.00000 ; HB2- CB- CG- HG
4904 4903 4906 4908 3 0.66944 2.00832 0.00000
-2.67776 0.00000 0.00000 ; HB2- CB- CG- CD1
4904 4903 4906 4912 3 0.66944 2.00832 0.00000
-2.67776 0.00000 0.00000 ; HB2- CB- CG- CD2
4905 4903 4901 4916 3 0.65084 1.95253 0.00000
-2.60338 0.00000 0.00000 ; HB3- CB- CA- C
4905 4903 4906 4907 3 0.62760 1.88280 0.00000
-2.51040 0.00000 0.00000 ; HB3- CB- CG- HG
4905 4903 4906 4908 3 0.66944 2.00832 0.00000
-2.67776 0.00000 0.00000 ; HB3- CB- CG- CD1
4905 4903 4906 4912 3 0.66944 2.00832 0.00000
-2.67776 0.00000 0.00000 ; HB3- CB- CG- CD2
4906 4903 4901 4916 3 10.33448 -1.14642 -11.54784
-2.40998 6.35968 0.00000 ; CG- CB- CA- C
4907 4906 4908 4909 3 0.62760 1.88280 0.00000
-2.51040 0.00000 0.00000 ; HG- CG- CD1- HD11
4907 4906 4908 4910 3 0.62760 1.88280 0.00000
-2.51040 0.00000 0.00000 ; HG- CG- CD1- HD12
4907 4906 4908 4911 3 0.62760 1.88280 0.00000
-2.51040 0.00000 0.00000 ; HG- CG- CD1- HD13
4907 4906 4912 4913 3 0.62760 1.88280 0.00000
-2.51040 0.00000 0.00000 ; HG- CG- CD2- HD21
4907 4906 4912 4914 3 0.62760 1.88280 0.00000
-2.51040 0.00000 0.00000 ; HG- CG- CD2- HD22
4907 4906 4912 4915 3 0.62760 1.88280 0.00000
-2.51040 0.00000 0.00000 ; HG- CG- CD2- HD23
4908 4906 4912 4913 3 0.66944 2.00832 0.00000
-2.67776 0.00000 0.00000 ; CD1- CG- CD2- HD21
4908 4906 4912 4914 3 0.66944 2.00832 0.00000
-2.67776 0.00000 0.00000 ; CD1- CG- CD2- HD22
4908 4906 4912 4915 3 0.66944 2.00832 0.00000
-2.67776 0.00000 0.00000 ; CD1- CG- CD2- HD23
4909 4908 4906 4912 3 0.66944 2.00832 0.00000
-2.67776 0.00000 0.00000 ; HD11- CD1- CG- CD2
4910 4908 4906 4912 3 0.66944 2.00832 0.00000
-2.67776 0.00000 0.00000 ; HD12- CD1- CG- CD2
4911 4908 4906 4912 3 0.66944 2.00832 0.00000
-2.67776 0.00000 0.00000 ; HD13- CD1- CG- CD2
4916 4918 4920 4921 3 0.00000 0.00000 0.00000
0.00000 0.00000 0.00000 ; C- N- CA- HA
4916 4918 4920 4922 3 11.71520 1.67360 15.06240 -
13.38880 0.00000 0.00000 ; C- N- CA- CB
4916 4918 4920 4930 3 1.75728 5.27184 2.25936
-7.02912 0.00000 0.00000 ; C- N- CA- C
4917 4916 4918 4919 3 29.28800 -8.36800 -20.92000
0.00000 0.00000 0.00000 ; O- C- N- H
4917 4916 4918 4920 3 20.92000 0.00000 -20.92000
0.00000 0.00000 0.00000 ; O- C- N- CA
4918 4920 4922 4923 3 0.65084 1.95253 0.00000
-2.60338 0.00000 0.00000 ; N- CA- CB- HB

4918 4920 4922 4924 3 3.83673 0.44769 -1.77402
-2.47693 -0.03347 0.00000 ; N- CA- CB- CG2
4918 4920 4922 4928 3 4.93294 1.13386 -3.12963
-5.27184 3.17984 0.00000 ; N- CA- CB- OG1
4918 4920 4930 4931 3 0.00000 0.00000 0.00000
0.00000 0.00000 0.00000 ; N- CA- C- O
4918 4920 4930 4932 3 17.40544 -5.02080 -13.22144
9.20480 0.00000 0.00000 ; N- CA- C- N
4919 4918 4920 4921 3 0.00000 0.00000 0.00000
0.00000 0.00000 0.00000 ; H- N- CA- HA
4919 4918 4920 4922 3 0.00000 0.00000 0.00000
0.00000 0.00000 0.00000 ; H- N- CA- CB
4919 4918 4920 4930 3 0.00000 0.00000 0.00000
0.00000 0.00000 0.00000 ; H- N- CA- C
4920 4922 4924 4925 3 0.66944 2.00832 0.00000
-2.67776 0.00000 0.00000 ; CA- CB- CG2- HG21
4920 4922 4924 4926 3 0.66944 2.00832 0.00000
-2.67776 0.00000 0.00000 ; CA- CB- CG2- HG22
4920 4922 4924 4927 3 0.66944 2.00832 0.00000
-2.67776 0.00000 0.00000 ; CA- CB- CG2- HG23
4920 4922 4928 4929 3 1.12131 2.98738 1.66523
-3.94970 0.43514 0.00000 ; CA- CB- OG1- HG1
4920 4930 4932 4933 3 20.92000 0.00000 -20.92000
0.00000 0.00000 0.00000 ; CA- C- N- H
4920 4930 4932 4934 3 20.92000 0.00000 -20.92000
0.00000 0.00000 0.00000 ; CA- C- N- CA
4921 4920 4922 4923 3 0.65084 1.95253 0.00000
-2.60338 0.00000 0.00000 ; HA- CA- CB- HB
4921 4920 4922 4924 3 0.65084 1.95253 0.00000
-2.60338 0.00000 0.00000 ; HA- CA- CB- CG2
4921 4920 4922 4928 3 1.04600 -1.04600 0.00000
0.00000 0.00000 0.00000 ; HA- CA- CB- OG1
4921 4920 4930 4931 3 3.68192 -4.35136 0.00000
1.33888 0.00000 0.00000 ; HA- CA- C- O
4921 4920 4930 4932 3 0.00000 0.00000 0.00000
0.00000 0.00000 0.00000 ; HA- CA- C- N
4922 4920 4930 4931 3 0.00000 0.00000 0.00000
0.00000 0.00000 0.00000 ; CB- CA- C- O
4922 4920 4930 4932 3 2.51040 4.18400 1.67360
-6.69440 0.00000 0.00000 ; CB- CA- C- N
4923 4922 4920 4930 3 0.65084 1.95253 0.00000
-2.60338 0.00000 0.00000 ; HB- CB- CA- C
4923 4922 4924 4925 3 0.65084 1.95253 0.00000
-2.60338 0.00000 0.00000 ; HB- CB- CG2- HG21
4923 4922 4924 4926 3 0.65084 1.95253 0.00000
-2.60338 0.00000 0.00000 ; HB- CB- CG2- HG22
4923 4922 4924 4927 3 0.65084 1.95253 0.00000
-2.60338 0.00000 0.00000 ; HB- CB- CG2- HG23
4923 4922 4928 4929 3 0.69733 2.09200 0.00000
-2.78933 0.00000 0.00000 ; HB- CB- OG1- HG1
4924 4922 4920 4930 3 5.67350 3.55640 -6.16722
-2.47693 3.74886 0.00000 ; CG2- CB- CA- C

4924 4922 4928 4929 3 4.74047 0.27196 -2.26773
-3.94970 1.60666 0.00000 ; CG2- CB- OG1- HG1
4925 4924 4922 4928 3 1.04600 -1.04600 0.00000
0.00000 0.00000 0.00000 ; HG21- CG2- CB- OG1
4926 4924 4922 4928 3 1.04600 -1.04600 0.00000
0.00000 0.00000 0.00000 ; HG22- CG2- CB- OG1
4927 4924 4922 4928 3 1.04600 -1.04600 0.00000
0.00000 0.00000 0.00000 ; HG23- CG2- CB- OG1
4928 4922 4920 4930 3 6.53541 6.87013 -6.21742
-5.27184 5.22163 0.00000 ; OG1- CB- CA- C
4930 4932 4934 4935 3 0.00000 0.00000 0.00000
0.00000 0.00000 0.00000 ; C- N- CA- HA
4930 4932 4934 4936 3 11.71520 1.67360 15.06240 -
13.38880 0.00000 0.00000 ; C- N- CA- CB
4930 4932 4934 4949 3 1.75728 5.27184 2.25936
-7.02912 0.00000 0.00000 ; C- N- CA- C
4931 4930 4932 4933 3 29.28800 -8.36800 -20.92000
0.00000 0.00000 0.00000 ; O- C- N- H
4931 4930 4932 4934 3 20.92000 0.00000 -20.92000
0.00000 0.00000 0.00000 ; O- C- N- CA
4932 4934 4936 4937 3 0.65084 1.95253 0.00000
-2.60338 0.00000 0.00000 ; N- CA- CB- HB
4932 4934 4936 4938 3 3.83673 0.44769 -1.77402
-2.47693 -0.03347 0.00000 ; N- CA- CB- CG2
4932 4934 4936 4942 3 2.97482 0.12134 2.04179
-1.89117 -3.24678 0.00000 ; N- CA- CB- CG1
4932 4934 4949 4950 3 0.00000 0.00000 0.00000
0.00000 0.00000 0.00000 ; N- CA- C- O
4932 4934 4949 4951 3 17.40544 -5.02080 -13.22144
9.20480 0.00000 0.00000 ; N- CA- C- N
4933 4932 4934 4935 3 0.00000 0.00000 0.00000
0.00000 0.00000 0.00000 ; H- N- CA- HA
4933 4932 4934 4936 3 0.00000 0.00000 0.00000
0.00000 0.00000 0.00000 ; H- N- CA- CB
4933 4932 4934 4949 3 0.00000 0.00000 0.00000
0.00000 0.00000 0.00000 ; H- N- CA- C
4934 4936 4938 4939 3 0.66944 2.00832 0.00000
-2.67776 0.00000 0.00000 ; CA- CB- CG2- HG21
4934 4936 4938 4940 3 0.66944 2.00832 0.00000
-2.67776 0.00000 0.00000 ; CA- CB- CG2- HG22
4934 4936 4938 4941 3 0.66944 2.00832 0.00000
-2.67776 0.00000 0.00000 ; CA- CB- CG2- HG23
4934 4936 4942 4943 3 0.66944 2.00832 0.00000
-2.67776 0.00000 0.00000 ; CA- CB- CG1- HG12
4934 4936 4942 4944 3 0.66944 2.00832 0.00000
-2.67776 0.00000 0.00000 ; CA- CB- CG1- HG13
4934 4936 4942 4945 3 4.24258 -0.52718 -7.25506
-1.79075 7.69856 0.00000 ; CA- CB- CG1- CD1
4934 4949 4951 4952 3 20.92000 0.00000 -20.92000
0.00000 0.00000 0.00000 ; CA- C- N- H
4934 4949 4951 4953 3 20.92000 0.00000 -20.92000
0.00000 0.00000 0.00000 ; CA- C- N- CA

4935 4934 4936 4937 3 0.65084 1.95253 0.00000
-2.60338 0.00000 0.00000 ; HA- CA- CB- HB
4935 4934 4936 4938 3 0.65084 1.95253 0.00000
-2.60338 0.00000 0.00000 ; HA- CA- CB- CG2
4935 4934 4936 4942 3 0.65084 1.95253 0.00000
-2.60338 0.00000 0.00000 ; HA- CA- CB- CG1
4935 4934 4949 4950 3 3.68192 -4.35136 0.00000
1.33888 0.00000 0.00000 ; HA- CA- C- O
4935 4934 4949 4951 3 0.00000 0.00000 0.00000
0.00000 0.00000 0.00000 ; HA- CA- C- N
4936 4934 4949 4950 3 0.00000 0.00000 0.00000
0.00000 0.00000 0.00000 ; CB- CA- C- O
4936 4934 4949 4951 3 2.51040 4.18400 1.67360
-6.69440 0.00000 0.00000 ; CB- CA- C- N
4936 4942 4945 4946 3 0.66944 2.00832 0.00000
-2.67776 0.00000 0.00000 ; CB- CG1- CD1- HD11
4936 4942 4945 4947 3 0.66944 2.00832 0.00000
-2.67776 0.00000 0.00000 ; CB- CG1- CD1- HD12
4936 4942 4945 4948 3 0.66944 2.00832 0.00000
-2.67776 0.00000 0.00000 ; CB- CG1- CD1- HD13
4937 4936 4934 4949 3 0.65084 1.95253 0.00000
-2.60338 0.00000 0.00000 ; HB- CB- CA- C
4937 4936 4938 4939 3 0.62760 1.88280 0.00000
-2.51040 0.00000 0.00000 ; HB- CB- CG2- HG21
4937 4936 4938 4940 3 0.62760 1.88280 0.00000
-2.51040 0.00000 0.00000 ; HB- CB- CG2- HG22
4937 4936 4938 4941 3 0.62760 1.88280 0.00000
-2.51040 0.00000 0.00000 ; HB- CB- CG2- HG23
4937 4936 4942 4943 3 0.62760 1.88280 0.00000
-2.51040 0.00000 0.00000 ; HB- CB- CG1- HG12
4937 4936 4942 4944 3 0.62760 1.88280 0.00000
-2.51040 0.00000 0.00000 ; HB- CB- CG1- HG13
4937 4936 4942 4945 3 0.66944 2.00832 0.00000
-2.67776 0.00000 0.00000 ; HB- CB- CG1- CD1
4938 4936 4934 4949 3 5.67350 3.55640 -6.16722
-2.47693 3.74886 0.00000 ; CG2- CB- CA- C
4938 4936 4942 4943 3 0.66944 2.00832 0.00000
-2.67776 0.00000 0.00000 ; CG2- CB- CG1- HG12
4938 4936 4942 4944 3 0.66944 2.00832 0.00000
-2.67776 0.00000 0.00000 ; CG2- CB- CG1- HG13
4938 4936 4942 4945 3 3.81162 0.49790 -8.14206
-1.79075 7.49773 0.00000 ; CG2- CB- CG1- CD1
4939 4938 4936 4942 3 0.66944 2.00832 0.00000
-2.67776 0.00000 0.00000 ; HG21- CG2- CB- CG1
4940 4938 4936 4942 3 0.66944 2.00832 0.00000
-2.67776 0.00000 0.00000 ; HG22- CG2- CB- CG1
4941 4938 4936 4942 3 0.66944 2.00832 0.00000
-2.67776 0.00000 0.00000 ; HG23- CG2- CB- CG1
4942 4936 4934 4949 3 8.26340 0.74057 -9.99976
-1.89117 3.84928 0.00000 ; CG1- CB- CA- C
4943 4942 4945 4946 3 0.62760 1.88280 0.00000
-2.51040 0.00000 0.00000 ; HG12- CG1- CD1- HD11

4943 4942 4945 4947 3 0.62760 1.88280 0.00000
-2.51040 0.00000 0.00000 ; HG12- CG1- CD1- HD12
4943 4942 4945 4948 3 0.62760 1.88280 0.00000
-2.51040 0.00000 0.00000 ; HG12- CG1- CD1- HD13
4944 4942 4945 4946 3 0.62760 1.88280 0.00000
-2.51040 0.00000 0.00000 ; HG13- CG1- CD1- HD11
4944 4942 4945 4947 3 0.62760 1.88280 0.00000
-2.51040 0.00000 0.00000 ; HG13- CG1- CD1- HD12
4944 4942 4945 4948 3 0.62760 1.88280 0.00000
-2.51040 0.00000 0.00000 ; HG13- CG1- CD1- HD13
4949 4951 4953 4954 3 0.00000 0.00000 0.00000
0.00000 0.00000 0.00000 ; C- N- CA- HA
4949 4951 4953 4955 3 11.71520 1.67360 15.06240 -
13.38880 0.00000 0.00000 ; C- N- CA- CB
4949 4951 4953 4968 3 1.75728 5.27184 2.25936
-7.02912 0.00000 0.00000 ; C- N- CA- C
4950 4949 4951 4952 3 29.28800 -8.36800 -20.92000
0.00000 0.00000 0.00000 ; O- C- N- H
4950 4949 4951 4953 3 20.92000 0.00000 -20.92000
0.00000 0.00000 0.00000 ; O- C- N- CA
4951 4953 4955 4956 3 0.65084 1.95253 0.00000
-2.60338 0.00000 0.00000 ; N- CA- CB- HB
4951 4953 4955 4957 3 3.83673 0.44769 -1.77402
-2.47693 -0.03347 0.00000 ; N- CA- CB- CG2
4951 4953 4955 4961 3 2.97482 0.12134 2.04179
-1.89117 -3.24678 0.00000 ; N- CA- CB- CG1
4951 4953 4968 4969 3 0.00000 0.00000 0.00000
0.00000 0.00000 0.00000 ; N- CA- C- O
4951 4953 4968 4970 3 17.40544 -5.02080 -13.22144
9.20480 0.00000 0.00000 ; N- CA- C- N
4952 4951 4953 4954 3 0.00000 0.00000 0.00000
0.00000 0.00000 0.00000 ; H- N- CA- HA
4952 4951 4953 4955 3 0.00000 0.00000 0.00000
0.00000 0.00000 0.00000 ; H- N- CA- CB
4952 4951 4953 4968 3 0.00000 0.00000 0.00000
0.00000 0.00000 0.00000 ; H- N- CA- C
4953 4955 4957 4958 3 0.66944 2.00832 0.00000
-2.67776 0.00000 0.00000 ; CA- CB- CG2- HG21
4953 4955 4957 4959 3 0.66944 2.00832 0.00000
-2.67776 0.00000 0.00000 ; CA- CB- CG2- HG22
4953 4955 4957 4960 3 0.66944 2.00832 0.00000
-2.67776 0.00000 0.00000 ; CA- CB- CG2- HG23
4953 4955 4961 4962 3 0.66944 2.00832 0.00000
-2.67776 0.00000 0.00000 ; CA- CB- CG1- HG12
4953 4955 4961 4963 3 0.66944 2.00832 0.00000
-2.67776 0.00000 0.00000 ; CA- CB- CG1- HG13
4953 4955 4961 4964 3 4.24258 -0.52718 -7.25506
-1.79075 7.69856 0.00000 ; CA- CB- CG1- CD1
4953 4968 4970 4971 3 20.92000 0.00000 -20.92000
0.00000 0.00000 0.00000 ; CA- C- N- H
4953 4968 4970 4972 3 20.92000 0.00000 -20.92000
0.00000 0.00000 0.00000 ; CA- C- N- CA

4954 4953 4955 4956 3 0.65084 1.95253 0.00000
-2.60338 0.00000 0.00000 ; HA- CA- CB- HB
4954 4953 4955 4957 3 0.65084 1.95253 0.00000
-2.60338 0.00000 0.00000 ; HA- CA- CB- CG2
4954 4953 4955 4961 3 0.65084 1.95253 0.00000
-2.60338 0.00000 0.00000 ; HA- CA- CB- CG1
4954 4953 4968 4969 3 3.68192 -4.35136 0.00000
1.33888 0.00000 0.00000 ; HA- CA- C- O
4954 4953 4968 4970 3 0.00000 0.00000 0.00000
0.00000 0.00000 0.00000 ; HA- CA- C- N
4955 4953 4968 4969 3 0.00000 0.00000 0.00000
0.00000 0.00000 0.00000 ; CB- CA- C- O
4955 4953 4968 4970 3 2.51040 4.18400 1.67360
-6.69440 0.00000 0.00000 ; CB- CA- C- N
4955 4961 4964 4965 3 0.66944 2.00832 0.00000
-2.67776 0.00000 0.00000 ; CB- CG1- CD1- HD11
4955 4961 4964 4966 3 0.66944 2.00832 0.00000
-2.67776 0.00000 0.00000 ; CB- CG1- CD1- HD12
4955 4961 4964 4967 3 0.66944 2.00832 0.00000
-2.67776 0.00000 0.00000 ; CB- CG1- CD1- HD13
4956 4955 4953 4968 3 0.65084 1.95253 0.00000
-2.60338 0.00000 0.00000 ; HB- CB- CA- C
4956 4955 4957 4958 3 0.62760 1.88280 0.00000
-2.51040 0.00000 0.00000 ; HB- CB- CG2- HG21
4956 4955 4957 4959 3 0.62760 1.88280 0.00000
-2.51040 0.00000 0.00000 ; HB- CB- CG2- HG22
4956 4955 4957 4960 3 0.62760 1.88280 0.00000
-2.51040 0.00000 0.00000 ; HB- CB- CG2- HG23
4956 4955 4961 4962 3 0.62760 1.88280 0.00000
-2.51040 0.00000 0.00000 ; HB- CB- CG1- HG12
4956 4955 4961 4963 3 0.62760 1.88280 0.00000
-2.51040 0.00000 0.00000 ; HB- CB- CG1- HG13
4956 4955 4961 4964 3 0.66944 2.00832 0.00000
-2.67776 0.00000 0.00000 ; HB- CB- CG1- CD1
4957 4955 4953 4968 3 5.67350 3.55640 -6.16722
-2.47693 3.74886 0.00000 ; CG2- CB- CA- C
4957 4955 4961 4962 3 0.66944 2.00832 0.00000
-2.67776 0.00000 0.00000 ; CG2- CB- CG1- HG12
4957 4955 4961 4963 3 0.66944 2.00832 0.00000
-2.67776 0.00000 0.00000 ; CG2- CB- CG1- HG13
4957 4955 4961 4964 3 3.81162 0.49790 -8.14206
-1.79075 7.49773 0.00000 ; CG2- CB- CG1- CD1
4958 4957 4955 4961 3 0.66944 2.00832 0.00000
-2.67776 0.00000 0.00000 ; HG21- CG2- CB- CG1
4959 4957 4955 4961 3 0.66944 2.00832 0.00000
-2.67776 0.00000 0.00000 ; HG22- CG2- CB- CG1
4960 4957 4955 4961 3 0.66944 2.00832 0.00000
-2.67776 0.00000 0.00000 ; HG23- CG2- CB- CG1
4961 4955 4953 4968 3 8.26340 0.74057 -9.99976
-1.89117 3.84928 0.00000 ; CG1- CB- CA- C
4962 4961 4964 4965 3 0.62760 1.88280 0.00000
-2.51040 0.00000 0.00000 ; HG12- CG1- CD1- HD11

```
  4962   4961   4964   4966       3    0.62760    1.88280    0.00000
-2.51040    0.00000    0.00000 ;   HG12-   CG1-   CD1-  HD12
  4962   4961   4964   4967       3    0.62760    1.88280    0.00000
-2.51040    0.00000    0.00000 ;   HG12-   CG1-   CD1-  HD13
  4963   4961   4964   4965       3    0.62760    1.88280    0.00000
-2.51040    0.00000    0.00000 ;   HG13-   CG1-   CD1-  HD11
  4963   4961   4964   4966       3    0.62760    1.88280    0.00000
-2.51040    0.00000    0.00000 ;   HG13-   CG1-   CD1-  HD12
  4963   4961   4964   4967       3    0.62760    1.88280    0.00000
-2.51040    0.00000    0.00000 ;   HG13-   CG1-   CD1-  HD13
  4968   4970   4972   4973       3    0.00000    0.00000    0.00000
0.00000    0.00000    0.00000 ;      C-      N-    CA-    HA
  4968   4970   4972   4974       3   11.71520    1.67360   15.06240  -
13.38880    0.00000    0.00000 ;      C-      N-    CA-    CB
  4968   4970   4972   4978       3    1.75728    5.27184    2.25936
-7.02912    0.00000    0.00000 ;      C-      N-    CA-     C
  4969   4968   4970   4971       3   29.28800   -8.36800  -20.92000
0.00000    0.00000    0.00000 ;      O-      C-      N-     H
  4969   4968   4970   4972       3   20.92000    0.00000  -20.92000
0.00000    0.00000    0.00000 ;      O-      C-      N-    CA
  4970   4972   4974   4975       3    0.65084    1.95253    0.00000
-2.60338    0.00000    0.00000 ;      N-    CA-    CB-   HB1
  4970   4972   4974   4976       3    0.65084    1.95253    0.00000
-2.60338    0.00000    0.00000 ;      N-    CA-    CB-   HB2
  4970   4972   4974   4977       3    0.65084    1.95253    0.00000
-2.60338    0.00000    0.00000 ;      N-    CA-    CB-   HB3
  4970   4972   4978   4979       3    0.00000    0.00000    0.00000
0.00000    0.00000    0.00000 ;      N-    CA-     C-     O
  4970   4972   4978   4980       3   17.40544   -5.02080  -13.22144
9.20480    0.00000    0.00000 ;      N-    CA-     C-     N
  4971   4970   4972   4973       3    0.00000    0.00000    0.00000
0.00000    0.00000    0.00000 ;      H-      N-    CA-    HA
  4971   4970   4972   4974       3    0.00000    0.00000    0.00000
0.00000    0.00000    0.00000 ;      H-      N-    CA-    CB
  4971   4970   4972   4978       3    0.00000    0.00000    0.00000
0.00000    0.00000    0.00000 ;      H-      N-    CA-     C
  4972   4978   4980   4981       3   20.92000    0.00000  -20.92000
0.00000    0.00000    0.00000 ;     CA-      C-      N-     H
  4972   4978   4980   4982       3   20.92000    0.00000  -20.92000
0.00000    0.00000    0.00000 ;     CA-      C-      N-    CA
  4973   4972   4974   4975       3    0.65084    1.95253    0.00000
-2.60338    0.00000    0.00000 ;     HA-    CA-    CB-   HB1
  4973   4972   4974   4976       3    0.65084    1.95253    0.00000
-2.60338    0.00000    0.00000 ;     HA-    CA-    CB-   HB2
  4973   4972   4974   4977       3    0.65084    1.95253    0.00000
-2.60338    0.00000    0.00000 ;     HA-    CA-    CB-   HB3
  4973   4972   4978   4979       3    3.68192   -4.35136    0.00000
1.33888    0.00000    0.00000 ;     HA-    CA-     C-     O
  4973   4972   4978   4980       3    0.00000    0.00000    0.00000
0.00000    0.00000    0.00000 ;     HA-    CA-     C-     N
  4974   4972   4978   4979       3    0.00000    0.00000    0.00000
0.00000    0.00000    0.00000 ;     CB-    CA-     C-     O
```

4974 4972 4978 4980 3 2.51040 4.18400 1.67360
-6.69440 0.00000 0.00000 ; CB- CA- C- N
4975 4974 4972 4978 3 0.65084 1.95253 0.00000
-2.60338 0.00000 0.00000 ; HB1- CB- CA- C
4976 4974 4972 4978 3 0.65084 1.95253 0.00000
-2.60338 0.00000 0.00000 ; HB2- CB- CA- C
4977 4974 4972 4978 3 0.65084 1.95253 0.00000
-2.60338 0.00000 0.00000 ; HB3- CB- CA- C
4978 4980 4982 4983 3 0.00000 0.00000 0.00000
0.00000 0.00000 0.00000 ; C- N- CA- HA
4978 4980 4982 4984 3 11.71520 1.67360 15.06240 -
13.38880 0.00000 0.00000 ; C- N- CA- CB
4978 4980 4982 5000 3 1.75728 5.27184 2.25936
-7.02912 0.00000 0.00000 ; C- N- CA- C
4979 4978 4980 4981 3 29.28800 -8.36800 -20.92000
0.00000 0.00000 0.00000 ; O- C- N- H
4979 4978 4980 4982 3 20.92000 0.00000 -20.92000
0.00000 0.00000 0.00000 ; O- C- N- CA
4980 4982 4984 4985 3 0.65084 1.95253 0.00000
-2.60338 0.00000 0.00000 ; N- CA- CB- HB2
4980 4982 4984 4986 3 0.65084 1.95253 0.00000
-2.60338 0.00000 0.00000 ; N- CA- CB- HB3
4980 4982 4984 4987 3 0.65084 1.95253 0.00000
-2.60338 0.00000 0.00000 ; N- CA- CB- CG
4980 4982 5000 5001 3 0.00000 0.00000 0.00000
0.00000 0.00000 0.00000 ; N- CA- C- O
4980 4982 5000 5002 3 17.40544 -5.02080 -13.22144
9.20480 0.00000 0.00000 ; N- CA- C- N
4981 4980 4982 4983 3 0.00000 0.00000 0.00000
0.00000 0.00000 0.00000 ; H- N- CA- HA
4981 4980 4982 4984 3 0.00000 0.00000 0.00000
0.00000 0.00000 0.00000 ; H- N- CA- CB
4981 4980 4982 5000 3 0.00000 0.00000 0.00000
0.00000 0.00000 0.00000 ; H- N- CA- C
4982 4984 4987 4988 3 0.66944 2.00832 0.00000
-2.67776 0.00000 0.00000 ; CA- CB- CG- HG2
4982 4984 4987 4989 3 0.66944 2.00832 0.00000
-2.67776 0.00000 0.00000 ; CA- CB- CG- HG3
4982 4984 4987 4990 3 3.68192 3.09616 -2.09200
-3.01248 0.00000 0.00000 ; CA- CB- CG- CD
4982 5000 5002 5003 3 20.92000 0.00000 -20.92000
0.00000 0.00000 0.00000 ; CA- C- N- H
4982 5000 5002 5004 3 20.92000 0.00000 -20.92000
0.00000 0.00000 0.00000 ; CA- C- N- CA
4983 4982 4984 4985 3 0.65084 1.95253 0.00000
-2.60338 0.00000 0.00000 ; HA- CA- CB- HB2
4983 4982 4984 4986 3 0.65084 1.95253 0.00000
-2.60338 0.00000 0.00000 ; HA- CA- CB- HB3
4983 4982 4984 4987 3 0.65084 1.95253 0.00000
-2.60338 0.00000 0.00000 ; HA- CA- CB- CG
4983 4982 5000 5001 3 3.68192 -4.35136 0.00000
1.33888 0.00000 0.00000 ; HA- CA- C- O

4983 4982 5000 5002 3 0.00000 0.00000 0.00000
0.00000 0.00000 0.00000 ; HA- CA- C- N
4984 4982 5000 5001 3 0.00000 0.00000 0.00000
0.00000 0.00000 0.00000 ; CB- CA- C- O
4984 4982 5000 5002 3 2.51040 4.18400 1.67360
-6.69440 0.00000 0.00000 ; CB- CA- C- N
4984 4987 4990 4991 3 0.66944 2.00832 0.00000
-2.67776 0.00000 0.00000 ; CB- CG- CD- HD2
4984 4987 4990 4992 3 0.66944 2.00832 0.00000
-2.67776 0.00000 0.00000 ; CB- CG- CD- HD3
4984 4987 4990 4993 3 3.68192 3.09616 -2.09200
-3.01248 0.00000 0.00000 ; CB- CG- CD- CE
4985 4984 4982 5000 3 0.65084 1.95253 0.00000
-2.60338 0.00000 0.00000 ; HB2- CB- CA- C
4985 4984 4987 4988 3 0.62760 1.88280 0.00000
-2.51040 0.00000 0.00000 ; HB2- CB- CG- HG2
4985 4984 4987 4989 3 0.62760 1.88280 0.00000
-2.51040 0.00000 0.00000 ; HB2- CB- CG- HG3
4985 4984 4987 4990 3 0.66944 2.00832 0.00000
-2.67776 0.00000 0.00000 ; HB2- CB- CG- CD
4986 4984 4982 5000 3 0.65084 1.95253 0.00000
-2.60338 0.00000 0.00000 ; HB3- CB- CA- C
4986 4984 4987 4988 3 0.62760 1.88280 0.00000
-2.51040 0.00000 0.00000 ; HB3- CB- CG- HG2
4986 4984 4987 4989 3 0.62760 1.88280 0.00000
-2.51040 0.00000 0.00000 ; HB3- CB- CG- HG3
4986 4984 4987 4990 3 0.66944 2.00832 0.00000
-2.67776 0.00000 0.00000 ; HB3- CB- CG- CD
4987 4984 4982 5000 3 0.65084 1.95253 0.00000
-2.60338 0.00000 0.00000 ; CG- CB- CA- C
4987 4990 4993 4994 3 0.65084 1.95253 0.00000
-2.60338 0.00000 0.00000 ; CG- CD- CE- HE2
4987 4990 4993 4995 3 0.65084 1.95253 0.00000
-2.60338 0.00000 0.00000 ; CG- CD- CE- HE3
4987 4990 4993 4996 3 0.65084 1.95253 0.00000
-2.60338 0.00000 0.00000 ; CG- CD- CE- NZ
4988 4987 4990 4991 3 0.62760 1.88280 0.00000
-2.51040 0.00000 0.00000 ; HG2- CG- CD- HD2
4988 4987 4990 4992 3 0.62760 1.88280 0.00000
-2.51040 0.00000 0.00000 ; HG2- CG- CD- HD3
4988 4987 4990 4993 3 0.66944 2.00832 0.00000
-2.67776 0.00000 0.00000 ; HG2- CG- CD- CE
4989 4987 4990 4991 3 0.62760 1.88280 0.00000
-2.51040 0.00000 0.00000 ; HG3- CG- CD- HD2
4989 4987 4990 4992 3 0.62760 1.88280 0.00000
-2.51040 0.00000 0.00000 ; HG3- CG- CD- HD3
4989 4987 4990 4993 3 0.66944 2.00832 0.00000
-2.67776 0.00000 0.00000 ; HG3- CG- CD- CE
4990 4993 4996 4997 3 0.65084 1.95253 0.00000
-2.60338 0.00000 0.00000 ; CD- CE- NZ- HZ1
4990 4993 4996 4998 3 0.65084 1.95253 0.00000
-2.60338 0.00000 0.00000 ; CD- CE- NZ- HZ2

4990 4993 4996 4999 3 0.65084 1.95253 0.00000
-2.60338 0.00000 0.00000 ; CD- CE- NZ- HZ3
4991 4990 4993 4994 3 0.65084 1.95253 0.00000
-2.60338 0.00000 0.00000 ; HD2- CD- CE- HE2
4991 4990 4993 4995 3 0.65084 1.95253 0.00000
-2.60338 0.00000 0.00000 ; HD2- CD- CE- HE3
4991 4990 4993 4996 3 0.65084 1.95253 0.00000
-2.60338 0.00000 0.00000 ; HD2- CD- CE- NZ
4992 4990 4993 4994 3 0.65084 1.95253 0.00000
-2.60338 0.00000 0.00000 ; HD3- CD- CE- HE2
4992 4990 4993 4995 3 0.65084 1.95253 0.00000
-2.60338 0.00000 0.00000 ; HD3- CD- CE- HE3
4992 4990 4993 4996 3 0.65084 1.95253 0.00000
-2.60338 0.00000 0.00000 ; HD3- CD- CE- NZ
4994 4993 4996 4997 3 0.65084 1.95253 0.00000
-2.60338 0.00000 0.00000 ; HE2- CE- NZ- HZ1
4994 4993 4996 4998 3 0.65084 1.95253 0.00000
-2.60338 0.00000 0.00000 ; HE2- CE- NZ- HZ2
4994 4993 4996 4999 3 0.65084 1.95253 0.00000
-2.60338 0.00000 0.00000 ; HE2- CE- NZ- HZ3
4995 4993 4996 4997 3 0.65084 1.95253 0.00000
-2.60338 0.00000 0.00000 ; HE3- CE- NZ- HZ1
4995 4993 4996 4998 3 0.65084 1.95253 0.00000
-2.60338 0.00000 0.00000 ; HE3- CE- NZ- HZ2
4995 4993 4996 4999 3 0.65084 1.95253 0.00000
-2.60338 0.00000 0.00000 ; HE3- CE- NZ- HZ3
5000 5002 5004 5005 3 0.00000 0.00000 0.00000
0.00000 0.00000 0.00000 ; C- N- CA- HA
5000 5002 5004 5006 3 11.71520 1.67360 15.06240 -
13.38880 0.00000 0.00000 ; C- N- CA- CB
5000 5002 5004 5018 3 1.75728 5.27184 2.25936
-7.02912 0.00000 0.00000 ; C- N- CA- C
5001 5000 5002 5003 3 29.28800 -8.36800 -20.92000
0.00000 0.00000 0.00000 ; O- C- N- H
5001 5000 5002 5004 3 20.92000 0.00000 -20.92000
0.00000 0.00000 0.00000 ; O- C- N- CA
5002 5004 5006 5007 3 0.65084 1.95253 0.00000
-2.60338 0.00000 0.00000 ; N- CA- CB- HB2
5002 5004 5006 5008 3 0.65084 1.95253 0.00000
-2.60338 0.00000 0.00000 ; N- CA- CB- HB3
5002 5004 5006 5009 3 4.79068 4.02919 -4.82834
-3.66518 2.97901 0.00000 ; N- CA- CB- CG
5002 5004 5018 5019 3 0.00000 0.00000 0.00000
0.00000 0.00000 0.00000 ; N- CA- C- O
5002 5004 5018 5020 3 17.40544 -5.02080 -13.22144
9.20480 0.00000 0.00000 ; N- CA- C- N
5003 5002 5004 5005 3 0.00000 0.00000 0.00000
0.00000 0.00000 0.00000 ; H- N- CA- HA
5003 5002 5004 5006 3 0.00000 0.00000 0.00000
0.00000 0.00000 0.00000 ; H- N- CA- CB
5003 5002 5004 5018 3 0.00000 0.00000 0.00000
0.00000 0.00000 0.00000 ; H- N- CA- C

5004 5006 5009 5010 3 6.81992 9.28011 -2.04179 -
11.48090 -1.23846 0.00000 ; CA- CB- CG- ND1
5004 5006 5009 5016 3 0.00000 0.00000 0.00000
0.00000 0.00000 0.00000 ; CA- CB- CG- CD2
5004 5018 5020 5021 3 20.92000 0.00000 -20.92000
0.00000 0.00000 0.00000 ; CA- C- N- H
5004 5018 5020 5022 3 20.92000 0.00000 -20.92000
0.00000 0.00000 0.00000 ; CA- C- N- CA
5005 5004 5006 5007 3 0.65084 1.95253 0.00000
-2.60338 0.00000 0.00000 ; HA- CA- CB- HB2
5005 5004 5006 5008 3 0.65084 1.95253 0.00000
-2.60338 0.00000 0.00000 ; HA- CA- CB- HB3
5005 5004 5006 5009 3 0.65084 1.95253 0.00000
-2.60338 0.00000 0.00000 ; HA- CA- CB- CG
5005 5004 5018 5019 3 3.68192 -4.35136 0.00000
1.33888 0.00000 0.00000 ; HA- CA- C- O
5005 5004 5018 5020 3 0.00000 0.00000 0.00000
0.00000 0.00000 0.00000 ; HA- CA- C- N
5006 5004 5018 5019 3 0.00000 0.00000 0.00000
0.00000 0.00000 0.00000 ; CB- CA- C- O
5006 5004 5018 5020 3 2.51040 4.18400 1.67360
-6.69440 0.00000 0.00000 ; CB- CA- C- N
5006 5009 5010 5011 3 11.71520 0.00000 -11.71520
0.00000 0.00000 0.00000 ; CB- CG- ND1- HD1
5006 5009 5010 5012 3 11.71520 0.00000 -11.71520
0.00000 0.00000 0.00000 ; CB- CG- ND1- CE1
5006 5009 5016 5014 3 44.97800 0.00000 -44.97800
0.00000 0.00000 0.00000 ; CB- CG- CD2- NE2
5006 5009 5016 5017 3 44.97800 0.00000 -44.97800
0.00000 0.00000 0.00000 ; CB- CG- CD2- HD2
5007 5006 5004 5018 3 0.65084 1.95253 0.00000
-2.60338 0.00000 0.00000 ; HB2- CB- CA- C
5007 5006 5009 5010 3 0.00000 0.00000 0.00000
0.00000 0.00000 0.00000 ; HB2- CB- CG- ND1
5007 5006 5009 5016 3 0.00000 0.00000 0.00000
0.00000 0.00000 0.00000 ; HB2- CB- CG- CD2
5008 5006 5004 5018 3 0.65084 1.95253 0.00000
-2.60338 0.00000 0.00000 ; HB3- CB- CA- C
5008 5006 5009 5010 3 0.00000 0.00000 0.00000
0.00000 0.00000 0.00000 ; HB3- CB- CG- ND1
5008 5006 5009 5016 3 0.00000 0.00000 0.00000
0.00000 0.00000 0.00000 ; HB3- CB- CG- CD2
5009 5006 5004 5018 3 3.76560 3.34720 -2.87859
-3.66518 0.83680 0.00000 ; CG- CB- CA- C
5009 5010 5012 5013 3 19.45560 0.00000 -19.45560
0.00000 0.00000 0.00000 ; CG- ND1- CE1- HE1
5009 5010 5012 5014 3 19.45560 0.00000 -19.45560
0.00000 0.00000 0.00000 ; CG- ND1- CE1- NE2
5009 5016 5014 5012 3 12.55200 0.00000 -12.55200
0.00000 0.00000 0.00000 ; CG- CD2- NE2- CE1
5009 5016 5014 5015 3 12.55200 0.00000 -12.55200
0.00000 0.00000 0.00000 ; CG- CD2- NE2- HE2

5010 5009 5016 5014 3 44.97800 0.00000 -44.97800
0.00000 0.00000 0.00000 ; ND1- CG- CD2- NE2
5010 5009 5016 5017 3 44.97800 0.00000 -44.97800
0.00000 0.00000 0.00000 ; ND1- CG- CD2- HD2
5010 5012 5014 5015 3 19.45560 0.00000 -19.45560
0.00000 0.00000 0.00000 ; ND1- CE1- NE2- HE2
5010 5012 5014 5016 3 19.45560 0.00000 -19.45560
0.00000 0.00000 0.00000 ; ND1- CE1- NE2- CD2
5011 5010 5009 5016 3 11.71520 0.00000 -11.71520
0.00000 0.00000 0.00000 ; HD1- ND1- CG- CD2
5011 5010 5012 5013 3 19.45560 0.00000 -19.45560
0.00000 0.00000 0.00000 ; HD1- ND1- CE1- HE1
5011 5010 5012 5014 3 19.45560 0.00000 -19.45560
0.00000 0.00000 0.00000 ; HD1- ND1- CE1- NE2
5012 5010 5009 5016 3 11.71520 0.00000 -11.71520
0.00000 0.00000 0.00000 ; CE1- ND1- CG- CD2
5012 5014 5016 5017 3 12.55200 0.00000 -12.55200
0.00000 0.00000 0.00000 ; CE1- NE2- CD2- HD2
5013 5012 5014 5015 3 19.45560 0.00000 -19.45560
0.00000 0.00000 0.00000 ; HE1- CE1- NE2- HE2
5013 5012 5014 5016 3 19.45560 0.00000 -19.45560
0.00000 0.00000 0.00000 ; HE1- CE1- NE2- CD2
5015 5014 5016 5017 3 12.55200 0.00000 -12.55200
0.00000 0.00000 0.00000 ; HE2- NE2- CD2- HD2
5018 5020 5022 5023 3 0.00000 0.00000 0.00000
0.00000 0.00000 0.00000 ; C- N- CA- HA
5018 5020 5022 5024 3 11.71520 1.67360 15.06240 -
13.38880 0.00000 0.00000 ; C- N- CA- CB
5018 5020 5022 5037 3 1.75728 5.27184 2.25936
-7.02912 0.00000 0.00000 ; C- N- CA- C
5019 5018 5020 5021 3 29.28800 -8.36800 -20.92000
0.00000 0.00000 0.00000 ; O- C- N- H
5019 5018 5020 5022 3 20.92000 0.00000 -20.92000
0.00000 0.00000 0.00000 ; O- C- N- CA
5020 5022 5024 5025 3 0.65084 1.95253 0.00000
-2.60338 0.00000 0.00000 ; N- CA- CB- HB2
5020 5022 5024 5026 3 0.65084 1.95253 0.00000
-2.60338 0.00000 0.00000 ; N- CA- CB- HB3
5020 5022 5024 5027 3 3.79070 1.39746 -4.61077
-2.40998 2.44346 0.00000 ; N- CA- CB- CG
5020 5022 5037 5038 3 0.00000 0.00000 0.00000
0.00000 0.00000 0.00000 ; N- CA- C- O
5020 5022 5037 5039 3 17.40544 -5.02080 -13.22144
9.20480 0.00000 0.00000 ; N- CA- C- N
5021 5020 5022 5023 3 0.00000 0.00000 0.00000
0.00000 0.00000 0.00000 ; H- N- CA- HA
5021 5020 5022 5024 3 0.00000 0.00000 0.00000
0.00000 0.00000 0.00000 ; H- N- CA- CB
5021 5020 5022 5037 3 0.00000 0.00000 0.00000
0.00000 0.00000 0.00000 ; H- N- CA- C
5022 5024 5027 5028 3 0.66944 2.00832 0.00000
-2.67776 0.00000 0.00000 ; CA- CB- CG- HG

```
  5022   5024   5027   5029      3    3.90367    0.19665   -6.21742
-2.37651    5.99149    0.00000 ;     CA-    CB-    CG-   CD1
  5022   5024   5027   5033      3    3.90367    0.19665   -6.21742
-2.37651    5.99149    0.00000 ;     CA-    CB-    CG-   CD2
  5022   5037   5039   5040      3   20.92000    0.00000  -20.92000
0.00000    0.00000    0.00000 ;     CA-     C-     N-     H
  5022   5037   5039   5041      3   20.92000    0.00000  -20.92000
0.00000    0.00000    0.00000 ;     CA-     C-     N-    CA
  5023   5022   5024   5025      3    0.65084    1.95253    0.00000
-2.60338    0.00000    0.00000 ;     HA-    CA-    CB-   HB2
  5023   5022   5024   5026      3    0.65084    1.95253    0.00000
-2.60338    0.00000    0.00000 ;     HA-    CA-    CB-   HB3
  5023   5022   5024   5027      3    0.65084    1.95253    0.00000
-2.60338    0.00000    0.00000 ;     HA-    CA-    CB-    CG
  5023   5022   5037   5038      3    3.68192   -4.35136    0.00000
1.33888    0.00000    0.00000 ;     HA-    CA-     C-     O
  5023   5022   5037   5039      3    0.00000    0.00000    0.00000
0.00000    0.00000    0.00000 ;     HA-    CA-     C-     N
  5024   5022   5037   5038      3    0.00000    0.00000    0.00000
0.00000    0.00000    0.00000 ;     CB-    CA-     C-     O
  5024   5022   5037   5039      3    2.51040    4.18400    1.67360
-6.69440    0.00000    0.00000 ;     CB-    CA-     C-     N
  5024   5027   5029   5030      3    0.66944    2.00832    0.00000
-2.67776    0.00000    0.00000 ;     CB-    CG-   CD1-  HD11
  5024   5027   5029   5031      3    0.66944    2.00832    0.00000
-2.67776    0.00000    0.00000 ;     CB-    CG-   CD1-  HD12
  5024   5027   5029   5032      3    0.66944    2.00832    0.00000
-2.67776    0.00000    0.00000 ;     CB-    CG-   CD1-  HD13
  5024   5027   5033   5034      3    0.66944    2.00832    0.00000
-2.67776    0.00000    0.00000 ;     CB-    CG-   CD2-  HD21
  5024   5027   5033   5035      3    0.66944    2.00832    0.00000
-2.67776    0.00000    0.00000 ;     CB-    CG-   CD2-  HD22
  5024   5027   5033   5036      3    0.66944    2.00832    0.00000
-2.67776    0.00000    0.00000 ;     CB-    CG-   CD2-  HD23
  5025   5024   5022   5037      3    0.65084    1.95253    0.00000
-2.60338    0.00000    0.00000 ;    HB2-    CB-    CA-     C
  5025   5024   5027   5028      3    0.62760    1.88280    0.00000
-2.51040    0.00000    0.00000 ;    HB2-    CB-    CG-    HG
  5025   5024   5027   5029      3    0.66944    2.00832    0.00000
-2.67776    0.00000    0.00000 ;    HB2-    CB-    CG-   CD1
  5025   5024   5027   5033      3    0.66944    2.00832    0.00000
-2.67776    0.00000    0.00000 ;    HB2-    CB-    CG-   CD2
  5026   5024   5022   5037      3    0.65084    1.95253    0.00000
-2.60338    0.00000    0.00000 ;    HB3-    CB-    CA-     C
  5026   5024   5027   5028      3    0.62760    1.88280    0.00000
-2.51040    0.00000    0.00000 ;    HB3-    CB-    CG-    HG
  5026   5024   5027   5029      3    0.66944    2.00832    0.00000
-2.67776    0.00000    0.00000 ;    HB3-    CB-    CG-   CD1
  5026   5024   5027   5033      3    0.66944    2.00832    0.00000
-2.67776    0.00000    0.00000 ;    HB3-    CB-    CG-   CD2
  5027   5024   5022   5037      3   10.33448   -1.14642  -11.54784
-2.40998    6.35968    0.00000 ;     CG-    CB-    CA-     C
```

5028 5027 5029 5030 3 0.62760 1.88280 0.00000
-2.51040 0.00000 0.00000 ; HG- CG- CD1- HD11
5028 5027 5029 5031 3 0.62760 1.88280 0.00000
-2.51040 0.00000 0.00000 ; HG- CG- CD1- HD12
5028 5027 5029 5032 3 0.62760 1.88280 0.00000
-2.51040 0.00000 0.00000 ; HG- CG- CD1- HD13
5028 5027 5033 5034 3 0.62760 1.88280 0.00000
-2.51040 0.00000 0.00000 ; HG- CG- CD2- HD21
5028 5027 5033 5035 3 0.62760 1.88280 0.00000
-2.51040 0.00000 0.00000 ; HG- CG- CD2- HD22
5028 5027 5033 5036 3 0.62760 1.88280 0.00000
-2.51040 0.00000 0.00000 ; HG- CG- CD2- HD23
5029 5027 5033 5034 3 0.66944 2.00832 0.00000
-2.67776 0.00000 0.00000 ; CD1- CG- CD2- HD21
5029 5027 5033 5035 3 0.66944 2.00832 0.00000
-2.67776 0.00000 0.00000 ; CD1- CG- CD2- HD22
5029 5027 5033 5036 3 0.66944 2.00832 0.00000
-2.67776 0.00000 0.00000 ; CD1- CG- CD2- HD23
5030 5029 5027 5033 3 0.66944 2.00832 0.00000
-2.67776 0.00000 0.00000 ; HD11- CD1- CG- CD2
5031 5029 5027 5033 3 0.66944 2.00832 0.00000
-2.67776 0.00000 0.00000 ; HD12- CD1- CG- CD2
5032 5029 5027 5033 3 0.66944 2.00832 0.00000
-2.67776 0.00000 0.00000 ; HD13- CD1- CG- CD2
5037 5039 5041 5042 3 0.00000 0.00000 0.00000
0.00000 0.00000 0.00000 ; C- N- CA- HA
5037 5039 5041 5043 3 11.71520 1.67360 15.06240 -
13.38880 0.00000 0.00000 ; C- N- CA- CB
5037 5039 5041 5048 3 1.75728 5.27184 2.25936
-7.02912 0.00000 0.00000 ; C- N- CA- C
5038 5037 5039 5040 3 29.28800 -8.36800 -20.92000
0.00000 0.00000 0.00000 ; O- C- N- H
5038 5037 5039 5041 3 20.92000 0.00000 -20.92000
0.00000 0.00000 0.00000 ; O- C- N- CA
5039 5041 5043 5044 3 0.65084 1.95253 0.00000
-2.60338 0.00000 0.00000 ; N- CA- CB- HB2
5039 5041 5043 5045 3 0.65084 1.95253 0.00000
-2.60338 0.00000 0.00000 ; N- CA- CB- HB3
5039 5041 5043 5046 3 6.03751 2.50622 -5.17142
-4.20074 1.10458 0.00000 ; N- CA- CB- SG
5039 5041 5048 5049 3 0.00000 0.00000 0.00000
0.00000 0.00000 0.00000 ; N- CA- C- O
5039 5041 5048 5050 3 17.40544 -5.02080 -13.22144
9.20480 0.00000 0.00000 ; N- CA- C- N
5040 5039 5041 5042 3 0.00000 0.00000 0.00000
0.00000 0.00000 0.00000 ; H- N- CA- HA
5040 5039 5041 5043 3 0.00000 0.00000 0.00000
0.00000 0.00000 0.00000 ; H- N- CA- CB
5040 5039 5041 5048 3 0.00000 0.00000 0.00000
0.00000 0.00000 0.00000 ; H- N- CA- C
5041 5043 5046 5047 3 1.69034 2.77818 4.11706
-4.21747 1.00416 0.00000 ; CA- CB- SG- HG

5041 5048 5050 5051 3 20.92000 0.00000 -20.92000
0.00000 0.00000 0.00000 ; CA- C- N- H
5041 5048 5050 5052 3 20.92000 0.00000 -20.92000
0.00000 0.00000 0.00000 ; CA- C- N- CA
5042 5041 5043 5044 3 0.65084 1.95253 0.00000
-2.60338 0.00000 0.00000 ; HA- CA- CB- HB2
5042 5041 5043 5045 3 0.65084 1.95253 0.00000
-2.60338 0.00000 0.00000 ; HA- CA- CB- HB3
5042 5041 5043 5046 3 0.65084 1.95253 0.00000
-2.60338 0.00000 0.00000 ; HA- CA- CB- SG
5042 5041 5048 5049 3 3.68192 -4.35136 0.00000
1.33888 0.00000 0.00000 ; HA- CA- C- O
5042 5041 5048 5050 3 0.00000 0.00000 0.00000
0.00000 0.00000 0.00000 ; HA- CA- C- N
5043 5041 5048 5049 3 0.00000 0.00000 0.00000
0.00000 0.00000 0.00000 ; CB- CA- C- O
5043 5041 5048 5050 3 2.51040 4.18400 1.67360
-6.69440 0.00000 0.00000 ; CB- CA- C- N
5044 5043 5041 5048 3 0.65084 1.95253 0.00000
-2.60338 0.00000 0.00000 ; HB2- CB- CA- C
5044 5043 5046 5047 3 1.04600 3.13800 0.00000
-4.18400 0.00000 0.00000 ; HB2- CB- SG- HG
5045 5043 5041 5048 3 0.65084 1.95253 0.00000
-2.60338 0.00000 0.00000 ; HB3- CB- CA- C
5045 5043 5046 5047 3 1.04600 3.13800 0.00000
-4.18400 0.00000 0.00000 ; HB3- CB- SG- HG
5046 5043 5041 5048 3 5.62330 4.27605 -5.33042
-4.20074 2.51040 0.00000 ; SG- CB- CA- C
5048 5050 5052 5053 3 0.00000 0.00000 0.00000
0.00000 0.00000 0.00000 ; C- N- CA- HA
5048 5050 5052 5054 3 11.71520 1.67360 15.06240 -
13.38880 0.00000 0.00000 ; C- N- CA- CB
5048 5050 5052 5060 3 1.75728 5.27184 2.25936
-7.02912 0.00000 0.00000 ; C- N- CA- C
5049 5048 5050 5051 3 29.28800 -8.36800 -20.92000
0.00000 0.00000 0.00000 ; O- C- N- H
5049 5048 5050 5052 3 20.92000 0.00000 -20.92000
0.00000 0.00000 0.00000 ; O- C- N- CA
5050 5052 5054 5055 3 0.65084 1.95253 0.00000
-2.60338 0.00000 0.00000 ; N- CA- CB- HB2
5050 5052 5054 5056 3 0.65084 1.95253 0.00000
-2.60338 0.00000 0.00000 ; N- CA- CB- HB3
5050 5052 5054 5057 3 15.41386 9.74035 -8.39310
-0.97069 2.97901 0.00000 ; N- CA- CB- CG
5050 5052 5060 5061 3 0.00000 0.00000 0.00000
0.00000 0.00000 0.00000 ; N- CA- C- O
5050 5052 5060 5062 3 17.40544 -5.02080 -13.22144
9.20480 0.00000 0.00000 ; N- CA- C- N
5051 5050 5052 5053 3 0.00000 0.00000 0.00000
0.00000 0.00000 0.00000 ; H- N- CA- HA
5051 5050 5052 5054 3 0.00000 0.00000 0.00000
0.00000 0.00000 0.00000 ; H- N- CA- CB

5051 5050 5052 5060 3 0.00000 0.00000 0.00000
0.00000 0.00000 0.00000 ; H- N- CA- C
5052 5054 5057 5058 3 6.43499 0.00000 -5.39736
0.00000 -1.03763 0.00000 ; CA- CB- CG- OD1
5052 5054 5057 5059 3 6.43499 0.00000 -5.39736
0.00000 -1.03763 0.00000 ; CA- CB- CG- OD2
5052 5060 5062 5063 3 20.92000 0.00000 -20.92000
0.00000 0.00000 0.00000 ; CA- C- N- H
5052 5060 5062 5064 3 20.92000 0.00000 -20.92000
0.00000 0.00000 0.00000 ; CA- C- N- CA
5053 5052 5054 5055 3 0.65084 1.95253 0.00000
-2.60338 0.00000 0.00000 ; HA- CA- CB- HB2
5053 5052 5054 5056 3 0.65084 1.95253 0.00000
-2.60338 0.00000 0.00000 ; HA- CA- CB- HB3
5053 5052 5054 5057 3 0.65084 1.95253 0.00000
-2.60338 0.00000 0.00000 ; HA- CA- CB- CG
5053 5052 5060 5061 3 3.68192 -4.35136 0.00000
1.33888 0.00000 0.00000 ; HA- CA- C- O
5053 5052 5060 5062 3 0.00000 0.00000 0.00000
0.00000 0.00000 0.00000 ; HA- CA- C- N
5054 5052 5060 5061 3 0.00000 0.00000 0.00000
0.00000 0.00000 0.00000 ; CB- CA- C- O
5054 5052 5060 5062 3 2.51040 4.18400 1.67360
-6.69440 0.00000 0.00000 ; CB- CA- C- N
5055 5054 5052 5060 3 0.65084 1.95253 0.00000
-2.60338 0.00000 0.00000 ; HB2- CB- CA- C
5055 5054 5057 5058 3 0.00000 0.00000 0.00000
0.00000 0.00000 0.00000 ; HB2- CB- CG- OD1
5055 5054 5057 5059 3 0.00000 0.00000 0.00000
0.00000 0.00000 0.00000 ; HB2- CB- CG- OD2
5056 5054 5052 5060 3 0.65084 1.95253 0.00000
-2.60338 0.00000 0.00000 ; HB3- CB- CA- C
5056 5054 5057 5058 3 0.00000 0.00000 0.00000
0.00000 0.00000 0.00000 ; HB3- CB- CG- OD1
5056 5054 5057 5059 3 0.00000 0.00000 0.00000
0.00000 0.00000 0.00000 ; HB3- CB- CG- OD2
5057 5054 5052 5060 3 7.14627 -1.04600 -8.99560
-0.97069 5.15469 0.00000 ; CG- CB- CA- C
5060 5062 5064 5065 3 0.00000 0.00000 0.00000
0.00000 0.00000 0.00000 ; C- N- CA- HA
5060 5062 5064 5066 3 11.71520 1.67360 15.06240 -
13.38880 0.00000 0.00000 ; C- N- CA- CB
5060 5062 5064 5076 3 1.75728 5.27184 2.25936
-7.02912 0.00000 0.00000 ; C- N- CA- C
5061 5060 5062 5063 3 29.28800 -8.36800 -20.92000
0.00000 0.00000 0.00000 ; O- C- N- H
5061 5060 5062 5064 3 20.92000 0.00000 -20.92000
0.00000 0.00000 0.00000 ; O- C- N- CA
5062 5064 5066 5067 3 0.65084 1.95253 0.00000
-2.60338 0.00000 0.00000 ; N- CA- CB- HB
5062 5064 5066 5068 3 3.83673 0.44769 -1.77402
-2.47693 -0.03347 0.00000 ; N- CA- CB- CG1

5062 5064 5066 5072 3 3.83673 0.44769 -1.77402
-2.47693 -0.03347 0.00000 ; N- CA- CB- CG2
5062 5064 5076 5077 3 0.00000 0.00000 0.00000
0.00000 0.00000 0.00000 ; N- CA- C- O
5062 5064 5076 5078 3 17.40544 -5.02080 -13.22144
9.20480 0.00000 0.00000 ; N- CA- C- N
5063 5062 5064 5065 3 0.00000 0.00000 0.00000
0.00000 0.00000 0.00000 ; H- N- CA- HA
5063 5062 5064 5066 3 0.00000 0.00000 0.00000
0.00000 0.00000 0.00000 ; H- N- CA- CB
5063 5062 5064 5076 3 0.00000 0.00000 0.00000
0.00000 0.00000 0.00000 ; H- N- CA- C
5064 5066 5068 5069 3 0.66944 2.00832 0.00000
-2.67776 0.00000 0.00000 ; CA- CB- CG1- HG11
5064 5066 5068 5070 3 0.66944 2.00832 0.00000
-2.67776 0.00000 0.00000 ; CA- CB- CG1- HG12
5064 5066 5068 5071 3 0.66944 2.00832 0.00000
-2.67776 0.00000 0.00000 ; CA- CB- CG1- HG13
5064 5066 5072 5073 3 0.66944 2.00832 0.00000
-2.67776 0.00000 0.00000 ; CA- CB- CG2- HG21
5064 5066 5072 5074 3 0.66944 2.00832 0.00000
-2.67776 0.00000 0.00000 ; CA- CB- CG2- HG22
5064 5066 5072 5075 3 0.66944 2.00832 0.00000
-2.67776 0.00000 0.00000 ; CA- CB- CG2- HG23
5064 5076 5078 5079 3 20.92000 0.00000 -20.92000
0.00000 0.00000 0.00000 ; CA- C- N- H
5064 5076 5078 5080 3 20.92000 0.00000 -20.92000
0.00000 0.00000 0.00000 ; CA- C- N- CA
5065 5064 5066 5067 3 0.65084 1.95253 0.00000
-2.60338 0.00000 0.00000 ; HA- CA- CB- HB
5065 5064 5066 5068 3 0.65084 1.95253 0.00000
-2.60338 0.00000 0.00000 ; HA- CA- CB- CG1
5065 5064 5066 5072 3 0.65084 1.95253 0.00000
-2.60338 0.00000 0.00000 ; HA- CA- CB- CG2
5065 5064 5076 5077 3 3.68192 -4.35136 0.00000
1.33888 0.00000 0.00000 ; HA- CA- C- O
5065 5064 5076 5078 3 0.00000 0.00000 0.00000
0.00000 0.00000 0.00000 ; HA- CA- C- N
5066 5064 5076 5077 3 0.00000 0.00000 0.00000
0.00000 0.00000 0.00000 ; CB- CA- C- O
5066 5064 5076 5078 3 2.51040 4.18400 1.67360
-6.69440 0.00000 0.00000 ; CB- CA- C- N
5067 5066 5064 5076 3 0.65084 1.95253 0.00000
-2.60338 0.00000 0.00000 ; HB- CB- CA- C
5067 5066 5068 5069 3 0.62760 1.88280 0.00000
-2.51040 0.00000 0.00000 ; HB- CB- CG1- HG11
5067 5066 5068 5070 3 0.62760 1.88280 0.00000
-2.51040 0.00000 0.00000 ; HB- CB- CG1- HG12
5067 5066 5068 5071 3 0.62760 1.88280 0.00000
-2.51040 0.00000 0.00000 ; HB- CB- CG1- HG13
5067 5066 5072 5073 3 0.62760 1.88280 0.00000
-2.51040 0.00000 0.00000 ; HB- CB- CG2- HG21

5067 5066 5072 5074 3 0.62760 1.88280 0.00000
-2.51040 0.00000 0.00000 ; HB- CB- CG2- HG22
5067 5066 5072 5075 3 0.62760 1.88280 0.00000
-2.51040 0.00000 0.00000 ; HB- CB- CG2- HG23
5068 5066 5064 5076 3 5.67350 3.55640 -6.16722
-2.47693 3.74886 0.00000 ; CG1- CB- CA- C
5068 5066 5072 5073 3 0.66944 2.00832 0.00000
-2.67776 0.00000 0.00000 ; CG1- CB- CG2- HG21
5068 5066 5072 5074 3 0.66944 2.00832 0.00000
-2.67776 0.00000 0.00000 ; CG1- CB- CG2- HG22
5068 5066 5072 5075 3 0.66944 2.00832 0.00000
-2.67776 0.00000 0.00000 ; CG1- CB- CG2- HG23
5069 5068 5066 5072 3 0.66944 2.00832 0.00000
-2.67776 0.00000 0.00000 ; HG11- CG1- CB- CG2
5070 5068 5066 5072 3 0.66944 2.00832 0.00000
-2.67776 0.00000 0.00000 ; HG12- CG1- CB- CG2
5071 5068 5066 5072 3 0.66944 2.00832 0.00000
-2.67776 0.00000 0.00000 ; HG13- CG1- CB- CG2
5072 5066 5064 5076 3 5.67350 3.55640 -6.16722
-2.47693 3.74886 0.00000 ; CG2- CB- CA- C
5076 5078 5080 5081 3 0.00000 0.00000 0.00000
0.00000 0.00000 0.00000 ; C- N- CA- HA
5076 5078 5080 5082 3 11.71520 1.67360 15.06240 -
13.38880 0.00000 0.00000 ; C- N- CA- CB
5076 5078 5080 5091 3 1.75728 5.27184 2.25936
-7.02912 0.00000 0.00000 ; C- N- CA- C
5077 5076 5078 5079 3 29.28800 -8.36800 -20.92000
0.00000 0.00000 0.00000 ; O- C- N- H
5077 5076 5078 5080 3 20.92000 0.00000 -20.92000
0.00000 0.00000 0.00000 ; O- C- N- CA
5078 5080 5082 5083 3 0.65084 1.95253 0.00000
-2.60338 0.00000 0.00000 ; N- CA- CB- HB2
5078 5080 5082 5084 3 0.65084 1.95253 0.00000
-2.60338 0.00000 0.00000 ; N- CA- CB- HB3
5078 5080 5082 5085 3 3.21331 2.22589 -4.15053
-2.40998 2.61082 0.00000 ; N- CA- CB- CG
5078 5080 5091 5092 3 0.00000 0.00000 0.00000
0.00000 0.00000 0.00000 ; N- CA- C- O
5078 5080 5091 5093 3 17.40544 -5.02080 -13.22144
9.20480 0.00000 0.00000 ; N- CA- C- N
5079 5078 5080 5081 3 0.00000 0.00000 0.00000
0.00000 0.00000 0.00000 ; H- N- CA- HA
5079 5078 5080 5082 3 0.00000 0.00000 0.00000
0.00000 0.00000 0.00000 ; H- N- CA- CB
5079 5078 5080 5091 3 0.00000 0.00000 0.00000
0.00000 0.00000 0.00000 ; H- N- CA- C
5080 5082 5085 5086 3 0.66944 2.00832 0.00000
-2.67776 0.00000 0.00000 ; CA- CB- CG- HG2
5080 5082 5085 5087 3 0.66944 2.00832 0.00000
-2.67776 0.00000 0.00000 ; CA- CB- CG- HG3
5080 5082 5085 5088 3 10.12110 -1.91209 0.01674
10.17549 -1.87443 0.00000 ; CA- CB- CG- CD

5080 5091 5093 5094 3 20.92000 0.00000 -20.92000
0.00000 0.00000 0.00000 ; CA- C- N- H
5080 5091 5093 5095 3 20.92000 0.00000 -20.92000
0.00000 0.00000 0.00000 ; CA- C- N- CA
5081 5080 5082 5083 3 0.65084 1.95253 0.00000
-2.60338 0.00000 0.00000 ; HA- CA- CB- HB2
5081 5080 5082 5084 3 0.65084 1.95253 0.00000
-2.60338 0.00000 0.00000 ; HA- CA- CB- HB3
5081 5080 5082 5085 3 0.65084 1.95253 0.00000
-2.60338 0.00000 0.00000 ; HA- CA- CB- CG
5081 5080 5091 5092 3 3.68192 -4.35136 0.00000
1.33888 0.00000 0.00000 ; HA- CA- C- O
5081 5080 5091 5093 3 0.00000 0.00000 0.00000
0.00000 0.00000 0.00000 ; HA- CA- C- N
5082 5080 5091 5092 3 0.00000 0.00000 0.00000
0.00000 0.00000 0.00000 ; CB- CA- C- O
5082 5080 5091 5093 3 2.51040 4.18400 1.67360
-6.69440 0.00000 0.00000 ; CB- CA- C- N
5082 5085 5088 5089 3 3.79907 0.00000 -5.40573
0.00000 2.14221 0.00000 ; CB- CG- CD- OE1
5082 5085 5088 5090 3 3.79907 0.00000 -5.40573
0.00000 2.14221 0.00000 ; CB- CG- CD- OE2
5083 5082 5080 5091 3 0.65084 1.95253 0.00000
-2.60338 0.00000 0.00000 ; HB2- CB- CA- C
5083 5082 5085 5086 3 0.62760 1.88280 0.00000
-2.51040 0.00000 0.00000 ; HB2- CB- CG- HG2
5083 5082 5085 5087 3 0.62760 1.88280 0.00000
-2.51040 0.00000 0.00000 ; HB2- CB- CG- HG3
5083 5082 5085 5088 3 0.65084 1.95253 0.00000
-2.60338 0.00000 0.00000 ; HB2- CB- CG- CD
5084 5082 5080 5091 3 0.65084 1.95253 0.00000
-2.60338 0.00000 0.00000 ; HB3- CB- CA- C
5084 5082 5085 5086 3 0.62760 1.88280 0.00000
-2.51040 0.00000 0.00000 ; HB3- CB- CG- HG2
5084 5082 5085 5087 3 0.62760 1.88280 0.00000
-2.51040 0.00000 0.00000 ; HB3- CB- CG- HG3
5084 5082 5085 5088 3 0.65084 1.95253 0.00000
-2.60338 0.00000 0.00000 ; HB3- CB- CG- CD
5085 5082 5080 5091 3 6.86594 3.56895 -8.14206
-2.40998 4.85344 0.00000 ; CG- CB- CA- C
5086 5085 5088 5089 3 0.00000 0.00000 0.00000
0.00000 0.00000 0.00000 ; HG2- CG- CD- OE1
5086 5085 5088 5090 3 0.00000 0.00000 0.00000
0.00000 0.00000 0.00000 ; HG2- CG- CD- OE2
5087 5085 5088 5089 3 0.00000 0.00000 0.00000
0.00000 0.00000 0.00000 ; HG3- CG- CD- OE1
5087 5085 5088 5090 3 0.00000 0.00000 0.00000
0.00000 0.00000 0.00000 ; HG3- CG- CD- OE2
5091 5093 5095 5096 3 0.00000 0.00000 0.00000
0.00000 0.00000 0.00000 ; C- N- CA- HA
5091 5093 5095 5097 3 11.71520 1.67360 15.06240 -
13.38880 0.00000 0.00000 ; C- N- CA- CB

5091 5093 5095 5110 3 1.75728 5.27184 2.25936
-7.02912 0.00000 0.00000 ; C- N- CA- C
5092 5091 5093 5094 3 29.28800 -8.36800 -20.92000
0.00000 0.00000 0.00000 ; O- C- N- H
5092 5091 5093 5095 3 20.92000 0.00000 -20.92000
0.00000 0.00000 0.00000 ; O- C- N- CA
5093 5095 5097 5098 3 0.65084 1.95253 0.00000
-2.60338 0.00000 0.00000 ; N- CA- CB- HB
5093 5095 5097 5099 3 3.83673 0.44769 -1.77402
-2.47693 -0.03347 0.00000 ; N- CA- CB- CG2
5093 5095 5097 5103 3 2.97482 0.12134 2.04179
-1.89117 -3.24678 0.00000 ; N- CA- CB- CG1
5093 5095 5110 5111 3 0.00000 0.00000 0.00000
0.00000 0.00000 0.00000 ; N- CA- C- O
5093 5095 5110 5112 3 17.40544 -5.02080 -13.22144
9.20480 0.00000 0.00000 ; N- CA- C- N
5094 5093 5095 5096 3 0.00000 0.00000 0.00000
0.00000 0.00000 0.00000 ; H- N- CA- HA
5094 5093 5095 5097 3 0.00000 0.00000 0.00000
0.00000 0.00000 0.00000 ; H- N- CA- CB
5094 5093 5095 5110 3 0.00000 0.00000 0.00000
0.00000 0.00000 0.00000 ; H- N- CA- C
5095 5097 5099 5100 3 0.66944 2.00832 0.00000
-2.67776 0.00000 0.00000 ; CA- CB- CG2- HG21
5095 5097 5099 5101 3 0.66944 2.00832 0.00000
-2.67776 0.00000 0.00000 ; CA- CB- CG2- HG22
5095 5097 5099 5102 3 0.66944 2.00832 0.00000
-2.67776 0.00000 0.00000 ; CA- CB- CG2- HG23
5095 5097 5103 5104 3 0.66944 2.00832 0.00000
-2.67776 0.00000 0.00000 ; CA- CB- CG1- HG12
5095 5097 5103 5105 3 0.66944 2.00832 0.00000
-2.67776 0.00000 0.00000 ; CA- CB- CG1- HG13
5095 5097 5103 5106 3 4.24258 -0.52718 -7.25506
-1.79075 7.69856 0.00000 ; CA- CB- CG1- CD1
5095 5110 5112 5113 3 20.92000 0.00000 -20.92000
0.00000 0.00000 0.00000 ; CA- C- N- H
5095 5110 5112 5114 3 20.92000 0.00000 -20.92000
0.00000 0.00000 0.00000 ; CA- C- N- CA
5096 5095 5097 5098 3 0.65084 1.95253 0.00000
-2.60338 0.00000 0.00000 ; HA- CA- CB- HB
5096 5095 5097 5099 3 0.65084 1.95253 0.00000
-2.60338 0.00000 0.00000 ; HA- CA- CB- CG2
5096 5095 5097 5103 3 0.65084 1.95253 0.00000
-2.60338 0.00000 0.00000 ; HA- CA- CB- CG1
5096 5095 5110 5111 3 3.68192 -4.35136 0.00000
1.33888 0.00000 0.00000 ; HA- CA- C- O
5096 5095 5110 5112 3 0.00000 0.00000 0.00000
0.00000 0.00000 0.00000 ; HA- CA- C- N
5097 5095 5110 5111 3 0.00000 0.00000 0.00000
0.00000 0.00000 0.00000 ; CB- CA- C- O
5097 5095 5110 5112 3 2.51040 4.18400 1.67360
-6.69440 0.00000 0.00000 ; CB- CA- C- N

5097 5103 5106 5107 3 0.66944 2.00832 0.00000
-2.67776 0.00000 0.00000 ; CB- CG1- CD1- HD11
5097 5103 5106 5108 3 0.66944 2.00832 0.00000
-2.67776 0.00000 0.00000 ; CB- CG1- CD1- HD12
5097 5103 5106 5109 3 0.66944 2.00832 0.00000
-2.67776 0.00000 0.00000 ; CB- CG1- CD1- HD13
5098 5097 5095 5110 3 0.65084 1.95253 0.00000
-2.60338 0.00000 0.00000 ; HB- CB- CA- C
5098 5097 5099 5100 3 0.62760 1.88280 0.00000
-2.51040 0.00000 0.00000 ; HB- CB- CG2- HG21
5098 5097 5099 5101 3 0.62760 1.88280 0.00000
-2.51040 0.00000 0.00000 ; HB- CB- CG2- HG22
5098 5097 5099 5102 3 0.62760 1.88280 0.00000
-2.51040 0.00000 0.00000 ; HB- CB- CG2- HG23
5098 5097 5103 5104 3 0.62760 1.88280 0.00000
-2.51040 0.00000 0.00000 ; HB- CB- CG1- HG12
5098 5097 5103 5105 3 0.62760 1.88280 0.00000
-2.51040 0.00000 0.00000 ; HB- CB- CG1- HG13
5098 5097 5103 5106 3 0.66944 2.00832 0.00000
-2.67776 0.00000 0.00000 ; HB- CB- CG1- CD1
5099 5097 5095 5110 3 5.67350 3.55640 -6.16722
-2.47693 3.74886 0.00000 ; CG2- CB- CA- C
5099 5097 5103 5104 3 0.66944 2.00832 0.00000
-2.67776 0.00000 0.00000 ; CG2- CB- CG1- HG12
5099 5097 5103 5105 3 0.66944 2.00832 0.00000
-2.67776 0.00000 0.00000 ; CG2- CB- CG1- HG13
5099 5097 5103 5106 3 3.81162 0.49790 -8.14206
-1.79075 7.49773 0.00000 ; CG2- CB- CG1- CD1
5100 5099 5097 5103 3 0.66944 2.00832 0.00000
-2.67776 0.00000 0.00000 ; HG21- CG2- CB- CG1
5101 5099 5097 5103 3 0.66944 2.00832 0.00000
-2.67776 0.00000 0.00000 ; HG22- CG2- CB- CG1
5102 5099 5097 5103 3 0.66944 2.00832 0.00000
-2.67776 0.00000 0.00000 ; HG23- CG2- CB- CG1
5103 5097 5095 5110 3 8.26340 0.74057 -9.99976
-1.89117 3.84928 0.00000 ; CG1- CB- CA- C
5104 5103 5106 5107 3 0.62760 1.88280 0.00000
-2.51040 0.00000 0.00000 ; HG12- CG1- CD1- HD11
5104 5103 5106 5108 3 0.62760 1.88280 0.00000
-2.51040 0.00000 0.00000 ; HG12- CG1- CD1- HD12
5104 5103 5106 5109 3 0.62760 1.88280 0.00000
-2.51040 0.00000 0.00000 ; HG12- CG1- CD1- HD13
5105 5103 5106 5107 3 0.62760 1.88280 0.00000
-2.51040 0.00000 0.00000 ; HG13- CG1- CD1- HD11
5105 5103 5106 5108 3 0.62760 1.88280 0.00000
-2.51040 0.00000 0.00000 ; HG13- CG1- CD1- HD12
5105 5103 5106 5109 3 0.62760 1.88280 0.00000
-2.51040 0.00000 0.00000 ; HG13- CG1- CD1- HD13
5110 5112 5114 5115 3 0.00000 0.00000 0.00000
0.00000 0.00000 0.00000 ; C- N- CA- HA
5110 5112 5114 5116 3 11.71520 1.67360 15.06240 -
13.38880 0.00000 0.00000 ; C- N- CA- CB

```
  5110   5112   5114   5124      3    1.75728    5.27184    2.25936
-7.02912    0.00000    0.00000 ;      C-     N-    CA-     C
  5111   5110   5112   5113      3   29.28800   -8.36800  -20.92000
0.00000    0.00000    0.00000 ;      O-     C-     N-     H
  5111   5110   5112   5114      3   20.92000    0.00000  -20.92000
0.00000    0.00000    0.00000 ;      O-     C-     N-    CA
  5112   5114   5116   5117      3    0.65084    1.95253    0.00000
-2.60338    0.00000    0.00000 ;      N-    CA-    CB-   HB2
  5112   5114   5116   5118      3    0.65084    1.95253    0.00000
-2.60338    0.00000    0.00000 ;      N-    CA-    CB-   HB3
  5112   5114   5116   5119      3    5.99567    3.29281   -4.46014
-0.55229    1.97485    0.00000 ;      N-    CA-    CB-    CG
  5112   5114   5124   5125      3    0.00000    0.00000    0.00000
0.00000    0.00000    0.00000 ;     N-    CA-     C-     O
  5112   5114   5124   5126      3   17.40544   -5.02080  -13.22144
9.20480    0.00000    0.00000 ;     N-    CA-     C-     N
  5113   5112   5114   5115      3    0.00000    0.00000    0.00000
0.00000    0.00000    0.00000 ;     H-     N-    CA-    HA
  5113   5112   5114   5116      3    0.00000    0.00000    0.00000
0.00000    0.00000    0.00000 ;     H-     N-    CA-    CB
  5113   5112   5114   5124      3    0.00000    0.00000    0.00000
0.00000    0.00000    0.00000 ;     H-     N-    CA-     C
  5114   5116   5119   5120      3    0.00000    0.00000    0.00000
0.00000    0.00000    0.00000 ;    CA-    CB-    CG-   OD1
  5114   5116   5119   5121      3    8.84916   -0.31380   -4.32626
5.03754    0.26778    0.00000 ;    CA-    CB-    CG-   ND2
  5114   5124   5126   5127      3   20.92000    0.00000  -20.92000
0.00000    0.00000    0.00000 ;    CA-     C-     N-     H
  5114   5124   5126   5128      3   20.92000    0.00000  -20.92000
0.00000    0.00000    0.00000 ;    CA-     C-     N-    CA
  5115   5114   5116   5117      3    0.65084    1.95253    0.00000
-2.60338    0.00000    0.00000 ;     HA-    CA-    CB-   HB2
  5115   5114   5116   5118      3    0.65084    1.95253    0.00000
-2.60338    0.00000    0.00000 ;     HA-    CA-    CB-   HB3
  5115   5114   5116   5119      3    0.65084    1.95253    0.00000
-2.60338    0.00000    0.00000 ;     HA-    CA-    CB-    CG
  5115   5114   5124   5125      3    3.68192   -4.35136    0.00000
1.33888    0.00000    0.00000 ;    HA-    CA-     C-     O
  5115   5114   5124   5126      3    0.00000    0.00000    0.00000
0.00000    0.00000    0.00000 ;    HA-    CA-     C-     N
  5116   5114   5124   5125      3    0.00000    0.00000    0.00000
0.00000    0.00000    0.00000 ;    CB-    CA-     C-     O
  5116   5114   5124   5126      3    2.51040    4.18400    1.67360
-6.69440    0.00000    0.00000 ;     CB-    CA-     C-     N
  5116   5119   5121   5122      3   20.92000    0.00000  -20.92000
0.00000    0.00000    0.00000 ;    CB-    CG-    ND2-  HD21
  5116   5119   5121   5123      3   20.92000    0.00000  -20.92000
0.00000    0.00000    0.00000 ;    CB-    CG-    ND2-  HD22
  5117   5116   5114   5124      3    0.65084    1.95253    0.00000
-2.60338    0.00000    0.00000 ;    HB2-    CB-    CA-     C
  5117   5116   5119   5120      3    3.68192   -4.35136    0.00000
1.33888    0.00000    0.00000 ;   HB2-    CB-    CG-   OD1
```

5117 5116 5119 5121 3 0.00000 0.00000 0.00000
0.00000 0.00000 0.00000 ; HB2- CB- CG- ND2
5118 5116 5114 5124 3 0.65084 1.95253 0.00000
-2.60338 0.00000 0.00000 ; HB3- CB- CA- C
5118 5116 5119 5120 3 3.68192 -4.35136 0.00000
1.33888 0.00000 0.00000 ; HB3- CB- CG- OD1
5118 5116 5119 5121 3 0.00000 0.00000 0.00000
0.00000 0.00000 0.00000 ; HB3- CB- CG- ND2
5119 5116 5114 5124 3 7.94542 -3.96225 -6.11701
-0.55229 3.58150 0.00000 ; CG- CB- CA- C
5120 5119 5121 5122 3 29.28800 -8.36800 -20.92000
0.00000 0.00000 0.00000 ; OD1- CG- ND2- HD21
5120 5119 5121 5123 3 29.28800 -8.36800 -20.92000
0.00000 0.00000 0.00000 ; OD1- CG- ND2- HD22
5124 5126 5128 5129 3 0.00000 0.00000 0.00000
0.00000 0.00000 0.00000 ; C- N- CA- HA
5124 5126 5128 5130 3 11.71520 1.67360 15.06240 -
13.38880 0.00000 0.00000 ; C- N- CA- CB
5124 5126 5128 5138 3 1.75728 5.27184 2.25936
-7.02912 0.00000 0.00000 ; C- N- CA- C
5125 5124 5126 5127 3 29.28800 -8.36800 -20.92000
0.00000 0.00000 0.00000 ; O- C- N- H
5125 5124 5126 5128 3 20.92000 0.00000 -20.92000
0.00000 0.00000 0.00000 ; O- C- N- CA
5126 5128 5130 5131 3 0.65084 1.95253 0.00000
-2.60338 0.00000 0.00000 ; N- CA- CB- HB2
5126 5128 5130 5132 3 0.65084 1.95253 0.00000
-2.60338 0.00000 0.00000 ; N- CA- CB- HB3
5126 5128 5130 5133 3 5.99567 3.29281 -4.46014
-0.55229 1.97485 0.00000 ; N- CA- CB- CG
5126 5128 5138 5139 3 0.00000 0.00000 0.00000
0.00000 0.00000 0.00000 ; N- CA- C- O
5126 5128 5138 5140 3 17.40544 -5.02080 -13.22144
9.20480 0.00000 0.00000 ; N- CA- C- N
5127 5126 5128 5129 3 0.00000 0.00000 0.00000
0.00000 0.00000 0.00000 ; H- N- CA- HA
5127 5126 5128 5130 3 0.00000 0.00000 0.00000
0.00000 0.00000 0.00000 ; H- N- CA- CB
5127 5126 5128 5138 3 0.00000 0.00000 0.00000
0.00000 0.00000 0.00000 ; H- N- CA- C
5128 5130 5133 5134 3 0.00000 0.00000 0.00000
0.00000 0.00000 0.00000 ; CA- CB- CG- OD1
5128 5130 5133 5135 3 8.84916 -0.31380 -4.32626
5.03754 0.26778 0.00000 ; CA- CB- CG- ND2
5128 5138 5140 5141 3 20.92000 0.00000 -20.92000
0.00000 0.00000 0.00000 ; CA- C- N- H
5128 5138 5140 5142 3 20.92000 0.00000 -20.92000
0.00000 0.00000 0.00000 ; CA- C- N- CA
5129 5128 5130 5131 3 0.65084 1.95253 0.00000
-2.60338 0.00000 0.00000 ; HA- CA- CB- HB2
5129 5128 5130 5132 3 0.65084 1.95253 0.00000
-2.60338 0.00000 0.00000 ; HA- CA- CB- HB3

5129 5128 5130 5133 3 0.65084 1.95253 0.00000
-2.60338 0.00000 0.00000 ; HA- CA- CB- CG
5129 5128 5138 5139 3 3.68192 -4.35136 0.00000
1.33888 0.00000 0.00000 ; HA- CA- C- O
5129 5128 5138 5140 3 0.00000 0.00000 0.00000
0.00000 0.00000 0.00000 ; HA- CA- C- N
5130 5128 5138 5139 3 0.00000 0.00000 0.00000
0.00000 0.00000 0.00000 ; CB- CA- C- O
5130 5128 5138 5140 3 2.51040 4.18400 1.67360
-6.69440 0.00000 0.00000 ; CB- CA- C- N
5130 5133 5135 5136 3 20.92000 0.00000 -20.92000
0.00000 0.00000 0.00000 ; CB- CG- ND2- HD21
5130 5133 5135 5137 3 20.92000 0.00000 -20.92000
0.00000 0.00000 0.00000 ; CB- CG- ND2- HD22
5131 5130 5128 5138 3 0.65084 1.95253 0.00000
-2.60338 0.00000 0.00000 ; HB2- CB- CA- C
5131 5130 5133 5134 3 3.68192 -4.35136 0.00000
1.33888 0.00000 0.00000 ; HB2- CB- CG- OD1
5131 5130 5133 5135 3 0.00000 0.00000 0.00000
0.00000 0.00000 0.00000 ; HB2- CB- CG- ND2
5132 5130 5128 5138 3 0.65084 1.95253 0.00000
-2.60338 0.00000 0.00000 ; HB3- CB- CA- C
5132 5130 5133 5134 3 3.68192 -4.35136 0.00000
1.33888 0.00000 0.00000 ; HB3- CB- CG- OD1
5132 5130 5133 5135 3 0.00000 0.00000 0.00000
0.00000 0.00000 0.00000 ; HB3- CB- CG- ND2
5133 5130 5128 5138 3 7.94542 -3.96225 -6.11701
-0.55229 3.58150 0.00000 ; CG- CB- CA- C
5134 5133 5135 5136 3 29.28800 -8.36800 -20.92000
0.00000 0.00000 0.00000 ; OD1- CG- ND2- HD21
5134 5133 5135 5137 3 29.28800 -8.36800 -20.92000
0.00000 0.00000 0.00000 ; OD1- CG- ND2- HD22
5138 5140 5142 5143 3 0.00000 0.00000 0.00000
0.00000 0.00000 0.00000 ; C- N- CA- HA
5138 5140 5142 5144 3 11.71520 1.67360 15.06240 -
13.38880 0.00000 0.00000 ; C- N- CA- CB
5138 5140 5142 5157 3 1.75728 5.27184 2.25936
-7.02912 0.00000 0.00000 ; C- N- CA- C
5139 5138 5140 5141 3 29.28800 -8.36800 -20.92000
0.00000 0.00000 0.00000 ; O- C- N- H
5139 5138 5140 5142 3 20.92000 0.00000 -20.92000
0.00000 0.00000 0.00000 ; O- C- N- CA
5140 5142 5144 5145 3 0.65084 1.95253 0.00000
-2.60338 0.00000 0.00000 ; N- CA- CB- HB2
5140 5142 5144 5146 3 0.65084 1.95253 0.00000
-2.60338 0.00000 0.00000 ; N- CA- CB- HB3
5140 5142 5144 5147 3 3.79070 1.39746 -4.61077
-2.40998 2.44346 0.00000 ; N- CA- CB- CG
5140 5142 5157 5158 3 0.00000 0.00000 0.00000
0.00000 0.00000 0.00000 ; N- CA- C- O
5140 5142 5157 5159 3 17.40544 -5.02080 -13.22144
9.20480 0.00000 0.00000 ; N- CA- C- N

5141 5140 5142 5143 3 0.00000 0.00000 0.00000
0.00000 0.00000 0.00000 ; H- N- CA- HA
5141 5140 5142 5144 3 0.00000 0.00000 0.00000
0.00000 0.00000 0.00000 ; H- N- CA- CB
5141 5140 5142 5157 3 0.00000 0.00000 0.00000
0.00000 0.00000 0.00000 ; H- N- CA- C
5142 5144 5147 5148 3 0.66944 2.00832 0.00000
-2.67776 0.00000 0.00000 ; CA- CB- CG- HG
5142 5144 5147 5149 3 3.90367 0.19665 -6.21742
-2.37651 5.99149 0.00000 ; CA- CB- CG- CD1
5142 5144 5147 5153 3 3.90367 0.19665 -6.21742
-2.37651 5.99149 0.00000 ; CA- CB- CG- CD2
5142 5157 5159 5160 3 20.92000 0.00000 -20.92000
0.00000 0.00000 0.00000 ; CA- C- N- H
5142 5157 5159 5161 3 20.92000 0.00000 -20.92000
0.00000 0.00000 0.00000 ; CA- C- N- CA
5143 5142 5144 5145 3 0.65084 1.95253 0.00000
-2.60338 0.00000 0.00000 ; HA- CA- CB- HB2
5143 5142 5144 5146 3 0.65084 1.95253 0.00000
-2.60338 0.00000 0.00000 ; HA- CA- CB- HB3
5143 5142 5144 5147 3 0.65084 1.95253 0.00000
-2.60338 0.00000 0.00000 ; HA- CA- CB- CG
5143 5142 5157 5158 3 3.68192 -4.35136 0.00000
1.33888 0.00000 0.00000 ; HA- CA- C- O
5143 5142 5157 5159 3 0.00000 0.00000 0.00000
0.00000 0.00000 0.00000 ; HA- CA- C- N
5144 5142 5157 5158 3 0.00000 0.00000 0.00000
0.00000 0.00000 0.00000 ; CB- CA- C- O
5144 5142 5157 5159 3 2.51040 4.18400 1.67360
-6.69440 0.00000 0.00000 ; CB- CA- C- N
5144 5147 5149 5150 3 0.66944 2.00832 0.00000
-2.67776 0.00000 0.00000 ; CB- CG- CD1- HD11
5144 5147 5149 5151 3 0.66944 2.00832 0.00000
-2.67776 0.00000 0.00000 ; CB- CG- CD1- HD12
5144 5147 5149 5152 3 0.66944 2.00832 0.00000
-2.67776 0.00000 0.00000 ; CB- CG- CD1- HD13
5144 5147 5153 5154 3 0.66944 2.00832 0.00000
-2.67776 0.00000 0.00000 ; CB- CG- CD2- HD21
5144 5147 5153 5155 3 0.66944 2.00832 0.00000
-2.67776 0.00000 0.00000 ; CB- CG- CD2- HD22
5144 5147 5153 5156 3 0.66944 2.00832 0.00000
-2.67776 0.00000 0.00000 ; CB- CG- CD2- HD23
5145 5144 5142 5157 3 0.65084 1.95253 0.00000
-2.60338 0.00000 0.00000 ; HB2- CB- CA- C
5145 5144 5147 5148 3 0.62760 1.88280 0.00000
-2.51040 0.00000 0.00000 ; HB2- CB- CG- HG
5145 5144 5147 5149 3 0.66944 2.00832 0.00000
-2.67776 0.00000 0.00000 ; HB2- CB- CG- CD1
5145 5144 5147 5153 3 0.66944 2.00832 0.00000
-2.67776 0.00000 0.00000 ; HB2- CB- CG- CD2
5146 5144 5142 5157 3 0.65084 1.95253 0.00000
-2.60338 0.00000 0.00000 ; HB3- CB- CA- C

5146 5144 5147 5148 3 0.62760 1.88280 0.00000
-2.51040 0.00000 0.00000 ; HB3- CB- CG- HG
5146 5144 5147 5149 3 0.66944 2.00832 0.00000
-2.67776 0.00000 0.00000 ; HB3- CB- CG- CD1
5146 5144 5147 5153 3 0.66944 2.00832 0.00000
-2.67776 0.00000 0.00000 ; HB3- CB- CG- CD2
5147 5144 5142 5157 3 10.33448 -1.14642 -11.54784
-2.40998 6.35968 0.00000 ; CG- CB- CA- C
5148 5147 5149 5150 3 0.62760 1.88280 0.00000
-2.51040 0.00000 0.00000 ; HG- CG- CD1- HD11
5148 5147 5149 5151 3 0.62760 1.88280 0.00000
-2.51040 0.00000 0.00000 ; HG- CG- CD1- HD12
5148 5147 5149 5152 3 0.62760 1.88280 0.00000
-2.51040 0.00000 0.00000 ; HG- CG- CD1- HD13
5148 5147 5153 5154 3 0.62760 1.88280 0.00000
-2.51040 0.00000 0.00000 ; HG- CG- CD2- HD21
5148 5147 5153 5155 3 0.62760 1.88280 0.00000
-2.51040 0.00000 0.00000 ; HG- CG- CD2- HD22
5148 5147 5153 5156 3 0.62760 1.88280 0.00000
-2.51040 0.00000 0.00000 ; HG- CG- CD2- HD23
5149 5147 5153 5154 3 0.66944 2.00832 0.00000
-2.67776 0.00000 0.00000 ; CD1- CG- CD2- HD21
5149 5147 5153 5155 3 0.66944 2.00832 0.00000
-2.67776 0.00000 0.00000 ; CD1- CG- CD2- HD22
5149 5147 5153 5156 3 0.66944 2.00832 0.00000
-2.67776 0.00000 0.00000 ; CD1- CG- CD2- HD23
5150 5149 5147 5153 3 0.66944 2.00832 0.00000
-2.67776 0.00000 0.00000 ; HD11- CD1- CG- CD2
5151 5149 5147 5153 3 0.66944 2.00832 0.00000
-2.67776 0.00000 0.00000 ; HD12- CD1- CG- CD2
5152 5149 5147 5153 3 0.66944 2.00832 0.00000
-2.67776 0.00000 0.00000 ; HD13- CD1- CG- CD2
5157 5159 5161 5162 3 0.00000 0.00000 0.00000
0.00000 0.00000 0.00000 ; C- N- CA- HA
5157 5159 5161 5163 3 11.71520 1.67360 15.06240 -
13.38880 0.00000 0.00000 ; C- N- CA- CB
5157 5159 5161 5181 3 1.75728 5.27184 2.25936
-7.02912 0.00000 0.00000 ; C- N- CA- C
5158 5157 5159 5160 3 29.28800 -8.36800 -20.92000
0.00000 0.00000 0.00000 ; O- C- N- H
5158 5157 5159 5161 3 20.92000 0.00000 -20.92000
0.00000 0.00000 0.00000 ; O- C- N- CA
5159 5161 5163 5164 3 0.65084 1.95253 0.00000
-2.60338 0.00000 0.00000 ; N- CA- CB- HB2
5159 5161 5163 5165 3 0.65084 1.95253 0.00000
-2.60338 0.00000 0.00000 ; N- CA- CB- HB3
5159 5161 5163 5166 3 0.65084 1.95253 0.00000
-2.60338 0.00000 0.00000 ; N- CA- CB- CG
5159 5161 5181 5182 3 0.00000 0.00000 0.00000
0.00000 0.00000 0.00000 ; N- CA- C- O
5159 5161 5181 5183 3 17.40544 -5.02080 -13.22144
9.20480 0.00000 0.00000 ; N- CA- C- N

```
  5160   5159   5161   5162       3    0.00000    0.00000    0.00000
0.00000    0.00000    0.00000 ;      H-     N-    CA-    HA
  5160   5159   5161   5163       3    0.00000    0.00000    0.00000
0.00000    0.00000    0.00000 ;      H-     N-    CA-    CB
  5160   5159   5161   5181       3    0.00000    0.00000    0.00000
0.00000    0.00000    0.00000 ;      H-     N-    CA-     C
  5161   5163   5166   5167       3    0.66944    2.00832    0.00000
-2.67776    0.00000    0.00000 ;     CA-    CB-    CG-   HG2
  5161   5163   5166   5168       3    0.66944    2.00832    0.00000
-2.67776    0.00000    0.00000 ;     CA-    CB-    CG-   HG3
  5161   5163   5166   5169       3    3.68192    3.09616   -2.09200
-3.01248    0.00000    0.00000 ;     CA-    CB-    CG-    CD
  5161   5181   5183   5184       3   20.92000    0.00000  -20.92000
0.00000    0.00000    0.00000 ;     CA-     C-     N-     H
  5161   5181   5183   5185       3   20.92000    0.00000  -20.92000
0.00000    0.00000    0.00000 ;     CA-     C-     N-    CA
  5162   5161   5163   5164       3    0.65084    1.95253    0.00000
-2.60338    0.00000    0.00000 ;     HA-    CA-    CB-   HB2
  5162   5161   5163   5165       3    0.65084    1.95253    0.00000
-2.60338    0.00000    0.00000 ;     HA-    CA-    CB-   HB3
  5162   5161   5163   5166       3    0.65084    1.95253    0.00000
-2.60338    0.00000    0.00000 ;     HA-    CA-    CB-    CG
  5162   5161   5181   5182       3    3.68192   -4.35136    0.00000
1.33888    0.00000    0.00000 ;     HA-    CA-     C-     O
  5162   5161   5181   5183       3    0.00000    0.00000    0.00000
0.00000    0.00000    0.00000 ;     HA-    CA-     C-     N
  5163   5161   5181   5182       3    0.00000    0.00000    0.00000
0.00000    0.00000    0.00000 ;     CB-    CA-     C-     O
  5163   5161   5181   5183       3    2.51040    4.18400    1.67360
-6.69440    0.00000    0.00000 ;     CB-    CA-     C-     N
  5163   5166   5169   5170       3    0.65084    1.95253    0.00000
-2.60338    0.00000    0.00000 ;     CB-    CG-    CD-   HD2
  5163   5166   5169   5171       3    0.65084    1.95253    0.00000
-2.60338    0.00000    0.00000 ;     CB-    CG-    CD-   HD3
  5163   5166   5169   5172       3    0.65084    1.95253    0.00000
-2.60338    0.00000    0.00000 ;     CB-    CG-    CD-    NE
  5164   5163   5161   5181       3    0.65084    1.95253    0.00000
-2.60338    0.00000    0.00000 ;    HB2-    CB-    CA-     C
  5164   5163   5166   5167       3    0.62760    1.88280    0.00000
-2.51040    0.00000    0.00000 ;    HB2-    CB-    CG-   HG2
  5164   5163   5166   5168       3    0.62760    1.88280    0.00000
-2.51040    0.00000    0.00000 ;    HB2-    CB-    CG-   HG3
  5164   5163   5166   5169       3    0.66944    2.00832    0.00000
-2.67776    0.00000    0.00000 ;    HB2-    CB-    CG-    CD
  5165   5163   5161   5181       3    0.65084    1.95253    0.00000
-2.60338    0.00000    0.00000 ;    HB3-    CB-    CA-     C
  5165   5163   5166   5167       3    0.62760    1.88280    0.00000
-2.51040    0.00000    0.00000 ;    HB3-    CB-    CG-   HG2
  5165   5163   5166   5168       3    0.62760    1.88280    0.00000
-2.51040    0.00000    0.00000 ;    HB3-    CB-    CG-   HG3
  5165   5163   5166   5169       3    0.66944    2.00832    0.00000
-2.67776    0.00000    0.00000 ;    HB3-    CB-    CG-    CD
```

5166 5163 5161 5181 3 0.65084 1.95253 0.00000
-2.60338 0.00000 0.00000 ; CG- CB- CA- C
5166 5169 5172 5173 3 0.00000 0.00000 0.00000
0.00000 0.00000 0.00000 ; CG- CD- NE- HE
5166 5169 5172 5174 3 0.00000 0.00000 0.00000
0.00000 0.00000 0.00000 ; CG- CD- NE- CZ
5167 5166 5169 5170 3 0.65084 1.95253 0.00000
-2.60338 0.00000 0.00000 ; HG2- CG- CD- HD2
5167 5166 5169 5171 3 0.65084 1.95253 0.00000
-2.60338 0.00000 0.00000 ; HG2- CG- CD- HD3
5167 5166 5169 5172 3 0.65084 1.95253 0.00000
-2.60338 0.00000 0.00000 ; HG2- CG- CD- NE
5168 5166 5169 5170 3 0.65084 1.95253 0.00000
-2.60338 0.00000 0.00000 ; HG3- CG- CD- HD2
5168 5166 5169 5171 3 0.65084 1.95253 0.00000
-2.60338 0.00000 0.00000 ; HG3- CG- CD- HD3
5168 5166 5169 5172 3 0.65084 1.95253 0.00000
-2.60338 0.00000 0.00000 ; HG3- CG- CD- NE
5169 5172 5174 5175 3 20.08320 0.00000 -20.08320
0.00000 0.00000 0.00000 ; CD- NE- CZ- NH1
5169 5172 5174 5178 3 20.08320 0.00000 -20.08320
0.00000 0.00000 0.00000 ; CD- NE- CZ- NH2
5170 5169 5172 5173 3 0.00000 0.00000 0.00000
0.00000 0.00000 0.00000 ; HD2- CD- NE- HE
5170 5169 5172 5174 3 0.00000 0.00000 0.00000
0.00000 0.00000 0.00000 ; HD2- CD- NE- CZ
5171 5169 5172 5173 3 0.00000 0.00000 0.00000
0.00000 0.00000 0.00000 ; HD3- CD- NE- HE
5171 5169 5172 5174 3 0.00000 0.00000 0.00000
0.00000 0.00000 0.00000 ; HD3- CD- NE- CZ
5172 5174 5175 5176 3 20.08320 0.00000 -20.08320
0.00000 0.00000 0.00000 ; NE- CZ- NH1- HH11
5172 5174 5175 5177 3 20.08320 0.00000 -20.08320
0.00000 0.00000 0.00000 ; NE- CZ- NH1- HH12
5172 5174 5178 5179 3 20.08320 0.00000 -20.08320
0.00000 0.00000 0.00000 ; NE- CZ- NH2- HH21
5172 5174 5178 5180 3 20.08320 0.00000 -20.08320
0.00000 0.00000 0.00000 ; NE- CZ- NH2- HH22
5173 5172 5174 5175 3 20.08320 0.00000 -20.08320
0.00000 0.00000 0.00000 ; HE- NE- CZ- NH1
5173 5172 5174 5178 3 20.08320 0.00000 -20.08320
0.00000 0.00000 0.00000 ; HE- NE- CZ- NH2
5175 5174 5178 5179 3 20.08320 0.00000 -20.08320
0.00000 0.00000 0.00000 ; NH1- CZ- NH2- HH21
5175 5174 5178 5180 3 20.08320 0.00000 -20.08320
0.00000 0.00000 0.00000 ; NH1- CZ- NH2- HH22
5176 5175 5174 5178 3 20.08320 0.00000 -20.08320
0.00000 0.00000 0.00000 ; HH11- NH1- CZ- NH2
5177 5175 5174 5178 3 20.08320 0.00000 -20.08320
0.00000 0.00000 0.00000 ; HH12- NH1- CZ- NH2
5181 5183 5185 5186 3 0.00000 0.00000 0.00000
0.00000 0.00000 0.00000 ; C- N- CA- HA

5181 5183 5185 5187 3 11.71520 1.67360 15.06240 -
13.38880 0.00000 0.00000 ; C- N- CA- CB
5181 5183 5185 5195 3 1.75728 5.27184 2.25936
-7.02912 0.00000 0.00000 ; C- N- CA- C
5182 5181 5183 5184 3 29.28800 -8.36800 -20.92000
0.00000 0.00000 0.00000 ; O- C- N- H
5182 5181 5183 5185 3 20.92000 0.00000 -20.92000
0.00000 0.00000 0.00000 ; O- C- N- CA
5183 5185 5187 5188 3 0.65084 1.95253 0.00000
-2.60338 0.00000 0.00000 ; N- CA- CB- HB2
5183 5185 5187 5189 3 0.65084 1.95253 0.00000
-2.60338 0.00000 0.00000 ; N- CA- CB- HB3
5183 5185 5187 5190 3 5.99567 3.29281 -4.46014
-0.55229 1.97485 0.00000 ; N- CA- CB- CG
5183 5185 5195 5196 3 0.00000 0.00000 0.00000
0.00000 0.00000 0.00000 ; N- CA- C- O
5183 5185 5195 5197 3 17.40544 -5.02080 -13.22144
9.20480 0.00000 0.00000 ; N- CA- C- N
5184 5183 5185 5186 3 0.00000 0.00000 0.00000
0.00000 0.00000 0.00000 ; H- N- CA- HA
5184 5183 5185 5187 3 0.00000 0.00000 0.00000
0.00000 0.00000 0.00000 ; H- N- CA- CB
5184 5183 5185 5195 3 0.00000 0.00000 0.00000
0.00000 0.00000 0.00000 ; H- N- CA- C
5185 5187 5190 5191 3 0.00000 0.00000 0.00000
0.00000 0.00000 0.00000 ; CA- CB- CG- OD1
5185 5187 5190 5192 3 8.84916 -0.31380 -4.32626
5.03754 0.26778 0.00000 ; CA- CB- CG- ND2
5185 5195 5197 5198 3 20.92000 0.00000 -20.92000
0.00000 0.00000 0.00000 ; CA- C- N- H
5185 5195 5197 5199 3 20.92000 0.00000 -20.92000
0.00000 0.00000 0.00000 ; CA- C- N- CA
5186 5185 5187 5188 3 0.65084 1.95253 0.00000
-2.60338 0.00000 0.00000 ; HA- CA- CB- HB2
5186 5185 5187 5189 3 0.65084 1.95253 0.00000
-2.60338 0.00000 0.00000 ; HA- CA- CB- HB3
5186 5185 5187 5190 3 0.65084 1.95253 0.00000
-2.60338 0.00000 0.00000 ; HA- CA- CB- CG
5186 5185 5195 5196 3 3.68192 -4.35136 0.00000
1.33888 0.00000 0.00000 ; HA- CA- C- O
5186 5185 5195 5197 3 0.00000 0.00000 0.00000
0.00000 0.00000 0.00000 ; HA- CA- C- N
5187 5185 5195 5196 3 0.00000 0.00000 0.00000
0.00000 0.00000 0.00000 ; CB- CA- C- O
5187 5185 5195 5197 3 2.51040 4.18400 1.67360
-6.69440 0.00000 0.00000 ; CB- CA- C- N
5187 5190 5192 5193 3 20.92000 0.00000 -20.92000
0.00000 0.00000 0.00000 ; CB- CG- ND2- HD21
5187 5190 5192 5194 3 20.92000 0.00000 -20.92000
0.00000 0.00000 0.00000 ; CB- CG- ND2- HD22
5188 5187 5185 5195 3 0.65084 1.95253 0.00000
-2.60338 0.00000 0.00000 ; HB2- CB- CA- C

5188 5187 5190 5191 3 3.68192 -4.35136 0.00000
1.33888 0.00000 0.00000 ; HB2- CB- CG- OD1
5188 5187 5190 5192 3 0.00000 0.00000 0.00000
0.00000 0.00000 0.00000 ; HB2- CB- CG- ND2
5189 5187 5185 5195 3 0.65084 1.95253 0.00000
-2.60338 0.00000 0.00000 ; HB3- CB- CA- C
5189 5187 5190 5191 3 3.68192 -4.35136 0.00000
1.33888 0.00000 0.00000 ; HB3- CB- CG- OD1
5189 5187 5190 5192 3 0.00000 0.00000 0.00000
0.00000 0.00000 0.00000 ; HB3- CB- CG- ND2
5190 5187 5185 5195 3 7.94542 -3.96225 -6.11701
-0.55229 3.58150 0.00000 ; CG- CB- CA- C
5191 5190 5192 5193 3 29.28800 -8.36800 -20.92000
0.00000 0.00000 0.00000 ; OD1- CG- ND2- HD21
5191 5190 5192 5194 3 29.28800 -8.36800 -20.92000
0.00000 0.00000 0.00000 ; OD1- CG- ND2- HD22
5195 5197 5199 5200 3 0.00000 0.00000 0.00000
0.00000 0.00000 0.00000 ; C- N- CA- HA
5195 5197 5199 5201 3 11.71520 1.67360 15.06240 -
13.38880 0.00000 0.00000 ; C- N- CA- CB
5195 5197 5199 5214 3 1.75728 5.27184 2.25936
-7.02912 0.00000 0.00000 ; C- N- CA- C
5196 5195 5197 5198 3 29.28800 -8.36800 -20.92000
0.00000 0.00000 0.00000 ; O- C- N- H
5196 5195 5197 5199 3 20.92000 0.00000 -20.92000
0.00000 0.00000 0.00000 ; O- C- N- CA
5197 5199 5201 5202 3 0.65084 1.95253 0.00000
-2.60338 0.00000 0.00000 ; N- CA- CB- HB
5197 5199 5201 5203 3 3.83673 0.44769 -1.77402
-2.47693 -0.03347 0.00000 ; N- CA- CB- CG2
5197 5199 5201 5207 3 2.97482 0.12134 2.04179
-1.89117 -3.24678 0.00000 ; N- CA- CB- CG1
5197 5199 5214 5215 3 0.00000 0.00000 0.00000
0.00000 0.00000 0.00000 ; N- CA- C- O
5197 5199 5214 5216 3 17.40544 -5.02080 -13.22144
9.20480 0.00000 0.00000 ; N- CA- C- N
5198 5197 5199 5200 3 0.00000 0.00000 0.00000
0.00000 0.00000 0.00000 ; H- N- CA- HA
5198 5197 5199 5201 3 0.00000 0.00000 0.00000
0.00000 0.00000 0.00000 ; H- N- CA- CB
5198 5197 5199 5214 3 0.00000 0.00000 0.00000
0.00000 0.00000 0.00000 ; H- N- CA- C
5199 5201 5203 5204 3 0.66944 2.00832 0.00000
-2.67776 0.00000 0.00000 ; CA- CB- CG2- HG21
5199 5201 5203 5205 3 0.66944 2.00832 0.00000
-2.67776 0.00000 0.00000 ; CA- CB- CG2- HG22
5199 5201 5203 5206 3 0.66944 2.00832 0.00000
-2.67776 0.00000 0.00000 ; CA- CB- CG2- HG23
5199 5201 5207 5208 3 0.66944 2.00832 0.00000
-2.67776 0.00000 0.00000 ; CA- CB- CG1- HG12
5199 5201 5207 5209 3 0.66944 2.00832 0.00000
-2.67776 0.00000 0.00000 ; CA- CB- CG1- HG13

5199 5201 5207 5210 3 4.24258 -0.52718 -7.25506
-1.79075 7.69856 0.00000 ; CA- CB- CG1- CD1
5199 5214 5216 5217 3 20.92000 0.00000 -20.92000
0.00000 0.00000 0.00000 ; CA- C- N- H
5199 5214 5216 5218 3 20.92000 0.00000 -20.92000
0.00000 0.00000 0.00000 ; CA- C- N- CA
5200 5199 5201 5202 3 0.65084 1.95253 0.00000
-2.60338 0.00000 0.00000 ; HA- CA- CB- HB
5200 5199 5201 5203 3 0.65084 1.95253 0.00000
-2.60338 0.00000 0.00000 ; HA- CA- CB- CG2
5200 5199 5201 5207 3 0.65084 1.95253 0.00000
-2.60338 0.00000 0.00000 ; HA- CA- CB- CG1
5200 5199 5214 5215 3 3.68192 -4.35136 0.00000
1.33888 0.00000 0.00000 ; HA- CA- C- O
5200 5199 5214 5216 3 0.00000 0.00000 0.00000
0.00000 0.00000 0.00000 ; HA- CA- C- N
5201 5199 5214 5215 3 0.00000 0.00000 0.00000
0.00000 0.00000 0.00000 ; CB- CA- C- O
5201 5199 5214 5216 3 2.51040 4.18400 1.67360
-6.69440 0.00000 0.00000 ; CB- CA- C- N
5201 5207 5210 5211 3 0.66944 2.00832 0.00000
-2.67776 0.00000 0.00000 ; CB- CG1- CD1- HD11
5201 5207 5210 5212 3 0.66944 2.00832 0.00000
-2.67776 0.00000 0.00000 ; CB- CG1- CD1- HD12
5201 5207 5210 5213 3 0.66944 2.00832 0.00000
-2.67776 0.00000 0.00000 ; CB- CG1- CD1- HD13
5202 5201 5199 5214 3 0.65084 1.95253 0.00000
-2.60338 0.00000 0.00000 ; HB- CB- CA- C
5202 5201 5203 5204 3 0.62760 1.88280 0.00000
-2.51040 0.00000 0.00000 ; HB- CB- CG2- HG21
5202 5201 5203 5205 3 0.62760 1.88280 0.00000
-2.51040 0.00000 0.00000 ; HB- CB- CG2- HG22
5202 5201 5203 5206 3 0.62760 1.88280 0.00000
-2.51040 0.00000 0.00000 ; HB- CB- CG2- HG23
5202 5201 5207 5208 3 0.62760 1.88280 0.00000
-2.51040 0.00000 0.00000 ; HB- CB- CG1- HG12
5202 5201 5207 5209 3 0.62760 1.88280 0.00000
-2.51040 0.00000 0.00000 ; HB- CB- CG1- HG13
5202 5201 5207 5210 3 0.66944 2.00832 0.00000
-2.67776 0.00000 0.00000 ; HB- CB- CG1- CD1
5203 5201 5199 5214 3 5.67350 3.55640 -6.16722
-2.47693 3.74886 0.00000 ; CG2- CB- CA- C
5203 5201 5207 5208 3 0.66944 2.00832 0.00000
-2.67776 0.00000 0.00000 ; CG2- CB- CG1- HG12
5203 5201 5207 5209 3 0.66944 2.00832 0.00000
-2.67776 0.00000 0.00000 ; CG2- CB- CG1- HG13
5203 5201 5207 5210 3 3.81162 0.49790 -8.14206
-1.79075 7.49773 0.00000 ; CG2- CB- CG1- CD1
5204 5203 5201 5207 3 0.66944 2.00832 0.00000
-2.67776 0.00000 0.00000 ; HG21- CG2- CB- CG1
5205 5203 5201 5207 3 0.66944 2.00832 0.00000
-2.67776 0.00000 0.00000 ; HG22- CG2- CB- CG1

5206 5203 5201 5207 3 0.66944 2.00832 0.00000
-2.67776 0.00000 0.00000 ; HG23- CG2- CB- CG1
5207 5201 5199 5214 3 8.26340 0.74057 -9.99976
-1.89117 3.84928 0.00000 ; CG1- CB- CA- C
5208 5207 5210 5211 3 0.62760 1.88280 0.00000
-2.51040 0.00000 0.00000 ; HG12- CG1- CD1- HD11
5208 5207 5210 5212 3 0.62760 1.88280 0.00000
-2.51040 0.00000 0.00000 ; HG12- CG1- CD1- HD12
5208 5207 5210 5213 3 0.62760 1.88280 0.00000
-2.51040 0.00000 0.00000 ; HG12- CG1- CD1- HD13
5209 5207 5210 5211 3 0.62760 1.88280 0.00000
-2.51040 0.00000 0.00000 ; HG13- CG1- CD1- HD11
5209 5207 5210 5212 3 0.62760 1.88280 0.00000
-2.51040 0.00000 0.00000 ; HG13- CG1- CD1- HD12
5209 5207 5210 5213 3 0.62760 1.88280 0.00000
-2.51040 0.00000 0.00000 ; HG13- CG1- CD1- HD13
5214 5216 5218 5219 3 0.00000 0.00000 0.00000
0.00000 0.00000 0.00000 ; C- N- CA- HA
5214 5216 5218 5220 3 11.71520 1.67360 15.06240 -
13.38880 0.00000 0.00000 ; C- N- CA- CB
5214 5216 5218 5234 3 1.75728 5.27184 2.25936
-7.02912 0.00000 0.00000 ; C- N- CA- C
5215 5214 5216 5217 3 29.28800 -8.36800 -20.92000
0.00000 0.00000 0.00000 ; O- C- N- H
5215 5214 5216 5218 3 20.92000 0.00000 -20.92000
0.00000 0.00000 0.00000 ; O- C- N- CA
5216 5218 5220 5221 3 0.65084 1.95253 0.00000
-2.60338 0.00000 0.00000 ; N- CA- CB- HB2
5216 5218 5220 5222 3 0.65084 1.95253 0.00000
-2.60338 0.00000 0.00000 ; N- CA- CB- HB3
5216 5218 5220 5223 3 3.28026 2.46019 -2.19242
-3.21331 -0.23430 0.00000 ; N- CA- CB- CG
5216 5218 5234 5235 3 0.00000 0.00000 0.00000
0.00000 0.00000 0.00000 ; N- CA- C- O
5216 5218 5234 5236 3 17.40544 -5.02080 -13.22144
9.20480 0.00000 0.00000 ; N- CA- C- N
5217 5216 5218 5219 3 0.00000 0.00000 0.00000
0.00000 0.00000 0.00000 ; H- N- CA- HA
5217 5216 5218 5220 3 0.00000 0.00000 0.00000
0.00000 0.00000 0.00000 ; H- N- CA- CB
5217 5216 5218 5234 3 0.00000 0.00000 0.00000
0.00000 0.00000 0.00000 ; H- N- CA- C
5218 5220 5223 5224 3 0.57739 0.00000 1.02926
0.00000 -1.60666 0.00000 ; CA- CB- CG- CD1
5218 5220 5223 5232 3 0.57739 0.00000 1.02926
0.00000 -1.60666 0.00000 ; CA- CB- CG- CD2
5218 5234 5236 5237 3 20.92000 0.00000 -20.92000
0.00000 0.00000 0.00000 ; CA- C- N- H
5218 5234 5236 5238 3 20.92000 0.00000 -20.92000
0.00000 0.00000 0.00000 ; CA- C- N- CA
5219 5218 5220 5221 3 0.65084 1.95253 0.00000
-2.60338 0.00000 0.00000 ; HA- CA- CB- HB2

5219 5218 5220 5222 3 0.65084 1.95253 0.00000
-2.60338 0.00000 0.00000 ; HA- CA- CB- HB3
5219 5218 5220 5223 3 0.65084 1.95253 0.00000
-2.60338 0.00000 0.00000 ; HA- CA- CB- CG
5219 5218 5234 5235 3 3.68192 -4.35136 0.00000
1.33888 0.00000 0.00000 ; HA- CA- C- O
5219 5218 5234 5236 3 0.00000 0.00000 0.00000
0.00000 0.00000 0.00000 ; HA- CA- C- N
5220 5218 5234 5235 3 0.00000 0.00000 0.00000
0.00000 0.00000 0.00000 ; CB- CA- C- O
5220 5218 5234 5236 3 2.51040 4.18400 1.67360
-6.69440 0.00000 0.00000 ; CB- CA- C- N
5220 5223 5224 5225 3 30.33400 0.00000 -30.33400
0.00000 0.00000 0.00000 ; CB- CG- CD1- HD1
5220 5223 5224 5226 3 30.33400 0.00000 -30.33400
0.00000 0.00000 0.00000 ; CB- CG- CD1- CE1
5220 5223 5232 5230 3 30.33400 0.00000 -30.33400
0.00000 0.00000 0.00000 ; CB- CG- CD2- CE2
5220 5223 5232 5233 3 30.33400 0.00000 -30.33400
0.00000 0.00000 0.00000 ; CB- CG- CD2- HD2
5221 5220 5218 5234 3 0.65084 1.95253 0.00000
-2.60338 0.00000 0.00000 ; HB2- CB- CA- C
5221 5220 5223 5224 3 0.00000 0.00000 0.00000
0.00000 0.00000 0.00000 ; HB2- CB- CG- CD1
5221 5220 5223 5232 3 0.00000 0.00000 0.00000
0.00000 0.00000 0.00000 ; HB2- CB- CG- CD2
5222 5220 5218 5234 3 0.65084 1.95253 0.00000
-2.60338 0.00000 0.00000 ; HB3- CB- CA- C
5222 5220 5223 5224 3 0.00000 0.00000 0.00000
0.00000 0.00000 0.00000 ; HB3- CB- CG- CD1
5222 5220 5223 5232 3 0.00000 0.00000 0.00000
0.00000 0.00000 0.00000 ; HB3- CB- CG- CD2
5223 5220 5218 5234 3 4.95804 2.17986 -3.52293
-3.21331 -0.40166 0.00000 ; CG- CB- CA- C
5223 5224 5226 5227 3 30.33400 0.00000 -30.33400
0.00000 0.00000 0.00000 ; CG- CD1- CE1- HE1
5223 5224 5226 5228 3 30.33400 0.00000 -30.33400
0.00000 0.00000 0.00000 ; CG- CD1- CE1- CZ
5223 5232 5230 5228 3 30.33400 0.00000 -30.33400
0.00000 0.00000 0.00000 ; CG- CD2- CE2- CZ
5223 5232 5230 5231 3 30.33400 0.00000 -30.33400
0.00000 0.00000 0.00000 ; CG- CD2- CE2- HE2
5224 5223 5232 5230 3 30.33400 0.00000 -30.33400
0.00000 0.00000 0.00000 ; CD1- CG- CD2- CE2
5224 5223 5232 5233 3 30.33400 0.00000 -30.33400
0.00000 0.00000 0.00000 ; CD1- CG- CD2- HD2
5224 5226 5228 5229 3 30.33400 0.00000 -30.33400
0.00000 0.00000 0.00000 ; CD1- CE1- CZ- HZ
5224 5226 5228 5230 3 30.33400 0.00000 -30.33400
0.00000 0.00000 0.00000 ; CD1- CE1- CZ- CE2
5225 5224 5223 5232 3 30.33400 0.00000 -30.33400
0.00000 0.00000 0.00000 ; HD1- CD1- CG- CD2

5225 5224 5226 5227 3 30.33400 0.00000 -30.33400
0.00000 0.00000 0.00000 ; HD1- CD1- CE1- HE1
5225 5224 5226 5228 3 30.33400 0.00000 -30.33400
0.00000 0.00000 0.00000 ; HD1- CD1- CE1- CZ
5226 5224 5223 5232 3 30.33400 0.00000 -30.33400
0.00000 0.00000 0.00000 ; CE1- CD1- CG- CD2
5226 5228 5230 5231 3 30.33400 0.00000 -30.33400
0.00000 0.00000 0.00000 ; CE1- CZ- CE2- HE2
5226 5228 5230 5232 3 30.33400 0.00000 -30.33400
0.00000 0.00000 0.00000 ; CE1- CZ- CE2- CD2
5227 5226 5228 5229 3 30.33400 0.00000 -30.33400
0.00000 0.00000 0.00000 ; HE1- CE1- CZ- HZ
5227 5226 5228 5230 3 30.33400 0.00000 -30.33400
0.00000 0.00000 0.00000 ; HE1- CE1- CZ- CE2
5228 5230 5232 5233 3 30.33400 0.00000 -30.33400
0.00000 0.00000 0.00000 ; CZ- CE2- CD2- HD2
5229 5228 5230 5231 3 30.33400 0.00000 -30.33400
0.00000 0.00000 0.00000 ; HZ- CZ- CE2- HE2
5229 5228 5230 5232 3 30.33400 0.00000 -30.33400
0.00000 0.00000 0.00000 ; HZ- CZ- CE2- CD2
5231 5230 5232 5233 3 30.33400 0.00000 -30.33400
0.00000 0.00000 0.00000 ; HE2- CE2- CD2- HD2
5234 5236 5238 5239 3 0.00000 0.00000 0.00000
0.00000 0.00000 0.00000 ; C- N- CA- HA
5234 5236 5238 5240 3 11.71520 1.67360 15.06240 -
13.38880 0.00000 0.00000 ; C- N- CA- CB
5234 5236 5238 5253 3 1.75728 5.27184 2.25936
-7.02912 0.00000 0.00000 ; C- N- CA- C
5235 5234 5236 5237 3 29.28800 -8.36800 -20.92000
0.00000 0.00000 0.00000 ; O- C- N- H
5235 5234 5236 5238 3 20.92000 0.00000 -20.92000
0.00000 0.00000 0.00000 ; O- C- N- CA
5236 5238 5240 5241 3 0.65084 1.95253 0.00000
-2.60338 0.00000 0.00000 ; N- CA- CB- HB
5236 5238 5240 5242 3 3.83673 0.44769 -1.77402
-2.47693 -0.03347 0.00000 ; N- CA- CB- CG2
5236 5238 5240 5246 3 2.97482 0.12134 2.04179
-1.89117 -3.24678 0.00000 ; N- CA- CB- CG1
5236 5238 5253 5254 3 0.00000 0.00000 0.00000
0.00000 0.00000 0.00000 ; N- CA- C- O
5236 5238 5253 5255 3 17.40544 -5.02080 -13.22144
9.20480 0.00000 0.00000 ; N- CA- C- N
5237 5236 5238 5239 3 0.00000 0.00000 0.00000
0.00000 0.00000 0.00000 ; H- N- CA- HA
5237 5236 5238 5240 3 0.00000 0.00000 0.00000
0.00000 0.00000 0.00000 ; H- N- CA- CB
5237 5236 5238 5253 3 0.00000 0.00000 0.00000
0.00000 0.00000 0.00000 ; H- N- CA- C
5238 5240 5242 5243 3 0.66944 2.00832 0.00000
-2.67776 0.00000 0.00000 ; CA- CB- CG2- HG21
5238 5240 5242 5244 3 0.66944 2.00832 0.00000
-2.67776 0.00000 0.00000 ; CA- CB- CG2- HG22

```
  5238   5240   5242   5245      3    0.66944    2.00832    0.00000
-2.67776    0.00000    0.00000 ;     CA-    CB-   CG2-  HG23
  5238   5240   5246   5247      3    0.66944    2.00832    0.00000
-2.67776    0.00000    0.00000 ;     CA-    CB-   CG1-  HG12
  5238   5240   5246   5248      3    0.66944    2.00832    0.00000
-2.67776    0.00000    0.00000 ;     CA-    CB-   CG1-  HG13
  5238   5240   5246   5249      3    4.24258   -0.52718   -7.25506
-1.79075    7.69856    0.00000 ;     CA-    CB-   CG1-   CD1
  5238   5253   5255   5256      3   20.92000    0.00000  -20.92000
0.00000    0.00000    0.00000 ;     CA-     C-     N-     H
  5238   5253   5255   5257      3   20.92000    0.00000  -20.92000
0.00000    0.00000    0.00000 ;     CA-     C-     N-    CA
  5239   5238   5240   5241      3    0.65084    1.95253    0.00000
-2.60338    0.00000    0.00000 ;     HA-    CA-    CB-    HB
  5239   5238   5240   5242      3    0.65084    1.95253    0.00000
-2.60338    0.00000    0.00000 ;     HA-    CA-    CB-   CG2
  5239   5238   5240   5246      3    0.65084    1.95253    0.00000
-2.60338    0.00000    0.00000 ;     HA-    CA-    CB-   CG1
  5239   5238   5253   5254      3    3.68192   -4.35136    0.00000
1.33888    0.00000    0.00000 ;     HA-    CA-     C-     O
  5239   5238   5253   5255      3    0.00000    0.00000    0.00000
0.00000    0.00000    0.00000 ;     HA-    CA-     C-     N
  5240   5238   5253   5254      3    0.00000    0.00000    0.00000
0.00000    0.00000    0.00000 ;     CB-    CA-     C-     O
  5240   5238   5253   5255      3    2.51040    4.18400    1.67360
-6.69440    0.00000    0.00000 ;     CB-    CA-     C-     N
  5240   5246   5249   5250      3    0.66944    2.00832    0.00000
-2.67776    0.00000    0.00000 ;     CB-   CG1-   CD1-  HD11
  5240   5246   5249   5251      3    0.66944    2.00832    0.00000
-2.67776    0.00000    0.00000 ;     CB-   CG1-   CD1-  HD12
  5240   5246   5249   5252      3    0.66944    2.00832    0.00000
-2.67776    0.00000    0.00000 ;     CB-   CG1-   CD1-  HD13
  5241   5240   5238   5253      3    0.65084    1.95253    0.00000
-2.60338    0.00000    0.00000 ;     HB-    CB-    CA-     C
  5241   5240   5242   5243      3    0.62760    1.88280    0.00000
-2.51040    0.00000    0.00000 ;     HB-    CB-   CG2-  HG21
  5241   5240   5242   5244      3    0.62760    1.88280    0.00000
-2.51040    0.00000    0.00000 ;     HB-    CB-   CG2-  HG22
  5241   5240   5242   5245      3    0.62760    1.88280    0.00000
-2.51040    0.00000    0.00000 ;     HB-    CB-   CG2-  HG23
  5241   5240   5246   5247      3    0.62760    1.88280    0.00000
-2.51040    0.00000    0.00000 ;     HB-    CB-   CG1-  HG12
  5241   5240   5246   5248      3    0.62760    1.88280    0.00000
-2.51040    0.00000    0.00000 ;     HB-    CB-   CG1-  HG13
  5241   5240   5246   5249      3    0.66944    2.00832    0.00000
-2.67776    0.00000    0.00000 ;     HB-    CB-   CG1-   CD1
  5242   5240   5238   5253      3    5.67350    3.55640   -6.16722
-2.47693    3.74886    0.00000 ;    CG2-    CB-    CA-     C
  5242   5240   5246   5247      3    0.66944    2.00832    0.00000
-2.67776    0.00000    0.00000 ;    CG2-    CB-   CG1-  HG12
  5242   5240   5246   5248      3    0.66944    2.00832    0.00000
-2.67776    0.00000    0.00000 ;    CG2-    CB-   CG1-  HG13
```

5242 5240 5246 5249 3 3.81162 0.49790 -8.14206
-1.79075 7.49773 0.00000 ; CG2- CB- CG1- CD1
5243 5242 5240 5246 3 0.66944 2.00832 0.00000
-2.67776 0.00000 0.00000 ; HG21- CG2- CB- CG1
5244 5242 5240 5246 3 0.66944 2.00832 0.00000
-2.67776 0.00000 0.00000 ; HG22- CG2- CB- CG1
5245 5242 5240 5246 3 0.66944 2.00832 0.00000
-2.67776 0.00000 0.00000 ; HG23- CG2- CB- CG1
5246 5240 5238 5253 3 8.26340 0.74057 -9.99976
-1.89117 3.84928 0.00000 ; CG1- CB- CA- C
5247 5246 5249 5250 3 0.62760 1.88280 0.00000
-2.51040 0.00000 0.00000 ; HG12- CG1- CD1- HD11
5247 5246 5249 5251 3 0.62760 1.88280 0.00000
-2.51040 0.00000 0.00000 ; HG12- CG1- CD1- HD12
5247 5246 5249 5252 3 0.62760 1.88280 0.00000
-2.51040 0.00000 0.00000 ; HG12- CG1- CD1- HD13
5248 5246 5249 5250 3 0.62760 1.88280 0.00000
-2.51040 0.00000 0.00000 ; HG13- CG1- CD1- HD11
5248 5246 5249 5251 3 0.62760 1.88280 0.00000
-2.51040 0.00000 0.00000 ; HG13- CG1- CD1- HD12
5248 5246 5249 5252 3 0.62760 1.88280 0.00000
-2.51040 0.00000 0.00000 ; HG13- CG1- CD1- HD13
5253 5255 5257 5258 3 0.00000 0.00000 0.00000
0.00000 0.00000 0.00000 ; C- N- CA- HA
5253 5255 5257 5259 3 11.71520 1.67360 15.06240 -
13.38880 0.00000 0.00000 ; C- N- CA- CB
5253 5255 5257 5263 3 1.75728 5.27184 2.25936
-7.02912 0.00000 0.00000 ; C- N- CA- C
5254 5253 5255 5256 3 29.28800 -8.36800 -20.92000
0.00000 0.00000 0.00000 ; O- C- N- H
5254 5253 5255 5257 3 20.92000 0.00000 -20.92000
0.00000 0.00000 0.00000 ; O- C- N- CA
5255 5257 5259 5260 3 0.65084 1.95253 0.00000
-2.60338 0.00000 0.00000 ; N- CA- CB- HB1
5255 5257 5259 5261 3 0.65084 1.95253 0.00000
-2.60338 0.00000 0.00000 ; N- CA- CB- HB2
5255 5257 5259 5262 3 0.65084 1.95253 0.00000
-2.60338 0.00000 0.00000 ; N- CA- CB- HB3
5255 5257 5263 5264 3 0.00000 0.00000 0.00000
0.00000 0.00000 0.00000 ; N- CA- C- O
5255 5257 5263 5265 3 17.40544 -5.02080 -13.22144
9.20480 0.00000 0.00000 ; N- CA- C- N
5256 5255 5257 5258 3 0.00000 0.00000 0.00000
0.00000 0.00000 0.00000 ; H- N- CA- HA
5256 5255 5257 5259 3 0.00000 0.00000 0.00000
0.00000 0.00000 0.00000 ; H- N- CA- CB
5256 5255 5257 5263 3 0.00000 0.00000 0.00000
0.00000 0.00000 0.00000 ; H- N- CA- C
5257 5263 5265 5266 3 20.92000 0.00000 -20.92000
0.00000 0.00000 0.00000 ; CA- C- N- H
5257 5263 5265 5267 3 20.92000 0.00000 -20.92000
0.00000 0.00000 0.00000 ; CA- C- N- CA

5258 5257 5259 5260 3 0.65084 1.95253 0.00000
-2.60338 0.00000 0.00000 ; HA- CA- CB- HB1
5258 5257 5259 5261 3 0.65084 1.95253 0.00000
-2.60338 0.00000 0.00000 ; HA- CA- CB- HB2
5258 5257 5259 5262 3 0.65084 1.95253 0.00000
-2.60338 0.00000 0.00000 ; HA- CA- CB- HB3
5258 5257 5263 5264 3 3.68192 -4.35136 0.00000
1.33888 0.00000 0.00000 ; HA- CA- C- O
5258 5257 5263 5265 3 0.00000 0.00000 0.00000
0.00000 0.00000 0.00000 ; HA- CA- C- N
5259 5257 5263 5264 3 0.00000 0.00000 0.00000
0.00000 0.00000 0.00000 ; CB- CA- C- O
5259 5257 5263 5265 3 2.51040 4.18400 1.67360
-6.69440 0.00000 0.00000 ; CB- CA- C- N
5260 5259 5257 5263 3 0.65084 1.95253 0.00000
-2.60338 0.00000 0.00000 ; HB1- CB- CA- C
5261 5259 5257 5263 3 0.65084 1.95253 0.00000
-2.60338 0.00000 0.00000 ; HB2- CB- CA- C
5262 5259 5257 5263 3 0.65084 1.95253 0.00000
-2.60338 0.00000 0.00000 ; HB3- CB- CA- C
5263 5265 5267 5268 3 0.00000 0.00000 0.00000
0.00000 0.00000 0.00000 ; C- N- CA- HA
5263 5265 5267 5269 3 11.71520 1.67360 15.06240 -
13.38880 0.00000 0.00000 ; C- N- CA- CB
5263 5265 5267 5282 3 1.75728 5.27184 2.25936
-7.02912 0.00000 0.00000 ; C- N- CA- C
5264 5263 5265 5266 3 29.28800 -8.36800 -20.92000
0.00000 0.00000 0.00000 ; O- C- N- H
5264 5263 5265 5267 3 20.92000 0.00000 -20.92000
0.00000 0.00000 0.00000 ; O- C- N- CA
5265 5267 5269 5270 3 0.65084 1.95253 0.00000
-2.60338 0.00000 0.00000 ; N- CA- CB- HB2
5265 5267 5269 5271 3 0.65084 1.95253 0.00000
-2.60338 0.00000 0.00000 ; N- CA- CB- HB3
5265 5267 5269 5272 3 3.79070 1.39746 -4.61077
-2.40998 2.44346 0.00000 ; N- CA- CB- CG
5265 5267 5282 5283 3 0.00000 0.00000 0.00000
0.00000 0.00000 0.00000 ; N- CA- C- O
5265 5267 5282 5284 3 17.40544 -5.02080 -13.22144
9.20480 0.00000 0.00000 ; N- CA- C- N
5266 5265 5267 5268 3 0.00000 0.00000 0.00000
0.00000 0.00000 0.00000 ; H- N- CA- HA
5266 5265 5267 5269 3 0.00000 0.00000 0.00000
0.00000 0.00000 0.00000 ; H- N- CA- CB
5266 5265 5267 5282 3 0.00000 0.00000 0.00000
0.00000 0.00000 0.00000 ; H- N- CA- C
5267 5269 5272 5273 3 0.66944 2.00832 0.00000
-2.67776 0.00000 0.00000 ; CA- CB- CG- HG
5267 5269 5272 5274 3 3.90367 0.19665 -6.21742
-2.37651 5.99149 0.00000 ; CA- CB- CG- CD1
5267 5269 5272 5278 3 3.90367 0.19665 -6.21742
-2.37651 5.99149 0.00000 ; CA- CB- CG- CD2

5267 5282 5284 5285 3 20.92000 0.00000 -20.92000
0.00000 0.00000 0.00000 ; CA- C- N- H
5267 5282 5284 5286 3 20.92000 0.00000 -20.92000
0.00000 0.00000 0.00000 ; CA- C- N- CA
5268 5267 5269 5270 3 0.65084 1.95253 0.00000
-2.60338 0.00000 0.00000 ; HA- CA- CB- HB2
5268 5267 5269 5271 3 0.65084 1.95253 0.00000
-2.60338 0.00000 0.00000 ; HA- CA- CB- HB3
5268 5267 5269 5272 3 0.65084 1.95253 0.00000
-2.60338 0.00000 0.00000 ; HA- CA- CB- CG
5268 5267 5282 5283 3 3.68192 -4.35136 0.00000
1.33888 0.00000 0.00000 ; HA- CA- C- O
5268 5267 5282 5284 3 0.00000 0.00000 0.00000
0.00000 0.00000 0.00000 ; HA- CA- C- N
5269 5267 5282 5283 3 0.00000 0.00000 0.00000
0.00000 0.00000 0.00000 ; CB- CA- C- O
5269 5267 5282 5284 3 2.51040 4.18400 1.67360
-6.69440 0.00000 0.00000 ; CB- CA- C- N
5269 5272 5274 5275 3 0.66944 2.00832 0.00000
-2.67776 0.00000 0.00000 ; CB- CG- CD1- HD11
5269 5272 5274 5276 3 0.66944 2.00832 0.00000
-2.67776 0.00000 0.00000 ; CB- CG- CD1- HD12
5269 5272 5274 5277 3 0.66944 2.00832 0.00000
-2.67776 0.00000 0.00000 ; CB- CG- CD1- HD13
5269 5272 5278 5279 3 0.66944 2.00832 0.00000
-2.67776 0.00000 0.00000 ; CB- CG- CD2- HD21
5269 5272 5278 5280 3 0.66944 2.00832 0.00000
-2.67776 0.00000 0.00000 ; CB- CG- CD2- HD22
5269 5272 5278 5281 3 0.66944 2.00832 0.00000
-2.67776 0.00000 0.00000 ; CB- CG- CD2- HD23
5270 5269 5267 5282 3 0.65084 1.95253 0.00000
-2.60338 0.00000 0.00000 ; HB2- CB- CA- C
5270 5269 5272 5273 3 0.62760 1.88280 0.00000
-2.51040 0.00000 0.00000 ; HB2- CB- CG- HG
5270 5269 5272 5274 3 0.66944 2.00832 0.00000
-2.67776 0.00000 0.00000 ; HB2- CB- CG- CD1
5270 5269 5272 5278 3 0.66944 2.00832 0.00000
-2.67776 0.00000 0.00000 ; HB2- CB- CG- CD2
5271 5269 5267 5282 3 0.65084 1.95253 0.00000
-2.60338 0.00000 0.00000 ; HB3- CB- CA- C
5271 5269 5272 5273 3 0.62760 1.88280 0.00000
-2.51040 0.00000 0.00000 ; HB3- CB- CG- HG
5271 5269 5272 5274 3 0.66944 2.00832 0.00000
-2.67776 0.00000 0.00000 ; HB3- CB- CG- CD1
5271 5269 5272 5278 3 0.66944 2.00832 0.00000
-2.67776 0.00000 0.00000 ; HB3- CB- CG- CD2
5272 5269 5267 5282 3 10.33448 -1.14642 -11.54784
-2.40998 6.35968 0.00000 ; CG- CB- CA- C
5273 5272 5274 5275 3 0.62760 1.88280 0.00000
-2.51040 0.00000 0.00000 ; HG- CG- CD1- HD11
5273 5272 5274 5276 3 0.62760 1.88280 0.00000
-2.51040 0.00000 0.00000 ; HG- CG- CD1- HD12

5273 5272 5274 5277 3 0.62760 1.88280 0.00000
-2.51040 0.00000 0.00000 ; HG- CG- CD1- HD13
5273 5272 5278 5279 3 0.62760 1.88280 0.00000
-2.51040 0.00000 0.00000 ; HG- CG- CD2- HD21
5273 5272 5278 5280 3 0.62760 1.88280 0.00000
-2.51040 0.00000 0.00000 ; HG- CG- CD2- HD22
5273 5272 5278 5281 3 0.62760 1.88280 0.00000
-2.51040 0.00000 0.00000 ; HG- CG- CD2- HD23
5274 5272 5278 5279 3 0.66944 2.00832 0.00000
-2.67776 0.00000 0.00000 ; CD1- CG- CD2- HD21
5274 5272 5278 5280 3 0.66944 2.00832 0.00000
-2.67776 0.00000 0.00000 ; CD1- CG- CD2- HD22
5274 5272 5278 5281 3 0.66944 2.00832 0.00000
-2.67776 0.00000 0.00000 ; CD1- CG- CD2- HD23
5275 5274 5272 5278 3 0.66944 2.00832 0.00000
-2.67776 0.00000 0.00000 ; HD11- CD1- CG- CD2
5276 5274 5272 5278 3 0.66944 2.00832 0.00000
-2.67776 0.00000 0.00000 ; HD12- CD1- CG- CD2
5277 5274 5272 5278 3 0.66944 2.00832 0.00000
-2.67776 0.00000 0.00000 ; HD13- CD1- CG- CD2
5282 5284 5286 5287 3 0.00000 0.00000 0.00000
0.00000 0.00000 0.00000 ; C- N- CA- HA
5282 5284 5286 5288 3 11.71520 1.67360 15.06240 -
13.38880 0.00000 0.00000 ; C- N- CA- CB
5282 5284 5286 5294 3 1.75728 5.27184 2.25936
-7.02912 0.00000 0.00000 ; C- N- CA- C
5283 5282 5284 5285 3 29.28800 -8.36800 -20.92000
0.00000 0.00000 0.00000 ; O- C- N- H
5283 5282 5284 5286 3 20.92000 0.00000 -20.92000
0.00000 0.00000 0.00000 ; O- C- N- CA
5284 5286 5288 5289 3 0.65084 1.95253 0.00000
-2.60338 0.00000 0.00000 ; N- CA- CB- HB2
5284 5286 5288 5290 3 0.65084 1.95253 0.00000
-2.60338 0.00000 0.00000 ; N- CA- CB- HB3
5284 5286 5288 5291 3 15.41386 9.74035 -8.39310
-0.97069 2.97901 0.00000 ; N- CA- CB- CG
5284 5286 5294 5295 3 0.00000 0.00000 0.00000
0.00000 0.00000 0.00000 ; N- CA- C- O
5284 5286 5294 5296 3 17.40544 -5.02080 -13.22144
9.20480 0.00000 0.00000 ; N- CA- C- N
5285 5284 5286 5287 3 0.00000 0.00000 0.00000
0.00000 0.00000 0.00000 ; H- N- CA- HA
5285 5284 5286 5288 3 0.00000 0.00000 0.00000
0.00000 0.00000 0.00000 ; H- N- CA- CB
5285 5284 5286 5294 3 0.00000 0.00000 0.00000
0.00000 0.00000 0.00000 ; H- N- CA- C
5286 5288 5291 5292 3 6.43499 0.00000 -5.39736
0.00000 -1.03763 0.00000 ; CA- CB- CG- OD1
5286 5288 5291 5293 3 6.43499 0.00000 -5.39736
0.00000 -1.03763 0.00000 ; CA- CB- CG- OD2
5286 5294 5296 5297 3 20.92000 0.00000 -20.92000
0.00000 0.00000 0.00000 ; CA- C- N- H

5286 5294 5296 5298 3 20.92000 0.00000 -20.92000
0.00000 0.00000 0.00000 ; CA- C- N- CA
5287 5286 5288 5289 3 0.65084 1.95253 0.00000
-2.60338 0.00000 0.00000 ; HA- CA- CB- HB2
5287 5286 5288 5290 3 0.65084 1.95253 0.00000
-2.60338 0.00000 0.00000 ; HA- CA- CB- HB3
5287 5286 5288 5291 3 0.65084 1.95253 0.00000
-2.60338 0.00000 0.00000 ; HA- CA- CB- CG
5287 5286 5294 5295 3 3.68192 -4.35136 0.00000
1.33888 0.00000 0.00000 ; HA- CA- C- O
5287 5286 5294 5296 3 0.00000 0.00000 0.00000
0.00000 0.00000 0.00000 ; HA- CA- C- N
5288 5286 5294 5295 3 0.00000 0.00000 0.00000
0.00000 0.00000 0.00000 ; CB- CA- C- O
5288 5286 5294 5296 3 2.51040 4.18400 1.67360
-6.69440 0.00000 0.00000 ; CB- CA- C- N
5289 5288 5286 5294 3 0.65084 1.95253 0.00000
-2.60338 0.00000 0.00000 ; HB2- CB- CA- C
5289 5288 5291 5292 3 0.00000 0.00000 0.00000
0.00000 0.00000 0.00000 ; HB2- CB- CG- OD1
5289 5288 5291 5293 3 0.00000 0.00000 0.00000
0.00000 0.00000 0.00000 ; HB2- CB- CG- OD2
5290 5288 5286 5294 3 0.65084 1.95253 0.00000
-2.60338 0.00000 0.00000 ; HB3- CB- CA- C
5290 5288 5291 5292 3 0.00000 0.00000 0.00000
0.00000 0.00000 0.00000 ; HB3- CB- CG- OD1
5290 5288 5291 5293 3 0.00000 0.00000 0.00000
0.00000 0.00000 0.00000 ; HB3- CB- CG- OD2
5291 5288 5286 5294 3 7.14627 -1.04600 -8.99560
-0.97069 5.15469 0.00000 ; CG- CB- CA- C
5294 5296 5298 5299 3 0.00000 0.00000 0.00000
0.00000 0.00000 0.00000 ; C- N- CA- HA
5294 5296 5298 5300 3 11.71520 1.67360 15.06240 -
13.38880 0.00000 0.00000 ; C- N- CA- CB
5294 5296 5298 5310 3 1.75728 5.27184 2.25936
-7.02912 0.00000 0.00000 ; C- N- CA- C
5295 5294 5296 5297 3 29.28800 -8.36800 -20.92000
0.00000 0.00000 0.00000 ; O- C- N- H
5295 5294 5296 5298 3 20.92000 0.00000 -20.92000
0.00000 0.00000 0.00000 ; O- C- N- CA
5296 5298 5300 5301 3 0.65084 1.95253 0.00000
-2.60338 0.00000 0.00000 ; N- CA- CB- HB
5296 5298 5300 5302 3 3.83673 0.44769 -1.77402
-2.47693 -0.03347 0.00000 ; N- CA- CB- CG1
5296 5298 5300 5306 3 3.83673 0.44769 -1.77402
-2.47693 -0.03347 0.00000 ; N- CA- CB- CG2
5296 5298 5310 5311 3 0.00000 0.00000 0.00000
0.00000 0.00000 0.00000 ; N- CA- C- O
5296 5298 5310 5312 3 17.40544 -5.02080 -13.22144
9.20480 0.00000 0.00000 ; N- CA- C- N
5297 5296 5298 5299 3 0.00000 0.00000 0.00000
0.00000 0.00000 0.00000 ; H- N- CA- HA

5297 5296 5298 5300 3 0.00000 0.00000 0.00000
0.00000 0.00000 0.00000 ; H- N- CA- CB
5297 5296 5298 5310 3 0.00000 0.00000 0.00000
0.00000 0.00000 0.00000 ; H- N- CA- C
5298 5300 5302 5303 3 0.66944 2.00832 0.00000
-2.67776 0.00000 0.00000 ; CA- CB- CG1- HG11
5298 5300 5302 5304 3 0.66944 2.00832 0.00000
-2.67776 0.00000 0.00000 ; CA- CB- CG1- HG12
5298 5300 5302 5305 3 0.66944 2.00832 0.00000
-2.67776 0.00000 0.00000 ; CA- CB- CG1- HG13
5298 5300 5306 5307 3 0.66944 2.00832 0.00000
-2.67776 0.00000 0.00000 ; CA- CB- CG2- HG21
5298 5300 5306 5308 3 0.66944 2.00832 0.00000
-2.67776 0.00000 0.00000 ; CA- CB- CG2- HG22
5298 5300 5306 5309 3 0.66944 2.00832 0.00000
-2.67776 0.00000 0.00000 ; CA- CB- CG2- HG23
5298 5310 5312 5313 3 20.92000 0.00000 -20.92000
0.00000 0.00000 0.00000 ; CA- C- N- H
5298 5310 5312 5314 3 20.92000 0.00000 -20.92000
0.00000 0.00000 0.00000 ; CA- C- N- CA
5299 5298 5300 5301 3 0.65084 1.95253 0.00000
-2.60338 0.00000 0.00000 ; HA- CA- CB- HB
5299 5298 5300 5302 3 0.65084 1.95253 0.00000
-2.60338 0.00000 0.00000 ; HA- CA- CB- CG1
5299 5298 5300 5306 3 0.65084 1.95253 0.00000
-2.60338 0.00000 0.00000 ; HA- CA- CB- CG2
5299 5298 5310 5311 3 3.68192 -4.35136 0.00000
1.33888 0.00000 0.00000 ; HA- CA- C- O
5299 5298 5310 5312 3 0.00000 0.00000 0.00000
0.00000 0.00000 0.00000 ; HA- CA- C- N
5300 5298 5310 5311 3 0.00000 0.00000 0.00000
0.00000 0.00000 0.00000 ; CB- CA- C- O
5300 5298 5310 5312 3 2.51040 4.18400 1.67360
-6.69440 0.00000 0.00000 ; CB- CA- C- N
5301 5300 5298 5310 3 0.65084 1.95253 0.00000
-2.60338 0.00000 0.00000 ; HB- CB- CA- C
5301 5300 5302 5303 3 0.62760 1.88280 0.00000
-2.51040 0.00000 0.00000 ; HB- CB- CG1- HG11
5301 5300 5302 5304 3 0.62760 1.88280 0.00000
-2.51040 0.00000 0.00000 ; HB- CB- CG1- HG12
5301 5300 5302 5305 3 0.62760 1.88280 0.00000
-2.51040 0.00000 0.00000 ; HB- CB- CG1- HG13
5301 5300 5306 5307 3 0.62760 1.88280 0.00000
-2.51040 0.00000 0.00000 ; HB- CB- CG2- HG21
5301 5300 5306 5308 3 0.62760 1.88280 0.00000
-2.51040 0.00000 0.00000 ; HB- CB- CG2- HG22
5301 5300 5306 5309 3 0.62760 1.88280 0.00000
-2.51040 0.00000 0.00000 ; HB- CB- CG2- HG23
5302 5300 5298 5310 3 5.67350 3.55640 -6.16722
-2.47693 3.74886 0.00000 ; CG1- CB- CA- C
5302 5300 5306 5307 3 0.66944 2.00832 0.00000
-2.67776 0.00000 0.00000 ; CG1- CB- CG2- HG21

5302 5300 5306 5308 3 0.66944 2.00832 0.00000
-2.67776 0.00000 0.00000 ; CG1- CB- CG2- HG22
5302 5300 5306 5309 3 0.66944 2.00832 0.00000
-2.67776 0.00000 0.00000 ; CG1- CB- CG2- HG23
5303 5302 5300 5306 3 0.66944 2.00832 0.00000
-2.67776 0.00000 0.00000 ; HG11- CG1- CB- CG2
5304 5302 5300 5306 3 0.66944 2.00832 0.00000
-2.67776 0.00000 0.00000 ; HG12- CG1- CB- CG2
5305 5302 5300 5306 3 0.66944 2.00832 0.00000
-2.67776 0.00000 0.00000 ; HG13- CG1- CB- CG2
5306 5300 5298 5310 3 5.67350 3.55640 -6.16722
-2.47693 3.74886 0.00000 ; CG2- CB- CA- C
5310 5312 5314 5315 3 0.00000 0.00000 0.00000
0.00000 0.00000 0.00000 ; C- N- CA- HA
5310 5312 5314 5316 3 11.71520 1.67360 15.06240 -
13.38880 0.00000 0.00000 ; C- N- CA- CB
5310 5312 5314 5322 3 1.75728 5.27184 2.25936
-7.02912 0.00000 0.00000 ; C- N- CA- C
5311 5310 5312 5313 3 29.28800 -8.36800 -20.92000
0.00000 0.00000 0.00000 ; O- C- N- H
5311 5310 5312 5314 3 20.92000 0.00000 -20.92000
0.00000 0.00000 0.00000 ; O- C- N- CA
5312 5314 5316 5317 3 0.65084 1.95253 0.00000
-2.60338 0.00000 0.00000 ; N- CA- CB- HB2
5312 5314 5316 5318 3 0.65084 1.95253 0.00000
-2.60338 0.00000 0.00000 ; N- CA- CB- HB3
5312 5314 5316 5319 3 15.41386 9.74035 -8.39310
-0.97069 2.97901 0.00000 ; N- CA- CB- CG
5312 5314 5322 5323 3 0.00000 0.00000 0.00000
0.00000 0.00000 0.00000 ; N- CA- C- O
5312 5314 5322 5324 3 17.40544 -5.02080 -13.22144
9.20480 0.00000 0.00000 ; N- CA- C- N
5313 5312 5314 5315 3 0.00000 0.00000 0.00000
0.00000 0.00000 0.00000 ; H- N- CA- HA
5313 5312 5314 5316 3 0.00000 0.00000 0.00000
0.00000 0.00000 0.00000 ; H- N- CA- CB
5313 5312 5314 5322 3 0.00000 0.00000 0.00000
0.00000 0.00000 0.00000 ; H- N- CA- C
5314 5316 5319 5320 3 6.43499 0.00000 -5.39736
0.00000 -1.03763 0.00000 ; CA- CB- CG- OD1
5314 5316 5319 5321 3 6.43499 0.00000 -5.39736
0.00000 -1.03763 0.00000 ; CA- CB- CG- OD2
5314 5322 5324 5325 3 20.92000 0.00000 -20.92000
0.00000 0.00000 0.00000 ; CA- C- N- H
5314 5322 5324 5326 3 20.92000 0.00000 -20.92000
0.00000 0.00000 0.00000 ; CA- C- N- CA
5315 5314 5316 5317 3 0.65084 1.95253 0.00000
-2.60338 0.00000 0.00000 ; HA- CA- CB- HB2
5315 5314 5316 5318 3 0.65084 1.95253 0.00000
-2.60338 0.00000 0.00000 ; HA- CA- CB- HB3
5315 5314 5316 5319 3 0.65084 1.95253 0.00000
-2.60338 0.00000 0.00000 ; HA- CA- CB- CG

5315 5314 5322 5323 3 3.68192 -4.35136 0.00000
1.33888 0.00000 0.00000 ; HA- CA- C- O
5315 5314 5322 5324 3 0.00000 0.00000 0.00000
0.00000 0.00000 0.00000 ; HA- CA- C- N
5316 5314 5322 5323 3 0.00000 0.00000 0.00000
0.00000 0.00000 0.00000 ; CB- CA- C- O
5316 5314 5322 5324 3 2.51040 4.18400 1.67360
-6.69440 0.00000 0.00000 ; CB- CA- C- N
5317 5316 5314 5322 3 0.65084 1.95253 0.00000
-2.60338 0.00000 0.00000 ; HB2- CB- CA- C
5317 5316 5319 5320 3 0.00000 0.00000 0.00000
0.00000 0.00000 0.00000 ; HB2- CB- CG- OD1
5317 5316 5319 5321 3 0.00000 0.00000 0.00000
0.00000 0.00000 0.00000 ; HB2- CB- CG- OD2
5318 5316 5314 5322 3 0.65084 1.95253 0.00000
-2.60338 0.00000 0.00000 ; HB3- CB- CA- C
5318 5316 5319 5320 3 0.00000 0.00000 0.00000
0.00000 0.00000 0.00000 ; HB3- CB- CG- OD1
5318 5316 5319 5321 3 0.00000 0.00000 0.00000
0.00000 0.00000 0.00000 ; HB3- CB- CG- OD2
5319 5316 5314 5322 3 7.14627 -1.04600 -8.99560
-0.97069 5.15469 0.00000 ; CG- CB- CA- C
5322 5324 5326 5327 3 0.00000 0.00000 0.00000
0.00000 0.00000 0.00000 ; C- N- CA- HA
5322 5324 5326 5328 3 11.71520 1.67360 15.06240 -
13.38880 0.00000 0.00000 ; C- N- CA- CB
5322 5324 5326 5336 3 1.75728 5.27184 2.25936
-7.02912 0.00000 0.00000 ; C- N- CA- C
5323 5322 5324 5325 3 29.28800 -8.36800 -20.92000
0.00000 0.00000 0.00000 ; O- C- N- H
5323 5322 5324 5326 3 20.92000 0.00000 -20.92000
0.00000 0.00000 0.00000 ; O- C- N- CA
5324 5326 5328 5329 3 0.65084 1.95253 0.00000
-2.60338 0.00000 0.00000 ; N- CA- CB- HB2
5324 5326 5328 5330 3 0.65084 1.95253 0.00000
-2.60338 0.00000 0.00000 ; N- CA- CB- HB3
5324 5326 5328 5331 3 5.99567 3.29281 -4.46014
-0.55229 1.97485 0.00000 ; N- CA- CB- CG
5324 5326 5336 5337 3 0.00000 0.00000 0.00000
0.00000 0.00000 0.00000 ; N- CA- C- O
5324 5326 5336 5338 3 17.40544 -5.02080 -13.22144
9.20480 0.00000 0.00000 ; N- CA- C- N
5325 5324 5326 5327 3 0.00000 0.00000 0.00000
0.00000 0.00000 0.00000 ; H- N- CA- HA
5325 5324 5326 5328 3 0.00000 0.00000 0.00000
0.00000 0.00000 0.00000 ; H- N- CA- CB
5325 5324 5326 5336 3 0.00000 0.00000 0.00000
0.00000 0.00000 0.00000 ; H- N- CA- C
5326 5328 5331 5332 3 0.00000 0.00000 0.00000
0.00000 0.00000 0.00000 ; CA- CB- CG- OD1
5326 5328 5331 5333 3 8.84916 -0.31380 -4.32626
5.03754 0.26778 0.00000 ; CA- CB- CG- ND2

```
  5326   5336   5338   5339      3   20.92000    0.00000  -20.92000
0.00000    0.00000    0.00000 ;     CA-     C-     N-     H
  5326   5336   5338   5340      3   20.92000    0.00000  -20.92000
0.00000    0.00000    0.00000 ;     CA-     C-     N-    CA
  5327   5326   5328   5329      3    0.65084    1.95253    0.00000
-2.60338    0.00000    0.00000 ;     HA-    CA-    CB-   HB2
  5327   5326   5328   5330      3    0.65084    1.95253    0.00000
-2.60338    0.00000    0.00000 ;     HA-    CA-    CB-   HB3
  5327   5326   5328   5331      3    0.65084    1.95253    0.00000
-2.60338    0.00000    0.00000 ;     HA-    CA-    CB-    CG
  5327   5326   5336   5337      3    3.68192   -4.35136    0.00000
1.33888    0.00000    0.00000 ;     HA-    CA-     C-     O
  5327   5326   5336   5338      3    0.00000    0.00000    0.00000
0.00000    0.00000    0.00000 ;     HA-    CA-     C-     N
  5328   5326   5336   5337      3    0.00000    0.00000    0.00000
0.00000    0.00000    0.00000 ;     CB-    CA-     C-     O
  5328   5326   5336   5338      3    2.51040    4.18400    1.67360
-6.69440    0.00000    0.00000 ;     CB-    CA-     C-     N
  5328   5331   5333   5334      3   20.92000    0.00000  -20.92000
0.00000    0.00000    0.00000 ;     CB-    CG-   ND2-  HD21
  5328   5331   5333   5335      3   20.92000    0.00000  -20.92000
0.00000    0.00000    0.00000 ;     CB-    CG-   ND2-  HD22
  5329   5328   5326   5336      3    0.65084    1.95253    0.00000
-2.60338    0.00000    0.00000 ;    HB2-    CB-    CA-     C
  5329   5328   5331   5332      3    3.68192   -4.35136    0.00000
1.33888    0.00000    0.00000 ;    HB2-    CB-    CG-   OD1
  5329   5328   5331   5333      3    0.00000    0.00000    0.00000
0.00000    0.00000    0.00000 ;    HB2-    CB-    CG-   ND2
  5330   5328   5326   5336      3    0.65084    1.95253    0.00000
-2.60338    0.00000    0.00000 ;    HB3-    CB-    CA-     C
  5330   5328   5331   5332      3    3.68192   -4.35136    0.00000
1.33888    0.00000    0.00000 ;    HB3-    CB-    CG-   OD1
  5330   5328   5331   5333      3    0.00000    0.00000    0.00000
0.00000    0.00000    0.00000 ;    HB3-    CB-    CG-   ND2
  5331   5328   5326   5336      3    7.94542   -3.96225   -6.11701
-0.55229    3.58150    0.00000 ;     CG-    CB-    CA-     C
  5332   5331   5333   5334      3   29.28800   -8.36800  -20.92000
0.00000    0.00000    0.00000 ;    OD1-    CG-   ND2-  HD21
  5332   5331   5333   5335      3   29.28800   -8.36800  -20.92000
0.00000    0.00000    0.00000 ;    OD1-    CG-   ND2-  HD22
  5336   5338   5340   5341      3    0.00000    0.00000    0.00000
0.00000    0.00000    0.00000 ;      C-     N-    CA-    HA
  5336   5338   5340   5342      3   11.71520    1.67360   15.06240  -
13.38880    0.00000    0.00000 ;      C-     N-    CA-    CB
  5336   5338   5340   5347      3    1.75728    5.27184    2.25936
-7.02912    0.00000    0.00000 ;      C-     N-    CA-     C
  5337   5336   5338   5339      3   29.28800   -8.36800  -20.92000
0.00000    0.00000    0.00000 ;      O-     C-     N-     H
  5337   5336   5338   5340      3   20.92000    0.00000  -20.92000
0.00000    0.00000    0.00000 ;      O-     C-     N-    CA
  5338   5340   5342   5343      3    0.65084    1.95253    0.00000
-2.60338    0.00000    0.00000 ;      N-    CA-    CB-   HB2
```

5338 5340 5342 5344 3 0.65084 1.95253 0.00000
-2.60338 0.00000 0.00000 ; N- CA- CB- HB3
5338 5340 5342 5345 3 7.86174 2.24681 -7.41405
-6.71114 5.35552 0.00000 ; N- CA- CB- OG
5338 5340 5347 5348 3 0.00000 0.00000 0.00000
0.00000 0.00000 0.00000 ; N- CA- C- O
5338 5340 5347 5349 3 17.40544 -5.02080 -13.22144
9.20480 0.00000 0.00000 ; N- CA- C- N
5339 5338 5340 5341 3 0.00000 0.00000 0.00000
0.00000 0.00000 0.00000 ; H- N- CA- HA
5339 5338 5340 5342 3 0.00000 0.00000 0.00000
0.00000 0.00000 0.00000 ; H- N- CA- CB
5339 5338 5340 5347 3 0.00000 0.00000 0.00000
0.00000 0.00000 0.00000 ; H- N- CA- C
5340 5342 5345 5346 3 2.05853 2.46856 3.48109
-4.46851 0.23430 0.00000 ; CA- CB- OG- HG
5340 5347 5349 5350 3 20.92000 0.00000 -20.92000
0.00000 0.00000 0.00000 ; CA- C- N- H
5340 5347 5349 5351 3 20.92000 0.00000 -20.92000
0.00000 0.00000 0.00000 ; CA- C- N- CA
5341 5340 5342 5343 3 0.65084 1.95253 0.00000
-2.60338 0.00000 0.00000 ; HA- CA- CB- HB2
5341 5340 5342 5344 3 0.65084 1.95253 0.00000
-2.60338 0.00000 0.00000 ; HA- CA- CB- HB3
5341 5340 5342 5345 3 1.04600 -1.04600 0.00000
0.00000 0.00000 0.00000 ; HA- CA- CB- OG
5341 5340 5347 5348 3 3.68192 -4.35136 0.00000
1.33888 0.00000 0.00000 ; HA- CA- C- O
5341 5340 5347 5349 3 0.00000 0.00000 0.00000
0.00000 0.00000 0.00000 ; HA- CA- C- N
5342 5340 5347 5348 3 0.00000 0.00000 0.00000
0.00000 0.00000 0.00000 ; CB- CA- C- O
5342 5340 5347 5349 3 2.51040 4.18400 1.67360
-6.69440 0.00000 0.00000 ; CB- CA- C- N
5343 5342 5340 5347 3 0.65084 1.95253 0.00000
-2.60338 0.00000 0.00000 ; HB2- CB- CA- C
5343 5342 5345 5346 3 0.69733 2.09200 0.00000
-2.78933 0.00000 0.00000 ; HB2- CB- OG- HG
5344 5342 5340 5347 3 0.65084 1.95253 0.00000
-2.60338 0.00000 0.00000 ; HB3- CB- CA- C
5344 5342 5345 5346 3 0.69733 2.09200 0.00000
-2.78933 0.00000 0.00000 ; HB3- CB- OG- HG
5345 5342 5340 5347 3 7.34710 7.79898 -6.14211
-6.71114 4.31789 0.00000 ; OG- CB- CA- C
5347 5349 5351 5352 3 0.00000 0.00000 0.00000
0.00000 0.00000 0.00000 ; C- N- CA- HA2
5347 5349 5351 5353 3 0.00000 0.00000 0.00000
0.00000 0.00000 0.00000 ; C- N- CA- HA3
5347 5349 5351 5354 3 1.75728 5.27184 2.25936
-7.02912 0.00000 0.00000 ; C- N- CA- C
5348 5347 5349 5350 3 29.28800 -8.36800 -20.92000
0.00000 0.00000 0.00000 ; O- C- N- H

5348 5347 5349 5351 3 20.92000 0.00000 -20.92000
0.00000 0.00000 0.00000 ; O- C- N- CA
5349 5351 5354 5355 3 0.00000 0.00000 0.00000
0.00000 0.00000 0.00000 ; N- CA- C- O
5349 5351 5354 5356 3 17.40544 -5.02080 -13.22144
9.20480 0.00000 0.00000 ; N- CA- C- N
5350 5349 5351 5352 3 0.00000 0.00000 0.00000
0.00000 0.00000 0.00000 ; H- N- CA- HA2
5350 5349 5351 5353 3 0.00000 0.00000 0.00000
0.00000 0.00000 0.00000 ; H- N- CA- HA3
5350 5349 5351 5354 3 0.00000 0.00000 0.00000
0.00000 0.00000 0.00000 ; H- N- CA- C
5351 5354 5356 5357 3 20.92000 0.00000 -20.92000
0.00000 0.00000 0.00000 ; CA- C- N- H
5351 5354 5356 5358 3 20.92000 0.00000 -20.92000
0.00000 0.00000 0.00000 ; CA- C- N- CA
5352 5351 5354 5355 3 3.68192 -4.35136 0.00000
1.33888 0.00000 0.00000 ; HA2- CA- C- O
5352 5351 5354 5356 3 0.00000 0.00000 0.00000
0.00000 0.00000 0.00000 ; HA2- CA- C- N
5353 5351 5354 5355 3 3.68192 -4.35136 0.00000
1.33888 0.00000 0.00000 ; HA3- CA- C- O
5353 5351 5354 5356 3 0.00000 0.00000 0.00000
0.00000 0.00000 0.00000 ; HA3- CA- C- N
5354 5356 5358 5359 3 0.00000 0.00000 0.00000
0.00000 0.00000 0.00000 ; C- N- CA- HA
5354 5356 5358 5360 3 11.71520 1.67360 15.06240 -
13.38880 0.00000 0.00000 ; C- N- CA- CB
5354 5356 5358 5368 3 1.75728 5.27184 2.25936
-7.02912 0.00000 0.00000 ; C- N- CA- C
5355 5354 5356 5357 3 29.28800 -8.36800 -20.92000
0.00000 0.00000 0.00000 ; O- C- N- H
5355 5354 5356 5358 3 20.92000 0.00000 -20.92000
0.00000 0.00000 0.00000 ; O- C- N- CA
5356 5358 5360 5361 3 0.65084 1.95253 0.00000
-2.60338 0.00000 0.00000 ; N- CA- CB- HB
5356 5358 5360 5362 3 3.83673 0.44769 -1.77402
-2.47693 -0.03347 0.00000 ; N- CA- CB- CG2
5356 5358 5360 5366 3 4.93294 1.13386 -3.12963
-5.27184 3.17984 0.00000 ; N- CA- CB- OG1
5356 5358 5368 5369 3 0.00000 0.00000 0.00000
0.00000 0.00000 0.00000 ; N- CA- C- O
5356 5358 5368 5370 3 17.40544 -5.02080 -13.22144
9.20480 0.00000 0.00000 ; N- CA- C- N
5357 5356 5358 5359 3 0.00000 0.00000 0.00000
0.00000 0.00000 0.00000 ; H- N- CA- HA
5357 5356 5358 5360 3 0.00000 0.00000 0.00000
0.00000 0.00000 0.00000 ; H- N- CA- CB
5357 5356 5358 5368 3 0.00000 0.00000 0.00000
0.00000 0.00000 0.00000 ; H- N- CA- C
5358 5360 5362 5363 3 0.66944 2.00832 0.00000
-2.67776 0.00000 0.00000 ; CA- CB- CG2- HG21

5358 5360 5362 5364 3 0.66944 2.00832 0.00000
-2.67776 0.00000 0.00000 ; CA- CB- CG2- HG22
5358 5360 5362 5365 3 0.66944 2.00832 0.00000
-2.67776 0.00000 0.00000 ; CA- CB- CG2- HG23
5358 5360 5366 5367 3 1.12131 2.98738 1.66523
-3.94970 0.43514 0.00000 ; CA- CB- OG1- HG1
5358 5368 5370 5371 3 20.92000 0.00000 -20.92000
0.00000 0.00000 0.00000 ; CA- C- N- H
5358 5368 5370 5372 3 20.92000 0.00000 -20.92000
0.00000 0.00000 0.00000 ; CA- C- N- CA
5359 5358 5360 5361 3 0.65084 1.95253 0.00000
-2.60338 0.00000 0.00000 ; HA- CA- CB- HB
5359 5358 5360 5362 3 0.65084 1.95253 0.00000
-2.60338 0.00000 0.00000 ; HA- CA- CB- CG2
5359 5358 5360 5366 3 1.04600 -1.04600 0.00000
0.00000 0.00000 0.00000 ; HA- CA- CB- OG1
5359 5358 5368 5369 3 3.68192 -4.35136 0.00000
1.33888 0.00000 0.00000 ; HA- CA- C- O
5359 5358 5368 5370 3 0.00000 0.00000 0.00000
0.00000 0.00000 0.00000 ; HA- CA- C- N
5360 5358 5368 5369 3 0.00000 0.00000 0.00000
0.00000 0.00000 0.00000 ; CB- CA- C- O
5360 5358 5368 5370 3 2.51040 4.18400 1.67360
-6.69440 0.00000 0.00000 ; CB- CA- C- N
5361 5360 5358 5368 3 0.65084 1.95253 0.00000
-2.60338 0.00000 0.00000 ; HB- CB- CA- C
5361 5360 5362 5363 3 0.65084 1.95253 0.00000
-2.60338 0.00000 0.00000 ; HB- CB- CG2- HG21
5361 5360 5362 5364 3 0.65084 1.95253 0.00000
-2.60338 0.00000 0.00000 ; HB- CB- CG2- HG22
5361 5360 5362 5365 3 0.65084 1.95253 0.00000
-2.60338 0.00000 0.00000 ; HB- CB- CG2- HG23
5361 5360 5366 5367 3 0.69733 2.09200 0.00000
-2.78933 0.00000 0.00000 ; HB- CB- OG1- HG1
5362 5360 5358 5368 3 5.67350 3.55640 -6.16722
-2.47693 3.74886 0.00000 ; CG2- CB- CA- C
5362 5360 5366 5367 3 4.74047 0.27196 -2.26773
-3.94970 1.60666 0.00000 ; CG2- CB- OG1- HG1
5363 5362 5360 5366 3 1.04600 -1.04600 0.00000
0.00000 0.00000 0.00000 ; HG21- CG2- CB- OG1
5364 5362 5360 5366 3 1.04600 -1.04600 0.00000
0.00000 0.00000 0.00000 ; HG22- CG2- CB- OG1
5365 5362 5360 5366 3 1.04600 -1.04600 0.00000
0.00000 0.00000 0.00000 ; HG23- CG2- CB- OG1
5366 5360 5358 5368 3 6.53541 6.87013 -6.21742
-5.27184 5.22163 0.00000 ; OG1- CB- CA- C
5368 5370 5372 5373 3 0.00000 0.00000 0.00000
0.00000 0.00000 0.00000 ; C- N- CA- HA
5368 5370 5372 5374 3 11.71520 1.67360 15.06240 -
13.38880 0.00000 0.00000 ; C- N- CA- CB
5368 5370 5372 5387 3 1.75728 5.27184 2.25936
-7.02912 0.00000 0.00000 ; C- N- CA- C

5369 5368 5370 5371 3 29.28800 -8.36800 -20.92000
0.00000 0.00000 0.00000 ; O- C- N- H
5369 5368 5370 5372 3 20.92000 0.00000 -20.92000
0.00000 0.00000 0.00000 ; O- C- N- CA
5370 5372 5374 5375 3 0.65084 1.95253 0.00000
-2.60338 0.00000 0.00000 ; N- CA- CB- HB2
5370 5372 5374 5376 3 0.65084 1.95253 0.00000
-2.60338 0.00000 0.00000 ; N- CA- CB- HB3
5370 5372 5374 5377 3 3.79070 1.39746 -4.61077
-2.40998 2.44346 0.00000 ; N- CA- CB- CG
5370 5372 5387 5388 3 0.00000 0.00000 0.00000
0.00000 0.00000 0.00000 ; N- CA- C- O
5370 5372 5387 5389 3 17.40544 -5.02080 -13.22144
9.20480 0.00000 0.00000 ; N- CA- C- N
5371 5370 5372 5373 3 0.00000 0.00000 0.00000
0.00000 0.00000 0.00000 ; H- N- CA- HA
5371 5370 5372 5374 3 0.00000 0.00000 0.00000
0.00000 0.00000 0.00000 ; H- N- CA- CB
5371 5370 5372 5387 3 0.00000 0.00000 0.00000
0.00000 0.00000 0.00000 ; H- N- CA- C
5372 5374 5377 5378 3 0.66944 2.00832 0.00000
-2.67776 0.00000 0.00000 ; CA- CB- CG- HG
5372 5374 5377 5379 3 3.90367 0.19665 -6.21742
-2.37651 5.99149 0.00000 ; CA- CB- CG- CD1
5372 5374 5377 5383 3 3.90367 0.19665 -6.21742
-2.37651 5.99149 0.00000 ; CA- CB- CG- CD2
5372 5387 5389 5390 3 20.92000 0.00000 -20.92000
0.00000 0.00000 0.00000 ; CA- C- N- H
5372 5387 5389 5391 3 20.92000 0.00000 -20.92000
0.00000 0.00000 0.00000 ; CA- C- N- CA
5373 5372 5374 5375 3 0.65084 1.95253 0.00000
-2.60338 0.00000 0.00000 ; HA- CA- CB- HB2
5373 5372 5374 5376 3 0.65084 1.95253 0.00000
-2.60338 0.00000 0.00000 ; HA- CA- CB- HB3
5373 5372 5374 5377 3 0.65084 1.95253 0.00000
-2.60338 0.00000 0.00000 ; HA- CA- CB- CG
5373 5372 5387 5388 3 3.68192 -4.35136 0.00000
1.33888 0.00000 0.00000 ; HA- CA- C- O
5373 5372 5387 5389 3 0.00000 0.00000 0.00000
0.00000 0.00000 0.00000 ; HA- CA- C- N
5374 5372 5387 5388 3 0.00000 0.00000 0.00000
0.00000 0.00000 0.00000 ; CB- CA- C- O
5374 5372 5387 5389 3 2.51040 4.18400 1.67360
-6.69440 0.00000 0.00000 ; CB- CA- C- N
5374 5377 5379 5380 3 0.66944 2.00832 0.00000
-2.67776 0.00000 0.00000 ; CB- CG- CD1- HD11
5374 5377 5379 5381 3 0.66944 2.00832 0.00000
-2.67776 0.00000 0.00000 ; CB- CG- CD1- HD12
5374 5377 5379 5382 3 0.66944 2.00832 0.00000
-2.67776 0.00000 0.00000 ; CB- CG- CD1- HD13
5374 5377 5383 5384 3 0.66944 2.00832 0.00000
-2.67776 0.00000 0.00000 ; CB- CG- CD2- HD21

5374 5377 5383 5385 3 0.66944 2.00832 0.00000
-2.67776 0.00000 0.00000 ; CB- CG- CD2- HD22
5374 5377 5383 5386 3 0.66944 2.00832 0.00000
-2.67776 0.00000 0.00000 ; CB- CG- CD2- HD23
5375 5374 5372 5387 3 0.65084 1.95253 0.00000
-2.60338 0.00000 0.00000 ; HB2- CB- CA- C
5375 5374 5377 5378 3 0.62760 1.88280 0.00000
-2.51040 0.00000 0.00000 ; HB2- CB- CG- HG
5375 5374 5377 5379 3 0.66944 2.00832 0.00000
-2.67776 0.00000 0.00000 ; HB2- CB- CG- CD1
5375 5374 5377 5383 3 0.66944 2.00832 0.00000
-2.67776 0.00000 0.00000 ; HB2- CB- CG- CD2
5376 5374 5372 5387 3 0.65084 1.95253 0.00000
-2.60338 0.00000 0.00000 ; HB3- CB- CA- C
5376 5374 5377 5378 3 0.62760 1.88280 0.00000
-2.51040 0.00000 0.00000 ; HB3- CB- CG- HG
5376 5374 5377 5379 3 0.66944 2.00832 0.00000
-2.67776 0.00000 0.00000 ; HB3- CB- CG- CD1
5376 5374 5377 5383 3 0.66944 2.00832 0.00000
-2.67776 0.00000 0.00000 ; HB3- CB- CG- CD2
5377 5374 5372 5387 3 10.33448 -1.14642 -11.54784
-2.40998 6.35968 0.00000 ; CG- CB- CA- C
5378 5377 5379 5380 3 0.62760 1.88280 0.00000
-2.51040 0.00000 0.00000 ; HG- CG- CD1- HD11
5378 5377 5379 5381 3 0.62760 1.88280 0.00000
-2.51040 0.00000 0.00000 ; HG- CG- CD1- HD12
5378 5377 5379 5382 3 0.62760 1.88280 0.00000
-2.51040 0.00000 0.00000 ; HG- CG- CD1- HD13
5378 5377 5383 5384 3 0.62760 1.88280 0.00000
-2.51040 0.00000 0.00000 ; HG- CG- CD2- HD21
5378 5377 5383 5385 3 0.62760 1.88280 0.00000
-2.51040 0.00000 0.00000 ; HG- CG- CD2- HD22
5378 5377 5383 5386 3 0.62760 1.88280 0.00000
-2.51040 0.00000 0.00000 ; HG- CG- CD2- HD23
5379 5377 5383 5384 3 0.66944 2.00832 0.00000
-2.67776 0.00000 0.00000 ; CD1- CG- CD2- HD21
5379 5377 5383 5385 3 0.66944 2.00832 0.00000
-2.67776 0.00000 0.00000 ; CD1- CG- CD2- HD22
5379 5377 5383 5386 3 0.66944 2.00832 0.00000
-2.67776 0.00000 0.00000 ; CD1- CG- CD2- HD23
5380 5379 5377 5383 3 0.66944 2.00832 0.00000
-2.67776 0.00000 0.00000 ; HD11- CD1- CG- CD2
5381 5379 5377 5383 3 0.66944 2.00832 0.00000
-2.67776 0.00000 0.00000 ; HD12- CD1- CG- CD2
5382 5379 5377 5383 3 0.66944 2.00832 0.00000
-2.67776 0.00000 0.00000 ; HD13- CD1- CG- CD2
5387 5389 5391 5392 3 0.00000 0.00000 0.00000
0.00000 0.00000 0.00000 ; C- N- CA- HA
5387 5389 5391 5393 3 11.71520 1.67360 15.06240 -
13.38880 0.00000 0.00000 ; C- N- CA- CB
5387 5389 5391 5398 3 1.75728 5.27184 2.25936
-7.02912 0.00000 0.00000 ; C- N- CA- C

5388 5387 5389 5390 3 29.28800 -8.36800 -20.92000
0.00000 0.00000 0.00000 ; O- C- N- H
5388 5387 5389 5391 3 20.92000 0.00000 -20.92000
0.00000 0.00000 0.00000 ; O- C- N- CA
5389 5391 5393 5394 3 0.65084 1.95253 0.00000
-2.60338 0.00000 0.00000 ; N- CA- CB- HB2
5389 5391 5393 5395 3 0.65084 1.95253 0.00000
-2.60338 0.00000 0.00000 ; N- CA- CB- HB3
5389 5391 5393 5396 3 7.86174 2.24681 -7.41405
-6.71114 5.35552 0.00000 ; N- CA- CB- OG
5389 5391 5398 5399 3 0.00000 0.00000 0.00000
0.00000 0.00000 0.00000 ; N- CA- C- O
5389 5391 5398 5400 3 17.40544 -5.02080 -13.22144
9.20480 0.00000 0.00000 ; N- CA- C- N
5390 5389 5391 5392 3 0.00000 0.00000 0.00000
0.00000 0.00000 0.00000 ; H- N- CA- HA
5390 5389 5391 5393 3 0.00000 0.00000 0.00000
0.00000 0.00000 0.00000 ; H- N- CA- CB
5390 5389 5391 5398 3 0.00000 0.00000 0.00000
0.00000 0.00000 0.00000 ; H- N- CA- C
5391 5393 5396 5397 3 2.05853 2.46856 3.48109
-4.46851 0.23430 0.00000 ; CA- CB- OG- HG
5391 5398 5400 5401 3 20.92000 0.00000 -20.92000
0.00000 0.00000 0.00000 ; CA- C- N- H
5391 5398 5400 5402 3 20.92000 0.00000 -20.92000
0.00000 0.00000 0.00000 ; CA- C- N- CA
5392 5391 5393 5394 3 0.65084 1.95253 0.00000
-2.60338 0.00000 0.00000 ; HA- CA- CB- HB2
5392 5391 5393 5395 3 0.65084 1.95253 0.00000
-2.60338 0.00000 0.00000 ; HA- CA- CB- HB3
5392 5391 5393 5396 3 1.04600 -1.04600 0.00000
0.00000 0.00000 0.00000 ; HA- CA- CB- OG
5392 5391 5398 5399 3 3.68192 -4.35136 0.00000
1.33888 0.00000 0.00000 ; HA- CA- C- O
5392 5391 5398 5400 3 0.00000 0.00000 0.00000
0.00000 0.00000 0.00000 ; HA- CA- C- N
5393 5391 5398 5399 3 0.00000 0.00000 0.00000
0.00000 0.00000 0.00000 ; CB- CA- C- O
5393 5391 5398 5400 3 2.51040 4.18400 1.67360
-6.69440 0.00000 0.00000 ; CB- CA- C- N
5394 5393 5391 5398 3 0.65084 1.95253 0.00000
-2.60338 0.00000 0.00000 ; HB2- CB- CA- C
5394 5393 5396 5397 3 0.69733 2.09200 0.00000
-2.78933 0.00000 0.00000 ; HB2- CB- OG- HG
5395 5393 5391 5398 3 0.65084 1.95253 0.00000
-2.60338 0.00000 0.00000 ; HB3- CB- CA- C
5395 5393 5396 5397 3 0.69733 2.09200 0.00000
-2.78933 0.00000 0.00000 ; HB3- CB- OG- HG
5396 5393 5391 5398 3 7.34710 7.79898 -6.14211
-6.71114 4.31789 0.00000 ; OG- CB- CA- C
5398 5400 5402 5403 3 0.00000 0.00000 0.00000
0.00000 0.00000 0.00000 ; C- N- CA- HA

5398 5400 5402 5404 3 11.71520 1.67360 15.06240 -
13.38880 0.00000 0.00000 ; C- N- CA- CB
5398 5400 5402 5409 3 1.75728 5.27184 2.25936
-7.02912 0.00000 0.00000 ; C- N- CA- C
5399 5398 5400 5401 3 29.28800 -8.36800 -20.92000
0.00000 0.00000 0.00000 ; O- C- N- H
5399 5398 5400 5402 3 20.92000 0.00000 -20.92000
0.00000 0.00000 0.00000 ; O- C- N- CA
5400 5402 5404 5405 3 0.65084 1.95253 0.00000
-2.60338 0.00000 0.00000 ; N- CA- CB- HB2
5400 5402 5404 5406 3 0.65084 1.95253 0.00000
-2.60338 0.00000 0.00000 ; N- CA- CB- HB3
5400 5402 5404 5407 3 7.86174 2.24681 -7.41405
-6.71114 5.35552 0.00000 ; N- CA- CB- OG
5400 5402 5409 5410 3 0.00000 0.00000 0.00000
0.00000 0.00000 0.00000 ; N- CA- C- O
5400 5402 5409 5411 3 17.40544 -5.02080 -13.22144
9.20480 0.00000 0.00000 ; N- CA- C- N
5401 5400 5402 5403 3 0.00000 0.00000 0.00000
0.00000 0.00000 0.00000 ; H- N- CA- HA
5401 5400 5402 5404 3 0.00000 0.00000 0.00000
0.00000 0.00000 0.00000 ; H- N- CA- CB
5401 5400 5402 5409 3 0.00000 0.00000 0.00000
0.00000 0.00000 0.00000 ; H- N- CA- C
5402 5404 5407 5408 3 2.05853 2.46856 3.48109
-4.46851 0.23430 0.00000 ; CA- CB- OG- HG
5402 5409 5411 5412 3 20.92000 0.00000 -20.92000
0.00000 0.00000 0.00000 ; CA- C- N- H
5402 5409 5411 5413 3 20.92000 0.00000 -20.92000
0.00000 0.00000 0.00000 ; CA- C- N- CA
5403 5402 5404 5405 3 0.65084 1.95253 0.00000
-2.60338 0.00000 0.00000 ; HA- CA- CB- HB2
5403 5402 5404 5406 3 0.65084 1.95253 0.00000
-2.60338 0.00000 0.00000 ; HA- CA- CB- HB3
5403 5402 5404 5407 3 1.04600 -1.04600 0.00000
0.00000 0.00000 0.00000 ; HA- CA- CB- OG
5403 5402 5409 5410 3 3.68192 -4.35136 0.00000
1.33888 0.00000 0.00000 ; HA- CA- C- O
5403 5402 5409 5411 3 0.00000 0.00000 0.00000
0.00000 0.00000 0.00000 ; HA- CA- C- N
5404 5402 5409 5410 3 0.00000 0.00000 0.00000
0.00000 0.00000 0.00000 ; CB- CA- C- O
5404 5402 5409 5411 3 2.51040 4.18400 1.67360
-6.69440 0.00000 0.00000 ; CB- CA- C- N
5405 5404 5402 5409 3 0.65084 1.95253 0.00000
-2.60338 0.00000 0.00000 ; HB2- CB- CA- C
5405 5404 5407 5408 3 0.69733 2.09200 0.00000
-2.78933 0.00000 0.00000 ; HB2- CB- OG- HG
5406 5404 5402 5409 3 0.65084 1.95253 0.00000
-2.60338 0.00000 0.00000 ; HB3- CB- CA- C
5406 5404 5407 5408 3 0.69733 2.09200 0.00000
-2.78933 0.00000 0.00000 ; HB3- CB- OG- HG

5407 5404 5402 5409 3 7.34710 7.79898 -6.14211
-6.71114 4.31789 0.00000 ; OG- CB- CA- C
5409 5411 5413 5414 3 0.00000 0.00000 0.00000
0.00000 0.00000 0.00000 ; C- N- CA- HA
5409 5411 5413 5415 3 11.71520 1.67360 15.06240 -
13.38880 0.00000 0.00000 ; C- N- CA- CB
5409 5411 5413 5426 3 1.75728 5.27184 2.25936
-7.02912 0.00000 0.00000 ; C- N- CA- C
5410 5409 5411 5412 3 29.28800 -8.36800 -20.92000
0.00000 0.00000 0.00000 ; O- C- N- H
5410 5409 5411 5413 3 20.92000 0.00000 -20.92000
0.00000 0.00000 0.00000 ; O- C- N- CA
5411 5413 5415 5416 3 0.65084 1.95253 0.00000
-2.60338 0.00000 0.00000 ; N- CA- CB- HB2
5411 5413 5415 5417 3 0.65084 1.95253 0.00000
-2.60338 0.00000 0.00000 ; N- CA- CB- HB3
5411 5413 5415 5418 3 3.21331 2.22589 -4.15053
-2.40998 2.61082 0.00000 ; N- CA- CB- CG
5411 5413 5426 5427 3 0.00000 0.00000 0.00000
0.00000 0.00000 0.00000 ; N- CA- C- O
5411 5413 5426 5428 3 17.40544 -5.02080 -13.22144
9.20480 0.00000 0.00000 ; N- CA- C- N
5412 5411 5413 5414 3 0.00000 0.00000 0.00000
0.00000 0.00000 0.00000 ; H- N- CA- HA
5412 5411 5413 5415 3 0.00000 0.00000 0.00000
0.00000 0.00000 0.00000 ; H- N- CA- CB
5412 5411 5413 5426 3 0.00000 0.00000 0.00000
0.00000 0.00000 0.00000 ; H- N- CA- C
5413 5415 5418 5419 3 0.66944 2.00832 0.00000
-2.67776 0.00000 0.00000 ; CA- CB- CG- HG2
5413 5415 5418 5420 3 0.66944 2.00832 0.00000
-2.67776 0.00000 0.00000 ; CA- CB- CG- HG3
5413 5415 5418 5421 3 3.69866 -4.35136 -3.92459
6.89523 4.61914 0.00000 ; CA- CB- CG- CD
5413 5426 5428 5429 3 20.92000 0.00000 -20.92000
0.00000 0.00000 0.00000 ; CA- C- N- H
5413 5426 5428 5430 3 20.92000 0.00000 -20.92000
0.00000 0.00000 0.00000 ; CA- C- N- CA
5414 5413 5415 5416 3 0.65084 1.95253 0.00000
-2.60338 0.00000 0.00000 ; HA- CA- CB- HB2
5414 5413 5415 5417 3 0.65084 1.95253 0.00000
-2.60338 0.00000 0.00000 ; HA- CA- CB- HB3
5414 5413 5415 5418 3 0.65084 1.95253 0.00000
-2.60338 0.00000 0.00000 ; HA- CA- CB- CG
5414 5413 5426 5427 3 3.68192 -4.35136 0.00000
1.33888 0.00000 0.00000 ; HA- CA- C- O
5414 5413 5426 5428 3 0.00000 0.00000 0.00000
0.00000 0.00000 0.00000 ; HA- CA- C- N
5415 5413 5426 5427 3 0.00000 0.00000 0.00000
0.00000 0.00000 0.00000 ; CB- CA- C- O
5415 5413 5426 5428 3 2.51040 4.18400 1.67360
-6.69440 0.00000 0.00000 ; CB- CA- C- N

5415 5418 5421 5422 3 0.00000 0.00000 0.00000
0.00000 0.00000 0.00000 ; CB- CG- CD- OE1
5415 5418 5421 5423 3 10.32611 1.48114 -8.47678
1.42256 1.40582 0.00000 ; CB- CG- CD- NE2
5416 5415 5413 5426 3 0.65084 1.95253 0.00000
-2.60338 0.00000 0.00000 ; HB2- CB- CA- C
5416 5415 5418 5419 3 0.62760 1.88280 0.00000
-2.51040 0.00000 0.00000 ; HB2- CB- CG- HG2
5416 5415 5418 5420 3 0.62760 1.88280 0.00000
-2.51040 0.00000 0.00000 ; HB2- CB- CG- HG3
5416 5415 5418 5421 3 0.65084 1.95253 0.00000
-2.60338 0.00000 0.00000 ; HB2- CB- CG- CD
5417 5415 5413 5426 3 0.65084 1.95253 0.00000
-2.60338 0.00000 0.00000 ; HB3- CB- CA- C
5417 5415 5418 5419 3 0.62760 1.88280 0.00000
-2.51040 0.00000 0.00000 ; HB3- CB- CG- HG2
5417 5415 5418 5420 3 0.62760 1.88280 0.00000
-2.51040 0.00000 0.00000 ; HB3- CB- CG- HG3
5417 5415 5418 5421 3 0.65084 1.95253 0.00000
-2.60338 0.00000 0.00000 ; HB3- CB- CG- CD
5418 5415 5413 5426 3 6.86594 3.56895 -8.14206
-2.40998 4.85344 0.00000 ; CG- CB- CA- C
5418 5421 5423 5424 3 20.92000 0.00000 -20.92000
0.00000 0.00000 0.00000 ; CG- CD- NE2- HE21
5418 5421 5423 5425 3 20.92000 0.00000 -20.92000
0.00000 0.00000 0.00000 ; CG- CD- NE2- HE22
5419 5418 5421 5422 3 3.68192 -4.35136 0.00000
1.33888 0.00000 0.00000 ; HG2- CG- CD- OE1
5419 5418 5421 5423 3 0.00000 0.00000 0.00000
0.00000 0.00000 0.00000 ; HG2- CG- CD- NE2
5420 5418 5421 5422 3 3.68192 -4.35136 0.00000
1.33888 0.00000 0.00000 ; HG3- CG- CD- OE1
5420 5418 5421 5423 3 0.00000 0.00000 0.00000
0.00000 0.00000 0.00000 ; HG3- CG- CD- NE2
5422 5421 5423 5424 3 29.28800 -8.36800 -20.92000
0.00000 0.00000 0.00000 ; OE1- CD- NE2- HE21
5422 5421 5423 5425 3 29.28800 -8.36800 -20.92000
0.00000 0.00000 0.00000 ; OE1- CD- NE2- HE22
5426 5428 5430 5431 3 0.00000 0.00000 0.00000
0.00000 0.00000 0.00000 ; C- N- CA- HA
5426 5428 5430 5432 3 11.71520 1.67360 15.06240 -
13.38880 0.00000 0.00000 ; C- N- CA- CB
5426 5428 5430 5442 3 1.75728 5.27184 2.25936
-7.02912 0.00000 0.00000 ; C- N- CA- C
5427 5426 5428 5429 3 29.28800 -8.36800 -20.92000
0.00000 0.00000 0.00000 ; O- C- N- H
5427 5426 5428 5430 3 20.92000 0.00000 -20.92000
0.00000 0.00000 0.00000 ; O- C- N- CA
5428 5430 5432 5433 3 0.65084 1.95253 0.00000
-2.60338 0.00000 0.00000 ; N- CA- CB- HB2
5428 5430 5432 5434 3 0.65084 1.95253 0.00000
-2.60338 0.00000 0.00000 ; N- CA- CB- HB3

5428 5430 5432 5435 3 3.21331 2.22589 -4.15053
-2.40998 2.61082 0.00000 ; N- CA- CB- CG
5428 5430 5442 5443 3 0.00000 0.00000 0.00000
0.00000 0.00000 0.00000 ; N- CA- C- O
5428 5430 5442 5444 3 17.40544 -5.02080 -13.22144
9.20480 0.00000 0.00000 ; N- CA- C- N
5429 5428 5430 5431 3 0.00000 0.00000 0.00000
0.00000 0.00000 0.00000 ; H- N- CA- HA
5429 5428 5430 5432 3 0.00000 0.00000 0.00000
0.00000 0.00000 0.00000 ; H- N- CA- CB
5429 5428 5430 5442 3 0.00000 0.00000 0.00000
0.00000 0.00000 0.00000 ; H- N- CA- C
5430 5432 5435 5436 3 0.66944 2.00832 0.00000
-2.67776 0.00000 0.00000 ; CA- CB- CG- HG2
5430 5432 5435 5437 3 0.66944 2.00832 0.00000
-2.67776 0.00000 0.00000 ; CA- CB- CG- HG3
5430 5432 5435 5438 3 3.69866 -4.35136 -3.92459
6.89523 4.61914 0.00000 ; CA- CB- CG- CD
5430 5442 5444 5445 3 20.92000 0.00000 -20.92000
0.00000 0.00000 0.00000 ; CA- C- N- H
5430 5442 5444 5446 3 20.92000 0.00000 -20.92000
0.00000 0.00000 0.00000 ; CA- C- N- CA
5431 5430 5432 5433 3 0.65084 1.95253 0.00000
-2.60338 0.00000 0.00000 ; HA- CA- CB- HB2
5431 5430 5432 5434 3 0.65084 1.95253 0.00000
-2.60338 0.00000 0.00000 ; HA- CA- CB- HB3
5431 5430 5432 5435 3 0.65084 1.95253 0.00000
-2.60338 0.00000 0.00000 ; HA- CA- CB- CG
5431 5430 5442 5443 3 3.68192 -4.35136 0.00000
1.33888 0.00000 0.00000 ; HA- CA- C- O
5431 5430 5442 5444 3 0.00000 0.00000 0.00000
0.00000 0.00000 0.00000 ; HA- CA- C- N
5432 5430 5442 5443 3 0.00000 0.00000 0.00000
0.00000 0.00000 0.00000 ; CB- CA- C- O
5432 5430 5442 5444 3 2.51040 4.18400 1.67360
-6.69440 0.00000 0.00000 ; CB- CA- C- N
5432 5435 5438 5439 3 0.00000 0.00000 0.00000
0.00000 0.00000 0.00000 ; CB- CG- CD- OE1
5432 5435 5438 5440 3 12.79049 3.13382 -7.02912
0.41840 -2.20915 0.00000 ; CB- CG- CD- OE2
5433 5432 5430 5442 3 0.65084 1.95253 0.00000
-2.60338 0.00000 0.00000 ; HB2- CB- CA- C
5433 5432 5435 5436 3 0.62760 1.88280 0.00000
-2.51040 0.00000 0.00000 ; HB2- CB- CG- HG2
5433 5432 5435 5437 3 0.62760 1.88280 0.00000
-2.51040 0.00000 0.00000 ; HB2- CB- CG- HG3
5433 5432 5435 5438 3 0.65084 1.95253 0.00000
-2.60338 0.00000 0.00000 ; HB2- CB- CG- CD
5434 5432 5430 5442 3 0.65084 1.95253 0.00000
-2.60338 0.00000 0.00000 ; HB3- CB- CA- C
5434 5432 5435 5436 3 0.62760 1.88280 0.00000
-2.51040 0.00000 0.00000 ; HB3- CB- CG- HG2

5434 5432 5435 5437 3 0.62760 1.88280 0.00000
-2.51040 0.00000 0.00000 ; HB3- CB- CG- HG3
5434 5432 5435 5438 3 0.65084 1.95253 0.00000
-2.60338 0.00000 0.00000 ; HB3- CB- CG- CD
5435 5432 5430 5442 3 6.86594 3.56895 -8.14206
-2.40998 4.85344 0.00000 ; CG- CB- CA- C
5435 5438 5440 5441 3 27.46796 7.88684 -26.42614
-8.01654 3.78234 0.00000 ; CG- CD- OE2- HE2
5436 5435 5438 5439 3 3.68192 -4.35136 0.00000
1.33888 0.00000 0.00000 ; HG2- CG- CD- OE1
5436 5435 5438 5440 3 0.00000 0.00000 0.00000
0.00000 0.00000 0.00000 ; HG2- CG- CD- OE2
5437 5435 5438 5439 3 3.68192 -4.35136 0.00000
1.33888 0.00000 0.00000 ; HG3- CG- CD- OE1
5437 5435 5438 5440 3 0.00000 0.00000 0.00000
0.00000 0.00000 0.00000 ; HG3- CG- CD- OE2
5439 5438 5440 5441 3 27.19600 -7.94960 -19.24640
0.00000 0.00000 0.00000 ; OE1- CD- OE2- HE2
5442 5444 5446 5447 3 0.00000 0.00000 0.00000
0.00000 0.00000 0.00000 ; C- N- CA- HA
5442 5444 5446 5448 3 11.71520 1.67360 15.06240 -
13.38880 0.00000 0.00000 ; C- N- CA- CB
5442 5444 5446 5461 3 1.75728 5.27184 2.25936
-7.02912 0.00000 0.00000 ; C- N- CA- C
5443 5442 5444 5445 3 29.28800 -8.36800 -20.92000
0.00000 0.00000 0.00000 ; O- C- N- H
5443 5442 5444 5446 3 20.92000 0.00000 -20.92000
0.00000 0.00000 0.00000 ; O- C- N- CA
5444 5446 5448 5449 3 0.65084 1.95253 0.00000
-2.60338 0.00000 0.00000 ; N- CA- CB- HB
5444 5446 5448 5450 3 3.83673 0.44769 -1.77402
-2.47693 -0.03347 0.00000 ; N- CA- CB- CG2
5444 5446 5448 5454 3 2.97482 0.12134 2.04179
-1.89117 -3.24678 0.00000 ; N- CA- CB- CG1
5444 5446 5461 5462 3 0.00000 0.00000 0.00000
0.00000 0.00000 0.00000 ; N- CA- C- O
5444 5446 5461 5463 3 17.40544 -5.02080 -13.22144
9.20480 0.00000 0.00000 ; N- CA- C- N
5445 5444 5446 5447 3 0.00000 0.00000 0.00000
0.00000 0.00000 0.00000 ; H- N- CA- HA
5445 5444 5446 5448 3 0.00000 0.00000 0.00000
0.00000 0.00000 0.00000 ; H- N- CA- CB
5445 5444 5446 5461 3 0.00000 0.00000 0.00000
0.00000 0.00000 0.00000 ; H- N- CA- C
5446 5448 5450 5451 3 0.66944 2.00832 0.00000
-2.67776 0.00000 0.00000 ; CA- CB- CG2- HG21
5446 5448 5450 5452 3 0.66944 2.00832 0.00000
-2.67776 0.00000 0.00000 ; CA- CB- CG2- HG22
5446 5448 5450 5453 3 0.66944 2.00832 0.00000
-2.67776 0.00000 0.00000 ; CA- CB- CG2- HG23
5446 5448 5454 5455 3 0.66944 2.00832 0.00000
-2.67776 0.00000 0.00000 ; CA- CB- CG1- HG12

```
  5446   5448   5454   5456      3    0.66944    2.00832    0.00000
-2.67776    0.00000    0.00000 ;     CA-    CB-   CG1-  HG13
  5446   5448   5454   5457      3    4.24258   -0.52718   -7.25506
-1.79075    7.69856    0.00000 ;     CA-    CB-   CG1-   CD1
  5446   5461   5463   5464      3   20.92000    0.00000  -20.92000
0.00000    0.00000    0.00000 ;     CA-     C-     N-     H
  5446   5461   5463   5465      3   20.92000    0.00000  -20.92000
0.00000    0.00000    0.00000 ;     CA-     C-     N-    CA
  5447   5446   5448   5449      3    0.65084    1.95253    0.00000
-2.60338    0.00000    0.00000 ;     HA-    CA-    CB-    HB
  5447   5446   5448   5450      3    0.65084    1.95253    0.00000
-2.60338    0.00000    0.00000 ;     HA-    CA-    CB-   CG2
  5447   5446   5448   5454      3    0.65084    1.95253    0.00000
-2.60338    0.00000    0.00000 ;     HA-    CA-    CB-   CG1
  5447   5446   5461   5462      3    3.68192   -4.35136    0.00000
1.33888    0.00000    0.00000 ;     HA-    CA-     C-     O
  5447   5446   5461   5463      3    0.00000    0.00000    0.00000
0.00000    0.00000    0.00000 ;     HA-    CA-     C-     N
  5448   5446   5461   5462      3    0.00000    0.00000    0.00000
0.00000    0.00000    0.00000 ;     CB-    CA-     C-     O
  5448   5446   5461   5463      3    2.51040    4.18400    1.67360
-6.69440    0.00000    0.00000 ;     CB-    CA-     C-     N
  5448   5454   5457   5458      3    0.66944    2.00832    0.00000
-2.67776    0.00000    0.00000 ;     CB-   CG1-   CD1-  HD11
  5448   5454   5457   5459      3    0.66944    2.00832    0.00000
-2.67776    0.00000    0.00000 ;     CB-   CG1-   CD1-  HD12
  5448   5454   5457   5460      3    0.66944    2.00832    0.00000
-2.67776    0.00000    0.00000 ;     CB-   CG1-   CD1-  HD13
  5449   5448   5446   5461      3    0.65084    1.95253    0.00000
-2.60338    0.00000    0.00000 ;     HB-    CB-    CA-     C
  5449   5448   5450   5451      3    0.62760    1.88280    0.00000
-2.51040    0.00000    0.00000 ;     HB-    CB-   CG2-  HG21
  5449   5448   5450   5452      3    0.62760    1.88280    0.00000
-2.51040    0.00000    0.00000 ;     HB-    CB-   CG2-  HG22
  5449   5448   5450   5453      3    0.62760    1.88280    0.00000
-2.51040    0.00000    0.00000 ;     HB-    CB-   CG2-  HG23
  5449   5448   5454   5455      3    0.62760    1.88280    0.00000
-2.51040    0.00000    0.00000 ;     HB-    CB-   CG1-  HG12
  5449   5448   5454   5456      3    0.62760    1.88280    0.00000
-2.51040    0.00000    0.00000 ;     HB-    CB-   CG1-  HG13
  5449   5448   5454   5457      3    0.66944    2.00832    0.00000
-2.67776    0.00000    0.00000 ;     HB-    CB-   CG1-   CD1
  5450   5448   5446   5461      3    5.67350    3.55640   -6.16722
-2.47693    3.74886    0.00000 ;    CG2-    CB-    CA-     C
  5450   5448   5454   5455      3    0.66944    2.00832    0.00000
-2.67776    0.00000    0.00000 ;    CG2-    CB-   CG1-  HG12
  5450   5448   5454   5456      3    0.66944    2.00832    0.00000
-2.67776    0.00000    0.00000 ;    CG2-    CB-   CG1-  HG13
  5450   5448   5454   5457      3    3.81162    0.49790   -8.14206
-1.79075    7.49773    0.00000 ;    CG2-    CB-   CG1-   CD1
  5451   5450   5448   5454      3    0.66944    2.00832    0.00000
-2.67776    0.00000    0.00000 ;   HG21-   CG2-    CB-   CG1
```

5452 5450 5448 5454 3 0.66944 2.00832 0.00000
-2.67776 0.00000 0.00000 ; HG22- CG2- CB- CG1
5453 5450 5448 5454 3 0.66944 2.00832 0.00000
-2.67776 0.00000 0.00000 ; HG23- CG2- CB- CG1
5454 5448 5446 5461 3 8.26340 0.74057 -9.99976
-1.89117 3.84928 0.00000 ; CG1- CB- CA- C
5455 5454 5457 5458 3 0.62760 1.88280 0.00000
-2.51040 0.00000 0.00000 ; HG12- CG1- CD1- HD11
5455 5454 5457 5459 3 0.62760 1.88280 0.00000
-2.51040 0.00000 0.00000 ; HG12- CG1- CD1- HD12
5455 5454 5457 5460 3 0.62760 1.88280 0.00000
-2.51040 0.00000 0.00000 ; HG12- CG1- CD1- HD13
5456 5454 5457 5458 3 0.62760 1.88280 0.00000
-2.51040 0.00000 0.00000 ; HG13- CG1- CD1- HD11
5456 5454 5457 5459 3 0.62760 1.88280 0.00000
-2.51040 0.00000 0.00000 ; HG13- CG1- CD1- HD12
5456 5454 5457 5460 3 0.62760 1.88280 0.00000
-2.51040 0.00000 0.00000 ; HG13- CG1- CD1- HD13
5461 5463 5465 5466 3 0.00000 0.00000 0.00000
0.00000 0.00000 0.00000 ; C- N- CA- HA
5461 5463 5465 5467 3 11.71520 1.67360 15.06240 -
13.38880 0.00000 0.00000 ; C- N- CA- CB
5461 5463 5465 5480 3 1.75728 5.27184 2.25936
-7.02912 0.00000 0.00000 ; C- N- CA- C
5462 5461 5463 5464 3 29.28800 -8.36800 -20.92000
0.00000 0.00000 0.00000 ; O- C- N- H
5462 5461 5463 5465 3 20.92000 0.00000 -20.92000
0.00000 0.00000 0.00000 ; O- C- N- CA
5463 5465 5467 5468 3 0.65084 1.95253 0.00000
-2.60338 0.00000 0.00000 ; N- CA- CB- HB2
5463 5465 5467 5469 3 0.65084 1.95253 0.00000
-2.60338 0.00000 0.00000 ; N- CA- CB- HB3
5463 5465 5467 5470 3 3.79070 1.39746 -4.61077
-2.40998 2.44346 0.00000 ; N- CA- CB- CG
5463 5465 5480 5481 3 0.00000 0.00000 0.00000
0.00000 0.00000 0.00000 ; N- CA- C- O
5463 5465 5480 5482 3 17.40544 -5.02080 -13.22144
9.20480 0.00000 0.00000 ; N- CA- C- N
5464 5463 5465 5466 3 0.00000 0.00000 0.00000
0.00000 0.00000 0.00000 ; H- N- CA- HA
5464 5463 5465 5467 3 0.00000 0.00000 0.00000
0.00000 0.00000 0.00000 ; H- N- CA- CB
5464 5463 5465 5480 3 0.00000 0.00000 0.00000
0.00000 0.00000 0.00000 ; H- N- CA- C
5465 5467 5470 5471 3 0.66944 2.00832 0.00000
-2.67776 0.00000 0.00000 ; CA- CB- CG- HG
5465 5467 5470 5472 3 3.90367 0.19665 -6.21742
-2.37651 5.99149 0.00000 ; CA- CB- CG- CD1
5465 5467 5470 5476 3 3.90367 0.19665 -6.21742
-2.37651 5.99149 0.00000 ; CA- CB- CG- CD2
5465 5480 5482 5483 3 20.92000 0.00000 -20.92000
0.00000 0.00000 0.00000 ; CA- C- N- H

5465 5480 5482 5484 3 20.92000 0.00000 -20.92000
0.00000 0.00000 0.00000 ; CA- C- N- CA
5466 5465 5467 5468 3 0.65084 1.95253 0.00000
-2.60338 0.00000 0.00000 ; HA- CA- CB- HB2
5466 5465 5467 5469 3 0.65084 1.95253 0.00000
-2.60338 0.00000 0.00000 ; HA- CA- CB- HB3
5466 5465 5467 5470 3 0.65084 1.95253 0.00000
-2.60338 0.00000 0.00000 ; HA- CA- CB- CG
5466 5465 5480 5481 3 3.68192 -4.35136 0.00000
1.33888 0.00000 0.00000 ; HA- CA- C- O
5466 5465 5480 5482 3 0.00000 0.00000 0.00000
0.00000 0.00000 0.00000 ; HA- CA- C- N
5467 5465 5480 5481 3 0.00000 0.00000 0.00000
0.00000 0.00000 0.00000 ; CB- CA- C- O
5467 5465 5480 5482 3 2.51040 4.18400 1.67360
-6.69440 0.00000 0.00000 ; CB- CA- C- N
5467 5470 5472 5473 3 0.66944 2.00832 0.00000
-2.67776 0.00000 0.00000 ; CB- CG- CD1- HD11
5467 5470 5472 5474 3 0.66944 2.00832 0.00000
-2.67776 0.00000 0.00000 ; CB- CG- CD1- HD12
5467 5470 5472 5475 3 0.66944 2.00832 0.00000
-2.67776 0.00000 0.00000 ; CB- CG- CD1- HD13
5467 5470 5476 5477 3 0.66944 2.00832 0.00000
-2.67776 0.00000 0.00000 ; CB- CG- CD2- HD21
5467 5470 5476 5478 3 0.66944 2.00832 0.00000
-2.67776 0.00000 0.00000 ; CB- CG- CD2- HD22
5467 5470 5476 5479 3 0.66944 2.00832 0.00000
-2.67776 0.00000 0.00000 ; CB- CG- CD2- HD23
5468 5467 5465 5480 3 0.65084 1.95253 0.00000
-2.60338 0.00000 0.00000 ; HB2- CB- CA- C
5468 5467 5470 5471 3 0.62760 1.88280 0.00000
-2.51040 0.00000 0.00000 ; HB2- CB- CG- HG
5468 5467 5470 5472 3 0.66944 2.00832 0.00000
-2.67776 0.00000 0.00000 ; HB2- CB- CG- CD1
5468 5467 5470 5476 3 0.66944 2.00832 0.00000
-2.67776 0.00000 0.00000 ; HB2- CB- CG- CD2
5469 5467 5465 5480 3 0.65084 1.95253 0.00000
-2.60338 0.00000 0.00000 ; HB3- CB- CA- C
5469 5467 5470 5471 3 0.62760 1.88280 0.00000
-2.51040 0.00000 0.00000 ; HB3- CB- CG- HG
5469 5467 5470 5472 3 0.66944 2.00832 0.00000
-2.67776 0.00000 0.00000 ; HB3- CB- CG- CD1
5469 5467 5470 5476 3 0.66944 2.00832 0.00000
-2.67776 0.00000 0.00000 ; HB3- CB- CG- CD2
5470 5467 5465 5480 3 10.33448 -1.14642 -11.54784
-2.40998 6.35968 0.00000 ; CG- CB- CA- C
5471 5470 5472 5473 3 0.62760 1.88280 0.00000
-2.51040 0.00000 0.00000 ; HG- CG- CD1- HD11
5471 5470 5472 5474 3 0.62760 1.88280 0.00000
-2.51040 0.00000 0.00000 ; HG- CG- CD1- HD12
5471 5470 5472 5475 3 0.62760 1.88280 0.00000
-2.51040 0.00000 0.00000 ; HG- CG- CD1- HD13

5471 5470 5476 5477 3 0.62760 1.88280 0.00000
-2.51040 0.00000 0.00000 ; HG- CG- CD2- HD21
5471 5470 5476 5478 3 0.62760 1.88280 0.00000
-2.51040 0.00000 0.00000 ; HG- CG- CD2- HD22
5471 5470 5476 5479 3 0.62760 1.88280 0.00000
-2.51040 0.00000 0.00000 ; HG- CG- CD2- HD23
5472 5470 5476 5477 3 0.66944 2.00832 0.00000
-2.67776 0.00000 0.00000 ; CD1- CG- CD2- HD21
5472 5470 5476 5478 3 0.66944 2.00832 0.00000
-2.67776 0.00000 0.00000 ; CD1- CG- CD2- HD22
5472 5470 5476 5479 3 0.66944 2.00832 0.00000
-2.67776 0.00000 0.00000 ; CD1- CG- CD2- HD23
5473 5472 5470 5476 3 0.66944 2.00832 0.00000
-2.67776 0.00000 0.00000 ; HD11- CD1- CG- CD2
5474 5472 5470 5476 3 0.66944 2.00832 0.00000
-2.67776 0.00000 0.00000 ; HD12- CD1- CG- CD2
5475 5472 5470 5476 3 0.66944 2.00832 0.00000
-2.67776 0.00000 0.00000 ; HD13- CD1- CG- CD2
5480 5482 5484 5485 3 0.00000 0.00000 0.00000
0.00000 0.00000 0.00000 ; C- N- CA- HA
5480 5482 5484 5486 3 11.71520 1.67360 15.06240 -
13.38880 0.00000 0.00000 ; C- N- CA- CB
5480 5482 5484 5492 3 1.75728 5.27184 2.25936
-7.02912 0.00000 0.00000 ; C- N- CA- C
5481 5480 5482 5483 3 29.28800 -8.36800 -20.92000
0.00000 0.00000 0.00000 ; O- C- N- H
5481 5480 5482 5484 3 20.92000 0.00000 -20.92000
0.00000 0.00000 0.00000 ; O- C- N- CA
5482 5484 5486 5487 3 0.65084 1.95253 0.00000
-2.60338 0.00000 0.00000 ; N- CA- CB- HB2
5482 5484 5486 5488 3 0.65084 1.95253 0.00000
-2.60338 0.00000 0.00000 ; N- CA- CB- HB3
5482 5484 5486 5489 3 15.41386 9.74035 -8.39310
-0.97069 2.97901 0.00000 ; N- CA- CB- CG
5482 5484 5492 5493 3 0.00000 0.00000 0.00000
0.00000 0.00000 0.00000 ; N- CA- C- O
5482 5484 5492 5494 3 17.40544 -5.02080 -13.22144
9.20480 0.00000 0.00000 ; N- CA- C- N
5483 5482 5484 5485 3 0.00000 0.00000 0.00000
0.00000 0.00000 0.00000 ; H- N- CA- HA
5483 5482 5484 5486 3 0.00000 0.00000 0.00000
0.00000 0.00000 0.00000 ; H- N- CA- CB
5483 5482 5484 5492 3 0.00000 0.00000 0.00000
0.00000 0.00000 0.00000 ; H- N- CA- C
5484 5486 5489 5490 3 6.43499 0.00000 -5.39736
0.00000 -1.03763 0.00000 ; CA- CB- CG- OD1
5484 5486 5489 5491 3 6.43499 0.00000 -5.39736
0.00000 -1.03763 0.00000 ; CA- CB- CG- OD2
5484 5492 5494 5495 3 20.92000 0.00000 -20.92000
0.00000 0.00000 0.00000 ; CA- C- N- H
5484 5492 5494 5496 3 20.92000 0.00000 -20.92000
0.00000 0.00000 0.00000 ; CA- C- N- CA

5485 5484 5486 5487 3 0.65084 1.95253 0.00000
-2.60338 0.00000 0.00000 ; HA- CA- CB- HB2
5485 5484 5486 5488 3 0.65084 1.95253 0.00000
-2.60338 0.00000 0.00000 ; HA- CA- CB- HB3
5485 5484 5486 5489 3 0.65084 1.95253 0.00000
-2.60338 0.00000 0.00000 ; HA- CA- CB- CG
5485 5484 5492 5493 3 3.68192 -4.35136 0.00000
1.33888 0.00000 0.00000 ; HA- CA- C- O
5485 5484 5492 5494 3 0.00000 0.00000 0.00000
0.00000 0.00000 0.00000 ; HA- CA- C- N
5486 5484 5492 5493 3 0.00000 0.00000 0.00000
0.00000 0.00000 0.00000 ; CB- CA- C- O
5486 5484 5492 5494 3 2.51040 4.18400 1.67360
-6.69440 0.00000 0.00000 ; CB- CA- C- N
5487 5486 5484 5492 3 0.65084 1.95253 0.00000
-2.60338 0.00000 0.00000 ; HB2- CB- CA- C
5487 5486 5489 5490 3 0.00000 0.00000 0.00000
0.00000 0.00000 0.00000 ; HB2- CB- CG- OD1
5487 5486 5489 5491 3 0.00000 0.00000 0.00000
0.00000 0.00000 0.00000 ; HB2- CB- CG- OD2
5488 5486 5484 5492 3 0.65084 1.95253 0.00000
-2.60338 0.00000 0.00000 ; HB3- CB- CA- C
5488 5486 5489 5490 3 0.00000 0.00000 0.00000
0.00000 0.00000 0.00000 ; HB3- CB- CG- OD1
5488 5486 5489 5491 3 0.00000 0.00000 0.00000
0.00000 0.00000 0.00000 ; HB3- CB- CG- OD2
5489 5486 5484 5492 3 7.14627 -1.04600 -8.99560
-0.97069 5.15469 0.00000 ; CG- CB- CA- C
5492 5494 5496 5497 3 0.00000 0.00000 0.00000
0.00000 0.00000 0.00000 ; C- N- CA- HA2
5492 5494 5496 5498 3 0.00000 0.00000 0.00000
0.00000 0.00000 0.00000 ; C- N- CA- HA3
5492 5494 5496 5499 3 1.75728 5.27184 2.25936
-7.02912 0.00000 0.00000 ; C- N- CA- C
5493 5492 5494 5495 3 29.28800 -8.36800 -20.92000
0.00000 0.00000 0.00000 ; O- C- N- H
5493 5492 5494 5496 3 20.92000 0.00000 -20.92000
0.00000 0.00000 0.00000 ; O- C- N- CA
5494 5496 5499 5500 3 0.00000 0.00000 0.00000
0.00000 0.00000 0.00000 ; N- CA- C- O
5494 5496 5499 5501 3 17.40544 -5.02080 -13.22144
9.20480 0.00000 0.00000 ; N- CA- C- N
5495 5494 5496 5497 3 0.00000 0.00000 0.00000
0.00000 0.00000 0.00000 ; H- N- CA- HA2
5495 5494 5496 5498 3 0.00000 0.00000 0.00000
0.00000 0.00000 0.00000 ; H- N- CA- HA3
5495 5494 5496 5499 3 0.00000 0.00000 0.00000
0.00000 0.00000 0.00000 ; H- N- CA- C
5496 5499 5501 5502 3 20.92000 0.00000 -20.92000
0.00000 0.00000 0.00000 ; CA- C- N- H
5496 5499 5501 5503 3 20.92000 0.00000 -20.92000
0.00000 0.00000 0.00000 ; CA- C- N- CA

5497 5496 5499 5500 3 3.68192 -4.35136 0.00000
1.33888 0.00000 0.00000 ; HA2- CA- C- O
5497 5496 5499 5501 3 0.00000 0.00000 0.00000
0.00000 0.00000 0.00000 ; HA2- CA- C- N
5498 5496 5499 5500 3 3.68192 -4.35136 0.00000
1.33888 0.00000 0.00000 ; HA3- CA- C- O
5498 5496 5499 5501 3 0.00000 0.00000 0.00000
0.00000 0.00000 0.00000 ; HA3- CA- C- N
5499 5501 5503 5504 3 0.00000 0.00000 0.00000
0.00000 0.00000 0.00000 ; C- N- CA- HA
5499 5501 5503 5505 3 11.71520 1.67360 15.06240 -
13.38880 0.00000 0.00000 ; C- N- CA- CB
5499 5501 5503 5518 3 1.75728 5.27184 2.25936
-7.02912 0.00000 0.00000 ; C- N- CA- C
5500 5499 5501 5502 3 29.28800 -8.36800 -20.92000
0.00000 0.00000 0.00000 ; O- C- N- H
5500 5499 5501 5503 3 20.92000 0.00000 -20.92000
0.00000 0.00000 0.00000 ; O- C- N- CA
5501 5503 5505 5506 3 0.65084 1.95253 0.00000
-2.60338 0.00000 0.00000 ; N- CA- CB- HB2
5501 5503 5505 5507 3 0.65084 1.95253 0.00000
-2.60338 0.00000 0.00000 ; N- CA- CB- HB3
5501 5503 5505 5508 3 3.79070 1.39746 -4.61077
-2.40998 2.44346 0.00000 ; N- CA- CB- CG
5501 5503 5518 5519 3 0.00000 0.00000 0.00000
0.00000 0.00000 0.00000 ; N- CA- C- O
5501 5503 5518 5520 3 17.40544 -5.02080 -13.22144
9.20480 0.00000 0.00000 ; N- CA- C- N
5502 5501 5503 5504 3 0.00000 0.00000 0.00000
0.00000 0.00000 0.00000 ; H- N- CA- HA
5502 5501 5503 5505 3 0.00000 0.00000 0.00000
0.00000 0.00000 0.00000 ; H- N- CA- CB
5502 5501 5503 5518 3 0.00000 0.00000 0.00000
0.00000 0.00000 0.00000 ; H- N- CA- C
5503 5505 5508 5509 3 0.66944 2.00832 0.00000
-2.67776 0.00000 0.00000 ; CA- CB- CG- HG
5503 5505 5508 5510 3 3.90367 0.19665 -6.21742
-2.37651 5.99149 0.00000 ; CA- CB- CG- CD1
5503 5505 5508 5514 3 3.90367 0.19665 -6.21742
-2.37651 5.99149 0.00000 ; CA- CB- CG- CD2
5503 5518 5520 5521 3 20.92000 0.00000 -20.92000
0.00000 0.00000 0.00000 ; CA- C- N- H
5503 5518 5520 5522 3 20.92000 0.00000 -20.92000
0.00000 0.00000 0.00000 ; CA- C- N- CA
5504 5503 5505 5506 3 0.65084 1.95253 0.00000
-2.60338 0.00000 0.00000 ; HA- CA- CB- HB2
5504 5503 5505 5507 3 0.65084 1.95253 0.00000
-2.60338 0.00000 0.00000 ; HA- CA- CB- HB3
5504 5503 5505 5508 3 0.65084 1.95253 0.00000
-2.60338 0.00000 0.00000 ; HA- CA- CB- CG
5504 5503 5518 5519 3 3.68192 -4.35136 0.00000
1.33888 0.00000 0.00000 ; HA- CA- C- O

5504 5503 5518 5520 3 0.00000 0.00000 0.00000
0.00000 0.00000 0.00000 ; HA- CA- C- N
5505 5503 5518 5519 3 0.00000 0.00000 0.00000
0.00000 0.00000 0.00000 ; CB- CA- C- O
5505 5503 5518 5520 3 2.51040 4.18400 1.67360
-6.69440 0.00000 0.00000 ; CB- CA- C- N
5505 5508 5510 5511 3 0.66944 2.00832 0.00000
-2.67776 0.00000 0.00000 ; CB- CG- CD1- HD11
5505 5508 5510 5512 3 0.66944 2.00832 0.00000
-2.67776 0.00000 0.00000 ; CB- CG- CD1- HD12
5505 5508 5510 5513 3 0.66944 2.00832 0.00000
-2.67776 0.00000 0.00000 ; CB- CG- CD1- HD13
5505 5508 5514 5515 3 0.66944 2.00832 0.00000
-2.67776 0.00000 0.00000 ; CB- CG- CD2- HD21
5505 5508 5514 5516 3 0.66944 2.00832 0.00000
-2.67776 0.00000 0.00000 ; CB- CG- CD2- HD22
5505 5508 5514 5517 3 0.66944 2.00832 0.00000
-2.67776 0.00000 0.00000 ; CB- CG- CD2- HD23
5506 5505 5503 5518 3 0.65084 1.95253 0.00000
-2.60338 0.00000 0.00000 ; HB2- CB- CA- C
5506 5505 5508 5509 3 0.62760 1.88280 0.00000
-2.51040 0.00000 0.00000 ; HB2- CB- CG- HG
5506 5505 5508 5510 3 0.66944 2.00832 0.00000
-2.67776 0.00000 0.00000 ; HB2- CB- CG- CD1
5506 5505 5508 5514 3 0.66944 2.00832 0.00000
-2.67776 0.00000 0.00000 ; HB2- CB- CG- CD2
5507 5505 5503 5518 3 0.65084 1.95253 0.00000
-2.60338 0.00000 0.00000 ; HB3- CB- CA- C
5507 5505 5508 5509 3 0.62760 1.88280 0.00000
-2.51040 0.00000 0.00000 ; HB3- CB- CG- HG
5507 5505 5508 5510 3 0.66944 2.00832 0.00000
-2.67776 0.00000 0.00000 ; HB3- CB- CG- CD1
5507 5505 5508 5514 3 0.66944 2.00832 0.00000
-2.67776 0.00000 0.00000 ; HB3- CB- CG- CD2
5508 5505 5503 5518 3 10.33448 -1.14642 -11.54784
-2.40998 6.35968 0.00000 ; CG- CB- CA- C
5509 5508 5510 5511 3 0.62760 1.88280 0.00000
-2.51040 0.00000 0.00000 ; HG- CG- CD1- HD11
5509 5508 5510 5512 3 0.62760 1.88280 0.00000
-2.51040 0.00000 0.00000 ; HG- CG- CD1- HD12
5509 5508 5510 5513 3 0.62760 1.88280 0.00000
-2.51040 0.00000 0.00000 ; HG- CG- CD1- HD13
5509 5508 5514 5515 3 0.62760 1.88280 0.00000
-2.51040 0.00000 0.00000 ; HG- CG- CD2- HD21
5509 5508 5514 5516 3 0.62760 1.88280 0.00000
-2.51040 0.00000 0.00000 ; HG- CG- CD2- HD22
5509 5508 5514 5517 3 0.62760 1.88280 0.00000
-2.51040 0.00000 0.00000 ; HG- CG- CD2- HD23
5510 5508 5514 5515 3 0.66944 2.00832 0.00000
-2.67776 0.00000 0.00000 ; CD1- CG- CD2- HD21
5510 5508 5514 5516 3 0.66944 2.00832 0.00000
-2.67776 0.00000 0.00000 ; CD1- CG- CD2- HD22

5510 5508 5514 5517 3 0.66944 2.00832 0.00000
-2.67776 0.00000 0.00000 ; CD1- CG- CD2- HD23
5511 5510 5508 5514 3 0.66944 2.00832 0.00000
-2.67776 0.00000 0.00000 ; HD11- CD1- CG- CD2
5512 5510 5508 5514 3 0.66944 2.00832 0.00000
-2.67776 0.00000 0.00000 ; HD12- CD1- CG- CD2
5513 5510 5508 5514 3 0.66944 2.00832 0.00000
-2.67776 0.00000 0.00000 ; HD13- CD1- CG- CD2
5518 5520 5522 5523 3 0.00000 0.00000 0.00000
0.00000 0.00000 0.00000 ; C- N- CA- HA
5518 5520 5522 5524 3 11.71520 1.67360 15.06240 -
13.38880 0.00000 0.00000 ; C- N- CA- CB
5518 5520 5522 5540 3 1.75728 5.27184 2.25936
-7.02912 0.00000 0.00000 ; C- N- CA- C
5519 5518 5520 5521 3 29.28800 -8.36800 -20.92000
0.00000 0.00000 0.00000 ; O- C- N- H
5519 5518 5520 5522 3 20.92000 0.00000 -20.92000
0.00000 0.00000 0.00000 ; O- C- N- CA
5520 5522 5524 5525 3 0.65084 1.95253 0.00000
-2.60338 0.00000 0.00000 ; N- CA- CB- HB2
5520 5522 5524 5526 3 0.65084 1.95253 0.00000
-2.60338 0.00000 0.00000 ; N- CA- CB- HB3
5520 5522 5524 5527 3 0.65084 1.95253 0.00000
-2.60338 0.00000 0.00000 ; N- CA- CB- CG
5520 5522 5540 5541 3 0.00000 0.00000 0.00000
0.00000 0.00000 0.00000 ; N- CA- C- O
5520 5522 5540 5542 3 17.40544 -5.02080 -13.22144
9.20480 0.00000 0.00000 ; N- CA- C- N
5521 5520 5522 5523 3 0.00000 0.00000 0.00000
0.00000 0.00000 0.00000 ; H- N- CA- HA
5521 5520 5522 5524 3 0.00000 0.00000 0.00000
0.00000 0.00000 0.00000 ; H- N- CA- CB
5521 5520 5522 5540 3 0.00000 0.00000 0.00000
0.00000 0.00000 0.00000 ; H- N- CA- C
5522 5524 5527 5528 3 0.66944 2.00832 0.00000
-2.67776 0.00000 0.00000 ; CA- CB- CG- HG2
5522 5524 5527 5529 3 0.66944 2.00832 0.00000
-2.67776 0.00000 0.00000 ; CA- CB- CG- HG3
5522 5524 5527 5530 3 3.68192 3.09616 -2.09200
-3.01248 0.00000 0.00000 ; CA- CB- CG- CD
5522 5540 5542 5543 3 20.92000 0.00000 -20.92000
0.00000 0.00000 0.00000 ; CA- C- N- H
5522 5540 5542 5544 3 20.92000 0.00000 -20.92000
0.00000 0.00000 0.00000 ; CA- C- N- CA
5523 5522 5524 5525 3 0.65084 1.95253 0.00000
-2.60338 0.00000 0.00000 ; HA- CA- CB- HB2
5523 5522 5524 5526 3 0.65084 1.95253 0.00000
-2.60338 0.00000 0.00000 ; HA- CA- CB- HB3
5523 5522 5524 5527 3 0.65084 1.95253 0.00000
-2.60338 0.00000 0.00000 ; HA- CA- CB- CG
5523 5522 5540 5541 3 3.68192 -4.35136 0.00000
1.33888 0.00000 0.00000 ; HA- CA- C- O

```
  5523   5522   5540   5542      3    0.00000    0.00000    0.00000
0.00000    0.00000    0.00000 ;     HA-    CA-     C-     N
  5524   5522   5540   5541      3    0.00000    0.00000    0.00000
0.00000    0.00000    0.00000 ;     CB-    CA-     C-     O
  5524   5522   5540   5542      3    2.51040    4.18400    1.67360
-6.69440    0.00000    0.00000 ;     CB-    CA-     C-     N
  5524   5527   5530   5531      3    0.66944    2.00832    0.00000
-2.67776    0.00000    0.00000 ;     CB-    CG-    CD-   HD2
  5524   5527   5530   5532      3    0.66944    2.00832    0.00000
-2.67776    0.00000    0.00000 ;     CB-    CG-    CD-   HD3
  5524   5527   5530   5533      3    3.68192    3.09616   -2.09200
-3.01248    0.00000    0.00000 ;     CB-    CG-    CD-    CE
  5525   5524   5522   5540      3    0.65084    1.95253    0.00000
-2.60338    0.00000    0.00000 ;    HB2-    CB-    CA-     C
  5525   5524   5527   5528      3    0.62760    1.88280    0.00000
-2.51040    0.00000    0.00000 ;    HB2-    CB-    CG-   HG2
  5525   5524   5527   5529      3    0.62760    1.88280    0.00000
-2.51040    0.00000    0.00000 ;    HB2-    CB-    CG-   HG3
  5525   5524   5527   5530      3    0.66944    2.00832    0.00000
-2.67776    0.00000    0.00000 ;    HB2-    CB-    CG-    CD
  5526   5524   5522   5540      3    0.65084    1.95253    0.00000
-2.60338    0.00000    0.00000 ;    HB3-    CB-    CA-     C
  5526   5524   5527   5528      3    0.62760    1.88280    0.00000
-2.51040    0.00000    0.00000 ;    HB3-    CB-    CG-   HG2
  5526   5524   5527   5529      3    0.62760    1.88280    0.00000
-2.51040    0.00000    0.00000 ;    HB3-    CB-    CG-   HG3
  5526   5524   5527   5530      3    0.66944    2.00832    0.00000
-2.67776    0.00000    0.00000 ;    HB3-    CB-    CG-    CD
  5527   5524   5522   5540      3    0.65084    1.95253    0.00000
-2.60338    0.00000    0.00000 ;     CG-    CB-    CA-     C
  5527   5530   5533   5534      3    0.65084    1.95253    0.00000
-2.60338    0.00000    0.00000 ;     CG-    CD-    CE-   HE2
  5527   5530   5533   5535      3    0.65084    1.95253    0.00000
-2.60338    0.00000    0.00000 ;     CG-    CD-    CE-   HE3
  5527   5530   5533   5536      3    0.65084    1.95253    0.00000
-2.60338    0.00000    0.00000 ;     CG-    CD-    CE-    NZ
  5528   5527   5530   5531      3    0.62760    1.88280    0.00000
-2.51040    0.00000    0.00000 ;    HG2-    CG-    CD-   HD2
  5528   5527   5530   5532      3    0.62760    1.88280    0.00000
-2.51040    0.00000    0.00000 ;    HG2-    CG-    CD-   HD3
  5528   5527   5530   5533      3    0.66944    2.00832    0.00000
-2.67776    0.00000    0.00000 ;    HG2-    CG-    CD-    CE
  5529   5527   5530   5531      3    0.62760    1.88280    0.00000
-2.51040    0.00000    0.00000 ;    HG3-    CG-    CD-   HD2
  5529   5527   5530   5532      3    0.62760    1.88280    0.00000
-2.51040    0.00000    0.00000 ;    HG3-    CG-    CD-   HD3
  5529   5527   5530   5533      3    0.66944    2.00832    0.00000
-2.67776    0.00000    0.00000 ;    HG3-    CG-    CD-    CE
  5530   5533   5536   5537      3    0.65084    1.95253    0.00000
-2.60338    0.00000    0.00000 ;     CD-    CE-    NZ-   HZ1
  5530   5533   5536   5538      3    0.65084    1.95253    0.00000
-2.60338    0.00000    0.00000 ;     CD-    CE-    NZ-   HZ2
```

5530 5533 5536 5539 3 0.65084 1.95253 0.00000
-2.60338 0.00000 0.00000 ; CD- CE- NZ- HZ3
5531 5530 5533 5534 3 0.65084 1.95253 0.00000
-2.60338 0.00000 0.00000 ; HD2- CD- CE- HE2
5531 5530 5533 5535 3 0.65084 1.95253 0.00000
-2.60338 0.00000 0.00000 ; HD2- CD- CE- HE3
5531 5530 5533 5536 3 0.65084 1.95253 0.00000
-2.60338 0.00000 0.00000 ; HD2- CD- CE- NZ
5532 5530 5533 5534 3 0.65084 1.95253 0.00000
-2.60338 0.00000 0.00000 ; HD3- CD- CE- HE2
5532 5530 5533 5535 3 0.65084 1.95253 0.00000
-2.60338 0.00000 0.00000 ; HD3- CD- CE- HE3
5532 5530 5533 5536 3 0.65084 1.95253 0.00000
-2.60338 0.00000 0.00000 ; HD3- CD- CE- NZ
5534 5533 5536 5537 3 0.65084 1.95253 0.00000
-2.60338 0.00000 0.00000 ; HE2- CE- NZ- HZ1
5534 5533 5536 5538 3 0.65084 1.95253 0.00000
-2.60338 0.00000 0.00000 ; HE2- CE- NZ- HZ2
5534 5533 5536 5539 3 0.65084 1.95253 0.00000
-2.60338 0.00000 0.00000 ; HE2- CE- NZ- HZ3
5535 5533 5536 5537 3 0.65084 1.95253 0.00000
-2.60338 0.00000 0.00000 ; HE3- CE- NZ- HZ1
5535 5533 5536 5538 3 0.65084 1.95253 0.00000
-2.60338 0.00000 0.00000 ; HE3- CE- NZ- HZ2
5535 5533 5536 5539 3 0.65084 1.95253 0.00000
-2.60338 0.00000 0.00000 ; HE3- CE- NZ- HZ3
5540 5542 5544 5545 3 0.00000 0.00000 0.00000
0.00000 0.00000 0.00000 ; C- N- CA- HA
5540 5542 5544 5546 3 11.71520 1.67360 15.06240 -
13.38880 0.00000 0.00000 ; C- N- CA- CB
5540 5542 5544 5562 3 1.75728 5.27184 2.25936
-7.02912 0.00000 0.00000 ; C- N- CA- C
5541 5540 5542 5543 3 29.28800 -8.36800 -20.92000
0.00000 0.00000 0.00000 ; O- C- N- H
5541 5540 5542 5544 3 20.92000 0.00000 -20.92000
0.00000 0.00000 0.00000 ; O- C- N- CA
5542 5544 5546 5547 3 0.65084 1.95253 0.00000
-2.60338 0.00000 0.00000 ; N- CA- CB- HB2
5542 5544 5546 5548 3 0.65084 1.95253 0.00000
-2.60338 0.00000 0.00000 ; N- CA- CB- HB3
5542 5544 5546 5549 3 0.65084 1.95253 0.00000
-2.60338 0.00000 0.00000 ; N- CA- CB- CG
5542 5544 5562 5563 3 0.00000 0.00000 0.00000
0.00000 0.00000 0.00000 ; N- CA- C- O
5542 5544 5562 5564 3 17.40544 -5.02080 -13.22144
9.20480 0.00000 0.00000 ; N- CA- C- N
5543 5542 5544 5545 3 0.00000 0.00000 0.00000
0.00000 0.00000 0.00000 ; H- N- CA- HA
5543 5542 5544 5546 3 0.00000 0.00000 0.00000
0.00000 0.00000 0.00000 ; H- N- CA- CB
5543 5542 5544 5562 3 0.00000 0.00000 0.00000
0.00000 0.00000 0.00000 ; H- N- CA- C

5544 5546 5549 5550 3 0.66944 2.00832 0.00000
-2.67776 0.00000 0.00000 ; CA- CB- CG- HG2
5544 5546 5549 5551 3 0.66944 2.00832 0.00000
-2.67776 0.00000 0.00000 ; CA- CB- CG- HG3
5544 5546 5549 5552 3 3.68192 3.09616 -2.09200
-3.01248 0.00000 0.00000 ; CA- CB- CG- CD
5544 5562 5564 5565 3 20.92000 0.00000 -20.92000
0.00000 0.00000 0.00000 ; CA- C- N- H
5544 5562 5564 5566 3 20.92000 0.00000 -20.92000
0.00000 0.00000 0.00000 ; CA- C- N- CA
5545 5544 5546 5547 3 0.65084 1.95253 0.00000
-2.60338 0.00000 0.00000 ; HA- CA- CB- HB2
5545 5544 5546 5548 3 0.65084 1.95253 0.00000
-2.60338 0.00000 0.00000 ; HA- CA- CB- HB3
5545 5544 5546 5549 3 0.65084 1.95253 0.00000
-2.60338 0.00000 0.00000 ; HA- CA- CB- CG
5545 5544 5562 5563 3 3.68192 -4.35136 0.00000
1.33888 0.00000 0.00000 ; HA- CA- C- O
5545 5544 5562 5564 3 0.00000 0.00000 0.00000
0.00000 0.00000 0.00000 ; HA- CA- C- N
5546 5544 5562 5563 3 0.00000 0.00000 0.00000
0.00000 0.00000 0.00000 ; CB- CA- C- O
5546 5544 5562 5564 3 2.51040 4.18400 1.67360
-6.69440 0.00000 0.00000 ; CB- CA- C- N
5546 5549 5552 5553 3 0.66944 2.00832 0.00000
-2.67776 0.00000 0.00000 ; CB- CG- CD- HD2
5546 5549 5552 5554 3 0.66944 2.00832 0.00000
-2.67776 0.00000 0.00000 ; CB- CG- CD- HD3
5546 5549 5552 5555 3 3.68192 3.09616 -2.09200
-3.01248 0.00000 0.00000 ; CB- CG- CD- CE
5547 5546 5544 5562 3 0.65084 1.95253 0.00000
-2.60338 0.00000 0.00000 ; HB2- CB- CA- C
5547 5546 5549 5550 3 0.62760 1.88280 0.00000
-2.51040 0.00000 0.00000 ; HB2- CB- CG- HG2
5547 5546 5549 5551 3 0.62760 1.88280 0.00000
-2.51040 0.00000 0.00000 ; HB2- CB- CG- HG3
5547 5546 5549 5552 3 0.66944 2.00832 0.00000
-2.67776 0.00000 0.00000 ; HB2- CB- CG- CD
5548 5546 5544 5562 3 0.65084 1.95253 0.00000
-2.60338 0.00000 0.00000 ; HB3- CB- CA- C
5548 5546 5549 5550 3 0.62760 1.88280 0.00000
-2.51040 0.00000 0.00000 ; HB3- CB- CG- HG2
5548 5546 5549 5551 3 0.62760 1.88280 0.00000
-2.51040 0.00000 0.00000 ; HB3- CB- CG- HG3
5548 5546 5549 5552 3 0.66944 2.00832 0.00000
-2.67776 0.00000 0.00000 ; HB3- CB- CG- CD
5549 5546 5544 5562 3 0.65084 1.95253 0.00000
-2.60338 0.00000 0.00000 ; CG- CB- CA- C
5549 5552 5555 5556 3 0.65084 1.95253 0.00000
-2.60338 0.00000 0.00000 ; CG- CD- CE- HE2
5549 5552 5555 5557 3 0.65084 1.95253 0.00000
-2.60338 0.00000 0.00000 ; CG- CD- CE- HE3

5549 5552 5555 5558 3 0.65084 1.95253 0.00000
-2.60338 0.00000 0.00000 ; CG- CD- CE- NZ
5550 5549 5552 5553 3 0.62760 1.88280 0.00000
-2.51040 0.00000 0.00000 ; HG2- CG- CD- HD2
5550 5549 5552 5554 3 0.62760 1.88280 0.00000
-2.51040 0.00000 0.00000 ; HG2- CG- CD- HD3
5550 5549 5552 5555 3 0.66944 2.00832 0.00000
-2.67776 0.00000 0.00000 ; HG2- CG- CD- CE
5551 5549 5552 5553 3 0.62760 1.88280 0.00000
-2.51040 0.00000 0.00000 ; HG3- CG- CD- HD2
5551 5549 5552 5554 3 0.62760 1.88280 0.00000
-2.51040 0.00000 0.00000 ; HG3- CG- CD- HD3
5551 5549 5552 5555 3 0.66944 2.00832 0.00000
-2.67776 0.00000 0.00000 ; HG3- CG- CD- CE
5552 5555 5558 5559 3 0.65084 1.95253 0.00000
-2.60338 0.00000 0.00000 ; CD- CE- NZ- HZ1
5552 5555 5558 5560 3 0.65084 1.95253 0.00000
-2.60338 0.00000 0.00000 ; CD- CE- NZ- HZ2
5552 5555 5558 5561 3 0.65084 1.95253 0.00000
-2.60338 0.00000 0.00000 ; CD- CE- NZ- HZ3
5553 5552 5555 5556 3 0.65084 1.95253 0.00000
-2.60338 0.00000 0.00000 ; HD2- CD- CE- HE2
5553 5552 5555 5557 3 0.65084 1.95253 0.00000
-2.60338 0.00000 0.00000 ; HD2- CD- CE- HE3
5553 5552 5555 5558 3 0.65084 1.95253 0.00000
-2.60338 0.00000 0.00000 ; HD2- CD- CE- NZ
5554 5552 5555 5556 3 0.65084 1.95253 0.00000
-2.60338 0.00000 0.00000 ; HD3- CD- CE- HE2
5554 5552 5555 5557 3 0.65084 1.95253 0.00000
-2.60338 0.00000 0.00000 ; HD3- CD- CE- HE3
5554 5552 5555 5558 3 0.65084 1.95253 0.00000
-2.60338 0.00000 0.00000 ; HD3- CD- CE- NZ
5556 5555 5558 5559 3 0.65084 1.95253 0.00000
-2.60338 0.00000 0.00000 ; HE2- CE- NZ- HZ1
5556 5555 5558 5560 3 0.65084 1.95253 0.00000
-2.60338 0.00000 0.00000 ; HE2- CE- NZ- HZ2
5556 5555 5558 5561 3 0.65084 1.95253 0.00000
-2.60338 0.00000 0.00000 ; HE2- CE- NZ- HZ3
5557 5555 5558 5559 3 0.65084 1.95253 0.00000
-2.60338 0.00000 0.00000 ; HE3- CE- NZ- HZ1
5557 5555 5558 5560 3 0.65084 1.95253 0.00000
-2.60338 0.00000 0.00000 ; HE3- CE- NZ- HZ2
5557 5555 5558 5561 3 0.65084 1.95253 0.00000
-2.60338 0.00000 0.00000 ; HE3- CE- NZ- HZ3
5562 5564 5566 5567 3 0.00000 0.00000 0.00000
0.00000 0.00000 0.00000 ; C- N- CA- HA
5562 5564 5566 5568 3 11.71520 1.67360 15.06240 -
13.38880 0.00000 0.00000 ; C- N- CA- CB
5562 5564 5566 5581 3 1.75728 5.27184 2.25936
-7.02912 0.00000 0.00000 ; C- N- CA- C
5563 5562 5564 5565 3 29.28800 -8.36800 -20.92000
0.00000 0.00000 0.00000 ; O- C- N- H

5563 5562 5564 5566 3 20.92000 0.00000 -20.92000
0.00000 0.00000 0.00000 ; O- C- N- CA
5564 5566 5568 5569 3 0.65084 1.95253 0.00000
-2.60338 0.00000 0.00000 ; N- CA- CB- HB
5564 5566 5568 5570 3 3.83673 0.44769 -1.77402
-2.47693 -0.03347 0.00000 ; N- CA- CB- CG2
5564 5566 5568 5574 3 2.97482 0.12134 2.04179
-1.89117 -3.24678 0.00000 ; N- CA- CB- CG1
5564 5566 5581 5582 3 0.00000 0.00000 0.00000
0.00000 0.00000 0.00000 ; N- CA- C- O
5564 5566 5581 5583 3 17.40544 -5.02080 -13.22144
9.20480 0.00000 0.00000 ; N- CA- C- N
5565 5564 5566 5567 3 0.00000 0.00000 0.00000
0.00000 0.00000 0.00000 ; H- N- CA- HA
5565 5564 5566 5568 3 0.00000 0.00000 0.00000
0.00000 0.00000 0.00000 ; H- N- CA- CB
5565 5564 5566 5581 3 0.00000 0.00000 0.00000
0.00000 0.00000 0.00000 ; H- N- CA- C
5566 5568 5570 5571 3 0.66944 2.00832 0.00000
-2.67776 0.00000 0.00000 ; CA- CB- CG2- HG21
5566 5568 5570 5572 3 0.66944 2.00832 0.00000
-2.67776 0.00000 0.00000 ; CA- CB- CG2- HG22
5566 5568 5570 5573 3 0.66944 2.00832 0.00000
-2.67776 0.00000 0.00000 ; CA- CB- CG2- HG23
5566 5568 5574 5575 3 0.66944 2.00832 0.00000
-2.67776 0.00000 0.00000 ; CA- CB- CG1- HG12
5566 5568 5574 5576 3 0.66944 2.00832 0.00000
-2.67776 0.00000 0.00000 ; CA- CB- CG1- HG13
5566 5568 5574 5577 3 4.24258 -0.52718 -7.25506
-1.79075 7.69856 0.00000 ; CA- CB- CG1- CD1
5566 5581 5583 5584 3 20.92000 0.00000 -20.92000
0.00000 0.00000 0.00000 ; CA- C- N- H
5566 5581 5583 5585 3 20.92000 0.00000 -20.92000
0.00000 0.00000 0.00000 ; CA- C- N- CA
5567 5566 5568 5569 3 0.65084 1.95253 0.00000
-2.60338 0.00000 0.00000 ; HA- CA- CB- HB
5567 5566 5568 5570 3 0.65084 1.95253 0.00000
-2.60338 0.00000 0.00000 ; HA- CA- CB- CG2
5567 5566 5568 5574 3 0.65084 1.95253 0.00000
-2.60338 0.00000 0.00000 ; HA- CA- CB- CG1
5567 5566 5581 5582 3 3.68192 -4.35136 0.00000
1.33888 0.00000 0.00000 ; HA- CA- C- O
5567 5566 5581 5583 3 0.00000 0.00000 0.00000
0.00000 0.00000 0.00000 ; HA- CA- C- N
5568 5566 5581 5582 3 0.00000 0.00000 0.00000
0.00000 0.00000 0.00000 ; CB- CA- C- O
5568 5566 5581 5583 3 2.51040 4.18400 1.67360
-6.69440 0.00000 0.00000 ; CB- CA- C- N
5568 5574 5577 5578 3 0.66944 2.00832 0.00000
-2.67776 0.00000 0.00000 ; CB- CG1- CD1- HD11
5568 5574 5577 5579 3 0.66944 2.00832 0.00000
-2.67776 0.00000 0.00000 ; CB- CG1- CD1- HD12

5568 5574 5577 5580 3 0.66944 2.00832 0.00000
-2.67776 0.00000 0.00000 ; CB- CG1- CD1- HD13
5569 5568 5566 5581 3 0.65084 1.95253 0.00000
-2.60338 0.00000 0.00000 ; HB- CB- CA- C
5569 5568 5570 5571 3 0.62760 1.88280 0.00000
-2.51040 0.00000 0.00000 ; HB- CB- CG2- HG21
5569 5568 5570 5572 3 0.62760 1.88280 0.00000
-2.51040 0.00000 0.00000 ; HB- CB- CG2- HG22
5569 5568 5570 5573 3 0.62760 1.88280 0.00000
-2.51040 0.00000 0.00000 ; HB- CB- CG2- HG23
5569 5568 5574 5575 3 0.62760 1.88280 0.00000
-2.51040 0.00000 0.00000 ; HB- CB- CG1- HG12
5569 5568 5574 5576 3 0.62760 1.88280 0.00000
-2.51040 0.00000 0.00000 ; HB- CB- CG1- HG13
5569 5568 5574 5577 3 0.66944 2.00832 0.00000
-2.67776 0.00000 0.00000 ; HB- CB- CG1- CD1
5570 5568 5566 5581 3 5.67350 3.55640 -6.16722
-2.47693 3.74886 0.00000 ; CG2- CB- CA- C
5570 5568 5574 5575 3 0.66944 2.00832 0.00000
-2.67776 0.00000 0.00000 ; CG2- CB- CG1- HG12
5570 5568 5574 5576 3 0.66944 2.00832 0.00000
-2.67776 0.00000 0.00000 ; CG2- CB- CG1- HG13
5570 5568 5574 5577 3 3.81162 0.49790 -8.14206
-1.79075 7.49773 0.00000 ; CG2- CB- CG1- CD1
5571 5570 5568 5574 3 0.66944 2.00832 0.00000
-2.67776 0.00000 0.00000 ; HG21- CG2- CB- CG1
5572 5570 5568 5574 3 0.66944 2.00832 0.00000
-2.67776 0.00000 0.00000 ; HG22- CG2- CB- CG1
5573 5570 5568 5574 3 0.66944 2.00832 0.00000
-2.67776 0.00000 0.00000 ; HG23- CG2- CB- CG1
5574 5568 5566 5581 3 8.26340 0.74057 -9.99976
-1.89117 3.84928 0.00000 ; CG1- CB- CA- C
5575 5574 5577 5578 3 0.62760 1.88280 0.00000
-2.51040 0.00000 0.00000 ; HG12- CG1- CD1- HD11
5575 5574 5577 5579 3 0.62760 1.88280 0.00000
-2.51040 0.00000 0.00000 ; HG12- CG1- CD1- HD12
5575 5574 5577 5580 3 0.62760 1.88280 0.00000
-2.51040 0.00000 0.00000 ; HG12- CG1- CD1- HD13
5576 5574 5577 5578 3 0.62760 1.88280 0.00000
-2.51040 0.00000 0.00000 ; HG13- CG1- CD1- HD11
5576 5574 5577 5579 3 0.62760 1.88280 0.00000
-2.51040 0.00000 0.00000 ; HG13- CG1- CD1- HD12
5576 5574 5577 5580 3 0.62760 1.88280 0.00000
-2.51040 0.00000 0.00000 ; HG13- CG1- CD1- HD13
5581 5583 5585 5586 3 0.00000 0.00000 0.00000
0.00000 0.00000 0.00000 ; C- N- CA- HA2
5581 5583 5585 5587 3 0.00000 0.00000 0.00000
0.00000 0.00000 0.00000 ; C- N- CA- HA3
5581 5583 5585 5588 3 1.75728 5.27184 2.25936
-7.02912 0.00000 0.00000 ; C- N- CA- C
5582 5581 5583 5584 3 29.28800 -8.36800 -20.92000
0.00000 0.00000 0.00000 ; O- C- N- H

5582 5581 5583 5585 3 20.92000 0.00000 -20.92000
0.00000 0.00000 0.00000 ; O- C- N- CA
5583 5585 5588 5589 3 0.00000 0.00000 0.00000
0.00000 0.00000 0.00000 ; N- CA- C- O
5583 5585 5588 5590 3 17.40544 -5.02080 -13.22144
9.20480 0.00000 0.00000 ; N- CA- C- N
5584 5583 5585 5586 3 0.00000 0.00000 0.00000
0.00000 0.00000 0.00000 ; H- N- CA- HA2
5584 5583 5585 5587 3 0.00000 0.00000 0.00000
0.00000 0.00000 0.00000 ; H- N- CA- HA3
5584 5583 5585 5588 3 0.00000 0.00000 0.00000
0.00000 0.00000 0.00000 ; H- N- CA- C
5585 5588 5590 5591 3 20.92000 0.00000 -20.92000
0.00000 0.00000 0.00000 ; CA- C- N- H
5585 5588 5590 5592 3 20.92000 0.00000 -20.92000
0.00000 0.00000 0.00000 ; CA- C- N- CA
5586 5585 5588 5589 3 3.68192 -4.35136 0.00000
1.33888 0.00000 0.00000 ; HA2- CA- C- O
5586 5585 5588 5590 3 0.00000 0.00000 0.00000
0.00000 0.00000 0.00000 ; HA2- CA- C- N
5587 5585 5588 5589 3 3.68192 -4.35136 0.00000
1.33888 0.00000 0.00000 ; HA3- CA- C- O
5587 5585 5588 5590 3 0.00000 0.00000 0.00000
0.00000 0.00000 0.00000 ; HA3- CA- C- N
5588 5590 5592 5593 3 0.00000 0.00000 0.00000
0.00000 0.00000 0.00000 ; C- N- CA- HA
5588 5590 5592 5594 3 11.71520 1.67360 15.06240 -
13.38880 0.00000 0.00000 ; C- N- CA- CB
5588 5590 5592 5609 3 1.75728 5.27184 2.25936
-7.02912 0.00000 0.00000 ; C- N- CA- C
5589 5588 5590 5591 3 29.28800 -8.36800 -20.92000
0.00000 0.00000 0.00000 ; O- C- N- H
5589 5588 5590 5592 3 20.92000 0.00000 -20.92000
0.00000 0.00000 0.00000 ; O- C- N- CA
5590 5592 5594 5595 3 0.65084 1.95253 0.00000
-2.60338 0.00000 0.00000 ; N- CA- CB- HB2
5590 5592 5594 5596 3 0.65084 1.95253 0.00000
-2.60338 0.00000 0.00000 ; N- CA- CB- HB3
5590 5592 5594 5597 3 3.28026 2.46019 -2.19242
-3.21331 -0.23430 0.00000 ; N- CA- CB- CG
5590 5592 5609 5610 3 0.00000 0.00000 0.00000
0.00000 0.00000 0.00000 ; N- CA- C- O
5590 5592 5609 5611 3 17.40544 -5.02080 -13.22144
9.20480 0.00000 0.00000 ; N- CA- C- N
5591 5590 5592 5593 3 0.00000 0.00000 0.00000
0.00000 0.00000 0.00000 ; H- N- CA- HA
5591 5590 5592 5594 3 0.00000 0.00000 0.00000
0.00000 0.00000 0.00000 ; H- N- CA- CB
5591 5590 5592 5609 3 0.00000 0.00000 0.00000
0.00000 0.00000 0.00000 ; H- N- CA- C
5592 5594 5597 5598 3 0.57739 0.00000 1.02926
0.00000 -1.60666 0.00000 ; CA- CB- CG- CD1

5592 5594 5597 5607 3 0.57739 0.00000 1.02926
0.00000 -1.60666 0.00000 ; CA- CB- CG- CD2
5592 5609 5611 5612 3 20.92000 0.00000 -20.92000
0.00000 0.00000 0.00000 ; CA- C- N- H
5592 5609 5611 5613 3 20.92000 0.00000 -20.92000
0.00000 0.00000 0.00000 ; CA- C- N- CA
5593 5592 5594 5595 3 0.65084 1.95253 0.00000
-2.60338 0.00000 0.00000 ; HA- CA- CB- HB2
5593 5592 5594 5596 3 0.65084 1.95253 0.00000
-2.60338 0.00000 0.00000 ; HA- CA- CB- HB3
5593 5592 5594 5597 3 0.65084 1.95253 0.00000
-2.60338 0.00000 0.00000 ; HA- CA- CB- CG
5593 5592 5609 5610 3 3.68192 -4.35136 0.00000
1.33888 0.00000 0.00000 ; HA- CA- C- O
5593 5592 5609 5611 3 0.00000 0.00000 0.00000
0.00000 0.00000 0.00000 ; HA- CA- C- N
5594 5592 5609 5610 3 0.00000 0.00000 0.00000
0.00000 0.00000 0.00000 ; CB- CA- C- O
5594 5592 5609 5611 3 2.51040 4.18400 1.67360
-6.69440 0.00000 0.00000 ; CB- CA- C- N
5594 5597 5598 5599 3 30.33400 0.00000 -30.33400
0.00000 0.00000 0.00000 ; CB- CG- CD1- HD1
5594 5597 5598 5600 3 30.33400 0.00000 -30.33400
0.00000 0.00000 0.00000 ; CB- CG- CD1- CE1
5594 5597 5607 5605 3 30.33400 0.00000 -30.33400
0.00000 0.00000 0.00000 ; CB- CG- CD2- CE2
5594 5597 5607 5608 3 30.33400 0.00000 -30.33400
0.00000 0.00000 0.00000 ; CB- CG- CD2- HD2
5595 5594 5592 5609 3 0.65084 1.95253 0.00000
-2.60338 0.00000 0.00000 ; HB2- CB- CA- C
5595 5594 5597 5598 3 0.00000 0.00000 0.00000
0.00000 0.00000 0.00000 ; HB2- CB- CG- CD1
5595 5594 5597 5607 3 0.00000 0.00000 0.00000
0.00000 0.00000 0.00000 ; HB2- CB- CG- CD2
5596 5594 5592 5609 3 0.65084 1.95253 0.00000
-2.60338 0.00000 0.00000 ; HB3- CB- CA- C
5596 5594 5597 5598 3 0.00000 0.00000 0.00000
0.00000 0.00000 0.00000 ; HB3- CB- CG- CD1
5596 5594 5597 5607 3 0.00000 0.00000 0.00000
0.00000 0.00000 0.00000 ; HB3- CB- CG- CD2
5597 5594 5592 5609 3 4.95804 2.17986 -3.52293
-3.21331 -0.40166 0.00000 ; CG- CB- CA- C
5597 5598 5600 5601 3 30.33400 0.00000 -30.33400
0.00000 0.00000 0.00000 ; CG- CD1- CE1- HE1
5597 5598 5600 5602 3 30.33400 0.00000 -30.33400
0.00000 0.00000 0.00000 ; CG- CD1- CE1- CZ
5597 5607 5605 5602 3 30.33400 0.00000 -30.33400
0.00000 0.00000 0.00000 ; CG- CD2- CE2- CZ
5597 5607 5605 5606 3 30.33400 0.00000 -30.33400
0.00000 0.00000 0.00000 ; CG- CD2- CE2- HE2
5598 5597 5607 5605 3 30.33400 0.00000 -30.33400
0.00000 0.00000 0.00000 ; CD1- CG- CD2- CE2

5598 5597 5607 5608 3 30.33400 0.00000 -30.33400
0.00000 0.00000 0.00000 ; CD1- CG- CD2- HD2
5598 5600 5602 5603 3 30.33400 0.00000 -30.33400
0.00000 0.00000 0.00000 ; CD1- CE1- CZ- OH
5598 5600 5602 5605 3 30.33400 0.00000 -30.33400
0.00000 0.00000 0.00000 ; CD1- CE1- CZ- CE2
5599 5598 5597 5607 3 30.33400 0.00000 -30.33400
0.00000 0.00000 0.00000 ; HD1- CD1- CG- CD2
5599 5598 5600 5601 3 30.33400 0.00000 -30.33400
0.00000 0.00000 0.00000 ; HD1- CD1- CE1- HE1
5599 5598 5600 5602 3 30.33400 0.00000 -30.33400
0.00000 0.00000 0.00000 ; HD1- CD1- CE1- CZ
5600 5598 5597 5607 3 30.33400 0.00000 -30.33400
0.00000 0.00000 0.00000 ; CE1- CD1- CG- CD2
5600 5602 5603 5604 3 7.93286 0.00000 -9.56462
0.00000 2.17568 0.00000 ; CE1- CZ- OH- HH
5600 5602 5605 5606 3 30.33400 0.00000 -30.33400
0.00000 0.00000 0.00000 ; CE1- CZ- CE2- HE2
5600 5602 5605 5607 3 30.33400 0.00000 -30.33400
0.00000 0.00000 0.00000 ; CE1- CZ- CE2- CD2
5601 5600 5602 5603 3 30.33400 0.00000 -30.33400
0.00000 0.00000 0.00000 ; HE1- CE1- CZ- OH
5601 5600 5602 5605 3 30.33400 0.00000 -30.33400
0.00000 0.00000 0.00000 ; HE1- CE1- CZ- CE2
5602 5605 5607 5608 3 30.33400 0.00000 -30.33400
0.00000 0.00000 0.00000 ; CZ- CE2- CD2- HD2
5603 5602 5605 5606 3 30.33400 0.00000 -30.33400
0.00000 0.00000 0.00000 ; OH- CZ- CE2- HE2
5603 5602 5605 5607 3 30.33400 0.00000 -30.33400
0.00000 0.00000 0.00000 ; OH- CZ- CE2- CD2
5604 5603 5602 5605 3 7.93286 0.00000 -9.56462
0.00000 2.17568 0.00000 ; HH- OH- CZ- CE2
5606 5605 5607 5608 3 30.33400 0.00000 -30.33400
0.00000 0.00000 0.00000 ; HE2- CE2- CD2- HD2
5609 5611 5613 5614 3 0.00000 0.00000 0.00000
0.00000 0.00000 0.00000 ; C- N- CA- HA
5609 5611 5613 5615 3 11.71520 1.67360 15.06240 -
13.38880 0.00000 0.00000 ; C- N- CA- CB
5609 5611 5613 5626 3 1.75728 5.27184 2.25936
-7.02912 0.00000 0.00000 ; C- N- CA- C
5610 5609 5611 5612 3 29.28800 -8.36800 -20.92000
0.00000 0.00000 0.00000 ; O- C- N- H
5610 5609 5611 5613 3 20.92000 0.00000 -20.92000
0.00000 0.00000 0.00000 ; O- C- N- CA
5611 5613 5615 5616 3 0.65084 1.95253 0.00000
-2.60338 0.00000 0.00000 ; N- CA- CB- HB2
5611 5613 5615 5617 3 0.65084 1.95253 0.00000
-2.60338 0.00000 0.00000 ; N- CA- CB- HB3
5611 5613 5615 5618 3 3.21331 2.22589 -4.15053
-2.40998 2.61082 0.00000 ; N- CA- CB- CG
5611 5613 5626 5627 3 0.00000 0.00000 0.00000
0.00000 0.00000 0.00000 ; N- CA- C- O

5611 5613 5626 5628 3 17.40544 -5.02080 -13.22144
9.20480 0.00000 0.00000 ; N- CA- C- N
5612 5611 5613 5614 3 0.00000 0.00000 0.00000
0.00000 0.00000 0.00000 ; H- N- CA- HA
5612 5611 5613 5615 3 0.00000 0.00000 0.00000
0.00000 0.00000 0.00000 ; H- N- CA- CB
5612 5611 5613 5626 3 0.00000 0.00000 0.00000
0.00000 0.00000 0.00000 ; H- N- CA- C
5613 5615 5618 5619 3 0.66944 2.00832 0.00000
-2.67776 0.00000 0.00000 ; CA- CB- CG- HG2
5613 5615 5618 5620 3 0.66944 2.00832 0.00000
-2.67776 0.00000 0.00000 ; CA- CB- CG- HG3
5613 5615 5618 5621 3 3.69866 -4.35136 -3.92459
6.89523 4.61914 0.00000 ; CA- CB- CG- CD
5613 5626 5628 5629 3 20.92000 0.00000 -20.92000
0.00000 0.00000 0.00000 ; CA- C- N- H
5613 5626 5628 5630 3 20.92000 0.00000 -20.92000
0.00000 0.00000 0.00000 ; CA- C- N- CA
5614 5613 5615 5616 3 0.65084 1.95253 0.00000
-2.60338 0.00000 0.00000 ; HA- CA- CB- HB2
5614 5613 5615 5617 3 0.65084 1.95253 0.00000
-2.60338 0.00000 0.00000 ; HA- CA- CB- HB3
5614 5613 5615 5618 3 0.65084 1.95253 0.00000
-2.60338 0.00000 0.00000 ; HA- CA- CB- CG
5614 5613 5626 5627 3 3.68192 -4.35136 0.00000
1.33888 0.00000 0.00000 ; HA- CA- C- O
5614 5613 5626 5628 3 0.00000 0.00000 0.00000
0.00000 0.00000 0.00000 ; HA- CA- C- N
5615 5613 5626 5627 3 0.00000 0.00000 0.00000
0.00000 0.00000 0.00000 ; CB- CA- C- O
5615 5613 5626 5628 3 2.51040 4.18400 1.67360
-6.69440 0.00000 0.00000 ; CB- CA- C- N
5615 5618 5621 5622 3 0.00000 0.00000 0.00000
0.00000 0.00000 0.00000 ; CB- CG- CD- OE1
5615 5618 5621 5623 3 10.32611 1.48114 -8.47678
1.42256 1.40582 0.00000 ; CB- CG- CD- NE2
5616 5615 5613 5626 3 0.65084 1.95253 0.00000
-2.60338 0.00000 0.00000 ; HB2- CB- CA- C
5616 5615 5618 5619 3 0.62760 1.88280 0.00000
-2.51040 0.00000 0.00000 ; HB2- CB- CG- HG2
5616 5615 5618 5620 3 0.62760 1.88280 0.00000
-2.51040 0.00000 0.00000 ; HB2- CB- CG- HG3
5616 5615 5618 5621 3 0.65084 1.95253 0.00000
-2.60338 0.00000 0.00000 ; HB2- CB- CG- CD
5617 5615 5613 5626 3 0.65084 1.95253 0.00000
-2.60338 0.00000 0.00000 ; HB3- CB- CA- C
5617 5615 5618 5619 3 0.62760 1.88280 0.00000
-2.51040 0.00000 0.00000 ; HB3- CB- CG- HG2
5617 5615 5618 5620 3 0.62760 1.88280 0.00000
-2.51040 0.00000 0.00000 ; HB3- CB- CG- HG3
5617 5615 5618 5621 3 0.65084 1.95253 0.00000
-2.60338 0.00000 0.00000 ; HB3- CB- CG- CD

5618 5615 5613 5626 3 6.86594 3.56895 -8.14206
-2.40998 4.85344 0.00000 ; CG- CB- CA- C
5618 5621 5623 5624 3 20.92000 0.00000 -20.92000
0.00000 0.00000 0.00000 ; CG- CD- NE2- HE21
5618 5621 5623 5625 3 20.92000 0.00000 -20.92000
0.00000 0.00000 0.00000 ; CG- CD- NE2- HE22
5619 5618 5621 5622 3 3.68192 -4.35136 0.00000
1.33888 0.00000 0.00000 ; HG2- CG- CD- OE1
5619 5618 5621 5623 3 0.00000 0.00000 0.00000
0.00000 0.00000 0.00000 ; HG2- CG- CD- NE2
5620 5618 5621 5622 3 3.68192 -4.35136 0.00000
1.33888 0.00000 0.00000 ; HG3- CG- CD- OE1
5620 5618 5621 5623 3 0.00000 0.00000 0.00000
0.00000 0.00000 0.00000 ; HG3- CG- CD- NE2
5622 5621 5623 5624 3 29.28800 -8.36800 -20.92000
0.00000 0.00000 0.00000 ; OE1- CD- NE2- HE21
5622 5621 5623 5625 3 29.28800 -8.36800 -20.92000
0.00000 0.00000 0.00000 ; OE1- CD- NE2- HE22
5626 5628 5630 5631 3 0.00000 0.00000 0.00000
0.00000 0.00000 0.00000 ; C- N- CA- HA
5626 5628 5630 5632 3 11.71520 1.67360 15.06240 -
13.38880 0.00000 0.00000 ; C- N- CA- CB
5626 5628 5630 5648 3 1.75728 5.27184 2.25936
-7.02912 0.00000 0.00000 ; C- N- CA- C
5627 5626 5628 5629 3 29.28800 -8.36800 -20.92000
0.00000 0.00000 0.00000 ; O- C- N- H
5627 5626 5628 5630 3 20.92000 0.00000 -20.92000
0.00000 0.00000 0.00000 ; O- C- N- CA
5628 5630 5632 5633 3 0.65084 1.95253 0.00000
-2.60338 0.00000 0.00000 ; N- CA- CB- HB2
5628 5630 5632 5634 3 0.65084 1.95253 0.00000
-2.60338 0.00000 0.00000 ; N- CA- CB- HB3
5628 5630 5632 5635 3 0.65084 1.95253 0.00000
-2.60338 0.00000 0.00000 ; N- CA- CB- CG
5628 5630 5648 5649 3 0.00000 0.00000 0.00000
0.00000 0.00000 0.00000 ; N- CA- C- O
5628 5630 5648 5650 3 17.40544 -5.02080 -13.22144
9.20480 0.00000 0.00000 ; N- CA- C- N
5629 5628 5630 5631 3 0.00000 0.00000 0.00000
0.00000 0.00000 0.00000 ; H- N- CA- HA
5629 5628 5630 5632 3 0.00000 0.00000 0.00000
0.00000 0.00000 0.00000 ; H- N- CA- CB
5629 5628 5630 5648 3 0.00000 0.00000 0.00000
0.00000 0.00000 0.00000 ; H- N- CA- C
5630 5632 5635 5636 3 0.66944 2.00832 0.00000
-2.67776 0.00000 0.00000 ; CA- CB- CG- HG2
5630 5632 5635 5637 3 0.66944 2.00832 0.00000
-2.67776 0.00000 0.00000 ; CA- CB- CG- HG3
5630 5632 5635 5638 3 3.68192 3.09616 -2.09200
-3.01248 0.00000 0.00000 ; CA- CB- CG- CD
5630 5648 5650 5651 3 20.92000 0.00000 -20.92000
0.00000 0.00000 0.00000 ; CA- C- N- H

5630 5648 5650 5652 3 20.92000 0.00000 -20.92000
0.00000 0.00000 0.00000 ; CA- C- N- CA
5631 5630 5632 5633 3 0.65084 1.95253 0.00000
-2.60338 0.00000 0.00000 ; HA- CA- CB- HB2
5631 5630 5632 5634 3 0.65084 1.95253 0.00000
-2.60338 0.00000 0.00000 ; HA- CA- CB- HB3
5631 5630 5632 5635 3 0.65084 1.95253 0.00000
-2.60338 0.00000 0.00000 ; HA- CA- CB- CG
5631 5630 5648 5649 3 3.68192 -4.35136 0.00000
1.33888 0.00000 0.00000 ; HA- CA- C- O
5631 5630 5648 5650 3 0.00000 0.00000 0.00000
0.00000 0.00000 0.00000 ; HA- CA- C- N
5632 5630 5648 5649 3 0.00000 0.00000 0.00000
0.00000 0.00000 0.00000 ; CB- CA- C- O
5632 5630 5648 5650 3 2.51040 4.18400 1.67360
-6.69440 0.00000 0.00000 ; CB- CA- C- N
5632 5635 5638 5639 3 0.66944 2.00832 0.00000
-2.67776 0.00000 0.00000 ; CB- CG- CD- HD2
5632 5635 5638 5640 3 0.66944 2.00832 0.00000
-2.67776 0.00000 0.00000 ; CB- CG- CD- HD3
5632 5635 5638 5641 3 3.68192 3.09616 -2.09200
-3.01248 0.00000 0.00000 ; CB- CG- CD- CE
5633 5632 5630 5648 3 0.65084 1.95253 0.00000
-2.60338 0.00000 0.00000 ; HB2- CB- CA- C
5633 5632 5635 5636 3 0.62760 1.88280 0.00000
-2.51040 0.00000 0.00000 ; HB2- CB- CG- HG2
5633 5632 5635 5637 3 0.62760 1.88280 0.00000
-2.51040 0.00000 0.00000 ; HB2- CB- CG- HG3
5633 5632 5635 5638 3 0.66944 2.00832 0.00000
-2.67776 0.00000 0.00000 ; HB2- CB- CG- CD
5634 5632 5630 5648 3 0.65084 1.95253 0.00000
-2.60338 0.00000 0.00000 ; HB3- CB- CA- C
5634 5632 5635 5636 3 0.62760 1.88280 0.00000
-2.51040 0.00000 0.00000 ; HB3- CB- CG- HG2
5634 5632 5635 5637 3 0.62760 1.88280 0.00000
-2.51040 0.00000 0.00000 ; HB3- CB- CG- HG3
5634 5632 5635 5638 3 0.66944 2.00832 0.00000
-2.67776 0.00000 0.00000 ; HB3- CB- CG- CD
5635 5632 5630 5648 3 0.65084 1.95253 0.00000
-2.60338 0.00000 0.00000 ; CG- CB- CA- C
5635 5638 5641 5642 3 0.65084 1.95253 0.00000
-2.60338 0.00000 0.00000 ; CG- CD- CE- HE2
5635 5638 5641 5643 3 0.65084 1.95253 0.00000
-2.60338 0.00000 0.00000 ; CG- CD- CE- HE3
5635 5638 5641 5644 3 0.65084 1.95253 0.00000
-2.60338 0.00000 0.00000 ; CG- CD- CE- NZ
5636 5635 5638 5639 3 0.62760 1.88280 0.00000
-2.51040 0.00000 0.00000 ; HG2- CG- CD- HD2
5636 5635 5638 5640 3 0.62760 1.88280 0.00000
-2.51040 0.00000 0.00000 ; HG2- CG- CD- HD3
5636 5635 5638 5641 3 0.66944 2.00832 0.00000
-2.67776 0.00000 0.00000 ; HG2- CG- CD- CE

5637 5635 5638 5639 3 0.62760 1.88280 0.00000
-2.51040 0.00000 0.00000 ; HG3- CG- CD- HD2
5637 5635 5638 5640 3 0.62760 1.88280 0.00000
-2.51040 0.00000 0.00000 ; HG3- CG- CD- HD3
5637 5635 5638 5641 3 0.66944 2.00832 0.00000
-2.67776 0.00000 0.00000 ; HG3- CG- CD- CE
5638 5641 5644 5645 3 0.65084 1.95253 0.00000
-2.60338 0.00000 0.00000 ; CD- CE- NZ- HZ1
5638 5641 5644 5646 3 0.65084 1.95253 0.00000
-2.60338 0.00000 0.00000 ; CD- CE- NZ- HZ2
5638 5641 5644 5647 3 0.65084 1.95253 0.00000
-2.60338 0.00000 0.00000 ; CD- CE- NZ- HZ3
5639 5638 5641 5642 3 0.65084 1.95253 0.00000
-2.60338 0.00000 0.00000 ; HD2- CD- CE- HE2
5639 5638 5641 5643 3 0.65084 1.95253 0.00000
-2.60338 0.00000 0.00000 ; HD2- CD- CE- HE3
5639 5638 5641 5644 3 0.65084 1.95253 0.00000
-2.60338 0.00000 0.00000 ; HD2- CD- CE- NZ
5640 5638 5641 5642 3 0.65084 1.95253 0.00000
-2.60338 0.00000 0.00000 ; HD3- CD- CE- HE2
5640 5638 5641 5643 3 0.65084 1.95253 0.00000
-2.60338 0.00000 0.00000 ; HD3- CD- CE- HE3
5640 5638 5641 5644 3 0.65084 1.95253 0.00000
-2.60338 0.00000 0.00000 ; HD3- CD- CE- NZ
5642 5641 5644 5645 3 0.65084 1.95253 0.00000
-2.60338 0.00000 0.00000 ; HE2- CE- NZ- HZ1
5642 5641 5644 5646 3 0.65084 1.95253 0.00000
-2.60338 0.00000 0.00000 ; HE2- CE- NZ- HZ2
5642 5641 5644 5647 3 0.65084 1.95253 0.00000
-2.60338 0.00000 0.00000 ; HE2- CE- NZ- HZ3
5643 5641 5644 5645 3 0.65084 1.95253 0.00000
-2.60338 0.00000 0.00000 ; HE3- CE- NZ- HZ1
5643 5641 5644 5646 3 0.65084 1.95253 0.00000
-2.60338 0.00000 0.00000 ; HE3- CE- NZ- HZ2
5643 5641 5644 5647 3 0.65084 1.95253 0.00000
-2.60338 0.00000 0.00000 ; HE3- CE- NZ- HZ3
5648 5650 5652 5653 3 0.00000 0.00000 0.00000
0.00000 0.00000 0.00000 ; C- N- CA- HA
5648 5650 5652 5654 3 11.71520 1.67360 15.06240 -
13.38880 0.00000 0.00000 ; C- N- CA- CB
5648 5650 5652 5667 3 1.75728 5.27184 2.25936
-7.02912 0.00000 0.00000 ; C- N- CA- C
5649 5648 5650 5651 3 29.28800 -8.36800 -20.92000
0.00000 0.00000 0.00000 ; O- C- N- H
5649 5648 5650 5652 3 20.92000 0.00000 -20.92000
0.00000 0.00000 0.00000 ; O- C- N- CA
5650 5652 5654 5655 3 0.65084 1.95253 0.00000
-2.60338 0.00000 0.00000 ; N- CA- CB- HB
5650 5652 5654 5656 3 3.83673 0.44769 -1.77402
-2.47693 -0.03347 0.00000 ; N- CA- CB- CG2
5650 5652 5654 5660 3 2.97482 0.12134 2.04179
-1.89117 -3.24678 0.00000 ; N- CA- CB- CG1

5650 5652 5667 5668 3 0.00000 0.00000 0.00000
0.00000 0.00000 0.00000 ; N- CA- C- O
5650 5652 5667 5669 3 17.40544 -5.02080 -13.22144
9.20480 0.00000 0.00000 ; N- CA- C- N
5651 5650 5652 5653 3 0.00000 0.00000 0.00000
0.00000 0.00000 0.00000 ; H- N- CA- HA
5651 5650 5652 5654 3 0.00000 0.00000 0.00000
0.00000 0.00000 0.00000 ; H- N- CA- CB
5651 5650 5652 5667 3 0.00000 0.00000 0.00000
0.00000 0.00000 0.00000 ; H- N- CA- C
5652 5654 5656 5657 3 0.66944 2.00832 0.00000
-2.67776 0.00000 0.00000 ; CA- CB- CG2- HG21
5652 5654 5656 5658 3 0.66944 2.00832 0.00000
-2.67776 0.00000 0.00000 ; CA- CB- CG2- HG22
5652 5654 5656 5659 3 0.66944 2.00832 0.00000
-2.67776 0.00000 0.00000 ; CA- CB- CG2- HG23
5652 5654 5660 5661 3 0.66944 2.00832 0.00000
-2.67776 0.00000 0.00000 ; CA- CB- CG1- HG12
5652 5654 5660 5662 3 0.66944 2.00832 0.00000
-2.67776 0.00000 0.00000 ; CA- CB- CG1- HG13
5652 5654 5660 5663 3 4.24258 -0.52718 -7.25506
-1.79075 7.69856 0.00000 ; CA- CB- CG1- CD1
5652 5667 5669 5670 3 20.92000 0.00000 -20.92000
0.00000 0.00000 0.00000 ; CA- C- N- CD
5652 5667 5669 5679 3 20.92000 0.00000 -20.92000
0.00000 0.00000 0.00000 ; CA- C- N- CA
5653 5652 5654 5655 3 0.65084 1.95253 0.00000
-2.60338 0.00000 0.00000 ; HA- CA- CB- HB
5653 5652 5654 5656 3 0.65084 1.95253 0.00000
-2.60338 0.00000 0.00000 ; HA- CA- CB- CG2
5653 5652 5654 5660 3 0.65084 1.95253 0.00000
-2.60338 0.00000 0.00000 ; HA- CA- CB- CG1
5653 5652 5667 5668 3 3.68192 -4.35136 0.00000
1.33888 0.00000 0.00000 ; HA- CA- C- O
5653 5652 5667 5669 3 0.00000 0.00000 0.00000
0.00000 0.00000 0.00000 ; HA- CA- C- N
5654 5652 5667 5668 3 0.00000 0.00000 0.00000
0.00000 0.00000 0.00000 ; CB- CA- C- O
5654 5652 5667 5669 3 2.51040 4.18400 1.67360
-6.69440 0.00000 0.00000 ; CB- CA- C- N
5654 5660 5663 5664 3 0.66944 2.00832 0.00000
-2.67776 0.00000 0.00000 ; CB- CG1- CD1- HD11
5654 5660 5663 5665 3 0.66944 2.00832 0.00000
-2.67776 0.00000 0.00000 ; CB- CG1- CD1- HD12
5654 5660 5663 5666 3 0.66944 2.00832 0.00000
-2.67776 0.00000 0.00000 ; CB- CG1- CD1- HD13
5655 5654 5652 5667 3 0.65084 1.95253 0.00000
-2.60338 0.00000 0.00000 ; HB- CB- CA- C
5655 5654 5656 5657 3 0.62760 1.88280 0.00000
-2.51040 0.00000 0.00000 ; HB- CB- CG2- HG21
5655 5654 5656 5658 3 0.62760 1.88280 0.00000
-2.51040 0.00000 0.00000 ; HB- CB- CG2- HG22

5655 5654 5656 5659 3 0.62760 1.88280 0.00000
-2.51040 0.00000 0.00000 ; HB- CB- CG2- HG23
5655 5654 5660 5661 3 0.62760 1.88280 0.00000
-2.51040 0.00000 0.00000 ; HB- CB- CG1- HG12
5655 5654 5660 5662 3 0.62760 1.88280 0.00000
-2.51040 0.00000 0.00000 ; HB- CB- CG1- HG13
5655 5654 5660 5663 3 0.66944 2.00832 0.00000
-2.67776 0.00000 0.00000 ; HB- CB- CG1- CD1
5656 5654 5652 5667 3 5.67350 3.55640 -6.16722
-2.47693 3.74886 0.00000 ; CG2- CB- CA- C
5656 5654 5660 5661 3 0.66944 2.00832 0.00000
-2.67776 0.00000 0.00000 ; CG2- CB- CG1- HG12
5656 5654 5660 5662 3 0.66944 2.00832 0.00000
-2.67776 0.00000 0.00000 ; CG2- CB- CG1- HG13
5656 5654 5660 5663 3 3.81162 0.49790 -8.14206
-1.79075 7.49773 0.00000 ; CG2- CB- CG1- CD1
5657 5656 5654 5660 3 0.66944 2.00832 0.00000
-2.67776 0.00000 0.00000 ; HG21- CG2- CB- CG1
5658 5656 5654 5660 3 0.66944 2.00832 0.00000
-2.67776 0.00000 0.00000 ; HG22- CG2- CB- CG1
5659 5656 5654 5660 3 0.66944 2.00832 0.00000
-2.67776 0.00000 0.00000 ; HG23- CG2- CB- CG1
5660 5654 5652 5667 3 8.26340 0.74057 -9.99976
-1.89117 3.84928 0.00000 ; CG1- CB- CA- C
5661 5660 5663 5664 3 0.62760 1.88280 0.00000
-2.51040 0.00000 0.00000 ; HG12- CG1- CD1- HD11
5661 5660 5663 5665 3 0.62760 1.88280 0.00000
-2.51040 0.00000 0.00000 ; HG12- CG1- CD1- HD12
5661 5660 5663 5666 3 0.62760 1.88280 0.00000
-2.51040 0.00000 0.00000 ; HG12- CG1- CD1- HD13
5662 5660 5663 5664 3 0.62760 1.88280 0.00000
-2.51040 0.00000 0.00000 ; HG13- CG1- CD1- HD11
5662 5660 5663 5665 3 0.62760 1.88280 0.00000
-2.51040 0.00000 0.00000 ; HG13- CG1- CD1- HD12
5662 5660 5663 5666 3 0.62760 1.88280 0.00000
-2.51040 0.00000 0.00000 ; HG13- CG1- CD1- HD13
5667 5669 5670 5671 3 0.00000 0.00000 0.00000
0.00000 0.00000 0.00000 ; C- N- CD- HD2
5667 5669 5670 5672 3 0.00000 0.00000 0.00000
0.00000 0.00000 0.00000 ; C- N- CD- HD3
5667 5669 5670 5673 3 10.04160 -3.34720 16.73600
-6.69440 0.00000 0.00000 ; C- N- CD- CG
5667 5669 5679 5676 3 11.71520 1.67360 15.06240 -
13.38880 0.00000 0.00000 ; C- N- CA- CB
5667 5669 5679 5680 3 0.00000 0.00000 0.00000
0.00000 0.00000 0.00000 ; C- N- CA- HA
5667 5669 5679 5681 3 1.75728 5.27184 2.25936
-7.02912 0.00000 0.00000 ; C- N- CA- C
5668 5667 5669 5670 3 20.92000 0.00000 -20.92000
0.00000 0.00000 0.00000 ; O- C- N- CD
5668 5667 5669 5679 3 20.92000 0.00000 -20.92000
0.00000 0.00000 0.00000 ; O- C- N- CA

5669 5670 5673 5674 3 0.65084 1.95253 0.00000
-2.60338 0.00000 0.00000 ; N- CD- CG- HG2
5669 5670 5673 5675 3 0.65084 1.95253 0.00000
-2.60338 0.00000 0.00000 ; N- CD- CG- HG3
5669 5670 5673 5676 3 0.65084 1.95253 0.00000
-2.60338 0.00000 0.00000 ; N- CD- CG- CB
5669 5679 5676 5673 3 0.65084 1.95253 0.00000
-2.60338 0.00000 0.00000 ; N- CA- CB- CG
5669 5679 5676 5677 3 0.65084 1.95253 0.00000
-2.60338 0.00000 0.00000 ; N- CA- CB- HB2
5669 5679 5676 5678 3 0.65084 1.95253 0.00000
-2.60338 0.00000 0.00000 ; N- CA- CB- HB3
5669 5679 5681 5682 3 0.00000 0.00000 0.00000
0.00000 0.00000 0.00000 ; N- CA- C- O
5669 5679 5681 5683 3 17.40544 -5.02080 -13.22144
9.20480 0.00000 0.00000 ; N- CA- C- N
5670 5669 5679 5676 3 0.00000 0.00000 0.00000
0.00000 0.00000 0.00000 ; CD- N- CA- CB
5670 5669 5679 5680 3 0.00000 0.00000 0.00000
0.00000 0.00000 0.00000 ; CD- N- CA- HA
5670 5669 5679 5681 3 0.00000 0.00000 0.00000
0.00000 0.00000 0.00000 ; CD- N- CA- C
5670 5673 5676 5677 3 0.66944 2.00832 0.00000
-2.67776 0.00000 0.00000 ; CD- CG- CB- HB2
5670 5673 5676 5678 3 0.66944 2.00832 0.00000
-2.67776 0.00000 0.00000 ; CD- CG- CB- HB3
5670 5673 5676 5679 3 3.68192 3.09616 -2.09200
-3.01248 0.00000 0.00000 ; CD- CG- CB- CA
5671 5670 5669 5679 3 0.00000 0.00000 0.00000
0.00000 0.00000 0.00000 ; HD2- CD- N- CA
5671 5670 5673 5674 3 0.65084 1.95253 0.00000
-2.60338 0.00000 0.00000 ; HD2- CD- CG- HG2
5671 5670 5673 5675 3 0.65084 1.95253 0.00000
-2.60338 0.00000 0.00000 ; HD2- CD- CG- HG3
5671 5670 5673 5676 3 0.65084 1.95253 0.00000
-2.60338 0.00000 0.00000 ; HD2- CD- CG- CB
5672 5670 5669 5679 3 0.00000 0.00000 0.00000
0.00000 0.00000 0.00000 ; HD3- CD- N- CA
5672 5670 5673 5674 3 0.65084 1.95253 0.00000
-2.60338 0.00000 0.00000 ; HD3- CD- CG- HG2
5672 5670 5673 5675 3 0.65084 1.95253 0.00000
-2.60338 0.00000 0.00000 ; HD3- CD- CG- HG3
5672 5670 5673 5676 3 0.65084 1.95253 0.00000
-2.60338 0.00000 0.00000 ; HD3- CD- CG- CB
5673 5670 5669 5679 3 0.00000 0.00000 0.00000
0.00000 0.00000 0.00000 ; CG- CD- N- CA
5673 5676 5679 5680 3 0.65084 1.95253 0.00000
-2.60338 0.00000 0.00000 ; CG- CB- CA- HA
5673 5676 5679 5681 3 0.65084 1.95253 0.00000
-2.60338 0.00000 0.00000 ; CG- CB- CA- C
5674 5673 5676 5677 3 0.62760 1.88280 0.00000
-2.51040 0.00000 0.00000 ; HG2- CG- CB- HB2

5674 5673 5676 5678 3 0.62760 1.88280 0.00000
-2.51040 0.00000 0.00000 ; HG2- CG- CB- HB3
5674 5673 5676 5679 3 0.66944 2.00832 0.00000
-2.67776 0.00000 0.00000 ; HG2- CG- CB- CA
5675 5673 5676 5677 3 0.62760 1.88280 0.00000
-2.51040 0.00000 0.00000 ; HG3- CG- CB- HB2
5675 5673 5676 5678 3 0.62760 1.88280 0.00000
-2.51040 0.00000 0.00000 ; HG3- CG- CB- HB3
5675 5673 5676 5679 3 0.66944 2.00832 0.00000
-2.67776 0.00000 0.00000 ; HG3- CG- CB- CA
5676 5679 5681 5682 3 0.00000 0.00000 0.00000
0.00000 0.00000 0.00000 ; CB- CA- C- O
5676 5679 5681 5683 3 2.51040 4.18400 1.67360
-6.69440 0.00000 0.00000 ; CB- CA- C- N
5677 5676 5679 5680 3 0.65084 1.95253 0.00000
-2.60338 0.00000 0.00000 ; HB2- CB- CA- HA
5677 5676 5679 5681 3 0.65084 1.95253 0.00000
-2.60338 0.00000 0.00000 ; HB2- CB- CA- C
5678 5676 5679 5680 3 0.65084 1.95253 0.00000
-2.60338 0.00000 0.00000 ; HB3- CB- CA- HA
5678 5676 5679 5681 3 0.65084 1.95253 0.00000
-2.60338 0.00000 0.00000 ; HB3- CB- CA- C
5679 5681 5683 5684 3 20.92000 0.00000 -20.92000
0.00000 0.00000 0.00000 ; CA- C- N- CD
5679 5681 5683 5693 3 20.92000 0.00000 -20.92000
0.00000 0.00000 0.00000 ; CA- C- N- CA
5680 5679 5681 5682 3 3.68192 -4.35136 0.00000
1.33888 0.00000 0.00000 ; HA- CA- C- O
5680 5679 5681 5683 3 0.00000 0.00000 0.00000
0.00000 0.00000 0.00000 ; HA- CA- C- N
5681 5683 5684 5685 3 0.00000 0.00000 0.00000
0.00000 0.00000 0.00000 ; C- N- CD- HD2
5681 5683 5684 5686 3 0.00000 0.00000 0.00000
0.00000 0.00000 0.00000 ; C- N- CD- HD3
5681 5683 5684 5687 3 10.04160 -3.34720 16.73600
-6.69440 0.00000 0.00000 ; C- N- CD- CG
5681 5683 5693 5690 3 11.71520 1.67360 15.06240 -
13.38880 0.00000 0.00000 ; C- N- CA- CB
5681 5683 5693 5694 3 0.00000 0.00000 0.00000
0.00000 0.00000 0.00000 ; C- N- CA- HA
5681 5683 5693 5695 3 1.75728 5.27184 2.25936
-7.02912 0.00000 0.00000 ; C- N- CA- C
5682 5681 5683 5684 3 20.92000 0.00000 -20.92000
0.00000 0.00000 0.00000 ; O- C- N- CD
5682 5681 5683 5693 3 20.92000 0.00000 -20.92000
0.00000 0.00000 0.00000 ; O- C- N- CA
5683 5684 5687 5688 3 0.65084 1.95253 0.00000
-2.60338 0.00000 0.00000 ; N- CD- CG- HG2
5683 5684 5687 5689 3 0.65084 1.95253 0.00000
-2.60338 0.00000 0.00000 ; N- CD- CG- HG3
5683 5684 5687 5690 3 0.65084 1.95253 0.00000
-2.60338 0.00000 0.00000 ; N- CD- CG- CB

5683 5693 5690 5687 3 0.65084 1.95253 0.00000
-2.60338 0.00000 0.00000 ; N- CA- CB- CG
5683 5693 5690 5691 3 0.65084 1.95253 0.00000
-2.60338 0.00000 0.00000 ; N- CA- CB- HB2
5683 5693 5690 5692 3 0.65084 1.95253 0.00000
-2.60338 0.00000 0.00000 ; N- CA- CB- HB3
5683 5693 5695 5696 3 0.00000 0.00000 0.00000
0.00000 0.00000 0.00000 ; N- CA- C- O
5683 5693 5695 5697 3 17.40544 -5.02080 -13.22144
9.20480 0.00000 0.00000 ; N- CA- C- N
5684 5683 5693 5690 3 0.00000 0.00000 0.00000
0.00000 0.00000 0.00000 ; CD- N- CA- CB
5684 5683 5693 5694 3 0.00000 0.00000 0.00000
0.00000 0.00000 0.00000 ; CD- N- CA- HA
5684 5683 5693 5695 3 0.00000 0.00000 0.00000
0.00000 0.00000 0.00000 ; CD- N- CA- C
5684 5687 5690 5691 3 0.66944 2.00832 0.00000
-2.67776 0.00000 0.00000 ; CD- CG- CB- HB2
5684 5687 5690 5692 3 0.66944 2.00832 0.00000
-2.67776 0.00000 0.00000 ; CD- CG- CB- HB3
5684 5687 5690 5693 3 3.68192 3.09616 -2.09200
-3.01248 0.00000 0.00000 ; CD- CG- CB- CA
5685 5684 5683 5693 3 0.00000 0.00000 0.00000
0.00000 0.00000 0.00000 ; HD2- CD- N- CA
5685 5684 5687 5688 3 0.65084 1.95253 0.00000
-2.60338 0.00000 0.00000 ; HD2- CD- CG- HG2
5685 5684 5687 5689 3 0.65084 1.95253 0.00000
-2.60338 0.00000 0.00000 ; HD2- CD- CG- HG3
5685 5684 5687 5690 3 0.65084 1.95253 0.00000
-2.60338 0.00000 0.00000 ; HD2- CD- CG- CB
5686 5684 5683 5693 3 0.00000 0.00000 0.00000
0.00000 0.00000 0.00000 ; HD3- CD- N- CA
5686 5684 5687 5688 3 0.65084 1.95253 0.00000
-2.60338 0.00000 0.00000 ; HD3- CD- CG- HG2
5686 5684 5687 5689 3 0.65084 1.95253 0.00000
-2.60338 0.00000 0.00000 ; HD3- CD- CG- HG3
5686 5684 5687 5690 3 0.65084 1.95253 0.00000
-2.60338 0.00000 0.00000 ; HD3- CD- CG- CB
5687 5684 5683 5693 3 0.00000 0.00000 0.00000
0.00000 0.00000 0.00000 ; CG- CD- N- CA
5687 5690 5693 5694 3 0.65084 1.95253 0.00000
-2.60338 0.00000 0.00000 ; CG- CB- CA- HA
5687 5690 5693 5695 3 0.65084 1.95253 0.00000
-2.60338 0.00000 0.00000 ; CG- CB- CA- C
5688 5687 5690 5691 3 0.62760 1.88280 0.00000
-2.51040 0.00000 0.00000 ; HG2- CG- CB- HB2
5688 5687 5690 5692 3 0.62760 1.88280 0.00000
-2.51040 0.00000 0.00000 ; HG2- CG- CB- HB3
5688 5687 5690 5693 3 0.66944 2.00832 0.00000
-2.67776 0.00000 0.00000 ; HG2- CG- CB- CA
5689 5687 5690 5691 3 0.62760 1.88280 0.00000
-2.51040 0.00000 0.00000 ; HG3- CG- CB- HB2

5689 5687 5690 5692 3 0.62760 1.88280 0.00000
-2.51040 0.00000 0.00000 ; HG3- CG- CB- HB3
5689 5687 5690 5693 3 0.66944 2.00832 0.00000
-2.67776 0.00000 0.00000 ; HG3- CG- CB- CA
5690 5693 5695 5696 3 0.00000 0.00000 0.00000
0.00000 0.00000 0.00000 ; CB- CA- C- O
5690 5693 5695 5697 3 2.51040 4.18400 1.67360
-6.69440 0.00000 0.00000 ; CB- CA- C- N
5691 5690 5693 5694 3 0.65084 1.95253 0.00000
-2.60338 0.00000 0.00000 ; HB2- CB- CA- HA
5691 5690 5693 5695 3 0.65084 1.95253 0.00000
-2.60338 0.00000 0.00000 ; HB2- CB- CA- C
5692 5690 5693 5694 3 0.65084 1.95253 0.00000
-2.60338 0.00000 0.00000 ; HB3- CB- CA- HA
5692 5690 5693 5695 3 0.65084 1.95253 0.00000
-2.60338 0.00000 0.00000 ; HB3- CB- CA- C
5693 5695 5697 5698 3 20.92000 0.00000 -20.92000
0.00000 0.00000 0.00000 ; CA- C- N- H
5693 5695 5697 5699 3 20.92000 0.00000 -20.92000
0.00000 0.00000 0.00000 ; CA- C- N- CA
5694 5693 5695 5696 3 3.68192 -4.35136 0.00000
1.33888 0.00000 0.00000 ; HA- CA- C- O
5694 5693 5695 5697 3 0.00000 0.00000 0.00000
0.00000 0.00000 0.00000 ; HA- CA- C- N
5695 5697 5699 5700 3 0.00000 0.00000 0.00000
0.00000 0.00000 0.00000 ; C- N- CA- HA
5695 5697 5699 5701 3 11.71520 1.67360 15.06240 -
13.38880 0.00000 0.00000 ; C- N- CA- CB
5695 5697 5699 5707 3 1.75728 5.27184 2.25936
-7.02912 0.00000 0.00000 ; C- N- CA- C
5696 5695 5697 5698 3 29.28800 -8.36800 -20.92000
0.00000 0.00000 0.00000 ; O- C- N- H
5696 5695 5697 5699 3 20.92000 0.00000 -20.92000
0.00000 0.00000 0.00000 ; O- C- N- CA
5697 5699 5701 5702 3 0.65084 1.95253 0.00000
-2.60338 0.00000 0.00000 ; N- CA- CB- HB2
5697 5699 5701 5703 3 0.65084 1.95253 0.00000
-2.60338 0.00000 0.00000 ; N- CA- CB- HB3
5697 5699 5701 5704 3 15.41386 9.74035 -8.39310
-0.97069 2.97901 0.00000 ; N- CA- CB- CG
5697 5699 5707 5708 3 0.00000 0.00000 0.00000
0.00000 0.00000 0.00000 ; N- CA- C- O
5697 5699 5707 5709 3 17.40544 -5.02080 -13.22144
9.20480 0.00000 0.00000 ; N- CA- C- N
5698 5697 5699 5700 3 0.00000 0.00000 0.00000
0.00000 0.00000 0.00000 ; H- N- CA- HA
5698 5697 5699 5701 3 0.00000 0.00000 0.00000
0.00000 0.00000 0.00000 ; H- N- CA- CB
5698 5697 5699 5707 3 0.00000 0.00000 0.00000
0.00000 0.00000 0.00000 ; H- N- CA- C
5699 5701 5704 5705 3 6.43499 0.00000 -5.39736
0.00000 -1.03763 0.00000 ; CA- CB- CG- OD1

5699 5701 5704 5706 3 6.43499 0.00000 -5.39736
0.00000 -1.03763 0.00000 ; CA- CB- CG- OD2
5699 5707 5709 5710 3 20.92000 0.00000 -20.92000
0.00000 0.00000 0.00000 ; CA- C- N- H
5699 5707 5709 5711 3 20.92000 0.00000 -20.92000
0.00000 0.00000 0.00000 ; CA- C- N- CA
5700 5699 5701 5702 3 0.65084 1.95253 0.00000
-2.60338 0.00000 0.00000 ; HA- CA- CB- HB2
5700 5699 5701 5703 3 0.65084 1.95253 0.00000
-2.60338 0.00000 0.00000 ; HA- CA- CB- HB3
5700 5699 5701 5704 3 0.65084 1.95253 0.00000
-2.60338 0.00000 0.00000 ; HA- CA- CB- CG
5700 5699 5707 5708 3 3.68192 -4.35136 0.00000
1.33888 0.00000 0.00000 ; HA- CA- C- O
5700 5699 5707 5709 3 0.00000 0.00000 0.00000
0.00000 0.00000 0.00000 ; HA- CA- C- N
5701 5699 5707 5708 3 0.00000 0.00000 0.00000
0.00000 0.00000 0.00000 ; CB- CA- C- O
5701 5699 5707 5709 3 2.51040 4.18400 1.67360
-6.69440 0.00000 0.00000 ; CB- CA- C- N
5702 5701 5699 5707 3 0.65084 1.95253 0.00000
-2.60338 0.00000 0.00000 ; HB2- CB- CA- C
5702 5701 5704 5705 3 0.00000 0.00000 0.00000
0.00000 0.00000 0.00000 ; HB2- CB- CG- OD1
5702 5701 5704 5706 3 0.00000 0.00000 0.00000
0.00000 0.00000 0.00000 ; HB2- CB- CG- OD2
5703 5701 5699 5707 3 0.65084 1.95253 0.00000
-2.60338 0.00000 0.00000 ; HB3- CB- CA- C
5703 5701 5704 5705 3 0.00000 0.00000 0.00000
0.00000 0.00000 0.00000 ; HB3- CB- CG- OD1
5703 5701 5704 5706 3 0.00000 0.00000 0.00000
0.00000 0.00000 0.00000 ; HB3- CB- CG- OD2
5704 5701 5699 5707 3 7.14627 -1.04600 -8.99560
-0.97069 5.15469 0.00000 ; CG- CB- CA- C
5707 5709 5711 5712 3 0.00000 0.00000 0.00000
0.00000 0.00000 0.00000 ; C- N- CA- HA
5707 5709 5711 5713 3 11.71520 1.67360 15.06240 -
13.38880 0.00000 0.00000 ; C- N- CA- CB
5707 5709 5711 5726 3 1.75728 5.27184 2.25936
-7.02912 0.00000 0.00000 ; C- N- CA- C
5708 5707 5709 5710 3 29.28800 -8.36800 -20.92000
0.00000 0.00000 0.00000 ; O- C- N- H
5708 5707 5709 5711 3 20.92000 0.00000 -20.92000
0.00000 0.00000 0.00000 ; O- C- N- CA
5709 5711 5713 5714 3 0.65084 1.95253 0.00000
-2.60338 0.00000 0.00000 ; N- CA- CB- HB
5709 5711 5713 5715 3 3.83673 0.44769 -1.77402
-2.47693 -0.03347 0.00000 ; N- CA- CB- CG2
5709 5711 5713 5719 3 2.97482 0.12134 2.04179
-1.89117 -3.24678 0.00000 ; N- CA- CB- CG1
5709 5711 5726 5727 3 0.00000 0.00000 0.00000
0.00000 0.00000 0.00000 ; N- CA- C- O

5709 5711 5726 5728 3 17.40544 -5.02080 -13.22144
9.20480 0.00000 0.00000 ; N- CA- C- N
5710 5709 5711 5712 3 0.00000 0.00000 0.00000
0.00000 0.00000 0.00000 ; H- N- CA- HA
5710 5709 5711 5713 3 0.00000 0.00000 0.00000
0.00000 0.00000 0.00000 ; H- N- CA- CB
5710 5709 5711 5726 3 0.00000 0.00000 0.00000
0.00000 0.00000 0.00000 ; H- N- CA- C
5711 5713 5715 5716 3 0.66944 2.00832 0.00000
-2.67776 0.00000 0.00000 ; CA- CB- CG2- HG21
5711 5713 5715 5717 3 0.66944 2.00832 0.00000
-2.67776 0.00000 0.00000 ; CA- CB- CG2- HG22
5711 5713 5715 5718 3 0.66944 2.00832 0.00000
-2.67776 0.00000 0.00000 ; CA- CB- CG2- HG23
5711 5713 5719 5720 3 0.66944 2.00832 0.00000
-2.67776 0.00000 0.00000 ; CA- CB- CG1- HG12
5711 5713 5719 5721 3 0.66944 2.00832 0.00000
-2.67776 0.00000 0.00000 ; CA- CB- CG1- HG13
5711 5713 5719 5722 3 4.24258 -0.52718 -7.25506
-1.79075 7.69856 0.00000 ; CA- CB- CG1- CD1
5711 5726 5728 5729 3 20.92000 0.00000 -20.92000
0.00000 0.00000 0.00000 ; CA- C- N- H
5711 5726 5728 5730 3 20.92000 0.00000 -20.92000
0.00000 0.00000 0.00000 ; CA- C- N- CA
5712 5711 5713 5714 3 0.65084 1.95253 0.00000
-2.60338 0.00000 0.00000 ; HA- CA- CB- HB
5712 5711 5713 5715 3 0.65084 1.95253 0.00000
-2.60338 0.00000 0.00000 ; HA- CA- CB- CG2
5712 5711 5713 5719 3 0.65084 1.95253 0.00000
-2.60338 0.00000 0.00000 ; HA- CA- CB- CG1
5712 5711 5726 5727 3 3.68192 -4.35136 0.00000
1.33888 0.00000 0.00000 ; HA- CA- C- O
5712 5711 5726 5728 3 0.00000 0.00000 0.00000
0.00000 0.00000 0.00000 ; HA- CA- C- N
5713 5711 5726 5727 3 0.00000 0.00000 0.00000
0.00000 0.00000 0.00000 ; CB- CA- C- O
5713 5711 5726 5728 3 2.51040 4.18400 1.67360
-6.69440 0.00000 0.00000 ; CB- CA- C- N
5713 5719 5722 5723 3 0.66944 2.00832 0.00000
-2.67776 0.00000 0.00000 ; CB- CG1- CD1- HD11
5713 5719 5722 5724 3 0.66944 2.00832 0.00000
-2.67776 0.00000 0.00000 ; CB- CG1- CD1- HD12
5713 5719 5722 5725 3 0.66944 2.00832 0.00000
-2.67776 0.00000 0.00000 ; CB- CG1- CD1- HD13
5714 5713 5711 5726 3 0.65084 1.95253 0.00000
-2.60338 0.00000 0.00000 ; HB- CB- CA- C
5714 5713 5715 5716 3 0.62760 1.88280 0.00000
-2.51040 0.00000 0.00000 ; HB- CB- CG2- HG21
5714 5713 5715 5717 3 0.62760 1.88280 0.00000
-2.51040 0.00000 0.00000 ; HB- CB- CG2- HG22
5714 5713 5715 5718 3 0.62760 1.88280 0.00000
-2.51040 0.00000 0.00000 ; HB- CB- CG2- HG23

5714 5713 5719 5720 3 0.62760 1.88280 0.00000
-2.51040 0.00000 0.00000 ; HB- CB- CG1- HG12
5714 5713 5719 5721 3 0.62760 1.88280 0.00000
-2.51040 0.00000 0.00000 ; HB- CB- CG1- HG13
5714 5713 5719 5722 3 0.66944 2.00832 0.00000
-2.67776 0.00000 0.00000 ; HB- CB- CG1- CD1
5715 5713 5711 5726 3 5.67350 3.55640 -6.16722
-2.47693 3.74886 0.00000 ; CG2- CB- CA- C
5715 5713 5719 5720 3 0.66944 2.00832 0.00000
-2.67776 0.00000 0.00000 ; CG2- CB- CG1- HG12
5715 5713 5719 5721 3 0.66944 2.00832 0.00000
-2.67776 0.00000 0.00000 ; CG2- CB- CG1- HG13
5715 5713 5719 5722 3 3.81162 0.49790 -8.14206
-1.79075 7.49773 0.00000 ; CG2- CB- CG1- CD1
5716 5715 5713 5719 3 0.66944 2.00832 0.00000
-2.67776 0.00000 0.00000 ; HG21- CG2- CB- CG1
5717 5715 5713 5719 3 0.66944 2.00832 0.00000
-2.67776 0.00000 0.00000 ; HG22- CG2- CB- CG1
5718 5715 5713 5719 3 0.66944 2.00832 0.00000
-2.67776 0.00000 0.00000 ; HG23- CG2- CB- CG1
5719 5713 5711 5726 3 8.26340 0.74057 -9.99976
-1.89117 3.84928 0.00000 ; CG1- CB- CA- C
5720 5719 5722 5723 3 0.62760 1.88280 0.00000
-2.51040 0.00000 0.00000 ; HG12- CG1- CD1- HD11
5720 5719 5722 5724 3 0.62760 1.88280 0.00000
-2.51040 0.00000 0.00000 ; HG12- CG1- CD1- HD12
5720 5719 5722 5725 3 0.62760 1.88280 0.00000
-2.51040 0.00000 0.00000 ; HG12- CG1- CD1- HD13
5721 5719 5722 5723 3 0.62760 1.88280 0.00000
-2.51040 0.00000 0.00000 ; HG13- CG1- CD1- HD11
5721 5719 5722 5724 3 0.62760 1.88280 0.00000
-2.51040 0.00000 0.00000 ; HG13- CG1- CD1- HD12
5721 5719 5722 5725 3 0.62760 1.88280 0.00000
-2.51040 0.00000 0.00000 ; HG13- CG1- CD1- HD13
5726 5728 5730 5731 3 0.00000 0.00000 0.00000
0.00000 0.00000 0.00000 ; C- N- CA- HA
5726 5728 5730 5732 3 11.71520 1.67360 15.06240 -
13.38880 0.00000 0.00000 ; C- N- CA- CB
5726 5728 5730 5743 3 1.75728 5.27184 2.25936
-7.02912 0.00000 0.00000 ; C- N- CA- C
5727 5726 5728 5729 3 29.28800 -8.36800 -20.92000
0.00000 0.00000 0.00000 ; O- C- N- H
5727 5726 5728 5730 3 20.92000 0.00000 -20.92000
0.00000 0.00000 0.00000 ; O- C- N- CA
5728 5730 5732 5733 3 0.65084 1.95253 0.00000
-2.60338 0.00000 0.00000 ; N- CA- CB- HB2
5728 5730 5732 5734 3 0.65084 1.95253 0.00000
-2.60338 0.00000 0.00000 ; N- CA- CB- HB3
5728 5730 5732 5735 3 4.79068 4.02919 -4.82834
-3.66518 2.97901 0.00000 ; N- CA- CB- CG
5728 5730 5743 5744 3 0.00000 0.00000 0.00000
0.00000 0.00000 0.00000 ; N- CA- C- O

5728 5730 5743 5745 3 17.40544 -5.02080 -13.22144
9.20480 0.00000 0.00000 ; N- CA- C- N
5729 5728 5730 5731 3 0.00000 0.00000 0.00000
0.00000 0.00000 0.00000 ; H- N- CA- HA
5729 5728 5730 5732 3 0.00000 0.00000 0.00000
0.00000 0.00000 0.00000 ; H- N- CA- CB
5729 5728 5730 5743 3 0.00000 0.00000 0.00000
0.00000 0.00000 0.00000 ; H- N- CA- C
5730 5732 5735 5736 3 5.98312 6.40152 3.28026 -
12.38464 -1.57318 0.00000 ; CA- CB- CG- ND1
5730 5732 5735 5741 3 0.00000 0.00000 0.00000
0.00000 0.00000 0.00000 ; CA- CB- CG- CD2
5730 5743 5745 5746 3 20.92000 0.00000 -20.92000
0.00000 0.00000 0.00000 ; CA- C- N- H
5730 5743 5745 5747 3 20.92000 0.00000 -20.92000
0.00000 0.00000 0.00000 ; CA- C- N- CA
5731 5730 5732 5733 3 0.65084 1.95253 0.00000
-2.60338 0.00000 0.00000 ; HA- CA- CB- HB2
5731 5730 5732 5734 3 0.65084 1.95253 0.00000
-2.60338 0.00000 0.00000 ; HA- CA- CB- HB3
5731 5730 5732 5735 3 0.65084 1.95253 0.00000
-2.60338 0.00000 0.00000 ; HA- CA- CB- CG
5731 5730 5743 5744 3 3.68192 -4.35136 0.00000
1.33888 0.00000 0.00000 ; HA- CA- C- O
5731 5730 5743 5745 3 0.00000 0.00000 0.00000
0.00000 0.00000 0.00000 ; HA- CA- C- N
5732 5730 5743 5744 3 0.00000 0.00000 0.00000
0.00000 0.00000 0.00000 ; CB- CA- C- O
5732 5730 5743 5745 3 2.51040 4.18400 1.67360
-6.69440 0.00000 0.00000 ; CB- CA- C- N
5732 5735 5736 5737 3 20.08320 0.00000 -20.08320
0.00000 0.00000 0.00000 ; CB- CG- ND1- CE1
5732 5735 5741 5739 3 44.97800 0.00000 -44.97800
0.00000 0.00000 0.00000 ; CB- CG- CD2- NE2
5732 5735 5741 5742 3 44.97800 0.00000 -44.97800
0.00000 0.00000 0.00000 ; CB- CG- CD2- HD2
5733 5732 5730 5743 3 0.65084 1.95253 0.00000
-2.60338 0.00000 0.00000 ; HB2- CB- CA- C
5733 5732 5735 5736 3 0.00000 0.00000 0.00000
0.00000 0.00000 0.00000 ; HB2- CB- CG- ND1
5733 5732 5735 5741 3 0.00000 0.00000 0.00000
0.00000 0.00000 0.00000 ; HB2- CB- CG- CD2
5734 5732 5730 5743 3 0.65084 1.95253 0.00000
-2.60338 0.00000 0.00000 ; HB3- CB- CA- C
5734 5732 5735 5736 3 0.00000 0.00000 0.00000
0.00000 0.00000 0.00000 ; HB3- CB- CG- ND1
5734 5732 5735 5741 3 0.00000 0.00000 0.00000
0.00000 0.00000 0.00000 ; HB3- CB- CG- CD2
5735 5732 5730 5743 3 3.76560 3.34720 -2.87859
-3.66518 0.83680 0.00000 ; CG- CB- CA- C
5735 5736 5737 5738 3 41.84000 0.00000 -41.84000
0.00000 0.00000 0.00000 ; CG- ND1- CE1- HE1

5735 5736 5737 5739 3 41.84000 0.00000 -41.84000
0.00000 0.00000 0.00000 ; CG- ND1- CE1- NE2
5735 5741 5739 5737 3 12.55200 0.00000 -12.55200
0.00000 0.00000 0.00000 ; CG- CD2- NE2- CE1
5735 5741 5739 5740 3 12.55200 0.00000 -12.55200
0.00000 0.00000 0.00000 ; CG- CD2- NE2- HE2
5736 5735 5741 5739 3 44.97800 0.00000 -44.97800
0.00000 0.00000 0.00000 ; ND1- CG- CD2- NE2
5736 5735 5741 5742 3 44.97800 0.00000 -44.97800
0.00000 0.00000 0.00000 ; ND1- CG- CD2- HD2
5736 5737 5739 5740 3 19.45560 0.00000 -19.45560
0.00000 0.00000 0.00000 ; ND1- CE1- NE2- HE2
5736 5737 5739 5741 3 19.45560 0.00000 -19.45560
0.00000 0.00000 0.00000 ; ND1- CE1- NE2- CD2
5737 5736 5735 5741 3 20.08320 0.00000 -20.08320
0.00000 0.00000 0.00000 ; CE1- ND1- CG- CD2
5737 5739 5741 5742 3 12.55200 0.00000 -12.55200
0.00000 0.00000 0.00000 ; CE1- NE2- CD2- HD2
5738 5737 5739 5740 3 19.45560 0.00000 -19.45560
0.00000 0.00000 0.00000 ; HE1- CE1- NE2- HE2
5738 5737 5739 5741 3 19.45560 0.00000 -19.45560
0.00000 0.00000 0.00000 ; HE1- CE1- NE2- CD2
5740 5739 5741 5742 3 12.55200 0.00000 -12.55200
0.00000 0.00000 0.00000 ; HE2- NE2- CD2- HD2
5743 5745 5747 5748 3 0.00000 0.00000 0.00000
0.00000 0.00000 0.00000 ; C- N- CA- HA
5743 5745 5747 5749 3 11.71520 1.67360 15.06240 -
13.38880 0.00000 0.00000 ; C- N- CA- CB
5743 5745 5747 5760 3 1.75728 5.27184 2.25936
-7.02912 0.00000 0.00000 ; C- N- CA- C
5744 5743 5745 5746 3 29.28800 -8.36800 -20.92000
0.00000 0.00000 0.00000 ; O- C- N- H
5744 5743 5745 5747 3 20.92000 0.00000 -20.92000
0.00000 0.00000 0.00000 ; O- C- N- CA
5745 5747 5749 5750 3 0.65084 1.95253 0.00000
-2.60338 0.00000 0.00000 ; N- CA- CB- HB2
5745 5747 5749 5751 3 0.65084 1.95253 0.00000
-2.60338 0.00000 0.00000 ; N- CA- CB- HB3
5745 5747 5749 5752 3 3.21331 2.22589 -4.15053
-2.40998 2.61082 0.00000 ; N- CA- CB- CG
5745 5747 5760 5761 3 0.00000 0.00000 0.00000
0.00000 0.00000 0.00000 ; N- CA- C- O
5745 5747 5760 5762 3 17.40544 -5.02080 -13.22144
9.20480 0.00000 0.00000 ; N- CA- C- N
5746 5745 5747 5748 3 0.00000 0.00000 0.00000
0.00000 0.00000 0.00000 ; H- N- CA- HA
5746 5745 5747 5749 3 0.00000 0.00000 0.00000
0.00000 0.00000 0.00000 ; H- N- CA- CB
5746 5745 5747 5760 3 0.00000 0.00000 0.00000
0.00000 0.00000 0.00000 ; H- N- CA- C
5747 5749 5752 5753 3 0.66944 2.00832 0.00000
-2.67776 0.00000 0.00000 ; CA- CB- CG- HG2

5747 5749 5752 5754 3 0.66944 2.00832 0.00000
-2.67776 0.00000 0.00000 ; CA- CB- CG- HG3
5747 5749 5752 5755 3 3.69866 -4.35136 -3.92459
6.89523 4.61914 0.00000 ; CA- CB- CG- CD
5747 5760 5762 5763 3 20.92000 0.00000 -20.92000
0.00000 0.00000 0.00000 ; CA- C- N- H
5747 5760 5762 5764 3 20.92000 0.00000 -20.92000
0.00000 0.00000 0.00000 ; CA- C- N- CA
5748 5747 5749 5750 3 0.65084 1.95253 0.00000
-2.60338 0.00000 0.00000 ; HA- CA- CB- HB2
5748 5747 5749 5751 3 0.65084 1.95253 0.00000
-2.60338 0.00000 0.00000 ; HA- CA- CB- HB3
5748 5747 5749 5752 3 0.65084 1.95253 0.00000
-2.60338 0.00000 0.00000 ; HA- CA- CB- CG
5748 5747 5760 5761 3 3.68192 -4.35136 0.00000
1.33888 0.00000 0.00000 ; HA- CA- C- O
5748 5747 5760 5762 3 0.00000 0.00000 0.00000
0.00000 0.00000 0.00000 ; HA- CA- C- N
5749 5747 5760 5761 3 0.00000 0.00000 0.00000
0.00000 0.00000 0.00000 ; CB- CA- C- O
5749 5747 5760 5762 3 2.51040 4.18400 1.67360
-6.69440 0.00000 0.00000 ; CB- CA- C- N
5749 5752 5755 5756 3 0.00000 0.00000 0.00000
0.00000 0.00000 0.00000 ; CB- CG- CD- OE1
5749 5752 5755 5757 3 10.32611 1.48114 -8.47678
1.42256 1.40582 0.00000 ; CB- CG- CD- NE2
5750 5749 5747 5760 3 0.65084 1.95253 0.00000
-2.60338 0.00000 0.00000 ; HB2- CB- CA- C
5750 5749 5752 5753 3 0.62760 1.88280 0.00000
-2.51040 0.00000 0.00000 ; HB2- CB- CG- HG2
5750 5749 5752 5754 3 0.62760 1.88280 0.00000
-2.51040 0.00000 0.00000 ; HB2- CB- CG- HG3
5750 5749 5752 5755 3 0.65084 1.95253 0.00000
-2.60338 0.00000 0.00000 ; HB2- CB- CG- CD
5751 5749 5747 5760 3 0.65084 1.95253 0.00000
-2.60338 0.00000 0.00000 ; HB3- CB- CA- C
5751 5749 5752 5753 3 0.62760 1.88280 0.00000
-2.51040 0.00000 0.00000 ; HB3- CB- CG- HG2
5751 5749 5752 5754 3 0.62760 1.88280 0.00000
-2.51040 0.00000 0.00000 ; HB3- CB- CG- HG3
5751 5749 5752 5755 3 0.65084 1.95253 0.00000
-2.60338 0.00000 0.00000 ; HB3- CB- CG- CD
5752 5749 5747 5760 3 6.86594 3.56895 -8.14206
-2.40998 4.85344 0.00000 ; CG- CB- CA- C
5752 5755 5757 5758 3 20.92000 0.00000 -20.92000
0.00000 0.00000 0.00000 ; CG- CD- NE2- HE21
5752 5755 5757 5759 3 20.92000 0.00000 -20.92000
0.00000 0.00000 0.00000 ; CG- CD- NE2- HE22
5753 5752 5755 5756 3 3.68192 -4.35136 0.00000
1.33888 0.00000 0.00000 ; HG2- CG- CD- OE1
5753 5752 5755 5757 3 0.00000 0.00000 0.00000
0.00000 0.00000 0.00000 ; HG2- CG- CD- NE2

5754 5752 5755 5756 3 3.68192 -4.35136 0.00000
1.33888 0.00000 0.00000 ; HG3- CG- CD- OE1
5754 5752 5755 5757 3 0.00000 0.00000 0.00000
0.00000 0.00000 0.00000 ; HG3- CG- CD- NE2
5756 5755 5757 5758 3 29.28800 -8.36800 -20.92000
0.00000 0.00000 0.00000 ; OE1- CD- NE2- HE21
5756 5755 5757 5759 3 29.28800 -8.36800 -20.92000
0.00000 0.00000 0.00000 ; OE1- CD- NE2- HE22
5760 5762 5764 5765 3 0.00000 0.00000 0.00000
0.00000 0.00000 0.00000 ; C- N- CA- HA
5760 5762 5764 5766 3 11.71520 1.67360 15.06240 -
13.38880 0.00000 0.00000 ; C- N- CA- CB
5760 5762 5764 5776 3 1.75728 5.27184 2.25936
-7.02912 0.00000 0.00000 ; C- N- CA- C
5761 5760 5762 5763 3 29.28800 -8.36800 -20.92000
0.00000 0.00000 0.00000 ; O- C- N- H
5761 5760 5762 5764 3 20.92000 0.00000 -20.92000
0.00000 0.00000 0.00000 ; O- C- N- CA
5762 5764 5766 5767 3 0.65084 1.95253 0.00000
-2.60338 0.00000 0.00000 ; N- CA- CB- HB
5762 5764 5766 5768 3 3.83673 0.44769 -1.77402
-2.47693 -0.03347 0.00000 ; N- CA- CB- CG1
5762 5764 5766 5772 3 3.83673 0.44769 -1.77402
-2.47693 -0.03347 0.00000 ; N- CA- CB- CG2
5762 5764 5776 5777 3 0.00000 0.00000 0.00000
0.00000 0.00000 0.00000 ; N- CA- C- O
5762 5764 5776 5778 3 17.40544 -5.02080 -13.22144
9.20480 0.00000 0.00000 ; N- CA- C- N
5763 5762 5764 5765 3 0.00000 0.00000 0.00000
0.00000 0.00000 0.00000 ; H- N- CA- HA
5763 5762 5764 5766 3 0.00000 0.00000 0.00000
0.00000 0.00000 0.00000 ; H- N- CA- CB
5763 5762 5764 5776 3 0.00000 0.00000 0.00000
0.00000 0.00000 0.00000 ; H- N- CA- C
5764 5766 5768 5769 3 0.66944 2.00832 0.00000
-2.67776 0.00000 0.00000 ; CA- CB- CG1- HG11
5764 5766 5768 5770 3 0.66944 2.00832 0.00000
-2.67776 0.00000 0.00000 ; CA- CB- CG1- HG12
5764 5766 5768 5771 3 0.66944 2.00832 0.00000
-2.67776 0.00000 0.00000 ; CA- CB- CG1- HG13
5764 5766 5772 5773 3 0.66944 2.00832 0.00000
-2.67776 0.00000 0.00000 ; CA- CB- CG2- HG21
5764 5766 5772 5774 3 0.66944 2.00832 0.00000
-2.67776 0.00000 0.00000 ; CA- CB- CG2- HG22
5764 5766 5772 5775 3 0.66944 2.00832 0.00000
-2.67776 0.00000 0.00000 ; CA- CB- CG2- HG23
5764 5776 5778 5779 3 20.92000 0.00000 -20.92000
0.00000 0.00000 0.00000 ; CA- C- N- H
5764 5776 5778 5780 3 20.92000 0.00000 -20.92000
0.00000 0.00000 0.00000 ; CA- C- N- CA
5765 5764 5766 5767 3 0.65084 1.95253 0.00000
-2.60338 0.00000 0.00000 ; HA- CA- CB- HB

5765 5764 5766 5768 3 0.65084 1.95253 0.00000
-2.60338 0.00000 0.00000 ; HA- CA- CB- CG1
5765 5764 5766 5772 3 0.65084 1.95253 0.00000
-2.60338 0.00000 0.00000 ; HA- CA- CB- CG2
5765 5764 5776 5777 3 3.68192 -4.35136 0.00000
1.33888 0.00000 0.00000 ; HA- CA- C- O
5765 5764 5776 5778 3 0.00000 0.00000 0.00000
0.00000 0.00000 0.00000 ; HA- CA- C- N
5766 5764 5776 5777 3 0.00000 0.00000 0.00000
0.00000 0.00000 0.00000 ; CB- CA- C- O
5766 5764 5776 5778 3 2.51040 4.18400 1.67360
-6.69440 0.00000 0.00000 ; CB- CA- C- N
5767 5766 5764 5776 3 0.65084 1.95253 0.00000
-2.60338 0.00000 0.00000 ; HB- CB- CA- C
5767 5766 5768 5769 3 0.62760 1.88280 0.00000
-2.51040 0.00000 0.00000 ; HB- CB- CG1- HG11
5767 5766 5768 5770 3 0.62760 1.88280 0.00000
-2.51040 0.00000 0.00000 ; HB- CB- CG1- HG12
5767 5766 5768 5771 3 0.62760 1.88280 0.00000
-2.51040 0.00000 0.00000 ; HB- CB- CG1- HG13
5767 5766 5772 5773 3 0.62760 1.88280 0.00000
-2.51040 0.00000 0.00000 ; HB- CB- CG2- HG21
5767 5766 5772 5774 3 0.62760 1.88280 0.00000
-2.51040 0.00000 0.00000 ; HB- CB- CG2- HG22
5767 5766 5772 5775 3 0.62760 1.88280 0.00000
-2.51040 0.00000 0.00000 ; HB- CB- CG2- HG23
5768 5766 5764 5776 3 5.67350 3.55640 -6.16722
-2.47693 3.74886 0.00000 ; CG1- CB- CA- C
5768 5766 5772 5773 3 0.66944 2.00832 0.00000
-2.67776 0.00000 0.00000 ; CG1- CB- CG2- HG21
5768 5766 5772 5774 3 0.66944 2.00832 0.00000
-2.67776 0.00000 0.00000 ; CG1- CB- CG2- HG22
5768 5766 5772 5775 3 0.66944 2.00832 0.00000
-2.67776 0.00000 0.00000 ; CG1- CB- CG2- HG23
5769 5768 5766 5772 3 0.66944 2.00832 0.00000
-2.67776 0.00000 0.00000 ; HG11- CG1- CB- CG2
5770 5768 5766 5772 3 0.66944 2.00832 0.00000
-2.67776 0.00000 0.00000 ; HG12- CG1- CB- CG2
5771 5768 5766 5772 3 0.66944 2.00832 0.00000
-2.67776 0.00000 0.00000 ; HG13- CG1- CB- CG2
5772 5766 5764 5776 3 5.67350 3.55640 -6.16722
-2.47693 3.74886 0.00000 ; CG2- CB- CA- C
5776 5778 5780 5781 3 0.00000 0.00000 0.00000
0.00000 0.00000 0.00000 ; C- N- CA- HA
5776 5778 5780 5782 3 11.71520 1.67360 15.06240 -
13.38880 0.00000 0.00000 ; C- N- CA- CB
5776 5778 5780 5795 3 1.75728 5.27184 2.25936
-7.02912 0.00000 0.00000 ; C- N- CA- C
5777 5776 5778 5779 3 29.28800 -8.36800 -20.92000
0.00000 0.00000 0.00000 ; O- C- N- H
5777 5776 5778 5780 3 20.92000 0.00000 -20.92000
0.00000 0.00000 0.00000 ; O- C- N- CA

5778 5780 5782 5783 3 0.65084 1.95253 0.00000
-2.60338 0.00000 0.00000 ; N- CA- CB- HB2
5778 5780 5782 5784 3 0.65084 1.95253 0.00000
-2.60338 0.00000 0.00000 ; N- CA- CB- HB3
5778 5780 5782 5785 3 3.79070 1.39746 -4.61077
-2.40998 2.44346 0.00000 ; N- CA- CB- CG
5778 5780 5795 5796 3 0.00000 0.00000 0.00000
0.00000 0.00000 0.00000 ; N- CA- C- O
5778 5780 5795 5797 3 17.40544 -5.02080 -13.22144
9.20480 0.00000 0.00000 ; N- CA- C- N
5779 5778 5780 5781 3 0.00000 0.00000 0.00000
0.00000 0.00000 0.00000 ; H- N- CA- HA
5779 5778 5780 5782 3 0.00000 0.00000 0.00000
0.00000 0.00000 0.00000 ; H- N- CA- CB
5779 5778 5780 5795 3 0.00000 0.00000 0.00000
0.00000 0.00000 0.00000 ; H- N- CA- C
5780 5782 5785 5786 3 0.66944 2.00832 0.00000
-2.67776 0.00000 0.00000 ; CA- CB- CG- HG
5780 5782 5785 5787 3 3.90367 0.19665 -6.21742
-2.37651 5.99149 0.00000 ; CA- CB- CG- CD1
5780 5782 5785 5791 3 3.90367 0.19665 -6.21742
-2.37651 5.99149 0.00000 ; CA- CB- CG- CD2
5780 5795 5797 5798 3 20.92000 0.00000 -20.92000
0.00000 0.00000 0.00000 ; CA- C- N- H
5780 5795 5797 5799 3 20.92000 0.00000 -20.92000
0.00000 0.00000 0.00000 ; CA- C- N- CA
5781 5780 5782 5783 3 0.65084 1.95253 0.00000
-2.60338 0.00000 0.00000 ; HA- CA- CB- HB2
5781 5780 5782 5784 3 0.65084 1.95253 0.00000
-2.60338 0.00000 0.00000 ; HA- CA- CB- HB3
5781 5780 5782 5785 3 0.65084 1.95253 0.00000
-2.60338 0.00000 0.00000 ; HA- CA- CB- CG
5781 5780 5795 5796 3 3.68192 -4.35136 0.00000
1.33888 0.00000 0.00000 ; HA- CA- C- O
5781 5780 5795 5797 3 0.00000 0.00000 0.00000
0.00000 0.00000 0.00000 ; HA- CA- C- N
5782 5780 5795 5796 3 0.00000 0.00000 0.00000
0.00000 0.00000 0.00000 ; CB- CA- C- O
5782 5780 5795 5797 3 2.51040 4.18400 1.67360
-6.69440 0.00000 0.00000 ; CB- CA- C- N
5782 5785 5787 5788 3 0.66944 2.00832 0.00000
-2.67776 0.00000 0.00000 ; CB- CG- CD1- HD11
5782 5785 5787 5789 3 0.66944 2.00832 0.00000
-2.67776 0.00000 0.00000 ; CB- CG- CD1- HD12
5782 5785 5787 5790 3 0.66944 2.00832 0.00000
-2.67776 0.00000 0.00000 ; CB- CG- CD1- HD13
5782 5785 5791 5792 3 0.66944 2.00832 0.00000
-2.67776 0.00000 0.00000 ; CB- CG- CD2- HD21
5782 5785 5791 5793 3 0.66944 2.00832 0.00000
-2.67776 0.00000 0.00000 ; CB- CG- CD2- HD22
5782 5785 5791 5794 3 0.66944 2.00832 0.00000
-2.67776 0.00000 0.00000 ; CB- CG- CD2- HD23

5783 5782 5780 5795 3 0.65084 1.95253 0.00000
-2.60338 0.00000 0.00000 ; HB2- CB- CA- C
5783 5782 5785 5786 3 0.62760 1.88280 0.00000
-2.51040 0.00000 0.00000 ; HB2- CB- CG- HG
5783 5782 5785 5787 3 0.66944 2.00832 0.00000
-2.67776 0.00000 0.00000 ; HB2- CB- CG- CD1
5783 5782 5785 5791 3 0.66944 2.00832 0.00000
-2.67776 0.00000 0.00000 ; HB2- CB- CG- CD2
5784 5782 5780 5795 3 0.65084 1.95253 0.00000
-2.60338 0.00000 0.00000 ; HB3- CB- CA- C
5784 5782 5785 5786 3 0.62760 1.88280 0.00000
-2.51040 0.00000 0.00000 ; HB3- CB- CG- HG
5784 5782 5785 5787 3 0.66944 2.00832 0.00000
-2.67776 0.00000 0.00000 ; HB3- CB- CG- CD1
5784 5782 5785 5791 3 0.66944 2.00832 0.00000
-2.67776 0.00000 0.00000 ; HB3- CB- CG- CD2
5785 5782 5780 5795 3 10.33448 -1.14642 -11.54784
-2.40998 6.35968 0.00000 ; CG- CB- CA- C
5786 5785 5787 5788 3 0.62760 1.88280 0.00000
-2.51040 0.00000 0.00000 ; HG- CG- CD1- HD11
5786 5785 5787 5789 3 0.62760 1.88280 0.00000
-2.51040 0.00000 0.00000 ; HG- CG- CD1- HD12
5786 5785 5787 5790 3 0.62760 1.88280 0.00000
-2.51040 0.00000 0.00000 ; HG- CG- CD1- HD13
5786 5785 5791 5792 3 0.62760 1.88280 0.00000
-2.51040 0.00000 0.00000 ; HG- CG- CD2- HD21
5786 5785 5791 5793 3 0.62760 1.88280 0.00000
-2.51040 0.00000 0.00000 ; HG- CG- CD2- HD22
5786 5785 5791 5794 3 0.62760 1.88280 0.00000
-2.51040 0.00000 0.00000 ; HG- CG- CD2- HD23
5787 5785 5791 5792 3 0.66944 2.00832 0.00000
-2.67776 0.00000 0.00000 ; CD1- CG- CD2- HD21
5787 5785 5791 5793 3 0.66944 2.00832 0.00000
-2.67776 0.00000 0.00000 ; CD1- CG- CD2- HD22
5787 5785 5791 5794 3 0.66944 2.00832 0.00000
-2.67776 0.00000 0.00000 ; CD1- CG- CD2- HD23
5788 5787 5785 5791 3 0.66944 2.00832 0.00000
-2.67776 0.00000 0.00000 ; HD11- CD1- CG- CD2
5789 5787 5785 5791 3 0.66944 2.00832 0.00000
-2.67776 0.00000 0.00000 ; HD12- CD1- CG- CD2
5790 5787 5785 5791 3 0.66944 2.00832 0.00000
-2.67776 0.00000 0.00000 ; HD13- CD1- CG- CD2
5795 5797 5799 5800 3 0.00000 0.00000 0.00000
0.00000 0.00000 0.00000 ; C- N- CA- HA
5795 5797 5799 5801 3 11.71520 1.67360 15.06240 -
13.38880 0.00000 0.00000 ; C- N- CA- CB
5795 5797 5799 5819 3 1.75728 5.27184 2.25936
-7.02912 0.00000 0.00000 ; C- N- CA- C
5796 5795 5797 5798 3 29.28800 -8.36800 -20.92000
0.00000 0.00000 0.00000 ; O- C- N- H
5796 5795 5797 5799 3 20.92000 0.00000 -20.92000
0.00000 0.00000 0.00000 ; O- C- N- CA

5797 5799 5801 5802 3 0.65084 1.95253 0.00000
-2.60338 0.00000 0.00000 ; N- CA- CB- HB2
5797 5799 5801 5803 3 0.65084 1.95253 0.00000
-2.60338 0.00000 0.00000 ; N- CA- CB- HB3
5797 5799 5801 5804 3 0.65084 1.95253 0.00000
-2.60338 0.00000 0.00000 ; N- CA- CB- CG
5797 5799 5819 5820 3 0.00000 0.00000 0.00000
0.00000 0.00000 0.00000 ; N- CA- C- O
5797 5799 5819 5821 3 17.40544 -5.02080 -13.22144
9.20480 0.00000 0.00000 ; N- CA- C- N
5798 5797 5799 5800 3 0.00000 0.00000 0.00000
0.00000 0.00000 0.00000 ; H- N- CA- HA
5798 5797 5799 5801 3 0.00000 0.00000 0.00000
0.00000 0.00000 0.00000 ; H- N- CA- CB
5798 5797 5799 5819 3 0.00000 0.00000 0.00000
0.00000 0.00000 0.00000 ; H- N- CA- C
5799 5801 5804 5805 3 0.66944 2.00832 0.00000
-2.67776 0.00000 0.00000 ; CA- CB- CG- HG2
5799 5801 5804 5806 3 0.66944 2.00832 0.00000
-2.67776 0.00000 0.00000 ; CA- CB- CG- HG3
5799 5801 5804 5807 3 3.68192 3.09616 -2.09200
-3.01248 0.00000 0.00000 ; CA- CB- CG- CD
5799 5819 5821 5822 3 20.92000 0.00000 -20.92000
0.00000 0.00000 0.00000 ; CA- C- N- H
5799 5819 5821 5823 3 20.92000 0.00000 -20.92000
0.00000 0.00000 0.00000 ; CA- C- N- CA
5800 5799 5801 5802 3 0.65084 1.95253 0.00000
-2.60338 0.00000 0.00000 ; HA- CA- CB- HB2
5800 5799 5801 5803 3 0.65084 1.95253 0.00000
-2.60338 0.00000 0.00000 ; HA- CA- CB- HB3
5800 5799 5801 5804 3 0.65084 1.95253 0.00000
-2.60338 0.00000 0.00000 ; HA- CA- CB- CG
5800 5799 5819 5820 3 3.68192 -4.35136 0.00000
1.33888 0.00000 0.00000 ; HA- CA- C- O
5800 5799 5819 5821 3 0.00000 0.00000 0.00000
0.00000 0.00000 0.00000 ; HA- CA- C- N
5801 5799 5819 5820 3 0.00000 0.00000 0.00000
0.00000 0.00000 0.00000 ; CB- CA- C- O
5801 5799 5819 5821 3 2.51040 4.18400 1.67360
-6.69440 0.00000 0.00000 ; CB- CA- C- N
5801 5804 5807 5808 3 0.65084 1.95253 0.00000
-2.60338 0.00000 0.00000 ; CB- CG- CD- HD2
5801 5804 5807 5809 3 0.65084 1.95253 0.00000
-2.60338 0.00000 0.00000 ; CB- CG- CD- HD3
5801 5804 5807 5810 3 0.65084 1.95253 0.00000
-2.60338 0.00000 0.00000 ; CB- CG- CD- NE
5802 5801 5799 5819 3 0.65084 1.95253 0.00000
-2.60338 0.00000 0.00000 ; HB2- CB- CA- C
5802 5801 5804 5805 3 0.62760 1.88280 0.00000
-2.51040 0.00000 0.00000 ; HB2- CB- CG- HG2
5802 5801 5804 5806 3 0.62760 1.88280 0.00000
-2.51040 0.00000 0.00000 ; HB2- CB- CG- HG3

```
  5802   5801   5804   5807       3    0.66944    2.00832    0.00000
-2.67776    0.00000    0.00000 ;    HB2-   CB-    CG-    CD
  5803   5801   5799   5819       3    0.65084    1.95253    0.00000
-2.60338    0.00000    0.00000 ;    HB3-   CB-    CA-     C
  5803   5801   5804   5805       3    0.62760    1.88280    0.00000
-2.51040    0.00000    0.00000 ;    HB3-   CB-    CG-   HG2
  5803   5801   5804   5806       3    0.62760    1.88280    0.00000
-2.51040    0.00000    0.00000 ;    HB3-   CB-    CG-   HG3
  5803   5801   5804   5807       3    0.66944    2.00832    0.00000
-2.67776    0.00000    0.00000 ;    HB3-   CB-    CG-    CD
  5804   5801   5799   5819       3    0.65084    1.95253    0.00000
-2.60338    0.00000    0.00000 ;     CG-   CB-    CA-     C
  5804   5807   5810   5811       3    0.00000    0.00000    0.00000
0.00000    0.00000    0.00000 ;     CG-   CD-    NE-    HE
  5804   5807   5810   5812       3    0.00000    0.00000    0.00000
0.00000    0.00000    0.00000 ;     CG-   CD-    NE-    CZ
  5805   5804   5807   5808       3    0.65084    1.95253    0.00000
-2.60338    0.00000    0.00000 ;    HG2-   CG-    CD-   HD2
  5805   5804   5807   5809       3    0.65084    1.95253    0.00000
-2.60338    0.00000    0.00000 ;    HG2-   CG-    CD-   HD3
  5805   5804   5807   5810       3    0.65084    1.95253    0.00000
-2.60338    0.00000    0.00000 ;    HG2-   CG-    CD-    NE
  5806   5804   5807   5808       3    0.65084    1.95253    0.00000
-2.60338    0.00000    0.00000 ;    HG3-   CG-    CD-   HD2
  5806   5804   5807   5809       3    0.65084    1.95253    0.00000
-2.60338    0.00000    0.00000 ;    HG3-   CG-    CD-   HD3
  5806   5804   5807   5810       3    0.65084    1.95253    0.00000
-2.60338    0.00000    0.00000 ;    HG3-   CG-    CD-    NE
  5807   5810   5812   5813       3   20.08320    0.00000  -20.08320
0.00000    0.00000    0.00000 ;     CD-   NE-    CZ-   NH1
  5807   5810   5812   5816       3   20.08320    0.00000  -20.08320
0.00000    0.00000    0.00000 ;     CD-   NE-    CZ-   NH2
  5808   5807   5810   5811       3    0.00000    0.00000    0.00000
0.00000    0.00000    0.00000 ;    HD2-   CD-    NE-    HE
  5808   5807   5810   5812       3    0.00000    0.00000    0.00000
0.00000    0.00000    0.00000 ;    HD2-   CD-    NE-    CZ
  5809   5807   5810   5811       3    0.00000    0.00000    0.00000
0.00000    0.00000    0.00000 ;    HD3-   CD-    NE-    HE
  5809   5807   5810   5812       3    0.00000    0.00000    0.00000
0.00000    0.00000    0.00000 ;    HD3-   CD-    NE-    CZ
  5810   5812   5813   5814       3   20.08320    0.00000  -20.08320
0.00000    0.00000    0.00000 ;     NE-   CZ-   NH1-  HH11
  5810   5812   5813   5815       3   20.08320    0.00000  -20.08320
0.00000    0.00000    0.00000 ;     NE-   CZ-   NH1-  HH12
  5810   5812   5816   5817       3   20.08320    0.00000  -20.08320
0.00000    0.00000    0.00000 ;     NE-   CZ-   NH2-  HH21
  5810   5812   5816   5818       3   20.08320    0.00000  -20.08320
0.00000    0.00000    0.00000 ;     NE-   CZ-   NH2-  HH22
  5811   5810   5812   5813       3   20.08320    0.00000  -20.08320
0.00000    0.00000    0.00000 ;     HE-   NE-    CZ-   NH1
  5811   5810   5812   5816       3   20.08320    0.00000  -20.08320
0.00000    0.00000    0.00000 ;     HE-   NE-    CZ-   NH2
```

5813 5812 5816 5817 3 20.08320 0.00000 -20.08320
0.00000 0.00000 0.00000 ; NH1- CZ- NH2- HH21
5813 5812 5816 5818 3 20.08320 0.00000 -20.08320
0.00000 0.00000 0.00000 ; NH1- CZ- NH2- HH22
5814 5813 5812 5816 3 20.08320 0.00000 -20.08320
0.00000 0.00000 0.00000 ; HH11- NH1- CZ- NH2
5815 5813 5812 5816 3 20.08320 0.00000 -20.08320
0.00000 0.00000 0.00000 ; HH12- NH1- CZ- NH2
5819 5821 5823 5824 3 0.00000 0.00000 0.00000
0.00000 0.00000 0.00000 ; C- N- CA- HA
5819 5821 5823 5825 3 11.71520 1.67360 15.06240 -
13.38880 0.00000 0.00000 ; C- N- CA- CB
5819 5821 5823 5831 3 1.75728 5.27184 2.25936
-7.02912 0.00000 0.00000 ; C- N- CA- C
5820 5819 5821 5822 3 29.28800 -8.36800 -20.92000
0.00000 0.00000 0.00000 ; O- C- N- H
5820 5819 5821 5823 3 20.92000 0.00000 -20.92000
0.00000 0.00000 0.00000 ; O- C- N- CA
5821 5823 5825 5826 3 0.65084 1.95253 0.00000
-2.60338 0.00000 0.00000 ; N- CA- CB- HB2
5821 5823 5825 5827 3 0.65084 1.95253 0.00000
-2.60338 0.00000 0.00000 ; N- CA- CB- HB3
5821 5823 5825 5828 3 15.41386 9.74035 -8.39310
-0.97069 2.97901 0.00000 ; N- CA- CB- CG
5821 5823 5831 5832 3 0.00000 0.00000 0.00000
0.00000 0.00000 0.00000 ; N- CA- C- O
5821 5823 5831 5833 3 17.40544 -5.02080 -13.22144
9.20480 0.00000 0.00000 ; N- CA- C- N
5822 5821 5823 5824 3 0.00000 0.00000 0.00000
0.00000 0.00000 0.00000 ; H- N- CA- HA
5822 5821 5823 5825 3 0.00000 0.00000 0.00000
0.00000 0.00000 0.00000 ; H- N- CA- CB
5822 5821 5823 5831 3 0.00000 0.00000 0.00000
0.00000 0.00000 0.00000 ; H- N- CA- C
5823 5825 5828 5829 3 6.43499 0.00000 -5.39736
0.00000 -1.03763 0.00000 ; CA- CB- CG- OD1
5823 5825 5828 5830 3 6.43499 0.00000 -5.39736
0.00000 -1.03763 0.00000 ; CA- CB- CG- OD2
5823 5831 5833 5834 3 20.92000 0.00000 -20.92000
0.00000 0.00000 0.00000 ; CA- C- N- H
5823 5831 5833 5835 3 20.92000 0.00000 -20.92000
0.00000 0.00000 0.00000 ; CA- C- N- CA
5824 5823 5825 5826 3 0.65084 1.95253 0.00000
-2.60338 0.00000 0.00000 ; HA- CA- CB- HB2
5824 5823 5825 5827 3 0.65084 1.95253 0.00000
-2.60338 0.00000 0.00000 ; HA- CA- CB- HB3
5824 5823 5825 5828 3 0.65084 1.95253 0.00000
-2.60338 0.00000 0.00000 ; HA- CA- CB- CG
5824 5823 5831 5832 3 3.68192 -4.35136 0.00000
1.33888 0.00000 0.00000 ; HA- CA- C- O
5824 5823 5831 5833 3 0.00000 0.00000 0.00000
0.00000 0.00000 0.00000 ; HA- CA- C- N

5825 5823 5831 5832 3 0.00000 0.00000 0.00000
0.00000 0.00000 0.00000 ; CB- CA- C- O
5825 5823 5831 5833 3 2.51040 4.18400 1.67360
-6.69440 0.00000 0.00000 ; CB- CA- C- N
5826 5825 5823 5831 3 0.65084 1.95253 0.00000
-2.60338 0.00000 0.00000 ; HB2- CB- CA- C
5826 5825 5828 5829 3 0.00000 0.00000 0.00000
0.00000 0.00000 0.00000 ; HB2- CB- CG- OD1
5826 5825 5828 5830 3 0.00000 0.00000 0.00000
0.00000 0.00000 0.00000 ; HB2- CB- CG- OD2
5827 5825 5823 5831 3 0.65084 1.95253 0.00000
-2.60338 0.00000 0.00000 ; HB3- CB- CA- C
5827 5825 5828 5829 3 0.00000 0.00000 0.00000
0.00000 0.00000 0.00000 ; HB3- CB- CG- OD1
5827 5825 5828 5830 3 0.00000 0.00000 0.00000
0.00000 0.00000 0.00000 ; HB3- CB- CG- OD2
5828 5825 5823 5831 3 7.14627 -1.04600 -8.99560
-0.97069 5.15469 0.00000 ; CG- CB- CA- C
5831 5833 5835 5836 3 0.00000 0.00000 0.00000
0.00000 0.00000 0.00000 ; C- N- CA- HA
5831 5833 5835 5837 3 11.71520 1.67360 15.06240 -
13.38880 0.00000 0.00000 ; C- N- CA- CB
5831 5833 5835 5850 3 1.75728 5.27184 2.25936
-7.02912 0.00000 0.00000 ; C- N- CA- C
5832 5831 5833 5834 3 29.28800 -8.36800 -20.92000
0.00000 0.00000 0.00000 ; O- C- N- H
5832 5831 5833 5835 3 20.92000 0.00000 -20.92000
0.00000 0.00000 0.00000 ; O- C- N- CA
5833 5835 5837 5838 3 0.65084 1.95253 0.00000
-2.60338 0.00000 0.00000 ; N- CA- CB- HB
5833 5835 5837 5839 3 3.83673 0.44769 -1.77402
-2.47693 -0.03347 0.00000 ; N- CA- CB- CG2
5833 5835 5837 5843 3 2.97482 0.12134 2.04179
-1.89117 -3.24678 0.00000 ; N- CA- CB- CG1
5833 5835 5850 5851 3 0.00000 0.00000 0.00000
0.00000 0.00000 0.00000 ; N- CA- C- O
5833 5835 5850 5852 3 17.40544 -5.02080 -13.22144
9.20480 0.00000 0.00000 ; N- CA- C- N
5834 5833 5835 5836 3 0.00000 0.00000 0.00000
0.00000 0.00000 0.00000 ; H- N- CA- HA
5834 5833 5835 5837 3 0.00000 0.00000 0.00000
0.00000 0.00000 0.00000 ; H- N- CA- CB
5834 5833 5835 5850 3 0.00000 0.00000 0.00000
0.00000 0.00000 0.00000 ; H- N- CA- C
5835 5837 5839 5840 3 0.66944 2.00832 0.00000
-2.67776 0.00000 0.00000 ; CA- CB- CG2- HG21
5835 5837 5839 5841 3 0.66944 2.00832 0.00000
-2.67776 0.00000 0.00000 ; CA- CB- CG2- HG22
5835 5837 5839 5842 3 0.66944 2.00832 0.00000
-2.67776 0.00000 0.00000 ; CA- CB- CG2- HG23
5835 5837 5843 5844 3 0.66944 2.00832 0.00000
-2.67776 0.00000 0.00000 ; CA- CB- CG1- HG12

```
  5835   5837   5843   5845      3    0.66944    2.00832    0.00000
-2.67776    0.00000    0.00000 ;      CA-    CB-   CG1-  HG13
  5835   5837   5843   5846      3    4.24258   -0.52718   -7.25506
-1.79075    7.69856    0.00000 ;      CA-    CB-   CG1-   CD1
  5835   5850   5852   5853      3   20.92000    0.00000  -20.92000
0.00000    0.00000    0.00000 ;      CA-     C-     N-     H
  5835   5850   5852   5854      3   20.92000    0.00000  -20.92000
0.00000    0.00000    0.00000 ;      CA-     C-     N-    CA
  5836   5835   5837   5838      3    0.65084    1.95253    0.00000
-2.60338    0.00000    0.00000 ;      HA-    CA-    CB-    HB
  5836   5835   5837   5839      3    0.65084    1.95253    0.00000
-2.60338    0.00000    0.00000 ;      HA-    CA-    CB-   CG2
  5836   5835   5837   5843      3    0.65084    1.95253    0.00000
-2.60338    0.00000    0.00000 ;      HA-    CA-    CB-   CG1
  5836   5835   5850   5851      3    3.68192   -4.35136    0.00000
1.33888    0.00000    0.00000 ;      HA-    CA-     C-     O
  5836   5835   5850   5852      3    0.00000    0.00000    0.00000
0.00000    0.00000    0.00000 ;      HA-    CA-     C-     N
  5837   5835   5850   5851      3    0.00000    0.00000    0.00000
0.00000    0.00000    0.00000 ;      CB-    CA-     C-     O
  5837   5835   5850   5852      3    2.51040    4.18400    1.67360
-6.69440    0.00000    0.00000 ;      CB-    CA-     C-     N
  5837   5843   5846   5847      3    0.66944    2.00832    0.00000
-2.67776    0.00000    0.00000 ;      CB-   CG1-   CD1-  HD11
  5837   5843   5846   5848      3    0.66944    2.00832    0.00000
-2.67776    0.00000    0.00000 ;      CB-   CG1-   CD1-  HD12
  5837   5843   5846   5849      3    0.66944    2.00832    0.00000
-2.67776    0.00000    0.00000 ;      CB-   CG1-   CD1-  HD13
  5838   5837   5835   5850      3    0.65084    1.95253    0.00000
-2.60338    0.00000    0.00000 ;      HB-    CB-    CA-     C
  5838   5837   5839   5840      3    0.62760    1.88280    0.00000
-2.51040    0.00000    0.00000 ;      HB-    CB-   CG2-  HG21
  5838   5837   5839   5841      3    0.62760    1.88280    0.00000
-2.51040    0.00000    0.00000 ;      HB-    CB-   CG2-  HG22
  5838   5837   5839   5842      3    0.62760    1.88280    0.00000
-2.51040    0.00000    0.00000 ;      HB-    CB-   CG2-  HG23
  5838   5837   5843   5844      3    0.62760    1.88280    0.00000
-2.51040    0.00000    0.00000 ;      HB-    CB-   CG1-  HG12
  5838   5837   5843   5845      3    0.62760    1.88280    0.00000
-2.51040    0.00000    0.00000 ;      HB-    CB-   CG1-  HG13
  5838   5837   5843   5846      3    0.66944    2.00832    0.00000
-2.67776    0.00000    0.00000 ;      HB-    CB-   CG1-   CD1
  5839   5837   5835   5850      3    5.67350    3.55640   -6.16722
-2.47693    3.74886    0.00000 ;     CG2-    CB-    CA-     C
  5839   5837   5843   5844      3    0.66944    2.00832    0.00000
-2.67776    0.00000    0.00000 ;     CG2-    CB-   CG1-  HG12
  5839   5837   5843   5845      3    0.66944    2.00832    0.00000
-2.67776    0.00000    0.00000 ;     CG2-    CB-   CG1-  HG13
  5839   5837   5843   5846      3    3.81162    0.49790   -8.14206
-1.79075    7.49773    0.00000 ;     CG2-    CB-   CG1-   CD1
  5840   5839   5837   5843      3    0.66944    2.00832    0.00000
-2.67776    0.00000    0.00000 ;    HG21-   CG2-    CB-   CG1
```

5841 5839 5837 5843 3 0.66944 2.00832 0.00000
-2.67776 0.00000 0.00000 ; HG22- CG2- CB- CG1
5842 5839 5837 5843 3 0.66944 2.00832 0.00000
-2.67776 0.00000 0.00000 ; HG23- CG2- CB- CG1
5843 5837 5835 5850 3 8.26340 0.74057 -9.99976
-1.89117 3.84928 0.00000 ; CG1- CB- CA- C
5844 5843 5846 5847 3 0.62760 1.88280 0.00000
-2.51040 0.00000 0.00000 ; HG12- CG1- CD1- HD11
5844 5843 5846 5848 3 0.62760 1.88280 0.00000
-2.51040 0.00000 0.00000 ; HG12- CG1- CD1- HD12
5844 5843 5846 5849 3 0.62760 1.88280 0.00000
-2.51040 0.00000 0.00000 ; HG12- CG1- CD1- HD13
5845 5843 5846 5847 3 0.62760 1.88280 0.00000
-2.51040 0.00000 0.00000 ; HG13- CG1- CD1- HD11
5845 5843 5846 5848 3 0.62760 1.88280 0.00000
-2.51040 0.00000 0.00000 ; HG13- CG1- CD1- HD12
5845 5843 5846 5849 3 0.62760 1.88280 0.00000
-2.51040 0.00000 0.00000 ; HG13- CG1- CD1- HD13
5850 5852 5854 5855 3 0.00000 0.00000 0.00000
0.00000 0.00000 0.00000 ; C- N- CA- HA
5850 5852 5854 5856 3 11.71520 1.67360 15.06240 -
13.38880 0.00000 0.00000 ; C- N- CA- CB
5850 5852 5854 5862 3 1.75728 5.27184 2.25936
-7.02912 0.00000 0.00000 ; C- N- CA- C
5851 5850 5852 5853 3 29.28800 -8.36800 -20.92000
0.00000 0.00000 0.00000 ; O- C- N- H
5851 5850 5852 5854 3 20.92000 0.00000 -20.92000
0.00000 0.00000 0.00000 ; O- C- N- CA
5852 5854 5856 5857 3 0.65084 1.95253 0.00000
-2.60338 0.00000 0.00000 ; N- CA- CB- HB2
5852 5854 5856 5858 3 0.65084 1.95253 0.00000
-2.60338 0.00000 0.00000 ; N- CA- CB- HB3
5852 5854 5856 5859 3 15.41386 9.74035 -8.39310
-0.97069 2.97901 0.00000 ; N- CA- CB- CG
5852 5854 5862 5863 3 0.00000 0.00000 0.00000
0.00000 0.00000 0.00000 ; N- CA- C- O
5852 5854 5862 5864 3 17.40544 -5.02080 -13.22144
9.20480 0.00000 0.00000 ; N- CA- C- N
5853 5852 5854 5855 3 0.00000 0.00000 0.00000
0.00000 0.00000 0.00000 ; H- N- CA- HA
5853 5852 5854 5856 3 0.00000 0.00000 0.00000
0.00000 0.00000 0.00000 ; H- N- CA- CB
5853 5852 5854 5862 3 0.00000 0.00000 0.00000
0.00000 0.00000 0.00000 ; H- N- CA- C
5854 5856 5859 5860 3 6.43499 0.00000 -5.39736
0.00000 -1.03763 0.00000 ; CA- CB- CG- OD1
5854 5856 5859 5861 3 6.43499 0.00000 -5.39736
0.00000 -1.03763 0.00000 ; CA- CB- CG- OD2
5854 5862 5864 5865 3 20.92000 0.00000 -20.92000
0.00000 0.00000 0.00000 ; CA- C- N- H
5854 5862 5864 5866 3 20.92000 0.00000 -20.92000
0.00000 0.00000 0.00000 ; CA- C- N- CA

5855 5854 5856 5857 3 0.65084 1.95253 0.00000
-2.60338 0.00000 0.00000 ; HA- CA- CB- HB2
5855 5854 5856 5858 3 0.65084 1.95253 0.00000
-2.60338 0.00000 0.00000 ; HA- CA- CB- HB3
5855 5854 5856 5859 3 0.65084 1.95253 0.00000
-2.60338 0.00000 0.00000 ; HA- CA- CB- CG
5855 5854 5862 5863 3 3.68192 -4.35136 0.00000
1.33888 0.00000 0.00000 ; HA- CA- C- O
5855 5854 5862 5864 3 0.00000 0.00000 0.00000
0.00000 0.00000 0.00000 ; HA- CA- C- N
5856 5854 5862 5863 3 0.00000 0.00000 0.00000
0.00000 0.00000 0.00000 ; CB- CA- C- O
5856 5854 5862 5864 3 2.51040 4.18400 1.67360
-6.69440 0.00000 0.00000 ; CB- CA- C- N
5857 5856 5854 5862 3 0.65084 1.95253 0.00000
-2.60338 0.00000 0.00000 ; HB2- CB- CA- C
5857 5856 5859 5860 3 0.00000 0.00000 0.00000
0.00000 0.00000 0.00000 ; HB2- CB- CG- OD1
5857 5856 5859 5861 3 0.00000 0.00000 0.00000
0.00000 0.00000 0.00000 ; HB2- CB- CG- OD2
5858 5856 5854 5862 3 0.65084 1.95253 0.00000
-2.60338 0.00000 0.00000 ; HB3- CB- CA- C
5858 5856 5859 5860 3 0.00000 0.00000 0.00000
0.00000 0.00000 0.00000 ; HB3- CB- CG- OD1
5858 5856 5859 5861 3 0.00000 0.00000 0.00000
0.00000 0.00000 0.00000 ; HB3- CB- CG- OD2
5859 5856 5854 5862 3 7.14627 -1.04600 -8.99560
-0.97069 5.15469 0.00000 ; CG- CB- CA- C
5862 5864 5866 5867 3 0.00000 0.00000 0.00000
0.00000 0.00000 0.00000 ; C- N- CA- HA
5862 5864 5866 5868 3 11.71520 1.67360 15.06240 -
13.38880 0.00000 0.00000 ; C- N- CA- CB
5862 5864 5866 5873 3 1.75728 5.27184 2.25936
-7.02912 0.00000 0.00000 ; C- N- CA- C
5863 5862 5864 5865 3 29.28800 -8.36800 -20.92000
0.00000 0.00000 0.00000 ; O- C- N- H
5863 5862 5864 5866 3 20.92000 0.00000 -20.92000
0.00000 0.00000 0.00000 ; O- C- N- CA
5864 5866 5868 5869 3 0.65084 1.95253 0.00000
-2.60338 0.00000 0.00000 ; N- CA- CB- HB2
5864 5866 5868 5870 3 0.65084 1.95253 0.00000
-2.60338 0.00000 0.00000 ; N- CA- CB- HB3
5864 5866 5868 5871 3 7.86174 2.24681 -7.41405
-6.71114 5.35552 0.00000 ; N- CA- CB- OG
5864 5866 5873 5874 3 0.00000 0.00000 0.00000
0.00000 0.00000 0.00000 ; N- CA- C- O
5864 5866 5873 5875 3 17.40544 -5.02080 -13.22144
9.20480 0.00000 0.00000 ; N- CA- C- N
5865 5864 5866 5867 3 0.00000 0.00000 0.00000
0.00000 0.00000 0.00000 ; H- N- CA- HA
5865 5864 5866 5868 3 0.00000 0.00000 0.00000
0.00000 0.00000 0.00000 ; H- N- CA- CB

5865 5864 5866 5873 3 0.00000 0.00000 0.00000
0.00000 0.00000 0.00000 ; H- N- CA- C
5866 5868 5871 5872 3 2.05853 2.46856 3.48109
-4.46851 0.23430 0.00000 ; CA- CB- OG- HG
5866 5873 5875 5876 3 20.92000 0.00000 -20.92000
0.00000 0.00000 0.00000 ; CA- C- N- H
5866 5873 5875 5877 3 20.92000 0.00000 -20.92000
0.00000 0.00000 0.00000 ; CA- C- N- CA
5867 5866 5868 5869 3 0.65084 1.95253 0.00000
-2.60338 0.00000 0.00000 ; HA- CA- CB- HB2
5867 5866 5868 5870 3 0.65084 1.95253 0.00000
-2.60338 0.00000 0.00000 ; HA- CA- CB- HB3
5867 5866 5868 5871 3 1.04600 -1.04600 0.00000
0.00000 0.00000 0.00000 ; HA- CA- CB- OG
5867 5866 5873 5874 3 3.68192 -4.35136 0.00000
1.33888 0.00000 0.00000 ; HA- CA- C- O
5867 5866 5873 5875 3 0.00000 0.00000 0.00000
0.00000 0.00000 0.00000 ; HA- CA- C- N
5868 5866 5873 5874 3 0.00000 0.00000 0.00000
0.00000 0.00000 0.00000 ; CB- CA- C- O
5868 5866 5873 5875 3 2.51040 4.18400 1.67360
-6.69440 0.00000 0.00000 ; CB- CA- C- N
5869 5868 5866 5873 3 0.65084 1.95253 0.00000
-2.60338 0.00000 0.00000 ; HB2- CB- CA- C
5869 5868 5871 5872 3 0.69733 2.09200 0.00000
-2.78933 0.00000 0.00000 ; HB2- CB- OG- HG
5870 5868 5866 5873 3 0.65084 1.95253 0.00000
-2.60338 0.00000 0.00000 ; HB3- CB- CA- C
5870 5868 5871 5872 3 0.69733 2.09200 0.00000
-2.78933 0.00000 0.00000 ; HB3- CB- OG- HG
5871 5868 5866 5873 3 7.34710 7.79898 -6.14211
-6.71114 4.31789 0.00000 ; OG- CB- CA- C
5873 5875 5877 5878 3 0.00000 0.00000 0.00000
0.00000 0.00000 0.00000 ; C- N- CA- HA
5873 5875 5877 5879 3 11.71520 1.67360 15.06240 -
13.38880 0.00000 0.00000 ; C- N- CA- CB
5873 5875 5877 5887 3 1.75728 5.27184 2.25936
-7.02912 0.00000 0.00000 ; C- N- CA- C
5874 5873 5875 5876 3 29.28800 -8.36800 -20.92000
0.00000 0.00000 0.00000 ; O- C- N- H
5874 5873 5875 5877 3 20.92000 0.00000 -20.92000
0.00000 0.00000 0.00000 ; O- C- N- CA
5875 5877 5879 5880 3 0.65084 1.95253 0.00000
-2.60338 0.00000 0.00000 ; N- CA- CB- HB2
5875 5877 5879 5881 3 0.65084 1.95253 0.00000
-2.60338 0.00000 0.00000 ; N- CA- CB- HB3
5875 5877 5879 5882 3 5.99567 3.29281 -4.46014
-0.55229 1.97485 0.00000 ; N- CA- CB- CG
5875 5877 5887 5888 3 0.00000 0.00000 0.00000
0.00000 0.00000 0.00000 ; N- CA- C- O
5875 5877 5887 5889 3 17.40544 -5.02080 -13.22144
9.20480 0.00000 0.00000 ; N- CA- C- N

```
  5876   5875   5877   5878      3    0.00000    0.00000    0.00000
0.00000    0.00000    0.00000 ;      H-     N-    CA-    HA
  5876   5875   5877   5879      3    0.00000    0.00000    0.00000
0.00000    0.00000    0.00000 ;      H-     N-    CA-    CB
  5876   5875   5877   5887      3    0.00000    0.00000    0.00000
0.00000    0.00000    0.00000 ;      H-     N-    CA-     C
  5877   5879   5882   5883      3    0.00000    0.00000    0.00000
0.00000    0.00000    0.00000 ;     CA-    CB-    CG-   OD1
  5877   5879   5882   5884      3    8.84916   -0.31380   -4.32626
5.03754    0.26778    0.00000 ;     CA-    CB-    CG-   ND2
  5877   5887   5889   5890      3   20.92000    0.00000  -20.92000
0.00000    0.00000    0.00000 ;     CA-     C-     N-     H
  5877   5887   5889   5891      3   20.92000    0.00000  -20.92000
0.00000    0.00000    0.00000 ;     CA-     C-     N-    CA
  5878   5877   5879   5880      3    0.65084    1.95253    0.00000
-2.60338    0.00000    0.00000 ;     HA-    CA-    CB-   HB2
  5878   5877   5879   5881      3    0.65084    1.95253    0.00000
-2.60338    0.00000    0.00000 ;     HA-    CA-    CB-   HB3
  5878   5877   5879   5882      3    0.65084    1.95253    0.00000
-2.60338    0.00000    0.00000 ;     HA-    CA-    CB-    CG
  5878   5877   5887   5888      3    3.68192   -4.35136    0.00000
1.33888    0.00000    0.00000 ;     HA-    CA-     C-     O
  5878   5877   5887   5889      3    0.00000    0.00000    0.00000
0.00000    0.00000    0.00000 ;     HA-    CA-     C-     N
  5879   5877   5887   5888      3    0.00000    0.00000    0.00000
0.00000    0.00000    0.00000 ;     CB-    CA-     C-     O
  5879   5877   5887   5889      3    2.51040    4.18400    1.67360
-6.69440    0.00000    0.00000 ;     CB-    CA-     C-     N
  5879   5882   5884   5885      3   20.92000    0.00000  -20.92000
0.00000    0.00000    0.00000 ;     CB-    CG-   ND2-  HD21
  5879   5882   5884   5886      3   20.92000    0.00000  -20.92000
0.00000    0.00000    0.00000 ;     CB-    CG-   ND2-  HD22
  5880   5879   5877   5887      3    0.65084    1.95253    0.00000
-2.60338    0.00000    0.00000 ;    HB2-    CB-    CA-     C
  5880   5879   5882   5883      3    3.68192   -4.35136    0.00000
1.33888    0.00000    0.00000 ;    HB2-    CB-    CG-   OD1
  5880   5879   5882   5884      3    0.00000    0.00000    0.00000
0.00000    0.00000    0.00000 ;    HB2-    CB-    CG-   ND2
  5881   5879   5877   5887      3    0.65084    1.95253    0.00000
-2.60338    0.00000    0.00000 ;    HB3-    CB-    CA-     C
  5881   5879   5882   5883      3    3.68192   -4.35136    0.00000
1.33888    0.00000    0.00000 ;    HB3-    CB-    CG-   OD1
  5881   5879   5882   5884      3    0.00000    0.00000    0.00000
0.00000    0.00000    0.00000 ;    HB3-    CB-    CG-   ND2
  5882   5879   5877   5887      3    7.94542   -3.96225   -6.11701
-0.55229    3.58150    0.00000 ;     CG-    CB-    CA-     C
  5883   5882   5884   5885      3   29.28800   -8.36800  -20.92000
0.00000    0.00000    0.00000 ;    OD1-    CG-   ND2-  HD21
  5883   5882   5884   5886      3   29.28800   -8.36800  -20.92000
0.00000    0.00000    0.00000 ;    OD1-    CG-   ND2-  HD22
  5887   5889   5891   5892      3    0.00000    0.00000    0.00000
0.00000    0.00000    0.00000 ;      C-     N-    CA-    HA
```

5887 5889 5891 5893 3 11.71520 1.67360 15.06240 -
13.38880 0.00000 0.00000 ; C- N- CA- CB
5887 5889 5891 5897 3 1.75728 5.27184 2.25936
-7.02912 0.00000 0.00000 ; C- N- CA- C
5888 5887 5889 5890 3 29.28800 -8.36800 -20.92000
0.00000 0.00000 0.00000 ; O- C- N- H
5888 5887 5889 5891 3 20.92000 0.00000 -20.92000
0.00000 0.00000 0.00000 ; O- C- N- CA
5889 5891 5893 5894 3 0.65084 1.95253 0.00000
-2.60338 0.00000 0.00000 ; N- CA- CB- HB1
5889 5891 5893 5895 3 0.65084 1.95253 0.00000
-2.60338 0.00000 0.00000 ; N- CA- CB- HB2
5889 5891 5893 5896 3 0.65084 1.95253 0.00000
-2.60338 0.00000 0.00000 ; N- CA- CB- HB3
5889 5891 5897 5898 3 0.00000 0.00000 0.00000
0.00000 0.00000 0.00000 ; N- CA- C- O
5889 5891 5897 5899 3 17.40544 -5.02080 -13.22144
9.20480 0.00000 0.00000 ; N- CA- C- N
5890 5889 5891 5892 3 0.00000 0.00000 0.00000
0.00000 0.00000 0.00000 ; H- N- CA- HA
5890 5889 5891 5893 3 0.00000 0.00000 0.00000
0.00000 0.00000 0.00000 ; H- N- CA- CB
5890 5889 5891 5897 3 0.00000 0.00000 0.00000
0.00000 0.00000 0.00000 ; H- N- CA- C
5891 5897 5899 5900 3 20.92000 0.00000 -20.92000
0.00000 0.00000 0.00000 ; CA- C- N- H
5891 5897 5899 5901 3 20.92000 0.00000 -20.92000
0.00000 0.00000 0.00000 ; CA- C- N- CA
5892 5891 5893 5894 3 0.65084 1.95253 0.00000
-2.60338 0.00000 0.00000 ; HA- CA- CB- HB1
5892 5891 5893 5895 3 0.65084 1.95253 0.00000
-2.60338 0.00000 0.00000 ; HA- CA- CB- HB2
5892 5891 5893 5896 3 0.65084 1.95253 0.00000
-2.60338 0.00000 0.00000 ; HA- CA- CB- HB3
5892 5891 5897 5898 3 3.68192 -4.35136 0.00000
1.33888 0.00000 0.00000 ; HA- CA- C- O
5892 5891 5897 5899 3 0.00000 0.00000 0.00000
0.00000 0.00000 0.00000 ; HA- CA- C- N
5893 5891 5897 5898 3 0.00000 0.00000 0.00000
0.00000 0.00000 0.00000 ; CB- CA- C- O
5893 5891 5897 5899 3 2.51040 4.18400 1.67360
-6.69440 0.00000 0.00000 ; CB- CA- C- N
5894 5893 5891 5897 3 0.65084 1.95253 0.00000
-2.60338 0.00000 0.00000 ; HB1- CB- CA- C
5895 5893 5891 5897 3 0.65084 1.95253 0.00000
-2.60338 0.00000 0.00000 ; HB2- CB- CA- C
5896 5893 5891 5897 3 0.65084 1.95253 0.00000
-2.60338 0.00000 0.00000 ; HB3- CB- CA- C
5897 5899 5901 5902 3 0.00000 0.00000 0.00000
0.00000 0.00000 0.00000 ; C- N- CA- HA
5897 5899 5901 5903 3 11.71520 1.67360 15.06240 -
13.38880 0.00000 0.00000 ; C- N- CA- CB

5897 5899 5901 5908 3 1.75728 5.27184 2.25936
-7.02912 0.00000 0.00000 ; C- N- CA- C
5898 5897 5899 5900 3 29.28800 -8.36800 -20.92000
0.00000 0.00000 0.00000 ; O- C- N- H
5898 5897 5899 5901 3 20.92000 0.00000 -20.92000
0.00000 0.00000 0.00000 ; O- C- N- CA
5899 5901 5903 5904 3 0.65084 1.95253 0.00000
-2.60338 0.00000 0.00000 ; N- CA- CB- HB2
5899 5901 5903 5905 3 0.65084 1.95253 0.00000
-2.60338 0.00000 0.00000 ; N- CA- CB- HB3
5899 5901 5903 5906 3 7.86174 2.24681 -7.41405
-6.71114 5.35552 0.00000 ; N- CA- CB- OG
5899 5901 5908 5909 3 0.00000 0.00000 0.00000
0.00000 0.00000 0.00000 ; N- CA- C- O
5899 5901 5908 5910 3 17.40544 -5.02080 -13.22144
9.20480 0.00000 0.00000 ; N- CA- C- N
5900 5899 5901 5902 3 0.00000 0.00000 0.00000
0.00000 0.00000 0.00000 ; H- N- CA- HA
5900 5899 5901 5903 3 0.00000 0.00000 0.00000
0.00000 0.00000 0.00000 ; H- N- CA- CB
5900 5899 5901 5908 3 0.00000 0.00000 0.00000
0.00000 0.00000 0.00000 ; H- N- CA- C
5901 5903 5906 5907 3 2.05853 2.46856 3.48109
-4.46851 0.23430 0.00000 ; CA- CB- OG- HG
5901 5908 5910 5911 3 20.92000 0.00000 -20.92000
0.00000 0.00000 0.00000 ; CA- C- N- H
5901 5908 5910 5912 3 20.92000 0.00000 -20.92000
0.00000 0.00000 0.00000 ; CA- C- N- CA
5902 5901 5903 5904 3 0.65084 1.95253 0.00000
-2.60338 0.00000 0.00000 ; HA- CA- CB- HB2
5902 5901 5903 5905 3 0.65084 1.95253 0.00000
-2.60338 0.00000 0.00000 ; HA- CA- CB- HB3
5902 5901 5903 5906 3 1.04600 -1.04600 0.00000
0.00000 0.00000 0.00000 ; HA- CA- CB- OG
5902 5901 5908 5909 3 3.68192 -4.35136 0.00000
1.33888 0.00000 0.00000 ; HA- CA- C- O
5902 5901 5908 5910 3 0.00000 0.00000 0.00000
0.00000 0.00000 0.00000 ; HA- CA- C- N
5903 5901 5908 5909 3 0.00000 0.00000 0.00000
0.00000 0.00000 0.00000 ; CB- CA- C- O
5903 5901 5908 5910 3 2.51040 4.18400 1.67360
-6.69440 0.00000 0.00000 ; CB- CA- C- N
5904 5903 5901 5908 3 0.65084 1.95253 0.00000
-2.60338 0.00000 0.00000 ; HB2- CB- CA- C
5904 5903 5906 5907 3 0.69733 2.09200 0.00000
-2.78933 0.00000 0.00000 ; HB2- CB- OG- HG
5905 5903 5901 5908 3 0.65084 1.95253 0.00000
-2.60338 0.00000 0.00000 ; HB3- CB- CA- C
5905 5903 5906 5907 3 0.69733 2.09200 0.00000
-2.78933 0.00000 0.00000 ; HB3- CB- OG- HG
5906 5903 5901 5908 3 7.34710 7.79898 -6.14211
-6.71114 4.31789 0.00000 ; OG- CB- CA- C

5908 5910 5912 5913 3 0.00000 0.00000 0.00000
0.00000 0.00000 0.00000 ; C- N- CA- HA2
5908 5910 5912 5914 3 0.00000 0.00000 0.00000
0.00000 0.00000 0.00000 ; C- N- CA- HA3
5908 5910 5912 5915 3 1.75728 5.27184 2.25936
-7.02912 0.00000 0.00000 ; C- N- CA- C
5909 5908 5910 5911 3 29.28800 -8.36800 -20.92000
0.00000 0.00000 0.00000 ; O- C- N- H
5909 5908 5910 5912 3 20.92000 0.00000 -20.92000
0.00000 0.00000 0.00000 ; O- C- N- CA
5910 5912 5915 5916 3 0.00000 0.00000 0.00000
0.00000 0.00000 0.00000 ; N- CA- C- O
5910 5912 5915 5917 3 17.40544 -5.02080 -13.22144
9.20480 0.00000 0.00000 ; N- CA- C- N
5911 5910 5912 5913 3 0.00000 0.00000 0.00000
0.00000 0.00000 0.00000 ; H- N- CA- HA2
5911 5910 5912 5914 3 0.00000 0.00000 0.00000
0.00000 0.00000 0.00000 ; H- N- CA- HA3
5911 5910 5912 5915 3 0.00000 0.00000 0.00000
0.00000 0.00000 0.00000 ; H- N- CA- C
5912 5915 5917 5918 3 20.92000 0.00000 -20.92000
0.00000 0.00000 0.00000 ; CA- C- N- H
5912 5915 5917 5919 3 20.92000 0.00000 -20.92000
0.00000 0.00000 0.00000 ; CA- C- N- CA
5913 5912 5915 5916 3 3.68192 -4.35136 0.00000
1.33888 0.00000 0.00000 ; HA2- CA- C- O
5913 5912 5915 5917 3 0.00000 0.00000 0.00000
0.00000 0.00000 0.00000 ; HA2- CA- C- N
5914 5912 5915 5916 3 3.68192 -4.35136 0.00000
1.33888 0.00000 0.00000 ; HA3- CA- C- O
5914 5912 5915 5917 3 0.00000 0.00000 0.00000
0.00000 0.00000 0.00000 ; HA3- CA- C- N
5915 5917 5919 5920 3 0.00000 0.00000 0.00000
0.00000 0.00000 0.00000 ; C- N- CA- HA
5915 5917 5919 5921 3 11.71520 1.67360 15.06240 -
13.38880 0.00000 0.00000 ; C- N- CA- CB
5915 5917 5919 5932 3 1.75728 5.27184 2.25936
-7.02912 0.00000 0.00000 ; C- N- CA- C
5916 5915 5917 5918 3 29.28800 -8.36800 -20.92000
0.00000 0.00000 0.00000 ; O- C- N- H
5916 5915 5917 5919 3 20.92000 0.00000 -20.92000
0.00000 0.00000 0.00000 ; O- C- N- CA
5917 5919 5921 5922 3 0.65084 1.95253 0.00000
-2.60338 0.00000 0.00000 ; N- CA- CB- HB2
5917 5919 5921 5923 3 0.65084 1.95253 0.00000
-2.60338 0.00000 0.00000 ; N- CA- CB- HB3
5917 5919 5921 5924 3 3.21331 2.22589 -4.15053
-2.40998 2.61082 0.00000 ; N- CA- CB- CG
5917 5919 5932 5933 3 0.00000 0.00000 0.00000
0.00000 0.00000 0.00000 ; N- CA- C- O
5917 5919 5932 5934 3 17.40544 -5.02080 -13.22144
9.20480 0.00000 0.00000 ; N- CA- C- N

5918 5917 5919 5920 3 0.00000 0.00000 0.00000
0.00000 0.00000 0.00000 ; H- N- CA- HA
5918 5917 5919 5921 3 0.00000 0.00000 0.00000
0.00000 0.00000 0.00000 ; H- N- CA- CB
5918 5917 5919 5932 3 0.00000 0.00000 0.00000
0.00000 0.00000 0.00000 ; H- N- CA- C
5919 5921 5924 5925 3 0.66944 2.00832 0.00000
-2.67776 0.00000 0.00000 ; CA- CB- CG- HG2
5919 5921 5924 5926 3 0.66944 2.00832 0.00000
-2.67776 0.00000 0.00000 ; CA- CB- CG- HG3
5919 5921 5924 5927 3 3.69866 -4.35136 -3.92459
6.89523 4.61914 0.00000 ; CA- CB- CG- CD
5919 5932 5934 5935 3 20.92000 0.00000 -20.92000
0.00000 0.00000 0.00000 ; CA- C- N- H
5919 5932 5934 5936 3 20.92000 0.00000 -20.92000
0.00000 0.00000 0.00000 ; CA- C- N- CA
5920 5919 5921 5922 3 0.65084 1.95253 0.00000
-2.60338 0.00000 0.00000 ; HA- CA- CB- HB2
5920 5919 5921 5923 3 0.65084 1.95253 0.00000
-2.60338 0.00000 0.00000 ; HA- CA- CB- HB3
5920 5919 5921 5924 3 0.65084 1.95253 0.00000
-2.60338 0.00000 0.00000 ; HA- CA- CB- CG
5920 5919 5932 5933 3 3.68192 -4.35136 0.00000
1.33888 0.00000 0.00000 ; HA- CA- C- O
5920 5919 5932 5934 3 0.00000 0.00000 0.00000
0.00000 0.00000 0.00000 ; HA- CA- C- N
5921 5919 5932 5933 3 0.00000 0.00000 0.00000
0.00000 0.00000 0.00000 ; CB- CA- C- O
5921 5919 5932 5934 3 2.51040 4.18400 1.67360
-6.69440 0.00000 0.00000 ; CB- CA- C- N
5921 5924 5927 5928 3 0.00000 0.00000 0.00000
0.00000 0.00000 0.00000 ; CB- CG- CD- OE1
5921 5924 5927 5929 3 10.32611 1.48114 -8.47678
1.42256 1.40582 0.00000 ; CB- CG- CD- NE2
5922 5921 5919 5932 3 0.65084 1.95253 0.00000
-2.60338 0.00000 0.00000 ; HB2- CB- CA- C
5922 5921 5924 5925 3 0.62760 1.88280 0.00000
-2.51040 0.00000 0.00000 ; HB2- CB- CG- HG2
5922 5921 5924 5926 3 0.62760 1.88280 0.00000
-2.51040 0.00000 0.00000 ; HB2- CB- CG- HG3
5922 5921 5924 5927 3 0.65084 1.95253 0.00000
-2.60338 0.00000 0.00000 ; HB2- CB- CG- CD
5923 5921 5919 5932 3 0.65084 1.95253 0.00000
-2.60338 0.00000 0.00000 ; HB3- CB- CA- C
5923 5921 5924 5925 3 0.62760 1.88280 0.00000
-2.51040 0.00000 0.00000 ; HB3- CB- CG- HG2
5923 5921 5924 5926 3 0.62760 1.88280 0.00000
-2.51040 0.00000 0.00000 ; HB3- CB- CG- HG3
5923 5921 5924 5927 3 0.65084 1.95253 0.00000
-2.60338 0.00000 0.00000 ; HB3- CB- CG- CD
5924 5921 5919 5932 3 6.86594 3.56895 -8.14206
-2.40998 4.85344 0.00000 ; CG- CB- CA- C

5924 5927 5929 5930 3 20.92000 0.00000 -20.92000
0.00000 0.00000 0.00000 ; CG- CD- NE2- HE21
5924 5927 5929 5931 3 20.92000 0.00000 -20.92000
0.00000 0.00000 0.00000 ; CG- CD- NE2- HE22
5925 5924 5927 5928 3 3.68192 -4.35136 0.00000
1.33888 0.00000 0.00000 ; HG2- CG- CD- OE1
5925 5924 5927 5929 3 0.00000 0.00000 0.00000
0.00000 0.00000 0.00000 ; HG2- CG- CD- NE2
5926 5924 5927 5928 3 3.68192 -4.35136 0.00000
1.33888 0.00000 0.00000 ; HG3- CG- CD- OE1
5926 5924 5927 5929 3 0.00000 0.00000 0.00000
0.00000 0.00000 0.00000 ; HG3- CG- CD- NE2
5928 5927 5929 5930 3 29.28800 -8.36800 -20.92000
0.00000 0.00000 0.00000 ; OE1- CD- NE2- HE21
5928 5927 5929 5931 3 29.28800 -8.36800 -20.92000
0.00000 0.00000 0.00000 ; OE1- CD- NE2- HE22
5932 5934 5936 5937 3 0.00000 0.00000 0.00000
0.00000 0.00000 0.00000 ; C- N- CA- HA
5932 5934 5936 5938 3 11.71520 1.67360 15.06240 -
13.38880 0.00000 0.00000 ; C- N- CA- CB
5932 5934 5936 5951 3 1.75728 5.27184 2.25936
-7.02912 0.00000 0.00000 ; C- N- CA- C
5933 5932 5934 5935 3 29.28800 -8.36800 -20.92000
0.00000 0.00000 0.00000 ; O- C- N- H
5933 5932 5934 5936 3 20.92000 0.00000 -20.92000
0.00000 0.00000 0.00000 ; O- C- N- CA
5934 5936 5938 5939 3 0.65084 1.95253 0.00000
-2.60338 0.00000 0.00000 ; N- CA- CB- HB
5934 5936 5938 5940 3 3.83673 0.44769 -1.77402
-2.47693 -0.03347 0.00000 ; N- CA- CB- CG2
5934 5936 5938 5944 3 2.97482 0.12134 2.04179
-1.89117 -3.24678 0.00000 ; N- CA- CB- CG1
5934 5936 5951 5952 3 0.00000 0.00000 0.00000
0.00000 0.00000 0.00000 ; N- CA- C- O
5934 5936 5951 5953 3 17.40544 -5.02080 -13.22144
9.20480 0.00000 0.00000 ; N- CA- C- N
5935 5934 5936 5937 3 0.00000 0.00000 0.00000
0.00000 0.00000 0.00000 ; H- N- CA- HA
5935 5934 5936 5938 3 0.00000 0.00000 0.00000
0.00000 0.00000 0.00000 ; H- N- CA- CB
5935 5934 5936 5951 3 0.00000 0.00000 0.00000
0.00000 0.00000 0.00000 ; H- N- CA- C
5936 5938 5940 5941 3 0.66944 2.00832 0.00000
-2.67776 0.00000 0.00000 ; CA- CB- CG2- HG21
5936 5938 5940 5942 3 0.66944 2.00832 0.00000
-2.67776 0.00000 0.00000 ; CA- CB- CG2- HG22
5936 5938 5940 5943 3 0.66944 2.00832 0.00000
-2.67776 0.00000 0.00000 ; CA- CB- CG2- HG23
5936 5938 5944 5945 3 0.66944 2.00832 0.00000
-2.67776 0.00000 0.00000 ; CA- CB- CG1- HG12
5936 5938 5944 5946 3 0.66944 2.00832 0.00000
-2.67776 0.00000 0.00000 ; CA- CB- CG1- HG13

5936 5938 5944 5947 3 4.24258 -0.52718 -7.25506
-1.79075 7.69856 0.00000 ; CA- CB- CG1- CD1
5936 5951 5953 5954 3 20.92000 0.00000 -20.92000
0.00000 0.00000 0.00000 ; CA- C- N- H
5936 5951 5953 5955 3 20.92000 0.00000 -20.92000
0.00000 0.00000 0.00000 ; CA- C- N- CA
5937 5936 5938 5939 3 0.65084 1.95253 0.00000
-2.60338 0.00000 0.00000 ; HA- CA- CB- HB
5937 5936 5938 5940 3 0.65084 1.95253 0.00000
-2.60338 0.00000 0.00000 ; HA- CA- CB- CG2
5937 5936 5938 5944 3 0.65084 1.95253 0.00000
-2.60338 0.00000 0.00000 ; HA- CA- CB- CG1
5937 5936 5951 5952 3 3.68192 -4.35136 0.00000
1.33888 0.00000 0.00000 ; HA- CA- C- O
5937 5936 5951 5953 3 0.00000 0.00000 0.00000
0.00000 0.00000 0.00000 ; HA- CA- C- N
5938 5936 5951 5952 3 0.00000 0.00000 0.00000
0.00000 0.00000 0.00000 ; CB- CA- C- O
5938 5936 5951 5953 3 2.51040 4.18400 1.67360
-6.69440 0.00000 0.00000 ; CB- CA- C- N
5938 5944 5947 5948 3 0.66944 2.00832 0.00000
-2.67776 0.00000 0.00000 ; CB- CG1- CD1- HD11
5938 5944 5947 5949 3 0.66944 2.00832 0.00000
-2.67776 0.00000 0.00000 ; CB- CG1- CD1- HD12
5938 5944 5947 5950 3 0.66944 2.00832 0.00000
-2.67776 0.00000 0.00000 ; CB- CG1- CD1- HD13
5939 5938 5936 5951 3 0.65084 1.95253 0.00000
-2.60338 0.00000 0.00000 ; HB- CB- CA- C
5939 5938 5940 5941 3 0.62760 1.88280 0.00000
-2.51040 0.00000 0.00000 ; HB- CB- CG2- HG21
5939 5938 5940 5942 3 0.62760 1.88280 0.00000
-2.51040 0.00000 0.00000 ; HB- CB- CG2- HG22
5939 5938 5940 5943 3 0.62760 1.88280 0.00000
-2.51040 0.00000 0.00000 ; HB- CB- CG2- HG23
5939 5938 5944 5945 3 0.62760 1.88280 0.00000
-2.51040 0.00000 0.00000 ; HB- CB- CG1- HG12
5939 5938 5944 5946 3 0.62760 1.88280 0.00000
-2.51040 0.00000 0.00000 ; HB- CB- CG1- HG13
5939 5938 5944 5947 3 0.66944 2.00832 0.00000
-2.67776 0.00000 0.00000 ; HB- CB- CG1- CD1
5940 5938 5936 5951 3 5.67350 3.55640 -6.16722
-2.47693 3.74886 0.00000 ; CG2- CB- CA- C
5940 5938 5944 5945 3 0.66944 2.00832 0.00000
-2.67776 0.00000 0.00000 ; CG2- CB- CG1- HG12
5940 5938 5944 5946 3 0.66944 2.00832 0.00000
-2.67776 0.00000 0.00000 ; CG2- CB- CG1- HG13
5940 5938 5944 5947 3 3.81162 0.49790 -8.14206
-1.79075 7.49773 0.00000 ; CG2- CB- CG1- CD1
5941 5940 5938 5944 3 0.66944 2.00832 0.00000
-2.67776 0.00000 0.00000 ; HG21- CG2- CB- CG1
5942 5940 5938 5944 3 0.66944 2.00832 0.00000
-2.67776 0.00000 0.00000 ; HG22- CG2- CB- CG1

5943 5940 5938 5944 3 0.66944 2.00832 0.00000
-2.67776 0.00000 0.00000 ; HG23- CG2- CB- CG1
5944 5938 5936 5951 3 8.26340 0.74057 -9.99976
-1.89117 3.84928 0.00000 ; CG1- CB- CA- C
5945 5944 5947 5948 3 0.62760 1.88280 0.00000
-2.51040 0.00000 0.00000 ; HG12- CG1- CD1- HD11
5945 5944 5947 5949 3 0.62760 1.88280 0.00000
-2.51040 0.00000 0.00000 ; HG12- CG1- CD1- HD12
5945 5944 5947 5950 3 0.62760 1.88280 0.00000
-2.51040 0.00000 0.00000 ; HG12- CG1- CD1- HD13
5946 5944 5947 5948 3 0.62760 1.88280 0.00000
-2.51040 0.00000 0.00000 ; HG13- CG1- CD1- HD11
5946 5944 5947 5949 3 0.62760 1.88280 0.00000
-2.51040 0.00000 0.00000 ; HG13- CG1- CD1- HD12
5946 5944 5947 5950 3 0.62760 1.88280 0.00000
-2.51040 0.00000 0.00000 ; HG13- CG1- CD1- HD13
5951 5953 5955 5956 3 0.00000 0.00000 0.00000
0.00000 0.00000 0.00000 ; C- N- CA- HA
5951 5953 5955 5957 3 11.71520 1.67360 15.06240 -
13.38880 0.00000 0.00000 ; C- N- CA- CB
5951 5953 5955 5969 3 1.75728 5.27184 2.25936
-7.02912 0.00000 0.00000 ; C- N- CA- C
5952 5951 5953 5954 3 29.28800 -8.36800 -20.92000
0.00000 0.00000 0.00000 ; O- C- N- H
5952 5951 5953 5955 3 20.92000 0.00000 -20.92000
0.00000 0.00000 0.00000 ; O- C- N- CA
5953 5955 5957 5958 3 0.65084 1.95253 0.00000
-2.60338 0.00000 0.00000 ; N- CA- CB- HB2
5953 5955 5957 5959 3 0.65084 1.95253 0.00000
-2.60338 0.00000 0.00000 ; N- CA- CB- HB3
5953 5955 5957 5960 3 4.79068 4.02919 -4.82834
-3.66518 2.97901 0.00000 ; N- CA- CB- CG
5953 5955 5969 5970 3 0.00000 0.00000 0.00000
0.00000 0.00000 0.00000 ; N- CA- C- O
5953 5955 5969 5971 3 17.40544 -5.02080 -13.22144
9.20480 0.00000 0.00000 ; N- CA- C- N
5954 5953 5955 5956 3 0.00000 0.00000 0.00000
0.00000 0.00000 0.00000 ; H- N- CA- HA
5954 5953 5955 5957 3 0.00000 0.00000 0.00000
0.00000 0.00000 0.00000 ; H- N- CA- CB
5954 5953 5955 5969 3 0.00000 0.00000 0.00000
0.00000 0.00000 0.00000 ; H- N- CA- C
5955 5957 5960 5961 3 6.81992 9.28011 -2.04179 -
11.48090 -1.23846 0.00000 ; CA- CB- CG- ND1
5955 5957 5960 5967 3 0.00000 0.00000 0.00000
0.00000 0.00000 0.00000 ; CA- CB- CG- CD2
5955 5969 5971 5972 3 20.92000 0.00000 -20.92000
0.00000 0.00000 0.00000 ; CA- C- N- H
5955 5969 5971 5973 3 20.92000 0.00000 -20.92000
0.00000 0.00000 0.00000 ; CA- C- N- CA
5956 5955 5957 5958 3 0.65084 1.95253 0.00000
-2.60338 0.00000 0.00000 ; HA- CA- CB- HB2

5956 5955 5957 5959 3 0.65084 1.95253 0.00000
-2.60338 0.00000 0.00000 ; HA- CA- CB- HB3
5956 5955 5957 5960 3 0.65084 1.95253 0.00000
-2.60338 0.00000 0.00000 ; HA- CA- CB- CG
5956 5955 5969 5970 3 3.68192 -4.35136 0.00000
1.33888 0.00000 0.00000 ; HA- CA- C- O
5956 5955 5969 5971 3 0.00000 0.00000 0.00000
0.00000 0.00000 0.00000 ; HA- CA- C- N
5957 5955 5969 5970 3 0.00000 0.00000 0.00000
0.00000 0.00000 0.00000 ; CB- CA- C- O
5957 5955 5969 5971 3 2.51040 4.18400 1.67360
-6.69440 0.00000 0.00000 ; CB- CA- C- N
5957 5960 5961 5962 3 11.71520 0.00000 -11.71520
0.00000 0.00000 0.00000 ; CB- CG- ND1- HD1
5957 5960 5961 5963 3 11.71520 0.00000 -11.71520
0.00000 0.00000 0.00000 ; CB- CG- ND1- CE1
5957 5960 5967 5965 3 44.97800 0.00000 -44.97800
0.00000 0.00000 0.00000 ; CB- CG- CD2- NE2
5957 5960 5967 5968 3 44.97800 0.00000 -44.97800
0.00000 0.00000 0.00000 ; CB- CG- CD2- HD2
5958 5957 5955 5969 3 0.65084 1.95253 0.00000
-2.60338 0.00000 0.00000 ; HB2- CB- CA- C
5958 5957 5960 5961 3 0.00000 0.00000 0.00000
0.00000 0.00000 0.00000 ; HB2- CB- CG- ND1
5958 5957 5960 5967 3 0.00000 0.00000 0.00000
0.00000 0.00000 0.00000 ; HB2- CB- CG- CD2
5959 5957 5955 5969 3 0.65084 1.95253 0.00000
-2.60338 0.00000 0.00000 ; HB3- CB- CA- C
5959 5957 5960 5961 3 0.00000 0.00000 0.00000
0.00000 0.00000 0.00000 ; HB3- CB- CG- ND1
5959 5957 5960 5967 3 0.00000 0.00000 0.00000
0.00000 0.00000 0.00000 ; HB3- CB- CG- CD2
5960 5957 5955 5969 3 3.76560 3.34720 -2.87859
-3.66518 0.83680 0.00000 ; CG- CB- CA- C
5960 5961 5963 5964 3 19.45560 0.00000 -19.45560
0.00000 0.00000 0.00000 ; CG- ND1- CE1- HE1
5960 5961 5963 5965 3 19.45560 0.00000 -19.45560
0.00000 0.00000 0.00000 ; CG- ND1- CE1- NE2
5960 5967 5965 5963 3 12.55200 0.00000 -12.55200
0.00000 0.00000 0.00000 ; CG- CD2- NE2- CE1
5960 5967 5965 5966 3 12.55200 0.00000 -12.55200
0.00000 0.00000 0.00000 ; CG- CD2- NE2- HE2
5961 5960 5967 5965 3 44.97800 0.00000 -44.97800
0.00000 0.00000 0.00000 ; ND1- CG- CD2- NE2
5961 5960 5967 5968 3 44.97800 0.00000 -44.97800
0.00000 0.00000 0.00000 ; ND1- CG- CD2- HD2
5961 5963 5965 5966 3 19.45560 0.00000 -19.45560
0.00000 0.00000 0.00000 ; ND1- CE1- NE2- HE2
5961 5963 5965 5967 3 19.45560 0.00000 -19.45560
0.00000 0.00000 0.00000 ; ND1- CE1- NE2- CD2
5962 5961 5960 5967 3 11.71520 0.00000 -11.71520
0.00000 0.00000 0.00000 ; HD1- ND1- CG- CD2

5962 5961 5963 5964 3 19.45560 0.00000 -19.45560
0.00000 0.00000 0.00000 ; HD1- ND1- CE1- HE1
5962 5961 5963 5965 3 19.45560 0.00000 -19.45560
0.00000 0.00000 0.00000 ; HD1- ND1- CE1- NE2
5963 5961 5960 5967 3 11.71520 0.00000 -11.71520
0.00000 0.00000 0.00000 ; CE1- ND1- CG- CD2
5963 5965 5967 5968 3 12.55200 0.00000 -12.55200
0.00000 0.00000 0.00000 ; CE1- NE2- CD2- HD2
5964 5963 5965 5966 3 19.45560 0.00000 -19.45560
0.00000 0.00000 0.00000 ; HE1- CE1- NE2- HE2
5964 5963 5965 5967 3 19.45560 0.00000 -19.45560
0.00000 0.00000 0.00000 ; HE1- CE1- NE2- CD2
5966 5965 5967 5968 3 12.55200 0.00000 -12.55200
0.00000 0.00000 0.00000 ; HE2- NE2- CD2- HD2
5969 5971 5973 5974 3 0.00000 0.00000 0.00000
0.00000 0.00000 0.00000 ; C- N- CA- HA
5969 5971 5973 5975 3 11.71520 1.67360 15.06240 -
13.38880 0.00000 0.00000 ; C- N- CA- CB
5969 5971 5973 5990 3 1.75728 5.27184 2.25936
-7.02912 0.00000 0.00000 ; C- N- CA- C
5970 5969 5971 5972 3 29.28800 -8.36800 -20.92000
0.00000 0.00000 0.00000 ; O- C- N- H
5970 5969 5971 5973 3 20.92000 0.00000 -20.92000
0.00000 0.00000 0.00000 ; O- C- N- CA
5971 5973 5975 5976 3 0.65084 1.95253 0.00000
-2.60338 0.00000 0.00000 ; N- CA- CB- HB2
5971 5973 5975 5977 3 0.65084 1.95253 0.00000
-2.60338 0.00000 0.00000 ; N- CA- CB- HB3
5971 5973 5975 5978 3 3.28026 2.46019 -2.19242
-3.21331 -0.23430 0.00000 ; N- CA- CB- CG
5971 5973 5990 5991 3 0.00000 0.00000 0.00000
0.00000 0.00000 0.00000 ; N- CA- C- O
5971 5973 5990 5992 3 17.40544 -5.02080 -13.22144
9.20480 0.00000 0.00000 ; N- CA- C- N
5972 5971 5973 5974 3 0.00000 0.00000 0.00000
0.00000 0.00000 0.00000 ; H- N- CA- HA
5972 5971 5973 5975 3 0.00000 0.00000 0.00000
0.00000 0.00000 0.00000 ; H- N- CA- CB
5972 5971 5973 5990 3 0.00000 0.00000 0.00000
0.00000 0.00000 0.00000 ; H- N- CA- C
5973 5975 5978 5979 3 0.57739 0.00000 1.02926
0.00000 -1.60666 0.00000 ; CA- CB- CG- CD1
5973 5975 5978 5988 3 0.57739 0.00000 1.02926
0.00000 -1.60666 0.00000 ; CA- CB- CG- CD2
5973 5990 5992 5993 3 20.92000 0.00000 -20.92000
0.00000 0.00000 0.00000 ; CA- C- N- H
5973 5990 5992 5994 3 20.92000 0.00000 -20.92000
0.00000 0.00000 0.00000 ; CA- C- N- CA
5974 5973 5975 5976 3 0.65084 1.95253 0.00000
-2.60338 0.00000 0.00000 ; HA- CA- CB- HB2
5974 5973 5975 5977 3 0.65084 1.95253 0.00000
-2.60338 0.00000 0.00000 ; HA- CA- CB- HB3

5974 5973 5975 5978 3 0.65084 1.95253 0.00000
-2.60338 0.00000 0.00000 ; HA- CA- CB- CG
5974 5973 5990 5991 3 3.68192 -4.35136 0.00000
1.33888 0.00000 0.00000 ; HA- CA- C- O
5974 5973 5990 5992 3 0.00000 0.00000 0.00000
0.00000 0.00000 0.00000 ; HA- CA- C- N
5975 5973 5990 5991 3 0.00000 0.00000 0.00000
0.00000 0.00000 0.00000 ; CB- CA- C- O
5975 5973 5990 5992 3 2.51040 4.18400 1.67360
-6.69440 0.00000 0.00000 ; CB- CA- C- N
5975 5978 5979 5980 3 30.33400 0.00000 -30.33400
0.00000 0.00000 0.00000 ; CB- CG- CD1- HD1
5975 5978 5979 5981 3 30.33400 0.00000 -30.33400
0.00000 0.00000 0.00000 ; CB- CG- CD1- CE1
5975 5978 5988 5986 3 30.33400 0.00000 -30.33400
0.00000 0.00000 0.00000 ; CB- CG- CD2- CE2
5975 5978 5988 5989 3 30.33400 0.00000 -30.33400
0.00000 0.00000 0.00000 ; CB- CG- CD2- HD2
5976 5975 5973 5990 3 0.65084 1.95253 0.00000
-2.60338 0.00000 0.00000 ; HB2- CB- CA- C
5976 5975 5978 5979 3 0.00000 0.00000 0.00000
0.00000 0.00000 0.00000 ; HB2- CB- CG- CD1
5976 5975 5978 5988 3 0.00000 0.00000 0.00000
0.00000 0.00000 0.00000 ; HB2- CB- CG- CD2
5977 5975 5973 5990 3 0.65084 1.95253 0.00000
-2.60338 0.00000 0.00000 ; HB3- CB- CA- C
5977 5975 5978 5979 3 0.00000 0.00000 0.00000
0.00000 0.00000 0.00000 ; HB3- CB- CG- CD1
5977 5975 5978 5988 3 0.00000 0.00000 0.00000
0.00000 0.00000 0.00000 ; HB3- CB- CG- CD2
5978 5975 5973 5990 3 4.95804 2.17986 -3.52293
-3.21331 -0.40166 0.00000 ; CG- CB- CA- C
5978 5979 5981 5982 3 30.33400 0.00000 -30.33400
0.00000 0.00000 0.00000 ; CG- CD1- CE1- HE1
5978 5979 5981 5983 3 30.33400 0.00000 -30.33400
0.00000 0.00000 0.00000 ; CG- CD1- CE1- CZ
5978 5988 5986 5983 3 30.33400 0.00000 -30.33400
0.00000 0.00000 0.00000 ; CG- CD2- CE2- CZ
5978 5988 5986 5987 3 30.33400 0.00000 -30.33400
0.00000 0.00000 0.00000 ; CG- CD2- CE2- HE2
5979 5978 5988 5986 3 30.33400 0.00000 -30.33400
0.00000 0.00000 0.00000 ; CD1- CG- CD2- CE2
5979 5978 5988 5989 3 30.33400 0.00000 -30.33400
0.00000 0.00000 0.00000 ; CD1- CG- CD2- HD2
5979 5981 5983 5984 3 30.33400 0.00000 -30.33400
0.00000 0.00000 0.00000 ; CD1- CE1- CZ- OH
5979 5981 5983 5986 3 30.33400 0.00000 -30.33400
0.00000 0.00000 0.00000 ; CD1- CE1- CZ- CE2
5980 5979 5978 5988 3 30.33400 0.00000 -30.33400
0.00000 0.00000 0.00000 ; HD1- CD1- CG- CD2
5980 5979 5981 5982 3 30.33400 0.00000 -30.33400
0.00000 0.00000 0.00000 ; HD1- CD1- CE1- HE1

5980 5979 5981 5983 3 30.33400 0.00000 -30.33400
0.00000 0.00000 0.00000 ; HD1- CD1- CE1- CZ
5981 5979 5978 5988 3 30.33400 0.00000 -30.33400
0.00000 0.00000 0.00000 ; CE1- CD1- CG- CD2
5981 5983 5984 5985 3 7.93286 0.00000 -9.56462
0.00000 2.17568 0.00000 ; CE1- CZ- OH- HH
5981 5983 5986 5987 3 30.33400 0.00000 -30.33400
0.00000 0.00000 0.00000 ; CE1- CZ- CE2- HE2
5981 5983 5986 5988 3 30.33400 0.00000 -30.33400
0.00000 0.00000 0.00000 ; CE1- CZ- CE2- CD2
5982 5981 5983 5984 3 30.33400 0.00000 -30.33400
0.00000 0.00000 0.00000 ; HE1- CE1- CZ- OH
5982 5981 5983 5986 3 30.33400 0.00000 -30.33400
0.00000 0.00000 0.00000 ; HE1- CE1- CZ- CE2
5983 5986 5988 5989 3 30.33400 0.00000 -30.33400
0.00000 0.00000 0.00000 ; CZ- CE2- CD2- HD2
5984 5983 5986 5987 3 30.33400 0.00000 -30.33400
0.00000 0.00000 0.00000 ; OH- CZ- CE2- HE2
5984 5983 5986 5988 3 30.33400 0.00000 -30.33400
0.00000 0.00000 0.00000 ; OH- CZ- CE2- CD2
5985 5984 5983 5986 3 7.93286 0.00000 -9.56462
0.00000 2.17568 0.00000 ; HH- OH- CZ- CE2
5987 5986 5988 5989 3 30.33400 0.00000 -30.33400
0.00000 0.00000 0.00000 ; HE2- CE2- CD2- HD2
5990 5992 5994 5995 3 0.00000 0.00000 0.00000
0.00000 0.00000 0.00000 ; C- N- CA- HA
5990 5992 5994 5996 3 11.71520 1.67360 15.06240 -
13.38880 0.00000 0.00000 ; C- N- CA- CB
5990 5992 5994 6004 3 1.75728 5.27184 2.25936
-7.02912 0.00000 0.00000 ; C- N- CA- C
5991 5990 5992 5993 3 29.28800 -8.36800 -20.92000
0.00000 0.00000 0.00000 ; O- C- N- H
5991 5990 5992 5994 3 20.92000 0.00000 -20.92000
0.00000 0.00000 0.00000 ; O- C- N- CA
5992 5994 5996 5997 3 0.65084 1.95253 0.00000
-2.60338 0.00000 0.00000 ; N- CA- CB- HB
5992 5994 5996 5998 3 3.83673 0.44769 -1.77402
-2.47693 -0.03347 0.00000 ; N- CA- CB- CG2
5992 5994 5996 6002 3 4.93294 1.13386 -3.12963
-5.27184 3.17984 0.00000 ; N- CA- CB- OG1
5992 5994 6004 6005 3 0.00000 0.00000 0.00000
0.00000 0.00000 0.00000 ; N- CA- C- O
5992 5994 6004 6006 3 17.40544 -5.02080 -13.22144
9.20480 0.00000 0.00000 ; N- CA- C- N
5993 5992 5994 5995 3 0.00000 0.00000 0.00000
0.00000 0.00000 0.00000 ; H- N- CA- HA
5993 5992 5994 5996 3 0.00000 0.00000 0.00000
0.00000 0.00000 0.00000 ; H- N- CA- CB
5993 5992 5994 6004 3 0.00000 0.00000 0.00000
0.00000 0.00000 0.00000 ; H- N- CA- C
5994 5996 5998 5999 3 0.66944 2.00832 0.00000
-2.67776 0.00000 0.00000 ; CA- CB- CG2- HG21

5994 5996 5998 6000 3 0.66944 2.00832 0.00000
-2.67776 0.00000 0.00000 ; CA- CB- CG2- HG22
5994 5996 5998 6001 3 0.66944 2.00832 0.00000
-2.67776 0.00000 0.00000 ; CA- CB- CG2- HG23
5994 5996 6002 6003 3 1.12131 2.98738 1.66523
-3.94970 0.43514 0.00000 ; CA- CB- OG1- HG1
5994 6004 6006 6007 3 20.92000 0.00000 -20.92000
0.00000 0.00000 0.00000 ; CA- C- N- H
5994 6004 6006 6008 3 20.92000 0.00000 -20.92000
0.00000 0.00000 0.00000 ; CA- C- N- CA
5995 5994 5996 5997 3 0.65084 1.95253 0.00000
-2.60338 0.00000 0.00000 ; HA- CA- CB- HB
5995 5994 5996 5998 3 0.65084 1.95253 0.00000
-2.60338 0.00000 0.00000 ; HA- CA- CB- CG2
5995 5994 5996 6002 3 1.04600 -1.04600 0.00000
0.00000 0.00000 0.00000 ; HA- CA- CB- OG1
5995 5994 6004 6005 3 3.68192 -4.35136 0.00000
1.33888 0.00000 0.00000 ; HA- CA- C- O
5995 5994 6004 6006 3 0.00000 0.00000 0.00000
0.00000 0.00000 0.00000 ; HA- CA- C- N
5996 5994 6004 6005 3 0.00000 0.00000 0.00000
0.00000 0.00000 0.00000 ; CB- CA- C- O
5996 5994 6004 6006 3 2.51040 4.18400 1.67360
-6.69440 0.00000 0.00000 ; CB- CA- C- N
5997 5996 5994 6004 3 0.65084 1.95253 0.00000
-2.60338 0.00000 0.00000 ; HB- CB- CA- C
5997 5996 5998 5999 3 0.65084 1.95253 0.00000
-2.60338 0.00000 0.00000 ; HB- CB- CG2- HG21
5997 5996 5998 6000 3 0.65084 1.95253 0.00000
-2.60338 0.00000 0.00000 ; HB- CB- CG2- HG22
5997 5996 5998 6001 3 0.65084 1.95253 0.00000
-2.60338 0.00000 0.00000 ; HB- CB- CG2- HG23
5997 5996 6002 6003 3 0.69733 2.09200 0.00000
-2.78933 0.00000 0.00000 ; HB- CB- OG1- HG1
5998 5996 5994 6004 3 5.67350 3.55640 -6.16722
-2.47693 3.74886 0.00000 ; CG2- CB- CA- C
5998 5996 6002 6003 3 4.74047 0.27196 -2.26773
-3.94970 1.60666 0.00000 ; CG2- CB- OG1- HG1
5999 5998 5996 6002 3 1.04600 -1.04600 0.00000
0.00000 0.00000 0.00000 ; HG21- CG2- CB- OG1
6000 5998 5996 6002 3 1.04600 -1.04600 0.00000
0.00000 0.00000 0.00000 ; HG22- CG2- CB- OG1
6001 5998 5996 6002 3 1.04600 -1.04600 0.00000
0.00000 0.00000 0.00000 ; HG23- CG2- CB- OG1
6002 5996 5994 6004 3 6.53541 6.87013 -6.21742
-5.27184 5.22163 0.00000 ; OG1- CB- CA- C
6004 6006 6008 6009 3 0.00000 0.00000 0.00000
0.00000 0.00000 0.00000 ; C- N- CA- HA
6004 6006 6008 6010 3 11.71520 1.67360 15.06240 -
13.38880 0.00000 0.00000 ; C- N- CA- CB
6004 6006 6008 6016 3 1.75728 5.27184 2.25936
-7.02912 0.00000 0.00000 ; C- N- CA- C

6005 6004 6006 6007 3 29.28800 -8.36800 -20.92000
0.00000 0.00000 0.00000 ; O- C- N- H
6005 6004 6006 6008 3 20.92000 0.00000 -20.92000
0.00000 0.00000 0.00000 ; O- C- N- CA
6006 6008 6010 6011 3 0.65084 1.95253 0.00000
-2.60338 0.00000 0.00000 ; N- CA- CB- HB2
6006 6008 6010 6012 3 0.65084 1.95253 0.00000
-2.60338 0.00000 0.00000 ; N- CA- CB- HB3
6006 6008 6010 6013 3 15.41386 9.74035 -8.39310
-0.97069 2.97901 0.00000 ; N- CA- CB- CG
6006 6008 6016 6017 3 0.00000 0.00000 0.00000
0.00000 0.00000 0.00000 ; N- CA- C- O
6006 6008 6016 6018 3 17.40544 -5.02080 -13.22144
9.20480 0.00000 0.00000 ; N- CA- C- N
6007 6006 6008 6009 3 0.00000 0.00000 0.00000
0.00000 0.00000 0.00000 ; H- N- CA- HA
6007 6006 6008 6010 3 0.00000 0.00000 0.00000
0.00000 0.00000 0.00000 ; H- N- CA- CB
6007 6006 6008 6016 3 0.00000 0.00000 0.00000
0.00000 0.00000 0.00000 ; H- N- CA- C
6008 6010 6013 6014 3 6.43499 0.00000 -5.39736
0.00000 -1.03763 0.00000 ; CA- CB- CG- OD1
6008 6010 6013 6015 3 6.43499 0.00000 -5.39736
0.00000 -1.03763 0.00000 ; CA- CB- CG- OD2
6008 6016 6018 6019 3 20.92000 0.00000 -20.92000
0.00000 0.00000 0.00000 ; CA- C- N- H
6008 6016 6018 6020 3 20.92000 0.00000 -20.92000
0.00000 0.00000 0.00000 ; CA- C- N- CA
6009 6008 6010 6011 3 0.65084 1.95253 0.00000
-2.60338 0.00000 0.00000 ; HA- CA- CB- HB2
6009 6008 6010 6012 3 0.65084 1.95253 0.00000
-2.60338 0.00000 0.00000 ; HA- CA- CB- HB3
6009 6008 6010 6013 3 0.65084 1.95253 0.00000
-2.60338 0.00000 0.00000 ; HA- CA- CB- CG
6009 6008 6016 6017 3 3.68192 -4.35136 0.00000
1.33888 0.00000 0.00000 ; HA- CA- C- O
6009 6008 6016 6018 3 0.00000 0.00000 0.00000
0.00000 0.00000 0.00000 ; HA- CA- C- N
6010 6008 6016 6017 3 0.00000 0.00000 0.00000
0.00000 0.00000 0.00000 ; CB- CA- C- O
6010 6008 6016 6018 3 2.51040 4.18400 1.67360
-6.69440 0.00000 0.00000 ; CB- CA- C- N
6011 6010 6008 6016 3 0.65084 1.95253 0.00000
-2.60338 0.00000 0.00000 ; HB2- CB- CA- C
6011 6010 6013 6014 3 0.00000 0.00000 0.00000
0.00000 0.00000 0.00000 ; HB2- CB- CG- OD1
6011 6010 6013 6015 3 0.00000 0.00000 0.00000
0.00000 0.00000 0.00000 ; HB2- CB- CG- OD2
6012 6010 6008 6016 3 0.65084 1.95253 0.00000
-2.60338 0.00000 0.00000 ; HB3- CB- CA- C
6012 6010 6013 6014 3 0.00000 0.00000 0.00000
0.00000 0.00000 0.00000 ; HB3- CB- CG- OD1

6012 6010 6013 6015 3 0.00000 0.00000 0.00000
0.00000 0.00000 0.00000 ; HB3- CB- CG- OD2
6013 6010 6008 6016 3 7.14627 -1.04600 -8.99560
-0.97069 5.15469 0.00000 ; CG- CB- CA- C
6016 6018 6020 6021 3 0.00000 0.00000 0.00000
0.00000 0.00000 0.00000 ; C- N- CA- HA
6016 6018 6020 6022 3 11.71520 1.67360 15.06240 -
13.38880 0.00000 0.00000 ; C- N- CA- CB
6016 6018 6020 6036 3 1.75728 5.27184 2.25936
-7.02912 0.00000 0.00000 ; C- N- CA- C
6017 6016 6018 6019 3 29.28800 -8.36800 -20.92000
0.00000 0.00000 0.00000 ; O- C- N- H
6017 6016 6018 6020 3 20.92000 0.00000 -20.92000
0.00000 0.00000 0.00000 ; O- C- N- CA
6018 6020 6022 6023 3 0.65084 1.95253 0.00000
-2.60338 0.00000 0.00000 ; N- CA- CB- HB2
6018 6020 6022 6024 3 0.65084 1.95253 0.00000
-2.60338 0.00000 0.00000 ; N- CA- CB- HB3
6018 6020 6022 6025 3 3.28026 2.46019 -2.19242
-3.21331 -0.23430 0.00000 ; N- CA- CB- CG
6018 6020 6036 6037 3 0.00000 0.00000 0.00000
0.00000 0.00000 0.00000 ; N- CA- C- O
6018 6020 6036 6038 3 17.40544 -5.02080 -13.22144
9.20480 0.00000 0.00000 ; N- CA- C- N
6019 6018 6020 6021 3 0.00000 0.00000 0.00000
0.00000 0.00000 0.00000 ; H- N- CA- HA
6019 6018 6020 6022 3 0.00000 0.00000 0.00000
0.00000 0.00000 0.00000 ; H- N- CA- CB
6019 6018 6020 6036 3 0.00000 0.00000 0.00000
0.00000 0.00000 0.00000 ; H- N- CA- C
6020 6022 6025 6026 3 0.57739 0.00000 1.02926
0.00000 -1.60666 0.00000 ; CA- CB- CG- CD1
6020 6022 6025 6034 3 0.57739 0.00000 1.02926
0.00000 -1.60666 0.00000 ; CA- CB- CG- CD2
6020 6036 6038 6039 3 20.92000 0.00000 -20.92000
0.00000 0.00000 0.00000 ; CA- C- N- H
6020 6036 6038 6040 3 20.92000 0.00000 -20.92000
0.00000 0.00000 0.00000 ; CA- C- N- CA
6021 6020 6022 6023 3 0.65084 1.95253 0.00000
-2.60338 0.00000 0.00000 ; HA- CA- CB- HB2
6021 6020 6022 6024 3 0.65084 1.95253 0.00000
-2.60338 0.00000 0.00000 ; HA- CA- CB- HB3
6021 6020 6022 6025 3 0.65084 1.95253 0.00000
-2.60338 0.00000 0.00000 ; HA- CA- CB- CG
6021 6020 6036 6037 3 3.68192 -4.35136 0.00000
1.33888 0.00000 0.00000 ; HA- CA- C- O
6021 6020 6036 6038 3 0.00000 0.00000 0.00000
0.00000 0.00000 0.00000 ; HA- CA- C- N
6022 6020 6036 6037 3 0.00000 0.00000 0.00000
0.00000 0.00000 0.00000 ; CB- CA- C- O
6022 6020 6036 6038 3 2.51040 4.18400 1.67360
-6.69440 0.00000 0.00000 ; CB- CA- C- N

6022 6025 6026 6027 3 30.33400 0.00000 -30.33400
0.00000 0.00000 0.00000 ; CB- CG- CD1- HD1
6022 6025 6026 6028 3 30.33400 0.00000 -30.33400
0.00000 0.00000 0.00000 ; CB- CG- CD1- CE1
6022 6025 6034 6032 3 30.33400 0.00000 -30.33400
0.00000 0.00000 0.00000 ; CB- CG- CD2- CE2
6022 6025 6034 6035 3 30.33400 0.00000 -30.33400
0.00000 0.00000 0.00000 ; CB- CG- CD2- HD2
6023 6022 6020 6036 3 0.65084 1.95253 0.00000
-2.60338 0.00000 0.00000 ; HB2- CB- CA- C
6023 6022 6025 6026 3 0.00000 0.00000 0.00000
0.00000 0.00000 0.00000 ; HB2- CB- CG- CD1
6023 6022 6025 6034 3 0.00000 0.00000 0.00000
0.00000 0.00000 0.00000 ; HB2- CB- CG- CD2
6024 6022 6020 6036 3 0.65084 1.95253 0.00000
-2.60338 0.00000 0.00000 ; HB3- CB- CA- C
6024 6022 6025 6026 3 0.00000 0.00000 0.00000
0.00000 0.00000 0.00000 ; HB3- CB- CG- CD1
6024 6022 6025 6034 3 0.00000 0.00000 0.00000
0.00000 0.00000 0.00000 ; HB3- CB- CG- CD2
6025 6022 6020 6036 3 4.95804 2.17986 -3.52293
-3.21331 -0.40166 0.00000 ; CG- CB- CA- C
6025 6026 6028 6029 3 30.33400 0.00000 -30.33400
0.00000 0.00000 0.00000 ; CG- CD1- CE1- HE1
6025 6026 6028 6030 3 30.33400 0.00000 -30.33400
0.00000 0.00000 0.00000 ; CG- CD1- CE1- CZ
6025 6034 6032 6030 3 30.33400 0.00000 -30.33400
0.00000 0.00000 0.00000 ; CG- CD2- CE2- CZ
6025 6034 6032 6033 3 30.33400 0.00000 -30.33400
0.00000 0.00000 0.00000 ; CG- CD2- CE2- HE2
6026 6025 6034 6032 3 30.33400 0.00000 -30.33400
0.00000 0.00000 0.00000 ; CD1- CG- CD2- CE2
6026 6025 6034 6035 3 30.33400 0.00000 -30.33400
0.00000 0.00000 0.00000 ; CD1- CG- CD2- HD2
6026 6028 6030 6031 3 30.33400 0.00000 -30.33400
0.00000 0.00000 0.00000 ; CD1- CE1- CZ- HZ
6026 6028 6030 6032 3 30.33400 0.00000 -30.33400
0.00000 0.00000 0.00000 ; CD1- CE1- CZ- CE2
6027 6026 6025 6034 3 30.33400 0.00000 -30.33400
0.00000 0.00000 0.00000 ; HD1- CD1- CG- CD2
6027 6026 6028 6029 3 30.33400 0.00000 -30.33400
0.00000 0.00000 0.00000 ; HD1- CD1- CE1- HE1
6027 6026 6028 6030 3 30.33400 0.00000 -30.33400
0.00000 0.00000 0.00000 ; HD1- CD1- CE1- CZ
6028 6026 6025 6034 3 30.33400 0.00000 -30.33400
0.00000 0.00000 0.00000 ; CE1- CD1- CG- CD2
6028 6030 6032 6033 3 30.33400 0.00000 -30.33400
0.00000 0.00000 0.00000 ; CE1- CZ- CE2- HE2
6028 6030 6032 6034 3 30.33400 0.00000 -30.33400
0.00000 0.00000 0.00000 ; CE1- CZ- CE2- CD2
6029 6028 6030 6031 3 30.33400 0.00000 -30.33400
0.00000 0.00000 0.00000 ; HE1- CE1- CZ- HZ

6029 6028 6030 6032 3 30.33400 0.00000 -30.33400
0.00000 0.00000 0.00000 ; HE1- CE1- CZ- CE2
6030 6032 6034 6035 3 30.33400 0.00000 -30.33400
0.00000 0.00000 0.00000 ; CZ- CE2- CD2- HD2
6031 6030 6032 6033 3 30.33400 0.00000 -30.33400
0.00000 0.00000 0.00000 ; HZ- CZ- CE2- HE2
6031 6030 6032 6034 3 30.33400 0.00000 -30.33400
0.00000 0.00000 0.00000 ; HZ- CZ- CE2- CD2
6033 6032 6034 6035 3 30.33400 0.00000 -30.33400
0.00000 0.00000 0.00000 ; HE2- CE2- CD2- HD2
6036 6038 6040 6041 3 0.00000 0.00000 0.00000
0.00000 0.00000 0.00000 ; C- N- CA- HA
6036 6038 6040 6042 3 11.71520 1.67360 15.06240 -
13.38880 0.00000 0.00000 ; C- N- CA- CB
6036 6038 6040 6055 3 1.75728 5.27184 2.25936
-7.02912 0.00000 0.00000 ; C- N- CA- C
6037 6036 6038 6039 3 29.28800 -8.36800 -20.92000
0.00000 0.00000 0.00000 ; O- C- N- H
6037 6036 6038 6040 3 20.92000 0.00000 -20.92000
0.00000 0.00000 0.00000 ; O- C- N- CA
6038 6040 6042 6043 3 0.65084 1.95253 0.00000
-2.60338 0.00000 0.00000 ; N- CA- CB- HB2
6038 6040 6042 6044 3 0.65084 1.95253 0.00000
-2.60338 0.00000 0.00000 ; N- CA- CB- HB3
6038 6040 6042 6045 3 3.79070 1.39746 -4.61077
-2.40998 2.44346 0.00000 ; N- CA- CB- CG
6038 6040 6055 6056 3 0.00000 0.00000 0.00000
0.00000 0.00000 0.00000 ; N- CA- C- O
6038 6040 6055 6057 3 17.40544 -5.02080 -13.22144
9.20480 0.00000 0.00000 ; N- CA- C- N
6039 6038 6040 6041 3 0.00000 0.00000 0.00000
0.00000 0.00000 0.00000 ; H- N- CA- HA
6039 6038 6040 6042 3 0.00000 0.00000 0.00000
0.00000 0.00000 0.00000 ; H- N- CA- CB
6039 6038 6040 6055 3 0.00000 0.00000 0.00000
0.00000 0.00000 0.00000 ; H- N- CA- C
6040 6042 6045 6046 3 0.66944 2.00832 0.00000
-2.67776 0.00000 0.00000 ; CA- CB- CG- HG
6040 6042 6045 6047 3 3.90367 0.19665 -6.21742
-2.37651 5.99149 0.00000 ; CA- CB- CG- CD1
6040 6042 6045 6051 3 3.90367 0.19665 -6.21742
-2.37651 5.99149 0.00000 ; CA- CB- CG- CD2
6040 6055 6057 6058 3 20.92000 0.00000 -20.92000
0.00000 0.00000 0.00000 ; CA- C- N- H
6040 6055 6057 6059 3 20.92000 0.00000 -20.92000
0.00000 0.00000 0.00000 ; CA- C- N- CA
6041 6040 6042 6043 3 0.65084 1.95253 0.00000
-2.60338 0.00000 0.00000 ; HA- CA- CB- HB2
6041 6040 6042 6044 3 0.65084 1.95253 0.00000
-2.60338 0.00000 0.00000 ; HA- CA- CB- HB3
6041 6040 6042 6045 3 0.65084 1.95253 0.00000
-2.60338 0.00000 0.00000 ; HA- CA- CB- CG

6041 6040 6055 6056 3 3.68192 -4.35136 0.00000
1.33888 0.00000 0.00000 ; HA- CA- C- O
6041 6040 6055 6057 3 0.00000 0.00000 0.00000
0.00000 0.00000 0.00000 ; HA- CA- C- N
6042 6040 6055 6056 3 0.00000 0.00000 0.00000
0.00000 0.00000 0.00000 ; CB- CA- C- O
6042 6040 6055 6057 3 2.51040 4.18400 1.67360
-6.69440 0.00000 0.00000 ; CB- CA- C- N
6042 6045 6047 6048 3 0.66944 2.00832 0.00000
-2.67776 0.00000 0.00000 ; CB- CG- CD1- HD11
6042 6045 6047 6049 3 0.66944 2.00832 0.00000
-2.67776 0.00000 0.00000 ; CB- CG- CD1- HD12
6042 6045 6047 6050 3 0.66944 2.00832 0.00000
-2.67776 0.00000 0.00000 ; CB- CG- CD1- HD13
6042 6045 6051 6052 3 0.66944 2.00832 0.00000
-2.67776 0.00000 0.00000 ; CB- CG- CD2- HD21
6042 6045 6051 6053 3 0.66944 2.00832 0.00000
-2.67776 0.00000 0.00000 ; CB- CG- CD2- HD22
6042 6045 6051 6054 3 0.66944 2.00832 0.00000
-2.67776 0.00000 0.00000 ; CB- CG- CD2- HD23
6043 6042 6040 6055 3 0.65084 1.95253 0.00000
-2.60338 0.00000 0.00000 ; HB2- CB- CA- C
6043 6042 6045 6046 3 0.62760 1.88280 0.00000
-2.51040 0.00000 0.00000 ; HB2- CB- CG- HG
6043 6042 6045 6047 3 0.66944 2.00832 0.00000
-2.67776 0.00000 0.00000 ; HB2- CB- CG- CD1
6043 6042 6045 6051 3 0.66944 2.00832 0.00000
-2.67776 0.00000 0.00000 ; HB2- CB- CG- CD2
6044 6042 6040 6055 3 0.65084 1.95253 0.00000
-2.60338 0.00000 0.00000 ; HB3- CB- CA- C
6044 6042 6045 6046 3 0.62760 1.88280 0.00000
-2.51040 0.00000 0.00000 ; HB3- CB- CG- HG
6044 6042 6045 6047 3 0.66944 2.00832 0.00000
-2.67776 0.00000 0.00000 ; HB3- CB- CG- CD1
6044 6042 6045 6051 3 0.66944 2.00832 0.00000
-2.67776 0.00000 0.00000 ; HB3- CB- CG- CD2
6045 6042 6040 6055 3 10.33448 -1.14642 -11.54784
-2.40998 6.35968 0.00000 ; CG- CB- CA- C
6046 6045 6047 6048 3 0.62760 1.88280 0.00000
-2.51040 0.00000 0.00000 ; HG- CG- CD1- HD11
6046 6045 6047 6049 3 0.62760 1.88280 0.00000
-2.51040 0.00000 0.00000 ; HG- CG- CD1- HD12
6046 6045 6047 6050 3 0.62760 1.88280 0.00000
-2.51040 0.00000 0.00000 ; HG- CG- CD1- HD13
6046 6045 6051 6052 3 0.62760 1.88280 0.00000
-2.51040 0.00000 0.00000 ; HG- CG- CD2- HD21
6046 6045 6051 6053 3 0.62760 1.88280 0.00000
-2.51040 0.00000 0.00000 ; HG- CG- CD2- HD22
6046 6045 6051 6054 3 0.62760 1.88280 0.00000
-2.51040 0.00000 0.00000 ; HG- CG- CD2- HD23
6047 6045 6051 6052 3 0.66944 2.00832 0.00000
-2.67776 0.00000 0.00000 ; CD1- CG- CD2- HD21

```
  6047   6045   6051   6053       3    0.66944    2.00832    0.00000
-2.67776    0.00000    0.00000 ;    CD1-    CG-   CD2-  HD22
  6047   6045   6051   6054       3    0.66944    2.00832    0.00000
-2.67776    0.00000    0.00000 ;    CD1-    CG-   CD2-  HD23
  6048   6047   6045   6051       3    0.66944    2.00832    0.00000
-2.67776    0.00000    0.00000 ;   HD11-   CD1-    CG-   CD2
  6049   6047   6045   6051       3    0.66944    2.00832    0.00000
-2.67776    0.00000    0.00000 ;   HD12-   CD1-    CG-   CD2
  6050   6047   6045   6051       3    0.66944    2.00832    0.00000
-2.67776    0.00000    0.00000 ;   HD13-   CD1-    CG-   CD2
  6055   6057   6059   6060       3    0.00000    0.00000    0.00000
0.00000    0.00000    0.00000 ;      C-     N-    CA-    HA
  6055   6057   6059   6061       3   11.71520    1.67360   15.06240  -
13.38880    0.00000    0.00000 ;      C-     N-    CA-    CB
  6055   6057   6059   6065       3    1.75728    5.27184    2.25936
-7.02912    0.00000    0.00000 ;      C-     N-    CA-     C
  6056   6055   6057   6058       3   29.28800   -8.36800  -20.92000
0.00000    0.00000    0.00000 ;      O-     C-     N-     H
  6056   6055   6057   6059       3   20.92000    0.00000  -20.92000
0.00000    0.00000    0.00000 ;      O-     C-     N-    CA
  6057   6059   6061   6062       3    0.65084    1.95253    0.00000
-2.60338    0.00000    0.00000 ;      N-    CA-    CB-   HB1
  6057   6059   6061   6063       3    0.65084    1.95253    0.00000
-2.60338    0.00000    0.00000 ;      N-    CA-    CB-   HB2
  6057   6059   6061   6064       3    0.65084    1.95253    0.00000
-2.60338    0.00000    0.00000 ;      N-    CA-    CB-   HB3
  6057   6059   6065   6066       3    0.00000    0.00000    0.00000
0.00000    0.00000    0.00000 ;      N-    CA-     C-     O
  6057   6059   6065   6067       3   17.40544   -5.02080  -13.22144
9.20480    0.00000    0.00000 ;      N-    CA-     C-     N
  6058   6057   6059   6060       3    0.00000    0.00000    0.00000
0.00000    0.00000    0.00000 ;      H-     N-    CA-    HA
  6058   6057   6059   6061       3    0.00000    0.00000    0.00000
0.00000    0.00000    0.00000 ;      H-     N-    CA-    CB
  6058   6057   6059   6065       3    0.00000    0.00000    0.00000
0.00000    0.00000    0.00000 ;      H-     N-    CA-     C
  6059   6065   6067   6068       3   20.92000    0.00000  -20.92000
0.00000    0.00000    0.00000 ;     CA-     C-     N-     H
  6059   6065   6067   6069       3   20.92000    0.00000  -20.92000
0.00000    0.00000    0.00000 ;     CA-     C-     N-    CA
  6060   6059   6061   6062       3    0.65084    1.95253    0.00000
-2.60338    0.00000    0.00000 ;     HA-    CA-    CB-   HB1
  6060   6059   6061   6063       3    0.65084    1.95253    0.00000
-2.60338    0.00000    0.00000 ;     HA-    CA-    CB-   HB2
  6060   6059   6061   6064       3    0.65084    1.95253    0.00000
-2.60338    0.00000    0.00000 ;     HA-    CA-    CB-   HB3
  6060   6059   6065   6066       3    3.68192   -4.35136    0.00000
1.33888    0.00000    0.00000 ;     HA-    CA-     C-     O
  6060   6059   6065   6067       3    0.00000    0.00000    0.00000
0.00000    0.00000    0.00000 ;     HA-    CA-     C-     N
  6061   6059   6065   6066       3    0.00000    0.00000    0.00000
0.00000    0.00000    0.00000 ;     CB-    CA-     C-     O
```

6061 6059 6065 6067 3 2.51040 4.18400 1.67360
-6.69440 0.00000 0.00000 ; CB- CA- C- N
6062 6061 6059 6065 3 0.65084 1.95253 0.00000
-2.60338 0.00000 0.00000 ; HB1- CB- CA- C
6063 6061 6059 6065 3 0.65084 1.95253 0.00000
-2.60338 0.00000 0.00000 ; HB2- CB- CA- C
6064 6061 6059 6065 3 0.65084 1.95253 0.00000
-2.60338 0.00000 0.00000 ; HB3- CB- CA- C
6065 6067 6069 6070 3 0.00000 0.00000 0.00000
0.00000 0.00000 0.00000 ; C- N- CA- HA
6065 6067 6069 6071 3 11.71520 1.67360 15.06240 -
13.38880 0.00000 0.00000 ; C- N- CA- CB
6065 6067 6069 6075 3 1.75728 5.27184 2.25936
-7.02912 0.00000 0.00000 ; C- N- CA- C
6066 6065 6067 6068 3 29.28800 -8.36800 -20.92000
0.00000 0.00000 0.00000 ; O- C- N- H
6066 6065 6067 6069 3 20.92000 0.00000 -20.92000
0.00000 0.00000 0.00000 ; O- C- N- CA
6067 6069 6071 6072 3 0.65084 1.95253 0.00000
-2.60338 0.00000 0.00000 ; N- CA- CB- HB1
6067 6069 6071 6073 3 0.65084 1.95253 0.00000
-2.60338 0.00000 0.00000 ; N- CA- CB- HB2
6067 6069 6071 6074 3 0.65084 1.95253 0.00000
-2.60338 0.00000 0.00000 ; N- CA- CB- HB3
6067 6069 6075 6076 3 0.00000 0.00000 0.00000
0.00000 0.00000 0.00000 ; N- CA- C- O
6067 6069 6075 6077 3 17.40544 -5.02080 -13.22144
9.20480 0.00000 0.00000 ; N- CA- C- N
6068 6067 6069 6070 3 0.00000 0.00000 0.00000
0.00000 0.00000 0.00000 ; H- N- CA- HA
6068 6067 6069 6071 3 0.00000 0.00000 0.00000
0.00000 0.00000 0.00000 ; H- N- CA- CB
6068 6067 6069 6075 3 0.00000 0.00000 0.00000
0.00000 0.00000 0.00000 ; H- N- CA- C
6069 6075 6077 6078 3 20.92000 0.00000 -20.92000
0.00000 0.00000 0.00000 ; CA- C- N- H
6069 6075 6077 6079 3 20.92000 0.00000 -20.92000
0.00000 0.00000 0.00000 ; CA- C- N- CA
6070 6069 6071 6072 3 0.65084 1.95253 0.00000
-2.60338 0.00000 0.00000 ; HA- CA- CB- HB1
6070 6069 6071 6073 3 0.65084 1.95253 0.00000
-2.60338 0.00000 0.00000 ; HA- CA- CB- HB2
6070 6069 6071 6074 3 0.65084 1.95253 0.00000
-2.60338 0.00000 0.00000 ; HA- CA- CB- HB3
6070 6069 6075 6076 3 3.68192 -4.35136 0.00000
1.33888 0.00000 0.00000 ; HA- CA- C- O
6070 6069 6075 6077 3 0.00000 0.00000 0.00000
0.00000 0.00000 0.00000 ; HA- CA- C- N
6071 6069 6075 6076 3 0.00000 0.00000 0.00000
0.00000 0.00000 0.00000 ; CB- CA- C- O
6071 6069 6075 6077 3 2.51040 4.18400 1.67360
-6.69440 0.00000 0.00000 ; CB- CA- C- N

6072 6071 6069 6075 3 0.65084 1.95253 0.00000
-2.60338 0.00000 0.00000 ; HB1- CB- CA- C
6073 6071 6069 6075 3 0.65084 1.95253 0.00000
-2.60338 0.00000 0.00000 ; HB2- CB- CA- C
6074 6071 6069 6075 3 0.65084 1.95253 0.00000
-2.60338 0.00000 0.00000 ; HB3- CB- CA- C
6075 6077 6079 6080 3 0.00000 0.00000 0.00000
0.00000 0.00000 0.00000 ; C- N- CA- HA
6075 6077 6079 6081 3 11.71520 1.67360 15.06240 -
13.38880 0.00000 0.00000 ; C- N- CA- CB
6075 6077 6079 6089 3 1.75728 5.27184 2.25936
-7.02912 0.00000 0.00000 ; C- N- CA- C
6076 6075 6077 6078 3 29.28800 -8.36800 -20.92000
0.00000 0.00000 0.00000 ; O- C- N- H
6076 6075 6077 6079 3 20.92000 0.00000 -20.92000
0.00000 0.00000 0.00000 ; O- C- N- CA
6077 6079 6081 6082 3 0.65084 1.95253 0.00000
-2.60338 0.00000 0.00000 ; N- CA- CB- HB
6077 6079 6081 6083 3 3.83673 0.44769 -1.77402
-2.47693 -0.03347 0.00000 ; N- CA- CB- CG2
6077 6079 6081 6087 3 4.93294 1.13386 -3.12963
-5.27184 3.17984 0.00000 ; N- CA- CB- OG1
6077 6079 6089 6090 3 0.00000 0.00000 0.00000
0.00000 0.00000 0.00000 ; N- CA- C- O
6077 6079 6089 6091 3 17.40544 -5.02080 -13.22144
9.20480 0.00000 0.00000 ; N- CA- C- N
6078 6077 6079 6080 3 0.00000 0.00000 0.00000
0.00000 0.00000 0.00000 ; H- N- CA- HA
6078 6077 6079 6081 3 0.00000 0.00000 0.00000
0.00000 0.00000 0.00000 ; H- N- CA- CB
6078 6077 6079 6089 3 0.00000 0.00000 0.00000
0.00000 0.00000 0.00000 ; H- N- CA- C
6079 6081 6083 6084 3 0.66944 2.00832 0.00000
-2.67776 0.00000 0.00000 ; CA- CB- CG2- HG21
6079 6081 6083 6085 3 0.66944 2.00832 0.00000
-2.67776 0.00000 0.00000 ; CA- CB- CG2- HG22
6079 6081 6083 6086 3 0.66944 2.00832 0.00000
-2.67776 0.00000 0.00000 ; CA- CB- CG2- HG23
6079 6081 6087 6088 3 1.12131 2.98738 1.66523
-3.94970 0.43514 0.00000 ; CA- CB- OG1- HG1
6079 6089 6091 6092 3 20.92000 0.00000 -20.92000
0.00000 0.00000 0.00000 ; CA- C- N- H
6079 6089 6091 6093 3 20.92000 0.00000 -20.92000
0.00000 0.00000 0.00000 ; CA- C- N- CA
6080 6079 6081 6082 3 0.65084 1.95253 0.00000
-2.60338 0.00000 0.00000 ; HA- CA- CB- HB
6080 6079 6081 6083 3 0.65084 1.95253 0.00000
-2.60338 0.00000 0.00000 ; HA- CA- CB- CG2
6080 6079 6081 6087 3 1.04600 -1.04600 0.00000
0.00000 0.00000 0.00000 ; HA- CA- CB- OG1
6080 6079 6089 6090 3 3.68192 -4.35136 0.00000
1.33888 0.00000 0.00000 ; HA- CA- C- O

6080 6079 6089 6091 3 0.00000 0.00000 0.00000
0.00000 0.00000 0.00000 ; HA- CA- C- N
6081 6079 6089 6090 3 0.00000 0.00000 0.00000
0.00000 0.00000 0.00000 ; CB- CA- C- O
6081 6079 6089 6091 3 2.51040 4.18400 1.67360
-6.69440 0.00000 0.00000 ; CB- CA- C- N
6082 6081 6079 6089 3 0.65084 1.95253 0.00000
-2.60338 0.00000 0.00000 ; HB- CB- CA- C
6082 6081 6083 6084 3 0.65084 1.95253 0.00000
-2.60338 0.00000 0.00000 ; HB- CB- CG2- HG21
6082 6081 6083 6085 3 0.65084 1.95253 0.00000
-2.60338 0.00000 0.00000 ; HB- CB- CG2- HG22
6082 6081 6083 6086 3 0.65084 1.95253 0.00000
-2.60338 0.00000 0.00000 ; HB- CB- CG2- HG23
6082 6081 6087 6088 3 0.69733 2.09200 0.00000
-2.78933 0.00000 0.00000 ; HB- CB- OG1- HG1
6083 6081 6079 6089 3 5.67350 3.55640 -6.16722
-2.47693 3.74886 0.00000 ; CG2- CB- CA- C
6083 6081 6087 6088 3 4.74047 0.27196 -2.26773
-3.94970 1.60666 0.00000 ; CG2- CB- OG1- HG1
6084 6083 6081 6087 3 1.04600 -1.04600 0.00000
0.00000 0.00000 0.00000 ; HG21- CG2- CB- OG1
6085 6083 6081 6087 3 1.04600 -1.04600 0.00000
0.00000 0.00000 0.00000 ; HG22- CG2- CB- OG1
6086 6083 6081 6087 3 1.04600 -1.04600 0.00000
0.00000 0.00000 0.00000 ; HG23- CG2- CB- OG1
6087 6081 6079 6089 3 6.53541 6.87013 -6.21742
-5.27184 5.22163 0.00000 ; OG1- CB- CA- C
6089 6091 6093 6094 3 0.00000 0.00000 0.00000
0.00000 0.00000 0.00000 ; C- N- CA- HA
6089 6091 6093 6095 3 11.71520 1.67360 15.06240 -
13.38880 0.00000 0.00000 ; C- N- CA- CB
6089 6091 6093 6108 3 1.75728 5.27184 2.25936
-7.02912 0.00000 0.00000 ; C- N- CA- C
6090 6089 6091 6092 3 29.28800 -8.36800 -20.92000
0.00000 0.00000 0.00000 ; O- C- N- H
6090 6089 6091 6093 3 20.92000 0.00000 -20.92000
0.00000 0.00000 0.00000 ; O- C- N- CA
6091 6093 6095 6096 3 0.65084 1.95253 0.00000
-2.60338 0.00000 0.00000 ; N- CA- CB- HB
6091 6093 6095 6097 3 3.83673 0.44769 -1.77402
-2.47693 -0.03347 0.00000 ; N- CA- CB- CG2
6091 6093 6095 6101 3 2.97482 0.12134 2.04179
-1.89117 -3.24678 0.00000 ; N- CA- CB- CG1
6091 6093 6108 6109 3 0.00000 0.00000 0.00000
0.00000 0.00000 0.00000 ; N- CA- C- O
6091 6093 6108 6110 3 17.40544 -5.02080 -13.22144
9.20480 0.00000 0.00000 ; N- CA- C- N
6092 6091 6093 6094 3 0.00000 0.00000 0.00000
0.00000 0.00000 0.00000 ; H- N- CA- HA
6092 6091 6093 6095 3 0.00000 0.00000 0.00000
0.00000 0.00000 0.00000 ; H- N- CA- CB

6092 6091 6093 6108 3 0.00000 0.00000 0.00000
0.00000 0.00000 0.00000 ; H- N- CA- C
6093 6095 6097 6098 3 0.66944 2.00832 0.00000
-2.67776 0.00000 0.00000 ; CA- CB- CG2- HG21
6093 6095 6097 6099 3 0.66944 2.00832 0.00000
-2.67776 0.00000 0.00000 ; CA- CB- CG2- HG22
6093 6095 6097 6100 3 0.66944 2.00832 0.00000
-2.67776 0.00000 0.00000 ; CA- CB- CG2- HG23
6093 6095 6101 6102 3 0.66944 2.00832 0.00000
-2.67776 0.00000 0.00000 ; CA- CB- CG1- HG12
6093 6095 6101 6103 3 0.66944 2.00832 0.00000
-2.67776 0.00000 0.00000 ; CA- CB- CG1- HG13
6093 6095 6101 6104 3 4.24258 -0.52718 -7.25506
-1.79075 7.69856 0.00000 ; CA- CB- CG1- CD1
6093 6108 6110 6111 3 20.92000 0.00000 -20.92000
0.00000 0.00000 0.00000 ; CA- C- N- H
6093 6108 6110 6112 3 20.92000 0.00000 -20.92000
0.00000 0.00000 0.00000 ; CA- C- N- CA
6094 6093 6095 6096 3 0.65084 1.95253 0.00000
-2.60338 0.00000 0.00000 ; HA- CA- CB- HB
6094 6093 6095 6097 3 0.65084 1.95253 0.00000
-2.60338 0.00000 0.00000 ; HA- CA- CB- CG2
6094 6093 6095 6101 3 0.65084 1.95253 0.00000
-2.60338 0.00000 0.00000 ; HA- CA- CB- CG1
6094 6093 6108 6109 3 3.68192 -4.35136 0.00000
1.33888 0.00000 0.00000 ; HA- CA- C- O
6094 6093 6108 6110 3 0.00000 0.00000 0.00000
0.00000 0.00000 0.00000 ; HA- CA- C- N
6095 6093 6108 6109 3 0.00000 0.00000 0.00000
0.00000 0.00000 0.00000 ; CB- CA- C- O
6095 6093 6108 6110 3 2.51040 4.18400 1.67360
-6.69440 0.00000 0.00000 ; CB- CA- C- N
6095 6101 6104 6105 3 0.66944 2.00832 0.00000
-2.67776 0.00000 0.00000 ; CB- CG1- CD1- HD11
6095 6101 6104 6106 3 0.66944 2.00832 0.00000
-2.67776 0.00000 0.00000 ; CB- CG1- CD1- HD12
6095 6101 6104 6107 3 0.66944 2.00832 0.00000
-2.67776 0.00000 0.00000 ; CB- CG1- CD1- HD13
6096 6095 6093 6108 3 0.65084 1.95253 0.00000
-2.60338 0.00000 0.00000 ; HB- CB- CA- C
6096 6095 6097 6098 3 0.62760 1.88280 0.00000
-2.51040 0.00000 0.00000 ; HB- CB- CG2- HG21
6096 6095 6097 6099 3 0.62760 1.88280 0.00000
-2.51040 0.00000 0.00000 ; HB- CB- CG2- HG22
6096 6095 6097 6100 3 0.62760 1.88280 0.00000
-2.51040 0.00000 0.00000 ; HB- CB- CG2- HG23
6096 6095 6101 6102 3 0.62760 1.88280 0.00000
-2.51040 0.00000 0.00000 ; HB- CB- CG1- HG12
6096 6095 6101 6103 3 0.62760 1.88280 0.00000
-2.51040 0.00000 0.00000 ; HB- CB- CG1- HG13
6096 6095 6101 6104 3 0.66944 2.00832 0.00000
-2.67776 0.00000 0.00000 ; HB- CB- CG1- CD1

6097 6095 6093 6108 3 5.67350 3.55640 -6.16722
-2.47693 3.74886 0.00000 ; CG2- CB- CA- C
6097 6095 6101 6102 3 0.66944 2.00832 0.00000
-2.67776 0.00000 0.00000 ; CG2- CB- CG1- HG12
6097 6095 6101 6103 3 0.66944 2.00832 0.00000
-2.67776 0.00000 0.00000 ; CG2- CB- CG1- HG13
6097 6095 6101 6104 3 3.81162 0.49790 -8.14206
-1.79075 7.49773 0.00000 ; CG2- CB- CG1- CD1
6098 6097 6095 6101 3 0.66944 2.00832 0.00000
-2.67776 0.00000 0.00000 ; HG21- CG2- CB- CG1
6099 6097 6095 6101 3 0.66944 2.00832 0.00000
-2.67776 0.00000 0.00000 ; HG22- CG2- CB- CG1
6100 6097 6095 6101 3 0.66944 2.00832 0.00000
-2.67776 0.00000 0.00000 ; HG23- CG2- CB- CG1
6101 6095 6093 6108 3 8.26340 0.74057 -9.99976
-1.89117 3.84928 0.00000 ; CG1- CB- CA- C
6102 6101 6104 6105 3 0.62760 1.88280 0.00000
-2.51040 0.00000 0.00000 ; HG12- CG1- CD1- HD11
6102 6101 6104 6106 3 0.62760 1.88280 0.00000
-2.51040 0.00000 0.00000 ; HG12- CG1- CD1- HD12
6102 6101 6104 6107 3 0.62760 1.88280 0.00000
-2.51040 0.00000 0.00000 ; HG12- CG1- CD1- HD13
6103 6101 6104 6105 3 0.62760 1.88280 0.00000
-2.51040 0.00000 0.00000 ; HG13- CG1- CD1- HD11
6103 6101 6104 6106 3 0.62760 1.88280 0.00000
-2.51040 0.00000 0.00000 ; HG13- CG1- CD1- HD12
6103 6101 6104 6107 3 0.62760 1.88280 0.00000
-2.51040 0.00000 0.00000 ; HG13- CG1- CD1- HD13
6108 6110 6112 6113 3 0.00000 0.00000 0.00000
0.00000 0.00000 0.00000 ; C- N- CA- HA
6108 6110 6112 6114 3 11.71520 1.67360 15.06240 -
13.38880 0.00000 0.00000 ; C- N- CA- CB
6108 6110 6112 6120 3 1.75728 5.27184 2.25936
-7.02912 0.00000 0.00000 ; C- N- CA- C
6109 6108 6110 6111 3 29.28800 -8.36800 -20.92000
0.00000 0.00000 0.00000 ; O- C- N- H
6109 6108 6110 6112 3 20.92000 0.00000 -20.92000
0.00000 0.00000 0.00000 ; O- C- N- CA
6110 6112 6114 6115 3 0.65084 1.95253 0.00000
-2.60338 0.00000 0.00000 ; N- CA- CB- HB2
6110 6112 6114 6116 3 0.65084 1.95253 0.00000
-2.60338 0.00000 0.00000 ; N- CA- CB- HB3
6110 6112 6114 6117 3 15.41386 9.74035 -8.39310
-0.97069 2.97901 0.00000 ; N- CA- CB- CG
6110 6112 6120 6121 3 0.00000 0.00000 0.00000
0.00000 0.00000 0.00000 ; N- CA- C- O
6110 6112 6120 6122 3 17.40544 -5.02080 -13.22144
9.20480 0.00000 0.00000 ; N- CA- C- N
6111 6110 6112 6113 3 0.00000 0.00000 0.00000
0.00000 0.00000 0.00000 ; H- N- CA- HA
6111 6110 6112 6114 3 0.00000 0.00000 0.00000
0.00000 0.00000 0.00000 ; H- N- CA- CB

6111 6110 6112 6120 3 0.00000 0.00000 0.00000
0.00000 0.00000 0.00000 ; H- N- CA- C
6112 6114 6117 6118 3 6.43499 0.00000 -5.39736
0.00000 -1.03763 0.00000 ; CA- CB- CG- OD1
6112 6114 6117 6119 3 6.43499 0.00000 -5.39736
0.00000 -1.03763 0.00000 ; CA- CB- CG- OD2
6112 6120 6122 6123 3 20.92000 0.00000 -20.92000
0.00000 0.00000 0.00000 ; CA- C- N- H
6112 6120 6122 6124 3 20.92000 0.00000 -20.92000
0.00000 0.00000 0.00000 ; CA- C- N- CA
6113 6112 6114 6115 3 0.65084 1.95253 0.00000
-2.60338 0.00000 0.00000 ; HA- CA- CB- HB2
6113 6112 6114 6116 3 0.65084 1.95253 0.00000
-2.60338 0.00000 0.00000 ; HA- CA- CB- HB3
6113 6112 6114 6117 3 0.65084 1.95253 0.00000
-2.60338 0.00000 0.00000 ; HA- CA- CB- CG
6113 6112 6120 6121 3 3.68192 -4.35136 0.00000
1.33888 0.00000 0.00000 ; HA- CA- C- O
6113 6112 6120 6122 3 0.00000 0.00000 0.00000
0.00000 0.00000 0.00000 ; HA- CA- C- N
6114 6112 6120 6121 3 0.00000 0.00000 0.00000
0.00000 0.00000 0.00000 ; CB- CA- C- O
6114 6112 6120 6122 3 2.51040 4.18400 1.67360
-6.69440 0.00000 0.00000 ; CB- CA- C- N
6115 6114 6112 6120 3 0.65084 1.95253 0.00000
-2.60338 0.00000 0.00000 ; HB2- CB- CA- C
6115 6114 6117 6118 3 0.00000 0.00000 0.00000
0.00000 0.00000 0.00000 ; HB2- CB- CG- OD1
6115 6114 6117 6119 3 0.00000 0.00000 0.00000
0.00000 0.00000 0.00000 ; HB2- CB- CG- OD2
6116 6114 6112 6120 3 0.65084 1.95253 0.00000
-2.60338 0.00000 0.00000 ; HB3- CB- CA- C
6116 6114 6117 6118 3 0.00000 0.00000 0.00000
0.00000 0.00000 0.00000 ; HB3- CB- CG- OD1
6116 6114 6117 6119 3 0.00000 0.00000 0.00000
0.00000 0.00000 0.00000 ; HB3- CB- CG- OD2
6117 6114 6112 6120 3 7.14627 -1.04600 -8.99560
-0.97069 5.15469 0.00000 ; CG- CB- CA- C
6120 6122 6124 6125 3 0.00000 0.00000 0.00000
0.00000 0.00000 0.00000 ; C- N- CA- HA
6120 6122 6124 6126 3 11.71520 1.67360 15.06240 -
13.38880 0.00000 0.00000 ; C- N- CA- CB
6120 6122 6124 6142 3 1.75728 5.27184 2.25936
-7.02912 0.00000 0.00000 ; C- N- CA- C
6121 6120 6122 6123 3 29.28800 -8.36800 -20.92000
0.00000 0.00000 0.00000 ; O- C- N- H
6121 6120 6122 6124 3 20.92000 0.00000 -20.92000
0.00000 0.00000 0.00000 ; O- C- N- CA
6122 6124 6126 6127 3 0.65084 1.95253 0.00000
-2.60338 0.00000 0.00000 ; N- CA- CB- HB2
6122 6124 6126 6128 3 0.65084 1.95253 0.00000
-2.60338 0.00000 0.00000 ; N- CA- CB- HB3

6122 6124 6126 6129 3 0.65084 1.95253 0.00000
-2.60338 0.00000 0.00000 ; N- CA- CB- CG
6122 6124 6142 6143 3 0.00000 0.00000 0.00000
0.00000 0.00000 0.00000 ; N- CA- C- O
6122 6124 6142 6144 3 17.40544 -5.02080 -13.22144
9.20480 0.00000 0.00000 ; N- CA- C- N
6123 6122 6124 6125 3 0.00000 0.00000 0.00000
0.00000 0.00000 0.00000 ; H- N- CA- HA
6123 6122 6124 6126 3 0.00000 0.00000 0.00000
0.00000 0.00000 0.00000 ; H- N- CA- CB
6123 6122 6124 6142 3 0.00000 0.00000 0.00000
0.00000 0.00000 0.00000 ; H- N- CA- C
6124 6126 6129 6130 3 0.66944 2.00832 0.00000
-2.67776 0.00000 0.00000 ; CA- CB- CG- HG2
6124 6126 6129 6131 3 0.66944 2.00832 0.00000
-2.67776 0.00000 0.00000 ; CA- CB- CG- HG3
6124 6126 6129 6132 3 3.68192 3.09616 -2.09200
-3.01248 0.00000 0.00000 ; CA- CB- CG- CD
6124 6142 6144 6145 3 20.92000 0.00000 -20.92000
0.00000 0.00000 0.00000 ; CA- C- N- H
6124 6142 6144 6146 3 20.92000 0.00000 -20.92000
0.00000 0.00000 0.00000 ; CA- C- N- CA
6125 6124 6126 6127 3 0.65084 1.95253 0.00000
-2.60338 0.00000 0.00000 ; HA- CA- CB- HB2
6125 6124 6126 6128 3 0.65084 1.95253 0.00000
-2.60338 0.00000 0.00000 ; HA- CA- CB- HB3
6125 6124 6126 6129 3 0.65084 1.95253 0.00000
-2.60338 0.00000 0.00000 ; HA- CA- CB- CG
6125 6124 6142 6143 3 3.68192 -4.35136 0.00000
1.33888 0.00000 0.00000 ; HA- CA- C- O
6125 6124 6142 6144 3 0.00000 0.00000 0.00000
0.00000 0.00000 0.00000 ; HA- CA- C- N
6126 6124 6142 6143 3 0.00000 0.00000 0.00000
0.00000 0.00000 0.00000 ; CB- CA- C- O
6126 6124 6142 6144 3 2.51040 4.18400 1.67360
-6.69440 0.00000 0.00000 ; CB- CA- C- N
6126 6129 6132 6133 3 0.66944 2.00832 0.00000
-2.67776 0.00000 0.00000 ; CB- CG- CD- HD2
6126 6129 6132 6134 3 0.66944 2.00832 0.00000
-2.67776 0.00000 0.00000 ; CB- CG- CD- HD3
6126 6129 6132 6135 3 3.68192 3.09616 -2.09200
-3.01248 0.00000 0.00000 ; CB- CG- CD- CE
6127 6126 6124 6142 3 0.65084 1.95253 0.00000
-2.60338 0.00000 0.00000 ; HB2- CB- CA- C
6127 6126 6129 6130 3 0.62760 1.88280 0.00000
-2.51040 0.00000 0.00000 ; HB2- CB- CG- HG2
6127 6126 6129 6131 3 0.62760 1.88280 0.00000
-2.51040 0.00000 0.00000 ; HB2- CB- CG- HG3
6127 6126 6129 6132 3 0.66944 2.00832 0.00000
-2.67776 0.00000 0.00000 ; HB2- CB- CG- CD
6128 6126 6124 6142 3 0.65084 1.95253 0.00000
-2.60338 0.00000 0.00000 ; HB3- CB- CA- C

```
  6128   6126   6129   6130      3    0.62760    1.88280    0.00000
-2.51040    0.00000    0.00000 ;    HB3-    CB-    CG-   HG2
  6128   6126   6129   6131      3    0.62760    1.88280    0.00000
-2.51040    0.00000    0.00000 ;    HB3-    CB-    CG-   HG3
  6128   6126   6129   6132      3    0.66944    2.00832    0.00000
-2.67776    0.00000    0.00000 ;    HB3-    CB-    CG-    CD
  6129   6126   6124   6142      3    0.65084    1.95253    0.00000
-2.60338    0.00000    0.00000 ;     CG-    CB-    CA-     C
  6129   6132   6135   6136      3    0.65084    1.95253    0.00000
-2.60338    0.00000    0.00000 ;     CG-    CD-    CE-   HE2
  6129   6132   6135   6137      3    0.65084    1.95253    0.00000
-2.60338    0.00000    0.00000 ;     CG-    CD-    CE-   HE3
  6129   6132   6135   6138      3    0.65084    1.95253    0.00000
-2.60338    0.00000    0.00000 ;     CG-    CD-    CE-    NZ
  6130   6129   6132   6133      3    0.62760    1.88280    0.00000
-2.51040    0.00000    0.00000 ;    HG2-    CG-    CD-   HD2
  6130   6129   6132   6134      3    0.62760    1.88280    0.00000
-2.51040    0.00000    0.00000 ;    HG2-    CG-    CD-   HD3
  6130   6129   6132   6135      3    0.66944    2.00832    0.00000
-2.67776    0.00000    0.00000 ;    HG2-    CG-    CD-    CE
  6131   6129   6132   6133      3    0.62760    1.88280    0.00000
-2.51040    0.00000    0.00000 ;    HG3-    CG-    CD-   HD2
  6131   6129   6132   6134      3    0.62760    1.88280    0.00000
-2.51040    0.00000    0.00000 ;    HG3-    CG-    CD-   HD3
  6131   6129   6132   6135      3    0.66944    2.00832    0.00000
-2.67776    0.00000    0.00000 ;    HG3-    CG-    CD-    CE
  6132   6135   6138   6139      3    0.65084    1.95253    0.00000
-2.60338    0.00000    0.00000 ;     CD-    CE-    NZ-   HZ1
  6132   6135   6138   6140      3    0.65084    1.95253    0.00000
-2.60338    0.00000    0.00000 ;     CD-    CE-    NZ-   HZ2
  6132   6135   6138   6141      3    0.65084    1.95253    0.00000
-2.60338    0.00000    0.00000 ;     CD-    CE-    NZ-   HZ3
  6133   6132   6135   6136      3    0.65084    1.95253    0.00000
-2.60338    0.00000    0.00000 ;    HD2-    CD-    CE-   HE2
  6133   6132   6135   6137      3    0.65084    1.95253    0.00000
-2.60338    0.00000    0.00000 ;    HD2-    CD-    CE-   HE3
  6133   6132   6135   6138      3    0.65084    1.95253    0.00000
-2.60338    0.00000    0.00000 ;    HD2-    CD-    CE-    NZ
  6134   6132   6135   6136      3    0.65084    1.95253    0.00000
-2.60338    0.00000    0.00000 ;    HD3-    CD-    CE-   HE2
  6134   6132   6135   6137      3    0.65084    1.95253    0.00000
-2.60338    0.00000    0.00000 ;    HD3-    CD-    CE-   HE3
  6134   6132   6135   6138      3    0.65084    1.95253    0.00000
-2.60338    0.00000    0.00000 ;    HD3-    CD-    CE-    NZ
  6136   6135   6138   6139      3    0.65084    1.95253    0.00000
-2.60338    0.00000    0.00000 ;    HE2-    CE-    NZ-   HZ1
  6136   6135   6138   6140      3    0.65084    1.95253    0.00000
-2.60338    0.00000    0.00000 ;    HE2-    CE-    NZ-   HZ2
  6136   6135   6138   6141      3    0.65084    1.95253    0.00000
-2.60338    0.00000    0.00000 ;    HE2-    CE-    NZ-   HZ3
  6137   6135   6138   6139      3    0.65084    1.95253    0.00000
-2.60338    0.00000    0.00000 ;    HE3-    CE-    NZ-   HZ1
```

6137 6135 6138 6140 3 0.65084 1.95253 0.00000
-2.60338 0.00000 0.00000 ; HE3- CE- NZ- HZ2
6137 6135 6138 6141 3 0.65084 1.95253 0.00000
-2.60338 0.00000 0.00000 ; HE3- CE- NZ- HZ3
6142 6144 6146 6147 3 0.00000 0.00000 0.00000
0.00000 0.00000 0.00000 ; C- N- CA- HA
6142 6144 6146 6148 3 11.71520 1.67360 15.06240 -
13.38880 0.00000 0.00000 ; C- N- CA- CB
6142 6144 6146 6159 3 1.75728 5.27184 2.25936
-7.02912 0.00000 0.00000 ; C- N- CA- C
6143 6142 6144 6145 3 29.28800 -8.36800 -20.92000
0.00000 0.00000 0.00000 ; O- C- N- H
6143 6142 6144 6146 3 20.92000 0.00000 -20.92000
0.00000 0.00000 0.00000 ; O- C- N- CA
6144 6146 6148 6149 3 0.65084 1.95253 0.00000
-2.60338 0.00000 0.00000 ; N- CA- CB- HB2
6144 6146 6148 6150 3 0.65084 1.95253 0.00000
-2.60338 0.00000 0.00000 ; N- CA- CB- HB3
6144 6146 6148 6151 3 3.21331 2.22589 -4.15053
-2.40998 2.61082 0.00000 ; N- CA- CB- CG
6144 6146 6159 6160 3 0.00000 0.00000 0.00000
0.00000 0.00000 0.00000 ; N- CA- C- O
6144 6146 6159 6161 3 17.40544 -5.02080 -13.22144
9.20480 0.00000 0.00000 ; N- CA- C- N
6145 6144 6146 6147 3 0.00000 0.00000 0.00000
0.00000 0.00000 0.00000 ; H- N- CA- HA
6145 6144 6146 6148 3 0.00000 0.00000 0.00000
0.00000 0.00000 0.00000 ; H- N- CA- CB
6145 6144 6146 6159 3 0.00000 0.00000 0.00000
0.00000 0.00000 0.00000 ; H- N- CA- C
6146 6148 6151 6152 3 0.66944 2.00832 0.00000
-2.67776 0.00000 0.00000 ; CA- CB- CG- HG2
6146 6148 6151 6153 3 0.66944 2.00832 0.00000
-2.67776 0.00000 0.00000 ; CA- CB- CG- HG3
6146 6148 6151 6154 3 3.69866 -4.35136 -3.92459
6.89523 4.61914 0.00000 ; CA- CB- CG- CD
6146 6159 6161 6162 3 20.92000 0.00000 -20.92000
0.00000 0.00000 0.00000 ; CA- C- N- H
6146 6159 6161 6163 3 20.92000 0.00000 -20.92000
0.00000 0.00000 0.00000 ; CA- C- N- CA
6147 6146 6148 6149 3 0.65084 1.95253 0.00000
-2.60338 0.00000 0.00000 ; HA- CA- CB- HB2
6147 6146 6148 6150 3 0.65084 1.95253 0.00000
-2.60338 0.00000 0.00000 ; HA- CA- CB- HB3
6147 6146 6148 6151 3 0.65084 1.95253 0.00000
-2.60338 0.00000 0.00000 ; HA- CA- CB- CG
6147 6146 6159 6160 3 3.68192 -4.35136 0.00000
1.33888 0.00000 0.00000 ; HA- CA- C- O
6147 6146 6159 6161 3 0.00000 0.00000 0.00000
0.00000 0.00000 0.00000 ; HA- CA- C- N
6148 6146 6159 6160 3 0.00000 0.00000 0.00000
0.00000 0.00000 0.00000 ; CB- CA- C- O

6148 6146 6159 6161 3 2.51040 4.18400 1.67360
-6.69440 0.00000 0.00000 ; CB- CA- C- N
6148 6151 6154 6155 3 0.00000 0.00000 0.00000
0.00000 0.00000 0.00000 ; CB- CG- CD- OE1
6148 6151 6154 6156 3 10.32611 1.48114 -8.47678
1.42256 1.40582 0.00000 ; CB- CG- CD- NE2
6149 6148 6146 6159 3 0.65084 1.95253 0.00000
-2.60338 0.00000 0.00000 ; HB2- CB- CA- C
6149 6148 6151 6152 3 0.62760 1.88280 0.00000
-2.51040 0.00000 0.00000 ; HB2- CB- CG- HG2
6149 6148 6151 6153 3 0.62760 1.88280 0.00000
-2.51040 0.00000 0.00000 ; HB2- CB- CG- HG3
6149 6148 6151 6154 3 0.65084 1.95253 0.00000
-2.60338 0.00000 0.00000 ; HB2- CB- CG- CD
6150 6148 6146 6159 3 0.65084 1.95253 0.00000
-2.60338 0.00000 0.00000 ; HB3- CB- CA- C
6150 6148 6151 6152 3 0.62760 1.88280 0.00000
-2.51040 0.00000 0.00000 ; HB3- CB- CG- HG2
6150 6148 6151 6153 3 0.62760 1.88280 0.00000
-2.51040 0.00000 0.00000 ; HB3- CB- CG- HG3
6150 6148 6151 6154 3 0.65084 1.95253 0.00000
-2.60338 0.00000 0.00000 ; HB3- CB- CG- CD
6151 6148 6146 6159 3 6.86594 3.56895 -8.14206
-2.40998 4.85344 0.00000 ; CG- CB- CA- C
6151 6154 6156 6157 3 20.92000 0.00000 -20.92000
0.00000 0.00000 0.00000 ; CG- CD- NE2- HE21
6151 6154 6156 6158 3 20.92000 0.00000 -20.92000
0.00000 0.00000 0.00000 ; CG- CD- NE2- HE22
6152 6151 6154 6155 3 3.68192 -4.35136 0.00000
1.33888 0.00000 0.00000 ; HG2- CG- CD- OE1
6152 6151 6154 6156 3 0.00000 0.00000 0.00000
0.00000 0.00000 0.00000 ; HG2- CG- CD- NE2
6153 6151 6154 6155 3 3.68192 -4.35136 0.00000
1.33888 0.00000 0.00000 ; HG3- CG- CD- OE1
6153 6151 6154 6156 3 0.00000 0.00000 0.00000
0.00000 0.00000 0.00000 ; HG3- CG- CD- NE2
6155 6154 6156 6157 3 29.28800 -8.36800 -20.92000
0.00000 0.00000 0.00000 ; OE1- CD- NE2- HE21
6155 6154 6156 6158 3 29.28800 -8.36800 -20.92000
0.00000 0.00000 0.00000 ; OE1- CD- NE2- HE22
6159 6161 6163 6164 3 0.00000 0.00000 0.00000
0.00000 0.00000 0.00000 ; C- N- CA- HA
6159 6161 6163 6165 3 11.71520 1.67360 15.06240 -
13.38880 0.00000 0.00000 ; C- N- CA- CB
6159 6161 6163 6173 3 1.75728 5.27184 2.25936
-7.02912 0.00000 0.00000 ; C- N- CA- C
6160 6159 6161 6162 3 29.28800 -8.36800 -20.92000
0.00000 0.00000 0.00000 ; O- C- N- H
6160 6159 6161 6163 3 20.92000 0.00000 -20.92000
0.00000 0.00000 0.00000 ; O- C- N- CA
6161 6163 6165 6166 3 0.65084 1.95253 0.00000
-2.60338 0.00000 0.00000 ; N- CA- CB- HB

6161 6163 6165 6167 3 3.83673 0.44769 -1.77402
-2.47693 -0.03347 0.00000 ; N- CA- CB- CG2
6161 6163 6165 6171 3 4.93294 1.13386 -3.12963
-5.27184 3.17984 0.00000 ; N- CA- CB- OG1
6161 6163 6173 6174 3 0.00000 0.00000 0.00000
0.00000 0.00000 0.00000 ; N- CA- C- O
6161 6163 6173 6175 3 17.40544 -5.02080 -13.22144
9.20480 0.00000 0.00000 ; N- CA- C- N
6162 6161 6163 6164 3 0.00000 0.00000 0.00000
0.00000 0.00000 0.00000 ; H- N- CA- HA
6162 6161 6163 6165 3 0.00000 0.00000 0.00000
0.00000 0.00000 0.00000 ; H- N- CA- CB
6162 6161 6163 6173 3 0.00000 0.00000 0.00000
0.00000 0.00000 0.00000 ; H- N- CA- C
6163 6165 6167 6168 3 0.66944 2.00832 0.00000
-2.67776 0.00000 0.00000 ; CA- CB- CG2- HG21
6163 6165 6167 6169 3 0.66944 2.00832 0.00000
-2.67776 0.00000 0.00000 ; CA- CB- CG2- HG22
6163 6165 6167 6170 3 0.66944 2.00832 0.00000
-2.67776 0.00000 0.00000 ; CA- CB- CG2- HG23
6163 6165 6171 6172 3 1.12131 2.98738 1.66523
-3.94970 0.43514 0.00000 ; CA- CB- OG1- HG1
6163 6173 6175 6176 3 20.92000 0.00000 -20.92000
0.00000 0.00000 0.00000 ; CA- C- N- H
6163 6173 6175 6177 3 20.92000 0.00000 -20.92000
0.00000 0.00000 0.00000 ; CA- C- N- CA
6164 6163 6165 6166 3 0.65084 1.95253 0.00000
-2.60338 0.00000 0.00000 ; HA- CA- CB- HB
6164 6163 6165 6167 3 0.65084 1.95253 0.00000
-2.60338 0.00000 0.00000 ; HA- CA- CB- CG2
6164 6163 6165 6171 3 1.04600 -1.04600 0.00000
0.00000 0.00000 0.00000 ; HA- CA- CB- OG1
6164 6163 6173 6174 3 3.68192 -4.35136 0.00000
1.33888 0.00000 0.00000 ; HA- CA- C- O
6164 6163 6173 6175 3 0.00000 0.00000 0.00000
0.00000 0.00000 0.00000 ; HA- CA- C- N
6165 6163 6173 6174 3 0.00000 0.00000 0.00000
0.00000 0.00000 0.00000 ; CB- CA- C- O
6165 6163 6173 6175 3 2.51040 4.18400 1.67360
-6.69440 0.00000 0.00000 ; CB- CA- C- N
6166 6165 6163 6173 3 0.65084 1.95253 0.00000
-2.60338 0.00000 0.00000 ; HB- CB- CA- C
6166 6165 6167 6168 3 0.65084 1.95253 0.00000
-2.60338 0.00000 0.00000 ; HB- CB- CG2- HG21
6166 6165 6167 6169 3 0.65084 1.95253 0.00000
-2.60338 0.00000 0.00000 ; HB- CB- CG2- HG22
6166 6165 6167 6170 3 0.65084 1.95253 0.00000
-2.60338 0.00000 0.00000 ; HB- CB- CG2- HG23
6166 6165 6171 6172 3 0.69733 2.09200 0.00000
-2.78933 0.00000 0.00000 ; HB- CB- OG1- HG1
6167 6165 6163 6173 3 5.67350 3.55640 -6.16722
-2.47693 3.74886 0.00000 ; CG2- CB- CA- C

6167 6165 6171 6172 3 4.74047 0.27196 -2.26773
-3.94970 1.60666 0.00000 ; CG2- CB- OG1- HG1
6168 6167 6165 6171 3 1.04600 -1.04600 0.00000
0.00000 0.00000 0.00000 ; HG21- CG2- CB- OG1
6169 6167 6165 6171 3 1.04600 -1.04600 0.00000
0.00000 0.00000 0.00000 ; HG22- CG2- CB- OG1
6170 6167 6165 6171 3 1.04600 -1.04600 0.00000
0.00000 0.00000 0.00000 ; HG23- CG2- CB- OG1
6171 6165 6163 6173 3 6.53541 6.87013 -6.21742
-5.27184 5.22163 0.00000 ; OG1- CB- CA- C
6173 6175 6177 6178 3 0.00000 0.00000 0.00000
0.00000 0.00000 0.00000 ; C- N- CA- HA
6173 6175 6177 6179 3 11.71520 1.67360 15.06240 -
13.38880 0.00000 0.00000 ; C- N- CA- CB
6173 6175 6177 6194 3 1.75728 5.27184 2.25936
-7.02912 0.00000 0.00000 ; C- N- CA- C
6174 6173 6175 6176 3 29.28800 -8.36800 -20.92000
0.00000 0.00000 0.00000 ; O- C- N- H
6174 6173 6175 6177 3 20.92000 0.00000 -20.92000
0.00000 0.00000 0.00000 ; O- C- N- CA
6175 6177 6179 6180 3 0.65084 1.95253 0.00000
-2.60338 0.00000 0.00000 ; N- CA- CB- HB2
6175 6177 6179 6181 3 0.65084 1.95253 0.00000
-2.60338 0.00000 0.00000 ; N- CA- CB- HB3
6175 6177 6179 6182 3 3.28026 2.46019 -2.19242
-3.21331 -0.23430 0.00000 ; N- CA- CB- CG
6175 6177 6194 6195 3 0.00000 0.00000 0.00000
0.00000 0.00000 0.00000 ; N- CA- C- O
6175 6177 6194 6196 3 17.40544 -5.02080 -13.22144
9.20480 0.00000 0.00000 ; N- CA- C- N
6176 6175 6177 6178 3 0.00000 0.00000 0.00000
0.00000 0.00000 0.00000 ; H- N- CA- HA
6176 6175 6177 6179 3 0.00000 0.00000 0.00000
0.00000 0.00000 0.00000 ; H- N- CA- CB
6176 6175 6177 6194 3 0.00000 0.00000 0.00000
0.00000 0.00000 0.00000 ; H- N- CA- C
6177 6179 6182 6183 3 0.57739 0.00000 1.02926
0.00000 -1.60666 0.00000 ; CA- CB- CG- CD1
6177 6179 6182 6192 3 0.57739 0.00000 1.02926
0.00000 -1.60666 0.00000 ; CA- CB- CG- CD2
6177 6194 6196 6197 3 20.92000 0.00000 -20.92000
0.00000 0.00000 0.00000 ; CA- C- N- H
6177 6194 6196 6198 3 20.92000 0.00000 -20.92000
0.00000 0.00000 0.00000 ; CA- C- N- CA
6178 6177 6179 6180 3 0.65084 1.95253 0.00000
-2.60338 0.00000 0.00000 ; HA- CA- CB- HB2
6178 6177 6179 6181 3 0.65084 1.95253 0.00000
-2.60338 0.00000 0.00000 ; HA- CA- CB- HB3
6178 6177 6179 6182 3 0.65084 1.95253 0.00000
-2.60338 0.00000 0.00000 ; HA- CA- CB- CG
6178 6177 6194 6195 3 3.68192 -4.35136 0.00000
1.33888 0.00000 0.00000 ; HA- CA- C- O

6178 6177 6194 6196 3 0.00000 0.00000 0.00000
0.00000 0.00000 0.00000 ; HA- CA- C- N
6179 6177 6194 6195 3 0.00000 0.00000 0.00000
0.00000 0.00000 0.00000 ; CB- CA- C- O
6179 6177 6194 6196 3 2.51040 4.18400 1.67360
-6.69440 0.00000 0.00000 ; CB- CA- C- N
6179 6182 6183 6184 3 30.33400 0.00000 -30.33400
0.00000 0.00000 0.00000 ; CB- CG- CD1- HD1
6179 6182 6183 6185 3 30.33400 0.00000 -30.33400
0.00000 0.00000 0.00000 ; CB- CG- CD1- CE1
6179 6182 6192 6190 3 30.33400 0.00000 -30.33400
0.00000 0.00000 0.00000 ; CB- CG- CD2- CE2
6179 6182 6192 6193 3 30.33400 0.00000 -30.33400
0.00000 0.00000 0.00000 ; CB- CG- CD2- HD2
6180 6179 6177 6194 3 0.65084 1.95253 0.00000
-2.60338 0.00000 0.00000 ; HB2- CB- CA- C
6180 6179 6182 6183 3 0.00000 0.00000 0.00000
0.00000 0.00000 0.00000 ; HB2- CB- CG- CD1
6180 6179 6182 6192 3 0.00000 0.00000 0.00000
0.00000 0.00000 0.00000 ; HB2- CB- CG- CD2
6181 6179 6177 6194 3 0.65084 1.95253 0.00000
-2.60338 0.00000 0.00000 ; HB3- CB- CA- C
6181 6179 6182 6183 3 0.00000 0.00000 0.00000
0.00000 0.00000 0.00000 ; HB3- CB- CG- CD1
6181 6179 6182 6192 3 0.00000 0.00000 0.00000
0.00000 0.00000 0.00000 ; HB3- CB- CG- CD2
6182 6179 6177 6194 3 4.95804 2.17986 -3.52293
-3.21331 -0.40166 0.00000 ; CG- CB- CA- C
6182 6183 6185 6186 3 30.33400 0.00000 -30.33400
0.00000 0.00000 0.00000 ; CG- CD1- CE1- HE1
6182 6183 6185 6187 3 30.33400 0.00000 -30.33400
0.00000 0.00000 0.00000 ; CG- CD1- CE1- CZ
6182 6192 6190 6187 3 30.33400 0.00000 -30.33400
0.00000 0.00000 0.00000 ; CG- CD2- CE2- CZ
6182 6192 6190 6191 3 30.33400 0.00000 -30.33400
0.00000 0.00000 0.00000 ; CG- CD2- CE2- HE2
6183 6182 6192 6190 3 30.33400 0.00000 -30.33400
0.00000 0.00000 0.00000 ; CD1- CG- CD2- CE2
6183 6182 6192 6193 3 30.33400 0.00000 -30.33400
0.00000 0.00000 0.00000 ; CD1- CG- CD2- HD2
6183 6185 6187 6188 3 30.33400 0.00000 -30.33400
0.00000 0.00000 0.00000 ; CD1- CE1- CZ- OH
6183 6185 6187 6190 3 30.33400 0.00000 -30.33400
0.00000 0.00000 0.00000 ; CD1- CE1- CZ- CE2
6184 6183 6182 6192 3 30.33400 0.00000 -30.33400
0.00000 0.00000 0.00000 ; HD1- CD1- CG- CD2
6184 6183 6185 6186 3 30.33400 0.00000 -30.33400
0.00000 0.00000 0.00000 ; HD1- CD1- CE1- HE1
6184 6183 6185 6187 3 30.33400 0.00000 -30.33400
0.00000 0.00000 0.00000 ; HD1- CD1- CE1- CZ
6185 6183 6182 6192 3 30.33400 0.00000 -30.33400
0.00000 0.00000 0.00000 ; CE1- CD1- CG- CD2

6185 6187 6188 6189 3 7.93286 0.00000 -9.56462
0.00000 2.17568 0.00000 ; CE1- CZ- OH- HH
6185 6187 6190 6191 3 30.33400 0.00000 -30.33400
0.00000 0.00000 0.00000 ; CE1- CZ- CE2- HE2
6185 6187 6190 6192 3 30.33400 0.00000 -30.33400
0.00000 0.00000 0.00000 ; CE1- CZ- CE2- CD2
6186 6185 6187 6188 3 30.33400 0.00000 -30.33400
0.00000 0.00000 0.00000 ; HE1- CE1- CZ- OH
6186 6185 6187 6190 3 30.33400 0.00000 -30.33400
0.00000 0.00000 0.00000 ; HE1- CE1- CZ- CE2
6187 6190 6192 6193 3 30.33400 0.00000 -30.33400
0.00000 0.00000 0.00000 ; CZ- CE2- CD2- HD2
6188 6187 6190 6191 3 30.33400 0.00000 -30.33400
0.00000 0.00000 0.00000 ; OH- CZ- CE2- HE2
6188 6187 6190 6192 3 30.33400 0.00000 -30.33400
0.00000 0.00000 0.00000 ; OH- CZ- CE2- CD2
6189 6188 6187 6190 3 7.93286 0.00000 -9.56462
0.00000 2.17568 0.00000 ; HH- OH- CZ- CE2
6191 6190 6192 6193 3 30.33400 0.00000 -30.33400
0.00000 0.00000 0.00000 ; HE2- CE2- CD2- HD2
6194 6196 6198 6199 3 0.00000 0.00000 0.00000
0.00000 0.00000 0.00000 ; C- N- CA- HA
6194 6196 6198 6200 3 11.71520 1.67360 15.06240 -
13.38880 0.00000 0.00000 ; C- N- CA- CB
6194 6196 6198 6213 3 1.75728 5.27184 2.25936
-7.02912 0.00000 0.00000 ; C- N- CA- C
6195 6194 6196 6197 3 29.28800 -8.36800 -20.92000
0.00000 0.00000 0.00000 ; O- C- N- H
6195 6194 6196 6198 3 20.92000 0.00000 -20.92000
0.00000 0.00000 0.00000 ; O- C- N- CA
6196 6198 6200 6201 3 0.65084 1.95253 0.00000
-2.60338 0.00000 0.00000 ; N- CA- CB- HB2
6196 6198 6200 6202 3 0.65084 1.95253 0.00000
-2.60338 0.00000 0.00000 ; N- CA- CB- HB3
6196 6198 6200 6203 3 3.79070 1.39746 -4.61077
-2.40998 2.44346 0.00000 ; N- CA- CB- CG
6196 6198 6213 6214 3 0.00000 0.00000 0.00000
0.00000 0.00000 0.00000 ; N- CA- C- O
6196 6198 6213 6215 3 17.40544 -5.02080 -13.22144
9.20480 0.00000 0.00000 ; N- CA- C- N
6197 6196 6198 6199 3 0.00000 0.00000 0.00000
0.00000 0.00000 0.00000 ; H- N- CA- HA
6197 6196 6198 6200 3 0.00000 0.00000 0.00000
0.00000 0.00000 0.00000 ; H- N- CA- CB
6197 6196 6198 6213 3 0.00000 0.00000 0.00000
0.00000 0.00000 0.00000 ; H- N- CA- C
6198 6200 6203 6204 3 0.66944 2.00832 0.00000
-2.67776 0.00000 0.00000 ; CA- CB- CG- HG
6198 6200 6203 6205 3 3.90367 0.19665 -6.21742
-2.37651 5.99149 0.00000 ; CA- CB- CG- CD1
6198 6200 6203 6209 3 3.90367 0.19665 -6.21742
-2.37651 5.99149 0.00000 ; CA- CB- CG- CD2

6198 6213 6215 6216 3 20.92000 0.00000 -20.92000
0.00000 0.00000 0.00000 ; CA- C- N- H
6198 6213 6215 6217 3 20.92000 0.00000 -20.92000
0.00000 0.00000 0.00000 ; CA- C- N- CA
6199 6198 6200 6201 3 0.65084 1.95253 0.00000
-2.60338 0.00000 0.00000 ; HA- CA- CB- HB2
6199 6198 6200 6202 3 0.65084 1.95253 0.00000
-2.60338 0.00000 0.00000 ; HA- CA- CB- HB3
6199 6198 6200 6203 3 0.65084 1.95253 0.00000
-2.60338 0.00000 0.00000 ; HA- CA- CB- CG
6199 6198 6213 6214 3 3.68192 -4.35136 0.00000
1.33888 0.00000 0.00000 ; HA- CA- C- O
6199 6198 6213 6215 3 0.00000 0.00000 0.00000
0.00000 0.00000 0.00000 ; HA- CA- C- N
6200 6198 6213 6214 3 0.00000 0.00000 0.00000
0.00000 0.00000 0.00000 ; CB- CA- C- O
6200 6198 6213 6215 3 2.51040 4.18400 1.67360
-6.69440 0.00000 0.00000 ; CB- CA- C- N
6200 6203 6205 6206 3 0.66944 2.00832 0.00000
-2.67776 0.00000 0.00000 ; CB- CG- CD1- HD11
6200 6203 6205 6207 3 0.66944 2.00832 0.00000
-2.67776 0.00000 0.00000 ; CB- CG- CD1- HD12
6200 6203 6205 6208 3 0.66944 2.00832 0.00000
-2.67776 0.00000 0.00000 ; CB- CG- CD1- HD13
6200 6203 6209 6210 3 0.66944 2.00832 0.00000
-2.67776 0.00000 0.00000 ; CB- CG- CD2- HD21
6200 6203 6209 6211 3 0.66944 2.00832 0.00000
-2.67776 0.00000 0.00000 ; CB- CG- CD2- HD22
6200 6203 6209 6212 3 0.66944 2.00832 0.00000
-2.67776 0.00000 0.00000 ; CB- CG- CD2- HD23
6201 6200 6198 6213 3 0.65084 1.95253 0.00000
-2.60338 0.00000 0.00000 ; HB2- CB- CA- C
6201 6200 6203 6204 3 0.62760 1.88280 0.00000
-2.51040 0.00000 0.00000 ; HB2- CB- CG- HG
6201 6200 6203 6205 3 0.66944 2.00832 0.00000
-2.67776 0.00000 0.00000 ; HB2- CB- CG- CD1
6201 6200 6203 6209 3 0.66944 2.00832 0.00000
-2.67776 0.00000 0.00000 ; HB2- CB- CG- CD2
6202 6200 6198 6213 3 0.65084 1.95253 0.00000
-2.60338 0.00000 0.00000 ; HB3- CB- CA- C
6202 6200 6203 6204 3 0.62760 1.88280 0.00000
-2.51040 0.00000 0.00000 ; HB3- CB- CG- HG
6202 6200 6203 6205 3 0.66944 2.00832 0.00000
-2.67776 0.00000 0.00000 ; HB3- CB- CG- CD1
6202 6200 6203 6209 3 0.66944 2.00832 0.00000
-2.67776 0.00000 0.00000 ; HB3- CB- CG- CD2
6203 6200 6198 6213 3 10.33448 -1.14642 -11.54784
-2.40998 6.35968 0.00000 ; CG- CB- CA- C
6204 6203 6205 6206 3 0.62760 1.88280 0.00000
-2.51040 0.00000 0.00000 ; HG- CG- CD1- HD11
6204 6203 6205 6207 3 0.62760 1.88280 0.00000
-2.51040 0.00000 0.00000 ; HG- CG- CD1- HD12

6204 6203 6205 6208 3 0.62760 1.88280 0.00000
-2.51040 0.00000 0.00000 ; HG- CG- CD1- HD13
6204 6203 6209 6210 3 0.62760 1.88280 0.00000
-2.51040 0.00000 0.00000 ; HG- CG- CD2- HD21
6204 6203 6209 6211 3 0.62760 1.88280 0.00000
-2.51040 0.00000 0.00000 ; HG- CG- CD2- HD22
6204 6203 6209 6212 3 0.62760 1.88280 0.00000
-2.51040 0.00000 0.00000 ; HG- CG- CD2- HD23
6205 6203 6209 6210 3 0.66944 2.00832 0.00000
-2.67776 0.00000 0.00000 ; CD1- CG- CD2- HD21
6205 6203 6209 6211 3 0.66944 2.00832 0.00000
-2.67776 0.00000 0.00000 ; CD1- CG- CD2- HD22
6205 6203 6209 6212 3 0.66944 2.00832 0.00000
-2.67776 0.00000 0.00000 ; CD1- CG- CD2- HD23
6206 6205 6203 6209 3 0.66944 2.00832 0.00000
-2.67776 0.00000 0.00000 ; HD11- CD1- CG- CD2
6207 6205 6203 6209 3 0.66944 2.00832 0.00000
-2.67776 0.00000 0.00000 ; HD12- CD1- CG- CD2
6208 6205 6203 6209 3 0.66944 2.00832 0.00000
-2.67776 0.00000 0.00000 ; HD13- CD1- CG- CD2
6213 6215 6217 6218 3 0.00000 0.00000 0.00000
0.00000 0.00000 0.00000 ; C- N- CA- HA
6213 6215 6217 6219 3 11.71520 1.67360 15.06240 -
13.38880 0.00000 0.00000 ; C- N- CA- CB
6213 6215 6217 6235 3 1.75728 5.27184 2.25936
-7.02912 0.00000 0.00000 ; C- N- CA- C
6214 6213 6215 6216 3 29.28800 -8.36800 -20.92000
0.00000 0.00000 0.00000 ; O- C- N- H
6214 6213 6215 6217 3 20.92000 0.00000 -20.92000
0.00000 0.00000 0.00000 ; O- C- N- CA
6215 6217 6219 6220 3 0.65084 1.95253 0.00000
-2.60338 0.00000 0.00000 ; N- CA- CB- HB2
6215 6217 6219 6221 3 0.65084 1.95253 0.00000
-2.60338 0.00000 0.00000 ; N- CA- CB- HB3
6215 6217 6219 6222 3 0.65084 1.95253 0.00000
-2.60338 0.00000 0.00000 ; N- CA- CB- CG
6215 6217 6235 6236 3 0.00000 0.00000 0.00000
0.00000 0.00000 0.00000 ; N- CA- C- O
6215 6217 6235 6237 3 17.40544 -5.02080 -13.22144
9.20480 0.00000 0.00000 ; N- CA- C- N
6216 6215 6217 6218 3 0.00000 0.00000 0.00000
0.00000 0.00000 0.00000 ; H- N- CA- HA
6216 6215 6217 6219 3 0.00000 0.00000 0.00000
0.00000 0.00000 0.00000 ; H- N- CA- CB
6216 6215 6217 6235 3 0.00000 0.00000 0.00000
0.00000 0.00000 0.00000 ; H- N- CA- C
6217 6219 6222 6223 3 0.66944 2.00832 0.00000
-2.67776 0.00000 0.00000 ; CA- CB- CG- HG2
6217 6219 6222 6224 3 0.66944 2.00832 0.00000
-2.67776 0.00000 0.00000 ; CA- CB- CG- HG3
6217 6219 6222 6225 3 3.68192 3.09616 -2.09200
-3.01248 0.00000 0.00000 ; CA- CB- CG- CD

```
  6217   6235   6237   6238      3   20.92000    0.00000  -20.92000
0.00000    0.00000    0.00000 ;     CA-     C-     N-     H
  6217   6235   6237   6239      3   20.92000    0.00000  -20.92000
0.00000    0.00000    0.00000 ;     CA-     C-     N-    CA
  6218   6217   6219   6220      3    0.65084    1.95253    0.00000
-2.60338    0.00000    0.00000 ;     HA-    CA-    CB-   HB2
  6218   6217   6219   6221      3    0.65084    1.95253    0.00000
-2.60338    0.00000    0.00000 ;     HA-    CA-    CB-   HB3
  6218   6217   6219   6222      3    0.65084    1.95253    0.00000
-2.60338    0.00000    0.00000 ;     HA-    CA-    CB-    CG
  6218   6217   6235   6236      3    3.68192   -4.35136    0.00000
1.33888    0.00000    0.00000 ;     HA-    CA-     C-     O
  6218   6217   6235   6237      3    0.00000    0.00000    0.00000
0.00000    0.00000    0.00000 ;     HA-    CA-     C-     N
  6219   6217   6235   6236      3    0.00000    0.00000    0.00000
0.00000    0.00000    0.00000 ;     CB-    CA-     C-     O
  6219   6217   6235   6237      3    2.51040    4.18400    1.67360
-6.69440    0.00000    0.00000 ;     CB-    CA-     C-     N
  6219   6222   6225   6226      3    0.66944    2.00832    0.00000
-2.67776    0.00000    0.00000 ;     CB-    CG-    CD-   HD2
  6219   6222   6225   6227      3    0.66944    2.00832    0.00000
-2.67776    0.00000    0.00000 ;     CB-    CG-    CD-   HD3
  6219   6222   6225   6228      3    3.68192    3.09616   -2.09200
-3.01248    0.00000    0.00000 ;     CB-    CG-    CD-    CE
  6220   6219   6217   6235      3    0.65084    1.95253    0.00000
-2.60338    0.00000    0.00000 ;    HB2-    CB-    CA-     C
  6220   6219   6222   6223      3    0.62760    1.88280    0.00000
-2.51040    0.00000    0.00000 ;    HB2-    CB-    CG-   HG2
  6220   6219   6222   6224      3    0.62760    1.88280    0.00000
-2.51040    0.00000    0.00000 ;    HB2-    CB-    CG-   HG3
  6220   6219   6222   6225      3    0.66944    2.00832    0.00000
-2.67776    0.00000    0.00000 ;    HB2-    CB-    CG-    CD
  6221   6219   6217   6235      3    0.65084    1.95253    0.00000
-2.60338    0.00000    0.00000 ;    HB3-    CB-    CA-     C
  6221   6219   6222   6223      3    0.62760    1.88280    0.00000
-2.51040    0.00000    0.00000 ;    HB3-    CB-    CG-   HG2
  6221   6219   6222   6224      3    0.62760    1.88280    0.00000
-2.51040    0.00000    0.00000 ;    HB3-    CB-    CG-   HG3
  6221   6219   6222   6225      3    0.66944    2.00832    0.00000
-2.67776    0.00000    0.00000 ;    HB3-    CB-    CG-    CD
  6222   6219   6217   6235      3    0.65084    1.95253    0.00000
-2.60338    0.00000    0.00000 ;     CG-    CB-    CA-     C
  6222   6225   6228   6229      3    0.65084    1.95253    0.00000
-2.60338    0.00000    0.00000 ;     CG-    CD-    CE-   HE2
  6222   6225   6228   6230      3    0.65084    1.95253    0.00000
-2.60338    0.00000    0.00000 ;     CG-    CD-    CE-   HE3
  6222   6225   6228   6231      3    0.65084    1.95253    0.00000
-2.60338    0.00000    0.00000 ;     CG-    CD-    CE-    NZ
  6223   6222   6225   6226      3    0.62760    1.88280    0.00000
-2.51040    0.00000    0.00000 ;    HG2-    CG-    CD-   HD2
  6223   6222   6225   6227      3    0.62760    1.88280    0.00000
-2.51040    0.00000    0.00000 ;    HG2-    CG-    CD-   HD3
```

6223 6222 6225 6228 3 0.66944 2.00832 0.00000
-2.67776 0.00000 0.00000 ; HG2- CG- CD- CE
6224 6222 6225 6226 3 0.62760 1.88280 0.00000
-2.51040 0.00000 0.00000 ; HG3- CG- CD- HD2
6224 6222 6225 6227 3 0.62760 1.88280 0.00000
-2.51040 0.00000 0.00000 ; HG3- CG- CD- HD3
6224 6222 6225 6228 3 0.66944 2.00832 0.00000
-2.67776 0.00000 0.00000 ; HG3- CG- CD- CE
6225 6228 6231 6232 3 0.65084 1.95253 0.00000
-2.60338 0.00000 0.00000 ; CD- CE- NZ- HZ1
6225 6228 6231 6233 3 0.65084 1.95253 0.00000
-2.60338 0.00000 0.00000 ; CD- CE- NZ- HZ2
6225 6228 6231 6234 3 0.65084 1.95253 0.00000
-2.60338 0.00000 0.00000 ; CD- CE- NZ- HZ3
6226 6225 6228 6229 3 0.65084 1.95253 0.00000
-2.60338 0.00000 0.00000 ; HD2- CD- CE- HE2
6226 6225 6228 6230 3 0.65084 1.95253 0.00000
-2.60338 0.00000 0.00000 ; HD2- CD- CE- HE3
6226 6225 6228 6231 3 0.65084 1.95253 0.00000
-2.60338 0.00000 0.00000 ; HD2- CD- CE- NZ
6227 6225 6228 6229 3 0.65084 1.95253 0.00000
-2.60338 0.00000 0.00000 ; HD3- CD- CE- HE2
6227 6225 6228 6230 3 0.65084 1.95253 0.00000
-2.60338 0.00000 0.00000 ; HD3- CD- CE- HE3
6227 6225 6228 6231 3 0.65084 1.95253 0.00000
-2.60338 0.00000 0.00000 ; HD3- CD- CE- NZ
6229 6228 6231 6232 3 0.65084 1.95253 0.00000
-2.60338 0.00000 0.00000 ; HE2- CE- NZ- HZ1
6229 6228 6231 6233 3 0.65084 1.95253 0.00000
-2.60338 0.00000 0.00000 ; HE2- CE- NZ- HZ2
6229 6228 6231 6234 3 0.65084 1.95253 0.00000
-2.60338 0.00000 0.00000 ; HE2- CE- NZ- HZ3
6230 6228 6231 6232 3 0.65084 1.95253 0.00000
-2.60338 0.00000 0.00000 ; HE3- CE- NZ- HZ1
6230 6228 6231 6233 3 0.65084 1.95253 0.00000
-2.60338 0.00000 0.00000 ; HE3- CE- NZ- HZ2
6230 6228 6231 6234 3 0.65084 1.95253 0.00000
-2.60338 0.00000 0.00000 ; HE3- CE- NZ- HZ3
6235 6237 6239 6240 3 0.00000 0.00000 0.00000
0.00000 0.00000 0.00000 ; C- N- CA- HA
6235 6237 6239 6241 3 11.71520 1.67360 15.06240 -
13.38880 0.00000 0.00000 ; C- N- CA- CB
6235 6237 6239 6257 3 1.75728 5.27184 2.25936
-7.02912 0.00000 0.00000 ; C- N- CA- C
6236 6235 6237 6238 3 29.28800 -8.36800 -20.92000
0.00000 0.00000 0.00000 ; O- C- N- H
6236 6235 6237 6239 3 20.92000 0.00000 -20.92000
0.00000 0.00000 0.00000 ; O- C- N- CA
6237 6239 6241 6242 3 0.65084 1.95253 0.00000
-2.60338 0.00000 0.00000 ; N- CA- CB- HB2
6237 6239 6241 6243 3 0.65084 1.95253 0.00000
-2.60338 0.00000 0.00000 ; N- CA- CB- HB3

6237 6239 6241 6244 3 0.65084 1.95253 0.00000
-2.60338 0.00000 0.00000 ; N- CA- CB- CG
6237 6239 6257 6258 3 0.00000 0.00000 0.00000
0.00000 0.00000 0.00000 ; N- CA- C- O
6237 6239 6257 6259 3 17.40544 -5.02080 -13.22144
9.20480 0.00000 0.00000 ; N- CA- C- N
6238 6237 6239 6240 3 0.00000 0.00000 0.00000
0.00000 0.00000 0.00000 ; H- N- CA- HA
6238 6237 6239 6241 3 0.00000 0.00000 0.00000
0.00000 0.00000 0.00000 ; H- N- CA- CB
6238 6237 6239 6257 3 0.00000 0.00000 0.00000
0.00000 0.00000 0.00000 ; H- N- CA- C
6239 6241 6244 6245 3 0.66944 2.00832 0.00000
-2.67776 0.00000 0.00000 ; CA- CB- CG- HG2
6239 6241 6244 6246 3 0.66944 2.00832 0.00000
-2.67776 0.00000 0.00000 ; CA- CB- CG- HG3
6239 6241 6244 6247 3 3.68192 3.09616 -2.09200
-3.01248 0.00000 0.00000 ; CA- CB- CG- CD
6239 6257 6259 6260 3 20.92000 0.00000 -20.92000
0.00000 0.00000 0.00000 ; CA- C- N- H
6239 6257 6259 6261 3 20.92000 0.00000 -20.92000
0.00000 0.00000 0.00000 ; CA- C- N- CA
6240 6239 6241 6242 3 0.65084 1.95253 0.00000
-2.60338 0.00000 0.00000 ; HA- CA- CB- HB2
6240 6239 6241 6243 3 0.65084 1.95253 0.00000
-2.60338 0.00000 0.00000 ; HA- CA- CB- HB3
6240 6239 6241 6244 3 0.65084 1.95253 0.00000
-2.60338 0.00000 0.00000 ; HA- CA- CB- CG
6240 6239 6257 6258 3 3.68192 -4.35136 0.00000
1.33888 0.00000 0.00000 ; HA- CA- C- O
6240 6239 6257 6259 3 0.00000 0.00000 0.00000
0.00000 0.00000 0.00000 ; HA- CA- C- N
6241 6239 6257 6258 3 0.00000 0.00000 0.00000
0.00000 0.00000 0.00000 ; CB- CA- C- O
6241 6239 6257 6259 3 2.51040 4.18400 1.67360
-6.69440 0.00000 0.00000 ; CB- CA- C- N
6241 6244 6247 6248 3 0.66944 2.00832 0.00000
-2.67776 0.00000 0.00000 ; CB- CG- CD- HD2
6241 6244 6247 6249 3 0.66944 2.00832 0.00000
-2.67776 0.00000 0.00000 ; CB- CG- CD- HD3
6241 6244 6247 6250 3 3.68192 3.09616 -2.09200
-3.01248 0.00000 0.00000 ; CB- CG- CD- CE
6242 6241 6239 6257 3 0.65084 1.95253 0.00000
-2.60338 0.00000 0.00000 ; HB2- CB- CA- C
6242 6241 6244 6245 3 0.62760 1.88280 0.00000
-2.51040 0.00000 0.00000 ; HB2- CB- CG- HG2
6242 6241 6244 6246 3 0.62760 1.88280 0.00000
-2.51040 0.00000 0.00000 ; HB2- CB- CG- HG3
6242 6241 6244 6247 3 0.66944 2.00832 0.00000
-2.67776 0.00000 0.00000 ; HB2- CB- CG- CD
6243 6241 6239 6257 3 0.65084 1.95253 0.00000
-2.60338 0.00000 0.00000 ; HB3- CB- CA- C

```
  6243   6241   6244   6245       3    0.62760    1.88280    0.00000
-2.51040    0.00000    0.00000 ;    HB3-    CB-    CG-   HG2
  6243   6241   6244   6246       3    0.62760    1.88280    0.00000
-2.51040    0.00000    0.00000 ;    HB3-    CB-    CG-   HG3
  6243   6241   6244   6247       3    0.66944    2.00832    0.00000
-2.67776    0.00000    0.00000 ;    HB3-    CB-    CG-    CD
  6244   6241   6239   6257       3    0.65084    1.95253    0.00000
-2.60338    0.00000    0.00000 ;     CG-    CB-    CA-     C
  6244   6247   6250   6251       3    0.65084    1.95253    0.00000
-2.60338    0.00000    0.00000 ;     CG-    CD-    CE-   HE2
  6244   6247   6250   6252       3    0.65084    1.95253    0.00000
-2.60338    0.00000    0.00000 ;     CG-    CD-    CE-   HE3
  6244   6247   6250   6253       3    0.65084    1.95253    0.00000
-2.60338    0.00000    0.00000 ;     CG-    CD-    CE-    NZ
  6245   6244   6247   6248       3    0.62760    1.88280    0.00000
-2.51040    0.00000    0.00000 ;    HG2-    CG-    CD-   HD2
  6245   6244   6247   6249       3    0.62760    1.88280    0.00000
-2.51040    0.00000    0.00000 ;    HG2-    CG-    CD-   HD3
  6245   6244   6247   6250       3    0.66944    2.00832    0.00000
-2.67776    0.00000    0.00000 ;    HG2-    CG-    CD-    CE
  6246   6244   6247   6248       3    0.62760    1.88280    0.00000
-2.51040    0.00000    0.00000 ;    HG3-    CG-    CD-   HD2
  6246   6244   6247   6249       3    0.62760    1.88280    0.00000
-2.51040    0.00000    0.00000 ;    HG3-    CG-    CD-   HD3
  6246   6244   6247   6250       3    0.66944    2.00832    0.00000
-2.67776    0.00000    0.00000 ;    HG3-    CG-    CD-    CE
  6247   6250   6253   6254       3    0.65084    1.95253    0.00000
-2.60338    0.00000    0.00000 ;     CD-    CE-    NZ-   HZ1
  6247   6250   6253   6255       3    0.65084    1.95253    0.00000
-2.60338    0.00000    0.00000 ;     CD-    CE-    NZ-   HZ2
  6247   6250   6253   6256       3    0.65084    1.95253    0.00000
-2.60338    0.00000    0.00000 ;     CD-    CE-    NZ-   HZ3
  6248   6247   6250   6251       3    0.65084    1.95253    0.00000
-2.60338    0.00000    0.00000 ;    HD2-    CD-    CE-   HE2
  6248   6247   6250   6252       3    0.65084    1.95253    0.00000
-2.60338    0.00000    0.00000 ;    HD2-    CD-    CE-   HE3
  6248   6247   6250   6253       3    0.65084    1.95253    0.00000
-2.60338    0.00000    0.00000 ;    HD2-    CD-    CE-    NZ
  6249   6247   6250   6251       3    0.65084    1.95253    0.00000
-2.60338    0.00000    0.00000 ;    HD3-    CD-    CE-   HE2
  6249   6247   6250   6252       3    0.65084    1.95253    0.00000
-2.60338    0.00000    0.00000 ;    HD3-    CD-    CE-   HE3
  6249   6247   6250   6253       3    0.65084    1.95253    0.00000
-2.60338    0.00000    0.00000 ;    HD3-    CD-    CE-    NZ
  6251   6250   6253   6254       3    0.65084    1.95253    0.00000
-2.60338    0.00000    0.00000 ;    HE2-    CE-    NZ-   HZ1
  6251   6250   6253   6255       3    0.65084    1.95253    0.00000
-2.60338    0.00000    0.00000 ;    HE2-    CE-    NZ-   HZ2
  6251   6250   6253   6256       3    0.65084    1.95253    0.00000
-2.60338    0.00000    0.00000 ;    HE2-    CE-    NZ-   HZ3
  6252   6250   6253   6254       3    0.65084    1.95253    0.00000
-2.60338    0.00000    0.00000 ;    HE3-    CE-    NZ-   HZ1
```

6252 6250 6253 6255 3 0.65084 1.95253 0.00000
-2.60338 0.00000 0.00000 ; HE3- CE- NZ- HZ2
6252 6250 6253 6256 3 0.65084 1.95253 0.00000
-2.60338 0.00000 0.00000 ; HE3- CE- NZ- HZ3
6257 6259 6261 6262 3 0.00000 0.00000 0.00000
0.00000 0.00000 0.00000 ; C- N- CA- HA
6257 6259 6261 6263 3 11.71520 1.67360 15.06240 -
13.38880 0.00000 0.00000 ; C- N- CA- CB
6257 6259 6261 6272 3 1.75728 5.27184 2.25936
-7.02912 0.00000 0.00000 ; C- N- CA- C
6258 6257 6259 6260 3 29.28800 -8.36800 -20.92000
0.00000 0.00000 0.00000 ; O- C- N- H
6258 6257 6259 6261 3 20.92000 0.00000 -20.92000
0.00000 0.00000 0.00000 ; O- C- N- CA
6259 6261 6263 6264 3 0.65084 1.95253 0.00000
-2.60338 0.00000 0.00000 ; N- CA- CB- HB2
6259 6261 6263 6265 3 0.65084 1.95253 0.00000
-2.60338 0.00000 0.00000 ; N- CA- CB- HB3
6259 6261 6263 6266 3 3.21331 2.22589 -4.15053
-2.40998 2.61082 0.00000 ; N- CA- CB- CG
6259 6261 6272 6273 3 0.00000 0.00000 0.00000
0.00000 0.00000 0.00000 ; N- CA- C- O
6259 6261 6272 6274 3 17.40544 -5.02080 -13.22144
9.20480 0.00000 0.00000 ; N- CA- C- N
6260 6259 6261 6262 3 0.00000 0.00000 0.00000
0.00000 0.00000 0.00000 ; H- N- CA- HA
6260 6259 6261 6263 3 0.00000 0.00000 0.00000
0.00000 0.00000 0.00000 ; H- N- CA- CB
6260 6259 6261 6272 3 0.00000 0.00000 0.00000
0.00000 0.00000 0.00000 ; H- N- CA- C
6261 6263 6266 6267 3 0.66944 2.00832 0.00000
-2.67776 0.00000 0.00000 ; CA- CB- CG- HG2
6261 6263 6266 6268 3 0.66944 2.00832 0.00000
-2.67776 0.00000 0.00000 ; CA- CB- CG- HG3
6261 6263 6266 6269 3 10.12110 -1.91209 0.01674
10.17549 -1.87443 0.00000 ; CA- CB- CG- CD
6261 6272 6274 6275 3 20.92000 0.00000 -20.92000
0.00000 0.00000 0.00000 ; CA- C- N- H
6261 6272 6274 6276 3 20.92000 0.00000 -20.92000
0.00000 0.00000 0.00000 ; CA- C- N- CA
6262 6261 6263 6264 3 0.65084 1.95253 0.00000
-2.60338 0.00000 0.00000 ; HA- CA- CB- HB2
6262 6261 6263 6265 3 0.65084 1.95253 0.00000
-2.60338 0.00000 0.00000 ; HA- CA- CB- HB3
6262 6261 6263 6266 3 0.65084 1.95253 0.00000
-2.60338 0.00000 0.00000 ; HA- CA- CB- CG
6262 6261 6272 6273 3 3.68192 -4.35136 0.00000
1.33888 0.00000 0.00000 ; HA- CA- C- O
6262 6261 6272 6274 3 0.00000 0.00000 0.00000
0.00000 0.00000 0.00000 ; HA- CA- C- N
6263 6261 6272 6273 3 0.00000 0.00000 0.00000
0.00000 0.00000 0.00000 ; CB- CA- C- O

6263 6261 6272 6274 3 2.51040 4.18400 1.67360
-6.69440 0.00000 0.00000 ; CB- CA- C- N
6263 6266 6269 6270 3 3.79907 0.00000 -5.40573
0.00000 2.14221 0.00000 ; CB- CG- CD- OE1
6263 6266 6269 6271 3 3.79907 0.00000 -5.40573
0.00000 2.14221 0.00000 ; CB- CG- CD- OE2
6264 6263 6261 6272 3 0.65084 1.95253 0.00000
-2.60338 0.00000 0.00000 ; HB2- CB- CA- C
6264 6263 6266 6267 3 0.62760 1.88280 0.00000
-2.51040 0.00000 0.00000 ; HB2- CB- CG- HG2
6264 6263 6266 6268 3 0.62760 1.88280 0.00000
-2.51040 0.00000 0.00000 ; HB2- CB- CG- HG3
6264 6263 6266 6269 3 0.65084 1.95253 0.00000
-2.60338 0.00000 0.00000 ; HB2- CB- CG- CD
6265 6263 6261 6272 3 0.65084 1.95253 0.00000
-2.60338 0.00000 0.00000 ; HB3- CB- CA- C
6265 6263 6266 6267 3 0.62760 1.88280 0.00000
-2.51040 0.00000 0.00000 ; HB3- CB- CG- HG2
6265 6263 6266 6268 3 0.62760 1.88280 0.00000
-2.51040 0.00000 0.00000 ; HB3- CB- CG- HG3
6265 6263 6266 6269 3 0.65084 1.95253 0.00000
-2.60338 0.00000 0.00000 ; HB3- CB- CG- CD
6266 6263 6261 6272 3 6.86594 3.56895 -8.14206
-2.40998 4.85344 0.00000 ; CG- CB- CA- C
6267 6266 6269 6270 3 0.00000 0.00000 0.00000
0.00000 0.00000 0.00000 ; HG2- CG- CD- OE1
6267 6266 6269 6271 3 0.00000 0.00000 0.00000
0.00000 0.00000 0.00000 ; HG2- CG- CD- OE2
6268 6266 6269 6270 3 0.00000 0.00000 0.00000
0.00000 0.00000 0.00000 ; HG3- CG- CD- OE1
6268 6266 6269 6271 3 0.00000 0.00000 0.00000
0.00000 0.00000 0.00000 ; HG3- CG- CD- OE2
6272 6274 6276 6277 3 0.00000 0.00000 0.00000
0.00000 0.00000 0.00000 ; C- N- CA- HA
6272 6274 6276 6278 3 11.71520 1.67360 15.06240 -
13.38880 0.00000 0.00000 ; C- N- CA- CB
6272 6274 6276 6288 3 1.75728 5.27184 2.25936
-7.02912 0.00000 0.00000 ; C- N- CA- C
6273 6272 6274 6275 3 29.28800 -8.36800 -20.92000
0.00000 0.00000 0.00000 ; O- C- N- H
6273 6272 6274 6276 3 20.92000 0.00000 -20.92000
0.00000 0.00000 0.00000 ; O- C- N- CA
6274 6276 6278 6279 3 0.65084 1.95253 0.00000
-2.60338 0.00000 0.00000 ; N- CA- CB- HB
6274 6276 6278 6280 3 3.83673 0.44769 -1.77402
-2.47693 -0.03347 0.00000 ; N- CA- CB- CG1
6274 6276 6278 6284 3 3.83673 0.44769 -1.77402
-2.47693 -0.03347 0.00000 ; N- CA- CB- CG2
6274 6276 6288 6289 3 0.00000 0.00000 0.00000
0.00000 0.00000 0.00000 ; N- CA- C- O
6274 6276 6288 6290 3 17.40544 -5.02080 -13.22144
9.20480 0.00000 0.00000 ; N- CA- C- N

6275 6274 6276 6277 3 0.00000 0.00000 0.00000
0.00000 0.00000 0.00000 ; H- N- CA- HA
6275 6274 6276 6278 3 0.00000 0.00000 0.00000
0.00000 0.00000 0.00000 ; H- N- CA- CB
6275 6274 6276 6288 3 0.00000 0.00000 0.00000
0.00000 0.00000 0.00000 ; H- N- CA- C
6276 6278 6280 6281 3 0.66944 2.00832 0.00000
-2.67776 0.00000 0.00000 ; CA- CB- CG1- HG11
6276 6278 6280 6282 3 0.66944 2.00832 0.00000
-2.67776 0.00000 0.00000 ; CA- CB- CG1- HG12
6276 6278 6280 6283 3 0.66944 2.00832 0.00000
-2.67776 0.00000 0.00000 ; CA- CB- CG1- HG13
6276 6278 6284 6285 3 0.66944 2.00832 0.00000
-2.67776 0.00000 0.00000 ; CA- CB- CG2- HG21
6276 6278 6284 6286 3 0.66944 2.00832 0.00000
-2.67776 0.00000 0.00000 ; CA- CB- CG2- HG22
6276 6278 6284 6287 3 0.66944 2.00832 0.00000
-2.67776 0.00000 0.00000 ; CA- CB- CG2- HG23
6276 6288 6290 6291 3 20.92000 0.00000 -20.92000
0.00000 0.00000 0.00000 ; CA- C- N- H
6276 6288 6290 6292 3 20.92000 0.00000 -20.92000
0.00000 0.00000 0.00000 ; CA- C- N- CA
6277 6276 6278 6279 3 0.65084 1.95253 0.00000
-2.60338 0.00000 0.00000 ; HA- CA- CB- HB
6277 6276 6278 6280 3 0.65084 1.95253 0.00000
-2.60338 0.00000 0.00000 ; HA- CA- CB- CG1
6277 6276 6278 6284 3 0.65084 1.95253 0.00000
-2.60338 0.00000 0.00000 ; HA- CA- CB- CG2
6277 6276 6288 6289 3 3.68192 -4.35136 0.00000
1.33888 0.00000 0.00000 ; HA- CA- C- O
6277 6276 6288 6290 3 0.00000 0.00000 0.00000
0.00000 0.00000 0.00000 ; HA- CA- C- N
6278 6276 6288 6289 3 0.00000 0.00000 0.00000
0.00000 0.00000 0.00000 ; CB- CA- C- O
6278 6276 6288 6290 3 2.51040 4.18400 1.67360
-6.69440 0.00000 0.00000 ; CB- CA- C- N
6279 6278 6276 6288 3 0.65084 1.95253 0.00000
-2.60338 0.00000 0.00000 ; HB- CB- CA- C
6279 6278 6280 6281 3 0.62760 1.88280 0.00000
-2.51040 0.00000 0.00000 ; HB- CB- CG1- HG11
6279 6278 6280 6282 3 0.62760 1.88280 0.00000
-2.51040 0.00000 0.00000 ; HB- CB- CG1- HG12
6279 6278 6280 6283 3 0.62760 1.88280 0.00000
-2.51040 0.00000 0.00000 ; HB- CB- CG1- HG13
6279 6278 6284 6285 3 0.62760 1.88280 0.00000
-2.51040 0.00000 0.00000 ; HB- CB- CG2- HG21
6279 6278 6284 6286 3 0.62760 1.88280 0.00000
-2.51040 0.00000 0.00000 ; HB- CB- CG2- HG22
6279 6278 6284 6287 3 0.62760 1.88280 0.00000
-2.51040 0.00000 0.00000 ; HB- CB- CG2- HG23
6280 6278 6276 6288 3 5.67350 3.55640 -6.16722
-2.47693 3.74886 0.00000 ; CG1- CB- CA- C

6280 6278 6284 6285 3 0.66944 2.00832 0.00000
-2.67776 0.00000 0.00000 ; CG1- CB- CG2- HG21
6280 6278 6284 6286 3 0.66944 2.00832 0.00000
-2.67776 0.00000 0.00000 ; CG1- CB- CG2- HG22
6280 6278 6284 6287 3 0.66944 2.00832 0.00000
-2.67776 0.00000 0.00000 ; CG1- CB- CG2- HG23
6281 6280 6278 6284 3 0.66944 2.00832 0.00000
-2.67776 0.00000 0.00000 ; HG11- CG1- CB- CG2
6282 6280 6278 6284 3 0.66944 2.00832 0.00000
-2.67776 0.00000 0.00000 ; HG12- CG1- CB- CG2
6283 6280 6278 6284 3 0.66944 2.00832 0.00000
-2.67776 0.00000 0.00000 ; HG13- CG1- CB- CG2
6284 6278 6276 6288 3 5.67350 3.55640 -6.16722
-2.47693 3.74886 0.00000 ; CG2- CB- CA- C
6288 6290 6292 6293 3 0.00000 0.00000 0.00000
0.00000 0.00000 0.00000 ; C- N- CA- HA
6288 6290 6292 6294 3 11.71520 1.67360 15.06240 -
13.38880 0.00000 0.00000 ; C- N- CA- CB
6288 6290 6292 6299 3 1.75728 5.27184 2.25936
-7.02912 0.00000 0.00000 ; C- N- CA- C
6289 6288 6290 6291 3 29.28800 -8.36800 -20.92000
0.00000 0.00000 0.00000 ; O- C- N- H
6289 6288 6290 6292 3 20.92000 0.00000 -20.92000
0.00000 0.00000 0.00000 ; O- C- N- CA
6290 6292 6294 6295 3 0.65084 1.95253 0.00000
-2.60338 0.00000 0.00000 ; N- CA- CB- HB2
6290 6292 6294 6296 3 0.65084 1.95253 0.00000
-2.60338 0.00000 0.00000 ; N- CA- CB- HB3
6290 6292 6294 6297 3 6.03751 2.50622 -5.17142
-4.20074 1.10458 0.00000 ; N- CA- CB- SG
6290 6292 6299 6300 3 0.00000 0.00000 0.00000
0.00000 0.00000 0.00000 ; N- CA- C- O
6290 6292 6299 6301 3 17.40544 -5.02080 -13.22144
9.20480 0.00000 0.00000 ; N- CA- C- N
6291 6290 6292 6293 3 0.00000 0.00000 0.00000
0.00000 0.00000 0.00000 ; H- N- CA- HA
6291 6290 6292 6294 3 0.00000 0.00000 0.00000
0.00000 0.00000 0.00000 ; H- N- CA- CB
6291 6290 6292 6299 3 0.00000 0.00000 0.00000
0.00000 0.00000 0.00000 ; H- N- CA- C
6292 6294 6297 6298 3 1.69034 2.77818 4.11706
-4.21747 1.00416 0.00000 ; CA- CB- SG- HG
6292 6299 6301 6302 3 20.92000 0.00000 -20.92000
0.00000 0.00000 0.00000 ; CA- C- N- H
6292 6299 6301 6303 3 20.92000 0.00000 -20.92000
0.00000 0.00000 0.00000 ; CA- C- N- CA
6293 6292 6294 6295 3 0.65084 1.95253 0.00000
-2.60338 0.00000 0.00000 ; HA- CA- CB- HB2
6293 6292 6294 6296 3 0.65084 1.95253 0.00000
-2.60338 0.00000 0.00000 ; HA- CA- CB- HB3
6293 6292 6294 6297 3 0.65084 1.95253 0.00000
-2.60338 0.00000 0.00000 ; HA- CA- CB- SG

6293 6292 6299 6300 3 3.68192 -4.35136 0.00000
1.33888 0.00000 0.00000 ; HA- CA- C- O
6293 6292 6299 6301 3 0.00000 0.00000 0.00000
0.00000 0.00000 0.00000 ; HA- CA- C- N
6294 6292 6299 6300 3 0.00000 0.00000 0.00000
0.00000 0.00000 0.00000 ; CB- CA- C- O
6294 6292 6299 6301 3 2.51040 4.18400 1.67360
-6.69440 0.00000 0.00000 ; CB- CA- C- N
6295 6294 6292 6299 3 0.65084 1.95253 0.00000
-2.60338 0.00000 0.00000 ; HB2- CB- CA- C
6295 6294 6297 6298 3 1.04600 3.13800 0.00000
-4.18400 0.00000 0.00000 ; HB2- CB- SG- HG
6296 6294 6292 6299 3 0.65084 1.95253 0.00000
-2.60338 0.00000 0.00000 ; HB3- CB- CA- C
6296 6294 6297 6298 3 1.04600 3.13800 0.00000
-4.18400 0.00000 0.00000 ; HB3- CB- SG- HG
6297 6294 6292 6299 3 5.62330 4.27605 -5.33042
-4.20074 2.51040 0.00000 ; SG- CB- CA- C
6299 6301 6303 6304 3 0.00000 0.00000 0.00000
0.00000 0.00000 0.00000 ; C- N- CA- HA
6299 6301 6303 6305 3 11.71520 1.67360 15.06240 -
13.38880 0.00000 0.00000 ; C- N- CA- CB
6299 6301 6303 6318 3 1.75728 5.27184 2.25936
-7.02912 0.00000 0.00000 ; C- N- CA- C
6300 6299 6301 6302 3 29.28800 -8.36800 -20.92000
0.00000 0.00000 0.00000 ; O- C- N- H
6300 6299 6301 6303 3 20.92000 0.00000 -20.92000
0.00000 0.00000 0.00000 ; O- C- N- CA
6301 6303 6305 6306 3 0.65084 1.95253 0.00000
-2.60338 0.00000 0.00000 ; N- CA- CB- HB2
6301 6303 6305 6307 3 0.65084 1.95253 0.00000
-2.60338 0.00000 0.00000 ; N- CA- CB- HB3
6301 6303 6305 6308 3 3.79070 1.39746 -4.61077
-2.40998 2.44346 0.00000 ; N- CA- CB- CG
6301 6303 6318 6319 3 0.00000 0.00000 0.00000
0.00000 0.00000 0.00000 ; N- CA- C- O
6301 6303 6318 6320 3 17.40544 -5.02080 -13.22144
9.20480 0.00000 0.00000 ; N- CA- C- N
6302 6301 6303 6304 3 0.00000 0.00000 0.00000
0.00000 0.00000 0.00000 ; H- N- CA- HA
6302 6301 6303 6305 3 0.00000 0.00000 0.00000
0.00000 0.00000 0.00000 ; H- N- CA- CB
6302 6301 6303 6318 3 0.00000 0.00000 0.00000
0.00000 0.00000 0.00000 ; H- N- CA- C
6303 6305 6308 6309 3 0.66944 2.00832 0.00000
-2.67776 0.00000 0.00000 ; CA- CB- CG- HG
6303 6305 6308 6310 3 3.90367 0.19665 -6.21742
-2.37651 5.99149 0.00000 ; CA- CB- CG- CD1
6303 6305 6308 6314 3 3.90367 0.19665 -6.21742
-2.37651 5.99149 0.00000 ; CA- CB- CG- CD2
6303 6318 6320 6321 3 20.92000 0.00000 -20.92000
0.00000 0.00000 0.00000 ; CA- C- N- H

6303 6318 6320 6322 3 20.92000 0.00000 -20.92000
0.00000 0.00000 0.00000 ; CA- C- N- CA
6304 6303 6305 6306 3 0.65084 1.95253 0.00000
-2.60338 0.00000 0.00000 ; HA- CA- CB- HB2
6304 6303 6305 6307 3 0.65084 1.95253 0.00000
-2.60338 0.00000 0.00000 ; HA- CA- CB- HB3
6304 6303 6305 6308 3 0.65084 1.95253 0.00000
-2.60338 0.00000 0.00000 ; HA- CA- CB- CG
6304 6303 6318 6319 3 3.68192 -4.35136 0.00000
1.33888 0.00000 0.00000 ; HA- CA- C- O
6304 6303 6318 6320 3 0.00000 0.00000 0.00000
0.00000 0.00000 0.00000 ; HA- CA- C- N
6305 6303 6318 6319 3 0.00000 0.00000 0.00000
0.00000 0.00000 0.00000 ; CB- CA- C- O
6305 6303 6318 6320 3 2.51040 4.18400 1.67360
-6.69440 0.00000 0.00000 ; CB- CA- C- N
6305 6308 6310 6311 3 0.66944 2.00832 0.00000
-2.67776 0.00000 0.00000 ; CB- CG- CD1- HD11
6305 6308 6310 6312 3 0.66944 2.00832 0.00000
-2.67776 0.00000 0.00000 ; CB- CG- CD1- HD12
6305 6308 6310 6313 3 0.66944 2.00832 0.00000
-2.67776 0.00000 0.00000 ; CB- CG- CD1- HD13
6305 6308 6314 6315 3 0.66944 2.00832 0.00000
-2.67776 0.00000 0.00000 ; CB- CG- CD2- HD21
6305 6308 6314 6316 3 0.66944 2.00832 0.00000
-2.67776 0.00000 0.00000 ; CB- CG- CD2- HD22
6305 6308 6314 6317 3 0.66944 2.00832 0.00000
-2.67776 0.00000 0.00000 ; CB- CG- CD2- HD23
6306 6305 6303 6318 3 0.65084 1.95253 0.00000
-2.60338 0.00000 0.00000 ; HB2- CB- CA- C
6306 6305 6308 6309 3 0.62760 1.88280 0.00000
-2.51040 0.00000 0.00000 ; HB2- CB- CG- HG
6306 6305 6308 6310 3 0.66944 2.00832 0.00000
-2.67776 0.00000 0.00000 ; HB2- CB- CG- CD1
6306 6305 6308 6314 3 0.66944 2.00832 0.00000
-2.67776 0.00000 0.00000 ; HB2- CB- CG- CD2
6307 6305 6303 6318 3 0.65084 1.95253 0.00000
-2.60338 0.00000 0.00000 ; HB3- CB- CA- C
6307 6305 6308 6309 3 0.62760 1.88280 0.00000
-2.51040 0.00000 0.00000 ; HB3- CB- CG- HG
6307 6305 6308 6310 3 0.66944 2.00832 0.00000
-2.67776 0.00000 0.00000 ; HB3- CB- CG- CD1
6307 6305 6308 6314 3 0.66944 2.00832 0.00000
-2.67776 0.00000 0.00000 ; HB3- CB- CG- CD2
6308 6305 6303 6318 3 10.33448 -1.14642 -11.54784
-2.40998 6.35968 0.00000 ; CG- CB- CA- C
6309 6308 6310 6311 3 0.62760 1.88280 0.00000
-2.51040 0.00000 0.00000 ; HG- CG- CD1- HD11
6309 6308 6310 6312 3 0.62760 1.88280 0.00000
-2.51040 0.00000 0.00000 ; HG- CG- CD1- HD12
6309 6308 6310 6313 3 0.62760 1.88280 0.00000
-2.51040 0.00000 0.00000 ; HG- CG- CD1- HD13

6309 6308 6314 6315 3 0.62760 1.88280 0.00000
-2.51040 0.00000 0.00000 ; HG- CG- CD2- HD21
6309 6308 6314 6316 3 0.62760 1.88280 0.00000
-2.51040 0.00000 0.00000 ; HG- CG- CD2- HD22
6309 6308 6314 6317 3 0.62760 1.88280 0.00000
-2.51040 0.00000 0.00000 ; HG- CG- CD2- HD23
6310 6308 6314 6315 3 0.66944 2.00832 0.00000
-2.67776 0.00000 0.00000 ; CD1- CG- CD2- HD21
6310 6308 6314 6316 3 0.66944 2.00832 0.00000
-2.67776 0.00000 0.00000 ; CD1- CG- CD2- HD22
6310 6308 6314 6317 3 0.66944 2.00832 0.00000
-2.67776 0.00000 0.00000 ; CD1- CG- CD2- HD23
6311 6310 6308 6314 3 0.66944 2.00832 0.00000
-2.67776 0.00000 0.00000 ; HD11- CD1- CG- CD2
6312 6310 6308 6314 3 0.66944 2.00832 0.00000
-2.67776 0.00000 0.00000 ; HD12- CD1- CG- CD2
6313 6310 6308 6314 3 0.66944 2.00832 0.00000
-2.67776 0.00000 0.00000 ; HD13- CD1- CG- CD2
6318 6320 6322 6323 3 0.00000 0.00000 0.00000
0.00000 0.00000 0.00000 ; C- N- CA- HA
6318 6320 6322 6324 3 11.71520 1.67360 15.06240 -
13.38880 0.00000 0.00000 ; C- N- CA- CB
6318 6320 6322 6337 3 1.75728 5.27184 2.25936
-7.02912 0.00000 0.00000 ; C- N- CA- C
6319 6318 6320 6321 3 29.28800 -8.36800 -20.92000
0.00000 0.00000 0.00000 ; O- C- N- H
6319 6318 6320 6322 3 20.92000 0.00000 -20.92000
0.00000 0.00000 0.00000 ; O- C- N- CA
6320 6322 6324 6325 3 0.65084 1.95253 0.00000
-2.60338 0.00000 0.00000 ; N- CA- CB- HB
6320 6322 6324 6326 3 3.83673 0.44769 -1.77402
-2.47693 -0.03347 0.00000 ; N- CA- CB- CG2
6320 6322 6324 6330 3 2.97482 0.12134 2.04179
-1.89117 -3.24678 0.00000 ; N- CA- CB- CG1
6320 6322 6337 6338 3 0.00000 0.00000 0.00000
0.00000 0.00000 0.00000 ; N- CA- C- O
6320 6322 6337 6339 3 17.40544 -5.02080 -13.22144
9.20480 0.00000 0.00000 ; N- CA- C- N
6321 6320 6322 6323 3 0.00000 0.00000 0.00000
0.00000 0.00000 0.00000 ; H- N- CA- HA
6321 6320 6322 6324 3 0.00000 0.00000 0.00000
0.00000 0.00000 0.00000 ; H- N- CA- CB
6321 6320 6322 6337 3 0.00000 0.00000 0.00000
0.00000 0.00000 0.00000 ; H- N- CA- C
6322 6324 6326 6327 3 0.66944 2.00832 0.00000
-2.67776 0.00000 0.00000 ; CA- CB- CG2- HG21
6322 6324 6326 6328 3 0.66944 2.00832 0.00000
-2.67776 0.00000 0.00000 ; CA- CB- CG2- HG22
6322 6324 6326 6329 3 0.66944 2.00832 0.00000
-2.67776 0.00000 0.00000 ; CA- CB- CG2- HG23
6322 6324 6330 6331 3 0.66944 2.00832 0.00000
-2.67776 0.00000 0.00000 ; CA- CB- CG1- HG12

6322 6324 6330 6332 3 0.66944 2.00832 0.00000
-2.67776 0.00000 0.00000 ; CA- CB- CG1- HG13
6322 6324 6330 6333 3 4.24258 -0.52718 -7.25506
-1.79075 7.69856 0.00000 ; CA- CB- CG1- CD1
6322 6337 6339 6340 3 20.92000 0.00000 -20.92000
0.00000 0.00000 0.00000 ; CA- C- N- CD
6322 6337 6339 6349 3 20.92000 0.00000 -20.92000
0.00000 0.00000 0.00000 ; CA- C- N- CA
6323 6322 6324 6325 3 0.65084 1.95253 0.00000
-2.60338 0.00000 0.00000 ; HA- CA- CB- HB
6323 6322 6324 6326 3 0.65084 1.95253 0.00000
-2.60338 0.00000 0.00000 ; HA- CA- CB- CG2
6323 6322 6324 6330 3 0.65084 1.95253 0.00000
-2.60338 0.00000 0.00000 ; HA- CA- CB- CG1
6323 6322 6337 6338 3 3.68192 -4.35136 0.00000
1.33888 0.00000 0.00000 ; HA- CA- C- O
6323 6322 6337 6339 3 0.00000 0.00000 0.00000
0.00000 0.00000 0.00000 ; HA- CA- C- N
6324 6322 6337 6338 3 0.00000 0.00000 0.00000
0.00000 0.00000 0.00000 ; CB- CA- C- O
6324 6322 6337 6339 3 2.51040 4.18400 1.67360
-6.69440 0.00000 0.00000 ; CB- CA- C- N
6324 6330 6333 6334 3 0.66944 2.00832 0.00000
-2.67776 0.00000 0.00000 ; CB- CG1- CD1- HD11
6324 6330 6333 6335 3 0.66944 2.00832 0.00000
-2.67776 0.00000 0.00000 ; CB- CG1- CD1- HD12
6324 6330 6333 6336 3 0.66944 2.00832 0.00000
-2.67776 0.00000 0.00000 ; CB- CG1- CD1- HD13
6325 6324 6322 6337 3 0.65084 1.95253 0.00000
-2.60338 0.00000 0.00000 ; HB- CB- CA- C
6325 6324 6326 6327 3 0.62760 1.88280 0.00000
-2.51040 0.00000 0.00000 ; HB- CB- CG2- HG21
6325 6324 6326 6328 3 0.62760 1.88280 0.00000
-2.51040 0.00000 0.00000 ; HB- CB- CG2- HG22
6325 6324 6326 6329 3 0.62760 1.88280 0.00000
-2.51040 0.00000 0.00000 ; HB- CB- CG2- HG23
6325 6324 6330 6331 3 0.62760 1.88280 0.00000
-2.51040 0.00000 0.00000 ; HB- CB- CG1- HG12
6325 6324 6330 6332 3 0.62760 1.88280 0.00000
-2.51040 0.00000 0.00000 ; HB- CB- CG1- HG13
6325 6324 6330 6333 3 0.66944 2.00832 0.00000
-2.67776 0.00000 0.00000 ; HB- CB- CG1- CD1
6326 6324 6322 6337 3 5.67350 3.55640 -6.16722
-2.47693 3.74886 0.00000 ; CG2- CB- CA- C
6326 6324 6330 6331 3 0.66944 2.00832 0.00000
-2.67776 0.00000 0.00000 ; CG2- CB- CG1- HG12
6326 6324 6330 6332 3 0.66944 2.00832 0.00000
-2.67776 0.00000 0.00000 ; CG2- CB- CG1- HG13
6326 6324 6330 6333 3 3.81162 0.49790 -8.14206
-1.79075 7.49773 0.00000 ; CG2- CB- CG1- CD1
6327 6326 6324 6330 3 0.66944 2.00832 0.00000
-2.67776 0.00000 0.00000 ; HG21- CG2- CB- CG1

6328 6326 6324 6330 3 0.66944 2.00832 0.00000
-2.67776 0.00000 0.00000 ; HG22- CG2- CB- CG1
6329 6326 6324 6330 3 0.66944 2.00832 0.00000
-2.67776 0.00000 0.00000 ; HG23- CG2- CB- CG1
6330 6324 6322 6337 3 8.26340 0.74057 -9.99976
-1.89117 3.84928 0.00000 ; CG1- CB- CA- C
6331 6330 6333 6334 3 0.62760 1.88280 0.00000
-2.51040 0.00000 0.00000 ; HG12- CG1- CD1- HD11
6331 6330 6333 6335 3 0.62760 1.88280 0.00000
-2.51040 0.00000 0.00000 ; HG12- CG1- CD1- HD12
6331 6330 6333 6336 3 0.62760 1.88280 0.00000
-2.51040 0.00000 0.00000 ; HG12- CG1- CD1- HD13
6332 6330 6333 6334 3 0.62760 1.88280 0.00000
-2.51040 0.00000 0.00000 ; HG13- CG1- CD1- HD11
6332 6330 6333 6335 3 0.62760 1.88280 0.00000
-2.51040 0.00000 0.00000 ; HG13- CG1- CD1- HD12
6332 6330 6333 6336 3 0.62760 1.88280 0.00000
-2.51040 0.00000 0.00000 ; HG13- CG1- CD1- HD13
6337 6339 6340 6341 3 0.00000 0.00000 0.00000
0.00000 0.00000 0.00000 ; C- N- CD- HD2
6337 6339 6340 6342 3 0.00000 0.00000 0.00000
0.00000 0.00000 0.00000 ; C- N- CD- HD3
6337 6339 6340 6343 3 10.04160 -3.34720 16.73600
-6.69440 0.00000 0.00000 ; C- N- CD- CG
6337 6339 6349 6346 3 11.71520 1.67360 15.06240 -
13.38880 0.00000 0.00000 ; C- N- CA- CB
6337 6339 6349 6350 3 0.00000 0.00000 0.00000
0.00000 0.00000 0.00000 ; C- N- CA- HA
6337 6339 6349 6351 3 1.75728 5.27184 2.25936
-7.02912 0.00000 0.00000 ; C- N- CA- C
6338 6337 6339 6340 3 20.92000 0.00000 -20.92000
0.00000 0.00000 0.00000 ; O- C- N- CD
6338 6337 6339 6349 3 20.92000 0.00000 -20.92000
0.00000 0.00000 0.00000 ; O- C- N- CA
6339 6340 6343 6344 3 0.65084 1.95253 0.00000
-2.60338 0.00000 0.00000 ; N- CD- CG- HG2
6339 6340 6343 6345 3 0.65084 1.95253 0.00000
-2.60338 0.00000 0.00000 ; N- CD- CG- HG3
6339 6340 6343 6346 3 0.65084 1.95253 0.00000
-2.60338 0.00000 0.00000 ; N- CD- CG- CB
6339 6349 6346 6343 3 0.65084 1.95253 0.00000
-2.60338 0.00000 0.00000 ; N- CA- CB- CG
6339 6349 6346 6347 3 0.65084 1.95253 0.00000
-2.60338 0.00000 0.00000 ; N- CA- CB- HB2
6339 6349 6346 6348 3 0.65084 1.95253 0.00000
-2.60338 0.00000 0.00000 ; N- CA- CB- HB3
6339 6349 6351 6352 3 0.00000 0.00000 0.00000
0.00000 0.00000 0.00000 ; N- CA- C- O
6339 6349 6351 6353 3 17.40544 -5.02080 -13.22144
9.20480 0.00000 0.00000 ; N- CA- C- N
6340 6339 6349 6346 3 0.00000 0.00000 0.00000
0.00000 0.00000 0.00000 ; CD- N- CA- CB

```
  6340   6339   6349   6350       3    0.00000    0.00000    0.00000
0.00000    0.00000    0.00000 ;     CD-     N-    CA-    HA
  6340   6339   6349   6351       3    0.00000    0.00000    0.00000
0.00000    0.00000    0.00000 ;     CD-     N-    CA-     C
  6340   6343   6346   6347       3    0.66944    2.00832    0.00000
-2.67776    0.00000    0.00000 ;     CD-    CG-    CB-   HB2
  6340   6343   6346   6348       3    0.66944    2.00832    0.00000
-2.67776    0.00000    0.00000 ;     CD-    CG-    CB-   HB3
  6340   6343   6346   6349       3    3.68192    3.09616   -2.09200
-3.01248    0.00000    0.00000 ;     CD-    CG-    CB-    CA
  6341   6340   6339   6349       3    0.00000    0.00000    0.00000
0.00000    0.00000    0.00000 ;    HD2-    CD-     N-    CA
  6341   6340   6343   6344       3    0.65084    1.95253    0.00000
-2.60338    0.00000    0.00000 ;    HD2-    CD-    CG-   HG2
  6341   6340   6343   6345       3    0.65084    1.95253    0.00000
-2.60338    0.00000    0.00000 ;    HD2-    CD-    CG-   HG3
  6341   6340   6343   6346       3    0.65084    1.95253    0.00000
-2.60338    0.00000    0.00000 ;    HD2-    CD-    CG-    CB
  6342   6340   6339   6349       3    0.00000    0.00000    0.00000
0.00000    0.00000    0.00000 ;    HD3-    CD-     N-    CA
  6342   6340   6343   6344       3    0.65084    1.95253    0.00000
-2.60338    0.00000    0.00000 ;    HD3-    CD-    CG-   HG2
  6342   6340   6343   6345       3    0.65084    1.95253    0.00000
-2.60338    0.00000    0.00000 ;    HD3-    CD-    CG-   HG3
  6342   6340   6343   6346       3    0.65084    1.95253    0.00000
-2.60338    0.00000    0.00000 ;    HD3-    CD-    CG-    CB
  6343   6340   6339   6349       3    0.00000    0.00000    0.00000
0.00000    0.00000    0.00000 ;     CG-    CD-     N-    CA
  6343   6346   6349   6350       3    0.65084    1.95253    0.00000
-2.60338    0.00000    0.00000 ;     CG-    CB-    CA-    HA
  6343   6346   6349   6351       3    0.65084    1.95253    0.00000
-2.60338    0.00000    0.00000 ;     CG-    CB-    CA-     C
  6344   6343   6346   6347       3    0.62760    1.88280    0.00000
-2.51040    0.00000    0.00000 ;    HG2-    CG-    CB-   HB2
  6344   6343   6346   6348       3    0.62760    1.88280    0.00000
-2.51040    0.00000    0.00000 ;    HG2-    CG-    CB-   HB3
  6344   6343   6346   6349       3    0.66944    2.00832    0.00000
-2.67776    0.00000    0.00000 ;    HG2-    CG-    CB-    CA
  6345   6343   6346   6347       3    0.62760    1.88280    0.00000
-2.51040    0.00000    0.00000 ;    HG3-    CG-    CB-   HB2
  6345   6343   6346   6348       3    0.62760    1.88280    0.00000
-2.51040    0.00000    0.00000 ;    HG3-    CG-    CB-   HB3
  6345   6343   6346   6349       3    0.66944    2.00832    0.00000
-2.67776    0.00000    0.00000 ;    HG3-    CG-    CB-    CA
  6346   6349   6351   6352       3    0.00000    0.00000    0.00000
0.00000    0.00000    0.00000 ;     CB-    CA-     C-     O
  6346   6349   6351   6353       3    2.51040    4.18400    1.67360
-6.69440    0.00000    0.00000 ;     CB-    CA-     C-     N
  6347   6346   6349   6350       3    0.65084    1.95253    0.00000
-2.60338    0.00000    0.00000 ;    HB2-    CB-    CA-    HA
  6347   6346   6349   6351       3    0.65084    1.95253    0.00000
-2.60338    0.00000    0.00000 ;    HB2-    CB-    CA-     C
```

6348 6346 6349 6350 3 0.65084 1.95253 0.00000
-2.60338 0.00000 0.00000 ; HB3- CB- CA- HA
6348 6346 6349 6351 3 0.65084 1.95253 0.00000
-2.60338 0.00000 0.00000 ; HB3- CB- CA- C
6349 6351 6353 6354 3 20.92000 0.00000 -20.92000
0.00000 0.00000 0.00000 ; CA- C- N- H
6349 6351 6353 6355 3 20.92000 0.00000 -20.92000
0.00000 0.00000 0.00000 ; CA- C- N- CA
6350 6349 6351 6352 3 3.68192 -4.35136 0.00000
1.33888 0.00000 0.00000 ; HA- CA- C- O
6350 6349 6351 6353 3 0.00000 0.00000 0.00000
0.00000 0.00000 0.00000 ; HA- CA- C- N
6351 6353 6355 6356 3 0.00000 0.00000 0.00000
0.00000 0.00000 0.00000 ; C- N- CA- HA
6351 6353 6355 6357 3 11.71520 1.67360 15.06240 -
13.38880 0.00000 0.00000 ; C- N- CA- CB
6351 6353 6355 6371 3 1.75728 5.27184 2.25936
-7.02912 0.00000 0.00000 ; C- N- CA- C
6352 6351 6353 6354 3 29.28800 -8.36800 -20.92000
0.00000 0.00000 0.00000 ; O- C- N- H
6352 6351 6353 6355 3 20.92000 0.00000 -20.92000
0.00000 0.00000 0.00000 ; O- C- N- CA
6353 6355 6357 6358 3 0.65084 1.95253 0.00000
-2.60338 0.00000 0.00000 ; N- CA- CB- HB2
6353 6355 6357 6359 3 0.65084 1.95253 0.00000
-2.60338 0.00000 0.00000 ; N- CA- CB- HB3
6353 6355 6357 6360 3 3.28026 2.46019 -2.19242
-3.21331 -0.23430 0.00000 ; N- CA- CB- CG
6353 6355 6371 6372 3 0.00000 0.00000 0.00000
0.00000 0.00000 0.00000 ; N- CA- C- O
6353 6355 6371 6373 3 17.40544 -5.02080 -13.22144
9.20480 0.00000 0.00000 ; N- CA- C- N
6354 6353 6355 6356 3 0.00000 0.00000 0.00000
0.00000 0.00000 0.00000 ; H- N- CA- HA
6354 6353 6355 6357 3 0.00000 0.00000 0.00000
0.00000 0.00000 0.00000 ; H- N- CA- CB
6354 6353 6355 6371 3 0.00000 0.00000 0.00000
0.00000 0.00000 0.00000 ; H- N- CA- C
6355 6357 6360 6361 3 0.57739 0.00000 1.02926
0.00000 -1.60666 0.00000 ; CA- CB- CG- CD1
6355 6357 6360 6369 3 0.57739 0.00000 1.02926
0.00000 -1.60666 0.00000 ; CA- CB- CG- CD2
6355 6371 6373 6374 3 20.92000 0.00000 -20.92000
0.00000 0.00000 0.00000 ; CA- C- N- H
6355 6371 6373 6375 3 20.92000 0.00000 -20.92000
0.00000 0.00000 0.00000 ; CA- C- N- CA
6356 6355 6357 6358 3 0.65084 1.95253 0.00000
-2.60338 0.00000 0.00000 ; HA- CA- CB- HB2
6356 6355 6357 6359 3 0.65084 1.95253 0.00000
-2.60338 0.00000 0.00000 ; HA- CA- CB- HB3
6356 6355 6357 6360 3 0.65084 1.95253 0.00000
-2.60338 0.00000 0.00000 ; HA- CA- CB- CG

6356 6355 6371 6372 3 3.68192 -4.35136 0.00000
1.33888 0.00000 0.00000 ; HA- CA- C- O
6356 6355 6371 6373 3 0.00000 0.00000 0.00000
0.00000 0.00000 0.00000 ; HA- CA- C- N
6357 6355 6371 6372 3 0.00000 0.00000 0.00000
0.00000 0.00000 0.00000 ; CB- CA- C- O
6357 6355 6371 6373 3 2.51040 4.18400 1.67360
-6.69440 0.00000 0.00000 ; CB- CA- C- N
6357 6360 6361 6362 3 30.33400 0.00000 -30.33400
0.00000 0.00000 0.00000 ; CB- CG- CD1- HD1
6357 6360 6361 6363 3 30.33400 0.00000 -30.33400
0.00000 0.00000 0.00000 ; CB- CG- CD1- CE1
6357 6360 6369 6367 3 30.33400 0.00000 -30.33400
0.00000 0.00000 0.00000 ; CB- CG- CD2- CE2
6357 6360 6369 6370 3 30.33400 0.00000 -30.33400
0.00000 0.00000 0.00000 ; CB- CG- CD2- HD2
6358 6357 6355 6371 3 0.65084 1.95253 0.00000
-2.60338 0.00000 0.00000 ; HB2- CB- CA- C
6358 6357 6360 6361 3 0.00000 0.00000 0.00000
0.00000 0.00000 0.00000 ; HB2- CB- CG- CD1
6358 6357 6360 6369 3 0.00000 0.00000 0.00000
0.00000 0.00000 0.00000 ; HB2- CB- CG- CD2
6359 6357 6355 6371 3 0.65084 1.95253 0.00000
-2.60338 0.00000 0.00000 ; HB3- CB- CA- C
6359 6357 6360 6361 3 0.00000 0.00000 0.00000
0.00000 0.00000 0.00000 ; HB3- CB- CG- CD1
6359 6357 6360 6369 3 0.00000 0.00000 0.00000
0.00000 0.00000 0.00000 ; HB3- CB- CG- CD2
6360 6357 6355 6371 3 4.95804 2.17986 -3.52293
-3.21331 -0.40166 0.00000 ; CG- CB- CA- C
6360 6361 6363 6364 3 30.33400 0.00000 -30.33400
0.00000 0.00000 0.00000 ; CG- CD1- CE1- HE1
6360 6361 6363 6365 3 30.33400 0.00000 -30.33400
0.00000 0.00000 0.00000 ; CG- CD1- CE1- CZ
6360 6369 6367 6365 3 30.33400 0.00000 -30.33400
0.00000 0.00000 0.00000 ; CG- CD2- CE2- CZ
6360 6369 6367 6368 3 30.33400 0.00000 -30.33400
0.00000 0.00000 0.00000 ; CG- CD2- CE2- HE2
6361 6360 6369 6367 3 30.33400 0.00000 -30.33400
0.00000 0.00000 0.00000 ; CD1- CG- CD2- CE2
6361 6360 6369 6370 3 30.33400 0.00000 -30.33400
0.00000 0.00000 0.00000 ; CD1- CG- CD2- HD2
6361 6363 6365 6366 3 30.33400 0.00000 -30.33400
0.00000 0.00000 0.00000 ; CD1- CE1- CZ- HZ
6361 6363 6365 6367 3 30.33400 0.00000 -30.33400
0.00000 0.00000 0.00000 ; CD1- CE1- CZ- CE2
6362 6361 6360 6369 3 30.33400 0.00000 -30.33400
0.00000 0.00000 0.00000 ; HD1- CD1- CG- CD2
6362 6361 6363 6364 3 30.33400 0.00000 -30.33400
0.00000 0.00000 0.00000 ; HD1- CD1- CE1- HE1
6362 6361 6363 6365 3 30.33400 0.00000 -30.33400
0.00000 0.00000 0.00000 ; HD1- CD1- CE1- CZ

6363 6361 6360 6369 3 30.33400 0.00000 -30.33400
0.00000 0.00000 0.00000 ; CE1- CD1- CG- CD2
6363 6365 6367 6368 3 30.33400 0.00000 -30.33400
0.00000 0.00000 0.00000 ; CE1- CZ- CE2- HE2
6363 6365 6367 6369 3 30.33400 0.00000 -30.33400
0.00000 0.00000 0.00000 ; CE1- CZ- CE2- CD2
6364 6363 6365 6366 3 30.33400 0.00000 -30.33400
0.00000 0.00000 0.00000 ; HE1- CE1- CZ- HZ
6364 6363 6365 6367 3 30.33400 0.00000 -30.33400
0.00000 0.00000 0.00000 ; HE1- CE1- CZ- CE2
6365 6367 6369 6370 3 30.33400 0.00000 -30.33400
0.00000 0.00000 0.00000 ; CZ- CE2- CD2- HD2
6366 6365 6367 6368 3 30.33400 0.00000 -30.33400
0.00000 0.00000 0.00000 ; HZ- CZ- CE2- HE2
6366 6365 6367 6369 3 30.33400 0.00000 -30.33400
0.00000 0.00000 0.00000 ; HZ- CZ- CE2- CD2
6368 6367 6369 6370 3 30.33400 0.00000 -30.33400
0.00000 0.00000 0.00000 ; HE2- CE2- CD2- HD2
6371 6373 6375 6376 3 0.00000 0.00000 0.00000
0.00000 0.00000 0.00000 ; C- N- CA- HA
6371 6373 6375 6377 3 11.71520 1.67360 15.06240 -
13.38880 0.00000 0.00000 ; C- N- CA- CB
6371 6373 6375 6393 3 1.75728 5.27184 2.25936
-7.02912 0.00000 0.00000 ; C- N- CA- C
6372 6371 6373 6374 3 29.28800 -8.36800 -20.92000
0.00000 0.00000 0.00000 ; O- C- N- H
6372 6371 6373 6375 3 20.92000 0.00000 -20.92000
0.00000 0.00000 0.00000 ; O- C- N- CA
6373 6375 6377 6378 3 0.65084 1.95253 0.00000
-2.60338 0.00000 0.00000 ; N- CA- CB- HB2
6373 6375 6377 6379 3 0.65084 1.95253 0.00000
-2.60338 0.00000 0.00000 ; N- CA- CB- HB3
6373 6375 6377 6380 3 0.65084 1.95253 0.00000
-2.60338 0.00000 0.00000 ; N- CA- CB- CG
6373 6375 6393 6394 3 0.00000 0.00000 0.00000
0.00000 0.00000 0.00000 ; N- CA- C- O
6373 6375 6393 6395 3 17.40544 -5.02080 -13.22144
9.20480 0.00000 0.00000 ; N- CA- C- N
6374 6373 6375 6376 3 0.00000 0.00000 0.00000
0.00000 0.00000 0.00000 ; H- N- CA- HA
6374 6373 6375 6377 3 0.00000 0.00000 0.00000
0.00000 0.00000 0.00000 ; H- N- CA- CB
6374 6373 6375 6393 3 0.00000 0.00000 0.00000
0.00000 0.00000 0.00000 ; H- N- CA- C
6375 6377 6380 6381 3 0.66944 2.00832 0.00000
-2.67776 0.00000 0.00000 ; CA- CB- CG- HG2
6375 6377 6380 6382 3 0.66944 2.00832 0.00000
-2.67776 0.00000 0.00000 ; CA- CB- CG- HG3
6375 6377 6380 6383 3 3.68192 3.09616 -2.09200
-3.01248 0.00000 0.00000 ; CA- CB- CG- CD
6375 6393 6395 6396 3 20.92000 0.00000 -20.92000
0.00000 0.00000 0.00000 ; CA- C- N- H

6375 6393 6395 6397 3 20.92000 0.00000 -20.92000
0.00000 0.00000 0.00000 ; CA- C- N- CA
6376 6375 6377 6378 3 0.65084 1.95253 0.00000
-2.60338 0.00000 0.00000 ; HA- CA- CB- HB2
6376 6375 6377 6379 3 0.65084 1.95253 0.00000
-2.60338 0.00000 0.00000 ; HA- CA- CB- HB3
6376 6375 6377 6380 3 0.65084 1.95253 0.00000
-2.60338 0.00000 0.00000 ; HA- CA- CB- CG
6376 6375 6393 6394 3 3.68192 -4.35136 0.00000
1.33888 0.00000 0.00000 ; HA- CA- C- O
6376 6375 6393 6395 3 0.00000 0.00000 0.00000
0.00000 0.00000 0.00000 ; HA- CA- C- N
6377 6375 6393 6394 3 0.00000 0.00000 0.00000
0.00000 0.00000 0.00000 ; CB- CA- C- O
6377 6375 6393 6395 3 2.51040 4.18400 1.67360
-6.69440 0.00000 0.00000 ; CB- CA- C- N
6377 6380 6383 6384 3 0.66944 2.00832 0.00000
-2.67776 0.00000 0.00000 ; CB- CG- CD- HD2
6377 6380 6383 6385 3 0.66944 2.00832 0.00000
-2.67776 0.00000 0.00000 ; CB- CG- CD- HD3
6377 6380 6383 6386 3 3.68192 3.09616 -2.09200
-3.01248 0.00000 0.00000 ; CB- CG- CD- CE
6378 6377 6375 6393 3 0.65084 1.95253 0.00000
-2.60338 0.00000 0.00000 ; HB2- CB- CA- C
6378 6377 6380 6381 3 0.62760 1.88280 0.00000
-2.51040 0.00000 0.00000 ; HB2- CB- CG- HG2
6378 6377 6380 6382 3 0.62760 1.88280 0.00000
-2.51040 0.00000 0.00000 ; HB2- CB- CG- HG3
6378 6377 6380 6383 3 0.66944 2.00832 0.00000
-2.67776 0.00000 0.00000 ; HB2- CB- CG- CD
6379 6377 6375 6393 3 0.65084 1.95253 0.00000
-2.60338 0.00000 0.00000 ; HB3- CB- CA- C
6379 6377 6380 6381 3 0.62760 1.88280 0.00000
-2.51040 0.00000 0.00000 ; HB3- CB- CG- HG2
6379 6377 6380 6382 3 0.62760 1.88280 0.00000
-2.51040 0.00000 0.00000 ; HB3- CB- CG- HG3
6379 6377 6380 6383 3 0.66944 2.00832 0.00000
-2.67776 0.00000 0.00000 ; HB3- CB- CG- CD
6380 6377 6375 6393 3 0.65084 1.95253 0.00000
-2.60338 0.00000 0.00000 ; CG- CB- CA- C
6380 6383 6386 6387 3 0.65084 1.95253 0.00000
-2.60338 0.00000 0.00000 ; CG- CD- CE- HE2
6380 6383 6386 6388 3 0.65084 1.95253 0.00000
-2.60338 0.00000 0.00000 ; CG- CD- CE- HE3
6380 6383 6386 6389 3 0.65084 1.95253 0.00000
-2.60338 0.00000 0.00000 ; CG- CD- CE- NZ
6381 6380 6383 6384 3 0.62760 1.88280 0.00000
-2.51040 0.00000 0.00000 ; HG2- CG- CD- HD2
6381 6380 6383 6385 3 0.62760 1.88280 0.00000
-2.51040 0.00000 0.00000 ; HG2- CG- CD- HD3
6381 6380 6383 6386 3 0.66944 2.00832 0.00000
-2.67776 0.00000 0.00000 ; HG2- CG- CD- CE

6382 6380 6383 6384 3 0.62760 1.88280 0.00000
-2.51040 0.00000 0.00000 ; HG3- CG- CD- HD2
6382 6380 6383 6385 3 0.62760 1.88280 0.00000
-2.51040 0.00000 0.00000 ; HG3- CG- CD- HD3
6382 6380 6383 6386 3 0.66944 2.00832 0.00000
-2.67776 0.00000 0.00000 ; HG3- CG- CD- CE
6383 6386 6389 6390 3 0.65084 1.95253 0.00000
-2.60338 0.00000 0.00000 ; CD- CE- NZ- HZ1
6383 6386 6389 6391 3 0.65084 1.95253 0.00000
-2.60338 0.00000 0.00000 ; CD- CE- NZ- HZ2
6383 6386 6389 6392 3 0.65084 1.95253 0.00000
-2.60338 0.00000 0.00000 ; CD- CE- NZ- HZ3
6384 6383 6386 6387 3 0.65084 1.95253 0.00000
-2.60338 0.00000 0.00000 ; HD2- CD- CE- HE2
6384 6383 6386 6388 3 0.65084 1.95253 0.00000
-2.60338 0.00000 0.00000 ; HD2- CD- CE- HE3
6384 6383 6386 6389 3 0.65084 1.95253 0.00000
-2.60338 0.00000 0.00000 ; HD2- CD- CE- NZ
6385 6383 6386 6387 3 0.65084 1.95253 0.00000
-2.60338 0.00000 0.00000 ; HD3- CD- CE- HE2
6385 6383 6386 6388 3 0.65084 1.95253 0.00000
-2.60338 0.00000 0.00000 ; HD3- CD- CE- HE3
6385 6383 6386 6389 3 0.65084 1.95253 0.00000
-2.60338 0.00000 0.00000 ; HD3- CD- CE- NZ
6387 6386 6389 6390 3 0.65084 1.95253 0.00000
-2.60338 0.00000 0.00000 ; HE2- CE- NZ- HZ1
6387 6386 6389 6391 3 0.65084 1.95253 0.00000
-2.60338 0.00000 0.00000 ; HE2- CE- NZ- HZ2
6387 6386 6389 6392 3 0.65084 1.95253 0.00000
-2.60338 0.00000 0.00000 ; HE2- CE- NZ- HZ3
6388 6386 6389 6390 3 0.65084 1.95253 0.00000
-2.60338 0.00000 0.00000 ; HE3- CE- NZ- HZ1
6388 6386 6389 6391 3 0.65084 1.95253 0.00000
-2.60338 0.00000 0.00000 ; HE3- CE- NZ- HZ2
6388 6386 6389 6392 3 0.65084 1.95253 0.00000
-2.60338 0.00000 0.00000 ; HE3- CE- NZ- HZ3
6393 6395 6397 6398 3 0.00000 0.00000 0.00000
0.00000 0.00000 0.00000 ; C- N- CA- HA
6393 6395 6397 6399 3 11.71520 1.67360 15.06240 -
13.38880 0.00000 0.00000 ; C- N- CA- CB
6393 6395 6397 6413 3 1.75728 5.27184 2.25936
-7.02912 0.00000 0.00000 ; C- N- CA- C
6394 6393 6395 6396 3 29.28800 -8.36800 -20.92000
0.00000 0.00000 0.00000 ; O- C- N- H
6394 6393 6395 6397 3 20.92000 0.00000 -20.92000
0.00000 0.00000 0.00000 ; O- C- N- CA
6395 6397 6399 6400 3 0.65084 1.95253 0.00000
-2.60338 0.00000 0.00000 ; N- CA- CB- HB2
6395 6397 6399 6401 3 0.65084 1.95253 0.00000
-2.60338 0.00000 0.00000 ; N- CA- CB- HB3
6395 6397 6399 6402 3 3.28026 2.46019 -2.19242
-3.21331 -0.23430 0.00000 ; N- CA- CB- CG

6395 6397 6413 6414 3 0.00000 0.00000 0.00000
0.00000 0.00000 0.00000 ; N- CA- C- O
6395 6397 6413 6415 3 17.40544 -5.02080 -13.22144
9.20480 0.00000 0.00000 ; N- CA- C- N
6396 6395 6397 6398 3 0.00000 0.00000 0.00000
0.00000 0.00000 0.00000 ; H- N- CA- HA
6396 6395 6397 6399 3 0.00000 0.00000 0.00000
0.00000 0.00000 0.00000 ; H- N- CA- CB
6396 6395 6397 6413 3 0.00000 0.00000 0.00000
0.00000 0.00000 0.00000 ; H- N- CA- C
6397 6399 6402 6403 3 0.57739 0.00000 1.02926
0.00000 -1.60666 0.00000 ; CA- CB- CG- CD1
6397 6399 6402 6411 3 0.57739 0.00000 1.02926
0.00000 -1.60666 0.00000 ; CA- CB- CG- CD2
6397 6413 6415 6416 3 20.92000 0.00000 -20.92000
0.00000 0.00000 0.00000 ; CA- C- N- H
6397 6413 6415 6417 3 20.92000 0.00000 -20.92000
0.00000 0.00000 0.00000 ; CA- C- N- CA
6398 6397 6399 6400 3 0.65084 1.95253 0.00000
-2.60338 0.00000 0.00000 ; HA- CA- CB- HB2
6398 6397 6399 6401 3 0.65084 1.95253 0.00000
-2.60338 0.00000 0.00000 ; HA- CA- CB- HB3
6398 6397 6399 6402 3 0.65084 1.95253 0.00000
-2.60338 0.00000 0.00000 ; HA- CA- CB- CG
6398 6397 6413 6414 3 3.68192 -4.35136 0.00000
1.33888 0.00000 0.00000 ; HA- CA- C- O
6398 6397 6413 6415 3 0.00000 0.00000 0.00000
0.00000 0.00000 0.00000 ; HA- CA- C- N
6399 6397 6413 6414 3 0.00000 0.00000 0.00000
0.00000 0.00000 0.00000 ; CB- CA- C- O
6399 6397 6413 6415 3 2.51040 4.18400 1.67360
-6.69440 0.00000 0.00000 ; CB- CA- C- N
6399 6402 6403 6404 3 30.33400 0.00000 -30.33400
0.00000 0.00000 0.00000 ; CB- CG- CD1- HD1
6399 6402 6403 6405 3 30.33400 0.00000 -30.33400
0.00000 0.00000 0.00000 ; CB- CG- CD1- CE1
6399 6402 6411 6409 3 30.33400 0.00000 -30.33400
0.00000 0.00000 0.00000 ; CB- CG- CD2- CE2
6399 6402 6411 6412 3 30.33400 0.00000 -30.33400
0.00000 0.00000 0.00000 ; CB- CG- CD2- HD2
6400 6399 6397 6413 3 0.65084 1.95253 0.00000
-2.60338 0.00000 0.00000 ; HB2- CB- CA- C
6400 6399 6402 6403 3 0.00000 0.00000 0.00000
0.00000 0.00000 0.00000 ; HB2- CB- CG- CD1
6400 6399 6402 6411 3 0.00000 0.00000 0.00000
0.00000 0.00000 0.00000 ; HB2- CB- CG- CD2
6401 6399 6397 6413 3 0.65084 1.95253 0.00000
-2.60338 0.00000 0.00000 ; HB3- CB- CA- C
6401 6399 6402 6403 3 0.00000 0.00000 0.00000
0.00000 0.00000 0.00000 ; HB3- CB- CG- CD1
6401 6399 6402 6411 3 0.00000 0.00000 0.00000
0.00000 0.00000 0.00000 ; HB3- CB- CG- CD2

6402 6399 6397 6413 3 4.95804 2.17986 -3.52293
-3.21331 -0.40166 0.00000 ; CG- CB- CA- C
6402 6403 6405 6406 3 30.33400 0.00000 -30.33400
0.00000 0.00000 0.00000 ; CG- CD1- CE1- HE1
6402 6403 6405 6407 3 30.33400 0.00000 -30.33400
0.00000 0.00000 0.00000 ; CG- CD1- CE1- CZ
6402 6411 6409 6407 3 30.33400 0.00000 -30.33400
0.00000 0.00000 0.00000 ; CG- CD2- CE2- CZ
6402 6411 6409 6410 3 30.33400 0.00000 -30.33400
0.00000 0.00000 0.00000 ; CG- CD2- CE2- HE2
6403 6402 6411 6409 3 30.33400 0.00000 -30.33400
0.00000 0.00000 0.00000 ; CD1- CG- CD2- CE2
6403 6402 6411 6412 3 30.33400 0.00000 -30.33400
0.00000 0.00000 0.00000 ; CD1- CG- CD2- HD2
6403 6405 6407 6408 3 30.33400 0.00000 -30.33400
0.00000 0.00000 0.00000 ; CD1- CE1- CZ- HZ
6403 6405 6407 6409 3 30.33400 0.00000 -30.33400
0.00000 0.00000 0.00000 ; CD1- CE1- CZ- CE2
6404 6403 6402 6411 3 30.33400 0.00000 -30.33400
0.00000 0.00000 0.00000 ; HD1- CD1- CG- CD2
6404 6403 6405 6406 3 30.33400 0.00000 -30.33400
0.00000 0.00000 0.00000 ; HD1- CD1- CE1- HE1
6404 6403 6405 6407 3 30.33400 0.00000 -30.33400
0.00000 0.00000 0.00000 ; HD1- CD1- CE1- CZ
6405 6403 6402 6411 3 30.33400 0.00000 -30.33400
0.00000 0.00000 0.00000 ; CE1- CD1- CG- CD2
6405 6407 6409 6410 3 30.33400 0.00000 -30.33400
0.00000 0.00000 0.00000 ; CE1- CZ- CE2- HE2
6405 6407 6409 6411 3 30.33400 0.00000 -30.33400
0.00000 0.00000 0.00000 ; CE1- CZ- CE2- CD2
6406 6405 6407 6408 3 30.33400 0.00000 -30.33400
0.00000 0.00000 0.00000 ; HE1- CE1- CZ- HZ
6406 6405 6407 6409 3 30.33400 0.00000 -30.33400
0.00000 0.00000 0.00000 ; HE1- CE1- CZ- CE2
6407 6409 6411 6412 3 30.33400 0.00000 -30.33400
0.00000 0.00000 0.00000 ; CZ- CE2- CD2- HD2
6408 6407 6409 6410 3 30.33400 0.00000 -30.33400
0.00000 0.00000 0.00000 ; HZ- CZ- CE2- HE2
6408 6407 6409 6411 3 30.33400 0.00000 -30.33400
0.00000 0.00000 0.00000 ; HZ- CZ- CE2- CD2
6410 6409 6411 6412 3 30.33400 0.00000 -30.33400
0.00000 0.00000 0.00000 ; HE2- CE2- CD2- HD2
6413 6415 6417 6418 3 0.00000 0.00000 0.00000
0.00000 0.00000 0.00000 ; C- N- CA- HA
6413 6415 6417 6419 3 11.71520 1.67360 15.06240 -
13.38880 0.00000 0.00000 ; C- N- CA- CB
6413 6415 6417 6433 3 1.75728 5.27184 2.25936
-7.02912 0.00000 0.00000 ; C- N- CA- C
6414 6413 6415 6416 3 29.28800 -8.36800 -20.92000
0.00000 0.00000 0.00000 ; O- C- N- H
6414 6413 6415 6417 3 20.92000 0.00000 -20.92000
0.00000 0.00000 0.00000 ; O- C- N- CA

6415 6417 6419 6420 3 0.65084 1.95253 0.00000
-2.60338 0.00000 0.00000 ; N- CA- CB- HB2
6415 6417 6419 6421 3 0.65084 1.95253 0.00000
-2.60338 0.00000 0.00000 ; N- CA- CB- HB3
6415 6417 6419 6422 3 3.28026 2.46019 -2.19242
-3.21331 -0.23430 0.00000 ; N- CA- CB- CG
6415 6417 6433 6434 3 0.00000 0.00000 0.00000
0.00000 0.00000 0.00000 ; N- CA- C- O
6415 6417 6433 6435 3 17.40544 -5.02080 -13.22144
9.20480 0.00000 0.00000 ; N- CA- C- N
6416 6415 6417 6418 3 0.00000 0.00000 0.00000
0.00000 0.00000 0.00000 ; H- N- CA- HA
6416 6415 6417 6419 3 0.00000 0.00000 0.00000
0.00000 0.00000 0.00000 ; H- N- CA- CB
6416 6415 6417 6433 3 0.00000 0.00000 0.00000
0.00000 0.00000 0.00000 ; H- N- CA- C
6417 6419 6422 6423 3 0.57739 0.00000 1.02926
0.00000 -1.60666 0.00000 ; CA- CB- CG- CD1
6417 6419 6422 6431 3 0.57739 0.00000 1.02926
0.00000 -1.60666 0.00000 ; CA- CB- CG- CD2
6417 6433 6435 6436 3 20.92000 0.00000 -20.92000
0.00000 0.00000 0.00000 ; CA- C- N- H
6417 6433 6435 6437 3 20.92000 0.00000 -20.92000
0.00000 0.00000 0.00000 ; CA- C- N- CA
6418 6417 6419 6420 3 0.65084 1.95253 0.00000
-2.60338 0.00000 0.00000 ; HA- CA- CB- HB2
6418 6417 6419 6421 3 0.65084 1.95253 0.00000
-2.60338 0.00000 0.00000 ; HA- CA- CB- HB3
6418 6417 6419 6422 3 0.65084 1.95253 0.00000
-2.60338 0.00000 0.00000 ; HA- CA- CB- CG
6418 6417 6433 6434 3 3.68192 -4.35136 0.00000
1.33888 0.00000 0.00000 ; HA- CA- C- O
6418 6417 6433 6435 3 0.00000 0.00000 0.00000
0.00000 0.00000 0.00000 ; HA- CA- C- N
6419 6417 6433 6434 3 0.00000 0.00000 0.00000
0.00000 0.00000 0.00000 ; CB- CA- C- O
6419 6417 6433 6435 3 2.51040 4.18400 1.67360
-6.69440 0.00000 0.00000 ; CB- CA- C- N
6419 6422 6423 6424 3 30.33400 0.00000 -30.33400
0.00000 0.00000 0.00000 ; CB- CG- CD1- HD1
6419 6422 6423 6425 3 30.33400 0.00000 -30.33400
0.00000 0.00000 0.00000 ; CB- CG- CD1- CE1
6419 6422 6431 6429 3 30.33400 0.00000 -30.33400
0.00000 0.00000 0.00000 ; CB- CG- CD2- CE2
6419 6422 6431 6432 3 30.33400 0.00000 -30.33400
0.00000 0.00000 0.00000 ; CB- CG- CD2- HD2
6420 6419 6417 6433 3 0.65084 1.95253 0.00000
-2.60338 0.00000 0.00000 ; HB2- CB- CA- C
6420 6419 6422 6423 3 0.00000 0.00000 0.00000
0.00000 0.00000 0.00000 ; HB2- CB- CG- CD1
6420 6419 6422 6431 3 0.00000 0.00000 0.00000
0.00000 0.00000 0.00000 ; HB2- CB- CG- CD2

6421 6419 6417 6433 3 0.65084 1.95253 0.00000
-2.60338 0.00000 0.00000 ; HB3- CB- CA- C
6421 6419 6422 6423 3 0.00000 0.00000 0.00000
0.00000 0.00000 0.00000 ; HB3- CB- CG- CD1
6421 6419 6422 6431 3 0.00000 0.00000 0.00000
0.00000 0.00000 0.00000 ; HB3- CB- CG- CD2
6422 6419 6417 6433 3 4.95804 2.17986 -3.52293
-3.21331 -0.40166 0.00000 ; CG- CB- CA- C
6422 6423 6425 6426 3 30.33400 0.00000 -30.33400
0.00000 0.00000 0.00000 ; CG- CD1- CE1- HE1
6422 6423 6425 6427 3 30.33400 0.00000 -30.33400
0.00000 0.00000 0.00000 ; CG- CD1- CE1- CZ
6422 6431 6429 6427 3 30.33400 0.00000 -30.33400
0.00000 0.00000 0.00000 ; CG- CD2- CE2- CZ
6422 6431 6429 6430 3 30.33400 0.00000 -30.33400
0.00000 0.00000 0.00000 ; CG- CD2- CE2- HE2
6423 6422 6431 6429 3 30.33400 0.00000 -30.33400
0.00000 0.00000 0.00000 ; CD1- CG- CD2- CE2
6423 6422 6431 6432 3 30.33400 0.00000 -30.33400
0.00000 0.00000 0.00000 ; CD1- CG- CD2- HD2
6423 6425 6427 6428 3 30.33400 0.00000 -30.33400
0.00000 0.00000 0.00000 ; CD1- CE1- CZ- HZ
6423 6425 6427 6429 3 30.33400 0.00000 -30.33400
0.00000 0.00000 0.00000 ; CD1- CE1- CZ- CE2
6424 6423 6422 6431 3 30.33400 0.00000 -30.33400
0.00000 0.00000 0.00000 ; HD1- CD1- CG- CD2
6424 6423 6425 6426 3 30.33400 0.00000 -30.33400
0.00000 0.00000 0.00000 ; HD1- CD1- CE1- HE1
6424 6423 6425 6427 3 30.33400 0.00000 -30.33400
0.00000 0.00000 0.00000 ; HD1- CD1- CE1- CZ
6425 6423 6422 6431 3 30.33400 0.00000 -30.33400
0.00000 0.00000 0.00000 ; CE1- CD1- CG- CD2
6425 6427 6429 6430 3 30.33400 0.00000 -30.33400
0.00000 0.00000 0.00000 ; CE1- CZ- CE2- HE2
6425 6427 6429 6431 3 30.33400 0.00000 -30.33400
0.00000 0.00000 0.00000 ; CE1- CZ- CE2- CD2
6426 6425 6427 6428 3 30.33400 0.00000 -30.33400
0.00000 0.00000 0.00000 ; HE1- CE1- CZ- HZ
6426 6425 6427 6429 3 30.33400 0.00000 -30.33400
0.00000 0.00000 0.00000 ; HE1- CE1- CZ- CE2
6427 6429 6431 6432 3 30.33400 0.00000 -30.33400
0.00000 0.00000 0.00000 ; CZ- CE2- CD2- HD2
6428 6427 6429 6430 3 30.33400 0.00000 -30.33400
0.00000 0.00000 0.00000 ; HZ- CZ- CE2- HE2
6428 6427 6429 6431 3 30.33400 0.00000 -30.33400
0.00000 0.00000 0.00000 ; HZ- CZ- CE2- CD2
6430 6429 6431 6432 3 30.33400 0.00000 -30.33400
0.00000 0.00000 0.00000 ; HE2- CE2- CD2- HD2
6433 6435 6437 6438 3 0.00000 0.00000 0.00000
0.00000 0.00000 0.00000 ; C- N- CA- HA
6433 6435 6437 6439 3 11.71520 1.67360 15.06240 -
13.38880 0.00000 0.00000 ; C- N- CA- CB

```
  6433   6435   6437   6445      3    1.75728    5.27184    2.25936
-7.02912    0.00000    0.00000 ;      C-     N-    CA-     C
  6434   6433   6435   6436      3   29.28800   -8.36800  -20.92000
0.00000    0.00000    0.00000 ;      O-     C-     N-     H
  6434   6433   6435   6437      3   20.92000    0.00000  -20.92000
0.00000    0.00000    0.00000 ;      O-     C-     N-    CA
  6435   6437   6439   6440      3    0.65084    1.95253    0.00000
-2.60338    0.00000    0.00000 ;      N-    CA-    CB-   HB2
  6435   6437   6439   6441      3    0.65084    1.95253    0.00000
-2.60338    0.00000    0.00000 ;      N-    CA-    CB-   HB3
  6435   6437   6439   6442      3   15.41386    9.74035   -8.39310
-0.97069    2.97901    0.00000 ;      N-    CA-    CB-    CG
  6435   6437   6445   6446      3    0.00000    0.00000    0.00000
0.00000    0.00000    0.00000 ;      N-    CA-     C-     O
  6435   6437   6445   6447      3   17.40544   -5.02080  -13.22144
9.20480    0.00000    0.00000 ;      N-    CA-     C-     N
  6436   6435   6437   6438      3    0.00000    0.00000    0.00000
0.00000    0.00000    0.00000 ;      H-     N-    CA-    HA
  6436   6435   6437   6439      3    0.00000    0.00000    0.00000
0.00000    0.00000    0.00000 ;      H-     N-    CA-    CB
  6436   6435   6437   6445      3    0.00000    0.00000    0.00000
0.00000    0.00000    0.00000 ;      H-     N-    CA-     C
  6437   6439   6442   6443      3    6.43499    0.00000   -5.39736
0.00000   -1.03763    0.00000 ;     CA-    CB-    CG-   OD1
  6437   6439   6442   6444      3    6.43499    0.00000   -5.39736
0.00000   -1.03763    0.00000 ;     CA-    CB-    CG-   OD2
  6437   6445   6447   6448      3   20.92000    0.00000  -20.92000
0.00000    0.00000    0.00000 ;     CA-     C-     N-     H
  6437   6445   6447   6449      3   20.92000    0.00000  -20.92000
0.00000    0.00000    0.00000 ;     CA-     C-     N-    CA
  6438   6437   6439   6440      3    0.65084    1.95253    0.00000
-2.60338    0.00000    0.00000 ;     HA-    CA-    CB-   HB2
  6438   6437   6439   6441      3    0.65084    1.95253    0.00000
-2.60338    0.00000    0.00000 ;     HA-    CA-    CB-   HB3
  6438   6437   6439   6442      3    0.65084    1.95253    0.00000
-2.60338    0.00000    0.00000 ;     HA-    CA-    CB-    CG
  6438   6437   6445   6446      3    3.68192   -4.35136    0.00000
1.33888    0.00000    0.00000 ;     HA-    CA-     C-     O
  6438   6437   6445   6447      3    0.00000    0.00000    0.00000
0.00000    0.00000    0.00000 ;     HA-    CA-     C-     N
  6439   6437   6445   6446      3    0.00000    0.00000    0.00000
0.00000    0.00000    0.00000 ;     CB-    CA-     C-     O
  6439   6437   6445   6447      3    2.51040    4.18400    1.67360
-6.69440    0.00000    0.00000 ;     CB-    CA-     C-     N
  6440   6439   6437   6445      3    0.65084    1.95253    0.00000
-2.60338    0.00000    0.00000 ;    HB2-    CB-    CA-     C
  6440   6439   6442   6443      3    0.00000    0.00000    0.00000
0.00000    0.00000    0.00000 ;    HB2-    CB-    CG-   OD1
  6440   6439   6442   6444      3    0.00000    0.00000    0.00000
0.00000    0.00000    0.00000 ;    HB2-    CB-    CG-   OD2
  6441   6439   6437   6445      3    0.65084    1.95253    0.00000
-2.60338    0.00000    0.00000 ;    HB3-    CB-    CA-     C
```

6441 6439 6442 6443 3 0.00000 0.00000 0.00000
0.00000 0.00000 0.00000 ; HB3- CB- CG- OD1
6441 6439 6442 6444 3 0.00000 0.00000 0.00000
0.00000 0.00000 0.00000 ; HB3- CB- CG- OD2
6442 6439 6437 6445 3 7.14627 -1.04600 -8.99560
-0.97069 5.15469 0.00000 ; CG- CB- CA- C
6445 6447 6449 6450 3 0.00000 0.00000 0.00000
0.00000 0.00000 0.00000 ; C- N- CA- HA
6445 6447 6449 6451 3 11.71520 1.67360 15.06240 -
13.38880 0.00000 0.00000 ; C- N- CA- CB
6445 6447 6449 6464 3 1.75728 5.27184 2.25936
-7.02912 0.00000 0.00000 ; C- N- CA- C
6446 6445 6447 6448 3 29.28800 -8.36800 -20.92000
0.00000 0.00000 0.00000 ; O- C- N- H
6446 6445 6447 6449 3 20.92000 0.00000 -20.92000
0.00000 0.00000 0.00000 ; O- C- N- CA
6447 6449 6451 6452 3 0.65084 1.95253 0.00000
-2.60338 0.00000 0.00000 ; N- CA- CB- HB
6447 6449 6451 6453 3 3.83673 0.44769 -1.77402
-2.47693 -0.03347 0.00000 ; N- CA- CB- CG2
6447 6449 6451 6457 3 2.97482 0.12134 2.04179
-1.89117 -3.24678 0.00000 ; N- CA- CB- CG1
6447 6449 6464 6465 3 0.00000 0.00000 0.00000
0.00000 0.00000 0.00000 ; N- CA- C- O
6447 6449 6464 6466 3 17.40544 -5.02080 -13.22144
9.20480 0.00000 0.00000 ; N- CA- C- N
6448 6447 6449 6450 3 0.00000 0.00000 0.00000
0.00000 0.00000 0.00000 ; H- N- CA- HA
6448 6447 6449 6451 3 0.00000 0.00000 0.00000
0.00000 0.00000 0.00000 ; H- N- CA- CB
6448 6447 6449 6464 3 0.00000 0.00000 0.00000
0.00000 0.00000 0.00000 ; H- N- CA- C
6449 6451 6453 6454 3 0.66944 2.00832 0.00000
-2.67776 0.00000 0.00000 ; CA- CB- CG2- HG21
6449 6451 6453 6455 3 0.66944 2.00832 0.00000
-2.67776 0.00000 0.00000 ; CA- CB- CG2- HG22
6449 6451 6453 6456 3 0.66944 2.00832 0.00000
-2.67776 0.00000 0.00000 ; CA- CB- CG2- HG23
6449 6451 6457 6458 3 0.66944 2.00832 0.00000
-2.67776 0.00000 0.00000 ; CA- CB- CG1- HG12
6449 6451 6457 6459 3 0.66944 2.00832 0.00000
-2.67776 0.00000 0.00000 ; CA- CB- CG1- HG13
6449 6451 6457 6460 3 4.24258 -0.52718 -7.25506
-1.79075 7.69856 0.00000 ; CA- CB- CG1- CD1
6449 6464 6466 6467 3 20.92000 0.00000 -20.92000
0.00000 0.00000 0.00000 ; CA- C- N- H
6449 6464 6466 6468 3 20.92000 0.00000 -20.92000
0.00000 0.00000 0.00000 ; CA- C- N- CA
6450 6449 6451 6452 3 0.65084 1.95253 0.00000
-2.60338 0.00000 0.00000 ; HA- CA- CB- HB
6450 6449 6451 6453 3 0.65084 1.95253 0.00000
-2.60338 0.00000 0.00000 ; HA- CA- CB- CG2

6450 6449 6451 6457 3 0.65084 1.95253 0.00000
-2.60338 0.00000 0.00000 ; HA- CA- CB- CG1
6450 6449 6464 6465 3 3.68192 -4.35136 0.00000
1.33888 0.00000 0.00000 ; HA- CA- C- O
6450 6449 6464 6466 3 0.00000 0.00000 0.00000
0.00000 0.00000 0.00000 ; HA- CA- C- N
6451 6449 6464 6465 3 0.00000 0.00000 0.00000
0.00000 0.00000 0.00000 ; CB- CA- C- O
6451 6449 6464 6466 3 2.51040 4.18400 1.67360
-6.69440 0.00000 0.00000 ; CB- CA- C- N
6451 6457 6460 6461 3 0.66944 2.00832 0.00000
-2.67776 0.00000 0.00000 ; CB- CG1- CD1- HD11
6451 6457 6460 6462 3 0.66944 2.00832 0.00000
-2.67776 0.00000 0.00000 ; CB- CG1- CD1- HD12
6451 6457 6460 6463 3 0.66944 2.00832 0.00000
-2.67776 0.00000 0.00000 ; CB- CG1- CD1- HD13
6452 6451 6449 6464 3 0.65084 1.95253 0.00000
-2.60338 0.00000 0.00000 ; HB- CB- CA- C
6452 6451 6453 6454 3 0.62760 1.88280 0.00000
-2.51040 0.00000 0.00000 ; HB- CB- CG2- HG21
6452 6451 6453 6455 3 0.62760 1.88280 0.00000
-2.51040 0.00000 0.00000 ; HB- CB- CG2- HG22
6452 6451 6453 6456 3 0.62760 1.88280 0.00000
-2.51040 0.00000 0.00000 ; HB- CB- CG2- HG23
6452 6451 6457 6458 3 0.62760 1.88280 0.00000
-2.51040 0.00000 0.00000 ; HB- CB- CG1- HG12
6452 6451 6457 6459 3 0.62760 1.88280 0.00000
-2.51040 0.00000 0.00000 ; HB- CB- CG1- HG13
6452 6451 6457 6460 3 0.66944 2.00832 0.00000
-2.67776 0.00000 0.00000 ; HB- CB- CG1- CD1
6453 6451 6449 6464 3 5.67350 3.55640 -6.16722
-2.47693 3.74886 0.00000 ; CG2- CB- CA- C
6453 6451 6457 6458 3 0.66944 2.00832 0.00000
-2.67776 0.00000 0.00000 ; CG2- CB- CG1- HG12
6453 6451 6457 6459 3 0.66944 2.00832 0.00000
-2.67776 0.00000 0.00000 ; CG2- CB- CG1- HG13
6453 6451 6457 6460 3 3.81162 0.49790 -8.14206
-1.79075 7.49773 0.00000 ; CG2- CB- CG1- CD1
6454 6453 6451 6457 3 0.66944 2.00832 0.00000
-2.67776 0.00000 0.00000 ; HG21- CG2- CB- CG1
6455 6453 6451 6457 3 0.66944 2.00832 0.00000
-2.67776 0.00000 0.00000 ; HG22- CG2- CB- CG1
6456 6453 6451 6457 3 0.66944 2.00832 0.00000
-2.67776 0.00000 0.00000 ; HG23- CG2- CB- CG1
6457 6451 6449 6464 3 8.26340 0.74057 -9.99976
-1.89117 3.84928 0.00000 ; CG1- CB- CA- C
6458 6457 6460 6461 3 0.62760 1.88280 0.00000
-2.51040 0.00000 0.00000 ; HG12- CG1- CD1- HD11
6458 6457 6460 6462 3 0.62760 1.88280 0.00000
-2.51040 0.00000 0.00000 ; HG12- CG1- CD1- HD12
6458 6457 6460 6463 3 0.62760 1.88280 0.00000
-2.51040 0.00000 0.00000 ; HG12- CG1- CD1- HD13

6459 6457 6460 6461 3 0.62760 1.88280 0.00000
-2.51040 0.00000 0.00000 ; HG13- CG1- CD1- HD11
6459 6457 6460 6462 3 0.62760 1.88280 0.00000
-2.51040 0.00000 0.00000 ; HG13- CG1- CD1- HD12
6459 6457 6460 6463 3 0.62760 1.88280 0.00000
-2.51040 0.00000 0.00000 ; HG13- CG1- CD1- HD13
6464 6466 6468 6469 3 0.00000 0.00000 0.00000
0.00000 0.00000 0.00000 ; C- N- CA- HA
6464 6466 6468 6470 3 11.71520 1.67360 15.06240 -
13.38880 0.00000 0.00000 ; C- N- CA- CB
6464 6466 6468 6476 3 1.75728 5.27184 2.25936
-7.02912 0.00000 0.00000 ; C- N- CA- C
6465 6464 6466 6467 3 29.28800 -8.36800 -20.92000
0.00000 0.00000 0.00000 ; O- C- N- H
6465 6464 6466 6468 3 20.92000 0.00000 -20.92000
0.00000 0.00000 0.00000 ; O- C- N- CA
6466 6468 6470 6471 3 0.65084 1.95253 0.00000
-2.60338 0.00000 0.00000 ; N- CA- CB- HB2
6466 6468 6470 6472 3 0.65084 1.95253 0.00000
-2.60338 0.00000 0.00000 ; N- CA- CB- HB3
6466 6468 6470 6473 3 15.41386 9.74035 -8.39310
-0.97069 2.97901 0.00000 ; N- CA- CB- CG
6466 6468 6476 6477 3 0.00000 0.00000 0.00000
0.00000 0.00000 0.00000 ; N- CA- C- O
6466 6468 6476 6478 3 17.40544 -5.02080 -13.22144
9.20480 0.00000 0.00000 ; N- CA- C- N
6467 6466 6468 6469 3 0.00000 0.00000 0.00000
0.00000 0.00000 0.00000 ; H- N- CA- HA
6467 6466 6468 6470 3 0.00000 0.00000 0.00000
0.00000 0.00000 0.00000 ; H- N- CA- CB
6467 6466 6468 6476 3 0.00000 0.00000 0.00000
0.00000 0.00000 0.00000 ; H- N- CA- C
6468 6470 6473 6474 3 6.43499 0.00000 -5.39736
0.00000 -1.03763 0.00000 ; CA- CB- CG- OD1
6468 6470 6473 6475 3 6.43499 0.00000 -5.39736
0.00000 -1.03763 0.00000 ; CA- CB- CG- OD2
6468 6476 6478 6479 3 20.92000 0.00000 -20.92000
0.00000 0.00000 0.00000 ; CA- C- N- H
6468 6476 6478 6480 3 20.92000 0.00000 -20.92000
0.00000 0.00000 0.00000 ; CA- C- N- CA
6469 6468 6470 6471 3 0.65084 1.95253 0.00000
-2.60338 0.00000 0.00000 ; HA- CA- CB- HB2
6469 6468 6470 6472 3 0.65084 1.95253 0.00000
-2.60338 0.00000 0.00000 ; HA- CA- CB- HB3
6469 6468 6470 6473 3 0.65084 1.95253 0.00000
-2.60338 0.00000 0.00000 ; HA- CA- CB- CG
6469 6468 6476 6477 3 3.68192 -4.35136 0.00000
1.33888 0.00000 0.00000 ; HA- CA- C- O
6469 6468 6476 6478 3 0.00000 0.00000 0.00000
0.00000 0.00000 0.00000 ; HA- CA- C- N
6470 6468 6476 6477 3 0.00000 0.00000 0.00000
0.00000 0.00000 0.00000 ; CB- CA- C- O

6470 6468 6476 6478 3 2.51040 4.18400 1.67360
-6.69440 0.00000 0.00000 ; CB- CA- C- N
6471 6470 6468 6476 3 0.65084 1.95253 0.00000
-2.60338 0.00000 0.00000 ; HB2- CB- CA- C
6471 6470 6473 6474 3 0.00000 0.00000 0.00000
0.00000 0.00000 0.00000 ; HB2- CB- CG- OD1
6471 6470 6473 6475 3 0.00000 0.00000 0.00000
0.00000 0.00000 0.00000 ; HB2- CB- CG- OD2
6472 6470 6468 6476 3 0.65084 1.95253 0.00000
-2.60338 0.00000 0.00000 ; HB3- CB- CA- C
6472 6470 6473 6474 3 0.00000 0.00000 0.00000
0.00000 0.00000 0.00000 ; HB3- CB- CG- OD1
6472 6470 6473 6475 3 0.00000 0.00000 0.00000
0.00000 0.00000 0.00000 ; HB3- CB- CG- OD2
6473 6470 6468 6476 3 7.14627 -1.04600 -8.99560
-0.97069 5.15469 0.00000 ; CG- CB- CA- C
6476 6478 6480 6481 3 0.00000 0.00000 0.00000
0.00000 0.00000 0.00000 ; C- N- CA- HA2
6476 6478 6480 6482 3 0.00000 0.00000 0.00000
0.00000 0.00000 0.00000 ; C- N- CA- HA3
6476 6478 6480 6483 3 1.75728 5.27184 2.25936
-7.02912 0.00000 0.00000 ; C- N- CA- C
6477 6476 6478 6479 3 29.28800 -8.36800 -20.92000
0.00000 0.00000 0.00000 ; O- C- N- H
6477 6476 6478 6480 3 20.92000 0.00000 -20.92000
0.00000 0.00000 0.00000 ; O- C- N- CA
6478 6480 6483 6484 3 0.00000 0.00000 0.00000
0.00000 0.00000 0.00000 ; N- CA- C- O
6478 6480 6483 6485 3 17.40544 -5.02080 -13.22144
9.20480 0.00000 0.00000 ; N- CA- C- N
6479 6478 6480 6481 3 0.00000 0.00000 0.00000
0.00000 0.00000 0.00000 ; H- N- CA- HA2
6479 6478 6480 6482 3 0.00000 0.00000 0.00000
0.00000 0.00000 0.00000 ; H- N- CA- HA3
6479 6478 6480 6483 3 0.00000 0.00000 0.00000
0.00000 0.00000 0.00000 ; H- N- CA- C
6480 6483 6485 6486 3 20.92000 0.00000 -20.92000
0.00000 0.00000 0.00000 ; CA- C- N- H
6480 6483 6485 6487 3 20.92000 0.00000 -20.92000
0.00000 0.00000 0.00000 ; CA- C- N- CA
6481 6480 6483 6484 3 3.68192 -4.35136 0.00000
1.33888 0.00000 0.00000 ; HA2- CA- C- O
6481 6480 6483 6485 3 0.00000 0.00000 0.00000
0.00000 0.00000 0.00000 ; HA2- CA- C- N
6482 6480 6483 6484 3 3.68192 -4.35136 0.00000
1.33888 0.00000 0.00000 ; HA3- CA- C- O
6482 6480 6483 6485 3 0.00000 0.00000 0.00000
0.00000 0.00000 0.00000 ; HA3- CA- C- N
6483 6485 6487 6488 3 0.00000 0.00000 0.00000
0.00000 0.00000 0.00000 ; C- N- CA- HA
6483 6485 6487 6489 3 11.71520 1.67360 15.06240 -
13.38880 0.00000 0.00000 ; C- N- CA- CB

6483 6485 6487 6497 3 1.75728 5.27184 2.25936
-7.02912 0.00000 0.00000 ; C- N- CA- C
6484 6483 6485 6486 3 29.28800 -8.36800 -20.92000
0.00000 0.00000 0.00000 ; O- C- N- H
6484 6483 6485 6487 3 20.92000 0.00000 -20.92000
0.00000 0.00000 0.00000 ; O- C- N- CA
6485 6487 6489 6490 3 0.65084 1.95253 0.00000
-2.60338 0.00000 0.00000 ; N- CA- CB- HB2
6485 6487 6489 6491 3 0.65084 1.95253 0.00000
-2.60338 0.00000 0.00000 ; N- CA- CB- HB3
6485 6487 6489 6492 3 5.99567 3.29281 -4.46014
-0.55229 1.97485 0.00000 ; N- CA- CB- CG
6485 6487 6497 6498 3 0.00000 0.00000 0.00000
0.00000 0.00000 0.00000 ; N- CA- C- O
6485 6487 6497 6499 3 17.40544 -5.02080 -13.22144
9.20480 0.00000 0.00000 ; N- CA- C- N
6486 6485 6487 6488 3 0.00000 0.00000 0.00000
0.00000 0.00000 0.00000 ; H- N- CA- HA
6486 6485 6487 6489 3 0.00000 0.00000 0.00000
0.00000 0.00000 0.00000 ; H- N- CA- CB
6486 6485 6487 6497 3 0.00000 0.00000 0.00000
0.00000 0.00000 0.00000 ; H- N- CA- C
6487 6489 6492 6493 3 0.00000 0.00000 0.00000
0.00000 0.00000 0.00000 ; CA- CB- CG- OD1
6487 6489 6492 6494 3 8.84916 -0.31380 -4.32626
5.03754 0.26778 0.00000 ; CA- CB- CG- ND2
6487 6497 6499 6500 3 20.92000 0.00000 -20.92000
0.00000 0.00000 0.00000 ; CA- C- N- H
6487 6497 6499 6501 3 20.92000 0.00000 -20.92000
0.00000 0.00000 0.00000 ; CA- C- N- CA
6488 6487 6489 6490 3 0.65084 1.95253 0.00000
-2.60338 0.00000 0.00000 ; HA- CA- CB- HB2
6488 6487 6489 6491 3 0.65084 1.95253 0.00000
-2.60338 0.00000 0.00000 ; HA- CA- CB- HB3
6488 6487 6489 6492 3 0.65084 1.95253 0.00000
-2.60338 0.00000 0.00000 ; HA- CA- CB- CG
6488 6487 6497 6498 3 3.68192 -4.35136 0.00000
1.33888 0.00000 0.00000 ; HA- CA- C- O
6488 6487 6497 6499 3 0.00000 0.00000 0.00000
0.00000 0.00000 0.00000 ; HA- CA- C- N
6489 6487 6497 6498 3 0.00000 0.00000 0.00000
0.00000 0.00000 0.00000 ; CB- CA- C- O
6489 6487 6497 6499 3 2.51040 4.18400 1.67360
-6.69440 0.00000 0.00000 ; CB- CA- C- N
6489 6492 6494 6495 3 20.92000 0.00000 -20.92000
0.00000 0.00000 0.00000 ; CB- CG- ND2- HD21
6489 6492 6494 6496 3 20.92000 0.00000 -20.92000
0.00000 0.00000 0.00000 ; CB- CG- ND2- HD22
6490 6489 6487 6497 3 0.65084 1.95253 0.00000
-2.60338 0.00000 0.00000 ; HB2- CB- CA- C
6490 6489 6492 6493 3 3.68192 -4.35136 0.00000
1.33888 0.00000 0.00000 ; HB2- CB- CG- OD1

6490 6489 6492 6494 3 0.00000 0.00000 0.00000
0.00000 0.00000 0.00000 ; HB2- CB- CG- ND2
6491 6489 6487 6497 3 0.65084 1.95253 0.00000
-2.60338 0.00000 0.00000 ; HB3- CB- CA- C
6491 6489 6492 6493 3 3.68192 -4.35136 0.00000
1.33888 0.00000 0.00000 ; HB3- CB- CG- OD1
6491 6489 6492 6494 3 0.00000 0.00000 0.00000
0.00000 0.00000 0.00000 ; HB3- CB- CG- ND2
6492 6489 6487 6497 3 7.94542 -3.96225 -6.11701
-0.55229 3.58150 0.00000 ; CG- CB- CA- C
6493 6492 6494 6495 3 29.28800 -8.36800 -20.92000
0.00000 0.00000 0.00000 ; OD1- CG- ND2- HD21
6493 6492 6494 6496 3 29.28800 -8.36800 -20.92000
0.00000 0.00000 0.00000 ; OD1- CG- ND2- HD22
6497 6499 6501 6502 3 0.00000 0.00000 0.00000
0.00000 0.00000 0.00000 ; C- N- CA- HA2
6497 6499 6501 6503 3 0.00000 0.00000 0.00000
0.00000 0.00000 0.00000 ; C- N- CA- HA3
6497 6499 6501 6504 3 1.75728 5.27184 2.25936
-7.02912 0.00000 0.00000 ; C- N- CA- C
6498 6497 6499 6500 3 29.28800 -8.36800 -20.92000
0.00000 0.00000 0.00000 ; O- C- N- H
6498 6497 6499 6501 3 20.92000 0.00000 -20.92000
0.00000 0.00000 0.00000 ; O- C- N- CA
6499 6501 6504 6505 3 0.00000 0.00000 0.00000
0.00000 0.00000 0.00000 ; N- CA- C- O
6499 6501 6504 6506 3 17.40544 -5.02080 -13.22144
9.20480 0.00000 0.00000 ; N- CA- C- N
6500 6499 6501 6502 3 0.00000 0.00000 0.00000
0.00000 0.00000 0.00000 ; H- N- CA- HA2
6500 6499 6501 6503 3 0.00000 0.00000 0.00000
0.00000 0.00000 0.00000 ; H- N- CA- HA3
6500 6499 6501 6504 3 0.00000 0.00000 0.00000
0.00000 0.00000 0.00000 ; H- N- CA- C
6501 6504 6506 6507 3 20.92000 0.00000 -20.92000
0.00000 0.00000 0.00000 ; CA- C- N- H
6501 6504 6506 6508 3 20.92000 0.00000 -20.92000
0.00000 0.00000 0.00000 ; CA- C- N- CA
6502 6501 6504 6505 3 3.68192 -4.35136 0.00000
1.33888 0.00000 0.00000 ; HA2- CA- C- O
6502 6501 6504 6506 3 0.00000 0.00000 0.00000
0.00000 0.00000 0.00000 ; HA2- CA- C- N
6503 6501 6504 6505 3 3.68192 -4.35136 0.00000
1.33888 0.00000 0.00000 ; HA3- CA- C- O
6503 6501 6504 6506 3 0.00000 0.00000 0.00000
0.00000 0.00000 0.00000 ; HA3- CA- C- N
6504 6506 6508 6509 3 0.00000 0.00000 0.00000
0.00000 0.00000 0.00000 ; C- N- CA- HA
6504 6506 6508 6510 3 11.71520 1.67360 15.06240 -
13.38880 0.00000 0.00000 ; C- N- CA- CB
6504 6506 6508 6526 3 1.75728 5.27184 2.25936
-7.02912 0.00000 0.00000 ; C- N- CA- C

6505 6504 6506 6507 3 29.28800 -8.36800 -20.92000
0.00000 0.00000 0.00000 ; O- C- N- H
6505 6504 6506 6508 3 20.92000 0.00000 -20.92000
0.00000 0.00000 0.00000 ; O- C- N- CA
6506 6508 6510 6511 3 0.65084 1.95253 0.00000
-2.60338 0.00000 0.00000 ; N- CA- CB- HB2
6506 6508 6510 6512 3 0.65084 1.95253 0.00000
-2.60338 0.00000 0.00000 ; N- CA- CB- HB3
6506 6508 6510 6513 3 0.65084 1.95253 0.00000
-2.60338 0.00000 0.00000 ; N- CA- CB- CG
6506 6508 6526 6527 3 0.00000 0.00000 0.00000
0.00000 0.00000 0.00000 ; N- CA- C- O
6506 6508 6526 6528 3 17.40544 -5.02080 -13.22144
9.20480 0.00000 0.00000 ; N- CA- C- N
6507 6506 6508 6509 3 0.00000 0.00000 0.00000
0.00000 0.00000 0.00000 ; H- N- CA- HA
6507 6506 6508 6510 3 0.00000 0.00000 0.00000
0.00000 0.00000 0.00000 ; H- N- CA- CB
6507 6506 6508 6526 3 0.00000 0.00000 0.00000
0.00000 0.00000 0.00000 ; H- N- CA- C
6508 6510 6513 6514 3 0.66944 2.00832 0.00000
-2.67776 0.00000 0.00000 ; CA- CB- CG- HG2
6508 6510 6513 6515 3 0.66944 2.00832 0.00000
-2.67776 0.00000 0.00000 ; CA- CB- CG- HG3
6508 6510 6513 6516 3 3.68192 3.09616 -2.09200
-3.01248 0.00000 0.00000 ; CA- CB- CG- CD
6508 6526 6528 6529 3 20.92000 0.00000 -20.92000
0.00000 0.00000 0.00000 ; CA- C- N- H
6508 6526 6528 6530 3 20.92000 0.00000 -20.92000
0.00000 0.00000 0.00000 ; CA- C- N- CA
6509 6508 6510 6511 3 0.65084 1.95253 0.00000
-2.60338 0.00000 0.00000 ; HA- CA- CB- HB2
6509 6508 6510 6512 3 0.65084 1.95253 0.00000
-2.60338 0.00000 0.00000 ; HA- CA- CB- HB3
6509 6508 6510 6513 3 0.65084 1.95253 0.00000
-2.60338 0.00000 0.00000 ; HA- CA- CB- CG
6509 6508 6526 6527 3 3.68192 -4.35136 0.00000
1.33888 0.00000 0.00000 ; HA- CA- C- O
6509 6508 6526 6528 3 0.00000 0.00000 0.00000
0.00000 0.00000 0.00000 ; HA- CA- C- N
6510 6508 6526 6527 3 0.00000 0.00000 0.00000
0.00000 0.00000 0.00000 ; CB- CA- C- O
6510 6508 6526 6528 3 2.51040 4.18400 1.67360
-6.69440 0.00000 0.00000 ; CB- CA- C- N
6510 6513 6516 6517 3 0.66944 2.00832 0.00000
-2.67776 0.00000 0.00000 ; CB- CG- CD- HD2
6510 6513 6516 6518 3 0.66944 2.00832 0.00000
-2.67776 0.00000 0.00000 ; CB- CG- CD- HD3
6510 6513 6516 6519 3 3.68192 3.09616 -2.09200
-3.01248 0.00000 0.00000 ; CB- CG- CD- CE
6511 6510 6508 6526 3 0.65084 1.95253 0.00000
-2.60338 0.00000 0.00000 ; HB2- CB- CA- C

```
  6511   6510   6513   6514      3    0.62760    1.88280    0.00000
-2.51040    0.00000    0.00000 ;    HB2-    CB-    CG-   HG2
  6511   6510   6513   6515      3    0.62760    1.88280    0.00000
-2.51040    0.00000    0.00000 ;    HB2-    CB-    CG-   HG3
  6511   6510   6513   6516      3    0.66944    2.00832    0.00000
-2.67776    0.00000    0.00000 ;    HB2-    CB-    CG-    CD
  6512   6510   6508   6526      3    0.65084    1.95253    0.00000
-2.60338    0.00000    0.00000 ;    HB3-    CB-    CA-     C
  6512   6510   6513   6514      3    0.62760    1.88280    0.00000
-2.51040    0.00000    0.00000 ;    HB3-    CB-    CG-   HG2
  6512   6510   6513   6515      3    0.62760    1.88280    0.00000
-2.51040    0.00000    0.00000 ;    HB3-    CB-    CG-   HG3
  6512   6510   6513   6516      3    0.66944    2.00832    0.00000
-2.67776    0.00000    0.00000 ;    HB3-    CB-    CG-    CD
  6513   6510   6508   6526      3    0.65084    1.95253    0.00000
-2.60338    0.00000    0.00000 ;     CG-    CB-    CA-     C
  6513   6516   6519   6520      3    0.65084    1.95253    0.00000
-2.60338    0.00000    0.00000 ;     CG-    CD-    CE-   HE2
  6513   6516   6519   6521      3    0.65084    1.95253    0.00000
-2.60338    0.00000    0.00000 ;     CG-    CD-    CE-   HE3
  6513   6516   6519   6522      3    0.65084    1.95253    0.00000
-2.60338    0.00000    0.00000 ;     CG-    CD-    CE-    NZ
  6514   6513   6516   6517      3    0.62760    1.88280    0.00000
-2.51040    0.00000    0.00000 ;    HG2-    CG-    CD-   HD2
  6514   6513   6516   6518      3    0.62760    1.88280    0.00000
-2.51040    0.00000    0.00000 ;    HG2-    CG-    CD-   HD3
  6514   6513   6516   6519      3    0.66944    2.00832    0.00000
-2.67776    0.00000    0.00000 ;    HG2-    CG-    CD-    CE
  6515   6513   6516   6517      3    0.62760    1.88280    0.00000
-2.51040    0.00000    0.00000 ;    HG3-    CG-    CD-   HD2
  6515   6513   6516   6518      3    0.62760    1.88280    0.00000
-2.51040    0.00000    0.00000 ;    HG3-    CG-    CD-   HD3
  6515   6513   6516   6519      3    0.66944    2.00832    0.00000
-2.67776    0.00000    0.00000 ;    HG3-    CG-    CD-    CE
  6516   6519   6522   6523      3    0.65084    1.95253    0.00000
-2.60338    0.00000    0.00000 ;     CD-    CE-    NZ-   HZ1
  6516   6519   6522   6524      3    0.65084    1.95253    0.00000
-2.60338    0.00000    0.00000 ;     CD-    CE-    NZ-   HZ2
  6516   6519   6522   6525      3    0.65084    1.95253    0.00000
-2.60338    0.00000    0.00000 ;     CD-    CE-    NZ-   HZ3
  6517   6516   6519   6520      3    0.65084    1.95253    0.00000
-2.60338    0.00000    0.00000 ;    HD2-    CD-    CE-   HE2
  6517   6516   6519   6521      3    0.65084    1.95253    0.00000
-2.60338    0.00000    0.00000 ;    HD2-    CD-    CE-   HE3
  6517   6516   6519   6522      3    0.65084    1.95253    0.00000
-2.60338    0.00000    0.00000 ;    HD2-    CD-    CE-    NZ
  6518   6516   6519   6520      3    0.65084    1.95253    0.00000
-2.60338    0.00000    0.00000 ;    HD3-    CD-    CE-   HE2
  6518   6516   6519   6521      3    0.65084    1.95253    0.00000
-2.60338    0.00000    0.00000 ;    HD3-    CD-    CE-   HE3
  6518   6516   6519   6522      3    0.65084    1.95253    0.00000
-2.60338    0.00000    0.00000 ;    HD3-    CD-    CE-    NZ
```

6520 6519 6522 6523 3 0.65084 1.95253 0.00000
-2.60338 0.00000 0.00000 ; HE2- CE- NZ- HZ1
6520 6519 6522 6524 3 0.65084 1.95253 0.00000
-2.60338 0.00000 0.00000 ; HE2- CE- NZ- HZ2
6520 6519 6522 6525 3 0.65084 1.95253 0.00000
-2.60338 0.00000 0.00000 ; HE2- CE- NZ- HZ3
6521 6519 6522 6523 3 0.65084 1.95253 0.00000
-2.60338 0.00000 0.00000 ; HE3- CE- NZ- HZ1
6521 6519 6522 6524 3 0.65084 1.95253 0.00000
-2.60338 0.00000 0.00000 ; HE3- CE- NZ- HZ2
6521 6519 6522 6525 3 0.65084 1.95253 0.00000
-2.60338 0.00000 0.00000 ; HE3- CE- NZ- HZ3
6526 6528 6530 6531 3 0.00000 0.00000 0.00000
0.00000 0.00000 0.00000 ; C- N- CA- HA
6526 6528 6530 6532 3 11.71520 1.67360 15.06240 -
13.38880 0.00000 0.00000 ; C- N- CA- CB
6526 6528 6530 6545 3 1.75728 5.27184 2.25936
-7.02912 0.00000 0.00000 ; C- N- CA- C
6527 6526 6528 6529 3 29.28800 -8.36800 -20.92000
0.00000 0.00000 0.00000 ; O- C- N- H
6527 6526 6528 6530 3 20.92000 0.00000 -20.92000
0.00000 0.00000 0.00000 ; O- C- N- CA
6528 6530 6532 6533 3 0.65084 1.95253 0.00000
-2.60338 0.00000 0.00000 ; N- CA- CB- HB
6528 6530 6532 6534 3 3.83673 0.44769 -1.77402
-2.47693 -0.03347 0.00000 ; N- CA- CB- CG2
6528 6530 6532 6538 3 2.97482 0.12134 2.04179
-1.89117 -3.24678 0.00000 ; N- CA- CB- CG1
6528 6530 6545 6546 3 0.00000 0.00000 0.00000
0.00000 0.00000 0.00000 ; N- CA- C- O
6528 6530 6545 6547 3 17.40544 -5.02080 -13.22144
9.20480 0.00000 0.00000 ; N- CA- C- N
6529 6528 6530 6531 3 0.00000 0.00000 0.00000
0.00000 0.00000 0.00000 ; H- N- CA- HA
6529 6528 6530 6532 3 0.00000 0.00000 0.00000
0.00000 0.00000 0.00000 ; H- N- CA- CB
6529 6528 6530 6545 3 0.00000 0.00000 0.00000
0.00000 0.00000 0.00000 ; H- N- CA- C
6530 6532 6534 6535 3 0.66944 2.00832 0.00000
-2.67776 0.00000 0.00000 ; CA- CB- CG2- HG21
6530 6532 6534 6536 3 0.66944 2.00832 0.00000
-2.67776 0.00000 0.00000 ; CA- CB- CG2- HG22
6530 6532 6534 6537 3 0.66944 2.00832 0.00000
-2.67776 0.00000 0.00000 ; CA- CB- CG2- HG23
6530 6532 6538 6539 3 0.66944 2.00832 0.00000
-2.67776 0.00000 0.00000 ; CA- CB- CG1- HG12
6530 6532 6538 6540 3 0.66944 2.00832 0.00000
-2.67776 0.00000 0.00000 ; CA- CB- CG1- HG13
6530 6532 6538 6541 3 4.24258 -0.52718 -7.25506
-1.79075 7.69856 0.00000 ; CA- CB- CG1- CD1
6530 6545 6547 6548 3 20.92000 0.00000 -20.92000
0.00000 0.00000 0.00000 ; CA- C- N- H

6530 6545 6547 6549 3 20.92000 0.00000 -20.92000
0.00000 0.00000 0.00000 ; CA- C- N- CA
6531 6530 6532 6533 3 0.65084 1.95253 0.00000
-2.60338 0.00000 0.00000 ; HA- CA- CB- HB
6531 6530 6532 6534 3 0.65084 1.95253 0.00000
-2.60338 0.00000 0.00000 ; HA- CA- CB- CG2
6531 6530 6532 6538 3 0.65084 1.95253 0.00000
-2.60338 0.00000 0.00000 ; HA- CA- CB- CG1
6531 6530 6545 6546 3 3.68192 -4.35136 0.00000
1.33888 0.00000 0.00000 ; HA- CA- C- O
6531 6530 6545 6547 3 0.00000 0.00000 0.00000
0.00000 0.00000 0.00000 ; HA- CA- C- N
6532 6530 6545 6546 3 0.00000 0.00000 0.00000
0.00000 0.00000 0.00000 ; CB- CA- C- O
6532 6530 6545 6547 3 2.51040 4.18400 1.67360
-6.69440 0.00000 0.00000 ; CB- CA- C- N
6532 6538 6541 6542 3 0.66944 2.00832 0.00000
-2.67776 0.00000 0.00000 ; CB- CG1- CD1- HD11
6532 6538 6541 6543 3 0.66944 2.00832 0.00000
-2.67776 0.00000 0.00000 ; CB- CG1- CD1- HD12
6532 6538 6541 6544 3 0.66944 2.00832 0.00000
-2.67776 0.00000 0.00000 ; CB- CG1- CD1- HD13
6533 6532 6530 6545 3 0.65084 1.95253 0.00000
-2.60338 0.00000 0.00000 ; HB- CB- CA- C
6533 6532 6534 6535 3 0.62760 1.88280 0.00000
-2.51040 0.00000 0.00000 ; HB- CB- CG2- HG21
6533 6532 6534 6536 3 0.62760 1.88280 0.00000
-2.51040 0.00000 0.00000 ; HB- CB- CG2- HG22
6533 6532 6534 6537 3 0.62760 1.88280 0.00000
-2.51040 0.00000 0.00000 ; HB- CB- CG2- HG23
6533 6532 6538 6539 3 0.62760 1.88280 0.00000
-2.51040 0.00000 0.00000 ; HB- CB- CG1- HG12
6533 6532 6538 6540 3 0.62760 1.88280 0.00000
-2.51040 0.00000 0.00000 ; HB- CB- CG1- HG13
6533 6532 6538 6541 3 0.66944 2.00832 0.00000
-2.67776 0.00000 0.00000 ; HB- CB- CG1- CD1
6534 6532 6530 6545 3 5.67350 3.55640 -6.16722
-2.47693 3.74886 0.00000 ; CG2- CB- CA- C
6534 6532 6538 6539 3 0.66944 2.00832 0.00000
-2.67776 0.00000 0.00000 ; CG2- CB- CG1- HG12
6534 6532 6538 6540 3 0.66944 2.00832 0.00000
-2.67776 0.00000 0.00000 ; CG2- CB- CG1- HG13
6534 6532 6538 6541 3 3.81162 0.49790 -8.14206
-1.79075 7.49773 0.00000 ; CG2- CB- CG1- CD1
6535 6534 6532 6538 3 0.66944 2.00832 0.00000
-2.67776 0.00000 0.00000 ; HG21- CG2- CB- CG1
6536 6534 6532 6538 3 0.66944 2.00832 0.00000
-2.67776 0.00000 0.00000 ; HG22- CG2- CB- CG1
6537 6534 6532 6538 3 0.66944 2.00832 0.00000
-2.67776 0.00000 0.00000 ; HG23- CG2- CB- CG1
6538 6532 6530 6545 3 8.26340 0.74057 -9.99976
-1.89117 3.84928 0.00000 ; CG1- CB- CA- C

6539 6538 6541 6542 3 0.62760 1.88280 0.00000
-2.51040 0.00000 0.00000 ; HG12- CG1- CD1- HD11
6539 6538 6541 6543 3 0.62760 1.88280 0.00000
-2.51040 0.00000 0.00000 ; HG12- CG1- CD1- HD12
6539 6538 6541 6544 3 0.62760 1.88280 0.00000
-2.51040 0.00000 0.00000 ; HG12- CG1- CD1- HD13
6540 6538 6541 6542 3 0.62760 1.88280 0.00000
-2.51040 0.00000 0.00000 ; HG13- CG1- CD1- HD11
6540 6538 6541 6543 3 0.62760 1.88280 0.00000
-2.51040 0.00000 0.00000 ; HG13- CG1- CD1- HD12
6540 6538 6541 6544 3 0.62760 1.88280 0.00000
-2.51040 0.00000 0.00000 ; HG13- CG1- CD1- HD13
6545 6547 6549 6550 3 0.00000 0.00000 0.00000
0.00000 0.00000 0.00000 ; C- N- CA- HA
6545 6547 6549 6551 3 11.71520 1.67360 15.06240 -
13.38880 0.00000 0.00000 ; C- N- CA- CB
6545 6547 6549 6556 3 1.75728 5.27184 2.25936
-7.02912 0.00000 0.00000 ; C- N- CA- C
6546 6545 6547 6548 3 29.28800 -8.36800 -20.92000
0.00000 0.00000 0.00000 ; O- C- N- H
6546 6545 6547 6549 3 20.92000 0.00000 -20.92000
0.00000 0.00000 0.00000 ; O- C- N- CA
6547 6549 6551 6552 3 0.65084 1.95253 0.00000
-2.60338 0.00000 0.00000 ; N- CA- CB- HB2
6547 6549 6551 6553 3 0.65084 1.95253 0.00000
-2.60338 0.00000 0.00000 ; N- CA- CB- HB3
6547 6549 6551 6554 3 7.86174 2.24681 -7.41405
-6.71114 5.35552 0.00000 ; N- CA- CB- OG
6547 6549 6556 6557 3 0.00000 0.00000 0.00000
0.00000 0.00000 0.00000 ; N- CA- C- O
6547 6549 6556 6558 3 17.40544 -5.02080 -13.22144
9.20480 0.00000 0.00000 ; N- CA- C- N
6548 6547 6549 6550 3 0.00000 0.00000 0.00000
0.00000 0.00000 0.00000 ; H- N- CA- HA
6548 6547 6549 6551 3 0.00000 0.00000 0.00000
0.00000 0.00000 0.00000 ; H- N- CA- CB
6548 6547 6549 6556 3 0.00000 0.00000 0.00000
0.00000 0.00000 0.00000 ; H- N- CA- C
6549 6551 6554 6555 3 2.05853 2.46856 3.48109
-4.46851 0.23430 0.00000 ; CA- CB- OG- HG
6549 6556 6558 6559 3 20.92000 0.00000 -20.92000
0.00000 0.00000 0.00000 ; CA- C- N- H
6549 6556 6558 6560 3 20.92000 0.00000 -20.92000
0.00000 0.00000 0.00000 ; CA- C- N- CA
6550 6549 6551 6552 3 0.65084 1.95253 0.00000
-2.60338 0.00000 0.00000 ; HA- CA- CB- HB2
6550 6549 6551 6553 3 0.65084 1.95253 0.00000
-2.60338 0.00000 0.00000 ; HA- CA- CB- HB3
6550 6549 6551 6554 3 1.04600 -1.04600 0.00000
0.00000 0.00000 0.00000 ; HA- CA- CB- OG
6550 6549 6556 6557 3 3.68192 -4.35136 0.00000
1.33888 0.00000 0.00000 ; HA- CA- C- O

6550 6549 6556 6558 3 0.00000 0.00000 0.00000
0.00000 0.00000 0.00000 ; HA- CA- C- N
6551 6549 6556 6557 3 0.00000 0.00000 0.00000
0.00000 0.00000 0.00000 ; CB- CA- C- O
6551 6549 6556 6558 3 2.51040 4.18400 1.67360
-6.69440 0.00000 0.00000 ; CB- CA- C- N
6552 6551 6549 6556 3 0.65084 1.95253 0.00000
-2.60338 0.00000 0.00000 ; HB2- CB- CA- C
6552 6551 6554 6555 3 0.69733 2.09200 0.00000
-2.78933 0.00000 0.00000 ; HB2- CB- OG- HG
6553 6551 6549 6556 3 0.65084 1.95253 0.00000
-2.60338 0.00000 0.00000 ; HB3- CB- CA- C
6553 6551 6554 6555 3 0.69733 2.09200 0.00000
-2.78933 0.00000 0.00000 ; HB3- CB- OG- HG
6554 6551 6549 6556 3 7.34710 7.79898 -6.14211
-6.71114 4.31789 0.00000 ; OG- CB- CA- C
6556 6558 6560 6561 3 0.00000 0.00000 0.00000
0.00000 0.00000 0.00000 ; C- N- CA- HA
6556 6558 6560 6562 3 11.71520 1.67360 15.06240 -
13.38880 0.00000 0.00000 ; C- N- CA- CB
6556 6558 6560 6572 3 1.75728 5.27184 2.25936
-7.02912 0.00000 0.00000 ; C- N- CA- C
6557 6556 6558 6559 3 29.28800 -8.36800 -20.92000
0.00000 0.00000 0.00000 ; O- C- N- H
6557 6556 6558 6560 3 20.92000 0.00000 -20.92000
0.00000 0.00000 0.00000 ; O- C- N- CA
6558 6560 6562 6563 3 0.65084 1.95253 0.00000
-2.60338 0.00000 0.00000 ; N- CA- CB- HB
6558 6560 6562 6564 3 3.83673 0.44769 -1.77402
-2.47693 -0.03347 0.00000 ; N- CA- CB- CG1
6558 6560 6562 6568 3 3.83673 0.44769 -1.77402
-2.47693 -0.03347 0.00000 ; N- CA- CB- CG2
6558 6560 6572 6573 3 0.00000 0.00000 0.00000
0.00000 0.00000 0.00000 ; N- CA- C- O
6558 6560 6572 6574 3 17.40544 -5.02080 -13.22144
9.20480 0.00000 0.00000 ; N- CA- C- N
6559 6558 6560 6561 3 0.00000 0.00000 0.00000
0.00000 0.00000 0.00000 ; H- N- CA- HA
6559 6558 6560 6562 3 0.00000 0.00000 0.00000
0.00000 0.00000 0.00000 ; H- N- CA- CB
6559 6558 6560 6572 3 0.00000 0.00000 0.00000
0.00000 0.00000 0.00000 ; H- N- CA- C
6560 6562 6564 6565 3 0.66944 2.00832 0.00000
-2.67776 0.00000 0.00000 ; CA- CB- CG1- HG11
6560 6562 6564 6566 3 0.66944 2.00832 0.00000
-2.67776 0.00000 0.00000 ; CA- CB- CG1- HG12
6560 6562 6564 6567 3 0.66944 2.00832 0.00000
-2.67776 0.00000 0.00000 ; CA- CB- CG1- HG13
6560 6562 6568 6569 3 0.66944 2.00832 0.00000
-2.67776 0.00000 0.00000 ; CA- CB- CG2- HG21
6560 6562 6568 6570 3 0.66944 2.00832 0.00000
-2.67776 0.00000 0.00000 ; CA- CB- CG2- HG22

6560 6562 6568 6571 3 0.66944 2.00832 0.00000
-2.67776 0.00000 0.00000 ; CA- CB- CG2- HG23
6560 6572 6574 6575 3 20.92000 0.00000 -20.92000
0.00000 0.00000 0.00000 ; CA- C- N- H
6560 6572 6574 6576 3 20.92000 0.00000 -20.92000
0.00000 0.00000 0.00000 ; CA- C- N- CA
6561 6560 6562 6563 3 0.65084 1.95253 0.00000
-2.60338 0.00000 0.00000 ; HA- CA- CB- HB
6561 6560 6562 6564 3 0.65084 1.95253 0.00000
-2.60338 0.00000 0.00000 ; HA- CA- CB- CG1
6561 6560 6562 6568 3 0.65084 1.95253 0.00000
-2.60338 0.00000 0.00000 ; HA- CA- CB- CG2
6561 6560 6572 6573 3 3.68192 -4.35136 0.00000
1.33888 0.00000 0.00000 ; HA- CA- C- O
6561 6560 6572 6574 3 0.00000 0.00000 0.00000
0.00000 0.00000 0.00000 ; HA- CA- C- N
6562 6560 6572 6573 3 0.00000 0.00000 0.00000
0.00000 0.00000 0.00000 ; CB- CA- C- O
6562 6560 6572 6574 3 2.51040 4.18400 1.67360
-6.69440 0.00000 0.00000 ; CB- CA- C- N
6563 6562 6560 6572 3 0.65084 1.95253 0.00000
-2.60338 0.00000 0.00000 ; HB- CB- CA- C
6563 6562 6564 6565 3 0.62760 1.88280 0.00000
-2.51040 0.00000 0.00000 ; HB- CB- CG1- HG11
6563 6562 6564 6566 3 0.62760 1.88280 0.00000
-2.51040 0.00000 0.00000 ; HB- CB- CG1- HG12
6563 6562 6564 6567 3 0.62760 1.88280 0.00000
-2.51040 0.00000 0.00000 ; HB- CB- CG1- HG13
6563 6562 6568 6569 3 0.62760 1.88280 0.00000
-2.51040 0.00000 0.00000 ; HB- CB- CG2- HG21
6563 6562 6568 6570 3 0.62760 1.88280 0.00000
-2.51040 0.00000 0.00000 ; HB- CB- CG2- HG22
6563 6562 6568 6571 3 0.62760 1.88280 0.00000
-2.51040 0.00000 0.00000 ; HB- CB- CG2- HG23
6564 6562 6560 6572 3 5.67350 3.55640 -6.16722
-2.47693 3.74886 0.00000 ; CG1- CB- CA- C
6564 6562 6568 6569 3 0.66944 2.00832 0.00000
-2.67776 0.00000 0.00000 ; CG1- CB- CG2- HG21
6564 6562 6568 6570 3 0.66944 2.00832 0.00000
-2.67776 0.00000 0.00000 ; CG1- CB- CG2- HG22
6564 6562 6568 6571 3 0.66944 2.00832 0.00000
-2.67776 0.00000 0.00000 ; CG1- CB- CG2- HG23
6565 6564 6562 6568 3 0.66944 2.00832 0.00000
-2.67776 0.00000 0.00000 ; HG11- CG1- CB- CG2
6566 6564 6562 6568 3 0.66944 2.00832 0.00000
-2.67776 0.00000 0.00000 ; HG12- CG1- CB- CG2
6567 6564 6562 6568 3 0.66944 2.00832 0.00000
-2.67776 0.00000 0.00000 ; HG13- CG1- CB- CG2
6568 6562 6560 6572 3 5.67350 3.55640 -6.16722
-2.47693 3.74886 0.00000 ; CG2- CB- CA- C
6572 6574 6576 6577 3 0.00000 0.00000 0.00000
0.00000 0.00000 0.00000 ; C- N- CA- HA

6572 6574 6576 6578 3 11.71520 1.67360 15.06240 -
13.38880 0.00000 0.00000 ; C- N- CA- CB
6572 6574 6576 6587 3 1.75728 5.27184 2.25936
-7.02912 0.00000 0.00000 ; C- N- CA- C
6573 6572 6574 6575 3 29.28800 -8.36800 -20.92000
0.00000 0.00000 0.00000 ; O- C- N- H
6573 6572 6574 6576 3 20.92000 0.00000 -20.92000
0.00000 0.00000 0.00000 ; O- C- N- CA
6574 6576 6578 6579 3 0.65084 1.95253 0.00000
-2.60338 0.00000 0.00000 ; N- CA- CB- HB2
6574 6576 6578 6580 3 0.65084 1.95253 0.00000
-2.60338 0.00000 0.00000 ; N- CA- CB- HB3
6574 6576 6578 6581 3 3.21331 2.22589 -4.15053
-2.40998 2.61082 0.00000 ; N- CA- CB- CG
6574 6576 6587 6588 3 0.00000 0.00000 0.00000
0.00000 0.00000 0.00000 ; N- CA- C- O
6574 6576 6587 6589 3 17.40544 -5.02080 -13.22144
9.20480 0.00000 0.00000 ; N- CA- C- N
6575 6574 6576 6577 3 0.00000 0.00000 0.00000
0.00000 0.00000 0.00000 ; H- N- CA- HA
6575 6574 6576 6578 3 0.00000 0.00000 0.00000
0.00000 0.00000 0.00000 ; H- N- CA- CB
6575 6574 6576 6587 3 0.00000 0.00000 0.00000
0.00000 0.00000 0.00000 ; H- N- CA- C
6576 6578 6581 6582 3 0.66944 2.00832 0.00000
-2.67776 0.00000 0.00000 ; CA- CB- CG- HG2
6576 6578 6581 6583 3 0.66944 2.00832 0.00000
-2.67776 0.00000 0.00000 ; CA- CB- CG- HG3
6576 6578 6581 6584 3 10.12110 -1.91209 0.01674
10.17549 -1.87443 0.00000 ; CA- CB- CG- CD
6576 6587 6589 6590 3 20.92000 0.00000 -20.92000
0.00000 0.00000 0.00000 ; CA- C- N- H
6576 6587 6589 6591 3 20.92000 0.00000 -20.92000
0.00000 0.00000 0.00000 ; CA- C- N- CA
6577 6576 6578 6579 3 0.65084 1.95253 0.00000
-2.60338 0.00000 0.00000 ; HA- CA- CB- HB2
6577 6576 6578 6580 3 0.65084 1.95253 0.00000
-2.60338 0.00000 0.00000 ; HA- CA- CB- HB3
6577 6576 6578 6581 3 0.65084 1.95253 0.00000
-2.60338 0.00000 0.00000 ; HA- CA- CB- CG
6577 6576 6587 6588 3 3.68192 -4.35136 0.00000
1.33888 0.00000 0.00000 ; HA- CA- C- O
6577 6576 6587 6589 3 0.00000 0.00000 0.00000
0.00000 0.00000 0.00000 ; HA- CA- C- N
6578 6576 6587 6588 3 0.00000 0.00000 0.00000
0.00000 0.00000 0.00000 ; CB- CA- C- O
6578 6576 6587 6589 3 2.51040 4.18400 1.67360
-6.69440 0.00000 0.00000 ; CB- CA- C- N
6578 6581 6584 6585 3 3.79907 0.00000 -5.40573
0.00000 2.14221 0.00000 ; CB- CG- CD- OE1
6578 6581 6584 6586 3 3.79907 0.00000 -5.40573
0.00000 2.14221 0.00000 ; CB- CG- CD- OE2

```
  6579   6578   6576   6587      3    0.65084    1.95253    0.00000
-2.60338    0.00000    0.00000 ;    HB2-    CB-    CA-     C
  6579   6578   6581   6582      3    0.62760    1.88280    0.00000
-2.51040    0.00000    0.00000 ;    HB2-    CB-    CG-   HG2
  6579   6578   6581   6583      3    0.62760    1.88280    0.00000
-2.51040    0.00000    0.00000 ;    HB2-    CB-    CG-   HG3
  6579   6578   6581   6584      3    0.65084    1.95253    0.00000
-2.60338    0.00000    0.00000 ;    HB2-    CB-    CG-    CD
  6580   6578   6576   6587      3    0.65084    1.95253    0.00000
-2.60338    0.00000    0.00000 ;    HB3-    CB-    CA-     C
  6580   6578   6581   6582      3    0.62760    1.88280    0.00000
-2.51040    0.00000    0.00000 ;    HB3-    CB-    CG-   HG2
  6580   6578   6581   6583      3    0.62760    1.88280    0.00000
-2.51040    0.00000    0.00000 ;    HB3-    CB-    CG-   HG3
  6580   6578   6581   6584      3    0.65084    1.95253    0.00000
-2.60338    0.00000    0.00000 ;    HB3-    CB-    CG-    CD
  6581   6578   6576   6587      3    6.86594    3.56895   -8.14206
-2.40998    4.85344    0.00000 ;     CG-    CB-    CA-     C
  6582   6581   6584   6585      3    0.00000    0.00000    0.00000
0.00000    0.00000    0.00000 ;    HG2-    CG-    CD-   OE1
  6582   6581   6584   6586      3    0.00000    0.00000    0.00000
0.00000    0.00000    0.00000 ;    HG2-    CG-    CD-   OE2
  6583   6581   6584   6585      3    0.00000    0.00000    0.00000
0.00000    0.00000    0.00000 ;    HG3-    CG-    CD-   OE1
  6583   6581   6584   6586      3    0.00000    0.00000    0.00000
0.00000    0.00000    0.00000 ;    HG3-    CG-    CD-   OE2
  6587   6589   6591   6592      3    0.00000    0.00000    0.00000
0.00000    0.00000    0.00000 ;      C-     N-    CA-    HA
  6587   6589   6591   6593      3   11.71520    1.67360   15.06240  -
13.38880    0.00000    0.00000 ;      C-     N-    CA-    CB
  6587   6589   6591   6603      3    1.75728    5.27184    2.25936
-7.02912    0.00000    0.00000 ;      C-     N-    CA-     C
  6588   6587   6589   6590      3   29.28800   -8.36800  -20.92000
0.00000    0.00000    0.00000 ;      O-     C-     N-     H
  6588   6587   6589   6591      3   20.92000    0.00000  -20.92000
0.00000    0.00000    0.00000 ;      O-     C-     N-    CA
  6589   6591   6593   6594      3    0.65084    1.95253    0.00000
-2.60338    0.00000    0.00000 ;      N-    CA-    CB-   HB2
  6589   6591   6593   6595      3    0.65084    1.95253    0.00000
-2.60338    0.00000    0.00000 ;      N-    CA-    CB-   HB3
  6589   6591   6593   6596      3    3.21331    2.22589   -4.15053
-2.40998    2.61082    0.00000 ;      N-    CA-    CB-    CG
  6589   6591   6603   6604      3    0.00000    0.00000    0.00000
0.00000    0.00000    0.00000 ;      N-    CA-     C-     O
  6589   6591   6603   6605      3   17.40544   -5.02080  -13.22144
9.20480    0.00000    0.00000 ;      N-    CA-     C-     N
  6590   6589   6591   6592      3    0.00000    0.00000    0.00000
0.00000    0.00000    0.00000 ;      H-     N-    CA-    HA
  6590   6589   6591   6593      3    0.00000    0.00000    0.00000
0.00000    0.00000    0.00000 ;      H-     N-    CA-    CB
  6590   6589   6591   6603      3    0.00000    0.00000    0.00000
0.00000    0.00000    0.00000 ;      H-     N-    CA-     C
```

6591 6593 6596 6597 3 0.66944 2.00832 0.00000
-2.67776 0.00000 0.00000 ; CA- CB- CG- HG2
6591 6593 6596 6598 3 0.66944 2.00832 0.00000
-2.67776 0.00000 0.00000 ; CA- CB- CG- HG3
6591 6593 6596 6599 3 3.69866 -4.35136 -3.92459
6.89523 4.61914 0.00000 ; CA- CB- CG- CD
6591 6603 6605 6606 3 20.92000 0.00000 -20.92000
0.00000 0.00000 0.00000 ; CA- C- N- H
6591 6603 6605 6607 3 20.92000 0.00000 -20.92000
0.00000 0.00000 0.00000 ; CA- C- N- CA
6592 6591 6593 6594 3 0.65084 1.95253 0.00000
-2.60338 0.00000 0.00000 ; HA- CA- CB- HB2
6592 6591 6593 6595 3 0.65084 1.95253 0.00000
-2.60338 0.00000 0.00000 ; HA- CA- CB- HB3
6592 6591 6593 6596 3 0.65084 1.95253 0.00000
-2.60338 0.00000 0.00000 ; HA- CA- CB- CG
6592 6591 6603 6604 3 3.68192 -4.35136 0.00000
1.33888 0.00000 0.00000 ; HA- CA- C- O
6592 6591 6603 6605 3 0.00000 0.00000 0.00000
0.00000 0.00000 0.00000 ; HA- CA- C- N
6593 6591 6603 6604 3 0.00000 0.00000 0.00000
0.00000 0.00000 0.00000 ; CB- CA- C- O
6593 6591 6603 6605 3 2.51040 4.18400 1.67360
-6.69440 0.00000 0.00000 ; CB- CA- C- N
6593 6596 6599 6600 3 0.00000 0.00000 0.00000
0.00000 0.00000 0.00000 ; CB- CG- CD- OE1
6593 6596 6599 6601 3 12.79049 3.13382 -7.02912
0.41840 -2.20915 0.00000 ; CB- CG- CD- OE2
6594 6593 6591 6603 3 0.65084 1.95253 0.00000
-2.60338 0.00000 0.00000 ; HB2- CB- CA- C
6594 6593 6596 6597 3 0.62760 1.88280 0.00000
-2.51040 0.00000 0.00000 ; HB2- CB- CG- HG2
6594 6593 6596 6598 3 0.62760 1.88280 0.00000
-2.51040 0.00000 0.00000 ; HB2- CB- CG- HG3
6594 6593 6596 6599 3 0.65084 1.95253 0.00000
-2.60338 0.00000 0.00000 ; HB2- CB- CG- CD
6595 6593 6591 6603 3 0.65084 1.95253 0.00000
-2.60338 0.00000 0.00000 ; HB3- CB- CA- C
6595 6593 6596 6597 3 0.62760 1.88280 0.00000
-2.51040 0.00000 0.00000 ; HB3- CB- CG- HG2
6595 6593 6596 6598 3 0.62760 1.88280 0.00000
-2.51040 0.00000 0.00000 ; HB3- CB- CG- HG3
6595 6593 6596 6599 3 0.65084 1.95253 0.00000
-2.60338 0.00000 0.00000 ; HB3- CB- CG- CD
6596 6593 6591 6603 3 6.86594 3.56895 -8.14206
-2.40998 4.85344 0.00000 ; CG- CB- CA- C
6596 6599 6601 6602 3 27.46796 7.88684 -26.42614
-8.01654 3.78234 0.00000 ; CG- CD- OE2- HE2
6597 6596 6599 6600 3 3.68192 -4.35136 0.00000
1.33888 0.00000 0.00000 ; HG2- CG- CD- OE1
6597 6596 6599 6601 3 0.00000 0.00000 0.00000
0.00000 0.00000 0.00000 ; HG2- CG- CD- OE2

6598 6596 6599 6600 3 3.68192 -4.35136 0.00000
1.33888 0.00000 0.00000 ; HG3- CG- CD- OE1
6598 6596 6599 6601 3 0.00000 0.00000 0.00000
0.00000 0.00000 0.00000 ; HG3- CG- CD- OE2
6600 6599 6601 6602 3 27.19600 -7.94960 -19.24640
0.00000 0.00000 0.00000 ; OE1- CD- OE2- HE2
6603 6605 6607 6608 3 0.00000 0.00000 0.00000
0.00000 0.00000 0.00000 ; C- N- CA- HA
6603 6605 6607 6609 3 11.71520 1.67360 15.06240 -
13.38880 0.00000 0.00000 ; C- N- CA- CB
6603 6605 6607 6622 3 1.75728 5.27184 2.25936
-7.02912 0.00000 0.00000 ; C- N- CA- C
6604 6603 6605 6606 3 29.28800 -8.36800 -20.92000
0.00000 0.00000 0.00000 ; O- C- N- H
6604 6603 6605 6607 3 20.92000 0.00000 -20.92000
0.00000 0.00000 0.00000 ; O- C- N- CA
6605 6607 6609 6610 3 0.65084 1.95253 0.00000
-2.60338 0.00000 0.00000 ; N- CA- CB- HB2
6605 6607 6609 6611 3 0.65084 1.95253 0.00000
-2.60338 0.00000 0.00000 ; N- CA- CB- HB3
6605 6607 6609 6612 3 3.79070 1.39746 -4.61077
-2.40998 2.44346 0.00000 ; N- CA- CB- CG
6605 6607 6622 6623 3 0.00000 0.00000 0.00000
0.00000 0.00000 0.00000 ; N- CA- C- O
6605 6607 6622 6624 3 17.40544 -5.02080 -13.22144
9.20480 0.00000 0.00000 ; N- CA- C- N
6606 6605 6607 6608 3 0.00000 0.00000 0.00000
0.00000 0.00000 0.00000 ; H- N- CA- HA
6606 6605 6607 6609 3 0.00000 0.00000 0.00000
0.00000 0.00000 0.00000 ; H- N- CA- CB
6606 6605 6607 6622 3 0.00000 0.00000 0.00000
0.00000 0.00000 0.00000 ; H- N- CA- C
6607 6609 6612 6613 3 0.66944 2.00832 0.00000
-2.67776 0.00000 0.00000 ; CA- CB- CG- HG
6607 6609 6612 6614 3 3.90367 0.19665 -6.21742
-2.37651 5.99149 0.00000 ; CA- CB- CG- CD1
6607 6609 6612 6618 3 3.90367 0.19665 -6.21742
-2.37651 5.99149 0.00000 ; CA- CB- CG- CD2
6607 6622 6624 6625 3 20.92000 0.00000 -20.92000
0.00000 0.00000 0.00000 ; CA- C- N- H
6607 6622 6624 6626 3 20.92000 0.00000 -20.92000
0.00000 0.00000 0.00000 ; CA- C- N- CA
6608 6607 6609 6610 3 0.65084 1.95253 0.00000
-2.60338 0.00000 0.00000 ; HA- CA- CB- HB2
6608 6607 6609 6611 3 0.65084 1.95253 0.00000
-2.60338 0.00000 0.00000 ; HA- CA- CB- HB3
6608 6607 6609 6612 3 0.65084 1.95253 0.00000
-2.60338 0.00000 0.00000 ; HA- CA- CB- CG
6608 6607 6622 6623 3 3.68192 -4.35136 0.00000
1.33888 0.00000 0.00000 ; HA- CA- C- O
6608 6607 6622 6624 3 0.00000 0.00000 0.00000
0.00000 0.00000 0.00000 ; HA- CA- C- N

```
  6609   6607   6622   6623      3    0.00000    0.00000    0.00000
0.00000    0.00000    0.00000 ;     CB-    CA-     C-     O
  6609   6607   6622   6624      3    2.51040    4.18400    1.67360
-6.69440    0.00000    0.00000 ;     CB-    CA-     C-     N
  6609   6612   6614   6615      3    0.66944    2.00832    0.00000
-2.67776    0.00000    0.00000 ;     CB-    CG-   CD1-  HD11
  6609   6612   6614   6616      3    0.66944    2.00832    0.00000
-2.67776    0.00000    0.00000 ;     CB-    CG-   CD1-  HD12
  6609   6612   6614   6617      3    0.66944    2.00832    0.00000
-2.67776    0.00000    0.00000 ;     CB-    CG-   CD1-  HD13
  6609   6612   6618   6619      3    0.66944    2.00832    0.00000
-2.67776    0.00000    0.00000 ;     CB-    CG-   CD2-  HD21
  6609   6612   6618   6620      3    0.66944    2.00832    0.00000
-2.67776    0.00000    0.00000 ;     CB-    CG-   CD2-  HD22
  6609   6612   6618   6621      3    0.66944    2.00832    0.00000
-2.67776    0.00000    0.00000 ;     CB-    CG-   CD2-  HD23
  6610   6609   6607   6622      3    0.65084    1.95253    0.00000
-2.60338    0.00000    0.00000 ;    HB2-    CB-    CA-     C
  6610   6609   6612   6613      3    0.62760    1.88280    0.00000
-2.51040    0.00000    0.00000 ;    HB2-    CB-    CG-    HG
  6610   6609   6612   6614      3    0.66944    2.00832    0.00000
-2.67776    0.00000    0.00000 ;    HB2-    CB-    CG-   CD1
  6610   6609   6612   6618      3    0.66944    2.00832    0.00000
-2.67776    0.00000    0.00000 ;    HB2-    CB-    CG-   CD2
  6611   6609   6607   6622      3    0.65084    1.95253    0.00000
-2.60338    0.00000    0.00000 ;    HB3-    CB-    CA-     C
  6611   6609   6612   6613      3    0.62760    1.88280    0.00000
-2.51040    0.00000    0.00000 ;    HB3-    CB-    CG-    HG
  6611   6609   6612   6614      3    0.66944    2.00832    0.00000
-2.67776    0.00000    0.00000 ;    HB3-    CB-    CG-   CD1
  6611   6609   6612   6618      3    0.66944    2.00832    0.00000
-2.67776    0.00000    0.00000 ;    HB3-    CB-    CG-   CD2
  6612   6609   6607   6622      3   10.33448   -1.14642  -11.54784
-2.40998    6.35968    0.00000 ;     CG-    CB-    CA-     C
  6613   6612   6614   6615      3    0.62760    1.88280    0.00000
-2.51040    0.00000    0.00000 ;     HG-    CG-   CD1-  HD11
  6613   6612   6614   6616      3    0.62760    1.88280    0.00000
-2.51040    0.00000    0.00000 ;     HG-    CG-   CD1-  HD12
  6613   6612   6614   6617      3    0.62760    1.88280    0.00000
-2.51040    0.00000    0.00000 ;     HG-    CG-   CD1-  HD13
  6613   6612   6618   6619      3    0.62760    1.88280    0.00000
-2.51040    0.00000    0.00000 ;     HG-    CG-   CD2-  HD21
  6613   6612   6618   6620      3    0.62760    1.88280    0.00000
-2.51040    0.00000    0.00000 ;     HG-    CG-   CD2-  HD22
  6613   6612   6618   6621      3    0.62760    1.88280    0.00000
-2.51040    0.00000    0.00000 ;     HG-    CG-   CD2-  HD23
  6614   6612   6618   6619      3    0.66944    2.00832    0.00000
-2.67776    0.00000    0.00000 ;    CD1-    CG-   CD2-  HD21
  6614   6612   6618   6620      3    0.66944    2.00832    0.00000
-2.67776    0.00000    0.00000 ;    CD1-    CG-   CD2-  HD22
  6614   6612   6618   6621      3    0.66944    2.00832    0.00000
-2.67776    0.00000    0.00000 ;    CD1-    CG-   CD2-  HD23
```

6615 6614 6612 6618 3 0.66944 2.00832 0.00000
-2.67776 0.00000 0.00000 ; HD11- CD1- CG- CD2
6616 6614 6612 6618 3 0.66944 2.00832 0.00000
-2.67776 0.00000 0.00000 ; HD12- CD1- CG- CD2
6617 6614 6612 6618 3 0.66944 2.00832 0.00000
-2.67776 0.00000 0.00000 ; HD13- CD1- CG- CD2
6622 6624 6626 6627 3 0.00000 0.00000 0.00000
0.00000 0.00000 0.00000 ; C- N- CA- HA
6622 6624 6626 6628 3 11.71520 1.67360 15.06240 -
13.38880 0.00000 0.00000 ; C- N- CA- CB
6622 6624 6626 6644 3 1.75728 5.27184 2.25936
-7.02912 0.00000 0.00000 ; C- N- CA- C
6623 6622 6624 6625 3 29.28800 -8.36800 -20.92000
0.00000 0.00000 0.00000 ; O- C- N- H
6623 6622 6624 6626 3 20.92000 0.00000 -20.92000
0.00000 0.00000 0.00000 ; O- C- N- CA
6624 6626 6628 6629 3 0.65084 1.95253 0.00000
-2.60338 0.00000 0.00000 ; N- CA- CB- HB2
6624 6626 6628 6630 3 0.65084 1.95253 0.00000
-2.60338 0.00000 0.00000 ; N- CA- CB- HB3
6624 6626 6628 6631 3 0.65084 1.95253 0.00000
-2.60338 0.00000 0.00000 ; N- CA- CB- CG
6624 6626 6644 6645 3 0.00000 0.00000 0.00000
0.00000 0.00000 0.00000 ; N- CA- C- O
6624 6626 6644 6646 3 17.40544 -5.02080 -13.22144
9.20480 0.00000 0.00000 ; N- CA- C- N
6625 6624 6626 6627 3 0.00000 0.00000 0.00000
0.00000 0.00000 0.00000 ; H- N- CA- HA
6625 6624 6626 6628 3 0.00000 0.00000 0.00000
0.00000 0.00000 0.00000 ; H- N- CA- CB
6625 6624 6626 6644 3 0.00000 0.00000 0.00000
0.00000 0.00000 0.00000 ; H- N- CA- C
6626 6628 6631 6632 3 0.66944 2.00832 0.00000
-2.67776 0.00000 0.00000 ; CA- CB- CG- HG2
6626 6628 6631 6633 3 0.66944 2.00832 0.00000
-2.67776 0.00000 0.00000 ; CA- CB- CG- HG3
6626 6628 6631 6634 3 3.68192 3.09616 -2.09200
-3.01248 0.00000 0.00000 ; CA- CB- CG- CD
6626 6644 6646 6647 3 20.92000 0.00000 -20.92000
0.00000 0.00000 0.00000 ; CA- C- N- H
6626 6644 6646 6648 3 20.92000 0.00000 -20.92000
0.00000 0.00000 0.00000 ; CA- C- N- CA
6627 6626 6628 6629 3 0.65084 1.95253 0.00000
-2.60338 0.00000 0.00000 ; HA- CA- CB- HB2
6627 6626 6628 6630 3 0.65084 1.95253 0.00000
-2.60338 0.00000 0.00000 ; HA- CA- CB- HB3
6627 6626 6628 6631 3 0.65084 1.95253 0.00000
-2.60338 0.00000 0.00000 ; HA- CA- CB- CG
6627 6626 6644 6645 3 3.68192 -4.35136 0.00000
1.33888 0.00000 0.00000 ; HA- CA- C- O
6627 6626 6644 6646 3 0.00000 0.00000 0.00000
0.00000 0.00000 0.00000 ; HA- CA- C- N

6628 6626 6644 6645 3 0.00000 0.00000 0.00000
0.00000 0.00000 0.00000 ; CB- CA- C- O
6628 6626 6644 6646 3 2.51040 4.18400 1.67360
-6.69440 0.00000 0.00000 ; CB- CA- C- N
6628 6631 6634 6635 3 0.66944 2.00832 0.00000
-2.67776 0.00000 0.00000 ; CB- CG- CD- HD2
6628 6631 6634 6636 3 0.66944 2.00832 0.00000
-2.67776 0.00000 0.00000 ; CB- CG- CD- HD3
6628 6631 6634 6637 3 3.68192 3.09616 -2.09200
-3.01248 0.00000 0.00000 ; CB- CG- CD- CE
6629 6628 6626 6644 3 0.65084 1.95253 0.00000
-2.60338 0.00000 0.00000 ; HB2- CB- CA- C
6629 6628 6631 6632 3 0.62760 1.88280 0.00000
-2.51040 0.00000 0.00000 ; HB2- CB- CG- HG2
6629 6628 6631 6633 3 0.62760 1.88280 0.00000
-2.51040 0.00000 0.00000 ; HB2- CB- CG- HG3
6629 6628 6631 6634 3 0.66944 2.00832 0.00000
-2.67776 0.00000 0.00000 ; HB2- CB- CG- CD
6630 6628 6626 6644 3 0.65084 1.95253 0.00000
-2.60338 0.00000 0.00000 ; HB3- CB- CA- C
6630 6628 6631 6632 3 0.62760 1.88280 0.00000
-2.51040 0.00000 0.00000 ; HB3- CB- CG- HG2
6630 6628 6631 6633 3 0.62760 1.88280 0.00000
-2.51040 0.00000 0.00000 ; HB3- CB- CG- HG3
6630 6628 6631 6634 3 0.66944 2.00832 0.00000
-2.67776 0.00000 0.00000 ; HB3- CB- CG- CD
6631 6628 6626 6644 3 0.65084 1.95253 0.00000
-2.60338 0.00000 0.00000 ; CG- CB- CA- C
6631 6634 6637 6638 3 0.65084 1.95253 0.00000
-2.60338 0.00000 0.00000 ; CG- CD- CE- HE2
6631 6634 6637 6639 3 0.65084 1.95253 0.00000
-2.60338 0.00000 0.00000 ; CG- CD- CE- HE3
6631 6634 6637 6640 3 0.65084 1.95253 0.00000
-2.60338 0.00000 0.00000 ; CG- CD- CE- NZ
6632 6631 6634 6635 3 0.62760 1.88280 0.00000
-2.51040 0.00000 0.00000 ; HG2- CG- CD- HD2
6632 6631 6634 6636 3 0.62760 1.88280 0.00000
-2.51040 0.00000 0.00000 ; HG2- CG- CD- HD3
6632 6631 6634 6637 3 0.66944 2.00832 0.00000
-2.67776 0.00000 0.00000 ; HG2- CG- CD- CE
6633 6631 6634 6635 3 0.62760 1.88280 0.00000
-2.51040 0.00000 0.00000 ; HG3- CG- CD- HD2
6633 6631 6634 6636 3 0.62760 1.88280 0.00000
-2.51040 0.00000 0.00000 ; HG3- CG- CD- HD3
6633 6631 6634 6637 3 0.66944 2.00832 0.00000
-2.67776 0.00000 0.00000 ; HG3- CG- CD- CE
6634 6637 6640 6641 3 0.65084 1.95253 0.00000
-2.60338 0.00000 0.00000 ; CD- CE- NZ- HZ1
6634 6637 6640 6642 3 0.65084 1.95253 0.00000
-2.60338 0.00000 0.00000 ; CD- CE- NZ- HZ2
6634 6637 6640 6643 3 0.65084 1.95253 0.00000
-2.60338 0.00000 0.00000 ; CD- CE- NZ- HZ3

6635 6634 6637 6638 3 0.65084 1.95253 0.00000
-2.60338 0.00000 0.00000 ; HD2- CD- CE- HE2
6635 6634 6637 6639 3 0.65084 1.95253 0.00000
-2.60338 0.00000 0.00000 ; HD2- CD- CE- HE3
6635 6634 6637 6640 3 0.65084 1.95253 0.00000
-2.60338 0.00000 0.00000 ; HD2- CD- CE- NZ
6636 6634 6637 6638 3 0.65084 1.95253 0.00000
-2.60338 0.00000 0.00000 ; HD3- CD- CE- HE2
6636 6634 6637 6639 3 0.65084 1.95253 0.00000
-2.60338 0.00000 0.00000 ; HD3- CD- CE- HE3
6636 6634 6637 6640 3 0.65084 1.95253 0.00000
-2.60338 0.00000 0.00000 ; HD3- CD- CE- NZ
6638 6637 6640 6641 3 0.65084 1.95253 0.00000
-2.60338 0.00000 0.00000 ; HE2- CE- NZ- HZ1
6638 6637 6640 6642 3 0.65084 1.95253 0.00000
-2.60338 0.00000 0.00000 ; HE2- CE- NZ- HZ2
6638 6637 6640 6643 3 0.65084 1.95253 0.00000
-2.60338 0.00000 0.00000 ; HE2- CE- NZ- HZ3
6639 6637 6640 6641 3 0.65084 1.95253 0.00000
-2.60338 0.00000 0.00000 ; HE3- CE- NZ- HZ1
6639 6637 6640 6642 3 0.65084 1.95253 0.00000
-2.60338 0.00000 0.00000 ; HE3- CE- NZ- HZ2
6639 6637 6640 6643 3 0.65084 1.95253 0.00000
-2.60338 0.00000 0.00000 ; HE3- CE- NZ- HZ3
6644 6646 6648 6649 3 0.00000 0.00000 0.00000
0.00000 0.00000 0.00000 ; C- N- CA- HA
6644 6646 6648 6650 3 11.71520 1.67360 15.06240 -
13.38880 0.00000 0.00000 ; C- N- CA- CB
6644 6646 6648 6668 3 1.75728 5.27184 2.25936
-7.02912 0.00000 0.00000 ; C- N- CA- C
6645 6644 6646 6647 3 29.28800 -8.36800 -20.92000
0.00000 0.00000 0.00000 ; O- C- N- H
6645 6644 6646 6648 3 20.92000 0.00000 -20.92000
0.00000 0.00000 0.00000 ; O- C- N- CA
6646 6648 6650 6651 3 0.65084 1.95253 0.00000
-2.60338 0.00000 0.00000 ; N- CA- CB- HB2
6646 6648 6650 6652 3 0.65084 1.95253 0.00000
-2.60338 0.00000 0.00000 ; N- CA- CB- HB3
6646 6648 6650 6653 3 0.65084 1.95253 0.00000
-2.60338 0.00000 0.00000 ; N- CA- CB- CG
6646 6648 6668 6669 3 0.00000 0.00000 0.00000
0.00000 0.00000 0.00000 ; N- CA- C- O
6646 6648 6668 6670 3 17.40544 -5.02080 -13.22144
9.20480 0.00000 0.00000 ; N- CA- C- N
6647 6646 6648 6649 3 0.00000 0.00000 0.00000
0.00000 0.00000 0.00000 ; H- N- CA- HA
6647 6646 6648 6650 3 0.00000 0.00000 0.00000
0.00000 0.00000 0.00000 ; H- N- CA- CB
6647 6646 6648 6668 3 0.00000 0.00000 0.00000
0.00000 0.00000 0.00000 ; H- N- CA- C
6648 6650 6653 6654 3 0.66944 2.00832 0.00000
-2.67776 0.00000 0.00000 ; CA- CB- CG- HG2

6648 6650 6653 6655 3 0.66944 2.00832 0.00000
-2.67776 0.00000 0.00000 ; CA- CB- CG- HG3
6648 6650 6653 6656 3 3.68192 3.09616 -2.09200
-3.01248 0.00000 0.00000 ; CA- CB- CG- CD
6648 6668 6670 6671 3 20.92000 0.00000 -20.92000
0.00000 0.00000 0.00000 ; CA- C- N- H
6648 6668 6670 6672 3 20.92000 0.00000 -20.92000
0.00000 0.00000 0.00000 ; CA- C- N- CA
6649 6648 6650 6651 3 0.65084 1.95253 0.00000
-2.60338 0.00000 0.00000 ; HA- CA- CB- HB2
6649 6648 6650 6652 3 0.65084 1.95253 0.00000
-2.60338 0.00000 0.00000 ; HA- CA- CB- HB3
6649 6648 6650 6653 3 0.65084 1.95253 0.00000
-2.60338 0.00000 0.00000 ; HA- CA- CB- CG
6649 6648 6668 6669 3 3.68192 -4.35136 0.00000
1.33888 0.00000 0.00000 ; HA- CA- C- O
6649 6648 6668 6670 3 0.00000 0.00000 0.00000
0.00000 0.00000 0.00000 ; HA- CA- C- N
6650 6648 6668 6669 3 0.00000 0.00000 0.00000
0.00000 0.00000 0.00000 ; CB- CA- C- O
6650 6648 6668 6670 3 2.51040 4.18400 1.67360
-6.69440 0.00000 0.00000 ; CB- CA- C- N
6650 6653 6656 6657 3 0.65084 1.95253 0.00000
-2.60338 0.00000 0.00000 ; CB- CG- CD- HD2
6650 6653 6656 6658 3 0.65084 1.95253 0.00000
-2.60338 0.00000 0.00000 ; CB- CG- CD- HD3
6650 6653 6656 6659 3 0.65084 1.95253 0.00000
-2.60338 0.00000 0.00000 ; CB- CG- CD- NE
6651 6650 6648 6668 3 0.65084 1.95253 0.00000
-2.60338 0.00000 0.00000 ; HB2- CB- CA- C
6651 6650 6653 6654 3 0.62760 1.88280 0.00000
-2.51040 0.00000 0.00000 ; HB2- CB- CG- HG2
6651 6650 6653 6655 3 0.62760 1.88280 0.00000
-2.51040 0.00000 0.00000 ; HB2- CB- CG- HG3
6651 6650 6653 6656 3 0.66944 2.00832 0.00000
-2.67776 0.00000 0.00000 ; HB2- CB- CG- CD
6652 6650 6648 6668 3 0.65084 1.95253 0.00000
-2.60338 0.00000 0.00000 ; HB3- CB- CA- C
6652 6650 6653 6654 3 0.62760 1.88280 0.00000
-2.51040 0.00000 0.00000 ; HB3- CB- CG- HG2
6652 6650 6653 6655 3 0.62760 1.88280 0.00000
-2.51040 0.00000 0.00000 ; HB3- CB- CG- HG3
6652 6650 6653 6656 3 0.66944 2.00832 0.00000
-2.67776 0.00000 0.00000 ; HB3- CB- CG- CD
6653 6650 6648 6668 3 0.65084 1.95253 0.00000
-2.60338 0.00000 0.00000 ; CG- CB- CA- C
6653 6656 6659 6660 3 0.00000 0.00000 0.00000
0.00000 0.00000 0.00000 ; CG- CD- NE- HE
6653 6656 6659 6661 3 0.00000 0.00000 0.00000
0.00000 0.00000 0.00000 ; CG- CD- NE- CZ
6654 6653 6656 6657 3 0.65084 1.95253 0.00000
-2.60338 0.00000 0.00000 ; HG2- CG- CD- HD2

6654 6653 6656 6658 3 0.65084 1.95253 0.00000
-2.60338 0.00000 0.00000 ; HG2- CG- CD- HD3
6654 6653 6656 6659 3 0.65084 1.95253 0.00000
-2.60338 0.00000 0.00000 ; HG2- CG- CD- NE
6655 6653 6656 6657 3 0.65084 1.95253 0.00000
-2.60338 0.00000 0.00000 ; HG3- CG- CD- HD2
6655 6653 6656 6658 3 0.65084 1.95253 0.00000
-2.60338 0.00000 0.00000 ; HG3- CG- CD- HD3
6655 6653 6656 6659 3 0.65084 1.95253 0.00000
-2.60338 0.00000 0.00000 ; HG3- CG- CD- NE
6656 6659 6661 6662 3 20.08320 0.00000 -20.08320
0.00000 0.00000 0.00000 ; CD- NE- CZ- NH1
6656 6659 6661 6665 3 20.08320 0.00000 -20.08320
0.00000 0.00000 0.00000 ; CD- NE- CZ- NH2
6657 6656 6659 6660 3 0.00000 0.00000 0.00000
0.00000 0.00000 0.00000 ; HD2- CD- NE- HE
6657 6656 6659 6661 3 0.00000 0.00000 0.00000
0.00000 0.00000 0.00000 ; HD2- CD- NE- CZ
6658 6656 6659 6660 3 0.00000 0.00000 0.00000
0.00000 0.00000 0.00000 ; HD3- CD- NE- HE
6658 6656 6659 6661 3 0.00000 0.00000 0.00000
0.00000 0.00000 0.00000 ; HD3- CD- NE- CZ
6659 6661 6662 6663 3 20.08320 0.00000 -20.08320
0.00000 0.00000 0.00000 ; NE- CZ- NH1- HH11
6659 6661 6662 6664 3 20.08320 0.00000 -20.08320
0.00000 0.00000 0.00000 ; NE- CZ- NH1- HH12
6659 6661 6665 6666 3 20.08320 0.00000 -20.08320
0.00000 0.00000 0.00000 ; NE- CZ- NH2- HH21
6659 6661 6665 6667 3 20.08320 0.00000 -20.08320
0.00000 0.00000 0.00000 ; NE- CZ- NH2- HH22
6660 6659 6661 6662 3 20.08320 0.00000 -20.08320
0.00000 0.00000 0.00000 ; HE- NE- CZ- NH1
6660 6659 6661 6665 3 20.08320 0.00000 -20.08320
0.00000 0.00000 0.00000 ; HE- NE- CZ- NH2
6662 6661 6665 6666 3 20.08320 0.00000 -20.08320
0.00000 0.00000 0.00000 ; NH1- CZ- NH2- HH21
6662 6661 6665 6667 3 20.08320 0.00000 -20.08320
0.00000 0.00000 0.00000 ; NH1- CZ- NH2- HH22
6663 6662 6661 6665 3 20.08320 0.00000 -20.08320
0.00000 0.00000 0.00000 ; HH11- NH1- CZ- NH2
6664 6662 6661 6665 3 20.08320 0.00000 -20.08320
0.00000 0.00000 0.00000 ; HH12- NH1- CZ- NH2
6668 6670 6672 6673 3 0.00000 0.00000 0.00000
0.00000 0.00000 0.00000 ; C- N- CA- HA
6668 6670 6672 6674 3 11.71520 1.67360 15.06240 -
13.38880 0.00000 0.00000 ; C- N- CA- CB
6668 6670 6672 6687 3 1.75728 5.27184 2.25936
-7.02912 0.00000 0.00000 ; C- N- CA- C
6669 6668 6670 6671 3 29.28800 -8.36800 -20.92000
0.00000 0.00000 0.00000 ; O- C- N- H
6669 6668 6670 6672 3 20.92000 0.00000 -20.92000
0.00000 0.00000 0.00000 ; O- C- N- CA

```
  6670   6672   6674   6675      3    0.65084    1.95253    0.00000
-2.60338    0.00000    0.00000 ;      N-    CA-    CB-    HB
  6670   6672   6674   6676      3    3.83673    0.44769   -1.77402
-2.47693   -0.03347    0.00000 ;      N-    CA-    CB-   CG2
  6670   6672   6674   6680      3    2.97482    0.12134    2.04179
-1.89117   -3.24678    0.00000 ;      N-    CA-    CB-   CG1
  6670   6672   6687   6688      3    0.00000    0.00000    0.00000
0.00000    0.00000    0.00000 ;      N-    CA-     C-     O
  6670   6672   6687   6689      3   17.40544   -5.02080  -13.22144
9.20480    0.00000    0.00000 ;      N-    CA-     C-     N
  6671   6670   6672   6673      3    0.00000    0.00000    0.00000
0.00000    0.00000    0.00000 ;      H-     N-    CA-    HA
  6671   6670   6672   6674      3    0.00000    0.00000    0.00000
0.00000    0.00000    0.00000 ;      H-     N-    CA-    CB
  6671   6670   6672   6687      3    0.00000    0.00000    0.00000
0.00000    0.00000    0.00000 ;      H-     N-    CA-     C
  6672   6674   6676   6677      3    0.66944    2.00832    0.00000
-2.67776    0.00000    0.00000 ;     CA-    CB-   CG2-  HG21
  6672   6674   6676   6678      3    0.66944    2.00832    0.00000
-2.67776    0.00000    0.00000 ;     CA-    CB-   CG2-  HG22
  6672   6674   6676   6679      3    0.66944    2.00832    0.00000
-2.67776    0.00000    0.00000 ;     CA-    CB-   CG2-  HG23
  6672   6674   6680   6681      3    0.66944    2.00832    0.00000
-2.67776    0.00000    0.00000 ;     CA-    CB-   CG1-  HG12
  6672   6674   6680   6682      3    0.66944    2.00832    0.00000
-2.67776    0.00000    0.00000 ;     CA-    CB-   CG1-  HG13
  6672   6674   6680   6683      3    4.24258   -0.52718   -7.25506
-1.79075    7.69856    0.00000 ;     CA-    CB-   CG1-   CD1
  6672   6687   6689   6690      3   20.92000    0.00000  -20.92000
0.00000    0.00000    0.00000 ;     CA-     C-     N-     H
  6672   6687   6689   6691      3   20.92000    0.00000  -20.92000
0.00000    0.00000    0.00000 ;     CA-     C-     N-    CA
  6673   6672   6674   6675      3    0.65084    1.95253    0.00000
-2.60338    0.00000    0.00000 ;     HA-    CA-    CB-    HB
  6673   6672   6674   6676      3    0.65084    1.95253    0.00000
-2.60338    0.00000    0.00000 ;     HA-    CA-    CB-   CG2
  6673   6672   6674   6680      3    0.65084    1.95253    0.00000
-2.60338    0.00000    0.00000 ;     HA-    CA-    CB-   CG1
  6673   6672   6687   6688      3    3.68192   -4.35136    0.00000
1.33888    0.00000    0.00000 ;     HA-    CA-     C-     O
  6673   6672   6687   6689      3    0.00000    0.00000    0.00000
0.00000    0.00000    0.00000 ;     HA-    CA-     C-     N
  6674   6672   6687   6688      3    0.00000    0.00000    0.00000
0.00000    0.00000    0.00000 ;     CB-    CA-     C-     O
  6674   6672   6687   6689      3    2.51040    4.18400    1.67360
-6.69440    0.00000    0.00000 ;     CB-    CA-     C-     N
  6674   6680   6683   6684      3    0.66944    2.00832    0.00000
-2.67776    0.00000    0.00000 ;     CB-   CG1-   CD1-  HD11
  6674   6680   6683   6685      3    0.66944    2.00832    0.00000
-2.67776    0.00000    0.00000 ;     CB-   CG1-   CD1-  HD12
  6674   6680   6683   6686      3    0.66944    2.00832    0.00000
-2.67776    0.00000    0.00000 ;     CB-   CG1-   CD1-  HD13
```

```
  6675   6674   6672   6687       3    0.65084    1.95253    0.00000
-2.60338    0.00000    0.00000 ;      HB-    CB-    CA-     C
  6675   6674   6676   6677       3    0.62760    1.88280    0.00000
-2.51040    0.00000    0.00000 ;      HB-    CB-   CG2-  HG21
  6675   6674   6676   6678       3    0.62760    1.88280    0.00000
-2.51040    0.00000    0.00000 ;      HB-    CB-   CG2-  HG22
  6675   6674   6676   6679       3    0.62760    1.88280    0.00000
-2.51040    0.00000    0.00000 ;      HB-    CB-   CG2-  HG23
  6675   6674   6680   6681       3    0.62760    1.88280    0.00000
-2.51040    0.00000    0.00000 ;      HB-    CB-   CG1-  HG12
  6675   6674   6680   6682       3    0.62760    1.88280    0.00000
-2.51040    0.00000    0.00000 ;      HB-    CB-   CG1-  HG13
  6675   6674   6680   6683       3    0.66944    2.00832    0.00000
-2.67776    0.00000    0.00000 ;      HB-    CB-   CG1-   CD1
  6676   6674   6672   6687       3    5.67350    3.55640   -6.16722
-2.47693    3.74886    0.00000 ;     CG2-    CB-    CA-     C
  6676   6674   6680   6681       3    0.66944    2.00832    0.00000
-2.67776    0.00000    0.00000 ;     CG2-    CB-   CG1-  HG12
  6676   6674   6680   6682       3    0.66944    2.00832    0.00000
-2.67776    0.00000    0.00000 ;     CG2-    CB-   CG1-  HG13
  6676   6674   6680   6683       3    3.81162    0.49790   -8.14206
-1.79075    7.49773    0.00000 ;     CG2-    CB-   CG1-   CD1
  6677   6676   6674   6680       3    0.66944    2.00832    0.00000
-2.67776    0.00000    0.00000 ;    HG21-   CG2-    CB-   CG1
  6678   6676   6674   6680       3    0.66944    2.00832    0.00000
-2.67776    0.00000    0.00000 ;    HG22-   CG2-    CB-   CG1
  6679   6676   6674   6680       3    0.66944    2.00832    0.00000
-2.67776    0.00000    0.00000 ;    HG23-   CG2-    CB-   CG1
  6680   6674   6672   6687       3    8.26340    0.74057   -9.99976
-1.89117    3.84928    0.00000 ;     CG1-    CB-    CA-     C
  6681   6680   6683   6684       3    0.62760    1.88280    0.00000
-2.51040    0.00000    0.00000 ;    HG12-   CG1-   CD1-  HD11
  6681   6680   6683   6685       3    0.62760    1.88280    0.00000
-2.51040    0.00000    0.00000 ;    HG12-   CG1-   CD1-  HD12
  6681   6680   6683   6686       3    0.62760    1.88280    0.00000
-2.51040    0.00000    0.00000 ;    HG12-   CG1-   CD1-  HD13
  6682   6680   6683   6684       3    0.62760    1.88280    0.00000
-2.51040    0.00000    0.00000 ;    HG13-   CG1-   CD1-  HD11
  6682   6680   6683   6685       3    0.62760    1.88280    0.00000
-2.51040    0.00000    0.00000 ;    HG13-   CG1-   CD1-  HD12
  6682   6680   6683   6686       3    0.62760    1.88280    0.00000
-2.51040    0.00000    0.00000 ;    HG13-   CG1-   CD1-  HD13
  6687   6689   6691   6692       3    0.00000    0.00000    0.00000
0.00000    0.00000    0.00000 ;      C-     N-    CA-    HA
  6687   6689   6691   6693       3   11.71520    1.67360   15.06240  -
13.38880    0.00000    0.00000 ;      C-     N-    CA-    CB
  6687   6689   6691   6707       3    1.75728    5.27184    2.25936
-7.02912    0.00000    0.00000 ;      C-     N-    CA-     C
  6688   6687   6689   6690       3   29.28800   -8.36800  -20.92000
0.00000    0.00000    0.00000 ;      O-     C-     N-     H
  6688   6687   6689   6691       3   20.92000    0.00000  -20.92000
0.00000    0.00000    0.00000 ;      O-     C-     N-    CA
```

6689 6691 6693 6694 3 0.65084 1.95253 0.00000
-2.60338 0.00000 0.00000 ; N- CA- CB- HB2
6689 6691 6693 6695 3 0.65084 1.95253 0.00000
-2.60338 0.00000 0.00000 ; N- CA- CB- HB3
6689 6691 6693 6696 3 3.28026 2.46019 -2.19242
-3.21331 -0.23430 0.00000 ; N- CA- CB- CG
6689 6691 6707 6708 3 0.00000 0.00000 0.00000
0.00000 0.00000 0.00000 ; N- CA- C- O
6689 6691 6707 6709 3 17.40544 -5.02080 -13.22144
9.20480 0.00000 0.00000 ; N- CA- C- N
6690 6689 6691 6692 3 0.00000 0.00000 0.00000
0.00000 0.00000 0.00000 ; H- N- CA- HA
6690 6689 6691 6693 3 0.00000 0.00000 0.00000
0.00000 0.00000 0.00000 ; H- N- CA- CB
6690 6689 6691 6707 3 0.00000 0.00000 0.00000
0.00000 0.00000 0.00000 ; H- N- CA- C
6691 6693 6696 6697 3 0.57739 0.00000 1.02926
0.00000 -1.60666 0.00000 ; CA- CB- CG- CD1
6691 6693 6696 6705 3 0.57739 0.00000 1.02926
0.00000 -1.60666 0.00000 ; CA- CB- CG- CD2
6691 6707 6709 6710 3 20.92000 0.00000 -20.92000
0.00000 0.00000 0.00000 ; CA- C- N- H
6691 6707 6709 6711 3 20.92000 0.00000 -20.92000
0.00000 0.00000 0.00000 ; CA- C- N- CA
6692 6691 6693 6694 3 0.65084 1.95253 0.00000
-2.60338 0.00000 0.00000 ; HA- CA- CB- HB2
6692 6691 6693 6695 3 0.65084 1.95253 0.00000
-2.60338 0.00000 0.00000 ; HA- CA- CB- HB3
6692 6691 6693 6696 3 0.65084 1.95253 0.00000
-2.60338 0.00000 0.00000 ; HA- CA- CB- CG
6692 6691 6707 6708 3 3.68192 -4.35136 0.00000
1.33888 0.00000 0.00000 ; HA- CA- C- O
6692 6691 6707 6709 3 0.00000 0.00000 0.00000
0.00000 0.00000 0.00000 ; HA- CA- C- N
6693 6691 6707 6708 3 0.00000 0.00000 0.00000
0.00000 0.00000 0.00000 ; CB- CA- C- O
6693 6691 6707 6709 3 2.51040 4.18400 1.67360
-6.69440 0.00000 0.00000 ; CB- CA- C- N
6693 6696 6697 6698 3 30.33400 0.00000 -30.33400
0.00000 0.00000 0.00000 ; CB- CG- CD1- HD1
6693 6696 6697 6699 3 30.33400 0.00000 -30.33400
0.00000 0.00000 0.00000 ; CB- CG- CD1- CE1
6693 6696 6705 6703 3 30.33400 0.00000 -30.33400
0.00000 0.00000 0.00000 ; CB- CG- CD2- CE2
6693 6696 6705 6706 3 30.33400 0.00000 -30.33400
0.00000 0.00000 0.00000 ; CB- CG- CD2- HD2
6694 6693 6691 6707 3 0.65084 1.95253 0.00000
-2.60338 0.00000 0.00000 ; HB2- CB- CA- C
6694 6693 6696 6697 3 0.00000 0.00000 0.00000
0.00000 0.00000 0.00000 ; HB2- CB- CG- CD1
6694 6693 6696 6705 3 0.00000 0.00000 0.00000
0.00000 0.00000 0.00000 ; HB2- CB- CG- CD2

6695 6693 6691 6707 3 0.65084 1.95253 0.00000
-2.60338 0.00000 0.00000 ; HB3- CB- CA- C
6695 6693 6696 6697 3 0.00000 0.00000 0.00000
0.00000 0.00000 0.00000 ; HB3- CB- CG- CD1
6695 6693 6696 6705 3 0.00000 0.00000 0.00000
0.00000 0.00000 0.00000 ; HB3- CB- CG- CD2
6696 6693 6691 6707 3 4.95804 2.17986 -3.52293
-3.21331 -0.40166 0.00000 ; CG- CB- CA- C
6696 6697 6699 6700 3 30.33400 0.00000 -30.33400
0.00000 0.00000 0.00000 ; CG- CD1- CE1- HE1
6696 6697 6699 6701 3 30.33400 0.00000 -30.33400
0.00000 0.00000 0.00000 ; CG- CD1- CE1- CZ
6696 6705 6703 6701 3 30.33400 0.00000 -30.33400
0.00000 0.00000 0.00000 ; CG- CD2- CE2- CZ
6696 6705 6703 6704 3 30.33400 0.00000 -30.33400
0.00000 0.00000 0.00000 ; CG- CD2- CE2- HE2
6697 6696 6705 6703 3 30.33400 0.00000 -30.33400
0.00000 0.00000 0.00000 ; CD1- CG- CD2- CE2
6697 6696 6705 6706 3 30.33400 0.00000 -30.33400
0.00000 0.00000 0.00000 ; CD1- CG- CD2- HD2
6697 6699 6701 6702 3 30.33400 0.00000 -30.33400
0.00000 0.00000 0.00000 ; CD1- CE1- CZ- HZ
6697 6699 6701 6703 3 30.33400 0.00000 -30.33400
0.00000 0.00000 0.00000 ; CD1- CE1- CZ- CE2
6698 6697 6696 6705 3 30.33400 0.00000 -30.33400
0.00000 0.00000 0.00000 ; HD1- CD1- CG- CD2
6698 6697 6699 6700 3 30.33400 0.00000 -30.33400
0.00000 0.00000 0.00000 ; HD1- CD1- CE1- HE1
6698 6697 6699 6701 3 30.33400 0.00000 -30.33400
0.00000 0.00000 0.00000 ; HD1- CD1- CE1- CZ
6699 6697 6696 6705 3 30.33400 0.00000 -30.33400
0.00000 0.00000 0.00000 ; CE1- CD1- CG- CD2
6699 6701 6703 6704 3 30.33400 0.00000 -30.33400
0.00000 0.00000 0.00000 ; CE1- CZ- CE2- HE2
6699 6701 6703 6705 3 30.33400 0.00000 -30.33400
0.00000 0.00000 0.00000 ; CE1- CZ- CE2- CD2
6700 6699 6701 6702 3 30.33400 0.00000 -30.33400
0.00000 0.00000 0.00000 ; HE1- CE1- CZ- HZ
6700 6699 6701 6703 3 30.33400 0.00000 -30.33400
0.00000 0.00000 0.00000 ; HE1- CE1- CZ- CE2
6701 6703 6705 6706 3 30.33400 0.00000 -30.33400
0.00000 0.00000 0.00000 ; CZ- CE2- CD2- HD2
6702 6701 6703 6704 3 30.33400 0.00000 -30.33400
0.00000 0.00000 0.00000 ; HZ- CZ- CE2- HE2
6702 6701 6703 6705 3 30.33400 0.00000 -30.33400
0.00000 0.00000 0.00000 ; HZ- CZ- CE2- CD2
6704 6703 6705 6706 3 30.33400 0.00000 -30.33400
0.00000 0.00000 0.00000 ; HE2- CE2- CD2- HD2
6707 6709 6711 6712 3 0.00000 0.00000 0.00000
0.00000 0.00000 0.00000 ; C- N- CA- HA2
6707 6709 6711 6713 3 0.00000 0.00000 0.00000
0.00000 0.00000 0.00000 ; C- N- CA- HA3

6707 6709 6711 6714 3 1.75728 5.27184 2.25936
-7.02912 0.00000 0.00000 ; C- N- CA- C
6708 6707 6709 6710 3 29.28800 -8.36800 -20.92000
0.00000 0.00000 0.00000 ; O- C- N- H
6708 6707 6709 6711 3 20.92000 0.00000 -20.92000
0.00000 0.00000 0.00000 ; O- C- N- CA
6709 6711 6714 6715 3 0.00000 0.00000 0.00000
0.00000 0.00000 0.00000 ; N- CA- C- O
6709 6711 6714 6716 3 17.40544 -5.02080 -13.22144
9.20480 0.00000 0.00000 ; N- CA- C- N
6710 6709 6711 6712 3 0.00000 0.00000 0.00000
0.00000 0.00000 0.00000 ; H- N- CA- HA2
6710 6709 6711 6713 3 0.00000 0.00000 0.00000
0.00000 0.00000 0.00000 ; H- N- CA- HA3
6710 6709 6711 6714 3 0.00000 0.00000 0.00000
0.00000 0.00000 0.00000 ; H- N- CA- C
6711 6714 6716 6717 3 20.92000 0.00000 -20.92000
0.00000 0.00000 0.00000 ; CA- C- N- H
6711 6714 6716 6718 3 20.92000 0.00000 -20.92000
0.00000 0.00000 0.00000 ; CA- C- N- CA
6712 6711 6714 6715 3 3.68192 -4.35136 0.00000
1.33888 0.00000 0.00000 ; HA2- CA- C- O
6712 6711 6714 6716 3 0.00000 0.00000 0.00000
0.00000 0.00000 0.00000 ; HA2- CA- C- N
6713 6711 6714 6715 3 3.68192 -4.35136 0.00000
1.33888 0.00000 0.00000 ; HA3- CA- C- O
6713 6711 6714 6716 3 0.00000 0.00000 0.00000
0.00000 0.00000 0.00000 ; HA3- CA- C- N
6714 6716 6718 6719 3 0.00000 0.00000 0.00000
0.00000 0.00000 0.00000 ; C- N- CA- HA
6714 6716 6718 6720 3 11.71520 1.67360 15.06240 -
13.38880 0.00000 0.00000 ; C- N- CA- CB
6714 6716 6718 6738 3 1.75728 5.27184 2.25936
-7.02912 0.00000 0.00000 ; C- N- CA- C
6715 6714 6716 6717 3 29.28800 -8.36800 -20.92000
0.00000 0.00000 0.00000 ; O- C- N- H
6715 6714 6716 6718 3 20.92000 0.00000 -20.92000
0.00000 0.00000 0.00000 ; O- C- N- CA
6716 6718 6720 6721 3 0.65084 1.95253 0.00000
-2.60338 0.00000 0.00000 ; N- CA- CB- HB2
6716 6718 6720 6722 3 0.65084 1.95253 0.00000
-2.60338 0.00000 0.00000 ; N- CA- CB- HB3
6716 6718 6720 6723 3 0.65084 1.95253 0.00000
-2.60338 0.00000 0.00000 ; N- CA- CB- CG
6716 6718 6738 6739 3 0.00000 0.00000 0.00000
0.00000 0.00000 0.00000 ; N- CA- C- O
6716 6718 6738 6740 3 17.40544 -5.02080 -13.22144
9.20480 0.00000 0.00000 ; N- CA- C- N
6717 6716 6718 6719 3 0.00000 0.00000 0.00000
0.00000 0.00000 0.00000 ; H- N- CA- HA
6717 6716 6718 6720 3 0.00000 0.00000 0.00000
0.00000 0.00000 0.00000 ; H- N- CA- CB

6717 6716 6718 6738 3 0.00000 0.00000 0.00000
0.00000 0.00000 0.00000 ; H- N- CA- C
6718 6720 6723 6724 3 0.66944 2.00832 0.00000
-2.67776 0.00000 0.00000 ; CA- CB- CG- HG2
6718 6720 6723 6725 3 0.66944 2.00832 0.00000
-2.67776 0.00000 0.00000 ; CA- CB- CG- HG3
6718 6720 6723 6726 3 3.68192 3.09616 -2.09200
-3.01248 0.00000 0.00000 ; CA- CB- CG- CD
6718 6738 6740 6741 3 20.92000 0.00000 -20.92000
0.00000 0.00000 0.00000 ; CA- C- N- H
6718 6738 6740 6742 3 20.92000 0.00000 -20.92000
0.00000 0.00000 0.00000 ; CA- C- N- CA
6719 6718 6720 6721 3 0.65084 1.95253 0.00000
-2.60338 0.00000 0.00000 ; HA- CA- CB- HB2
6719 6718 6720 6722 3 0.65084 1.95253 0.00000
-2.60338 0.00000 0.00000 ; HA- CA- CB- HB3
6719 6718 6720 6723 3 0.65084 1.95253 0.00000
-2.60338 0.00000 0.00000 ; HA- CA- CB- CG
6719 6718 6738 6739 3 3.68192 -4.35136 0.00000
1.33888 0.00000 0.00000 ; HA- CA- C- O
6719 6718 6738 6740 3 0.00000 0.00000 0.00000
0.00000 0.00000 0.00000 ; HA- CA- C- N
6720 6718 6738 6739 3 0.00000 0.00000 0.00000
0.00000 0.00000 0.00000 ; CB- CA- C- O
6720 6718 6738 6740 3 2.51040 4.18400 1.67360
-6.69440 0.00000 0.00000 ; CB- CA- C- N
6720 6723 6726 6727 3 0.65084 1.95253 0.00000
-2.60338 0.00000 0.00000 ; CB- CG- CD- HD2
6720 6723 6726 6728 3 0.65084 1.95253 0.00000
-2.60338 0.00000 0.00000 ; CB- CG- CD- HD3
6720 6723 6726 6729 3 0.65084 1.95253 0.00000
-2.60338 0.00000 0.00000 ; CB- CG- CD- NE
6721 6720 6718 6738 3 0.65084 1.95253 0.00000
-2.60338 0.00000 0.00000 ; HB2- CB- CA- C
6721 6720 6723 6724 3 0.62760 1.88280 0.00000
-2.51040 0.00000 0.00000 ; HB2- CB- CG- HG2
6721 6720 6723 6725 3 0.62760 1.88280 0.00000
-2.51040 0.00000 0.00000 ; HB2- CB- CG- HG3
6721 6720 6723 6726 3 0.66944 2.00832 0.00000
-2.67776 0.00000 0.00000 ; HB2- CB- CG- CD
6722 6720 6718 6738 3 0.65084 1.95253 0.00000
-2.60338 0.00000 0.00000 ; HB3- CB- CA- C
6722 6720 6723 6724 3 0.62760 1.88280 0.00000
-2.51040 0.00000 0.00000 ; HB3- CB- CG- HG2
6722 6720 6723 6725 3 0.62760 1.88280 0.00000
-2.51040 0.00000 0.00000 ; HB3- CB- CG- HG3
6722 6720 6723 6726 3 0.66944 2.00832 0.00000
-2.67776 0.00000 0.00000 ; HB3- CB- CG- CD
6723 6720 6718 6738 3 0.65084 1.95253 0.00000
-2.60338 0.00000 0.00000 ; CG- CB- CA- C
6723 6726 6729 6730 3 0.00000 0.00000 0.00000
0.00000 0.00000 0.00000 ; CG- CD- NE- HE

6723 6726 6729 6731 3 0.00000 0.00000 0.00000
0.00000 0.00000 0.00000 ; CG- CD- NE- CZ
6724 6723 6726 6727 3 0.65084 1.95253 0.00000
-2.60338 0.00000 0.00000 ; HG2- CG- CD- HD2
6724 6723 6726 6728 3 0.65084 1.95253 0.00000
-2.60338 0.00000 0.00000 ; HG2- CG- CD- HD3
6724 6723 6726 6729 3 0.65084 1.95253 0.00000
-2.60338 0.00000 0.00000 ; HG2- CG- CD- NE
6725 6723 6726 6727 3 0.65084 1.95253 0.00000
-2.60338 0.00000 0.00000 ; HG3- CG- CD- HD2
6725 6723 6726 6728 3 0.65084 1.95253 0.00000
-2.60338 0.00000 0.00000 ; HG3- CG- CD- HD3
6725 6723 6726 6729 3 0.65084 1.95253 0.00000
-2.60338 0.00000 0.00000 ; HG3- CG- CD- NE
6726 6729 6731 6732 3 20.08320 0.00000 -20.08320
0.00000 0.00000 0.00000 ; CD- NE- CZ- NH1
6726 6729 6731 6735 3 20.08320 0.00000 -20.08320
0.00000 0.00000 0.00000 ; CD- NE- CZ- NH2
6727 6726 6729 6730 3 0.00000 0.00000 0.00000
0.00000 0.00000 0.00000 ; HD2- CD- NE- HE
6727 6726 6729 6731 3 0.00000 0.00000 0.00000
0.00000 0.00000 0.00000 ; HD2- CD- NE- CZ
6728 6726 6729 6730 3 0.00000 0.00000 0.00000
0.00000 0.00000 0.00000 ; HD3- CD- NE- HE
6728 6726 6729 6731 3 0.00000 0.00000 0.00000
0.00000 0.00000 0.00000 ; HD3- CD- NE- CZ
6729 6731 6732 6733 3 20.08320 0.00000 -20.08320
0.00000 0.00000 0.00000 ; NE- CZ- NH1- HH11
6729 6731 6732 6734 3 20.08320 0.00000 -20.08320
0.00000 0.00000 0.00000 ; NE- CZ- NH1- HH12
6729 6731 6735 6736 3 20.08320 0.00000 -20.08320
0.00000 0.00000 0.00000 ; NE- CZ- NH2- HH21
6729 6731 6735 6737 3 20.08320 0.00000 -20.08320
0.00000 0.00000 0.00000 ; NE- CZ- NH2- HH22
6730 6729 6731 6732 3 20.08320 0.00000 -20.08320
0.00000 0.00000 0.00000 ; HE- NE- CZ- NH1
6730 6729 6731 6735 3 20.08320 0.00000 -20.08320
0.00000 0.00000 0.00000 ; HE- NE- CZ- NH2
6732 6731 6735 6736 3 20.08320 0.00000 -20.08320
0.00000 0.00000 0.00000 ; NH1- CZ- NH2- HH21
6732 6731 6735 6737 3 20.08320 0.00000 -20.08320
0.00000 0.00000 0.00000 ; NH1- CZ- NH2- HH22
6733 6732 6731 6735 3 20.08320 0.00000 -20.08320
0.00000 0.00000 0.00000 ; HH11- NH1- CZ- NH2
6734 6732 6731 6735 3 20.08320 0.00000 -20.08320
0.00000 0.00000 0.00000 ; HH12- NH1- CZ- NH2
6738 6740 6742 6743 3 0.00000 0.00000 0.00000
0.00000 0.00000 0.00000 ; C- N- CA- HA
6738 6740 6742 6744 3 11.71520 1.67360 15.06240 -
13.38880 0.00000 0.00000 ; C- N- CA- CB
6738 6740 6742 6750 3 1.75728 5.27184 2.25936
-7.02912 0.00000 0.00000 ; C- N- CA- C

6739 6738 6740 6741 3 29.28800 -8.36800 -20.92000
0.00000 0.00000 0.00000 ; O- C- N- H
6739 6738 6740 6742 3 20.92000 0.00000 -20.92000
0.00000 0.00000 0.00000 ; O- C- N- CA
6740 6742 6744 6745 3 0.65084 1.95253 0.00000
-2.60338 0.00000 0.00000 ; N- CA- CB- HB2
6740 6742 6744 6746 3 0.65084 1.95253 0.00000
-2.60338 0.00000 0.00000 ; N- CA- CB- HB3
6740 6742 6744 6747 3 15.41386 9.74035 -8.39310
-0.97069 2.97901 0.00000 ; N- CA- CB- CG
6740 6742 6750 6751 3 0.00000 0.00000 0.00000
0.00000 0.00000 0.00000 ; N- CA- C- O
6740 6742 6750 6752 3 17.40544 -5.02080 -13.22144
9.20480 0.00000 0.00000 ; N- CA- C- N
6741 6740 6742 6743 3 0.00000 0.00000 0.00000
0.00000 0.00000 0.00000 ; H- N- CA- HA
6741 6740 6742 6744 3 0.00000 0.00000 0.00000
0.00000 0.00000 0.00000 ; H- N- CA- CB
6741 6740 6742 6750 3 0.00000 0.00000 0.00000
0.00000 0.00000 0.00000 ; H- N- CA- C
6742 6744 6747 6748 3 6.43499 0.00000 -5.39736
0.00000 -1.03763 0.00000 ; CA- CB- CG- OD1
6742 6744 6747 6749 3 6.43499 0.00000 -5.39736
0.00000 -1.03763 0.00000 ; CA- CB- CG- OD2
6742 6750 6752 6753 3 20.92000 0.00000 -20.92000
0.00000 0.00000 0.00000 ; CA- C- N- H
6742 6750 6752 6754 3 20.92000 0.00000 -20.92000
0.00000 0.00000 0.00000 ; CA- C- N- CA
6743 6742 6744 6745 3 0.65084 1.95253 0.00000
-2.60338 0.00000 0.00000 ; HA- CA- CB- HB2
6743 6742 6744 6746 3 0.65084 1.95253 0.00000
-2.60338 0.00000 0.00000 ; HA- CA- CB- HB3
6743 6742 6744 6747 3 0.65084 1.95253 0.00000
-2.60338 0.00000 0.00000 ; HA- CA- CB- CG
6743 6742 6750 6751 3 3.68192 -4.35136 0.00000
1.33888 0.00000 0.00000 ; HA- CA- C- O
6743 6742 6750 6752 3 0.00000 0.00000 0.00000
0.00000 0.00000 0.00000 ; HA- CA- C- N
6744 6742 6750 6751 3 0.00000 0.00000 0.00000
0.00000 0.00000 0.00000 ; CB- CA- C- O
6744 6742 6750 6752 3 2.51040 4.18400 1.67360
-6.69440 0.00000 0.00000 ; CB- CA- C- N
6745 6744 6742 6750 3 0.65084 1.95253 0.00000
-2.60338 0.00000 0.00000 ; HB2- CB- CA- C
6745 6744 6747 6748 3 0.00000 0.00000 0.00000
0.00000 0.00000 0.00000 ; HB2- CB- CG- OD1
6745 6744 6747 6749 3 0.00000 0.00000 0.00000
0.00000 0.00000 0.00000 ; HB2- CB- CG- OD2
6746 6744 6742 6750 3 0.65084 1.95253 0.00000
-2.60338 0.00000 0.00000 ; HB3- CB- CA- C
6746 6744 6747 6748 3 0.00000 0.00000 0.00000
0.00000 0.00000 0.00000 ; HB3- CB- CG- OD1

6746 6744 6747 6749 3 0.00000 0.00000 0.00000
0.00000 0.00000 0.00000 ; HB3- CB- CG- OD2
6747 6744 6742 6750 3 7.14627 -1.04600 -8.99560
-0.97069 5.15469 0.00000 ; CG- CB- CA- C
6750 6752 6754 6755 3 0.00000 0.00000 0.00000
0.00000 0.00000 0.00000 ; C- N- CA- HA
6750 6752 6754 6756 3 11.71520 1.67360 15.06240 -
13.38880 0.00000 0.00000 ; C- N- CA- CB
6750 6752 6754 6762 3 1.75728 5.27184 2.25936
-7.02912 0.00000 0.00000 ; C- N- CA- C
6751 6750 6752 6753 3 29.28800 -8.36800 -20.92000
0.00000 0.00000 0.00000 ; O- C- N- H
6751 6750 6752 6754 3 20.92000 0.00000 -20.92000
0.00000 0.00000 0.00000 ; O- C- N- CA
6752 6754 6756 6757 3 0.65084 1.95253 0.00000
-2.60338 0.00000 0.00000 ; N- CA- CB- HB2
6752 6754 6756 6758 3 0.65084 1.95253 0.00000
-2.60338 0.00000 0.00000 ; N- CA- CB- HB3
6752 6754 6756 6759 3 15.41386 9.74035 -8.39310
-0.97069 2.97901 0.00000 ; N- CA- CB- CG
6752 6754 6762 6763 3 0.00000 0.00000 0.00000
0.00000 0.00000 0.00000 ; N- CA- C- O
6752 6754 6762 6764 3 17.40544 -5.02080 -13.22144
9.20480 0.00000 0.00000 ; N- CA- C- N
6753 6752 6754 6755 3 0.00000 0.00000 0.00000
0.00000 0.00000 0.00000 ; H- N- CA- HA
6753 6752 6754 6756 3 0.00000 0.00000 0.00000
0.00000 0.00000 0.00000 ; H- N- CA- CB
6753 6752 6754 6762 3 0.00000 0.00000 0.00000
0.00000 0.00000 0.00000 ; H- N- CA- C
6754 6756 6759 6760 3 6.43499 0.00000 -5.39736
0.00000 -1.03763 0.00000 ; CA- CB- CG- OD1
6754 6756 6759 6761 3 6.43499 0.00000 -5.39736
0.00000 -1.03763 0.00000 ; CA- CB- CG- OD2
6754 6762 6764 6765 3 20.92000 0.00000 -20.92000
0.00000 0.00000 0.00000 ; CA- C- N- H
6754 6762 6764 6766 3 20.92000 0.00000 -20.92000
0.00000 0.00000 0.00000 ; CA- C- N- CA
6755 6754 6756 6757 3 0.65084 1.95253 0.00000
-2.60338 0.00000 0.00000 ; HA- CA- CB- HB2
6755 6754 6756 6758 3 0.65084 1.95253 0.00000
-2.60338 0.00000 0.00000 ; HA- CA- CB- HB3
6755 6754 6756 6759 3 0.65084 1.95253 0.00000
-2.60338 0.00000 0.00000 ; HA- CA- CB- CG
6755 6754 6762 6763 3 3.68192 -4.35136 0.00000
1.33888 0.00000 0.00000 ; HA- CA- C- O
6755 6754 6762 6764 3 0.00000 0.00000 0.00000
0.00000 0.00000 0.00000 ; HA- CA- C- N
6756 6754 6762 6763 3 0.00000 0.00000 0.00000
0.00000 0.00000 0.00000 ; CB- CA- C- O
6756 6754 6762 6764 3 2.51040 4.18400 1.67360
-6.69440 0.00000 0.00000 ; CB- CA- C- N

6757 6756 6754 6762 3 0.65084 1.95253 0.00000
-2.60338 0.00000 0.00000 ; HB2- CB- CA- C
6757 6756 6759 6760 3 0.00000 0.00000 0.00000
0.00000 0.00000 0.00000 ; HB2- CB- CG- OD1
6757 6756 6759 6761 3 0.00000 0.00000 0.00000
0.00000 0.00000 0.00000 ; HB2- CB- CG- OD2
6758 6756 6754 6762 3 0.65084 1.95253 0.00000
-2.60338 0.00000 0.00000 ; HB3- CB- CA- C
6758 6756 6759 6760 3 0.00000 0.00000 0.00000
0.00000 0.00000 0.00000 ; HB3- CB- CG- OD1
6758 6756 6759 6761 3 0.00000 0.00000 0.00000
0.00000 0.00000 0.00000 ; HB3- CB- CG- OD2
6759 6756 6754 6762 3 7.14627 -1.04600 -8.99560
-0.97069 5.15469 0.00000 ; CG- CB- CA- C
6762 6764 6766 6767 3 0.00000 0.00000 0.00000
0.00000 0.00000 0.00000 ; C- N- CA- HA
6762 6764 6766 6768 3 11.71520 1.67360 15.06240 -
13.38880 0.00000 0.00000 ; C- N- CA- CB
6762 6764 6766 6781 3 1.75728 5.27184 2.25936
-7.02912 0.00000 0.00000 ; C- N- CA- C
6763 6762 6764 6765 3 29.28800 -8.36800 -20.92000
0.00000 0.00000 0.00000 ; O- C- N- H
6763 6762 6764 6766 3 20.92000 0.00000 -20.92000
0.00000 0.00000 0.00000 ; O- C- N- CA
6764 6766 6768 6769 3 0.65084 1.95253 0.00000
-2.60338 0.00000 0.00000 ; N- CA- CB- HB
6764 6766 6768 6770 3 3.83673 0.44769 -1.77402
-2.47693 -0.03347 0.00000 ; N- CA- CB- CG2
6764 6766 6768 6774 3 2.97482 0.12134 2.04179
-1.89117 -3.24678 0.00000 ; N- CA- CB- CG1
6764 6766 6781 6782 3 0.00000 0.00000 0.00000
0.00000 0.00000 0.00000 ; N- CA- C- O
6764 6766 6781 6783 3 17.40544 -5.02080 -13.22144
9.20480 0.00000 0.00000 ; N- CA- C- N
6765 6764 6766 6767 3 0.00000 0.00000 0.00000
0.00000 0.00000 0.00000 ; H- N- CA- HA
6765 6764 6766 6768 3 0.00000 0.00000 0.00000
0.00000 0.00000 0.00000 ; H- N- CA- CB
6765 6764 6766 6781 3 0.00000 0.00000 0.00000
0.00000 0.00000 0.00000 ; H- N- CA- C
6766 6768 6770 6771 3 0.66944 2.00832 0.00000
-2.67776 0.00000 0.00000 ; CA- CB- CG2- HG21
6766 6768 6770 6772 3 0.66944 2.00832 0.00000
-2.67776 0.00000 0.00000 ; CA- CB- CG2- HG22
6766 6768 6770 6773 3 0.66944 2.00832 0.00000
-2.67776 0.00000 0.00000 ; CA- CB- CG2- HG23
6766 6768 6774 6775 3 0.66944 2.00832 0.00000
-2.67776 0.00000 0.00000 ; CA- CB- CG1- HG12
6766 6768 6774 6776 3 0.66944 2.00832 0.00000
-2.67776 0.00000 0.00000 ; CA- CB- CG1- HG13
6766 6768 6774 6777 3 4.24258 -0.52718 -7.25506
-1.79075 7.69856 0.00000 ; CA- CB- CG1- CD1

```
  6766   6781   6783   6784       3   20.92000    0.00000  -20.92000
0.00000    0.00000    0.00000 ;      CA-     C-     N-     H
  6766   6781   6783   6785       3   20.92000    0.00000  -20.92000
0.00000    0.00000    0.00000 ;      CA-     C-     N-    CA
  6767   6766   6768   6769       3    0.65084    1.95253    0.00000
-2.60338    0.00000    0.00000 ;      HA-    CA-    CB-    HB
  6767   6766   6768   6770       3    0.65084    1.95253    0.00000
-2.60338    0.00000    0.00000 ;      HA-    CA-    CB-   CG2
  6767   6766   6768   6774       3    0.65084    1.95253    0.00000
-2.60338    0.00000    0.00000 ;      HA-    CA-    CB-   CG1
  6767   6766   6781   6782       3    3.68192   -4.35136    0.00000
1.33888    0.00000    0.00000 ;      HA-    CA-     C-     O
  6767   6766   6781   6783       3    0.00000    0.00000    0.00000
0.00000    0.00000    0.00000 ;      HA-    CA-     C-     N
  6768   6766   6781   6782       3    0.00000    0.00000    0.00000
0.00000    0.00000    0.00000 ;      CB-    CA-     C-     O
  6768   6766   6781   6783       3    2.51040    4.18400    1.67360
-6.69440    0.00000    0.00000 ;      CB-    CA-     C-     N
  6768   6774   6777   6778       3    0.66944    2.00832    0.00000
-2.67776    0.00000    0.00000 ;      CB-   CG1-   CD1-  HD11
  6768   6774   6777   6779       3    0.66944    2.00832    0.00000
-2.67776    0.00000    0.00000 ;      CB-   CG1-   CD1-  HD12
  6768   6774   6777   6780       3    0.66944    2.00832    0.00000
-2.67776    0.00000    0.00000 ;      CB-   CG1-   CD1-  HD13
  6769   6768   6766   6781       3    0.65084    1.95253    0.00000
-2.60338    0.00000    0.00000 ;      HB-    CB-    CA-     C
  6769   6768   6770   6771       3    0.62760    1.88280    0.00000
-2.51040    0.00000    0.00000 ;      HB-    CB-   CG2-  HG21
  6769   6768   6770   6772       3    0.62760    1.88280    0.00000
-2.51040    0.00000    0.00000 ;      HB-    CB-   CG2-  HG22
  6769   6768   6770   6773       3    0.62760    1.88280    0.00000
-2.51040    0.00000    0.00000 ;      HB-    CB-   CG2-  HG23
  6769   6768   6774   6775       3    0.62760    1.88280    0.00000
-2.51040    0.00000    0.00000 ;      HB-    CB-   CG1-  HG12
  6769   6768   6774   6776       3    0.62760    1.88280    0.00000
-2.51040    0.00000    0.00000 ;      HB-    CB-   CG1-  HG13
  6769   6768   6774   6777       3    0.66944    2.00832    0.00000
-2.67776    0.00000    0.00000 ;      HB-    CB-   CG1-   CD1
  6770   6768   6766   6781       3    5.67350    3.55640   -6.16722
-2.47693    3.74886    0.00000 ;     CG2-    CB-    CA-     C
  6770   6768   6774   6775       3    0.66944    2.00832    0.00000
-2.67776    0.00000    0.00000 ;     CG2-    CB-   CG1-  HG12
  6770   6768   6774   6776       3    0.66944    2.00832    0.00000
-2.67776    0.00000    0.00000 ;     CG2-    CB-   CG1-  HG13
  6770   6768   6774   6777       3    3.81162    0.49790   -8.14206
-1.79075    7.49773    0.00000 ;     CG2-    CB-   CG1-   CD1
  6771   6770   6768   6774       3    0.66944    2.00832    0.00000
-2.67776    0.00000    0.00000 ;    HG21-   CG2-    CB-   CG1
  6772   6770   6768   6774       3    0.66944    2.00832    0.00000
-2.67776    0.00000    0.00000 ;    HG22-   CG2-    CB-   CG1
  6773   6770   6768   6774       3    0.66944    2.00832    0.00000
-2.67776    0.00000    0.00000 ;    HG23-   CG2-    CB-   CG1
```

6774 6768 6766 6781 3 8.26340 0.74057 -9.99976
-1.89117 3.84928 0.00000 ; CG1- CB- CA- C
6775 6774 6777 6778 3 0.62760 1.88280 0.00000
-2.51040 0.00000 0.00000 ; HG12- CG1- CD1- HD11
6775 6774 6777 6779 3 0.62760 1.88280 0.00000
-2.51040 0.00000 0.00000 ; HG12- CG1- CD1- HD12
6775 6774 6777 6780 3 0.62760 1.88280 0.00000
-2.51040 0.00000 0.00000 ; HG12- CG1- CD1- HD13
6776 6774 6777 6778 3 0.62760 1.88280 0.00000
-2.51040 0.00000 0.00000 ; HG13- CG1- CD1- HD11
6776 6774 6777 6779 3 0.62760 1.88280 0.00000
-2.51040 0.00000 0.00000 ; HG13- CG1- CD1- HD12
6776 6774 6777 6780 3 0.62760 1.88280 0.00000
-2.51040 0.00000 0.00000 ; HG13- CG1- CD1- HD13
6781 6783 6785 6786 3 0.00000 0.00000 0.00000
0.00000 0.00000 0.00000 ; C- N- CA- HA
6781 6783 6785 6787 3 11.71520 1.67360 15.06240 -
13.38880 0.00000 0.00000 ; C- N- CA- CB
6781 6783 6785 6796 3 1.75728 5.27184 2.25936
-7.02912 0.00000 0.00000 ; C- N- CA- C
6782 6781 6783 6784 3 29.28800 -8.36800 -20.92000
0.00000 0.00000 0.00000 ; O- C- N- H
6782 6781 6783 6785 3 20.92000 0.00000 -20.92000
0.00000 0.00000 0.00000 ; O- C- N- CA
6783 6785 6787 6788 3 0.65084 1.95253 0.00000
-2.60338 0.00000 0.00000 ; N- CA- CB- HB2
6783 6785 6787 6789 3 0.65084 1.95253 0.00000
-2.60338 0.00000 0.00000 ; N- CA- CB- HB3
6783 6785 6787 6790 3 3.21331 2.22589 -4.15053
-2.40998 2.61082 0.00000 ; N- CA- CB- CG
6783 6785 6796 6797 3 0.00000 0.00000 0.00000
0.00000 0.00000 0.00000 ; N- CA- C- O
6783 6785 6796 6798 3 17.40544 -5.02080 -13.22144
9.20480 0.00000 0.00000 ; N- CA- C- N
6784 6783 6785 6786 3 0.00000 0.00000 0.00000
0.00000 0.00000 0.00000 ; H- N- CA- HA
6784 6783 6785 6787 3 0.00000 0.00000 0.00000
0.00000 0.00000 0.00000 ; H- N- CA- CB
6784 6783 6785 6796 3 0.00000 0.00000 0.00000
0.00000 0.00000 0.00000 ; H- N- CA- C
6785 6787 6790 6791 3 0.66944 2.00832 0.00000
-2.67776 0.00000 0.00000 ; CA- CB- CG- HG2
6785 6787 6790 6792 3 0.66944 2.00832 0.00000
-2.67776 0.00000 0.00000 ; CA- CB- CG- HG3
6785 6787 6790 6793 3 10.12110 -1.91209 0.01674
10.17549 -1.87443 0.00000 ; CA- CB- CG- CD
6785 6796 6798 6799 3 20.92000 0.00000 -20.92000
0.00000 0.00000 0.00000 ; CA- C- N- H
6785 6796 6798 6800 3 20.92000 0.00000 -20.92000
0.00000 0.00000 0.00000 ; CA- C- N- CA
6786 6785 6787 6788 3 0.65084 1.95253 0.00000
-2.60338 0.00000 0.00000 ; HA- CA- CB- HB2

6786 6785 6787 6789 3 0.65084 1.95253 0.00000
-2.60338 0.00000 0.00000 ; HA- CA- CB- HB3
6786 6785 6787 6790 3 0.65084 1.95253 0.00000
-2.60338 0.00000 0.00000 ; HA- CA- CB- CG
6786 6785 6796 6797 3 3.68192 -4.35136 0.00000
1.33888 0.00000 0.00000 ; HA- CA- C- O
6786 6785 6796 6798 3 0.00000 0.00000 0.00000
0.00000 0.00000 0.00000 ; HA- CA- C- N
6787 6785 6796 6797 3 0.00000 0.00000 0.00000
0.00000 0.00000 0.00000 ; CB- CA- C- O
6787 6785 6796 6798 3 2.51040 4.18400 1.67360
-6.69440 0.00000 0.00000 ; CB- CA- C- N
6787 6790 6793 6794 3 3.79907 0.00000 -5.40573
0.00000 2.14221 0.00000 ; CB- CG- CD- OE1
6787 6790 6793 6795 3 3.79907 0.00000 -5.40573
0.00000 2.14221 0.00000 ; CB- CG- CD- OE2
6788 6787 6785 6796 3 0.65084 1.95253 0.00000
-2.60338 0.00000 0.00000 ; HB2- CB- CA- C
6788 6787 6790 6791 3 0.62760 1.88280 0.00000
-2.51040 0.00000 0.00000 ; HB2- CB- CG- HG2
6788 6787 6790 6792 3 0.62760 1.88280 0.00000
-2.51040 0.00000 0.00000 ; HB2- CB- CG- HG3
6788 6787 6790 6793 3 0.65084 1.95253 0.00000
-2.60338 0.00000 0.00000 ; HB2- CB- CG- CD
6789 6787 6785 6796 3 0.65084 1.95253 0.00000
-2.60338 0.00000 0.00000 ; HB3- CB- CA- C
6789 6787 6790 6791 3 0.62760 1.88280 0.00000
-2.51040 0.00000 0.00000 ; HB3- CB- CG- HG2
6789 6787 6790 6792 3 0.62760 1.88280 0.00000
-2.51040 0.00000 0.00000 ; HB3- CB- CG- HG3
6789 6787 6790 6793 3 0.65084 1.95253 0.00000
-2.60338 0.00000 0.00000 ; HB3- CB- CG- CD
6790 6787 6785 6796 3 6.86594 3.56895 -8.14206
-2.40998 4.85344 0.00000 ; CG- CB- CA- C
6791 6790 6793 6794 3 0.00000 0.00000 0.00000
0.00000 0.00000 0.00000 ; HG2- CG- CD- OE1
6791 6790 6793 6795 3 0.00000 0.00000 0.00000
0.00000 0.00000 0.00000 ; HG2- CG- CD- OE2
6792 6790 6793 6794 3 0.00000 0.00000 0.00000
0.00000 0.00000 0.00000 ; HG3- CG- CD- OE1
6792 6790 6793 6795 3 0.00000 0.00000 0.00000
0.00000 0.00000 0.00000 ; HG3- CG- CD- OE2
6796 6798 6800 6801 3 0.00000 0.00000 0.00000
0.00000 0.00000 0.00000 ; C- N- CA- HA
6796 6798 6800 6802 3 11.71520 1.67360 15.06240 -
13.38880 0.00000 0.00000 ; C- N- CA- CB
6796 6798 6800 6810 3 1.75728 5.27184 2.25936
-7.02912 0.00000 0.00000 ; C- N- CA- C
6797 6796 6798 6799 3 29.28800 -8.36800 -20.92000
0.00000 0.00000 0.00000 ; O- C- N- H
6797 6796 6798 6800 3 20.92000 0.00000 -20.92000
0.00000 0.00000 0.00000 ; O- C- N- CA

6798 6800 6802 6803 3 0.65084 1.95253 0.00000
-2.60338 0.00000 0.00000 ; N- CA- CB- HB2
6798 6800 6802 6804 3 0.65084 1.95253 0.00000
-2.60338 0.00000 0.00000 ; N- CA- CB- HB3
6798 6800 6802 6805 3 5.99567 3.29281 -4.46014
-0.55229 1.97485 0.00000 ; N- CA- CB- CG
6798 6800 6810 6811 3 0.00000 0.00000 0.00000
0.00000 0.00000 0.00000 ; N- CA- C- O
6798 6800 6810 6812 3 17.40544 -5.02080 -13.22144
9.20480 0.00000 0.00000 ; N- CA- C- N
6799 6798 6800 6801 3 0.00000 0.00000 0.00000
0.00000 0.00000 0.00000 ; H- N- CA- HA
6799 6798 6800 6802 3 0.00000 0.00000 0.00000
0.00000 0.00000 0.00000 ; H- N- CA- CB
6799 6798 6800 6810 3 0.00000 0.00000 0.00000
0.00000 0.00000 0.00000 ; H- N- CA- C
6800 6802 6805 6806 3 0.00000 0.00000 0.00000
0.00000 0.00000 0.00000 ; CA- CB- CG- OD1
6800 6802 6805 6807 3 8.84916 -0.31380 -4.32626
5.03754 0.26778 0.00000 ; CA- CB- CG- ND2
6800 6810 6812 6813 3 20.92000 0.00000 -20.92000
0.00000 0.00000 0.00000 ; CA- C- N- CD
6800 6810 6812 6822 3 20.92000 0.00000 -20.92000
0.00000 0.00000 0.00000 ; CA- C- N- CA
6801 6800 6802 6803 3 0.65084 1.95253 0.00000
-2.60338 0.00000 0.00000 ; HA- CA- CB- HB2
6801 6800 6802 6804 3 0.65084 1.95253 0.00000
-2.60338 0.00000 0.00000 ; HA- CA- CB- HB3
6801 6800 6802 6805 3 0.65084 1.95253 0.00000
-2.60338 0.00000 0.00000 ; HA- CA- CB- CG
6801 6800 6810 6811 3 3.68192 -4.35136 0.00000
1.33888 0.00000 0.00000 ; HA- CA- C- O
6801 6800 6810 6812 3 0.00000 0.00000 0.00000
0.00000 0.00000 0.00000 ; HA- CA- C- N
6802 6800 6810 6811 3 0.00000 0.00000 0.00000
0.00000 0.00000 0.00000 ; CB- CA- C- O
6802 6800 6810 6812 3 2.51040 4.18400 1.67360
-6.69440 0.00000 0.00000 ; CB- CA- C- N
6802 6805 6807 6808 3 20.92000 0.00000 -20.92000
0.00000 0.00000 0.00000 ; CB- CG- ND2- HD21
6802 6805 6807 6809 3 20.92000 0.00000 -20.92000
0.00000 0.00000 0.00000 ; CB- CG- ND2- HD22
6803 6802 6800 6810 3 0.65084 1.95253 0.00000
-2.60338 0.00000 0.00000 ; HB2- CB- CA- C
6803 6802 6805 6806 3 3.68192 -4.35136 0.00000
1.33888 0.00000 0.00000 ; HB2- CB- CG- OD1
6803 6802 6805 6807 3 0.00000 0.00000 0.00000
0.00000 0.00000 0.00000 ; HB2- CB- CG- ND2
6804 6802 6800 6810 3 0.65084 1.95253 0.00000
-2.60338 0.00000 0.00000 ; HB3- CB- CA- C
6804 6802 6805 6806 3 3.68192 -4.35136 0.00000
1.33888 0.00000 0.00000 ; HB3- CB- CG- OD1

```
  6804   6802   6805   6807      3    0.00000    0.00000    0.00000
0.00000    0.00000    0.00000 ;    HB3-    CB-    CG-   ND2
  6805   6802   6800   6810      3    7.94542   -3.96225   -6.11701
-0.55229    3.58150    0.00000 ;     CG-    CB-    CA-     C
  6806   6805   6807   6808      3   29.28800   -8.36800  -20.92000
0.00000    0.00000    0.00000 ;    OD1-    CG-   ND2-  HD21
  6806   6805   6807   6809      3   29.28800   -8.36800  -20.92000
0.00000    0.00000    0.00000 ;    OD1-    CG-   ND2-  HD22
  6810   6812   6813   6814      3    0.00000    0.00000    0.00000
0.00000    0.00000    0.00000 ;     C-     N-    CD-   HD2
  6810   6812   6813   6815      3    0.00000    0.00000    0.00000
0.00000    0.00000    0.00000 ;     C-     N-    CD-   HD3
  6810   6812   6813   6816      3   10.04160   -3.34720   16.73600
-6.69440    0.00000    0.00000 ;     C-     N-    CD-    CG
  6810   6812   6822   6819      3   11.71520    1.67360   15.06240  -
13.38880    0.00000    0.00000 ;     C-     N-    CA-    CB
  6810   6812   6822   6823      3    0.00000    0.00000    0.00000
0.00000    0.00000    0.00000 ;     C-     N-    CA-    HA
  6810   6812   6822   6824      3    1.75728    5.27184    2.25936
-7.02912    0.00000    0.00000 ;     C-     N-    CA-     C
  6811   6810   6812   6813      3   20.92000    0.00000  -20.92000
0.00000    0.00000    0.00000 ;     O-     C-     N-    CD
  6811   6810   6812   6822      3   20.92000    0.00000  -20.92000
0.00000    0.00000    0.00000 ;     O-     C-     N-    CA
  6812   6813   6816   6817      3    0.65084    1.95253    0.00000
-2.60338    0.00000    0.00000 ;     N-    CD-    CG-   HG2
  6812   6813   6816   6818      3    0.65084    1.95253    0.00000
-2.60338    0.00000    0.00000 ;     N-    CD-    CG-   HG3
  6812   6813   6816   6819      3    0.65084    1.95253    0.00000
-2.60338    0.00000    0.00000 ;     N-    CD-    CG-    CB
  6812   6822   6819   6816      3    0.65084    1.95253    0.00000
-2.60338    0.00000    0.00000 ;     N-    CA-    CB-    CG
  6812   6822   6819   6820      3    0.65084    1.95253    0.00000
-2.60338    0.00000    0.00000 ;     N-    CA-    CB-   HB2
  6812   6822   6819   6821      3    0.65084    1.95253    0.00000
-2.60338    0.00000    0.00000 ;     N-    CA-    CB-   HB3
  6812   6822   6824   6825      3    0.00000    0.00000    0.00000
0.00000    0.00000    0.00000 ;     N-    CA-     C-     O
  6812   6822   6824   6826      3   17.40544   -5.02080  -13.22144
9.20480    0.00000    0.00000 ;     N-    CA-     C-     N
  6813   6812   6822   6819      3    0.00000    0.00000    0.00000
0.00000    0.00000    0.00000 ;    CD-     N-    CA-    CB
  6813   6812   6822   6823      3    0.00000    0.00000    0.00000
0.00000    0.00000    0.00000 ;    CD-     N-    CA-    HA
  6813   6812   6822   6824      3    0.00000    0.00000    0.00000
0.00000    0.00000    0.00000 ;    CD-     N-    CA-     C
  6813   6816   6819   6820      3    0.66944    2.00832    0.00000
-2.67776    0.00000    0.00000 ;    CD-    CG-    CB-   HB2
  6813   6816   6819   6821      3    0.66944    2.00832    0.00000
-2.67776    0.00000    0.00000 ;    CD-    CG-    CB-   HB3
  6813   6816   6819   6822      3    3.68192    3.09616   -2.09200
-3.01248    0.00000    0.00000 ;    CD-    CG-    CB-    CA
```

```
  6814   6813   6812   6822       3    0.00000    0.00000    0.00000
0.00000    0.00000    0.00000 ;    HD2-    CD-     N-    CA
  6814   6813   6816   6817       3    0.65084    1.95253    0.00000
-2.60338    0.00000    0.00000 ;    HD2-    CD-    CG-   HG2
  6814   6813   6816   6818       3    0.65084    1.95253    0.00000
-2.60338    0.00000    0.00000 ;    HD2-    CD-    CG-   HG3
  6814   6813   6816   6819       3    0.65084    1.95253    0.00000
-2.60338    0.00000    0.00000 ;    HD2-    CD-    CG-    CB
  6815   6813   6812   6822       3    0.00000    0.00000    0.00000
0.00000    0.00000    0.00000 ;    HD3-    CD-     N-    CA
  6815   6813   6816   6817       3    0.65084    1.95253    0.00000
-2.60338    0.00000    0.00000 ;    HD3-    CD-    CG-   HG2
  6815   6813   6816   6818       3    0.65084    1.95253    0.00000
-2.60338    0.00000    0.00000 ;    HD3-    CD-    CG-   HG3
  6815   6813   6816   6819       3    0.65084    1.95253    0.00000
-2.60338    0.00000    0.00000 ;    HD3-    CD-    CG-    CB
  6816   6813   6812   6822       3    0.00000    0.00000    0.00000
0.00000    0.00000    0.00000 ;     CG-    CD-     N-    CA
  6816   6819   6822   6823       3    0.65084    1.95253    0.00000
-2.60338    0.00000    0.00000 ;     CG-    CB-    CA-    HA
  6816   6819   6822   6824       3    0.65084    1.95253    0.00000
-2.60338    0.00000    0.00000 ;     CG-    CB-    CA-     C
  6817   6816   6819   6820       3    0.62760    1.88280    0.00000
-2.51040    0.00000    0.00000 ;    HG2-    CG-    CB-   HB2
  6817   6816   6819   6821       3    0.62760    1.88280    0.00000
-2.51040    0.00000    0.00000 ;    HG2-    CG-    CB-   HB3
  6817   6816   6819   6822       3    0.66944    2.00832    0.00000
-2.67776    0.00000    0.00000 ;    HG2-    CG-    CB-    CA
  6818   6816   6819   6820       3    0.62760    1.88280    0.00000
-2.51040    0.00000    0.00000 ;    HG3-    CG-    CB-   HB2
  6818   6816   6819   6821       3    0.62760    1.88280    0.00000
-2.51040    0.00000    0.00000 ;    HG3-    CG-    CB-   HB3
  6818   6816   6819   6822       3    0.66944    2.00832    0.00000
-2.67776    0.00000    0.00000 ;    HG3-    CG-    CB-    CA
  6819   6822   6824   6825       3    0.00000    0.00000    0.00000
0.00000    0.00000    0.00000 ;     CB-    CA-     C-     O
  6819   6822   6824   6826       3    2.51040    4.18400    1.67360
-6.69440    0.00000    0.00000 ;     CB-    CA-     C-     N
  6820   6819   6822   6823       3    0.65084    1.95253    0.00000
-2.60338    0.00000    0.00000 ;    HB2-    CB-    CA-    HA
  6820   6819   6822   6824       3    0.65084    1.95253    0.00000
-2.60338    0.00000    0.00000 ;    HB2-    CB-    CA-     C
  6821   6819   6822   6823       3    0.65084    1.95253    0.00000
-2.60338    0.00000    0.00000 ;    HB3-    CB-    CA-    HA
  6821   6819   6822   6824       3    0.65084    1.95253    0.00000
-2.60338    0.00000    0.00000 ;    HB3-    CB-    CA-     C
  6822   6824   6826   6827       3   20.92000    0.00000  -20.92000
0.00000    0.00000    0.00000 ;     CA-     C-     N-     H
  6822   6824   6826   6828       3   20.92000    0.00000  -20.92000
0.00000    0.00000    0.00000 ;     CA-     C-     N-    CA
  6823   6822   6824   6825       3    3.68192   -4.35136    0.00000
1.33888    0.00000    0.00000 ;     HA-    CA-     C-     O
```

6823 6822 6824 6826 3 0.00000 0.00000 0.00000
0.00000 0.00000 0.00000 ; HA- CA- C- N
6824 6826 6828 6829 3 0.00000 0.00000 0.00000
0.00000 0.00000 0.00000 ; C- N- CA- HA
6824 6826 6828 6830 3 11.71520 1.67360 15.06240 -
13.38880 0.00000 0.00000 ; C- N- CA- CB
6824 6826 6828 6843 3 1.75728 5.27184 2.25936
-7.02912 0.00000 0.00000 ; C- N- CA- C
6825 6824 6826 6827 3 29.28800 -8.36800 -20.92000
0.00000 0.00000 0.00000 ; O- C- N- H
6825 6824 6826 6828 3 20.92000 0.00000 -20.92000
0.00000 0.00000 0.00000 ; O- C- N- CA
6826 6828 6830 6831 3 0.65084 1.95253 0.00000
-2.60338 0.00000 0.00000 ; N- CA- CB- HB2
6826 6828 6830 6832 3 0.65084 1.95253 0.00000
-2.60338 0.00000 0.00000 ; N- CA- CB- HB3
6826 6828 6830 6833 3 3.79070 1.39746 -4.61077
-2.40998 2.44346 0.00000 ; N- CA- CB- CG
6826 6828 6843 6844 3 0.00000 0.00000 0.00000
0.00000 0.00000 0.00000 ; N- CA- C- O
6826 6828 6843 6845 3 17.40544 -5.02080 -13.22144
9.20480 0.00000 0.00000 ; N- CA- C- N
6827 6826 6828 6829 3 0.00000 0.00000 0.00000
0.00000 0.00000 0.00000 ; H- N- CA- HA
6827 6826 6828 6830 3 0.00000 0.00000 0.00000
0.00000 0.00000 0.00000 ; H- N- CA- CB
6827 6826 6828 6843 3 0.00000 0.00000 0.00000
0.00000 0.00000 0.00000 ; H- N- CA- C
6828 6830 6833 6834 3 0.66944 2.00832 0.00000
-2.67776 0.00000 0.00000 ; CA- CB- CG- HG
6828 6830 6833 6835 3 3.90367 0.19665 -6.21742
-2.37651 5.99149 0.00000 ; CA- CB- CG- CD1
6828 6830 6833 6839 3 3.90367 0.19665 -6.21742
-2.37651 5.99149 0.00000 ; CA- CB- CG- CD2
6828 6843 6845 6846 3 20.92000 0.00000 -20.92000
0.00000 0.00000 0.00000 ; CA- C- N- H
6828 6843 6845 6847 3 20.92000 0.00000 -20.92000
0.00000 0.00000 0.00000 ; CA- C- N- CA
6829 6828 6830 6831 3 0.65084 1.95253 0.00000
-2.60338 0.00000 0.00000 ; HA- CA- CB- HB2
6829 6828 6830 6832 3 0.65084 1.95253 0.00000
-2.60338 0.00000 0.00000 ; HA- CA- CB- HB3
6829 6828 6830 6833 3 0.65084 1.95253 0.00000
-2.60338 0.00000 0.00000 ; HA- CA- CB- CG
6829 6828 6843 6844 3 3.68192 -4.35136 0.00000
1.33888 0.00000 0.00000 ; HA- CA- C- O
6829 6828 6843 6845 3 0.00000 0.00000 0.00000
0.00000 0.00000 0.00000 ; HA- CA- C- N
6830 6828 6843 6844 3 0.00000 0.00000 0.00000
0.00000 0.00000 0.00000 ; CB- CA- C- O
6830 6828 6843 6845 3 2.51040 4.18400 1.67360
-6.69440 0.00000 0.00000 ; CB- CA- C- N

```
  6830   6833   6835   6836      3    0.66944    2.00832    0.00000
-2.67776    0.00000    0.00000 ;     CB-    CG-   CD1-  HD11
  6830   6833   6835   6837      3    0.66944    2.00832    0.00000
-2.67776    0.00000    0.00000 ;     CB-    CG-   CD1-  HD12
  6830   6833   6835   6838      3    0.66944    2.00832    0.00000
-2.67776    0.00000    0.00000 ;     CB-    CG-   CD1-  HD13
  6830   6833   6839   6840      3    0.66944    2.00832    0.00000
-2.67776    0.00000    0.00000 ;     CB-    CG-   CD2-  HD21
  6830   6833   6839   6841      3    0.66944    2.00832    0.00000
-2.67776    0.00000    0.00000 ;     CB-    CG-   CD2-  HD22
  6830   6833   6839   6842      3    0.66944    2.00832    0.00000
-2.67776    0.00000    0.00000 ;     CB-    CG-   CD2-  HD23
  6831   6830   6828   6843      3    0.65084    1.95253    0.00000
-2.60338    0.00000    0.00000 ;    HB2-    CB-    CA-     C
  6831   6830   6833   6834      3    0.62760    1.88280    0.00000
-2.51040    0.00000    0.00000 ;    HB2-    CB-    CG-    HG
  6831   6830   6833   6835      3    0.66944    2.00832    0.00000
-2.67776    0.00000    0.00000 ;    HB2-    CB-    CG-   CD1
  6831   6830   6833   6839      3    0.66944    2.00832    0.00000
-2.67776    0.00000    0.00000 ;    HB2-    CB-    CG-   CD2
  6832   6830   6828   6843      3    0.65084    1.95253    0.00000
-2.60338    0.00000    0.00000 ;    HB3-    CB-    CA-     C
  6832   6830   6833   6834      3    0.62760    1.88280    0.00000
-2.51040    0.00000    0.00000 ;    HB3-    CB-    CG-    HG
  6832   6830   6833   6835      3    0.66944    2.00832    0.00000
-2.67776    0.00000    0.00000 ;    HB3-    CB-    CG-   CD1
  6832   6830   6833   6839      3    0.66944    2.00832    0.00000
-2.67776    0.00000    0.00000 ;    HB3-    CB-    CG-   CD2
  6833   6830   6828   6843      3   10.33448   -1.14642  -11.54784
-2.40998    6.35968    0.00000 ;     CG-    CB-    CA-     C
  6834   6833   6835   6836      3    0.62760    1.88280    0.00000
-2.51040    0.00000    0.00000 ;     HG-    CG-   CD1-  HD11
  6834   6833   6835   6837      3    0.62760    1.88280    0.00000
-2.51040    0.00000    0.00000 ;     HG-    CG-   CD1-  HD12
  6834   6833   6835   6838      3    0.62760    1.88280    0.00000
-2.51040    0.00000    0.00000 ;     HG-    CG-   CD1-  HD13
  6834   6833   6839   6840      3    0.62760    1.88280    0.00000
-2.51040    0.00000    0.00000 ;     HG-    CG-   CD2-  HD21
  6834   6833   6839   6841      3    0.62760    1.88280    0.00000
-2.51040    0.00000    0.00000 ;     HG-    CG-   CD2-  HD22
  6834   6833   6839   6842      3    0.62760    1.88280    0.00000
-2.51040    0.00000    0.00000 ;     HG-    CG-   CD2-  HD23
  6835   6833   6839   6840      3    0.66944    2.00832    0.00000
-2.67776    0.00000    0.00000 ;    CD1-    CG-   CD2-  HD21
  6835   6833   6839   6841      3    0.66944    2.00832    0.00000
-2.67776    0.00000    0.00000 ;    CD1-    CG-   CD2-  HD22
  6835   6833   6839   6842      3    0.66944    2.00832    0.00000
-2.67776    0.00000    0.00000 ;    CD1-    CG-   CD2-  HD23
  6836   6835   6833   6839      3    0.66944    2.00832    0.00000
-2.67776    0.00000    0.00000 ;   HD11-   CD1-    CG-   CD2
  6837   6835   6833   6839      3    0.66944    2.00832    0.00000
-2.67776    0.00000    0.00000 ;   HD12-   CD1-    CG-   CD2
```

```
  6838   6835   6833   6839       3    0.66944    2.00832    0.00000
-2.67776    0.00000    0.00000 ;   HD13-   CD1-    CG-   CD2
  6843   6845   6847   6848       3    0.00000    0.00000    0.00000
0.00000    0.00000    0.00000 ;      C-     N-    CA-    HA
  6843   6845   6847   6849       3   11.71520    1.67360   15.06240  -
13.38880    0.00000    0.00000 ;      C-     N-    CA-    CB
  6843   6845   6847   6862       3    1.75728    5.27184    2.25936
-7.02912    0.00000    0.00000 ;      C-     N-    CA-     C
  6844   6843   6845   6846       3   29.28800   -8.36800  -20.92000
0.00000    0.00000    0.00000 ;      O-     C-     N-     H
  6844   6843   6845   6847       3   20.92000    0.00000  -20.92000
0.00000    0.00000    0.00000 ;      O-     C-     N-    CA
  6845   6847   6849   6850       3    0.65084    1.95253    0.00000
-2.60338    0.00000    0.00000 ;      N-    CA-    CB-    HB
  6845   6847   6849   6851       3    3.83673    0.44769   -1.77402
-2.47693   -0.03347    0.00000 ;      N-    CA-    CB-   CG2
  6845   6847   6849   6855       3    2.97482    0.12134    2.04179
-1.89117   -3.24678    0.00000 ;      N-    CA-    CB-   CG1
  6845   6847   6862   6863       3    0.00000    0.00000    0.00000
0.00000    0.00000    0.00000 ;      N-    CA-     C-     O
  6845   6847   6862   6864       3   17.40544   -5.02080  -13.22144
9.20480    0.00000    0.00000 ;      N-    CA-     C-     N
  6846   6845   6847   6848       3    0.00000    0.00000    0.00000
0.00000    0.00000    0.00000 ;      H-     N-    CA-    HA
  6846   6845   6847   6849       3    0.00000    0.00000    0.00000
0.00000    0.00000    0.00000 ;      H-     N-    CA-    CB
  6846   6845   6847   6862       3    0.00000    0.00000    0.00000
0.00000    0.00000    0.00000 ;      H-     N-    CA-     C
  6847   6849   6851   6852       3    0.66944    2.00832    0.00000
-2.67776    0.00000    0.00000 ;     CA-    CB-   CG2-  HG21
  6847   6849   6851   6853       3    0.66944    2.00832    0.00000
-2.67776    0.00000    0.00000 ;     CA-    CB-   CG2-  HG22
  6847   6849   6851   6854       3    0.66944    2.00832    0.00000
-2.67776    0.00000    0.00000 ;     CA-    CB-   CG2-  HG23
  6847   6849   6855   6856       3    0.66944    2.00832    0.00000
-2.67776    0.00000    0.00000 ;     CA-    CB-   CG1-  HG12
  6847   6849   6855   6857       3    0.66944    2.00832    0.00000
-2.67776    0.00000    0.00000 ;     CA-    CB-   CG1-  HG13
  6847   6849   6855   6858       3    4.24258   -0.52718   -7.25506
-1.79075    7.69856    0.00000 ;     CA-    CB-   CG1-   CD1
  6847   6862   6864   6865       3   20.92000    0.00000  -20.92000
0.00000    0.00000    0.00000 ;     CA-     C-     N-     H
  6847   6862   6864   6866       3   20.92000    0.00000  -20.92000
0.00000    0.00000    0.00000 ;     CA-     C-     N-    CA
  6848   6847   6849   6850       3    0.65084    1.95253    0.00000
-2.60338    0.00000    0.00000 ;     HA-    CA-    CB-    HB
  6848   6847   6849   6851       3    0.65084    1.95253    0.00000
-2.60338    0.00000    0.00000 ;     HA-    CA-    CB-   CG2
  6848   6847   6849   6855       3    0.65084    1.95253    0.00000
-2.60338    0.00000    0.00000 ;     HA-    CA-    CB-   CG1
  6848   6847   6862   6863       3    3.68192   -4.35136    0.00000
1.33888    0.00000    0.00000 ;     HA-    CA-     C-     O
```

6848 6847 6862 6864 3 0.00000 0.00000 0.00000
0.00000 0.00000 0.00000 ; HA- CA- C- N
6849 6847 6862 6863 3 0.00000 0.00000 0.00000
0.00000 0.00000 0.00000 ; CB- CA- C- O
6849 6847 6862 6864 3 2.51040 4.18400 1.67360
-6.69440 0.00000 0.00000 ; CB- CA- C- N
6849 6855 6858 6859 3 0.66944 2.00832 0.00000
-2.67776 0.00000 0.00000 ; CB- CG1- CD1- HD11
6849 6855 6858 6860 3 0.66944 2.00832 0.00000
-2.67776 0.00000 0.00000 ; CB- CG1- CD1- HD12
6849 6855 6858 6861 3 0.66944 2.00832 0.00000
-2.67776 0.00000 0.00000 ; CB- CG1- CD1- HD13
6850 6849 6847 6862 3 0.65084 1.95253 0.00000
-2.60338 0.00000 0.00000 ; HB- CB- CA- C
6850 6849 6851 6852 3 0.62760 1.88280 0.00000
-2.51040 0.00000 0.00000 ; HB- CB- CG2- HG21
6850 6849 6851 6853 3 0.62760 1.88280 0.00000
-2.51040 0.00000 0.00000 ; HB- CB- CG2- HG22
6850 6849 6851 6854 3 0.62760 1.88280 0.00000
-2.51040 0.00000 0.00000 ; HB- CB- CG2- HG23
6850 6849 6855 6856 3 0.62760 1.88280 0.00000
-2.51040 0.00000 0.00000 ; HB- CB- CG1- HG12
6850 6849 6855 6857 3 0.62760 1.88280 0.00000
-2.51040 0.00000 0.00000 ; HB- CB- CG1- HG13
6850 6849 6855 6858 3 0.66944 2.00832 0.00000
-2.67776 0.00000 0.00000 ; HB- CB- CG1- CD1
6851 6849 6847 6862 3 5.67350 3.55640 -6.16722
-2.47693 3.74886 0.00000 ; CG2- CB- CA- C
6851 6849 6855 6856 3 0.66944 2.00832 0.00000
-2.67776 0.00000 0.00000 ; CG2- CB- CG1- HG12
6851 6849 6855 6857 3 0.66944 2.00832 0.00000
-2.67776 0.00000 0.00000 ; CG2- CB- CG1- HG13
6851 6849 6855 6858 3 3.81162 0.49790 -8.14206
-1.79075 7.49773 0.00000 ; CG2- CB- CG1- CD1
6852 6851 6849 6855 3 0.66944 2.00832 0.00000
-2.67776 0.00000 0.00000 ; HG21- CG2- CB- CG1
6853 6851 6849 6855 3 0.66944 2.00832 0.00000
-2.67776 0.00000 0.00000 ; HG22- CG2- CB- CG1
6854 6851 6849 6855 3 0.66944 2.00832 0.00000
-2.67776 0.00000 0.00000 ; HG23- CG2- CB- CG1
6855 6849 6847 6862 3 8.26340 0.74057 -9.99976
-1.89117 3.84928 0.00000 ; CG1- CB- CA- C
6856 6855 6858 6859 3 0.62760 1.88280 0.00000
-2.51040 0.00000 0.00000 ; HG12- CG1- CD1- HD11
6856 6855 6858 6860 3 0.62760 1.88280 0.00000
-2.51040 0.00000 0.00000 ; HG12- CG1- CD1- HD12
6856 6855 6858 6861 3 0.62760 1.88280 0.00000
-2.51040 0.00000 0.00000 ; HG12- CG1- CD1- HD13
6857 6855 6858 6859 3 0.62760 1.88280 0.00000
-2.51040 0.00000 0.00000 ; HG13- CG1- CD1- HD11
6857 6855 6858 6860 3 0.62760 1.88280 0.00000
-2.51040 0.00000 0.00000 ; HG13- CG1- CD1- HD12

6857 6855 6858 6861 3 0.62760 1.88280 0.00000
-2.51040 0.00000 0.00000 ; HG13- CG1- CD1- HD13
6862 6864 6866 6867 3 0.00000 0.00000 0.00000
0.00000 0.00000 0.00000 ; C- N- CA- HA
6862 6864 6866 6868 3 11.71520 1.67360 15.06240 -
13.38880 0.00000 0.00000 ; C- N- CA- CB
6862 6864 6866 6874 3 1.75728 5.27184 2.25936
-7.02912 0.00000 0.00000 ; C- N- CA- C
6863 6862 6864 6865 3 29.28800 -8.36800 -20.92000
0.00000 0.00000 0.00000 ; O- C- N- H
6863 6862 6864 6866 3 20.92000 0.00000 -20.92000
0.00000 0.00000 0.00000 ; O- C- N- CA
6864 6866 6868 6869 3 0.65084 1.95253 0.00000
-2.60338 0.00000 0.00000 ; N- CA- CB- HB2
6864 6866 6868 6870 3 0.65084 1.95253 0.00000
-2.60338 0.00000 0.00000 ; N- CA- CB- HB3
6864 6866 6868 6871 3 15.41386 9.74035 -8.39310
-0.97069 2.97901 0.00000 ; N- CA- CB- CG
6864 6866 6874 6875 3 0.00000 0.00000 0.00000
0.00000 0.00000 0.00000 ; N- CA- C- O
6864 6866 6874 6876 3 17.40544 -5.02080 -13.22144
9.20480 0.00000 0.00000 ; N- CA- C- N
6865 6864 6866 6867 3 0.00000 0.00000 0.00000
0.00000 0.00000 0.00000 ; H- N- CA- HA
6865 6864 6866 6868 3 0.00000 0.00000 0.00000
0.00000 0.00000 0.00000 ; H- N- CA- CB
6865 6864 6866 6874 3 0.00000 0.00000 0.00000
0.00000 0.00000 0.00000 ; H- N- CA- C
6866 6868 6871 6872 3 6.43499 0.00000 -5.39736
0.00000 -1.03763 0.00000 ; CA- CB- CG- OD1
6866 6868 6871 6873 3 6.43499 0.00000 -5.39736
0.00000 -1.03763 0.00000 ; CA- CB- CG- OD2
6866 6874 6876 6877 3 20.92000 0.00000 -20.92000
0.00000 0.00000 0.00000 ; CA- C- N- H
6866 6874 6876 6878 3 20.92000 0.00000 -20.92000
0.00000 0.00000 0.00000 ; CA- C- N- CA
6867 6866 6868 6869 3 0.65084 1.95253 0.00000
-2.60338 0.00000 0.00000 ; HA- CA- CB- HB2
6867 6866 6868 6870 3 0.65084 1.95253 0.00000
-2.60338 0.00000 0.00000 ; HA- CA- CB- HB3
6867 6866 6868 6871 3 0.65084 1.95253 0.00000
-2.60338 0.00000 0.00000 ; HA- CA- CB- CG
6867 6866 6874 6875 3 3.68192 -4.35136 0.00000
1.33888 0.00000 0.00000 ; HA- CA- C- O
6867 6866 6874 6876 3 0.00000 0.00000 0.00000
0.00000 0.00000 0.00000 ; HA- CA- C- N
6868 6866 6874 6875 3 0.00000 0.00000 0.00000
0.00000 0.00000 0.00000 ; CB- CA- C- O
6868 6866 6874 6876 3 2.51040 4.18400 1.67360
-6.69440 0.00000 0.00000 ; CB- CA- C- N
6869 6868 6866 6874 3 0.65084 1.95253 0.00000
-2.60338 0.00000 0.00000 ; HB2- CB- CA- C

6869 6868 6871 6872 3 0.00000 0.00000 0.00000
0.00000 0.00000 0.00000 ; HB2- CB- CG- OD1
6869 6868 6871 6873 3 0.00000 0.00000 0.00000
0.00000 0.00000 0.00000 ; HB2- CB- CG- OD2
6870 6868 6866 6874 3 0.65084 1.95253 0.00000
-2.60338 0.00000 0.00000 ; HB3- CB- CA- C
6870 6868 6871 6872 3 0.00000 0.00000 0.00000
0.00000 0.00000 0.00000 ; HB3- CB- CG- OD1
6870 6868 6871 6873 3 0.00000 0.00000 0.00000
0.00000 0.00000 0.00000 ; HB3- CB- CG- OD2
6871 6868 6866 6874 3 7.14627 -1.04600 -8.99560
-0.97069 5.15469 0.00000 ; CG- CB- CA- C
6874 6876 6878 6879 3 0.00000 0.00000 0.00000
0.00000 0.00000 0.00000 ; C- N- CA- HA
6874 6876 6878 6880 3 11.71520 1.67360 15.06240 -
13.38880 0.00000 0.00000 ; C- N- CA- CB
6874 6876 6878 6896 3 1.75728 5.27184 2.25936
-7.02912 0.00000 0.00000 ; C- N- CA- C
6875 6874 6876 6877 3 29.28800 -8.36800 -20.92000
0.00000 0.00000 0.00000 ; O- C- N- H
6875 6874 6876 6878 3 20.92000 0.00000 -20.92000
0.00000 0.00000 0.00000 ; O- C- N- CA
6876 6878 6880 6881 3 0.65084 1.95253 0.00000
-2.60338 0.00000 0.00000 ; N- CA- CB- HB2
6876 6878 6880 6882 3 0.65084 1.95253 0.00000
-2.60338 0.00000 0.00000 ; N- CA- CB- HB3
6876 6878 6880 6883 3 0.65084 1.95253 0.00000
-2.60338 0.00000 0.00000 ; N- CA- CB- CG
6876 6878 6896 6897 3 0.00000 0.00000 0.00000
0.00000 0.00000 0.00000 ; N- CA- C- O
6876 6878 6896 6898 3 17.40544 -5.02080 -13.22144
9.20480 0.00000 0.00000 ; N- CA- C- N
6877 6876 6878 6879 3 0.00000 0.00000 0.00000
0.00000 0.00000 0.00000 ; H- N- CA- HA
6877 6876 6878 6880 3 0.00000 0.00000 0.00000
0.00000 0.00000 0.00000 ; H- N- CA- CB
6877 6876 6878 6896 3 0.00000 0.00000 0.00000
0.00000 0.00000 0.00000 ; H- N- CA- C
6878 6880 6883 6884 3 0.66944 2.00832 0.00000
-2.67776 0.00000 0.00000 ; CA- CB- CG- HG2
6878 6880 6883 6885 3 0.66944 2.00832 0.00000
-2.67776 0.00000 0.00000 ; CA- CB- CG- HG3
6878 6880 6883 6886 3 3.68192 3.09616 -2.09200
-3.01248 0.00000 0.00000 ; CA- CB- CG- CD
6878 6896 6898 6899 3 20.92000 0.00000 -20.92000
0.00000 0.00000 0.00000 ; CA- C- N- H
6878 6896 6898 6900 3 20.92000 0.00000 -20.92000
0.00000 0.00000 0.00000 ; CA- C- N- CA
6879 6878 6880 6881 3 0.65084 1.95253 0.00000
-2.60338 0.00000 0.00000 ; HA- CA- CB- HB2
6879 6878 6880 6882 3 0.65084 1.95253 0.00000
-2.60338 0.00000 0.00000 ; HA- CA- CB- HB3

6879 6878 6880 6883 3 0.65084 1.95253 0.00000
-2.60338 0.00000 0.00000 ; HA- CA- CB- CG
6879 6878 6896 6897 3 3.68192 -4.35136 0.00000
1.33888 0.00000 0.00000 ; HA- CA- C- O
6879 6878 6896 6898 3 0.00000 0.00000 0.00000
0.00000 0.00000 0.00000 ; HA- CA- C- N
6880 6878 6896 6897 3 0.00000 0.00000 0.00000
0.00000 0.00000 0.00000 ; CB- CA- C- O
6880 6878 6896 6898 3 2.51040 4.18400 1.67360
-6.69440 0.00000 0.00000 ; CB- CA- C- N
6880 6883 6886 6887 3 0.66944 2.00832 0.00000
-2.67776 0.00000 0.00000 ; CB- CG- CD- HD2
6880 6883 6886 6888 3 0.66944 2.00832 0.00000
-2.67776 0.00000 0.00000 ; CB- CG- CD- HD3
6880 6883 6886 6889 3 3.68192 3.09616 -2.09200
-3.01248 0.00000 0.00000 ; CB- CG- CD- CE
6881 6880 6878 6896 3 0.65084 1.95253 0.00000
-2.60338 0.00000 0.00000 ; HB2- CB- CA- C
6881 6880 6883 6884 3 0.62760 1.88280 0.00000
-2.51040 0.00000 0.00000 ; HB2- CB- CG- HG2
6881 6880 6883 6885 3 0.62760 1.88280 0.00000
-2.51040 0.00000 0.00000 ; HB2- CB- CG- HG3
6881 6880 6883 6886 3 0.66944 2.00832 0.00000
-2.67776 0.00000 0.00000 ; HB2- CB- CG- CD
6882 6880 6878 6896 3 0.65084 1.95253 0.00000
-2.60338 0.00000 0.00000 ; HB3- CB- CA- C
6882 6880 6883 6884 3 0.62760 1.88280 0.00000
-2.51040 0.00000 0.00000 ; HB3- CB- CG- HG2
6882 6880 6883 6885 3 0.62760 1.88280 0.00000
-2.51040 0.00000 0.00000 ; HB3- CB- CG- HG3
6882 6880 6883 6886 3 0.66944 2.00832 0.00000
-2.67776 0.00000 0.00000 ; HB3- CB- CG- CD
6883 6880 6878 6896 3 0.65084 1.95253 0.00000
-2.60338 0.00000 0.00000 ; CG- CB- CA- C
6883 6886 6889 6890 3 0.65084 1.95253 0.00000
-2.60338 0.00000 0.00000 ; CG- CD- CE- HE2
6883 6886 6889 6891 3 0.65084 1.95253 0.00000
-2.60338 0.00000 0.00000 ; CG- CD- CE- HE3
6883 6886 6889 6892 3 0.65084 1.95253 0.00000
-2.60338 0.00000 0.00000 ; CG- CD- CE- NZ
6884 6883 6886 6887 3 0.62760 1.88280 0.00000
-2.51040 0.00000 0.00000 ; HG2- CG- CD- HD2
6884 6883 6886 6888 3 0.62760 1.88280 0.00000
-2.51040 0.00000 0.00000 ; HG2- CG- CD- HD3
6884 6883 6886 6889 3 0.66944 2.00832 0.00000
-2.67776 0.00000 0.00000 ; HG2- CG- CD- CE
6885 6883 6886 6887 3 0.62760 1.88280 0.00000
-2.51040 0.00000 0.00000 ; HG3- CG- CD- HD2
6885 6883 6886 6888 3 0.62760 1.88280 0.00000
-2.51040 0.00000 0.00000 ; HG3- CG- CD- HD3
6885 6883 6886 6889 3 0.66944 2.00832 0.00000
-2.67776 0.00000 0.00000 ; HG3- CG- CD- CE

6886 6889 6892 6893 3 0.65084 1.95253 0.00000
-2.60338 0.00000 0.00000 ; CD- CE- NZ- HZ1
6886 6889 6892 6894 3 0.65084 1.95253 0.00000
-2.60338 0.00000 0.00000 ; CD- CE- NZ- HZ2
6886 6889 6892 6895 3 0.65084 1.95253 0.00000
-2.60338 0.00000 0.00000 ; CD- CE- NZ- HZ3
6887 6886 6889 6890 3 0.65084 1.95253 0.00000
-2.60338 0.00000 0.00000 ; HD2- CD- CE- HE2
6887 6886 6889 6891 3 0.65084 1.95253 0.00000
-2.60338 0.00000 0.00000 ; HD2- CD- CE- HE3
6887 6886 6889 6892 3 0.65084 1.95253 0.00000
-2.60338 0.00000 0.00000 ; HD2- CD- CE- NZ
6888 6886 6889 6890 3 0.65084 1.95253 0.00000
-2.60338 0.00000 0.00000 ; HD3- CD- CE- HE2
6888 6886 6889 6891 3 0.65084 1.95253 0.00000
-2.60338 0.00000 0.00000 ; HD3- CD- CE- HE3
6888 6886 6889 6892 3 0.65084 1.95253 0.00000
-2.60338 0.00000 0.00000 ; HD3- CD- CE- NZ
6890 6889 6892 6893 3 0.65084 1.95253 0.00000
-2.60338 0.00000 0.00000 ; HE2- CE- NZ- HZ1
6890 6889 6892 6894 3 0.65084 1.95253 0.00000
-2.60338 0.00000 0.00000 ; HE2- CE- NZ- HZ2
6890 6889 6892 6895 3 0.65084 1.95253 0.00000
-2.60338 0.00000 0.00000 ; HE2- CE- NZ- HZ3
6891 6889 6892 6893 3 0.65084 1.95253 0.00000
-2.60338 0.00000 0.00000 ; HE3- CE- NZ- HZ1
6891 6889 6892 6894 3 0.65084 1.95253 0.00000
-2.60338 0.00000 0.00000 ; HE3- CE- NZ- HZ2
6891 6889 6892 6895 3 0.65084 1.95253 0.00000
-2.60338 0.00000 0.00000 ; HE3- CE- NZ- HZ3
6896 6898 6900 6901 3 0.00000 0.00000 0.00000
0.00000 0.00000 0.00000 ; C- N- CA- HA
6896 6898 6900 6902 3 11.71520 1.67360 15.06240 -
13.38880 0.00000 0.00000 ; C- N- CA- CB
6896 6898 6900 6906 3 1.75728 5.27184 2.25936
-7.02912 0.00000 0.00000 ; C- N- CA- C
6897 6896 6898 6899 3 29.28800 -8.36800 -20.92000
0.00000 0.00000 0.00000 ; O- C- N- H
6897 6896 6898 6900 3 20.92000 0.00000 -20.92000
0.00000 0.00000 0.00000 ; O- C- N- CA
6898 6900 6902 6903 3 0.65084 1.95253 0.00000
-2.60338 0.00000 0.00000 ; N- CA- CB- HB1
6898 6900 6902 6904 3 0.65084 1.95253 0.00000
-2.60338 0.00000 0.00000 ; N- CA- CB- HB2
6898 6900 6902 6905 3 0.65084 1.95253 0.00000
-2.60338 0.00000 0.00000 ; N- CA- CB- HB3
6898 6900 6906 6907 3 0.00000 0.00000 0.00000
0.00000 0.00000 0.00000 ; N- CA- C- O
6898 6900 6906 6908 3 17.40544 -5.02080 -13.22144
9.20480 0.00000 0.00000 ; N- CA- C- N
6899 6898 6900 6901 3 0.00000 0.00000 0.00000
0.00000 0.00000 0.00000 ; H- N- CA- HA

6899 6898 6900 6902 3 0.00000 0.00000 0.00000
0.00000 0.00000 0.00000 ; H- N- CA- CB
6899 6898 6900 6906 3 0.00000 0.00000 0.00000
0.00000 0.00000 0.00000 ; H- N- CA- C
6900 6906 6908 6909 3 20.92000 0.00000 -20.92000
0.00000 0.00000 0.00000 ; CA- C- N- H
6900 6906 6908 6910 3 20.92000 0.00000 -20.92000
0.00000 0.00000 0.00000 ; CA- C- N- CA
6901 6900 6902 6903 3 0.65084 1.95253 0.00000
-2.60338 0.00000 0.00000 ; HA- CA- CB- HB1
6901 6900 6902 6904 3 0.65084 1.95253 0.00000
-2.60338 0.00000 0.00000 ; HA- CA- CB- HB2
6901 6900 6902 6905 3 0.65084 1.95253 0.00000
-2.60338 0.00000 0.00000 ; HA- CA- CB- HB3
6901 6900 6906 6907 3 3.68192 -4.35136 0.00000
1.33888 0.00000 0.00000 ; HA- CA- C- O
6901 6900 6906 6908 3 0.00000 0.00000 0.00000
0.00000 0.00000 0.00000 ; HA- CA- C- N
6902 6900 6906 6907 3 0.00000 0.00000 0.00000
0.00000 0.00000 0.00000 ; CB- CA- C- O
6902 6900 6906 6908 3 2.51040 4.18400 1.67360
-6.69440 0.00000 0.00000 ; CB- CA- C- N
6903 6902 6900 6906 3 0.65084 1.95253 0.00000
-2.60338 0.00000 0.00000 ; HB1- CB- CA- C
6904 6902 6900 6906 3 0.65084 1.95253 0.00000
-2.60338 0.00000 0.00000 ; HB2- CB- CA- C
6905 6902 6900 6906 3 0.65084 1.95253 0.00000
-2.60338 0.00000 0.00000 ; HB3- CB- CA- C
6906 6908 6910 6911 3 0.00000 0.00000 0.00000
0.00000 0.00000 0.00000 ; C- N- CA- HA
6906 6908 6910 6912 3 11.71520 1.67360 15.06240 -
13.38880 0.00000 0.00000 ; C- N- CA- CB
6906 6908 6910 6925 3 1.75728 5.27184 2.25936
-7.02912 0.00000 0.00000 ; C- N- CA- C
6907 6906 6908 6909 3 29.28800 -8.36800 -20.92000
0.00000 0.00000 0.00000 ; O- C- N- H
6907 6906 6908 6910 3 20.92000 0.00000 -20.92000
0.00000 0.00000 0.00000 ; O- C- N- CA
6908 6910 6912 6913 3 0.65084 1.95253 0.00000
-2.60338 0.00000 0.00000 ; N- CA- CB- HB
6908 6910 6912 6914 3 3.83673 0.44769 -1.77402
-2.47693 -0.03347 0.00000 ; N- CA- CB- CG2
6908 6910 6912 6918 3 2.97482 0.12134 2.04179
-1.89117 -3.24678 0.00000 ; N- CA- CB- CG1
6908 6910 6925 6926 3 0.00000 0.00000 0.00000
0.00000 0.00000 0.00000 ; N- CA- C- O
6908 6910 6925 6927 3 17.40544 -5.02080 -13.22144
9.20480 0.00000 0.00000 ; N- CA- C- N
6909 6908 6910 6911 3 0.00000 0.00000 0.00000
0.00000 0.00000 0.00000 ; H- N- CA- HA
6909 6908 6910 6912 3 0.00000 0.00000 0.00000
0.00000 0.00000 0.00000 ; H- N- CA- CB

6909 6908 6910 6925 3 0.00000 0.00000 0.00000
0.00000 0.00000 0.00000 ; H- N- CA- C
6910 6912 6914 6915 3 0.66944 2.00832 0.00000
-2.67776 0.00000 0.00000 ; CA- CB- CG2- HG21
6910 6912 6914 6916 3 0.66944 2.00832 0.00000
-2.67776 0.00000 0.00000 ; CA- CB- CG2- HG22
6910 6912 6914 6917 3 0.66944 2.00832 0.00000
-2.67776 0.00000 0.00000 ; CA- CB- CG2- HG23
6910 6912 6918 6919 3 0.66944 2.00832 0.00000
-2.67776 0.00000 0.00000 ; CA- CB- CG1- HG12
6910 6912 6918 6920 3 0.66944 2.00832 0.00000
-2.67776 0.00000 0.00000 ; CA- CB- CG1- HG13
6910 6912 6918 6921 3 4.24258 -0.52718 -7.25506
-1.79075 7.69856 0.00000 ; CA- CB- CG1- CD1
6910 6925 6927 6928 3 20.92000 0.00000 -20.92000
0.00000 0.00000 0.00000 ; CA- C- N- H
6910 6925 6927 6929 3 20.92000 0.00000 -20.92000
0.00000 0.00000 0.00000 ; CA- C- N- CA
6911 6910 6912 6913 3 0.65084 1.95253 0.00000
-2.60338 0.00000 0.00000 ; HA- CA- CB- HB
6911 6910 6912 6914 3 0.65084 1.95253 0.00000
-2.60338 0.00000 0.00000 ; HA- CA- CB- CG2
6911 6910 6912 6918 3 0.65084 1.95253 0.00000
-2.60338 0.00000 0.00000 ; HA- CA- CB- CG1
6911 6910 6925 6926 3 3.68192 -4.35136 0.00000
1.33888 0.00000 0.00000 ; HA- CA- C- O
6911 6910 6925 6927 3 0.00000 0.00000 0.00000
0.00000 0.00000 0.00000 ; HA- CA- C- N
6912 6910 6925 6926 3 0.00000 0.00000 0.00000
0.00000 0.00000 0.00000 ; CB- CA- C- O
6912 6910 6925 6927 3 2.51040 4.18400 1.67360
-6.69440 0.00000 0.00000 ; CB- CA- C- N
6912 6918 6921 6922 3 0.66944 2.00832 0.00000
-2.67776 0.00000 0.00000 ; CB- CG1- CD1- HD11
6912 6918 6921 6923 3 0.66944 2.00832 0.00000
-2.67776 0.00000 0.00000 ; CB- CG1- CD1- HD12
6912 6918 6921 6924 3 0.66944 2.00832 0.00000
-2.67776 0.00000 0.00000 ; CB- CG1- CD1- HD13
6913 6912 6910 6925 3 0.65084 1.95253 0.00000
-2.60338 0.00000 0.00000 ; HB- CB- CA- C
6913 6912 6914 6915 3 0.62760 1.88280 0.00000
-2.51040 0.00000 0.00000 ; HB- CB- CG2- HG21
6913 6912 6914 6916 3 0.62760 1.88280 0.00000
-2.51040 0.00000 0.00000 ; HB- CB- CG2- HG22
6913 6912 6914 6917 3 0.62760 1.88280 0.00000
-2.51040 0.00000 0.00000 ; HB- CB- CG2- HG23
6913 6912 6918 6919 3 0.62760 1.88280 0.00000
-2.51040 0.00000 0.00000 ; HB- CB- CG1- HG12
6913 6912 6918 6920 3 0.62760 1.88280 0.00000
-2.51040 0.00000 0.00000 ; HB- CB- CG1- HG13
6913 6912 6918 6921 3 0.66944 2.00832 0.00000
-2.67776 0.00000 0.00000 ; HB- CB- CG1- CD1

6914 6912 6910 6925 3 5.67350 3.55640 -6.16722
-2.47693 3.74886 0.00000 ; CG2- CB- CA- C
6914 6912 6918 6919 3 0.66944 2.00832 0.00000
-2.67776 0.00000 0.00000 ; CG2- CB- CG1- HG12
6914 6912 6918 6920 3 0.66944 2.00832 0.00000
-2.67776 0.00000 0.00000 ; CG2- CB- CG1- HG13
6914 6912 6918 6921 3 3.81162 0.49790 -8.14206
-1.79075 7.49773 0.00000 ; CG2- CB- CG1- CD1
6915 6914 6912 6918 3 0.66944 2.00832 0.00000
-2.67776 0.00000 0.00000 ; HG21- CG2- CB- CG1
6916 6914 6912 6918 3 0.66944 2.00832 0.00000
-2.67776 0.00000 0.00000 ; HG22- CG2- CB- CG1
6917 6914 6912 6918 3 0.66944 2.00832 0.00000
-2.67776 0.00000 0.00000 ; HG23- CG2- CB- CG1
6918 6912 6910 6925 3 8.26340 0.74057 -9.99976
-1.89117 3.84928 0.00000 ; CG1- CB- CA- C
6919 6918 6921 6922 3 0.62760 1.88280 0.00000
-2.51040 0.00000 0.00000 ; HG12- CG1- CD1- HD11
6919 6918 6921 6923 3 0.62760 1.88280 0.00000
-2.51040 0.00000 0.00000 ; HG12- CG1- CD1- HD12
6919 6918 6921 6924 3 0.62760 1.88280 0.00000
-2.51040 0.00000 0.00000 ; HG12- CG1- CD1- HD13
6920 6918 6921 6922 3 0.62760 1.88280 0.00000
-2.51040 0.00000 0.00000 ; HG13- CG1- CD1- HD11
6920 6918 6921 6923 3 0.62760 1.88280 0.00000
-2.51040 0.00000 0.00000 ; HG13- CG1- CD1- HD12
6920 6918 6921 6924 3 0.62760 1.88280 0.00000
-2.51040 0.00000 0.00000 ; HG13- CG1- CD1- HD13
6925 6927 6929 6930 3 0.00000 0.00000 0.00000
0.00000 0.00000 0.00000 ; C- N- CA- HA
6925 6927 6929 6931 3 11.71520 1.67360 15.06240 -
13.38880 0.00000 0.00000 ; C- N- CA- CB
6925 6927 6929 6937 3 1.75728 5.27184 2.25936
-7.02912 0.00000 0.00000 ; C- N- CA- C
6926 6925 6927 6928 3 29.28800 -8.36800 -20.92000
0.00000 0.00000 0.00000 ; O- C- N- H
6926 6925 6927 6929 3 20.92000 0.00000 -20.92000
0.00000 0.00000 0.00000 ; O- C- N- CA
6927 6929 6931 6932 3 0.65084 1.95253 0.00000
-2.60338 0.00000 0.00000 ; N- CA- CB- HB2
6927 6929 6931 6933 3 0.65084 1.95253 0.00000
-2.60338 0.00000 0.00000 ; N- CA- CB- HB3
6927 6929 6931 6934 3 15.41386 9.74035 -8.39310
-0.97069 2.97901 0.00000 ; N- CA- CB- CG
6927 6929 6937 6938 3 0.00000 0.00000 0.00000
0.00000 0.00000 0.00000 ; N- CA- C- O
6927 6929 6937 6939 3 17.40544 -5.02080 -13.22144
9.20480 0.00000 0.00000 ; N- CA- C- N
6928 6927 6929 6930 3 0.00000 0.00000 0.00000
0.00000 0.00000 0.00000 ; H- N- CA- HA
6928 6927 6929 6931 3 0.00000 0.00000 0.00000
0.00000 0.00000 0.00000 ; H- N- CA- CB

6928 6927 6929 6937 3 0.00000 0.00000 0.00000
0.00000 0.00000 0.00000 ; H- N- CA- C
6929 6931 6934 6935 3 6.43499 0.00000 -5.39736
0.00000 -1.03763 0.00000 ; CA- CB- CG- OD1
6929 6931 6934 6936 3 6.43499 0.00000 -5.39736
0.00000 -1.03763 0.00000 ; CA- CB- CG- OD2
6929 6937 6939 6940 3 20.92000 0.00000 -20.92000
0.00000 0.00000 0.00000 ; CA- C- N- H
6929 6937 6939 6941 3 20.92000 0.00000 -20.92000
0.00000 0.00000 0.00000 ; CA- C- N- CA
6930 6929 6931 6932 3 0.65084 1.95253 0.00000
-2.60338 0.00000 0.00000 ; HA- CA- CB- HB2
6930 6929 6931 6933 3 0.65084 1.95253 0.00000
-2.60338 0.00000 0.00000 ; HA- CA- CB- HB3
6930 6929 6931 6934 3 0.65084 1.95253 0.00000
-2.60338 0.00000 0.00000 ; HA- CA- CB- CG
6930 6929 6937 6938 3 3.68192 -4.35136 0.00000
1.33888 0.00000 0.00000 ; HA- CA- C- O
6930 6929 6937 6939 3 0.00000 0.00000 0.00000
0.00000 0.00000 0.00000 ; HA- CA- C- N
6931 6929 6937 6938 3 0.00000 0.00000 0.00000
0.00000 0.00000 0.00000 ; CB- CA- C- O
6931 6929 6937 6939 3 2.51040 4.18400 1.67360
-6.69440 0.00000 0.00000 ; CB- CA- C- N
6932 6931 6929 6937 3 0.65084 1.95253 0.00000
-2.60338 0.00000 0.00000 ; HB2- CB- CA- C
6932 6931 6934 6935 3 0.00000 0.00000 0.00000
0.00000 0.00000 0.00000 ; HB2- CB- CG- OD1
6932 6931 6934 6936 3 0.00000 0.00000 0.00000
0.00000 0.00000 0.00000 ; HB2- CB- CG- OD2
6933 6931 6929 6937 3 0.65084 1.95253 0.00000
-2.60338 0.00000 0.00000 ; HB3- CB- CA- C
6933 6931 6934 6935 3 0.00000 0.00000 0.00000
0.00000 0.00000 0.00000 ; HB3- CB- CG- OD1
6933 6931 6934 6936 3 0.00000 0.00000 0.00000
0.00000 0.00000 0.00000 ; HB3- CB- CG- OD2
6934 6931 6929 6937 3 7.14627 -1.04600 -8.99560
-0.97069 5.15469 0.00000 ; CG- CB- CA- C
6937 6939 6941 6942 3 0.00000 0.00000 0.00000
0.00000 0.00000 0.00000 ; C- N- CA- HA
6937 6939 6941 6943 3 11.71520 1.67360 15.06240 -
13.38880 0.00000 0.00000 ; C- N- CA- CB
6937 6939 6941 6948 3 1.75728 5.27184 2.25936
-7.02912 0.00000 0.00000 ; C- N- CA- C
6938 6937 6939 6940 3 29.28800 -8.36800 -20.92000
0.00000 0.00000 0.00000 ; O- C- N- H
6938 6937 6939 6941 3 20.92000 0.00000 -20.92000
0.00000 0.00000 0.00000 ; O- C- N- CA
6939 6941 6943 6944 3 0.65084 1.95253 0.00000
-2.60338 0.00000 0.00000 ; N- CA- CB- HB2
6939 6941 6943 6945 3 0.65084 1.95253 0.00000
-2.60338 0.00000 0.00000 ; N- CA- CB- HB3

6939 6941 6943 6946 3 7.86174 2.24681 -7.41405
-6.71114 5.35552 0.00000 ; N- CA- CB- OG
6939 6941 6948 6949 3 0.00000 0.00000 0.00000
0.00000 0.00000 0.00000 ; N- CA- C- O
6939 6941 6948 6950 3 17.40544 -5.02080 -13.22144
9.20480 0.00000 0.00000 ; N- CA- C- N
6940 6939 6941 6942 3 0.00000 0.00000 0.00000
0.00000 0.00000 0.00000 ; H- N- CA- HA
6940 6939 6941 6943 3 0.00000 0.00000 0.00000
0.00000 0.00000 0.00000 ; H- N- CA- CB
6940 6939 6941 6948 3 0.00000 0.00000 0.00000
0.00000 0.00000 0.00000 ; H- N- CA- C
6941 6943 6946 6947 3 2.05853 2.46856 3.48109
-4.46851 0.23430 0.00000 ; CA- CB- OG- HG
6941 6948 6950 6951 3 20.92000 0.00000 -20.92000
0.00000 0.00000 0.00000 ; CA- C- N- H
6941 6948 6950 6952 3 20.92000 0.00000 -20.92000
0.00000 0.00000 0.00000 ; CA- C- N- CA
6942 6941 6943 6944 3 0.65084 1.95253 0.00000
-2.60338 0.00000 0.00000 ; HA- CA- CB- HB2
6942 6941 6943 6945 3 0.65084 1.95253 0.00000
-2.60338 0.00000 0.00000 ; HA- CA- CB- HB3
6942 6941 6943 6946 3 1.04600 -1.04600 0.00000
0.00000 0.00000 0.00000 ; HA- CA- CB- OG
6942 6941 6948 6949 3 3.68192 -4.35136 0.00000
1.33888 0.00000 0.00000 ; HA- CA- C- O
6942 6941 6948 6950 3 0.00000 0.00000 0.00000
0.00000 0.00000 0.00000 ; HA- CA- C- N
6943 6941 6948 6949 3 0.00000 0.00000 0.00000
0.00000 0.00000 0.00000 ; CB- CA- C- O
6943 6941 6948 6950 3 2.51040 4.18400 1.67360
-6.69440 0.00000 0.00000 ; CB- CA- C- N
6944 6943 6941 6948 3 0.65084 1.95253 0.00000
-2.60338 0.00000 0.00000 ; HB2- CB- CA- C
6944 6943 6946 6947 3 0.69733 2.09200 0.00000
-2.78933 0.00000 0.00000 ; HB2- CB- OG- HG
6945 6943 6941 6948 3 0.65084 1.95253 0.00000
-2.60338 0.00000 0.00000 ; HB3- CB- CA- C
6945 6943 6946 6947 3 0.69733 2.09200 0.00000
-2.78933 0.00000 0.00000 ; HB3- CB- OG- HG
6946 6943 6941 6948 3 7.34710 7.79898 -6.14211
-6.71114 4.31789 0.00000 ; OG- CB- CA- C
6948 6950 6952 6953 3 0.00000 0.00000 0.00000
0.00000 0.00000 0.00000 ; C- N- CA- HA
6948 6950 6952 6954 3 11.71520 1.67360 15.06240 -
13.38880 0.00000 0.00000 ; C- N- CA- CB
6948 6950 6952 6967 3 1.75728 5.27184 2.25936
-7.02912 0.00000 0.00000 ; C- N- CA- C
6949 6948 6950 6951 3 29.28800 -8.36800 -20.92000
0.00000 0.00000 0.00000 ; O- C- N- H
6949 6948 6950 6952 3 20.92000 0.00000 -20.92000
0.00000 0.00000 0.00000 ; O- C- N- CA

```
  6950   6952   6954   6955       3    0.65084    1.95253    0.00000
-2.60338    0.00000    0.00000 ;       N-    CA-    CB-   HB2
  6950   6952   6954   6956       3    0.65084    1.95253    0.00000
-2.60338    0.00000    0.00000 ;       N-    CA-    CB-   HB3
  6950   6952   6954   6957       3    3.79070    1.39746   -4.61077
-2.40998    2.44346    0.00000 ;       N-    CA-    CB-    CG
  6950   6952   6967   6968       3    0.00000    0.00000    0.00000
0.00000    0.00000    0.00000 ;       N-    CA-     C-     O
  6950   6952   6967   6969       3   17.40544   -5.02080  -13.22144
9.20480    0.00000    0.00000 ;       N-    CA-     C-     N
  6951   6950   6952   6953       3    0.00000    0.00000    0.00000
0.00000    0.00000    0.00000 ;       H-     N-    CA-    HA
  6951   6950   6952   6954       3    0.00000    0.00000    0.00000
0.00000    0.00000    0.00000 ;       H-     N-    CA-    CB
  6951   6950   6952   6967       3    0.00000    0.00000    0.00000
0.00000    0.00000    0.00000 ;       H-     N-    CA-     C
  6952   6954   6957   6958       3    0.66944    2.00832    0.00000
-2.67776    0.00000    0.00000 ;      CA-    CB-    CG-    HG
  6952   6954   6957   6959       3    3.90367    0.19665   -6.21742
-2.37651    5.99149    0.00000 ;      CA-    CB-    CG-   CD1
  6952   6954   6957   6963       3    3.90367    0.19665   -6.21742
-2.37651    5.99149    0.00000 ;      CA-    CB-    CG-   CD2
  6952   6967   6969   6970       3   20.92000    0.00000  -20.92000
0.00000    0.00000    0.00000 ;      CA-     C-     N-     H
  6952   6967   6969   6971       3   20.92000    0.00000  -20.92000
0.00000    0.00000    0.00000 ;      CA-     C-     N-    CA
  6953   6952   6954   6955       3    0.65084    1.95253    0.00000
-2.60338    0.00000    0.00000 ;      HA-    CA-    CB-   HB2
  6953   6952   6954   6956       3    0.65084    1.95253    0.00000
-2.60338    0.00000    0.00000 ;      HA-    CA-    CB-   HB3
  6953   6952   6954   6957       3    0.65084    1.95253    0.00000
-2.60338    0.00000    0.00000 ;      HA-    CA-    CB-    CG
  6953   6952   6967   6968       3    3.68192   -4.35136    0.00000
1.33888    0.00000    0.00000 ;      HA-    CA-     C-     O
  6953   6952   6967   6969       3    0.00000    0.00000    0.00000
0.00000    0.00000    0.00000 ;      HA-    CA-     C-     N
  6954   6952   6967   6968       3    0.00000    0.00000    0.00000
0.00000    0.00000    0.00000 ;      CB-    CA-     C-     O
  6954   6952   6967   6969       3    2.51040    4.18400    1.67360
-6.69440    0.00000    0.00000 ;      CB-    CA-     C-     N
  6954   6957   6959   6960       3    0.66944    2.00832    0.00000
-2.67776    0.00000    0.00000 ;      CB-    CG-   CD1-  HD11
  6954   6957   6959   6961       3    0.66944    2.00832    0.00000
-2.67776    0.00000    0.00000 ;      CB-    CG-   CD1-  HD12
  6954   6957   6959   6962       3    0.66944    2.00832    0.00000
-2.67776    0.00000    0.00000 ;      CB-    CG-   CD1-  HD13
  6954   6957   6963   6964       3    0.66944    2.00832    0.00000
-2.67776    0.00000    0.00000 ;      CB-    CG-   CD2-  HD21
  6954   6957   6963   6965       3    0.66944    2.00832    0.00000
-2.67776    0.00000    0.00000 ;      CB-    CG-   CD2-  HD22
  6954   6957   6963   6966       3    0.66944    2.00832    0.00000
-2.67776    0.00000    0.00000 ;      CB-    CG-   CD2-  HD23
```

6955 6954 6952 6967 3 0.65084 1.95253 0.00000
-2.60338 0.00000 0.00000 ; HB2- CB- CA- C
6955 6954 6957 6958 3 0.62760 1.88280 0.00000
-2.51040 0.00000 0.00000 ; HB2- CB- CG- HG
6955 6954 6957 6959 3 0.66944 2.00832 0.00000
-2.67776 0.00000 0.00000 ; HB2- CB- CG- CD1
6955 6954 6957 6963 3 0.66944 2.00832 0.00000
-2.67776 0.00000 0.00000 ; HB2- CB- CG- CD2
6956 6954 6952 6967 3 0.65084 1.95253 0.00000
-2.60338 0.00000 0.00000 ; HB3- CB- CA- C
6956 6954 6957 6958 3 0.62760 1.88280 0.00000
-2.51040 0.00000 0.00000 ; HB3- CB- CG- HG
6956 6954 6957 6959 3 0.66944 2.00832 0.00000
-2.67776 0.00000 0.00000 ; HB3- CB- CG- CD1
6956 6954 6957 6963 3 0.66944 2.00832 0.00000
-2.67776 0.00000 0.00000 ; HB3- CB- CG- CD2
6957 6954 6952 6967 3 10.33448 -1.14642 -11.54784
-2.40998 6.35968 0.00000 ; CG- CB- CA- C
6958 6957 6959 6960 3 0.62760 1.88280 0.00000
-2.51040 0.00000 0.00000 ; HG- CG- CD1- HD11
6958 6957 6959 6961 3 0.62760 1.88280 0.00000
-2.51040 0.00000 0.00000 ; HG- CG- CD1- HD12
6958 6957 6959 6962 3 0.62760 1.88280 0.00000
-2.51040 0.00000 0.00000 ; HG- CG- CD1- HD13
6958 6957 6963 6964 3 0.62760 1.88280 0.00000
-2.51040 0.00000 0.00000 ; HG- CG- CD2- HD21
6958 6957 6963 6965 3 0.62760 1.88280 0.00000
-2.51040 0.00000 0.00000 ; HG- CG- CD2- HD22
6958 6957 6963 6966 3 0.62760 1.88280 0.00000
-2.51040 0.00000 0.00000 ; HG- CG- CD2- HD23
6959 6957 6963 6964 3 0.66944 2.00832 0.00000
-2.67776 0.00000 0.00000 ; CD1- CG- CD2- HD21
6959 6957 6963 6965 3 0.66944 2.00832 0.00000
-2.67776 0.00000 0.00000 ; CD1- CG- CD2- HD22
6959 6957 6963 6966 3 0.66944 2.00832 0.00000
-2.67776 0.00000 0.00000 ; CD1- CG- CD2- HD23
6960 6959 6957 6963 3 0.66944 2.00832 0.00000
-2.67776 0.00000 0.00000 ; HD11- CD1- CG- CD2
6961 6959 6957 6963 3 0.66944 2.00832 0.00000
-2.67776 0.00000 0.00000 ; HD12- CD1- CG- CD2
6962 6959 6957 6963 3 0.66944 2.00832 0.00000
-2.67776 0.00000 0.00000 ; HD13- CD1- CG- CD2
6967 6969 6971 6972 3 0.00000 0.00000 0.00000
0.00000 0.00000 0.00000 ; C- N- CA- HA
6967 6969 6971 6973 3 11.71520 1.67360 15.06240 -
13.38880 0.00000 0.00000 ; C- N- CA- CB
6967 6969 6971 6986 3 1.75728 5.27184 2.25936
-7.02912 0.00000 0.00000 ; C- N- CA- C
6968 6967 6969 6970 3 29.28800 -8.36800 -20.92000
0.00000 0.00000 0.00000 ; O- C- N- H
6968 6967 6969 6971 3 20.92000 0.00000 -20.92000
0.00000 0.00000 0.00000 ; O- C- N- CA

6969 6971 6973 6974 3 0.65084 1.95253 0.00000
-2.60338 0.00000 0.00000 ; N- CA- CB- HB2
6969 6971 6973 6975 3 0.65084 1.95253 0.00000
-2.60338 0.00000 0.00000 ; N- CA- CB- HB3
6969 6971 6973 6976 3 3.79070 1.39746 -4.61077
-2.40998 2.44346 0.00000 ; N- CA- CB- CG
6969 6971 6986 6987 3 0.00000 0.00000 0.00000
0.00000 0.00000 0.00000 ; N- CA- C- O
6969 6971 6986 6988 3 17.40544 -5.02080 -13.22144
9.20480 0.00000 0.00000 ; N- CA- C- N
6970 6969 6971 6972 3 0.00000 0.00000 0.00000
0.00000 0.00000 0.00000 ; H- N- CA- HA
6970 6969 6971 6973 3 0.00000 0.00000 0.00000
0.00000 0.00000 0.00000 ; H- N- CA- CB
6970 6969 6971 6986 3 0.00000 0.00000 0.00000
0.00000 0.00000 0.00000 ; H- N- CA- C
6971 6973 6976 6977 3 0.66944 2.00832 0.00000
-2.67776 0.00000 0.00000 ; CA- CB- CG- HG
6971 6973 6976 6978 3 3.90367 0.19665 -6.21742
-2.37651 5.99149 0.00000 ; CA- CB- CG- CD1
6971 6973 6976 6982 3 3.90367 0.19665 -6.21742
-2.37651 5.99149 0.00000 ; CA- CB- CG- CD2
6971 6986 6988 6989 3 20.92000 0.00000 -20.92000
0.00000 0.00000 0.00000 ; CA- C- N- H
6971 6986 6988 6990 3 20.92000 0.00000 -20.92000
0.00000 0.00000 0.00000 ; CA- C- N- CA
6972 6971 6973 6974 3 0.65084 1.95253 0.00000
-2.60338 0.00000 0.00000 ; HA- CA- CB- HB2
6972 6971 6973 6975 3 0.65084 1.95253 0.00000
-2.60338 0.00000 0.00000 ; HA- CA- CB- HB3
6972 6971 6973 6976 3 0.65084 1.95253 0.00000
-2.60338 0.00000 0.00000 ; HA- CA- CB- CG
6972 6971 6986 6987 3 3.68192 -4.35136 0.00000
1.33888 0.00000 0.00000 ; HA- CA- C- O
6972 6971 6986 6988 3 0.00000 0.00000 0.00000
0.00000 0.00000 0.00000 ; HA- CA- C- N
6973 6971 6986 6987 3 0.00000 0.00000 0.00000
0.00000 0.00000 0.00000 ; CB- CA- C- O
6973 6971 6986 6988 3 2.51040 4.18400 1.67360
-6.69440 0.00000 0.00000 ; CB- CA- C- N
6973 6976 6978 6979 3 0.66944 2.00832 0.00000
-2.67776 0.00000 0.00000 ; CB- CG- CD1- HD11
6973 6976 6978 6980 3 0.66944 2.00832 0.00000
-2.67776 0.00000 0.00000 ; CB- CG- CD1- HD12
6973 6976 6978 6981 3 0.66944 2.00832 0.00000
-2.67776 0.00000 0.00000 ; CB- CG- CD1- HD13
6973 6976 6982 6983 3 0.66944 2.00832 0.00000
-2.67776 0.00000 0.00000 ; CB- CG- CD2- HD21
6973 6976 6982 6984 3 0.66944 2.00832 0.00000
-2.67776 0.00000 0.00000 ; CB- CG- CD2- HD22
6973 6976 6982 6985 3 0.66944 2.00832 0.00000
-2.67776 0.00000 0.00000 ; CB- CG- CD2- HD23

```
  6974   6973   6971   6986       3    0.65084    1.95253    0.00000
-2.60338    0.00000    0.00000 ;    HB2-    CB-    CA-     C
  6974   6973   6976   6977       3    0.62760    1.88280    0.00000
-2.51040    0.00000    0.00000 ;    HB2-    CB-    CG-    HG
  6974   6973   6976   6978       3    0.66944    2.00832    0.00000
-2.67776    0.00000    0.00000 ;    HB2-    CB-    CG-   CD1
  6974   6973   6976   6982       3    0.66944    2.00832    0.00000
-2.67776    0.00000    0.00000 ;    HB2-    CB-    CG-   CD2
  6975   6973   6971   6986       3    0.65084    1.95253    0.00000
-2.60338    0.00000    0.00000 ;    HB3-    CB-    CA-     C
  6975   6973   6976   6977       3    0.62760    1.88280    0.00000
-2.51040    0.00000    0.00000 ;    HB3-    CB-    CG-    HG
  6975   6973   6976   6978       3    0.66944    2.00832    0.00000
-2.67776    0.00000    0.00000 ;    HB3-    CB-    CG-   CD1
  6975   6973   6976   6982       3    0.66944    2.00832    0.00000
-2.67776    0.00000    0.00000 ;    HB3-    CB-    CG-   CD2
  6976   6973   6971   6986       3   10.33448   -1.14642  -11.54784
-2.40998    6.35968    0.00000 ;     CG-    CB-    CA-     C
  6977   6976   6978   6979       3    0.62760    1.88280    0.00000
-2.51040    0.00000    0.00000 ;     HG-    CG-   CD1-  HD11
  6977   6976   6978   6980       3    0.62760    1.88280    0.00000
-2.51040    0.00000    0.00000 ;     HG-    CG-   CD1-  HD12
  6977   6976   6978   6981       3    0.62760    1.88280    0.00000
-2.51040    0.00000    0.00000 ;     HG-    CG-   CD1-  HD13
  6977   6976   6982   6983       3    0.62760    1.88280    0.00000
-2.51040    0.00000    0.00000 ;     HG-    CG-   CD2-  HD21
  6977   6976   6982   6984       3    0.62760    1.88280    0.00000
-2.51040    0.00000    0.00000 ;     HG-    CG-   CD2-  HD22
  6977   6976   6982   6985       3    0.62760    1.88280    0.00000
-2.51040    0.00000    0.00000 ;     HG-    CG-   CD2-  HD23
  6978   6976   6982   6983       3    0.66944    2.00832    0.00000
-2.67776    0.00000    0.00000 ;    CD1-    CG-   CD2-  HD21
  6978   6976   6982   6984       3    0.66944    2.00832    0.00000
-2.67776    0.00000    0.00000 ;    CD1-    CG-   CD2-  HD22
  6978   6976   6982   6985       3    0.66944    2.00832    0.00000
-2.67776    0.00000    0.00000 ;    CD1-    CG-   CD2-  HD23
  6979   6978   6976   6982       3    0.66944    2.00832    0.00000
-2.67776    0.00000    0.00000 ;   HD11-   CD1-    CG-   CD2
  6980   6978   6976   6982       3    0.66944    2.00832    0.00000
-2.67776    0.00000    0.00000 ;   HD12-   CD1-    CG-   CD2
  6981   6978   6976   6982       3    0.66944    2.00832    0.00000
-2.67776    0.00000    0.00000 ;   HD13-   CD1-    CG-   CD2
  6986   6988   6990   6991       3    0.00000    0.00000    0.00000
0.00000    0.00000    0.00000 ;      C-     N-    CA-    HA
  6986   6988   6990   6992       3   11.71520    1.67360   15.06240  -
13.38880    0.00000    0.00000 ;      C-     N-    CA-    CB
  6986   6988   6990   7003       3    1.75728    5.27184    2.25936
-7.02912    0.00000    0.00000 ;      C-     N-    CA-     C
  6987   6986   6988   6989       3   29.28800   -8.36800  -20.92000
0.00000    0.00000    0.00000 ;      O-     C-     N-     H
  6987   6986   6988   6990       3   20.92000    0.00000  -20.92000
0.00000    0.00000    0.00000 ;      O-     C-     N-    CA
```

6988 6990 6992 6993 3 0.65084 1.95253 0.00000
-2.60338 0.00000 0.00000 ; N- CA- CB- HB2
6988 6990 6992 6994 3 0.65084 1.95253 0.00000
-2.60338 0.00000 0.00000 ; N- CA- CB- HB3
6988 6990 6992 6995 3 3.21331 2.22589 -4.15053
-2.40998 2.61082 0.00000 ; N- CA- CB- CG
6988 6990 7003 7004 3 0.00000 0.00000 0.00000
0.00000 0.00000 0.00000 ; N- CA- C- O
6988 6990 7003 7005 3 17.40544 -5.02080 -13.22144
9.20480 0.00000 0.00000 ; N- CA- C- N
6989 6988 6990 6991 3 0.00000 0.00000 0.00000
0.00000 0.00000 0.00000 ; H- N- CA- HA
6989 6988 6990 6992 3 0.00000 0.00000 0.00000
0.00000 0.00000 0.00000 ; H- N- CA- CB
6989 6988 6990 7003 3 0.00000 0.00000 0.00000
0.00000 0.00000 0.00000 ; H- N- CA- C
6990 6992 6995 6996 3 0.66944 2.00832 0.00000
-2.67776 0.00000 0.00000 ; CA- CB- CG- HG2
6990 6992 6995 6997 3 0.66944 2.00832 0.00000
-2.67776 0.00000 0.00000 ; CA- CB- CG- HG3
6990 6992 6995 6998 3 3.69866 -4.35136 -3.92459
6.89523 4.61914 0.00000 ; CA- CB- CG- CD
6990 7003 7005 7006 3 20.92000 0.00000 -20.92000
0.00000 0.00000 0.00000 ; CA- C- N- H
6990 7003 7005 7007 3 20.92000 0.00000 -20.92000
0.00000 0.00000 0.00000 ; CA- C- N- CA
6991 6990 6992 6993 3 0.65084 1.95253 0.00000
-2.60338 0.00000 0.00000 ; HA- CA- CB- HB2
6991 6990 6992 6994 3 0.65084 1.95253 0.00000
-2.60338 0.00000 0.00000 ; HA- CA- CB- HB3
6991 6990 6992 6995 3 0.65084 1.95253 0.00000
-2.60338 0.00000 0.00000 ; HA- CA- CB- CG
6991 6990 7003 7004 3 3.68192 -4.35136 0.00000
1.33888 0.00000 0.00000 ; HA- CA- C- O
6991 6990 7003 7005 3 0.00000 0.00000 0.00000
0.00000 0.00000 0.00000 ; HA- CA- C- N
6992 6990 7003 7004 3 0.00000 0.00000 0.00000
0.00000 0.00000 0.00000 ; CB- CA- C- O
6992 6990 7003 7005 3 2.51040 4.18400 1.67360
-6.69440 0.00000 0.00000 ; CB- CA- C- N
6992 6995 6998 6999 3 0.00000 0.00000 0.00000
0.00000 0.00000 0.00000 ; CB- CG- CD- OE1
6992 6995 6998 7000 3 10.32611 1.48114 -8.47678
1.42256 1.40582 0.00000 ; CB- CG- CD- NE2
6993 6992 6990 7003 3 0.65084 1.95253 0.00000
-2.60338 0.00000 0.00000 ; HB2- CB- CA- C
6993 6992 6995 6996 3 0.62760 1.88280 0.00000
-2.51040 0.00000 0.00000 ; HB2- CB- CG- HG2
6993 6992 6995 6997 3 0.62760 1.88280 0.00000
-2.51040 0.00000 0.00000 ; HB2- CB- CG- HG3
6993 6992 6995 6998 3 0.65084 1.95253 0.00000
-2.60338 0.00000 0.00000 ; HB2- CB- CG- CD

6994 6992 6990 7003 3 0.65084 1.95253 0.00000
-2.60338 0.00000 0.00000 ; HB3- CB- CA- C
6994 6992 6995 6996 3 0.62760 1.88280 0.00000
-2.51040 0.00000 0.00000 ; HB3- CB- CG- HG2
6994 6992 6995 6997 3 0.62760 1.88280 0.00000
-2.51040 0.00000 0.00000 ; HB3- CB- CG- HG3
6994 6992 6995 6998 3 0.65084 1.95253 0.00000
-2.60338 0.00000 0.00000 ; HB3- CB- CG- CD
6995 6992 6990 7003 3 6.86594 3.56895 -8.14206
-2.40998 4.85344 0.00000 ; CG- CB- CA- C
6995 6998 7000 7001 3 20.92000 0.00000 -20.92000
0.00000 0.00000 0.00000 ; CG- CD- NE2- HE21
6995 6998 7000 7002 3 20.92000 0.00000 -20.92000
0.00000 0.00000 0.00000 ; CG- CD- NE2- HE22
6996 6995 6998 6999 3 3.68192 -4.35136 0.00000
1.33888 0.00000 0.00000 ; HG2- CG- CD- OE1
6996 6995 6998 7000 3 0.00000 0.00000 0.00000
0.00000 0.00000 0.00000 ; HG2- CG- CD- NE2
6997 6995 6998 6999 3 3.68192 -4.35136 0.00000
1.33888 0.00000 0.00000 ; HG3- CG- CD- OE1
6997 6995 6998 7000 3 0.00000 0.00000 0.00000
0.00000 0.00000 0.00000 ; HG3- CG- CD- NE2
6999 6998 7000 7001 3 29.28800 -8.36800 -20.92000
0.00000 0.00000 0.00000 ; OE1- CD- NE2- HE21
6999 6998 7000 7002 3 29.28800 -8.36800 -20.92000
0.00000 0.00000 0.00000 ; OE1- CD- NE2- HE22
7003 7005 7007 7008 3 0.00000 0.00000 0.00000
0.00000 0.00000 0.00000 ; C- N- CA- HA
7003 7005 7007 7009 3 11.71520 1.67360 15.06240 -
13.38880 0.00000 0.00000 ; C- N- CA- CB
7003 7005 7007 7018 3 1.75728 5.27184 2.25936
-7.02912 0.00000 0.00000 ; C- N- CA- C
7004 7003 7005 7006 3 29.28800 -8.36800 -20.92000
0.00000 0.00000 0.00000 ; O- C- N- H
7004 7003 7005 7007 3 20.92000 0.00000 -20.92000
0.00000 0.00000 0.00000 ; O- C- N- CA
7005 7007 7009 7010 3 0.65084 1.95253 0.00000
-2.60338 0.00000 0.00000 ; N- CA- CB- HB2
7005 7007 7009 7011 3 0.65084 1.95253 0.00000
-2.60338 0.00000 0.00000 ; N- CA- CB- HB3
7005 7007 7009 7012 3 3.21331 2.22589 -4.15053
-2.40998 2.61082 0.00000 ; N- CA- CB- CG
7005 7007 7018 7019 3 0.00000 0.00000 0.00000
0.00000 0.00000 0.00000 ; N- CA- C- O
7005 7007 7018 7020 3 17.40544 -5.02080 -13.22144
9.20480 0.00000 0.00000 ; N- CA- C- N
7006 7005 7007 7008 3 0.00000 0.00000 0.00000
0.00000 0.00000 0.00000 ; H- N- CA- HA
7006 7005 7007 7009 3 0.00000 0.00000 0.00000
0.00000 0.00000 0.00000 ; H- N- CA- CB
7006 7005 7007 7018 3 0.00000 0.00000 0.00000
0.00000 0.00000 0.00000 ; H- N- CA- C

7007 7009 7012 7013 3 0.66944 2.00832 0.00000
-2.67776 0.00000 0.00000 ; CA- CB- CG- HG2
7007 7009 7012 7014 3 0.66944 2.00832 0.00000
-2.67776 0.00000 0.00000 ; CA- CB- CG- HG3
7007 7009 7012 7015 3 10.12110 -1.91209 0.01674
10.17549 -1.87443 0.00000 ; CA- CB- CG- CD
7007 7018 7020 7021 3 20.92000 0.00000 -20.92000
0.00000 0.00000 0.00000 ; CA- C- N- H
7007 7018 7020 7022 3 20.92000 0.00000 -20.92000
0.00000 0.00000 0.00000 ; CA- C- N- CA
7008 7007 7009 7010 3 0.65084 1.95253 0.00000
-2.60338 0.00000 0.00000 ; HA- CA- CB- HB2
7008 7007 7009 7011 3 0.65084 1.95253 0.00000
-2.60338 0.00000 0.00000 ; HA- CA- CB- HB3
7008 7007 7009 7012 3 0.65084 1.95253 0.00000
-2.60338 0.00000 0.00000 ; HA- CA- CB- CG
7008 7007 7018 7019 3 3.68192 -4.35136 0.00000
1.33888 0.00000 0.00000 ; HA- CA- C- O
7008 7007 7018 7020 3 0.00000 0.00000 0.00000
0.00000 0.00000 0.00000 ; HA- CA- C- N
7009 7007 7018 7019 3 0.00000 0.00000 0.00000
0.00000 0.00000 0.00000 ; CB- CA- C- O
7009 7007 7018 7020 3 2.51040 4.18400 1.67360
-6.69440 0.00000 0.00000 ; CB- CA- C- N
7009 7012 7015 7016 3 3.79907 0.00000 -5.40573
0.00000 2.14221 0.00000 ; CB- CG- CD- OE1
7009 7012 7015 7017 3 3.79907 0.00000 -5.40573
0.00000 2.14221 0.00000 ; CB- CG- CD- OE2
7010 7009 7007 7018 3 0.65084 1.95253 0.00000
-2.60338 0.00000 0.00000 ; HB2- CB- CA- C
7010 7009 7012 7013 3 0.62760 1.88280 0.00000
-2.51040 0.00000 0.00000 ; HB2- CB- CG- HG2
7010 7009 7012 7014 3 0.62760 1.88280 0.00000
-2.51040 0.00000 0.00000 ; HB2- CB- CG- HG3
7010 7009 7012 7015 3 0.65084 1.95253 0.00000
-2.60338 0.00000 0.00000 ; HB2- CB- CG- CD
7011 7009 7007 7018 3 0.65084 1.95253 0.00000
-2.60338 0.00000 0.00000 ; HB3- CB- CA- C
7011 7009 7012 7013 3 0.62760 1.88280 0.00000
-2.51040 0.00000 0.00000 ; HB3- CB- CG- HG2
7011 7009 7012 7014 3 0.62760 1.88280 0.00000
-2.51040 0.00000 0.00000 ; HB3- CB- CG- HG3
7011 7009 7012 7015 3 0.65084 1.95253 0.00000
-2.60338 0.00000 0.00000 ; HB3- CB- CG- CD
7012 7009 7007 7018 3 6.86594 3.56895 -8.14206
-2.40998 4.85344 0.00000 ; CG- CB- CA- C
7013 7012 7015 7016 3 0.00000 0.00000 0.00000
0.00000 0.00000 0.00000 ; HG2- CG- CD- OE1
7013 7012 7015 7017 3 0.00000 0.00000 0.00000
0.00000 0.00000 0.00000 ; HG2- CG- CD- OE2
7014 7012 7015 7016 3 0.00000 0.00000 0.00000
0.00000 0.00000 0.00000 ; HG3- CG- CD- OE1

7014 7012 7015 7017 3 0.00000 0.00000 0.00000
0.00000 0.00000 0.00000 ; HG3- CG- CD- OE2
7018 7020 7022 7023 3 0.00000 0.00000 0.00000
0.00000 0.00000 0.00000 ; C- N- CA- HA
7018 7020 7022 7024 3 11.71520 1.67360 15.06240 -
13.38880 0.00000 0.00000 ; C- N- CA- CB
7018 7020 7022 7034 3 1.75728 5.27184 2.25936
-7.02912 0.00000 0.00000 ; C- N- CA- C
7019 7018 7020 7021 3 29.28800 -8.36800 -20.92000
0.00000 0.00000 0.00000 ; O- C- N- H
7019 7018 7020 7022 3 20.92000 0.00000 -20.92000
0.00000 0.00000 0.00000 ; O- C- N- CA
7020 7022 7024 7025 3 0.65084 1.95253 0.00000
-2.60338 0.00000 0.00000 ; N- CA- CB- HB
7020 7022 7024 7026 3 3.83673 0.44769 -1.77402
-2.47693 -0.03347 0.00000 ; N- CA- CB- CG1
7020 7022 7024 7030 3 3.83673 0.44769 -1.77402
-2.47693 -0.03347 0.00000 ; N- CA- CB- CG2
7020 7022 7034 7035 3 0.00000 0.00000 0.00000
0.00000 0.00000 0.00000 ; N- CA- C- O
7020 7022 7034 7036 3 17.40544 -5.02080 -13.22144
9.20480 0.00000 0.00000 ; N- CA- C- N
7021 7020 7022 7023 3 0.00000 0.00000 0.00000
0.00000 0.00000 0.00000 ; H- N- CA- HA
7021 7020 7022 7024 3 0.00000 0.00000 0.00000
0.00000 0.00000 0.00000 ; H- N- CA- CB
7021 7020 7022 7034 3 0.00000 0.00000 0.00000
0.00000 0.00000 0.00000 ; H- N- CA- C
7022 7024 7026 7027 3 0.66944 2.00832 0.00000
-2.67776 0.00000 0.00000 ; CA- CB- CG1- HG11
7022 7024 7026 7028 3 0.66944 2.00832 0.00000
-2.67776 0.00000 0.00000 ; CA- CB- CG1- HG12
7022 7024 7026 7029 3 0.66944 2.00832 0.00000
-2.67776 0.00000 0.00000 ; CA- CB- CG1- HG13
7022 7024 7030 7031 3 0.66944 2.00832 0.00000
-2.67776 0.00000 0.00000 ; CA- CB- CG2- HG21
7022 7024 7030 7032 3 0.66944 2.00832 0.00000
-2.67776 0.00000 0.00000 ; CA- CB- CG2- HG22
7022 7024 7030 7033 3 0.66944 2.00832 0.00000
-2.67776 0.00000 0.00000 ; CA- CB- CG2- HG23
7022 7034 7036 7037 3 20.92000 0.00000 -20.92000
0.00000 0.00000 0.00000 ; CA- C- N- H
7022 7034 7036 7038 3 20.92000 0.00000 -20.92000
0.00000 0.00000 0.00000 ; CA- C- N- CA
7023 7022 7024 7025 3 0.65084 1.95253 0.00000
-2.60338 0.00000 0.00000 ; HA- CA- CB- HB
7023 7022 7024 7026 3 0.65084 1.95253 0.00000
-2.60338 0.00000 0.00000 ; HA- CA- CB- CG1
7023 7022 7024 7030 3 0.65084 1.95253 0.00000
-2.60338 0.00000 0.00000 ; HA- CA- CB- CG2
7023 7022 7034 7035 3 3.68192 -4.35136 0.00000
1.33888 0.00000 0.00000 ; HA- CA- C- O

7023 7022 7034 7036 3 0.00000 0.00000 0.00000
0.00000 0.00000 0.00000 ; HA- CA- C- N
7024 7022 7034 7035 3 0.00000 0.00000 0.00000
0.00000 0.00000 0.00000 ; CB- CA- C- O
7024 7022 7034 7036 3 2.51040 4.18400 1.67360
-6.69440 0.00000 0.00000 ; CB- CA- C- N
7025 7024 7022 7034 3 0.65084 1.95253 0.00000
-2.60338 0.00000 0.00000 ; HB- CB- CA- C
7025 7024 7026 7027 3 0.62760 1.88280 0.00000
-2.51040 0.00000 0.00000 ; HB- CB- CG1- HG11
7025 7024 7026 7028 3 0.62760 1.88280 0.00000
-2.51040 0.00000 0.00000 ; HB- CB- CG1- HG12
7025 7024 7026 7029 3 0.62760 1.88280 0.00000
-2.51040 0.00000 0.00000 ; HB- CB- CG1- HG13
7025 7024 7030 7031 3 0.62760 1.88280 0.00000
-2.51040 0.00000 0.00000 ; HB- CB- CG2- HG21
7025 7024 7030 7032 3 0.62760 1.88280 0.00000
-2.51040 0.00000 0.00000 ; HB- CB- CG2- HG22
7025 7024 7030 7033 3 0.62760 1.88280 0.00000
-2.51040 0.00000 0.00000 ; HB- CB- CG2- HG23
7026 7024 7022 7034 3 5.67350 3.55640 -6.16722
-2.47693 3.74886 0.00000 ; CG1- CB- CA- C
7026 7024 7030 7031 3 0.66944 2.00832 0.00000
-2.67776 0.00000 0.00000 ; CG1- CB- CG2- HG21
7026 7024 7030 7032 3 0.66944 2.00832 0.00000
-2.67776 0.00000 0.00000 ; CG1- CB- CG2- HG22
7026 7024 7030 7033 3 0.66944 2.00832 0.00000
-2.67776 0.00000 0.00000 ; CG1- CB- CG2- HG23
7027 7026 7024 7030 3 0.66944 2.00832 0.00000
-2.67776 0.00000 0.00000 ; HG11- CG1- CB- CG2
7028 7026 7024 7030 3 0.66944 2.00832 0.00000
-2.67776 0.00000 0.00000 ; HG12- CG1- CB- CG2
7029 7026 7024 7030 3 0.66944 2.00832 0.00000
-2.67776 0.00000 0.00000 ; HG13- CG1- CB- CG2
7030 7024 7022 7034 3 5.67350 3.55640 -6.16722
-2.47693 3.74886 0.00000 ; CG2- CB- CA- C
7034 7036 7038 7039 3 0.00000 0.00000 0.00000
0.00000 0.00000 0.00000 ; C- N- CA- HA
7034 7036 7038 7040 3 11.71520 1.67360 15.06240 -
13.38880 0.00000 0.00000 ; C- N- CA- CB
7034 7036 7038 7046 3 1.75728 5.27184 2.25936
-7.02912 0.00000 0.00000 ; C- N- CA- C
7035 7034 7036 7037 3 29.28800 -8.36800 -20.92000
0.00000 0.00000 0.00000 ; O- C- N- H
7035 7034 7036 7038 3 20.92000 0.00000 -20.92000
0.00000 0.00000 0.00000 ; O- C- N- CA
7036 7038 7040 7041 3 0.65084 1.95253 0.00000
-2.60338 0.00000 0.00000 ; N- CA- CB- HB2
7036 7038 7040 7042 3 0.65084 1.95253 0.00000
-2.60338 0.00000 0.00000 ; N- CA- CB- HB3
7036 7038 7040 7043 3 15.41386 9.74035 -8.39310
-0.97069 2.97901 0.00000 ; N- CA- CB- CG

7036 7038 7046 7047 3 0.00000 0.00000 0.00000
0.00000 0.00000 0.00000 ; N- CA- C- O
7036 7038 7046 7048 3 17.40544 -5.02080 -13.22144
9.20480 0.00000 0.00000 ; N- CA- C- N
7037 7036 7038 7039 3 0.00000 0.00000 0.00000
0.00000 0.00000 0.00000 ; H- N- CA- HA
7037 7036 7038 7040 3 0.00000 0.00000 0.00000
0.00000 0.00000 0.00000 ; H- N- CA- CB
7037 7036 7038 7046 3 0.00000 0.00000 0.00000
0.00000 0.00000 0.00000 ; H- N- CA- C
7038 7040 7043 7044 3 6.43499 0.00000 -5.39736
0.00000 -1.03763 0.00000 ; CA- CB- CG- OD1
7038 7040 7043 7045 3 6.43499 0.00000 -5.39736
0.00000 -1.03763 0.00000 ; CA- CB- CG- OD2
7038 7046 7048 7049 3 20.92000 0.00000 -20.92000
0.00000 0.00000 0.00000 ; CA- C- N- H
7038 7046 7048 7050 3 20.92000 0.00000 -20.92000
0.00000 0.00000 0.00000 ; CA- C- N- CA
7039 7038 7040 7041 3 0.65084 1.95253 0.00000
-2.60338 0.00000 0.00000 ; HA- CA- CB- HB2
7039 7038 7040 7042 3 0.65084 1.95253 0.00000
-2.60338 0.00000 0.00000 ; HA- CA- CB- HB3
7039 7038 7040 7043 3 0.65084 1.95253 0.00000
-2.60338 0.00000 0.00000 ; HA- CA- CB- CG
7039 7038 7046 7047 3 3.68192 -4.35136 0.00000
1.33888 0.00000 0.00000 ; HA- CA- C- O
7039 7038 7046 7048 3 0.00000 0.00000 0.00000
0.00000 0.00000 0.00000 ; HA- CA- C- N
7040 7038 7046 7047 3 0.00000 0.00000 0.00000
0.00000 0.00000 0.00000 ; CB- CA- C- O
7040 7038 7046 7048 3 2.51040 4.18400 1.67360
-6.69440 0.00000 0.00000 ; CB- CA- C- N
7041 7040 7038 7046 3 0.65084 1.95253 0.00000
-2.60338 0.00000 0.00000 ; HB2- CB- CA- C
7041 7040 7043 7044 3 0.00000 0.00000 0.00000
0.00000 0.00000 0.00000 ; HB2- CB- CG- OD1
7041 7040 7043 7045 3 0.00000 0.00000 0.00000
0.00000 0.00000 0.00000 ; HB2- CB- CG- OD2
7042 7040 7038 7046 3 0.65084 1.95253 0.00000
-2.60338 0.00000 0.00000 ; HB3- CB- CA- C
7042 7040 7043 7044 3 0.00000 0.00000 0.00000
0.00000 0.00000 0.00000 ; HB3- CB- CG- OD1
7042 7040 7043 7045 3 0.00000 0.00000 0.00000
0.00000 0.00000 0.00000 ; HB3- CB- CG- OD2
7043 7040 7038 7046 3 7.14627 -1.04600 -8.99560
-0.97069 5.15469 0.00000 ; CG- CB- CA- C
7046 7048 7050 7051 3 0.00000 0.00000 0.00000
0.00000 0.00000 0.00000 ; C- N- CA- HA
7046 7048 7050 7052 3 11.71520 1.67360 15.06240 -
13.38880 0.00000 0.00000 ; C- N- CA- CB
7046 7048 7050 7065 3 1.75728 5.27184 2.25936
-7.02912 0.00000 0.00000 ; C- N- CA- C

7047 7046 7048 7049 3 29.28800 -8.36800 -20.92000
0.00000 0.00000 0.00000 ; O- C- N- H
7047 7046 7048 7050 3 20.92000 0.00000 -20.92000
0.00000 0.00000 0.00000 ; O- C- N- CA
7048 7050 7052 7053 3 0.65084 1.95253 0.00000
-2.60338 0.00000 0.00000 ; N- CA- CB- HB2
7048 7050 7052 7054 3 0.65084 1.95253 0.00000
-2.60338 0.00000 0.00000 ; N- CA- CB- HB3
7048 7050 7052 7055 3 3.79070 1.39746 -4.61077
-2.40998 2.44346 0.00000 ; N- CA- CB- CG
7048 7050 7065 7066 3 0.00000 0.00000 0.00000
0.00000 0.00000 0.00000 ; N- CA- C- O
7048 7050 7065 7067 3 17.40544 -5.02080 -13.22144
9.20480 0.00000 0.00000 ; N- CA- C- N
7049 7048 7050 7051 3 0.00000 0.00000 0.00000
0.00000 0.00000 0.00000 ; H- N- CA- HA
7049 7048 7050 7052 3 0.00000 0.00000 0.00000
0.00000 0.00000 0.00000 ; H- N- CA- CB
7049 7048 7050 7065 3 0.00000 0.00000 0.00000
0.00000 0.00000 0.00000 ; H- N- CA- C
7050 7052 7055 7056 3 0.66944 2.00832 0.00000
-2.67776 0.00000 0.00000 ; CA- CB- CG- HG
7050 7052 7055 7057 3 3.90367 0.19665 -6.21742
-2.37651 5.99149 0.00000 ; CA- CB- CG- CD1
7050 7052 7055 7061 3 3.90367 0.19665 -6.21742
-2.37651 5.99149 0.00000 ; CA- CB- CG- CD2
7050 7065 7067 7068 3 20.92000 0.00000 -20.92000
0.00000 0.00000 0.00000 ; CA- C- N- H
7050 7065 7067 7069 3 20.92000 0.00000 -20.92000
0.00000 0.00000 0.00000 ; CA- C- N- CA
7051 7050 7052 7053 3 0.65084 1.95253 0.00000
-2.60338 0.00000 0.00000 ; HA- CA- CB- HB2
7051 7050 7052 7054 3 0.65084 1.95253 0.00000
-2.60338 0.00000 0.00000 ; HA- CA- CB- HB3
7051 7050 7052 7055 3 0.65084 1.95253 0.00000
-2.60338 0.00000 0.00000 ; HA- CA- CB- CG
7051 7050 7065 7066 3 3.68192 -4.35136 0.00000
1.33888 0.00000 0.00000 ; HA- CA- C- O
7051 7050 7065 7067 3 0.00000 0.00000 0.00000
0.00000 0.00000 0.00000 ; HA- CA- C- N
7052 7050 7065 7066 3 0.00000 0.00000 0.00000
0.00000 0.00000 0.00000 ; CB- CA- C- O
7052 7050 7065 7067 3 2.51040 4.18400 1.67360
-6.69440 0.00000 0.00000 ; CB- CA- C- N
7052 7055 7057 7058 3 0.66944 2.00832 0.00000
-2.67776 0.00000 0.00000 ; CB- CG- CD1- HD11
7052 7055 7057 7059 3 0.66944 2.00832 0.00000
-2.67776 0.00000 0.00000 ; CB- CG- CD1- HD12
7052 7055 7057 7060 3 0.66944 2.00832 0.00000
-2.67776 0.00000 0.00000 ; CB- CG- CD1- HD13
7052 7055 7061 7062 3 0.66944 2.00832 0.00000
-2.67776 0.00000 0.00000 ; CB- CG- CD2- HD21

7052 7055 7061 7063 3 0.66944 2.00832 0.00000
-2.67776 0.00000 0.00000 ; CB- CG- CD2- HD22
7052 7055 7061 7064 3 0.66944 2.00832 0.00000
-2.67776 0.00000 0.00000 ; CB- CG- CD2- HD23
7053 7052 7050 7065 3 0.65084 1.95253 0.00000
-2.60338 0.00000 0.00000 ; HB2- CB- CA- C
7053 7052 7055 7056 3 0.62760 1.88280 0.00000
-2.51040 0.00000 0.00000 ; HB2- CB- CG- HG
7053 7052 7055 7057 3 0.66944 2.00832 0.00000
-2.67776 0.00000 0.00000 ; HB2- CB- CG- CD1
7053 7052 7055 7061 3 0.66944 2.00832 0.00000
-2.67776 0.00000 0.00000 ; HB2- CB- CG- CD2
7054 7052 7050 7065 3 0.65084 1.95253 0.00000
-2.60338 0.00000 0.00000 ; HB3- CB- CA- C
7054 7052 7055 7056 3 0.62760 1.88280 0.00000
-2.51040 0.00000 0.00000 ; HB3- CB- CG- HG
7054 7052 7055 7057 3 0.66944 2.00832 0.00000
-2.67776 0.00000 0.00000 ; HB3- CB- CG- CD1
7054 7052 7055 7061 3 0.66944 2.00832 0.00000
-2.67776 0.00000 0.00000 ; HB3- CB- CG- CD2
7055 7052 7050 7065 3 10.33448 -1.14642 -11.54784
-2.40998 6.35968 0.00000 ; CG- CB- CA- C
7056 7055 7057 7058 3 0.62760 1.88280 0.00000
-2.51040 0.00000 0.00000 ; HG- CG- CD1- HD11
7056 7055 7057 7059 3 0.62760 1.88280 0.00000
-2.51040 0.00000 0.00000 ; HG- CG- CD1- HD12
7056 7055 7057 7060 3 0.62760 1.88280 0.00000
-2.51040 0.00000 0.00000 ; HG- CG- CD1- HD13
7056 7055 7061 7062 3 0.62760 1.88280 0.00000
-2.51040 0.00000 0.00000 ; HG- CG- CD2- HD21
7056 7055 7061 7063 3 0.62760 1.88280 0.00000
-2.51040 0.00000 0.00000 ; HG- CG- CD2- HD22
7056 7055 7061 7064 3 0.62760 1.88280 0.00000
-2.51040 0.00000 0.00000 ; HG- CG- CD2- HD23
7057 7055 7061 7062 3 0.66944 2.00832 0.00000
-2.67776 0.00000 0.00000 ; CD1- CG- CD2- HD21
7057 7055 7061 7063 3 0.66944 2.00832 0.00000
-2.67776 0.00000 0.00000 ; CD1- CG- CD2- HD22
7057 7055 7061 7064 3 0.66944 2.00832 0.00000
-2.67776 0.00000 0.00000 ; CD1- CG- CD2- HD23
7058 7057 7055 7061 3 0.66944 2.00832 0.00000
-2.67776 0.00000 0.00000 ; HD11- CD1- CG- CD2
7059 7057 7055 7061 3 0.66944 2.00832 0.00000
-2.67776 0.00000 0.00000 ; HD12- CD1- CG- CD2
7060 7057 7055 7061 3 0.66944 2.00832 0.00000
-2.67776 0.00000 0.00000 ; HD13- CD1- CG- CD2
7065 7067 7069 7070 3 0.00000 0.00000 0.00000
0.00000 0.00000 0.00000 ; C- N- CA- HA
7065 7067 7069 7071 3 11.71520 1.67360 15.06240 -
13.38880 0.00000 0.00000 ; C- N- CA- CB
7065 7067 7069 7079 3 1.75728 5.27184 2.25936
-7.02912 0.00000 0.00000 ; C- N- CA- C

7066 7065 7067 7068 3 29.28800 -8.36800 -20.92000
0.00000 0.00000 0.00000 ; O- C- N- H
7066 7065 7067 7069 3 20.92000 0.00000 -20.92000
0.00000 0.00000 0.00000 ; O- C- N- CA
7067 7069 7071 7072 3 0.65084 1.95253 0.00000
-2.60338 0.00000 0.00000 ; N- CA- CB- HB2
7067 7069 7071 7073 3 0.65084 1.95253 0.00000
-2.60338 0.00000 0.00000 ; N- CA- CB- HB3
7067 7069 7071 7074 3 5.99567 3.29281 -4.46014
-0.55229 1.97485 0.00000 ; N- CA- CB- CG
7067 7069 7079 7080 3 0.00000 0.00000 0.00000
0.00000 0.00000 0.00000 ; N- CA- C- O
7067 7069 7079 7081 3 17.40544 -5.02080 -13.22144
9.20480 0.00000 0.00000 ; N- CA- C- N
7068 7067 7069 7070 3 0.00000 0.00000 0.00000
0.00000 0.00000 0.00000 ; H- N- CA- HA
7068 7067 7069 7071 3 0.00000 0.00000 0.00000
0.00000 0.00000 0.00000 ; H- N- CA- CB
7068 7067 7069 7079 3 0.00000 0.00000 0.00000
0.00000 0.00000 0.00000 ; H- N- CA- C
7069 7071 7074 7075 3 0.00000 0.00000 0.00000
0.00000 0.00000 0.00000 ; CA- CB- CG- OD1
7069 7071 7074 7076 3 8.84916 -0.31380 -4.32626
5.03754 0.26778 0.00000 ; CA- CB- CG- ND2
7069 7079 7081 7082 3 20.92000 0.00000 -20.92000
0.00000 0.00000 0.00000 ; CA- C- N- H
7069 7079 7081 7083 3 20.92000 0.00000 -20.92000
0.00000 0.00000 0.00000 ; CA- C- N- CA
7070 7069 7071 7072 3 0.65084 1.95253 0.00000
-2.60338 0.00000 0.00000 ; HA- CA- CB- HB2
7070 7069 7071 7073 3 0.65084 1.95253 0.00000
-2.60338 0.00000 0.00000 ; HA- CA- CB- HB3
7070 7069 7071 7074 3 0.65084 1.95253 0.00000
-2.60338 0.00000 0.00000 ; HA- CA- CB- CG
7070 7069 7079 7080 3 3.68192 -4.35136 0.00000
1.33888 0.00000 0.00000 ; HA- CA- C- O
7070 7069 7079 7081 3 0.00000 0.00000 0.00000
0.00000 0.00000 0.00000 ; HA- CA- C- N
7071 7069 7079 7080 3 0.00000 0.00000 0.00000
0.00000 0.00000 0.00000 ; CB- CA- C- O
7071 7069 7079 7081 3 2.51040 4.18400 1.67360
-6.69440 0.00000 0.00000 ; CB- CA- C- N
7071 7074 7076 7077 3 20.92000 0.00000 -20.92000
0.00000 0.00000 0.00000 ; CB- CG- ND2- HD21
7071 7074 7076 7078 3 20.92000 0.00000 -20.92000
0.00000 0.00000 0.00000 ; CB- CG- ND2- HD22
7072 7071 7069 7079 3 0.65084 1.95253 0.00000
-2.60338 0.00000 0.00000 ; HB2- CB- CA- C
7072 7071 7074 7075 3 3.68192 -4.35136 0.00000
1.33888 0.00000 0.00000 ; HB2- CB- CG- OD1
7072 7071 7074 7076 3 0.00000 0.00000 0.00000
0.00000 0.00000 0.00000 ; HB2- CB- CG- ND2

7073 7071 7069 7079 3 0.65084 1.95253 0.00000
-2.60338 0.00000 0.00000 ; HB3- CB- CA- C
7073 7071 7074 7075 3 3.68192 -4.35136 0.00000
1.33888 0.00000 0.00000 ; HB3- CB- CG- OD1
7073 7071 7074 7076 3 0.00000 0.00000 0.00000
0.00000 0.00000 0.00000 ; HB3- CB- CG- ND2
7074 7071 7069 7079 3 7.94542 -3.96225 -6.11701
-0.55229 3.58150 0.00000 ; CG- CB- CA- C
7075 7074 7076 7077 3 29.28800 -8.36800 -20.92000
0.00000 0.00000 0.00000 ; OD1- CG- ND2- HD21
7075 7074 7076 7078 3 29.28800 -8.36800 -20.92000
0.00000 0.00000 0.00000 ; OD1- CG- ND2- HD22
7079 7081 7083 7084 3 0.00000 0.00000 0.00000
0.00000 0.00000 0.00000 ; C- N- CA- HA2
7079 7081 7083 7085 3 0.00000 0.00000 0.00000
0.00000 0.00000 0.00000 ; C- N- CA- HA3
7079 7081 7083 7086 3 1.75728 5.27184 2.25936
-7.02912 0.00000 0.00000 ; C- N- CA- C
7080 7079 7081 7082 3 29.28800 -8.36800 -20.92000
0.00000 0.00000 0.00000 ; O- C- N- H
7080 7079 7081 7083 3 20.92000 0.00000 -20.92000
0.00000 0.00000 0.00000 ; O- C- N- CA
7081 7083 7086 7087 3 0.00000 0.00000 0.00000
0.00000 0.00000 0.00000 ; N- CA- C- O
7081 7083 7086 7088 3 17.40544 -5.02080 -13.22144
9.20480 0.00000 0.00000 ; N- CA- C- N
7082 7081 7083 7084 3 0.00000 0.00000 0.00000
0.00000 0.00000 0.00000 ; H- N- CA- HA2
7082 7081 7083 7085 3 0.00000 0.00000 0.00000
0.00000 0.00000 0.00000 ; H- N- CA- HA3
7082 7081 7083 7086 3 0.00000 0.00000 0.00000
0.00000 0.00000 0.00000 ; H- N- CA- C
7083 7086 7088 7089 3 20.92000 0.00000 -20.92000
0.00000 0.00000 0.00000 ; CA- C- N- H
7083 7086 7088 7090 3 20.92000 0.00000 -20.92000
0.00000 0.00000 0.00000 ; CA- C- N- CA
7084 7083 7086 7087 3 3.68192 -4.35136 0.00000
1.33888 0.00000 0.00000 ; HA2- CA- C- O
7084 7083 7086 7088 3 0.00000 0.00000 0.00000
0.00000 0.00000 0.00000 ; HA2- CA- C- N
7085 7083 7086 7087 3 3.68192 -4.35136 0.00000
1.33888 0.00000 0.00000 ; HA3- CA- C- O
7085 7083 7086 7088 3 0.00000 0.00000 0.00000
0.00000 0.00000 0.00000 ; HA3- CA- C- N
7086 7088 7090 7091 3 0.00000 0.00000 0.00000
0.00000 0.00000 0.00000 ; C- N- CA- HA
7086 7088 7090 7092 3 11.71520 1.67360 15.06240 -
13.38880 0.00000 0.00000 ; C- N- CA- CB
7086 7088 7090 7099 3 1.75728 5.27184 2.25936
-7.02912 0.00000 0.00000 ; C- N- CA- C
7087 7086 7088 7089 3 29.28800 -8.36800 -20.92000
0.00000 0.00000 0.00000 ; O- C- N- H

7087 7086 7088 7090 3 20.92000 0.00000 -20.92000
0.00000 0.00000 0.00000 ; O- C- N- CA
7088 7090 7092 7093 3 0.65084 1.95253 0.00000
-2.60338 0.00000 0.00000 ; N- CA- CB- HB2
7088 7090 7092 7094 3 0.65084 1.95253 0.00000
-2.60338 0.00000 0.00000 ; N- CA- CB- HB3
7088 7090 7092 7095 3 5.99567 3.29281 -4.46014
-0.55229 1.97485 0.00000 ; N- CA- CB- CG
7088 7090 7099 7100 3 0.00000 0.00000 0.00000
0.00000 0.00000 0.00000 ; N- CA- C- O
7088 7090 7099 7101 3 17.40544 -5.02080 -13.22144
9.20480 0.00000 0.00000 ; N- CA- C- N
7089 7088 7090 7091 3 0.00000 0.00000 0.00000
0.00000 0.00000 0.00000 ; H- N- CA- HA
7089 7088 7090 7092 3 0.00000 0.00000 0.00000
0.00000 0.00000 0.00000 ; H- N- CA- CB
7089 7088 7090 7099 3 0.00000 0.00000 0.00000
0.00000 0.00000 0.00000 ; H- N- CA- C
7090 7092 7095 7096 3 0.00000 0.00000 0.00000
0.00000 0.00000 0.00000 ; CA- CB- CG- OD1
7090 7092 7095 7097 3 9.86169 5.11703 1.84933
-0.13389 -6.66093 0.00000 ; CA- CB- CG- OD2
7090 7099 7101 7102 3 20.92000 0.00000 -20.92000
0.00000 0.00000 0.00000 ; CA- C- N- H
7090 7099 7101 7103 3 20.92000 0.00000 -20.92000
0.00000 0.00000 0.00000 ; CA- C- N- CA
7091 7090 7092 7093 3 0.65084 1.95253 0.00000
-2.60338 0.00000 0.00000 ; HA- CA- CB- HB2
7091 7090 7092 7094 3 0.65084 1.95253 0.00000
-2.60338 0.00000 0.00000 ; HA- CA- CB- HB3
7091 7090 7092 7095 3 0.65084 1.95253 0.00000
-2.60338 0.00000 0.00000 ; HA- CA- CB- CG
7091 7090 7099 7100 3 3.68192 -4.35136 0.00000
1.33888 0.00000 0.00000 ; HA- CA- C- O
7091 7090 7099 7101 3 0.00000 0.00000 0.00000
0.00000 0.00000 0.00000 ; HA- CA- C- N
7092 7090 7099 7100 3 0.00000 0.00000 0.00000
0.00000 0.00000 0.00000 ; CB- CA- C- O
7092 7090 7099 7101 3 2.51040 4.18400 1.67360
-6.69440 0.00000 0.00000 ; CB- CA- C- N
7092 7095 7097 7098 3 27.46796 7.88684 -26.42614
-8.01654 3.78234 0.00000 ; CB- CG- OD2- HD2
7093 7092 7090 7099 3 0.65084 1.95253 0.00000
-2.60338 0.00000 0.00000 ; HB2- CB- CA- C
7093 7092 7095 7096 3 3.68192 -4.35136 0.00000
1.33888 0.00000 0.00000 ; HB2- CB- CG- OD1
7093 7092 7095 7097 3 0.00000 0.00000 0.00000
0.00000 0.00000 0.00000 ; HB2- CB- CG- OD2
7094 7092 7090 7099 3 0.65084 1.95253 0.00000
-2.60338 0.00000 0.00000 ; HB3- CB- CA- C
7094 7092 7095 7096 3 3.68192 -4.35136 0.00000
1.33888 0.00000 0.00000 ; HB3- CB- CG- OD1

7094 7092 7095 7097 3 0.00000 0.00000 0.00000
0.00000 0.00000 0.00000 ; HB3- CB- CG- OD2
7095 7092 7090 7099 3 7.94542 -3.96225 -6.11701
-0.55229 3.58150 0.00000 ; CG- CB- CA- C
7096 7095 7097 7098 3 27.19600 -7.94960 -19.24640
0.00000 0.00000 0.00000 ; OD1- CG- OD2- HD2
7099 7101 7103 7104 3 0.00000 0.00000 0.00000
0.00000 0.00000 0.00000 ; C- N- CA- HA2
7099 7101 7103 7105 3 0.00000 0.00000 0.00000
0.00000 0.00000 0.00000 ; C- N- CA- HA3
7099 7101 7103 7106 3 1.75728 5.27184 2.25936
-7.02912 0.00000 0.00000 ; C- N- CA- C
7100 7099 7101 7102 3 29.28800 -8.36800 -20.92000
0.00000 0.00000 0.00000 ; O- C- N- H
7100 7099 7101 7103 3 20.92000 0.00000 -20.92000
0.00000 0.00000 0.00000 ; O- C- N- CA
7101 7103 7106 7107 3 0.00000 0.00000 0.00000
0.00000 0.00000 0.00000 ; N- CA- C- O
7101 7103 7106 7108 3 17.40544 -5.02080 -13.22144
9.20480 0.00000 0.00000 ; N- CA- C- N
7102 7101 7103 7104 3 0.00000 0.00000 0.00000
0.00000 0.00000 0.00000 ; H- N- CA- HA2
7102 7101 7103 7105 3 0.00000 0.00000 0.00000
0.00000 0.00000 0.00000 ; H- N- CA- HA3
7102 7101 7103 7106 3 0.00000 0.00000 0.00000
0.00000 0.00000 0.00000 ; H- N- CA- C
7103 7106 7108 7109 3 20.92000 0.00000 -20.92000
0.00000 0.00000 0.00000 ; CA- C- N- H
7103 7106 7108 7110 3 20.92000 0.00000 -20.92000
0.00000 0.00000 0.00000 ; CA- C- N- CA
7104 7103 7106 7107 3 3.68192 -4.35136 0.00000
1.33888 0.00000 0.00000 ; HA2- CA- C- O
7104 7103 7106 7108 3 0.00000 0.00000 0.00000
0.00000 0.00000 0.00000 ; HA2- CA- C- N
7105 7103 7106 7107 3 3.68192 -4.35136 0.00000
1.33888 0.00000 0.00000 ; HA3- CA- C- O
7105 7103 7106 7108 3 0.00000 0.00000 0.00000
0.00000 0.00000 0.00000 ; HA3- CA- C- N
7106 7108 7110 7111 3 0.00000 0.00000 0.00000
0.00000 0.00000 0.00000 ; C- N- CA- HA
7106 7108 7110 7112 3 11.71520 1.67360 15.06240 -
13.38880 0.00000 0.00000 ; C- N- CA- CB
7106 7108 7110 7121 3 1.75728 5.27184 2.25936
-7.02912 0.00000 0.00000 ; C- N- CA- C
7107 7106 7108 7109 3 29.28800 -8.36800 -20.92000
0.00000 0.00000 0.00000 ; O- C- N- H
7107 7106 7108 7110 3 20.92000 0.00000 -20.92000
0.00000 0.00000 0.00000 ; O- C- N- CA
7108 7110 7112 7113 3 0.65084 1.95253 0.00000
-2.60338 0.00000 0.00000 ; N- CA- CB- HB2
7108 7110 7112 7114 3 0.65084 1.95253 0.00000
-2.60338 0.00000 0.00000 ; N- CA- CB- HB3

```
  7108   7110   7112   7115      3    3.21331    2.22589   -4.15053
-2.40998    2.61082    0.00000 ;      N-    CA-    CB-    CG
  7108   7110   7121   7122      3    0.00000    0.00000    0.00000
0.00000    0.00000    0.00000 ;      N-    CA-     C-     O
  7108   7110   7121   7123      3   17.40544   -5.02080  -13.22144
9.20480    0.00000    0.00000 ;      N-    CA-     C-     N
  7109   7108   7110   7111      3    0.00000    0.00000    0.00000
0.00000    0.00000    0.00000 ;      H-     N-    CA-    HA
  7109   7108   7110   7112      3    0.00000    0.00000    0.00000
0.00000    0.00000    0.00000 ;      H-     N-    CA-    CB
  7109   7108   7110   7121      3    0.00000    0.00000    0.00000
0.00000    0.00000    0.00000 ;      H-     N-    CA-     C
  7110   7112   7115   7116      3    0.66944    2.00832    0.00000
-2.67776    0.00000    0.00000 ;     CA-    CB-    CG-   HG2
  7110   7112   7115   7117      3    0.66944    2.00832    0.00000
-2.67776    0.00000    0.00000 ;     CA-    CB-    CG-   HG3
  7110   7112   7115   7118      3   10.12110   -1.91209    0.01674
10.17549   -1.87443    0.00000 ;     CA-    CB-    CG-    CD
  7110   7121   7123   7124      3   20.92000    0.00000  -20.92000
0.00000    0.00000    0.00000 ;     CA-     C-     N-     H
  7110   7121   7123   7125      3   20.92000    0.00000  -20.92000
0.00000    0.00000    0.00000 ;     CA-     C-     N-    CA
  7111   7110   7112   7113      3    0.65084    1.95253    0.00000
-2.60338    0.00000    0.00000 ;     HA-    CA-    CB-   HB2
  7111   7110   7112   7114      3    0.65084    1.95253    0.00000
-2.60338    0.00000    0.00000 ;     HA-    CA-    CB-   HB3
  7111   7110   7112   7115      3    0.65084    1.95253    0.00000
-2.60338    0.00000    0.00000 ;     HA-    CA-    CB-    CG
  7111   7110   7121   7122      3    3.68192   -4.35136    0.00000
1.33888    0.00000    0.00000 ;     HA-    CA-     C-     O
  7111   7110   7121   7123      3    0.00000    0.00000    0.00000
0.00000    0.00000    0.00000 ;     HA-    CA-     C-     N
  7112   7110   7121   7122      3    0.00000    0.00000    0.00000
0.00000    0.00000    0.00000 ;     CB-    CA-     C-     O
  7112   7110   7121   7123      3    2.51040    4.18400    1.67360
-6.69440    0.00000    0.00000 ;     CB-    CA-     C-     N
  7112   7115   7118   7119      3    3.79907    0.00000   -5.40573
0.00000    2.14221    0.00000 ;     CB-    CG-    CD-   OE1
  7112   7115   7118   7120      3    3.79907    0.00000   -5.40573
0.00000    2.14221    0.00000 ;     CB-    CG-    CD-   OE2
  7113   7112   7110   7121      3    0.65084    1.95253    0.00000
-2.60338    0.00000    0.00000 ;    HB2-    CB-    CA-     C
  7113   7112   7115   7116      3    0.62760    1.88280    0.00000
-2.51040    0.00000    0.00000 ;    HB2-    CB-    CG-   HG2
  7113   7112   7115   7117      3    0.62760    1.88280    0.00000
-2.51040    0.00000    0.00000 ;    HB2-    CB-    CG-   HG3
  7113   7112   7115   7118      3    0.65084    1.95253    0.00000
-2.60338    0.00000    0.00000 ;    HB2-    CB-    CG-    CD
  7114   7112   7110   7121      3    0.65084    1.95253    0.00000
-2.60338    0.00000    0.00000 ;    HB3-    CB-    CA-     C
  7114   7112   7115   7116      3    0.62760    1.88280    0.00000
-2.51040    0.00000    0.00000 ;    HB3-    CB-    CG-   HG2
```

7114 7112 7115 7117 3 0.62760 1.88280 0.00000
-2.51040 0.00000 0.00000 ; HB3- CB- CG- HG3
7114 7112 7115 7118 3 0.65084 1.95253 0.00000
-2.60338 0.00000 0.00000 ; HB3- CB- CG- CD
7115 7112 7110 7121 3 6.86594 3.56895 -8.14206
-2.40998 4.85344 0.00000 ; CG- CB- CA- C
7116 7115 7118 7119 3 0.00000 0.00000 0.00000
0.00000 0.00000 0.00000 ; HG2- CG- CD- OE1
7116 7115 7118 7120 3 0.00000 0.00000 0.00000
0.00000 0.00000 0.00000 ; HG2- CG- CD- OE2
7117 7115 7118 7119 3 0.00000 0.00000 0.00000
0.00000 0.00000 0.00000 ; HG3- CG- CD- OE1
7117 7115 7118 7120 3 0.00000 0.00000 0.00000
0.00000 0.00000 0.00000 ; HG3- CG- CD- OE2
7121 7123 7125 7126 3 0.00000 0.00000 0.00000
0.00000 0.00000 0.00000 ; C- N- CA- HA
7121 7123 7125 7127 3 11.71520 1.67360 15.06240 -
13.38880 0.00000 0.00000 ; C- N- CA- CB
7121 7123 7125 7140 3 1.75728 5.27184 2.25936
-7.02912 0.00000 0.00000 ; C- N- CA- C
7122 7121 7123 7124 3 29.28800 -8.36800 -20.92000
0.00000 0.00000 0.00000 ; O- C- N- H
7122 7121 7123 7125 3 20.92000 0.00000 -20.92000
0.00000 0.00000 0.00000 ; O- C- N- CA
7123 7125 7127 7128 3 0.65084 1.95253 0.00000
-2.60338 0.00000 0.00000 ; N- CA- CB- HB
7123 7125 7127 7129 3 3.83673 0.44769 -1.77402
-2.47693 -0.03347 0.00000 ; N- CA- CB- CG2
7123 7125 7127 7133 3 2.97482 0.12134 2.04179
-1.89117 -3.24678 0.00000 ; N- CA- CB- CG1
7123 7125 7140 7141 3 0.00000 0.00000 0.00000
0.00000 0.00000 0.00000 ; N- CA- C- O
7123 7125 7140 7142 3 17.40544 -5.02080 -13.22144
9.20480 0.00000 0.00000 ; N- CA- C- N
7124 7123 7125 7126 3 0.00000 0.00000 0.00000
0.00000 0.00000 0.00000 ; H- N- CA- HA
7124 7123 7125 7127 3 0.00000 0.00000 0.00000
0.00000 0.00000 0.00000 ; H- N- CA- CB
7124 7123 7125 7140 3 0.00000 0.00000 0.00000
0.00000 0.00000 0.00000 ; H- N- CA- C
7125 7127 7129 7130 3 0.66944 2.00832 0.00000
-2.67776 0.00000 0.00000 ; CA- CB- CG2- HG21
7125 7127 7129 7131 3 0.66944 2.00832 0.00000
-2.67776 0.00000 0.00000 ; CA- CB- CG2- HG22
7125 7127 7129 7132 3 0.66944 2.00832 0.00000
-2.67776 0.00000 0.00000 ; CA- CB- CG2- HG23
7125 7127 7133 7134 3 0.66944 2.00832 0.00000
-2.67776 0.00000 0.00000 ; CA- CB- CG1- HG12
7125 7127 7133 7135 3 0.66944 2.00832 0.00000
-2.67776 0.00000 0.00000 ; CA- CB- CG1- HG13
7125 7127 7133 7136 3 4.24258 -0.52718 -7.25506
-1.79075 7.69856 0.00000 ; CA- CB- CG1- CD1

7125 7140 7142 7143 3 20.92000 0.00000 -20.92000
0.00000 0.00000 0.00000 ; CA- C- N- H
7125 7140 7142 7144 3 20.92000 0.00000 -20.92000
0.00000 0.00000 0.00000 ; CA- C- N- CA
7126 7125 7127 7128 3 0.65084 1.95253 0.00000
-2.60338 0.00000 0.00000 ; HA- CA- CB- HB
7126 7125 7127 7129 3 0.65084 1.95253 0.00000
-2.60338 0.00000 0.00000 ; HA- CA- CB- CG2
7126 7125 7127 7133 3 0.65084 1.95253 0.00000
-2.60338 0.00000 0.00000 ; HA- CA- CB- CG1
7126 7125 7140 7141 3 3.68192 -4.35136 0.00000
1.33888 0.00000 0.00000 ; HA- CA- C- O
7126 7125 7140 7142 3 0.00000 0.00000 0.00000
0.00000 0.00000 0.00000 ; HA- CA- C- N
7127 7125 7140 7141 3 0.00000 0.00000 0.00000
0.00000 0.00000 0.00000 ; CB- CA- C- O
7127 7125 7140 7142 3 2.51040 4.18400 1.67360
-6.69440 0.00000 0.00000 ; CB- CA- C- N
7127 7133 7136 7137 3 0.66944 2.00832 0.00000
-2.67776 0.00000 0.00000 ; CB- CG1- CD1- HD11
7127 7133 7136 7138 3 0.66944 2.00832 0.00000
-2.67776 0.00000 0.00000 ; CB- CG1- CD1- HD12
7127 7133 7136 7139 3 0.66944 2.00832 0.00000
-2.67776 0.00000 0.00000 ; CB- CG1- CD1- HD13
7128 7127 7125 7140 3 0.65084 1.95253 0.00000
-2.60338 0.00000 0.00000 ; HB- CB- CA- C
7128 7127 7129 7130 3 0.62760 1.88280 0.00000
-2.51040 0.00000 0.00000 ; HB- CB- CG2- HG21
7128 7127 7129 7131 3 0.62760 1.88280 0.00000
-2.51040 0.00000 0.00000 ; HB- CB- CG2- HG22
7128 7127 7129 7132 3 0.62760 1.88280 0.00000
-2.51040 0.00000 0.00000 ; HB- CB- CG2- HG23
7128 7127 7133 7134 3 0.62760 1.88280 0.00000
-2.51040 0.00000 0.00000 ; HB- CB- CG1- HG12
7128 7127 7133 7135 3 0.62760 1.88280 0.00000
-2.51040 0.00000 0.00000 ; HB- CB- CG1- HG13
7128 7127 7133 7136 3 0.66944 2.00832 0.00000
-2.67776 0.00000 0.00000 ; HB- CB- CG1- CD1
7129 7127 7125 7140 3 5.67350 3.55640 -6.16722
-2.47693 3.74886 0.00000 ; CG2- CB- CA- C
7129 7127 7133 7134 3 0.66944 2.00832 0.00000
-2.67776 0.00000 0.00000 ; CG2- CB- CG1- HG12
7129 7127 7133 7135 3 0.66944 2.00832 0.00000
-2.67776 0.00000 0.00000 ; CG2- CB- CG1- HG13
7129 7127 7133 7136 3 3.81162 0.49790 -8.14206
-1.79075 7.49773 0.00000 ; CG2- CB- CG1- CD1
7130 7129 7127 7133 3 0.66944 2.00832 0.00000
-2.67776 0.00000 0.00000 ; HG21- CG2- CB- CG1
7131 7129 7127 7133 3 0.66944 2.00832 0.00000
-2.67776 0.00000 0.00000 ; HG22- CG2- CB- CG1
7132 7129 7127 7133 3 0.66944 2.00832 0.00000
-2.67776 0.00000 0.00000 ; HG23- CG2- CB- CG1

```
  7133   7127   7125   7140      3    8.26340    0.74057   -9.99976
-1.89117    3.84928    0.00000 ;   CG1-   CB-   CA-    C
  7134   7133   7136   7137      3    0.62760    1.88280    0.00000
-2.51040    0.00000    0.00000 ;  HG12-  CG1-  CD1-  HD11
  7134   7133   7136   7138      3    0.62760    1.88280    0.00000
-2.51040    0.00000    0.00000 ;  HG12-  CG1-  CD1-  HD12
  7134   7133   7136   7139      3    0.62760    1.88280    0.00000
-2.51040    0.00000    0.00000 ;  HG12-  CG1-  CD1-  HD13
  7135   7133   7136   7137      3    0.62760    1.88280    0.00000
-2.51040    0.00000    0.00000 ;  HG13-  CG1-  CD1-  HD11
  7135   7133   7136   7138      3    0.62760    1.88280    0.00000
-2.51040    0.00000    0.00000 ;  HG13-  CG1-  CD1-  HD12
  7135   7133   7136   7139      3    0.62760    1.88280    0.00000
-2.51040    0.00000    0.00000 ;  HG13-  CG1-  CD1-  HD13
  7140   7142   7144   7145      3    0.00000    0.00000    0.00000
0.00000    0.00000    0.00000 ;     C-    N-   CA-   HA
  7140   7142   7144   7146      3   11.71520    1.67360   15.06240  -
13.38880    0.00000    0.00000 ;     C-    N-   CA-   CB
  7140   7142   7144   7152      3    1.75728    5.27184    2.25936
-7.02912    0.00000    0.00000 ;     C-    N-   CA-    C
  7141   7140   7142   7143      3   29.28800   -8.36800  -20.92000
0.00000    0.00000    0.00000 ;     O-    C-    N-    H
  7141   7140   7142   7144      3   20.92000    0.00000  -20.92000
0.00000    0.00000    0.00000 ;     O-    C-    N-   CA
  7142   7144   7146   7147      3    0.65084    1.95253    0.00000
-2.60338    0.00000    0.00000 ;     N-   CA-   CB-   HB2
  7142   7144   7146   7148      3    0.65084    1.95253    0.00000
-2.60338    0.00000    0.00000 ;     N-   CA-   CB-   HB3
  7142   7144   7146   7149      3   15.41386    9.74035   -8.39310
-0.97069    2.97901    0.00000 ;     N-   CA-   CB-   CG
  7142   7144   7152   7153      3    0.00000    0.00000    0.00000
0.00000    0.00000    0.00000 ;     N-   CA-    C-    O
  7142   7144   7152   7154      3   17.40544   -5.02080  -13.22144
9.20480    0.00000    0.00000 ;     N-   CA-    C-    N
  7143   7142   7144   7145      3    0.00000    0.00000    0.00000
0.00000    0.00000    0.00000 ;     H-    N-   CA-   HA
  7143   7142   7144   7146      3    0.00000    0.00000    0.00000
0.00000    0.00000    0.00000 ;     H-    N-   CA-   CB
  7143   7142   7144   7152      3    0.00000    0.00000    0.00000
0.00000    0.00000    0.00000 ;     H-    N-   CA-    C
  7144   7146   7149   7150      3    6.43499    0.00000   -5.39736
0.00000   -1.03763    0.00000 ;    CA-   CB-   CG-  OD1
  7144   7146   7149   7151      3    6.43499    0.00000   -5.39736
0.00000   -1.03763    0.00000 ;    CA-   CB-   CG-  OD2
  7144   7152   7154   7155      3   20.92000    0.00000  -20.92000
0.00000    0.00000    0.00000 ;    CA-    C-    N-    H
  7144   7152   7154   7156      3   20.92000    0.00000  -20.92000
0.00000    0.00000    0.00000 ;    CA-    C-    N-   CA
  7145   7144   7146   7147      3    0.65084    1.95253    0.00000
-2.60338    0.00000    0.00000 ;    HA-   CA-   CB-   HB2
  7145   7144   7146   7148      3    0.65084    1.95253    0.00000
-2.60338    0.00000    0.00000 ;    HA-   CA-   CB-   HB3
```

7145 7144 7146 7149 3 0.65084 1.95253 0.00000
-2.60338 0.00000 0.00000 ; HA- CA- CB- CG
7145 7144 7152 7153 3 3.68192 -4.35136 0.00000
1.33888 0.00000 0.00000 ; HA- CA- C- O
7145 7144 7152 7154 3 0.00000 0.00000 0.00000
0.00000 0.00000 0.00000 ; HA- CA- C- N
7146 7144 7152 7153 3 0.00000 0.00000 0.00000
0.00000 0.00000 0.00000 ; CB- CA- C- O
7146 7144 7152 7154 3 2.51040 4.18400 1.67360
-6.69440 0.00000 0.00000 ; CB- CA- C- N
7147 7146 7144 7152 3 0.65084 1.95253 0.00000
-2.60338 0.00000 0.00000 ; HB2- CB- CA- C
7147 7146 7149 7150 3 0.00000 0.00000 0.00000
0.00000 0.00000 0.00000 ; HB2- CB- CG- OD1
7147 7146 7149 7151 3 0.00000 0.00000 0.00000
0.00000 0.00000 0.00000 ; HB2- CB- CG- OD2
7148 7146 7144 7152 3 0.65084 1.95253 0.00000
-2.60338 0.00000 0.00000 ; HB3- CB- CA- C
7148 7146 7149 7150 3 0.00000 0.00000 0.00000
0.00000 0.00000 0.00000 ; HB3- CB- CG- OD1
7148 7146 7149 7151 3 0.00000 0.00000 0.00000
0.00000 0.00000 0.00000 ; HB3- CB- CG- OD2
7149 7146 7144 7152 3 7.14627 -1.04600 -8.99560
-0.97069 5.15469 0.00000 ; CG- CB- CA- C
7152 7154 7156 7157 3 0.00000 0.00000 0.00000
0.00000 0.00000 0.00000 ; C- N- CA- HA
7152 7154 7156 7158 3 11.71520 1.67360 15.06240 -
13.38880 0.00000 0.00000 ; C- N- CA- CB
7152 7154 7156 7172 3 1.75728 5.27184 2.25936
-7.02912 0.00000 0.00000 ; C- N- CA- C
7153 7152 7154 7155 3 29.28800 -8.36800 -20.92000
0.00000 0.00000 0.00000 ; O- C- N- H
7153 7152 7154 7156 3 20.92000 0.00000 -20.92000
0.00000 0.00000 0.00000 ; O- C- N- CA
7154 7156 7158 7159 3 0.65084 1.95253 0.00000
-2.60338 0.00000 0.00000 ; N- CA- CB- HB2
7154 7156 7158 7160 3 0.65084 1.95253 0.00000
-2.60338 0.00000 0.00000 ; N- CA- CB- HB3
7154 7156 7158 7161 3 3.28026 2.46019 -2.19242
-3.21331 -0.23430 0.00000 ; N- CA- CB- CG
7154 7156 7172 7173 3 0.00000 0.00000 0.00000
0.00000 0.00000 0.00000 ; N- CA- C- O
7154 7156 7172 7174 3 17.40544 -5.02080 -13.22144
9.20480 0.00000 0.00000 ; N- CA- C- N
7155 7154 7156 7157 3 0.00000 0.00000 0.00000
0.00000 0.00000 0.00000 ; H- N- CA- HA
7155 7154 7156 7158 3 0.00000 0.00000 0.00000
0.00000 0.00000 0.00000 ; H- N- CA- CB
7155 7154 7156 7172 3 0.00000 0.00000 0.00000
0.00000 0.00000 0.00000 ; H- N- CA- C
7156 7158 7161 7162 3 0.57739 0.00000 1.02926
0.00000 -1.60666 0.00000 ; CA- CB- CG- CD1

7156 7158 7161 7170 3 0.57739 0.00000 1.02926
0.00000 -1.60666 0.00000 ; CA- CB- CG- CD2
7156 7172 7174 7175 3 20.92000 0.00000 -20.92000
0.00000 0.00000 0.00000 ; CA- C- N- H
7156 7172 7174 7176 3 20.92000 0.00000 -20.92000
0.00000 0.00000 0.00000 ; CA- C- N- CA
7157 7156 7158 7159 3 0.65084 1.95253 0.00000
-2.60338 0.00000 0.00000 ; HA- CA- CB- HB2
7157 7156 7158 7160 3 0.65084 1.95253 0.00000
-2.60338 0.00000 0.00000 ; HA- CA- CB- HB3
7157 7156 7158 7161 3 0.65084 1.95253 0.00000
-2.60338 0.00000 0.00000 ; HA- CA- CB- CG
7157 7156 7172 7173 3 3.68192 -4.35136 0.00000
1.33888 0.00000 0.00000 ; HA- CA- C- O
7157 7156 7172 7174 3 0.00000 0.00000 0.00000
0.00000 0.00000 0.00000 ; HA- CA- C- N
7158 7156 7172 7173 3 0.00000 0.00000 0.00000
0.00000 0.00000 0.00000 ; CB- CA- C- O
7158 7156 7172 7174 3 2.51040 4.18400 1.67360
-6.69440 0.00000 0.00000 ; CB- CA- C- N
7158 7161 7162 7163 3 30.33400 0.00000 -30.33400
0.00000 0.00000 0.00000 ; CB- CG- CD1- HD1
7158 7161 7162 7164 3 30.33400 0.00000 -30.33400
0.00000 0.00000 0.00000 ; CB- CG- CD1- CE1
7158 7161 7170 7168 3 30.33400 0.00000 -30.33400
0.00000 0.00000 0.00000 ; CB- CG- CD2- CE2
7158 7161 7170 7171 3 30.33400 0.00000 -30.33400
0.00000 0.00000 0.00000 ; CB- CG- CD2- HD2
7159 7158 7156 7172 3 0.65084 1.95253 0.00000
-2.60338 0.00000 0.00000 ; HB2- CB- CA- C
7159 7158 7161 7162 3 0.00000 0.00000 0.00000
0.00000 0.00000 0.00000 ; HB2- CB- CG- CD1
7159 7158 7161 7170 3 0.00000 0.00000 0.00000
0.00000 0.00000 0.00000 ; HB2- CB- CG- CD2
7160 7158 7156 7172 3 0.65084 1.95253 0.00000
-2.60338 0.00000 0.00000 ; HB3- CB- CA- C
7160 7158 7161 7162 3 0.00000 0.00000 0.00000
0.00000 0.00000 0.00000 ; HB3- CB- CG- CD1
7160 7158 7161 7170 3 0.00000 0.00000 0.00000
0.00000 0.00000 0.00000 ; HB3- CB- CG- CD2
7161 7158 7156 7172 3 4.95804 2.17986 -3.52293
-3.21331 -0.40166 0.00000 ; CG- CB- CA- C
7161 7162 7164 7165 3 30.33400 0.00000 -30.33400
0.00000 0.00000 0.00000 ; CG- CD1- CE1- HE1
7161 7162 7164 7166 3 30.33400 0.00000 -30.33400
0.00000 0.00000 0.00000 ; CG- CD1- CE1- CZ
7161 7170 7168 7166 3 30.33400 0.00000 -30.33400
0.00000 0.00000 0.00000 ; CG- CD2- CE2- CZ
7161 7170 7168 7169 3 30.33400 0.00000 -30.33400
0.00000 0.00000 0.00000 ; CG- CD2- CE2- HE2
7162 7161 7170 7168 3 30.33400 0.00000 -30.33400
0.00000 0.00000 0.00000 ; CD1- CG- CD2- CE2

7162 7161 7170 7171 3 30.33400 0.00000 -30.33400
0.00000 0.00000 0.00000 ; CD1- CG- CD2- HD2
7162 7164 7166 7167 3 30.33400 0.00000 -30.33400
0.00000 0.00000 0.00000 ; CD1- CE1- CZ- HZ
7162 7164 7166 7168 3 30.33400 0.00000 -30.33400
0.00000 0.00000 0.00000 ; CD1- CE1- CZ- CE2
7163 7162 7161 7170 3 30.33400 0.00000 -30.33400
0.00000 0.00000 0.00000 ; HD1- CD1- CG- CD2
7163 7162 7164 7165 3 30.33400 0.00000 -30.33400
0.00000 0.00000 0.00000 ; HD1- CD1- CE1- HE1
7163 7162 7164 7166 3 30.33400 0.00000 -30.33400
0.00000 0.00000 0.00000 ; HD1- CD1- CE1- CZ
7164 7162 7161 7170 3 30.33400 0.00000 -30.33400
0.00000 0.00000 0.00000 ; CE1- CD1- CG- CD2
7164 7166 7168 7169 3 30.33400 0.00000 -30.33400
0.00000 0.00000 0.00000 ; CE1- CZ- CE2- HE2
7164 7166 7168 7170 3 30.33400 0.00000 -30.33400
0.00000 0.00000 0.00000 ; CE1- CZ- CE2- CD2
7165 7164 7166 7167 3 30.33400 0.00000 -30.33400
0.00000 0.00000 0.00000 ; HE1- CE1- CZ- HZ
7165 7164 7166 7168 3 30.33400 0.00000 -30.33400
0.00000 0.00000 0.00000 ; HE1- CE1- CZ- CE2
7166 7168 7170 7171 3 30.33400 0.00000 -30.33400
0.00000 0.00000 0.00000 ; CZ- CE2- CD2- HD2
7167 7166 7168 7169 3 30.33400 0.00000 -30.33400
0.00000 0.00000 0.00000 ; HZ- CZ- CE2- HE2
7167 7166 7168 7170 3 30.33400 0.00000 -30.33400
0.00000 0.00000 0.00000 ; HZ- CZ- CE2- CD2
7169 7168 7170 7171 3 30.33400 0.00000 -30.33400
0.00000 0.00000 0.00000 ; HE2- CE2- CD2- HD2
7172 7174 7176 7177 3 0.00000 0.00000 0.00000
0.00000 0.00000 0.00000 ; C- N- CA- HA
7172 7174 7176 7178 3 11.71520 1.67360 15.06240 -
13.38880 0.00000 0.00000 ; C- N- CA- CB
7172 7174 7176 7189 3 1.75728 5.27184 2.25936
-7.02912 0.00000 0.00000 ; C- N- CA- C
7173 7172 7174 7175 3 29.28800 -8.36800 -20.92000
0.00000 0.00000 0.00000 ; O- C- N- H
7173 7172 7174 7176 3 20.92000 0.00000 -20.92000
0.00000 0.00000 0.00000 ; O- C- N- CA
7174 7176 7178 7179 3 0.65084 1.95253 0.00000
-2.60338 0.00000 0.00000 ; N- CA- CB- HB2
7174 7176 7178 7180 3 0.65084 1.95253 0.00000
-2.60338 0.00000 0.00000 ; N- CA- CB- HB3
7174 7176 7178 7181 3 4.79068 4.02919 -4.82834
-3.66518 2.97901 0.00000 ; N- CA- CB- CG
7174 7176 7189 7190 3 0.00000 0.00000 0.00000
0.00000 0.00000 0.00000 ; N- CA- C- O
7174 7176 7189 7191 3 17.40544 -5.02080 -13.22144
9.20480 0.00000 0.00000 ; N- CA- C- N
7175 7174 7176 7177 3 0.00000 0.00000 0.00000
0.00000 0.00000 0.00000 ; H- N- CA- HA

7175 7174 7176 7178 3 0.00000 0.00000 0.00000
0.00000 0.00000 0.00000 ; H- N- CA- CB
7175 7174 7176 7189 3 0.00000 0.00000 0.00000
0.00000 0.00000 0.00000 ; H- N- CA- C
7176 7178 7181 7182 3 5.98312 6.40152 3.28026 -
12.38464 -1.57318 0.00000 ; CA- CB- CG- ND1
7176 7178 7181 7187 3 0.00000 0.00000 0.00000
0.00000 0.00000 0.00000 ; CA- CB- CG- CD2
7176 7189 7191 7192 3 20.92000 0.00000 -20.92000
0.00000 0.00000 0.00000 ; CA- C- N- H
7176 7189 7191 7193 3 20.92000 0.00000 -20.92000
0.00000 0.00000 0.00000 ; CA- C- N- CA
7177 7176 7178 7179 3 0.65084 1.95253 0.00000
-2.60338 0.00000 0.00000 ; HA- CA- CB- HB2
7177 7176 7178 7180 3 0.65084 1.95253 0.00000
-2.60338 0.00000 0.00000 ; HA- CA- CB- HB3
7177 7176 7178 7181 3 0.65084 1.95253 0.00000
-2.60338 0.00000 0.00000 ; HA- CA- CB- CG
7177 7176 7189 7190 3 3.68192 -4.35136 0.00000
1.33888 0.00000 0.00000 ; HA- CA- C- O
7177 7176 7189 7191 3 0.00000 0.00000 0.00000
0.00000 0.00000 0.00000 ; HA- CA- C- N
7178 7176 7189 7190 3 0.00000 0.00000 0.00000
0.00000 0.00000 0.00000 ; CB- CA- C- O
7178 7176 7189 7191 3 2.51040 4.18400 1.67360
-6.69440 0.00000 0.00000 ; CB- CA- C- N
7178 7181 7182 7183 3 20.08320 0.00000 -20.08320
0.00000 0.00000 0.00000 ; CB- CG- ND1- CE1
7178 7181 7187 7185 3 44.97800 0.00000 -44.97800
0.00000 0.00000 0.00000 ; CB- CG- CD2- NE2
7178 7181 7187 7188 3 44.97800 0.00000 -44.97800
0.00000 0.00000 0.00000 ; CB- CG- CD2- HD2
7179 7178 7176 7189 3 0.65084 1.95253 0.00000
-2.60338 0.00000 0.00000 ; HB2- CB- CA- C
7179 7178 7181 7182 3 0.00000 0.00000 0.00000
0.00000 0.00000 0.00000 ; HB2- CB- CG- ND1
7179 7178 7181 7187 3 0.00000 0.00000 0.00000
0.00000 0.00000 0.00000 ; HB2- CB- CG- CD2
7180 7178 7176 7189 3 0.65084 1.95253 0.00000
-2.60338 0.00000 0.00000 ; HB3- CB- CA- C
7180 7178 7181 7182 3 0.00000 0.00000 0.00000
0.00000 0.00000 0.00000 ; HB3- CB- CG- ND1
7180 7178 7181 7187 3 0.00000 0.00000 0.00000
0.00000 0.00000 0.00000 ; HB3- CB- CG- CD2
7181 7178 7176 7189 3 3.76560 3.34720 -2.87859
-3.66518 0.83680 0.00000 ; CG- CB- CA- C
7181 7182 7183 7184 3 41.84000 0.00000 -41.84000
0.00000 0.00000 0.00000 ; CG- ND1- CE1- HE1
7181 7182 7183 7185 3 41.84000 0.00000 -41.84000
0.00000 0.00000 0.00000 ; CG- ND1- CE1- NE2
7181 7187 7185 7183 3 12.55200 0.00000 -12.55200
0.00000 0.00000 0.00000 ; CG- CD2- NE2- CE1

7181 7187 7185 7186 3 12.55200 0.00000 -12.55200
0.00000 0.00000 0.00000 ; CG- CD2- NE2- HE2
7182 7181 7187 7185 3 44.97800 0.00000 -44.97800
0.00000 0.00000 0.00000 ; ND1- CG- CD2- NE2
7182 7181 7187 7188 3 44.97800 0.00000 -44.97800
0.00000 0.00000 0.00000 ; ND1- CG- CD2- HD2
7182 7183 7185 7186 3 19.45560 0.00000 -19.45560
0.00000 0.00000 0.00000 ; ND1- CE1- NE2- HE2
7182 7183 7185 7187 3 19.45560 0.00000 -19.45560
0.00000 0.00000 0.00000 ; ND1- CE1- NE2- CD2
7183 7182 7181 7187 3 20.08320 0.00000 -20.08320
0.00000 0.00000 0.00000 ; CE1- ND1- CG- CD2
7183 7185 7187 7188 3 12.55200 0.00000 -12.55200
0.00000 0.00000 0.00000 ; CE1- NE2- CD2- HD2
7184 7183 7185 7186 3 19.45560 0.00000 -19.45560
0.00000 0.00000 0.00000 ; HE1- CE1- NE2- HE2
7184 7183 7185 7187 3 19.45560 0.00000 -19.45560
0.00000 0.00000 0.00000 ; HE1- CE1- NE2- CD2
7186 7185 7187 7188 3 12.55200 0.00000 -12.55200
0.00000 0.00000 0.00000 ; HE2- NE2- CD2- HD2
7189 7191 7193 7194 3 0.00000 0.00000 0.00000
0.00000 0.00000 0.00000 ; C- N- CA- HA
7189 7191 7193 7195 3 11.71520 1.67360 15.06240 -
13.38880 0.00000 0.00000 ; C- N- CA- CB
7189 7191 7193 7205 3 1.75728 5.27184 2.25936
-7.02912 0.00000 0.00000 ; C- N- CA- C
7190 7189 7191 7192 3 29.28800 -8.36800 -20.92000
0.00000 0.00000 0.00000 ; O- C- N- H
7190 7189 7191 7193 3 20.92000 0.00000 -20.92000
0.00000 0.00000 0.00000 ; O- C- N- CA
7191 7193 7195 7196 3 0.65084 1.95253 0.00000
-2.60338 0.00000 0.00000 ; N- CA- CB- HB2
7191 7193 7195 7197 3 0.65084 1.95253 0.00000
-2.60338 0.00000 0.00000 ; N- CA- CB- HB3
7191 7193 7195 7198 3 3.21331 2.22589 -4.15053
-2.40998 2.61082 0.00000 ; N- CA- CB- CG
7191 7193 7205 7206 3 0.00000 0.00000 0.00000
0.00000 0.00000 0.00000 ; N- CA- C- O
7191 7193 7205 7207 3 17.40544 -5.02080 -13.22144
9.20480 0.00000 0.00000 ; N- CA- C- N
7192 7191 7193 7194 3 0.00000 0.00000 0.00000
0.00000 0.00000 0.00000 ; H- N- CA- HA
7192 7191 7193 7195 3 0.00000 0.00000 0.00000
0.00000 0.00000 0.00000 ; H- N- CA- CB
7192 7191 7193 7205 3 0.00000 0.00000 0.00000
0.00000 0.00000 0.00000 ; H- N- CA- C
7193 7195 7198 7199 3 0.66944 2.00832 0.00000
-2.67776 0.00000 0.00000 ; CA- CB- CG- HG2
7193 7195 7198 7200 3 0.66944 2.00832 0.00000
-2.67776 0.00000 0.00000 ; CA- CB- CG- HG3
7193 7195 7198 7201 3 3.69866 -4.35136 -3.92459
6.89523 4.61914 0.00000 ; CA- CB- CG- CD

```
  7193   7205   7207   7208      3   20.92000    0.00000  -20.92000
0.00000    0.00000    0.00000 ;     CA-     C-     N-     H
  7193   7205   7207   7209      3   20.92000    0.00000  -20.92000
0.00000    0.00000    0.00000 ;     CA-     C-     N-    CA
  7194   7193   7195   7196      3    0.65084    1.95253    0.00000
-2.60338    0.00000    0.00000 ;     HA-    CA-    CB-   HB2
  7194   7193   7195   7197      3    0.65084    1.95253    0.00000
-2.60338    0.00000    0.00000 ;     HA-    CA-    CB-   HB3
  7194   7193   7195   7198      3    0.65084    1.95253    0.00000
-2.60338    0.00000    0.00000 ;     HA-    CA-    CB-    CG
  7194   7193   7205   7206      3    3.68192   -4.35136    0.00000
1.33888    0.00000    0.00000 ;     HA-    CA-     C-     O
  7194   7193   7205   7207      3    0.00000    0.00000    0.00000
0.00000    0.00000    0.00000 ;     HA-    CA-     C-     N
  7195   7193   7205   7206      3    0.00000    0.00000    0.00000
0.00000    0.00000    0.00000 ;     CB-    CA-     C-     O
  7195   7193   7205   7207      3    2.51040    4.18400    1.67360
-6.69440    0.00000    0.00000 ;     CB-    CA-     C-     N
  7195   7198   7201   7202      3    0.00000    0.00000    0.00000
0.00000    0.00000    0.00000 ;     CB-    CG-    CD-   OE1
  7195   7198   7201   7203      3   12.79049    3.13382   -7.02912
0.41840   -2.20915    0.00000 ;     CB-    CG-    CD-   OE2
  7196   7195   7193   7205      3    0.65084    1.95253    0.00000
-2.60338    0.00000    0.00000 ;    HB2-    CB-    CA-     C
  7196   7195   7198   7199      3    0.62760    1.88280    0.00000
-2.51040    0.00000    0.00000 ;    HB2-    CB-    CG-   HG2
  7196   7195   7198   7200      3    0.62760    1.88280    0.00000
-2.51040    0.00000    0.00000 ;    HB2-    CB-    CG-   HG3
  7196   7195   7198   7201      3    0.65084    1.95253    0.00000
-2.60338    0.00000    0.00000 ;    HB2-    CB-    CG-    CD
  7197   7195   7193   7205      3    0.65084    1.95253    0.00000
-2.60338    0.00000    0.00000 ;    HB3-    CB-    CA-     C
  7197   7195   7198   7199      3    0.62760    1.88280    0.00000
-2.51040    0.00000    0.00000 ;    HB3-    CB-    CG-   HG2
  7197   7195   7198   7200      3    0.62760    1.88280    0.00000
-2.51040    0.00000    0.00000 ;    HB3-    CB-    CG-   HG3
  7197   7195   7198   7201      3    0.65084    1.95253    0.00000
-2.60338    0.00000    0.00000 ;    HB3-    CB-    CG-    CD
  7198   7195   7193   7205      3    6.86594    3.56895   -8.14206
-2.40998    4.85344    0.00000 ;     CG-    CB-    CA-     C
  7198   7201   7203   7204      3   27.46796    7.88684  -26.42614
-8.01654    3.78234    0.00000 ;     CG-    CD-   OE2-   HE2
  7199   7198   7201   7202      3    3.68192   -4.35136    0.00000
1.33888    0.00000    0.00000 ;    HG2-    CG-    CD-   OE1
  7199   7198   7201   7203      3    0.00000    0.00000    0.00000
0.00000    0.00000    0.00000 ;    HG2-    CG-    CD-   OE2
  7200   7198   7201   7202      3    3.68192   -4.35136    0.00000
1.33888    0.00000    0.00000 ;    HG3-    CG-    CD-   OE1
  7200   7198   7201   7203      3    0.00000    0.00000    0.00000
0.00000    0.00000    0.00000 ;    HG3-    CG-    CD-   OE2
  7202   7201   7203   7204      3   27.19600   -7.94960  -19.24640
0.00000    0.00000    0.00000 ;    OE1-    CD-   OE2-   HE2
```

7205 7207 7209 7210 3 0.00000 0.00000 0.00000
0.00000 0.00000 0.00000 ; C- N- CA- HA
7205 7207 7209 7211 3 11.71520 1.67360 15.06240 -
13.38880 0.00000 0.00000 ; C- N- CA- CB
7205 7207 7209 7225 3 1.75728 5.27184 2.25936
-7.02912 0.00000 0.00000 ; C- N- CA- C
7206 7205 7207 7208 3 29.28800 -8.36800 -20.92000
0.00000 0.00000 0.00000 ; O- C- N- H
7206 7205 7207 7209 3 20.92000 0.00000 -20.92000
0.00000 0.00000 0.00000 ; O- C- N- CA
7207 7209 7211 7212 3 0.65084 1.95253 0.00000
-2.60338 0.00000 0.00000 ; N- CA- CB- HB2
7207 7209 7211 7213 3 0.65084 1.95253 0.00000
-2.60338 0.00000 0.00000 ; N- CA- CB- HB3
7207 7209 7211 7214 3 3.28026 2.46019 -2.19242
-3.21331 -0.23430 0.00000 ; N- CA- CB- CG
7207 7209 7225 7226 3 0.00000 0.00000 0.00000
0.00000 0.00000 0.00000 ; N- CA- C- O
7207 7209 7225 7227 3 17.40544 -5.02080 -13.22144
9.20480 0.00000 0.00000 ; N- CA- C- N
7208 7207 7209 7210 3 0.00000 0.00000 0.00000
0.00000 0.00000 0.00000 ; H- N- CA- HA
7208 7207 7209 7211 3 0.00000 0.00000 0.00000
0.00000 0.00000 0.00000 ; H- N- CA- CB
7208 7207 7209 7225 3 0.00000 0.00000 0.00000
0.00000 0.00000 0.00000 ; H- N- CA- C
7209 7211 7214 7215 3 0.57739 0.00000 1.02926
0.00000 -1.60666 0.00000 ; CA- CB- CG- CD1
7209 7211 7214 7223 3 0.57739 0.00000 1.02926
0.00000 -1.60666 0.00000 ; CA- CB- CG- CD2
7209 7225 7227 7228 3 20.92000 0.00000 -20.92000
0.00000 0.00000 0.00000 ; CA- C- N- H
7209 7225 7227 7229 3 20.92000 0.00000 -20.92000
0.00000 0.00000 0.00000 ; CA- C- N- CA
7210 7209 7211 7212 3 0.65084 1.95253 0.00000
-2.60338 0.00000 0.00000 ; HA- CA- CB- HB2
7210 7209 7211 7213 3 0.65084 1.95253 0.00000
-2.60338 0.00000 0.00000 ; HA- CA- CB- HB3
7210 7209 7211 7214 3 0.65084 1.95253 0.00000
-2.60338 0.00000 0.00000 ; HA- CA- CB- CG
7210 7209 7225 7226 3 3.68192 -4.35136 0.00000
1.33888 0.00000 0.00000 ; HA- CA- C- O
7210 7209 7225 7227 3 0.00000 0.00000 0.00000
0.00000 0.00000 0.00000 ; HA- CA- C- N
7211 7209 7225 7226 3 0.00000 0.00000 0.00000
0.00000 0.00000 0.00000 ; CB- CA- C- O
7211 7209 7225 7227 3 2.51040 4.18400 1.67360
-6.69440 0.00000 0.00000 ; CB- CA- C- N
7211 7214 7215 7216 3 30.33400 0.00000 -30.33400
0.00000 0.00000 0.00000 ; CB- CG- CD1- HD1
7211 7214 7215 7217 3 30.33400 0.00000 -30.33400
0.00000 0.00000 0.00000 ; CB- CG- CD1- CE1

7211 7214 7223 7221 3 30.33400 0.00000 -30.33400
0.00000 0.00000 0.00000 ; CB- CG- CD2- CE2
7211 7214 7223 7224 3 30.33400 0.00000 -30.33400
0.00000 0.00000 0.00000 ; CB- CG- CD2- HD2
7212 7211 7209 7225 3 0.65084 1.95253 0.00000
-2.60338 0.00000 0.00000 ; HB2- CB- CA- C
7212 7211 7214 7215 3 0.00000 0.00000 0.00000
0.00000 0.00000 0.00000 ; HB2- CB- CG- CD1
7212 7211 7214 7223 3 0.00000 0.00000 0.00000
0.00000 0.00000 0.00000 ; HB2- CB- CG- CD2
7213 7211 7209 7225 3 0.65084 1.95253 0.00000
-2.60338 0.00000 0.00000 ; HB3- CB- CA- C
7213 7211 7214 7215 3 0.00000 0.00000 0.00000
0.00000 0.00000 0.00000 ; HB3- CB- CG- CD1
7213 7211 7214 7223 3 0.00000 0.00000 0.00000
0.00000 0.00000 0.00000 ; HB3- CB- CG- CD2
7214 7211 7209 7225 3 4.95804 2.17986 -3.52293
-3.21331 -0.40166 0.00000 ; CG- CB- CA- C
7214 7215 7217 7218 3 30.33400 0.00000 -30.33400
0.00000 0.00000 0.00000 ; CG- CD1- CE1- HE1
7214 7215 7217 7219 3 30.33400 0.00000 -30.33400
0.00000 0.00000 0.00000 ; CG- CD1- CE1- CZ
7214 7223 7221 7219 3 30.33400 0.00000 -30.33400
0.00000 0.00000 0.00000 ; CG- CD2- CE2- CZ
7214 7223 7221 7222 3 30.33400 0.00000 -30.33400
0.00000 0.00000 0.00000 ; CG- CD2- CE2- HE2
7215 7214 7223 7221 3 30.33400 0.00000 -30.33400
0.00000 0.00000 0.00000 ; CD1- CG- CD2- CE2
7215 7214 7223 7224 3 30.33400 0.00000 -30.33400
0.00000 0.00000 0.00000 ; CD1- CG- CD2- HD2
7215 7217 7219 7220 3 30.33400 0.00000 -30.33400
0.00000 0.00000 0.00000 ; CD1- CE1- CZ- HZ
7215 7217 7219 7221 3 30.33400 0.00000 -30.33400
0.00000 0.00000 0.00000 ; CD1- CE1- CZ- CE2
7216 7215 7214 7223 3 30.33400 0.00000 -30.33400
0.00000 0.00000 0.00000 ; HD1- CD1- CG- CD2
7216 7215 7217 7218 3 30.33400 0.00000 -30.33400
0.00000 0.00000 0.00000 ; HD1- CD1- CE1- HE1
7216 7215 7217 7219 3 30.33400 0.00000 -30.33400
0.00000 0.00000 0.00000 ; HD1- CD1- CE1- CZ
7217 7215 7214 7223 3 30.33400 0.00000 -30.33400
0.00000 0.00000 0.00000 ; CE1- CD1- CG- CD2
7217 7219 7221 7222 3 30.33400 0.00000 -30.33400
0.00000 0.00000 0.00000 ; CE1- CZ- CE2- HE2
7217 7219 7221 7223 3 30.33400 0.00000 -30.33400
0.00000 0.00000 0.00000 ; CE1- CZ- CE2- CD2
7218 7217 7219 7220 3 30.33400 0.00000 -30.33400
0.00000 0.00000 0.00000 ; HE1- CE1- CZ- HZ
7218 7217 7219 7221 3 30.33400 0.00000 -30.33400
0.00000 0.00000 0.00000 ; HE1- CE1- CZ- CE2
7219 7221 7223 7224 3 30.33400 0.00000 -30.33400
0.00000 0.00000 0.00000 ; CZ- CE2- CD2- HD2

7220 7219 7221 7222 3 30.33400 0.00000 -30.33400
0.00000 0.00000 0.00000 ; HZ- CZ- CE2- HE2
7220 7219 7221 7223 3 30.33400 0.00000 -30.33400
0.00000 0.00000 0.00000 ; HZ- CZ- CE2- CD2
7222 7221 7223 7224 3 30.33400 0.00000 -30.33400
0.00000 0.00000 0.00000 ; HE2- CE2- CD2- HD2
7225 7227 7229 7230 3 0.00000 0.00000 0.00000
0.00000 0.00000 0.00000 ; C- N- CA- HA
7225 7227 7229 7231 3 11.71520 1.67360 15.06240 -
13.38880 0.00000 0.00000 ; C- N- CA- CB
7225 7227 7229 7242 3 1.75728 5.27184 2.25936
-7.02912 0.00000 0.00000 ; C- N- CA- C
7226 7225 7227 7228 3 29.28800 -8.36800 -20.92000
0.00000 0.00000 0.00000 ; O- C- N- H
7226 7225 7227 7229 3 20.92000 0.00000 -20.92000
0.00000 0.00000 0.00000 ; O- C- N- CA
7227 7229 7231 7232 3 0.65084 1.95253 0.00000
-2.60338 0.00000 0.00000 ; N- CA- CB- HB2
7227 7229 7231 7233 3 0.65084 1.95253 0.00000
-2.60338 0.00000 0.00000 ; N- CA- CB- HB3
7227 7229 7231 7234 3 3.21331 2.22589 -4.15053
-2.40998 2.61082 0.00000 ; N- CA- CB- CG
7227 7229 7242 7243 3 0.00000 0.00000 0.00000
0.00000 0.00000 0.00000 ; N- CA- C- O
7227 7229 7242 7244 3 17.40544 -5.02080 -13.22144
9.20480 0.00000 0.00000 ; N- CA- C- N
7228 7227 7229 7230 3 0.00000 0.00000 0.00000
0.00000 0.00000 0.00000 ; H- N- CA- HA
7228 7227 7229 7231 3 0.00000 0.00000 0.00000
0.00000 0.00000 0.00000 ; H- N- CA- CB
7228 7227 7229 7242 3 0.00000 0.00000 0.00000
0.00000 0.00000 0.00000 ; H- N- CA- C
7229 7231 7234 7235 3 0.65084 1.95253 0.00000
-2.60338 0.00000 0.00000 ; CA- CB- CG- HG2
7229 7231 7234 7236 3 0.65084 1.95253 0.00000
-2.60338 0.00000 0.00000 ; CA- CB- CG- HG3
7229 7231 7234 7237 3 2.04598 -1.54390 1.11294
-0.26778 0.93722 0.00000 ; CA- CB- CG- SD
7229 7242 7244 7245 3 20.92000 0.00000 -20.92000
0.00000 0.00000 0.00000 ; CA- C- N- H
7229 7242 7244 7246 3 20.92000 0.00000 -20.92000
0.00000 0.00000 0.00000 ; CA- C- N- CA
7230 7229 7231 7232 3 0.65084 1.95253 0.00000
-2.60338 0.00000 0.00000 ; HA- CA- CB- HB2
7230 7229 7231 7233 3 0.65084 1.95253 0.00000
-2.60338 0.00000 0.00000 ; HA- CA- CB- HB3
7230 7229 7231 7234 3 0.65084 1.95253 0.00000
-2.60338 0.00000 0.00000 ; HA- CA- CB- CG
7230 7229 7242 7243 3 3.68192 -4.35136 0.00000
1.33888 0.00000 0.00000 ; HA- CA- C- O
7230 7229 7242 7244 3 0.00000 0.00000 0.00000
0.00000 0.00000 0.00000 ; HA- CA- C- N

7231 7229 7242 7243 3 0.00000 0.00000 0.00000
0.00000 0.00000 0.00000 ; CB- CA- C- O
7231 7229 7242 7244 3 2.51040 4.18400 1.67360
-6.69440 0.00000 0.00000 ; CB- CA- C- N
7231 7234 7237 7238 3 3.24260 6.22998 1.79075
-6.92870 1.90790 0.00000 ; CB- CG- SD- CE
7232 7231 7229 7242 3 0.65084 1.95253 0.00000
-2.60338 0.00000 0.00000 ; HB2- CB- CA- C
7232 7231 7234 7235 3 0.65084 1.95253 0.00000
-2.60338 0.00000 0.00000 ; HB2- CB- CG- HG2
7232 7231 7234 7236 3 0.65084 1.95253 0.00000
-2.60338 0.00000 0.00000 ; HB2- CB- CG- HG3
7232 7231 7234 7237 3 0.65084 1.95253 0.00000
-2.60338 0.00000 0.00000 ; HB2- CB- CG- SD
7233 7231 7229 7242 3 0.65084 1.95253 0.00000
-2.60338 0.00000 0.00000 ; HB3- CB- CA- C
7233 7231 7234 7235 3 0.65084 1.95253 0.00000
-2.60338 0.00000 0.00000 ; HB3- CB- CG- HG2
7233 7231 7234 7236 3 0.65084 1.95253 0.00000
-2.60338 0.00000 0.00000 ; HB3- CB- CG- HG3
7233 7231 7234 7237 3 0.65084 1.95253 0.00000
-2.60338 0.00000 0.00000 ; HB3- CB- CG- SD
7234 7231 7229 7242 3 6.86594 3.56895 -8.14206
-2.40998 4.85344 0.00000 ; CG- CB- CA- C
7234 7237 7238 7239 3 1.39467 4.18400 0.00000
-5.57867 0.00000 0.00000 ; CG- SD- CE- HE1
7234 7237 7238 7240 3 1.39467 4.18400 0.00000
-5.57867 0.00000 0.00000 ; CG- SD- CE- HE2
7234 7237 7238 7241 3 1.39467 4.18400 0.00000
-5.57867 0.00000 0.00000 ; CG- SD- CE- HE3
7235 7234 7237 7238 3 1.39467 4.18400 0.00000
-5.57867 0.00000 0.00000 ; HG2- CG- SD- CE
7236 7234 7237 7238 3 1.39467 4.18400 0.00000
-5.57867 0.00000 0.00000 ; HG3- CG- SD- CE
7242 7244 7246 7247 3 0.00000 0.00000 0.00000
0.00000 0.00000 0.00000 ; C- N- CA- HA
7242 7244 7246 7248 3 11.71520 1.67360 15.06240 -
13.38880 0.00000 0.00000 ; C- N- CA- CB
7242 7244 7246 7261 3 1.75728 5.27184 2.25936
-7.02912 0.00000 0.00000 ; C- N- CA- C
7243 7242 7244 7245 3 29.28800 -8.36800 -20.92000
0.00000 0.00000 0.00000 ; O- C- N- H
7243 7242 7244 7246 3 20.92000 0.00000 -20.92000
0.00000 0.00000 0.00000 ; O- C- N- CA
7244 7246 7248 7249 3 0.65084 1.95253 0.00000
-2.60338 0.00000 0.00000 ; N- CA- CB- HB2
7244 7246 7248 7250 3 0.65084 1.95253 0.00000
-2.60338 0.00000 0.00000 ; N- CA- CB- HB3
7244 7246 7248 7251 3 3.79070 1.39746 -4.61077
-2.40998 2.44346 0.00000 ; N- CA- CB- CG
7244 7246 7261 7262 3 0.00000 0.00000 0.00000
0.00000 0.00000 0.00000 ; N- CA- C- O

7244 7246 7261 7263 3 17.40544 -5.02080 -13.22144
9.20480 0.00000 0.00000 ; N- CA- C- N
7245 7244 7246 7247 3 0.00000 0.00000 0.00000
0.00000 0.00000 0.00000 ; H- N- CA- HA
7245 7244 7246 7248 3 0.00000 0.00000 0.00000
0.00000 0.00000 0.00000 ; H- N- CA- CB
7245 7244 7246 7261 3 0.00000 0.00000 0.00000
0.00000 0.00000 0.00000 ; H- N- CA- C
7246 7248 7251 7252 3 0.66944 2.00832 0.00000
-2.67776 0.00000 0.00000 ; CA- CB- CG- HG
7246 7248 7251 7253 3 3.90367 0.19665 -6.21742
-2.37651 5.99149 0.00000 ; CA- CB- CG- CD1
7246 7248 7251 7257 3 3.90367 0.19665 -6.21742
-2.37651 5.99149 0.00000 ; CA- CB- CG- CD2
7246 7261 7263 7264 3 20.92000 0.00000 -20.92000
0.00000 0.00000 0.00000 ; CA- C- N- H
7246 7261 7263 7265 3 20.92000 0.00000 -20.92000
0.00000 0.00000 0.00000 ; CA- C- N- CA
7247 7246 7248 7249 3 0.65084 1.95253 0.00000
-2.60338 0.00000 0.00000 ; HA- CA- CB- HB2
7247 7246 7248 7250 3 0.65084 1.95253 0.00000
-2.60338 0.00000 0.00000 ; HA- CA- CB- HB3
7247 7246 7248 7251 3 0.65084 1.95253 0.00000
-2.60338 0.00000 0.00000 ; HA- CA- CB- CG
7247 7246 7261 7262 3 3.68192 -4.35136 0.00000
1.33888 0.00000 0.00000 ; HA- CA- C- O
7247 7246 7261 7263 3 0.00000 0.00000 0.00000
0.00000 0.00000 0.00000 ; HA- CA- C- N
7248 7246 7261 7262 3 0.00000 0.00000 0.00000
0.00000 0.00000 0.00000 ; CB- CA- C- O
7248 7246 7261 7263 3 2.51040 4.18400 1.67360
-6.69440 0.00000 0.00000 ; CB- CA- C- N
7248 7251 7253 7254 3 0.66944 2.00832 0.00000
-2.67776 0.00000 0.00000 ; CB- CG- CD1- HD11
7248 7251 7253 7255 3 0.66944 2.00832 0.00000
-2.67776 0.00000 0.00000 ; CB- CG- CD1- HD12
7248 7251 7253 7256 3 0.66944 2.00832 0.00000
-2.67776 0.00000 0.00000 ; CB- CG- CD1- HD13
7248 7251 7257 7258 3 0.66944 2.00832 0.00000
-2.67776 0.00000 0.00000 ; CB- CG- CD2- HD21
7248 7251 7257 7259 3 0.66944 2.00832 0.00000
-2.67776 0.00000 0.00000 ; CB- CG- CD2- HD22
7248 7251 7257 7260 3 0.66944 2.00832 0.00000
-2.67776 0.00000 0.00000 ; CB- CG- CD2- HD23
7249 7248 7246 7261 3 0.65084 1.95253 0.00000
-2.60338 0.00000 0.00000 ; HB2- CB- CA- C
7249 7248 7251 7252 3 0.62760 1.88280 0.00000
-2.51040 0.00000 0.00000 ; HB2- CB- CG- HG
7249 7248 7251 7253 3 0.66944 2.00832 0.00000
-2.67776 0.00000 0.00000 ; HB2- CB- CG- CD1
7249 7248 7251 7257 3 0.66944 2.00832 0.00000
-2.67776 0.00000 0.00000 ; HB2- CB- CG- CD2

7250 7248 7246 7261 3 0.65084 1.95253 0.00000
-2.60338 0.00000 0.00000 ; HB3- CB- CA- C
7250 7248 7251 7252 3 0.62760 1.88280 0.00000
-2.51040 0.00000 0.00000 ; HB3- CB- CG- HG
7250 7248 7251 7253 3 0.66944 2.00832 0.00000
-2.67776 0.00000 0.00000 ; HB3- CB- CG- CD1
7250 7248 7251 7257 3 0.66944 2.00832 0.00000
-2.67776 0.00000 0.00000 ; HB3- CB- CG- CD2
7251 7248 7246 7261 3 10.33448 -1.14642 -11.54784
-2.40998 6.35968 0.00000 ; CG- CB- CA- C
7252 7251 7253 7254 3 0.62760 1.88280 0.00000
-2.51040 0.00000 0.00000 ; HG- CG- CD1- HD11
7252 7251 7253 7255 3 0.62760 1.88280 0.00000
-2.51040 0.00000 0.00000 ; HG- CG- CD1- HD12
7252 7251 7253 7256 3 0.62760 1.88280 0.00000
-2.51040 0.00000 0.00000 ; HG- CG- CD1- HD13
7252 7251 7257 7258 3 0.62760 1.88280 0.00000
-2.51040 0.00000 0.00000 ; HG- CG- CD2- HD21
7252 7251 7257 7259 3 0.62760 1.88280 0.00000
-2.51040 0.00000 0.00000 ; HG- CG- CD2- HD22
7252 7251 7257 7260 3 0.62760 1.88280 0.00000
-2.51040 0.00000 0.00000 ; HG- CG- CD2- HD23
7253 7251 7257 7258 3 0.66944 2.00832 0.00000
-2.67776 0.00000 0.00000 ; CD1- CG- CD2- HD21
7253 7251 7257 7259 3 0.66944 2.00832 0.00000
-2.67776 0.00000 0.00000 ; CD1- CG- CD2- HD22
7253 7251 7257 7260 3 0.66944 2.00832 0.00000
-2.67776 0.00000 0.00000 ; CD1- CG- CD2- HD23
7254 7253 7251 7257 3 0.66944 2.00832 0.00000
-2.67776 0.00000 0.00000 ; HD11- CD1- CG- CD2
7255 7253 7251 7257 3 0.66944 2.00832 0.00000
-2.67776 0.00000 0.00000 ; HD12- CD1- CG- CD2
7256 7253 7251 7257 3 0.66944 2.00832 0.00000
-2.67776 0.00000 0.00000 ; HD13- CD1- CG- CD2
7261 7263 7265 7266 3 0.00000 0.00000 0.00000
0.00000 0.00000 0.00000 ; C- N- CA- HA
7261 7263 7265 7267 3 11.71520 1.67360 15.06240 -
13.38880 0.00000 0.00000 ; C- N- CA- CB
7261 7263 7265 7278 3 1.75728 5.27184 2.25936
-7.02912 0.00000 0.00000 ; C- N- CA- C
7262 7261 7263 7264 3 29.28800 -8.36800 -20.92000
0.00000 0.00000 0.00000 ; O- C- N- H
7262 7261 7263 7265 3 20.92000 0.00000 -20.92000
0.00000 0.00000 0.00000 ; O- C- N- CA
7263 7265 7267 7268 3 0.65084 1.95253 0.00000
-2.60338 0.00000 0.00000 ; N- CA- CB- HB2
7263 7265 7267 7269 3 0.65084 1.95253 0.00000
-2.60338 0.00000 0.00000 ; N- CA- CB- HB3
7263 7265 7267 7270 3 3.21331 2.22589 -4.15053
-2.40998 2.61082 0.00000 ; N- CA- CB- CG
7263 7265 7278 7279 3 0.00000 0.00000 0.00000
0.00000 0.00000 0.00000 ; N- CA- C- O

7263 7265 7278 7280 3 17.40544 -5.02080 -13.22144
9.20480 0.00000 0.00000 ; N- CA- C- N
7264 7263 7265 7266 3 0.00000 0.00000 0.00000
0.00000 0.00000 0.00000 ; H- N- CA- HA
7264 7263 7265 7267 3 0.00000 0.00000 0.00000
0.00000 0.00000 0.00000 ; H- N- CA- CB
7264 7263 7265 7278 3 0.00000 0.00000 0.00000
0.00000 0.00000 0.00000 ; H- N- CA- C
7265 7267 7270 7271 3 0.65084 1.95253 0.00000
-2.60338 0.00000 0.00000 ; CA- CB- CG- HG2
7265 7267 7270 7272 3 0.65084 1.95253 0.00000
-2.60338 0.00000 0.00000 ; CA- CB- CG- HG3
7265 7267 7270 7273 3 2.04598 -1.54390 1.11294
-0.26778 0.93722 0.00000 ; CA- CB- CG- SD
7265 7278 7280 7281 3 20.92000 0.00000 -20.92000
0.00000 0.00000 0.00000 ; CA- C- N- H
7265 7278 7280 7282 3 20.92000 0.00000 -20.92000
0.00000 0.00000 0.00000 ; CA- C- N- CA
7266 7265 7267 7268 3 0.65084 1.95253 0.00000
-2.60338 0.00000 0.00000 ; HA- CA- CB- HB2
7266 7265 7267 7269 3 0.65084 1.95253 0.00000
-2.60338 0.00000 0.00000 ; HA- CA- CB- HB3
7266 7265 7267 7270 3 0.65084 1.95253 0.00000
-2.60338 0.00000 0.00000 ; HA- CA- CB- CG
7266 7265 7278 7279 3 3.68192 -4.35136 0.00000
1.33888 0.00000 0.00000 ; HA- CA- C- O
7266 7265 7278 7280 3 0.00000 0.00000 0.00000
0.00000 0.00000 0.00000 ; HA- CA- C- N
7267 7265 7278 7279 3 0.00000 0.00000 0.00000
0.00000 0.00000 0.00000 ; CB- CA- C- O
7267 7265 7278 7280 3 2.51040 4.18400 1.67360
-6.69440 0.00000 0.00000 ; CB- CA- C- N
7267 7270 7273 7274 3 3.24260 6.22998 1.79075
-6.92870 1.90790 0.00000 ; CB- CG- SD- CE
7268 7267 7265 7278 3 0.65084 1.95253 0.00000
-2.60338 0.00000 0.00000 ; HB2- CB- CA- C
7268 7267 7270 7271 3 0.65084 1.95253 0.00000
-2.60338 0.00000 0.00000 ; HB2- CB- CG- HG2
7268 7267 7270 7272 3 0.65084 1.95253 0.00000
-2.60338 0.00000 0.00000 ; HB2- CB- CG- HG3
7268 7267 7270 7273 3 0.65084 1.95253 0.00000
-2.60338 0.00000 0.00000 ; HB2- CB- CG- SD
7269 7267 7265 7278 3 0.65084 1.95253 0.00000
-2.60338 0.00000 0.00000 ; HB3- CB- CA- C
7269 7267 7270 7271 3 0.65084 1.95253 0.00000
-2.60338 0.00000 0.00000 ; HB3- CB- CG- HG2
7269 7267 7270 7272 3 0.65084 1.95253 0.00000
-2.60338 0.00000 0.00000 ; HB3- CB- CG- HG3
7269 7267 7270 7273 3 0.65084 1.95253 0.00000
-2.60338 0.00000 0.00000 ; HB3- CB- CG- SD
7270 7267 7265 7278 3 6.86594 3.56895 -8.14206
-2.40998 4.85344 0.00000 ; CG- CB- CA- C

7270 7273 7274 7275 3 1.39467 4.18400 0.00000
-5.57867 0.00000 0.00000 ; CG- SD- CE- HE1
7270 7273 7274 7276 3 1.39467 4.18400 0.00000
-5.57867 0.00000 0.00000 ; CG- SD- CE- HE2
7270 7273 7274 7277 3 1.39467 4.18400 0.00000
-5.57867 0.00000 0.00000 ; CG- SD- CE- HE3
7271 7270 7273 7274 3 1.39467 4.18400 0.00000
-5.57867 0.00000 0.00000 ; HG2- CG- SD- CE
7272 7270 7273 7274 3 1.39467 4.18400 0.00000
-5.57867 0.00000 0.00000 ; HG3- CG- SD- CE
7278 7280 7282 7283 3 0.00000 0.00000 0.00000
0.00000 0.00000 0.00000 ; C- N- CA- HA
7278 7280 7282 7284 3 11.71520 1.67360 15.06240 -
13.38880 0.00000 0.00000 ; C- N- CA- CB
7278 7280 7282 7295 3 1.75728 5.27184 2.25936
-7.02912 0.00000 0.00000 ; C- N- CA- C
7279 7278 7280 7281 3 29.28800 -8.36800 -20.92000
0.00000 0.00000 0.00000 ; O- C- N- H
7279 7278 7280 7282 3 20.92000 0.00000 -20.92000
0.00000 0.00000 0.00000 ; O- C- N- CA
7280 7282 7284 7285 3 0.65084 1.95253 0.00000
-2.60338 0.00000 0.00000 ; N- CA- CB- HB2
7280 7282 7284 7286 3 0.65084 1.95253 0.00000
-2.60338 0.00000 0.00000 ; N- CA- CB- HB3
7280 7282 7284 7287 3 3.21331 2.22589 -4.15053
-2.40998 2.61082 0.00000 ; N- CA- CB- CG
7280 7282 7295 7296 3 0.00000 0.00000 0.00000
0.00000 0.00000 0.00000 ; N- CA- C- O
7280 7282 7295 7297 3 17.40544 -5.02080 -13.22144
9.20480 0.00000 0.00000 ; N- CA- C- N
7281 7280 7282 7283 3 0.00000 0.00000 0.00000
0.00000 0.00000 0.00000 ; H- N- CA- HA
7281 7280 7282 7284 3 0.00000 0.00000 0.00000
0.00000 0.00000 0.00000 ; H- N- CA- CB
7281 7280 7282 7295 3 0.00000 0.00000 0.00000
0.00000 0.00000 0.00000 ; H- N- CA- C
7282 7284 7287 7288 3 0.65084 1.95253 0.00000
-2.60338 0.00000 0.00000 ; CA- CB- CG- HG2
7282 7284 7287 7289 3 0.65084 1.95253 0.00000
-2.60338 0.00000 0.00000 ; CA- CB- CG- HG3
7282 7284 7287 7290 3 2.04598 -1.54390 1.11294
-0.26778 0.93722 0.00000 ; CA- CB- CG- SD
7282 7295 7297 7298 3 20.92000 0.00000 -20.92000
0.00000 0.00000 0.00000 ; CA- C- N- H
7282 7295 7297 7299 3 20.92000 0.00000 -20.92000
0.00000 0.00000 0.00000 ; CA- C- N- CA
7283 7282 7284 7285 3 0.65084 1.95253 0.00000
-2.60338 0.00000 0.00000 ; HA- CA- CB- HB2
7283 7282 7284 7286 3 0.65084 1.95253 0.00000
-2.60338 0.00000 0.00000 ; HA- CA- CB- HB3
7283 7282 7284 7287 3 0.65084 1.95253 0.00000
-2.60338 0.00000 0.00000 ; HA- CA- CB- CG

7283 7282 7295 7296 3 3.68192 -4.35136 0.00000
1.33888 0.00000 0.00000 ; HA- CA- C- O
7283 7282 7295 7297 3 0.00000 0.00000 0.00000
0.00000 0.00000 0.00000 ; HA- CA- C- N
7284 7282 7295 7296 3 0.00000 0.00000 0.00000
0.00000 0.00000 0.00000 ; CB- CA- C- O
7284 7282 7295 7297 3 2.51040 4.18400 1.67360
-6.69440 0.00000 0.00000 ; CB- CA- C- N
7284 7287 7290 7291 3 3.24260 6.22998 1.79075
-6.92870 1.90790 0.00000 ; CB- CG- SD- CE
7285 7284 7282 7295 3 0.65084 1.95253 0.00000
-2.60338 0.00000 0.00000 ; HB2- CB- CA- C
7285 7284 7287 7288 3 0.65084 1.95253 0.00000
-2.60338 0.00000 0.00000 ; HB2- CB- CG- HG2
7285 7284 7287 7289 3 0.65084 1.95253 0.00000
-2.60338 0.00000 0.00000 ; HB2- CB- CG- HG3
7285 7284 7287 7290 3 0.65084 1.95253 0.00000
-2.60338 0.00000 0.00000 ; HB2- CB- CG- SD
7286 7284 7282 7295 3 0.65084 1.95253 0.00000
-2.60338 0.00000 0.00000 ; HB3- CB- CA- C
7286 7284 7287 7288 3 0.65084 1.95253 0.00000
-2.60338 0.00000 0.00000 ; HB3- CB- CG- HG2
7286 7284 7287 7289 3 0.65084 1.95253 0.00000
-2.60338 0.00000 0.00000 ; HB3- CB- CG- HG3
7286 7284 7287 7290 3 0.65084 1.95253 0.00000
-2.60338 0.00000 0.00000 ; HB3- CB- CG- SD
7287 7284 7282 7295 3 6.86594 3.56895 -8.14206
-2.40998 4.85344 0.00000 ; CG- CB- CA- C
7287 7290 7291 7292 3 1.39467 4.18400 0.00000
-5.57867 0.00000 0.00000 ; CG- SD- CE- HE1
7287 7290 7291 7293 3 1.39467 4.18400 0.00000
-5.57867 0.00000 0.00000 ; CG- SD- CE- HE2
7287 7290 7291 7294 3 1.39467 4.18400 0.00000
-5.57867 0.00000 0.00000 ; CG- SD- CE- HE3
7288 7287 7290 7291 3 1.39467 4.18400 0.00000
-5.57867 0.00000 0.00000 ; HG2- CG- SD- CE
7289 7287 7290 7291 3 1.39467 4.18400 0.00000
-5.57867 0.00000 0.00000 ; HG3- CG- SD- CE
7295 7297 7299 7300 3 0.00000 0.00000 0.00000
0.00000 0.00000 0.00000 ; C- N- CA- HA
7295 7297 7299 7301 3 11.71520 1.67360 15.06240 -
13.38880 0.00000 0.00000 ; C- N- CA- CB
7295 7297 7299 7306 3 1.75728 5.27184 2.25936
-7.02912 0.00000 0.00000 ; C- N- CA- C
7296 7295 7297 7298 3 29.28800 -8.36800 -20.92000
0.00000 0.00000 0.00000 ; O- C- N- H
7296 7295 7297 7299 3 20.92000 0.00000 -20.92000
0.00000 0.00000 0.00000 ; O- C- N- CA
7297 7299 7301 7302 3 0.65084 1.95253 0.00000
-2.60338 0.00000 0.00000 ; N- CA- CB- HB2
7297 7299 7301 7303 3 0.65084 1.95253 0.00000
-2.60338 0.00000 0.00000 ; N- CA- CB- HB3

7297 7299 7301 7304 3 7.86174 2.24681 -7.41405
-6.71114 5.35552 0.00000 ; N- CA- CB- OG
7297 7299 7306 7307 3 0.00000 0.00000 0.00000
0.00000 0.00000 0.00000 ; N- CA- C- O
7297 7299 7306 7308 3 17.40544 -5.02080 -13.22144
9.20480 0.00000 0.00000 ; N- CA- C- N
7298 7297 7299 7300 3 0.00000 0.00000 0.00000
0.00000 0.00000 0.00000 ; H- N- CA- HA
7298 7297 7299 7301 3 0.00000 0.00000 0.00000
0.00000 0.00000 0.00000 ; H- N- CA- CB
7298 7297 7299 7306 3 0.00000 0.00000 0.00000
0.00000 0.00000 0.00000 ; H- N- CA- C
7299 7301 7304 7305 3 2.05853 2.46856 3.48109
-4.46851 0.23430 0.00000 ; CA- CB- OG- HG
7299 7306 7308 7309 3 20.92000 0.00000 -20.92000
0.00000 0.00000 0.00000 ; CA- C- N- H
7299 7306 7308 7310 3 20.92000 0.00000 -20.92000
0.00000 0.00000 0.00000 ; CA- C- N- CA
7300 7299 7301 7302 3 0.65084 1.95253 0.00000
-2.60338 0.00000 0.00000 ; HA- CA- CB- HB2
7300 7299 7301 7303 3 0.65084 1.95253 0.00000
-2.60338 0.00000 0.00000 ; HA- CA- CB- HB3
7300 7299 7301 7304 3 1.04600 -1.04600 0.00000
0.00000 0.00000 0.00000 ; HA- CA- CB- OG
7300 7299 7306 7307 3 3.68192 -4.35136 0.00000
1.33888 0.00000 0.00000 ; HA- CA- C- O
7300 7299 7306 7308 3 0.00000 0.00000 0.00000
0.00000 0.00000 0.00000 ; HA- CA- C- N
7301 7299 7306 7307 3 0.00000 0.00000 0.00000
0.00000 0.00000 0.00000 ; CB- CA- C- O
7301 7299 7306 7308 3 2.51040 4.18400 1.67360
-6.69440 0.00000 0.00000 ; CB- CA- C- N
7302 7301 7299 7306 3 0.65084 1.95253 0.00000
-2.60338 0.00000 0.00000 ; HB2- CB- CA- C
7302 7301 7304 7305 3 0.69733 2.09200 0.00000
-2.78933 0.00000 0.00000 ; HB2- CB- OG- HG
7303 7301 7299 7306 3 0.65084 1.95253 0.00000
-2.60338 0.00000 0.00000 ; HB3- CB- CA- C
7303 7301 7304 7305 3 0.69733 2.09200 0.00000
-2.78933 0.00000 0.00000 ; HB3- CB- OG- HG
7304 7301 7299 7306 3 7.34710 7.79898 -6.14211
-6.71114 4.31789 0.00000 ; OG- CB- CA- C
7306 7308 7310 7311 3 0.00000 0.00000 0.00000
0.00000 0.00000 0.00000 ; C- N- CA- HA
7306 7308 7310 7312 3 11.71520 1.67360 15.06240 -
13.38880 0.00000 0.00000 ; C- N- CA- CB
7306 7308 7310 7328 3 1.75728 5.27184 2.25936
-7.02912 0.00000 0.00000 ; C- N- CA- C
7307 7306 7308 7309 3 29.28800 -8.36800 -20.92000
0.00000 0.00000 0.00000 ; O- C- N- H
7307 7306 7308 7310 3 20.92000 0.00000 -20.92000
0.00000 0.00000 0.00000 ; O- C- N- CA

```
  7308   7310   7312   7313      3    0.65084    1.95253    0.00000
-2.60338    0.00000    0.00000 ;      N-    CA-    CB-   HB2
  7308   7310   7312   7314      3    0.65084    1.95253    0.00000
-2.60338    0.00000    0.00000 ;      N-    CA-    CB-   HB3
  7308   7310   7312   7315      3    0.65084    1.95253    0.00000
-2.60338    0.00000    0.00000 ;      N-    CA-    CB-    CG
  7308   7310   7328   7329      3    0.00000    0.00000    0.00000
0.00000    0.00000    0.00000 ;      N-    CA-     C-     O
  7308   7310   7328   7330      3   17.40544   -5.02080  -13.22144
9.20480    0.00000    0.00000 ;      N-    CA-     C-     N
  7309   7308   7310   7311      3    0.00000    0.00000    0.00000
0.00000    0.00000    0.00000 ;      H-     N-    CA-    HA
  7309   7308   7310   7312      3    0.00000    0.00000    0.00000
0.00000    0.00000    0.00000 ;      H-     N-    CA-    CB
  7309   7308   7310   7328      3    0.00000    0.00000    0.00000
0.00000    0.00000    0.00000 ;      H-     N-    CA-     C
  7310   7312   7315   7316      3    0.66944    2.00832    0.00000
-2.67776    0.00000    0.00000 ;     CA-    CB-    CG-   HG2
  7310   7312   7315   7317      3    0.66944    2.00832    0.00000
-2.67776    0.00000    0.00000 ;     CA-    CB-    CG-   HG3
  7310   7312   7315   7318      3    3.68192    3.09616   -2.09200
-3.01248    0.00000    0.00000 ;     CA-    CB-    CG-    CD
  7310   7328   7330   7331      3   20.92000    0.00000  -20.92000
0.00000    0.00000    0.00000 ;     CA-     C-     N-     H
  7310   7328   7330   7332      3   20.92000    0.00000  -20.92000
0.00000    0.00000    0.00000 ;     CA-     C-     N-    CA
  7311   7310   7312   7313      3    0.65084    1.95253    0.00000
-2.60338    0.00000    0.00000 ;     HA-    CA-    CB-   HB2
  7311   7310   7312   7314      3    0.65084    1.95253    0.00000
-2.60338    0.00000    0.00000 ;     HA-    CA-    CB-   HB3
  7311   7310   7312   7315      3    0.65084    1.95253    0.00000
-2.60338    0.00000    0.00000 ;     HA-    CA-    CB-    CG
  7311   7310   7328   7329      3    3.68192   -4.35136    0.00000
1.33888    0.00000    0.00000 ;     HA-    CA-     C-     O
  7311   7310   7328   7330      3    0.00000    0.00000    0.00000
0.00000    0.00000    0.00000 ;     HA-    CA-     C-     N
  7312   7310   7328   7329      3    0.00000    0.00000    0.00000
0.00000    0.00000    0.00000 ;     CB-    CA-     C-     O
  7312   7310   7328   7330      3    2.51040    4.18400    1.67360
-6.69440    0.00000    0.00000 ;     CB-    CA-     C-     N
  7312   7315   7318   7319      3    0.66944    2.00832    0.00000
-2.67776    0.00000    0.00000 ;     CB-    CG-    CD-   HD2
  7312   7315   7318   7320      3    0.66944    2.00832    0.00000
-2.67776    0.00000    0.00000 ;     CB-    CG-    CD-   HD3
  7312   7315   7318   7321      3    3.68192    3.09616   -2.09200
-3.01248    0.00000    0.00000 ;     CB-    CG-    CD-    CE
  7313   7312   7310   7328      3    0.65084    1.95253    0.00000
-2.60338    0.00000    0.00000 ;    HB2-    CB-    CA-     C
  7313   7312   7315   7316      3    0.62760    1.88280    0.00000
-2.51040    0.00000    0.00000 ;    HB2-    CB-    CG-   HG2
  7313   7312   7315   7317      3    0.62760    1.88280    0.00000
-2.51040    0.00000    0.00000 ;    HB2-    CB-    CG-   HG3
```

```
  7313   7312   7315   7318       3    0.66944    2.00832    0.00000
-2.67776    0.00000    0.00000 ;     HB2-    CB-    CG-    CD
  7314   7312   7310   7328       3    0.65084    1.95253    0.00000
-2.60338    0.00000    0.00000 ;     HB3-    CB-    CA-     C
  7314   7312   7315   7316       3    0.62760    1.88280    0.00000
-2.51040    0.00000    0.00000 ;     HB3-    CB-    CG-   HG2
  7314   7312   7315   7317       3    0.62760    1.88280    0.00000
-2.51040    0.00000    0.00000 ;     HB3-    CB-    CG-   HG3
  7314   7312   7315   7318       3    0.66944    2.00832    0.00000
-2.67776    0.00000    0.00000 ;     HB3-    CB-    CG-    CD
  7315   7312   7310   7328       3    0.65084    1.95253    0.00000
-2.60338    0.00000    0.00000 ;      CG-    CB-    CA-     C
  7315   7318   7321   7322       3    0.65084    1.95253    0.00000
-2.60338    0.00000    0.00000 ;      CG-    CD-    CE-   HE2
  7315   7318   7321   7323       3    0.65084    1.95253    0.00000
-2.60338    0.00000    0.00000 ;      CG-    CD-    CE-   HE3
  7315   7318   7321   7324       3    0.65084    1.95253    0.00000
-2.60338    0.00000    0.00000 ;      CG-    CD-    CE-    NZ
  7316   7315   7318   7319       3    0.62760    1.88280    0.00000
-2.51040    0.00000    0.00000 ;     HG2-    CG-    CD-   HD2
  7316   7315   7318   7320       3    0.62760    1.88280    0.00000
-2.51040    0.00000    0.00000 ;     HG2-    CG-    CD-   HD3
  7316   7315   7318   7321       3    0.66944    2.00832    0.00000
-2.67776    0.00000    0.00000 ;     HG2-    CG-    CD-    CE
  7317   7315   7318   7319       3    0.62760    1.88280    0.00000
-2.51040    0.00000    0.00000 ;     HG3-    CG-    CD-   HD2
  7317   7315   7318   7320       3    0.62760    1.88280    0.00000
-2.51040    0.00000    0.00000 ;     HG3-    CG-    CD-   HD3
  7317   7315   7318   7321       3    0.66944    2.00832    0.00000
-2.67776    0.00000    0.00000 ;     HG3-    CG-    CD-    CE
  7318   7321   7324   7325       3    0.65084    1.95253    0.00000
-2.60338    0.00000    0.00000 ;      CD-    CE-    NZ-   HZ1
  7318   7321   7324   7326       3    0.65084    1.95253    0.00000
-2.60338    0.00000    0.00000 ;      CD-    CE-    NZ-   HZ2
  7318   7321   7324   7327       3    0.65084    1.95253    0.00000
-2.60338    0.00000    0.00000 ;      CD-    CE-    NZ-   HZ3
  7319   7318   7321   7322       3    0.65084    1.95253    0.00000
-2.60338    0.00000    0.00000 ;     HD2-    CD-    CE-   HE2
  7319   7318   7321   7323       3    0.65084    1.95253    0.00000
-2.60338    0.00000    0.00000 ;     HD2-    CD-    CE-   HE3
  7319   7318   7321   7324       3    0.65084    1.95253    0.00000
-2.60338    0.00000    0.00000 ;     HD2-    CD-    CE-    NZ
  7320   7318   7321   7322       3    0.65084    1.95253    0.00000
-2.60338    0.00000    0.00000 ;     HD3-    CD-    CE-   HE2
  7320   7318   7321   7323       3    0.65084    1.95253    0.00000
-2.60338    0.00000    0.00000 ;     HD3-    CD-    CE-   HE3
  7320   7318   7321   7324       3    0.65084    1.95253    0.00000
-2.60338    0.00000    0.00000 ;     HD3-    CD-    CE-    NZ
  7322   7321   7324   7325       3    0.65084    1.95253    0.00000
-2.60338    0.00000    0.00000 ;     HE2-    CE-    NZ-   HZ1
  7322   7321   7324   7326       3    0.65084    1.95253    0.00000
-2.60338    0.00000    0.00000 ;     HE2-    CE-    NZ-   HZ2
```

7322 7321 7324 7327 3 0.65084 1.95253 0.00000
-2.60338 0.00000 0.00000 ; HE2- CE- NZ- HZ3
7323 7321 7324 7325 3 0.65084 1.95253 0.00000
-2.60338 0.00000 0.00000 ; HE3- CE- NZ- HZ1
7323 7321 7324 7326 3 0.65084 1.95253 0.00000
-2.60338 0.00000 0.00000 ; HE3- CE- NZ- HZ2
7323 7321 7324 7327 3 0.65084 1.95253 0.00000
-2.60338 0.00000 0.00000 ; HE3- CE- NZ- HZ3
7328 7330 7332 7333 3 0.00000 0.00000 0.00000
0.00000 0.00000 0.00000 ; C- N- CA- HA
7328 7330 7332 7334 3 11.71520 1.67360 15.06240 -
13.38880 0.00000 0.00000 ; C- N- CA- CB
7328 7330 7332 7350 3 1.75728 5.27184 2.25936
-7.02912 0.00000 0.00000 ; C- N- CA- C
7329 7328 7330 7331 3 29.28800 -8.36800 -20.92000
0.00000 0.00000 0.00000 ; O- C- N- H
7329 7328 7330 7332 3 20.92000 0.00000 -20.92000
0.00000 0.00000 0.00000 ; O- C- N- CA
7330 7332 7334 7335 3 0.65084 1.95253 0.00000
-2.60338 0.00000 0.00000 ; N- CA- CB- HB2
7330 7332 7334 7336 3 0.65084 1.95253 0.00000
-2.60338 0.00000 0.00000 ; N- CA- CB- HB3
7330 7332 7334 7337 3 0.65084 1.95253 0.00000
-2.60338 0.00000 0.00000 ; N- CA- CB- CG
7330 7332 7350 7351 3 0.00000 0.00000 0.00000
0.00000 0.00000 0.00000 ; N- CA- C- O
7330 7332 7350 7352 3 17.40544 -5.02080 -13.22144
9.20480 0.00000 0.00000 ; N- CA- C- N
7331 7330 7332 7333 3 0.00000 0.00000 0.00000
0.00000 0.00000 0.00000 ; H- N- CA- HA
7331 7330 7332 7334 3 0.00000 0.00000 0.00000
0.00000 0.00000 0.00000 ; H- N- CA- CB
7331 7330 7332 7350 3 0.00000 0.00000 0.00000
0.00000 0.00000 0.00000 ; H- N- CA- C
7332 7334 7337 7338 3 0.66944 2.00832 0.00000
-2.67776 0.00000 0.00000 ; CA- CB- CG- HG2
7332 7334 7337 7339 3 0.66944 2.00832 0.00000
-2.67776 0.00000 0.00000 ; CA- CB- CG- HG3
7332 7334 7337 7340 3 3.68192 3.09616 -2.09200
-3.01248 0.00000 0.00000 ; CA- CB- CG- CD
7332 7350 7352 7353 3 20.92000 0.00000 -20.92000
0.00000 0.00000 0.00000 ; CA- C- N- H
7332 7350 7352 7354 3 20.92000 0.00000 -20.92000
0.00000 0.00000 0.00000 ; CA- C- N- CA
7333 7332 7334 7335 3 0.65084 1.95253 0.00000
-2.60338 0.00000 0.00000 ; HA- CA- CB- HB2
7333 7332 7334 7336 3 0.65084 1.95253 0.00000
-2.60338 0.00000 0.00000 ; HA- CA- CB- HB3
7333 7332 7334 7337 3 0.65084 1.95253 0.00000
-2.60338 0.00000 0.00000 ; HA- CA- CB- CG
7333 7332 7350 7351 3 3.68192 -4.35136 0.00000
1.33888 0.00000 0.00000 ; HA- CA- C- O

7333 7332 7350 7352 3 0.00000 0.00000 0.00000
0.00000 0.00000 0.00000 ; HA- CA- C- N
7334 7332 7350 7351 3 0.00000 0.00000 0.00000
0.00000 0.00000 0.00000 ; CB- CA- C- O
7334 7332 7350 7352 3 2.51040 4.18400 1.67360
-6.69440 0.00000 0.00000 ; CB- CA- C- N
7334 7337 7340 7341 3 0.66944 2.00832 0.00000
-2.67776 0.00000 0.00000 ; CB- CG- CD- HD2
7334 7337 7340 7342 3 0.66944 2.00832 0.00000
-2.67776 0.00000 0.00000 ; CB- CG- CD- HD3
7334 7337 7340 7343 3 3.68192 3.09616 -2.09200
-3.01248 0.00000 0.00000 ; CB- CG- CD- CE
7335 7334 7332 7350 3 0.65084 1.95253 0.00000
-2.60338 0.00000 0.00000 ; HB2- CB- CA- C
7335 7334 7337 7338 3 0.62760 1.88280 0.00000
-2.51040 0.00000 0.00000 ; HB2- CB- CG- HG2
7335 7334 7337 7339 3 0.62760 1.88280 0.00000
-2.51040 0.00000 0.00000 ; HB2- CB- CG- HG3
7335 7334 7337 7340 3 0.66944 2.00832 0.00000
-2.67776 0.00000 0.00000 ; HB2- CB- CG- CD
7336 7334 7332 7350 3 0.65084 1.95253 0.00000
-2.60338 0.00000 0.00000 ; HB3- CB- CA- C
7336 7334 7337 7338 3 0.62760 1.88280 0.00000
-2.51040 0.00000 0.00000 ; HB3- CB- CG- HG2
7336 7334 7337 7339 3 0.62760 1.88280 0.00000
-2.51040 0.00000 0.00000 ; HB3- CB- CG- HG3
7336 7334 7337 7340 3 0.66944 2.00832 0.00000
-2.67776 0.00000 0.00000 ; HB3- CB- CG- CD
7337 7334 7332 7350 3 0.65084 1.95253 0.00000
-2.60338 0.00000 0.00000 ; CG- CB- CA- C
7337 7340 7343 7344 3 0.65084 1.95253 0.00000
-2.60338 0.00000 0.00000 ; CG- CD- CE- HE2
7337 7340 7343 7345 3 0.65084 1.95253 0.00000
-2.60338 0.00000 0.00000 ; CG- CD- CE- HE3
7337 7340 7343 7346 3 0.65084 1.95253 0.00000
-2.60338 0.00000 0.00000 ; CG- CD- CE- NZ
7338 7337 7340 7341 3 0.62760 1.88280 0.00000
-2.51040 0.00000 0.00000 ; HG2- CG- CD- HD2
7338 7337 7340 7342 3 0.62760 1.88280 0.00000
-2.51040 0.00000 0.00000 ; HG2- CG- CD- HD3
7338 7337 7340 7343 3 0.66944 2.00832 0.00000
-2.67776 0.00000 0.00000 ; HG2- CG- CD- CE
7339 7337 7340 7341 3 0.62760 1.88280 0.00000
-2.51040 0.00000 0.00000 ; HG3- CG- CD- HD2
7339 7337 7340 7342 3 0.62760 1.88280 0.00000
-2.51040 0.00000 0.00000 ; HG3- CG- CD- HD3
7339 7337 7340 7343 3 0.66944 2.00832 0.00000
-2.67776 0.00000 0.00000 ; HG3- CG- CD- CE
7340 7343 7346 7347 3 0.65084 1.95253 0.00000
-2.60338 0.00000 0.00000 ; CD- CE- NZ- HZ1
7340 7343 7346 7348 3 0.65084 1.95253 0.00000
-2.60338 0.00000 0.00000 ; CD- CE- NZ- HZ2

7340 7343 7346 7349 3 0.65084 1.95253 0.00000
-2.60338 0.00000 0.00000 ; CD- CE- NZ- HZ3
7341 7340 7343 7344 3 0.65084 1.95253 0.00000
-2.60338 0.00000 0.00000 ; HD2- CD- CE- HE2
7341 7340 7343 7345 3 0.65084 1.95253 0.00000
-2.60338 0.00000 0.00000 ; HD2- CD- CE- HE3
7341 7340 7343 7346 3 0.65084 1.95253 0.00000
-2.60338 0.00000 0.00000 ; HD2- CD- CE- NZ
7342 7340 7343 7344 3 0.65084 1.95253 0.00000
-2.60338 0.00000 0.00000 ; HD3- CD- CE- HE2
7342 7340 7343 7345 3 0.65084 1.95253 0.00000
-2.60338 0.00000 0.00000 ; HD3- CD- CE- HE3
7342 7340 7343 7346 3 0.65084 1.95253 0.00000
-2.60338 0.00000 0.00000 ; HD3- CD- CE- NZ
7344 7343 7346 7347 3 0.65084 1.95253 0.00000
-2.60338 0.00000 0.00000 ; HE2- CE- NZ- HZ1
7344 7343 7346 7348 3 0.65084 1.95253 0.00000
-2.60338 0.00000 0.00000 ; HE2- CE- NZ- HZ2
7344 7343 7346 7349 3 0.65084 1.95253 0.00000
-2.60338 0.00000 0.00000 ; HE2- CE- NZ- HZ3
7345 7343 7346 7347 3 0.65084 1.95253 0.00000
-2.60338 0.00000 0.00000 ; HE3- CE- NZ- HZ1
7345 7343 7346 7348 3 0.65084 1.95253 0.00000
-2.60338 0.00000 0.00000 ; HE3- CE- NZ- HZ2
7345 7343 7346 7349 3 0.65084 1.95253 0.00000
-2.60338 0.00000 0.00000 ; HE3- CE- NZ- HZ3
7350 7352 7354 7355 3 0.00000 0.00000 0.00000
0.00000 0.00000 0.00000 ; C- N- CA- HA
7350 7352 7354 7356 3 11.71520 1.67360 15.06240 -
13.38880 0.00000 0.00000 ; C- N- CA- CB
7350 7352 7354 7372 3 1.75728 5.27184 2.25936
-7.02912 0.00000 0.00000 ; C- N- CA- C
7351 7350 7352 7353 3 29.28800 -8.36800 -20.92000
0.00000 0.00000 0.00000 ; O- C- N- H
7351 7350 7352 7354 3 20.92000 0.00000 -20.92000
0.00000 0.00000 0.00000 ; O- C- N- CA
7352 7354 7356 7357 3 0.65084 1.95253 0.00000
-2.60338 0.00000 0.00000 ; N- CA- CB- HB2
7352 7354 7356 7358 3 0.65084 1.95253 0.00000
-2.60338 0.00000 0.00000 ; N- CA- CB- HB3
7352 7354 7356 7359 3 0.65084 1.95253 0.00000
-2.60338 0.00000 0.00000 ; N- CA- CB- CG
7352 7354 7372 7373 3 0.00000 0.00000 0.00000
0.00000 0.00000 0.00000 ; N- CA- C- O
7352 7354 7372 7374 3 0.00000 0.00000 0.00000
0.00000 0.00000 0.00000 ; N- CA- C- OXT
7353 7352 7354 7355 3 0.00000 0.00000 0.00000
0.00000 0.00000 0.00000 ; H- N- CA- HA
7353 7352 7354 7356 3 0.00000 0.00000 0.00000
0.00000 0.00000 0.00000 ; H- N- CA- CB
7353 7352 7354 7372 3 0.00000 0.00000 0.00000
0.00000 0.00000 0.00000 ; H- N- CA- C

7354 7356 7359 7360 3 0.66944 2.00832 0.00000
-2.67776 0.00000 0.00000 ; CA- CB- CG- HG2
7354 7356 7359 7361 3 0.66944 2.00832 0.00000
-2.67776 0.00000 0.00000 ; CA- CB- CG- HG3
7354 7356 7359 7362 3 3.68192 3.09616 -2.09200
-3.01248 0.00000 0.00000 ; CA- CB- CG- CD
7355 7354 7356 7357 3 0.65084 1.95253 0.00000
-2.60338 0.00000 0.00000 ; HA- CA- CB- HB2
7355 7354 7356 7358 3 0.65084 1.95253 0.00000
-2.60338 0.00000 0.00000 ; HA- CA- CB- HB3
7355 7354 7356 7359 3 0.65084 1.95253 0.00000
-2.60338 0.00000 0.00000 ; HA- CA- CB- CG
7355 7354 7372 7373 3 0.00000 0.00000 0.00000
0.00000 0.00000 0.00000 ; HA- CA- C- O
7355 7354 7372 7374 3 0.00000 0.00000 0.00000
0.00000 0.00000 0.00000 ; HA- CA- C- OXT
7356 7354 7372 7373 3 0.00000 0.00000 0.00000
0.00000 0.00000 0.00000 ; CB- CA- C- O
7356 7354 7372 7374 3 0.00000 0.00000 0.00000
0.00000 0.00000 0.00000 ; CB- CA- C- OXT
7356 7359 7362 7363 3 0.66944 2.00832 0.00000
-2.67776 0.00000 0.00000 ; CB- CG- CD- HD2
7356 7359 7362 7364 3 0.66944 2.00832 0.00000
-2.67776 0.00000 0.00000 ; CB- CG- CD- HD3
7356 7359 7362 7365 3 3.68192 3.09616 -2.09200
-3.01248 0.00000 0.00000 ; CB- CG- CD- CE
7357 7356 7354 7372 3 0.65084 1.95253 0.00000
-2.60338 0.00000 0.00000 ; HB2- CB- CA- C
7357 7356 7359 7360 3 0.62760 1.88280 0.00000
-2.51040 0.00000 0.00000 ; HB2- CB- CG- HG2
7357 7356 7359 7361 3 0.62760 1.88280 0.00000
-2.51040 0.00000 0.00000 ; HB2- CB- CG- HG3
7357 7356 7359 7362 3 0.66944 2.00832 0.00000
-2.67776 0.00000 0.00000 ; HB2- CB- CG- CD
7358 7356 7354 7372 3 0.65084 1.95253 0.00000
-2.60338 0.00000 0.00000 ; HB3- CB- CA- C
7358 7356 7359 7360 3 0.62760 1.88280 0.00000
-2.51040 0.00000 0.00000 ; HB3- CB- CG- HG2
7358 7356 7359 7361 3 0.62760 1.88280 0.00000
-2.51040 0.00000 0.00000 ; HB3- CB- CG- HG3
7358 7356 7359 7362 3 0.66944 2.00832 0.00000
-2.67776 0.00000 0.00000 ; HB3- CB- CG- CD
7359 7356 7354 7372 3 0.65084 1.95253 0.00000
-2.60338 0.00000 0.00000 ; CG- CB- CA- C
7359 7362 7365 7366 3 0.65084 1.95253 0.00000
-2.60338 0.00000 0.00000 ; CG- CD- CE- HE2
7359 7362 7365 7367 3 0.65084 1.95253 0.00000
-2.60338 0.00000 0.00000 ; CG- CD- CE- HE3
7359 7362 7365 7368 3 0.65084 1.95253 0.00000
-2.60338 0.00000 0.00000 ; CG- CD- CE- NZ
7360 7359 7362 7363 3 0.62760 1.88280 0.00000
-2.51040 0.00000 0.00000 ; HG2- CG- CD- HD2

```
  7360   7359   7362   7364      3    0.62760    1.88280    0.00000
-2.51040    0.00000    0.00000 ;    HG2-   CG-    CD-   HD3
  7360   7359   7362   7365      3    0.66944    2.00832    0.00000
-2.67776    0.00000    0.00000 ;    HG2-   CG-    CD-    CE
  7361   7359   7362   7363      3    0.62760    1.88280    0.00000
-2.51040    0.00000    0.00000 ;    HG3-   CG-    CD-   HD2
  7361   7359   7362   7364      3    0.62760    1.88280    0.00000
-2.51040    0.00000    0.00000 ;    HG3-   CG-    CD-   HD3
  7361   7359   7362   7365      3    0.66944    2.00832    0.00000
-2.67776    0.00000    0.00000 ;    HG3-   CG-    CD-    CE
  7362   7365   7368   7369      3    0.65084    1.95253    0.00000
-2.60338    0.00000    0.00000 ;     CD-   CE-    NZ-   HZ1
  7362   7365   7368   7370      3    0.65084    1.95253    0.00000
-2.60338    0.00000    0.00000 ;     CD-   CE-    NZ-   HZ2
  7362   7365   7368   7371      3    0.65084    1.95253    0.00000
-2.60338    0.00000    0.00000 ;     CD-   CE-    NZ-   HZ3
  7363   7362   7365   7366      3    0.65084    1.95253    0.00000
-2.60338    0.00000    0.00000 ;    HD2-   CD-    CE-   HE2
  7363   7362   7365   7367      3    0.65084    1.95253    0.00000
-2.60338    0.00000    0.00000 ;    HD2-   CD-    CE-   HE3
  7363   7362   7365   7368      3    0.65084    1.95253    0.00000
-2.60338    0.00000    0.00000 ;    HD2-   CD-    CE-    NZ
  7364   7362   7365   7366      3    0.65084    1.95253    0.00000
-2.60338    0.00000    0.00000 ;    HD3-   CD-    CE-   HE2
  7364   7362   7365   7367      3    0.65084    1.95253    0.00000
-2.60338    0.00000    0.00000 ;    HD3-   CD-    CE-   HE3
  7364   7362   7365   7368      3    0.65084    1.95253    0.00000
-2.60338    0.00000    0.00000 ;    HD3-   CD-    CE-    NZ
  7366   7365   7368   7369      3    0.65084    1.95253    0.00000
-2.60338    0.00000    0.00000 ;    HE2-   CE-    NZ-   HZ1
  7366   7365   7368   7370      3    0.65084    1.95253    0.00000
-2.60338    0.00000    0.00000 ;    HE2-   CE-    NZ-   HZ2
  7366   7365   7368   7371      3    0.65084    1.95253    0.00000
-2.60338    0.00000    0.00000 ;    HE2-   CE-    NZ-   HZ3
  7367   7365   7368   7369      3    0.65084    1.95253    0.00000
-2.60338    0.00000    0.00000 ;    HE3-   CE-    NZ-   HZ1
  7367   7365   7368   7370      3    0.65084    1.95253    0.00000
-2.60338    0.00000    0.00000 ;    HE3-   CE-    NZ-   HZ2
  7367   7365   7368   7371      3    0.65084    1.95253    0.00000
-2.60338    0.00000    0.00000 ;    HE3-   CE-    NZ-   HZ3
  7375   7376   7377   7378      3   55.64720    0.00000  -55.64720
0.00000    0.00000    0.00000 ;     F2-   C38-   C39-   N23
  7375   7376   7377   7423      3   55.64720    0.00000  -55.64720
0.00000    0.00000    0.00000 ;     F2-   C38-   C39-   H39
  7375   7376   7383   7384      3   55.64720    0.00000  -55.64720
0.00000    0.00000    0.00000 ;     F2-   C38-   C37-   C36
  7375   7376   7383   7424      3   55.64720    0.00000  -55.64720
0.00000    0.00000    0.00000 ;     F2-   C38-   C37-   H37
  7376   7377   7378   7385      3   34.72720    0.00000  -34.72720
0.00000    0.00000    0.00000 ;    C38-   C39-   N23-   C40
  7376   7377   7378   7433      3    0.00000    0.00000    0.00000
0.00000    0.00000    0.00000 ;    C38-   C39-   N23-    RU
```

```
  7376   7383   7384   7385      3    5.85760    0.00000   -5.85760
0.00000   0.00000   0.00000 ;   C38-  C37-   C36-  C40
  7376   7383   7384   7386      3    5.85760    0.00000   -5.85760
0.00000   0.00000   0.00000 ;   C38-  C37-   C36-  C35
  7377   7376   7383   7384      3   55.64720    0.00000  -55.64720
0.00000   0.00000   0.00000 ;   C39-  C38-   C37-  C36
  7377   7376   7383   7424      3   55.64720    0.00000  -55.64720
0.00000   0.00000   0.00000 ;   C39-  C38-   C37-  H37
  7377   7378   7385   7384      3    0.00000    0.00000    0.00000
0.00000   0.00000   0.00000 ;   C39-  N23-   C40-  C36
  7377   7378   7385   7395      3    0.00000    0.00000    0.00000
0.00000   0.00000   0.00000 ;   C39-  N23-   C40-  C24
  7377   7378   7433   7379      3    0.00000    0.00000    0.00000
0.00000   0.00000   0.00000 ;   C39-  N23-    RU-  N21
  7377   7378   7433   7380      3    0.00000    0.00000    0.00000
0.00000   0.00000   0.00000 ;   C39-  N23-    RU-  C13
  7377   7378   7433   7382      3    0.00000    0.00000    0.00000
0.00000   0.00000   0.00000 ;   C39-  N23-    RU- CL17
  7377   7378   7433   7402      3    0.00000    0.00000    0.00000
0.00000   0.00000   0.00000 ;   C39-  N23-    RU-  N15
  7378   7377   7376   7383      3   55.64720    0.00000  -55.64720
0.00000   0.00000   0.00000 ;   N23-  C39-   C38-  C37
  7378   7385   7384   7383      3   30.33400    0.00000  -30.33400
0.00000   0.00000   0.00000 ;   N23-  C40-   C36-  C37
  7378   7385   7384   7386      3   30.33400    0.00000  -30.33400
0.00000   0.00000   0.00000 ;   N23-  C40-   C36-  C35
  7378   7385   7395   7379      3   30.33400    0.00000  -30.33400
0.00000   0.00000   0.00000 ;   N23-  C40-   C24-  N21
  7378   7385   7395   7394      3   30.33400    0.00000  -30.33400
0.00000   0.00000   0.00000 ;   N23-  C40-   C24-  C31
  7378   7433   7379   7395      3    0.00000    0.00000    0.00000
0.00000   0.00000   0.00000 ;   N23-   RU-   N21-  C24
  7378   7433   7379   7397      3    0.00000    0.00000    0.00000
0.00000   0.00000   0.00000 ;   N23-   RU-   N21-  C25
  7378   7433   7380   7381      3    0.00000    0.00000    0.00000
0.00000   0.00000   0.00000 ;   N23-   RU-   C13-  O14
  7378   7433   7402   7403      3    0.00000    0.00000    0.00000
0.00000   0.00000   0.00000 ;   N23-   RU-   N15-   C7
  7378   7433   7402   7404      3    0.00000    0.00000    0.00000
0.00000   0.00000   0.00000 ;   N23-   RU-   N15-  C18
  7379   7395   7385   7384      3   30.33400    0.00000  -30.33400
0.00000   0.00000   0.00000 ;   N21-  C24-   C40-  C36
  7379   7395   7394   7391      3   30.33400    0.00000  -30.33400
0.00000   0.00000   0.00000 ;   N21-  C24-   C31-  C32
  7379   7395   7394   7396      3   30.33400    0.00000  -30.33400
0.00000   0.00000   0.00000 ;   N21-  C24-   C31-  C30
  7379   7397   7396   7394      3   30.33400    0.00000  -30.33400
0.00000   0.00000   0.00000 ;   N21-  C25-   C30-  C31
  7379   7397   7396   7398      3   30.33400    0.00000  -30.33400
0.00000   0.00000   0.00000 ;   N21-  C25-   C30-  C29
  7379   7397   7401   7400      3   30.33400    0.00000  -30.33400
0.00000   0.00000   0.00000 ;   N21-  C25-   C26-  C27
```

7379 7397 7401 7428 3 30.33400 0.00000 -30.33400
0.00000 0.00000 0.00000 ; N21- C25- C26- H26
7379 7433 7378 7385 3 0.00000 0.00000 0.00000
0.00000 0.00000 0.00000 ; N21- RU- N23- C40
7379 7433 7380 7381 3 0.00000 0.00000 0.00000
0.00000 0.00000 0.00000 ; N21- RU- C13- O14
7379 7433 7402 7403 3 0.00000 0.00000 0.00000
0.00000 0.00000 0.00000 ; N21- RU- N15- C7
7379 7433 7402 7404 3 0.00000 0.00000 0.00000
0.00000 0.00000 0.00000 ; N21- RU- N15- C18
7380 7433 7378 7385 3 0.00000 0.00000 0.00000
0.00000 0.00000 0.00000 ; C13- RU- N23- C40
7380 7433 7379 7395 3 0.00000 0.00000 0.00000
0.00000 0.00000 0.00000 ; C13- RU- N21- C24
7380 7433 7379 7397 3 0.00000 0.00000 0.00000
0.00000 0.00000 0.00000 ; C13- RU- N21- C25
7380 7433 7402 7403 3 0.00000 0.00000 0.00000
0.00000 0.00000 0.00000 ; C13- RU- N15- C7
7380 7433 7402 7404 3 0.00000 0.00000 0.00000
0.00000 0.00000 0.00000 ; C13- RU- N15- C18
7381 7380 7433 7382 3 0.00000 0.00000 0.00000
0.00000 0.00000 0.00000 ; O14- C13- RU- CL17
7381 7380 7433 7402 3 0.00000 0.00000 0.00000
0.00000 0.00000 0.00000 ; O14- C13- RU- N15
7382 7433 7378 7385 3 0.00000 0.00000 0.00000
0.00000 0.00000 0.00000 ; CL17- RU- N23- C40
7382 7433 7379 7395 3 0.00000 0.00000 0.00000
0.00000 0.00000 0.00000 ; CL17- RU- N21- C24
7382 7433 7379 7397 3 0.00000 0.00000 0.00000
0.00000 0.00000 0.00000 ; CL17- RU- N21- C25
7382 7433 7402 7403 3 0.00000 0.00000 0.00000
0.00000 0.00000 0.00000 ; CL17- RU- N15- C7
7382 7433 7402 7404 3 0.00000 0.00000 0.00000
0.00000 0.00000 0.00000 ; CL17- RU- N15- C18
7383 7376 7377 7423 3 55.64720 0.00000 -55.64720
0.00000 0.00000 0.00000 ; C37- C38- C39- H39
7383 7384 7385 7395 3 30.33400 0.00000 -30.33400
0.00000 0.00000 0.00000 ; C37- C36- C40- C24
7383 7384 7386 7387 3 30.33400 0.00000 -30.33400
0.00000 0.00000 0.00000 ; C37- C36- C35- C34
7383 7384 7386 7391 3 30.33400 0.00000 -30.33400
0.00000 0.00000 0.00000 ; C37- C36- C35- C32
7384 7385 7378 7433 3 0.00000 0.00000 0.00000
0.00000 0.00000 0.00000 ; C36- C40- N23- RU
7384 7385 7395 7394 3 30.33400 0.00000 -30.33400
0.00000 0.00000 0.00000 ; C36- C40- C24- C31
7384 7386 7387 7388 3 5.85760 0.00000 -5.85760
0.00000 0.00000 0.00000 ; C36- C35- C34- N22
7384 7386 7387 7390 3 5.85760 0.00000 -5.85760
0.00000 0.00000 0.00000 ; C36- C35- C34- O42
7384 7386 7391 7392 3 30.33400 0.00000 -30.33400
0.00000 0.00000 0.00000 ; C36- C35- C32- C33

7384 7386 7391 7394 3 30.33400 0.00000 -30.33400
0.00000 0.00000 0.00000 ; C36- C35- C32- C31
7385 7378 7377 7423 3 34.72720 0.00000 -34.72720
0.00000 0.00000 0.00000 ; C40- N23- C39- H39
7385 7378 7433 7402 3 0.00000 0.00000 0.00000
0.00000 0.00000 0.00000 ; C40- N23- RU- N15
7385 7384 7383 7424 3 5.85760 0.00000 -5.85760
0.00000 0.00000 0.00000 ; C40- C36- C37- H37
7385 7384 7386 7387 3 30.33400 0.00000 -30.33400
0.00000 0.00000 0.00000 ; C40- C36- C35- C34
7385 7384 7386 7391 3 30.33400 0.00000 -30.33400
0.00000 0.00000 0.00000 ; C40- C36- C35- C32
7385 7395 7379 7397 3 0.00000 0.00000 0.00000
0.00000 0.00000 0.00000 ; C40- C24- N21- C25
7385 7395 7379 7433 3 0.00000 0.00000 0.00000
0.00000 0.00000 0.00000 ; C40- C24- N21- RU
7385 7395 7394 7391 3 30.33400 0.00000 -30.33400
0.00000 0.00000 0.00000 ; C40- C24- C31- C32
7385 7395 7394 7396 3 30.33400 0.00000 -30.33400
0.00000 0.00000 0.00000 ; C40- C24- C31- C30
7386 7384 7383 7424 3 5.85760 0.00000 -5.85760
0.00000 0.00000 0.00000 ; C35- C36- C37- H37
7386 7384 7385 7395 3 30.33400 0.00000 -30.33400
0.00000 0.00000 0.00000 ; C35- C36- C40- C24
7386 7387 7388 7389 3 5.43920 0.00000 -5.43920
0.00000 0.00000 0.00000 ; C35- C34- N22- H22
7386 7387 7388 7392 3 5.43920 0.00000 -5.43920
0.00000 0.00000 0.00000 ; C35- C34- N22- C33
7386 7391 7392 7388 3 5.85760 0.00000 -5.85760
0.00000 0.00000 0.00000 ; C35- C32- C33- N22
7386 7391 7392 7393 3 5.85760 0.00000 -5.85760
0.00000 0.00000 0.00000 ; C35- C32- C33- O41
7386 7391 7394 7395 3 30.33400 0.00000 -30.33400
0.00000 0.00000 0.00000 ; C35- C32- C31- C24
7386 7391 7394 7396 3 30.33400 0.00000 -30.33400
0.00000 0.00000 0.00000 ; C35- C32- C31- C30
7387 7386 7391 7392 3 30.33400 0.00000 -30.33400
0.00000 0.00000 0.00000 ; C34- C35- C32- C33
7387 7386 7391 7394 3 30.33400 0.00000 -30.33400
0.00000 0.00000 0.00000 ; C34- C35- C32- C31
7387 7388 7392 7391 3 5.43920 0.00000 -5.43920
0.00000 0.00000 0.00000 ; C34- N22- C33- C32
7387 7388 7392 7393 3 5.43920 0.00000 -5.43920
0.00000 0.00000 0.00000 ; C34- N22- C33- O41
7388 7387 7386 7391 3 5.85760 0.00000 -5.85760
0.00000 0.00000 0.00000 ; N22- C34- C35- C32
7388 7392 7391 7394 3 5.85760 0.00000 -5.85760
0.00000 0.00000 0.00000 ; N22- C33- C32- C31
7389 7388 7387 7390 3 5.43920 0.00000 -5.43920
0.00000 0.00000 0.00000 ; H22- N22- C34- O42
7389 7388 7392 7391 3 5.43920 0.00000 -5.43920
0.00000 0.00000 0.00000 ; H22- N22- C33- C32

7389 7388 7392 7393 3 5.43920 0.00000 -5.43920
0.00000 0.00000 0.00000 ; H22- N22- C33- O41
7390 7387 7386 7391 3 5.85760 0.00000 -5.85760
0.00000 0.00000 0.00000 ; O42- C34- C35- C32
7390 7387 7388 7392 3 5.43920 0.00000 -5.43920
0.00000 0.00000 0.00000 ; O42- C34- N22- C33
7391 7394 7396 7397 3 8.36800 0.00000 -8.36800
0.00000 0.00000 0.00000 ; C32- C31- C30- C25
7391 7394 7396 7398 3 8.36800 0.00000 -8.36800
0.00000 0.00000 0.00000 ; C32- C31- C30- C29
7392 7391 7394 7395 3 30.33400 0.00000 -30.33400
0.00000 0.00000 0.00000 ; C33- C32- C31- C24
7392 7391 7394 7396 3 30.33400 0.00000 -30.33400
0.00000 0.00000 0.00000 ; C33- C32- C31- C30
7393 7392 7391 7394 3 5.85760 0.00000 -5.85760
0.00000 0.00000 0.00000 ; O41- C33- C32- C31
7394 7395 7379 7397 3 0.00000 0.00000 0.00000
0.00000 0.00000 0.00000 ; C31- C24- N21- C25
7394 7395 7379 7433 3 0.00000 0.00000 0.00000
0.00000 0.00000 0.00000 ; C31- C24- N21- RU
7394 7396 7397 7401 3 30.33400 0.00000 -30.33400
0.00000 0.00000 0.00000 ; C31- C30- C25- C26
7394 7396 7398 7399 3 30.33400 0.00000 -30.33400
0.00000 0.00000 0.00000 ; C31- C30- C29- C28
7394 7396 7398 7426 3 30.33400 0.00000 -30.33400
0.00000 0.00000 0.00000 ; C31- C30- C29- H29
7395 7379 7397 7396 3 0.00000 0.00000 0.00000
0.00000 0.00000 0.00000 ; C24- N21- C25- C30
7395 7379 7397 7401 3 0.00000 0.00000 0.00000
0.00000 0.00000 0.00000 ; C24- N21- C25- C26
7395 7379 7433 7402 3 0.00000 0.00000 0.00000
0.00000 0.00000 0.00000 ; C24- N21- RU- N15
7395 7385 7378 7433 3 0.00000 0.00000 0.00000
0.00000 0.00000 0.00000 ; C24- C40- N23- RU
7395 7394 7396 7397 3 8.36800 0.00000 -8.36800
0.00000 0.00000 0.00000 ; C24- C31- C30- C25
7395 7394 7396 7398 3 8.36800 0.00000 -8.36800
0.00000 0.00000 0.00000 ; C24- C31- C30- C29
7396 7397 7379 7433 3 0.00000 0.00000 0.00000
0.00000 0.00000 0.00000 ; C30- C25- N21- RU
7396 7397 7401 7400 3 30.33400 0.00000 -30.33400
0.00000 0.00000 0.00000 ; C30- C25- C26- C27
7396 7397 7401 7428 3 30.33400 0.00000 -30.33400
0.00000 0.00000 0.00000 ; C30- C25- C26- H26
7396 7398 7399 7400 3 30.33400 0.00000 -30.33400
0.00000 0.00000 0.00000 ; C30- C29- C28- C27
7396 7398 7399 7416 3 30.33400 0.00000 -30.33400
0.00000 0.00000 0.00000 ; C30- C29- C28- O1
7397 7379 7433 7402 3 0.00000 0.00000 0.00000
0.00000 0.00000 0.00000 ; C25- N21- RU- N15
7397 7396 7398 7399 3 30.33400 0.00000 -30.33400
0.00000 0.00000 0.00000 ; C25- C30- C29- C28

```
  7397   7396   7398   7426      3   30.33400     0.00000  -30.33400
0.00000   0.00000   0.00000 ;    C25-   C30-   C29-   H29
  7397   7401   7400   7399      3   30.33400     0.00000  -30.33400
0.00000   0.00000   0.00000 ;    C25-   C26-   C27-   C28
  7397   7401   7400   7427      3   30.33400     0.00000  -30.33400
0.00000   0.00000   0.00000 ;    C25-   C26-   C27-   H27
  7398   7396   7397   7401      3   30.33400     0.00000  -30.33400
0.00000   0.00000   0.00000 ;    C29-   C30-   C25-   C26
  7398   7399   7400   7401      3   30.33400     0.00000  -30.33400
0.00000   0.00000   0.00000 ;    C29-   C28-   C27-   C26
  7398   7399   7400   7427      3   30.33400     0.00000  -30.33400
0.00000   0.00000   0.00000 ;    C29-   C28-   C27-   H27
  7398   7399   7416   7417      3    7.53120     0.00000   -7.53120
0.00000   0.00000   0.00000 ;    C29-   C28-    O1-    H1
  7399   7400   7401   7428      3   30.33400     0.00000  -30.33400
0.00000   0.00000   0.00000 ;    C28-   C27-   C26-   H26
  7400   7399   7398   7426      3   30.33400     0.00000  -30.33400
0.00000   0.00000   0.00000 ;    C27-   C28-   C29-   H29
  7400   7399   7416   7417      3    7.53120     0.00000   -7.53120
0.00000   0.00000   0.00000 ;    C27-   C28-    O1-    H1
  7401   7397   7379   7433      3    0.00000     0.00000    0.00000
0.00000   0.00000   0.00000 ;    C26-   C25-   N21-    RU
  7401   7400   7399   7416      3   30.33400     0.00000  -30.33400
0.00000   0.00000   0.00000 ;    C26-   C27-   C28-    O1
  7402   7403   7410   7411      3   55.64720     0.00000  -55.64720
0.00000   0.00000   0.00000 ;    N15-    C7-    C8-    C9
  7402   7403   7410   7415      3   55.64720     0.00000  -55.64720
0.00000   0.00000   0.00000 ;    N15-    C7-    C8-   N16
  7402   7404   7405   7406      3   30.33400     0.00000  -30.33400
0.00000   0.00000   0.00000 ;    N15-   C18-    C5-    C6
  7402   7404   7405   7421      3   30.33400     0.00000  -30.33400
0.00000   0.00000   0.00000 ;    N15-   C18-    C5-    H5
  7402   7404   7409   7408      3   30.33400     0.00000  -30.33400
0.00000   0.00000   0.00000 ;    N15-   C18-   C19-    C3
  7402   7404   7409   7429      3   30.33400     0.00000  -30.33400
0.00000   0.00000   0.00000 ;    N15-   C18-   C19-   H19
  7403   7402   7404   7405      3    0.00000     0.00000    0.00000
0.00000   0.00000   0.00000 ;     C7-   N15-   C18-    C5
  7403   7402   7404   7409      3    0.00000     0.00000    0.00000
0.00000   0.00000   0.00000 ;     C7-   N15-   C18-   C19
  7403   7410   7411   7412      3   55.64720     0.00000  -55.64720
0.00000   0.00000   0.00000 ;     C7-    C8-    C9-   C10
  7403   7410   7411   7418      3   55.64720     0.00000  -55.64720
0.00000   0.00000   0.00000 ;     C7-    C8-    C9-    H9
  7403   7410   7415   7414      3   34.72720     0.00000  -34.72720
0.00000   0.00000   0.00000 ;     C7-    C8-   N16-   C11
  7404   7402   7403   7410      3   34.72720     0.00000  -34.72720
0.00000   0.00000   0.00000 ;    C18-   N15-    C7-    C8
  7404   7402   7403   7419      3   34.72720     0.00000  -34.72720
0.00000   0.00000   0.00000 ;    C18-   N15-    C7-   H7A
  7404   7405   7406   7407      3   30.33400     0.00000  -30.33400
0.00000   0.00000   0.00000 ;    C18-    C5-    C6-    C4
```

```
  7404   7405   7406   7420       3   30.33400     0.00000  -30.33400
0.00000    0.00000    0.00000 ;    C18-   C5-    C6-    H6
  7404   7409   7408   7407       3   30.33400     0.00000  -30.33400
0.00000    0.00000    0.00000 ;    C18-   C19-    C3-    C4
  7404   7409   7408   7425       3   30.33400     0.00000  -30.33400
0.00000    0.00000    0.00000 ;    C18-   C19-    C3-    H3
  7405   7404   7402   7433       3    0.00000     0.00000    0.00000
0.00000    0.00000    0.00000 ;     C5-   C18-   N15-    RU
  7405   7404   7409   7408       3   30.33400     0.00000  -30.33400
0.00000    0.00000    0.00000 ;     C5-   C18-   C19-    C3
  7405   7404   7409   7429       3   30.33400     0.00000  -30.33400
0.00000    0.00000    0.00000 ;     C5-   C18-   C19-   H19
  7405   7406   7407   7408       3   30.33400     0.00000  -30.33400
0.00000    0.00000    0.00000 ;     C5-    C6-    C4-    C3
  7405   7406   7407   7422       3   30.33400     0.00000  -30.33400
0.00000    0.00000    0.00000 ;     C5-    C6-    C4-    H4
  7406   7405   7404   7409       3   30.33400     0.00000  -30.33400
0.00000    0.00000    0.00000 ;     C6-    C5-   C18-   C19
  7406   7407   7408   7409       3   30.33400     0.00000  -30.33400
0.00000    0.00000    0.00000 ;     C6-    C4-    C3-   C19
  7406   7407   7408   7425       3   30.33400     0.00000  -30.33400
0.00000    0.00000    0.00000 ;     C6-    C4-    C3-    H3
  7407   7406   7405   7421       3   30.33400     0.00000  -30.33400
0.00000    0.00000    0.00000 ;     C4-    C6-    C5-    H5
  7407   7408   7409   7429       3   30.33400     0.00000  -30.33400
0.00000    0.00000    0.00000 ;     C4-    C3-   C19-   H19
  7408   7407   7406   7420       3   30.33400     0.00000  -30.33400
0.00000    0.00000    0.00000 ;     C3-    C4-    C6-    H6
  7409   7404   7402   7433       3    0.00000     0.00000    0.00000
0.00000    0.00000    0.00000 ;    C19-   C18-   N15-    RU
  7409   7404   7405   7421       3   30.33400     0.00000  -30.33400
0.00000    0.00000    0.00000 ;    C19-   C18-    C5-    H5
  7409   7408   7407   7422       3   30.33400     0.00000  -30.33400
0.00000    0.00000    0.00000 ;    C19-    C3-    C4-    H4
  7410   7403   7402   7433       3    0.00000     0.00000    0.00000
0.00000    0.00000    0.00000 ;     C8-    C7-   N15-    RU
  7410   7411   7412   7413       3   55.64720     0.00000  -55.64720
0.00000    0.00000    0.00000 ;     C8-    C9-   C10-   C12
  7410   7411   7412   7432       3   55.64720     0.00000  -55.64720
0.00000    0.00000    0.00000 ;     C8-    C9-   C10-   H10
  7410   7415   7414   7413       3   34.72720     0.00000  -34.72720
0.00000    0.00000    0.00000 ;     C8-   N16-   C11-   C12
  7410   7415   7414   7431       3   34.72720     0.00000  -34.72720
0.00000    0.00000    0.00000 ;     C8-   N16-   C11-   H11
  7411   7410   7403   7419       3   55.64720     0.00000  -55.64720
0.00000    0.00000    0.00000 ;     C9-    C8-    C7-   H7A
  7411   7410   7415   7414       3   34.72720     0.00000  -34.72720
0.00000    0.00000    0.00000 ;     C9-    C8-   N16-   C11
  7411   7412   7413   7414       3   55.64720     0.00000  -55.64720
0.00000    0.00000    0.00000 ;     C9-   C10-   C12-   C11
  7411   7412   7413   7430       3   55.64720     0.00000  -55.64720
0.00000    0.00000    0.00000 ;     C9-   C10-   C12-   H12
```

```
  7412   7411   7410   7415      3   55.64720    0.00000  -55.64720
0.00000    0.00000    0.00000 ;    C10-   C9-    C8-   N16
  7412   7413   7414   7415      3   55.64720    0.00000  -55.64720
0.00000    0.00000    0.00000 ;    C10-   C12-   C11-   N16
  7412   7413   7414   7431      3   55.64720    0.00000  -55.64720
0.00000    0.00000    0.00000 ;    C10-   C12-   C11-   H11
  7413   7412   7411   7418      3   55.64720    0.00000  -55.64720
0.00000    0.00000    0.00000 ;    C12-   C10-    C9-    H9
  7414   7413   7412   7432      3   55.64720    0.00000  -55.64720
0.00000    0.00000    0.00000 ;    C11-   C12-   C10-   H10
  7415   7410   7403   7419      3   55.64720    0.00000  -55.64720
0.00000    0.00000    0.00000 ;    N16-    C8-    C7-   H7A
  7415   7410   7411   7418      3   55.64720    0.00000  -55.64720
0.00000    0.00000    0.00000 ;    N16-    C8-    C9-    H9
  7415   7414   7413   7430      3   55.64720    0.00000  -55.64720
0.00000    0.00000    0.00000 ;    N16-   C11-   C12-   H12
  7416   7399   7398   7426      3   30.33400    0.00000  -30.33400
0.00000    0.00000    0.00000 ;     O1-   C28-   C29-   H29
  7416   7399   7400   7427      3   30.33400    0.00000  -30.33400
0.00000    0.00000    0.00000 ;     O1-   C28-   C27-   H27
  7418   7411   7412   7432      3   55.64720    0.00000  -55.64720
0.00000    0.00000    0.00000 ;     H9-    C9-   C10-   H10
  7419   7403   7402   7433      3    0.00000    0.00000    0.00000
0.00000    0.00000    0.00000 ;    H7A-    C7-   N15-    RU
  7420   7406   7405   7421      3   30.33400    0.00000  -30.33400
0.00000    0.00000    0.00000 ;     H6-    C6-    C5-    H5
  7420   7406   7407   7422      3   30.33400    0.00000  -30.33400
0.00000    0.00000    0.00000 ;     H6-    C6-    C4-    H4
  7422   7407   7408   7425      3   30.33400    0.00000  -30.33400
0.00000    0.00000    0.00000 ;     H4-    C4-    C3-    H3
  7423   7377   7378   7433      3    0.00000    0.00000    0.00000
0.00000    0.00000    0.00000 ;    H39-   C39-   N23-    RU
  7425   7408   7409   7429      3   30.33400    0.00000  -30.33400
0.00000    0.00000    0.00000 ;     H3-    C3-   C19-   H19
  7427   7400   7401   7428      3   30.33400    0.00000  -30.33400
0.00000    0.00000    0.00000 ;    H27-   C27-   C26-   H26
  7430   7413   7412   7432      3   55.64720    0.00000  -55.64720
0.00000    0.00000    0.00000 ;    H12-   C12-   C10-   H10
  7430   7413   7414   7431      3   55.64720    0.00000  -55.64720
0.00000    0.00000    0.00000 ;    H12-   C12-   C11-   H11

[ dihedrals ] ; impropers
; treated as propers in GROMACS to use correct AMBER analytical
function
;    i      j      k      l   func   phase     kd      pn
     5     22     20     21      1   180.00  43.93200   2 ;     CA-
N-     C-     O
    20     24     22     23      1   180.00   4.60240   2 ;      C-
CA-     N-     H
    24     41     39     40      1   180.00  43.93200   2 ;     CA-
N-     C-     O
```

39 43 41 42 1 180.00 4.60240 2 ; C-
CA- N- H
43 60 58 59 1 180.00 43.93200 2 ; CA-
N- C- O
58 62 60 61 1 180.00 4.60240 2 ; C-
CA- N- H
62 71 69 70 1 180.00 43.93200 2 ; CA-
N- C- O
69 73 71 72 1 180.00 4.60240 2 ; C-
CA- N- H
73 88 86 87 1 180.00 43.93200 2 ; CA-
N- C- O
75 84 78 79 1 180.00 4.60240 2 ; CB-
CD2- CG- ND1
78 82 84 85 1 180.00 4.60240 2 ; CG-
NE2- CD2- HD2
79 82 80 81 1 180.00 4.60240 2 ; ND1-
NE2- CE1- HE1
80 84 82 83 1 180.00 4.18400 2 ; CE1-
CD2- NE2- HE2
86 90 88 89 1 180.00 4.60240 2 ; C-
CA- N- H
90 99 97 98 1 180.00 43.93200 2 ; CA-
N- C- O
97 101 99 100 1 180.00 4.60240 2 ; C-
CA- N- H
101 106 104 105 1 180.00 43.93200 2 ; CA-
N- C- O
104 108 106 107 1 180.00 4.60240 2 ; C-
CA- N- H
108 128 126 127 1 180.00 43.93200 2 ; CA-
N- C- O
126 130 128 129 1 180.00 4.60240 2 ; C-
CA- N- H
130 147 145 146 1 180.00 43.93200 2 ; CA-
N- C- O
145 149 147 148 1 180.00 4.60240 2 ; C-
CA- N- H
149 162 160 161 1 180.00 43.93200 2 ; CA-
N- C- O
154 158 157 159 1 180.00 43.93200 2 ; CG-
OE1- CD- OE2
160 164 162 163 1 180.00 4.60240 2 ; C-
CA- N- H
164 174 172 173 1 180.00 43.93200 2 ; CA-
N- C- O
166 170 169 171 1 180.00 43.93200 2 ; CB-
OD1- CG- OD2
172 176 174 175 1 180.00 4.60240 2 ; C-
CA- N- H
176 196 194 195 1 180.00 43.93200 2 ; CA-
N- C- O

194 198 196 197 1 180.00 4.60240 2 ; C-
CA- N- H
198 217 215 216 1 180.00 43.93200 2 ; CA-
N- C- O
203 206 204 205 1 180.00 4.60240 2 ; CG-
CE1- CD1- HD1
203 211 213 214 1 180.00 4.60240 2 ; CG-
CE2- CD2- HD2
204 208 206 207 1 180.00 4.60240 2 ; CD1-
CZ- CE1- HE1
204 213 203 200 1 180.00 4.60240 2 ; CD1-
CD2- CG- CB
206 211 208 209 1 180.00 4.60240 2 ; CE1-
CE2- CZ- OH
208 213 211 212 1 180.00 4.60240 2 ; CZ-
CD2- CE2- HE2
215 219 217 218 1 180.00 4.60240 2 ; C-
CA- N- H
219 236 234 235 1 180.00 43.93200 2 ; CA-
N- C- O
234 238 236 237 1 180.00 4.60240 2 ; C-
CA- N- H
238 255 253 254 1 180.00 43.93200 2 ; CA-
N- C- O
253 257 255 256 1 180.00 4.60240 2 ; C-
CA- N- H
257 267 265 266 1 180.00 43.93200 2 ; CA-
N- C- O
259 263 262 264 1 180.00 43.93200 2 ; CB-
OD1- CG- OD2
265 269 267 268 1 180.00 4.60240 2 ; C-
CA- N- H
269 282 280 281 1 180.00 43.93200 2 ; CA-
N- C- O
274 278 277 279 1 180.00 43.93200 2 ; CG-
OE1- CD- OE2
280 284 282 283 1 180.00 4.60240 2 ; C-
CA- N- H
284 304 302 303 1 180.00 43.93200 2 ; CA-
N- C- O
302 306 304 305 1 180.00 4.60240 2 ; C-
CA- N- H
306 323 321 322 1 180.00 43.93200 2 ; CA-
N- C- O
321 325 323 324 1 180.00 4.60240 2 ; C-
CA- N- H
325 330 328 329 1 180.00 43.93200 2 ; CA-
N- C- O
328 332 330 331 1 180.00 4.60240 2 ; C-
CA- N- H
332 347 345 346 1 180.00 43.93200 2 ; CA-
N- C- O

337 342 340 341 1 180.00 43.93200 2 ; CG-
NE2- CD- OE1
340 343 342 344 1 180.00 4.18400 2 ; CD-
HE21- NE2- HE22
345 349 347 348 1 180.00 4.60240 2 ; C-
CA- N- H
349 354 352 353 1 180.00 43.93200 2 ; CA-
N- C- O
352 356 354 355 1 180.00 4.60240 2 ; C-
CA- N- H
356 368 366 367 1 180.00 43.93200 2 ; CA-
N- C- O
366 370 368 369 1 180.00 4.60240 2 ; C-
CA- N- H
370 389 387 388 1 180.00 43.93200 2 ; CA-
N- C- O
375 378 376 377 1 180.00 4.60240 2 ; CG-
CE1- CD1- HD1
375 383 385 386 1 180.00 4.60240 2 ; CG-
CE2- CD2- HD2
376 380 378 379 1 180.00 4.60240 2 ; CD1-
CZ- CE1- HE1
376 385 375 372 1 180.00 4.60240 2 ; CD1-
CD2- CG- CB
378 383 380 381 1 180.00 4.60240 2 ; CE1-
CE2- CZ- OH
380 385 383 384 1 180.00 4.60240 2 ; CZ-
CD2- CE2- HE2
387 391 389 390 1 180.00 4.60240 2 ; C-
CA- N- H
391 396 394 395 1 180.00 43.93200 2 ; CA-
N- C- O
394 398 396 397 1 180.00 4.60240 2 ; C-
CA- N- H
398 407 405 406 1 180.00 43.93200 2 ; CA-
N- C- O
405 409 407 408 1 180.00 4.60240 2 ; C-
CA- N- H
409 423 421 422 1 180.00 43.93200 2 ; CA-
N- C- O
421 425 423 424 1 180.00 4.60240 2 ; C-
CA- N- H
425 444 442 443 1 180.00 43.93200 2 ; CA-
N- C- O
430 433 431 432 1 180.00 4.60240 2 ; CG-
CE1- CD1- HD1
430 438 440 441 1 180.00 4.60240 2 ; CG-
CE2- CD2- HD2
431 435 433 434 1 180.00 4.60240 2 ; CD1-
CZ- CE1- HE1
431 440 430 427 1 180.00 4.60240 2 ; CD1-
CD2- CG- CB

433 438 435 436 1 180.00 4.60240 2 ; CE1-
CE2- CZ- OH
435 440 438 439 1 180.00 4.60240 2 ; CZ-
CD2- CE2- HE2
442 446 444 445 1 180.00 4.60240 2 ; C-
CA- N- H
446 466 464 465 1 180.00 43.93200 2 ; CA-
N- C- O
464 468 466 467 1 180.00 4.60240 2 ; C-
CA- N- H
468 473 471 472 1 180.00 43.93200 2 ; CA-
N- C- O
471 475 473 474 1 180.00 4.60240 2 ; C-
CA- N- H
475 492 490 491 1 180.00 43.93200 2 ; CA-
N- C- O
490 494 492 493 1 180.00 4.60240 2 ; C-
CA- N- H
494 504 502 503 1 180.00 43.93200 2 ; CA-
N- C- O
496 500 499 501 1 180.00 43.93200 2 ; CB-
OD1- CG- OD2
502 506 504 505 1 180.00 4.60240 2 ; C-
CA- N- H
506 526 524 525 1 180.00 43.93200 2 ; CA-
N- C- O
524 528 526 527 1 180.00 4.60240 2 ; C-
CA- N- H
528 542 540 541 1 180.00 43.93200 2 ; CA-
N- C- O
540 544 542 543 1 180.00 4.60240 2 ; C-
CA- N- H
544 556 554 555 1 180.00 43.93200 2 ; CA-
N- C- O
554 558 556 557 1 180.00 4.60240 2 ; C-
CA- N- H
558 570 568 569 1 180.00 43.93200 2 ; CA-
N- C- O
560 565 563 564 1 180.00 43.93200 2 ; CB-
ND2- CG- OD1
563 566 565 567 1 180.00 4.18400 2 ; CG-
HD21- ND2- HD22
568 572 570 571 1 180.00 4.60240 2 ; C-
CA- N- H
572 587 585 586 1 180.00 43.93200 2 ; CA-
N- C- O
577 582 580 581 1 180.00 43.93200 2 ; CG-
NE2- CD- OE1
580 583 582 584 1 180.00 4.18400 2 ; CD-
HE21- NE2- HE22
585 589 587 588 1 180.00 4.60240 2 ; C-
CA- N- H

```
   589    606    604    605       1   180.00  43.93200   2 ;      CA-
N-     C-     O
   604    608    606    607       1   180.00   4.60240   2 ;       C-
CA-     N-     H
   608    627    625    626       1   180.00  43.93200   2 ;      CA-
N-     C-     O
   613    616    614    615       1   180.00   4.60240   2 ;      CG-
CE1-   CD1-   HD1
   613    621    623    624       1   180.00   4.60240   2 ;      CG-
CE2-   CD2-   HD2
   614    618    616    617       1   180.00   4.60240   2 ;     CD1-
CZ-   CE1-   HE1
   614    623    613    610       1   180.00   4.60240   2 ;     CD1-
CD2-    CG-    CB
   616    621    618    619       1   180.00   4.60240   2 ;     CE1-
CE2-    CZ-    OH
   618    623    621    622       1   180.00   4.60240   2 ;      CZ-
CD2-   CE2-   HE2
   625    629    627    628       1   180.00   4.60240   2 ;       C-
CA-     N-     H
   629    637    635    636       1   180.00  43.93200   2 ;      CA-
N-     C-     O
   635    639    637    638       1   180.00   4.60240   2 ;       C-
CA-     N-     H
   639    656    654    655       1   180.00  43.93200   2 ;      CA-
N-     C-     O
   654    658    656    657       1   180.00   4.60240   2 ;       C-
CA-     N-     H
   658    678    676    677       1   180.00  43.93200   2 ;      CA-
N-     C-     O
   676    680    678    679       1   180.00   4.60240   2 ;       C-
CA-     N-     H
   680    693    691    692       1   180.00  43.93200   2 ;      CA-
N-     C-     O
   685    689    688    690       1   180.00  43.93200   2 ;      CG-
OE1-    CD-   OE2
   691    695    693    694       1   180.00   4.60240   2 ;       C-
CA-     N-     H
   695    709    707    708       1   180.00  43.93200   2 ;      CA-
N-     C-     O
   700    704    703    705       1   180.00  43.93200   2 ;      CG-
OE1-    CD-   OE2
   707    711    709    710       1   180.00   4.60240   2 ;       C-
CA-     N-     H
   711    731    729    730       1   180.00  43.93200   2 ;      CA-
N-     C-     O
   729    733    731    732       1   180.00   4.60240   2 ;       C-
CA-     N-     H
   733    753    751    752       1   180.00  43.93200   2 ;      CA-
N-     C-     O
   751    755    753    754       1   180.00   4.60240   2 ;       C-
CA-     N-     H
```

755 765 763 764 1 180.00 43.93200 2 ; CA-
N- C- O
757 761 760 762 1 180.00 43.93200 2 ; CB-
OD1- CG- OD2
763 767 765 766 1 180.00 4.60240 2 ; C-
CA- N- H
767 789 787 788 1 180.00 43.93200 2 ; CA-
N- C- O
775 780 778 779 1 180.00 4.18400 2 ; CD-
CZ- NE- HE
778 781 780 784 1 180.00 43.93200 2 ; NE-
NH1- CZ- NH2
780 782 781 783 1 180.00 4.18400 2 ; CZ-
HH11- NH1- HH12
780 785 784 786 1 180.00 4.18400 2 ; CZ-
HH21- NH2- HH22
787 791 789 790 1 180.00 4.60240 2 ; C-
CA- N- H
791 808 806 807 1 180.00 43.93200 2 ; CA-
N- C- O
806 810 808 809 1 180.00 4.60240 2 ; C-
CA- N- H
810 830 828 829 1 180.00 43.93200 2 ; CA-
N- C- O
828 832 830 831 1 180.00 4.60240 2 ; C-
CA- N- H
832 844 842 843 1 180.00 43.93200 2 ; CA-
N- C- O
834 839 837 838 1 180.00 43.93200 2 ; CB-
ND2- CG- OD1
837 840 839 841 1 180.00 4.18400 2 ; CG-
HD21- ND2- HD22
842 846 844 845 1 180.00 4.60240 2 ; C-
CA- N- H
846 863 861 862 1 180.00 43.93200 2 ; CA-
N- C- O
861 865 863 864 1 180.00 4.60240 2 ; C-
CA- N- H
865 877 875 876 1 180.00 43.93200 2 ; CA-
N- C- O
867 872 870 871 1 180.00 43.93200 2 ; CB-
ND2- CG- OD1
870 873 872 874 1 180.00 4.18400 2 ; CG-
HD21- ND2- HD22
875 879 877 878 1 180.00 4.60240 2 ; C-
CA- N- H
879 901 899 900 1 180.00 43.93200 2 ; CA-
N- C- O
887 892 890 891 1 180.00 4.18400 2 ; CD-
CZ- NE- HE
890 893 892 896 1 180.00 43.93200 2 ; NE-
NH1- CZ- NH2

```
   892    894    893    895       1   180.00   4.18400   2 ;      CZ-
HH11-   NH1-  HH12
   892    897    896    898       1   180.00   4.18400   2 ;      CZ-
HH21-   NH2-  HH22
   899    903    901    902       1   180.00   4.60240   2 ;       C-
CA-     N-     H
   903    921    919    920       1   180.00  43.93200   2 ;      CA-
N-     C-      O
   908    911    909    910       1   180.00   4.60240   2 ;      CG-
CE1-   CD1-   HD1
   908    915    917    918       1   180.00   4.60240   2 ;      CG-
CE2-   CD2-   HD2
   909    913    911    912       1   180.00   4.60240   2 ;     CD1-
CZ-   CE1-   HE1
   909    917    908    905       1   180.00   4.60240   2 ;     CD1-
CD2-    CG-    CB
   911    915    913    914       1   180.00   4.60240   2 ;     CE1-
CE2-    CZ-    HZ
   913    917    915    916       1   180.00   4.60240   2 ;      CZ-
CD2-   CE2-   HE2
   919    923    921    922       1   180.00   4.60240   2 ;       C-
CA-     N-     H
   923    941    939    940       1   180.00  43.93200   2 ;      CA-
N-     C-      O
   928    931    929    930       1   180.00   4.60240   2 ;      CG-
CE1-   CD1-   HD1
   928    935    937    938       1   180.00   4.60240   2 ;      CG-
CE2-   CD2-   HD2
   929    933    931    932       1   180.00   4.60240   2 ;     CD1-
CZ-   CE1-   HE1
   929    937    928    925       1   180.00   4.60240   2 ;     CD1-
CD2-    CG-    CB
   931    935    933    934       1   180.00   4.60240   2 ;     CE1-
CE2-    CZ-    HZ
   933    937    935    936       1   180.00   4.60240   2 ;      CZ-
CD2-   CE2-   HE2
   939    943    941    942       1   180.00   4.60240   2 ;       C-
CA-     N-     H
   943    958    956    957       1   180.00  43.93200   2 ;      CA-
N-     C-      O
   948    953    951    952       1   180.00  43.93200   2 ;      CG-
NE2-    CD-   OE1
   951    954    953    955       1   180.00   4.18400   2 ;      CD-
HE21-   NE2-  HE22
   956    960    958    959       1   180.00   4.60240   2 ;       C-
CA-     N-     H
   960    973    971    972       1   180.00  43.93200   2 ;      CA-
N-     C-      O
   965    969    968    970       1   180.00  43.93200   2 ;      CG-
OE1-    CD-   OE2
   971    975    973    974       1   180.00   4.60240   2 ;       C-
CA-     N-     H
```

975 992 990 991 1 180.00 43.93200 2 ; CA-
N- C- O
990 994 992 993 1 180.00 4.60240 2 ; C-
CA- N- H
994 1007 1005 1006 1 180.00 43.93200 2 ; CA-
N- C- O
999 1003 1002 1004 1 180.00 43.93200 2 ; CG-
OE1- CD- OE2
1005 1009 1007 1008 1 180.00 4.60240 2 ; C-
CA- N- H
1009 1026 1024 1025 1 180.00 43.93200 2 ; CA-
N- C- O
1024 1028 1026 1027 1 180.00 4.60240 2 ; C-
CA- N- H
1028 1043 1041 1042 1 180.00 43.93200 2 ; CA-
N- C- O
1041 1045 1043 1044 1 180.00 4.60240 2 ; C-
CA- N- H
1045 1065 1063 1064 1 180.00 43.93200 2 ; CA-
N- C- O
1063 1067 1065 1066 1 180.00 4.60240 2 ; C-
CA- N- H
1067 1087 1085 1086 1 180.00 43.93200 2 ; CA-
N- C- O
1085 1089 1087 1088 1 180.00 4.60240 2 ; C-
CA- N- H
1089 1106 1104 1105 1 180.00 43.93200 2 ; CA-
N- C- O
1104 1108 1106 1107 1 180.00 4.60240 2 ; C-
CA- N- H
1108 1118 1116 1117 1 180.00 43.93200 2 ; CA-
N- C- O
1110 1114 1113 1115 1 180.00 43.93200 2 ; CB-
OD1- CG- OD2
1116 1120 1118 1119 1 180.00 4.60240 2 ; C-
CA- N- H
1120 1135 1133 1134 1 180.00 43.93200 2 ; CA-
N- C- O
1122 1131 1125 1126 1 180.00 4.60240 2 ; CB-
CD2- CG- ND1
1125 1129 1131 1132 1 180.00 4.60240 2 ; CG-
NE2- CD2- HD2
1126 1129 1127 1128 1 180.00 4.60240 2 ; ND1-
NE2- CE1- HE1
1127 1131 1129 1130 1 180.00 4.18400 2 ; CE1-
CD2- NE2- HE2
1133 1136 1135 1145 1 180.00 4.18400 2 ; C-
CD- N- CA
1145 1149 1147 1148 1 180.00 43.93200 2 ; CA-
N- C- O
1147 1151 1149 1150 1 180.00 4.60240 2 ; C-
CA- N- H

1151 1163 1161 1162 1 180.00 43.93200 2 ; CA-
N- C- O
1153 1158 1156 1157 1 180.00 43.93200 2 ; CB-
ND2- CG- OD1
1156 1159 1158 1160 1 180.00 4.18400 2 ; CG-
HD21- ND2- HD22
1161 1165 1163 1164 1 180.00 4.60240 2 ; C-
CA- N- H
1165 1182 1180 1181 1 180.00 43.93200 2 ; CA-
N- C- O
1180 1184 1182 1183 1 180.00 4.60240 2 ; C-
CA- N- H
1184 1198 1196 1197 1 180.00 43.93200 2 ; CA-
N- C- O
1196 1200 1198 1199 1 180.00 4.60240 2 ; C-
CA- N- H
1200 1220 1218 1219 1 180.00 43.93200 2 ; CA-
N- C- O
1218 1222 1220 1221 1 180.00 4.60240 2 ; C-
CA- N- H
1222 1239 1237 1238 1 180.00 43.93200 2 ; CA-
N- C- O
1237 1241 1239 1240 1 180.00 4.60240 2 ; C-
CA- N- H
1241 1260 1258 1259 1 180.00 43.93200 2 ; CA-
N- C- O
1246 1249 1247 1248 1 180.00 4.60240 2 ; CG-
CE1- CD1- HD1
1246 1254 1256 1257 1 180.00 4.60240 2 ; CG-
CE2- CD2- HD2
1247 1251 1249 1250 1 180.00 4.60240 2 ; CD1-
CZ- CE1- HE1
1247 1256 1246 1243 1 180.00 4.60240 2 ; CD1-
CD2- CG- CB
1249 1254 1251 1252 1 180.00 4.60240 2 ; CE1-
CE2- CZ- OH
1251 1256 1254 1255 1 180.00 4.60240 2 ; CZ-
CD2- CE2- HE2
1258 1262 1260 1261 1 180.00 4.60240 2 ; C-
CA- N- H
1262 1275 1273 1274 1 180.00 43.93200 2 ; CA-
N- C- O
1267 1271 1270 1272 1 180.00 43.93200 2 ; CG-
OE1- CD- OE2
1273 1277 1275 1276 1 180.00 4.60240 2 ; C-
CA- N- H
1277 1289 1287 1288 1 180.00 43.93200 2 ; CA-
N- C- O
1287 1291 1289 1290 1 180.00 4.60240 2 ; C-
CA- N- H
1291 1310 1308 1309 1 180.00 43.93200 2 ; CA-
N- C- O

1296 1299 1297 1298 1 180.00 4.60240 2 ; CG-
CE1- CD1- HD1
1296 1304 1306 1307 1 180.00 4.60240 2 ; CG-
CE2- CD2- HD2
1297 1301 1299 1300 1 180.00 4.60240 2 ; CD1-
CZ- CE1- HE1
1297 1306 1296 1293 1 180.00 4.60240 2 ; CD1-
CD2- CG- CB
1299 1304 1301 1302 1 180.00 4.60240 2 ; CE1-
CE2- CZ- OH
1301 1306 1304 1305 1 180.00 4.60240 2 ; CZ-
CD2- CE2- HE2
1308 1312 1310 1311 1 180.00 4.60240 2 ; C-
CA- N- H
1312 1325 1323 1324 1 180.00 43.93200 2 ; CA-
N- C- O
1317 1321 1320 1322 1 180.00 43.93200 2 ; CG-
OE1- CD- OE2
1323 1327 1325 1326 1 180.00 4.60240 2 ; C-
CA- N- H
1327 1339 1337 1338 1 180.00 43.93200 2 ; CA-
N- C- O
1329 1334 1332 1333 1 180.00 43.93200 2 ; CB-
ND2- CG- OD1
1332 1335 1334 1336 1 180.00 4.18400 2 ; CG-
HD21- ND2- HD22
1337 1341 1339 1340 1 180.00 4.60240 2 ; C-
CA- N- H
1341 1351 1349 1350 1 180.00 43.93200 2 ; CA-
N- C- O
1343 1347 1346 1348 1 180.00 43.93200 2 ; CB-
OD1- CG- OD2
1349 1353 1351 1352 1 180.00 4.60240 2 ; C-
CA- N- H
1353 1365 1363 1364 1 180.00 43.93200 2 ; CA-
N- C- O
1355 1360 1358 1359 1 180.00 43.93200 2 ; CB-
ND2- CG- OD1
1358 1361 1360 1362 1 180.00 4.18400 2 ; CG-
HD21- ND2- HD22
1363 1367 1365 1366 1 180.00 4.60240 2 ; C-
CA- N- H
1367 1386 1384 1385 1 180.00 43.93200 2 ; CA-
N- C- O
1372 1375 1373 1374 1 180.00 4.60240 2 ; CG-
CE1- CD1- HD1
1372 1380 1382 1383 1 180.00 4.60240 2 ; CG-
CE2- CD2- HD2
1373 1377 1375 1376 1 180.00 4.60240 2 ; CD1-
CZ- CE1- HE1
1373 1382 1372 1369 1 180.00 4.60240 2 ; CD1-
CD2- CG- CB

1375 1380 1377 1378 1 180.00 4.60240 2 ; CE1-
CE2- CZ- OH
1377 1382 1380 1381 1 180.00 4.60240 2 ; CZ-
CD2- CE2- HE2
1384 1388 1386 1387 1 180.00 4.60240 2 ; C-
CA- N- H
1388 1405 1403 1404 1 180.00 43.93200 2 ; CA-
N- C- O
1403 1407 1405 1406 1 180.00 4.60240 2 ; C-
CA- N- H
1407 1426 1424 1425 1 180.00 43.93200 2 ; CA-
N- C- O
1412 1415 1413 1414 1 180.00 4.60240 2 ; CG-
CE1- CD1- HD1
1412 1420 1422 1423 1 180.00 4.60240 2 ; CG-
CE2- CD2- HD2
1413 1417 1415 1416 1 180.00 4.60240 2 ; CD1-
CZ- CE1- HE1
1413 1422 1412 1409 1 180.00 4.60240 2 ; CD1-
CD2- CG- CB
1415 1420 1417 1418 1 180.00 4.60240 2 ; CE1-
CE2- CZ- OH
1417 1422 1420 1421 1 180.00 4.60240 2 ; CZ-
CD2- CE2- HE2
1424 1428 1426 1427 1 180.00 4.60240 2 ; C-
CA- N- H
1428 1445 1443 1444 1 180.00 43.93200 2 ; CA-
N- C- O
1443 1447 1445 1446 1 180.00 4.60240 2 ; C-
CA- N- H
1447 1464 1462 1463 1 180.00 43.93200 2 ; CA-
N- C- O
1462 1466 1464 1465 1 180.00 4.60240 2 ; C-
CA- N- H
1466 1481 1479 1480 1 180.00 43.93200 2 ; CA-
N- C- O
1479 1483 1481 1482 1 180.00 4.60240 2 ; C-
CA- N- H
1483 1496 1494 1495 1 180.00 43.93200 2 ; CA-
N- C- O
1488 1492 1491 1493 1 180.00 43.93200 2 ; CG-
OE1- CD- OE2
1494 1498 1496 1497 1 180.00 4.60240 2 ; C-
CA- N- H
1498 1515 1513 1514 1 180.00 43.93200 2 ; CA-
N- C- O
1513 1517 1515 1516 1 180.00 4.60240 2 ; C-
CA- N- H
1517 1526 1524 1525 1 180.00 43.93200 2 ; CA-
N- C- O
1524 1528 1526 1527 1 180.00 4.60240 2 ; C-
CA- N- H

1528 1537 1535 1536 1 180.00 43.93200 2 ; CA-
N- C- O
1535 1539 1537 1538 1 180.00 4.60240 2 ; C-
CA- N- H
1539 1544 1542 1543 1 180.00 43.93200 2 ; CA-
N- C- O
1542 1546 1544 1545 1 180.00 4.60240 2 ; C-
CA- N- H
1546 1568 1566 1567 1 180.00 43.93200 2 ; CA-
N- C- O
1554 1559 1557 1558 1 180.00 4.18400 2 ; CD-
CZ- NE- HE
1557 1560 1559 1563 1 180.00 43.93200 2 ; NE-
NH1- CZ- NH2
1559 1561 1560 1562 1 180.00 4.18400 2 ; CZ-
HH11- NH1- HH12
1559 1564 1563 1565 1 180.00 4.18400 2 ; CZ-
HH21- NH2- HH22
1566 1570 1568 1569 1 180.00 4.60240 2 ; C-
CA- N- H
1570 1583 1581 1582 1 180.00 43.93200 2 ; CA-
N- C- O
1575 1579 1578 1580 1 180.00 43.93200 2 ; CG-
OE1- CD- OE2
1581 1585 1583 1584 1 180.00 4.60240 2 ; C-
CA- N- H
1585 1602 1600 1601 1 180.00 43.93200 2 ; CA-
N- C- O
1600 1604 1602 1603 1 180.00 4.60240 2 ; C-
CA- N- H
1604 1622 1620 1621 1 180.00 43.93200 2 ; CA-
N- C- O
1609 1612 1610 1611 1 180.00 4.60240 2 ; CG-
CE1- CD1- HD1
1609 1616 1618 1619 1 180.00 4.60240 2 ; CG-
CE2- CD2- HD2
1610 1614 1612 1613 1 180.00 4.60240 2 ; CD1-
CZ- CE1- HE1
1610 1618 1609 1606 1 180.00 4.60240 2 ; CD1-
CD2- CG- CB
1612 1616 1614 1615 1 180.00 4.60240 2 ; CE1-
CE2- CZ- HZ
1614 1618 1616 1617 1 180.00 4.60240 2 ; CZ-
CD2- CE2- HE2
1620 1624 1622 1623 1 180.00 4.60240 2 ; C-
CA- N- H
1624 1634 1632 1633 1 180.00 43.93200 2 ; CA-
N- C- O
1626 1630 1629 1631 1 180.00 43.93200 2 ; CB-
OD1- CG- OD2
1632 1636 1634 1635 1 180.00 4.60240 2 ; C-
CA- N- H

```
  1636   1645   1643   1644       1   180.00  43.93200   2 ;      CA-
N-     C-     O
  1643   1647   1645   1646       1   180.00   4.60240   2 ;       C-
CA-     N-     H
  1647   1664   1662   1663       1   180.00  43.93200   2 ;      CA-
N-     C-     O
  1662   1666   1664   1665       1   180.00   4.60240   2 ;       C-
CA-     N-     H
  1666   1683   1681   1682       1   180.00  43.93200   2 ;      CA-
N-     C-     O
  1681   1685   1683   1684       1   180.00   4.60240   2 ;       C-
CA-     N-     H
  1685   1698   1696   1697       1   180.00  43.93200   2 ;      CA-
N-     C-     O
  1690   1694   1693   1695       1   180.00  43.93200   2 ;      CG-
OE1-    CD-   OE2
  1696   1700   1698   1699       1   180.00   4.60240   2 ;       C-
CA-     N-     H
  1700   1712   1710   1711       1   180.00  43.93200   2 ;      CA-
N-     C-     O
  1702   1707   1705   1706       1   180.00  43.93200   2 ;      CB-
ND2-    CG-   OD1
  1705   1708   1707   1709       1   180.00   4.18400   2 ;      CG-
HD21-   ND2-  HD22
  1710   1714   1712   1713       1   180.00   4.60240   2 ;       C-
CA-     N-     H
  1714   1719   1717   1718       1   180.00  43.93200   2 ;      CA-
N-     C-     O
  1717   1721   1719   1720       1   180.00   4.60240   2 ;       C-
CA-     N-     H
  1721   1730   1728   1729       1   180.00  43.93200   2 ;      CA-
N-     C-     O
  1728   1732   1730   1731       1   180.00   4.60240   2 ;       C-
CA-     N-     H
  1732   1750   1748   1749       1   180.00  43.93200   2 ;      CA-
N-     C-     O
  1737   1740   1738   1739       1   180.00   4.60240   2 ;      CG-
CE1-   CD1-   HD1
  1737   1744   1746   1747       1   180.00   4.60240   2 ;      CG-
CE2-   CD2-   HD2
  1738   1742   1740   1741       1   180.00   4.60240   2 ;     CD1-
CZ-   CE1-   HE1
  1738   1746   1737   1734       1   180.00   4.60240   2 ;     CD1-
CD2-    CG-    CB
  1740   1744   1742   1743       1   180.00   4.60240   2 ;     CE1-
CE2-    CZ-    HZ
  1742   1746   1744   1745       1   180.00   4.60240   2 ;      CZ-
CD2-   CE2-   HE2
  1748   1752   1750   1751       1   180.00   4.60240   2 ;       C-
CA-     N-     H
  1752   1764   1762   1763       1   180.00  43.93200   2 ;      CA-
N-     C-     O
```

1762 1766 1764 1765 1 180.00 4.60240 2 ; C-
CA- N- H
1766 1779 1777 1778 1 180.00 43.93200 2 ; CA-
N- C- O
1771 1775 1774 1776 1 180.00 43.93200 2 ; CG-
OE1- CD- OE2
1777 1781 1779 1780 1 180.00 4.60240 2 ; C-
CA- N- H
1781 1801 1799 1800 1 180.00 43.93200 2 ; CA-
N- C- O
1799 1803 1801 1802 1 180.00 4.60240 2 ; C-
CA- N- H
1803 1815 1813 1814 1 180.00 43.93200 2 ; CA-
N- C- O
1805 1810 1808 1809 1 180.00 43.93200 2 ; CB-
ND2- CG- OD1
1808 1811 1810 1812 1 180.00 4.18400 2 ; CG-
HD21- ND2- HD22
1813 1817 1815 1816 1 180.00 4.60240 2 ; C-
CA- N- H
1817 1825 1823 1824 1 180.00 43.93200 2 ; CA-
N- C- O
1823 1827 1825 1826 1 180.00 4.60240 2 ; C-
CA- N- H
1827 1835 1833 1834 1 180.00 43.93200 2 ; CA-
N- C- O
1833 1837 1835 1836 1 180.00 4.60240 2 ; C-
CA- N- H
1837 1849 1847 1848 1 180.00 43.93200 2 ; CA-
N- C- O
1847 1851 1849 1850 1 180.00 4.60240 2 ; C-
CA- N- H
1851 1868 1866 1867 1 180.00 43.93200 2 ; CA-
N- C- O
1866 1870 1868 1869 1 180.00 4.60240 2 ; C-
CA- N- H
1870 1885 1883 1884 1 180.00 43.93200 2 ; CA-
N- C- O
1883 1887 1885 1886 1 180.00 4.60240 2 ; C-
CA- N- H
1887 1907 1905 1906 1 180.00 43.93200 2 ; CA-
N- C- O
1905 1909 1907 1908 1 180.00 4.60240 2 ; C-
CA- N- H
1909 1924 1922 1923 1 180.00 43.93200 2 ; CA-
N- C- O
1914 1919 1917 1918 1 180.00 43.93200 2 ; CG-
NE2- CD- OE1
1917 1920 1919 1921 1 180.00 4.18400 2 ; CD-
HE21- NE2- HE22
1922 1926 1924 1925 1 180.00 4.60240 2 ; C-
CA- N- H

1926 1943 1941 1942 1 180.00 43.93200 2 ; CA-
N- C- O
1941 1945 1943 1944 1 180.00 4.60240 2 ; C-
CA- N- H
1945 1963 1961 1962 1 180.00 43.93200 2 ; CA-
N- C- O
1950 1953 1951 1952 1 180.00 4.60240 2 ; CG-
CE1- CD1- HD1
1950 1957 1959 1960 1 180.00 4.60240 2 ; CG-
CE2- CD2- HD2
1951 1955 1953 1954 1 180.00 4.60240 2 ; CD1-
CZ- CE1- HE1
1951 1959 1950 1947 1 180.00 4.60240 2 ; CD1-
CD2- CG- CB
1953 1957 1955 1956 1 180.00 4.60240 2 ; CE1-
CE2- CZ- HZ
1955 1959 1957 1958 1 180.00 4.60240 2 ; CZ-
CD2- CE2- HE2
1961 1965 1963 1964 1 180.00 4.60240 2 ; C-
CA- N- H
1965 1974 1972 1973 1 180.00 43.93200 2 ; CA-
N- C- O
1972 1976 1974 1975 1 180.00 4.60240 2 ; C-
CA- N- H
1976 1984 1982 1983 1 180.00 43.93200 2 ; CA-
N- C- O
1982 1986 1984 1985 1 180.00 4.60240 2 ; C-
CA- N- H
1986 2003 2001 2002 1 180.00 43.93200 2 ; CA-
N- C- O
2001 2005 2003 2004 1 180.00 4.60240 2 ; C-
CA- N- H
2005 2023 2021 2022 1 180.00 43.93200 2 ; CA-
N- C- O
2010 2013 2011 2012 1 180.00 4.60240 2 ; CG-
CE1- CD1- HD1
2010 2017 2019 2020 1 180.00 4.60240 2 ; CG-
CE2- CD2- HD2
2011 2015 2013 2014 1 180.00 4.60240 2 ; CD1-
CZ- CE1- HE1
2011 2019 2010 2007 1 180.00 4.60240 2 ; CD1-
CD2- CG- CB
2013 2017 2015 2016 1 180.00 4.60240 2 ; CE1-
CE2- CZ- HZ
2015 2019 2017 2018 1 180.00 4.60240 2 ; CZ-
CD2- CE2- HE2
2021 2025 2023 2024 1 180.00 4.60240 2 ; C-
CA- N- H
2025 2044 2042 2043 1 180.00 43.93200 2 ; CA-
N- C- O
2030 2033 2031 2032 1 180.00 4.60240 2 ; CG-
CE1- CD1- HD1

2030 2038 2040 2041 1 180.00 4.60240 2 ; CG-
CE2- CD2- HD2
2031 2035 2033 2034 1 180.00 4.60240 2 ; CD1-
CZ- CE1- HE1
2031 2040 2030 2027 1 180.00 4.60240 2 ; CD1-
CD2- CG- CB
2033 2038 2035 2036 1 180.00 4.60240 2 ; CE1-
CE2- CZ- OH
2035 2040 2038 2039 1 180.00 4.60240 2 ; CZ-
CD2- CE2- HE2
2042 2046 2044 2045 1 180.00 4.60240 2 ; C-
CA- N- H
2046 2063 2061 2062 1 180.00 43.93200 2 ; CA-
N- C- O
2061 2065 2063 2064 1 180.00 4.60240 2 ; C-
CA- N- H
2065 2081 2079 2080 1 180.00 43.93200 2 ; CA-
N- C- O
2067 2077 2070 2071 1 180.00 4.60240 2 ; CB-
CD2- CG- ND1
2070 2073 2071 2072 1 180.00 4.18400 2 ; CG-
CE1- ND1- HD1
2070 2075 2077 2078 1 180.00 4.60240 2 ; CG-
NE2- CD2- HD2
2071 2075 2073 2074 1 180.00 4.60240 2 ; ND1-
NE2- CE1- HE1
2073 2077 2075 2076 1 180.00 4.18400 2 ; CE1-
CD2- NE2- HE2
2079 2083 2081 2082 1 180.00 4.60240 2 ; C-
CA- N- H
2083 2092 2090 2091 1 180.00 43.93200 2 ; CA-
N- C- O
2090 2094 2092 2093 1 180.00 4.60240 2 ; C-
CA- N- H
2094 2111 2109 2110 1 180.00 43.93200 2 ; CA-
N- C- O
2109 2113 2111 2112 1 180.00 4.60240 2 ; C-
CA- N- H
2113 2125 2123 2124 1 180.00 43.93200 2 ; CA-
N- C- O
2115 2120 2118 2119 1 180.00 43.93200 2 ; CB-
ND2- CG- OD1
2118 2121 2120 2122 1 180.00 4.18400 2 ; CG-
HD21- ND2- HD22
2123 2127 2125 2126 1 180.00 4.60240 2 ; C-
CA- N- H
2127 2144 2142 2143 1 180.00 43.93200 2 ; CA-
N- C- O
2142 2146 2144 2145 1 180.00 4.60240 2 ; C-
CA- N- H
2146 2160 2158 2159 1 180.00 43.93200 2 ; CA-
N- C- O

```
  2158   2162   2160   2161       1   180.00   4.60240   2 ;      C-
CA-     N-     H
  2162   2177   2175   2176       1   180.00  43.93200   2 ;     CA-
N-      C-     O
  2164   2173   2167   2168       1   180.00   4.60240   2 ;     CB-
CD2-    CG-   ND1
  2167   2171   2173   2174       1   180.00   4.60240   2 ;     CG-
NE2-   CD2-   HD2
  2168   2171   2169   2170       1   180.00   4.60240   2 ;    ND1-
NE2-   CE1-   HE1
  2169   2173   2171   2172       1   180.00   4.18400   2 ;    CE1-
CD2-   NE2-   HE2
  2175   2179   2177   2178       1   180.00   4.60240   2 ;      C-
CA-     N-     H
  2179   2201   2199   2200       1   180.00  43.93200   2 ;     CA-
N-      C-     O
  2187   2192   2190   2191       1   180.00   4.18400   2 ;     CD-
CZ-    NE-    HE
  2190   2193   2192   2196       1   180.00  43.93200   2 ;     NE-
NH1-    CZ-   NH2
  2192   2194   2193   2195       1   180.00   4.18400   2 ;     CZ-
HH11-   NH1-  HH12
  2192   2197   2196   2198       1   180.00   4.18400   2 ;     CZ-
HH21-   NH2-  HH22
  2199   2203   2201   2202       1   180.00   4.60240   2 ;      C-
CA-     N-     H
  2203   2213   2211   2212       1   180.00  43.93200   2 ;     CA-
N-      C-     O
  2205   2209   2208   2210       1   180.00  43.93200   2 ;     CB-
OD1-    CG-   OD2
  2211   2215   2213   2214       1   180.00   4.60240   2 ;      C-
CA-     N-     H
  2215   2232   2230   2231       1   180.00  43.93200   2 ;     CA-
N-      C-     O
  2230   2234   2232   2233       1   180.00   4.60240   2 ;      C-
CA-     N-     H
  2234   2254   2252   2253       1   180.00  43.93200   2 ;     CA-
N-      C-     O
  2252   2255   2254   2264       1   180.00   4.18400   2 ;      C-
CD-     N-    CA
  2264   2268   2266   2267       1   180.00  43.93200   2 ;     CA-
N-      C-     O
  2266   2270   2268   2269       1   180.00   4.60240   2 ;      C-
CA-     N-     H
  2270   2283   2281   2282       1   180.00  43.93200   2 ;     CA-
N-      C-     O
  2275   2279   2278   2280       1   180.00  43.93200   2 ;     CG-
OE1-    CD-   OE2
  2281   2285   2283   2284       1   180.00   4.60240   2 ;      C-
CA-     N-     H
  2285   2297   2295   2296       1   180.00  43.93200   2 ;     CA-
N-      C-     O
```

2287 2292 2290 2291 1 180.00 43.93200 2 ; CB-
ND2- CG- OD1
2290 2293 2292 2294 1 180.00 4.18400 2 ; CG-
HD21- ND2- HD22
2295 2299 2297 2298 1 180.00 4.60240 2 ; C-
CA- N- H
2299 2317 2315 2316 1 180.00 43.93200 2 ; CA-
N- C- O
2304 2307 2305 2306 1 180.00 4.60240 2 ; CG-
CE1- CD1- HD1
2304 2311 2313 2314 1 180.00 4.60240 2 ; CG-
CE2- CD2- HD2
2305 2309 2307 2308 1 180.00 4.60240 2 ; CD1-
CZ- CE1- HE1
2305 2313 2304 2301 1 180.00 4.60240 2 ; CD1-
CD2- CG- CB
2307 2311 2309 2310 1 180.00 4.60240 2 ; CE1-
CE2- CZ- HZ
2309 2313 2311 2312 1 180.00 4.60240 2 ; CZ-
CD2- CE2- HE2
2315 2319 2317 2318 1 180.00 4.60240 2 ; C-
CA- N- H
2319 2336 2334 2335 1 180.00 43.93200 2 ; CA-
N- C- O
2334 2338 2336 2337 1 180.00 4.60240 2 ; C-
CA- N- H
2338 2356 2354 2355 1 180.00 43.93200 2 ; CA-
N- C- O
2343 2346 2344 2345 1 180.00 4.60240 2 ; CG-
CE1- CD1- HD1
2343 2350 2352 2353 1 180.00 4.60240 2 ; CG-
CE2- CD2- HD2
2344 2348 2346 2347 1 180.00 4.60240 2 ; CD1-
CZ- CE1- HE1
2344 2352 2343 2340 1 180.00 4.60240 2 ; CD1-
CD2- CG- CB
2346 2350 2348 2349 1 180.00 4.60240 2 ; CE1-
CE2- CZ- HZ
2348 2352 2350 2351 1 180.00 4.60240 2 ; CZ-
CD2- CE2- HE2
2354 2358 2356 2357 1 180.00 4.60240 2 ; C-
CA- N- H
2358 2373 2371 2372 1 180.00 43.93200 2 ; CA-
N- C- O
2363 2368 2366 2367 1 180.00 43.93200 2 ; CG-
NE2- CD- OE1
2366 2369 2368 2370 1 180.00 4.18400 2 ; CD-
HE21- NE2- HE22
2371 2375 2373 2374 1 180.00 4.60240 2 ; C-
CA- N- H
2375 2384 2382 2383 1 180.00 43.93200 2 ; CA-
N- C- O

2382 2386 2384 2385 1 180.00 4.60240 2 ; C-
CA- N- H
2386 2399 2397 2398 1 180.00 43.93200 2 ; CA-
N- C- O
2391 2395 2394 2396 1 180.00 43.93200 2 ; CG-
OE1- CD- OE2
2397 2401 2399 2400 1 180.00 4.60240 2 ; C-
CA- N- H
2401 2413 2411 2412 1 180.00 43.93200 2 ; CA-
N- C- O
2403 2408 2406 2407 1 180.00 43.93200 2 ; CB-
ND2- CG- OD1
2406 2409 2408 2410 1 180.00 4.18400 2 ; CG-
HD21- ND2- HD22
2411 2415 2413 2414 1 180.00 4.60240 2 ; C-
CA- N- H
2415 2435 2433 2434 1 180.00 43.93200 2 ; CA-
N- C- O
2433 2437 2435 2436 1 180.00 4.60240 2 ; C-
CA- N- H
2437 2447 2445 2446 1 180.00 43.93200 2 ; CA-
N- C- O
2439 2443 2442 2444 1 180.00 43.93200 2 ; CB-
OD1- CG- OD2
2445 2449 2447 2448 1 180.00 4.60240 2 ; C-
CA- N- H
2449 2458 2456 2457 1 180.00 43.93200 2 ; CA-
N- C- O
2456 2460 2458 2459 1 180.00 4.60240 2 ; C-
CA- N- H
2460 2477 2475 2476 1 180.00 43.93200 2 ; CA-
N- C- O
2475 2479 2477 2478 1 180.00 4.60240 2 ; C-
CA- N- H
2479 2496 2494 2495 1 180.00 43.93200 2 ; CA-
N- C- O
2494 2498 2496 2497 1 180.00 4.60240 2 ; C-
CA- N- H
2498 2518 2516 2517 1 180.00 43.93200 2 ; CA-
N- C- O
2516 2520 2518 2519 1 180.00 4.60240 2 ; C-
CA- N- H
2520 2537 2535 2536 1 180.00 43.93200 2 ; CA-
N- C- O
2535 2539 2537 2538 1 180.00 4.60240 2 ; C-
CA- N- H
2539 2556 2554 2555 1 180.00 43.93200 2 ; CA-
N- C- O
2554 2558 2556 2557 1 180.00 4.60240 2 ; C-
CA- N- H
2558 2568 2566 2567 1 180.00 43.93200 2 ; CA-
N- C- O

2560 2564 2563 2565 1 180.00 43.93200 2 ; CB-
OD1- CG- OD2
2566 2570 2568 2569 1 180.00 4.60240 2 ; C-
CA- N- H
2570 2588 2586 2587 1 180.00 43.93200 2 ; CA-
N- C- O
2575 2578 2576 2577 1 180.00 4.60240 2 ; CG-
CE1- CD1- HD1
2575 2582 2584 2585 1 180.00 4.60240 2 ; CG-
CE2- CD2- HD2
2576 2580 2578 2579 1 180.00 4.60240 2 ; CD1-
CZ- CE1- HE1
2576 2584 2575 2572 1 180.00 4.60240 2 ; CD1-
CD2- CG- CB
2578 2582 2580 2581 1 180.00 4.60240 2 ; CE1-
CE2- CZ- HZ
2580 2584 2582 2583 1 180.00 4.60240 2 ; CZ-
CD2- CE2- HE2
2586 2590 2588 2589 1 180.00 4.60240 2 ; C-
CA- N- H
2590 2595 2593 2594 1 180.00 43.93200 2 ; CA-
N- C- O
2593 2597 2595 2596 1 180.00 4.60240 2 ; C-
CA- N- H
2597 2614 2612 2613 1 180.00 43.93200 2 ; CA-
N- C- O
2612 2616 2614 2615 1 180.00 4.60240 2 ; C-
CA- N- H
2616 2625 2623 2624 1 180.00 43.93200 2 ; CA-
N- C- O
2623 2627 2625 2626 1 180.00 4.60240 2 ; C-
CA- N- H
2627 2647 2645 2646 1 180.00 43.93200 2 ; CA-
N- C- O
2645 2649 2647 2648 1 180.00 4.60240 2 ; C-
CA- N- H
2649 2661 2659 2660 1 180.00 43.93200 2 ; CA-
N- C- O
2651 2656 2654 2655 1 180.00 43.93200 2 ; CB-
ND2- CG- OD1
2654 2657 2656 2658 1 180.00 4.18400 2 ; CG-
HD21- ND2- HD22
2659 2663 2661 2662 1 180.00 4.60240 2 ; C-
CA- N- H
2663 2680 2678 2679 1 180.00 43.93200 2 ; CA-
N- C- O
2678 2682 2680 2681 1 180.00 4.60240 2 ; C-
CA- N- H
2682 2687 2685 2686 1 180.00 43.93200 2 ; CA-
N- C- O
2685 2689 2687 2688 1 180.00 4.60240 2 ; C-
CA- N- H

2689 2701 2699 2700 1 180.00 43.93200 2 ; CA-
N- C- O
2699 2703 2701 2702 1 180.00 4.60240 2 ; C-
CA- N- H
2703 2708 2706 2707 1 180.00 43.93200 2 ; CA-
N- C- O
2706 2710 2708 2709 1 180.00 4.60240 2 ; C-
CA- N- H
2710 2723 2721 2722 1 180.00 43.93200 2 ; CA-
N- C- O
2715 2719 2718 2720 1 180.00 43.93200 2 ; CG-
OE1- CD- OE2
2721 2725 2723 2724 1 180.00 4.60240 2 ; C-
CA- N- H
2725 2743 2741 2742 1 180.00 43.93200 2 ; CA-
N- C- O
2730 2733 2731 2732 1 180.00 4.60240 2 ; CG-
CE1- CD1- HD1
2730 2737 2739 2740 1 180.00 4.60240 2 ; CG-
CE2- CD2- HD2
2731 2735 2733 2734 1 180.00 4.60240 2 ; CD1-
CZ- CE1- HE1
2731 2739 2730 2727 1 180.00 4.60240 2 ; CD1-
CD2- CG- CB
2733 2737 2735 2736 1 180.00 4.60240 2 ; CE1-
CE2- CZ- HZ
2735 2739 2737 2738 1 180.00 4.60240 2 ; CZ-
CD2- CE2- HE2
2741 2745 2743 2744 1 180.00 4.60240 2 ; C-
CA- N- H
2745 2757 2755 2756 1 180.00 43.93200 2 ; CA-
N- C- O
2755 2759 2757 2758 1 180.00 4.60240 2 ; C-
CA- N- H
2759 2771 2769 2770 1 180.00 43.93200 2 ; CA-
N- C- O
2769 2773 2771 2772 1 180.00 4.60240 2 ; C-
CA- N- H
2773 2785 2783 2784 1 180.00 43.93200 2 ; CA-
N- C- O
2783 2787 2785 2786 1 180.00 4.60240 2 ; C-
CA- N- H
2787 2807 2805 2806 1 180.00 43.93200 2 ; CA-
N- C- O
2805 2809 2807 2808 1 180.00 4.60240 2 ; C-
CA- N- H
2809 2817 2815 2816 1 180.00 43.93200 2 ; CA-
N- C- O
2815 2819 2817 2818 1 180.00 4.60240 2 ; C-
CA- N- H
2819 2824 2822 2823 1 180.00 43.93200 2 ; CA-
N- C- O

2822 2826 2824 2825 1 180.00 4.60240 2 ; C-
CA- N- H
2826 2838 2836 2837 1 180.00 43.93200 2 ; CA-
N- C- O
2836 2839 2838 2848 1 180.00 4.18400 2 ; C-
CD- N- CA
2848 2852 2850 2851 1 180.00 43.93200 2 ; CA-
N- C- O
2850 2854 2852 2853 1 180.00 4.60240 2 ; C-
CA- N- H
2854 2873 2871 2872 1 180.00 43.93200 2 ; CA-
N- C- O
2859 2862 2860 2861 1 180.00 4.60240 2 ; CG-
CE1- CD1- HD1
2859 2867 2869 2870 1 180.00 4.60240 2 ; CG-
CE2- CD2- HD2
2860 2864 2862 2863 1 180.00 4.60240 2 ; CD1-
CZ- CE1- HE1
2860 2869 2859 2856 1 180.00 4.60240 2 ; CD1-
CD2- CG- CB
2862 2867 2864 2865 1 180.00 4.60240 2 ; CE1-
CE2- CZ- OH
2864 2869 2867 2868 1 180.00 4.60240 2 ; CZ-
CD2- CE2- HE2
2871 2875 2873 2874 1 180.00 4.60240 2 ; C-
CA- N- H
2875 2894 2892 2893 1 180.00 43.93200 2 ; CA-
N- C- O
2880 2883 2881 2882 1 180.00 4.60240 2 ; CG-
CE1- CD1- HD1
2880 2888 2890 2891 1 180.00 4.60240 2 ; CG-
CE2- CD2- HD2
2881 2885 2883 2884 1 180.00 4.60240 2 ; CD1-
CZ- CE1- HE1
2881 2890 2880 2877 1 180.00 4.60240 2 ; CD1-
CD2- CG- CB
2883 2888 2885 2886 1 180.00 4.60240 2 ; CE1-
CE2- CZ- OH
2885 2890 2888 2889 1 180.00 4.60240 2 ; CZ-
CD2- CE2- HE2
2892 2896 2894 2895 1 180.00 4.60240 2 ; C-
CA- N- H
2896 2910 2908 2909 1 180.00 43.93200 2 ; CA-
N- C- O
2908 2912 2910 2911 1 180.00 4.60240 2 ; C-
CA- N- H
2912 2920 2918 2919 1 180.00 43.93200 2 ; CA-
N- C- O
2918 2921 2920 2930 1 180.00 4.18400 2 ; C-
CD- N- CA
2930 2934 2932 2933 1 180.00 43.93200 2 ; CA-
N- C- O

2932 2936 2934 2935 1 180.00 4.60240 2 ; C-
CA- N- H
2936 2951 2949 2950 1 180.00 43.93200 2 ; CA-
N- C- O
2941 2946 2944 2945 1 180.00 43.93200 2 ; CG-
NE2- CD- OE1
2944 2947 2946 2948 1 180.00 4.18400 2 ; CD-
HE21- NE2- HE22
2949 2953 2951 2952 1 180.00 4.60240 2 ; C-
CA- N- H
2953 2967 2965 2966 1 180.00 43.93200 2 ; CA-
N- C- O
2965 2969 2967 2968 1 180.00 4.60240 2 ; C-
CA- N- H
2969 2986 2984 2985 1 180.00 43.93200 2 ; CA-
N- C- O
2984 2988 2986 2987 1 180.00 4.60240 2 ; C-
CA- N- H
2988 2998 2996 2997 1 180.00 43.93200 2 ; CA-
N- C- O
2990 2994 2993 2995 1 180.00 43.93200 2 ; CB-
OD1- CG- OD2
2996 3000 2998 2999 1 180.00 4.60240 2 ; C-
CA- N- H
3000 3005 3003 3004 1 180.00 43.93200 2 ; CA-
N- C- O
3003 3007 3005 3006 1 180.00 4.60240 2 ; C-
CA- N- H
3007 3027 3025 3026 1 180.00 43.93200 2 ; CA-
N- C- O
3025 3029 3027 3028 1 180.00 4.60240 2 ; C-
CA- N- H
3029 3048 3046 3047 1 180.00 43.93200 2 ; CA-
N- C- O
3034 3037 3035 3036 1 180.00 4.60240 2 ; CG-
CE1- CD1- HD1
3034 3042 3044 3045 1 180.00 4.60240 2 ; CG-
CE2- CD2- HD2
3035 3039 3037 3038 1 180.00 4.60240 2 ; CD1-
CZ- CE1- HE1
3035 3044 3034 3031 1 180.00 4.60240 2 ; CD1-
CD2- CG- CB
3037 3042 3039 3040 1 180.00 4.60240 2 ; CE1-
CE2- CZ- OH
3039 3044 3042 3043 1 180.00 4.60240 2 ; CZ-
CD2- CE2- HE2
3046 3050 3048 3049 1 180.00 4.60240 2 ; C-
CA- N- H
3050 3060 3058 3059 1 180.00 43.93200 2 ; CA-
N- C- O
3052 3056 3055 3057 1 180.00 43.93200 2 ; CB-
OD1- CG- OD2

3058 3062 3060 3061 1 180.00 4.60240 2 ; C-
CA- N- H
3062 3082 3080 3081 1 180.00 43.93200 2 ; CA-
N- C- O
3080 3084 3082 3083 1 180.00 4.60240 2 ; C-
CA- N- H
3084 3104 3102 3103 1 180.00 43.93200 2 ; CA-
N- C- O
3102 3106 3104 3105 1 180.00 4.60240 2 ; C-
CA- N- H
3106 3115 3113 3114 1 180.00 43.93200 2 ; CA-
N- C- O
3113 3117 3115 3116 1 180.00 4.60240 2 ; C-
CA- N- H
3117 3127 3125 3126 1 180.00 43.93200 2 ; CA-
N- C- O
3119 3123 3122 3124 1 180.00 43.93200 2 ; CB-
OD1- CG- OD2
3125 3129 3127 3128 1 180.00 4.60240 2 ; C-
CA- N- H
3129 3146 3144 3145 1 180.00 43.93200 2 ; CA-
N- C- O
3144 3148 3146 3147 1 180.00 4.60240 2 ; C-
CA- N- H
3148 3170 3168 3169 1 180.00 43.93200 2 ; CA-
N- C- O
3153 3156 3154 3155 1 180.00 4.60240 2 ; CG-
NE1- CD1- HD1
3154 3158 3156 3157 1 180.00 4.18400 2 ; CD1-
CE2- NE1- HE1
3158 3161 3159 3160 1 180.00 4.60240 2 ; CE2-
CH2- CZ2- HZ2
3159 3163 3161 3162 1 180.00 4.60240 2 ; CZ2-
CZ3- CH2- HH2
3161 3165 3163 3164 1 180.00 4.60240 2 ; CH2-
CE3- CZ3- HZ3
3163 3167 3165 3166 1 180.00 4.60240 2 ; CZ3-
CD2- CE3- HE3
3167 3150 3153 3154 1 180.00 4.60240 2 ; CD2-
CB- CG- CD1
3168 3172 3170 3171 1 180.00 4.60240 2 ; C-
CA- N- H
3172 3181 3179 3180 1 180.00 43.93200 2 ; CA-
N- C- O
3179 3183 3181 3182 1 180.00 4.60240 2 ; C-
CA- N- H
3183 3192 3190 3191 1 180.00 43.93200 2 ; CA-
N- C- O
3190 3194 3192 3193 1 180.00 4.60240 2 ; C-
CA- N- H
3194 3199 3197 3198 1 180.00 43.93200 2 ; CA-
N- C- O

3197 3201 3199 3200 1 180.00 4.60240 2 ; C-
CA- N- H
3201 3215 3213 3214 1 180.00 43.93200 2 ; CA-
N- C- O
3213 3217 3215 3216 1 180.00 4.60240 2 ; C-
CA- N- H
3217 3234 3232 3233 1 180.00 43.93200 2 ; CA-
N- C- O
3232 3236 3234 3235 1 180.00 4.60240 2 ; C-
CA- N- H
3236 3251 3249 3250 1 180.00 43.93200 2 ; CA-
N- C- O
3249 3253 3251 3252 1 180.00 4.60240 2 ; C-
CA- N- H
3253 3272 3270 3271 1 180.00 43.93200 2 ; CA-
N- C- O
3258 3261 3259 3260 1 180.00 4.60240 2 ; CG-
CE1- CD1- HD1
3258 3266 3268 3269 1 180.00 4.60240 2 ; CG-
CE2- CD2- HD2
3259 3263 3261 3262 1 180.00 4.60240 2 ; CD1-
CZ- CE1- HE1
3259 3268 3258 3255 1 180.00 4.60240 2 ; CD1-
CD2- CG- CB
3261 3266 3263 3264 1 180.00 4.60240 2 ; CE1-
CE2- CZ- OH
3263 3268 3266 3267 1 180.00 4.60240 2 ; CZ-
CD2- CE2- HE2
3270 3274 3272 3273 1 180.00 4.60240 2 ; C-
CA- N- H
3274 3286 3284 3285 1 180.00 43.93200 2 ; CA-
N- C- O
3284 3288 3286 3287 1 180.00 4.60240 2 ; C-
CA- N- H
3288 3305 3303 3304 1 180.00 43.93200 2 ; CA-
N- C- O
3303 3307 3305 3306 1 180.00 4.60240 2 ; C-
CA- N- H
3307 3324 3322 3323 1 180.00 43.93200 2 ; CA-
N- C- O
3322 3326 3324 3325 1 180.00 4.60240 2 ; C-
CA- N- H
3326 3335 3333 3334 1 180.00 43.93200 2 ; CA-
N- C- O
3333 3337 3335 3336 1 180.00 4.60240 2 ; C-
CA- N- H
3337 3342 3340 3341 1 180.00 43.93200 2 ; CA-
N- C- O
3340 3344 3342 3343 1 180.00 4.60240 2 ; C-
CA- N- H
3344 3363 3361 3362 1 180.00 43.93200 2 ; CA-
N- C- O

3349 3352 3350 3351 1 180.00 4.60240 2 ; CG-
CE1- CD1- HD1
3349 3357 3359 3360 1 180.00 4.60240 2 ; CG-
CE2- CD2- HD2
3350 3354 3352 3353 1 180.00 4.60240 2 ; CD1-
CZ- CE1- HE1
3350 3359 3349 3346 1 180.00 4.60240 2 ; CD1-
CD2- CG- CB
3352 3357 3354 3355 1 180.00 4.60240 2 ; CE1-
CE2- CZ- OH
3354 3359 3357 3358 1 180.00 4.60240 2 ; CZ-
CD2- CE2- HE2
3361 3364 3363 3373 1 180.00 4.18400 2 ; C-
CD- N- CA
3373 3377 3375 3376 1 180.00 43.93200 2 ; CA-
N- C- O
3375 3378 3377 3387 1 180.00 4.18400 2 ; C-
CD- N- CA
3387 3391 3389 3390 1 180.00 43.93200 2 ; CA-
N- C- O
3389 3393 3391 3392 1 180.00 4.60240 2 ; C-
CA- N- H
3393 3411 3409 3410 1 180.00 43.93200 2 ; CA-
N- C- O
3398 3401 3399 3400 1 180.00 4.60240 2 ; CG-
CE1- CD1- HD1
3398 3405 3407 3408 1 180.00 4.60240 2 ; CG-
CE2- CD2- HD2
3399 3403 3401 3402 1 180.00 4.60240 2 ; CD1-
CZ- CE1- HE1
3399 3407 3398 3395 1 180.00 4.60240 2 ; CD1-
CD2- CG- CB
3401 3405 3403 3404 1 180.00 4.60240 2 ; CE1-
CE2- CZ- HZ
3403 3407 3405 3406 1 180.00 4.60240 2 ; CZ-
CD2- CE2- HE2
3409 3413 3411 3412 1 180.00 4.60240 2 ; C-
CA- N- H
3413 3432 3430 3431 1 180.00 43.93200 2 ; CA-
N- C- O
3418 3421 3419 3420 1 180.00 4.60240 2 ; CG-
CE1- CD1- HD1
3418 3426 3428 3429 1 180.00 4.60240 2 ; CG-
CE2- CD2- HD2
3419 3423 3421 3422 1 180.00 4.60240 2 ; CD1-
CZ- CE1- HE1
3419 3428 3418 3415 1 180.00 4.60240 2 ; CD1-
CD2- CG- CB
3421 3426 3423 3424 1 180.00 4.60240 2 ; CE1-
CE2- CZ- OH
3423 3428 3426 3427 1 180.00 4.60240 2 ; CZ-
CD2- CE2- HE2

```
  3430   3434   3432   3433       1   180.00   4.60240   2 ;      C-
CA-     N-      H
  3434   3439   3437   3438       1   180.00  43.93200   2 ;     CA-
N-     C-      O
  3437   3441   3439   3440       1   180.00   4.60240   2 ;      C-
CA-     N-      H
  3441   3451   3449   3450       1   180.00  43.93200   2 ;     CA-
N-     C-      O
  3443   3447   3446   3448       1   180.00  43.93200   2 ;     CB-
OD1-    CG-   OD2
  3449   3453   3451   3452       1   180.00   4.60240   2 ;      C-
CA-     N-      H
  3453   3465   3463   3464       1   180.00  43.93200   2 ;     CA-
N-     C-      O
  3463   3467   3465   3466       1   180.00   4.60240   2 ;      C-
CA-     N-      H
  3467   3477   3475   3476       1   180.00  43.93200   2 ;     CA-
N-     C-      O
  3469   3473   3472   3474       1   180.00  43.93200   2 ;     CB-
OD1-    CG-   OD2
  3475   3479   3477   3478       1   180.00   4.60240   2 ;      C-
CA-     N-      H
  3479   3491   3489   3490       1   180.00  43.93200   2 ;     CA-
N-     C-      O
  3481   3486   3484   3485       1   180.00  43.93200   2 ;     CB-
ND2-    CG-   OD1
  3484   3487   3486   3488       1   180.00   4.18400   2 ;     CG-
HD21-   ND2-  HD22
  3489   3493   3491   3492       1   180.00   4.60240   2 ;      C-
CA-     N-      H
  3493   3506   3504   3505       1   180.00  43.93200   2 ;     CA-
N-     C-      O
  3498   3502   3501   3503       1   180.00  43.93200   2 ;     CG-
OE1-    CD-   OE2
  3504   3508   3506   3507       1   180.00   4.60240   2 ;      C-
CA-     N-      H
  3508   3522   3520   3521       1   180.00  43.93200   2 ;     CA-
N-     C-      O
  3520   3524   3522   3523       1   180.00   4.60240   2 ;      C-
CA-     N-      H
  3524   3541   3539   3540       1   180.00  43.93200   2 ;     CA-
N-     C-      O
  3539   3543   3541   3542       1   180.00   4.60240   2 ;      C-
CA-     N-      H
  3543   3563   3561   3562       1   180.00  43.93200   2 ;     CA-
N-     C-      O
  3561   3565   3563   3564       1   180.00   4.60240   2 ;      C-
CA-     N-      H
  3565   3585   3583   3584       1   180.00  43.93200   2 ;     CA-
N-     C-      O
  3583   3587   3585   3586       1   180.00   4.60240   2 ;      C-
CA-     N-      H
```

3587 3601 3599 3600 1 180.00 43.93200 2 ; CA-
N- C- O
3599 3603 3601 3602 1 180.00 4.60240 2 ; C-
CA- N- H
3603 3623 3621 3622 1 180.00 43.93200 2 ; CA-
N- C- O
3621 3625 3623 3624 1 180.00 4.60240 2 ; C-
CA- N- H
3625 3645 3643 3644 1 180.00 43.93200 2 ; CA-
N- C- O
3643 3647 3645 3646 1 180.00 4.60240 2 ; C-
CA- N- H
3647 3652 3650 3651 1 180.00 43.93200 2 ; CA-
N- C- O
3650 3654 3652 3653 1 180.00 4.60240 2 ; C-
CA- N- H
3654 3667 3665 3666 1 180.00 43.93200 2 ; CA-
N- C- O
3659 3663 3662 3664 1 180.00 43.93200 2 ; CG-
OE1- CD- OE2
3665 3669 3667 3668 1 180.00 4.60240 2 ; C-
CA- N- H
3669 3687 3685 3686 1 180.00 43.93200 2 ; CA-
N- C- O
3674 3677 3675 3676 1 180.00 4.60240 2 ; CG-
CE1- CD1- HD1
3674 3681 3683 3684 1 180.00 4.60240 2 ; CG-
CE2- CD2- HD2
3675 3679 3677 3678 1 180.00 4.60240 2 ; CD1-
CZ- CE1- HE1
3675 3683 3674 3671 1 180.00 4.60240 2 ; CD1-
CD2- CG- CB
3677 3681 3679 3680 1 180.00 4.60240 2 ; CE1-
CE2- CZ- HZ
3679 3683 3681 3682 1 180.00 4.60240 2 ; CZ-
CD2- CE2- HE2
3685 3689 3687 3688 1 180.00 4.60240 2 ; C-
CA- N- H
3689 3698 3696 3697 1 180.00 43.93200 2 ; CA-
N- C- O
3696 3700 3698 3699 1 180.00 4.60240 2 ; C-
CA- N- H
3700 3718 3716 3717 1 180.00 43.93200 2 ; CA-
N- C- O
3705 3708 3706 3707 1 180.00 4.60240 2 ; CG-
CE1- CD1- HD1
3705 3712 3714 3715 1 180.00 4.60240 2 ; CG-
CE2- CD2- HD2
3706 3710 3708 3709 1 180.00 4.60240 2 ; CD1-
CZ- CE1- HE1
3706 3714 3705 3702 1 180.00 4.60240 2 ; CD1-
CD2- CG- CB

3708 3712 3710 3711 1 180.00 4.60240 2 ; CE1-
CE2- CZ- HZ
3710 3714 3712 3713 1 180.00 4.60240 2 ; CZ-
CD2- CE2- HE2
3716 3720 3718 3719 1 180.00 4.60240 2 ; C-
CA- N- H
3720 3739 3737 3738 1 180.00 43.93200 2 ; CA-
N- C- O
3725 3728 3726 3727 1 180.00 4.60240 2 ; CG-
CE1- CD1- HD1
3725 3733 3735 3736 1 180.00 4.60240 2 ; CG-
CE2- CD2- HD2
3726 3730 3728 3729 1 180.00 4.60240 2 ; CD1-
CZ- CE1- HE1
3726 3735 3725 3722 1 180.00 4.60240 2 ; CD1-
CD2- CG- CB
3728 3733 3730 3731 1 180.00 4.60240 2 ; CE1-
CE2- CZ- OH
3730 3735 3733 3734 1 180.00 4.60240 2 ; CZ-
CD2- CE2- HE2
3737 3741 3739 3740 1 180.00 4.60240 2 ; C-
CA- N- H
3741 3754 3752 3753 1 180.00 43.93200 2 ; CA-
N- C- O
3746 3750 3749 3751 1 180.00 43.93200 2 ; CG-
OE1- CD- OE2
3752 3756 3754 3755 1 180.00 4.60240 2 ; C-
CA- N- H
3756 3768 3766 3767 1 180.00 43.93200 2 ; CA-
N- C- O
3758 3763 3761 3762 1 180.00 43.93200 2 ; CB-
ND2- CG- OD1
3761 3764 3763 3765 1 180.00 4.18400 2 ; CG-
HD21- ND2- HD22
3766 3770 3768 3769 1 180.00 4.60240 2 ; C-
CA- N- H
3770 3780 3778 3779 1 180.00 43.93200 2 ; CA-
N- C- O
3772 3776 3775 3777 1 180.00 43.93200 2 ; CB-
OD1- CG- OD2
3778 3782 3780 3781 1 180.00 4.60240 2 ; C-
CA- N- H
3782 3804 3802 3803 1 180.00 43.93200 2 ; CA-
N- C- O
3787 3790 3788 3789 1 180.00 4.60240 2 ; CG-
NE1- CD1- HD1
3788 3792 3790 3791 1 180.00 4.18400 2 ; CD1-
CE2- NE1- HE1
3792 3795 3793 3794 1 180.00 4.60240 2 ; CE2-
CH2- CZ2- HZ2
3793 3797 3795 3796 1 180.00 4.60240 2 ; CZ2-
CZ3- CH2- HH2

3795 3799 3797 3798 1 180.00 4.60240 2 ; CH2-
CE3- CZ3- HZ3
3797 3801 3799 3800 1 180.00 4.60240 2 ; CZ3-
CD2- CE3- HE3
3801 3784 3787 3788 1 180.00 4.60240 2 ; CD2-
CB- CG- CD1
3802 3806 3804 3805 1 180.00 4.60240 2 ; C-
CA- N- H
3806 3811 3809 3810 1 180.00 43.93200 2 ; CA-
N- C- O
3809 3813 3811 3812 1 180.00 4.60240 2 ; C-
CA- N- H
3813 3822 3820 3821 1 180.00 43.93200 2 ; CA-
N- C- O
3820 3824 3822 3823 1 180.00 4.60240 2 ; C-
CA- N- H
3824 3841 3839 3840 1 180.00 43.93200 2 ; CA-
N- C- O
3839 3843 3841 3842 1 180.00 4.60240 2 ; C-
CA- N- H
3843 3852 3850 3851 1 180.00 43.93200 2 ; CA-
N- C- O
3850 3854 3852 3853 1 180.00 4.60240 2 ; C-
CA- N- H
3854 3863 3861 3862 1 180.00 43.93200 2 ; CA-
N- C- O
3861 3865 3863 3864 1 180.00 4.60240 2 ; C-
CA- N- H
3865 3875 3873 3874 1 180.00 43.93200 2 ; CA-
N- C- O
3867 3871 3870 3872 1 180.00 43.93200 2 ; CB-
OD1- CG- OD2
3873 3877 3875 3876 1 180.00 4.60240 2 ; C-
CA- N- H
3877 3885 3883 3884 1 180.00 43.93200 2 ; CA-
N- C- O
3883 3887 3885 3886 1 180.00 4.60240 2 ; C-
CA- N- H
3887 3907 3905 3906 1 180.00 43.93200 2 ; CA-
N- C- O
3905 3909 3907 3908 1 180.00 4.60240 2 ; C-
CA- N- H
3909 3921 3919 3920 1 180.00 43.93200 2 ; CA-
N- C- O
3911 3916 3914 3915 1 180.00 43.93200 2 ; CB-
ND2- CG- OD1
3914 3917 3916 3918 1 180.00 4.18400 2 ; CG-
HD21- ND2- HD22
3919 3923 3921 3922 1 180.00 4.60240 2 ; C-
CA- N- H
3923 3940 3938 3939 1 180.00 43.93200 2 ; CA-
N- C- O

3938 3942 3940 3941 1 180.00 4.60240 2 ; C-
CA- N- H
3942 3959 3957 3958 1 180.00 43.93200 2 ; CA-
N- C- O
3957 3961 3959 3960 1 180.00 4.60240 2 ; C-
CA- N- H
3961 3973 3971 3972 1 180.00 43.93200 2 ; CA-
N- C- O
3971 3975 3973 3974 1 180.00 4.60240 2 ; C-
CA- N- H
3975 3995 3993 3994 1 180.00 43.93200 2 ; CA-
N- C- O
3993 3997 3995 3996 1 180.00 4.60240 2 ; C-
CA- N- H
3997 4014 4012 4013 1 180.00 43.93200 2 ; CA-
N- C- O
4012 4016 4014 4015 1 180.00 4.60240 2 ; C-
CA- N- H
4016 4033 4031 4032 1 180.00 43.93200 2 ; CA-
N- C- O
4031 4035 4033 4034 1 180.00 4.60240 2 ; C-
CA- N- H
4035 4047 4045 4046 1 180.00 43.93200 2 ; CA-
N- C- O
4045 4049 4047 4048 1 180.00 4.60240 2 ; C-
CA- N- H
4049 4068 4066 4067 1 180.00 43.93200 2 ; CA-
N- C- O
4054 4057 4055 4056 1 180.00 4.60240 2 ; CG-
CE1- CD1- HD1
4054 4062 4064 4065 1 180.00 4.60240 2 ; CG-
CE2- CD2- HD2
4055 4059 4057 4058 1 180.00 4.60240 2 ; CD1-
CZ- CE1- HE1
4055 4064 4054 4051 1 180.00 4.60240 2 ; CD1-
CD2- CG- CB
4057 4062 4059 4060 1 180.00 4.60240 2 ; CE1-
CE2- CZ- OH
4059 4064 4062 4063 1 180.00 4.60240 2 ; CZ-
CD2- CE2- HE2
4066 4070 4068 4069 1 180.00 4.60240 2 ; C-
CA- N- H
4070 4082 4080 4081 1 180.00 43.93200 2 ; CA-
N- C- O
4072 4077 4075 4076 1 180.00 43.93200 2 ; CB-
ND2- CG- OD1
4075 4078 4077 4079 1 180.00 4.18400 2 ; CG-
HD21- ND2- HD22
4080 4083 4082 4092 1 180.00 4.18400 2 ; C-
CD- N- CA
4092 4096 4094 4095 1 180.00 43.93200 2 ; CA-
N- C- O

```
  4094   4098   4096   4097      1   180.00   4.60240   2 ;      C-
CA-     N-     H
  4098   4110   4108   4109      1   180.00  43.93200   2 ;     CA-
N-     C-     O
  4100   4105   4103   4104      1   180.00  43.93200   2 ;     CB-
ND2-    CG-   OD1
  4103   4106   4105   4107      1   180.00   4.18400   2 ;     CG-
HD21-   ND2-  HD22
  4108   4112   4110   4111      1   180.00   4.60240   2 ;      C-
CA-     N-     H
  4112   4125   4123   4124      1   180.00  43.93200   2 ;     CA-
N-     C-     O
  4117   4121   4120   4122      1   180.00  43.93200   2 ;     CG-
OE1-    CD-   OE2
  4123   4127   4125   4126      1   180.00   4.60240   2 ;      C-
CA-     N-     H
  4127   4149   4147   4148      1   180.00  43.93200   2 ;     CA-
N-     C-     O
  4135   4140   4138   4139      1   180.00   4.18400   2 ;     CD-
CZ-    NE-    HE
  4138   4141   4140   4144      1   180.00  43.93200   2 ;     NE-
NH1-    CZ-   NH2
  4140   4142   4141   4143      1   180.00   4.18400   2 ;     CZ-
HH11-   NH1-  HH12
  4140   4145   4144   4146      1   180.00   4.18400   2 ;     CZ-
HH21-   NH2-  HH22
  4147   4151   4149   4150      1   180.00   4.60240   2 ;      C-
CA-     N-     H
  4151   4160   4158   4159      1   180.00  43.93200   2 ;     CA-
N-     C-     O
  4158   4162   4160   4161      1   180.00   4.60240   2 ;      C-
CA-     N-     H
  4162   4174   4172   4173      1   180.00  43.93200   2 ;     CA-
N-     C-     O
  4172   4176   4174   4175      1   180.00   4.60240   2 ;      C-
CA-     N-     H
  4176   4193   4191   4192      1   180.00  43.93200   2 ;     CA-
N-     C-     O
  4191   4195   4193   4194      1   180.00   4.60240   2 ;      C-
CA-     N-     H
  4195   4208   4206   4207      1   180.00  43.93200   2 ;     CA-
N-     C-     O
  4200   4204   4203   4205      1   180.00  43.93200   2 ;     CG-
OE1-    CD-   OE2
  4206   4210   4208   4209      1   180.00   4.60240   2 ;      C-
CA-     N-     H
  4210   4223   4221   4222      1   180.00  43.93200   2 ;     CA-
N-     C-     O
  4215   4219   4218   4220      1   180.00  43.93200   2 ;     CG-
OE1-    CD-   OE2
  4221   4225   4223   4224      1   180.00   4.60240   2 ;      C-
CA-     N-     H
```

4225 4233 4231 4232 1 180.00 43.93200 2 ; CA-
N- C- O
4231 4235 4233 4234 1 180.00 4.60240 2 ; C-
CA- N- H
4235 4252 4250 4251 1 180.00 43.93200 2 ; CA-
N- C- O
4250 4254 4252 4253 1 180.00 4.60240 2 ; C-
CA- N- H
4254 4266 4264 4265 1 180.00 43.93200 2 ; CA-
N- C- O
4256 4261 4259 4260 1 180.00 43.93200 2 ; CB-
ND2- CG- OD1
4259 4262 4261 4263 1 180.00 4.18400 2 ; CG-
HD21- ND2- HD22
4264 4268 4266 4267 1 180.00 4.60240 2 ; C-
CA- N- H
4268 4283 4281 4282 1 180.00 43.93200 2 ; CA-
N- C- O
4270 4279 4273 4274 1 180.00 4.60240 2 ; CB-
CD2- CG- ND1
4273 4277 4279 4280 1 180.00 4.60240 2 ; CG-
NE2- CD2- HD2
4274 4277 4275 4276 1 180.00 4.60240 2 ; ND1-
NE2- CE1- HE1
4275 4279 4277 4278 1 180.00 4.18400 2 ; CE1-
CD2- NE2- HE2
4281 4284 4283 4293 1 180.00 4.18400 2 ; C-
CD- N- CA
4293 4297 4295 4296 1 180.00 43.93200 2 ; CA-
N- C- O
4295 4299 4297 4298 1 180.00 4.60240 2 ; C-
CA- N- H
4299 4321 4319 4320 1 180.00 43.93200 2 ; CA-
N- C- O
4304 4307 4305 4306 1 180.00 4.60240 2 ; CG-
NE1- CD1- HD1
4305 4309 4307 4308 1 180.00 4.18400 2 ; CD1-
CE2- NE1- HE1
4309 4312 4310 4311 1 180.00 4.60240 2 ; CE2-
CH2- CZ2- HZ2
4310 4314 4312 4313 1 180.00 4.60240 2 ; CZ2-
CZ3- CH2- HH2
4312 4316 4314 4315 1 180.00 4.60240 2 ; CH2-
CE3- CZ3- HZ3
4314 4318 4316 4317 1 180.00 4.60240 2 ; CZ3-
CD2- CE3- HE3
4318 4301 4304 4305 1 180.00 4.60240 2 ; CD2-
CB- CG- CD1
4319 4323 4321 4322 1 180.00 4.60240 2 ; C-
CA- N- H
4323 4340 4338 4339 1 180.00 43.93200 2 ; CA-
N- C- O

4338 4342 4340 4341 1 180.00 4.60240 2 ; C-
CA- N- H
4342 4354 4352 4353 1 180.00 43.93200 2 ; CA-
N- C- O
4352 4356 4354 4355 1 180.00 4.60240 2 ; C-
CA- N- H
4356 4371 4369 4370 1 180.00 43.93200 2 ; CA-
N- C- O
4361 4366 4364 4365 1 180.00 43.93200 2 ; CG-
NE2- CD- OE1
4364 4367 4366 4368 1 180.00 4.18400 2 ; CD-
HE21- NE2- HE22
4369 4373 4371 4372 1 180.00 4.60240 2 ; C-
CA- N- H
4373 4388 4386 4387 1 180.00 43.93200 2 ; CA-
N- C- O
4386 4390 4388 4389 1 180.00 4.60240 2 ; C-
CA- N- H
4390 4402 4400 4401 1 180.00 43.93200 2 ; CA-
N- C- O
4400 4404 4402 4403 1 180.00 4.60240 2 ; C-
CA- N- H
4404 4424 4422 4423 1 180.00 43.93200 2 ; CA-
N- C- O
4422 4426 4424 4425 1 180.00 4.60240 2 ; C-
CA- N- H
4426 4435 4433 4434 1 180.00 43.93200 2 ; CA-
N- C- O
4433 4437 4435 4436 1 180.00 4.60240 2 ; C-
CA- N- H
4437 4452 4450 4451 1 180.00 43.93200 2 ; CA-
N- C- O
4439 4448 4442 4443 1 180.00 4.60240 2 ; CB-
CD2- CG- ND1
4442 4446 4448 4449 1 180.00 4.60240 2 ; CG-
NE2- CD2- HD2
4443 4446 4444 4445 1 180.00 4.60240 2 ; ND1-
NE2- CE1- HE1
4444 4448 4446 4447 1 180.00 4.18400 2 ; CE1-
CD2- NE2- HE2
4450 4454 4452 4453 1 180.00 4.60240 2 ; C-
CA- N- H
4454 4467 4465 4466 1 180.00 43.93200 2 ; CA-
N- C- O
4459 4463 4462 4464 1 180.00 43.93200 2 ; CG-
OE1- CD- OE2
4465 4469 4467 4468 1 180.00 4.60240 2 ; C-
CA- N- H
4469 4484 4482 4483 1 180.00 43.93200 2 ; CA-
N- C- O
4471 4480 4474 4475 1 180.00 4.60240 2 ; CB-
CD2- CG- ND1

4474 4478 4480 4481 1 180.00 4.60240 2 ; CG-
NE2- CD2- HD2
4475 4478 4476 4477 1 180.00 4.60240 2 ; ND1-
NE2- CE1- HE1
4476 4480 4478 4479 1 180.00 4.18400 2 ; CE1-
CD2- NE2- HE2
4482 4486 4484 4485 1 180.00 4.60240 2 ; C-
CA- N- H
4486 4500 4498 4499 1 180.00 43.93200 2 ; CA-
N- C- O
4498 4502 4500 4501 1 180.00 4.60240 2 ; C-
CA- N- H
4502 4515 4513 4514 1 180.00 43.93200 2 ; CA-
N- C- O
4507 4511 4510 4512 1 180.00 43.93200 2 ; CG-
OE1- CD- OE2
4513 4517 4515 4516 1 180.00 4.60240 2 ; C-
CA- N- H
4517 4534 4532 4533 1 180.00 43.93200 2 ; CA-
N- C- O
4532 4536 4534 4535 1 180.00 4.60240 2 ; C-
CA- N- H
4536 4545 4543 4544 1 180.00 43.93200 2 ; CA-
N- C- O
4543 4547 4545 4546 1 180.00 4.60240 2 ; C-
CA- N- H
4547 4556 4554 4555 1 180.00 43.93200 2 ; CA-
N- C- O
4554 4558 4556 4557 1 180.00 4.60240 2 ; C-
CA- N- H
4558 4570 4568 4569 1 180.00 43.93200 2 ; CA-
N- C- O
4568 4572 4570 4571 1 180.00 4.60240 2 ; C-
CA- N- H
4572 4589 4587 4588 1 180.00 43.93200 2 ; CA-
N- C- O
4587 4591 4589 4590 1 180.00 4.60240 2 ; C-
CA- N- H
4591 4608 4606 4607 1 180.00 43.93200 2 ; CA-
N- C- O
4606 4610 4608 4609 1 180.00 4.60240 2 ; C-
CA- N- H
4610 4630 4628 4629 1 180.00 43.93200 2 ; CA-
N- C- O
4628 4632 4630 4631 1 180.00 4.60240 2 ; C-
CA- N- H
4632 4644 4642 4643 1 180.00 43.93200 2 ; CA-
N- C- O
4634 4639 4637 4638 1 180.00 43.93200 2 ; CB-
ND2- CG- OD1
4637 4640 4639 4641 1 180.00 4.18400 2 ; CG-
HD21- ND2- HD22

```
  4642   4646   4644   4645       1   180.00   4.60240   2 ;      C-
CA-     N-     H
  4646   4663   4661   4662       1   180.00  43.93200   2 ;     CA-
N-     C-     O
  4661   4665   4663   4664       1   180.00   4.60240   2 ;      C-
CA-     N-     H
  4665   4685   4683   4684       1   180.00  43.93200   2 ;     CA-
N-     C-     O
  4683   4687   4685   4686       1   180.00   4.60240   2 ;      C-
CA-     N-     H
  4687   4699   4697   4698       1   180.00  43.93200   2 ;     CA-
N-     C-     O
  4689   4694   4692   4693       1   180.00  43.93200   2 ;     CB-
ND2-    CG-   OD1
  4692   4695   4694   4696       1   180.00   4.18400   2 ;     CG-
HD21-   ND2-  HD22
  4697   4701   4699   4700       1   180.00   4.60240   2 ;      C-
CA-     N-     H
  4701   4719   4717   4718       1   180.00  43.93200   2 ;     CA-
N-     C-     O
  4706   4709   4707   4708       1   180.00   4.60240   2 ;     CG-
CE1-   CD1-   HD1
  4706   4713   4715   4716       1   180.00   4.60240   2 ;     CG-
CE2-   CD2-   HD2
  4707   4711   4709   4710       1   180.00   4.60240   2 ;    CD1-
CZ-   CE1-   HE1
  4707   4715   4706   4703       1   180.00   4.60240   2 ;    CD1-
CD2-    CG-    CB
  4709   4713   4711   4712       1   180.00   4.60240   2 ;    CE1-
CE2-    CZ-    HZ
  4711   4715   4713   4714       1   180.00   4.60240   2 ;     CZ-
CD2-   CE2-   HE2
  4717   4721   4719   4720       1   180.00   4.60240   2 ;      C-
CA-     N-     H
  4721   4741   4739   4740       1   180.00  43.93200   2 ;     CA-
N-     C-     O
  4739   4743   4741   4742       1   180.00   4.60240   2 ;      C-
CA-     N-     H
  4743   4763   4761   4762       1   180.00  43.93200   2 ;     CA-
N-     C-     O
  4761   4765   4763   4764       1   180.00   4.60240   2 ;      C-
CA-     N-     H
  4765   4778   4776   4777       1   180.00  43.93200   2 ;     CA-
N-     C-     O
  4770   4774   4773   4775       1   180.00  43.93200   2 ;     CG-
OE1-    CD-   OE2
  4776   4780   4778   4779       1   180.00   4.60240   2 ;      C-
CA-     N-     H
  4780   4792   4790   4791       1   180.00  43.93200   2 ;     CA-
N-     C-     O
  4782   4787   4785   4786       1   180.00  43.93200   2 ;     CB-
ND2-    CG-   OD1
```

4785 4788 4787 4789 1 180.00 4.18400 2 ; CG-
HD21- ND2- HD22
4790 4794 4792 4793 1 180.00 4.60240 2 ; C-
CA- N- H
4794 4807 4805 4806 1 180.00 43.93200 2 ; CA-
N- C- O
4799 4803 4802 4804 1 180.00 43.93200 2 ; CG-
OE1- CD- OE2
4805 4809 4807 4808 1 180.00 4.60240 2 ; C-
CA- N- H
4809 4826 4824 4825 1 180.00 43.93200 2 ; CA-
N- C- O
4824 4828 4826 4827 1 180.00 4.60240 2 ; C-
CA- N- H
4828 4848 4846 4847 1 180.00 43.93200 2 ; CA-
N- C- O
4846 4850 4848 4849 1 180.00 4.60240 2 ; C-
CA- N- H
4850 4870 4868 4869 1 180.00 43.93200 2 ; CA-
N- C- O
4868 4872 4870 4871 1 180.00 4.60240 2 ; C-
CA- N- H
4872 4889 4887 4888 1 180.00 43.93200 2 ; CA-
N- C- O
4887 4891 4889 4890 1 180.00 4.60240 2 ; C-
CA- N- H
4891 4899 4897 4898 1 180.00 43.93200 2 ; CA-
N- C- O
4897 4901 4899 4900 1 180.00 4.60240 2 ; C-
CA- N- H
4901 4918 4916 4917 1 180.00 43.93200 2 ; CA-
N- C- O
4916 4920 4918 4919 1 180.00 4.60240 2 ; C-
CA- N- H
4920 4932 4930 4931 1 180.00 43.93200 2 ; CA-
N- C- O
4930 4934 4932 4933 1 180.00 4.60240 2 ; C-
CA- N- H
4934 4951 4949 4950 1 180.00 43.93200 2 ; CA-
N- C- O
4949 4953 4951 4952 1 180.00 4.60240 2 ; C-
CA- N- H
4953 4970 4968 4969 1 180.00 43.93200 2 ; CA-
N- C- O
4968 4972 4970 4971 1 180.00 4.60240 2 ; C-
CA- N- H
4972 4980 4978 4979 1 180.00 43.93200 2 ; CA-
N- C- O
4978 4982 4980 4981 1 180.00 4.60240 2 ; C-
CA- N- H
4982 5002 5000 5001 1 180.00 43.93200 2 ; CA-
N- C- O

```
  5000   5004   5002   5003       1   180.00   4.60240   2 ;      C-
CA-     N-      H
  5004   5020   5018   5019       1   180.00  43.93200   2 ;     CA-
N-     C-      O
  5006   5016   5009   5010       1   180.00   4.60240   2 ;     CB-
CD2-    CG-   ND1
  5009   5012   5010   5011       1   180.00   4.18400   2 ;     CG-
CE1-   ND1-   HD1
  5009   5014   5016   5017       1   180.00   4.60240   2 ;     CG-
NE2-   CD2-   HD2
  5010   5014   5012   5013       1   180.00   4.60240   2 ;    ND1-
NE2-   CE1-   HE1
  5012   5016   5014   5015       1   180.00   4.18400   2 ;    CE1-
CD2-   NE2-   HE2
  5018   5022   5020   5021       1   180.00   4.60240   2 ;      C-
CA-     N-      H
  5022   5039   5037   5038       1   180.00  43.93200   2 ;     CA-
N-     C-      O
  5037   5041   5039   5040       1   180.00   4.60240   2 ;      C-
CA-     N-      H
  5041   5050   5048   5049       1   180.00  43.93200   2 ;     CA-
N-     C-      O
  5048   5052   5050   5051       1   180.00   4.60240   2 ;      C-
CA-     N-      H
  5052   5062   5060   5061       1   180.00  43.93200   2 ;     CA-
N-     C-      O
  5054   5058   5057   5059       1   180.00  43.93200   2 ;     CB-
OD1-    CG-   OD2
  5060   5064   5062   5063       1   180.00   4.60240   2 ;      C-
CA-     N-      H
  5064   5078   5076   5077       1   180.00  43.93200   2 ;     CA-
N-     C-      O
  5076   5080   5078   5079       1   180.00   4.60240   2 ;      C-
CA-     N-      H
  5080   5093   5091   5092       1   180.00  43.93200   2 ;     CA-
N-     C-      O
  5085   5089   5088   5090       1   180.00  43.93200   2 ;     CG-
OE1-    CD-   OE2
  5091   5095   5093   5094       1   180.00   4.60240   2 ;      C-
CA-     N-      H
  5095   5112   5110   5111       1   180.00  43.93200   2 ;     CA-
N-     C-      O
  5110   5114   5112   5113       1   180.00   4.60240   2 ;      C-
CA-     N-      H
  5114   5126   5124   5125       1   180.00  43.93200   2 ;     CA-
N-     C-      O
  5116   5121   5119   5120       1   180.00  43.93200   2 ;     CB-
ND2-    CG-   OD1
  5119   5122   5121   5123       1   180.00   4.18400   2 ;     CG-
HD21-   ND2-  HD22
  5124   5128   5126   5127       1   180.00   4.60240   2 ;      C-
CA-     N-      H
```

5128 5140 5138 5139 1 180.00 43.93200 2 ; CA-
N- C- O
5130 5135 5133 5134 1 180.00 43.93200 2 ; CB-
ND2- CG- OD1
5133 5136 5135 5137 1 180.00 4.18400 2 ; CG-
HD21- ND2- HD22
5138 5142 5140 5141 1 180.00 4.60240 2 ; C-
CA- N- H
5142 5159 5157 5158 1 180.00 43.93200 2 ; CA-
N- C- O
5157 5161 5159 5160 1 180.00 4.60240 2 ; C-
CA- N- H
5161 5183 5181 5182 1 180.00 43.93200 2 ; CA-
N- C- O
5169 5174 5172 5173 1 180.00 4.18400 2 ; CD-
CZ- NE- HE
5172 5175 5174 5178 1 180.00 43.93200 2 ; NE-
NH1- CZ- NH2
5174 5176 5175 5177 1 180.00 4.18400 2 ; CZ-
HH11- NH1- HH12
5174 5179 5178 5180 1 180.00 4.18400 2 ; CZ-
HH21- NH2- HH22
5181 5185 5183 5184 1 180.00 4.60240 2 ; C-
CA- N- H
5185 5197 5195 5196 1 180.00 43.93200 2 ; CA-
N- C- O
5187 5192 5190 5191 1 180.00 43.93200 2 ; CB-
ND2- CG- OD1
5190 5193 5192 5194 1 180.00 4.18400 2 ; CG-
HD21- ND2- HD22
5195 5199 5197 5198 1 180.00 4.60240 2 ; C-
CA- N- H
5199 5216 5214 5215 1 180.00 43.93200 2 ; CA-
N- C- O
5214 5218 5216 5217 1 180.00 4.60240 2 ; C-
CA- N- H
5218 5236 5234 5235 1 180.00 43.93200 2 ; CA-
N- C- O
5223 5226 5224 5225 1 180.00 4.60240 2 ; CG-
CE1- CD1- HD1
5223 5230 5232 5233 1 180.00 4.60240 2 ; CG-
CE2- CD2- HD2
5224 5228 5226 5227 1 180.00 4.60240 2 ; CD1-
CZ- CE1- HE1
5224 5232 5223 5220 1 180.00 4.60240 2 ; CD1-
CD2- CG- CB
5226 5230 5228 5229 1 180.00 4.60240 2 ; CE1-
CE2- CZ- HZ
5228 5232 5230 5231 1 180.00 4.60240 2 ; CZ-
CD2- CE2- HE2
5234 5238 5236 5237 1 180.00 4.60240 2 ; C-
CA- N- H

```
  5238   5255   5253   5254       1   180.00  43.93200   2 ;      CA-
N-     C-     O
  5253   5257   5255   5256       1   180.00   4.60240   2 ;       C-
CA-     N-     H
  5257   5265   5263   5264       1   180.00  43.93200   2 ;      CA-
N-     C-     O
  5263   5267   5265   5266       1   180.00   4.60240   2 ;       C-
CA-     N-     H
  5267   5284   5282   5283       1   180.00  43.93200   2 ;      CA-
N-     C-     O
  5282   5286   5284   5285       1   180.00   4.60240   2 ;       C-
CA-     N-     H
  5286   5296   5294   5295       1   180.00  43.93200   2 ;      CA-
N-     C-     O
  5288   5292   5291   5293       1   180.00  43.93200   2 ;      CB-
OD1-    CG-   OD2
  5294   5298   5296   5297       1   180.00   4.60240   2 ;       C-
CA-     N-     H
  5298   5312   5310   5311       1   180.00  43.93200   2 ;      CA-
N-     C-     O
  5310   5314   5312   5313       1   180.00   4.60240   2 ;       C-
CA-     N-     H
  5314   5324   5322   5323       1   180.00  43.93200   2 ;      CA-
N-     C-     O
  5316   5320   5319   5321       1   180.00  43.93200   2 ;      CB-
OD1-    CG-   OD2
  5322   5326   5324   5325       1   180.00   4.60240   2 ;       C-
CA-     N-     H
  5326   5338   5336   5337       1   180.00  43.93200   2 ;      CA-
N-     C-     O
  5328   5333   5331   5332       1   180.00  43.93200   2 ;      CB-
ND2-    CG-   OD1
  5331   5334   5333   5335       1   180.00   4.18400   2 ;      CG-
HD21-   ND2-  HD22
  5336   5340   5338   5339       1   180.00   4.60240   2 ;       C-
CA-     N-     H
  5340   5349   5347   5348       1   180.00  43.93200   2 ;      CA-
N-     C-     O
  5347   5351   5349   5350       1   180.00   4.60240   2 ;       C-
CA-     N-     H
  5351   5356   5354   5355       1   180.00  43.93200   2 ;      CA-
N-     C-     O
  5354   5358   5356   5357       1   180.00   4.60240   2 ;       C-
CA-     N-     H
  5358   5370   5368   5369       1   180.00  43.93200   2 ;      CA-
N-     C-     O
  5368   5372   5370   5371       1   180.00   4.60240   2 ;       C-
CA-     N-     H
  5372   5389   5387   5388       1   180.00  43.93200   2 ;      CA-
N-     C-     O
  5387   5391   5389   5390       1   180.00   4.60240   2 ;       C-
CA-     N-     H
```

5391 5400 5398 5399 1 180.00 43.93200 2 ; CA-
N- C- O
5398 5402 5400 5401 1 180.00 4.60240 2 ; C-
CA- N- H
5402 5411 5409 5410 1 180.00 43.93200 2 ; CA-
N- C- O
5409 5413 5411 5412 1 180.00 4.60240 2 ; C-
CA- N- H
5413 5428 5426 5427 1 180.00 43.93200 2 ; CA-
N- C- O
5418 5423 5421 5422 1 180.00 43.93200 2 ; CG-
NE2- CD- OE1
5421 5424 5423 5425 1 180.00 4.18400 2 ; CD-
HE21- NE2- HE22
5426 5430 5428 5429 1 180.00 4.60240 2 ; C-
CA- N- H
5430 5444 5442 5443 1 180.00 43.93200 2 ; CA-
N- C- O
5435 5439 5438 5440 1 180.00 43.93200 2 ; CG-
OE1- CD- OE2
5442 5446 5444 5445 1 180.00 4.60240 2 ; C-
CA- N- H
5446 5463 5461 5462 1 180.00 43.93200 2 ; CA-
N- C- O
5461 5465 5463 5464 1 180.00 4.60240 2 ; C-
CA- N- H
5465 5482 5480 5481 1 180.00 43.93200 2 ; CA-
N- C- O
5480 5484 5482 5483 1 180.00 4.60240 2 ; C-
CA- N- H
5484 5494 5492 5493 1 180.00 43.93200 2 ; CA-
N- C- O
5486 5490 5489 5491 1 180.00 43.93200 2 ; CB-
OD1- CG- OD2
5492 5496 5494 5495 1 180.00 4.60240 2 ; C-
CA- N- H
5496 5501 5499 5500 1 180.00 43.93200 2 ; CA-
N- C- O
5499 5503 5501 5502 1 180.00 4.60240 2 ; C-
CA- N- H
5503 5520 5518 5519 1 180.00 43.93200 2 ; CA-
N- C- O
5518 5522 5520 5521 1 180.00 4.60240 2 ; C-
CA- N- H
5522 5542 5540 5541 1 180.00 43.93200 2 ; CA-
N- C- O
5540 5544 5542 5543 1 180.00 4.60240 2 ; C-
CA- N- H
5544 5564 5562 5563 1 180.00 43.93200 2 ; CA-
N- C- O
5562 5566 5564 5565 1 180.00 4.60240 2 ; C-
CA- N- H

```
  5566   5583   5581   5582       1   180.00  43.93200   2 ;      CA-
N-     C-      O
  5581   5585   5583   5584       1   180.00   4.60240   2 ;       C-
CA-     N-     H
  5585   5590   5588   5589       1   180.00  43.93200   2 ;      CA-
N-     C-      O
  5588   5592   5590   5591       1   180.00   4.60240   2 ;       C-
CA-     N-     H
  5592   5611   5609   5610       1   180.00  43.93200   2 ;      CA-
N-     C-      O
  5597   5600   5598   5599       1   180.00   4.60240   2 ;      CG-
CE1-   CD1-   HD1
  5597   5605   5607   5608       1   180.00   4.60240   2 ;      CG-
CE2-   CD2-   HD2
  5598   5602   5600   5601       1   180.00   4.60240   2 ;     CD1-
CZ-   CE1-   HE1
  5598   5607   5597   5594       1   180.00   4.60240   2 ;     CD1-
CD2-    CG-    CB
  5600   5605   5602   5603       1   180.00   4.60240   2 ;     CE1-
CE2-    CZ-    OH
  5602   5607   5605   5606       1   180.00   4.60240   2 ;      CZ-
CD2-   CE2-   HE2
  5609   5613   5611   5612       1   180.00   4.60240   2 ;       C-
CA-     N-     H
  5613   5628   5626   5627       1   180.00  43.93200   2 ;      CA-
N-     C-      O
  5618   5623   5621   5622       1   180.00  43.93200   2 ;      CG-
NE2-    CD-   OE1
  5621   5624   5623   5625       1   180.00   4.18400   2 ;      CD-
HE21-   NE2-  HE22
  5626   5630   5628   5629       1   180.00   4.60240   2 ;       C-
CA-     N-     H
  5630   5650   5648   5649       1   180.00  43.93200   2 ;      CA-
N-     C-      O
  5648   5652   5650   5651       1   180.00   4.60240   2 ;       C-
CA-     N-     H
  5652   5669   5667   5668       1   180.00  43.93200   2 ;      CA-
N-     C-      O
  5667   5670   5669   5679       1   180.00   4.18400   2 ;       C-
CD-     N-    CA
  5679   5683   5681   5682       1   180.00  43.93200   2 ;      CA-
N-     C-      O
  5681   5684   5683   5693       1   180.00   4.18400   2 ;       C-
CD-     N-    CA
  5693   5697   5695   5696       1   180.00  43.93200   2 ;      CA-
N-     C-      O
  5695   5699   5697   5698       1   180.00   4.60240   2 ;       C-
CA-     N-     H
  5699   5709   5707   5708       1   180.00  43.93200   2 ;      CA-
N-     C-      O
  5701   5705   5704   5706       1   180.00  43.93200   2 ;      CB-
OD1-    CG-   OD2
```

5707 5711 5709 5710 1 180.00 4.60240 2 ; C-
CA- N- H
5711 5728 5726 5727 1 180.00 43.93200 2 ; CA-
N- C- O
5726 5730 5728 5729 1 180.00 4.60240 2 ; C-
CA- N- H
5730 5745 5743 5744 1 180.00 43.93200 2 ; CA-
N- C- O
5732 5741 5735 5736 1 180.00 4.60240 2 ; CB-
CD2- CG- ND1
5735 5739 5741 5742 1 180.00 4.60240 2 ; CG-
NE2- CD2- HD2
5736 5739 5737 5738 1 180.00 4.60240 2 ; ND1-
NE2- CE1- HE1
5737 5741 5739 5740 1 180.00 4.18400 2 ; CE1-
CD2- NE2- HE2
5743 5747 5745 5746 1 180.00 4.60240 2 ; C-
CA- N- H
5747 5762 5760 5761 1 180.00 43.93200 2 ; CA-
N- C- O
5752 5757 5755 5756 1 180.00 43.93200 2 ; CG-
NE2- CD- OE1
5755 5758 5757 5759 1 180.00 4.18400 2 ; CD-
HE21- NE2- HE22
5760 5764 5762 5763 1 180.00 4.60240 2 ; C-
CA- N- H
5764 5778 5776 5777 1 180.00 43.93200 2 ; CA-
N- C- O
5776 5780 5778 5779 1 180.00 4.60240 2 ; C-
CA- N- H
5780 5797 5795 5796 1 180.00 43.93200 2 ; CA-
N- C- O
5795 5799 5797 5798 1 180.00 4.60240 2 ; C-
CA- N- H
5799 5821 5819 5820 1 180.00 43.93200 2 ; CA-
N- C- O
5807 5812 5810 5811 1 180.00 4.18400 2 ; CD-
CZ- NE- HE
5810 5813 5812 5816 1 180.00 43.93200 2 ; NE-
NH1- CZ- NH2
5812 5814 5813 5815 1 180.00 4.18400 2 ; CZ-
HH11- NH1- HH12
5812 5817 5816 5818 1 180.00 4.18400 2 ; CZ-
HH21- NH2- HH22
5819 5823 5821 5822 1 180.00 4.60240 2 ; C-
CA- N- H
5823 5833 5831 5832 1 180.00 43.93200 2 ; CA-
N- C- O
5825 5829 5828 5830 1 180.00 43.93200 2 ; CB-
OD1- CG- OD2
5831 5835 5833 5834 1 180.00 4.60240 2 ; C-
CA- N- H

5835   5852   5850   5851       1   180.00  43.93200   2 ;      CA-
N-     C-     O
5850   5854   5852   5853       1   180.00   4.60240   2 ;       C-
CA-     N-     H
5854   5864   5862   5863       1   180.00  43.93200   2 ;      CA-
N-     C-     O
5856   5860   5859   5861       1   180.00  43.93200   2 ;      CB-
OD1-    CG-   OD2
5862   5866   5864   5865       1   180.00   4.60240   2 ;       C-
CA-     N-     H
5866   5875   5873   5874       1   180.00  43.93200   2 ;      CA-
N-     C-     O
5873   5877   5875   5876       1   180.00   4.60240   2 ;       C-
CA-     N-     H
5877   5889   5887   5888       1   180.00  43.93200   2 ;      CA-
N-     C-     O
5879   5884   5882   5883       1   180.00  43.93200   2 ;      CB-
ND2-    CG-   OD1
5882   5885   5884   5886       1   180.00   4.18400   2 ;      CG-
HD21-   ND2-  HD22
5887   5891   5889   5890       1   180.00   4.60240   2 ;       C-
CA-     N-     H
5891   5899   5897   5898       1   180.00  43.93200   2 ;      CA-
N-     C-     O
5897   5901   5899   5900       1   180.00   4.60240   2 ;       C-
CA-     N-     H
5901   5910   5908   5909       1   180.00  43.93200   2 ;      CA-
N-     C-     O
5908   5912   5910   5911       1   180.00   4.60240   2 ;       C-
CA-     N-     H
5912   5917   5915   5916       1   180.00  43.93200   2 ;      CA-
N-     C-     O
5915   5919   5917   5918       1   180.00   4.60240   2 ;       C-
CA-     N-     H
5919   5934   5932   5933       1   180.00  43.93200   2 ;      CA-
N-     C-     O
5924   5929   5927   5928       1   180.00  43.93200   2 ;      CG-
NE2-    CD-   OE1
5927   5930   5929   5931       1   180.00   4.18400   2 ;      CD-
HE21-   NE2-  HE22
5932   5936   5934   5935       1   180.00   4.60240   2 ;       C-
CA-     N-     H
5936   5953   5951   5952       1   180.00  43.93200   2 ;      CA-
N-     C-     O
5951   5955   5953   5954       1   180.00   4.60240   2 ;       C-
CA-     N-     H
5955   5971   5969   5970       1   180.00  43.93200   2 ;      CA-
N-     C-     O
5957   5967   5960   5961       1   180.00   4.60240   2 ;      CB-
CD2-    CG-   ND1
5960   5963   5961   5962       1   180.00   4.18400   2 ;      CG-
CE1-   ND1-   HD1

```
  5960   5965   5967   5968       1   180.00   4.60240   2 ;      CG-
NE2-   CD2-   HD2
  5961   5965   5963   5964       1   180.00   4.60240   2 ;     ND1-
NE2-   CE1-   HE1
  5963   5967   5965   5966       1   180.00   4.18400   2 ;     CE1-
CD2-   NE2-   HE2
  5969   5973   5971   5972       1   180.00   4.60240   2 ;       C-
CA-     N-     H
  5973   5992   5990   5991       1   180.00  43.93200   2 ;      CA-
N-     C-     O
  5978   5981   5979   5980       1   180.00   4.60240   2 ;      CG-
CE1-   CD1-   HD1
  5978   5986   5988   5989       1   180.00   4.60240   2 ;      CG-
CE2-   CD2-   HD2
  5979   5983   5981   5982       1   180.00   4.60240   2 ;     CD1-
CZ-   CE1-   HE1
  5979   5988   5978   5975       1   180.00   4.60240   2 ;     CD1-
CD2-    CG-    CB
  5981   5986   5983   5984       1   180.00   4.60240   2 ;     CE1-
CE2-    CZ-    OH
  5983   5988   5986   5987       1   180.00   4.60240   2 ;      CZ-
CD2-   CE2-   HE2
  5990   5994   5992   5993       1   180.00   4.60240   2 ;       C-
CA-     N-     H
  5994   6006   6004   6005       1   180.00  43.93200   2 ;      CA-
N-     C-     O
  6004   6008   6006   6007       1   180.00   4.60240   2 ;       C-
CA-     N-     H
  6008   6018   6016   6017       1   180.00  43.93200   2 ;      CA-
N-     C-     O
  6010   6014   6013   6015       1   180.00  43.93200   2 ;      CB-
OD1-    CG-   OD2
  6016   6020   6018   6019       1   180.00   4.60240   2 ;       C-
CA-     N-     H
  6020   6038   6036   6037       1   180.00  43.93200   2 ;      CA-
N-     C-     O
  6025   6028   6026   6027       1   180.00   4.60240   2 ;      CG-
CE1-   CD1-   HD1
  6025   6032   6034   6035       1   180.00   4.60240   2 ;      CG-
CE2-   CD2-   HD2
  6026   6030   6028   6029       1   180.00   4.60240   2 ;     CD1-
CZ-   CE1-   HE1
  6026   6034   6025   6022       1   180.00   4.60240   2 ;     CD1-
CD2-    CG-    CB
  6028   6032   6030   6031       1   180.00   4.60240   2 ;     CE1-
CE2-    CZ-    HZ
  6030   6034   6032   6033       1   180.00   4.60240   2 ;      CZ-
CD2-   CE2-   HE2
  6036   6040   6038   6039       1   180.00   4.60240   2 ;       C-
CA-     N-     H
  6040   6057   6055   6056       1   180.00  43.93200   2 ;      CA-
N-     C-     O
```

```
  6055   6059   6057   6058       1   180.00   4.60240   2 ;      C-
CA-     N-     H
  6059   6067   6065   6066       1   180.00  43.93200   2 ;     CA-
N-     C-     O
  6065   6069   6067   6068       1   180.00   4.60240   2 ;      C-
CA-     N-     H
  6069   6077   6075   6076       1   180.00  43.93200   2 ;     CA-
N-     C-     O
  6075   6079   6077   6078       1   180.00   4.60240   2 ;      C-
CA-     N-     H
  6079   6091   6089   6090       1   180.00  43.93200   2 ;     CA-
N-     C-     O
  6089   6093   6091   6092       1   180.00   4.60240   2 ;      C-
CA-     N-     H
  6093   6110   6108   6109       1   180.00  43.93200   2 ;     CA-
N-     C-     O
  6108   6112   6110   6111       1   180.00   4.60240   2 ;      C-
CA-     N-     H
  6112   6122   6120   6121       1   180.00  43.93200   2 ;     CA-
N-     C-     O
  6114   6118   6117   6119       1   180.00  43.93200   2 ;     CB-
OD1-    CG-   OD2
  6120   6124   6122   6123       1   180.00   4.60240   2 ;      C-
CA-     N-     H
  6124   6144   6142   6143       1   180.00  43.93200   2 ;     CA-
N-     C-     O
  6142   6146   6144   6145       1   180.00   4.60240   2 ;      C-
CA-     N-     H
  6146   6161   6159   6160       1   180.00  43.93200   2 ;     CA-
N-     C-     O
  6151   6156   6154   6155       1   180.00  43.93200   2 ;     CG-
NE2-    CD-   OE1
  6154   6157   6156   6158       1   180.00   4.18400   2 ;     CD-
HE21-   NE2-  HE22
  6159   6163   6161   6162       1   180.00   4.60240   2 ;      C-
CA-     N-     H
  6163   6175   6173   6174       1   180.00  43.93200   2 ;     CA-
N-     C-     O
  6173   6177   6175   6176       1   180.00   4.60240   2 ;      C-
CA-     N-     H
  6177   6196   6194   6195       1   180.00  43.93200   2 ;     CA-
N-     C-     O
  6182   6185   6183   6184       1   180.00   4.60240   2 ;     CG-
CE1-   CD1-   HD1
  6182   6190   6192   6193       1   180.00   4.60240   2 ;     CG-
CE2-   CD2-   HD2
  6183   6187   6185   6186       1   180.00   4.60240   2 ;    CD1-
CZ-   CE1-   HE1
  6183   6192   6182   6179       1   180.00   4.60240   2 ;    CD1-
CD2-    CG-    CB
  6185   6190   6187   6188       1   180.00   4.60240   2 ;    CE1-
CE2-    CZ-    OH
```

6187 6192 6190 6191 1 180.00 4.60240 2 ; CZ-
CD2- CE2- HE2
6194 6198 6196 6197 1 180.00 4.60240 2 ; C-
CA- N- H
6198 6215 6213 6214 1 180.00 43.93200 2 ; CA-
N- C- O
6213 6217 6215 6216 1 180.00 4.60240 2 ; C-
CA- N- H
6217 6237 6235 6236 1 180.00 43.93200 2 ; CA-
N- C- O
6235 6239 6237 6238 1 180.00 4.60240 2 ; C-
CA- N- H
6239 6259 6257 6258 1 180.00 43.93200 2 ; CA-
N- C- O
6257 6261 6259 6260 1 180.00 4.60240 2 ; C-
CA- N- H
6261 6274 6272 6273 1 180.00 43.93200 2 ; CA-
N- C- O
6266 6270 6269 6271 1 180.00 43.93200 2 ; CG-
OE1- CD- OE2
6272 6276 6274 6275 1 180.00 4.60240 2 ; C-
CA- N- H
6276 6290 6288 6289 1 180.00 43.93200 2 ; CA-
N- C- O
6288 6292 6290 6291 1 180.00 4.60240 2 ; C-
CA- N- H
6292 6301 6299 6300 1 180.00 43.93200 2 ; CA-
N- C- O
6299 6303 6301 6302 1 180.00 4.60240 2 ; C-
CA- N- H
6303 6320 6318 6319 1 180.00 43.93200 2 ; CA-
N- C- O
6318 6322 6320 6321 1 180.00 4.60240 2 ; C-
CA- N- H
6322 6339 6337 6338 1 180.00 43.93200 2 ; CA-
N- C- O
6337 6340 6339 6349 1 180.00 4.18400 2 ; C-
CD- N- CA
6349 6353 6351 6352 1 180.00 43.93200 2 ; CA-
N- C- O
6351 6355 6353 6354 1 180.00 4.60240 2 ; C-
CA- N- H
6355 6373 6371 6372 1 180.00 43.93200 2 ; CA-
N- C- O
6360 6363 6361 6362 1 180.00 4.60240 2 ; CG-
CE1- CD1- HD1
6360 6367 6369 6370 1 180.00 4.60240 2 ; CG-
CE2- CD2- HD2
6361 6365 6363 6364 1 180.00 4.60240 2 ; CD1-
CZ- CE1- HE1
6361 6369 6360 6357 1 180.00 4.60240 2 ; CD1-
CD2- CG- CB

6363 6367 6365 6366 1 180.00 4.60240 2 ; CE1-
CE2- CZ- HZ
6365 6369 6367 6368 1 180.00 4.60240 2 ; CZ-
CD2- CE2- HE2
6371 6375 6373 6374 1 180.00 4.60240 2 ; C-
CA- N- H
6375 6395 6393 6394 1 180.00 43.93200 2 ; CA-
N- C- O
6393 6397 6395 6396 1 180.00 4.60240 2 ; C-
CA- N- H
6397 6415 6413 6414 1 180.00 43.93200 2 ; CA-
N- C- O
6402 6405 6403 6404 1 180.00 4.60240 2 ; CG-
CE1- CD1- HD1
6402 6409 6411 6412 1 180.00 4.60240 2 ; CG-
CE2- CD2- HD2
6403 6407 6405 6406 1 180.00 4.60240 2 ; CD1-
CZ- CE1- HE1
6403 6411 6402 6399 1 180.00 4.60240 2 ; CD1-
CD2- CG- CB
6405 6409 6407 6408 1 180.00 4.60240 2 ; CE1-
CE2- CZ- HZ
6407 6411 6409 6410 1 180.00 4.60240 2 ; CZ-
CD2- CE2- HE2
6413 6417 6415 6416 1 180.00 4.60240 2 ; C-
CA- N- H
6417 6435 6433 6434 1 180.00 43.93200 2 ; CA-
N- C- O
6422 6425 6423 6424 1 180.00 4.60240 2 ; CG-
CE1- CD1- HD1
6422 6429 6431 6432 1 180.00 4.60240 2 ; CG-
CE2- CD2- HD2
6423 6427 6425 6426 1 180.00 4.60240 2 ; CD1-
CZ- CE1- HE1
6423 6431 6422 6419 1 180.00 4.60240 2 ; CD1-
CD2- CG- CB
6425 6429 6427 6428 1 180.00 4.60240 2 ; CE1-
CE2- CZ- HZ
6427 6431 6429 6430 1 180.00 4.60240 2 ; CZ-
CD2- CE2- HE2
6433 6437 6435 6436 1 180.00 4.60240 2 ; C-
CA- N- H
6437 6447 6445 6446 1 180.00 43.93200 2 ; CA-
N- C- O
6439 6443 6442 6444 1 180.00 43.93200 2 ; CB-
OD1- CG- OD2
6445 6449 6447 6448 1 180.00 4.60240 2 ; C-
CA- N- H
6449 6466 6464 6465 1 180.00 43.93200 2 ; CA-
N- C- O
6464 6468 6466 6467 1 180.00 4.60240 2 ; C-
CA- N- H

6468 6478 6476 6477 1 180.00 43.93200 2 ; CA-
N- C- O
6470 6474 6473 6475 1 180.00 43.93200 2 ; CB-
OD1- CG- OD2
6476 6480 6478 6479 1 180.00 4.60240 2 ; C-
CA- N- H
6480 6485 6483 6484 1 180.00 43.93200 2 ; CA-
N- C- O
6483 6487 6485 6486 1 180.00 4.60240 2 ; C-
CA- N- H
6487 6499 6497 6498 1 180.00 43.93200 2 ; CA-
N- C- O
6489 6494 6492 6493 1 180.00 43.93200 2 ; CB-
ND2- CG- OD1
6492 6495 6494 6496 1 180.00 4.18400 2 ; CG-
HD21- ND2- HD22
6497 6501 6499 6500 1 180.00 4.60240 2 ; C-
CA- N- H
6501 6506 6504 6505 1 180.00 43.93200 2 ; CA-
N- C- O
6504 6508 6506 6507 1 180.00 4.60240 2 ; C-
CA- N- H
6508 6528 6526 6527 1 180.00 43.93200 2 ; CA-
N- C- O
6526 6530 6528 6529 1 180.00 4.60240 2 ; C-
CA- N- H
6530 6547 6545 6546 1 180.00 43.93200 2 ; CA-
N- C- O
6545 6549 6547 6548 1 180.00 4.60240 2 ; C-
CA- N- H
6549 6558 6556 6557 1 180.00 43.93200 2 ; CA-
N- C- O
6556 6560 6558 6559 1 180.00 4.60240 2 ; C-
CA- N- H
6560 6574 6572 6573 1 180.00 43.93200 2 ; CA-
N- C- O
6572 6576 6574 6575 1 180.00 4.60240 2 ; C-
CA- N- H
6576 6589 6587 6588 1 180.00 43.93200 2 ; CA-
N- C- O
6581 6585 6584 6586 1 180.00 43.93200 2 ; CG-
OE1- CD- OE2
6587 6591 6589 6590 1 180.00 4.60240 2 ; C-
CA- N- H
6591 6605 6603 6604 1 180.00 43.93200 2 ; CA-
N- C- O
6596 6600 6599 6601 1 180.00 43.93200 2 ; CG-
OE1- CD- OE2
6603 6607 6605 6606 1 180.00 4.60240 2 ; C-
CA- N- H
6607 6624 6622 6623 1 180.00 43.93200 2 ; CA-
N- C- O

6622 6626 6624 6625 1 180.00 4.60240 2 ; C-
CA- N- H
6626 6646 6644 6645 1 180.00 43.93200 2 ; CA-
N- C- O
6644 6648 6646 6647 1 180.00 4.60240 2 ; C-
CA- N- H
6648 6670 6668 6669 1 180.00 43.93200 2 ; CA-
N- C- O
6656 6661 6659 6660 1 180.00 4.18400 2 ; CD-
CZ- NE- HE
6659 6662 6661 6665 1 180.00 43.93200 2 ; NE-
NH1- CZ- NH2
6661 6663 6662 6664 1 180.00 4.18400 2 ; CZ-
HH11- NH1- HH12
6661 6666 6665 6667 1 180.00 4.18400 2 ; CZ-
HH21- NH2- HH22
6668 6672 6670 6671 1 180.00 4.60240 2 ; C-
CA- N- H
6672 6689 6687 6688 1 180.00 43.93200 2 ; CA-
N- C- O
6687 6691 6689 6690 1 180.00 4.60240 2 ; C-
CA- N- H
6691 6709 6707 6708 1 180.00 43.93200 2 ; CA-
N- C- O
6696 6699 6697 6698 1 180.00 4.60240 2 ; CG-
CE1- CD1- HD1
6696 6703 6705 6706 1 180.00 4.60240 2 ; CG-
CE2- CD2- HD2
6697 6701 6699 6700 1 180.00 4.60240 2 ; CD1-
CZ- CE1- HE1
6697 6705 6696 6693 1 180.00 4.60240 2 ; CD1-
CD2- CG- CB
6699 6703 6701 6702 1 180.00 4.60240 2 ; CE1-
CE2- CZ- HZ
6701 6705 6703 6704 1 180.00 4.60240 2 ; CZ-
CD2- CE2- HE2
6707 6711 6709 6710 1 180.00 4.60240 2 ; C-
CA- N- H
6711 6716 6714 6715 1 180.00 43.93200 2 ; CA-
N- C- O
6714 6718 6716 6717 1 180.00 4.60240 2 ; C-
CA- N- H
6718 6740 6738 6739 1 180.00 43.93200 2 ; CA-
N- C- O
6726 6731 6729 6730 1 180.00 4.18400 2 ; CD-
CZ- NE- HE
6729 6732 6731 6735 1 180.00 43.93200 2 ; NE-
NH1- CZ- NH2
6731 6733 6732 6734 1 180.00 4.18400 2 ; CZ-
HH11- NH1- HH12
6731 6736 6735 6737 1 180.00 4.18400 2 ; CZ-
HH21- NH2- HH22

6738 6742 6740 6741 1 180.00 4.60240 2 ; C-
CA- N- H
6742 6752 6750 6751 1 180.00 43.93200 2 ; CA-
N- C- O
6744 6748 6747 6749 1 180.00 43.93200 2 ; CB-
OD1- CG- OD2
6750 6754 6752 6753 1 180.00 4.60240 2 ; C-
CA- N- H
6754 6764 6762 6763 1 180.00 43.93200 2 ; CA-
N- C- O
6756 6760 6759 6761 1 180.00 43.93200 2 ; CB-
OD1- CG- OD2
6762 6766 6764 6765 1 180.00 4.60240 2 ; C-
CA- N- H
6766 6783 6781 6782 1 180.00 43.93200 2 ; CA-
N- C- O
6781 6785 6783 6784 1 180.00 4.60240 2 ; C-
CA- N- H
6785 6798 6796 6797 1 180.00 43.93200 2 ; CA-
N- C- O
6790 6794 6793 6795 1 180.00 43.93200 2 ; CG-
OE1- CD- OE2
6796 6800 6798 6799 1 180.00 4.60240 2 ; C-
CA- N- H
6800 6812 6810 6811 1 180.00 43.93200 2 ; CA-
N- C- O
6802 6807 6805 6806 1 180.00 43.93200 2 ; CB-
ND2- CG- OD1
6805 6808 6807 6809 1 180.00 4.18400 2 ; CG-
HD21- ND2- HD22
6810 6813 6812 6822 1 180.00 4.18400 2 ; C-
CD- N- CA
6822 6826 6824 6825 1 180.00 43.93200 2 ; CA-
N- C- O
6824 6828 6826 6827 1 180.00 4.60240 2 ; C-
CA- N- H
6828 6845 6843 6844 1 180.00 43.93200 2 ; CA-
N- C- O
6843 6847 6845 6846 1 180.00 4.60240 2 ; C-
CA- N- H
6847 6864 6862 6863 1 180.00 43.93200 2 ; CA-
N- C- O
6862 6866 6864 6865 1 180.00 4.60240 2 ; C-
CA- N- H
6866 6876 6874 6875 1 180.00 43.93200 2 ; CA-
N- C- O
6868 6872 6871 6873 1 180.00 43.93200 2 ; CB-
OD1- CG- OD2
6874 6878 6876 6877 1 180.00 4.60240 2 ; C-
CA- N- H
6878 6898 6896 6897 1 180.00 43.93200 2 ; CA-
N- C- O

6896 6900 6898 6899 1 180.00 4.60240 2 ; C-
CA- N- H
6900 6908 6906 6907 1 180.00 43.93200 2 ; CA-
N- C- O
6906 6910 6908 6909 1 180.00 4.60240 2 ; C-
CA- N- H
6910 6927 6925 6926 1 180.00 43.93200 2 ; CA-
N- C- O
6925 6929 6927 6928 1 180.00 4.60240 2 ; C-
CA- N- H
6929 6939 6937 6938 1 180.00 43.93200 2 ; CA-
N- C- O
6931 6935 6934 6936 1 180.00 43.93200 2 ; CB-
OD1- CG- OD2
6937 6941 6939 6940 1 180.00 4.60240 2 ; C-
CA- N- H
6941 6950 6948 6949 1 180.00 43.93200 2 ; CA-
N- C- O
6948 6952 6950 6951 1 180.00 4.60240 2 ; C-
CA- N- H
6952 6969 6967 6968 1 180.00 43.93200 2 ; CA-
N- C- O
6967 6971 6969 6970 1 180.00 4.60240 2 ; C-
CA- N- H
6971 6988 6986 6987 1 180.00 43.93200 2 ; CA-
N- C- O
6986 6990 6988 6989 1 180.00 4.60240 2 ; C-
CA- N- H
6990 7005 7003 7004 1 180.00 43.93200 2 ; CA-
N- C- O
6995 7000 6998 6999 1 180.00 43.93200 2 ; CG-
NE2- CD- OE1
6998 7001 7000 7002 1 180.00 4.18400 2 ; CD-
HE21- NE2- HE22
7003 7007 7005 7006 1 180.00 4.60240 2 ; C-
CA- N- H
7007 7020 7018 7019 1 180.00 43.93200 2 ; CA-
N- C- O
7012 7016 7015 7017 1 180.00 43.93200 2 ; CG-
OE1- CD- OE2
7018 7022 7020 7021 1 180.00 4.60240 2 ; C-
CA- N- H
7022 7036 7034 7035 1 180.00 43.93200 2 ; CA-
N- C- O
7034 7038 7036 7037 1 180.00 4.60240 2 ; C-
CA- N- H
7038 7048 7046 7047 1 180.00 43.93200 2 ; CA-
N- C- O
7040 7044 7043 7045 1 180.00 43.93200 2 ; CB-
OD1- CG- OD2
7046 7050 7048 7049 1 180.00 4.60240 2 ; C-
CA- N- H

7050 7067 7065 7066 1 180.00 43.93200 2 ; CA-
N- C- O
7065 7069 7067 7068 1 180.00 4.60240 2 ; C-
CA- N- H
7069 7081 7079 7080 1 180.00 43.93200 2 ; CA-
N- C- O
7071 7076 7074 7075 1 180.00 43.93200 2 ; CB-
ND2- CG- OD1
7074 7077 7076 7078 1 180.00 4.18400 2 ; CG-
HD21- ND2- HD22
7079 7083 7081 7082 1 180.00 4.60240 2 ; C-
CA- N- H
7083 7088 7086 7087 1 180.00 43.93200 2 ; CA-
N- C- O
7086 7090 7088 7089 1 180.00 4.60240 2 ; C-
CA- N- H
7090 7101 7099 7100 1 180.00 43.93200 2 ; CA-
N- C- O
7092 7096 7095 7097 1 180.00 43.93200 2 ; CB-
OD1- CG- OD2
7099 7103 7101 7102 1 180.00 4.60240 2 ; C-
CA- N- H
7103 7108 7106 7107 1 180.00 43.93200 2 ; CA-
N- C- O
7106 7110 7108 7109 1 180.00 4.60240 2 ; C-
CA- N- H
7110 7123 7121 7122 1 180.00 43.93200 2 ; CA-
N- C- O
7115 7119 7118 7120 1 180.00 43.93200 2 ; CG-
OE1- CD- OE2
7121 7125 7123 7124 1 180.00 4.60240 2 ; C-
CA- N- H
7125 7142 7140 7141 1 180.00 43.93200 2 ; CA-
N- C- O
7140 7144 7142 7143 1 180.00 4.60240 2 ; C-
CA- N- H
7144 7154 7152 7153 1 180.00 43.93200 2 ; CA-
N- C- O
7146 7150 7149 7151 1 180.00 43.93200 2 ; CB-
OD1- CG- OD2
7152 7156 7154 7155 1 180.00 4.60240 2 ; C-
CA- N- H
7156 7174 7172 7173 1 180.00 43.93200 2 ; CA-
N- C- O
7161 7164 7162 7163 1 180.00 4.60240 2 ; CG-
CE1- CD1- HD1
7161 7168 7170 7171 1 180.00 4.60240 2 ; CG-
CE2- CD2- HD2
7162 7166 7164 7165 1 180.00 4.60240 2 ; CD1-
CZ- CE1- HE1
7162 7170 7161 7158 1 180.00 4.60240 2 ; CD1-
CD2- CG- CB

7164 7168 7166 7167 1 180.00 4.60240 2 ; CE1-
CE2- CZ- HZ
7166 7170 7168 7169 1 180.00 4.60240 2 ; CZ-
CD2- CE2- HE2
7172 7176 7174 7175 1 180.00 4.60240 2 ; C-
CA- N- H
7176 7191 7189 7190 1 180.00 43.93200 2 ; CA-
N- C- O
7178 7187 7181 7182 1 180.00 4.60240 2 ; CB-
CD2- CG- ND1
7181 7185 7187 7188 1 180.00 4.60240 2 ; CG-
NE2- CD2- HD2
7182 7185 7183 7184 1 180.00 4.60240 2 ; ND1-
NE2- CE1- HE1
7183 7187 7185 7186 1 180.00 4.18400 2 ; CE1-
CD2- NE2- HE2
7189 7193 7191 7192 1 180.00 4.60240 2 ; C-
CA- N- H
7193 7207 7205 7206 1 180.00 43.93200 2 ; CA-
N- C- O
7198 7202 7201 7203 1 180.00 43.93200 2 ; CG-
OE1- CD- OE2
7205 7209 7207 7208 1 180.00 4.60240 2 ; C-
CA- N- H
7209 7227 7225 7226 1 180.00 43.93200 2 ; CA-
N- C- O
7214 7217 7215 7216 1 180.00 4.60240 2 ; CG-
CE1- CD1- HD1
7214 7221 7223 7224 1 180.00 4.60240 2 ; CG-
CE2- CD2- HD2
7215 7219 7217 7218 1 180.00 4.60240 2 ; CD1-
CZ- CE1- HE1
7215 7223 7214 7211 1 180.00 4.60240 2 ; CD1-
CD2- CG- CB
7217 7221 7219 7220 1 180.00 4.60240 2 ; CE1-
CE2- CZ- HZ
7219 7223 7221 7222 1 180.00 4.60240 2 ; CZ-
CD2- CE2- HE2
7225 7229 7227 7228 1 180.00 4.60240 2 ; C-
CA- N- H
7229 7244 7242 7243 1 180.00 43.93200 2 ; CA-
N- C- O
7242 7246 7244 7245 1 180.00 4.60240 2 ; C-
CA- N- H
7246 7263 7261 7262 1 180.00 43.93200 2 ; CA-
N- C- O
7261 7265 7263 7264 1 180.00 4.60240 2 ; C-
CA- N- H
7265 7280 7278 7279 1 180.00 43.93200 2 ; CA-
N- C- O
7278 7282 7280 7281 1 180.00 4.60240 2 ; C-
CA- N- H

7282 7297 7295 7296 1 180.00 43.93200 2 ; CA-
N- C- O
7295 7299 7297 7298 1 180.00 4.60240 2 ; C-
CA- N- H
7299 7308 7306 7307 1 180.00 43.93200 2 ; CA-
N- C- O
7306 7310 7308 7309 1 180.00 4.60240 2 ; C-
CA- N- H
7310 7330 7328 7329 1 180.00 43.93200 2 ; CA-
N- C- O
7328 7332 7330 7331 1 180.00 4.60240 2 ; C-
CA- N- H
7332 7352 7350 7351 1 180.00 43.93200 2 ; CA-
N- C- O
7350 7354 7352 7353 1 180.00 4.60240 2 ; C-
CA- N- H
7354 7373 7372 7374 1 180.00 43.93200 2 ; CA-
O- C- OXT
7376 7384 7383 7424 1 180.00 4.60240 2 ; C38-
C36- C37- H37
7377 7383 7376 7375 1 180.00 4.60240 2 ; C39-
C37- C38- F2
7378 7376 7377 7423 1 180.00 4.60240 2 ; N23-
C38- C39- H39
7378 7384 7385 7395 1 180.00 4.60240 2 ; N23-
C36- C40- C24
7379 7385 7395 7394 1 180.00 4.60240 2 ; N21-
C40- C24- C31
7379 7401 7397 7396 1 180.00 4.60240 2 ; N21-
C26- C25- C30
7383 7385 7384 7386 1 180.00 4.60240 2 ; C37-
C40- C36- C35
7384 7391 7386 7387 1 180.00 4.60240 2 ; C36-
C32- C35- C34
7386 7388 7387 7390 1 180.00 4.60240 2 ; C35-
N22- C34- O42
7387 7392 7388 7389 1 180.00 4.60240 2 ; C34-
C33- N22- H22
7391 7388 7392 7393 1 180.00 4.60240 2 ; C32-
N22- C33- O41
7391 7395 7394 7396 1 180.00 4.60240 2 ; C32-
C24- C31- C30
7392 7386 7391 7394 1 180.00 4.60240 2 ; C33-
C35- C32- C31
7397 7398 7396 7394 1 180.00 4.60240 2 ; C25-
C29- C30- C31
7397 7400 7401 7428 1 180.00 4.60240 2 ; C25-
C27- C26- H26
7398 7400 7399 7416 1 180.00 4.60240 2 ; C29-
C27- C28- O1
7399 7396 7398 7426 1 180.00 4.60240 2 ; C28-
C30- C29- H29

```
  7399   7401   7400   7427      1   180.00   4.60240   2 ;    C28-
C26-   C27-   H27
  7402   7405   7404   7409      1   180.00   4.60240   2 ;    N15-
C5-   C18-   C19
  7402   7410   7403   7419      1   180.00   4.60240   2 ;    N15-
C8-    C7-   H7A
  7403   7411   7410   7415      1   180.00   4.60240   2 ;     C7-
C9-    C8-   N16
  7404   7406   7405   7421      1   180.00   4.60240   2 ;    C18-
C6-    C5-    H5
  7404   7408   7409   7429      1   180.00   4.60240   2 ;    C18-
C3-   C19-   H19
  7405   7407   7406   7420      1   180.00   4.60240   2 ;     C5-
C4-    C6-    H6
  7406   7408   7407   7422      1   180.00   4.60240   2 ;     C6-
C3-    C4-    H4
  7407   7409   7408   7425      1   180.00   4.60240   2 ;     C4-
C19-    C3-    H3
  7410   7412   7411   7418      1   180.00   4.60240   2 ;     C8-
C10-    C9-    H9
  7411   7413   7412   7432      1   180.00   4.60240   2 ;     C9-
C12-   C10-   H10
  7412   7414   7413   7430      1   180.00   4.60240   2 ;    C10-
C11-   C12-   H12
  7413   7431   7414   7415      1   180.00   4.60240   2 ;    C12-
H11-   C11-   N16

[ system ]
 4MVF_dry

[ molecules ]
; Compound        nmols
 4MVF_dry         1
```
