## Supplementary material for "Exploring Ruthenium-based organometallic inhibitors against Plasmodium Calcium Dependent Kinase 2 (PfCDPK2): a combined ensemble docking, QM paramterization and molecular dynamics study": Table S1

**Title: Topology file for 4MVF with E52 ligand prepared after docking and MCPB.py calculation using Gaussian**

```
; 4MVF_E52_dry_GMX.top created by acpype (Rev: 0) on Thu Dec  6
01:21:21 2018

[ defaults ]
; nbfunc        comb-rule       gen-pairs       fudgeLJ fudgeQQ
1               2               yes             0.5     0.8333

[ atomtypes ]
;name   bond_type     mass     charge   ptype   sigma         epsilon
Amb
 N3       N3          0.00000  0.00000   A     3.25000e-01   7.11280e-
01 ; 1.82  0.1700
 H        H           0.00000  0.00000   A     1.06908e-01   6.56888e-
02 ; 0.60  0.0157
 CX       CX          0.00000  0.00000   A     3.39967e-01   4.57730e-
01 ; 1.91  0.1094
 HP       HP          0.00000  0.00000   A     1.95998e-01   6.56888e-
02 ; 1.10  0.0157
 A2C      A2C         0.00000  0.00000   A     3.39967e-01   4.57730e-
01 ; 1.91  0.1094
 HC       HC          0.00000  0.00000   A     2.64953e-01   6.56888e-
02 ; 1.49  0.0157
 A3C      A3C         0.00000  0.00000   A     3.39967e-01   4.57730e-
01 ; 1.91  0.1094
 CT       CT          0.00000  0.00000   A     3.39967e-01   4.57730e-
01 ; 1.91  0.1094
 C        C           0.00000  0.00000   A     3.39967e-01   3.59824e-
01 ; 1.91  0.0860
 O        O           0.00000  0.00000   A     2.95992e-01   8.78640e-
01 ; 1.66  0.2100
 N        N           0.00000  0.00000   A     3.25000e-01   7.11280e-
01 ; 1.82  0.1700
 H1       H1          0.00000  0.00000   A     2.47135e-01   6.56888e-
02 ; 1.39  0.0157
 SH       SH          0.00000  0.00000   A     3.56359e-01
1.04600e+00 ; 2.00  0.2500
 HS       HS          0.00000  0.00000   A     1.06908e-01   6.56888e-
02 ; 0.60  0.0157
 CC       CC          0.00000  0.00000   A     3.39967e-01   3.59824e-
01 ; 1.91  0.0860
 NA       NA          0.00000  0.00000   A     3.25000e-01   7.11280e-
01 ; 1.82  0.1700
 CR       CR          0.00000  0.00000   A     3.39967e-01   3.59824e-
01 ; 1.91  0.0860
 H5       H5          0.00000  0.00000   A     2.42146e-01   6.27600e-
02 ; 1.36  0.0150
 CW       CW          0.00000  0.00000   A     3.39967e-01   3.59824e-
01 ; 1.91  0.0860
```

```
 H4       H4          0.00000  0.00000   A     2.51055e-01   6.27600e-
02 ; 1.41  0.0150
 OH       OH          0.00000  0.00000   A     3.06647e-01   8.80314e-
01 ; 1.72  0.2104
 HO       HO          0.00000  0.00000   A     0.00000e+00
0.00000e+00 ; 0.00  0.0000
 C8       C8          0.00000  0.00000   A     3.39967e-01   4.57730e-
01 ; 1.91  0.1094
 CO       CO          0.00000  0.00000   A     3.39967e-01   3.59824e-
01 ; 1.91  0.0860
 O2       O2          0.00000  0.00000   A     2.95992e-01   8.78640e-
01 ; 1.66  0.2100
 CA       CA          0.00000  0.00000   A     3.39967e-01   3.59824e-
01 ; 1.91  0.0860
 HA       HA          0.00000  0.00000   A     2.59964e-01   6.27600e-
02 ; 1.46  0.0150
 N2       N2          0.00000  0.00000   A     3.25000e-01   7.11280e-
01 ; 1.82  0.1700
 S        S           0.00000  0.00000   A     3.56359e-01
1.04600e+00 ; 2.00  0.2500
 NB       NB          0.00000  0.00000   A     3.25000e-01   7.11280e-
01 ; 1.82  0.1700
 C*       C*          0.00000  0.00000   A     3.39967e-01   3.59824e-
01 ; 1.91  0.0860
 CN       CN          0.00000  0.00000   A     3.39967e-01   3.59824e-
01 ; 1.91  0.0860
 CB       CB          0.00000  0.00000   A     3.39967e-01   3.59824e-
01 ; 1.91  0.0860
 cp       cp          0.00000  0.00000   A     3.39967e-01   3.59824e-
01 ; 1.91  0.0860
 cq       cq          0.00000  0.00000   A     3.39967e-01   3.59824e-
01 ; 1.91  0.0860
 c2       c2          0.00000  0.00000   A     3.39967e-01   3.59824e-
01 ; 1.91  0.0860
 f        f           0.00000  0.00000   A     3.11815e-01   2.55224e-
01 ; 1.75  0.0610
 Y1       Y1          0.00000  0.00000   A     3.25000e-01   7.11280e-
01 ; 1.82  0.1700
 Y2       Y2          0.00000  0.00000   A     3.25000e-01   7.11280e-
01 ; 1.82  0.1700
 Y3       Y3          0.00000  0.00000   A     3.39967e-01   3.59824e-
01 ; 1.91  0.0860
 Y4       Y4          0.00000  0.00000   A     3.39967e-01   3.59824e-
01 ; 1.91  0.0860
 Y5       Y5          0.00000  0.00000   A     3.39967e-01   3.59824e-
01 ; 1.91  0.0860
 Y6       Y6          0.00000  0.00000   A     3.39967e-01   3.59824e-
01 ; 1.91  0.0860
 Y7       Y7          0.00000  0.00000   A     3.39967e-01   3.59824e-
01 ; 1.91  0.0860
 Y8       Y8          0.00000  0.00000   A     3.39967e-01   8.78640e-
01 ; 1.91  0.2100
```

```
 c3       c3          0.00000  0.00000   A     3.39967e-01   4.57730e-
01 ; 1.91  0.1094
 hn       hn          0.00000  0.00000   A     1.06908e-01   6.56888e-
02 ; 0.60  0.0157
 M1       M1          0.00000  0.00000   A     2.63973e-01   2.34304e-
01 ; 1.48  0.0560

[ moleculetype ]
;name            nrexcl
 4MVF_E52_dry     3

[ atoms ]
;   nr  type  resi  res  atom  cgnr     charge      mass       ; qtot
bond_type
     1   N3     1   LEU     N    1     0.101000     14.01000 ; qtot
0.101
     2    H     1   LEU    H1    2     0.214800      1.00800 ; qtot
0.316
     3    H     1   LEU    H2    3     0.214800      1.00800 ; qtot
0.531
     4    H     1   LEU    H3    4     0.214800      1.00800 ; qtot
0.745
     5   CX     1   LEU    CA    5     0.010400     12.01000 ; qtot
0.756
     6   HP     1   LEU    HA    6     0.105300      1.00800 ; qtot
0.861
     7  A2C     1   LEU    CB    7    -0.024400     12.01000 ; qtot
0.837
     8   HC     1   LEU   HB2    8     0.025600      1.00800 ; qtot
0.862
     9   HC     1   LEU   HB3    9     0.025600      1.00800 ; qtot
0.888
    10  A3C     1   LEU    CG   10     0.342100     12.01000 ; qtot
1.230
    11   HC     1   LEU    HG   11    -0.038000      1.00800 ; qtot
1.192
    12   CT     1   LEU   CD1   12    -0.410600     12.01000 ; qtot
0.781
    13   HC     1   LEU  HD11   13     0.098000      1.00800 ; qtot
0.879
    14   HC     1   LEU  HD12   14     0.098000      1.00800 ; qtot
0.977
    15   HC     1   LEU  HD13   15     0.098000      1.00800 ; qtot
1.075
    16   CT     1   LEU   CD2   16    -0.410400     12.01000 ; qtot
0.665
    17   HC     1   LEU  HD21   17     0.098000      1.00800 ; qtot
0.763
    18   HC     1   LEU  HD22   18     0.098000      1.00800 ; qtot
0.861
    19   HC     1   LEU  HD23   19     0.098000      1.00800 ; qtot
```

```
0.786
    46   HC      3   LEU   HB2    46      0.045700      1.00800 ; qtot
0.832
    47   HC      3   LEU   HB3    47      0.045700      1.00800 ; qtot
0.878
    48  A3C      3   LEU    CG    48      0.353100     12.01000 ; qtot
1.231
    49   HC      3   LEU    HG    49     -0.036100      1.00800 ; qtot
1.195
    50   CT      3   LEU   CD1    50     -0.412100     12.01000 ; qtot
0.783
    51   HC      3   LEU  HD11    51      0.100000      1.00800 ; qtot
0.883
    52   HC      3   LEU  HD12    52      0.100000      1.00800 ; qtot
0.983
    53   HC      3   LEU  HD13    53      0.100000      1.00800 ; qtot
1.083
    54   CT      3   LEU   CD2    54     -0.412100     12.01000 ; qtot
0.671
    55   HC      3   LEU  HD21    55      0.100000      1.00800 ; qtot
0.771
    56   HC      3   LEU  HD22    56      0.100000      1.00800 ; qtot
0.871
    57   HC      3   LEU  HD23    57      0.100000      1.00800 ; qtot
0.971
    58    C      3   LEU     C    58      0.597301     12.01000 ; qtot
1.568
    59    O      3   LEU     O    59     -0.567901     16.00000 ; qtot
1.000
    60    N      4   CYS     N    60     -0.415700     14.01000 ; qtot
0.584
    61    H      4   CYS     H    61      0.271900      1.00800 ; qtot
0.856
    62   CX      4   CYS    CA    62      0.021300     12.01000 ; qtot
0.878
    63   H1      4   CYS    HA    63      0.112400      1.00800 ; qtot
0.990
    64  A2C      4   CYS    CB    64     -0.123100     12.01000 ; qtot
0.867
    65   H1      4   CYS   HB2    65      0.111200      1.00800 ; qtot
0.978
    66   H1      4   CYS   HB3    66      0.111200      1.00800 ; qtot
1.089
    67   SH      4   CYS    SG    67     -0.311900     32.06000 ; qtot
0.777
    68   HS      4   CYS    HG    68      0.193300      1.00800 ; qtot
0.971
    69    C      4   CYS     C    69      0.597301     12.01000 ; qtot
1.568
    70    O      4   CYS     O    70     -0.567901     16.00000 ; qtot
1.000
    71    N      5   HIP     N    71     -0.347900     14.01000 ; qtot
```

```
0.652
    72    H      5   HIP      H   72     0.274700      1.00800 ; qtot
0.927
    73   CX      5   HIP     CA   73    -0.135400     12.01000 ; qtot
0.791
    74   H1      5   HIP     HA   74     0.121200      1.00800 ; qtot
0.913
    75   CT      5   HIP     CB   75    -0.041400     12.01000 ; qtot
0.871
    76   HC      5   HIP    HB2   76     0.081000      1.00800 ; qtot
0.952
    77   HC      5   HIP    HB3   77     0.081000      1.00800 ; qtot
1.033
    78   CC      5   HIP     CG   78    -0.001200     12.01000 ; qtot
1.032
    79   NA      5   HIP    ND1   79    -0.151300     14.01000 ; qtot
0.881
    80    H      5   HIP    HD1   80     0.386600      1.00800 ; qtot
1.267
    81   CR      5   HIP    CE1   81    -0.017000     12.01000 ; qtot
1.250
    82   H5      5   HIP    HE1   82     0.268100      1.00800 ; qtot
1.518
    83   NA      5   HIP    NE2   83    -0.171800     14.01000 ; qtot
1.347
    84    H      5   HIP    HE2   84     0.391100      1.00800 ; qtot
1.738
    85   CW      5   HIP    CD2   85    -0.114100     12.01000 ; qtot
1.624
    86   H4      5   HIP    HD2   86     0.231700      1.00800 ; qtot
1.855
    87    C      5   HIP      C   87     0.734101     12.01000 ; qtot
2.589
    88    O      5   HIP      O   88    -0.589401     16.00000 ; qtot
2.000
    89    N      6   SER      N   89    -0.415700     14.01000 ; qtot
1.584
    90    H      6   SER      H   90     0.271900      1.00800 ; qtot
1.856
    91   CX      6   SER     CA   91    -0.024900     12.01000 ; qtot
1.831
    92   H1      6   SER     HA   92     0.084300      1.00800 ; qtot
1.916
    93  A2C      6   SER     CB   93     0.211700     12.01000 ; qtot
2.127
    94   H1      6   SER    HB2   94     0.035200      1.00800 ; qtot
2.163
    95   H1      6   SER    HB3   95     0.035200      1.00800 ; qtot
2.198
    96   OH      6   SER     OG   96    -0.654601     16.00000 ; qtot
1.543
    97   HO      6   SER     HG   97     0.427500      1.00800 ; qtot
```

```
1.971
    98    C     6   SER     C   98     0.597301     12.01000 ; qtot
2.568
    99    O     6   SER     O   99    -0.567901     16.00000 ; qtot
2.000
   100    N     7   GLY     N  100    -0.415700     14.01000 ; qtot
1.584
   101    H     7   GLY     H  101     0.271900      1.00800 ; qtot
1.856
   102   CX     7   GLY    CA  102    -0.025200     12.01000 ; qtot
1.831
   103   H1     7   GLY   HA2  103     0.069800      1.00800 ; qtot
1.901
   104   H1     7   GLY   HA3  104     0.069800      1.00800 ; qtot
1.971
   105    C     7   GLY     C  105     0.597301     12.01000 ; qtot
2.568
   106    O     7   GLY     O  106    -0.567901     16.00000 ; qtot
2.000
   107    N     8   LYS     N  107    -0.347900     14.01000 ; qtot
1.652
   108    H     8   LYS     H  108     0.274700      1.00800 ; qtot
1.927
   109   CX     8   LYS    CA  109    -0.240000     12.01000 ; qtot
1.687
   110   H1     8   LYS    HA  110     0.142600      1.00800 ; qtot
1.829
   111   C8     8   LYS    CB  111    -0.009400     12.01000 ; qtot
1.820
   112   HC     8   LYS   HB2  112     0.036200      1.00800 ; qtot
1.856
   113   HC     8   LYS   HB3  113     0.036200      1.00800 ; qtot
1.892
   114   C8     8   LYS    CG  114     0.018700     12.01000 ; qtot
1.911
   115   HC     8   LYS   HG2  115     0.010300      1.00800 ; qtot
1.921
   116   HC     8   LYS   HG3  116     0.010300      1.00800 ; qtot
1.932
   117   C8     8   LYS    CD  117    -0.047900     12.01000 ; qtot
1.884
   118   HC     8   LYS   HD2  118     0.062100      1.00800 ; qtot
1.946
   119   HC     8   LYS   HD3  119     0.062100      1.00800 ; qtot
2.008
   120   C8     8   LYS    CE  120    -0.014300     12.01000 ; qtot
1.994
   121   HP     8   LYS   HE2  121     0.113500      1.00800 ; qtot
2.107
   122   HP     8   LYS   HE3  122     0.113500      1.00800 ; qtot
2.221
   123   N3     8   LYS    NZ  123    -0.385400     14.01000 ; qtot
```

```
1.835
   124    H     8   LYS   HZ1  124     0.340000      1.00800 ; qtot
2.175
   125    H     8   LYS   HZ2  125     0.340000      1.00800 ; qtot
2.515
   126    H     8   LYS   HZ3  126     0.340000      1.00800 ; qtot
2.855
   127    C     8   LYS     C  127     0.734101     12.01000 ; qtot
3.589
   128    O     8   LYS     O  128    -0.589401     16.00000 ; qtot
3.000
   129    N     9   LEU     N  129    -0.415700     14.01000 ; qtot
2.584
   130    H     9   LEU     H  130     0.271900      1.00800 ; qtot
2.856
   131   CX     9   LEU    CA  131    -0.051800     12.01000 ; qtot
2.804
   132   H1     9   LEU    HA  132     0.092200      1.00800 ; qtot
2.897
   133  A2C     9   LEU    CB  133    -0.110200     12.01000 ; qtot
2.786
   134   HC     9   LEU   HB2  134     0.045700      1.00800 ; qtot
2.832
   135   HC     9   LEU   HB3  135     0.045700      1.00800 ; qtot
2.878
   136  A3C     9   LEU    CG  136     0.353100     12.01000 ; qtot
3.231
   137   HC     9   LEU    HG  137    -0.036100      1.00800 ; qtot
3.195
   138   CT     9   LEU   CD1  138    -0.412100     12.01000 ; qtot
2.783
   139   HC     9   LEU  HD11  139     0.100000      1.00800 ; qtot
2.883
   140   HC     9   LEU  HD12  140     0.100000      1.00800 ; qtot
2.983
   141   HC     9   LEU  HD13  141     0.100000      1.00800 ; qtot
3.083
   142   CT     9   LEU   CD2  142    -0.412100     12.01000 ; qtot
2.671
   143   HC     9   LEU  HD21  143     0.100000      1.00800 ; qtot
2.771
   144   HC     9   LEU  HD22  144     0.100000      1.00800 ; qtot
2.871
   145   HC     9   LEU  HD23  145     0.100000      1.00800 ; qtot
2.971
   146    C     9   LEU     C  146     0.597301     12.01000 ; qtot
3.568
   147    O     9   LEU     O  147    -0.567901     16.00000 ; qtot
3.000
   148    N    10   GLU     N  148    -0.516301     14.01000 ; qtot
2.484
   149    H    10   GLU     H  149     0.293600      1.00800 ; qtot
```

2.777
   150   CX    10   GLU    CA  150     0.039700     12.01000 ; qtot
2.817
   151   H1    10   GLU    HA  151     0.110500      1.00800 ; qtot
2.928
   152  A2C    10   GLU    CB  152     0.056000     12.01000 ; qtot
2.984
   153   HC    10   GLU   HB2  153    -0.017300      1.00800 ; qtot
2.966
   154   HC    10   GLU   HB3  154    -0.017300      1.00800 ; qtot
2.949
   155  A2C    10   GLU    CG  155     0.013600     12.01000 ; qtot
2.963
   156   HC    10   GLU   HG2  156    -0.042500      1.00800 ; qtot
2.920
   157   HC    10   GLU   HG3  157    -0.042500      1.00800 ; qtot
2.878
   158   CO    10   GLU    CD  158     0.805401     12.01000 ; qtot
3.683
   159   O2    10   GLU   OE1  159    -0.818801     16.00000 ; qtot
2.864
   160   O2    10   GLU   OE2  160    -0.818801     16.00000 ; qtot
2.045
   161    C    10   GLU     C  161     0.536601     12.01000 ; qtot
2.582
   162    O    10   GLU     O  162    -0.581901     16.00000 ; qtot
2.000
   163    N    11   ASP     N  163    -0.516301     14.01000 ; qtot
1.484
   164    H    11   ASP     H  164     0.293600      1.00800 ; qtot
1.777
   165   CX    11   ASP    CA  165     0.038100     12.01000 ; qtot
1.815
   166   H1    11   ASP    HA  166     0.088000      1.00800 ; qtot
1.903
   167  A2C    11   ASP    CB  167    -0.030300     12.01000 ; qtot
1.873
   168   HC    11   ASP   HB2  168    -0.012200      1.00800 ; qtot
1.861
   169   HC    11   ASP   HB3  169    -0.012200      1.00800 ; qtot
1.849
   170   CO    11   ASP    CG  170     0.799401     12.01000 ; qtot
2.648
   171   O2    11   ASP   OD1  171    -0.801401     16.00000 ; qtot
1.847
   172   O2    11   ASP   OD2  172    -0.801401     16.00000 ; qtot
1.045
   173    C    11   ASP     C  173     0.536601     12.01000 ; qtot
1.582
   174    O    11   ASP     O  174    -0.581901     16.00000 ; qtot
1.000
   175    N    12   LYS     N  175    -0.347900     14.01000 ; qtot

```
0.652
   176    H     12   LYS     H  176     0.274700      1.00800 ; qtot
0.927
   177   CX     12   LYS    CA  177    -0.240000     12.01000 ; qtot
0.687
   178   H1     12   LYS    HA  178     0.142600      1.00800 ; qtot
0.829
   179   C8     12   LYS    CB  179    -0.009400     12.01000 ; qtot
0.820
   180   HC     12   LYS   HB2  180     0.036200      1.00800 ; qtot
0.856
   181   HC     12   LYS   HB3  181     0.036200      1.00800 ; qtot
0.892
   182   C8     12   LYS    CG  182     0.018700     12.01000 ; qtot
0.911
   183   HC     12   LYS   HG2  183     0.010300      1.00800 ; qtot
0.921
   184   HC     12   LYS   HG3  184     0.010300      1.00800 ; qtot
0.932
   185   C8     12   LYS    CD  185    -0.047900     12.01000 ; qtot
0.884
   186   HC     12   LYS   HD2  186     0.062100      1.00800 ; qtot
0.946
   187   HC     12   LYS   HD3  187     0.062100      1.00800 ; qtot
1.008
   188   C8     12   LYS    CE  188    -0.014300     12.01000 ; qtot
0.994
   189   HP     12   LYS   HE2  189     0.113500      1.00800 ; qtot
1.107
   190   HP     12   LYS   HE3  190     0.113500      1.00800 ; qtot
1.221
   191   N3     12   LYS    NZ  191    -0.385400     14.01000 ; qtot
0.835
   192    H     12   LYS   HZ1  192     0.340000      1.00800 ; qtot
1.175
   193    H     12   LYS   HZ2  193     0.340000      1.00800 ; qtot
1.515
   194    H     12   LYS   HZ3  194     0.340000      1.00800 ; qtot
1.855
   195    C     12   LYS     C  195     0.734101     12.01000 ; qtot
2.589
   196    O     12   LYS     O  196    -0.589401     16.00000 ; qtot
2.000
   197    N     13   TYR     N  197    -0.415700     14.01000 ; qtot
1.584
   198    H     13   TYR     H  198     0.271900      1.00800 ; qtot
1.856
   199   CX     13   TYR    CA  199    -0.001400     12.01000 ; qtot
1.855
   200   H1     13   TYR    HA  200     0.087600      1.00800 ; qtot
1.942
   201   CT     13   TYR    CB  201    -0.015200     12.01000 ; qtot
```

1.927
202 HC 13 TYR HB2 202 0.029500 1.00800 ; qtot
1.957
203 HC 13 TYR HB3 203 0.029500 1.00800 ; qtot
1.986
204 CA 13 TYR CG 204 -0.001100 12.01000 ; qtot
1.985
205 CA 13 TYR CD1 205 -0.190600 12.01000 ; qtot
1.795
206 HA 13 TYR HD1 206 0.169900 1.00800 ; qtot
1.964
207 CA 13 TYR CE1 207 -0.234100 12.01000 ; qtot
1.730
208 HA 13 TYR HE1 208 0.165600 1.00800 ; qtot
1.896
209 C 13 TYR CZ 209 0.322600 12.01000 ; qtot
2.219
210 OH 13 TYR OH 210 -0.557901 16.00000 ; qtot
1.661
211 HO 13 TYR HH 211 0.399200 1.00800 ; qtot
2.060
212 CA 13 TYR CE2 212 -0.234100 12.01000 ; qtot
1.826
213 HA 13 TYR HE2 213 0.165600 1.00800 ; qtot
1.991
214 CA 13 TYR CD2 214 -0.190600 12.01000 ; qtot
1.801
215 HA 13 TYR HD2 215 0.169900 1.00800 ; qtot
1.971
216 C 13 TYR C 216 0.597301 12.01000 ; qtot
2.568
217 O 13 TYR O 217 -0.567901 16.00000 ; qtot
2.000
218 N 14 ILE N 218 -0.415700 14.01000 ; qtot
1.584
219 H 14 ILE H 219 0.271900 1.00800 ; qtot
1.856
220 CX 14 ILE CA 220 -0.059700 12.01000 ; qtot
1.797
221 H1 14 ILE HA 221 0.086900 1.00800 ; qtot
1.883
222 A3C 14 ILE CB 222 0.130300 12.01000 ; qtot
2.014
223 HC 14 ILE HB 223 0.018700 1.00800 ; qtot
2.032
224 CT 14 ILE CG2 224 -0.320400 12.01000 ; qtot
1.712
225 HC 14 ILE HG21 225 0.088200 1.00800 ; qtot
1.800
226 HC 14 ILE HG22 226 0.088200 1.00800 ; qtot
1.888
227 HC 14 ILE HG23 227 0.088200 1.00800 ; qtot

```
1.977
   228  A2C    14   ILE   CG1  228    -0.043000     12.01000 ; qtot
1.934
   229   HC    14   ILE  HG12  229     0.023600      1.00800 ; qtot
1.957
   230   HC    14   ILE  HG13  230     0.023600      1.00800 ; qtot
1.981
   231   CT    14   ILE   CD1  231    -0.066000     12.01000 ; qtot
1.915
   232   HC    14   ILE  HD11  232     0.018600      1.00800 ; qtot
1.933
   233   HC    14   ILE  HD12  233     0.018600      1.00800 ; qtot
1.952
   234   HC    14   ILE  HD13  234     0.018600      1.00800 ; qtot
1.971
   235    C    14   ILE     C  235     0.597301     12.01000 ; qtot
2.568
   236    O    14   ILE     O  236    -0.567901     16.00000 ; qtot
2.000
   237    N    15   ILE     N  237    -0.415700     14.01000 ; qtot
1.584
   238    H    15   ILE     H  238     0.271900      1.00800 ; qtot
1.856
   239   CX    15   ILE    CA  239    -0.059700     12.01000 ; qtot
1.797
   240   H1    15   ILE    HA  240     0.086900      1.00800 ; qtot
1.883
   241  A3C    15   ILE    CB  241     0.130300     12.01000 ; qtot
2.014
   242   HC    15   ILE    HB  242     0.018700      1.00800 ; qtot
2.032
   243   CT    15   ILE   CG2  243    -0.320400     12.01000 ; qtot
1.712
   244   HC    15   ILE  HG21  244     0.088200      1.00800 ; qtot
1.800
   245   HC    15   ILE  HG22  245     0.088200      1.00800 ; qtot
1.888
   246   HC    15   ILE  HG23  246     0.088200      1.00800 ; qtot
1.977
   247  A2C    15   ILE   CG1  247    -0.043000     12.01000 ; qtot
1.934
   248   HC    15   ILE  HG12  248     0.023600      1.00800 ; qtot
1.957
   249   HC    15   ILE  HG13  249     0.023600      1.00800 ; qtot
1.981
   250   CT    15   ILE   CD1  250    -0.066000     12.01000 ; qtot
1.915
   251   HC    15   ILE  HD11  251     0.018600      1.00800 ; qtot
1.933
   252   HC    15   ILE  HD12  252     0.018600      1.00800 ; qtot
1.952
   253   HC    15   ILE  HD13  253     0.018600      1.00800 ; qtot
```

```
1.971
   254    C    15   ILE     C  254     0.597301     12.01000 ; qtot
2.568
   255    O    15   ILE     O  255    -0.567901     16.00000 ; qtot
2.000
   256    N    16   ASP     N  256    -0.516301     14.01000 ; qtot
1.484
   257    H    16   ASP     H  257     0.293600      1.00800 ; qtot
1.777
   258   CX    16   ASP    CA  258     0.038100     12.01000 ; qtot
1.815
   259   H1    16   ASP    HA  259     0.088000      1.00800 ; qtot
1.903
   260  A2C    16   ASP    CB  260    -0.030300     12.01000 ; qtot
1.873
   261   HC    16   ASP   HB2  261    -0.012200      1.00800 ; qtot
1.861
   262   HC    16   ASP   HB3  262    -0.012200      1.00800 ; qtot
1.849
   263   CO    16   ASP    CG  263     0.799401     12.01000 ; qtot
2.648
   264   O2    16   ASP   OD1  264    -0.801401     16.00000 ; qtot
1.847
   265   O2    16   ASP   OD2  265    -0.801401     16.00000 ; qtot
1.045
   266    C    16   ASP     C  266     0.536601     12.01000 ; qtot
1.582
   267    O    16   ASP     O  267    -0.581901     16.00000 ; qtot
1.000
   268    N    17   GLU     N  268    -0.516301     14.01000 ; qtot
0.484
   269    H    17   GLU     H  269     0.293600      1.00800 ; qtot
0.777
   270   CX    17   GLU    CA  270     0.039700     12.01000 ; qtot
0.817
   271   H1    17   GLU    HA  271     0.110500      1.00800 ; qtot
0.928
   272  A2C    17   GLU    CB  272     0.056000     12.01000 ; qtot
0.984
   273   HC    17   GLU   HB2  273    -0.017300      1.00800 ; qtot
0.966
   274   HC    17   GLU   HB3  274    -0.017300      1.00800 ; qtot
0.949
   275  A2C    17   GLU    CG  275     0.013600     12.01000 ; qtot
0.963
   276   HC    17   GLU   HG2  276    -0.042500      1.00800 ; qtot
0.920
   277   HC    17   GLU   HG3  277    -0.042500      1.00800 ; qtot
0.878
   278   CO    17   GLU    CD  278     0.805401     12.01000 ; qtot
1.683
   279   O2    17   GLU   OE1  279    -0.818801     16.00000 ; qtot
```

0.864
280 O2 17 GLU OE2 280 -0.818801 16.00000 ; qtot
0.045
281 C 17 GLU C 281 0.536601 12.01000 ; qtot
0.582
282 O 17 GLU O 282 -0.581901 16.00000 ; qtot -
0.000
283 N 18 LYS N 283 -0.347900 14.01000 ; qtot -
0.348
284 H 18 LYS H 284 0.274700 1.00800 ; qtot -
0.073
285 CX 18 LYS CA 285 -0.240000 12.01000 ; qtot -
0.313
286 H1 18 LYS HA 286 0.142600 1.00800 ; qtot -
0.171
287 C8 18 LYS CB 287 -0.009400 12.01000 ; qtot -
0.180
288 HC 18 LYS HB2 288 0.036200 1.00800 ; qtot -
0.144
289 HC 18 LYS HB3 289 0.036200 1.00800 ; qtot -
0.108
290 C8 18 LYS CG 290 0.018700 12.01000 ; qtot -
0.089
291 HC 18 LYS HG2 291 0.010300 1.00800 ; qtot -
0.079
292 HC 18 LYS HG3 292 0.010300 1.00800 ; qtot -
0.068
293 C8 18 LYS CD 293 -0.047900 12.01000 ; qtot -
0.116
294 HC 18 LYS HD2 294 0.062100 1.00800 ; qtot -
0.054
295 HC 18 LYS HD3 295 0.062100 1.00800 ; qtot
0.008
296 C8 18 LYS CE 296 -0.014300 12.01000 ; qtot -
0.006
297 HP 18 LYS HE2 297 0.113500 1.00800 ; qtot
0.107
298 HP 18 LYS HE3 298 0.113500 1.00800 ; qtot
0.221
299 N3 18 LYS NZ 299 -0.385400 14.01000 ; qtot -
0.165
300 H 18 LYS HZ1 300 0.340000 1.00800 ; qtot
0.175
301 H 18 LYS HZ2 301 0.340000 1.00800 ; qtot
0.515
302 H 18 LYS HZ3 302 0.340000 1.00800 ; qtot
0.855
303 C 18 LYS C 303 0.734101 12.01000 ; qtot
1.589
304 O 18 LYS O 304 -0.589401 16.00000 ; qtot
1.000
305 N 19 LEU N 305 -0.415700 14.01000 ; qtot

```
0.584
   306    H    19   LEU     H  306     0.271900      1.00800 ; qtot
0.856
   307   CX    19   LEU    CA  307    -0.051800     12.01000 ; qtot
0.804
   308   H1    19   LEU    HA  308     0.092200      1.00800 ; qtot
0.897
   309  A2C    19   LEU    CB  309    -0.110200     12.01000 ; qtot
0.786
   310   HC    19   LEU   HB2  310     0.045700      1.00800 ; qtot
0.832
   311   HC    19   LEU   HB3  311     0.045700      1.00800 ; qtot
0.878
   312  A3C    19   LEU    CG  312     0.353100     12.01000 ; qtot
1.231
   313   HC    19   LEU    HG  313    -0.036100      1.00800 ; qtot
1.195
   314   CT    19   LEU   CD1  314    -0.412100     12.01000 ; qtot
0.783
   315   HC    19   LEU  HD11  315     0.100000      1.00800 ; qtot
0.883
   316   HC    19   LEU  HD12  316     0.100000      1.00800 ; qtot
0.983
   317   HC    19   LEU  HD13  317     0.100000      1.00800 ; qtot
1.083
   318   CT    19   LEU   CD2  318    -0.412100     12.01000 ; qtot
0.671
   319   HC    19   LEU  HD21  319     0.100000      1.00800 ; qtot
0.771
   320   HC    19   LEU  HD22  320     0.100000      1.00800 ; qtot
0.871
   321   HC    19   LEU  HD23  321     0.100000      1.00800 ; qtot
0.971
   322    C    19   LEU     C  322     0.597301     12.01000 ; qtot
1.568
   323    O    19   LEU     O  323    -0.567901     16.00000 ; qtot
1.000
   324    N    20   GLY     N  324    -0.415700     14.01000 ; qtot
0.584
   325    H    20   GLY     H  325     0.271900      1.00800 ; qtot
0.856
   326   CX    20   GLY    CA  326    -0.025200     12.01000 ; qtot
0.831
   327   H1    20   GLY   HA2  327     0.069800      1.00800 ; qtot
0.901
   328   H1    20   GLY   HA3  328     0.069800      1.00800 ; qtot
0.971
   329    C    20   GLY     C  329     0.597301     12.01000 ; qtot
1.568
   330    O    20   GLY     O  330    -0.567901     16.00000 ; qtot
1.000
   331    N    21   GLN     N  331    -0.415700     14.01000 ; qtot
```

```
0.584
   332    H    21   GLN      H  332      0.271900       1.00800 ; qtot
0.856
   333   CX    21   GLN     CA  333     -0.003100      12.01000 ; qtot
0.853
   334   H1    21   GLN     HA  334      0.085000       1.00800 ; qtot
0.938
   335  A2C    21   GLN     CB  335     -0.003600      12.01000 ; qtot
0.935
   336   HC    21   GLN    HB2  336      0.017100       1.00800 ; qtot
0.952
   337   HC    21   GLN    HB3  337      0.017100       1.00800 ; qtot
0.969
   338  A2C    21   GLN     CG  338     -0.064500      12.01000 ; qtot
0.904
   339   HC    21   GLN    HG2  339      0.035200       1.00800 ; qtot
0.939
   340   HC    21   GLN    HG3  340      0.035200       1.00800 ; qtot
0.975
   341    C    21   GLN     CD  341      0.695101      12.01000 ; qtot
1.670
   342    O    21   GLN    OE1  342     -0.608601      16.00000 ; qtot
1.061
   343    N    21   GLN    NE2  343     -0.940701      14.01000 ; qtot
0.120
   344    H    21   GLN   HE21  344      0.425100       1.00800 ; qtot
0.546
   345    H    21   GLN   HE22  345      0.425100       1.00800 ; qtot
0.971
   346    C    21   GLN      C  346      0.597301      12.01000 ; qtot
1.568
   347    O    21   GLN      O  347     -0.567901      16.00000 ; qtot
1.000
   348    N    22   GLY      N  348     -0.415700      14.01000 ; qtot
0.584
   349    H    22   GLY      H  349      0.271900       1.00800 ; qtot
0.856
   350   CX    22   GLY     CA  350     -0.025200      12.01000 ; qtot
0.831
   351   H1    22   GLY    HA2  351      0.069800       1.00800 ; qtot
0.901
   352   H1    22   GLY    HA3  352      0.069800       1.00800 ; qtot
0.971
   353    C    22   GLY      C  353      0.597301      12.01000 ; qtot
1.568
   354    O    22   GLY      O  354     -0.567901      16.00000 ; qtot
1.000
   355    N    23   THR      N  355     -0.415700      14.01000 ; qtot
0.584
   356    H    23   THR      H  356      0.271900       1.00800 ; qtot
0.856
   357   CX    23   THR     CA  357     -0.038900      12.01000 ; qtot
```

```
0.817
   358   H1    23   THR    HA  358     0.100700      1.00800 ; qtot
0.918
   359  A3C    23   THR    CB  359     0.365400     12.01000 ; qtot
1.283
   360   H1    23   THR    HB  360     0.004300      1.00800 ; qtot
1.288
   361   CT    23   THR   CG2  361    -0.243800     12.01000 ; qtot
1.044
   362   HC    23   THR  HG21  362     0.064200      1.00800 ; qtot
1.108
   363   HC    23   THR  HG22  363     0.064200      1.00800 ; qtot
1.172
   364   HC    23   THR  HG23  364     0.064200      1.00800 ; qtot
1.237
   365   OH    23   THR   OG1  365    -0.676101     16.00000 ; qtot
0.560
   366   HO    23   THR   HG1  366     0.410200      1.00800 ; qtot
0.971
   367    C    23   THR     C  367     0.597301     12.01000 ; qtot
1.568
   368    O    23   THR     O  368    -0.567901     16.00000 ; qtot
1.000
   369    N    24   TYR     N  369    -0.415700     14.01000 ; qtot
0.584
   370    H    24   TYR     H  370     0.271900      1.00800 ; qtot
0.856
   371   CX    24   TYR    CA  371    -0.001400     12.01000 ; qtot
0.855
   372   H1    24   TYR    HA  372     0.087600      1.00800 ; qtot
0.942
   373   CT    24   TYR    CB  373    -0.015200     12.01000 ; qtot
0.927
   374   HC    24   TYR   HB2  374     0.029500      1.00800 ; qtot
0.957
   375   HC    24   TYR   HB3  375     0.029500      1.00800 ; qtot
0.986
   376   CA    24   TYR    CG  376    -0.001100     12.01000 ; qtot
0.985
   377   CA    24   TYR   CD1  377    -0.190600     12.01000 ; qtot
0.795
   378   HA    24   TYR   HD1  378     0.169900      1.00800 ; qtot
0.964
   379   CA    24   TYR   CE1  379    -0.234100     12.01000 ; qtot
0.730
   380   HA    24   TYR   HE1  380     0.165600      1.00800 ; qtot
0.896
   381    C    24   TYR    CZ  381     0.322600     12.01000 ; qtot
1.219
   382   OH    24   TYR    OH  382    -0.557901     16.00000 ; qtot
0.661
   383   HO    24   TYR    HH  383     0.399200      1.00800 ; qtot
```

1.060
384 CA 24 TYR CE2 384 -0.234100 12.01000 ; qtot
0.826
385 HA 24 TYR HE2 385 0.165600 1.00800 ; qtot
0.991
386 CA 24 TYR CD2 386 -0.190600 12.01000 ; qtot
0.801
387 HA 24 TYR HD2 387 0.169900 1.00800 ; qtot
0.971
388 C 24 TYR C 388 0.597301 12.01000 ; qtot
1.568
389 O 24 TYR O 389 -0.567901 16.00000 ; qtot
1.000
390 N 25 GLY N 390 -0.415700 14.01000 ; qtot
0.584
391 H 25 GLY H 391 0.271900 1.00800 ; qtot
0.856
392 CX 25 GLY CA 392 -0.025200 12.01000 ; qtot
0.831
393 H1 25 GLY HA2 393 0.069800 1.00800 ; qtot
0.901
394 H1 25 GLY HA3 394 0.069800 1.00800 ; qtot
0.971
395 C 25 GLY C 395 0.597301 12.01000 ; qtot
1.568
396 O 25 GLY O 396 -0.567901 16.00000 ; qtot
1.000
397 N 26 CYS N 397 -0.415700 14.01000 ; qtot
0.584
398 H 26 CYS H 398 0.271900 1.00800 ; qtot
0.856
399 CX 26 CYS CA 399 0.021300 12.01000 ; qtot
0.878
400 H1 26 CYS HA 400 0.112400 1.00800 ; qtot
0.990
401 A2C 26 CYS CB 401 -0.123100 12.01000 ; qtot
0.867
402 H1 26 CYS HB2 402 0.111200 1.00800 ; qtot
0.978
403 H1 26 CYS HB3 403 0.111200 1.00800 ; qtot
1.089
404 SH 26 CYS SG 404 -0.311900 32.06000 ; qtot
0.777
405 HS 26 CYS HG 405 0.193300 1.00800 ; qtot
0.971
406 C 26 CYS C 406 0.597301 12.01000 ; qtot
1.568
407 O 26 CYS O 407 -0.567901 16.00000 ; qtot
1.000
408 N 27 VAL N 408 -0.415700 14.01000 ; qtot
0.584
409 H 27 VAL H 409 0.271900 1.00800 ; qtot

0.856
   410   CX    27   VAL     CA  410    -0.087500     12.01000 ; qtot
0.769
   411   H1    27   VAL     HA  411     0.096900      1.00800 ; qtot
0.866
   412  A3C    27   VAL     CB  412     0.298500     12.01000 ; qtot
1.164
   413   HC    27   VAL     HB  413    -0.029700      1.00800 ; qtot
1.134
   414   CT    27   VAL    CG1  414    -0.319200     12.01000 ; qtot
0.815
   415   HC    27   VAL   HG11  415     0.079100      1.00800 ; qtot
0.894
   416   HC    27   VAL   HG12  416     0.079100      1.00800 ; qtot
0.973
   417   HC    27   VAL   HG13  417     0.079100      1.00800 ; qtot
1.053
   418   CT    27   VAL    CG2  418    -0.319200     12.01000 ; qtot
0.733
   419   HC    27   VAL   HG21  419     0.079100      1.00800 ; qtot
0.812
   420   HC    27   VAL   HG22  420     0.079100      1.00800 ; qtot
0.892
   421   HC    27   VAL   HG23  421     0.079100      1.00800 ; qtot
0.971
   422    C    27   VAL      C  422     0.597301     12.01000 ; qtot
1.568
   423    O    27   VAL      O  423    -0.567901     16.00000 ; qtot
1.000
   424    N    28   TYR      N  424    -0.415700     14.01000 ; qtot
0.584
   425    H    28   TYR      H  425     0.271900      1.00800 ; qtot
0.856
   426   CX    28   TYR     CA  426    -0.001400     12.01000 ; qtot
0.855
   427   H1    28   TYR     HA  427     0.087600      1.00800 ; qtot
0.942
   428   CT    28   TYR     CB  428    -0.015200     12.01000 ; qtot
0.927
   429   HC    28   TYR    HB2  429     0.029500      1.00800 ; qtot
0.957
   430   HC    28   TYR    HB3  430     0.029500      1.00800 ; qtot
0.986
   431   CA    28   TYR     CG  431    -0.001100     12.01000 ; qtot
0.985
   432   CA    28   TYR    CD1  432    -0.190600     12.01000 ; qtot
0.795
   433   HA    28   TYR    HD1  433     0.169900      1.00800 ; qtot
0.964
   434   CA    28   TYR    CE1  434    -0.234100     12.01000 ; qtot
0.730
   435   HA    28   TYR    HE1  435     0.165600      1.00800 ; qtot

0.896
436 C 28 TYR CZ 436 0.322600 12.01000 ; qtot
1.219
437 OH 28 TYR OH 437 -0.557901 16.00000 ; qtot
0.661
438 HO 28 TYR HH 438 0.399200 1.00800 ; qtot
1.060
439 CA 28 TYR CE2 439 -0.234100 12.01000 ; qtot
0.826
440 HA 28 TYR HE2 440 0.165600 1.00800 ; qtot
0.991
441 CA 28 TYR CD2 441 -0.190600 12.01000 ; qtot
0.801
442 HA 28 TYR HD2 442 0.169900 1.00800 ; qtot
0.971
443 C 28 TYR C 443 0.597301 12.01000 ; qtot
1.568
444 O 28 TYR O 444 -0.567901 16.00000 ; qtot
1.000
445 N 29 LYS N 445 -0.347900 14.01000 ; qtot
0.652
446 H 29 LYS H 446 0.274700 1.00800 ; qtot
0.927
447 CX 29 LYS CA 447 -0.240000 12.01000 ; qtot
0.687
448 H1 29 LYS HA 448 0.142600 1.00800 ; qtot
0.829
449 C8 29 LYS CB 449 -0.009400 12.01000 ; qtot
0.820
450 HC 29 LYS HB2 450 0.036200 1.00800 ; qtot
0.856
451 HC 29 LYS HB3 451 0.036200 1.00800 ; qtot
0.892
452 C8 29 LYS CG 452 0.018700 12.01000 ; qtot
0.911
453 HC 29 LYS HG2 453 0.010300 1.00800 ; qtot
0.921
454 HC 29 LYS HG3 454 0.010300 1.00800 ; qtot
0.932
455 C8 29 LYS CD 455 -0.047900 12.01000 ; qtot
0.884
456 HC 29 LYS HD2 456 0.062100 1.00800 ; qtot
0.946
457 HC 29 LYS HD3 457 0.062100 1.00800 ; qtot
1.008
458 C8 29 LYS CE 458 -0.014300 12.01000 ; qtot
0.994
459 HP 29 LYS HE2 459 0.113500 1.00800 ; qtot
1.107
460 HP 29 LYS HE3 460 0.113500 1.00800 ; qtot
1.221
461 N3 29 LYS NZ 461 -0.385400 14.01000 ; qtot

0.835
   462    H    29   LYS   HZ1  462    0.340000     1.00800 ; qtot
1.175
   463    H    29   LYS   HZ2  463    0.340000     1.00800 ; qtot
1.515
   464    H    29   LYS   HZ3  464    0.340000     1.00800 ; qtot
1.855
   465    C    29   LYS     C  465    0.734101    12.01000 ; qtot
2.589
   466    O    29   LYS     O  466   -0.589401    16.00000 ; qtot
2.000
   467    N    30   GLY     N  467   -0.415700    14.01000 ; qtot
1.584
   468    H    30   GLY     H  468    0.271900     1.00800 ; qtot
1.856
   469   CX    30   GLY    CA  469   -0.025200    12.01000 ; qtot
1.831
   470   H1    30   GLY   HA2  470    0.069800     1.00800 ; qtot
1.901
   471   H1    30   GLY   HA3  471    0.069800     1.00800 ; qtot
1.971
   472    C    30   GLY     C  472    0.597301    12.01000 ; qtot
2.568
   473    O    30   GLY     O  473   -0.567901    16.00000 ; qtot
2.000
   474    N    31   ILE     N  474   -0.415700    14.01000 ; qtot
1.584
   475    H    31   ILE     H  475    0.271900     1.00800 ; qtot
1.856
   476   CX    31   ILE    CA  476   -0.059700    12.01000 ; qtot
1.797
   477   H1    31   ILE    HA  477    0.086900     1.00800 ; qtot
1.883
   478  A3C    31   ILE    CB  478    0.130300    12.01000 ; qtot
2.014
   479   HC    31   ILE    HB  479    0.018700     1.00800 ; qtot
2.032
   480   CT    31   ILE   CG2  480   -0.320400    12.01000 ; qtot
1.712
   481   HC    31   ILE  HG21  481    0.088200     1.00800 ; qtot
1.800
   482   HC    31   ILE  HG22  482    0.088200     1.00800 ; qtot
1.888
   483   HC    31   ILE  HG23  483    0.088200     1.00800 ; qtot
1.977
   484  A2C    31   ILE   CG1  484   -0.043000    12.01000 ; qtot
1.934
   485   HC    31   ILE  HG12  485    0.023600     1.00800 ; qtot
1.957
   486   HC    31   ILE  HG13  486    0.023600     1.00800 ; qtot
1.981
   487   CT    31   ILE   CD1  487   -0.066000    12.01000 ; qtot

1.915
488 HC 31 ILE HD11 488 0.018600 1.00800 ; qtot
1.933
489 HC 31 ILE HD12 489 0.018600 1.00800 ; qtot
1.952
490 HC 31 ILE HD13 490 0.018600 1.00800 ; qtot
1.971
491 C 31 ILE C 491 0.597301 12.01000 ; qtot
2.568
492 O 31 ILE O 492 -0.567901 16.00000 ; qtot
2.000
493 N 32 ASP N 493 -0.516301 14.01000 ; qtot
1.484
494 H 32 ASP H 494 0.293600 1.00800 ; qtot
1.777
495 CX 32 ASP CA 495 0.038100 12.01000 ; qtot
1.815
496 H1 32 ASP HA 496 0.088000 1.00800 ; qtot
1.903
497 A2C 32 ASP CB 497 -0.030300 12.01000 ; qtot
1.873
498 HC 32 ASP HB2 498 -0.012200 1.00800 ; qtot
1.861
499 HC 32 ASP HB3 499 -0.012200 1.00800 ; qtot
1.849
500 CO 32 ASP CG 500 0.799401 12.01000 ; qtot
2.648
501 O2 32 ASP OD1 501 -0.801401 16.00000 ; qtot
1.847
502 O2 32 ASP OD2 502 -0.801401 16.00000 ; qtot
1.045
503 C 32 ASP C 503 0.536601 12.01000 ; qtot
1.582
504 O 32 ASP O 504 -0.581901 16.00000 ; qtot
1.000
505 N 33 LYS N 505 -0.347900 14.01000 ; qtot
0.652
506 H 33 LYS H 506 0.274700 1.00800 ; qtot
0.927
507 CX 33 LYS CA 507 -0.240000 12.01000 ; qtot
0.687
508 H1 33 LYS HA 508 0.142600 1.00800 ; qtot
0.829
509 C8 33 LYS CB 509 -0.009400 12.01000 ; qtot
0.820
510 HC 33 LYS HB2 510 0.036200 1.00800 ; qtot
0.856
511 HC 33 LYS HB3 511 0.036200 1.00800 ; qtot
0.892
512 C8 33 LYS CG 512 0.018700 12.01000 ; qtot
0.911
513 HC 33 LYS HG2 513 0.010300 1.00800 ; qtot

```
0.921
   514   HC     33   LYS   HG3  514     0.010300      1.00800 ; qtot
0.932
   515   C8     33   LYS    CD  515    -0.047900     12.01000 ; qtot
0.884
   516   HC     33   LYS   HD2  516     0.062100      1.00800 ; qtot
0.946
   517   HC     33   LYS   HD3  517     0.062100      1.00800 ; qtot
1.008
   518   C8     33   LYS    CE  518    -0.014300     12.01000 ; qtot
0.994
   519   HP     33   LYS   HE2  519     0.113500      1.00800 ; qtot
1.107
   520   HP     33   LYS   HE3  520     0.113500      1.00800 ; qtot
1.221
   521   N3     33   LYS    NZ  521    -0.385400     14.01000 ; qtot
0.835
   522    H     33   LYS   HZ1  522     0.340000      1.00800 ; qtot
1.175
   523    H     33   LYS   HZ2  523     0.340000      1.00800 ; qtot
1.515
   524    H     33   LYS   HZ3  524     0.340000      1.00800 ; qtot
1.855
   525    C     33   LYS     C  525     0.734101     12.01000 ; qtot
2.589
   526    O     33   LYS     O  526    -0.589401     16.00000 ; qtot
2.000
   527    N     34   VAL     N  527    -0.415700     14.01000 ; qtot
1.584
   528    H     34   VAL     H  528     0.271900      1.00800 ; qtot
1.856
   529   CX     34   VAL    CA  529    -0.087500     12.01000 ; qtot
1.769
   530   H1     34   VAL    HA  530     0.096900      1.00800 ; qtot
1.866
   531  A3C     34   VAL    CB  531     0.298500     12.01000 ; qtot
2.164
   532   HC     34   VAL    HB  532    -0.029700      1.00800 ; qtot
2.134
   533   CT     34   VAL   CG1  533    -0.319200     12.01000 ; qtot
1.815
   534   HC     34   VAL  HG11  534     0.079100      1.00800 ; qtot
1.894
   535   HC     34   VAL  HG12  535     0.079100      1.00800 ; qtot
1.973
   536   HC     34   VAL  HG13  536     0.079100      1.00800 ; qtot
2.053
   537   CT     34   VAL   CG2  537    -0.319200     12.01000 ; qtot
1.733
   538   HC     34   VAL  HG21  538     0.079100      1.00800 ; qtot
1.812
   539   HC     34   VAL  HG22  539     0.079100      1.00800 ; qtot
```

1.892
540 HC 34 VAL HG23 540 0.079100 1.00800 ; qtot
1.971
541 C 34 VAL C 541 0.597301 12.01000 ; qtot
2.568
542 O 34 VAL O 542 -0.567901 16.00000 ; qtot
2.000
543 N 35 THR N 543 -0.415700 14.01000 ; qtot
1.584
544 H 35 THR H 544 0.271900 1.00800 ; qtot
1.856
545 CX 35 THR CA 545 -0.038900 12.01000 ; qtot
1.817
546 H1 35 THR HA 546 0.100700 1.00800 ; qtot
1.918
547 A3C 35 THR CB 547 0.365400 12.01000 ; qtot
2.283
548 H1 35 THR HB 548 0.004300 1.00800 ; qtot
2.288
549 CT 35 THR CG2 549 -0.243800 12.01000 ; qtot
2.044
550 HC 35 THR HG21 550 0.064200 1.00800 ; qtot
2.108
551 HC 35 THR HG22 551 0.064200 1.00800 ; qtot
2.172
552 HC 35 THR HG23 552 0.064200 1.00800 ; qtot
2.237
553 OH 35 THR OG1 553 -0.676101 16.00000 ; qtot
1.560
554 HO 35 THR HG1 554 0.410200 1.00800 ; qtot
1.971
555 C 35 THR C 555 0.597301 12.01000 ; qtot
2.568
556 O 35 THR O 556 -0.567901 16.00000 ; qtot
2.000
557 N 36 ASN N 557 -0.415700 14.01000 ; qtot
1.584
558 H 36 ASN H 558 0.271900 1.00800 ; qtot
1.856
559 CX 36 ASN CA 559 0.014300 12.01000 ; qtot
1.871
560 H1 36 ASN HA 560 0.104800 1.00800 ; qtot
1.975
561 A2C 36 ASN CB 561 -0.204100 12.01000 ; qtot
1.771
562 HC 36 ASN HB2 562 0.079700 1.00800 ; qtot
1.851
563 HC 36 ASN HB3 563 0.079700 1.00800 ; qtot
1.931
564 C 36 ASN CG 564 0.713001 12.01000 ; qtot
2.644
565 O 36 ASN OD1 565 -0.593101 16.00000 ; qtot

```
2.051
   566    N    36   ASN   ND2  566    -0.919101     14.01000 ; qtot
1.131
   567    H    36   ASN  HD21  567     0.419600      1.00800 ; qtot
1.551
   568    H    36   ASN  HD22  568     0.419600      1.00800 ; qtot
1.971
   569    C    36   ASN     C  569     0.597301     12.01000 ; qtot
2.568
   570    O    36   ASN     O  570    -0.567901     16.00000 ; qtot
2.000
   571    N    37   GLN     N  571    -0.415700     14.01000 ; qtot
1.584
   572    H    37   GLN     H  572     0.271900      1.00800 ; qtot
1.856
   573   CX    37   GLN    CA  573    -0.003100     12.01000 ; qtot
1.853
   574   H1    37   GLN    HA  574     0.085000      1.00800 ; qtot
1.938
   575  A2C    37   GLN    CB  575    -0.003600     12.01000 ; qtot
1.935
   576   HC    37   GLN   HB2  576     0.017100      1.00800 ; qtot
1.952
   577   HC    37   GLN   HB3  577     0.017100      1.00800 ; qtot
1.969
   578  A2C    37   GLN    CG  578    -0.064500     12.01000 ; qtot
1.904
   579   HC    37   GLN   HG2  579     0.035200      1.00800 ; qtot
1.939
   580   HC    37   GLN   HG3  580     0.035200      1.00800 ; qtot
1.975
   581    C    37   GLN    CD  581     0.695101     12.01000 ; qtot
2.670
   582    O    37   GLN   OE1  582    -0.608601     16.00000 ; qtot
2.061
   583    N    37   GLN   NE2  583    -0.940701     14.01000 ; qtot
1.120
   584    H    37   GLN  HE21  584     0.425100      1.00800 ; qtot
1.546
   585    H    37   GLN  HE22  585     0.425100      1.00800 ; qtot
1.971
   586    C    37   GLN     C  586     0.597301     12.01000 ; qtot
2.568
   587    O    37   GLN     O  587    -0.567901     16.00000 ; qtot
2.000
   588    N    38   LEU     N  588    -0.415700     14.01000 ; qtot
1.584
   589    H    38   LEU     H  589     0.271900      1.00800 ; qtot
1.856
   590   CX    38   LEU    CA  590    -0.051800     12.01000 ; qtot
1.804
   591   H1    38   LEU    HA  591     0.092200      1.00800 ; qtot
```

1.897
592 A2C 38 LEU CB 592 -0.110200 12.01000 ; qtot
1.786
593 HC 38 LEU HB2 593 0.045700 1.00800 ; qtot
1.832
594 HC 38 LEU HB3 594 0.045700 1.00800 ; qtot
1.878
595 A3C 38 LEU CG 595 0.353100 12.01000 ; qtot
2.231
596 HC 38 LEU HG 596 -0.036100 1.00800 ; qtot
2.195
597 CT 38 LEU CD1 597 -0.412100 12.01000 ; qtot
1.783
598 HC 38 LEU HD11 598 0.100000 1.00800 ; qtot
1.883
599 HC 38 LEU HD12 599 0.100000 1.00800 ; qtot
1.983
600 HC 38 LEU HD13 600 0.100000 1.00800 ; qtot
2.083
601 CT 38 LEU CD2 601 -0.412100 12.01000 ; qtot
1.671
602 HC 38 LEU HD21 602 0.100000 1.00800 ; qtot
1.771
603 HC 38 LEU HD22 603 0.100000 1.00800 ; qtot
1.871
604 HC 38 LEU HD23 604 0.100000 1.00800 ; qtot
1.971
605 C 38 LEU C 605 0.597301 12.01000 ; qtot
2.568
606 O 38 LEU O 606 -0.567901 16.00000 ; qtot
2.000
607 N 39 TYR N 607 -0.415700 14.01000 ; qtot
1.584
608 H 39 TYR H 608 0.271900 1.00800 ; qtot
1.856
609 CX 39 TYR CA 609 -0.001400 12.01000 ; qtot
1.855
610 H1 39 TYR HA 610 0.087600 1.00800 ; qtot
1.942
611 CT 39 TYR CB 611 -0.015200 12.01000 ; qtot
1.927
612 HC 39 TYR HB2 612 0.029500 1.00800 ; qtot
1.957
613 HC 39 TYR HB3 613 0.029500 1.00800 ; qtot
1.986
614 CA 39 TYR CG 614 -0.001100 12.01000 ; qtot
1.985
615 CA 39 TYR CD1 615 -0.190600 12.01000 ; qtot
1.795
616 HA 39 TYR HD1 616 0.169900 1.00800 ; qtot
1.964
617 CA 39 TYR CE1 617 -0.234100 12.01000 ; qtot

1.730
618 HA 39 TYR HE1 618 0.165600 1.00800 ; qtot
1.896
619 C 39 TYR CZ 619 0.322600 12.01000 ; qtot
2.219
620 OH 39 TYR OH 620 -0.557901 16.00000 ; qtot
1.661
621 HO 39 TYR HH 621 0.399200 1.00800 ; qtot
2.060
622 CA 39 TYR CE2 622 -0.234100 12.01000 ; qtot
1.826
623 HA 39 TYR HE2 623 0.165600 1.00800 ; qtot
1.991
624 CA 39 TYR CD2 624 -0.190600 12.01000 ; qtot
1.801
625 HA 39 TYR HD2 625 0.169900 1.00800 ; qtot
1.971
626 C 39 TYR C 626 0.597301 12.01000 ; qtot
2.568
627 O 39 TYR O 627 -0.567901 16.00000 ; qtot
2.000
628 N 40 ALA N 628 -0.415700 14.01000 ; qtot
1.584
629 H 40 ALA H 629 0.271900 1.00800 ; qtot
1.856
630 CX 40 ALA CA 630 0.033700 12.01000 ; qtot
1.890
631 H1 40 ALA HA 631 0.082300 1.00800 ; qtot
1.972
632 CT 40 ALA CB 632 -0.182500 12.01000 ; qtot
1.790
633 HC 40 ALA HB1 633 0.060300 1.00800 ; qtot
1.850
634 HC 40 ALA HB2 634 0.060300 1.00800 ; qtot
1.910
635 HC 40 ALA HB3 635 0.060300 1.00800 ; qtot
1.971
636 C 40 ALA C 636 0.597301 12.01000 ; qtot
2.568
637 O 40 ALA O 637 -0.567901 16.00000 ; qtot
2.000
638 N 41 ILE N 638 -0.415700 14.01000 ; qtot
1.584
639 H 41 ILE H 639 0.271900 1.00800 ; qtot
1.856
640 CX 41 ILE CA 640 -0.059700 12.01000 ; qtot
1.797
641 H1 41 ILE HA 641 0.086900 1.00800 ; qtot
1.883
642 A3C 41 ILE CB 642 0.130300 12.01000 ; qtot
2.014
643 HC 41 ILE HB 643 0.018700 1.00800 ; qtot

```
2.032
   644   CT    41   ILE   CG2  644    -0.320400     12.01000 ; qtot
1.712
   645   HC    41   ILE  HG21  645     0.088200      1.00800 ; qtot
1.800
   646   HC    41   ILE  HG22  646     0.088200      1.00800 ; qtot
1.888
   647   HC    41   ILE  HG23  647     0.088200      1.00800 ; qtot
1.977
   648  A2C    41   ILE   CG1  648    -0.043000     12.01000 ; qtot
1.934
   649   HC    41   ILE  HG12  649     0.023600      1.00800 ; qtot
1.957
   650   HC    41   ILE  HG13  650     0.023600      1.00800 ; qtot
1.981
   651   CT    41   ILE   CD1  651    -0.066000     12.01000 ; qtot
1.915
   652   HC    41   ILE  HD11  652     0.018600      1.00800 ; qtot
1.933
   653   HC    41   ILE  HD12  653     0.018600      1.00800 ; qtot
1.952
   654   HC    41   ILE  HD13  654     0.018600      1.00800 ; qtot
1.971
   655    C    41   ILE     C  655     0.597301     12.01000 ; qtot
2.568
   656    O    41   ILE     O  656    -0.567901     16.00000 ; qtot
2.000
   657    N    42   LYS     N  657    -0.347900     14.01000 ; qtot
1.652
   658    H    42   LYS     H  658     0.274700      1.00800 ; qtot
1.927
   659   CX    42   LYS    CA  659    -0.240000     12.01000 ; qtot
1.687
   660   H1    42   LYS    HA  660     0.142600      1.00800 ; qtot
1.829
   661   C8    42   LYS    CB  661    -0.009400     12.01000 ; qtot
1.820
   662   HC    42   LYS   HB2  662     0.036200      1.00800 ; qtot
1.856
   663   HC    42   LYS   HB3  663     0.036200      1.00800 ; qtot
1.892
   664   C8    42   LYS    CG  664     0.018700     12.01000 ; qtot
1.911
   665   HC    42   LYS   HG2  665     0.010300      1.00800 ; qtot
1.921
   666   HC    42   LYS   HG3  666     0.010300      1.00800 ; qtot
1.932
   667   C8    42   LYS    CD  667    -0.047900     12.01000 ; qtot
1.884
   668   HC    42   LYS   HD2  668     0.062100      1.00800 ; qtot
1.946
   669   HC    42   LYS   HD3  669     0.062100      1.00800 ; qtot
```

```
2.008
   670   C8    42   LYS    CE  670    -0.014300     12.01000 ; qtot
1.994
   671   HP    42   LYS   HE2  671     0.113500      1.00800 ; qtot
2.107
   672   HP    42   LYS   HE3  672     0.113500      1.00800 ; qtot
2.221
   673   N3    42   LYS    NZ  673    -0.385400     14.01000 ; qtot
1.835
   674    H    42   LYS   HZ1  674     0.340000      1.00800 ; qtot
2.175
   675    H    42   LYS   HZ2  675     0.340000      1.00800 ; qtot
2.515
   676    H    42   LYS   HZ3  676     0.340000      1.00800 ; qtot
2.855
   677    C    42   LYS     C  677     0.734101     12.01000 ; qtot
3.589
   678    O    42   LYS     O  678    -0.589401     16.00000 ; qtot
3.000
   679    N    43   GLU     N  679    -0.516301     14.01000 ; qtot
2.484
   680    H    43   GLU     H  680     0.293600      1.00800 ; qtot
2.777
   681   CX    43   GLU    CA  681     0.039700     12.01000 ; qtot
2.817
   682   H1    43   GLU    HA  682     0.110500      1.00800 ; qtot
2.928
   683  A2C    43   GLU    CB  683     0.056000     12.01000 ; qtot
2.984
   684   HC    43   GLU   HB2  684    -0.017300      1.00800 ; qtot
2.966
   685   HC    43   GLU   HB3  685    -0.017300      1.00800 ; qtot
2.949
   686  A2C    43   GLU    CG  686     0.013600     12.01000 ; qtot
2.963
   687   HC    43   GLU   HG2  687    -0.042500      1.00800 ; qtot
2.920
   688   HC    43   GLU   HG3  688    -0.042500      1.00800 ; qtot
2.878
   689   CO    43   GLU    CD  689     0.805401     12.01000 ; qtot
3.683
   690   O2    43   GLU   OE1  690    -0.818801     16.00000 ; qtot
2.864
   691   O2    43   GLU   OE2  691    -0.818801     16.00000 ; qtot
2.045
   692    C    43   GLU     C  692     0.536601     12.01000 ; qtot
2.582
   693    O    43   GLU     O  693    -0.581901     16.00000 ; qtot
2.000
   694    N    44   GLU     N  694    -0.516301     14.01000 ; qtot
1.484
   695    H    44   GLU     H  695     0.293600      1.00800 ; qtot
```

```
1.777
   696   CX    44   GLU    CA  696     0.039700     12.01000 ; qtot
1.817
   697   H1    44   GLU    HA  697     0.110500      1.00800 ; qtot
1.928
   698  A2C    44   GLU    CB  698     0.056000     12.01000 ; qtot
1.984
   699   HC    44   GLU   HB2  699    -0.017300      1.00800 ; qtot
1.966
   700   HC    44   GLU   HB3  700    -0.017300      1.00800 ; qtot
1.949
   701  A2C    44   GLU    CG  701     0.013600     12.01000 ; qtot
1.963
   702   HC    44   GLU   HG2  702    -0.042500      1.00800 ; qtot
1.920
   703   HC    44   GLU   HG3  703    -0.042500      1.00800 ; qtot
1.878
   704   CO    44   GLU    CD  704     0.805401     12.01000 ; qtot
2.683
   705   O2    44   GLU   OE1  705    -0.818801     16.00000 ; qtot
1.864
   706   O2    44   GLU   OE2  706    -0.818801     16.00000 ; qtot
1.045
   707    C    44   GLU     C  707     0.536601     12.01000 ; qtot
1.582
   708    O    44   GLU     O  708    -0.581901     16.00000 ; qtot
1.000
   709    N    45   LYS     N  709    -0.347900     14.01000 ; qtot
0.652
   710    H    45   LYS     H  710     0.274700      1.00800 ; qtot
0.927
   711   CX    45   LYS    CA  711    -0.240000     12.01000 ; qtot
0.687
   712   H1    45   LYS    HA  712     0.142600      1.00800 ; qtot
0.829
   713   C8    45   LYS    CB  713    -0.009400     12.01000 ; qtot
0.820
   714   HC    45   LYS   HB2  714     0.036200      1.00800 ; qtot
0.856
   715   HC    45   LYS   HB3  715     0.036200      1.00800 ; qtot
0.892
   716   C8    45   LYS    CG  716     0.018700     12.01000 ; qtot
0.911
   717   HC    45   LYS   HG2  717     0.010300      1.00800 ; qtot
0.921
   718   HC    45   LYS   HG3  718     0.010300      1.00800 ; qtot
0.932
   719   C8    45   LYS    CD  719    -0.047900     12.01000 ; qtot
0.884
   720   HC    45   LYS   HD2  720     0.062100      1.00800 ; qtot
0.946
   721   HC    45   LYS   HD3  721     0.062100      1.00800 ; qtot
```

```
0.045
  1632    C    97   ASP      C 1632     0.536601     12.01000 ; qtot
0.582
  1633    O    97   ASP      O 1633    -0.581901     16.00000 ; qtot -
0.000
  1634    N    98   SER      N 1634    -0.415700     14.01000 ; qtot -
0.416
  1635    H    98   SER      H 1635     0.271900      1.00800 ; qtot -
0.144
  1636   CX    98   SER     CA 1636    -0.024900     12.01000 ; qtot -
0.169
  1637   H1    98   SER     HA 1637     0.084300      1.00800 ; qtot -
0.084
  1638  A2C    98   SER     CB 1638     0.211700     12.01000 ; qtot
0.127
  1639   H1    98   SER    HB2 1639     0.035200      1.00800 ; qtot
0.163
  1640   H1    98   SER    HB3 1640     0.035200      1.00800 ; qtot
0.198
  1641   OH    98   SER     OG 1641    -0.654601     16.00000 ; qtot -
0.457
  1642   HO    98   SER     HG 1642     0.427500      1.00800 ; qtot -
0.029
  1643    C    98   SER      C 1643     0.597301     12.01000 ; qtot
0.568
  1644    O    98   SER      O 1644    -0.567901     16.00000 ; qtot -
0.000
  1645    N    99   ILE      N 1645    -0.415700     14.01000 ; qtot -
0.416
  1646    H    99   ILE      H 1646     0.271900      1.00800 ; qtot -
0.144
  1647   CX    99   ILE     CA 1647    -0.059700     12.01000 ; qtot -
0.204
  1648   H1    99   ILE     HA 1648     0.086900      1.00800 ; qtot -
0.117
  1649  A3C    99   ILE     CB 1649     0.130300     12.01000 ; qtot
0.014
  1650   HC    99   ILE     HB 1650     0.018700      1.00800 ; qtot
0.032
  1651   CT    99   ILE    CG2 1651    -0.320400     12.01000 ; qtot -
0.288
  1652   HC    99   ILE   HG21 1652     0.088200      1.00800 ; qtot -
0.200
  1653   HC    99   ILE   HG22 1653     0.088200      1.00800 ; qtot -
0.112
  1654   HC    99   ILE   HG23 1654     0.088200      1.00800 ; qtot -
0.023
  1655  A2C    99   ILE    CG1 1655    -0.043000     12.01000 ; qtot -
0.066
  1656   HC    99   ILE   HG12 1656     0.023600      1.00800 ; qtot -
0.043
  1657   HC    99   ILE   HG13 1657     0.023600      1.00800 ; qtot -
```

```
0.019
  1658   CT    99   ILE   CD1 1658    -0.066000     12.01000 ; qtot -
0.085
  1659   HC    99   ILE  HD11 1659     0.018600      1.00800 ; qtot -
0.067
  1660   HC    99   ILE  HD12 1660     0.018600      1.00800 ; qtot -
0.048
  1661   HC    99   ILE  HD13 1661     0.018600      1.00800 ; qtot -
0.029
  1662    C    99   ILE     C 1662     0.597301     12.01000 ; qtot
0.568
  1663    O    99   ILE     O 1663    -0.567901     16.00000 ; qtot -
0.000
  1664    N   100   ILE     N 1664    -0.415700     14.01000 ; qtot -
0.416
  1665    H   100   ILE     H 1665     0.271900      1.00800 ; qtot -
0.144
  1666   CX   100   ILE    CA 1666    -0.059700     12.01000 ; qtot -
0.204
  1667   H1   100   ILE    HA 1667     0.086900      1.00800 ; qtot -
0.117
  1668  A3C   100   ILE    CB 1668     0.130300     12.01000 ; qtot
0.014
  1669   HC   100   ILE    HB 1669     0.018700      1.00800 ; qtot
0.032
  1670   CT   100   ILE   CG2 1670    -0.320400     12.01000 ; qtot -
0.288
  1671   HC   100   ILE  HG21 1671     0.088200      1.00800 ; qtot -
0.200
  1672   HC   100   ILE  HG22 1672     0.088200      1.00800 ; qtot -
0.112
  1673   HC   100   ILE  HG23 1673     0.088200      1.00800 ; qtot -
0.023
  1674  A2C   100   ILE   CG1 1674    -0.043000     12.01000 ; qtot -
0.066
  1675   HC   100   ILE  HG12 1675     0.023600      1.00800 ; qtot -
0.043
  1676   HC   100   ILE  HG13 1676     0.023600      1.00800 ; qtot -
0.019
  1677   CT   100   ILE   CD1 1677    -0.066000     12.01000 ; qtot -
0.085
  1678   HC   100   ILE  HD11 1678     0.018600      1.00800 ; qtot -
0.067
  1679   HC   100   ILE  HD12 1679     0.018600      1.00800 ; qtot -
0.048
  1680   HC   100   ILE  HD13 1680     0.018600      1.00800 ; qtot -
0.029
  1681    C   100   ILE     C 1681     0.597301     12.01000 ; qtot
0.568
  1682    O   100   ILE     O 1682    -0.567901     16.00000 ; qtot -
0.000
  1683    N   101   GLU     N 1683    -0.516301     14.01000 ; qtot -
```

```
0.198
  1970   OH   119   SER    OG 1970    -0.654601     16.00000 ; qtot -
0.457
  1971   HO   119   SER    HG 1971     0.427500      1.00800 ; qtot -
0.029
  1972    C   119   SER     C 1972     0.597301     12.01000 ; qtot
0.568
  1973    O   119   SER     O 1973    -0.567901     16.00000 ; qtot -
0.000
  1974    N   120   ALA     N 1974    -0.415700     14.01000 ; qtot -
0.416
  1975    H   120   ALA     H 1975     0.271900      1.00800 ; qtot -
0.144
  1976   CX   120   ALA    CA 1976     0.033700     12.01000 ; qtot -
0.110
  1977   H1   120   ALA    HA 1977     0.082300      1.00800 ; qtot -
0.028
  1978   CT   120   ALA    CB 1978    -0.182500     12.01000 ; qtot -
0.210
  1979   HC   120   ALA   HB1 1979     0.060300      1.00800 ; qtot -
0.150
  1980   HC   120   ALA   HB2 1980     0.060300      1.00800 ; qtot -
0.090
  1981   HC   120   ALA   HB3 1981     0.060300      1.00800 ; qtot -
0.029
  1982    C   120   ALA     C 1982     0.597301     12.01000 ; qtot
0.568
  1983    O   120   ALA     O 1983    -0.567901     16.00000 ; qtot -
0.000
  1984    N   121   ILE     N 1984    -0.415700     14.01000 ; qtot -
0.416
  1985    H   121   ILE     H 1985     0.271900      1.00800 ; qtot -
0.144
  1986   CX   121   ILE    CA 1986    -0.059700     12.01000 ; qtot -
0.204
  1987   H1   121   ILE    HA 1987     0.086900      1.00800 ; qtot -
0.117
  1988  A3C   121   ILE    CB 1988     0.130300     12.01000 ; qtot
0.014
  1989   HC   121   ILE    HB 1989     0.018700      1.00800 ; qtot
0.032
  1990   CT   121   ILE   CG2 1990    -0.320400     12.01000 ; qtot -
0.288
  1991   HC   121   ILE  HG21 1991     0.088200      1.00800 ; qtot -
0.200
  1992   HC   121   ILE  HG22 1992     0.088200      1.00800 ; qtot -
0.112
  1993   HC   121   ILE  HG23 1993     0.088200      1.00800 ; qtot -
0.023
  1994  A2C   121   ILE   CG1 1994    -0.043000     12.01000 ; qtot -
0.066
  1995   HC   121   ILE  HG12 1995     0.023600      1.00800 ; qtot -
```

```
0.043
  1996   HC   121   ILE  HG13 1996     0.023600      1.00800 ; qtot -
0.019
  1997   CT   121   ILE   CD1 1997    -0.066000     12.01000 ; qtot -
0.085
  1998   HC   121   ILE  HD11 1998     0.018600      1.00800 ; qtot -
0.067
  1999   HC   121   ILE  HD12 1999     0.018600      1.00800 ; qtot -
0.048
  2000   HC   121   ILE  HD13 2000     0.018600      1.00800 ; qtot -
0.029
  2001    C   121   ILE     C 2001     0.597301     12.01000 ; qtot
0.568
  2002    O   121   ILE     O 2002    -0.567901     16.00000 ; qtot -
0.000
  2003    N   122   PHE     N 2003    -0.415700     14.01000 ; qtot -
0.416
  2004    H   122   PHE     H 2004     0.271900      1.00800 ; qtot -
0.144
  2005   CX   122   PHE    CA 2005    -0.002400     12.01000 ; qtot -
0.146
  2006   H1   122   PHE    HA 2006     0.097800      1.00800 ; qtot -
0.048
  2007   CT   122   PHE    CB 2007    -0.034300     12.01000 ; qtot -
0.083
  2008   HC   122   PHE   HB2 2008     0.029500      1.00800 ; qtot -
0.053
  2009   HC   122   PHE   HB3 2009     0.029500      1.00800 ; qtot -
0.024
  2010   CA   122   PHE    CG 2010     0.011800     12.01000 ; qtot -
0.012
  2011   CA   122   PHE   CD1 2011    -0.125600     12.01000 ; qtot -
0.138
  2012   HA   122   PHE   HD1 2012     0.133000      1.00800 ; qtot -
0.005
  2013   CA   122   PHE   CE1 2013    -0.170400     12.01000 ; qtot -
0.175
  2014   HA   122   PHE   HE1 2014     0.143000      1.00800 ; qtot -
0.032
  2015   CA   122   PHE    CZ 2015    -0.107200     12.01000 ; qtot -
0.139
  2016   HA   122   PHE    HZ 2016     0.129700      1.00800 ; qtot -
0.009
  2017   CA   122   PHE   CE2 2017    -0.170400     12.01000 ; qtot -
0.180
  2018   HA   122   PHE   HE2 2018     0.143000      1.00800 ; qtot -
0.037
  2019   CA   122   PHE   CD2 2019    -0.125600     12.01000 ; qtot -
0.162
  2020   HA   122   PHE   HD2 2020     0.133000      1.00800 ; qtot -
0.029
  2021    C   122   PHE     C 2021     0.597301     12.01000 ; qtot
```

0.568
2022 O 122 PHE O 2022 -0.567901 16.00000 ; qtot -
0.000
2023 N 123 TYR N 2023 -0.415700 14.01000 ; qtot -
0.416
2024 H 123 TYR H 2024 0.271900 1.00800 ; qtot -
0.144
2025 CX 123 TYR CA 2025 -0.001400 12.01000 ; qtot -
0.145
2026 H1 123 TYR HA 2026 0.087600 1.00800 ; qtot -
0.058
2027 CT 123 TYR CB 2027 -0.015200 12.01000 ; qtot -
0.073
2028 HC 123 TYR HB2 2028 0.029500 1.00800 ; qtot -
0.043
2029 HC 123 TYR HB3 2029 0.029500 1.00800 ; qtot -
0.014
2030 CA 123 TYR CG 2030 -0.001100 12.01000 ; qtot -
0.015
2031 CA 123 TYR CD1 2031 -0.190600 12.01000 ; qtot -
0.206
2032 HA 123 TYR HD1 2032 0.169900 1.00800 ; qtot -
0.036
2033 CA 123 TYR CE1 2033 -0.234100 12.01000 ; qtot -
0.270
2034 HA 123 TYR HE1 2034 0.165600 1.00800 ; qtot -
0.104
2035 C 123 TYR CZ 2035 0.322600 12.01000 ; qtot
0.219
2036 OH 123 TYR OH 2036 -0.557901 16.00000 ; qtot -
0.339
2037 HO 123 TYR HH 2037 0.399200 1.00800 ; qtot
0.060
2038 CA 123 TYR CE2 2038 -0.234100 12.01000 ; qtot -
0.174
2039 HA 123 TYR HE2 2039 0.165600 1.00800 ; qtot -
0.009
2040 CA 123 TYR CD2 2040 -0.190600 12.01000 ; qtot -
0.199
2041 HA 123 TYR HD2 2041 0.169900 1.00800 ; qtot -
0.029
2042 C 123 TYR C 2042 0.597301 12.01000 ; qtot
0.568
2043 O 123 TYR O 2043 -0.567901 16.00000 ; qtot -
0.000
2044 N 124 LEU N 2044 -0.415700 14.01000 ; qtot -
0.416
2045 H 124 LEU H 2045 0.271900 1.00800 ; qtot -
0.144
2046 CX 124 LEU CA 2046 -0.051800 12.01000 ; qtot -
0.196
2047 H1 124 LEU HA 2047 0.092200 1.00800 ; qtot -

0.103
  2048  A2C   124   LEU     CB 2048    -0.110200     12.01000 ; qtot -
0.214
  2049   HC   124   LEU    HB2 2049     0.045700      1.00800 ; qtot -
0.168
  2050   HC   124   LEU    HB3 2050     0.045700      1.00800 ; qtot -
0.122
  2051  A3C   124   LEU     CG 2051     0.353100     12.01000 ; qtot
0.231
  2052   HC   124   LEU     HG 2052    -0.036100      1.00800 ; qtot
0.195
  2053   CT   124   LEU    CD1 2053    -0.412100     12.01000 ; qtot -
0.217
  2054   HC   124   LEU   HD11 2054     0.100000      1.00800 ; qtot -
0.117
  2055   HC   124   LEU   HD12 2055     0.100000      1.00800 ; qtot -
0.017
  2056   HC   124   LEU   HD13 2056     0.100000      1.00800 ; qtot
0.083
  2057   CT   124   LEU    CD2 2057    -0.412100     12.01000 ; qtot -
0.329
  2058   HC   124   LEU   HD21 2058     0.100000      1.00800 ; qtot -
0.229
  2059   HC   124   LEU   HD22 2059     0.100000      1.00800 ; qtot -
0.129
  2060   HC   124   LEU   HD23 2060     0.100000      1.00800 ; qtot -
0.029
  2061    C   124   LEU      C 2061     0.597301     12.01000 ; qtot
0.568
  2062    O   124   LEU      O 2062    -0.567901     16.00000 ; qtot -
0.000
  2063    N   125   HIE      N 2063    -0.415700     14.01000 ; qtot -
0.416
  2064    H   125   HIE      H 2064     0.271900      1.00800 ; qtot -
0.144
  2065   CX   125   HIE     CA 2065    -0.058100     12.01000 ; qtot -
0.202
  2066   H1   125   HIE     HA 2066     0.136000      1.00800 ; qtot -
0.066
  2067   CT   125   HIE     CB 2067    -0.007400     12.01000 ; qtot -
0.073
  2068   HC   125   HIE    HB2 2068     0.036700      1.00800 ; qtot -
0.037
  2069   HC   125   HIE    HB3 2069     0.036700      1.00800 ; qtot
0.000
  2070   CC   125   HIE     CG 2070     0.186800     12.01000 ; qtot
0.187
  2071   NB   125   HIE    ND1 2071    -0.543201     14.01000 ; qtot -
0.356
  2072   CR   125   HIE    CE1 2072     0.163500     12.01000 ; qtot -
0.193
  2073   H5   125   HIE    HE1 2073     0.143500      1.00800 ; qtot -

0.049
  2074   NA   125   HIE   NE2 2074    -0.279500     14.01000 ; qtot -
0.329
  2075    H   125   HIE   HE2 2075     0.333900      1.00800 ; qtot
0.005
  2076   CW   125   HIE   CD2 2076    -0.220700     12.01000 ; qtot -
0.216
  2077   H4   125   HIE   HD2 2077     0.186200      1.00800 ; qtot -
0.029
  2078    C   125   HIE     C 2078     0.597301     12.01000 ; qtot
0.568
  2079    O   125   HIE     O 2079    -0.567901     16.00000 ; qtot -
0.000
  2080    N   126   SER     N 2080    -0.415700     14.01000 ; qtot -
0.416
  2081    H   126   SER     H 2081     0.271900      1.00800 ; qtot -
0.144
  2082   CX   126   SER    CA 2082    -0.024900     12.01000 ; qtot -
0.169
  2083   H1   126   SER    HA 2083     0.084300      1.00800 ; qtot -
0.084
  2084  A2C   126   SER    CB 2084     0.211700     12.01000 ; qtot
0.127
  2085   H1   126   SER   HB2 2085     0.035200      1.00800 ; qtot
0.163
  2086   H1   126   SER   HB3 2086     0.035200      1.00800 ; qtot
0.198
  2087   OH   126   SER    OG 2087    -0.654601     16.00000 ; qtot -
0.457
  2088   HO   126   SER    HG 2088     0.427500      1.00800 ; qtot -
0.029
  2089    C   126   SER     C 2089     0.597301     12.01000 ; qtot
0.568
  2090    O   126   SER     O 2090    -0.567901     16.00000 ; qtot -
0.000
  2091    N   127   LEU     N 2091    -0.415700     14.01000 ; qtot -
0.416
  2092    H   127   LEU     H 2092     0.271900      1.00800 ; qtot -
0.144
  2093   CX   127   LEU    CA 2093    -0.051800     12.01000 ; qtot -
0.196
  2094   H1   127   LEU    HA 2094     0.092200      1.00800 ; qtot -
0.103
  2095  A2C   127   LEU    CB 2095    -0.110200     12.01000 ; qtot -
0.214
  2096   HC   127   LEU   HB2 2096     0.045700      1.00800 ; qtot -
0.168
  2097   HC   127   LEU   HB3 2097     0.045700      1.00800 ; qtot -
0.122
  2098  A3C   127   LEU    CG 2098     0.353100     12.01000 ; qtot
0.231
  2099   HC   127   LEU    HG 2099    -0.036100      1.00800 ; qtot

0.195
  2100   CT   127   LEU   CD1 2100    -0.412100     12.01000 ; qtot -
0.217
  2101   HC   127   LEU  HD11 2101     0.100000      1.00800 ; qtot -
0.117
  2102   HC   127   LEU  HD12 2102     0.100000      1.00800 ; qtot -
0.017
  2103   HC   127   LEU  HD13 2103     0.100000      1.00800 ; qtot
0.083
  2104   CT   127   LEU   CD2 2104    -0.412100     12.01000 ; qtot -
0.329
  2105   HC   127   LEU  HD21 2105     0.100000      1.00800 ; qtot -
0.229
  2106   HC   127   LEU  HD22 2106     0.100000      1.00800 ; qtot -
0.129
  2107   HC   127   LEU  HD23 2107     0.100000      1.00800 ; qtot -
0.029
  2108    C   127   LEU     C 2108     0.597301     12.01000 ; qtot
0.568
  2109    O   127   LEU     O 2109    -0.567901     16.00000 ; qtot -
0.000
  2110    N   128   ASN     N 2110    -0.415700     14.01000 ; qtot -
0.416
  2111    H   128   ASN     H 2111     0.271900      1.00800 ; qtot -
0.144
  2112   CX   128   ASN    CA 2112     0.014300     12.01000 ; qtot -
0.130
  2113   H1   128   ASN    HA 2113     0.104800      1.00800 ; qtot -
0.025
  2114  A2C   128   ASN    CB 2114    -0.204100     12.01000 ; qtot -
0.229
  2115   HC   128   ASN   HB2 2115     0.079700      1.00800 ; qtot -
0.149
  2116   HC   128   ASN   HB3 2116     0.079700      1.00800 ; qtot -
0.069
  2117    C   128   ASN    CG 2117     0.713001     12.01000 ; qtot
0.644
  2118    O   128   ASN   OD1 2118    -0.593101     16.00000 ; qtot
0.050
  2119    N   128   ASN   ND2 2119    -0.919101     14.01000 ; qtot -
0.869
  2120    H   128   ASN  HD21 2120     0.419600      1.00800 ; qtot -
0.449
  2121    H   128   ASN  HD22 2121     0.419600      1.00800 ; qtot -
0.029
  2122    C   128   ASN     C 2122     0.597301     12.01000 ; qtot
0.568
  2123    O   128   ASN     O 2123    -0.567901     16.00000 ; qtot -
0.000
  2124    N   129   ILE     N 2124    -0.415700     14.01000 ; qtot -
0.416
  2125    H   129   ILE     H 2125     0.271900      1.00800 ; qtot -

0.144
  2126   CX   129   ILE     CA 2126    -0.059700     12.01000 ; qtot -
0.204
  2127   H1   129   ILE     HA 2127     0.086900      1.00800 ; qtot -
0.117
  2128  A3C   129   ILE     CB 2128     0.130300     12.01000 ; qtot
0.014
  2129   HC   129   ILE     HB 2129     0.018700      1.00800 ; qtot
0.032
  2130   CT   129   ILE    CG2 2130    -0.320400     12.01000 ; qtot -
0.288
  2131   HC   129   ILE   HG21 2131     0.088200      1.00800 ; qtot -
0.200
  2132   HC   129   ILE   HG22 2132     0.088200      1.00800 ; qtot -
0.112
  2133   HC   129   ILE   HG23 2133     0.088200      1.00800 ; qtot -
0.023
  2134  A2C   129   ILE    CG1 2134    -0.043000     12.01000 ; qtot -
0.066
  2135   HC   129   ILE   HG12 2135     0.023600      1.00800 ; qtot -
0.043
  2136   HC   129   ILE   HG13 2136     0.023600      1.00800 ; qtot -
0.019
  2137   CT   129   ILE    CD1 2137    -0.066000     12.01000 ; qtot -
0.085
  2138   HC   129   ILE   HD11 2138     0.018600      1.00800 ; qtot -
0.067
  2139   HC   129   ILE   HD12 2139     0.018600      1.00800 ; qtot -
0.048
  2140   HC   129   ILE   HD13 2140     0.018600      1.00800 ; qtot -
0.029
  2141    C   129   ILE      C 2141     0.597301     12.01000 ; qtot
0.568
  2142    O   129   ILE      O 2142    -0.567901     16.00000 ; qtot -
0.000
  2143    N   130   VAL      N 2143    -0.415700     14.01000 ; qtot -
0.416
  2144    H   130   VAL      H 2144     0.271900      1.00800 ; qtot -
0.144
  2145   CX   130   VAL     CA 2145    -0.087500     12.01000 ; qtot -
0.231
  2146   H1   130   VAL     HA 2146     0.096900      1.00800 ; qtot -
0.134
  2147  A3C   130   VAL     CB 2147     0.298500     12.01000 ; qtot
0.164
  2148   HC   130   VAL     HB 2148    -0.029700      1.00800 ; qtot
0.134
  2149   CT   130   VAL    CG1 2149    -0.319200     12.01000 ; qtot -
0.185
  2150   HC   130   VAL   HG11 2150     0.079100      1.00800 ; qtot -
0.106
  2151   HC   130   VAL   HG12 2151     0.079100      1.00800 ; qtot -

```
0.027
  2152   HC   130   VAL  HG13 2152     0.079100      1.00800 ; qtot
0.052
  2153   CT   130   VAL   CG2 2153    -0.319200     12.01000 ; qtot -
0.267
  2154   HC   130   VAL  HG21 2154     0.079100      1.00800 ; qtot -
0.188
  2155   HC   130   VAL  HG22 2155     0.079100      1.00800 ; qtot -
0.109
  2156   HC   130   VAL  HG23 2156     0.079100      1.00800 ; qtot -
0.029
  2157    C   130   VAL     C 2157     0.597301     12.01000 ; qtot
0.568
  2158    O   130   VAL     O 2158    -0.567901     16.00000 ; qtot -
0.000
  2159    N   131   HIE     N 2159    -0.415700     14.01000 ; qtot -
0.416
  2160    H   131   HIE     H 2160     0.271900      1.00800 ; qtot -
0.144
  2161   CX   131   HIE    CA 2161    -0.058100     12.01000 ; qtot -
0.202
  2162   H1   131   HIE    HA 2162     0.136000      1.00800 ; qtot -
0.066
  2163   CT   131   HIE    CB 2163    -0.007400     12.01000 ; qtot -
0.073
  2164   HC   131   HIE   HB2 2164     0.036700      1.00800 ; qtot -
0.037
  2165   HC   131   HIE   HB3 2165     0.036700      1.00800 ; qtot
0.000
  2166   CC   131   HIE    CG 2166     0.186800     12.01000 ; qtot
0.187
  2167   NB   131   HIE   ND1 2167    -0.543201     14.01000 ; qtot -
0.356
  2168   CR   131   HIE   CE1 2168     0.163500     12.01000 ; qtot -
0.193
  2169   H5   131   HIE   HE1 2169     0.143500      1.00800 ; qtot -
0.049
  2170   NA   131   HIE   NE2 2170    -0.279500     14.01000 ; qtot -
0.329
  2171    H   131   HIE   HE2 2171     0.333900      1.00800 ; qtot
0.005
  2172   CW   131   HIE   CD2 2172    -0.220700     12.01000 ; qtot -
0.216
  2173   H4   131   HIE   HD2 2173     0.186200      1.00800 ; qtot -
0.029
  2174    C   131   HIE     C 2174     0.597301     12.01000 ; qtot
0.568
  2175    O   131   HIE     O 2175    -0.567901     16.00000 ; qtot -
0.000
  2176    N   132   ARG     N 2176    -0.347900     14.01000 ; qtot -
0.348
  2177    H   132   ARG     H 2177     0.274700      1.00800 ; qtot -
```

```
0.073
  2178   CX   132   ARG    CA 2178    -0.263700     12.01000 ; qtot -
0.337
  2179   H1   132   ARG    HA 2179     0.156000      1.00800 ; qtot -
0.181
  2180   C8   132   ARG    CB 2180    -0.000700     12.01000 ; qtot -
0.182
  2181   HC   132   ARG   HB2 2181     0.032700      1.00800 ; qtot -
0.149
  2182   HC   132   ARG   HB3 2182     0.032700      1.00800 ; qtot -
0.116
  2183   C8   132   ARG    CG 2183     0.039000     12.01000 ; qtot -
0.077
  2184   HC   132   ARG   HG2 2184     0.028500      1.00800 ; qtot -
0.049
  2185   HC   132   ARG   HG3 2185     0.028500      1.00800 ; qtot -
0.020
  2186   C8   132   ARG    CD 2186     0.048600     12.01000 ; qtot
0.028
  2187   H1   132   ARG   HD2 2187     0.068700      1.00800 ; qtot
0.097
  2188   H1   132   ARG   HD3 2188     0.068700      1.00800 ; qtot
0.166
  2189   N2   132   ARG    NE 2189    -0.529501     14.01000 ; qtot -
0.364
  2190    H   132   ARG    HE 2190     0.345600      1.00800 ; qtot -
0.018
  2191   CA   132   ARG    CZ 2191     0.807601     12.01000 ; qtot
0.790
  2192   N2   132   ARG   NH1 2192    -0.862701     14.01000 ; qtot -
0.073
  2193    H   132   ARG  HH11 2193     0.447800      1.00800 ; qtot
0.375
  2194    H   132   ARG  HH12 2194     0.447800      1.00800 ; qtot
0.822
  2195   N2   132   ARG   NH2 2195    -0.862701     14.01000 ; qtot -
0.040
  2196    H   132   ARG  HH21 2196     0.447800      1.00800 ; qtot
0.408
  2197    H   132   ARG  HH22 2197     0.447800      1.00800 ; qtot
0.855
  2198    C   132   ARG     C 2198     0.734101     12.01000 ; qtot
1.589
  2199    O   132   ARG     O 2199    -0.589401     16.00000 ; qtot
1.000
  2200    N   133   ASP     N 2200    -0.516301     14.01000 ; qtot
0.484
  2201    H   133   ASP     H 2201     0.293600      1.00800 ; qtot
0.777
  2202   CX   133   ASP    CA 2202     0.038100     12.01000 ; qtot
0.815
  2203   H1   133   ASP    HA 2203     0.088000      1.00800 ; qtot
```

```
0.903
  2204  A2C   133   ASP    CB 2204    -0.030300     12.01000 ; qtot
0.873
  2205   HC   133   ASP   HB2 2205    -0.012200      1.00800 ; qtot
0.861
  2206   HC   133   ASP   HB3 2206    -0.012200      1.00800 ; qtot
0.849
  2207   CO   133   ASP    CG 2207     0.799401     12.01000 ; qtot
1.648
  2208   O2   133   ASP   OD1 2208    -0.801401     16.00000 ; qtot
0.847
  2209   O2   133   ASP   OD2 2209    -0.801401     16.00000 ; qtot
0.045
  2210    C   133   ASP     C 2210     0.536601     12.01000 ; qtot
0.582
  2211    O   133   ASP     O 2211    -0.581901     16.00000 ; qtot -
0.000
  2212    N   134   LEU     N 2212    -0.415700     14.01000 ; qtot -
0.416
  2213    H   134   LEU     H 2213     0.271900      1.00800 ; qtot -
0.144
  2214   CX   134   LEU    CA 2214    -0.051800     12.01000 ; qtot -
0.196
  2215   H1   134   LEU    HA 2215     0.092200      1.00800 ; qtot -
0.103
  2216  A2C   134   LEU    CB 2216    -0.110200     12.01000 ; qtot -
0.214
  2217   HC   134   LEU   HB2 2217     0.045700      1.00800 ; qtot -
0.168
  2218   HC   134   LEU   HB3 2218     0.045700      1.00800 ; qtot -
0.122
  2219  A3C   134   LEU    CG 2219     0.353100     12.01000 ; qtot
0.231
  2220   HC   134   LEU    HG 2220    -0.036100      1.00800 ; qtot
0.195
  2221   CT   134   LEU   CD1 2221    -0.412100     12.01000 ; qtot -
0.217
  2222   HC   134   LEU  HD11 2222     0.100000      1.00800 ; qtot -
0.117
  2223   HC   134   LEU  HD12 2223     0.100000      1.00800 ; qtot -
0.017
  2224   HC   134   LEU  HD13 2224     0.100000      1.00800 ; qtot
0.083
  2225   CT   134   LEU   CD2 2225    -0.412100     12.01000 ; qtot -
0.329
  2226   HC   134   LEU  HD21 2226     0.100000      1.00800 ; qtot -
0.229
  2227   HC   134   LEU  HD22 2227     0.100000      1.00800 ; qtot -
0.129
  2228   HC   134   LEU  HD23 2228     0.100000      1.00800 ; qtot -
0.029
  2229    C   134   LEU     C 2229     0.597301     12.01000 ; qtot
```

0.568
  2230    O   134   LEU      O 2230    -0.567901     16.00000 ; qtot -
0.000
  2231    N   135   LYS      N 2231    -0.347900     14.01000 ; qtot -
0.348
  2232    H   135   LYS      H 2232     0.274700      1.00800 ; qtot -
0.073
  2233   CX   135   LYS     CA 2233    -0.240000     12.01000 ; qtot -
0.313
  2234   H1   135   LYS     HA 2234     0.142600      1.00800 ; qtot -
0.171
  2235   C8   135   LYS     CB 2235    -0.009400     12.01000 ; qtot -
0.180
  2236   HC   135   LYS    HB2 2236     0.036200      1.00800 ; qtot -
0.144
  2237   HC   135   LYS    HB3 2237     0.036200      1.00800 ; qtot -
0.108
  2238   C8   135   LYS     CG 2238     0.018700     12.01000 ; qtot -
0.089
  2239   HC   135   LYS    HG2 2239     0.010300      1.00800 ; qtot -
0.079
  2240   HC   135   LYS    HG3 2240     0.010300      1.00800 ; qtot -
0.068
  2241   C8   135   LYS     CD 2241    -0.047900     12.01000 ; qtot -
0.116
  2242   HC   135   LYS    HD2 2242     0.062100      1.00800 ; qtot -
0.054
  2243   HC   135   LYS    HD3 2243     0.062100      1.00800 ; qtot
0.008
  2244   C8   135   LYS     CE 2244    -0.014300     12.01000 ; qtot -
0.006
  2245   HP   135   LYS    HE2 2245     0.113500      1.00800 ; qtot
0.107
  2246   HP   135   LYS    HE3 2246     0.113500      1.00800 ; qtot
0.221
  2247   N3   135   LYS     NZ 2247    -0.385400     14.01000 ; qtot -
0.165
  2248    H   135   LYS    HZ1 2248     0.340000      1.00800 ; qtot
0.175
  2249    H   135   LYS    HZ2 2249     0.340000      1.00800 ; qtot
0.515
  2250    H   135   LYS    HZ3 2250     0.340000      1.00800 ; qtot
0.855
  2251    C   135   LYS      C 2251     0.734101     12.01000 ; qtot
1.589
  2252    O   135   LYS      O 2252    -0.589401     16.00000 ; qtot
1.000
  2253    N   136   PRO      N 2253    -0.254800     14.01000 ; qtot
0.745
  2254   CT   136   PRO     CD 2254     0.019200     12.01000 ; qtot
0.764
  2255   H1   136   PRO    HD2 2255     0.039100      1.00800 ; qtot

0.804
2256 H1 136 PRO HD3 2256 0.039100 1.00800 ; qtot
0.843
2257 CT 136 PRO CG 2257 0.018900 12.01000 ; qtot
0.862
2258 HC 136 PRO HG2 2258 0.021300 1.00800 ; qtot
0.883
2259 HC 136 PRO HG3 2259 0.021300 1.00800 ; qtot
0.904
2260 CT 136 PRO CB 2260 -0.007000 12.01000 ; qtot
0.897
2261 HC 136 PRO HB2 2261 0.025300 1.00800 ; qtot
0.922
2262 HC 136 PRO HB3 2262 0.025300 1.00800 ; qtot
0.948
2263 CX 136 PRO CA 2263 -0.026600 12.01000 ; qtot
0.921
2264 H1 136 PRO HA 2264 0.064100 1.00800 ; qtot
0.985
2265 C 136 PRO C 2265 0.589601 12.01000 ; qtot
1.575
2266 O 136 PRO O 2266 -0.574801 16.00000 ; qtot
1.000
2267 N 137 GLU N 2267 -0.516301 14.01000 ; qtot
0.484
2268 H 137 GLU H 2268 0.293600 1.00800 ; qtot
0.777
2269 CX 137 GLU CA 2269 0.039700 12.01000 ; qtot
0.817
2270 H1 137 GLU HA 2270 0.110500 1.00800 ; qtot
0.928
2271 A2C 137 GLU CB 2271 0.056000 12.01000 ; qtot
0.984
2272 HC 137 GLU HB2 2272 -0.017300 1.00800 ; qtot
0.966
2273 HC 137 GLU HB3 2273 -0.017300 1.00800 ; qtot
0.949
2274 A2C 137 GLU CG 2274 0.013600 12.01000 ; qtot
0.963
2275 HC 137 GLU HG2 2275 -0.042500 1.00800 ; qtot
0.920
2276 HC 137 GLU HG3 2276 -0.042500 1.00800 ; qtot
0.878
2277 CO 137 GLU CD 2277 0.805401 12.01000 ; qtot
1.683
2278 O2 137 GLU OE1 2278 -0.818801 16.00000 ; qtot
0.864
2279 O2 137 GLU OE2 2279 -0.818801 16.00000 ; qtot
0.045
2280 C 137 GLU C 2280 0.536601 12.01000 ; qtot
0.582
2281 O 137 GLU O 2281 -0.581901 16.00000 ; qtot -

0.000
2282 N 138 ASN N 2282 -0.415700 14.01000 ; qtot -
0.416
2283 H 138 ASN H 2283 0.271900 1.00800 ; qtot -
0.144
2284 CX 138 ASN CA 2284 0.014300 12.01000 ; qtot -
0.130
2285 H1 138 ASN HA 2285 0.104800 1.00800 ; qtot -
0.025
2286 A2C 138 ASN CB 2286 -0.204100 12.01000 ; qtot -
0.229
2287 HC 138 ASN HB2 2287 0.079700 1.00800 ; qtot -
0.149
2288 HC 138 ASN HB3 2288 0.079700 1.00800 ; qtot -
0.069
2289 C 138 ASN CG 2289 0.713001 12.01000 ; qtot
0.644
2290 O 138 ASN OD1 2290 -0.593101 16.00000 ; qtot
0.050
2291 N 138 ASN ND2 2291 -0.919101 14.01000 ; qtot -
0.869
2292 H 138 ASN HD21 2292 0.419600 1.00800 ; qtot -
0.449
2293 H 138 ASN HD22 2293 0.419600 1.00800 ; qtot -
0.029
2294 C 138 ASN C 2294 0.597301 12.01000 ; qtot
0.568
2295 O 138 ASN O 2295 -0.567901 16.00000 ; qtot -
0.000
2296 N 139 PHE N 2296 -0.415700 14.01000 ; qtot -
0.416
2297 H 139 PHE H 2297 0.271900 1.00800 ; qtot -
0.144
2298 CX 139 PHE CA 2298 -0.002400 12.01000 ; qtot -
0.146
2299 H1 139 PHE HA 2299 0.097800 1.00800 ; qtot -
0.048
2300 CT 139 PHE CB 2300 -0.034300 12.01000 ; qtot -
0.083
2301 HC 139 PHE HB2 2301 0.029500 1.00800 ; qtot -
0.053
2302 HC 139 PHE HB3 2302 0.029500 1.00800 ; qtot -
0.024
2303 CA 139 PHE CG 2303 0.011800 12.01000 ; qtot -
0.012
2304 CA 139 PHE CD1 2304 -0.125600 12.01000 ; qtot -
0.138
2305 HA 139 PHE HD1 2305 0.133000 1.00800 ; qtot -
0.005
2306 CA 139 PHE CE1 2306 -0.170400 12.01000 ; qtot -
0.175
2307 HA 139 PHE HE1 2307 0.143000 1.00800 ; qtot -

0.032
  2308   CA   139   PHE     CZ 2308    -0.107200     12.01000 ; qtot -
0.139
  2309   HA   139   PHE     HZ 2309     0.129700      1.00800 ; qtot -
0.009
  2310   CA   139   PHE    CE2 2310    -0.170400     12.01000 ; qtot -
0.180
  2311   HA   139   PHE    HE2 2311     0.143000      1.00800 ; qtot -
0.037
  2312   CA   139   PHE    CD2 2312    -0.125600     12.01000 ; qtot -
0.162
  2313   HA   139   PHE    HD2 2313     0.133000      1.00800 ; qtot -
0.029
  2314    C   139   PHE      C 2314     0.597301     12.01000 ; qtot
0.568
  2315    O   139   PHE      O 2315    -0.567901     16.00000 ; qtot -
0.000
  2316    N   140   LEU      N 2316    -0.415700     14.01000 ; qtot -
0.416
  2317    H   140   LEU      H 2317     0.271900      1.00800 ; qtot -
0.144
  2318   CX   140   LEU     CA 2318    -0.051800     12.01000 ; qtot -
0.196
  2319   H1   140   LEU     HA 2319     0.092200      1.00800 ; qtot -
0.103
  2320  A2C   140   LEU     CB 2320    -0.110200     12.01000 ; qtot -
0.214
  2321   HC   140   LEU    HB2 2321     0.045700      1.00800 ; qtot -
0.168
  2322   HC   140   LEU    HB3 2322     0.045700      1.00800 ; qtot -
0.122
  2323  A3C   140   LEU     CG 2323     0.353100     12.01000 ; qtot
0.231
  2324   HC   140   LEU     HG 2324    -0.036100      1.00800 ; qtot
0.195
  2325   CT   140   LEU    CD1 2325    -0.412100     12.01000 ; qtot -
0.217
  2326   HC   140   LEU   HD11 2326     0.100000      1.00800 ; qtot -
0.117
  2327   HC   140   LEU   HD12 2327     0.100000      1.00800 ; qtot -
0.017
  2328   HC   140   LEU   HD13 2328     0.100000      1.00800 ; qtot
0.083
  2329   CT   140   LEU    CD2 2329    -0.412100     12.01000 ; qtot -
0.329
  2330   HC   140   LEU   HD21 2330     0.100000      1.00800 ; qtot -
0.229
  2331   HC   140   LEU   HD22 2331     0.100000      1.00800 ; qtot -
0.129
  2332   HC   140   LEU   HD23 2332     0.100000      1.00800 ; qtot -
0.029
  2333    C   140   LEU      C 2333     0.597301     12.01000 ; qtot

0.568
2334 O 140 LEU O 2334 -0.567901 16.00000 ; qtot -
0.000
2335 N 141 PHE N 2335 -0.415700 14.01000 ; qtot -
0.416
2336 H 141 PHE H 2336 0.271900 1.00800 ; qtot -
0.144
2337 CX 141 PHE CA 2337 -0.002400 12.01000 ; qtot -
0.146
2338 H1 141 PHE HA 2338 0.097800 1.00800 ; qtot -
0.048
2339 CT 141 PHE CB 2339 -0.034300 12.01000 ; qtot -
0.083
2340 HC 141 PHE HB2 2340 0.029500 1.00800 ; qtot -
0.053
2341 HC 141 PHE HB3 2341 0.029500 1.00800 ; qtot -
0.024
2342 CA 141 PHE CG 2342 0.011800 12.01000 ; qtot -
0.012
2343 CA 141 PHE CD1 2343 -0.125600 12.01000 ; qtot -
0.138
2344 HA 141 PHE HD1 2344 0.133000 1.00800 ; qtot -
0.005
2345 CA 141 PHE CE1 2345 -0.170400 12.01000 ; qtot -
0.175
2346 HA 141 PHE HE1 2346 0.143000 1.00800 ; qtot -
0.032
2347 CA 141 PHE CZ 2347 -0.107200 12.01000 ; qtot -
0.139
2348 HA 141 PHE HZ 2348 0.129700 1.00800 ; qtot -
0.009
2349 CA 141 PHE CE2 2349 -0.170400 12.01000 ; qtot -
0.180
2350 HA 141 PHE HE2 2350 0.143000 1.00800 ; qtot -
0.037
2351 CA 141 PHE CD2 2351 -0.125600 12.01000 ; qtot -
0.162
2352 HA 141 PHE HD2 2352 0.133000 1.00800 ; qtot -
0.029
2353 C 141 PHE C 2353 0.597301 12.01000 ; qtot
0.568
2354 O 141 PHE O 2354 -0.567901 16.00000 ; qtot -
0.000
2355 N 142 GLN N 2355 -0.415700 14.01000 ; qtot -
0.416
2356 H 142 GLN H 2356 0.271900 1.00800 ; qtot -
0.144
2357 CX 142 GLN CA 2357 -0.003100 12.01000 ; qtot -
0.147
2358 H1 142 GLN HA 2358 0.085000 1.00800 ; qtot -
0.062
2359 A2C 142 GLN CB 2359 -0.003600 12.01000 ; qtot -

0.066
  2360   HC   142   GLN   HB2 2360     0.017100      1.00800 ; qtot -
0.048
  2361   HC   142   GLN   HB3 2361     0.017100      1.00800 ; qtot -
0.031
  2362  A2C   142   GLN    CG 2362    -0.064500     12.01000 ; qtot -
0.096
  2363   HC   142   GLN   HG2 2363     0.035200      1.00800 ; qtot -
0.061
  2364   HC   142   GLN   HG3 2364     0.035200      1.00800 ; qtot -
0.025
  2365    C   142   GLN    CD 2365     0.695101     12.01000 ; qtot
0.670
  2366    O   142   GLN   OE1 2366    -0.608601     16.00000 ; qtot
0.061
  2367    N   142   GLN   NE2 2367    -0.940701     14.01000 ; qtot -
0.880
  2368    H   142   GLN  HE21 2368     0.425100      1.00800 ; qtot -
0.455
  2369    H   142   GLN  HE22 2369     0.425100      1.00800 ; qtot -
0.029
  2370    C   142   GLN     C 2370     0.597301     12.01000 ; qtot
0.568
  2371    O   142   GLN     O 2371    -0.567901     16.00000 ; qtot -
0.000
  2372    N   143   SER     N 2372    -0.415700     14.01000 ; qtot -
0.416
  2373    H   143   SER     H 2373     0.271900      1.00800 ; qtot -
0.144
  2374   CX   143   SER    CA 2374    -0.024900     12.01000 ; qtot -
0.169
  2375   H1   143   SER    HA 2375     0.084300      1.00800 ; qtot -
0.084
  2376  A2C   143   SER    CB 2376     0.211700     12.01000 ; qtot
0.127
  2377   H1   143   SER   HB2 2377     0.035200      1.00800 ; qtot
0.163
  2378   H1   143   SER   HB3 2378     0.035200      1.00800 ; qtot
0.198
  2379   OH   143   SER    OG 2379    -0.654601     16.00000 ; qtot -
0.457
  2380   HO   143   SER    HG 2380     0.427500      1.00800 ; qtot -
0.029
  2381    C   143   SER     C 2381     0.597301     12.01000 ; qtot
0.568
  2382    O   143   SER     O 2382    -0.567901     16.00000 ; qtot -
0.000
  2383    N   144   GLU     N 2383    -0.516301     14.01000 ; qtot -
0.516
  2384    H   144   GLU     H 2384     0.293600      1.00800 ; qtot -
0.223
  2385   CX   144   GLU    CA 2385     0.039700     12.01000 ; qtot -

0.183
 2386 H1 144 GLU HA 2386 0.110500 1.00800 ; qtot -
0.073
 2387 A2C 144 GLU CB 2387 0.056000 12.01000 ; qtot -
0.017
 2388 HC 144 GLU HB2 2388 -0.017300 1.00800 ; qtot -
0.034
 2389 HC 144 GLU HB3 2389 -0.017300 1.00800 ; qtot -
0.051
 2390 A2C 144 GLU CG 2390 0.013600 12.01000 ; qtot -
0.038
 2391 HC 144 GLU HG2 2391 -0.042500 1.00800 ; qtot -
0.080
 2392 HC 144 GLU HG3 2392 -0.042500 1.00800 ; qtot -
0.123
 2393 CO 144 GLU CD 2393 0.805401 12.01000 ; qtot
0.683
 2394 O2 144 GLU OE1 2394 -0.818801 16.00000 ; qtot -
0.136
 2395 O2 144 GLU OE2 2395 -0.818801 16.00000 ; qtot -
0.955
 2396 C 144 GLU C 2396 0.536601 12.01000 ; qtot -
0.418
 2397 O 144 GLU O 2397 -0.581901 16.00000 ; qtot -
1.000
 2398 N 145 ASN N 2398 -0.415700 14.01000 ; qtot -
1.416
 2399 H 145 ASN H 2399 0.271900 1.00800 ; qtot -
1.144
 2400 CX 145 ASN CA 2400 0.014300 12.01000 ; qtot -
1.130
 2401 H1 145 ASN HA 2401 0.104800 1.00800 ; qtot -
1.025
 2402 A2C 145 ASN CB 2402 -0.204100 12.01000 ; qtot -
1.229
 2403 HC 145 ASN HB2 2403 0.079700 1.00800 ; qtot -
1.149
 2404 HC 145 ASN HB3 2404 0.079700 1.00800 ; qtot -
1.069
 2405 C 145 ASN CG 2405 0.713001 12.01000 ; qtot -
0.356
 2406 O 145 ASN OD1 2406 -0.593101 16.00000 ; qtot -
0.950
 2407 N 145 ASN ND2 2407 -0.919101 14.01000 ; qtot -
1.869
 2408 H 145 ASN HD21 2408 0.419600 1.00800 ; qtot -
1.449
 2409 H 145 ASN HD22 2409 0.419600 1.00800 ; qtot -
1.029
 2410 C 145 ASN C 2410 0.597301 12.01000 ; qtot -
0.432
 2411 O 145 ASN O 2411 -0.567901 16.00000 ; qtot -

1.000
  2412    N   146   LYS      N 2412    -0.347900     14.01000 ; qtot -
1.348
  2413    H   146   LYS      H 2413     0.274700      1.00800 ; qtot -
1.073
  2414   CX   146   LYS     CA 2414    -0.240000     12.01000 ; qtot -
1.313
  2415   H1   146   LYS     HA 2415     0.142600      1.00800 ; qtot -
1.171
  2416   C8   146   LYS     CB 2416    -0.009400     12.01000 ; qtot -
1.180
  2417   HC   146   LYS    HB2 2417     0.036200      1.00800 ; qtot -
1.144
  2418   HC   146   LYS    HB3 2418     0.036200      1.00800 ; qtot -
1.108
  2419   C8   146   LYS     CG 2419     0.018700     12.01000 ; qtot -
1.089
  2420   HC   146   LYS    HG2 2420     0.010300      1.00800 ; qtot -
1.079
  2421   HC   146   LYS    HG3 2421     0.010300      1.00800 ; qtot -
1.068
  2422   C8   146   LYS     CD 2422    -0.047900     12.01000 ; qtot -
1.116
  2423   HC   146   LYS    HD2 2423     0.062100      1.00800 ; qtot -
1.054
  2424   HC   146   LYS    HD3 2424     0.062100      1.00800 ; qtot -
0.992
  2425   C8   146   LYS     CE 2425    -0.014300     12.01000 ; qtot -
1.006
  2426   HP   146   LYS    HE2 2426     0.113500      1.00800 ; qtot -
0.893
  2427   HP   146   LYS    HE3 2427     0.113500      1.00800 ; qtot -
0.779
  2428   N3   146   LYS     NZ 2428    -0.385400     14.01000 ; qtot -
1.165
  2429    H   146   LYS    HZ1 2429     0.340000      1.00800 ; qtot -
0.825
  2430    H   146   LYS    HZ2 2430     0.340000      1.00800 ; qtot -
0.485
  2431    H   146   LYS    HZ3 2431     0.340000      1.00800 ; qtot -
0.145
  2432    C   146   LYS      C 2432     0.734101     12.01000 ; qtot
0.589
  2433    O   146   LYS      O 2433    -0.589401     16.00000 ; qtot -
0.000
  2434    N   147   ASP      N 2434    -0.516301     14.01000 ; qtot -
0.516
  2435    H   147   ASP      H 2435     0.293600      1.00800 ; qtot -
0.223
  2436   CX   147   ASP     CA 2436     0.038100     12.01000 ; qtot -
0.185
  2437   H1   147   ASP     HA 2437     0.088000      1.00800 ; qtot -

0.097
  2438  A2C   147   ASP    CB 2438    -0.030300     12.01000 ; qtot -
0.127
  2439   HC   147   ASP   HB2 2439    -0.012200      1.00800 ; qtot -
0.139
  2440   HC   147   ASP   HB3 2440    -0.012200      1.00800 ; qtot -
0.151
  2441   CO   147   ASP    CG 2441     0.799401     12.01000 ; qtot
0.648
  2442   O2   147   ASP   OD1 2442    -0.801401     16.00000 ; qtot -
0.153
  2443   O2   147   ASP   OD2 2443    -0.801401     16.00000 ; qtot -
0.955
  2444    C   147   ASP     C 2444     0.536601     12.01000 ; qtot -
0.418
  2445    O   147   ASP     O 2445    -0.581901     16.00000 ; qtot -
1.000
  2446    N   148   SER     N 2446    -0.415700     14.01000 ; qtot -
1.416
  2447    H   148   SER     H 2447     0.271900      1.00800 ; qtot -
1.144
  2448   CX   148   SER    CA 2448    -0.024900     12.01000 ; qtot -
1.169
  2449   H1   148   SER    HA 2449     0.084300      1.00800 ; qtot -
1.084
  2450  A2C   148   SER    CB 2450     0.211700     12.01000 ; qtot -
0.873
  2451   H1   148   SER   HB2 2451     0.035200      1.00800 ; qtot -
0.838
  2452   H1   148   SER   HB3 2452     0.035200      1.00800 ; qtot -
0.802
  2453   OH   148   SER    OG 2453    -0.654601     16.00000 ; qtot -
1.457
  2454   HO   148   SER    HG 2454     0.427500      1.00800 ; qtot -
1.029
  2455    C   148   SER     C 2455     0.597301     12.01000 ; qtot -
0.432
  2456    O   148   SER     O 2456    -0.567901     16.00000 ; qtot -
1.000
  2457    N   149   LEU     N 2457    -0.415700     14.01000 ; qtot -
1.416
  2458    H   149   LEU     H 2458     0.271900      1.00800 ; qtot -
1.144
  2459   CX   149   LEU    CA 2459    -0.051800     12.01000 ; qtot -
1.196
  2460   H1   149   LEU    HA 2460     0.092200      1.00800 ; qtot -
1.103
  2461  A2C   149   LEU    CB 2461    -0.110200     12.01000 ; qtot -
1.214
  2462   HC   149   LEU   HB2 2462     0.045700      1.00800 ; qtot -
1.168
  2463   HC   149   LEU   HB3 2463     0.045700      1.00800 ; qtot -

1.122
2464 A3C 149 LEU CG 2464 0.353100 12.01000 ; qtot -
0.769
2465 HC 149 LEU HG 2465 -0.036100 1.00800 ; qtot -
0.805
2466 CT 149 LEU CD1 2466 -0.412100 12.01000 ; qtot -
1.217
2467 HC 149 LEU HD11 2467 0.100000 1.00800 ; qtot -
1.117
2468 HC 149 LEU HD12 2468 0.100000 1.00800 ; qtot -
1.017
2469 HC 149 LEU HD13 2469 0.100000 1.00800 ; qtot -
0.917
2470 CT 149 LEU CD2 2470 -0.412100 12.01000 ; qtot -
1.329
2471 HC 149 LEU HD21 2471 0.100000 1.00800 ; qtot -
1.229
2472 HC 149 LEU HD22 2472 0.100000 1.00800 ; qtot -
1.129
2473 HC 149 LEU HD23 2473 0.100000 1.00800 ; qtot -
1.029
2474 C 149 LEU C 2474 0.597301 12.01000 ; qtot -
0.432
2475 O 149 LEU O 2475 -0.567901 16.00000 ; qtot -
1.000
2476 N 150 LEU N 2476 -0.415700 14.01000 ; qtot -
1.416
2477 H 150 LEU H 2477 0.271900 1.00800 ; qtot -
1.144
2478 CX 150 LEU CA 2478 -0.051800 12.01000 ; qtot -
1.196
2479 H1 150 LEU HA 2479 0.092200 1.00800 ; qtot -
1.103
2480 A2C 150 LEU CB 2480 -0.110200 12.01000 ; qtot -
1.214
2481 HC 150 LEU HB2 2481 0.045700 1.00800 ; qtot -
1.168
2482 HC 150 LEU HB3 2482 0.045700 1.00800 ; qtot -
1.122
2483 A3C 150 LEU CG 2483 0.353100 12.01000 ; qtot -
0.769
2484 HC 150 LEU HG 2484 -0.036100 1.00800 ; qtot -
0.805
2485 CT 150 LEU CD1 2485 -0.412100 12.01000 ; qtot -
1.217
2486 HC 150 LEU HD11 2486 0.100000 1.00800 ; qtot -
1.117
2487 HC 150 LEU HD12 2487 0.100000 1.00800 ; qtot -
1.017
2488 HC 150 LEU HD13 2488 0.100000 1.00800 ; qtot -
0.917
2489 CT 150 LEU CD2 2489 -0.412100 12.01000 ; qtot -

1.329
2490 HC 150 LEU HD21 2490 0.100000 1.00800 ; qtot -
1.229
2491 HC 150 LEU HD22 2491 0.100000 1.00800 ; qtot -
1.129
2492 HC 150 LEU HD23 2492 0.100000 1.00800 ; qtot -
1.029
2493 C 150 LEU C 2493 0.597301 12.01000 ; qtot -
0.432
2494 O 150 LEU O 2494 -0.567901 16.00000 ; qtot -
1.000
2495 N 151 LYS N 2495 -0.347900 14.01000 ; qtot -
1.348
2496 H 151 LYS H 2496 0.274700 1.00800 ; qtot -
1.073
2497 CX 151 LYS CA 2497 -0.240000 12.01000 ; qtot -
1.313
2498 H1 151 LYS HA 2498 0.142600 1.00800 ; qtot -
1.171
2499 C8 151 LYS CB 2499 -0.009400 12.01000 ; qtot -
1.180
2500 HC 151 LYS HB2 2500 0.036200 1.00800 ; qtot -
1.144
2501 HC 151 LYS HB3 2501 0.036200 1.00800 ; qtot -
1.108
2502 C8 151 LYS CG 2502 0.018700 12.01000 ; qtot -
1.089
2503 HC 151 LYS HG2 2503 0.010300 1.00800 ; qtot -
1.079
2504 HC 151 LYS HG3 2504 0.010300 1.00800 ; qtot -
1.068
2505 C8 151 LYS CD 2505 -0.047900 12.01000 ; qtot -
1.116
2506 HC 151 LYS HD2 2506 0.062100 1.00800 ; qtot -
1.054
2507 HC 151 LYS HD3 2507 0.062100 1.00800 ; qtot -
0.992
2508 C8 151 LYS CE 2508 -0.014300 12.01000 ; qtot -
1.006
2509 HP 151 LYS HE2 2509 0.113500 1.00800 ; qtot -
0.893
2510 HP 151 LYS HE3 2510 0.113500 1.00800 ; qtot -
0.779
2511 N3 151 LYS NZ 2511 -0.385400 14.01000 ; qtot -
1.165
2512 H 151 LYS HZ1 2512 0.340000 1.00800 ; qtot -
0.825
2513 H 151 LYS HZ2 2513 0.340000 1.00800 ; qtot -
0.485
2514 H 151 LYS HZ3 2514 0.340000 1.00800 ; qtot -
0.145
2515 C 151 LYS C 2515 0.734101 12.01000 ; qtot

0.589
  2516    O   151   LYS      O 2516    -0.589401     16.00000 ; qtot -
0.000
  2517    N   152   ILE      N 2517    -0.415700     14.01000 ; qtot -
0.416
  2518    H   152   ILE      H 2518     0.271900      1.00800 ; qtot -
0.144
  2519   CX   152   ILE     CA 2519    -0.059700     12.01000 ; qtot -
0.204
  2520   H1   152   ILE     HA 2520     0.086900      1.00800 ; qtot -
0.117
  2521  A3C   152   ILE     CB 2521     0.130300     12.01000 ; qtot
0.014
  2522   HC   152   ILE     HB 2522     0.018700      1.00800 ; qtot
0.032
  2523   CT   152   ILE    CG2 2523    -0.320400     12.01000 ; qtot -
0.288
  2524   HC   152   ILE   HG21 2524     0.088200      1.00800 ; qtot -
0.200
  2525   HC   152   ILE   HG22 2525     0.088200      1.00800 ; qtot -
0.112
  2526   HC   152   ILE   HG23 2526     0.088200      1.00800 ; qtot -
0.023
  2527  A2C   152   ILE    CG1 2527    -0.043000     12.01000 ; qtot -
0.066
  2528   HC   152   ILE   HG12 2528     0.023600      1.00800 ; qtot -
0.043
  2529   HC   152   ILE   HG13 2529     0.023600      1.00800 ; qtot -
0.019
  2530   CT   152   ILE    CD1 2530    -0.066000     12.01000 ; qtot -
0.085
  2531   HC   152   ILE   HD11 2531     0.018600      1.00800 ; qtot -
0.067
  2532   HC   152   ILE   HD12 2532     0.018600      1.00800 ; qtot -
0.048
  2533   HC   152   ILE   HD13 2533     0.018600      1.00800 ; qtot -
0.029
  2534    C   152   ILE      C 2534     0.597301     12.01000 ; qtot
0.568
  2535    O   152   ILE      O 2535    -0.567901     16.00000 ; qtot -
0.000
  2536    N   153   ILE      N 2536    -0.415700     14.01000 ; qtot -
0.416
  2537    H   153   ILE      H 2537     0.271900      1.00800 ; qtot -
0.144
  2538   CX   153   ILE     CA 2538    -0.059700     12.01000 ; qtot -
0.204
  2539   H1   153   ILE     HA 2539     0.086900      1.00800 ; qtot -
0.117
  2540  A3C   153   ILE     CB 2540     0.130300     12.01000 ; qtot
0.014
  2541   HC   153   ILE     HB 2541     0.018700      1.00800 ; qtot

0.032
  2542   CT   153   ILE   CG2 2542    -0.320400     12.01000 ; qtot -
0.288
  2543   HC   153   ILE  HG21 2543     0.088200      1.00800 ; qtot -
0.200
  2544   HC   153   ILE  HG22 2544     0.088200      1.00800 ; qtot -
0.112
  2545   HC   153   ILE  HG23 2545     0.088200      1.00800 ; qtot -
0.023
  2546  A2C   153   ILE   CG1 2546    -0.043000     12.01000 ; qtot -
0.066
  2547   HC   153   ILE  HG12 2547     0.023600      1.00800 ; qtot -
0.043
  2548   HC   153   ILE  HG13 2548     0.023600      1.00800 ; qtot -
0.019
  2549   CT   153   ILE   CD1 2549    -0.066000     12.01000 ; qtot -
0.085
  2550   HC   153   ILE  HD11 2550     0.018600      1.00800 ; qtot -
0.067
  2551   HC   153   ILE  HD12 2551     0.018600      1.00800 ; qtot -
0.048
  2552   HC   153   ILE  HD13 2552     0.018600      1.00800 ; qtot -
0.029
  2553    C   153   ILE     C 2553     0.597301     12.01000 ; qtot
0.568
  2554    O   153   ILE     O 2554    -0.567901     16.00000 ; qtot -
0.000
  2555    N   154   ASP     N 2555    -0.516301     14.01000 ; qtot -
0.516
  2556    H   154   ASP     H 2556     0.293600      1.00800 ; qtot -
0.223
  2557   CX   154   ASP    CA 2557     0.038100     12.01000 ; qtot -
0.185
  2558   H1   154   ASP    HA 2558     0.088000      1.00800 ; qtot -
0.097
  2559  A2C   154   ASP    CB 2559    -0.030300     12.01000 ; qtot -
0.127
  2560   HC   154   ASP   HB2 2560    -0.012200      1.00800 ; qtot -
0.139
  2561   HC   154   ASP   HB3 2561    -0.012200      1.00800 ; qtot -
0.151
  2562   CO   154   ASP    CG 2562     0.799401     12.01000 ; qtot
0.648
  2563   O2   154   ASP   OD1 2563    -0.801401     16.00000 ; qtot -
0.153
  2564   O2   154   ASP   OD2 2564    -0.801401     16.00000 ; qtot -
0.955
  2565    C   154   ASP     C 2565     0.536601     12.01000 ; qtot -
0.418
  2566    O   154   ASP     O 2566    -0.581901     16.00000 ; qtot -
1.000
  2567    N   155   PHE     N 2567    -0.415700     14.01000 ; qtot -

```
1.416
  2568    H   155   PHE     H 2568     0.271900      1.00800 ; qtot -
1.144
  2569   CX   155   PHE    CA 2569    -0.002400     12.01000 ; qtot -
1.146
  2570   H1   155   PHE    HA 2570     0.097800      1.00800 ; qtot -
1.048
  2571   CT   155   PHE    CB 2571    -0.034300     12.01000 ; qtot -
1.083
  2572   HC   155   PHE   HB2 2572     0.029500      1.00800 ; qtot -
1.053
  2573   HC   155   PHE   HB3 2573     0.029500      1.00800 ; qtot -
1.024
  2574   CA   155   PHE    CG 2574     0.011800     12.01000 ; qtot -
1.012
  2575   CA   155   PHE   CD1 2575    -0.125600     12.01000 ; qtot -
1.138
  2576   HA   155   PHE   HD1 2576     0.133000      1.00800 ; qtot -
1.005
  2577   CA   155   PHE   CE1 2577    -0.170400     12.01000 ; qtot -
1.175
  2578   HA   155   PHE   HE1 2578     0.143000      1.00800 ; qtot -
1.032
  2579   CA   155   PHE    CZ 2579    -0.107200     12.01000 ; qtot -
1.139
  2580   HA   155   PHE    HZ 2580     0.129700      1.00800 ; qtot -
1.009
  2581   CA   155   PHE   CE2 2581    -0.170400     12.01000 ; qtot -
1.180
  2582   HA   155   PHE   HE2 2582     0.143000      1.00800 ; qtot -
1.037
  2583   CA   155   PHE   CD2 2583    -0.125600     12.01000 ; qtot -
1.162
  2584   HA   155   PHE   HD2 2584     0.133000      1.00800 ; qtot -
1.029
  2585    C   155   PHE     C 2585     0.597301     12.01000 ; qtot -
0.432
  2586    O   155   PHE     O 2586    -0.567901     16.00000 ; qtot -
1.000
  2587    N   156   GLY     N 2587    -0.415700     14.01000 ; qtot -
1.416
  2588    H   156   GLY     H 2588     0.271900      1.00800 ; qtot -
1.144
  2589   CX   156   GLY    CA 2589    -0.025200     12.01000 ; qtot -
1.169
  2590   H1   156   GLY   HA2 2590     0.069800      1.00800 ; qtot -
1.099
  2591   H1   156   GLY   HA3 2591     0.069800      1.00800 ; qtot -
1.029
  2592    C   156   GLY     C 2592     0.597301     12.01000 ; qtot -
0.432
  2593    O   156   GLY     O 2593    -0.567901     16.00000 ; qtot -
```

```
1.000
  2594    N   157   LEU      N 2594    -0.415700     14.01000 ; qtot -
1.416
  2595    H   157   LEU      H 2595     0.271900      1.00800 ; qtot -
1.144
  2596   CX   157   LEU     CA 2596    -0.051800     12.01000 ; qtot -
1.196
  2597   H1   157   LEU     HA 2597     0.092200      1.00800 ; qtot -
1.103
  2598  A2C   157   LEU     CB 2598    -0.110200     12.01000 ; qtot -
1.214
  2599   HC   157   LEU    HB2 2599     0.045700      1.00800 ; qtot -
1.168
  2600   HC   157   LEU    HB3 2600     0.045700      1.00800 ; qtot -
1.122
  2601  A3C   157   LEU     CG 2601     0.353100     12.01000 ; qtot -
0.769
  2602   HC   157   LEU     HG 2602    -0.036100      1.00800 ; qtot -
0.805
  2603   CT   157   LEU    CD1 2603    -0.412100     12.01000 ; qtot -
1.217
  2604   HC   157   LEU   HD11 2604     0.100000      1.00800 ; qtot -
1.117
  2605   HC   157   LEU   HD12 2605     0.100000      1.00800 ; qtot -
1.017
  2606   HC   157   LEU   HD13 2606     0.100000      1.00800 ; qtot -
0.917
  2607   CT   157   LEU    CD2 2607    -0.412100     12.01000 ; qtot -
1.329
  2608   HC   157   LEU   HD21 2608     0.100000      1.00800 ; qtot -
1.229
  2609   HC   157   LEU   HD22 2609     0.100000      1.00800 ; qtot -
1.129
  2610   HC   157   LEU   HD23 2610     0.100000      1.00800 ; qtot -
1.029
  2611    C   157   LEU      C 2611     0.597301     12.01000 ; qtot -
0.432
  2612    O   157   LEU      O 2612    -0.567901     16.00000 ; qtot -
1.000
  2613    N   158   SER      N 2613    -0.415700     14.01000 ; qtot -
1.416
  2614    H   158   SER      H 2614     0.271900      1.00800 ; qtot -
1.144
  2615   CX   158   SER     CA 2615    -0.024900     12.01000 ; qtot -
1.169
  2616   H1   158   SER     HA 2616     0.084300      1.00800 ; qtot -
1.084
  2617  A2C   158   SER     CB 2617     0.211700     12.01000 ; qtot -
0.873
  2618   H1   158   SER    HB2 2618     0.035200      1.00800 ; qtot -
0.838
  2619   H1   158   SER    HB3 2619     0.035200      1.00800 ; qtot -
```

```
0.802
  2620   OH   158   SER    OG 2620    -0.654601     16.00000 ; qtot -
1.457
  2621   HO   158   SER    HG 2621     0.427500      1.00800 ; qtot -
1.029
  2622    C   158   SER     C 2622     0.597301     12.01000 ; qtot -
0.432
  2623    O   158   SER     O 2623    -0.567901     16.00000 ; qtot -
1.000
  2624    N   159   LYS     N 2624    -0.347900     14.01000 ; qtot -
1.348
  2625    H   159   LYS     H 2625     0.274700      1.00800 ; qtot -
1.073
  2626   CX   159   LYS    CA 2626    -0.240000     12.01000 ; qtot -
1.313
  2627   H1   159   LYS    HA 2627     0.142600      1.00800 ; qtot -
1.171
  2628   C8   159   LYS    CB 2628    -0.009400     12.01000 ; qtot -
1.180
  2629   HC   159   LYS   HB2 2629     0.036200      1.00800 ; qtot -
1.144
  2630   HC   159   LYS   HB3 2630     0.036200      1.00800 ; qtot -
1.108
  2631   C8   159   LYS    CG 2631     0.018700     12.01000 ; qtot -
1.089
  2632   HC   159   LYS   HG2 2632     0.010300      1.00800 ; qtot -
1.079
  2633   HC   159   LYS   HG3 2633     0.010300      1.00800 ; qtot -
1.068
  2634   C8   159   LYS    CD 2634    -0.047900     12.01000 ; qtot -
1.116
  2635   HC   159   LYS   HD2 2635     0.062100      1.00800 ; qtot -
1.054
  2636   HC   159   LYS   HD3 2636     0.062100      1.00800 ; qtot -
0.992
  2637   C8   159   LYS    CE 2637    -0.014300     12.01000 ; qtot -
1.006
  2638   HP   159   LYS   HE2 2638     0.113500      1.00800 ; qtot -
0.893
  2639   HP   159   LYS   HE3 2639     0.113500      1.00800 ; qtot -
0.779
  2640   N3   159   LYS    NZ 2640    -0.385400     14.01000 ; qtot -
1.165
  2641    H   159   LYS   HZ1 2641     0.340000      1.00800 ; qtot -
0.825
  2642    H   159   LYS   HZ2 2642     0.340000      1.00800 ; qtot -
0.485
  2643    H   159   LYS   HZ3 2643     0.340000      1.00800 ; qtot -
0.145
  2644    C   159   LYS     C 2644     0.734101     12.01000 ; qtot
0.589
  2645    O   159   LYS     O 2645    -0.589401     16.00000 ; qtot -
```

```
0.000
  2646    N   160   ASN     N 2646    -0.415700     14.01000 ; qtot -
0.416
  2647    H   160   ASN     H 2647     0.271900      1.00800 ; qtot -
0.144
  2648   CX   160   ASN    CA 2648     0.014300     12.01000 ; qtot -
0.130
  2649   H1   160   ASN    HA 2649     0.104800      1.00800 ; qtot -
0.025
  2650  A2C   160   ASN    CB 2650    -0.204100     12.01000 ; qtot -
0.229
  2651   HC   160   ASN   HB2 2651     0.079700      1.00800 ; qtot -
0.149
  2652   HC   160   ASN   HB3 2652     0.079700      1.00800 ; qtot -
0.069
  2653    C   160   ASN    CG 2653     0.713001     12.01000 ; qtot
0.644
  2654    O   160   ASN   OD1 2654    -0.593101     16.00000 ; qtot
0.050
  2655    N   160   ASN   ND2 2655    -0.919101     14.01000 ; qtot -
0.869
  2656    H   160   ASN  HD21 2656     0.419600      1.00800 ; qtot -
0.449
  2657    H   160   ASN  HD22 2657     0.419600      1.00800 ; qtot -
0.029
  2658    C   160   ASN     C 2658     0.597301     12.01000 ; qtot
0.568
  2659    O   160   ASN     O 2659    -0.567901     16.00000 ; qtot -
0.000
  2660    N   161   LEU     N 2660    -0.415700     14.01000 ; qtot -
0.416
  2661    H   161   LEU     H 2661     0.271900      1.00800 ; qtot -
0.144
  2662   CX   161   LEU    CA 2662    -0.051800     12.01000 ; qtot -
0.196
  2663   H1   161   LEU    HA 2663     0.092200      1.00800 ; qtot -
0.103
  2664  A2C   161   LEU    CB 2664    -0.110200     12.01000 ; qtot -
0.214
  2665   HC   161   LEU   HB2 2665     0.045700      1.00800 ; qtot -
0.168
  2666   HC   161   LEU   HB3 2666     0.045700      1.00800 ; qtot -
0.122
  2667  A3C   161   LEU    CG 2667     0.353100     12.01000 ; qtot
0.231
  2668   HC   161   LEU    HG 2668    -0.036100      1.00800 ; qtot
0.195
  2669   CT   161   LEU   CD1 2669    -0.412100     12.01000 ; qtot -
0.217
  2670   HC   161   LEU  HD11 2670     0.100000      1.00800 ; qtot -
0.117
  2671   HC   161   LEU  HD12 2671     0.100000      1.00800 ; qtot -
```

0.017
  2672   HC   161   LEU  HD13 2672     0.100000      1.00800 ; qtot
0.083
  2673   CT   161   LEU   CD2 2673    -0.412100     12.01000 ; qtot -
0.329
  2674   HC   161   LEU  HD21 2674     0.100000      1.00800 ; qtot -
0.229
  2675   HC   161   LEU  HD22 2675     0.100000      1.00800 ; qtot -
0.129
  2676   HC   161   LEU  HD23 2676     0.100000      1.00800 ; qtot -
0.029
  2677    C   161   LEU     C 2677     0.597301     12.01000 ; qtot
0.568
  2678    O   161   LEU     O 2678    -0.567901     16.00000 ; qtot -
0.000
  2679    N   162   GLY     N 2679    -0.415700     14.01000 ; qtot -
0.416
  2680    H   162   GLY     H 2680     0.271900      1.00800 ; qtot -
0.144
  2681   CX   162   GLY    CA 2681    -0.025200     12.01000 ; qtot -
0.169
  2682   H1   162   GLY   HA2 2682     0.069800      1.00800 ; qtot -
0.099
  2683   H1   162   GLY   HA3 2683     0.069800      1.00800 ; qtot -
0.029
  2684    C   162   GLY     C 2684     0.597301     12.01000 ; qtot
0.568
  2685    O   162   GLY     O 2685    -0.567901     16.00000 ; qtot -
0.000
  2686    N   163   THR     N 2686    -0.415700     14.01000 ; qtot -
0.416
  2687    H   163   THR     H 2687     0.271900      1.00800 ; qtot -
0.144
  2688   CX   163   THR    CA 2688    -0.038900     12.01000 ; qtot -
0.183
  2689   H1   163   THR    HA 2689     0.100700      1.00800 ; qtot -
0.082
  2690  A3C   163   THR    CB 2690     0.365400     12.01000 ; qtot
0.283
  2691   H1   163   THR    HB 2691     0.004300      1.00800 ; qtot
0.288
  2692   CT   163   THR   CG2 2692    -0.243800     12.01000 ; qtot
0.044
  2693   HC   163   THR  HG21 2693     0.064200      1.00800 ; qtot
0.108
  2694   HC   163   THR  HG22 2694     0.064200      1.00800 ; qtot
0.172
  2695   HC   163   THR  HG23 2695     0.064200      1.00800 ; qtot
0.237
  2696   OH   163   THR   OG1 2696    -0.676101     16.00000 ; qtot -
0.440
  2697   HO   163   THR   HG1 2697     0.410200      1.00800 ; qtot -

```
0.029
  2698    C   163   THR      C 2698     0.597301     12.01000 ; qtot
0.568
  2699    O   163   THR      O 2699    -0.567901     16.00000 ; qtot -
0.000
  2700    N   164   GLY      N 2700    -0.415700     14.01000 ; qtot -
0.416
  2701    H   164   GLY      H 2701     0.271900      1.00800 ; qtot -
0.144
  2702   CX   164   GLY     CA 2702    -0.025200     12.01000 ; qtot -
0.169
  2703   H1   164   GLY    HA2 2703     0.069800      1.00800 ; qtot -
0.099
  2704   H1   164   GLY    HA3 2704     0.069800      1.00800 ; qtot -
0.029
  2705    C   164   GLY      C 2705     0.597301     12.01000 ; qtot
0.568
  2706    O   164   GLY      O 2706    -0.567901     16.00000 ; qtot -
0.000
  2707    N   165   GLU      N 2707    -0.516301     14.01000 ; qtot -
0.516
  2708    H   165   GLU      H 2708     0.293600      1.00800 ; qtot -
0.223
  2709   CX   165   GLU     CA 2709     0.039700     12.01000 ; qtot -
0.183
  2710   H1   165   GLU     HA 2710     0.110500      1.00800 ; qtot -
0.073
  2711  A2C   165   GLU     CB 2711     0.056000     12.01000 ; qtot -
0.017
  2712   HC   165   GLU    HB2 2712    -0.017300      1.00800 ; qtot -
0.034
  2713   HC   165   GLU    HB3 2713    -0.017300      1.00800 ; qtot -
0.051
  2714  A2C   165   GLU     CG 2714     0.013600     12.01000 ; qtot -
0.038
  2715   HC   165   GLU    HG2 2715    -0.042500      1.00800 ; qtot -
0.080
  2716   HC   165   GLU    HG3 2716    -0.042500      1.00800 ; qtot -
0.123
  2717   CO   165   GLU     CD 2717     0.805401     12.01000 ; qtot
0.683
  2718   O2   165   GLU    OE1 2718    -0.818801     16.00000 ; qtot -
0.136
  2719   O2   165   GLU    OE2 2719    -0.818801     16.00000 ; qtot -
0.955
  2720    C   165   GLU      C 2720     0.536601     12.01000 ; qtot -
0.418
  2721    O   165   GLU      O 2721    -0.581901     16.00000 ; qtot -
1.000
  2722    N   166   PHE      N 2722    -0.415700     14.01000 ; qtot -
1.416
  2723    H   166   PHE      H 2723     0.271900      1.00800 ; qtot -
```

1.144
2724 CX 166 PHE CA 2724 -0.002400 12.01000 ; qtot -
1.146
2725 H1 166 PHE HA 2725 0.097800 1.00800 ; qtot -
1.048
2726 CT 166 PHE CB 2726 -0.034300 12.01000 ; qtot -
1.083
2727 HC 166 PHE HB2 2727 0.029500 1.00800 ; qtot -
1.053
2728 HC 166 PHE HB3 2728 0.029500 1.00800 ; qtot -
1.024
2729 CA 166 PHE CG 2729 0.011800 12.01000 ; qtot -
1.012
2730 CA 166 PHE CD1 2730 -0.125600 12.01000 ; qtot -
1.138
2731 HA 166 PHE HD1 2731 0.133000 1.00800 ; qtot -
1.005
2732 CA 166 PHE CE1 2732 -0.170400 12.01000 ; qtot -
1.175
2733 HA 166 PHE HE1 2733 0.143000 1.00800 ; qtot -
1.032
2734 CA 166 PHE CZ 2734 -0.107200 12.01000 ; qtot -
1.139
2735 HA 166 PHE HZ 2735 0.129700 1.00800 ; qtot -
1.009
2736 CA 166 PHE CE2 2736 -0.170400 12.01000 ; qtot -
1.180
2737 HA 166 PHE HE2 2737 0.143000 1.00800 ; qtot -
1.037
2738 CA 166 PHE CD2 2738 -0.125600 12.01000 ; qtot -
1.162
2739 HA 166 PHE HD2 2739 0.133000 1.00800 ; qtot -
1.029
2740 C 166 PHE C 2740 0.597301 12.01000 ; qtot -
0.432
2741 O 166 PHE O 2741 -0.567901 16.00000 ; qtot -
1.000
2742 N 167 THR N 2742 -0.415700 14.01000 ; qtot -
1.416
2743 H 167 THR H 2743 0.271900 1.00800 ; qtot -
1.144
2744 CX 167 THR CA 2744 -0.038900 12.01000 ; qtot -
1.183
2745 H1 167 THR HA 2745 0.100700 1.00800 ; qtot -
1.082
2746 A3C 167 THR CB 2746 0.365400 12.01000 ; qtot -
0.717
2747 H1 167 THR HB 2747 0.004300 1.00800 ; qtot -
0.712
2748 CT 167 THR CG2 2748 -0.243800 12.01000 ; qtot -
0.956
2749 HC 167 THR HG21 2749 0.064200 1.00800 ; qtot -

0.892
  2750   HC   167   THR  HG22 2750     0.064200      1.00800 ; qtot -
0.828
  2751   HC   167   THR  HG23 2751     0.064200      1.00800 ; qtot -
0.764
  2752   OH   167   THR   OG1 2752    -0.676101     16.00000 ; qtot -
1.440
  2753   HO   167   THR   HG1 2753     0.410200      1.00800 ; qtot -
1.029
  2754    C   167   THR     C 2754     0.597301     12.01000 ; qtot -
0.432
  2755    O   167   THR     O 2755    -0.567901     16.00000 ; qtot -
1.000
  2756    N   168   THR     N 2756    -0.415700     14.01000 ; qtot -
1.416
  2757    H   168   THR     H 2757     0.271900      1.00800 ; qtot -
1.144
  2758   CX   168   THR    CA 2758    -0.038900     12.01000 ; qtot -
1.183
  2759   H1   168   THR    HA 2759     0.100700      1.00800 ; qtot -
1.082
  2760  A3C   168   THR    CB 2760     0.365400     12.01000 ; qtot -
0.717
  2761   H1   168   THR    HB 2761     0.004300      1.00800 ; qtot -
0.712
  2762   CT   168   THR   CG2 2762    -0.243800     12.01000 ; qtot -
0.956
  2763   HC   168   THR  HG21 2763     0.064200      1.00800 ; qtot -
0.892
  2764   HC   168   THR  HG22 2764     0.064200      1.00800 ; qtot -
0.828
  2765   HC   168   THR  HG23 2765     0.064200      1.00800 ; qtot -
0.764
  2766   OH   168   THR   OG1 2766    -0.676101     16.00000 ; qtot -
1.440
  2767   HO   168   THR   HG1 2767     0.410200      1.00800 ; qtot -
1.029
  2768    C   168   THR     C 2768     0.597301     12.01000 ; qtot -
0.432
  2769    O   168   THR     O 2769    -0.567901     16.00000 ; qtot -
1.000
  2770    N   169   THR     N 2770    -0.415700     14.01000 ; qtot -
1.416
  2771    H   169   THR     H 2771     0.271900      1.00800 ; qtot -
1.144
  2772   CX   169   THR    CA 2772    -0.038900     12.01000 ; qtot -
1.183
  2773   H1   169   THR    HA 2773     0.100700      1.00800 ; qtot -
1.082
  2774  A3C   169   THR    CB 2774     0.365400     12.01000 ; qtot -
0.717
  2775   H1   169   THR    HB 2775     0.004300      1.00800 ; qtot -

0.712
  2776   CT   169   THR   CG2 2776    -0.243800     12.01000 ; qtot -
0.956
  2777   HC   169   THR  HG21 2777     0.064200      1.00800 ; qtot -
0.892
  2778   HC   169   THR  HG22 2778     0.064200      1.00800 ; qtot -
0.828
  2779   HC   169   THR  HG23 2779     0.064200      1.00800 ; qtot -
0.764
  2780   OH   169   THR   OG1 2780    -0.676101     16.00000 ; qtot -
1.440
  2781   HO   169   THR   HG1 2781     0.410200      1.00800 ; qtot -
1.029
  2782    C   169   THR     C 2782     0.597301     12.01000 ; qtot -
0.432
  2783    O   169   THR     O 2783    -0.567901     16.00000 ; qtot -
1.000
  2784    N   170   LYS     N 2784    -0.347900     14.01000 ; qtot -
1.348
  2785    H   170   LYS     H 2785     0.274700      1.00800 ; qtot -
1.073
  2786   CX   170   LYS    CA 2786    -0.240000     12.01000 ; qtot -
1.313
  2787   H1   170   LYS    HA 2787     0.142600      1.00800 ; qtot -
1.171
  2788   C8   170   LYS    CB 2788    -0.009400     12.01000 ; qtot -
1.180
  2789   HC   170   LYS   HB2 2789     0.036200      1.00800 ; qtot -
1.144
  2790   HC   170   LYS   HB3 2790     0.036200      1.00800 ; qtot -
1.108
  2791   C8   170   LYS    CG 2791     0.018700     12.01000 ; qtot -
1.089
  2792   HC   170   LYS   HG2 2792     0.010300      1.00800 ; qtot -
1.079
  2793   HC   170   LYS   HG3 2793     0.010300      1.00800 ; qtot -
1.068
  2794   C8   170   LYS    CD 2794    -0.047900     12.01000 ; qtot -
1.116
  2795   HC   170   LYS   HD2 2795     0.062100      1.00800 ; qtot -
1.054
  2796   HC   170   LYS   HD3 2796     0.062100      1.00800 ; qtot -
0.992
  2797   C8   170   LYS    CE 2797    -0.014300     12.01000 ; qtot -
1.006
  2798   HP   170   LYS   HE2 2798     0.113500      1.00800 ; qtot -
0.893
  2799   HP   170   LYS   HE3 2799     0.113500      1.00800 ; qtot -
0.779
  2800   N3   170   LYS    NZ 2800    -0.385400     14.01000 ; qtot -
1.165
  2801    H   170   LYS   HZ1 2801     0.340000      1.00800 ; qtot -

0.825
2802 H 170 LYS HZ2 2802 0.340000 1.00800 ; qtot -
0.485
2803 H 170 LYS HZ3 2803 0.340000 1.00800 ; qtot -
0.145
2804 C 170 LYS C 2804 0.734101 12.01000 ; qtot
0.589
2805 O 170 LYS O 2805 -0.589401 16.00000 ; qtot -
0.000
2806 N 171 ALA N 2806 -0.415700 14.01000 ; qtot -
0.416
2807 H 171 ALA H 2807 0.271900 1.00800 ; qtot -
0.144
2808 CX 171 ALA CA 2808 0.033700 12.01000 ; qtot -
0.110
2809 H1 171 ALA HA 2809 0.082300 1.00800 ; qtot -
0.028
2810 CT 171 ALA CB 2810 -0.182500 12.01000 ; qtot -
0.210
2811 HC 171 ALA HB1 2811 0.060300 1.00800 ; qtot -
0.150
2812 HC 171 ALA HB2 2812 0.060300 1.00800 ; qtot -
0.090
2813 HC 171 ALA HB3 2813 0.060300 1.00800 ; qtot -
0.029
2814 C 171 ALA C 2814 0.597301 12.01000 ; qtot
0.568
2815 O 171 ALA O 2815 -0.567901 16.00000 ; qtot -
0.000
2816 N 172 GLY N 2816 -0.415700 14.01000 ; qtot -
0.416
2817 H 172 GLY H 2817 0.271900 1.00800 ; qtot -
0.144
2818 CX 172 GLY CA 2818 -0.025200 12.01000 ; qtot -
0.169
2819 H1 172 GLY HA2 2819 0.069800 1.00800 ; qtot -
0.099
2820 H1 172 GLY HA3 2820 0.069800 1.00800 ; qtot -
0.029
2821 C 172 GLY C 2821 0.597301 12.01000 ; qtot
0.568
2822 O 172 GLY O 2822 -0.567901 16.00000 ; qtot -
0.000
2823 N 173 THR N 2823 -0.415700 14.01000 ; qtot -
0.416
2824 H 173 THR H 2824 0.271900 1.00800 ; qtot -
0.144
2825 CX 173 THR CA 2825 -0.038900 12.01000 ; qtot -
0.183
2826 H1 173 THR HA 2826 0.100700 1.00800 ; qtot -
0.082
2827 A3C 173 THR CB 2827 0.365400 12.01000 ; qtot

0.283
  2828   H1   173   THR    HB 2828     0.004300      1.00800 ; qtot
0.288
  2829   CT   173   THR   CG2 2829    -0.243800     12.01000 ; qtot
0.044
  2830   HC   173   THR  HG21 2830     0.064200      1.00800 ; qtot
0.108
  2831   HC   173   THR  HG22 2831     0.064200      1.00800 ; qtot
0.172
  2832   HC   173   THR  HG23 2832     0.064200      1.00800 ; qtot
0.237
  2833   OH   173   THR   OG1 2833    -0.676101     16.00000 ; qtot -
0.440
  2834   HO   173   THR   HG1 2834     0.410200      1.00800 ; qtot -
0.029
  2835    C   173   THR     C 2835     0.597301     12.01000 ; qtot
0.568
  2836    O   173   THR     O 2836    -0.567901     16.00000 ; qtot -
0.000
  2837    N   174   PRO     N 2837    -0.254800     14.01000 ; qtot -
0.255
  2838   CT   174   PRO    CD 2838     0.019200     12.01000 ; qtot -
0.236
  2839   H1   174   PRO   HD2 2839     0.039100      1.00800 ; qtot -
0.197
  2840   H1   174   PRO   HD3 2840     0.039100      1.00800 ; qtot -
0.157
  2841   CT   174   PRO    CG 2841     0.018900     12.01000 ; qtot -
0.139
  2842   HC   174   PRO   HG2 2842     0.021300      1.00800 ; qtot -
0.117
  2843   HC   174   PRO   HG3 2843     0.021300      1.00800 ; qtot -
0.096
  2844   CT   174   PRO    CB 2844    -0.007000     12.01000 ; qtot -
0.103
  2845   HC   174   PRO   HB2 2845     0.025300      1.00800 ; qtot -
0.078
  2846   HC   174   PRO   HB3 2846     0.025300      1.00800 ; qtot -
0.052
  2847   CX   174   PRO    CA 2847    -0.026600     12.01000 ; qtot -
0.079
  2848   H1   174   PRO    HA 2848     0.064100      1.00800 ; qtot -
0.015
  2849    C   174   PRO     C 2849     0.589601     12.01000 ; qtot
0.575
  2850    O   174   PRO     O 2850    -0.574801     16.00000 ; qtot -
0.000
  2851    N   175   TYR     N 2851    -0.415700     14.01000 ; qtot -
0.416
  2852    H   175   TYR     H 2852     0.271900      1.00800 ; qtot -
0.144
  2853   CX   175   TYR    CA 2853    -0.001400     12.01000 ; qtot -

```
0.145
  2854   H1   175   TYR    HA 2854     0.087600      1.00800 ; qtot -
0.058
  2855   CT   175   TYR    CB 2855    -0.015200     12.01000 ; qtot -
0.073
  2856   HC   175   TYR   HB2 2856     0.029500      1.00800 ; qtot -
0.043
  2857   HC   175   TYR   HB3 2857     0.029500      1.00800 ; qtot -
0.014
  2858   CA   175   TYR    CG 2858    -0.001100     12.01000 ; qtot -
0.015
  2859   CA   175   TYR   CD1 2859    -0.190600     12.01000 ; qtot -
0.206
  2860   HA   175   TYR   HD1 2860     0.169900      1.00800 ; qtot -
0.036
  2861   CA   175   TYR   CE1 2861    -0.234100     12.01000 ; qtot -
0.270
  2862   HA   175   TYR   HE1 2862     0.165600      1.00800 ; qtot -
0.104
  2863    C   175   TYR    CZ 2863     0.322600     12.01000 ; qtot
0.219
  2864   OH   175   TYR    OH 2864    -0.557901     16.00000 ; qtot -
0.339
  2865   HO   175   TYR    HH 2865     0.399200      1.00800 ; qtot
0.060
  2866   CA   175   TYR   CE2 2866    -0.234100     12.01000 ; qtot -
0.174
  2867   HA   175   TYR   HE2 2867     0.165600      1.00800 ; qtot -
0.009
  2868   CA   175   TYR   CD2 2868    -0.190600     12.01000 ; qtot -
0.199
  2869   HA   175   TYR   HD2 2869     0.169900      1.00800 ; qtot -
0.029
  2870    C   175   TYR     C 2870     0.597301     12.01000 ; qtot
0.568
  2871    O   175   TYR     O 2871    -0.567901     16.00000 ; qtot -
0.000
  2872    N   176   TYR     N 2872    -0.415700     14.01000 ; qtot -
0.416
  2873    H   176   TYR     H 2873     0.271900      1.00800 ; qtot -
0.144
  2874   CX   176   TYR    CA 2874    -0.001400     12.01000 ; qtot -
0.145
  2875   H1   176   TYR    HA 2875     0.087600      1.00800 ; qtot -
0.058
  2876   CT   176   TYR    CB 2876    -0.015200     12.01000 ; qtot -
0.073
  2877   HC   176   TYR   HB2 2877     0.029500      1.00800 ; qtot -
0.043
  2878   HC   176   TYR   HB3 2878     0.029500      1.00800 ; qtot -
0.014
  2879   CA   176   TYR    CG 2879    -0.001100     12.01000 ; qtot -
```

0.015
2880 CA 176 TYR CD1 2880 -0.190600 12.01000 ; qtot -
0.206
2881 HA 176 TYR HD1 2881 0.169900 1.00800 ; qtot -
0.036
2882 CA 176 TYR CE1 2882 -0.234100 12.01000 ; qtot -
0.270
2883 HA 176 TYR HE1 2883 0.165600 1.00800 ; qtot -
0.104
2884 C 176 TYR CZ 2884 0.322600 12.01000 ; qtot
0.219
2885 OH 176 TYR OH 2885 -0.557901 16.00000 ; qtot -
0.339
2886 HO 176 TYR HH 2886 0.399200 1.00800 ; qtot
0.060
2887 CA 176 TYR CE2 2887 -0.234100 12.01000 ; qtot -
0.174
2888 HA 176 TYR HE2 2888 0.165600 1.00800 ; qtot -
0.009
2889 CA 176 TYR CD2 2889 -0.190600 12.01000 ; qtot -
0.199
2890 HA 176 TYR HD2 2890 0.169900 1.00800 ; qtot -
0.029
2891 C 176 TYR C 2891 0.597301 12.01000 ; qtot
0.568
2892 O 176 TYR O 2892 -0.567901 16.00000 ; qtot -
0.000
2893 N 177 VAL N 2893 -0.415700 14.01000 ; qtot -
0.416
2894 H 177 VAL H 2894 0.271900 1.00800 ; qtot -
0.144
2895 CX 177 VAL CA 2895 -0.087500 12.01000 ; qtot -
0.231
2896 H1 177 VAL HA 2896 0.096900 1.00800 ; qtot -
0.134
2897 A3C 177 VAL CB 2897 0.298500 12.01000 ; qtot
0.164
2898 HC 177 VAL HB 2898 -0.029700 1.00800 ; qtot
0.134
2899 CT 177 VAL CG1 2899 -0.319200 12.01000 ; qtot -
0.185
2900 HC 177 VAL HG11 2900 0.079100 1.00800 ; qtot -
0.106
2901 HC 177 VAL HG12 2901 0.079100 1.00800 ; qtot -
0.027
2902 HC 177 VAL HG13 2902 0.079100 1.00800 ; qtot
0.052
2903 CT 177 VAL CG2 2903 -0.319200 12.01000 ; qtot -
0.267
2904 HC 177 VAL HG21 2904 0.079100 1.00800 ; qtot -
0.188
2905 HC 177 VAL HG22 2905 0.079100 1.00800 ; qtot -

0.109
  2906   HC   177   VAL  HG23 2906     0.079100      1.00800 ; qtot -
0.029
  2907    C   177   VAL     C 2907     0.597301     12.01000 ; qtot
0.568
  2908    O   177   VAL     O 2908    -0.567901     16.00000 ; qtot -
0.000
  2909    N   178   ALA     N 2909    -0.415700     14.01000 ; qtot -
0.416
  2910    H   178   ALA     H 2910     0.271900      1.00800 ; qtot -
0.144
  2911   CX   178   ALA    CA 2911     0.033700     12.01000 ; qtot -
0.110
  2912   H1   178   ALA    HA 2912     0.082300      1.00800 ; qtot -
0.028
  2913   CT   178   ALA    CB 2913    -0.182500     12.01000 ; qtot -
0.210
  2914   HC   178   ALA   HB1 2914     0.060300      1.00800 ; qtot -
0.150
  2915   HC   178   ALA   HB2 2915     0.060300      1.00800 ; qtot -
0.090
  2916   HC   178   ALA   HB3 2916     0.060300      1.00800 ; qtot -
0.029
  2917    C   178   ALA     C 2917     0.597301     12.01000 ; qtot
0.568
  2918    O   178   ALA     O 2918    -0.567901     16.00000 ; qtot -
0.000
  2919    N   179   PRO     N 2919    -0.254800     14.01000 ; qtot -
0.255
  2920   CT   179   PRO    CD 2920     0.019200     12.01000 ; qtot -
0.236
  2921   H1   179   PRO   HD2 2921     0.039100      1.00800 ; qtot -
0.197
  2922   H1   179   PRO   HD3 2922     0.039100      1.00800 ; qtot -
0.157
  2923   CT   179   PRO    CG 2923     0.018900     12.01000 ; qtot -
0.139
  2924   HC   179   PRO   HG2 2924     0.021300      1.00800 ; qtot -
0.117
  2925   HC   179   PRO   HG3 2925     0.021300      1.00800 ; qtot -
0.096
  2926   CT   179   PRO    CB 2926    -0.007000     12.01000 ; qtot -
0.103
  2927   HC   179   PRO   HB2 2927     0.025300      1.00800 ; qtot -
0.078
  2928   HC   179   PRO   HB3 2928     0.025300      1.00800 ; qtot -
0.052
  2929   CX   179   PRO    CA 2929    -0.026600     12.01000 ; qtot -
0.079
  2930   H1   179   PRO    HA 2930     0.064100      1.00800 ; qtot -
0.015
  2931    C   179   PRO     C 2931     0.589601     12.01000 ; qtot

0.575
  2932    O   179   PRO      O 2932    -0.574801     16.00000 ; qtot -
0.000
  2933    N   180   GLN      N 2933    -0.415700     14.01000 ; qtot -
0.416
  2934    H   180   GLN      H 2934     0.271900      1.00800 ; qtot -
0.144
  2935   CX   180   GLN     CA 2935    -0.003100     12.01000 ; qtot -
0.147
  2936   H1   180   GLN     HA 2936     0.085000      1.00800 ; qtot -
0.062
  2937  A2C   180   GLN     CB 2937    -0.003600     12.01000 ; qtot -
0.066
  2938   HC   180   GLN    HB2 2938     0.017100      1.00800 ; qtot -
0.048
  2939   HC   180   GLN    HB3 2939     0.017100      1.00800 ; qtot -
0.031
  2940  A2C   180   GLN     CG 2940    -0.064500     12.01000 ; qtot -
0.096
  2941   HC   180   GLN    HG2 2941     0.035200      1.00800 ; qtot -
0.061
  2942   HC   180   GLN    HG3 2942     0.035200      1.00800 ; qtot -
0.025
  2943    C   180   GLN     CD 2943     0.695101     12.01000 ; qtot
0.670
  2944    O   180   GLN    OE1 2944    -0.608601     16.00000 ; qtot
0.061
  2945    N   180   GLN    NE2 2945    -0.940701     14.01000 ; qtot -
0.880
  2946    H   180   GLN   HE21 2946     0.425100      1.00800 ; qtot -
0.455
  2947    H   180   GLN   HE22 2947     0.425100      1.00800 ; qtot -
0.029
  2948    C   180   GLN      C 2948     0.597301     12.01000 ; qtot
0.568
  2949    O   180   GLN      O 2949    -0.567901     16.00000 ; qtot -
0.000
  2950    N   181   VAL      N 2950    -0.415700     14.01000 ; qtot -
0.416
  2951    H   181   VAL      H 2951     0.271900      1.00800 ; qtot -
0.144
  2952   CX   181   VAL     CA 2952    -0.087500     12.01000 ; qtot -
0.231
  2953   H1   181   VAL     HA 2953     0.096900      1.00800 ; qtot -
0.134
  2954  A3C   181   VAL     CB 2954     0.298500     12.01000 ; qtot
0.164
  2955   HC   181   VAL     HB 2955    -0.029700      1.00800 ; qtot
0.134
  2956   CT   181   VAL    CG1 2956    -0.319200     12.01000 ; qtot -
0.185
  2957   HC   181   VAL   HG11 2957     0.079100      1.00800 ; qtot -

0.106
  2958   HC   181   VAL  HG12 2958     0.079100      1.00800 ; qtot -
0.027
  2959   HC   181   VAL  HG13 2959     0.079100      1.00800 ; qtot
0.052
  2960   CT   181   VAL   CG2 2960    -0.319200     12.01000 ; qtot -
0.267
  2961   HC   181   VAL  HG21 2961     0.079100      1.00800 ; qtot -
0.188
  2962   HC   181   VAL  HG22 2962     0.079100      1.00800 ; qtot -
0.109
  2963   HC   181   VAL  HG23 2963     0.079100      1.00800 ; qtot -
0.029
  2964    C   181   VAL     C 2964     0.597301     12.01000 ; qtot
0.568
  2965    O   181   VAL     O 2965    -0.567901     16.00000 ; qtot -
0.000
  2966    N   182   LEU     N 2966    -0.415700     14.01000 ; qtot -
0.416
  2967    H   182   LEU     H 2967     0.271900      1.00800 ; qtot -
0.144
  2968   CX   182   LEU    CA 2968    -0.051800     12.01000 ; qtot -
0.196
  2969   H1   182   LEU    HA 2969     0.092200      1.00800 ; qtot -
0.103
  2970  A2C   182   LEU    CB 2970    -0.110200     12.01000 ; qtot -
0.214
  2971   HC   182   LEU   HB2 2971     0.045700      1.00800 ; qtot -
0.168
  2972   HC   182   LEU   HB3 2972     0.045700      1.00800 ; qtot -
0.122
  2973  A3C   182   LEU    CG 2973     0.353100     12.01000 ; qtot
0.231
  2974   HC   182   LEU    HG 2974    -0.036100      1.00800 ; qtot
0.195
  2975   CT   182   LEU   CD1 2975    -0.412100     12.01000 ; qtot -
0.217
  2976   HC   182   LEU  HD11 2976     0.100000      1.00800 ; qtot -
0.117
  2977   HC   182   LEU  HD12 2977     0.100000      1.00800 ; qtot -
0.017
  2978   HC   182   LEU  HD13 2978     0.100000      1.00800 ; qtot
0.083
  2979   CT   182   LEU   CD2 2979    -0.412100     12.01000 ; qtot -
0.329
  2980   HC   182   LEU  HD21 2980     0.100000      1.00800 ; qtot -
0.229
  2981   HC   182   LEU  HD22 2981     0.100000      1.00800 ; qtot -
0.129
  2982   HC   182   LEU  HD23 2982     0.100000      1.00800 ; qtot -
0.029
  2983    C   182   LEU     C 2983     0.597301     12.01000 ; qtot

0.568
2984 O 182 LEU O 2984 -0.567901 16.00000 ; qtot -
0.000
2985 N 183 ASP N 2985 -0.516301 14.01000 ; qtot -
0.516
2986 H 183 ASP H 2986 0.293600 1.00800 ; qtot -
0.223
2987 CX 183 ASP CA 2987 0.038100 12.01000 ; qtot -
0.185
2988 H1 183 ASP HA 2988 0.088000 1.00800 ; qtot -
0.097
2989 A2C 183 ASP CB 2989 -0.030300 12.01000 ; qtot -
0.127
2990 HC 183 ASP HB2 2990 -0.012200 1.00800 ; qtot -
0.139
2991 HC 183 ASP HB3 2991 -0.012200 1.00800 ; qtot -
0.151
2992 CO 183 ASP CG 2992 0.799401 12.01000 ; qtot
0.648
2993 O2 183 ASP OD1 2993 -0.801401 16.00000 ; qtot -
0.153
2994 O2 183 ASP OD2 2994 -0.801401 16.00000 ; qtot -
0.955
2995 C 183 ASP C 2995 0.536601 12.01000 ; qtot -
0.418
2996 O 183 ASP O 2996 -0.581901 16.00000 ; qtot -
1.000
2997 N 184 GLY N 2997 -0.415700 14.01000 ; qtot -
1.416
2998 H 184 GLY H 2998 0.271900 1.00800 ; qtot -
1.144
2999 CX 184 GLY CA 2999 -0.025200 12.01000 ; qtot -
1.169
3000 H1 184 GLY HA2 3000 0.069800 1.00800 ; qtot -
1.099
3001 H1 184 GLY HA3 3001 0.069800 1.00800 ; qtot -
1.029
3002 C 184 GLY C 3002 0.597301 12.01000 ; qtot -
0.432
3003 O 184 GLY O 3003 -0.567901 16.00000 ; qtot -
1.000
3004 N 185 LYS N 3004 -0.347900 14.01000 ; qtot -
1.348
3005 H 185 LYS H 3005 0.274700 1.00800 ; qtot -
1.073
3006 CX 185 LYS CA 3006 -0.240000 12.01000 ; qtot -
1.313
3007 H1 185 LYS HA 3007 0.142600 1.00800 ; qtot -
1.171
3008 C8 185 LYS CB 3008 -0.009400 12.01000 ; qtot -
1.180
3009 HC 185 LYS HB2 3009 0.036200 1.00800 ; qtot -

1.144
  3010   HC   185   LYS   HB3 3010     0.036200      1.00800 ; qtot -
1.108
  3011   C8   185   LYS    CG 3011     0.018700     12.01000 ; qtot -
1.089
  3012   HC   185   LYS   HG2 3012     0.010300      1.00800 ; qtot -
1.079
  3013   HC   185   LYS   HG3 3013     0.010300      1.00800 ; qtot -
1.068
  3014   C8   185   LYS    CD 3014    -0.047900     12.01000 ; qtot -
1.116
  3015   HC   185   LYS   HD2 3015     0.062100      1.00800 ; qtot -
1.054
  3016   HC   185   LYS   HD3 3016     0.062100      1.00800 ; qtot -
0.992
  3017   C8   185   LYS    CE 3017    -0.014300     12.01000 ; qtot -
1.006
  3018   HP   185   LYS   HE2 3018     0.113500      1.00800 ; qtot -
0.893
  3019   HP   185   LYS   HE3 3019     0.113500      1.00800 ; qtot -
0.779
  3020   N3   185   LYS    NZ 3020    -0.385400     14.01000 ; qtot -
1.165
  3021    H   185   LYS   HZ1 3021     0.340000      1.00800 ; qtot -
0.825
  3022    H   185   LYS   HZ2 3022     0.340000      1.00800 ; qtot -
0.485
  3023    H   185   LYS   HZ3 3023     0.340000      1.00800 ; qtot -
0.145
  3024    C   185   LYS     C 3024     0.734101     12.01000 ; qtot
0.589
  3025    O   185   LYS     O 3025    -0.589401     16.00000 ; qtot -
0.000
  3026    N   186   TYR     N 3026    -0.415700     14.01000 ; qtot -
0.416
  3027    H   186   TYR     H 3027     0.271900      1.00800 ; qtot -
0.144
  3028   CX   186   TYR    CA 3028    -0.001400     12.01000 ; qtot -
0.145
  3029   H1   186   TYR    HA 3029     0.087600      1.00800 ; qtot -
0.058
  3030   CT   186   TYR    CB 3030    -0.015200     12.01000 ; qtot -
0.073
  3031   HC   186   TYR   HB2 3031     0.029500      1.00800 ; qtot -
0.043
  3032   HC   186   TYR   HB3 3032     0.029500      1.00800 ; qtot -
0.014
  3033   CA   186   TYR    CG 3033    -0.001100     12.01000 ; qtot -
0.015
  3034   CA   186   TYR   CD1 3034    -0.190600     12.01000 ; qtot -
0.206
  3035   HA   186   TYR   HD1 3035     0.169900      1.00800 ; qtot -

0.036
  3036   CA   186   TYR   CE1 3036    -0.234100     12.01000 ; qtot -
0.270
  3037   HA   186   TYR   HE1 3037     0.165600      1.00800 ; qtot -
0.104
  3038    C   186   TYR    CZ 3038     0.322600     12.01000 ; qtot
0.219
  3039   OH   186   TYR    OH 3039    -0.557901     16.00000 ; qtot -
0.339
  3040   HO   186   TYR    HH 3040     0.399200      1.00800 ; qtot
0.060
  3041   CA   186   TYR   CE2 3041    -0.234100     12.01000 ; qtot -
0.174
  3042   HA   186   TYR   HE2 3042     0.165600      1.00800 ; qtot -
0.009
  3043   CA   186   TYR   CD2 3043    -0.190600     12.01000 ; qtot -
0.199
  3044   HA   186   TYR   HD2 3044     0.169900      1.00800 ; qtot -
0.029
  3045    C   186   TYR     C 3045     0.597301     12.01000 ; qtot
0.568
  3046    O   186   TYR     O 3046    -0.567901     16.00000 ; qtot -
0.000
  3047    N   187   ASP     N 3047    -0.516301     14.01000 ; qtot -
0.516
  3048    H   187   ASP     H 3048     0.293600      1.00800 ; qtot -
0.223
  3049   CX   187   ASP    CA 3049     0.038100     12.01000 ; qtot -
0.185
  3050   H1   187   ASP    HA 3050     0.088000      1.00800 ; qtot -
0.097
  3051  A2C   187   ASP    CB 3051    -0.030300     12.01000 ; qtot -
0.127
  3052   HC   187   ASP   HB2 3052    -0.012200      1.00800 ; qtot -
0.139
  3053   HC   187   ASP   HB3 3053    -0.012200      1.00800 ; qtot -
0.151
  3054   CO   187   ASP    CG 3054     0.799401     12.01000 ; qtot
0.648
  3055   O2   187   ASP   OD1 3055    -0.801401     16.00000 ; qtot -
0.153
  3056   O2   187   ASP   OD2 3056    -0.801401     16.00000 ; qtot -
0.955
  3057    C   187   ASP     C 3057     0.536601     12.01000 ; qtot -
0.418
  3058    O   187   ASP     O 3058    -0.581901     16.00000 ; qtot -
1.000
  3059    N   188   LYS     N 3059    -0.347900     14.01000 ; qtot -
1.348
  3060    H   188   LYS     H 3060     0.274700      1.00800 ; qtot -
1.073
  3061   CX   188   LYS    CA 3061    -0.240000     12.01000 ; qtot -

1.313
3062 H1 188 LYS HA 3062 0.142600 1.00800 ; qtot -
1.171
3063 C8 188 LYS CB 3063 -0.009400 12.01000 ; qtot -
1.180
3064 HC 188 LYS HB2 3064 0.036200 1.00800 ; qtot -
1.144
3065 HC 188 LYS HB3 3065 0.036200 1.00800 ; qtot -
1.108
3066 C8 188 LYS CG 3066 0.018700 12.01000 ; qtot -
1.089
3067 HC 188 LYS HG2 3067 0.010300 1.00800 ; qtot -
1.079
3068 HC 188 LYS HG3 3068 0.010300 1.00800 ; qtot -
1.068
3069 C8 188 LYS CD 3069 -0.047900 12.01000 ; qtot -
1.116
3070 HC 188 LYS HD2 3070 0.062100 1.00800 ; qtot -
1.054
3071 HC 188 LYS HD3 3071 0.062100 1.00800 ; qtot -
0.992
3072 C8 188 LYS CE 3072 -0.014300 12.01000 ; qtot -
1.006
3073 HP 188 LYS HE2 3073 0.113500 1.00800 ; qtot -
0.893
3074 HP 188 LYS HE3 3074 0.113500 1.00800 ; qtot -
0.779
3075 N3 188 LYS NZ 3075 -0.385400 14.01000 ; qtot -
1.165
3076 H 188 LYS HZ1 3076 0.340000 1.00800 ; qtot -
0.825
3077 H 188 LYS HZ2 3077 0.340000 1.00800 ; qtot -
0.485
3078 H 188 LYS HZ3 3078 0.340000 1.00800 ; qtot -
0.145
3079 C 188 LYS C 3079 0.734101 12.01000 ; qtot
0.589
3080 O 188 LYS O 3080 -0.589401 16.00000 ; qtot -
0.000
3081 N 189 LYN N 3081 -0.415700 14.01000 ; qtot -
0.416
3082 H 189 LYN H 3082 0.271900 1.00800 ; qtot -
0.144
3083 CX 189 LYN CA 3083 -0.072060 12.01000 ; qtot -
0.216
3084 H1 189 LYN HA 3084 0.099400 1.00800 ; qtot -
0.116
3085 CT 189 LYN CB 3085 -0.048450 12.01000 ; qtot -
0.165
3086 HC 189 LYN HB2 3086 0.034000 1.00800 ; qtot -
0.131
3087 HC 189 LYN HB3 3087 0.034000 1.00800 ; qtot -

```
0.097
  3088   CT   189   LYN    CG 3088     0.066120     12.01000 ; qtot -
0.031
  3089   HC   189   LYN   HG2 3089     0.010410      1.00800 ; qtot -
0.020
  3090   HC   189   LYN   HG3 3090     0.010410      1.00800 ; qtot -
0.010
  3091   CT   189   LYN    CD 3091    -0.037680     12.01000 ; qtot -
0.048
  3092   HC   189   LYN   HD2 3092     0.011550      1.00800 ; qtot -
0.036
  3093   HC   189   LYN   HD3 3093     0.011550      1.00800 ; qtot -
0.025
  3094   CT   189   LYN    CE 3094     0.326040     12.01000 ; qtot
0.301
  3095   HP   189   LYN   HE2 3095    -0.033580      1.00800 ; qtot
0.268
  3096   HP   189   LYN   HE3 3096    -0.033580      1.00800 ; qtot
0.234
  3097   N3   189   LYN    NZ 3097    -1.035805     14.01000 ; qtot -
0.801
  3098    H   189   LYN   HZ2 3098     0.386040      1.00800 ; qtot -
0.415
  3099    H   189   LYN   HZ3 3099     0.386040      1.00800 ; qtot -
0.029
  3100    C   189   LYN     C 3100     0.597301     12.01000 ; qtot
0.568
  3101    O   189   LYN     O 3101    -0.567901     16.00000 ; qtot
0.000
  3102    N   190   CYS     N 3102    -0.415700     14.01000 ; qtot -
0.416
  3103    H   190   CYS     H 3103     0.271900      1.00800 ; qtot -
0.144
  3104   CX   190   CYS    CA 3104     0.021300     12.01000 ; qtot -
0.122
  3105   H1   190   CYS    HA 3105     0.112400      1.00800 ; qtot -
0.010
  3106  A2C   190   CYS    CB 3106    -0.123100     12.01000 ; qtot -
0.133
  3107   H1   190   CYS   HB2 3107     0.111200      1.00800 ; qtot -
0.022
  3108   H1   190   CYS   HB3 3108     0.111200      1.00800 ; qtot
0.089
  3109   SH   190   CYS    SG 3109    -0.311900     32.06000 ; qtot -
0.223
  3110   HS   190   CYS    HG 3110     0.193300      1.00800 ; qtot -
0.029
  3111    C   190   CYS     C 3111     0.597301     12.01000 ; qtot
0.568
  3112    O   190   CYS     O 3112    -0.567901     16.00000 ; qtot
0.000
  3113    N   191   ASP     N 3113    -0.516301     14.01000 ; qtot -
```

0.516
  3114    H   191   ASP      H 3114     0.293600      1.00800 ; qtot -
0.223
  3115   CX   191   ASP     CA 3115     0.038100     12.01000 ; qtot -
0.185
  3116   H1   191   ASP     HA 3116     0.088000      1.00800 ; qtot -
0.097
  3117  A2C   191   ASP     CB 3117    -0.030300     12.01000 ; qtot -
0.127
  3118   HC   191   ASP    HB2 3118    -0.012200      1.00800 ; qtot -
0.139
  3119   HC   191   ASP    HB3 3119    -0.012200      1.00800 ; qtot -
0.151
  3120   CO   191   ASP     CG 3120     0.799401     12.01000 ; qtot
0.648
  3121   O2   191   ASP    OD1 3121    -0.801401     16.00000 ; qtot -
0.153
  3122   O2   191   ASP    OD2 3122    -0.801401     16.00000 ; qtot -
0.955
  3123    C   191   ASP      C 3123     0.536601     12.01000 ; qtot -
0.418
  3124    O   191   ASP      O 3124    -0.581901     16.00000 ; qtot -
1.000
  3125    N   192   ILE      N 3125    -0.415700     14.01000 ; qtot -
1.416
  3126    H   192   ILE      H 3126     0.271900      1.00800 ; qtot -
1.144
  3127   CX   192   ILE     CA 3127    -0.059700     12.01000 ; qtot -
1.203
  3128   H1   192   ILE     HA 3128     0.086900      1.00800 ; qtot -
1.117
  3129  A3C   192   ILE     CB 3129     0.130300     12.01000 ; qtot -
0.986
  3130   HC   192   ILE     HB 3130     0.018700      1.00800 ; qtot -
0.968
  3131   CT   192   ILE    CG2 3131    -0.320400     12.01000 ; qtot -
1.288
  3132   HC   192   ILE   HG21 3132     0.088200      1.00800 ; qtot -
1.200
  3133   HC   192   ILE   HG22 3133     0.088200      1.00800 ; qtot -
1.112
  3134   HC   192   ILE   HG23 3134     0.088200      1.00800 ; qtot -
1.023
  3135  A2C   192   ILE    CG1 3135    -0.043000     12.01000 ; qtot -
1.066
  3136   HC   192   ILE   HG12 3136     0.023600      1.00800 ; qtot -
1.043
  3137   HC   192   ILE   HG13 3137     0.023600      1.00800 ; qtot -
1.019
  3138   CT   192   ILE    CD1 3138    -0.066000     12.01000 ; qtot -
1.085
  3139   HC   192   ILE   HD11 3139     0.018600      1.00800 ; qtot -

1.067
  3140   HC   192   ILE  HD12 3140     0.018600      1.00800 ; qtot -
1.048
  3141   HC   192   ILE  HD13 3141     0.018600      1.00800 ; qtot -
1.029
  3142    C   192   ILE     C 3142     0.597301     12.01000 ; qtot -
0.432
  3143    O   192   ILE     O 3143    -0.567901     16.00000 ; qtot -
1.000
  3144    N   193   TRP     N 3144    -0.415700     14.01000 ; qtot -
1.416
  3145    H   193   TRP     H 3145     0.271900      1.00800 ; qtot -
1.144
  3146   CX   193   TRP    CA 3146    -0.027500     12.01000 ; qtot -
1.171
  3147   H1   193   TRP    HA 3147     0.112300      1.00800 ; qtot -
1.059
  3148   CT   193   TRP    CB 3148    -0.005000     12.01000 ; qtot -
1.064
  3149   HC   193   TRP   HB2 3149     0.033900      1.00800 ; qtot -
1.030
  3150   HC   193   TRP   HB3 3150     0.033900      1.00800 ; qtot -
0.996
  3151   C*   193   TRP    CG 3151    -0.141500     12.01000 ; qtot -
1.138
  3152   CW   193   TRP   CD1 3152    -0.163800     12.01000 ; qtot -
1.301
  3153   H4   193   TRP   HD1 3153     0.206200      1.00800 ; qtot -
1.095
  3154   NA   193   TRP   NE1 3154    -0.341800     14.01000 ; qtot -
1.437
  3155    H   193   TRP   HE1 3155     0.341200      1.00800 ; qtot -
1.096
  3156   CN   193   TRP   CE2 3156     0.138000     12.01000 ; qtot -
0.958
  3157   CA   193   TRP   CZ2 3157    -0.260100     12.01000 ; qtot -
1.218
  3158   HA   193   TRP   HZ2 3158     0.157200      1.00800 ; qtot -
1.061
  3159   CA   193   TRP   CH2 3159    -0.113400     12.01000 ; qtot -
1.174
  3160   HA   193   TRP   HH2 3160     0.141700      1.00800 ; qtot -
1.032
  3161   CA   193   TRP   CZ3 3161    -0.197200     12.01000 ; qtot -
1.230
  3162   HA   193   TRP   HZ3 3162     0.144700      1.00800 ; qtot -
1.085
  3163   CA   193   TRP   CE3 3163    -0.238700     12.01000 ; qtot -
1.324
  3164   HA   193   TRP   HE3 3164     0.170000      1.00800 ; qtot -
1.154
  3165   CB   193   TRP   CD2 3165     0.124300     12.01000 ; qtot -

```
1.029
  3166    C   193   TRP      C 3166     0.597301     12.01000 ; qtot -
0.432
  3167    O   193   TRP      O 3167    -0.567901     16.00000 ; qtot -
1.000
  3168    N   194   SER      N 3168    -0.415700     14.01000 ; qtot -
1.416
  3169    H   194   SER      H 3169     0.271900      1.00800 ; qtot -
1.144
  3170   CX   194   SER     CA 3170    -0.024900     12.01000 ; qtot -
1.169
  3171   H1   194   SER     HA 3171     0.084300      1.00800 ; qtot -
1.084
  3172  A2C   194   SER     CB 3172     0.211700     12.01000 ; qtot -
0.873
  3173   H1   194   SER    HB2 3173     0.035200      1.00800 ; qtot -
0.837
  3174   H1   194   SER    HB3 3174     0.035200      1.00800 ; qtot -
0.802
  3175   OH   194   SER     OG 3175    -0.654601     16.00000 ; qtot -
1.457
  3176   HO   194   SER     HG 3176     0.427500      1.00800 ; qtot -
1.029
  3177    C   194   SER      C 3177     0.597301     12.01000 ; qtot -
0.432
  3178    O   194   SER      O 3178    -0.567901     16.00000 ; qtot -
1.000
  3179    N   195   SER      N 3179    -0.415700     14.01000 ; qtot -
1.416
  3180    H   195   SER      H 3180     0.271900      1.00800 ; qtot -
1.144
  3181   CX   195   SER     CA 3181    -0.024900     12.01000 ; qtot -
1.169
  3182   H1   195   SER     HA 3182     0.084300      1.00800 ; qtot -
1.084
  3183  A2C   195   SER     CB 3183     0.211700     12.01000 ; qtot -
0.873
  3184   H1   195   SER    HB2 3184     0.035200      1.00800 ; qtot -
0.837
  3185   H1   195   SER    HB3 3185     0.035200      1.00800 ; qtot -
0.802
  3186   OH   195   SER     OG 3186    -0.654601     16.00000 ; qtot -
1.457
  3187   HO   195   SER     HG 3187     0.427500      1.00800 ; qtot -
1.029
  3188    C   195   SER      C 3188     0.597301     12.01000 ; qtot -
0.432
  3189    O   195   SER      O 3189    -0.567901     16.00000 ; qtot -
1.000
  3190    N   196   GLY      N 3190    -0.415700     14.01000 ; qtot -
1.416
  3191    H   196   GLY      H 3191     0.271900      1.00800 ; qtot -
```

1.144
3192 CX 196 GLY CA 3192 -0.025200 12.01000 ; qtot -
1.169
3193 H1 196 GLY HA2 3193 0.069800 1.00800 ; qtot -
1.099
3194 H1 196 GLY HA3 3194 0.069800 1.00800 ; qtot -
1.029
3195 C 196 GLY C 3195 0.597301 12.01000 ; qtot -
0.432
3196 O 196 GLY O 3196 -0.567901 16.00000 ; qtot -
1.000
3197 N 197 VAL N 3197 -0.415700 14.01000 ; qtot -
1.416
3198 H 197 VAL H 3198 0.271900 1.00800 ; qtot -
1.144
3199 CX 197 VAL CA 3199 -0.087500 12.01000 ; qtot -
1.231
3200 H1 197 VAL HA 3200 0.096900 1.00800 ; qtot -
1.134
3201 A3C 197 VAL CB 3201 0.298500 12.01000 ; qtot -
0.836
3202 HC 197 VAL HB 3202 -0.029700 1.00800 ; qtot -
0.866
3203 CT 197 VAL CG1 3203 -0.319200 12.01000 ; qtot -
1.185
3204 HC 197 VAL HG11 3204 0.079100 1.00800 ; qtot -
1.106
3205 HC 197 VAL HG12 3205 0.079100 1.00800 ; qtot -
1.027
3206 HC 197 VAL HG13 3206 0.079100 1.00800 ; qtot -
0.947
3207 CT 197 VAL CG2 3207 -0.319200 12.01000 ; qtot -
1.267
3208 HC 197 VAL HG21 3208 0.079100 1.00800 ; qtot -
1.188
3209 HC 197 VAL HG22 3209 0.079100 1.00800 ; qtot -
1.108
3210 HC 197 VAL HG23 3210 0.079100 1.00800 ; qtot -
1.029
3211 C 197 VAL C 3211 0.597301 12.01000 ; qtot -
0.432
3212 O 197 VAL O 3212 -0.567901 16.00000 ; qtot -
1.000
3213 N 198 ILE N 3213 -0.415700 14.01000 ; qtot -
1.416
3214 H 198 ILE H 3214 0.271900 1.00800 ; qtot -
1.144
3215 CX 198 ILE CA 3215 -0.059700 12.01000 ; qtot -
1.203
3216 H1 198 ILE HA 3216 0.086900 1.00800 ; qtot -
1.117
3217 A3C 198 ILE CB 3217 0.130300 12.01000 ; qtot -

0.986
3218 HC 198 ILE HB 3218 0.018700 1.00800 ; qtot -
0.968
3219 CT 198 ILE CG2 3219 -0.320400 12.01000 ; qtot -
1.288
3220 HC 198 ILE HG21 3220 0.088200 1.00800 ; qtot -
1.200
3221 HC 198 ILE HG22 3221 0.088200 1.00800 ; qtot -
1.112
3222 HC 198 ILE HG23 3222 0.088200 1.00800 ; qtot -
1.023
3223 A2C 198 ILE CG1 3223 -0.043000 12.01000 ; qtot -
1.066
3224 HC 198 ILE HG12 3224 0.023600 1.00800 ; qtot -
1.043
3225 HC 198 ILE HG13 3225 0.023600 1.00800 ; qtot -
1.019
3226 CT 198 ILE CD1 3226 -0.066000 12.01000 ; qtot -
1.085
3227 HC 198 ILE HD11 3227 0.018600 1.00800 ; qtot -
1.067
3228 HC 198 ILE HD12 3228 0.018600 1.00800 ; qtot -
1.048
3229 HC 198 ILE HD13 3229 0.018600 1.00800 ; qtot -
1.029
3230 C 198 ILE C 3230 0.597301 12.01000 ; qtot -
0.432
3231 O 198 ILE O 3231 -0.567901 16.00000 ; qtot -
1.000
3232 N 199 MET N 3232 -0.415700 14.01000 ; qtot -
1.416
3233 H 199 MET H 3233 0.271900 1.00800 ; qtot -
1.144
3234 CX 199 MET CA 3234 -0.023700 12.01000 ; qtot -
1.167
3235 H1 199 MET HA 3235 0.088000 1.00800 ; qtot -
1.079
3236 A2C 199 MET CB 3236 0.034200 12.01000 ; qtot -
1.045
3237 HC 199 MET HB2 3237 0.024100 1.00800 ; qtot -
1.021
3238 HC 199 MET HB3 3238 0.024100 1.00800 ; qtot -
0.997
3239 A2C 199 MET CG 3239 0.001800 12.01000 ; qtot -
0.995
3240 H1 199 MET HG2 3240 0.044000 1.00800 ; qtot -
0.951
3241 H1 199 MET HG3 3241 0.044000 1.00800 ; qtot -
0.907
3242 S 199 MET SD 3242 -0.273700 32.06000 ; qtot -
1.181
3243 CT 199 MET CE 3243 -0.053600 12.01000 ; qtot -

1.235
  3244   H1   199   MET   HE1 3244     0.068400      1.00800 ; qtot -
1.166
  3245   H1   199   MET   HE2 3245     0.068400      1.00800 ; qtot -
1.098
  3246   H1   199   MET   HE3 3246     0.068400      1.00800 ; qtot -
1.029
  3247    C   199   MET     C 3247     0.597301     12.01000 ; qtot -
0.432
  3248    O   199   MET     O 3248    -0.567901     16.00000 ; qtot -
1.000
  3249    N   200   TYR     N 3249    -0.415700     14.01000 ; qtot -
1.416
  3250    H   200   TYR     H 3250     0.271900      1.00800 ; qtot -
1.144
  3251   CX   200   TYR    CA 3251    -0.001400     12.01000 ; qtot -
1.145
  3252   H1   200   TYR    HA 3252     0.087600      1.00800 ; qtot -
1.058
  3253   CT   200   TYR    CB 3253    -0.015200     12.01000 ; qtot -
1.073
  3254   HC   200   TYR   HB2 3254     0.029500      1.00800 ; qtot -
1.043
  3255   HC   200   TYR   HB3 3255     0.029500      1.00800 ; qtot -
1.014
  3256   CA   200   TYR    CG 3256    -0.001100     12.01000 ; qtot -
1.015
  3257   CA   200   TYR   CD1 3257    -0.190600     12.01000 ; qtot -
1.205
  3258   HA   200   TYR   HD1 3258     0.169900      1.00800 ; qtot -
1.036
  3259   CA   200   TYR   CE1 3259    -0.234100     12.01000 ; qtot -
1.270
  3260   HA   200   TYR   HE1 3260     0.165600      1.00800 ; qtot -
1.104
  3261    C   200   TYR    CZ 3261     0.322600     12.01000 ; qtot -
0.781
  3262   OH   200   TYR    OH 3262    -0.557901     16.00000 ; qtot -
1.339
  3263   HO   200   TYR    HH 3263     0.399200      1.00800 ; qtot -
0.940
  3264   CA   200   TYR   CE2 3264    -0.234100     12.01000 ; qtot -
1.174
  3265   HA   200   TYR   HE2 3265     0.165600      1.00800 ; qtot -
1.009
  3266   CA   200   TYR   CD2 3266    -0.190600     12.01000 ; qtot -
1.199
  3267   HA   200   TYR   HD2 3267     0.169900      1.00800 ; qtot -
1.029
  3268    C   200   TYR     C 3268     0.597301     12.01000 ; qtot -
0.432
  3269    O   200   TYR     O 3269    -0.567901     16.00000 ; qtot -

1.000
  3270    N   201   THR      N 3270    -0.415700     14.01000 ; qtot -
1.416
  3271    H   201   THR      H 3271     0.271900      1.00800 ; qtot -
1.144
  3272   CX   201   THR     CA 3272    -0.038900     12.01000 ; qtot -
1.183
  3273   H1   201   THR     HA 3273     0.100700      1.00800 ; qtot -
1.082
  3274  A3C   201   THR     CB 3274     0.365400     12.01000 ; qtot -
0.717
  3275   H1   201   THR     HB 3275     0.004300      1.00800 ; qtot -
0.712
  3276   CT   201   THR    CG2 3276    -0.243800     12.01000 ; qtot -
0.956
  3277   HC   201   THR   HG21 3277     0.064200      1.00800 ; qtot -
0.892
  3278   HC   201   THR   HG22 3278     0.064200      1.00800 ; qtot -
0.828
  3279   HC   201   THR   HG23 3279     0.064200      1.00800 ; qtot -
0.763
  3280   OH   201   THR    OG1 3280    -0.676101     16.00000 ; qtot -
1.440
  3281   HO   201   THR    HG1 3281     0.410200      1.00800 ; qtot -
1.029
  3282    C   201   THR      C 3282     0.597301     12.01000 ; qtot -
0.432
  3283    O   201   THR      O 3283    -0.567901     16.00000 ; qtot -
1.000
  3284    N   202   LEU      N 3284    -0.415700     14.01000 ; qtot -
1.416
  3285    H   202   LEU      H 3285     0.271900      1.00800 ; qtot -
1.144
  3286   CX   202   LEU     CA 3286    -0.051800     12.01000 ; qtot -
1.196
  3287   H1   202   LEU     HA 3287     0.092200      1.00800 ; qtot -
1.103
  3288  A2C   202   LEU     CB 3288    -0.110200     12.01000 ; qtot -
1.214
  3289   HC   202   LEU    HB2 3289     0.045700      1.00800 ; qtot -
1.168
  3290   HC   202   LEU    HB3 3290     0.045700      1.00800 ; qtot -
1.122
  3291  A3C   202   LEU     CG 3291     0.353100     12.01000 ; qtot -
0.769
  3292   HC   202   LEU     HG 3292    -0.036100      1.00800 ; qtot -
0.805
  3293   CT   202   LEU    CD1 3293    -0.412100     12.01000 ; qtot -
1.217
  3294   HC   202   LEU   HD11 3294     0.100000      1.00800 ; qtot -
1.117
  3295   HC   202   LEU   HD12 3295     0.100000      1.00800 ; qtot -

1.017
3296 HC 202 LEU HD13 3296 0.100000 1.00800 ; qtot -
0.917
3297 CT 202 LEU CD2 3297 -0.412100 12.01000 ; qtot -
1.329
3298 HC 202 LEU HD21 3298 0.100000 1.00800 ; qtot -
1.229
3299 HC 202 LEU HD22 3299 0.100000 1.00800 ; qtot -
1.129
3300 HC 202 LEU HD23 3300 0.100000 1.00800 ; qtot -
1.029
3301 C 202 LEU C 3301 0.597301 12.01000 ; qtot -
0.432
3302 O 202 LEU O 3302 -0.567901 16.00000 ; qtot -
1.000
3303 N 203 LEU N 3303 -0.415700 14.01000 ; qtot -
1.416
3304 H 203 LEU H 3304 0.271900 1.00800 ; qtot -
1.144
3305 CX 203 LEU CA 3305 -0.051800 12.01000 ; qtot -
1.196
3306 H1 203 LEU HA 3306 0.092200 1.00800 ; qtot -
1.103
3307 A2C 203 LEU CB 3307 -0.110200 12.01000 ; qtot -
1.214
3308 HC 203 LEU HB2 3308 0.045700 1.00800 ; qtot -
1.168
3309 HC 203 LEU HB3 3309 0.045700 1.00800 ; qtot -
1.122
3310 A3C 203 LEU CG 3310 0.353100 12.01000 ; qtot -
0.769
3311 HC 203 LEU HG 3311 -0.036100 1.00800 ; qtot -
0.805
3312 CT 203 LEU CD1 3312 -0.412100 12.01000 ; qtot -
1.217
3313 HC 203 LEU HD11 3313 0.100000 1.00800 ; qtot -
1.117
3314 HC 203 LEU HD12 3314 0.100000 1.00800 ; qtot -
1.017
3315 HC 203 LEU HD13 3315 0.100000 1.00800 ; qtot -
0.917
3316 CT 203 LEU CD2 3316 -0.412100 12.01000 ; qtot -
1.329
3317 HC 203 LEU HD21 3317 0.100000 1.00800 ; qtot -
1.229
3318 HC 203 LEU HD22 3318 0.100000 1.00800 ; qtot -
1.129
3319 HC 203 LEU HD23 3319 0.100000 1.00800 ; qtot -
1.029
3320 C 203 LEU C 3320 0.597301 12.01000 ; qtot -
0.432
3321 O 203 LEU O 3321 -0.567901 16.00000 ; qtot -

1.000
3322 N 204 CYS N 3322 -0.415700 14.01000 ; qtot -
1.416
3323 H 204 CYS H 3323 0.271900 1.00800 ; qtot -
1.144
3324 CX 204 CYS CA 3324 0.021300 12.01000 ; qtot -
1.122
3325 H1 204 CYS HA 3325 0.112400 1.00800 ; qtot -
1.010
3326 A2C 204 CYS CB 3326 -0.123100 12.01000 ; qtot -
1.133
3327 H1 204 CYS HB2 3327 0.111200 1.00800 ; qtot -
1.022
3328 H1 204 CYS HB3 3328 0.111200 1.00800 ; qtot -
0.911
3329 SH 204 CYS SG 3329 -0.311900 32.06000 ; qtot -
1.223
3330 HS 204 CYS HG 3330 0.193300 1.00800 ; qtot -
1.029
3331 C 204 CYS C 3331 0.597301 12.01000 ; qtot -
0.432
3332 O 204 CYS O 3332 -0.567901 16.00000 ; qtot -
1.000
3333 N 205 GLY N 3333 -0.415700 14.01000 ; qtot -
1.416
3334 H 205 GLY H 3334 0.271900 1.00800 ; qtot -
1.144
3335 CX 205 GLY CA 3335 -0.025200 12.01000 ; qtot -
1.169
3336 H1 205 GLY HA2 3336 0.069800 1.00800 ; qtot -
1.099
3337 H1 205 GLY HA3 3337 0.069800 1.00800 ; qtot -
1.029
3338 C 205 GLY C 3338 0.597301 12.01000 ; qtot -
0.432
3339 O 205 GLY O 3339 -0.567901 16.00000 ; qtot -
1.000
3340 N 206 TYR N 3340 -0.415700 14.01000 ; qtot -
1.416
3341 H 206 TYR H 3341 0.271900 1.00800 ; qtot -
1.144
3342 CX 206 TYR CA 3342 -0.001400 12.01000 ; qtot -
1.145
3343 H1 206 TYR HA 3343 0.087600 1.00800 ; qtot -
1.058
3344 CT 206 TYR CB 3344 -0.015200 12.01000 ; qtot -
1.073
3345 HC 206 TYR HB2 3345 0.029500 1.00800 ; qtot -
1.043
3346 HC 206 TYR HB3 3346 0.029500 1.00800 ; qtot -
1.014
3347 CA 206 TYR CG 3347 -0.001100 12.01000 ; qtot -

1.015
  3348   CA   206   TYR   CD1 3348    -0.190600     12.01000 ; qtot -
1.205
  3349   HA   206   TYR   HD1 3349     0.169900      1.00800 ; qtot -
1.036
  3350   CA   206   TYR   CE1 3350    -0.234100     12.01000 ; qtot -
1.270
  3351   HA   206   TYR   HE1 3351     0.165600      1.00800 ; qtot -
1.104
  3352    C   206   TYR    CZ 3352     0.322600     12.01000 ; qtot -
0.781
  3353   OH   206   TYR    OH 3353    -0.557901     16.00000 ; qtot -
1.339
  3354   HO   206   TYR    HH 3354     0.399200      1.00800 ; qtot -
0.940
  3355   CA   206   TYR   CE2 3355    -0.234100     12.01000 ; qtot -
1.174
  3356   HA   206   TYR   HE2 3356     0.165600      1.00800 ; qtot -
1.009
  3357   CA   206   TYR   CD2 3357    -0.190600     12.01000 ; qtot -
1.199
  3358   HA   206   TYR   HD2 3358     0.169900      1.00800 ; qtot -
1.029
  3359    C   206   TYR     C 3359     0.597301     12.01000 ; qtot -
0.432
  3360    O   206   TYR     O 3360    -0.567901     16.00000 ; qtot -
1.000
  3361    N   207   PRO     N 3361    -0.254800     14.01000 ; qtot -
1.255
  3362   CT   207   PRO    CD 3362     0.019200     12.01000 ; qtot -
1.236
  3363   H1   207   PRO   HD2 3363     0.039100      1.00800 ; qtot -
1.196
  3364   H1   207   PRO   HD3 3364     0.039100      1.00800 ; qtot -
1.157
  3365   CT   207   PRO    CG 3365     0.018900     12.01000 ; qtot -
1.138
  3366   HC   207   PRO   HG2 3366     0.021300      1.00800 ; qtot -
1.117
  3367   HC   207   PRO   HG3 3367     0.021300      1.00800 ; qtot -
1.096
  3368   CT   207   PRO    CB 3368    -0.007000     12.01000 ; qtot -
1.103
  3369   HC   207   PRO   HB2 3369     0.025300      1.00800 ; qtot -
1.078
  3370   HC   207   PRO   HB3 3370     0.025300      1.00800 ; qtot -
1.052
  3371   CX   207   PRO    CA 3371    -0.026600     12.01000 ; qtot -
1.079
  3372   H1   207   PRO    HA 3372     0.064100      1.00800 ; qtot -
1.015
  3373    C   207   PRO     C 3373     0.589601     12.01000 ; qtot -

```
0.425
  3374    O   207   PRO      O 3374    -0.574801     16.00000 ; qtot -
1.000
  3375    N   208   PRO      N 3375    -0.254800     14.01000 ; qtot -
1.255
  3376   CT   208   PRO     CD 3376     0.019200     12.01000 ; qtot -
1.236
  3377   H1   208   PRO    HD2 3377     0.039100      1.00800 ; qtot -
1.196
  3378   H1   208   PRO    HD3 3378     0.039100      1.00800 ; qtot -
1.157
  3379   CT   208   PRO     CG 3379     0.018900     12.01000 ; qtot -
1.138
  3380   HC   208   PRO    HG2 3380     0.021300      1.00800 ; qtot -
1.117
  3381   HC   208   PRO    HG3 3381     0.021300      1.00800 ; qtot -
1.096
  3382   CT   208   PRO     CB 3382    -0.007000     12.01000 ; qtot -
1.103
  3383   HC   208   PRO    HB2 3383     0.025300      1.00800 ; qtot -
1.078
  3384   HC   208   PRO    HB3 3384     0.025300      1.00800 ; qtot -
1.052
  3385   CX   208   PRO     CA 3385    -0.026600     12.01000 ; qtot -
1.079
  3386   H1   208   PRO     HA 3386     0.064100      1.00800 ; qtot -
1.015
  3387    C   208   PRO      C 3387     0.589601     12.01000 ; qtot -
0.425
  3388    O   208   PRO      O 3388    -0.574801     16.00000 ; qtot -
1.000
  3389    N   209   PHE      N 3389    -0.415700     14.01000 ; qtot -
1.416
  3390    H   209   PHE      H 3390     0.271900      1.00800 ; qtot -
1.144
  3391   CX   209   PHE     CA 3391    -0.002400     12.01000 ; qtot -
1.146
  3392   H1   209   PHE     HA 3392     0.097800      1.00800 ; qtot -
1.048
  3393   CT   209   PHE     CB 3393    -0.034300     12.01000 ; qtot -
1.083
  3394   HC   209   PHE    HB2 3394     0.029500      1.00800 ; qtot -
1.053
  3395   HC   209   PHE    HB3 3395     0.029500      1.00800 ; qtot -
1.024
  3396   CA   209   PHE     CG 3396     0.011800     12.01000 ; qtot -
1.012
  3397   CA   209   PHE    CD1 3397    -0.125600     12.01000 ; qtot -
1.137
  3398   HA   209   PHE    HD1 3398     0.133000      1.00800 ; qtot -
1.004
  3399   CA   209   PHE    CE1 3399    -0.170400     12.01000 ; qtot -
```

1.175
3400 HA 209 PHE HE1 3400 0.143000 1.00800 ; qtot -
1.032
3401 CA 209 PHE CZ 3401 -0.107200 12.01000 ; qtot -
1.139
3402 HA 209 PHE HZ 3402 0.129700 1.00800 ; qtot -
1.009
3403 CA 209 PHE CE2 3403 -0.170400 12.01000 ; qtot -
1.180
3404 HA 209 PHE HE2 3404 0.143000 1.00800 ; qtot -
1.037
3405 CA 209 PHE CD2 3405 -0.125600 12.01000 ; qtot -
1.162
3406 HA 209 PHE HD2 3406 0.133000 1.00800 ; qtot -
1.029
3407 C 209 PHE C 3407 0.597301 12.01000 ; qtot -
0.432
3408 O 209 PHE O 3408 -0.567901 16.00000 ; qtot -
1.000
3409 N 210 TYR N 3409 -0.415700 14.01000 ; qtot -
1.416
3410 H 210 TYR H 3410 0.271900 1.00800 ; qtot -
1.144
3411 CX 210 TYR CA 3411 -0.001400 12.01000 ; qtot -
1.145
3412 H1 210 TYR HA 3412 0.087600 1.00800 ; qtot -
1.058
3413 CT 210 TYR CB 3413 -0.015200 12.01000 ; qtot -
1.073
3414 HC 210 TYR HB2 3414 0.029500 1.00800 ; qtot -
1.043
3415 HC 210 TYR HB3 3415 0.029500 1.00800 ; qtot -
1.014
3416 CA 210 TYR CG 3416 -0.001100 12.01000 ; qtot -
1.015
3417 CA 210 TYR CD1 3417 -0.190600 12.01000 ; qtot -
1.205
3418 HA 210 TYR HD1 3418 0.169900 1.00800 ; qtot -
1.036
3419 CA 210 TYR CE1 3419 -0.234100 12.01000 ; qtot -
1.270
3420 HA 210 TYR HE1 3420 0.165600 1.00800 ; qtot -
1.104
3421 C 210 TYR CZ 3421 0.322600 12.01000 ; qtot -
0.781
3422 OH 210 TYR OH 3422 -0.557901 16.00000 ; qtot -
1.339
3423 HO 210 TYR HH 3423 0.399200 1.00800 ; qtot -
0.940
3424 CA 210 TYR CE2 3424 -0.234100 12.01000 ; qtot -
1.174
3425 HA 210 TYR HE2 3425 0.165600 1.00800 ; qtot -

```
1.009
  3426   CA   210   TYR   CD2 3426    -0.190600     12.01000 ; qtot -
1.199
  3427   HA   210   TYR   HD2 3427     0.169900      1.00800 ; qtot -
1.029
  3428    C   210   TYR     C 3428     0.597301     12.01000 ; qtot -
0.432
  3429    O   210   TYR     O 3429    -0.567901     16.00000 ; qtot -
1.000
  3430    N   211   GLY     N 3430    -0.415700     14.01000 ; qtot -
1.416
  3431    H   211   GLY     H 3431     0.271900      1.00800 ; qtot -
1.144
  3432   CX   211   GLY    CA 3432    -0.025200     12.01000 ; qtot -
1.169
  3433   H1   211   GLY   HA2 3433     0.069800      1.00800 ; qtot -
1.099
  3434   H1   211   GLY   HA3 3434     0.069800      1.00800 ; qtot -
1.029
  3435    C   211   GLY     C 3435     0.597301     12.01000 ; qtot -
0.432
  3436    O   211   GLY     O 3436    -0.567901     16.00000 ; qtot -
1.000
  3437    N   212   ASP     N 3437    -0.516301     14.01000 ; qtot -
1.516
  3438    H   212   ASP     H 3438     0.293600      1.00800 ; qtot -
1.223
  3439   CX   212   ASP    CA 3439     0.038100     12.01000 ; qtot -
1.185
  3440   H1   212   ASP    HA 3440     0.088000      1.00800 ; qtot -
1.097
  3441  A2C   212   ASP    CB 3441    -0.030300     12.01000 ; qtot -
1.127
  3442   HC   212   ASP   HB2 3442    -0.012200      1.00800 ; qtot -
1.139
  3443   HC   212   ASP   HB3 3443    -0.012200      1.00800 ; qtot -
1.151
  3444   CO   212   ASP    CG 3444     0.799401     12.01000 ; qtot -
0.352
  3445   O2   212   ASP   OD1 3445    -0.801401     16.00000 ; qtot -
1.153
  3446   O2   212   ASP   OD2 3446    -0.801401     16.00000 ; qtot -
1.955
  3447    C   212   ASP     C 3447     0.536601     12.01000 ; qtot -
1.418
  3448    O   212   ASP     O 3448    -0.581901     16.00000 ; qtot -
2.000
  3449    N   213   THR     N 3449    -0.415700     14.01000 ; qtot -
2.416
  3450    H   213   THR     H 3450     0.271900      1.00800 ; qtot -
2.144
  3451   CX   213   THR    CA 3451    -0.038900     12.01000 ; qtot -
```

```
2.183
  3452   H1   213   THR    HA 3452     0.100700      1.00800 ; qtot -
2.082
  3453  A3C   213   THR    CB 3453     0.365400     12.01000 ; qtot -
1.717
  3454   H1   213   THR    HB 3454     0.004300      1.00800 ; qtot -
1.712
  3455   CT   213   THR   CG2 3455    -0.243800     12.01000 ; qtot -
1.956
  3456   HC   213   THR  HG21 3456     0.064200      1.00800 ; qtot -
1.892
  3457   HC   213   THR  HG22 3457     0.064200      1.00800 ; qtot -
1.828
  3458   HC   213   THR  HG23 3458     0.064200      1.00800 ; qtot -
1.763
  3459   OH   213   THR   OG1 3459    -0.676101     16.00000 ; qtot -
2.440
  3460   HO   213   THR   HG1 3460     0.410200      1.00800 ; qtot -
2.029
  3461    C   213   THR     C 3461     0.597301     12.01000 ; qtot -
1.432
  3462    O   213   THR     O 3462    -0.567901     16.00000 ; qtot -
2.000
  3463    N   214   ASP     N 3463    -0.516301     14.01000 ; qtot -
2.516
  3464    H   214   ASP     H 3464     0.293600      1.00800 ; qtot -
2.223
  3465   CX   214   ASP    CA 3465     0.038100     12.01000 ; qtot -
2.185
  3466   H1   214   ASP    HA 3466     0.088000      1.00800 ; qtot -
2.097
  3467  A2C   214   ASP    CB 3467    -0.030300     12.01000 ; qtot -
2.127
  3468   HC   214   ASP   HB2 3468    -0.012200      1.00800 ; qtot -
2.139
  3469   HC   214   ASP   HB3 3469    -0.012200      1.00800 ; qtot -
2.151
  3470   CO   214   ASP    CG 3470     0.799401     12.01000 ; qtot -
1.352
  3471   O2   214   ASP   OD1 3471    -0.801401     16.00000 ; qtot -
2.153
  3472   O2   214   ASP   OD2 3472    -0.801401     16.00000 ; qtot -
2.955
  3473    C   214   ASP     C 3473     0.536601     12.01000 ; qtot -
2.418
  3474    O   214   ASP     O 3474    -0.581901     16.00000 ; qtot -
3.000
  3475    N   215   ASN     N 3475    -0.415700     14.01000 ; qtot -
3.416
  3476    H   215   ASN     H 3476     0.271900      1.00800 ; qtot -
3.144
  3477   CX   215   ASN    CA 3477     0.014300     12.01000 ; qtot -
```

```
3.129
  3478   H1   215   ASN    HA 3478     0.104800      1.00800 ; qtot -
3.025
  3479  A2C   215   ASN    CB 3479    -0.204100     12.01000 ; qtot -
3.229
  3480   HC   215   ASN   HB2 3480     0.079700      1.00800 ; qtot -
3.149
  3481   HC   215   ASN   HB3 3481     0.079700      1.00800 ; qtot -
3.069
  3482    C   215   ASN    CG 3482     0.713001     12.01000 ; qtot -
2.356
  3483    O   215   ASN   OD1 3483    -0.593101     16.00000 ; qtot -
2.949
  3484    N   215   ASN   ND2 3484    -0.919101     14.01000 ; qtot -
3.869
  3485    H   215   ASN  HD21 3485     0.419600      1.00800 ; qtot -
3.449
  3486    H   215   ASN  HD22 3486     0.419600      1.00800 ; qtot -
3.029
  3487    C   215   ASN     C 3487     0.597301     12.01000 ; qtot -
2.432
  3488    O   215   ASN     O 3488    -0.567901     16.00000 ; qtot -
3.000
  3489    N   216   GLU     N 3489    -0.516301     14.01000 ; qtot -
3.516
  3490    H   216   GLU     H 3490     0.293600      1.00800 ; qtot -
3.223
  3491   CX   216   GLU    CA 3491     0.039700     12.01000 ; qtot -
3.183
  3492   H1   216   GLU    HA 3492     0.110500      1.00800 ; qtot -
3.072
  3493  A2C   216   GLU    CB 3493     0.056000     12.01000 ; qtot -
3.016
  3494   HC   216   GLU   HB2 3494    -0.017300      1.00800 ; qtot -
3.034
  3495   HC   216   GLU   HB3 3495    -0.017300      1.00800 ; qtot -
3.051
  3496  A2C   216   GLU    CG 3496     0.013600     12.01000 ; qtot -
3.037
  3497   HC   216   GLU   HG2 3497    -0.042500      1.00800 ; qtot -
3.080
  3498   HC   216   GLU   HG3 3498    -0.042500      1.00800 ; qtot -
3.122
  3499   CO   216   GLU    CD 3499     0.805401     12.01000 ; qtot -
2.317
  3500   O2   216   GLU   OE1 3500    -0.818801     16.00000 ; qtot -
3.136
  3501   O2   216   GLU   OE2 3501    -0.818801     16.00000 ; qtot -
3.955
  3502    C   216   GLU     C 3502     0.536601     12.01000 ; qtot -
3.418
  3503    O   216   GLU     O 3503    -0.581901     16.00000 ; qtot -
```

```
4.000
  3504    N   217   VAL      N 3504    -0.415700     14.01000 ; qtot -
4.416
  3505    H   217   VAL      H 3505     0.271900      1.00800 ; qtot -
4.144
  3506   CX   217   VAL     CA 3506    -0.087500     12.01000 ; qtot -
4.231
  3507   H1   217   VAL     HA 3507     0.096900      1.00800 ; qtot -
4.134
  3508  A3C   217   VAL     CB 3508     0.298500     12.01000 ; qtot -
3.836
  3509   HC   217   VAL     HB 3509    -0.029700      1.00800 ; qtot -
3.866
  3510   CT   217   VAL    CG1 3510    -0.319200     12.01000 ; qtot -
4.185
  3511   HC   217   VAL   HG11 3511     0.079100      1.00800 ; qtot -
4.106
  3512   HC   217   VAL   HG12 3512     0.079100      1.00800 ; qtot -
4.027
  3513   HC   217   VAL   HG13 3513     0.079100      1.00800 ; qtot -
3.947
  3514   CT   217   VAL    CG2 3514    -0.319200     12.01000 ; qtot -
4.267
  3515   HC   217   VAL   HG21 3515     0.079100      1.00800 ; qtot -
4.188
  3516   HC   217   VAL   HG22 3516     0.079100      1.00800 ; qtot -
4.108
  3517   HC   217   VAL   HG23 3517     0.079100      1.00800 ; qtot -
4.029
  3518    C   217   VAL      C 3518     0.597301     12.01000 ; qtot -
3.432
  3519    O   217   VAL      O 3519    -0.567901     16.00000 ; qtot -
4.000
  3520    N   218   LEU      N 3520    -0.415700     14.01000 ; qtot -
4.416
  3521    H   218   LEU      H 3521     0.271900      1.00800 ; qtot -
4.144
  3522   CX   218   LEU     CA 3522    -0.051800     12.01000 ; qtot -
4.196
  3523   H1   218   LEU     HA 3523     0.092200      1.00800 ; qtot -
4.103
  3524  A2C   218   LEU     CB 3524    -0.110200     12.01000 ; qtot -
4.214
  3525   HC   218   LEU    HB2 3525     0.045700      1.00800 ; qtot -
4.168
  3526   HC   218   LEU    HB3 3526     0.045700      1.00800 ; qtot -
4.122
  3527  A3C   218   LEU     CG 3527     0.353100     12.01000 ; qtot -
3.769
  3528   HC   218   LEU     HG 3528    -0.036100      1.00800 ; qtot -
3.805
  3529   CT   218   LEU    CD1 3529    -0.412100     12.01000 ; qtot -
```

```
4.217
  3530   HC   218   LEU  HD11 3530     0.100000      1.00800 ; qtot -
4.117
  3531   HC   218   LEU  HD12 3531     0.100000      1.00800 ; qtot -
4.017
  3532   HC   218   LEU  HD13 3532     0.100000      1.00800 ; qtot -
3.917
  3533   CT   218   LEU   CD2 3533    -0.412100     12.01000 ; qtot -
4.329
  3534   HC   218   LEU  HD21 3534     0.100000      1.00800 ; qtot -
4.229
  3535   HC   218   LEU  HD22 3535     0.100000      1.00800 ; qtot -
4.129
  3536   HC   218   LEU  HD23 3536     0.100000      1.00800 ; qtot -
4.029
  3537    C   218   LEU     C 3537     0.597301     12.01000 ; qtot -
3.432
  3538    O   218   LEU     O 3538    -0.567901     16.00000 ; qtot -
4.000
  3539    N   219   LYS     N 3539    -0.347900     14.01000 ; qtot -
4.348
  3540    H   219   LYS     H 3540     0.274700      1.00800 ; qtot -
4.073
  3541   CX   219   LYS    CA 3541    -0.240000     12.01000 ; qtot -
4.313
  3542   H1   219   LYS    HA 3542     0.142600      1.00800 ; qtot -
4.171
  3543   C8   219   LYS    CB 3543    -0.009400     12.01000 ; qtot -
4.180
  3544   HC   219   LYS   HB2 3544     0.036200      1.00800 ; qtot -
4.144
  3545   HC   219   LYS   HB3 3545     0.036200      1.00800 ; qtot -
4.108
  3546   C8   219   LYS    CG 3546     0.018700     12.01000 ; qtot -
4.089
  3547   HC   219   LYS   HG2 3547     0.010300      1.00800 ; qtot -
4.079
  3548   HC   219   LYS   HG3 3548     0.010300      1.00800 ; qtot -
4.068
  3549   C8   219   LYS    CD 3549    -0.047900     12.01000 ; qtot -
4.116
  3550   HC   219   LYS   HD2 3550     0.062100      1.00800 ; qtot -
4.054
  3551   HC   219   LYS   HD3 3551     0.062100      1.00800 ; qtot -
3.992
  3552   C8   219   LYS    CE 3552    -0.014300     12.01000 ; qtot -
4.006
  3553   HP   219   LYS   HE2 3553     0.113500      1.00800 ; qtot -
3.893
  3554   HP   219   LYS   HE3 3554     0.113500      1.00800 ; qtot -
3.779
  3555   N3   219   LYS    NZ 3555    -0.385400     14.01000 ; qtot -
```

4.165
  3556    H   219   LYS   HZ1 3556     0.340000      1.00800 ; qtot -
3.825
  3557    H   219   LYS   HZ2 3557     0.340000      1.00800 ; qtot -
3.485
  3558    H   219   LYS   HZ3 3558     0.340000      1.00800 ; qtot -
3.145
  3559    C   219   LYS     C 3559     0.734101     12.01000 ; qtot -
2.411
  3560    O   219   LYS     O 3560    -0.589401     16.00000 ; qtot -
3.000
  3561    N   220   LYS     N 3561    -0.347900     14.01000 ; qtot -
3.348
  3562    H   220   LYS     H 3562     0.274700      1.00800 ; qtot -
3.073
  3563   CX   220   LYS    CA 3563    -0.240000     12.01000 ; qtot -
3.313
  3564   H1   220   LYS    HA 3564     0.142600      1.00800 ; qtot -
3.171
  3565   C8   220   LYS    CB 3565    -0.009400     12.01000 ; qtot -
3.180
  3566   HC   220   LYS   HB2 3566     0.036200      1.00800 ; qtot -
3.144
  3567   HC   220   LYS   HB3 3567     0.036200      1.00800 ; qtot -
3.108
  3568   C8   220   LYS    CG 3568     0.018700     12.01000 ; qtot -
3.089
  3569   HC   220   LYS   HG2 3569     0.010300      1.00800 ; qtot -
3.079
  3570   HC   220   LYS   HG3 3570     0.010300      1.00800 ; qtot -
3.068
  3571   C8   220   LYS    CD 3571    -0.047900     12.01000 ; qtot -
3.116
  3572   HC   220   LYS   HD2 3572     0.062100      1.00800 ; qtot -
3.054
  3573   HC   220   LYS   HD3 3573     0.062100      1.00800 ; qtot -
2.992
  3574   C8   220   LYS    CE 3574    -0.014300     12.01000 ; qtot -
3.006
  3575   HP   220   LYS   HE2 3575     0.113500      1.00800 ; qtot -
2.893
  3576   HP   220   LYS   HE3 3576     0.113500      1.00800 ; qtot -
2.779
  3577   N3   220   LYS    NZ 3577    -0.385400     14.01000 ; qtot -
3.165
  3578    H   220   LYS   HZ1 3578     0.340000      1.00800 ; qtot -
2.825
  3579    H   220   LYS   HZ2 3579     0.340000      1.00800 ; qtot -
2.485
  3580    H   220   LYS   HZ3 3580     0.340000      1.00800 ; qtot -
2.145
  3581    C   220   LYS     C 3581     0.734101     12.01000 ; qtot -

```
1.411
  3582    O   220   LYS      O 3582    -0.589401     16.00000 ; qtot -
2.000
  3583    N   221   VAL      N 3583    -0.415700     14.01000 ; qtot -
2.416
  3584    H   221   VAL      H 3584     0.271900      1.00800 ; qtot -
2.144
  3585   CX   221   VAL     CA 3585    -0.087500     12.01000 ; qtot -
2.231
  3586   H1   221   VAL     HA 3586     0.096900      1.00800 ; qtot -
2.134
  3587  A3C   221   VAL     CB 3587     0.298500     12.01000 ; qtot -
1.836
  3588   HC   221   VAL     HB 3588    -0.029700      1.00800 ; qtot -
1.866
  3589   CT   221   VAL    CG1 3589    -0.319200     12.01000 ; qtot -
2.185
  3590   HC   221   VAL   HG11 3590     0.079100      1.00800 ; qtot -
2.106
  3591   HC   221   VAL   HG12 3591     0.079100      1.00800 ; qtot -
2.027
  3592   HC   221   VAL   HG13 3592     0.079100      1.00800 ; qtot -
1.947
  3593   CT   221   VAL    CG2 3593    -0.319200     12.01000 ; qtot -
2.267
  3594   HC   221   VAL   HG21 3594     0.079100      1.00800 ; qtot -
2.188
  3595   HC   221   VAL   HG22 3595     0.079100      1.00800 ; qtot -
2.108
  3596   HC   221   VAL   HG23 3596     0.079100      1.00800 ; qtot -
2.029
  3597    C   221   VAL      C 3597     0.597301     12.01000 ; qtot -
1.432
  3598    O   221   VAL      O 3598    -0.567901     16.00000 ; qtot -
2.000
  3599    N   222   LYS      N 3599    -0.347900     14.01000 ; qtot -
2.348
  3600    H   222   LYS      H 3600     0.274700      1.00800 ; qtot -
2.073
  3601   CX   222   LYS     CA 3601    -0.240000     12.01000 ; qtot -
2.313
  3602   H1   222   LYS     HA 3602     0.142600      1.00800 ; qtot -
2.171
  3603   C8   222   LYS     CB 3603    -0.009400     12.01000 ; qtot -
2.180
  3604   HC   222   LYS    HB2 3604     0.036200      1.00800 ; qtot -
2.144
  3605   HC   222   LYS    HB3 3605     0.036200      1.00800 ; qtot -
2.108
  3606   C8   222   LYS     CG 3606     0.018700     12.01000 ; qtot -
2.089
  3607   HC   222   LYS    HG2 3607     0.010300      1.00800 ; qtot -
```

2.079
  3608   HC   222   LYS   HG3 3608     0.010300      1.00800 ; qtot -
2.068
  3609   C8   222   LYS    CD 3609    -0.047900     12.01000 ; qtot -
2.116
  3610   HC   222   LYS   HD2 3610     0.062100      1.00800 ; qtot -
2.054
  3611   HC   222   LYS   HD3 3611     0.062100      1.00800 ; qtot -
1.992
  3612   C8   222   LYS    CE 3612    -0.014300     12.01000 ; qtot -
2.006
  3613   HP   222   LYS   HE2 3613     0.113500      1.00800 ; qtot -
1.893
  3614   HP   222   LYS   HE3 3614     0.113500      1.00800 ; qtot -
1.779
  3615   N3   222   LYS    NZ 3615    -0.385400     14.01000 ; qtot -
2.165
  3616    H   222   LYS   HZ1 3616     0.340000      1.00800 ; qtot -
1.825
  3617    H   222   LYS   HZ2 3617     0.340000      1.00800 ; qtot -
1.485
  3618    H   222   LYS   HZ3 3618     0.340000      1.00800 ; qtot -
1.145
  3619    C   222   LYS     C 3619     0.734101     12.01000 ; qtot -
0.411
  3620    O   222   LYS     O 3620    -0.589401     16.00000 ; qtot -
1.000
  3621    N   223   LYS     N 3621    -0.347900     14.01000 ; qtot -
1.348
  3622    H   223   LYS     H 3622     0.274700      1.00800 ; qtot -
1.073
  3623   CX   223   LYS    CA 3623    -0.240000     12.01000 ; qtot -
1.313
  3624   H1   223   LYS    HA 3624     0.142600      1.00800 ; qtot -
1.171
  3625   C8   223   LYS    CB 3625    -0.009400     12.01000 ; qtot -
1.180
  3626   HC   223   LYS   HB2 3626     0.036200      1.00800 ; qtot -
1.144
  3627   HC   223   LYS   HB3 3627     0.036200      1.00800 ; qtot -
1.108
  3628   C8   223   LYS    CG 3628     0.018700     12.01000 ; qtot -
1.089
  3629   HC   223   LYS   HG2 3629     0.010300      1.00800 ; qtot -
1.079
  3630   HC   223   LYS   HG3 3630     0.010300      1.00800 ; qtot -
1.068
  3631   C8   223   LYS    CD 3631    -0.047900     12.01000 ; qtot -
1.116
  3632   HC   223   LYS   HD2 3632     0.062100      1.00800 ; qtot -
1.054
  3633   HC   223   LYS   HD3 3633     0.062100      1.00800 ; qtot -

0.992
3634 C8 223 LYS CE 3634 -0.014300 12.01000 ; qtot -
1.006
3635 HP 223 LYS HE2 3635 0.113500 1.00800 ; qtot -
0.893
3636 HP 223 LYS HE3 3636 0.113500 1.00800 ; qtot -
0.779
3637 N3 223 LYS NZ 3637 -0.385400 14.01000 ; qtot -
1.165
3638 H 223 LYS HZ1 3638 0.340000 1.00800 ; qtot -
0.825
3639 H 223 LYS HZ2 3639 0.340000 1.00800 ; qtot -
0.485
3640 H 223 LYS HZ3 3640 0.340000 1.00800 ; qtot -
0.145
3641 C 223 LYS C 3641 0.734101 12.01000 ; qtot
0.589
3642 O 223 LYS O 3642 -0.589401 16.00000 ; qtot
0.000
3643 N 224 GLY N 3643 -0.415700 14.01000 ; qtot -
0.416
3644 H 224 GLY H 3644 0.271900 1.00800 ; qtot -
0.144
3645 CX 224 GLY CA 3645 -0.025200 12.01000 ; qtot -
0.169
3646 H1 224 GLY HA2 3646 0.069800 1.00800 ; qtot -
0.099
3647 H1 224 GLY HA3 3647 0.069800 1.00800 ; qtot -
0.029
3648 C 224 GLY C 3648 0.597301 12.01000 ; qtot
0.568
3649 O 224 GLY O 3649 -0.567901 16.00000 ; qtot
0.000
3650 N 225 GLU N 3650 -0.516301 14.01000 ; qtot -
0.516
3651 H 225 GLU H 3651 0.293600 1.00800 ; qtot -
0.223
3652 CX 225 GLU CA 3652 0.039700 12.01000 ; qtot -
0.183
3653 H1 225 GLU HA 3653 0.110500 1.00800 ; qtot -
0.072
3654 A2C 225 GLU CB 3654 0.056000 12.01000 ; qtot -
0.016
3655 HC 225 GLU HB2 3655 -0.017300 1.00800 ; qtot -
0.034
3656 HC 225 GLU HB3 3656 -0.017300 1.00800 ; qtot -
0.051
3657 A2C 225 GLU CG 3657 0.013600 12.01000 ; qtot -
0.037
3658 HC 225 GLU HG2 3658 -0.042500 1.00800 ; qtot -
0.080
3659 HC 225 GLU HG3 3659 -0.042500 1.00800 ; qtot -

0.122
  3660   CO   225   GLU    CD 3660     0.805401     12.01000 ; qtot
0.683
  3661   O2   225   GLU   OE1 3661    -0.818801     16.00000 ; qtot -
0.136
  3662   O2   225   GLU   OE2 3662    -0.818801     16.00000 ; qtot -
0.955
  3663    C   225   GLU     C 3663     0.536601     12.01000 ; qtot -
0.418
  3664    O   225   GLU     O 3664    -0.581901     16.00000 ; qtot -
1.000
  3665    N   226   PHE     N 3665    -0.415700     14.01000 ; qtot -
1.416
  3666    H   226   PHE     H 3666     0.271900      1.00800 ; qtot -
1.144
  3667   CX   226   PHE    CA 3667    -0.002400     12.01000 ; qtot -
1.146
  3668   H1   226   PHE    HA 3668     0.097800      1.00800 ; qtot -
1.048
  3669   CT   226   PHE    CB 3669    -0.034300     12.01000 ; qtot -
1.083
  3670   HC   226   PHE   HB2 3670     0.029500      1.00800 ; qtot -
1.053
  3671   HC   226   PHE   HB3 3671     0.029500      1.00800 ; qtot -
1.024
  3672   CA   226   PHE    CG 3672     0.011800     12.01000 ; qtot -
1.012
  3673   CA   226   PHE   CD1 3673    -0.125600     12.01000 ; qtot -
1.137
  3674   HA   226   PHE   HD1 3674     0.133000      1.00800 ; qtot -
1.004
  3675   CA   226   PHE   CE1 3675    -0.170400     12.01000 ; qtot -
1.175
  3676   HA   226   PHE   HE1 3676     0.143000      1.00800 ; qtot -
1.032
  3677   CA   226   PHE    CZ 3677    -0.107200     12.01000 ; qtot -
1.139
  3678   HA   226   PHE    HZ 3678     0.129700      1.00800 ; qtot -
1.009
  3679   CA   226   PHE   CE2 3679    -0.170400     12.01000 ; qtot -
1.180
  3680   HA   226   PHE   HE2 3680     0.143000      1.00800 ; qtot -
1.037
  3681   CA   226   PHE   CD2 3681    -0.125600     12.01000 ; qtot -
1.162
  3682   HA   226   PHE   HD2 3682     0.133000      1.00800 ; qtot -
1.029
  3683    C   226   PHE     C 3683     0.597301     12.01000 ; qtot -
0.432
  3684    O   226   PHE     O 3684    -0.567901     16.00000 ; qtot -
1.000
  3685    N   227   CYS     N 3685    -0.415700     14.01000 ; qtot -

1.416
  3686    H   227   CYS     H 3686     0.271900      1.00800 ; qtot -
1.144
  3687   CX   227   CYS    CA 3687     0.021300     12.01000 ; qtot -
1.122
  3688   H1   227   CYS    HA 3688     0.112400      1.00800 ; qtot -
1.010
  3689  A2C   227   CYS    CB 3689    -0.123100     12.01000 ; qtot -
1.133
  3690   H1   227   CYS   HB2 3690     0.111200      1.00800 ; qtot -
1.022
  3691   H1   227   CYS   HB3 3691     0.111200      1.00800 ; qtot -
0.911
  3692   SH   227   CYS    SG 3692    -0.311900     32.06000 ; qtot -
1.223
  3693   HS   227   CYS    HG 3693     0.193300      1.00800 ; qtot -
1.029
  3694    C   227   CYS     C 3694     0.597301     12.01000 ; qtot -
0.432
  3695    O   227   CYS     O 3695    -0.567901     16.00000 ; qtot -
1.000
  3696    N   228   PHE     N 3696    -0.415700     14.01000 ; qtot -
1.416
  3697    H   228   PHE     H 3697     0.271900      1.00800 ; qtot -
1.144
  3698   CX   228   PHE    CA 3698    -0.002400     12.01000 ; qtot -
1.146
  3699   H1   228   PHE    HA 3699     0.097800      1.00800 ; qtot -
1.048
  3700   CT   228   PHE    CB 3700    -0.034300     12.01000 ; qtot -
1.083
  3701   HC   228   PHE   HB2 3701     0.029500      1.00800 ; qtot -
1.053
  3702   HC   228   PHE   HB3 3702     0.029500      1.00800 ; qtot -
1.024
  3703   CA   228   PHE    CG 3703     0.011800     12.01000 ; qtot -
1.012
  3704   CA   228   PHE   CD1 3704    -0.125600     12.01000 ; qtot -
1.137
  3705   HA   228   PHE   HD1 3705     0.133000      1.00800 ; qtot -
1.004
  3706   CA   228   PHE   CE1 3706    -0.170400     12.01000 ; qtot -
1.175
  3707   HA   228   PHE   HE1 3707     0.143000      1.00800 ; qtot -
1.032
  3708   CA   228   PHE    CZ 3708    -0.107200     12.01000 ; qtot -
1.139
  3709   HA   228   PHE    HZ 3709     0.129700      1.00800 ; qtot -
1.009
  3710   CA   228   PHE   CE2 3710    -0.170400     12.01000 ; qtot -
1.180
  3711   HA   228   PHE   HE2 3711     0.143000      1.00800 ; qtot -

1.037
3712 CA 228 PHE CD2 3712 -0.125600 12.01000 ; qtot -
1.162
3713 HA 228 PHE HD2 3713 0.133000 1.00800 ; qtot -
1.029
3714 C 228 PHE C 3714 0.597301 12.01000 ; qtot -
0.432
3715 O 228 PHE O 3715 -0.567901 16.00000 ; qtot -
1.000
3716 N 229 TYR N 3716 -0.415700 14.01000 ; qtot -
1.416
3717 H 229 TYR H 3717 0.271900 1.00800 ; qtot -
1.144
3718 CX 229 TYR CA 3718 -0.001400 12.01000 ; qtot -
1.145
3719 H1 229 TYR HA 3719 0.087600 1.00800 ; qtot -
1.058
3720 CT 229 TYR CB 3720 -0.015200 12.01000 ; qtot -
1.073
3721 HC 229 TYR HB2 3721 0.029500 1.00800 ; qtot -
1.043
3722 HC 229 TYR HB3 3722 0.029500 1.00800 ; qtot -
1.014
3723 CA 229 TYR CG 3723 -0.001100 12.01000 ; qtot -
1.015
3724 CA 229 TYR CD1 3724 -0.190600 12.01000 ; qtot -
1.205
3725 HA 229 TYR HD1 3725 0.169900 1.00800 ; qtot -
1.036
3726 CA 229 TYR CE1 3726 -0.234100 12.01000 ; qtot -
1.270
3727 HA 229 TYR HE1 3727 0.165600 1.00800 ; qtot -
1.104
3728 C 229 TYR CZ 3728 0.322600 12.01000 ; qtot -
0.781
3729 OH 229 TYR OH 3729 -0.557901 16.00000 ; qtot -
1.339
3730 HO 229 TYR HH 3730 0.399200 1.00800 ; qtot -
0.940
3731 CA 229 TYR CE2 3731 -0.234100 12.01000 ; qtot -
1.174
3732 HA 229 TYR HE2 3732 0.165600 1.00800 ; qtot -
1.009
3733 CA 229 TYR CD2 3733 -0.190600 12.01000 ; qtot -
1.199
3734 HA 229 TYR HD2 3734 0.169900 1.00800 ; qtot -
1.029
3735 C 229 TYR C 3735 0.597301 12.01000 ; qtot -
0.432
3736 O 229 TYR O 3736 -0.567901 16.00000 ; qtot -
1.000
3737 N 230 GLU N 3737 -0.516301 14.01000 ; qtot -

1.516
  3738    H   230   GLU     H 3738     0.293600     1.00800 ; qtot -
1.223
  3739   CX   230   GLU    CA 3739     0.039700    12.01000 ; qtot -
1.183
  3740   H1   230   GLU    HA 3740     0.110500     1.00800 ; qtot -
1.072
  3741  A2C   230   GLU    CB 3741     0.056000    12.01000 ; qtot -
1.016
  3742   HC   230   GLU   HB2 3742    -0.017300     1.00800 ; qtot -
1.034
  3743   HC   230   GLU   HB3 3743    -0.017300     1.00800 ; qtot -
1.051
  3744  A2C   230   GLU    CG 3744     0.013600    12.01000 ; qtot -
1.037
  3745   HC   230   GLU   HG2 3745    -0.042500     1.00800 ; qtot -
1.080
  3746   HC   230   GLU   HG3 3746    -0.042500     1.00800 ; qtot -
1.122
  3747   CO   230   GLU    CD 3747     0.805401    12.01000 ; qtot -
0.317
  3748   O2   230   GLU   OE1 3748    -0.818801    16.00000 ; qtot -
1.136
  3749   O2   230   GLU   OE2 3749    -0.818801    16.00000 ; qtot -
1.955
  3750    C   230   GLU     C 3750     0.536601    12.01000 ; qtot -
1.418
  3751    O   230   GLU     O 3751    -0.581901    16.00000 ; qtot -
2.000
  3752    N   231   ASN     N 3752    -0.415700    14.01000 ; qtot -
2.416
  3753    H   231   ASN     H 3753     0.271900     1.00800 ; qtot -
2.144
  3754   CX   231   ASN    CA 3754     0.014300    12.01000 ; qtot -
2.129
  3755   H1   231   ASN    HA 3755     0.104800     1.00800 ; qtot -
2.025
  3756  A2C   231   ASN    CB 3756    -0.204100    12.01000 ; qtot -
2.229
  3757   HC   231   ASN   HB2 3757     0.079700     1.00800 ; qtot -
2.149
  3758   HC   231   ASN   HB3 3758     0.079700     1.00800 ; qtot -
2.069
  3759    C   231   ASN    CG 3759     0.713001    12.01000 ; qtot -
1.356
  3760    O   231   ASN   OD1 3760    -0.593101    16.00000 ; qtot -
1.949
  3761    N   231   ASN   ND2 3761    -0.919101    14.01000 ; qtot -
2.869
  3762    H   231   ASN  HD21 3762     0.419600     1.00800 ; qtot -
2.449
  3763    H   231   ASN  HD22 3763     0.419600     1.00800 ; qtot -

2.029
3764 C 231 ASN C 3764 0.597301 12.01000 ; qtot -
1.432
3765 O 231 ASN O 3765 -0.567901 16.00000 ; qtot -
2.000
3766 N 232 ASP N 3766 -0.516301 14.01000 ; qtot -
2.516
3767 H 232 ASP H 3767 0.293600 1.00800 ; qtot -
2.223
3768 CX 232 ASP CA 3768 0.038100 12.01000 ; qtot -
2.185
3769 H1 232 ASP HA 3769 0.088000 1.00800 ; qtot -
2.097
3770 A2C 232 ASP CB 3770 -0.030300 12.01000 ; qtot -
2.127
3771 HC 232 ASP HB2 3771 -0.012200 1.00800 ; qtot -
2.139
3772 HC 232 ASP HB3 3772 -0.012200 1.00800 ; qtot -
2.151
3773 CO 232 ASP CG 3773 0.799401 12.01000 ; qtot -
1.352
3774 O2 232 ASP OD1 3774 -0.801401 16.00000 ; qtot -
2.153
3775 O2 232 ASP OD2 3775 -0.801401 16.00000 ; qtot -
2.955
3776 C 232 ASP C 3776 0.536601 12.01000 ; qtot -
2.418
3777 O 232 ASP O 3777 -0.581901 16.00000 ; qtot -
3.000
3778 N 233 TRP N 3778 -0.415700 14.01000 ; qtot -
3.416
3779 H 233 TRP H 3779 0.271900 1.00800 ; qtot -
3.144
3780 CX 233 TRP CA 3780 -0.027500 12.01000 ; qtot -
3.171
3781 H1 233 TRP HA 3781 0.112300 1.00800 ; qtot -
3.059
3782 CT 233 TRP CB 3782 -0.005000 12.01000 ; qtot -
3.064
3783 HC 233 TRP HB2 3783 0.033900 1.00800 ; qtot -
3.030
3784 HC 233 TRP HB3 3784 0.033900 1.00800 ; qtot -
2.996
3785 C* 233 TRP CG 3785 -0.141500 12.01000 ; qtot -
3.138
3786 CW 233 TRP CD1 3786 -0.163800 12.01000 ; qtot -
3.301
3787 H4 233 TRP HD1 3787 0.206200 1.00800 ; qtot -
3.095
3788 NA 233 TRP NE1 3788 -0.341800 14.01000 ; qtot -
3.437
3789 H 233 TRP HE1 3789 0.341200 1.00800 ; qtot -

3.096
  3790   CN   233   TRP   CE2 3790     0.138000     12.01000 ; qtot -
2.958
  3791   CA   233   TRP   CZ2 3791    -0.260100     12.01000 ; qtot -
3.218
  3792   HA   233   TRP   HZ2 3792     0.157200      1.00800 ; qtot -
3.061
  3793   CA   233   TRP   CH2 3793    -0.113400     12.01000 ; qtot -
3.174
  3794   HA   233   TRP   HH2 3794     0.141700      1.00800 ; qtot -
3.032
  3795   CA   233   TRP   CZ3 3795    -0.197200     12.01000 ; qtot -
3.230
  3796   HA   233   TRP   HZ3 3796     0.144700      1.00800 ; qtot -
3.085
  3797   CA   233   TRP   CE3 3797    -0.238700     12.01000 ; qtot -
3.324
  3798   HA   233   TRP   HE3 3798     0.170000      1.00800 ; qtot -
3.154
  3799   CB   233   TRP   CD2 3799     0.124300     12.01000 ; qtot -
3.029
  3800    C   233   TRP     C 3800     0.597301     12.01000 ; qtot -
2.432
  3801    O   233   TRP     O 3801    -0.567901     16.00000 ; qtot -
3.000
  3802    N   234   GLY     N 3802    -0.415700     14.01000 ; qtot -
3.416
  3803    H   234   GLY     H 3803     0.271900      1.00800 ; qtot -
3.144
  3804   CX   234   GLY    CA 3804    -0.025200     12.01000 ; qtot -
3.169
  3805   H1   234   GLY   HA2 3805     0.069800      1.00800 ; qtot -
3.099
  3806   H1   234   GLY   HA3 3806     0.069800      1.00800 ; qtot -
3.029
  3807    C   234   GLY     C 3807     0.597301     12.01000 ; qtot -
2.432
  3808    O   234   GLY     O 3808    -0.567901     16.00000 ; qtot -
3.000
  3809    N   235   SER     N 3809    -0.415700     14.01000 ; qtot -
3.416
  3810    H   235   SER     H 3810     0.271900      1.00800 ; qtot -
3.144
  3811   CX   235   SER    CA 3811    -0.024900     12.01000 ; qtot -
3.169
  3812   H1   235   SER    HA 3812     0.084300      1.00800 ; qtot -
3.084
  3813  A2C   235   SER    CB 3813     0.211700     12.01000 ; qtot -
2.873
  3814   H1   235   SER   HB2 3814     0.035200      1.00800 ; qtot -
2.837
  3815   H1   235   SER   HB3 3815     0.035200      1.00800 ; qtot -

```
2.802
  3816   OH   235   SER    OG 3816    -0.654601     16.00000 ; qtot -
3.457
  3817   HO   235   SER    HG 3817     0.427500      1.00800 ; qtot -
3.029
  3818    C   235   SER     C 3818     0.597301     12.01000 ; qtot -
2.432
  3819    O   235   SER     O 3819    -0.567901     16.00000 ; qtot -
3.000
  3820    N   236   ILE     N 3820    -0.415700     14.01000 ; qtot -
3.416
  3821    H   236   ILE     H 3821     0.271900      1.00800 ; qtot -
3.144
  3822   CX   236   ILE    CA 3822    -0.059700     12.01000 ; qtot -
3.203
  3823   H1   236   ILE    HA 3823     0.086900      1.00800 ; qtot -
3.117
  3824  A3C   236   ILE    CB 3824     0.130300     12.01000 ; qtot -
2.986
  3825   HC   236   ILE    HB 3825     0.018700      1.00800 ; qtot -
2.968
  3826   CT   236   ILE   CG2 3826    -0.320400     12.01000 ; qtot -
3.288
  3827   HC   236   ILE  HG21 3827     0.088200      1.00800 ; qtot -
3.200
  3828   HC   236   ILE  HG22 3828     0.088200      1.00800 ; qtot -
3.112
  3829   HC   236   ILE  HG23 3829     0.088200      1.00800 ; qtot -
3.023
  3830  A2C   236   ILE   CG1 3830    -0.043000     12.01000 ; qtot -
3.066
  3831   HC   236   ILE  HG12 3831     0.023600      1.00800 ; qtot -
3.043
  3832   HC   236   ILE  HG13 3832     0.023600      1.00800 ; qtot -
3.019
  3833   CT   236   ILE   CD1 3833    -0.066000     12.01000 ; qtot -
3.085
  3834   HC   236   ILE  HD11 3834     0.018600      1.00800 ; qtot -
3.067
  3835   HC   236   ILE  HD12 3835     0.018600      1.00800 ; qtot -
3.048
  3836   HC   236   ILE  HD13 3836     0.018600      1.00800 ; qtot -
3.029
  3837    C   236   ILE     C 3837     0.597301     12.01000 ; qtot -
2.432
  3838    O   236   ILE     O 3838    -0.567901     16.00000 ; qtot -
3.000
  3839    N   237   SER     N 3839    -0.415700     14.01000 ; qtot -
3.416
  3840    H   237   SER     H 3840     0.271900      1.00800 ; qtot -
3.144
  3841   CX   237   SER    CA 3841    -0.024900     12.01000 ; qtot -
```

3.169
 3842 H1 237 SER HA 3842 0.084300 1.00800 ; qtot -
3.084
 3843 A2C 237 SER CB 3843 0.211700 12.01000 ; qtot -
2.873
 3844 H1 237 SER HB2 3844 0.035200 1.00800 ; qtot -
2.837
 3845 H1 237 SER HB3 3845 0.035200 1.00800 ; qtot -
2.802
 3846 OH 237 SER OG 3846 -0.654601 16.00000 ; qtot -
3.457
 3847 HO 237 SER HG 3847 0.427500 1.00800 ; qtot -
3.029
 3848 C 237 SER C 3848 0.597301 12.01000 ; qtot -
2.432
 3849 O 237 SER O 3849 -0.567901 16.00000 ; qtot -
3.000
 3850 N 238 SER N 3850 -0.415700 14.01000 ; qtot -
3.416
 3851 H 238 SER H 3851 0.271900 1.00800 ; qtot -
3.144
 3852 CX 238 SER CA 3852 -0.024900 12.01000 ; qtot -
3.169
 3853 H1 238 SER HA 3853 0.084300 1.00800 ; qtot -
3.084
 3854 A2C 238 SER CB 3854 0.211700 12.01000 ; qtot -
2.873
 3855 H1 238 SER HB2 3855 0.035200 1.00800 ; qtot -
2.837
 3856 H1 238 SER HB3 3856 0.035200 1.00800 ; qtot -
2.802
 3857 OH 238 SER OG 3857 -0.654601 16.00000 ; qtot -
3.457
 3858 HO 238 SER HG 3858 0.427500 1.00800 ; qtot -
3.029
 3859 C 238 SER C 3859 0.597301 12.01000 ; qtot -
2.432
 3860 O 238 SER O 3860 -0.567901 16.00000 ; qtot -
3.000
 3861 N 239 ASP N 3861 -0.516301 14.01000 ; qtot -
3.516
 3862 H 239 ASP H 3862 0.293600 1.00800 ; qtot -
3.223
 3863 CX 239 ASP CA 3863 0.038100 12.01000 ; qtot -
3.185
 3864 H1 239 ASP HA 3864 0.088000 1.00800 ; qtot -
3.097
 3865 A2C 239 ASP CB 3865 -0.030300 12.01000 ; qtot -
3.127
 3866 HC 239 ASP HB2 3866 -0.012200 1.00800 ; qtot -
3.139
 3867 HC 239 ASP HB3 3867 -0.012200 1.00800 ; qtot -

3.151
  3868   CO   239   ASP    CG 3868     0.799401     12.01000 ; qtot -
2.352
  3869   O2   239   ASP   OD1 3869    -0.801401     16.00000 ; qtot -
3.153
  3870   O2   239   ASP   OD2 3870    -0.801401     16.00000 ; qtot -
3.955
  3871    C   239   ASP     C 3871     0.536601     12.01000 ; qtot -
3.418
  3872    O   239   ASP     O 3872    -0.581901     16.00000 ; qtot -
4.000
  3873    N   240   ALA     N 3873    -0.415700     14.01000 ; qtot -
4.416
  3874    H   240   ALA     H 3874     0.271900      1.00800 ; qtot -
4.144
  3875   CX   240   ALA    CA 3875     0.033700     12.01000 ; qtot -
4.110
  3876   H1   240   ALA    HA 3876     0.082300      1.00800 ; qtot -
4.028
  3877   CT   240   ALA    CB 3877    -0.182500     12.01000 ; qtot -
4.210
  3878   HC   240   ALA   HB1 3878     0.060300      1.00800 ; qtot -
4.150
  3879   HC   240   ALA   HB2 3879     0.060300      1.00800 ; qtot -
4.090
  3880   HC   240   ALA   HB3 3880     0.060300      1.00800 ; qtot -
4.029
  3881    C   240   ALA     C 3881     0.597301     12.01000 ; qtot -
3.432
  3882    O   240   ALA     O 3882    -0.567901     16.00000 ; qtot -
4.000
  3883    N   241   LYS     N 3883    -0.347900     14.01000 ; qtot -
4.348
  3884    H   241   LYS     H 3884     0.274700      1.00800 ; qtot -
4.073
  3885   CX   241   LYS    CA 3885    -0.240000     12.01000 ; qtot -
4.313
  3886   H1   241   LYS    HA 3886     0.142600      1.00800 ; qtot -
4.171
  3887   C8   241   LYS    CB 3887    -0.009400     12.01000 ; qtot -
4.180
  3888   HC   241   LYS   HB2 3888     0.036200      1.00800 ; qtot -
4.144
  3889   HC   241   LYS   HB3 3889     0.036200      1.00800 ; qtot -
4.108
  3890   C8   241   LYS    CG 3890     0.018700     12.01000 ; qtot -
4.089
  3891   HC   241   LYS   HG2 3891     0.010300      1.00800 ; qtot -
4.079
  3892   HC   241   LYS   HG3 3892     0.010300      1.00800 ; qtot -
4.068
  3893   C8   241   LYS    CD 3893    -0.047900     12.01000 ; qtot -

4.116
3894 HC 241 LYS HD2 3894 0.062100 1.00800 ; qtot -
4.054
3895 HC 241 LYS HD3 3895 0.062100 1.00800 ; qtot -
3.992
3896 C8 241 LYS CE 3896 -0.014300 12.01000 ; qtot -
4.006
3897 HP 241 LYS HE2 3897 0.113500 1.00800 ; qtot -
3.893
3898 HP 241 LYS HE3 3898 0.113500 1.00800 ; qtot -
3.779
3899 N3 241 LYS NZ 3899 -0.385400 14.01000 ; qtot -
4.165
3900 H 241 LYS HZ1 3900 0.340000 1.00800 ; qtot -
3.825
3901 H 241 LYS HZ2 3901 0.340000 1.00800 ; qtot -
3.485
3902 H 241 LYS HZ3 3902 0.340000 1.00800 ; qtot -
3.145
3903 C 241 LYS C 3903 0.734101 12.01000 ; qtot -
2.411
3904 O 241 LYS O 3904 -0.589401 16.00000 ; qtot -
3.000
3905 N 242 ASN N 3905 -0.415700 14.01000 ; qtot -
3.416
3906 H 242 ASN H 3906 0.271900 1.00800 ; qtot -
3.144
3907 CX 242 ASN CA 3907 0.014300 12.01000 ; qtot -
3.129
3908 H1 242 ASN HA 3908 0.104800 1.00800 ; qtot -
3.025
3909 A2C 242 ASN CB 3909 -0.204100 12.01000 ; qtot -
3.229
3910 HC 242 ASN HB2 3910 0.079700 1.00800 ; qtot -
3.149
3911 HC 242 ASN HB3 3911 0.079700 1.00800 ; qtot -
3.069
3912 C 242 ASN CG 3912 0.713001 12.01000 ; qtot -
2.356
3913 O 242 ASN OD1 3913 -0.593101 16.00000 ; qtot -
2.949
3914 N 242 ASN ND2 3914 -0.919101 14.01000 ; qtot -
3.869
3915 H 242 ASN HD21 3915 0.419600 1.00800 ; qtot -
3.449
3916 H 242 ASN HD22 3916 0.419600 1.00800 ; qtot -
3.029
3917 C 242 ASN C 3917 0.597301 12.01000 ; qtot -
2.432
3918 O 242 ASN O 3918 -0.567901 16.00000 ; qtot -
3.000
3919 N 243 LEU N 3919 -0.415700 14.01000 ; qtot -

3.416
  3920    H   243   LEU      H 3920     0.271900      1.00800 ; qtot -
3.144
  3921   CX   243   LEU     CA 3921    -0.051800     12.01000 ; qtot -
3.196
  3922   H1   243   LEU     HA 3922     0.092200      1.00800 ; qtot -
3.103
  3923  A2C   243   LEU     CB 3923    -0.110200     12.01000 ; qtot -
3.214
  3924   HC   243   LEU    HB2 3924     0.045700      1.00800 ; qtot -
3.168
  3925   HC   243   LEU    HB3 3925     0.045700      1.00800 ; qtot -
3.122
  3926  A3C   243   LEU     CG 3926     0.353100     12.01000 ; qtot -
2.769
  3927   HC   243   LEU     HG 3927    -0.036100      1.00800 ; qtot -
2.805
  3928   CT   243   LEU    CD1 3928    -0.412100     12.01000 ; qtot -
3.217
  3929   HC   243   LEU   HD11 3929     0.100000      1.00800 ; qtot -
3.117
  3930   HC   243   LEU   HD12 3930     0.100000      1.00800 ; qtot -
3.017
  3931   HC   243   LEU   HD13 3931     0.100000      1.00800 ; qtot -
2.917
  3932   CT   243   LEU    CD2 3932    -0.412100     12.01000 ; qtot -
3.329
  3933   HC   243   LEU   HD21 3933     0.100000      1.00800 ; qtot -
3.229
  3934   HC   243   LEU   HD22 3934     0.100000      1.00800 ; qtot -
3.129
  3935   HC   243   LEU   HD23 3935     0.100000      1.00800 ; qtot -
3.029
  3936    C   243   LEU      C 3936     0.597301     12.01000 ; qtot -
2.432
  3937    O   243   LEU      O 3937    -0.567901     16.00000 ; qtot -
3.000
  3938    N   244   ILE      N 3938    -0.415700     14.01000 ; qtot -
3.416
  3939    H   244   ILE      H 3939     0.271900      1.00800 ; qtot -
3.144
  3940   CX   244   ILE     CA 3940    -0.059700     12.01000 ; qtot -
3.203
  3941   H1   244   ILE     HA 3941     0.086900      1.00800 ; qtot -
3.117
  3942  A3C   244   ILE     CB 3942     0.130300     12.01000 ; qtot -
2.986
  3943   HC   244   ILE     HB 3943     0.018700      1.00800 ; qtot -
2.968
  3944   CT   244   ILE    CG2 3944    -0.320400     12.01000 ; qtot -
3.288
  3945   HC   244   ILE   HG21 3945     0.088200      1.00800 ; qtot -

3.200
  3946   HC   244   ILE  HG22 3946     0.088200      1.00800 ; qtot -
3.112
  3947   HC   244   ILE  HG23 3947     0.088200      1.00800 ; qtot -
3.023
  3948  A2C   244   ILE   CG1 3948    -0.043000     12.01000 ; qtot -
3.066
  3949   HC   244   ILE  HG12 3949     0.023600      1.00800 ; qtot -
3.043
  3950   HC   244   ILE  HG13 3950     0.023600      1.00800 ; qtot -
3.019
  3951   CT   244   ILE   CD1 3951    -0.066000     12.01000 ; qtot -
3.085
  3952   HC   244   ILE  HD11 3952     0.018600      1.00800 ; qtot -
3.067
  3953   HC   244   ILE  HD12 3953     0.018600      1.00800 ; qtot -
3.048
  3954   HC   244   ILE  HD13 3954     0.018600      1.00800 ; qtot -
3.029
  3955    C   244   ILE     C 3955     0.597301     12.01000 ; qtot -
2.432
  3956    O   244   ILE     O 3956    -0.567901     16.00000 ; qtot -
3.000
  3957    N   245   THR     N 3957    -0.415700     14.01000 ; qtot -
3.416
  3958    H   245   THR     H 3958     0.271900      1.00800 ; qtot -
3.144
  3959   CX   245   THR    CA 3959    -0.038900     12.01000 ; qtot -
3.183
  3960   H1   245   THR    HA 3960     0.100700      1.00800 ; qtot -
3.082
  3961  A3C   245   THR    CB 3961     0.365400     12.01000 ; qtot -
2.717
  3962   H1   245   THR    HB 3962     0.004300      1.00800 ; qtot -
2.712
  3963   CT   245   THR   CG2 3963    -0.243800     12.01000 ; qtot -
2.956
  3964   HC   245   THR  HG21 3964     0.064200      1.00800 ; qtot -
2.892
  3965   HC   245   THR  HG22 3965     0.064200      1.00800 ; qtot -
2.828
  3966   HC   245   THR  HG23 3966     0.064200      1.00800 ; qtot -
2.763
  3967   OH   245   THR   OG1 3967    -0.676101     16.00000 ; qtot -
3.440
  3968   HO   245   THR   HG1 3968     0.410200      1.00800 ; qtot -
3.029
  3969    C   245   THR     C 3969     0.597301     12.01000 ; qtot -
2.432
  3970    O   245   THR     O 3970    -0.567901     16.00000 ; qtot -
3.000
  3971    N   246   LYS     N 3971    -0.347900     14.01000 ; qtot -

3.348
  3972    H   246   LYS      H 3972      0.274700       1.00800 ; qtot -
3.073
  3973   CX   246   LYS     CA 3973     -0.240000      12.01000 ; qtot -
3.313
  3974   H1   246   LYS     HA 3974      0.142600       1.00800 ; qtot -
3.171
  3975   C8   246   LYS     CB 3975     -0.009400      12.01000 ; qtot -
3.180
  3976   HC   246   LYS    HB2 3976      0.036200       1.00800 ; qtot -
3.144
  3977   HC   246   LYS    HB3 3977      0.036200       1.00800 ; qtot -
3.108
  3978   C8   246   LYS     CG 3978      0.018700      12.01000 ; qtot -
3.089
  3979   HC   246   LYS    HG2 3979      0.010300       1.00800 ; qtot -
3.079
  3980   HC   246   LYS    HG3 3980      0.010300       1.00800 ; qtot -
3.068
  3981   C8   246   LYS     CD 3981     -0.047900      12.01000 ; qtot -
3.116
  3982   HC   246   LYS    HD2 3982      0.062100       1.00800 ; qtot -
3.054
  3983   HC   246   LYS    HD3 3983      0.062100       1.00800 ; qtot -
2.992
  3984   C8   246   LYS     CE 3984     -0.014300      12.01000 ; qtot -
3.006
  3985   HP   246   LYS    HE2 3985      0.113500       1.00800 ; qtot -
2.893
  3986   HP   246   LYS    HE3 3986      0.113500       1.00800 ; qtot -
2.779
  3987   N3   246   LYS     NZ 3987     -0.385400      14.01000 ; qtot -
3.165
  3988    H   246   LYS    HZ1 3988      0.340000       1.00800 ; qtot -
2.825
  3989    H   246   LYS    HZ2 3989      0.340000       1.00800 ; qtot -
2.485
  3990    H   246   LYS    HZ3 3990      0.340000       1.00800 ; qtot -
2.145
  3991    C   246   LYS      C 3991      0.734101      12.01000 ; qtot -
1.411
  3992    O   246   LYS      O 3992     -0.589401      16.00000 ; qtot -
2.000
  3993    N   247   LEU      N 3993     -0.415700      14.01000 ; qtot -
2.416
  3994    H   247   LEU      H 3994      0.271900       1.00800 ; qtot -
2.144
  3995   CX   247   LEU     CA 3995     -0.051800      12.01000 ; qtot -
2.196
  3996   H1   247   LEU     HA 3996      0.092200       1.00800 ; qtot -
2.103
  3997  A2C   247   LEU     CB 3997     -0.110200      12.01000 ; qtot -

2.214
  3998   HC   247   LEU   HB2 3998      0.045700      1.00800 ; qtot -
2.168
  3999   HC   247   LEU   HB3 3999      0.045700      1.00800 ; qtot -
2.122
  4000  A3C   247   LEU    CG 4000      0.353100     12.01000 ; qtot -
1.769
  4001   HC   247   LEU    HG 4001     -0.036100      1.00800 ; qtot -
1.805
  4002   CT   247   LEU   CD1 4002     -0.412100     12.01000 ; qtot -
2.217
  4003   HC   247   LEU  HD11 4003      0.100000      1.00800 ; qtot -
2.117
  4004   HC   247   LEU  HD12 4004      0.100000      1.00800 ; qtot -
2.017
  4005   HC   247   LEU  HD13 4005      0.100000      1.00800 ; qtot -
1.917
  4006   CT   247   LEU   CD2 4006     -0.412100     12.01000 ; qtot -
2.329
  4007   HC   247   LEU  HD21 4007      0.100000      1.00800 ; qtot -
2.229
  4008   HC   247   LEU  HD22 4008      0.100000      1.00800 ; qtot -
2.129
  4009   HC   247   LEU  HD23 4009      0.100000      1.00800 ; qtot -
2.029
  4010    C   247   LEU     C 4010      0.597301     12.01000 ; qtot -
1.432
  4011    O   247   LEU     O 4011     -0.567901     16.00000 ; qtot -
2.000
  4012    N   248   LEU     N 4012     -0.415700     14.01000 ; qtot -
2.416
  4013    H   248   LEU     H 4013      0.271900      1.00800 ; qtot -
2.144
  4014   CX   248   LEU    CA 4014     -0.051800     12.01000 ; qtot -
2.196
  4015   H1   248   LEU    HA 4015      0.092200      1.00800 ; qtot -
2.103
  4016  A2C   248   LEU    CB 4016     -0.110200     12.01000 ; qtot -
2.214
  4017   HC   248   LEU   HB2 4017      0.045700      1.00800 ; qtot -
2.168
  4018   HC   248   LEU   HB3 4018      0.045700      1.00800 ; qtot -
2.122
  4019  A3C   248   LEU    CG 4019      0.353100     12.01000 ; qtot -
1.769
  4020   HC   248   LEU    HG 4020     -0.036100      1.00800 ; qtot -
1.805
  4021   CT   248   LEU   CD1 4021     -0.412100     12.01000 ; qtot -
2.217
  4022   HC   248   LEU  HD11 4022      0.100000      1.00800 ; qtot -
2.117
  4023   HC   248   LEU  HD12 4023      0.100000      1.00800 ; qtot -

2.017
  4024   HC   248   LEU  HD13 4024     0.100000      1.00800 ; qtot -
1.917
  4025   CT   248   LEU   CD2 4025    -0.412100     12.01000 ; qtot -
2.329
  4026   HC   248   LEU  HD21 4026     0.100000      1.00800 ; qtot -
2.229
  4027   HC   248   LEU  HD22 4027     0.100000      1.00800 ; qtot -
2.129
  4028   HC   248   LEU  HD23 4028     0.100000      1.00800 ; qtot -
2.029
  4029    C   248   LEU     C 4029     0.597301     12.01000 ; qtot -
1.432
  4030    O   248   LEU     O 4030    -0.567901     16.00000 ; qtot -
2.000
  4031    N   249   THR     N 4031    -0.415700     14.01000 ; qtot -
2.416
  4032    H   249   THR     H 4032     0.271900      1.00800 ; qtot -
2.144
  4033   CX   249   THR    CA 4033    -0.038900     12.01000 ; qtot -
2.183
  4034   H1   249   THR    HA 4034     0.100700      1.00800 ; qtot -
2.082
  4035  A3C   249   THR    CB 4035     0.365400     12.01000 ; qtot -
1.717
  4036   H1   249   THR    HB 4036     0.004300      1.00800 ; qtot -
1.712
  4037   CT   249   THR   CG2 4037    -0.243800     12.01000 ; qtot -
1.956
  4038   HC   249   THR  HG21 4038     0.064200      1.00800 ; qtot -
1.892
  4039   HC   249   THR  HG22 4039     0.064200      1.00800 ; qtot -
1.828
  4040   HC   249   THR  HG23 4040     0.064200      1.00800 ; qtot -
1.763
  4041   OH   249   THR   OG1 4041    -0.676101     16.00000 ; qtot -
2.440
  4042   HO   249   THR   HG1 4042     0.410200      1.00800 ; qtot -
2.029
  4043    C   249   THR     C 4043     0.597301     12.01000 ; qtot -
1.432
  4044    O   249   THR     O 4044    -0.567901     16.00000 ; qtot -
2.000
  4045    N   250   TYR     N 4045    -0.415700     14.01000 ; qtot -
2.416
  4046    H   250   TYR     H 4046     0.271900      1.00800 ; qtot -
2.144
  4047   CX   250   TYR    CA 4047    -0.001400     12.01000 ; qtot -
2.145
  4048   H1   250   TYR    HA 4048     0.087600      1.00800 ; qtot -
2.058
  4049   CT   250   TYR    CB 4049    -0.015200     12.01000 ; qtot -

2.073
  4050   HC   250   TYR   HB2 4050     0.029500      1.00800 ; qtot -
2.043
  4051   HC   250   TYR   HB3 4051     0.029500      1.00800 ; qtot -
2.014
  4052   CA   250   TYR    CG 4052    -0.001100     12.01000 ; qtot -
2.015
  4053   CA   250   TYR   CD1 4053    -0.190600     12.01000 ; qtot -
2.205
  4054   HA   250   TYR   HD1 4054     0.169900      1.00800 ; qtot -
2.036
  4055   CA   250   TYR   CE1 4055    -0.234100     12.01000 ; qtot -
2.270
  4056   HA   250   TYR   HE1 4056     0.165600      1.00800 ; qtot -
2.104
  4057    C   250   TYR    CZ 4057     0.322600     12.01000 ; qtot -
1.781
  4058   OH   250   TYR    OH 4058    -0.557901     16.00000 ; qtot -
2.339
  4059   HO   250   TYR    HH 4059     0.399200      1.00800 ; qtot -
1.940
  4060   CA   250   TYR   CE2 4060    -0.234100     12.01000 ; qtot -
2.174
  4061   HA   250   TYR   HE2 4061     0.165600      1.00800 ; qtot -
2.009
  4062   CA   250   TYR   CD2 4062    -0.190600     12.01000 ; qtot -
2.199
  4063   HA   250   TYR   HD2 4063     0.169900      1.00800 ; qtot -
2.029
  4064    C   250   TYR     C 4064     0.597301     12.01000 ; qtot -
1.432
  4065    O   250   TYR     O 4065    -0.567901     16.00000 ; qtot -
2.000
  4066    N   251   ASN     N 4066    -0.415700     14.01000 ; qtot -
2.416
  4067    H   251   ASN     H 4067     0.271900      1.00800 ; qtot -
2.144
  4068   CX   251   ASN    CA 4068     0.014300     12.01000 ; qtot -
2.129
  4069   H1   251   ASN    HA 4069     0.104800      1.00800 ; qtot -
2.025
  4070  A2C   251   ASN    CB 4070    -0.204100     12.01000 ; qtot -
2.229
  4071   HC   251   ASN   HB2 4071     0.079700      1.00800 ; qtot -
2.149
  4072   HC   251   ASN   HB3 4072     0.079700      1.00800 ; qtot -
2.069
  4073    C   251   ASN    CG 4073     0.713001     12.01000 ; qtot -
1.356
  4074    O   251   ASN   OD1 4074    -0.593101     16.00000 ; qtot -
1.949
  4075    N   251   ASN   ND2 4075    -0.919101     14.01000 ; qtot -

2.869
  4076    H   251   ASN  HD21 4076     0.419600      1.00800 ; qtot -
2.449
  4077    H   251   ASN  HD22 4077     0.419600      1.00800 ; qtot -
2.029
  4078    C   251   ASN     C 4078     0.597301     12.01000 ; qtot -
1.432
  4079    O   251   ASN     O 4079    -0.567901     16.00000 ; qtot -
2.000
  4080    N   252   PRO     N 4080    -0.254800     14.01000 ; qtot -
2.255
  4081   CT   252   PRO    CD 4081     0.019200     12.01000 ; qtot -
2.236
  4082   H1   252   PRO   HD2 4082     0.039100      1.00800 ; qtot -
2.196
  4083   H1   252   PRO   HD3 4083     0.039100      1.00800 ; qtot -
2.157
  4084   CT   252   PRO    CG 4084     0.018900     12.01000 ; qtot -
2.138
  4085   HC   252   PRO   HG2 4085     0.021300      1.00800 ; qtot -
2.117
  4086   HC   252   PRO   HG3 4086     0.021300      1.00800 ; qtot -
2.096
  4087   CT   252   PRO    CB 4087    -0.007000     12.01000 ; qtot -
2.103
  4088   HC   252   PRO   HB2 4088     0.025300      1.00800 ; qtot -
2.078
  4089   HC   252   PRO   HB3 4089     0.025300      1.00800 ; qtot -
2.052
  4090   CX   252   PRO    CA 4090    -0.026600     12.01000 ; qtot -
2.079
  4091   H1   252   PRO    HA 4091     0.064100      1.00800 ; qtot -
2.015
  4092    C   252   PRO     C 4092     0.589601     12.01000 ; qtot -
1.425
  4093    O   252   PRO     O 4093    -0.574801     16.00000 ; qtot -
2.000
  4094    N   253   ASN     N 4094    -0.415700     14.01000 ; qtot -
2.416
  4095    H   253   ASN     H 4095     0.271900      1.00800 ; qtot -
2.144
  4096   CX   253   ASN    CA 4096     0.014300     12.01000 ; qtot -
2.129
  4097   H1   253   ASN    HA 4097     0.104800      1.00800 ; qtot -
2.025
  4098  A2C   253   ASN    CB 4098    -0.204100     12.01000 ; qtot -
2.229
  4099   HC   253   ASN   HB2 4099     0.079700      1.00800 ; qtot -
2.149
  4100   HC   253   ASN   HB3 4100     0.079700      1.00800 ; qtot -
2.069
  4101    C   253   ASN    CG 4101     0.713001     12.01000 ; qtot -

1.356
  4102    O   253   ASN   OD1 4102    -0.593101     16.00000 ; qtot -
1.949
  4103    N   253   ASN   ND2 4103    -0.919101     14.01000 ; qtot -
2.869
  4104    H   253   ASN  HD21 4104     0.419600      1.00800 ; qtot -
2.449
  4105    H   253   ASN  HD22 4105     0.419600      1.00800 ; qtot -
2.029
  4106    C   253   ASN     C 4106     0.597301     12.01000 ; qtot -
1.432
  4107    O   253   ASN     O 4107    -0.567901     16.00000 ; qtot -
2.000
  4108    N   254   GLU     N 4108    -0.516301     14.01000 ; qtot -
2.516
  4109    H   254   GLU     H 4109     0.293600      1.00800 ; qtot -
2.223
  4110   CX   254   GLU    CA 4110     0.039700     12.01000 ; qtot -
2.183
  4111   H1   254   GLU    HA 4111     0.110500      1.00800 ; qtot -
2.072
  4112  A2C   254   GLU    CB 4112     0.056000     12.01000 ; qtot -
2.016
  4113   HC   254   GLU   HB2 4113    -0.017300      1.00800 ; qtot -
2.034
  4114   HC   254   GLU   HB3 4114    -0.017300      1.00800 ; qtot -
2.051
  4115  A2C   254   GLU    CG 4115     0.013600     12.01000 ; qtot -
2.037
  4116   HC   254   GLU   HG2 4116    -0.042500      1.00800 ; qtot -
2.080
  4117   HC   254   GLU   HG3 4117    -0.042500      1.00800 ; qtot -
2.122
  4118   CO   254   GLU    CD 4118     0.805401     12.01000 ; qtot -
1.317
  4119   O2   254   GLU   OE1 4119    -0.818801     16.00000 ; qtot -
2.136
  4120   O2   254   GLU   OE2 4120    -0.818801     16.00000 ; qtot -
2.955
  4121    C   254   GLU     C 4121     0.536601     12.01000 ; qtot -
2.418
  4122    O   254   GLU     O 4122    -0.581901     16.00000 ; qtot -
3.000
  4123    N   255   ARG     N 4123    -0.347900     14.01000 ; qtot -
3.348
  4124    H   255   ARG     H 4124     0.274700      1.00800 ; qtot -
3.073
  4125   CX   255   ARG    CA 4125    -0.263700     12.01000 ; qtot -
3.337
  4126   H1   255   ARG    HA 4126     0.156000      1.00800 ; qtot -
3.181
  4127   C8   255   ARG    CB 4127    -0.000700     12.01000 ; qtot -

3.182
4128 HC 255 ARG HB2 4128 0.032700 1.00800 ; qtot -
3.149
4129 HC 255 ARG HB3 4129 0.032700 1.00800 ; qtot -
3.116
4130 C8 255 ARG CG 4130 0.039000 12.01000 ; qtot -
3.077
4131 HC 255 ARG HG2 4131 0.028500 1.00800 ; qtot -
3.049
4132 HC 255 ARG HG3 4132 0.028500 1.00800 ; qtot -
3.020
4133 C8 255 ARG CD 4133 0.048600 12.01000 ; qtot -
2.972
4134 H1 255 ARG HD2 4134 0.068700 1.00800 ; qtot -
2.903
4135 H1 255 ARG HD3 4135 0.068700 1.00800 ; qtot -
2.834
4136 N2 255 ARG NE 4136 -0.529501 14.01000 ; qtot -
3.364
4137 H 255 ARG HE 4137 0.345600 1.00800 ; qtot -
3.018
4138 CA 255 ARG CZ 4138 0.807601 12.01000 ; qtot -
2.210
4139 N2 255 ARG NH1 4139 -0.862701 14.01000 ; qtot -
3.073
4140 H 255 ARG HH11 4140 0.447800 1.00800 ; qtot -
2.625
4141 H 255 ARG HH12 4141 0.447800 1.00800 ; qtot -
2.178
4142 N2 255 ARG NH2 4142 -0.862701 14.01000 ; qtot -
3.040
4143 H 255 ARG HH21 4143 0.447800 1.00800 ; qtot -
2.592
4144 H 255 ARG HH22 4144 0.447800 1.00800 ; qtot -
2.145
4145 C 255 ARG C 4145 0.734101 12.01000 ; qtot -
1.411
4146 O 255 ARG O 4146 -0.589401 16.00000 ; qtot -
2.000
4147 N 256 CYS N 4147 -0.415700 14.01000 ; qtot -
2.416
4148 H 256 CYS H 4148 0.271900 1.00800 ; qtot -
2.144
4149 CX 256 CYS CA 4149 0.021300 12.01000 ; qtot -
2.122
4150 H1 256 CYS HA 4150 0.112400 1.00800 ; qtot -
2.010
4151 A2C 256 CYS CB 4151 -0.123100 12.01000 ; qtot -
2.133
4152 H1 256 CYS HB2 4152 0.111200 1.00800 ; qtot -
2.022
4153 H1 256 CYS HB3 4153 0.111200 1.00800 ; qtot -

1.911
  4154   SH   256   CYS    SG 4154    -0.311900     32.06000 ; qtot -
2.223
  4155   HS   256   CYS    HG 4155     0.193300      1.00800 ; qtot -
2.029
  4156    C   256   CYS     C 4156     0.597301     12.01000 ; qtot -
1.432
  4157    O   256   CYS     O 4157    -0.567901     16.00000 ; qtot -
2.000
  4158    N   257   THR     N 4158    -0.415700     14.01000 ; qtot -
2.416
  4159    H   257   THR     H 4159     0.271900      1.00800 ; qtot -
2.144
  4160   CX   257   THR    CA 4160    -0.038900     12.01000 ; qtot -
2.183
  4161   H1   257   THR    HA 4161     0.100700      1.00800 ; qtot -
2.082
  4162  A3C   257   THR    CB 4162     0.365400     12.01000 ; qtot -
1.717
  4163   H1   257   THR    HB 4163     0.004300      1.00800 ; qtot -
1.712
  4164   CT   257   THR   CG2 4164    -0.243800     12.01000 ; qtot -
1.956
  4165   HC   257   THR  HG21 4165     0.064200      1.00800 ; qtot -
1.892
  4166   HC   257   THR  HG22 4166     0.064200      1.00800 ; qtot -
1.828
  4167   HC   257   THR  HG23 4167     0.064200      1.00800 ; qtot -
1.763
  4168   OH   257   THR   OG1 4168    -0.676101     16.00000 ; qtot -
2.440
  4169   HO   257   THR   HG1 4169     0.410200      1.00800 ; qtot -
2.029
  4170    C   257   THR     C 4170     0.597301     12.01000 ; qtot -
1.432
  4171    O   257   THR     O 4171    -0.567901     16.00000 ; qtot -
2.000
  4172    N   258   ILE     N 4172    -0.415700     14.01000 ; qtot -
2.416
  4173    H   258   ILE     H 4173     0.271900      1.00800 ; qtot -
2.144
  4174   CX   258   ILE    CA 4174    -0.059700     12.01000 ; qtot -
2.203
  4175   H1   258   ILE    HA 4175     0.086900      1.00800 ; qtot -
2.117
  4176  A3C   258   ILE    CB 4176     0.130300     12.01000 ; qtot -
1.986
  4177   HC   258   ILE    HB 4177     0.018700      1.00800 ; qtot -
1.968
  4178   CT   258   ILE   CG2 4178    -0.320400     12.01000 ; qtot -
2.288
  4179   HC   258   ILE  HG21 4179     0.088200      1.00800 ; qtot -

2.200
4180 HC 258 ILE HG22 4180 0.088200 1.00800 ; qtot -2.112
4181 HC 258 ILE HG23 4181 0.088200 1.00800 ; qtot -2.023
4182 A2C 258 ILE CG1 4182 -0.043000 12.01000 ; qtot -2.066
4183 HC 258 ILE HG12 4183 0.023600 1.00800 ; qtot -2.043
4184 HC 258 ILE HG13 4184 0.023600 1.00800 ; qtot -2.019
4185 CT 258 ILE CD1 4185 -0.066000 12.01000 ; qtot -2.085
4186 HC 258 ILE HD11 4186 0.018600 1.00800 ; qtot -2.067
4187 HC 258 ILE HD12 4187 0.018600 1.00800 ; qtot -2.048
4188 HC 258 ILE HD13 4188 0.018600 1.00800 ; qtot -2.029
4189 C 258 ILE C 4189 0.597301 12.01000 ; qtot -1.432
4190 O 258 ILE O 4190 -0.567901 16.00000 ; qtot -2.000
4191 N 259 GLU N 4191 -0.516301 14.01000 ; qtot -2.516
4192 H 259 GLU H 4192 0.293600 1.00800 ; qtot -2.223
4193 CX 259 GLU CA 4193 0.039700 12.01000 ; qtot -2.183
4194 H1 259 GLU HA 4194 0.110500 1.00800 ; qtot -2.072
4195 A2C 259 GLU CB 4195 0.056000 12.01000 ; qtot -2.016
4196 HC 259 GLU HB2 4196 -0.017300 1.00800 ; qtot -2.034
4197 HC 259 GLU HB3 4197 -0.017300 1.00800 ; qtot -2.051
4198 A2C 259 GLU CG 4198 0.013600 12.01000 ; qtot -2.037
4199 HC 259 GLU HG2 4199 -0.042500 1.00800 ; qtot -2.080
4200 HC 259 GLU HG3 4200 -0.042500 1.00800 ; qtot -2.122
4201 CO 259 GLU CD 4201 0.805401 12.01000 ; qtot -1.317
4202 O2 259 GLU OE1 4202 -0.818801 16.00000 ; qtot -2.136
4203 O2 259 GLU OE2 4203 -0.818801 16.00000 ; qtot -2.955
4204 C 259 GLU C 4204 0.536601 12.01000 ; qtot -2.418
4205 O 259 GLU O 4205 -0.581901 16.00000 ; qtot -

3.000
  4206    N   260   GLU     N 4206    -0.516301     14.01000 ; qtot -
3.516
  4207    H   260   GLU     H 4207     0.293600      1.00800 ; qtot -
3.223
  4208   CX   260   GLU    CA 4208     0.039700     12.01000 ; qtot -
3.183
  4209   H1   260   GLU    HA 4209     0.110500      1.00800 ; qtot -
3.072
  4210  A2C   260   GLU    CB 4210     0.056000     12.01000 ; qtot -
3.016
  4211   HC   260   GLU   HB2 4211    -0.017300      1.00800 ; qtot -
3.034
  4212   HC   260   GLU   HB3 4212    -0.017300      1.00800 ; qtot -
3.051
  4213  A2C   260   GLU    CG 4213     0.013600     12.01000 ; qtot -
3.037
  4214   HC   260   GLU   HG2 4214    -0.042500      1.00800 ; qtot -
3.080
  4215   HC   260   GLU   HG3 4215    -0.042500      1.00800 ; qtot -
3.122
  4216   CO   260   GLU    CD 4216     0.805401     12.01000 ; qtot -
2.317
  4217   O2   260   GLU   OE1 4217    -0.818801     16.00000 ; qtot -
3.136
  4218   O2   260   GLU   OE2 4218    -0.818801     16.00000 ; qtot -
3.955
  4219    C   260   GLU     C 4219     0.536601     12.01000 ; qtot -
3.418
  4220    O   260   GLU     O 4220    -0.581901     16.00000 ; qtot -
4.000
  4221    N   261   ALA     N 4221    -0.415700     14.01000 ; qtot -
4.416
  4222    H   261   ALA     H 4222     0.271900      1.00800 ; qtot -
4.144
  4223   CX   261   ALA    CA 4223     0.033700     12.01000 ; qtot -
4.110
  4224   H1   261   ALA    HA 4224     0.082300      1.00800 ; qtot -
4.028
  4225   CT   261   ALA    CB 4225    -0.182500     12.01000 ; qtot -
4.210
  4226   HC   261   ALA   HB1 4226     0.060300      1.00800 ; qtot -
4.150
  4227   HC   261   ALA   HB2 4227     0.060300      1.00800 ; qtot -
4.090
  4228   HC   261   ALA   HB3 4228     0.060300      1.00800 ; qtot -
4.029
  4229    C   261   ALA     C 4229     0.597301     12.01000 ; qtot -
3.432
  4230    O   261   ALA     O 4230    -0.567901     16.00000 ; qtot -
4.000
  4231    N   262   LEU     N 4231    -0.415700     14.01000 ; qtot -

4.416
  4232    H   262   LEU      H 4232      0.271900      1.00800 ; qtot -
4.144
  4233   CX   262   LEU     CA 4233     -0.051800     12.01000 ; qtot -
4.196
  4234   H1   262   LEU     HA 4234      0.092200      1.00800 ; qtot -
4.103
  4235  A2C   262   LEU     CB 4235     -0.110200     12.01000 ; qtot -
4.214
  4236   HC   262   LEU    HB2 4236      0.045700      1.00800 ; qtot -
4.168
  4237   HC   262   LEU    HB3 4237      0.045700      1.00800 ; qtot -
4.122
  4238  A3C   262   LEU     CG 4238      0.353100     12.01000 ; qtot -
3.769
  4239   HC   262   LEU     HG 4239     -0.036100      1.00800 ; qtot -
3.805
  4240   CT   262   LEU    CD1 4240     -0.412100     12.01000 ; qtot -
4.217
  4241   HC   262   LEU   HD11 4241      0.100000      1.00800 ; qtot -
4.117
  4242   HC   262   LEU   HD12 4242      0.100000      1.00800 ; qtot -
4.017
  4243   HC   262   LEU   HD13 4243      0.100000      1.00800 ; qtot -
3.917
  4244   CT   262   LEU    CD2 4244     -0.412100     12.01000 ; qtot -
4.329
  4245   HC   262   LEU   HD21 4245      0.100000      1.00800 ; qtot -
4.229
  4246   HC   262   LEU   HD22 4246      0.100000      1.00800 ; qtot -
4.129
  4247   HC   262   LEU   HD23 4247      0.100000      1.00800 ; qtot -
4.029
  4248    C   262   LEU      C 4248      0.597301     12.01000 ; qtot -
3.432
  4249    O   262   LEU      O 4249     -0.567901     16.00000 ; qtot -
4.000
  4250    N   263   ASN      N 4250     -0.415700     14.01000 ; qtot -
4.416
  4251    H   263   ASN      H 4251      0.271900      1.00800 ; qtot -
4.144
  4252   CX   263   ASN     CA 4252      0.014300     12.01000 ; qtot -
4.129
  4253   H1   263   ASN     HA 4253      0.104800      1.00800 ; qtot -
4.025
  4254  A2C   263   ASN     CB 4254     -0.204100     12.01000 ; qtot -
4.229
  4255   HC   263   ASN    HB2 4255      0.079700      1.00800 ; qtot -
4.149
  4256   HC   263   ASN    HB3 4256      0.079700      1.00800 ; qtot -
4.069
  4257    C   263   ASN     CG 4257      0.713001     12.01000 ; qtot -

3.356
  4258    O   263   ASN   OD1 4258    -0.593101     16.00000 ; qtot -
3.949
  4259    N   263   ASN   ND2 4259    -0.919101     14.01000 ; qtot -
4.869
  4260    H   263   ASN  HD21 4260     0.419600      1.00800 ; qtot -
4.449
  4261    H   263   ASN  HD22 4261     0.419600      1.00800 ; qtot -
4.029
  4262    C   263   ASN     C 4262     0.597301     12.01000 ; qtot -
3.432
  4263    O   263   ASN     O 4263    -0.567901     16.00000 ; qtot -
4.000
  4264    N   264   HIP     N 4264    -0.347900     14.01000 ; qtot -
4.348
  4265    H   264   HIP     H 4265     0.274700      1.00800 ; qtot -
4.073
  4266   CX   264   HIP    CA 4266    -0.135400     12.01000 ; qtot -
4.209
  4267   H1   264   HIP    HA 4267     0.121200      1.00800 ; qtot -
4.087
  4268   CT   264   HIP    CB 4268    -0.041400     12.01000 ; qtot -
4.129
  4269   HC   264   HIP   HB2 4269     0.081000      1.00800 ; qtot -
4.048
  4270   HC   264   HIP   HB3 4270     0.081000      1.00800 ; qtot -
3.967
  4271   CC   264   HIP    CG 4271    -0.001200     12.01000 ; qtot -
3.968
  4272   NA   264   HIP   ND1 4272    -0.151300     14.01000 ; qtot -
4.119
  4273    H   264   HIP   HD1 4273     0.386600      1.00800 ; qtot -
3.733
  4274   CR   264   HIP   CE1 4274    -0.017000     12.01000 ; qtot -
3.750
  4275   H5   264   HIP   HE1 4275     0.268100      1.00800 ; qtot -
3.482
  4276   NA   264   HIP   NE2 4276    -0.171800     14.01000 ; qtot -
3.653
  4277    H   264   HIP   HE2 4277     0.391100      1.00800 ; qtot -
3.262
  4278   CW   264   HIP   CD2 4278    -0.114100     12.01000 ; qtot -
3.376
  4279   H4   264   HIP   HD2 4279     0.231700      1.00800 ; qtot -
3.145
  4280    C   264   HIP     C 4280     0.734101     12.01000 ; qtot -
2.411
  4281    O   264   HIP     O 4281    -0.589401     16.00000 ; qtot -
3.000
  4282    N   265   PRO     N 4282    -0.254800     14.01000 ; qtot -
3.255
  4283   CT   265   PRO    CD 4283     0.019200     12.01000 ; qtot -

```
3.236
  4284   H1   265   PRO   HD2 4284     0.039100      1.00800 ; qtot -
3.196
  4285   H1   265   PRO   HD3 4285     0.039100      1.00800 ; qtot -
3.157
  4286   CT   265   PRO    CG 4286     0.018900     12.01000 ; qtot -
3.138
  4287   HC   265   PRO   HG2 4287     0.021300      1.00800 ; qtot -
3.117
  4288   HC   265   PRO   HG3 4288     0.021300      1.00800 ; qtot -
3.096
  4289   CT   265   PRO    CB 4289    -0.007000     12.01000 ; qtot -
3.103
  4290   HC   265   PRO   HB2 4290     0.025300      1.00800 ; qtot -
3.078
  4291   HC   265   PRO   HB3 4291     0.025300      1.00800 ; qtot -
3.052
  4292   CX   265   PRO    CA 4292    -0.026600     12.01000 ; qtot -
3.079
  4293   H1   265   PRO    HA 4293     0.064100      1.00800 ; qtot -
3.015
  4294    C   265   PRO     C 4294     0.589601     12.01000 ; qtot -
2.425
  4295    O   265   PRO     O 4295    -0.574801     16.00000 ; qtot -
3.000
  4296    N   266   TRP     N 4296    -0.415700     14.01000 ; qtot -
3.416
  4297    H   266   TRP     H 4297     0.271900      1.00800 ; qtot -
3.144
  4298   CX   266   TRP    CA 4298    -0.027500     12.01000 ; qtot -
3.171
  4299   H1   266   TRP    HA 4299     0.112300      1.00800 ; qtot -
3.059
  4300   CT   266   TRP    CB 4300    -0.005000     12.01000 ; qtot -
3.064
  4301   HC   266   TRP   HB2 4301     0.033900      1.00800 ; qtot -
3.030
  4302   HC   266   TRP   HB3 4302     0.033900      1.00800 ; qtot -
2.996
  4303   C*   266   TRP    CG 4303    -0.141500     12.01000 ; qtot -
3.138
  4304   CW   266   TRP   CD1 4304    -0.163800     12.01000 ; qtot -
3.301
  4305   H4   266   TRP   HD1 4305     0.206200      1.00800 ; qtot -
3.095
  4306   NA   266   TRP   NE1 4306    -0.341800     14.01000 ; qtot -
3.437
  4307    H   266   TRP   HE1 4307     0.341200      1.00800 ; qtot -
3.096
  4308   CN   266   TRP   CE2 4308     0.138000     12.01000 ; qtot -
2.958
  4309   CA   266   TRP   CZ2 4309    -0.260100     12.01000 ; qtot -
```

3.218
  4310    HA   266   TRP    HZ2 4310      0.157200      1.00800 ; qtot -
3.061
  4311    CA   266   TRP    CH2 4311     -0.113400     12.01000 ; qtot -
3.174
  4312    HA   266   TRP    HH2 4312      0.141700      1.00800 ; qtot -
3.032
  4313    CA   266   TRP    CZ3 4313     -0.197200     12.01000 ; qtot -
3.230
  4314    HA   266   TRP    HZ3 4314      0.144700      1.00800 ; qtot -
3.085
  4315    CA   266   TRP    CE3 4315     -0.238700     12.01000 ; qtot -
3.324
  4316    HA   266   TRP    HE3 4316      0.170000      1.00800 ; qtot -
3.154
  4317    CB   266   TRP    CD2 4317      0.124300     12.01000 ; qtot -
3.029
  4318     C   266   TRP      C 4318      0.597301     12.01000 ; qtot -
2.432
  4319     O   266   TRP      O 4319     -0.567901     16.00000 ; qtot -
3.000
  4320     N   267   ILE      N 4320     -0.415700     14.01000 ; qtot -
3.416
  4321     H   267   ILE      H 4321      0.271900      1.00800 ; qtot -
3.144
  4322    CX   267   ILE     CA 4322     -0.059700     12.01000 ; qtot -
3.203
  4323    H1   267   ILE     HA 4323      0.086900      1.00800 ; qtot -
3.117
  4324   A3C   267   ILE     CB 4324      0.130300     12.01000 ; qtot -
2.986
  4325    HC   267   ILE     HB 4325      0.018700      1.00800 ; qtot -
2.968
  4326    CT   267   ILE    CG2 4326     -0.320400     12.01000 ; qtot -
3.288
  4327    HC   267   ILE   HG21 4327      0.088200      1.00800 ; qtot -
3.200
  4328    HC   267   ILE   HG22 4328      0.088200      1.00800 ; qtot -
3.112
  4329    HC   267   ILE   HG23 4329      0.088200      1.00800 ; qtot -
3.023
  4330   A2C   267   ILE    CG1 4330     -0.043000     12.01000 ; qtot -
3.066
  4331    HC   267   ILE   HG12 4331      0.023600      1.00800 ; qtot -
3.043
  4332    HC   267   ILE   HG13 4332      0.023600      1.00800 ; qtot -
3.019
  4333    CT   267   ILE    CD1 4333     -0.066000     12.01000 ; qtot -
3.085
  4334    HC   267   ILE   HD11 4334      0.018600      1.00800 ; qtot -
3.067
  4335    HC   267   ILE   HD12 4335      0.018600      1.00800 ; qtot -

3.048
4336 HC 267 ILE HD13 4336 0.018600 1.00800 ; qtot -3.029
4337 C 267 ILE C 4337 0.597301 12.01000 ; qtot -2.432
4338 O 267 ILE O 4338 -0.567901 16.00000 ; qtot -3.000
4339 N 268 THR N 4339 -0.415700 14.01000 ; qtot -3.416
4340 H 268 THR H 4340 0.271900 1.00800 ; qtot -3.144
4341 CX 268 THR CA 4341 -0.038900 12.01000 ; qtot -3.183
4342 H1 268 THR HA 4342 0.100700 1.00800 ; qtot -3.082
4343 A3C 268 THR CB 4343 0.365400 12.01000 ; qtot -2.717
4344 H1 268 THR HB 4344 0.004300 1.00800 ; qtot -2.712
4345 CT 268 THR CG2 4345 -0.243800 12.01000 ; qtot -2.956
4346 HC 268 THR HG21 4346 0.064200 1.00800 ; qtot -2.892
4347 HC 268 THR HG22 4347 0.064200 1.00800 ; qtot -2.828
4348 HC 268 THR HG23 4348 0.064200 1.00800 ; qtot -2.763
4349 OH 268 THR OG1 4349 -0.676101 16.00000 ; qtot -3.440
4350 HO 268 THR HG1 4350 0.410200 1.00800 ; qtot -3.029
4351 C 268 THR C 4351 0.597301 12.01000 ; qtot -2.432
4352 O 268 THR O 4352 -0.567901 16.00000 ; qtot -3.000
4353 N 269 GLN N 4353 -0.415700 14.01000 ; qtot -3.416
4354 H 269 GLN H 4354 0.271900 1.00800 ; qtot -3.144
4355 CX 269 GLN CA 4355 -0.003100 12.01000 ; qtot -3.147
4356 H1 269 GLN HA 4356 0.085000 1.00800 ; qtot -3.062
4357 A2C 269 GLN CB 4357 -0.003600 12.01000 ; qtot -3.065
4358 HC 269 GLN HB2 4358 0.017100 1.00800 ; qtot -3.048
4359 HC 269 GLN HB3 4359 0.017100 1.00800 ; qtot -3.031
4360 A2C 269 GLN CG 4360 -0.064500 12.01000 ; qtot -3.096
4361 HC 269 GLN HG2 4361 0.035200 1.00800 ; qtot -

3.061
4362 HC 269 GLN HG3 4362 0.035200 1.00800 ; qtot -3.025
4363 C 269 GLN CD 4363 0.695101 12.01000 ; qtot -2.330
4364 O 269 GLN OE1 4364 -0.608601 16.00000 ; qtot -2.939
4365 N 269 GLN NE2 4365 -0.940701 14.01000 ; qtot -3.880
4366 H 269 GLN HE21 4366 0.425100 1.00800 ; qtot -3.454
4367 H 269 GLN HE22 4367 0.425100 1.00800 ; qtot -3.029
4368 C 269 GLN C 4368 0.597301 12.01000 ; qtot -2.432
4369 O 269 GLN O 4369 -0.567901 16.00000 ; qtot -3.000
4370 N 270 MET N 4370 -0.415700 14.01000 ; qtot -3.416
4371 H 270 MET H 4371 0.271900 1.00800 ; qtot -3.144
4372 CX 270 MET CA 4372 -0.023700 12.01000 ; qtot -3.167
4373 H1 270 MET HA 4373 0.088000 1.00800 ; qtot -3.079
4374 A2C 270 MET CB 4374 0.034200 12.01000 ; qtot -3.045
4375 HC 270 MET HB2 4375 0.024100 1.00800 ; qtot -3.021
4376 HC 270 MET HB3 4376 0.024100 1.00800 ; qtot -2.997
4377 A2C 270 MET CG 4377 0.001800 12.01000 ; qtot -2.995
4378 H1 270 MET HG2 4378 0.044000 1.00800 ; qtot -2.951
4379 H1 270 MET HG3 4379 0.044000 1.00800 ; qtot -2.907
4380 S 270 MET SD 4380 -0.273700 32.06000 ; qtot -3.181
4381 CT 270 MET CE 4381 -0.053600 12.01000 ; qtot -3.235
4382 H1 270 MET HE1 4382 0.068400 1.00800 ; qtot -3.166
4383 H1 270 MET HE2 4383 0.068400 1.00800 ; qtot -3.098
4384 H1 270 MET HE3 4384 0.068400 1.00800 ; qtot -3.029
4385 C 270 MET C 4385 0.597301 12.01000 ; qtot -2.432
4386 O 270 MET O 4386 -0.567901 16.00000 ; qtot -3.000
4387 N 271 THR N 4387 -0.415700 14.01000 ; qtot -

```
3.416
  4388    H   271   THR      H 4388     0.271900      1.00800 ; qtot -
3.144
  4389   CX   271   THR     CA 4389    -0.038900     12.01000 ; qtot -
3.183
  4390   H1   271   THR     HA 4390     0.100700      1.00800 ; qtot -
3.082
  4391  A3C   271   THR     CB 4391     0.365400     12.01000 ; qtot -
2.717
  4392   H1   271   THR     HB 4392     0.004300      1.00800 ; qtot -
2.712
  4393   CT   271   THR    CG2 4393    -0.243800     12.01000 ; qtot -
2.956
  4394   HC   271   THR   HG21 4394     0.064200      1.00800 ; qtot -
2.892
  4395   HC   271   THR   HG22 4395     0.064200      1.00800 ; qtot -
2.828
  4396   HC   271   THR   HG23 4396     0.064200      1.00800 ; qtot -
2.763
  4397   OH   271   THR    OG1 4397    -0.676101     16.00000 ; qtot -
3.440
  4398   HO   271   THR    HG1 4398     0.410200      1.00800 ; qtot -
3.029
  4399    C   271   THR      C 4399     0.597301     12.01000 ; qtot -
2.432
  4400    O   271   THR      O 4400    -0.567901     16.00000 ; qtot -
3.000
  4401    N   272   LYS      N 4401    -0.347900     14.01000 ; qtot -
3.348
  4402    H   272   LYS      H 4402     0.274700      1.00800 ; qtot -
3.073
  4403   CX   272   LYS     CA 4403    -0.240000     12.01000 ; qtot -
3.313
  4404   H1   272   LYS     HA 4404     0.142600      1.00800 ; qtot -
3.171
  4405   C8   272   LYS     CB 4405    -0.009400     12.01000 ; qtot -
3.180
  4406   HC   272   LYS    HB2 4406     0.036200      1.00800 ; qtot -
3.144
  4407   HC   272   LYS    HB3 4407     0.036200      1.00800 ; qtot -
3.108
  4408   C8   272   LYS     CG 4408     0.018700     12.01000 ; qtot -
3.089
  4409   HC   272   LYS    HG2 4409     0.010300      1.00800 ; qtot -
3.079
  4410   HC   272   LYS    HG3 4410     0.010300      1.00800 ; qtot -
3.068
  4411   C8   272   LYS     CD 4411    -0.047900     12.01000 ; qtot -
3.116
  4412   HC   272   LYS    HD2 4412     0.062100      1.00800 ; qtot -
3.054
  4413   HC   272   LYS    HD3 4413     0.062100      1.00800 ; qtot -
```

```
2.992
  4414   C8   272   LYS    CE 4414    -0.014300     12.01000 ; qtot -
3.006
  4415   HP   272   LYS   HE2 4415     0.113500      1.00800 ; qtot -
2.893
  4416   HP   272   LYS   HE3 4416     0.113500      1.00800 ; qtot -
2.779
  4417   N3   272   LYS    NZ 4417    -0.385400     14.01000 ; qtot -
3.165
  4418    H   272   LYS   HZ1 4418     0.340000      1.00800 ; qtot -
2.825
  4419    H   272   LYS   HZ2 4419     0.340000      1.00800 ; qtot -
2.485
  4420    H   272   LYS   HZ3 4420     0.340000      1.00800 ; qtot -
2.145
  4421    C   272   LYS     C 4421     0.734101     12.01000 ; qtot -
1.411
  4422    O   272   LYS     O 4422    -0.589401     16.00000 ; qtot -
2.000
  4423    N   273   SER     N 4423    -0.415700     14.01000 ; qtot -
2.416
  4424    H   273   SER     H 4424     0.271900      1.00800 ; qtot -
2.144
  4425   CX   273   SER    CA 4425    -0.024900     12.01000 ; qtot -
2.169
  4426   H1   273   SER    HA 4426     0.084300      1.00800 ; qtot -
2.084
  4427  A2C   273   SER    CB 4427     0.211700     12.01000 ; qtot -
1.873
  4428   H1   273   SER   HB2 4428     0.035200      1.00800 ; qtot -
1.837
  4429   H1   273   SER   HB3 4429     0.035200      1.00800 ; qtot -
1.802
  4430   OH   273   SER    OG 4430    -0.654601     16.00000 ; qtot -
2.457
  4431   HO   273   SER    HG 4431     0.427500      1.00800 ; qtot -
2.029
  4432    C   273   SER     C 4432     0.597301     12.01000 ; qtot -
1.432
  4433    O   273   SER     O 4433    -0.567901     16.00000 ; qtot -
2.000
  4434    N   274   HIE     N 4434    -0.415700     14.01000 ; qtot -
2.416
  4435    H   274   HIE     H 4435     0.271900      1.00800 ; qtot -
2.144
  4436   CX   274   HIE    CA 4436    -0.058100     12.01000 ; qtot -
2.202
  4437   H1   274   HIE    HA 4437     0.136000      1.00800 ; qtot -
2.066
  4438   CT   274   HIE    CB 4438    -0.007400     12.01000 ; qtot -
2.073
  4439   HC   274   HIE   HB2 4439     0.036700      1.00800 ; qtot -
```

2.037
4440 HC 274 HIE HB3 4440 0.036700 1.00800 ; qtot -
2.000
4441 CC 274 HIE CG 4441 0.186800 12.01000 ; qtot -
1.813
4442 NB 274 HIE ND1 4442 -0.543201 14.01000 ; qtot -
2.356
4443 CR 274 HIE CE1 4443 0.163500 12.01000 ; qtot -
2.193
4444 H5 274 HIE HE1 4444 0.143500 1.00800 ; qtot -
2.049
4445 NA 274 HIE NE2 4445 -0.279500 14.01000 ; qtot -
2.329
4446 H 274 HIE HE2 4446 0.333900 1.00800 ; qtot -
1.995
4447 CW 274 HIE CD2 4447 -0.220700 12.01000 ; qtot -
2.216
4448 H4 274 HIE HD2 4448 0.186200 1.00800 ; qtot -
2.029
4449 C 274 HIE C 4449 0.597301 12.01000 ; qtot -
1.432
4450 O 274 HIE O 4450 -0.567901 16.00000 ; qtot -
2.000
4451 N 275 GLU N 4451 -0.516301 14.01000 ; qtot -
2.516
4452 H 275 GLU H 4452 0.293600 1.00800 ; qtot -
2.223
4453 CX 275 GLU CA 4453 0.039700 12.01000 ; qtot -
2.183
4454 H1 275 GLU HA 4454 0.110500 1.00800 ; qtot -
2.072
4455 A2C 275 GLU CB 4455 0.056000 12.01000 ; qtot -
2.016
4456 HC 275 GLU HB2 4456 -0.017300 1.00800 ; qtot -
2.034
4457 HC 275 GLU HB3 4457 -0.017300 1.00800 ; qtot -
2.051
4458 A2C 275 GLU CG 4458 0.013600 12.01000 ; qtot -
2.037
4459 HC 275 GLU HG2 4459 -0.042500 1.00800 ; qtot -
2.080
4460 HC 275 GLU HG3 4460 -0.042500 1.00800 ; qtot -
2.122
4461 CO 275 GLU CD 4461 0.805401 12.01000 ; qtot -
1.317
4462 O2 275 GLU OE1 4462 -0.818801 16.00000 ; qtot -
2.136
4463 O2 275 GLU OE2 4463 -0.818801 16.00000 ; qtot -
2.955
4464 C 275 GLU C 4464 0.536601 12.01000 ; qtot -
2.418
4465 O 275 GLU O 4465 -0.581901 16.00000 ; qtot -

3.000
  4466    N   276   HIE      N 4466    -0.415700     14.01000 ; qtot -
3.416
  4467    H   276   HIE      H 4467     0.271900      1.00800 ; qtot -
3.144
  4468   CX   276   HIE     CA 4468    -0.058100     12.01000 ; qtot -
3.202
  4469   H1   276   HIE     HA 4469     0.136000      1.00800 ; qtot -
3.066
  4470   CT   276   HIE     CB 4470    -0.007400     12.01000 ; qtot -
3.073
  4471   HC   276   HIE    HB2 4471     0.036700      1.00800 ; qtot -
3.037
  4472   HC   276   HIE    HB3 4472     0.036700      1.00800 ; qtot -
3.000
  4473   CC   276   HIE     CG 4473     0.186800     12.01000 ; qtot -
2.813
  4474   NB   276   HIE    ND1 4474    -0.543201     14.01000 ; qtot -
3.356
  4475   CR   276   HIE    CE1 4475     0.163500     12.01000 ; qtot -
3.193
  4476   H5   276   HIE    HE1 4476     0.143500      1.00800 ; qtot -
3.049
  4477   NA   276   HIE    NE2 4477    -0.279500     14.01000 ; qtot -
3.329
  4478    H   276   HIE    HE2 4478     0.333900      1.00800 ; qtot -
2.995
  4479   CW   276   HIE    CD2 4479    -0.220700     12.01000 ; qtot -
3.216
  4480   H4   276   HIE    HD2 4480     0.186200      1.00800 ; qtot -
3.029
  4481    C   276   HIE      C 4481     0.597301     12.01000 ; qtot -
2.432
  4482    O   276   HIE      O 4482    -0.567901     16.00000 ; qtot -
3.000
  4483    N   277   VAL      N 4483    -0.415700     14.01000 ; qtot -
3.416
  4484    H   277   VAL      H 4484     0.271900      1.00800 ; qtot -
3.144
  4485   CX   277   VAL     CA 4485    -0.087500     12.01000 ; qtot -
3.231
  4486   H1   277   VAL     HA 4486     0.096900      1.00800 ; qtot -
3.134
  4487  A3C   277   VAL     CB 4487     0.298500     12.01000 ; qtot -
2.836
  4488   HC   277   VAL     HB 4488    -0.029700      1.00800 ; qtot -
2.866
  4489   CT   277   VAL    CG1 4489    -0.319200     12.01000 ; qtot -
3.185
  4490   HC   277   VAL   HG11 4490     0.079100      1.00800 ; qtot -
3.106
  4491   HC   277   VAL   HG12 4491     0.079100      1.00800 ; qtot -

3.027
 4492 HC 277 VAL HG13 4492 0.079100 1.00800 ; qtot -
2.947
 4493 CT 277 VAL CG2 4493 -0.319200 12.01000 ; qtot -
3.267
 4494 HC 277 VAL HG21 4494 0.079100 1.00800 ; qtot -
3.188
 4495 HC 277 VAL HG22 4495 0.079100 1.00800 ; qtot -
3.108
 4496 HC 277 VAL HG23 4496 0.079100 1.00800 ; qtot -
3.029
 4497 C 277 VAL C 4497 0.597301 12.01000 ; qtot -
2.432
 4498 O 277 VAL O 4498 -0.567901 16.00000 ; qtot -
3.000
 4499 N 278 GLU N 4499 -0.516301 14.01000 ; qtot -
3.516
 4500 H 278 GLU H 4500 0.293600 1.00800 ; qtot -
3.223
 4501 CX 278 GLU CA 4501 0.039700 12.01000 ; qtot -
3.183
 4502 H1 278 GLU HA 4502 0.110500 1.00800 ; qtot -
3.072
 4503 A2C 278 GLU CB 4503 0.056000 12.01000 ; qtot -
3.016
 4504 HC 278 GLU HB2 4504 -0.017300 1.00800 ; qtot -
3.034
 4505 HC 278 GLU HB3 4505 -0.017300 1.00800 ; qtot -
3.051
 4506 A2C 278 GLU CG 4506 0.013600 12.01000 ; qtot -
3.037
 4507 HC 278 GLU HG2 4507 -0.042500 1.00800 ; qtot -
3.080
 4508 HC 278 GLU HG3 4508 -0.042500 1.00800 ; qtot -
3.122
 4509 CO 278 GLU CD 4509 0.805401 12.01000 ; qtot -
2.317
 4510 O2 278 GLU OE1 4510 -0.818801 16.00000 ; qtot -
3.136
 4511 O2 278 GLU OE2 4511 -0.818801 16.00000 ; qtot -
3.955
 4512 C 278 GLU C 4512 0.536601 12.01000 ; qtot -
3.418
 4513 O 278 GLU O 4513 -0.581901 16.00000 ; qtot -
4.000
 4514 N 279 LEU N 4514 -0.415700 14.01000 ; qtot -
4.416
 4515 H 279 LEU H 4515 0.271900 1.00800 ; qtot -
4.144
 4516 CX 279 LEU CA 4516 -0.051800 12.01000 ; qtot -
4.196
 4517 H1 279 LEU HA 4517 0.092200 1.00800 ; qtot -

4.103
4518 A2C 279 LEU CB 4518 -0.110200 12.01000 ; qtot -
4.214
4519 HC 279 LEU HB2 4519 0.045700 1.00800 ; qtot -
4.168
4520 HC 279 LEU HB3 4520 0.045700 1.00800 ; qtot -
4.122
4521 A3C 279 LEU CG 4521 0.353100 12.01000 ; qtot -
3.769
4522 HC 279 LEU HG 4522 -0.036100 1.00800 ; qtot -
3.805
4523 CT 279 LEU CD1 4523 -0.412100 12.01000 ; qtot -
4.217
4524 HC 279 LEU HD11 4524 0.100000 1.00800 ; qtot -
4.117
4525 HC 279 LEU HD12 4525 0.100000 1.00800 ; qtot -
4.017
4526 HC 279 LEU HD13 4526 0.100000 1.00800 ; qtot -
3.917
4527 CT 279 LEU CD2 4527 -0.412100 12.01000 ; qtot -
4.329
4528 HC 279 LEU HD21 4528 0.100000 1.00800 ; qtot -
4.229
4529 HC 279 LEU HD22 4529 0.100000 1.00800 ; qtot -
4.129
4530 HC 279 LEU HD23 4530 0.100000 1.00800 ; qtot -
4.029
4531 C 279 LEU C 4531 0.597301 12.01000 ; qtot -
3.432
4532 O 279 LEU O 4532 -0.567901 16.00000 ; qtot -
4.000
4533 N 280 SER N 4533 -0.415700 14.01000 ; qtot -
4.416
4534 H 280 SER H 4534 0.271900 1.00800 ; qtot -
4.144
4535 CX 280 SER CA 4535 -0.024900 12.01000 ; qtot -
4.169
4536 H1 280 SER HA 4536 0.084300 1.00800 ; qtot -
4.084
4537 A2C 280 SER CB 4537 0.211700 12.01000 ; qtot -
3.873
4538 H1 280 SER HB2 4538 0.035200 1.00800 ; qtot -
3.837
4539 H1 280 SER HB3 4539 0.035200 1.00800 ; qtot -
3.802
4540 OH 280 SER OG 4540 -0.654601 16.00000 ; qtot -
4.457
4541 HO 280 SER HG 4541 0.427500 1.00800 ; qtot -
4.029
4542 C 280 SER C 4542 0.597301 12.01000 ; qtot -
3.432
4543 O 280 SER O 4543 -0.567901 16.00000 ; qtot -

4.000
  4544    N   281   SER      N 4544    -0.415700     14.01000 ; qtot -
4.416
  4545    H   281   SER      H 4545     0.271900      1.00800 ; qtot -
4.144
  4546   CX   281   SER     CA 4546    -0.024900     12.01000 ; qtot -
4.169
  4547   H1   281   SER     HA 4547     0.084300      1.00800 ; qtot -
4.084
  4548  A2C   281   SER     CB 4548     0.211700     12.01000 ; qtot -
3.873
  4549   H1   281   SER    HB2 4549     0.035200      1.00800 ; qtot -
3.837
  4550   H1   281   SER    HB3 4550     0.035200      1.00800 ; qtot -
3.802
  4551   OH   281   SER     OG 4551    -0.654601     16.00000 ; qtot -
4.457
  4552   HO   281   SER     HG 4552     0.427500      1.00800 ; qtot -
4.029
  4553    C   281   SER      C 4553     0.597301     12.01000 ; qtot -
3.432
  4554    O   281   SER      O 4554    -0.567901     16.00000 ; qtot -
4.000
  4555    N   282   THR      N 4555    -0.415700     14.01000 ; qtot -
4.416
  4556    H   282   THR      H 4556     0.271900      1.00800 ; qtot -
4.144
  4557   CX   282   THR     CA 4557    -0.038900     12.01000 ; qtot -
4.183
  4558   H1   282   THR     HA 4558     0.100700      1.00800 ; qtot -
4.082
  4559  A3C   282   THR     CB 4559     0.365400     12.01000 ; qtot -
3.717
  4560   H1   282   THR     HB 4560     0.004300      1.00800 ; qtot -
3.712
  4561   CT   282   THR    CG2 4561    -0.243800     12.01000 ; qtot -
3.956
  4562   HC   282   THR   HG21 4562     0.064200      1.00800 ; qtot -
3.892
  4563   HC   282   THR   HG22 4563     0.064200      1.00800 ; qtot -
3.828
  4564   HC   282   THR   HG23 4564     0.064200      1.00800 ; qtot -
3.763
  4565   OH   282   THR    OG1 4565    -0.676101     16.00000 ; qtot -
4.440
  4566   HO   282   THR    HG1 4566     0.410200      1.00800 ; qtot -
4.029
  4567    C   282   THR      C 4567     0.597301     12.01000 ; qtot -
3.432
  4568    O   282   THR      O 4568    -0.567901     16.00000 ; qtot -
4.000
  4569    N   283   LEU      N 4569    -0.415700     14.01000 ; qtot -

4.416
  4570    H   283   LEU     H 4570     0.271900     1.00800 ; qtot -
4.144
  4571   CX   283   LEU    CA 4571    -0.051800    12.01000 ; qtot -
4.196
  4572   H1   283   LEU    HA 4572     0.092200     1.00800 ; qtot -
4.103
  4573  A2C   283   LEU    CB 4573    -0.110200    12.01000 ; qtot -
4.214
  4574   HC   283   LEU   HB2 4574     0.045700     1.00800 ; qtot -
4.168
  4575   HC   283   LEU   HB3 4575     0.045700     1.00800 ; qtot -
4.122
  4576  A3C   283   LEU    CG 4576     0.353100    12.01000 ; qtot -
3.769
  4577   HC   283   LEU    HG 4577    -0.036100     1.00800 ; qtot -
3.805
  4578   CT   283   LEU   CD1 4578    -0.412100    12.01000 ; qtot -
4.217
  4579   HC   283   LEU  HD11 4579     0.100000     1.00800 ; qtot -
4.117
  4580   HC   283   LEU  HD12 4580     0.100000     1.00800 ; qtot -
4.017
  4581   HC   283   LEU  HD13 4581     0.100000     1.00800 ; qtot -
3.917
  4582   CT   283   LEU   CD2 4582    -0.412100    12.01000 ; qtot -
4.329
  4583   HC   283   LEU  HD21 4583     0.100000     1.00800 ; qtot -
4.229
  4584   HC   283   LEU  HD22 4584     0.100000     1.00800 ; qtot -
4.129
  4585   HC   283   LEU  HD23 4585     0.100000     1.00800 ; qtot -
4.029
  4586    C   283   LEU     C 4586     0.597301    12.01000 ; qtot -
3.432
  4587    O   283   LEU     O 4587    -0.567901    16.00000 ; qtot -
4.000
  4588    N   284   LEU     N 4588    -0.415700    14.01000 ; qtot -
4.416
  4589    H   284   LEU     H 4589     0.271900     1.00800 ; qtot -
4.144
  4590   CX   284   LEU    CA 4590    -0.051800    12.01000 ; qtot -
4.196
  4591   H1   284   LEU    HA 4591     0.092200     1.00800 ; qtot -
4.103
  4592  A2C   284   LEU    CB 4592    -0.110200    12.01000 ; qtot -
4.214
  4593   HC   284   LEU   HB2 4593     0.045700     1.00800 ; qtot -
4.168
  4594   HC   284   LEU   HB3 4594     0.045700     1.00800 ; qtot -
4.122
  4595  A3C   284   LEU    CG 4595     0.353100    12.01000 ; qtot -

3.769
  4596   HC   284   LEU    HG 4596    -0.036100      1.00800 ; qtot -
3.805
  4597   CT   284   LEU   CD1 4597    -0.412100     12.01000 ; qtot -
4.217
  4598   HC   284   LEU  HD11 4598     0.100000      1.00800 ; qtot -
4.117
  4599   HC   284   LEU  HD12 4599     0.100000      1.00800 ; qtot -
4.017
  4600   HC   284   LEU  HD13 4600     0.100000      1.00800 ; qtot -
3.917
  4601   CT   284   LEU   CD2 4601    -0.412100     12.01000 ; qtot -
4.329
  4602   HC   284   LEU  HD21 4602     0.100000      1.00800 ; qtot -
4.229
  4603   HC   284   LEU  HD22 4603     0.100000      1.00800 ; qtot -
4.129
  4604   HC   284   LEU  HD23 4604     0.100000      1.00800 ; qtot -
4.029
  4605    C   284   LEU     C 4605     0.597301     12.01000 ; qtot -
3.432
  4606    O   284   LEU     O 4606    -0.567901     16.00000 ; qtot -
4.000
  4607    N   285   LYS     N 4607    -0.347900     14.01000 ; qtot -
4.348
  4608    H   285   LYS     H 4608     0.274700      1.00800 ; qtot -
4.073
  4609   CX   285   LYS    CA 4609    -0.240000     12.01000 ; qtot -
4.313
  4610   H1   285   LYS    HA 4610     0.142600      1.00800 ; qtot -
4.171
  4611   C8   285   LYS    CB 4611    -0.009400     12.01000 ; qtot -
4.180
  4612   HC   285   LYS   HB2 4612     0.036200      1.00800 ; qtot -
4.144
  4613   HC   285   LYS   HB3 4613     0.036200      1.00800 ; qtot -
4.108
  4614   C8   285   LYS    CG 4614     0.018700     12.01000 ; qtot -
4.089
  4615   HC   285   LYS   HG2 4615     0.010300      1.00800 ; qtot -
4.079
  4616   HC   285   LYS   HG3 4616     0.010300      1.00800 ; qtot -
4.068
  4617   C8   285   LYS    CD 4617    -0.047900     12.01000 ; qtot -
4.116
  4618   HC   285   LYS   HD2 4618     0.062100      1.00800 ; qtot -
4.054
  4619   HC   285   LYS   HD3 4619     0.062100      1.00800 ; qtot -
3.992
  4620   C8   285   LYS    CE 4620    -0.014300     12.01000 ; qtot -
4.006
  4621   HP   285   LYS   HE2 4621     0.113500      1.00800 ; qtot -

3.893
  4622   HP   285   LYS   HE3 4622     0.113500      1.00800 ; qtot -
3.779
  4623   N3   285   LYS    NZ 4623    -0.385400     14.01000 ; qtot -
4.165
  4624    H   285   LYS   HZ1 4624     0.340000      1.00800 ; qtot -
3.825
  4625    H   285   LYS   HZ2 4625     0.340000      1.00800 ; qtot -
3.485
  4626    H   285   LYS   HZ3 4626     0.340000      1.00800 ; qtot -
3.145
  4627    C   285   LYS     C 4627     0.734101     12.01000 ; qtot -
2.411
  4628    O   285   LYS     O 4628    -0.589401     16.00000 ; qtot -
3.000
  4629    N   286   ASN     N 4629    -0.415700     14.01000 ; qtot -
3.416
  4630    H   286   ASN     H 4630     0.271900      1.00800 ; qtot -
3.144
  4631   CX   286   ASN    CA 4631     0.014300     12.01000 ; qtot -
3.129
  4632   H1   286   ASN    HA 4632     0.104800      1.00800 ; qtot -
3.025
  4633  A2C   286   ASN    CB 4633    -0.204100     12.01000 ; qtot -
3.229
  4634   HC   286   ASN   HB2 4634     0.079700      1.00800 ; qtot -
3.149
  4635   HC   286   ASN   HB3 4635     0.079700      1.00800 ; qtot -
3.069
  4636    C   286   ASN    CG 4636     0.713001     12.01000 ; qtot -
2.356
  4637    O   286   ASN   OD1 4637    -0.593101     16.00000 ; qtot -
2.949
  4638    N   286   ASN   ND2 4638    -0.919101     14.01000 ; qtot -
3.869
  4639    H   286   ASN  HD21 4639     0.419600      1.00800 ; qtot -
3.449
  4640    H   286   ASN  HD22 4640     0.419600      1.00800 ; qtot -
3.029
  4641    C   286   ASN     C 4641     0.597301     12.01000 ; qtot -
2.432
  4642    O   286   ASN     O 4642    -0.567901     16.00000 ; qtot -
3.000
  4643    N   287   LEU     N 4643    -0.415700     14.01000 ; qtot -
3.416
  4644    H   287   LEU     H 4644     0.271900      1.00800 ; qtot -
3.144
  4645   CX   287   LEU    CA 4645    -0.051800     12.01000 ; qtot -
3.196
  4646   H1   287   LEU    HA 4646     0.092200      1.00800 ; qtot -
3.103
  4647  A2C   287   LEU    CB 4647    -0.110200     12.01000 ; qtot -

3.214
  4648   HC   287   LEU   HB2 4648     0.045700      1.00800 ; qtot -
3.168
  4649   HC   287   LEU   HB3 4649     0.045700      1.00800 ; qtot -
3.122
  4650  A3C   287   LEU    CG 4650     0.353100     12.01000 ; qtot -
2.769
  4651   HC   287   LEU    HG 4651    -0.036100      1.00800 ; qtot -
2.805
  4652   CT   287   LEU   CD1 4652    -0.412100     12.01000 ; qtot -
3.217
  4653   HC   287   LEU  HD11 4653     0.100000      1.00800 ; qtot -
3.117
  4654   HC   287   LEU  HD12 4654     0.100000      1.00800 ; qtot -
3.017
  4655   HC   287   LEU  HD13 4655     0.100000      1.00800 ; qtot -
2.917
  4656   CT   287   LEU   CD2 4656    -0.412100     12.01000 ; qtot -
3.329
  4657   HC   287   LEU  HD21 4657     0.100000      1.00800 ; qtot -
3.229
  4658   HC   287   LEU  HD22 4658     0.100000      1.00800 ; qtot -
3.129
  4659   HC   287   LEU  HD23 4659     0.100000      1.00800 ; qtot -
3.029
  4660    C   287   LEU     C 4660     0.597301     12.01000 ; qtot -
2.432
  4661    O   287   LEU     O 4661    -0.567901     16.00000 ; qtot -
3.000
  4662    N   288   LYS     N 4662    -0.347900     14.01000 ; qtot -
3.348
  4663    H   288   LYS     H 4663     0.274700      1.00800 ; qtot -
3.073
  4664   CX   288   LYS    CA 4664    -0.240000     12.01000 ; qtot -
3.313
  4665   H1   288   LYS    HA 4665     0.142600      1.00800 ; qtot -
3.171
  4666   C8   288   LYS    CB 4666    -0.009400     12.01000 ; qtot -
3.180
  4667   HC   288   LYS   HB2 4667     0.036200      1.00800 ; qtot -
3.144
  4668   HC   288   LYS   HB3 4668     0.036200      1.00800 ; qtot -
3.108
  4669   C8   288   LYS    CG 4669     0.018700     12.01000 ; qtot -
3.089
  4670   HC   288   LYS   HG2 4670     0.010300      1.00800 ; qtot -
3.079
  4671   HC   288   LYS   HG3 4671     0.010300      1.00800 ; qtot -
3.068
  4672   C8   288   LYS    CD 4672    -0.047900     12.01000 ; qtot -
3.116
  4673   HC   288   LYS   HD2 4673     0.062100      1.00800 ; qtot -

3.054
  4674   HC   288   LYS   HD3 4674     0.062100      1.00800 ; qtot -
2.992
  4675   C8   288   LYS    CE 4675    -0.014300     12.01000 ; qtot -
3.006
  4676   HP   288   LYS   HE2 4676     0.113500      1.00800 ; qtot -
2.893
  4677   HP   288   LYS   HE3 4677     0.113500      1.00800 ; qtot -
2.779
  4678   N3   288   LYS    NZ 4678    -0.385400     14.01000 ; qtot -
3.165
  4679    H   288   LYS   HZ1 4679     0.340000      1.00800 ; qtot -
2.825
  4680    H   288   LYS   HZ2 4680     0.340000      1.00800 ; qtot -
2.485
  4681    H   288   LYS   HZ3 4681     0.340000      1.00800 ; qtot -
2.145
  4682    C   288   LYS     C 4682     0.734101     12.01000 ; qtot -
1.411
  4683    O   288   LYS     O 4683    -0.589401     16.00000 ; qtot -
2.000
  4684    N   289   ASN     N 4684    -0.415700     14.01000 ; qtot -
2.416
  4685    H   289   ASN     H 4685     0.271900      1.00800 ; qtot -
2.144
  4686   CX   289   ASN    CA 4686     0.014300     12.01000 ; qtot -
2.129
  4687   H1   289   ASN    HA 4687     0.104800      1.00800 ; qtot -
2.025
  4688  A2C   289   ASN    CB 4688    -0.204100     12.01000 ; qtot -
2.229
  4689   HC   289   ASN   HB2 4689     0.079700      1.00800 ; qtot -
2.149
  4690   HC   289   ASN   HB3 4690     0.079700      1.00800 ; qtot -
2.069
  4691    C   289   ASN    CG 4691     0.713001     12.01000 ; qtot -
1.356
  4692    O   289   ASN   OD1 4692    -0.593101     16.00000 ; qtot -
1.949
  4693    N   289   ASN   ND2 4693    -0.919101     14.01000 ; qtot -
2.869
  4694    H   289   ASN  HD21 4694     0.419600      1.00800 ; qtot -
2.449
  4695    H   289   ASN  HD22 4695     0.419600      1.00800 ; qtot -
2.029
  4696    C   289   ASN     C 4696     0.597301     12.01000 ; qtot -
1.432
  4697    O   289   ASN     O 4697    -0.567901     16.00000 ; qtot -
2.000
  4698    N   290   PHE     N 4698    -0.415700     14.01000 ; qtot -
2.416
  4699    H   290   PHE     H 4699     0.271900      1.00800 ; qtot -

2.144
 4700 CX 290 PHE CA 4700 -0.002400 12.01000 ; qtot -
2.146
 4701 H1 290 PHE HA 4701 0.097800 1.00800 ; qtot -
2.048
 4702 CT 290 PHE CB 4702 -0.034300 12.01000 ; qtot -
2.083
 4703 HC 290 PHE HB2 4703 0.029500 1.00800 ; qtot -
2.053
 4704 HC 290 PHE HB3 4704 0.029500 1.00800 ; qtot -
2.024
 4705 CA 290 PHE CG 4705 0.011800 12.01000 ; qtot -
2.012
 4706 CA 290 PHE CD1 4706 -0.125600 12.01000 ; qtot -
2.137
 4707 HA 290 PHE HD1 4707 0.133000 1.00800 ; qtot -
2.004
 4708 CA 290 PHE CE1 4708 -0.170400 12.01000 ; qtot -
2.175
 4709 HA 290 PHE HE1 4709 0.143000 1.00800 ; qtot -
2.032
 4710 CA 290 PHE CZ 4710 -0.107200 12.01000 ; qtot -
2.139
 4711 HA 290 PHE HZ 4711 0.129700 1.00800 ; qtot -
2.009
 4712 CA 290 PHE CE2 4712 -0.170400 12.01000 ; qtot -
2.180
 4713 HA 290 PHE HE2 4713 0.143000 1.00800 ; qtot -
2.037
 4714 CA 290 PHE CD2 4714 -0.125600 12.01000 ; qtot -
2.162
 4715 HA 290 PHE HD2 4715 0.133000 1.00800 ; qtot -
2.029
 4716 C 290 PHE C 4716 0.597301 12.01000 ; qtot -
1.432
 4717 O 290 PHE O 4717 -0.567901 16.00000 ; qtot -
2.000
 4718 N 291 LYS N 4718 -0.347900 14.01000 ; qtot -
2.348
 4719 H 291 LYS H 4719 0.274700 1.00800 ; qtot -
2.073
 4720 CX 291 LYS CA 4720 -0.240000 12.01000 ; qtot -
2.313
 4721 H1 291 LYS HA 4721 0.142600 1.00800 ; qtot -
2.171
 4722 C8 291 LYS CB 4722 -0.009400 12.01000 ; qtot -
2.180
 4723 HC 291 LYS HB2 4723 0.036200 1.00800 ; qtot -
2.144
 4724 HC 291 LYS HB3 4724 0.036200 1.00800 ; qtot -
2.108
 4725 C8 291 LYS CG 4725 0.018700 12.01000 ; qtot -

2.089
4726 HC 291 LYS HG2 4726 0.010300 1.00800 ; qtot -
2.079
4727 HC 291 LYS HG3 4727 0.010300 1.00800 ; qtot -
2.068
4728 C8 291 LYS CD 4728 -0.047900 12.01000 ; qtot -
2.116
4729 HC 291 LYS HD2 4729 0.062100 1.00800 ; qtot -
2.054
4730 HC 291 LYS HD3 4730 0.062100 1.00800 ; qtot -
1.992
4731 C8 291 LYS CE 4731 -0.014300 12.01000 ; qtot -
2.006
4732 HP 291 LYS HE2 4732 0.113500 1.00800 ; qtot -
1.893
4733 HP 291 LYS HE3 4733 0.113500 1.00800 ; qtot -
1.779
4734 N3 291 LYS NZ 4734 -0.385400 14.01000 ; qtot -
2.165
4735 H 291 LYS HZ1 4735 0.340000 1.00800 ; qtot -
1.825
4736 H 291 LYS HZ2 4736 0.340000 1.00800 ; qtot -
1.485
4737 H 291 LYS HZ3 4737 0.340000 1.00800 ; qtot -
1.145
4738 C 291 LYS C 4738 0.734101 12.01000 ; qtot -
0.411
4739 O 291 LYS O 4739 -0.589401 16.00000 ; qtot -
1.000
4740 N 292 LYS N 4740 -0.347900 14.01000 ; qtot -
1.348
4741 H 292 LYS H 4741 0.274700 1.00800 ; qtot -
1.073
4742 CX 292 LYS CA 4742 -0.240000 12.01000 ; qtot -
1.313
4743 H1 292 LYS HA 4743 0.142600 1.00800 ; qtot -
1.171
4744 C8 292 LYS CB 4744 -0.009400 12.01000 ; qtot -
1.180
4745 HC 292 LYS HB2 4745 0.036200 1.00800 ; qtot -
1.144
4746 HC 292 LYS HB3 4746 0.036200 1.00800 ; qtot -
1.108
4747 C8 292 LYS CG 4747 0.018700 12.01000 ; qtot -
1.089
4748 HC 292 LYS HG2 4748 0.010300 1.00800 ; qtot -
1.079
4749 HC 292 LYS HG3 4749 0.010300 1.00800 ; qtot -
1.068
4750 C8 292 LYS CD 4750 -0.047900 12.01000 ; qtot -
1.116
4751 HC 292 LYS HD2 4751 0.062100 1.00800 ; qtot -

1.054
4752 HC 292 LYS HD3 4752 0.062100 1.00800 ; qtot -
0.992
4753 C8 292 LYS CE 4753 -0.014300 12.01000 ; qtot -
1.006
4754 HP 292 LYS HE2 4754 0.113500 1.00800 ; qtot -
0.893
4755 HP 292 LYS HE3 4755 0.113500 1.00800 ; qtot -
0.779
4756 N3 292 LYS NZ 4756 -0.385400 14.01000 ; qtot -
1.165
4757 H 292 LYS HZ1 4757 0.340000 1.00800 ; qtot -
0.825
4758 H 292 LYS HZ2 4758 0.340000 1.00800 ; qtot -
0.485
4759 H 292 LYS HZ3 4759 0.340000 1.00800 ; qtot -
0.145
4760 C 292 LYS C 4760 0.734101 12.01000 ; qtot
0.589
4761 O 292 LYS O 4761 -0.589401 16.00000 ; qtot
0.000
4762 N 293 GLU N 4762 -0.516301 14.01000 ; qtot -
0.516
4763 H 293 GLU H 4763 0.293600 1.00800 ; qtot -
0.223
4764 CX 293 GLU CA 4764 0.039700 12.01000 ; qtot -
0.183
4765 H1 293 GLU HA 4765 0.110500 1.00800 ; qtot -
0.072
4766 A2C 293 GLU CB 4766 0.056000 12.01000 ; qtot -
0.016
4767 HC 293 GLU HB2 4767 -0.017300 1.00800 ; qtot -
0.034
4768 HC 293 GLU HB3 4768 -0.017300 1.00800 ; qtot -
0.051
4769 A2C 293 GLU CG 4769 0.013600 12.01000 ; qtot -
0.037
4770 HC 293 GLU HG2 4770 -0.042500 1.00800 ; qtot -
0.080
4771 HC 293 GLU HG3 4771 -0.042500 1.00800 ; qtot -
0.122
4772 CO 293 GLU CD 4772 0.805401 12.01000 ; qtot
0.683
4773 O2 293 GLU OE1 4773 -0.818801 16.00000 ; qtot -
0.136
4774 O2 293 GLU OE2 4774 -0.818801 16.00000 ; qtot -
0.955
4775 C 293 GLU C 4775 0.536601 12.01000 ; qtot -
0.418
4776 O 293 GLU O 4776 -0.581901 16.00000 ; qtot -
1.000
4777 N 294 ASN N 4777 -0.415700 14.01000 ; qtot -

1.416
 4778 H 294 ASN H 4778 0.271900 1.00800 ; qtot -
1.144
 4779 CX 294 ASN CA 4779 0.014300 12.01000 ; qtot -
1.129
 4780 H1 294 ASN HA 4780 0.104800 1.00800 ; qtot -
1.025
 4781 A2C 294 ASN CB 4781 -0.204100 12.01000 ; qtot -
1.229
 4782 HC 294 ASN HB2 4782 0.079700 1.00800 ; qtot -
1.149
 4783 HC 294 ASN HB3 4783 0.079700 1.00800 ; qtot -
1.069
 4784 C 294 ASN CG 4784 0.713001 12.01000 ; qtot -
0.356
 4785 O 294 ASN OD1 4785 -0.593101 16.00000 ; qtot -
0.949
 4786 N 294 ASN ND2 4786 -0.919101 14.01000 ; qtot -
1.869
 4787 H 294 ASN HD21 4787 0.419600 1.00800 ; qtot -
1.449
 4788 H 294 ASN HD22 4788 0.419600 1.00800 ; qtot -
1.029
 4789 C 294 ASN C 4789 0.597301 12.01000 ; qtot -
0.432
 4790 O 294 ASN O 4790 -0.567901 16.00000 ; qtot -
1.000
 4791 N 295 GLU N 4791 -0.516301 14.01000 ; qtot -
1.516
 4792 H 295 GLU H 4792 0.293600 1.00800 ; qtot -
1.223
 4793 CX 295 GLU CA 4793 0.039700 12.01000 ; qtot -
1.183
 4794 H1 295 GLU HA 4794 0.110500 1.00800 ; qtot -
1.072
 4795 A2C 295 GLU CB 4795 0.056000 12.01000 ; qtot -
1.016
 4796 HC 295 GLU HB2 4796 -0.017300 1.00800 ; qtot -
1.034
 4797 HC 295 GLU HB3 4797 -0.017300 1.00800 ; qtot -
1.051
 4798 A2C 295 GLU CG 4798 0.013600 12.01000 ; qtot -
1.037
 4799 HC 295 GLU HG2 4799 -0.042500 1.00800 ; qtot -
1.080
 4800 HC 295 GLU HG3 4800 -0.042500 1.00800 ; qtot -
1.122
 4801 CO 295 GLU CD 4801 0.805401 12.01000 ; qtot -
0.317
 4802 O2 295 GLU OE1 4802 -0.818801 16.00000 ; qtot -
1.136
 4803 O2 295 GLU OE2 4803 -0.818801 16.00000 ; qtot -

1.955
  4804    C   295   GLU      C 4804     0.536601     12.01000 ; qtot -
1.418
  4805    O   295   GLU      O 4805    -0.581901     16.00000 ; qtot -
2.000
  4806    N   296   LEU      N 4806    -0.415700     14.01000 ; qtot -
2.416
  4807    H   296   LEU      H 4807     0.271900      1.00800 ; qtot -
2.144
  4808   CX   296   LEU     CA 4808    -0.051800     12.01000 ; qtot -
2.196
  4809   H1   296   LEU     HA 4809     0.092200      1.00800 ; qtot -
2.103
  4810  A2C   296   LEU     CB 4810    -0.110200     12.01000 ; qtot -
2.214
  4811   HC   296   LEU    HB2 4811     0.045700      1.00800 ; qtot -
2.168
  4812   HC   296   LEU    HB3 4812     0.045700      1.00800 ; qtot -
2.122
  4813  A3C   296   LEU     CG 4813     0.353100     12.01000 ; qtot -
1.769
  4814   HC   296   LEU     HG 4814    -0.036100      1.00800 ; qtot -
1.805
  4815   CT   296   LEU    CD1 4815    -0.412100     12.01000 ; qtot -
2.217
  4816   HC   296   LEU   HD11 4816     0.100000      1.00800 ; qtot -
2.117
  4817   HC   296   LEU   HD12 4817     0.100000      1.00800 ; qtot -
2.017
  4818   HC   296   LEU   HD13 4818     0.100000      1.00800 ; qtot -
1.917
  4819   CT   296   LEU    CD2 4819    -0.412100     12.01000 ; qtot -
2.329
  4820   HC   296   LEU   HD21 4820     0.100000      1.00800 ; qtot -
2.229
  4821   HC   296   LEU   HD22 4821     0.100000      1.00800 ; qtot -
2.129
  4822   HC   296   LEU   HD23 4822     0.100000      1.00800 ; qtot -
2.029
  4823    C   296   LEU      C 4823     0.597301     12.01000 ; qtot -
1.432
  4824    O   296   LEU      O 4824    -0.567901     16.00000 ; qtot -
2.000
  4825    N   297   LYS      N 4825    -0.347900     14.01000 ; qtot -
2.348
  4826    H   297   LYS      H 4826     0.274700      1.00800 ; qtot -
2.073
  4827   CX   297   LYS     CA 4827    -0.240000     12.01000 ; qtot -
2.313
  4828   H1   297   LYS     HA 4828     0.142600      1.00800 ; qtot -
2.171
  4829   C8   297   LYS     CB 4829    -0.009400     12.01000 ; qtot -

2.180
4830 HC 297 LYS HB2 4830 0.036200 1.00800 ; qtot -
2.144
4831 HC 297 LYS HB3 4831 0.036200 1.00800 ; qtot -
2.108
4832 C8 297 LYS CG 4832 0.018700 12.01000 ; qtot -
2.089
4833 HC 297 LYS HG2 4833 0.010300 1.00800 ; qtot -
2.079
4834 HC 297 LYS HG3 4834 0.010300 1.00800 ; qtot -
2.068
4835 C8 297 LYS CD 4835 -0.047900 12.01000 ; qtot -
2.116
4836 HC 297 LYS HD2 4836 0.062100 1.00800 ; qtot -
2.054
4837 HC 297 LYS HD3 4837 0.062100 1.00800 ; qtot -
1.992
4838 C8 297 LYS CE 4838 -0.014300 12.01000 ; qtot -
2.006
4839 HP 297 LYS HE2 4839 0.113500 1.00800 ; qtot -
1.893
4840 HP 297 LYS HE3 4840 0.113500 1.00800 ; qtot -
1.779
4841 N3 297 LYS NZ 4841 -0.385400 14.01000 ; qtot -
2.165
4842 H 297 LYS HZ1 4842 0.340000 1.00800 ; qtot -
1.825
4843 H 297 LYS HZ2 4843 0.340000 1.00800 ; qtot -
1.485
4844 H 297 LYS HZ3 4844 0.340000 1.00800 ; qtot -
1.145
4845 C 297 LYS C 4845 0.734101 12.01000 ; qtot -
0.411
4846 O 297 LYS O 4846 -0.589401 16.00000 ; qtot -
1.000
4847 N 298 LYS N 4847 -0.347900 14.01000 ; qtot -
1.348
4848 H 298 LYS H 4848 0.274700 1.00800 ; qtot -
1.073
4849 CX 298 LYS CA 4849 -0.240000 12.01000 ; qtot -
1.313
4850 H1 298 LYS HA 4850 0.142600 1.00800 ; qtot -
1.171
4851 C8 298 LYS CB 4851 -0.009400 12.01000 ; qtot -
1.180
4852 HC 298 LYS HB2 4852 0.036200 1.00800 ; qtot -
1.144
4853 HC 298 LYS HB3 4853 0.036200 1.00800 ; qtot -
1.108
4854 C8 298 LYS CG 4854 0.018700 12.01000 ; qtot -
1.089
4855 HC 298 LYS HG2 4855 0.010300 1.00800 ; qtot -

1.079
4856 HC 298 LYS HG3 4856 0.010300 1.00800 ; qtot -
1.068
4857 C8 298 LYS CD 4857 -0.047900 12.01000 ; qtot -
1.116
4858 HC 298 LYS HD2 4858 0.062100 1.00800 ; qtot -
1.054
4859 HC 298 LYS HD3 4859 0.062100 1.00800 ; qtot -
0.992
4860 C8 298 LYS CE 4860 -0.014300 12.01000 ; qtot -
1.006
4861 HP 298 LYS HE2 4861 0.113500 1.00800 ; qtot -
0.893
4862 HP 298 LYS HE3 4862 0.113500 1.00800 ; qtot -
0.779
4863 N3 298 LYS NZ 4863 -0.385400 14.01000 ; qtot -
1.165
4864 H 298 LYS HZ1 4864 0.340000 1.00800 ; qtot -
0.825
4865 H 298 LYS HZ2 4865 0.340000 1.00800 ; qtot -
0.485
4866 H 298 LYS HZ3 4866 0.340000 1.00800 ; qtot -
0.145
4867 C 298 LYS C 4867 0.734101 12.01000 ; qtot
0.589
4868 O 298 LYS O 4868 -0.589401 16.00000 ; qtot
0.000
4869 N 299 ILE N 4869 -0.415700 14.01000 ; qtot -
0.416
4870 H 299 ILE H 4870 0.271900 1.00800 ; qtot -
0.144
4871 CX 299 ILE CA 4871 -0.059700 12.01000 ; qtot -
0.203
4872 H1 299 ILE HA 4872 0.086900 1.00800 ; qtot -
0.117
4873 A3C 299 ILE CB 4873 0.130300 12.01000 ; qtot
0.014
4874 HC 299 ILE HB 4874 0.018700 1.00800 ; qtot
0.032
4875 CT 299 ILE CG2 4875 -0.320400 12.01000 ; qtot -
0.288
4876 HC 299 ILE HG21 4876 0.088200 1.00800 ; qtot -
0.200
4877 HC 299 ILE HG22 4877 0.088200 1.00800 ; qtot -
0.112
4878 HC 299 ILE HG23 4878 0.088200 1.00800 ; qtot -
0.023
4879 A2C 299 ILE CG1 4879 -0.043000 12.01000 ; qtot -
0.066
4880 HC 299 ILE HG12 4880 0.023600 1.00800 ; qtot -
0.043
4881 HC 299 ILE HG13 4881 0.023600 1.00800 ; qtot -

0.019
  4882   CT   299   ILE   CD1 4882    -0.066000     12.01000 ; qtot -
0.085
  4883   HC   299   ILE  HD11 4883     0.018600      1.00800 ; qtot -
0.067
  4884   HC   299   ILE  HD12 4884     0.018600      1.00800 ; qtot -
0.048
  4885   HC   299   ILE  HD13 4885     0.018600      1.00800 ; qtot -
0.029
  4886    C   299   ILE     C 4886     0.597301     12.01000 ; qtot
0.568
  4887    O   299   ILE     O 4887    -0.567901     16.00000 ; qtot
0.000
  4888    N   300   ALA     N 4888    -0.415700     14.01000 ; qtot -
0.416
  4889    H   300   ALA     H 4889     0.271900      1.00800 ; qtot -
0.144
  4890   CX   300   ALA    CA 4890     0.033700     12.01000 ; qtot -
0.110
  4891   H1   300   ALA    HA 4891     0.082300      1.00800 ; qtot -
0.028
  4892   CT   300   ALA    CB 4892    -0.182500     12.01000 ; qtot -
0.210
  4893   HC   300   ALA   HB1 4893     0.060300      1.00800 ; qtot -
0.150
  4894   HC   300   ALA   HB2 4894     0.060300      1.00800 ; qtot -
0.090
  4895   HC   300   ALA   HB3 4895     0.060300      1.00800 ; qtot -
0.029
  4896    C   300   ALA     C 4896     0.597301     12.01000 ; qtot
0.568
  4897    O   300   ALA     O 4897    -0.567901     16.00000 ; qtot
0.000
  4898    N   301   LEU     N 4898    -0.415700     14.01000 ; qtot -
0.416
  4899    H   301   LEU     H 4899     0.271900      1.00800 ; qtot -
0.144
  4900   CX   301   LEU    CA 4900    -0.051800     12.01000 ; qtot -
0.196
  4901   H1   301   LEU    HA 4901     0.092200      1.00800 ; qtot -
0.103
  4902  A2C   301   LEU    CB 4902    -0.110200     12.01000 ; qtot -
0.214
  4903   HC   301   LEU   HB2 4903     0.045700      1.00800 ; qtot -
0.168
  4904   HC   301   LEU   HB3 4904     0.045700      1.00800 ; qtot -
0.122
  4905  A3C   301   LEU    CG 4905     0.353100     12.01000 ; qtot
0.231
  4906   HC   301   LEU    HG 4906    -0.036100      1.00800 ; qtot
0.195
  4907   CT   301   LEU   CD1 4907    -0.412100     12.01000 ; qtot -

0.217
  4908   HC   301   LEU  HD11 4908     0.100000      1.00800 ; qtot -
0.117
  4909   HC   301   LEU  HD12 4909     0.100000      1.00800 ; qtot -
0.017
  4910   HC   301   LEU  HD13 4910     0.100000      1.00800 ; qtot
0.083
  4911   CT   301   LEU   CD2 4911    -0.412100     12.01000 ; qtot -
0.329
  4912   HC   301   LEU  HD21 4912     0.100000      1.00800 ; qtot -
0.229
  4913   HC   301   LEU  HD22 4913     0.100000      1.00800 ; qtot -
0.129
  4914   HC   301   LEU  HD23 4914     0.100000      1.00800 ; qtot -
0.029
  4915    C   301   LEU     C 4915     0.597301     12.01000 ; qtot
0.568
  4916    O   301   LEU     O 4916    -0.567901     16.00000 ; qtot
0.000
  4917    N   302   THR     N 4917    -0.415700     14.01000 ; qtot -
0.416
  4918    H   302   THR     H 4918     0.271900      1.00800 ; qtot -
0.144
  4919   CX   302   THR    CA 4919    -0.038900     12.01000 ; qtot -
0.183
  4920   H1   302   THR    HA 4920     0.100700      1.00800 ; qtot -
0.082
  4921  A3C   302   THR    CB 4921     0.365400     12.01000 ; qtot
0.283
  4922   H1   302   THR    HB 4922     0.004300      1.00800 ; qtot
0.288
  4923   CT   302   THR   CG2 4923    -0.243800     12.01000 ; qtot
0.044
  4924   HC   302   THR  HG21 4924     0.064200      1.00800 ; qtot
0.108
  4925   HC   302   THR  HG22 4925     0.064200      1.00800 ; qtot
0.172
  4926   HC   302   THR  HG23 4926     0.064200      1.00800 ; qtot
0.237
  4927   OH   302   THR   OG1 4927    -0.676101     16.00000 ; qtot -
0.440
  4928   HO   302   THR   HG1 4928     0.410200      1.00800 ; qtot -
0.029
  4929    C   302   THR     C 4929     0.597301     12.01000 ; qtot
0.568
  4930    O   302   THR     O 4930    -0.567901     16.00000 ; qtot
0.000
  4931    N   303   ILE     N 4931    -0.415700     14.01000 ; qtot -
0.416
  4932    H   303   ILE     H 4932     0.271900      1.00800 ; qtot -
0.144
  4933   CX   303   ILE    CA 4933    -0.059700     12.01000 ; qtot -

0.203
4934 H1 303 ILE HA 4934 0.086900 1.00800 ; qtot -
0.117
4935 A3C 303 ILE CB 4935 0.130300 12.01000 ; qtot
0.014
4936 HC 303 ILE HB 4936 0.018700 1.00800 ; qtot
0.032
4937 CT 303 ILE CG2 4937 -0.320400 12.01000 ; qtot -
0.288
4938 HC 303 ILE HG21 4938 0.088200 1.00800 ; qtot -
0.200
4939 HC 303 ILE HG22 4939 0.088200 1.00800 ; qtot -
0.112
4940 HC 303 ILE HG23 4940 0.088200 1.00800 ; qtot -
0.023
4941 A2C 303 ILE CG1 4941 -0.043000 12.01000 ; qtot -
0.066
4942 HC 303 ILE HG12 4942 0.023600 1.00800 ; qtot -
0.043
4943 HC 303 ILE HG13 4943 0.023600 1.00800 ; qtot -
0.019
4944 CT 303 ILE CD1 4944 -0.066000 12.01000 ; qtot -
0.085
4945 HC 303 ILE HD11 4945 0.018600 1.00800 ; qtot -
0.067
4946 HC 303 ILE HD12 4946 0.018600 1.00800 ; qtot -
0.048
4947 HC 303 ILE HD13 4947 0.018600 1.00800 ; qtot -
0.029
4948 C 303 ILE C 4948 0.597301 12.01000 ; qtot
0.568
4949 O 303 ILE O 4949 -0.567901 16.00000 ; qtot
0.000
4950 N 304 ILE N 4950 -0.415700 14.01000 ; qtot -
0.416
4951 H 304 ILE H 4951 0.271900 1.00800 ; qtot -
0.144
4952 CX 304 ILE CA 4952 -0.059700 12.01000 ; qtot -
0.203
4953 H1 304 ILE HA 4953 0.086900 1.00800 ; qtot -
0.117
4954 A3C 304 ILE CB 4954 0.130300 12.01000 ; qtot
0.014
4955 HC 304 ILE HB 4955 0.018700 1.00800 ; qtot
0.032
4956 CT 304 ILE CG2 4956 -0.320400 12.01000 ; qtot -
0.288
4957 HC 304 ILE HG21 4957 0.088200 1.00800 ; qtot -
0.200
4958 HC 304 ILE HG22 4958 0.088200 1.00800 ; qtot -
0.112
4959 HC 304 ILE HG23 4959 0.088200 1.00800 ; qtot -

0.023
4960 A2C 304 ILE CG1 4960 -0.043000 12.01000 ; qtot -
0.066
4961 HC 304 ILE HG12 4961 0.023600 1.00800 ; qtot -
0.043
4962 HC 304 ILE HG13 4962 0.023600 1.00800 ; qtot -
0.019
4963 CT 304 ILE CD1 4963 -0.066000 12.01000 ; qtot -
0.085
4964 HC 304 ILE HD11 4964 0.018600 1.00800 ; qtot -
0.067
4965 HC 304 ILE HD12 4965 0.018600 1.00800 ; qtot -
0.048
4966 HC 304 ILE HD13 4966 0.018600 1.00800 ; qtot -
0.029
4967 C 304 ILE C 4967 0.597301 12.01000 ; qtot
0.568
4968 O 304 ILE O 4968 -0.567901 16.00000 ; qtot
0.000
4969 N 305 ALA N 4969 -0.415700 14.01000 ; qtot -
0.416
4970 H 305 ALA H 4970 0.271900 1.00800 ; qtot -
0.144
4971 CX 305 ALA CA 4971 0.033700 12.01000 ; qtot -
0.110
4972 H1 305 ALA HA 4972 0.082300 1.00800 ; qtot -
0.028
4973 CT 305 ALA CB 4973 -0.182500 12.01000 ; qtot -
0.210
4974 HC 305 ALA HB1 4974 0.060300 1.00800 ; qtot -
0.150
4975 HC 305 ALA HB2 4975 0.060300 1.00800 ; qtot -
0.090
4976 HC 305 ALA HB3 4976 0.060300 1.00800 ; qtot -
0.029
4977 C 305 ALA C 4977 0.597301 12.01000 ; qtot
0.568
4978 O 305 ALA O 4978 -0.567901 16.00000 ; qtot
0.000
4979 N 306 LYS N 4979 -0.347900 14.01000 ; qtot -
0.348
4980 H 306 LYS H 4980 0.274700 1.00800 ; qtot -
0.073
4981 CX 306 LYS CA 4981 -0.240000 12.01000 ; qtot -
0.313
4982 H1 306 LYS HA 4982 0.142600 1.00800 ; qtot -
0.171
4983 C8 306 LYS CB 4983 -0.009400 12.01000 ; qtot -
0.180
4984 HC 306 LYS HB2 4984 0.036200 1.00800 ; qtot -
0.144
4985 HC 306 LYS HB3 4985 0.036200 1.00800 ; qtot -

0.108
  4986   C8   306   LYS    CG 4986     0.018700     12.01000 ; qtot -
0.089
  4987   HC   306   LYS   HG2 4987     0.010300      1.00800 ; qtot -
0.079
  4988   HC   306   LYS   HG3 4988     0.010300      1.00800 ; qtot -
0.068
  4989   C8   306   LYS    CD 4989    -0.047900     12.01000 ; qtot -
0.116
  4990   HC   306   LYS   HD2 4990     0.062100      1.00800 ; qtot -
0.054
  4991   HC   306   LYS   HD3 4991     0.062100      1.00800 ; qtot
0.008
  4992   C8   306   LYS    CE 4992    -0.014300     12.01000 ; qtot -
0.006
  4993   HP   306   LYS   HE2 4993     0.113500      1.00800 ; qtot
0.107
  4994   HP   306   LYS   HE3 4994     0.113500      1.00800 ; qtot
0.221
  4995   N3   306   LYS    NZ 4995    -0.385400     14.01000 ; qtot -
0.165
  4996    H   306   LYS   HZ1 4996     0.340000      1.00800 ; qtot
0.175
  4997    H   306   LYS   HZ2 4997     0.340000      1.00800 ; qtot
0.515
  4998    H   306   LYS   HZ3 4998     0.340000      1.00800 ; qtot
0.855
  4999    C   306   LYS     C 4999     0.734101     12.01000 ; qtot
1.589
  5000    O   306   LYS     O 5000    -0.589401     16.00000 ; qtot
1.000
  5001    N   307   HIE     N 5001    -0.415700     14.01000 ; qtot
0.584
  5002    H   307   HIE     H 5002     0.271900      1.00800 ; qtot
0.856
  5003   CX   307   HIE    CA 5003    -0.058100     12.01000 ; qtot
0.798
  5004   H1   307   HIE    HA 5004     0.136000      1.00800 ; qtot
0.934
  5005   CT   307   HIE    CB 5005    -0.007400     12.01000 ; qtot
0.927
  5006   HC   307   HIE   HB2 5006     0.036700      1.00800 ; qtot
0.963
  5007   HC   307   HIE   HB3 5007     0.036700      1.00800 ; qtot
1.000
  5008   CC   307   HIE    CG 5008     0.186800     12.01000 ; qtot
1.187
  5009   NB   307   HIE   ND1 5009    -0.543201     14.01000 ; qtot
0.644
  5010   CR   307   HIE   CE1 5010     0.163500     12.01000 ; qtot
0.807
  5011   H5   307   HIE   HE1 5011     0.143500      1.00800 ; qtot

0.951
  5012   NA   307   HIE   NE2 5012    -0.279500     14.01000 ; qtot
0.671
  5013    H   307   HIE   HE2 5013     0.333900      1.00800 ; qtot
1.005
  5014   CW   307   HIE   CD2 5014    -0.220700     12.01000 ; qtot
0.784
  5015   H4   307   HIE   HD2 5015     0.186200      1.00800 ; qtot
0.971
  5016    C   307   HIE     C 5016     0.597301     12.01000 ; qtot
1.568
  5017    O   307   HIE     O 5017    -0.567901     16.00000 ; qtot
1.000
  5018    N   308   LEU     N 5018    -0.415700     14.01000 ; qtot
0.584
  5019    H   308   LEU     H 5019     0.271900      1.00800 ; qtot
0.856
  5020   CX   308   LEU    CA 5020    -0.051800     12.01000 ; qtot
0.804
  5021   H1   308   LEU    HA 5021     0.092200      1.00800 ; qtot
0.897
  5022  A2C   308   LEU    CB 5022    -0.110200     12.01000 ; qtot
0.786
  5023   HC   308   LEU   HB2 5023     0.045700      1.00800 ; qtot
0.832
  5024   HC   308   LEU   HB3 5024     0.045700      1.00800 ; qtot
0.878
  5025  A3C   308   LEU    CG 5025     0.353100     12.01000 ; qtot
1.231
  5026   HC   308   LEU    HG 5026    -0.036100      1.00800 ; qtot
1.195
  5027   CT   308   LEU   CD1 5027    -0.412100     12.01000 ; qtot
0.783
  5028   HC   308   LEU  HD11 5028     0.100000      1.00800 ; qtot
0.883
  5029   HC   308   LEU  HD12 5029     0.100000      1.00800 ; qtot
0.983
  5030   HC   308   LEU  HD13 5030     0.100000      1.00800 ; qtot
1.083
  5031   CT   308   LEU   CD2 5031    -0.412100     12.01000 ; qtot
0.671
  5032   HC   308   LEU  HD21 5032     0.100000      1.00800 ; qtot
0.771
  5033   HC   308   LEU  HD22 5033     0.100000      1.00800 ; qtot
0.871
  5034   HC   308   LEU  HD23 5034     0.100000      1.00800 ; qtot
0.971
  5035    C   308   LEU     C 5035     0.597301     12.01000 ; qtot
1.568
  5036    O   308   LEU     O 5036    -0.567901     16.00000 ; qtot
1.000
  5037    N   309   CYS     N 5037    -0.415700     14.01000 ; qtot

0.584
  5038    H   309   CYS     H 5038     0.271900      1.00800 ; qtot
0.856
  5039   CX   309   CYS    CA 5039     0.021300     12.01000 ; qtot
0.878
  5040   H1   309   CYS    HA 5040     0.112400      1.00800 ; qtot
0.990
  5041  A2C   309   CYS    CB 5041    -0.123100     12.01000 ; qtot
0.867
  5042   H1   309   CYS   HB2 5042     0.111200      1.00800 ; qtot
0.978
  5043   H1   309   CYS   HB3 5043     0.111200      1.00800 ; qtot
1.089
  5044   SH   309   CYS    SG 5044    -0.311900     32.06000 ; qtot
0.777
  5045   HS   309   CYS    HG 5045     0.193300      1.00800 ; qtot
0.971
  5046    C   309   CYS     C 5046     0.597301     12.01000 ; qtot
1.568
  5047    O   309   CYS     O 5047    -0.567901     16.00000 ; qtot
1.000
  5048    N   310   ASP     N 5048    -0.516301     14.01000 ; qtot
0.484
  5049    H   310   ASP     H 5049     0.293600      1.00800 ; qtot
0.777
  5050   CX   310   ASP    CA 5050     0.038100     12.01000 ; qtot
0.815
  5051   H1   310   ASP    HA 5051     0.088000      1.00800 ; qtot
0.903
  5052  A2C   310   ASP    CB 5052    -0.030300     12.01000 ; qtot
0.873
  5053   HC   310   ASP   HB2 5053    -0.012200      1.00800 ; qtot
0.861
  5054   HC   310   ASP   HB3 5054    -0.012200      1.00800 ; qtot
0.849
  5055   CO   310   ASP    CG 5055     0.799401     12.01000 ; qtot
1.648
  5056   O2   310   ASP   OD1 5056    -0.801401     16.00000 ; qtot
0.847
  5057   O2   310   ASP   OD2 5057    -0.801401     16.00000 ; qtot
0.045
  5058    C   310   ASP     C 5058     0.536601     12.01000 ; qtot
0.582
  5059    O   310   ASP     O 5059    -0.581901     16.00000 ; qtot
0.000
  5060    N   311   VAL     N 5060    -0.415700     14.01000 ; qtot -
0.416
  5061    H   311   VAL     H 5061     0.271900      1.00800 ; qtot -
0.144
  5062   CX   311   VAL    CA 5062    -0.087500     12.01000 ; qtot -
0.231
  5063   H1   311   VAL    HA 5063     0.096900      1.00800 ; qtot -

0.134
  5064  A3C   311   VAL     CB 5064      0.298500     12.01000 ; qtot
0.164
  5065   HC   311   VAL     HB 5065    -0.029700      1.00800 ; qtot
0.134
  5066   CT   311   VAL    CG1 5066    -0.319200     12.01000 ; qtot -
0.185
  5067   HC   311   VAL   HG11 5067      0.079100      1.00800 ; qtot -
0.106
  5068   HC   311   VAL   HG12 5068      0.079100      1.00800 ; qtot -
0.027
  5069   HC   311   VAL   HG13 5069      0.079100      1.00800 ; qtot
0.053
  5070   CT   311   VAL    CG2 5070    -0.319200     12.01000 ; qtot -
0.267
  5071   HC   311   VAL   HG21 5071      0.079100      1.00800 ; qtot -
0.188
  5072   HC   311   VAL   HG22 5072      0.079100      1.00800 ; qtot -
0.108
  5073   HC   311   VAL   HG23 5073      0.079100      1.00800 ; qtot -
0.029
  5074    C   311   VAL      C 5074      0.597301     12.01000 ; qtot
0.568
  5075    O   311   VAL      O 5075    -0.567901     16.00000 ; qtot
0.000
  5076    N   312   GLU      N 5076    -0.516301     14.01000 ; qtot -
0.516
  5077    H   312   GLU      H 5077      0.293600      1.00800 ; qtot -
0.223
  5078   CX   312   GLU     CA 5078      0.039700     12.01000 ; qtot -
0.183
  5079   H1   312   GLU     HA 5079      0.110500      1.00800 ; qtot -
0.072
  5080  A2C   312   GLU     CB 5080      0.056000     12.01000 ; qtot -
0.016
  5081   HC   312   GLU    HB2 5081    -0.017300      1.00800 ; qtot -
0.034
  5082   HC   312   GLU    HB3 5082    -0.017300      1.00800 ; qtot -
0.051
  5083  A2C   312   GLU     CG 5083      0.013600     12.01000 ; qtot -
0.037
  5084   HC   312   GLU    HG2 5084    -0.042500      1.00800 ; qtot -
0.080
  5085   HC   312   GLU    HG3 5085    -0.042500      1.00800 ; qtot -
0.122
  5086   CO   312   GLU     CD 5086      0.805401     12.01000 ; qtot
0.683
  5087   O2   312   GLU    OE1 5087    -0.818801     16.00000 ; qtot -
0.136
  5088   O2   312   GLU    OE2 5088    -0.818801     16.00000 ; qtot -
0.955
  5089    C   312   GLU      C 5089      0.536601     12.01000 ; qtot -

0.418
  5090    O   312   GLU      O 5090    -0.581901     16.00000 ; qtot -1.000
  5091    N   313   ILE      N 5091    -0.415700     14.01000 ; qtot -1.416
  5092    H   313   ILE      H 5092     0.271900      1.00800 ; qtot -1.144
  5093   CX   313   ILE     CA 5093    -0.059700     12.01000 ; qtot -1.203
  5094   H1   313   ILE     HA 5094     0.086900      1.00800 ; qtot -1.117
  5095  A3C   313   ILE     CB 5095     0.130300     12.01000 ; qtot -0.986
  5096   HC   313   ILE     HB 5096     0.018700      1.00800 ; qtot -0.968
  5097   CT   313   ILE    CG2 5097    -0.320400     12.01000 ; qtot -1.288
  5098   HC   313   ILE   HG21 5098     0.088200      1.00800 ; qtot -1.200
  5099   HC   313   ILE   HG22 5099     0.088200      1.00800 ; qtot -1.112
  5100   HC   313   ILE   HG23 5100     0.088200      1.00800 ; qtot -1.023
  5101  A2C   313   ILE    CG1 5101    -0.043000     12.01000 ; qtot -1.066
  5102   HC   313   ILE   HG12 5102     0.023600      1.00800 ; qtot -1.043
  5103   HC   313   ILE   HG13 5103     0.023600      1.00800 ; qtot -1.019
  5104   CT   313   ILE    CD1 5104    -0.066000     12.01000 ; qtot -1.085
  5105   HC   313   ILE   HD11 5105     0.018600      1.00800 ; qtot -1.067
  5106   HC   313   ILE   HD12 5106     0.018600      1.00800 ; qtot -1.048
  5107   HC   313   ILE   HD13 5107     0.018600      1.00800 ; qtot -1.029
  5108    C   313   ILE      C 5108     0.597301     12.01000 ; qtot -0.432
  5109    O   313   ILE      O 5109    -0.567901     16.00000 ; qtot -1.000
  5110    N   314   ASN      N 5110    -0.415700     14.01000 ; qtot -1.416
  5111    H   314   ASN      H 5111     0.271900      1.00800 ; qtot -1.144
  5112   CX   314   ASN     CA 5112     0.014300     12.01000 ; qtot -1.129
  5113   H1   314   ASN     HA 5113     0.104800      1.00800 ; qtot -1.025
  5114  A2C   314   ASN     CB 5114    -0.204100     12.01000 ; qtot -1.229
  5115   HC   314   ASN    HB2 5115     0.079700      1.00800 ; qtot -

```
1.149
  5116   HC   314   ASN   HB3 5116     0.079700      1.00800 ; qtot -
1.069
  5117    C   314   ASN    CG 5117     0.713001     12.01000 ; qtot -
0.356
  5118    O   314   ASN   OD1 5118    -0.593101     16.00000 ; qtot -
0.949
  5119    N   314   ASN   ND2 5119    -0.919101     14.01000 ; qtot -
1.869
  5120    H   314   ASN  HD21 5120     0.419600      1.00800 ; qtot -
1.449
  5121    H   314   ASN  HD22 5121     0.419600      1.00800 ; qtot -
1.029
  5122    C   314   ASN     C 5122     0.597301     12.01000 ; qtot -
0.432
  5123    O   314   ASN     O 5123    -0.567901     16.00000 ; qtot -
1.000
  5124    N   315   ASN     N 5124    -0.415700     14.01000 ; qtot -
1.416
  5125    H   315   ASN     H 5125     0.271900      1.00800 ; qtot -
1.144
  5126   CX   315   ASN    CA 5126     0.014300     12.01000 ; qtot -
1.129
  5127   H1   315   ASN    HA 5127     0.104800      1.00800 ; qtot -
1.025
  5128  A2C   315   ASN    CB 5128    -0.204100     12.01000 ; qtot -
1.229
  5129   HC   315   ASN   HB2 5129     0.079700      1.00800 ; qtot -
1.149
  5130   HC   315   ASN   HB3 5130     0.079700      1.00800 ; qtot -
1.069
  5131    C   315   ASN    CG 5131     0.713001     12.01000 ; qtot -
0.356
  5132    O   315   ASN   OD1 5132    -0.593101     16.00000 ; qtot -
0.949
  5133    N   315   ASN   ND2 5133    -0.919101     14.01000 ; qtot -
1.869
  5134    H   315   ASN  HD21 5134     0.419600      1.00800 ; qtot -
1.449
  5135    H   315   ASN  HD22 5135     0.419600      1.00800 ; qtot -
1.029
  5136    C   315   ASN     C 5136     0.597301     12.01000 ; qtot -
0.432
  5137    O   315   ASN     O 5137    -0.567901     16.00000 ; qtot -
1.000
  5138    N   316   LEU     N 5138    -0.415700     14.01000 ; qtot -
1.416
  5139    H   316   LEU     H 5139     0.271900      1.00800 ; qtot -
1.144
  5140   CX   316   LEU    CA 5140    -0.051800     12.01000 ; qtot -
1.196
  5141   H1   316   LEU    HA 5141     0.092200      1.00800 ; qtot -
```

```
1.103
  5142  A2C   316   LEU    CB 5142    -0.110200     12.01000 ; qtot -
1.214
  5143   HC   316   LEU   HB2 5143     0.045700      1.00800 ; qtot -
1.168
  5144   HC   316   LEU   HB3 5144     0.045700      1.00800 ; qtot -
1.122
  5145  A3C   316   LEU    CG 5145     0.353100     12.01000 ; qtot -
0.769
  5146   HC   316   LEU    HG 5146    -0.036100      1.00800 ; qtot -
0.805
  5147   CT   316   LEU   CD1 5147    -0.412100     12.01000 ; qtot -
1.217
  5148   HC   316   LEU  HD11 5148     0.100000      1.00800 ; qtot -
1.117
  5149   HC   316   LEU  HD12 5149     0.100000      1.00800 ; qtot -
1.017
  5150   HC   316   LEU  HD13 5150     0.100000      1.00800 ; qtot -
0.917
  5151   CT   316   LEU   CD2 5151    -0.412100     12.01000 ; qtot -
1.329
  5152   HC   316   LEU  HD21 5152     0.100000      1.00800 ; qtot -
1.229
  5153   HC   316   LEU  HD22 5153     0.100000      1.00800 ; qtot -
1.129
  5154   HC   316   LEU  HD23 5154     0.100000      1.00800 ; qtot -
1.029
  5155    C   316   LEU     C 5155     0.597301     12.01000 ; qtot -
0.432
  5156    O   316   LEU     O 5156    -0.567901     16.00000 ; qtot -
1.000
  5157    N   317   ARG     N 5157    -0.347900     14.01000 ; qtot -
1.348
  5158    H   317   ARG     H 5158     0.274700      1.00800 ; qtot -
1.073
  5159   CX   317   ARG    CA 5159    -0.263700     12.01000 ; qtot -
1.337
  5160   H1   317   ARG    HA 5160     0.156000      1.00800 ; qtot -
1.181
  5161   C8   317   ARG    CB 5161    -0.000700     12.01000 ; qtot -
1.182
  5162   HC   317   ARG   HB2 5162     0.032700      1.00800 ; qtot -
1.149
  5163   HC   317   ARG   HB3 5163     0.032700      1.00800 ; qtot -
1.116
  5164   C8   317   ARG    CG 5164     0.039000     12.01000 ; qtot -
1.077
  5165   HC   317   ARG   HG2 5165     0.028500      1.00800 ; qtot -
1.049
  5166   HC   317   ARG   HG3 5166     0.028500      1.00800 ; qtot -
1.020
  5167   C8   317   ARG    CD 5167     0.048600     12.01000 ; qtot -
```

0.972
5168 H1 317 ARG HD2 5168 0.068700 1.00800 ; qtot -0.903
5169 H1 317 ARG HD3 5169 0.068700 1.00800 ; qtot -0.834
5170 N2 317 ARG NE 5170 -0.529501 14.01000 ; qtot -1.364
5171 H 317 ARG HE 5171 0.345600 1.00800 ; qtot -1.018
5172 CA 317 ARG CZ 5172 0.807601 12.01000 ; qtot -0.210
5173 N2 317 ARG NH1 5173 -0.862701 14.01000 ; qtot -1.073
5174 H 317 ARG HH11 5174 0.447800 1.00800 ; qtot -0.625
5175 H 317 ARG HH12 5175 0.447800 1.00800 ; qtot -0.178
5176 N2 317 ARG NH2 5176 -0.862701 14.01000 ; qtot -1.040
5177 H 317 ARG HH21 5177 0.447800 1.00800 ; qtot -0.592
5178 H 317 ARG HH22 5178 0.447800 1.00800 ; qtot -0.145
5179 C 317 ARG C 5179 0.734101 12.01000 ; qtot 0.589
5180 O 317 ARG O 5180 -0.589401 16.00000 ; qtot 0.000
5181 N 318 ASN N 5181 -0.415700 14.01000 ; qtot -0.416
5182 H 318 ASN H 5182 0.271900 1.00800 ; qtot -0.144
5183 CX 318 ASN CA 5183 0.014300 12.01000 ; qtot -0.129
5184 H1 318 ASN HA 5184 0.104800 1.00800 ; qtot -0.025
5185 A2C 318 ASN CB 5185 -0.204100 12.01000 ; qtot -0.229
5186 HC 318 ASN HB2 5186 0.079700 1.00800 ; qtot -0.149
5187 HC 318 ASN HB3 5187 0.079700 1.00800 ; qtot -0.069
5188 C 318 ASN CG 5188 0.713001 12.01000 ; qtot 0.644
5189 O 318 ASN OD1 5189 -0.593101 16.00000 ; qtot 0.051
5190 N 318 ASN ND2 5190 -0.919101 14.01000 ; qtot -0.869
5191 H 318 ASN HD21 5191 0.419600 1.00800 ; qtot -0.449
5192 H 318 ASN HD22 5192 0.419600 1.00800 ; qtot -0.029
5193 C 318 ASN C 5193 0.597301 12.01000 ; qtot

0.568
  5194    O   318   ASN      O 5194    -0.567901     16.00000 ; qtot
0.000
  5195    N   319   ILE      N 5195    -0.415700     14.01000 ; qtot -
0.416
  5196    H   319   ILE      H 5196     0.271900      1.00800 ; qtot -
0.144
  5197   CX   319   ILE     CA 5197    -0.059700     12.01000 ; qtot -
0.203
  5198   H1   319   ILE     HA 5198     0.086900      1.00800 ; qtot -
0.117
  5199  A3C   319   ILE     CB 5199     0.130300     12.01000 ; qtot
0.014
  5200   HC   319   ILE     HB 5200     0.018700      1.00800 ; qtot
0.032
  5201   CT   319   ILE    CG2 5201    -0.320400     12.01000 ; qtot -
0.288
  5202   HC   319   ILE   HG21 5202     0.088200      1.00800 ; qtot -
0.200
  5203   HC   319   ILE   HG22 5203     0.088200      1.00800 ; qtot -
0.112
  5204   HC   319   ILE   HG23 5204     0.088200      1.00800 ; qtot -
0.023
  5205  A2C   319   ILE    CG1 5205    -0.043000     12.01000 ; qtot -
0.066
  5206   HC   319   ILE   HG12 5206     0.023600      1.00800 ; qtot -
0.043
  5207   HC   319   ILE   HG13 5207     0.023600      1.00800 ; qtot -
0.019
  5208   CT   319   ILE    CD1 5208    -0.066000     12.01000 ; qtot -
0.085
  5209   HC   319   ILE   HD11 5209     0.018600      1.00800 ; qtot -
0.067
  5210   HC   319   ILE   HD12 5210     0.018600      1.00800 ; qtot -
0.048
  5211   HC   319   ILE   HD13 5211     0.018600      1.00800 ; qtot -
0.029
  5212    C   319   ILE      C 5212     0.597301     12.01000 ; qtot
0.568
  5213    O   319   ILE      O 5213    -0.567901     16.00000 ; qtot
0.000
  5214    N   320   PHE      N 5214    -0.415700     14.01000 ; qtot -
0.416
  5215    H   320   PHE      H 5215     0.271900      1.00800 ; qtot -
0.144
  5216   CX   320   PHE     CA 5216    -0.002400     12.01000 ; qtot -
0.146
  5217   H1   320   PHE     HA 5217     0.097800      1.00800 ; qtot -
0.048
  5218   CT   320   PHE     CB 5218    -0.034300     12.01000 ; qtot -
0.083
  5219   HC   320   PHE    HB2 5219     0.029500      1.00800 ; qtot -

0.053
  5220   HC   320   PHE   HB3 5220     0.029500      1.00800 ; qtot -
0.024
  5221   CA   320   PHE    CG 5221     0.011800     12.01000 ; qtot -
0.012
  5222   CA   320   PHE   CD1 5222    -0.125600     12.01000 ; qtot -
0.137
  5223   HA   320   PHE   HD1 5223     0.133000      1.00800 ; qtot -
0.004
  5224   CA   320   PHE   CE1 5224    -0.170400     12.01000 ; qtot -
0.175
  5225   HA   320   PHE   HE1 5225     0.143000      1.00800 ; qtot -
0.032
  5226   CA   320   PHE    CZ 5226    -0.107200     12.01000 ; qtot -
0.139
  5227   HA   320   PHE    HZ 5227     0.129700      1.00800 ; qtot -
0.009
  5228   CA   320   PHE   CE2 5228    -0.170400     12.01000 ; qtot -
0.180
  5229   HA   320   PHE   HE2 5229     0.143000      1.00800 ; qtot -
0.037
  5230   CA   320   PHE   CD2 5230    -0.125600     12.01000 ; qtot -
0.162
  5231   HA   320   PHE   HD2 5231     0.133000      1.00800 ; qtot -
0.029
  5232    C   320   PHE     C 5232     0.597301     12.01000 ; qtot
0.568
  5233    O   320   PHE     O 5233    -0.567901     16.00000 ; qtot
0.000
  5234    N   321   ILE     N 5234    -0.415700     14.01000 ; qtot -
0.416
  5235    H   321   ILE     H 5235     0.271900      1.00800 ; qtot -
0.144
  5236   CX   321   ILE    CA 5236    -0.059700     12.01000 ; qtot -
0.203
  5237   H1   321   ILE    HA 5237     0.086900      1.00800 ; qtot -
0.117
  5238  A3C   321   ILE    CB 5238     0.130300     12.01000 ; qtot
0.014
  5239   HC   321   ILE    HB 5239     0.018700      1.00800 ; qtot
0.032
  5240   CT   321   ILE   CG2 5240    -0.320400     12.01000 ; qtot -
0.288
  5241   HC   321   ILE  HG21 5241     0.088200      1.00800 ; qtot -
0.200
  5242   HC   321   ILE  HG22 5242     0.088200      1.00800 ; qtot -
0.112
  5243   HC   321   ILE  HG23 5243     0.088200      1.00800 ; qtot -
0.023
  5244  A2C   321   ILE   CG1 5244    -0.043000     12.01000 ; qtot -
0.066
  5245   HC   321   ILE  HG12 5245     0.023600      1.00800 ; qtot -

0.043
5246 HC 321 ILE HG13 5246 0.023600 1.00800 ; qtot -
0.019
5247 CT 321 ILE CD1 5247 -0.066000 12.01000 ; qtot -
0.085
5248 HC 321 ILE HD11 5248 0.018600 1.00800 ; qtot -
0.067
5249 HC 321 ILE HD12 5249 0.018600 1.00800 ; qtot -
0.048
5250 HC 321 ILE HD13 5250 0.018600 1.00800 ; qtot -
0.029
5251 C 321 ILE C 5251 0.597301 12.01000 ; qtot
0.568
5252 O 321 ILE O 5252 -0.567901 16.00000 ; qtot
0.000
5253 N 322 ALA N 5253 -0.415700 14.01000 ; qtot -
0.416
5254 H 322 ALA H 5254 0.271900 1.00800 ; qtot -
0.144
5255 CX 322 ALA CA 5255 0.033700 12.01000 ; qtot -
0.110
5256 H1 322 ALA HA 5256 0.082300 1.00800 ; qtot -
0.028
5257 CT 322 ALA CB 5257 -0.182500 12.01000 ; qtot -
0.210
5258 HC 322 ALA HB1 5258 0.060300 1.00800 ; qtot -
0.150
5259 HC 322 ALA HB2 5259 0.060300 1.00800 ; qtot -
0.090
5260 HC 322 ALA HB3 5260 0.060300 1.00800 ; qtot -
0.029
5261 C 322 ALA C 5261 0.597301 12.01000 ; qtot
0.568
5262 O 322 ALA O 5262 -0.567901 16.00000 ; qtot
0.000
5263 N 323 LEU N 5263 -0.415700 14.01000 ; qtot -
0.416
5264 H 323 LEU H 5264 0.271900 1.00800 ; qtot -
0.144
5265 CX 323 LEU CA 5265 -0.051800 12.01000 ; qtot -
0.196
5266 H1 323 LEU HA 5266 0.092200 1.00800 ; qtot -
0.103
5267 A2C 323 LEU CB 5267 -0.110200 12.01000 ; qtot -
0.214
5268 HC 323 LEU HB2 5268 0.045700 1.00800 ; qtot -
0.168
5269 HC 323 LEU HB3 5269 0.045700 1.00800 ; qtot -
0.122
5270 A3C 323 LEU CG 5270 0.353100 12.01000 ; qtot
0.231
5271 HC 323 LEU HG 5271 -0.036100 1.00800 ; qtot

0.195
5272 CT 323 LEU CD1 5272 -0.412100 12.01000 ; qtot -
0.217
5273 HC 323 LEU HD11 5273 0.100000 1.00800 ; qtot -
0.117
5274 HC 323 LEU HD12 5274 0.100000 1.00800 ; qtot -
0.017
5275 HC 323 LEU HD13 5275 0.100000 1.00800 ; qtot
0.083
5276 CT 323 LEU CD2 5276 -0.412100 12.01000 ; qtot -
0.329
5277 HC 323 LEU HD21 5277 0.100000 1.00800 ; qtot -
0.229
5278 HC 323 LEU HD22 5278 0.100000 1.00800 ; qtot -
0.129
5279 HC 323 LEU HD23 5279 0.100000 1.00800 ; qtot -
0.029
5280 C 323 LEU C 5280 0.597301 12.01000 ; qtot
0.568
5281 O 323 LEU O 5281 -0.567901 16.00000 ; qtot
0.000
5282 N 324 ASP N 5282 -0.516301 14.01000 ; qtot -
0.516
5283 H 324 ASP H 5283 0.293600 1.00800 ; qtot -
0.223
5284 CX 324 ASP CA 5284 0.038100 12.01000 ; qtot -
0.185
5285 H1 324 ASP HA 5285 0.088000 1.00800 ; qtot -
0.097
5286 A2C 324 ASP CB 5286 -0.030300 12.01000 ; qtot -
0.127
5287 HC 324 ASP HB2 5287 -0.012200 1.00800 ; qtot -
0.139
5288 HC 324 ASP HB3 5288 -0.012200 1.00800 ; qtot -
0.151
5289 CO 324 ASP CG 5289 0.799401 12.01000 ; qtot
0.648
5290 O2 324 ASP OD1 5290 -0.801401 16.00000 ; qtot -
0.153
5291 O2 324 ASP OD2 5291 -0.801401 16.00000 ; qtot -
0.955
5292 C 324 ASP C 5292 0.536601 12.01000 ; qtot -
0.418
5293 O 324 ASP O 5293 -0.581901 16.00000 ; qtot -
1.000
5294 N 325 VAL N 5294 -0.415700 14.01000 ; qtot -
1.416
5295 H 325 VAL H 5295 0.271900 1.00800 ; qtot -
1.144
5296 CX 325 VAL CA 5296 -0.087500 12.01000 ; qtot -
1.231
5297 H1 325 VAL HA 5297 0.096900 1.00800 ; qtot -

1.134
  5298  A3C   325   VAL     CB 5298      0.298500     12.01000 ; qtot -
0.836
  5299   HC   325   VAL     HB 5299     -0.029700      1.00800 ; qtot -
0.866
  5300   CT   325   VAL    CG1 5300     -0.319200     12.01000 ; qtot -
1.185
  5301   HC   325   VAL   HG11 5301      0.079100      1.00800 ; qtot -
1.106
  5302   HC   325   VAL   HG12 5302      0.079100      1.00800 ; qtot -
1.027
  5303   HC   325   VAL   HG13 5303      0.079100      1.00800 ; qtot -
0.947
  5304   CT   325   VAL    CG2 5304     -0.319200     12.01000 ; qtot -
1.267
  5305   HC   325   VAL   HG21 5305      0.079100      1.00800 ; qtot -
1.188
  5306   HC   325   VAL   HG22 5306      0.079100      1.00800 ; qtot -
1.108
  5307   HC   325   VAL   HG23 5307      0.079100      1.00800 ; qtot -
1.029
  5308    C   325   VAL      C 5308      0.597301     12.01000 ; qtot -
0.432
  5309    O   325   VAL      O 5309     -0.567901     16.00000 ; qtot -
1.000
  5310    N   326   ASP      N 5310     -0.516301     14.01000 ; qtot -
1.516
  5311    H   326   ASP      H 5311      0.293600      1.00800 ; qtot -
1.223
  5312   CX   326   ASP     CA 5312      0.038100     12.01000 ; qtot -
1.185
  5313   H1   326   ASP     HA 5313      0.088000      1.00800 ; qtot -
1.097
  5314  A2C   326   ASP     CB 5314     -0.030300     12.01000 ; qtot -
1.127
  5315   HC   326   ASP    HB2 5315     -0.012200      1.00800 ; qtot -
1.139
  5316   HC   326   ASP    HB3 5316     -0.012200      1.00800 ; qtot -
1.151
  5317   CO   326   ASP     CG 5317      0.799401     12.01000 ; qtot -
0.352
  5318   O2   326   ASP    OD1 5318     -0.801401     16.00000 ; qtot -
1.153
  5319   O2   326   ASP    OD2 5319     -0.801401     16.00000 ; qtot -
1.955
  5320    C   326   ASP      C 5320      0.536601     12.01000 ; qtot -
1.418
  5321    O   326   ASP      O 5321     -0.581901     16.00000 ; qtot -
2.000
  5322    N   327   ASN      N 5322     -0.415700     14.01000 ; qtot -
2.416
  5323    H   327   ASN      H 5323      0.271900      1.00800 ; qtot -

2.144
5324 CX 327 ASN CA 5324 0.014300 12.01000 ; qtot -
2.129
5325 H1 327 ASN HA 5325 0.104800 1.00800 ; qtot -
2.025
5326 A2C 327 ASN CB 5326 -0.204100 12.01000 ; qtot -
2.229
5327 HC 327 ASN HB2 5327 0.079700 1.00800 ; qtot -
2.149
5328 HC 327 ASN HB3 5328 0.079700 1.00800 ; qtot -
2.069
5329 C 327 ASN CG 5329 0.713001 12.01000 ; qtot -
1.356
5330 O 327 ASN OD1 5330 -0.593101 16.00000 ; qtot -
1.949
5331 N 327 ASN ND2 5331 -0.919101 14.01000 ; qtot -
2.869
5332 H 327 ASN HD21 5332 0.419600 1.00800 ; qtot -
2.449
5333 H 327 ASN HD22 5333 0.419600 1.00800 ; qtot -
2.029
5334 C 327 ASN C 5334 0.597301 12.01000 ; qtot -
1.432
5335 O 327 ASN O 5335 -0.567901 16.00000 ; qtot -
2.000
5336 N 328 SER N 5336 -0.415700 14.01000 ; qtot -
2.416
5337 H 328 SER H 5337 0.271900 1.00800 ; qtot -
2.144
5338 CX 328 SER CA 5338 -0.024900 12.01000 ; qtot -
2.169
5339 H1 328 SER HA 5339 0.084300 1.00800 ; qtot -
2.084
5340 A2C 328 SER CB 5340 0.211700 12.01000 ; qtot -
1.873
5341 H1 328 SER HB2 5341 0.035200 1.00800 ; qtot -
1.837
5342 H1 328 SER HB3 5342 0.035200 1.00800 ; qtot -
1.802
5343 OH 328 SER OG 5343 -0.654601 16.00000 ; qtot -
2.457
5344 HO 328 SER HG 5344 0.427500 1.00800 ; qtot -
2.029
5345 C 328 SER C 5345 0.597301 12.01000 ; qtot -
1.432
5346 O 328 SER O 5346 -0.567901 16.00000 ; qtot -
2.000
5347 N 329 GLY N 5347 -0.415700 14.01000 ; qtot -
2.416
5348 H 329 GLY H 5348 0.271900 1.00800 ; qtot -
2.144
5349 CX 329 GLY CA 5349 -0.025200 12.01000 ; qtot -

2.169
  5350   H1   329   GLY   HA2 5350     0.069800      1.00800 ; qtot -
2.099
  5351   H1   329   GLY   HA3 5351     0.069800      1.00800 ; qtot -
2.029
  5352    C   329   GLY     C 5352     0.597301     12.01000 ; qtot -
1.432
  5353    O   329   GLY     O 5353    -0.567901     16.00000 ; qtot -
2.000
  5354    N   330   THR     N 5354    -0.415700     14.01000 ; qtot -
2.416
  5355    H   330   THR     H 5355     0.271900      1.00800 ; qtot -
2.144
  5356   CX   330   THR    CA 5356    -0.038900     12.01000 ; qtot -
2.183
  5357   H1   330   THR    HA 5357     0.100700      1.00800 ; qtot -
2.082
  5358  A3C   330   THR    CB 5358     0.365400     12.01000 ; qtot -
1.717
  5359   H1   330   THR    HB 5359     0.004300      1.00800 ; qtot -
1.712
  5360   CT   330   THR   CG2 5360    -0.243800     12.01000 ; qtot -
1.956
  5361   HC   330   THR  HG21 5361     0.064200      1.00800 ; qtot -
1.892
  5362   HC   330   THR  HG22 5362     0.064200      1.00800 ; qtot -
1.828
  5363   HC   330   THR  HG23 5363     0.064200      1.00800 ; qtot -
1.763
  5364   OH   330   THR   OG1 5364    -0.676101     16.00000 ; qtot -
2.440
  5365   HO   330   THR   HG1 5365     0.410200      1.00800 ; qtot -
2.029
  5366    C   330   THR     C 5366     0.597301     12.01000 ; qtot -
1.432
  5367    O   330   THR     O 5367    -0.567901     16.00000 ; qtot -
2.000
  5368    N   331   LEU     N 5368    -0.415700     14.01000 ; qtot -
2.416
  5369    H   331   LEU     H 5369     0.271900      1.00800 ; qtot -
2.144
  5370   CX   331   LEU    CA 5370    -0.051800     12.01000 ; qtot -
2.196
  5371   H1   331   LEU    HA 5371     0.092200      1.00800 ; qtot -
2.103
  5372  A2C   331   LEU    CB 5372    -0.110200     12.01000 ; qtot -
2.214
  5373   HC   331   LEU   HB2 5373     0.045700      1.00800 ; qtot -
2.168
  5374   HC   331   LEU   HB3 5374     0.045700      1.00800 ; qtot -
2.122
  5375  A3C   331   LEU    CG 5375     0.353100     12.01000 ; qtot -

1.769
5376 HC 331 LEU HG 5376 -0.036100 1.00800 ; qtot -
1.805
5377 CT 331 LEU CD1 5377 -0.412100 12.01000 ; qtot -
2.217
5378 HC 331 LEU HD11 5378 0.100000 1.00800 ; qtot -
2.117
5379 HC 331 LEU HD12 5379 0.100000 1.00800 ; qtot -
2.017
5380 HC 331 LEU HD13 5380 0.100000 1.00800 ; qtot -
1.917
5381 CT 331 LEU CD2 5381 -0.412100 12.01000 ; qtot -
2.329
5382 HC 331 LEU HD21 5382 0.100000 1.00800 ; qtot -
2.229
5383 HC 331 LEU HD22 5383 0.100000 1.00800 ; qtot -
2.129
5384 HC 331 LEU HD23 5384 0.100000 1.00800 ; qtot -
2.029
5385 C 331 LEU C 5385 0.597301 12.01000 ; qtot -
1.432
5386 O 331 LEU O 5386 -0.567901 16.00000 ; qtot -
2.000
5387 N 332 SER N 5387 -0.415700 14.01000 ; qtot -
2.416
5388 H 332 SER H 5388 0.271900 1.00800 ; qtot -
2.144
5389 CX 332 SER CA 5389 -0.024900 12.01000 ; qtot -
2.169
5390 H1 332 SER HA 5390 0.084300 1.00800 ; qtot -
2.084
5391 A2C 332 SER CB 5391 0.211700 12.01000 ; qtot -
1.873
5392 H1 332 SER HB2 5392 0.035200 1.00800 ; qtot -
1.837
5393 H1 332 SER HB3 5393 0.035200 1.00800 ; qtot -
1.802
5394 OH 332 SER OG 5394 -0.654601 16.00000 ; qtot -
2.457
5395 HO 332 SER HG 5395 0.427500 1.00800 ; qtot -
2.029
5396 C 332 SER C 5396 0.597301 12.01000 ; qtot -
1.432
5397 O 332 SER O 5397 -0.567901 16.00000 ; qtot -
2.000
5398 N 333 SER N 5398 -0.415700 14.01000 ; qtot -
2.416
5399 H 333 SER H 5399 0.271900 1.00800 ; qtot -
2.144
5400 CX 333 SER CA 5400 -0.024900 12.01000 ; qtot -
2.169
5401 H1 333 SER HA 5401 0.084300 1.00800 ; qtot -

2.084
 5402 A2C 333 SER CB 5402 0.211700 12.01000 ; qtot -
1.873
 5403 H1 333 SER HB2 5403 0.035200 1.00800 ; qtot -
1.837
 5404 H1 333 SER HB3 5404 0.035200 1.00800 ; qtot -
1.802
 5405 OH 333 SER OG 5405 -0.654601 16.00000 ; qtot -
2.457
 5406 HO 333 SER HG 5406 0.427500 1.00800 ; qtot -
2.029
 5407 C 333 SER C 5407 0.597301 12.01000 ; qtot -
1.432
 5408 O 333 SER O 5408 -0.567901 16.00000 ; qtot -
2.000
 5409 N 334 GLN N 5409 -0.415700 14.01000 ; qtot -
2.416
 5410 H 334 GLN H 5410 0.271900 1.00800 ; qtot -
2.144
 5411 CX 334 GLN CA 5411 -0.003100 12.01000 ; qtot -
2.147
 5412 H1 334 GLN HA 5412 0.085000 1.00800 ; qtot -
2.062
 5413 A2C 334 GLN CB 5413 -0.003600 12.01000 ; qtot -
2.065
 5414 HC 334 GLN HB2 5414 0.017100 1.00800 ; qtot -
2.048
 5415 HC 334 GLN HB3 5415 0.017100 1.00800 ; qtot -
2.031
 5416 A2C 334 GLN CG 5416 -0.064500 12.01000 ; qtot -
2.096
 5417 HC 334 GLN HG2 5417 0.035200 1.00800 ; qtot -
2.061
 5418 HC 334 GLN HG3 5418 0.035200 1.00800 ; qtot -
2.025
 5419 C 334 GLN CD 5419 0.695101 12.01000 ; qtot -
1.330
 5420 O 334 GLN OE1 5420 -0.608601 16.00000 ; qtot -
1.939
 5421 N 334 GLN NE2 5421 -0.940701 14.01000 ; qtot -
2.880
 5422 H 334 GLN HE21 5422 0.425100 1.00800 ; qtot -
2.454
 5423 H 334 GLN HE22 5423 0.425100 1.00800 ; qtot -
2.029
 5424 C 334 GLN C 5424 0.597301 12.01000 ; qtot -
1.432
 5425 O 334 GLN O 5425 -0.567901 16.00000 ; qtot -
2.000
 5426 N 335 GLH N 5426 -0.415700 14.01000 ; qtot -
2.416
 5427 H 335 GLH H 5427 0.271900 1.00800 ; qtot -

2.144
5428 CX 335 GLH CA 5428 0.014500 12.01000 ; qtot -
2.129
5429 H1 335 GLH HA 5429 0.077900 1.00800 ; qtot -
2.051
5430 A2C 335 GLH CB 5430 -0.007100 12.01000 ; qtot -
2.058
5431 HC 335 GLH HB2 5431 0.025600 1.00800 ; qtot -
2.033
5432 HC 335 GLH HB3 5432 0.025600 1.00800 ; qtot -
2.007
5433 A2C 335 GLH CG 5433 -0.017400 12.01000 ; qtot -
2.025
5434 HC 335 GLH HG2 5434 0.043000 1.00800 ; qtot -
1.982
5435 HC 335 GLH HG3 5435 0.043000 1.00800 ; qtot -
1.939
5436 C 335 GLH CD 5436 0.680101 12.01000 ; qtot -
1.259
5437 O 335 GLH OE1 5437 -0.583801 16.00000 ; qtot -
1.842
5438 OH 335 GLH OE2 5438 -0.651101 16.00000 ; qtot -
2.493
5439 HO 335 GLH HE2 5439 0.464100 1.00800 ; qtot -
2.029
5440 C 335 GLH C 5440 0.597301 12.01000 ; qtot -
1.432
5441 O 335 GLH O 5441 -0.567901 16.00000 ; qtot -
2.000
5442 N 336 ILE N 5442 -0.415700 14.01000 ; qtot -
2.416
5443 H 336 ILE H 5443 0.271900 1.00800 ; qtot -
2.144
5444 CX 336 ILE CA 5444 -0.059700 12.01000 ; qtot -
2.203
5445 H1 336 ILE HA 5445 0.086900 1.00800 ; qtot -
2.117
5446 A3C 336 ILE CB 5446 0.130300 12.01000 ; qtot -
1.986
5447 HC 336 ILE HB 5447 0.018700 1.00800 ; qtot -
1.968
5448 CT 336 ILE CG2 5448 -0.320400 12.01000 ; qtot -
2.288
5449 HC 336 ILE HG21 5449 0.088200 1.00800 ; qtot -
2.200
5450 HC 336 ILE HG22 5450 0.088200 1.00800 ; qtot -
2.112
5451 HC 336 ILE HG23 5451 0.088200 1.00800 ; qtot -
2.023
5452 A2C 336 ILE CG1 5452 -0.043000 12.01000 ; qtot -
2.066
5453 HC 336 ILE HG12 5453 0.023600 1.00800 ; qtot -

2.043
  5454   HC   336   ILE  HG13 5454     0.023600      1.00800 ; qtot -
2.019
  5455   CT   336   ILE   CD1 5455    -0.066000     12.01000 ; qtot -
2.085
  5456   HC   336   ILE  HD11 5456     0.018600      1.00800 ; qtot -
2.067
  5457   HC   336   ILE  HD12 5457     0.018600      1.00800 ; qtot -
2.048
  5458   HC   336   ILE  HD13 5458     0.018600      1.00800 ; qtot -
2.029
  5459    C   336   ILE     C 5459     0.597301     12.01000 ; qtot -
1.432
  5460    O   336   ILE     O 5460    -0.567901     16.00000 ; qtot -
2.000
  5461    N   337   LEU     N 5461    -0.415700     14.01000 ; qtot -
2.416
  5462    H   337   LEU     H 5462     0.271900      1.00800 ; qtot -
2.144
  5463   CX   337   LEU    CA 5463    -0.051800     12.01000 ; qtot -
2.196
  5464   H1   337   LEU    HA 5464     0.092200      1.00800 ; qtot -
2.103
  5465  A2C   337   LEU    CB 5465    -0.110200     12.01000 ; qtot -
2.214
  5466   HC   337   LEU   HB2 5466     0.045700      1.00800 ; qtot -
2.168
  5467   HC   337   LEU   HB3 5467     0.045700      1.00800 ; qtot -
2.122
  5468  A3C   337   LEU    CG 5468     0.353100     12.01000 ; qtot -
1.769
  5469   HC   337   LEU    HG 5469    -0.036100      1.00800 ; qtot -
1.805
  5470   CT   337   LEU   CD1 5470    -0.412100     12.01000 ; qtot -
2.217
  5471   HC   337   LEU  HD11 5471     0.100000      1.00800 ; qtot -
2.117
  5472   HC   337   LEU  HD12 5472     0.100000      1.00800 ; qtot -
2.017
  5473   HC   337   LEU  HD13 5473     0.100000      1.00800 ; qtot -
1.917
  5474   CT   337   LEU   CD2 5474    -0.412100     12.01000 ; qtot -
2.329
  5475   HC   337   LEU  HD21 5475     0.100000      1.00800 ; qtot -
2.229
  5476   HC   337   LEU  HD22 5476     0.100000      1.00800 ; qtot -
2.129
  5477   HC   337   LEU  HD23 5477     0.100000      1.00800 ; qtot -
2.029
  5478    C   337   LEU     C 5478     0.597301     12.01000 ; qtot -
1.432
  5479    O   337   LEU     O 5479    -0.567901     16.00000 ; qtot -

2.000
  5480    N   338   ASP      N 5480    -0.516301     14.01000 ; qtot -
2.516
  5481    H   338   ASP      H 5481     0.293600      1.00800 ; qtot -
2.223
  5482   CX   338   ASP     CA 5482     0.038100     12.01000 ; qtot -
2.185
  5483   H1   338   ASP     HA 5483     0.088000      1.00800 ; qtot -
2.097
  5484  A2C   338   ASP     CB 5484    -0.030300     12.01000 ; qtot -
2.127
  5485   HC   338   ASP    HB2 5485    -0.012200      1.00800 ; qtot -
2.139
  5486   HC   338   ASP    HB3 5486    -0.012200      1.00800 ; qtot -
2.151
  5487   CO   338   ASP     CG 5487     0.799401     12.01000 ; qtot -
1.352
  5488   O2   338   ASP    OD1 5488    -0.801401     16.00000 ; qtot -
2.153
  5489   O2   338   ASP    OD2 5489    -0.801401     16.00000 ; qtot -
2.955
  5490    C   338   ASP      C 5490     0.536601     12.01000 ; qtot -
2.418
  5491    O   338   ASP      O 5491    -0.581901     16.00000 ; qtot -
3.000
  5492    N   339   GLY      N 5492    -0.415700     14.01000 ; qtot -
3.416
  5493    H   339   GLY      H 5493     0.271900      1.00800 ; qtot -
3.144
  5494   CX   339   GLY     CA 5494    -0.025200     12.01000 ; qtot -
3.169
  5495   H1   339   GLY    HA2 5495     0.069800      1.00800 ; qtot -
3.099
  5496   H1   339   GLY    HA3 5496     0.069800      1.00800 ; qtot -
3.029
  5497    C   339   GLY      C 5497     0.597301     12.01000 ; qtot -
2.432
  5498    O   339   GLY      O 5498    -0.567901     16.00000 ; qtot -
3.000
  5499    N   340   LEU      N 5499    -0.415700     14.01000 ; qtot -
3.416
  5500    H   340   LEU      H 5500     0.271900      1.00800 ; qtot -
3.144
  5501   CX   340   LEU     CA 5501    -0.051800     12.01000 ; qtot -
3.196
  5502   H1   340   LEU     HA 5502     0.092200      1.00800 ; qtot -
3.103
  5503  A2C   340   LEU     CB 5503    -0.110200     12.01000 ; qtot -
3.214
  5504   HC   340   LEU    HB2 5504     0.045700      1.00800 ; qtot -
3.168
  5505   HC   340   LEU    HB3 5505     0.045700      1.00800 ; qtot -

```
3.122
  5506  A3C   340   LEU    CG 5506     0.353100     12.01000 ; qtot -
2.769
  5507   HC   340   LEU    HG 5507    -0.036100      1.00800 ; qtot -
2.805
  5508   CT   340   LEU   CD1 5508    -0.412100     12.01000 ; qtot -
3.217
  5509   HC   340   LEU  HD11 5509     0.100000      1.00800 ; qtot -
3.117
  5510   HC   340   LEU  HD12 5510     0.100000      1.00800 ; qtot -
3.017
  5511   HC   340   LEU  HD13 5511     0.100000      1.00800 ; qtot -
2.917
  5512   CT   340   LEU   CD2 5512    -0.412100     12.01000 ; qtot -
3.329
  5513   HC   340   LEU  HD21 5513     0.100000      1.00800 ; qtot -
3.229
  5514   HC   340   LEU  HD22 5514     0.100000      1.00800 ; qtot -
3.129
  5515   HC   340   LEU  HD23 5515     0.100000      1.00800 ; qtot -
3.029
  5516    C   340   LEU     C 5516     0.597301     12.01000 ; qtot -
2.432
  5517    O   340   LEU     O 5517    -0.567901     16.00000 ; qtot -
3.000
  5518    N   341   LYS     N 5518    -0.347900     14.01000 ; qtot -
3.348
  5519    H   341   LYS     H 5519     0.274700      1.00800 ; qtot -
3.073
  5520   CX   341   LYS    CA 5520    -0.240000     12.01000 ; qtot -
3.313
  5521   H1   341   LYS    HA 5521     0.142600      1.00800 ; qtot -
3.171
  5522   C8   341   LYS    CB 5522    -0.009400     12.01000 ; qtot -
3.180
  5523   HC   341   LYS   HB2 5523     0.036200      1.00800 ; qtot -
3.144
  5524   HC   341   LYS   HB3 5524     0.036200      1.00800 ; qtot -
3.108
  5525   C8   341   LYS    CG 5525     0.018700     12.01000 ; qtot -
3.089
  5526   HC   341   LYS   HG2 5526     0.010300      1.00800 ; qtot -
3.079
  5527   HC   341   LYS   HG3 5527     0.010300      1.00800 ; qtot -
3.068
  5528   C8   341   LYS    CD 5528    -0.047900     12.01000 ; qtot -
3.116
  5529   HC   341   LYS   HD2 5529     0.062100      1.00800 ; qtot -
3.054
  5530   HC   341   LYS   HD3 5530     0.062100      1.00800 ; qtot -
2.992
  5531   C8   341   LYS    CE 5531    -0.014300     12.01000 ; qtot -
```

3.006
 5532 HP 341 LYS HE2 5532 0.113500 1.00800 ; qtot -
2.893
 5533 HP 341 LYS HE3 5533 0.113500 1.00800 ; qtot -
2.779
 5534 N3 341 LYS NZ 5534 -0.385400 14.01000 ; qtot -
3.165
 5535 H 341 LYS HZ1 5535 0.340000 1.00800 ; qtot -
2.825
 5536 H 341 LYS HZ2 5536 0.340000 1.00800 ; qtot -
2.485
 5537 H 341 LYS HZ3 5537 0.340000 1.00800 ; qtot -
2.145
 5538 C 341 LYS C 5538 0.734101 12.01000 ; qtot -
1.411
 5539 O 341 LYS O 5539 -0.589401 16.00000 ; qtot -
2.000
 5540 N 342 LYS N 5540 -0.347900 14.01000 ; qtot -
2.348
 5541 H 342 LYS H 5541 0.274700 1.00800 ; qtot -
2.073
 5542 CX 342 LYS CA 5542 -0.240000 12.01000 ; qtot -
2.313
 5543 H1 342 LYS HA 5543 0.142600 1.00800 ; qtot -
2.171
 5544 C8 342 LYS CB 5544 -0.009400 12.01000 ; qtot -
2.180
 5545 HC 342 LYS HB2 5545 0.036200 1.00800 ; qtot -
2.144
 5546 HC 342 LYS HB3 5546 0.036200 1.00800 ; qtot -
2.108
 5547 C8 342 LYS CG 5547 0.018700 12.01000 ; qtot -
2.089
 5548 HC 342 LYS HG2 5548 0.010300 1.00800 ; qtot -
2.079
 5549 HC 342 LYS HG3 5549 0.010300 1.00800 ; qtot -
2.068
 5550 C8 342 LYS CD 5550 -0.047900 12.01000 ; qtot -
2.116
 5551 HC 342 LYS HD2 5551 0.062100 1.00800 ; qtot -
2.054
 5552 HC 342 LYS HD3 5552 0.062100 1.00800 ; qtot -
1.992
 5553 C8 342 LYS CE 5553 -0.014300 12.01000 ; qtot -
2.006
 5554 HP 342 LYS HE2 5554 0.113500 1.00800 ; qtot -
1.893
 5555 HP 342 LYS HE3 5555 0.113500 1.00800 ; qtot -
1.779
 5556 N3 342 LYS NZ 5556 -0.385400 14.01000 ; qtot -
2.165
 5557 H 342 LYS HZ1 5557 0.340000 1.00800 ; qtot -

1.825
 5558 H 342 LYS HZ2 5558 0.340000 1.00800 ; qtot -
1.485
 5559 H 342 LYS HZ3 5559 0.340000 1.00800 ; qtot -
1.145
 5560 C 342 LYS C 5560 0.734101 12.01000 ; qtot -
0.411
 5561 O 342 LYS O 5561 -0.589401 16.00000 ; qtot -
1.000
 5562 N 343 ILE N 5562 -0.415700 14.01000 ; qtot -
1.416
 5563 H 343 ILE H 5563 0.271900 1.00800 ; qtot -
1.144
 5564 CX 343 ILE CA 5564 -0.059700 12.01000 ; qtot -
1.203
 5565 H1 343 ILE HA 5565 0.086900 1.00800 ; qtot -
1.117
 5566 A3C 343 ILE CB 5566 0.130300 12.01000 ; qtot -
0.986
 5567 HC 343 ILE HB 5567 0.018700 1.00800 ; qtot -
0.968
 5568 CT 343 ILE CG2 5568 -0.320400 12.01000 ; qtot -
1.288
 5569 HC 343 ILE HG21 5569 0.088200 1.00800 ; qtot -
1.200
 5570 HC 343 ILE HG22 5570 0.088200 1.00800 ; qtot -
1.112
 5571 HC 343 ILE HG23 5571 0.088200 1.00800 ; qtot -
1.023
 5572 A2C 343 ILE CG1 5572 -0.043000 12.01000 ; qtot -
1.066
 5573 HC 343 ILE HG12 5573 0.023600 1.00800 ; qtot -
1.043
 5574 HC 343 ILE HG13 5574 0.023600 1.00800 ; qtot -
1.019
 5575 CT 343 ILE CD1 5575 -0.066000 12.01000 ; qtot -
1.085
 5576 HC 343 ILE HD11 5576 0.018600 1.00800 ; qtot -
1.067
 5577 HC 343 ILE HD12 5577 0.018600 1.00800 ; qtot -
1.048
 5578 HC 343 ILE HD13 5578 0.018600 1.00800 ; qtot -
1.029
 5579 C 343 ILE C 5579 0.597301 12.01000 ; qtot -
0.432
 5580 O 343 ILE O 5580 -0.567901 16.00000 ; qtot -
1.000
 5581 N 344 GLY N 5581 -0.415700 14.01000 ; qtot -
1.416
 5582 H 344 GLY H 5582 0.271900 1.00800 ; qtot -
1.144
 5583 CX 344 GLY CA 5583 -0.025200 12.01000 ; qtot -

```
1.169
  5584   H1   344   GLY   HA2 5584     0.069800      1.00800 ; qtot -
1.099
  5585   H1   344   GLY   HA3 5585     0.069800      1.00800 ; qtot -
1.029
  5586    C   344   GLY     C 5586     0.597301     12.01000 ; qtot -
0.432
  5587    O   344   GLY     O 5587    -0.567901     16.00000 ; qtot -
1.000
  5588    N   345   TYR     N 5588    -0.415700     14.01000 ; qtot -
1.416
  5589    H   345   TYR     H 5589     0.271900      1.00800 ; qtot -
1.144
  5590   CX   345   TYR    CA 5590    -0.001400     12.01000 ; qtot -
1.145
  5591   H1   345   TYR    HA 5591     0.087600      1.00800 ; qtot -
1.058
  5592   CT   345   TYR    CB 5592    -0.015200     12.01000 ; qtot -
1.073
  5593   HC   345   TYR   HB2 5593     0.029500      1.00800 ; qtot -
1.043
  5594   HC   345   TYR   HB3 5594     0.029500      1.00800 ; qtot -
1.014
  5595   CA   345   TYR    CG 5595    -0.001100     12.01000 ; qtot -
1.015
  5596   CA   345   TYR   CD1 5596    -0.190600     12.01000 ; qtot -
1.205
  5597   HA   345   TYR   HD1 5597     0.169900      1.00800 ; qtot -
1.036
  5598   CA   345   TYR   CE1 5598    -0.234100     12.01000 ; qtot -
1.270
  5599   HA   345   TYR   HE1 5599     0.165600      1.00800 ; qtot -
1.104
  5600    C   345   TYR    CZ 5600     0.322600     12.01000 ; qtot -
0.781
  5601   OH   345   TYR    OH 5601    -0.557901     16.00000 ; qtot -
1.339
  5602   HO   345   TYR    HH 5602     0.399200      1.00800 ; qtot -
0.940
  5603   CA   345   TYR   CE2 5603    -0.234100     12.01000 ; qtot -
1.174
  5604   HA   345   TYR   HE2 5604     0.165600      1.00800 ; qtot -
1.009
  5605   CA   345   TYR   CD2 5605    -0.190600     12.01000 ; qtot -
1.199
  5606   HA   345   TYR   HD2 5606     0.169900      1.00800 ; qtot -
1.029
  5607    C   345   TYR     C 5607     0.597301     12.01000 ; qtot -
0.432
  5608    O   345   TYR     O 5608    -0.567901     16.00000 ; qtot -
1.000
  5609    N   346   GLN     N 5609    -0.415700     14.01000 ; qtot -
```

```
1.416
  5610    H   346   GLN     H 5610     0.271900      1.00800 ; qtot -
1.144
  5611   CX   346   GLN    CA 5611    -0.003100     12.01000 ; qtot -
1.147
  5612   H1   346   GLN    HA 5612     0.085000      1.00800 ; qtot -
1.062
  5613  A2C   346   GLN    CB 5613    -0.003600     12.01000 ; qtot -
1.065
  5614   HC   346   GLN   HB2 5614     0.017100      1.00800 ; qtot -
1.048
  5615   HC   346   GLN   HB3 5615     0.017100      1.00800 ; qtot -
1.031
  5616  A2C   346   GLN    CG 5616    -0.064500     12.01000 ; qtot -
1.096
  5617   HC   346   GLN   HG2 5617     0.035200      1.00800 ; qtot -
1.061
  5618   HC   346   GLN   HG3 5618     0.035200      1.00800 ; qtot -
1.025
  5619    C   346   GLN    CD 5619     0.695101     12.01000 ; qtot -
0.330
  5620    O   346   GLN   OE1 5620    -0.608601     16.00000 ; qtot -
0.939
  5621    N   346   GLN   NE2 5621    -0.940701     14.01000 ; qtot -
1.880
  5622    H   346   GLN  HE21 5622     0.425100      1.00800 ; qtot -
1.454
  5623    H   346   GLN  HE22 5623     0.425100      1.00800 ; qtot -
1.029
  5624    C   346   GLN     C 5624     0.597301     12.01000 ; qtot -
0.432
  5625    O   346   GLN     O 5625    -0.567901     16.00000 ; qtot -
1.000
  5626    N   347   LYS     N 5626    -0.347900     14.01000 ; qtot -
1.348
  5627    H   347   LYS     H 5627     0.274700      1.00800 ; qtot -
1.073
  5628   CX   347   LYS    CA 5628    -0.240000     12.01000 ; qtot -
1.313
  5629   H1   347   LYS    HA 5629     0.142600      1.00800 ; qtot -
1.171
  5630   C8   347   LYS    CB 5630    -0.009400     12.01000 ; qtot -
1.180
  5631   HC   347   LYS   HB2 5631     0.036200      1.00800 ; qtot -
1.144
  5632   HC   347   LYS   HB3 5632     0.036200      1.00800 ; qtot -
1.108
  5633   C8   347   LYS    CG 5633     0.018700     12.01000 ; qtot -
1.089
  5634   HC   347   LYS   HG2 5634     0.010300      1.00800 ; qtot -
1.079
  5635   HC   347   LYS   HG3 5635     0.010300      1.00800 ; qtot -
```

1.068
  5636   C8   347   LYS    CD 5636    -0.047900     12.01000 ; qtot -
1.116
  5637   HC   347   LYS   HD2 5637     0.062100      1.00800 ; qtot -
1.054
  5638   HC   347   LYS   HD3 5638     0.062100      1.00800 ; qtot -
0.992
  5639   C8   347   LYS    CE 5639    -0.014300     12.01000 ; qtot -
1.006
  5640   HP   347   LYS   HE2 5640     0.113500      1.00800 ; qtot -
0.893
  5641   HP   347   LYS   HE3 5641     0.113500      1.00800 ; qtot -
0.779
  5642   N3   347   LYS    NZ 5642    -0.385400     14.01000 ; qtot -
1.165
  5643    H   347   LYS   HZ1 5643     0.340000      1.00800 ; qtot -
0.825
  5644    H   347   LYS   HZ2 5644     0.340000      1.00800 ; qtot -
0.485
  5645    H   347   LYS   HZ3 5645     0.340000      1.00800 ; qtot -
0.145
  5646    C   347   LYS     C 5646     0.734101     12.01000 ; qtot
0.589
  5647    O   347   LYS     O 5647    -0.589401     16.00000 ; qtot
0.000
  5648    N   348   ILE     N 5648    -0.415700     14.01000 ; qtot -
0.416
  5649    H   348   ILE     H 5649     0.271900      1.00800 ; qtot -
0.144
  5650   CX   348   ILE    CA 5650    -0.059700     12.01000 ; qtot -
0.203
  5651   H1   348   ILE    HA 5651     0.086900      1.00800 ; qtot -
0.117
  5652  A3C   348   ILE    CB 5652     0.130300     12.01000 ; qtot
0.014
  5653   HC   348   ILE    HB 5653     0.018700      1.00800 ; qtot
0.032
  5654   CT   348   ILE   CG2 5654    -0.320400     12.01000 ; qtot -
0.288
  5655   HC   348   ILE  HG21 5655     0.088200      1.00800 ; qtot -
0.200
  5656   HC   348   ILE  HG22 5656     0.088200      1.00800 ; qtot -
0.112
  5657   HC   348   ILE  HG23 5657     0.088200      1.00800 ; qtot -
0.023
  5658  A2C   348   ILE   CG1 5658    -0.043000     12.01000 ; qtot -
0.066
  5659   HC   348   ILE  HG12 5659     0.023600      1.00800 ; qtot -
0.043
  5660   HC   348   ILE  HG13 5660     0.023600      1.00800 ; qtot -
0.019
  5661   CT   348   ILE   CD1 5661    -0.066000     12.01000 ; qtot -

0.085
  5662   HC   348   ILE  HD11 5662     0.018600      1.00800 ; qtot -
0.067
  5663   HC   348   ILE  HD12 5663     0.018600      1.00800 ; qtot -
0.048
  5664   HC   348   ILE  HD13 5664     0.018600      1.00800 ; qtot -
0.029
  5665    C   348   ILE     C 5665     0.597301     12.01000 ; qtot
0.568
  5666    O   348   ILE     O 5666    -0.567901     16.00000 ; qtot
0.000
  5667    N   349   PRO     N 5667    -0.254800     14.01000 ; qtot -
0.255
  5668   CT   349   PRO    CD 5668     0.019200     12.01000 ; qtot -
0.236
  5669   H1   349   PRO   HD2 5669     0.039100      1.00800 ; qtot -
0.196
  5670   H1   349   PRO   HD3 5670     0.039100      1.00800 ; qtot -
0.157
  5671   CT   349   PRO    CG 5671     0.018900     12.01000 ; qtot -
0.138
  5672   HC   349   PRO   HG2 5672     0.021300      1.00800 ; qtot -
0.117
  5673   HC   349   PRO   HG3 5673     0.021300      1.00800 ; qtot -
0.096
  5674   CT   349   PRO    CB 5674    -0.007000     12.01000 ; qtot -
0.103
  5675   HC   349   PRO   HB2 5675     0.025300      1.00800 ; qtot -
0.078
  5676   HC   349   PRO   HB3 5676     0.025300      1.00800 ; qtot -
0.052
  5677   CX   349   PRO    CA 5677    -0.026600     12.01000 ; qtot -
0.079
  5678   H1   349   PRO    HA 5678     0.064100      1.00800 ; qtot -
0.015
  5679    C   349   PRO     C 5679     0.589601     12.01000 ; qtot
0.575
  5680    O   349   PRO     O 5680    -0.574801     16.00000 ; qtot
0.000
  5681    N   350   PRO     N 5681    -0.254800     14.01000 ; qtot -
0.255
  5682   CT   350   PRO    CD 5682     0.019200     12.01000 ; qtot -
0.236
  5683   H1   350   PRO   HD2 5683     0.039100      1.00800 ; qtot -
0.196
  5684   H1   350   PRO   HD3 5684     0.039100      1.00800 ; qtot -
0.157
  5685   CT   350   PRO    CG 5685     0.018900     12.01000 ; qtot -
0.138
  5686   HC   350   PRO   HG2 5686     0.021300      1.00800 ; qtot -
0.117
  5687   HC   350   PRO   HG3 5687     0.021300      1.00800 ; qtot -

```
0.096
  5688   CT   350   PRO    CB 5688    -0.007000     12.01000 ; qtot -
0.103
  5689   HC   350   PRO   HB2 5689     0.025300      1.00800 ; qtot -
0.078
  5690   HC   350   PRO   HB3 5690     0.025300      1.00800 ; qtot -
0.052
  5691   CX   350   PRO    CA 5691    -0.026600     12.01000 ; qtot -
0.079
  5692   H1   350   PRO    HA 5692     0.064100      1.00800 ; qtot -
0.015
  5693    C   350   PRO     C 5693     0.589601     12.01000 ; qtot
0.575
  5694    O   350   PRO     O 5694    -0.574801     16.00000 ; qtot
0.000
  5695    N   351   ASP     N 5695    -0.516301     14.01000 ; qtot -
0.516
  5696    H   351   ASP     H 5696     0.293600      1.00800 ; qtot -
0.223
  5697   CX   351   ASP    CA 5697     0.038100     12.01000 ; qtot -
0.185
  5698   H1   351   ASP    HA 5698     0.088000      1.00800 ; qtot -
0.097
  5699  A2C   351   ASP    CB 5699    -0.030300     12.01000 ; qtot -
0.127
  5700   HC   351   ASP   HB2 5700    -0.012200      1.00800 ; qtot -
0.139
  5701   HC   351   ASP   HB3 5701    -0.012200      1.00800 ; qtot -
0.151
  5702   CO   351   ASP    CG 5702     0.799401     12.01000 ; qtot
0.648
  5703   O2   351   ASP   OD1 5703    -0.801401     16.00000 ; qtot -
0.153
  5704   O2   351   ASP   OD2 5704    -0.801401     16.00000 ; qtot -
0.955
  5705    C   351   ASP     C 5705     0.536601     12.01000 ; qtot -
0.418
  5706    O   351   ASP     O 5706    -0.581901     16.00000 ; qtot -
1.000
  5707    N   352   ILE     N 5707    -0.415700     14.01000 ; qtot -
1.416
  5708    H   352   ILE     H 5708     0.271900      1.00800 ; qtot -
1.144
  5709   CX   352   ILE    CA 5709    -0.059700     12.01000 ; qtot -
1.203
  5710   H1   352   ILE    HA 5710     0.086900      1.00800 ; qtot -
1.117
  5711  A3C   352   ILE    CB 5711     0.130300     12.01000 ; qtot -
0.986
  5712   HC   352   ILE    HB 5712     0.018700      1.00800 ; qtot -
0.968
  5713   CT   352   ILE   CG2 5713    -0.320400     12.01000 ; qtot -
```

1.288
5714 HC 352 ILE HG21 5714 0.088200 1.00800 ; qtot -
1.200
5715 HC 352 ILE HG22 5715 0.088200 1.00800 ; qtot -
1.112
5716 HC 352 ILE HG23 5716 0.088200 1.00800 ; qtot -
1.023
5717 A2C 352 ILE CG1 5717 -0.043000 12.01000 ; qtot -
1.066
5718 HC 352 ILE HG12 5718 0.023600 1.00800 ; qtot -
1.043
5719 HC 352 ILE HG13 5719 0.023600 1.00800 ; qtot -
1.019
5720 CT 352 ILE CD1 5720 -0.066000 12.01000 ; qtot -
1.085
5721 HC 352 ILE HD11 5721 0.018600 1.00800 ; qtot -
1.067
5722 HC 352 ILE HD12 5722 0.018600 1.00800 ; qtot -
1.048
5723 HC 352 ILE HD13 5723 0.018600 1.00800 ; qtot -
1.029
5724 C 352 ILE C 5724 0.597301 12.01000 ; qtot -
0.432
5725 O 352 ILE O 5725 -0.567901 16.00000 ; qtot -
1.000
5726 N 353 HIE N 5726 -0.415700 14.01000 ; qtot -
1.416
5727 H 353 HIE H 5727 0.271900 1.00800 ; qtot -
1.144
5728 CX 353 HIE CA 5728 -0.058100 12.01000 ; qtot -
1.202
5729 H1 353 HIE HA 5729 0.136000 1.00800 ; qtot -
1.066
5730 CT 353 HIE CB 5730 -0.007400 12.01000 ; qtot -
1.073
5731 HC 353 HIE HB2 5731 0.036700 1.00800 ; qtot -
1.037
5732 HC 353 HIE HB3 5732 0.036700 1.00800 ; qtot -
1.000
5733 CC 353 HIE CG 5733 0.186800 12.01000 ; qtot -
0.813
5734 NB 353 HIE ND1 5734 -0.543201 14.01000 ; qtot -
1.356
5735 CR 353 HIE CE1 5735 0.163500 12.01000 ; qtot -
1.193
5736 H5 353 HIE HE1 5736 0.143500 1.00800 ; qtot -
1.049
5737 NA 353 HIE NE2 5737 -0.279500 14.01000 ; qtot -
1.329
5738 H 353 HIE HE2 5738 0.333900 1.00800 ; qtot -
0.995
5739 CW 353 HIE CD2 5739 -0.220700 12.01000 ; qtot -

1.216
  5740   H4   353   HIE   HD2 5740     0.186200      1.00800 ; qtot -
1.029
  5741    C   353   HIE     C 5741     0.597301     12.01000 ; qtot -
0.432
  5742    O   353   HIE     O 5742    -0.567901     16.00000 ; qtot -
1.000
  5743    N   354   GLN     N 5743    -0.415700     14.01000 ; qtot -
1.416
  5744    H   354   GLN     H 5744     0.271900      1.00800 ; qtot -
1.144
  5745   CX   354   GLN    CA 5745    -0.003100     12.01000 ; qtot -
1.147
  5746   H1   354   GLN    HA 5746     0.085000      1.00800 ; qtot -
1.062
  5747  A2C   354   GLN    CB 5747    -0.003600     12.01000 ; qtot -
1.065
  5748   HC   354   GLN   HB2 5748     0.017100      1.00800 ; qtot -
1.048
  5749   HC   354   GLN   HB3 5749     0.017100      1.00800 ; qtot -
1.031
  5750  A2C   354   GLN    CG 5750    -0.064500     12.01000 ; qtot -
1.096
  5751   HC   354   GLN   HG2 5751     0.035200      1.00800 ; qtot -
1.061
  5752   HC   354   GLN   HG3 5752     0.035200      1.00800 ; qtot -
1.025
  5753    C   354   GLN    CD 5753     0.695101     12.01000 ; qtot -
0.330
  5754    O   354   GLN   OE1 5754    -0.608601     16.00000 ; qtot -
0.939
  5755    N   354   GLN   NE2 5755    -0.940701     14.01000 ; qtot -
1.880
  5756    H   354   GLN  HE21 5756     0.425100      1.00800 ; qtot -
1.454
  5757    H   354   GLN  HE22 5757     0.425100      1.00800 ; qtot -
1.029
  5758    C   354   GLN     C 5758     0.597301     12.01000 ; qtot -
0.432
  5759    O   354   GLN     O 5759    -0.567901     16.00000 ; qtot -
1.000
  5760    N   355   VAL     N 5760    -0.415700     14.01000 ; qtot -
1.416
  5761    H   355   VAL     H 5761     0.271900      1.00800 ; qtot -
1.144
  5762   CX   355   VAL    CA 5762    -0.087500     12.01000 ; qtot -
1.231
  5763   H1   355   VAL    HA 5763     0.096900      1.00800 ; qtot -
1.134
  5764  A3C   355   VAL    CB 5764     0.298500     12.01000 ; qtot -
0.836
  5765   HC   355   VAL    HB 5765    -0.029700      1.00800 ; qtot -

```
0.866
  5766   CT   355   VAL   CG1 5766    -0.319200     12.01000 ; qtot -
1.185
  5767   HC   355   VAL  HG11 5767     0.079100      1.00800 ; qtot -
1.106
  5768   HC   355   VAL  HG12 5768     0.079100      1.00800 ; qtot -
1.027
  5769   HC   355   VAL  HG13 5769     0.079100      1.00800 ; qtot -
0.947
  5770   CT   355   VAL   CG2 5770    -0.319200     12.01000 ; qtot -
1.267
  5771   HC   355   VAL  HG21 5771     0.079100      1.00800 ; qtot -
1.188
  5772   HC   355   VAL  HG22 5772     0.079100      1.00800 ; qtot -
1.108
  5773   HC   355   VAL  HG23 5773     0.079100      1.00800 ; qtot -
1.029
  5774    C   355   VAL     C 5774     0.597301     12.01000 ; qtot -
0.432
  5775    O   355   VAL     O 5775    -0.567901     16.00000 ; qtot -
1.000
  5776    N   356   LEU     N 5776    -0.415700     14.01000 ; qtot -
1.416
  5777    H   356   LEU     H 5777     0.271900      1.00800 ; qtot -
1.144
  5778   CX   356   LEU    CA 5778    -0.051800     12.01000 ; qtot -
1.196
  5779   H1   356   LEU    HA 5779     0.092200      1.00800 ; qtot -
1.103
  5780  A2C   356   LEU    CB 5780    -0.110200     12.01000 ; qtot -
1.214
  5781   HC   356   LEU   HB2 5781     0.045700      1.00800 ; qtot -
1.168
  5782   HC   356   LEU   HB3 5782     0.045700      1.00800 ; qtot -
1.122
  5783  A3C   356   LEU    CG 5783     0.353100     12.01000 ; qtot -
0.769
  5784   HC   356   LEU    HG 5784    -0.036100      1.00800 ; qtot -
0.805
  5785   CT   356   LEU   CD1 5785    -0.412100     12.01000 ; qtot -
1.217
  5786   HC   356   LEU  HD11 5786     0.100000      1.00800 ; qtot -
1.117
  5787   HC   356   LEU  HD12 5787     0.100000      1.00800 ; qtot -
1.017
  5788   HC   356   LEU  HD13 5788     0.100000      1.00800 ; qtot -
0.917
  5789   CT   356   LEU   CD2 5789    -0.412100     12.01000 ; qtot -
1.329
  5790   HC   356   LEU  HD21 5790     0.100000      1.00800 ; qtot -
1.229
  5791   HC   356   LEU  HD22 5791     0.100000      1.00800 ; qtot -
```

1.129
  5792   HC   356   LEU  HD23 5792      0.100000      1.00800 ; qtot -
1.029
  5793    C   356   LEU     C 5793      0.597301     12.01000 ; qtot -
0.432
  5794    O   356   LEU     O 5794     -0.567901     16.00000 ; qtot -
1.000
  5795    N   357   ARG     N 5795     -0.347900     14.01000 ; qtot -
1.348
  5796    H   357   ARG     H 5796      0.274700      1.00800 ; qtot -
1.073
  5797   CX   357   ARG    CA 5797     -0.263700     12.01000 ; qtot -
1.337
  5798   H1   357   ARG    HA 5798      0.156000      1.00800 ; qtot -
1.181
  5799   C8   357   ARG    CB 5799     -0.000700     12.01000 ; qtot -
1.182
  5800   HC   357   ARG   HB2 5800      0.032700      1.00800 ; qtot -
1.149
  5801   HC   357   ARG   HB3 5801      0.032700      1.00800 ; qtot -
1.116
  5802   C8   357   ARG    CG 5802      0.039000     12.01000 ; qtot -
1.077
  5803   HC   357   ARG   HG2 5803      0.028500      1.00800 ; qtot -
1.049
  5804   HC   357   ARG   HG3 5804      0.028500      1.00800 ; qtot -
1.020
  5805   C8   357   ARG    CD 5805      0.048600     12.01000 ; qtot -
0.972
  5806   H1   357   ARG   HD2 5806      0.068700      1.00800 ; qtot -
0.903
  5807   H1   357   ARG   HD3 5807      0.068700      1.00800 ; qtot -
0.834
  5808   N2   357   ARG    NE 5808     -0.529501     14.01000 ; qtot -
1.364
  5809    H   357   ARG    HE 5809      0.345600      1.00800 ; qtot -
1.018
  5810   CA   357   ARG    CZ 5810      0.807601     12.01000 ; qtot -
0.210
  5811   N2   357   ARG   NH1 5811     -0.862701     14.01000 ; qtot -
1.073
  5812    H   357   ARG  HH11 5812      0.447800      1.00800 ; qtot -
0.625
  5813    H   357   ARG  HH12 5813      0.447800      1.00800 ; qtot -
0.178
  5814   N2   357   ARG   NH2 5814     -0.862701     14.01000 ; qtot -
1.040
  5815    H   357   ARG  HH21 5815      0.447800      1.00800 ; qtot -
0.592
  5816    H   357   ARG  HH22 5816      0.447800      1.00800 ; qtot -
0.145
  5817    C   357   ARG     C 5817      0.734101     12.01000 ; qtot

```
0.589
  5818    O   357   ARG      O 5818    -0.589401     16.00000 ; qtot 
0.000
  5819    N   358   ASP      N 5819    -0.516301     14.01000 ; qtot -
0.516
  5820    H   358   ASP      H 5820     0.293600      1.00800 ; qtot -
0.223
  5821   CX   358   ASP     CA 5821     0.038100     12.01000 ; qtot -
0.185
  5822   H1   358   ASP     HA 5822     0.088000      1.00800 ; qtot -
0.097
  5823  A2C   358   ASP     CB 5823    -0.030300     12.01000 ; qtot -
0.127
  5824   HC   358   ASP    HB2 5824    -0.012200      1.00800 ; qtot -
0.139
  5825   HC   358   ASP    HB3 5825    -0.012200      1.00800 ; qtot -
0.151
  5826   CO   358   ASP     CG 5826     0.799401     12.01000 ; qtot 
0.648
  5827   O2   358   ASP    OD1 5827    -0.801401     16.00000 ; qtot -
0.153
  5828   O2   358   ASP    OD2 5828    -0.801401     16.00000 ; qtot -
0.955
  5829    C   358   ASP      C 5829     0.536601     12.01000 ; qtot -
0.418
  5830    O   358   ASP      O 5830    -0.581901     16.00000 ; qtot -
1.000
  5831    N   359   ILE      N 5831    -0.415700     14.01000 ; qtot -
1.416
  5832    H   359   ILE      H 5832     0.271900      1.00800 ; qtot -
1.144
  5833   CX   359   ILE     CA 5833    -0.059700     12.01000 ; qtot -
1.203
  5834   H1   359   ILE     HA 5834     0.086900      1.00800 ; qtot -
1.117
  5835  A3C   359   ILE     CB 5835     0.130300     12.01000 ; qtot -
0.986
  5836   HC   359   ILE     HB 5836     0.018700      1.00800 ; qtot -
0.968
  5837   CT   359   ILE    CG2 5837    -0.320400     12.01000 ; qtot -
1.288
  5838   HC   359   ILE   HG21 5838     0.088200      1.00800 ; qtot -
1.200
  5839   HC   359   ILE   HG22 5839     0.088200      1.00800 ; qtot -
1.112
  5840   HC   359   ILE   HG23 5840     0.088200      1.00800 ; qtot -
1.023
  5841  A2C   359   ILE    CG1 5841    -0.043000     12.01000 ; qtot -
1.066
  5842   HC   359   ILE   HG12 5842     0.023600      1.00800 ; qtot -
1.043
  5843   HC   359   ILE   HG13 5843     0.023600      1.00800 ; qtot -
```

1.019
5844 CT 359 ILE CD1 5844 -0.066000 12.01000 ; qtot -
1.085
5845 HC 359 ILE HD11 5845 0.018600 1.00800 ; qtot -
1.067
5846 HC 359 ILE HD12 5846 0.018600 1.00800 ; qtot -
1.048
5847 HC 359 ILE HD13 5847 0.018600 1.00800 ; qtot -
1.029
5848 C 359 ILE C 5848 0.597301 12.01000 ; qtot -
0.432
5849 O 359 ILE O 5849 -0.567901 16.00000 ; qtot -
1.000
5850 N 360 ASP N 5850 -0.516301 14.01000 ; qtot -
1.516
5851 H 360 ASP H 5851 0.293600 1.00800 ; qtot -
1.223
5852 CX 360 ASP CA 5852 0.038100 12.01000 ; qtot -
1.185
5853 H1 360 ASP HA 5853 0.088000 1.00800 ; qtot -
1.097
5854 A2C 360 ASP CB 5854 -0.030300 12.01000 ; qtot -
1.127
5855 HC 360 ASP HB2 5855 -0.012200 1.00800 ; qtot -
1.139
5856 HC 360 ASP HB3 5856 -0.012200 1.00800 ; qtot -
1.151
5857 CO 360 ASP CG 5857 0.799401 12.01000 ; qtot -
0.352
5858 O2 360 ASP OD1 5858 -0.801401 16.00000 ; qtot -
1.153
5859 O2 360 ASP OD2 5859 -0.801401 16.00000 ; qtot -
1.955
5860 C 360 ASP C 5860 0.536601 12.01000 ; qtot -
1.418
5861 O 360 ASP O 5861 -0.581901 16.00000 ; qtot -
2.000
5862 N 361 SER N 5862 -0.415700 14.01000 ; qtot -
2.416
5863 H 361 SER H 5863 0.271900 1.00800 ; qtot -
2.144
5864 CX 361 SER CA 5864 -0.024900 12.01000 ; qtot -
2.169
5865 H1 361 SER HA 5865 0.084300 1.00800 ; qtot -
2.084
5866 A2C 361 SER CB 5866 0.211700 12.01000 ; qtot -
1.873
5867 H1 361 SER HB2 5867 0.035200 1.00800 ; qtot -
1.837
5868 H1 361 SER HB3 5868 0.035200 1.00800 ; qtot -
1.802
5869 OH 361 SER OG 5869 -0.654601 16.00000 ; qtot -

2.457
  5870   HO   361   SER     HG 5870     0.427500      1.00800 ; qtot -
2.029
  5871    C   361   SER      C 5871     0.597301     12.01000 ; qtot -
1.432
  5872    O   361   SER      O 5872    -0.567901     16.00000 ; qtot -
2.000
  5873    N   362   ASN      N 5873    -0.415700     14.01000 ; qtot -
2.416
  5874    H   362   ASN      H 5874     0.271900      1.00800 ; qtot -
2.144
  5875   CX   362   ASN     CA 5875     0.014300     12.01000 ; qtot -
2.129
  5876   H1   362   ASN     HA 5876     0.104800      1.00800 ; qtot -
2.025
  5877  A2C   362   ASN     CB 5877    -0.204100     12.01000 ; qtot -
2.229
  5878   HC   362   ASN    HB2 5878     0.079700      1.00800 ; qtot -
2.149
  5879   HC   362   ASN    HB3 5879     0.079700      1.00800 ; qtot -
2.069
  5880    C   362   ASN     CG 5880     0.713001     12.01000 ; qtot -
1.356
  5881    O   362   ASN    OD1 5881    -0.593101     16.00000 ; qtot -
1.949
  5882    N   362   ASN    ND2 5882    -0.919101     14.01000 ; qtot -
2.869
  5883    H   362   ASN   HD21 5883     0.419600      1.00800 ; qtot -
2.449
  5884    H   362   ASN   HD22 5884     0.419600      1.00800 ; qtot -
2.029
  5885    C   362   ASN      C 5885     0.597301     12.01000 ; qtot -
1.432
  5886    O   362   ASN      O 5886    -0.567901     16.00000 ; qtot -
2.000
  5887    N   363   ALA      N 5887    -0.415700     14.01000 ; qtot -
2.416
  5888    H   363   ALA      H 5888     0.271900      1.00800 ; qtot -
2.144
  5889   CX   363   ALA     CA 5889     0.033700     12.01000 ; qtot -
2.110
  5890   H1   363   ALA     HA 5890     0.082300      1.00800 ; qtot -
2.028
  5891   CT   363   ALA     CB 5891    -0.182500     12.01000 ; qtot -
2.210
  5892   HC   363   ALA    HB1 5892     0.060300      1.00800 ; qtot -
2.150
  5893   HC   363   ALA    HB2 5893     0.060300      1.00800 ; qtot -
2.090
  5894   HC   363   ALA    HB3 5894     0.060300      1.00800 ; qtot -
2.029
  5895    C   363   ALA      C 5895     0.597301     12.01000 ; qtot -

1.432
  5896     O   363   ALA      O 5896    -0.567901     16.00000 ; qtot -
2.000
  5897     N   364   SER      N 5897    -0.415700     14.01000 ; qtot -
2.416
  5898     H   364   SER      H 5898     0.271900      1.00800 ; qtot -
2.144
  5899    CX   364   SER     CA 5899    -0.024900     12.01000 ; qtot -
2.169
  5900    H1   364   SER     HA 5900     0.084300      1.00800 ; qtot -
2.084
  5901   A2C   364   SER     CB 5901     0.211700     12.01000 ; qtot -
1.873
  5902    H1   364   SER    HB2 5902     0.035200      1.00800 ; qtot -
1.837
  5903    H1   364   SER    HB3 5903     0.035200      1.00800 ; qtot -
1.802
  5904    OH   364   SER     OG 5904    -0.654601     16.00000 ; qtot -
2.457
  5905    HO   364   SER     HG 5905     0.427500      1.00800 ; qtot -
2.029
  5906     C   364   SER      C 5906     0.597301     12.01000 ; qtot -
1.432
  5907     O   364   SER      O 5907    -0.567901     16.00000 ; qtot -
2.000
  5908     N   365   GLY      N 5908    -0.415700     14.01000 ; qtot -
2.416
  5909     H   365   GLY      H 5909     0.271900      1.00800 ; qtot -
2.144
  5910    CX   365   GLY     CA 5910    -0.025200     12.01000 ; qtot -
2.169
  5911    H1   365   GLY    HA2 5911     0.069800      1.00800 ; qtot -
2.099
  5912    H1   365   GLY    HA3 5912     0.069800      1.00800 ; qtot -
2.029
  5913     C   365   GLY      C 5913     0.597301     12.01000 ; qtot -
1.432
  5914     O   365   GLY      O 5914    -0.567901     16.00000 ; qtot -
2.000
  5915     N   366   GLN      N 5915    -0.415700     14.01000 ; qtot -
2.416
  5916     H   366   GLN      H 5916     0.271900      1.00800 ; qtot -
2.144
  5917    CX   366   GLN     CA 5917    -0.003100     12.01000 ; qtot -
2.147
  5918    H1   366   GLN     HA 5918     0.085000      1.00800 ; qtot -
2.062
  5919   A2C   366   GLN     CB 5919    -0.003600     12.01000 ; qtot -
2.065
  5920    HC   366   GLN    HB2 5920     0.017100      1.00800 ; qtot -
2.048
  5921    HC   366   GLN    HB3 5921     0.017100      1.00800 ; qtot -

2.031
  5922  A2C   366   GLN    CG 5922    -0.064500     12.01000 ; qtot -
2.096
  5923   HC   366   GLN   HG2 5923     0.035200      1.00800 ; qtot -
2.061
  5924   HC   366   GLN   HG3 5924     0.035200      1.00800 ; qtot -
2.025
  5925    C   366   GLN    CD 5925     0.695101     12.01000 ; qtot -
1.330
  5926    O   366   GLN   OE1 5926    -0.608601     16.00000 ; qtot -
1.939
  5927    N   366   GLN   NE2 5927    -0.940701     14.01000 ; qtot -
2.880
  5928    H   366   GLN  HE21 5928     0.425100      1.00800 ; qtot -
2.454
  5929    H   366   GLN  HE22 5929     0.425100      1.00800 ; qtot -
2.029
  5930    C   366   GLN     C 5930     0.597301     12.01000 ; qtot -
1.432
  5931    O   366   GLN     O 5931    -0.567901     16.00000 ; qtot -
2.000
  5932    N   367   ILE     N 5932    -0.415700     14.01000 ; qtot -
2.416
  5933    H   367   ILE     H 5933     0.271900      1.00800 ; qtot -
2.144
  5934   CX   367   ILE    CA 5934    -0.059700     12.01000 ; qtot -
2.203
  5935   H1   367   ILE    HA 5935     0.086900      1.00800 ; qtot -
2.117
  5936  A3C   367   ILE    CB 5936     0.130300     12.01000 ; qtot -
1.986
  5937   HC   367   ILE    HB 5937     0.018700      1.00800 ; qtot -
1.968
  5938   CT   367   ILE   CG2 5938    -0.320400     12.01000 ; qtot -
2.288
  5939   HC   367   ILE  HG21 5939     0.088200      1.00800 ; qtot -
2.200
  5940   HC   367   ILE  HG22 5940     0.088200      1.00800 ; qtot -
2.112
  5941   HC   367   ILE  HG23 5941     0.088200      1.00800 ; qtot -
2.023
  5942  A2C   367   ILE   CG1 5942    -0.043000     12.01000 ; qtot -
2.066
  5943   HC   367   ILE  HG12 5943     0.023600      1.00800 ; qtot -
2.043
  5944   HC   367   ILE  HG13 5944     0.023600      1.00800 ; qtot -
2.019
  5945   CT   367   ILE   CD1 5945    -0.066000     12.01000 ; qtot -
2.085
  5946   HC   367   ILE  HD11 5946     0.018600      1.00800 ; qtot -
2.067
  5947   HC   367   ILE  HD12 5947     0.018600      1.00800 ; qtot -

2.048
5948 HC 367 ILE HD13 5948 0.018600 1.00800 ; qtot -2.029
5949 C 367 ILE C 5949 0.597301 12.01000 ; qtot -1.432
5950 O 367 ILE O 5950 -0.567901 16.00000 ; qtot -2.000
5951 N 368 HIE N 5951 -0.415700 14.01000 ; qtot -2.416
5952 H 368 HIE H 5952 0.271900 1.00800 ; qtot -2.144
5953 CX 368 HIE CA 5953 -0.058100 12.01000 ; qtot -2.202
5954 H1 368 HIE HA 5954 0.136000 1.00800 ; qtot -2.066
5955 CT 368 HIE CB 5955 -0.007400 12.01000 ; qtot -2.073
5956 HC 368 HIE HB2 5956 0.036700 1.00800 ; qtot -2.037
5957 HC 368 HIE HB3 5957 0.036700 1.00800 ; qtot -2.000
5958 CC 368 HIE CG 5958 0.186800 12.01000 ; qtot -1.813
5959 NB 368 HIE ND1 5959 -0.543201 14.01000 ; qtot -2.356
5960 CR 368 HIE CE1 5960 0.163500 12.01000 ; qtot -2.193
5961 H5 368 HIE HE1 5961 0.143500 1.00800 ; qtot -2.049
5962 NA 368 HIE NE2 5962 -0.279500 14.01000 ; qtot -2.329
5963 H 368 HIE HE2 5963 0.333900 1.00800 ; qtot -1.995
5964 CW 368 HIE CD2 5964 -0.220700 12.01000 ; qtot -2.216
5965 H4 368 HIE HD2 5965 0.186200 1.00800 ; qtot -2.029
5966 C 368 HIE C 5966 0.597301 12.01000 ; qtot -1.432
5967 O 368 HIE O 5967 -0.567901 16.00000 ; qtot -2.000
5968 N 369 TYR N 5968 -0.415700 14.01000 ; qtot -2.416
5969 H 369 TYR H 5969 0.271900 1.00800 ; qtot -2.144
5970 CX 369 TYR CA 5970 -0.001400 12.01000 ; qtot -2.145
5971 H1 369 TYR HA 5971 0.087600 1.00800 ; qtot -2.058
5972 CT 369 TYR CB 5972 -0.015200 12.01000 ; qtot -2.073
5973 HC 369 TYR HB2 5973 0.029500 1.00800 ; qtot -

2.043
  5974   HC   369   TYR   HB3 5974     0.029500      1.00800 ; qtot -
2.014
  5975   CA   369   TYR    CG 5975    -0.001100     12.01000 ; qtot -
2.015
  5976   CA   369   TYR   CD1 5976    -0.190600     12.01000 ; qtot -
2.205
  5977   HA   369   TYR   HD1 5977     0.169900      1.00800 ; qtot -
2.036
  5978   CA   369   TYR   CE1 5978    -0.234100     12.01000 ; qtot -
2.270
  5979   HA   369   TYR   HE1 5979     0.165600      1.00800 ; qtot -
2.104
  5980    C   369   TYR    CZ 5980     0.322600     12.01000 ; qtot -
1.781
  5981   OH   369   TYR    OH 5981    -0.557901     16.00000 ; qtot -
2.339
  5982   HO   369   TYR    HH 5982     0.399200      1.00800 ; qtot -
1.940
  5983   CA   369   TYR   CE2 5983    -0.234100     12.01000 ; qtot -
2.174
  5984   HA   369   TYR   HE2 5984     0.165600      1.00800 ; qtot -
2.009
  5985   CA   369   TYR   CD2 5985    -0.190600     12.01000 ; qtot -
2.199
  5986   HA   369   TYR   HD2 5986     0.169900      1.00800 ; qtot -
2.029
  5987    C   369   TYR     C 5987     0.597301     12.01000 ; qtot -
1.432
  5988    O   369   TYR     O 5988    -0.567901     16.00000 ; qtot -
2.000
  5989    N   370   THR     N 5989    -0.415700     14.01000 ; qtot -
2.416
  5990    H   370   THR     H 5990     0.271900      1.00800 ; qtot -
2.144
  5991   CX   370   THR    CA 5991    -0.038900     12.01000 ; qtot -
2.183
  5992   H1   370   THR    HA 5992     0.100700      1.00800 ; qtot -
2.082
  5993  A3C   370   THR    CB 5993     0.365400     12.01000 ; qtot -
1.717
  5994   H1   370   THR    HB 5994     0.004300      1.00800 ; qtot -
1.712
  5995   CT   370   THR   CG2 5995    -0.243800     12.01000 ; qtot -
1.956
  5996   HC   370   THR  HG21 5996     0.064200      1.00800 ; qtot -
1.892
  5997   HC   370   THR  HG22 5997     0.064200      1.00800 ; qtot -
1.828
  5998   HC   370   THR  HG23 5998     0.064200      1.00800 ; qtot -
1.763
  5999   OH   370   THR   OG1 5999    -0.676101     16.00000 ; qtot -

2.440
  6000   HO   370   THR   HG1 6000     0.410200      1.00800 ; qtot -
2.029
  6001    C   370   THR     C 6001     0.597301     12.01000 ; qtot -
1.432
  6002    O   370   THR     O 6002    -0.567901     16.00000 ; qtot -
2.000
  6003    N   371   ASP     N 6003    -0.516301     14.01000 ; qtot -
2.516
  6004    H   371   ASP     H 6004     0.293600      1.00800 ; qtot -
2.223
  6005   CX   371   ASP    CA 6005     0.038100     12.01000 ; qtot -
2.185
  6006   H1   371   ASP    HA 6006     0.088000      1.00800 ; qtot -
2.097
  6007  A2C   371   ASP    CB 6007    -0.030300     12.01000 ; qtot -
2.127
  6008   HC   371   ASP   HB2 6008    -0.012200      1.00800 ; qtot -
2.139
  6009   HC   371   ASP   HB3 6009    -0.012200      1.00800 ; qtot -
2.151
  6010   CO   371   ASP    CG 6010     0.799401     12.01000 ; qtot -
1.352
  6011   O2   371   ASP   OD1 6011    -0.801401     16.00000 ; qtot -
2.153
  6012   O2   371   ASP   OD2 6012    -0.801401     16.00000 ; qtot -
2.955
  6013    C   371   ASP     C 6013     0.536601     12.01000 ; qtot -
2.418
  6014    O   371   ASP     O 6014    -0.581901     16.00000 ; qtot -
3.000
  6015    N   372   PHE     N 6015    -0.415700     14.01000 ; qtot -
3.416
  6016    H   372   PHE     H 6016     0.271900      1.00800 ; qtot -
3.144
  6017   CX   372   PHE    CA 6017    -0.002400     12.01000 ; qtot -
3.146
  6018   H1   372   PHE    HA 6018     0.097800      1.00800 ; qtot -
3.048
  6019   CT   372   PHE    CB 6019    -0.034300     12.01000 ; qtot -
3.083
  6020   HC   372   PHE   HB2 6020     0.029500      1.00800 ; qtot -
3.053
  6021   HC   372   PHE   HB3 6021     0.029500      1.00800 ; qtot -
3.024
  6022   CA   372   PHE    CG 6022     0.011800     12.01000 ; qtot -
3.012
  6023   CA   372   PHE   CD1 6023    -0.125600     12.01000 ; qtot -
3.137
  6024   HA   372   PHE   HD1 6024     0.133000      1.00800 ; qtot -
3.004
  6025   CA   372   PHE   CE1 6025    -0.170400     12.01000 ; qtot -

```
3.175
  6026   HA   372   PHE   HE1 6026     0.143000      1.00800 ; qtot -
3.032
  6027   CA   372   PHE    CZ 6027    -0.107200     12.01000 ; qtot -
3.139
  6028   HA   372   PHE    HZ 6028     0.129700      1.00800 ; qtot -
3.009
  6029   CA   372   PHE   CE2 6029    -0.170400     12.01000 ; qtot -
3.180
  6030   HA   372   PHE   HE2 6030     0.143000      1.00800 ; qtot -
3.037
  6031   CA   372   PHE   CD2 6031    -0.125600     12.01000 ; qtot -
3.162
  6032   HA   372   PHE   HD2 6032     0.133000      1.00800 ; qtot -
3.029
  6033    C   372   PHE     C 6033     0.597301     12.01000 ; qtot -
2.432
  6034    O   372   PHE     O 6034    -0.567901     16.00000 ; qtot -
3.000
  6035    N   373   LEU     N 6035    -0.415700     14.01000 ; qtot -
3.416
  6036    H   373   LEU     H 6036     0.271900      1.00800 ; qtot -
3.144
  6037   CX   373   LEU    CA 6037    -0.051800     12.01000 ; qtot -
3.196
  6038   H1   373   LEU    HA 6038     0.092200      1.00800 ; qtot -
3.103
  6039  A2C   373   LEU    CB 6039    -0.110200     12.01000 ; qtot -
3.214
  6040   HC   373   LEU   HB2 6040     0.045700      1.00800 ; qtot -
3.168
  6041   HC   373   LEU   HB3 6041     0.045700      1.00800 ; qtot -
3.122
  6042  A3C   373   LEU    CG 6042     0.353100     12.01000 ; qtot -
2.769
  6043   HC   373   LEU    HG 6043    -0.036100      1.00800 ; qtot -
2.805
  6044   CT   373   LEU   CD1 6044    -0.412100     12.01000 ; qtot -
3.217
  6045   HC   373   LEU  HD11 6045     0.100000      1.00800 ; qtot -
3.117
  6046   HC   373   LEU  HD12 6046     0.100000      1.00800 ; qtot -
3.017
  6047   HC   373   LEU  HD13 6047     0.100000      1.00800 ; qtot -
2.917
  6048   CT   373   LEU   CD2 6048    -0.412100     12.01000 ; qtot -
3.329
  6049   HC   373   LEU  HD21 6049     0.100000      1.00800 ; qtot -
3.229
  6050   HC   373   LEU  HD22 6050     0.100000      1.00800 ; qtot -
3.129
  6051   HC   373   LEU  HD23 6051     0.100000      1.00800 ; qtot -
```

```
3.029
  6052    C   373   LEU      C 6052     0.597301     12.01000 ; qtot -
2.432
  6053    O   373   LEU      O 6053    -0.567901     16.00000 ; qtot -
3.000
  6054    N   374   ALA      N 6054    -0.415700     14.01000 ; qtot -
3.416
  6055    H   374   ALA      H 6055     0.271900      1.00800 ; qtot -
3.144
  6056   CX   374   ALA     CA 6056     0.033700     12.01000 ; qtot -
3.110
  6057   H1   374   ALA     HA 6057     0.082300      1.00800 ; qtot -
3.028
  6058   CT   374   ALA     CB 6058    -0.182500     12.01000 ; qtot -
3.210
  6059   HC   374   ALA    HB1 6059     0.060300      1.00800 ; qtot -
3.150
  6060   HC   374   ALA    HB2 6060     0.060300      1.00800 ; qtot -
3.090
  6061   HC   374   ALA    HB3 6061     0.060300      1.00800 ; qtot -
3.029
  6062    C   374   ALA      C 6062     0.597301     12.01000 ; qtot -
2.432
  6063    O   374   ALA      O 6063    -0.567901     16.00000 ; qtot -
3.000
  6064    N   375   ALA      N 6064    -0.415700     14.01000 ; qtot -
3.416
  6065    H   375   ALA      H 6065     0.271900      1.00800 ; qtot -
3.144
  6066   CX   375   ALA     CA 6066     0.033700     12.01000 ; qtot -
3.110
  6067   H1   375   ALA     HA 6067     0.082300      1.00800 ; qtot -
3.028
  6068   CT   375   ALA     CB 6068    -0.182500     12.01000 ; qtot -
3.210
  6069   HC   375   ALA    HB1 6069     0.060300      1.00800 ; qtot -
3.150
  6070   HC   375   ALA    HB2 6070     0.060300      1.00800 ; qtot -
3.090
  6071   HC   375   ALA    HB3 6071     0.060300      1.00800 ; qtot -
3.029
  6072    C   375   ALA      C 6072     0.597301     12.01000 ; qtot -
2.432
  6073    O   375   ALA      O 6073    -0.567901     16.00000 ; qtot -
3.000
  6074    N   376   THR      N 6074    -0.415700     14.01000 ; qtot -
3.416
  6075    H   376   THR      H 6075     0.271900      1.00800 ; qtot -
3.144
  6076   CX   376   THR     CA 6076    -0.038900     12.01000 ; qtot -
3.183
  6077   H1   376   THR     HA 6077     0.100700      1.00800 ; qtot -
```

3.082
6078 A3C 376 THR CB 6078 0.365400 12.01000 ; qtot -
2.717
6079 H1 376 THR HB 6079 0.004300 1.00800 ; qtot -
2.712
6080 CT 376 THR CG2 6080 -0.243800 12.01000 ; qtot -
2.956
6081 HC 376 THR HG21 6081 0.064200 1.00800 ; qtot -
2.892
6082 HC 376 THR HG22 6082 0.064200 1.00800 ; qtot -
2.828
6083 HC 376 THR HG23 6083 0.064200 1.00800 ; qtot -
2.763
6084 OH 376 THR OG1 6084 -0.676101 16.00000 ; qtot -
3.440
6085 HO 376 THR HG1 6085 0.410200 1.00800 ; qtot -
3.029
6086 C 376 THR C 6086 0.597301 12.01000 ; qtot -
2.432
6087 O 376 THR O 6087 -0.567901 16.00000 ; qtot -
3.000
6088 N 377 ILE N 6088 -0.415700 14.01000 ; qtot -
3.416
6089 H 377 ILE H 6089 0.271900 1.00800 ; qtot -
3.144
6090 CX 377 ILE CA 6090 -0.059700 12.01000 ; qtot -
3.203
6091 H1 377 ILE HA 6091 0.086900 1.00800 ; qtot -
3.117
6092 A3C 377 ILE CB 6092 0.130300 12.01000 ; qtot -
2.986
6093 HC 377 ILE HB 6093 0.018700 1.00800 ; qtot -
2.968
6094 CT 377 ILE CG2 6094 -0.320400 12.01000 ; qtot -
3.288
6095 HC 377 ILE HG21 6095 0.088200 1.00800 ; qtot -
3.200
6096 HC 377 ILE HG22 6096 0.088200 1.00800 ; qtot -
3.112
6097 HC 377 ILE HG23 6097 0.088200 1.00800 ; qtot -
3.023
6098 A2C 377 ILE CG1 6098 -0.043000 12.01000 ; qtot -
3.066
6099 HC 377 ILE HG12 6099 0.023600 1.00800 ; qtot -
3.043
6100 HC 377 ILE HG13 6100 0.023600 1.00800 ; qtot -
3.019
6101 CT 377 ILE CD1 6101 -0.066000 12.01000 ; qtot -
3.085
6102 HC 377 ILE HD11 6102 0.018600 1.00800 ; qtot -
3.067
6103 HC 377 ILE HD12 6103 0.018600 1.00800 ; qtot -

```
3.048
  6104   HC   377   ILE  HD13 6104     0.018600      1.00800 ; qtot -
3.029
  6105    C   377   ILE     C 6105     0.597301     12.01000 ; qtot -
2.432
  6106    O   377   ILE     O 6106    -0.567901     16.00000 ; qtot -
3.000
  6107    N   378   ASP     N 6107    -0.516301     14.01000 ; qtot -
3.516
  6108    H   378   ASP     H 6108     0.293600      1.00800 ; qtot -
3.223
  6109   CX   378   ASP    CA 6109     0.038100     12.01000 ; qtot -
3.185
  6110   H1   378   ASP    HA 6110     0.088000      1.00800 ; qtot -
3.097
  6111  A2C   378   ASP    CB 6111    -0.030300     12.01000 ; qtot -
3.127
  6112   HC   378   ASP   HB2 6112    -0.012200      1.00800 ; qtot -
3.139
  6113   HC   378   ASP   HB3 6113    -0.012200      1.00800 ; qtot -
3.151
  6114   CO   378   ASP    CG 6114     0.799401     12.01000 ; qtot -
2.352
  6115   O2   378   ASP   OD1 6115    -0.801401     16.00000 ; qtot -
3.153
  6116   O2   378   ASP   OD2 6116    -0.801401     16.00000 ; qtot -
3.955
  6117    C   378   ASP     C 6117     0.536601     12.01000 ; qtot -
3.418
  6118    O   378   ASP     O 6118    -0.581901     16.00000 ; qtot -
4.000
  6119    N   379   LYS     N 6119    -0.347900     14.01000 ; qtot -
4.348
  6120    H   379   LYS     H 6120     0.274700      1.00800 ; qtot -
4.073
  6121   CX   379   LYS    CA 6121    -0.240000     12.01000 ; qtot -
4.313
  6122   H1   379   LYS    HA 6122     0.142600      1.00800 ; qtot -
4.171
  6123   C8   379   LYS    CB 6123    -0.009400     12.01000 ; qtot -
4.180
  6124   HC   379   LYS   HB2 6124     0.036200      1.00800 ; qtot -
4.144
  6125   HC   379   LYS   HB3 6125     0.036200      1.00800 ; qtot -
4.108
  6126   C8   379   LYS    CG 6126     0.018700     12.01000 ; qtot -
4.089
  6127   HC   379   LYS   HG2 6127     0.010300      1.00800 ; qtot -
4.079
  6128   HC   379   LYS   HG3 6128     0.010300      1.00800 ; qtot -
4.068
  6129   C8   379   LYS    CD 6129    -0.047900     12.01000 ; qtot -
```

4.116
  6130   HC   379   LYS   HD2 6130     0.062100      1.00800 ; qtot -
4.054
  6131   HC   379   LYS   HD3 6131     0.062100      1.00800 ; qtot -
3.992
  6132   C8   379   LYS    CE 6132    -0.014300     12.01000 ; qtot -
4.006
  6133   HP   379   LYS   HE2 6133     0.113500      1.00800 ; qtot -
3.893
  6134   HP   379   LYS   HE3 6134     0.113500      1.00800 ; qtot -
3.779
  6135   N3   379   LYS    NZ 6135    -0.385400     14.01000 ; qtot -
4.165
  6136    H   379   LYS   HZ1 6136     0.340000      1.00800 ; qtot -
3.825
  6137    H   379   LYS   HZ2 6137     0.340000      1.00800 ; qtot -
3.485
  6138    H   379   LYS   HZ3 6138     0.340000      1.00800 ; qtot -
3.145
  6139    C   379   LYS     C 6139     0.734101     12.01000 ; qtot -
2.411
  6140    O   379   LYS     O 6140    -0.589401     16.00000 ; qtot -
3.000
  6141    N   380   GLN     N 6141    -0.415700     14.01000 ; qtot -
3.416
  6142    H   380   GLN     H 6142     0.271900      1.00800 ; qtot -
3.144
  6143   CX   380   GLN    CA 6143    -0.003100     12.01000 ; qtot -
3.147
  6144   H1   380   GLN    HA 6144     0.085000      1.00800 ; qtot -
3.062
  6145  A2C   380   GLN    CB 6145    -0.003600     12.01000 ; qtot -
3.065
  6146   HC   380   GLN   HB2 6146     0.017100      1.00800 ; qtot -
3.048
  6147   HC   380   GLN   HB3 6147     0.017100      1.00800 ; qtot -
3.031
  6148  A2C   380   GLN    CG 6148    -0.064500     12.01000 ; qtot -
3.096
  6149   HC   380   GLN   HG2 6149     0.035200      1.00800 ; qtot -
3.061
  6150   HC   380   GLN   HG3 6150     0.035200      1.00800 ; qtot -
3.025
  6151    C   380   GLN    CD 6151     0.695101     12.01000 ; qtot -
2.330
  6152    O   380   GLN   OE1 6152    -0.608601     16.00000 ; qtot -
2.939
  6153    N   380   GLN   NE2 6153    -0.940701     14.01000 ; qtot -
3.880
  6154    H   380   GLN  HE21 6154     0.425100      1.00800 ; qtot -
3.454
  6155    H   380   GLN  HE22 6155     0.425100      1.00800 ; qtot -

```
3.029
  6156    C   380   GLN      C 6156     0.597301     12.01000 ; qtot -
2.432
  6157    O   380   GLN      O 6157    -0.567901     16.00000 ; qtot -
3.000
  6158    N   381   THR      N 6158    -0.415700     14.01000 ; qtot -
3.416
  6159    H   381   THR      H 6159     0.271900      1.00800 ; qtot -
3.144
  6160   CX   381   THR     CA 6160    -0.038900     12.01000 ; qtot -
3.183
  6161   H1   381   THR     HA 6161     0.100700      1.00800 ; qtot -
3.082
  6162  A3C   381   THR     CB 6162     0.365400     12.01000 ; qtot -
2.717
  6163   H1   381   THR     HB 6163     0.004300      1.00800 ; qtot -
2.712
  6164   CT   381   THR    CG2 6164    -0.243800     12.01000 ; qtot -
2.956
  6165   HC   381   THR   HG21 6165     0.064200      1.00800 ; qtot -
2.892
  6166   HC   381   THR   HG22 6166     0.064200      1.00800 ; qtot -
2.828
  6167   HC   381   THR   HG23 6167     0.064200      1.00800 ; qtot -
2.763
  6168   OH   381   THR    OG1 6168    -0.676101     16.00000 ; qtot -
3.440
  6169   HO   381   THR    HG1 6169     0.410200      1.00800 ; qtot -
3.029
  6170    C   381   THR      C 6170     0.597301     12.01000 ; qtot -
2.432
  6171    O   381   THR      O 6171    -0.567901     16.00000 ; qtot -
3.000
  6172    N   382   TYR      N 6172    -0.415700     14.01000 ; qtot -
3.416
  6173    H   382   TYR      H 6173     0.271900      1.00800 ; qtot -
3.144
  6174   CX   382   TYR     CA 6174    -0.001400     12.01000 ; qtot -
3.145
  6175   H1   382   TYR     HA 6175     0.087600      1.00800 ; qtot -
3.058
  6176   CT   382   TYR     CB 6176    -0.015200     12.01000 ; qtot -
3.073
  6177   HC   382   TYR    HB2 6177     0.029500      1.00800 ; qtot -
3.043
  6178   HC   382   TYR    HB3 6178     0.029500      1.00800 ; qtot -
3.014
  6179   CA   382   TYR     CG 6179    -0.001100     12.01000 ; qtot -
3.015
  6180   CA   382   TYR    CD1 6180    -0.190600     12.01000 ; qtot -
3.205
  6181   HA   382   TYR    HD1 6181     0.169900      1.00800 ; qtot -
```

```
3.036
  6182   CA   382   TYR   CE1 6182    -0.234100     12.01000 ; qtot -
3.270
  6183   HA   382   TYR   HE1 6183     0.165600      1.00800 ; qtot -
3.104
  6184    C   382   TYR    CZ 6184     0.322600     12.01000 ; qtot -
2.781
  6185   OH   382   TYR    OH 6185    -0.557901     16.00000 ; qtot -
3.339
  6186   HO   382   TYR    HH 6186     0.399200      1.00800 ; qtot -
2.940
  6187   CA   382   TYR   CE2 6187    -0.234100     12.01000 ; qtot -
3.174
  6188   HA   382   TYR   HE2 6188     0.165600      1.00800 ; qtot -
3.009
  6189   CA   382   TYR   CD2 6189    -0.190600     12.01000 ; qtot -
3.199
  6190   HA   382   TYR   HD2 6190     0.169900      1.00800 ; qtot -
3.029
  6191    C   382   TYR     C 6191     0.597301     12.01000 ; qtot -
2.432
  6192    O   382   TYR     O 6192    -0.567901     16.00000 ; qtot -
3.000
  6193    N   383   LEU     N 6193    -0.415700     14.01000 ; qtot -
3.416
  6194    H   383   LEU     H 6194     0.271900      1.00800 ; qtot -
3.144
  6195   CX   383   LEU    CA 6195    -0.051800     12.01000 ; qtot -
3.196
  6196   H1   383   LEU    HA 6196     0.092200      1.00800 ; qtot -
3.103
  6197  A2C   383   LEU    CB 6197    -0.110200     12.01000 ; qtot -
3.214
  6198   HC   383   LEU   HB2 6198     0.045700      1.00800 ; qtot -
3.168
  6199   HC   383   LEU   HB3 6199     0.045700      1.00800 ; qtot -
3.122
  6200  A3C   383   LEU    CG 6200     0.353100     12.01000 ; qtot -
2.769
  6201   HC   383   LEU    HG 6201    -0.036100      1.00800 ; qtot -
2.805
  6202   CT   383   LEU   CD1 6202    -0.412100     12.01000 ; qtot -
3.217
  6203   HC   383   LEU  HD11 6203     0.100000      1.00800 ; qtot -
3.117
  6204   HC   383   LEU  HD12 6204     0.100000      1.00800 ; qtot -
3.017
  6205   HC   383   LEU  HD13 6205     0.100000      1.00800 ; qtot -
2.917
  6206   CT   383   LEU   CD2 6206    -0.412100     12.01000 ; qtot -
3.329
  6207   HC   383   LEU  HD21 6207     0.100000      1.00800 ; qtot -
```

3.229
6208 HC 383 LEU HD22 6208 0.100000 1.00800 ; qtot -
3.129
6209 HC 383 LEU HD23 6209 0.100000 1.00800 ; qtot -
3.029
6210 C 383 LEU C 6210 0.597301 12.01000 ; qtot -
2.432
6211 O 383 LEU O 6211 -0.567901 16.00000 ; qtot -
3.000
6212 N 384 LYS N 6212 -0.347900 14.01000 ; qtot -
3.348
6213 H 384 LYS H 6213 0.274700 1.00800 ; qtot -
3.073
6214 CX 384 LYS CA 6214 -0.240000 12.01000 ; qtot -
3.313
6215 H1 384 LYS HA 6215 0.142600 1.00800 ; qtot -
3.171
6216 C8 384 LYS CB 6216 -0.009400 12.01000 ; qtot -
3.180
6217 HC 384 LYS HB2 6217 0.036200 1.00800 ; qtot -
3.144
6218 HC 384 LYS HB3 6218 0.036200 1.00800 ; qtot -
3.108
6219 C8 384 LYS CG 6219 0.018700 12.01000 ; qtot -
3.089
6220 HC 384 LYS HG2 6220 0.010300 1.00800 ; qtot -
3.079
6221 HC 384 LYS HG3 6221 0.010300 1.00800 ; qtot -
3.068
6222 C8 384 LYS CD 6222 -0.047900 12.01000 ; qtot -
3.116
6223 HC 384 LYS HD2 6223 0.062100 1.00800 ; qtot -
3.054
6224 HC 384 LYS HD3 6224 0.062100 1.00800 ; qtot -
2.992
6225 C8 384 LYS CE 6225 -0.014300 12.01000 ; qtot -
3.006
6226 HP 384 LYS HE2 6226 0.113500 1.00800 ; qtot -
2.893
6227 HP 384 LYS HE3 6227 0.113500 1.00800 ; qtot -
2.779
6228 N3 384 LYS NZ 6228 -0.385400 14.01000 ; qtot -
3.165
6229 H 384 LYS HZ1 6229 0.340000 1.00800 ; qtot -
2.825
6230 H 384 LYS HZ2 6230 0.340000 1.00800 ; qtot -
2.485
6231 H 384 LYS HZ3 6231 0.340000 1.00800 ; qtot -
2.145
6232 C 384 LYS C 6232 0.734101 12.01000 ; qtot -
1.411
6233 O 384 LYS O 6233 -0.589401 16.00000 ; qtot -

2.000
6234 N 385 LYS N 6234 -0.347900 14.01000 ; qtot -
2.348
6235 H 385 LYS H 6235 0.274700 1.00800 ; qtot -
2.073
6236 CX 385 LYS CA 6236 -0.240000 12.01000 ; qtot -
2.313
6237 H1 385 LYS HA 6237 0.142600 1.00800 ; qtot -
2.171
6238 C8 385 LYS CB 6238 -0.009400 12.01000 ; qtot -
2.180
6239 HC 385 LYS HB2 6239 0.036200 1.00800 ; qtot -
2.144
6240 HC 385 LYS HB3 6240 0.036200 1.00800 ; qtot -
2.108
6241 C8 385 LYS CG 6241 0.018700 12.01000 ; qtot -
2.089
6242 HC 385 LYS HG2 6242 0.010300 1.00800 ; qtot -
2.079
6243 HC 385 LYS HG3 6243 0.010300 1.00800 ; qtot -
2.068
6244 C8 385 LYS CD 6244 -0.047900 12.01000 ; qtot -
2.116
6245 HC 385 LYS HD2 6245 0.062100 1.00800 ; qtot -
2.054
6246 HC 385 LYS HD3 6246 0.062100 1.00800 ; qtot -
1.992
6247 C8 385 LYS CE 6247 -0.014300 12.01000 ; qtot -
2.006
6248 HP 385 LYS HE2 6248 0.113500 1.00800 ; qtot -
1.893
6249 HP 385 LYS HE3 6249 0.113500 1.00800 ; qtot -
1.779
6250 N3 385 LYS NZ 6250 -0.385400 14.01000 ; qtot -
2.165
6251 H 385 LYS HZ1 6251 0.340000 1.00800 ; qtot -
1.825
6252 H 385 LYS HZ2 6252 0.340000 1.00800 ; qtot -
1.485
6253 H 385 LYS HZ3 6253 0.340000 1.00800 ; qtot -
1.145
6254 C 385 LYS C 6254 0.734101 12.01000 ; qtot -
0.411
6255 O 385 LYS O 6255 -0.589401 16.00000 ; qtot -
1.000
6256 N 386 GLU N 6256 -0.516301 14.01000 ; qtot -
1.516
6257 H 386 GLU H 6257 0.293600 1.00800 ; qtot -
1.223
6258 CX 386 GLU CA 6258 0.039700 12.01000 ; qtot -
1.183
6259 H1 386 GLU HA 6259 0.110500 1.00800 ; qtot -

1.072
  6260  A2C   386   GLU    CB 6260     0.056000     12.01000 ; qtot -
1.016
  6261   HC   386   GLU   HB2 6261    -0.017300      1.00800 ; qtot -
1.034
  6262   HC   386   GLU   HB3 6262    -0.017300      1.00800 ; qtot -
1.051
  6263  A2C   386   GLU    CG 6263     0.013600     12.01000 ; qtot -
1.037
  6264   HC   386   GLU   HG2 6264    -0.042500      1.00800 ; qtot -
1.080
  6265   HC   386   GLU   HG3 6265    -0.042500      1.00800 ; qtot -
1.122
  6266   CO   386   GLU    CD 6266     0.805401     12.01000 ; qtot -
0.317
  6267   O2   386   GLU   OE1 6267    -0.818801     16.00000 ; qtot -
1.136
  6268   O2   386   GLU   OE2 6268    -0.818801     16.00000 ; qtot -
1.955
  6269    C   386   GLU     C 6269     0.536601     12.01000 ; qtot -
1.418
  6270    O   386   GLU     O 6270    -0.581901     16.00000 ; qtot -
2.000
  6271    N   387   VAL     N 6271    -0.415700     14.01000 ; qtot -
2.416
  6272    H   387   VAL     H 6272     0.271900      1.00800 ; qtot -
2.144
  6273   CX   387   VAL    CA 6273    -0.087500     12.01000 ; qtot -
2.231
  6274   H1   387   VAL    HA 6274     0.096900      1.00800 ; qtot -
2.134
  6275  A3C   387   VAL    CB 6275     0.298500     12.01000 ; qtot -
1.836
  6276   HC   387   VAL    HB 6276    -0.029700      1.00800 ; qtot -
1.866
  6277   CT   387   VAL   CG1 6277    -0.319200     12.01000 ; qtot -
2.185
  6278   HC   387   VAL  HG11 6278     0.079100      1.00800 ; qtot -
2.106
  6279   HC   387   VAL  HG12 6279     0.079100      1.00800 ; qtot -
2.027
  6280   HC   387   VAL  HG13 6280     0.079100      1.00800 ; qtot -
1.947
  6281   CT   387   VAL   CG2 6281    -0.319200     12.01000 ; qtot -
2.267
  6282   HC   387   VAL  HG21 6282     0.079100      1.00800 ; qtot -
2.188
  6283   HC   387   VAL  HG22 6283     0.079100      1.00800 ; qtot -
2.108
  6284   HC   387   VAL  HG23 6284     0.079100      1.00800 ; qtot -
2.029
  6285    C   387   VAL     C 6285     0.597301     12.01000 ; qtot -

```
1.432
   6286    O   387   VAL      O 6286    -0.567901     16.00000 ; qtot -
2.000
   6287    N   388   CYS      N 6287    -0.415700     14.01000 ; qtot -
2.416
   6288    H   388   CYS      H 6288     0.271900      1.00800 ; qtot -
2.144
   6289   CX   388   CYS     CA 6289     0.021300     12.01000 ; qtot -
2.122
   6290   H1   388   CYS     HA 6290     0.112400      1.00800 ; qtot -
2.010
   6291  A2C   388   CYS     CB 6291    -0.123100     12.01000 ; qtot -
2.133
   6292   H1   388   CYS    HB2 6292     0.111200      1.00800 ; qtot -
2.022
   6293   H1   388   CYS    HB3 6293     0.111200      1.00800 ; qtot -
1.911
   6294   SH   388   CYS     SG 6294    -0.311900     32.06000 ; qtot -
2.223
   6295   HS   388   CYS     HG 6295     0.193300      1.00800 ; qtot -
2.029
   6296    C   388   CYS      C 6296     0.597301     12.01000 ; qtot -
1.432
   6297    O   388   CYS      O 6297    -0.567901     16.00000 ; qtot -
2.000
   6298    N   389   LEU      N 6298    -0.415700     14.01000 ; qtot -
2.416
   6299    H   389   LEU      H 6299     0.271900      1.00800 ; qtot -
2.144
   6300   CX   389   LEU     CA 6300    -0.051800     12.01000 ; qtot -
2.196
   6301   H1   389   LEU     HA 6301     0.092200      1.00800 ; qtot -
2.103
   6302  A2C   389   LEU     CB 6302    -0.110200     12.01000 ; qtot -
2.214
   6303   HC   389   LEU    HB2 6303     0.045700      1.00800 ; qtot -
2.168
   6304   HC   389   LEU    HB3 6304     0.045700      1.00800 ; qtot -
2.122
   6305  A3C   389   LEU     CG 6305     0.353100     12.01000 ; qtot -
1.769
   6306   HC   389   LEU     HG 6306    -0.036100      1.00800 ; qtot -
1.805
   6307   CT   389   LEU    CD1 6307    -0.412100     12.01000 ; qtot -
2.217
   6308   HC   389   LEU   HD11 6308     0.100000      1.00800 ; qtot -
2.117
   6309   HC   389   LEU   HD12 6309     0.100000      1.00800 ; qtot -
2.017
   6310   HC   389   LEU   HD13 6310     0.100000      1.00800 ; qtot -
1.917
   6311   CT   389   LEU    CD2 6311    -0.412100     12.01000 ; qtot -
```

2.329
6312 HC 389 LEU HD21 6312 0.100000 1.00800 ; qtot -
2.229
6313 HC 389 LEU HD22 6313 0.100000 1.00800 ; qtot -
2.129
6314 HC 389 LEU HD23 6314 0.100000 1.00800 ; qtot -
2.029
6315 C 389 LEU C 6315 0.597301 12.01000 ; qtot -
1.432
6316 O 389 LEU O 6316 -0.567901 16.00000 ; qtot -
2.000
6317 N 390 ILE N 6317 -0.415700 14.01000 ; qtot -
2.416
6318 H 390 ILE H 6318 0.271900 1.00800 ; qtot -
2.144
6319 CX 390 ILE CA 6319 -0.059700 12.01000 ; qtot -
2.203
6320 H1 390 ILE HA 6320 0.086900 1.00800 ; qtot -
2.117
6321 A3C 390 ILE CB 6321 0.130300 12.01000 ; qtot -
1.986
6322 HC 390 ILE HB 6322 0.018700 1.00800 ; qtot -
1.968
6323 CT 390 ILE CG2 6323 -0.320400 12.01000 ; qtot -
2.288
6324 HC 390 ILE HG21 6324 0.088200 1.00800 ; qtot -
2.200
6325 HC 390 ILE HG22 6325 0.088200 1.00800 ; qtot -
2.112
6326 HC 390 ILE HG23 6326 0.088200 1.00800 ; qtot -
2.023
6327 A2C 390 ILE CG1 6327 -0.043000 12.01000 ; qtot -
2.066
6328 HC 390 ILE HG12 6328 0.023600 1.00800 ; qtot -
2.043
6329 HC 390 ILE HG13 6329 0.023600 1.00800 ; qtot -
2.019
6330 CT 390 ILE CD1 6330 -0.066000 12.01000 ; qtot -
2.085
6331 HC 390 ILE HD11 6331 0.018600 1.00800 ; qtot -
2.067
6332 HC 390 ILE HD12 6332 0.018600 1.00800 ; qtot -
2.048
6333 HC 390 ILE HD13 6333 0.018600 1.00800 ; qtot -
2.029
6334 C 390 ILE C 6334 0.597301 12.01000 ; qtot -
1.432
6335 O 390 ILE O 6335 -0.567901 16.00000 ; qtot -
2.000
6336 N 391 PRO N 6336 -0.254800 14.01000 ; qtot -
2.255
6337 CT 391 PRO CD 6337 0.019200 12.01000 ; qtot -

2.236
  6338   H1   391   PRO   HD2 6338     0.039100      1.00800 ; qtot -
2.196
  6339   H1   391   PRO   HD3 6339     0.039100      1.00800 ; qtot -
2.157
  6340   CT   391   PRO    CG 6340     0.018900     12.01000 ; qtot -
2.138
  6341   HC   391   PRO   HG2 6341     0.021300      1.00800 ; qtot -
2.117
  6342   HC   391   PRO   HG3 6342     0.021300      1.00800 ; qtot -
2.096
  6343   CT   391   PRO    CB 6343    -0.007000     12.01000 ; qtot -
2.103
  6344   HC   391   PRO   HB2 6344     0.025300      1.00800 ; qtot -
2.078
  6345   HC   391   PRO   HB3 6345     0.025300      1.00800 ; qtot -
2.052
  6346   CX   391   PRO    CA 6346    -0.026600     12.01000 ; qtot -
2.079
  6347   H1   391   PRO    HA 6347     0.064100      1.00800 ; qtot -
2.015
  6348    C   391   PRO     C 6348     0.589601     12.01000 ; qtot -
1.425
  6349    O   391   PRO     O 6349    -0.574801     16.00000 ; qtot -
2.000
  6350    N   392   PHE     N 6350    -0.415700     14.01000 ; qtot -
2.416
  6351    H   392   PHE     H 6351     0.271900      1.00800 ; qtot -
2.144
  6352   CX   392   PHE    CA 6352    -0.002400     12.01000 ; qtot -
2.146
  6353   H1   392   PHE    HA 6353     0.097800      1.00800 ; qtot -
2.048
  6354   CT   392   PHE    CB 6354    -0.034300     12.01000 ; qtot -
2.083
  6355   HC   392   PHE   HB2 6355     0.029500      1.00800 ; qtot -
2.053
  6356   HC   392   PHE   HB3 6356     0.029500      1.00800 ; qtot -
2.024
  6357   CA   392   PHE    CG 6357     0.011800     12.01000 ; qtot -
2.012
  6358   CA   392   PHE   CD1 6358    -0.125600     12.01000 ; qtot -
2.137
  6359   HA   392   PHE   HD1 6359     0.133000      1.00800 ; qtot -
2.004
  6360   CA   392   PHE   CE1 6360    -0.170400     12.01000 ; qtot -
2.175
  6361   HA   392   PHE   HE1 6361     0.143000      1.00800 ; qtot -
2.032
  6362   CA   392   PHE    CZ 6362    -0.107200     12.01000 ; qtot -
2.139
  6363   HA   392   PHE    HZ 6363     0.129700      1.00800 ; qtot -

2.009
  6364   CA   392   PHE   CE2 6364    -0.170400     12.01000 ; qtot -
2.180
  6365   HA   392   PHE   HE2 6365     0.143000      1.00800 ; qtot -
2.037
  6366   CA   392   PHE   CD2 6366    -0.125600     12.01000 ; qtot -
2.162
  6367   HA   392   PHE   HD2 6367     0.133000      1.00800 ; qtot -
2.029
  6368    C   392   PHE     C 6368     0.597301     12.01000 ; qtot -
1.432
  6369    O   392   PHE     O 6369    -0.567901     16.00000 ; qtot -
2.000
  6370    N   393   LYS     N 6370    -0.347900     14.01000 ; qtot -
2.348
  6371    H   393   LYS     H 6371     0.274700      1.00800 ; qtot -
2.073
  6372   CX   393   LYS    CA 6372    -0.240000     12.01000 ; qtot -
2.313
  6373   H1   393   LYS    HA 6373     0.142600      1.00800 ; qtot -
2.171
  6374   C8   393   LYS    CB 6374    -0.009400     12.01000 ; qtot -
2.180
  6375   HC   393   LYS   HB2 6375     0.036200      1.00800 ; qtot -
2.144
  6376   HC   393   LYS   HB3 6376     0.036200      1.00800 ; qtot -
2.108
  6377   C8   393   LYS    CG 6377     0.018700     12.01000 ; qtot -
2.089
  6378   HC   393   LYS   HG2 6378     0.010300      1.00800 ; qtot -
2.079
  6379   HC   393   LYS   HG3 6379     0.010300      1.00800 ; qtot -
2.068
  6380   C8   393   LYS    CD 6380    -0.047900     12.01000 ; qtot -
2.116
  6381   HC   393   LYS   HD2 6381     0.062100      1.00800 ; qtot -
2.054
  6382   HC   393   LYS   HD3 6382     0.062100      1.00800 ; qtot -
1.992
  6383   C8   393   LYS    CE 6383    -0.014300     12.01000 ; qtot -
2.006
  6384   HP   393   LYS   HE2 6384     0.113500      1.00800 ; qtot -
1.893
  6385   HP   393   LYS   HE3 6385     0.113500      1.00800 ; qtot -
1.779
  6386   N3   393   LYS    NZ 6386    -0.385400     14.01000 ; qtot -
2.165
  6387    H   393   LYS   HZ1 6387     0.340000      1.00800 ; qtot -
1.825
  6388    H   393   LYS   HZ2 6388     0.340000      1.00800 ; qtot -
1.485
  6389    H   393   LYS   HZ3 6389     0.340000      1.00800 ; qtot -

1.145
6390 C 393 LYS C 6390 0.734101 12.01000 ; qtot -
0.411
6391 O 393 LYS O 6391 -0.589401 16.00000 ; qtot -
1.000
6392 N 394 PHE N 6392 -0.415700 14.01000 ; qtot -
1.416
6393 H 394 PHE H 6393 0.271900 1.00800 ; qtot -
1.144
6394 CX 394 PHE CA 6394 -0.002400 12.01000 ; qtot -
1.146
6395 H1 394 PHE HA 6395 0.097800 1.00800 ; qtot -
1.048
6396 CT 394 PHE CB 6396 -0.034300 12.01000 ; qtot -
1.083
6397 HC 394 PHE HB2 6397 0.029500 1.00800 ; qtot -
1.053
6398 HC 394 PHE HB3 6398 0.029500 1.00800 ; qtot -
1.024
6399 CA 394 PHE CG 6399 0.011800 12.01000 ; qtot -
1.012
6400 CA 394 PHE CD1 6400 -0.125600 12.01000 ; qtot -
1.137
6401 HA 394 PHE HD1 6401 0.133000 1.00800 ; qtot -
1.004
6402 CA 394 PHE CE1 6402 -0.170400 12.01000 ; qtot -
1.175
6403 HA 394 PHE HE1 6403 0.143000 1.00800 ; qtot -
1.032
6404 CA 394 PHE CZ 6404 -0.107200 12.01000 ; qtot -
1.139
6405 HA 394 PHE HZ 6405 0.129700 1.00800 ; qtot -
1.009
6406 CA 394 PHE CE2 6406 -0.170400 12.01000 ; qtot -
1.180
6407 HA 394 PHE HE2 6407 0.143000 1.00800 ; qtot -
1.037
6408 CA 394 PHE CD2 6408 -0.125600 12.01000 ; qtot -
1.162
6409 HA 394 PHE HD2 6409 0.133000 1.00800 ; qtot -
1.029
6410 C 394 PHE C 6410 0.597301 12.01000 ; qtot -
0.432
6411 O 394 PHE O 6411 -0.567901 16.00000 ; qtot -
1.000
6412 N 395 PHE N 6412 -0.415700 14.01000 ; qtot -
1.416
6413 H 395 PHE H 6413 0.271900 1.00800 ; qtot -
1.144
6414 CX 395 PHE CA 6414 -0.002400 12.01000 ; qtot -
1.146
6415 H1 395 PHE HA 6415 0.097800 1.00800 ; qtot -

1.048
6416 CT 395 PHE CB 6416 -0.034300 12.01000 ; qtot -1.083
6417 HC 395 PHE HB2 6417 0.029500 1.00800 ; qtot -1.053
6418 HC 395 PHE HB3 6418 0.029500 1.00800 ; qtot -1.024
6419 CA 395 PHE CG 6419 0.011800 12.01000 ; qtot -1.012
6420 CA 395 PHE CD1 6420 -0.125600 12.01000 ; qtot -1.137
6421 HA 395 PHE HD1 6421 0.133000 1.00800 ; qtot -1.004
6422 CA 395 PHE CE1 6422 -0.170400 12.01000 ; qtot -1.175
6423 HA 395 PHE HE1 6423 0.143000 1.00800 ; qtot -1.032
6424 CA 395 PHE CZ 6424 -0.107200 12.01000 ; qtot -1.139
6425 HA 395 PHE HZ 6425 0.129700 1.00800 ; qtot -1.009
6426 CA 395 PHE CE2 6426 -0.170400 12.01000 ; qtot -1.180
6427 HA 395 PHE HE2 6427 0.143000 1.00800 ; qtot -1.037
6428 CA 395 PHE CD2 6428 -0.125600 12.01000 ; qtot -1.162
6429 HA 395 PHE HD2 6429 0.133000 1.00800 ; qtot -1.029
6430 C 395 PHE C 6430 0.597301 12.01000 ; qtot -0.432
6431 O 395 PHE O 6431 -0.567901 16.00000 ; qtot -1.000
6432 N 396 ASP N 6432 -0.516301 14.01000 ; qtot -1.516
6433 H 396 ASP H 6433 0.293600 1.00800 ; qtot -1.223
6434 CX 396 ASP CA 6434 0.038100 12.01000 ; qtot -1.185
6435 H1 396 ASP HA 6435 0.088000 1.00800 ; qtot -1.097
6436 A2C 396 ASP CB 6436 -0.030300 12.01000 ; qtot -1.127
6437 HC 396 ASP HB2 6437 -0.012200 1.00800 ; qtot -1.139
6438 HC 396 ASP HB3 6438 -0.012200 1.00800 ; qtot -1.151
6439 CO 396 ASP CG 6439 0.799401 12.01000 ; qtot -0.352
6440 O2 396 ASP OD1 6440 -0.801401 16.00000 ; qtot -1.153
6441 O2 396 ASP OD2 6441 -0.801401 16.00000 ; qtot -

```
1.955
  6442    C   396   ASP     C 6442     0.536601     12.01000 ; qtot -
1.418
  6443    O   396   ASP     O 6443    -0.581901     16.00000 ; qtot -
2.000
  6444    N   397   ILE     N 6444    -0.415700     14.01000 ; qtot -
2.416
  6445    H   397   ILE     H 6445     0.271900      1.00800 ; qtot -
2.144
  6446   CX   397   ILE    CA 6446    -0.059700     12.01000 ; qtot -
2.203
  6447   H1   397   ILE    HA 6447     0.086900      1.00800 ; qtot -
2.117
  6448  A3C   397   ILE    CB 6448     0.130300     12.01000 ; qtot -
1.986
  6449   HC   397   ILE    HB 6449     0.018700      1.00800 ; qtot -
1.968
  6450   CT   397   ILE   CG2 6450    -0.320400     12.01000 ; qtot -
2.288
  6451   HC   397   ILE  HG21 6451     0.088200      1.00800 ; qtot -
2.200
  6452   HC   397   ILE  HG22 6452     0.088200      1.00800 ; qtot -
2.112
  6453   HC   397   ILE  HG23 6453     0.088200      1.00800 ; qtot -
2.023
  6454  A2C   397   ILE   CG1 6454    -0.043000     12.01000 ; qtot -
2.066
  6455   HC   397   ILE  HG12 6455     0.023600      1.00800 ; qtot -
2.043
  6456   HC   397   ILE  HG13 6456     0.023600      1.00800 ; qtot -
2.019
  6457   CT   397   ILE   CD1 6457    -0.066000     12.01000 ; qtot -
2.085
  6458   HC   397   ILE  HD11 6458     0.018600      1.00800 ; qtot -
2.067
  6459   HC   397   ILE  HD12 6459     0.018600      1.00800 ; qtot -
2.048
  6460   HC   397   ILE  HD13 6460     0.018600      1.00800 ; qtot -
2.029
  6461    C   397   ILE     C 6461     0.597301     12.01000 ; qtot -
1.432
  6462    O   397   ILE     O 6462    -0.567901     16.00000 ; qtot -
2.000
  6463    N   398   ASP     N 6463    -0.516301     14.01000 ; qtot -
2.516
  6464    H   398   ASP     H 6464     0.293600      1.00800 ; qtot -
2.223
  6465   CX   398   ASP    CA 6465     0.038100     12.01000 ; qtot -
2.185
  6466   H1   398   ASP    HA 6466     0.088000      1.00800 ; qtot -
2.097
  6467  A2C   398   ASP    CB 6467    -0.030300     12.01000 ; qtot -
```

2.127
 6468 HC 398 ASP HB2 6468 -0.012200 1.00800 ; qtot -
2.139
 6469 HC 398 ASP HB3 6469 -0.012200 1.00800 ; qtot -
2.151
 6470 CO 398 ASP CG 6470 0.799401 12.01000 ; qtot -
1.352
 6471 O2 398 ASP OD1 6471 -0.801401 16.00000 ; qtot -
2.153
 6472 O2 398 ASP OD2 6472 -0.801401 16.00000 ; qtot -
2.955
 6473 C 398 ASP C 6473 0.536601 12.01000 ; qtot -
2.418
 6474 O 398 ASP O 6474 -0.581901 16.00000 ; qtot -
3.000
 6475 N 399 GLY N 6475 -0.415700 14.01000 ; qtot -
3.416
 6476 H 399 GLY H 6476 0.271900 1.00800 ; qtot -
3.144
 6477 CX 399 GLY CA 6477 -0.025200 12.01000 ; qtot -
3.169
 6478 H1 399 GLY HA2 6478 0.069800 1.00800 ; qtot -
3.099
 6479 H1 399 GLY HA3 6479 0.069800 1.00800 ; qtot -
3.029
 6480 C 399 GLY C 6480 0.597301 12.01000 ; qtot -
2.432
 6481 O 399 GLY O 6481 -0.567901 16.00000 ; qtot -
3.000
 6482 N 400 ASN N 6482 -0.415700 14.01000 ; qtot -
3.416
 6483 H 400 ASN H 6483 0.271900 1.00800 ; qtot -
3.144
 6484 CX 400 ASN CA 6484 0.014300 12.01000 ; qtot -
3.129
 6485 H1 400 ASN HA 6485 0.104800 1.00800 ; qtot -
3.025
 6486 A2C 400 ASN CB 6486 -0.204100 12.01000 ; qtot -
3.229
 6487 HC 400 ASN HB2 6487 0.079700 1.00800 ; qtot -
3.149
 6488 HC 400 ASN HB3 6488 0.079700 1.00800 ; qtot -
3.069
 6489 C 400 ASN CG 6489 0.713001 12.01000 ; qtot -
2.356
 6490 O 400 ASN OD1 6490 -0.593101 16.00000 ; qtot -
2.949
 6491 N 400 ASN ND2 6491 -0.919101 14.01000 ; qtot -
3.869
 6492 H 400 ASN HD21 6492 0.419600 1.00800 ; qtot -
3.449
 6493 H 400 ASN HD22 6493 0.419600 1.00800 ; qtot -

```
3.029
  6494    C   400   ASN      C 6494     0.597301     12.01000 ; qtot -
2.432
  6495    O   400   ASN      O 6495    -0.567901     16.00000 ; qtot -
3.000
  6496    N   401   GLY      N 6496    -0.415700     14.01000 ; qtot -
3.416
  6497    H   401   GLY      H 6497     0.271900      1.00800 ; qtot -
3.144
  6498   CX   401   GLY     CA 6498    -0.025200     12.01000 ; qtot -
3.169
  6499   H1   401   GLY    HA2 6499     0.069800      1.00800 ; qtot -
3.099
  6500   H1   401   GLY    HA3 6500     0.069800      1.00800 ; qtot -
3.029
  6501    C   401   GLY      C 6501     0.597301     12.01000 ; qtot -
2.432
  6502    O   401   GLY      O 6502    -0.567901     16.00000 ; qtot -
3.000
  6503    N   402   LYS      N 6503    -0.347900     14.01000 ; qtot -
3.348
  6504    H   402   LYS      H 6504     0.274700      1.00800 ; qtot -
3.073
  6505   CX   402   LYS     CA 6505    -0.240000     12.01000 ; qtot -
3.313
  6506   H1   402   LYS     HA 6506     0.142600      1.00800 ; qtot -
3.171
  6507   C8   402   LYS     CB 6507    -0.009400     12.01000 ; qtot -
3.180
  6508   HC   402   LYS    HB2 6508     0.036200      1.00800 ; qtot -
3.144
  6509   HC   402   LYS    HB3 6509     0.036200      1.00800 ; qtot -
3.108
  6510   C8   402   LYS     CG 6510     0.018700     12.01000 ; qtot -
3.089
  6511   HC   402   LYS    HG2 6511     0.010300      1.00800 ; qtot -
3.079
  6512   HC   402   LYS    HG3 6512     0.010300      1.00800 ; qtot -
3.068
  6513   C8   402   LYS     CD 6513    -0.047900     12.01000 ; qtot -
3.116
  6514   HC   402   LYS    HD2 6514     0.062100      1.00800 ; qtot -
3.054
  6515   HC   402   LYS    HD3 6515     0.062100      1.00800 ; qtot -
2.992
  6516   C8   402   LYS     CE 6516    -0.014300     12.01000 ; qtot -
3.006
  6517   HP   402   LYS    HE2 6517     0.113500      1.00800 ; qtot -
2.893
  6518   HP   402   LYS    HE3 6518     0.113500      1.00800 ; qtot -
2.779
  6519   N3   402   LYS     NZ 6519    -0.385400     14.01000 ; qtot -
```

```
3.165
  6520    H   402   LYS   HZ1 6520     0.340000      1.00800 ; qtot -
2.825
  6521    H   402   LYS   HZ2 6521     0.340000      1.00800 ; qtot -
2.485
  6522    H   402   LYS   HZ3 6522     0.340000      1.00800 ; qtot -
2.145
  6523    C   402   LYS     C 6523     0.734101     12.01000 ; qtot -
1.411
  6524    O   402   LYS     O 6524    -0.589401     16.00000 ; qtot -
2.000
  6525    N   403   ILE     N 6525    -0.415700     14.01000 ; qtot -
2.416
  6526    H   403   ILE     H 6526     0.271900      1.00800 ; qtot -
2.144
  6527   CX   403   ILE    CA 6527    -0.059700     12.01000 ; qtot -
2.203
  6528   H1   403   ILE    HA 6528     0.086900      1.00800 ; qtot -
2.117
  6529  A3C   403   ILE    CB 6529     0.130300     12.01000 ; qtot -
1.986
  6530   HC   403   ILE    HB 6530     0.018700      1.00800 ; qtot -
1.968
  6531   CT   403   ILE   CG2 6531    -0.320400     12.01000 ; qtot -
2.288
  6532   HC   403   ILE  HG21 6532     0.088200      1.00800 ; qtot -
2.200
  6533   HC   403   ILE  HG22 6533     0.088200      1.00800 ; qtot -
2.112
  6534   HC   403   ILE  HG23 6534     0.088200      1.00800 ; qtot -
2.023
  6535  A2C   403   ILE   CG1 6535    -0.043000     12.01000 ; qtot -
2.066
  6536   HC   403   ILE  HG12 6536     0.023600      1.00800 ; qtot -
2.043
  6537   HC   403   ILE  HG13 6537     0.023600      1.00800 ; qtot -
2.019
  6538   CT   403   ILE   CD1 6538    -0.066000     12.01000 ; qtot -
2.085
  6539   HC   403   ILE  HD11 6539     0.018600      1.00800 ; qtot -
2.067
  6540   HC   403   ILE  HD12 6540     0.018600      1.00800 ; qtot -
2.048
  6541   HC   403   ILE  HD13 6541     0.018600      1.00800 ; qtot -
2.029
  6542    C   403   ILE     C 6542     0.597301     12.01000 ; qtot -
1.432
  6543    O   403   ILE     O 6543    -0.567901     16.00000 ; qtot -
2.000
  6544    N   404   SER     N 6544    -0.415700     14.01000 ; qtot -
2.416
  6545    H   404   SER     H 6545     0.271900      1.00800 ; qtot -
```

2.144
6546 CX 404 SER CA 6546 -0.024900 12.01000 ; qtot -
2.169
6547 H1 404 SER HA 6547 0.084300 1.00800 ; qtot -
2.084
6548 A2C 404 SER CB 6548 0.211700 12.01000 ; qtot -
1.873
6549 H1 404 SER HB2 6549 0.035200 1.00800 ; qtot -
1.837
6550 H1 404 SER HB3 6550 0.035200 1.00800 ; qtot -
1.802
6551 OH 404 SER OG 6551 -0.654601 16.00000 ; qtot -
2.457
6552 HO 404 SER HG 6552 0.427500 1.00800 ; qtot -
2.029
6553 C 404 SER C 6553 0.597301 12.01000 ; qtot -
1.432
6554 O 404 SER O 6554 -0.567901 16.00000 ; qtot -
2.000
6555 N 405 VAL N 6555 -0.415700 14.01000 ; qtot -
2.416
6556 H 405 VAL H 6556 0.271900 1.00800 ; qtot -
2.144
6557 CX 405 VAL CA 6557 -0.087500 12.01000 ; qtot -
2.231
6558 H1 405 VAL HA 6558 0.096900 1.00800 ; qtot -
2.134
6559 A3C 405 VAL CB 6559 0.298500 12.01000 ; qtot -
1.836
6560 HC 405 VAL HB 6560 -0.029700 1.00800 ; qtot -
1.866
6561 CT 405 VAL CG1 6561 -0.319200 12.01000 ; qtot -
2.185
6562 HC 405 VAL HG11 6562 0.079100 1.00800 ; qtot -
2.106
6563 HC 405 VAL HG12 6563 0.079100 1.00800 ; qtot -
2.027
6564 HC 405 VAL HG13 6564 0.079100 1.00800 ; qtot -
1.947
6565 CT 405 VAL CG2 6565 -0.319200 12.01000 ; qtot -
2.267
6566 HC 405 VAL HG21 6566 0.079100 1.00800 ; qtot -
2.188
6567 HC 405 VAL HG22 6567 0.079100 1.00800 ; qtot -
2.108
6568 HC 405 VAL HG23 6568 0.079100 1.00800 ; qtot -
2.029
6569 C 405 VAL C 6569 0.597301 12.01000 ; qtot -
1.432
6570 O 405 VAL O 6570 -0.567901 16.00000 ; qtot -
2.000
6571 N 406 GLU N 6571 -0.516301 14.01000 ; qtot -

```
2.516
  6572    H   406   GLU     H 6572     0.293600      1.00800 ; qtot -
2.223
  6573   CX   406   GLU    CA 6573     0.039700     12.01000 ; qtot -
2.183
  6574   H1   406   GLU    HA 6574     0.110500      1.00800 ; qtot -
2.072
  6575  A2C   406   GLU    CB 6575     0.056000     12.01000 ; qtot -
2.016
  6576   HC   406   GLU   HB2 6576    -0.017300      1.00800 ; qtot -
2.034
  6577   HC   406   GLU   HB3 6577    -0.017300      1.00800 ; qtot -
2.051
  6578  A2C   406   GLU    CG 6578     0.013600     12.01000 ; qtot -
2.037
  6579   HC   406   GLU   HG2 6579    -0.042500      1.00800 ; qtot -
2.080
  6580   HC   406   GLU   HG3 6580    -0.042500      1.00800 ; qtot -
2.122
  6581   CO   406   GLU    CD 6581     0.805401     12.01000 ; qtot -
1.317
  6582   O2   406   GLU   OE1 6582    -0.818801     16.00000 ; qtot -
2.136
  6583   O2   406   GLU   OE2 6583    -0.818801     16.00000 ; qtot -
2.955
  6584    C   406   GLU     C 6584     0.536601     12.01000 ; qtot -
2.418
  6585    O   406   GLU     O 6585    -0.581901     16.00000 ; qtot -
3.000
  6586    N   407   GLH     N 6586    -0.415700     14.01000 ; qtot -
3.416
  6587    H   407   GLH     H 6587     0.271900      1.00800 ; qtot -
3.144
  6588   CX   407   GLH    CA 6588     0.014500     12.01000 ; qtot -
3.129
  6589   H1   407   GLH    HA 6589     0.077900      1.00800 ; qtot -
3.051
  6590  A2C   407   GLH    CB 6590    -0.007100     12.01000 ; qtot -
3.058
  6591   HC   407   GLH   HB2 6591     0.025600      1.00800 ; qtot -
3.033
  6592   HC   407   GLH   HB3 6592     0.025600      1.00800 ; qtot -
3.007
  6593  A2C   407   GLH    CG 6593    -0.017400     12.01000 ; qtot -
3.025
  6594   HC   407   GLH   HG2 6594     0.043000      1.00800 ; qtot -
2.982
  6595   HC   407   GLH   HG3 6595     0.043000      1.00800 ; qtot -
2.939
  6596    C   407   GLH    CD 6596     0.680101     12.01000 ; qtot -
2.259
  6597    O   407   GLH   OE1 6597    -0.583801     16.00000 ; qtot -
```

2.842
6598 OH 407 GLH OE2 6598 -0.651101 16.00000 ; qtot -
3.493
6599 HO 407 GLH HE2 6599 0.464100 1.00800 ; qtot -
3.029
6600 C 407 GLH C 6600 0.597301 12.01000 ; qtot -
2.432
6601 O 407 GLH O 6601 -0.567901 16.00000 ; qtot -
3.000
6602 N 408 LEU N 6602 -0.415700 14.01000 ; qtot -
3.416
6603 H 408 LEU H 6603 0.271900 1.00800 ; qtot -
3.144
6604 CX 408 LEU CA 6604 -0.051800 12.01000 ; qtot -
3.196
6605 H1 408 LEU HA 6605 0.092200 1.00800 ; qtot -
3.103
6606 A2C 408 LEU CB 6606 -0.110200 12.01000 ; qtot -
3.214
6607 HC 408 LEU HB2 6607 0.045700 1.00800 ; qtot -
3.168
6608 HC 408 LEU HB3 6608 0.045700 1.00800 ; qtot -
3.122
6609 A3C 408 LEU CG 6609 0.353100 12.01000 ; qtot -
2.769
6610 HC 408 LEU HG 6610 -0.036100 1.00800 ; qtot -
2.805
6611 CT 408 LEU CD1 6611 -0.412100 12.01000 ; qtot -
3.217
6612 HC 408 LEU HD11 6612 0.100000 1.00800 ; qtot -
3.117
6613 HC 408 LEU HD12 6613 0.100000 1.00800 ; qtot -
3.017
6614 HC 408 LEU HD13 6614 0.100000 1.00800 ; qtot -
2.917
6615 CT 408 LEU CD2 6615 -0.412100 12.01000 ; qtot -
3.329
6616 HC 408 LEU HD21 6616 0.100000 1.00800 ; qtot -
3.229
6617 HC 408 LEU HD22 6617 0.100000 1.00800 ; qtot -
3.129
6618 HC 408 LEU HD23 6618 0.100000 1.00800 ; qtot -
3.029
6619 C 408 LEU C 6619 0.597301 12.01000 ; qtot -
2.432
6620 O 408 LEU O 6620 -0.567901 16.00000 ; qtot -
3.000
6621 N 409 LYS N 6621 -0.347900 14.01000 ; qtot -
3.348
6622 H 409 LYS H 6622 0.274700 1.00800 ; qtot -
3.073
6623 CX 409 LYS CA 6623 -0.240000 12.01000 ; qtot -

3.313
  6624   H1   409   LYS    HA 6624     0.142600      1.00800 ; qtot -
3.171
  6625   C8   409   LYS    CB 6625    -0.009400     12.01000 ; qtot -
3.180
  6626   HC   409   LYS   HB2 6626     0.036200      1.00800 ; qtot -
3.144
  6627   HC   409   LYS   HB3 6627     0.036200      1.00800 ; qtot -
3.108
  6628   C8   409   LYS    CG 6628     0.018700     12.01000 ; qtot -
3.089
  6629   HC   409   LYS   HG2 6629     0.010300      1.00800 ; qtot -
3.079
  6630   HC   409   LYS   HG3 6630     0.010300      1.00800 ; qtot -
3.068
  6631   C8   409   LYS    CD 6631    -0.047900     12.01000 ; qtot -
3.116
  6632   HC   409   LYS   HD2 6632     0.062100      1.00800 ; qtot -
3.054
  6633   HC   409   LYS   HD3 6633     0.062100      1.00800 ; qtot -
2.992
  6634   C8   409   LYS    CE 6634    -0.014300     12.01000 ; qtot -
3.006
  6635   HP   409   LYS   HE2 6635     0.113500      1.00800 ; qtot -
2.893
  6636   HP   409   LYS   HE3 6636     0.113500      1.00800 ; qtot -
2.779
  6637   N3   409   LYS    NZ 6637    -0.385400     14.01000 ; qtot -
3.165
  6638    H   409   LYS   HZ1 6638     0.340000      1.00800 ; qtot -
2.825
  6639    H   409   LYS   HZ2 6639     0.340000      1.00800 ; qtot -
2.485
  6640    H   409   LYS   HZ3 6640     0.340000      1.00800 ; qtot -
2.145
  6641    C   409   LYS     C 6641     0.734101     12.01000 ; qtot -
1.411
  6642    O   409   LYS     O 6642    -0.589401     16.00000 ; qtot -
2.000
  6643    N   410   ARG     N 6643    -0.347900     14.01000 ; qtot -
2.348
  6644    H   410   ARG     H 6644     0.274700      1.00800 ; qtot -
2.073
  6645   CX   410   ARG    CA 6645    -0.263700     12.01000 ; qtot -
2.337
  6646   H1   410   ARG    HA 6646     0.156000      1.00800 ; qtot -
2.181
  6647   C8   410   ARG    CB 6647    -0.000700     12.01000 ; qtot -
2.182
  6648   HC   410   ARG   HB2 6648     0.032700      1.00800 ; qtot -
2.149
  6649   HC   410   ARG   HB3 6649     0.032700      1.00800 ; qtot -

2.116
  6650   C8   410   ARG    CG 6650     0.039000     12.01000 ; qtot -
2.077
  6651   HC   410   ARG   HG2 6651     0.028500      1.00800 ; qtot -
2.049
  6652   HC   410   ARG   HG3 6652     0.028500      1.00800 ; qtot -
2.020
  6653   C8   410   ARG    CD 6653     0.048600     12.01000 ; qtot -
1.972
  6654   H1   410   ARG   HD2 6654     0.068700      1.00800 ; qtot -
1.903
  6655   H1   410   ARG   HD3 6655     0.068700      1.00800 ; qtot -
1.834
  6656   N2   410   ARG    NE 6656    -0.529501     14.01000 ; qtot -
2.364
  6657    H   410   ARG    HE 6657     0.345600      1.00800 ; qtot -
2.018
  6658   CA   410   ARG    CZ 6658     0.807601     12.01000 ; qtot -
1.210
  6659   N2   410   ARG   NH1 6659    -0.862701     14.01000 ; qtot -
2.073
  6660    H   410   ARG  HH11 6660     0.447800      1.00800 ; qtot -
1.625
  6661    H   410   ARG  HH12 6661     0.447800      1.00800 ; qtot -
1.178
  6662   N2   410   ARG   NH2 6662    -0.862701     14.01000 ; qtot -
2.040
  6663    H   410   ARG  HH21 6663     0.447800      1.00800 ; qtot -
1.592
  6664    H   410   ARG  HH22 6664     0.447800      1.00800 ; qtot -
1.145
  6665    C   410   ARG     C 6665     0.734101     12.01000 ; qtot -
0.411
  6666    O   410   ARG     O 6666    -0.589401     16.00000 ; qtot -
1.000
  6667    N   411   ILE     N 6667    -0.415700     14.01000 ; qtot -
1.416
  6668    H   411   ILE     H 6668     0.271900      1.00800 ; qtot -
1.144
  6669   CX   411   ILE    CA 6669    -0.059700     12.01000 ; qtot -
1.203
  6670   H1   411   ILE    HA 6670     0.086900      1.00800 ; qtot -
1.117
  6671  A3C   411   ILE    CB 6671     0.130300     12.01000 ; qtot -
0.986
  6672   HC   411   ILE    HB 6672     0.018700      1.00800 ; qtot -
0.968
  6673   CT   411   ILE   CG2 6673    -0.320400     12.01000 ; qtot -
1.288
  6674   HC   411   ILE  HG21 6674     0.088200      1.00800 ; qtot -
1.200
  6675   HC   411   ILE  HG22 6675     0.088200      1.00800 ; qtot -

1.112
 6676 HC 411 ILE HG23 6676 0.088200 1.00800 ; qtot -
1.023
 6677 A2C 411 ILE CG1 6677 -0.043000 12.01000 ; qtot -
1.066
 6678 HC 411 ILE HG12 6678 0.023600 1.00800 ; qtot -
1.043
 6679 HC 411 ILE HG13 6679 0.023600 1.00800 ; qtot -
1.019
 6680 CT 411 ILE CD1 6680 -0.066000 12.01000 ; qtot -
1.085
 6681 HC 411 ILE HD11 6681 0.018600 1.00800 ; qtot -
1.067
 6682 HC 411 ILE HD12 6682 0.018600 1.00800 ; qtot -
1.048
 6683 HC 411 ILE HD13 6683 0.018600 1.00800 ; qtot -
1.029
 6684 C 411 ILE C 6684 0.597301 12.01000 ; qtot -
0.432
 6685 O 411 ILE O 6685 -0.567901 16.00000 ; qtot -
1.000
 6686 N 412 PHE N 6686 -0.415700 14.01000 ; qtot -
1.416
 6687 H 412 PHE H 6687 0.271900 1.00800 ; qtot -
1.144
 6688 CX 412 PHE CA 6688 -0.002400 12.01000 ; qtot -
1.146
 6689 H1 412 PHE HA 6689 0.097800 1.00800 ; qtot -
1.048
 6690 CT 412 PHE CB 6690 -0.034300 12.01000 ; qtot -
1.083
 6691 HC 412 PHE HB2 6691 0.029500 1.00800 ; qtot -
1.053
 6692 HC 412 PHE HB3 6692 0.029500 1.00800 ; qtot -
1.024
 6693 CA 412 PHE CG 6693 0.011800 12.01000 ; qtot -
1.012
 6694 CA 412 PHE CD1 6694 -0.125600 12.01000 ; qtot -
1.137
 6695 HA 412 PHE HD1 6695 0.133000 1.00800 ; qtot -
1.004
 6696 CA 412 PHE CE1 6696 -0.170400 12.01000 ; qtot -
1.175
 6697 HA 412 PHE HE1 6697 0.143000 1.00800 ; qtot -
1.032
 6698 CA 412 PHE CZ 6698 -0.107200 12.01000 ; qtot -
1.139
 6699 HA 412 PHE HZ 6699 0.129700 1.00800 ; qtot -
1.009
 6700 CA 412 PHE CE2 6700 -0.170400 12.01000 ; qtot -
1.180
 6701 HA 412 PHE HE2 6701 0.143000 1.00800 ; qtot -

1.037
  6702   CA   412   PHE   CD2 6702    -0.125600     12.01000 ; qtot -
1.162
  6703   HA   412   PHE   HD2 6703     0.133000      1.00800 ; qtot -
1.029
  6704    C   412   PHE     C 6704     0.597301     12.01000 ; qtot -
0.432
  6705    O   412   PHE     O 6705    -0.567901     16.00000 ; qtot -
1.000
  6706    N   413   GLY     N 6706    -0.415700     14.01000 ; qtot -
1.416
  6707    H   413   GLY     H 6707     0.271900      1.00800 ; qtot -
1.144
  6708   CX   413   GLY    CA 6708    -0.025200     12.01000 ; qtot -
1.169
  6709   H1   413   GLY   HA2 6709     0.069800      1.00800 ; qtot -
1.099
  6710   H1   413   GLY   HA3 6710     0.069800      1.00800 ; qtot -
1.029
  6711    C   413   GLY     C 6711     0.597301     12.01000 ; qtot -
0.432
  6712    O   413   GLY     O 6712    -0.567901     16.00000 ; qtot -
1.000
  6713    N   414   ARG     N 6713    -0.347900     14.01000 ; qtot -
1.348
  6714    H   414   ARG     H 6714     0.274700      1.00800 ; qtot -
1.073
  6715   CX   414   ARG    CA 6715    -0.263700     12.01000 ; qtot -
1.337
  6716   H1   414   ARG    HA 6716     0.156000      1.00800 ; qtot -
1.181
  6717   C8   414   ARG    CB 6717    -0.000700     12.01000 ; qtot -
1.182
  6718   HC   414   ARG   HB2 6718     0.032700      1.00800 ; qtot -
1.149
  6719   HC   414   ARG   HB3 6719     0.032700      1.00800 ; qtot -
1.116
  6720   C8   414   ARG    CG 6720     0.039000     12.01000 ; qtot -
1.077
  6721   HC   414   ARG   HG2 6721     0.028500      1.00800 ; qtot -
1.049
  6722   HC   414   ARG   HG3 6722     0.028500      1.00800 ; qtot -
1.020
  6723   C8   414   ARG    CD 6723     0.048600     12.01000 ; qtot -
0.972
  6724   H1   414   ARG   HD2 6724     0.068700      1.00800 ; qtot -
0.903
  6725   H1   414   ARG   HD3 6725     0.068700      1.00800 ; qtot -
0.834
  6726   N2   414   ARG    NE 6726    -0.529501     14.01000 ; qtot -
1.364
  6727    H   414   ARG    HE 6727     0.345600      1.00800 ; qtot -

1.018
  6728   CA   414   ARG    CZ 6728     0.807601     12.01000 ; qtot -
0.210
  6729   N2   414   ARG   NH1 6729    -0.862701     14.01000 ; qtot -
1.073
  6730    H   414   ARG  HH11 6730     0.447800      1.00800 ; qtot -
0.625
  6731    H   414   ARG  HH12 6731     0.447800      1.00800 ; qtot -
0.178
  6732   N2   414   ARG   NH2 6732    -0.862701     14.01000 ; qtot -
1.040
  6733    H   414   ARG  HH21 6733     0.447800      1.00800 ; qtot -
0.592
  6734    H   414   ARG  HH22 6734     0.447800      1.00800 ; qtot -
0.145
  6735    C   414   ARG     C 6735     0.734101     12.01000 ; qtot
0.589
  6736    O   414   ARG     O 6736    -0.589401     16.00000 ; qtot
0.000
  6737    N   415   ASP     N 6737    -0.516301     14.01000 ; qtot -
0.516
  6738    H   415   ASP     H 6738     0.293600      1.00800 ; qtot -
0.223
  6739   CX   415   ASP    CA 6739     0.038100     12.01000 ; qtot -
0.185
  6740   H1   415   ASP    HA 6740     0.088000      1.00800 ; qtot -
0.097
  6741  A2C   415   ASP    CB 6741    -0.030300     12.01000 ; qtot -
0.127
  6742   HC   415   ASP   HB2 6742    -0.012200      1.00800 ; qtot -
0.139
  6743   HC   415   ASP   HB3 6743    -0.012200      1.00800 ; qtot -
0.151
  6744   CO   415   ASP    CG 6744     0.799401     12.01000 ; qtot
0.648
  6745   O2   415   ASP   OD1 6745    -0.801401     16.00000 ; qtot -
0.153
  6746   O2   415   ASP   OD2 6746    -0.801401     16.00000 ; qtot -
0.955
  6747    C   415   ASP     C 6747     0.536601     12.01000 ; qtot -
0.418
  6748    O   415   ASP     O 6748    -0.581901     16.00000 ; qtot -
1.000
  6749    N   416   ASP     N 6749    -0.516301     14.01000 ; qtot -
1.516
  6750    H   416   ASP     H 6750     0.293600      1.00800 ; qtot -
1.223
  6751   CX   416   ASP    CA 6751     0.038100     12.01000 ; qtot -
1.185
  6752   H1   416   ASP    HA 6752     0.088000      1.00800 ; qtot -
1.097
  6753  A2C   416   ASP    CB 6753    -0.030300     12.01000 ; qtot -

1.127
6754 HC 416 ASP HB2 6754 -0.012200 1.00800 ; qtot -1.139
6755 HC 416 ASP HB3 6755 -0.012200 1.00800 ; qtot -1.151
6756 CO 416 ASP CG 6756 0.799401 12.01000 ; qtot -0.352
6757 O2 416 ASP OD1 6757 -0.801401 16.00000 ; qtot -1.153
6758 O2 416 ASP OD2 6758 -0.801401 16.00000 ; qtot -1.955
6759 C 416 ASP C 6759 0.536601 12.01000 ; qtot -1.418
6760 O 416 ASP O 6760 -0.581901 16.00000 ; qtot -2.000
6761 N 417 ILE N 6761 -0.415700 14.01000 ; qtot -2.416
6762 H 417 ILE H 6762 0.271900 1.00800 ; qtot -2.144
6763 CX 417 ILE CA 6763 -0.059700 12.01000 ; qtot -2.203
6764 H1 417 ILE HA 6764 0.086900 1.00800 ; qtot -2.117
6765 A3C 417 ILE CB 6765 0.130300 12.01000 ; qtot -1.986
6766 HC 417 ILE HB 6766 0.018700 1.00800 ; qtot -1.968
6767 CT 417 ILE CG2 6767 -0.320400 12.01000 ; qtot -2.288
6768 HC 417 ILE HG21 6768 0.088200 1.00800 ; qtot -2.200
6769 HC 417 ILE HG22 6769 0.088200 1.00800 ; qtot -2.112
6770 HC 417 ILE HG23 6770 0.088200 1.00800 ; qtot -2.023
6771 A2C 417 ILE CG1 6771 -0.043000 12.01000 ; qtot -2.066
6772 HC 417 ILE HG12 6772 0.023600 1.00800 ; qtot -2.043
6773 HC 417 ILE HG13 6773 0.023600 1.00800 ; qtot -2.019
6774 CT 417 ILE CD1 6774 -0.066000 12.01000 ; qtot -2.085
6775 HC 417 ILE HD11 6775 0.018600 1.00800 ; qtot -2.067
6776 HC 417 ILE HD12 6776 0.018600 1.00800 ; qtot -2.048
6777 HC 417 ILE HD13 6777 0.018600 1.00800 ; qtot -2.029
6778 C 417 ILE C 6778 0.597301 12.01000 ; qtot -1.432
6779 O 417 ILE O 6779 -0.567901 16.00000 ; qtot -

2.000
6780 N 418 GLU N 6780 -0.516301 14.01000 ; qtot -2.516
6781 H 418 GLU H 6781 0.293600 1.00800 ; qtot -2.223
6782 CX 418 GLU CA 6782 0.039700 12.01000 ; qtot -2.183
6783 H1 418 GLU HA 6783 0.110500 1.00800 ; qtot -2.072
6784 A2C 418 GLU CB 6784 0.056000 12.01000 ; qtot -2.016
6785 HC 418 GLU HB2 6785 -0.017300 1.00800 ; qtot -2.034
6786 HC 418 GLU HB3 6786 -0.017300 1.00800 ; qtot -2.051
6787 A2C 418 GLU CG 6787 0.013600 12.01000 ; qtot -2.037
6788 HC 418 GLU HG2 6788 -0.042500 1.00800 ; qtot -2.080
6789 HC 418 GLU HG3 6789 -0.042500 1.00800 ; qtot -2.122
6790 CO 418 GLU CD 6790 0.805401 12.01000 ; qtot -1.317
6791 O2 418 GLU OE1 6791 -0.818801 16.00000 ; qtot -2.136
6792 O2 418 GLU OE2 6792 -0.818801 16.00000 ; qtot -2.955
6793 C 418 GLU C 6793 0.536601 12.01000 ; qtot -2.418
6794 O 418 GLU O 6794 -0.581901 16.00000 ; qtot -3.000
6795 N 419 ASN N 6795 -0.415700 14.01000 ; qtot -3.416
6796 H 419 ASN H 6796 0.271900 1.00800 ; qtot -3.144
6797 CX 419 ASN CA 6797 0.014300 12.01000 ; qtot -3.129
6798 H1 419 ASN HA 6798 0.104800 1.00800 ; qtot -3.025
6799 A2C 419 ASN CB 6799 -0.204100 12.01000 ; qtot -3.229
6800 HC 419 ASN HB2 6800 0.079700 1.00800 ; qtot -3.149
6801 HC 419 ASN HB3 6801 0.079700 1.00800 ; qtot -3.069
6802 C 419 ASN CG 6802 0.713001 12.01000 ; qtot -2.356
6803 O 419 ASN OD1 6803 -0.593101 16.00000 ; qtot -2.949
6804 N 419 ASN ND2 6804 -0.919101 14.01000 ; qtot -3.869
6805 H 419 ASN HD21 6805 0.419600 1.00800 ; qtot -

3.449
6806 H 419 ASN HD22 6806 0.419600 1.00800 ; qtot -
3.029
6807 C 419 ASN C 6807 0.597301 12.01000 ; qtot -
2.432
6808 O 419 ASN O 6808 -0.567901 16.00000 ; qtot -
3.000
6809 N 420 PRO N 6809 -0.254800 14.01000 ; qtot -
3.255
6810 CT 420 PRO CD 6810 0.019200 12.01000 ; qtot -
3.236
6811 H1 420 PRO HD2 6811 0.039100 1.00800 ; qtot -
3.196
6812 H1 420 PRO HD3 6812 0.039100 1.00800 ; qtot -
3.157
6813 CT 420 PRO CG 6813 0.018900 12.01000 ; qtot -
3.138
6814 HC 420 PRO HG2 6814 0.021300 1.00800 ; qtot -
3.117
6815 HC 420 PRO HG3 6815 0.021300 1.00800 ; qtot -
3.096
6816 CT 420 PRO CB 6816 -0.007000 12.01000 ; qtot -
3.103
6817 HC 420 PRO HB2 6817 0.025300 1.00800 ; qtot -
3.078
6818 HC 420 PRO HB3 6818 0.025300 1.00800 ; qtot -
3.052
6819 CX 420 PRO CA 6819 -0.026600 12.01000 ; qtot -
3.079
6820 H1 420 PRO HA 6820 0.064100 1.00800 ; qtot -
3.015
6821 C 420 PRO C 6821 0.589601 12.01000 ; qtot -
2.425
6822 O 420 PRO O 6822 -0.574801 16.00000 ; qtot -
3.000
6823 N 421 LEU N 6823 -0.415700 14.01000 ; qtot -
3.416
6824 H 421 LEU H 6824 0.271900 1.00800 ; qtot -
3.144
6825 CX 421 LEU CA 6825 -0.051800 12.01000 ; qtot -
3.196
6826 H1 421 LEU HA 6826 0.092200 1.00800 ; qtot -
3.103
6827 A2C 421 LEU CB 6827 -0.110200 12.01000 ; qtot -
3.214
6828 HC 421 LEU HB2 6828 0.045700 1.00800 ; qtot -
3.168
6829 HC 421 LEU HB3 6829 0.045700 1.00800 ; qtot -
3.122
6830 A3C 421 LEU CG 6830 0.353100 12.01000 ; qtot -
2.769
6831 HC 421 LEU HG 6831 -0.036100 1.00800 ; qtot -

2.805
  6832   CT   421   LEU   CD1 6832    -0.412100     12.01000 ; qtot -
3.217
  6833   HC   421   LEU  HD11 6833     0.100000      1.00800 ; qtot -
3.117
  6834   HC   421   LEU  HD12 6834     0.100000      1.00800 ; qtot -
3.017
  6835   HC   421   LEU  HD13 6835     0.100000      1.00800 ; qtot -
2.917
  6836   CT   421   LEU   CD2 6836    -0.412100     12.01000 ; qtot -
3.329
  6837   HC   421   LEU  HD21 6837     0.100000      1.00800 ; qtot -
3.229
  6838   HC   421   LEU  HD22 6838     0.100000      1.00800 ; qtot -
3.129
  6839   HC   421   LEU  HD23 6839     0.100000      1.00800 ; qtot -
3.029
  6840    C   421   LEU     C 6840     0.597301     12.01000 ; qtot -
2.432
  6841    O   421   LEU     O 6841    -0.567901     16.00000 ; qtot -
3.000
  6842    N   422   ILE     N 6842    -0.415700     14.01000 ; qtot -
3.416
  6843    H   422   ILE     H 6843     0.271900      1.00800 ; qtot -
3.144
  6844   CX   422   ILE    CA 6844    -0.059700     12.01000 ; qtot -
3.203
  6845   H1   422   ILE    HA 6845     0.086900      1.00800 ; qtot -
3.117
  6846  A3C   422   ILE    CB 6846     0.130300     12.01000 ; qtot -
2.986
  6847   HC   422   ILE    HB 6847     0.018700      1.00800 ; qtot -
2.968
  6848   CT   422   ILE   CG2 6848    -0.320400     12.01000 ; qtot -
3.288
  6849   HC   422   ILE  HG21 6849     0.088200      1.00800 ; qtot -
3.200
  6850   HC   422   ILE  HG22 6850     0.088200      1.00800 ; qtot -
3.112
  6851   HC   422   ILE  HG23 6851     0.088200      1.00800 ; qtot -
3.023
  6852  A2C   422   ILE   CG1 6852    -0.043000     12.01000 ; qtot -
3.066
  6853   HC   422   ILE  HG12 6853     0.023600      1.00800 ; qtot -
3.043
  6854   HC   422   ILE  HG13 6854     0.023600      1.00800 ; qtot -
3.019
  6855   CT   422   ILE   CD1 6855    -0.066000     12.01000 ; qtot -
3.085
  6856   HC   422   ILE  HD11 6856     0.018600      1.00800 ; qtot -
3.067
  6857   HC   422   ILE  HD12 6857     0.018600      1.00800 ; qtot -

```
3.048
  6858   HC   422   ILE  HD13 6858     0.018600      1.00800 ; qtot -
3.029
  6859    C   422   ILE     C 6859     0.597301     12.01000 ; qtot -
2.432
  6860    O   422   ILE     O 6860    -0.567901     16.00000 ; qtot -
3.000
  6861    N   423   ASP     N 6861    -0.516301     14.01000 ; qtot -
3.516
  6862    H   423   ASP     H 6862     0.293600      1.00800 ; qtot -
3.223
  6863   CX   423   ASP    CA 6863     0.038100     12.01000 ; qtot -
3.185
  6864   H1   423   ASP    HA 6864     0.088000      1.00800 ; qtot -
3.097
  6865  A2C   423   ASP    CB 6865    -0.030300     12.01000 ; qtot -
3.127
  6866   HC   423   ASP   HB2 6866    -0.012200      1.00800 ; qtot -
3.139
  6867   HC   423   ASP   HB3 6867    -0.012200      1.00800 ; qtot -
3.151
  6868   CO   423   ASP    CG 6868     0.799401     12.01000 ; qtot -
2.352
  6869   O2   423   ASP   OD1 6869    -0.801401     16.00000 ; qtot -
3.153
  6870   O2   423   ASP   OD2 6870    -0.801401     16.00000 ; qtot -
3.955
  6871    C   423   ASP     C 6871     0.536601     12.01000 ; qtot -
3.418
  6872    O   423   ASP     O 6872    -0.581901     16.00000 ; qtot -
4.000
  6873    N   424   LYS     N 6873    -0.347900     14.01000 ; qtot -
4.348
  6874    H   424   LYS     H 6874     0.274700      1.00800 ; qtot -
4.073
  6875   CX   424   LYS    CA 6875    -0.240000     12.01000 ; qtot -
4.313
  6876   H1   424   LYS    HA 6876     0.142600      1.00800 ; qtot -
4.171
  6877   C8   424   LYS    CB 6877    -0.009400     12.01000 ; qtot -
4.180
  6878   HC   424   LYS   HB2 6878     0.036200      1.00800 ; qtot -
4.144
  6879   HC   424   LYS   HB3 6879     0.036200      1.00800 ; qtot -
4.108
  6880   C8   424   LYS    CG 6880     0.018700     12.01000 ; qtot -
4.089
  6881   HC   424   LYS   HG2 6881     0.010300      1.00800 ; qtot -
4.079
  6882   HC   424   LYS   HG3 6882     0.010300      1.00800 ; qtot -
4.068
  6883   C8   424   LYS    CD 6883    -0.047900     12.01000 ; qtot -
```

4.116
6884 HC 424 LYS HD2 6884 0.062100 1.00800 ; qtot -
4.054
6885 HC 424 LYS HD3 6885 0.062100 1.00800 ; qtot -
3.992
6886 C8 424 LYS CE 6886 -0.014300 12.01000 ; qtot -
4.006
6887 HP 424 LYS HE2 6887 0.113500 1.00800 ; qtot -
3.893
6888 HP 424 LYS HE3 6888 0.113500 1.00800 ; qtot -
3.779
6889 N3 424 LYS NZ 6889 -0.385400 14.01000 ; qtot -
4.165
6890 H 424 LYS HZ1 6890 0.340000 1.00800 ; qtot -
3.825
6891 H 424 LYS HZ2 6891 0.340000 1.00800 ; qtot -
3.485
6892 H 424 LYS HZ3 6892 0.340000 1.00800 ; qtot -
3.145
6893 C 424 LYS C 6893 0.734101 12.01000 ; qtot -
2.411
6894 O 424 LYS O 6894 -0.589401 16.00000 ; qtot -
3.000
6895 N 425 ALA N 6895 -0.415700 14.01000 ; qtot -
3.416
6896 H 425 ALA H 6896 0.271900 1.00800 ; qtot -
3.144
6897 CX 425 ALA CA 6897 0.033700 12.01000 ; qtot -
3.110
6898 H1 425 ALA HA 6898 0.082300 1.00800 ; qtot -
3.028
6899 CT 425 ALA CB 6899 -0.182500 12.01000 ; qtot -
3.210
6900 HC 425 ALA HB1 6900 0.060300 1.00800 ; qtot -
3.150
6901 HC 425 ALA HB2 6901 0.060300 1.00800 ; qtot -
3.090
6902 HC 425 ALA HB3 6902 0.060300 1.00800 ; qtot -
3.029
6903 C 425 ALA C 6903 0.597301 12.01000 ; qtot -
2.432
6904 O 425 ALA O 6904 -0.567901 16.00000 ; qtot -
3.000
6905 N 426 ILE N 6905 -0.415700 14.01000 ; qtot -
3.416
6906 H 426 ILE H 6906 0.271900 1.00800 ; qtot -
3.144
6907 CX 426 ILE CA 6907 -0.059700 12.01000 ; qtot -
3.203
6908 H1 426 ILE HA 6908 0.086900 1.00800 ; qtot -
3.117
6909 A3C 426 ILE CB 6909 0.130300 12.01000 ; qtot -

2.986
6910 HC 426 ILE HB 6910 0.018700 1.00800 ; qtot -
2.968
6911 CT 426 ILE CG2 6911 -0.320400 12.01000 ; qtot -
3.288
6912 HC 426 ILE HG21 6912 0.088200 1.00800 ; qtot -
3.200
6913 HC 426 ILE HG22 6913 0.088200 1.00800 ; qtot -
3.112
6914 HC 426 ILE HG23 6914 0.088200 1.00800 ; qtot -
3.023
6915 A2C 426 ILE CG1 6915 -0.043000 12.01000 ; qtot -
3.066
6916 HC 426 ILE HG12 6916 0.023600 1.00800 ; qtot -
3.043
6917 HC 426 ILE HG13 6917 0.023600 1.00800 ; qtot -
3.019
6918 CT 426 ILE CD1 6918 -0.066000 12.01000 ; qtot -
3.085
6919 HC 426 ILE HD11 6919 0.018600 1.00800 ; qtot -
3.067
6920 HC 426 ILE HD12 6920 0.018600 1.00800 ; qtot -
3.048
6921 HC 426 ILE HD13 6921 0.018600 1.00800 ; qtot -
3.029
6922 C 426 ILE C 6922 0.597301 12.01000 ; qtot -
2.432
6923 O 426 ILE O 6923 -0.567901 16.00000 ; qtot -
3.000
6924 N 427 ASP N 6924 -0.516301 14.01000 ; qtot -
3.516
6925 H 427 ASP H 6925 0.293600 1.00800 ; qtot -
3.223
6926 CX 427 ASP CA 6926 0.038100 12.01000 ; qtot -
3.185
6927 H1 427 ASP HA 6927 0.088000 1.00800 ; qtot -
3.097
6928 A2C 427 ASP CB 6928 -0.030300 12.01000 ; qtot -
3.127
6929 HC 427 ASP HB2 6929 -0.012200 1.00800 ; qtot -
3.139
6930 HC 427 ASP HB3 6930 -0.012200 1.00800 ; qtot -
3.151
6931 CO 427 ASP CG 6931 0.799401 12.01000 ; qtot -
2.352
6932 O2 427 ASP OD1 6932 -0.801401 16.00000 ; qtot -
3.153
6933 O2 427 ASP OD2 6933 -0.801401 16.00000 ; qtot -
3.955
6934 C 427 ASP C 6934 0.536601 12.01000 ; qtot -
3.418
6935 O 427 ASP O 6935 -0.581901 16.00000 ; qtot -

4.000
 6936 N 428 SER N 6936 -0.415700 14.01000 ; qtot -
4.416
 6937 H 428 SER H 6937 0.271900 1.00800 ; qtot -
4.144
 6938 CX 428 SER CA 6938 -0.024900 12.01000 ; qtot -
4.169
 6939 H1 428 SER HA 6939 0.084300 1.00800 ; qtot -
4.084
 6940 A2C 428 SER CB 6940 0.211700 12.01000 ; qtot -
3.873
 6941 H1 428 SER HB2 6941 0.035200 1.00800 ; qtot -
3.837
 6942 H1 428 SER HB3 6942 0.035200 1.00800 ; qtot -
3.802
 6943 OH 428 SER OG 6943 -0.654601 16.00000 ; qtot -
4.457
 6944 HO 428 SER HG 6944 0.427500 1.00800 ; qtot -
4.029
 6945 C 428 SER C 6945 0.597301 12.01000 ; qtot -
3.432
 6946 O 428 SER O 6946 -0.567901 16.00000 ; qtot -
4.000
 6947 N 429 LEU N 6947 -0.415700 14.01000 ; qtot -
4.416
 6948 H 429 LEU H 6948 0.271900 1.00800 ; qtot -
4.144
 6949 CX 429 LEU CA 6949 -0.051800 12.01000 ; qtot -
4.196
 6950 H1 429 LEU HA 6950 0.092200 1.00800 ; qtot -
4.103
 6951 A2C 429 LEU CB 6951 -0.110200 12.01000 ; qtot -
4.214
 6952 HC 429 LEU HB2 6952 0.045700 1.00800 ; qtot -
4.168
 6953 HC 429 LEU HB3 6953 0.045700 1.00800 ; qtot -
4.122
 6954 A3C 429 LEU CG 6954 0.353100 12.01000 ; qtot -
3.769
 6955 HC 429 LEU HG 6955 -0.036100 1.00800 ; qtot -
3.805
 6956 CT 429 LEU CD1 6956 -0.412100 12.01000 ; qtot -
4.217
 6957 HC 429 LEU HD11 6957 0.100000 1.00800 ; qtot -
4.117
 6958 HC 429 LEU HD12 6958 0.100000 1.00800 ; qtot -
4.017
 6959 HC 429 LEU HD13 6959 0.100000 1.00800 ; qtot -
3.917
 6960 CT 429 LEU CD2 6960 -0.412100 12.01000 ; qtot -
4.329
 6961 HC 429 LEU HD21 6961 0.100000 1.00800 ; qtot -

4.229
6962 HC 429 LEU HD22 6962 0.100000 1.00800 ; qtot -
4.129
6963 HC 429 LEU HD23 6963 0.100000 1.00800 ; qtot -
4.029
6964 C 429 LEU C 6964 0.597301 12.01000 ; qtot -
3.432
6965 O 429 LEU O 6965 -0.567901 16.00000 ; qtot -
4.000
6966 N 430 LEU N 6966 -0.415700 14.01000 ; qtot -
4.416
6967 H 430 LEU H 6967 0.271900 1.00800 ; qtot -
4.144
6968 CX 430 LEU CA 6968 -0.051800 12.01000 ; qtot -
4.196
6969 H1 430 LEU HA 6969 0.092200 1.00800 ; qtot -
4.103
6970 A2C 430 LEU CB 6970 -0.110200 12.01000 ; qtot -
4.214
6971 HC 430 LEU HB2 6971 0.045700 1.00800 ; qtot -
4.168
6972 HC 430 LEU HB3 6972 0.045700 1.00800 ; qtot -
4.122
6973 A3C 430 LEU CG 6973 0.353100 12.01000 ; qtot -
3.769
6974 HC 430 LEU HG 6974 -0.036100 1.00800 ; qtot -
3.805
6975 CT 430 LEU CD1 6975 -0.412100 12.01000 ; qtot -
4.217
6976 HC 430 LEU HD11 6976 0.100000 1.00800 ; qtot -
4.117
6977 HC 430 LEU HD12 6977 0.100000 1.00800 ; qtot -
4.017
6978 HC 430 LEU HD13 6978 0.100000 1.00800 ; qtot -
3.917
6979 CT 430 LEU CD2 6979 -0.412100 12.01000 ; qtot -
4.329
6980 HC 430 LEU HD21 6980 0.100000 1.00800 ; qtot -
4.229
6981 HC 430 LEU HD22 6981 0.100000 1.00800 ; qtot -
4.129
6982 HC 430 LEU HD23 6982 0.100000 1.00800 ; qtot -
4.029
6983 C 430 LEU C 6983 0.597301 12.01000 ; qtot -
3.432
6984 O 430 LEU O 6984 -0.567901 16.00000 ; qtot -
4.000
6985 N 431 GLN N 6985 -0.415700 14.01000 ; qtot -
4.416
6986 H 431 GLN H 6986 0.271900 1.00800 ; qtot -
4.144
6987 CX 431 GLN CA 6987 -0.003100 12.01000 ; qtot -

4.147
 6988 H1 431 GLN HA 6988 0.085000 1.00800 ; qtot -
4.062
 6989 A2C 431 GLN CB 6989 -0.003600 12.01000 ; qtot -
4.065
 6990 HC 431 GLN HB2 6990 0.017100 1.00800 ; qtot -
4.048
 6991 HC 431 GLN HB3 6991 0.017100 1.00800 ; qtot -
4.031
 6992 A2C 431 GLN CG 6992 -0.064500 12.01000 ; qtot -
4.096
 6993 HC 431 GLN HG2 6993 0.035200 1.00800 ; qtot -
4.061
 6994 HC 431 GLN HG3 6994 0.035200 1.00800 ; qtot -
4.025
 6995 C 431 GLN CD 6995 0.695101 12.01000 ; qtot -
3.330
 6996 O 431 GLN OE1 6996 -0.608601 16.00000 ; qtot -
3.939
 6997 N 431 GLN NE2 6997 -0.940701 14.01000 ; qtot -
4.880
 6998 H 431 GLN HE21 6998 0.425100 1.00800 ; qtot -
4.454
 6999 H 431 GLN HE22 6999 0.425100 1.00800 ; qtot -
4.029
 7000 C 431 GLN C 7000 0.597301 12.01000 ; qtot -
3.432
 7001 O 431 GLN O 7001 -0.567901 16.00000 ; qtot -
4.000
 7002 N 432 GLU N 7002 -0.516301 14.01000 ; qtot -
4.516
 7003 H 432 GLU H 7003 0.293600 1.00800 ; qtot -
4.223
 7004 CX 432 GLU CA 7004 0.039700 12.01000 ; qtot -
4.183
 7005 H1 432 GLU HA 7005 0.110500 1.00800 ; qtot -
4.072
 7006 A2C 432 GLU CB 7006 0.056000 12.01000 ; qtot -
4.016
 7007 HC 432 GLU HB2 7007 -0.017300 1.00800 ; qtot -
4.034
 7008 HC 432 GLU HB3 7008 -0.017300 1.00800 ; qtot -
4.051
 7009 A2C 432 GLU CG 7009 0.013600 12.01000 ; qtot -
4.037
 7010 HC 432 GLU HG2 7010 -0.042500 1.00800 ; qtot -
4.080
 7011 HC 432 GLU HG3 7011 -0.042500 1.00800 ; qtot -
4.122
 7012 CO 432 GLU CD 7012 0.805401 12.01000 ; qtot -
3.317
 7013 O2 432 GLU OE1 7013 -0.818801 16.00000 ; qtot -

```
4.136
  7014   O2   432   GLU   OE2 7014    -0.818801     16.00000 ; qtot -
4.955
  7015    C   432   GLU     C 7015     0.536601     12.01000 ; qtot -
4.418
  7016    O   432   GLU     O 7016    -0.581901     16.00000 ; qtot -
5.000
  7017    N   433   VAL     N 7017    -0.415700     14.01000 ; qtot -
5.416
  7018    H   433   VAL     H 7018     0.271900      1.00800 ; qtot -
5.144
  7019   CX   433   VAL    CA 7019    -0.087500     12.01000 ; qtot -
5.231
  7020   H1   433   VAL    HA 7020     0.096900      1.00800 ; qtot -
5.134
  7021  A3C   433   VAL    CB 7021     0.298500     12.01000 ; qtot -
4.836
  7022   HC   433   VAL    HB 7022    -0.029700      1.00800 ; qtot -
4.866
  7023   CT   433   VAL   CG1 7023    -0.319200     12.01000 ; qtot -
5.185
  7024   HC   433   VAL  HG11 7024     0.079100      1.00800 ; qtot -
5.106
  7025   HC   433   VAL  HG12 7025     0.079100      1.00800 ; qtot -
5.027
  7026   HC   433   VAL  HG13 7026     0.079100      1.00800 ; qtot -
4.947
  7027   CT   433   VAL   CG2 7027    -0.319200     12.01000 ; qtot -
5.267
  7028   HC   433   VAL  HG21 7028     0.079100      1.00800 ; qtot -
5.188
  7029   HC   433   VAL  HG22 7029     0.079100      1.00800 ; qtot -
5.108
  7030   HC   433   VAL  HG23 7030     0.079100      1.00800 ; qtot -
5.029
  7031    C   433   VAL     C 7031     0.597301     12.01000 ; qtot -
4.432
  7032    O   433   VAL     O 7032    -0.567901     16.00000 ; qtot -
5.000
  7033    N   434   ASP     N 7033    -0.516301     14.01000 ; qtot -
5.516
  7034    H   434   ASP     H 7034     0.293600      1.00800 ; qtot -
5.223
  7035   CX   434   ASP    CA 7035     0.038100     12.01000 ; qtot -
5.185
  7036   H1   434   ASP    HA 7036     0.088000      1.00800 ; qtot -
5.097
  7037  A2C   434   ASP    CB 7037    -0.030300     12.01000 ; qtot -
5.127
  7038   HC   434   ASP   HB2 7038    -0.012200      1.00800 ; qtot -
5.139
  7039   HC   434   ASP   HB3 7039    -0.012200      1.00800 ; qtot -
```

5.151
  7040   CO   434   ASP    CG 7040     0.799401     12.01000 ; qtot -
4.352
  7041   O2   434   ASP   OD1 7041    -0.801401     16.00000 ; qtot -
5.153
  7042   O2   434   ASP   OD2 7042    -0.801401     16.00000 ; qtot -
5.955
  7043    C   434   ASP     C 7043     0.536601     12.01000 ; qtot -
5.418
  7044    O   434   ASP     O 7044    -0.581901     16.00000 ; qtot -
6.000
  7045    N   435   LEU     N 7045    -0.415700     14.01000 ; qtot -
6.416
  7046    H   435   LEU     H 7046     0.271900      1.00800 ; qtot -
6.144
  7047   CX   435   LEU    CA 7047    -0.051800     12.01000 ; qtot -
6.196
  7048   H1   435   LEU    HA 7048     0.092200      1.00800 ; qtot -
6.103
  7049  A2C   435   LEU    CB 7049    -0.110200     12.01000 ; qtot -
6.214
  7050   HC   435   LEU   HB2 7050     0.045700      1.00800 ; qtot -
6.168
  7051   HC   435   LEU   HB3 7051     0.045700      1.00800 ; qtot -
6.122
  7052  A3C   435   LEU    CG 7052     0.353100     12.01000 ; qtot -
5.769
  7053   HC   435   LEU    HG 7053    -0.036100      1.00800 ; qtot -
5.805
  7054   CT   435   LEU   CD1 7054    -0.412100     12.01000 ; qtot -
6.217
  7055   HC   435   LEU  HD11 7055     0.100000      1.00800 ; qtot -
6.117
  7056   HC   435   LEU  HD12 7056     0.100000      1.00800 ; qtot -
6.017
  7057   HC   435   LEU  HD13 7057     0.100000      1.00800 ; qtot -
5.917
  7058   CT   435   LEU   CD2 7058    -0.412100     12.01000 ; qtot -
6.329
  7059   HC   435   LEU  HD21 7059     0.100000      1.00800 ; qtot -
6.229
  7060   HC   435   LEU  HD22 7060     0.100000      1.00800 ; qtot -
6.129
  7061   HC   435   LEU  HD23 7061     0.100000      1.00800 ; qtot -
6.029
  7062    C   435   LEU     C 7062     0.597301     12.01000 ; qtot -
5.432
  7063    O   435   LEU     O 7063    -0.567901     16.00000 ; qtot -
6.000
  7064    N   436   ASN     N 7064    -0.415700     14.01000 ; qtot -
6.416
  7065    H   436   ASN     H 7065     0.271900      1.00800 ; qtot -

6.144
  7066   CX   436   ASN     CA 7066     0.014300     12.01000 ; qtot -
6.130
  7067   H1   436   ASN     HA 7067     0.104800      1.00800 ; qtot -
6.025
  7068  A2C   436   ASN     CB 7068    -0.204100     12.01000 ; qtot -
6.229
  7069   HC   436   ASN    HB2 7069     0.079700      1.00800 ; qtot -
6.149
  7070   HC   436   ASN    HB3 7070     0.079700      1.00800 ; qtot -
6.069
  7071    C   436   ASN     CG 7071     0.713001     12.01000 ; qtot -
5.356
  7072    O   436   ASN    OD1 7072    -0.593101     16.00000 ; qtot -
5.949
  7073    N   436   ASN    ND2 7073    -0.919101     14.01000 ; qtot -
6.869
  7074    H   436   ASN   HD21 7074     0.419600      1.00800 ; qtot -
6.449
  7075    H   436   ASN   HD22 7075     0.419600      1.00800 ; qtot -
6.029
  7076    C   436   ASN      C 7076     0.597301     12.01000 ; qtot -
5.432
  7077    O   436   ASN      O 7077    -0.567901     16.00000 ; qtot -
6.000
  7078    N   437   GLY      N 7078    -0.415700     14.01000 ; qtot -
6.416
  7079    H   437   GLY      H 7079     0.271900      1.00800 ; qtot -
6.144
  7080   CX   437   GLY     CA 7080    -0.025200     12.01000 ; qtot -
6.169
  7081   H1   437   GLY    HA2 7081     0.069800      1.00800 ; qtot -
6.099
  7082   H1   437   GLY    HA3 7082     0.069800      1.00800 ; qtot -
6.029
  7083    C   437   GLY      C 7083     0.597301     12.01000 ; qtot -
5.432
  7084    O   437   GLY      O 7084    -0.567901     16.00000 ; qtot -
6.000
  7085    N   438   ASP      N 7085    -0.516301     14.01000 ; qtot -
6.516
  7086    H   438   ASP      H 7086     0.293600      1.00800 ; qtot -
6.223
  7087   CX   438   ASP     CA 7087     0.038100     12.01000 ; qtot -
6.185
  7088   H1   438   ASP     HA 7088     0.088000      1.00800 ; qtot -
6.097
  7089  A2C   438   ASP     CB 7089    -0.030300     12.01000 ; qtot -
6.127
  7090   HC   438   ASP    HB2 7090    -0.012200      1.00800 ; qtot -
6.139
  7091   HC   438   ASP    HB3 7091    -0.012200      1.00800 ; qtot -

6.151
  7092   CO   438   ASP    CG 7092     0.799401     12.01000 ; qtot -
5.352
  7093   O2   438   ASP   OD1 7093    -0.801401     16.00000 ; qtot -
6.153
  7094   O2   438   ASP   OD2 7094    -0.801401     16.00000 ; qtot -
6.955
  7095    C   438   ASP     C 7095     0.536601     12.01000 ; qtot -
6.418
  7096    O   438   ASP     O 7096    -0.581901     16.00000 ; qtot -
7.000
  7097    N   439   GLY     N 7097    -0.415700     14.01000 ; qtot -
7.416
  7098    H   439   GLY     H 7098     0.271900      1.00800 ; qtot -
7.144
  7099   CX   439   GLY    CA 7099    -0.025200     12.01000 ; qtot -
7.169
  7100   H1   439   GLY   HA2 7100     0.069800      1.00800 ; qtot -
7.099
  7101   H1   439   GLY   HA3 7101     0.069800      1.00800 ; qtot -
7.029
  7102    C   439   GLY     C 7102     0.597301     12.01000 ; qtot -
6.432
  7103    O   439   GLY     O 7103    -0.567901     16.00000 ; qtot -
7.000
  7104    N   440   GLU     N 7104    -0.516301     14.01000 ; qtot -
7.516
  7105    H   440   GLU     H 7105     0.293600      1.00800 ; qtot -
7.223
  7106   CX   440   GLU    CA 7106     0.039700     12.01000 ; qtot -
7.183
  7107   H1   440   GLU    HA 7107     0.110500      1.00800 ; qtot -
7.073
  7108  A2C   440   GLU    CB 7108     0.056000     12.01000 ; qtot -
7.017
  7109   HC   440   GLU   HB2 7109    -0.017300      1.00800 ; qtot -
7.034
  7110   HC   440   GLU   HB3 7110    -0.017300      1.00800 ; qtot -
7.051
  7111  A2C   440   GLU    CG 7111     0.013600     12.01000 ; qtot -
7.038
  7112   HC   440   GLU   HG2 7112    -0.042500      1.00800 ; qtot -
7.080
  7113   HC   440   GLU   HG3 7113    -0.042500      1.00800 ; qtot -
7.123
  7114   CO   440   GLU    CD 7114     0.805401     12.01000 ; qtot -
6.317
  7115   O2   440   GLU   OE1 7115    -0.818801     16.00000 ; qtot -
7.136
  7116   O2   440   GLU   OE2 7116    -0.818801     16.00000 ; qtot -
7.955
  7117    C   440   GLU     C 7117     0.536601     12.01000 ; qtot -

```
7.418
  7118    O   440   GLU      O 7118    -0.581901     16.00000 ; qtot -
8.000
  7119    N   441   ILE      N 7119    -0.415700     14.01000 ; qtot -
8.416
  7120    H   441   ILE      H 7120     0.271900      1.00800 ; qtot -
8.144
  7121   CX   441   ILE     CA 7121    -0.059700     12.01000 ; qtot -
8.204
  7122   H1   441   ILE     HA 7122     0.086900      1.00800 ; qtot -
8.117
  7123  A3C   441   ILE     CB 7123     0.130300     12.01000 ; qtot -
7.986
  7124   HC   441   ILE     HB 7124     0.018700      1.00800 ; qtot -
7.968
  7125   CT   441   ILE    CG2 7125    -0.320400     12.01000 ; qtot -
8.288
  7126   HC   441   ILE   HG21 7126     0.088200      1.00800 ; qtot -
8.200
  7127   HC   441   ILE   HG22 7127     0.088200      1.00800 ; qtot -
8.112
  7128   HC   441   ILE   HG23 7128     0.088200      1.00800 ; qtot -
8.023
  7129  A2C   441   ILE    CG1 7129    -0.043000     12.01000 ; qtot -
8.066
  7130   HC   441   ILE   HG12 7130     0.023600      1.00800 ; qtot -
8.043
  7131   HC   441   ILE   HG13 7131     0.023600      1.00800 ; qtot -
8.019
  7132   CT   441   ILE    CD1 7132    -0.066000     12.01000 ; qtot -
8.085
  7133   HC   441   ILE   HD11 7133     0.018600      1.00800 ; qtot -
8.067
  7134   HC   441   ILE   HD12 7134     0.018600      1.00800 ; qtot -
8.048
  7135   HC   441   ILE   HD13 7135     0.018600      1.00800 ; qtot -
8.029
  7136    C   441   ILE      C 7136     0.597301     12.01000 ; qtot -
7.432
  7137    O   441   ILE      O 7137    -0.567901     16.00000 ; qtot -
8.000
  7138    N   442   ASP      N 7138    -0.516301     14.01000 ; qtot -
8.516
  7139    H   442   ASP      H 7139     0.293600      1.00800 ; qtot -
8.223
  7140   CX   442   ASP     CA 7140     0.038100     12.01000 ; qtot -
8.185
  7141   H1   442   ASP     HA 7141     0.088000      1.00800 ; qtot -
8.097
  7142  A2C   442   ASP     CB 7142    -0.030300     12.01000 ; qtot -
8.127
  7143   HC   442   ASP    HB2 7143    -0.012200      1.00800 ; qtot -
```

8.139
7144 HC 442 ASP HB3 7144 -0.012200 1.00800 ; qtot -8.151
7145 CO 442 ASP CG 7145 0.799401 12.01000 ; qtot -7.352
7146 O2 442 ASP OD1 7146 -0.801401 16.00000 ; qtot -8.153
7147 O2 442 ASP OD2 7147 -0.801401 16.00000 ; qtot -8.955
7148 C 442 ASP C 7148 0.536601 12.01000 ; qtot -8.418
7149 O 442 ASP O 7149 -0.581901 16.00000 ; qtot -9.000
7150 N 443 PHE N 7150 -0.415700 14.01000 ; qtot -9.416
7151 H 443 PHE H 7151 0.271900 1.00800 ; qtot -9.144
7152 CX 443 PHE CA 7152 -0.002400 12.01000 ; qtot -9.146
7153 H1 443 PHE HA 7153 0.097800 1.00800 ; qtot -9.048
7154 CT 443 PHE CB 7154 -0.034300 12.01000 ; qtot -9.083
7155 HC 443 PHE HB2 7155 0.029500 1.00800 ; qtot -9.053
7156 HC 443 PHE HB3 7156 0.029500 1.00800 ; qtot -9.024
7157 CA 443 PHE CG 7157 0.011800 12.01000 ; qtot -9.012
7158 CA 443 PHE CD1 7158 -0.125600 12.01000 ; qtot -9.138
7159 HA 443 PHE HD1 7159 0.133000 1.00800 ; qtot -9.005
7160 CA 443 PHE CE1 7160 -0.170400 12.01000 ; qtot -9.175
7161 HA 443 PHE HE1 7161 0.143000 1.00800 ; qtot -9.032
7162 CA 443 PHE CZ 7162 -0.107200 12.01000 ; qtot -9.139
7163 HA 443 PHE HZ 7163 0.129700 1.00800 ; qtot -9.009
7164 CA 443 PHE CE2 7164 -0.170400 12.01000 ; qtot -9.180
7165 HA 443 PHE HE2 7165 0.143000 1.00800 ; qtot -9.037
7166 CA 443 PHE CD2 7166 -0.125600 12.01000 ; qtot -9.162
7167 HA 443 PHE HD2 7167 0.133000 1.00800 ; qtot -9.029
7168 C 443 PHE C 7168 0.597301 12.01000 ; qtot -8.432
7169 O 443 PHE O 7169 -0.567901 16.00000 ; qtot -

```
9.000
  7170    N   444   HIP     N 7170    -0.347900     14.01000 ; qtot -
9.348
  7171    H   444   HIP     H 7171     0.274700      1.00800 ; qtot -
9.073
  7172   CX   444   HIP    CA 7172    -0.135400     12.01000 ; qtot -
9.209
  7173   H1   444   HIP    HA 7173     0.121200      1.00800 ; qtot -
9.087
  7174   CT   444   HIP    CB 7174    -0.041400     12.01000 ; qtot -
9.129
  7175   HC   444   HIP   HB2 7175     0.081000      1.00800 ; qtot -
9.048
  7176   HC   444   HIP   HB3 7176     0.081000      1.00800 ; qtot -
8.967
  7177   CC   444   HIP    CG 7177    -0.001200     12.01000 ; qtot -
8.968
  7178   NA   444   HIP   ND1 7178    -0.151300     14.01000 ; qtot -
9.119
  7179    H   444   HIP   HD1 7179     0.386600      1.00800 ; qtot -
8.733
  7180   CR   444   HIP   CE1 7180    -0.017000     12.01000 ; qtot -
8.750
  7181   H5   444   HIP   HE1 7181     0.268100      1.00800 ; qtot -
8.482
  7182   NA   444   HIP   NE2 7182    -0.171800     14.01000 ; qtot -
8.653
  7183    H   444   HIP   HE2 7183     0.391100      1.00800 ; qtot -
8.262
  7184   CW   444   HIP   CD2 7184    -0.114100     12.01000 ; qtot -
8.376
  7185   H4   444   HIP   HD2 7185     0.231700      1.00800 ; qtot -
8.145
  7186    C   444   HIP     C 7186     0.734101     12.01000 ; qtot -
7.411
  7187    O   444   HIP     O 7187    -0.589401     16.00000 ; qtot -
8.000
  7188    N   445   GLH     N 7188    -0.415700     14.01000 ; qtot -
8.416
  7189    H   445   GLH     H 7189     0.271900      1.00800 ; qtot -
8.144
  7190   CX   445   GLH    CA 7190     0.014500     12.01000 ; qtot -
8.129
  7191   H1   445   GLH    HA 7191     0.077900      1.00800 ; qtot -
8.051
  7192  A2C   445   GLH    CB 7192    -0.007100     12.01000 ; qtot -
8.059
  7193   HC   445   GLH   HB2 7193     0.025600      1.00800 ; qtot -
8.033
  7194   HC   445   GLH   HB3 7194     0.025600      1.00800 ; qtot -
8.007
  7195  A2C   445   GLH    CG 7195    -0.017400     12.01000 ; qtot -
```

8.025
  7196   HC   445   GLH   HG2 7196     0.043000      1.00800 ; qtot -
7.982
  7197   HC   445   GLH   HG3 7197     0.043000      1.00800 ; qtot -
7.939
  7198    C   445   GLH    CD 7198     0.680101     12.01000 ; qtot -
7.259
  7199    O   445   GLH   OE1 7199    -0.583801     16.00000 ; qtot -
7.842
  7200   OH   445   GLH   OE2 7200    -0.651101     16.00000 ; qtot -
8.494
  7201   HO   445   GLH   HE2 7201     0.464100      1.00800 ; qtot -
8.029
  7202    C   445   GLH     C 7202     0.597301     12.01000 ; qtot -
7.432
  7203    O   445   GLH     O 7203    -0.567901     16.00000 ; qtot -
8.000
  7204    N   446   PHE     N 7204    -0.415700     14.01000 ; qtot -
8.416
  7205    H   446   PHE     H 7205     0.271900      1.00800 ; qtot -
8.144
  7206   CX   446   PHE    CA 7206    -0.002400     12.01000 ; qtot -
8.146
  7207   H1   446   PHE    HA 7207     0.097800      1.00800 ; qtot -
8.048
  7208   CT   446   PHE    CB 7208    -0.034300     12.01000 ; qtot -
8.083
  7209   HC   446   PHE   HB2 7209     0.029500      1.00800 ; qtot -
8.053
  7210   HC   446   PHE   HB3 7210     0.029500      1.00800 ; qtot -
8.024
  7211   CA   446   PHE    CG 7211     0.011800     12.01000 ; qtot -
8.012
  7212   CA   446   PHE   CD1 7212    -0.125600     12.01000 ; qtot -
8.138
  7213   HA   446   PHE   HD1 7213     0.133000      1.00800 ; qtot -
8.005
  7214   CA   446   PHE   CE1 7214    -0.170400     12.01000 ; qtot -
8.175
  7215   HA   446   PHE   HE1 7215     0.143000      1.00800 ; qtot -
8.032
  7216   CA   446   PHE    CZ 7216    -0.107200     12.01000 ; qtot -
8.139
  7217   HA   446   PHE    HZ 7217     0.129700      1.00800 ; qtot -
8.009
  7218   CA   446   PHE   CE2 7218    -0.170400     12.01000 ; qtot -
8.180
  7219   HA   446   PHE   HE2 7219     0.143000      1.00800 ; qtot -
8.037
  7220   CA   446   PHE   CD2 7220    -0.125600     12.01000 ; qtot -
8.162
  7221   HA   446   PHE   HD2 7221     0.133000      1.00800 ; qtot -

8.029
  7222    C   446   PHE      C 7222     0.597301     12.01000 ; qtot -
7.432
  7223    O   446   PHE      O 7223    -0.567901     16.00000 ; qtot -
8.000
  7224    N   447   MET      N 7224    -0.415700     14.01000 ; qtot -
8.416
  7225    H   447   MET      H 7225     0.271900      1.00800 ; qtot -
8.144
  7226   CX   447   MET     CA 7226    -0.023700     12.01000 ; qtot -
8.168
  7227   H1   447   MET     HA 7227     0.088000      1.00800 ; qtot -
8.080
  7228  A2C   447   MET     CB 7228     0.034200     12.01000 ; qtot -
8.045
  7229   HC   447   MET    HB2 7229     0.024100      1.00800 ; qtot -
8.021
  7230   HC   447   MET    HB3 7230     0.024100      1.00800 ; qtot -
7.997
  7231  A2C   447   MET     CG 7231     0.001800     12.01000 ; qtot -
7.995
  7232   H1   447   MET    HG2 7232     0.044000      1.00800 ; qtot -
7.951
  7233   H1   447   MET    HG3 7233     0.044000      1.00800 ; qtot -
7.907
  7234    S   447   MET     SD 7234    -0.273700     32.06000 ; qtot -
8.181
  7235   CT   447   MET     CE 7235    -0.053600     12.01000 ; qtot -
8.235
  7236   H1   447   MET    HE1 7236     0.068400      1.00800 ; qtot -
8.166
  7237   H1   447   MET    HE2 7237     0.068400      1.00800 ; qtot -
8.098
  7238   H1   447   MET    HE3 7238     0.068400      1.00800 ; qtot -
8.029
  7239    C   447   MET      C 7239     0.597301     12.01000 ; qtot -
7.432
  7240    O   447   MET      O 7240    -0.567901     16.00000 ; qtot -
8.000
  7241    N   448   LEU      N 7241    -0.415700     14.01000 ; qtot -
8.416
  7242    H   448   LEU      H 7242     0.271900      1.00800 ; qtot -
8.144
  7243   CX   448   LEU     CA 7243    -0.051800     12.01000 ; qtot -
8.196
  7244   H1   448   LEU     HA 7244     0.092200      1.00800 ; qtot -
8.103
  7245  A2C   448   LEU     CB 7245    -0.110200     12.01000 ; qtot -
8.214
  7246   HC   448   LEU    HB2 7246     0.045700      1.00800 ; qtot -
8.168
  7247   HC   448   LEU    HB3 7247     0.045700      1.00800 ; qtot -

8.122
  7248  A3C   448   LEU    CG 7248     0.353100     12.01000 ; qtot -
7.769
  7249   HC   448   LEU    HG 7249    -0.036100      1.00800 ; qtot -
7.805
  7250   CT   448   LEU   CD1 7250    -0.412100     12.01000 ; qtot -
8.217
  7251   HC   448   LEU  HD11 7251     0.100000      1.00800 ; qtot -
8.117
  7252   HC   448   LEU  HD12 7252     0.100000      1.00800 ; qtot -
8.017
  7253   HC   448   LEU  HD13 7253     0.100000      1.00800 ; qtot -
7.917
  7254   CT   448   LEU   CD2 7254    -0.412100     12.01000 ; qtot -
8.329
  7255   HC   448   LEU  HD21 7255     0.100000      1.00800 ; qtot -
8.229
  7256   HC   448   LEU  HD22 7256     0.100000      1.00800 ; qtot -
8.129
  7257   HC   448   LEU  HD23 7257     0.100000      1.00800 ; qtot -
8.029
  7258    C   448   LEU     C 7258     0.597301     12.01000 ; qtot -
7.432
  7259    O   448   LEU     O 7259    -0.567901     16.00000 ; qtot -
8.000
  7260    N   449   MET     N 7260    -0.415700     14.01000 ; qtot -
8.416
  7261    H   449   MET     H 7261     0.271900      1.00800 ; qtot -
8.144
  7262   CX   449   MET    CA 7262    -0.023700     12.01000 ; qtot -
8.168
  7263   H1   449   MET    HA 7263     0.088000      1.00800 ; qtot -
8.080
  7264  A2C   449   MET    CB 7264     0.034200     12.01000 ; qtot -
8.045
  7265   HC   449   MET   HB2 7265     0.024100      1.00800 ; qtot -
8.021
  7266   HC   449   MET   HB3 7266     0.024100      1.00800 ; qtot -
7.997
  7267  A2C   449   MET    CG 7267     0.001800     12.01000 ; qtot -
7.995
  7268   H1   449   MET   HG2 7268     0.044000      1.00800 ; qtot -
7.951
  7269   H1   449   MET   HG3 7269     0.044000      1.00800 ; qtot -
7.907
  7270    S   449   MET    SD 7270    -0.273700     32.06000 ; qtot -
8.181
  7271   CT   449   MET    CE 7271    -0.053600     12.01000 ; qtot -
8.235
  7272   H1   449   MET   HE1 7272     0.068400      1.00800 ; qtot -
8.166
  7273   H1   449   MET   HE2 7273     0.068400      1.00800 ; qtot -

8.098
7274 H1 449 MET HE3 7274 0.068400 1.00800 ; qtot -
8.029
7275 C 449 MET C 7275 0.597301 12.01000 ; qtot -
7.432
7276 O 449 MET O 7276 -0.567901 16.00000 ; qtot -
8.000
7277 N 450 MET N 7277 -0.415700 14.01000 ; qtot -
8.416
7278 H 450 MET H 7278 0.271900 1.00800 ; qtot -
8.144
7279 CX 450 MET CA 7279 -0.023700 12.01000 ; qtot -
8.168
7280 H1 450 MET HA 7280 0.088000 1.00800 ; qtot -
8.080
7281 A2C 450 MET CB 7281 0.034200 12.01000 ; qtot -
8.045
7282 HC 450 MET HB2 7282 0.024100 1.00800 ; qtot -
8.021
7283 HC 450 MET HB3 7283 0.024100 1.00800 ; qtot -
7.997
7284 A2C 450 MET CG 7284 0.001800 12.01000 ; qtot -
7.995
7285 H1 450 MET HG2 7285 0.044000 1.00800 ; qtot -
7.951
7286 H1 450 MET HG3 7286 0.044000 1.00800 ; qtot -
7.907
7287 S 450 MET SD 7287 -0.273700 32.06000 ; qtot -
8.181
7288 CT 450 MET CE 7288 -0.053600 12.01000 ; qtot -
8.235
7289 H1 450 MET HE1 7289 0.068400 1.00800 ; qtot -
8.166
7290 H1 450 MET HE2 7290 0.068400 1.00800 ; qtot -
8.098
7291 H1 450 MET HE3 7291 0.068400 1.00800 ; qtot -
8.029
7292 C 450 MET C 7292 0.597301 12.01000 ; qtot -
7.432
7293 O 450 MET O 7293 -0.567901 16.00000 ; qtot -
8.000
7294 N 451 SER N 7294 -0.415700 14.01000 ; qtot -
8.416
7295 H 451 SER H 7295 0.271900 1.00800 ; qtot -
8.144
7296 CX 451 SER CA 7296 -0.024900 12.01000 ; qtot -
8.169
7297 H1 451 SER HA 7297 0.084300 1.00800 ; qtot -
8.084
7298 A2C 451 SER CB 7298 0.211700 12.01000 ; qtot -
7.873
7299 H1 451 SER HB2 7299 0.035200 1.00800 ; qtot -

7.838
7300 H1 451 SER HB3 7300 0.035200 1.00800 ; qtot -
7.802
7301 OH 451 SER OG 7301 -0.654601 16.00000 ; qtot -
8.457
7302 HO 451 SER HG 7302 0.427500 1.00800 ; qtot -
8.029
7303 C 451 SER C 7303 0.597301 12.01000 ; qtot -
7.432
7304 O 451 SER O 7304 -0.567901 16.00000 ; qtot -
8.000
7305 N 452 LYS N 7305 -0.347900 14.01000 ; qtot -
8.348
7306 H 452 LYS H 7306 0.274700 1.00800 ; qtot -
8.073
7307 CX 452 LYS CA 7307 -0.240000 12.01000 ; qtot -
8.313
7308 H1 452 LYS HA 7308 0.142600 1.00800 ; qtot -
8.171
7309 C8 452 LYS CB 7309 -0.009400 12.01000 ; qtot -
8.180
7310 HC 452 LYS HB2 7310 0.036200 1.00800 ; qtot -
8.144
7311 HC 452 LYS HB3 7311 0.036200 1.00800 ; qtot -
8.108
7312 C8 452 LYS CG 7312 0.018700 12.01000 ; qtot -
8.089
7313 HC 452 LYS HG2 7313 0.010300 1.00800 ; qtot -
8.079
7314 HC 452 LYS HG3 7314 0.010300 1.00800 ; qtot -
8.068
7315 C8 452 LYS CD 7315 -0.047900 12.01000 ; qtot -
8.116
7316 HC 452 LYS HD2 7316 0.062100 1.00800 ; qtot -
8.054
7317 HC 452 LYS HD3 7317 0.062100 1.00800 ; qtot -
7.992
7318 C8 452 LYS CE 7318 -0.014300 12.01000 ; qtot -
8.006
7319 HP 452 LYS HE2 7319 0.113500 1.00800 ; qtot -
7.893
7320 HP 452 LYS HE3 7320 0.113500 1.00800 ; qtot -
7.779
7321 N3 452 LYS NZ 7321 -0.385400 14.01000 ; qtot -
8.165
7322 H 452 LYS HZ1 7322 0.340000 1.00800 ; qtot -
7.825
7323 H 452 LYS HZ2 7323 0.340000 1.00800 ; qtot -
7.485
7324 H 452 LYS HZ3 7324 0.340000 1.00800 ; qtot -
7.145
7325 C 452 LYS C 7325 0.734101 12.01000 ; qtot -

6.411
  7326    O   452   LYS     O 7326    -0.589401     16.00000 ; qtot -
7.000
  7327    N   453   LYS     N 7327    -0.347900     14.01000 ; qtot -
7.348
  7328    H   453   LYS     H 7328     0.274700      1.00800 ; qtot -
7.073
  7329   CX   453   LYS    CA 7329    -0.240000     12.01000 ; qtot -
7.313
  7330   H1   453   LYS    HA 7330     0.142600      1.00800 ; qtot -
7.171
  7331   C8   453   LYS    CB 7331    -0.009400     12.01000 ; qtot -
7.180
  7332   HC   453   LYS   HB2 7332     0.036200      1.00800 ; qtot -
7.144
  7333   HC   453   LYS   HB3 7333     0.036200      1.00800 ; qtot -
7.108
  7334   C8   453   LYS    CG 7334     0.018700     12.01000 ; qtot -
7.089
  7335   HC   453   LYS   HG2 7335     0.010300      1.00800 ; qtot -
7.079
  7336   HC   453   LYS   HG3 7336     0.010300      1.00800 ; qtot -
7.068
  7337   C8   453   LYS    CD 7337    -0.047900     12.01000 ; qtot -
7.116
  7338   HC   453   LYS   HD2 7338     0.062100      1.00800 ; qtot -
7.054
  7339   HC   453   LYS   HD3 7339     0.062100      1.00800 ; qtot -
6.992
  7340   C8   453   LYS    CE 7340    -0.014300     12.01000 ; qtot -
7.006
  7341   HP   453   LYS   HE2 7341     0.113500      1.00800 ; qtot -
6.893
  7342   HP   453   LYS   HE3 7342     0.113500      1.00800 ; qtot -
6.779
  7343   N3   453   LYS    NZ 7343    -0.385400     14.01000 ; qtot -
7.165
  7344    H   453   LYS   HZ1 7344     0.340000      1.00800 ; qtot -
6.825
  7345    H   453   LYS   HZ2 7345     0.340000      1.00800 ; qtot -
6.485
  7346    H   453   LYS   HZ3 7346     0.340000      1.00800 ; qtot -
6.145
  7347    C   453   LYS     C 7347     0.734101     12.01000 ; qtot -
5.411
  7348    O   453   LYS     O 7348    -0.589401     16.00000 ; qtot -
6.000
  7349    N   454   LYS     N 7349    -0.348100     14.01000 ; qtot -
6.348
  7350    H   454   LYS     H 7350     0.276400      1.00800 ; qtot -
6.072
  7351   CX   454   LYS    CA 7351    -0.290300     12.01000 ; qtot -

6.362
  7352   H1   454   LYS    HA 7352     0.143800      1.00800 ; qtot -
6.218
  7353   C8   454   LYS    CB 7353    -0.053800     12.01000 ; qtot -
6.272
  7354   HC   454   LYS   HB2 7354     0.048200      1.00800 ; qtot -
6.224
  7355   HC   454   LYS   HB3 7355     0.048200      1.00800 ; qtot -
6.176
  7356   C8   454   LYS    CG 7356     0.022700     12.01000 ; qtot -
6.153
  7357   HC   454   LYS   HG2 7357     0.013400      1.00800 ; qtot -
6.140
  7358   HC   454   LYS   HG3 7358     0.013400      1.00800 ; qtot -
6.126
  7359   C8   454   LYS    CD 7359    -0.039200     12.01000 ; qtot -
6.165
  7360   HC   454   LYS   HD2 7360     0.061100      1.00800 ; qtot -
6.104
  7361   HC   454   LYS   HD3 7361     0.061100      1.00800 ; qtot -
6.043
  7362   C8   454   LYS    CE 7362    -0.017600     12.01000 ; qtot -
6.061
  7363   HP   454   LYS   HE2 7363     0.112100      1.00800 ; qtot -
5.949
  7364   HP   454   LYS   HE3 7364     0.112100      1.00800 ; qtot -
5.836
  7365   N3   454   LYS    NZ 7365    -0.374100     14.01000 ; qtot -
6.211
  7366    H   454   LYS   HZ1 7366     0.337400      1.00800 ; qtot -
5.873
  7367    H   454   LYS   HZ2 7367     0.337400      1.00800 ; qtot -
5.536
  7368    H   454   LYS   HZ3 7368     0.337400      1.00800 ; qtot -
5.198
  7369    C   454   LYS     C 7369     0.848801     12.01000 ; qtot -
4.350
  7370   O2   454   LYS     O 7370    -0.825201     16.00000 ; qtot -
5.175
  7371   O2   454   LYS   OXT 7371    -0.825201     16.00000 ; qtot -
6.000
  7372   CA   455   E21    C1 7372     0.229575     12.01000 ; qtot -
5.770
  7373   CA   455   E21    C2 7373    -0.215845     12.01000 ; qtot -
5.986
  7374   CA   455   E21    C3 7374    -0.223493     12.01000 ; qtot -
6.210
  7375   CA   455   E21    C4 7375     0.034966     12.01000 ; qtot -
6.175
  7376   cp   455   E21    C5 7376    -0.101855     12.01000 ; qtot -
6.277
  7377   CA   455   E21    C6 7377    -0.183343     12.01000 ; qtot -

6.460
  7378   CA   455   E21    C7 7378     0.029538     12.01000 ; qtot -
6.430
  7379   cq   455   E21    C8 7379    -0.005720     12.01000 ; qtot -
6.436
  7380   CA   455   E21    C9 7380     0.047186     12.01000 ; qtot -
6.389
  7381   CA   455   E21   C10 7381    -0.025281     12.01000 ; qtot -
6.414
  7382   c2   455   E21   C11 7382    -0.025445     12.01000 ; qtot -
6.440
  7383   c2   455   E21   C12 7383     0.190979     12.01000 ; qtot -
6.249
  7384   c2   455   E21   C13 7384    -0.142239     12.01000 ; qtot -
6.391
  7385    f   455   E21   F14 7385    -0.119942     19.00000 ; qtot -
6.511
  7386   Y1   455   E21   N15 7386    -0.080082     14.01000 ; qtot -
6.591
  7387   Y2   455   E21   N16 7387     0.003613     14.01000 ; qtot -
6.587
  7388   OH   455   E21   O17 7388    -0.552817     16.00000 ; qtot -
7.140
  7389   Y3   455   E21   C18 7389    -0.140021     12.01000 ; qtot -
7.280
  7390   Y4   455   E21   C19 7390     0.006309     12.01000 ; qtot -
7.274
  7391   Y5   455   E21   C20 7391    -0.258138     12.01000 ; qtot -
7.532
  7392   Y6   455   E21   C21 7392    -0.064007     12.01000 ; qtot -
7.596
  7393   Y7   455   E21   C22 7393    -0.229683     12.01000 ; qtot -
7.826
  7394   Y8   455   E21   C23 7394     0.245906     12.01000 ; qtot -
7.580
  7395    O   455   E21   O24 7395    -0.199654     16.00000 ; qtot -
7.779
  7396   c2   455   E21   C25 7396     0.315263     12.01000 ; qtot -
7.464
  7397    N   455   E21   N26 7397     0.047093     14.01000 ; qtot -
7.417
  7398    O   455   E21   O27 7398    -0.398729     16.00000 ; qtot -
7.816
  7399   c3   455   E21   C28 7399     0.108116     12.01000 ; qtot -
7.708
  7400   c3   455   E21   C29 7400    -0.113698     12.01000 ; qtot -
7.821
  7401   c3   455   E21   C30 7401     0.035550     12.01000 ; qtot -
7.786
  7402   OH   455   E21   O31 7402    -0.576089     16.00000 ; qtot -
8.362
  7403   c3   455   E21   C32 7403     0.066802     12.01000 ; qtot -

8.295
 7404 OH 455 E21 O33 7404 -0.593303 16.00000 ; qtot -
8.888
 7405 CA 455 E21 C34 7405 -0.120089 12.01000 ; qtot -
9.009
 7406 c2 455 E21 C35 7406 0.455528 12.01000 ; qtot -
8.553
 7407 N 455 E21 N36 7407 -0.097051 14.01000 ; qtot -
8.650
 7408 O 455 E21 O37 7408 -0.404709 16.00000 ; qtot -
9.055
 7409 c3 455 E21 C38 7409 -0.196725 12.01000 ; qtot -
9.252
 7410 c2 455 E21 C39 7410 0.502612 12.01000 ; qtot -
8.749
 7411 O 455 E21 O40 7411 -0.424252 16.00000 ; qtot -
9.173
 7412 CA 455 E21 C41 7412 -0.199105 12.01000 ; qtot -
9.372
 7413 HA 455 E21 H42 7413 0.151373 1.00800 ; qtot -
9.221
 7414 HA 455 E21 H43 7414 0.164316 1.00800 ; qtot -
9.057
 7415 HA 455 E21 H44 7415 0.167656 1.00800 ; qtot -
8.889
 7416 H4 455 E21 H45 7416 -0.070672 1.00800 ; qtot -
8.960
 7417 HA 455 E21 H46 7417 0.160147 1.00800 ; qtot -
8.799
 7418 HA 455 E21 H47 7418 0.193602 1.00800 ; qtot -
8.606
 7419 HA 455 E21 H48 7419 0.174693 1.00800 ; qtot -
8.431
 7420 HA 455 E21 H49 7420 0.180458 1.00800 ; qtot -
8.251
 7421 H1 455 E21 H50 7421 0.059671 1.00800 ; qtot -
8.191
 7422 H1 455 E21 H51 7422 0.059671 1.00800 ; qtot -
8.131
 7423 H1 455 E21 H52 7423 0.067003 1.00800 ; qtot -
8.064
 7424 H1 455 E21 H53 7424 0.067003 1.00800 ; qtot -
7.997
 7425 HO 455 E21 H54 7425 0.412888 1.00800 ; qtot -
7.584
 7426 H1 455 E21 H55 7426 0.099097 1.00800 ; qtot -
7.485
 7427 H1 455 E21 H56 7427 0.099097 1.00800 ; qtot -
7.386
 7428 H1 455 E21 H57 7428 0.099097 1.00800 ; qtot -
7.287
 7429 HO 455 E21 H58 7429 0.407133 1.00800 ; qtot -

```
6.880
  7430   HO   455   E21   H59 7430     0.413326      1.00800 ; qtot -
6.467
  7431   HC   455   E21   H60 7431     0.033376      1.00800 ; qtot -
6.433
  7432   HC   455   E21   H61 7432     0.033376      1.00800 ; qtot -
6.400
  7433   HC   455   E21   H62 7433     0.033376      1.00800 ; qtot -
6.367
  7434   hn   455   E21   H63 7434     0.192377      1.00800 ; qtot -
6.174
  7435   HA   455   E21   H64 7435     0.208916      1.00800 ; qtot -
5.965
  7436   M1   456   RU1    RU 7436    -0.034703    101.07000 ; qtot -
6.000

[ bonds ]
;   ai     aj funct   r             k
     1      2   1    1.0100e-01    3.6317e+05 ;      N - H1
     1      3   1    1.0100e-01    3.6317e+05 ;      N - H2
     1      4   1    1.0100e-01    3.6317e+05 ;      N - H3
     1      5   1    1.4710e-01    3.0711e+05 ;      N - CA
     5      6   1    1.0900e-01    2.8451e+05 ;     CA - HA
     5      7   1    1.5260e-01    2.5941e+05 ;     CA - CB
     5     20   1    1.5220e-01    2.6527e+05 ;     CA - C
     7      8   1    1.0900e-01    2.8451e+05 ;     CB - HB2
     7      9   1    1.0900e-01    2.8451e+05 ;     CB - HB3
     7     10   1    1.5260e-01    2.5941e+05 ;     CB - CG
    10     11   1    1.0900e-01    2.8451e+05 ;     CG - HG
    10     12   1    1.5260e-01    2.5941e+05 ;     CG - CD1
    10     16   1    1.5260e-01    2.5941e+05 ;     CG - CD2
    12     13   1    1.0900e-01    2.8451e+05 ;    CD1 - HD11
    12     14   1    1.0900e-01    2.8451e+05 ;    CD1 - HD12
    12     15   1    1.0900e-01    2.8451e+05 ;    CD1 - HD13
    16     17   1    1.0900e-01    2.8451e+05 ;    CD2 - HD21
    16     18   1    1.0900e-01    2.8451e+05 ;    CD2 - HD22
    16     19   1    1.0900e-01    2.8451e+05 ;    CD2 - HD23
    20     21   1    1.2290e-01    4.7698e+05 ;      C - O
    20     22   1    1.3350e-01    4.1003e+05 ;      C - N
    22     23   1    1.0100e-01    3.6317e+05 ;      N - H
    22     24   1    1.4490e-01    2.8200e+05 ;      N - CA
    24     25   1    1.0900e-01    2.8451e+05 ;     CA - HA
    24     26   1    1.5260e-01    2.5941e+05 ;     CA - CB
    24     39   1    1.5220e-01    2.6527e+05 ;     CA - C
    26     27   1    1.0900e-01    2.8451e+05 ;     CB - HB
    26     28   1    1.5260e-01    2.5941e+05 ;     CB - CG2
    26     32   1    1.5260e-01    2.5941e+05 ;     CB - CG1
    28     29   1    1.0900e-01    2.8451e+05 ;    CG2 - HG21
    28     30   1    1.0900e-01    2.8451e+05 ;    CG2 - HG22
    28     31   1    1.0900e-01    2.8451e+05 ;    CG2 - HG23
    32     33   1    1.0900e-01    2.8451e+05 ;    CG1 - HG12
```

| | | | | | | | | |
|---|---|---|---|---|---|---|---|---|
| 83 | 85 | 1 | 1.3810e-01 | 3.5731e+05 | ; | NE2 | - | CD2 |
| 85 | 86 | 1 | 1.0800e-01 | 3.0711e+05 | ; | CD2 | - | HD2 |
| 87 | 88 | 1 | 1.2290e-01 | 4.7698e+05 | ; | C | - | O |
| 87 | 89 | 1 | 1.3350e-01 | 4.1003e+05 | ; | C | - | N |
| 89 | 90 | 1 | 1.0100e-01 | 3.6317e+05 | ; | N | - | H |
| 89 | 91 | 1 | 1.4490e-01 | 2.8200e+05 | ; | N | - | CA |
| 91 | 92 | 1 | 1.0900e-01 | 2.8451e+05 | ; | CA | - | HA |
| 91 | 93 | 1 | 1.5260e-01 | 2.5941e+05 | ; | CA | - | CB |
| 91 | 98 | 1 | 1.5220e-01 | 2.6527e+05 | ; | CA | - | C |
| 93 | 94 | 1 | 1.0900e-01 | 2.8451e+05 | ; | CB | - | HB2 |
| 93 | 95 | 1 | 1.0900e-01 | 2.8451e+05 | ; | CB | - | HB3 |
| 93 | 96 | 1 | 1.4100e-01 | 2.6778e+05 | ; | CB | - | OG |
| 96 | 97 | 1 | 9.6000e-02 | 4.6275e+05 | ; | OG | - | HG |
| 98 | 99 | 1 | 1.2290e-01 | 4.7698e+05 | ; | C | - | O |
| 98 | 100 | 1 | 1.3350e-01 | 4.1003e+05 | ; | C | - | N |
| 100 | 101 | 1 | 1.0100e-01 | 3.6317e+05 | ; | N | - | H |
| 100 | 102 | 1 | 1.4490e-01 | 2.8200e+05 | ; | N | - | CA |
| 102 | 103 | 1 | 1.0900e-01 | 2.8451e+05 | ; | CA | - | HA2 |
| 102 | 104 | 1 | 1.0900e-01 | 2.8451e+05 | ; | CA | - | HA3 |
| 102 | 105 | 1 | 1.5220e-01 | 2.6527e+05 | ; | CA | - | C |
| 105 | 106 | 1 | 1.2290e-01 | 4.7698e+05 | ; | C | - | O |
| 105 | 107 | 1 | 1.3350e-01 | 4.1003e+05 | ; | C | - | N |
| 107 | 108 | 1 | 1.0100e-01 | 3.6317e+05 | ; | N | - | H |
| 107 | 109 | 1 | 1.4490e-01 | 2.8200e+05 | ; | N | - | CA |
| 109 | 110 | 1 | 1.0900e-01 | 2.8451e+05 | ; | CA | - | HA |
| 109 | 111 | 1 | 1.5260e-01 | 2.5941e+05 | ; | CA | - | CB |
| 109 | 127 | 1 | 1.5220e-01 | 2.6527e+05 | ; | CA | - | C |
| 111 | 112 | 1 | 1.0900e-01 | 2.8451e+05 | ; | CB | - | HB2 |
| 111 | 113 | 1 | 1.0900e-01 | 2.8451e+05 | ; | CB | - | HB3 |
| 111 | 114 | 1 | 1.5260e-01 | 2.5941e+05 | ; | CB | - | CG |
| 114 | 115 | 1 | 1.0900e-01 | 2.8451e+05 | ; | CG | - | HG2 |
| 114 | 116 | 1 | 1.0900e-01 | 2.8451e+05 | ; | CG | - | HG3 |
| 114 | 117 | 1 | 1.5260e-01 | 2.5941e+05 | ; | CG | - | CD |
| 117 | 118 | 1 | 1.0900e-01 | 2.8451e+05 | ; | CD | - | HD2 |
| 117 | 119 | 1 | 1.0900e-01 | 2.8451e+05 | ; | CD | - | HD3 |
| 117 | 120 | 1 | 1.5260e-01 | 2.5941e+05 | ; | CD | - | CE |
| 120 | 121 | 1 | 1.0900e-01 | 2.8451e+05 | ; | CE | - | HE2 |
| 120 | 122 | 1 | 1.0900e-01 | 2.8451e+05 | ; | CE | - | HE3 |
| 120 | 123 | 1 | 1.4710e-01 | 3.0711e+05 | ; | CE | - | NZ |
| 123 | 124 | 1 | 1.0100e-01 | 3.6317e+05 | ; | NZ | - | HZ1 |
| 123 | 125 | 1 | 1.0100e-01 | 3.6317e+05 | ; | NZ | - | HZ2 |
| 123 | 126 | 1 | 1.0100e-01 | 3.6317e+05 | ; | NZ | - | HZ3 |
| 127 | 128 | 1 | 1.2290e-01 | 4.7698e+05 | ; | C | - | O |
| 127 | 129 | 1 | 1.3350e-01 | 4.1003e+05 | ; | C | - | N |
| 129 | 130 | 1 | 1.0100e-01 | 3.6317e+05 | ; | N | - | H |
| 129 | 131 | 1 | 1.4490e-01 | 2.8200e+05 | ; | N | - | CA |
| 131 | 132 | 1 | 1.0900e-01 | 2.8451e+05 | ; | CA | - | HA |
| 131 | 133 | 1 | 1.5260e-01 | 2.5941e+05 | ; | CA | - | CB |
| 131 | 146 | 1 | 1.5220e-01 | 2.6527e+05 | ; | CA | - | C |
| 133 | 134 | 1 | 1.0900e-01 | 2.8451e+05 | ; | CB | - | HB2 |
| 133 | 135 | 1 | 1.0900e-01 | 2.8451e+05 | ; | CB | - | HB3 |
| 133 | 136 | 1 | 1.5260e-01 | 2.5941e+05 | ; | CB | - | CG |

| | | | | | | |
|---|---|---|---|---|---|---|
| 136 | 137 | 1 | 1.0900e-01 | 2.8451e+05 | ; | CG - HG |
| 136 | 138 | 1 | 1.5260e-01 | 2.5941e+05 | ; | CG - CD1 |
| 136 | 142 | 1 | 1.5260e-01 | 2.5941e+05 | ; | CG - CD2 |
| 138 | 139 | 1 | 1.0900e-01 | 2.8451e+05 | ; | CD1 - HD11 |
| 138 | 140 | 1 | 1.0900e-01 | 2.8451e+05 | ; | CD1 - HD12 |
| 138 | 141 | 1 | 1.0900e-01 | 2.8451e+05 | ; | CD1 - HD13 |
| 142 | 143 | 1 | 1.0900e-01 | 2.8451e+05 | ; | CD2 - HD21 |
| 142 | 144 | 1 | 1.0900e-01 | 2.8451e+05 | ; | CD2 - HD22 |
| 142 | 145 | 1 | 1.0900e-01 | 2.8451e+05 | ; | CD2 - HD23 |
| 146 | 147 | 1 | 1.2290e-01 | 4.7698e+05 | ; | C - O |
| 146 | 148 | 1 | 1.3350e-01 | 4.1003e+05 | ; | C - N |
| 148 | 149 | 1 | 1.0100e-01 | 3.6317e+05 | ; | N - H |
| 148 | 150 | 1 | 1.4490e-01 | 2.8200e+05 | ; | N - CA |
| 150 | 151 | 1 | 1.0900e-01 | 2.8451e+05 | ; | CA - HA |
| 150 | 152 | 1 | 1.5260e-01 | 2.5941e+05 | ; | CA - CB |
| 150 | 161 | 1 | 1.5220e-01 | 2.6527e+05 | ; | CA - C |
| 152 | 153 | 1 | 1.0900e-01 | 2.8451e+05 | ; | CB - HB2 |
| 152 | 154 | 1 | 1.0900e-01 | 2.8451e+05 | ; | CB - HB3 |
| 152 | 155 | 1 | 1.5260e-01 | 2.5941e+05 | ; | CB - CG |
| 155 | 156 | 1 | 1.0900e-01 | 2.8451e+05 | ; | CG - HG2 |
| 155 | 157 | 1 | 1.0900e-01 | 2.8451e+05 | ; | CG - HG3 |
| 155 | 158 | 1 | 1.5220e-01 | 2.6527e+05 | ; | CG - CD |
| 158 | 159 | 1 | 1.2500e-01 | 5.4894e+05 | ; | CD - OE1 |
| 158 | 160 | 1 | 1.2500e-01 | 5.4894e+05 | ; | CD - OE2 |
| 161 | 162 | 1 | 1.2290e-01 | 4.7698e+05 | ; | C - O |
| 161 | 163 | 1 | 1.3350e-01 | 4.1003e+05 | ; | C - N |
| 163 | 164 | 1 | 1.0100e-01 | 3.6317e+05 | ; | N - H |
| 163 | 165 | 1 | 1.4490e-01 | 2.8200e+05 | ; | N - CA |
| 165 | 166 | 1 | 1.0900e-01 | 2.8451e+05 | ; | CA - HA |
| 165 | 167 | 1 | 1.5260e-01 | 2.5941e+05 | ; | CA - CB |
| 165 | 173 | 1 | 1.5220e-01 | 2.6527e+05 | ; | CA - C |
| 167 | 168 | 1 | 1.0900e-01 | 2.8451e+05 | ; | CB - HB2 |
| 167 | 169 | 1 | 1.0900e-01 | 2.8451e+05 | ; | CB - HB3 |
| 167 | 170 | 1 | 1.5220e-01 | 2.6527e+05 | ; | CB - CG |
| 170 | 171 | 1 | 1.2500e-01 | 5.4894e+05 | ; | CG - OD1 |
| 170 | 172 | 1 | 1.2500e-01 | 5.4894e+05 | ; | CG - OD2 |
| 173 | 174 | 1 | 1.2290e-01 | 4.7698e+05 | ; | C - O |
| 173 | 175 | 1 | 1.3350e-01 | 4.1003e+05 | ; | C - N |
| 175 | 176 | 1 | 1.0100e-01 | 3.6317e+05 | ; | N - H |
| 175 | 177 | 1 | 1.4490e-01 | 2.8200e+05 | ; | N - CA |
| 177 | 178 | 1 | 1.0900e-01 | 2.8451e+05 | ; | CA - HA |
| 177 | 179 | 1 | 1.5260e-01 | 2.5941e+05 | ; | CA - CB |
| 177 | 195 | 1 | 1.5220e-01 | 2.6527e+05 | ; | CA - C |
| 179 | 180 | 1 | 1.0900e-01 | 2.8451e+05 | ; | CB - HB2 |
| 179 | 181 | 1 | 1.0900e-01 | 2.8451e+05 | ; | CB - HB3 |
| 179 | 182 | 1 | 1.5260e-01 | 2.5941e+05 | ; | CB - CG |
| 182 | 183 | 1 | 1.0900e-01 | 2.8451e+05 | ; | CG - HG2 |
| 182 | 184 | 1 | 1.0900e-01 | 2.8451e+05 | ; | CG - HG3 |
| 182 | 185 | 1 | 1.5260e-01 | 2.5941e+05 | ; | CG - CD |
| 185 | 186 | 1 | 1.0900e-01 | 2.8451e+05 | ; | CD - HD2 |
| 185 | 187 | 1 | 1.0900e-01 | 2.8451e+05 | ; | CD - HD3 |
| 185 | 188 | 1 | 1.5260e-01 | 2.5941e+05 | ; | CD - CE |

| | | | | | | |
|---|---|---|---|---|---|---|
| 188 | 189 | 1 | 1.0900e-01 | 2.8451e+05 | ; | CE - HE2 |
| 188 | 190 | 1 | 1.0900e-01 | 2.8451e+05 | ; | CE - HE3 |
| 188 | 191 | 1 | 1.4710e-01 | 3.0711e+05 | ; | CE - NZ |
| 191 | 192 | 1 | 1.0100e-01 | 3.6317e+05 | ; | NZ - HZ1 |
| 191 | 193 | 1 | 1.0100e-01 | 3.6317e+05 | ; | NZ - HZ2 |
| 191 | 194 | 1 | 1.0100e-01 | 3.6317e+05 | ; | NZ - HZ3 |
| 195 | 196 | 1 | 1.2290e-01 | 4.7698e+05 | ; | C - O |
| 195 | 197 | 1 | 1.3350e-01 | 4.1003e+05 | ; | C - N |
| 197 | 198 | 1 | 1.0100e-01 | 3.6317e+05 | ; | N - H |
| 197 | 199 | 1 | 1.4490e-01 | 2.8200e+05 | ; | N - CA |
| 199 | 200 | 1 | 1.0900e-01 | 2.8451e+05 | ; | CA - HA |
| 199 | 201 | 1 | 1.5260e-01 | 2.5941e+05 | ; | CA - CB |
| 199 | 216 | 1 | 1.5220e-01 | 2.6527e+05 | ; | CA - C |
| 201 | 202 | 1 | 1.0900e-01 | 2.8451e+05 | ; | CB - HB2 |
| 201 | 203 | 1 | 1.0900e-01 | 2.8451e+05 | ; | CB - HB3 |
| 201 | 204 | 1 | 1.5100e-01 | 2.6527e+05 | ; | CB - CG |
| 204 | 205 | 1 | 1.4000e-01 | 3.9246e+05 | ; | CG - CD1 |
| 204 | 214 | 1 | 1.4000e-01 | 3.9246e+05 | ; | CG - CD2 |
| 205 | 206 | 1 | 1.0800e-01 | 3.0711e+05 | ; | CD1 - HD1 |
| 205 | 207 | 1 | 1.4000e-01 | 3.9246e+05 | ; | CD1 - CE1 |
| 207 | 208 | 1 | 1.0800e-01 | 3.0711e+05 | ; | CE1 - HE1 |
| 207 | 209 | 1 | 1.4090e-01 | 3.9246e+05 | ; | CE1 - CZ |
| 209 | 210 | 1 | 1.3640e-01 | 3.7656e+05 | ; | CZ - OH |
| 209 | 212 | 1 | 1.4090e-01 | 3.9246e+05 | ; | CZ - CE2 |
| 210 | 211 | 1 | 9.6000e-02 | 4.6275e+05 | ; | OH - HH |
| 212 | 213 | 1 | 1.0800e-01 | 3.0711e+05 | ; | CE2 - HE2 |
| 212 | 214 | 1 | 1.4000e-01 | 3.9246e+05 | ; | CE2 - CD2 |
| 214 | 215 | 1 | 1.0800e-01 | 3.0711e+05 | ; | CD2 - HD2 |
| 216 | 217 | 1 | 1.2290e-01 | 4.7698e+05 | ; | C - O |
| 216 | 218 | 1 | 1.3350e-01 | 4.1003e+05 | ; | C - N |
| 218 | 219 | 1 | 1.0100e-01 | 3.6317e+05 | ; | N - H |
| 218 | 220 | 1 | 1.4490e-01 | 2.8200e+05 | ; | N - CA |
| 220 | 221 | 1 | 1.0900e-01 | 2.8451e+05 | ; | CA - HA |
| 220 | 222 | 1 | 1.5260e-01 | 2.5941e+05 | ; | CA - CB |
| 220 | 235 | 1 | 1.5220e-01 | 2.6527e+05 | ; | CA - C |
| 222 | 223 | 1 | 1.0900e-01 | 2.8451e+05 | ; | CB - HB |
| 222 | 224 | 1 | 1.5260e-01 | 2.5941e+05 | ; | CB - CG2 |
| 222 | 228 | 1 | 1.5260e-01 | 2.5941e+05 | ; | CB - CG1 |
| 224 | 225 | 1 | 1.0900e-01 | 2.8451e+05 | ; | CG2 - HG21 |
| 224 | 226 | 1 | 1.0900e-01 | 2.8451e+05 | ; | CG2 - HG22 |
| 224 | 227 | 1 | 1.0900e-01 | 2.8451e+05 | ; | CG2 - HG23 |
| 228 | 229 | 1 | 1.0900e-01 | 2.8451e+05 | ; | CG1 - HG12 |
| 228 | 230 | 1 | 1.0900e-01 | 2.8451e+05 | ; | CG1 - HG13 |
| 228 | 231 | 1 | 1.5260e-01 | 2.5941e+05 | ; | CG1 - CD1 |
| 231 | 232 | 1 | 1.0900e-01 | 2.8451e+05 | ; | CD1 - HD11 |
| 231 | 233 | 1 | 1.0900e-01 | 2.8451e+05 | ; | CD1 - HD12 |
| 231 | 234 | 1 | 1.0900e-01 | 2.8451e+05 | ; | CD1 - HD13 |
| 235 | 236 | 1 | 1.2290e-01 | 4.7698e+05 | ; | C - O |
| 235 | 237 | 1 | 1.3350e-01 | 4.1003e+05 | ; | C - N |
| 237 | 238 | 1 | 1.0100e-01 | 3.6317e+05 | ; | N - H |
| 237 | 239 | 1 | 1.4490e-01 | 2.8200e+05 | ; | N - CA |
| 239 | 240 | 1 | 1.0900e-01 | 2.8451e+05 | ; | CA - HA |

| | | | | | | | | |
|---|---|---|---|---|---|---|---|---|
| 239 | 241 | 1 | 1.5260e-01 | 2.5941e+05 | ; | CA | - | CB |
| 239 | 254 | 1 | 1.5220e-01 | 2.6527e+05 | ; | CA | - | C |
| 241 | 242 | 1 | 1.0900e-01 | 2.8451e+05 | ; | CB | - | HB |
| 241 | 243 | 1 | 1.5260e-01 | 2.5941e+05 | ; | CB | - | CG2 |
| 241 | 247 | 1 | 1.5260e-01 | 2.5941e+05 | ; | CB | - | CG1 |
| 243 | 244 | 1 | 1.0900e-01 | 2.8451e+05 | ; | CG2 | - | HG21 |
| 243 | 245 | 1 | 1.0900e-01 | 2.8451e+05 | ; | CG2 | - | HG22 |
| 243 | 246 | 1 | 1.0900e-01 | 2.8451e+05 | ; | CG2 | - | HG23 |
| 247 | 248 | 1 | 1.0900e-01 | 2.8451e+05 | ; | CG1 | - | HG12 |
| 247 | 249 | 1 | 1.0900e-01 | 2.8451e+05 | ; | CG1 | - | HG13 |
| 247 | 250 | 1 | 1.5260e-01 | 2.5941e+05 | ; | CG1 | - | CD1 |
| 250 | 251 | 1 | 1.0900e-01 | 2.8451e+05 | ; | CD1 | - | HD11 |
| 250 | 252 | 1 | 1.0900e-01 | 2.8451e+05 | ; | CD1 | - | HD12 |
| 250 | 253 | 1 | 1.0900e-01 | 2.8451e+05 | ; | CD1 | - | HD13 |
| 254 | 255 | 1 | 1.2290e-01 | 4.7698e+05 | ; | C | - | O |
| 254 | 256 | 1 | 1.3350e-01 | 4.1003e+05 | ; | C | - | N |
| 256 | 257 | 1 | 1.0100e-01 | 3.6317e+05 | ; | N | - | H |
| 256 | 258 | 1 | 1.4490e-01 | 2.8200e+05 | ; | N | - | CA |
| 258 | 259 | 1 | 1.0900e-01 | 2.8451e+05 | ; | CA | - | HA |
| 258 | 260 | 1 | 1.5260e-01 | 2.5941e+05 | ; | CA | - | CB |
| 258 | 266 | 1 | 1.5220e-01 | 2.6527e+05 | ; | CA | - | C |
| 260 | 261 | 1 | 1.0900e-01 | 2.8451e+05 | ; | CB | - | HB2 |
| 260 | 262 | 1 | 1.0900e-01 | 2.8451e+05 | ; | CB | - | HB3 |
| 260 | 263 | 1 | 1.5220e-01 | 2.6527e+05 | ; | CB | - | CG |
| 263 | 264 | 1 | 1.2500e-01 | 5.4894e+05 | ; | CG | - | OD1 |
| 263 | 265 | 1 | 1.2500e-01 | 5.4894e+05 | ; | CG | - | OD2 |
| 266 | 267 | 1 | 1.2290e-01 | 4.7698e+05 | ; | C | - | O |
| 266 | 268 | 1 | 1.3350e-01 | 4.1003e+05 | ; | C | - | N |
| 268 | 269 | 1 | 1.0100e-01 | 3.6317e+05 | ; | N | - | H |
| 268 | 270 | 1 | 1.4490e-01 | 2.8200e+05 | ; | N | - | CA |
| 270 | 271 | 1 | 1.0900e-01 | 2.8451e+05 | ; | CA | - | HA |
| 270 | 272 | 1 | 1.5260e-01 | 2.5941e+05 | ; | CA | - | CB |
| 270 | 281 | 1 | 1.5220e-01 | 2.6527e+05 | ; | CA | - | C |
| 272 | 273 | 1 | 1.0900e-01 | 2.8451e+05 | ; | CB | - | HB2 |
| 272 | 274 | 1 | 1.0900e-01 | 2.8451e+05 | ; | CB | - | HB3 |
| 272 | 275 | 1 | 1.5260e-01 | 2.5941e+05 | ; | CB | - | CG |
| 275 | 276 | 1 | 1.0900e-01 | 2.8451e+05 | ; | CG | - | HG2 |
| 275 | 277 | 1 | 1.0900e-01 | 2.8451e+05 | ; | CG | - | HG3 |
| 275 | 278 | 1 | 1.5220e-01 | 2.6527e+05 | ; | CG | - | CD |
| 278 | 279 | 1 | 1.2500e-01 | 5.4894e+05 | ; | CD | - | OE1 |
| 278 | 280 | 1 | 1.2500e-01 | 5.4894e+05 | ; | CD | - | OE2 |
| 281 | 282 | 1 | 1.2290e-01 | 4.7698e+05 | ; | C | - | O |
| 281 | 283 | 1 | 1.3350e-01 | 4.1003e+05 | ; | C | - | N |
| 283 | 284 | 1 | 1.0100e-01 | 3.6317e+05 | ; | N | - | H |
| 283 | 285 | 1 | 1.4490e-01 | 2.8200e+05 | ; | N | - | CA |
| 285 | 286 | 1 | 1.0900e-01 | 2.8451e+05 | ; | CA | - | HA |
| 285 | 287 | 1 | 1.5260e-01 | 2.5941e+05 | ; | CA | - | CB |
| 285 | 303 | 1 | 1.5220e-01 | 2.6527e+05 | ; | CA | - | C |
| 287 | 288 | 1 | 1.0900e-01 | 2.8451e+05 | ; | CB | - | HB2 |
| 287 | 289 | 1 | 1.0900e-01 | 2.8451e+05 | ; | CB | - | HB3 |
| 287 | 290 | 1 | 1.5260e-01 | 2.5941e+05 | ; | CB | - | CG |
| 290 | 291 | 1 | 1.0900e-01 | 2.8451e+05 | ; | CG | - | HG2 |

| | | | | | | |
|---|---|---|---|---|---|---|
| 290 | 292 | 1 | 1.0900e-01 | 2.8451e+05 | ; | CG - HG3 |
| 290 | 293 | 1 | 1.5260e-01 | 2.5941e+05 | ; | CG - CD |
| 293 | 294 | 1 | 1.0900e-01 | 2.8451e+05 | ; | CD - HD2 |
| 293 | 295 | 1 | 1.0900e-01 | 2.8451e+05 | ; | CD - HD3 |
| 293 | 296 | 1 | 1.5260e-01 | 2.5941e+05 | ; | CD - CE |
| 296 | 297 | 1 | 1.0900e-01 | 2.8451e+05 | ; | CE - HE2 |
| 296 | 298 | 1 | 1.0900e-01 | 2.8451e+05 | ; | CE - HE3 |
| 296 | 299 | 1 | 1.4710e-01 | 3.0711e+05 | ; | CE - NZ |
| 299 | 300 | 1 | 1.0100e-01 | 3.6317e+05 | ; | NZ - HZ1 |
| 299 | 301 | 1 | 1.0100e-01 | 3.6317e+05 | ; | NZ - HZ2 |
| 299 | 302 | 1 | 1.0100e-01 | 3.6317e+05 | ; | NZ - HZ3 |
| 303 | 304 | 1 | 1.2290e-01 | 4.7698e+05 | ; | C - O |
| 303 | 305 | 1 | 1.3350e-01 | 4.1003e+05 | ; | C - N |
| 305 | 306 | 1 | 1.0100e-01 | 3.6317e+05 | ; | N - H |
| 305 | 307 | 1 | 1.4490e-01 | 2.8200e+05 | ; | N - CA |
| 307 | 308 | 1 | 1.0900e-01 | 2.8451e+05 | ; | CA - HA |
| 307 | 309 | 1 | 1.5260e-01 | 2.5941e+05 | ; | CA - CB |
| 307 | 322 | 1 | 1.5220e-01 | 2.6527e+05 | ; | CA - C |
| 309 | 310 | 1 | 1.0900e-01 | 2.8451e+05 | ; | CB - HB2 |
| 309 | 311 | 1 | 1.0900e-01 | 2.8451e+05 | ; | CB - HB3 |
| 309 | 312 | 1 | 1.5260e-01 | 2.5941e+05 | ; | CB - CG |
| 312 | 313 | 1 | 1.0900e-01 | 2.8451e+05 | ; | CG - HG |
| 312 | 314 | 1 | 1.5260e-01 | 2.5941e+05 | ; | CG - CD1 |
| 312 | 318 | 1 | 1.5260e-01 | 2.5941e+05 | ; | CG - CD2 |
| 314 | 315 | 1 | 1.0900e-01 | 2.8451e+05 | ; | CD1 - HD11 |
| 314 | 316 | 1 | 1.0900e-01 | 2.8451e+05 | ; | CD1 - HD12 |
| 314 | 317 | 1 | 1.0900e-01 | 2.8451e+05 | ; | CD1 - HD13 |
| 318 | 319 | 1 | 1.0900e-01 | 2.8451e+05 | ; | CD2 - HD21 |
| 318 | 320 | 1 | 1.0900e-01 | 2.8451e+05 | ; | CD2 - HD22 |
| 318 | 321 | 1 | 1.0900e-01 | 2.8451e+05 | ; | CD2 - HD23 |
| 322 | 323 | 1 | 1.2290e-01 | 4.7698e+05 | ; | C - O |
| 322 | 324 | 1 | 1.3350e-01 | 4.1003e+05 | ; | C - N |
| 324 | 325 | 1 | 1.0100e-01 | 3.6317e+05 | ; | N - H |
| 324 | 326 | 1 | 1.4490e-01 | 2.8200e+05 | ; | N - CA |
| 326 | 327 | 1 | 1.0900e-01 | 2.8451e+05 | ; | CA - HA2 |
| 326 | 328 | 1 | 1.0900e-01 | 2.8451e+05 | ; | CA - HA3 |
| 326 | 329 | 1 | 1.5220e-01 | 2.6527e+05 | ; | CA - C |
| 329 | 330 | 1 | 1.2290e-01 | 4.7698e+05 | ; | C - O |
| 329 | 331 | 1 | 1.3350e-01 | 4.1003e+05 | ; | C - N |
| 331 | 332 | 1 | 1.0100e-01 | 3.6317e+05 | ; | N - H |
| 331 | 333 | 1 | 1.4490e-01 | 2.8200e+05 | ; | N - CA |
| 333 | 334 | 1 | 1.0900e-01 | 2.8451e+05 | ; | CA - HA |
| 333 | 335 | 1 | 1.5260e-01 | 2.5941e+05 | ; | CA - CB |
| 333 | 346 | 1 | 1.5220e-01 | 2.6527e+05 | ; | CA - C |
| 335 | 336 | 1 | 1.0900e-01 | 2.8451e+05 | ; | CB - HB2 |
| 335 | 337 | 1 | 1.0900e-01 | 2.8451e+05 | ; | CB - HB3 |
| 335 | 338 | 1 | 1.5260e-01 | 2.5941e+05 | ; | CB - CG |
| 338 | 339 | 1 | 1.0900e-01 | 2.8451e+05 | ; | CG - HG2 |
| 338 | 340 | 1 | 1.0900e-01 | 2.8451e+05 | ; | CG - HG3 |
| 338 | 341 | 1 | 1.5220e-01 | 2.6527e+05 | ; | CG - CD |
| 341 | 342 | 1 | 1.2290e-01 | 4.7698e+05 | ; | CD - OE1 |
| 341 | 343 | 1 | 1.3350e-01 | 4.1003e+05 | ; | CD - NE2 |

| | | | | | | | | |
|---|---|---|---|---|---|---|---|---|
| 343 | 344 | 1 | 1.0100e-01 | 3.6317e+05 | ; | NE2 | - | HE21 |
| 343 | 345 | 1 | 1.0100e-01 | 3.6317e+05 | ; | NE2 | - | HE22 |
| 346 | 347 | 1 | 1.2290e-01 | 4.7698e+05 | ; | C | - | O |
| 346 | 348 | 1 | 1.3350e-01 | 4.1003e+05 | ; | C | - | N |
| 348 | 349 | 1 | 1.0100e-01 | 3.6317e+05 | ; | N | - | H |
| 348 | 350 | 1 | 1.4490e-01 | 2.8200e+05 | ; | N | - | CA |
| 350 | 351 | 1 | 1.0900e-01 | 2.8451e+05 | ; | CA | - | HA2 |
| 350 | 352 | 1 | 1.0900e-01 | 2.8451e+05 | ; | CA | - | HA3 |
| 350 | 353 | 1 | 1.5220e-01 | 2.6527e+05 | ; | CA | - | C |
| 353 | 354 | 1 | 1.2290e-01 | 4.7698e+05 | ; | C | - | O |
| 353 | 355 | 1 | 1.3350e-01 | 4.1003e+05 | ; | C | - | N |
| 355 | 356 | 1 | 1.0100e-01 | 3.6317e+05 | ; | N | - | H |
| 355 | 357 | 1 | 1.4490e-01 | 2.8200e+05 | ; | N | - | CA |
| 357 | 358 | 1 | 1.0900e-01 | 2.8451e+05 | ; | CA | - | HA |
| 357 | 359 | 1 | 1.5260e-01 | 2.5941e+05 | ; | CA | - | CB |
| 357 | 367 | 1 | 1.5220e-01 | 2.6527e+05 | ; | CA | - | C |
| 359 | 360 | 1 | 1.0900e-01 | 2.8451e+05 | ; | CB | - | HB |
| 359 | 361 | 1 | 1.5260e-01 | 2.5941e+05 | ; | CB | - | CG2 |
| 359 | 365 | 1 | 1.4100e-01 | 2.6778e+05 | ; | CB | - | OG1 |
| 361 | 362 | 1 | 1.0900e-01 | 2.8451e+05 | ; | CG2 | - | HG21 |
| 361 | 363 | 1 | 1.0900e-01 | 2.8451e+05 | ; | CG2 | - | HG22 |
| 361 | 364 | 1 | 1.0900e-01 | 2.8451e+05 | ; | CG2 | - | HG23 |
| 365 | 366 | 1 | 9.6000e-02 | 4.6275e+05 | ; | OG1 | - | HG1 |
| 367 | 368 | 1 | 1.2290e-01 | 4.7698e+05 | ; | C | - | O |
| 367 | 369 | 1 | 1.3350e-01 | 4.1003e+05 | ; | C | - | N |
| 369 | 370 | 1 | 1.0100e-01 | 3.6317e+05 | ; | N | - | H |
| 369 | 371 | 1 | 1.4490e-01 | 2.8200e+05 | ; | N | - | CA |
| 371 | 372 | 1 | 1.0900e-01 | 2.8451e+05 | ; | CA | - | HA |
| 371 | 373 | 1 | 1.5260e-01 | 2.5941e+05 | ; | CA | - | CB |
| 371 | 388 | 1 | 1.5220e-01 | 2.6527e+05 | ; | CA | - | C |
| 373 | 374 | 1 | 1.0900e-01 | 2.8451e+05 | ; | CB | - | HB2 |
| 373 | 375 | 1 | 1.0900e-01 | 2.8451e+05 | ; | CB | - | HB3 |
| 373 | 376 | 1 | 1.5100e-01 | 2.6527e+05 | ; | CB | - | CG |
| 376 | 377 | 1 | 1.4000e-01 | 3.9246e+05 | ; | CG | - | CD1 |
| 376 | 386 | 1 | 1.4000e-01 | 3.9246e+05 | ; | CG | - | CD2 |
| 377 | 378 | 1 | 1.0800e-01 | 3.0711e+05 | ; | CD1 | - | HD1 |
| 377 | 379 | 1 | 1.4000e-01 | 3.9246e+05 | ; | CD1 | - | CE1 |
| 379 | 380 | 1 | 1.0800e-01 | 3.0711e+05 | ; | CE1 | - | HE1 |
| 379 | 381 | 1 | 1.4090e-01 | 3.9246e+05 | ; | CE1 | - | CZ |
| 381 | 382 | 1 | 1.3640e-01 | 3.7656e+05 | ; | CZ | - | OH |
| 381 | 384 | 1 | 1.4090e-01 | 3.9246e+05 | ; | CZ | - | CE2 |
| 382 | 383 | 1 | 9.6000e-02 | 4.6275e+05 | ; | OH | - | HH |
| 384 | 385 | 1 | 1.0800e-01 | 3.0711e+05 | ; | CE2 | - | HE2 |
| 384 | 386 | 1 | 1.4000e-01 | 3.9246e+05 | ; | CE2 | - | CD2 |
| 386 | 387 | 1 | 1.0800e-01 | 3.0711e+05 | ; | CD2 | - | HD2 |
| 388 | 389 | 1 | 1.2290e-01 | 4.7698e+05 | ; | C | - | O |
| 388 | 390 | 1 | 1.3350e-01 | 4.1003e+05 | ; | C | - | N |
| 390 | 391 | 1 | 1.0100e-01 | 3.6317e+05 | ; | N | - | H |
| 390 | 392 | 1 | 1.4490e-01 | 2.8200e+05 | ; | N | - | CA |
| 392 | 393 | 1 | 1.0900e-01 | 2.8451e+05 | ; | CA | - | HA2 |
| 392 | 394 | 1 | 1.0900e-01 | 2.8451e+05 | ; | CA | - | HA3 |
| 392 | 395 | 1 | 1.5220e-01 | 2.6527e+05 | ; | CA | - | C |

| | | | | | | | |
|---|---|---|---|---|---|---|---|
| 395 | 396 | 1 | 1.2290e-01 | 4.7698e+05 ; | C | - | O |
| 395 | 397 | 1 | 1.3350e-01 | 4.1003e+05 ; | C | - | N |
| 397 | 398 | 1 | 1.0100e-01 | 3.6317e+05 ; | N | - | H |
| 397 | 399 | 1 | 1.4490e-01 | 2.8200e+05 ; | N | - | CA |
| 399 | 400 | 1 | 1.0900e-01 | 2.8451e+05 ; | CA | - | HA |
| 399 | 401 | 1 | 1.5260e-01 | 2.5941e+05 ; | CA | - | CB |
| 399 | 406 | 1 | 1.5220e-01 | 2.6527e+05 ; | CA | - | C |
| 401 | 402 | 1 | 1.0900e-01 | 2.8451e+05 ; | CB | - | HB2 |
| 401 | 403 | 1 | 1.0900e-01 | 2.8451e+05 ; | CB | - | HB3 |
| 401 | 404 | 1 | 1.8100e-01 | 1.9832e+05 ; | CB | - | SG |
| 404 | 405 | 1 | 1.3360e-01 | 2.2928e+05 ; | SG | - | HG |
| 406 | 407 | 1 | 1.2290e-01 | 4.7698e+05 ; | C | - | O |
| 406 | 408 | 1 | 1.3350e-01 | 4.1003e+05 ; | C | - | N |
| 408 | 409 | 1 | 1.0100e-01 | 3.6317e+05 ; | N | - | H |
| 408 | 410 | 1 | 1.4490e-01 | 2.8200e+05 ; | N | - | CA |
| 410 | 411 | 1 | 1.0900e-01 | 2.8451e+05 ; | CA | - | HA |
| 410 | 412 | 1 | 1.5260e-01 | 2.5941e+05 ; | CA | - | CB |
| 410 | 422 | 1 | 1.5220e-01 | 2.6527e+05 ; | CA | - | C |
| 412 | 413 | 1 | 1.0900e-01 | 2.8451e+05 ; | CB | - | HB |
| 412 | 414 | 1 | 1.5260e-01 | 2.5941e+05 ; | CB | - | CG1 |
| 412 | 418 | 1 | 1.5260e-01 | 2.5941e+05 ; | CB | - | CG2 |
| 414 | 415 | 1 | 1.0900e-01 | 2.8451e+05 ; | CG1 | - | HG11 |
| 414 | 416 | 1 | 1.0900e-01 | 2.8451e+05 ; | CG1 | - | HG12 |
| 414 | 417 | 1 | 1.0900e-01 | 2.8451e+05 ; | CG1 | - | HG13 |
| 418 | 419 | 1 | 1.0900e-01 | 2.8451e+05 ; | CG2 | - | HG21 |
| 418 | 420 | 1 | 1.0900e-01 | 2.8451e+05 ; | CG2 | - | HG22 |
| 418 | 421 | 1 | 1.0900e-01 | 2.8451e+05 ; | CG2 | - | HG23 |
| 422 | 423 | 1 | 1.2290e-01 | 4.7698e+05 ; | C | - | O |
| 422 | 424 | 1 | 1.3350e-01 | 4.1003e+05 ; | C | - | N |
| 424 | 425 | 1 | 1.0100e-01 | 3.6317e+05 ; | N | - | H |
| 424 | 426 | 1 | 1.4490e-01 | 2.8200e+05 ; | N | - | CA |
| 426 | 427 | 1 | 1.0900e-01 | 2.8451e+05 ; | CA | - | HA |
| 426 | 428 | 1 | 1.5260e-01 | 2.5941e+05 ; | CA | - | CB |
| 426 | 443 | 1 | 1.5220e-01 | 2.6527e+05 ; | CA | - | C |
| 428 | 429 | 1 | 1.0900e-01 | 2.8451e+05 ; | CB | - | HB2 |
| 428 | 430 | 1 | 1.0900e-01 | 2.8451e+05 ; | CB | - | HB3 |
| 428 | 431 | 1 | 1.5100e-01 | 2.6527e+05 ; | CB | - | CG |
| 431 | 432 | 1 | 1.4000e-01 | 3.9246e+05 ; | CG | - | CD1 |
| 431 | 441 | 1 | 1.4000e-01 | 3.9246e+05 ; | CG | - | CD2 |
| 432 | 433 | 1 | 1.0800e-01 | 3.0711e+05 ; | CD1 | - | HD1 |
| 432 | 434 | 1 | 1.4000e-01 | 3.9246e+05 ; | CD1 | - | CE1 |
| 434 | 435 | 1 | 1.0800e-01 | 3.0711e+05 ; | CE1 | - | HE1 |
| 434 | 436 | 1 | 1.4090e-01 | 3.9246e+05 ; | CE1 | - | CZ |
| 436 | 437 | 1 | 1.3640e-01 | 3.7656e+05 ; | CZ | - | OH |
| 436 | 439 | 1 | 1.4090e-01 | 3.9246e+05 ; | CZ | - | CE2 |
| 437 | 438 | 1 | 9.6000e-02 | 4.6275e+05 ; | OH | - | HH |
| 439 | 440 | 1 | 1.0800e-01 | 3.0711e+05 ; | CE2 | - | HE2 |
| 439 | 441 | 1 | 1.4000e-01 | 3.9246e+05 ; | CE2 | - | CD2 |
| 441 | 442 | 1 | 1.0800e-01 | 3.0711e+05 ; | CD2 | - | HD2 |
| 443 | 444 | 1 | 1.2290e-01 | 4.7698e+05 ; | C | - | O |
| 443 | 445 | 1 | 1.3350e-01 | 4.1003e+05 ; | C | - | N |
| 445 | 446 | 1 | 1.0100e-01 | 3.6317e+05 ; | N | - | H |

| | | | | | | |
|---|---|---|---|---|---|---|
| 445 | 447 | 1 | 1.4490e-01 | 2.8200e+05 | ; | N - CA |
| 447 | 448 | 1 | 1.0900e-01 | 2.8451e+05 | ; | CA - HA |
| 447 | 449 | 1 | 1.5260e-01 | 2.5941e+05 | ; | CA - CB |
| 447 | 465 | 1 | 1.5220e-01 | 2.6527e+05 | ; | CA - C |
| 449 | 450 | 1 | 1.0900e-01 | 2.8451e+05 | ; | CB - HB2 |
| 449 | 451 | 1 | 1.0900e-01 | 2.8451e+05 | ; | CB - HB3 |
| 449 | 452 | 1 | 1.5260e-01 | 2.5941e+05 | ; | CB - CG |
| 452 | 453 | 1 | 1.0900e-01 | 2.8451e+05 | ; | CG - HG2 |
| 452 | 454 | 1 | 1.0900e-01 | 2.8451e+05 | ; | CG - HG3 |
| 452 | 455 | 1 | 1.5260e-01 | 2.5941e+05 | ; | CG - CD |
| 455 | 456 | 1 | 1.0900e-01 | 2.8451e+05 | ; | CD - HD2 |
| 455 | 457 | 1 | 1.0900e-01 | 2.8451e+05 | ; | CD - HD3 |
| 455 | 458 | 1 | 1.5260e-01 | 2.5941e+05 | ; | CD - CE |
| 458 | 459 | 1 | 1.0900e-01 | 2.8451e+05 | ; | CE - HE2 |
| 458 | 460 | 1 | 1.0900e-01 | 2.8451e+05 | ; | CE - HE3 |
| 458 | 461 | 1 | 1.4710e-01 | 3.0711e+05 | ; | CE - NZ |
| 461 | 462 | 1 | 1.0100e-01 | 3.6317e+05 | ; | NZ - HZ1 |
| 461 | 463 | 1 | 1.0100e-01 | 3.6317e+05 | ; | NZ - HZ2 |
| 461 | 464 | 1 | 1.0100e-01 | 3.6317e+05 | ; | NZ - HZ3 |
| 465 | 466 | 1 | 1.2290e-01 | 4.7698e+05 | ; | C - O |
| 465 | 467 | 1 | 1.3350e-01 | 4.1003e+05 | ; | C - N |
| 467 | 468 | 1 | 1.0100e-01 | 3.6317e+05 | ; | N - H |
| 467 | 469 | 1 | 1.4490e-01 | 2.8200e+05 | ; | N - CA |
| 469 | 470 | 1 | 1.0900e-01 | 2.8451e+05 | ; | CA - HA2 |
| 469 | 471 | 1 | 1.0900e-01 | 2.8451e+05 | ; | CA - HA3 |
| 469 | 472 | 1 | 1.5220e-01 | 2.6527e+05 | ; | CA - C |
| 472 | 473 | 1 | 1.2290e-01 | 4.7698e+05 | ; | C - O |
| 472 | 474 | 1 | 1.3350e-01 | 4.1003e+05 | ; | C - N |
| 474 | 475 | 1 | 1.0100e-01 | 3.6317e+05 | ; | N - H |
| 474 | 476 | 1 | 1.4490e-01 | 2.8200e+05 | ; | N - CA |
| 476 | 477 | 1 | 1.0900e-01 | 2.8451e+05 | ; | CA - HA |
| 476 | 478 | 1 | 1.5260e-01 | 2.5941e+05 | ; | CA - CB |
| 476 | 491 | 1 | 1.5220e-01 | 2.6527e+05 | ; | CA - C |
| 478 | 479 | 1 | 1.0900e-01 | 2.8451e+05 | ; | CB - HB |
| 478 | 480 | 1 | 1.5260e-01 | 2.5941e+05 | ; | CB - CG2 |
| 478 | 484 | 1 | 1.5260e-01 | 2.5941e+05 | ; | CB - CG1 |
| 480 | 481 | 1 | 1.0900e-01 | 2.8451e+05 | ; | CG2 - HG21 |
| 480 | 482 | 1 | 1.0900e-01 | 2.8451e+05 | ; | CG2 - HG22 |
| 480 | 483 | 1 | 1.0900e-01 | 2.8451e+05 | ; | CG2 - HG23 |
| 484 | 485 | 1 | 1.0900e-01 | 2.8451e+05 | ; | CG1 - HG12 |
| 484 | 486 | 1 | 1.0900e-01 | 2.8451e+05 | ; | CG1 - HG13 |
| 484 | 487 | 1 | 1.5260e-01 | 2.5941e+05 | ; | CG1 - CD1 |
| 487 | 488 | 1 | 1.0900e-01 | 2.8451e+05 | ; | CD1 - HD11 |
| 487 | 489 | 1 | 1.0900e-01 | 2.8451e+05 | ; | CD1 - HD12 |
| 487 | 490 | 1 | 1.0900e-01 | 2.8451e+05 | ; | CD1 - HD13 |
| 491 | 492 | 1 | 1.2290e-01 | 4.7698e+05 | ; | C - O |
| 491 | 493 | 1 | 1.3350e-01 | 4.1003e+05 | ; | C - N |
| 493 | 494 | 1 | 1.0100e-01 | 3.6317e+05 | ; | N - H |
| 493 | 495 | 1 | 1.4490e-01 | 2.8200e+05 | ; | N - CA |
| 495 | 496 | 1 | 1.0900e-01 | 2.8451e+05 | ; | CA - HA |
| 495 | 497 | 1 | 1.5260e-01 | 2.5941e+05 | ; | CA - CB |
| 495 | 503 | 1 | 1.5220e-01 | 2.6527e+05 | ; | CA - C |

| | | | | | |
|---|---|---|---|---|---|
| 497 | 498 | 1 | 1.0900e-01 | 2.8451e+05 ; | CB - HB2 |
| 497 | 499 | 1 | 1.0900e-01 | 2.8451e+05 ; | CB - HB3 |
| 497 | 500 | 1 | 1.5220e-01 | 2.6527e+05 ; | CB - CG |
| 500 | 501 | 1 | 1.2500e-01 | 5.4894e+05 ; | CG - OD1 |
| 500 | 502 | 1 | 1.2500e-01 | 5.4894e+05 ; | CG - OD2 |
| 503 | 504 | 1 | 1.2290e-01 | 4.7698e+05 ; | C - O |
| 503 | 505 | 1 | 1.3350e-01 | 4.1003e+05 ; | C - N |
| 505 | 506 | 1 | 1.0100e-01 | 3.6317e+05 ; | N - H |
| 505 | 507 | 1 | 1.4490e-01 | 2.8200e+05 ; | N - CA |
| 507 | 508 | 1 | 1.0900e-01 | 2.8451e+05 ; | CA - HA |
| 507 | 509 | 1 | 1.5260e-01 | 2.5941e+05 ; | CA - CB |
| 507 | 525 | 1 | 1.5220e-01 | 2.6527e+05 ; | CA - C |
| 509 | 510 | 1 | 1.0900e-01 | 2.8451e+05 ; | CB - HB2 |
| 509 | 511 | 1 | 1.0900e-01 | 2.8451e+05 ; | CB - HB3 |
| 509 | 512 | 1 | 1.5260e-01 | 2.5941e+05 ; | CB - CG |
| 512 | 513 | 1 | 1.0900e-01 | 2.8451e+05 ; | CG - HG2 |
| 512 | 514 | 1 | 1.0900e-01 | 2.8451e+05 ; | CG - HG3 |
| 512 | 515 | 1 | 1.5260e-01 | 2.5941e+05 ; | CG - CD |
| 515 | 516 | 1 | 1.0900e-01 | 2.8451e+05 ; | CD - HD2 |
| 515 | 517 | 1 | 1.0900e-01 | 2.8451e+05 ; | CD - HD3 |
| 515 | 518 | 1 | 1.5260e-01 | 2.5941e+05 ; | CD - CE |
| 518 | 519 | 1 | 1.0900e-01 | 2.8451e+05 ; | CE - HE2 |
| 518 | 520 | 1 | 1.0900e-01 | 2.8451e+05 ; | CE - HE3 |
| 518 | 521 | 1 | 1.4710e-01 | 3.0711e+05 ; | CE - NZ |
| 521 | 522 | 1 | 1.0100e-01 | 3.6317e+05 ; | NZ - HZ1 |
| 521 | 523 | 1 | 1.0100e-01 | 3.6317e+05 ; | NZ - HZ2 |
| 521 | 524 | 1 | 1.0100e-01 | 3.6317e+05 ; | NZ - HZ3 |
| 525 | 526 | 1 | 1.2290e-01 | 4.7698e+05 ; | C - O |
| 525 | 527 | 1 | 1.3350e-01 | 4.1003e+05 ; | C - N |
| 527 | 528 | 1 | 1.0100e-01 | 3.6317e+05 ; | N - H |
| 527 | 529 | 1 | 1.4490e-01 | 2.8200e+05 ; | N - CA |
| 529 | 530 | 1 | 1.0900e-01 | 2.8451e+05 ; | CA - HA |
| 529 | 531 | 1 | 1.5260e-01 | 2.5941e+05 ; | CA - CB |
| 529 | 541 | 1 | 1.5220e-01 | 2.6527e+05 ; | CA - C |
| 531 | 532 | 1 | 1.0900e-01 | 2.8451e+05 ; | CB - HB |
| 531 | 533 | 1 | 1.5260e-01 | 2.5941e+05 ; | CB - CG1 |
| 531 | 537 | 1 | 1.5260e-01 | 2.5941e+05 ; | CB - CG2 |
| 533 | 534 | 1 | 1.0900e-01 | 2.8451e+05 ; | CG1 - HG11 |
| 533 | 535 | 1 | 1.0900e-01 | 2.8451e+05 ; | CG1 - HG12 |
| 533 | 536 | 1 | 1.0900e-01 | 2.8451e+05 ; | CG1 - HG13 |
| 537 | 538 | 1 | 1.0900e-01 | 2.8451e+05 ; | CG2 - HG21 |
| 537 | 539 | 1 | 1.0900e-01 | 2.8451e+05 ; | CG2 - HG22 |
| 537 | 540 | 1 | 1.0900e-01 | 2.8451e+05 ; | CG2 - HG23 |
| 541 | 542 | 1 | 1.2290e-01 | 4.7698e+05 ; | C - O |
| 541 | 543 | 1 | 1.3350e-01 | 4.1003e+05 ; | C - N |
| 543 | 544 | 1 | 1.0100e-01 | 3.6317e+05 ; | N - H |
| 543 | 545 | 1 | 1.4490e-01 | 2.8200e+05 ; | N - CA |
| 545 | 546 | 1 | 1.0900e-01 | 2.8451e+05 ; | CA - HA |
| 545 | 547 | 1 | 1.5260e-01 | 2.5941e+05 ; | CA - CB |
| 545 | 555 | 1 | 1.5220e-01 | 2.6527e+05 ; | CA - C |
| 547 | 548 | 1 | 1.0900e-01 | 2.8451e+05 ; | CB - HB |
| 547 | 549 | 1 | 1.5260e-01 | 2.5941e+05 ; | CB - CG2 |

| | | | | | |
|---|---|---|---|---|---|
| 547 | 553 | 1 | 1.4100e-01 | 2.6778e+05 ; | CB - OG1 |
| 549 | 550 | 1 | 1.0900e-01 | 2.8451e+05 ; | CG2 - HG21 |
| 549 | 551 | 1 | 1.0900e-01 | 2.8451e+05 ; | CG2 - HG22 |
| 549 | 552 | 1 | 1.0900e-01 | 2.8451e+05 ; | CG2 - HG23 |
| 553 | 554 | 1 | 9.6000e-02 | 4.6275e+05 ; | OG1 - HG1 |
| 555 | 556 | 1 | 1.2290e-01 | 4.7698e+05 ; | C - O |
| 555 | 557 | 1 | 1.3350e-01 | 4.1003e+05 ; | C - N |
| 557 | 558 | 1 | 1.0100e-01 | 3.6317e+05 ; | N - H |
| 557 | 559 | 1 | 1.4490e-01 | 2.8200e+05 ; | N - CA |
| 559 | 560 | 1 | 1.0900e-01 | 2.8451e+05 ; | CA - HA |
| 559 | 561 | 1 | 1.5260e-01 | 2.5941e+05 ; | CA - CB |
| 559 | 569 | 1 | 1.5220e-01 | 2.6527e+05 ; | CA - C |
| 561 | 562 | 1 | 1.0900e-01 | 2.8451e+05 ; | CB - HB2 |
| 561 | 563 | 1 | 1.0900e-01 | 2.8451e+05 ; | CB - HB3 |
| 561 | 564 | 1 | 1.5220e-01 | 2.6527e+05 ; | CB - CG |
| 564 | 565 | 1 | 1.2290e-01 | 4.7698e+05 ; | CG - OD1 |
| 564 | 566 | 1 | 1.3350e-01 | 4.1003e+05 ; | CG - ND2 |
| 566 | 567 | 1 | 1.0100e-01 | 3.6317e+05 ; | ND2 - HD21 |
| 566 | 568 | 1 | 1.0100e-01 | 3.6317e+05 ; | ND2 - HD22 |
| 569 | 570 | 1 | 1.2290e-01 | 4.7698e+05 ; | C - O |
| 569 | 571 | 1 | 1.3350e-01 | 4.1003e+05 ; | C - N |
| 571 | 572 | 1 | 1.0100e-01 | 3.6317e+05 ; | N - H |
| 571 | 573 | 1 | 1.4490e-01 | 2.8200e+05 ; | N - CA |
| 573 | 574 | 1 | 1.0900e-01 | 2.8451e+05 ; | CA - HA |
| 573 | 575 | 1 | 1.5260e-01 | 2.5941e+05 ; | CA - CB |
| 573 | 586 | 1 | 1.5220e-01 | 2.6527e+05 ; | CA - C |
| 575 | 576 | 1 | 1.0900e-01 | 2.8451e+05 ; | CB - HB2 |
| 575 | 577 | 1 | 1.0900e-01 | 2.8451e+05 ; | CB - HB3 |
| 575 | 578 | 1 | 1.5260e-01 | 2.5941e+05 ; | CB - CG |
| 578 | 579 | 1 | 1.0900e-01 | 2.8451e+05 ; | CG - HG2 |
| 578 | 580 | 1 | 1.0900e-01 | 2.8451e+05 ; | CG - HG3 |
| 578 | 581 | 1 | 1.5220e-01 | 2.6527e+05 ; | CG - CD |
| 581 | 582 | 1 | 1.2290e-01 | 4.7698e+05 ; | CD - OE1 |
| 581 | 583 | 1 | 1.3350e-01 | 4.1003e+05 ; | CD - NE2 |
| 583 | 584 | 1 | 1.0100e-01 | 3.6317e+05 ; | NE2 - HE21 |
| 583 | 585 | 1 | 1.0100e-01 | 3.6317e+05 ; | NE2 - HE22 |
| 586 | 587 | 1 | 1.2290e-01 | 4.7698e+05 ; | C - O |
| 586 | 588 | 1 | 1.3350e-01 | 4.1003e+05 ; | C - N |
| 588 | 589 | 1 | 1.0100e-01 | 3.6317e+05 ; | N - H |
| 588 | 590 | 1 | 1.4490e-01 | 2.8200e+05 ; | N - CA |
| 590 | 591 | 1 | 1.0900e-01 | 2.8451e+05 ; | CA - HA |
| 590 | 592 | 1 | 1.5260e-01 | 2.5941e+05 ; | CA - CB |
| 590 | 605 | 1 | 1.5220e-01 | 2.6527e+05 ; | CA - C |
| 592 | 593 | 1 | 1.0900e-01 | 2.8451e+05 ; | CB - HB2 |
| 592 | 594 | 1 | 1.0900e-01 | 2.8451e+05 ; | CB - HB3 |
| 592 | 595 | 1 | 1.5260e-01 | 2.5941e+05 ; | CB - CG |
| 595 | 596 | 1 | 1.0900e-01 | 2.8451e+05 ; | CG - HG |
| 595 | 597 | 1 | 1.5260e-01 | 2.5941e+05 ; | CG - CD1 |
| 595 | 601 | 1 | 1.5260e-01 | 2.5941e+05 ; | CG - CD2 |
| 597 | 598 | 1 | 1.0900e-01 | 2.8451e+05 ; | CD1 - HD11 |
| 597 | 599 | 1 | 1.0900e-01 | 2.8451e+05 ; | CD1 - HD12 |
| 597 | 600 | 1 | 1.0900e-01 | 2.8451e+05 ; | CD1 - HD13 |

| | | | | | | |
|---|---|---|---|---|---|---|
| 601 | 602 | 1 | 1.0900e-01 | 2.8451e+05 | ; | CD2 - HD21 |
| 601 | 603 | 1 | 1.0900e-01 | 2.8451e+05 | ; | CD2 - HD22 |
| 601 | 604 | 1 | 1.0900e-01 | 2.8451e+05 | ; | CD2 - HD23 |
| 605 | 606 | 1 | 1.2290e-01 | 4.7698e+05 | ; | C - O |
| 605 | 607 | 1 | 1.3350e-01 | 4.1003e+05 | ; | C - N |
| 607 | 608 | 1 | 1.0100e-01 | 3.6317e+05 | ; | N - H |
| 607 | 609 | 1 | 1.4490e-01 | 2.8200e+05 | ; | N - CA |
| 609 | 610 | 1 | 1.0900e-01 | 2.8451e+05 | ; | CA - HA |
| 609 | 611 | 1 | 1.5260e-01 | 2.5941e+05 | ; | CA - CB |
| 609 | 626 | 1 | 1.5220e-01 | 2.6527e+05 | ; | CA - C |
| 611 | 612 | 1 | 1.0900e-01 | 2.8451e+05 | ; | CB - HB2 |
| 611 | 613 | 1 | 1.0900e-01 | 2.8451e+05 | ; | CB - HB3 |
| 611 | 614 | 1 | 1.5100e-01 | 2.6527e+05 | ; | CB - CG |
| 614 | 615 | 1 | 1.4000e-01 | 3.9246e+05 | ; | CG - CD1 |
| 614 | 624 | 1 | 1.4000e-01 | 3.9246e+05 | ; | CG - CD2 |
| 615 | 616 | 1 | 1.0800e-01 | 3.0711e+05 | ; | CD1 - HD1 |
| 615 | 617 | 1 | 1.4000e-01 | 3.9246e+05 | ; | CD1 - CE1 |
| 617 | 618 | 1 | 1.0800e-01 | 3.0711e+05 | ; | CE1 - HE1 |
| 617 | 619 | 1 | 1.4090e-01 | 3.9246e+05 | ; | CE1 - CZ |
| 619 | 620 | 1 | 1.3640e-01 | 3.7656e+05 | ; | CZ - OH |
| 619 | 622 | 1 | 1.4090e-01 | 3.9246e+05 | ; | CZ - CE2 |
| 620 | 621 | 1 | 9.6000e-02 | 4.6275e+05 | ; | OH - HH |
| 622 | 623 | 1 | 1.0800e-01 | 3.0711e+05 | ; | CE2 - HE2 |
| 622 | 624 | 1 | 1.4000e-01 | 3.9246e+05 | ; | CE2 - CD2 |
| 624 | 625 | 1 | 1.0800e-01 | 3.0711e+05 | ; | CD2 - HD2 |
| 626 | 627 | 1 | 1.2290e-01 | 4.7698e+05 | ; | C - O |
| 626 | 628 | 1 | 1.3350e-01 | 4.1003e+05 | ; | C - N |
| 628 | 629 | 1 | 1.0100e-01 | 3.6317e+05 | ; | N - H |
| 628 | 630 | 1 | 1.4490e-01 | 2.8200e+05 | ; | N - CA |
| 630 | 631 | 1 | 1.0900e-01 | 2.8451e+05 | ; | CA - HA |
| 630 | 632 | 1 | 1.5260e-01 | 2.5941e+05 | ; | CA - CB |
| 630 | 636 | 1 | 1.5220e-01 | 2.6527e+05 | ; | CA - C |
| 632 | 633 | 1 | 1.0900e-01 | 2.8451e+05 | ; | CB - HB1 |
| 632 | 634 | 1 | 1.0900e-01 | 2.8451e+05 | ; | CB - HB2 |
| 632 | 635 | 1 | 1.0900e-01 | 2.8451e+05 | ; | CB - HB3 |
| 636 | 637 | 1 | 1.2290e-01 | 4.7698e+05 | ; | C - O |
| 636 | 638 | 1 | 1.3350e-01 | 4.1003e+05 | ; | C - N |
| 638 | 639 | 1 | 1.0100e-01 | 3.6317e+05 | ; | N - H |
| 638 | 640 | 1 | 1.4490e-01 | 2.8200e+05 | ; | N - CA |
| 640 | 641 | 1 | 1.0900e-01 | 2.8451e+05 | ; | CA - HA |
| 640 | 642 | 1 | 1.5260e-01 | 2.5941e+05 | ; | CA - CB |
| 640 | 655 | 1 | 1.5220e-01 | 2.6527e+05 | ; | CA - C |
| 642 | 643 | 1 | 1.0900e-01 | 2.8451e+05 | ; | CB - HB |
| 642 | 644 | 1 | 1.5260e-01 | 2.5941e+05 | ; | CB - CG2 |
| 642 | 648 | 1 | 1.5260e-01 | 2.5941e+05 | ; | CB - CG1 |
| 644 | 645 | 1 | 1.0900e-01 | 2.8451e+05 | ; | CG2 - HG21 |
| 644 | 646 | 1 | 1.0900e-01 | 2.8451e+05 | ; | CG2 - HG22 |
| 644 | 647 | 1 | 1.0900e-01 | 2.8451e+05 | ; | CG2 - HG23 |
| 648 | 649 | 1 | 1.0900e-01 | 2.8451e+05 | ; | CG1 - HG12 |
| 648 | 650 | 1 | 1.0900e-01 | 2.8451e+05 | ; | CG1 - HG13 |
| 648 | 651 | 1 | 1.5260e-01 | 2.5941e+05 | ; | CG1 - CD1 |
| 651 | 652 | 1 | 1.0900e-01 | 2.8451e+05 | ; | CD1 - HD11 |

| | | | | | |
|---|---|---|---|---|---|
| 651 | 653 | 1 | 1.0900e-01 | 2.8451e+05 ; | CD1 - HD12 |
| 651 | 654 | 1 | 1.0900e-01 | 2.8451e+05 ; | CD1 - HD13 |
| 655 | 656 | 1 | 1.2290e-01 | 4.7698e+05 ; | C - O |
| 655 | 657 | 1 | 1.3350e-01 | 4.1003e+05 ; | C - N |
| 657 | 658 | 1 | 1.0100e-01 | 3.6317e+05 ; | N - H |
| 657 | 659 | 1 | 1.4490e-01 | 2.8200e+05 ; | N - CA |
| 659 | 660 | 1 | 1.0900e-01 | 2.8451e+05 ; | CA - HA |
| 659 | 661 | 1 | 1.5260e-01 | 2.5941e+05 ; | CA - CB |
| 659 | 677 | 1 | 1.5220e-01 | 2.6527e+05 ; | CA - C |
| 661 | 662 | 1 | 1.0900e-01 | 2.8451e+05 ; | CB - HB2 |
| 661 | 663 | 1 | 1.0900e-01 | 2.8451e+05 ; | CB - HB3 |
| 661 | 664 | 1 | 1.5260e-01 | 2.5941e+05 ; | CB - CG |
| 664 | 665 | 1 | 1.0900e-01 | 2.8451e+05 ; | CG - HG2 |
| 664 | 666 | 1 | 1.0900e-01 | 2.8451e+05 ; | CG - HG3 |
| 664 | 667 | 1 | 1.5260e-01 | 2.5941e+05 ; | CG - CD |
| 667 | 668 | 1 | 1.0900e-01 | 2.8451e+05 ; | CD - HD2 |
| 667 | 669 | 1 | 1.0900e-01 | 2.8451e+05 ; | CD - HD3 |
| 667 | 670 | 1 | 1.5260e-01 | 2.5941e+05 ; | CD - CE |
| 670 | 671 | 1 | 1.0900e-01 | 2.8451e+05 ; | CE - HE2 |
| 670 | 672 | 1 | 1.0900e-01 | 2.8451e+05 ; | CE - HE3 |
| 670 | 673 | 1 | 1.4710e-01 | 3.0711e+05 ; | CE - NZ |
| 673 | 674 | 1 | 1.0100e-01 | 3.6317e+05 ; | NZ - HZ1 |
| 673 | 675 | 1 | 1.0100e-01 | 3.6317e+05 ; | NZ - HZ2 |
| 673 | 676 | 1 | 1.0100e-01 | 3.6317e+05 ; | NZ - HZ3 |
| 677 | 678 | 1 | 1.2290e-01 | 4.7698e+05 ; | C - O |
| 677 | 679 | 1 | 1.3350e-01 | 4.1003e+05 ; | C - N |
| 679 | 680 | 1 | 1.0100e-01 | 3.6317e+05 ; | N - H |
| 679 | 681 | 1 | 1.4490e-01 | 2.8200e+05 ; | N - CA |
| 681 | 682 | 1 | 1.0900e-01 | 2.8451e+05 ; | CA - HA |
| 681 | 683 | 1 | 1.5260e-01 | 2.5941e+05 ; | CA - CB |
| 681 | 692 | 1 | 1.5220e-01 | 2.6527e+05 ; | CA - C |
| 683 | 684 | 1 | 1.0900e-01 | 2.8451e+05 ; | CB - HB2 |
| 683 | 685 | 1 | 1.0900e-01 | 2.8451e+05 ; | CB - HB3 |
| 683 | 686 | 1 | 1.5260e-01 | 2.5941e+05 ; | CB - CG |
| 686 | 687 | 1 | 1.0900e-01 | 2.8451e+05 ; | CG - HG2 |
| 686 | 688 | 1 | 1.0900e-01 | 2.8451e+05 ; | CG - HG3 |
| 686 | 689 | 1 | 1.5220e-01 | 2.6527e+05 ; | CG - CD |
| 689 | 690 | 1 | 1.2500e-01 | 5.4894e+05 ; | CD - OE1 |
| 689 | 691 | 1 | 1.2500e-01 | 5.4894e+05 ; | CD - OE2 |
| 692 | 693 | 1 | 1.2290e-01 | 4.7698e+05 ; | C - O |
| 692 | 694 | 1 | 1.3350e-01 | 4.1003e+05 ; | C - N |
| 694 | 695 | 1 | 1.0100e-01 | 3.6317e+05 ; | N - H |
| 694 | 696 | 1 | 1.4490e-01 | 2.8200e+05 ; | N - CA |
| 696 | 697 | 1 | 1.0900e-01 | 2.8451e+05 ; | CA - HA |
| 696 | 698 | 1 | 1.5260e-01 | 2.5941e+05 ; | CA - CB |
| 696 | 707 | 1 | 1.5220e-01 | 2.6527e+05 ; | CA - C |
| 698 | 699 | 1 | 1.0900e-01 | 2.8451e+05 ; | CB - HB2 |
| 698 | 700 | 1 | 1.0900e-01 | 2.8451e+05 ; | CB - HB3 |
| 698 | 701 | 1 | 1.5260e-01 | 2.5941e+05 ; | CB - CG |
| 701 | 702 | 1 | 1.0900e-01 | 2.8451e+05 ; | CG - HG2 |
| 701 | 703 | 1 | 1.0900e-01 | 2.8451e+05 ; | CG - HG3 |
| 701 | 704 | 1 | 1.5220e-01 | 2.6527e+05 ; | CG - CD |

| | | | | | | |
|---|---|---|---|---|---|---|
| 2044 | 2045 | 1 | 1.0100e-01 | 3.6317e+05 | ; | N - H |
| 2044 | 2046 | 1 | 1.4490e-01 | 2.8200e+05 | ; | N - CA |
| 2046 | 2047 | 1 | 1.0900e-01 | 2.8451e+05 | ; | CA - HA |
| 2046 | 2048 | 1 | 1.5260e-01 | 2.5941e+05 | ; | CA - CB |
| 2046 | 2061 | 1 | 1.5220e-01 | 2.6527e+05 | ; | CA - C |
| 2048 | 2049 | 1 | 1.0900e-01 | 2.8451e+05 | ; | CB - HB2 |
| 2048 | 2050 | 1 | 1.0900e-01 | 2.8451e+05 | ; | CB - HB3 |
| 2048 | 2051 | 1 | 1.5260e-01 | 2.5941e+05 | ; | CB - CG |
| 2051 | 2052 | 1 | 1.0900e-01 | 2.8451e+05 | ; | CG - HG |
| 2051 | 2053 | 1 | 1.5260e-01 | 2.5941e+05 | ; | CG - CD1 |
| 2051 | 2057 | 1 | 1.5260e-01 | 2.5941e+05 | ; | CG - CD2 |
| 2053 | 2054 | 1 | 1.0900e-01 | 2.8451e+05 | ; | CD1 - HD11 |
| 2053 | 2055 | 1 | 1.0900e-01 | 2.8451e+05 | ; | CD1 - HD12 |
| 2053 | 2056 | 1 | 1.0900e-01 | 2.8451e+05 | ; | CD1 - HD13 |
| 2057 | 2058 | 1 | 1.0900e-01 | 2.8451e+05 | ; | CD2 - HD21 |
| 2057 | 2059 | 1 | 1.0900e-01 | 2.8451e+05 | ; | CD2 - HD22 |
| 2057 | 2060 | 1 | 1.0900e-01 | 2.8451e+05 | ; | CD2 - HD23 |
| 2061 | 2062 | 1 | 1.2290e-01 | 4.7698e+05 | ; | C - O |
| 2061 | 2063 | 1 | 1.3350e-01 | 4.1003e+05 | ; | C - N |
| 2063 | 2064 | 1 | 1.0100e-01 | 3.6317e+05 | ; | N - H |
| 2063 | 2065 | 1 | 1.4490e-01 | 2.8200e+05 | ; | N - CA |
| 2065 | 2066 | 1 | 1.0900e-01 | 2.8451e+05 | ; | CA - HA |
| 2065 | 2067 | 1 | 1.5260e-01 | 2.5941e+05 | ; | CA - CB |
| 2065 | 2078 | 1 | 1.5220e-01 | 2.6527e+05 | ; | CA - C |
| 2067 | 2068 | 1 | 1.0900e-01 | 2.8451e+05 | ; | CB - HB2 |
| 2067 | 2069 | 1 | 1.0900e-01 | 2.8451e+05 | ; | CB - HB3 |
| 2067 | 2070 | 1 | 1.5040e-01 | 2.6527e+05 | ; | CB - CG |
| 2070 | 2071 | 1 | 1.3940e-01 | 3.4309e+05 | ; | CG - ND1 |
| 2070 | 2076 | 1 | 1.3710e-01 | 4.3346e+05 | ; | CG - CD2 |
| 2071 | 2072 | 1 | 1.3350e-01 | 4.0836e+05 | ; | ND1 - CE1 |
| 2072 | 2073 | 1 | 1.0800e-01 | 3.0711e+05 | ; | CE1 - HE1 |
| 2072 | 2074 | 1 | 1.3430e-01 | 3.9915e+05 | ; | CE1 - NE2 |
| 2074 | 2075 | 1 | 1.0100e-01 | 3.6317e+05 | ; | NE2 - HE2 |
| 2074 | 2076 | 1 | 1.3810e-01 | 3.5731e+05 | ; | NE2 - CD2 |
| 2076 | 2077 | 1 | 1.0800e-01 | 3.0711e+05 | ; | CD2 - HD2 |
| 2078 | 2079 | 1 | 1.2290e-01 | 4.7698e+05 | ; | C - O |
| 2078 | 2080 | 1 | 1.3350e-01 | 4.1003e+05 | ; | C - N |
| 2080 | 2081 | 1 | 1.0100e-01 | 3.6317e+05 | ; | N - H |
| 2080 | 2082 | 1 | 1.4490e-01 | 2.8200e+05 | ; | N - CA |
| 2082 | 2083 | 1 | 1.0900e-01 | 2.8451e+05 | ; | CA - HA |
| 2082 | 2084 | 1 | 1.5260e-01 | 2.5941e+05 | ; | CA - CB |
| 2082 | 2089 | 1 | 1.5220e-01 | 2.6527e+05 | ; | CA - C |
| 2084 | 2085 | 1 | 1.0900e-01 | 2.8451e+05 | ; | CB - HB2 |
| 2084 | 2086 | 1 | 1.0900e-01 | 2.8451e+05 | ; | CB - HB3 |
| 2084 | 2087 | 1 | 1.4100e-01 | 2.6778e+05 | ; | CB - OG |
| 2087 | 2088 | 1 | 9.6000e-02 | 4.6275e+05 | ; | OG - HG |
| 2089 | 2090 | 1 | 1.2290e-01 | 4.7698e+05 | ; | C - O |
| 2089 | 2091 | 1 | 1.3350e-01 | 4.1003e+05 | ; | C - N |
| 2091 | 2092 | 1 | 1.0100e-01 | 3.6317e+05 | ; | N - H |
| 2091 | 2093 | 1 | 1.4490e-01 | 2.8200e+05 | ; | N - CA |
| 2093 | 2094 | 1 | 1.0900e-01 | 2.8451e+05 | ; | CA - HA |
| 2093 | 2095 | 1 | 1.5260e-01 | 2.5941e+05 | ; | CA - CB |

| | | | | | | |
|---|---|---|---|---|---|---|
| 2093 | 2108 | 1 | 1.5220e-01 | 2.6527e+05 | ; | CA - C |
| 2095 | 2096 | 1 | 1.0900e-01 | 2.8451e+05 | ; | CB - HB2 |
| 2095 | 2097 | 1 | 1.0900e-01 | 2.8451e+05 | ; | CB - HB3 |
| 2095 | 2098 | 1 | 1.5260e-01 | 2.5941e+05 | ; | CB - CG |
| 2098 | 2099 | 1 | 1.0900e-01 | 2.8451e+05 | ; | CG - HG |
| 2098 | 2100 | 1 | 1.5260e-01 | 2.5941e+05 | ; | CG - CD1 |
| 2098 | 2104 | 1 | 1.5260e-01 | 2.5941e+05 | ; | CG - CD2 |
| 2100 | 2101 | 1 | 1.0900e-01 | 2.8451e+05 | ; | CD1 - HD11 |
| 2100 | 2102 | 1 | 1.0900e-01 | 2.8451e+05 | ; | CD1 - HD12 |
| 2100 | 2103 | 1 | 1.0900e-01 | 2.8451e+05 | ; | CD1 - HD13 |
| 2104 | 2105 | 1 | 1.0900e-01 | 2.8451e+05 | ; | CD2 - HD21 |
| 2104 | 2106 | 1 | 1.0900e-01 | 2.8451e+05 | ; | CD2 - HD22 |
| 2104 | 2107 | 1 | 1.0900e-01 | 2.8451e+05 | ; | CD2 - HD23 |
| 2108 | 2109 | 1 | 1.2290e-01 | 4.7698e+05 | ; | C - O |
| 2108 | 2110 | 1 | 1.3350e-01 | 4.1003e+05 | ; | C - N |
| 2110 | 2111 | 1 | 1.0100e-01 | 3.6317e+05 | ; | N - H |
| 2110 | 2112 | 1 | 1.4490e-01 | 2.8200e+05 | ; | N - CA |
| 2112 | 2113 | 1 | 1.0900e-01 | 2.8451e+05 | ; | CA - HA |
| 2112 | 2114 | 1 | 1.5260e-01 | 2.5941e+05 | ; | CA - CB |
| 2112 | 2122 | 1 | 1.5220e-01 | 2.6527e+05 | ; | CA - C |
| 2114 | 2115 | 1 | 1.0900e-01 | 2.8451e+05 | ; | CB - HB2 |
| 2114 | 2116 | 1 | 1.0900e-01 | 2.8451e+05 | ; | CB - HB3 |
| 2114 | 2117 | 1 | 1.5220e-01 | 2.6527e+05 | ; | CB - CG |
| 2117 | 2118 | 1 | 1.2290e-01 | 4.7698e+05 | ; | CG - OD1 |
| 2117 | 2119 | 1 | 1.3350e-01 | 4.1003e+05 | ; | CG - ND2 |
| 2119 | 2120 | 1 | 1.0100e-01 | 3.6317e+05 | ; | ND2 - HD21 |
| 2119 | 2121 | 1 | 1.0100e-01 | 3.6317e+05 | ; | ND2 - HD22 |
| 2122 | 2123 | 1 | 1.2290e-01 | 4.7698e+05 | ; | C - O |
| 2122 | 2124 | 1 | 1.3350e-01 | 4.1003e+05 | ; | C - N |
| 2124 | 2125 | 1 | 1.0100e-01 | 3.6317e+05 | ; | N - H |
| 2124 | 2126 | 1 | 1.4490e-01 | 2.8200e+05 | ; | N - CA |
| 2126 | 2127 | 1 | 1.0900e-01 | 2.8451e+05 | ; | CA - HA |
| 2126 | 2128 | 1 | 1.5260e-01 | 2.5941e+05 | ; | CA - CB |
| 2126 | 2141 | 1 | 1.5220e-01 | 2.6527e+05 | ; | CA - C |
| 2128 | 2129 | 1 | 1.0900e-01 | 2.8451e+05 | ; | CB - HB |
| 2128 | 2130 | 1 | 1.5260e-01 | 2.5941e+05 | ; | CB - CG2 |
| 2128 | 2134 | 1 | 1.5260e-01 | 2.5941e+05 | ; | CB - CG1 |
| 2130 | 2131 | 1 | 1.0900e-01 | 2.8451e+05 | ; | CG2 - HG21 |
| 2130 | 2132 | 1 | 1.0900e-01 | 2.8451e+05 | ; | CG2 - HG22 |
| 2130 | 2133 | 1 | 1.0900e-01 | 2.8451e+05 | ; | CG2 - HG23 |
| 2134 | 2135 | 1 | 1.0900e-01 | 2.8451e+05 | ; | CG1 - HG12 |
| 2134 | 2136 | 1 | 1.0900e-01 | 2.8451e+05 | ; | CG1 - HG13 |
| 2134 | 2137 | 1 | 1.5260e-01 | 2.5941e+05 | ; | CG1 - CD1 |
| 2137 | 2138 | 1 | 1.0900e-01 | 2.8451e+05 | ; | CD1 - HD11 |
| 2137 | 2139 | 1 | 1.0900e-01 | 2.8451e+05 | ; | CD1 - HD12 |
| 2137 | 2140 | 1 | 1.0900e-01 | 2.8451e+05 | ; | CD1 - HD13 |
| 2141 | 2142 | 1 | 1.2290e-01 | 4.7698e+05 | ; | C - O |
| 2141 | 2143 | 1 | 1.3350e-01 | 4.1003e+05 | ; | C - N |
| 2143 | 2144 | 1 | 1.0100e-01 | 3.6317e+05 | ; | N - H |
| 2143 | 2145 | 1 | 1.4490e-01 | 2.8200e+05 | ; | N - CA |
| 2145 | 2146 | 1 | 1.0900e-01 | 2.8451e+05 | ; | CA - HA |
| 2145 | 2147 | 1 | 1.5260e-01 | 2.5941e+05 | ; | CA - CB |

| | | | | | |
|---|---|---|---|---|---|
| 2145 | 2157 | 1 | 1.5220e-01 | 2.6527e+05 ; | CA - C |
| 2147 | 2148 | 1 | 1.0900e-01 | 2.8451e+05 ; | CB - HB |
| 2147 | 2149 | 1 | 1.5260e-01 | 2.5941e+05 ; | CB - CG1 |
| 2147 | 2153 | 1 | 1.5260e-01 | 2.5941e+05 ; | CB - CG2 |
| 2149 | 2150 | 1 | 1.0900e-01 | 2.8451e+05 ; | CG1 - HG11 |
| 2149 | 2151 | 1 | 1.0900e-01 | 2.8451e+05 ; | CG1 - HG12 |
| 2149 | 2152 | 1 | 1.0900e-01 | 2.8451e+05 ; | CG1 - HG13 |
| 2153 | 2154 | 1 | 1.0900e-01 | 2.8451e+05 ; | CG2 - HG21 |
| 2153 | 2155 | 1 | 1.0900e-01 | 2.8451e+05 ; | CG2 - HG22 |
| 2153 | 2156 | 1 | 1.0900e-01 | 2.8451e+05 ; | CG2 - HG23 |
| 2157 | 2158 | 1 | 1.2290e-01 | 4.7698e+05 ; | C - O |
| 2157 | 2159 | 1 | 1.3350e-01 | 4.1003e+05 ; | C - N |
| 2159 | 2160 | 1 | 1.0100e-01 | 3.6317e+05 ; | N - H |
| 2159 | 2161 | 1 | 1.4490e-01 | 2.8200e+05 ; | N - CA |
| 2161 | 2162 | 1 | 1.0900e-01 | 2.8451e+05 ; | CA - HA |
| 2161 | 2163 | 1 | 1.5260e-01 | 2.5941e+05 ; | CA - CB |
| 2161 | 2174 | 1 | 1.5220e-01 | 2.6527e+05 ; | CA - C |
| 2163 | 2164 | 1 | 1.0900e-01 | 2.8451e+05 ; | CB - HB2 |
| 2163 | 2165 | 1 | 1.0900e-01 | 2.8451e+05 ; | CB - HB3 |
| 2163 | 2166 | 1 | 1.5040e-01 | 2.6527e+05 ; | CB - CG |
| 2166 | 2167 | 1 | 1.3940e-01 | 3.4309e+05 ; | CG - ND1 |
| 2166 | 2172 | 1 | 1.3710e-01 | 4.3346e+05 ; | CG - CD2 |
| 2167 | 2168 | 1 | 1.3350e-01 | 4.0836e+05 ; | ND1 - CE1 |
| 2168 | 2169 | 1 | 1.0800e-01 | 3.0711e+05 ; | CE1 - HE1 |
| 2168 | 2170 | 1 | 1.3430e-01 | 3.9915e+05 ; | CE1 - NE2 |
| 2170 | 2171 | 1 | 1.0100e-01 | 3.6317e+05 ; | NE2 - HE2 |
| 2170 | 2172 | 1 | 1.3810e-01 | 3.5731e+05 ; | NE2 - CD2 |
| 2172 | 2173 | 1 | 1.0800e-01 | 3.0711e+05 ; | CD2 - HD2 |
| 2174 | 2175 | 1 | 1.2290e-01 | 4.7698e+05 ; | C - O |
| 2174 | 2176 | 1 | 1.3350e-01 | 4.1003e+05 ; | C - N |
| 2176 | 2177 | 1 | 1.0100e-01 | 3.6317e+05 ; | N - H |
| 2176 | 2178 | 1 | 1.4490e-01 | 2.8200e+05 ; | N - CA |
| 2178 | 2179 | 1 | 1.0900e-01 | 2.8451e+05 ; | CA - HA |
| 2178 | 2180 | 1 | 1.5260e-01 | 2.5941e+05 ; | CA - CB |
| 2178 | 2198 | 1 | 1.5220e-01 | 2.6527e+05 ; | CA - C |
| 2180 | 2181 | 1 | 1.0900e-01 | 2.8451e+05 ; | CB - HB2 |
| 2180 | 2182 | 1 | 1.0900e-01 | 2.8451e+05 ; | CB - HB3 |
| 2180 | 2183 | 1 | 1.5260e-01 | 2.5941e+05 ; | CB - CG |
| 2183 | 2184 | 1 | 1.0900e-01 | 2.8451e+05 ; | CG - HG2 |
| 2183 | 2185 | 1 | 1.0900e-01 | 2.8451e+05 ; | CG - HG3 |
| 2183 | 2186 | 1 | 1.5260e-01 | 2.5941e+05 ; | CG - CD |
| 2186 | 2187 | 1 | 1.0900e-01 | 2.8451e+05 ; | CD - HD2 |
| 2186 | 2188 | 1 | 1.0900e-01 | 2.8451e+05 ; | CD - HD3 |
| 2186 | 2189 | 1 | 1.4630e-01 | 2.8200e+05 ; | CD - NE |
| 2189 | 2190 | 1 | 1.0100e-01 | 3.6317e+05 ; | NE - HE |
| 2189 | 2191 | 1 | 1.3400e-01 | 4.0250e+05 ; | NE - CZ |
| 2191 | 2192 | 1 | 1.3400e-01 | 4.0250e+05 ; | CZ - NH1 |
| 2191 | 2195 | 1 | 1.3400e-01 | 4.0250e+05 ; | CZ - NH2 |
| 2192 | 2193 | 1 | 1.0100e-01 | 3.6317e+05 ; | NH1 - HH11 |
| 2192 | 2194 | 1 | 1.0100e-01 | 3.6317e+05 ; | NH1 - HH12 |
| 2195 | 2196 | 1 | 1.0100e-01 | 3.6317e+05 ; | NH2 - HH21 |
| 2195 | 2197 | 1 | 1.0100e-01 | 3.6317e+05 ; | NH2 - HH22 |

| | | | | | | |
|---|---|---|---|---|---|---|
| 2198 | 2199 | 1 | 1.2290e-01 | 4.7698e+05 | ; | C - O |
| 2198 | 2200 | 1 | 1.3350e-01 | 4.1003e+05 | ; | C - N |
| 2200 | 2201 | 1 | 1.0100e-01 | 3.6317e+05 | ; | N - H |
| 2200 | 2202 | 1 | 1.4490e-01 | 2.8200e+05 | ; | N - CA |
| 2202 | 2203 | 1 | 1.0900e-01 | 2.8451e+05 | ; | CA - HA |
| 2202 | 2204 | 1 | 1.5260e-01 | 2.5941e+05 | ; | CA - CB |
| 2202 | 2210 | 1 | 1.5220e-01 | 2.6527e+05 | ; | CA - C |
| 2204 | 2205 | 1 | 1.0900e-01 | 2.8451e+05 | ; | CB - HB2 |
| 2204 | 2206 | 1 | 1.0900e-01 | 2.8451e+05 | ; | CB - HB3 |
| 2204 | 2207 | 1 | 1.5220e-01 | 2.6527e+05 | ; | CB - CG |
| 2207 | 2208 | 1 | 1.2500e-01 | 5.4894e+05 | ; | CG - OD1 |
| 2207 | 2209 | 1 | 1.2500e-01 | 5.4894e+05 | ; | CG - OD2 |
| 2210 | 2211 | 1 | 1.2290e-01 | 4.7698e+05 | ; | C - O |
| 2210 | 2212 | 1 | 1.3350e-01 | 4.1003e+05 | ; | C - N |
| 2212 | 2213 | 1 | 1.0100e-01 | 3.6317e+05 | ; | N - H |
| 2212 | 2214 | 1 | 1.4490e-01 | 2.8200e+05 | ; | N - CA |
| 2214 | 2215 | 1 | 1.0900e-01 | 2.8451e+05 | ; | CA - HA |
| 2214 | 2216 | 1 | 1.5260e-01 | 2.5941e+05 | ; | CA - CB |
| 2214 | 2229 | 1 | 1.5220e-01 | 2.6527e+05 | ; | CA - C |
| 2216 | 2217 | 1 | 1.0900e-01 | 2.8451e+05 | ; | CB - HB2 |
| 2216 | 2218 | 1 | 1.0900e-01 | 2.8451e+05 | ; | CB - HB3 |
| 2216 | 2219 | 1 | 1.5260e-01 | 2.5941e+05 | ; | CB - CG |
| 2219 | 2220 | 1 | 1.0900e-01 | 2.8451e+05 | ; | CG - HG |
| 2219 | 2221 | 1 | 1.5260e-01 | 2.5941e+05 | ; | CG - CD1 |
| 2219 | 2225 | 1 | 1.5260e-01 | 2.5941e+05 | ; | CG - CD2 |
| 2221 | 2222 | 1 | 1.0900e-01 | 2.8451e+05 | ; | CD1 - HD11 |
| 2221 | 2223 | 1 | 1.0900e-01 | 2.8451e+05 | ; | CD1 - HD12 |
| 2221 | 2224 | 1 | 1.0900e-01 | 2.8451e+05 | ; | CD1 - HD13 |
| 2225 | 2226 | 1 | 1.0900e-01 | 2.8451e+05 | ; | CD2 - HD21 |
| 2225 | 2227 | 1 | 1.0900e-01 | 2.8451e+05 | ; | CD2 - HD22 |
| 2225 | 2228 | 1 | 1.0900e-01 | 2.8451e+05 | ; | CD2 - HD23 |
| 2229 | 2230 | 1 | 1.2290e-01 | 4.7698e+05 | ; | C - O |
| 2229 | 2231 | 1 | 1.3350e-01 | 4.1003e+05 | ; | C - N |
| 2231 | 2232 | 1 | 1.0100e-01 | 3.6317e+05 | ; | N - H |
| 2231 | 2233 | 1 | 1.4490e-01 | 2.8200e+05 | ; | N - CA |
| 2233 | 2234 | 1 | 1.0900e-01 | 2.8451e+05 | ; | CA - HA |
| 2233 | 2235 | 1 | 1.5260e-01 | 2.5941e+05 | ; | CA - CB |
| 2233 | 2251 | 1 | 1.5220e-01 | 2.6527e+05 | ; | CA - C |
| 2235 | 2236 | 1 | 1.0900e-01 | 2.8451e+05 | ; | CB - HB2 |
| 2235 | 2237 | 1 | 1.0900e-01 | 2.8451e+05 | ; | CB - HB3 |
| 2235 | 2238 | 1 | 1.5260e-01 | 2.5941e+05 | ; | CB - CG |
| 2238 | 2239 | 1 | 1.0900e-01 | 2.8451e+05 | ; | CG - HG2 |
| 2238 | 2240 | 1 | 1.0900e-01 | 2.8451e+05 | ; | CG - HG3 |
| 2238 | 2241 | 1 | 1.5260e-01 | 2.5941e+05 | ; | CG - CD |
| 2241 | 2242 | 1 | 1.0900e-01 | 2.8451e+05 | ; | CD - HD2 |
| 2241 | 2243 | 1 | 1.0900e-01 | 2.8451e+05 | ; | CD - HD3 |
| 2241 | 2244 | 1 | 1.5260e-01 | 2.5941e+05 | ; | CD - CE |
| 2244 | 2245 | 1 | 1.0900e-01 | 2.8451e+05 | ; | CE - HE2 |
| 2244 | 2246 | 1 | 1.0900e-01 | 2.8451e+05 | ; | CE - HE3 |
| 2244 | 2247 | 1 | 1.4710e-01 | 3.0711e+05 | ; | CE - NZ |
| 2247 | 2248 | 1 | 1.0100e-01 | 3.6317e+05 | ; | NZ - HZ1 |
| 2247 | 2249 | 1 | 1.0100e-01 | 3.6317e+05 | ; | NZ - HZ2 |

| | | | | | |
|---|---|---|---|---|---|
| 2247 | 2250 | 1 | 1.0100e-01 | 3.6317e+05 ; | NZ - HZ3 |
| 2251 | 2252 | 1 | 1.2290e-01 | 4.7698e+05 ; | C - O |
| 2251 | 2253 | 1 | 1.3350e-01 | 4.1003e+05 ; | C - N |
| 2253 | 2254 | 1 | 1.4490e-01 | 2.8200e+05 ; | N - CD |
| 2253 | 2263 | 1 | 1.4490e-01 | 2.8200e+05 ; | N - CA |
| 2254 | 2255 | 1 | 1.0900e-01 | 2.8451e+05 ; | CD - HD2 |
| 2254 | 2256 | 1 | 1.0900e-01 | 2.8451e+05 ; | CD - HD3 |
| 2254 | 2257 | 1 | 1.5260e-01 | 2.5941e+05 ; | CD - CG |
| 2257 | 2258 | 1 | 1.0900e-01 | 2.8451e+05 ; | CG - HG2 |
| 2257 | 2259 | 1 | 1.0900e-01 | 2.8451e+05 ; | CG - HG3 |
| 2257 | 2260 | 1 | 1.5260e-01 | 2.5941e+05 ; | CG - CB |
| 2260 | 2261 | 1 | 1.0900e-01 | 2.8451e+05 ; | CB - HB2 |
| 2260 | 2262 | 1 | 1.0900e-01 | 2.8451e+05 ; | CB - HB3 |
| 2260 | 2263 | 1 | 1.5260e-01 | 2.5941e+05 ; | CB - CA |
| 2263 | 2264 | 1 | 1.0900e-01 | 2.8451e+05 ; | CA - HA |
| 2263 | 2265 | 1 | 1.5220e-01 | 2.6527e+05 ; | CA - C |
| 2265 | 2266 | 1 | 1.2290e-01 | 4.7698e+05 ; | C - O |
| 2265 | 2267 | 1 | 1.3350e-01 | 4.1003e+05 ; | C - N |
| 2267 | 2268 | 1 | 1.0100e-01 | 3.6317e+05 ; | N - H |
| 2267 | 2269 | 1 | 1.4490e-01 | 2.8200e+05 ; | N - CA |
| 2269 | 2270 | 1 | 1.0900e-01 | 2.8451e+05 ; | CA - HA |
| 2269 | 2271 | 1 | 1.5260e-01 | 2.5941e+05 ; | CA - CB |
| 2269 | 2280 | 1 | 1.5220e-01 | 2.6527e+05 ; | CA - C |
| 2271 | 2272 | 1 | 1.0900e-01 | 2.8451e+05 ; | CB - HB2 |
| 2271 | 2273 | 1 | 1.0900e-01 | 2.8451e+05 ; | CB - HB3 |
| 2271 | 2274 | 1 | 1.5260e-01 | 2.5941e+05 ; | CB - CG |
| 2274 | 2275 | 1 | 1.0900e-01 | 2.8451e+05 ; | CG - HG2 |
| 2274 | 2276 | 1 | 1.0900e-01 | 2.8451e+05 ; | CG - HG3 |
| 2274 | 2277 | 1 | 1.5220e-01 | 2.6527e+05 ; | CG - CD |
| 2277 | 2278 | 1 | 1.2500e-01 | 5.4894e+05 ; | CD - OE1 |
| 2277 | 2279 | 1 | 1.2500e-01 | 5.4894e+05 ; | CD - OE2 |
| 2280 | 2281 | 1 | 1.2290e-01 | 4.7698e+05 ; | C - O |
| 2280 | 2282 | 1 | 1.3350e-01 | 4.1003e+05 ; | C - N |
| 2282 | 2283 | 1 | 1.0100e-01 | 3.6317e+05 ; | N - H |
| 2282 | 2284 | 1 | 1.4490e-01 | 2.8200e+05 ; | N - CA |
| 2284 | 2285 | 1 | 1.0900e-01 | 2.8451e+05 ; | CA - HA |
| 2284 | 2286 | 1 | 1.5260e-01 | 2.5941e+05 ; | CA - CB |
| 2284 | 2294 | 1 | 1.5220e-01 | 2.6527e+05 ; | CA - C |
| 2286 | 2287 | 1 | 1.0900e-01 | 2.8451e+05 ; | CB - HB2 |
| 2286 | 2288 | 1 | 1.0900e-01 | 2.8451e+05 ; | CB - HB3 |
| 2286 | 2289 | 1 | 1.5220e-01 | 2.6527e+05 ; | CB - CG |
| 2289 | 2290 | 1 | 1.2290e-01 | 4.7698e+05 ; | CG - OD1 |
| 2289 | 2291 | 1 | 1.3350e-01 | 4.1003e+05 ; | CG - ND2 |
| 2291 | 2292 | 1 | 1.0100e-01 | 3.6317e+05 ; | ND2 - HD21 |
| 2291 | 2293 | 1 | 1.0100e-01 | 3.6317e+05 ; | ND2 - HD22 |
| 2294 | 2295 | 1 | 1.2290e-01 | 4.7698e+05 ; | C - O |
| 2294 | 2296 | 1 | 1.3350e-01 | 4.1003e+05 ; | C - N |
| 2296 | 2297 | 1 | 1.0100e-01 | 3.6317e+05 ; | N - H |
| 2296 | 2298 | 1 | 1.4490e-01 | 2.8200e+05 ; | N - CA |
| 2298 | 2299 | 1 | 1.0900e-01 | 2.8451e+05 ; | CA - HA |
| 2298 | 2300 | 1 | 1.5260e-01 | 2.5941e+05 ; | CA - CB |
| 2298 | 2314 | 1 | 1.5220e-01 | 2.6527e+05 ; | CA - C |

| | | | | | |
|---|---|---|---|---|---|
| 2300 | 2301 | 1 | 1.0900e-01 | 2.8451e+05 ; | CB - HB2 |
| 2300 | 2302 | 1 | 1.0900e-01 | 2.8451e+05 ; | CB - HB3 |
| 2300 | 2303 | 1 | 1.5100e-01 | 2.6527e+05 ; | CB - CG |
| 2303 | 2304 | 1 | 1.4000e-01 | 3.9246e+05 ; | CG - CD1 |
| 2303 | 2312 | 1 | 1.4000e-01 | 3.9246e+05 ; | CG - CD2 |
| 2304 | 2305 | 1 | 1.0800e-01 | 3.0711e+05 ; | CD1 - HD1 |
| 2304 | 2306 | 1 | 1.4000e-01 | 3.9246e+05 ; | CD1 - CE1 |
| 2306 | 2307 | 1 | 1.0800e-01 | 3.0711e+05 ; | CE1 - HE1 |
| 2306 | 2308 | 1 | 1.4000e-01 | 3.9246e+05 ; | CE1 - CZ |
| 2308 | 2309 | 1 | 1.0800e-01 | 3.0711e+05 ; | CZ - HZ |
| 2308 | 2310 | 1 | 1.4000e-01 | 3.9246e+05 ; | CZ - CE2 |
| 2310 | 2311 | 1 | 1.0800e-01 | 3.0711e+05 ; | CE2 - HE2 |
| 2310 | 2312 | 1 | 1.4000e-01 | 3.9246e+05 ; | CE2 - CD2 |
| 2312 | 2313 | 1 | 1.0800e-01 | 3.0711e+05 ; | CD2 - HD2 |
| 2314 | 2315 | 1 | 1.2290e-01 | 4.7698e+05 ; | C - O |
| 2314 | 2316 | 1 | 1.3350e-01 | 4.1003e+05 ; | C - N |
| 2316 | 2317 | 1 | 1.0100e-01 | 3.6317e+05 ; | N - H |
| 2316 | 2318 | 1 | 1.4490e-01 | 2.8200e+05 ; | N - CA |
| 2318 | 2319 | 1 | 1.0900e-01 | 2.8451e+05 ; | CA - HA |
| 2318 | 2320 | 1 | 1.5260e-01 | 2.5941e+05 ; | CA - CB |
| 2318 | 2333 | 1 | 1.5220e-01 | 2.6527e+05 ; | CA - C |
| 2320 | 2321 | 1 | 1.0900e-01 | 2.8451e+05 ; | CB - HB2 |
| 2320 | 2322 | 1 | 1.0900e-01 | 2.8451e+05 ; | CB - HB3 |
| 2320 | 2323 | 1 | 1.5260e-01 | 2.5941e+05 ; | CB - CG |
| 2323 | 2324 | 1 | 1.0900e-01 | 2.8451e+05 ; | CG - HG |
| 2323 | 2325 | 1 | 1.5260e-01 | 2.5941e+05 ; | CG - CD1 |
| 2323 | 2329 | 1 | 1.5260e-01 | 2.5941e+05 ; | CG - CD2 |
| 2325 | 2326 | 1 | 1.0900e-01 | 2.8451e+05 ; | CD1 - HD11 |
| 2325 | 2327 | 1 | 1.0900e-01 | 2.8451e+05 ; | CD1 - HD12 |
| 2325 | 2328 | 1 | 1.0900e-01 | 2.8451e+05 ; | CD1 - HD13 |
| 2329 | 2330 | 1 | 1.0900e-01 | 2.8451e+05 ; | CD2 - HD21 |
| 2329 | 2331 | 1 | 1.0900e-01 | 2.8451e+05 ; | CD2 - HD22 |
| 2329 | 2332 | 1 | 1.0900e-01 | 2.8451e+05 ; | CD2 - HD23 |
| 2333 | 2334 | 1 | 1.2290e-01 | 4.7698e+05 ; | C - O |
| 2333 | 2335 | 1 | 1.3350e-01 | 4.1003e+05 ; | C - N |
| 2335 | 2336 | 1 | 1.0100e-01 | 3.6317e+05 ; | N - H |
| 2335 | 2337 | 1 | 1.4490e-01 | 2.8200e+05 ; | N - CA |
| 2337 | 2338 | 1 | 1.0900e-01 | 2.8451e+05 ; | CA - HA |
| 2337 | 2339 | 1 | 1.5260e-01 | 2.5941e+05 ; | CA - CB |
| 2337 | 2353 | 1 | 1.5220e-01 | 2.6527e+05 ; | CA - C |
| 2339 | 2340 | 1 | 1.0900e-01 | 2.8451e+05 ; | CB - HB2 |
| 2339 | 2341 | 1 | 1.0900e-01 | 2.8451e+05 ; | CB - HB3 |
| 2339 | 2342 | 1 | 1.5100e-01 | 2.6527e+05 ; | CB - CG |
| 2342 | 2343 | 1 | 1.4000e-01 | 3.9246e+05 ; | CG - CD1 |
| 2342 | 2351 | 1 | 1.4000e-01 | 3.9246e+05 ; | CG - CD2 |
| 2343 | 2344 | 1 | 1.0800e-01 | 3.0711e+05 ; | CD1 - HD1 |
| 2343 | 2345 | 1 | 1.4000e-01 | 3.9246e+05 ; | CD1 - CE1 |
| 2345 | 2346 | 1 | 1.0800e-01 | 3.0711e+05 ; | CE1 - HE1 |
| 2345 | 2347 | 1 | 1.4000e-01 | 3.9246e+05 ; | CE1 - CZ |
| 2347 | 2348 | 1 | 1.0800e-01 | 3.0711e+05 ; | CZ - HZ |
| 2347 | 2349 | 1 | 1.4000e-01 | 3.9246e+05 ; | CZ - CE2 |
| 2349 | 2350 | 1 | 1.0800e-01 | 3.0711e+05 ; | CE2 - HE2 |

| | | | | | |
|---|---|---|---|---|---|
| 2349 | 2351 | 1 | 1.4000e-01 | 3.9246e+05 ; | CE2 - CD2 |
| 2351 | 2352 | 1 | 1.0800e-01 | 3.0711e+05 ; | CD2 - HD2 |
| 2353 | 2354 | 1 | 1.2290e-01 | 4.7698e+05 ; | C - O |
| 2353 | 2355 | 1 | 1.3350e-01 | 4.1003e+05 ; | C - N |
| 2355 | 2356 | 1 | 1.0100e-01 | 3.6317e+05 ; | N - H |
| 2355 | 2357 | 1 | 1.4490e-01 | 2.8200e+05 ; | N - CA |
| 2357 | 2358 | 1 | 1.0900e-01 | 2.8451e+05 ; | CA - HA |
| 2357 | 2359 | 1 | 1.5260e-01 | 2.5941e+05 ; | CA - CB |
| 2357 | 2370 | 1 | 1.5220e-01 | 2.6527e+05 ; | CA - C |
| 2359 | 2360 | 1 | 1.0900e-01 | 2.8451e+05 ; | CB - HB2 |
| 2359 | 2361 | 1 | 1.0900e-01 | 2.8451e+05 ; | CB - HB3 |
| 2359 | 2362 | 1 | 1.5260e-01 | 2.5941e+05 ; | CB - CG |
| 2362 | 2363 | 1 | 1.0900e-01 | 2.8451e+05 ; | CG - HG2 |
| 2362 | 2364 | 1 | 1.0900e-01 | 2.8451e+05 ; | CG - HG3 |
| 2362 | 2365 | 1 | 1.5220e-01 | 2.6527e+05 ; | CG - CD |
| 2365 | 2366 | 1 | 1.2290e-01 | 4.7698e+05 ; | CD - OE1 |
| 2365 | 2367 | 1 | 1.3350e-01 | 4.1003e+05 ; | CD - NE2 |
| 2367 | 2368 | 1 | 1.0100e-01 | 3.6317e+05 ; | NE2 - HE21 |
| 2367 | 2369 | 1 | 1.0100e-01 | 3.6317e+05 ; | NE2 - HE22 |
| 2370 | 2371 | 1 | 1.2290e-01 | 4.7698e+05 ; | C - O |
| 2370 | 2372 | 1 | 1.3350e-01 | 4.1003e+05 ; | C - N |
| 2372 | 2373 | 1 | 1.0100e-01 | 3.6317e+05 ; | N - H |
| 2372 | 2374 | 1 | 1.4490e-01 | 2.8200e+05 ; | N - CA |
| 2374 | 2375 | 1 | 1.0900e-01 | 2.8451e+05 ; | CA - HA |
| 2374 | 2376 | 1 | 1.5260e-01 | 2.5941e+05 ; | CA - CB |
| 2374 | 2381 | 1 | 1.5220e-01 | 2.6527e+05 ; | CA - C |
| 2376 | 2377 | 1 | 1.0900e-01 | 2.8451e+05 ; | CB - HB2 |
| 2376 | 2378 | 1 | 1.0900e-01 | 2.8451e+05 ; | CB - HB3 |
| 2376 | 2379 | 1 | 1.4100e-01 | 2.6778e+05 ; | CB - OG |
| 2379 | 2380 | 1 | 9.6000e-02 | 4.6275e+05 ; | OG - HG |
| 2381 | 2382 | 1 | 1.2290e-01 | 4.7698e+05 ; | C - O |
| 2381 | 2383 | 1 | 1.3350e-01 | 4.1003e+05 ; | C - N |
| 2383 | 2384 | 1 | 1.0100e-01 | 3.6317e+05 ; | N - H |
| 2383 | 2385 | 1 | 1.4490e-01 | 2.8200e+05 ; | N - CA |
| 2385 | 2386 | 1 | 1.0900e-01 | 2.8451e+05 ; | CA - HA |
| 2385 | 2387 | 1 | 1.5260e-01 | 2.5941e+05 ; | CA - CB |
| 2385 | 2396 | 1 | 1.5220e-01 | 2.6527e+05 ; | CA - C |
| 2387 | 2388 | 1 | 1.0900e-01 | 2.8451e+05 ; | CB - HB2 |
| 2387 | 2389 | 1 | 1.0900e-01 | 2.8451e+05 ; | CB - HB3 |
| 2387 | 2390 | 1 | 1.5260e-01 | 2.5941e+05 ; | CB - CG |
| 2390 | 2391 | 1 | 1.0900e-01 | 2.8451e+05 ; | CG - HG2 |
| 2390 | 2392 | 1 | 1.0900e-01 | 2.8451e+05 ; | CG - HG3 |
| 2390 | 2393 | 1 | 1.5220e-01 | 2.6527e+05 ; | CG - CD |
| 2393 | 2394 | 1 | 1.2500e-01 | 5.4894e+05 ; | CD - OE1 |
| 2393 | 2395 | 1 | 1.2500e-01 | 5.4894e+05 ; | CD - OE2 |
| 2396 | 2397 | 1 | 1.2290e-01 | 4.7698e+05 ; | C - O |
| 2396 | 2398 | 1 | 1.3350e-01 | 4.1003e+05 ; | C - N |
| 2398 | 2399 | 1 | 1.0100e-01 | 3.6317e+05 ; | N - H |
| 2398 | 2400 | 1 | 1.4490e-01 | 2.8200e+05 ; | N - CA |
| 2400 | 2401 | 1 | 1.0900e-01 | 2.8451e+05 ; | CA - HA |
| 2400 | 2402 | 1 | 1.5260e-01 | 2.5941e+05 ; | CA - CB |
| 2400 | 2410 | 1 | 1.5220e-01 | 2.6527e+05 ; | CA - C |

| | | | | | | |
|---|---|---|---|---|---|---|
| 2402 | 2403 | 1 | 1.0900e-01 | 2.8451e+05 | ; | CB - HB2 |
| 2402 | 2404 | 1 | 1.0900e-01 | 2.8451e+05 | ; | CB - HB3 |
| 2402 | 2405 | 1 | 1.5220e-01 | 2.6527e+05 | ; | CB - CG |
| 2405 | 2406 | 1 | 1.2290e-01 | 4.7698e+05 | ; | CG - OD1 |
| 2405 | 2407 | 1 | 1.3350e-01 | 4.1003e+05 | ; | CG - ND2 |
| 2407 | 2408 | 1 | 1.0100e-01 | 3.6317e+05 | ; | ND2 - HD21 |
| 2407 | 2409 | 1 | 1.0100e-01 | 3.6317e+05 | ; | ND2 - HD22 |
| 2410 | 2411 | 1 | 1.2290e-01 | 4.7698e+05 | ; | C - O |
| 2410 | 2412 | 1 | 1.3350e-01 | 4.1003e+05 | ; | C - N |
| 2412 | 2413 | 1 | 1.0100e-01 | 3.6317e+05 | ; | N - H |
| 2412 | 2414 | 1 | 1.4490e-01 | 2.8200e+05 | ; | N - CA |
| 2414 | 2415 | 1 | 1.0900e-01 | 2.8451e+05 | ; | CA - HA |
| 2414 | 2416 | 1 | 1.5260e-01 | 2.5941e+05 | ; | CA - CB |
| 2414 | 2432 | 1 | 1.5220e-01 | 2.6527e+05 | ; | CA - C |
| 2416 | 2417 | 1 | 1.0900e-01 | 2.8451e+05 | ; | CB - HB2 |
| 2416 | 2418 | 1 | 1.0900e-01 | 2.8451e+05 | ; | CB - HB3 |
| 2416 | 2419 | 1 | 1.5260e-01 | 2.5941e+05 | ; | CB - CG |
| 2419 | 2420 | 1 | 1.0900e-01 | 2.8451e+05 | ; | CG - HG2 |
| 2419 | 2421 | 1 | 1.0900e-01 | 2.8451e+05 | ; | CG - HG3 |
| 2419 | 2422 | 1 | 1.5260e-01 | 2.5941e+05 | ; | CG - CD |
| 2422 | 2423 | 1 | 1.0900e-01 | 2.8451e+05 | ; | CD - HD2 |
| 2422 | 2424 | 1 | 1.0900e-01 | 2.8451e+05 | ; | CD - HD3 |
| 2422 | 2425 | 1 | 1.5260e-01 | 2.5941e+05 | ; | CD - CE |
| 2425 | 2426 | 1 | 1.0900e-01 | 2.8451e+05 | ; | CE - HE2 |
| 2425 | 2427 | 1 | 1.0900e-01 | 2.8451e+05 | ; | CE - HE3 |
| 2425 | 2428 | 1 | 1.4710e-01 | 3.0711e+05 | ; | CE - NZ |
| 2428 | 2429 | 1 | 1.0100e-01 | 3.6317e+05 | ; | NZ - HZ1 |
| 2428 | 2430 | 1 | 1.0100e-01 | 3.6317e+05 | ; | NZ - HZ2 |
| 2428 | 2431 | 1 | 1.0100e-01 | 3.6317e+05 | ; | NZ - HZ3 |
| 2432 | 2433 | 1 | 1.2290e-01 | 4.7698e+05 | ; | C - O |
| 2432 | 2434 | 1 | 1.3350e-01 | 4.1003e+05 | ; | C - N |
| 2434 | 2435 | 1 | 1.0100e-01 | 3.6317e+05 | ; | N - H |
| 2434 | 2436 | 1 | 1.4490e-01 | 2.8200e+05 | ; | N - CA |
| 2436 | 2437 | 1 | 1.0900e-01 | 2.8451e+05 | ; | CA - HA |
| 2436 | 2438 | 1 | 1.5260e-01 | 2.5941e+05 | ; | CA - CB |
| 2436 | 2444 | 1 | 1.5220e-01 | 2.6527e+05 | ; | CA - C |
| 2438 | 2439 | 1 | 1.0900e-01 | 2.8451e+05 | ; | CB - HB2 |
| 2438 | 2440 | 1 | 1.0900e-01 | 2.8451e+05 | ; | CB - HB3 |
| 2438 | 2441 | 1 | 1.5220e-01 | 2.6527e+05 | ; | CB - CG |
| 2441 | 2442 | 1 | 1.2500e-01 | 5.4894e+05 | ; | CG - OD1 |
| 2441 | 2443 | 1 | 1.2500e-01 | 5.4894e+05 | ; | CG - OD2 |
| 2444 | 2445 | 1 | 1.2290e-01 | 4.7698e+05 | ; | C - O |
| 2444 | 2446 | 1 | 1.3350e-01 | 4.1003e+05 | ; | C - N |
| 2446 | 2447 | 1 | 1.0100e-01 | 3.6317e+05 | ; | N - H |
| 2446 | 2448 | 1 | 1.4490e-01 | 2.8200e+05 | ; | N - CA |
| 2448 | 2449 | 1 | 1.0900e-01 | 2.8451e+05 | ; | CA - HA |
| 2448 | 2450 | 1 | 1.5260e-01 | 2.5941e+05 | ; | CA - CB |
| 2448 | 2455 | 1 | 1.5220e-01 | 2.6527e+05 | ; | CA - C |
| 2450 | 2451 | 1 | 1.0900e-01 | 2.8451e+05 | ; | CB - HB2 |
| 2450 | 2452 | 1 | 1.0900e-01 | 2.8451e+05 | ; | CB - HB3 |
| 2450 | 2453 | 1 | 1.4100e-01 | 2.6778e+05 | ; | CB - OG |
| 2453 | 2454 | 1 | 9.6000e-02 | 4.6275e+05 | ; | OG - HG |

| | | | | | |
|---|---|---|---|---|---|
| 2455 | 2456 | 1 | 1.2290e-01 | 4.7698e+05 ; | C - O |
| 2455 | 2457 | 1 | 1.3350e-01 | 4.1003e+05 ; | C - N |
| 2457 | 2458 | 1 | 1.0100e-01 | 3.6317e+05 ; | N - H |
| 2457 | 2459 | 1 | 1.4490e-01 | 2.8200e+05 ; | N - CA |
| 2459 | 2460 | 1 | 1.0900e-01 | 2.8451e+05 ; | CA - HA |
| 2459 | 2461 | 1 | 1.5260e-01 | 2.5941e+05 ; | CA - CB |
| 2459 | 2474 | 1 | 1.5220e-01 | 2.6527e+05 ; | CA - C |
| 2461 | 2462 | 1 | 1.0900e-01 | 2.8451e+05 ; | CB - HB2 |
| 2461 | 2463 | 1 | 1.0900e-01 | 2.8451e+05 ; | CB - HB3 |
| 2461 | 2464 | 1 | 1.5260e-01 | 2.5941e+05 ; | CB - CG |
| 2464 | 2465 | 1 | 1.0900e-01 | 2.8451e+05 ; | CG - HG |
| 2464 | 2466 | 1 | 1.5260e-01 | 2.5941e+05 ; | CG - CD1 |
| 2464 | 2470 | 1 | 1.5260e-01 | 2.5941e+05 ; | CG - CD2 |
| 2466 | 2467 | 1 | 1.0900e-01 | 2.8451e+05 ; | CD1 - HD11 |
| 2466 | 2468 | 1 | 1.0900e-01 | 2.8451e+05 ; | CD1 - HD12 |
| 2466 | 2469 | 1 | 1.0900e-01 | 2.8451e+05 ; | CD1 - HD13 |
| 2470 | 2471 | 1 | 1.0900e-01 | 2.8451e+05 ; | CD2 - HD21 |
| 2470 | 2472 | 1 | 1.0900e-01 | 2.8451e+05 ; | CD2 - HD22 |
| 2470 | 2473 | 1 | 1.0900e-01 | 2.8451e+05 ; | CD2 - HD23 |
| 2474 | 2475 | 1 | 1.2290e-01 | 4.7698e+05 ; | C - O |
| 2474 | 2476 | 1 | 1.3350e-01 | 4.1003e+05 ; | C - N |
| 2476 | 2477 | 1 | 1.0100e-01 | 3.6317e+05 ; | N - H |
| 2476 | 2478 | 1 | 1.4490e-01 | 2.8200e+05 ; | N - CA |
| 2478 | 2479 | 1 | 1.0900e-01 | 2.8451e+05 ; | CA - HA |
| 2478 | 2480 | 1 | 1.5260e-01 | 2.5941e+05 ; | CA - CB |
| 2478 | 2493 | 1 | 1.5220e-01 | 2.6527e+05 ; | CA - C |
| 2480 | 2481 | 1 | 1.0900e-01 | 2.8451e+05 ; | CB - HB2 |
| 2480 | 2482 | 1 | 1.0900e-01 | 2.8451e+05 ; | CB - HB3 |
| 2480 | 2483 | 1 | 1.5260e-01 | 2.5941e+05 ; | CB - CG |
| 2483 | 2484 | 1 | 1.0900e-01 | 2.8451e+05 ; | CG - HG |
| 2483 | 2485 | 1 | 1.5260e-01 | 2.5941e+05 ; | CG - CD1 |
| 2483 | 2489 | 1 | 1.5260e-01 | 2.5941e+05 ; | CG - CD2 |
| 2485 | 2486 | 1 | 1.0900e-01 | 2.8451e+05 ; | CD1 - HD11 |
| 2485 | 2487 | 1 | 1.0900e-01 | 2.8451e+05 ; | CD1 - HD12 |
| 2485 | 2488 | 1 | 1.0900e-01 | 2.8451e+05 ; | CD1 - HD13 |
| 2489 | 2490 | 1 | 1.0900e-01 | 2.8451e+05 ; | CD2 - HD21 |
| 2489 | 2491 | 1 | 1.0900e-01 | 2.8451e+05 ; | CD2 - HD22 |
| 2489 | 2492 | 1 | 1.0900e-01 | 2.8451e+05 ; | CD2 - HD23 |
| 2493 | 2494 | 1 | 1.2290e-01 | 4.7698e+05 ; | C - O |
| 2493 | 2495 | 1 | 1.3350e-01 | 4.1003e+05 ; | C - N |
| 2495 | 2496 | 1 | 1.0100e-01 | 3.6317e+05 ; | N - H |
| 2495 | 2497 | 1 | 1.4490e-01 | 2.8200e+05 ; | N - CA |
| 2497 | 2498 | 1 | 1.0900e-01 | 2.8451e+05 ; | CA - HA |
| 2497 | 2499 | 1 | 1.5260e-01 | 2.5941e+05 ; | CA - CB |
| 2497 | 2515 | 1 | 1.5220e-01 | 2.6527e+05 ; | CA - C |
| 2499 | 2500 | 1 | 1.0900e-01 | 2.8451e+05 ; | CB - HB2 |
| 2499 | 2501 | 1 | 1.0900e-01 | 2.8451e+05 ; | CB - HB3 |
| 2499 | 2502 | 1 | 1.5260e-01 | 2.5941e+05 ; | CB - CG |
| 2502 | 2503 | 1 | 1.0900e-01 | 2.8451e+05 ; | CG - HG2 |
| 2502 | 2504 | 1 | 1.0900e-01 | 2.8451e+05 ; | CG - HG3 |
| 2502 | 2505 | 1 | 1.5260e-01 | 2.5941e+05 ; | CG - CD |
| 2505 | 2506 | 1 | 1.0900e-01 | 2.8451e+05 ; | CD - HD2 |

| | | | | | | |
|---|---|---|---|---|---|---|
| 2505 | 2507 | 1 | 1.0900e-01 | 2.8451e+05 | ; | CD - HD3 |
| 2505 | 2508 | 1 | 1.5260e-01 | 2.5941e+05 | ; | CD - CE |
| 2508 | 2509 | 1 | 1.0900e-01 | 2.8451e+05 | ; | CE - HE2 |
| 2508 | 2510 | 1 | 1.0900e-01 | 2.8451e+05 | ; | CE - HE3 |
| 2508 | 2511 | 1 | 1.4710e-01 | 3.0711e+05 | ; | CE - NZ |
| 2511 | 2512 | 1 | 1.0100e-01 | 3.6317e+05 | ; | NZ - HZ1 |
| 2511 | 2513 | 1 | 1.0100e-01 | 3.6317e+05 | ; | NZ - HZ2 |
| 2511 | 2514 | 1 | 1.0100e-01 | 3.6317e+05 | ; | NZ - HZ3 |
| 2515 | 2516 | 1 | 1.2290e-01 | 4.7698e+05 | ; | C - O |
| 2515 | 2517 | 1 | 1.3350e-01 | 4.1003e+05 | ; | C - N |
| 2517 | 2518 | 1 | 1.0100e-01 | 3.6317e+05 | ; | N - H |
| 2517 | 2519 | 1 | 1.4490e-01 | 2.8200e+05 | ; | N - CA |
| 2519 | 2520 | 1 | 1.0900e-01 | 2.8451e+05 | ; | CA - HA |
| 2519 | 2521 | 1 | 1.5260e-01 | 2.5941e+05 | ; | CA - CB |
| 2519 | 2534 | 1 | 1.5220e-01 | 2.6527e+05 | ; | CA - C |
| 2521 | 2522 | 1 | 1.0900e-01 | 2.8451e+05 | ; | CB - HB |
| 2521 | 2523 | 1 | 1.5260e-01 | 2.5941e+05 | ; | CB - CG2 |
| 2521 | 2527 | 1 | 1.5260e-01 | 2.5941e+05 | ; | CB - CG1 |
| 2523 | 2524 | 1 | 1.0900e-01 | 2.8451e+05 | ; | CG2 - HG21 |
| 2523 | 2525 | 1 | 1.0900e-01 | 2.8451e+05 | ; | CG2 - HG22 |
| 2523 | 2526 | 1 | 1.0900e-01 | 2.8451e+05 | ; | CG2 - HG23 |
| 2527 | 2528 | 1 | 1.0900e-01 | 2.8451e+05 | ; | CG1 - HG12 |
| 2527 | 2529 | 1 | 1.0900e-01 | 2.8451e+05 | ; | CG1 - HG13 |
| 2527 | 2530 | 1 | 1.5260e-01 | 2.5941e+05 | ; | CG1 - CD1 |
| 2530 | 2531 | 1 | 1.0900e-01 | 2.8451e+05 | ; | CD1 - HD11 |
| 2530 | 2532 | 1 | 1.0900e-01 | 2.8451e+05 | ; | CD1 - HD12 |
| 2530 | 2533 | 1 | 1.0900e-01 | 2.8451e+05 | ; | CD1 - HD13 |
| 2534 | 2535 | 1 | 1.2290e-01 | 4.7698e+05 | ; | C - O |
| 2534 | 2536 | 1 | 1.3350e-01 | 4.1003e+05 | ; | C - N |
| 2536 | 2537 | 1 | 1.0100e-01 | 3.6317e+05 | ; | N - H |
| 2536 | 2538 | 1 | 1.4490e-01 | 2.8200e+05 | ; | N - CA |
| 2538 | 2539 | 1 | 1.0900e-01 | 2.8451e+05 | ; | CA - HA |
| 2538 | 2540 | 1 | 1.5260e-01 | 2.5941e+05 | ; | CA - CB |
| 2538 | 2553 | 1 | 1.5220e-01 | 2.6527e+05 | ; | CA - C |
| 2540 | 2541 | 1 | 1.0900e-01 | 2.8451e+05 | ; | CB - HB |
| 2540 | 2542 | 1 | 1.5260e-01 | 2.5941e+05 | ; | CB - CG2 |
| 2540 | 2546 | 1 | 1.5260e-01 | 2.5941e+05 | ; | CB - CG1 |
| 2542 | 2543 | 1 | 1.0900e-01 | 2.8451e+05 | ; | CG2 - HG21 |
| 2542 | 2544 | 1 | 1.0900e-01 | 2.8451e+05 | ; | CG2 - HG22 |
| 2542 | 2545 | 1 | 1.0900e-01 | 2.8451e+05 | ; | CG2 - HG23 |
| 2546 | 2547 | 1 | 1.0900e-01 | 2.8451e+05 | ; | CG1 - HG12 |
| 2546 | 2548 | 1 | 1.0900e-01 | 2.8451e+05 | ; | CG1 - HG13 |
| 2546 | 2549 | 1 | 1.5260e-01 | 2.5941e+05 | ; | CG1 - CD1 |
| 2549 | 2550 | 1 | 1.0900e-01 | 2.8451e+05 | ; | CD1 - HD11 |
| 2549 | 2551 | 1 | 1.0900e-01 | 2.8451e+05 | ; | CD1 - HD12 |
| 2549 | 2552 | 1 | 1.0900e-01 | 2.8451e+05 | ; | CD1 - HD13 |
| 2553 | 2554 | 1 | 1.2290e-01 | 4.7698e+05 | ; | C - O |
| 2553 | 2555 | 1 | 1.3350e-01 | 4.1003e+05 | ; | C - N |
| 2555 | 2556 | 1 | 1.0100e-01 | 3.6317e+05 | ; | N - H |
| 2555 | 2557 | 1 | 1.4490e-01 | 2.8200e+05 | ; | N - CA |
| 2557 | 2558 | 1 | 1.0900e-01 | 2.8451e+05 | ; | CA - HA |
| 2557 | 2559 | 1 | 1.5260e-01 | 2.5941e+05 | ; | CA - CB |

| | | | | | | |
|---|---|---|---|---|---|---|
| 2557 | 2565 | 1 | 1.5220e-01 | 2.6527e+05 | ; | CA - C |
| 2559 | 2560 | 1 | 1.0900e-01 | 2.8451e+05 | ; | CB - HB2 |
| 2559 | 2561 | 1 | 1.0900e-01 | 2.8451e+05 | ; | CB - HB3 |
| 2559 | 2562 | 1 | 1.5220e-01 | 2.6527e+05 | ; | CB - CG |
| 2562 | 2563 | 1 | 1.2500e-01 | 5.4894e+05 | ; | CG - OD1 |
| 2562 | 2564 | 1 | 1.2500e-01 | 5.4894e+05 | ; | CG - OD2 |
| 2565 | 2566 | 1 | 1.2290e-01 | 4.7698e+05 | ; | C - O |
| 2565 | 2567 | 1 | 1.3350e-01 | 4.1003e+05 | ; | C - N |
| 2567 | 2568 | 1 | 1.0100e-01 | 3.6317e+05 | ; | N - H |
| 2567 | 2569 | 1 | 1.4490e-01 | 2.8200e+05 | ; | N - CA |
| 2569 | 2570 | 1 | 1.0900e-01 | 2.8451e+05 | ; | CA - HA |
| 2569 | 2571 | 1 | 1.5260e-01 | 2.5941e+05 | ; | CA - CB |
| 2569 | 2585 | 1 | 1.5220e-01 | 2.6527e+05 | ; | CA - C |
| 2571 | 2572 | 1 | 1.0900e-01 | 2.8451e+05 | ; | CB - HB2 |
| 2571 | 2573 | 1 | 1.0900e-01 | 2.8451e+05 | ; | CB - HB3 |
| 2571 | 2574 | 1 | 1.5100e-01 | 2.6527e+05 | ; | CB - CG |
| 2574 | 2575 | 1 | 1.4000e-01 | 3.9246e+05 | ; | CG - CD1 |
| 2574 | 2583 | 1 | 1.4000e-01 | 3.9246e+05 | ; | CG - CD2 |
| 2575 | 2576 | 1 | 1.0800e-01 | 3.0711e+05 | ; | CD1 - HD1 |
| 2575 | 2577 | 1 | 1.4000e-01 | 3.9246e+05 | ; | CD1 - CE1 |
| 2577 | 2578 | 1 | 1.0800e-01 | 3.0711e+05 | ; | CE1 - HE1 |
| 2577 | 2579 | 1 | 1.4000e-01 | 3.9246e+05 | ; | CE1 - CZ |
| 2579 | 2580 | 1 | 1.0800e-01 | 3.0711e+05 | ; | CZ - HZ |
| 2579 | 2581 | 1 | 1.4000e-01 | 3.9246e+05 | ; | CZ - CE2 |
| 2581 | 2582 | 1 | 1.0800e-01 | 3.0711e+05 | ; | CE2 - HE2 |
| 2581 | 2583 | 1 | 1.4000e-01 | 3.9246e+05 | ; | CE2 - CD2 |
| 2583 | 2584 | 1 | 1.0800e-01 | 3.0711e+05 | ; | CD2 - HD2 |
| 2585 | 2586 | 1 | 1.2290e-01 | 4.7698e+05 | ; | C - O |
| 2585 | 2587 | 1 | 1.3350e-01 | 4.1003e+05 | ; | C - N |
| 2587 | 2588 | 1 | 1.0100e-01 | 3.6317e+05 | ; | N - H |
| 2587 | 2589 | 1 | 1.4490e-01 | 2.8200e+05 | ; | N - CA |
| 2589 | 2590 | 1 | 1.0900e-01 | 2.8451e+05 | ; | CA - HA2 |
| 2589 | 2591 | 1 | 1.0900e-01 | 2.8451e+05 | ; | CA - HA3 |
| 2589 | 2592 | 1 | 1.5220e-01 | 2.6527e+05 | ; | CA - C |
| 2592 | 2593 | 1 | 1.2290e-01 | 4.7698e+05 | ; | C - O |
| 2592 | 2594 | 1 | 1.3350e-01 | 4.1003e+05 | ; | C - N |
| 2594 | 2595 | 1 | 1.0100e-01 | 3.6317e+05 | ; | N - H |
| 2594 | 2596 | 1 | 1.4490e-01 | 2.8200e+05 | ; | N - CA |
| 2596 | 2597 | 1 | 1.0900e-01 | 2.8451e+05 | ; | CA - HA |
| 2596 | 2598 | 1 | 1.5260e-01 | 2.5941e+05 | ; | CA - CB |
| 2596 | 2611 | 1 | 1.5220e-01 | 2.6527e+05 | ; | CA - C |
| 2598 | 2599 | 1 | 1.0900e-01 | 2.8451e+05 | ; | CB - HB2 |
| 2598 | 2600 | 1 | 1.0900e-01 | 2.8451e+05 | ; | CB - HB3 |
| 2598 | 2601 | 1 | 1.5260e-01 | 2.5941e+05 | ; | CB - CG |
| 2601 | 2602 | 1 | 1.0900e-01 | 2.8451e+05 | ; | CG - HG |
| 2601 | 2603 | 1 | 1.5260e-01 | 2.5941e+05 | ; | CG - CD1 |
| 2601 | 2607 | 1 | 1.5260e-01 | 2.5941e+05 | ; | CG - CD2 |
| 2603 | 2604 | 1 | 1.0900e-01 | 2.8451e+05 | ; | CD1 - HD11 |
| 2603 | 2605 | 1 | 1.0900e-01 | 2.8451e+05 | ; | CD1 - HD12 |
| 2603 | 2606 | 1 | 1.0900e-01 | 2.8451e+05 | ; | CD1 - HD13 |
| 2607 | 2608 | 1 | 1.0900e-01 | 2.8451e+05 | ; | CD2 - HD21 |
| 2607 | 2609 | 1 | 1.0900e-01 | 2.8451e+05 | ; | CD2 - HD22 |

| | | | | | | |
|---|---|---|---|---|---|---|
| 2607 | 2610 | 1 | 1.0900e-01 | 2.8451e+05 | ; | CD2 - HD23 |
| 2611 | 2612 | 1 | 1.2290e-01 | 4.7698e+05 | ; | C - O |
| 2611 | 2613 | 1 | 1.3350e-01 | 4.1003e+05 | ; | C - N |
| 2613 | 2614 | 1 | 1.0100e-01 | 3.6317e+05 | ; | N - H |
| 2613 | 2615 | 1 | 1.4490e-01 | 2.8200e+05 | ; | N - CA |
| 2615 | 2616 | 1 | 1.0900e-01 | 2.8451e+05 | ; | CA - HA |
| 2615 | 2617 | 1 | 1.5260e-01 | 2.5941e+05 | ; | CA - CB |
| 2615 | 2622 | 1 | 1.5220e-01 | 2.6527e+05 | ; | CA - C |
| 2617 | 2618 | 1 | 1.0900e-01 | 2.8451e+05 | ; | CB - HB2 |
| 2617 | 2619 | 1 | 1.0900e-01 | 2.8451e+05 | ; | CB - HB3 |
| 2617 | 2620 | 1 | 1.4100e-01 | 2.6778e+05 | ; | CB - OG |
| 2620 | 2621 | 1 | 9.6000e-02 | 4.6275e+05 | ; | OG - HG |
| 2622 | 2623 | 1 | 1.2290e-01 | 4.7698e+05 | ; | C - O |
| 2622 | 2624 | 1 | 1.3350e-01 | 4.1003e+05 | ; | C - N |
| 2624 | 2625 | 1 | 1.0100e-01 | 3.6317e+05 | ; | N - H |
| 2624 | 2626 | 1 | 1.4490e-01 | 2.8200e+05 | ; | N - CA |
| 2626 | 2627 | 1 | 1.0900e-01 | 2.8451e+05 | ; | CA - HA |
| 2626 | 2628 | 1 | 1.5260e-01 | 2.5941e+05 | ; | CA - CB |
| 2626 | 2644 | 1 | 1.5220e-01 | 2.6527e+05 | ; | CA - C |
| 2628 | 2629 | 1 | 1.0900e-01 | 2.8451e+05 | ; | CB - HB2 |
| 2628 | 2630 | 1 | 1.0900e-01 | 2.8451e+05 | ; | CB - HB3 |
| 2628 | 2631 | 1 | 1.5260e-01 | 2.5941e+05 | ; | CB - CG |
| 2631 | 2632 | 1 | 1.0900e-01 | 2.8451e+05 | ; | CG - HG2 |
| 2631 | 2633 | 1 | 1.0900e-01 | 2.8451e+05 | ; | CG - HG3 |
| 2631 | 2634 | 1 | 1.5260e-01 | 2.5941e+05 | ; | CG - CD |
| 2634 | 2635 | 1 | 1.0900e-01 | 2.8451e+05 | ; | CD - HD2 |
| 2634 | 2636 | 1 | 1.0900e-01 | 2.8451e+05 | ; | CD - HD3 |
| 2634 | 2637 | 1 | 1.5260e-01 | 2.5941e+05 | ; | CD - CE |
| 2637 | 2638 | 1 | 1.0900e-01 | 2.8451e+05 | ; | CE - HE2 |
| 2637 | 2639 | 1 | 1.0900e-01 | 2.8451e+05 | ; | CE - HE3 |
| 2637 | 2640 | 1 | 1.4710e-01 | 3.0711e+05 | ; | CE - NZ |
| 2640 | 2641 | 1 | 1.0100e-01 | 3.6317e+05 | ; | NZ - HZ1 |
| 2640 | 2642 | 1 | 1.0100e-01 | 3.6317e+05 | ; | NZ - HZ2 |
| 2640 | 2643 | 1 | 1.0100e-01 | 3.6317e+05 | ; | NZ - HZ3 |
| 2644 | 2645 | 1 | 1.2290e-01 | 4.7698e+05 | ; | C - O |
| 2644 | 2646 | 1 | 1.3350e-01 | 4.1003e+05 | ; | C - N |
| 2646 | 2647 | 1 | 1.0100e-01 | 3.6317e+05 | ; | N - H |
| 2646 | 2648 | 1 | 1.4490e-01 | 2.8200e+05 | ; | N - CA |
| 2648 | 2649 | 1 | 1.0900e-01 | 2.8451e+05 | ; | CA - HA |
| 2648 | 2650 | 1 | 1.5260e-01 | 2.5941e+05 | ; | CA - CB |
| 2648 | 2658 | 1 | 1.5220e-01 | 2.6527e+05 | ; | CA - C |
| 2650 | 2651 | 1 | 1.0900e-01 | 2.8451e+05 | ; | CB - HB2 |
| 2650 | 2652 | 1 | 1.0900e-01 | 2.8451e+05 | ; | CB - HB3 |
| 2650 | 2653 | 1 | 1.5220e-01 | 2.6527e+05 | ; | CB - CG |
| 2653 | 2654 | 1 | 1.2290e-01 | 4.7698e+05 | ; | CG - OD1 |
| 2653 | 2655 | 1 | 1.3350e-01 | 4.1003e+05 | ; | CG - ND2 |
| 2655 | 2656 | 1 | 1.0100e-01 | 3.6317e+05 | ; | ND2 - HD21 |
| 2655 | 2657 | 1 | 1.0100e-01 | 3.6317e+05 | ; | ND2 - HD22 |
| 2658 | 2659 | 1 | 1.2290e-01 | 4.7698e+05 | ; | C - O |
| 2658 | 2660 | 1 | 1.3350e-01 | 4.1003e+05 | ; | C - N |
| 2660 | 2661 | 1 | 1.0100e-01 | 3.6317e+05 | ; | N - H |
| 2660 | 2662 | 1 | 1.4490e-01 | 2.8200e+05 | ; | N - CA |

| | | | | | | | | |
|---|---|---|---|---|---|---|---|---|
| 2662 | 2663 | 1 | 1.0900e-01 | 2.8451e+05 | ; | CA | - | HA |
| 2662 | 2664 | 1 | 1.5260e-01 | 2.5941e+05 | ; | CA | - | CB |
| 2662 | 2677 | 1 | 1.5220e-01 | 2.6527e+05 | ; | CA | - | C |
| 2664 | 2665 | 1 | 1.0900e-01 | 2.8451e+05 | ; | CB | - | HB2 |
| 2664 | 2666 | 1 | 1.0900e-01 | 2.8451e+05 | ; | CB | - | HB3 |
| 2664 | 2667 | 1 | 1.5260e-01 | 2.5941e+05 | ; | CB | - | CG |
| 2667 | 2668 | 1 | 1.0900e-01 | 2.8451e+05 | ; | CG | - | HG |
| 2667 | 2669 | 1 | 1.5260e-01 | 2.5941e+05 | ; | CG | - | CD1 |
| 2667 | 2673 | 1 | 1.5260e-01 | 2.5941e+05 | ; | CG | - | CD2 |
| 2669 | 2670 | 1 | 1.0900e-01 | 2.8451e+05 | ; | CD1 | - | HD11 |
| 2669 | 2671 | 1 | 1.0900e-01 | 2.8451e+05 | ; | CD1 | - | HD12 |
| 2669 | 2672 | 1 | 1.0900e-01 | 2.8451e+05 | ; | CD1 | - | HD13 |
| 2673 | 2674 | 1 | 1.0900e-01 | 2.8451e+05 | ; | CD2 | - | HD21 |
| 2673 | 2675 | 1 | 1.0900e-01 | 2.8451e+05 | ; | CD2 | - | HD22 |
| 2673 | 2676 | 1 | 1.0900e-01 | 2.8451e+05 | ; | CD2 | - | HD23 |
| 2677 | 2678 | 1 | 1.2290e-01 | 4.7698e+05 | ; | C | - | O |
| 2677 | 2679 | 1 | 1.3350e-01 | 4.1003e+05 | ; | C | - | N |
| 2679 | 2680 | 1 | 1.0100e-01 | 3.6317e+05 | ; | N | - | H |
| 2679 | 2681 | 1 | 1.4490e-01 | 2.8200e+05 | ; | N | - | CA |
| 2681 | 2682 | 1 | 1.0900e-01 | 2.8451e+05 | ; | CA | - | HA2 |
| 2681 | 2683 | 1 | 1.0900e-01 | 2.8451e+05 | ; | CA | - | HA3 |
| 2681 | 2684 | 1 | 1.5220e-01 | 2.6527e+05 | ; | CA | - | C |
| 2684 | 2685 | 1 | 1.2290e-01 | 4.7698e+05 | ; | C | - | O |
| 2684 | 2686 | 1 | 1.3350e-01 | 4.1003e+05 | ; | C | - | N |
| 2686 | 2687 | 1 | 1.0100e-01 | 3.6317e+05 | ; | N | - | H |
| 2686 | 2688 | 1 | 1.4490e-01 | 2.8200e+05 | ; | N | - | CA |
| 2688 | 2689 | 1 | 1.0900e-01 | 2.8451e+05 | ; | CA | - | HA |
| 2688 | 2690 | 1 | 1.5260e-01 | 2.5941e+05 | ; | CA | - | CB |
| 2688 | 2698 | 1 | 1.5220e-01 | 2.6527e+05 | ; | CA | - | C |
| 2690 | 2691 | 1 | 1.0900e-01 | 2.8451e+05 | ; | CB | - | HB |
| 2690 | 2692 | 1 | 1.5260e-01 | 2.5941e+05 | ; | CB | - | CG2 |
| 2690 | 2696 | 1 | 1.4100e-01 | 2.6778e+05 | ; | CB | - | OG1 |
| 2692 | 2693 | 1 | 1.0900e-01 | 2.8451e+05 | ; | CG2 | - | HG21 |
| 2692 | 2694 | 1 | 1.0900e-01 | 2.8451e+05 | ; | CG2 | - | HG22 |
| 2692 | 2695 | 1 | 1.0900e-01 | 2.8451e+05 | ; | CG2 | - | HG23 |
| 2696 | 2697 | 1 | 9.6000e-02 | 4.6275e+05 | ; | OG1 | - | HG1 |
| 2698 | 2699 | 1 | 1.2290e-01 | 4.7698e+05 | ; | C | - | O |
| 2698 | 2700 | 1 | 1.3350e-01 | 4.1003e+05 | ; | C | - | N |
| 2700 | 2701 | 1 | 1.0100e-01 | 3.6317e+05 | ; | N | - | H |
| 2700 | 2702 | 1 | 1.4490e-01 | 2.8200e+05 | ; | N | - | CA |
| 2702 | 2703 | 1 | 1.0900e-01 | 2.8451e+05 | ; | CA | - | HA2 |
| 2702 | 2704 | 1 | 1.0900e-01 | 2.8451e+05 | ; | CA | - | HA3 |
| 2702 | 2705 | 1 | 1.5220e-01 | 2.6527e+05 | ; | CA | - | C |
| 2705 | 2706 | 1 | 1.2290e-01 | 4.7698e+05 | ; | C | - | O |
| 2705 | 2707 | 1 | 1.3350e-01 | 4.1003e+05 | ; | C | - | N |
| 2707 | 2708 | 1 | 1.0100e-01 | 3.6317e+05 | ; | N | - | H |
| 2707 | 2709 | 1 | 1.4490e-01 | 2.8200e+05 | ; | N | - | CA |
| 2709 | 2710 | 1 | 1.0900e-01 | 2.8451e+05 | ; | CA | - | HA |
| 2709 | 2711 | 1 | 1.5260e-01 | 2.5941e+05 | ; | CA | - | CB |
| 2709 | 2720 | 1 | 1.5220e-01 | 2.6527e+05 | ; | CA | - | C |
| 2711 | 2712 | 1 | 1.0900e-01 | 2.8451e+05 | ; | CB | - | HB2 |
| 2711 | 2713 | 1 | 1.0900e-01 | 2.8451e+05 | ; | CB | - | HB3 |

| | | | | | |
|---|---|---|---|---|---|
| 2711 | 2714 | 1 | 1.5260e-01 | 2.5941e+05 ; | CB - CG |
| 2714 | 2715 | 1 | 1.0900e-01 | 2.8451e+05 ; | CG - HG2 |
| 2714 | 2716 | 1 | 1.0900e-01 | 2.8451e+05 ; | CG - HG3 |
| 2714 | 2717 | 1 | 1.5220e-01 | 2.6527e+05 ; | CG - CD |
| 2717 | 2718 | 1 | 1.2500e-01 | 5.4894e+05 ; | CD - OE1 |
| 2717 | 2719 | 1 | 1.2500e-01 | 5.4894e+05 ; | CD - OE2 |
| 2720 | 2721 | 1 | 1.2290e-01 | 4.7698e+05 ; | C - O |
| 2720 | 2722 | 1 | 1.3350e-01 | 4.1003e+05 ; | C - N |
| 2722 | 2723 | 1 | 1.0100e-01 | 3.6317e+05 ; | N - H |
| 2722 | 2724 | 1 | 1.4490e-01 | 2.8200e+05 ; | N - CA |
| 2724 | 2725 | 1 | 1.0900e-01 | 2.8451e+05 ; | CA - HA |
| 2724 | 2726 | 1 | 1.5260e-01 | 2.5941e+05 ; | CA - CB |
| 2724 | 2740 | 1 | 1.5220e-01 | 2.6527e+05 ; | CA - C |
| 2726 | 2727 | 1 | 1.0900e-01 | 2.8451e+05 ; | CB - HB2 |
| 2726 | 2728 | 1 | 1.0900e-01 | 2.8451e+05 ; | CB - HB3 |
| 2726 | 2729 | 1 | 1.5100e-01 | 2.6527e+05 ; | CB - CG |
| 2729 | 2730 | 1 | 1.4000e-01 | 3.9246e+05 ; | CG - CD1 |
| 2729 | 2738 | 1 | 1.4000e-01 | 3.9246e+05 ; | CG - CD2 |
| 2730 | 2731 | 1 | 1.0800e-01 | 3.0711e+05 ; | CD1 - HD1 |
| 2730 | 2732 | 1 | 1.4000e-01 | 3.9246e+05 ; | CD1 - CE1 |
| 2732 | 2733 | 1 | 1.0800e-01 | 3.0711e+05 ; | CE1 - HE1 |
| 2732 | 2734 | 1 | 1.4000e-01 | 3.9246e+05 ; | CE1 - CZ |
| 2734 | 2735 | 1 | 1.0800e-01 | 3.0711e+05 ; | CZ - HZ |
| 2734 | 2736 | 1 | 1.4000e-01 | 3.9246e+05 ; | CZ - CE2 |
| 2736 | 2737 | 1 | 1.0800e-01 | 3.0711e+05 ; | CE2 - HE2 |
| 2736 | 2738 | 1 | 1.4000e-01 | 3.9246e+05 ; | CE2 - CD2 |
| 2738 | 2739 | 1 | 1.0800e-01 | 3.0711e+05 ; | CD2 - HD2 |
| 2740 | 2741 | 1 | 1.2290e-01 | 4.7698e+05 ; | C - O |
| 2740 | 2742 | 1 | 1.3350e-01 | 4.1003e+05 ; | C - N |
| 2742 | 2743 | 1 | 1.0100e-01 | 3.6317e+05 ; | N - H |
| 2742 | 2744 | 1 | 1.4490e-01 | 2.8200e+05 ; | N - CA |
| 2744 | 2745 | 1 | 1.0900e-01 | 2.8451e+05 ; | CA - HA |
| 2744 | 2746 | 1 | 1.5260e-01 | 2.5941e+05 ; | CA - CB |
| 2744 | 2754 | 1 | 1.5220e-01 | 2.6527e+05 ; | CA - C |
| 2746 | 2747 | 1 | 1.0900e-01 | 2.8451e+05 ; | CB - HB |
| 2746 | 2748 | 1 | 1.5260e-01 | 2.5941e+05 ; | CB - CG2 |
| 2746 | 2752 | 1 | 1.4100e-01 | 2.6778e+05 ; | CB - OG1 |
| 2748 | 2749 | 1 | 1.0900e-01 | 2.8451e+05 ; | CG2 - HG21 |
| 2748 | 2750 | 1 | 1.0900e-01 | 2.8451e+05 ; | CG2 - HG22 |
| 2748 | 2751 | 1 | 1.0900e-01 | 2.8451e+05 ; | CG2 - HG23 |
| 2752 | 2753 | 1 | 9.6000e-02 | 4.6275e+05 ; | OG1 - HG1 |
| 2754 | 2755 | 1 | 1.2290e-01 | 4.7698e+05 ; | C - O |
| 2754 | 2756 | 1 | 1.3350e-01 | 4.1003e+05 ; | C - N |
| 2756 | 2757 | 1 | 1.0100e-01 | 3.6317e+05 ; | N - H |
| 2756 | 2758 | 1 | 1.4490e-01 | 2.8200e+05 ; | N - CA |
| 2758 | 2759 | 1 | 1.0900e-01 | 2.8451e+05 ; | CA - HA |
| 2758 | 2760 | 1 | 1.5260e-01 | 2.5941e+05 ; | CA - CB |
| 2758 | 2768 | 1 | 1.5220e-01 | 2.6527e+05 ; | CA - C |
| 2760 | 2761 | 1 | 1.0900e-01 | 2.8451e+05 ; | CB - HB |
| 2760 | 2762 | 1 | 1.5260e-01 | 2.5941e+05 ; | CB - CG2 |
| 2760 | 2766 | 1 | 1.4100e-01 | 2.6778e+05 ; | CB - OG1 |
| 2762 | 2763 | 1 | 1.0900e-01 | 2.8451e+05 ; | CG2 - HG21 |

| | | | | | |
|---|---|---|---|---|---|
| 2762 | 2764 | 1 | 1.0900e-01 | 2.8451e+05 ; | CG2 - HG22 |
| 2762 | 2765 | 1 | 1.0900e-01 | 2.8451e+05 ; | CG2 - HG23 |
| 2766 | 2767 | 1 | 9.6000e-02 | 4.6275e+05 ; | OG1 - HG1 |
| 2768 | 2769 | 1 | 1.2290e-01 | 4.7698e+05 ; | C - O |
| 2768 | 2770 | 1 | 1.3350e-01 | 4.1003e+05 ; | C - N |
| 2770 | 2771 | 1 | 1.0100e-01 | 3.6317e+05 ; | N - H |
| 2770 | 2772 | 1 | 1.4490e-01 | 2.8200e+05 ; | N - CA |
| 2772 | 2773 | 1 | 1.0900e-01 | 2.8451e+05 ; | CA - HA |
| 2772 | 2774 | 1 | 1.5260e-01 | 2.5941e+05 ; | CA - CB |
| 2772 | 2782 | 1 | 1.5220e-01 | 2.6527e+05 ; | CA - C |
| 2774 | 2775 | 1 | 1.0900e-01 | 2.8451e+05 ; | CB - HB |
| 2774 | 2776 | 1 | 1.5260e-01 | 2.5941e+05 ; | CB - CG2 |
| 2774 | 2780 | 1 | 1.4100e-01 | 2.6778e+05 ; | CB - OG1 |
| 2776 | 2777 | 1 | 1.0900e-01 | 2.8451e+05 ; | CG2 - HG21 |
| 2776 | 2778 | 1 | 1.0900e-01 | 2.8451e+05 ; | CG2 - HG22 |
| 2776 | 2779 | 1 | 1.0900e-01 | 2.8451e+05 ; | CG2 - HG23 |
| 2780 | 2781 | 1 | 9.6000e-02 | 4.6275e+05 ; | OG1 - HG1 |
| 2782 | 2783 | 1 | 1.2290e-01 | 4.7698e+05 ; | C - O |
| 2782 | 2784 | 1 | 1.3350e-01 | 4.1003e+05 ; | C - N |
| 2784 | 2785 | 1 | 1.0100e-01 | 3.6317e+05 ; | N - H |
| 2784 | 2786 | 1 | 1.4490e-01 | 2.8200e+05 ; | N - CA |
| 2786 | 2787 | 1 | 1.0900e-01 | 2.8451e+05 ; | CA - HA |
| 2786 | 2788 | 1 | 1.5260e-01 | 2.5941e+05 ; | CA - CB |
| 2786 | 2804 | 1 | 1.5220e-01 | 2.6527e+05 ; | CA - C |
| 2788 | 2789 | 1 | 1.0900e-01 | 2.8451e+05 ; | CB - HB2 |
| 2788 | 2790 | 1 | 1.0900e-01 | 2.8451e+05 ; | CB - HB3 |
| 2788 | 2791 | 1 | 1.5260e-01 | 2.5941e+05 ; | CB - CG |
| 2791 | 2792 | 1 | 1.0900e-01 | 2.8451e+05 ; | CG - HG2 |
| 2791 | 2793 | 1 | 1.0900e-01 | 2.8451e+05 ; | CG - HG3 |
| 2791 | 2794 | 1 | 1.5260e-01 | 2.5941e+05 ; | CG - CD |
| 2794 | 2795 | 1 | 1.0900e-01 | 2.8451e+05 ; | CD - HD2 |
| 2794 | 2796 | 1 | 1.0900e-01 | 2.8451e+05 ; | CD - HD3 |
| 2794 | 2797 | 1 | 1.5260e-01 | 2.5941e+05 ; | CD - CE |
| 2797 | 2798 | 1 | 1.0900e-01 | 2.8451e+05 ; | CE - HE2 |
| 2797 | 2799 | 1 | 1.0900e-01 | 2.8451e+05 ; | CE - HE3 |
| 2797 | 2800 | 1 | 1.4710e-01 | 3.0711e+05 ; | CE - NZ |
| 2800 | 2801 | 1 | 1.0100e-01 | 3.6317e+05 ; | NZ - HZ1 |
| 2800 | 2802 | 1 | 1.0100e-01 | 3.6317e+05 ; | NZ - HZ2 |
| 2800 | 2803 | 1 | 1.0100e-01 | 3.6317e+05 ; | NZ - HZ3 |
| 2804 | 2805 | 1 | 1.2290e-01 | 4.7698e+05 ; | C - O |
| 2804 | 2806 | 1 | 1.3350e-01 | 4.1003e+05 ; | C - N |
| 2806 | 2807 | 1 | 1.0100e-01 | 3.6317e+05 ; | N - H |
| 2806 | 2808 | 1 | 1.4490e-01 | 2.8200e+05 ; | N - CA |
| 2808 | 2809 | 1 | 1.0900e-01 | 2.8451e+05 ; | CA - HA |
| 2808 | 2810 | 1 | 1.5260e-01 | 2.5941e+05 ; | CA - CB |
| 2808 | 2814 | 1 | 1.5220e-01 | 2.6527e+05 ; | CA - C |
| 2810 | 2811 | 1 | 1.0900e-01 | 2.8451e+05 ; | CB - HB1 |
| 2810 | 2812 | 1 | 1.0900e-01 | 2.8451e+05 ; | CB - HB2 |
| 2810 | 2813 | 1 | 1.0900e-01 | 2.8451e+05 ; | CB - HB3 |
| 2814 | 2815 | 1 | 1.2290e-01 | 4.7698e+05 ; | C - O |
| 2814 | 2816 | 1 | 1.3350e-01 | 4.1003e+05 ; | C - N |
| 2816 | 2817 | 1 | 1.0100e-01 | 3.6317e+05 ; | N - H |

| | | | | | |
|---|---|---|---|---|---|
| 2816 | 2818 | 1 | 1.4490e-01 | 2.8200e+05 ; | N - CA |
| 2818 | 2819 | 1 | 1.0900e-01 | 2.8451e+05 ; | CA - HA2 |
| 2818 | 2820 | 1 | 1.0900e-01 | 2.8451e+05 ; | CA - HA3 |
| 2818 | 2821 | 1 | 1.5220e-01 | 2.6527e+05 ; | CA - C |
| 2821 | 2822 | 1 | 1.2290e-01 | 4.7698e+05 ; | C - O |
| 2821 | 2823 | 1 | 1.3350e-01 | 4.1003e+05 ; | C - N |
| 2823 | 2824 | 1 | 1.0100e-01 | 3.6317e+05 ; | N - H |
| 2823 | 2825 | 1 | 1.4490e-01 | 2.8200e+05 ; | N - CA |
| 2825 | 2826 | 1 | 1.0900e-01 | 2.8451e+05 ; | CA - HA |
| 2825 | 2827 | 1 | 1.5260e-01 | 2.5941e+05 ; | CA - CB |
| 2825 | 2835 | 1 | 1.5220e-01 | 2.6527e+05 ; | CA - C |
| 2827 | 2828 | 1 | 1.0900e-01 | 2.8451e+05 ; | CB - HB |
| 2827 | 2829 | 1 | 1.5260e-01 | 2.5941e+05 ; | CB - CG2 |
| 2827 | 2833 | 1 | 1.4100e-01 | 2.6778e+05 ; | CB - OG1 |
| 2829 | 2830 | 1 | 1.0900e-01 | 2.8451e+05 ; | CG2 - HG21 |
| 2829 | 2831 | 1 | 1.0900e-01 | 2.8451e+05 ; | CG2 - HG22 |
| 2829 | 2832 | 1 | 1.0900e-01 | 2.8451e+05 ; | CG2 - HG23 |
| 2833 | 2834 | 1 | 9.6000e-02 | 4.6275e+05 ; | OG1 - HG1 |
| 2835 | 2836 | 1 | 1.2290e-01 | 4.7698e+05 ; | C - O |
| 2835 | 2837 | 1 | 1.3350e-01 | 4.1003e+05 ; | C - N |
| 2837 | 2838 | 1 | 1.4490e-01 | 2.8200e+05 ; | N - CD |
| 2837 | 2847 | 1 | 1.4490e-01 | 2.8200e+05 ; | N - CA |
| 2838 | 2839 | 1 | 1.0900e-01 | 2.8451e+05 ; | CD - HD2 |
| 2838 | 2840 | 1 | 1.0900e-01 | 2.8451e+05 ; | CD - HD3 |
| 2838 | 2841 | 1 | 1.5260e-01 | 2.5941e+05 ; | CD - CG |
| 2841 | 2842 | 1 | 1.0900e-01 | 2.8451e+05 ; | CG - HG2 |
| 2841 | 2843 | 1 | 1.0900e-01 | 2.8451e+05 ; | CG - HG3 |
| 2841 | 2844 | 1 | 1.5260e-01 | 2.5941e+05 ; | CG - CB |
| 2844 | 2845 | 1 | 1.0900e-01 | 2.8451e+05 ; | CB - HB2 |
| 2844 | 2846 | 1 | 1.0900e-01 | 2.8451e+05 ; | CB - HB3 |
| 2844 | 2847 | 1 | 1.5260e-01 | 2.5941e+05 ; | CB - CA |
| 2847 | 2848 | 1 | 1.0900e-01 | 2.8451e+05 ; | CA - HA |
| 2847 | 2849 | 1 | 1.5220e-01 | 2.6527e+05 ; | CA - C |
| 2849 | 2850 | 1 | 1.2290e-01 | 4.7698e+05 ; | C - O |
| 2849 | 2851 | 1 | 1.3350e-01 | 4.1003e+05 ; | C - N |
| 2851 | 2852 | 1 | 1.0100e-01 | 3.6317e+05 ; | N - H |
| 2851 | 2853 | 1 | 1.4490e-01 | 2.8200e+05 ; | N - CA |
| 2853 | 2854 | 1 | 1.0900e-01 | 2.8451e+05 ; | CA - HA |
| 2853 | 2855 | 1 | 1.5260e-01 | 2.5941e+05 ; | CA - CB |
| 2853 | 2870 | 1 | 1.5220e-01 | 2.6527e+05 ; | CA - C |
| 2855 | 2856 | 1 | 1.0900e-01 | 2.8451e+05 ; | CB - HB2 |
| 2855 | 2857 | 1 | 1.0900e-01 | 2.8451e+05 ; | CB - HB3 |
| 2855 | 2858 | 1 | 1.5100e-01 | 2.6527e+05 ; | CB - CG |
| 2858 | 2859 | 1 | 1.4000e-01 | 3.9246e+05 ; | CG - CD1 |
| 2858 | 2868 | 1 | 1.4000e-01 | 3.9246e+05 ; | CG - CD2 |
| 2859 | 2860 | 1 | 1.0800e-01 | 3.0711e+05 ; | CD1 - HD1 |
| 2859 | 2861 | 1 | 1.4000e-01 | 3.9246e+05 ; | CD1 - CE1 |
| 2861 | 2862 | 1 | 1.0800e-01 | 3.0711e+05 ; | CE1 - HE1 |
| 2861 | 2863 | 1 | 1.4090e-01 | 3.9246e+05 ; | CE1 - CZ |
| 2863 | 2864 | 1 | 1.3640e-01 | 3.7656e+05 ; | CZ - OH |
| 2863 | 2866 | 1 | 1.4090e-01 | 3.9246e+05 ; | CZ - CE2 |
| 2864 | 2865 | 1 | 9.6000e-02 | 4.6275e+05 ; | OH - HH |

| | | | | | | |
|---|---|---|---|---|---|---|
| 2866 | 2867 | 1 | 1.0800e-01 | 3.0711e+05 | ; | CE2 - HE2 |
| 2866 | 2868 | 1 | 1.4000e-01 | 3.9246e+05 | ; | CE2 - CD2 |
| 2868 | 2869 | 1 | 1.0800e-01 | 3.0711e+05 | ; | CD2 - HD2 |
| 2870 | 2871 | 1 | 1.2290e-01 | 4.7698e+05 | ; | C - O |
| 2870 | 2872 | 1 | 1.3350e-01 | 4.1003e+05 | ; | C - N |
| 2872 | 2873 | 1 | 1.0100e-01 | 3.6317e+05 | ; | N - H |
| 2872 | 2874 | 1 | 1.4490e-01 | 2.8200e+05 | ; | N - CA |
| 2874 | 2875 | 1 | 1.0900e-01 | 2.8451e+05 | ; | CA - HA |
| 2874 | 2876 | 1 | 1.5260e-01 | 2.5941e+05 | ; | CA - CB |
| 2874 | 2891 | 1 | 1.5220e-01 | 2.6527e+05 | ; | CA - C |
| 2876 | 2877 | 1 | 1.0900e-01 | 2.8451e+05 | ; | CB - HB2 |
| 2876 | 2878 | 1 | 1.0900e-01 | 2.8451e+05 | ; | CB - HB3 |
| 2876 | 2879 | 1 | 1.5100e-01 | 2.6527e+05 | ; | CB - CG |
| 2879 | 2880 | 1 | 1.4000e-01 | 3.9246e+05 | ; | CG - CD1 |
| 2879 | 2889 | 1 | 1.4000e-01 | 3.9246e+05 | ; | CG - CD2 |
| 2880 | 2881 | 1 | 1.0800e-01 | 3.0711e+05 | ; | CD1 - HD1 |
| 2880 | 2882 | 1 | 1.4000e-01 | 3.9246e+05 | ; | CD1 - CE1 |
| 2882 | 2883 | 1 | 1.0800e-01 | 3.0711e+05 | ; | CE1 - HE1 |
| 2882 | 2884 | 1 | 1.4090e-01 | 3.9246e+05 | ; | CE1 - CZ |
| 2884 | 2885 | 1 | 1.3640e-01 | 3.7656e+05 | ; | CZ - OH |
| 2884 | 2887 | 1 | 1.4090e-01 | 3.9246e+05 | ; | CZ - CE2 |
| 2885 | 2886 | 1 | 9.6000e-02 | 4.6275e+05 | ; | OH - HH |
| 2887 | 2888 | 1 | 1.0800e-01 | 3.0711e+05 | ; | CE2 - HE2 |
| 2887 | 2889 | 1 | 1.4000e-01 | 3.9246e+05 | ; | CE2 - CD2 |
| 2889 | 2890 | 1 | 1.0800e-01 | 3.0711e+05 | ; | CD2 - HD2 |
| 2891 | 2892 | 1 | 1.2290e-01 | 4.7698e+05 | ; | C - O |
| 2891 | 2893 | 1 | 1.3350e-01 | 4.1003e+05 | ; | C - N |
| 2893 | 2894 | 1 | 1.0100e-01 | 3.6317e+05 | ; | N - H |
| 2893 | 2895 | 1 | 1.4490e-01 | 2.8200e+05 | ; | N - CA |
| 2895 | 2896 | 1 | 1.0900e-01 | 2.8451e+05 | ; | CA - HA |
| 2895 | 2897 | 1 | 1.5260e-01 | 2.5941e+05 | ; | CA - CB |
| 2895 | 2907 | 1 | 1.5220e-01 | 2.6527e+05 | ; | CA - C |
| 2897 | 2898 | 1 | 1.0900e-01 | 2.8451e+05 | ; | CB - HB |
| 2897 | 2899 | 1 | 1.5260e-01 | 2.5941e+05 | ; | CB - CG1 |
| 2897 | 2903 | 1 | 1.5260e-01 | 2.5941e+05 | ; | CB - CG2 |
| 2899 | 2900 | 1 | 1.0900e-01 | 2.8451e+05 | ; | CG1 - HG11 |
| 2899 | 2901 | 1 | 1.0900e-01 | 2.8451e+05 | ; | CG1 - HG12 |
| 2899 | 2902 | 1 | 1.0900e-01 | 2.8451e+05 | ; | CG1 - HG13 |
| 2903 | 2904 | 1 | 1.0900e-01 | 2.8451e+05 | ; | CG2 - HG21 |
| 2903 | 2905 | 1 | 1.0900e-01 | 2.8451e+05 | ; | CG2 - HG22 |
| 2903 | 2906 | 1 | 1.0900e-01 | 2.8451e+05 | ; | CG2 - HG23 |
| 2907 | 2908 | 1 | 1.2290e-01 | 4.7698e+05 | ; | C - O |
| 2907 | 2909 | 1 | 1.3350e-01 | 4.1003e+05 | ; | C - N |
| 2909 | 2910 | 1 | 1.0100e-01 | 3.6317e+05 | ; | N - H |
| 2909 | 2911 | 1 | 1.4490e-01 | 2.8200e+05 | ; | N - CA |
| 2911 | 2912 | 1 | 1.0900e-01 | 2.8451e+05 | ; | CA - HA |
| 2911 | 2913 | 1 | 1.5260e-01 | 2.5941e+05 | ; | CA - CB |
| 2911 | 2917 | 1 | 1.5220e-01 | 2.6527e+05 | ; | CA - C |
| 2913 | 2914 | 1 | 1.0900e-01 | 2.8451e+05 | ; | CB - HB1 |
| 2913 | 2915 | 1 | 1.0900e-01 | 2.8451e+05 | ; | CB - HB2 |
| 2913 | 2916 | 1 | 1.0900e-01 | 2.8451e+05 | ; | CB - HB3 |
| 2917 | 2918 | 1 | 1.2290e-01 | 4.7698e+05 | ; | C - O |

| | | | | | | | | |
|---|---|---|---|---|---|---|---|---|
| 2917 | 2919 | 1 | 1.3350e-01 | 4.1003e+05 | ; | C | - | N |
| 2919 | 2920 | 1 | 1.4490e-01 | 2.8200e+05 | ; | N | - | CD |
| 2919 | 2929 | 1 | 1.4490e-01 | 2.8200e+05 | ; | N | - | CA |
| 2920 | 2921 | 1 | 1.0900e-01 | 2.8451e+05 | ; | CD | - | HD2 |
| 2920 | 2922 | 1 | 1.0900e-01 | 2.8451e+05 | ; | CD | - | HD3 |
| 2920 | 2923 | 1 | 1.5260e-01 | 2.5941e+05 | ; | CD | - | CG |
| 2923 | 2924 | 1 | 1.0900e-01 | 2.8451e+05 | ; | CG | - | HG2 |
| 2923 | 2925 | 1 | 1.0900e-01 | 2.8451e+05 | ; | CG | - | HG3 |
| 2923 | 2926 | 1 | 1.5260e-01 | 2.5941e+05 | ; | CG | - | CB |
| 2926 | 2927 | 1 | 1.0900e-01 | 2.8451e+05 | ; | CB | - | HB2 |
| 2926 | 2928 | 1 | 1.0900e-01 | 2.8451e+05 | ; | CB | - | HB3 |
| 2926 | 2929 | 1 | 1.5260e-01 | 2.5941e+05 | ; | CB | - | CA |
| 2929 | 2930 | 1 | 1.0900e-01 | 2.8451e+05 | ; | CA | - | HA |
| 2929 | 2931 | 1 | 1.5220e-01 | 2.6527e+05 | ; | CA | - | C |
| 2931 | 2932 | 1 | 1.2290e-01 | 4.7698e+05 | ; | C | - | O |
| 2931 | 2933 | 1 | 1.3350e-01 | 4.1003e+05 | ; | C | - | N |
| 2933 | 2934 | 1 | 1.0100e-01 | 3.6317e+05 | ; | N | - | H |
| 2933 | 2935 | 1 | 1.4490e-01 | 2.8200e+05 | ; | N | - | CA |
| 2935 | 2936 | 1 | 1.0900e-01 | 2.8451e+05 | ; | CA | - | HA |
| 2935 | 2937 | 1 | 1.5260e-01 | 2.5941e+05 | ; | CA | - | CB |
| 2935 | 2948 | 1 | 1.5220e-01 | 2.6527e+05 | ; | CA | - | C |
| 2937 | 2938 | 1 | 1.0900e-01 | 2.8451e+05 | ; | CB | - | HB2 |
| 2937 | 2939 | 1 | 1.0900e-01 | 2.8451e+05 | ; | CB | - | HB3 |
| 2937 | 2940 | 1 | 1.5260e-01 | 2.5941e+05 | ; | CB | - | CG |
| 2940 | 2941 | 1 | 1.0900e-01 | 2.8451e+05 | ; | CG | - | HG2 |
| 2940 | 2942 | 1 | 1.0900e-01 | 2.8451e+05 | ; | CG | - | HG3 |
| 2940 | 2943 | 1 | 1.5220e-01 | 2.6527e+05 | ; | CG | - | CD |
| 2943 | 2944 | 1 | 1.2290e-01 | 4.7698e+05 | ; | CD | - | OE1 |
| 2943 | 2945 | 1 | 1.3350e-01 | 4.1003e+05 | ; | CD | - | NE2 |
| 2945 | 2946 | 1 | 1.0100e-01 | 3.6317e+05 | ; | NE2 | - | HE21 |
| 2945 | 2947 | 1 | 1.0100e-01 | 3.6317e+05 | ; | NE2 | - | HE22 |
| 2948 | 2949 | 1 | 1.2290e-01 | 4.7698e+05 | ; | C | - | O |
| 2948 | 2950 | 1 | 1.3350e-01 | 4.1003e+05 | ; | C | - | N |
| 2950 | 2951 | 1 | 1.0100e-01 | 3.6317e+05 | ; | N | - | H |
| 2950 | 2952 | 1 | 1.4490e-01 | 2.8200e+05 | ; | N | - | CA |
| 2952 | 2953 | 1 | 1.0900e-01 | 2.8451e+05 | ; | CA | - | HA |
| 2952 | 2954 | 1 | 1.5260e-01 | 2.5941e+05 | ; | CA | - | CB |
| 2952 | 2964 | 1 | 1.5220e-01 | 2.6527e+05 | ; | CA | - | C |
| 2954 | 2955 | 1 | 1.0900e-01 | 2.8451e+05 | ; | CB | - | HB |
| 2954 | 2956 | 1 | 1.5260e-01 | 2.5941e+05 | ; | CB | - | CG1 |
| 2954 | 2960 | 1 | 1.5260e-01 | 2.5941e+05 | ; | CB | - | CG2 |
| 2956 | 2957 | 1 | 1.0900e-01 | 2.8451e+05 | ; | CG1 | - | HG11 |
| 2956 | 2958 | 1 | 1.0900e-01 | 2.8451e+05 | ; | CG1 | - | HG12 |
| 2956 | 2959 | 1 | 1.0900e-01 | 2.8451e+05 | ; | CG1 | - | HG13 |
| 2960 | 2961 | 1 | 1.0900e-01 | 2.8451e+05 | ; | CG2 | - | HG21 |
| 2960 | 2962 | 1 | 1.0900e-01 | 2.8451e+05 | ; | CG2 | - | HG22 |
| 2960 | 2963 | 1 | 1.0900e-01 | 2.8451e+05 | ; | CG2 | - | HG23 |
| 2964 | 2965 | 1 | 1.2290e-01 | 4.7698e+05 | ; | C | - | O |
| 2964 | 2966 | 1 | 1.3350e-01 | 4.1003e+05 | ; | C | - | N |
| 2966 | 2967 | 1 | 1.0100e-01 | 3.6317e+05 | ; | N | - | H |
| 2966 | 2968 | 1 | 1.4490e-01 | 2.8200e+05 | ; | N | - | CA |
| 2968 | 2969 | 1 | 1.0900e-01 | 2.8451e+05 | ; | CA | - | HA |

| | | | | | | | | |
|---|---|---|---|---|---|---|---|---|
| 2968 | 2970 | 1 | 1.5260e-01 | 2.5941e+05 | ; | CA | - | CB |
| 2968 | 2983 | 1 | 1.5220e-01 | 2.6527e+05 | ; | CA | - | C |
| 2970 | 2971 | 1 | 1.0900e-01 | 2.8451e+05 | ; | CB | - | HB2 |
| 2970 | 2972 | 1 | 1.0900e-01 | 2.8451e+05 | ; | CB | - | HB3 |
| 2970 | 2973 | 1 | 1.5260e-01 | 2.5941e+05 | ; | CB | - | CG |
| 2973 | 2974 | 1 | 1.0900e-01 | 2.8451e+05 | ; | CG | - | HG |
| 2973 | 2975 | 1 | 1.5260e-01 | 2.5941e+05 | ; | CG | - | CD1 |
| 2973 | 2979 | 1 | 1.5260e-01 | 2.5941e+05 | ; | CG | - | CD2 |
| 2975 | 2976 | 1 | 1.0900e-01 | 2.8451e+05 | ; | CD1 | - | HD11 |
| 2975 | 2977 | 1 | 1.0900e-01 | 2.8451e+05 | ; | CD1 | - | HD12 |
| 2975 | 2978 | 1 | 1.0900e-01 | 2.8451e+05 | ; | CD1 | - | HD13 |
| 2979 | 2980 | 1 | 1.0900e-01 | 2.8451e+05 | ; | CD2 | - | HD21 |
| 2979 | 2981 | 1 | 1.0900e-01 | 2.8451e+05 | ; | CD2 | - | HD22 |
| 2979 | 2982 | 1 | 1.0900e-01 | 2.8451e+05 | ; | CD2 | - | HD23 |
| 2983 | 2984 | 1 | 1.2290e-01 | 4.7698e+05 | ; | C | - | O |
| 2983 | 2985 | 1 | 1.3350e-01 | 4.1003e+05 | ; | C | - | N |
| 2985 | 2986 | 1 | 1.0100e-01 | 3.6317e+05 | ; | N | - | H |
| 2985 | 2987 | 1 | 1.4490e-01 | 2.8200e+05 | ; | N | - | CA |
| 2987 | 2988 | 1 | 1.0900e-01 | 2.8451e+05 | ; | CA | - | HA |
| 2987 | 2989 | 1 | 1.5260e-01 | 2.5941e+05 | ; | CA | - | CB |
| 2987 | 2995 | 1 | 1.5220e-01 | 2.6527e+05 | ; | CA | - | C |
| 2989 | 2990 | 1 | 1.0900e-01 | 2.8451e+05 | ; | CB | - | HB2 |
| 2989 | 2991 | 1 | 1.0900e-01 | 2.8451e+05 | ; | CB | - | HB3 |
| 2989 | 2992 | 1 | 1.5220e-01 | 2.6527e+05 | ; | CB | - | CG |
| 2992 | 2993 | 1 | 1.2500e-01 | 5.4894e+05 | ; | CG | - | OD1 |
| 2992 | 2994 | 1 | 1.2500e-01 | 5.4894e+05 | ; | CG | - | OD2 |
| 2995 | 2996 | 1 | 1.2290e-01 | 4.7698e+05 | ; | C | - | O |
| 2995 | 2997 | 1 | 1.3350e-01 | 4.1003e+05 | ; | C | - | N |
| 2997 | 2998 | 1 | 1.0100e-01 | 3.6317e+05 | ; | N | - | H |
| 2997 | 2999 | 1 | 1.4490e-01 | 2.8200e+05 | ; | N | - | CA |
| 2999 | 3000 | 1 | 1.0900e-01 | 2.8451e+05 | ; | CA | - | HA2 |
| 2999 | 3001 | 1 | 1.0900e-01 | 2.8451e+05 | ; | CA | - | HA3 |
| 2999 | 3002 | 1 | 1.5220e-01 | 2.6527e+05 | ; | CA | - | C |
| 3002 | 3003 | 1 | 1.2290e-01 | 4.7698e+05 | ; | C | - | O |
| 3002 | 3004 | 1 | 1.3350e-01 | 4.1003e+05 | ; | C | - | N |
| 3004 | 3005 | 1 | 1.0100e-01 | 3.6317e+05 | ; | N | - | H |
| 3004 | 3006 | 1 | 1.4490e-01 | 2.8200e+05 | ; | N | - | CA |
| 3006 | 3007 | 1 | 1.0900e-01 | 2.8451e+05 | ; | CA | - | HA |
| 3006 | 3008 | 1 | 1.5260e-01 | 2.5941e+05 | ; | CA | - | CB |
| 3006 | 3024 | 1 | 1.5220e-01 | 2.6527e+05 | ; | CA | - | C |
| 3008 | 3009 | 1 | 1.0900e-01 | 2.8451e+05 | ; | CB | - | HB2 |
| 3008 | 3010 | 1 | 1.0900e-01 | 2.8451e+05 | ; | CB | - | HB3 |
| 3008 | 3011 | 1 | 1.5260e-01 | 2.5941e+05 | ; | CB | - | CG |
| 3011 | 3012 | 1 | 1.0900e-01 | 2.8451e+05 | ; | CG | - | HG2 |
| 3011 | 3013 | 1 | 1.0900e-01 | 2.8451e+05 | ; | CG | - | HG3 |
| 3011 | 3014 | 1 | 1.5260e-01 | 2.5941e+05 | ; | CG | - | CD |
| 3014 | 3015 | 1 | 1.0900e-01 | 2.8451e+05 | ; | CD | - | HD2 |
| 3014 | 3016 | 1 | 1.0900e-01 | 2.8451e+05 | ; | CD | - | HD3 |
| 3014 | 3017 | 1 | 1.5260e-01 | 2.5941e+05 | ; | CD | - | CE |
| 3017 | 3018 | 1 | 1.0900e-01 | 2.8451e+05 | ; | CE | - | HE2 |
| 3017 | 3019 | 1 | 1.0900e-01 | 2.8451e+05 | ; | CE | - | HE3 |
| 3017 | 3020 | 1 | 1.4710e-01 | 3.0711e+05 | ; | CE | - | NZ |

| | | | | | | | | |
|---|---|---|---|---|---|---|---|---|
| 3020 | 3021 | 1 | 1.0100e-01 | 3.6317e+05 | ; | NZ | - | HZ1 |
| 3020 | 3022 | 1 | 1.0100e-01 | 3.6317e+05 | ; | NZ | - | HZ2 |
| 3020 | 3023 | 1 | 1.0100e-01 | 3.6317e+05 | ; | NZ | - | HZ3 |
| 3024 | 3025 | 1 | 1.2290e-01 | 4.7698e+05 | ; | C | - | O |
| 3024 | 3026 | 1 | 1.3350e-01 | 4.1003e+05 | ; | C | - | N |
| 3026 | 3027 | 1 | 1.0100e-01 | 3.6317e+05 | ; | N | - | H |
| 3026 | 3028 | 1 | 1.4490e-01 | 2.8200e+05 | ; | N | - | CA |
| 3028 | 3029 | 1 | 1.0900e-01 | 2.8451e+05 | ; | CA | - | HA |
| 3028 | 3030 | 1 | 1.5260e-01 | 2.5941e+05 | ; | CA | - | CB |
| 3028 | 3045 | 1 | 1.5220e-01 | 2.6527e+05 | ; | CA | - | C |
| 3030 | 3031 | 1 | 1.0900e-01 | 2.8451e+05 | ; | CB | - | HB2 |
| 3030 | 3032 | 1 | 1.0900e-01 | 2.8451e+05 | ; | CB | - | HB3 |
| 3030 | 3033 | 1 | 1.5100e-01 | 2.6527e+05 | ; | CB | - | CG |
| 3033 | 3034 | 1 | 1.4000e-01 | 3.9246e+05 | ; | CG | - | CD1 |
| 3033 | 3043 | 1 | 1.4000e-01 | 3.9246e+05 | ; | CG | - | CD2 |
| 3034 | 3035 | 1 | 1.0800e-01 | 3.0711e+05 | ; | CD1 | - | HD1 |
| 3034 | 3036 | 1 | 1.4000e-01 | 3.9246e+05 | ; | CD1 | - | CE1 |
| 3036 | 3037 | 1 | 1.0800e-01 | 3.0711e+05 | ; | CE1 | - | HE1 |
| 3036 | 3038 | 1 | 1.4090e-01 | 3.9246e+05 | ; | CE1 | - | CZ |
| 3038 | 3039 | 1 | 1.3640e-01 | 3.7656e+05 | ; | CZ | - | OH |
| 3038 | 3041 | 1 | 1.4090e-01 | 3.9246e+05 | ; | CZ | - | CE2 |
| 3039 | 3040 | 1 | 9.6000e-02 | 4.6275e+05 | ; | OH | - | HH |
| 3041 | 3042 | 1 | 1.0800e-01 | 3.0711e+05 | ; | CE2 | - | HE2 |
| 3041 | 3043 | 1 | 1.4000e-01 | 3.9246e+05 | ; | CE2 | - | CD2 |
| 3043 | 3044 | 1 | 1.0800e-01 | 3.0711e+05 | ; | CD2 | - | HD2 |
| 3045 | 3046 | 1 | 1.2290e-01 | 4.7698e+05 | ; | C | - | O |
| 3045 | 3047 | 1 | 1.3350e-01 | 4.1003e+05 | ; | C | - | N |
| 3047 | 3048 | 1 | 1.0100e-01 | 3.6317e+05 | ; | N | - | H |
| 3047 | 3049 | 1 | 1.4490e-01 | 2.8200e+05 | ; | N | - | CA |
| 3049 | 3050 | 1 | 1.0900e-01 | 2.8451e+05 | ; | CA | - | HA |
| 3049 | 3051 | 1 | 1.5260e-01 | 2.5941e+05 | ; | CA | - | CB |
| 3049 | 3057 | 1 | 1.5220e-01 | 2.6527e+05 | ; | CA | - | C |
| 3051 | 3052 | 1 | 1.0900e-01 | 2.8451e+05 | ; | CB | - | HB2 |
| 3051 | 3053 | 1 | 1.0900e-01 | 2.8451e+05 | ; | CB | - | HB3 |
| 3051 | 3054 | 1 | 1.5220e-01 | 2.6527e+05 | ; | CB | - | CG |
| 3054 | 3055 | 1 | 1.2500e-01 | 5.4894e+05 | ; | CG | - | OD1 |
| 3054 | 3056 | 1 | 1.2500e-01 | 5.4894e+05 | ; | CG | - | OD2 |
| 3057 | 3058 | 1 | 1.2290e-01 | 4.7698e+05 | ; | C | - | O |
| 3057 | 3059 | 1 | 1.3350e-01 | 4.1003e+05 | ; | C | - | N |
| 3059 | 3060 | 1 | 1.0100e-01 | 3.6317e+05 | ; | N | - | H |
| 3059 | 3061 | 1 | 1.4490e-01 | 2.8200e+05 | ; | N | - | CA |
| 3061 | 3062 | 1 | 1.0900e-01 | 2.8451e+05 | ; | CA | - | HA |
| 3061 | 3063 | 1 | 1.5260e-01 | 2.5941e+05 | ; | CA | - | CB |
| 3061 | 3079 | 1 | 1.5220e-01 | 2.6527e+05 | ; | CA | - | C |
| 3063 | 3064 | 1 | 1.0900e-01 | 2.8451e+05 | ; | CB | - | HB2 |
| 3063 | 3065 | 1 | 1.0900e-01 | 2.8451e+05 | ; | CB | - | HB3 |
| 3063 | 3066 | 1 | 1.5260e-01 | 2.5941e+05 | ; | CB | - | CG |
| 3066 | 3067 | 1 | 1.0900e-01 | 2.8451e+05 | ; | CG | - | HG2 |
| 3066 | 3068 | 1 | 1.0900e-01 | 2.8451e+05 | ; | CG | - | HG3 |
| 3066 | 3069 | 1 | 1.5260e-01 | 2.5941e+05 | ; | CG | - | CD |
| 3069 | 3070 | 1 | 1.0900e-01 | 2.8451e+05 | ; | CD | - | HD2 |
| 3069 | 3071 | 1 | 1.0900e-01 | 2.8451e+05 | ; | CD | - | HD3 |

| | | | | | | |
|---|---|---|---|---|---|---|
| 3069 | 3072 | 1 | 1.5260e-01 | 2.5941e+05 | ; | CD - CE |
| 3072 | 3073 | 1 | 1.0900e-01 | 2.8451e+05 | ; | CE - HE2 |
| 3072 | 3074 | 1 | 1.0900e-01 | 2.8451e+05 | ; | CE - HE3 |
| 3072 | 3075 | 1 | 1.4710e-01 | 3.0711e+05 | ; | CE - NZ |
| 3075 | 3076 | 1 | 1.0100e-01 | 3.6317e+05 | ; | NZ - HZ1 |
| 3075 | 3077 | 1 | 1.0100e-01 | 3.6317e+05 | ; | NZ - HZ2 |
| 3075 | 3078 | 1 | 1.0100e-01 | 3.6317e+05 | ; | NZ - HZ3 |
| 3079 | 3080 | 1 | 1.2290e-01 | 4.7698e+05 | ; | C - O |
| 3079 | 3081 | 1 | 1.3350e-01 | 4.1003e+05 | ; | C - N |
| 3081 | 3082 | 1 | 1.0100e-01 | 3.6317e+05 | ; | N - H |
| 3081 | 3083 | 1 | 1.4490e-01 | 2.8200e+05 | ; | N - CA |
| 3083 | 3084 | 1 | 1.0900e-01 | 2.8451e+05 | ; | CA - HA |
| 3083 | 3085 | 1 | 1.5260e-01 | 2.5941e+05 | ; | CA - CB |
| 3083 | 3100 | 1 | 1.5220e-01 | 2.6527e+05 | ; | CA - C |
| 3085 | 3086 | 1 | 1.0900e-01 | 2.8451e+05 | ; | CB - HB2 |
| 3085 | 3087 | 1 | 1.0900e-01 | 2.8451e+05 | ; | CB - HB3 |
| 3085 | 3088 | 1 | 1.5260e-01 | 2.5941e+05 | ; | CB - CG |
| 3088 | 3089 | 1 | 1.0900e-01 | 2.8451e+05 | ; | CG - HG2 |
| 3088 | 3090 | 1 | 1.0900e-01 | 2.8451e+05 | ; | CG - HG3 |
| 3088 | 3091 | 1 | 1.5260e-01 | 2.5941e+05 | ; | CG - CD |
| 3091 | 3092 | 1 | 1.0900e-01 | 2.8451e+05 | ; | CD - HD2 |
| 3091 | 3093 | 1 | 1.0900e-01 | 2.8451e+05 | ; | CD - HD3 |
| 3091 | 3094 | 1 | 1.5260e-01 | 2.5941e+05 | ; | CD - CE |
| 3094 | 3095 | 1 | 1.0900e-01 | 2.8451e+05 | ; | CE - HE2 |
| 3094 | 3096 | 1 | 1.0900e-01 | 2.8451e+05 | ; | CE - HE3 |
| 3094 | 3097 | 1 | 1.4710e-01 | 3.0711e+05 | ; | CE - NZ |
| 3097 | 3098 | 1 | 1.0100e-01 | 3.6317e+05 | ; | NZ - HZ2 |
| 3097 | 3099 | 1 | 1.0100e-01 | 3.6317e+05 | ; | NZ - HZ3 |
| 3100 | 3101 | 1 | 1.2290e-01 | 4.7698e+05 | ; | C - O |
| 3100 | 3102 | 1 | 1.3350e-01 | 4.1003e+05 | ; | C - N |
| 3102 | 3103 | 1 | 1.0100e-01 | 3.6317e+05 | ; | N - H |
| 3102 | 3104 | 1 | 1.4490e-01 | 2.8200e+05 | ; | N - CA |
| 3104 | 3105 | 1 | 1.0900e-01 | 2.8451e+05 | ; | CA - HA |
| 3104 | 3106 | 1 | 1.5260e-01 | 2.5941e+05 | ; | CA - CB |
| 3104 | 3111 | 1 | 1.5220e-01 | 2.6527e+05 | ; | CA - C |
| 3106 | 3107 | 1 | 1.0900e-01 | 2.8451e+05 | ; | CB - HB2 |
| 3106 | 3108 | 1 | 1.0900e-01 | 2.8451e+05 | ; | CB - HB3 |
| 3106 | 3109 | 1 | 1.8100e-01 | 1.9832e+05 | ; | CB - SG |
| 3109 | 3110 | 1 | 1.3360e-01 | 2.2928e+05 | ; | SG - HG |
| 3111 | 3112 | 1 | 1.2290e-01 | 4.7698e+05 | ; | C - O |
| 3111 | 3113 | 1 | 1.3350e-01 | 4.1003e+05 | ; | C - N |
| 3113 | 3114 | 1 | 1.0100e-01 | 3.6317e+05 | ; | N - H |
| 3113 | 3115 | 1 | 1.4490e-01 | 2.8200e+05 | ; | N - CA |
| 3115 | 3116 | 1 | 1.0900e-01 | 2.8451e+05 | ; | CA - HA |
| 3115 | 3117 | 1 | 1.5260e-01 | 2.5941e+05 | ; | CA - CB |
| 3115 | 3123 | 1 | 1.5220e-01 | 2.6527e+05 | ; | CA - C |
| 3117 | 3118 | 1 | 1.0900e-01 | 2.8451e+05 | ; | CB - HB2 |
| 3117 | 3119 | 1 | 1.0900e-01 | 2.8451e+05 | ; | CB - HB3 |
| 3117 | 3120 | 1 | 1.5220e-01 | 2.6527e+05 | ; | CB - CG |
| 3120 | 3121 | 1 | 1.2500e-01 | 5.4894e+05 | ; | CG - OD1 |
| 3120 | 3122 | 1 | 1.2500e-01 | 5.4894e+05 | ; | CG - OD2 |
| 3123 | 3124 | 1 | 1.2290e-01 | 4.7698e+05 | ; | C - O |

| | | | | | | |
|---|---|---|---|---|---|---|
| 3123 | 3125 | 1 | 1.3350e-01 | 4.1003e+05 | ; | C - N |
| 3125 | 3126 | 1 | 1.0100e-01 | 3.6317e+05 | ; | N - H |
| 3125 | 3127 | 1 | 1.4490e-01 | 2.8200e+05 | ; | N - CA |
| 3127 | 3128 | 1 | 1.0900e-01 | 2.8451e+05 | ; | CA - HA |
| 3127 | 3129 | 1 | 1.5260e-01 | 2.5941e+05 | ; | CA - CB |
| 3127 | 3142 | 1 | 1.5220e-01 | 2.6527e+05 | ; | CA - C |
| 3129 | 3130 | 1 | 1.0900e-01 | 2.8451e+05 | ; | CB - HB |
| 3129 | 3131 | 1 | 1.5260e-01 | 2.5941e+05 | ; | CB - CG2 |
| 3129 | 3135 | 1 | 1.5260e-01 | 2.5941e+05 | ; | CB - CG1 |
| 3131 | 3132 | 1 | 1.0900e-01 | 2.8451e+05 | ; | CG2 - HG21 |
| 3131 | 3133 | 1 | 1.0900e-01 | 2.8451e+05 | ; | CG2 - HG22 |
| 3131 | 3134 | 1 | 1.0900e-01 | 2.8451e+05 | ; | CG2 - HG23 |
| 3135 | 3136 | 1 | 1.0900e-01 | 2.8451e+05 | ; | CG1 - HG12 |
| 3135 | 3137 | 1 | 1.0900e-01 | 2.8451e+05 | ; | CG1 - HG13 |
| 3135 | 3138 | 1 | 1.5260e-01 | 2.5941e+05 | ; | CG1 - CD1 |
| 3138 | 3139 | 1 | 1.0900e-01 | 2.8451e+05 | ; | CD1 - HD11 |
| 3138 | 3140 | 1 | 1.0900e-01 | 2.8451e+05 | ; | CD1 - HD12 |
| 3138 | 3141 | 1 | 1.0900e-01 | 2.8451e+05 | ; | CD1 - HD13 |
| 3142 | 3143 | 1 | 1.2290e-01 | 4.7698e+05 | ; | C - O |
| 3142 | 3144 | 1 | 1.3350e-01 | 4.1003e+05 | ; | C - N |
| 3144 | 3145 | 1 | 1.0100e-01 | 3.6317e+05 | ; | N - H |
| 3144 | 3146 | 1 | 1.4490e-01 | 2.8200e+05 | ; | N - CA |
| 3146 | 3147 | 1 | 1.0900e-01 | 2.8451e+05 | ; | CA - HA |
| 3146 | 3148 | 1 | 1.5260e-01 | 2.5941e+05 | ; | CA - CB |
| 3146 | 3166 | 1 | 1.5220e-01 | 2.6527e+05 | ; | CA - C |
| 3148 | 3149 | 1 | 1.0900e-01 | 2.8451e+05 | ; | CB - HB2 |
| 3148 | 3150 | 1 | 1.0900e-01 | 2.8451e+05 | ; | CB - HB3 |
| 3148 | 3151 | 1 | 1.4950e-01 | 2.6527e+05 | ; | CB - CG |
| 3151 | 3152 | 1 | 1.3520e-01 | 4.5689e+05 | ; | CG - CD1 |
| 3151 | 3165 | 1 | 1.4590e-01 | 3.2468e+05 | ; | CG - CD2 |
| 3152 | 3153 | 1 | 1.0800e-01 | 3.0711e+05 | ; | CD1 - HD1 |
| 3152 | 3154 | 1 | 1.3810e-01 | 3.5731e+05 | ; | CD1 - NE1 |
| 3154 | 3155 | 1 | 1.0100e-01 | 3.6317e+05 | ; | NE1 - HE1 |
| 3154 | 3156 | 1 | 1.3800e-01 | 3.5815e+05 | ; | NE1 - CE2 |
| 3156 | 3157 | 1 | 1.4000e-01 | 3.9246e+05 | ; | CE2 - CZ2 |
| 3156 | 3165 | 1 | 1.4190e-01 | 3.7405e+05 | ; | CE2 - CD2 |
| 3157 | 3158 | 1 | 1.0800e-01 | 3.0711e+05 | ; | CZ2 - HZ2 |
| 3157 | 3159 | 1 | 1.4000e-01 | 3.9246e+05 | ; | CZ2 - CH2 |
| 3159 | 3160 | 1 | 1.0800e-01 | 3.0711e+05 | ; | CH2 - HH2 |
| 3159 | 3161 | 1 | 1.4000e-01 | 3.9246e+05 | ; | CH2 - CZ3 |
| 3161 | 3162 | 1 | 1.0800e-01 | 3.0711e+05 | ; | CZ3 - HZ3 |
| 3161 | 3163 | 1 | 1.4000e-01 | 3.9246e+05 | ; | CZ3 - CE3 |
| 3163 | 3164 | 1 | 1.0800e-01 | 3.0711e+05 | ; | CE3 - HE3 |
| 3163 | 3165 | 1 | 1.4040e-01 | 3.9246e+05 | ; | CE3 - CD2 |
| 3166 | 3167 | 1 | 1.2290e-01 | 4.7698e+05 | ; | C - O |
| 3166 | 3168 | 1 | 1.3350e-01 | 4.1003e+05 | ; | C - N |
| 3168 | 3169 | 1 | 1.0100e-01 | 3.6317e+05 | ; | N - H |
| 3168 | 3170 | 1 | 1.4490e-01 | 2.8200e+05 | ; | N - CA |
| 3170 | 3171 | 1 | 1.0900e-01 | 2.8451e+05 | ; | CA - HA |
| 3170 | 3172 | 1 | 1.5260e-01 | 2.5941e+05 | ; | CA - CB |
| 3170 | 3177 | 1 | 1.5220e-01 | 2.6527e+05 | ; | CA - C |
| 3172 | 3173 | 1 | 1.0900e-01 | 2.8451e+05 | ; | CB - HB2 |

| | | | | | | |
|---|---|---|---|---|---|---|
| 3172 | 3174 | 1 | 1.0900e-01 | 2.8451e+05 | ; | CB - HB3 |
| 3172 | 3175 | 1 | 1.4100e-01 | 2.6778e+05 | ; | CB - OG |
| 3175 | 3176 | 1 | 9.6000e-02 | 4.6275e+05 | ; | OG - HG |
| 3177 | 3178 | 1 | 1.2290e-01 | 4.7698e+05 | ; | C - O |
| 3177 | 3179 | 1 | 1.3350e-01 | 4.1003e+05 | ; | C - N |
| 3179 | 3180 | 1 | 1.0100e-01 | 3.6317e+05 | ; | N - H |
| 3179 | 3181 | 1 | 1.4490e-01 | 2.8200e+05 | ; | N - CA |
| 3181 | 3182 | 1 | 1.0900e-01 | 2.8451e+05 | ; | CA - HA |
| 3181 | 3183 | 1 | 1.5260e-01 | 2.5941e+05 | ; | CA - CB |
| 3181 | 3188 | 1 | 1.5220e-01 | 2.6527e+05 | ; | CA - C |
| 3183 | 3184 | 1 | 1.0900e-01 | 2.8451e+05 | ; | CB - HB2 |
| 3183 | 3185 | 1 | 1.0900e-01 | 2.8451e+05 | ; | CB - HB3 |
| 3183 | 3186 | 1 | 1.4100e-01 | 2.6778e+05 | ; | CB - OG |
| 3186 | 3187 | 1 | 9.6000e-02 | 4.6275e+05 | ; | OG - HG |
| 3188 | 3189 | 1 | 1.2290e-01 | 4.7698e+05 | ; | C - O |
| 3188 | 3190 | 1 | 1.3350e-01 | 4.1003e+05 | ; | C - N |
| 3190 | 3191 | 1 | 1.0100e-01 | 3.6317e+05 | ; | N - H |
| 3190 | 3192 | 1 | 1.4490e-01 | 2.8200e+05 | ; | N - CA |
| 3192 | 3193 | 1 | 1.0900e-01 | 2.8451e+05 | ; | CA - HA2 |
| 3192 | 3194 | 1 | 1.0900e-01 | 2.8451e+05 | ; | CA - HA3 |
| 3192 | 3195 | 1 | 1.5220e-01 | 2.6527e+05 | ; | CA - C |
| 3195 | 3196 | 1 | 1.2290e-01 | 4.7698e+05 | ; | C - O |
| 3195 | 3197 | 1 | 1.3350e-01 | 4.1003e+05 | ; | C - N |
| 3197 | 3198 | 1 | 1.0100e-01 | 3.6317e+05 | ; | N - H |
| 3197 | 3199 | 1 | 1.4490e-01 | 2.8200e+05 | ; | N - CA |
| 3199 | 3200 | 1 | 1.0900e-01 | 2.8451e+05 | ; | CA - HA |
| 3199 | 3201 | 1 | 1.5260e-01 | 2.5941e+05 | ; | CA - CB |
| 3199 | 3211 | 1 | 1.5220e-01 | 2.6527e+05 | ; | CA - C |
| 3201 | 3202 | 1 | 1.0900e-01 | 2.8451e+05 | ; | CB - HB |
| 3201 | 3203 | 1 | 1.5260e-01 | 2.5941e+05 | ; | CB - CG1 |
| 3201 | 3207 | 1 | 1.5260e-01 | 2.5941e+05 | ; | CB - CG2 |
| 3203 | 3204 | 1 | 1.0900e-01 | 2.8451e+05 | ; | CG1 - HG11 |
| 3203 | 3205 | 1 | 1.0900e-01 | 2.8451e+05 | ; | CG1 - HG12 |
| 3203 | 3206 | 1 | 1.0900e-01 | 2.8451e+05 | ; | CG1 - HG13 |
| 3207 | 3208 | 1 | 1.0900e-01 | 2.8451e+05 | ; | CG2 - HG21 |
| 3207 | 3209 | 1 | 1.0900e-01 | 2.8451e+05 | ; | CG2 - HG22 |
| 3207 | 3210 | 1 | 1.0900e-01 | 2.8451e+05 | ; | CG2 - HG23 |
| 3211 | 3212 | 1 | 1.2290e-01 | 4.7698e+05 | ; | C - O |
| 3211 | 3213 | 1 | 1.3350e-01 | 4.1003e+05 | ; | C - N |
| 3213 | 3214 | 1 | 1.0100e-01 | 3.6317e+05 | ; | N - H |
| 3213 | 3215 | 1 | 1.4490e-01 | 2.8200e+05 | ; | N - CA |
| 3215 | 3216 | 1 | 1.0900e-01 | 2.8451e+05 | ; | CA - HA |
| 3215 | 3217 | 1 | 1.5260e-01 | 2.5941e+05 | ; | CA - CB |
| 3215 | 3230 | 1 | 1.5220e-01 | 2.6527e+05 | ; | CA - C |
| 3217 | 3218 | 1 | 1.0900e-01 | 2.8451e+05 | ; | CB - HB |
| 3217 | 3219 | 1 | 1.5260e-01 | 2.5941e+05 | ; | CB - CG2 |
| 3217 | 3223 | 1 | 1.5260e-01 | 2.5941e+05 | ; | CB - CG1 |
| 3219 | 3220 | 1 | 1.0900e-01 | 2.8451e+05 | ; | CG2 - HG21 |
| 3219 | 3221 | 1 | 1.0900e-01 | 2.8451e+05 | ; | CG2 - HG22 |
| 3219 | 3222 | 1 | 1.0900e-01 | 2.8451e+05 | ; | CG2 - HG23 |
| 3223 | 3224 | 1 | 1.0900e-01 | 2.8451e+05 | ; | CG1 - HG12 |
| 3223 | 3225 | 1 | 1.0900e-01 | 2.8451e+05 | ; | CG1 - HG13 |

| | | | | | |
|---|---|---|---|---|---|
| 3223 | 3226 | 1 | 1.5260e-01 | 2.5941e+05 ; | CG1 - CD1 |
| 3226 | 3227 | 1 | 1.0900e-01 | 2.8451e+05 ; | CD1 - HD11 |
| 3226 | 3228 | 1 | 1.0900e-01 | 2.8451e+05 ; | CD1 - HD12 |
| 3226 | 3229 | 1 | 1.0900e-01 | 2.8451e+05 ; | CD1 - HD13 |
| 3230 | 3231 | 1 | 1.2290e-01 | 4.7698e+05 ; | C - O |
| 3230 | 3232 | 1 | 1.3350e-01 | 4.1003e+05 ; | C - N |
| 3232 | 3233 | 1 | 1.0100e-01 | 3.6317e+05 ; | N - H |
| 3232 | 3234 | 1 | 1.4490e-01 | 2.8200e+05 ; | N - CA |
| 3234 | 3235 | 1 | 1.0900e-01 | 2.8451e+05 ; | CA - HA |
| 3234 | 3236 | 1 | 1.5260e-01 | 2.5941e+05 ; | CA - CB |
| 3234 | 3247 | 1 | 1.5220e-01 | 2.6527e+05 ; | CA - C |
| 3236 | 3237 | 1 | 1.0900e-01 | 2.8451e+05 ; | CB - HB2 |
| 3236 | 3238 | 1 | 1.0900e-01 | 2.8451e+05 ; | CB - HB3 |
| 3236 | 3239 | 1 | 1.5260e-01 | 2.5941e+05 ; | CB - CG |
| 3239 | 3240 | 1 | 1.0900e-01 | 2.8451e+05 ; | CG - HG2 |
| 3239 | 3241 | 1 | 1.0900e-01 | 2.8451e+05 ; | CG - HG3 |
| 3239 | 3242 | 1 | 1.8100e-01 | 1.8995e+05 ; | CG - SD |
| 3242 | 3243 | 1 | 1.8100e-01 | 1.8995e+05 ; | SD - CE |
| 3243 | 3244 | 1 | 1.0900e-01 | 2.8451e+05 ; | CE - HE1 |
| 3243 | 3245 | 1 | 1.0900e-01 | 2.8451e+05 ; | CE - HE2 |
| 3243 | 3246 | 1 | 1.0900e-01 | 2.8451e+05 ; | CE - HE3 |
| 3247 | 3248 | 1 | 1.2290e-01 | 4.7698e+05 ; | C - O |
| 3247 | 3249 | 1 | 1.3350e-01 | 4.1003e+05 ; | C - N |
| 3249 | 3250 | 1 | 1.0100e-01 | 3.6317e+05 ; | N - H |
| 3249 | 3251 | 1 | 1.4490e-01 | 2.8200e+05 ; | N - CA |
| 3251 | 3252 | 1 | 1.0900e-01 | 2.8451e+05 ; | CA - HA |
| 3251 | 3253 | 1 | 1.5260e-01 | 2.5941e+05 ; | CA - CB |
| 3251 | 3268 | 1 | 1.5220e-01 | 2.6527e+05 ; | CA - C |
| 3253 | 3254 | 1 | 1.0900e-01 | 2.8451e+05 ; | CB - HB2 |
| 3253 | 3255 | 1 | 1.0900e-01 | 2.8451e+05 ; | CB - HB3 |
| 3253 | 3256 | 1 | 1.5100e-01 | 2.6527e+05 ; | CB - CG |
| 3256 | 3257 | 1 | 1.4000e-01 | 3.9246e+05 ; | CG - CD1 |
| 3256 | 3266 | 1 | 1.4000e-01 | 3.9246e+05 ; | CG - CD2 |
| 3257 | 3258 | 1 | 1.0800e-01 | 3.0711e+05 ; | CD1 - HD1 |
| 3257 | 3259 | 1 | 1.4000e-01 | 3.9246e+05 ; | CD1 - CE1 |
| 3259 | 3260 | 1 | 1.0800e-01 | 3.0711e+05 ; | CE1 - HE1 |
| 3259 | 3261 | 1 | 1.4090e-01 | 3.9246e+05 ; | CE1 - CZ |
| 3261 | 3262 | 1 | 1.3640e-01 | 3.7656e+05 ; | CZ - OH |
| 3261 | 3264 | 1 | 1.4090e-01 | 3.9246e+05 ; | CZ - CE2 |
| 3262 | 3263 | 1 | 9.6000e-02 | 4.6275e+05 ; | OH - HH |
| 3264 | 3265 | 1 | 1.0800e-01 | 3.0711e+05 ; | CE2 - HE2 |
| 3264 | 3266 | 1 | 1.4000e-01 | 3.9246e+05 ; | CE2 - CD2 |
| 3266 | 3267 | 1 | 1.0800e-01 | 3.0711e+05 ; | CD2 - HD2 |
| 3268 | 3269 | 1 | 1.2290e-01 | 4.7698e+05 ; | C - O |
| 3268 | 3270 | 1 | 1.3350e-01 | 4.1003e+05 ; | C - N |
| 3270 | 3271 | 1 | 1.0100e-01 | 3.6317e+05 ; | N - H |
| 3270 | 3272 | 1 | 1.4490e-01 | 2.8200e+05 ; | N - CA |
| 3272 | 3273 | 1 | 1.0900e-01 | 2.8451e+05 ; | CA - HA |
| 3272 | 3274 | 1 | 1.5260e-01 | 2.5941e+05 ; | CA - CB |
| 3272 | 3282 | 1 | 1.5220e-01 | 2.6527e+05 ; | CA - C |
| 3274 | 3275 | 1 | 1.0900e-01 | 2.8451e+05 ; | CB - HB |
| 3274 | 3276 | 1 | 1.5260e-01 | 2.5941e+05 ; | CB - CG2 |

| | | | | | | |
|---|---|---|---|---|---|---|
| 3274 | 3280 | 1 | 1.4100e-01 | 2.6778e+05 | ; | CB - OG1 |
| 3276 | 3277 | 1 | 1.0900e-01 | 2.8451e+05 | ; | CG2 - HG21 |
| 3276 | 3278 | 1 | 1.0900e-01 | 2.8451e+05 | ; | CG2 - HG22 |
| 3276 | 3279 | 1 | 1.0900e-01 | 2.8451e+05 | ; | CG2 - HG23 |
| 3280 | 3281 | 1 | 9.6000e-02 | 4.6275e+05 | ; | OG1 - HG1 |
| 3282 | 3283 | 1 | 1.2290e-01 | 4.7698e+05 | ; | C - O |
| 3282 | 3284 | 1 | 1.3350e-01 | 4.1003e+05 | ; | C - N |
| 3284 | 3285 | 1 | 1.0100e-01 | 3.6317e+05 | ; | N - H |
| 3284 | 3286 | 1 | 1.4490e-01 | 2.8200e+05 | ; | N - CA |
| 3286 | 3287 | 1 | 1.0900e-01 | 2.8451e+05 | ; | CA - HA |
| 3286 | 3288 | 1 | 1.5260e-01 | 2.5941e+05 | ; | CA - CB |
| 3286 | 3301 | 1 | 1.5220e-01 | 2.6527e+05 | ; | CA - C |
| 3288 | 3289 | 1 | 1.0900e-01 | 2.8451e+05 | ; | CB - HB2 |
| 3288 | 3290 | 1 | 1.0900e-01 | 2.8451e+05 | ; | CB - HB3 |
| 3288 | 3291 | 1 | 1.5260e-01 | 2.5941e+05 | ; | CB - CG |
| 3291 | 3292 | 1 | 1.0900e-01 | 2.8451e+05 | ; | CG - HG |
| 3291 | 3293 | 1 | 1.5260e-01 | 2.5941e+05 | ; | CG - CD1 |
| 3291 | 3297 | 1 | 1.5260e-01 | 2.5941e+05 | ; | CG - CD2 |
| 3293 | 3294 | 1 | 1.0900e-01 | 2.8451e+05 | ; | CD1 - HD11 |
| 3293 | 3295 | 1 | 1.0900e-01 | 2.8451e+05 | ; | CD1 - HD12 |
| 3293 | 3296 | 1 | 1.0900e-01 | 2.8451e+05 | ; | CD1 - HD13 |
| 3297 | 3298 | 1 | 1.0900e-01 | 2.8451e+05 | ; | CD2 - HD21 |
| 3297 | 3299 | 1 | 1.0900e-01 | 2.8451e+05 | ; | CD2 - HD22 |
| 3297 | 3300 | 1 | 1.0900e-01 | 2.8451e+05 | ; | CD2 - HD23 |
| 3301 | 3302 | 1 | 1.2290e-01 | 4.7698e+05 | ; | C - O |
| 3301 | 3303 | 1 | 1.3350e-01 | 4.1003e+05 | ; | C - N |
| 3303 | 3304 | 1 | 1.0100e-01 | 3.6317e+05 | ; | N - H |
| 3303 | 3305 | 1 | 1.4490e-01 | 2.8200e+05 | ; | N - CA |
| 3305 | 3306 | 1 | 1.0900e-01 | 2.8451e+05 | ; | CA - HA |
| 3305 | 3307 | 1 | 1.5260e-01 | 2.5941e+05 | ; | CA - CB |
| 3305 | 3320 | 1 | 1.5220e-01 | 2.6527e+05 | ; | CA - C |
| 3307 | 3308 | 1 | 1.0900e-01 | 2.8451e+05 | ; | CB - HB2 |
| 3307 | 3309 | 1 | 1.0900e-01 | 2.8451e+05 | ; | CB - HB3 |
| 3307 | 3310 | 1 | 1.5260e-01 | 2.5941e+05 | ; | CB - CG |
| 3310 | 3311 | 1 | 1.0900e-01 | 2.8451e+05 | ; | CG - HG |
| 3310 | 3312 | 1 | 1.5260e-01 | 2.5941e+05 | ; | CG - CD1 |
| 3310 | 3316 | 1 | 1.5260e-01 | 2.5941e+05 | ; | CG - CD2 |
| 3312 | 3313 | 1 | 1.0900e-01 | 2.8451e+05 | ; | CD1 - HD11 |
| 3312 | 3314 | 1 | 1.0900e-01 | 2.8451e+05 | ; | CD1 - HD12 |
| 3312 | 3315 | 1 | 1.0900e-01 | 2.8451e+05 | ; | CD1 - HD13 |
| 3316 | 3317 | 1 | 1.0900e-01 | 2.8451e+05 | ; | CD2 - HD21 |
| 3316 | 3318 | 1 | 1.0900e-01 | 2.8451e+05 | ; | CD2 - HD22 |
| 3316 | 3319 | 1 | 1.0900e-01 | 2.8451e+05 | ; | CD2 - HD23 |
| 3320 | 3321 | 1 | 1.2290e-01 | 4.7698e+05 | ; | C - O |
| 3320 | 3322 | 1 | 1.3350e-01 | 4.1003e+05 | ; | C - N |
| 3322 | 3323 | 1 | 1.0100e-01 | 3.6317e+05 | ; | N - H |
| 3322 | 3324 | 1 | 1.4490e-01 | 2.8200e+05 | ; | N - CA |
| 3324 | 3325 | 1 | 1.0900e-01 | 2.8451e+05 | ; | CA - HA |
| 3324 | 3326 | 1 | 1.5260e-01 | 2.5941e+05 | ; | CA - CB |
| 3324 | 3331 | 1 | 1.5220e-01 | 2.6527e+05 | ; | CA - C |
| 3326 | 3327 | 1 | 1.0900e-01 | 2.8451e+05 | ; | CB - HB2 |
| 3326 | 3328 | 1 | 1.0900e-01 | 2.8451e+05 | ; | CB - HB3 |

| | | | | | | |
|---|---|---|---|---|---|---|
| 3326 | 3329 | 1 | 1.8100e-01 | 1.9832e+05 | ; | CB - SG |
| 3329 | 3330 | 1 | 1.3360e-01 | 2.2928e+05 | ; | SG - HG |
| 3331 | 3332 | 1 | 1.2290e-01 | 4.7698e+05 | ; | C - O |
| 3331 | 3333 | 1 | 1.3350e-01 | 4.1003e+05 | ; | C - N |
| 3333 | 3334 | 1 | 1.0100e-01 | 3.6317e+05 | ; | N - H |
| 3333 | 3335 | 1 | 1.4490e-01 | 2.8200e+05 | ; | N - CA |
| 3335 | 3336 | 1 | 1.0900e-01 | 2.8451e+05 | ; | CA - HA2 |
| 3335 | 3337 | 1 | 1.0900e-01 | 2.8451e+05 | ; | CA - HA3 |
| 3335 | 3338 | 1 | 1.5220e-01 | 2.6527e+05 | ; | CA - C |
| 3338 | 3339 | 1 | 1.2290e-01 | 4.7698e+05 | ; | C - O |
| 3338 | 3340 | 1 | 1.3350e-01 | 4.1003e+05 | ; | C - N |
| 3340 | 3341 | 1 | 1.0100e-01 | 3.6317e+05 | ; | N - H |
| 3340 | 3342 | 1 | 1.4490e-01 | 2.8200e+05 | ; | N - CA |
| 3342 | 3343 | 1 | 1.0900e-01 | 2.8451e+05 | ; | CA - HA |
| 3342 | 3344 | 1 | 1.5260e-01 | 2.5941e+05 | ; | CA - CB |
| 3342 | 3359 | 1 | 1.5220e-01 | 2.6527e+05 | ; | CA - C |
| 3344 | 3345 | 1 | 1.0900e-01 | 2.8451e+05 | ; | CB - HB2 |
| 3344 | 3346 | 1 | 1.0900e-01 | 2.8451e+05 | ; | CB - HB3 |
| 3344 | 3347 | 1 | 1.5100e-01 | 2.6527e+05 | ; | CB - CG |
| 3347 | 3348 | 1 | 1.4000e-01 | 3.9246e+05 | ; | CG - CD1 |
| 3347 | 3357 | 1 | 1.4000e-01 | 3.9246e+05 | ; | CG - CD2 |
| 3348 | 3349 | 1 | 1.0800e-01 | 3.0711e+05 | ; | CD1 - HD1 |
| 3348 | 3350 | 1 | 1.4000e-01 | 3.9246e+05 | ; | CD1 - CE1 |
| 3350 | 3351 | 1 | 1.0800e-01 | 3.0711e+05 | ; | CE1 - HE1 |
| 3350 | 3352 | 1 | 1.4090e-01 | 3.9246e+05 | ; | CE1 - CZ |
| 3352 | 3353 | 1 | 1.3640e-01 | 3.7656e+05 | ; | CZ - OH |
| 3352 | 3355 | 1 | 1.4090e-01 | 3.9246e+05 | ; | CZ - CE2 |
| 3353 | 3354 | 1 | 9.6000e-02 | 4.6275e+05 | ; | OH - HH |
| 3355 | 3356 | 1 | 1.0800e-01 | 3.0711e+05 | ; | CE2 - HE2 |
| 3355 | 3357 | 1 | 1.4000e-01 | 3.9246e+05 | ; | CE2 - CD2 |
| 3357 | 3358 | 1 | 1.0800e-01 | 3.0711e+05 | ; | CD2 - HD2 |
| 3359 | 3360 | 1 | 1.2290e-01 | 4.7698e+05 | ; | C - O |
| 3359 | 3361 | 1 | 1.3350e-01 | 4.1003e+05 | ; | C - N |
| 3361 | 3362 | 1 | 1.4490e-01 | 2.8200e+05 | ; | N - CD |
| 3361 | 3371 | 1 | 1.4490e-01 | 2.8200e+05 | ; | N - CA |
| 3362 | 3363 | 1 | 1.0900e-01 | 2.8451e+05 | ; | CD - HD2 |
| 3362 | 3364 | 1 | 1.0900e-01 | 2.8451e+05 | ; | CD - HD3 |
| 3362 | 3365 | 1 | 1.5260e-01 | 2.5941e+05 | ; | CD - CG |
| 3365 | 3366 | 1 | 1.0900e-01 | 2.8451e+05 | ; | CG - HG2 |
| 3365 | 3367 | 1 | 1.0900e-01 | 2.8451e+05 | ; | CG - HG3 |
| 3365 | 3368 | 1 | 1.5260e-01 | 2.5941e+05 | ; | CG - CB |
| 3368 | 3369 | 1 | 1.0900e-01 | 2.8451e+05 | ; | CB - HB2 |
| 3368 | 3370 | 1 | 1.0900e-01 | 2.8451e+05 | ; | CB - HB3 |
| 3368 | 3371 | 1 | 1.5260e-01 | 2.5941e+05 | ; | CB - CA |
| 3371 | 3372 | 1 | 1.0900e-01 | 2.8451e+05 | ; | CA - HA |
| 3371 | 3373 | 1 | 1.5220e-01 | 2.6527e+05 | ; | CA - C |
| 3373 | 3374 | 1 | 1.2290e-01 | 4.7698e+05 | ; | C - O |
| 3373 | 3375 | 1 | 1.3350e-01 | 4.1003e+05 | ; | C - N |
| 3375 | 3376 | 1 | 1.4490e-01 | 2.8200e+05 | ; | N - CD |
| 3375 | 3385 | 1 | 1.4490e-01 | 2.8200e+05 | ; | N - CA |
| 3376 | 3377 | 1 | 1.0900e-01 | 2.8451e+05 | ; | CD - HD2 |
| 3376 | 3378 | 1 | 1.0900e-01 | 2.8451e+05 | ; | CD - HD3 |

| | | | | | | | | |
|---|---|---|---|---|---|---|---|---|
| 3376 | 3379 | 1 | 1.5260e-01 | 2.5941e+05 | ; | CD | - | CG |
| 3379 | 3380 | 1 | 1.0900e-01 | 2.8451e+05 | ; | CG | - | HG2 |
| 3379 | 3381 | 1 | 1.0900e-01 | 2.8451e+05 | ; | CG | - | HG3 |
| 3379 | 3382 | 1 | 1.5260e-01 | 2.5941e+05 | ; | CG | - | CB |
| 3382 | 3383 | 1 | 1.0900e-01 | 2.8451e+05 | ; | CB | - | HB2 |
| 3382 | 3384 | 1 | 1.0900e-01 | 2.8451e+05 | ; | CB | - | HB3 |
| 3382 | 3385 | 1 | 1.5260e-01 | 2.5941e+05 | ; | CB | - | CA |
| 3385 | 3386 | 1 | 1.0900e-01 | 2.8451e+05 | ; | CA | - | HA |
| 3385 | 3387 | 1 | 1.5220e-01 | 2.6527e+05 | ; | CA | - | C |
| 3387 | 3388 | 1 | 1.2290e-01 | 4.7698e+05 | ; | C | - | O |
| 3387 | 3389 | 1 | 1.3350e-01 | 4.1003e+05 | ; | C | - | N |
| 3389 | 3390 | 1 | 1.0100e-01 | 3.6317e+05 | ; | N | - | H |
| 3389 | 3391 | 1 | 1.4490e-01 | 2.8200e+05 | ; | N | - | CA |
| 3391 | 3392 | 1 | 1.0900e-01 | 2.8451e+05 | ; | CA | - | HA |
| 3391 | 3393 | 1 | 1.5260e-01 | 2.5941e+05 | ; | CA | - | CB |
| 3391 | 3407 | 1 | 1.5220e-01 | 2.6527e+05 | ; | CA | - | C |
| 3393 | 3394 | 1 | 1.0900e-01 | 2.8451e+05 | ; | CB | - | HB2 |
| 3393 | 3395 | 1 | 1.0900e-01 | 2.8451e+05 | ; | CB | - | HB3 |
| 3393 | 3396 | 1 | 1.5100e-01 | 2.6527e+05 | ; | CB | - | CG |
| 3396 | 3397 | 1 | 1.4000e-01 | 3.9246e+05 | ; | CG | - | CD1 |
| 3396 | 3405 | 1 | 1.4000e-01 | 3.9246e+05 | ; | CG | - | CD2 |
| 3397 | 3398 | 1 | 1.0800e-01 | 3.0711e+05 | ; | CD1 | - | HD1 |
| 3397 | 3399 | 1 | 1.4000e-01 | 3.9246e+05 | ; | CD1 | - | CE1 |
| 3399 | 3400 | 1 | 1.0800e-01 | 3.0711e+05 | ; | CE1 | - | HE1 |
| 3399 | 3401 | 1 | 1.4000e-01 | 3.9246e+05 | ; | CE1 | - | CZ |
| 3401 | 3402 | 1 | 1.0800e-01 | 3.0711e+05 | ; | CZ | - | HZ |
| 3401 | 3403 | 1 | 1.4000e-01 | 3.9246e+05 | ; | CZ | - | CE2 |
| 3403 | 3404 | 1 | 1.0800e-01 | 3.0711e+05 | ; | CE2 | - | HE2 |
| 3403 | 3405 | 1 | 1.4000e-01 | 3.9246e+05 | ; | CE2 | - | CD2 |
| 3405 | 3406 | 1 | 1.0800e-01 | 3.0711e+05 | ; | CD2 | - | HD2 |
| 3407 | 3408 | 1 | 1.2290e-01 | 4.7698e+05 | ; | C | - | O |
| 3407 | 3409 | 1 | 1.3350e-01 | 4.1003e+05 | ; | C | - | N |
| 3409 | 3410 | 1 | 1.0100e-01 | 3.6317e+05 | ; | N | - | H |
| 3409 | 3411 | 1 | 1.4490e-01 | 2.8200e+05 | ; | N | - | CA |
| 3411 | 3412 | 1 | 1.0900e-01 | 2.8451e+05 | ; | CA | - | HA |
| 3411 | 3413 | 1 | 1.5260e-01 | 2.5941e+05 | ; | CA | - | CB |
| 3411 | 3428 | 1 | 1.5220e-01 | 2.6527e+05 | ; | CA | - | C |
| 3413 | 3414 | 1 | 1.0900e-01 | 2.8451e+05 | ; | CB | - | HB2 |
| 3413 | 3415 | 1 | 1.0900e-01 | 2.8451e+05 | ; | CB | - | HB3 |
| 3413 | 3416 | 1 | 1.5100e-01 | 2.6527e+05 | ; | CB | - | CG |
| 3416 | 3417 | 1 | 1.4000e-01 | 3.9246e+05 | ; | CG | - | CD1 |
| 3416 | 3426 | 1 | 1.4000e-01 | 3.9246e+05 | ; | CG | - | CD2 |
| 3417 | 3418 | 1 | 1.0800e-01 | 3.0711e+05 | ; | CD1 | - | HD1 |
| 3417 | 3419 | 1 | 1.4000e-01 | 3.9246e+05 | ; | CD1 | - | CE1 |
| 3419 | 3420 | 1 | 1.0800e-01 | 3.0711e+05 | ; | CE1 | - | HE1 |
| 3419 | 3421 | 1 | 1.4090e-01 | 3.9246e+05 | ; | CE1 | - | CZ |
| 3421 | 3422 | 1 | 1.3640e-01 | 3.7656e+05 | ; | CZ | - | OH |
| 3421 | 3424 | 1 | 1.4090e-01 | 3.9246e+05 | ; | CZ | - | CE2 |
| 3422 | 3423 | 1 | 9.6000e-02 | 4.6275e+05 | ; | OH | - | HH |
| 3424 | 3425 | 1 | 1.0800e-01 | 3.0711e+05 | ; | CE2 | - | HE2 |
| 3424 | 3426 | 1 | 1.4000e-01 | 3.9246e+05 | ; | CE2 | - | CD2 |
| 3426 | 3427 | 1 | 1.0800e-01 | 3.0711e+05 | ; | CD2 | - | HD2 |

| | | | | | | |
|---|---|---|---|---|---|---|
| 3428 | 3429 | 1 | 1.2290e-01 | 4.7698e+05 | ; | C - O |
| 3428 | 3430 | 1 | 1.3350e-01 | 4.1003e+05 | ; | C - N |
| 3430 | 3431 | 1 | 1.0100e-01 | 3.6317e+05 | ; | N - H |
| 3430 | 3432 | 1 | 1.4490e-01 | 2.8200e+05 | ; | N - CA |
| 3432 | 3433 | 1 | 1.0900e-01 | 2.8451e+05 | ; | CA - HA2 |
| 3432 | 3434 | 1 | 1.0900e-01 | 2.8451e+05 | ; | CA - HA3 |
| 3432 | 3435 | 1 | 1.5220e-01 | 2.6527e+05 | ; | CA - C |
| 3435 | 3436 | 1 | 1.2290e-01 | 4.7698e+05 | ; | C - O |
| 3435 | 3437 | 1 | 1.3350e-01 | 4.1003e+05 | ; | C - N |
| 3437 | 3438 | 1 | 1.0100e-01 | 3.6317e+05 | ; | N - H |
| 3437 | 3439 | 1 | 1.4490e-01 | 2.8200e+05 | ; | N - CA |
| 3439 | 3440 | 1 | 1.0900e-01 | 2.8451e+05 | ; | CA - HA |
| 3439 | 3441 | 1 | 1.5260e-01 | 2.5941e+05 | ; | CA - CB |
| 3439 | 3447 | 1 | 1.5220e-01 | 2.6527e+05 | ; | CA - C |
| 3441 | 3442 | 1 | 1.0900e-01 | 2.8451e+05 | ; | CB - HB2 |
| 3441 | 3443 | 1 | 1.0900e-01 | 2.8451e+05 | ; | CB - HB3 |
| 3441 | 3444 | 1 | 1.5220e-01 | 2.6527e+05 | ; | CB - CG |
| 3444 | 3445 | 1 | 1.2500e-01 | 5.4894e+05 | ; | CG - OD1 |
| 3444 | 3446 | 1 | 1.2500e-01 | 5.4894e+05 | ; | CG - OD2 |
| 3447 | 3448 | 1 | 1.2290e-01 | 4.7698e+05 | ; | C - O |
| 3447 | 3449 | 1 | 1.3350e-01 | 4.1003e+05 | ; | C - N |
| 3449 | 3450 | 1 | 1.0100e-01 | 3.6317e+05 | ; | N - H |
| 3449 | 3451 | 1 | 1.4490e-01 | 2.8200e+05 | ; | N - CA |
| 3451 | 3452 | 1 | 1.0900e-01 | 2.8451e+05 | ; | CA - HA |
| 3451 | 3453 | 1 | 1.5260e-01 | 2.5941e+05 | ; | CA - CB |
| 3451 | 3461 | 1 | 1.5220e-01 | 2.6527e+05 | ; | CA - C |
| 3453 | 3454 | 1 | 1.0900e-01 | 2.8451e+05 | ; | CB - HB |
| 3453 | 3455 | 1 | 1.5260e-01 | 2.5941e+05 | ; | CB - CG2 |
| 3453 | 3459 | 1 | 1.4100e-01 | 2.6778e+05 | ; | CB - OG1 |
| 3455 | 3456 | 1 | 1.0900e-01 | 2.8451e+05 | ; | CG2 - HG21 |
| 3455 | 3457 | 1 | 1.0900e-01 | 2.8451e+05 | ; | CG2 - HG22 |
| 3455 | 3458 | 1 | 1.0900e-01 | 2.8451e+05 | ; | CG2 - HG23 |
| 3459 | 3460 | 1 | 9.6000e-02 | 4.6275e+05 | ; | OG1 - HG1 |
| 3461 | 3462 | 1 | 1.2290e-01 | 4.7698e+05 | ; | C - O |
| 3461 | 3463 | 1 | 1.3350e-01 | 4.1003e+05 | ; | C - N |
| 3463 | 3464 | 1 | 1.0100e-01 | 3.6317e+05 | ; | N - H |
| 3463 | 3465 | 1 | 1.4490e-01 | 2.8200e+05 | ; | N - CA |
| 3465 | 3466 | 1 | 1.0900e-01 | 2.8451e+05 | ; | CA - HA |
| 3465 | 3467 | 1 | 1.5260e-01 | 2.5941e+05 | ; | CA - CB |
| 3465 | 3473 | 1 | 1.5220e-01 | 2.6527e+05 | ; | CA - C |
| 3467 | 3468 | 1 | 1.0900e-01 | 2.8451e+05 | ; | CB - HB2 |
| 3467 | 3469 | 1 | 1.0900e-01 | 2.8451e+05 | ; | CB - HB3 |
| 3467 | 3470 | 1 | 1.5220e-01 | 2.6527e+05 | ; | CB - CG |
| 3470 | 3471 | 1 | 1.2500e-01 | 5.4894e+05 | ; | CG - OD1 |
| 3470 | 3472 | 1 | 1.2500e-01 | 5.4894e+05 | ; | CG - OD2 |
| 3473 | 3474 | 1 | 1.2290e-01 | 4.7698e+05 | ; | C - O |
| 3473 | 3475 | 1 | 1.3350e-01 | 4.1003e+05 | ; | C - N |
| 3475 | 3476 | 1 | 1.0100e-01 | 3.6317e+05 | ; | N - H |
| 3475 | 3477 | 1 | 1.4490e-01 | 2.8200e+05 | ; | N - CA |
| 3477 | 3478 | 1 | 1.0900e-01 | 2.8451e+05 | ; | CA - HA |
| 3477 | 3479 | 1 | 1.5260e-01 | 2.5941e+05 | ; | CA - CB |
| 3477 | 3487 | 1 | 1.5220e-01 | 2.6527e+05 | ; | CA - C |

| | | | | | |
|---|---|---|---|---|---|
| 3479 | 3480 | 1 | 1.0900e-01 | 2.8451e+05 ; | CB - HB2 |
| 3479 | 3481 | 1 | 1.0900e-01 | 2.8451e+05 ; | CB - HB3 |
| 3479 | 3482 | 1 | 1.5220e-01 | 2.6527e+05 ; | CB - CG |
| 3482 | 3483 | 1 | 1.2290e-01 | 4.7698e+05 ; | CG - OD1 |
| 3482 | 3484 | 1 | 1.3350e-01 | 4.1003e+05 ; | CG - ND2 |
| 3484 | 3485 | 1 | 1.0100e-01 | 3.6317e+05 ; | ND2 - HD21 |
| 3484 | 3486 | 1 | 1.0100e-01 | 3.6317e+05 ; | ND2 - HD22 |
| 3487 | 3488 | 1 | 1.2290e-01 | 4.7698e+05 ; | C - O |
| 3487 | 3489 | 1 | 1.3350e-01 | 4.1003e+05 ; | C - N |
| 3489 | 3490 | 1 | 1.0100e-01 | 3.6317e+05 ; | N - H |
| 3489 | 3491 | 1 | 1.4490e-01 | 2.8200e+05 ; | N - CA |
| 3491 | 3492 | 1 | 1.0900e-01 | 2.8451e+05 ; | CA - HA |
| 3491 | 3493 | 1 | 1.5260e-01 | 2.5941e+05 ; | CA - CB |
| 3491 | 3502 | 1 | 1.5220e-01 | 2.6527e+05 ; | CA - C |
| 3493 | 3494 | 1 | 1.0900e-01 | 2.8451e+05 ; | CB - HB2 |
| 3493 | 3495 | 1 | 1.0900e-01 | 2.8451e+05 ; | CB - HB3 |
| 3493 | 3496 | 1 | 1.5260e-01 | 2.5941e+05 ; | CB - CG |
| 3496 | 3497 | 1 | 1.0900e-01 | 2.8451e+05 ; | CG - HG2 |
| 3496 | 3498 | 1 | 1.0900e-01 | 2.8451e+05 ; | CG - HG3 |
| 3496 | 3499 | 1 | 1.5220e-01 | 2.6527e+05 ; | CG - CD |
| 3499 | 3500 | 1 | 1.2500e-01 | 5.4894e+05 ; | CD - OE1 |
| 3499 | 3501 | 1 | 1.2500e-01 | 5.4894e+05 ; | CD - OE2 |
| 3502 | 3503 | 1 | 1.2290e-01 | 4.7698e+05 ; | C - O |
| 3502 | 3504 | 1 | 1.3350e-01 | 4.1003e+05 ; | C - N |
| 3504 | 3505 | 1 | 1.0100e-01 | 3.6317e+05 ; | N - H |
| 3504 | 3506 | 1 | 1.4490e-01 | 2.8200e+05 ; | N - CA |
| 3506 | 3507 | 1 | 1.0900e-01 | 2.8451e+05 ; | CA - HA |
| 3506 | 3508 | 1 | 1.5260e-01 | 2.5941e+05 ; | CA - CB |
| 3506 | 3518 | 1 | 1.5220e-01 | 2.6527e+05 ; | CA - C |
| 3508 | 3509 | 1 | 1.0900e-01 | 2.8451e+05 ; | CB - HB |
| 3508 | 3510 | 1 | 1.5260e-01 | 2.5941e+05 ; | CB - CG1 |
| 3508 | 3514 | 1 | 1.5260e-01 | 2.5941e+05 ; | CB - CG2 |
| 3510 | 3511 | 1 | 1.0900e-01 | 2.8451e+05 ; | CG1 - HG11 |
| 3510 | 3512 | 1 | 1.0900e-01 | 2.8451e+05 ; | CG1 - HG12 |
| 3510 | 3513 | 1 | 1.0900e-01 | 2.8451e+05 ; | CG1 - HG13 |
| 3514 | 3515 | 1 | 1.0900e-01 | 2.8451e+05 ; | CG2 - HG21 |
| 3514 | 3516 | 1 | 1.0900e-01 | 2.8451e+05 ; | CG2 - HG22 |
| 3514 | 3517 | 1 | 1.0900e-01 | 2.8451e+05 ; | CG2 - HG23 |
| 3518 | 3519 | 1 | 1.2290e-01 | 4.7698e+05 ; | C - O |
| 3518 | 3520 | 1 | 1.3350e-01 | 4.1003e+05 ; | C - N |
| 3520 | 3521 | 1 | 1.0100e-01 | 3.6317e+05 ; | N - H |
| 3520 | 3522 | 1 | 1.4490e-01 | 2.8200e+05 ; | N - CA |
| 3522 | 3523 | 1 | 1.0900e-01 | 2.8451e+05 ; | CA - HA |
| 3522 | 3524 | 1 | 1.5260e-01 | 2.5941e+05 ; | CA - CB |
| 3522 | 3537 | 1 | 1.5220e-01 | 2.6527e+05 ; | CA - C |
| 3524 | 3525 | 1 | 1.0900e-01 | 2.8451e+05 ; | CB - HB2 |
| 3524 | 3526 | 1 | 1.0900e-01 | 2.8451e+05 ; | CB - HB3 |
| 3524 | 3527 | 1 | 1.5260e-01 | 2.5941e+05 ; | CB - CG |
| 3527 | 3528 | 1 | 1.0900e-01 | 2.8451e+05 ; | CG - HG |
| 3527 | 3529 | 1 | 1.5260e-01 | 2.5941e+05 ; | CG - CD1 |
| 3527 | 3533 | 1 | 1.5260e-01 | 2.5941e+05 ; | CG - CD2 |
| 3529 | 3530 | 1 | 1.0900e-01 | 2.8451e+05 ; | CD1 - HD11 |

| | | | | | | | | |
|---|---|---|---|---|---|---|---|---|
| 3529 | 3531 | 1 | 1.0900e-01 | 2.8451e+05 | ; | CD1 | - | HD12 |
| 3529 | 3532 | 1 | 1.0900e-01 | 2.8451e+05 | ; | CD1 | - | HD13 |
| 3533 | 3534 | 1 | 1.0900e-01 | 2.8451e+05 | ; | CD2 | - | HD21 |
| 3533 | 3535 | 1 | 1.0900e-01 | 2.8451e+05 | ; | CD2 | - | HD22 |
| 3533 | 3536 | 1 | 1.0900e-01 | 2.8451e+05 | ; | CD2 | - | HD23 |
| 3537 | 3538 | 1 | 1.2290e-01 | 4.7698e+05 | ; | C | - | O |
| 3537 | 3539 | 1 | 1.3350e-01 | 4.1003e+05 | ; | C | - | N |
| 3539 | 3540 | 1 | 1.0100e-01 | 3.6317e+05 | ; | N | - | H |
| 3539 | 3541 | 1 | 1.4490e-01 | 2.8200e+05 | ; | N | - | CA |
| 3541 | 3542 | 1 | 1.0900e-01 | 2.8451e+05 | ; | CA | - | HA |
| 3541 | 3543 | 1 | 1.5260e-01 | 2.5941e+05 | ; | CA | - | CB |
| 3541 | 3559 | 1 | 1.5220e-01 | 2.6527e+05 | ; | CA | - | C |
| 3543 | 3544 | 1 | 1.0900e-01 | 2.8451e+05 | ; | CB | - | HB2 |
| 3543 | 3545 | 1 | 1.0900e-01 | 2.8451e+05 | ; | CB | - | HB3 |
| 3543 | 3546 | 1 | 1.5260e-01 | 2.5941e+05 | ; | CB | - | CG |
| 3546 | 3547 | 1 | 1.0900e-01 | 2.8451e+05 | ; | CG | - | HG2 |
| 3546 | 3548 | 1 | 1.0900e-01 | 2.8451e+05 | ; | CG | - | HG3 |
| 3546 | 3549 | 1 | 1.5260e-01 | 2.5941e+05 | ; | CG | - | CD |
| 3549 | 3550 | 1 | 1.0900e-01 | 2.8451e+05 | ; | CD | - | HD2 |
| 3549 | 3551 | 1 | 1.0900e-01 | 2.8451e+05 | ; | CD | - | HD3 |
| 3549 | 3552 | 1 | 1.5260e-01 | 2.5941e+05 | ; | CD | - | CE |
| 3552 | 3553 | 1 | 1.0900e-01 | 2.8451e+05 | ; | CE | - | HE2 |
| 3552 | 3554 | 1 | 1.0900e-01 | 2.8451e+05 | ; | CE | - | HE3 |
| 3552 | 3555 | 1 | 1.4710e-01 | 3.0711e+05 | ; | CE | - | NZ |
| 3555 | 3556 | 1 | 1.0100e-01 | 3.6317e+05 | ; | NZ | - | HZ1 |
| 3555 | 3557 | 1 | 1.0100e-01 | 3.6317e+05 | ; | NZ | - | HZ2 |
| 3555 | 3558 | 1 | 1.0100e-01 | 3.6317e+05 | ; | NZ | - | HZ3 |
| 3559 | 3560 | 1 | 1.2290e-01 | 4.7698e+05 | ; | C | - | O |
| 3559 | 3561 | 1 | 1.3350e-01 | 4.1003e+05 | ; | C | - | N |
| 3561 | 3562 | 1 | 1.0100e-01 | 3.6317e+05 | ; | N | - | H |
| 3561 | 3563 | 1 | 1.4490e-01 | 2.8200e+05 | ; | N | - | CA |
| 3563 | 3564 | 1 | 1.0900e-01 | 2.8451e+05 | ; | CA | - | HA |
| 3563 | 3565 | 1 | 1.5260e-01 | 2.5941e+05 | ; | CA | - | CB |
| 3563 | 3581 | 1 | 1.5220e-01 | 2.6527e+05 | ; | CA | - | C |
| 3565 | 3566 | 1 | 1.0900e-01 | 2.8451e+05 | ; | CB | - | HB2 |
| 3565 | 3567 | 1 | 1.0900e-01 | 2.8451e+05 | ; | CB | - | HB3 |
| 3565 | 3568 | 1 | 1.5260e-01 | 2.5941e+05 | ; | CB | - | CG |
| 3568 | 3569 | 1 | 1.0900e-01 | 2.8451e+05 | ; | CG | - | HG2 |
| 3568 | 3570 | 1 | 1.0900e-01 | 2.8451e+05 | ; | CG | - | HG3 |
| 3568 | 3571 | 1 | 1.5260e-01 | 2.5941e+05 | ; | CG | - | CD |
| 3571 | 3572 | 1 | 1.0900e-01 | 2.8451e+05 | ; | CD | - | HD2 |
| 3571 | 3573 | 1 | 1.0900e-01 | 2.8451e+05 | ; | CD | - | HD3 |
| 3571 | 3574 | 1 | 1.5260e-01 | 2.5941e+05 | ; | CD | - | CE |
| 3574 | 3575 | 1 | 1.0900e-01 | 2.8451e+05 | ; | CE | - | HE2 |
| 3574 | 3576 | 1 | 1.0900e-01 | 2.8451e+05 | ; | CE | - | HE3 |
| 3574 | 3577 | 1 | 1.4710e-01 | 3.0711e+05 | ; | CE | - | NZ |
| 3577 | 3578 | 1 | 1.0100e-01 | 3.6317e+05 | ; | NZ | - | HZ1 |
| 3577 | 3579 | 1 | 1.0100e-01 | 3.6317e+05 | ; | NZ | - | HZ2 |
| 3577 | 3580 | 1 | 1.0100e-01 | 3.6317e+05 | ; | NZ | - | HZ3 |
| 3581 | 3582 | 1 | 1.2290e-01 | 4.7698e+05 | ; | C | - | O |
| 3581 | 3583 | 1 | 1.3350e-01 | 4.1003e+05 | ; | C | - | N |
| 3583 | 3584 | 1 | 1.0100e-01 | 3.6317e+05 | ; | N | - | H |

| | | | | | | | | |
|---|---|---|---|---|---|---|---|---|
| 3583 | 3585 | 1 | 1.4490e-01 | 2.8200e+05 | ; | N | - | CA |
| 3585 | 3586 | 1 | 1.0900e-01 | 2.8451e+05 | ; | CA | - | HA |
| 3585 | 3587 | 1 | 1.5260e-01 | 2.5941e+05 | ; | CA | - | CB |
| 3585 | 3597 | 1 | 1.5220e-01 | 2.6527e+05 | ; | CA | - | C |
| 3587 | 3588 | 1 | 1.0900e-01 | 2.8451e+05 | ; | CB | - | HB |
| 3587 | 3589 | 1 | 1.5260e-01 | 2.5941e+05 | ; | CB | - | CG1 |
| 3587 | 3593 | 1 | 1.5260e-01 | 2.5941e+05 | ; | CB | - | CG2 |
| 3589 | 3590 | 1 | 1.0900e-01 | 2.8451e+05 | ; | CG1 | - | HG11 |
| 3589 | 3591 | 1 | 1.0900e-01 | 2.8451e+05 | ; | CG1 | - | HG12 |
| 3589 | 3592 | 1 | 1.0900e-01 | 2.8451e+05 | ; | CG1 | - | HG13 |
| 3593 | 3594 | 1 | 1.0900e-01 | 2.8451e+05 | ; | CG2 | - | HG21 |
| 3593 | 3595 | 1 | 1.0900e-01 | 2.8451e+05 | ; | CG2 | - | HG22 |
| 3593 | 3596 | 1 | 1.0900e-01 | 2.8451e+05 | ; | CG2 | - | HG23 |
| 3597 | 3598 | 1 | 1.2290e-01 | 4.7698e+05 | ; | C | - | O |
| 3597 | 3599 | 1 | 1.3350e-01 | 4.1003e+05 | ; | C | - | N |
| 3599 | 3600 | 1 | 1.0100e-01 | 3.6317e+05 | ; | N | - | H |
| 3599 | 3601 | 1 | 1.4490e-01 | 2.8200e+05 | ; | N | - | CA |
| 3601 | 3602 | 1 | 1.0900e-01 | 2.8451e+05 | ; | CA | - | HA |
| 3601 | 3603 | 1 | 1.5260e-01 | 2.5941e+05 | ; | CA | - | CB |
| 3601 | 3619 | 1 | 1.5220e-01 | 2.6527e+05 | ; | CA | - | C |
| 3603 | 3604 | 1 | 1.0900e-01 | 2.8451e+05 | ; | CB | - | HB2 |
| 3603 | 3605 | 1 | 1.0900e-01 | 2.8451e+05 | ; | CB | - | HB3 |
| 3603 | 3606 | 1 | 1.5260e-01 | 2.5941e+05 | ; | CB | - | CG |
| 3606 | 3607 | 1 | 1.0900e-01 | 2.8451e+05 | ; | CG | - | HG2 |
| 3606 | 3608 | 1 | 1.0900e-01 | 2.8451e+05 | ; | CG | - | HG3 |
| 3606 | 3609 | 1 | 1.5260e-01 | 2.5941e+05 | ; | CG | - | CD |
| 3609 | 3610 | 1 | 1.0900e-01 | 2.8451e+05 | ; | CD | - | HD2 |
| 3609 | 3611 | 1 | 1.0900e-01 | 2.8451e+05 | ; | CD | - | HD3 |
| 3609 | 3612 | 1 | 1.5260e-01 | 2.5941e+05 | ; | CD | - | CE |
| 3612 | 3613 | 1 | 1.0900e-01 | 2.8451e+05 | ; | CE | - | HE2 |
| 3612 | 3614 | 1 | 1.0900e-01 | 2.8451e+05 | ; | CE | - | HE3 |
| 3612 | 3615 | 1 | 1.4710e-01 | 3.0711e+05 | ; | CE | - | NZ |
| 3615 | 3616 | 1 | 1.0100e-01 | 3.6317e+05 | ; | NZ | - | HZ1 |
| 3615 | 3617 | 1 | 1.0100e-01 | 3.6317e+05 | ; | NZ | - | HZ2 |
| 3615 | 3618 | 1 | 1.0100e-01 | 3.6317e+05 | ; | NZ | - | HZ3 |
| 3619 | 3620 | 1 | 1.2290e-01 | 4.7698e+05 | ; | C | - | O |
| 3619 | 3621 | 1 | 1.3350e-01 | 4.1003e+05 | ; | C | - | N |
| 3621 | 3622 | 1 | 1.0100e-01 | 3.6317e+05 | ; | N | - | H |
| 3621 | 3623 | 1 | 1.4490e-01 | 2.8200e+05 | ; | N | - | CA |
| 3623 | 3624 | 1 | 1.0900e-01 | 2.8451e+05 | ; | CA | - | HA |
| 3623 | 3625 | 1 | 1.5260e-01 | 2.5941e+05 | ; | CA | - | CB |
| 3623 | 3641 | 1 | 1.5220e-01 | 2.6527e+05 | ; | CA | - | C |
| 3625 | 3626 | 1 | 1.0900e-01 | 2.8451e+05 | ; | CB | - | HB2 |
| 3625 | 3627 | 1 | 1.0900e-01 | 2.8451e+05 | ; | CB | - | HB3 |
| 3625 | 3628 | 1 | 1.5260e-01 | 2.5941e+05 | ; | CB | - | CG |
| 3628 | 3629 | 1 | 1.0900e-01 | 2.8451e+05 | ; | CG | - | HG2 |
| 3628 | 3630 | 1 | 1.0900e-01 | 2.8451e+05 | ; | CG | - | HG3 |
| 3628 | 3631 | 1 | 1.5260e-01 | 2.5941e+05 | ; | CG | - | CD |
| 3631 | 3632 | 1 | 1.0900e-01 | 2.8451e+05 | ; | CD | - | HD2 |
| 3631 | 3633 | 1 | 1.0900e-01 | 2.8451e+05 | ; | CD | - | HD3 |
| 3631 | 3634 | 1 | 1.5260e-01 | 2.5941e+05 | ; | CD | - | CE |
| 3634 | 3635 | 1 | 1.0900e-01 | 2.8451e+05 | ; | CE | - | HE2 |

| | | | | | | |
|---|---|---|---|---|---|---|
| 3634 | 3636 | 1 | 1.0900e-01 | 2.8451e+05 | ; | CE - HE3 |
| 3634 | 3637 | 1 | 1.4710e-01 | 3.0711e+05 | ; | CE - NZ |
| 3637 | 3638 | 1 | 1.0100e-01 | 3.6317e+05 | ; | NZ - HZ1 |
| 3637 | 3639 | 1 | 1.0100e-01 | 3.6317e+05 | ; | NZ - HZ2 |
| 3637 | 3640 | 1 | 1.0100e-01 | 3.6317e+05 | ; | NZ - HZ3 |
| 3641 | 3642 | 1 | 1.2290e-01 | 4.7698e+05 | ; | C - O |
| 3641 | 3643 | 1 | 1.3350e-01 | 4.1003e+05 | ; | C - N |
| 3643 | 3644 | 1 | 1.0100e-01 | 3.6317e+05 | ; | N - H |
| 3643 | 3645 | 1 | 1.4490e-01 | 2.8200e+05 | ; | N - CA |
| 3645 | 3646 | 1 | 1.0900e-01 | 2.8451e+05 | ; | CA - HA2 |
| 3645 | 3647 | 1 | 1.0900e-01 | 2.8451e+05 | ; | CA - HA3 |
| 3645 | 3648 | 1 | 1.5220e-01 | 2.6527e+05 | ; | CA - C |
| 3648 | 3649 | 1 | 1.2290e-01 | 4.7698e+05 | ; | C - O |
| 3648 | 3650 | 1 | 1.3350e-01 | 4.1003e+05 | ; | C - N |
| 3650 | 3651 | 1 | 1.0100e-01 | 3.6317e+05 | ; | N - H |
| 3650 | 3652 | 1 | 1.4490e-01 | 2.8200e+05 | ; | N - CA |
| 3652 | 3653 | 1 | 1.0900e-01 | 2.8451e+05 | ; | CA - HA |
| 3652 | 3654 | 1 | 1.5260e-01 | 2.5941e+05 | ; | CA - CB |
| 3652 | 3663 | 1 | 1.5220e-01 | 2.6527e+05 | ; | CA - C |
| 3654 | 3655 | 1 | 1.0900e-01 | 2.8451e+05 | ; | CB - HB2 |
| 3654 | 3656 | 1 | 1.0900e-01 | 2.8451e+05 | ; | CB - HB3 |
| 3654 | 3657 | 1 | 1.5260e-01 | 2.5941e+05 | ; | CB - CG |
| 3657 | 3658 | 1 | 1.0900e-01 | 2.8451e+05 | ; | CG - HG2 |
| 3657 | 3659 | 1 | 1.0900e-01 | 2.8451e+05 | ; | CG - HG3 |
| 3657 | 3660 | 1 | 1.5220e-01 | 2.6527e+05 | ; | CG - CD |
| 3660 | 3661 | 1 | 1.2500e-01 | 5.4894e+05 | ; | CD - OE1 |
| 3660 | 3662 | 1 | 1.2500e-01 | 5.4894e+05 | ; | CD - OE2 |
| 3663 | 3664 | 1 | 1.2290e-01 | 4.7698e+05 | ; | C - O |
| 3663 | 3665 | 1 | 1.3350e-01 | 4.1003e+05 | ; | C - N |
| 3665 | 3666 | 1 | 1.0100e-01 | 3.6317e+05 | ; | N - H |
| 3665 | 3667 | 1 | 1.4490e-01 | 2.8200e+05 | ; | N - CA |
| 3667 | 3668 | 1 | 1.0900e-01 | 2.8451e+05 | ; | CA - HA |
| 3667 | 3669 | 1 | 1.5260e-01 | 2.5941e+05 | ; | CA - CB |
| 3667 | 3683 | 1 | 1.5220e-01 | 2.6527e+05 | ; | CA - C |
| 3669 | 3670 | 1 | 1.0900e-01 | 2.8451e+05 | ; | CB - HB2 |
| 3669 | 3671 | 1 | 1.0900e-01 | 2.8451e+05 | ; | CB - HB3 |
| 3669 | 3672 | 1 | 1.5100e-01 | 2.6527e+05 | ; | CB - CG |
| 3672 | 3673 | 1 | 1.4000e-01 | 3.9246e+05 | ; | CG - CD1 |
| 3672 | 3681 | 1 | 1.4000e-01 | 3.9246e+05 | ; | CG - CD2 |
| 3673 | 3674 | 1 | 1.0800e-01 | 3.0711e+05 | ; | CD1 - HD1 |
| 3673 | 3675 | 1 | 1.4000e-01 | 3.9246e+05 | ; | CD1 - CE1 |
| 3675 | 3676 | 1 | 1.0800e-01 | 3.0711e+05 | ; | CE1 - HE1 |
| 3675 | 3677 | 1 | 1.4000e-01 | 3.9246e+05 | ; | CE1 - CZ |
| 3677 | 3678 | 1 | 1.0800e-01 | 3.0711e+05 | ; | CZ - HZ |
| 3677 | 3679 | 1 | 1.4000e-01 | 3.9246e+05 | ; | CZ - CE2 |
| 3679 | 3680 | 1 | 1.0800e-01 | 3.0711e+05 | ; | CE2 - HE2 |
| 3679 | 3681 | 1 | 1.4000e-01 | 3.9246e+05 | ; | CE2 - CD2 |
| 3681 | 3682 | 1 | 1.0800e-01 | 3.0711e+05 | ; | CD2 - HD2 |
| 3683 | 3684 | 1 | 1.2290e-01 | 4.7698e+05 | ; | C - O |
| 3683 | 3685 | 1 | 1.3350e-01 | 4.1003e+05 | ; | C - N |
| 3685 | 3686 | 1 | 1.0100e-01 | 3.6317e+05 | ; | N - H |
| 3685 | 3687 | 1 | 1.4490e-01 | 2.8200e+05 | ; | N - CA |

| | | | | | | | | |
|---|---|---|---|---|---|---|---|---|
| 3687 | 3688 | 1 | 1.0900e-01 | 2.8451e+05 | ; | CA | - | HA |
| 3687 | 3689 | 1 | 1.5260e-01 | 2.5941e+05 | ; | CA | - | CB |
| 3687 | 3694 | 1 | 1.5220e-01 | 2.6527e+05 | ; | CA | - | C |
| 3689 | 3690 | 1 | 1.0900e-01 | 2.8451e+05 | ; | CB | - | HB2 |
| 3689 | 3691 | 1 | 1.0900e-01 | 2.8451e+05 | ; | CB | - | HB3 |
| 3689 | 3692 | 1 | 1.8100e-01 | 1.9832e+05 | ; | CB | - | SG |
| 3692 | 3693 | 1 | 1.3360e-01 | 2.2928e+05 | ; | SG | - | HG |
| 3694 | 3695 | 1 | 1.2290e-01 | 4.7698e+05 | ; | C | - | O |
| 3694 | 3696 | 1 | 1.3350e-01 | 4.1003e+05 | ; | C | - | N |
| 3696 | 3697 | 1 | 1.0100e-01 | 3.6317e+05 | ; | N | - | H |
| 3696 | 3698 | 1 | 1.4490e-01 | 2.8200e+05 | ; | N | - | CA |
| 3698 | 3699 | 1 | 1.0900e-01 | 2.8451e+05 | ; | CA | - | HA |
| 3698 | 3700 | 1 | 1.5260e-01 | 2.5941e+05 | ; | CA | - | CB |
| 3698 | 3714 | 1 | 1.5220e-01 | 2.6527e+05 | ; | CA | - | C |
| 3700 | 3701 | 1 | 1.0900e-01 | 2.8451e+05 | ; | CB | - | HB2 |
| 3700 | 3702 | 1 | 1.0900e-01 | 2.8451e+05 | ; | CB | - | HB3 |
| 3700 | 3703 | 1 | 1.5100e-01 | 2.6527e+05 | ; | CB | - | CG |
| 3703 | 3704 | 1 | 1.4000e-01 | 3.9246e+05 | ; | CG | - | CD1 |
| 3703 | 3712 | 1 | 1.4000e-01 | 3.9246e+05 | ; | CG | - | CD2 |
| 3704 | 3705 | 1 | 1.0800e-01 | 3.0711e+05 | ; | CD1 | - | HD1 |
| 3704 | 3706 | 1 | 1.4000e-01 | 3.9246e+05 | ; | CD1 | - | CE1 |
| 3706 | 3707 | 1 | 1.0800e-01 | 3.0711e+05 | ; | CE1 | - | HE1 |
| 3706 | 3708 | 1 | 1.4000e-01 | 3.9246e+05 | ; | CE1 | - | CZ |
| 3708 | 3709 | 1 | 1.0800e-01 | 3.0711e+05 | ; | CZ | - | HZ |
| 3708 | 3710 | 1 | 1.4000e-01 | 3.9246e+05 | ; | CZ | - | CE2 |
| 3710 | 3711 | 1 | 1.0800e-01 | 3.0711e+05 | ; | CE2 | - | HE2 |
| 3710 | 3712 | 1 | 1.4000e-01 | 3.9246e+05 | ; | CE2 | - | CD2 |
| 3712 | 3713 | 1 | 1.0800e-01 | 3.0711e+05 | ; | CD2 | - | HD2 |
| 3714 | 3715 | 1 | 1.2290e-01 | 4.7698e+05 | ; | C | - | O |
| 3714 | 3716 | 1 | 1.3350e-01 | 4.1003e+05 | ; | C | - | N |
| 3716 | 3717 | 1 | 1.0100e-01 | 3.6317e+05 | ; | N | - | H |
| 3716 | 3718 | 1 | 1.4490e-01 | 2.8200e+05 | ; | N | - | CA |
| 3718 | 3719 | 1 | 1.0900e-01 | 2.8451e+05 | ; | CA | - | HA |
| 3718 | 3720 | 1 | 1.5260e-01 | 2.5941e+05 | ; | CA | - | CB |
| 3718 | 3735 | 1 | 1.5220e-01 | 2.6527e+05 | ; | CA | - | C |
| 3720 | 3721 | 1 | 1.0900e-01 | 2.8451e+05 | ; | CB | - | HB2 |
| 3720 | 3722 | 1 | 1.0900e-01 | 2.8451e+05 | ; | CB | - | HB3 |
| 3720 | 3723 | 1 | 1.5100e-01 | 2.6527e+05 | ; | CB | - | CG |
| 3723 | 3724 | 1 | 1.4000e-01 | 3.9246e+05 | ; | CG | - | CD1 |
| 3723 | 3733 | 1 | 1.4000e-01 | 3.9246e+05 | ; | CG | - | CD2 |
| 3724 | 3725 | 1 | 1.0800e-01 | 3.0711e+05 | ; | CD1 | - | HD1 |
| 3724 | 3726 | 1 | 1.4000e-01 | 3.9246e+05 | ; | CD1 | - | CE1 |
| 3726 | 3727 | 1 | 1.0800e-01 | 3.0711e+05 | ; | CE1 | - | HE1 |
| 3726 | 3728 | 1 | 1.4090e-01 | 3.9246e+05 | ; | CE1 | - | CZ |
| 3728 | 3729 | 1 | 1.3640e-01 | 3.7656e+05 | ; | CZ | - | OH |
| 3728 | 3731 | 1 | 1.4090e-01 | 3.9246e+05 | ; | CZ | - | CE2 |
| 3729 | 3730 | 1 | 9.6000e-02 | 4.6275e+05 | ; | OH | - | HH |
| 3731 | 3732 | 1 | 1.0800e-01 | 3.0711e+05 | ; | CE2 | - | HE2 |
| 3731 | 3733 | 1 | 1.4000e-01 | 3.9246e+05 | ; | CE2 | - | CD2 |
| 3733 | 3734 | 1 | 1.0800e-01 | 3.0711e+05 | ; | CD2 | - | HD2 |
| 3735 | 3736 | 1 | 1.2290e-01 | 4.7698e+05 | ; | C | - | O |
| 3735 | 3737 | 1 | 1.3350e-01 | 4.1003e+05 | ; | C | - | N |

| | | | | | | | | |
|---|---|---|---|---|---|---|---|---|
| 3737 | 3738 | 1 | 1.0100e-01 | 3.6317e+05 | ; | N | - | H |
| 3737 | 3739 | 1 | 1.4490e-01 | 2.8200e+05 | ; | N | - | CA |
| 3739 | 3740 | 1 | 1.0900e-01 | 2.8451e+05 | ; | CA | - | HA |
| 3739 | 3741 | 1 | 1.5260e-01 | 2.5941e+05 | ; | CA | - | CB |
| 3739 | 3750 | 1 | 1.5220e-01 | 2.6527e+05 | ; | CA | - | C |
| 3741 | 3742 | 1 | 1.0900e-01 | 2.8451e+05 | ; | CB | - | HB2 |
| 3741 | 3743 | 1 | 1.0900e-01 | 2.8451e+05 | ; | CB | - | HB3 |
| 3741 | 3744 | 1 | 1.5260e-01 | 2.5941e+05 | ; | CB | - | CG |
| 3744 | 3745 | 1 | 1.0900e-01 | 2.8451e+05 | ; | CG | - | HG2 |
| 3744 | 3746 | 1 | 1.0900e-01 | 2.8451e+05 | ; | CG | - | HG3 |
| 3744 | 3747 | 1 | 1.5220e-01 | 2.6527e+05 | ; | CG | - | CD |
| 3747 | 3748 | 1 | 1.2500e-01 | 5.4894e+05 | ; | CD | - | OE1 |
| 3747 | 3749 | 1 | 1.2500e-01 | 5.4894e+05 | ; | CD | - | OE2 |
| 3750 | 3751 | 1 | 1.2290e-01 | 4.7698e+05 | ; | C | - | O |
| 3750 | 3752 | 1 | 1.3350e-01 | 4.1003e+05 | ; | C | - | N |
| 3752 | 3753 | 1 | 1.0100e-01 | 3.6317e+05 | ; | N | - | H |
| 3752 | 3754 | 1 | 1.4490e-01 | 2.8200e+05 | ; | N | - | CA |
| 3754 | 3755 | 1 | 1.0900e-01 | 2.8451e+05 | ; | CA | - | HA |
| 3754 | 3756 | 1 | 1.5260e-01 | 2.5941e+05 | ; | CA | - | CB |
| 3754 | 3764 | 1 | 1.5220e-01 | 2.6527e+05 | ; | CA | - | C |
| 3756 | 3757 | 1 | 1.0900e-01 | 2.8451e+05 | ; | CB | - | HB2 |
| 3756 | 3758 | 1 | 1.0900e-01 | 2.8451e+05 | ; | CB | - | HB3 |
| 3756 | 3759 | 1 | 1.5220e-01 | 2.6527e+05 | ; | CB | - | CG |
| 3759 | 3760 | 1 | 1.2290e-01 | 4.7698e+05 | ; | CG | - | OD1 |
| 3759 | 3761 | 1 | 1.3350e-01 | 4.1003e+05 | ; | CG | - | ND2 |
| 3761 | 3762 | 1 | 1.0100e-01 | 3.6317e+05 | ; | ND2 | - | HD21 |
| 3761 | 3763 | 1 | 1.0100e-01 | 3.6317e+05 | ; | ND2 | - | HD22 |
| 3764 | 3765 | 1 | 1.2290e-01 | 4.7698e+05 | ; | C | - | O |
| 3764 | 3766 | 1 | 1.3350e-01 | 4.1003e+05 | ; | C | - | N |
| 3766 | 3767 | 1 | 1.0100e-01 | 3.6317e+05 | ; | N | - | H |
| 3766 | 3768 | 1 | 1.4490e-01 | 2.8200e+05 | ; | N | - | CA |
| 3768 | 3769 | 1 | 1.0900e-01 | 2.8451e+05 | ; | CA | - | HA |
| 3768 | 3770 | 1 | 1.5260e-01 | 2.5941e+05 | ; | CA | - | CB |
| 3768 | 3776 | 1 | 1.5220e-01 | 2.6527e+05 | ; | CA | - | C |
| 3770 | 3771 | 1 | 1.0900e-01 | 2.8451e+05 | ; | CB | - | HB2 |
| 3770 | 3772 | 1 | 1.0900e-01 | 2.8451e+05 | ; | CB | - | HB3 |
| 3770 | 3773 | 1 | 1.5220e-01 | 2.6527e+05 | ; | CB | - | CG |
| 3773 | 3774 | 1 | 1.2500e-01 | 5.4894e+05 | ; | CG | - | OD1 |
| 3773 | 3775 | 1 | 1.2500e-01 | 5.4894e+05 | ; | CG | - | OD2 |
| 3776 | 3777 | 1 | 1.2290e-01 | 4.7698e+05 | ; | C | - | O |
| 3776 | 3778 | 1 | 1.3350e-01 | 4.1003e+05 | ; | C | - | N |
| 3778 | 3779 | 1 | 1.0100e-01 | 3.6317e+05 | ; | N | - | H |
| 3778 | 3780 | 1 | 1.4490e-01 | 2.8200e+05 | ; | N | - | CA |
| 3780 | 3781 | 1 | 1.0900e-01 | 2.8451e+05 | ; | CA | - | HA |
| 3780 | 3782 | 1 | 1.5260e-01 | 2.5941e+05 | ; | CA | - | CB |
| 3780 | 3800 | 1 | 1.5220e-01 | 2.6527e+05 | ; | CA | - | C |
| 3782 | 3783 | 1 | 1.0900e-01 | 2.8451e+05 | ; | CB | - | HB2 |
| 3782 | 3784 | 1 | 1.0900e-01 | 2.8451e+05 | ; | CB | - | HB3 |
| 3782 | 3785 | 1 | 1.4950e-01 | 2.6527e+05 | ; | CB | - | CG |
| 3785 | 3786 | 1 | 1.3520e-01 | 4.5689e+05 | ; | CG | - | CD1 |
| 3785 | 3799 | 1 | 1.4590e-01 | 3.2468e+05 | ; | CG | - | CD2 |
| 3786 | 3787 | 1 | 1.0800e-01 | 3.0711e+05 | ; | CD1 | - | HD1 |

| | | | | | | |
|---|---|---|---|---|---|---|
| 3786 | 3788 | 1 | 1.3810e-01 | 3.5731e+05 | ; | CD1 - NE1 |
| 3788 | 3789 | 1 | 1.0100e-01 | 3.6317e+05 | ; | NE1 - HE1 |
| 3788 | 3790 | 1 | 1.3800e-01 | 3.5815e+05 | ; | NE1 - CE2 |
| 3790 | 3791 | 1 | 1.4000e-01 | 3.9246e+05 | ; | CE2 - CZ2 |
| 3790 | 3799 | 1 | 1.4190e-01 | 3.7405e+05 | ; | CE2 - CD2 |
| 3791 | 3792 | 1 | 1.0800e-01 | 3.0711e+05 | ; | CZ2 - HZ2 |
| 3791 | 3793 | 1 | 1.4000e-01 | 3.9246e+05 | ; | CZ2 - CH2 |
| 3793 | 3794 | 1 | 1.0800e-01 | 3.0711e+05 | ; | CH2 - HH2 |
| 3793 | 3795 | 1 | 1.4000e-01 | 3.9246e+05 | ; | CH2 - CZ3 |
| 3795 | 3796 | 1 | 1.0800e-01 | 3.0711e+05 | ; | CZ3 - HZ3 |
| 3795 | 3797 | 1 | 1.4000e-01 | 3.9246e+05 | ; | CZ3 - CE3 |
| 3797 | 3798 | 1 | 1.0800e-01 | 3.0711e+05 | ; | CE3 - HE3 |
| 3797 | 3799 | 1 | 1.4040e-01 | 3.9246e+05 | ; | CE3 - CD2 |
| 3800 | 3801 | 1 | 1.2290e-01 | 4.7698e+05 | ; | C - O |
| 3800 | 3802 | 1 | 1.3350e-01 | 4.1003e+05 | ; | C - N |
| 3802 | 3803 | 1 | 1.0100e-01 | 3.6317e+05 | ; | N - H |
| 3802 | 3804 | 1 | 1.4490e-01 | 2.8200e+05 | ; | N - CA |
| 3804 | 3805 | 1 | 1.0900e-01 | 2.8451e+05 | ; | CA - HA2 |
| 3804 | 3806 | 1 | 1.0900e-01 | 2.8451e+05 | ; | CA - HA3 |
| 3804 | 3807 | 1 | 1.5220e-01 | 2.6527e+05 | ; | CA - C |
| 3807 | 3808 | 1 | 1.2290e-01 | 4.7698e+05 | ; | C - O |
| 3807 | 3809 | 1 | 1.3350e-01 | 4.1003e+05 | ; | C - N |
| 3809 | 3810 | 1 | 1.0100e-01 | 3.6317e+05 | ; | N - H |
| 3809 | 3811 | 1 | 1.4490e-01 | 2.8200e+05 | ; | N - CA |
| 3811 | 3812 | 1 | 1.0900e-01 | 2.8451e+05 | ; | CA - HA |
| 3811 | 3813 | 1 | 1.5260e-01 | 2.5941e+05 | ; | CA - CB |
| 3811 | 3818 | 1 | 1.5220e-01 | 2.6527e+05 | ; | CA - C |
| 3813 | 3814 | 1 | 1.0900e-01 | 2.8451e+05 | ; | CB - HB2 |
| 3813 | 3815 | 1 | 1.0900e-01 | 2.8451e+05 | ; | CB - HB3 |
| 3813 | 3816 | 1 | 1.4100e-01 | 2.6778e+05 | ; | CB - OG |
| 3816 | 3817 | 1 | 9.6000e-02 | 4.6275e+05 | ; | OG - HG |
| 3818 | 3819 | 1 | 1.2290e-01 | 4.7698e+05 | ; | C - O |
| 3818 | 3820 | 1 | 1.3350e-01 | 4.1003e+05 | ; | C - N |
| 3820 | 3821 | 1 | 1.0100e-01 | 3.6317e+05 | ; | N - H |
| 3820 | 3822 | 1 | 1.4490e-01 | 2.8200e+05 | ; | N - CA |
| 3822 | 3823 | 1 | 1.0900e-01 | 2.8451e+05 | ; | CA - HA |
| 3822 | 3824 | 1 | 1.5260e-01 | 2.5941e+05 | ; | CA - CB |
| 3822 | 3837 | 1 | 1.5220e-01 | 2.6527e+05 | ; | CA - C |
| 3824 | 3825 | 1 | 1.0900e-01 | 2.8451e+05 | ; | CB - HB |
| 3824 | 3826 | 1 | 1.5260e-01 | 2.5941e+05 | ; | CB - CG2 |
| 3824 | 3830 | 1 | 1.5260e-01 | 2.5941e+05 | ; | CB - CG1 |
| 3826 | 3827 | 1 | 1.0900e-01 | 2.8451e+05 | ; | CG2 - HG21 |
| 3826 | 3828 | 1 | 1.0900e-01 | 2.8451e+05 | ; | CG2 - HG22 |
| 3826 | 3829 | 1 | 1.0900e-01 | 2.8451e+05 | ; | CG2 - HG23 |
| 3830 | 3831 | 1 | 1.0900e-01 | 2.8451e+05 | ; | CG1 - HG12 |
| 3830 | 3832 | 1 | 1.0900e-01 | 2.8451e+05 | ; | CG1 - HG13 |
| 3830 | 3833 | 1 | 1.5260e-01 | 2.5941e+05 | ; | CG1 - CD1 |
| 3833 | 3834 | 1 | 1.0900e-01 | 2.8451e+05 | ; | CD1 - HD11 |
| 3833 | 3835 | 1 | 1.0900e-01 | 2.8451e+05 | ; | CD1 - HD12 |
| 3833 | 3836 | 1 | 1.0900e-01 | 2.8451e+05 | ; | CD1 - HD13 |
| 3837 | 3838 | 1 | 1.2290e-01 | 4.7698e+05 | ; | C - O |
| 3837 | 3839 | 1 | 1.3350e-01 | 4.1003e+05 | ; | C - N |

| | | | | | | |
|---|---|---|---|---|---|---|
| 3839 | 3840 | 1 | 1.0100e-01 | 3.6317e+05 | ; | N - H |
| 3839 | 3841 | 1 | 1.4490e-01 | 2.8200e+05 | ; | N - CA |
| 3841 | 3842 | 1 | 1.0900e-01 | 2.8451e+05 | ; | CA - HA |
| 3841 | 3843 | 1 | 1.5260e-01 | 2.5941e+05 | ; | CA - CB |
| 3841 | 3848 | 1 | 1.5220e-01 | 2.6527e+05 | ; | CA - C |
| 3843 | 3844 | 1 | 1.0900e-01 | 2.8451e+05 | ; | CB - HB2 |
| 3843 | 3845 | 1 | 1.0900e-01 | 2.8451e+05 | ; | CB - HB3 |
| 3843 | 3846 | 1 | 1.4100e-01 | 2.6778e+05 | ; | CB - OG |
| 3846 | 3847 | 1 | 9.6000e-02 | 4.6275e+05 | ; | OG - HG |
| 3848 | 3849 | 1 | 1.2290e-01 | 4.7698e+05 | ; | C - O |
| 3848 | 3850 | 1 | 1.3350e-01 | 4.1003e+05 | ; | C - N |
| 3850 | 3851 | 1 | 1.0100e-01 | 3.6317e+05 | ; | N - H |
| 3850 | 3852 | 1 | 1.4490e-01 | 2.8200e+05 | ; | N - CA |
| 3852 | 3853 | 1 | 1.0900e-01 | 2.8451e+05 | ; | CA - HA |
| 3852 | 3854 | 1 | 1.5260e-01 | 2.5941e+05 | ; | CA - CB |
| 3852 | 3859 | 1 | 1.5220e-01 | 2.6527e+05 | ; | CA - C |
| 3854 | 3855 | 1 | 1.0900e-01 | 2.8451e+05 | ; | CB - HB2 |
| 3854 | 3856 | 1 | 1.0900e-01 | 2.8451e+05 | ; | CB - HB3 |
| 3854 | 3857 | 1 | 1.4100e-01 | 2.6778e+05 | ; | CB - OG |
| 3857 | 3858 | 1 | 9.6000e-02 | 4.6275e+05 | ; | OG - HG |
| 3859 | 3860 | 1 | 1.2290e-01 | 4.7698e+05 | ; | C - O |
| 3859 | 3861 | 1 | 1.3350e-01 | 4.1003e+05 | ; | C - N |
| 3861 | 3862 | 1 | 1.0100e-01 | 3.6317e+05 | ; | N - H |
| 3861 | 3863 | 1 | 1.4490e-01 | 2.8200e+05 | ; | N - CA |
| 3863 | 3864 | 1 | 1.0900e-01 | 2.8451e+05 | ; | CA - HA |
| 3863 | 3865 | 1 | 1.5260e-01 | 2.5941e+05 | ; | CA - CB |
| 3863 | 3871 | 1 | 1.5220e-01 | 2.6527e+05 | ; | CA - C |
| 3865 | 3866 | 1 | 1.0900e-01 | 2.8451e+05 | ; | CB - HB2 |
| 3865 | 3867 | 1 | 1.0900e-01 | 2.8451e+05 | ; | CB - HB3 |
| 3865 | 3868 | 1 | 1.5220e-01 | 2.6527e+05 | ; | CB - CG |
| 3868 | 3869 | 1 | 1.2500e-01 | 5.4894e+05 | ; | CG - OD1 |
| 3868 | 3870 | 1 | 1.2500e-01 | 5.4894e+05 | ; | CG - OD2 |
| 3871 | 3872 | 1 | 1.2290e-01 | 4.7698e+05 | ; | C - O |
| 3871 | 3873 | 1 | 1.3350e-01 | 4.1003e+05 | ; | C - N |
| 3873 | 3874 | 1 | 1.0100e-01 | 3.6317e+05 | ; | N - H |
| 3873 | 3875 | 1 | 1.4490e-01 | 2.8200e+05 | ; | N - CA |
| 3875 | 3876 | 1 | 1.0900e-01 | 2.8451e+05 | ; | CA - HA |
| 3875 | 3877 | 1 | 1.5260e-01 | 2.5941e+05 | ; | CA - CB |
| 3875 | 3881 | 1 | 1.5220e-01 | 2.6527e+05 | ; | CA - C |
| 3877 | 3878 | 1 | 1.0900e-01 | 2.8451e+05 | ; | CB - HB1 |
| 3877 | 3879 | 1 | 1.0900e-01 | 2.8451e+05 | ; | CB - HB2 |
| 3877 | 3880 | 1 | 1.0900e-01 | 2.8451e+05 | ; | CB - HB3 |
| 3881 | 3882 | 1 | 1.2290e-01 | 4.7698e+05 | ; | C - O |
| 3881 | 3883 | 1 | 1.3350e-01 | 4.1003e+05 | ; | C - N |
| 3883 | 3884 | 1 | 1.0100e-01 | 3.6317e+05 | ; | N - H |
| 3883 | 3885 | 1 | 1.4490e-01 | 2.8200e+05 | ; | N - CA |
| 3885 | 3886 | 1 | 1.0900e-01 | 2.8451e+05 | ; | CA - HA |
| 3885 | 3887 | 1 | 1.5260e-01 | 2.5941e+05 | ; | CA - CB |
| 3885 | 3903 | 1 | 1.5220e-01 | 2.6527e+05 | ; | CA - C |
| 3887 | 3888 | 1 | 1.0900e-01 | 2.8451e+05 | ; | CB - HB2 |
| 3887 | 3889 | 1 | 1.0900e-01 | 2.8451e+05 | ; | CB - HB3 |
| 3887 | 3890 | 1 | 1.5260e-01 | 2.5941e+05 | ; | CB - CG |

| | | | | | |
|---|---|---|---|---|---|
| 3890 | 3891 | 1 | 1.0900e-01 | 2.8451e+05 ; | CG - HG2 |
| 3890 | 3892 | 1 | 1.0900e-01 | 2.8451e+05 ; | CG - HG3 |
| 3890 | 3893 | 1 | 1.5260e-01 | 2.5941e+05 ; | CG - CD |
| 3893 | 3894 | 1 | 1.0900e-01 | 2.8451e+05 ; | CD - HD2 |
| 3893 | 3895 | 1 | 1.0900e-01 | 2.8451e+05 ; | CD - HD3 |
| 3893 | 3896 | 1 | 1.5260e-01 | 2.5941e+05 ; | CD - CE |
| 3896 | 3897 | 1 | 1.0900e-01 | 2.8451e+05 ; | CE - HE2 |
| 3896 | 3898 | 1 | 1.0900e-01 | 2.8451e+05 ; | CE - HE3 |
| 3896 | 3899 | 1 | 1.4710e-01 | 3.0711e+05 ; | CE - NZ |
| 3899 | 3900 | 1 | 1.0100e-01 | 3.6317e+05 ; | NZ - HZ1 |
| 3899 | 3901 | 1 | 1.0100e-01 | 3.6317e+05 ; | NZ - HZ2 |
| 3899 | 3902 | 1 | 1.0100e-01 | 3.6317e+05 ; | NZ - HZ3 |
| 3903 | 3904 | 1 | 1.2290e-01 | 4.7698e+05 ; | C - O |
| 3903 | 3905 | 1 | 1.3350e-01 | 4.1003e+05 ; | C - N |
| 3905 | 3906 | 1 | 1.0100e-01 | 3.6317e+05 ; | N - H |
| 3905 | 3907 | 1 | 1.4490e-01 | 2.8200e+05 ; | N - CA |
| 3907 | 3908 | 1 | 1.0900e-01 | 2.8451e+05 ; | CA - HA |
| 3907 | 3909 | 1 | 1.5260e-01 | 2.5941e+05 ; | CA - CB |
| 3907 | 3917 | 1 | 1.5220e-01 | 2.6527e+05 ; | CA - C |
| 3909 | 3910 | 1 | 1.0900e-01 | 2.8451e+05 ; | CB - HB2 |
| 3909 | 3911 | 1 | 1.0900e-01 | 2.8451e+05 ; | CB - HB3 |
| 3909 | 3912 | 1 | 1.5220e-01 | 2.6527e+05 ; | CB - CG |
| 3912 | 3913 | 1 | 1.2290e-01 | 4.7698e+05 ; | CG - OD1 |
| 3912 | 3914 | 1 | 1.3350e-01 | 4.1003e+05 ; | CG - ND2 |
| 3914 | 3915 | 1 | 1.0100e-01 | 3.6317e+05 ; | ND2 - HD21 |
| 3914 | 3916 | 1 | 1.0100e-01 | 3.6317e+05 ; | ND2 - HD22 |
| 3917 | 3918 | 1 | 1.2290e-01 | 4.7698e+05 ; | C - O |
| 3917 | 3919 | 1 | 1.3350e-01 | 4.1003e+05 ; | C - N |
| 3919 | 3920 | 1 | 1.0100e-01 | 3.6317e+05 ; | N - H |
| 3919 | 3921 | 1 | 1.4490e-01 | 2.8200e+05 ; | N - CA |
| 3921 | 3922 | 1 | 1.0900e-01 | 2.8451e+05 ; | CA - HA |
| 3921 | 3923 | 1 | 1.5260e-01 | 2.5941e+05 ; | CA - CB |
| 3921 | 3936 | 1 | 1.5220e-01 | 2.6527e+05 ; | CA - C |
| 3923 | 3924 | 1 | 1.0900e-01 | 2.8451e+05 ; | CB - HB2 |
| 3923 | 3925 | 1 | 1.0900e-01 | 2.8451e+05 ; | CB - HB3 |
| 3923 | 3926 | 1 | 1.5260e-01 | 2.5941e+05 ; | CB - CG |
| 3926 | 3927 | 1 | 1.0900e-01 | 2.8451e+05 ; | CG - HG |
| 3926 | 3928 | 1 | 1.5260e-01 | 2.5941e+05 ; | CG - CD1 |
| 3926 | 3932 | 1 | 1.5260e-01 | 2.5941e+05 ; | CG - CD2 |
| 3928 | 3929 | 1 | 1.0900e-01 | 2.8451e+05 ; | CD1 - HD11 |
| 3928 | 3930 | 1 | 1.0900e-01 | 2.8451e+05 ; | CD1 - HD12 |
| 3928 | 3931 | 1 | 1.0900e-01 | 2.8451e+05 ; | CD1 - HD13 |
| 3932 | 3933 | 1 | 1.0900e-01 | 2.8451e+05 ; | CD2 - HD21 |
| 3932 | 3934 | 1 | 1.0900e-01 | 2.8451e+05 ; | CD2 - HD22 |
| 3932 | 3935 | 1 | 1.0900e-01 | 2.8451e+05 ; | CD2 - HD23 |
| 3936 | 3937 | 1 | 1.2290e-01 | 4.7698e+05 ; | C - O |
| 3936 | 3938 | 1 | 1.3350e-01 | 4.1003e+05 ; | C - N |
| 3938 | 3939 | 1 | 1.0100e-01 | 3.6317e+05 ; | N - H |
| 3938 | 3940 | 1 | 1.4490e-01 | 2.8200e+05 ; | N - CA |
| 3940 | 3941 | 1 | 1.0900e-01 | 2.8451e+05 ; | CA - HA |
| 3940 | 3942 | 1 | 1.5260e-01 | 2.5941e+05 ; | CA - CB |
| 3940 | 3955 | 1 | 1.5220e-01 | 2.6527e+05 ; | CA - C |

| | | | | | |
|---|---|---|---|---|---|
| 3942 | 3943 | 1 | 1.0900e-01 | 2.8451e+05 ; | CB - HB |
| 3942 | 3944 | 1 | 1.5260e-01 | 2.5941e+05 ; | CB - CG2 |
| 3942 | 3948 | 1 | 1.5260e-01 | 2.5941e+05 ; | CB - CG1 |
| 3944 | 3945 | 1 | 1.0900e-01 | 2.8451e+05 ; | CG2 - HG21 |
| 3944 | 3946 | 1 | 1.0900e-01 | 2.8451e+05 ; | CG2 - HG22 |
| 3944 | 3947 | 1 | 1.0900e-01 | 2.8451e+05 ; | CG2 - HG23 |
| 3948 | 3949 | 1 | 1.0900e-01 | 2.8451e+05 ; | CG1 - HG12 |
| 3948 | 3950 | 1 | 1.0900e-01 | 2.8451e+05 ; | CG1 - HG13 |
| 3948 | 3951 | 1 | 1.5260e-01 | 2.5941e+05 ; | CG1 - CD1 |
| 3951 | 3952 | 1 | 1.0900e-01 | 2.8451e+05 ; | CD1 - HD11 |
| 3951 | 3953 | 1 | 1.0900e-01 | 2.8451e+05 ; | CD1 - HD12 |
| 3951 | 3954 | 1 | 1.0900e-01 | 2.8451e+05 ; | CD1 - HD13 |
| 3955 | 3956 | 1 | 1.2290e-01 | 4.7698e+05 ; | C - O |
| 3955 | 3957 | 1 | 1.3350e-01 | 4.1003e+05 ; | C - N |
| 3957 | 3958 | 1 | 1.0100e-01 | 3.6317e+05 ; | N - H |
| 3957 | 3959 | 1 | 1.4490e-01 | 2.8200e+05 ; | N - CA |
| 3959 | 3960 | 1 | 1.0900e-01 | 2.8451e+05 ; | CA - HA |
| 3959 | 3961 | 1 | 1.5260e-01 | 2.5941e+05 ; | CA - CB |
| 3959 | 3969 | 1 | 1.5220e-01 | 2.6527e+05 ; | CA - C |
| 3961 | 3962 | 1 | 1.0900e-01 | 2.8451e+05 ; | CB - HB |
| 3961 | 3963 | 1 | 1.5260e-01 | 2.5941e+05 ; | CB - CG2 |
| 3961 | 3967 | 1 | 1.4100e-01 | 2.6778e+05 ; | CB - OG1 |
| 3963 | 3964 | 1 | 1.0900e-01 | 2.8451e+05 ; | CG2 - HG21 |
| 3963 | 3965 | 1 | 1.0900e-01 | 2.8451e+05 ; | CG2 - HG22 |
| 3963 | 3966 | 1 | 1.0900e-01 | 2.8451e+05 ; | CG2 - HG23 |
| 3967 | 3968 | 1 | 9.6000e-02 | 4.6275e+05 ; | OG1 - HG1 |
| 3969 | 3970 | 1 | 1.2290e-01 | 4.7698e+05 ; | C - O |
| 3969 | 3971 | 1 | 1.3350e-01 | 4.1003e+05 ; | C - N |
| 3971 | 3972 | 1 | 1.0100e-01 | 3.6317e+05 ; | N - H |
| 3971 | 3973 | 1 | 1.4490e-01 | 2.8200e+05 ; | N - CA |
| 3973 | 3974 | 1 | 1.0900e-01 | 2.8451e+05 ; | CA - HA |
| 3973 | 3975 | 1 | 1.5260e-01 | 2.5941e+05 ; | CA - CB |
| 3973 | 3991 | 1 | 1.5220e-01 | 2.6527e+05 ; | CA - C |
| 3975 | 3976 | 1 | 1.0900e-01 | 2.8451e+05 ; | CB - HB2 |
| 3975 | 3977 | 1 | 1.0900e-01 | 2.8451e+05 ; | CB - HB3 |
| 3975 | 3978 | 1 | 1.5260e-01 | 2.5941e+05 ; | CB - CG |
| 3978 | 3979 | 1 | 1.0900e-01 | 2.8451e+05 ; | CG - HG2 |
| 3978 | 3980 | 1 | 1.0900e-01 | 2.8451e+05 ; | CG - HG3 |
| 3978 | 3981 | 1 | 1.5260e-01 | 2.5941e+05 ; | CG - CD |
| 3981 | 3982 | 1 | 1.0900e-01 | 2.8451e+05 ; | CD - HD2 |
| 3981 | 3983 | 1 | 1.0900e-01 | 2.8451e+05 ; | CD - HD3 |
| 3981 | 3984 | 1 | 1.5260e-01 | 2.5941e+05 ; | CD - CE |
| 3984 | 3985 | 1 | 1.0900e-01 | 2.8451e+05 ; | CE - HE2 |
| 3984 | 3986 | 1 | 1.0900e-01 | 2.8451e+05 ; | CE - HE3 |
| 3984 | 3987 | 1 | 1.4710e-01 | 3.0711e+05 ; | CE - NZ |
| 3987 | 3988 | 1 | 1.0100e-01 | 3.6317e+05 ; | NZ - HZ1 |
| 3987 | 3989 | 1 | 1.0100e-01 | 3.6317e+05 ; | NZ - HZ2 |
| 3987 | 3990 | 1 | 1.0100e-01 | 3.6317e+05 ; | NZ - HZ3 |
| 3991 | 3992 | 1 | 1.2290e-01 | 4.7698e+05 ; | C - O |
| 3991 | 3993 | 1 | 1.3350e-01 | 4.1003e+05 ; | C - N |
| 3993 | 3994 | 1 | 1.0100e-01 | 3.6317e+05 ; | N - H |
| 3993 | 3995 | 1 | 1.4490e-01 | 2.8200e+05 ; | N - CA |

| | | | | | | |
|---|---|---|---|---|---|---|
| 3995 | 3996 | 1 | 1.0900e-01 | 2.8451e+05 | ; | CA - HA |
| 3995 | 3997 | 1 | 1.5260e-01 | 2.5941e+05 | ; | CA - CB |
| 3995 | 4010 | 1 | 1.5220e-01 | 2.6527e+05 | ; | CA - C |
| 3997 | 3998 | 1 | 1.0900e-01 | 2.8451e+05 | ; | CB - HB2 |
| 3997 | 3999 | 1 | 1.0900e-01 | 2.8451e+05 | ; | CB - HB3 |
| 3997 | 4000 | 1 | 1.5260e-01 | 2.5941e+05 | ; | CB - CG |
| 4000 | 4001 | 1 | 1.0900e-01 | 2.8451e+05 | ; | CG - HG |
| 4000 | 4002 | 1 | 1.5260e-01 | 2.5941e+05 | ; | CG - CD1 |
| 4000 | 4006 | 1 | 1.5260e-01 | 2.5941e+05 | ; | CG - CD2 |
| 4002 | 4003 | 1 | 1.0900e-01 | 2.8451e+05 | ; | CD1 - HD11 |
| 4002 | 4004 | 1 | 1.0900e-01 | 2.8451e+05 | ; | CD1 - HD12 |
| 4002 | 4005 | 1 | 1.0900e-01 | 2.8451e+05 | ; | CD1 - HD13 |
| 4006 | 4007 | 1 | 1.0900e-01 | 2.8451e+05 | ; | CD2 - HD21 |
| 4006 | 4008 | 1 | 1.0900e-01 | 2.8451e+05 | ; | CD2 - HD22 |
| 4006 | 4009 | 1 | 1.0900e-01 | 2.8451e+05 | ; | CD2 - HD23 |
| 4010 | 4011 | 1 | 1.2290e-01 | 4.7698e+05 | ; | C - O |
| 4010 | 4012 | 1 | 1.3350e-01 | 4.1003e+05 | ; | C - N |
| 4012 | 4013 | 1 | 1.0100e-01 | 3.6317e+05 | ; | N - H |
| 4012 | 4014 | 1 | 1.4490e-01 | 2.8200e+05 | ; | N - CA |
| 4014 | 4015 | 1 | 1.0900e-01 | 2.8451e+05 | ; | CA - HA |
| 4014 | 4016 | 1 | 1.5260e-01 | 2.5941e+05 | ; | CA - CB |
| 4014 | 4029 | 1 | 1.5220e-01 | 2.6527e+05 | ; | CA - C |
| 4016 | 4017 | 1 | 1.0900e-01 | 2.8451e+05 | ; | CB - HB2 |
| 4016 | 4018 | 1 | 1.0900e-01 | 2.8451e+05 | ; | CB - HB3 |
| 4016 | 4019 | 1 | 1.5260e-01 | 2.5941e+05 | ; | CB - CG |
| 4019 | 4020 | 1 | 1.0900e-01 | 2.8451e+05 | ; | CG - HG |
| 4019 | 4021 | 1 | 1.5260e-01 | 2.5941e+05 | ; | CG - CD1 |
| 4019 | 4025 | 1 | 1.5260e-01 | 2.5941e+05 | ; | CG - CD2 |
| 4021 | 4022 | 1 | 1.0900e-01 | 2.8451e+05 | ; | CD1 - HD11 |
| 4021 | 4023 | 1 | 1.0900e-01 | 2.8451e+05 | ; | CD1 - HD12 |
| 4021 | 4024 | 1 | 1.0900e-01 | 2.8451e+05 | ; | CD1 - HD13 |
| 4025 | 4026 | 1 | 1.0900e-01 | 2.8451e+05 | ; | CD2 - HD21 |
| 4025 | 4027 | 1 | 1.0900e-01 | 2.8451e+05 | ; | CD2 - HD22 |
| 4025 | 4028 | 1 | 1.0900e-01 | 2.8451e+05 | ; | CD2 - HD23 |
| 4029 | 4030 | 1 | 1.2290e-01 | 4.7698e+05 | ; | C - O |
| 4029 | 4031 | 1 | 1.3350e-01 | 4.1003e+05 | ; | C - N |
| 4031 | 4032 | 1 | 1.0100e-01 | 3.6317e+05 | ; | N - H |
| 4031 | 4033 | 1 | 1.4490e-01 | 2.8200e+05 | ; | N - CA |
| 4033 | 4034 | 1 | 1.0900e-01 | 2.8451e+05 | ; | CA - HA |
| 4033 | 4035 | 1 | 1.5260e-01 | 2.5941e+05 | ; | CA - CB |
| 4033 | 4043 | 1 | 1.5220e-01 | 2.6527e+05 | ; | CA - C |
| 4035 | 4036 | 1 | 1.0900e-01 | 2.8451e+05 | ; | CB - HB |
| 4035 | 4037 | 1 | 1.5260e-01 | 2.5941e+05 | ; | CB - CG2 |
| 4035 | 4041 | 1 | 1.4100e-01 | 2.6778e+05 | ; | CB - OG1 |
| 4037 | 4038 | 1 | 1.0900e-01 | 2.8451e+05 | ; | CG2 - HG21 |
| 4037 | 4039 | 1 | 1.0900e-01 | 2.8451e+05 | ; | CG2 - HG22 |
| 4037 | 4040 | 1 | 1.0900e-01 | 2.8451e+05 | ; | CG2 - HG23 |
| 4041 | 4042 | 1 | 9.6000e-02 | 4.6275e+05 | ; | OG1 - HG1 |
| 4043 | 4044 | 1 | 1.2290e-01 | 4.7698e+05 | ; | C - O |
| 4043 | 4045 | 1 | 1.3350e-01 | 4.1003e+05 | ; | C - N |
| 4045 | 4046 | 1 | 1.0100e-01 | 3.6317e+05 | ; | N - H |
| 4045 | 4047 | 1 | 1.4490e-01 | 2.8200e+05 | ; | N - CA |

| | | | | | | | | |
|---|---|---|---|---|---|---|---|---|
| 4047 | 4048 | 1 | 1.0900e-01 | 2.8451e+05 | ; | CA | - | HA |
| 4047 | 4049 | 1 | 1.5260e-01 | 2.5941e+05 | ; | CA | - | CB |
| 4047 | 4064 | 1 | 1.5220e-01 | 2.6527e+05 | ; | CA | - | C |
| 4049 | 4050 | 1 | 1.0900e-01 | 2.8451e+05 | ; | CB | - | HB2 |
| 4049 | 4051 | 1 | 1.0900e-01 | 2.8451e+05 | ; | CB | - | HB3 |
| 4049 | 4052 | 1 | 1.5100e-01 | 2.6527e+05 | ; | CB | - | CG |
| 4052 | 4053 | 1 | 1.4000e-01 | 3.9246e+05 | ; | CG | - | CD1 |
| 4052 | 4062 | 1 | 1.4000e-01 | 3.9246e+05 | ; | CG | - | CD2 |
| 4053 | 4054 | 1 | 1.0800e-01 | 3.0711e+05 | ; | CD1 | - | HD1 |
| 4053 | 4055 | 1 | 1.4000e-01 | 3.9246e+05 | ; | CD1 | - | CE1 |
| 4055 | 4056 | 1 | 1.0800e-01 | 3.0711e+05 | ; | CE1 | - | HE1 |
| 4055 | 4057 | 1 | 1.4090e-01 | 3.9246e+05 | ; | CE1 | - | CZ |
| 4057 | 4058 | 1 | 1.3640e-01 | 3.7656e+05 | ; | CZ | - | OH |
| 4057 | 4060 | 1 | 1.4090e-01 | 3.9246e+05 | ; | CZ | - | CE2 |
| 4058 | 4059 | 1 | 9.6000e-02 | 4.6275e+05 | ; | OH | - | HH |
| 4060 | 4061 | 1 | 1.0800e-01 | 3.0711e+05 | ; | CE2 | - | HE2 |
| 4060 | 4062 | 1 | 1.4000e-01 | 3.9246e+05 | ; | CE2 | - | CD2 |
| 4062 | 4063 | 1 | 1.0800e-01 | 3.0711e+05 | ; | CD2 | - | HD2 |
| 4064 | 4065 | 1 | 1.2290e-01 | 4.7698e+05 | ; | C | - | O |
| 4064 | 4066 | 1 | 1.3350e-01 | 4.1003e+05 | ; | C | - | N |
| 4066 | 4067 | 1 | 1.0100e-01 | 3.6317e+05 | ; | N | - | H |
| 4066 | 4068 | 1 | 1.4490e-01 | 2.8200e+05 | ; | N | - | CA |
| 4068 | 4069 | 1 | 1.0900e-01 | 2.8451e+05 | ; | CA | - | HA |
| 4068 | 4070 | 1 | 1.5260e-01 | 2.5941e+05 | ; | CA | - | CB |
| 4068 | 4078 | 1 | 1.5220e-01 | 2.6527e+05 | ; | CA | - | C |
| 4070 | 4071 | 1 | 1.0900e-01 | 2.8451e+05 | ; | CB | - | HB2 |
| 4070 | 4072 | 1 | 1.0900e-01 | 2.8451e+05 | ; | CB | - | HB3 |
| 4070 | 4073 | 1 | 1.5220e-01 | 2.6527e+05 | ; | CB | - | CG |
| 4073 | 4074 | 1 | 1.2290e-01 | 4.7698e+05 | ; | CG | - | OD1 |
| 4073 | 4075 | 1 | 1.3350e-01 | 4.1003e+05 | ; | CG | - | ND2 |
| 4075 | 4076 | 1 | 1.0100e-01 | 3.6317e+05 | ; | ND2 | - | HD21 |
| 4075 | 4077 | 1 | 1.0100e-01 | 3.6317e+05 | ; | ND2 | - | HD22 |
| 4078 | 4079 | 1 | 1.2290e-01 | 4.7698e+05 | ; | C | - | O |
| 4078 | 4080 | 1 | 1.3350e-01 | 4.1003e+05 | ; | C | - | N |
| 4080 | 4081 | 1 | 1.4490e-01 | 2.8200e+05 | ; | N | - | CD |
| 4080 | 4090 | 1 | 1.4490e-01 | 2.8200e+05 | ; | N | - | CA |
| 4081 | 4082 | 1 | 1.0900e-01 | 2.8451e+05 | ; | CD | - | HD2 |
| 4081 | 4083 | 1 | 1.0900e-01 | 2.8451e+05 | ; | CD | - | HD3 |
| 4081 | 4084 | 1 | 1.5260e-01 | 2.5941e+05 | ; | CD | - | CG |
| 4084 | 4085 | 1 | 1.0900e-01 | 2.8451e+05 | ; | CG | - | HG2 |
| 4084 | 4086 | 1 | 1.0900e-01 | 2.8451e+05 | ; | CG | - | HG3 |
| 4084 | 4087 | 1 | 1.5260e-01 | 2.5941e+05 | ; | CG | - | CB |
| 4087 | 4088 | 1 | 1.0900e-01 | 2.8451e+05 | ; | CB | - | HB2 |
| 4087 | 4089 | 1 | 1.0900e-01 | 2.8451e+05 | ; | CB | - | HB3 |
| 4087 | 4090 | 1 | 1.5260e-01 | 2.5941e+05 | ; | CB | - | CA |
| 4090 | 4091 | 1 | 1.0900e-01 | 2.8451e+05 | ; | CA | - | HA |
| 4090 | 4092 | 1 | 1.5220e-01 | 2.6527e+05 | ; | CA | - | C |
| 4092 | 4093 | 1 | 1.2290e-01 | 4.7698e+05 | ; | C | - | O |
| 4092 | 4094 | 1 | 1.3350e-01 | 4.1003e+05 | ; | C | - | N |
| 4094 | 4095 | 1 | 1.0100e-01 | 3.6317e+05 | ; | N | - | H |
| 4094 | 4096 | 1 | 1.4490e-01 | 2.8200e+05 | ; | N | - | CA |
| 4096 | 4097 | 1 | 1.0900e-01 | 2.8451e+05 | ; | CA | - | HA |

| | | | | | |
|---|---|---|---|---|---|
| 4096 | 4098 | 1 | 1.5260e-01 | 2.5941e+05 ; | CA - CB |
| 4096 | 4106 | 1 | 1.5220e-01 | 2.6527e+05 ; | CA - C |
| 4098 | 4099 | 1 | 1.0900e-01 | 2.8451e+05 ; | CB - HB2 |
| 4098 | 4100 | 1 | 1.0900e-01 | 2.8451e+05 ; | CB - HB3 |
| 4098 | 4101 | 1 | 1.5220e-01 | 2.6527e+05 ; | CB - CG |
| 4101 | 4102 | 1 | 1.2290e-01 | 4.7698e+05 ; | CG - OD1 |
| 4101 | 4103 | 1 | 1.3350e-01 | 4.1003e+05 ; | CG - ND2 |
| 4103 | 4104 | 1 | 1.0100e-01 | 3.6317e+05 ; | ND2 - HD21 |
| 4103 | 4105 | 1 | 1.0100e-01 | 3.6317e+05 ; | ND2 - HD22 |
| 4106 | 4107 | 1 | 1.2290e-01 | 4.7698e+05 ; | C - O |
| 4106 | 4108 | 1 | 1.3350e-01 | 4.1003e+05 ; | C - N |
| 4108 | 4109 | 1 | 1.0100e-01 | 3.6317e+05 ; | N - H |
| 4108 | 4110 | 1 | 1.4490e-01 | 2.8200e+05 ; | N - CA |
| 4110 | 4111 | 1 | 1.0900e-01 | 2.8451e+05 ; | CA - HA |
| 4110 | 4112 | 1 | 1.5260e-01 | 2.5941e+05 ; | CA - CB |
| 4110 | 4121 | 1 | 1.5220e-01 | 2.6527e+05 ; | CA - C |
| 4112 | 4113 | 1 | 1.0900e-01 | 2.8451e+05 ; | CB - HB2 |
| 4112 | 4114 | 1 | 1.0900e-01 | 2.8451e+05 ; | CB - HB3 |
| 4112 | 4115 | 1 | 1.5260e-01 | 2.5941e+05 ; | CB - CG |
| 4115 | 4116 | 1 | 1.0900e-01 | 2.8451e+05 ; | CG - HG2 |
| 4115 | 4117 | 1 | 1.0900e-01 | 2.8451e+05 ; | CG - HG3 |
| 4115 | 4118 | 1 | 1.5220e-01 | 2.6527e+05 ; | CG - CD |
| 4118 | 4119 | 1 | 1.2500e-01 | 5.4894e+05 ; | CD - OE1 |
| 4118 | 4120 | 1 | 1.2500e-01 | 5.4894e+05 ; | CD - OE2 |
| 4121 | 4122 | 1 | 1.2290e-01 | 4.7698e+05 ; | C - O |
| 4121 | 4123 | 1 | 1.3350e-01 | 4.1003e+05 ; | C - N |
| 4123 | 4124 | 1 | 1.0100e-01 | 3.6317e+05 ; | N - H |
| 4123 | 4125 | 1 | 1.4490e-01 | 2.8200e+05 ; | N - CA |
| 4125 | 4126 | 1 | 1.0900e-01 | 2.8451e+05 ; | CA - HA |
| 4125 | 4127 | 1 | 1.5260e-01 | 2.5941e+05 ; | CA - CB |
| 4125 | 4145 | 1 | 1.5220e-01 | 2.6527e+05 ; | CA - C |
| 4127 | 4128 | 1 | 1.0900e-01 | 2.8451e+05 ; | CB - HB2 |
| 4127 | 4129 | 1 | 1.0900e-01 | 2.8451e+05 ; | CB - HB3 |
| 4127 | 4130 | 1 | 1.5260e-01 | 2.5941e+05 ; | CB - CG |
| 4130 | 4131 | 1 | 1.0900e-01 | 2.8451e+05 ; | CG - HG2 |
| 4130 | 4132 | 1 | 1.0900e-01 | 2.8451e+05 ; | CG - HG3 |
| 4130 | 4133 | 1 | 1.5260e-01 | 2.5941e+05 ; | CG - CD |
| 4133 | 4134 | 1 | 1.0900e-01 | 2.8451e+05 ; | CD - HD2 |
| 4133 | 4135 | 1 | 1.0900e-01 | 2.8451e+05 ; | CD - HD3 |
| 4133 | 4136 | 1 | 1.4630e-01 | 2.8200e+05 ; | CD - NE |
| 4136 | 4137 | 1 | 1.0100e-01 | 3.6317e+05 ; | NE - HE |
| 4136 | 4138 | 1 | 1.3400e-01 | 4.0250e+05 ; | NE - CZ |
| 4138 | 4139 | 1 | 1.3400e-01 | 4.0250e+05 ; | CZ - NH1 |
| 4138 | 4142 | 1 | 1.3400e-01 | 4.0250e+05 ; | CZ - NH2 |
| 4139 | 4140 | 1 | 1.0100e-01 | 3.6317e+05 ; | NH1 - HH11 |
| 4139 | 4141 | 1 | 1.0100e-01 | 3.6317e+05 ; | NH1 - HH12 |
| 4142 | 4143 | 1 | 1.0100e-01 | 3.6317e+05 ; | NH2 - HH21 |
| 4142 | 4144 | 1 | 1.0100e-01 | 3.6317e+05 ; | NH2 - HH22 |
| 4145 | 4146 | 1 | 1.2290e-01 | 4.7698e+05 ; | C - O |
| 4145 | 4147 | 1 | 1.3350e-01 | 4.1003e+05 ; | C - N |
| 4147 | 4148 | 1 | 1.0100e-01 | 3.6317e+05 ; | N - H |
| 4147 | 4149 | 1 | 1.4490e-01 | 2.8200e+05 ; | N - CA |

| | | | | | | |
|---|---|---|---|---|---|---|
| 4149 | 4150 | 1 | 1.0900e-01 | 2.8451e+05 | ; | CA - HA |
| 4149 | 4151 | 1 | 1.5260e-01 | 2.5941e+05 | ; | CA - CB |
| 4149 | 4156 | 1 | 1.5220e-01 | 2.6527e+05 | ; | CA - C |
| 4151 | 4152 | 1 | 1.0900e-01 | 2.8451e+05 | ; | CB - HB2 |
| 4151 | 4153 | 1 | 1.0900e-01 | 2.8451e+05 | ; | CB - HB3 |
| 4151 | 4154 | 1 | 1.8100e-01 | 1.9832e+05 | ; | CB - SG |
| 4154 | 4155 | 1 | 1.3360e-01 | 2.2928e+05 | ; | SG - HG |
| 4156 | 4157 | 1 | 1.2290e-01 | 4.7698e+05 | ; | C - O |
| 4156 | 4158 | 1 | 1.3350e-01 | 4.1003e+05 | ; | C - N |
| 4158 | 4159 | 1 | 1.0100e-01 | 3.6317e+05 | ; | N - H |
| 4158 | 4160 | 1 | 1.4490e-01 | 2.8200e+05 | ; | N - CA |
| 4160 | 4161 | 1 | 1.0900e-01 | 2.8451e+05 | ; | CA - HA |
| 4160 | 4162 | 1 | 1.5260e-01 | 2.5941e+05 | ; | CA - CB |
| 4160 | 4170 | 1 | 1.5220e-01 | 2.6527e+05 | ; | CA - C |
| 4162 | 4163 | 1 | 1.0900e-01 | 2.8451e+05 | ; | CB - HB |
| 4162 | 4164 | 1 | 1.5260e-01 | 2.5941e+05 | ; | CB - CG2 |
| 4162 | 4168 | 1 | 1.4100e-01 | 2.6778e+05 | ; | CB - OG1 |
| 4164 | 4165 | 1 | 1.0900e-01 | 2.8451e+05 | ; | CG2 - HG21 |
| 4164 | 4166 | 1 | 1.0900e-01 | 2.8451e+05 | ; | CG2 - HG22 |
| 4164 | 4167 | 1 | 1.0900e-01 | 2.8451e+05 | ; | CG2 - HG23 |
| 4168 | 4169 | 1 | 9.6000e-02 | 4.6275e+05 | ; | OG1 - HG1 |
| 4170 | 4171 | 1 | 1.2290e-01 | 4.7698e+05 | ; | C - O |
| 4170 | 4172 | 1 | 1.3350e-01 | 4.1003e+05 | ; | C - N |
| 4172 | 4173 | 1 | 1.0100e-01 | 3.6317e+05 | ; | N - H |
| 4172 | 4174 | 1 | 1.4490e-01 | 2.8200e+05 | ; | N - CA |
| 4174 | 4175 | 1 | 1.0900e-01 | 2.8451e+05 | ; | CA - HA |
| 4174 | 4176 | 1 | 1.5260e-01 | 2.5941e+05 | ; | CA - CB |
| 4174 | 4189 | 1 | 1.5220e-01 | 2.6527e+05 | ; | CA - C |
| 4176 | 4177 | 1 | 1.0900e-01 | 2.8451e+05 | ; | CB - HB |
| 4176 | 4178 | 1 | 1.5260e-01 | 2.5941e+05 | ; | CB - CG2 |
| 4176 | 4182 | 1 | 1.5260e-01 | 2.5941e+05 | ; | CB - CG1 |
| 4178 | 4179 | 1 | 1.0900e-01 | 2.8451e+05 | ; | CG2 - HG21 |
| 4178 | 4180 | 1 | 1.0900e-01 | 2.8451e+05 | ; | CG2 - HG22 |
| 4178 | 4181 | 1 | 1.0900e-01 | 2.8451e+05 | ; | CG2 - HG23 |
| 4182 | 4183 | 1 | 1.0900e-01 | 2.8451e+05 | ; | CG1 - HG12 |
| 4182 | 4184 | 1 | 1.0900e-01 | 2.8451e+05 | ; | CG1 - HG13 |
| 4182 | 4185 | 1 | 1.5260e-01 | 2.5941e+05 | ; | CG1 - CD1 |
| 4185 | 4186 | 1 | 1.0900e-01 | 2.8451e+05 | ; | CD1 - HD11 |
| 4185 | 4187 | 1 | 1.0900e-01 | 2.8451e+05 | ; | CD1 - HD12 |
| 4185 | 4188 | 1 | 1.0900e-01 | 2.8451e+05 | ; | CD1 - HD13 |
| 4189 | 4190 | 1 | 1.2290e-01 | 4.7698e+05 | ; | C - O |
| 4189 | 4191 | 1 | 1.3350e-01 | 4.1003e+05 | ; | C - N |
| 4191 | 4192 | 1 | 1.0100e-01 | 3.6317e+05 | ; | N - H |
| 4191 | 4193 | 1 | 1.4490e-01 | 2.8200e+05 | ; | N - CA |
| 4193 | 4194 | 1 | 1.0900e-01 | 2.8451e+05 | ; | CA - HA |
| 4193 | 4195 | 1 | 1.5260e-01 | 2.5941e+05 | ; | CA - CB |
| 4193 | 4204 | 1 | 1.5220e-01 | 2.6527e+05 | ; | CA - C |
| 4195 | 4196 | 1 | 1.0900e-01 | 2.8451e+05 | ; | CB - HB2 |
| 4195 | 4197 | 1 | 1.0900e-01 | 2.8451e+05 | ; | CB - HB3 |
| 4195 | 4198 | 1 | 1.5260e-01 | 2.5941e+05 | ; | CB - CG |
| 4198 | 4199 | 1 | 1.0900e-01 | 2.8451e+05 | ; | CG - HG2 |
| 4198 | 4200 | 1 | 1.0900e-01 | 2.8451e+05 | ; | CG - HG3 |

| | | | | | | |
|---|---|---|---|---|---|---|
| 4198 | 4201 | 1 | 1.5220e-01 | 2.6527e+05 | ; | CG - CD |
| 4201 | 4202 | 1 | 1.2500e-01 | 5.4894e+05 | ; | CD - OE1 |
| 4201 | 4203 | 1 | 1.2500e-01 | 5.4894e+05 | ; | CD - OE2 |
| 4204 | 4205 | 1 | 1.2290e-01 | 4.7698e+05 | ; | C - O |
| 4204 | 4206 | 1 | 1.3350e-01 | 4.1003e+05 | ; | C - N |
| 4206 | 4207 | 1 | 1.0100e-01 | 3.6317e+05 | ; | N - H |
| 4206 | 4208 | 1 | 1.4490e-01 | 2.8200e+05 | ; | N - CA |
| 4208 | 4209 | 1 | 1.0900e-01 | 2.8451e+05 | ; | CA - HA |
| 4208 | 4210 | 1 | 1.5260e-01 | 2.5941e+05 | ; | CA - CB |
| 4208 | 4219 | 1 | 1.5220e-01 | 2.6527e+05 | ; | CA - C |
| 4210 | 4211 | 1 | 1.0900e-01 | 2.8451e+05 | ; | CB - HB2 |
| 4210 | 4212 | 1 | 1.0900e-01 | 2.8451e+05 | ; | CB - HB3 |
| 4210 | 4213 | 1 | 1.5260e-01 | 2.5941e+05 | ; | CB - CG |
| 4213 | 4214 | 1 | 1.0900e-01 | 2.8451e+05 | ; | CG - HG2 |
| 4213 | 4215 | 1 | 1.0900e-01 | 2.8451e+05 | ; | CG - HG3 |
| 4213 | 4216 | 1 | 1.5220e-01 | 2.6527e+05 | ; | CG - CD |
| 4216 | 4217 | 1 | 1.2500e-01 | 5.4894e+05 | ; | CD - OE1 |
| 4216 | 4218 | 1 | 1.2500e-01 | 5.4894e+05 | ; | CD - OE2 |
| 4219 | 4220 | 1 | 1.2290e-01 | 4.7698e+05 | ; | C - O |
| 4219 | 4221 | 1 | 1.3350e-01 | 4.1003e+05 | ; | C - N |
| 4221 | 4222 | 1 | 1.0100e-01 | 3.6317e+05 | ; | N - H |
| 4221 | 4223 | 1 | 1.4490e-01 | 2.8200e+05 | ; | N - CA |
| 4223 | 4224 | 1 | 1.0900e-01 | 2.8451e+05 | ; | CA - HA |
| 4223 | 4225 | 1 | 1.5260e-01 | 2.5941e+05 | ; | CA - CB |
| 4223 | 4229 | 1 | 1.5220e-01 | 2.6527e+05 | ; | CA - C |
| 4225 | 4226 | 1 | 1.0900e-01 | 2.8451e+05 | ; | CB - HB1 |
| 4225 | 4227 | 1 | 1.0900e-01 | 2.8451e+05 | ; | CB - HB2 |
| 4225 | 4228 | 1 | 1.0900e-01 | 2.8451e+05 | ; | CB - HB3 |
| 4229 | 4230 | 1 | 1.2290e-01 | 4.7698e+05 | ; | C - O |
| 4229 | 4231 | 1 | 1.3350e-01 | 4.1003e+05 | ; | C - N |
| 4231 | 4232 | 1 | 1.0100e-01 | 3.6317e+05 | ; | N - H |
| 4231 | 4233 | 1 | 1.4490e-01 | 2.8200e+05 | ; | N - CA |
| 4233 | 4234 | 1 | 1.0900e-01 | 2.8451e+05 | ; | CA - HA |
| 4233 | 4235 | 1 | 1.5260e-01 | 2.5941e+05 | ; | CA - CB |
| 4233 | 4248 | 1 | 1.5220e-01 | 2.6527e+05 | ; | CA - C |
| 4235 | 4236 | 1 | 1.0900e-01 | 2.8451e+05 | ; | CB - HB2 |
| 4235 | 4237 | 1 | 1.0900e-01 | 2.8451e+05 | ; | CB - HB3 |
| 4235 | 4238 | 1 | 1.5260e-01 | 2.5941e+05 | ; | CB - CG |
| 4238 | 4239 | 1 | 1.0900e-01 | 2.8451e+05 | ; | CG - HG |
| 4238 | 4240 | 1 | 1.5260e-01 | 2.5941e+05 | ; | CG - CD1 |
| 4238 | 4244 | 1 | 1.5260e-01 | 2.5941e+05 | ; | CG - CD2 |
| 4240 | 4241 | 1 | 1.0900e-01 | 2.8451e+05 | ; | CD1 - HD11 |
| 4240 | 4242 | 1 | 1.0900e-01 | 2.8451e+05 | ; | CD1 - HD12 |
| 4240 | 4243 | 1 | 1.0900e-01 | 2.8451e+05 | ; | CD1 - HD13 |
| 4244 | 4245 | 1 | 1.0900e-01 | 2.8451e+05 | ; | CD2 - HD21 |
| 4244 | 4246 | 1 | 1.0900e-01 | 2.8451e+05 | ; | CD2 - HD22 |
| 4244 | 4247 | 1 | 1.0900e-01 | 2.8451e+05 | ; | CD2 - HD23 |
| 4248 | 4249 | 1 | 1.2290e-01 | 4.7698e+05 | ; | C - O |
| 4248 | 4250 | 1 | 1.3350e-01 | 4.1003e+05 | ; | C - N |
| 4250 | 4251 | 1 | 1.0100e-01 | 3.6317e+05 | ; | N - H |
| 4250 | 4252 | 1 | 1.4490e-01 | 2.8200e+05 | ; | N - CA |
| 4252 | 4253 | 1 | 1.0900e-01 | 2.8451e+05 | ; | CA - HA |

| | | | | | |
|---|---|---|---|---|---|
| 4252 | 4254 | 1 | 1.5260e-01 | 2.5941e+05 ; | CA - CB |
| 4252 | 4262 | 1 | 1.5220e-01 | 2.6527e+05 ; | CA - C |
| 4254 | 4255 | 1 | 1.0900e-01 | 2.8451e+05 ; | CB - HB2 |
| 4254 | 4256 | 1 | 1.0900e-01 | 2.8451e+05 ; | CB - HB3 |
| 4254 | 4257 | 1 | 1.5220e-01 | 2.6527e+05 ; | CB - CG |
| 4257 | 4258 | 1 | 1.2290e-01 | 4.7698e+05 ; | CG - OD1 |
| 4257 | 4259 | 1 | 1.3350e-01 | 4.1003e+05 ; | CG - ND2 |
| 4259 | 4260 | 1 | 1.0100e-01 | 3.6317e+05 ; | ND2 - HD21 |
| 4259 | 4261 | 1 | 1.0100e-01 | 3.6317e+05 ; | ND2 - HD22 |
| 4262 | 4263 | 1 | 1.2290e-01 | 4.7698e+05 ; | C - O |
| 4262 | 4264 | 1 | 1.3350e-01 | 4.1003e+05 ; | C - N |
| 4264 | 4265 | 1 | 1.0100e-01 | 3.6317e+05 ; | N - H |
| 4264 | 4266 | 1 | 1.4490e-01 | 2.8200e+05 ; | N - CA |
| 4266 | 4267 | 1 | 1.0900e-01 | 2.8451e+05 ; | CA - HA |
| 4266 | 4268 | 1 | 1.5260e-01 | 2.5941e+05 ; | CA - CB |
| 4266 | 4280 | 1 | 1.5220e-01 | 2.6527e+05 ; | CA - C |
| 4268 | 4269 | 1 | 1.0900e-01 | 2.8451e+05 ; | CB - HB2 |
| 4268 | 4270 | 1 | 1.0900e-01 | 2.8451e+05 ; | CB - HB3 |
| 4268 | 4271 | 1 | 1.5040e-01 | 2.6527e+05 ; | CB - CG |
| 4271 | 4272 | 1 | 1.3850e-01 | 3.5313e+05 ; | CG - ND1 |
| 4271 | 4278 | 1 | 1.3710e-01 | 4.3346e+05 ; | CG - CD2 |
| 4272 | 4273 | 1 | 1.0100e-01 | 3.6317e+05 ; | ND1 - HD1 |
| 4272 | 4274 | 1 | 1.3430e-01 | 3.9915e+05 ; | ND1 - CE1 |
| 4274 | 4275 | 1 | 1.0800e-01 | 3.0711e+05 ; | CE1 - HE1 |
| 4274 | 4276 | 1 | 1.3430e-01 | 3.9915e+05 ; | CE1 - NE2 |
| 4276 | 4277 | 1 | 1.0100e-01 | 3.6317e+05 ; | NE2 - HE2 |
| 4276 | 4278 | 1 | 1.3810e-01 | 3.5731e+05 ; | NE2 - CD2 |
| 4278 | 4279 | 1 | 1.0800e-01 | 3.0711e+05 ; | CD2 - HD2 |
| 4280 | 4281 | 1 | 1.2290e-01 | 4.7698e+05 ; | C - O |
| 4280 | 4282 | 1 | 1.3350e-01 | 4.1003e+05 ; | C - N |
| 4282 | 4283 | 1 | 1.4490e-01 | 2.8200e+05 ; | N - CD |
| 4282 | 4292 | 1 | 1.4490e-01 | 2.8200e+05 ; | N - CA |
| 4283 | 4284 | 1 | 1.0900e-01 | 2.8451e+05 ; | CD - HD2 |
| 4283 | 4285 | 1 | 1.0900e-01 | 2.8451e+05 ; | CD - HD3 |
| 4283 | 4286 | 1 | 1.5260e-01 | 2.5941e+05 ; | CD - CG |
| 4286 | 4287 | 1 | 1.0900e-01 | 2.8451e+05 ; | CG - HG2 |
| 4286 | 4288 | 1 | 1.0900e-01 | 2.8451e+05 ; | CG - HG3 |
| 4286 | 4289 | 1 | 1.5260e-01 | 2.5941e+05 ; | CG - CB |
| 4289 | 4290 | 1 | 1.0900e-01 | 2.8451e+05 ; | CB - HB2 |
| 4289 | 4291 | 1 | 1.0900e-01 | 2.8451e+05 ; | CB - HB3 |
| 4289 | 4292 | 1 | 1.5260e-01 | 2.5941e+05 ; | CB - CA |
| 4292 | 4293 | 1 | 1.0900e-01 | 2.8451e+05 ; | CA - HA |
| 4292 | 4294 | 1 | 1.5220e-01 | 2.6527e+05 ; | CA - C |
| 4294 | 4295 | 1 | 1.2290e-01 | 4.7698e+05 ; | C - O |
| 4294 | 4296 | 1 | 1.3350e-01 | 4.1003e+05 ; | C - N |
| 4296 | 4297 | 1 | 1.0100e-01 | 3.6317e+05 ; | N - H |
| 4296 | 4298 | 1 | 1.4490e-01 | 2.8200e+05 ; | N - CA |
| 4298 | 4299 | 1 | 1.0900e-01 | 2.8451e+05 ; | CA - HA |
| 4298 | 4300 | 1 | 1.5260e-01 | 2.5941e+05 ; | CA - CB |
| 4298 | 4318 | 1 | 1.5220e-01 | 2.6527e+05 ; | CA - C |
| 4300 | 4301 | 1 | 1.0900e-01 | 2.8451e+05 ; | CB - HB2 |
| 4300 | 4302 | 1 | 1.0900e-01 | 2.8451e+05 ; | CB - HB3 |

| | | | | | |
|---|---|---|---|---|---|
| 4300 | 4303 | 1 | 1.4950e-01 | 2.6527e+05 ; | CB - CG |
| 4303 | 4304 | 1 | 1.3520e-01 | 4.5689e+05 ; | CG - CD1 |
| 4303 | 4317 | 1 | 1.4590e-01 | 3.2468e+05 ; | CG - CD2 |
| 4304 | 4305 | 1 | 1.0800e-01 | 3.0711e+05 ; | CD1 - HD1 |
| 4304 | 4306 | 1 | 1.3810e-01 | 3.5731e+05 ; | CD1 - NE1 |
| 4306 | 4307 | 1 | 1.0100e-01 | 3.6317e+05 ; | NE1 - HE1 |
| 4306 | 4308 | 1 | 1.3800e-01 | 3.5815e+05 ; | NE1 - CE2 |
| 4308 | 4309 | 1 | 1.4000e-01 | 3.9246e+05 ; | CE2 - CZ2 |
| 4308 | 4317 | 1 | 1.4190e-01 | 3.7405e+05 ; | CE2 - CD2 |
| 4309 | 4310 | 1 | 1.0800e-01 | 3.0711e+05 ; | CZ2 - HZ2 |
| 4309 | 4311 | 1 | 1.4000e-01 | 3.9246e+05 ; | CZ2 - CH2 |
| 4311 | 4312 | 1 | 1.0800e-01 | 3.0711e+05 ; | CH2 - HH2 |
| 4311 | 4313 | 1 | 1.4000e-01 | 3.9246e+05 ; | CH2 - CZ3 |
| 4313 | 4314 | 1 | 1.0800e-01 | 3.0711e+05 ; | CZ3 - HZ3 |
| 4313 | 4315 | 1 | 1.4000e-01 | 3.9246e+05 ; | CZ3 - CE3 |
| 4315 | 4316 | 1 | 1.0800e-01 | 3.0711e+05 ; | CE3 - HE3 |
| 4315 | 4317 | 1 | 1.4040e-01 | 3.9246e+05 ; | CE3 - CD2 |
| 4318 | 4319 | 1 | 1.2290e-01 | 4.7698e+05 ; | C - O |
| 4318 | 4320 | 1 | 1.3350e-01 | 4.1003e+05 ; | C - N |
| 4320 | 4321 | 1 | 1.0100e-01 | 3.6317e+05 ; | N - H |
| 4320 | 4322 | 1 | 1.4490e-01 | 2.8200e+05 ; | N - CA |
| 4322 | 4323 | 1 | 1.0900e-01 | 2.8451e+05 ; | CA - HA |
| 4322 | 4324 | 1 | 1.5260e-01 | 2.5941e+05 ; | CA - CB |
| 4322 | 4337 | 1 | 1.5220e-01 | 2.6527e+05 ; | CA - C |
| 4324 | 4325 | 1 | 1.0900e-01 | 2.8451e+05 ; | CB - HB |
| 4324 | 4326 | 1 | 1.5260e-01 | 2.5941e+05 ; | CB - CG2 |
| 4324 | 4330 | 1 | 1.5260e-01 | 2.5941e+05 ; | CB - CG1 |
| 4326 | 4327 | 1 | 1.0900e-01 | 2.8451e+05 ; | CG2 - HG21 |
| 4326 | 4328 | 1 | 1.0900e-01 | 2.8451e+05 ; | CG2 - HG22 |
| 4326 | 4329 | 1 | 1.0900e-01 | 2.8451e+05 ; | CG2 - HG23 |
| 4330 | 4331 | 1 | 1.0900e-01 | 2.8451e+05 ; | CG1 - HG12 |
| 4330 | 4332 | 1 | 1.0900e-01 | 2.8451e+05 ; | CG1 - HG13 |
| 4330 | 4333 | 1 | 1.5260e-01 | 2.5941e+05 ; | CG1 - CD1 |
| 4333 | 4334 | 1 | 1.0900e-01 | 2.8451e+05 ; | CD1 - HD11 |
| 4333 | 4335 | 1 | 1.0900e-01 | 2.8451e+05 ; | CD1 - HD12 |
| 4333 | 4336 | 1 | 1.0900e-01 | 2.8451e+05 ; | CD1 - HD13 |
| 4337 | 4338 | 1 | 1.2290e-01 | 4.7698e+05 ; | C - O |
| 4337 | 4339 | 1 | 1.3350e-01 | 4.1003e+05 ; | C - N |
| 4339 | 4340 | 1 | 1.0100e-01 | 3.6317e+05 ; | N - H |
| 4339 | 4341 | 1 | 1.4490e-01 | 2.8200e+05 ; | N - CA |
| 4341 | 4342 | 1 | 1.0900e-01 | 2.8451e+05 ; | CA - HA |
| 4341 | 4343 | 1 | 1.5260e-01 | 2.5941e+05 ; | CA - CB |
| 4341 | 4351 | 1 | 1.5220e-01 | 2.6527e+05 ; | CA - C |
| 4343 | 4344 | 1 | 1.0900e-01 | 2.8451e+05 ; | CB - HB |
| 4343 | 4345 | 1 | 1.5260e-01 | 2.5941e+05 ; | CB - CG2 |
| 4343 | 4349 | 1 | 1.4100e-01 | 2.6778e+05 ; | CB - OG1 |
| 4345 | 4346 | 1 | 1.0900e-01 | 2.8451e+05 ; | CG2 - HG21 |
| 4345 | 4347 | 1 | 1.0900e-01 | 2.8451e+05 ; | CG2 - HG22 |
| 4345 | 4348 | 1 | 1.0900e-01 | 2.8451e+05 ; | CG2 - HG23 |
| 4349 | 4350 | 1 | 9.6000e-02 | 4.6275e+05 ; | OG1 - HG1 |
| 4351 | 4352 | 1 | 1.2290e-01 | 4.7698e+05 ; | C - O |
| 4351 | 4353 | 1 | 1.3350e-01 | 4.1003e+05 ; | C - N |

| | | | | | |
|---|---|---|---|---|---|
| 4353 | 4354 | 1 | 1.0100e-01 | 3.6317e+05 ; | N - H |
| 4353 | 4355 | 1 | 1.4490e-01 | 2.8200e+05 ; | N - CA |
| 4355 | 4356 | 1 | 1.0900e-01 | 2.8451e+05 ; | CA - HA |
| 4355 | 4357 | 1 | 1.5260e-01 | 2.5941e+05 ; | CA - CB |
| 4355 | 4368 | 1 | 1.5220e-01 | 2.6527e+05 ; | CA - C |
| 4357 | 4358 | 1 | 1.0900e-01 | 2.8451e+05 ; | CB - HB2 |
| 4357 | 4359 | 1 | 1.0900e-01 | 2.8451e+05 ; | CB - HB3 |
| 4357 | 4360 | 1 | 1.5260e-01 | 2.5941e+05 ; | CB - CG |
| 4360 | 4361 | 1 | 1.0900e-01 | 2.8451e+05 ; | CG - HG2 |
| 4360 | 4362 | 1 | 1.0900e-01 | 2.8451e+05 ; | CG - HG3 |
| 4360 | 4363 | 1 | 1.5220e-01 | 2.6527e+05 ; | CG - CD |
| 4363 | 4364 | 1 | 1.2290e-01 | 4.7698e+05 ; | CD - OE1 |
| 4363 | 4365 | 1 | 1.3350e-01 | 4.1003e+05 ; | CD - NE2 |
| 4365 | 4366 | 1 | 1.0100e-01 | 3.6317e+05 ; | NE2 - HE21 |
| 4365 | 4367 | 1 | 1.0100e-01 | 3.6317e+05 ; | NE2 - HE22 |
| 4368 | 4369 | 1 | 1.2290e-01 | 4.7698e+05 ; | C - O |
| 4368 | 4370 | 1 | 1.3350e-01 | 4.1003e+05 ; | C - N |
| 4370 | 4371 | 1 | 1.0100e-01 | 3.6317e+05 ; | N - H |
| 4370 | 4372 | 1 | 1.4490e-01 | 2.8200e+05 ; | N - CA |
| 4372 | 4373 | 1 | 1.0900e-01 | 2.8451e+05 ; | CA - HA |
| 4372 | 4374 | 1 | 1.5260e-01 | 2.5941e+05 ; | CA - CB |
| 4372 | 4385 | 1 | 1.5220e-01 | 2.6527e+05 ; | CA - C |
| 4374 | 4375 | 1 | 1.0900e-01 | 2.8451e+05 ; | CB - HB2 |
| 4374 | 4376 | 1 | 1.0900e-01 | 2.8451e+05 ; | CB - HB3 |
| 4374 | 4377 | 1 | 1.5260e-01 | 2.5941e+05 ; | CB - CG |
| 4377 | 4378 | 1 | 1.0900e-01 | 2.8451e+05 ; | CG - HG2 |
| 4377 | 4379 | 1 | 1.0900e-01 | 2.8451e+05 ; | CG - HG3 |
| 4377 | 4380 | 1 | 1.8100e-01 | 1.8995e+05 ; | CG - SD |
| 4380 | 4381 | 1 | 1.8100e-01 | 1.8995e+05 ; | SD - CE |
| 4381 | 4382 | 1 | 1.0900e-01 | 2.8451e+05 ; | CE - HE1 |
| 4381 | 4383 | 1 | 1.0900e-01 | 2.8451e+05 ; | CE - HE2 |
| 4381 | 4384 | 1 | 1.0900e-01 | 2.8451e+05 ; | CE - HE3 |
| 4385 | 4386 | 1 | 1.2290e-01 | 4.7698e+05 ; | C - O |
| 4385 | 4387 | 1 | 1.3350e-01 | 4.1003e+05 ; | C - N |
| 4387 | 4388 | 1 | 1.0100e-01 | 3.6317e+05 ; | N - H |
| 4387 | 4389 | 1 | 1.4490e-01 | 2.8200e+05 ; | N - CA |
| 4389 | 4390 | 1 | 1.0900e-01 | 2.8451e+05 ; | CA - HA |
| 4389 | 4391 | 1 | 1.5260e-01 | 2.5941e+05 ; | CA - CB |
| 4389 | 4399 | 1 | 1.5220e-01 | 2.6527e+05 ; | CA - C |
| 4391 | 4392 | 1 | 1.0900e-01 | 2.8451e+05 ; | CB - HB |
| 4391 | 4393 | 1 | 1.5260e-01 | 2.5941e+05 ; | CB - CG2 |
| 4391 | 4397 | 1 | 1.4100e-01 | 2.6778e+05 ; | CB - OG1 |
| 4393 | 4394 | 1 | 1.0900e-01 | 2.8451e+05 ; | CG2 - HG21 |
| 4393 | 4395 | 1 | 1.0900e-01 | 2.8451e+05 ; | CG2 - HG22 |
| 4393 | 4396 | 1 | 1.0900e-01 | 2.8451e+05 ; | CG2 - HG23 |
| 4397 | 4398 | 1 | 9.6000e-02 | 4.6275e+05 ; | OG1 - HG1 |
| 4399 | 4400 | 1 | 1.2290e-01 | 4.7698e+05 ; | C - O |
| 4399 | 4401 | 1 | 1.3350e-01 | 4.1003e+05 ; | C - N |
| 4401 | 4402 | 1 | 1.0100e-01 | 3.6317e+05 ; | N - H |
| 4401 | 4403 | 1 | 1.4490e-01 | 2.8200e+05 ; | N - CA |
| 4403 | 4404 | 1 | 1.0900e-01 | 2.8451e+05 ; | CA - HA |
| 4403 | 4405 | 1 | 1.5260e-01 | 2.5941e+05 ; | CA - CB |

| | | | | | | |
|---|---|---|---|---|---|---|
| 4403 | 4421 | 1 | 1.5220e-01 | 2.6527e+05 | ; | CA - C |
| 4405 | 4406 | 1 | 1.0900e-01 | 2.8451e+05 | ; | CB - HB2 |
| 4405 | 4407 | 1 | 1.0900e-01 | 2.8451e+05 | ; | CB - HB3 |
| 4405 | 4408 | 1 | 1.5260e-01 | 2.5941e+05 | ; | CB - CG |
| 4408 | 4409 | 1 | 1.0900e-01 | 2.8451e+05 | ; | CG - HG2 |
| 4408 | 4410 | 1 | 1.0900e-01 | 2.8451e+05 | ; | CG - HG3 |
| 4408 | 4411 | 1 | 1.5260e-01 | 2.5941e+05 | ; | CG - CD |
| 4411 | 4412 | 1 | 1.0900e-01 | 2.8451e+05 | ; | CD - HD2 |
| 4411 | 4413 | 1 | 1.0900e-01 | 2.8451e+05 | ; | CD - HD3 |
| 4411 | 4414 | 1 | 1.5260e-01 | 2.5941e+05 | ; | CD - CE |
| 4414 | 4415 | 1 | 1.0900e-01 | 2.8451e+05 | ; | CE - HE2 |
| 4414 | 4416 | 1 | 1.0900e-01 | 2.8451e+05 | ; | CE - HE3 |
| 4414 | 4417 | 1 | 1.4710e-01 | 3.0711e+05 | ; | CE - NZ |
| 4417 | 4418 | 1 | 1.0100e-01 | 3.6317e+05 | ; | NZ - HZ1 |
| 4417 | 4419 | 1 | 1.0100e-01 | 3.6317e+05 | ; | NZ - HZ2 |
| 4417 | 4420 | 1 | 1.0100e-01 | 3.6317e+05 | ; | NZ - HZ3 |
| 4421 | 4422 | 1 | 1.2290e-01 | 4.7698e+05 | ; | C - O |
| 4421 | 4423 | 1 | 1.3350e-01 | 4.1003e+05 | ; | C - N |
| 4423 | 4424 | 1 | 1.0100e-01 | 3.6317e+05 | ; | N - H |
| 4423 | 4425 | 1 | 1.4490e-01 | 2.8200e+05 | ; | N - CA |
| 4425 | 4426 | 1 | 1.0900e-01 | 2.8451e+05 | ; | CA - HA |
| 4425 | 4427 | 1 | 1.5260e-01 | 2.5941e+05 | ; | CA - CB |
| 4425 | 4432 | 1 | 1.5220e-01 | 2.6527e+05 | ; | CA - C |
| 4427 | 4428 | 1 | 1.0900e-01 | 2.8451e+05 | ; | CB - HB2 |
| 4427 | 4429 | 1 | 1.0900e-01 | 2.8451e+05 | ; | CB - HB3 |
| 4427 | 4430 | 1 | 1.4100e-01 | 2.6778e+05 | ; | CB - OG |
| 4430 | 4431 | 1 | 9.6000e-02 | 4.6275e+05 | ; | OG - HG |
| 4432 | 4433 | 1 | 1.2290e-01 | 4.7698e+05 | ; | C - O |
| 4432 | 4434 | 1 | 1.3350e-01 | 4.1003e+05 | ; | C - N |
| 4434 | 4435 | 1 | 1.0100e-01 | 3.6317e+05 | ; | N - H |
| 4434 | 4436 | 1 | 1.4490e-01 | 2.8200e+05 | ; | N - CA |
| 4436 | 4437 | 1 | 1.0900e-01 | 2.8451e+05 | ; | CA - HA |
| 4436 | 4438 | 1 | 1.5260e-01 | 2.5941e+05 | ; | CA - CB |
| 4436 | 4449 | 1 | 1.5220e-01 | 2.6527e+05 | ; | CA - C |
| 4438 | 4439 | 1 | 1.0900e-01 | 2.8451e+05 | ; | CB - HB2 |
| 4438 | 4440 | 1 | 1.0900e-01 | 2.8451e+05 | ; | CB - HB3 |
| 4438 | 4441 | 1 | 1.5040e-01 | 2.6527e+05 | ; | CB - CG |
| 4441 | 4442 | 1 | 1.3940e-01 | 3.4309e+05 | ; | CG - ND1 |
| 4441 | 4447 | 1 | 1.3710e-01 | 4.3346e+05 | ; | CG - CD2 |
| 4442 | 4443 | 1 | 1.3350e-01 | 4.0836e+05 | ; | ND1 - CE1 |
| 4443 | 4444 | 1 | 1.0800e-01 | 3.0711e+05 | ; | CE1 - HE1 |
| 4443 | 4445 | 1 | 1.3430e-01 | 3.9915e+05 | ; | CE1 - NE2 |
| 4445 | 4446 | 1 | 1.0100e-01 | 3.6317e+05 | ; | NE2 - HE2 |
| 4445 | 4447 | 1 | 1.3810e-01 | 3.5731e+05 | ; | NE2 - CD2 |
| 4447 | 4448 | 1 | 1.0800e-01 | 3.0711e+05 | ; | CD2 - HD2 |
| 4449 | 4450 | 1 | 1.2290e-01 | 4.7698e+05 | ; | C - O |
| 4449 | 4451 | 1 | 1.3350e-01 | 4.1003e+05 | ; | C - N |
| 4451 | 4452 | 1 | 1.0100e-01 | 3.6317e+05 | ; | N - H |
| 4451 | 4453 | 1 | 1.4490e-01 | 2.8200e+05 | ; | N - CA |
| 4453 | 4454 | 1 | 1.0900e-01 | 2.8451e+05 | ; | CA - HA |
| 4453 | 4455 | 1 | 1.5260e-01 | 2.5941e+05 | ; | CA - CB |
| 4453 | 4464 | 1 | 1.5220e-01 | 2.6527e+05 | ; | CA - C |

| | | | | | |
|---|---|---|---|---|---|
| 4455 | 4456 | 1 | 1.0900e-01 | 2.8451e+05 ; | CB - HB2 |
| 4455 | 4457 | 1 | 1.0900e-01 | 2.8451e+05 ; | CB - HB3 |
| 4455 | 4458 | 1 | 1.5260e-01 | 2.5941e+05 ; | CB - CG |
| 4458 | 4459 | 1 | 1.0900e-01 | 2.8451e+05 ; | CG - HG2 |
| 4458 | 4460 | 1 | 1.0900e-01 | 2.8451e+05 ; | CG - HG3 |
| 4458 | 4461 | 1 | 1.5220e-01 | 2.6527e+05 ; | CG - CD |
| 4461 | 4462 | 1 | 1.2500e-01 | 5.4894e+05 ; | CD - OE1 |
| 4461 | 4463 | 1 | 1.2500e-01 | 5.4894e+05 ; | CD - OE2 |
| 4464 | 4465 | 1 | 1.2290e-01 | 4.7698e+05 ; | C - O |
| 4464 | 4466 | 1 | 1.3350e-01 | 4.1003e+05 ; | C - N |
| 4466 | 4467 | 1 | 1.0100e-01 | 3.6317e+05 ; | N - H |
| 4466 | 4468 | 1 | 1.4490e-01 | 2.8200e+05 ; | N - CA |
| 4468 | 4469 | 1 | 1.0900e-01 | 2.8451e+05 ; | CA - HA |
| 4468 | 4470 | 1 | 1.5260e-01 | 2.5941e+05 ; | CA - CB |
| 4468 | 4481 | 1 | 1.5220e-01 | 2.6527e+05 ; | CA - C |
| 4470 | 4471 | 1 | 1.0900e-01 | 2.8451e+05 ; | CB - HB2 |
| 4470 | 4472 | 1 | 1.0900e-01 | 2.8451e+05 ; | CB - HB3 |
| 4470 | 4473 | 1 | 1.5040e-01 | 2.6527e+05 ; | CB - CG |
| 4473 | 4474 | 1 | 1.3940e-01 | 3.4309e+05 ; | CG - ND1 |
| 4473 | 4479 | 1 | 1.3710e-01 | 4.3346e+05 ; | CG - CD2 |
| 4474 | 4475 | 1 | 1.3350e-01 | 4.0836e+05 ; | ND1 - CE1 |
| 4475 | 4476 | 1 | 1.0800e-01 | 3.0711e+05 ; | CE1 - HE1 |
| 4475 | 4477 | 1 | 1.3430e-01 | 3.9915e+05 ; | CE1 - NE2 |
| 4477 | 4478 | 1 | 1.0100e-01 | 3.6317e+05 ; | NE2 - HE2 |
| 4477 | 4479 | 1 | 1.3810e-01 | 3.5731e+05 ; | NE2 - CD2 |
| 4479 | 4480 | 1 | 1.0800e-01 | 3.0711e+05 ; | CD2 - HD2 |
| 4481 | 4482 | 1 | 1.2290e-01 | 4.7698e+05 ; | C - O |
| 4481 | 4483 | 1 | 1.3350e-01 | 4.1003e+05 ; | C - N |
| 4483 | 4484 | 1 | 1.0100e-01 | 3.6317e+05 ; | N - H |
| 4483 | 4485 | 1 | 1.4490e-01 | 2.8200e+05 ; | N - CA |
| 4485 | 4486 | 1 | 1.0900e-01 | 2.8451e+05 ; | CA - HA |
| 4485 | 4487 | 1 | 1.5260e-01 | 2.5941e+05 ; | CA - CB |
| 4485 | 4497 | 1 | 1.5220e-01 | 2.6527e+05 ; | CA - C |
| 4487 | 4488 | 1 | 1.0900e-01 | 2.8451e+05 ; | CB - HB |
| 4487 | 4489 | 1 | 1.5260e-01 | 2.5941e+05 ; | CB - CG1 |
| 4487 | 4493 | 1 | 1.5260e-01 | 2.5941e+05 ; | CB - CG2 |
| 4489 | 4490 | 1 | 1.0900e-01 | 2.8451e+05 ; | CG1 - HG11 |
| 4489 | 4491 | 1 | 1.0900e-01 | 2.8451e+05 ; | CG1 - HG12 |
| 4489 | 4492 | 1 | 1.0900e-01 | 2.8451e+05 ; | CG1 - HG13 |
| 4493 | 4494 | 1 | 1.0900e-01 | 2.8451e+05 ; | CG2 - HG21 |
| 4493 | 4495 | 1 | 1.0900e-01 | 2.8451e+05 ; | CG2 - HG22 |
| 4493 | 4496 | 1 | 1.0900e-01 | 2.8451e+05 ; | CG2 - HG23 |
| 4497 | 4498 | 1 | 1.2290e-01 | 4.7698e+05 ; | C - O |
| 4497 | 4499 | 1 | 1.3350e-01 | 4.1003e+05 ; | C - N |
| 4499 | 4500 | 1 | 1.0100e-01 | 3.6317e+05 ; | N - H |
| 4499 | 4501 | 1 | 1.4490e-01 | 2.8200e+05 ; | N - CA |
| 4501 | 4502 | 1 | 1.0900e-01 | 2.8451e+05 ; | CA - HA |
| 4501 | 4503 | 1 | 1.5260e-01 | 2.5941e+05 ; | CA - CB |
| 4501 | 4512 | 1 | 1.5220e-01 | 2.6527e+05 ; | CA - C |
| 4503 | 4504 | 1 | 1.0900e-01 | 2.8451e+05 ; | CB - HB2 |
| 4503 | 4505 | 1 | 1.0900e-01 | 2.8451e+05 ; | CB - HB3 |
| 4503 | 4506 | 1 | 1.5260e-01 | 2.5941e+05 ; | CB - CG |

| | | | | | |
|---|---|---|---|---|---|
| 4506 | 4507 | 1 | 1.0900e-01 | 2.8451e+05 ; | CG - HG2 |
| 4506 | 4508 | 1 | 1.0900e-01 | 2.8451e+05 ; | CG - HG3 |
| 4506 | 4509 | 1 | 1.5220e-01 | 2.6527e+05 ; | CG - CD |
| 4509 | 4510 | 1 | 1.2500e-01 | 5.4894e+05 ; | CD - OE1 |
| 4509 | 4511 | 1 | 1.2500e-01 | 5.4894e+05 ; | CD - OE2 |
| 4512 | 4513 | 1 | 1.2290e-01 | 4.7698e+05 ; | C - O |
| 4512 | 4514 | 1 | 1.3350e-01 | 4.1003e+05 ; | C - N |
| 4514 | 4515 | 1 | 1.0100e-01 | 3.6317e+05 ; | N - H |
| 4514 | 4516 | 1 | 1.4490e-01 | 2.8200e+05 ; | N - CA |
| 4516 | 4517 | 1 | 1.0900e-01 | 2.8451e+05 ; | CA - HA |
| 4516 | 4518 | 1 | 1.5260e-01 | 2.5941e+05 ; | CA - CB |
| 4516 | 4531 | 1 | 1.5220e-01 | 2.6527e+05 ; | CA - C |
| 4518 | 4519 | 1 | 1.0900e-01 | 2.8451e+05 ; | CB - HB2 |
| 4518 | 4520 | 1 | 1.0900e-01 | 2.8451e+05 ; | CB - HB3 |
| 4518 | 4521 | 1 | 1.5260e-01 | 2.5941e+05 ; | CB - CG |
| 4521 | 4522 | 1 | 1.0900e-01 | 2.8451e+05 ; | CG - HG |
| 4521 | 4523 | 1 | 1.5260e-01 | 2.5941e+05 ; | CG - CD1 |
| 4521 | 4527 | 1 | 1.5260e-01 | 2.5941e+05 ; | CG - CD2 |
| 4523 | 4524 | 1 | 1.0900e-01 | 2.8451e+05 ; | CD1 - HD11 |
| 4523 | 4525 | 1 | 1.0900e-01 | 2.8451e+05 ; | CD1 - HD12 |
| 4523 | 4526 | 1 | 1.0900e-01 | 2.8451e+05 ; | CD1 - HD13 |
| 4527 | 4528 | 1 | 1.0900e-01 | 2.8451e+05 ; | CD2 - HD21 |
| 4527 | 4529 | 1 | 1.0900e-01 | 2.8451e+05 ; | CD2 - HD22 |
| 4527 | 4530 | 1 | 1.0900e-01 | 2.8451e+05 ; | CD2 - HD23 |
| 4531 | 4532 | 1 | 1.2290e-01 | 4.7698e+05 ; | C - O |
| 4531 | 4533 | 1 | 1.3350e-01 | 4.1003e+05 ; | C - N |
| 4533 | 4534 | 1 | 1.0100e-01 | 3.6317e+05 ; | N - H |
| 4533 | 4535 | 1 | 1.4490e-01 | 2.8200e+05 ; | N - CA |
| 4535 | 4536 | 1 | 1.0900e-01 | 2.8451e+05 ; | CA - HA |
| 4535 | 4537 | 1 | 1.5260e-01 | 2.5941e+05 ; | CA - CB |
| 4535 | 4542 | 1 | 1.5220e-01 | 2.6527e+05 ; | CA - C |
| 4537 | 4538 | 1 | 1.0900e-01 | 2.8451e+05 ; | CB - HB2 |
| 4537 | 4539 | 1 | 1.0900e-01 | 2.8451e+05 ; | CB - HB3 |
| 4537 | 4540 | 1 | 1.4100e-01 | 2.6778e+05 ; | CB - OG |
| 4540 | 4541 | 1 | 9.6000e-02 | 4.6275e+05 ; | OG - HG |
| 4542 | 4543 | 1 | 1.2290e-01 | 4.7698e+05 ; | C - O |
| 4542 | 4544 | 1 | 1.3350e-01 | 4.1003e+05 ; | C - N |
| 4544 | 4545 | 1 | 1.0100e-01 | 3.6317e+05 ; | N - H |
| 4544 | 4546 | 1 | 1.4490e-01 | 2.8200e+05 ; | N - CA |
| 4546 | 4547 | 1 | 1.0900e-01 | 2.8451e+05 ; | CA - HA |
| 4546 | 4548 | 1 | 1.5260e-01 | 2.5941e+05 ; | CA - CB |
| 4546 | 4553 | 1 | 1.5220e-01 | 2.6527e+05 ; | CA - C |
| 4548 | 4549 | 1 | 1.0900e-01 | 2.8451e+05 ; | CB - HB2 |
| 4548 | 4550 | 1 | 1.0900e-01 | 2.8451e+05 ; | CB - HB3 |
| 4548 | 4551 | 1 | 1.4100e-01 | 2.6778e+05 ; | CB - OG |
| 4551 | 4552 | 1 | 9.6000e-02 | 4.6275e+05 ; | OG - HG |
| 4553 | 4554 | 1 | 1.2290e-01 | 4.7698e+05 ; | C - O |
| 4553 | 4555 | 1 | 1.3350e-01 | 4.1003e+05 ; | C - N |
| 4555 | 4556 | 1 | 1.0100e-01 | 3.6317e+05 ; | N - H |
| 4555 | 4557 | 1 | 1.4490e-01 | 2.8200e+05 ; | N - CA |
| 4557 | 4558 | 1 | 1.0900e-01 | 2.8451e+05 ; | CA - HA |
| 4557 | 4559 | 1 | 1.5260e-01 | 2.5941e+05 ; | CA - CB |

| | | | | | | |
|---|---|---|---|---|---|---|
| 4557 | 4567 | 1 | 1.5220e-01 | 2.6527e+05 | ; | CA - C |
| 4559 | 4560 | 1 | 1.0900e-01 | 2.8451e+05 | ; | CB - HB |
| 4559 | 4561 | 1 | 1.5260e-01 | 2.5941e+05 | ; | CB - CG2 |
| 4559 | 4565 | 1 | 1.4100e-01 | 2.6778e+05 | ; | CB - OG1 |
| 4561 | 4562 | 1 | 1.0900e-01 | 2.8451e+05 | ; | CG2 - HG21 |
| 4561 | 4563 | 1 | 1.0900e-01 | 2.8451e+05 | ; | CG2 - HG22 |
| 4561 | 4564 | 1 | 1.0900e-01 | 2.8451e+05 | ; | CG2 - HG23 |
| 4565 | 4566 | 1 | 9.6000e-02 | 4.6275e+05 | ; | OG1 - HG1 |
| 4567 | 4568 | 1 | 1.2290e-01 | 4.7698e+05 | ; | C - O |
| 4567 | 4569 | 1 | 1.3350e-01 | 4.1003e+05 | ; | C - N |
| 4569 | 4570 | 1 | 1.0100e-01 | 3.6317e+05 | ; | N - H |
| 4569 | 4571 | 1 | 1.4490e-01 | 2.8200e+05 | ; | N - CA |
| 4571 | 4572 | 1 | 1.0900e-01 | 2.8451e+05 | ; | CA - HA |
| 4571 | 4573 | 1 | 1.5260e-01 | 2.5941e+05 | ; | CA - CB |
| 4571 | 4586 | 1 | 1.5220e-01 | 2.6527e+05 | ; | CA - C |
| 4573 | 4574 | 1 | 1.0900e-01 | 2.8451e+05 | ; | CB - HB2 |
| 4573 | 4575 | 1 | 1.0900e-01 | 2.8451e+05 | ; | CB - HB3 |
| 4573 | 4576 | 1 | 1.5260e-01 | 2.5941e+05 | ; | CB - CG |
| 4576 | 4577 | 1 | 1.0900e-01 | 2.8451e+05 | ; | CG - HG |
| 4576 | 4578 | 1 | 1.5260e-01 | 2.5941e+05 | ; | CG - CD1 |
| 4576 | 4582 | 1 | 1.5260e-01 | 2.5941e+05 | ; | CG - CD2 |
| 4578 | 4579 | 1 | 1.0900e-01 | 2.8451e+05 | ; | CD1 - HD11 |
| 4578 | 4580 | 1 | 1.0900e-01 | 2.8451e+05 | ; | CD1 - HD12 |
| 4578 | 4581 | 1 | 1.0900e-01 | 2.8451e+05 | ; | CD1 - HD13 |
| 4582 | 4583 | 1 | 1.0900e-01 | 2.8451e+05 | ; | CD2 - HD21 |
| 4582 | 4584 | 1 | 1.0900e-01 | 2.8451e+05 | ; | CD2 - HD22 |
| 4582 | 4585 | 1 | 1.0900e-01 | 2.8451e+05 | ; | CD2 - HD23 |
| 4586 | 4587 | 1 | 1.2290e-01 | 4.7698e+05 | ; | C - O |
| 4586 | 4588 | 1 | 1.3350e-01 | 4.1003e+05 | ; | C - N |
| 4588 | 4589 | 1 | 1.0100e-01 | 3.6317e+05 | ; | N - H |
| 4588 | 4590 | 1 | 1.4490e-01 | 2.8200e+05 | ; | N - CA |
| 4590 | 4591 | 1 | 1.0900e-01 | 2.8451e+05 | ; | CA - HA |
| 4590 | 4592 | 1 | 1.5260e-01 | 2.5941e+05 | ; | CA - CB |
| 4590 | 4605 | 1 | 1.5220e-01 | 2.6527e+05 | ; | CA - C |
| 4592 | 4593 | 1 | 1.0900e-01 | 2.8451e+05 | ; | CB - HB2 |
| 4592 | 4594 | 1 | 1.0900e-01 | 2.8451e+05 | ; | CB - HB3 |
| 4592 | 4595 | 1 | 1.5260e-01 | 2.5941e+05 | ; | CB - CG |
| 4595 | 4596 | 1 | 1.0900e-01 | 2.8451e+05 | ; | CG - HG |
| 4595 | 4597 | 1 | 1.5260e-01 | 2.5941e+05 | ; | CG - CD1 |
| 4595 | 4601 | 1 | 1.5260e-01 | 2.5941e+05 | ; | CG - CD2 |
| 4597 | 4598 | 1 | 1.0900e-01 | 2.8451e+05 | ; | CD1 - HD11 |
| 4597 | 4599 | 1 | 1.0900e-01 | 2.8451e+05 | ; | CD1 - HD12 |
| 4597 | 4600 | 1 | 1.0900e-01 | 2.8451e+05 | ; | CD1 - HD13 |
| 4601 | 4602 | 1 | 1.0900e-01 | 2.8451e+05 | ; | CD2 - HD21 |
| 4601 | 4603 | 1 | 1.0900e-01 | 2.8451e+05 | ; | CD2 - HD22 |
| 4601 | 4604 | 1 | 1.0900e-01 | 2.8451e+05 | ; | CD2 - HD23 |
| 4605 | 4606 | 1 | 1.2290e-01 | 4.7698e+05 | ; | C - O |
| 4605 | 4607 | 1 | 1.3350e-01 | 4.1003e+05 | ; | C - N |
| 4607 | 4608 | 1 | 1.0100e-01 | 3.6317e+05 | ; | N - H |
| 4607 | 4609 | 1 | 1.4490e-01 | 2.8200e+05 | ; | N - CA |
| 4609 | 4610 | 1 | 1.0900e-01 | 2.8451e+05 | ; | CA - HA |
| 4609 | 4611 | 1 | 1.5260e-01 | 2.5941e+05 | ; | CA - CB |

| | | | | | | |
|---|---|---|---|---|---|---|
| 4609 | 4627 | 1 | 1.5220e-01 | 2.6527e+05 | ; | CA - C |
| 4611 | 4612 | 1 | 1.0900e-01 | 2.8451e+05 | ; | CB - HB2 |
| 4611 | 4613 | 1 | 1.0900e-01 | 2.8451e+05 | ; | CB - HB3 |
| 4611 | 4614 | 1 | 1.5260e-01 | 2.5941e+05 | ; | CB - CG |
| 4614 | 4615 | 1 | 1.0900e-01 | 2.8451e+05 | ; | CG - HG2 |
| 4614 | 4616 | 1 | 1.0900e-01 | 2.8451e+05 | ; | CG - HG3 |
| 4614 | 4617 | 1 | 1.5260e-01 | 2.5941e+05 | ; | CG - CD |
| 4617 | 4618 | 1 | 1.0900e-01 | 2.8451e+05 | ; | CD - HD2 |
| 4617 | 4619 | 1 | 1.0900e-01 | 2.8451e+05 | ; | CD - HD3 |
| 4617 | 4620 | 1 | 1.5260e-01 | 2.5941e+05 | ; | CD - CE |
| 4620 | 4621 | 1 | 1.0900e-01 | 2.8451e+05 | ; | CE - HE2 |
| 4620 | 4622 | 1 | 1.0900e-01 | 2.8451e+05 | ; | CE - HE3 |
| 4620 | 4623 | 1 | 1.4710e-01 | 3.0711e+05 | ; | CE - NZ |
| 4623 | 4624 | 1 | 1.0100e-01 | 3.6317e+05 | ; | NZ - HZ1 |
| 4623 | 4625 | 1 | 1.0100e-01 | 3.6317e+05 | ; | NZ - HZ2 |
| 4623 | 4626 | 1 | 1.0100e-01 | 3.6317e+05 | ; | NZ - HZ3 |
| 4627 | 4628 | 1 | 1.2290e-01 | 4.7698e+05 | ; | C - O |
| 4627 | 4629 | 1 | 1.3350e-01 | 4.1003e+05 | ; | C - N |
| 4629 | 4630 | 1 | 1.0100e-01 | 3.6317e+05 | ; | N - H |
| 4629 | 4631 | 1 | 1.4490e-01 | 2.8200e+05 | ; | N - CA |
| 4631 | 4632 | 1 | 1.0900e-01 | 2.8451e+05 | ; | CA - HA |
| 4631 | 4633 | 1 | 1.5260e-01 | 2.5941e+05 | ; | CA - CB |
| 4631 | 4641 | 1 | 1.5220e-01 | 2.6527e+05 | ; | CA - C |
| 4633 | 4634 | 1 | 1.0900e-01 | 2.8451e+05 | ; | CB - HB2 |
| 4633 | 4635 | 1 | 1.0900e-01 | 2.8451e+05 | ; | CB - HB3 |
| 4633 | 4636 | 1 | 1.5220e-01 | 2.6527e+05 | ; | CB - CG |
| 4636 | 4637 | 1 | 1.2290e-01 | 4.7698e+05 | ; | CG - OD1 |
| 4636 | 4638 | 1 | 1.3350e-01 | 4.1003e+05 | ; | CG - ND2 |
| 4638 | 4639 | 1 | 1.0100e-01 | 3.6317e+05 | ; | ND2 - HD21 |
| 4638 | 4640 | 1 | 1.0100e-01 | 3.6317e+05 | ; | ND2 - HD22 |
| 4641 | 4642 | 1 | 1.2290e-01 | 4.7698e+05 | ; | C - O |
| 4641 | 4643 | 1 | 1.3350e-01 | 4.1003e+05 | ; | C - N |
| 4643 | 4644 | 1 | 1.0100e-01 | 3.6317e+05 | ; | N - H |
| 4643 | 4645 | 1 | 1.4490e-01 | 2.8200e+05 | ; | N - CA |
| 4645 | 4646 | 1 | 1.0900e-01 | 2.8451e+05 | ; | CA - HA |
| 4645 | 4647 | 1 | 1.5260e-01 | 2.5941e+05 | ; | CA - CB |
| 4645 | 4660 | 1 | 1.5220e-01 | 2.6527e+05 | ; | CA - C |
| 4647 | 4648 | 1 | 1.0900e-01 | 2.8451e+05 | ; | CB - HB2 |
| 4647 | 4649 | 1 | 1.0900e-01 | 2.8451e+05 | ; | CB - HB3 |
| 4647 | 4650 | 1 | 1.5260e-01 | 2.5941e+05 | ; | CB - CG |
| 4650 | 4651 | 1 | 1.0900e-01 | 2.8451e+05 | ; | CG - HG |
| 4650 | 4652 | 1 | 1.5260e-01 | 2.5941e+05 | ; | CG - CD1 |
| 4650 | 4656 | 1 | 1.5260e-01 | 2.5941e+05 | ; | CG - CD2 |
| 4652 | 4653 | 1 | 1.0900e-01 | 2.8451e+05 | ; | CD1 - HD11 |
| 4652 | 4654 | 1 | 1.0900e-01 | 2.8451e+05 | ; | CD1 - HD12 |
| 4652 | 4655 | 1 | 1.0900e-01 | 2.8451e+05 | ; | CD1 - HD13 |
| 4656 | 4657 | 1 | 1.0900e-01 | 2.8451e+05 | ; | CD2 - HD21 |
| 4656 | 4658 | 1 | 1.0900e-01 | 2.8451e+05 | ; | CD2 - HD22 |
| 4656 | 4659 | 1 | 1.0900e-01 | 2.8451e+05 | ; | CD2 - HD23 |
| 4660 | 4661 | 1 | 1.2290e-01 | 4.7698e+05 | ; | C - O |
| 4660 | 4662 | 1 | 1.3350e-01 | 4.1003e+05 | ; | C - N |
| 4662 | 4663 | 1 | 1.0100e-01 | 3.6317e+05 | ; | N - H |

| | | | | | | | | |
|---|---|---|---|---|---|---|---|---|
| 4662 | 4664 | 1 | 1.4490e-01 | 2.8200e+05 | ; | N | - | CA |
| 4664 | 4665 | 1 | 1.0900e-01 | 2.8451e+05 | ; | CA | - | HA |
| 4664 | 4666 | 1 | 1.5260e-01 | 2.5941e+05 | ; | CA | - | CB |
| 4664 | 4682 | 1 | 1.5220e-01 | 2.6527e+05 | ; | CA | - | C |
| 4666 | 4667 | 1 | 1.0900e-01 | 2.8451e+05 | ; | CB | - | HB2 |
| 4666 | 4668 | 1 | 1.0900e-01 | 2.8451e+05 | ; | CB | - | HB3 |
| 4666 | 4669 | 1 | 1.5260e-01 | 2.5941e+05 | ; | CB | - | CG |
| 4669 | 4670 | 1 | 1.0900e-01 | 2.8451e+05 | ; | CG | - | HG2 |
| 4669 | 4671 | 1 | 1.0900e-01 | 2.8451e+05 | ; | CG | - | HG3 |
| 4669 | 4672 | 1 | 1.5260e-01 | 2.5941e+05 | ; | CG | - | CD |
| 4672 | 4673 | 1 | 1.0900e-01 | 2.8451e+05 | ; | CD | - | HD2 |
| 4672 | 4674 | 1 | 1.0900e-01 | 2.8451e+05 | ; | CD | - | HD3 |
| 4672 | 4675 | 1 | 1.5260e-01 | 2.5941e+05 | ; | CD | - | CE |
| 4675 | 4676 | 1 | 1.0900e-01 | 2.8451e+05 | ; | CE | - | HE2 |
| 4675 | 4677 | 1 | 1.0900e-01 | 2.8451e+05 | ; | CE | - | HE3 |
| 4675 | 4678 | 1 | 1.4710e-01 | 3.0711e+05 | ; | CE | - | NZ |
| 4678 | 4679 | 1 | 1.0100e-01 | 3.6317e+05 | ; | NZ | - | HZ1 |
| 4678 | 4680 | 1 | 1.0100e-01 | 3.6317e+05 | ; | NZ | - | HZ2 |
| 4678 | 4681 | 1 | 1.0100e-01 | 3.6317e+05 | ; | NZ | - | HZ3 |
| 4682 | 4683 | 1 | 1.2290e-01 | 4.7698e+05 | ; | C | - | O |
| 4682 | 4684 | 1 | 1.3350e-01 | 4.1003e+05 | ; | C | - | N |
| 4684 | 4685 | 1 | 1.0100e-01 | 3.6317e+05 | ; | N | - | H |
| 4684 | 4686 | 1 | 1.4490e-01 | 2.8200e+05 | ; | N | - | CA |
| 4686 | 4687 | 1 | 1.0900e-01 | 2.8451e+05 | ; | CA | - | HA |
| 4686 | 4688 | 1 | 1.5260e-01 | 2.5941e+05 | ; | CA | - | CB |
| 4686 | 4696 | 1 | 1.5220e-01 | 2.6527e+05 | ; | CA | - | C |
| 4688 | 4689 | 1 | 1.0900e-01 | 2.8451e+05 | ; | CB | - | HB2 |
| 4688 | 4690 | 1 | 1.0900e-01 | 2.8451e+05 | ; | CB | - | HB3 |
| 4688 | 4691 | 1 | 1.5220e-01 | 2.6527e+05 | ; | CB | - | CG |
| 4691 | 4692 | 1 | 1.2290e-01 | 4.7698e+05 | ; | CG | - | OD1 |
| 4691 | 4693 | 1 | 1.3350e-01 | 4.1003e+05 | ; | CG | - | ND2 |
| 4693 | 4694 | 1 | 1.0100e-01 | 3.6317e+05 | ; | ND2 | - | HD21 |
| 4693 | 4695 | 1 | 1.0100e-01 | 3.6317e+05 | ; | ND2 | - | HD22 |
| 4696 | 4697 | 1 | 1.2290e-01 | 4.7698e+05 | ; | C | - | O |
| 4696 | 4698 | 1 | 1.3350e-01 | 4.1003e+05 | ; | C | - | N |
| 4698 | 4699 | 1 | 1.0100e-01 | 3.6317e+05 | ; | N | - | H |
| 4698 | 4700 | 1 | 1.4490e-01 | 2.8200e+05 | ; | N | - | CA |
| 4700 | 4701 | 1 | 1.0900e-01 | 2.8451e+05 | ; | CA | - | HA |
| 4700 | 4702 | 1 | 1.5260e-01 | 2.5941e+05 | ; | CA | - | CB |
| 4700 | 4716 | 1 | 1.5220e-01 | 2.6527e+05 | ; | CA | - | C |
| 4702 | 4703 | 1 | 1.0900e-01 | 2.8451e+05 | ; | CB | - | HB2 |
| 4702 | 4704 | 1 | 1.0900e-01 | 2.8451e+05 | ; | CB | - | HB3 |
| 4702 | 4705 | 1 | 1.5100e-01 | 2.6527e+05 | ; | CB | - | CG |
| 4705 | 4706 | 1 | 1.4000e-01 | 3.9246e+05 | ; | CG | - | CD1 |
| 4705 | 4714 | 1 | 1.4000e-01 | 3.9246e+05 | ; | CG | - | CD2 |
| 4706 | 4707 | 1 | 1.0800e-01 | 3.0711e+05 | ; | CD1 | - | HD1 |
| 4706 | 4708 | 1 | 1.4000e-01 | 3.9246e+05 | ; | CD1 | - | CE1 |
| 4708 | 4709 | 1 | 1.0800e-01 | 3.0711e+05 | ; | CE1 | - | HE1 |
| 4708 | 4710 | 1 | 1.4000e-01 | 3.9246e+05 | ; | CE1 | - | CZ |
| 4710 | 4711 | 1 | 1.0800e-01 | 3.0711e+05 | ; | CZ | - | HZ |
| 4710 | 4712 | 1 | 1.4000e-01 | 3.9246e+05 | ; | CZ | - | CE2 |
| 4712 | 4713 | 1 | 1.0800e-01 | 3.0711e+05 | ; | CE2 | - | HE2 |

| | | | | | |
|---|---|---|---|---|---|
| 4712 | 4714 | 1 | 1.4000e-01 | 3.9246e+05 ; | CE2 - CD2 |
| 4714 | 4715 | 1 | 1.0800e-01 | 3.0711e+05 ; | CD2 - HD2 |
| 4716 | 4717 | 1 | 1.2290e-01 | 4.7698e+05 ; | C - O |
| 4716 | 4718 | 1 | 1.3350e-01 | 4.1003e+05 ; | C - N |
| 4718 | 4719 | 1 | 1.0100e-01 | 3.6317e+05 ; | N - H |
| 4718 | 4720 | 1 | 1.4490e-01 | 2.8200e+05 ; | N - CA |
| 4720 | 4721 | 1 | 1.0900e-01 | 2.8451e+05 ; | CA - HA |
| 4720 | 4722 | 1 | 1.5260e-01 | 2.5941e+05 ; | CA - CB |
| 4720 | 4738 | 1 | 1.5220e-01 | 2.6527e+05 ; | CA - C |
| 4722 | 4723 | 1 | 1.0900e-01 | 2.8451e+05 ; | CB - HB2 |
| 4722 | 4724 | 1 | 1.0900e-01 | 2.8451e+05 ; | CB - HB3 |
| 4722 | 4725 | 1 | 1.5260e-01 | 2.5941e+05 ; | CB - CG |
| 4725 | 4726 | 1 | 1.0900e-01 | 2.8451e+05 ; | CG - HG2 |
| 4725 | 4727 | 1 | 1.0900e-01 | 2.8451e+05 ; | CG - HG3 |
| 4725 | 4728 | 1 | 1.5260e-01 | 2.5941e+05 ; | CG - CD |
| 4728 | 4729 | 1 | 1.0900e-01 | 2.8451e+05 ; | CD - HD2 |
| 4728 | 4730 | 1 | 1.0900e-01 | 2.8451e+05 ; | CD - HD3 |
| 4728 | 4731 | 1 | 1.5260e-01 | 2.5941e+05 ; | CD - CE |
| 4731 | 4732 | 1 | 1.0900e-01 | 2.8451e+05 ; | CE - HE2 |
| 4731 | 4733 | 1 | 1.0900e-01 | 2.8451e+05 ; | CE - HE3 |
| 4731 | 4734 | 1 | 1.4710e-01 | 3.0711e+05 ; | CE - NZ |
| 4734 | 4735 | 1 | 1.0100e-01 | 3.6317e+05 ; | NZ - HZ1 |
| 4734 | 4736 | 1 | 1.0100e-01 | 3.6317e+05 ; | NZ - HZ2 |
| 4734 | 4737 | 1 | 1.0100e-01 | 3.6317e+05 ; | NZ - HZ3 |
| 4738 | 4739 | 1 | 1.2290e-01 | 4.7698e+05 ; | C - O |
| 4738 | 4740 | 1 | 1.3350e-01 | 4.1003e+05 ; | C - N |
| 4740 | 4741 | 1 | 1.0100e-01 | 3.6317e+05 ; | N - H |
| 4740 | 4742 | 1 | 1.4490e-01 | 2.8200e+05 ; | N - CA |
| 4742 | 4743 | 1 | 1.0900e-01 | 2.8451e+05 ; | CA - HA |
| 4742 | 4744 | 1 | 1.5260e-01 | 2.5941e+05 ; | CA - CB |
| 4742 | 4760 | 1 | 1.5220e-01 | 2.6527e+05 ; | CA - C |
| 4744 | 4745 | 1 | 1.0900e-01 | 2.8451e+05 ; | CB - HB2 |
| 4744 | 4746 | 1 | 1.0900e-01 | 2.8451e+05 ; | CB - HB3 |
| 4744 | 4747 | 1 | 1.5260e-01 | 2.5941e+05 ; | CB - CG |
| 4747 | 4748 | 1 | 1.0900e-01 | 2.8451e+05 ; | CG - HG2 |
| 4747 | 4749 | 1 | 1.0900e-01 | 2.8451e+05 ; | CG - HG3 |
| 4747 | 4750 | 1 | 1.5260e-01 | 2.5941e+05 ; | CG - CD |
| 4750 | 4751 | 1 | 1.0900e-01 | 2.8451e+05 ; | CD - HD2 |
| 4750 | 4752 | 1 | 1.0900e-01 | 2.8451e+05 ; | CD - HD3 |
| 4750 | 4753 | 1 | 1.5260e-01 | 2.5941e+05 ; | CD - CE |
| 4753 | 4754 | 1 | 1.0900e-01 | 2.8451e+05 ; | CE - HE2 |
| 4753 | 4755 | 1 | 1.0900e-01 | 2.8451e+05 ; | CE - HE3 |
| 4753 | 4756 | 1 | 1.4710e-01 | 3.0711e+05 ; | CE - NZ |
| 4756 | 4757 | 1 | 1.0100e-01 | 3.6317e+05 ; | NZ - HZ1 |
| 4756 | 4758 | 1 | 1.0100e-01 | 3.6317e+05 ; | NZ - HZ2 |
| 4756 | 4759 | 1 | 1.0100e-01 | 3.6317e+05 ; | NZ - HZ3 |
| 4760 | 4761 | 1 | 1.2290e-01 | 4.7698e+05 ; | C - O |
| 4760 | 4762 | 1 | 1.3350e-01 | 4.1003e+05 ; | C - N |
| 4762 | 4763 | 1 | 1.0100e-01 | 3.6317e+05 ; | N - H |
| 4762 | 4764 | 1 | 1.4490e-01 | 2.8200e+05 ; | N - CA |
| 4764 | 4765 | 1 | 1.0900e-01 | 2.8451e+05 ; | CA - HA |
| 4764 | 4766 | 1 | 1.5260e-01 | 2.5941e+05 ; | CA - CB |

| | | | | | |
|---|---|---|---|---|---|
| 4764 | 4775 | 1 | 1.5220e-01 | 2.6527e+05 ; | CA - C |
| 4766 | 4767 | 1 | 1.0900e-01 | 2.8451e+05 ; | CB - HB2 |
| 4766 | 4768 | 1 | 1.0900e-01 | 2.8451e+05 ; | CB - HB3 |
| 4766 | 4769 | 1 | 1.5260e-01 | 2.5941e+05 ; | CB - CG |
| 4769 | 4770 | 1 | 1.0900e-01 | 2.8451e+05 ; | CG - HG2 |
| 4769 | 4771 | 1 | 1.0900e-01 | 2.8451e+05 ; | CG - HG3 |
| 4769 | 4772 | 1 | 1.5220e-01 | 2.6527e+05 ; | CG - CD |
| 4772 | 4773 | 1 | 1.2500e-01 | 5.4894e+05 ; | CD - OE1 |
| 4772 | 4774 | 1 | 1.2500e-01 | 5.4894e+05 ; | CD - OE2 |
| 4775 | 4776 | 1 | 1.2290e-01 | 4.7698e+05 ; | C - O |
| 4775 | 4777 | 1 | 1.3350e-01 | 4.1003e+05 ; | C - N |
| 4777 | 4778 | 1 | 1.0100e-01 | 3.6317e+05 ; | N - H |
| 4777 | 4779 | 1 | 1.4490e-01 | 2.8200e+05 ; | N - CA |
| 4779 | 4780 | 1 | 1.0900e-01 | 2.8451e+05 ; | CA - HA |
| 4779 | 4781 | 1 | 1.5260e-01 | 2.5941e+05 ; | CA - CB |
| 4779 | 4789 | 1 | 1.5220e-01 | 2.6527e+05 ; | CA - C |
| 4781 | 4782 | 1 | 1.0900e-01 | 2.8451e+05 ; | CB - HB2 |
| 4781 | 4783 | 1 | 1.0900e-01 | 2.8451e+05 ; | CB - HB3 |
| 4781 | 4784 | 1 | 1.5220e-01 | 2.6527e+05 ; | CB - CG |
| 4784 | 4785 | 1 | 1.2290e-01 | 4.7698e+05 ; | CG - OD1 |
| 4784 | 4786 | 1 | 1.3350e-01 | 4.1003e+05 ; | CG - ND2 |
| 4786 | 4787 | 1 | 1.0100e-01 | 3.6317e+05 ; | ND2 - HD21 |
| 4786 | 4788 | 1 | 1.0100e-01 | 3.6317e+05 ; | ND2 - HD22 |
| 4789 | 4790 | 1 | 1.2290e-01 | 4.7698e+05 ; | C - O |
| 4789 | 4791 | 1 | 1.3350e-01 | 4.1003e+05 ; | C - N |
| 4791 | 4792 | 1 | 1.0100e-01 | 3.6317e+05 ; | N - H |
| 4791 | 4793 | 1 | 1.4490e-01 | 2.8200e+05 ; | N - CA |
| 4793 | 4794 | 1 | 1.0900e-01 | 2.8451e+05 ; | CA - HA |
| 4793 | 4795 | 1 | 1.5260e-01 | 2.5941e+05 ; | CA - CB |
| 4793 | 4804 | 1 | 1.5220e-01 | 2.6527e+05 ; | CA - C |
| 4795 | 4796 | 1 | 1.0900e-01 | 2.8451e+05 ; | CB - HB2 |
| 4795 | 4797 | 1 | 1.0900e-01 | 2.8451e+05 ; | CB - HB3 |
| 4795 | 4798 | 1 | 1.5260e-01 | 2.5941e+05 ; | CB - CG |
| 4798 | 4799 | 1 | 1.0900e-01 | 2.8451e+05 ; | CG - HG2 |
| 4798 | 4800 | 1 | 1.0900e-01 | 2.8451e+05 ; | CG - HG3 |
| 4798 | 4801 | 1 | 1.5220e-01 | 2.6527e+05 ; | CG - CD |
| 4801 | 4802 | 1 | 1.2500e-01 | 5.4894e+05 ; | CD - OE1 |
| 4801 | 4803 | 1 | 1.2500e-01 | 5.4894e+05 ; | CD - OE2 |
| 4804 | 4805 | 1 | 1.2290e-01 | 4.7698e+05 ; | C - O |
| 4804 | 4806 | 1 | 1.3350e-01 | 4.1003e+05 ; | C - N |
| 4806 | 4807 | 1 | 1.0100e-01 | 3.6317e+05 ; | N - H |
| 4806 | 4808 | 1 | 1.4490e-01 | 2.8200e+05 ; | N - CA |
| 4808 | 4809 | 1 | 1.0900e-01 | 2.8451e+05 ; | CA - HA |
| 4808 | 4810 | 1 | 1.5260e-01 | 2.5941e+05 ; | CA - CB |
| 4808 | 4823 | 1 | 1.5220e-01 | 2.6527e+05 ; | CA - C |
| 4810 | 4811 | 1 | 1.0900e-01 | 2.8451e+05 ; | CB - HB2 |
| 4810 | 4812 | 1 | 1.0900e-01 | 2.8451e+05 ; | CB - HB3 |
| 4810 | 4813 | 1 | 1.5260e-01 | 2.5941e+05 ; | CB - CG |
| 4813 | 4814 | 1 | 1.0900e-01 | 2.8451e+05 ; | CG - HG |
| 4813 | 4815 | 1 | 1.5260e-01 | 2.5941e+05 ; | CG - CD1 |
| 4813 | 4819 | 1 | 1.5260e-01 | 2.5941e+05 ; | CG - CD2 |
| 4815 | 4816 | 1 | 1.0900e-01 | 2.8451e+05 ; | CD1 - HD11 |

| | | | | | | |
|---|---|---|---|---|---|---|
| 4815 | 4817 | 1 | 1.0900e-01 | 2.8451e+05 | ; | CD1 - HD12 |
| 4815 | 4818 | 1 | 1.0900e-01 | 2.8451e+05 | ; | CD1 - HD13 |
| 4819 | 4820 | 1 | 1.0900e-01 | 2.8451e+05 | ; | CD2 - HD21 |
| 4819 | 4821 | 1 | 1.0900e-01 | 2.8451e+05 | ; | CD2 - HD22 |
| 4819 | 4822 | 1 | 1.0900e-01 | 2.8451e+05 | ; | CD2 - HD23 |
| 4823 | 4824 | 1 | 1.2290e-01 | 4.7698e+05 | ; | C - O |
| 4823 | 4825 | 1 | 1.3350e-01 | 4.1003e+05 | ; | C - N |
| 4825 | 4826 | 1 | 1.0100e-01 | 3.6317e+05 | ; | N - H |
| 4825 | 4827 | 1 | 1.4490e-01 | 2.8200e+05 | ; | N - CA |
| 4827 | 4828 | 1 | 1.0900e-01 | 2.8451e+05 | ; | CA - HA |
| 4827 | 4829 | 1 | 1.5260e-01 | 2.5941e+05 | ; | CA - CB |
| 4827 | 4845 | 1 | 1.5220e-01 | 2.6527e+05 | ; | CA - C |
| 4829 | 4830 | 1 | 1.0900e-01 | 2.8451e+05 | ; | CB - HB2 |
| 4829 | 4831 | 1 | 1.0900e-01 | 2.8451e+05 | ; | CB - HB3 |
| 4829 | 4832 | 1 | 1.5260e-01 | 2.5941e+05 | ; | CB - CG |
| 4832 | 4833 | 1 | 1.0900e-01 | 2.8451e+05 | ; | CG - HG2 |
| 4832 | 4834 | 1 | 1.0900e-01 | 2.8451e+05 | ; | CG - HG3 |
| 4832 | 4835 | 1 | 1.5260e-01 | 2.5941e+05 | ; | CG - CD |
| 4835 | 4836 | 1 | 1.0900e-01 | 2.8451e+05 | ; | CD - HD2 |
| 4835 | 4837 | 1 | 1.0900e-01 | 2.8451e+05 | ; | CD - HD3 |
| 4835 | 4838 | 1 | 1.5260e-01 | 2.5941e+05 | ; | CD - CE |
| 4838 | 4839 | 1 | 1.0900e-01 | 2.8451e+05 | ; | CE - HE2 |
| 4838 | 4840 | 1 | 1.0900e-01 | 2.8451e+05 | ; | CE - HE3 |
| 4838 | 4841 | 1 | 1.4710e-01 | 3.0711e+05 | ; | CE - NZ |
| 4841 | 4842 | 1 | 1.0100e-01 | 3.6317e+05 | ; | NZ - HZ1 |
| 4841 | 4843 | 1 | 1.0100e-01 | 3.6317e+05 | ; | NZ - HZ2 |
| 4841 | 4844 | 1 | 1.0100e-01 | 3.6317e+05 | ; | NZ - HZ3 |
| 4845 | 4846 | 1 | 1.2290e-01 | 4.7698e+05 | ; | C - O |
| 4845 | 4847 | 1 | 1.3350e-01 | 4.1003e+05 | ; | C - N |
| 4847 | 4848 | 1 | 1.0100e-01 | 3.6317e+05 | ; | N - H |
| 4847 | 4849 | 1 | 1.4490e-01 | 2.8200e+05 | ; | N - CA |
| 4849 | 4850 | 1 | 1.0900e-01 | 2.8451e+05 | ; | CA - HA |
| 4849 | 4851 | 1 | 1.5260e-01 | 2.5941e+05 | ; | CA - CB |
| 4849 | 4867 | 1 | 1.5220e-01 | 2.6527e+05 | ; | CA - C |
| 4851 | 4852 | 1 | 1.0900e-01 | 2.8451e+05 | ; | CB - HB2 |
| 4851 | 4853 | 1 | 1.0900e-01 | 2.8451e+05 | ; | CB - HB3 |
| 4851 | 4854 | 1 | 1.5260e-01 | 2.5941e+05 | ; | CB - CG |
| 4854 | 4855 | 1 | 1.0900e-01 | 2.8451e+05 | ; | CG - HG2 |
| 4854 | 4856 | 1 | 1.0900e-01 | 2.8451e+05 | ; | CG - HG3 |
| 4854 | 4857 | 1 | 1.5260e-01 | 2.5941e+05 | ; | CG - CD |
| 4857 | 4858 | 1 | 1.0900e-01 | 2.8451e+05 | ; | CD - HD2 |
| 4857 | 4859 | 1 | 1.0900e-01 | 2.8451e+05 | ; | CD - HD3 |
| 4857 | 4860 | 1 | 1.5260e-01 | 2.5941e+05 | ; | CD - CE |
| 4860 | 4861 | 1 | 1.0900e-01 | 2.8451e+05 | ; | CE - HE2 |
| 4860 | 4862 | 1 | 1.0900e-01 | 2.8451e+05 | ; | CE - HE3 |
| 4860 | 4863 | 1 | 1.4710e-01 | 3.0711e+05 | ; | CE - NZ |
| 4863 | 4864 | 1 | 1.0100e-01 | 3.6317e+05 | ; | NZ - HZ1 |
| 4863 | 4865 | 1 | 1.0100e-01 | 3.6317e+05 | ; | NZ - HZ2 |
| 4863 | 4866 | 1 | 1.0100e-01 | 3.6317e+05 | ; | NZ - HZ3 |
| 4867 | 4868 | 1 | 1.2290e-01 | 4.7698e+05 | ; | C - O |
| 4867 | 4869 | 1 | 1.3350e-01 | 4.1003e+05 | ; | C - N |
| 4869 | 4870 | 1 | 1.0100e-01 | 3.6317e+05 | ; | N - H |

```
4869   4871   1    1.4490e-01    2.8200e+05 ;       N - CA
4871   4872   1    1.0900e-01    2.8451e+05 ;      CA - HA
4871   4873   1    1.5260e-01    2.5941e+05 ;      CA - CB
4871   4886   1    1.5220e-01    2.6527e+05 ;      CA - C
4873   4874   1    1.0900e-01    2.8451e+05 ;      CB - HB
4873   4875   1    1.5260e-01    2.5941e+05 ;      CB - CG2
4873   4879   1    1.5260e-01    2.5941e+05 ;      CB - CG1
4875   4876   1    1.0900e-01    2.8451e+05 ;     CG2 - HG21
4875   4877   1    1.0900e-01    2.8451e+05 ;     CG2 - HG22
4875   4878   1    1.0900e-01    2.8451e+05 ;     CG2 - HG23
4879   4880   1    1.0900e-01    2.8451e+05 ;     CG1 - HG12
4879   4881   1    1.0900e-01    2.8451e+05 ;     CG1 - HG13
4879   4882   1    1.5260e-01    2.5941e+05 ;     CG1 - CD1
4882   4883   1    1.0900e-01    2.8451e+05 ;     CD1 - HD11
4882   4884   1    1.0900e-01    2.8451e+05 ;     CD1 - HD12
4882   4885   1    1.0900e-01    2.8451e+05 ;     CD1 - HD13
4886   4887   1    1.2290e-01    4.7698e+05 ;       C - O
4886   4888   1    1.3350e-01    4.1003e+05 ;       C - N
4888   4889   1    1.0100e-01    3.6317e+05 ;       N - H
4888   4890   1    1.4490e-01    2.8200e+05 ;       N - CA
4890   4891   1    1.0900e-01    2.8451e+05 ;      CA - HA
4890   4892   1    1.5260e-01    2.5941e+05 ;      CA - CB
4890   4896   1    1.5220e-01    2.6527e+05 ;      CA - C
4892   4893   1    1.0900e-01    2.8451e+05 ;      CB - HB1
4892   4894   1    1.0900e-01    2.8451e+05 ;      CB - HB2
4892   4895   1    1.0900e-01    2.8451e+05 ;      CB - HB3
4896   4897   1    1.2290e-01    4.7698e+05 ;       C - O
4896   4898   1    1.3350e-01    4.1003e+05 ;       C - N
4898   4899   1    1.0100e-01    3.6317e+05 ;       N - H
4898   4900   1    1.4490e-01    2.8200e+05 ;       N - CA
4900   4901   1    1.0900e-01    2.8451e+05 ;      CA - HA
4900   4902   1    1.5260e-01    2.5941e+05 ;      CA - CB
4900   4915   1    1.5220e-01    2.6527e+05 ;      CA - C
4902   4903   1    1.0900e-01    2.8451e+05 ;      CB - HB2
4902   4904   1    1.0900e-01    2.8451e+05 ;      CB - HB3
4902   4905   1    1.5260e-01    2.5941e+05 ;      CB - CG
4905   4906   1    1.0900e-01    2.8451e+05 ;      CG - HG
4905   4907   1    1.5260e-01    2.5941e+05 ;      CG - CD1
4905   4911   1    1.5260e-01    2.5941e+05 ;      CG - CD2
4907   4908   1    1.0900e-01    2.8451e+05 ;     CD1 - HD11
4907   4909   1    1.0900e-01    2.8451e+05 ;     CD1 - HD12
4907   4910   1    1.0900e-01    2.8451e+05 ;     CD1 - HD13
4911   4912   1    1.0900e-01    2.8451e+05 ;     CD2 - HD21
4911   4913   1    1.0900e-01    2.8451e+05 ;     CD2 - HD22
4911   4914   1    1.0900e-01    2.8451e+05 ;     CD2 - HD23
4915   4916   1    1.2290e-01    4.7698e+05 ;       C - O
4915   4917   1    1.3350e-01    4.1003e+05 ;       C - N
4917   4918   1    1.0100e-01    3.6317e+05 ;       N - H
4917   4919   1    1.4490e-01    2.8200e+05 ;       N - CA
4919   4920   1    1.0900e-01    2.8451e+05 ;      CA - HA
4919   4921   1    1.5260e-01    2.5941e+05 ;      CA - CB
4919   4929   1    1.5220e-01    2.6527e+05 ;      CA - C
```

| | | | | | | |
|---|---|---|---|---|---|---|
| 4921 | 4922 | 1 | 1.0900e-01 | 2.8451e+05 | ; | CB - HB |
| 4921 | 4923 | 1 | 1.5260e-01 | 2.5941e+05 | ; | CB - CG2 |
| 4921 | 4927 | 1 | 1.4100e-01 | 2.6778e+05 | ; | CB - OG1 |
| 4923 | 4924 | 1 | 1.0900e-01 | 2.8451e+05 | ; | CG2 - HG21 |
| 4923 | 4925 | 1 | 1.0900e-01 | 2.8451e+05 | ; | CG2 - HG22 |
| 4923 | 4926 | 1 | 1.0900e-01 | 2.8451e+05 | ; | CG2 - HG23 |
| 4927 | 4928 | 1 | 9.6000e-02 | 4.6275e+05 | ; | OG1 - HG1 |
| 4929 | 4930 | 1 | 1.2290e-01 | 4.7698e+05 | ; | C - O |
| 4929 | 4931 | 1 | 1.3350e-01 | 4.1003e+05 | ; | C - N |
| 4931 | 4932 | 1 | 1.0100e-01 | 3.6317e+05 | ; | N - H |
| 4931 | 4933 | 1 | 1.4490e-01 | 2.8200e+05 | ; | N - CA |
| 4933 | 4934 | 1 | 1.0900e-01 | 2.8451e+05 | ; | CA - HA |
| 4933 | 4935 | 1 | 1.5260e-01 | 2.5941e+05 | ; | CA - CB |
| 4933 | 4948 | 1 | 1.5220e-01 | 2.6527e+05 | ; | CA - C |
| 4935 | 4936 | 1 | 1.0900e-01 | 2.8451e+05 | ; | CB - HB |
| 4935 | 4937 | 1 | 1.5260e-01 | 2.5941e+05 | ; | CB - CG2 |
| 4935 | 4941 | 1 | 1.5260e-01 | 2.5941e+05 | ; | CB - CG1 |
| 4937 | 4938 | 1 | 1.0900e-01 | 2.8451e+05 | ; | CG2 - HG21 |
| 4937 | 4939 | 1 | 1.0900e-01 | 2.8451e+05 | ; | CG2 - HG22 |
| 4937 | 4940 | 1 | 1.0900e-01 | 2.8451e+05 | ; | CG2 - HG23 |
| 4941 | 4942 | 1 | 1.0900e-01 | 2.8451e+05 | ; | CG1 - HG12 |
| 4941 | 4943 | 1 | 1.0900e-01 | 2.8451e+05 | ; | CG1 - HG13 |
| 4941 | 4944 | 1 | 1.5260e-01 | 2.5941e+05 | ; | CG1 - CD1 |
| 4944 | 4945 | 1 | 1.0900e-01 | 2.8451e+05 | ; | CD1 - HD11 |
| 4944 | 4946 | 1 | 1.0900e-01 | 2.8451e+05 | ; | CD1 - HD12 |
| 4944 | 4947 | 1 | 1.0900e-01 | 2.8451e+05 | ; | CD1 - HD13 |
| 4948 | 4949 | 1 | 1.2290e-01 | 4.7698e+05 | ; | C - O |
| 4948 | 4950 | 1 | 1.3350e-01 | 4.1003e+05 | ; | C - N |
| 4950 | 4951 | 1 | 1.0100e-01 | 3.6317e+05 | ; | N - H |
| 4950 | 4952 | 1 | 1.4490e-01 | 2.8200e+05 | ; | N - CA |
| 4952 | 4953 | 1 | 1.0900e-01 | 2.8451e+05 | ; | CA - HA |
| 4952 | 4954 | 1 | 1.5260e-01 | 2.5941e+05 | ; | CA - CB |
| 4952 | 4967 | 1 | 1.5220e-01 | 2.6527e+05 | ; | CA - C |
| 4954 | 4955 | 1 | 1.0900e-01 | 2.8451e+05 | ; | CB - HB |
| 4954 | 4956 | 1 | 1.5260e-01 | 2.5941e+05 | ; | CB - CG2 |
| 4954 | 4960 | 1 | 1.5260e-01 | 2.5941e+05 | ; | CB - CG1 |
| 4956 | 4957 | 1 | 1.0900e-01 | 2.8451e+05 | ; | CG2 - HG21 |
| 4956 | 4958 | 1 | 1.0900e-01 | 2.8451e+05 | ; | CG2 - HG22 |
| 4956 | 4959 | 1 | 1.0900e-01 | 2.8451e+05 | ; | CG2 - HG23 |
| 4960 | 4961 | 1 | 1.0900e-01 | 2.8451e+05 | ; | CG1 - HG12 |
| 4960 | 4962 | 1 | 1.0900e-01 | 2.8451e+05 | ; | CG1 - HG13 |
| 4960 | 4963 | 1 | 1.5260e-01 | 2.5941e+05 | ; | CG1 - CD1 |
| 4963 | 4964 | 1 | 1.0900e-01 | 2.8451e+05 | ; | CD1 - HD11 |
| 4963 | 4965 | 1 | 1.0900e-01 | 2.8451e+05 | ; | CD1 - HD12 |
| 4963 | 4966 | 1 | 1.0900e-01 | 2.8451e+05 | ; | CD1 - HD13 |
| 4967 | 4968 | 1 | 1.2290e-01 | 4.7698e+05 | ; | C - O |
| 4967 | 4969 | 1 | 1.3350e-01 | 4.1003e+05 | ; | C - N |
| 4969 | 4970 | 1 | 1.0100e-01 | 3.6317e+05 | ; | N - H |
| 4969 | 4971 | 1 | 1.4490e-01 | 2.8200e+05 | ; | N - CA |
| 4971 | 4972 | 1 | 1.0900e-01 | 2.8451e+05 | ; | CA - HA |
| 4971 | 4973 | 1 | 1.5260e-01 | 2.5941e+05 | ; | CA - CB |
| 4971 | 4977 | 1 | 1.5220e-01 | 2.6527e+05 | ; | CA - C |

| | | | | | | | | |
|---|---|---|---|---|---|---|---|---|
| 4973 | 4974 | 1 | 1.0900e-01 | 2.8451e+05 | ; | CB | - | HB1 |
| 4973 | 4975 | 1 | 1.0900e-01 | 2.8451e+05 | ; | CB | - | HB2 |
| 4973 | 4976 | 1 | 1.0900e-01 | 2.8451e+05 | ; | CB | - | HB3 |
| 4977 | 4978 | 1 | 1.2290e-01 | 4.7698e+05 | ; | C | - | O |
| 4977 | 4979 | 1 | 1.3350e-01 | 4.1003e+05 | ; | C | - | N |
| 4979 | 4980 | 1 | 1.0100e-01 | 3.6317e+05 | ; | N | - | H |
| 4979 | 4981 | 1 | 1.4490e-01 | 2.8200e+05 | ; | N | - | CA |
| 4981 | 4982 | 1 | 1.0900e-01 | 2.8451e+05 | ; | CA | - | HA |
| 4981 | 4983 | 1 | 1.5260e-01 | 2.5941e+05 | ; | CA | - | CB |
| 4981 | 4999 | 1 | 1.5220e-01 | 2.6527e+05 | ; | CA | - | C |
| 4983 | 4984 | 1 | 1.0900e-01 | 2.8451e+05 | ; | CB | - | HB2 |
| 4983 | 4985 | 1 | 1.0900e-01 | 2.8451e+05 | ; | CB | - | HB3 |
| 4983 | 4986 | 1 | 1.5260e-01 | 2.5941e+05 | ; | CB | - | CG |
| 4986 | 4987 | 1 | 1.0900e-01 | 2.8451e+05 | ; | CG | - | HG2 |
| 4986 | 4988 | 1 | 1.0900e-01 | 2.8451e+05 | ; | CG | - | HG3 |
| 4986 | 4989 | 1 | 1.5260e-01 | 2.5941e+05 | ; | CG | - | CD |
| 4989 | 4990 | 1 | 1.0900e-01 | 2.8451e+05 | ; | CD | - | HD2 |
| 4989 | 4991 | 1 | 1.0900e-01 | 2.8451e+05 | ; | CD | - | HD3 |
| 4989 | 4992 | 1 | 1.5260e-01 | 2.5941e+05 | ; | CD | - | CE |
| 4992 | 4993 | 1 | 1.0900e-01 | 2.8451e+05 | ; | CE | - | HE2 |
| 4992 | 4994 | 1 | 1.0900e-01 | 2.8451e+05 | ; | CE | - | HE3 |
| 4992 | 4995 | 1 | 1.4710e-01 | 3.0711e+05 | ; | CE | - | NZ |
| 4995 | 4996 | 1 | 1.0100e-01 | 3.6317e+05 | ; | NZ | - | HZ1 |
| 4995 | 4997 | 1 | 1.0100e-01 | 3.6317e+05 | ; | NZ | - | HZ2 |
| 4995 | 4998 | 1 | 1.0100e-01 | 3.6317e+05 | ; | NZ | - | HZ3 |
| 4999 | 5000 | 1 | 1.2290e-01 | 4.7698e+05 | ; | C | - | O |
| 4999 | 5001 | 1 | 1.3350e-01 | 4.1003e+05 | ; | C | - | N |
| 5001 | 5002 | 1 | 1.0100e-01 | 3.6317e+05 | ; | N | - | H |
| 5001 | 5003 | 1 | 1.4490e-01 | 2.8200e+05 | ; | N | - | CA |
| 5003 | 5004 | 1 | 1.0900e-01 | 2.8451e+05 | ; | CA | - | HA |
| 5003 | 5005 | 1 | 1.5260e-01 | 2.5941e+05 | ; | CA | - | CB |
| 5003 | 5016 | 1 | 1.5220e-01 | 2.6527e+05 | ; | CA | - | C |
| 5005 | 5006 | 1 | 1.0900e-01 | 2.8451e+05 | ; | CB | - | HB2 |
| 5005 | 5007 | 1 | 1.0900e-01 | 2.8451e+05 | ; | CB | - | HB3 |
| 5005 | 5008 | 1 | 1.5040e-01 | 2.6527e+05 | ; | CB | - | CG |
| 5008 | 5009 | 1 | 1.3940e-01 | 3.4309e+05 | ; | CG | - | ND1 |
| 5008 | 5014 | 1 | 1.3710e-01 | 4.3346e+05 | ; | CG | - | CD2 |
| 5009 | 5010 | 1 | 1.3350e-01 | 4.0836e+05 | ; | ND1 | - | CE1 |
| 5010 | 5011 | 1 | 1.0800e-01 | 3.0711e+05 | ; | CE1 | - | HE1 |
| 5010 | 5012 | 1 | 1.3430e-01 | 3.9915e+05 | ; | CE1 | - | NE2 |
| 5012 | 5013 | 1 | 1.0100e-01 | 3.6317e+05 | ; | NE2 | - | HE2 |
| 5012 | 5014 | 1 | 1.3810e-01 | 3.5731e+05 | ; | NE2 | - | CD2 |
| 5014 | 5015 | 1 | 1.0800e-01 | 3.0711e+05 | ; | CD2 | - | HD2 |
| 5016 | 5017 | 1 | 1.2290e-01 | 4.7698e+05 | ; | C | - | O |
| 5016 | 5018 | 1 | 1.3350e-01 | 4.1003e+05 | ; | C | - | N |
| 5018 | 5019 | 1 | 1.0100e-01 | 3.6317e+05 | ; | N | - | H |
| 5018 | 5020 | 1 | 1.4490e-01 | 2.8200e+05 | ; | N | - | CA |
| 5020 | 5021 | 1 | 1.0900e-01 | 2.8451e+05 | ; | CA | - | HA |
| 5020 | 5022 | 1 | 1.5260e-01 | 2.5941e+05 | ; | CA | - | CB |
| 5020 | 5035 | 1 | 1.5220e-01 | 2.6527e+05 | ; | CA | - | C |
| 5022 | 5023 | 1 | 1.0900e-01 | 2.8451e+05 | ; | CB | - | HB2 |
| 5022 | 5024 | 1 | 1.0900e-01 | 2.8451e+05 | ; | CB | - | HB3 |

| | | | | | |
|---|---|---|---|---|---|
| 5022 | 5025 | 1 | 1.5260e-01 | 2.5941e+05 ; | CB - CG |
| 5025 | 5026 | 1 | 1.0900e-01 | 2.8451e+05 ; | CG - HG |
| 5025 | 5027 | 1 | 1.5260e-01 | 2.5941e+05 ; | CG - CD1 |
| 5025 | 5031 | 1 | 1.5260e-01 | 2.5941e+05 ; | CG - CD2 |
| 5027 | 5028 | 1 | 1.0900e-01 | 2.8451e+05 ; | CD1 - HD11 |
| 5027 | 5029 | 1 | 1.0900e-01 | 2.8451e+05 ; | CD1 - HD12 |
| 5027 | 5030 | 1 | 1.0900e-01 | 2.8451e+05 ; | CD1 - HD13 |
| 5031 | 5032 | 1 | 1.0900e-01 | 2.8451e+05 ; | CD2 - HD21 |
| 5031 | 5033 | 1 | 1.0900e-01 | 2.8451e+05 ; | CD2 - HD22 |
| 5031 | 5034 | 1 | 1.0900e-01 | 2.8451e+05 ; | CD2 - HD23 |
| 5035 | 5036 | 1 | 1.2290e-01 | 4.7698e+05 ; | C - O |
| 5035 | 5037 | 1 | 1.3350e-01 | 4.1003e+05 ; | C - N |
| 5037 | 5038 | 1 | 1.0100e-01 | 3.6317e+05 ; | N - H |
| 5037 | 5039 | 1 | 1.4490e-01 | 2.8200e+05 ; | N - CA |
| 5039 | 5040 | 1 | 1.0900e-01 | 2.8451e+05 ; | CA - HA |
| 5039 | 5041 | 1 | 1.5260e-01 | 2.5941e+05 ; | CA - CB |
| 5039 | 5046 | 1 | 1.5220e-01 | 2.6527e+05 ; | CA - C |
| 5041 | 5042 | 1 | 1.0900e-01 | 2.8451e+05 ; | CB - HB2 |
| 5041 | 5043 | 1 | 1.0900e-01 | 2.8451e+05 ; | CB - HB3 |
| 5041 | 5044 | 1 | 1.8100e-01 | 1.9832e+05 ; | CB - SG |
| 5044 | 5045 | 1 | 1.3360e-01 | 2.2928e+05 ; | SG - HG |
| 5046 | 5047 | 1 | 1.2290e-01 | 4.7698e+05 ; | C - O |
| 5046 | 5048 | 1 | 1.3350e-01 | 4.1003e+05 ; | C - N |
| 5048 | 5049 | 1 | 1.0100e-01 | 3.6317e+05 ; | N - H |
| 5048 | 5050 | 1 | 1.4490e-01 | 2.8200e+05 ; | N - CA |
| 5050 | 5051 | 1 | 1.0900e-01 | 2.8451e+05 ; | CA - HA |
| 5050 | 5052 | 1 | 1.5260e-01 | 2.5941e+05 ; | CA - CB |
| 5050 | 5058 | 1 | 1.5220e-01 | 2.6527e+05 ; | CA - C |
| 5052 | 5053 | 1 | 1.0900e-01 | 2.8451e+05 ; | CB - HB2 |
| 5052 | 5054 | 1 | 1.0900e-01 | 2.8451e+05 ; | CB - HB3 |
| 5052 | 5055 | 1 | 1.5220e-01 | 2.6527e+05 ; | CB - CG |
| 5055 | 5056 | 1 | 1.2500e-01 | 5.4894e+05 ; | CG - OD1 |
| 5055 | 5057 | 1 | 1.2500e-01 | 5.4894e+05 ; | CG - OD2 |
| 5058 | 5059 | 1 | 1.2290e-01 | 4.7698e+05 ; | C - O |
| 5058 | 5060 | 1 | 1.3350e-01 | 4.1003e+05 ; | C - N |
| 5060 | 5061 | 1 | 1.0100e-01 | 3.6317e+05 ; | N - H |
| 5060 | 5062 | 1 | 1.4490e-01 | 2.8200e+05 ; | N - CA |
| 5062 | 5063 | 1 | 1.0900e-01 | 2.8451e+05 ; | CA - HA |
| 5062 | 5064 | 1 | 1.5260e-01 | 2.5941e+05 ; | CA - CB |
| 5062 | 5074 | 1 | 1.5220e-01 | 2.6527e+05 ; | CA - C |
| 5064 | 5065 | 1 | 1.0900e-01 | 2.8451e+05 ; | CB - HB |
| 5064 | 5066 | 1 | 1.5260e-01 | 2.5941e+05 ; | CB - CG1 |
| 5064 | 5070 | 1 | 1.5260e-01 | 2.5941e+05 ; | CB - CG2 |
| 5066 | 5067 | 1 | 1.0900e-01 | 2.8451e+05 ; | CG1 - HG11 |
| 5066 | 5068 | 1 | 1.0900e-01 | 2.8451e+05 ; | CG1 - HG12 |
| 5066 | 5069 | 1 | 1.0900e-01 | 2.8451e+05 ; | CG1 - HG13 |
| 5070 | 5071 | 1 | 1.0900e-01 | 2.8451e+05 ; | CG2 - HG21 |
| 5070 | 5072 | 1 | 1.0900e-01 | 2.8451e+05 ; | CG2 - HG22 |
| 5070 | 5073 | 1 | 1.0900e-01 | 2.8451e+05 ; | CG2 - HG23 |
| 5074 | 5075 | 1 | 1.2290e-01 | 4.7698e+05 ; | C - O |
| 5074 | 5076 | 1 | 1.3350e-01 | 4.1003e+05 ; | C - N |
| 5076 | 5077 | 1 | 1.0100e-01 | 3.6317e+05 ; | N - H |

| | | | | | | |
|---|---|---|---|---|---|---|
| 5076 | 5078 | 1 | 1.4490e-01 | 2.8200e+05 | ; | N - CA |
| 5078 | 5079 | 1 | 1.0900e-01 | 2.8451e+05 | ; | CA - HA |
| 5078 | 5080 | 1 | 1.5260e-01 | 2.5941e+05 | ; | CA - CB |
| 5078 | 5089 | 1 | 1.5220e-01 | 2.6527e+05 | ; | CA - C |
| 5080 | 5081 | 1 | 1.0900e-01 | 2.8451e+05 | ; | CB - HB2 |
| 5080 | 5082 | 1 | 1.0900e-01 | 2.8451e+05 | ; | CB - HB3 |
| 5080 | 5083 | 1 | 1.5260e-01 | 2.5941e+05 | ; | CB - CG |
| 5083 | 5084 | 1 | 1.0900e-01 | 2.8451e+05 | ; | CG - HG2 |
| 5083 | 5085 | 1 | 1.0900e-01 | 2.8451e+05 | ; | CG - HG3 |
| 5083 | 5086 | 1 | 1.5220e-01 | 2.6527e+05 | ; | CG - CD |
| 5086 | 5087 | 1 | 1.2500e-01 | 5.4894e+05 | ; | CD - OE1 |
| 5086 | 5088 | 1 | 1.2500e-01 | 5.4894e+05 | ; | CD - OE2 |
| 5089 | 5090 | 1 | 1.2290e-01 | 4.7698e+05 | ; | C - O |
| 5089 | 5091 | 1 | 1.3350e-01 | 4.1003e+05 | ; | C - N |
| 5091 | 5092 | 1 | 1.0100e-01 | 3.6317e+05 | ; | N - H |
| 5091 | 5093 | 1 | 1.4490e-01 | 2.8200e+05 | ; | N - CA |
| 5093 | 5094 | 1 | 1.0900e-01 | 2.8451e+05 | ; | CA - HA |
| 5093 | 5095 | 1 | 1.5260e-01 | 2.5941e+05 | ; | CA - CB |
| 5093 | 5108 | 1 | 1.5220e-01 | 2.6527e+05 | ; | CA - C |
| 5095 | 5096 | 1 | 1.0900e-01 | 2.8451e+05 | ; | CB - HB |
| 5095 | 5097 | 1 | 1.5260e-01 | 2.5941e+05 | ; | CB - CG2 |
| 5095 | 5101 | 1 | 1.5260e-01 | 2.5941e+05 | ; | CB - CG1 |
| 5097 | 5098 | 1 | 1.0900e-01 | 2.8451e+05 | ; | CG2 - HG21 |
| 5097 | 5099 | 1 | 1.0900e-01 | 2.8451e+05 | ; | CG2 - HG22 |
| 5097 | 5100 | 1 | 1.0900e-01 | 2.8451e+05 | ; | CG2 - HG23 |
| 5101 | 5102 | 1 | 1.0900e-01 | 2.8451e+05 | ; | CG1 - HG12 |
| 5101 | 5103 | 1 | 1.0900e-01 | 2.8451e+05 | ; | CG1 - HG13 |
| 5101 | 5104 | 1 | 1.5260e-01 | 2.5941e+05 | ; | CG1 - CD1 |
| 5104 | 5105 | 1 | 1.0900e-01 | 2.8451e+05 | ; | CD1 - HD11 |
| 5104 | 5106 | 1 | 1.0900e-01 | 2.8451e+05 | ; | CD1 - HD12 |
| 5104 | 5107 | 1 | 1.0900e-01 | 2.8451e+05 | ; | CD1 - HD13 |
| 5108 | 5109 | 1 | 1.2290e-01 | 4.7698e+05 | ; | C - O |
| 5108 | 5110 | 1 | 1.3350e-01 | 4.1003e+05 | ; | C - N |
| 5110 | 5111 | 1 | 1.0100e-01 | 3.6317e+05 | ; | N - H |
| 5110 | 5112 | 1 | 1.4490e-01 | 2.8200e+05 | ; | N - CA |
| 5112 | 5113 | 1 | 1.0900e-01 | 2.8451e+05 | ; | CA - HA |
| 5112 | 5114 | 1 | 1.5260e-01 | 2.5941e+05 | ; | CA - CB |
| 5112 | 5122 | 1 | 1.5220e-01 | 2.6527e+05 | ; | CA - C |
| 5114 | 5115 | 1 | 1.0900e-01 | 2.8451e+05 | ; | CB - HB2 |
| 5114 | 5116 | 1 | 1.0900e-01 | 2.8451e+05 | ; | CB - HB3 |
| 5114 | 5117 | 1 | 1.5220e-01 | 2.6527e+05 | ; | CB - CG |
| 5117 | 5118 | 1 | 1.2290e-01 | 4.7698e+05 | ; | CG - OD1 |
| 5117 | 5119 | 1 | 1.3350e-01 | 4.1003e+05 | ; | CG - ND2 |
| 5119 | 5120 | 1 | 1.0100e-01 | 3.6317e+05 | ; | ND2 - HD21 |
| 5119 | 5121 | 1 | 1.0100e-01 | 3.6317e+05 | ; | ND2 - HD22 |
| 5122 | 5123 | 1 | 1.2290e-01 | 4.7698e+05 | ; | C - O |
| 5122 | 5124 | 1 | 1.3350e-01 | 4.1003e+05 | ; | C - N |
| 5124 | 5125 | 1 | 1.0100e-01 | 3.6317e+05 | ; | N - H |
| 5124 | 5126 | 1 | 1.4490e-01 | 2.8200e+05 | ; | N - CA |
| 5126 | 5127 | 1 | 1.0900e-01 | 2.8451e+05 | ; | CA - HA |
| 5126 | 5128 | 1 | 1.5260e-01 | 2.5941e+05 | ; | CA - CB |
| 5126 | 5136 | 1 | 1.5220e-01 | 2.6527e+05 | ; | CA - C |

| | | | | | | | | |
|---|---|---|---|---|---|---|---|---|
| 5128 | 5129 | 1 | 1.0900e-01 | 2.8451e+05 | ; | CB | - | HB2 |
| 5128 | 5130 | 1 | 1.0900e-01 | 2.8451e+05 | ; | CB | - | HB3 |
| 5128 | 5131 | 1 | 1.5220e-01 | 2.6527e+05 | ; | CB | - | CG |
| 5131 | 5132 | 1 | 1.2290e-01 | 4.7698e+05 | ; | CG | - | OD1 |
| 5131 | 5133 | 1 | 1.3350e-01 | 4.1003e+05 | ; | CG | - | ND2 |
| 5133 | 5134 | 1 | 1.0100e-01 | 3.6317e+05 | ; | ND2 | - | HD21 |
| 5133 | 5135 | 1 | 1.0100e-01 | 3.6317e+05 | ; | ND2 | - | HD22 |
| 5136 | 5137 | 1 | 1.2290e-01 | 4.7698e+05 | ; | C | - | O |
| 5136 | 5138 | 1 | 1.3350e-01 | 4.1003e+05 | ; | C | - | N |
| 5138 | 5139 | 1 | 1.0100e-01 | 3.6317e+05 | ; | N | - | H |
| 5138 | 5140 | 1 | 1.4490e-01 | 2.8200e+05 | ; | N | - | CA |
| 5140 | 5141 | 1 | 1.0900e-01 | 2.8451e+05 | ; | CA | - | HA |
| 5140 | 5142 | 1 | 1.5260e-01 | 2.5941e+05 | ; | CA | - | CB |
| 5140 | 5155 | 1 | 1.5220e-01 | 2.6527e+05 | ; | CA | - | C |
| 5142 | 5143 | 1 | 1.0900e-01 | 2.8451e+05 | ; | CB | - | HB2 |
| 5142 | 5144 | 1 | 1.0900e-01 | 2.8451e+05 | ; | CB | - | HB3 |
| 5142 | 5145 | 1 | 1.5260e-01 | 2.5941e+05 | ; | CB | - | CG |
| 5145 | 5146 | 1 | 1.0900e-01 | 2.8451e+05 | ; | CG | - | HG |
| 5145 | 5147 | 1 | 1.5260e-01 | 2.5941e+05 | ; | CG | - | CD1 |
| 5145 | 5151 | 1 | 1.5260e-01 | 2.5941e+05 | ; | CG | - | CD2 |
| 5147 | 5148 | 1 | 1.0900e-01 | 2.8451e+05 | ; | CD1 | - | HD11 |
| 5147 | 5149 | 1 | 1.0900e-01 | 2.8451e+05 | ; | CD1 | - | HD12 |
| 5147 | 5150 | 1 | 1.0900e-01 | 2.8451e+05 | ; | CD1 | - | HD13 |
| 5151 | 5152 | 1 | 1.0900e-01 | 2.8451e+05 | ; | CD2 | - | HD21 |
| 5151 | 5153 | 1 | 1.0900e-01 | 2.8451e+05 | ; | CD2 | - | HD22 |
| 5151 | 5154 | 1 | 1.0900e-01 | 2.8451e+05 | ; | CD2 | - | HD23 |
| 5155 | 5156 | 1 | 1.2290e-01 | 4.7698e+05 | ; | C | - | O |
| 5155 | 5157 | 1 | 1.3350e-01 | 4.1003e+05 | ; | C | - | N |
| 5157 | 5158 | 1 | 1.0100e-01 | 3.6317e+05 | ; | N | - | H |
| 5157 | 5159 | 1 | 1.4490e-01 | 2.8200e+05 | ; | N | - | CA |
| 5159 | 5160 | 1 | 1.0900e-01 | 2.8451e+05 | ; | CA | - | HA |
| 5159 | 5161 | 1 | 1.5260e-01 | 2.5941e+05 | ; | CA | - | CB |
| 5159 | 5179 | 1 | 1.5220e-01 | 2.6527e+05 | ; | CA | - | C |
| 5161 | 5162 | 1 | 1.0900e-01 | 2.8451e+05 | ; | CB | - | HB2 |
| 5161 | 5163 | 1 | 1.0900e-01 | 2.8451e+05 | ; | CB | - | HB3 |
| 5161 | 5164 | 1 | 1.5260e-01 | 2.5941e+05 | ; | CB | - | CG |
| 5164 | 5165 | 1 | 1.0900e-01 | 2.8451e+05 | ; | CG | - | HG2 |
| 5164 | 5166 | 1 | 1.0900e-01 | 2.8451e+05 | ; | CG | - | HG3 |
| 5164 | 5167 | 1 | 1.5260e-01 | 2.5941e+05 | ; | CG | - | CD |
| 5167 | 5168 | 1 | 1.0900e-01 | 2.8451e+05 | ; | CD | - | HD2 |
| 5167 | 5169 | 1 | 1.0900e-01 | 2.8451e+05 | ; | CD | - | HD3 |
| 5167 | 5170 | 1 | 1.4630e-01 | 2.8200e+05 | ; | CD | - | NE |
| 5170 | 5171 | 1 | 1.0100e-01 | 3.6317e+05 | ; | NE | - | HE |
| 5170 | 5172 | 1 | 1.3400e-01 | 4.0250e+05 | ; | NE | - | CZ |
| 5172 | 5173 | 1 | 1.3400e-01 | 4.0250e+05 | ; | CZ | - | NH1 |
| 5172 | 5176 | 1 | 1.3400e-01 | 4.0250e+05 | ; | CZ | - | NH2 |
| 5173 | 5174 | 1 | 1.0100e-01 | 3.6317e+05 | ; | NH1 | - | HH11 |
| 5173 | 5175 | 1 | 1.0100e-01 | 3.6317e+05 | ; | NH1 | - | HH12 |
| 5176 | 5177 | 1 | 1.0100e-01 | 3.6317e+05 | ; | NH2 | - | HH21 |
| 5176 | 5178 | 1 | 1.0100e-01 | 3.6317e+05 | ; | NH2 | - | HH22 |
| 5179 | 5180 | 1 | 1.2290e-01 | 4.7698e+05 | ; | C | - | O |
| 5179 | 5181 | 1 | 1.3350e-01 | 4.1003e+05 | ; | C | - | N |

| | | | | | |
|---|---|---|---|---|---|
| 5181 | 5182 | 1 | 1.0100e-01 | 3.6317e+05 ; | N - H |
| 5181 | 5183 | 1 | 1.4490e-01 | 2.8200e+05 ; | N - CA |
| 5183 | 5184 | 1 | 1.0900e-01 | 2.8451e+05 ; | CA - HA |
| 5183 | 5185 | 1 | 1.5260e-01 | 2.5941e+05 ; | CA - CB |
| 5183 | 5193 | 1 | 1.5220e-01 | 2.6527e+05 ; | CA - C |
| 5185 | 5186 | 1 | 1.0900e-01 | 2.8451e+05 ; | CB - HB2 |
| 5185 | 5187 | 1 | 1.0900e-01 | 2.8451e+05 ; | CB - HB3 |
| 5185 | 5188 | 1 | 1.5220e-01 | 2.6527e+05 ; | CB - CG |
| 5188 | 5189 | 1 | 1.2290e-01 | 4.7698e+05 ; | CG - OD1 |
| 5188 | 5190 | 1 | 1.3350e-01 | 4.1003e+05 ; | CG - ND2 |
| 5190 | 5191 | 1 | 1.0100e-01 | 3.6317e+05 ; | ND2 - HD21 |
| 5190 | 5192 | 1 | 1.0100e-01 | 3.6317e+05 ; | ND2 - HD22 |
| 5193 | 5194 | 1 | 1.2290e-01 | 4.7698e+05 ; | C - O |
| 5193 | 5195 | 1 | 1.3350e-01 | 4.1003e+05 ; | C - N |
| 5195 | 5196 | 1 | 1.0100e-01 | 3.6317e+05 ; | N - H |
| 5195 | 5197 | 1 | 1.4490e-01 | 2.8200e+05 ; | N - CA |
| 5197 | 5198 | 1 | 1.0900e-01 | 2.8451e+05 ; | CA - HA |
| 5197 | 5199 | 1 | 1.5260e-01 | 2.5941e+05 ; | CA - CB |
| 5197 | 5212 | 1 | 1.5220e-01 | 2.6527e+05 ; | CA - C |
| 5199 | 5200 | 1 | 1.0900e-01 | 2.8451e+05 ; | CB - HB |
| 5199 | 5201 | 1 | 1.5260e-01 | 2.5941e+05 ; | CB - CG2 |
| 5199 | 5205 | 1 | 1.5260e-01 | 2.5941e+05 ; | CB - CG1 |
| 5201 | 5202 | 1 | 1.0900e-01 | 2.8451e+05 ; | CG2 - HG21 |
| 5201 | 5203 | 1 | 1.0900e-01 | 2.8451e+05 ; | CG2 - HG22 |
| 5201 | 5204 | 1 | 1.0900e-01 | 2.8451e+05 ; | CG2 - HG23 |
| 5205 | 5206 | 1 | 1.0900e-01 | 2.8451e+05 ; | CG1 - HG12 |
| 5205 | 5207 | 1 | 1.0900e-01 | 2.8451e+05 ; | CG1 - HG13 |
| 5205 | 5208 | 1 | 1.5260e-01 | 2.5941e+05 ; | CG1 - CD1 |
| 5208 | 5209 | 1 | 1.0900e-01 | 2.8451e+05 ; | CD1 - HD11 |
| 5208 | 5210 | 1 | 1.0900e-01 | 2.8451e+05 ; | CD1 - HD12 |
| 5208 | 5211 | 1 | 1.0900e-01 | 2.8451e+05 ; | CD1 - HD13 |
| 5212 | 5213 | 1 | 1.2290e-01 | 4.7698e+05 ; | C - O |
| 5212 | 5214 | 1 | 1.3350e-01 | 4.1003e+05 ; | C - N |
| 5214 | 5215 | 1 | 1.0100e-01 | 3.6317e+05 ; | N - H |
| 5214 | 5216 | 1 | 1.4490e-01 | 2.8200e+05 ; | N - CA |
| 5216 | 5217 | 1 | 1.0900e-01 | 2.8451e+05 ; | CA - HA |
| 5216 | 5218 | 1 | 1.5260e-01 | 2.5941e+05 ; | CA - CB |
| 5216 | 5232 | 1 | 1.5220e-01 | 2.6527e+05 ; | CA - C |
| 5218 | 5219 | 1 | 1.0900e-01 | 2.8451e+05 ; | CB - HB2 |
| 5218 | 5220 | 1 | 1.0900e-01 | 2.8451e+05 ; | CB - HB3 |
| 5218 | 5221 | 1 | 1.5100e-01 | 2.6527e+05 ; | CB - CG |
| 5221 | 5222 | 1 | 1.4000e-01 | 3.9246e+05 ; | CG - CD1 |
| 5221 | 5230 | 1 | 1.4000e-01 | 3.9246e+05 ; | CG - CD2 |
| 5222 | 5223 | 1 | 1.0800e-01 | 3.0711e+05 ; | CD1 - HD1 |
| 5222 | 5224 | 1 | 1.4000e-01 | 3.9246e+05 ; | CD1 - CE1 |
| 5224 | 5225 | 1 | 1.0800e-01 | 3.0711e+05 ; | CE1 - HE1 |
| 5224 | 5226 | 1 | 1.4000e-01 | 3.9246e+05 ; | CE1 - CZ |
| 5226 | 5227 | 1 | 1.0800e-01 | 3.0711e+05 ; | CZ - HZ |
| 5226 | 5228 | 1 | 1.4000e-01 | 3.9246e+05 ; | CZ - CE2 |
| 5228 | 5229 | 1 | 1.0800e-01 | 3.0711e+05 ; | CE2 - HE2 |
| 5228 | 5230 | 1 | 1.4000e-01 | 3.9246e+05 ; | CE2 - CD2 |
| 5230 | 5231 | 1 | 1.0800e-01 | 3.0711e+05 ; | CD2 - HD2 |

| | | | | | |
|---|---|---|---|---|---|
| 5232 | 5233 | 1 | 1.2290e-01 | 4.7698e+05 ; | C - O |
| 5232 | 5234 | 1 | 1.3350e-01 | 4.1003e+05 ; | C - N |
| 5234 | 5235 | 1 | 1.0100e-01 | 3.6317e+05 ; | N - H |
| 5234 | 5236 | 1 | 1.4490e-01 | 2.8200e+05 ; | N - CA |
| 5236 | 5237 | 1 | 1.0900e-01 | 2.8451e+05 ; | CA - HA |
| 5236 | 5238 | 1 | 1.5260e-01 | 2.5941e+05 ; | CA - CB |
| 5236 | 5251 | 1 | 1.5220e-01 | 2.6527e+05 ; | CA - C |
| 5238 | 5239 | 1 | 1.0900e-01 | 2.8451e+05 ; | CB - HB |
| 5238 | 5240 | 1 | 1.5260e-01 | 2.5941e+05 ; | CB - CG2 |
| 5238 | 5244 | 1 | 1.5260e-01 | 2.5941e+05 ; | CB - CG1 |
| 5240 | 5241 | 1 | 1.0900e-01 | 2.8451e+05 ; | CG2 - HG21 |
| 5240 | 5242 | 1 | 1.0900e-01 | 2.8451e+05 ; | CG2 - HG22 |
| 5240 | 5243 | 1 | 1.0900e-01 | 2.8451e+05 ; | CG2 - HG23 |
| 5244 | 5245 | 1 | 1.0900e-01 | 2.8451e+05 ; | CG1 - HG12 |
| 5244 | 5246 | 1 | 1.0900e-01 | 2.8451e+05 ; | CG1 - HG13 |
| 5244 | 5247 | 1 | 1.5260e-01 | 2.5941e+05 ; | CG1 - CD1 |
| 5247 | 5248 | 1 | 1.0900e-01 | 2.8451e+05 ; | CD1 - HD11 |
| 5247 | 5249 | 1 | 1.0900e-01 | 2.8451e+05 ; | CD1 - HD12 |
| 5247 | 5250 | 1 | 1.0900e-01 | 2.8451e+05 ; | CD1 - HD13 |
| 5251 | 5252 | 1 | 1.2290e-01 | 4.7698e+05 ; | C - O |
| 5251 | 5253 | 1 | 1.3350e-01 | 4.1003e+05 ; | C - N |
| 5253 | 5254 | 1 | 1.0100e-01 | 3.6317e+05 ; | N - H |
| 5253 | 5255 | 1 | 1.4490e-01 | 2.8200e+05 ; | N - CA |
| 5255 | 5256 | 1 | 1.0900e-01 | 2.8451e+05 ; | CA - HA |
| 5255 | 5257 | 1 | 1.5260e-01 | 2.5941e+05 ; | CA - CB |
| 5255 | 5261 | 1 | 1.5220e-01 | 2.6527e+05 ; | CA - C |
| 5257 | 5258 | 1 | 1.0900e-01 | 2.8451e+05 ; | CB - HB1 |
| 5257 | 5259 | 1 | 1.0900e-01 | 2.8451e+05 ; | CB - HB2 |
| 5257 | 5260 | 1 | 1.0900e-01 | 2.8451e+05 ; | CB - HB3 |
| 5261 | 5262 | 1 | 1.2290e-01 | 4.7698e+05 ; | C - O |
| 5261 | 5263 | 1 | 1.3350e-01 | 4.1003e+05 ; | C - N |
| 5263 | 5264 | 1 | 1.0100e-01 | 3.6317e+05 ; | N - H |
| 5263 | 5265 | 1 | 1.4490e-01 | 2.8200e+05 ; | N - CA |
| 5265 | 5266 | 1 | 1.0900e-01 | 2.8451e+05 ; | CA - HA |
| 5265 | 5267 | 1 | 1.5260e-01 | 2.5941e+05 ; | CA - CB |
| 5265 | 5280 | 1 | 1.5220e-01 | 2.6527e+05 ; | CA - C |
| 5267 | 5268 | 1 | 1.0900e-01 | 2.8451e+05 ; | CB - HB2 |
| 5267 | 5269 | 1 | 1.0900e-01 | 2.8451e+05 ; | CB - HB3 |
| 5267 | 5270 | 1 | 1.5260e-01 | 2.5941e+05 ; | CB - CG |
| 5270 | 5271 | 1 | 1.0900e-01 | 2.8451e+05 ; | CG - HG |
| 5270 | 5272 | 1 | 1.5260e-01 | 2.5941e+05 ; | CG - CD1 |
| 5270 | 5276 | 1 | 1.5260e-01 | 2.5941e+05 ; | CG - CD2 |
| 5272 | 5273 | 1 | 1.0900e-01 | 2.8451e+05 ; | CD1 - HD11 |
| 5272 | 5274 | 1 | 1.0900e-01 | 2.8451e+05 ; | CD1 - HD12 |
| 5272 | 5275 | 1 | 1.0900e-01 | 2.8451e+05 ; | CD1 - HD13 |
| 5276 | 5277 | 1 | 1.0900e-01 | 2.8451e+05 ; | CD2 - HD21 |
| 5276 | 5278 | 1 | 1.0900e-01 | 2.8451e+05 ; | CD2 - HD22 |
| 5276 | 5279 | 1 | 1.0900e-01 | 2.8451e+05 ; | CD2 - HD23 |
| 5280 | 5281 | 1 | 1.2290e-01 | 4.7698e+05 ; | C - O |
| 5280 | 5282 | 1 | 1.3350e-01 | 4.1003e+05 ; | C - N |
| 5282 | 5283 | 1 | 1.0100e-01 | 3.6317e+05 ; | N - H |
| 5282 | 5284 | 1 | 1.4490e-01 | 2.8200e+05 ; | N - CA |

| | | | | | | |
|---|---|---|---|---|---|---|
| 5284 | 5285 | 1 | 1.0900e-01 | 2.8451e+05 | ; | CA - HA |
| 5284 | 5286 | 1 | 1.5260e-01 | 2.5941e+05 | ; | CA - CB |
| 5284 | 5292 | 1 | 1.5220e-01 | 2.6527e+05 | ; | CA - C |
| 5286 | 5287 | 1 | 1.0900e-01 | 2.8451e+05 | ; | CB - HB2 |
| 5286 | 5288 | 1 | 1.0900e-01 | 2.8451e+05 | ; | CB - HB3 |
| 5286 | 5289 | 1 | 1.5220e-01 | 2.6527e+05 | ; | CB - CG |
| 5289 | 5290 | 1 | 1.2500e-01 | 5.4894e+05 | ; | CG - OD1 |
| 5289 | 5291 | 1 | 1.2500e-01 | 5.4894e+05 | ; | CG - OD2 |
| 5292 | 5293 | 1 | 1.2290e-01 | 4.7698e+05 | ; | C - O |
| 5292 | 5294 | 1 | 1.3350e-01 | 4.1003e+05 | ; | C - N |
| 5294 | 5295 | 1 | 1.0100e-01 | 3.6317e+05 | ; | N - H |
| 5294 | 5296 | 1 | 1.4490e-01 | 2.8200e+05 | ; | N - CA |
| 5296 | 5297 | 1 | 1.0900e-01 | 2.8451e+05 | ; | CA - HA |
| 5296 | 5298 | 1 | 1.5260e-01 | 2.5941e+05 | ; | CA - CB |
| 5296 | 5308 | 1 | 1.5220e-01 | 2.6527e+05 | ; | CA - C |
| 5298 | 5299 | 1 | 1.0900e-01 | 2.8451e+05 | ; | CB - HB |
| 5298 | 5300 | 1 | 1.5260e-01 | 2.5941e+05 | ; | CB - CG1 |
| 5298 | 5304 | 1 | 1.5260e-01 | 2.5941e+05 | ; | CB - CG2 |
| 5300 | 5301 | 1 | 1.0900e-01 | 2.8451e+05 | ; | CG1 - HG11 |
| 5300 | 5302 | 1 | 1.0900e-01 | 2.8451e+05 | ; | CG1 - HG12 |
| 5300 | 5303 | 1 | 1.0900e-01 | 2.8451e+05 | ; | CG1 - HG13 |
| 5304 | 5305 | 1 | 1.0900e-01 | 2.8451e+05 | ; | CG2 - HG21 |
| 5304 | 5306 | 1 | 1.0900e-01 | 2.8451e+05 | ; | CG2 - HG22 |
| 5304 | 5307 | 1 | 1.0900e-01 | 2.8451e+05 | ; | CG2 - HG23 |
| 5308 | 5309 | 1 | 1.2290e-01 | 4.7698e+05 | ; | C - O |
| 5308 | 5310 | 1 | 1.3350e-01 | 4.1003e+05 | ; | C - N |
| 5310 | 5311 | 1 | 1.0100e-01 | 3.6317e+05 | ; | N - H |
| 5310 | 5312 | 1 | 1.4490e-01 | 2.8200e+05 | ; | N - CA |
| 5312 | 5313 | 1 | 1.0900e-01 | 2.8451e+05 | ; | CA - HA |
| 5312 | 5314 | 1 | 1.5260e-01 | 2.5941e+05 | ; | CA - CB |
| 5312 | 5320 | 1 | 1.5220e-01 | 2.6527e+05 | ; | CA - C |
| 5314 | 5315 | 1 | 1.0900e-01 | 2.8451e+05 | ; | CB - HB2 |
| 5314 | 5316 | 1 | 1.0900e-01 | 2.8451e+05 | ; | CB - HB3 |
| 5314 | 5317 | 1 | 1.5220e-01 | 2.6527e+05 | ; | CB - CG |
| 5317 | 5318 | 1 | 1.2500e-01 | 5.4894e+05 | ; | CG - OD1 |
| 5317 | 5319 | 1 | 1.2500e-01 | 5.4894e+05 | ; | CG - OD2 |
| 5320 | 5321 | 1 | 1.2290e-01 | 4.7698e+05 | ; | C - O |
| 5320 | 5322 | 1 | 1.3350e-01 | 4.1003e+05 | ; | C - N |
| 5322 | 5323 | 1 | 1.0100e-01 | 3.6317e+05 | ; | N - H |
| 5322 | 5324 | 1 | 1.4490e-01 | 2.8200e+05 | ; | N - CA |
| 5324 | 5325 | 1 | 1.0900e-01 | 2.8451e+05 | ; | CA - HA |
| 5324 | 5326 | 1 | 1.5260e-01 | 2.5941e+05 | ; | CA - CB |
| 5324 | 5334 | 1 | 1.5220e-01 | 2.6527e+05 | ; | CA - C |
| 5326 | 5327 | 1 | 1.0900e-01 | 2.8451e+05 | ; | CB - HB2 |
| 5326 | 5328 | 1 | 1.0900e-01 | 2.8451e+05 | ; | CB - HB3 |
| 5326 | 5329 | 1 | 1.5220e-01 | 2.6527e+05 | ; | CB - CG |
| 5329 | 5330 | 1 | 1.2290e-01 | 4.7698e+05 | ; | CG - OD1 |
| 5329 | 5331 | 1 | 1.3350e-01 | 4.1003e+05 | ; | CG - ND2 |
| 5331 | 5332 | 1 | 1.0100e-01 | 3.6317e+05 | ; | ND2 - HD21 |
| 5331 | 5333 | 1 | 1.0100e-01 | 3.6317e+05 | ; | ND2 - HD22 |
| 5334 | 5335 | 1 | 1.2290e-01 | 4.7698e+05 | ; | C - O |
| 5334 | 5336 | 1 | 1.3350e-01 | 4.1003e+05 | ; | C - N |

| | | | | | |
|---|---|---|---|---|---|
| 5336 | 5337 | 1 | 1.0100e-01 | 3.6317e+05 ; | N - H |
| 5336 | 5338 | 1 | 1.4490e-01 | 2.8200e+05 ; | N - CA |
| 5338 | 5339 | 1 | 1.0900e-01 | 2.8451e+05 ; | CA - HA |
| 5338 | 5340 | 1 | 1.5260e-01 | 2.5941e+05 ; | CA - CB |
| 5338 | 5345 | 1 | 1.5220e-01 | 2.6527e+05 ; | CA - C |
| 5340 | 5341 | 1 | 1.0900e-01 | 2.8451e+05 ; | CB - HB2 |
| 5340 | 5342 | 1 | 1.0900e-01 | 2.8451e+05 ; | CB - HB3 |
| 5340 | 5343 | 1 | 1.4100e-01 | 2.6778e+05 ; | CB - OG |
| 5343 | 5344 | 1 | 9.6000e-02 | 4.6275e+05 ; | OG - HG |
| 5345 | 5346 | 1 | 1.2290e-01 | 4.7698e+05 ; | C - O |
| 5345 | 5347 | 1 | 1.3350e-01 | 4.1003e+05 ; | C - N |
| 5347 | 5348 | 1 | 1.0100e-01 | 3.6317e+05 ; | N - H |
| 5347 | 5349 | 1 | 1.4490e-01 | 2.8200e+05 ; | N - CA |
| 5349 | 5350 | 1 | 1.0900e-01 | 2.8451e+05 ; | CA - HA2 |
| 5349 | 5351 | 1 | 1.0900e-01 | 2.8451e+05 ; | CA - HA3 |
| 5349 | 5352 | 1 | 1.5220e-01 | 2.6527e+05 ; | CA - C |
| 5352 | 5353 | 1 | 1.2290e-01 | 4.7698e+05 ; | C - O |
| 5352 | 5354 | 1 | 1.3350e-01 | 4.1003e+05 ; | C - N |
| 5354 | 5355 | 1 | 1.0100e-01 | 3.6317e+05 ; | N - H |
| 5354 | 5356 | 1 | 1.4490e-01 | 2.8200e+05 ; | N - CA |
| 5356 | 5357 | 1 | 1.0900e-01 | 2.8451e+05 ; | CA - HA |
| 5356 | 5358 | 1 | 1.5260e-01 | 2.5941e+05 ; | CA - CB |
| 5356 | 5366 | 1 | 1.5220e-01 | 2.6527e+05 ; | CA - C |
| 5358 | 5359 | 1 | 1.0900e-01 | 2.8451e+05 ; | CB - HB |
| 5358 | 5360 | 1 | 1.5260e-01 | 2.5941e+05 ; | CB - CG2 |
| 5358 | 5364 | 1 | 1.4100e-01 | 2.6778e+05 ; | CB - OG1 |
| 5360 | 5361 | 1 | 1.0900e-01 | 2.8451e+05 ; | CG2 - HG21 |
| 5360 | 5362 | 1 | 1.0900e-01 | 2.8451e+05 ; | CG2 - HG22 |
| 5360 | 5363 | 1 | 1.0900e-01 | 2.8451e+05 ; | CG2 - HG23 |
| 5364 | 5365 | 1 | 9.6000e-02 | 4.6275e+05 ; | OG1 - HG1 |
| 5366 | 5367 | 1 | 1.2290e-01 | 4.7698e+05 ; | C - O |
| 5366 | 5368 | 1 | 1.3350e-01 | 4.1003e+05 ; | C - N |
| 5368 | 5369 | 1 | 1.0100e-01 | 3.6317e+05 ; | N - H |
| 5368 | 5370 | 1 | 1.4490e-01 | 2.8200e+05 ; | N - CA |
| 5370 | 5371 | 1 | 1.0900e-01 | 2.8451e+05 ; | CA - HA |
| 5370 | 5372 | 1 | 1.5260e-01 | 2.5941e+05 ; | CA - CB |
| 5370 | 5385 | 1 | 1.5220e-01 | 2.6527e+05 ; | CA - C |
| 5372 | 5373 | 1 | 1.0900e-01 | 2.8451e+05 ; | CB - HB2 |
| 5372 | 5374 | 1 | 1.0900e-01 | 2.8451e+05 ; | CB - HB3 |
| 5372 | 5375 | 1 | 1.5260e-01 | 2.5941e+05 ; | CB - CG |
| 5375 | 5376 | 1 | 1.0900e-01 | 2.8451e+05 ; | CG - HG |
| 5375 | 5377 | 1 | 1.5260e-01 | 2.5941e+05 ; | CG - CD1 |
| 5375 | 5381 | 1 | 1.5260e-01 | 2.5941e+05 ; | CG - CD2 |
| 5377 | 5378 | 1 | 1.0900e-01 | 2.8451e+05 ; | CD1 - HD11 |
| 5377 | 5379 | 1 | 1.0900e-01 | 2.8451e+05 ; | CD1 - HD12 |
| 5377 | 5380 | 1 | 1.0900e-01 | 2.8451e+05 ; | CD1 - HD13 |
| 5381 | 5382 | 1 | 1.0900e-01 | 2.8451e+05 ; | CD2 - HD21 |
| 5381 | 5383 | 1 | 1.0900e-01 | 2.8451e+05 ; | CD2 - HD22 |
| 5381 | 5384 | 1 | 1.0900e-01 | 2.8451e+05 ; | CD2 - HD23 |
| 5385 | 5386 | 1 | 1.2290e-01 | 4.7698e+05 ; | C - O |
| 5385 | 5387 | 1 | 1.3350e-01 | 4.1003e+05 ; | C - N |
| 5387 | 5388 | 1 | 1.0100e-01 | 3.6317e+05 ; | N - H |

| | | | | | | |
|---|---|---|---|---|---|---|
| 5387 | 5389 | 1 | 1.4490e-01 | 2.8200e+05 | ; | N - CA |
| 5389 | 5390 | 1 | 1.0900e-01 | 2.8451e+05 | ; | CA - HA |
| 5389 | 5391 | 1 | 1.5260e-01 | 2.5941e+05 | ; | CA - CB |
| 5389 | 5396 | 1 | 1.5220e-01 | 2.6527e+05 | ; | CA - C |
| 5391 | 5392 | 1 | 1.0900e-01 | 2.8451e+05 | ; | CB - HB2 |
| 5391 | 5393 | 1 | 1.0900e-01 | 2.8451e+05 | ; | CB - HB3 |
| 5391 | 5394 | 1 | 1.4100e-01 | 2.6778e+05 | ; | CB - OG |
| 5394 | 5395 | 1 | 9.6000e-02 | 4.6275e+05 | ; | OG - HG |
| 5396 | 5397 | 1 | 1.2290e-01 | 4.7698e+05 | ; | C - O |
| 5396 | 5398 | 1 | 1.3350e-01 | 4.1003e+05 | ; | C - N |
| 5398 | 5399 | 1 | 1.0100e-01 | 3.6317e+05 | ; | N - H |
| 5398 | 5400 | 1 | 1.4490e-01 | 2.8200e+05 | ; | N - CA |
| 5400 | 5401 | 1 | 1.0900e-01 | 2.8451e+05 | ; | CA - HA |
| 5400 | 5402 | 1 | 1.5260e-01 | 2.5941e+05 | ; | CA - CB |
| 5400 | 5407 | 1 | 1.5220e-01 | 2.6527e+05 | ; | CA - C |
| 5402 | 5403 | 1 | 1.0900e-01 | 2.8451e+05 | ; | CB - HB2 |
| 5402 | 5404 | 1 | 1.0900e-01 | 2.8451e+05 | ; | CB - HB3 |
| 5402 | 5405 | 1 | 1.4100e-01 | 2.6778e+05 | ; | CB - OG |
| 5405 | 5406 | 1 | 9.6000e-02 | 4.6275e+05 | ; | OG - HG |
| 5407 | 5408 | 1 | 1.2290e-01 | 4.7698e+05 | ; | C - O |
| 5407 | 5409 | 1 | 1.3350e-01 | 4.1003e+05 | ; | C - N |
| 5409 | 5410 | 1 | 1.0100e-01 | 3.6317e+05 | ; | N - H |
| 5409 | 5411 | 1 | 1.4490e-01 | 2.8200e+05 | ; | N - CA |
| 5411 | 5412 | 1 | 1.0900e-01 | 2.8451e+05 | ; | CA - HA |
| 5411 | 5413 | 1 | 1.5260e-01 | 2.5941e+05 | ; | CA - CB |
| 5411 | 5424 | 1 | 1.5220e-01 | 2.6527e+05 | ; | CA - C |
| 5413 | 5414 | 1 | 1.0900e-01 | 2.8451e+05 | ; | CB - HB2 |
| 5413 | 5415 | 1 | 1.0900e-01 | 2.8451e+05 | ; | CB - HB3 |
| 5413 | 5416 | 1 | 1.5260e-01 | 2.5941e+05 | ; | CB - CG |
| 5416 | 5417 | 1 | 1.0900e-01 | 2.8451e+05 | ; | CG - HG2 |
| 5416 | 5418 | 1 | 1.0900e-01 | 2.8451e+05 | ; | CG - HG3 |
| 5416 | 5419 | 1 | 1.5220e-01 | 2.6527e+05 | ; | CG - CD |
| 5419 | 5420 | 1 | 1.2290e-01 | 4.7698e+05 | ; | CD - OE1 |
| 5419 | 5421 | 1 | 1.3350e-01 | 4.1003e+05 | ; | CD - NE2 |
| 5421 | 5422 | 1 | 1.0100e-01 | 3.6317e+05 | ; | NE2 - HE21 |
| 5421 | 5423 | 1 | 1.0100e-01 | 3.6317e+05 | ; | NE2 - HE22 |
| 5424 | 5425 | 1 | 1.2290e-01 | 4.7698e+05 | ; | C - O |
| 5424 | 5426 | 1 | 1.3350e-01 | 4.1003e+05 | ; | C - N |
| 5426 | 5427 | 1 | 1.0100e-01 | 3.6317e+05 | ; | N - H |
| 5426 | 5428 | 1 | 1.4490e-01 | 2.8200e+05 | ; | N - CA |
| 5428 | 5429 | 1 | 1.0900e-01 | 2.8451e+05 | ; | CA - HA |
| 5428 | 5430 | 1 | 1.5260e-01 | 2.5941e+05 | ; | CA - CB |
| 5428 | 5440 | 1 | 1.5220e-01 | 2.6527e+05 | ; | CA - C |
| 5430 | 5431 | 1 | 1.0900e-01 | 2.8451e+05 | ; | CB - HB2 |
| 5430 | 5432 | 1 | 1.0900e-01 | 2.8451e+05 | ; | CB - HB3 |
| 5430 | 5433 | 1 | 1.5260e-01 | 2.5941e+05 | ; | CB - CG |
| 5433 | 5434 | 1 | 1.0900e-01 | 2.8451e+05 | ; | CG - HG2 |
| 5433 | 5435 | 1 | 1.0900e-01 | 2.8451e+05 | ; | CG - HG3 |
| 5433 | 5436 | 1 | 1.5220e-01 | 2.6527e+05 | ; | CG - CD |
| 5436 | 5437 | 1 | 1.2290e-01 | 4.7698e+05 | ; | CD - OE1 |
| 5436 | 5438 | 1 | 1.3640e-01 | 3.7656e+05 | ; | CD - OE2 |
| 5438 | 5439 | 1 | 9.6000e-02 | 4.6275e+05 | ; | OE2 - HE2 |

| | | | | | |
|---|---|---|---|---|---|
| 5440 | 5441 | 1 | 1.2290e-01 | 4.7698e+05 ; | C - O |
| 5440 | 5442 | 1 | 1.3350e-01 | 4.1003e+05 ; | C - N |
| 5442 | 5443 | 1 | 1.0100e-01 | 3.6317e+05 ; | N - H |
| 5442 | 5444 | 1 | 1.4490e-01 | 2.8200e+05 ; | N - CA |
| 5444 | 5445 | 1 | 1.0900e-01 | 2.8451e+05 ; | CA - HA |
| 5444 | 5446 | 1 | 1.5260e-01 | 2.5941e+05 ; | CA - CB |
| 5444 | 5459 | 1 | 1.5220e-01 | 2.6527e+05 ; | CA - C |
| 5446 | 5447 | 1 | 1.0900e-01 | 2.8451e+05 ; | CB - HB |
| 5446 | 5448 | 1 | 1.5260e-01 | 2.5941e+05 ; | CB - CG2 |
| 5446 | 5452 | 1 | 1.5260e-01 | 2.5941e+05 ; | CB - CG1 |
| 5448 | 5449 | 1 | 1.0900e-01 | 2.8451e+05 ; | CG2 - HG21 |
| 5448 | 5450 | 1 | 1.0900e-01 | 2.8451e+05 ; | CG2 - HG22 |
| 5448 | 5451 | 1 | 1.0900e-01 | 2.8451e+05 ; | CG2 - HG23 |
| 5452 | 5453 | 1 | 1.0900e-01 | 2.8451e+05 ; | CG1 - HG12 |
| 5452 | 5454 | 1 | 1.0900e-01 | 2.8451e+05 ; | CG1 - HG13 |
| 5452 | 5455 | 1 | 1.5260e-01 | 2.5941e+05 ; | CG1 - CD1 |
| 5455 | 5456 | 1 | 1.0900e-01 | 2.8451e+05 ; | CD1 - HD11 |
| 5455 | 5457 | 1 | 1.0900e-01 | 2.8451e+05 ; | CD1 - HD12 |
| 5455 | 5458 | 1 | 1.0900e-01 | 2.8451e+05 ; | CD1 - HD13 |
| 5459 | 5460 | 1 | 1.2290e-01 | 4.7698e+05 ; | C - O |
| 5459 | 5461 | 1 | 1.3350e-01 | 4.1003e+05 ; | C - N |
| 5461 | 5462 | 1 | 1.0100e-01 | 3.6317e+05 ; | N - H |
| 5461 | 5463 | 1 | 1.4490e-01 | 2.8200e+05 ; | N - CA |
| 5463 | 5464 | 1 | 1.0900e-01 | 2.8451e+05 ; | CA - HA |
| 5463 | 5465 | 1 | 1.5260e-01 | 2.5941e+05 ; | CA - CB |
| 5463 | 5478 | 1 | 1.5220e-01 | 2.6527e+05 ; | CA - C |
| 5465 | 5466 | 1 | 1.0900e-01 | 2.8451e+05 ; | CB - HB2 |
| 5465 | 5467 | 1 | 1.0900e-01 | 2.8451e+05 ; | CB - HB3 |
| 5465 | 5468 | 1 | 1.5260e-01 | 2.5941e+05 ; | CB - CG |
| 5468 | 5469 | 1 | 1.0900e-01 | 2.8451e+05 ; | CG - HG |
| 5468 | 5470 | 1 | 1.5260e-01 | 2.5941e+05 ; | CG - CD1 |
| 5468 | 5474 | 1 | 1.5260e-01 | 2.5941e+05 ; | CG - CD2 |
| 5470 | 5471 | 1 | 1.0900e-01 | 2.8451e+05 ; | CD1 - HD11 |
| 5470 | 5472 | 1 | 1.0900e-01 | 2.8451e+05 ; | CD1 - HD12 |
| 5470 | 5473 | 1 | 1.0900e-01 | 2.8451e+05 ; | CD1 - HD13 |
| 5474 | 5475 | 1 | 1.0900e-01 | 2.8451e+05 ; | CD2 - HD21 |
| 5474 | 5476 | 1 | 1.0900e-01 | 2.8451e+05 ; | CD2 - HD22 |
| 5474 | 5477 | 1 | 1.0900e-01 | 2.8451e+05 ; | CD2 - HD23 |
| 5478 | 5479 | 1 | 1.2290e-01 | 4.7698e+05 ; | C - O |
| 5478 | 5480 | 1 | 1.3350e-01 | 4.1003e+05 ; | C - N |
| 5480 | 5481 | 1 | 1.0100e-01 | 3.6317e+05 ; | N - H |
| 5480 | 5482 | 1 | 1.4490e-01 | 2.8200e+05 ; | N - CA |
| 5482 | 5483 | 1 | 1.0900e-01 | 2.8451e+05 ; | CA - HA |
| 5482 | 5484 | 1 | 1.5260e-01 | 2.5941e+05 ; | CA - CB |
| 5482 | 5490 | 1 | 1.5220e-01 | 2.6527e+05 ; | CA - C |
| 5484 | 5485 | 1 | 1.0900e-01 | 2.8451e+05 ; | CB - HB2 |
| 5484 | 5486 | 1 | 1.0900e-01 | 2.8451e+05 ; | CB - HB3 |
| 5484 | 5487 | 1 | 1.5220e-01 | 2.6527e+05 ; | CB - CG |
| 5487 | 5488 | 1 | 1.2500e-01 | 5.4894e+05 ; | CG - OD1 |
| 5487 | 5489 | 1 | 1.2500e-01 | 5.4894e+05 ; | CG - OD2 |
| 5490 | 5491 | 1 | 1.2290e-01 | 4.7698e+05 ; | C - O |
| 5490 | 5492 | 1 | 1.3350e-01 | 4.1003e+05 ; | C - N |

| | | | | | | |
|---|---|---|---|---|---|---|
| 5492 | 5493 | 1 | 1.0100e-01 | 3.6317e+05 | ; | N - H |
| 5492 | 5494 | 1 | 1.4490e-01 | 2.8200e+05 | ; | N - CA |
| 5494 | 5495 | 1 | 1.0900e-01 | 2.8451e+05 | ; | CA - HA2 |
| 5494 | 5496 | 1 | 1.0900e-01 | 2.8451e+05 | ; | CA - HA3 |
| 5494 | 5497 | 1 | 1.5220e-01 | 2.6527e+05 | ; | CA - C |
| 5497 | 5498 | 1 | 1.2290e-01 | 4.7698e+05 | ; | C - O |
| 5497 | 5499 | 1 | 1.3350e-01 | 4.1003e+05 | ; | C - N |
| 5499 | 5500 | 1 | 1.0100e-01 | 3.6317e+05 | ; | N - H |
| 5499 | 5501 | 1 | 1.4490e-01 | 2.8200e+05 | ; | N - CA |
| 5501 | 5502 | 1 | 1.0900e-01 | 2.8451e+05 | ; | CA - HA |
| 5501 | 5503 | 1 | 1.5260e-01 | 2.5941e+05 | ; | CA - CB |
| 5501 | 5516 | 1 | 1.5220e-01 | 2.6527e+05 | ; | CA - C |
| 5503 | 5504 | 1 | 1.0900e-01 | 2.8451e+05 | ; | CB - HB2 |
| 5503 | 5505 | 1 | 1.0900e-01 | 2.8451e+05 | ; | CB - HB3 |
| 5503 | 5506 | 1 | 1.5260e-01 | 2.5941e+05 | ; | CB - CG |
| 5506 | 5507 | 1 | 1.0900e-01 | 2.8451e+05 | ; | CG - HG |
| 5506 | 5508 | 1 | 1.5260e-01 | 2.5941e+05 | ; | CG - CD1 |
| 5506 | 5512 | 1 | 1.5260e-01 | 2.5941e+05 | ; | CG - CD2 |
| 5508 | 5509 | 1 | 1.0900e-01 | 2.8451e+05 | ; | CD1 - HD11 |
| 5508 | 5510 | 1 | 1.0900e-01 | 2.8451e+05 | ; | CD1 - HD12 |
| 5508 | 5511 | 1 | 1.0900e-01 | 2.8451e+05 | ; | CD1 - HD13 |
| 5512 | 5513 | 1 | 1.0900e-01 | 2.8451e+05 | ; | CD2 - HD21 |
| 5512 | 5514 | 1 | 1.0900e-01 | 2.8451e+05 | ; | CD2 - HD22 |
| 5512 | 5515 | 1 | 1.0900e-01 | 2.8451e+05 | ; | CD2 - HD23 |
| 5516 | 5517 | 1 | 1.2290e-01 | 4.7698e+05 | ; | C - O |
| 5516 | 5518 | 1 | 1.3350e-01 | 4.1003e+05 | ; | C - N |
| 5518 | 5519 | 1 | 1.0100e-01 | 3.6317e+05 | ; | N - H |
| 5518 | 5520 | 1 | 1.4490e-01 | 2.8200e+05 | ; | N - CA |
| 5520 | 5521 | 1 | 1.0900e-01 | 2.8451e+05 | ; | CA - HA |
| 5520 | 5522 | 1 | 1.5260e-01 | 2.5941e+05 | ; | CA - CB |
| 5520 | 5538 | 1 | 1.5220e-01 | 2.6527e+05 | ; | CA - C |
| 5522 | 5523 | 1 | 1.0900e-01 | 2.8451e+05 | ; | CB - HB2 |
| 5522 | 5524 | 1 | 1.0900e-01 | 2.8451e+05 | ; | CB - HB3 |
| 5522 | 5525 | 1 | 1.5260e-01 | 2.5941e+05 | ; | CB - CG |
| 5525 | 5526 | 1 | 1.0900e-01 | 2.8451e+05 | ; | CG - HG2 |
| 5525 | 5527 | 1 | 1.0900e-01 | 2.8451e+05 | ; | CG - HG3 |
| 5525 | 5528 | 1 | 1.5260e-01 | 2.5941e+05 | ; | CG - CD |
| 5528 | 5529 | 1 | 1.0900e-01 | 2.8451e+05 | ; | CD - HD2 |
| 5528 | 5530 | 1 | 1.0900e-01 | 2.8451e+05 | ; | CD - HD3 |
| 5528 | 5531 | 1 | 1.5260e-01 | 2.5941e+05 | ; | CD - CE |
| 5531 | 5532 | 1 | 1.0900e-01 | 2.8451e+05 | ; | CE - HE2 |
| 5531 | 5533 | 1 | 1.0900e-01 | 2.8451e+05 | ; | CE - HE3 |
| 5531 | 5534 | 1 | 1.4710e-01 | 3.0711e+05 | ; | CE - NZ |
| 5534 | 5535 | 1 | 1.0100e-01 | 3.6317e+05 | ; | NZ - HZ1 |
| 5534 | 5536 | 1 | 1.0100e-01 | 3.6317e+05 | ; | NZ - HZ2 |
| 5534 | 5537 | 1 | 1.0100e-01 | 3.6317e+05 | ; | NZ - HZ3 |
| 5538 | 5539 | 1 | 1.2290e-01 | 4.7698e+05 | ; | C - O |
| 5538 | 5540 | 1 | 1.3350e-01 | 4.1003e+05 | ; | C - N |
| 5540 | 5541 | 1 | 1.0100e-01 | 3.6317e+05 | ; | N - H |
| 5540 | 5542 | 1 | 1.4490e-01 | 2.8200e+05 | ; | N - CA |
| 5542 | 5543 | 1 | 1.0900e-01 | 2.8451e+05 | ; | CA - HA |
| 5542 | 5544 | 1 | 1.5260e-01 | 2.5941e+05 | ; | CA - CB |

| | | | | | | |
|---|---|---|---|---|---|---|
| 5542 | 5560 | 1 | 1.5220e-01 | 2.6527e+05 | ; | CA - C |
| 5544 | 5545 | 1 | 1.0900e-01 | 2.8451e+05 | ; | CB - HB2 |
| 5544 | 5546 | 1 | 1.0900e-01 | 2.8451e+05 | ; | CB - HB3 |
| 5544 | 5547 | 1 | 1.5260e-01 | 2.5941e+05 | ; | CB - CG |
| 5547 | 5548 | 1 | 1.0900e-01 | 2.8451e+05 | ; | CG - HG2 |
| 5547 | 5549 | 1 | 1.0900e-01 | 2.8451e+05 | ; | CG - HG3 |
| 5547 | 5550 | 1 | 1.5260e-01 | 2.5941e+05 | ; | CG - CD |
| 5550 | 5551 | 1 | 1.0900e-01 | 2.8451e+05 | ; | CD - HD2 |
| 5550 | 5552 | 1 | 1.0900e-01 | 2.8451e+05 | ; | CD - HD3 |
| 5550 | 5553 | 1 | 1.5260e-01 | 2.5941e+05 | ; | CD - CE |
| 5553 | 5554 | 1 | 1.0900e-01 | 2.8451e+05 | ; | CE - HE2 |
| 5553 | 5555 | 1 | 1.0900e-01 | 2.8451e+05 | ; | CE - HE3 |
| 5553 | 5556 | 1 | 1.4710e-01 | 3.0711e+05 | ; | CE - NZ |
| 5556 | 5557 | 1 | 1.0100e-01 | 3.6317e+05 | ; | NZ - HZ1 |
| 5556 | 5558 | 1 | 1.0100e-01 | 3.6317e+05 | ; | NZ - HZ2 |
| 5556 | 5559 | 1 | 1.0100e-01 | 3.6317e+05 | ; | NZ - HZ3 |
| 5560 | 5561 | 1 | 1.2290e-01 | 4.7698e+05 | ; | C - O |
| 5560 | 5562 | 1 | 1.3350e-01 | 4.1003e+05 | ; | C - N |
| 5562 | 5563 | 1 | 1.0100e-01 | 3.6317e+05 | ; | N - H |
| 5562 | 5564 | 1 | 1.4490e-01 | 2.8200e+05 | ; | N - CA |
| 5564 | 5565 | 1 | 1.0900e-01 | 2.8451e+05 | ; | CA - HA |
| 5564 | 5566 | 1 | 1.5260e-01 | 2.5941e+05 | ; | CA - CB |
| 5564 | 5579 | 1 | 1.5220e-01 | 2.6527e+05 | ; | CA - C |
| 5566 | 5567 | 1 | 1.0900e-01 | 2.8451e+05 | ; | CB - HB |
| 5566 | 5568 | 1 | 1.5260e-01 | 2.5941e+05 | ; | CB - CG2 |
| 5566 | 5572 | 1 | 1.5260e-01 | 2.5941e+05 | ; | CB - CG1 |
| 5568 | 5569 | 1 | 1.0900e-01 | 2.8451e+05 | ; | CG2 - HG21 |
| 5568 | 5570 | 1 | 1.0900e-01 | 2.8451e+05 | ; | CG2 - HG22 |
| 5568 | 5571 | 1 | 1.0900e-01 | 2.8451e+05 | ; | CG2 - HG23 |
| 5572 | 5573 | 1 | 1.0900e-01 | 2.8451e+05 | ; | CG1 - HG12 |
| 5572 | 5574 | 1 | 1.0900e-01 | 2.8451e+05 | ; | CG1 - HG13 |
| 5572 | 5575 | 1 | 1.5260e-01 | 2.5941e+05 | ; | CG1 - CD1 |
| 5575 | 5576 | 1 | 1.0900e-01 | 2.8451e+05 | ; | CD1 - HD11 |
| 5575 | 5577 | 1 | 1.0900e-01 | 2.8451e+05 | ; | CD1 - HD12 |
| 5575 | 5578 | 1 | 1.0900e-01 | 2.8451e+05 | ; | CD1 - HD13 |
| 5579 | 5580 | 1 | 1.2290e-01 | 4.7698e+05 | ; | C - O |
| 5579 | 5581 | 1 | 1.3350e-01 | 4.1003e+05 | ; | C - N |
| 5581 | 5582 | 1 | 1.0100e-01 | 3.6317e+05 | ; | N - H |
| 5581 | 5583 | 1 | 1.4490e-01 | 2.8200e+05 | ; | N - CA |
| 5583 | 5584 | 1 | 1.0900e-01 | 2.8451e+05 | ; | CA - HA2 |
| 5583 | 5585 | 1 | 1.0900e-01 | 2.8451e+05 | ; | CA - HA3 |
| 5583 | 5586 | 1 | 1.5220e-01 | 2.6527e+05 | ; | CA - C |
| 5586 | 5587 | 1 | 1.2290e-01 | 4.7698e+05 | ; | C - O |
| 5586 | 5588 | 1 | 1.3350e-01 | 4.1003e+05 | ; | C - N |
| 5588 | 5589 | 1 | 1.0100e-01 | 3.6317e+05 | ; | N - H |
| 5588 | 5590 | 1 | 1.4490e-01 | 2.8200e+05 | ; | N - CA |
| 5590 | 5591 | 1 | 1.0900e-01 | 2.8451e+05 | ; | CA - HA |
| 5590 | 5592 | 1 | 1.5260e-01 | 2.5941e+05 | ; | CA - CB |
| 5590 | 5607 | 1 | 1.5220e-01 | 2.6527e+05 | ; | CA - C |
| 5592 | 5593 | 1 | 1.0900e-01 | 2.8451e+05 | ; | CB - HB2 |
| 5592 | 5594 | 1 | 1.0900e-01 | 2.8451e+05 | ; | CB - HB3 |
| 5592 | 5595 | 1 | 1.5100e-01 | 2.6527e+05 | ; | CB - CG |

| | | | | | | | | |
|---|---|---|---|---|---|---|---|---|
| 5595 | 5596 | 1 | 1.4000e-01 | 3.9246e+05 | ; | CG | - | CD1 |
| 5595 | 5605 | 1 | 1.4000e-01 | 3.9246e+05 | ; | CG | - | CD2 |
| 5596 | 5597 | 1 | 1.0800e-01 | 3.0711e+05 | ; | CD1 | - | HD1 |
| 5596 | 5598 | 1 | 1.4000e-01 | 3.9246e+05 | ; | CD1 | - | CE1 |
| 5598 | 5599 | 1 | 1.0800e-01 | 3.0711e+05 | ; | CE1 | - | HE1 |
| 5598 | 5600 | 1 | 1.4090e-01 | 3.9246e+05 | ; | CE1 | - | CZ |
| 5600 | 5601 | 1 | 1.3640e-01 | 3.7656e+05 | ; | CZ | - | OH |
| 5600 | 5603 | 1 | 1.4090e-01 | 3.9246e+05 | ; | CZ | - | CE2 |
| 5601 | 5602 | 1 | 9.6000e-02 | 4.6275e+05 | ; | OH | - | HH |
| 5603 | 5604 | 1 | 1.0800e-01 | 3.0711e+05 | ; | CE2 | - | HE2 |
| 5603 | 5605 | 1 | 1.4000e-01 | 3.9246e+05 | ; | CE2 | - | CD2 |
| 5605 | 5606 | 1 | 1.0800e-01 | 3.0711e+05 | ; | CD2 | - | HD2 |
| 5607 | 5608 | 1 | 1.2290e-01 | 4.7698e+05 | ; | C | - | O |
| 5607 | 5609 | 1 | 1.3350e-01 | 4.1003e+05 | ; | C | - | N |
| 5609 | 5610 | 1 | 1.0100e-01 | 3.6317e+05 | ; | N | - | H |
| 5609 | 5611 | 1 | 1.4490e-01 | 2.8200e+05 | ; | N | - | CA |
| 5611 | 5612 | 1 | 1.0900e-01 | 2.8451e+05 | ; | CA | - | HA |
| 5611 | 5613 | 1 | 1.5260e-01 | 2.5941e+05 | ; | CA | - | CB |
| 5611 | 5624 | 1 | 1.5220e-01 | 2.6527e+05 | ; | CA | - | C |
| 5613 | 5614 | 1 | 1.0900e-01 | 2.8451e+05 | ; | CB | - | HB2 |
| 5613 | 5615 | 1 | 1.0900e-01 | 2.8451e+05 | ; | CB | - | HB3 |
| 5613 | 5616 | 1 | 1.5260e-01 | 2.5941e+05 | ; | CB | - | CG |
| 5616 | 5617 | 1 | 1.0900e-01 | 2.8451e+05 | ; | CG | - | HG2 |
| 5616 | 5618 | 1 | 1.0900e-01 | 2.8451e+05 | ; | CG | - | HG3 |
| 5616 | 5619 | 1 | 1.5220e-01 | 2.6527e+05 | ; | CG | - | CD |
| 5619 | 5620 | 1 | 1.2290e-01 | 4.7698e+05 | ; | CD | - | OE1 |
| 5619 | 5621 | 1 | 1.3350e-01 | 4.1003e+05 | ; | CD | - | NE2 |
| 5621 | 5622 | 1 | 1.0100e-01 | 3.6317e+05 | ; | NE2 | - | HE21 |
| 5621 | 5623 | 1 | 1.0100e-01 | 3.6317e+05 | ; | NE2 | - | HE22 |
| 5624 | 5625 | 1 | 1.2290e-01 | 4.7698e+05 | ; | C | - | O |
| 5624 | 5626 | 1 | 1.3350e-01 | 4.1003e+05 | ; | C | - | N |
| 5626 | 5627 | 1 | 1.0100e-01 | 3.6317e+05 | ; | N | - | H |
| 5626 | 5628 | 1 | 1.4490e-01 | 2.8200e+05 | ; | N | - | CA |
| 5628 | 5629 | 1 | 1.0900e-01 | 2.8451e+05 | ; | CA | - | HA |
| 5628 | 5630 | 1 | 1.5260e-01 | 2.5941e+05 | ; | CA | - | CB |
| 5628 | 5646 | 1 | 1.5220e-01 | 2.6527e+05 | ; | CA | - | C |
| 5630 | 5631 | 1 | 1.0900e-01 | 2.8451e+05 | ; | CB | - | HB2 |
| 5630 | 5632 | 1 | 1.0900e-01 | 2.8451e+05 | ; | CB | - | HB3 |
| 5630 | 5633 | 1 | 1.5260e-01 | 2.5941e+05 | ; | CB | - | CG |
| 5633 | 5634 | 1 | 1.0900e-01 | 2.8451e+05 | ; | CG | - | HG2 |
| 5633 | 5635 | 1 | 1.0900e-01 | 2.8451e+05 | ; | CG | - | HG3 |
| 5633 | 5636 | 1 | 1.5260e-01 | 2.5941e+05 | ; | CG | - | CD |
| 5636 | 5637 | 1 | 1.0900e-01 | 2.8451e+05 | ; | CD | - | HD2 |
| 5636 | 5638 | 1 | 1.0900e-01 | 2.8451e+05 | ; | CD | - | HD3 |
| 5636 | 5639 | 1 | 1.5260e-01 | 2.5941e+05 | ; | CD | - | CE |
| 5639 | 5640 | 1 | 1.0900e-01 | 2.8451e+05 | ; | CE | - | HE2 |
| 5639 | 5641 | 1 | 1.0900e-01 | 2.8451e+05 | ; | CE | - | HE3 |
| 5639 | 5642 | 1 | 1.4710e-01 | 3.0711e+05 | ; | CE | - | NZ |
| 5642 | 5643 | 1 | 1.0100e-01 | 3.6317e+05 | ; | NZ | - | HZ1 |
| 5642 | 5644 | 1 | 1.0100e-01 | 3.6317e+05 | ; | NZ | - | HZ2 |
| 5642 | 5645 | 1 | 1.0100e-01 | 3.6317e+05 | ; | NZ | - | HZ3 |
| 5646 | 5647 | 1 | 1.2290e-01 | 4.7698e+05 | ; | C | - | O |

| | | | | | | |
|---|---|---|---|---|---|---|
| 5646 | 5648 | 1 | 1.3350e-01 | 4.1003e+05 | ; | C - N |
| 5648 | 5649 | 1 | 1.0100e-01 | 3.6317e+05 | ; | N - H |
| 5648 | 5650 | 1 | 1.4490e-01 | 2.8200e+05 | ; | N - CA |
| 5650 | 5651 | 1 | 1.0900e-01 | 2.8451e+05 | ; | CA - HA |
| 5650 | 5652 | 1 | 1.5260e-01 | 2.5941e+05 | ; | CA - CB |
| 5650 | 5665 | 1 | 1.5220e-01 | 2.6527e+05 | ; | CA - C |
| 5652 | 5653 | 1 | 1.0900e-01 | 2.8451e+05 | ; | CB - HB |
| 5652 | 5654 | 1 | 1.5260e-01 | 2.5941e+05 | ; | CB - CG2 |
| 5652 | 5658 | 1 | 1.5260e-01 | 2.5941e+05 | ; | CB - CG1 |
| 5654 | 5655 | 1 | 1.0900e-01 | 2.8451e+05 | ; | CG2 - HG21 |
| 5654 | 5656 | 1 | 1.0900e-01 | 2.8451e+05 | ; | CG2 - HG22 |
| 5654 | 5657 | 1 | 1.0900e-01 | 2.8451e+05 | ; | CG2 - HG23 |
| 5658 | 5659 | 1 | 1.0900e-01 | 2.8451e+05 | ; | CG1 - HG12 |
| 5658 | 5660 | 1 | 1.0900e-01 | 2.8451e+05 | ; | CG1 - HG13 |
| 5658 | 5661 | 1 | 1.5260e-01 | 2.5941e+05 | ; | CG1 - CD1 |
| 5661 | 5662 | 1 | 1.0900e-01 | 2.8451e+05 | ; | CD1 - HD11 |
| 5661 | 5663 | 1 | 1.0900e-01 | 2.8451e+05 | ; | CD1 - HD12 |
| 5661 | 5664 | 1 | 1.0900e-01 | 2.8451e+05 | ; | CD1 - HD13 |
| 5665 | 5666 | 1 | 1.2290e-01 | 4.7698e+05 | ; | C - O |
| 5665 | 5667 | 1 | 1.3350e-01 | 4.1003e+05 | ; | C - N |
| 5667 | 5668 | 1 | 1.4490e-01 | 2.8200e+05 | ; | N - CD |
| 5667 | 5677 | 1 | 1.4490e-01 | 2.8200e+05 | ; | N - CA |
| 5668 | 5669 | 1 | 1.0900e-01 | 2.8451e+05 | ; | CD - HD2 |
| 5668 | 5670 | 1 | 1.0900e-01 | 2.8451e+05 | ; | CD - HD3 |
| 5668 | 5671 | 1 | 1.5260e-01 | 2.5941e+05 | ; | CD - CG |
| 5671 | 5672 | 1 | 1.0900e-01 | 2.8451e+05 | ; | CG - HG2 |
| 5671 | 5673 | 1 | 1.0900e-01 | 2.8451e+05 | ; | CG - HG3 |
| 5671 | 5674 | 1 | 1.5260e-01 | 2.5941e+05 | ; | CG - CB |
| 5674 | 5675 | 1 | 1.0900e-01 | 2.8451e+05 | ; | CB - HB2 |
| 5674 | 5676 | 1 | 1.0900e-01 | 2.8451e+05 | ; | CB - HB3 |
| 5674 | 5677 | 1 | 1.5260e-01 | 2.5941e+05 | ; | CB - CA |
| 5677 | 5678 | 1 | 1.0900e-01 | 2.8451e+05 | ; | CA - HA |
| 5677 | 5679 | 1 | 1.5220e-01 | 2.6527e+05 | ; | CA - C |
| 5679 | 5680 | 1 | 1.2290e-01 | 4.7698e+05 | ; | C - O |
| 5679 | 5681 | 1 | 1.3350e-01 | 4.1003e+05 | ; | C - N |
| 5681 | 5682 | 1 | 1.4490e-01 | 2.8200e+05 | ; | N - CD |
| 5681 | 5691 | 1 | 1.4490e-01 | 2.8200e+05 | ; | N - CA |
| 5682 | 5683 | 1 | 1.0900e-01 | 2.8451e+05 | ; | CD - HD2 |
| 5682 | 5684 | 1 | 1.0900e-01 | 2.8451e+05 | ; | CD - HD3 |
| 5682 | 5685 | 1 | 1.5260e-01 | 2.5941e+05 | ; | CD - CG |
| 5685 | 5686 | 1 | 1.0900e-01 | 2.8451e+05 | ; | CG - HG2 |
| 5685 | 5687 | 1 | 1.0900e-01 | 2.8451e+05 | ; | CG - HG3 |
| 5685 | 5688 | 1 | 1.5260e-01 | 2.5941e+05 | ; | CG - CB |
| 5688 | 5689 | 1 | 1.0900e-01 | 2.8451e+05 | ; | CB - HB2 |
| 5688 | 5690 | 1 | 1.0900e-01 | 2.8451e+05 | ; | CB - HB3 |
| 5688 | 5691 | 1 | 1.5260e-01 | 2.5941e+05 | ; | CB - CA |
| 5691 | 5692 | 1 | 1.0900e-01 | 2.8451e+05 | ; | CA - HA |
| 5691 | 5693 | 1 | 1.5220e-01 | 2.6527e+05 | ; | CA - C |
| 5693 | 5694 | 1 | 1.2290e-01 | 4.7698e+05 | ; | C - O |
| 5693 | 5695 | 1 | 1.3350e-01 | 4.1003e+05 | ; | C - N |
| 5695 | 5696 | 1 | 1.0100e-01 | 3.6317e+05 | ; | N - H |
| 5695 | 5697 | 1 | 1.4490e-01 | 2.8200e+05 | ; | N - CA |

| | | | | | |
|---|---|---|---|---|---|
| 5697 | 5698 | 1 | 1.0900e-01 | 2.8451e+05 ; | CA - HA |
| 5697 | 5699 | 1 | 1.5260e-01 | 2.5941e+05 ; | CA - CB |
| 5697 | 5705 | 1 | 1.5220e-01 | 2.6527e+05 ; | CA - C |
| 5699 | 5700 | 1 | 1.0900e-01 | 2.8451e+05 ; | CB - HB2 |
| 5699 | 5701 | 1 | 1.0900e-01 | 2.8451e+05 ; | CB - HB3 |
| 5699 | 5702 | 1 | 1.5220e-01 | 2.6527e+05 ; | CB - CG |
| 5702 | 5703 | 1 | 1.2500e-01 | 5.4894e+05 ; | CG - OD1 |
| 5702 | 5704 | 1 | 1.2500e-01 | 5.4894e+05 ; | CG - OD2 |
| 5705 | 5706 | 1 | 1.2290e-01 | 4.7698e+05 ; | C - O |
| 5705 | 5707 | 1 | 1.3350e-01 | 4.1003e+05 ; | C - N |
| 5707 | 5708 | 1 | 1.0100e-01 | 3.6317e+05 ; | N - H |
| 5707 | 5709 | 1 | 1.4490e-01 | 2.8200e+05 ; | N - CA |
| 5709 | 5710 | 1 | 1.0900e-01 | 2.8451e+05 ; | CA - HA |
| 5709 | 5711 | 1 | 1.5260e-01 | 2.5941e+05 ; | CA - CB |
| 5709 | 5724 | 1 | 1.5220e-01 | 2.6527e+05 ; | CA - C |
| 5711 | 5712 | 1 | 1.0900e-01 | 2.8451e+05 ; | CB - HB |
| 5711 | 5713 | 1 | 1.5260e-01 | 2.5941e+05 ; | CB - CG2 |
| 5711 | 5717 | 1 | 1.5260e-01 | 2.5941e+05 ; | CB - CG1 |
| 5713 | 5714 | 1 | 1.0900e-01 | 2.8451e+05 ; | CG2 - HG21 |
| 5713 | 5715 | 1 | 1.0900e-01 | 2.8451e+05 ; | CG2 - HG22 |
| 5713 | 5716 | 1 | 1.0900e-01 | 2.8451e+05 ; | CG2 - HG23 |
| 5717 | 5718 | 1 | 1.0900e-01 | 2.8451e+05 ; | CG1 - HG12 |
| 5717 | 5719 | 1 | 1.0900e-01 | 2.8451e+05 ; | CG1 - HG13 |
| 5717 | 5720 | 1 | 1.5260e-01 | 2.5941e+05 ; | CG1 - CD1 |
| 5720 | 5721 | 1 | 1.0900e-01 | 2.8451e+05 ; | CD1 - HD11 |
| 5720 | 5722 | 1 | 1.0900e-01 | 2.8451e+05 ; | CD1 - HD12 |
| 5720 | 5723 | 1 | 1.0900e-01 | 2.8451e+05 ; | CD1 - HD13 |
| 5724 | 5725 | 1 | 1.2290e-01 | 4.7698e+05 ; | C - O |
| 5724 | 5726 | 1 | 1.3350e-01 | 4.1003e+05 ; | C - N |
| 5726 | 5727 | 1 | 1.0100e-01 | 3.6317e+05 ; | N - H |
| 5726 | 5728 | 1 | 1.4490e-01 | 2.8200e+05 ; | N - CA |
| 5728 | 5729 | 1 | 1.0900e-01 | 2.8451e+05 ; | CA - HA |
| 5728 | 5730 | 1 | 1.5260e-01 | 2.5941e+05 ; | CA - CB |
| 5728 | 5741 | 1 | 1.5220e-01 | 2.6527e+05 ; | CA - C |
| 5730 | 5731 | 1 | 1.0900e-01 | 2.8451e+05 ; | CB - HB2 |
| 5730 | 5732 | 1 | 1.0900e-01 | 2.8451e+05 ; | CB - HB3 |
| 5730 | 5733 | 1 | 1.5040e-01 | 2.6527e+05 ; | CB - CG |
| 5733 | 5734 | 1 | 1.3940e-01 | 3.4309e+05 ; | CG - ND1 |
| 5733 | 5739 | 1 | 1.3710e-01 | 4.3346e+05 ; | CG - CD2 |
| 5734 | 5735 | 1 | 1.3350e-01 | 4.0836e+05 ; | ND1 - CE1 |
| 5735 | 5736 | 1 | 1.0800e-01 | 3.0711e+05 ; | CE1 - HE1 |
| 5735 | 5737 | 1 | 1.3430e-01 | 3.9915e+05 ; | CE1 - NE2 |
| 5737 | 5738 | 1 | 1.0100e-01 | 3.6317e+05 ; | NE2 - HE2 |
| 5737 | 5739 | 1 | 1.3810e-01 | 3.5731e+05 ; | NE2 - CD2 |
| 5739 | 5740 | 1 | 1.0800e-01 | 3.0711e+05 ; | CD2 - HD2 |
| 5741 | 5742 | 1 | 1.2290e-01 | 4.7698e+05 ; | C - O |
| 5741 | 5743 | 1 | 1.3350e-01 | 4.1003e+05 ; | C - N |
| 5743 | 5744 | 1 | 1.0100e-01 | 3.6317e+05 ; | N - H |
| 5743 | 5745 | 1 | 1.4490e-01 | 2.8200e+05 ; | N - CA |
| 5745 | 5746 | 1 | 1.0900e-01 | 2.8451e+05 ; | CA - HA |
| 5745 | 5747 | 1 | 1.5260e-01 | 2.5941e+05 ; | CA - CB |
| 5745 | 5758 | 1 | 1.5220e-01 | 2.6527e+05 ; | CA - C |

| | | | | | |
|---|---|---|---|---|---|
| 5747 | 5748 | 1 | 1.0900e-01 | 2.8451e+05 ; | CB - HB2 |
| 5747 | 5749 | 1 | 1.0900e-01 | 2.8451e+05 ; | CB - HB3 |
| 5747 | 5750 | 1 | 1.5260e-01 | 2.5941e+05 ; | CB - CG |
| 5750 | 5751 | 1 | 1.0900e-01 | 2.8451e+05 ; | CG - HG2 |
| 5750 | 5752 | 1 | 1.0900e-01 | 2.8451e+05 ; | CG - HG3 |
| 5750 | 5753 | 1 | 1.5220e-01 | 2.6527e+05 ; | CG - CD |
| 5753 | 5754 | 1 | 1.2290e-01 | 4.7698e+05 ; | CD - OE1 |
| 5753 | 5755 | 1 | 1.3350e-01 | 4.1003e+05 ; | CD - NE2 |
| 5755 | 5756 | 1 | 1.0100e-01 | 3.6317e+05 ; | NE2 - HE21 |
| 5755 | 5757 | 1 | 1.0100e-01 | 3.6317e+05 ; | NE2 - HE22 |
| 5758 | 5759 | 1 | 1.2290e-01 | 4.7698e+05 ; | C - O |
| 5758 | 5760 | 1 | 1.3350e-01 | 4.1003e+05 ; | C - N |
| 5760 | 5761 | 1 | 1.0100e-01 | 3.6317e+05 ; | N - H |
| 5760 | 5762 | 1 | 1.4490e-01 | 2.8200e+05 ; | N - CA |
| 5762 | 5763 | 1 | 1.0900e-01 | 2.8451e+05 ; | CA - HA |
| 5762 | 5764 | 1 | 1.5260e-01 | 2.5941e+05 ; | CA - CB |
| 5762 | 5774 | 1 | 1.5220e-01 | 2.6527e+05 ; | CA - C |
| 5764 | 5765 | 1 | 1.0900e-01 | 2.8451e+05 ; | CB - HB |
| 5764 | 5766 | 1 | 1.5260e-01 | 2.5941e+05 ; | CB - CG1 |
| 5764 | 5770 | 1 | 1.5260e-01 | 2.5941e+05 ; | CB - CG2 |
| 5766 | 5767 | 1 | 1.0900e-01 | 2.8451e+05 ; | CG1 - HG11 |
| 5766 | 5768 | 1 | 1.0900e-01 | 2.8451e+05 ; | CG1 - HG12 |
| 5766 | 5769 | 1 | 1.0900e-01 | 2.8451e+05 ; | CG1 - HG13 |
| 5770 | 5771 | 1 | 1.0900e-01 | 2.8451e+05 ; | CG2 - HG21 |
| 5770 | 5772 | 1 | 1.0900e-01 | 2.8451e+05 ; | CG2 - HG22 |
| 5770 | 5773 | 1 | 1.0900e-01 | 2.8451e+05 ; | CG2 - HG23 |
| 5774 | 5775 | 1 | 1.2290e-01 | 4.7698e+05 ; | C - O |
| 5774 | 5776 | 1 | 1.3350e-01 | 4.1003e+05 ; | C - N |
| 5776 | 5777 | 1 | 1.0100e-01 | 3.6317e+05 ; | N - H |
| 5776 | 5778 | 1 | 1.4490e-01 | 2.8200e+05 ; | N - CA |
| 5778 | 5779 | 1 | 1.0900e-01 | 2.8451e+05 ; | CA - HA |
| 5778 | 5780 | 1 | 1.5260e-01 | 2.5941e+05 ; | CA - CB |
| 5778 | 5793 | 1 | 1.5220e-01 | 2.6527e+05 ; | CA - C |
| 5780 | 5781 | 1 | 1.0900e-01 | 2.8451e+05 ; | CB - HB2 |
| 5780 | 5782 | 1 | 1.0900e-01 | 2.8451e+05 ; | CB - HB3 |
| 5780 | 5783 | 1 | 1.5260e-01 | 2.5941e+05 ; | CB - CG |
| 5783 | 5784 | 1 | 1.0900e-01 | 2.8451e+05 ; | CG - HG |
| 5783 | 5785 | 1 | 1.5260e-01 | 2.5941e+05 ; | CG - CD1 |
| 5783 | 5789 | 1 | 1.5260e-01 | 2.5941e+05 ; | CG - CD2 |
| 5785 | 5786 | 1 | 1.0900e-01 | 2.8451e+05 ; | CD1 - HD11 |
| 5785 | 5787 | 1 | 1.0900e-01 | 2.8451e+05 ; | CD1 - HD12 |
| 5785 | 5788 | 1 | 1.0900e-01 | 2.8451e+05 ; | CD1 - HD13 |
| 5789 | 5790 | 1 | 1.0900e-01 | 2.8451e+05 ; | CD2 - HD21 |
| 5789 | 5791 | 1 | 1.0900e-01 | 2.8451e+05 ; | CD2 - HD22 |
| 5789 | 5792 | 1 | 1.0900e-01 | 2.8451e+05 ; | CD2 - HD23 |
| 5793 | 5794 | 1 | 1.2290e-01 | 4.7698e+05 ; | C - O |
| 5793 | 5795 | 1 | 1.3350e-01 | 4.1003e+05 ; | C - N |
| 5795 | 5796 | 1 | 1.0100e-01 | 3.6317e+05 ; | N - H |
| 5795 | 5797 | 1 | 1.4490e-01 | 2.8200e+05 ; | N - CA |
| 5797 | 5798 | 1 | 1.0900e-01 | 2.8451e+05 ; | CA - HA |
| 5797 | 5799 | 1 | 1.5260e-01 | 2.5941e+05 ; | CA - CB |
| 5797 | 5817 | 1 | 1.5220e-01 | 2.6527e+05 ; | CA - C |

| | | | | | | | | |
|---|---|---|---|---|---|---|---|---|
| 5799 | 5800 | 1 | 1.0900e-01 | 2.8451e+05 | ; | CB | - | HB2 |
| 5799 | 5801 | 1 | 1.0900e-01 | 2.8451e+05 | ; | CB | - | HB3 |
| 5799 | 5802 | 1 | 1.5260e-01 | 2.5941e+05 | ; | CB | - | CG |
| 5802 | 5803 | 1 | 1.0900e-01 | 2.8451e+05 | ; | CG | - | HG2 |
| 5802 | 5804 | 1 | 1.0900e-01 | 2.8451e+05 | ; | CG | - | HG3 |
| 5802 | 5805 | 1 | 1.5260e-01 | 2.5941e+05 | ; | CG | - | CD |
| 5805 | 5806 | 1 | 1.0900e-01 | 2.8451e+05 | ; | CD | - | HD2 |
| 5805 | 5807 | 1 | 1.0900e-01 | 2.8451e+05 | ; | CD | - | HD3 |
| 5805 | 5808 | 1 | 1.4630e-01 | 2.8200e+05 | ; | CD | - | NE |
| 5808 | 5809 | 1 | 1.0100e-01 | 3.6317e+05 | ; | NE | - | HE |
| 5808 | 5810 | 1 | 1.3400e-01 | 4.0250e+05 | ; | NE | - | CZ |
| 5810 | 5811 | 1 | 1.3400e-01 | 4.0250e+05 | ; | CZ | - | NH1 |
| 5810 | 5814 | 1 | 1.3400e-01 | 4.0250e+05 | ; | CZ | - | NH2 |
| 5811 | 5812 | 1 | 1.0100e-01 | 3.6317e+05 | ; | NH1 | - | HH11 |
| 5811 | 5813 | 1 | 1.0100e-01 | 3.6317e+05 | ; | NH1 | - | HH12 |
| 5814 | 5815 | 1 | 1.0100e-01 | 3.6317e+05 | ; | NH2 | - | HH21 |
| 5814 | 5816 | 1 | 1.0100e-01 | 3.6317e+05 | ; | NH2 | - | HH22 |
| 5817 | 5818 | 1 | 1.2290e-01 | 4.7698e+05 | ; | C | - | O |
| 5817 | 5819 | 1 | 1.3350e-01 | 4.1003e+05 | ; | C | - | N |
| 5819 | 5820 | 1 | 1.0100e-01 | 3.6317e+05 | ; | N | - | H |
| 5819 | 5821 | 1 | 1.4490e-01 | 2.8200e+05 | ; | N | - | CA |
| 5821 | 5822 | 1 | 1.0900e-01 | 2.8451e+05 | ; | CA | - | HA |
| 5821 | 5823 | 1 | 1.5260e-01 | 2.5941e+05 | ; | CA | - | CB |
| 5821 | 5829 | 1 | 1.5220e-01 | 2.6527e+05 | ; | CA | - | C |
| 5823 | 5824 | 1 | 1.0900e-01 | 2.8451e+05 | ; | CB | - | HB2 |
| 5823 | 5825 | 1 | 1.0900e-01 | 2.8451e+05 | ; | CB | - | HB3 |
| 5823 | 5826 | 1 | 1.5220e-01 | 2.6527e+05 | ; | CB | - | CG |
| 5826 | 5827 | 1 | 1.2500e-01 | 5.4894e+05 | ; | CG | - | OD1 |
| 5826 | 5828 | 1 | 1.2500e-01 | 5.4894e+05 | ; | CG | - | OD2 |
| 5829 | 5830 | 1 | 1.2290e-01 | 4.7698e+05 | ; | C | - | O |
| 5829 | 5831 | 1 | 1.3350e-01 | 4.1003e+05 | ; | C | - | N |
| 5831 | 5832 | 1 | 1.0100e-01 | 3.6317e+05 | ; | N | - | H |
| 5831 | 5833 | 1 | 1.4490e-01 | 2.8200e+05 | ; | N | - | CA |
| 5833 | 5834 | 1 | 1.0900e-01 | 2.8451e+05 | ; | CA | - | HA |
| 5833 | 5835 | 1 | 1.5260e-01 | 2.5941e+05 | ; | CA | - | CB |
| 5833 | 5848 | 1 | 1.5220e-01 | 2.6527e+05 | ; | CA | - | C |
| 5835 | 5836 | 1 | 1.0900e-01 | 2.8451e+05 | ; | CB | - | HB |
| 5835 | 5837 | 1 | 1.5260e-01 | 2.5941e+05 | ; | CB | - | CG2 |
| 5835 | 5841 | 1 | 1.5260e-01 | 2.5941e+05 | ; | CB | - | CG1 |
| 5837 | 5838 | 1 | 1.0900e-01 | 2.8451e+05 | ; | CG2 | - | HG21 |
| 5837 | 5839 | 1 | 1.0900e-01 | 2.8451e+05 | ; | CG2 | - | HG22 |
| 5837 | 5840 | 1 | 1.0900e-01 | 2.8451e+05 | ; | CG2 | - | HG23 |
| 5841 | 5842 | 1 | 1.0900e-01 | 2.8451e+05 | ; | CG1 | - | HG12 |
| 5841 | 5843 | 1 | 1.0900e-01 | 2.8451e+05 | ; | CG1 | - | HG13 |
| 5841 | 5844 | 1 | 1.5260e-01 | 2.5941e+05 | ; | CG1 | - | CD1 |
| 5844 | 5845 | 1 | 1.0900e-01 | 2.8451e+05 | ; | CD1 | - | HD11 |
| 5844 | 5846 | 1 | 1.0900e-01 | 2.8451e+05 | ; | CD1 | - | HD12 |
| 5844 | 5847 | 1 | 1.0900e-01 | 2.8451e+05 | ; | CD1 | - | HD13 |
| 5848 | 5849 | 1 | 1.2290e-01 | 4.7698e+05 | ; | C | - | O |
| 5848 | 5850 | 1 | 1.3350e-01 | 4.1003e+05 | ; | C | - | N |
| 5850 | 5851 | 1 | 1.0100e-01 | 3.6317e+05 | ; | N | - | H |
| 5850 | 5852 | 1 | 1.4490e-01 | 2.8200e+05 | ; | N | - | CA |

| | | | | | |
|---|---|---|---|---|---|
| 5852 | 5853 | 1 | 1.0900e-01 | 2.8451e+05 ; | CA - HA |
| 5852 | 5854 | 1 | 1.5260e-01 | 2.5941e+05 ; | CA - CB |
| 5852 | 5860 | 1 | 1.5220e-01 | 2.6527e+05 ; | CA - C |
| 5854 | 5855 | 1 | 1.0900e-01 | 2.8451e+05 ; | CB - HB2 |
| 5854 | 5856 | 1 | 1.0900e-01 | 2.8451e+05 ; | CB - HB3 |
| 5854 | 5857 | 1 | 1.5220e-01 | 2.6527e+05 ; | CB - CG |
| 5857 | 5858 | 1 | 1.2500e-01 | 5.4894e+05 ; | CG - OD1 |
| 5857 | 5859 | 1 | 1.2500e-01 | 5.4894e+05 ; | CG - OD2 |
| 5860 | 5861 | 1 | 1.2290e-01 | 4.7698e+05 ; | C - O |
| 5860 | 5862 | 1 | 1.3350e-01 | 4.1003e+05 ; | C - N |
| 5862 | 5863 | 1 | 1.0100e-01 | 3.6317e+05 ; | N - H |
| 5862 | 5864 | 1 | 1.4490e-01 | 2.8200e+05 ; | N - CA |
| 5864 | 5865 | 1 | 1.0900e-01 | 2.8451e+05 ; | CA - HA |
| 5864 | 5866 | 1 | 1.5260e-01 | 2.5941e+05 ; | CA - CB |
| 5864 | 5871 | 1 | 1.5220e-01 | 2.6527e+05 ; | CA - C |
| 5866 | 5867 | 1 | 1.0900e-01 | 2.8451e+05 ; | CB - HB2 |
| 5866 | 5868 | 1 | 1.0900e-01 | 2.8451e+05 ; | CB - HB3 |
| 5866 | 5869 | 1 | 1.4100e-01 | 2.6778e+05 ; | CB - OG |
| 5869 | 5870 | 1 | 9.6000e-02 | 4.6275e+05 ; | OG - HG |
| 5871 | 5872 | 1 | 1.2290e-01 | 4.7698e+05 ; | C - O |
| 5871 | 5873 | 1 | 1.3350e-01 | 4.1003e+05 ; | C - N |
| 5873 | 5874 | 1 | 1.0100e-01 | 3.6317e+05 ; | N - H |
| 5873 | 5875 | 1 | 1.4490e-01 | 2.8200e+05 ; | N - CA |
| 5875 | 5876 | 1 | 1.0900e-01 | 2.8451e+05 ; | CA - HA |
| 5875 | 5877 | 1 | 1.5260e-01 | 2.5941e+05 ; | CA - CB |
| 5875 | 5885 | 1 | 1.5220e-01 | 2.6527e+05 ; | CA - C |
| 5877 | 5878 | 1 | 1.0900e-01 | 2.8451e+05 ; | CB - HB2 |
| 5877 | 5879 | 1 | 1.0900e-01 | 2.8451e+05 ; | CB - HB3 |
| 5877 | 5880 | 1 | 1.5220e-01 | 2.6527e+05 ; | CB - CG |
| 5880 | 5881 | 1 | 1.2290e-01 | 4.7698e+05 ; | CG - OD1 |
| 5880 | 5882 | 1 | 1.3350e-01 | 4.1003e+05 ; | CG - ND2 |
| 5882 | 5883 | 1 | 1.0100e-01 | 3.6317e+05 ; | ND2 - HD21 |
| 5882 | 5884 | 1 | 1.0100e-01 | 3.6317e+05 ; | ND2 - HD22 |
| 5885 | 5886 | 1 | 1.2290e-01 | 4.7698e+05 ; | C - O |
| 5885 | 5887 | 1 | 1.3350e-01 | 4.1003e+05 ; | C - N |
| 5887 | 5888 | 1 | 1.0100e-01 | 3.6317e+05 ; | N - H |
| 5887 | 5889 | 1 | 1.4490e-01 | 2.8200e+05 ; | N - CA |
| 5889 | 5890 | 1 | 1.0900e-01 | 2.8451e+05 ; | CA - HA |
| 5889 | 5891 | 1 | 1.5260e-01 | 2.5941e+05 ; | CA - CB |
| 5889 | 5895 | 1 | 1.5220e-01 | 2.6527e+05 ; | CA - C |
| 5891 | 5892 | 1 | 1.0900e-01 | 2.8451e+05 ; | CB - HB1 |
| 5891 | 5893 | 1 | 1.0900e-01 | 2.8451e+05 ; | CB - HB2 |
| 5891 | 5894 | 1 | 1.0900e-01 | 2.8451e+05 ; | CB - HB3 |
| 5895 | 5896 | 1 | 1.2290e-01 | 4.7698e+05 ; | C - O |
| 5895 | 5897 | 1 | 1.3350e-01 | 4.1003e+05 ; | C - N |
| 5897 | 5898 | 1 | 1.0100e-01 | 3.6317e+05 ; | N - H |
| 5897 | 5899 | 1 | 1.4490e-01 | 2.8200e+05 ; | N - CA |
| 5899 | 5900 | 1 | 1.0900e-01 | 2.8451e+05 ; | CA - HA |
| 5899 | 5901 | 1 | 1.5260e-01 | 2.5941e+05 ; | CA - CB |
| 5899 | 5906 | 1 | 1.5220e-01 | 2.6527e+05 ; | CA - C |
| 5901 | 5902 | 1 | 1.0900e-01 | 2.8451e+05 ; | CB - HB2 |
| 5901 | 5903 | 1 | 1.0900e-01 | 2.8451e+05 ; | CB - HB3 |

| | | | | | | | | |
|---|---|---|---|---|---|---|---|---|
| 5901 | 5904 | 1 | 1.4100e-01 | 2.6778e+05 | ; | CB | - | OG |
| 5904 | 5905 | 1 | 9.6000e-02 | 4.6275e+05 | ; | OG | - | HG |
| 5906 | 5907 | 1 | 1.2290e-01 | 4.7698e+05 | ; | C | - | O |
| 5906 | 5908 | 1 | 1.3350e-01 | 4.1003e+05 | ; | C | - | N |
| 5908 | 5909 | 1 | 1.0100e-01 | 3.6317e+05 | ; | N | - | H |
| 5908 | 5910 | 1 | 1.4490e-01 | 2.8200e+05 | ; | N | - | CA |
| 5910 | 5911 | 1 | 1.0900e-01 | 2.8451e+05 | ; | CA | - | HA2 |
| 5910 | 5912 | 1 | 1.0900e-01 | 2.8451e+05 | ; | CA | - | HA3 |
| 5910 | 5913 | 1 | 1.5220e-01 | 2.6527e+05 | ; | CA | - | C |
| 5913 | 5914 | 1 | 1.2290e-01 | 4.7698e+05 | ; | C | - | O |
| 5913 | 5915 | 1 | 1.3350e-01 | 4.1003e+05 | ; | C | - | N |
| 5915 | 5916 | 1 | 1.0100e-01 | 3.6317e+05 | ; | N | - | H |
| 5915 | 5917 | 1 | 1.4490e-01 | 2.8200e+05 | ; | N | - | CA |
| 5917 | 5918 | 1 | 1.0900e-01 | 2.8451e+05 | ; | CA | - | HA |
| 5917 | 5919 | 1 | 1.5260e-01 | 2.5941e+05 | ; | CA | - | CB |
| 5917 | 5930 | 1 | 1.5220e-01 | 2.6527e+05 | ; | CA | - | C |
| 5919 | 5920 | 1 | 1.0900e-01 | 2.8451e+05 | ; | CB | - | HB2 |
| 5919 | 5921 | 1 | 1.0900e-01 | 2.8451e+05 | ; | CB | - | HB3 |
| 5919 | 5922 | 1 | 1.5260e-01 | 2.5941e+05 | ; | CB | - | CG |
| 5922 | 5923 | 1 | 1.0900e-01 | 2.8451e+05 | ; | CG | - | HG2 |
| 5922 | 5924 | 1 | 1.0900e-01 | 2.8451e+05 | ; | CG | - | HG3 |
| 5922 | 5925 | 1 | 1.5220e-01 | 2.6527e+05 | ; | CG | - | CD |
| 5925 | 5926 | 1 | 1.2290e-01 | 4.7698e+05 | ; | CD | - | OE1 |
| 5925 | 5927 | 1 | 1.3350e-01 | 4.1003e+05 | ; | CD | - | NE2 |
| 5927 | 5928 | 1 | 1.0100e-01 | 3.6317e+05 | ; | NE2 | - | HE21 |
| 5927 | 5929 | 1 | 1.0100e-01 | 3.6317e+05 | ; | NE2 | - | HE22 |
| 5930 | 5931 | 1 | 1.2290e-01 | 4.7698e+05 | ; | C | - | O |
| 5930 | 5932 | 1 | 1.3350e-01 | 4.1003e+05 | ; | C | - | N |
| 5932 | 5933 | 1 | 1.0100e-01 | 3.6317e+05 | ; | N | - | H |
| 5932 | 5934 | 1 | 1.4490e-01 | 2.8200e+05 | ; | N | - | CA |
| 5934 | 5935 | 1 | 1.0900e-01 | 2.8451e+05 | ; | CA | - | HA |
| 5934 | 5936 | 1 | 1.5260e-01 | 2.5941e+05 | ; | CA | - | CB |
| 5934 | 5949 | 1 | 1.5220e-01 | 2.6527e+05 | ; | CA | - | C |
| 5936 | 5937 | 1 | 1.0900e-01 | 2.8451e+05 | ; | CB | - | HB |
| 5936 | 5938 | 1 | 1.5260e-01 | 2.5941e+05 | ; | CB | - | CG2 |
| 5936 | 5942 | 1 | 1.5260e-01 | 2.5941e+05 | ; | CB | - | CG1 |
| 5938 | 5939 | 1 | 1.0900e-01 | 2.8451e+05 | ; | CG2 | - | HG21 |
| 5938 | 5940 | 1 | 1.0900e-01 | 2.8451e+05 | ; | CG2 | - | HG22 |
| 5938 | 5941 | 1 | 1.0900e-01 | 2.8451e+05 | ; | CG2 | - | HG23 |
| 5942 | 5943 | 1 | 1.0900e-01 | 2.8451e+05 | ; | CG1 | - | HG12 |
| 5942 | 5944 | 1 | 1.0900e-01 | 2.8451e+05 | ; | CG1 | - | HG13 |
| 5942 | 5945 | 1 | 1.5260e-01 | 2.5941e+05 | ; | CG1 | - | CD1 |
| 5945 | 5946 | 1 | 1.0900e-01 | 2.8451e+05 | ; | CD1 | - | HD11 |
| 5945 | 5947 | 1 | 1.0900e-01 | 2.8451e+05 | ; | CD1 | - | HD12 |
| 5945 | 5948 | 1 | 1.0900e-01 | 2.8451e+05 | ; | CD1 | - | HD13 |
| 5949 | 5950 | 1 | 1.2290e-01 | 4.7698e+05 | ; | C | - | O |
| 5949 | 5951 | 1 | 1.3350e-01 | 4.1003e+05 | ; | C | - | N |
| 5951 | 5952 | 1 | 1.0100e-01 | 3.6317e+05 | ; | N | - | H |
| 5951 | 5953 | 1 | 1.4490e-01 | 2.8200e+05 | ; | N | - | CA |
| 5953 | 5954 | 1 | 1.0900e-01 | 2.8451e+05 | ; | CA | - | HA |
| 5953 | 5955 | 1 | 1.5260e-01 | 2.5941e+05 | ; | CA | - | CB |
| 5953 | 5966 | 1 | 1.5220e-01 | 2.6527e+05 | ; | CA | - | C |

| | | | | | | | | |
|---|---|---|---|---|---|---|---|---|
| 5955 | 5956 | 1 | 1.0900e-01 | 2.8451e+05 | ; | CB | - | HB2 |
| 5955 | 5957 | 1 | 1.0900e-01 | 2.8451e+05 | ; | CB | - | HB3 |
| 5955 | 5958 | 1 | 1.5040e-01 | 2.6527e+05 | ; | CB | - | CG |
| 5958 | 5959 | 1 | 1.3940e-01 | 3.4309e+05 | ; | CG | - | ND1 |
| 5958 | 5964 | 1 | 1.3710e-01 | 4.3346e+05 | ; | CG | - | CD2 |
| 5959 | 5960 | 1 | 1.3350e-01 | 4.0836e+05 | ; | ND1 | - | CE1 |
| 5960 | 5961 | 1 | 1.0800e-01 | 3.0711e+05 | ; | CE1 | - | HE1 |
| 5960 | 5962 | 1 | 1.3430e-01 | 3.9915e+05 | ; | CE1 | - | NE2 |
| 5962 | 5963 | 1 | 1.0100e-01 | 3.6317e+05 | ; | NE2 | - | HE2 |
| 5962 | 5964 | 1 | 1.3810e-01 | 3.5731e+05 | ; | NE2 | - | CD2 |
| 5964 | 5965 | 1 | 1.0800e-01 | 3.0711e+05 | ; | CD2 | - | HD2 |
| 5966 | 5967 | 1 | 1.2290e-01 | 4.7698e+05 | ; | C | - | O |
| 5966 | 5968 | 1 | 1.3350e-01 | 4.1003e+05 | ; | C | - | N |
| 5968 | 5969 | 1 | 1.0100e-01 | 3.6317e+05 | ; | N | - | H |
| 5968 | 5970 | 1 | 1.4490e-01 | 2.8200e+05 | ; | N | - | CA |
| 5970 | 5971 | 1 | 1.0900e-01 | 2.8451e+05 | ; | CA | - | HA |
| 5970 | 5972 | 1 | 1.5260e-01 | 2.5941e+05 | ; | CA | - | CB |
| 5970 | 5987 | 1 | 1.5220e-01 | 2.6527e+05 | ; | CA | - | C |
| 5972 | 5973 | 1 | 1.0900e-01 | 2.8451e+05 | ; | CB | - | HB2 |
| 5972 | 5974 | 1 | 1.0900e-01 | 2.8451e+05 | ; | CB | - | HB3 |
| 5972 | 5975 | 1 | 1.5100e-01 | 2.6527e+05 | ; | CB | - | CG |
| 5975 | 5976 | 1 | 1.4000e-01 | 3.9246e+05 | ; | CG | - | CD1 |
| 5975 | 5985 | 1 | 1.4000e-01 | 3.9246e+05 | ; | CG | - | CD2 |
| 5976 | 5977 | 1 | 1.0800e-01 | 3.0711e+05 | ; | CD1 | - | HD1 |
| 5976 | 5978 | 1 | 1.4000e-01 | 3.9246e+05 | ; | CD1 | - | CE1 |
| 5978 | 5979 | 1 | 1.0800e-01 | 3.0711e+05 | ; | CE1 | - | HE1 |
| 5978 | 5980 | 1 | 1.4090e-01 | 3.9246e+05 | ; | CE1 | - | CZ |
| 5980 | 5981 | 1 | 1.3640e-01 | 3.7656e+05 | ; | CZ | - | OH |
| 5980 | 5983 | 1 | 1.4090e-01 | 3.9246e+05 | ; | CZ | - | CE2 |
| 5981 | 5982 | 1 | 9.6000e-02 | 4.6275e+05 | ; | OH | - | HH |
| 5983 | 5984 | 1 | 1.0800e-01 | 3.0711e+05 | ; | CE2 | - | HE2 |
| 5983 | 5985 | 1 | 1.4000e-01 | 3.9246e+05 | ; | CE2 | - | CD2 |
| 5985 | 5986 | 1 | 1.0800e-01 | 3.0711e+05 | ; | CD2 | - | HD2 |
| 5987 | 5988 | 1 | 1.2290e-01 | 4.7698e+05 | ; | C | - | O |
| 5987 | 5989 | 1 | 1.3350e-01 | 4.1003e+05 | ; | C | - | N |
| 5989 | 5990 | 1 | 1.0100e-01 | 3.6317e+05 | ; | N | - | H |
| 5989 | 5991 | 1 | 1.4490e-01 | 2.8200e+05 | ; | N | - | CA |
| 5991 | 5992 | 1 | 1.0900e-01 | 2.8451e+05 | ; | CA | - | HA |
| 5991 | 5993 | 1 | 1.5260e-01 | 2.5941e+05 | ; | CA | - | CB |
| 5991 | 6001 | 1 | 1.5220e-01 | 2.6527e+05 | ; | CA | - | C |
| 5993 | 5994 | 1 | 1.0900e-01 | 2.8451e+05 | ; | CB | - | HB |
| 5993 | 5995 | 1 | 1.5260e-01 | 2.5941e+05 | ; | CB | - | CG2 |
| 5993 | 5999 | 1 | 1.4100e-01 | 2.6778e+05 | ; | CB | - | OG1 |
| 5995 | 5996 | 1 | 1.0900e-01 | 2.8451e+05 | ; | CG2 | - | HG21 |
| 5995 | 5997 | 1 | 1.0900e-01 | 2.8451e+05 | ; | CG2 | - | HG22 |
| 5995 | 5998 | 1 | 1.0900e-01 | 2.8451e+05 | ; | CG2 | - | HG23 |
| 5999 | 6000 | 1 | 9.6000e-02 | 4.6275e+05 | ; | OG1 | - | HG1 |
| 6001 | 6002 | 1 | 1.2290e-01 | 4.7698e+05 | ; | C | - | O |
| 6001 | 6003 | 1 | 1.3350e-01 | 4.1003e+05 | ; | C | - | N |
| 6003 | 6004 | 1 | 1.0100e-01 | 3.6317e+05 | ; | N | - | H |
| 6003 | 6005 | 1 | 1.4490e-01 | 2.8200e+05 | ; | N | - | CA |
| 6005 | 6006 | 1 | 1.0900e-01 | 2.8451e+05 | ; | CA | - | HA |

| | | | | | |
|---|---|---|---|---|---|
| 6005 | 6007 | 1 | 1.5260e-01 | 2.5941e+05 ; | CA - CB |
| 6005 | 6013 | 1 | 1.5220e-01 | 2.6527e+05 ; | CA - C |
| 6007 | 6008 | 1 | 1.0900e-01 | 2.8451e+05 ; | CB - HB2 |
| 6007 | 6009 | 1 | 1.0900e-01 | 2.8451e+05 ; | CB - HB3 |
| 6007 | 6010 | 1 | 1.5220e-01 | 2.6527e+05 ; | CB - CG |
| 6010 | 6011 | 1 | 1.2500e-01 | 5.4894e+05 ; | CG - OD1 |
| 6010 | 6012 | 1 | 1.2500e-01 | 5.4894e+05 ; | CG - OD2 |
| 6013 | 6014 | 1 | 1.2290e-01 | 4.7698e+05 ; | C - O |
| 6013 | 6015 | 1 | 1.3350e-01 | 4.1003e+05 ; | C - N |
| 6015 | 6016 | 1 | 1.0100e-01 | 3.6317e+05 ; | N - H |
| 6015 | 6017 | 1 | 1.4490e-01 | 2.8200e+05 ; | N - CA |
| 6017 | 6018 | 1 | 1.0900e-01 | 2.8451e+05 ; | CA - HA |
| 6017 | 6019 | 1 | 1.5260e-01 | 2.5941e+05 ; | CA - CB |
| 6017 | 6033 | 1 | 1.5220e-01 | 2.6527e+05 ; | CA - C |
| 6019 | 6020 | 1 | 1.0900e-01 | 2.8451e+05 ; | CB - HB2 |
| 6019 | 6021 | 1 | 1.0900e-01 | 2.8451e+05 ; | CB - HB3 |
| 6019 | 6022 | 1 | 1.5100e-01 | 2.6527e+05 ; | CB - CG |
| 6022 | 6023 | 1 | 1.4000e-01 | 3.9246e+05 ; | CG - CD1 |
| 6022 | 6031 | 1 | 1.4000e-01 | 3.9246e+05 ; | CG - CD2 |
| 6023 | 6024 | 1 | 1.0800e-01 | 3.0711e+05 ; | CD1 - HD1 |
| 6023 | 6025 | 1 | 1.4000e-01 | 3.9246e+05 ; | CD1 - CE1 |
| 6025 | 6026 | 1 | 1.0800e-01 | 3.0711e+05 ; | CE1 - HE1 |
| 6025 | 6027 | 1 | 1.4000e-01 | 3.9246e+05 ; | CE1 - CZ |
| 6027 | 6028 | 1 | 1.0800e-01 | 3.0711e+05 ; | CZ - HZ |
| 6027 | 6029 | 1 | 1.4000e-01 | 3.9246e+05 ; | CZ - CE2 |
| 6029 | 6030 | 1 | 1.0800e-01 | 3.0711e+05 ; | CE2 - HE2 |
| 6029 | 6031 | 1 | 1.4000e-01 | 3.9246e+05 ; | CE2 - CD2 |
| 6031 | 6032 | 1 | 1.0800e-01 | 3.0711e+05 ; | CD2 - HD2 |
| 6033 | 6034 | 1 | 1.2290e-01 | 4.7698e+05 ; | C - O |
| 6033 | 6035 | 1 | 1.3350e-01 | 4.1003e+05 ; | C - N |
| 6035 | 6036 | 1 | 1.0100e-01 | 3.6317e+05 ; | N - H |
| 6035 | 6037 | 1 | 1.4490e-01 | 2.8200e+05 ; | N - CA |
| 6037 | 6038 | 1 | 1.0900e-01 | 2.8451e+05 ; | CA - HA |
| 6037 | 6039 | 1 | 1.5260e-01 | 2.5941e+05 ; | CA - CB |
| 6037 | 6052 | 1 | 1.5220e-01 | 2.6527e+05 ; | CA - C |
| 6039 | 6040 | 1 | 1.0900e-01 | 2.8451e+05 ; | CB - HB2 |
| 6039 | 6041 | 1 | 1.0900e-01 | 2.8451e+05 ; | CB - HB3 |
| 6039 | 6042 | 1 | 1.5260e-01 | 2.5941e+05 ; | CB - CG |
| 6042 | 6043 | 1 | 1.0900e-01 | 2.8451e+05 ; | CG - HG |
| 6042 | 6044 | 1 | 1.5260e-01 | 2.5941e+05 ; | CG - CD1 |
| 6042 | 6048 | 1 | 1.5260e-01 | 2.5941e+05 ; | CG - CD2 |
| 6044 | 6045 | 1 | 1.0900e-01 | 2.8451e+05 ; | CD1 - HD11 |
| 6044 | 6046 | 1 | 1.0900e-01 | 2.8451e+05 ; | CD1 - HD12 |
| 6044 | 6047 | 1 | 1.0900e-01 | 2.8451e+05 ; | CD1 - HD13 |
| 6048 | 6049 | 1 | 1.0900e-01 | 2.8451e+05 ; | CD2 - HD21 |
| 6048 | 6050 | 1 | 1.0900e-01 | 2.8451e+05 ; | CD2 - HD22 |
| 6048 | 6051 | 1 | 1.0900e-01 | 2.8451e+05 ; | CD2 - HD23 |
| 6052 | 6053 | 1 | 1.2290e-01 | 4.7698e+05 ; | C - O |
| 6052 | 6054 | 1 | 1.3350e-01 | 4.1003e+05 ; | C - N |
| 6054 | 6055 | 1 | 1.0100e-01 | 3.6317e+05 ; | N - H |
| 6054 | 6056 | 1 | 1.4490e-01 | 2.8200e+05 ; | N - CA |
| 6056 | 6057 | 1 | 1.0900e-01 | 2.8451e+05 ; | CA - HA |

| | | | | | | | | |
|---|---|---|---|---|---|---|---|---|
| 6056 | 6058 | 1 | 1.5260e-01 | 2.5941e+05 | ; | CA | - | CB |
| 6056 | 6062 | 1 | 1.5220e-01 | 2.6527e+05 | ; | CA | - | C |
| 6058 | 6059 | 1 | 1.0900e-01 | 2.8451e+05 | ; | CB | - | HB1 |
| 6058 | 6060 | 1 | 1.0900e-01 | 2.8451e+05 | ; | CB | - | HB2 |
| 6058 | 6061 | 1 | 1.0900e-01 | 2.8451e+05 | ; | CB | - | HB3 |
| 6062 | 6063 | 1 | 1.2290e-01 | 4.7698e+05 | ; | C | - | O |
| 6062 | 6064 | 1 | 1.3350e-01 | 4.1003e+05 | ; | C | - | N |
| 6064 | 6065 | 1 | 1.0100e-01 | 3.6317e+05 | ; | N | - | H |
| 6064 | 6066 | 1 | 1.4490e-01 | 2.8200e+05 | ; | N | - | CA |
| 6066 | 6067 | 1 | 1.0900e-01 | 2.8451e+05 | ; | CA | - | HA |
| 6066 | 6068 | 1 | 1.5260e-01 | 2.5941e+05 | ; | CA | - | CB |
| 6066 | 6072 | 1 | 1.5220e-01 | 2.6527e+05 | ; | CA | - | C |
| 6068 | 6069 | 1 | 1.0900e-01 | 2.8451e+05 | ; | CB | - | HB1 |
| 6068 | 6070 | 1 | 1.0900e-01 | 2.8451e+05 | ; | CB | - | HB2 |
| 6068 | 6071 | 1 | 1.0900e-01 | 2.8451e+05 | ; | CB | - | HB3 |
| 6072 | 6073 | 1 | 1.2290e-01 | 4.7698e+05 | ; | C | - | O |
| 6072 | 6074 | 1 | 1.3350e-01 | 4.1003e+05 | ; | C | - | N |
| 6074 | 6075 | 1 | 1.0100e-01 | 3.6317e+05 | ; | N | - | H |
| 6074 | 6076 | 1 | 1.4490e-01 | 2.8200e+05 | ; | N | - | CA |
| 6076 | 6077 | 1 | 1.0900e-01 | 2.8451e+05 | ; | CA | - | HA |
| 6076 | 6078 | 1 | 1.5260e-01 | 2.5941e+05 | ; | CA | - | CB |
| 6076 | 6086 | 1 | 1.5220e-01 | 2.6527e+05 | ; | CA | - | C |
| 6078 | 6079 | 1 | 1.0900e-01 | 2.8451e+05 | ; | CB | - | HB |
| 6078 | 6080 | 1 | 1.5260e-01 | 2.5941e+05 | ; | CB | - | CG2 |
| 6078 | 6084 | 1 | 1.4100e-01 | 2.6778e+05 | ; | CB | - | OG1 |
| 6080 | 6081 | 1 | 1.0900e-01 | 2.8451e+05 | ; | CG2 | - | HG21 |
| 6080 | 6082 | 1 | 1.0900e-01 | 2.8451e+05 | ; | CG2 | - | HG22 |
| 6080 | 6083 | 1 | 1.0900e-01 | 2.8451e+05 | ; | CG2 | - | HG23 |
| 6084 | 6085 | 1 | 9.6000e-02 | 4.6275e+05 | ; | OG1 | - | HG1 |
| 6086 | 6087 | 1 | 1.2290e-01 | 4.7698e+05 | ; | C | - | O |
| 6086 | 6088 | 1 | 1.3350e-01 | 4.1003e+05 | ; | C | - | N |
| 6088 | 6089 | 1 | 1.0100e-01 | 3.6317e+05 | ; | N | - | H |
| 6088 | 6090 | 1 | 1.4490e-01 | 2.8200e+05 | ; | N | - | CA |
| 6090 | 6091 | 1 | 1.0900e-01 | 2.8451e+05 | ; | CA | - | HA |
| 6090 | 6092 | 1 | 1.5260e-01 | 2.5941e+05 | ; | CA | - | CB |
| 6090 | 6105 | 1 | 1.5220e-01 | 2.6527e+05 | ; | CA | - | C |
| 6092 | 6093 | 1 | 1.0900e-01 | 2.8451e+05 | ; | CB | - | HB |
| 6092 | 6094 | 1 | 1.5260e-01 | 2.5941e+05 | ; | CB | - | CG2 |
| 6092 | 6098 | 1 | 1.5260e-01 | 2.5941e+05 | ; | CB | - | CG1 |
| 6094 | 6095 | 1 | 1.0900e-01 | 2.8451e+05 | ; | CG2 | - | HG21 |
| 6094 | 6096 | 1 | 1.0900e-01 | 2.8451e+05 | ; | CG2 | - | HG22 |
| 6094 | 6097 | 1 | 1.0900e-01 | 2.8451e+05 | ; | CG2 | - | HG23 |
| 6098 | 6099 | 1 | 1.0900e-01 | 2.8451e+05 | ; | CG1 | - | HG12 |
| 6098 | 6100 | 1 | 1.0900e-01 | 2.8451e+05 | ; | CG1 | - | HG13 |
| 6098 | 6101 | 1 | 1.5260e-01 | 2.5941e+05 | ; | CG1 | - | CD1 |
| 6101 | 6102 | 1 | 1.0900e-01 | 2.8451e+05 | ; | CD1 | - | HD11 |
| 6101 | 6103 | 1 | 1.0900e-01 | 2.8451e+05 | ; | CD1 | - | HD12 |
| 6101 | 6104 | 1 | 1.0900e-01 | 2.8451e+05 | ; | CD1 | - | HD13 |
| 6105 | 6106 | 1 | 1.2290e-01 | 4.7698e+05 | ; | C | - | O |
| 6105 | 6107 | 1 | 1.3350e-01 | 4.1003e+05 | ; | C | - | N |
| 6107 | 6108 | 1 | 1.0100e-01 | 3.6317e+05 | ; | N | - | H |
| 6107 | 6109 | 1 | 1.4490e-01 | 2.8200e+05 | ; | N | - | CA |

| | | | | | |
|---|---|---|---|---|---|
| 6109 | 6110 | 1 | 1.0900e-01 | 2.8451e+05 ; | CA - HA |
| 6109 | 6111 | 1 | 1.5260e-01 | 2.5941e+05 ; | CA - CB |
| 6109 | 6117 | 1 | 1.5220e-01 | 2.6527e+05 ; | CA - C |
| 6111 | 6112 | 1 | 1.0900e-01 | 2.8451e+05 ; | CB - HB2 |
| 6111 | 6113 | 1 | 1.0900e-01 | 2.8451e+05 ; | CB - HB3 |
| 6111 | 6114 | 1 | 1.5220e-01 | 2.6527e+05 ; | CB - CG |
| 6114 | 6115 | 1 | 1.2500e-01 | 5.4894e+05 ; | CG - OD1 |
| 6114 | 6116 | 1 | 1.2500e-01 | 5.4894e+05 ; | CG - OD2 |
| 6117 | 6118 | 1 | 1.2290e-01 | 4.7698e+05 ; | C - O |
| 6117 | 6119 | 1 | 1.3350e-01 | 4.1003e+05 ; | C - N |
| 6119 | 6120 | 1 | 1.0100e-01 | 3.6317e+05 ; | N - H |
| 6119 | 6121 | 1 | 1.4490e-01 | 2.8200e+05 ; | N - CA |
| 6121 | 6122 | 1 | 1.0900e-01 | 2.8451e+05 ; | CA - HA |
| 6121 | 6123 | 1 | 1.5260e-01 | 2.5941e+05 ; | CA - CB |
| 6121 | 6139 | 1 | 1.5220e-01 | 2.6527e+05 ; | CA - C |
| 6123 | 6124 | 1 | 1.0900e-01 | 2.8451e+05 ; | CB - HB2 |
| 6123 | 6125 | 1 | 1.0900e-01 | 2.8451e+05 ; | CB - HB3 |
| 6123 | 6126 | 1 | 1.5260e-01 | 2.5941e+05 ; | CB - CG |
| 6126 | 6127 | 1 | 1.0900e-01 | 2.8451e+05 ; | CG - HG2 |
| 6126 | 6128 | 1 | 1.0900e-01 | 2.8451e+05 ; | CG - HG3 |
| 6126 | 6129 | 1 | 1.5260e-01 | 2.5941e+05 ; | CG - CD |
| 6129 | 6130 | 1 | 1.0900e-01 | 2.8451e+05 ; | CD - HD2 |
| 6129 | 6131 | 1 | 1.0900e-01 | 2.8451e+05 ; | CD - HD3 |
| 6129 | 6132 | 1 | 1.5260e-01 | 2.5941e+05 ; | CD - CE |
| 6132 | 6133 | 1 | 1.0900e-01 | 2.8451e+05 ; | CE - HE2 |
| 6132 | 6134 | 1 | 1.0900e-01 | 2.8451e+05 ; | CE - HE3 |
| 6132 | 6135 | 1 | 1.4710e-01 | 3.0711e+05 ; | CE - NZ |
| 6135 | 6136 | 1 | 1.0100e-01 | 3.6317e+05 ; | NZ - HZ1 |
| 6135 | 6137 | 1 | 1.0100e-01 | 3.6317e+05 ; | NZ - HZ2 |
| 6135 | 6138 | 1 | 1.0100e-01 | 3.6317e+05 ; | NZ - HZ3 |
| 6139 | 6140 | 1 | 1.2290e-01 | 4.7698e+05 ; | C - O |
| 6139 | 6141 | 1 | 1.3350e-01 | 4.1003e+05 ; | C - N |
| 6141 | 6142 | 1 | 1.0100e-01 | 3.6317e+05 ; | N - H |
| 6141 | 6143 | 1 | 1.4490e-01 | 2.8200e+05 ; | N - CA |
| 6143 | 6144 | 1 | 1.0900e-01 | 2.8451e+05 ; | CA - HA |
| 6143 | 6145 | 1 | 1.5260e-01 | 2.5941e+05 ; | CA - CB |
| 6143 | 6156 | 1 | 1.5220e-01 | 2.6527e+05 ; | CA - C |
| 6145 | 6146 | 1 | 1.0900e-01 | 2.8451e+05 ; | CB - HB2 |
| 6145 | 6147 | 1 | 1.0900e-01 | 2.8451e+05 ; | CB - HB3 |
| 6145 | 6148 | 1 | 1.5260e-01 | 2.5941e+05 ; | CB - CG |
| 6148 | 6149 | 1 | 1.0900e-01 | 2.8451e+05 ; | CG - HG2 |
| 6148 | 6150 | 1 | 1.0900e-01 | 2.8451e+05 ; | CG - HG3 |
| 6148 | 6151 | 1 | 1.5220e-01 | 2.6527e+05 ; | CG - CD |
| 6151 | 6152 | 1 | 1.2290e-01 | 4.7698e+05 ; | CD - OE1 |
| 6151 | 6153 | 1 | 1.3350e-01 | 4.1003e+05 ; | CD - NE2 |
| 6153 | 6154 | 1 | 1.0100e-01 | 3.6317e+05 ; | NE2 - HE21 |
| 6153 | 6155 | 1 | 1.0100e-01 | 3.6317e+05 ; | NE2 - HE22 |
| 6156 | 6157 | 1 | 1.2290e-01 | 4.7698e+05 ; | C - O |
| 6156 | 6158 | 1 | 1.3350e-01 | 4.1003e+05 ; | C - N |
| 6158 | 6159 | 1 | 1.0100e-01 | 3.6317e+05 ; | N - H |
| 6158 | 6160 | 1 | 1.4490e-01 | 2.8200e+05 ; | N - CA |
| 6160 | 6161 | 1 | 1.0900e-01 | 2.8451e+05 ; | CA - HA |

| | | | | | | | | |
|---|---|---|---|---|---|---|---|---|
| 6160 | 6162 | 1 | 1.5260e-01 | 2.5941e+05 | ; | CA | - | CB |
| 6160 | 6170 | 1 | 1.5220e-01 | 2.6527e+05 | ; | CA | - | C |
| 6162 | 6163 | 1 | 1.0900e-01 | 2.8451e+05 | ; | CB | - | HB |
| 6162 | 6164 | 1 | 1.5260e-01 | 2.5941e+05 | ; | CB | - | CG2 |
| 6162 | 6168 | 1 | 1.4100e-01 | 2.6778e+05 | ; | CB | - | OG1 |
| 6164 | 6165 | 1 | 1.0900e-01 | 2.8451e+05 | ; | CG2 | - | HG21 |
| 6164 | 6166 | 1 | 1.0900e-01 | 2.8451e+05 | ; | CG2 | - | HG22 |
| 6164 | 6167 | 1 | 1.0900e-01 | 2.8451e+05 | ; | CG2 | - | HG23 |
| 6168 | 6169 | 1 | 9.6000e-02 | 4.6275e+05 | ; | OG1 | - | HG1 |
| 6170 | 6171 | 1 | 1.2290e-01 | 4.7698e+05 | ; | C | - | O |
| 6170 | 6172 | 1 | 1.3350e-01 | 4.1003e+05 | ; | C | - | N |
| 6172 | 6173 | 1 | 1.0100e-01 | 3.6317e+05 | ; | N | - | H |
| 6172 | 6174 | 1 | 1.4490e-01 | 2.8200e+05 | ; | N | - | CA |
| 6174 | 6175 | 1 | 1.0900e-01 | 2.8451e+05 | ; | CA | - | HA |
| 6174 | 6176 | 1 | 1.5260e-01 | 2.5941e+05 | ; | CA | - | CB |
| 6174 | 6191 | 1 | 1.5220e-01 | 2.6527e+05 | ; | CA | - | C |
| 6176 | 6177 | 1 | 1.0900e-01 | 2.8451e+05 | ; | CB | - | HB2 |
| 6176 | 6178 | 1 | 1.0900e-01 | 2.8451e+05 | ; | CB | - | HB3 |
| 6176 | 6179 | 1 | 1.5100e-01 | 2.6527e+05 | ; | CB | - | CG |
| 6179 | 6180 | 1 | 1.4000e-01 | 3.9246e+05 | ; | CG | - | CD1 |
| 6179 | 6189 | 1 | 1.4000e-01 | 3.9246e+05 | ; | CG | - | CD2 |
| 6180 | 6181 | 1 | 1.0800e-01 | 3.0711e+05 | ; | CD1 | - | HD1 |
| 6180 | 6182 | 1 | 1.4000e-01 | 3.9246e+05 | ; | CD1 | - | CE1 |
| 6182 | 6183 | 1 | 1.0800e-01 | 3.0711e+05 | ; | CE1 | - | HE1 |
| 6182 | 6184 | 1 | 1.4090e-01 | 3.9246e+05 | ; | CE1 | - | CZ |
| 6184 | 6185 | 1 | 1.3640e-01 | 3.7656e+05 | ; | CZ | - | OH |
| 6184 | 6187 | 1 | 1.4090e-01 | 3.9246e+05 | ; | CZ | - | CE2 |
| 6185 | 6186 | 1 | 9.6000e-02 | 4.6275e+05 | ; | OH | - | HH |
| 6187 | 6188 | 1 | 1.0800e-01 | 3.0711e+05 | ; | CE2 | - | HE2 |
| 6187 | 6189 | 1 | 1.4000e-01 | 3.9246e+05 | ; | CE2 | - | CD2 |
| 6189 | 6190 | 1 | 1.0800e-01 | 3.0711e+05 | ; | CD2 | - | HD2 |
| 6191 | 6192 | 1 | 1.2290e-01 | 4.7698e+05 | ; | C | - | O |
| 6191 | 6193 | 1 | 1.3350e-01 | 4.1003e+05 | ; | C | - | N |
| 6193 | 6194 | 1 | 1.0100e-01 | 3.6317e+05 | ; | N | - | H |
| 6193 | 6195 | 1 | 1.4490e-01 | 2.8200e+05 | ; | N | - | CA |
| 6195 | 6196 | 1 | 1.0900e-01 | 2.8451e+05 | ; | CA | - | HA |
| 6195 | 6197 | 1 | 1.5260e-01 | 2.5941e+05 | ; | CA | - | CB |
| 6195 | 6210 | 1 | 1.5220e-01 | 2.6527e+05 | ; | CA | - | C |
| 6197 | 6198 | 1 | 1.0900e-01 | 2.8451e+05 | ; | CB | - | HB2 |
| 6197 | 6199 | 1 | 1.0900e-01 | 2.8451e+05 | ; | CB | - | HB3 |
| 6197 | 6200 | 1 | 1.5260e-01 | 2.5941e+05 | ; | CB | - | CG |
| 6200 | 6201 | 1 | 1.0900e-01 | 2.8451e+05 | ; | CG | - | HG |
| 6200 | 6202 | 1 | 1.5260e-01 | 2.5941e+05 | ; | CG | - | CD1 |
| 6200 | 6206 | 1 | 1.5260e-01 | 2.5941e+05 | ; | CG | - | CD2 |
| 6202 | 6203 | 1 | 1.0900e-01 | 2.8451e+05 | ; | CD1 | - | HD11 |
| 6202 | 6204 | 1 | 1.0900e-01 | 2.8451e+05 | ; | CD1 | - | HD12 |
| 6202 | 6205 | 1 | 1.0900e-01 | 2.8451e+05 | ; | CD1 | - | HD13 |
| 6206 | 6207 | 1 | 1.0900e-01 | 2.8451e+05 | ; | CD2 | - | HD21 |
| 6206 | 6208 | 1 | 1.0900e-01 | 2.8451e+05 | ; | CD2 | - | HD22 |
| 6206 | 6209 | 1 | 1.0900e-01 | 2.8451e+05 | ; | CD2 | - | HD23 |
| 6210 | 6211 | 1 | 1.2290e-01 | 4.7698e+05 | ; | C | - | O |
| 6210 | 6212 | 1 | 1.3350e-01 | 4.1003e+05 | ; | C | - | N |

| | | | | | |
|---|---|---|---|---|---|
| 6212 | 6213 | 1 | 1.0100e-01 | 3.6317e+05 ; | N - H |
| 6212 | 6214 | 1 | 1.4490e-01 | 2.8200e+05 ; | N - CA |
| 6214 | 6215 | 1 | 1.0900e-01 | 2.8451e+05 ; | CA - HA |
| 6214 | 6216 | 1 | 1.5260e-01 | 2.5941e+05 ; | CA - CB |
| 6214 | 6232 | 1 | 1.5220e-01 | 2.6527e+05 ; | CA - C |
| 6216 | 6217 | 1 | 1.0900e-01 | 2.8451e+05 ; | CB - HB2 |
| 6216 | 6218 | 1 | 1.0900e-01 | 2.8451e+05 ; | CB - HB3 |
| 6216 | 6219 | 1 | 1.5260e-01 | 2.5941e+05 ; | CB - CG |
| 6219 | 6220 | 1 | 1.0900e-01 | 2.8451e+05 ; | CG - HG2 |
| 6219 | 6221 | 1 | 1.0900e-01 | 2.8451e+05 ; | CG - HG3 |
| 6219 | 6222 | 1 | 1.5260e-01 | 2.5941e+05 ; | CG - CD |
| 6222 | 6223 | 1 | 1.0900e-01 | 2.8451e+05 ; | CD - HD2 |
| 6222 | 6224 | 1 | 1.0900e-01 | 2.8451e+05 ; | CD - HD3 |
| 6222 | 6225 | 1 | 1.5260e-01 | 2.5941e+05 ; | CD - CE |
| 6225 | 6226 | 1 | 1.0900e-01 | 2.8451e+05 ; | CE - HE2 |
| 6225 | 6227 | 1 | 1.0900e-01 | 2.8451e+05 ; | CE - HE3 |
| 6225 | 6228 | 1 | 1.4710e-01 | 3.0711e+05 ; | CE - NZ |
| 6228 | 6229 | 1 | 1.0100e-01 | 3.6317e+05 ; | NZ - HZ1 |
| 6228 | 6230 | 1 | 1.0100e-01 | 3.6317e+05 ; | NZ - HZ2 |
| 6228 | 6231 | 1 | 1.0100e-01 | 3.6317e+05 ; | NZ - HZ3 |
| 6232 | 6233 | 1 | 1.2290e-01 | 4.7698e+05 ; | C - O |
| 6232 | 6234 | 1 | 1.3350e-01 | 4.1003e+05 ; | C - N |
| 6234 | 6235 | 1 | 1.0100e-01 | 3.6317e+05 ; | N - H |
| 6234 | 6236 | 1 | 1.4490e-01 | 2.8200e+05 ; | N - CA |
| 6236 | 6237 | 1 | 1.0900e-01 | 2.8451e+05 ; | CA - HA |
| 6236 | 6238 | 1 | 1.5260e-01 | 2.5941e+05 ; | CA - CB |
| 6236 | 6254 | 1 | 1.5220e-01 | 2.6527e+05 ; | CA - C |
| 6238 | 6239 | 1 | 1.0900e-01 | 2.8451e+05 ; | CB - HB2 |
| 6238 | 6240 | 1 | 1.0900e-01 | 2.8451e+05 ; | CB - HB3 |
| 6238 | 6241 | 1 | 1.5260e-01 | 2.5941e+05 ; | CB - CG |
| 6241 | 6242 | 1 | 1.0900e-01 | 2.8451e+05 ; | CG - HG2 |
| 6241 | 6243 | 1 | 1.0900e-01 | 2.8451e+05 ; | CG - HG3 |
| 6241 | 6244 | 1 | 1.5260e-01 | 2.5941e+05 ; | CG - CD |
| 6244 | 6245 | 1 | 1.0900e-01 | 2.8451e+05 ; | CD - HD2 |
| 6244 | 6246 | 1 | 1.0900e-01 | 2.8451e+05 ; | CD - HD3 |
| 6244 | 6247 | 1 | 1.5260e-01 | 2.5941e+05 ; | CD - CE |
| 6247 | 6248 | 1 | 1.0900e-01 | 2.8451e+05 ; | CE - HE2 |
| 6247 | 6249 | 1 | 1.0900e-01 | 2.8451e+05 ; | CE - HE3 |
| 6247 | 6250 | 1 | 1.4710e-01 | 3.0711e+05 ; | CE - NZ |
| 6250 | 6251 | 1 | 1.0100e-01 | 3.6317e+05 ; | NZ - HZ1 |
| 6250 | 6252 | 1 | 1.0100e-01 | 3.6317e+05 ; | NZ - HZ2 |
| 6250 | 6253 | 1 | 1.0100e-01 | 3.6317e+05 ; | NZ - HZ3 |
| 6254 | 6255 | 1 | 1.2290e-01 | 4.7698e+05 ; | C - O |
| 6254 | 6256 | 1 | 1.3350e-01 | 4.1003e+05 ; | C - N |
| 6256 | 6257 | 1 | 1.0100e-01 | 3.6317e+05 ; | N - H |
| 6256 | 6258 | 1 | 1.4490e-01 | 2.8200e+05 ; | N - CA |
| 6258 | 6259 | 1 | 1.0900e-01 | 2.8451e+05 ; | CA - HA |
| 6258 | 6260 | 1 | 1.5260e-01 | 2.5941e+05 ; | CA - CB |
| 6258 | 6269 | 1 | 1.5220e-01 | 2.6527e+05 ; | CA - C |
| 6260 | 6261 | 1 | 1.0900e-01 | 2.8451e+05 ; | CB - HB2 |
| 6260 | 6262 | 1 | 1.0900e-01 | 2.8451e+05 ; | CB - HB3 |
| 6260 | 6263 | 1 | 1.5260e-01 | 2.5941e+05 ; | CB - CG |

```
6263   6264   1    1.0900e-01    2.8451e+05 ;     CG - HG2
6263   6265   1    1.0900e-01    2.8451e+05 ;     CG - HG3
6263   6266   1    1.5220e-01    2.6527e+05 ;     CG - CD
6266   6267   1    1.2500e-01    5.4894e+05 ;     CD - OE1
6266   6268   1    1.2500e-01    5.4894e+05 ;     CD - OE2
6269   6270   1    1.2290e-01    4.7698e+05 ;      C - O
6269   6271   1    1.3350e-01    4.1003e+05 ;      C - N
6271   6272   1    1.0100e-01    3.6317e+05 ;      N - H
6271   6273   1    1.4490e-01    2.8200e+05 ;      N - CA
6273   6274   1    1.0900e-01    2.8451e+05 ;     CA - HA
6273   6275   1    1.5260e-01    2.5941e+05 ;     CA - CB
6273   6285   1    1.5220e-01    2.6527e+05 ;     CA - C
6275   6276   1    1.0900e-01    2.8451e+05 ;     CB - HB
6275   6277   1    1.5260e-01    2.5941e+05 ;     CB - CG1
6275   6281   1    1.5260e-01    2.5941e+05 ;     CB - CG2
6277   6278   1    1.0900e-01    2.8451e+05 ;    CG1 - HG11
6277   6279   1    1.0900e-01    2.8451e+05 ;    CG1 - HG12
6277   6280   1    1.0900e-01    2.8451e+05 ;    CG1 - HG13
6281   6282   1    1.0900e-01    2.8451e+05 ;    CG2 - HG21
6281   6283   1    1.0900e-01    2.8451e+05 ;    CG2 - HG22
6281   6284   1    1.0900e-01    2.8451e+05 ;    CG2 - HG23
6285   6286   1    1.2290e-01    4.7698e+05 ;      C - O
6285   6287   1    1.3350e-01    4.1003e+05 ;      C - N
6287   6288   1    1.0100e-01    3.6317e+05 ;      N - H
6287   6289   1    1.4490e-01    2.8200e+05 ;      N - CA
6289   6290   1    1.0900e-01    2.8451e+05 ;     CA - HA
6289   6291   1    1.5260e-01    2.5941e+05 ;     CA - CB
6289   6296   1    1.5220e-01    2.6527e+05 ;     CA - C
6291   6292   1    1.0900e-01    2.8451e+05 ;     CB - HB2
6291   6293   1    1.0900e-01    2.8451e+05 ;     CB - HB3
6291   6294   1    1.8100e-01    1.9832e+05 ;     CB - SG
6294   6295   1    1.3360e-01    2.2928e+05 ;     SG - HG
6296   6297   1    1.2290e-01    4.7698e+05 ;      C - O
6296   6298   1    1.3350e-01    4.1003e+05 ;      C - N
6298   6299   1    1.0100e-01    3.6317e+05 ;      N - H
6298   6300   1    1.4490e-01    2.8200e+05 ;      N - CA
6300   6301   1    1.0900e-01    2.8451e+05 ;     CA - HA
6300   6302   1    1.5260e-01    2.5941e+05 ;     CA - CB
6300   6315   1    1.5220e-01    2.6527e+05 ;     CA - C
6302   6303   1    1.0900e-01    2.8451e+05 ;     CB - HB2
6302   6304   1    1.0900e-01    2.8451e+05 ;     CB - HB3
6302   6305   1    1.5260e-01    2.5941e+05 ;     CB - CG
6305   6306   1    1.0900e-01    2.8451e+05 ;     CG - HG
6305   6307   1    1.5260e-01    2.5941e+05 ;     CG - CD1
6305   6311   1    1.5260e-01    2.5941e+05 ;     CG - CD2
6307   6308   1    1.0900e-01    2.8451e+05 ;    CD1 - HD11
6307   6309   1    1.0900e-01    2.8451e+05 ;    CD1 - HD12
6307   6310   1    1.0900e-01    2.8451e+05 ;    CD1 - HD13
6311   6312   1    1.0900e-01    2.8451e+05 ;    CD2 - HD21
6311   6313   1    1.0900e-01    2.8451e+05 ;    CD2 - HD22
6311   6314   1    1.0900e-01    2.8451e+05 ;    CD2 - HD23
6315   6316   1    1.2290e-01    4.7698e+05 ;      C - O
```

| | | | | | | | | |
|---|---|---|---|---|---|---|---|---|
| 6315 | 6317 | 1 | 1.3350e-01 | 4.1003e+05 | ; | C | - | N |
| 6317 | 6318 | 1 | 1.0100e-01 | 3.6317e+05 | ; | N | - | H |
| 6317 | 6319 | 1 | 1.4490e-01 | 2.8200e+05 | ; | N | - | CA |
| 6319 | 6320 | 1 | 1.0900e-01 | 2.8451e+05 | ; | CA | - | HA |
| 6319 | 6321 | 1 | 1.5260e-01 | 2.5941e+05 | ; | CA | - | CB |
| 6319 | 6334 | 1 | 1.5220e-01 | 2.6527e+05 | ; | CA | - | C |
| 6321 | 6322 | 1 | 1.0900e-01 | 2.8451e+05 | ; | CB | - | HB |
| 6321 | 6323 | 1 | 1.5260e-01 | 2.5941e+05 | ; | CB | - | CG2 |
| 6321 | 6327 | 1 | 1.5260e-01 | 2.5941e+05 | ; | CB | - | CG1 |
| 6323 | 6324 | 1 | 1.0900e-01 | 2.8451e+05 | ; | CG2 | - | HG21 |
| 6323 | 6325 | 1 | 1.0900e-01 | 2.8451e+05 | ; | CG2 | - | HG22 |
| 6323 | 6326 | 1 | 1.0900e-01 | 2.8451e+05 | ; | CG2 | - | HG23 |
| 6327 | 6328 | 1 | 1.0900e-01 | 2.8451e+05 | ; | CG1 | - | HG12 |
| 6327 | 6329 | 1 | 1.0900e-01 | 2.8451e+05 | ; | CG1 | - | HG13 |
| 6327 | 6330 | 1 | 1.5260e-01 | 2.5941e+05 | ; | CG1 | - | CD1 |
| 6330 | 6331 | 1 | 1.0900e-01 | 2.8451e+05 | ; | CD1 | - | HD11 |
| 6330 | 6332 | 1 | 1.0900e-01 | 2.8451e+05 | ; | CD1 | - | HD12 |
| 6330 | 6333 | 1 | 1.0900e-01 | 2.8451e+05 | ; | CD1 | - | HD13 |
| 6334 | 6335 | 1 | 1.2290e-01 | 4.7698e+05 | ; | C | - | O |
| 6334 | 6336 | 1 | 1.3350e-01 | 4.1003e+05 | ; | C | - | N |
| 6336 | 6337 | 1 | 1.4490e-01 | 2.8200e+05 | ; | N | - | CD |
| 6336 | 6346 | 1 | 1.4490e-01 | 2.8200e+05 | ; | N | - | CA |
| 6337 | 6338 | 1 | 1.0900e-01 | 2.8451e+05 | ; | CD | - | HD2 |
| 6337 | 6339 | 1 | 1.0900e-01 | 2.8451e+05 | ; | CD | - | HD3 |
| 6337 | 6340 | 1 | 1.5260e-01 | 2.5941e+05 | ; | CD | - | CG |
| 6340 | 6341 | 1 | 1.0900e-01 | 2.8451e+05 | ; | CG | - | HG2 |
| 6340 | 6342 | 1 | 1.0900e-01 | 2.8451e+05 | ; | CG | - | HG3 |
| 6340 | 6343 | 1 | 1.5260e-01 | 2.5941e+05 | ; | CG | - | CB |
| 6343 | 6344 | 1 | 1.0900e-01 | 2.8451e+05 | ; | CB | - | HB2 |
| 6343 | 6345 | 1 | 1.0900e-01 | 2.8451e+05 | ; | CB | - | HB3 |
| 6343 | 6346 | 1 | 1.5260e-01 | 2.5941e+05 | ; | CB | - | CA |
| 6346 | 6347 | 1 | 1.0900e-01 | 2.8451e+05 | ; | CA | - | HA |
| 6346 | 6348 | 1 | 1.5220e-01 | 2.6527e+05 | ; | CA | - | C |
| 6348 | 6349 | 1 | 1.2290e-01 | 4.7698e+05 | ; | C | - | O |
| 6348 | 6350 | 1 | 1.3350e-01 | 4.1003e+05 | ; | C | - | N |
| 6350 | 6351 | 1 | 1.0100e-01 | 3.6317e+05 | ; | N | - | H |
| 6350 | 6352 | 1 | 1.4490e-01 | 2.8200e+05 | ; | N | - | CA |
| 6352 | 6353 | 1 | 1.0900e-01 | 2.8451e+05 | ; | CA | - | HA |
| 6352 | 6354 | 1 | 1.5260e-01 | 2.5941e+05 | ; | CA | - | CB |
| 6352 | 6368 | 1 | 1.5220e-01 | 2.6527e+05 | ; | CA | - | C |
| 6354 | 6355 | 1 | 1.0900e-01 | 2.8451e+05 | ; | CB | - | HB2 |
| 6354 | 6356 | 1 | 1.0900e-01 | 2.8451e+05 | ; | CB | - | HB3 |
| 6354 | 6357 | 1 | 1.5100e-01 | 2.6527e+05 | ; | CB | - | CG |
| 6357 | 6358 | 1 | 1.4000e-01 | 3.9246e+05 | ; | CG | - | CD1 |
| 6357 | 6366 | 1 | 1.4000e-01 | 3.9246e+05 | ; | CG | - | CD2 |
| 6358 | 6359 | 1 | 1.0800e-01 | 3.0711e+05 | ; | CD1 | - | HD1 |
| 6358 | 6360 | 1 | 1.4000e-01 | 3.9246e+05 | ; | CD1 | - | CE1 |
| 6360 | 6361 | 1 | 1.0800e-01 | 3.0711e+05 | ; | CE1 | - | HE1 |
| 6360 | 6362 | 1 | 1.4000e-01 | 3.9246e+05 | ; | CE1 | - | CZ |
| 6362 | 6363 | 1 | 1.0800e-01 | 3.0711e+05 | ; | CZ | - | HZ |
| 6362 | 6364 | 1 | 1.4000e-01 | 3.9246e+05 | ; | CZ | - | CE2 |
| 6364 | 6365 | 1 | 1.0800e-01 | 3.0711e+05 | ; | CE2 | - | HE2 |

| | | | | | | |
|---|---|---|---|---|---|---|
| 6364 | 6366 | 1 | 1.4000e-01 | 3.9246e+05 | ; | CE2 - CD2 |
| 6366 | 6367 | 1 | 1.0800e-01 | 3.0711e+05 | ; | CD2 - HD2 |
| 6368 | 6369 | 1 | 1.2290e-01 | 4.7698e+05 | ; | C - O |
| 6368 | 6370 | 1 | 1.3350e-01 | 4.1003e+05 | ; | C - N |
| 6370 | 6371 | 1 | 1.0100e-01 | 3.6317e+05 | ; | N - H |
| 6370 | 6372 | 1 | 1.4490e-01 | 2.8200e+05 | ; | N - CA |
| 6372 | 6373 | 1 | 1.0900e-01 | 2.8451e+05 | ; | CA - HA |
| 6372 | 6374 | 1 | 1.5260e-01 | 2.5941e+05 | ; | CA - CB |
| 6372 | 6390 | 1 | 1.5220e-01 | 2.6527e+05 | ; | CA - C |
| 6374 | 6375 | 1 | 1.0900e-01 | 2.8451e+05 | ; | CB - HB2 |
| 6374 | 6376 | 1 | 1.0900e-01 | 2.8451e+05 | ; | CB - HB3 |
| 6374 | 6377 | 1 | 1.5260e-01 | 2.5941e+05 | ; | CB - CG |
| 6377 | 6378 | 1 | 1.0900e-01 | 2.8451e+05 | ; | CG - HG2 |
| 6377 | 6379 | 1 | 1.0900e-01 | 2.8451e+05 | ; | CG - HG3 |
| 6377 | 6380 | 1 | 1.5260e-01 | 2.5941e+05 | ; | CG - CD |
| 6380 | 6381 | 1 | 1.0900e-01 | 2.8451e+05 | ; | CD - HD2 |
| 6380 | 6382 | 1 | 1.0900e-01 | 2.8451e+05 | ; | CD - HD3 |
| 6380 | 6383 | 1 | 1.5260e-01 | 2.5941e+05 | ; | CD - CE |
| 6383 | 6384 | 1 | 1.0900e-01 | 2.8451e+05 | ; | CE - HE2 |
| 6383 | 6385 | 1 | 1.0900e-01 | 2.8451e+05 | ; | CE - HE3 |
| 6383 | 6386 | 1 | 1.4710e-01 | 3.0711e+05 | ; | CE - NZ |
| 6386 | 6387 | 1 | 1.0100e-01 | 3.6317e+05 | ; | NZ - HZ1 |
| 6386 | 6388 | 1 | 1.0100e-01 | 3.6317e+05 | ; | NZ - HZ2 |
| 6386 | 6389 | 1 | 1.0100e-01 | 3.6317e+05 | ; | NZ - HZ3 |
| 6390 | 6391 | 1 | 1.2290e-01 | 4.7698e+05 | ; | C - O |
| 6390 | 6392 | 1 | 1.3350e-01 | 4.1003e+05 | ; | C - N |
| 6392 | 6393 | 1 | 1.0100e-01 | 3.6317e+05 | ; | N - H |
| 6392 | 6394 | 1 | 1.4490e-01 | 2.8200e+05 | ; | N - CA |
| 6394 | 6395 | 1 | 1.0900e-01 | 2.8451e+05 | ; | CA - HA |
| 6394 | 6396 | 1 | 1.5260e-01 | 2.5941e+05 | ; | CA - CB |
| 6394 | 6410 | 1 | 1.5220e-01 | 2.6527e+05 | ; | CA - C |
| 6396 | 6397 | 1 | 1.0900e-01 | 2.8451e+05 | ; | CB - HB2 |
| 6396 | 6398 | 1 | 1.0900e-01 | 2.8451e+05 | ; | CB - HB3 |
| 6396 | 6399 | 1 | 1.5100e-01 | 2.6527e+05 | ; | CB - CG |
| 6399 | 6400 | 1 | 1.4000e-01 | 3.9246e+05 | ; | CG - CD1 |
| 6399 | 6408 | 1 | 1.4000e-01 | 3.9246e+05 | ; | CG - CD2 |
| 6400 | 6401 | 1 | 1.0800e-01 | 3.0711e+05 | ; | CD1 - HD1 |
| 6400 | 6402 | 1 | 1.4000e-01 | 3.9246e+05 | ; | CD1 - CE1 |
| 6402 | 6403 | 1 | 1.0800e-01 | 3.0711e+05 | ; | CE1 - HE1 |
| 6402 | 6404 | 1 | 1.4000e-01 | 3.9246e+05 | ; | CE1 - CZ |
| 6404 | 6405 | 1 | 1.0800e-01 | 3.0711e+05 | ; | CZ - HZ |
| 6404 | 6406 | 1 | 1.4000e-01 | 3.9246e+05 | ; | CZ - CE2 |
| 6406 | 6407 | 1 | 1.0800e-01 | 3.0711e+05 | ; | CE2 - HE2 |
| 6406 | 6408 | 1 | 1.4000e-01 | 3.9246e+05 | ; | CE2 - CD2 |
| 6408 | 6409 | 1 | 1.0800e-01 | 3.0711e+05 | ; | CD2 - HD2 |
| 6410 | 6411 | 1 | 1.2290e-01 | 4.7698e+05 | ; | C - O |
| 6410 | 6412 | 1 | 1.3350e-01 | 4.1003e+05 | ; | C - N |
| 6412 | 6413 | 1 | 1.0100e-01 | 3.6317e+05 | ; | N - H |
| 6412 | 6414 | 1 | 1.4490e-01 | 2.8200e+05 | ; | N - CA |
| 6414 | 6415 | 1 | 1.0900e-01 | 2.8451e+05 | ; | CA - HA |
| 6414 | 6416 | 1 | 1.5260e-01 | 2.5941e+05 | ; | CA - CB |
| 6414 | 6430 | 1 | 1.5220e-01 | 2.6527e+05 | ; | CA - C |

| | | | | | | | | |
|---|---|---|---|---|---|---|---|---|
| 6416 | 6417 | 1 | 1.0900e-01 | 2.8451e+05 | ; | CB | - | HB2 |
| 6416 | 6418 | 1 | 1.0900e-01 | 2.8451e+05 | ; | CB | - | HB3 |
| 6416 | 6419 | 1 | 1.5100e-01 | 2.6527e+05 | ; | CB | - | CG |
| 6419 | 6420 | 1 | 1.4000e-01 | 3.9246e+05 | ; | CG | - | CD1 |
| 6419 | 6428 | 1 | 1.4000e-01 | 3.9246e+05 | ; | CG | - | CD2 |
| 6420 | 6421 | 1 | 1.0800e-01 | 3.0711e+05 | ; | CD1 | - | HD1 |
| 6420 | 6422 | 1 | 1.4000e-01 | 3.9246e+05 | ; | CD1 | - | CE1 |
| 6422 | 6423 | 1 | 1.0800e-01 | 3.0711e+05 | ; | CE1 | - | HE1 |
| 6422 | 6424 | 1 | 1.4000e-01 | 3.9246e+05 | ; | CE1 | - | CZ |
| 6424 | 6425 | 1 | 1.0800e-01 | 3.0711e+05 | ; | CZ | - | HZ |
| 6424 | 6426 | 1 | 1.4000e-01 | 3.9246e+05 | ; | CZ | - | CE2 |
| 6426 | 6427 | 1 | 1.0800e-01 | 3.0711e+05 | ; | CE2 | - | HE2 |
| 6426 | 6428 | 1 | 1.4000e-01 | 3.9246e+05 | ; | CE2 | - | CD2 |
| 6428 | 6429 | 1 | 1.0800e-01 | 3.0711e+05 | ; | CD2 | - | HD2 |
| 6430 | 6431 | 1 | 1.2290e-01 | 4.7698e+05 | ; | C | - | O |
| 6430 | 6432 | 1 | 1.3350e-01 | 4.1003e+05 | ; | C | - | N |
| 6432 | 6433 | 1 | 1.0100e-01 | 3.6317e+05 | ; | N | - | H |
| 6432 | 6434 | 1 | 1.4490e-01 | 2.8200e+05 | ; | N | - | CA |
| 6434 | 6435 | 1 | 1.0900e-01 | 2.8451e+05 | ; | CA | - | HA |
| 6434 | 6436 | 1 | 1.5260e-01 | 2.5941e+05 | ; | CA | - | CB |
| 6434 | 6442 | 1 | 1.5220e-01 | 2.6527e+05 | ; | CA | - | C |
| 6436 | 6437 | 1 | 1.0900e-01 | 2.8451e+05 | ; | CB | - | HB2 |
| 6436 | 6438 | 1 | 1.0900e-01 | 2.8451e+05 | ; | CB | - | HB3 |
| 6436 | 6439 | 1 | 1.5220e-01 | 2.6527e+05 | ; | CB | - | CG |
| 6439 | 6440 | 1 | 1.2500e-01 | 5.4894e+05 | ; | CG | - | OD1 |
| 6439 | 6441 | 1 | 1.2500e-01 | 5.4894e+05 | ; | CG | - | OD2 |
| 6442 | 6443 | 1 | 1.2290e-01 | 4.7698e+05 | ; | C | - | O |
| 6442 | 6444 | 1 | 1.3350e-01 | 4.1003e+05 | ; | C | - | N |
| 6444 | 6445 | 1 | 1.0100e-01 | 3.6317e+05 | ; | N | - | H |
| 6444 | 6446 | 1 | 1.4490e-01 | 2.8200e+05 | ; | N | - | CA |
| 6446 | 6447 | 1 | 1.0900e-01 | 2.8451e+05 | ; | CA | - | HA |
| 6446 | 6448 | 1 | 1.5260e-01 | 2.5941e+05 | ; | CA | - | CB |
| 6446 | 6461 | 1 | 1.5220e-01 | 2.6527e+05 | ; | CA | - | C |
| 6448 | 6449 | 1 | 1.0900e-01 | 2.8451e+05 | ; | CB | - | HB |
| 6448 | 6450 | 1 | 1.5260e-01 | 2.5941e+05 | ; | CB | - | CG2 |
| 6448 | 6454 | 1 | 1.5260e-01 | 2.5941e+05 | ; | CB | - | CG1 |
| 6450 | 6451 | 1 | 1.0900e-01 | 2.8451e+05 | ; | CG2 | - | HG21 |
| 6450 | 6452 | 1 | 1.0900e-01 | 2.8451e+05 | ; | CG2 | - | HG22 |
| 6450 | 6453 | 1 | 1.0900e-01 | 2.8451e+05 | ; | CG2 | - | HG23 |
| 6454 | 6455 | 1 | 1.0900e-01 | 2.8451e+05 | ; | CG1 | - | HG12 |
| 6454 | 6456 | 1 | 1.0900e-01 | 2.8451e+05 | ; | CG1 | - | HG13 |
| 6454 | 6457 | 1 | 1.5260e-01 | 2.5941e+05 | ; | CG1 | - | CD1 |
| 6457 | 6458 | 1 | 1.0900e-01 | 2.8451e+05 | ; | CD1 | - | HD11 |
| 6457 | 6459 | 1 | 1.0900e-01 | 2.8451e+05 | ; | CD1 | - | HD12 |
| 6457 | 6460 | 1 | 1.0900e-01 | 2.8451e+05 | ; | CD1 | - | HD13 |
| 6461 | 6462 | 1 | 1.2290e-01 | 4.7698e+05 | ; | C | - | O |
| 6461 | 6463 | 1 | 1.3350e-01 | 4.1003e+05 | ; | C | - | N |
| 6463 | 6464 | 1 | 1.0100e-01 | 3.6317e+05 | ; | N | - | H |
| 6463 | 6465 | 1 | 1.4490e-01 | 2.8200e+05 | ; | N | - | CA |
| 6465 | 6466 | 1 | 1.0900e-01 | 2.8451e+05 | ; | CA | - | HA |
| 6465 | 6467 | 1 | 1.5260e-01 | 2.5941e+05 | ; | CA | - | CB |
| 6465 | 6473 | 1 | 1.5220e-01 | 2.6527e+05 | ; | CA | - | C |

| | | | | | | | | |
|---|---|---|---|---|---|---|---|---|
| 6467 | 6468 | 1 | 1.0900e-01 | 2.8451e+05 | ; | CB | - | HB2 |
| 6467 | 6469 | 1 | 1.0900e-01 | 2.8451e+05 | ; | CB | - | HB3 |
| 6467 | 6470 | 1 | 1.5220e-01 | 2.6527e+05 | ; | CB | - | CG |
| 6470 | 6471 | 1 | 1.2500e-01 | 5.4894e+05 | ; | CG | - | OD1 |
| 6470 | 6472 | 1 | 1.2500e-01 | 5.4894e+05 | ; | CG | - | OD2 |
| 6473 | 6474 | 1 | 1.2290e-01 | 4.7698e+05 | ; | C | - | O |
| 6473 | 6475 | 1 | 1.3350e-01 | 4.1003e+05 | ; | C | - | N |
| 6475 | 6476 | 1 | 1.0100e-01 | 3.6317e+05 | ; | N | - | H |
| 6475 | 6477 | 1 | 1.4490e-01 | 2.8200e+05 | ; | N | - | CA |
| 6477 | 6478 | 1 | 1.0900e-01 | 2.8451e+05 | ; | CA | - | HA2 |
| 6477 | 6479 | 1 | 1.0900e-01 | 2.8451e+05 | ; | CA | - | HA3 |
| 6477 | 6480 | 1 | 1.5220e-01 | 2.6527e+05 | ; | CA | - | C |
| 6480 | 6481 | 1 | 1.2290e-01 | 4.7698e+05 | ; | C | - | O |
| 6480 | 6482 | 1 | 1.3350e-01 | 4.1003e+05 | ; | C | - | N |
| 6482 | 6483 | 1 | 1.0100e-01 | 3.6317e+05 | ; | N | - | H |
| 6482 | 6484 | 1 | 1.4490e-01 | 2.8200e+05 | ; | N | - | CA |
| 6484 | 6485 | 1 | 1.0900e-01 | 2.8451e+05 | ; | CA | - | HA |
| 6484 | 6486 | 1 | 1.5260e-01 | 2.5941e+05 | ; | CA | - | CB |
| 6484 | 6494 | 1 | 1.5220e-01 | 2.6527e+05 | ; | CA | - | C |
| 6486 | 6487 | 1 | 1.0900e-01 | 2.8451e+05 | ; | CB | - | HB2 |
| 6486 | 6488 | 1 | 1.0900e-01 | 2.8451e+05 | ; | CB | - | HB3 |
| 6486 | 6489 | 1 | 1.5220e-01 | 2.6527e+05 | ; | CB | - | CG |
| 6489 | 6490 | 1 | 1.2290e-01 | 4.7698e+05 | ; | CG | - | OD1 |
| 6489 | 6491 | 1 | 1.3350e-01 | 4.1003e+05 | ; | CG | - | ND2 |
| 6491 | 6492 | 1 | 1.0100e-01 | 3.6317e+05 | ; | ND2 | - | HD21 |
| 6491 | 6493 | 1 | 1.0100e-01 | 3.6317e+05 | ; | ND2 | - | HD22 |
| 6494 | 6495 | 1 | 1.2290e-01 | 4.7698e+05 | ; | C | - | O |
| 6494 | 6496 | 1 | 1.3350e-01 | 4.1003e+05 | ; | C | - | N |
| 6496 | 6497 | 1 | 1.0100e-01 | 3.6317e+05 | ; | N | - | H |
| 6496 | 6498 | 1 | 1.4490e-01 | 2.8200e+05 | ; | N | - | CA |
| 6498 | 6499 | 1 | 1.0900e-01 | 2.8451e+05 | ; | CA | - | HA2 |
| 6498 | 6500 | 1 | 1.0900e-01 | 2.8451e+05 | ; | CA | - | HA3 |
| 6498 | 6501 | 1 | 1.5220e-01 | 2.6527e+05 | ; | CA | - | C |
| 6501 | 6502 | 1 | 1.2290e-01 | 4.7698e+05 | ; | C | - | O |
| 6501 | 6503 | 1 | 1.3350e-01 | 4.1003e+05 | ; | C | - | N |
| 6503 | 6504 | 1 | 1.0100e-01 | 3.6317e+05 | ; | N | - | H |
| 6503 | 6505 | 1 | 1.4490e-01 | 2.8200e+05 | ; | N | - | CA |
| 6505 | 6506 | 1 | 1.0900e-01 | 2.8451e+05 | ; | CA | - | HA |
| 6505 | 6507 | 1 | 1.5260e-01 | 2.5941e+05 | ; | CA | - | CB |
| 6505 | 6523 | 1 | 1.5220e-01 | 2.6527e+05 | ; | CA | - | C |
| 6507 | 6508 | 1 | 1.0900e-01 | 2.8451e+05 | ; | CB | - | HB2 |
| 6507 | 6509 | 1 | 1.0900e-01 | 2.8451e+05 | ; | CB | - | HB3 |
| 6507 | 6510 | 1 | 1.5260e-01 | 2.5941e+05 | ; | CB | - | CG |
| 6510 | 6511 | 1 | 1.0900e-01 | 2.8451e+05 | ; | CG | - | HG2 |
| 6510 | 6512 | 1 | 1.0900e-01 | 2.8451e+05 | ; | CG | - | HG3 |
| 6510 | 6513 | 1 | 1.5260e-01 | 2.5941e+05 | ; | CG | - | CD |
| 6513 | 6514 | 1 | 1.0900e-01 | 2.8451e+05 | ; | CD | - | HD2 |
| 6513 | 6515 | 1 | 1.0900e-01 | 2.8451e+05 | ; | CD | - | HD3 |
| 6513 | 6516 | 1 | 1.5260e-01 | 2.5941e+05 | ; | CD | - | CE |
| 6516 | 6517 | 1 | 1.0900e-01 | 2.8451e+05 | ; | CE | - | HE2 |
| 6516 | 6518 | 1 | 1.0900e-01 | 2.8451e+05 | ; | CE | - | HE3 |
| 6516 | 6519 | 1 | 1.4710e-01 | 3.0711e+05 | ; | CE | - | NZ |

| | | | | | | |
|---|---|---|---|---|---|---|
| 6519 | 6520 | 1 | 1.0100e-01 | 3.6317e+05 | ; | NZ - HZ1 |
| 6519 | 6521 | 1 | 1.0100e-01 | 3.6317e+05 | ; | NZ - HZ2 |
| 6519 | 6522 | 1 | 1.0100e-01 | 3.6317e+05 | ; | NZ - HZ3 |
| 6523 | 6524 | 1 | 1.2290e-01 | 4.7698e+05 | ; | C - O |
| 6523 | 6525 | 1 | 1.3350e-01 | 4.1003e+05 | ; | C - N |
| 6525 | 6526 | 1 | 1.0100e-01 | 3.6317e+05 | ; | N - H |
| 6525 | 6527 | 1 | 1.4490e-01 | 2.8200e+05 | ; | N - CA |
| 6527 | 6528 | 1 | 1.0900e-01 | 2.8451e+05 | ; | CA - HA |
| 6527 | 6529 | 1 | 1.5260e-01 | 2.5941e+05 | ; | CA - CB |
| 6527 | 6542 | 1 | 1.5220e-01 | 2.6527e+05 | ; | CA - C |
| 6529 | 6530 | 1 | 1.0900e-01 | 2.8451e+05 | ; | CB - HB |
| 6529 | 6531 | 1 | 1.5260e-01 | 2.5941e+05 | ; | CB - CG2 |
| 6529 | 6535 | 1 | 1.5260e-01 | 2.5941e+05 | ; | CB - CG1 |
| 6531 | 6532 | 1 | 1.0900e-01 | 2.8451e+05 | ; | CG2 - HG21 |
| 6531 | 6533 | 1 | 1.0900e-01 | 2.8451e+05 | ; | CG2 - HG22 |
| 6531 | 6534 | 1 | 1.0900e-01 | 2.8451e+05 | ; | CG2 - HG23 |
| 6535 | 6536 | 1 | 1.0900e-01 | 2.8451e+05 | ; | CG1 - HG12 |
| 6535 | 6537 | 1 | 1.0900e-01 | 2.8451e+05 | ; | CG1 - HG13 |
| 6535 | 6538 | 1 | 1.5260e-01 | 2.5941e+05 | ; | CG1 - CD1 |
| 6538 | 6539 | 1 | 1.0900e-01 | 2.8451e+05 | ; | CD1 - HD11 |
| 6538 | 6540 | 1 | 1.0900e-01 | 2.8451e+05 | ; | CD1 - HD12 |
| 6538 | 6541 | 1 | 1.0900e-01 | 2.8451e+05 | ; | CD1 - HD13 |
| 6542 | 6543 | 1 | 1.2290e-01 | 4.7698e+05 | ; | C - O |
| 6542 | 6544 | 1 | 1.3350e-01 | 4.1003e+05 | ; | C - N |
| 6544 | 6545 | 1 | 1.0100e-01 | 3.6317e+05 | ; | N - H |
| 6544 | 6546 | 1 | 1.4490e-01 | 2.8200e+05 | ; | N - CA |
| 6546 | 6547 | 1 | 1.0900e-01 | 2.8451e+05 | ; | CA - HA |
| 6546 | 6548 | 1 | 1.5260e-01 | 2.5941e+05 | ; | CA - CB |
| 6546 | 6553 | 1 | 1.5220e-01 | 2.6527e+05 | ; | CA - C |
| 6548 | 6549 | 1 | 1.0900e-01 | 2.8451e+05 | ; | CB - HB2 |
| 6548 | 6550 | 1 | 1.0900e-01 | 2.8451e+05 | ; | CB - HB3 |
| 6548 | 6551 | 1 | 1.4100e-01 | 2.6778e+05 | ; | CB - OG |
| 6551 | 6552 | 1 | 9.6000e-02 | 4.6275e+05 | ; | OG - HG |
| 6553 | 6554 | 1 | 1.2290e-01 | 4.7698e+05 | ; | C - O |
| 6553 | 6555 | 1 | 1.3350e-01 | 4.1003e+05 | ; | C - N |
| 6555 | 6556 | 1 | 1.0100e-01 | 3.6317e+05 | ; | N - H |
| 6555 | 6557 | 1 | 1.4490e-01 | 2.8200e+05 | ; | N - CA |
| 6557 | 6558 | 1 | 1.0900e-01 | 2.8451e+05 | ; | CA - HA |
| 6557 | 6559 | 1 | 1.5260e-01 | 2.5941e+05 | ; | CA - CB |
| 6557 | 6569 | 1 | 1.5220e-01 | 2.6527e+05 | ; | CA - C |
| 6559 | 6560 | 1 | 1.0900e-01 | 2.8451e+05 | ; | CB - HB |
| 6559 | 6561 | 1 | 1.5260e-01 | 2.5941e+05 | ; | CB - CG1 |
| 6559 | 6565 | 1 | 1.5260e-01 | 2.5941e+05 | ; | CB - CG2 |
| 6561 | 6562 | 1 | 1.0900e-01 | 2.8451e+05 | ; | CG1 - HG11 |
| 6561 | 6563 | 1 | 1.0900e-01 | 2.8451e+05 | ; | CG1 - HG12 |
| 6561 | 6564 | 1 | 1.0900e-01 | 2.8451e+05 | ; | CG1 - HG13 |
| 6565 | 6566 | 1 | 1.0900e-01 | 2.8451e+05 | ; | CG2 - HG21 |
| 6565 | 6567 | 1 | 1.0900e-01 | 2.8451e+05 | ; | CG2 - HG22 |
| 6565 | 6568 | 1 | 1.0900e-01 | 2.8451e+05 | ; | CG2 - HG23 |
| 6569 | 6570 | 1 | 1.2290e-01 | 4.7698e+05 | ; | C - O |
| 6569 | 6571 | 1 | 1.3350e-01 | 4.1003e+05 | ; | C - N |
| 6571 | 6572 | 1 | 1.0100e-01 | 3.6317e+05 | ; | N - H |

| | | | | | | |
|---|---|---|---|---|---|---|
| 6571 | 6573 | 1 | 1.4490e-01 | 2.8200e+05 | ; | N - CA |
| 6573 | 6574 | 1 | 1.0900e-01 | 2.8451e+05 | ; | CA - HA |
| 6573 | 6575 | 1 | 1.5260e-01 | 2.5941e+05 | ; | CA - CB |
| 6573 | 6584 | 1 | 1.5220e-01 | 2.6527e+05 | ; | CA - C |
| 6575 | 6576 | 1 | 1.0900e-01 | 2.8451e+05 | ; | CB - HB2 |
| 6575 | 6577 | 1 | 1.0900e-01 | 2.8451e+05 | ; | CB - HB3 |
| 6575 | 6578 | 1 | 1.5260e-01 | 2.5941e+05 | ; | CB - CG |
| 6578 | 6579 | 1 | 1.0900e-01 | 2.8451e+05 | ; | CG - HG2 |
| 6578 | 6580 | 1 | 1.0900e-01 | 2.8451e+05 | ; | CG - HG3 |
| 6578 | 6581 | 1 | 1.5220e-01 | 2.6527e+05 | ; | CG - CD |
| 6581 | 6582 | 1 | 1.2500e-01 | 5.4894e+05 | ; | CD - OE1 |
| 6581 | 6583 | 1 | 1.2500e-01 | 5.4894e+05 | ; | CD - OE2 |
| 6584 | 6585 | 1 | 1.2290e-01 | 4.7698e+05 | ; | C - O |
| 6584 | 6586 | 1 | 1.3350e-01 | 4.1003e+05 | ; | C - N |
| 6586 | 6587 | 1 | 1.0100e-01 | 3.6317e+05 | ; | N - H |
| 6586 | 6588 | 1 | 1.4490e-01 | 2.8200e+05 | ; | N - CA |
| 6588 | 6589 | 1 | 1.0900e-01 | 2.8451e+05 | ; | CA - HA |
| 6588 | 6590 | 1 | 1.5260e-01 | 2.5941e+05 | ; | CA - CB |
| 6588 | 6600 | 1 | 1.5220e-01 | 2.6527e+05 | ; | CA - C |
| 6590 | 6591 | 1 | 1.0900e-01 | 2.8451e+05 | ; | CB - HB2 |
| 6590 | 6592 | 1 | 1.0900e-01 | 2.8451e+05 | ; | CB - HB3 |
| 6590 | 6593 | 1 | 1.5260e-01 | 2.5941e+05 | ; | CB - CG |
| 6593 | 6594 | 1 | 1.0900e-01 | 2.8451e+05 | ; | CG - HG2 |
| 6593 | 6595 | 1 | 1.0900e-01 | 2.8451e+05 | ; | CG - HG3 |
| 6593 | 6596 | 1 | 1.5220e-01 | 2.6527e+05 | ; | CG - CD |
| 6596 | 6597 | 1 | 1.2290e-01 | 4.7698e+05 | ; | CD - OE1 |
| 6596 | 6598 | 1 | 1.3640e-01 | 3.7656e+05 | ; | CD - OE2 |
| 6598 | 6599 | 1 | 9.6000e-02 | 4.6275e+05 | ; | OE2 - HE2 |
| 6600 | 6601 | 1 | 1.2290e-01 | 4.7698e+05 | ; | C - O |
| 6600 | 6602 | 1 | 1.3350e-01 | 4.1003e+05 | ; | C - N |
| 6602 | 6603 | 1 | 1.0100e-01 | 3.6317e+05 | ; | N - H |
| 6602 | 6604 | 1 | 1.4490e-01 | 2.8200e+05 | ; | N - CA |
| 6604 | 6605 | 1 | 1.0900e-01 | 2.8451e+05 | ; | CA - HA |
| 6604 | 6606 | 1 | 1.5260e-01 | 2.5941e+05 | ; | CA - CB |
| 6604 | 6619 | 1 | 1.5220e-01 | 2.6527e+05 | ; | CA - C |
| 6606 | 6607 | 1 | 1.0900e-01 | 2.8451e+05 | ; | CB - HB2 |
| 6606 | 6608 | 1 | 1.0900e-01 | 2.8451e+05 | ; | CB - HB3 |
| 6606 | 6609 | 1 | 1.5260e-01 | 2.5941e+05 | ; | CB - CG |
| 6609 | 6610 | 1 | 1.0900e-01 | 2.8451e+05 | ; | CG - HG |
| 6609 | 6611 | 1 | 1.5260e-01 | 2.5941e+05 | ; | CG - CD1 |
| 6609 | 6615 | 1 | 1.5260e-01 | 2.5941e+05 | ; | CG - CD2 |
| 6611 | 6612 | 1 | 1.0900e-01 | 2.8451e+05 | ; | CD1 - HD11 |
| 6611 | 6613 | 1 | 1.0900e-01 | 2.8451e+05 | ; | CD1 - HD12 |
| 6611 | 6614 | 1 | 1.0900e-01 | 2.8451e+05 | ; | CD1 - HD13 |
| 6615 | 6616 | 1 | 1.0900e-01 | 2.8451e+05 | ; | CD2 - HD21 |
| 6615 | 6617 | 1 | 1.0900e-01 | 2.8451e+05 | ; | CD2 - HD22 |
| 6615 | 6618 | 1 | 1.0900e-01 | 2.8451e+05 | ; | CD2 - HD23 |
| 6619 | 6620 | 1 | 1.2290e-01 | 4.7698e+05 | ; | C - O |
| 6619 | 6621 | 1 | 1.3350e-01 | 4.1003e+05 | ; | C - N |
| 6621 | 6622 | 1 | 1.0100e-01 | 3.6317e+05 | ; | N - H |
| 6621 | 6623 | 1 | 1.4490e-01 | 2.8200e+05 | ; | N - CA |
| 6623 | 6624 | 1 | 1.0900e-01 | 2.8451e+05 | ; | CA - HA |

| | | | | | | |
|---|---|---|---|---|---|---|
| 6623 | 6625 | 1 | 1.5260e-01 | 2.5941e+05 | ; | CA - CB |
| 6623 | 6641 | 1 | 1.5220e-01 | 2.6527e+05 | ; | CA - C |
| 6625 | 6626 | 1 | 1.0900e-01 | 2.8451e+05 | ; | CB - HB2 |
| 6625 | 6627 | 1 | 1.0900e-01 | 2.8451e+05 | ; | CB - HB3 |
| 6625 | 6628 | 1 | 1.5260e-01 | 2.5941e+05 | ; | CB - CG |
| 6628 | 6629 | 1 | 1.0900e-01 | 2.8451e+05 | ; | CG - HG2 |
| 6628 | 6630 | 1 | 1.0900e-01 | 2.8451e+05 | ; | CG - HG3 |
| 6628 | 6631 | 1 | 1.5260e-01 | 2.5941e+05 | ; | CG - CD |
| 6631 | 6632 | 1 | 1.0900e-01 | 2.8451e+05 | ; | CD - HD2 |
| 6631 | 6633 | 1 | 1.0900e-01 | 2.8451e+05 | ; | CD - HD3 |
| 6631 | 6634 | 1 | 1.5260e-01 | 2.5941e+05 | ; | CD - CE |
| 6634 | 6635 | 1 | 1.0900e-01 | 2.8451e+05 | ; | CE - HE2 |
| 6634 | 6636 | 1 | 1.0900e-01 | 2.8451e+05 | ; | CE - HE3 |
| 6634 | 6637 | 1 | 1.4710e-01 | 3.0711e+05 | ; | CE - NZ |
| 6637 | 6638 | 1 | 1.0100e-01 | 3.6317e+05 | ; | NZ - HZ1 |
| 6637 | 6639 | 1 | 1.0100e-01 | 3.6317e+05 | ; | NZ - HZ2 |
| 6637 | 6640 | 1 | 1.0100e-01 | 3.6317e+05 | ; | NZ - HZ3 |
| 6641 | 6642 | 1 | 1.2290e-01 | 4.7698e+05 | ; | C - O |
| 6641 | 6643 | 1 | 1.3350e-01 | 4.1003e+05 | ; | C - N |
| 6643 | 6644 | 1 | 1.0100e-01 | 3.6317e+05 | ; | N - H |
| 6643 | 6645 | 1 | 1.4490e-01 | 2.8200e+05 | ; | N - CA |
| 6645 | 6646 | 1 | 1.0900e-01 | 2.8451e+05 | ; | CA - HA |
| 6645 | 6647 | 1 | 1.5260e-01 | 2.5941e+05 | ; | CA - CB |
| 6645 | 6665 | 1 | 1.5220e-01 | 2.6527e+05 | ; | CA - C |
| 6647 | 6648 | 1 | 1.0900e-01 | 2.8451e+05 | ; | CB - HB2 |
| 6647 | 6649 | 1 | 1.0900e-01 | 2.8451e+05 | ; | CB - HB3 |
| 6647 | 6650 | 1 | 1.5260e-01 | 2.5941e+05 | ; | CB - CG |
| 6650 | 6651 | 1 | 1.0900e-01 | 2.8451e+05 | ; | CG - HG2 |
| 6650 | 6652 | 1 | 1.0900e-01 | 2.8451e+05 | ; | CG - HG3 |
| 6650 | 6653 | 1 | 1.5260e-01 | 2.5941e+05 | ; | CG - CD |
| 6653 | 6654 | 1 | 1.0900e-01 | 2.8451e+05 | ; | CD - HD2 |
| 6653 | 6655 | 1 | 1.0900e-01 | 2.8451e+05 | ; | CD - HD3 |
| 6653 | 6656 | 1 | 1.4630e-01 | 2.8200e+05 | ; | CD - NE |
| 6656 | 6657 | 1 | 1.0100e-01 | 3.6317e+05 | ; | NE - HE |
| 6656 | 6658 | 1 | 1.3400e-01 | 4.0250e+05 | ; | NE - CZ |
| 6658 | 6659 | 1 | 1.3400e-01 | 4.0250e+05 | ; | CZ - NH1 |
| 6658 | 6662 | 1 | 1.3400e-01 | 4.0250e+05 | ; | CZ - NH2 |
| 6659 | 6660 | 1 | 1.0100e-01 | 3.6317e+05 | ; | NH1 - HH11 |
| 6659 | 6661 | 1 | 1.0100e-01 | 3.6317e+05 | ; | NH1 - HH12 |
| 6662 | 6663 | 1 | 1.0100e-01 | 3.6317e+05 | ; | NH2 - HH21 |
| 6662 | 6664 | 1 | 1.0100e-01 | 3.6317e+05 | ; | NH2 - HH22 |
| 6665 | 6666 | 1 | 1.2290e-01 | 4.7698e+05 | ; | C - O |
| 6665 | 6667 | 1 | 1.3350e-01 | 4.1003e+05 | ; | C - N |
| 6667 | 6668 | 1 | 1.0100e-01 | 3.6317e+05 | ; | N - H |
| 6667 | 6669 | 1 | 1.4490e-01 | 2.8200e+05 | ; | N - CA |
| 6669 | 6670 | 1 | 1.0900e-01 | 2.8451e+05 | ; | CA - HA |
| 6669 | 6671 | 1 | 1.5260e-01 | 2.5941e+05 | ; | CA - CB |
| 6669 | 6684 | 1 | 1.5220e-01 | 2.6527e+05 | ; | CA - C |
| 6671 | 6672 | 1 | 1.0900e-01 | 2.8451e+05 | ; | CB - HB |
| 6671 | 6673 | 1 | 1.5260e-01 | 2.5941e+05 | ; | CB - CG2 |
| 6671 | 6677 | 1 | 1.5260e-01 | 2.5941e+05 | ; | CB - CG1 |
| 6673 | 6674 | 1 | 1.0900e-01 | 2.8451e+05 | ; | CG2 - HG21 |

| | | | | | | | | |
|---|---|---|---|---|---|---|---|---|
| 6673 | 6675 | 1 | 1.0900e-01 | 2.8451e+05 | ; | CG2 | - | HG22 |
| 6673 | 6676 | 1 | 1.0900e-01 | 2.8451e+05 | ; | CG2 | - | HG23 |
| 6677 | 6678 | 1 | 1.0900e-01 | 2.8451e+05 | ; | CG1 | - | HG12 |
| 6677 | 6679 | 1 | 1.0900e-01 | 2.8451e+05 | ; | CG1 | - | HG13 |
| 6677 | 6680 | 1 | 1.5260e-01 | 2.5941e+05 | ; | CG1 | - | CD1 |
| 6680 | 6681 | 1 | 1.0900e-01 | 2.8451e+05 | ; | CD1 | - | HD11 |
| 6680 | 6682 | 1 | 1.0900e-01 | 2.8451e+05 | ; | CD1 | - | HD12 |
| 6680 | 6683 | 1 | 1.0900e-01 | 2.8451e+05 | ; | CD1 | - | HD13 |
| 6684 | 6685 | 1 | 1.2290e-01 | 4.7698e+05 | ; | C | - | O |
| 6684 | 6686 | 1 | 1.3350e-01 | 4.1003e+05 | ; | C | - | N |
| 6686 | 6687 | 1 | 1.0100e-01 | 3.6317e+05 | ; | N | - | H |
| 6686 | 6688 | 1 | 1.4490e-01 | 2.8200e+05 | ; | N | - | CA |
| 6688 | 6689 | 1 | 1.0900e-01 | 2.8451e+05 | ; | CA | - | HA |
| 6688 | 6690 | 1 | 1.5260e-01 | 2.5941e+05 | ; | CA | - | CB |
| 6688 | 6704 | 1 | 1.5220e-01 | 2.6527e+05 | ; | CA | - | C |
| 6690 | 6691 | 1 | 1.0900e-01 | 2.8451e+05 | ; | CB | - | HB2 |
| 6690 | 6692 | 1 | 1.0900e-01 | 2.8451e+05 | ; | CB | - | HB3 |
| 6690 | 6693 | 1 | 1.5100e-01 | 2.6527e+05 | ; | CB | - | CG |
| 6693 | 6694 | 1 | 1.4000e-01 | 3.9246e+05 | ; | CG | - | CD1 |
| 6693 | 6702 | 1 | 1.4000e-01 | 3.9246e+05 | ; | CG | - | CD2 |
| 6694 | 6695 | 1 | 1.0800e-01 | 3.0711e+05 | ; | CD1 | - | HD1 |
| 6694 | 6696 | 1 | 1.4000e-01 | 3.9246e+05 | ; | CD1 | - | CE1 |
| 6696 | 6697 | 1 | 1.0800e-01 | 3.0711e+05 | ; | CE1 | - | HE1 |
| 6696 | 6698 | 1 | 1.4000e-01 | 3.9246e+05 | ; | CE1 | - | CZ |
| 6698 | 6699 | 1 | 1.0800e-01 | 3.0711e+05 | ; | CZ | - | HZ |
| 6698 | 6700 | 1 | 1.4000e-01 | 3.9246e+05 | ; | CZ | - | CE2 |
| 6700 | 6701 | 1 | 1.0800e-01 | 3.0711e+05 | ; | CE2 | - | HE2 |
| 6700 | 6702 | 1 | 1.4000e-01 | 3.9246e+05 | ; | CE2 | - | CD2 |
| 6702 | 6703 | 1 | 1.0800e-01 | 3.0711e+05 | ; | CD2 | - | HD2 |
| 6704 | 6705 | 1 | 1.2290e-01 | 4.7698e+05 | ; | C | - | O |
| 6704 | 6706 | 1 | 1.3350e-01 | 4.1003e+05 | ; | C | - | N |
| 6706 | 6707 | 1 | 1.0100e-01 | 3.6317e+05 | ; | N | - | H |
| 6706 | 6708 | 1 | 1.4490e-01 | 2.8200e+05 | ; | N | - | CA |
| 6708 | 6709 | 1 | 1.0900e-01 | 2.8451e+05 | ; | CA | - | HA2 |
| 6708 | 6710 | 1 | 1.0900e-01 | 2.8451e+05 | ; | CA | - | HA3 |
| 6708 | 6711 | 1 | 1.5220e-01 | 2.6527e+05 | ; | CA | - | C |
| 6711 | 6712 | 1 | 1.2290e-01 | 4.7698e+05 | ; | C | - | O |
| 6711 | 6713 | 1 | 1.3350e-01 | 4.1003e+05 | ; | C | - | N |
| 6713 | 6714 | 1 | 1.0100e-01 | 3.6317e+05 | ; | N | - | H |
| 6713 | 6715 | 1 | 1.4490e-01 | 2.8200e+05 | ; | N | - | CA |
| 6715 | 6716 | 1 | 1.0900e-01 | 2.8451e+05 | ; | CA | - | HA |
| 6715 | 6717 | 1 | 1.5260e-01 | 2.5941e+05 | ; | CA | - | CB |
| 6715 | 6735 | 1 | 1.5220e-01 | 2.6527e+05 | ; | CA | - | C |
| 6717 | 6718 | 1 | 1.0900e-01 | 2.8451e+05 | ; | CB | - | HB2 |
| 6717 | 6719 | 1 | 1.0900e-01 | 2.8451e+05 | ; | CB | - | HB3 |
| 6717 | 6720 | 1 | 1.5260e-01 | 2.5941e+05 | ; | CB | - | CG |
| 6720 | 6721 | 1 | 1.0900e-01 | 2.8451e+05 | ; | CG | - | HG2 |
| 6720 | 6722 | 1 | 1.0900e-01 | 2.8451e+05 | ; | CG | - | HG3 |
| 6720 | 6723 | 1 | 1.5260e-01 | 2.5941e+05 | ; | CG | - | CD |
| 6723 | 6724 | 1 | 1.0900e-01 | 2.8451e+05 | ; | CD | - | HD2 |
| 6723 | 6725 | 1 | 1.0900e-01 | 2.8451e+05 | ; | CD | - | HD3 |
| 6723 | 6726 | 1 | 1.4630e-01 | 2.8200e+05 | ; | CD | - | NE |

| | | | | | | | | |
|---|---|---|---|---|---|---|---|---|
| 6726 | 6727 | 1 | 1.0100e-01 | 3.6317e+05 | ; | NE | - | HE |
| 6726 | 6728 | 1 | 1.3400e-01 | 4.0250e+05 | ; | NE | - | CZ |
| 6728 | 6729 | 1 | 1.3400e-01 | 4.0250e+05 | ; | CZ | - | NH1 |
| 6728 | 6732 | 1 | 1.3400e-01 | 4.0250e+05 | ; | CZ | - | NH2 |
| 6729 | 6730 | 1 | 1.0100e-01 | 3.6317e+05 | ; | NH1 | - | HH11 |
| 6729 | 6731 | 1 | 1.0100e-01 | 3.6317e+05 | ; | NH1 | - | HH12 |
| 6732 | 6733 | 1 | 1.0100e-01 | 3.6317e+05 | ; | NH2 | - | HH21 |
| 6732 | 6734 | 1 | 1.0100e-01 | 3.6317e+05 | ; | NH2 | - | HH22 |
| 6735 | 6736 | 1 | 1.2290e-01 | 4.7698e+05 | ; | C | - | O |
| 6735 | 6737 | 1 | 1.3350e-01 | 4.1003e+05 | ; | C | - | N |
| 6737 | 6738 | 1 | 1.0100e-01 | 3.6317e+05 | ; | N | - | H |
| 6737 | 6739 | 1 | 1.4490e-01 | 2.8200e+05 | ; | N | - | CA |
| 6739 | 6740 | 1 | 1.0900e-01 | 2.8451e+05 | ; | CA | - | HA |
| 6739 | 6741 | 1 | 1.5260e-01 | 2.5941e+05 | ; | CA | - | CB |
| 6739 | 6747 | 1 | 1.5220e-01 | 2.6527e+05 | ; | CA | - | C |
| 6741 | 6742 | 1 | 1.0900e-01 | 2.8451e+05 | ; | CB | - | HB2 |
| 6741 | 6743 | 1 | 1.0900e-01 | 2.8451e+05 | ; | CB | - | HB3 |
| 6741 | 6744 | 1 | 1.5220e-01 | 2.6527e+05 | ; | CB | - | CG |
| 6744 | 6745 | 1 | 1.2500e-01 | 5.4894e+05 | ; | CG | - | OD1 |
| 6744 | 6746 | 1 | 1.2500e-01 | 5.4894e+05 | ; | CG | - | OD2 |
| 6747 | 6748 | 1 | 1.2290e-01 | 4.7698e+05 | ; | C | - | O |
| 6747 | 6749 | 1 | 1.3350e-01 | 4.1003e+05 | ; | C | - | N |
| 6749 | 6750 | 1 | 1.0100e-01 | 3.6317e+05 | ; | N | - | H |
| 6749 | 6751 | 1 | 1.4490e-01 | 2.8200e+05 | ; | N | - | CA |
| 6751 | 6752 | 1 | 1.0900e-01 | 2.8451e+05 | ; | CA | - | HA |
| 6751 | 6753 | 1 | 1.5260e-01 | 2.5941e+05 | ; | CA | - | CB |
| 6751 | 6759 | 1 | 1.5220e-01 | 2.6527e+05 | ; | CA | - | C |
| 6753 | 6754 | 1 | 1.0900e-01 | 2.8451e+05 | ; | CB | - | HB2 |
| 6753 | 6755 | 1 | 1.0900e-01 | 2.8451e+05 | ; | CB | - | HB3 |
| 6753 | 6756 | 1 | 1.5220e-01 | 2.6527e+05 | ; | CB | - | CG |
| 6756 | 6757 | 1 | 1.2500e-01 | 5.4894e+05 | ; | CG | - | OD1 |
| 6756 | 6758 | 1 | 1.2500e-01 | 5.4894e+05 | ; | CG | - | OD2 |
| 6759 | 6760 | 1 | 1.2290e-01 | 4.7698e+05 | ; | C | - | O |
| 6759 | 6761 | 1 | 1.3350e-01 | 4.1003e+05 | ; | C | - | N |
| 6761 | 6762 | 1 | 1.0100e-01 | 3.6317e+05 | ; | N | - | H |
| 6761 | 6763 | 1 | 1.4490e-01 | 2.8200e+05 | ; | N | - | CA |
| 6763 | 6764 | 1 | 1.0900e-01 | 2.8451e+05 | ; | CA | - | HA |
| 6763 | 6765 | 1 | 1.5260e-01 | 2.5941e+05 | ; | CA | - | CB |
| 6763 | 6778 | 1 | 1.5220e-01 | 2.6527e+05 | ; | CA | - | C |
| 6765 | 6766 | 1 | 1.0900e-01 | 2.8451e+05 | ; | CB | - | HB |
| 6765 | 6767 | 1 | 1.5260e-01 | 2.5941e+05 | ; | CB | - | CG2 |
| 6765 | 6771 | 1 | 1.5260e-01 | 2.5941e+05 | ; | CB | - | CG1 |
| 6767 | 6768 | 1 | 1.0900e-01 | 2.8451e+05 | ; | CG2 | - | HG21 |
| 6767 | 6769 | 1 | 1.0900e-01 | 2.8451e+05 | ; | CG2 | - | HG22 |
| 6767 | 6770 | 1 | 1.0900e-01 | 2.8451e+05 | ; | CG2 | - | HG23 |
| 6771 | 6772 | 1 | 1.0900e-01 | 2.8451e+05 | ; | CG1 | - | HG12 |
| 6771 | 6773 | 1 | 1.0900e-01 | 2.8451e+05 | ; | CG1 | - | HG13 |
| 6771 | 6774 | 1 | 1.5260e-01 | 2.5941e+05 | ; | CG1 | - | CD1 |
| 6774 | 6775 | 1 | 1.0900e-01 | 2.8451e+05 | ; | CD1 | - | HD11 |
| 6774 | 6776 | 1 | 1.0900e-01 | 2.8451e+05 | ; | CD1 | - | HD12 |
| 6774 | 6777 | 1 | 1.0900e-01 | 2.8451e+05 | ; | CD1 | - | HD13 |
| 6778 | 6779 | 1 | 1.2290e-01 | 4.7698e+05 | ; | C | - | O |

| | | | | | | | | |
|---|---|---|---|---|---|---|---|---|
| 6778 | 6780 | 1 | 1.3350e-01 | 4.1003e+05 | ; | C | - | N |
| 6780 | 6781 | 1 | 1.0100e-01 | 3.6317e+05 | ; | N | - | H |
| 6780 | 6782 | 1 | 1.4490e-01 | 2.8200e+05 | ; | N | - | CA |
| 6782 | 6783 | 1 | 1.0900e-01 | 2.8451e+05 | ; | CA | - | HA |
| 6782 | 6784 | 1 | 1.5260e-01 | 2.5941e+05 | ; | CA | - | CB |
| 6782 | 6793 | 1 | 1.5220e-01 | 2.6527e+05 | ; | CA | - | C |
| 6784 | 6785 | 1 | 1.0900e-01 | 2.8451e+05 | ; | CB | - | HB2 |
| 6784 | 6786 | 1 | 1.0900e-01 | 2.8451e+05 | ; | CB | - | HB3 |
| 6784 | 6787 | 1 | 1.5260e-01 | 2.5941e+05 | ; | CB | - | CG |
| 6787 | 6788 | 1 | 1.0900e-01 | 2.8451e+05 | ; | CG | - | HG2 |
| 6787 | 6789 | 1 | 1.0900e-01 | 2.8451e+05 | ; | CG | - | HG3 |
| 6787 | 6790 | 1 | 1.5220e-01 | 2.6527e+05 | ; | CG | - | CD |
| 6790 | 6791 | 1 | 1.2500e-01 | 5.4894e+05 | ; | CD | - | OE1 |
| 6790 | 6792 | 1 | 1.2500e-01 | 5.4894e+05 | ; | CD | - | OE2 |
| 6793 | 6794 | 1 | 1.2290e-01 | 4.7698e+05 | ; | C | - | O |
| 6793 | 6795 | 1 | 1.3350e-01 | 4.1003e+05 | ; | C | - | N |
| 6795 | 6796 | 1 | 1.0100e-01 | 3.6317e+05 | ; | N | - | H |
| 6795 | 6797 | 1 | 1.4490e-01 | 2.8200e+05 | ; | N | - | CA |
| 6797 | 6798 | 1 | 1.0900e-01 | 2.8451e+05 | ; | CA | - | HA |
| 6797 | 6799 | 1 | 1.5260e-01 | 2.5941e+05 | ; | CA | - | CB |
| 6797 | 6807 | 1 | 1.5220e-01 | 2.6527e+05 | ; | CA | - | C |
| 6799 | 6800 | 1 | 1.0900e-01 | 2.8451e+05 | ; | CB | - | HB2 |
| 6799 | 6801 | 1 | 1.0900e-01 | 2.8451e+05 | ; | CB | - | HB3 |
| 6799 | 6802 | 1 | 1.5220e-01 | 2.6527e+05 | ; | CB | - | CG |
| 6802 | 6803 | 1 | 1.2290e-01 | 4.7698e+05 | ; | CG | - | OD1 |
| 6802 | 6804 | 1 | 1.3350e-01 | 4.1003e+05 | ; | CG | - | ND2 |
| 6804 | 6805 | 1 | 1.0100e-01 | 3.6317e+05 | ; | ND2 | - | HD21 |
| 6804 | 6806 | 1 | 1.0100e-01 | 3.6317e+05 | ; | ND2 | - | HD22 |
| 6807 | 6808 | 1 | 1.2290e-01 | 4.7698e+05 | ; | C | - | O |
| 6807 | 6809 | 1 | 1.3350e-01 | 4.1003e+05 | ; | C | - | N |
| 6809 | 6810 | 1 | 1.4490e-01 | 2.8200e+05 | ; | N | - | CD |
| 6809 | 6819 | 1 | 1.4490e-01 | 2.8200e+05 | ; | N | - | CA |
| 6810 | 6811 | 1 | 1.0900e-01 | 2.8451e+05 | ; | CD | - | HD2 |
| 6810 | 6812 | 1 | 1.0900e-01 | 2.8451e+05 | ; | CD | - | HD3 |
| 6810 | 6813 | 1 | 1.5260e-01 | 2.5941e+05 | ; | CD | - | CG |
| 6813 | 6814 | 1 | 1.0900e-01 | 2.8451e+05 | ; | CG | - | HG2 |
| 6813 | 6815 | 1 | 1.0900e-01 | 2.8451e+05 | ; | CG | - | HG3 |
| 6813 | 6816 | 1 | 1.5260e-01 | 2.5941e+05 | ; | CG | - | CB |
| 6816 | 6817 | 1 | 1.0900e-01 | 2.8451e+05 | ; | CB | - | HB2 |
| 6816 | 6818 | 1 | 1.0900e-01 | 2.8451e+05 | ; | CB | - | HB3 |
| 6816 | 6819 | 1 | 1.5260e-01 | 2.5941e+05 | ; | CB | - | CA |
| 6819 | 6820 | 1 | 1.0900e-01 | 2.8451e+05 | ; | CA | - | HA |
| 6819 | 6821 | 1 | 1.5220e-01 | 2.6527e+05 | ; | CA | - | C |
| 6821 | 6822 | 1 | 1.2290e-01 | 4.7698e+05 | ; | C | - | O |
| 6821 | 6823 | 1 | 1.3350e-01 | 4.1003e+05 | ; | C | - | N |
| 6823 | 6824 | 1 | 1.0100e-01 | 3.6317e+05 | ; | N | - | H |
| 6823 | 6825 | 1 | 1.4490e-01 | 2.8200e+05 | ; | N | - | CA |
| 6825 | 6826 | 1 | 1.0900e-01 | 2.8451e+05 | ; | CA | - | HA |
| 6825 | 6827 | 1 | 1.5260e-01 | 2.5941e+05 | ; | CA | - | CB |
| 6825 | 6840 | 1 | 1.5220e-01 | 2.6527e+05 | ; | CA | - | C |
| 6827 | 6828 | 1 | 1.0900e-01 | 2.8451e+05 | ; | CB | - | HB2 |
| 6827 | 6829 | 1 | 1.0900e-01 | 2.8451e+05 | ; | CB | - | HB3 |

| | | | | | | |
|---|---|---|---|---|---|---|
| 6827 | 6830 | 1 | 1.5260e-01 | 2.5941e+05 | ; | CB - CG |
| 6830 | 6831 | 1 | 1.0900e-01 | 2.8451e+05 | ; | CG - HG |
| 6830 | 6832 | 1 | 1.5260e-01 | 2.5941e+05 | ; | CG - CD1 |
| 6830 | 6836 | 1 | 1.5260e-01 | 2.5941e+05 | ; | CG - CD2 |
| 6832 | 6833 | 1 | 1.0900e-01 | 2.8451e+05 | ; | CD1 - HD11 |
| 6832 | 6834 | 1 | 1.0900e-01 | 2.8451e+05 | ; | CD1 - HD12 |
| 6832 | 6835 | 1 | 1.0900e-01 | 2.8451e+05 | ; | CD1 - HD13 |
| 6836 | 6837 | 1 | 1.0900e-01 | 2.8451e+05 | ; | CD2 - HD21 |
| 6836 | 6838 | 1 | 1.0900e-01 | 2.8451e+05 | ; | CD2 - HD22 |
| 6836 | 6839 | 1 | 1.0900e-01 | 2.8451e+05 | ; | CD2 - HD23 |
| 6840 | 6841 | 1 | 1.2290e-01 | 4.7698e+05 | ; | C - O |
| 6840 | 6842 | 1 | 1.3350e-01 | 4.1003e+05 | ; | C - N |
| 6842 | 6843 | 1 | 1.0100e-01 | 3.6317e+05 | ; | N - H |
| 6842 | 6844 | 1 | 1.4490e-01 | 2.8200e+05 | ; | N - CA |
| 6844 | 6845 | 1 | 1.0900e-01 | 2.8451e+05 | ; | CA - HA |
| 6844 | 6846 | 1 | 1.5260e-01 | 2.5941e+05 | ; | CA - CB |
| 6844 | 6859 | 1 | 1.5220e-01 | 2.6527e+05 | ; | CA - C |
| 6846 | 6847 | 1 | 1.0900e-01 | 2.8451e+05 | ; | CB - HB |
| 6846 | 6848 | 1 | 1.5260e-01 | 2.5941e+05 | ; | CB - CG2 |
| 6846 | 6852 | 1 | 1.5260e-01 | 2.5941e+05 | ; | CB - CG1 |
| 6848 | 6849 | 1 | 1.0900e-01 | 2.8451e+05 | ; | CG2 - HG21 |
| 6848 | 6850 | 1 | 1.0900e-01 | 2.8451e+05 | ; | CG2 - HG22 |
| 6848 | 6851 | 1 | 1.0900e-01 | 2.8451e+05 | ; | CG2 - HG23 |
| 6852 | 6853 | 1 | 1.0900e-01 | 2.8451e+05 | ; | CG1 - HG12 |
| 6852 | 6854 | 1 | 1.0900e-01 | 2.8451e+05 | ; | CG1 - HG13 |
| 6852 | 6855 | 1 | 1.5260e-01 | 2.5941e+05 | ; | CG1 - CD1 |
| 6855 | 6856 | 1 | 1.0900e-01 | 2.8451e+05 | ; | CD1 - HD11 |
| 6855 | 6857 | 1 | 1.0900e-01 | 2.8451e+05 | ; | CD1 - HD12 |
| 6855 | 6858 | 1 | 1.0900e-01 | 2.8451e+05 | ; | CD1 - HD13 |
| 6859 | 6860 | 1 | 1.2290e-01 | 4.7698e+05 | ; | C - O |
| 6859 | 6861 | 1 | 1.3350e-01 | 4.1003e+05 | ; | C - N |
| 6861 | 6862 | 1 | 1.0100e-01 | 3.6317e+05 | ; | N - H |
| 6861 | 6863 | 1 | 1.4490e-01 | 2.8200e+05 | ; | N - CA |
| 6863 | 6864 | 1 | 1.0900e-01 | 2.8451e+05 | ; | CA - HA |
| 6863 | 6865 | 1 | 1.5260e-01 | 2.5941e+05 | ; | CA - CB |
| 6863 | 6871 | 1 | 1.5220e-01 | 2.6527e+05 | ; | CA - C |
| 6865 | 6866 | 1 | 1.0900e-01 | 2.8451e+05 | ; | CB - HB2 |
| 6865 | 6867 | 1 | 1.0900e-01 | 2.8451e+05 | ; | CB - HB3 |
| 6865 | 6868 | 1 | 1.5220e-01 | 2.6527e+05 | ; | CB - CG |
| 6868 | 6869 | 1 | 1.2500e-01 | 5.4894e+05 | ; | CG - OD1 |
| 6868 | 6870 | 1 | 1.2500e-01 | 5.4894e+05 | ; | CG - OD2 |
| 6871 | 6872 | 1 | 1.2290e-01 | 4.7698e+05 | ; | C - O |
| 6871 | 6873 | 1 | 1.3350e-01 | 4.1003e+05 | ; | C - N |
| 6873 | 6874 | 1 | 1.0100e-01 | 3.6317e+05 | ; | N - H |
| 6873 | 6875 | 1 | 1.4490e-01 | 2.8200e+05 | ; | N - CA |
| 6875 | 6876 | 1 | 1.0900e-01 | 2.8451e+05 | ; | CA - HA |
| 6875 | 6877 | 1 | 1.5260e-01 | 2.5941e+05 | ; | CA - CB |
| 6875 | 6893 | 1 | 1.5220e-01 | 2.6527e+05 | ; | CA - C |
| 6877 | 6878 | 1 | 1.0900e-01 | 2.8451e+05 | ; | CB - HB2 |
| 6877 | 6879 | 1 | 1.0900e-01 | 2.8451e+05 | ; | CB - HB3 |
| 6877 | 6880 | 1 | 1.5260e-01 | 2.5941e+05 | ; | CB - CG |
| 6880 | 6881 | 1 | 1.0900e-01 | 2.8451e+05 | ; | CG - HG2 |

| | | | | | | | | |
|---|---|---|---|---|---|---|---|---|
| 6880 | 6882 | 1 | 1.0900e-01 | 2.8451e+05 | ; | CG | - | HG3 |
| 6880 | 6883 | 1 | 1.5260e-01 | 2.5941e+05 | ; | CG | - | CD |
| 6883 | 6884 | 1 | 1.0900e-01 | 2.8451e+05 | ; | CD | - | HD2 |
| 6883 | 6885 | 1 | 1.0900e-01 | 2.8451e+05 | ; | CD | - | HD3 |
| 6883 | 6886 | 1 | 1.5260e-01 | 2.5941e+05 | ; | CD | - | CE |
| 6886 | 6887 | 1 | 1.0900e-01 | 2.8451e+05 | ; | CE | - | HE2 |
| 6886 | 6888 | 1 | 1.0900e-01 | 2.8451e+05 | ; | CE | - | HE3 |
| 6886 | 6889 | 1 | 1.4710e-01 | 3.0711e+05 | ; | CE | - | NZ |
| 6889 | 6890 | 1 | 1.0100e-01 | 3.6317e+05 | ; | NZ | - | HZ1 |
| 6889 | 6891 | 1 | 1.0100e-01 | 3.6317e+05 | ; | NZ | - | HZ2 |
| 6889 | 6892 | 1 | 1.0100e-01 | 3.6317e+05 | ; | NZ | - | HZ3 |
| 6893 | 6894 | 1 | 1.2290e-01 | 4.7698e+05 | ; | C | - | O |
| 6893 | 6895 | 1 | 1.3350e-01 | 4.1003e+05 | ; | C | - | N |
| 6895 | 6896 | 1 | 1.0100e-01 | 3.6317e+05 | ; | N | - | H |
| 6895 | 6897 | 1 | 1.4490e-01 | 2.8200e+05 | ; | N | - | CA |
| 6897 | 6898 | 1 | 1.0900e-01 | 2.8451e+05 | ; | CA | - | HA |
| 6897 | 6899 | 1 | 1.5260e-01 | 2.5941e+05 | ; | CA | - | CB |
| 6897 | 6903 | 1 | 1.5220e-01 | 2.6527e+05 | ; | CA | - | C |
| 6899 | 6900 | 1 | 1.0900e-01 | 2.8451e+05 | ; | CB | - | HB1 |
| 6899 | 6901 | 1 | 1.0900e-01 | 2.8451e+05 | ; | CB | - | HB2 |
| 6899 | 6902 | 1 | 1.0900e-01 | 2.8451e+05 | ; | CB | - | HB3 |
| 6903 | 6904 | 1 | 1.2290e-01 | 4.7698e+05 | ; | C | - | O |
| 6903 | 6905 | 1 | 1.3350e-01 | 4.1003e+05 | ; | C | - | N |
| 6905 | 6906 | 1 | 1.0100e-01 | 3.6317e+05 | ; | N | - | H |
| 6905 | 6907 | 1 | 1.4490e-01 | 2.8200e+05 | ; | N | - | CA |
| 6907 | 6908 | 1 | 1.0900e-01 | 2.8451e+05 | ; | CA | - | HA |
| 6907 | 6909 | 1 | 1.5260e-01 | 2.5941e+05 | ; | CA | - | CB |
| 6907 | 6922 | 1 | 1.5220e-01 | 2.6527e+05 | ; | CA | - | C |
| 6909 | 6910 | 1 | 1.0900e-01 | 2.8451e+05 | ; | CB | - | HB |
| 6909 | 6911 | 1 | 1.5260e-01 | 2.5941e+05 | ; | CB | - | CG2 |
| 6909 | 6915 | 1 | 1.5260e-01 | 2.5941e+05 | ; | CB | - | CG1 |
| 6911 | 6912 | 1 | 1.0900e-01 | 2.8451e+05 | ; | CG2 | - | HG21 |
| 6911 | 6913 | 1 | 1.0900e-01 | 2.8451e+05 | ; | CG2 | - | HG22 |
| 6911 | 6914 | 1 | 1.0900e-01 | 2.8451e+05 | ; | CG2 | - | HG23 |
| 6915 | 6916 | 1 | 1.0900e-01 | 2.8451e+05 | ; | CG1 | - | HG12 |
| 6915 | 6917 | 1 | 1.0900e-01 | 2.8451e+05 | ; | CG1 | - | HG13 |
| 6915 | 6918 | 1 | 1.5260e-01 | 2.5941e+05 | ; | CG1 | - | CD1 |
| 6918 | 6919 | 1 | 1.0900e-01 | 2.8451e+05 | ; | CD1 | - | HD11 |
| 6918 | 6920 | 1 | 1.0900e-01 | 2.8451e+05 | ; | CD1 | - | HD12 |
| 6918 | 6921 | 1 | 1.0900e-01 | 2.8451e+05 | ; | CD1 | - | HD13 |
| 6922 | 6923 | 1 | 1.2290e-01 | 4.7698e+05 | ; | C | - | O |
| 6922 | 6924 | 1 | 1.3350e-01 | 4.1003e+05 | ; | C | - | N |
| 6924 | 6925 | 1 | 1.0100e-01 | 3.6317e+05 | ; | N | - | H |
| 6924 | 6926 | 1 | 1.4490e-01 | 2.8200e+05 | ; | N | - | CA |
| 6926 | 6927 | 1 | 1.0900e-01 | 2.8451e+05 | ; | CA | - | HA |
| 6926 | 6928 | 1 | 1.5260e-01 | 2.5941e+05 | ; | CA | - | CB |
| 6926 | 6934 | 1 | 1.5220e-01 | 2.6527e+05 | ; | CA | - | C |
| 6928 | 6929 | 1 | 1.0900e-01 | 2.8451e+05 | ; | CB | - | HB2 |
| 6928 | 6930 | 1 | 1.0900e-01 | 2.8451e+05 | ; | CB | - | HB3 |
| 6928 | 6931 | 1 | 1.5220e-01 | 2.6527e+05 | ; | CB | - | CG |
| 6931 | 6932 | 1 | 1.2500e-01 | 5.4894e+05 | ; | CG | - | OD1 |
| 6931 | 6933 | 1 | 1.2500e-01 | 5.4894e+05 | ; | CG | - | OD2 |

| | | | | | |
|---|---|---|---|---|---|
| 6934 | 6935 | 1 | 1.2290e-01 | 4.7698e+05 ; | C - O |
| 6934 | 6936 | 1 | 1.3350e-01 | 4.1003e+05 ; | C - N |
| 6936 | 6937 | 1 | 1.0100e-01 | 3.6317e+05 ; | N - H |
| 6936 | 6938 | 1 | 1.4490e-01 | 2.8200e+05 ; | N - CA |
| 6938 | 6939 | 1 | 1.0900e-01 | 2.8451e+05 ; | CA - HA |
| 6938 | 6940 | 1 | 1.5260e-01 | 2.5941e+05 ; | CA - CB |
| 6938 | 6945 | 1 | 1.5220e-01 | 2.6527e+05 ; | CA - C |
| 6940 | 6941 | 1 | 1.0900e-01 | 2.8451e+05 ; | CB - HB2 |
| 6940 | 6942 | 1 | 1.0900e-01 | 2.8451e+05 ; | CB - HB3 |
| 6940 | 6943 | 1 | 1.4100e-01 | 2.6778e+05 ; | CB - OG |
| 6943 | 6944 | 1 | 9.6000e-02 | 4.6275e+05 ; | OG - HG |
| 6945 | 6946 | 1 | 1.2290e-01 | 4.7698e+05 ; | C - O |
| 6945 | 6947 | 1 | 1.3350e-01 | 4.1003e+05 ; | C - N |
| 6947 | 6948 | 1 | 1.0100e-01 | 3.6317e+05 ; | N - H |
| 6947 | 6949 | 1 | 1.4490e-01 | 2.8200e+05 ; | N - CA |
| 6949 | 6950 | 1 | 1.0900e-01 | 2.8451e+05 ; | CA - HA |
| 6949 | 6951 | 1 | 1.5260e-01 | 2.5941e+05 ; | CA - CB |
| 6949 | 6964 | 1 | 1.5220e-01 | 2.6527e+05 ; | CA - C |
| 6951 | 6952 | 1 | 1.0900e-01 | 2.8451e+05 ; | CB - HB2 |
| 6951 | 6953 | 1 | 1.0900e-01 | 2.8451e+05 ; | CB - HB3 |
| 6951 | 6954 | 1 | 1.5260e-01 | 2.5941e+05 ; | CB - CG |
| 6954 | 6955 | 1 | 1.0900e-01 | 2.8451e+05 ; | CG - HG |
| 6954 | 6956 | 1 | 1.5260e-01 | 2.5941e+05 ; | CG - CD1 |
| 6954 | 6960 | 1 | 1.5260e-01 | 2.5941e+05 ; | CG - CD2 |
| 6956 | 6957 | 1 | 1.0900e-01 | 2.8451e+05 ; | CD1 - HD11 |
| 6956 | 6958 | 1 | 1.0900e-01 | 2.8451e+05 ; | CD1 - HD12 |
| 6956 | 6959 | 1 | 1.0900e-01 | 2.8451e+05 ; | CD1 - HD13 |
| 6960 | 6961 | 1 | 1.0900e-01 | 2.8451e+05 ; | CD2 - HD21 |
| 6960 | 6962 | 1 | 1.0900e-01 | 2.8451e+05 ; | CD2 - HD22 |
| 6960 | 6963 | 1 | 1.0900e-01 | 2.8451e+05 ; | CD2 - HD23 |
| 6964 | 6965 | 1 | 1.2290e-01 | 4.7698e+05 ; | C - O |
| 6964 | 6966 | 1 | 1.3350e-01 | 4.1003e+05 ; | C - N |
| 6966 | 6967 | 1 | 1.0100e-01 | 3.6317e+05 ; | N - H |
| 6966 | 6968 | 1 | 1.4490e-01 | 2.8200e+05 ; | N - CA |
| 6968 | 6969 | 1 | 1.0900e-01 | 2.8451e+05 ; | CA - HA |
| 6968 | 6970 | 1 | 1.5260e-01 | 2.5941e+05 ; | CA - CB |
| 6968 | 6983 | 1 | 1.5220e-01 | 2.6527e+05 ; | CA - C |
| 6970 | 6971 | 1 | 1.0900e-01 | 2.8451e+05 ; | CB - HB2 |
| 6970 | 6972 | 1 | 1.0900e-01 | 2.8451e+05 ; | CB - HB3 |
| 6970 | 6973 | 1 | 1.5260e-01 | 2.5941e+05 ; | CB - CG |
| 6973 | 6974 | 1 | 1.0900e-01 | 2.8451e+05 ; | CG - HG |
| 6973 | 6975 | 1 | 1.5260e-01 | 2.5941e+05 ; | CG - CD1 |
| 6973 | 6979 | 1 | 1.5260e-01 | 2.5941e+05 ; | CG - CD2 |
| 6975 | 6976 | 1 | 1.0900e-01 | 2.8451e+05 ; | CD1 - HD11 |
| 6975 | 6977 | 1 | 1.0900e-01 | 2.8451e+05 ; | CD1 - HD12 |
| 6975 | 6978 | 1 | 1.0900e-01 | 2.8451e+05 ; | CD1 - HD13 |
| 6979 | 6980 | 1 | 1.0900e-01 | 2.8451e+05 ; | CD2 - HD21 |
| 6979 | 6981 | 1 | 1.0900e-01 | 2.8451e+05 ; | CD2 - HD22 |
| 6979 | 6982 | 1 | 1.0900e-01 | 2.8451e+05 ; | CD2 - HD23 |
| 6983 | 6984 | 1 | 1.2290e-01 | 4.7698e+05 ; | C - O |
| 6983 | 6985 | 1 | 1.3350e-01 | 4.1003e+05 ; | C - N |
| 6985 | 6986 | 1 | 1.0100e-01 | 3.6317e+05 ; | N - H |

```
6985   6987   1    1.4490e-01    2.8200e+05 ;      N - CA
6987   6988   1    1.0900e-01    2.8451e+05 ;     CA - HA
6987   6989   1    1.5260e-01    2.5941e+05 ;     CA - CB
6987   7000   1    1.5220e-01    2.6527e+05 ;     CA - C
6989   6990   1    1.0900e-01    2.8451e+05 ;     CB - HB2
6989   6991   1    1.0900e-01    2.8451e+05 ;     CB - HB3
6989   6992   1    1.5260e-01    2.5941e+05 ;     CB - CG
6992   6993   1    1.0900e-01    2.8451e+05 ;     CG - HG2
6992   6994   1    1.0900e-01    2.8451e+05 ;     CG - HG3
6992   6995   1    1.5220e-01    2.6527e+05 ;     CG - CD
6995   6996   1    1.2290e-01    4.7698e+05 ;     CD - OE1
6995   6997   1    1.3350e-01    4.1003e+05 ;     CD - NE2
6997   6998   1    1.0100e-01    3.6317e+05 ;    NE2 - HE21
6997   6999   1    1.0100e-01    3.6317e+05 ;    NE2 - HE22
7000   7001   1    1.2290e-01    4.7698e+05 ;      C - O
7000   7002   1    1.3350e-01    4.1003e+05 ;      C - N
7002   7003   1    1.0100e-01    3.6317e+05 ;      N - H
7002   7004   1    1.4490e-01    2.8200e+05 ;      N - CA
7004   7005   1    1.0900e-01    2.8451e+05 ;     CA - HA
7004   7006   1    1.5260e-01    2.5941e+05 ;     CA - CB
7004   7015   1    1.5220e-01    2.6527e+05 ;     CA - C
7006   7007   1    1.0900e-01    2.8451e+05 ;     CB - HB2
7006   7008   1    1.0900e-01    2.8451e+05 ;     CB - HB3
7006   7009   1    1.5260e-01    2.5941e+05 ;     CB - CG
7009   7010   1    1.0900e-01    2.8451e+05 ;     CG - HG2
7009   7011   1    1.0900e-01    2.8451e+05 ;     CG - HG3
7009   7012   1    1.5220e-01    2.6527e+05 ;     CG - CD
7012   7013   1    1.2500e-01    5.4894e+05 ;     CD - OE1
7012   7014   1    1.2500e-01    5.4894e+05 ;     CD - OE2
7015   7016   1    1.2290e-01    4.7698e+05 ;      C - O
7015   7017   1    1.3350e-01    4.1003e+05 ;      C - N
7017   7018   1    1.0100e-01    3.6317e+05 ;      N - H
7017   7019   1    1.4490e-01    2.8200e+05 ;      N - CA
7019   7020   1    1.0900e-01    2.8451e+05 ;     CA - HA
7019   7021   1    1.5260e-01    2.5941e+05 ;     CA - CB
7019   7031   1    1.5220e-01    2.6527e+05 ;     CA - C
7021   7022   1    1.0900e-01    2.8451e+05 ;     CB - HB
7021   7023   1    1.5260e-01    2.5941e+05 ;     CB - CG1
7021   7027   1    1.5260e-01    2.5941e+05 ;     CB - CG2
7023   7024   1    1.0900e-01    2.8451e+05 ;    CG1 - HG11
7023   7025   1    1.0900e-01    2.8451e+05 ;    CG1 - HG12
7023   7026   1    1.0900e-01    2.8451e+05 ;    CG1 - HG13
7027   7028   1    1.0900e-01    2.8451e+05 ;    CG2 - HG21
7027   7029   1    1.0900e-01    2.8451e+05 ;    CG2 - HG22
7027   7030   1    1.0900e-01    2.8451e+05 ;    CG2 - HG23
7031   7032   1    1.2290e-01    4.7698e+05 ;      C - O
7031   7033   1    1.3350e-01    4.1003e+05 ;      C - N
7033   7034   1    1.0100e-01    3.6317e+05 ;      N - H
7033   7035   1    1.4490e-01    2.8200e+05 ;      N - CA
7035   7036   1    1.0900e-01    2.8451e+05 ;     CA - HA
7035   7037   1    1.5260e-01    2.5941e+05 ;     CA - CB
7035   7043   1    1.5220e-01    2.6527e+05 ;     CA - C
```

| | | | | | | | | |
|---|---|---|---|---|---|---|---|---|
| 7037 | 7038 | 1 | 1.0900e-01 | 2.8451e+05 | ; | CB | - | HB2 |
| 7037 | 7039 | 1 | 1.0900e-01 | 2.8451e+05 | ; | CB | - | HB3 |
| 7037 | 7040 | 1 | 1.5220e-01 | 2.6527e+05 | ; | CB | - | CG |
| 7040 | 7041 | 1 | 1.2500e-01 | 5.4894e+05 | ; | CG | - | OD1 |
| 7040 | 7042 | 1 | 1.2500e-01 | 5.4894e+05 | ; | CG | - | OD2 |
| 7043 | 7044 | 1 | 1.2290e-01 | 4.7698e+05 | ; | C | - | O |
| 7043 | 7045 | 1 | 1.3350e-01 | 4.1003e+05 | ; | C | - | N |
| 7045 | 7046 | 1 | 1.0100e-01 | 3.6317e+05 | ; | N | - | H |
| 7045 | 7047 | 1 | 1.4490e-01 | 2.8200e+05 | ; | N | - | CA |
| 7047 | 7048 | 1 | 1.0900e-01 | 2.8451e+05 | ; | CA | - | HA |
| 7047 | 7049 | 1 | 1.5260e-01 | 2.5941e+05 | ; | CA | - | CB |
| 7047 | 7062 | 1 | 1.5220e-01 | 2.6527e+05 | ; | CA | - | C |
| 7049 | 7050 | 1 | 1.0900e-01 | 2.8451e+05 | ; | CB | - | HB2 |
| 7049 | 7051 | 1 | 1.0900e-01 | 2.8451e+05 | ; | CB | - | HB3 |
| 7049 | 7052 | 1 | 1.5260e-01 | 2.5941e+05 | ; | CB | - | CG |
| 7052 | 7053 | 1 | 1.0900e-01 | 2.8451e+05 | ; | CG | - | HG |
| 7052 | 7054 | 1 | 1.5260e-01 | 2.5941e+05 | ; | CG | - | CD1 |
| 7052 | 7058 | 1 | 1.5260e-01 | 2.5941e+05 | ; | CG | - | CD2 |
| 7054 | 7055 | 1 | 1.0900e-01 | 2.8451e+05 | ; | CD1 | - | HD11 |
| 7054 | 7056 | 1 | 1.0900e-01 | 2.8451e+05 | ; | CD1 | - | HD12 |
| 7054 | 7057 | 1 | 1.0900e-01 | 2.8451e+05 | ; | CD1 | - | HD13 |
| 7058 | 7059 | 1 | 1.0900e-01 | 2.8451e+05 | ; | CD2 | - | HD21 |
| 7058 | 7060 | 1 | 1.0900e-01 | 2.8451e+05 | ; | CD2 | - | HD22 |
| 7058 | 7061 | 1 | 1.0900e-01 | 2.8451e+05 | ; | CD2 | - | HD23 |
| 7062 | 7063 | 1 | 1.2290e-01 | 4.7698e+05 | ; | C | - | O |
| 7062 | 7064 | 1 | 1.3350e-01 | 4.1003e+05 | ; | C | - | N |
| 7064 | 7065 | 1 | 1.0100e-01 | 3.6317e+05 | ; | N | - | H |
| 7064 | 7066 | 1 | 1.4490e-01 | 2.8200e+05 | ; | N | - | CA |
| 7066 | 7067 | 1 | 1.0900e-01 | 2.8451e+05 | ; | CA | - | HA |
| 7066 | 7068 | 1 | 1.5260e-01 | 2.5941e+05 | ; | CA | - | CB |
| 7066 | 7076 | 1 | 1.5220e-01 | 2.6527e+05 | ; | CA | - | C |
| 7068 | 7069 | 1 | 1.0900e-01 | 2.8451e+05 | ; | CB | - | HB2 |
| 7068 | 7070 | 1 | 1.0900e-01 | 2.8451e+05 | ; | CB | - | HB3 |
| 7068 | 7071 | 1 | 1.5220e-01 | 2.6527e+05 | ; | CB | - | CG |
| 7071 | 7072 | 1 | 1.2290e-01 | 4.7698e+05 | ; | CG | - | OD1 |
| 7071 | 7073 | 1 | 1.3350e-01 | 4.1003e+05 | ; | CG | - | ND2 |
| 7073 | 7074 | 1 | 1.0100e-01 | 3.6317e+05 | ; | ND2 | - | HD21 |
| 7073 | 7075 | 1 | 1.0100e-01 | 3.6317e+05 | ; | ND2 | - | HD22 |
| 7076 | 7077 | 1 | 1.2290e-01 | 4.7698e+05 | ; | C | - | O |
| 7076 | 7078 | 1 | 1.3350e-01 | 4.1003e+05 | ; | C | - | N |
| 7078 | 7079 | 1 | 1.0100e-01 | 3.6317e+05 | ; | N | - | H |
| 7078 | 7080 | 1 | 1.4490e-01 | 2.8200e+05 | ; | N | - | CA |
| 7080 | 7081 | 1 | 1.0900e-01 | 2.8451e+05 | ; | CA | - | HA2 |
| 7080 | 7082 | 1 | 1.0900e-01 | 2.8451e+05 | ; | CA | - | HA3 |
| 7080 | 7083 | 1 | 1.5220e-01 | 2.6527e+05 | ; | CA | - | C |
| 7083 | 7084 | 1 | 1.2290e-01 | 4.7698e+05 | ; | C | - | O |
| 7083 | 7085 | 1 | 1.3350e-01 | 4.1003e+05 | ; | C | - | N |
| 7085 | 7086 | 1 | 1.0100e-01 | 3.6317e+05 | ; | N | - | H |
| 7085 | 7087 | 1 | 1.4490e-01 | 2.8200e+05 | ; | N | - | CA |
| 7087 | 7088 | 1 | 1.0900e-01 | 2.8451e+05 | ; | CA | - | HA |
| 7087 | 7089 | 1 | 1.5260e-01 | 2.5941e+05 | ; | CA | - | CB |
| 7087 | 7095 | 1 | 1.5220e-01 | 2.6527e+05 | ; | CA | - | C |

| | | | | | |
|---|---|---|---|---|---|
| 7089 | 7090 | 1 | 1.0900e-01 | 2.8451e+05 ; | CB - HB2 |
| 7089 | 7091 | 1 | 1.0900e-01 | 2.8451e+05 ; | CB - HB3 |
| 7089 | 7092 | 1 | 1.5220e-01 | 2.6527e+05 ; | CB - CG |
| 7092 | 7093 | 1 | 1.2500e-01 | 5.4894e+05 ; | CG - OD1 |
| 7092 | 7094 | 1 | 1.2500e-01 | 5.4894e+05 ; | CG - OD2 |
| 7095 | 7096 | 1 | 1.2290e-01 | 4.7698e+05 ; | C - O |
| 7095 | 7097 | 1 | 1.3350e-01 | 4.1003e+05 ; | C - N |
| 7097 | 7098 | 1 | 1.0100e-01 | 3.6317e+05 ; | N - H |
| 7097 | 7099 | 1 | 1.4490e-01 | 2.8200e+05 ; | N - CA |
| 7099 | 7100 | 1 | 1.0900e-01 | 2.8451e+05 ; | CA - HA2 |
| 7099 | 7101 | 1 | 1.0900e-01 | 2.8451e+05 ; | CA - HA3 |
| 7099 | 7102 | 1 | 1.5220e-01 | 2.6527e+05 ; | CA - C |
| 7102 | 7103 | 1 | 1.2290e-01 | 4.7698e+05 ; | C - O |
| 7102 | 7104 | 1 | 1.3350e-01 | 4.1003e+05 ; | C - N |
| 7104 | 7105 | 1 | 1.0100e-01 | 3.6317e+05 ; | N - H |
| 7104 | 7106 | 1 | 1.4490e-01 | 2.8200e+05 ; | N - CA |
| 7106 | 7107 | 1 | 1.0900e-01 | 2.8451e+05 ; | CA - HA |
| 7106 | 7108 | 1 | 1.5260e-01 | 2.5941e+05 ; | CA - CB |
| 7106 | 7117 | 1 | 1.5220e-01 | 2.6527e+05 ; | CA - C |
| 7108 | 7109 | 1 | 1.0900e-01 | 2.8451e+05 ; | CB - HB2 |
| 7108 | 7110 | 1 | 1.0900e-01 | 2.8451e+05 ; | CB - HB3 |
| 7108 | 7111 | 1 | 1.5260e-01 | 2.5941e+05 ; | CB - CG |
| 7111 | 7112 | 1 | 1.0900e-01 | 2.8451e+05 ; | CG - HG2 |
| 7111 | 7113 | 1 | 1.0900e-01 | 2.8451e+05 ; | CG - HG3 |
| 7111 | 7114 | 1 | 1.5220e-01 | 2.6527e+05 ; | CG - CD |
| 7114 | 7115 | 1 | 1.2500e-01 | 5.4894e+05 ; | CD - OE1 |
| 7114 | 7116 | 1 | 1.2500e-01 | 5.4894e+05 ; | CD - OE2 |
| 7117 | 7118 | 1 | 1.2290e-01 | 4.7698e+05 ; | C - O |
| 7117 | 7119 | 1 | 1.3350e-01 | 4.1003e+05 ; | C - N |
| 7119 | 7120 | 1 | 1.0100e-01 | 3.6317e+05 ; | N - H |
| 7119 | 7121 | 1 | 1.4490e-01 | 2.8200e+05 ; | N - CA |
| 7121 | 7122 | 1 | 1.0900e-01 | 2.8451e+05 ; | CA - HA |
| 7121 | 7123 | 1 | 1.5260e-01 | 2.5941e+05 ; | CA - CB |
| 7121 | 7136 | 1 | 1.5220e-01 | 2.6527e+05 ; | CA - C |
| 7123 | 7124 | 1 | 1.0900e-01 | 2.8451e+05 ; | CB - HB |
| 7123 | 7125 | 1 | 1.5260e-01 | 2.5941e+05 ; | CB - CG2 |
| 7123 | 7129 | 1 | 1.5260e-01 | 2.5941e+05 ; | CB - CG1 |
| 7125 | 7126 | 1 | 1.0900e-01 | 2.8451e+05 ; | CG2 - HG21 |
| 7125 | 7127 | 1 | 1.0900e-01 | 2.8451e+05 ; | CG2 - HG22 |
| 7125 | 7128 | 1 | 1.0900e-01 | 2.8451e+05 ; | CG2 - HG23 |
| 7129 | 7130 | 1 | 1.0900e-01 | 2.8451e+05 ; | CG1 - HG12 |
| 7129 | 7131 | 1 | 1.0900e-01 | 2.8451e+05 ; | CG1 - HG13 |
| 7129 | 7132 | 1 | 1.5260e-01 | 2.5941e+05 ; | CG1 - CD1 |
| 7132 | 7133 | 1 | 1.0900e-01 | 2.8451e+05 ; | CD1 - HD11 |
| 7132 | 7134 | 1 | 1.0900e-01 | 2.8451e+05 ; | CD1 - HD12 |
| 7132 | 7135 | 1 | 1.0900e-01 | 2.8451e+05 ; | CD1 - HD13 |
| 7136 | 7137 | 1 | 1.2290e-01 | 4.7698e+05 ; | C - O |
| 7136 | 7138 | 1 | 1.3350e-01 | 4.1003e+05 ; | C - N |
| 7138 | 7139 | 1 | 1.0100e-01 | 3.6317e+05 ; | N - H |
| 7138 | 7140 | 1 | 1.4490e-01 | 2.8200e+05 ; | N - CA |
| 7140 | 7141 | 1 | 1.0900e-01 | 2.8451e+05 ; | CA - HA |
| 7140 | 7142 | 1 | 1.5260e-01 | 2.5941e+05 ; | CA - CB |

| | | | | | | | | |
|---|---|---|---|---|---|---|---|---|
| 7140 | 7148 | 1 | 1.5220e-01 | 2.6527e+05 | ; | CA | - | C |
| 7142 | 7143 | 1 | 1.0900e-01 | 2.8451e+05 | ; | CB | - | HB2 |
| 7142 | 7144 | 1 | 1.0900e-01 | 2.8451e+05 | ; | CB | - | HB3 |
| 7142 | 7145 | 1 | 1.5220e-01 | 2.6527e+05 | ; | CB | - | CG |
| 7145 | 7146 | 1 | 1.2500e-01 | 5.4894e+05 | ; | CG | - | OD1 |
| 7145 | 7147 | 1 | 1.2500e-01 | 5.4894e+05 | ; | CG | - | OD2 |
| 7148 | 7149 | 1 | 1.2290e-01 | 4.7698e+05 | ; | C | - | O |
| 7148 | 7150 | 1 | 1.3350e-01 | 4.1003e+05 | ; | C | - | N |
| 7150 | 7151 | 1 | 1.0100e-01 | 3.6317e+05 | ; | N | - | H |
| 7150 | 7152 | 1 | 1.4490e-01 | 2.8200e+05 | ; | N | - | CA |
| 7152 | 7153 | 1 | 1.0900e-01 | 2.8451e+05 | ; | CA | - | HA |
| 7152 | 7154 | 1 | 1.5260e-01 | 2.5941e+05 | ; | CA | - | CB |
| 7152 | 7168 | 1 | 1.5220e-01 | 2.6527e+05 | ; | CA | - | C |
| 7154 | 7155 | 1 | 1.0900e-01 | 2.8451e+05 | ; | CB | - | HB2 |
| 7154 | 7156 | 1 | 1.0900e-01 | 2.8451e+05 | ; | CB | - | HB3 |
| 7154 | 7157 | 1 | 1.5100e-01 | 2.6527e+05 | ; | CB | - | CG |
| 7157 | 7158 | 1 | 1.4000e-01 | 3.9246e+05 | ; | CG | - | CD1 |
| 7157 | 7166 | 1 | 1.4000e-01 | 3.9246e+05 | ; | CG | - | CD2 |
| 7158 | 7159 | 1 | 1.0800e-01 | 3.0711e+05 | ; | CD1 | - | HD1 |
| 7158 | 7160 | 1 | 1.4000e-01 | 3.9246e+05 | ; | CD1 | - | CE1 |
| 7160 | 7161 | 1 | 1.0800e-01 | 3.0711e+05 | ; | CE1 | - | HE1 |
| 7160 | 7162 | 1 | 1.4000e-01 | 3.9246e+05 | ; | CE1 | - | CZ |
| 7162 | 7163 | 1 | 1.0800e-01 | 3.0711e+05 | ; | CZ | - | HZ |
| 7162 | 7164 | 1 | 1.4000e-01 | 3.9246e+05 | ; | CZ | - | CE2 |
| 7164 | 7165 | 1 | 1.0800e-01 | 3.0711e+05 | ; | CE2 | - | HE2 |
| 7164 | 7166 | 1 | 1.4000e-01 | 3.9246e+05 | ; | CE2 | - | CD2 |
| 7166 | 7167 | 1 | 1.0800e-01 | 3.0711e+05 | ; | CD2 | - | HD2 |
| 7168 | 7169 | 1 | 1.2290e-01 | 4.7698e+05 | ; | C | - | O |
| 7168 | 7170 | 1 | 1.3350e-01 | 4.1003e+05 | ; | C | - | N |
| 7170 | 7171 | 1 | 1.0100e-01 | 3.6317e+05 | ; | N | - | H |
| 7170 | 7172 | 1 | 1.4490e-01 | 2.8200e+05 | ; | N | - | CA |
| 7172 | 7173 | 1 | 1.0900e-01 | 2.8451e+05 | ; | CA | - | HA |
| 7172 | 7174 | 1 | 1.5260e-01 | 2.5941e+05 | ; | CA | - | CB |
| 7172 | 7186 | 1 | 1.5220e-01 | 2.6527e+05 | ; | CA | - | C |
| 7174 | 7175 | 1 | 1.0900e-01 | 2.8451e+05 | ; | CB | - | HB2 |
| 7174 | 7176 | 1 | 1.0900e-01 | 2.8451e+05 | ; | CB | - | HB3 |
| 7174 | 7177 | 1 | 1.5040e-01 | 2.6527e+05 | ; | CB | - | CG |
| 7177 | 7178 | 1 | 1.3850e-01 | 3.5313e+05 | ; | CG | - | ND1 |
| 7177 | 7184 | 1 | 1.3710e-01 | 4.3346e+05 | ; | CG | - | CD2 |
| 7178 | 7179 | 1 | 1.0100e-01 | 3.6317e+05 | ; | ND1 | - | HD1 |
| 7178 | 7180 | 1 | 1.3430e-01 | 3.9915e+05 | ; | ND1 | - | CE1 |
| 7180 | 7181 | 1 | 1.0800e-01 | 3.0711e+05 | ; | CE1 | - | HE1 |
| 7180 | 7182 | 1 | 1.3430e-01 | 3.9915e+05 | ; | CE1 | - | NE2 |
| 7182 | 7183 | 1 | 1.0100e-01 | 3.6317e+05 | ; | NE2 | - | HE2 |
| 7182 | 7184 | 1 | 1.3810e-01 | 3.5731e+05 | ; | NE2 | - | CD2 |
| 7184 | 7185 | 1 | 1.0800e-01 | 3.0711e+05 | ; | CD2 | - | HD2 |
| 7186 | 7187 | 1 | 1.2290e-01 | 4.7698e+05 | ; | C | - | O |
| 7186 | 7188 | 1 | 1.3350e-01 | 4.1003e+05 | ; | C | - | N |
| 7188 | 7189 | 1 | 1.0100e-01 | 3.6317e+05 | ; | N | - | H |
| 7188 | 7190 | 1 | 1.4490e-01 | 2.8200e+05 | ; | N | - | CA |
| 7190 | 7191 | 1 | 1.0900e-01 | 2.8451e+05 | ; | CA | - | HA |
| 7190 | 7192 | 1 | 1.5260e-01 | 2.5941e+05 | ; | CA | - | CB |

| | | | | | | | | |
|---|---|---|---|---|---|---|---|---|
| 7190 | 7202 | 1 | 1.5220e-01 | 2.6527e+05 | ; | CA | - | C |
| 7192 | 7193 | 1 | 1.0900e-01 | 2.8451e+05 | ; | CB | - | HB2 |
| 7192 | 7194 | 1 | 1.0900e-01 | 2.8451e+05 | ; | CB | - | HB3 |
| 7192 | 7195 | 1 | 1.5260e-01 | 2.5941e+05 | ; | CB | - | CG |
| 7195 | 7196 | 1 | 1.0900e-01 | 2.8451e+05 | ; | CG | - | HG2 |
| 7195 | 7197 | 1 | 1.0900e-01 | 2.8451e+05 | ; | CG | - | HG3 |
| 7195 | 7198 | 1 | 1.5220e-01 | 2.6527e+05 | ; | CG | - | CD |
| 7198 | 7199 | 1 | 1.2290e-01 | 4.7698e+05 | ; | CD | - | OE1 |
| 7198 | 7200 | 1 | 1.3640e-01 | 3.7656e+05 | ; | CD | - | OE2 |
| 7200 | 7201 | 1 | 9.6000e-02 | 4.6275e+05 | ; | OE2 | - | HE2 |
| 7202 | 7203 | 1 | 1.2290e-01 | 4.7698e+05 | ; | C | - | O |
| 7202 | 7204 | 1 | 1.3350e-01 | 4.1003e+05 | ; | C | - | N |
| 7204 | 7205 | 1 | 1.0100e-01 | 3.6317e+05 | ; | N | - | H |
| 7204 | 7206 | 1 | 1.4490e-01 | 2.8200e+05 | ; | N | - | CA |
| 7206 | 7207 | 1 | 1.0900e-01 | 2.8451e+05 | ; | CA | - | HA |
| 7206 | 7208 | 1 | 1.5260e-01 | 2.5941e+05 | ; | CA | - | CB |
| 7206 | 7222 | 1 | 1.5220e-01 | 2.6527e+05 | ; | CA | - | C |
| 7208 | 7209 | 1 | 1.0900e-01 | 2.8451e+05 | ; | CB | - | HB2 |
| 7208 | 7210 | 1 | 1.0900e-01 | 2.8451e+05 | ; | CB | - | HB3 |
| 7208 | 7211 | 1 | 1.5100e-01 | 2.6527e+05 | ; | CB | - | CG |
| 7211 | 7212 | 1 | 1.4000e-01 | 3.9246e+05 | ; | CG | - | CD1 |
| 7211 | 7220 | 1 | 1.4000e-01 | 3.9246e+05 | ; | CG | - | CD2 |
| 7212 | 7213 | 1 | 1.0800e-01 | 3.0711e+05 | ; | CD1 | - | HD1 |
| 7212 | 7214 | 1 | 1.4000e-01 | 3.9246e+05 | ; | CD1 | - | CE1 |
| 7214 | 7215 | 1 | 1.0800e-01 | 3.0711e+05 | ; | CE1 | - | HE1 |
| 7214 | 7216 | 1 | 1.4000e-01 | 3.9246e+05 | ; | CE1 | - | CZ |
| 7216 | 7217 | 1 | 1.0800e-01 | 3.0711e+05 | ; | CZ | - | HZ |
| 7216 | 7218 | 1 | 1.4000e-01 | 3.9246e+05 | ; | CZ | - | CE2 |
| 7218 | 7219 | 1 | 1.0800e-01 | 3.0711e+05 | ; | CE2 | - | HE2 |
| 7218 | 7220 | 1 | 1.4000e-01 | 3.9246e+05 | ; | CE2 | - | CD2 |
| 7220 | 7221 | 1 | 1.0800e-01 | 3.0711e+05 | ; | CD2 | - | HD2 |
| 7222 | 7223 | 1 | 1.2290e-01 | 4.7698e+05 | ; | C | - | O |
| 7222 | 7224 | 1 | 1.3350e-01 | 4.1003e+05 | ; | C | - | N |
| 7224 | 7225 | 1 | 1.0100e-01 | 3.6317e+05 | ; | N | - | H |
| 7224 | 7226 | 1 | 1.4490e-01 | 2.8200e+05 | ; | N | - | CA |
| 7226 | 7227 | 1 | 1.0900e-01 | 2.8451e+05 | ; | CA | - | HA |
| 7226 | 7228 | 1 | 1.5260e-01 | 2.5941e+05 | ; | CA | - | CB |
| 7226 | 7239 | 1 | 1.5220e-01 | 2.6527e+05 | ; | CA | - | C |
| 7228 | 7229 | 1 | 1.0900e-01 | 2.8451e+05 | ; | CB | - | HB2 |
| 7228 | 7230 | 1 | 1.0900e-01 | 2.8451e+05 | ; | CB | - | HB3 |
| 7228 | 7231 | 1 | 1.5260e-01 | 2.5941e+05 | ; | CB | - | CG |
| 7231 | 7232 | 1 | 1.0900e-01 | 2.8451e+05 | ; | CG | - | HG2 |
| 7231 | 7233 | 1 | 1.0900e-01 | 2.8451e+05 | ; | CG | - | HG3 |
| 7231 | 7234 | 1 | 1.8100e-01 | 1.8995e+05 | ; | CG | - | SD |
| 7234 | 7235 | 1 | 1.8100e-01 | 1.8995e+05 | ; | SD | - | CE |
| 7235 | 7236 | 1 | 1.0900e-01 | 2.8451e+05 | ; | CE | - | HE1 |
| 7235 | 7237 | 1 | 1.0900e-01 | 2.8451e+05 | ; | CE | - | HE2 |
| 7235 | 7238 | 1 | 1.0900e-01 | 2.8451e+05 | ; | CE | - | HE3 |
| 7239 | 7240 | 1 | 1.2290e-01 | 4.7698e+05 | ; | C | - | O |
| 7239 | 7241 | 1 | 1.3350e-01 | 4.1003e+05 | ; | C | - | N |
| 7241 | 7242 | 1 | 1.0100e-01 | 3.6317e+05 | ; | N | - | H |
| 7241 | 7243 | 1 | 1.4490e-01 | 2.8200e+05 | ; | N | - | CA |

| | | | | | | |
|---|---|---|---|---|---|---|
| 7243 | 7244 | 1 | 1.0900e-01 | 2.8451e+05 | ; | CA - HA |
| 7243 | 7245 | 1 | 1.5260e-01 | 2.5941e+05 | ; | CA - CB |
| 7243 | 7258 | 1 | 1.5220e-01 | 2.6527e+05 | ; | CA - C |
| 7245 | 7246 | 1 | 1.0900e-01 | 2.8451e+05 | ; | CB - HB2 |
| 7245 | 7247 | 1 | 1.0900e-01 | 2.8451e+05 | ; | CB - HB3 |
| 7245 | 7248 | 1 | 1.5260e-01 | 2.5941e+05 | ; | CB - CG |
| 7248 | 7249 | 1 | 1.0900e-01 | 2.8451e+05 | ; | CG - HG |
| 7248 | 7250 | 1 | 1.5260e-01 | 2.5941e+05 | ; | CG - CD1 |
| 7248 | 7254 | 1 | 1.5260e-01 | 2.5941e+05 | ; | CG - CD2 |
| 7250 | 7251 | 1 | 1.0900e-01 | 2.8451e+05 | ; | CD1 - HD11 |
| 7250 | 7252 | 1 | 1.0900e-01 | 2.8451e+05 | ; | CD1 - HD12 |
| 7250 | 7253 | 1 | 1.0900e-01 | 2.8451e+05 | ; | CD1 - HD13 |
| 7254 | 7255 | 1 | 1.0900e-01 | 2.8451e+05 | ; | CD2 - HD21 |
| 7254 | 7256 | 1 | 1.0900e-01 | 2.8451e+05 | ; | CD2 - HD22 |
| 7254 | 7257 | 1 | 1.0900e-01 | 2.8451e+05 | ; | CD2 - HD23 |
| 7258 | 7259 | 1 | 1.2290e-01 | 4.7698e+05 | ; | C - O |
| 7258 | 7260 | 1 | 1.3350e-01 | 4.1003e+05 | ; | C - N |
| 7260 | 7261 | 1 | 1.0100e-01 | 3.6317e+05 | ; | N - H |
| 7260 | 7262 | 1 | 1.4490e-01 | 2.8200e+05 | ; | N - CA |
| 7262 | 7263 | 1 | 1.0900e-01 | 2.8451e+05 | ; | CA - HA |
| 7262 | 7264 | 1 | 1.5260e-01 | 2.5941e+05 | ; | CA - CB |
| 7262 | 7275 | 1 | 1.5220e-01 | 2.6527e+05 | ; | CA - C |
| 7264 | 7265 | 1 | 1.0900e-01 | 2.8451e+05 | ; | CB - HB2 |
| 7264 | 7266 | 1 | 1.0900e-01 | 2.8451e+05 | ; | CB - HB3 |
| 7264 | 7267 | 1 | 1.5260e-01 | 2.5941e+05 | ; | CB - CG |
| 7267 | 7268 | 1 | 1.0900e-01 | 2.8451e+05 | ; | CG - HG2 |
| 7267 | 7269 | 1 | 1.0900e-01 | 2.8451e+05 | ; | CG - HG3 |
| 7267 | 7270 | 1 | 1.8100e-01 | 1.8995e+05 | ; | CG - SD |
| 7270 | 7271 | 1 | 1.8100e-01 | 1.8995e+05 | ; | SD - CE |
| 7271 | 7272 | 1 | 1.0900e-01 | 2.8451e+05 | ; | CE - HE1 |
| 7271 | 7273 | 1 | 1.0900e-01 | 2.8451e+05 | ; | CE - HE2 |
| 7271 | 7274 | 1 | 1.0900e-01 | 2.8451e+05 | ; | CE - HE3 |
| 7275 | 7276 | 1 | 1.2290e-01 | 4.7698e+05 | ; | C - O |
| 7275 | 7277 | 1 | 1.3350e-01 | 4.1003e+05 | ; | C - N |
| 7277 | 7278 | 1 | 1.0100e-01 | 3.6317e+05 | ; | N - H |
| 7277 | 7279 | 1 | 1.4490e-01 | 2.8200e+05 | ; | N - CA |
| 7279 | 7280 | 1 | 1.0900e-01 | 2.8451e+05 | ; | CA - HA |
| 7279 | 7281 | 1 | 1.5260e-01 | 2.5941e+05 | ; | CA - CB |
| 7279 | 7292 | 1 | 1.5220e-01 | 2.6527e+05 | ; | CA - C |
| 7281 | 7282 | 1 | 1.0900e-01 | 2.8451e+05 | ; | CB - HB2 |
| 7281 | 7283 | 1 | 1.0900e-01 | 2.8451e+05 | ; | CB - HB3 |
| 7281 | 7284 | 1 | 1.5260e-01 | 2.5941e+05 | ; | CB - CG |
| 7284 | 7285 | 1 | 1.0900e-01 | 2.8451e+05 | ; | CG - HG2 |
| 7284 | 7286 | 1 | 1.0900e-01 | 2.8451e+05 | ; | CG - HG3 |
| 7284 | 7287 | 1 | 1.8100e-01 | 1.8995e+05 | ; | CG - SD |
| 7287 | 7288 | 1 | 1.8100e-01 | 1.8995e+05 | ; | SD - CE |
| 7288 | 7289 | 1 | 1.0900e-01 | 2.8451e+05 | ; | CE - HE1 |
| 7288 | 7290 | 1 | 1.0900e-01 | 2.8451e+05 | ; | CE - HE2 |
| 7288 | 7291 | 1 | 1.0900e-01 | 2.8451e+05 | ; | CE - HE3 |
| 7292 | 7293 | 1 | 1.2290e-01 | 4.7698e+05 | ; | C - O |
| 7292 | 7294 | 1 | 1.3350e-01 | 4.1003e+05 | ; | C - N |
| 7294 | 7295 | 1 | 1.0100e-01 | 3.6317e+05 | ; | N - H |

| | | | | | | |
|---|---|---|---|---|---|---|
| 7294 | 7296 | 1 | 1.4490e-01 | 2.8200e+05 | ; | N - CA |
| 7296 | 7297 | 1 | 1.0900e-01 | 2.8451e+05 | ; | CA - HA |
| 7296 | 7298 | 1 | 1.5260e-01 | 2.5941e+05 | ; | CA - CB |
| 7296 | 7303 | 1 | 1.5220e-01 | 2.6527e+05 | ; | CA - C |
| 7298 | 7299 | 1 | 1.0900e-01 | 2.8451e+05 | ; | CB - HB2 |
| 7298 | 7300 | 1 | 1.0900e-01 | 2.8451e+05 | ; | CB - HB3 |
| 7298 | 7301 | 1 | 1.4100e-01 | 2.6778e+05 | ; | CB - OG |
| 7301 | 7302 | 1 | 9.6000e-02 | 4.6275e+05 | ; | OG - HG |
| 7303 | 7304 | 1 | 1.2290e-01 | 4.7698e+05 | ; | C - O |
| 7303 | 7305 | 1 | 1.3350e-01 | 4.1003e+05 | ; | C - N |
| 7305 | 7306 | 1 | 1.0100e-01 | 3.6317e+05 | ; | N - H |
| 7305 | 7307 | 1 | 1.4490e-01 | 2.8200e+05 | ; | N - CA |
| 7307 | 7308 | 1 | 1.0900e-01 | 2.8451e+05 | ; | CA - HA |
| 7307 | 7309 | 1 | 1.5260e-01 | 2.5941e+05 | ; | CA - CB |
| 7307 | 7325 | 1 | 1.5220e-01 | 2.6527e+05 | ; | CA - C |
| 7309 | 7310 | 1 | 1.0900e-01 | 2.8451e+05 | ; | CB - HB2 |
| 7309 | 7311 | 1 | 1.0900e-01 | 2.8451e+05 | ; | CB - HB3 |
| 7309 | 7312 | 1 | 1.5260e-01 | 2.5941e+05 | ; | CB - CG |
| 7312 | 7313 | 1 | 1.0900e-01 | 2.8451e+05 | ; | CG - HG2 |
| 7312 | 7314 | 1 | 1.0900e-01 | 2.8451e+05 | ; | CG - HG3 |
| 7312 | 7315 | 1 | 1.5260e-01 | 2.5941e+05 | ; | CG - CD |
| 7315 | 7316 | 1 | 1.0900e-01 | 2.8451e+05 | ; | CD - HD2 |
| 7315 | 7317 | 1 | 1.0900e-01 | 2.8451e+05 | ; | CD - HD3 |
| 7315 | 7318 | 1 | 1.5260e-01 | 2.5941e+05 | ; | CD - CE |
| 7318 | 7319 | 1 | 1.0900e-01 | 2.8451e+05 | ; | CE - HE2 |
| 7318 | 7320 | 1 | 1.0900e-01 | 2.8451e+05 | ; | CE - HE3 |
| 7318 | 7321 | 1 | 1.4710e-01 | 3.0711e+05 | ; | CE - NZ |
| 7321 | 7322 | 1 | 1.0100e-01 | 3.6317e+05 | ; | NZ - HZ1 |
| 7321 | 7323 | 1 | 1.0100e-01 | 3.6317e+05 | ; | NZ - HZ2 |
| 7321 | 7324 | 1 | 1.0100e-01 | 3.6317e+05 | ; | NZ - HZ3 |
| 7325 | 7326 | 1 | 1.2290e-01 | 4.7698e+05 | ; | C - O |
| 7325 | 7327 | 1 | 1.3350e-01 | 4.1003e+05 | ; | C - N |
| 7327 | 7328 | 1 | 1.0100e-01 | 3.6317e+05 | ; | N - H |
| 7327 | 7329 | 1 | 1.4490e-01 | 2.8200e+05 | ; | N - CA |
| 7329 | 7330 | 1 | 1.0900e-01 | 2.8451e+05 | ; | CA - HA |
| 7329 | 7331 | 1 | 1.5260e-01 | 2.5941e+05 | ; | CA - CB |
| 7329 | 7347 | 1 | 1.5220e-01 | 2.6527e+05 | ; | CA - C |
| 7331 | 7332 | 1 | 1.0900e-01 | 2.8451e+05 | ; | CB - HB2 |
| 7331 | 7333 | 1 | 1.0900e-01 | 2.8451e+05 | ; | CB - HB3 |
| 7331 | 7334 | 1 | 1.5260e-01 | 2.5941e+05 | ; | CB - CG |
| 7334 | 7335 | 1 | 1.0900e-01 | 2.8451e+05 | ; | CG - HG2 |
| 7334 | 7336 | 1 | 1.0900e-01 | 2.8451e+05 | ; | CG - HG3 |
| 7334 | 7337 | 1 | 1.5260e-01 | 2.5941e+05 | ; | CG - CD |
| 7337 | 7338 | 1 | 1.0900e-01 | 2.8451e+05 | ; | CD - HD2 |
| 7337 | 7339 | 1 | 1.0900e-01 | 2.8451e+05 | ; | CD - HD3 |
| 7337 | 7340 | 1 | 1.5260e-01 | 2.5941e+05 | ; | CD - CE |
| 7340 | 7341 | 1 | 1.0900e-01 | 2.8451e+05 | ; | CE - HE2 |
| 7340 | 7342 | 1 | 1.0900e-01 | 2.8451e+05 | ; | CE - HE3 |
| 7340 | 7343 | 1 | 1.4710e-01 | 3.0711e+05 | ; | CE - NZ |
| 7343 | 7344 | 1 | 1.0100e-01 | 3.6317e+05 | ; | NZ - HZ1 |
| 7343 | 7345 | 1 | 1.0100e-01 | 3.6317e+05 | ; | NZ - HZ2 |
| 7343 | 7346 | 1 | 1.0100e-01 | 3.6317e+05 | ; | NZ - HZ3 |

| | | | | | | | | |
|---|---|---|---|---|---|---|---|---|
| 7347 | 7348 | 1 | 1.2290e-01 | 4.7698e+05 | ; | C | - | O |
| 7347 | 7349 | 1 | 1.3350e-01 | 4.1003e+05 | ; | C | - | N |
| 7349 | 7350 | 1 | 1.0100e-01 | 3.6317e+05 | ; | N | - | H |
| 7349 | 7351 | 1 | 1.4490e-01 | 2.8200e+05 | ; | N | - | CA |
| 7351 | 7352 | 1 | 1.0900e-01 | 2.8451e+05 | ; | CA | - | HA |
| 7351 | 7353 | 1 | 1.5260e-01 | 2.5941e+05 | ; | CA | - | CB |
| 7351 | 7369 | 1 | 1.5220e-01 | 2.6527e+05 | ; | CA | - | C |
| 7353 | 7354 | 1 | 1.0900e-01 | 2.8451e+05 | ; | CB | - | HB2 |
| 7353 | 7355 | 1 | 1.0900e-01 | 2.8451e+05 | ; | CB | - | HB3 |
| 7353 | 7356 | 1 | 1.5260e-01 | 2.5941e+05 | ; | CB | - | CG |
| 7356 | 7357 | 1 | 1.0900e-01 | 2.8451e+05 | ; | CG | - | HG2 |
| 7356 | 7358 | 1 | 1.0900e-01 | 2.8451e+05 | ; | CG | - | HG3 |
| 7356 | 7359 | 1 | 1.5260e-01 | 2.5941e+05 | ; | CG | - | CD |
| 7359 | 7360 | 1 | 1.0900e-01 | 2.8451e+05 | ; | CD | - | HD2 |
| 7359 | 7361 | 1 | 1.0900e-01 | 2.8451e+05 | ; | CD | - | HD3 |
| 7359 | 7362 | 1 | 1.5260e-01 | 2.5941e+05 | ; | CD | - | CE |
| 7362 | 7363 | 1 | 1.0900e-01 | 2.8451e+05 | ; | CE | - | HE2 |
| 7362 | 7364 | 1 | 1.0900e-01 | 2.8451e+05 | ; | CE | - | HE3 |
| 7362 | 7365 | 1 | 1.4710e-01 | 3.0711e+05 | ; | CE | - | NZ |
| 7365 | 7366 | 1 | 1.0100e-01 | 3.6317e+05 | ; | NZ | - | HZ1 |
| 7365 | 7367 | 1 | 1.0100e-01 | 3.6317e+05 | ; | NZ | - | HZ2 |
| 7365 | 7368 | 1 | 1.0100e-01 | 3.6317e+05 | ; | NZ | - | HZ3 |
| 7369 | 7370 | 1 | 1.2500e-01 | 5.4894e+05 | ; | C | - | O |
| 7369 | 7371 | 1 | 1.2500e-01 | 5.4894e+05 | ; | C | - | OXT |
| 7372 | 7373 | 1 | 1.3984e-01 | 3.8585e+05 | ; | C1 | - | C2 |
| 7372 | 7377 | 1 | 1.3984e-01 | 3.8585e+05 | ; | C1 | - | C6 |
| 7372 | 7388 | 1 | 1.3637e-01 | 3.2133e+05 | ; | C1 | - | O17 |
| 7373 | 7374 | 1 | 1.3984e-01 | 3.8585e+05 | ; | C2 | - | C3 |
| 7373 | 7413 | 1 | 1.0860e-01 | 2.8937e+05 | ; | C2 | - | H42 |
| 7374 | 7375 | 1 | 1.3984e-01 | 3.8585e+05 | ; | C3 | - | C4 |
| 7374 | 7414 | 1 | 1.0860e-01 | 2.8937e+05 | ; | C3 | - | H43 |
| 7375 | 7376 | 1 | 1.4058e-01 | 3.7673e+05 | ; | C4 | - | C5 |
| 7375 | 7387 | 1 | 1.3030e-01 | 4.6158e+05 | ; | C4 | - | N16 |
| 7376 | 7377 | 1 | 1.4058e-01 | 3.7673e+05 | ; | C5 | - | C6 |
| 7376 | 7379 | 1 | 1.4542e-01 | 3.2351e+05 | ; | C5 | - | C8 |
| 7377 | 7415 | 1 | 1.0860e-01 | 2.8937e+05 | ; | C6 | - | H44 |
| 7378 | 7379 | 1 | 1.4058e-01 | 3.7673e+05 | ; | C7 | - | C8 |
| 7378 | 7380 | 1 | 1.3984e-01 | 3.8585e+05 | ; | C7 | - | C9 |
| 7378 | 7387 | 1 | 1.3030e-01 | 4.6158e+05 | ; | C7 | - | N16 |
| 7379 | 7405 | 1 | 1.4058e-01 | 3.7673e+05 | ; | C8 | - | C34 |
| 7380 | 7381 | 1 | 1.3984e-01 | 3.8585e+05 | ; | C9 | - | C10 |
| 7380 | 7386 | 1 | 1.3030e-01 | 4.6158e+05 | ; | C9 | - | N15 |
| 7381 | 7384 | 1 | 1.3846e-01 | 4.0342e+05 | ; | C10 | - | C13 |
| 7381 | 7412 | 1 | 1.3984e-01 | 3.8585e+05 | ; | C10 | - | C41 |
| 7382 | 7383 | 1 | 1.3343e-01 | 4.7647e+05 | ; | C11 | - | C12 |
| 7382 | 7386 | 1 | 1.2817e-01 | 4.9714e+05 | ; | C11 | - | N15 |
| 7382 | 7416 | 1 | 1.0868e-01 | 2.8836e+05 | ; | C11 | - | H45 |
| 7383 | 7384 | 1 | 1.3343e-01 | 4.7647e+05 | ; | C12 | - | C13 |
| 7383 | 7385 | 1 | 1.3385e-01 | 3.1012e+05 | ; | C12 | - | F14 |
| 7384 | 7417 | 1 | 1.0879e-01 | 2.8711e+05 | ; | C13 | - | H46 |
| 7386 | 7436 | 1 | 2.1490e-01 | 5.8827e+04 | ; | N15 | - | RU |
| 7387 | 7436 | 1 | 2.1112e-01 | 8.4349e+04 | ; | N16 | - | RU |

```
  7388   7430   1    9.7300e-02    3.1079e+05 ;    O17 - H59
  7389   7390   1    1.3343e-01    4.7647e+05 ;    C18 - C19
  7389   7393   1    1.3343e-01    4.7647e+05 ;    C18 - C22
  7389   7418   1    1.0879e-01    2.8711e+05 ;    C18 - H47
  7389   7436   1    2.2671e-01    4.4267e+04 ;    C18 - RU
  7390   7391   1    1.3343e-01    4.7647e+05 ;    C19 - C20
  7390   7396   1    1.3343e-01    4.7647e+05 ;    C19 - C25
  7390   7436   1    2.2930e-01    5.3388e+04 ;    C19 - RU
  7391   7392   1    1.3343e-01    4.7647e+05 ;    C20 - C21
  7391   7435   1    1.0879e-01    2.8711e+05 ;    C20 - H64
  7391   7436   1    2.3731e-01    2.6108e+04 ;    C20 - RU
  7392   7393   1    1.3343e-01    4.7647e+05 ;    C21 - C22
  7392   7419   1    1.0879e-01    2.8711e+05 ;    C21 - H48
  7392   7436   1    2.3799e-01    2.8200e+04 ;    C21 - RU
  7393   7420   1    1.0879e-01    2.8711e+05 ;    C22 - H49
  7393   7436   1    2.3012e-01    5.2049e+04 ;    C22 - RU
  7394   7395   1    1.1724e-01    6.3438e+05 ;    C23 - O24
  7394   7436   1    1.8797e-01    1.4895e+05 ;    C23 - RU
  7396   7397   1    1.3994e-01    3.3480e+05 ;    C25 - N26
  7396   7398   1    1.2247e-01    5.2124e+05 ;    C25 - O27
  7397   7399   1    1.4619e-01    2.7506e+05 ;    N26 - C28
  7397   7434   1    1.0129e-01    3.3740e+05 ;    N26 - H63
  7399   7400   1    1.5375e-01    2.5179e+05 ;    C28 - C29
  7399   7401   1    1.5375e-01    2.5179e+05 ;    C28 - C30
  7399   7403   1    1.5375e-01    2.5179e+05 ;    C28 - C32
  7400   7431   1    1.0969e-01    2.7665e+05 ;    C29 - H60
  7400   7432   1    1.0969e-01    2.7665e+05 ;    C29 - H61
  7400   7433   1    1.0969e-01    2.7665e+05 ;    C29 - H62
  7401   7402   1    1.4233e-01    2.6501e+05 ;    C30 - O31
  7401   7421   1    1.0969e-01    2.7665e+05 ;    C30 - H50
  7401   7422   1    1.0969e-01    2.7665e+05 ;    C30 - H51
  7402   7425   1    9.7300e-02    3.1079e+05 ;    O31 - H54
  7403   7404   1    1.4233e-01    2.6501e+05 ;    C32 - O33
  7403   7423   1    1.0969e-01    2.7665e+05 ;    C32 - H52
  7403   7424   1    1.0969e-01    2.7665e+05 ;    C32 - H53
  7404   7429   1    9.7300e-02    3.1079e+05 ;    O33 - H58
  7405   7406   1    1.3846e-01    4.0342e+05 ;    C34 - C35
  7405   7412   1    1.3984e-01    3.8585e+05 ;    C34 - C41
  7406   7407   1    1.3994e-01    3.3480e+05 ;    C35 - N36
  7406   7408   1    1.2247e-01    5.2124e+05 ;    C35 - O37
  7407   7409   1    1.4619e-01    2.7506e+05 ;    N36 - C38
  7407   7410   1    1.3994e-01    3.3480e+05 ;    N36 - C39
  7409   7426   1    1.0969e-01    2.7665e+05 ;    C38 - H55
  7409   7427   1    1.0969e-01    2.7665e+05 ;    C38 - H56
  7409   7428   1    1.0969e-01    2.7665e+05 ;    C38 - H57
  7410   7411   1    1.2247e-01    5.2124e+05 ;    C39 - O40
  7410   7412   1    1.3846e-01    4.0342e+05 ;    C39 - C41

[ pairs ]
;   ai     aj    funct
     1      8      1 ;      N - HB2
```

```
58      63       1 ;      C - HA
58      64       1 ;      C - CB
58      69       1 ;      C - C
59      61       1 ;      O - H
59      62       1 ;      O - CA
60      65       1 ;      N - HB2
60      66       1 ;      N - HB3
60      67       1 ;      N - SG
60      70       1 ;      N - O
60      71       1 ;      N - N
61      63       1 ;      H - HA
61      64       1 ;      H - CB
61      69       1 ;      H - C
62      68       1 ;     CA - HG
62      72       1 ;     CA - H
62      73       1 ;     CA - CA
63      65       1 ;     HA - HB2
63      66       1 ;     HA - HB3
63      67       1 ;     HA - SG
63      70       1 ;     HA - O
63      71       1 ;     HA - N
64      70       1 ;     CB - O
64      71       1 ;     CB - N
65      68       1 ;    HB2 - HG
65      69       1 ;    HB2 - C
66      68       1 ;    HB3 - HG
66      69       1 ;    HB3 - C
67      69       1 ;     SG - C
69      74       1 ;      C - HA
69      75       1 ;      C - CB
69      87       1 ;      C - C
70      72       1 ;      O - H
70      73       1 ;      O - CA
71      76       1 ;      N - HB2
71      77       1 ;      N - HB3
71      78       1 ;      N - CG
71      88       1 ;      N - O
71      89       1 ;      N - N
72      74       1 ;      H - HA
72      75       1 ;      H - CB
72      87       1 ;      H - C
73      79       1 ;     CA - ND1
73      85       1 ;     CA - CD2
73      90       1 ;     CA - H
73      91       1 ;     CA - CA
74      76       1 ;     HA - HB2
74      77       1 ;     HA - HB3
74      78       1 ;     HA - CG
74      88       1 ;     HA - O
74      89       1 ;     HA - N
75      80       1 ;     CB - HD1
75      81       1 ;     CB - CE1
```

```
    75     83       1 ;      CB - NE2
    75     86       1 ;      CB - HD2
    75     88       1 ;      CB - O
    75     89       1 ;      CB - N
    76     79       1 ;     HB2 - ND1
    76     85       1 ;     HB2 - CD2
    76     87       1 ;     HB2 - C
    77     79       1 ;     HB3 - ND1
    77     85       1 ;     HB3 - CD2
    77     87       1 ;     HB3 - C
    78     82       1 ;      CG - HE1
    78     84       1 ;      CG - HE2
    78     87       1 ;      CG - C
    79     84       1 ;     ND1 - HE2
    79     86       1 ;     ND1 - HD2
    80     82       1 ;     HD1 - HE1
    80     83       1 ;     HD1 - NE2
    80     85       1 ;     HD1 - CD2
    81     86       1 ;     CE1 - HD2
    82     84       1 ;     HE1 - HE2
    82     85       1 ;     HE1 - CD2
    84     86       1 ;     HE2 - HD2
    87     92       1 ;       C - HA
    87     93       1 ;       C - CB
    87     98       1 ;       C - C
    88     90       1 ;       O - H
    88     91       1 ;       O - CA
    89     94       1 ;       N - HB2
    89     95       1 ;       N - HB3
    89     96       1 ;       N - OG
    89     99       1 ;       N - O
    89    100       1 ;       N - N
    90     92       1 ;       H - HA
    90     93       1 ;       H - CB
    90     98       1 ;       H - C
    91     97       1 ;      CA - HG
    91    101       1 ;      CA - H
    91    102       1 ;      CA - CA
    92     94       1 ;      HA - HB2
    92     95       1 ;      HA - HB3
    92     96       1 ;      HA - OG
    92     99       1 ;      HA - O
    92    100       1 ;      HA - N
    93     99       1 ;      CB - O
    93    100       1 ;      CB - N
    94     97       1 ;     HB2 - HG
    94     98       1 ;     HB2 - C
    95     97       1 ;     HB3 - HG
    95     98       1 ;     HB3 - C
    96     98       1 ;      OG - C
    98    103       1 ;       C - HA2
    98    104       1 ;       C - HA3
```

```
 98    105       1 ;       C - C
 99    101       1 ;       O - H
 99    102       1 ;       O - CA
100    106       1 ;       N - O
100    107       1 ;       N - N
101    103       1 ;       H - HA2
101    104       1 ;       H - HA3
101    105       1 ;       H - C
102    108       1 ;      CA - H
102    109       1 ;      CA - CA
103    106       1 ;     HA2 - O
103    107       1 ;     HA2 - N
104    106       1 ;     HA3 - O
104    107       1 ;     HA3 - N
105    110       1 ;       C - HA
105    111       1 ;       C - CB
105    127       1 ;       C - C
106    108       1 ;       O - H
106    109       1 ;       O - CA
107    112       1 ;       N - HB2
107    113       1 ;       N - HB3
107    114       1 ;       N - CG
107    128       1 ;       N - O
107    129       1 ;       N - N
108    110       1 ;       H - HA
108    111       1 ;       H - CB
108    127       1 ;       H - C
109    115       1 ;      CA - HG2
109    116       1 ;      CA - HG3
109    117       1 ;      CA - CD
109    130       1 ;      CA - H
109    131       1 ;      CA - CA
110    112       1 ;      HA - HB2
110    113       1 ;      HA - HB3
110    114       1 ;      HA - CG
110    128       1 ;      HA - O
110    129       1 ;      HA - N
111    118       1 ;      CB - HD2
111    119       1 ;      CB - HD3
111    120       1 ;      CB - CE
111    128       1 ;      CB - O
111    129       1 ;      CB - N
112    115       1 ;     HB2 - HG2
112    116       1 ;     HB2 - HG3
112    117       1 ;     HB2 - CD
112    127       1 ;     HB2 - C
113    115       1 ;     HB3 - HG2
113    116       1 ;     HB3 - HG3
113    117       1 ;     HB3 - CD
113    127       1 ;     HB3 - C
114    121       1 ;      CG - HE2
114    122       1 ;      CG - HE3
```

```
114    123       1 ;      CG - NZ
114    127       1 ;      CG - C
115    118       1 ;     HG2 - HD2
115    119       1 ;     HG2 - HD3
115    120       1 ;     HG2 - CE
116    118       1 ;     HG3 - HD2
116    119       1 ;     HG3 - HD3
116    120       1 ;     HG3 - CE
117    124       1 ;      CD - HZ1
117    125       1 ;      CD - HZ2
117    126       1 ;      CD - HZ3
118    121       1 ;     HD2 - HE2
118    122       1 ;     HD2 - HE3
118    123       1 ;     HD2 - NZ
119    121       1 ;     HD3 - HE2
119    122       1 ;     HD3 - HE3
119    123       1 ;     HD3 - NZ
121    124       1 ;     HE2 - HZ1
121    125       1 ;     HE2 - HZ2
121    126       1 ;     HE2 - HZ3
122    124       1 ;     HE3 - HZ1
122    125       1 ;     HE3 - HZ2
122    126       1 ;     HE3 - HZ3
127    132       1 ;       C - HA
127    133       1 ;       C - CB
127    146       1 ;       C - C
128    130       1 ;       O - H
128    131       1 ;       O - CA
129    134       1 ;       N - HB2
129    135       1 ;       N - HB3
129    136       1 ;       N - CG
129    147       1 ;       N - O
129    148       1 ;       N - N
130    132       1 ;       H - HA
130    133       1 ;       H - CB
130    146       1 ;       H - C
131    137       1 ;      CA - HG
131    138       1 ;      CA - CD1
131    142       1 ;      CA - CD2
131    149       1 ;      CA - H
131    150       1 ;      CA - CA
132    134       1 ;      HA - HB2
132    135       1 ;      HA - HB3
132    136       1 ;      HA - CG
132    147       1 ;      HA - O
132    148       1 ;      HA - N
133    139       1 ;      CB - HD11
133    140       1 ;      CB - HD12
133    141       1 ;      CB - HD13
133    143       1 ;      CB - HD21
133    144       1 ;      CB - HD22
133    145       1 ;      CB - HD23
```

```
133    147       1 ;      CB - O
133    148       1 ;      CB - N
134    137       1 ;     HB2 - HG
134    138       1 ;     HB2 - CD1
134    142       1 ;     HB2 - CD2
134    146       1 ;     HB2 - C
135    137       1 ;     HB3 - HG
135    138       1 ;     HB3 - CD1
135    142       1 ;     HB3 - CD2
135    146       1 ;     HB3 - C
136    146       1 ;      CG - C
137    139       1 ;      HG - HD11
137    140       1 ;      HG - HD12
137    141       1 ;      HG - HD13
137    143       1 ;      HG - HD21
137    144       1 ;      HG - HD22
137    145       1 ;      HG - HD23
138    143       1 ;     CD1 - HD21
138    144       1 ;     CD1 - HD22
138    145       1 ;     CD1 - HD23
139    142       1 ;    HD11 - CD2
140    142       1 ;    HD12 - CD2
141    142       1 ;    HD13 - CD2
146    151       1 ;       C - HA
146    152       1 ;       C - CB
146    161       1 ;       C - C
147    149       1 ;       O - H
147    150       1 ;       O - CA
148    153       1 ;       N - HB2
148    154       1 ;       N - HB3
148    155       1 ;       N - CG
148    162       1 ;       N - O
148    163       1 ;       N - N
149    151       1 ;       H - HA
149    152       1 ;       H - CB
149    161       1 ;       H - C
150    156       1 ;      CA - HG2
150    157       1 ;      CA - HG3
150    158       1 ;      CA - CD
150    164       1 ;      CA - H
150    165       1 ;      CA - CA
151    153       1 ;      HA - HB2
151    154       1 ;      HA - HB3
151    155       1 ;      HA - CG
151    162       1 ;      HA - O
151    163       1 ;      HA - N
152    159       1 ;      CB - OE1
152    160       1 ;      CB - OE2
152    162       1 ;      CB - O
152    163       1 ;      CB - N
153    156       1 ;     HB2 - HG2
153    157       1 ;     HB2 - HG3
```

```
153    158      1 ;    HB2 - CD
153    161      1 ;    HB2 - C
154    156      1 ;    HB3 - HG2
154    157      1 ;    HB3 - HG3
154    158      1 ;    HB3 - CD
154    161      1 ;    HB3 - C
155    161      1 ;     CG - C
156    159      1 ;    HG2 - OE1
156    160      1 ;    HG2 - OE2
157    159      1 ;    HG3 - OE1
157    160      1 ;    HG3 - OE2
161    166      1 ;      C - HA
161    167      1 ;      C - CB
161    173      1 ;      C - C
162    164      1 ;      O - H
162    165      1 ;      O - CA
163    168      1 ;      N - HB2
163    169      1 ;      N - HB3
163    170      1 ;      N - CG
163    174      1 ;      N - O
163    175      1 ;      N - N
164    166      1 ;      H - HA
164    167      1 ;      H - CB
164    173      1 ;      H - C
165    171      1 ;     CA - OD1
165    172      1 ;     CA - OD2
165    176      1 ;     CA - H
165    177      1 ;     CA - CA
166    168      1 ;     HA - HB2
166    169      1 ;     HA - HB3
166    170      1 ;     HA - CG
166    174      1 ;     HA - O
166    175      1 ;     HA - N
167    174      1 ;     CB - O
167    175      1 ;     CB - N
168    171      1 ;    HB2 - OD1
168    172      1 ;    HB2 - OD2
168    173      1 ;    HB2 - C
169    171      1 ;    HB3 - OD1
169    172      1 ;    HB3 - OD2
169    173      1 ;    HB3 - C
170    173      1 ;     CG - C
173    178      1 ;      C - HA
173    179      1 ;      C - CB
173    195      1 ;      C - C
174    176      1 ;      O - H
174    177      1 ;      O - CA
175    180      1 ;      N - HB2
175    181      1 ;      N - HB3
175    182      1 ;      N - CG
175    196      1 ;      N - O
175    197      1 ;      N - N
```

```
176    178       1 ;       H - HA
176    179       1 ;       H - CB
176    195       1 ;       H - C
177    183       1 ;      CA - HG2
177    184       1 ;      CA - HG3
177    185       1 ;      CA - CD
177    198       1 ;      CA - H
177    199       1 ;      CA - CA
178    180       1 ;      HA - HB2
178    181       1 ;      HA - HB3
178    182       1 ;      HA - CG
178    196       1 ;      HA - O
178    197       1 ;      HA - N
179    186       1 ;      CB - HD2
179    187       1 ;      CB - HD3
179    188       1 ;      CB - CE
179    196       1 ;      CB - O
179    197       1 ;      CB - N
180    183       1 ;     HB2 - HG2
180    184       1 ;     HB2 - HG3
180    185       1 ;     HB2 - CD
180    195       1 ;     HB2 - C
181    183       1 ;     HB3 - HG2
181    184       1 ;     HB3 - HG3
181    185       1 ;     HB3 - CD
181    195       1 ;     HB3 - C
182    189       1 ;      CG - HE2
182    190       1 ;      CG - HE3
182    191       1 ;      CG - NZ
182    195       1 ;      CG - C
183    186       1 ;     HG2 - HD2
183    187       1 ;     HG2 - HD3
183    188       1 ;     HG2 - CE
184    186       1 ;     HG3 - HD2
184    187       1 ;     HG3 - HD3
184    188       1 ;     HG3 - CE
185    192       1 ;      CD - HZ1
185    193       1 ;      CD - HZ2
185    194       1 ;      CD - HZ3
186    189       1 ;     HD2 - HE2
186    190       1 ;     HD2 - HE3
186    191       1 ;     HD2 - NZ
187    189       1 ;     HD3 - HE2
187    190       1 ;     HD3 - HE3
187    191       1 ;     HD3 - NZ
189    192       1 ;     HE2 - HZ1
189    193       1 ;     HE2 - HZ2
189    194       1 ;     HE2 - HZ3
190    192       1 ;     HE3 - HZ1
190    193       1 ;     HE3 - HZ2
190    194       1 ;     HE3 - HZ3
195    200       1 ;       C - HA
```

| | | | |
|---|---|---|---|
| 195 | 201 | 1 ; | C - CB |
| 195 | 216 | 1 ; | C - C |
| 196 | 198 | 1 ; | O - H |
| 196 | 199 | 1 ; | O - CA |
| 197 | 202 | 1 ; | N - HB2 |
| 197 | 203 | 1 ; | N - HB3 |
| 197 | 204 | 1 ; | N - CG |
| 197 | 217 | 1 ; | N - O |
| 197 | 218 | 1 ; | N - N |
| 198 | 200 | 1 ; | H - HA |
| 198 | 201 | 1 ; | H - CB |
| 198 | 216 | 1 ; | H - C |
| 199 | 205 | 1 ; | CA - CD1 |
| 199 | 214 | 1 ; | CA - CD2 |
| 199 | 219 | 1 ; | CA - H |
| 199 | 220 | 1 ; | CA - CA |
| 200 | 202 | 1 ; | HA - HB2 |
| 200 | 203 | 1 ; | HA - HB3 |
| 200 | 204 | 1 ; | HA - CG |
| 200 | 217 | 1 ; | HA - O |
| 200 | 218 | 1 ; | HA - N |
| 201 | 206 | 1 ; | CB - HD1 |
| 201 | 207 | 1 ; | CB - CE1 |
| 201 | 212 | 1 ; | CB - CE2 |
| 201 | 215 | 1 ; | CB - HD2 |
| 201 | 217 | 1 ; | CB - O |
| 201 | 218 | 1 ; | CB - N |
| 202 | 205 | 1 ; | HB2 - CD1 |
| 202 | 214 | 1 ; | HB2 - CD2 |
| 202 | 216 | 1 ; | HB2 - C |
| 203 | 205 | 1 ; | HB3 - CD1 |
| 203 | 214 | 1 ; | HB3 - CD2 |
| 203 | 216 | 1 ; | HB3 - C |
| 204 | 208 | 1 ; | CG - HE1 |
| 204 | 209 | 1 ; | CG - CZ |
| 204 | 213 | 1 ; | CG - HE2 |
| 204 | 216 | 1 ; | CG - C |
| 205 | 210 | 1 ; | CD1 - OH |
| 205 | 212 | 1 ; | CD1 - CE2 |
| 205 | 215 | 1 ; | CD1 - HD2 |
| 206 | 208 | 1 ; | HD1 - HE1 |
| 206 | 209 | 1 ; | HD1 - CZ |
| 206 | 214 | 1 ; | HD1 - CD2 |
| 207 | 211 | 1 ; | CE1 - HH |
| 207 | 213 | 1 ; | CE1 - HE2 |
| 207 | 214 | 1 ; | CE1 - CD2 |
| 208 | 210 | 1 ; | HE1 - OH |
| 208 | 212 | 1 ; | HE1 - CE2 |
| 209 | 215 | 1 ; | CZ - HD2 |
| 210 | 213 | 1 ; | OH - HE2 |
| 210 | 214 | 1 ; | OH - CD2 |
| 211 | 212 | 1 ; | HH - CE2 |

```
 213    215       1 ;     HE2 - HD2
 216    221       1 ;       C - HA
 216    222       1 ;       C - CB
 216    235       1 ;       C - C
 217    219       1 ;       O - H
 217    220       1 ;       O - CA
 218    223       1 ;       N - HB
 218    224       1 ;       N - CG2
 218    228       1 ;       N - CG1
 218    236       1 ;       N - O
 218    237       1 ;       N - N
 219    221       1 ;       H - HA
 219    222       1 ;       H - CB
 219    235       1 ;       H - C
 220    225       1 ;      CA - HG21
 220    226       1 ;      CA - HG22
 220    227       1 ;      CA - HG23
 220    229       1 ;      CA - HG12
 220    230       1 ;      CA - HG13
 220    231       1 ;      CA - CD1
 220    238       1 ;      CA - H
 220    239       1 ;      CA - CA
 221    223       1 ;      HA - HB
 221    224       1 ;      HA - CG2
 221    228       1 ;      HA - CG1
 221    236       1 ;      HA - O
 221    237       1 ;      HA - N
 222    232       1 ;      CB - HD11
 222    233       1 ;      CB - HD12
 222    234       1 ;      CB - HD13
 222    236       1 ;      CB - O
 222    237       1 ;      CB - N
 223    225       1 ;      HB - HG21
 223    226       1 ;      HB - HG22
 223    227       1 ;      HB - HG23
 223    229       1 ;      HB - HG12
 223    230       1 ;      HB - HG13
 223    231       1 ;      HB - CD1
 223    235       1 ;      HB - C
 224    229       1 ;     CG2 - HG12
 224    230       1 ;     CG2 - HG13
 224    231       1 ;     CG2 - CD1
 224    235       1 ;     CG2 - C
 225    228       1 ;    HG21 - CG1
 226    228       1 ;    HG22 - CG1
 227    228       1 ;    HG23 - CG1
 228    235       1 ;     CG1 - C
 229    232       1 ;    HG12 - HD11
 229    233       1 ;    HG12 - HD12
 229    234       1 ;    HG12 - HD13
 230    232       1 ;    HG13 - HD11
 230    233       1 ;    HG13 - HD12
```

```
230    234      1 ;   HG13 - HD13
235    240      1 ;      C - HA
235    241      1 ;      C - CB
235    254      1 ;      C - C
236    238      1 ;      O - H
236    239      1 ;      O - CA
237    242      1 ;      N - HB
237    243      1 ;      N - CG2
237    247      1 ;      N - CG1
237    255      1 ;      N - O
237    256      1 ;      N - N
238    240      1 ;      H - HA
238    241      1 ;      H - CB
238    254      1 ;      H - C
239    244      1 ;     CA - HG21
239    245      1 ;     CA - HG22
239    246      1 ;     CA - HG23
239    248      1 ;     CA - HG12
239    249      1 ;     CA - HG13
239    250      1 ;     CA - CD1
239    257      1 ;     CA - H
239    258      1 ;     CA - CA
240    242      1 ;     HA - HB
240    243      1 ;     HA - CG2
240    247      1 ;     HA - CG1
240    255      1 ;     HA - O
240    256      1 ;     HA - N
241    251      1 ;     CB - HD11
241    252      1 ;     CB - HD12
241    253      1 ;     CB - HD13
241    255      1 ;     CB - O
241    256      1 ;     CB - N
242    244      1 ;     HB - HG21
242    245      1 ;     HB - HG22
242    246      1 ;     HB - HG23
242    248      1 ;     HB - HG12
242    249      1 ;     HB - HG13
242    250      1 ;     HB - CD1
242    254      1 ;     HB - C
243    248      1 ;    CG2 - HG12
243    249      1 ;    CG2 - HG13
243    250      1 ;    CG2 - CD1
243    254      1 ;    CG2 - C
244    247      1 ;   HG21 - CG1
245    247      1 ;   HG22 - CG1
246    247      1 ;   HG23 - CG1
247    254      1 ;    CG1 - C
248    251      1 ;   HG12 - HD11
248    252      1 ;   HG12 - HD12
248    253      1 ;   HG12 - HD13
249    251      1 ;   HG13 - HD11
249    252      1 ;   HG13 - HD12
```

| | | | |
|---|---|---|---|
| 249 | 253 | 1 ; | HG13 - HD13 |
| 254 | 259 | 1 ; | C - HA |
| 254 | 260 | 1 ; | C - CB |
| 254 | 266 | 1 ; | C - C |
| 255 | 257 | 1 ; | O - H |
| 255 | 258 | 1 ; | O - CA |
| 256 | 261 | 1 ; | N - HB2 |
| 256 | 262 | 1 ; | N - HB3 |
| 256 | 263 | 1 ; | N - CG |
| 256 | 267 | 1 ; | N - O |
| 256 | 268 | 1 ; | N - N |
| 257 | 259 | 1 ; | H - HA |
| 257 | 260 | 1 ; | H - CB |
| 257 | 266 | 1 ; | H - C |
| 258 | 264 | 1 ; | CA - OD1 |
| 258 | 265 | 1 ; | CA - OD2 |
| 258 | 269 | 1 ; | CA - H |
| 258 | 270 | 1 ; | CA - CA |
| 259 | 261 | 1 ; | HA - HB2 |
| 259 | 262 | 1 ; | HA - HB3 |
| 259 | 263 | 1 ; | HA - CG |
| 259 | 267 | 1 ; | HA - O |
| 259 | 268 | 1 ; | HA - N |
| 260 | 267 | 1 ; | CB - O |
| 260 | 268 | 1 ; | CB - N |
| 261 | 264 | 1 ; | HB2 - OD1 |
| 261 | 265 | 1 ; | HB2 - OD2 |
| 261 | 266 | 1 ; | HB2 - C |
| 262 | 264 | 1 ; | HB3 - OD1 |
| 262 | 265 | 1 ; | HB3 - OD2 |
| 262 | 266 | 1 ; | HB3 - C |
| 263 | 266 | 1 ; | CG - C |
| 266 | 271 | 1 ; | C - HA |
| 266 | 272 | 1 ; | C - CB |
| 266 | 281 | 1 ; | C - C |
| 267 | 269 | 1 ; | O - H |
| 267 | 270 | 1 ; | O - CA |
| 268 | 273 | 1 ; | N - HB2 |
| 268 | 274 | 1 ; | N - HB3 |
| 268 | 275 | 1 ; | N - CG |
| 268 | 282 | 1 ; | N - O |
| 268 | 283 | 1 ; | N - N |
| 269 | 271 | 1 ; | H - HA |
| 269 | 272 | 1 ; | H - CB |
| 269 | 281 | 1 ; | H - C |
| 270 | 276 | 1 ; | CA - HG2 |
| 270 | 277 | 1 ; | CA - HG3 |
| 270 | 278 | 1 ; | CA - CD |
| 270 | 284 | 1 ; | CA - H |
| 270 | 285 | 1 ; | CA - CA |
| 271 | 273 | 1 ; | HA - HB2 |
| 271 | 274 | 1 ; | HA - HB3 |

| | | | |
|---|---|---|---|
| 271 | 275 | 1 ; | HA - CG |
| 271 | 282 | 1 ; | HA - O |
| 271 | 283 | 1 ; | HA - N |
| 272 | 279 | 1 ; | CB - OE1 |
| 272 | 280 | 1 ; | CB - OE2 |
| 272 | 282 | 1 ; | CB - O |
| 272 | 283 | 1 ; | CB - N |
| 273 | 276 | 1 ; | HB2 - HG2 |
| 273 | 277 | 1 ; | HB2 - HG3 |
| 273 | 278 | 1 ; | HB2 - CD |
| 273 | 281 | 1 ; | HB2 - C |
| 274 | 276 | 1 ; | HB3 - HG2 |
| 274 | 277 | 1 ; | HB3 - HG3 |
| 274 | 278 | 1 ; | HB3 - CD |
| 274 | 281 | 1 ; | HB3 - C |
| 275 | 281 | 1 ; | CG - C |
| 276 | 279 | 1 ; | HG2 - OE1 |
| 276 | 280 | 1 ; | HG2 - OE2 |
| 277 | 279 | 1 ; | HG3 - OE1 |
| 277 | 280 | 1 ; | HG3 - OE2 |
| 281 | 286 | 1 ; | C - HA |
| 281 | 287 | 1 ; | C - CB |
| 281 | 303 | 1 ; | C - C |
| 282 | 284 | 1 ; | O - H |
| 282 | 285 | 1 ; | O - CA |
| 283 | 288 | 1 ; | N - HB2 |
| 283 | 289 | 1 ; | N - HB3 |
| 283 | 290 | 1 ; | N - CG |
| 283 | 304 | 1 ; | N - O |
| 283 | 305 | 1 ; | N - N |
| 284 | 286 | 1 ; | H - HA |
| 284 | 287 | 1 ; | H - CB |
| 284 | 303 | 1 ; | H - C |
| 285 | 291 | 1 ; | CA - HG2 |
| 285 | 292 | 1 ; | CA - HG3 |
| 285 | 293 | 1 ; | CA - CD |
| 285 | 306 | 1 ; | CA - H |
| 285 | 307 | 1 ; | CA - CA |
| 286 | 288 | 1 ; | HA - HB2 |
| 286 | 289 | 1 ; | HA - HB3 |
| 286 | 290 | 1 ; | HA - CG |
| 286 | 304 | 1 ; | HA - O |
| 286 | 305 | 1 ; | HA - N |
| 287 | 294 | 1 ; | CB - HD2 |
| 287 | 295 | 1 ; | CB - HD3 |
| 287 | 296 | 1 ; | CB - CE |
| 287 | 304 | 1 ; | CB - O |
| 287 | 305 | 1 ; | CB - N |
| 288 | 291 | 1 ; | HB2 - HG2 |
| 288 | 292 | 1 ; | HB2 - HG3 |
| 288 | 293 | 1 ; | HB2 - CD |
| 288 | 303 | 1 ; | HB2 - C |

```
289    291      1 ;    HB3 - HG2
289    292      1 ;    HB3 - HG3
289    293      1 ;    HB3 - CD
289    303      1 ;    HB3 - C
290    297      1 ;     CG - HE2
290    298      1 ;     CG - HE3
290    299      1 ;     CG - NZ
290    303      1 ;     CG - C
291    294      1 ;    HG2 - HD2
291    295      1 ;    HG2 - HD3
291    296      1 ;    HG2 - CE
292    294      1 ;    HG3 - HD2
292    295      1 ;    HG3 - HD3
292    296      1 ;    HG3 - CE
293    300      1 ;     CD - HZ1
293    301      1 ;     CD - HZ2
293    302      1 ;     CD - HZ3
294    297      1 ;    HD2 - HE2
294    298      1 ;    HD2 - HE3
294    299      1 ;    HD2 - NZ
295    297      1 ;    HD3 - HE2
295    298      1 ;    HD3 - HE3
295    299      1 ;    HD3 - NZ
297    300      1 ;    HE2 - HZ1
297    301      1 ;    HE2 - HZ2
297    302      1 ;    HE2 - HZ3
298    300      1 ;    HE3 - HZ1
298    301      1 ;    HE3 - HZ2
298    302      1 ;    HE3 - HZ3
303    308      1 ;      C - HA
303    309      1 ;      C - CB
303    322      1 ;      C - C
304    306      1 ;      O - H
304    307      1 ;      O - CA
305    310      1 ;      N - HB2
305    311      1 ;      N - HB3
305    312      1 ;      N - CG
305    323      1 ;      N - O
305    324      1 ;      N - N
306    308      1 ;      H - HA
306    309      1 ;      H - CB
306    322      1 ;      H - C
307    313      1 ;     CA - HG
307    314      1 ;     CA - CD1
307    318      1 ;     CA - CD2
307    325      1 ;     CA - H
307    326      1 ;     CA - CA
308    310      1 ;     HA - HB2
308    311      1 ;     HA - HB3
308    312      1 ;     HA - CG
308    323      1 ;     HA - O
308    324      1 ;     HA - N
```

```
  309   315     1 ;     CB - HD11
  309   316     1 ;     CB - HD12
  309   317     1 ;     CB - HD13
  309   319     1 ;     CB - HD21
  309   320     1 ;     CB - HD22
  309   321     1 ;     CB - HD23
  309   323     1 ;     CB - O
  309   324     1 ;     CB - N
  310   313     1 ;    HB2 - HG
  310   314     1 ;    HB2 - CD1
  310   318     1 ;    HB2 - CD2
  310   322     1 ;    HB2 - C
  311   313     1 ;    HB3 - HG
  311   314     1 ;    HB3 - CD1
  311   318     1 ;    HB3 - CD2
  311   322     1 ;    HB3 - C
  312   322     1 ;     CG - C
  313   315     1 ;     HG - HD11
  313   316     1 ;     HG - HD12
  313   317     1 ;     HG - HD13
  313   319     1 ;     HG - HD21
  313   320     1 ;     HG - HD22
  313   321     1 ;     HG - HD23
  314   319     1 ;    CD1 - HD21
  314   320     1 ;    CD1 - HD22
  314   321     1 ;    CD1 - HD23
  315   318     1 ;   HD11 - CD2
  316   318     1 ;   HD12 - CD2
  317   318     1 ;   HD13 - CD2
  322   327     1 ;      C - HA2
  322   328     1 ;      C - HA3
  322   329     1 ;      C - C
  323   325     1 ;      O - H
  323   326     1 ;      O - CA
  324   330     1 ;      N - O
  324   331     1 ;      N - N
  325   327     1 ;      H - HA2
  325   328     1 ;      H - HA3
  325   329     1 ;      H - C
  326   332     1 ;     CA - H
  326   333     1 ;     CA - CA
  327   330     1 ;    HA2 - O
  327   331     1 ;    HA2 - N
  328   330     1 ;    HA3 - O
  328   331     1 ;    HA3 - N
  329   334     1 ;      C - HA
  329   335     1 ;      C - CB
  329   346     1 ;      C - C
  330   332     1 ;      O - H
  330   333     1 ;      O - CA
  331   336     1 ;      N - HB2
  331   337     1 ;      N - HB3
```

```
331    338     1 ;      N - CG
331    347     1 ;      N - O
331    348     1 ;      N - N
332    334     1 ;      H - HA
332    335     1 ;      H - CB
332    346     1 ;      H - C
333    339     1 ;     CA - HG2
333    340     1 ;     CA - HG3
333    341     1 ;     CA - CD
333    349     1 ;     CA - H
333    350     1 ;     CA - CA
334    336     1 ;     HA - HB2
334    337     1 ;     HA - HB3
334    338     1 ;     HA - CG
334    347     1 ;     HA - O
334    348     1 ;     HA - N
335    342     1 ;     CB - OE1
335    343     1 ;     CB - NE2
335    347     1 ;     CB - O
335    348     1 ;     CB - N
336    339     1 ;    HB2 - HG2
336    340     1 ;    HB2 - HG3
336    341     1 ;    HB2 - CD
336    346     1 ;    HB2 - C
337    339     1 ;    HB3 - HG2
337    340     1 ;    HB3 - HG3
337    341     1 ;    HB3 - CD
337    346     1 ;    HB3 - C
338    344     1 ;     CG - HE21
338    345     1 ;     CG - HE22
338    346     1 ;     CG - C
339    342     1 ;    HG2 - OE1
339    343     1 ;    HG2 - NE2
340    342     1 ;    HG3 - OE1
340    343     1 ;    HG3 - NE2
342    344     1 ;    OE1 - HE21
342    345     1 ;    OE1 - HE22
346    351     1 ;      C - HA2
346    352     1 ;      C - HA3
346    353     1 ;      C - C
347    349     1 ;      O - H
347    350     1 ;      O - CA
348    354     1 ;      N - O
348    355     1 ;      N - N
349    351     1 ;      H - HA2
349    352     1 ;      H - HA3
349    353     1 ;      H - C
350    356     1 ;     CA - H
350    357     1 ;     CA - CA
351    354     1 ;    HA2 - O
351    355     1 ;    HA2 - N
352    354     1 ;    HA3 - O
```

```
352     355       1 ;    HA3 - N
353     358       1 ;      C - HA
353     359       1 ;      C - CB
353     367       1 ;      C - C
354     356       1 ;      O - H
354     357       1 ;      O - CA
355     360       1 ;      N - HB
355     361       1 ;      N - CG2
355     365       1 ;      N - OG1
355     368       1 ;      N - O
355     369       1 ;      N - N
356     358       1 ;      H - HA
356     359       1 ;      H - CB
356     367       1 ;      H - C
357     362       1 ;     CA - HG21
357     363       1 ;     CA - HG22
357     364       1 ;     CA - HG23
357     366       1 ;     CA - HG1
357     370       1 ;     CA - H
357     371       1 ;     CA - CA
358     360       1 ;     HA - HB
358     361       1 ;     HA - CG2
358     365       1 ;     HA - OG1
358     368       1 ;     HA - O
358     369       1 ;     HA - N
359     368       1 ;     CB - O
359     369       1 ;     CB - N
360     362       1 ;     HB - HG21
360     363       1 ;     HB - HG22
360     364       1 ;     HB - HG23
360     366       1 ;     HB - HG1
360     367       1 ;     HB - C
361     366       1 ;    CG2 - HG1
361     367       1 ;    CG2 - C
362     365       1 ;   HG21 - OG1
363     365       1 ;   HG22 - OG1
364     365       1 ;   HG23 - OG1
365     367       1 ;    OG1 - C
367     372       1 ;      C - HA
367     373       1 ;      C - CB
367     388       1 ;      C - C
368     370       1 ;      O - H
368     371       1 ;      O - CA
369     374       1 ;      N - HB2
369     375       1 ;      N - HB3
369     376       1 ;      N - CG
369     389       1 ;      N - O
369     390       1 ;      N - N
370     372       1 ;      H - HA
370     373       1 ;      H - CB
370     388       1 ;      H - C
371     377       1 ;     CA - CD1
```

```
371    386      1 ;     CA - CD2
371    391      1 ;     CA - H
371    392      1 ;     CA - CA
372    374      1 ;     HA - HB2
372    375      1 ;     HA - HB3
372    376      1 ;     HA - CG
372    389      1 ;     HA - O
372    390      1 ;     HA - N
373    378      1 ;     CB - HD1
373    379      1 ;     CB - CE1
373    384      1 ;     CB - CE2
373    387      1 ;     CB - HD2
373    389      1 ;     CB - O
373    390      1 ;     CB - N
374    377      1 ;    HB2 - CD1
374    386      1 ;    HB2 - CD2
374    388      1 ;    HB2 - C
375    377      1 ;    HB3 - CD1
375    386      1 ;    HB3 - CD2
375    388      1 ;    HB3 - C
376    380      1 ;     CG - HE1
376    381      1 ;     CG - CZ
376    385      1 ;     CG - HE2
376    388      1 ;     CG - C
377    382      1 ;    CD1 - OH
377    384      1 ;    CD1 - CE2
377    387      1 ;    CD1 - HD2
378    380      1 ;    HD1 - HE1
378    381      1 ;    HD1 - CZ
378    386      1 ;    HD1 - CD2
379    383      1 ;    CE1 - HH
379    385      1 ;    CE1 - HE2
379    386      1 ;    CE1 - CD2
380    382      1 ;    HE1 - OH
380    384      1 ;    HE1 - CE2
381    387      1 ;     CZ - HD2
382    385      1 ;     OH - HE2
382    386      1 ;     OH - CD2
383    384      1 ;     HH - CE2
385    387      1 ;    HE2 - HD2
388    393      1 ;      C - HA2
388    394      1 ;      C - HA3
388    395      1 ;      C - C
389    391      1 ;      O - H
389    392      1 ;      O - CA
390    396      1 ;      N - O
390    397      1 ;      N - N
391    393      1 ;      H - HA2
391    394      1 ;      H - HA3
391    395      1 ;      H - C
392    398      1 ;     CA - H
392    399      1 ;     CA - CA
```

```
393    396      1 ;    HA2 - O
393    397      1 ;    HA2 - N
394    396      1 ;    HA3 - O
394    397      1 ;    HA3 - N
395    400      1 ;      C - HA
395    401      1 ;      C - CB
395    406      1 ;      C - C
396    398      1 ;      O - H
396    399      1 ;      O - CA
397    402      1 ;      N - HB2
397    403      1 ;      N - HB3
397    404      1 ;      N - SG
397    407      1 ;      N - O
397    408      1 ;      N - N
398    400      1 ;      H - HA
398    401      1 ;      H - CB
398    406      1 ;      H - C
399    405      1 ;     CA - HG
399    409      1 ;     CA - H
399    410      1 ;     CA - CA
400    402      1 ;     HA - HB2
400    403      1 ;     HA - HB3
400    404      1 ;     HA - SG
400    407      1 ;     HA - O
400    408      1 ;     HA - N
401    407      1 ;     CB - O
401    408      1 ;     CB - N
402    405      1 ;    HB2 - HG
402    406      1 ;    HB2 - C
403    405      1 ;    HB3 - HG
403    406      1 ;    HB3 - C
404    406      1 ;     SG - C
406    411      1 ;      C - HA
406    412      1 ;      C - CB
406    422      1 ;      C - C
407    409      1 ;      O - H
407    410      1 ;      O - CA
408    413      1 ;      N - HB
408    414      1 ;      N - CG1
408    418      1 ;      N - CG2
408    423      1 ;      N - O
408    424      1 ;      N - N
409    411      1 ;      H - HA
409    412      1 ;      H - CB
409    422      1 ;      H - C
410    415      1 ;     CA - HG11
410    416      1 ;     CA - HG12
410    417      1 ;     CA - HG13
410    419      1 ;     CA - HG21
410    420      1 ;     CA - HG22
410    421      1 ;     CA - HG23
410    425      1 ;     CA - H
```

```
410    426      1 ;      CA - CA
411    413      1 ;      HA - HB
411    414      1 ;      HA - CG1
411    418      1 ;      HA - CG2
411    423      1 ;      HA - O
411    424      1 ;      HA - N
412    423      1 ;      CB - O
412    424      1 ;      CB - N
413    415      1 ;      HB - HG11
413    416      1 ;      HB - HG12
413    417      1 ;      HB - HG13
413    419      1 ;      HB - HG21
413    420      1 ;      HB - HG22
413    421      1 ;      HB - HG23
413    422      1 ;      HB - C
414    419      1 ;     CG1 - HG21
414    420      1 ;     CG1 - HG22
414    421      1 ;     CG1 - HG23
414    422      1 ;     CG1 - C
415    418      1 ;    HG11 - CG2
416    418      1 ;    HG12 - CG2
417    418      1 ;    HG13 - CG2
418    422      1 ;     CG2 - C
422    427      1 ;       C - HA
422    428      1 ;       C - CB
422    443      1 ;       C - C
423    425      1 ;       O - H
423    426      1 ;       O - CA
424    429      1 ;       N - HB2
424    430      1 ;       N - HB3
424    431      1 ;       N - CG
424    444      1 ;       N - O
424    445      1 ;       N - N
425    427      1 ;       H - HA
425    428      1 ;       H - CB
425    443      1 ;       H - C
426    432      1 ;      CA - CD1
426    441      1 ;      CA - CD2
426    446      1 ;      CA - H
426    447      1 ;      CA - CA
427    429      1 ;      HA - HB2
427    430      1 ;      HA - HB3
427    431      1 ;      HA - CG
427    444      1 ;      HA - O
427    445      1 ;      HA - N
428    433      1 ;      CB - HD1
428    434      1 ;      CB - CE1
428    439      1 ;      CB - CE2
428    442      1 ;      CB - HD2
428    444      1 ;      CB - O
428    445      1 ;      CB - N
429    432      1 ;     HB2 - CD1
```

| | | | |
|---|---|---|---|
| 429 | 441 | 1 ; | HB2 - CD2 |
| 429 | 443 | 1 ; | HB2 - C |
| 430 | 432 | 1 ; | HB3 - CD1 |
| 430 | 441 | 1 ; | HB3 - CD2 |
| 430 | 443 | 1 ; | HB3 - C |
| 431 | 435 | 1 ; | CG - HE1 |
| 431 | 436 | 1 ; | CG - CZ |
| 431 | 440 | 1 ; | CG - HE2 |
| 431 | 443 | 1 ; | CG - C |
| 432 | 437 | 1 ; | CD1 - OH |
| 432 | 439 | 1 ; | CD1 - CE2 |
| 432 | 442 | 1 ; | CD1 - HD2 |
| 433 | 435 | 1 ; | HD1 - HE1 |
| 433 | 436 | 1 ; | HD1 - CZ |
| 433 | 441 | 1 ; | HD1 - CD2 |
| 434 | 438 | 1 ; | CE1 - HH |
| 434 | 440 | 1 ; | CE1 - HE2 |
| 434 | 441 | 1 ; | CE1 - CD2 |
| 435 | 437 | 1 ; | HE1 - OH |
| 435 | 439 | 1 ; | HE1 - CE2 |
| 436 | 442 | 1 ; | CZ - HD2 |
| 437 | 440 | 1 ; | OH - HE2 |
| 437 | 441 | 1 ; | OH - CD2 |
| 438 | 439 | 1 ; | HH - CE2 |
| 440 | 442 | 1 ; | HE2 - HD2 |
| 443 | 448 | 1 ; | C - HA |
| 443 | 449 | 1 ; | C - CB |
| 443 | 465 | 1 ; | C - C |
| 444 | 446 | 1 ; | O - H |
| 444 | 447 | 1 ; | O - CA |
| 445 | 450 | 1 ; | N - HB2 |
| 445 | 451 | 1 ; | N - HB3 |
| 445 | 452 | 1 ; | N - CG |
| 445 | 466 | 1 ; | N - O |
| 445 | 467 | 1 ; | N - N |
| 446 | 448 | 1 ; | H - HA |
| 446 | 449 | 1 ; | H - CB |
| 446 | 465 | 1 ; | H - C |
| 447 | 453 | 1 ; | CA - HG2 |
| 447 | 454 | 1 ; | CA - HG3 |
| 447 | 455 | 1 ; | CA - CD |
| 447 | 468 | 1 ; | CA - H |
| 447 | 469 | 1 ; | CA - CA |
| 448 | 450 | 1 ; | HA - HB2 |
| 448 | 451 | 1 ; | HA - HB3 |
| 448 | 452 | 1 ; | HA - CG |
| 448 | 466 | 1 ; | HA - O |
| 448 | 467 | 1 ; | HA - N |
| 449 | 456 | 1 ; | CB - HD2 |
| 449 | 457 | 1 ; | CB - HD3 |
| 449 | 458 | 1 ; | CB - CE |
| 449 | 466 | 1 ; | CB - O |

```
  449    467       1 ;      CB - N
  450    453       1 ;     HB2 - HG2
  450    454       1 ;     HB2 - HG3
  450    455       1 ;     HB2 - CD
  450    465       1 ;     HB2 - C
  451    453       1 ;     HB3 - HG2
  451    454       1 ;     HB3 - HG3
  451    455       1 ;     HB3 - CD
  451    465       1 ;     HB3 - C
  452    459       1 ;      CG - HE2
  452    460       1 ;      CG - HE3
  452    461       1 ;      CG - NZ
  452    465       1 ;      CG - C
  453    456       1 ;     HG2 - HD2
  453    457       1 ;     HG2 - HD3
  453    458       1 ;     HG2 - CE
  454    456       1 ;     HG3 - HD2
  454    457       1 ;     HG3 - HD3
  454    458       1 ;     HG3 - CE
  455    462       1 ;      CD - HZ1
  455    463       1 ;      CD - HZ2
  455    464       1 ;      CD - HZ3
  456    459       1 ;     HD2 - HE2
  456    460       1 ;     HD2 - HE3
  456    461       1 ;     HD2 - NZ
  457    459       1 ;     HD3 - HE2
  457    460       1 ;     HD3 - HE3
  457    461       1 ;     HD3 - NZ
  459    462       1 ;     HE2 - HZ1
  459    463       1 ;     HE2 - HZ2
  459    464       1 ;     HE2 - HZ3
  460    462       1 ;     HE3 - HZ1
  460    463       1 ;     HE3 - HZ2
  460    464       1 ;     HE3 - HZ3
  465    470       1 ;       C - HA2
  465    471       1 ;       C - HA3
  465    472       1 ;       C - C
  466    468       1 ;       O - H
  466    469       1 ;       O - CA
  467    473       1 ;       N - O
  467    474       1 ;       N - N
  468    470       1 ;       H - HA2
  468    471       1 ;       H - HA3
  468    472       1 ;       H - C
  469    475       1 ;      CA - H
  469    476       1 ;      CA - CA
  470    473       1 ;     HA2 - O
  470    474       1 ;     HA2 - N
  471    473       1 ;     HA3 - O
  471    474       1 ;     HA3 - N
  472    477       1 ;       C - HA
  472    478       1 ;       C - CB
```

```
472    491       1 ;      C - C
473    475       1 ;      O - H
473    476       1 ;      O - CA
474    479       1 ;      N - HB
474    480       1 ;      N - CG2
474    484       1 ;      N - CG1
474    492       1 ;      N - O
474    493       1 ;      N - N
475    477       1 ;      H - HA
475    478       1 ;      H - CB
475    491       1 ;      H - C
476    481       1 ;     CA - HG21
476    482       1 ;     CA - HG22
476    483       1 ;     CA - HG23
476    485       1 ;     CA - HG12
476    486       1 ;     CA - HG13
476    487       1 ;     CA - CD1
476    494       1 ;     CA - H
476    495       1 ;     CA - CA
477    479       1 ;     HA - HB
477    480       1 ;     HA - CG2
477    484       1 ;     HA - CG1
477    492       1 ;     HA - O
477    493       1 ;     HA - N
478    488       1 ;     CB - HD11
478    489       1 ;     CB - HD12
478    490       1 ;     CB - HD13
478    492       1 ;     CB - O
478    493       1 ;     CB - N
479    481       1 ;     HB - HG21
479    482       1 ;     HB - HG22
479    483       1 ;     HB - HG23
479    485       1 ;     HB - HG12
479    486       1 ;     HB - HG13
479    487       1 ;     HB - CD1
479    491       1 ;     HB - C
480    485       1 ;    CG2 - HG12
480    486       1 ;    CG2 - HG13
480    487       1 ;    CG2 - CD1
480    491       1 ;    CG2 - C
481    484       1 ;   HG21 - CG1
482    484       1 ;   HG22 - CG1
483    484       1 ;   HG23 - CG1
484    491       1 ;    CG1 - C
485    488       1 ;   HG12 - HD11
485    489       1 ;   HG12 - HD12
485    490       1 ;   HG12 - HD13
486    488       1 ;   HG13 - HD11
486    489       1 ;   HG13 - HD12
486    490       1 ;   HG13 - HD13
491    496       1 ;      C - HA
491    497       1 ;      C - CB
```

```
491    503     1 ;     C - C
492    494     1 ;     O - H
492    495     1 ;     O - CA
493    498     1 ;     N - HB2
493    499     1 ;     N - HB3
493    500     1 ;     N - CG
493    504     1 ;     N - O
493    505     1 ;     N - N
494    496     1 ;     H - HA
494    497     1 ;     H - CB
494    503     1 ;     H - C
495    501     1 ;    CA - OD1
495    502     1 ;    CA - OD2
495    506     1 ;    CA - H
495    507     1 ;    CA - CA
496    498     1 ;    HA - HB2
496    499     1 ;    HA - HB3
496    500     1 ;    HA - CG
496    504     1 ;    HA - O
496    505     1 ;    HA - N
497    504     1 ;    CB - O
497    505     1 ;    CB - N
498    501     1 ;   HB2 - OD1
498    502     1 ;   HB2 - OD2
498    503     1 ;   HB2 - C
499    501     1 ;   HB3 - OD1
499    502     1 ;   HB3 - OD2
499    503     1 ;   HB3 - C
500    503     1 ;    CG - C
503    508     1 ;     C - HA
503    509     1 ;     C - CB
503    525     1 ;     C - C
504    506     1 ;     O - H
504    507     1 ;     O - CA
505    510     1 ;     N - HB2
505    511     1 ;     N - HB3
505    512     1 ;     N - CG
505    526     1 ;     N - O
505    527     1 ;     N - N
506    508     1 ;     H - HA
506    509     1 ;     H - CB
506    525     1 ;     H - C
507    513     1 ;    CA - HG2
507    514     1 ;    CA - HG3
507    515     1 ;    CA - CD
507    528     1 ;    CA - H
507    529     1 ;    CA - CA
508    510     1 ;    HA - HB2
508    511     1 ;    HA - HB3
508    512     1 ;    HA - CG
508    526     1 ;    HA - O
508    527     1 ;    HA - N
```

```
509    516       1 ;      CB - HD2
509    517       1 ;      CB - HD3
509    518       1 ;      CB - CE
509    526       1 ;      CB - O
509    527       1 ;      CB - N
510    513       1 ;     HB2 - HG2
510    514       1 ;     HB2 - HG3
510    515       1 ;     HB2 - CD
510    525       1 ;     HB2 - C
511    513       1 ;     HB3 - HG2
511    514       1 ;     HB3 - HG3
511    515       1 ;     HB3 - CD
511    525       1 ;     HB3 - C
512    519       1 ;      CG - HE2
512    520       1 ;      CG - HE3
512    521       1 ;      CG - NZ
512    525       1 ;      CG - C
513    516       1 ;     HG2 - HD2
513    517       1 ;     HG2 - HD3
513    518       1 ;     HG2 - CE
514    516       1 ;     HG3 - HD2
514    517       1 ;     HG3 - HD3
514    518       1 ;     HG3 - CE
515    522       1 ;      CD - HZ1
515    523       1 ;      CD - HZ2
515    524       1 ;      CD - HZ3
516    519       1 ;     HD2 - HE2
516    520       1 ;     HD2 - HE3
516    521       1 ;     HD2 - NZ
517    519       1 ;     HD3 - HE2
517    520       1 ;     HD3 - HE3
517    521       1 ;     HD3 - NZ
519    522       1 ;     HE2 - HZ1
519    523       1 ;     HE2 - HZ2
519    524       1 ;     HE2 - HZ3
520    522       1 ;     HE3 - HZ1
520    523       1 ;     HE3 - HZ2
520    524       1 ;     HE3 - HZ3
525    530       1 ;       C - HA
525    531       1 ;       C - CB
525    541       1 ;       C - C
526    528       1 ;       O - H
526    529       1 ;       O - CA
527    532       1 ;       N - HB
527    533       1 ;       N - CG1
527    537       1 ;       N - CG2
527    542       1 ;       N - O
527    543       1 ;       N - N
528    530       1 ;       H - HA
528    531       1 ;       H - CB
528    541       1 ;       H - C
529    534       1 ;      CA - HG11
```

| | | | | |
|---|---|---|---|---|
| 529 | 535 | 1 | ; | CA - HG12 |
| 529 | 536 | 1 | ; | CA - HG13 |
| 529 | 538 | 1 | ; | CA - HG21 |
| 529 | 539 | 1 | ; | CA - HG22 |
| 529 | 540 | 1 | ; | CA - HG23 |
| 529 | 544 | 1 | ; | CA - H |
| 529 | 545 | 1 | ; | CA - CA |
| 530 | 532 | 1 | ; | HA - HB |
| 530 | 533 | 1 | ; | HA - CG1 |
| 530 | 537 | 1 | ; | HA - CG2 |
| 530 | 542 | 1 | ; | HA - O |
| 530 | 543 | 1 | ; | HA - N |
| 531 | 542 | 1 | ; | CB - O |
| 531 | 543 | 1 | ; | CB - N |
| 532 | 534 | 1 | ; | HB - HG11 |
| 532 | 535 | 1 | ; | HB - HG12 |
| 532 | 536 | 1 | ; | HB - HG13 |
| 532 | 538 | 1 | ; | HB - HG21 |
| 532 | 539 | 1 | ; | HB - HG22 |
| 532 | 540 | 1 | ; | HB - HG23 |
| 532 | 541 | 1 | ; | HB - C |
| 533 | 538 | 1 | ; | CG1 - HG21 |
| 533 | 539 | 1 | ; | CG1 - HG22 |
| 533 | 540 | 1 | ; | CG1 - HG23 |
| 533 | 541 | 1 | ; | CG1 - C |
| 534 | 537 | 1 | ; | HG11 - CG2 |
| 535 | 537 | 1 | ; | HG12 - CG2 |
| 536 | 537 | 1 | ; | HG13 - CG2 |
| 537 | 541 | 1 | ; | CG2 - C |
| 541 | 546 | 1 | ; | C - HA |
| 541 | 547 | 1 | ; | C - CB |
| 541 | 555 | 1 | ; | C - C |
| 542 | 544 | 1 | ; | O - H |
| 542 | 545 | 1 | ; | O - CA |
| 543 | 548 | 1 | ; | N - HB |
| 543 | 549 | 1 | ; | N - CG2 |
| 543 | 553 | 1 | ; | N - OG1 |
| 543 | 556 | 1 | ; | N - O |
| 543 | 557 | 1 | ; | N - N |
| 544 | 546 | 1 | ; | H - HA |
| 544 | 547 | 1 | ; | H - CB |
| 544 | 555 | 1 | ; | H - C |
| 545 | 550 | 1 | ; | CA - HG21 |
| 545 | 551 | 1 | ; | CA - HG22 |
| 545 | 552 | 1 | ; | CA - HG23 |
| 545 | 554 | 1 | ; | CA - HG1 |
| 545 | 558 | 1 | ; | CA - H |
| 545 | 559 | 1 | ; | CA - CA |
| 546 | 548 | 1 | ; | HA - HB |
| 546 | 549 | 1 | ; | HA - CG2 |
| 546 | 553 | 1 | ; | HA - OG1 |
| 546 | 556 | 1 | ; | HA - O |

```
546    557      1 ;     HA - N
547    556      1 ;     CB - O
547    557      1 ;     CB - N
548    550      1 ;     HB - HG21
548    551      1 ;     HB - HG22
548    552      1 ;     HB - HG23
548    554      1 ;     HB - HG1
548    555      1 ;     HB - C
549    554      1 ;    CG2 - HG1
549    555      1 ;    CG2 - C
550    553      1 ;   HG21 - OG1
551    553      1 ;   HG22 - OG1
552    553      1 ;   HG23 - OG1
553    555      1 ;    OG1 - C
555    560      1 ;      C - HA
555    561      1 ;      C - CB
555    569      1 ;      C - C
556    558      1 ;      O - H
556    559      1 ;      O - CA
557    562      1 ;      N - HB2
557    563      1 ;      N - HB3
557    564      1 ;      N - CG
557    570      1 ;      N - O
557    571      1 ;      N - N
558    560      1 ;      H - HA
558    561      1 ;      H - CB
558    569      1 ;      H - C
559    565      1 ;     CA - OD1
559    566      1 ;     CA - ND2
559    572      1 ;     CA - H
559    573      1 ;     CA - CA
560    562      1 ;     HA - HB2
560    563      1 ;     HA - HB3
560    564      1 ;     HA - CG
560    570      1 ;     HA - O
560    571      1 ;     HA - N
561    567      1 ;     CB - HD21
561    568      1 ;     CB - HD22
561    570      1 ;     CB - O
561    571      1 ;     CB - N
562    565      1 ;    HB2 - OD1
562    566      1 ;    HB2 - ND2
562    569      1 ;    HB2 - C
563    565      1 ;    HB3 - OD1
563    566      1 ;    HB3 - ND2
563    569      1 ;    HB3 - C
564    569      1 ;     CG - C
565    567      1 ;    OD1 - HD21
565    568      1 ;    OD1 - HD22
569    574      1 ;      C - HA
569    575      1 ;      C - CB
569    586      1 ;      C - C
```

```
570    572    1 ;     O - H
570    573    1 ;     O - CA
571    576    1 ;     N - HB2
571    577    1 ;     N - HB3
571    578    1 ;     N - CG
571    587    1 ;     N - O
571    588    1 ;     N - N
572    574    1 ;     H - HA
572    575    1 ;     H - CB
572    586    1 ;     H - C
573    579    1 ;    CA - HG2
573    580    1 ;    CA - HG3
573    581    1 ;    CA - CD
573    589    1 ;    CA - H
573    590    1 ;    CA - CA
574    576    1 ;    HA - HB2
574    577    1 ;    HA - HB3
574    578    1 ;    HA - CG
574    587    1 ;    HA - O
574    588    1 ;    HA - N
575    582    1 ;    CB - OE1
575    583    1 ;    CB - NE2
575    587    1 ;    CB - O
575    588    1 ;    CB - N
576    579    1 ;   HB2 - HG2
576    580    1 ;   HB2 - HG3
576    581    1 ;   HB2 - CD
576    586    1 ;   HB2 - C
577    579    1 ;   HB3 - HG2
577    580    1 ;   HB3 - HG3
577    581    1 ;   HB3 - CD
577    586    1 ;   HB3 - C
578    584    1 ;    CG - HE21
578    585    1 ;    CG - HE22
578    586    1 ;    CG - C
579    582    1 ;   HG2 - OE1
579    583    1 ;   HG2 - NE2
580    582    1 ;   HG3 - OE1
580    583    1 ;   HG3 - NE2
582    584    1 ;   OE1 - HE21
582    585    1 ;   OE1 - HE22
586    591    1 ;     C - HA
586    592    1 ;     C - CB
586    605    1 ;     C - C
587    589    1 ;     O - H
587    590    1 ;     O - CA
588    593    1 ;     N - HB2
588    594    1 ;     N - HB3
588    595    1 ;     N - CG
588    606    1 ;     N - O
588    607    1 ;     N - N
589    591    1 ;     H - HA
```

```
589    592       1 ;       H - CB
589    605       1 ;       H - C
590    596       1 ;      CA - HG
590    597       1 ;      CA - CD1
590    601       1 ;      CA - CD2
590    608       1 ;      CA - H
590    609       1 ;      CA - CA
591    593       1 ;      HA - HB2
591    594       1 ;      HA - HB3
591    595       1 ;      HA - CG
591    606       1 ;      HA - O
591    607       1 ;      HA - N
592    598       1 ;      CB - HD11
592    599       1 ;      CB - HD12
592    600       1 ;      CB - HD13
592    602       1 ;      CB - HD21
592    603       1 ;      CB - HD22
592    604       1 ;      CB - HD23
592    606       1 ;      CB - O
592    607       1 ;      CB - N
593    596       1 ;     HB2 - HG
593    597       1 ;     HB2 - CD1
593    601       1 ;     HB2 - CD2
593    605       1 ;     HB2 - C
594    596       1 ;     HB3 - HG
594    597       1 ;     HB3 - CD1
594    601       1 ;     HB3 - CD2
594    605       1 ;     HB3 - C
595    605       1 ;      CG - C
596    598       1 ;      HG - HD11
596    599       1 ;      HG - HD12
596    600       1 ;      HG - HD13
596    602       1 ;      HG - HD21
596    603       1 ;      HG - HD22
596    604       1 ;      HG - HD23
597    602       1 ;     CD1 - HD21
597    603       1 ;     CD1 - HD22
597    604       1 ;     CD1 - HD23
598    601       1 ;    HD11 - CD2
599    601       1 ;    HD12 - CD2
600    601       1 ;    HD13 - CD2
605    610       1 ;       C - HA
605    611       1 ;       C - CB
605    626       1 ;       C - C
606    608       1 ;       O - H
606    609       1 ;       O - CA
607    612       1 ;       N - HB2
607    613       1 ;       N - HB3
607    614       1 ;       N - CG
607    627       1 ;       N - O
607    628       1 ;       N - N
608    610       1 ;       H - HA
```

```
  608    611      1 ;      H - CB
  608    626      1 ;      H - C
  609    615      1 ;     CA - CD1
  609    624      1 ;     CA - CD2
  609    629      1 ;     CA - H
  609    630      1 ;     CA - CA
  610    612      1 ;     HA - HB2
  610    613      1 ;     HA - HB3
  610    614      1 ;     HA - CG
  610    627      1 ;     HA - O
  610    628      1 ;     HA - N
  611    616      1 ;     CB - HD1
  611    617      1 ;     CB - CE1
  611    622      1 ;     CB - CE2
  611    625      1 ;     CB - HD2
  611    627      1 ;     CB - O
  611    628      1 ;     CB - N
  612    615      1 ;    HB2 - CD1
  612    624      1 ;    HB2 - CD2
  612    626      1 ;    HB2 - C
  613    615      1 ;    HB3 - CD1
  613    624      1 ;    HB3 - CD2
  613    626      1 ;    HB3 - C
  614    618      1 ;     CG - HE1
  614    619      1 ;     CG - CZ
  614    623      1 ;     CG - HE2
  614    626      1 ;     CG - C
  615    620      1 ;    CD1 - OH
  615    622      1 ;    CD1 - CE2
  615    625      1 ;    CD1 - HD2
  616    618      1 ;    HD1 - HE1
  616    619      1 ;    HD1 - CZ
  616    624      1 ;    HD1 - CD2
  617    621      1 ;    CE1 - HH
  617    623      1 ;    CE1 - HE2
  617    624      1 ;    CE1 - CD2
  618    620      1 ;    HE1 - OH
  618    622      1 ;    HE1 - CE2
  619    625      1 ;     CZ - HD2
  620    623      1 ;     OH - HE2
  620    624      1 ;     OH - CD2
  621    622      1 ;     HH - CE2
  623    625      1 ;    HE2 - HD2
  626    631      1 ;      C - HA
  626    632      1 ;      C - CB
  626    636      1 ;      C - C
  627    629      1 ;      O - H
  627    630      1 ;      O - CA
  628    633      1 ;      N - HB1
  628    634      1 ;      N - HB2
  628    635      1 ;      N - HB3
  628    637      1 ;      N - O
```

```
628    638       1 ;      N - N
629    631       1 ;      H - HA
629    632       1 ;      H - CB
629    636       1 ;      H - C
630    639       1 ;     CA - H
630    640       1 ;     CA - CA
631    633       1 ;     HA - HB1
631    634       1 ;     HA - HB2
631    635       1 ;     HA - HB3
631    637       1 ;     HA - O
631    638       1 ;     HA - N
632    637       1 ;     CB - O
632    638       1 ;     CB - N
633    636       1 ;    HB1 - C
634    636       1 ;    HB2 - C
635    636       1 ;    HB3 - C
636    641       1 ;      C - HA
636    642       1 ;      C - CB
636    655       1 ;      C - C
637    639       1 ;      O - H
637    640       1 ;      O - CA
638    643       1 ;      N - HB
638    644       1 ;      N - CG2
638    648       1 ;      N - CG1
638    656       1 ;      N - O
638    657       1 ;      N - N
639    641       1 ;      H - HA
639    642       1 ;      H - CB
639    655       1 ;      H - C
640    645       1 ;     CA - HG21
640    646       1 ;     CA - HG22
640    647       1 ;     CA - HG23
640    649       1 ;     CA - HG12
640    650       1 ;     CA - HG13
640    651       1 ;     CA - CD1
640    658       1 ;     CA - H
640    659       1 ;     CA - CA
641    643       1 ;     HA - HB
641    644       1 ;     HA - CG2
641    648       1 ;     HA - CG1
641    656       1 ;     HA - O
641    657       1 ;     HA - N
642    652       1 ;     CB - HD11
642    653       1 ;     CB - HD12
642    654       1 ;     CB - HD13
642    656       1 ;     CB - O
642    657       1 ;     CB - N
643    645       1 ;     HB - HG21
643    646       1 ;     HB - HG22
643    647       1 ;     HB - HG23
643    649       1 ;     HB - HG12
643    650       1 ;     HB - HG13
```

| | | | | |
|---|---|---|---|---|
| 643 | 651 | 1 ; | HB - | CD1 |
| 643 | 655 | 1 ; | HB - | C |
| 644 | 649 | 1 ; | CG2 - | HG12 |
| 644 | 650 | 1 ; | CG2 - | HG13 |
| 644 | 651 | 1 ; | CG2 - | CD1 |
| 644 | 655 | 1 ; | CG2 - | C |
| 645 | 648 | 1 ; | HG21 - | CG1 |
| 646 | 648 | 1 ; | HG22 - | CG1 |
| 647 | 648 | 1 ; | HG23 - | CG1 |
| 648 | 655 | 1 ; | CG1 - | C |
| 649 | 652 | 1 ; | HG12 - | HD11 |
| 649 | 653 | 1 ; | HG12 - | HD12 |
| 649 | 654 | 1 ; | HG12 - | HD13 |
| 650 | 652 | 1 ; | HG13 - | HD11 |
| 650 | 653 | 1 ; | HG13 - | HD12 |
| 650 | 654 | 1 ; | HG13 - | HD13 |
| 655 | 660 | 1 ; | C - | HA |
| 655 | 661 | 1 ; | C - | CB |
| 655 | 677 | 1 ; | C - | C |
| 656 | 658 | 1 ; | O - | H |
| 656 | 659 | 1 ; | O - | CA |
| 657 | 662 | 1 ; | N - | HB2 |
| 657 | 663 | 1 ; | N - | HB3 |
| 657 | 664 | 1 ; | N - | CG |
| 657 | 678 | 1 ; | N - | O |
| 657 | 679 | 1 ; | N - | N |
| 658 | 660 | 1 ; | H - | HA |
| 658 | 661 | 1 ; | H - | CB |
| 658 | 677 | 1 ; | H - | C |
| 659 | 665 | 1 ; | CA - | HG2 |
| 659 | 666 | 1 ; | CA - | HG3 |
| 659 | 667 | 1 ; | CA - | CD |
| 659 | 680 | 1 ; | CA - | H |
| 659 | 681 | 1 ; | CA - | CA |
| 660 | 662 | 1 ; | HA - | HB2 |
| 660 | 663 | 1 ; | HA - | HB3 |
| 660 | 664 | 1 ; | HA - | CG |
| 660 | 678 | 1 ; | HA - | O |
| 660 | 679 | 1 ; | HA - | N |
| 661 | 668 | 1 ; | CB - | HD2 |
| 661 | 669 | 1 ; | CB - | HD3 |
| 661 | 670 | 1 ; | CB - | CE |
| 661 | 678 | 1 ; | CB - | O |
| 661 | 679 | 1 ; | CB - | N |
| 662 | 665 | 1 ; | HB2 - | HG2 |
| 662 | 666 | 1 ; | HB2 - | HG3 |
| 662 | 667 | 1 ; | HB2 - | CD |
| 662 | 677 | 1 ; | HB2 - | C |
| 663 | 665 | 1 ; | HB3 - | HG2 |
| 663 | 666 | 1 ; | HB3 - | HG3 |
| 663 | 667 | 1 ; | HB3 - | CD |
| 663 | 677 | 1 ; | HB3 - | C |

```
664    671       1 ;      CG - HE2
664    672       1 ;      CG - HE3
664    673       1 ;      CG - NZ
664    677       1 ;      CG - C
665    668       1 ;     HG2 - HD2
665    669       1 ;     HG2 - HD3
665    670       1 ;     HG2 - CE
666    668       1 ;     HG3 - HD2
666    669       1 ;     HG3 - HD3
666    670       1 ;     HG3 - CE
667    674       1 ;      CD - HZ1
667    675       1 ;      CD - HZ2
667    676       1 ;      CD - HZ3
668    671       1 ;     HD2 - HE2
668    672       1 ;     HD2 - HE3
668    673       1 ;     HD2 - NZ
669    671       1 ;     HD3 - HE2
669    672       1 ;     HD3 - HE3
669    673       1 ;     HD3 - NZ
671    674       1 ;     HE2 - HZ1
671    675       1 ;     HE2 - HZ2
671    676       1 ;     HE2 - HZ3
672    674       1 ;     HE3 - HZ1
672    675       1 ;     HE3 - HZ2
672    676       1 ;     HE3 - HZ3
677    682       1 ;       C - HA
677    683       1 ;       C - CB
677    692       1 ;       C - C
678    680       1 ;       O - H
678    681       1 ;       O - CA
679    684       1 ;       N - HB2
679    685       1 ;       N - HB3
679    686       1 ;       N - CG
679    693       1 ;       N - O
679    694       1 ;       N - N
680    682       1 ;       H - HA
680    683       1 ;       H - CB
680    692       1 ;       H - C
681    687       1 ;      CA - HG2
681    688       1 ;      CA - HG3
681    689       1 ;      CA - CD
681    695       1 ;      CA - H
681    696       1 ;      CA - CA
682    684       1 ;      HA - HB2
682    685       1 ;      HA - HB3
682    686       1 ;      HA - CG
682    693       1 ;      HA - O
682    694       1 ;      HA - N
683    690       1 ;      CB - OE1
683    691       1 ;      CB - OE2
683    693       1 ;      CB - O
683    694       1 ;      CB - N
```

```
684    687       1 ;    HB2 - HG2
684    688       1 ;    HB2 - HG3
684    689       1 ;    HB2 - CD
684    692       1 ;    HB2 - C
685    687       1 ;    HB3 - HG2
685    688       1 ;    HB3 - HG3
685    689       1 ;    HB3 - CD
685    692       1 ;    HB3 - C
686    692       1 ;     CG - C
687    690       1 ;    HG2 - OE1
687    691       1 ;    HG2 - OE2
688    690       1 ;    HG3 - OE1
688    691       1 ;    HG3 - OE2
692    697       1 ;      C - HA
692    698       1 ;      C - CB
692    707       1 ;      C - C
693    695       1 ;      O - H
693    696       1 ;      O - CA
694    699       1 ;      N - HB2
694    700       1 ;      N - HB3
694    701       1 ;      N - CG
694    708       1 ;      N - O
694    709       1 ;      N - N
695    697       1 ;      H - HA
695    698       1 ;      H - CB
695    707       1 ;      H - C
696    702       1 ;     CA - HG2
696    703       1 ;     CA - HG3
696    704       1 ;     CA - CD
696    710       1 ;     CA - H
696    711       1 ;     CA - CA
697    699       1 ;     HA - HB2
697    700       1 ;     HA - HB3
697    701       1 ;     HA - CG
697    708       1 ;     HA - O
697    709       1 ;     HA - N
698    705       1 ;     CB - OE1
698    706       1 ;     CB - OE2
698    708       1 ;     CB - O
698    709       1 ;     CB - N
699    702       1 ;    HB2 - HG2
699    703       1 ;    HB2 - HG3
699    704       1 ;    HB2 - CD
699    707       1 ;    HB2 - C
700    702       1 ;    HB3 - HG2
700    703       1 ;    HB3 - HG3
700    704       1 ;    HB3 - CD
700    707       1 ;    HB3 - C
701    707       1 ;     CG - C
702    705       1 ;    HG2 - OE1
702    706       1 ;    HG2 - OE2
703    705       1 ;    HG3 - OE1
```

```
703    706     1 ;   HG3 - OE2
707    712     1 ;     C - HA
707    713     1 ;     C - CB
707    729     1 ;     C - C
708    710     1 ;     O - H
708    711     1 ;     O - CA
709    714     1 ;     N - HB2
709    715     1 ;     N - HB3
709    716     1 ;     N - CG
709    730     1 ;     N - O
709    731     1 ;     N - N
710    712     1 ;     H - HA
710    713     1 ;     H - CB
710    729     1 ;     H - C
711    717     1 ;    CA - HG2
711    718     1 ;    CA - HG3
711    719     1 ;    CA - CD
711    732     1 ;    CA - H
711    733     1 ;    CA - CA
712    714     1 ;    HA - HB2
712    715     1 ;    HA - HB3
712    716     1 ;    HA - CG
712    730     1 ;    HA - O
712    731     1 ;    HA - N
713    720     1 ;    CB - HD2
713    721     1 ;    CB - HD3
713    722     1 ;    CB - CE
713    730     1 ;    CB - O
713    731     1 ;    CB - N
714    717     1 ;   HB2 - HG2
714    718     1 ;   HB2 - HG3
714    719     1 ;   HB2 - CD
714    729     1 ;   HB2 - C
715    717     1 ;   HB3 - HG2
715    718     1 ;   HB3 - HG3
715    719     1 ;   HB3 - CD
715    729     1 ;   HB3 - C
716    723     1 ;    CG - HE2
716    724     1 ;    CG - HE3
716    725     1 ;    CG - NZ
716    729     1 ;    CG - C
717    720     1 ;   HG2 - HD2
717    721     1 ;   HG2 - HD3
717    722     1 ;   HG2 - CE
718    720     1 ;   HG3 - HD2
718    721     1 ;   HG3 - HD3
718    722     1 ;   HG3 - CE
719    726     1 ;    CD - HZ1
719    727     1 ;    CD - HZ2
719    728     1 ;    CD - HZ3
720    723     1 ;   HD2 - HE2
720    724     1 ;   HD2 - HE3
```

```
2046   2053      1 ;     CA - CD1
2046   2057      1 ;     CA - CD2
2046   2064      1 ;     CA - H
2046   2065      1 ;     CA - CA
2047   2049      1 ;     HA - HB2
2047   2050      1 ;     HA - HB3
2047   2051      1 ;     HA - CG
2047   2062      1 ;     HA - O
2047   2063      1 ;     HA - N
2048   2054      1 ;     CB - HD11
2048   2055      1 ;     CB - HD12
2048   2056      1 ;     CB - HD13
2048   2058      1 ;     CB - HD21
2048   2059      1 ;     CB - HD22
2048   2060      1 ;     CB - HD23
2048   2062      1 ;     CB - O
2048   2063      1 ;     CB - N
2049   2052      1 ;    HB2 - HG
2049   2053      1 ;    HB2 - CD1
2049   2057      1 ;    HB2 - CD2
2049   2061      1 ;    HB2 - C
2050   2052      1 ;    HB3 - HG
2050   2053      1 ;    HB3 - CD1
2050   2057      1 ;    HB3 - CD2
2050   2061      1 ;    HB3 - C
2051   2061      1 ;     CG - C
2052   2054      1 ;     HG - HD11
2052   2055      1 ;     HG - HD12
2052   2056      1 ;     HG - HD13
2052   2058      1 ;     HG - HD21
2052   2059      1 ;     HG - HD22
2052   2060      1 ;     HG - HD23
2053   2058      1 ;    CD1 - HD21
2053   2059      1 ;    CD1 - HD22
2053   2060      1 ;    CD1 - HD23
2054   2057      1 ;   HD11 - CD2
2055   2057      1 ;   HD12 - CD2
2056   2057      1 ;   HD13 - CD2
2061   2066      1 ;      C - HA
2061   2067      1 ;      C - CB
2061   2078      1 ;      C - C
2062   2064      1 ;      O - H
2062   2065      1 ;      O - CA
2063   2068      1 ;      N - HB2
2063   2069      1 ;      N - HB3
2063   2070      1 ;      N - CG
2063   2079      1 ;      N - O
2063   2080      1 ;      N - N
2064   2066      1 ;      H - HA
2064   2067      1 ;      H - CB
2064   2078      1 ;      H - C
2065   2071      1 ;     CA - ND1
```

```
2065   2076       1 ;      CA - CD2
2065   2081       1 ;      CA - H
2065   2082       1 ;      CA - CA
2066   2068       1 ;      HA - HB2
2066   2069       1 ;      HA - HB3
2066   2070       1 ;      HA - CG
2066   2079       1 ;      HA - O
2066   2080       1 ;      HA - N
2067   2072       1 ;      CB - CE1
2067   2074       1 ;      CB - NE2
2067   2077       1 ;      CB - HD2
2067   2079       1 ;      CB - O
2067   2080       1 ;      CB - N
2068   2071       1 ;     HB2 - ND1
2068   2076       1 ;     HB2 - CD2
2068   2078       1 ;     HB2 - C
2069   2071       1 ;     HB3 - ND1
2069   2076       1 ;     HB3 - CD2
2069   2078       1 ;     HB3 - C
2070   2073       1 ;      CG - HE1
2070   2075       1 ;      CG - HE2
2070   2078       1 ;      CG - C
2071   2075       1 ;     ND1 - HE2
2071   2077       1 ;     ND1 - HD2
2072   2077       1 ;     CE1 - HD2
2073   2075       1 ;     HE1 - HE2
2073   2076       1 ;     HE1 - CD2
2075   2077       1 ;     HE2 - HD2
2078   2083       1 ;       C - HA
2078   2084       1 ;       C - CB
2078   2089       1 ;       C - C
2079   2081       1 ;       O - H
2079   2082       1 ;       O - CA
2080   2085       1 ;       N - HB2
2080   2086       1 ;       N - HB3
2080   2087       1 ;       N - OG
2080   2090       1 ;       N - O
2080   2091       1 ;       N - N
2081   2083       1 ;       H - HA
2081   2084       1 ;       H - CB
2081   2089       1 ;       H - C
2082   2088       1 ;      CA - HG
2082   2092       1 ;      CA - H
2082   2093       1 ;      CA - CA
2083   2085       1 ;      HA - HB2
2083   2086       1 ;      HA - HB3
2083   2087       1 ;      HA - OG
2083   2090       1 ;      HA - O
2083   2091       1 ;      HA - N
2084   2090       1 ;      CB - O
2084   2091       1 ;      CB - N
2085   2088       1 ;     HB2 - HG
```

```
2085   2089      1 ;    HB2 - C
2086   2088      1 ;    HB3 - HG
2086   2089      1 ;    HB3 - C
2087   2089      1 ;     OG - C
2089   2094      1 ;      C - HA
2089   2095      1 ;      C - CB
2089   2108      1 ;      C - C
2090   2092      1 ;      O - H
2090   2093      1 ;      O - CA
2091   2096      1 ;      N - HB2
2091   2097      1 ;      N - HB3
2091   2098      1 ;      N - CG
2091   2109      1 ;      N - O
2091   2110      1 ;      N - N
2092   2094      1 ;      H - HA
2092   2095      1 ;      H - CB
2092   2108      1 ;      H - C
2093   2099      1 ;     CA - HG
2093   2100      1 ;     CA - CD1
2093   2104      1 ;     CA - CD2
2093   2111      1 ;     CA - H
2093   2112      1 ;     CA - CA
2094   2096      1 ;     HA - HB2
2094   2097      1 ;     HA - HB3
2094   2098      1 ;     HA - CG
2094   2109      1 ;     HA - O
2094   2110      1 ;     HA - N
2095   2101      1 ;     CB - HD11
2095   2102      1 ;     CB - HD12
2095   2103      1 ;     CB - HD13
2095   2105      1 ;     CB - HD21
2095   2106      1 ;     CB - HD22
2095   2107      1 ;     CB - HD23
2095   2109      1 ;     CB - O
2095   2110      1 ;     CB - N
2096   2099      1 ;    HB2 - HG
2096   2100      1 ;    HB2 - CD1
2096   2104      1 ;    HB2 - CD2
2096   2108      1 ;    HB2 - C
2097   2099      1 ;    HB3 - HG
2097   2100      1 ;    HB3 - CD1
2097   2104      1 ;    HB3 - CD2
2097   2108      1 ;    HB3 - C
2098   2108      1 ;     CG - C
2099   2101      1 ;     HG - HD11
2099   2102      1 ;     HG - HD12
2099   2103      1 ;     HG - HD13
2099   2105      1 ;     HG - HD21
2099   2106      1 ;     HG - HD22
2099   2107      1 ;     HG - HD23
2100   2105      1 ;    CD1 - HD21
2100   2106      1 ;    CD1 - HD22
```

| | | | | | |
|---|---|---|---|---|---|
| 2100 | 2107 | 1 ; | CD1 | - | HD23 |
| 2101 | 2104 | 1 ; | HD11 | - | CD2 |
| 2102 | 2104 | 1 ; | HD12 | - | CD2 |
| 2103 | 2104 | 1 ; | HD13 | - | CD2 |
| 2108 | 2113 | 1 ; | C | - | HA |
| 2108 | 2114 | 1 ; | C | - | CB |
| 2108 | 2122 | 1 ; | C | - | C |
| 2109 | 2111 | 1 ; | O | - | H |
| 2109 | 2112 | 1 ; | O | - | CA |
| 2110 | 2115 | 1 ; | N | - | HB2 |
| 2110 | 2116 | 1 ; | N | - | HB3 |
| 2110 | 2117 | 1 ; | N | - | CG |
| 2110 | 2123 | 1 ; | N | - | O |
| 2110 | 2124 | 1 ; | N | - | N |
| 2111 | 2113 | 1 ; | H | - | HA |
| 2111 | 2114 | 1 ; | H | - | CB |
| 2111 | 2122 | 1 ; | H | - | C |
| 2112 | 2118 | 1 ; | CA | - | OD1 |
| 2112 | 2119 | 1 ; | CA | - | ND2 |
| 2112 | 2125 | 1 ; | CA | - | H |
| 2112 | 2126 | 1 ; | CA | - | CA |
| 2113 | 2115 | 1 ; | HA | - | HB2 |
| 2113 | 2116 | 1 ; | HA | - | HB3 |
| 2113 | 2117 | 1 ; | HA | - | CG |
| 2113 | 2123 | 1 ; | HA | - | O |
| 2113 | 2124 | 1 ; | HA | - | N |
| 2114 | 2120 | 1 ; | CB | - | HD21 |
| 2114 | 2121 | 1 ; | CB | - | HD22 |
| 2114 | 2123 | 1 ; | CB | - | O |
| 2114 | 2124 | 1 ; | CB | - | N |
| 2115 | 2118 | 1 ; | HB2 | - | OD1 |
| 2115 | 2119 | 1 ; | HB2 | - | ND2 |
| 2115 | 2122 | 1 ; | HB2 | - | C |
| 2116 | 2118 | 1 ; | HB3 | - | OD1 |
| 2116 | 2119 | 1 ; | HB3 | - | ND2 |
| 2116 | 2122 | 1 ; | HB3 | - | C |
| 2117 | 2122 | 1 ; | CG | - | C |
| 2118 | 2120 | 1 ; | OD1 | - | HD21 |
| 2118 | 2121 | 1 ; | OD1 | - | HD22 |
| 2122 | 2127 | 1 ; | C | - | HA |
| 2122 | 2128 | 1 ; | C | - | CB |
| 2122 | 2141 | 1 ; | C | - | C |
| 2123 | 2125 | 1 ; | O | - | H |
| 2123 | 2126 | 1 ; | O | - | CA |
| 2124 | 2129 | 1 ; | N | - | HB |
| 2124 | 2130 | 1 ; | N | - | CG2 |
| 2124 | 2134 | 1 ; | N | - | CG1 |
| 2124 | 2142 | 1 ; | N | - | O |
| 2124 | 2143 | 1 ; | N | - | N |
| 2125 | 2127 | 1 ; | H | - | HA |
| 2125 | 2128 | 1 ; | H | - | CB |
| 2125 | 2141 | 1 ; | H | - | C |

| | | | | |
|---|---|---|---|---|
| 2126 | 2131 | 1 ; | CA - | HG21 |
| 2126 | 2132 | 1 ; | CA - | HG22 |
| 2126 | 2133 | 1 ; | CA - | HG23 |
| 2126 | 2135 | 1 ; | CA - | HG12 |
| 2126 | 2136 | 1 ; | CA - | HG13 |
| 2126 | 2137 | 1 ; | CA - | CD1 |
| 2126 | 2144 | 1 ; | CA - | H |
| 2126 | 2145 | 1 ; | CA - | CA |
| 2127 | 2129 | 1 ; | HA - | HB |
| 2127 | 2130 | 1 ; | HA - | CG2 |
| 2127 | 2134 | 1 ; | HA - | CG1 |
| 2127 | 2142 | 1 ; | HA - | O |
| 2127 | 2143 | 1 ; | HA - | N |
| 2128 | 2138 | 1 ; | CB - | HD11 |
| 2128 | 2139 | 1 ; | CB - | HD12 |
| 2128 | 2140 | 1 ; | CB - | HD13 |
| 2128 | 2142 | 1 ; | CB - | O |
| 2128 | 2143 | 1 ; | CB - | N |
| 2129 | 2131 | 1 ; | HB - | HG21 |
| 2129 | 2132 | 1 ; | HB - | HG22 |
| 2129 | 2133 | 1 ; | HB - | HG23 |
| 2129 | 2135 | 1 ; | HB - | HG12 |
| 2129 | 2136 | 1 ; | HB - | HG13 |
| 2129 | 2137 | 1 ; | HB - | CD1 |
| 2129 | 2141 | 1 ; | HB - | C |
| 2130 | 2135 | 1 ; | CG2 - | HG12 |
| 2130 | 2136 | 1 ; | CG2 - | HG13 |
| 2130 | 2137 | 1 ; | CG2 - | CD1 |
| 2130 | 2141 | 1 ; | CG2 - | C |
| 2131 | 2134 | 1 ; | HG21 - | CG1 |
| 2132 | 2134 | 1 ; | HG22 - | CG1 |
| 2133 | 2134 | 1 ; | HG23 - | CG1 |
| 2134 | 2141 | 1 ; | CG1 - | C |
| 2135 | 2138 | 1 ; | HG12 - | HD11 |
| 2135 | 2139 | 1 ; | HG12 - | HD12 |
| 2135 | 2140 | 1 ; | HG12 - | HD13 |
| 2136 | 2138 | 1 ; | HG13 - | HD11 |
| 2136 | 2139 | 1 ; | HG13 - | HD12 |
| 2136 | 2140 | 1 ; | HG13 - | HD13 |
| 2141 | 2146 | 1 ; | C - | HA |
| 2141 | 2147 | 1 ; | C - | CB |
| 2141 | 2157 | 1 ; | C - | C |
| 2142 | 2144 | 1 ; | O - | H |
| 2142 | 2145 | 1 ; | O - | CA |
| 2143 | 2148 | 1 ; | N - | HB |
| 2143 | 2149 | 1 ; | N - | CG1 |
| 2143 | 2153 | 1 ; | N - | CG2 |
| 2143 | 2158 | 1 ; | N - | O |
| 2143 | 2159 | 1 ; | N - | N |
| 2144 | 2146 | 1 ; | H - | HA |
| 2144 | 2147 | 1 ; | H - | CB |
| 2144 | 2157 | 1 ; | H - | C |

```
2145   2150       1 ;      CA - HG11
2145   2151       1 ;      CA - HG12
2145   2152       1 ;      CA - HG13
2145   2154       1 ;      CA - HG21
2145   2155       1 ;      CA - HG22
2145   2156       1 ;      CA - HG23
2145   2160       1 ;      CA - H
2145   2161       1 ;      CA - CA
2146   2148       1 ;      HA - HB
2146   2149       1 ;      HA - CG1
2146   2153       1 ;      HA - CG2
2146   2158       1 ;      HA - O
2146   2159       1 ;      HA - N
2147   2158       1 ;      CB - O
2147   2159       1 ;      CB - N
2148   2150       1 ;      HB - HG11
2148   2151       1 ;      HB - HG12
2148   2152       1 ;      HB - HG13
2148   2154       1 ;      HB - HG21
2148   2155       1 ;      HB - HG22
2148   2156       1 ;      HB - HG23
2148   2157       1 ;      HB - C
2149   2154       1 ;     CG1 - HG21
2149   2155       1 ;     CG1 - HG22
2149   2156       1 ;     CG1 - HG23
2149   2157       1 ;     CG1 - C
2150   2153       1 ;    HG11 - CG2
2151   2153       1 ;    HG12 - CG2
2152   2153       1 ;    HG13 - CG2
2153   2157       1 ;     CG2 - C
2157   2162       1 ;       C - HA
2157   2163       1 ;       C - CB
2157   2174       1 ;       C - C
2158   2160       1 ;       O - H
2158   2161       1 ;       O - CA
2159   2164       1 ;       N - HB2
2159   2165       1 ;       N - HB3
2159   2166       1 ;       N - CG
2159   2175       1 ;       N - O
2159   2176       1 ;       N - N
2160   2162       1 ;       H - HA
2160   2163       1 ;       H - CB
2160   2174       1 ;       H - C
2161   2167       1 ;      CA - ND1
2161   2172       1 ;      CA - CD2
2161   2177       1 ;      CA - H
2161   2178       1 ;      CA - CA
2162   2164       1 ;      HA - HB2
2162   2165       1 ;      HA - HB3
2162   2166       1 ;      HA - CG
2162   2175       1 ;      HA - O
2162   2176       1 ;      HA - N
```

| | | | |
|---|---|---|---|
| 2163 | 2168 | 1 ; | CB - CE1 |
| 2163 | 2170 | 1 ; | CB - NE2 |
| 2163 | 2173 | 1 ; | CB - HD2 |
| 2163 | 2175 | 1 ; | CB - O |
| 2163 | 2176 | 1 ; | CB - N |
| 2164 | 2167 | 1 ; | HB2 - ND1 |
| 2164 | 2172 | 1 ; | HB2 - CD2 |
| 2164 | 2174 | 1 ; | HB2 - C |
| 2165 | 2167 | 1 ; | HB3 - ND1 |
| 2165 | 2172 | 1 ; | HB3 - CD2 |
| 2165 | 2174 | 1 ; | HB3 - C |
| 2166 | 2169 | 1 ; | CG - HE1 |
| 2166 | 2171 | 1 ; | CG - HE2 |
| 2166 | 2174 | 1 ; | CG - C |
| 2167 | 2171 | 1 ; | ND1 - HE2 |
| 2167 | 2173 | 1 ; | ND1 - HD2 |
| 2168 | 2173 | 1 ; | CE1 - HD2 |
| 2169 | 2171 | 1 ; | HE1 - HE2 |
| 2169 | 2172 | 1 ; | HE1 - CD2 |
| 2171 | 2173 | 1 ; | HE2 - HD2 |
| 2174 | 2179 | 1 ; | C - HA |
| 2174 | 2180 | 1 ; | C - CB |
| 2174 | 2198 | 1 ; | C - C |
| 2175 | 2177 | 1 ; | O - H |
| 2175 | 2178 | 1 ; | O - CA |
| 2176 | 2181 | 1 ; | N - HB2 |
| 2176 | 2182 | 1 ; | N - HB3 |
| 2176 | 2183 | 1 ; | N - CG |
| 2176 | 2199 | 1 ; | N - O |
| 2176 | 2200 | 1 ; | N - N |
| 2177 | 2179 | 1 ; | H - HA |
| 2177 | 2180 | 1 ; | H - CB |
| 2177 | 2198 | 1 ; | H - C |
| 2178 | 2184 | 1 ; | CA - HG2 |
| 2178 | 2185 | 1 ; | CA - HG3 |
| 2178 | 2186 | 1 ; | CA - CD |
| 2178 | 2201 | 1 ; | CA - H |
| 2178 | 2202 | 1 ; | CA - CA |
| 2179 | 2181 | 1 ; | HA - HB2 |
| 2179 | 2182 | 1 ; | HA - HB3 |
| 2179 | 2183 | 1 ; | HA - CG |
| 2179 | 2199 | 1 ; | HA - O |
| 2179 | 2200 | 1 ; | HA - N |
| 2180 | 2187 | 1 ; | CB - HD2 |
| 2180 | 2188 | 1 ; | CB - HD3 |
| 2180 | 2189 | 1 ; | CB - NE |
| 2180 | 2199 | 1 ; | CB - O |
| 2180 | 2200 | 1 ; | CB - N |
| 2181 | 2184 | 1 ; | HB2 - HG2 |
| 2181 | 2185 | 1 ; | HB2 - HG3 |
| 2181 | 2186 | 1 ; | HB2 - CD |
| 2181 | 2198 | 1 ; | HB2 - C |

| | | | | | |
|---|---|---|---|---|---|
| 2182 | 2184 | 1 ; | HB3 | - | HG2 |
| 2182 | 2185 | 1 ; | HB3 | - | HG3 |
| 2182 | 2186 | 1 ; | HB3 | - | CD |
| 2182 | 2198 | 1 ; | HB3 | - | C |
| 2183 | 2190 | 1 ; | CG | - | HE |
| 2183 | 2191 | 1 ; | CG | - | CZ |
| 2183 | 2198 | 1 ; | CG | - | C |
| 2184 | 2187 | 1 ; | HG2 | - | HD2 |
| 2184 | 2188 | 1 ; | HG2 | - | HD3 |
| 2184 | 2189 | 1 ; | HG2 | - | NE |
| 2185 | 2187 | 1 ; | HG3 | - | HD2 |
| 2185 | 2188 | 1 ; | HG3 | - | HD3 |
| 2185 | 2189 | 1 ; | HG3 | - | NE |
| 2186 | 2192 | 1 ; | CD | - | NH1 |
| 2186 | 2195 | 1 ; | CD | - | NH2 |
| 2187 | 2190 | 1 ; | HD2 | - | HE |
| 2187 | 2191 | 1 ; | HD2 | - | CZ |
| 2188 | 2190 | 1 ; | HD3 | - | HE |
| 2188 | 2191 | 1 ; | HD3 | - | CZ |
| 2189 | 2193 | 1 ; | NE | - | HH11 |
| 2189 | 2194 | 1 ; | NE | - | HH12 |
| 2189 | 2196 | 1 ; | NE | - | HH21 |
| 2189 | 2197 | 1 ; | NE | - | HH22 |
| 2190 | 2192 | 1 ; | HE | - | NH1 |
| 2190 | 2195 | 1 ; | HE | - | NH2 |
| 2192 | 2196 | 1 ; | NH1 | - | HH21 |
| 2192 | 2197 | 1 ; | NH1 | - | HH22 |
| 2193 | 2195 | 1 ; | HH11 | - | NH2 |
| 2194 | 2195 | 1 ; | HH12 | - | NH2 |
| 2198 | 2203 | 1 ; | C | - | HA |
| 2198 | 2204 | 1 ; | C | - | CB |
| 2198 | 2210 | 1 ; | C | - | C |
| 2199 | 2201 | 1 ; | O | - | H |
| 2199 | 2202 | 1 ; | O | - | CA |
| 2200 | 2205 | 1 ; | N | - | HB2 |
| 2200 | 2206 | 1 ; | N | - | HB3 |
| 2200 | 2207 | 1 ; | N | - | CG |
| 2200 | 2211 | 1 ; | N | - | O |
| 2200 | 2212 | 1 ; | N | - | N |
| 2201 | 2203 | 1 ; | H | - | HA |
| 2201 | 2204 | 1 ; | H | - | CB |
| 2201 | 2210 | 1 ; | H | - | C |
| 2202 | 2208 | 1 ; | CA | - | OD1 |
| 2202 | 2209 | 1 ; | CA | - | OD2 |
| 2202 | 2213 | 1 ; | CA | - | H |
| 2202 | 2214 | 1 ; | CA | - | CA |
| 2203 | 2205 | 1 ; | HA | - | HB2 |
| 2203 | 2206 | 1 ; | HA | - | HB3 |
| 2203 | 2207 | 1 ; | HA | - | CG |
| 2203 | 2211 | 1 ; | HA | - | O |
| 2203 | 2212 | 1 ; | HA | - | N |
| 2204 | 2211 | 1 ; | CB | - | O |

| | | | | | | |
|---|---|---|---|---|---|---|
| 2204 | 2212 | 1 | ; | CB | - | N |
| 2205 | 2208 | 1 | ; | HB2 | - | OD1 |
| 2205 | 2209 | 1 | ; | HB2 | - | OD2 |
| 2205 | 2210 | 1 | ; | HB2 | - | C |
| 2206 | 2208 | 1 | ; | HB3 | - | OD1 |
| 2206 | 2209 | 1 | ; | HB3 | - | OD2 |
| 2206 | 2210 | 1 | ; | HB3 | - | C |
| 2207 | 2210 | 1 | ; | CG | - | C |
| 2210 | 2215 | 1 | ; | C | - | HA |
| 2210 | 2216 | 1 | ; | C | - | CB |
| 2210 | 2229 | 1 | ; | C | - | C |
| 2211 | 2213 | 1 | ; | O | - | H |
| 2211 | 2214 | 1 | ; | O | - | CA |
| 2212 | 2217 | 1 | ; | N | - | HB2 |
| 2212 | 2218 | 1 | ; | N | - | HB3 |
| 2212 | 2219 | 1 | ; | N | - | CG |
| 2212 | 2230 | 1 | ; | N | - | O |
| 2212 | 2231 | 1 | ; | N | - | N |
| 2213 | 2215 | 1 | ; | H | - | HA |
| 2213 | 2216 | 1 | ; | H | - | CB |
| 2213 | 2229 | 1 | ; | H | - | C |
| 2214 | 2220 | 1 | ; | CA | - | HG |
| 2214 | 2221 | 1 | ; | CA | - | CD1 |
| 2214 | 2225 | 1 | ; | CA | - | CD2 |
| 2214 | 2232 | 1 | ; | CA | - | H |
| 2214 | 2233 | 1 | ; | CA | - | CA |
| 2215 | 2217 | 1 | ; | HA | - | HB2 |
| 2215 | 2218 | 1 | ; | HA | - | HB3 |
| 2215 | 2219 | 1 | ; | HA | - | CG |
| 2215 | 2230 | 1 | ; | HA | - | O |
| 2215 | 2231 | 1 | ; | HA | - | N |
| 2216 | 2222 | 1 | ; | CB | - | HD11 |
| 2216 | 2223 | 1 | ; | CB | - | HD12 |
| 2216 | 2224 | 1 | ; | CB | - | HD13 |
| 2216 | 2226 | 1 | ; | CB | - | HD21 |
| 2216 | 2227 | 1 | ; | CB | - | HD22 |
| 2216 | 2228 | 1 | ; | CB | - | HD23 |
| 2216 | 2230 | 1 | ; | CB | - | O |
| 2216 | 2231 | 1 | ; | CB | - | N |
| 2217 | 2220 | 1 | ; | HB2 | - | HG |
| 2217 | 2221 | 1 | ; | HB2 | - | CD1 |
| 2217 | 2225 | 1 | ; | HB2 | - | CD2 |
| 2217 | 2229 | 1 | ; | HB2 | - | C |
| 2218 | 2220 | 1 | ; | HB3 | - | HG |
| 2218 | 2221 | 1 | ; | HB3 | - | CD1 |
| 2218 | 2225 | 1 | ; | HB3 | - | CD2 |
| 2218 | 2229 | 1 | ; | HB3 | - | C |
| 2219 | 2229 | 1 | ; | CG | - | C |
| 2220 | 2222 | 1 | ; | HG | - | HD11 |
| 2220 | 2223 | 1 | ; | HG | - | HD12 |
| 2220 | 2224 | 1 | ; | HG | - | HD13 |
| 2220 | 2226 | 1 | ; | HG | - | HD21 |

```
2220   2227      1 ;     HG - HD22
2220   2228      1 ;     HG - HD23
2221   2226      1 ;    CD1 - HD21
2221   2227      1 ;    CD1 - HD22
2221   2228      1 ;    CD1 - HD23
2222   2225      1 ;   HD11 - CD2
2223   2225      1 ;   HD12 - CD2
2224   2225      1 ;   HD13 - CD2
2229   2234      1 ;      C - HA
2229   2235      1 ;      C - CB
2229   2251      1 ;      C - C
2230   2232      1 ;      O - H
2230   2233      1 ;      O - CA
2231   2236      1 ;      N - HB2
2231   2237      1 ;      N - HB3
2231   2238      1 ;      N - CG
2231   2252      1 ;      N - O
2231   2253      1 ;      N - N
2232   2234      1 ;      H - HA
2232   2235      1 ;      H - CB
2232   2251      1 ;      H - C
2233   2239      1 ;     CA - HG2
2233   2240      1 ;     CA - HG3
2233   2241      1 ;     CA - CD
2233   2254      1 ;     CA - CD
2233   2263      1 ;     CA - CA
2234   2236      1 ;     HA - HB2
2234   2237      1 ;     HA - HB3
2234   2238      1 ;     HA - CG
2234   2252      1 ;     HA - O
2234   2253      1 ;     HA - N
2235   2242      1 ;     CB - HD2
2235   2243      1 ;     CB - HD3
2235   2244      1 ;     CB - CE
2235   2252      1 ;     CB - O
2235   2253      1 ;     CB - N
2236   2239      1 ;    HB2 - HG2
2236   2240      1 ;    HB2 - HG3
2236   2241      1 ;    HB2 - CD
2236   2251      1 ;    HB2 - C
2237   2239      1 ;    HB3 - HG2
2237   2240      1 ;    HB3 - HG3
2237   2241      1 ;    HB3 - CD
2237   2251      1 ;    HB3 - C
2238   2245      1 ;     CG - HE2
2238   2246      1 ;     CG - HE3
2238   2247      1 ;     CG - NZ
2238   2251      1 ;     CG - C
2239   2242      1 ;    HG2 - HD2
2239   2243      1 ;    HG2 - HD3
2239   2244      1 ;    HG2 - CE
2240   2242      1 ;    HG3 - HD2
```

```
2240   2243      1 ;    HG3 - HD3
2240   2244      1 ;    HG3 - CE
2241   2248      1 ;     CD - HZ1
2241   2249      1 ;     CD - HZ2
2241   2250      1 ;     CD - HZ3
2242   2245      1 ;    HD2 - HE2
2242   2246      1 ;    HD2 - HE3
2242   2247      1 ;    HD2 - NZ
2243   2245      1 ;    HD3 - HE2
2243   2246      1 ;    HD3 - HE3
2243   2247      1 ;    HD3 - NZ
2245   2248      1 ;    HE2 - HZ1
2245   2249      1 ;    HE2 - HZ2
2245   2250      1 ;    HE2 - HZ3
2246   2248      1 ;    HE3 - HZ1
2246   2249      1 ;    HE3 - HZ2
2246   2250      1 ;    HE3 - HZ3
2251   2255      1 ;      C - HD2
2251   2256      1 ;      C - HD3
2251   2257      1 ;      C - CG
2251   2260      1 ;      C - CB
2251   2264      1 ;      C - HA
2251   2265      1 ;      C - C
2252   2254      1 ;      O - CD
2252   2263      1 ;      O - CA
2253   2258      1 ;      N - HG2
2253   2259      1 ;      N - HG3
2253   2261      1 ;      N - HB2
2253   2262      1 ;      N - HB3
2253   2266      1 ;      N - O
2253   2267      1 ;      N - N
2254   2261      1 ;     CD - HB2
2254   2262      1 ;     CD - HB3
2254   2264      1 ;     CD - HA
2254   2265      1 ;     CD - C
2255   2258      1 ;    HD2 - HG2
2255   2259      1 ;    HD2 - HG3
2255   2260      1 ;    HD2 - CB
2255   2263      1 ;    HD2 - CA
2256   2258      1 ;    HD3 - HG2
2256   2259      1 ;    HD3 - HG3
2256   2260      1 ;    HD3 - CB
2256   2263      1 ;    HD3 - CA
2257   2264      1 ;     CG - HA
2257   2265      1 ;     CG - C
2258   2261      1 ;    HG2 - HB2
2258   2262      1 ;    HG2 - HB3
2258   2263      1 ;    HG2 - CA
2259   2261      1 ;    HG3 - HB2
2259   2262      1 ;    HG3 - HB3
2259   2263      1 ;    HG3 - CA
2260   2266      1 ;     CB - O
```

```
2260   2267       1 ;      CB - N
2261   2264       1 ;     HB2 - HA
2261   2265       1 ;     HB2 - C
2262   2264       1 ;     HB3 - HA
2262   2265       1 ;     HB3 - C
2263   2268       1 ;      CA - H
2263   2269       1 ;      CA - CA
2264   2266       1 ;      HA - O
2264   2267       1 ;      HA - N
2265   2270       1 ;       C - HA
2265   2271       1 ;       C - CB
2265   2280       1 ;       C - C
2266   2268       1 ;       O - H
2266   2269       1 ;       O - CA
2267   2272       1 ;       N - HB2
2267   2273       1 ;       N - HB3
2267   2274       1 ;       N - CG
2267   2281       1 ;       N - O
2267   2282       1 ;       N - N
2268   2270       1 ;       H - HA
2268   2271       1 ;       H - CB
2268   2280       1 ;       H - C
2269   2275       1 ;      CA - HG2
2269   2276       1 ;      CA - HG3
2269   2277       1 ;      CA - CD
2269   2283       1 ;      CA - H
2269   2284       1 ;      CA - CA
2270   2272       1 ;      HA - HB2
2270   2273       1 ;      HA - HB3
2270   2274       1 ;      HA - CG
2270   2281       1 ;      HA - O
2270   2282       1 ;      HA - N
2271   2278       1 ;      CB - OE1
2271   2279       1 ;      CB - OE2
2271   2281       1 ;      CB - O
2271   2282       1 ;      CB - N
2272   2275       1 ;     HB2 - HG2
2272   2276       1 ;     HB2 - HG3
2272   2277       1 ;     HB2 - CD
2272   2280       1 ;     HB2 - C
2273   2275       1 ;     HB3 - HG2
2273   2276       1 ;     HB3 - HG3
2273   2277       1 ;     HB3 - CD
2273   2280       1 ;     HB3 - C
2274   2280       1 ;      CG - C
2275   2278       1 ;     HG2 - OE1
2275   2279       1 ;     HG2 - OE2
2276   2278       1 ;     HG3 - OE1
2276   2279       1 ;     HG3 - OE2
2280   2285       1 ;       C - HA
2280   2286       1 ;       C - CB
2280   2294       1 ;       C - C
```

```
2281   2283       1 ;      O - H
2281   2284       1 ;      O - CA
2282   2287       1 ;      N - HB2
2282   2288       1 ;      N - HB3
2282   2289       1 ;      N - CG
2282   2295       1 ;      N - O
2282   2296       1 ;      N - N
2283   2285       1 ;      H - HA
2283   2286       1 ;      H - CB
2283   2294       1 ;      H - C
2284   2290       1 ;     CA - OD1
2284   2291       1 ;     CA - ND2
2284   2297       1 ;     CA - H
2284   2298       1 ;     CA - CA
2285   2287       1 ;     HA - HB2
2285   2288       1 ;     HA - HB3
2285   2289       1 ;     HA - CG
2285   2295       1 ;     HA - O
2285   2296       1 ;     HA - N
2286   2292       1 ;     CB - HD21
2286   2293       1 ;     CB - HD22
2286   2295       1 ;     CB - O
2286   2296       1 ;     CB - N
2287   2290       1 ;    HB2 - OD1
2287   2291       1 ;    HB2 - ND2
2287   2294       1 ;    HB2 - C
2288   2290       1 ;    HB3 - OD1
2288   2291       1 ;    HB3 - ND2
2288   2294       1 ;    HB3 - C
2289   2294       1 ;     CG - C
2290   2292       1 ;    OD1 - HD21
2290   2293       1 ;    OD1 - HD22
2294   2299       1 ;      C - HA
2294   2300       1 ;      C - CB
2294   2314       1 ;      C - C
2295   2297       1 ;      O - H
2295   2298       1 ;      O - CA
2296   2301       1 ;      N - HB2
2296   2302       1 ;      N - HB3
2296   2303       1 ;      N - CG
2296   2315       1 ;      N - O
2296   2316       1 ;      N - N
2297   2299       1 ;      H - HA
2297   2300       1 ;      H - CB
2297   2314       1 ;      H - C
2298   2304       1 ;     CA - CD1
2298   2312       1 ;     CA - CD2
2298   2317       1 ;     CA - H
2298   2318       1 ;     CA - CA
2299   2301       1 ;     HA - HB2
2299   2302       1 ;     HA - HB3
2299   2303       1 ;     HA - CG
```

```
2299   2315       1 ;      HA - O
2299   2316       1 ;      HA - N
2300   2305       1 ;      CB - HD1
2300   2306       1 ;      CB - CE1
2300   2310       1 ;      CB - CE2
2300   2313       1 ;      CB - HD2
2300   2315       1 ;      CB - O
2300   2316       1 ;      CB - N
2301   2304       1 ;     HB2 - CD1
2301   2312       1 ;     HB2 - CD2
2301   2314       1 ;     HB2 - C
2302   2304       1 ;     HB3 - CD1
2302   2312       1 ;     HB3 - CD2
2302   2314       1 ;     HB3 - C
2303   2307       1 ;      CG - HE1
2303   2308       1 ;      CG - CZ
2303   2311       1 ;      CG - HE2
2303   2314       1 ;      CG - C
2304   2309       1 ;     CD1 - HZ
2304   2310       1 ;     CD1 - CE2
2304   2313       1 ;     CD1 - HD2
2305   2307       1 ;     HD1 - HE1
2305   2308       1 ;     HD1 - CZ
2305   2312       1 ;     HD1 - CD2
2306   2311       1 ;     CE1 - HE2
2306   2312       1 ;     CE1 - CD2
2307   2309       1 ;     HE1 - HZ
2307   2310       1 ;     HE1 - CE2
2308   2313       1 ;      CZ - HD2
2309   2311       1 ;      HZ - HE2
2309   2312       1 ;      HZ - CD2
2311   2313       1 ;     HE2 - HD2
2314   2319       1 ;       C - HA
2314   2320       1 ;       C - CB
2314   2333       1 ;       C - C
2315   2317       1 ;       O - H
2315   2318       1 ;       O - CA
2316   2321       1 ;       N - HB2
2316   2322       1 ;       N - HB3
2316   2323       1 ;       N - CG
2316   2334       1 ;       N - O
2316   2335       1 ;       N - N
2317   2319       1 ;       H - HA
2317   2320       1 ;       H - CB
2317   2333       1 ;       H - C
2318   2324       1 ;      CA - HG
2318   2325       1 ;      CA - CD1
2318   2329       1 ;      CA - CD2
2318   2336       1 ;      CA - H
2318   2337       1 ;      CA - CA
2319   2321       1 ;      HA - HB2
2319   2322       1 ;      HA - HB3
```

| | | | | | |
|---|---|---|---|---|---|
| 2319 | 2323 | 1 ; | HA | - | CG |
| 2319 | 2334 | 1 ; | HA | - | O |
| 2319 | 2335 | 1 ; | HA | - | N |
| 2320 | 2326 | 1 ; | CB | - | HD11 |
| 2320 | 2327 | 1 ; | CB | - | HD12 |
| 2320 | 2328 | 1 ; | CB | - | HD13 |
| 2320 | 2330 | 1 ; | CB | - | HD21 |
| 2320 | 2331 | 1 ; | CB | - | HD22 |
| 2320 | 2332 | 1 ; | CB | - | HD23 |
| 2320 | 2334 | 1 ; | CB | - | O |
| 2320 | 2335 | 1 ; | CB | - | N |
| 2321 | 2324 | 1 ; | HB2 | - | HG |
| 2321 | 2325 | 1 ; | HB2 | - | CD1 |
| 2321 | 2329 | 1 ; | HB2 | - | CD2 |
| 2321 | 2333 | 1 ; | HB2 | - | C |
| 2322 | 2324 | 1 ; | HB3 | - | HG |
| 2322 | 2325 | 1 ; | HB3 | - | CD1 |
| 2322 | 2329 | 1 ; | HB3 | - | CD2 |
| 2322 | 2333 | 1 ; | HB3 | - | C |
| 2323 | 2333 | 1 ; | CG | - | C |
| 2324 | 2326 | 1 ; | HG | - | HD11 |
| 2324 | 2327 | 1 ; | HG | - | HD12 |
| 2324 | 2328 | 1 ; | HG | - | HD13 |
| 2324 | 2330 | 1 ; | HG | - | HD21 |
| 2324 | 2331 | 1 ; | HG | - | HD22 |
| 2324 | 2332 | 1 ; | HG | - | HD23 |
| 2325 | 2330 | 1 ; | CD1 | - | HD21 |
| 2325 | 2331 | 1 ; | CD1 | - | HD22 |
| 2325 | 2332 | 1 ; | CD1 | - | HD23 |
| 2326 | 2329 | 1 ; | HD11 | - | CD2 |
| 2327 | 2329 | 1 ; | HD12 | - | CD2 |
| 2328 | 2329 | 1 ; | HD13 | - | CD2 |
| 2333 | 2338 | 1 ; | C | - | HA |
| 2333 | 2339 | 1 ; | C | - | CB |
| 2333 | 2353 | 1 ; | C | - | C |
| 2334 | 2336 | 1 ; | O | - | H |
| 2334 | 2337 | 1 ; | O | - | CA |
| 2335 | 2340 | 1 ; | N | - | HB2 |
| 2335 | 2341 | 1 ; | N | - | HB3 |
| 2335 | 2342 | 1 ; | N | - | CG |
| 2335 | 2354 | 1 ; | N | - | O |
| 2335 | 2355 | 1 ; | N | - | N |
| 2336 | 2338 | 1 ; | H | - | HA |
| 2336 | 2339 | 1 ; | H | - | CB |
| 2336 | 2353 | 1 ; | H | - | C |
| 2337 | 2343 | 1 ; | CA | - | CD1 |
| 2337 | 2351 | 1 ; | CA | - | CD2 |
| 2337 | 2356 | 1 ; | CA | - | H |
| 2337 | 2357 | 1 ; | CA | - | CA |
| 2338 | 2340 | 1 ; | HA | - | HB2 |
| 2338 | 2341 | 1 ; | HA | - | HB3 |
| 2338 | 2342 | 1 ; | HA | - | CG |

| | | | | | | |
|---|---|---|---|---|---|---|
| 2338 | 2354 | 1 | ; | HA | - | O |
| 2338 | 2355 | 1 | ; | HA | - | N |
| 2339 | 2344 | 1 | ; | CB | - | HD1 |
| 2339 | 2345 | 1 | ; | CB | - | CE1 |
| 2339 | 2349 | 1 | ; | CB | - | CE2 |
| 2339 | 2352 | 1 | ; | CB | - | HD2 |
| 2339 | 2354 | 1 | ; | CB | - | O |
| 2339 | 2355 | 1 | ; | CB | - | N |
| 2340 | 2343 | 1 | ; | HB2 | - | CD1 |
| 2340 | 2351 | 1 | ; | HB2 | - | CD2 |
| 2340 | 2353 | 1 | ; | HB2 | - | C |
| 2341 | 2343 | 1 | ; | HB3 | - | CD1 |
| 2341 | 2351 | 1 | ; | HB3 | - | CD2 |
| 2341 | 2353 | 1 | ; | HB3 | - | C |
| 2342 | 2346 | 1 | ; | CG | - | HE1 |
| 2342 | 2347 | 1 | ; | CG | - | CZ |
| 2342 | 2350 | 1 | ; | CG | - | HE2 |
| 2342 | 2353 | 1 | ; | CG | - | C |
| 2343 | 2348 | 1 | ; | CD1 | - | HZ |
| 2343 | 2349 | 1 | ; | CD1 | - | CE2 |
| 2343 | 2352 | 1 | ; | CD1 | - | HD2 |
| 2344 | 2346 | 1 | ; | HD1 | - | HE1 |
| 2344 | 2347 | 1 | ; | HD1 | - | CZ |
| 2344 | 2351 | 1 | ; | HD1 | - | CD2 |
| 2345 | 2350 | 1 | ; | CE1 | - | HE2 |
| 2345 | 2351 | 1 | ; | CE1 | - | CD2 |
| 2346 | 2348 | 1 | ; | HE1 | - | HZ |
| 2346 | 2349 | 1 | ; | HE1 | - | CE2 |
| 2347 | 2352 | 1 | ; | CZ | - | HD2 |
| 2348 | 2350 | 1 | ; | HZ | - | HE2 |
| 2348 | 2351 | 1 | ; | HZ | - | CD2 |
| 2350 | 2352 | 1 | ; | HE2 | - | HD2 |
| 2353 | 2358 | 1 | ; | C | - | HA |
| 2353 | 2359 | 1 | ; | C | - | CB |
| 2353 | 2370 | 1 | ; | C | - | C |
| 2354 | 2356 | 1 | ; | O | - | H |
| 2354 | 2357 | 1 | ; | O | - | CA |
| 2355 | 2360 | 1 | ; | N | - | HB2 |
| 2355 | 2361 | 1 | ; | N | - | HB3 |
| 2355 | 2362 | 1 | ; | N | - | CG |
| 2355 | 2371 | 1 | ; | N | - | O |
| 2355 | 2372 | 1 | ; | N | - | N |
| 2356 | 2358 | 1 | ; | H | - | HA |
| 2356 | 2359 | 1 | ; | H | - | CB |
| 2356 | 2370 | 1 | ; | H | - | C |
| 2357 | 2363 | 1 | ; | CA | - | HG2 |
| 2357 | 2364 | 1 | ; | CA | - | HG3 |
| 2357 | 2365 | 1 | ; | CA | - | CD |
| 2357 | 2373 | 1 | ; | CA | - | H |
| 2357 | 2374 | 1 | ; | CA | - | CA |
| 2358 | 2360 | 1 | ; | HA | - | HB2 |
| 2358 | 2361 | 1 | ; | HA | - | HB3 |

| | | | | | | |
|---|---|---|---|---|---|---|
| 2358 | 2362 | 1 | ; | HA | - | CG |
| 2358 | 2371 | 1 | ; | HA | - | O |
| 2358 | 2372 | 1 | ; | HA | - | N |
| 2359 | 2366 | 1 | ; | CB | - | OE1 |
| 2359 | 2367 | 1 | ; | CB | - | NE2 |
| 2359 | 2371 | 1 | ; | CB | - | O |
| 2359 | 2372 | 1 | ; | CB | - | N |
| 2360 | 2363 | 1 | ; | HB2 | - | HG2 |
| 2360 | 2364 | 1 | ; | HB2 | - | HG3 |
| 2360 | 2365 | 1 | ; | HB2 | - | CD |
| 2360 | 2370 | 1 | ; | HB2 | - | C |
| 2361 | 2363 | 1 | ; | HB3 | - | HG2 |
| 2361 | 2364 | 1 | ; | HB3 | - | HG3 |
| 2361 | 2365 | 1 | ; | HB3 | - | CD |
| 2361 | 2370 | 1 | ; | HB3 | - | C |
| 2362 | 2368 | 1 | ; | CG | - | HE21 |
| 2362 | 2369 | 1 | ; | CG | - | HE22 |
| 2362 | 2370 | 1 | ; | CG | - | C |
| 2363 | 2366 | 1 | ; | HG2 | - | OE1 |
| 2363 | 2367 | 1 | ; | HG2 | - | NE2 |
| 2364 | 2366 | 1 | ; | HG3 | - | OE1 |
| 2364 | 2367 | 1 | ; | HG3 | - | NE2 |
| 2366 | 2368 | 1 | ; | OE1 | - | HE21 |
| 2366 | 2369 | 1 | ; | OE1 | - | HE22 |
| 2370 | 2375 | 1 | ; | C | - | HA |
| 2370 | 2376 | 1 | ; | C | - | CB |
| 2370 | 2381 | 1 | ; | C | - | C |
| 2371 | 2373 | 1 | ; | O | - | H |
| 2371 | 2374 | 1 | ; | O | - | CA |
| 2372 | 2377 | 1 | ; | N | - | HB2 |
| 2372 | 2378 | 1 | ; | N | - | HB3 |
| 2372 | 2379 | 1 | ; | N | - | OG |
| 2372 | 2382 | 1 | ; | N | - | O |
| 2372 | 2383 | 1 | ; | N | - | N |
| 2373 | 2375 | 1 | ; | H | - | HA |
| 2373 | 2376 | 1 | ; | H | - | CB |
| 2373 | 2381 | 1 | ; | H | - | C |
| 2374 | 2380 | 1 | ; | CA | - | HG |
| 2374 | 2384 | 1 | ; | CA | - | H |
| 2374 | 2385 | 1 | ; | CA | - | CA |
| 2375 | 2377 | 1 | ; | HA | - | HB2 |
| 2375 | 2378 | 1 | ; | HA | - | HB3 |
| 2375 | 2379 | 1 | ; | HA | - | OG |
| 2375 | 2382 | 1 | ; | HA | - | O |
| 2375 | 2383 | 1 | ; | HA | - | N |
| 2376 | 2382 | 1 | ; | CB | - | O |
| 2376 | 2383 | 1 | ; | CB | - | N |
| 2377 | 2380 | 1 | ; | HB2 | - | HG |
| 2377 | 2381 | 1 | ; | HB2 | - | C |
| 2378 | 2380 | 1 | ; | HB3 | - | HG |
| 2378 | 2381 | 1 | ; | HB3 | - | C |
| 2379 | 2381 | 1 | ; | OG | - | C |

2381 2386 1 ; C - HA
2381 2387 1 ; C - CB
2381 2396 1 ; C - C
2382 2384 1 ; O - H
2382 2385 1 ; O - CA
2383 2388 1 ; N - HB2
2383 2389 1 ; N - HB3
2383 2390 1 ; N - CG
2383 2397 1 ; N - O
2383 2398 1 ; N - N
2384 2386 1 ; H - HA
2384 2387 1 ; H - CB
2384 2396 1 ; H - C
2385 2391 1 ; CA - HG2
2385 2392 1 ; CA - HG3
2385 2393 1 ; CA - CD
2385 2399 1 ; CA - H
2385 2400 1 ; CA - CA
2386 2388 1 ; HA - HB2
2386 2389 1 ; HA - HB3
2386 2390 1 ; HA - CG
2386 2397 1 ; HA - O
2386 2398 1 ; HA - N
2387 2394 1 ; CB - OE1
2387 2395 1 ; CB - OE2
2387 2397 1 ; CB - O
2387 2398 1 ; CB - N
2388 2391 1 ; HB2 - HG2
2388 2392 1 ; HB2 - HG3
2388 2393 1 ; HB2 - CD
2388 2396 1 ; HB2 - C
2389 2391 1 ; HB3 - HG2
2389 2392 1 ; HB3 - HG3
2389 2393 1 ; HB3 - CD
2389 2396 1 ; HB3 - C
2390 2396 1 ; CG - C
2391 2394 1 ; HG2 - OE1
2391 2395 1 ; HG2 - OE2
2392 2394 1 ; HG3 - OE1
2392 2395 1 ; HG3 - OE2
2396 2401 1 ; C - HA
2396 2402 1 ; C - CB
2396 2410 1 ; C - C
2397 2399 1 ; O - H
2397 2400 1 ; O - CA
2398 2403 1 ; N - HB2
2398 2404 1 ; N - HB3
2398 2405 1 ; N - CG
2398 2411 1 ; N - O
2398 2412 1 ; N - N
2399 2401 1 ; H - HA
2399 2402 1 ; H - CB

```
2399   2410       1 ;      H - C
2400   2406       1 ;     CA - OD1
2400   2407       1 ;     CA - ND2
2400   2413       1 ;     CA - H
2400   2414       1 ;     CA - CA
2401   2403       1 ;     HA - HB2
2401   2404       1 ;     HA - HB3
2401   2405       1 ;     HA - CG
2401   2411       1 ;     HA - O
2401   2412       1 ;     HA - N
2402   2408       1 ;     CB - HD21
2402   2409       1 ;     CB - HD22
2402   2411       1 ;     CB - O
2402   2412       1 ;     CB - N
2403   2406       1 ;    HB2 - OD1
2403   2407       1 ;    HB2 - ND2
2403   2410       1 ;    HB2 - C
2404   2406       1 ;    HB3 - OD1
2404   2407       1 ;    HB3 - ND2
2404   2410       1 ;    HB3 - C
2405   2410       1 ;     CG - C
2406   2408       1 ;    OD1 - HD21
2406   2409       1 ;    OD1 - HD22
2410   2415       1 ;      C - HA
2410   2416       1 ;      C - CB
2410   2432       1 ;      C - C
2411   2413       1 ;      O - H
2411   2414       1 ;      O - CA
2412   2417       1 ;      N - HB2
2412   2418       1 ;      N - HB3
2412   2419       1 ;      N - CG
2412   2433       1 ;      N - O
2412   2434       1 ;      N - N
2413   2415       1 ;      H - HA
2413   2416       1 ;      H - CB
2413   2432       1 ;      H - C
2414   2420       1 ;     CA - HG2
2414   2421       1 ;     CA - HG3
2414   2422       1 ;     CA - CD
2414   2435       1 ;     CA - H
2414   2436       1 ;     CA - CA
2415   2417       1 ;     HA - HB2
2415   2418       1 ;     HA - HB3
2415   2419       1 ;     HA - CG
2415   2433       1 ;     HA - O
2415   2434       1 ;     HA - N
2416   2423       1 ;     CB - HD2
2416   2424       1 ;     CB - HD3
2416   2425       1 ;     CB - CE
2416   2433       1 ;     CB - O
2416   2434       1 ;     CB - N
2417   2420       1 ;    HB2 - HG2
```

```
2417   2421       1 ;    HB2 - HG3
2417   2422       1 ;    HB2 - CD
2417   2432       1 ;    HB2 - C
2418   2420       1 ;    HB3 - HG2
2418   2421       1 ;    HB3 - HG3
2418   2422       1 ;    HB3 - CD
2418   2432       1 ;    HB3 - C
2419   2426       1 ;     CG - HE2
2419   2427       1 ;     CG - HE3
2419   2428       1 ;     CG - NZ
2419   2432       1 ;     CG - C
2420   2423       1 ;    HG2 - HD2
2420   2424       1 ;    HG2 - HD3
2420   2425       1 ;    HG2 - CE
2421   2423       1 ;    HG3 - HD2
2421   2424       1 ;    HG3 - HD3
2421   2425       1 ;    HG3 - CE
2422   2429       1 ;     CD - HZ1
2422   2430       1 ;     CD - HZ2
2422   2431       1 ;     CD - HZ3
2423   2426       1 ;    HD2 - HE2
2423   2427       1 ;    HD2 - HE3
2423   2428       1 ;    HD2 - NZ
2424   2426       1 ;    HD3 - HE2
2424   2427       1 ;    HD3 - HE3
2424   2428       1 ;    HD3 - NZ
2426   2429       1 ;    HE2 - HZ1
2426   2430       1 ;    HE2 - HZ2
2426   2431       1 ;    HE2 - HZ3
2427   2429       1 ;    HE3 - HZ1
2427   2430       1 ;    HE3 - HZ2
2427   2431       1 ;    HE3 - HZ3
2432   2437       1 ;      C - HA
2432   2438       1 ;      C - CB
2432   2444       1 ;      C - C
2433   2435       1 ;      O - H
2433   2436       1 ;      O - CA
2434   2439       1 ;      N - HB2
2434   2440       1 ;      N - HB3
2434   2441       1 ;      N - CG
2434   2445       1 ;      N - O
2434   2446       1 ;      N - N
2435   2437       1 ;      H - HA
2435   2438       1 ;      H - CB
2435   2444       1 ;      H - C
2436   2442       1 ;     CA - OD1
2436   2443       1 ;     CA - OD2
2436   2447       1 ;     CA - H
2436   2448       1 ;     CA - CA
2437   2439       1 ;     HA - HB2
2437   2440       1 ;     HA - HB3
2437   2441       1 ;     HA - CG
```

| | | | | | |
|---|---|---|---|---|---|
| 2437 | 2445 | 1 ; | HA | - | O |
| 2437 | 2446 | 1 ; | HA | - | N |
| 2438 | 2445 | 1 ; | CB | - | O |
| 2438 | 2446 | 1 ; | CB | - | N |
| 2439 | 2442 | 1 ; | HB2 | - | OD1 |
| 2439 | 2443 | 1 ; | HB2 | - | OD2 |
| 2439 | 2444 | 1 ; | HB2 | - | C |
| 2440 | 2442 | 1 ; | HB3 | - | OD1 |
| 2440 | 2443 | 1 ; | HB3 | - | OD2 |
| 2440 | 2444 | 1 ; | HB3 | - | C |
| 2441 | 2444 | 1 ; | CG | - | C |
| 2444 | 2449 | 1 ; | C | - | HA |
| 2444 | 2450 | 1 ; | C | - | CB |
| 2444 | 2455 | 1 ; | C | - | C |
| 2445 | 2447 | 1 ; | O | - | H |
| 2445 | 2448 | 1 ; | O | - | CA |
| 2446 | 2451 | 1 ; | N | - | HB2 |
| 2446 | 2452 | 1 ; | N | - | HB3 |
| 2446 | 2453 | 1 ; | N | - | OG |
| 2446 | 2456 | 1 ; | N | - | O |
| 2446 | 2457 | 1 ; | N | - | N |
| 2447 | 2449 | 1 ; | H | - | HA |
| 2447 | 2450 | 1 ; | H | - | CB |
| 2447 | 2455 | 1 ; | H | - | C |
| 2448 | 2454 | 1 ; | CA | - | HG |
| 2448 | 2458 | 1 ; | CA | - | H |
| 2448 | 2459 | 1 ; | CA | - | CA |
| 2449 | 2451 | 1 ; | HA | - | HB2 |
| 2449 | 2452 | 1 ; | HA | - | HB3 |
| 2449 | 2453 | 1 ; | HA | - | OG |
| 2449 | 2456 | 1 ; | HA | - | O |
| 2449 | 2457 | 1 ; | HA | - | N |
| 2450 | 2456 | 1 ; | CB | - | O |
| 2450 | 2457 | 1 ; | CB | - | N |
| 2451 | 2454 | 1 ; | HB2 | - | HG |
| 2451 | 2455 | 1 ; | HB2 | - | C |
| 2452 | 2454 | 1 ; | HB3 | - | HG |
| 2452 | 2455 | 1 ; | HB3 | - | C |
| 2453 | 2455 | 1 ; | OG | - | C |
| 2455 | 2460 | 1 ; | C | - | HA |
| 2455 | 2461 | 1 ; | C | - | CB |
| 2455 | 2474 | 1 ; | C | - | C |
| 2456 | 2458 | 1 ; | O | - | H |
| 2456 | 2459 | 1 ; | O | - | CA |
| 2457 | 2462 | 1 ; | N | - | HB2 |
| 2457 | 2463 | 1 ; | N | - | HB3 |
| 2457 | 2464 | 1 ; | N | - | CG |
| 2457 | 2475 | 1 ; | N | - | O |
| 2457 | 2476 | 1 ; | N | - | N |
| 2458 | 2460 | 1 ; | H | - | HA |
| 2458 | 2461 | 1 ; | H | - | CB |
| 2458 | 2474 | 1 ; | H | - | C |

```
2459   2465      1 ;     CA - HG
2459   2466      1 ;     CA - CD1
2459   2470      1 ;     CA - CD2
2459   2477      1 ;     CA - H
2459   2478      1 ;     CA - CA
2460   2462      1 ;     HA - HB2
2460   2463      1 ;     HA - HB3
2460   2464      1 ;     HA - CG
2460   2475      1 ;     HA - O
2460   2476      1 ;     HA - N
2461   2467      1 ;     CB - HD11
2461   2468      1 ;     CB - HD12
2461   2469      1 ;     CB - HD13
2461   2471      1 ;     CB - HD21
2461   2472      1 ;     CB - HD22
2461   2473      1 ;     CB - HD23
2461   2475      1 ;     CB - O
2461   2476      1 ;     CB - N
2462   2465      1 ;    HB2 - HG
2462   2466      1 ;    HB2 - CD1
2462   2470      1 ;    HB2 - CD2
2462   2474      1 ;    HB2 - C
2463   2465      1 ;    HB3 - HG
2463   2466      1 ;    HB3 - CD1
2463   2470      1 ;    HB3 - CD2
2463   2474      1 ;    HB3 - C
2464   2474      1 ;     CG - C
2465   2467      1 ;     HG - HD11
2465   2468      1 ;     HG - HD12
2465   2469      1 ;     HG - HD13
2465   2471      1 ;     HG - HD21
2465   2472      1 ;     HG - HD22
2465   2473      1 ;     HG - HD23
2466   2471      1 ;    CD1 - HD21
2466   2472      1 ;    CD1 - HD22
2466   2473      1 ;    CD1 - HD23
2467   2470      1 ;   HD11 - CD2
2468   2470      1 ;   HD12 - CD2
2469   2470      1 ;   HD13 - CD2
2474   2479      1 ;      C - HA
2474   2480      1 ;      C - CB
2474   2493      1 ;      C - C
2475   2477      1 ;      O - H
2475   2478      1 ;      O - CA
2476   2481      1 ;      N - HB2
2476   2482      1 ;      N - HB3
2476   2483      1 ;      N - CG
2476   2494      1 ;      N - O
2476   2495      1 ;      N - N
2477   2479      1 ;      H - HA
2477   2480      1 ;      H - CB
2477   2493      1 ;      H - C
```

```
2478   2484       1 ;      CA - HG
2478   2485       1 ;      CA - CD1
2478   2489       1 ;      CA - CD2
2478   2496       1 ;      CA - H
2478   2497       1 ;      CA - CA
2479   2481       1 ;      HA - HB2
2479   2482       1 ;      HA - HB3
2479   2483       1 ;      HA - CG
2479   2494       1 ;      HA - O
2479   2495       1 ;      HA - N
2480   2486       1 ;      CB - HD11
2480   2487       1 ;      CB - HD12
2480   2488       1 ;      CB - HD13
2480   2490       1 ;      CB - HD21
2480   2491       1 ;      CB - HD22
2480   2492       1 ;      CB - HD23
2480   2494       1 ;      CB - O
2480   2495       1 ;      CB - N
2481   2484       1 ;     HB2 - HG
2481   2485       1 ;     HB2 - CD1
2481   2489       1 ;     HB2 - CD2
2481   2493       1 ;     HB2 - C
2482   2484       1 ;     HB3 - HG
2482   2485       1 ;     HB3 - CD1
2482   2489       1 ;     HB3 - CD2
2482   2493       1 ;     HB3 - C
2483   2493       1 ;      CG - C
2484   2486       1 ;      HG - HD11
2484   2487       1 ;      HG - HD12
2484   2488       1 ;      HG - HD13
2484   2490       1 ;      HG - HD21
2484   2491       1 ;      HG - HD22
2484   2492       1 ;      HG - HD23
2485   2490       1 ;     CD1 - HD21
2485   2491       1 ;     CD1 - HD22
2485   2492       1 ;     CD1 - HD23
2486   2489       1 ;    HD11 - CD2
2487   2489       1 ;    HD12 - CD2
2488   2489       1 ;    HD13 - CD2
2493   2498       1 ;       C - HA
2493   2499       1 ;       C - CB
2493   2515       1 ;       C - C
2494   2496       1 ;       O - H
2494   2497       1 ;       O - CA
2495   2500       1 ;       N - HB2
2495   2501       1 ;       N - HB3
2495   2502       1 ;       N - CG
2495   2516       1 ;       N - O
2495   2517       1 ;       N - N
2496   2498       1 ;       H - HA
2496   2499       1 ;       H - CB
2496   2515       1 ;       H - C
```

```
2497   2503      1 ;     CA - HG2
2497   2504      1 ;     CA - HG3
2497   2505      1 ;     CA - CD
2497   2518      1 ;     CA - H
2497   2519      1 ;     CA - CA
2498   2500      1 ;     HA - HB2
2498   2501      1 ;     HA - HB3
2498   2502      1 ;     HA - CG
2498   2516      1 ;     HA - O
2498   2517      1 ;     HA - N
2499   2506      1 ;     CB - HD2
2499   2507      1 ;     CB - HD3
2499   2508      1 ;     CB - CE
2499   2516      1 ;     CB - O
2499   2517      1 ;     CB - N
2500   2503      1 ;    HB2 - HG2
2500   2504      1 ;    HB2 - HG3
2500   2505      1 ;    HB2 - CD
2500   2515      1 ;    HB2 - C
2501   2503      1 ;    HB3 - HG2
2501   2504      1 ;    HB3 - HG3
2501   2505      1 ;    HB3 - CD
2501   2515      1 ;    HB3 - C
2502   2509      1 ;     CG - HE2
2502   2510      1 ;     CG - HE3
2502   2511      1 ;     CG - NZ
2502   2515      1 ;     CG - C
2503   2506      1 ;    HG2 - HD2
2503   2507      1 ;    HG2 - HD3
2503   2508      1 ;    HG2 - CE
2504   2506      1 ;    HG3 - HD2
2504   2507      1 ;    HG3 - HD3
2504   2508      1 ;    HG3 - CE
2505   2512      1 ;     CD - HZ1
2505   2513      1 ;     CD - HZ2
2505   2514      1 ;     CD - HZ3
2506   2509      1 ;    HD2 - HE2
2506   2510      1 ;    HD2 - HE3
2506   2511      1 ;    HD2 - NZ
2507   2509      1 ;    HD3 - HE2
2507   2510      1 ;    HD3 - HE3
2507   2511      1 ;    HD3 - NZ
2509   2512      1 ;    HE2 - HZ1
2509   2513      1 ;    HE2 - HZ2
2509   2514      1 ;    HE2 - HZ3
2510   2512      1 ;    HE3 - HZ1
2510   2513      1 ;    HE3 - HZ2
2510   2514      1 ;    HE3 - HZ3
2515   2520      1 ;      C - HA
2515   2521      1 ;      C - CB
2515   2534      1 ;      C - C
2516   2518      1 ;      O - H
```

```
2516   2519      1 ;      O - CA
2517   2522      1 ;      N - HB
2517   2523      1 ;      N - CG2
2517   2527      1 ;      N - CG1
2517   2535      1 ;      N - O
2517   2536      1 ;      N - N
2518   2520      1 ;      H - HA
2518   2521      1 ;      H - CB
2518   2534      1 ;      H - C
2519   2524      1 ;     CA - HG21
2519   2525      1 ;     CA - HG22
2519   2526      1 ;     CA - HG23
2519   2528      1 ;     CA - HG12
2519   2529      1 ;     CA - HG13
2519   2530      1 ;     CA - CD1
2519   2537      1 ;     CA - H
2519   2538      1 ;     CA - CA
2520   2522      1 ;     HA - HB
2520   2523      1 ;     HA - CG2
2520   2527      1 ;     HA - CG1
2520   2535      1 ;     HA - O
2520   2536      1 ;     HA - N
2521   2531      1 ;     CB - HD11
2521   2532      1 ;     CB - HD12
2521   2533      1 ;     CB - HD13
2521   2535      1 ;     CB - O
2521   2536      1 ;     CB - N
2522   2524      1 ;     HB - HG21
2522   2525      1 ;     HB - HG22
2522   2526      1 ;     HB - HG23
2522   2528      1 ;     HB - HG12
2522   2529      1 ;     HB - HG13
2522   2530      1 ;     HB - CD1
2522   2534      1 ;     HB - C
2523   2528      1 ;    CG2 - HG12
2523   2529      1 ;    CG2 - HG13
2523   2530      1 ;    CG2 - CD1
2523   2534      1 ;    CG2 - C
2524   2527      1 ;   HG21 - CG1
2525   2527      1 ;   HG22 - CG1
2526   2527      1 ;   HG23 - CG1
2527   2534      1 ;    CG1 - C
2528   2531      1 ;   HG12 - HD11
2528   2532      1 ;   HG12 - HD12
2528   2533      1 ;   HG12 - HD13
2529   2531      1 ;   HG13 - HD11
2529   2532      1 ;   HG13 - HD12
2529   2533      1 ;   HG13 - HD13
2534   2539      1 ;      C - HA
2534   2540      1 ;      C - CB
2534   2553      1 ;      C - C
2535   2537      1 ;      O - H
```

```
2535   2538       1 ;      O - CA
2536   2541       1 ;      N - HB
2536   2542       1 ;      N - CG2
2536   2546       1 ;      N - CG1
2536   2554       1 ;      N - O
2536   2555       1 ;      N - N
2537   2539       1 ;      H - HA
2537   2540       1 ;      H - CB
2537   2553       1 ;      H - C
2538   2543       1 ;     CA - HG21
2538   2544       1 ;     CA - HG22
2538   2545       1 ;     CA - HG23
2538   2547       1 ;     CA - HG12
2538   2548       1 ;     CA - HG13
2538   2549       1 ;     CA - CD1
2538   2556       1 ;     CA - H
2538   2557       1 ;     CA - CA
2539   2541       1 ;     HA - HB
2539   2542       1 ;     HA - CG2
2539   2546       1 ;     HA - CG1
2539   2554       1 ;     HA - O
2539   2555       1 ;     HA - N
2540   2550       1 ;     CB - HD11
2540   2551       1 ;     CB - HD12
2540   2552       1 ;     CB - HD13
2540   2554       1 ;     CB - O
2540   2555       1 ;     CB - N
2541   2543       1 ;     HB - HG21
2541   2544       1 ;     HB - HG22
2541   2545       1 ;     HB - HG23
2541   2547       1 ;     HB - HG12
2541   2548       1 ;     HB - HG13
2541   2549       1 ;     HB - CD1
2541   2553       1 ;     HB - C
2542   2547       1 ;    CG2 - HG12
2542   2548       1 ;    CG2 - HG13
2542   2549       1 ;    CG2 - CD1
2542   2553       1 ;    CG2 - C
2543   2546       1 ;   HG21 - CG1
2544   2546       1 ;   HG22 - CG1
2545   2546       1 ;   HG23 - CG1
2546   2553       1 ;    CG1 - C
2547   2550       1 ;   HG12 - HD11
2547   2551       1 ;   HG12 - HD12
2547   2552       1 ;   HG12 - HD13
2548   2550       1 ;   HG13 - HD11
2548   2551       1 ;   HG13 - HD12
2548   2552       1 ;   HG13 - HD13
2553   2558       1 ;      C - HA
2553   2559       1 ;      C - CB
2553   2565       1 ;      C - C
2554   2556       1 ;      O - H
```

| | | | |
|---|---|---|---|
| 2554 | 2557 | 1 ; | O - CA |
| 2555 | 2560 | 1 ; | N - HB2 |
| 2555 | 2561 | 1 ; | N - HB3 |
| 2555 | 2562 | 1 ; | N - CG |
| 2555 | 2566 | 1 ; | N - O |
| 2555 | 2567 | 1 ; | N - N |
| 2556 | 2558 | 1 ; | H - HA |
| 2556 | 2559 | 1 ; | H - CB |
| 2556 | 2565 | 1 ; | H - C |
| 2557 | 2563 | 1 ; | CA - OD1 |
| 2557 | 2564 | 1 ; | CA - OD2 |
| 2557 | 2568 | 1 ; | CA - H |
| 2557 | 2569 | 1 ; | CA - CA |
| 2558 | 2560 | 1 ; | HA - HB2 |
| 2558 | 2561 | 1 ; | HA - HB3 |
| 2558 | 2562 | 1 ; | HA - CG |
| 2558 | 2566 | 1 ; | HA - O |
| 2558 | 2567 | 1 ; | HA - N |
| 2559 | 2566 | 1 ; | CB - O |
| 2559 | 2567 | 1 ; | CB - N |
| 2560 | 2563 | 1 ; | HB2 - OD1 |
| 2560 | 2564 | 1 ; | HB2 - OD2 |
| 2560 | 2565 | 1 ; | HB2 - C |
| 2561 | 2563 | 1 ; | HB3 - OD1 |
| 2561 | 2564 | 1 ; | HB3 - OD2 |
| 2561 | 2565 | 1 ; | HB3 - C |
| 2562 | 2565 | 1 ; | CG - C |
| 2565 | 2570 | 1 ; | C - HA |
| 2565 | 2571 | 1 ; | C - CB |
| 2565 | 2585 | 1 ; | C - C |
| 2566 | 2568 | 1 ; | O - H |
| 2566 | 2569 | 1 ; | O - CA |
| 2567 | 2572 | 1 ; | N - HB2 |
| 2567 | 2573 | 1 ; | N - HB3 |
| 2567 | 2574 | 1 ; | N - CG |
| 2567 | 2586 | 1 ; | N - O |
| 2567 | 2587 | 1 ; | N - N |
| 2568 | 2570 | 1 ; | H - HA |
| 2568 | 2571 | 1 ; | H - CB |
| 2568 | 2585 | 1 ; | H - C |
| 2569 | 2575 | 1 ; | CA - CD1 |
| 2569 | 2583 | 1 ; | CA - CD2 |
| 2569 | 2588 | 1 ; | CA - H |
| 2569 | 2589 | 1 ; | CA - CA |
| 2570 | 2572 | 1 ; | HA - HB2 |
| 2570 | 2573 | 1 ; | HA - HB3 |
| 2570 | 2574 | 1 ; | HA - CG |
| 2570 | 2586 | 1 ; | HA - O |
| 2570 | 2587 | 1 ; | HA - N |
| 2571 | 2576 | 1 ; | CB - HD1 |
| 2571 | 2577 | 1 ; | CB - CE1 |
| 2571 | 2581 | 1 ; | CB - CE2 |

| | | | | | |
|---|---|---|---|---|---|
| 2571 | 2584 | 1 ; | CB | - | HD2 |
| 2571 | 2586 | 1 ; | CB | - | O |
| 2571 | 2587 | 1 ; | CB | - | N |
| 2572 | 2575 | 1 ; | HB2 | - | CD1 |
| 2572 | 2583 | 1 ; | HB2 | - | CD2 |
| 2572 | 2585 | 1 ; | HB2 | - | C |
| 2573 | 2575 | 1 ; | HB3 | - | CD1 |
| 2573 | 2583 | 1 ; | HB3 | - | CD2 |
| 2573 | 2585 | 1 ; | HB3 | - | C |
| 2574 | 2578 | 1 ; | CG | - | HE1 |
| 2574 | 2579 | 1 ; | CG | - | CZ |
| 2574 | 2582 | 1 ; | CG | - | HE2 |
| 2574 | 2585 | 1 ; | CG | - | C |
| 2575 | 2580 | 1 ; | CD1 | - | HZ |
| 2575 | 2581 | 1 ; | CD1 | - | CE2 |
| 2575 | 2584 | 1 ; | CD1 | - | HD2 |
| 2576 | 2578 | 1 ; | HD1 | - | HE1 |
| 2576 | 2579 | 1 ; | HD1 | - | CZ |
| 2576 | 2583 | 1 ; | HD1 | - | CD2 |
| 2577 | 2582 | 1 ; | CE1 | - | HE2 |
| 2577 | 2583 | 1 ; | CE1 | - | CD2 |
| 2578 | 2580 | 1 ; | HE1 | - | HZ |
| 2578 | 2581 | 1 ; | HE1 | - | CE2 |
| 2579 | 2584 | 1 ; | CZ | - | HD2 |
| 2580 | 2582 | 1 ; | HZ | - | HE2 |
| 2580 | 2583 | 1 ; | HZ | - | CD2 |
| 2582 | 2584 | 1 ; | HE2 | - | HD2 |
| 2585 | 2590 | 1 ; | C | - | HA2 |
| 2585 | 2591 | 1 ; | C | - | HA3 |
| 2585 | 2592 | 1 ; | C | - | C |
| 2586 | 2588 | 1 ; | O | - | H |
| 2586 | 2589 | 1 ; | O | - | CA |
| 2587 | 2593 | 1 ; | N | - | O |
| 2587 | 2594 | 1 ; | N | - | N |
| 2588 | 2590 | 1 ; | H | - | HA2 |
| 2588 | 2591 | 1 ; | H | - | HA3 |
| 2588 | 2592 | 1 ; | H | - | C |
| 2589 | 2595 | 1 ; | CA | - | H |
| 2589 | 2596 | 1 ; | CA | - | CA |
| 2590 | 2593 | 1 ; | HA2 | - | O |
| 2590 | 2594 | 1 ; | HA2 | - | N |
| 2591 | 2593 | 1 ; | HA3 | - | O |
| 2591 | 2594 | 1 ; | HA3 | - | N |
| 2592 | 2597 | 1 ; | C | - | HA |
| 2592 | 2598 | 1 ; | C | - | CB |
| 2592 | 2611 | 1 ; | C | - | C |
| 2593 | 2595 | 1 ; | O | - | H |
| 2593 | 2596 | 1 ; | O | - | CA |
| 2594 | 2599 | 1 ; | N | - | HB2 |
| 2594 | 2600 | 1 ; | N | - | HB3 |
| 2594 | 2601 | 1 ; | N | - | CG |
| 2594 | 2612 | 1 ; | N | - | O |

| | | | |
|---|---|---|---|
| 2594 | 2613 | 1 ; | N - N |
| 2595 | 2597 | 1 ; | H - HA |
| 2595 | 2598 | 1 ; | H - CB |
| 2595 | 2611 | 1 ; | H - C |
| 2596 | 2602 | 1 ; | CA - HG |
| 2596 | 2603 | 1 ; | CA - CD1 |
| 2596 | 2607 | 1 ; | CA - CD2 |
| 2596 | 2614 | 1 ; | CA - H |
| 2596 | 2615 | 1 ; | CA - CA |
| 2597 | 2599 | 1 ; | HA - HB2 |
| 2597 | 2600 | 1 ; | HA - HB3 |
| 2597 | 2601 | 1 ; | HA - CG |
| 2597 | 2612 | 1 ; | HA - O |
| 2597 | 2613 | 1 ; | HA - N |
| 2598 | 2604 | 1 ; | CB - HD11 |
| 2598 | 2605 | 1 ; | CB - HD12 |
| 2598 | 2606 | 1 ; | CB - HD13 |
| 2598 | 2608 | 1 ; | CB - HD21 |
| 2598 | 2609 | 1 ; | CB - HD22 |
| 2598 | 2610 | 1 ; | CB - HD23 |
| 2598 | 2612 | 1 ; | CB - O |
| 2598 | 2613 | 1 ; | CB - N |
| 2599 | 2602 | 1 ; | HB2 - HG |
| 2599 | 2603 | 1 ; | HB2 - CD1 |
| 2599 | 2607 | 1 ; | HB2 - CD2 |
| 2599 | 2611 | 1 ; | HB2 - C |
| 2600 | 2602 | 1 ; | HB3 - HG |
| 2600 | 2603 | 1 ; | HB3 - CD1 |
| 2600 | 2607 | 1 ; | HB3 - CD2 |
| 2600 | 2611 | 1 ; | HB3 - C |
| 2601 | 2611 | 1 ; | CG - C |
| 2602 | 2604 | 1 ; | HG - HD11 |
| 2602 | 2605 | 1 ; | HG - HD12 |
| 2602 | 2606 | 1 ; | HG - HD13 |
| 2602 | 2608 | 1 ; | HG - HD21 |
| 2602 | 2609 | 1 ; | HG - HD22 |
| 2602 | 2610 | 1 ; | HG - HD23 |
| 2603 | 2608 | 1 ; | CD1 - HD21 |
| 2603 | 2609 | 1 ; | CD1 - HD22 |
| 2603 | 2610 | 1 ; | CD1 - HD23 |
| 2604 | 2607 | 1 ; | HD11 - CD2 |
| 2605 | 2607 | 1 ; | HD12 - CD2 |
| 2606 | 2607 | 1 ; | HD13 - CD2 |
| 2611 | 2616 | 1 ; | C - HA |
| 2611 | 2617 | 1 ; | C - CB |
| 2611 | 2622 | 1 ; | C - C |
| 2612 | 2614 | 1 ; | O - H |
| 2612 | 2615 | 1 ; | O - CA |
| 2613 | 2618 | 1 ; | N - HB2 |
| 2613 | 2619 | 1 ; | N - HB3 |
| 2613 | 2620 | 1 ; | N - OG |
| 2613 | 2623 | 1 ; | N - O |

```
2613   2624       1 ;       N - N
2614   2616       1 ;       H - HA
2614   2617       1 ;       H - CB
2614   2622       1 ;       H - C
2615   2621       1 ;      CA - HG
2615   2625       1 ;      CA - H
2615   2626       1 ;      CA - CA
2616   2618       1 ;      HA - HB2
2616   2619       1 ;      HA - HB3
2616   2620       1 ;      HA - OG
2616   2623       1 ;      HA - O
2616   2624       1 ;      HA - N
2617   2623       1 ;      CB - O
2617   2624       1 ;      CB - N
2618   2621       1 ;     HB2 - HG
2618   2622       1 ;     HB2 - C
2619   2621       1 ;     HB3 - HG
2619   2622       1 ;     HB3 - C
2620   2622       1 ;      OG - C
2622   2627       1 ;       C - HA
2622   2628       1 ;       C - CB
2622   2644       1 ;       C - C
2623   2625       1 ;       O - H
2623   2626       1 ;       O - CA
2624   2629       1 ;       N - HB2
2624   2630       1 ;       N - HB3
2624   2631       1 ;       N - CG
2624   2645       1 ;       N - O
2624   2646       1 ;       N - N
2625   2627       1 ;       H - HA
2625   2628       1 ;       H - CB
2625   2644       1 ;       H - C
2626   2632       1 ;      CA - HG2
2626   2633       1 ;      CA - HG3
2626   2634       1 ;      CA - CD
2626   2647       1 ;      CA - H
2626   2648       1 ;      CA - CA
2627   2629       1 ;      HA - HB2
2627   2630       1 ;      HA - HB3
2627   2631       1 ;      HA - CG
2627   2645       1 ;      HA - O
2627   2646       1 ;      HA - N
2628   2635       1 ;      CB - HD2
2628   2636       1 ;      CB - HD3
2628   2637       1 ;      CB - CE
2628   2645       1 ;      CB - O
2628   2646       1 ;      CB - N
2629   2632       1 ;     HB2 - HG2
2629   2633       1 ;     HB2 - HG3
2629   2634       1 ;     HB2 - CD
2629   2644       1 ;     HB2 - C
2630   2632       1 ;     HB3 - HG2
```

```
2630   2633      1 ;    HB3 - HG3
2630   2634      1 ;    HB3 - CD
2630   2644      1 ;    HB3 - C
2631   2638      1 ;     CG - HE2
2631   2639      1 ;     CG - HE3
2631   2640      1 ;     CG - NZ
2631   2644      1 ;     CG - C
2632   2635      1 ;    HG2 - HD2
2632   2636      1 ;    HG2 - HD3
2632   2637      1 ;    HG2 - CE
2633   2635      1 ;    HG3 - HD2
2633   2636      1 ;    HG3 - HD3
2633   2637      1 ;    HG3 - CE
2634   2641      1 ;     CD - HZ1
2634   2642      1 ;     CD - HZ2
2634   2643      1 ;     CD - HZ3
2635   2638      1 ;    HD2 - HE2
2635   2639      1 ;    HD2 - HE3
2635   2640      1 ;    HD2 - NZ
2636   2638      1 ;    HD3 - HE2
2636   2639      1 ;    HD3 - HE3
2636   2640      1 ;    HD3 - NZ
2638   2641      1 ;    HE2 - HZ1
2638   2642      1 ;    HE2 - HZ2
2638   2643      1 ;    HE2 - HZ3
2639   2641      1 ;    HE3 - HZ1
2639   2642      1 ;    HE3 - HZ2
2639   2643      1 ;    HE3 - HZ3
2644   2649      1 ;      C - HA
2644   2650      1 ;      C - CB
2644   2658      1 ;      C - C
2645   2647      1 ;      O - H
2645   2648      1 ;      O - CA
2646   2651      1 ;      N - HB2
2646   2652      1 ;      N - HB3
2646   2653      1 ;      N - CG
2646   2659      1 ;      N - O
2646   2660      1 ;      N - N
2647   2649      1 ;      H - HA
2647   2650      1 ;      H - CB
2647   2658      1 ;      H - C
2648   2654      1 ;     CA - OD1
2648   2655      1 ;     CA - ND2
2648   2661      1 ;     CA - H
2648   2662      1 ;     CA - CA
2649   2651      1 ;     HA - HB2
2649   2652      1 ;     HA - HB3
2649   2653      1 ;     HA - CG
2649   2659      1 ;     HA - O
2649   2660      1 ;     HA - N
2650   2656      1 ;     CB - HD21
2650   2657      1 ;     CB - HD22
```

| | | | | | |
|---|---|---|---|---|---|
| 2650 | 2659 | 1 ; | CB | - | O |
| 2650 | 2660 | 1 ; | CB | - | N |
| 2651 | 2654 | 1 ; | HB2 | - | OD1 |
| 2651 | 2655 | 1 ; | HB2 | - | ND2 |
| 2651 | 2658 | 1 ; | HB2 | - | C |
| 2652 | 2654 | 1 ; | HB3 | - | OD1 |
| 2652 | 2655 | 1 ; | HB3 | - | ND2 |
| 2652 | 2658 | 1 ; | HB3 | - | C |
| 2653 | 2658 | 1 ; | CG | - | C |
| 2654 | 2656 | 1 ; | OD1 | - | HD21 |
| 2654 | 2657 | 1 ; | OD1 | - | HD22 |
| 2658 | 2663 | 1 ; | C | - | HA |
| 2658 | 2664 | 1 ; | C | - | CB |
| 2658 | 2677 | 1 ; | C | - | C |
| 2659 | 2661 | 1 ; | O | - | H |
| 2659 | 2662 | 1 ; | O | - | CA |
| 2660 | 2665 | 1 ; | N | - | HB2 |
| 2660 | 2666 | 1 ; | N | - | HB3 |
| 2660 | 2667 | 1 ; | N | - | CG |
| 2660 | 2678 | 1 ; | N | - | O |
| 2660 | 2679 | 1 ; | N | - | N |
| 2661 | 2663 | 1 ; | H | - | HA |
| 2661 | 2664 | 1 ; | H | - | CB |
| 2661 | 2677 | 1 ; | H | - | C |
| 2662 | 2668 | 1 ; | CA | - | HG |
| 2662 | 2669 | 1 ; | CA | - | CD1 |
| 2662 | 2673 | 1 ; | CA | - | CD2 |
| 2662 | 2680 | 1 ; | CA | - | H |
| 2662 | 2681 | 1 ; | CA | - | CA |
| 2663 | 2665 | 1 ; | HA | - | HB2 |
| 2663 | 2666 | 1 ; | HA | - | HB3 |
| 2663 | 2667 | 1 ; | HA | - | CG |
| 2663 | 2678 | 1 ; | HA | - | O |
| 2663 | 2679 | 1 ; | HA | - | N |
| 2664 | 2670 | 1 ; | CB | - | HD11 |
| 2664 | 2671 | 1 ; | CB | - | HD12 |
| 2664 | 2672 | 1 ; | CB | - | HD13 |
| 2664 | 2674 | 1 ; | CB | - | HD21 |
| 2664 | 2675 | 1 ; | CB | - | HD22 |
| 2664 | 2676 | 1 ; | CB | - | HD23 |
| 2664 | 2678 | 1 ; | CB | - | O |
| 2664 | 2679 | 1 ; | CB | - | N |
| 2665 | 2668 | 1 ; | HB2 | - | HG |
| 2665 | 2669 | 1 ; | HB2 | - | CD1 |
| 2665 | 2673 | 1 ; | HB2 | - | CD2 |
| 2665 | 2677 | 1 ; | HB2 | - | C |
| 2666 | 2668 | 1 ; | HB3 | - | HG |
| 2666 | 2669 | 1 ; | HB3 | - | CD1 |
| 2666 | 2673 | 1 ; | HB3 | - | CD2 |
| 2666 | 2677 | 1 ; | HB3 | - | C |
| 2667 | 2677 | 1 ; | CG | - | C |
| 2668 | 2670 | 1 ; | HG | - | HD11 |

```
2668   2671       1 ;      HG - HD12
2668   2672       1 ;      HG - HD13
2668   2674       1 ;      HG - HD21
2668   2675       1 ;      HG - HD22
2668   2676       1 ;      HG - HD23
2669   2674       1 ;     CD1 - HD21
2669   2675       1 ;     CD1 - HD22
2669   2676       1 ;     CD1 - HD23
2670   2673       1 ;    HD11 - CD2
2671   2673       1 ;    HD12 - CD2
2672   2673       1 ;    HD13 - CD2
2677   2682       1 ;       C - HA2
2677   2683       1 ;       C - HA3
2677   2684       1 ;       C - C
2678   2680       1 ;       O - H
2678   2681       1 ;       O - CA
2679   2685       1 ;       N - O
2679   2686       1 ;       N - N
2680   2682       1 ;       H - HA2
2680   2683       1 ;       H - HA3
2680   2684       1 ;       H - C
2681   2687       1 ;      CA - H
2681   2688       1 ;      CA - CA
2682   2685       1 ;     HA2 - O
2682   2686       1 ;     HA2 - N
2683   2685       1 ;     HA3 - O
2683   2686       1 ;     HA3 - N
2684   2689       1 ;       C - HA
2684   2690       1 ;       C - CB
2684   2698       1 ;       C - C
2685   2687       1 ;       O - H
2685   2688       1 ;       O - CA
2686   2691       1 ;       N - HB
2686   2692       1 ;       N - CG2
2686   2696       1 ;       N - OG1
2686   2699       1 ;       N - O
2686   2700       1 ;       N - N
2687   2689       1 ;       H - HA
2687   2690       1 ;       H - CB
2687   2698       1 ;       H - C
2688   2693       1 ;      CA - HG21
2688   2694       1 ;      CA - HG22
2688   2695       1 ;      CA - HG23
2688   2697       1 ;      CA - HG1
2688   2701       1 ;      CA - H
2688   2702       1 ;      CA - CA
2689   2691       1 ;      HA - HB
2689   2692       1 ;      HA - CG2
2689   2696       1 ;      HA - OG1
2689   2699       1 ;      HA - O
2689   2700       1 ;      HA - N
2690   2699       1 ;      CB - O
```

```
2690   2700      1 ;      CB - N
2691   2693      1 ;      HB - HG21
2691   2694      1 ;      HB - HG22
2691   2695      1 ;      HB - HG23
2691   2697      1 ;      HB - HG1
2691   2698      1 ;      HB - C
2692   2697      1 ;     CG2 - HG1
2692   2698      1 ;     CG2 - C
2693   2696      1 ;    HG21 - OG1
2694   2696      1 ;    HG22 - OG1
2695   2696      1 ;    HG23 - OG1
2696   2698      1 ;     OG1 - C
2698   2703      1 ;       C - HA2
2698   2704      1 ;       C - HA3
2698   2705      1 ;       C - C
2699   2701      1 ;       O - H
2699   2702      1 ;       O - CA
2700   2706      1 ;       N - O
2700   2707      1 ;       N - N
2701   2703      1 ;       H - HA2
2701   2704      1 ;       H - HA3
2701   2705      1 ;       H - C
2702   2708      1 ;      CA - H
2702   2709      1 ;      CA - CA
2703   2706      1 ;     HA2 - O
2703   2707      1 ;     HA2 - N
2704   2706      1 ;     HA3 - O
2704   2707      1 ;     HA3 - N
2705   2710      1 ;       C - HA
2705   2711      1 ;       C - CB
2705   2720      1 ;       C - C
2706   2708      1 ;       O - H
2706   2709      1 ;       O - CA
2707   2712      1 ;       N - HB2
2707   2713      1 ;       N - HB3
2707   2714      1 ;       N - CG
2707   2721      1 ;       N - O
2707   2722      1 ;       N - N
2708   2710      1 ;       H - HA
2708   2711      1 ;       H - CB
2708   2720      1 ;       H - C
2709   2715      1 ;      CA - HG2
2709   2716      1 ;      CA - HG3
2709   2717      1 ;      CA - CD
2709   2723      1 ;      CA - H
2709   2724      1 ;      CA - CA
2710   2712      1 ;      HA - HB2
2710   2713      1 ;      HA - HB3
2710   2714      1 ;      HA - CG
2710   2721      1 ;      HA - O
2710   2722      1 ;      HA - N
2711   2718      1 ;      CB - OE1
```

```
2711   2719       1 ;      CB - OE2
2711   2721       1 ;      CB - O
2711   2722       1 ;      CB - N
2712   2715       1 ;     HB2 - HG2
2712   2716       1 ;     HB2 - HG3
2712   2717       1 ;     HB2 - CD
2712   2720       1 ;     HB2 - C
2713   2715       1 ;     HB3 - HG2
2713   2716       1 ;     HB3 - HG3
2713   2717       1 ;     HB3 - CD
2713   2720       1 ;     HB3 - C
2714   2720       1 ;      CG - C
2715   2718       1 ;     HG2 - OE1
2715   2719       1 ;     HG2 - OE2
2716   2718       1 ;     HG3 - OE1
2716   2719       1 ;     HG3 - OE2
2720   2725       1 ;       C - HA
2720   2726       1 ;       C - CB
2720   2740       1 ;       C - C
2721   2723       1 ;       O - H
2721   2724       1 ;       O - CA
2722   2727       1 ;       N - HB2
2722   2728       1 ;       N - HB3
2722   2729       1 ;       N - CG
2722   2741       1 ;       N - O
2722   2742       1 ;       N - N
2723   2725       1 ;       H - HA
2723   2726       1 ;       H - CB
2723   2740       1 ;       H - C
2724   2730       1 ;      CA - CD1
2724   2738       1 ;      CA - CD2
2724   2743       1 ;      CA - H
2724   2744       1 ;      CA - CA
2725   2727       1 ;      HA - HB2
2725   2728       1 ;      HA - HB3
2725   2729       1 ;      HA - CG
2725   2741       1 ;      HA - O
2725   2742       1 ;      HA - N
2726   2731       1 ;      CB - HD1
2726   2732       1 ;      CB - CE1
2726   2736       1 ;      CB - CE2
2726   2739       1 ;      CB - HD2
2726   2741       1 ;      CB - O
2726   2742       1 ;      CB - N
2727   2730       1 ;     HB2 - CD1
2727   2738       1 ;     HB2 - CD2
2727   2740       1 ;     HB2 - C
2728   2730       1 ;     HB3 - CD1
2728   2738       1 ;     HB3 - CD2
2728   2740       1 ;     HB3 - C
2729   2733       1 ;      CG - HE1
2729   2734       1 ;      CG - CZ
```

```
2729   2737       1 ;      CG - HE2
2729   2740       1 ;      CG - C
2730   2735       1 ;     CD1 - HZ
2730   2736       1 ;     CD1 - CE2
2730   2739       1 ;     CD1 - HD2
2731   2733       1 ;     HD1 - HE1
2731   2734       1 ;     HD1 - CZ
2731   2738       1 ;     HD1 - CD2
2732   2737       1 ;     CE1 - HE2
2732   2738       1 ;     CE1 - CD2
2733   2735       1 ;     HE1 - HZ
2733   2736       1 ;     HE1 - CE2
2734   2739       1 ;      CZ - HD2
2735   2737       1 ;      HZ - HE2
2735   2738       1 ;      HZ - CD2
2737   2739       1 ;     HE2 - HD2
2740   2745       1 ;       C - HA
2740   2746       1 ;       C - CB
2740   2754       1 ;       C - C
2741   2743       1 ;       O - H
2741   2744       1 ;       O - CA
2742   2747       1 ;       N - HB
2742   2748       1 ;       N - CG2
2742   2752       1 ;       N - OG1
2742   2755       1 ;       N - O
2742   2756       1 ;       N - N
2743   2745       1 ;       H - HA
2743   2746       1 ;       H - CB
2743   2754       1 ;       H - C
2744   2749       1 ;      CA - HG21
2744   2750       1 ;      CA - HG22
2744   2751       1 ;      CA - HG23
2744   2753       1 ;      CA - HG1
2744   2757       1 ;      CA - H
2744   2758       1 ;      CA - CA
2745   2747       1 ;      HA - HB
2745   2748       1 ;      HA - CG2
2745   2752       1 ;      HA - OG1
2745   2755       1 ;      HA - O
2745   2756       1 ;      HA - N
2746   2755       1 ;      CB - O
2746   2756       1 ;      CB - N
2747   2749       1 ;      HB - HG21
2747   2750       1 ;      HB - HG22
2747   2751       1 ;      HB - HG23
2747   2753       1 ;      HB - HG1
2747   2754       1 ;      HB - C
2748   2753       1 ;     CG2 - HG1
2748   2754       1 ;     CG2 - C
2749   2752       1 ;    HG21 - OG1
2750   2752       1 ;    HG22 - OG1
2751   2752       1 ;    HG23 - OG1
```

```
2752   2754       1 ;    OG1 - C
2754   2759       1 ;      C - HA
2754   2760       1 ;      C - CB
2754   2768       1 ;      C - C
2755   2757       1 ;      O - H
2755   2758       1 ;      O - CA
2756   2761       1 ;      N - HB
2756   2762       1 ;      N - CG2
2756   2766       1 ;      N - OG1
2756   2769       1 ;      N - O
2756   2770       1 ;      N - N
2757   2759       1 ;      H - HA
2757   2760       1 ;      H - CB
2757   2768       1 ;      H - C
2758   2763       1 ;     CA - HG21
2758   2764       1 ;     CA - HG22
2758   2765       1 ;     CA - HG23
2758   2767       1 ;     CA - HG1
2758   2771       1 ;     CA - H
2758   2772       1 ;     CA - CA
2759   2761       1 ;     HA - HB
2759   2762       1 ;     HA - CG2
2759   2766       1 ;     HA - OG1
2759   2769       1 ;     HA - O
2759   2770       1 ;     HA - N
2760   2769       1 ;     CB - O
2760   2770       1 ;     CB - N
2761   2763       1 ;     HB - HG21
2761   2764       1 ;     HB - HG22
2761   2765       1 ;     HB - HG23
2761   2767       1 ;     HB - HG1
2761   2768       1 ;     HB - C
2762   2767       1 ;    CG2 - HG1
2762   2768       1 ;    CG2 - C
2763   2766       1 ;   HG21 - OG1
2764   2766       1 ;   HG22 - OG1
2765   2766       1 ;   HG23 - OG1
2766   2768       1 ;    OG1 - C
2768   2773       1 ;      C - HA
2768   2774       1 ;      C - CB
2768   2782       1 ;      C - C
2769   2771       1 ;      O - H
2769   2772       1 ;      O - CA
2770   2775       1 ;      N - HB
2770   2776       1 ;      N - CG2
2770   2780       1 ;      N - OG1
2770   2783       1 ;      N - O
2770   2784       1 ;      N - N
2771   2773       1 ;      H - HA
2771   2774       1 ;      H - CB
2771   2782       1 ;      H - C
2772   2777       1 ;     CA - HG21
```

```
2772   2778       1 ;      CA - HG22
2772   2779       1 ;      CA - HG23
2772   2781       1 ;      CA - HG1
2772   2785       1 ;      CA - H
2772   2786       1 ;      CA - CA
2773   2775       1 ;      HA - HB
2773   2776       1 ;      HA - CG2
2773   2780       1 ;      HA - OG1
2773   2783       1 ;      HA - O
2773   2784       1 ;      HA - N
2774   2783       1 ;      CB - O
2774   2784       1 ;      CB - N
2775   2777       1 ;      HB - HG21
2775   2778       1 ;      HB - HG22
2775   2779       1 ;      HB - HG23
2775   2781       1 ;      HB - HG1
2775   2782       1 ;      HB - C
2776   2781       1 ;     CG2 - HG1
2776   2782       1 ;     CG2 - C
2777   2780       1 ;    HG21 - OG1
2778   2780       1 ;    HG22 - OG1
2779   2780       1 ;    HG23 - OG1
2780   2782       1 ;     OG1 - C
2782   2787       1 ;       C - HA
2782   2788       1 ;       C - CB
2782   2804       1 ;       C - C
2783   2785       1 ;       O - H
2783   2786       1 ;       O - CA
2784   2789       1 ;       N - HB2
2784   2790       1 ;       N - HB3
2784   2791       1 ;       N - CG
2784   2805       1 ;       N - O
2784   2806       1 ;       N - N
2785   2787       1 ;       H - HA
2785   2788       1 ;       H - CB
2785   2804       1 ;       H - C
2786   2792       1 ;      CA - HG2
2786   2793       1 ;      CA - HG3
2786   2794       1 ;      CA - CD
2786   2807       1 ;      CA - H
2786   2808       1 ;      CA - CA
2787   2789       1 ;      HA - HB2
2787   2790       1 ;      HA - HB3
2787   2791       1 ;      HA - CG
2787   2805       1 ;      HA - O
2787   2806       1 ;      HA - N
2788   2795       1 ;      CB - HD2
2788   2796       1 ;      CB - HD3
2788   2797       1 ;      CB - CE
2788   2805       1 ;      CB - O
2788   2806       1 ;      CB - N
2789   2792       1 ;     HB2 - HG2
```

| | | | | | |
|---|---|---|---|---|---|
| 2789 | 2793 | 1 ; | HB2 | - | HG3 |
| 2789 | 2794 | 1 ; | HB2 | - | CD |
| 2789 | 2804 | 1 ; | HB2 | - | C |
| 2790 | 2792 | 1 ; | HB3 | - | HG2 |
| 2790 | 2793 | 1 ; | HB3 | - | HG3 |
| 2790 | 2794 | 1 ; | HB3 | - | CD |
| 2790 | 2804 | 1 ; | HB3 | - | C |
| 2791 | 2798 | 1 ; | CG | - | HE2 |
| 2791 | 2799 | 1 ; | CG | - | HE3 |
| 2791 | 2800 | 1 ; | CG | - | NZ |
| 2791 | 2804 | 1 ; | CG | - | C |
| 2792 | 2795 | 1 ; | HG2 | - | HD2 |
| 2792 | 2796 | 1 ; | HG2 | - | HD3 |
| 2792 | 2797 | 1 ; | HG2 | - | CE |
| 2793 | 2795 | 1 ; | HG3 | - | HD2 |
| 2793 | 2796 | 1 ; | HG3 | - | HD3 |
| 2793 | 2797 | 1 ; | HG3 | - | CE |
| 2794 | 2801 | 1 ; | CD | - | HZ1 |
| 2794 | 2802 | 1 ; | CD | - | HZ2 |
| 2794 | 2803 | 1 ; | CD | - | HZ3 |
| 2795 | 2798 | 1 ; | HD2 | - | HE2 |
| 2795 | 2799 | 1 ; | HD2 | - | HE3 |
| 2795 | 2800 | 1 ; | HD2 | - | NZ |
| 2796 | 2798 | 1 ; | HD3 | - | HE2 |
| 2796 | 2799 | 1 ; | HD3 | - | HE3 |
| 2796 | 2800 | 1 ; | HD3 | - | NZ |
| 2798 | 2801 | 1 ; | HE2 | - | HZ1 |
| 2798 | 2802 | 1 ; | HE2 | - | HZ2 |
| 2798 | 2803 | 1 ; | HE2 | - | HZ3 |
| 2799 | 2801 | 1 ; | HE3 | - | HZ1 |
| 2799 | 2802 | 1 ; | HE3 | - | HZ2 |
| 2799 | 2803 | 1 ; | HE3 | - | HZ3 |
| 2804 | 2809 | 1 ; | C | - | HA |
| 2804 | 2810 | 1 ; | C | - | CB |
| 2804 | 2814 | 1 ; | C | - | C |
| 2805 | 2807 | 1 ; | O | - | H |
| 2805 | 2808 | 1 ; | O | - | CA |
| 2806 | 2811 | 1 ; | N | - | HB1 |
| 2806 | 2812 | 1 ; | N | - | HB2 |
| 2806 | 2813 | 1 ; | N | - | HB3 |
| 2806 | 2815 | 1 ; | N | - | O |
| 2806 | 2816 | 1 ; | N | - | N |
| 2807 | 2809 | 1 ; | H | - | HA |
| 2807 | 2810 | 1 ; | H | - | CB |
| 2807 | 2814 | 1 ; | H | - | C |
| 2808 | 2817 | 1 ; | CA | - | H |
| 2808 | 2818 | 1 ; | CA | - | CA |
| 2809 | 2811 | 1 ; | HA | - | HB1 |
| 2809 | 2812 | 1 ; | HA | - | HB2 |
| 2809 | 2813 | 1 ; | HA | - | HB3 |
| 2809 | 2815 | 1 ; | HA | - | O |
| 2809 | 2816 | 1 ; | HA | - | N |

```
2810   2815       1 ;      CB - O
2810   2816       1 ;      CB - N
2811   2814       1 ;     HB1 - C
2812   2814       1 ;     HB2 - C
2813   2814       1 ;     HB3 - C
2814   2819       1 ;       C - HA2
2814   2820       1 ;       C - HA3
2814   2821       1 ;       C - C
2815   2817       1 ;       O - H
2815   2818       1 ;       O - CA
2816   2822       1 ;       N - O
2816   2823       1 ;       N - N
2817   2819       1 ;       H - HA2
2817   2820       1 ;       H - HA3
2817   2821       1 ;       H - C
2818   2824       1 ;      CA - H
2818   2825       1 ;      CA - CA
2819   2822       1 ;     HA2 - O
2819   2823       1 ;     HA2 - N
2820   2822       1 ;     HA3 - O
2820   2823       1 ;     HA3 - N
2821   2826       1 ;       C - HA
2821   2827       1 ;       C - CB
2821   2835       1 ;       C - C
2822   2824       1 ;       O - H
2822   2825       1 ;       O - CA
2823   2828       1 ;       N - HB
2823   2829       1 ;       N - CG2
2823   2833       1 ;       N - OG1
2823   2836       1 ;       N - O
2823   2837       1 ;       N - N
2824   2826       1 ;       H - HA
2824   2827       1 ;       H - CB
2824   2835       1 ;       H - C
2825   2830       1 ;      CA - HG21
2825   2831       1 ;      CA - HG22
2825   2832       1 ;      CA - HG23
2825   2834       1 ;      CA - HG1
2825   2838       1 ;      CA - CD
2825   2847       1 ;      CA - CA
2826   2828       1 ;      HA - HB
2826   2829       1 ;      HA - CG2
2826   2833       1 ;      HA - OG1
2826   2836       1 ;      HA - O
2826   2837       1 ;      HA - N
2827   2836       1 ;      CB - O
2827   2837       1 ;      CB - N
2828   2830       1 ;      HB - HG21
2828   2831       1 ;      HB - HG22
2828   2832       1 ;      HB - HG23
2828   2834       1 ;      HB - HG1
2828   2835       1 ;      HB - C
```

```
2829   2834      1 ;    CG2 - HG1
2829   2835      1 ;    CG2 - C
2830   2833      1 ;   HG21 - OG1
2831   2833      1 ;   HG22 - OG1
2832   2833      1 ;   HG23 - OG1
2833   2835      1 ;    OG1 - C
2835   2839      1 ;      C - HD2
2835   2840      1 ;      C - HD3
2835   2841      1 ;      C - CG
2835   2844      1 ;      C - CB
2835   2848      1 ;      C - HA
2835   2849      1 ;      C - C
2836   2838      1 ;      O - CD
2836   2847      1 ;      O - CA
2837   2842      1 ;      N - HG2
2837   2843      1 ;      N - HG3
2837   2845      1 ;      N - HB2
2837   2846      1 ;      N - HB3
2837   2850      1 ;      N - O
2837   2851      1 ;      N - N
2838   2845      1 ;     CD - HB2
2838   2846      1 ;     CD - HB3
2838   2848      1 ;     CD - HA
2838   2849      1 ;     CD - C
2839   2842      1 ;    HD2 - HG2
2839   2843      1 ;    HD2 - HG3
2839   2844      1 ;    HD2 - CB
2839   2847      1 ;    HD2 - CA
2840   2842      1 ;    HD3 - HG2
2840   2843      1 ;    HD3 - HG3
2840   2844      1 ;    HD3 - CB
2840   2847      1 ;    HD3 - CA
2841   2848      1 ;     CG - HA
2841   2849      1 ;     CG - C
2842   2845      1 ;    HG2 - HB2
2842   2846      1 ;    HG2 - HB3
2842   2847      1 ;    HG2 - CA
2843   2845      1 ;    HG3 - HB2
2843   2846      1 ;    HG3 - HB3
2843   2847      1 ;    HG3 - CA
2844   2850      1 ;     CB - O
2844   2851      1 ;     CB - N
2845   2848      1 ;    HB2 - HA
2845   2849      1 ;    HB2 - C
2846   2848      1 ;    HB3 - HA
2846   2849      1 ;    HB3 - C
2847   2852      1 ;     CA - H
2847   2853      1 ;     CA - CA
2848   2850      1 ;     HA - O
2848   2851      1 ;     HA - N
2849   2854      1 ;      C - HA
2849   2855      1 ;      C - CB
```

| | | | | | | |
|---|---|---|---|---|---|---|
| 2849 | 2870 | 1 | ; | C | - | C |
| 2850 | 2852 | 1 | ; | O | - | H |
| 2850 | 2853 | 1 | ; | O | - | CA |
| 2851 | 2856 | 1 | ; | N | - | HB2 |
| 2851 | 2857 | 1 | ; | N | - | HB3 |
| 2851 | 2858 | 1 | ; | N | - | CG |
| 2851 | 2871 | 1 | ; | N | - | O |
| 2851 | 2872 | 1 | ; | N | - | N |
| 2852 | 2854 | 1 | ; | H | - | HA |
| 2852 | 2855 | 1 | ; | H | - | CB |
| 2852 | 2870 | 1 | ; | H | - | C |
| 2853 | 2859 | 1 | ; | CA | - | CD1 |
| 2853 | 2868 | 1 | ; | CA | - | CD2 |
| 2853 | 2873 | 1 | ; | CA | - | H |
| 2853 | 2874 | 1 | ; | CA | - | CA |
| 2854 | 2856 | 1 | ; | HA | - | HB2 |
| 2854 | 2857 | 1 | ; | HA | - | HB3 |
| 2854 | 2858 | 1 | ; | HA | - | CG |
| 2854 | 2871 | 1 | ; | HA | - | O |
| 2854 | 2872 | 1 | ; | HA | - | N |
| 2855 | 2860 | 1 | ; | CB | - | HD1 |
| 2855 | 2861 | 1 | ; | CB | - | CE1 |
| 2855 | 2866 | 1 | ; | CB | - | CE2 |
| 2855 | 2869 | 1 | ; | CB | - | HD2 |
| 2855 | 2871 | 1 | ; | CB | - | O |
| 2855 | 2872 | 1 | ; | CB | - | N |
| 2856 | 2859 | 1 | ; | HB2 | - | CD1 |
| 2856 | 2868 | 1 | ; | HB2 | - | CD2 |
| 2856 | 2870 | 1 | ; | HB2 | - | C |
| 2857 | 2859 | 1 | ; | HB3 | - | CD1 |
| 2857 | 2868 | 1 | ; | HB3 | - | CD2 |
| 2857 | 2870 | 1 | ; | HB3 | - | C |
| 2858 | 2862 | 1 | ; | CG | - | HE1 |
| 2858 | 2863 | 1 | ; | CG | - | CZ |
| 2858 | 2867 | 1 | ; | CG | - | HE2 |
| 2858 | 2870 | 1 | ; | CG | - | C |
| 2859 | 2864 | 1 | ; | CD1 | - | OH |
| 2859 | 2866 | 1 | ; | CD1 | - | CE2 |
| 2859 | 2869 | 1 | ; | CD1 | - | HD2 |
| 2860 | 2862 | 1 | ; | HD1 | - | HE1 |
| 2860 | 2863 | 1 | ; | HD1 | - | CZ |
| 2860 | 2868 | 1 | ; | HD1 | - | CD2 |
| 2861 | 2865 | 1 | ; | CE1 | - | HH |
| 2861 | 2867 | 1 | ; | CE1 | - | HE2 |
| 2861 | 2868 | 1 | ; | CE1 | - | CD2 |
| 2862 | 2864 | 1 | ; | HE1 | - | OH |
| 2862 | 2866 | 1 | ; | HE1 | - | CE2 |
| 2863 | 2869 | 1 | ; | CZ | - | HD2 |
| 2864 | 2867 | 1 | ; | OH | - | HE2 |
| 2864 | 2868 | 1 | ; | OH | - | CD2 |
| 2865 | 2866 | 1 | ; | HH | - | CE2 |
| 2867 | 2869 | 1 | ; | HE2 | - | HD2 |

```
2870   2875     1 ;     C - HA
2870   2876     1 ;     C - CB
2870   2891     1 ;     C - C
2871   2873     1 ;     O - H
2871   2874     1 ;     O - CA
2872   2877     1 ;     N - HB2
2872   2878     1 ;     N - HB3
2872   2879     1 ;     N - CG
2872   2892     1 ;     N - O
2872   2893     1 ;     N - N
2873   2875     1 ;     H - HA
2873   2876     1 ;     H - CB
2873   2891     1 ;     H - C
2874   2880     1 ;    CA - CD1
2874   2889     1 ;    CA - CD2
2874   2894     1 ;    CA - H
2874   2895     1 ;    CA - CA
2875   2877     1 ;    HA - HB2
2875   2878     1 ;    HA - HB3
2875   2879     1 ;    HA - CG
2875   2892     1 ;    HA - O
2875   2893     1 ;    HA - N
2876   2881     1 ;    CB - HD1
2876   2882     1 ;    CB - CE1
2876   2887     1 ;    CB - CE2
2876   2890     1 ;    CB - HD2
2876   2892     1 ;    CB - O
2876   2893     1 ;    CB - N
2877   2880     1 ;   HB2 - CD1
2877   2889     1 ;   HB2 - CD2
2877   2891     1 ;   HB2 - C
2878   2880     1 ;   HB3 - CD1
2878   2889     1 ;   HB3 - CD2
2878   2891     1 ;   HB3 - C
2879   2883     1 ;    CG - HE1
2879   2884     1 ;    CG - CZ
2879   2888     1 ;    CG - HE2
2879   2891     1 ;    CG - C
2880   2885     1 ;   CD1 - OH
2880   2887     1 ;   CD1 - CE2
2880   2890     1 ;   CD1 - HD2
2881   2883     1 ;   HD1 - HE1
2881   2884     1 ;   HD1 - CZ
2881   2889     1 ;   HD1 - CD2
2882   2886     1 ;   CE1 - HH
2882   2888     1 ;   CE1 - HE2
2882   2889     1 ;   CE1 - CD2
2883   2885     1 ;   HE1 - OH
2883   2887     1 ;   HE1 - CE2
2884   2890     1 ;    CZ - HD2
2885   2888     1 ;    OH - HE2
2885   2889     1 ;    OH - CD2
```

| | | | | | |
|---|---|---|---|---|---|
| 2886 | 2887 | 1 ; | HH | - | CE2 |
| 2888 | 2890 | 1 ; | HE2 | - | HD2 |
| 2891 | 2896 | 1 ; | C | - | HA |
| 2891 | 2897 | 1 ; | C | - | CB |
| 2891 | 2907 | 1 ; | C | - | C |
| 2892 | 2894 | 1 ; | O | - | H |
| 2892 | 2895 | 1 ; | O | - | CA |
| 2893 | 2898 | 1 ; | N | - | HB |
| 2893 | 2899 | 1 ; | N | - | CG1 |
| 2893 | 2903 | 1 ; | N | - | CG2 |
| 2893 | 2908 | 1 ; | N | - | O |
| 2893 | 2909 | 1 ; | N | - | N |
| 2894 | 2896 | 1 ; | H | - | HA |
| 2894 | 2897 | 1 ; | H | - | CB |
| 2894 | 2907 | 1 ; | H | - | C |
| 2895 | 2900 | 1 ; | CA | - | HG11 |
| 2895 | 2901 | 1 ; | CA | - | HG12 |
| 2895 | 2902 | 1 ; | CA | - | HG13 |
| 2895 | 2904 | 1 ; | CA | - | HG21 |
| 2895 | 2905 | 1 ; | CA | - | HG22 |
| 2895 | 2906 | 1 ; | CA | - | HG23 |
| 2895 | 2910 | 1 ; | CA | - | H |
| 2895 | 2911 | 1 ; | CA | - | CA |
| 2896 | 2898 | 1 ; | HA | - | HB |
| 2896 | 2899 | 1 ; | HA | - | CG1 |
| 2896 | 2903 | 1 ; | HA | - | CG2 |
| 2896 | 2908 | 1 ; | HA | - | O |
| 2896 | 2909 | 1 ; | HA | - | N |
| 2897 | 2908 | 1 ; | CB | - | O |
| 2897 | 2909 | 1 ; | CB | - | N |
| 2898 | 2900 | 1 ; | HB | - | HG11 |
| 2898 | 2901 | 1 ; | HB | - | HG12 |
| 2898 | 2902 | 1 ; | HB | - | HG13 |
| 2898 | 2904 | 1 ; | HB | - | HG21 |
| 2898 | 2905 | 1 ; | HB | - | HG22 |
| 2898 | 2906 | 1 ; | HB | - | HG23 |
| 2898 | 2907 | 1 ; | HB | - | C |
| 2899 | 2904 | 1 ; | CG1 | - | HG21 |
| 2899 | 2905 | 1 ; | CG1 | - | HG22 |
| 2899 | 2906 | 1 ; | CG1 | - | HG23 |
| 2899 | 2907 | 1 ; | CG1 | - | C |
| 2900 | 2903 | 1 ; | HG11 | - | CG2 |
| 2901 | 2903 | 1 ; | HG12 | - | CG2 |
| 2902 | 2903 | 1 ; | HG13 | - | CG2 |
| 2903 | 2907 | 1 ; | CG2 | - | C |
| 2907 | 2912 | 1 ; | C | - | HA |
| 2907 | 2913 | 1 ; | C | - | CB |
| 2907 | 2917 | 1 ; | C | - | C |
| 2908 | 2910 | 1 ; | O | - | H |
| 2908 | 2911 | 1 ; | O | - | CA |
| 2909 | 2914 | 1 ; | N | - | HB1 |
| 2909 | 2915 | 1 ; | N | - | HB2 |

| | | | | | | |
|---|---|---|---|---|---|---|
| 2909 | 2916 | 1 | ; | N | - | HB3 |
| 2909 | 2918 | 1 | ; | N | - | O |
| 2909 | 2919 | 1 | ; | N | - | N |
| 2910 | 2912 | 1 | ; | H | - | HA |
| 2910 | 2913 | 1 | ; | H | - | CB |
| 2910 | 2917 | 1 | ; | H | - | C |
| 2911 | 2920 | 1 | ; | CA | - | CD |
| 2911 | 2929 | 1 | ; | CA | - | CA |
| 2912 | 2914 | 1 | ; | HA | - | HB1 |
| 2912 | 2915 | 1 | ; | HA | - | HB2 |
| 2912 | 2916 | 1 | ; | HA | - | HB3 |
| 2912 | 2918 | 1 | ; | HA | - | O |
| 2912 | 2919 | 1 | ; | HA | - | N |
| 2913 | 2918 | 1 | ; | CB | - | O |
| 2913 | 2919 | 1 | ; | CB | - | N |
| 2914 | 2917 | 1 | ; | HB1 | - | C |
| 2915 | 2917 | 1 | ; | HB2 | - | C |
| 2916 | 2917 | 1 | ; | HB3 | - | C |
| 2917 | 2921 | 1 | ; | C | - | HD2 |
| 2917 | 2922 | 1 | ; | C | - | HD3 |
| 2917 | 2923 | 1 | ; | C | - | CG |
| 2917 | 2926 | 1 | ; | C | - | CB |
| 2917 | 2930 | 1 | ; | C | - | HA |
| 2917 | 2931 | 1 | ; | C | - | C |
| 2918 | 2920 | 1 | ; | O | - | CD |
| 2918 | 2929 | 1 | ; | O | - | CA |
| 2919 | 2924 | 1 | ; | N | - | HG2 |
| 2919 | 2925 | 1 | ; | N | - | HG3 |
| 2919 | 2927 | 1 | ; | N | - | HB2 |
| 2919 | 2928 | 1 | ; | N | - | HB3 |
| 2919 | 2932 | 1 | ; | N | - | O |
| 2919 | 2933 | 1 | ; | N | - | N |
| 2920 | 2927 | 1 | ; | CD | - | HB2 |
| 2920 | 2928 | 1 | ; | CD | - | HB3 |
| 2920 | 2930 | 1 | ; | CD | - | HA |
| 2920 | 2931 | 1 | ; | CD | - | C |
| 2921 | 2924 | 1 | ; | HD2 | - | HG2 |
| 2921 | 2925 | 1 | ; | HD2 | - | HG3 |
| 2921 | 2926 | 1 | ; | HD2 | - | CB |
| 2921 | 2929 | 1 | ; | HD2 | - | CA |
| 2922 | 2924 | 1 | ; | HD3 | - | HG2 |
| 2922 | 2925 | 1 | ; | HD3 | - | HG3 |
| 2922 | 2926 | 1 | ; | HD3 | - | CB |
| 2922 | 2929 | 1 | ; | HD3 | - | CA |
| 2923 | 2930 | 1 | ; | CG | - | HA |
| 2923 | 2931 | 1 | ; | CG | - | C |
| 2924 | 2927 | 1 | ; | HG2 | - | HB2 |
| 2924 | 2928 | 1 | ; | HG2 | - | HB3 |
| 2924 | 2929 | 1 | ; | HG2 | - | CA |
| 2925 | 2927 | 1 | ; | HG3 | - | HB2 |
| 2925 | 2928 | 1 | ; | HG3 | - | HB3 |
| 2925 | 2929 | 1 | ; | HG3 | - | CA |

```
2926   2932       1 ;      CB - O
2926   2933       1 ;      CB - N
2927   2930       1 ;     HB2 - HA
2927   2931       1 ;     HB2 - C
2928   2930       1 ;     HB3 - HA
2928   2931       1 ;     HB3 - C
2929   2934       1 ;      CA - H
2929   2935       1 ;      CA - CA
2930   2932       1 ;      HA - O
2930   2933       1 ;      HA - N
2931   2936       1 ;       C - HA
2931   2937       1 ;       C - CB
2931   2948       1 ;       C - C
2932   2934       1 ;       O - H
2932   2935       1 ;       O - CA
2933   2938       1 ;       N - HB2
2933   2939       1 ;       N - HB3
2933   2940       1 ;       N - CG
2933   2949       1 ;       N - O
2933   2950       1 ;       N - N
2934   2936       1 ;       H - HA
2934   2937       1 ;       H - CB
2934   2948       1 ;       H - C
2935   2941       1 ;      CA - HG2
2935   2942       1 ;      CA - HG3
2935   2943       1 ;      CA - CD
2935   2951       1 ;      CA - H
2935   2952       1 ;      CA - CA
2936   2938       1 ;      HA - HB2
2936   2939       1 ;      HA - HB3
2936   2940       1 ;      HA - CG
2936   2949       1 ;      HA - O
2936   2950       1 ;      HA - N
2937   2944       1 ;      CB - OE1
2937   2945       1 ;      CB - NE2
2937   2949       1 ;      CB - O
2937   2950       1 ;      CB - N
2938   2941       1 ;     HB2 - HG2
2938   2942       1 ;     HB2 - HG3
2938   2943       1 ;     HB2 - CD
2938   2948       1 ;     HB2 - C
2939   2941       1 ;     HB3 - HG2
2939   2942       1 ;     HB3 - HG3
2939   2943       1 ;     HB3 - CD
2939   2948       1 ;     HB3 - C
2940   2946       1 ;      CG - HE21
2940   2947       1 ;      CG - HE22
2940   2948       1 ;      CG - C
2941   2944       1 ;     HG2 - OE1
2941   2945       1 ;     HG2 - NE2
2942   2944       1 ;     HG3 - OE1
2942   2945       1 ;     HG3 - NE2
```

```
2944   2946       1 ;     OE1 - HE21
2944   2947       1 ;     OE1 - HE22
2948   2953       1 ;       C - HA
2948   2954       1 ;       C - CB
2948   2964       1 ;       C - C
2949   2951       1 ;       O - H
2949   2952       1 ;       O - CA
2950   2955       1 ;       N - HB
2950   2956       1 ;       N - CG1
2950   2960       1 ;       N - CG2
2950   2965       1 ;       N - O
2950   2966       1 ;       N - N
2951   2953       1 ;       H - HA
2951   2954       1 ;       H - CB
2951   2964       1 ;       H - C
2952   2957       1 ;      CA - HG11
2952   2958       1 ;      CA - HG12
2952   2959       1 ;      CA - HG13
2952   2961       1 ;      CA - HG21
2952   2962       1 ;      CA - HG22
2952   2963       1 ;      CA - HG23
2952   2967       1 ;      CA - H
2952   2968       1 ;      CA - CA
2953   2955       1 ;      HA - HB
2953   2956       1 ;      HA - CG1
2953   2960       1 ;      HA - CG2
2953   2965       1 ;      HA - O
2953   2966       1 ;      HA - N
2954   2965       1 ;      CB - O
2954   2966       1 ;      CB - N
2955   2957       1 ;      HB - HG11
2955   2958       1 ;      HB - HG12
2955   2959       1 ;      HB - HG13
2955   2961       1 ;      HB - HG21
2955   2962       1 ;      HB - HG22
2955   2963       1 ;      HB - HG23
2955   2964       1 ;      HB - C
2956   2961       1 ;     CG1 - HG21
2956   2962       1 ;     CG1 - HG22
2956   2963       1 ;     CG1 - HG23
2956   2964       1 ;     CG1 - C
2957   2960       1 ;    HG11 - CG2
2958   2960       1 ;    HG12 - CG2
2959   2960       1 ;    HG13 - CG2
2960   2964       1 ;     CG2 - C
2964   2969       1 ;       C - HA
2964   2970       1 ;       C - CB
2964   2983       1 ;       C - C
2965   2967       1 ;       O - H
2965   2968       1 ;       O - CA
2966   2971       1 ;       N - HB2
2966   2972       1 ;       N - HB3
```

```
2966   2973       1 ;       N - CG
2966   2984       1 ;       N - O
2966   2985       1 ;       N - N
2967   2969       1 ;       H - HA
2967   2970       1 ;       H - CB
2967   2983       1 ;       H - C
2968   2974       1 ;      CA - HG
2968   2975       1 ;      CA - CD1
2968   2979       1 ;      CA - CD2
2968   2986       1 ;      CA - H
2968   2987       1 ;      CA - CA
2969   2971       1 ;      HA - HB2
2969   2972       1 ;      HA - HB3
2969   2973       1 ;      HA - CG
2969   2984       1 ;      HA - O
2969   2985       1 ;      HA - N
2970   2976       1 ;      CB - HD11
2970   2977       1 ;      CB - HD12
2970   2978       1 ;      CB - HD13
2970   2980       1 ;      CB - HD21
2970   2981       1 ;      CB - HD22
2970   2982       1 ;      CB - HD23
2970   2984       1 ;      CB - O
2970   2985       1 ;      CB - N
2971   2974       1 ;     HB2 - HG
2971   2975       1 ;     HB2 - CD1
2971   2979       1 ;     HB2 - CD2
2971   2983       1 ;     HB2 - C
2972   2974       1 ;     HB3 - HG
2972   2975       1 ;     HB3 - CD1
2972   2979       1 ;     HB3 - CD2
2972   2983       1 ;     HB3 - C
2973   2983       1 ;      CG - C
2974   2976       1 ;      HG - HD11
2974   2977       1 ;      HG - HD12
2974   2978       1 ;      HG - HD13
2974   2980       1 ;      HG - HD21
2974   2981       1 ;      HG - HD22
2974   2982       1 ;      HG - HD23
2975   2980       1 ;     CD1 - HD21
2975   2981       1 ;     CD1 - HD22
2975   2982       1 ;     CD1 - HD23
2976   2979       1 ;    HD11 - CD2
2977   2979       1 ;    HD12 - CD2
2978   2979       1 ;    HD13 - CD2
2983   2988       1 ;       C - HA
2983   2989       1 ;       C - CB
2983   2995       1 ;       C - C
2984   2986       1 ;       O - H
2984   2987       1 ;       O - CA
2985   2990       1 ;       N - HB2
2985   2991       1 ;       N - HB3
```

| | | | |
|---|---|---|---|
| 2985 | 2992 | 1 ; | N - CG |
| 2985 | 2996 | 1 ; | N - O |
| 2985 | 2997 | 1 ; | N - N |
| 2986 | 2988 | 1 ; | H - HA |
| 2986 | 2989 | 1 ; | H - CB |
| 2986 | 2995 | 1 ; | H - C |
| 2987 | 2993 | 1 ; | CA - OD1 |
| 2987 | 2994 | 1 ; | CA - OD2 |
| 2987 | 2998 | 1 ; | CA - H |
| 2987 | 2999 | 1 ; | CA - CA |
| 2988 | 2990 | 1 ; | HA - HB2 |
| 2988 | 2991 | 1 ; | HA - HB3 |
| 2988 | 2992 | 1 ; | HA - CG |
| 2988 | 2996 | 1 ; | HA - O |
| 2988 | 2997 | 1 ; | HA - N |
| 2989 | 2996 | 1 ; | CB - O |
| 2989 | 2997 | 1 ; | CB - N |
| 2990 | 2993 | 1 ; | HB2 - OD1 |
| 2990 | 2994 | 1 ; | HB2 - OD2 |
| 2990 | 2995 | 1 ; | HB2 - C |
| 2991 | 2993 | 1 ; | HB3 - OD1 |
| 2991 | 2994 | 1 ; | HB3 - OD2 |
| 2991 | 2995 | 1 ; | HB3 - C |
| 2992 | 2995 | 1 ; | CG - C |
| 2995 | 3000 | 1 ; | C - HA2 |
| 2995 | 3001 | 1 ; | C - HA3 |
| 2995 | 3002 | 1 ; | C - C |
| 2996 | 2998 | 1 ; | O - H |
| 2996 | 2999 | 1 ; | O - CA |
| 2997 | 3003 | 1 ; | N - O |
| 2997 | 3004 | 1 ; | N - N |
| 2998 | 3000 | 1 ; | H - HA2 |
| 2998 | 3001 | 1 ; | H - HA3 |
| 2998 | 3002 | 1 ; | H - C |
| 2999 | 3005 | 1 ; | CA - H |
| 2999 | 3006 | 1 ; | CA - CA |
| 3000 | 3003 | 1 ; | HA2 - O |
| 3000 | 3004 | 1 ; | HA2 - N |
| 3001 | 3003 | 1 ; | HA3 - O |
| 3001 | 3004 | 1 ; | HA3 - N |
| 3002 | 3007 | 1 ; | C - HA |
| 3002 | 3008 | 1 ; | C - CB |
| 3002 | 3024 | 1 ; | C - C |
| 3003 | 3005 | 1 ; | O - H |
| 3003 | 3006 | 1 ; | O - CA |
| 3004 | 3009 | 1 ; | N - HB2 |
| 3004 | 3010 | 1 ; | N - HB3 |
| 3004 | 3011 | 1 ; | N - CG |
| 3004 | 3025 | 1 ; | N - O |
| 3004 | 3026 | 1 ; | N - N |
| 3005 | 3007 | 1 ; | H - HA |
| 3005 | 3008 | 1 ; | H - CB |

```
3005   3024       1 ;       H - C
3006   3012       1 ;      CA - HG2
3006   3013       1 ;      CA - HG3
3006   3014       1 ;      CA - CD
3006   3027       1 ;      CA - H
3006   3028       1 ;      CA - CA
3007   3009       1 ;      HA - HB2
3007   3010       1 ;      HA - HB3
3007   3011       1 ;      HA - CG
3007   3025       1 ;      HA - O
3007   3026       1 ;      HA - N
3008   3015       1 ;      CB - HD2
3008   3016       1 ;      CB - HD3
3008   3017       1 ;      CB - CE
3008   3025       1 ;      CB - O
3008   3026       1 ;      CB - N
3009   3012       1 ;     HB2 - HG2
3009   3013       1 ;     HB2 - HG3
3009   3014       1 ;     HB2 - CD
3009   3024       1 ;     HB2 - C
3010   3012       1 ;     HB3 - HG2
3010   3013       1 ;     HB3 - HG3
3010   3014       1 ;     HB3 - CD
3010   3024       1 ;     HB3 - C
3011   3018       1 ;      CG - HE2
3011   3019       1 ;      CG - HE3
3011   3020       1 ;      CG - NZ
3011   3024       1 ;      CG - C
3012   3015       1 ;     HG2 - HD2
3012   3016       1 ;     HG2 - HD3
3012   3017       1 ;     HG2 - CE
3013   3015       1 ;     HG3 - HD2
3013   3016       1 ;     HG3 - HD3
3013   3017       1 ;     HG3 - CE
3014   3021       1 ;      CD - HZ1
3014   3022       1 ;      CD - HZ2
3014   3023       1 ;      CD - HZ3
3015   3018       1 ;     HD2 - HE2
3015   3019       1 ;     HD2 - HE3
3015   3020       1 ;     HD2 - NZ
3016   3018       1 ;     HD3 - HE2
3016   3019       1 ;     HD3 - HE3
3016   3020       1 ;     HD3 - NZ
3018   3021       1 ;     HE2 - HZ1
3018   3022       1 ;     HE2 - HZ2
3018   3023       1 ;     HE2 - HZ3
3019   3021       1 ;     HE3 - HZ1
3019   3022       1 ;     HE3 - HZ2
3019   3023       1 ;     HE3 - HZ3
3024   3029       1 ;       C - HA
3024   3030       1 ;       C - CB
3024   3045       1 ;       C - C
```

| | | | | | |
|---|---|---|---|---|---|
| 3025 | 3027 | 1 ; | O | - | H |
| 3025 | 3028 | 1 ; | O | - | CA |
| 3026 | 3031 | 1 ; | N | - | HB2 |
| 3026 | 3032 | 1 ; | N | - | HB3 |
| 3026 | 3033 | 1 ; | N | - | CG |
| 3026 | 3046 | 1 ; | N | - | O |
| 3026 | 3047 | 1 ; | N | - | N |
| 3027 | 3029 | 1 ; | H | - | HA |
| 3027 | 3030 | 1 ; | H | - | CB |
| 3027 | 3045 | 1 ; | H | - | C |
| 3028 | 3034 | 1 ; | CA | - | CD1 |
| 3028 | 3043 | 1 ; | CA | - | CD2 |
| 3028 | 3048 | 1 ; | CA | - | H |
| 3028 | 3049 | 1 ; | CA | - | CA |
| 3029 | 3031 | 1 ; | HA | - | HB2 |
| 3029 | 3032 | 1 ; | HA | - | HB3 |
| 3029 | 3033 | 1 ; | HA | - | CG |
| 3029 | 3046 | 1 ; | HA | - | O |
| 3029 | 3047 | 1 ; | HA | - | N |
| 3030 | 3035 | 1 ; | CB | - | HD1 |
| 3030 | 3036 | 1 ; | CB | - | CE1 |
| 3030 | 3041 | 1 ; | CB | - | CE2 |
| 3030 | 3044 | 1 ; | CB | - | HD2 |
| 3030 | 3046 | 1 ; | CB | - | O |
| 3030 | 3047 | 1 ; | CB | - | N |
| 3031 | 3034 | 1 ; | HB2 | - | CD1 |
| 3031 | 3043 | 1 ; | HB2 | - | CD2 |
| 3031 | 3045 | 1 ; | HB2 | - | C |
| 3032 | 3034 | 1 ; | HB3 | - | CD1 |
| 3032 | 3043 | 1 ; | HB3 | - | CD2 |
| 3032 | 3045 | 1 ; | HB3 | - | C |
| 3033 | 3037 | 1 ; | CG | - | HE1 |
| 3033 | 3038 | 1 ; | CG | - | CZ |
| 3033 | 3042 | 1 ; | CG | - | HE2 |
| 3033 | 3045 | 1 ; | CG | - | C |
| 3034 | 3039 | 1 ; | CD1 | - | OH |
| 3034 | 3041 | 1 ; | CD1 | - | CE2 |
| 3034 | 3044 | 1 ; | CD1 | - | HD2 |
| 3035 | 3037 | 1 ; | HD1 | - | HE1 |
| 3035 | 3038 | 1 ; | HD1 | - | CZ |
| 3035 | 3043 | 1 ; | HD1 | - | CD2 |
| 3036 | 3040 | 1 ; | CE1 | - | HH |
| 3036 | 3042 | 1 ; | CE1 | - | HE2 |
| 3036 | 3043 | 1 ; | CE1 | - | CD2 |
| 3037 | 3039 | 1 ; | HE1 | - | OH |
| 3037 | 3041 | 1 ; | HE1 | - | CE2 |
| 3038 | 3044 | 1 ; | CZ | - | HD2 |
| 3039 | 3042 | 1 ; | OH | - | HE2 |
| 3039 | 3043 | 1 ; | OH | - | CD2 |
| 3040 | 3041 | 1 ; | HH | - | CE2 |
| 3042 | 3044 | 1 ; | HE2 | - | HD2 |
| 3045 | 3050 | 1 ; | C | - | HA |

| | | | | |
|---|---|---|---|---|
| 3045 | 3051 | 1 ; | C - | CB |
| 3045 | 3057 | 1 ; | C - | C |
| 3046 | 3048 | 1 ; | O - | H |
| 3046 | 3049 | 1 ; | O - | CA |
| 3047 | 3052 | 1 ; | N - | HB2 |
| 3047 | 3053 | 1 ; | N - | HB3 |
| 3047 | 3054 | 1 ; | N - | CG |
| 3047 | 3058 | 1 ; | N - | O |
| 3047 | 3059 | 1 ; | N - | N |
| 3048 | 3050 | 1 ; | H - | HA |
| 3048 | 3051 | 1 ; | H - | CB |
| 3048 | 3057 | 1 ; | H - | C |
| 3049 | 3055 | 1 ; | CA - | OD1 |
| 3049 | 3056 | 1 ; | CA - | OD2 |
| 3049 | 3060 | 1 ; | CA - | H |
| 3049 | 3061 | 1 ; | CA - | CA |
| 3050 | 3052 | 1 ; | HA - | HB2 |
| 3050 | 3053 | 1 ; | HA - | HB3 |
| 3050 | 3054 | 1 ; | HA - | CG |
| 3050 | 3058 | 1 ; | HA - | O |
| 3050 | 3059 | 1 ; | HA - | N |
| 3051 | 3058 | 1 ; | CB - | O |
| 3051 | 3059 | 1 ; | CB - | N |
| 3052 | 3055 | 1 ; | HB2 - | OD1 |
| 3052 | 3056 | 1 ; | HB2 - | OD2 |
| 3052 | 3057 | 1 ; | HB2 - | C |
| 3053 | 3055 | 1 ; | HB3 - | OD1 |
| 3053 | 3056 | 1 ; | HB3 - | OD2 |
| 3053 | 3057 | 1 ; | HB3 - | C |
| 3054 | 3057 | 1 ; | CG - | C |
| 3057 | 3062 | 1 ; | C - | HA |
| 3057 | 3063 | 1 ; | C - | CB |
| 3057 | 3079 | 1 ; | C - | C |
| 3058 | 3060 | 1 ; | O - | H |
| 3058 | 3061 | 1 ; | O - | CA |
| 3059 | 3064 | 1 ; | N - | HB2 |
| 3059 | 3065 | 1 ; | N - | HB3 |
| 3059 | 3066 | 1 ; | N - | CG |
| 3059 | 3080 | 1 ; | N - | O |
| 3059 | 3081 | 1 ; | N - | N |
| 3060 | 3062 | 1 ; | H - | HA |
| 3060 | 3063 | 1 ; | H - | CB |
| 3060 | 3079 | 1 ; | H - | C |
| 3061 | 3067 | 1 ; | CA - | HG2 |
| 3061 | 3068 | 1 ; | CA - | HG3 |
| 3061 | 3069 | 1 ; | CA - | CD |
| 3061 | 3082 | 1 ; | CA - | H |
| 3061 | 3083 | 1 ; | CA - | CA |
| 3062 | 3064 | 1 ; | HA - | HB2 |
| 3062 | 3065 | 1 ; | HA - | HB3 |
| 3062 | 3066 | 1 ; | HA - | CG |
| 3062 | 3080 | 1 ; | HA - | O |

| | | | | | |
|---|---|---|---|---|---|
| 3062 | 3081 | 1 ; | HA | - | N |
| 3063 | 3070 | 1 ; | CB | - | HD2 |
| 3063 | 3071 | 1 ; | CB | - | HD3 |
| 3063 | 3072 | 1 ; | CB | - | CE |
| 3063 | 3080 | 1 ; | CB | - | O |
| 3063 | 3081 | 1 ; | CB | - | N |
| 3064 | 3067 | 1 ; | HB2 | - | HG2 |
| 3064 | 3068 | 1 ; | HB2 | - | HG3 |
| 3064 | 3069 | 1 ; | HB2 | - | CD |
| 3064 | 3079 | 1 ; | HB2 | - | C |
| 3065 | 3067 | 1 ; | HB3 | - | HG2 |
| 3065 | 3068 | 1 ; | HB3 | - | HG3 |
| 3065 | 3069 | 1 ; | HB3 | - | CD |
| 3065 | 3079 | 1 ; | HB3 | - | C |
| 3066 | 3073 | 1 ; | CG | - | HE2 |
| 3066 | 3074 | 1 ; | CG | - | HE3 |
| 3066 | 3075 | 1 ; | CG | - | NZ |
| 3066 | 3079 | 1 ; | CG | - | C |
| 3067 | 3070 | 1 ; | HG2 | - | HD2 |
| 3067 | 3071 | 1 ; | HG2 | - | HD3 |
| 3067 | 3072 | 1 ; | HG2 | - | CE |
| 3068 | 3070 | 1 ; | HG3 | - | HD2 |
| 3068 | 3071 | 1 ; | HG3 | - | HD3 |
| 3068 | 3072 | 1 ; | HG3 | - | CE |
| 3069 | 3076 | 1 ; | CD | - | HZ1 |
| 3069 | 3077 | 1 ; | CD | - | HZ2 |
| 3069 | 3078 | 1 ; | CD | - | HZ3 |
| 3070 | 3073 | 1 ; | HD2 | - | HE2 |
| 3070 | 3074 | 1 ; | HD2 | - | HE3 |
| 3070 | 3075 | 1 ; | HD2 | - | NZ |
| 3071 | 3073 | 1 ; | HD3 | - | HE2 |
| 3071 | 3074 | 1 ; | HD3 | - | HE3 |
| 3071 | 3075 | 1 ; | HD3 | - | NZ |
| 3073 | 3076 | 1 ; | HE2 | - | HZ1 |
| 3073 | 3077 | 1 ; | HE2 | - | HZ2 |
| 3073 | 3078 | 1 ; | HE2 | - | HZ3 |
| 3074 | 3076 | 1 ; | HE3 | - | HZ1 |
| 3074 | 3077 | 1 ; | HE3 | - | HZ2 |
| 3074 | 3078 | 1 ; | HE3 | - | HZ3 |
| 3079 | 3084 | 1 ; | C | - | HA |
| 3079 | 3085 | 1 ; | C | - | CB |
| 3079 | 3100 | 1 ; | C | - | C |
| 3080 | 3082 | 1 ; | O | - | H |
| 3080 | 3083 | 1 ; | O | - | CA |
| 3081 | 3086 | 1 ; | N | - | HB2 |
| 3081 | 3087 | 1 ; | N | - | HB3 |
| 3081 | 3088 | 1 ; | N | - | CG |
| 3081 | 3101 | 1 ; | N | - | O |
| 3081 | 3102 | 1 ; | N | - | N |
| 3082 | 3084 | 1 ; | H | - | HA |
| 3082 | 3085 | 1 ; | H | - | CB |
| 3082 | 3100 | 1 ; | H | - | C |

| | | | |
|---|---|---|---|
| 3083 | 3089 | 1 ; | CA - HG2 |
| 3083 | 3090 | 1 ; | CA - HG3 |
| 3083 | 3091 | 1 ; | CA - CD |
| 3083 | 3103 | 1 ; | CA - H |
| 3083 | 3104 | 1 ; | CA - CA |
| 3084 | 3086 | 1 ; | HA - HB2 |
| 3084 | 3087 | 1 ; | HA - HB3 |
| 3084 | 3088 | 1 ; | HA - CG |
| 3084 | 3101 | 1 ; | HA - O |
| 3084 | 3102 | 1 ; | HA - N |
| 3085 | 3092 | 1 ; | CB - HD2 |
| 3085 | 3093 | 1 ; | CB - HD3 |
| 3085 | 3094 | 1 ; | CB - CE |
| 3085 | 3101 | 1 ; | CB - O |
| 3085 | 3102 | 1 ; | CB - N |
| 3086 | 3089 | 1 ; | HB2 - HG2 |
| 3086 | 3090 | 1 ; | HB2 - HG3 |
| 3086 | 3091 | 1 ; | HB2 - CD |
| 3086 | 3100 | 1 ; | HB2 - C |
| 3087 | 3089 | 1 ; | HB3 - HG2 |
| 3087 | 3090 | 1 ; | HB3 - HG3 |
| 3087 | 3091 | 1 ; | HB3 - CD |
| 3087 | 3100 | 1 ; | HB3 - C |
| 3088 | 3095 | 1 ; | CG - HE2 |
| 3088 | 3096 | 1 ; | CG - HE3 |
| 3088 | 3097 | 1 ; | CG - NZ |
| 3088 | 3100 | 1 ; | CG - C |
| 3089 | 3092 | 1 ; | HG2 - HD2 |
| 3089 | 3093 | 1 ; | HG2 - HD3 |
| 3089 | 3094 | 1 ; | HG2 - CE |
| 3090 | 3092 | 1 ; | HG3 - HD2 |
| 3090 | 3093 | 1 ; | HG3 - HD3 |
| 3090 | 3094 | 1 ; | HG3 - CE |
| 3091 | 3098 | 1 ; | CD - HZ2 |
| 3091 | 3099 | 1 ; | CD - HZ3 |
| 3092 | 3095 | 1 ; | HD2 - HE2 |
| 3092 | 3096 | 1 ; | HD2 - HE3 |
| 3092 | 3097 | 1 ; | HD2 - NZ |
| 3093 | 3095 | 1 ; | HD3 - HE2 |
| 3093 | 3096 | 1 ; | HD3 - HE3 |
| 3093 | 3097 | 1 ; | HD3 - NZ |
| 3095 | 3098 | 1 ; | HE2 - HZ2 |
| 3095 | 3099 | 1 ; | HE2 - HZ3 |
| 3096 | 3098 | 1 ; | HE3 - HZ2 |
| 3096 | 3099 | 1 ; | HE3 - HZ3 |
| 3100 | 3105 | 1 ; | C - HA |
| 3100 | 3106 | 1 ; | C - CB |
| 3100 | 3111 | 1 ; | C - C |
| 3101 | 3103 | 1 ; | O - H |
| 3101 | 3104 | 1 ; | O - CA |
| 3102 | 3107 | 1 ; | N - HB2 |
| 3102 | 3108 | 1 ; | N - HB3 |

```
3102   3109       1 ;       N - SG
3102   3112       1 ;       N - O
3102   3113       1 ;       N - N
3103   3105       1 ;       H - HA
3103   3106       1 ;       H - CB
3103   3111       1 ;       H - C
3104   3110       1 ;      CA - HG
3104   3114       1 ;      CA - H
3104   3115       1 ;      CA - CA
3105   3107       1 ;      HA - HB2
3105   3108       1 ;      HA - HB3
3105   3109       1 ;      HA - SG
3105   3112       1 ;      HA - O
3105   3113       1 ;      HA - N
3106   3112       1 ;      CB - O
3106   3113       1 ;      CB - N
3107   3110       1 ;     HB2 - HG
3107   3111       1 ;     HB2 - C
3108   3110       1 ;     HB3 - HG
3108   3111       1 ;     HB3 - C
3109   3111       1 ;      SG - C
3111   3116       1 ;       C - HA
3111   3117       1 ;       C - CB
3111   3123       1 ;       C - C
3112   3114       1 ;       O - H
3112   3115       1 ;       O - CA
3113   3118       1 ;       N - HB2
3113   3119       1 ;       N - HB3
3113   3120       1 ;       N - CG
3113   3124       1 ;       N - O
3113   3125       1 ;       N - N
3114   3116       1 ;       H - HA
3114   3117       1 ;       H - CB
3114   3123       1 ;       H - C
3115   3121       1 ;      CA - OD1
3115   3122       1 ;      CA - OD2
3115   3126       1 ;      CA - H
3115   3127       1 ;      CA - CA
3116   3118       1 ;      HA - HB2
3116   3119       1 ;      HA - HB3
3116   3120       1 ;      HA - CG
3116   3124       1 ;      HA - O
3116   3125       1 ;      HA - N
3117   3124       1 ;      CB - O
3117   3125       1 ;      CB - N
3118   3121       1 ;     HB2 - OD1
3118   3122       1 ;     HB2 - OD2
3118   3123       1 ;     HB2 - C
3119   3121       1 ;     HB3 - OD1
3119   3122       1 ;     HB3 - OD2
3119   3123       1 ;     HB3 - C
3120   3123       1 ;      CG - C
```

```
3123   3128      1 ;      C - HA
3123   3129      1 ;      C - CB
3123   3142      1 ;      C - C
3124   3126      1 ;      O - H
3124   3127      1 ;      O - CA
3125   3130      1 ;      N - HB
3125   3131      1 ;      N - CG2
3125   3135      1 ;      N - CG1
3125   3143      1 ;      N - O
3125   3144      1 ;      N - N
3126   3128      1 ;      H - HA
3126   3129      1 ;      H - CB
3126   3142      1 ;      H - C
3127   3132      1 ;     CA - HG21
3127   3133      1 ;     CA - HG22
3127   3134      1 ;     CA - HG23
3127   3136      1 ;     CA - HG12
3127   3137      1 ;     CA - HG13
3127   3138      1 ;     CA - CD1
3127   3145      1 ;     CA - H
3127   3146      1 ;     CA - CA
3128   3130      1 ;     HA - HB
3128   3131      1 ;     HA - CG2
3128   3135      1 ;     HA - CG1
3128   3143      1 ;     HA - O
3128   3144      1 ;     HA - N
3129   3139      1 ;     CB - HD11
3129   3140      1 ;     CB - HD12
3129   3141      1 ;     CB - HD13
3129   3143      1 ;     CB - O
3129   3144      1 ;     CB - N
3130   3132      1 ;     HB - HG21
3130   3133      1 ;     HB - HG22
3130   3134      1 ;     HB - HG23
3130   3136      1 ;     HB - HG12
3130   3137      1 ;     HB - HG13
3130   3138      1 ;     HB - CD1
3130   3142      1 ;     HB - C
3131   3136      1 ;    CG2 - HG12
3131   3137      1 ;    CG2 - HG13
3131   3138      1 ;    CG2 - CD1
3131   3142      1 ;    CG2 - C
3132   3135      1 ;   HG21 - CG1
3133   3135      1 ;   HG22 - CG1
3134   3135      1 ;   HG23 - CG1
3135   3142      1 ;    CG1 - C
3136   3139      1 ;   HG12 - HD11
3136   3140      1 ;   HG12 - HD12
3136   3141      1 ;   HG12 - HD13
3137   3139      1 ;   HG13 - HD11
3137   3140      1 ;   HG13 - HD12
3137   3141      1 ;   HG13 - HD13
```

```
3142   3147      1 ;      C  -  HA
3142   3148      1 ;      C  -  CB
3142   3166      1 ;      C  -  C
3143   3145      1 ;      O  -  H
3143   3146      1 ;      O  -  CA
3144   3149      1 ;      N  -  HB2
3144   3150      1 ;      N  -  HB3
3144   3151      1 ;      N  -  CG
3144   3167      1 ;      N  -  O
3144   3168      1 ;      N  -  N
3145   3147      1 ;      H  -  HA
3145   3148      1 ;      H  -  CB
3145   3166      1 ;      H  -  C
3146   3152      1 ;     CA  -  CD1
3146   3165      1 ;     CA  -  CD2
3146   3169      1 ;     CA  -  H
3146   3170      1 ;     CA  -  CA
3147   3149      1 ;     HA  -  HB2
3147   3150      1 ;     HA  -  HB3
3147   3151      1 ;     HA  -  CG
3147   3167      1 ;     HA  -  O
3147   3168      1 ;     HA  -  N
3148   3153      1 ;     CB  -  HD1
3148   3154      1 ;     CB  -  NE1
3148   3156      1 ;     CB  -  CE2
3148   3163      1 ;     CB  -  CE3
3148   3167      1 ;     CB  -  O
3148   3168      1 ;     CB  -  N
3149   3152      1 ;    HB2  -  CD1
3149   3165      1 ;    HB2  -  CD2
3149   3166      1 ;    HB2  -  C
3150   3152      1 ;    HB3  -  CD1
3150   3165      1 ;    HB3  -  CD2
3150   3166      1 ;    HB3  -  C
3151   3155      1 ;     CG  -  HE1
3151   3157      1 ;     CG  -  CZ2
3151   3161      1 ;     CG  -  CZ3
3151   3164      1 ;     CG  -  HE3
3151   3166      1 ;     CG  -  C
3152   3157      1 ;    CD1  -  CZ2
3152   3163      1 ;    CD1  -  CE3
3153   3155      1 ;    HD1  -  HE1
3153   3156      1 ;    HD1  -  CE2
3153   3165      1 ;    HD1  -  CD2
3154   3158      1 ;    NE1  -  HZ2
3154   3159      1 ;    NE1  -  CH2
3154   3163      1 ;    NE1  -  CE3
3155   3157      1 ;    HE1  -  CZ2
3155   3165      1 ;    HE1  -  CD2
3156   3160      1 ;    CE2  -  HH2
3156   3161      1 ;    CE2  -  CZ3
3156   3164      1 ;    CE2  -  HE3
```

```
3157   3162       1 ;     CZ2 - HZ3
3157   3163       1 ;     CZ2 - CE3
3158   3160       1 ;     HZ2 - HH2
3158   3161       1 ;     HZ2 - CZ3
3158   3165       1 ;     HZ2 - CD2
3159   3164       1 ;     CH2 - HE3
3159   3165       1 ;     CH2 - CD2
3160   3162       1 ;     HH2 - HZ3
3160   3163       1 ;     HH2 - CE3
3162   3164       1 ;     HZ3 - HE3
3162   3165       1 ;     HZ3 - CD2
3166   3171       1 ;       C - HA
3166   3172       1 ;       C - CB
3166   3177       1 ;       C - C
3167   3169       1 ;       O - H
3167   3170       1 ;       O - CA
3168   3173       1 ;       N - HB2
3168   3174       1 ;       N - HB3
3168   3175       1 ;       N - OG
3168   3178       1 ;       N - O
3168   3179       1 ;       N - N
3169   3171       1 ;       H - HA
3169   3172       1 ;       H - CB
3169   3177       1 ;       H - C
3170   3176       1 ;      CA - HG
3170   3180       1 ;      CA - H
3170   3181       1 ;      CA - CA
3171   3173       1 ;      HA - HB2
3171   3174       1 ;      HA - HB3
3171   3175       1 ;      HA - OG
3171   3178       1 ;      HA - O
3171   3179       1 ;      HA - N
3172   3178       1 ;      CB - O
3172   3179       1 ;      CB - N
3173   3176       1 ;     HB2 - HG
3173   3177       1 ;     HB2 - C
3174   3176       1 ;     HB3 - HG
3174   3177       1 ;     HB3 - C
3175   3177       1 ;      OG - C
3177   3182       1 ;       C - HA
3177   3183       1 ;       C - CB
3177   3188       1 ;       C - C
3178   3180       1 ;       O - H
3178   3181       1 ;       O - CA
3179   3184       1 ;       N - HB2
3179   3185       1 ;       N - HB3
3179   3186       1 ;       N - OG
3179   3189       1 ;       N - O
3179   3190       1 ;       N - N
3180   3182       1 ;       H - HA
3180   3183       1 ;       H - CB
3180   3188       1 ;       H - C
```

```
3181   3187       1 ;      CA - HG
3181   3191       1 ;      CA - H
3181   3192       1 ;      CA - CA
3182   3184       1 ;      HA - HB2
3182   3185       1 ;      HA - HB3
3182   3186       1 ;      HA - OG
3182   3189       1 ;      HA - O
3182   3190       1 ;      HA - N
3183   3189       1 ;      CB - O
3183   3190       1 ;      CB - N
3184   3187       1 ;     HB2 - HG
3184   3188       1 ;     HB2 - C
3185   3187       1 ;     HB3 - HG
3185   3188       1 ;     HB3 - C
3186   3188       1 ;      OG - C
3188   3193       1 ;       C - HA2
3188   3194       1 ;       C - HA3
3188   3195       1 ;       C - C
3189   3191       1 ;       O - H
3189   3192       1 ;       O - CA
3190   3196       1 ;       N - O
3190   3197       1 ;       N - N
3191   3193       1 ;       H - HA2
3191   3194       1 ;       H - HA3
3191   3195       1 ;       H - C
3192   3198       1 ;      CA - H
3192   3199       1 ;      CA - CA
3193   3196       1 ;     HA2 - O
3193   3197       1 ;     HA2 - N
3194   3196       1 ;     HA3 - O
3194   3197       1 ;     HA3 - N
3195   3200       1 ;       C - HA
3195   3201       1 ;       C - CB
3195   3211       1 ;       C - C
3196   3198       1 ;       O - H
3196   3199       1 ;       O - CA
3197   3202       1 ;       N - HB
3197   3203       1 ;       N - CG1
3197   3207       1 ;       N - CG2
3197   3212       1 ;       N - O
3197   3213       1 ;       N - N
3198   3200       1 ;       H - HA
3198   3201       1 ;       H - CB
3198   3211       1 ;       H - C
3199   3204       1 ;      CA - HG11
3199   3205       1 ;      CA - HG12
3199   3206       1 ;      CA - HG13
3199   3208       1 ;      CA - HG21
3199   3209       1 ;      CA - HG22
3199   3210       1 ;      CA - HG23
3199   3214       1 ;      CA - H
3199   3215       1 ;      CA - CA
```

```
3200   3202       1 ;      HA - HB
3200   3203       1 ;      HA - CG1
3200   3207       1 ;      HA - CG2
3200   3212       1 ;      HA - O
3200   3213       1 ;      HA - N
3201   3212       1 ;      CB - O
3201   3213       1 ;      CB - N
3202   3204       1 ;      HB - HG11
3202   3205       1 ;      HB - HG12
3202   3206       1 ;      HB - HG13
3202   3208       1 ;      HB - HG21
3202   3209       1 ;      HB - HG22
3202   3210       1 ;      HB - HG23
3202   3211       1 ;      HB - C
3203   3208       1 ;     CG1 - HG21
3203   3209       1 ;     CG1 - HG22
3203   3210       1 ;     CG1 - HG23
3203   3211       1 ;     CG1 - C
3204   3207       1 ;    HG11 - CG2
3205   3207       1 ;    HG12 - CG2
3206   3207       1 ;    HG13 - CG2
3207   3211       1 ;     CG2 - C
3211   3216       1 ;       C - HA
3211   3217       1 ;       C - CB
3211   3230       1 ;       C - C
3212   3214       1 ;       O - H
3212   3215       1 ;       O - CA
3213   3218       1 ;       N - HB
3213   3219       1 ;       N - CG2
3213   3223       1 ;       N - CG1
3213   3231       1 ;       N - O
3213   3232       1 ;       N - N
3214   3216       1 ;       H - HA
3214   3217       1 ;       H - CB
3214   3230       1 ;       H - C
3215   3220       1 ;      CA - HG21
3215   3221       1 ;      CA - HG22
3215   3222       1 ;      CA - HG23
3215   3224       1 ;      CA - HG12
3215   3225       1 ;      CA - HG13
3215   3226       1 ;      CA - CD1
3215   3233       1 ;      CA - H
3215   3234       1 ;      CA - CA
3216   3218       1 ;      HA - HB
3216   3219       1 ;      HA - CG2
3216   3223       1 ;      HA - CG1
3216   3231       1 ;      HA - O
3216   3232       1 ;      HA - N
3217   3227       1 ;      CB - HD11
3217   3228       1 ;      CB - HD12
3217   3229       1 ;      CB - HD13
3217   3231       1 ;      CB - O
```

```
3217   3232       1 ;      CB - N
3218   3220       1 ;      HB - HG21
3218   3221       1 ;      HB - HG22
3218   3222       1 ;      HB - HG23
3218   3224       1 ;      HB - HG12
3218   3225       1 ;      HB - HG13
3218   3226       1 ;      HB - CD1
3218   3230       1 ;      HB - C
3219   3224       1 ;     CG2 - HG12
3219   3225       1 ;     CG2 - HG13
3219   3226       1 ;     CG2 - CD1
3219   3230       1 ;     CG2 - C
3220   3223       1 ;    HG21 - CG1
3221   3223       1 ;    HG22 - CG1
3222   3223       1 ;    HG23 - CG1
3223   3230       1 ;     CG1 - C
3224   3227       1 ;    HG12 - HD11
3224   3228       1 ;    HG12 - HD12
3224   3229       1 ;    HG12 - HD13
3225   3227       1 ;    HG13 - HD11
3225   3228       1 ;    HG13 - HD12
3225   3229       1 ;    HG13 - HD13
3230   3235       1 ;       C - HA
3230   3236       1 ;       C - CB
3230   3247       1 ;       C - C
3231   3233       1 ;       O - H
3231   3234       1 ;       O - CA
3232   3237       1 ;       N - HB2
3232   3238       1 ;       N - HB3
3232   3239       1 ;       N - CG
3232   3248       1 ;       N - O
3232   3249       1 ;       N - N
3233   3235       1 ;       H - HA
3233   3236       1 ;       H - CB
3233   3247       1 ;       H - C
3234   3240       1 ;      CA - HG2
3234   3241       1 ;      CA - HG3
3234   3242       1 ;      CA - SD
3234   3250       1 ;      CA - H
3234   3251       1 ;      CA - CA
3235   3237       1 ;      HA - HB2
3235   3238       1 ;      HA - HB3
3235   3239       1 ;      HA - CG
3235   3248       1 ;      HA - O
3235   3249       1 ;      HA - N
3236   3243       1 ;      CB - CE
3236   3248       1 ;      CB - O
3236   3249       1 ;      CB - N
3237   3240       1 ;     HB2 - HG2
3237   3241       1 ;     HB2 - HG3
3237   3242       1 ;     HB2 - SD
3237   3247       1 ;     HB2 - C
```

| | | | |
|---|---|---|---|
| 3238 | 3240 | 1 ; | HB3 - HG2 |
| 3238 | 3241 | 1 ; | HB3 - HG3 |
| 3238 | 3242 | 1 ; | HB3 - SD |
| 3238 | 3247 | 1 ; | HB3 - C |
| 3239 | 3244 | 1 ; | CG - HE1 |
| 3239 | 3245 | 1 ; | CG - HE2 |
| 3239 | 3246 | 1 ; | CG - HE3 |
| 3239 | 3247 | 1 ; | CG - C |
| 3240 | 3243 | 1 ; | HG2 - CE |
| 3241 | 3243 | 1 ; | HG3 - CE |
| 3247 | 3252 | 1 ; | C - HA |
| 3247 | 3253 | 1 ; | C - CB |
| 3247 | 3268 | 1 ; | C - C |
| 3248 | 3250 | 1 ; | O - H |
| 3248 | 3251 | 1 ; | O - CA |
| 3249 | 3254 | 1 ; | N - HB2 |
| 3249 | 3255 | 1 ; | N - HB3 |
| 3249 | 3256 | 1 ; | N - CG |
| 3249 | 3269 | 1 ; | N - O |
| 3249 | 3270 | 1 ; | N - N |
| 3250 | 3252 | 1 ; | H - HA |
| 3250 | 3253 | 1 ; | H - CB |
| 3250 | 3268 | 1 ; | H - C |
| 3251 | 3257 | 1 ; | CA - CD1 |
| 3251 | 3266 | 1 ; | CA - CD2 |
| 3251 | 3271 | 1 ; | CA - H |
| 3251 | 3272 | 1 ; | CA - CA |
| 3252 | 3254 | 1 ; | HA - HB2 |
| 3252 | 3255 | 1 ; | HA - HB3 |
| 3252 | 3256 | 1 ; | HA - CG |
| 3252 | 3269 | 1 ; | HA - O |
| 3252 | 3270 | 1 ; | HA - N |
| 3253 | 3258 | 1 ; | CB - HD1 |
| 3253 | 3259 | 1 ; | CB - CE1 |
| 3253 | 3264 | 1 ; | CB - CE2 |
| 3253 | 3267 | 1 ; | CB - HD2 |
| 3253 | 3269 | 1 ; | CB - O |
| 3253 | 3270 | 1 ; | CB - N |
| 3254 | 3257 | 1 ; | HB2 - CD1 |
| 3254 | 3266 | 1 ; | HB2 - CD2 |
| 3254 | 3268 | 1 ; | HB2 - C |
| 3255 | 3257 | 1 ; | HB3 - CD1 |
| 3255 | 3266 | 1 ; | HB3 - CD2 |
| 3255 | 3268 | 1 ; | HB3 - C |
| 3256 | 3260 | 1 ; | CG - HE1 |
| 3256 | 3261 | 1 ; | CG - CZ |
| 3256 | 3265 | 1 ; | CG - HE2 |
| 3256 | 3268 | 1 ; | CG - C |
| 3257 | 3262 | 1 ; | CD1 - OH |
| 3257 | 3264 | 1 ; | CD1 - CE2 |
| 3257 | 3267 | 1 ; | CD1 - HD2 |
| 3258 | 3260 | 1 ; | HD1 - HE1 |

```
3258   3261       1 ;    HD1 - CZ
3258   3266       1 ;    HD1 - CD2
3259   3263       1 ;    CE1 - HH
3259   3265       1 ;    CE1 - HE2
3259   3266       1 ;    CE1 - CD2
3260   3262       1 ;    HE1 - OH
3260   3264       1 ;    HE1 - CE2
3261   3267       1 ;     CZ - HD2
3262   3265       1 ;     OH - HE2
3262   3266       1 ;     OH - CD2
3263   3264       1 ;     HH - CE2
3265   3267       1 ;    HE2 - HD2
3268   3273       1 ;      C - HA
3268   3274       1 ;      C - CB
3268   3282       1 ;      C - C
3269   3271       1 ;      O - H
3269   3272       1 ;      O - CA
3270   3275       1 ;      N - HB
3270   3276       1 ;      N - CG2
3270   3280       1 ;      N - OG1
3270   3283       1 ;      N - O
3270   3284       1 ;      N - N
3271   3273       1 ;      H - HA
3271   3274       1 ;      H - CB
3271   3282       1 ;      H - C
3272   3277       1 ;     CA - HG21
3272   3278       1 ;     CA - HG22
3272   3279       1 ;     CA - HG23
3272   3281       1 ;     CA - HG1
3272   3285       1 ;     CA - H
3272   3286       1 ;     CA - CA
3273   3275       1 ;     HA - HB
3273   3276       1 ;     HA - CG2
3273   3280       1 ;     HA - OG1
3273   3283       1 ;     HA - O
3273   3284       1 ;     HA - N
3274   3283       1 ;     CB - O
3274   3284       1 ;     CB - N
3275   3277       1 ;     HB - HG21
3275   3278       1 ;     HB - HG22
3275   3279       1 ;     HB - HG23
3275   3281       1 ;     HB - HG1
3275   3282       1 ;     HB - C
3276   3281       1 ;    CG2 - HG1
3276   3282       1 ;    CG2 - C
3277   3280       1 ;   HG21 - OG1
3278   3280       1 ;   HG22 - OG1
3279   3280       1 ;   HG23 - OG1
3280   3282       1 ;    OG1 - C
3282   3287       1 ;      C - HA
3282   3288       1 ;      C - CB
3282   3301       1 ;      C - C
```

```
3283   3285      1 ;      O - H
3283   3286      1 ;      O - CA
3284   3289      1 ;      N - HB2
3284   3290      1 ;      N - HB3
3284   3291      1 ;      N - CG
3284   3302      1 ;      N - O
3284   3303      1 ;      N - N
3285   3287      1 ;      H - HA
3285   3288      1 ;      H - CB
3285   3301      1 ;      H - C
3286   3292      1 ;     CA - HG
3286   3293      1 ;     CA - CD1
3286   3297      1 ;     CA - CD2
3286   3304      1 ;     CA - H
3286   3305      1 ;     CA - CA
3287   3289      1 ;     HA - HB2
3287   3290      1 ;     HA - HB3
3287   3291      1 ;     HA - CG
3287   3302      1 ;     HA - O
3287   3303      1 ;     HA - N
3288   3294      1 ;     CB - HD11
3288   3295      1 ;     CB - HD12
3288   3296      1 ;     CB - HD13
3288   3298      1 ;     CB - HD21
3288   3299      1 ;     CB - HD22
3288   3300      1 ;     CB - HD23
3288   3302      1 ;     CB - O
3288   3303      1 ;     CB - N
3289   3292      1 ;    HB2 - HG
3289   3293      1 ;    HB2 - CD1
3289   3297      1 ;    HB2 - CD2
3289   3301      1 ;    HB2 - C
3290   3292      1 ;    HB3 - HG
3290   3293      1 ;    HB3 - CD1
3290   3297      1 ;    HB3 - CD2
3290   3301      1 ;    HB3 - C
3291   3301      1 ;     CG - C
3292   3294      1 ;     HG - HD11
3292   3295      1 ;     HG - HD12
3292   3296      1 ;     HG - HD13
3292   3298      1 ;     HG - HD21
3292   3299      1 ;     HG - HD22
3292   3300      1 ;     HG - HD23
3293   3298      1 ;    CD1 - HD21
3293   3299      1 ;    CD1 - HD22
3293   3300      1 ;    CD1 - HD23
3294   3297      1 ;   HD11 - CD2
3295   3297      1 ;   HD12 - CD2
3296   3297      1 ;   HD13 - CD2
3301   3306      1 ;      C - HA
3301   3307      1 ;      C - CB
3301   3320      1 ;      C - C
```

| | | | |
|---|---|---|---|
| 3302 | 3304 | 1 ; | O - H |
| 3302 | 3305 | 1 ; | O - CA |
| 3303 | 3308 | 1 ; | N - HB2 |
| 3303 | 3309 | 1 ; | N - HB3 |
| 3303 | 3310 | 1 ; | N - CG |
| 3303 | 3321 | 1 ; | N - O |
| 3303 | 3322 | 1 ; | N - N |
| 3304 | 3306 | 1 ; | H - HA |
| 3304 | 3307 | 1 ; | H - CB |
| 3304 | 3320 | 1 ; | H - C |
| 3305 | 3311 | 1 ; | CA - HG |
| 3305 | 3312 | 1 ; | CA - CD1 |
| 3305 | 3316 | 1 ; | CA - CD2 |
| 3305 | 3323 | 1 ; | CA - H |
| 3305 | 3324 | 1 ; | CA - CA |
| 3306 | 3308 | 1 ; | HA - HB2 |
| 3306 | 3309 | 1 ; | HA - HB3 |
| 3306 | 3310 | 1 ; | HA - CG |
| 3306 | 3321 | 1 ; | HA - O |
| 3306 | 3322 | 1 ; | HA - N |
| 3307 | 3313 | 1 ; | CB - HD11 |
| 3307 | 3314 | 1 ; | CB - HD12 |
| 3307 | 3315 | 1 ; | CB - HD13 |
| 3307 | 3317 | 1 ; | CB - HD21 |
| 3307 | 3318 | 1 ; | CB - HD22 |
| 3307 | 3319 | 1 ; | CB - HD23 |
| 3307 | 3321 | 1 ; | CB - O |
| 3307 | 3322 | 1 ; | CB - N |
| 3308 | 3311 | 1 ; | HB2 - HG |
| 3308 | 3312 | 1 ; | HB2 - CD1 |
| 3308 | 3316 | 1 ; | HB2 - CD2 |
| 3308 | 3320 | 1 ; | HB2 - C |
| 3309 | 3311 | 1 ; | HB3 - HG |
| 3309 | 3312 | 1 ; | HB3 - CD1 |
| 3309 | 3316 | 1 ; | HB3 - CD2 |
| 3309 | 3320 | 1 ; | HB3 - C |
| 3310 | 3320 | 1 ; | CG - C |
| 3311 | 3313 | 1 ; | HG - HD11 |
| 3311 | 3314 | 1 ; | HG - HD12 |
| 3311 | 3315 | 1 ; | HG - HD13 |
| 3311 | 3317 | 1 ; | HG - HD21 |
| 3311 | 3318 | 1 ; | HG - HD22 |
| 3311 | 3319 | 1 ; | HG - HD23 |
| 3312 | 3317 | 1 ; | CD1 - HD21 |
| 3312 | 3318 | 1 ; | CD1 - HD22 |
| 3312 | 3319 | 1 ; | CD1 - HD23 |
| 3313 | 3316 | 1 ; | HD11 - CD2 |
| 3314 | 3316 | 1 ; | HD12 - CD2 |
| 3315 | 3316 | 1 ; | HD13 - CD2 |
| 3320 | 3325 | 1 ; | C - HA |
| 3320 | 3326 | 1 ; | C - CB |
| 3320 | 3331 | 1 ; | C - C |

| | | | | | |
|---|---|---|---|---|---|
| 3321 | 3323 | 1 ; | O | - | H |
| 3321 | 3324 | 1 ; | O | - | CA |
| 3322 | 3327 | 1 ; | N | - | HB2 |
| 3322 | 3328 | 1 ; | N | - | HB3 |
| 3322 | 3329 | 1 ; | N | - | SG |
| 3322 | 3332 | 1 ; | N | - | O |
| 3322 | 3333 | 1 ; | N | - | N |
| 3323 | 3325 | 1 ; | H | - | HA |
| 3323 | 3326 | 1 ; | H | - | CB |
| 3323 | 3331 | 1 ; | H | - | C |
| 3324 | 3330 | 1 ; | CA | - | HG |
| 3324 | 3334 | 1 ; | CA | - | H |
| 3324 | 3335 | 1 ; | CA | - | CA |
| 3325 | 3327 | 1 ; | HA | - | HB2 |
| 3325 | 3328 | 1 ; | HA | - | HB3 |
| 3325 | 3329 | 1 ; | HA | - | SG |
| 3325 | 3332 | 1 ; | HA | - | O |
| 3325 | 3333 | 1 ; | HA | - | N |
| 3326 | 3332 | 1 ; | CB | - | O |
| 3326 | 3333 | 1 ; | CB | - | N |
| 3327 | 3330 | 1 ; | HB2 | - | HG |
| 3327 | 3331 | 1 ; | HB2 | - | C |
| 3328 | 3330 | 1 ; | HB3 | - | HG |
| 3328 | 3331 | 1 ; | HB3 | - | C |
| 3329 | 3331 | 1 ; | SG | - | C |
| 3331 | 3336 | 1 ; | C | - | HA2 |
| 3331 | 3337 | 1 ; | C | - | HA3 |
| 3331 | 3338 | 1 ; | C | - | C |
| 3332 | 3334 | 1 ; | O | - | H |
| 3332 | 3335 | 1 ; | O | - | CA |
| 3333 | 3339 | 1 ; | N | - | O |
| 3333 | 3340 | 1 ; | N | - | N |
| 3334 | 3336 | 1 ; | H | - | HA2 |
| 3334 | 3337 | 1 ; | H | - | HA3 |
| 3334 | 3338 | 1 ; | H | - | C |
| 3335 | 3341 | 1 ; | CA | - | H |
| 3335 | 3342 | 1 ; | CA | - | CA |
| 3336 | 3339 | 1 ; | HA2 | - | O |
| 3336 | 3340 | 1 ; | HA2 | - | N |
| 3337 | 3339 | 1 ; | HA3 | - | O |
| 3337 | 3340 | 1 ; | HA3 | - | N |
| 3338 | 3343 | 1 ; | C | - | HA |
| 3338 | 3344 | 1 ; | C | - | CB |
| 3338 | 3359 | 1 ; | C | - | C |
| 3339 | 3341 | 1 ; | O | - | H |
| 3339 | 3342 | 1 ; | O | - | CA |
| 3340 | 3345 | 1 ; | N | - | HB2 |
| 3340 | 3346 | 1 ; | N | - | HB3 |
| 3340 | 3347 | 1 ; | N | - | CG |
| 3340 | 3360 | 1 ; | N | - | O |
| 3340 | 3361 | 1 ; | N | - | N |
| 3341 | 3343 | 1 ; | H | - | HA |

```
3341   3344       1 ;      H - CB
3341   3359       1 ;      H - C
3342   3348       1 ;     CA - CD1
3342   3357       1 ;     CA - CD2
3342   3362       1 ;     CA - CD
3342   3371       1 ;     CA - CA
3343   3345       1 ;     HA - HB2
3343   3346       1 ;     HA - HB3
3343   3347       1 ;     HA - CG
3343   3360       1 ;     HA - O
3343   3361       1 ;     HA - N
3344   3349       1 ;     CB - HD1
3344   3350       1 ;     CB - CE1
3344   3355       1 ;     CB - CE2
3344   3358       1 ;     CB - HD2
3344   3360       1 ;     CB - O
3344   3361       1 ;     CB - N
3345   3348       1 ;    HB2 - CD1
3345   3357       1 ;    HB2 - CD2
3345   3359       1 ;    HB2 - C
3346   3348       1 ;    HB3 - CD1
3346   3357       1 ;    HB3 - CD2
3346   3359       1 ;    HB3 - C
3347   3351       1 ;     CG - HE1
3347   3352       1 ;     CG - CZ
3347   3356       1 ;     CG - HE2
3347   3359       1 ;     CG - C
3348   3353       1 ;    CD1 - OH
3348   3355       1 ;    CD1 - CE2
3348   3358       1 ;    CD1 - HD2
3349   3351       1 ;    HD1 - HE1
3349   3352       1 ;    HD1 - CZ
3349   3357       1 ;    HD1 - CD2
3350   3354       1 ;    CE1 - HH
3350   3356       1 ;    CE1 - HE2
3350   3357       1 ;    CE1 - CD2
3351   3353       1 ;    HE1 - OH
3351   3355       1 ;    HE1 - CE2
3352   3358       1 ;     CZ - HD2
3353   3356       1 ;     OH - HE2
3353   3357       1 ;     OH - CD2
3354   3355       1 ;     HH - CE2
3356   3358       1 ;    HE2 - HD2
3359   3363       1 ;      C - HD2
3359   3364       1 ;      C - HD3
3359   3365       1 ;      C - CG
3359   3368       1 ;      C - CB
3359   3372       1 ;      C - HA
3359   3373       1 ;      C - C
3360   3362       1 ;      O - CD
3360   3371       1 ;      O - CA
3361   3366       1 ;      N - HG2
```

```
3361   3367       1 ;       N - HG3
3361   3369       1 ;       N - HB2
3361   3370       1 ;       N - HB3
3361   3374       1 ;       N - O
3361   3375       1 ;       N - N
3362   3369       1 ;      CD - HB2
3362   3370       1 ;      CD - HB3
3362   3372       1 ;      CD - HA
3362   3373       1 ;      CD - C
3363   3366       1 ;     HD2 - HG2
3363   3367       1 ;     HD2 - HG3
3363   3368       1 ;     HD2 - CB
3363   3371       1 ;     HD2 - CA
3364   3366       1 ;     HD3 - HG2
3364   3367       1 ;     HD3 - HG3
3364   3368       1 ;     HD3 - CB
3364   3371       1 ;     HD3 - CA
3365   3372       1 ;      CG - HA
3365   3373       1 ;      CG - C
3366   3369       1 ;     HG2 - HB2
3366   3370       1 ;     HG2 - HB3
3366   3371       1 ;     HG2 - CA
3367   3369       1 ;     HG3 - HB2
3367   3370       1 ;     HG3 - HB3
3367   3371       1 ;     HG3 - CA
3368   3374       1 ;      CB - O
3368   3375       1 ;      CB - N
3369   3372       1 ;     HB2 - HA
3369   3373       1 ;     HB2 - C
3370   3372       1 ;     HB3 - HA
3370   3373       1 ;     HB3 - C
3371   3376       1 ;      CA - CD
3371   3385       1 ;      CA - CA
3372   3374       1 ;      HA - O
3372   3375       1 ;      HA - N
3373   3377       1 ;       C - HD2
3373   3378       1 ;       C - HD3
3373   3379       1 ;       C - CG
3373   3382       1 ;       C - CB
3373   3386       1 ;       C - HA
3373   3387       1 ;       C - C
3374   3376       1 ;       O - CD
3374   3385       1 ;       O - CA
3375   3380       1 ;       N - HG2
3375   3381       1 ;       N - HG3
3375   3383       1 ;       N - HB2
3375   3384       1 ;       N - HB3
3375   3388       1 ;       N - O
3375   3389       1 ;       N - N
3376   3383       1 ;      CD - HB2
3376   3384       1 ;      CD - HB3
3376   3386       1 ;      CD - HA
```

| | | | | |
|---|---|---|---|---|
| 3376 | 3387 | 1 ; | CD - | C |
| 3377 | 3380 | 1 ; | HD2 - | HG2 |
| 3377 | 3381 | 1 ; | HD2 - | HG3 |
| 3377 | 3382 | 1 ; | HD2 - | CB |
| 3377 | 3385 | 1 ; | HD2 - | CA |
| 3378 | 3380 | 1 ; | HD3 - | HG2 |
| 3378 | 3381 | 1 ; | HD3 - | HG3 |
| 3378 | 3382 | 1 ; | HD3 - | CB |
| 3378 | 3385 | 1 ; | HD3 - | CA |
| 3379 | 3386 | 1 ; | CG - | HA |
| 3379 | 3387 | 1 ; | CG - | C |
| 3380 | 3383 | 1 ; | HG2 - | HB2 |
| 3380 | 3384 | 1 ; | HG2 - | HB3 |
| 3380 | 3385 | 1 ; | HG2 - | CA |
| 3381 | 3383 | 1 ; | HG3 - | HB2 |
| 3381 | 3384 | 1 ; | HG3 - | HB3 |
| 3381 | 3385 | 1 ; | HG3 - | CA |
| 3382 | 3388 | 1 ; | CB - | O |
| 3382 | 3389 | 1 ; | CB - | N |
| 3383 | 3386 | 1 ; | HB2 - | HA |
| 3383 | 3387 | 1 ; | HB2 - | C |
| 3384 | 3386 | 1 ; | HB3 - | HA |
| 3384 | 3387 | 1 ; | HB3 - | C |
| 3385 | 3390 | 1 ; | CA - | H |
| 3385 | 3391 | 1 ; | CA - | CA |
| 3386 | 3388 | 1 ; | HA - | O |
| 3386 | 3389 | 1 ; | HA - | N |
| 3387 | 3392 | 1 ; | C - | HA |
| 3387 | 3393 | 1 ; | C - | CB |
| 3387 | 3407 | 1 ; | C - | C |
| 3388 | 3390 | 1 ; | O - | H |
| 3388 | 3391 | 1 ; | O - | CA |
| 3389 | 3394 | 1 ; | N - | HB2 |
| 3389 | 3395 | 1 ; | N - | HB3 |
| 3389 | 3396 | 1 ; | N - | CG |
| 3389 | 3408 | 1 ; | N - | O |
| 3389 | 3409 | 1 ; | N - | N |
| 3390 | 3392 | 1 ; | H - | HA |
| 3390 | 3393 | 1 ; | H - | CB |
| 3390 | 3407 | 1 ; | H - | C |
| 3391 | 3397 | 1 ; | CA - | CD1 |
| 3391 | 3405 | 1 ; | CA - | CD2 |
| 3391 | 3410 | 1 ; | CA - | H |
| 3391 | 3411 | 1 ; | CA - | CA |
| 3392 | 3394 | 1 ; | HA - | HB2 |
| 3392 | 3395 | 1 ; | HA - | HB3 |
| 3392 | 3396 | 1 ; | HA - | CG |
| 3392 | 3408 | 1 ; | HA - | O |
| 3392 | 3409 | 1 ; | HA - | N |
| 3393 | 3398 | 1 ; | CB - | HD1 |
| 3393 | 3399 | 1 ; | CB - | CE1 |
| 3393 | 3403 | 1 ; | CB - | CE2 |

| | | | | |
|---|---|---|---|---|
| 3393 | 3406 | 1 ; | CB - | HD2 |
| 3393 | 3408 | 1 ; | CB - | O |
| 3393 | 3409 | 1 ; | CB - | N |
| 3394 | 3397 | 1 ; | HB2 - | CD1 |
| 3394 | 3405 | 1 ; | HB2 - | CD2 |
| 3394 | 3407 | 1 ; | HB2 - | C |
| 3395 | 3397 | 1 ; | HB3 - | CD1 |
| 3395 | 3405 | 1 ; | HB3 - | CD2 |
| 3395 | 3407 | 1 ; | HB3 - | C |
| 3396 | 3400 | 1 ; | CG - | HE1 |
| 3396 | 3401 | 1 ; | CG - | CZ |
| 3396 | 3404 | 1 ; | CG - | HE2 |
| 3396 | 3407 | 1 ; | CG - | C |
| 3397 | 3402 | 1 ; | CD1 - | HZ |
| 3397 | 3403 | 1 ; | CD1 - | CE2 |
| 3397 | 3406 | 1 ; | CD1 - | HD2 |
| 3398 | 3400 | 1 ; | HD1 - | HE1 |
| 3398 | 3401 | 1 ; | HD1 - | CZ |
| 3398 | 3405 | 1 ; | HD1 - | CD2 |
| 3399 | 3404 | 1 ; | CE1 - | HE2 |
| 3399 | 3405 | 1 ; | CE1 - | CD2 |
| 3400 | 3402 | 1 ; | HE1 - | HZ |
| 3400 | 3403 | 1 ; | HE1 - | CE2 |
| 3401 | 3406 | 1 ; | CZ - | HD2 |
| 3402 | 3404 | 1 ; | HZ - | HE2 |
| 3402 | 3405 | 1 ; | HZ - | CD2 |
| 3404 | 3406 | 1 ; | HE2 - | HD2 |
| 3407 | 3412 | 1 ; | C - | HA |
| 3407 | 3413 | 1 ; | C - | CB |
| 3407 | 3428 | 1 ; | C - | C |
| 3408 | 3410 | 1 ; | O - | H |
| 3408 | 3411 | 1 ; | O - | CA |
| 3409 | 3414 | 1 ; | N - | HB2 |
| 3409 | 3415 | 1 ; | N - | HB3 |
| 3409 | 3416 | 1 ; | N - | CG |
| 3409 | 3429 | 1 ; | N - | O |
| 3409 | 3430 | 1 ; | N - | N |
| 3410 | 3412 | 1 ; | H - | HA |
| 3410 | 3413 | 1 ; | H - | CB |
| 3410 | 3428 | 1 ; | H - | C |
| 3411 | 3417 | 1 ; | CA - | CD1 |
| 3411 | 3426 | 1 ; | CA - | CD2 |
| 3411 | 3431 | 1 ; | CA - | H |
| 3411 | 3432 | 1 ; | CA - | CA |
| 3412 | 3414 | 1 ; | HA - | HB2 |
| 3412 | 3415 | 1 ; | HA - | HB3 |
| 3412 | 3416 | 1 ; | HA - | CG |
| 3412 | 3429 | 1 ; | HA - | O |
| 3412 | 3430 | 1 ; | HA - | N |
| 3413 | 3418 | 1 ; | CB - | HD1 |
| 3413 | 3419 | 1 ; | CB - | CE1 |
| 3413 | 3424 | 1 ; | CB - | CE2 |

| | | | | | |
|---|---|---|---|---|---|
| 3413 | 3427 | 1 ; | CB | - | HD2 |
| 3413 | 3429 | 1 ; | CB | - | O |
| 3413 | 3430 | 1 ; | CB | - | N |
| 3414 | 3417 | 1 ; | HB2 | - | CD1 |
| 3414 | 3426 | 1 ; | HB2 | - | CD2 |
| 3414 | 3428 | 1 ; | HB2 | - | C |
| 3415 | 3417 | 1 ; | HB3 | - | CD1 |
| 3415 | 3426 | 1 ; | HB3 | - | CD2 |
| 3415 | 3428 | 1 ; | HB3 | - | C |
| 3416 | 3420 | 1 ; | CG | - | HE1 |
| 3416 | 3421 | 1 ; | CG | - | CZ |
| 3416 | 3425 | 1 ; | CG | - | HE2 |
| 3416 | 3428 | 1 ; | CG | - | C |
| 3417 | 3422 | 1 ; | CD1 | - | OH |
| 3417 | 3424 | 1 ; | CD1 | - | CE2 |
| 3417 | 3427 | 1 ; | CD1 | - | HD2 |
| 3418 | 3420 | 1 ; | HD1 | - | HE1 |
| 3418 | 3421 | 1 ; | HD1 | - | CZ |
| 3418 | 3426 | 1 ; | HD1 | - | CD2 |
| 3419 | 3423 | 1 ; | CE1 | - | HH |
| 3419 | 3425 | 1 ; | CE1 | - | HE2 |
| 3419 | 3426 | 1 ; | CE1 | - | CD2 |
| 3420 | 3422 | 1 ; | HE1 | - | OH |
| 3420 | 3424 | 1 ; | HE1 | - | CE2 |
| 3421 | 3427 | 1 ; | CZ | - | HD2 |
| 3422 | 3425 | 1 ; | OH | - | HE2 |
| 3422 | 3426 | 1 ; | OH | - | CD2 |
| 3423 | 3424 | 1 ; | HH | - | CE2 |
| 3425 | 3427 | 1 ; | HE2 | - | HD2 |
| 3428 | 3433 | 1 ; | C | - | HA2 |
| 3428 | 3434 | 1 ; | C | - | HA3 |
| 3428 | 3435 | 1 ; | C | - | C |
| 3429 | 3431 | 1 ; | O | - | H |
| 3429 | 3432 | 1 ; | O | - | CA |
| 3430 | 3436 | 1 ; | N | - | O |
| 3430 | 3437 | 1 ; | N | - | N |
| 3431 | 3433 | 1 ; | H | - | HA2 |
| 3431 | 3434 | 1 ; | H | - | HA3 |
| 3431 | 3435 | 1 ; | H | - | C |
| 3432 | 3438 | 1 ; | CA | - | H |
| 3432 | 3439 | 1 ; | CA | - | CA |
| 3433 | 3436 | 1 ; | HA2 | - | O |
| 3433 | 3437 | 1 ; | HA2 | - | N |
| 3434 | 3436 | 1 ; | HA3 | - | O |
| 3434 | 3437 | 1 ; | HA3 | - | N |
| 3435 | 3440 | 1 ; | C | - | HA |
| 3435 | 3441 | 1 ; | C | - | CB |
| 3435 | 3447 | 1 ; | C | - | C |
| 3436 | 3438 | 1 ; | O | - | H |
| 3436 | 3439 | 1 ; | O | - | CA |
| 3437 | 3442 | 1 ; | N | - | HB2 |
| 3437 | 3443 | 1 ; | N | - | HB3 |

| | | | | | |
|---|---|---|---|---|---|
| 3437 | 3444 | 1 ; | N | - | CG |
| 3437 | 3448 | 1 ; | N | - | O |
| 3437 | 3449 | 1 ; | N | - | N |
| 3438 | 3440 | 1 ; | H | - | HA |
| 3438 | 3441 | 1 ; | H | - | CB |
| 3438 | 3447 | 1 ; | H | - | C |
| 3439 | 3445 | 1 ; | CA | - | OD1 |
| 3439 | 3446 | 1 ; | CA | - | OD2 |
| 3439 | 3450 | 1 ; | CA | - | H |
| 3439 | 3451 | 1 ; | CA | - | CA |
| 3440 | 3442 | 1 ; | HA | - | HB2 |
| 3440 | 3443 | 1 ; | HA | - | HB3 |
| 3440 | 3444 | 1 ; | HA | - | CG |
| 3440 | 3448 | 1 ; | HA | - | O |
| 3440 | 3449 | 1 ; | HA | - | N |
| 3441 | 3448 | 1 ; | CB | - | O |
| 3441 | 3449 | 1 ; | CB | - | N |
| 3442 | 3445 | 1 ; | HB2 | - | OD1 |
| 3442 | 3446 | 1 ; | HB2 | - | OD2 |
| 3442 | 3447 | 1 ; | HB2 | - | C |
| 3443 | 3445 | 1 ; | HB3 | - | OD1 |
| 3443 | 3446 | 1 ; | HB3 | - | OD2 |
| 3443 | 3447 | 1 ; | HB3 | - | C |
| 3444 | 3447 | 1 ; | CG | - | C |
| 3447 | 3452 | 1 ; | C | - | HA |
| 3447 | 3453 | 1 ; | C | - | CB |
| 3447 | 3461 | 1 ; | C | - | C |
| 3448 | 3450 | 1 ; | O | - | H |
| 3448 | 3451 | 1 ; | O | - | CA |
| 3449 | 3454 | 1 ; | N | - | HB |
| 3449 | 3455 | 1 ; | N | - | CG2 |
| 3449 | 3459 | 1 ; | N | - | OG1 |
| 3449 | 3462 | 1 ; | N | - | O |
| 3449 | 3463 | 1 ; | N | - | N |
| 3450 | 3452 | 1 ; | H | - | HA |
| 3450 | 3453 | 1 ; | H | - | CB |
| 3450 | 3461 | 1 ; | H | - | C |
| 3451 | 3456 | 1 ; | CA | - | HG21 |
| 3451 | 3457 | 1 ; | CA | - | HG22 |
| 3451 | 3458 | 1 ; | CA | - | HG23 |
| 3451 | 3460 | 1 ; | CA | - | HG1 |
| 3451 | 3464 | 1 ; | CA | - | H |
| 3451 | 3465 | 1 ; | CA | - | CA |
| 3452 | 3454 | 1 ; | HA | - | HB |
| 3452 | 3455 | 1 ; | HA | - | CG2 |
| 3452 | 3459 | 1 ; | HA | - | OG1 |
| 3452 | 3462 | 1 ; | HA | - | O |
| 3452 | 3463 | 1 ; | HA | - | N |
| 3453 | 3462 | 1 ; | CB | - | O |
| 3453 | 3463 | 1 ; | CB | - | N |
| 3454 | 3456 | 1 ; | HB | - | HG21 |
| 3454 | 3457 | 1 ; | HB | - | HG22 |

```
3454   3458      1 ;     HB - HG23
3454   3460      1 ;     HB - HG1
3454   3461      1 ;     HB - C
3455   3460      1 ;    CG2 - HG1
3455   3461      1 ;    CG2 - C
3456   3459      1 ;   HG21 - OG1
3457   3459      1 ;   HG22 - OG1
3458   3459      1 ;   HG23 - OG1
3459   3461      1 ;    OG1 - C
3461   3466      1 ;      C - HA
3461   3467      1 ;      C - CB
3461   3473      1 ;      C - C
3462   3464      1 ;      O - H
3462   3465      1 ;      O - CA
3463   3468      1 ;      N - HB2
3463   3469      1 ;      N - HB3
3463   3470      1 ;      N - CG
3463   3474      1 ;      N - O
3463   3475      1 ;      N - N
3464   3466      1 ;      H - HA
3464   3467      1 ;      H - CB
3464   3473      1 ;      H - C
3465   3471      1 ;     CA - OD1
3465   3472      1 ;     CA - OD2
3465   3476      1 ;     CA - H
3465   3477      1 ;     CA - CA
3466   3468      1 ;     HA - HB2
3466   3469      1 ;     HA - HB3
3466   3470      1 ;     HA - CG
3466   3474      1 ;     HA - O
3466   3475      1 ;     HA - N
3467   3474      1 ;     CB - O
3467   3475      1 ;     CB - N
3468   3471      1 ;    HB2 - OD1
3468   3472      1 ;    HB2 - OD2
3468   3473      1 ;    HB2 - C
3469   3471      1 ;    HB3 - OD1
3469   3472      1 ;    HB3 - OD2
3469   3473      1 ;    HB3 - C
3470   3473      1 ;     CG - C
3473   3478      1 ;      C - HA
3473   3479      1 ;      C - CB
3473   3487      1 ;      C - C
3474   3476      1 ;      O - H
3474   3477      1 ;      O - CA
3475   3480      1 ;      N - HB2
3475   3481      1 ;      N - HB3
3475   3482      1 ;      N - CG
3475   3488      1 ;      N - O
3475   3489      1 ;      N - N
3476   3478      1 ;      H - HA
3476   3479      1 ;      H - CB
```

| | | | | | | |
|---|---|---|---|---|---|---|
| 3476 | 3487 | 1 | ; | H | - | C |
| 3477 | 3483 | 1 | ; | CA | - | OD1 |
| 3477 | 3484 | 1 | ; | CA | - | ND2 |
| 3477 | 3490 | 1 | ; | CA | - | H |
| 3477 | 3491 | 1 | ; | CA | - | CA |
| 3478 | 3480 | 1 | ; | HA | - | HB2 |
| 3478 | 3481 | 1 | ; | HA | - | HB3 |
| 3478 | 3482 | 1 | ; | HA | - | CG |
| 3478 | 3488 | 1 | ; | HA | - | O |
| 3478 | 3489 | 1 | ; | HA | - | N |
| 3479 | 3485 | 1 | ; | CB | - | HD21 |
| 3479 | 3486 | 1 | ; | CB | - | HD22 |
| 3479 | 3488 | 1 | ; | CB | - | O |
| 3479 | 3489 | 1 | ; | CB | - | N |
| 3480 | 3483 | 1 | ; | HB2 | - | OD1 |
| 3480 | 3484 | 1 | ; | HB2 | - | ND2 |
| 3480 | 3487 | 1 | ; | HB2 | - | C |
| 3481 | 3483 | 1 | ; | HB3 | - | OD1 |
| 3481 | 3484 | 1 | ; | HB3 | - | ND2 |
| 3481 | 3487 | 1 | ; | HB3 | - | C |
| 3482 | 3487 | 1 | ; | CG | - | C |
| 3483 | 3485 | 1 | ; | OD1 | - | HD21 |
| 3483 | 3486 | 1 | ; | OD1 | - | HD22 |
| 3487 | 3492 | 1 | ; | C | - | HA |
| 3487 | 3493 | 1 | ; | C | - | CB |
| 3487 | 3502 | 1 | ; | C | - | C |
| 3488 | 3490 | 1 | ; | O | - | H |
| 3488 | 3491 | 1 | ; | O | - | CA |
| 3489 | 3494 | 1 | ; | N | - | HB2 |
| 3489 | 3495 | 1 | ; | N | - | HB3 |
| 3489 | 3496 | 1 | ; | N | - | CG |
| 3489 | 3503 | 1 | ; | N | - | O |
| 3489 | 3504 | 1 | ; | N | - | N |
| 3490 | 3492 | 1 | ; | H | - | HA |
| 3490 | 3493 | 1 | ; | H | - | CB |
| 3490 | 3502 | 1 | ; | H | - | C |
| 3491 | 3497 | 1 | ; | CA | - | HG2 |
| 3491 | 3498 | 1 | ; | CA | - | HG3 |
| 3491 | 3499 | 1 | ; | CA | - | CD |
| 3491 | 3505 | 1 | ; | CA | - | H |
| 3491 | 3506 | 1 | ; | CA | - | CA |
| 3492 | 3494 | 1 | ; | HA | - | HB2 |
| 3492 | 3495 | 1 | ; | HA | - | HB3 |
| 3492 | 3496 | 1 | ; | HA | - | CG |
| 3492 | 3503 | 1 | ; | HA | - | O |
| 3492 | 3504 | 1 | ; | HA | - | N |
| 3493 | 3500 | 1 | ; | CB | - | OE1 |
| 3493 | 3501 | 1 | ; | CB | - | OE2 |
| 3493 | 3503 | 1 | ; | CB | - | O |
| 3493 | 3504 | 1 | ; | CB | - | N |
| 3494 | 3497 | 1 | ; | HB2 | - | HG2 |
| 3494 | 3498 | 1 | ; | HB2 | - | HG3 |

```
3494   3499       1 ;    HB2 - CD
3494   3502       1 ;    HB2 - C
3495   3497       1 ;    HB3 - HG2
3495   3498       1 ;    HB3 - HG3
3495   3499       1 ;    HB3 - CD
3495   3502       1 ;    HB3 - C
3496   3502       1 ;     CG - C
3497   3500       1 ;    HG2 - OE1
3497   3501       1 ;    HG2 - OE2
3498   3500       1 ;    HG3 - OE1
3498   3501       1 ;    HG3 - OE2
3502   3507       1 ;      C - HA
3502   3508       1 ;      C - CB
3502   3518       1 ;      C - C
3503   3505       1 ;      O - H
3503   3506       1 ;      O - CA
3504   3509       1 ;      N - HB
3504   3510       1 ;      N - CG1
3504   3514       1 ;      N - CG2
3504   3519       1 ;      N - O
3504   3520       1 ;      N - N
3505   3507       1 ;      H - HA
3505   3508       1 ;      H - CB
3505   3518       1 ;      H - C
3506   3511       1 ;     CA - HG11
3506   3512       1 ;     CA - HG12
3506   3513       1 ;     CA - HG13
3506   3515       1 ;     CA - HG21
3506   3516       1 ;     CA - HG22
3506   3517       1 ;     CA - HG23
3506   3521       1 ;     CA - H
3506   3522       1 ;     CA - CA
3507   3509       1 ;     HA - HB
3507   3510       1 ;     HA - CG1
3507   3514       1 ;     HA - CG2
3507   3519       1 ;     HA - O
3507   3520       1 ;     HA - N
3508   3519       1 ;     CB - O
3508   3520       1 ;     CB - N
3509   3511       1 ;     HB - HG11
3509   3512       1 ;     HB - HG12
3509   3513       1 ;     HB - HG13
3509   3515       1 ;     HB - HG21
3509   3516       1 ;     HB - HG22
3509   3517       1 ;     HB - HG23
3509   3518       1 ;     HB - C
3510   3515       1 ;    CG1 - HG21
3510   3516       1 ;    CG1 - HG22
3510   3517       1 ;    CG1 - HG23
3510   3518       1 ;    CG1 - C
3511   3514       1 ;   HG11 - CG2
3512   3514       1 ;   HG12 - CG2
```

```
3513   3514       1 ;   HG13 - CG2
3514   3518       1 ;    CG2 - C
3518   3523       1 ;      C - HA
3518   3524       1 ;      C - CB
3518   3537       1 ;      C - C
3519   3521       1 ;      O - H
3519   3522       1 ;      O - CA
3520   3525       1 ;      N - HB2
3520   3526       1 ;      N - HB3
3520   3527       1 ;      N - CG
3520   3538       1 ;      N - O
3520   3539       1 ;      N - N
3521   3523       1 ;      H - HA
3521   3524       1 ;      H - CB
3521   3537       1 ;      H - C
3522   3528       1 ;     CA - HG
3522   3529       1 ;     CA - CD1
3522   3533       1 ;     CA - CD2
3522   3540       1 ;     CA - H
3522   3541       1 ;     CA - CA
3523   3525       1 ;     HA - HB2
3523   3526       1 ;     HA - HB3
3523   3527       1 ;     HA - CG
3523   3538       1 ;     HA - O
3523   3539       1 ;     HA - N
3524   3530       1 ;     CB - HD11
3524   3531       1 ;     CB - HD12
3524   3532       1 ;     CB - HD13
3524   3534       1 ;     CB - HD21
3524   3535       1 ;     CB - HD22
3524   3536       1 ;     CB - HD23
3524   3538       1 ;     CB - O
3524   3539       1 ;     CB - N
3525   3528       1 ;    HB2 - HG
3525   3529       1 ;    HB2 - CD1
3525   3533       1 ;    HB2 - CD2
3525   3537       1 ;    HB2 - C
3526   3528       1 ;    HB3 - HG
3526   3529       1 ;    HB3 - CD1
3526   3533       1 ;    HB3 - CD2
3526   3537       1 ;    HB3 - C
3527   3537       1 ;     CG - C
3528   3530       1 ;     HG - HD11
3528   3531       1 ;     HG - HD12
3528   3532       1 ;     HG - HD13
3528   3534       1 ;     HG - HD21
3528   3535       1 ;     HG - HD22
3528   3536       1 ;     HG - HD23
3529   3534       1 ;    CD1 - HD21
3529   3535       1 ;    CD1 - HD22
3529   3536       1 ;    CD1 - HD23
3530   3533       1 ;   HD11 - CD2
```

```
3531   3533       1 ;   HD12 - CD2
3532   3533       1 ;   HD13 - CD2
3537   3542       1 ;      C - HA
3537   3543       1 ;      C - CB
3537   3559       1 ;      C - C
3538   3540       1 ;      O - H
3538   3541       1 ;      O - CA
3539   3544       1 ;      N - HB2
3539   3545       1 ;      N - HB3
3539   3546       1 ;      N - CG
3539   3560       1 ;      N - O
3539   3561       1 ;      N - N
3540   3542       1 ;      H - HA
3540   3543       1 ;      H - CB
3540   3559       1 ;      H - C
3541   3547       1 ;     CA - HG2
3541   3548       1 ;     CA - HG3
3541   3549       1 ;     CA - CD
3541   3562       1 ;     CA - H
3541   3563       1 ;     CA - CA
3542   3544       1 ;     HA - HB2
3542   3545       1 ;     HA - HB3
3542   3546       1 ;     HA - CG
3542   3560       1 ;     HA - O
3542   3561       1 ;     HA - N
3543   3550       1 ;     CB - HD2
3543   3551       1 ;     CB - HD3
3543   3552       1 ;     CB - CE
3543   3560       1 ;     CB - O
3543   3561       1 ;     CB - N
3544   3547       1 ;    HB2 - HG2
3544   3548       1 ;    HB2 - HG3
3544   3549       1 ;    HB2 - CD
3544   3559       1 ;    HB2 - C
3545   3547       1 ;    HB3 - HG2
3545   3548       1 ;    HB3 - HG3
3545   3549       1 ;    HB3 - CD
3545   3559       1 ;    HB3 - C
3546   3553       1 ;     CG - HE2
3546   3554       1 ;     CG - HE3
3546   3555       1 ;     CG - NZ
3546   3559       1 ;     CG - C
3547   3550       1 ;    HG2 - HD2
3547   3551       1 ;    HG2 - HD3
3547   3552       1 ;    HG2 - CE
3548   3550       1 ;    HG3 - HD2
3548   3551       1 ;    HG3 - HD3
3548   3552       1 ;    HG3 - CE
3549   3556       1 ;     CD - HZ1
3549   3557       1 ;     CD - HZ2
3549   3558       1 ;     CD - HZ3
3550   3553       1 ;    HD2 - HE2
```

```
3550   3554       1 ;    HD2 - HE3
3550   3555       1 ;    HD2 - NZ
3551   3553       1 ;    HD3 - HE2
3551   3554       1 ;    HD3 - HE3
3551   3555       1 ;    HD3 - NZ
3553   3556       1 ;    HE2 - HZ1
3553   3557       1 ;    HE2 - HZ2
3553   3558       1 ;    HE2 - HZ3
3554   3556       1 ;    HE3 - HZ1
3554   3557       1 ;    HE3 - HZ2
3554   3558       1 ;    HE3 - HZ3
3559   3564       1 ;      C - HA
3559   3565       1 ;      C - CB
3559   3581       1 ;      C - C
3560   3562       1 ;      O - H
3560   3563       1 ;      O - CA
3561   3566       1 ;      N - HB2
3561   3567       1 ;      N - HB3
3561   3568       1 ;      N - CG
3561   3582       1 ;      N - O
3561   3583       1 ;      N - N
3562   3564       1 ;      H - HA
3562   3565       1 ;      H - CB
3562   3581       1 ;      H - C
3563   3569       1 ;     CA - HG2
3563   3570       1 ;     CA - HG3
3563   3571       1 ;     CA - CD
3563   3584       1 ;     CA - H
3563   3585       1 ;     CA - CA
3564   3566       1 ;     HA - HB2
3564   3567       1 ;     HA - HB3
3564   3568       1 ;     HA - CG
3564   3582       1 ;     HA - O
3564   3583       1 ;     HA - N
3565   3572       1 ;     CB - HD2
3565   3573       1 ;     CB - HD3
3565   3574       1 ;     CB - CE
3565   3582       1 ;     CB - O
3565   3583       1 ;     CB - N
3566   3569       1 ;    HB2 - HG2
3566   3570       1 ;    HB2 - HG3
3566   3571       1 ;    HB2 - CD
3566   3581       1 ;    HB2 - C
3567   3569       1 ;    HB3 - HG2
3567   3570       1 ;    HB3 - HG3
3567   3571       1 ;    HB3 - CD
3567   3581       1 ;    HB3 - C
3568   3575       1 ;     CG - HE2
3568   3576       1 ;     CG - HE3
3568   3577       1 ;     CG - NZ
3568   3581       1 ;     CG - C
3569   3572       1 ;    HG2 - HD2
```

```
3569   3573      1 ;    HG2 - HD3
3569   3574      1 ;    HG2 - CE
3570   3572      1 ;    HG3 - HD2
3570   3573      1 ;    HG3 - HD3
3570   3574      1 ;    HG3 - CE
3571   3578      1 ;     CD - HZ1
3571   3579      1 ;     CD - HZ2
3571   3580      1 ;     CD - HZ3
3572   3575      1 ;    HD2 - HE2
3572   3576      1 ;    HD2 - HE3
3572   3577      1 ;    HD2 - NZ
3573   3575      1 ;    HD3 - HE2
3573   3576      1 ;    HD3 - HE3
3573   3577      1 ;    HD3 - NZ
3575   3578      1 ;    HE2 - HZ1
3575   3579      1 ;    HE2 - HZ2
3575   3580      1 ;    HE2 - HZ3
3576   3578      1 ;    HE3 - HZ1
3576   3579      1 ;    HE3 - HZ2
3576   3580      1 ;    HE3 - HZ3
3581   3586      1 ;      C - HA
3581   3587      1 ;      C - CB
3581   3597      1 ;      C - C
3582   3584      1 ;      O - H
3582   3585      1 ;      O - CA
3583   3588      1 ;      N - HB
3583   3589      1 ;      N - CG1
3583   3593      1 ;      N - CG2
3583   3598      1 ;      N - O
3583   3599      1 ;      N - N
3584   3586      1 ;      H - HA
3584   3587      1 ;      H - CB
3584   3597      1 ;      H - C
3585   3590      1 ;     CA - HG11
3585   3591      1 ;     CA - HG12
3585   3592      1 ;     CA - HG13
3585   3594      1 ;     CA - HG21
3585   3595      1 ;     CA - HG22
3585   3596      1 ;     CA - HG23
3585   3600      1 ;     CA - H
3585   3601      1 ;     CA - CA
3586   3588      1 ;     HA - HB
3586   3589      1 ;     HA - CG1
3586   3593      1 ;     HA - CG2
3586   3598      1 ;     HA - O
3586   3599      1 ;     HA - N
3587   3598      1 ;     CB - O
3587   3599      1 ;     CB - N
3588   3590      1 ;     HB - HG11
3588   3591      1 ;     HB - HG12
3588   3592      1 ;     HB - HG13
3588   3594      1 ;     HB - HG21
```

```
3588   3595       1 ;      HB - HG22
3588   3596       1 ;      HB - HG23
3588   3597       1 ;      HB - C
3589   3594       1 ;     CG1 - HG21
3589   3595       1 ;     CG1 - HG22
3589   3596       1 ;     CG1 - HG23
3589   3597       1 ;     CG1 - C
3590   3593       1 ;    HG11 - CG2
3591   3593       1 ;    HG12 - CG2
3592   3593       1 ;    HG13 - CG2
3593   3597       1 ;     CG2 - C
3597   3602       1 ;       C - HA
3597   3603       1 ;       C - CB
3597   3619       1 ;       C - C
3598   3600       1 ;       O - H
3598   3601       1 ;       O - CA
3599   3604       1 ;       N - HB2
3599   3605       1 ;       N - HB3
3599   3606       1 ;       N - CG
3599   3620       1 ;       N - O
3599   3621       1 ;       N - N
3600   3602       1 ;       H - HA
3600   3603       1 ;       H - CB
3600   3619       1 ;       H - C
3601   3607       1 ;      CA - HG2
3601   3608       1 ;      CA - HG3
3601   3609       1 ;      CA - CD
3601   3622       1 ;      CA - H
3601   3623       1 ;      CA - CA
3602   3604       1 ;      HA - HB2
3602   3605       1 ;      HA - HB3
3602   3606       1 ;      HA - CG
3602   3620       1 ;      HA - O
3602   3621       1 ;      HA - N
3603   3610       1 ;      CB - HD2
3603   3611       1 ;      CB - HD3
3603   3612       1 ;      CB - CE
3603   3620       1 ;      CB - O
3603   3621       1 ;      CB - N
3604   3607       1 ;     HB2 - HG2
3604   3608       1 ;     HB2 - HG3
3604   3609       1 ;     HB2 - CD
3604   3619       1 ;     HB2 - C
3605   3607       1 ;     HB3 - HG2
3605   3608       1 ;     HB3 - HG3
3605   3609       1 ;     HB3 - CD
3605   3619       1 ;     HB3 - C
3606   3613       1 ;      CG - HE2
3606   3614       1 ;      CG - HE3
3606   3615       1 ;      CG - NZ
3606   3619       1 ;      CG - C
3607   3610       1 ;     HG2 - HD2
```

```
3607   3611       1 ;     HG2 - HD3
3607   3612       1 ;     HG2 - CE
3608   3610       1 ;     HG3 - HD2
3608   3611       1 ;     HG3 - HD3
3608   3612       1 ;     HG3 - CE
3609   3616       1 ;      CD - HZ1
3609   3617       1 ;      CD - HZ2
3609   3618       1 ;      CD - HZ3
3610   3613       1 ;     HD2 - HE2
3610   3614       1 ;     HD2 - HE3
3610   3615       1 ;     HD2 - NZ
3611   3613       1 ;     HD3 - HE2
3611   3614       1 ;     HD3 - HE3
3611   3615       1 ;     HD3 - NZ
3613   3616       1 ;     HE2 - HZ1
3613   3617       1 ;     HE2 - HZ2
3613   3618       1 ;     HE2 - HZ3
3614   3616       1 ;     HE3 - HZ1
3614   3617       1 ;     HE3 - HZ2
3614   3618       1 ;     HE3 - HZ3
3619   3624       1 ;       C - HA
3619   3625       1 ;       C - CB
3619   3641       1 ;       C - C
3620   3622       1 ;       O - H
3620   3623       1 ;       O - CA
3621   3626       1 ;       N - HB2
3621   3627       1 ;       N - HB3
3621   3628       1 ;       N - CG
3621   3642       1 ;       N - O
3621   3643       1 ;       N - N
3622   3624       1 ;       H - HA
3622   3625       1 ;       H - CB
3622   3641       1 ;       H - C
3623   3629       1 ;      CA - HG2
3623   3630       1 ;      CA - HG3
3623   3631       1 ;      CA - CD
3623   3644       1 ;      CA - H
3623   3645       1 ;      CA - CA
3624   3626       1 ;      HA - HB2
3624   3627       1 ;      HA - HB3
3624   3628       1 ;      HA - CG
3624   3642       1 ;      HA - O
3624   3643       1 ;      HA - N
3625   3632       1 ;      CB - HD2
3625   3633       1 ;      CB - HD3
3625   3634       1 ;      CB - CE
3625   3642       1 ;      CB - O
3625   3643       1 ;      CB - N
3626   3629       1 ;     HB2 - HG2
3626   3630       1 ;     HB2 - HG3
3626   3631       1 ;     HB2 - CD
3626   3641       1 ;     HB2 - C
```

| | | | | | |
|---|---|---|---|---|---|
| 3627 | 3629 | 1 ; | HB3 | - | HG2 |
| 3627 | 3630 | 1 ; | HB3 | - | HG3 |
| 3627 | 3631 | 1 ; | HB3 | - | CD |
| 3627 | 3641 | 1 ; | HB3 | - | C |
| 3628 | 3635 | 1 ; | CG | - | HE2 |
| 3628 | 3636 | 1 ; | CG | - | HE3 |
| 3628 | 3637 | 1 ; | CG | - | NZ |
| 3628 | 3641 | 1 ; | CG | - | C |
| 3629 | 3632 | 1 ; | HG2 | - | HD2 |
| 3629 | 3633 | 1 ; | HG2 | - | HD3 |
| 3629 | 3634 | 1 ; | HG2 | - | CE |
| 3630 | 3632 | 1 ; | HG3 | - | HD2 |
| 3630 | 3633 | 1 ; | HG3 | - | HD3 |
| 3630 | 3634 | 1 ; | HG3 | - | CE |
| 3631 | 3638 | 1 ; | CD | - | HZ1 |
| 3631 | 3639 | 1 ; | CD | - | HZ2 |
| 3631 | 3640 | 1 ; | CD | - | HZ3 |
| 3632 | 3635 | 1 ; | HD2 | - | HE2 |
| 3632 | 3636 | 1 ; | HD2 | - | HE3 |
| 3632 | 3637 | 1 ; | HD2 | - | NZ |
| 3633 | 3635 | 1 ; | HD3 | - | HE2 |
| 3633 | 3636 | 1 ; | HD3 | - | HE3 |
| 3633 | 3637 | 1 ; | HD3 | - | NZ |
| 3635 | 3638 | 1 ; | HE2 | - | HZ1 |
| 3635 | 3639 | 1 ; | HE2 | - | HZ2 |
| 3635 | 3640 | 1 ; | HE2 | - | HZ3 |
| 3636 | 3638 | 1 ; | HE3 | - | HZ1 |
| 3636 | 3639 | 1 ; | HE3 | - | HZ2 |
| 3636 | 3640 | 1 ; | HE3 | - | HZ3 |
| 3641 | 3646 | 1 ; | C | - | HA2 |
| 3641 | 3647 | 1 ; | C | - | HA3 |
| 3641 | 3648 | 1 ; | C | - | C |
| 3642 | 3644 | 1 ; | O | - | H |
| 3642 | 3645 | 1 ; | O | - | CA |
| 3643 | 3649 | 1 ; | N | - | O |
| 3643 | 3650 | 1 ; | N | - | N |
| 3644 | 3646 | 1 ; | H | - | HA2 |
| 3644 | 3647 | 1 ; | H | - | HA3 |
| 3644 | 3648 | 1 ; | H | - | C |
| 3645 | 3651 | 1 ; | CA | - | H |
| 3645 | 3652 | 1 ; | CA | - | CA |
| 3646 | 3649 | 1 ; | HA2 | - | O |
| 3646 | 3650 | 1 ; | HA2 | - | N |
| 3647 | 3649 | 1 ; | HA3 | - | O |
| 3647 | 3650 | 1 ; | HA3 | - | N |
| 3648 | 3653 | 1 ; | C | - | HA |
| 3648 | 3654 | 1 ; | C | - | CB |
| 3648 | 3663 | 1 ; | C | - | C |
| 3649 | 3651 | 1 ; | O | - | H |
| 3649 | 3652 | 1 ; | O | - | CA |
| 3650 | 3655 | 1 ; | N | - | HB2 |
| 3650 | 3656 | 1 ; | N | - | HB3 |

```
3650   3657       1 ;      N - CG
3650   3664       1 ;      N - O
3650   3665       1 ;      N - N
3651   3653       1 ;      H - HA
3651   3654       1 ;      H - CB
3651   3663       1 ;      H - C
3652   3658       1 ;     CA - HG2
3652   3659       1 ;     CA - HG3
3652   3660       1 ;     CA - CD
3652   3666       1 ;     CA - H
3652   3667       1 ;     CA - CA
3653   3655       1 ;     HA - HB2
3653   3656       1 ;     HA - HB3
3653   3657       1 ;     HA - CG
3653   3664       1 ;     HA - O
3653   3665       1 ;     HA - N
3654   3661       1 ;     CB - OE1
3654   3662       1 ;     CB - OE2
3654   3664       1 ;     CB - O
3654   3665       1 ;     CB - N
3655   3658       1 ;    HB2 - HG2
3655   3659       1 ;    HB2 - HG3
3655   3660       1 ;    HB2 - CD
3655   3663       1 ;    HB2 - C
3656   3658       1 ;    HB3 - HG2
3656   3659       1 ;    HB3 - HG3
3656   3660       1 ;    HB3 - CD
3656   3663       1 ;    HB3 - C
3657   3663       1 ;     CG - C
3658   3661       1 ;    HG2 - OE1
3658   3662       1 ;    HG2 - OE2
3659   3661       1 ;    HG3 - OE1
3659   3662       1 ;    HG3 - OE2
3663   3668       1 ;      C - HA
3663   3669       1 ;      C - CB
3663   3683       1 ;      C - C
3664   3666       1 ;      O - H
3664   3667       1 ;      O - CA
3665   3670       1 ;      N - HB2
3665   3671       1 ;      N - HB3
3665   3672       1 ;      N - CG
3665   3684       1 ;      N - O
3665   3685       1 ;      N - N
3666   3668       1 ;      H - HA
3666   3669       1 ;      H - CB
3666   3683       1 ;      H - C
3667   3673       1 ;     CA - CD1
3667   3681       1 ;     CA - CD2
3667   3686       1 ;     CA - H
3667   3687       1 ;     CA - CA
3668   3670       1 ;     HA - HB2
3668   3671       1 ;     HA - HB3
```

```
3668   3672      1 ;     HA - CG
3668   3684      1 ;     HA - O
3668   3685      1 ;     HA - N
3669   3674      1 ;     CB - HD1
3669   3675      1 ;     CB - CE1
3669   3679      1 ;     CB - CE2
3669   3682      1 ;     CB - HD2
3669   3684      1 ;     CB - O
3669   3685      1 ;     CB - N
3670   3673      1 ;    HB2 - CD1
3670   3681      1 ;    HB2 - CD2
3670   3683      1 ;    HB2 - C
3671   3673      1 ;    HB3 - CD1
3671   3681      1 ;    HB3 - CD2
3671   3683      1 ;    HB3 - C
3672   3676      1 ;     CG - HE1
3672   3677      1 ;     CG - CZ
3672   3680      1 ;     CG - HE2
3672   3683      1 ;     CG - C
3673   3678      1 ;    CD1 - HZ
3673   3679      1 ;    CD1 - CE2
3673   3682      1 ;    CD1 - HD2
3674   3676      1 ;    HD1 - HE1
3674   3677      1 ;    HD1 - CZ
3674   3681      1 ;    HD1 - CD2
3675   3680      1 ;    CE1 - HE2
3675   3681      1 ;    CE1 - CD2
3676   3678      1 ;    HE1 - HZ
3676   3679      1 ;    HE1 - CE2
3677   3682      1 ;     CZ - HD2
3678   3680      1 ;     HZ - HE2
3678   3681      1 ;     HZ - CD2
3680   3682      1 ;    HE2 - HD2
3683   3688      1 ;      C - HA
3683   3689      1 ;      C - CB
3683   3694      1 ;      C - C
3684   3686      1 ;      O - H
3684   3687      1 ;      O - CA
3685   3690      1 ;      N - HB2
3685   3691      1 ;      N - HB3
3685   3692      1 ;      N - SG
3685   3695      1 ;      N - O
3685   3696      1 ;      N - N
3686   3688      1 ;      H - HA
3686   3689      1 ;      H - CB
3686   3694      1 ;      H - C
3687   3693      1 ;     CA - HG
3687   3697      1 ;     CA - H
3687   3698      1 ;     CA - CA
3688   3690      1 ;     HA - HB2
3688   3691      1 ;     HA - HB3
3688   3692      1 ;     HA - SG
```

| | | | | | | |
|---|---|---|---|---|---|---|
| 3688 | 3695 | 1 | ; | HA | - | O |
| 3688 | 3696 | 1 | ; | HA | - | N |
| 3689 | 3695 | 1 | ; | CB | - | O |
| 3689 | 3696 | 1 | ; | CB | - | N |
| 3690 | 3693 | 1 | ; | HB2 | - | HG |
| 3690 | 3694 | 1 | ; | HB2 | - | C |
| 3691 | 3693 | 1 | ; | HB3 | - | HG |
| 3691 | 3694 | 1 | ; | HB3 | - | C |
| 3692 | 3694 | 1 | ; | SG | - | C |
| 3694 | 3699 | 1 | ; | C | - | HA |
| 3694 | 3700 | 1 | ; | C | - | CB |
| 3694 | 3714 | 1 | ; | C | - | C |
| 3695 | 3697 | 1 | ; | O | - | H |
| 3695 | 3698 | 1 | ; | O | - | CA |
| 3696 | 3701 | 1 | ; | N | - | HB2 |
| 3696 | 3702 | 1 | ; | N | - | HB3 |
| 3696 | 3703 | 1 | ; | N | - | CG |
| 3696 | 3715 | 1 | ; | N | - | O |
| 3696 | 3716 | 1 | ; | N | - | N |
| 3697 | 3699 | 1 | ; | H | - | HA |
| 3697 | 3700 | 1 | ; | H | - | CB |
| 3697 | 3714 | 1 | ; | H | - | C |
| 3698 | 3704 | 1 | ; | CA | - | CD1 |
| 3698 | 3712 | 1 | ; | CA | - | CD2 |
| 3698 | 3717 | 1 | ; | CA | - | H |
| 3698 | 3718 | 1 | ; | CA | - | CA |
| 3699 | 3701 | 1 | ; | HA | - | HB2 |
| 3699 | 3702 | 1 | ; | HA | - | HB3 |
| 3699 | 3703 | 1 | ; | HA | - | CG |
| 3699 | 3715 | 1 | ; | HA | - | O |
| 3699 | 3716 | 1 | ; | HA | - | N |
| 3700 | 3705 | 1 | ; | CB | - | HD1 |
| 3700 | 3706 | 1 | ; | CB | - | CE1 |
| 3700 | 3710 | 1 | ; | CB | - | CE2 |
| 3700 | 3713 | 1 | ; | CB | - | HD2 |
| 3700 | 3715 | 1 | ; | CB | - | O |
| 3700 | 3716 | 1 | ; | CB | - | N |
| 3701 | 3704 | 1 | ; | HB2 | - | CD1 |
| 3701 | 3712 | 1 | ; | HB2 | - | CD2 |
| 3701 | 3714 | 1 | ; | HB2 | - | C |
| 3702 | 3704 | 1 | ; | HB3 | - | CD1 |
| 3702 | 3712 | 1 | ; | HB3 | - | CD2 |
| 3702 | 3714 | 1 | ; | HB3 | - | C |
| 3703 | 3707 | 1 | ; | CG | - | HE1 |
| 3703 | 3708 | 1 | ; | CG | - | CZ |
| 3703 | 3711 | 1 | ; | CG | - | HE2 |
| 3703 | 3714 | 1 | ; | CG | - | C |
| 3704 | 3709 | 1 | ; | CD1 | - | HZ |
| 3704 | 3710 | 1 | ; | CD1 | - | CE2 |
| 3704 | 3713 | 1 | ; | CD1 | - | HD2 |
| 3705 | 3707 | 1 | ; | HD1 | - | HE1 |
| 3705 | 3708 | 1 | ; | HD1 | - | CZ |

| | | | |
|---|---|---|---|
| 3705 | 3712 | 1 ; | HD1 - CD2 |
| 3706 | 3711 | 1 ; | CE1 - HE2 |
| 3706 | 3712 | 1 ; | CE1 - CD2 |
| 3707 | 3709 | 1 ; | HE1 - HZ |
| 3707 | 3710 | 1 ; | HE1 - CE2 |
| 3708 | 3713 | 1 ; | CZ - HD2 |
| 3709 | 3711 | 1 ; | HZ - HE2 |
| 3709 | 3712 | 1 ; | HZ - CD2 |
| 3711 | 3713 | 1 ; | HE2 - HD2 |
| 3714 | 3719 | 1 ; | C - HA |
| 3714 | 3720 | 1 ; | C - CB |
| 3714 | 3735 | 1 ; | C - C |
| 3715 | 3717 | 1 ; | O - H |
| 3715 | 3718 | 1 ; | O - CA |
| 3716 | 3721 | 1 ; | N - HB2 |
| 3716 | 3722 | 1 ; | N - HB3 |
| 3716 | 3723 | 1 ; | N - CG |
| 3716 | 3736 | 1 ; | N - O |
| 3716 | 3737 | 1 ; | N - N |
| 3717 | 3719 | 1 ; | H - HA |
| 3717 | 3720 | 1 ; | H - CB |
| 3717 | 3735 | 1 ; | H - C |
| 3718 | 3724 | 1 ; | CA - CD1 |
| 3718 | 3733 | 1 ; | CA - CD2 |
| 3718 | 3738 | 1 ; | CA - H |
| 3718 | 3739 | 1 ; | CA - CA |
| 3719 | 3721 | 1 ; | HA - HB2 |
| 3719 | 3722 | 1 ; | HA - HB3 |
| 3719 | 3723 | 1 ; | HA - CG |
| 3719 | 3736 | 1 ; | HA - O |
| 3719 | 3737 | 1 ; | HA - N |
| 3720 | 3725 | 1 ; | CB - HD1 |
| 3720 | 3726 | 1 ; | CB - CE1 |
| 3720 | 3731 | 1 ; | CB - CE2 |
| 3720 | 3734 | 1 ; | CB - HD2 |
| 3720 | 3736 | 1 ; | CB - O |
| 3720 | 3737 | 1 ; | CB - N |
| 3721 | 3724 | 1 ; | HB2 - CD1 |
| 3721 | 3733 | 1 ; | HB2 - CD2 |
| 3721 | 3735 | 1 ; | HB2 - C |
| 3722 | 3724 | 1 ; | HB3 - CD1 |
| 3722 | 3733 | 1 ; | HB3 - CD2 |
| 3722 | 3735 | 1 ; | HB3 - C |
| 3723 | 3727 | 1 ; | CG - HE1 |
| 3723 | 3728 | 1 ; | CG - CZ |
| 3723 | 3732 | 1 ; | CG - HE2 |
| 3723 | 3735 | 1 ; | CG - C |
| 3724 | 3729 | 1 ; | CD1 - OH |
| 3724 | 3731 | 1 ; | CD1 - CE2 |
| 3724 | 3734 | 1 ; | CD1 - HD2 |
| 3725 | 3727 | 1 ; | HD1 - HE1 |
| 3725 | 3728 | 1 ; | HD1 - CZ |

```
3725   3733       1 ;    HD1 - CD2
3726   3730       1 ;    CE1 - HH
3726   3732       1 ;    CE1 - HE2
3726   3733       1 ;    CE1 - CD2
3727   3729       1 ;    HE1 - OH
3727   3731       1 ;    HE1 - CE2
3728   3734       1 ;     CZ - HD2
3729   3732       1 ;     OH - HE2
3729   3733       1 ;     OH - CD2
3730   3731       1 ;     HH - CE2
3732   3734       1 ;    HE2 - HD2
3735   3740       1 ;      C - HA
3735   3741       1 ;      C - CB
3735   3750       1 ;      C - C
3736   3738       1 ;      O - H
3736   3739       1 ;      O - CA
3737   3742       1 ;      N - HB2
3737   3743       1 ;      N - HB3
3737   3744       1 ;      N - CG
3737   3751       1 ;      N - O
3737   3752       1 ;      N - N
3738   3740       1 ;      H - HA
3738   3741       1 ;      H - CB
3738   3750       1 ;      H - C
3739   3745       1 ;     CA - HG2
3739   3746       1 ;     CA - HG3
3739   3747       1 ;     CA - CD
3739   3753       1 ;     CA - H
3739   3754       1 ;     CA - CA
3740   3742       1 ;     HA - HB2
3740   3743       1 ;     HA - HB3
3740   3744       1 ;     HA - CG
3740   3751       1 ;     HA - O
3740   3752       1 ;     HA - N
3741   3748       1 ;     CB - OE1
3741   3749       1 ;     CB - OE2
3741   3751       1 ;     CB - O
3741   3752       1 ;     CB - N
3742   3745       1 ;    HB2 - HG2
3742   3746       1 ;    HB2 - HG3
3742   3747       1 ;    HB2 - CD
3742   3750       1 ;    HB2 - C
3743   3745       1 ;    HB3 - HG2
3743   3746       1 ;    HB3 - HG3
3743   3747       1 ;    HB3 - CD
3743   3750       1 ;    HB3 - C
3744   3750       1 ;     CG - C
3745   3748       1 ;    HG2 - OE1
3745   3749       1 ;    HG2 - OE2
3746   3748       1 ;    HG3 - OE1
3746   3749       1 ;    HG3 - OE2
3750   3755       1 ;      C - HA
```

```
3750   3756       1 ;      C - CB
3750   3764       1 ;      C - C
3751   3753       1 ;      O - H
3751   3754       1 ;      O - CA
3752   3757       1 ;      N - HB2
3752   3758       1 ;      N - HB3
3752   3759       1 ;      N - CG
3752   3765       1 ;      N - O
3752   3766       1 ;      N - N
3753   3755       1 ;      H - HA
3753   3756       1 ;      H - CB
3753   3764       1 ;      H - C
3754   3760       1 ;     CA - OD1
3754   3761       1 ;     CA - ND2
3754   3767       1 ;     CA - H
3754   3768       1 ;     CA - CA
3755   3757       1 ;     HA - HB2
3755   3758       1 ;     HA - HB3
3755   3759       1 ;     HA - CG
3755   3765       1 ;     HA - O
3755   3766       1 ;     HA - N
3756   3762       1 ;     CB - HD21
3756   3763       1 ;     CB - HD22
3756   3765       1 ;     CB - O
3756   3766       1 ;     CB - N
3757   3760       1 ;    HB2 - OD1
3757   3761       1 ;    HB2 - ND2
3757   3764       1 ;    HB2 - C
3758   3760       1 ;    HB3 - OD1
3758   3761       1 ;    HB3 - ND2
3758   3764       1 ;    HB3 - C
3759   3764       1 ;     CG - C
3760   3762       1 ;    OD1 - HD21
3760   3763       1 ;    OD1 - HD22
3764   3769       1 ;      C - HA
3764   3770       1 ;      C - CB
3764   3776       1 ;      C - C
3765   3767       1 ;      O - H
3765   3768       1 ;      O - CA
3766   3771       1 ;      N - HB2
3766   3772       1 ;      N - HB3
3766   3773       1 ;      N - CG
3766   3777       1 ;      N - O
3766   3778       1 ;      N - N
3767   3769       1 ;      H - HA
3767   3770       1 ;      H - CB
3767   3776       1 ;      H - C
3768   3774       1 ;     CA - OD1
3768   3775       1 ;     CA - OD2
3768   3779       1 ;     CA - H
3768   3780       1 ;     CA - CA
3769   3771       1 ;     HA - HB2
```

```
3769   3772      1 ;     HA - HB3
3769   3773      1 ;     HA - CG
3769   3777      1 ;     HA - O
3769   3778      1 ;     HA - N
3770   3777      1 ;     CB - O
3770   3778      1 ;     CB - N
3771   3774      1 ;    HB2 - OD1
3771   3775      1 ;    HB2 - OD2
3771   3776      1 ;    HB2 - C
3772   3774      1 ;    HB3 - OD1
3772   3775      1 ;    HB3 - OD2
3772   3776      1 ;    HB3 - C
3773   3776      1 ;     CG - C
3776   3781      1 ;      C - HA
3776   3782      1 ;      C - CB
3776   3800      1 ;      C - C
3777   3779      1 ;      O - H
3777   3780      1 ;      O - CA
3778   3783      1 ;      N - HB2
3778   3784      1 ;      N - HB3
3778   3785      1 ;      N - CG
3778   3801      1 ;      N - O
3778   3802      1 ;      N - N
3779   3781      1 ;      H - HA
3779   3782      1 ;      H - CB
3779   3800      1 ;      H - C
3780   3786      1 ;     CA - CD1
3780   3799      1 ;     CA - CD2
3780   3803      1 ;     CA - H
3780   3804      1 ;     CA - CA
3781   3783      1 ;     HA - HB2
3781   3784      1 ;     HA - HB3
3781   3785      1 ;     HA - CG
3781   3801      1 ;     HA - O
3781   3802      1 ;     HA - N
3782   3787      1 ;     CB - HD1
3782   3788      1 ;     CB - NE1
3782   3790      1 ;     CB - CE2
3782   3797      1 ;     CB - CE3
3782   3801      1 ;     CB - O
3782   3802      1 ;     CB - N
3783   3786      1 ;    HB2 - CD1
3783   3799      1 ;    HB2 - CD2
3783   3800      1 ;    HB2 - C
3784   3786      1 ;    HB3 - CD1
3784   3799      1 ;    HB3 - CD2
3784   3800      1 ;    HB3 - C
3785   3789      1 ;     CG - HE1
3785   3791      1 ;     CG - CZ2
3785   3795      1 ;     CG - CZ3
3785   3798      1 ;     CG - HE3
3785   3800      1 ;     CG - C
```

```
3786   3791       1 ;    CD1 - CZ2
3786   3797       1 ;    CD1 - CE3
3787   3789       1 ;    HD1 - HE1
3787   3790       1 ;    HD1 - CE2
3787   3799       1 ;    HD1 - CD2
3788   3792       1 ;    NE1 - HZ2
3788   3793       1 ;    NE1 - CH2
3788   3797       1 ;    NE1 - CE3
3789   3791       1 ;    HE1 - CZ2
3789   3799       1 ;    HE1 - CD2
3790   3794       1 ;    CE2 - HH2
3790   3795       1 ;    CE2 - CZ3
3790   3798       1 ;    CE2 - HE3
3791   3796       1 ;    CZ2 - HZ3
3791   3797       1 ;    CZ2 - CE3
3792   3794       1 ;    HZ2 - HH2
3792   3795       1 ;    HZ2 - CZ3
3792   3799       1 ;    HZ2 - CD2
3793   3798       1 ;    CH2 - HE3
3793   3799       1 ;    CH2 - CD2
3794   3796       1 ;    HH2 - HZ3
3794   3797       1 ;    HH2 - CE3
3796   3798       1 ;    HZ3 - HE3
3796   3799       1 ;    HZ3 - CD2
3800   3805       1 ;      C - HA2
3800   3806       1 ;      C - HA3
3800   3807       1 ;      C - C
3801   3803       1 ;      O - H
3801   3804       1 ;      O - CA
3802   3808       1 ;      N - O
3802   3809       1 ;      N - N
3803   3805       1 ;      H - HA2
3803   3806       1 ;      H - HA3
3803   3807       1 ;      H - C
3804   3810       1 ;     CA - H
3804   3811       1 ;     CA - CA
3805   3808       1 ;    HA2 - O
3805   3809       1 ;    HA2 - N
3806   3808       1 ;    HA3 - O
3806   3809       1 ;    HA3 - N
3807   3812       1 ;      C - HA
3807   3813       1 ;      C - CB
3807   3818       1 ;      C - C
3808   3810       1 ;      O - H
3808   3811       1 ;      O - CA
3809   3814       1 ;      N - HB2
3809   3815       1 ;      N - HB3
3809   3816       1 ;      N - OG
3809   3819       1 ;      N - O
3809   3820       1 ;      N - N
3810   3812       1 ;      H - HA
3810   3813       1 ;      H - CB
```

```
3810   3818       1 ;       H - C
3811   3817       1 ;      CA - HG
3811   3821       1 ;      CA - H
3811   3822       1 ;      CA - CA
3812   3814       1 ;      HA - HB2
3812   3815       1 ;      HA - HB3
3812   3816       1 ;      HA - OG
3812   3819       1 ;      HA - O
3812   3820       1 ;      HA - N
3813   3819       1 ;      CB - O
3813   3820       1 ;      CB - N
3814   3817       1 ;     HB2 - HG
3814   3818       1 ;     HB2 - C
3815   3817       1 ;     HB3 - HG
3815   3818       1 ;     HB3 - C
3816   3818       1 ;      OG - C
3818   3823       1 ;       C - HA
3818   3824       1 ;       C - CB
3818   3837       1 ;       C - C
3819   3821       1 ;       O - H
3819   3822       1 ;       O - CA
3820   3825       1 ;       N - HB
3820   3826       1 ;       N - CG2
3820   3830       1 ;       N - CG1
3820   3838       1 ;       N - O
3820   3839       1 ;       N - N
3821   3823       1 ;       H - HA
3821   3824       1 ;       H - CB
3821   3837       1 ;       H - C
3822   3827       1 ;      CA - HG21
3822   3828       1 ;      CA - HG22
3822   3829       1 ;      CA - HG23
3822   3831       1 ;      CA - HG12
3822   3832       1 ;      CA - HG13
3822   3833       1 ;      CA - CD1
3822   3840       1 ;      CA - H
3822   3841       1 ;      CA - CA
3823   3825       1 ;      HA - HB
3823   3826       1 ;      HA - CG2
3823   3830       1 ;      HA - CG1
3823   3838       1 ;      HA - O
3823   3839       1 ;      HA - N
3824   3834       1 ;      CB - HD11
3824   3835       1 ;      CB - HD12
3824   3836       1 ;      CB - HD13
3824   3838       1 ;      CB - O
3824   3839       1 ;      CB - N
3825   3827       1 ;      HB - HG21
3825   3828       1 ;      HB - HG22
3825   3829       1 ;      HB - HG23
3825   3831       1 ;      HB - HG12
3825   3832       1 ;      HB - HG13
```

```
3825   3833       1 ;      HB - CD1
3825   3837       1 ;      HB - C
3826   3831       1 ;     CG2 - HG12
3826   3832       1 ;     CG2 - HG13
3826   3833       1 ;     CG2 - CD1
3826   3837       1 ;     CG2 - C
3827   3830       1 ;    HG21 - CG1
3828   3830       1 ;    HG22 - CG1
3829   3830       1 ;    HG23 - CG1
3830   3837       1 ;     CG1 - C
3831   3834       1 ;    HG12 - HD11
3831   3835       1 ;    HG12 - HD12
3831   3836       1 ;    HG12 - HD13
3832   3834       1 ;    HG13 - HD11
3832   3835       1 ;    HG13 - HD12
3832   3836       1 ;    HG13 - HD13
3837   3842       1 ;       C - HA
3837   3843       1 ;       C - CB
3837   3848       1 ;       C - C
3838   3840       1 ;       O - H
3838   3841       1 ;       O - CA
3839   3844       1 ;       N - HB2
3839   3845       1 ;       N - HB3
3839   3846       1 ;       N - OG
3839   3849       1 ;       N - O
3839   3850       1 ;       N - N
3840   3842       1 ;       H - HA
3840   3843       1 ;       H - CB
3840   3848       1 ;       H - C
3841   3847       1 ;      CA - HG
3841   3851       1 ;      CA - H
3841   3852       1 ;      CA - CA
3842   3844       1 ;      HA - HB2
3842   3845       1 ;      HA - HB3
3842   3846       1 ;      HA - OG
3842   3849       1 ;      HA - O
3842   3850       1 ;      HA - N
3843   3849       1 ;      CB - O
3843   3850       1 ;      CB - N
3844   3847       1 ;     HB2 - HG
3844   3848       1 ;     HB2 - C
3845   3847       1 ;     HB3 - HG
3845   3848       1 ;     HB3 - C
3846   3848       1 ;      OG - C
3848   3853       1 ;       C - HA
3848   3854       1 ;       C - CB
3848   3859       1 ;       C - C
3849   3851       1 ;       O - H
3849   3852       1 ;       O - CA
3850   3855       1 ;       N - HB2
3850   3856       1 ;       N - HB3
3850   3857       1 ;       N - OG
```

```
3850   3860      1 ;      N - O
3850   3861      1 ;      N - N
3851   3853      1 ;      H - HA
3851   3854      1 ;      H - CB
3851   3859      1 ;      H - C
3852   3858      1 ;     CA - HG
3852   3862      1 ;     CA - H
3852   3863      1 ;     CA - CA
3853   3855      1 ;     HA - HB2
3853   3856      1 ;     HA - HB3
3853   3857      1 ;     HA - OG
3853   3860      1 ;     HA - O
3853   3861      1 ;     HA - N
3854   3860      1 ;     CB - O
3854   3861      1 ;     CB - N
3855   3858      1 ;    HB2 - HG
3855   3859      1 ;    HB2 - C
3856   3858      1 ;    HB3 - HG
3856   3859      1 ;    HB3 - C
3857   3859      1 ;     OG - C
3859   3864      1 ;      C - HA
3859   3865      1 ;      C - CB
3859   3871      1 ;      C - C
3860   3862      1 ;      O - H
3860   3863      1 ;      O - CA
3861   3866      1 ;      N - HB2
3861   3867      1 ;      N - HB3
3861   3868      1 ;      N - CG
3861   3872      1 ;      N - O
3861   3873      1 ;      N - N
3862   3864      1 ;      H - HA
3862   3865      1 ;      H - CB
3862   3871      1 ;      H - C
3863   3869      1 ;     CA - OD1
3863   3870      1 ;     CA - OD2
3863   3874      1 ;     CA - H
3863   3875      1 ;     CA - CA
3864   3866      1 ;     HA - HB2
3864   3867      1 ;     HA - HB3
3864   3868      1 ;     HA - CG
3864   3872      1 ;     HA - O
3864   3873      1 ;     HA - N
3865   3872      1 ;     CB - O
3865   3873      1 ;     CB - N
3866   3869      1 ;    HB2 - OD1
3866   3870      1 ;    HB2 - OD2
3866   3871      1 ;    HB2 - C
3867   3869      1 ;    HB3 - OD1
3867   3870      1 ;    HB3 - OD2
3867   3871      1 ;    HB3 - C
3868   3871      1 ;     CG - C
3871   3876      1 ;      C - HA
```

| | | | | |
|---|---|---|---|---|
| 3871 | 3877 | 1 ; | C - | CB |
| 3871 | 3881 | 1 ; | C - | C |
| 3872 | 3874 | 1 ; | O - | H |
| 3872 | 3875 | 1 ; | O - | CA |
| 3873 | 3878 | 1 ; | N - | HB1 |
| 3873 | 3879 | 1 ; | N - | HB2 |
| 3873 | 3880 | 1 ; | N - | HB3 |
| 3873 | 3882 | 1 ; | N - | O |
| 3873 | 3883 | 1 ; | N - | N |
| 3874 | 3876 | 1 ; | H - | HA |
| 3874 | 3877 | 1 ; | H - | CB |
| 3874 | 3881 | 1 ; | H - | C |
| 3875 | 3884 | 1 ; | CA - | H |
| 3875 | 3885 | 1 ; | CA - | CA |
| 3876 | 3878 | 1 ; | HA - | HB1 |
| 3876 | 3879 | 1 ; | HA - | HB2 |
| 3876 | 3880 | 1 ; | HA - | HB3 |
| 3876 | 3882 | 1 ; | HA - | O |
| 3876 | 3883 | 1 ; | HA - | N |
| 3877 | 3882 | 1 ; | CB - | O |
| 3877 | 3883 | 1 ; | CB - | N |
| 3878 | 3881 | 1 ; | HB1 - | C |
| 3879 | 3881 | 1 ; | HB2 - | C |
| 3880 | 3881 | 1 ; | HB3 - | C |
| 3881 | 3886 | 1 ; | C - | HA |
| 3881 | 3887 | 1 ; | C - | CB |
| 3881 | 3903 | 1 ; | C - | C |
| 3882 | 3884 | 1 ; | O - | H |
| 3882 | 3885 | 1 ; | O - | CA |
| 3883 | 3888 | 1 ; | N - | HB2 |
| 3883 | 3889 | 1 ; | N - | HB3 |
| 3883 | 3890 | 1 ; | N - | CG |
| 3883 | 3904 | 1 ; | N - | O |
| 3883 | 3905 | 1 ; | N - | N |
| 3884 | 3886 | 1 ; | H - | HA |
| 3884 | 3887 | 1 ; | H - | CB |
| 3884 | 3903 | 1 ; | H - | C |
| 3885 | 3891 | 1 ; | CA - | HG2 |
| 3885 | 3892 | 1 ; | CA - | HG3 |
| 3885 | 3893 | 1 ; | CA - | CD |
| 3885 | 3906 | 1 ; | CA - | H |
| 3885 | 3907 | 1 ; | CA - | CA |
| 3886 | 3888 | 1 ; | HA - | HB2 |
| 3886 | 3889 | 1 ; | HA - | HB3 |
| 3886 | 3890 | 1 ; | HA - | CG |
| 3886 | 3904 | 1 ; | HA - | O |
| 3886 | 3905 | 1 ; | HA - | N |
| 3887 | 3894 | 1 ; | CB - | HD2 |
| 3887 | 3895 | 1 ; | CB - | HD3 |
| 3887 | 3896 | 1 ; | CB - | CE |
| 3887 | 3904 | 1 ; | CB - | O |
| 3887 | 3905 | 1 ; | CB - | N |

| | | | |
|---|---|---|---|
| 3888 | 3891 | 1 ; | HB2 - HG2 |
| 3888 | 3892 | 1 ; | HB2 - HG3 |
| 3888 | 3893 | 1 ; | HB2 - CD |
| 3888 | 3903 | 1 ; | HB2 - C |
| 3889 | 3891 | 1 ; | HB3 - HG2 |
| 3889 | 3892 | 1 ; | HB3 - HG3 |
| 3889 | 3893 | 1 ; | HB3 - CD |
| 3889 | 3903 | 1 ; | HB3 - C |
| 3890 | 3897 | 1 ; | CG - HE2 |
| 3890 | 3898 | 1 ; | CG - HE3 |
| 3890 | 3899 | 1 ; | CG - NZ |
| 3890 | 3903 | 1 ; | CG - C |
| 3891 | 3894 | 1 ; | HG2 - HD2 |
| 3891 | 3895 | 1 ; | HG2 - HD3 |
| 3891 | 3896 | 1 ; | HG2 - CE |
| 3892 | 3894 | 1 ; | HG3 - HD2 |
| 3892 | 3895 | 1 ; | HG3 - HD3 |
| 3892 | 3896 | 1 ; | HG3 - CE |
| 3893 | 3900 | 1 ; | CD - HZ1 |
| 3893 | 3901 | 1 ; | CD - HZ2 |
| 3893 | 3902 | 1 ; | CD - HZ3 |
| 3894 | 3897 | 1 ; | HD2 - HE2 |
| 3894 | 3898 | 1 ; | HD2 - HE3 |
| 3894 | 3899 | 1 ; | HD2 - NZ |
| 3895 | 3897 | 1 ; | HD3 - HE2 |
| 3895 | 3898 | 1 ; | HD3 - HE3 |
| 3895 | 3899 | 1 ; | HD3 - NZ |
| 3897 | 3900 | 1 ; | HE2 - HZ1 |
| 3897 | 3901 | 1 ; | HE2 - HZ2 |
| 3897 | 3902 | 1 ; | HE2 - HZ3 |
| 3898 | 3900 | 1 ; | HE3 - HZ1 |
| 3898 | 3901 | 1 ; | HE3 - HZ2 |
| 3898 | 3902 | 1 ; | HE3 - HZ3 |
| 3903 | 3908 | 1 ; | C - HA |
| 3903 | 3909 | 1 ; | C - CB |
| 3903 | 3917 | 1 ; | C - C |
| 3904 | 3906 | 1 ; | O - H |
| 3904 | 3907 | 1 ; | O - CA |
| 3905 | 3910 | 1 ; | N - HB2 |
| 3905 | 3911 | 1 ; | N - HB3 |
| 3905 | 3912 | 1 ; | N - CG |
| 3905 | 3918 | 1 ; | N - O |
| 3905 | 3919 | 1 ; | N - N |
| 3906 | 3908 | 1 ; | H - HA |
| 3906 | 3909 | 1 ; | H - CB |
| 3906 | 3917 | 1 ; | H - C |
| 3907 | 3913 | 1 ; | CA - OD1 |
| 3907 | 3914 | 1 ; | CA - ND2 |
| 3907 | 3920 | 1 ; | CA - H |
| 3907 | 3921 | 1 ; | CA - CA |
| 3908 | 3910 | 1 ; | HA - HB2 |
| 3908 | 3911 | 1 ; | HA - HB3 |

```
3908   3912       1 ;      HA - CG
3908   3918       1 ;      HA - O
3908   3919       1 ;      HA - N
3909   3915       1 ;      CB - HD21
3909   3916       1 ;      CB - HD22
3909   3918       1 ;      CB - O
3909   3919       1 ;      CB - N
3910   3913       1 ;     HB2 - OD1
3910   3914       1 ;     HB2 - ND2
3910   3917       1 ;     HB2 - C
3911   3913       1 ;     HB3 - OD1
3911   3914       1 ;     HB3 - ND2
3911   3917       1 ;     HB3 - C
3912   3917       1 ;      CG - C
3913   3915       1 ;     OD1 - HD21
3913   3916       1 ;     OD1 - HD22
3917   3922       1 ;       C - HA
3917   3923       1 ;       C - CB
3917   3936       1 ;       C - C
3918   3920       1 ;       O - H
3918   3921       1 ;       O - CA
3919   3924       1 ;       N - HB2
3919   3925       1 ;       N - HB3
3919   3926       1 ;       N - CG
3919   3937       1 ;       N - O
3919   3938       1 ;       N - N
3920   3922       1 ;       H - HA
3920   3923       1 ;       H - CB
3920   3936       1 ;       H - C
3921   3927       1 ;      CA - HG
3921   3928       1 ;      CA - CD1
3921   3932       1 ;      CA - CD2
3921   3939       1 ;      CA - H
3921   3940       1 ;      CA - CA
3922   3924       1 ;      HA - HB2
3922   3925       1 ;      HA - HB3
3922   3926       1 ;      HA - CG
3922   3937       1 ;      HA - O
3922   3938       1 ;      HA - N
3923   3929       1 ;      CB - HD11
3923   3930       1 ;      CB - HD12
3923   3931       1 ;      CB - HD13
3923   3933       1 ;      CB - HD21
3923   3934       1 ;      CB - HD22
3923   3935       1 ;      CB - HD23
3923   3937       1 ;      CB - O
3923   3938       1 ;      CB - N
3924   3927       1 ;     HB2 - HG
3924   3928       1 ;     HB2 - CD1
3924   3932       1 ;     HB2 - CD2
3924   3936       1 ;     HB2 - C
3925   3927       1 ;     HB3 - HG
```

```
3925   3928       1 ;     HB3 - CD1
3925   3932       1 ;     HB3 - CD2
3925   3936       1 ;     HB3 - C
3926   3936       1 ;      CG - C
3927   3929       1 ;      HG - HD11
3927   3930       1 ;      HG - HD12
3927   3931       1 ;      HG - HD13
3927   3933       1 ;      HG - HD21
3927   3934       1 ;      HG - HD22
3927   3935       1 ;      HG - HD23
3928   3933       1 ;     CD1 - HD21
3928   3934       1 ;     CD1 - HD22
3928   3935       1 ;     CD1 - HD23
3929   3932       1 ;    HD11 - CD2
3930   3932       1 ;    HD12 - CD2
3931   3932       1 ;    HD13 - CD2
3936   3941       1 ;       C - HA
3936   3942       1 ;       C - CB
3936   3955       1 ;       C - C
3937   3939       1 ;       O - H
3937   3940       1 ;       O - CA
3938   3943       1 ;       N - HB
3938   3944       1 ;       N - CG2
3938   3948       1 ;       N - CG1
3938   3956       1 ;       N - O
3938   3957       1 ;       N - N
3939   3941       1 ;       H - HA
3939   3942       1 ;       H - CB
3939   3955       1 ;       H - C
3940   3945       1 ;      CA - HG21
3940   3946       1 ;      CA - HG22
3940   3947       1 ;      CA - HG23
3940   3949       1 ;      CA - HG12
3940   3950       1 ;      CA - HG13
3940   3951       1 ;      CA - CD1
3940   3958       1 ;      CA - H
3940   3959       1 ;      CA - CA
3941   3943       1 ;      HA - HB
3941   3944       1 ;      HA - CG2
3941   3948       1 ;      HA - CG1
3941   3956       1 ;      HA - O
3941   3957       1 ;      HA - N
3942   3952       1 ;      CB - HD11
3942   3953       1 ;      CB - HD12
3942   3954       1 ;      CB - HD13
3942   3956       1 ;      CB - O
3942   3957       1 ;      CB - N
3943   3945       1 ;      HB - HG21
3943   3946       1 ;      HB - HG22
3943   3947       1 ;      HB - HG23
3943   3949       1 ;      HB - HG12
3943   3950       1 ;      HB - HG13
```

```
3943   3951       1 ;      HB - CD1
3943   3955       1 ;      HB - C
3944   3949       1 ;     CG2 - HG12
3944   3950       1 ;     CG2 - HG13
3944   3951       1 ;     CG2 - CD1
3944   3955       1 ;     CG2 - C
3945   3948       1 ;    HG21 - CG1
3946   3948       1 ;    HG22 - CG1
3947   3948       1 ;    HG23 - CG1
3948   3955       1 ;     CG1 - C
3949   3952       1 ;    HG12 - HD11
3949   3953       1 ;    HG12 - HD12
3949   3954       1 ;    HG12 - HD13
3950   3952       1 ;    HG13 - HD11
3950   3953       1 ;    HG13 - HD12
3950   3954       1 ;    HG13 - HD13
3955   3960       1 ;       C - HA
3955   3961       1 ;       C - CB
3955   3969       1 ;       C - C
3956   3958       1 ;       O - H
3956   3959       1 ;       O - CA
3957   3962       1 ;       N - HB
3957   3963       1 ;       N - CG2
3957   3967       1 ;       N - OG1
3957   3970       1 ;       N - O
3957   3971       1 ;       N - N
3958   3960       1 ;       H - HA
3958   3961       1 ;       H - CB
3958   3969       1 ;       H - C
3959   3964       1 ;      CA - HG21
3959   3965       1 ;      CA - HG22
3959   3966       1 ;      CA - HG23
3959   3968       1 ;      CA - HG1
3959   3972       1 ;      CA - H
3959   3973       1 ;      CA - CA
3960   3962       1 ;      HA - HB
3960   3963       1 ;      HA - CG2
3960   3967       1 ;      HA - OG1
3960   3970       1 ;      HA - O
3960   3971       1 ;      HA - N
3961   3970       1 ;      CB - O
3961   3971       1 ;      CB - N
3962   3964       1 ;      HB - HG21
3962   3965       1 ;      HB - HG22
3962   3966       1 ;      HB - HG23
3962   3968       1 ;      HB - HG1
3962   3969       1 ;      HB - C
3963   3968       1 ;     CG2 - HG1
3963   3969       1 ;     CG2 - C
3964   3967       1 ;    HG21 - OG1
3965   3967       1 ;    HG22 - OG1
3966   3967       1 ;    HG23 - OG1
```

```
3967   3969       1 ;     OG1 - C
3969   3974       1 ;       C - HA
3969   3975       1 ;       C - CB
3969   3991       1 ;       C - C
3970   3972       1 ;       O - H
3970   3973       1 ;       O - CA
3971   3976       1 ;       N - HB2
3971   3977       1 ;       N - HB3
3971   3978       1 ;       N - CG
3971   3992       1 ;       N - O
3971   3993       1 ;       N - N
3972   3974       1 ;       H - HA
3972   3975       1 ;       H - CB
3972   3991       1 ;       H - C
3973   3979       1 ;      CA - HG2
3973   3980       1 ;      CA - HG3
3973   3981       1 ;      CA - CD
3973   3994       1 ;      CA - H
3973   3995       1 ;      CA - CA
3974   3976       1 ;      HA - HB2
3974   3977       1 ;      HA - HB3
3974   3978       1 ;      HA - CG
3974   3992       1 ;      HA - O
3974   3993       1 ;      HA - N
3975   3982       1 ;      CB - HD2
3975   3983       1 ;      CB - HD3
3975   3984       1 ;      CB - CE
3975   3992       1 ;      CB - O
3975   3993       1 ;      CB - N
3976   3979       1 ;     HB2 - HG2
3976   3980       1 ;     HB2 - HG3
3976   3981       1 ;     HB2 - CD
3976   3991       1 ;     HB2 - C
3977   3979       1 ;     HB3 - HG2
3977   3980       1 ;     HB3 - HG3
3977   3981       1 ;     HB3 - CD
3977   3991       1 ;     HB3 - C
3978   3985       1 ;      CG - HE2
3978   3986       1 ;      CG - HE3
3978   3987       1 ;      CG - NZ
3978   3991       1 ;      CG - C
3979   3982       1 ;     HG2 - HD2
3979   3983       1 ;     HG2 - HD3
3979   3984       1 ;     HG2 - CE
3980   3982       1 ;     HG3 - HD2
3980   3983       1 ;     HG3 - HD3
3980   3984       1 ;     HG3 - CE
3981   3988       1 ;      CD - HZ1
3981   3989       1 ;      CD - HZ2
3981   3990       1 ;      CD - HZ3
3982   3985       1 ;     HD2 - HE2
3982   3986       1 ;     HD2 - HE3
```

| | | | | | | |
|---|---|---|---|---|---|---|
| 3982 | 3987 | 1 | ; | HD2 | - | NZ |
| 3983 | 3985 | 1 | ; | HD3 | - | HE2 |
| 3983 | 3986 | 1 | ; | HD3 | - | HE3 |
| 3983 | 3987 | 1 | ; | HD3 | - | NZ |
| 3985 | 3988 | 1 | ; | HE2 | - | HZ1 |
| 3985 | 3989 | 1 | ; | HE2 | - | HZ2 |
| 3985 | 3990 | 1 | ; | HE2 | - | HZ3 |
| 3986 | 3988 | 1 | ; | HE3 | - | HZ1 |
| 3986 | 3989 | 1 | ; | HE3 | - | HZ2 |
| 3986 | 3990 | 1 | ; | HE3 | - | HZ3 |
| 3991 | 3996 | 1 | ; | C | - | HA |
| 3991 | 3997 | 1 | ; | C | - | CB |
| 3991 | 4010 | 1 | ; | C | - | C |
| 3992 | 3994 | 1 | ; | O | - | H |
| 3992 | 3995 | 1 | ; | O | - | CA |
| 3993 | 3998 | 1 | ; | N | - | HB2 |
| 3993 | 3999 | 1 | ; | N | - | HB3 |
| 3993 | 4000 | 1 | ; | N | - | CG |
| 3993 | 4011 | 1 | ; | N | - | O |
| 3993 | 4012 | 1 | ; | N | - | N |
| 3994 | 3996 | 1 | ; | H | - | HA |
| 3994 | 3997 | 1 | ; | H | - | CB |
| 3994 | 4010 | 1 | ; | H | - | C |
| 3995 | 4001 | 1 | ; | CA | - | HG |
| 3995 | 4002 | 1 | ; | CA | - | CD1 |
| 3995 | 4006 | 1 | ; | CA | - | CD2 |
| 3995 | 4013 | 1 | ; | CA | - | H |
| 3995 | 4014 | 1 | ; | CA | - | CA |
| 3996 | 3998 | 1 | ; | HA | - | HB2 |
| 3996 | 3999 | 1 | ; | HA | - | HB3 |
| 3996 | 4000 | 1 | ; | HA | - | CG |
| 3996 | 4011 | 1 | ; | HA | - | O |
| 3996 | 4012 | 1 | ; | HA | - | N |
| 3997 | 4003 | 1 | ; | CB | - | HD11 |
| 3997 | 4004 | 1 | ; | CB | - | HD12 |
| 3997 | 4005 | 1 | ; | CB | - | HD13 |
| 3997 | 4007 | 1 | ; | CB | - | HD21 |
| 3997 | 4008 | 1 | ; | CB | - | HD22 |
| 3997 | 4009 | 1 | ; | CB | - | HD23 |
| 3997 | 4011 | 1 | ; | CB | - | O |
| 3997 | 4012 | 1 | ; | CB | - | N |
| 3998 | 4001 | 1 | ; | HB2 | - | HG |
| 3998 | 4002 | 1 | ; | HB2 | - | CD1 |
| 3998 | 4006 | 1 | ; | HB2 | - | CD2 |
| 3998 | 4010 | 1 | ; | HB2 | - | C |
| 3999 | 4001 | 1 | ; | HB3 | - | HG |
| 3999 | 4002 | 1 | ; | HB3 | - | CD1 |
| 3999 | 4006 | 1 | ; | HB3 | - | CD2 |
| 3999 | 4010 | 1 | ; | HB3 | - | C |
| 4000 | 4010 | 1 | ; | CG | - | C |
| 4001 | 4003 | 1 | ; | HG | - | HD11 |
| 4001 | 4004 | 1 | ; | HG | - | HD12 |

```
4001   4005      1 ;     HG - HD13
4001   4007      1 ;     HG - HD21
4001   4008      1 ;     HG - HD22
4001   4009      1 ;     HG - HD23
4002   4007      1 ;    CD1 - HD21
4002   4008      1 ;    CD1 - HD22
4002   4009      1 ;    CD1 - HD23
4003   4006      1 ;   HD11 - CD2
4004   4006      1 ;   HD12 - CD2
4005   4006      1 ;   HD13 - CD2
4010   4015      1 ;      C - HA
4010   4016      1 ;      C - CB
4010   4029      1 ;      C - C
4011   4013      1 ;      O - H
4011   4014      1 ;      O - CA
4012   4017      1 ;      N - HB2
4012   4018      1 ;      N - HB3
4012   4019      1 ;      N - CG
4012   4030      1 ;      N - O
4012   4031      1 ;      N - N
4013   4015      1 ;      H - HA
4013   4016      1 ;      H - CB
4013   4029      1 ;      H - C
4014   4020      1 ;     CA - HG
4014   4021      1 ;     CA - CD1
4014   4025      1 ;     CA - CD2
4014   4032      1 ;     CA - H
4014   4033      1 ;     CA - CA
4015   4017      1 ;     HA - HB2
4015   4018      1 ;     HA - HB3
4015   4019      1 ;     HA - CG
4015   4030      1 ;     HA - O
4015   4031      1 ;     HA - N
4016   4022      1 ;     CB - HD11
4016   4023      1 ;     CB - HD12
4016   4024      1 ;     CB - HD13
4016   4026      1 ;     CB - HD21
4016   4027      1 ;     CB - HD22
4016   4028      1 ;     CB - HD23
4016   4030      1 ;     CB - O
4016   4031      1 ;     CB - N
4017   4020      1 ;    HB2 - HG
4017   4021      1 ;    HB2 - CD1
4017   4025      1 ;    HB2 - CD2
4017   4029      1 ;    HB2 - C
4018   4020      1 ;    HB3 - HG
4018   4021      1 ;    HB3 - CD1
4018   4025      1 ;    HB3 - CD2
4018   4029      1 ;    HB3 - C
4019   4029      1 ;     CG - C
4020   4022      1 ;     HG - HD11
4020   4023      1 ;     HG - HD12
```

```
4020   4024       1 ;      HG - HD13
4020   4026       1 ;      HG - HD21
4020   4027       1 ;      HG - HD22
4020   4028       1 ;      HG - HD23
4021   4026       1 ;     CD1 - HD21
4021   4027       1 ;     CD1 - HD22
4021   4028       1 ;     CD1 - HD23
4022   4025       1 ;    HD11 - CD2
4023   4025       1 ;    HD12 - CD2
4024   4025       1 ;    HD13 - CD2
4029   4034       1 ;       C - HA
4029   4035       1 ;       C - CB
4029   4043       1 ;       C - C
4030   4032       1 ;       O - H
4030   4033       1 ;       O - CA
4031   4036       1 ;       N - HB
4031   4037       1 ;       N - CG2
4031   4041       1 ;       N - OG1
4031   4044       1 ;       N - O
4031   4045       1 ;       N - N
4032   4034       1 ;       H - HA
4032   4035       1 ;       H - CB
4032   4043       1 ;       H - C
4033   4038       1 ;      CA - HG21
4033   4039       1 ;      CA - HG22
4033   4040       1 ;      CA - HG23
4033   4042       1 ;      CA - HG1
4033   4046       1 ;      CA - H
4033   4047       1 ;      CA - CA
4034   4036       1 ;      HA - HB
4034   4037       1 ;      HA - CG2
4034   4041       1 ;      HA - OG1
4034   4044       1 ;      HA - O
4034   4045       1 ;      HA - N
4035   4044       1 ;      CB - O
4035   4045       1 ;      CB - N
4036   4038       1 ;      HB - HG21
4036   4039       1 ;      HB - HG22
4036   4040       1 ;      HB - HG23
4036   4042       1 ;      HB - HG1
4036   4043       1 ;      HB - C
4037   4042       1 ;     CG2 - HG1
4037   4043       1 ;     CG2 - C
4038   4041       1 ;    HG21 - OG1
4039   4041       1 ;    HG22 - OG1
4040   4041       1 ;    HG23 - OG1
4041   4043       1 ;     OG1 - C
4043   4048       1 ;       C - HA
4043   4049       1 ;       C - CB
4043   4064       1 ;       C - C
4044   4046       1 ;       O - H
4044   4047       1 ;       O - CA
```

```
4045   4050       1 ;       N - HB2
4045   4051       1 ;       N - HB3
4045   4052       1 ;       N - CG
4045   4065       1 ;       N - O
4045   4066       1 ;       N - N
4046   4048       1 ;       H - HA
4046   4049       1 ;       H - CB
4046   4064       1 ;       H - C
4047   4053       1 ;      CA - CD1
4047   4062       1 ;      CA - CD2
4047   4067       1 ;      CA - H
4047   4068       1 ;      CA - CA
4048   4050       1 ;      HA - HB2
4048   4051       1 ;      HA - HB3
4048   4052       1 ;      HA - CG
4048   4065       1 ;      HA - O
4048   4066       1 ;      HA - N
4049   4054       1 ;      CB - HD1
4049   4055       1 ;      CB - CE1
4049   4060       1 ;      CB - CE2
4049   4063       1 ;      CB - HD2
4049   4065       1 ;      CB - O
4049   4066       1 ;      CB - N
4050   4053       1 ;     HB2 - CD1
4050   4062       1 ;     HB2 - CD2
4050   4064       1 ;     HB2 - C
4051   4053       1 ;     HB3 - CD1
4051   4062       1 ;     HB3 - CD2
4051   4064       1 ;     HB3 - C
4052   4056       1 ;      CG - HE1
4052   4057       1 ;      CG - CZ
4052   4061       1 ;      CG - HE2
4052   4064       1 ;      CG - C
4053   4058       1 ;     CD1 - OH
4053   4060       1 ;     CD1 - CE2
4053   4063       1 ;     CD1 - HD2
4054   4056       1 ;     HD1 - HE1
4054   4057       1 ;     HD1 - CZ
4054   4062       1 ;     HD1 - CD2
4055   4059       1 ;     CE1 - HH
4055   4061       1 ;     CE1 - HE2
4055   4062       1 ;     CE1 - CD2
4056   4058       1 ;     HE1 - OH
4056   4060       1 ;     HE1 - CE2
4057   4063       1 ;      CZ - HD2
4058   4061       1 ;      OH - HE2
4058   4062       1 ;      OH - CD2
4059   4060       1 ;      HH - CE2
4061   4063       1 ;     HE2 - HD2
4064   4069       1 ;       C - HA
4064   4070       1 ;       C - CB
4064   4078       1 ;       C - C
```

```
4065   4067       1 ;       O - H
4065   4068       1 ;       O - CA
4066   4071       1 ;       N - HB2
4066   4072       1 ;       N - HB3
4066   4073       1 ;       N - CG
4066   4079       1 ;       N - O
4066   4080       1 ;       N - N
4067   4069       1 ;       H - HA
4067   4070       1 ;       H - CB
4067   4078       1 ;       H - C
4068   4074       1 ;      CA - OD1
4068   4075       1 ;      CA - ND2
4068   4081       1 ;      CA - CD
4068   4090       1 ;      CA - CA
4069   4071       1 ;      HA - HB2
4069   4072       1 ;      HA - HB3
4069   4073       1 ;      HA - CG
4069   4079       1 ;      HA - O
4069   4080       1 ;      HA - N
4070   4076       1 ;      CB - HD21
4070   4077       1 ;      CB - HD22
4070   4079       1 ;      CB - O
4070   4080       1 ;      CB - N
4071   4074       1 ;     HB2 - OD1
4071   4075       1 ;     HB2 - ND2
4071   4078       1 ;     HB2 - C
4072   4074       1 ;     HB3 - OD1
4072   4075       1 ;     HB3 - ND2
4072   4078       1 ;     HB3 - C
4073   4078       1 ;      CG - C
4074   4076       1 ;     OD1 - HD21
4074   4077       1 ;     OD1 - HD22
4078   4082       1 ;       C - HD2
4078   4083       1 ;       C - HD3
4078   4084       1 ;       C - CG
4078   4087       1 ;       C - CB
4078   4091       1 ;       C - HA
4078   4092       1 ;       C - C
4079   4081       1 ;       O - CD
4079   4090       1 ;       O - CA
4080   4085       1 ;       N - HG2
4080   4086       1 ;       N - HG3
4080   4088       1 ;       N - HB2
4080   4089       1 ;       N - HB3
4080   4093       1 ;       N - O
4080   4094       1 ;       N - N
4081   4088       1 ;      CD - HB2
4081   4089       1 ;      CD - HB3
4081   4091       1 ;      CD - HA
4081   4092       1 ;      CD - C
4082   4085       1 ;     HD2 - HG2
4082   4086       1 ;     HD2 - HG3
```

```
4082   4087       1 ;    HD2 - CB
4082   4090       1 ;    HD2 - CA
4083   4085       1 ;    HD3 - HG2
4083   4086       1 ;    HD3 - HG3
4083   4087       1 ;    HD3 - CB
4083   4090       1 ;    HD3 - CA
4084   4091       1 ;     CG - HA
4084   4092       1 ;     CG - C
4085   4088       1 ;    HG2 - HB2
4085   4089       1 ;    HG2 - HB3
4085   4090       1 ;    HG2 - CA
4086   4088       1 ;    HG3 - HB2
4086   4089       1 ;    HG3 - HB3
4086   4090       1 ;    HG3 - CA
4087   4093       1 ;     CB - O
4087   4094       1 ;     CB - N
4088   4091       1 ;    HB2 - HA
4088   4092       1 ;    HB2 - C
4089   4091       1 ;    HB3 - HA
4089   4092       1 ;    HB3 - C
4090   4095       1 ;     CA - H
4090   4096       1 ;     CA - CA
4091   4093       1 ;     HA - O
4091   4094       1 ;     HA - N
4092   4097       1 ;      C - HA
4092   4098       1 ;      C - CB
4092   4106       1 ;      C - C
4093   4095       1 ;      O - H
4093   4096       1 ;      O - CA
4094   4099       1 ;      N - HB2
4094   4100       1 ;      N - HB3
4094   4101       1 ;      N - CG
4094   4107       1 ;      N - O
4094   4108       1 ;      N - N
4095   4097       1 ;      H - HA
4095   4098       1 ;      H - CB
4095   4106       1 ;      H - C
4096   4102       1 ;     CA - OD1
4096   4103       1 ;     CA - ND2
4096   4109       1 ;     CA - H
4096   4110       1 ;     CA - CA
4097   4099       1 ;     HA - HB2
4097   4100       1 ;     HA - HB3
4097   4101       1 ;     HA - CG
4097   4107       1 ;     HA - O
4097   4108       1 ;     HA - N
4098   4104       1 ;     CB - HD21
4098   4105       1 ;     CB - HD22
4098   4107       1 ;     CB - O
4098   4108       1 ;     CB - N
4099   4102       1 ;    HB2 - OD1
4099   4103       1 ;    HB2 - ND2
```

```
4099   4106      1 ;    HB2 - C
4100   4102      1 ;    HB3 - OD1
4100   4103      1 ;    HB3 - ND2
4100   4106      1 ;    HB3 - C
4101   4106      1 ;     CG - C
4102   4104      1 ;    OD1 - HD21
4102   4105      1 ;    OD1 - HD22
4106   4111      1 ;      C - HA
4106   4112      1 ;      C - CB
4106   4121      1 ;      C - C
4107   4109      1 ;      O - H
4107   4110      1 ;      O - CA
4108   4113      1 ;      N - HB2
4108   4114      1 ;      N - HB3
4108   4115      1 ;      N - CG
4108   4122      1 ;      N - O
4108   4123      1 ;      N - N
4109   4111      1 ;      H - HA
4109   4112      1 ;      H - CB
4109   4121      1 ;      H - C
4110   4116      1 ;     CA - HG2
4110   4117      1 ;     CA - HG3
4110   4118      1 ;     CA - CD
4110   4124      1 ;     CA - H
4110   4125      1 ;     CA - CA
4111   4113      1 ;     HA - HB2
4111   4114      1 ;     HA - HB3
4111   4115      1 ;     HA - CG
4111   4122      1 ;     HA - O
4111   4123      1 ;     HA - N
4112   4119      1 ;     CB - OE1
4112   4120      1 ;     CB - OE2
4112   4122      1 ;     CB - O
4112   4123      1 ;     CB - N
4113   4116      1 ;    HB2 - HG2
4113   4117      1 ;    HB2 - HG3
4113   4118      1 ;    HB2 - CD
4113   4121      1 ;    HB2 - C
4114   4116      1 ;    HB3 - HG2
4114   4117      1 ;    HB3 - HG3
4114   4118      1 ;    HB3 - CD
4114   4121      1 ;    HB3 - C
4115   4121      1 ;     CG - C
4116   4119      1 ;    HG2 - OE1
4116   4120      1 ;    HG2 - OE2
4117   4119      1 ;    HG3 - OE1
4117   4120      1 ;    HG3 - OE2
4121   4126      1 ;      C - HA
4121   4127      1 ;      C - CB
4121   4145      1 ;      C - C
4122   4124      1 ;      O - H
4122   4125      1 ;      O - CA
```

```
4123   4128       1 ;       N - HB2
4123   4129       1 ;       N - HB3
4123   4130       1 ;       N - CG
4123   4146       1 ;       N - O
4123   4147       1 ;       N - N
4124   4126       1 ;       H - HA
4124   4127       1 ;       H - CB
4124   4145       1 ;       H - C
4125   4131       1 ;      CA - HG2
4125   4132       1 ;      CA - HG3
4125   4133       1 ;      CA - CD
4125   4148       1 ;      CA - H
4125   4149       1 ;      CA - CA
4126   4128       1 ;      HA - HB2
4126   4129       1 ;      HA - HB3
4126   4130       1 ;      HA - CG
4126   4146       1 ;      HA - O
4126   4147       1 ;      HA - N
4127   4134       1 ;      CB - HD2
4127   4135       1 ;      CB - HD3
4127   4136       1 ;      CB - NE
4127   4146       1 ;      CB - O
4127   4147       1 ;      CB - N
4128   4131       1 ;     HB2 - HG2
4128   4132       1 ;     HB2 - HG3
4128   4133       1 ;     HB2 - CD
4128   4145       1 ;     HB2 - C
4129   4131       1 ;     HB3 - HG2
4129   4132       1 ;     HB3 - HG3
4129   4133       1 ;     HB3 - CD
4129   4145       1 ;     HB3 - C
4130   4137       1 ;      CG - HE
4130   4138       1 ;      CG - CZ
4130   4145       1 ;      CG - C
4131   4134       1 ;     HG2 - HD2
4131   4135       1 ;     HG2 - HD3
4131   4136       1 ;     HG2 - NE
4132   4134       1 ;     HG3 - HD2
4132   4135       1 ;     HG3 - HD3
4132   4136       1 ;     HG3 - NE
4133   4139       1 ;      CD - NH1
4133   4142       1 ;      CD - NH2
4134   4137       1 ;     HD2 - HE
4134   4138       1 ;     HD2 - CZ
4135   4137       1 ;     HD3 - HE
4135   4138       1 ;     HD3 - CZ
4136   4140       1 ;      NE - HH11
4136   4141       1 ;      NE - HH12
4136   4143       1 ;      NE - HH21
4136   4144       1 ;      NE - HH22
4137   4139       1 ;      HE - NH1
4137   4142       1 ;      HE - NH2
```

```
4139   4143       1 ;    NH1 - HH21
4139   4144       1 ;    NH1 - HH22
4140   4142       1 ;   HH11 - NH2
4141   4142       1 ;   HH12 - NH2
4145   4150       1 ;      C - HA
4145   4151       1 ;      C - CB
4145   4156       1 ;      C - C
4146   4148       1 ;      O - H
4146   4149       1 ;      O - CA
4147   4152       1 ;      N - HB2
4147   4153       1 ;      N - HB3
4147   4154       1 ;      N - SG
4147   4157       1 ;      N - O
4147   4158       1 ;      N - N
4148   4150       1 ;      H - HA
4148   4151       1 ;      H - CB
4148   4156       1 ;      H - C
4149   4155       1 ;     CA - HG
4149   4159       1 ;     CA - H
4149   4160       1 ;     CA - CA
4150   4152       1 ;     HA - HB2
4150   4153       1 ;     HA - HB3
4150   4154       1 ;     HA - SG
4150   4157       1 ;     HA - O
4150   4158       1 ;     HA - N
4151   4157       1 ;     CB - O
4151   4158       1 ;     CB - N
4152   4155       1 ;    HB2 - HG
4152   4156       1 ;    HB2 - C
4153   4155       1 ;    HB3 - HG
4153   4156       1 ;    HB3 - C
4154   4156       1 ;     SG - C
4156   4161       1 ;      C - HA
4156   4162       1 ;      C - CB
4156   4170       1 ;      C - C
4157   4159       1 ;      O - H
4157   4160       1 ;      O - CA
4158   4163       1 ;      N - HB
4158   4164       1 ;      N - CG2
4158   4168       1 ;      N - OG1
4158   4171       1 ;      N - O
4158   4172       1 ;      N - N
4159   4161       1 ;      H - HA
4159   4162       1 ;      H - CB
4159   4170       1 ;      H - C
4160   4165       1 ;     CA - HG21
4160   4166       1 ;     CA - HG22
4160   4167       1 ;     CA - HG23
4160   4169       1 ;     CA - HG1
4160   4173       1 ;     CA - H
4160   4174       1 ;     CA - CA
4161   4163       1 ;     HA - HB
```

```
 4161   4164       1 ;      HA - CG2
 4161   4168       1 ;      HA - OG1
 4161   4171       1 ;      HA - O
 4161   4172       1 ;      HA - N
 4162   4171       1 ;      CB - O
 4162   4172       1 ;      CB - N
 4163   4165       1 ;      HB - HG21
 4163   4166       1 ;      HB - HG22
 4163   4167       1 ;      HB - HG23
 4163   4169       1 ;      HB - HG1
 4163   4170       1 ;      HB - C
 4164   4169       1 ;     CG2 - HG1
 4164   4170       1 ;     CG2 - C
 4165   4168       1 ;    HG21 - OG1
 4166   4168       1 ;    HG22 - OG1
 4167   4168       1 ;    HG23 - OG1
 4168   4170       1 ;     OG1 - C
 4170   4175       1 ;       C - HA
 4170   4176       1 ;       C - CB
 4170   4189       1 ;       C - C
 4171   4173       1 ;       O - H
 4171   4174       1 ;       O - CA
 4172   4177       1 ;       N - HB
 4172   4178       1 ;       N - CG2
 4172   4182       1 ;       N - CG1
 4172   4190       1 ;       N - O
 4172   4191       1 ;       N - N
 4173   4175       1 ;       H - HA
 4173   4176       1 ;       H - CB
 4173   4189       1 ;       H - C
 4174   4179       1 ;      CA - HG21
 4174   4180       1 ;      CA - HG22
 4174   4181       1 ;      CA - HG23
 4174   4183       1 ;      CA - HG12
 4174   4184       1 ;      CA - HG13
 4174   4185       1 ;      CA - CD1
 4174   4192       1 ;      CA - H
 4174   4193       1 ;      CA - CA
 4175   4177       1 ;      HA - HB
 4175   4178       1 ;      HA - CG2
 4175   4182       1 ;      HA - CG1
 4175   4190       1 ;      HA - O
 4175   4191       1 ;      HA - N
 4176   4186       1 ;      CB - HD11
 4176   4187       1 ;      CB - HD12
 4176   4188       1 ;      CB - HD13
 4176   4190       1 ;      CB - O
 4176   4191       1 ;      CB - N
 4177   4179       1 ;      HB - HG21
 4177   4180       1 ;      HB - HG22
 4177   4181       1 ;      HB - HG23
 4177   4183       1 ;      HB - HG12
```

| | | | | | | |
|---|---|---|---|---|---|---|
| 4177 | 4184 | 1 | ; | HB | - | HG13 |
| 4177 | 4185 | 1 | ; | HB | - | CD1 |
| 4177 | 4189 | 1 | ; | HB | - | C |
| 4178 | 4183 | 1 | ; | CG2 | - | HG12 |
| 4178 | 4184 | 1 | ; | CG2 | - | HG13 |
| 4178 | 4185 | 1 | ; | CG2 | - | CD1 |
| 4178 | 4189 | 1 | ; | CG2 | - | C |
| 4179 | 4182 | 1 | ; | HG21 | - | CG1 |
| 4180 | 4182 | 1 | ; | HG22 | - | CG1 |
| 4181 | 4182 | 1 | ; | HG23 | - | CG1 |
| 4182 | 4189 | 1 | ; | CG1 | - | C |
| 4183 | 4186 | 1 | ; | HG12 | - | HD11 |
| 4183 | 4187 | 1 | ; | HG12 | - | HD12 |
| 4183 | 4188 | 1 | ; | HG12 | - | HD13 |
| 4184 | 4186 | 1 | ; | HG13 | - | HD11 |
| 4184 | 4187 | 1 | ; | HG13 | - | HD12 |
| 4184 | 4188 | 1 | ; | HG13 | - | HD13 |
| 4189 | 4194 | 1 | ; | C | - | HA |
| 4189 | 4195 | 1 | ; | C | - | CB |
| 4189 | 4204 | 1 | ; | C | - | C |
| 4190 | 4192 | 1 | ; | O | - | H |
| 4190 | 4193 | 1 | ; | O | - | CA |
| 4191 | 4196 | 1 | ; | N | - | HB2 |
| 4191 | 4197 | 1 | ; | N | - | HB3 |
| 4191 | 4198 | 1 | ; | N | - | CG |
| 4191 | 4205 | 1 | ; | N | - | O |
| 4191 | 4206 | 1 | ; | N | - | N |
| 4192 | 4194 | 1 | ; | H | - | HA |
| 4192 | 4195 | 1 | ; | H | - | CB |
| 4192 | 4204 | 1 | ; | H | - | C |
| 4193 | 4199 | 1 | ; | CA | - | HG2 |
| 4193 | 4200 | 1 | ; | CA | - | HG3 |
| 4193 | 4201 | 1 | ; | CA | - | CD |
| 4193 | 4207 | 1 | ; | CA | - | H |
| 4193 | 4208 | 1 | ; | CA | - | CA |
| 4194 | 4196 | 1 | ; | HA | - | HB2 |
| 4194 | 4197 | 1 | ; | HA | - | HB3 |
| 4194 | 4198 | 1 | ; | HA | - | CG |
| 4194 | 4205 | 1 | ; | HA | - | O |
| 4194 | 4206 | 1 | ; | HA | - | N |
| 4195 | 4202 | 1 | ; | CB | - | OE1 |
| 4195 | 4203 | 1 | ; | CB | - | OE2 |
| 4195 | 4205 | 1 | ; | CB | - | O |
| 4195 | 4206 | 1 | ; | CB | - | N |
| 4196 | 4199 | 1 | ; | HB2 | - | HG2 |
| 4196 | 4200 | 1 | ; | HB2 | - | HG3 |
| 4196 | 4201 | 1 | ; | HB2 | - | CD |
| 4196 | 4204 | 1 | ; | HB2 | - | C |
| 4197 | 4199 | 1 | ; | HB3 | - | HG2 |
| 4197 | 4200 | 1 | ; | HB3 | - | HG3 |
| 4197 | 4201 | 1 | ; | HB3 | - | CD |
| 4197 | 4204 | 1 | ; | HB3 | - | C |

| | | | | | |
|---|---|---|---|---|---|
| 4198 | 4204 | 1 ; | CG | - | C |
| 4199 | 4202 | 1 ; | HG2 | - | OE1 |
| 4199 | 4203 | 1 ; | HG2 | - | OE2 |
| 4200 | 4202 | 1 ; | HG3 | - | OE1 |
| 4200 | 4203 | 1 ; | HG3 | - | OE2 |
| 4204 | 4209 | 1 ; | C | - | HA |
| 4204 | 4210 | 1 ; | C | - | CB |
| 4204 | 4219 | 1 ; | C | - | C |
| 4205 | 4207 | 1 ; | O | - | H |
| 4205 | 4208 | 1 ; | O | - | CA |
| 4206 | 4211 | 1 ; | N | - | HB2 |
| 4206 | 4212 | 1 ; | N | - | HB3 |
| 4206 | 4213 | 1 ; | N | - | CG |
| 4206 | 4220 | 1 ; | N | - | O |
| 4206 | 4221 | 1 ; | N | - | N |
| 4207 | 4209 | 1 ; | H | - | HA |
| 4207 | 4210 | 1 ; | H | - | CB |
| 4207 | 4219 | 1 ; | H | - | C |
| 4208 | 4214 | 1 ; | CA | - | HG2 |
| 4208 | 4215 | 1 ; | CA | - | HG3 |
| 4208 | 4216 | 1 ; | CA | - | CD |
| 4208 | 4222 | 1 ; | CA | - | H |
| 4208 | 4223 | 1 ; | CA | - | CA |
| 4209 | 4211 | 1 ; | HA | - | HB2 |
| 4209 | 4212 | 1 ; | HA | - | HB3 |
| 4209 | 4213 | 1 ; | HA | - | CG |
| 4209 | 4220 | 1 ; | HA | - | O |
| 4209 | 4221 | 1 ; | HA | - | N |
| 4210 | 4217 | 1 ; | CB | - | OE1 |
| 4210 | 4218 | 1 ; | CB | - | OE2 |
| 4210 | 4220 | 1 ; | CB | - | O |
| 4210 | 4221 | 1 ; | CB | - | N |
| 4211 | 4214 | 1 ; | HB2 | - | HG2 |
| 4211 | 4215 | 1 ; | HB2 | - | HG3 |
| 4211 | 4216 | 1 ; | HB2 | - | CD |
| 4211 | 4219 | 1 ; | HB2 | - | C |
| 4212 | 4214 | 1 ; | HB3 | - | HG2 |
| 4212 | 4215 | 1 ; | HB3 | - | HG3 |
| 4212 | 4216 | 1 ; | HB3 | - | CD |
| 4212 | 4219 | 1 ; | HB3 | - | C |
| 4213 | 4219 | 1 ; | CG | - | C |
| 4214 | 4217 | 1 ; | HG2 | - | OE1 |
| 4214 | 4218 | 1 ; | HG2 | - | OE2 |
| 4215 | 4217 | 1 ; | HG3 | - | OE1 |
| 4215 | 4218 | 1 ; | HG3 | - | OE2 |
| 4219 | 4224 | 1 ; | C | - | HA |
| 4219 | 4225 | 1 ; | C | - | CB |
| 4219 | 4229 | 1 ; | C | - | C |
| 4220 | 4222 | 1 ; | O | - | H |
| 4220 | 4223 | 1 ; | O | - | CA |
| 4221 | 4226 | 1 ; | N | - | HB1 |
| 4221 | 4227 | 1 ; | N | - | HB2 |

```
4221   4228       1 ;       N - HB3
4221   4230       1 ;       N - O
4221   4231       1 ;       N - N
4222   4224       1 ;       H - HA
4222   4225       1 ;       H - CB
4222   4229       1 ;       H - C
4223   4232       1 ;      CA - H
4223   4233       1 ;      CA - CA
4224   4226       1 ;      HA - HB1
4224   4227       1 ;      HA - HB2
4224   4228       1 ;      HA - HB3
4224   4230       1 ;      HA - O
4224   4231       1 ;      HA - N
4225   4230       1 ;      CB - O
4225   4231       1 ;      CB - N
4226   4229       1 ;     HB1 - C
4227   4229       1 ;     HB2 - C
4228   4229       1 ;     HB3 - C
4229   4234       1 ;       C - HA
4229   4235       1 ;       C - CB
4229   4248       1 ;       C - C
4230   4232       1 ;       O - H
4230   4233       1 ;       O - CA
4231   4236       1 ;       N - HB2
4231   4237       1 ;       N - HB3
4231   4238       1 ;       N - CG
4231   4249       1 ;       N - O
4231   4250       1 ;       N - N
4232   4234       1 ;       H - HA
4232   4235       1 ;       H - CB
4232   4248       1 ;       H - C
4233   4239       1 ;      CA - HG
4233   4240       1 ;      CA - CD1
4233   4244       1 ;      CA - CD2
4233   4251       1 ;      CA - H
4233   4252       1 ;      CA - CA
4234   4236       1 ;      HA - HB2
4234   4237       1 ;      HA - HB3
4234   4238       1 ;      HA - CG
4234   4249       1 ;      HA - O
4234   4250       1 ;      HA - N
4235   4241       1 ;      CB - HD11
4235   4242       1 ;      CB - HD12
4235   4243       1 ;      CB - HD13
4235   4245       1 ;      CB - HD21
4235   4246       1 ;      CB - HD22
4235   4247       1 ;      CB - HD23
4235   4249       1 ;      CB - O
4235   4250       1 ;      CB - N
4236   4239       1 ;     HB2 - HG
4236   4240       1 ;     HB2 - CD1
4236   4244       1 ;     HB2 - CD2
```

```
4236   4248      1 ;    HB2 - C
4237   4239      1 ;    HB3 - HG
4237   4240      1 ;    HB3 - CD1
4237   4244      1 ;    HB3 - CD2
4237   4248      1 ;    HB3 - C
4238   4248      1 ;     CG - C
4239   4241      1 ;     HG - HD11
4239   4242      1 ;     HG - HD12
4239   4243      1 ;     HG - HD13
4239   4245      1 ;     HG - HD21
4239   4246      1 ;     HG - HD22
4239   4247      1 ;     HG - HD23
4240   4245      1 ;    CD1 - HD21
4240   4246      1 ;    CD1 - HD22
4240   4247      1 ;    CD1 - HD23
4241   4244      1 ;   HD11 - CD2
4242   4244      1 ;   HD12 - CD2
4243   4244      1 ;   HD13 - CD2
4248   4253      1 ;      C - HA
4248   4254      1 ;      C - CB
4248   4262      1 ;      C - C
4249   4251      1 ;      O - H
4249   4252      1 ;      O - CA
4250   4255      1 ;      N - HB2
4250   4256      1 ;      N - HB3
4250   4257      1 ;      N - CG
4250   4263      1 ;      N - O
4250   4264      1 ;      N - N
4251   4253      1 ;      H - HA
4251   4254      1 ;      H - CB
4251   4262      1 ;      H - C
4252   4258      1 ;     CA - OD1
4252   4259      1 ;     CA - ND2
4252   4265      1 ;     CA - H
4252   4266      1 ;     CA - CA
4253   4255      1 ;     HA - HB2
4253   4256      1 ;     HA - HB3
4253   4257      1 ;     HA - CG
4253   4263      1 ;     HA - O
4253   4264      1 ;     HA - N
4254   4260      1 ;     CB - HD21
4254   4261      1 ;     CB - HD22
4254   4263      1 ;     CB - O
4254   4264      1 ;     CB - N
4255   4258      1 ;    HB2 - OD1
4255   4259      1 ;    HB2 - ND2
4255   4262      1 ;    HB2 - C
4256   4258      1 ;    HB3 - OD1
4256   4259      1 ;    HB3 - ND2
4256   4262      1 ;    HB3 - C
4257   4262      1 ;     CG - C
4258   4260      1 ;    OD1 - HD21
```

```
4258   4261       1 ;    OD1 - HD22
4262   4267       1 ;      C - HA
4262   4268       1 ;      C - CB
4262   4280       1 ;      C - C
4263   4265       1 ;      O - H
4263   4266       1 ;      O - CA
4264   4269       1 ;      N - HB2
4264   4270       1 ;      N - HB3
4264   4271       1 ;      N - CG
4264   4281       1 ;      N - O
4264   4282       1 ;      N - N
4265   4267       1 ;      H - HA
4265   4268       1 ;      H - CB
4265   4280       1 ;      H - C
4266   4272       1 ;     CA - ND1
4266   4278       1 ;     CA - CD2
4266   4283       1 ;     CA - CD
4266   4292       1 ;     CA - CA
4267   4269       1 ;     HA - HB2
4267   4270       1 ;     HA - HB3
4267   4271       1 ;     HA - CG
4267   4281       1 ;     HA - O
4267   4282       1 ;     HA - N
4268   4273       1 ;     CB - HD1
4268   4274       1 ;     CB - CE1
4268   4276       1 ;     CB - NE2
4268   4279       1 ;     CB - HD2
4268   4281       1 ;     CB - O
4268   4282       1 ;     CB - N
4269   4272       1 ;    HB2 - ND1
4269   4278       1 ;    HB2 - CD2
4269   4280       1 ;    HB2 - C
4270   4272       1 ;    HB3 - ND1
4270   4278       1 ;    HB3 - CD2
4270   4280       1 ;    HB3 - C
4271   4275       1 ;     CG - HE1
4271   4277       1 ;     CG - HE2
4271   4280       1 ;     CG - C
4272   4277       1 ;    ND1 - HE2
4272   4279       1 ;    ND1 - HD2
4273   4275       1 ;    HD1 - HE1
4273   4276       1 ;    HD1 - NE2
4273   4278       1 ;    HD1 - CD2
4274   4279       1 ;    CE1 - HD2
4275   4277       1 ;    HE1 - HE2
4275   4278       1 ;    HE1 - CD2
4277   4279       1 ;    HE2 - HD2
4280   4284       1 ;      C - HD2
4280   4285       1 ;      C - HD3
4280   4286       1 ;      C - CG
4280   4289       1 ;      C - CB
4280   4293       1 ;      C - HA
```

```
4280   4294       1 ;       C - C
4281   4283       1 ;       O - CD
4281   4292       1 ;       O - CA
4282   4287       1 ;       N - HG2
4282   4288       1 ;       N - HG3
4282   4290       1 ;       N - HB2
4282   4291       1 ;       N - HB3
4282   4295       1 ;       N - O
4282   4296       1 ;       N - N
4283   4290       1 ;      CD - HB2
4283   4291       1 ;      CD - HB3
4283   4293       1 ;      CD - HA
4283   4294       1 ;      CD - C
4284   4287       1 ;     HD2 - HG2
4284   4288       1 ;     HD2 - HG3
4284   4289       1 ;     HD2 - CB
4284   4292       1 ;     HD2 - CA
4285   4287       1 ;     HD3 - HG2
4285   4288       1 ;     HD3 - HG3
4285   4289       1 ;     HD3 - CB
4285   4292       1 ;     HD3 - CA
4286   4293       1 ;      CG - HA
4286   4294       1 ;      CG - C
4287   4290       1 ;     HG2 - HB2
4287   4291       1 ;     HG2 - HB3
4287   4292       1 ;     HG2 - CA
4288   4290       1 ;     HG3 - HB2
4288   4291       1 ;     HG3 - HB3
4288   4292       1 ;     HG3 - CA
4289   4295       1 ;      CB - O
4289   4296       1 ;      CB - N
4290   4293       1 ;     HB2 - HA
4290   4294       1 ;     HB2 - C
4291   4293       1 ;     HB3 - HA
4291   4294       1 ;     HB3 - C
4292   4297       1 ;      CA - H
4292   4298       1 ;      CA - CA
4293   4295       1 ;      HA - O
4293   4296       1 ;      HA - N
4294   4299       1 ;       C - HA
4294   4300       1 ;       C - CB
4294   4318       1 ;       C - C
4295   4297       1 ;       O - H
4295   4298       1 ;       O - CA
4296   4301       1 ;       N - HB2
4296   4302       1 ;       N - HB3
4296   4303       1 ;       N - CG
4296   4319       1 ;       N - O
4296   4320       1 ;       N - N
4297   4299       1 ;       H - HA
4297   4300       1 ;       H - CB
4297   4318       1 ;       H - C
```

| | | | | | | |
|---|---|---|---|---|---|---|
| 4298 | 4304 | 1 | ; | CA | - | CD1 |
| 4298 | 4317 | 1 | ; | CA | - | CD2 |
| 4298 | 4321 | 1 | ; | CA | - | H |
| 4298 | 4322 | 1 | ; | CA | - | CA |
| 4299 | 4301 | 1 | ; | HA | - | HB2 |
| 4299 | 4302 | 1 | ; | HA | - | HB3 |
| 4299 | 4303 | 1 | ; | HA | - | CG |
| 4299 | 4319 | 1 | ; | HA | - | O |
| 4299 | 4320 | 1 | ; | HA | - | N |
| 4300 | 4305 | 1 | ; | CB | - | HD1 |
| 4300 | 4306 | 1 | ; | CB | - | NE1 |
| 4300 | 4308 | 1 | ; | CB | - | CE2 |
| 4300 | 4315 | 1 | ; | CB | - | CE3 |
| 4300 | 4319 | 1 | ; | CB | - | O |
| 4300 | 4320 | 1 | ; | CB | - | N |
| 4301 | 4304 | 1 | ; | HB2 | - | CD1 |
| 4301 | 4317 | 1 | ; | HB2 | - | CD2 |
| 4301 | 4318 | 1 | ; | HB2 | - | C |
| 4302 | 4304 | 1 | ; | HB3 | - | CD1 |
| 4302 | 4317 | 1 | ; | HB3 | - | CD2 |
| 4302 | 4318 | 1 | ; | HB3 | - | C |
| 4303 | 4307 | 1 | ; | CG | - | HE1 |
| 4303 | 4309 | 1 | ; | CG | - | CZ2 |
| 4303 | 4313 | 1 | ; | CG | - | CZ3 |
| 4303 | 4316 | 1 | ; | CG | - | HE3 |
| 4303 | 4318 | 1 | ; | CG | - | C |
| 4304 | 4309 | 1 | ; | CD1 | - | CZ2 |
| 4304 | 4315 | 1 | ; | CD1 | - | CE3 |
| 4305 | 4307 | 1 | ; | HD1 | - | HE1 |
| 4305 | 4308 | 1 | ; | HD1 | - | CE2 |
| 4305 | 4317 | 1 | ; | HD1 | - | CD2 |
| 4306 | 4310 | 1 | ; | NE1 | - | HZ2 |
| 4306 | 4311 | 1 | ; | NE1 | - | CH2 |
| 4306 | 4315 | 1 | ; | NE1 | - | CE3 |
| 4307 | 4309 | 1 | ; | HE1 | - | CZ2 |
| 4307 | 4317 | 1 | ; | HE1 | - | CD2 |
| 4308 | 4312 | 1 | ; | CE2 | - | HH2 |
| 4308 | 4313 | 1 | ; | CE2 | - | CZ3 |
| 4308 | 4316 | 1 | ; | CE2 | - | HE3 |
| 4309 | 4314 | 1 | ; | CZ2 | - | HZ3 |
| 4309 | 4315 | 1 | ; | CZ2 | - | CE3 |
| 4310 | 4312 | 1 | ; | HZ2 | - | HH2 |
| 4310 | 4313 | 1 | ; | HZ2 | - | CZ3 |
| 4310 | 4317 | 1 | ; | HZ2 | - | CD2 |
| 4311 | 4316 | 1 | ; | CH2 | - | HE3 |
| 4311 | 4317 | 1 | ; | CH2 | - | CD2 |
| 4312 | 4314 | 1 | ; | HH2 | - | HZ3 |
| 4312 | 4315 | 1 | ; | HH2 | - | CE3 |
| 4314 | 4316 | 1 | ; | HZ3 | - | HE3 |
| 4314 | 4317 | 1 | ; | HZ3 | - | CD2 |
| 4318 | 4323 | 1 | ; | C | - | HA |
| 4318 | 4324 | 1 | ; | C | - | CB |

```
4318   4337       1 ;       C - C
4319   4321       1 ;       O - H
4319   4322       1 ;       O - CA
4320   4325       1 ;       N - HB
4320   4326       1 ;       N - CG2
4320   4330       1 ;       N - CG1
4320   4338       1 ;       N - O
4320   4339       1 ;       N - N
4321   4323       1 ;       H - HA
4321   4324       1 ;       H - CB
4321   4337       1 ;       H - C
4322   4327       1 ;      CA - HG21
4322   4328       1 ;      CA - HG22
4322   4329       1 ;      CA - HG23
4322   4331       1 ;      CA - HG12
4322   4332       1 ;      CA - HG13
4322   4333       1 ;      CA - CD1
4322   4340       1 ;      CA - H
4322   4341       1 ;      CA - CA
4323   4325       1 ;      HA - HB
4323   4326       1 ;      HA - CG2
4323   4330       1 ;      HA - CG1
4323   4338       1 ;      HA - O
4323   4339       1 ;      HA - N
4324   4334       1 ;      CB - HD11
4324   4335       1 ;      CB - HD12
4324   4336       1 ;      CB - HD13
4324   4338       1 ;      CB - O
4324   4339       1 ;      CB - N
4325   4327       1 ;      HB - HG21
4325   4328       1 ;      HB - HG22
4325   4329       1 ;      HB - HG23
4325   4331       1 ;      HB - HG12
4325   4332       1 ;      HB - HG13
4325   4333       1 ;      HB - CD1
4325   4337       1 ;      HB - C
4326   4331       1 ;     CG2 - HG12
4326   4332       1 ;     CG2 - HG13
4326   4333       1 ;     CG2 - CD1
4326   4337       1 ;     CG2 - C
4327   4330       1 ;    HG21 - CG1
4328   4330       1 ;    HG22 - CG1
4329   4330       1 ;    HG23 - CG1
4330   4337       1 ;     CG1 - C
4331   4334       1 ;    HG12 - HD11
4331   4335       1 ;    HG12 - HD12
4331   4336       1 ;    HG12 - HD13
4332   4334       1 ;    HG13 - HD11
4332   4335       1 ;    HG13 - HD12
4332   4336       1 ;    HG13 - HD13
4337   4342       1 ;       C - HA
4337   4343       1 ;       C - CB
```

```
4337   4351       1 ;       C - C
4338   4340       1 ;       O - H
4338   4341       1 ;       O - CA
4339   4344       1 ;       N - HB
4339   4345       1 ;       N - CG2
4339   4349       1 ;       N - OG1
4339   4352       1 ;       N - O
4339   4353       1 ;       N - N
4340   4342       1 ;       H - HA
4340   4343       1 ;       H - CB
4340   4351       1 ;       H - C
4341   4346       1 ;      CA - HG21
4341   4347       1 ;      CA - HG22
4341   4348       1 ;      CA - HG23
4341   4350       1 ;      CA - HG1
4341   4354       1 ;      CA - H
4341   4355       1 ;      CA - CA
4342   4344       1 ;      HA - HB
4342   4345       1 ;      HA - CG2
4342   4349       1 ;      HA - OG1
4342   4352       1 ;      HA - O
4342   4353       1 ;      HA - N
4343   4352       1 ;      CB - O
4343   4353       1 ;      CB - N
4344   4346       1 ;      HB - HG21
4344   4347       1 ;      HB - HG22
4344   4348       1 ;      HB - HG23
4344   4350       1 ;      HB - HG1
4344   4351       1 ;      HB - C
4345   4350       1 ;     CG2 - HG1
4345   4351       1 ;     CG2 - C
4346   4349       1 ;    HG21 - OG1
4347   4349       1 ;    HG22 - OG1
4348   4349       1 ;    HG23 - OG1
4349   4351       1 ;     OG1 - C
4351   4356       1 ;       C - HA
4351   4357       1 ;       C - CB
4351   4368       1 ;       C - C
4352   4354       1 ;       O - H
4352   4355       1 ;       O - CA
4353   4358       1 ;       N - HB2
4353   4359       1 ;       N - HB3
4353   4360       1 ;       N - CG
4353   4369       1 ;       N - O
4353   4370       1 ;       N - N
4354   4356       1 ;       H - HA
4354   4357       1 ;       H - CB
4354   4368       1 ;       H - C
4355   4361       1 ;      CA - HG2
4355   4362       1 ;      CA - HG3
4355   4363       1 ;      CA - CD
4355   4371       1 ;      CA - H
```

```
4355   4372       1 ;      CA - CA
4356   4358       1 ;      HA - HB2
4356   4359       1 ;      HA - HB3
4356   4360       1 ;      HA - CG
4356   4369       1 ;      HA - O
4356   4370       1 ;      HA - N
4357   4364       1 ;      CB - OE1
4357   4365       1 ;      CB - NE2
4357   4369       1 ;      CB - O
4357   4370       1 ;      CB - N
4358   4361       1 ;     HB2 - HG2
4358   4362       1 ;     HB2 - HG3
4358   4363       1 ;     HB2 - CD
4358   4368       1 ;     HB2 - C
4359   4361       1 ;     HB3 - HG2
4359   4362       1 ;     HB3 - HG3
4359   4363       1 ;     HB3 - CD
4359   4368       1 ;     HB3 - C
4360   4366       1 ;      CG - HE21
4360   4367       1 ;      CG - HE22
4360   4368       1 ;      CG - C
4361   4364       1 ;     HG2 - OE1
4361   4365       1 ;     HG2 - NE2
4362   4364       1 ;     HG3 - OE1
4362   4365       1 ;     HG3 - NE2
4364   4366       1 ;     OE1 - HE21
4364   4367       1 ;     OE1 - HE22
4368   4373       1 ;       C - HA
4368   4374       1 ;       C - CB
4368   4385       1 ;       C - C
4369   4371       1 ;       O - H
4369   4372       1 ;       O - CA
4370   4375       1 ;       N - HB2
4370   4376       1 ;       N - HB3
4370   4377       1 ;       N - CG
4370   4386       1 ;       N - O
4370   4387       1 ;       N - N
4371   4373       1 ;       H - HA
4371   4374       1 ;       H - CB
4371   4385       1 ;       H - C
4372   4378       1 ;      CA - HG2
4372   4379       1 ;      CA - HG3
4372   4380       1 ;      CA - SD
4372   4388       1 ;      CA - H
4372   4389       1 ;      CA - CA
4373   4375       1 ;      HA - HB2
4373   4376       1 ;      HA - HB3
4373   4377       1 ;      HA - CG
4373   4386       1 ;      HA - O
4373   4387       1 ;      HA - N
4374   4381       1 ;      CB - CE
4374   4386       1 ;      CB - O
```

```
4374   4387       1 ;      CB - N
4375   4378       1 ;     HB2 - HG2
4375   4379       1 ;     HB2 - HG3
4375   4380       1 ;     HB2 - SD
4375   4385       1 ;     HB2 - C
4376   4378       1 ;     HB3 - HG2
4376   4379       1 ;     HB3 - HG3
4376   4380       1 ;     HB3 - SD
4376   4385       1 ;     HB3 - C
4377   4382       1 ;      CG - HE1
4377   4383       1 ;      CG - HE2
4377   4384       1 ;      CG - HE3
4377   4385       1 ;      CG - C
4378   4381       1 ;     HG2 - CE
4379   4381       1 ;     HG3 - CE
4385   4390       1 ;       C - HA
4385   4391       1 ;       C - CB
4385   4399       1 ;       C - C
4386   4388       1 ;       O - H
4386   4389       1 ;       O - CA
4387   4392       1 ;       N - HB
4387   4393       1 ;       N - CG2
4387   4397       1 ;       N - OG1
4387   4400       1 ;       N - O
4387   4401       1 ;       N - N
4388   4390       1 ;       H - HA
4388   4391       1 ;       H - CB
4388   4399       1 ;       H - C
4389   4394       1 ;      CA - HG21
4389   4395       1 ;      CA - HG22
4389   4396       1 ;      CA - HG23
4389   4398       1 ;      CA - HG1
4389   4402       1 ;      CA - H
4389   4403       1 ;      CA - CA
4390   4392       1 ;      HA - HB
4390   4393       1 ;      HA - CG2
4390   4397       1 ;      HA - OG1
4390   4400       1 ;      HA - O
4390   4401       1 ;      HA - N
4391   4400       1 ;      CB - O
4391   4401       1 ;      CB - N
4392   4394       1 ;      HB - HG21
4392   4395       1 ;      HB - HG22
4392   4396       1 ;      HB - HG23
4392   4398       1 ;      HB - HG1
4392   4399       1 ;      HB - C
4393   4398       1 ;     CG2 - HG1
4393   4399       1 ;     CG2 - C
4394   4397       1 ;    HG21 - OG1
4395   4397       1 ;    HG22 - OG1
4396   4397       1 ;    HG23 - OG1
4397   4399       1 ;     OG1 - C
```

| | | | |
|---|---|---|---|
| 4399 | 4404 | 1 ; | C - HA |
| 4399 | 4405 | 1 ; | C - CB |
| 4399 | 4421 | 1 ; | C - C |
| 4400 | 4402 | 1 ; | O - H |
| 4400 | 4403 | 1 ; | O - CA |
| 4401 | 4406 | 1 ; | N - HB2 |
| 4401 | 4407 | 1 ; | N - HB3 |
| 4401 | 4408 | 1 ; | N - CG |
| 4401 | 4422 | 1 ; | N - O |
| 4401 | 4423 | 1 ; | N - N |
| 4402 | 4404 | 1 ; | H - HA |
| 4402 | 4405 | 1 ; | H - CB |
| 4402 | 4421 | 1 ; | H - C |
| 4403 | 4409 | 1 ; | CA - HG2 |
| 4403 | 4410 | 1 ; | CA - HG3 |
| 4403 | 4411 | 1 ; | CA - CD |
| 4403 | 4424 | 1 ; | CA - H |
| 4403 | 4425 | 1 ; | CA - CA |
| 4404 | 4406 | 1 ; | HA - HB2 |
| 4404 | 4407 | 1 ; | HA - HB3 |
| 4404 | 4408 | 1 ; | HA - CG |
| 4404 | 4422 | 1 ; | HA - O |
| 4404 | 4423 | 1 ; | HA - N |
| 4405 | 4412 | 1 ; | CB - HD2 |
| 4405 | 4413 | 1 ; | CB - HD3 |
| 4405 | 4414 | 1 ; | CB - CE |
| 4405 | 4422 | 1 ; | CB - O |
| 4405 | 4423 | 1 ; | CB - N |
| 4406 | 4409 | 1 ; | HB2 - HG2 |
| 4406 | 4410 | 1 ; | HB2 - HG3 |
| 4406 | 4411 | 1 ; | HB2 - CD |
| 4406 | 4421 | 1 ; | HB2 - C |
| 4407 | 4409 | 1 ; | HB3 - HG2 |
| 4407 | 4410 | 1 ; | HB3 - HG3 |
| 4407 | 4411 | 1 ; | HB3 - CD |
| 4407 | 4421 | 1 ; | HB3 - C |
| 4408 | 4415 | 1 ; | CG - HE2 |
| 4408 | 4416 | 1 ; | CG - HE3 |
| 4408 | 4417 | 1 ; | CG - NZ |
| 4408 | 4421 | 1 ; | CG - C |
| 4409 | 4412 | 1 ; | HG2 - HD2 |
| 4409 | 4413 | 1 ; | HG2 - HD3 |
| 4409 | 4414 | 1 ; | HG2 - CE |
| 4410 | 4412 | 1 ; | HG3 - HD2 |
| 4410 | 4413 | 1 ; | HG3 - HD3 |
| 4410 | 4414 | 1 ; | HG3 - CE |
| 4411 | 4418 | 1 ; | CD - HZ1 |
| 4411 | 4419 | 1 ; | CD - HZ2 |
| 4411 | 4420 | 1 ; | CD - HZ3 |
| 4412 | 4415 | 1 ; | HD2 - HE2 |
| 4412 | 4416 | 1 ; | HD2 - HE3 |
| 4412 | 4417 | 1 ; | HD2 - NZ |

| | | | |
|---|---|---|---|
| 4413 | 4415 | 1 ; | HD3 - HE2 |
| 4413 | 4416 | 1 ; | HD3 - HE3 |
| 4413 | 4417 | 1 ; | HD3 - NZ |
| 4415 | 4418 | 1 ; | HE2 - HZ1 |
| 4415 | 4419 | 1 ; | HE2 - HZ2 |
| 4415 | 4420 | 1 ; | HE2 - HZ3 |
| 4416 | 4418 | 1 ; | HE3 - HZ1 |
| 4416 | 4419 | 1 ; | HE3 - HZ2 |
| 4416 | 4420 | 1 ; | HE3 - HZ3 |
| 4421 | 4426 | 1 ; | C - HA |
| 4421 | 4427 | 1 ; | C - CB |
| 4421 | 4432 | 1 ; | C - C |
| 4422 | 4424 | 1 ; | O - H |
| 4422 | 4425 | 1 ; | O - CA |
| 4423 | 4428 | 1 ; | N - HB2 |
| 4423 | 4429 | 1 ; | N - HB3 |
| 4423 | 4430 | 1 ; | N - OG |
| 4423 | 4433 | 1 ; | N - O |
| 4423 | 4434 | 1 ; | N - N |
| 4424 | 4426 | 1 ; | H - HA |
| 4424 | 4427 | 1 ; | H - CB |
| 4424 | 4432 | 1 ; | H - C |
| 4425 | 4431 | 1 ; | CA - HG |
| 4425 | 4435 | 1 ; | CA - H |
| 4425 | 4436 | 1 ; | CA - CA |
| 4426 | 4428 | 1 ; | HA - HB2 |
| 4426 | 4429 | 1 ; | HA - HB3 |
| 4426 | 4430 | 1 ; | HA - OG |
| 4426 | 4433 | 1 ; | HA - O |
| 4426 | 4434 | 1 ; | HA - N |
| 4427 | 4433 | 1 ; | CB - O |
| 4427 | 4434 | 1 ; | CB - N |
| 4428 | 4431 | 1 ; | HB2 - HG |
| 4428 | 4432 | 1 ; | HB2 - C |
| 4429 | 4431 | 1 ; | HB3 - HG |
| 4429 | 4432 | 1 ; | HB3 - C |
| 4430 | 4432 | 1 ; | OG - C |
| 4432 | 4437 | 1 ; | C - HA |
| 4432 | 4438 | 1 ; | C - CB |
| 4432 | 4449 | 1 ; | C - C |
| 4433 | 4435 | 1 ; | O - H |
| 4433 | 4436 | 1 ; | O - CA |
| 4434 | 4439 | 1 ; | N - HB2 |
| 4434 | 4440 | 1 ; | N - HB3 |
| 4434 | 4441 | 1 ; | N - CG |
| 4434 | 4450 | 1 ; | N - O |
| 4434 | 4451 | 1 ; | N - N |
| 4435 | 4437 | 1 ; | H - HA |
| 4435 | 4438 | 1 ; | H - CB |
| 4435 | 4449 | 1 ; | H - C |
| 4436 | 4442 | 1 ; | CA - ND1 |
| 4436 | 4447 | 1 ; | CA - CD2 |

```
4436   4452       1 ;      CA - H
4436   4453       1 ;      CA - CA
4437   4439       1 ;      HA - HB2
4437   4440       1 ;      HA - HB3
4437   4441       1 ;      HA - CG
4437   4450       1 ;      HA - O
4437   4451       1 ;      HA - N
4438   4443       1 ;      CB - CE1
4438   4445       1 ;      CB - NE2
4438   4448       1 ;      CB - HD2
4438   4450       1 ;      CB - O
4438   4451       1 ;      CB - N
4439   4442       1 ;     HB2 - ND1
4439   4447       1 ;     HB2 - CD2
4439   4449       1 ;     HB2 - C
4440   4442       1 ;     HB3 - ND1
4440   4447       1 ;     HB3 - CD2
4440   4449       1 ;     HB3 - C
4441   4444       1 ;      CG - HE1
4441   4446       1 ;      CG - HE2
4441   4449       1 ;      CG - C
4442   4446       1 ;     ND1 - HE2
4442   4448       1 ;     ND1 - HD2
4443   4448       1 ;     CE1 - HD2
4444   4446       1 ;     HE1 - HE2
4444   4447       1 ;     HE1 - CD2
4446   4448       1 ;     HE2 - HD2
4449   4454       1 ;       C - HA
4449   4455       1 ;       C - CB
4449   4464       1 ;       C - C
4450   4452       1 ;       O - H
4450   4453       1 ;       O - CA
4451   4456       1 ;       N - HB2
4451   4457       1 ;       N - HB3
4451   4458       1 ;       N - CG
4451   4465       1 ;       N - O
4451   4466       1 ;       N - N
4452   4454       1 ;       H - HA
4452   4455       1 ;       H - CB
4452   4464       1 ;       H - C
4453   4459       1 ;      CA - HG2
4453   4460       1 ;      CA - HG3
4453   4461       1 ;      CA - CD
4453   4467       1 ;      CA - H
4453   4468       1 ;      CA - CA
4454   4456       1 ;      HA - HB2
4454   4457       1 ;      HA - HB3
4454   4458       1 ;      HA - CG
4454   4465       1 ;      HA - O
4454   4466       1 ;      HA - N
4455   4462       1 ;      CB - OE1
4455   4463       1 ;      CB - OE2
```

| | | | | |
|---|---|---|---|---|
| 4455 | 4465 | 1 ; | CB | - O |
| 4455 | 4466 | 1 ; | CB | - N |
| 4456 | 4459 | 1 ; | HB2 | - HG2 |
| 4456 | 4460 | 1 ; | HB2 | - HG3 |
| 4456 | 4461 | 1 ; | HB2 | - CD |
| 4456 | 4464 | 1 ; | HB2 | - C |
| 4457 | 4459 | 1 ; | HB3 | - HG2 |
| 4457 | 4460 | 1 ; | HB3 | - HG3 |
| 4457 | 4461 | 1 ; | HB3 | - CD |
| 4457 | 4464 | 1 ; | HB3 | - C |
| 4458 | 4464 | 1 ; | CG | - C |
| 4459 | 4462 | 1 ; | HG2 | - OE1 |
| 4459 | 4463 | 1 ; | HG2 | - OE2 |
| 4460 | 4462 | 1 ; | HG3 | - OE1 |
| 4460 | 4463 | 1 ; | HG3 | - OE2 |
| 4464 | 4469 | 1 ; | C | - HA |
| 4464 | 4470 | 1 ; | C | - CB |
| 4464 | 4481 | 1 ; | C | - C |
| 4465 | 4467 | 1 ; | O | - H |
| 4465 | 4468 | 1 ; | O | - CA |
| 4466 | 4471 | 1 ; | N | - HB2 |
| 4466 | 4472 | 1 ; | N | - HB3 |
| 4466 | 4473 | 1 ; | N | - CG |
| 4466 | 4482 | 1 ; | N | - O |
| 4466 | 4483 | 1 ; | N | - N |
| 4467 | 4469 | 1 ; | H | - HA |
| 4467 | 4470 | 1 ; | H | - CB |
| 4467 | 4481 | 1 ; | H | - C |
| 4468 | 4474 | 1 ; | CA | - ND1 |
| 4468 | 4479 | 1 ; | CA | - CD2 |
| 4468 | 4484 | 1 ; | CA | - H |
| 4468 | 4485 | 1 ; | CA | - CA |
| 4469 | 4471 | 1 ; | HA | - HB2 |
| 4469 | 4472 | 1 ; | HA | - HB3 |
| 4469 | 4473 | 1 ; | HA | - CG |
| 4469 | 4482 | 1 ; | HA | - O |
| 4469 | 4483 | 1 ; | HA | - N |
| 4470 | 4475 | 1 ; | CB | - CE1 |
| 4470 | 4477 | 1 ; | CB | - NE2 |
| 4470 | 4480 | 1 ; | CB | - HD2 |
| 4470 | 4482 | 1 ; | CB | - O |
| 4470 | 4483 | 1 ; | CB | - N |
| 4471 | 4474 | 1 ; | HB2 | - ND1 |
| 4471 | 4479 | 1 ; | HB2 | - CD2 |
| 4471 | 4481 | 1 ; | HB2 | - C |
| 4472 | 4474 | 1 ; | HB3 | - ND1 |
| 4472 | 4479 | 1 ; | HB3 | - CD2 |
| 4472 | 4481 | 1 ; | HB3 | - C |
| 4473 | 4476 | 1 ; | CG | - HE1 |
| 4473 | 4478 | 1 ; | CG | - HE2 |
| 4473 | 4481 | 1 ; | CG | - C |
| 4474 | 4478 | 1 ; | ND1 | - HE2 |

```
4474   4480       1 ;    ND1 - HD2
4475   4480       1 ;    CE1 - HD2
4476   4478       1 ;    HE1 - HE2
4476   4479       1 ;    HE1 - CD2
4478   4480       1 ;    HE2 - HD2
4481   4486       1 ;      C - HA
4481   4487       1 ;      C - CB
4481   4497       1 ;      C - C
4482   4484       1 ;      O - H
4482   4485       1 ;      O - CA
4483   4488       1 ;      N - HB
4483   4489       1 ;      N - CG1
4483   4493       1 ;      N - CG2
4483   4498       1 ;      N - O
4483   4499       1 ;      N - N
4484   4486       1 ;      H - HA
4484   4487       1 ;      H - CB
4484   4497       1 ;      H - C
4485   4490       1 ;     CA - HG11
4485   4491       1 ;     CA - HG12
4485   4492       1 ;     CA - HG13
4485   4494       1 ;     CA - HG21
4485   4495       1 ;     CA - HG22
4485   4496       1 ;     CA - HG23
4485   4500       1 ;     CA - H
4485   4501       1 ;     CA - CA
4486   4488       1 ;     HA - HB
4486   4489       1 ;     HA - CG1
4486   4493       1 ;     HA - CG2
4486   4498       1 ;     HA - O
4486   4499       1 ;     HA - N
4487   4498       1 ;     CB - O
4487   4499       1 ;     CB - N
4488   4490       1 ;     HB - HG11
4488   4491       1 ;     HB - HG12
4488   4492       1 ;     HB - HG13
4488   4494       1 ;     HB - HG21
4488   4495       1 ;     HB - HG22
4488   4496       1 ;     HB - HG23
4488   4497       1 ;     HB - C
4489   4494       1 ;    CG1 - HG21
4489   4495       1 ;    CG1 - HG22
4489   4496       1 ;    CG1 - HG23
4489   4497       1 ;    CG1 - C
4490   4493       1 ;   HG11 - CG2
4491   4493       1 ;   HG12 - CG2
4492   4493       1 ;   HG13 - CG2
4493   4497       1 ;    CG2 - C
4497   4502       1 ;      C - HA
4497   4503       1 ;      C - CB
4497   4512       1 ;      C - C
4498   4500       1 ;      O - H
```

| | | | | | |
|---|---|---|---|---|---|
| 4498 | 4501 | 1 ; | O | - | CA |
| 4499 | 4504 | 1 ; | N | - | HB2 |
| 4499 | 4505 | 1 ; | N | - | HB3 |
| 4499 | 4506 | 1 ; | N | - | CG |
| 4499 | 4513 | 1 ; | N | - | O |
| 4499 | 4514 | 1 ; | N | - | N |
| 4500 | 4502 | 1 ; | H | - | HA |
| 4500 | 4503 | 1 ; | H | - | CB |
| 4500 | 4512 | 1 ; | H | - | C |
| 4501 | 4507 | 1 ; | CA | - | HG2 |
| 4501 | 4508 | 1 ; | CA | - | HG3 |
| 4501 | 4509 | 1 ; | CA | - | CD |
| 4501 | 4515 | 1 ; | CA | - | H |
| 4501 | 4516 | 1 ; | CA | - | CA |
| 4502 | 4504 | 1 ; | HA | - | HB2 |
| 4502 | 4505 | 1 ; | HA | - | HB3 |
| 4502 | 4506 | 1 ; | HA | - | CG |
| 4502 | 4513 | 1 ; | HA | - | O |
| 4502 | 4514 | 1 ; | HA | - | N |
| 4503 | 4510 | 1 ; | CB | - | OE1 |
| 4503 | 4511 | 1 ; | CB | - | OE2 |
| 4503 | 4513 | 1 ; | CB | - | O |
| 4503 | 4514 | 1 ; | CB | - | N |
| 4504 | 4507 | 1 ; | HB2 | - | HG2 |
| 4504 | 4508 | 1 ; | HB2 | - | HG3 |
| 4504 | 4509 | 1 ; | HB2 | - | CD |
| 4504 | 4512 | 1 ; | HB2 | - | C |
| 4505 | 4507 | 1 ; | HB3 | - | HG2 |
| 4505 | 4508 | 1 ; | HB3 | - | HG3 |
| 4505 | 4509 | 1 ; | HB3 | - | CD |
| 4505 | 4512 | 1 ; | HB3 | - | C |
| 4506 | 4512 | 1 ; | CG | - | C |
| 4507 | 4510 | 1 ; | HG2 | - | OE1 |
| 4507 | 4511 | 1 ; | HG2 | - | OE2 |
| 4508 | 4510 | 1 ; | HG3 | - | OE1 |
| 4508 | 4511 | 1 ; | HG3 | - | OE2 |
| 4512 | 4517 | 1 ; | C | - | HA |
| 4512 | 4518 | 1 ; | C | - | CB |
| 4512 | 4531 | 1 ; | C | - | C |
| 4513 | 4515 | 1 ; | O | - | H |
| 4513 | 4516 | 1 ; | O | - | CA |
| 4514 | 4519 | 1 ; | N | - | HB2 |
| 4514 | 4520 | 1 ; | N | - | HB3 |
| 4514 | 4521 | 1 ; | N | - | CG |
| 4514 | 4532 | 1 ; | N | - | O |
| 4514 | 4533 | 1 ; | N | - | N |
| 4515 | 4517 | 1 ; | H | - | HA |
| 4515 | 4518 | 1 ; | H | - | CB |
| 4515 | 4531 | 1 ; | H | - | C |
| 4516 | 4522 | 1 ; | CA | - | HG |
| 4516 | 4523 | 1 ; | CA | - | CD1 |
| 4516 | 4527 | 1 ; | CA | - | CD2 |

```
4516   4534       1 ;      CA - H
4516   4535       1 ;      CA - CA
4517   4519       1 ;      HA - HB2
4517   4520       1 ;      HA - HB3
4517   4521       1 ;      HA - CG
4517   4532       1 ;      HA - O
4517   4533       1 ;      HA - N
4518   4524       1 ;      CB - HD11
4518   4525       1 ;      CB - HD12
4518   4526       1 ;      CB - HD13
4518   4528       1 ;      CB - HD21
4518   4529       1 ;      CB - HD22
4518   4530       1 ;      CB - HD23
4518   4532       1 ;      CB - O
4518   4533       1 ;      CB - N
4519   4522       1 ;     HB2 - HG
4519   4523       1 ;     HB2 - CD1
4519   4527       1 ;     HB2 - CD2
4519   4531       1 ;     HB2 - C
4520   4522       1 ;     HB3 - HG
4520   4523       1 ;     HB3 - CD1
4520   4527       1 ;     HB3 - CD2
4520   4531       1 ;     HB3 - C
4521   4531       1 ;      CG - C
4522   4524       1 ;      HG - HD11
4522   4525       1 ;      HG - HD12
4522   4526       1 ;      HG - HD13
4522   4528       1 ;      HG - HD21
4522   4529       1 ;      HG - HD22
4522   4530       1 ;      HG - HD23
4523   4528       1 ;     CD1 - HD21
4523   4529       1 ;     CD1 - HD22
4523   4530       1 ;     CD1 - HD23
4524   4527       1 ;    HD11 - CD2
4525   4527       1 ;    HD12 - CD2
4526   4527       1 ;    HD13 - CD2
4531   4536       1 ;       C - HA
4531   4537       1 ;       C - CB
4531   4542       1 ;       C - C
4532   4534       1 ;       O - H
4532   4535       1 ;       O - CA
4533   4538       1 ;       N - HB2
4533   4539       1 ;       N - HB3
4533   4540       1 ;       N - OG
4533   4543       1 ;       N - O
4533   4544       1 ;       N - N
4534   4536       1 ;       H - HA
4534   4537       1 ;       H - CB
4534   4542       1 ;       H - C
4535   4541       1 ;      CA - HG
4535   4545       1 ;      CA - H
4535   4546       1 ;      CA - CA
```

```
4536   4538       1 ;      HA - HB2
4536   4539       1 ;      HA - HB3
4536   4540       1 ;      HA - OG
4536   4543       1 ;      HA - O
4536   4544       1 ;      HA - N
4537   4543       1 ;      CB - O
4537   4544       1 ;      CB - N
4538   4541       1 ;     HB2 - HG
4538   4542       1 ;     HB2 - C
4539   4541       1 ;     HB3 - HG
4539   4542       1 ;     HB3 - C
4540   4542       1 ;      OG - C
4542   4547       1 ;       C - HA
4542   4548       1 ;       C - CB
4542   4553       1 ;       C - C
4543   4545       1 ;       O - H
4543   4546       1 ;       O - CA
4544   4549       1 ;       N - HB2
4544   4550       1 ;       N - HB3
4544   4551       1 ;       N - OG
4544   4554       1 ;       N - O
4544   4555       1 ;       N - N
4545   4547       1 ;       H - HA
4545   4548       1 ;       H - CB
4545   4553       1 ;       H - C
4546   4552       1 ;      CA - HG
4546   4556       1 ;      CA - H
4546   4557       1 ;      CA - CA
4547   4549       1 ;      HA - HB2
4547   4550       1 ;      HA - HB3
4547   4551       1 ;      HA - OG
4547   4554       1 ;      HA - O
4547   4555       1 ;      HA - N
4548   4554       1 ;      CB - O
4548   4555       1 ;      CB - N
4549   4552       1 ;     HB2 - HG
4549   4553       1 ;     HB2 - C
4550   4552       1 ;     HB3 - HG
4550   4553       1 ;     HB3 - C
4551   4553       1 ;      OG - C
4553   4558       1 ;       C - HA
4553   4559       1 ;       C - CB
4553   4567       1 ;       C - C
4554   4556       1 ;       O - H
4554   4557       1 ;       O - CA
4555   4560       1 ;       N - HB
4555   4561       1 ;       N - CG2
4555   4565       1 ;       N - OG1
4555   4568       1 ;       N - O
4555   4569       1 ;       N - N
4556   4558       1 ;       H - HA
4556   4559       1 ;       H - CB
```

```
4556   4567      1 ;      H - C
4557   4562      1 ;     CA - HG21
4557   4563      1 ;     CA - HG22
4557   4564      1 ;     CA - HG23
4557   4566      1 ;     CA - HG1
4557   4570      1 ;     CA - H
4557   4571      1 ;     CA - CA
4558   4560      1 ;     HA - HB
4558   4561      1 ;     HA - CG2
4558   4565      1 ;     HA - OG1
4558   4568      1 ;     HA - O
4558   4569      1 ;     HA - N
4559   4568      1 ;     CB - O
4559   4569      1 ;     CB - N
4560   4562      1 ;     HB - HG21
4560   4563      1 ;     HB - HG22
4560   4564      1 ;     HB - HG23
4560   4566      1 ;     HB - HG1
4560   4567      1 ;     HB - C
4561   4566      1 ;    CG2 - HG1
4561   4567      1 ;    CG2 - C
4562   4565      1 ;   HG21 - OG1
4563   4565      1 ;   HG22 - OG1
4564   4565      1 ;   HG23 - OG1
4565   4567      1 ;    OG1 - C
4567   4572      1 ;      C - HA
4567   4573      1 ;      C - CB
4567   4586      1 ;      C - C
4568   4570      1 ;      O - H
4568   4571      1 ;      O - CA
4569   4574      1 ;      N - HB2
4569   4575      1 ;      N - HB3
4569   4576      1 ;      N - CG
4569   4587      1 ;      N - O
4569   4588      1 ;      N - N
4570   4572      1 ;      H - HA
4570   4573      1 ;      H - CB
4570   4586      1 ;      H - C
4571   4577      1 ;     CA - HG
4571   4578      1 ;     CA - CD1
4571   4582      1 ;     CA - CD2
4571   4589      1 ;     CA - H
4571   4590      1 ;     CA - CA
4572   4574      1 ;     HA - HB2
4572   4575      1 ;     HA - HB3
4572   4576      1 ;     HA - CG
4572   4587      1 ;     HA - O
4572   4588      1 ;     HA - N
4573   4579      1 ;     CB - HD11
4573   4580      1 ;     CB - HD12
4573   4581      1 ;     CB - HD13
4573   4583      1 ;     CB - HD21
```

| | | | | | |
|---|---|---|---|---|---|
| 4573 | 4584 | 1 ; | CB | - | HD22 |
| 4573 | 4585 | 1 ; | CB | - | HD23 |
| 4573 | 4587 | 1 ; | CB | - | O |
| 4573 | 4588 | 1 ; | CB | - | N |
| 4574 | 4577 | 1 ; | HB2 | - | HG |
| 4574 | 4578 | 1 ; | HB2 | - | CD1 |
| 4574 | 4582 | 1 ; | HB2 | - | CD2 |
| 4574 | 4586 | 1 ; | HB2 | - | C |
| 4575 | 4577 | 1 ; | HB3 | - | HG |
| 4575 | 4578 | 1 ; | HB3 | - | CD1 |
| 4575 | 4582 | 1 ; | HB3 | - | CD2 |
| 4575 | 4586 | 1 ; | HB3 | - | C |
| 4576 | 4586 | 1 ; | CG | - | C |
| 4577 | 4579 | 1 ; | HG | - | HD11 |
| 4577 | 4580 | 1 ; | HG | - | HD12 |
| 4577 | 4581 | 1 ; | HG | - | HD13 |
| 4577 | 4583 | 1 ; | HG | - | HD21 |
| 4577 | 4584 | 1 ; | HG | - | HD22 |
| 4577 | 4585 | 1 ; | HG | - | HD23 |
| 4578 | 4583 | 1 ; | CD1 | - | HD21 |
| 4578 | 4584 | 1 ; | CD1 | - | HD22 |
| 4578 | 4585 | 1 ; | CD1 | - | HD23 |
| 4579 | 4582 | 1 ; | HD11 | - | CD2 |
| 4580 | 4582 | 1 ; | HD12 | - | CD2 |
| 4581 | 4582 | 1 ; | HD13 | - | CD2 |
| 4586 | 4591 | 1 ; | C | - | HA |
| 4586 | 4592 | 1 ; | C | - | CB |
| 4586 | 4605 | 1 ; | C | - | C |
| 4587 | 4589 | 1 ; | O | - | H |
| 4587 | 4590 | 1 ; | O | - | CA |
| 4588 | 4593 | 1 ; | N | - | HB2 |
| 4588 | 4594 | 1 ; | N | - | HB3 |
| 4588 | 4595 | 1 ; | N | - | CG |
| 4588 | 4606 | 1 ; | N | - | O |
| 4588 | 4607 | 1 ; | N | - | N |
| 4589 | 4591 | 1 ; | H | - | HA |
| 4589 | 4592 | 1 ; | H | - | CB |
| 4589 | 4605 | 1 ; | H | - | C |
| 4590 | 4596 | 1 ; | CA | - | HG |
| 4590 | 4597 | 1 ; | CA | - | CD1 |
| 4590 | 4601 | 1 ; | CA | - | CD2 |
| 4590 | 4608 | 1 ; | CA | - | H |
| 4590 | 4609 | 1 ; | CA | - | CA |
| 4591 | 4593 | 1 ; | HA | - | HB2 |
| 4591 | 4594 | 1 ; | HA | - | HB3 |
| 4591 | 4595 | 1 ; | HA | - | CG |
| 4591 | 4606 | 1 ; | HA | - | O |
| 4591 | 4607 | 1 ; | HA | - | N |
| 4592 | 4598 | 1 ; | CB | - | HD11 |
| 4592 | 4599 | 1 ; | CB | - | HD12 |
| 4592 | 4600 | 1 ; | CB | - | HD13 |
| 4592 | 4602 | 1 ; | CB | - | HD21 |

| | | | | |
|---|---|---|---|---|
| 4592 | 4603 | 1 ; | CB - | HD22 |
| 4592 | 4604 | 1 ; | CB - | HD23 |
| 4592 | 4606 | 1 ; | CB - | O |
| 4592 | 4607 | 1 ; | CB - | N |
| 4593 | 4596 | 1 ; | HB2 - | HG |
| 4593 | 4597 | 1 ; | HB2 - | CD1 |
| 4593 | 4601 | 1 ; | HB2 - | CD2 |
| 4593 | 4605 | 1 ; | HB2 - | C |
| 4594 | 4596 | 1 ; | HB3 - | HG |
| 4594 | 4597 | 1 ; | HB3 - | CD1 |
| 4594 | 4601 | 1 ; | HB3 - | CD2 |
| 4594 | 4605 | 1 ; | HB3 - | C |
| 4595 | 4605 | 1 ; | CG - | C |
| 4596 | 4598 | 1 ; | HG - | HD11 |
| 4596 | 4599 | 1 ; | HG - | HD12 |
| 4596 | 4600 | 1 ; | HG - | HD13 |
| 4596 | 4602 | 1 ; | HG - | HD21 |
| 4596 | 4603 | 1 ; | HG - | HD22 |
| 4596 | 4604 | 1 ; | HG - | HD23 |
| 4597 | 4602 | 1 ; | CD1 - | HD21 |
| 4597 | 4603 | 1 ; | CD1 - | HD22 |
| 4597 | 4604 | 1 ; | CD1 - | HD23 |
| 4598 | 4601 | 1 ; | HD11 - | CD2 |
| 4599 | 4601 | 1 ; | HD12 - | CD2 |
| 4600 | 4601 | 1 ; | HD13 - | CD2 |
| 4605 | 4610 | 1 ; | C - | HA |
| 4605 | 4611 | 1 ; | C - | CB |
| 4605 | 4627 | 1 ; | C - | C |
| 4606 | 4608 | 1 ; | O - | H |
| 4606 | 4609 | 1 ; | O - | CA |
| 4607 | 4612 | 1 ; | N - | HB2 |
| 4607 | 4613 | 1 ; | N - | HB3 |
| 4607 | 4614 | 1 ; | N - | CG |
| 4607 | 4628 | 1 ; | N - | O |
| 4607 | 4629 | 1 ; | N - | N |
| 4608 | 4610 | 1 ; | H - | HA |
| 4608 | 4611 | 1 ; | H - | CB |
| 4608 | 4627 | 1 ; | H - | C |
| 4609 | 4615 | 1 ; | CA - | HG2 |
| 4609 | 4616 | 1 ; | CA - | HG3 |
| 4609 | 4617 | 1 ; | CA - | CD |
| 4609 | 4630 | 1 ; | CA - | H |
| 4609 | 4631 | 1 ; | CA - | CA |
| 4610 | 4612 | 1 ; | HA - | HB2 |
| 4610 | 4613 | 1 ; | HA - | HB3 |
| 4610 | 4614 | 1 ; | HA - | CG |
| 4610 | 4628 | 1 ; | HA - | O |
| 4610 | 4629 | 1 ; | HA - | N |
| 4611 | 4618 | 1 ; | CB - | HD2 |
| 4611 | 4619 | 1 ; | CB - | HD3 |
| 4611 | 4620 | 1 ; | CB - | CE |
| 4611 | 4628 | 1 ; | CB - | O |

```
 4611   4629       1 ;      CB - N
 4612   4615       1 ;     HB2 - HG2
 4612   4616       1 ;     HB2 - HG3
 4612   4617       1 ;     HB2 - CD
 4612   4627       1 ;     HB2 - C
 4613   4615       1 ;     HB3 - HG2
 4613   4616       1 ;     HB3 - HG3
 4613   4617       1 ;     HB3 - CD
 4613   4627       1 ;     HB3 - C
 4614   4621       1 ;      CG - HE2
 4614   4622       1 ;      CG - HE3
 4614   4623       1 ;      CG - NZ
 4614   4627       1 ;      CG - C
 4615   4618       1 ;     HG2 - HD2
 4615   4619       1 ;     HG2 - HD3
 4615   4620       1 ;     HG2 - CE
 4616   4618       1 ;     HG3 - HD2
 4616   4619       1 ;     HG3 - HD3
 4616   4620       1 ;     HG3 - CE
 4617   4624       1 ;      CD - HZ1
 4617   4625       1 ;      CD - HZ2
 4617   4626       1 ;      CD - HZ3
 4618   4621       1 ;     HD2 - HE2
 4618   4622       1 ;     HD2 - HE3
 4618   4623       1 ;     HD2 - NZ
 4619   4621       1 ;     HD3 - HE2
 4619   4622       1 ;     HD3 - HE3
 4619   4623       1 ;     HD3 - NZ
 4621   4624       1 ;     HE2 - HZ1
 4621   4625       1 ;     HE2 - HZ2
 4621   4626       1 ;     HE2 - HZ3
 4622   4624       1 ;     HE3 - HZ1
 4622   4625       1 ;     HE3 - HZ2
 4622   4626       1 ;     HE3 - HZ3
 4627   4632       1 ;       C - HA
 4627   4633       1 ;       C - CB
 4627   4641       1 ;       C - C
 4628   4630       1 ;       O - H
 4628   4631       1 ;       O - CA
 4629   4634       1 ;       N - HB2
 4629   4635       1 ;       N - HB3
 4629   4636       1 ;       N - CG
 4629   4642       1 ;       N - O
 4629   4643       1 ;       N - N
 4630   4632       1 ;       H - HA
 4630   4633       1 ;       H - CB
 4630   4641       1 ;       H - C
 4631   4637       1 ;      CA - OD1
 4631   4638       1 ;      CA - ND2
 4631   4644       1 ;      CA - H
 4631   4645       1 ;      CA - CA
 4632   4634       1 ;      HA - HB2
```

```
4632   4635      1 ;     HA - HB3
4632   4636      1 ;     HA - CG
4632   4642      1 ;     HA - O
4632   4643      1 ;     HA - N
4633   4639      1 ;     CB - HD21
4633   4640      1 ;     CB - HD22
4633   4642      1 ;     CB - O
4633   4643      1 ;     CB - N
4634   4637      1 ;    HB2 - OD1
4634   4638      1 ;    HB2 - ND2
4634   4641      1 ;    HB2 - C
4635   4637      1 ;    HB3 - OD1
4635   4638      1 ;    HB3 - ND2
4635   4641      1 ;    HB3 - C
4636   4641      1 ;     CG - C
4637   4639      1 ;    OD1 - HD21
4637   4640      1 ;    OD1 - HD22
4641   4646      1 ;      C - HA
4641   4647      1 ;      C - CB
4641   4660      1 ;      C - C
4642   4644      1 ;      O - H
4642   4645      1 ;      O - CA
4643   4648      1 ;      N - HB2
4643   4649      1 ;      N - HB3
4643   4650      1 ;      N - CG
4643   4661      1 ;      N - O
4643   4662      1 ;      N - N
4644   4646      1 ;      H - HA
4644   4647      1 ;      H - CB
4644   4660      1 ;      H - C
4645   4651      1 ;     CA - HG
4645   4652      1 ;     CA - CD1
4645   4656      1 ;     CA - CD2
4645   4663      1 ;     CA - H
4645   4664      1 ;     CA - CA
4646   4648      1 ;     HA - HB2
4646   4649      1 ;     HA - HB3
4646   4650      1 ;     HA - CG
4646   4661      1 ;     HA - O
4646   4662      1 ;     HA - N
4647   4653      1 ;     CB - HD11
4647   4654      1 ;     CB - HD12
4647   4655      1 ;     CB - HD13
4647   4657      1 ;     CB - HD21
4647   4658      1 ;     CB - HD22
4647   4659      1 ;     CB - HD23
4647   4661      1 ;     CB - O
4647   4662      1 ;     CB - N
4648   4651      1 ;    HB2 - HG
4648   4652      1 ;    HB2 - CD1
4648   4656      1 ;    HB2 - CD2
4648   4660      1 ;    HB2 - C
```

```
4649   4651       1 ;    HB3 - HG
4649   4652       1 ;    HB3 - CD1
4649   4656       1 ;    HB3 - CD2
4649   4660       1 ;    HB3 - C
4650   4660       1 ;     CG - C
4651   4653       1 ;     HG - HD11
4651   4654       1 ;     HG - HD12
4651   4655       1 ;     HG - HD13
4651   4657       1 ;     HG - HD21
4651   4658       1 ;     HG - HD22
4651   4659       1 ;     HG - HD23
4652   4657       1 ;    CD1 - HD21
4652   4658       1 ;    CD1 - HD22
4652   4659       1 ;    CD1 - HD23
4653   4656       1 ;   HD11 - CD2
4654   4656       1 ;   HD12 - CD2
4655   4656       1 ;   HD13 - CD2
4660   4665       1 ;      C - HA
4660   4666       1 ;      C - CB
4660   4682       1 ;      C - C
4661   4663       1 ;      O - H
4661   4664       1 ;      O - CA
4662   4667       1 ;      N - HB2
4662   4668       1 ;      N - HB3
4662   4669       1 ;      N - CG
4662   4683       1 ;      N - O
4662   4684       1 ;      N - N
4663   4665       1 ;      H - HA
4663   4666       1 ;      H - CB
4663   4682       1 ;      H - C
4664   4670       1 ;     CA - HG2
4664   4671       1 ;     CA - HG3
4664   4672       1 ;     CA - CD
4664   4685       1 ;     CA - H
4664   4686       1 ;     CA - CA
4665   4667       1 ;     HA - HB2
4665   4668       1 ;     HA - HB3
4665   4669       1 ;     HA - CG
4665   4683       1 ;     HA - O
4665   4684       1 ;     HA - N
4666   4673       1 ;     CB - HD2
4666   4674       1 ;     CB - HD3
4666   4675       1 ;     CB - CE
4666   4683       1 ;     CB - O
4666   4684       1 ;     CB - N
4667   4670       1 ;    HB2 - HG2
4667   4671       1 ;    HB2 - HG3
4667   4672       1 ;    HB2 - CD
4667   4682       1 ;    HB2 - C
4668   4670       1 ;    HB3 - HG2
4668   4671       1 ;    HB3 - HG3
4668   4672       1 ;    HB3 - CD
```

```
4668   4682       1 ;    HB3 - C
4669   4676       1 ;     CG - HE2
4669   4677       1 ;     CG - HE3
4669   4678       1 ;     CG - NZ
4669   4682       1 ;     CG - C
4670   4673       1 ;    HG2 - HD2
4670   4674       1 ;    HG2 - HD3
4670   4675       1 ;    HG2 - CE
4671   4673       1 ;    HG3 - HD2
4671   4674       1 ;    HG3 - HD3
4671   4675       1 ;    HG3 - CE
4672   4679       1 ;     CD - HZ1
4672   4680       1 ;     CD - HZ2
4672   4681       1 ;     CD - HZ3
4673   4676       1 ;    HD2 - HE2
4673   4677       1 ;    HD2 - HE3
4673   4678       1 ;    HD2 - NZ
4674   4676       1 ;    HD3 - HE2
4674   4677       1 ;    HD3 - HE3
4674   4678       1 ;    HD3 - NZ
4676   4679       1 ;    HE2 - HZ1
4676   4680       1 ;    HE2 - HZ2
4676   4681       1 ;    HE2 - HZ3
4677   4679       1 ;    HE3 - HZ1
4677   4680       1 ;    HE3 - HZ2
4677   4681       1 ;    HE3 - HZ3
4682   4687       1 ;      C - HA
4682   4688       1 ;      C - CB
4682   4696       1 ;      C - C
4683   4685       1 ;      O - H
4683   4686       1 ;      O - CA
4684   4689       1 ;      N - HB2
4684   4690       1 ;      N - HB3
4684   4691       1 ;      N - CG
4684   4697       1 ;      N - O
4684   4698       1 ;      N - N
4685   4687       1 ;      H - HA
4685   4688       1 ;      H - CB
4685   4696       1 ;      H - C
4686   4692       1 ;     CA - OD1
4686   4693       1 ;     CA - ND2
4686   4699       1 ;     CA - H
4686   4700       1 ;     CA - CA
4687   4689       1 ;     HA - HB2
4687   4690       1 ;     HA - HB3
4687   4691       1 ;     HA - CG
4687   4697       1 ;     HA - O
4687   4698       1 ;     HA - N
4688   4694       1 ;     CB - HD21
4688   4695       1 ;     CB - HD22
4688   4697       1 ;     CB - O
4688   4698       1 ;     CB - N
```

```
4689   4692      1 ;    HB2 - OD1
4689   4693      1 ;    HB2 - ND2
4689   4696      1 ;    HB2 - C
4690   4692      1 ;    HB3 - OD1
4690   4693      1 ;    HB3 - ND2
4690   4696      1 ;    HB3 - C
4691   4696      1 ;     CG - C
4692   4694      1 ;    OD1 - HD21
4692   4695      1 ;    OD1 - HD22
4696   4701      1 ;      C - HA
4696   4702      1 ;      C - CB
4696   4716      1 ;      C - C
4697   4699      1 ;      O - H
4697   4700      1 ;      O - CA
4698   4703      1 ;      N - HB2
4698   4704      1 ;      N - HB3
4698   4705      1 ;      N - CG
4698   4717      1 ;      N - O
4698   4718      1 ;      N - N
4699   4701      1 ;      H - HA
4699   4702      1 ;      H - CB
4699   4716      1 ;      H - C
4700   4706      1 ;     CA - CD1
4700   4714      1 ;     CA - CD2
4700   4719      1 ;     CA - H
4700   4720      1 ;     CA - CA
4701   4703      1 ;     HA - HB2
4701   4704      1 ;     HA - HB3
4701   4705      1 ;     HA - CG
4701   4717      1 ;     HA - O
4701   4718      1 ;     HA - N
4702   4707      1 ;     CB - HD1
4702   4708      1 ;     CB - CE1
4702   4712      1 ;     CB - CE2
4702   4715      1 ;     CB - HD2
4702   4717      1 ;     CB - O
4702   4718      1 ;     CB - N
4703   4706      1 ;    HB2 - CD1
4703   4714      1 ;    HB2 - CD2
4703   4716      1 ;    HB2 - C
4704   4706      1 ;    HB3 - CD1
4704   4714      1 ;    HB3 - CD2
4704   4716      1 ;    HB3 - C
4705   4709      1 ;     CG - HE1
4705   4710      1 ;     CG - CZ
4705   4713      1 ;     CG - HE2
4705   4716      1 ;     CG - C
4706   4711      1 ;    CD1 - HZ
4706   4712      1 ;    CD1 - CE2
4706   4715      1 ;    CD1 - HD2
4707   4709      1 ;    HD1 - HE1
4707   4710      1 ;    HD1 - CZ
```

| | | | |
|---|---|---|---|
| 4707 | 4714 | 1 ; | HD1 - CD2 |
| 4708 | 4713 | 1 ; | CE1 - HE2 |
| 4708 | 4714 | 1 ; | CE1 - CD2 |
| 4709 | 4711 | 1 ; | HE1 - HZ |
| 4709 | 4712 | 1 ; | HE1 - CE2 |
| 4710 | 4715 | 1 ; | CZ - HD2 |
| 4711 | 4713 | 1 ; | HZ - HE2 |
| 4711 | 4714 | 1 ; | HZ - CD2 |
| 4713 | 4715 | 1 ; | HE2 - HD2 |
| 4716 | 4721 | 1 ; | C - HA |
| 4716 | 4722 | 1 ; | C - CB |
| 4716 | 4738 | 1 ; | C - C |
| 4717 | 4719 | 1 ; | O - H |
| 4717 | 4720 | 1 ; | O - CA |
| 4718 | 4723 | 1 ; | N - HB2 |
| 4718 | 4724 | 1 ; | N - HB3 |
| 4718 | 4725 | 1 ; | N - CG |
| 4718 | 4739 | 1 ; | N - O |
| 4718 | 4740 | 1 ; | N - N |
| 4719 | 4721 | 1 ; | H - HA |
| 4719 | 4722 | 1 ; | H - CB |
| 4719 | 4738 | 1 ; | H - C |
| 4720 | 4726 | 1 ; | CA - HG2 |
| 4720 | 4727 | 1 ; | CA - HG3 |
| 4720 | 4728 | 1 ; | CA - CD |
| 4720 | 4741 | 1 ; | CA - H |
| 4720 | 4742 | 1 ; | CA - CA |
| 4721 | 4723 | 1 ; | HA - HB2 |
| 4721 | 4724 | 1 ; | HA - HB3 |
| 4721 | 4725 | 1 ; | HA - CG |
| 4721 | 4739 | 1 ; | HA - O |
| 4721 | 4740 | 1 ; | HA - N |
| 4722 | 4729 | 1 ; | CB - HD2 |
| 4722 | 4730 | 1 ; | CB - HD3 |
| 4722 | 4731 | 1 ; | CB - CE |
| 4722 | 4739 | 1 ; | CB - O |
| 4722 | 4740 | 1 ; | CB - N |
| 4723 | 4726 | 1 ; | HB2 - HG2 |
| 4723 | 4727 | 1 ; | HB2 - HG3 |
| 4723 | 4728 | 1 ; | HB2 - CD |
| 4723 | 4738 | 1 ; | HB2 - C |
| 4724 | 4726 | 1 ; | HB3 - HG2 |
| 4724 | 4727 | 1 ; | HB3 - HG3 |
| 4724 | 4728 | 1 ; | HB3 - CD |
| 4724 | 4738 | 1 ; | HB3 - C |
| 4725 | 4732 | 1 ; | CG - HE2 |
| 4725 | 4733 | 1 ; | CG - HE3 |
| 4725 | 4734 | 1 ; | CG - NZ |
| 4725 | 4738 | 1 ; | CG - C |
| 4726 | 4729 | 1 ; | HG2 - HD2 |
| 4726 | 4730 | 1 ; | HG2 - HD3 |
| 4726 | 4731 | 1 ; | HG2 - CE |

| | | | | | |
|---|---|---|---|---|---|
| 4727 | 4729 | 1 ; | HG3 | - | HD2 |
| 4727 | 4730 | 1 ; | HG3 | - | HD3 |
| 4727 | 4731 | 1 ; | HG3 | - | CE |
| 4728 | 4735 | 1 ; | CD | - | HZ1 |
| 4728 | 4736 | 1 ; | CD | - | HZ2 |
| 4728 | 4737 | 1 ; | CD | - | HZ3 |
| 4729 | 4732 | 1 ; | HD2 | - | HE2 |
| 4729 | 4733 | 1 ; | HD2 | - | HE3 |
| 4729 | 4734 | 1 ; | HD2 | - | NZ |
| 4730 | 4732 | 1 ; | HD3 | - | HE2 |
| 4730 | 4733 | 1 ; | HD3 | - | HE3 |
| 4730 | 4734 | 1 ; | HD3 | - | NZ |
| 4732 | 4735 | 1 ; | HE2 | - | HZ1 |
| 4732 | 4736 | 1 ; | HE2 | - | HZ2 |
| 4732 | 4737 | 1 ; | HE2 | - | HZ3 |
| 4733 | 4735 | 1 ; | HE3 | - | HZ1 |
| 4733 | 4736 | 1 ; | HE3 | - | HZ2 |
| 4733 | 4737 | 1 ; | HE3 | - | HZ3 |
| 4738 | 4743 | 1 ; | C | - | HA |
| 4738 | 4744 | 1 ; | C | - | CB |
| 4738 | 4760 | 1 ; | C | - | C |
| 4739 | 4741 | 1 ; | O | - | H |
| 4739 | 4742 | 1 ; | O | - | CA |
| 4740 | 4745 | 1 ; | N | - | HB2 |
| 4740 | 4746 | 1 ; | N | - | HB3 |
| 4740 | 4747 | 1 ; | N | - | CG |
| 4740 | 4761 | 1 ; | N | - | O |
| 4740 | 4762 | 1 ; | N | - | N |
| 4741 | 4743 | 1 ; | H | - | HA |
| 4741 | 4744 | 1 ; | H | - | CB |
| 4741 | 4760 | 1 ; | H | - | C |
| 4742 | 4748 | 1 ; | CA | - | HG2 |
| 4742 | 4749 | 1 ; | CA | - | HG3 |
| 4742 | 4750 | 1 ; | CA | - | CD |
| 4742 | 4763 | 1 ; | CA | - | H |
| 4742 | 4764 | 1 ; | CA | - | CA |
| 4743 | 4745 | 1 ; | HA | - | HB2 |
| 4743 | 4746 | 1 ; | HA | - | HB3 |
| 4743 | 4747 | 1 ; | HA | - | CG |
| 4743 | 4761 | 1 ; | HA | - | O |
| 4743 | 4762 | 1 ; | HA | - | N |
| 4744 | 4751 | 1 ; | CB | - | HD2 |
| 4744 | 4752 | 1 ; | CB | - | HD3 |
| 4744 | 4753 | 1 ; | CB | - | CE |
| 4744 | 4761 | 1 ; | CB | - | O |
| 4744 | 4762 | 1 ; | CB | - | N |
| 4745 | 4748 | 1 ; | HB2 | - | HG2 |
| 4745 | 4749 | 1 ; | HB2 | - | HG3 |
| 4745 | 4750 | 1 ; | HB2 | - | CD |
| 4745 | 4760 | 1 ; | HB2 | - | C |
| 4746 | 4748 | 1 ; | HB3 | - | HG2 |
| 4746 | 4749 | 1 ; | HB3 | - | HG3 |

| | | | | | |
|---|---|---|---|---|---|
| 4746 | 4750 | 1 ; | HB3 | - | CD |
| 4746 | 4760 | 1 ; | HB3 | - | C |
| 4747 | 4754 | 1 ; | CG | - | HE2 |
| 4747 | 4755 | 1 ; | CG | - | HE3 |
| 4747 | 4756 | 1 ; | CG | - | NZ |
| 4747 | 4760 | 1 ; | CG | - | C |
| 4748 | 4751 | 1 ; | HG2 | - | HD2 |
| 4748 | 4752 | 1 ; | HG2 | - | HD3 |
| 4748 | 4753 | 1 ; | HG2 | - | CE |
| 4749 | 4751 | 1 ; | HG3 | - | HD2 |
| 4749 | 4752 | 1 ; | HG3 | - | HD3 |
| 4749 | 4753 | 1 ; | HG3 | - | CE |
| 4750 | 4757 | 1 ; | CD | - | HZ1 |
| 4750 | 4758 | 1 ; | CD | - | HZ2 |
| 4750 | 4759 | 1 ; | CD | - | HZ3 |
| 4751 | 4754 | 1 ; | HD2 | - | HE2 |
| 4751 | 4755 | 1 ; | HD2 | - | HE3 |
| 4751 | 4756 | 1 ; | HD2 | - | NZ |
| 4752 | 4754 | 1 ; | HD3 | - | HE2 |
| 4752 | 4755 | 1 ; | HD3 | - | HE3 |
| 4752 | 4756 | 1 ; | HD3 | - | NZ |
| 4754 | 4757 | 1 ; | HE2 | - | HZ1 |
| 4754 | 4758 | 1 ; | HE2 | - | HZ2 |
| 4754 | 4759 | 1 ; | HE2 | - | HZ3 |
| 4755 | 4757 | 1 ; | HE3 | - | HZ1 |
| 4755 | 4758 | 1 ; | HE3 | - | HZ2 |
| 4755 | 4759 | 1 ; | HE3 | - | HZ3 |
| 4760 | 4765 | 1 ; | C | - | HA |
| 4760 | 4766 | 1 ; | C | - | CB |
| 4760 | 4775 | 1 ; | C | - | C |
| 4761 | 4763 | 1 ; | O | - | H |
| 4761 | 4764 | 1 ; | O | - | CA |
| 4762 | 4767 | 1 ; | N | - | HB2 |
| 4762 | 4768 | 1 ; | N | - | HB3 |
| 4762 | 4769 | 1 ; | N | - | CG |
| 4762 | 4776 | 1 ; | N | - | O |
| 4762 | 4777 | 1 ; | N | - | N |
| 4763 | 4765 | 1 ; | H | - | HA |
| 4763 | 4766 | 1 ; | H | - | CB |
| 4763 | 4775 | 1 ; | H | - | C |
| 4764 | 4770 | 1 ; | CA | - | HG2 |
| 4764 | 4771 | 1 ; | CA | - | HG3 |
| 4764 | 4772 | 1 ; | CA | - | CD |
| 4764 | 4778 | 1 ; | CA | - | H |
| 4764 | 4779 | 1 ; | CA | - | CA |
| 4765 | 4767 | 1 ; | HA | - | HB2 |
| 4765 | 4768 | 1 ; | HA | - | HB3 |
| 4765 | 4769 | 1 ; | HA | - | CG |
| 4765 | 4776 | 1 ; | HA | - | O |
| 4765 | 4777 | 1 ; | HA | - | N |
| 4766 | 4773 | 1 ; | CB | - | OE1 |
| 4766 | 4774 | 1 ; | CB | - | OE2 |

| | | | |
|---|---|---|---|
| 4766 | 4776 | 1 ; | CB - O |
| 4766 | 4777 | 1 ; | CB - N |
| 4767 | 4770 | 1 ; | HB2 - HG2 |
| 4767 | 4771 | 1 ; | HB2 - HG3 |
| 4767 | 4772 | 1 ; | HB2 - CD |
| 4767 | 4775 | 1 ; | HB2 - C |
| 4768 | 4770 | 1 ; | HB3 - HG2 |
| 4768 | 4771 | 1 ; | HB3 - HG3 |
| 4768 | 4772 | 1 ; | HB3 - CD |
| 4768 | 4775 | 1 ; | HB3 - C |
| 4769 | 4775 | 1 ; | CG - C |
| 4770 | 4773 | 1 ; | HG2 - OE1 |
| 4770 | 4774 | 1 ; | HG2 - OE2 |
| 4771 | 4773 | 1 ; | HG3 - OE1 |
| 4771 | 4774 | 1 ; | HG3 - OE2 |
| 4775 | 4780 | 1 ; | C - HA |
| 4775 | 4781 | 1 ; | C - CB |
| 4775 | 4789 | 1 ; | C - C |
| 4776 | 4778 | 1 ; | O - H |
| 4776 | 4779 | 1 ; | O - CA |
| 4777 | 4782 | 1 ; | N - HB2 |
| 4777 | 4783 | 1 ; | N - HB3 |
| 4777 | 4784 | 1 ; | N - CG |
| 4777 | 4790 | 1 ; | N - O |
| 4777 | 4791 | 1 ; | N - N |
| 4778 | 4780 | 1 ; | H - HA |
| 4778 | 4781 | 1 ; | H - CB |
| 4778 | 4789 | 1 ; | H - C |
| 4779 | 4785 | 1 ; | CA - OD1 |
| 4779 | 4786 | 1 ; | CA - ND2 |
| 4779 | 4792 | 1 ; | CA - H |
| 4779 | 4793 | 1 ; | CA - CA |
| 4780 | 4782 | 1 ; | HA - HB2 |
| 4780 | 4783 | 1 ; | HA - HB3 |
| 4780 | 4784 | 1 ; | HA - CG |
| 4780 | 4790 | 1 ; | HA - O |
| 4780 | 4791 | 1 ; | HA - N |
| 4781 | 4787 | 1 ; | CB - HD21 |
| 4781 | 4788 | 1 ; | CB - HD22 |
| 4781 | 4790 | 1 ; | CB - O |
| 4781 | 4791 | 1 ; | CB - N |
| 4782 | 4785 | 1 ; | HB2 - OD1 |
| 4782 | 4786 | 1 ; | HB2 - ND2 |
| 4782 | 4789 | 1 ; | HB2 - C |
| 4783 | 4785 | 1 ; | HB3 - OD1 |
| 4783 | 4786 | 1 ; | HB3 - ND2 |
| 4783 | 4789 | 1 ; | HB3 - C |
| 4784 | 4789 | 1 ; | CG - C |
| 4785 | 4787 | 1 ; | OD1 - HD21 |
| 4785 | 4788 | 1 ; | OD1 - HD22 |
| 4789 | 4794 | 1 ; | C - HA |
| 4789 | 4795 | 1 ; | C - CB |

```
4789   4804       1 ;       C - C
4790   4792       1 ;       O - H
4790   4793       1 ;       O - CA
4791   4796       1 ;       N - HB2
4791   4797       1 ;       N - HB3
4791   4798       1 ;       N - CG
4791   4805       1 ;       N - O
4791   4806       1 ;       N - N
4792   4794       1 ;       H - HA
4792   4795       1 ;       H - CB
4792   4804       1 ;       H - C
4793   4799       1 ;      CA - HG2
4793   4800       1 ;      CA - HG3
4793   4801       1 ;      CA - CD
4793   4807       1 ;      CA - H
4793   4808       1 ;      CA - CA
4794   4796       1 ;      HA - HB2
4794   4797       1 ;      HA - HB3
4794   4798       1 ;      HA - CG
4794   4805       1 ;      HA - O
4794   4806       1 ;      HA - N
4795   4802       1 ;      CB - OE1
4795   4803       1 ;      CB - OE2
4795   4805       1 ;      CB - O
4795   4806       1 ;      CB - N
4796   4799       1 ;     HB2 - HG2
4796   4800       1 ;     HB2 - HG3
4796   4801       1 ;     HB2 - CD
4796   4804       1 ;     HB2 - C
4797   4799       1 ;     HB3 - HG2
4797   4800       1 ;     HB3 - HG3
4797   4801       1 ;     HB3 - CD
4797   4804       1 ;     HB3 - C
4798   4804       1 ;      CG - C
4799   4802       1 ;     HG2 - OE1
4799   4803       1 ;     HG2 - OE2
4800   4802       1 ;     HG3 - OE1
4800   4803       1 ;     HG3 - OE2
4804   4809       1 ;       C - HA
4804   4810       1 ;       C - CB
4804   4823       1 ;       C - C
4805   4807       1 ;       O - H
4805   4808       1 ;       O - CA
4806   4811       1 ;       N - HB2
4806   4812       1 ;       N - HB3
4806   4813       1 ;       N - CG
4806   4824       1 ;       N - O
4806   4825       1 ;       N - N
4807   4809       1 ;       H - HA
4807   4810       1 ;       H - CB
4807   4823       1 ;       H - C
4808   4814       1 ;      CA - HG
```

```
4808   4815      1 ;      CA - CD1
4808   4819      1 ;      CA - CD2
4808   4826      1 ;      CA - H
4808   4827      1 ;      CA - CA
4809   4811      1 ;      HA - HB2
4809   4812      1 ;      HA - HB3
4809   4813      1 ;      HA - CG
4809   4824      1 ;      HA - O
4809   4825      1 ;      HA - N
4810   4816      1 ;      CB - HD11
4810   4817      1 ;      CB - HD12
4810   4818      1 ;      CB - HD13
4810   4820      1 ;      CB - HD21
4810   4821      1 ;      CB - HD22
4810   4822      1 ;      CB - HD23
4810   4824      1 ;      CB - O
4810   4825      1 ;      CB - N
4811   4814      1 ;     HB2 - HG
4811   4815      1 ;     HB2 - CD1
4811   4819      1 ;     HB2 - CD2
4811   4823      1 ;     HB2 - C
4812   4814      1 ;     HB3 - HG
4812   4815      1 ;     HB3 - CD1
4812   4819      1 ;     HB3 - CD2
4812   4823      1 ;     HB3 - C
4813   4823      1 ;      CG - C
4814   4816      1 ;      HG - HD11
4814   4817      1 ;      HG - HD12
4814   4818      1 ;      HG - HD13
4814   4820      1 ;      HG - HD21
4814   4821      1 ;      HG - HD22
4814   4822      1 ;      HG - HD23
4815   4820      1 ;     CD1 - HD21
4815   4821      1 ;     CD1 - HD22
4815   4822      1 ;     CD1 - HD23
4816   4819      1 ;    HD11 - CD2
4817   4819      1 ;    HD12 - CD2
4818   4819      1 ;    HD13 - CD2
4823   4828      1 ;       C - HA
4823   4829      1 ;       C - CB
4823   4845      1 ;       C - C
4824   4826      1 ;       O - H
4824   4827      1 ;       O - CA
4825   4830      1 ;       N - HB2
4825   4831      1 ;       N - HB3
4825   4832      1 ;       N - CG
4825   4846      1 ;       N - O
4825   4847      1 ;       N - N
4826   4828      1 ;       H - HA
4826   4829      1 ;       H - CB
4826   4845      1 ;       H - C
4827   4833      1 ;      CA - HG2
```

```
 4827   4834       1 ;      CA - HG3
 4827   4835       1 ;      CA - CD
 4827   4848       1 ;      CA - H
 4827   4849       1 ;      CA - CA
 4828   4830       1 ;      HA - HB2
 4828   4831       1 ;      HA - HB3
 4828   4832       1 ;      HA - CG
 4828   4846       1 ;      HA - O
 4828   4847       1 ;      HA - N
 4829   4836       1 ;      CB - HD2
 4829   4837       1 ;      CB - HD3
 4829   4838       1 ;      CB - CE
 4829   4846       1 ;      CB - O
 4829   4847       1 ;      CB - N
 4830   4833       1 ;     HB2 - HG2
 4830   4834       1 ;     HB2 - HG3
 4830   4835       1 ;     HB2 - CD
 4830   4845       1 ;     HB2 - C
 4831   4833       1 ;     HB3 - HG2
 4831   4834       1 ;     HB3 - HG3
 4831   4835       1 ;     HB3 - CD
 4831   4845       1 ;     HB3 - C
 4832   4839       1 ;      CG - HE2
 4832   4840       1 ;      CG - HE3
 4832   4841       1 ;      CG - NZ
 4832   4845       1 ;      CG - C
 4833   4836       1 ;     HG2 - HD2
 4833   4837       1 ;     HG2 - HD3
 4833   4838       1 ;     HG2 - CE
 4834   4836       1 ;     HG3 - HD2
 4834   4837       1 ;     HG3 - HD3
 4834   4838       1 ;     HG3 - CE
 4835   4842       1 ;      CD - HZ1
 4835   4843       1 ;      CD - HZ2
 4835   4844       1 ;      CD - HZ3
 4836   4839       1 ;     HD2 - HE2
 4836   4840       1 ;     HD2 - HE3
 4836   4841       1 ;     HD2 - NZ
 4837   4839       1 ;     HD3 - HE2
 4837   4840       1 ;     HD3 - HE3
 4837   4841       1 ;     HD3 - NZ
 4839   4842       1 ;     HE2 - HZ1
 4839   4843       1 ;     HE2 - HZ2
 4839   4844       1 ;     HE2 - HZ3
 4840   4842       1 ;     HE3 - HZ1
 4840   4843       1 ;     HE3 - HZ2
 4840   4844       1 ;     HE3 - HZ3
 4845   4850       1 ;       C - HA
 4845   4851       1 ;       C - CB
 4845   4867       1 ;       C - C
 4846   4848       1 ;       O - H
 4846   4849       1 ;       O - CA
```

```
4847   4852       1 ;       N - HB2
4847   4853       1 ;       N - HB3
4847   4854       1 ;       N - CG
4847   4868       1 ;       N - O
4847   4869       1 ;       N - N
4848   4850       1 ;       H - HA
4848   4851       1 ;       H - CB
4848   4867       1 ;       H - C
4849   4855       1 ;      CA - HG2
4849   4856       1 ;      CA - HG3
4849   4857       1 ;      CA - CD
4849   4870       1 ;      CA - H
4849   4871       1 ;      CA - CA
4850   4852       1 ;      HA - HB2
4850   4853       1 ;      HA - HB3
4850   4854       1 ;      HA - CG
4850   4868       1 ;      HA - O
4850   4869       1 ;      HA - N
4851   4858       1 ;      CB - HD2
4851   4859       1 ;      CB - HD3
4851   4860       1 ;      CB - CE
4851   4868       1 ;      CB - O
4851   4869       1 ;      CB - N
4852   4855       1 ;     HB2 - HG2
4852   4856       1 ;     HB2 - HG3
4852   4857       1 ;     HB2 - CD
4852   4867       1 ;     HB2 - C
4853   4855       1 ;     HB3 - HG2
4853   4856       1 ;     HB3 - HG3
4853   4857       1 ;     HB3 - CD
4853   4867       1 ;     HB3 - C
4854   4861       1 ;      CG - HE2
4854   4862       1 ;      CG - HE3
4854   4863       1 ;      CG - NZ
4854   4867       1 ;      CG - C
4855   4858       1 ;     HG2 - HD2
4855   4859       1 ;     HG2 - HD3
4855   4860       1 ;     HG2 - CE
4856   4858       1 ;     HG3 - HD2
4856   4859       1 ;     HG3 - HD3
4856   4860       1 ;     HG3 - CE
4857   4864       1 ;      CD - HZ1
4857   4865       1 ;      CD - HZ2
4857   4866       1 ;      CD - HZ3
4858   4861       1 ;     HD2 - HE2
4858   4862       1 ;     HD2 - HE3
4858   4863       1 ;     HD2 - NZ
4859   4861       1 ;     HD3 - HE2
4859   4862       1 ;     HD3 - HE3
4859   4863       1 ;     HD3 - NZ
4861   4864       1 ;     HE2 - HZ1
4861   4865       1 ;     HE2 - HZ2
```

```
4861   4866      1 ;    HE2 - HZ3
4862   4864      1 ;    HE3 - HZ1
4862   4865      1 ;    HE3 - HZ2
4862   4866      1 ;    HE3 - HZ3
4867   4872      1 ;      C - HA
4867   4873      1 ;      C - CB
4867   4886      1 ;      C - C
4868   4870      1 ;      O - H
4868   4871      1 ;      O - CA
4869   4874      1 ;      N - HB
4869   4875      1 ;      N - CG2
4869   4879      1 ;      N - CG1
4869   4887      1 ;      N - O
4869   4888      1 ;      N - N
4870   4872      1 ;      H - HA
4870   4873      1 ;      H - CB
4870   4886      1 ;      H - C
4871   4876      1 ;     CA - HG21
4871   4877      1 ;     CA - HG22
4871   4878      1 ;     CA - HG23
4871   4880      1 ;     CA - HG12
4871   4881      1 ;     CA - HG13
4871   4882      1 ;     CA - CD1
4871   4889      1 ;     CA - H
4871   4890      1 ;     CA - CA
4872   4874      1 ;     HA - HB
4872   4875      1 ;     HA - CG2
4872   4879      1 ;     HA - CG1
4872   4887      1 ;     HA - O
4872   4888      1 ;     HA - N
4873   4883      1 ;     CB - HD11
4873   4884      1 ;     CB - HD12
4873   4885      1 ;     CB - HD13
4873   4887      1 ;     CB - O
4873   4888      1 ;     CB - N
4874   4876      1 ;     HB - HG21
4874   4877      1 ;     HB - HG22
4874   4878      1 ;     HB - HG23
4874   4880      1 ;     HB - HG12
4874   4881      1 ;     HB - HG13
4874   4882      1 ;     HB - CD1
4874   4886      1 ;     HB - C
4875   4880      1 ;    CG2 - HG12
4875   4881      1 ;    CG2 - HG13
4875   4882      1 ;    CG2 - CD1
4875   4886      1 ;    CG2 - C
4876   4879      1 ;   HG21 - CG1
4877   4879      1 ;   HG22 - CG1
4878   4879      1 ;   HG23 - CG1
4879   4886      1 ;    CG1 - C
4880   4883      1 ;   HG12 - HD11
4880   4884      1 ;   HG12 - HD12
```

| | | | | |
|---|---|---|---|---|
| 4880 | 4885 | 1 ; | HG12 - | HD13 |
| 4881 | 4883 | 1 ; | HG13 - | HD11 |
| 4881 | 4884 | 1 ; | HG13 - | HD12 |
| 4881 | 4885 | 1 ; | HG13 - | HD13 |
| 4886 | 4891 | 1 ; | C - | HA |
| 4886 | 4892 | 1 ; | C - | CB |
| 4886 | 4896 | 1 ; | C - | C |
| 4887 | 4889 | 1 ; | O - | H |
| 4887 | 4890 | 1 ; | O - | CA |
| 4888 | 4893 | 1 ; | N - | HB1 |
| 4888 | 4894 | 1 ; | N - | HB2 |
| 4888 | 4895 | 1 ; | N - | HB3 |
| 4888 | 4897 | 1 ; | N - | O |
| 4888 | 4898 | 1 ; | N - | N |
| 4889 | 4891 | 1 ; | H - | HA |
| 4889 | 4892 | 1 ; | H - | CB |
| 4889 | 4896 | 1 ; | H - | C |
| 4890 | 4899 | 1 ; | CA - | H |
| 4890 | 4900 | 1 ; | CA - | CA |
| 4891 | 4893 | 1 ; | HA - | HB1 |
| 4891 | 4894 | 1 ; | HA - | HB2 |
| 4891 | 4895 | 1 ; | HA - | HB3 |
| 4891 | 4897 | 1 ; | HA - | O |
| 4891 | 4898 | 1 ; | HA - | N |
| 4892 | 4897 | 1 ; | CB - | O |
| 4892 | 4898 | 1 ; | CB - | N |
| 4893 | 4896 | 1 ; | HB1 - | C |
| 4894 | 4896 | 1 ; | HB2 - | C |
| 4895 | 4896 | 1 ; | HB3 - | C |
| 4896 | 4901 | 1 ; | C - | HA |
| 4896 | 4902 | 1 ; | C - | CB |
| 4896 | 4915 | 1 ; | C - | C |
| 4897 | 4899 | 1 ; | O - | H |
| 4897 | 4900 | 1 ; | O - | CA |
| 4898 | 4903 | 1 ; | N - | HB2 |
| 4898 | 4904 | 1 ; | N - | HB3 |
| 4898 | 4905 | 1 ; | N - | CG |
| 4898 | 4916 | 1 ; | N - | O |
| 4898 | 4917 | 1 ; | N - | N |
| 4899 | 4901 | 1 ; | H - | HA |
| 4899 | 4902 | 1 ; | H - | CB |
| 4899 | 4915 | 1 ; | H - | C |
| 4900 | 4906 | 1 ; | CA - | HG |
| 4900 | 4907 | 1 ; | CA - | CD1 |
| 4900 | 4911 | 1 ; | CA - | CD2 |
| 4900 | 4918 | 1 ; | CA - | H |
| 4900 | 4919 | 1 ; | CA - | CA |
| 4901 | 4903 | 1 ; | HA - | HB2 |
| 4901 | 4904 | 1 ; | HA - | HB3 |
| 4901 | 4905 | 1 ; | HA - | CG |
| 4901 | 4916 | 1 ; | HA - | O |
| 4901 | 4917 | 1 ; | HA - | N |

| | | | | |
|---|---|---|---|---|
| 4902 | 4908 | 1 ; | CB - | HD11 |
| 4902 | 4909 | 1 ; | CB - | HD12 |
| 4902 | 4910 | 1 ; | CB - | HD13 |
| 4902 | 4912 | 1 ; | CB - | HD21 |
| 4902 | 4913 | 1 ; | CB - | HD22 |
| 4902 | 4914 | 1 ; | CB - | HD23 |
| 4902 | 4916 | 1 ; | CB - | O |
| 4902 | 4917 | 1 ; | CB - | N |
| 4903 | 4906 | 1 ; | HB2 - | HG |
| 4903 | 4907 | 1 ; | HB2 - | CD1 |
| 4903 | 4911 | 1 ; | HB2 - | CD2 |
| 4903 | 4915 | 1 ; | HB2 - | C |
| 4904 | 4906 | 1 ; | HB3 - | HG |
| 4904 | 4907 | 1 ; | HB3 - | CD1 |
| 4904 | 4911 | 1 ; | HB3 - | CD2 |
| 4904 | 4915 | 1 ; | HB3 - | C |
| 4905 | 4915 | 1 ; | CG - | C |
| 4906 | 4908 | 1 ; | HG - | HD11 |
| 4906 | 4909 | 1 ; | HG - | HD12 |
| 4906 | 4910 | 1 ; | HG - | HD13 |
| 4906 | 4912 | 1 ; | HG - | HD21 |
| 4906 | 4913 | 1 ; | HG - | HD22 |
| 4906 | 4914 | 1 ; | HG - | HD23 |
| 4907 | 4912 | 1 ; | CD1 - | HD21 |
| 4907 | 4913 | 1 ; | CD1 - | HD22 |
| 4907 | 4914 | 1 ; | CD1 - | HD23 |
| 4908 | 4911 | 1 ; | HD11 - | CD2 |
| 4909 | 4911 | 1 ; | HD12 - | CD2 |
| 4910 | 4911 | 1 ; | HD13 - | CD2 |
| 4915 | 4920 | 1 ; | C - | HA |
| 4915 | 4921 | 1 ; | C - | CB |
| 4915 | 4929 | 1 ; | C - | C |
| 4916 | 4918 | 1 ; | O - | H |
| 4916 | 4919 | 1 ; | O - | CA |
| 4917 | 4922 | 1 ; | N - | HB |
| 4917 | 4923 | 1 ; | N - | CG2 |
| 4917 | 4927 | 1 ; | N - | OG1 |
| 4917 | 4930 | 1 ; | N - | O |
| 4917 | 4931 | 1 ; | N - | N |
| 4918 | 4920 | 1 ; | H - | HA |
| 4918 | 4921 | 1 ; | H - | CB |
| 4918 | 4929 | 1 ; | H - | C |
| 4919 | 4924 | 1 ; | CA - | HG21 |
| 4919 | 4925 | 1 ; | CA - | HG22 |
| 4919 | 4926 | 1 ; | CA - | HG23 |
| 4919 | 4928 | 1 ; | CA - | HG1 |
| 4919 | 4932 | 1 ; | CA - | H |
| 4919 | 4933 | 1 ; | CA - | CA |
| 4920 | 4922 | 1 ; | HA - | HB |
| 4920 | 4923 | 1 ; | HA - | CG2 |
| 4920 | 4927 | 1 ; | HA - | OG1 |
| 4920 | 4930 | 1 ; | HA - | O |

```
4920   4931       1 ;      HA - N
4921   4930       1 ;      CB - O
4921   4931       1 ;      CB - N
4922   4924       1 ;      HB - HG21
4922   4925       1 ;      HB - HG22
4922   4926       1 ;      HB - HG23
4922   4928       1 ;      HB - HG1
4922   4929       1 ;      HB - C
4923   4928       1 ;     CG2 - HG1
4923   4929       1 ;     CG2 - C
4924   4927       1 ;    HG21 - OG1
4925   4927       1 ;    HG22 - OG1
4926   4927       1 ;    HG23 - OG1
4927   4929       1 ;     OG1 - C
4929   4934       1 ;       C - HA
4929   4935       1 ;       C - CB
4929   4948       1 ;       C - C
4930   4932       1 ;       O - H
4930   4933       1 ;       O - CA
4931   4936       1 ;       N - HB
4931   4937       1 ;       N - CG2
4931   4941       1 ;       N - CG1
4931   4949       1 ;       N - O
4931   4950       1 ;       N - N
4932   4934       1 ;       H - HA
4932   4935       1 ;       H - CB
4932   4948       1 ;       H - C
4933   4938       1 ;      CA - HG21
4933   4939       1 ;      CA - HG22
4933   4940       1 ;      CA - HG23
4933   4942       1 ;      CA - HG12
4933   4943       1 ;      CA - HG13
4933   4944       1 ;      CA - CD1
4933   4951       1 ;      CA - H
4933   4952       1 ;      CA - CA
4934   4936       1 ;      HA - HB
4934   4937       1 ;      HA - CG2
4934   4941       1 ;      HA - CG1
4934   4949       1 ;      HA - O
4934   4950       1 ;      HA - N
4935   4945       1 ;      CB - HD11
4935   4946       1 ;      CB - HD12
4935   4947       1 ;      CB - HD13
4935   4949       1 ;      CB - O
4935   4950       1 ;      CB - N
4936   4938       1 ;      HB - HG21
4936   4939       1 ;      HB - HG22
4936   4940       1 ;      HB - HG23
4936   4942       1 ;      HB - HG12
4936   4943       1 ;      HB - HG13
4936   4944       1 ;      HB - CD1
4936   4948       1 ;      HB - C
```

| | | | | | | |
|---|---|---|---|---|---|---|
| 4937 | 4942 | 1 | ; | CG2 | - | HG12 |
| 4937 | 4943 | 1 | ; | CG2 | - | HG13 |
| 4937 | 4944 | 1 | ; | CG2 | - | CD1 |
| 4937 | 4948 | 1 | ; | CG2 | - | C |
| 4938 | 4941 | 1 | ; | HG21 | - | CG1 |
| 4939 | 4941 | 1 | ; | HG22 | - | CG1 |
| 4940 | 4941 | 1 | ; | HG23 | - | CG1 |
| 4941 | 4948 | 1 | ; | CG1 | - | C |
| 4942 | 4945 | 1 | ; | HG12 | - | HD11 |
| 4942 | 4946 | 1 | ; | HG12 | - | HD12 |
| 4942 | 4947 | 1 | ; | HG12 | - | HD13 |
| 4943 | 4945 | 1 | ; | HG13 | - | HD11 |
| 4943 | 4946 | 1 | ; | HG13 | - | HD12 |
| 4943 | 4947 | 1 | ; | HG13 | - | HD13 |
| 4948 | 4953 | 1 | ; | C | - | HA |
| 4948 | 4954 | 1 | ; | C | - | CB |
| 4948 | 4967 | 1 | ; | C | - | C |
| 4949 | 4951 | 1 | ; | O | - | H |
| 4949 | 4952 | 1 | ; | O | - | CA |
| 4950 | 4955 | 1 | ; | N | - | HB |
| 4950 | 4956 | 1 | ; | N | - | CG2 |
| 4950 | 4960 | 1 | ; | N | - | CG1 |
| 4950 | 4968 | 1 | ; | N | - | O |
| 4950 | 4969 | 1 | ; | N | - | N |
| 4951 | 4953 | 1 | ; | H | - | HA |
| 4951 | 4954 | 1 | ; | H | - | CB |
| 4951 | 4967 | 1 | ; | H | - | C |
| 4952 | 4957 | 1 | ; | CA | - | HG21 |
| 4952 | 4958 | 1 | ; | CA | - | HG22 |
| 4952 | 4959 | 1 | ; | CA | - | HG23 |
| 4952 | 4961 | 1 | ; | CA | - | HG12 |
| 4952 | 4962 | 1 | ; | CA | - | HG13 |
| 4952 | 4963 | 1 | ; | CA | - | CD1 |
| 4952 | 4970 | 1 | ; | CA | - | H |
| 4952 | 4971 | 1 | ; | CA | - | CA |
| 4953 | 4955 | 1 | ; | HA | - | HB |
| 4953 | 4956 | 1 | ; | HA | - | CG2 |
| 4953 | 4960 | 1 | ; | HA | - | CG1 |
| 4953 | 4968 | 1 | ; | HA | - | O |
| 4953 | 4969 | 1 | ; | HA | - | N |
| 4954 | 4964 | 1 | ; | CB | - | HD11 |
| 4954 | 4965 | 1 | ; | CB | - | HD12 |
| 4954 | 4966 | 1 | ; | CB | - | HD13 |
| 4954 | 4968 | 1 | ; | CB | - | O |
| 4954 | 4969 | 1 | ; | CB | - | N |
| 4955 | 4957 | 1 | ; | HB | - | HG21 |
| 4955 | 4958 | 1 | ; | HB | - | HG22 |
| 4955 | 4959 | 1 | ; | HB | - | HG23 |
| 4955 | 4961 | 1 | ; | HB | - | HG12 |
| 4955 | 4962 | 1 | ; | HB | - | HG13 |
| 4955 | 4963 | 1 | ; | HB | - | CD1 |
| 4955 | 4967 | 1 | ; | HB | - | C |

```
4956   4961       1 ;    CG2 - HG12
4956   4962       1 ;    CG2 - HG13
4956   4963       1 ;    CG2 - CD1
4956   4967       1 ;    CG2 - C
4957   4960       1 ;   HG21 - CG1
4958   4960       1 ;   HG22 - CG1
4959   4960       1 ;   HG23 - CG1
4960   4967       1 ;    CG1 - C
4961   4964       1 ;   HG12 - HD11
4961   4965       1 ;   HG12 - HD12
4961   4966       1 ;   HG12 - HD13
4962   4964       1 ;   HG13 - HD11
4962   4965       1 ;   HG13 - HD12
4962   4966       1 ;   HG13 - HD13
4967   4972       1 ;      C - HA
4967   4973       1 ;      C - CB
4967   4977       1 ;      C - C
4968   4970       1 ;      O - H
4968   4971       1 ;      O - CA
4969   4974       1 ;      N - HB1
4969   4975       1 ;      N - HB2
4969   4976       1 ;      N - HB3
4969   4978       1 ;      N - O
4969   4979       1 ;      N - N
4970   4972       1 ;      H - HA
4970   4973       1 ;      H - CB
4970   4977       1 ;      H - C
4971   4980       1 ;     CA - H
4971   4981       1 ;     CA - CA
4972   4974       1 ;     HA - HB1
4972   4975       1 ;     HA - HB2
4972   4976       1 ;     HA - HB3
4972   4978       1 ;     HA - O
4972   4979       1 ;     HA - N
4973   4978       1 ;     CB - O
4973   4979       1 ;     CB - N
4974   4977       1 ;    HB1 - C
4975   4977       1 ;    HB2 - C
4976   4977       1 ;    HB3 - C
4977   4982       1 ;      C - HA
4977   4983       1 ;      C - CB
4977   4999       1 ;      C - C
4978   4980       1 ;      O - H
4978   4981       1 ;      O - CA
4979   4984       1 ;      N - HB2
4979   4985       1 ;      N - HB3
4979   4986       1 ;      N - CG
4979   5000       1 ;      N - O
4979   5001       1 ;      N - N
4980   4982       1 ;      H - HA
4980   4983       1 ;      H - CB
4980   4999       1 ;      H - C
```

| | | | |
|---|---|---|---|
| 4981 | 4987 | 1 ; | CA - HG2 |
| 4981 | 4988 | 1 ; | CA - HG3 |
| 4981 | 4989 | 1 ; | CA - CD |
| 4981 | 5002 | 1 ; | CA - H |
| 4981 | 5003 | 1 ; | CA - CA |
| 4982 | 4984 | 1 ; | HA - HB2 |
| 4982 | 4985 | 1 ; | HA - HB3 |
| 4982 | 4986 | 1 ; | HA - CG |
| 4982 | 5000 | 1 ; | HA - O |
| 4982 | 5001 | 1 ; | HA - N |
| 4983 | 4990 | 1 ; | CB - HD2 |
| 4983 | 4991 | 1 ; | CB - HD3 |
| 4983 | 4992 | 1 ; | CB - CE |
| 4983 | 5000 | 1 ; | CB - O |
| 4983 | 5001 | 1 ; | CB - N |
| 4984 | 4987 | 1 ; | HB2 - HG2 |
| 4984 | 4988 | 1 ; | HB2 - HG3 |
| 4984 | 4989 | 1 ; | HB2 - CD |
| 4984 | 4999 | 1 ; | HB2 - C |
| 4985 | 4987 | 1 ; | HB3 - HG2 |
| 4985 | 4988 | 1 ; | HB3 - HG3 |
| 4985 | 4989 | 1 ; | HB3 - CD |
| 4985 | 4999 | 1 ; | HB3 - C |
| 4986 | 4993 | 1 ; | CG - HE2 |
| 4986 | 4994 | 1 ; | CG - HE3 |
| 4986 | 4995 | 1 ; | CG - NZ |
| 4986 | 4999 | 1 ; | CG - C |
| 4987 | 4990 | 1 ; | HG2 - HD2 |
| 4987 | 4991 | 1 ; | HG2 - HD3 |
| 4987 | 4992 | 1 ; | HG2 - CE |
| 4988 | 4990 | 1 ; | HG3 - HD2 |
| 4988 | 4991 | 1 ; | HG3 - HD3 |
| 4988 | 4992 | 1 ; | HG3 - CE |
| 4989 | 4996 | 1 ; | CD - HZ1 |
| 4989 | 4997 | 1 ; | CD - HZ2 |
| 4989 | 4998 | 1 ; | CD - HZ3 |
| 4990 | 4993 | 1 ; | HD2 - HE2 |
| 4990 | 4994 | 1 ; | HD2 - HE3 |
| 4990 | 4995 | 1 ; | HD2 - NZ |
| 4991 | 4993 | 1 ; | HD3 - HE2 |
| 4991 | 4994 | 1 ; | HD3 - HE3 |
| 4991 | 4995 | 1 ; | HD3 - NZ |
| 4993 | 4996 | 1 ; | HE2 - HZ1 |
| 4993 | 4997 | 1 ; | HE2 - HZ2 |
| 4993 | 4998 | 1 ; | HE2 - HZ3 |
| 4994 | 4996 | 1 ; | HE3 - HZ1 |
| 4994 | 4997 | 1 ; | HE3 - HZ2 |
| 4994 | 4998 | 1 ; | HE3 - HZ3 |
| 4999 | 5004 | 1 ; | C - HA |
| 4999 | 5005 | 1 ; | C - CB |
| 4999 | 5016 | 1 ; | C - C |
| 5000 | 5002 | 1 ; | O - H |

```
5000   5003      1 ;      O - CA
5001   5006      1 ;      N - HB2
5001   5007      1 ;      N - HB3
5001   5008      1 ;      N - CG
5001   5017      1 ;      N - O
5001   5018      1 ;      N - N
5002   5004      1 ;      H - HA
5002   5005      1 ;      H - CB
5002   5016      1 ;      H - C
5003   5009      1 ;     CA - ND1
5003   5014      1 ;     CA - CD2
5003   5019      1 ;     CA - H
5003   5020      1 ;     CA - CA
5004   5006      1 ;     HA - HB2
5004   5007      1 ;     HA - HB3
5004   5008      1 ;     HA - CG
5004   5017      1 ;     HA - O
5004   5018      1 ;     HA - N
5005   5010      1 ;     CB - CE1
5005   5012      1 ;     CB - NE2
5005   5015      1 ;     CB - HD2
5005   5017      1 ;     CB - O
5005   5018      1 ;     CB - N
5006   5009      1 ;    HB2 - ND1
5006   5014      1 ;    HB2 - CD2
5006   5016      1 ;    HB2 - C
5007   5009      1 ;    HB3 - ND1
5007   5014      1 ;    HB3 - CD2
5007   5016      1 ;    HB3 - C
5008   5011      1 ;     CG - HE1
5008   5013      1 ;     CG - HE2
5008   5016      1 ;     CG - C
5009   5013      1 ;    ND1 - HE2
5009   5015      1 ;    ND1 - HD2
5010   5015      1 ;    CE1 - HD2
5011   5013      1 ;    HE1 - HE2
5011   5014      1 ;    HE1 - CD2
5013   5015      1 ;    HE2 - HD2
5016   5021      1 ;      C - HA
5016   5022      1 ;      C - CB
5016   5035      1 ;      C - C
5017   5019      1 ;      O - H
5017   5020      1 ;      O - CA
5018   5023      1 ;      N - HB2
5018   5024      1 ;      N - HB3
5018   5025      1 ;      N - CG
5018   5036      1 ;      N - O
5018   5037      1 ;      N - N
5019   5021      1 ;      H - HA
5019   5022      1 ;      H - CB
5019   5035      1 ;      H - C
5020   5026      1 ;     CA - HG
```

| | | | | |
|---|---|---|---|---|
| 5020 | 5027 | 1 ; | CA - | CD1 |
| 5020 | 5031 | 1 ; | CA - | CD2 |
| 5020 | 5038 | 1 ; | CA - | H |
| 5020 | 5039 | 1 ; | CA - | CA |
| 5021 | 5023 | 1 ; | HA - | HB2 |
| 5021 | 5024 | 1 ; | HA - | HB3 |
| 5021 | 5025 | 1 ; | HA - | CG |
| 5021 | 5036 | 1 ; | HA - | O |
| 5021 | 5037 | 1 ; | HA - | N |
| 5022 | 5028 | 1 ; | CB - | HD11 |
| 5022 | 5029 | 1 ; | CB - | HD12 |
| 5022 | 5030 | 1 ; | CB - | HD13 |
| 5022 | 5032 | 1 ; | CB - | HD21 |
| 5022 | 5033 | 1 ; | CB - | HD22 |
| 5022 | 5034 | 1 ; | CB - | HD23 |
| 5022 | 5036 | 1 ; | CB - | O |
| 5022 | 5037 | 1 ; | CB - | N |
| 5023 | 5026 | 1 ; | HB2 - | HG |
| 5023 | 5027 | 1 ; | HB2 - | CD1 |
| 5023 | 5031 | 1 ; | HB2 - | CD2 |
| 5023 | 5035 | 1 ; | HB2 - | C |
| 5024 | 5026 | 1 ; | HB3 - | HG |
| 5024 | 5027 | 1 ; | HB3 - | CD1 |
| 5024 | 5031 | 1 ; | HB3 - | CD2 |
| 5024 | 5035 | 1 ; | HB3 - | C |
| 5025 | 5035 | 1 ; | CG - | C |
| 5026 | 5028 | 1 ; | HG - | HD11 |
| 5026 | 5029 | 1 ; | HG - | HD12 |
| 5026 | 5030 | 1 ; | HG - | HD13 |
| 5026 | 5032 | 1 ; | HG - | HD21 |
| 5026 | 5033 | 1 ; | HG - | HD22 |
| 5026 | 5034 | 1 ; | HG - | HD23 |
| 5027 | 5032 | 1 ; | CD1 - | HD21 |
| 5027 | 5033 | 1 ; | CD1 - | HD22 |
| 5027 | 5034 | 1 ; | CD1 - | HD23 |
| 5028 | 5031 | 1 ; | HD11 - | CD2 |
| 5029 | 5031 | 1 ; | HD12 - | CD2 |
| 5030 | 5031 | 1 ; | HD13 - | CD2 |
| 5035 | 5040 | 1 ; | C - | HA |
| 5035 | 5041 | 1 ; | C - | CB |
| 5035 | 5046 | 1 ; | C - | C |
| 5036 | 5038 | 1 ; | O - | H |
| 5036 | 5039 | 1 ; | O - | CA |
| 5037 | 5042 | 1 ; | N - | HB2 |
| 5037 | 5043 | 1 ; | N - | HB3 |
| 5037 | 5044 | 1 ; | N - | SG |
| 5037 | 5047 | 1 ; | N - | O |
| 5037 | 5048 | 1 ; | N - | N |
| 5038 | 5040 | 1 ; | H - | HA |
| 5038 | 5041 | 1 ; | H - | CB |
| 5038 | 5046 | 1 ; | H - | C |
| 5039 | 5045 | 1 ; | CA - | HG |

| | | | |
|---|---|---|---|
| 5039 | 5049 | 1 ; | CA - H |
| 5039 | 5050 | 1 ; | CA - CA |
| 5040 | 5042 | 1 ; | HA - HB2 |
| 5040 | 5043 | 1 ; | HA - HB3 |
| 5040 | 5044 | 1 ; | HA - SG |
| 5040 | 5047 | 1 ; | HA - O |
| 5040 | 5048 | 1 ; | HA - N |
| 5041 | 5047 | 1 ; | CB - O |
| 5041 | 5048 | 1 ; | CB - N |
| 5042 | 5045 | 1 ; | HB2 - HG |
| 5042 | 5046 | 1 ; | HB2 - C |
| 5043 | 5045 | 1 ; | HB3 - HG |
| 5043 | 5046 | 1 ; | HB3 - C |
| 5044 | 5046 | 1 ; | SG - C |
| 5046 | 5051 | 1 ; | C - HA |
| 5046 | 5052 | 1 ; | C - CB |
| 5046 | 5058 | 1 ; | C - C |
| 5047 | 5049 | 1 ; | O - H |
| 5047 | 5050 | 1 ; | O - CA |
| 5048 | 5053 | 1 ; | N - HB2 |
| 5048 | 5054 | 1 ; | N - HB3 |
| 5048 | 5055 | 1 ; | N - CG |
| 5048 | 5059 | 1 ; | N - O |
| 5048 | 5060 | 1 ; | N - N |
| 5049 | 5051 | 1 ; | H - HA |
| 5049 | 5052 | 1 ; | H - CB |
| 5049 | 5058 | 1 ; | H - C |
| 5050 | 5056 | 1 ; | CA - OD1 |
| 5050 | 5057 | 1 ; | CA - OD2 |
| 5050 | 5061 | 1 ; | CA - H |
| 5050 | 5062 | 1 ; | CA - CA |
| 5051 | 5053 | 1 ; | HA - HB2 |
| 5051 | 5054 | 1 ; | HA - HB3 |
| 5051 | 5055 | 1 ; | HA - CG |
| 5051 | 5059 | 1 ; | HA - O |
| 5051 | 5060 | 1 ; | HA - N |
| 5052 | 5059 | 1 ; | CB - O |
| 5052 | 5060 | 1 ; | CB - N |
| 5053 | 5056 | 1 ; | HB2 - OD1 |
| 5053 | 5057 | 1 ; | HB2 - OD2 |
| 5053 | 5058 | 1 ; | HB2 - C |
| 5054 | 5056 | 1 ; | HB3 - OD1 |
| 5054 | 5057 | 1 ; | HB3 - OD2 |
| 5054 | 5058 | 1 ; | HB3 - C |
| 5055 | 5058 | 1 ; | CG - C |
| 5058 | 5063 | 1 ; | C - HA |
| 5058 | 5064 | 1 ; | C - CB |
| 5058 | 5074 | 1 ; | C - C |
| 5059 | 5061 | 1 ; | O - H |
| 5059 | 5062 | 1 ; | O - CA |
| 5060 | 5065 | 1 ; | N - HB |
| 5060 | 5066 | 1 ; | N - CG1 |

```
5060   5070       1 ;       N - CG2
5060   5075       1 ;       N - O
5060   5076       1 ;       N - N
5061   5063       1 ;       H - HA
5061   5064       1 ;       H - CB
5061   5074       1 ;       H - C
5062   5067       1 ;      CA - HG11
5062   5068       1 ;      CA - HG12
5062   5069       1 ;      CA - HG13
5062   5071       1 ;      CA - HG21
5062   5072       1 ;      CA - HG22
5062   5073       1 ;      CA - HG23
5062   5077       1 ;      CA - H
5062   5078       1 ;      CA - CA
5063   5065       1 ;      HA - HB
5063   5066       1 ;      HA - CG1
5063   5070       1 ;      HA - CG2
5063   5075       1 ;      HA - O
5063   5076       1 ;      HA - N
5064   5075       1 ;      CB - O
5064   5076       1 ;      CB - N
5065   5067       1 ;      HB - HG11
5065   5068       1 ;      HB - HG12
5065   5069       1 ;      HB - HG13
5065   5071       1 ;      HB - HG21
5065   5072       1 ;      HB - HG22
5065   5073       1 ;      HB - HG23
5065   5074       1 ;      HB - C
5066   5071       1 ;     CG1 - HG21
5066   5072       1 ;     CG1 - HG22
5066   5073       1 ;     CG1 - HG23
5066   5074       1 ;     CG1 - C
5067   5070       1 ;    HG11 - CG2
5068   5070       1 ;    HG12 - CG2
5069   5070       1 ;    HG13 - CG2
5070   5074       1 ;     CG2 - C
5074   5079       1 ;       C - HA
5074   5080       1 ;       C - CB
5074   5089       1 ;       C - C
5075   5077       1 ;       O - H
5075   5078       1 ;       O - CA
5076   5081       1 ;       N - HB2
5076   5082       1 ;       N - HB3
5076   5083       1 ;       N - CG
5076   5090       1 ;       N - O
5076   5091       1 ;       N - N
5077   5079       1 ;       H - HA
5077   5080       1 ;       H - CB
5077   5089       1 ;       H - C
5078   5084       1 ;      CA - HG2
5078   5085       1 ;      CA - HG3
5078   5086       1 ;      CA - CD
```

```
5078   5092       1 ;      CA - H
5078   5093       1 ;      CA - CA
5079   5081       1 ;      HA - HB2
5079   5082       1 ;      HA - HB3
5079   5083       1 ;      HA - CG
5079   5090       1 ;      HA - O
5079   5091       1 ;      HA - N
5080   5087       1 ;      CB - OE1
5080   5088       1 ;      CB - OE2
5080   5090       1 ;      CB - O
5080   5091       1 ;      CB - N
5081   5084       1 ;     HB2 - HG2
5081   5085       1 ;     HB2 - HG3
5081   5086       1 ;     HB2 - CD
5081   5089       1 ;     HB2 - C
5082   5084       1 ;     HB3 - HG2
5082   5085       1 ;     HB3 - HG3
5082   5086       1 ;     HB3 - CD
5082   5089       1 ;     HB3 - C
5083   5089       1 ;      CG - C
5084   5087       1 ;     HG2 - OE1
5084   5088       1 ;     HG2 - OE2
5085   5087       1 ;     HG3 - OE1
5085   5088       1 ;     HG3 - OE2
5089   5094       1 ;       C - HA
5089   5095       1 ;       C - CB
5089   5108       1 ;       C - C
5090   5092       1 ;       O - H
5090   5093       1 ;       O - CA
5091   5096       1 ;       N - HB
5091   5097       1 ;       N - CG2
5091   5101       1 ;       N - CG1
5091   5109       1 ;       N - O
5091   5110       1 ;       N - N
5092   5094       1 ;       H - HA
5092   5095       1 ;       H - CB
5092   5108       1 ;       H - C
5093   5098       1 ;      CA - HG21
5093   5099       1 ;      CA - HG22
5093   5100       1 ;      CA - HG23
5093   5102       1 ;      CA - HG12
5093   5103       1 ;      CA - HG13
5093   5104       1 ;      CA - CD1
5093   5111       1 ;      CA - H
5093   5112       1 ;      CA - CA
5094   5096       1 ;      HA - HB
5094   5097       1 ;      HA - CG2
5094   5101       1 ;      HA - CG1
5094   5109       1 ;      HA - O
5094   5110       1 ;      HA - N
5095   5105       1 ;      CB - HD11
5095   5106       1 ;      CB - HD12
```

| | | | |
|---|---|---|---|
| 5095 | 5107 | 1 ; | CB - HD13 |
| 5095 | 5109 | 1 ; | CB - O |
| 5095 | 5110 | 1 ; | CB - N |
| 5096 | 5098 | 1 ; | HB - HG21 |
| 5096 | 5099 | 1 ; | HB - HG22 |
| 5096 | 5100 | 1 ; | HB - HG23 |
| 5096 | 5102 | 1 ; | HB - HG12 |
| 5096 | 5103 | 1 ; | HB - HG13 |
| 5096 | 5104 | 1 ; | HB - CD1 |
| 5096 | 5108 | 1 ; | HB - C |
| 5097 | 5102 | 1 ; | CG2 - HG12 |
| 5097 | 5103 | 1 ; | CG2 - HG13 |
| 5097 | 5104 | 1 ; | CG2 - CD1 |
| 5097 | 5108 | 1 ; | CG2 - C |
| 5098 | 5101 | 1 ; | HG21 - CG1 |
| 5099 | 5101 | 1 ; | HG22 - CG1 |
| 5100 | 5101 | 1 ; | HG23 - CG1 |
| 5101 | 5108 | 1 ; | CG1 - C |
| 5102 | 5105 | 1 ; | HG12 - HD11 |
| 5102 | 5106 | 1 ; | HG12 - HD12 |
| 5102 | 5107 | 1 ; | HG12 - HD13 |
| 5103 | 5105 | 1 ; | HG13 - HD11 |
| 5103 | 5106 | 1 ; | HG13 - HD12 |
| 5103 | 5107 | 1 ; | HG13 - HD13 |
| 5108 | 5113 | 1 ; | C - HA |
| 5108 | 5114 | 1 ; | C - CB |
| 5108 | 5122 | 1 ; | C - C |
| 5109 | 5111 | 1 ; | O - H |
| 5109 | 5112 | 1 ; | O - CA |
| 5110 | 5115 | 1 ; | N - HB2 |
| 5110 | 5116 | 1 ; | N - HB3 |
| 5110 | 5117 | 1 ; | N - CG |
| 5110 | 5123 | 1 ; | N - O |
| 5110 | 5124 | 1 ; | N - N |
| 5111 | 5113 | 1 ; | H - HA |
| 5111 | 5114 | 1 ; | H - CB |
| 5111 | 5122 | 1 ; | H - C |
| 5112 | 5118 | 1 ; | CA - OD1 |
| 5112 | 5119 | 1 ; | CA - ND2 |
| 5112 | 5125 | 1 ; | CA - H |
| 5112 | 5126 | 1 ; | CA - CA |
| 5113 | 5115 | 1 ; | HA - HB2 |
| 5113 | 5116 | 1 ; | HA - HB3 |
| 5113 | 5117 | 1 ; | HA - CG |
| 5113 | 5123 | 1 ; | HA - O |
| 5113 | 5124 | 1 ; | HA - N |
| 5114 | 5120 | 1 ; | CB - HD21 |
| 5114 | 5121 | 1 ; | CB - HD22 |
| 5114 | 5123 | 1 ; | CB - O |
| 5114 | 5124 | 1 ; | CB - N |
| 5115 | 5118 | 1 ; | HB2 - OD1 |
| 5115 | 5119 | 1 ; | HB2 - ND2 |

```
5115   5122       1 ;     HB2 - C
5116   5118       1 ;     HB3 - OD1
5116   5119       1 ;     HB3 - ND2
5116   5122       1 ;     HB3 - C
5117   5122       1 ;      CG - C
5118   5120       1 ;     OD1 - HD21
5118   5121       1 ;     OD1 - HD22
5122   5127       1 ;       C - HA
5122   5128       1 ;       C - CB
5122   5136       1 ;       C - C
5123   5125       1 ;       O - H
5123   5126       1 ;       O - CA
5124   5129       1 ;       N - HB2
5124   5130       1 ;       N - HB3
5124   5131       1 ;       N - CG
5124   5137       1 ;       N - O
5124   5138       1 ;       N - N
5125   5127       1 ;       H - HA
5125   5128       1 ;       H - CB
5125   5136       1 ;       H - C
5126   5132       1 ;      CA - OD1
5126   5133       1 ;      CA - ND2
5126   5139       1 ;      CA - H
5126   5140       1 ;      CA - CA
5127   5129       1 ;      HA - HB2
5127   5130       1 ;      HA - HB3
5127   5131       1 ;      HA - CG
5127   5137       1 ;      HA - O
5127   5138       1 ;      HA - N
5128   5134       1 ;      CB - HD21
5128   5135       1 ;      CB - HD22
5128   5137       1 ;      CB - O
5128   5138       1 ;      CB - N
5129   5132       1 ;     HB2 - OD1
5129   5133       1 ;     HB2 - ND2
5129   5136       1 ;     HB2 - C
5130   5132       1 ;     HB3 - OD1
5130   5133       1 ;     HB3 - ND2
5130   5136       1 ;     HB3 - C
5131   5136       1 ;      CG - C
5132   5134       1 ;     OD1 - HD21
5132   5135       1 ;     OD1 - HD22
5136   5141       1 ;       C - HA
5136   5142       1 ;       C - CB
5136   5155       1 ;       C - C
5137   5139       1 ;       O - H
5137   5140       1 ;       O - CA
5138   5143       1 ;       N - HB2
5138   5144       1 ;       N - HB3
5138   5145       1 ;       N - CG
5138   5156       1 ;       N - O
5138   5157       1 ;       N - N
```

```
5139   5141       1 ;       H - HA
5139   5142       1 ;       H - CB
5139   5155       1 ;       H - C
5140   5146       1 ;      CA - HG
5140   5147       1 ;      CA - CD1
5140   5151       1 ;      CA - CD2
5140   5158       1 ;      CA - H
5140   5159       1 ;      CA - CA
5141   5143       1 ;      HA - HB2
5141   5144       1 ;      HA - HB3
5141   5145       1 ;      HA - CG
5141   5156       1 ;      HA - O
5141   5157       1 ;      HA - N
5142   5148       1 ;      CB - HD11
5142   5149       1 ;      CB - HD12
5142   5150       1 ;      CB - HD13
5142   5152       1 ;      CB - HD21
5142   5153       1 ;      CB - HD22
5142   5154       1 ;      CB - HD23
5142   5156       1 ;      CB - O
5142   5157       1 ;      CB - N
5143   5146       1 ;     HB2 - HG
5143   5147       1 ;     HB2 - CD1
5143   5151       1 ;     HB2 - CD2
5143   5155       1 ;     HB2 - C
5144   5146       1 ;     HB3 - HG
5144   5147       1 ;     HB3 - CD1
5144   5151       1 ;     HB3 - CD2
5144   5155       1 ;     HB3 - C
5145   5155       1 ;      CG - C
5146   5148       1 ;      HG - HD11
5146   5149       1 ;      HG - HD12
5146   5150       1 ;      HG - HD13
5146   5152       1 ;      HG - HD21
5146   5153       1 ;      HG - HD22
5146   5154       1 ;      HG - HD23
5147   5152       1 ;     CD1 - HD21
5147   5153       1 ;     CD1 - HD22
5147   5154       1 ;     CD1 - HD23
5148   5151       1 ;    HD11 - CD2
5149   5151       1 ;    HD12 - CD2
5150   5151       1 ;    HD13 - CD2
5155   5160       1 ;       C - HA
5155   5161       1 ;       C - CB
5155   5179       1 ;       C - C
5156   5158       1 ;       O - H
5156   5159       1 ;       O - CA
5157   5162       1 ;       N - HB2
5157   5163       1 ;       N - HB3
5157   5164       1 ;       N - CG
5157   5180       1 ;       N - O
5157   5181       1 ;       N - N
```

```
5158   5160      1 ;      H - HA
5158   5161      1 ;      H - CB
5158   5179      1 ;      H - C
5159   5165      1 ;     CA - HG2
5159   5166      1 ;     CA - HG3
5159   5167      1 ;     CA - CD
5159   5182      1 ;     CA - H
5159   5183      1 ;     CA - CA
5160   5162      1 ;     HA - HB2
5160   5163      1 ;     HA - HB3
5160   5164      1 ;     HA - CG
5160   5180      1 ;     HA - O
5160   5181      1 ;     HA - N
5161   5168      1 ;     CB - HD2
5161   5169      1 ;     CB - HD3
5161   5170      1 ;     CB - NE
5161   5180      1 ;     CB - O
5161   5181      1 ;     CB - N
5162   5165      1 ;    HB2 - HG2
5162   5166      1 ;    HB2 - HG3
5162   5167      1 ;    HB2 - CD
5162   5179      1 ;    HB2 - C
5163   5165      1 ;    HB3 - HG2
5163   5166      1 ;    HB3 - HG3
5163   5167      1 ;    HB3 - CD
5163   5179      1 ;    HB3 - C
5164   5171      1 ;     CG - HE
5164   5172      1 ;     CG - CZ
5164   5179      1 ;     CG - C
5165   5168      1 ;    HG2 - HD2
5165   5169      1 ;    HG2 - HD3
5165   5170      1 ;    HG2 - NE
5166   5168      1 ;    HG3 - HD2
5166   5169      1 ;    HG3 - HD3
5166   5170      1 ;    HG3 - NE
5167   5173      1 ;     CD - NH1
5167   5176      1 ;     CD - NH2
5168   5171      1 ;    HD2 - HE
5168   5172      1 ;    HD2 - CZ
5169   5171      1 ;    HD3 - HE
5169   5172      1 ;    HD3 - CZ
5170   5174      1 ;     NE - HH11
5170   5175      1 ;     NE - HH12
5170   5177      1 ;     NE - HH21
5170   5178      1 ;     NE - HH22
5171   5173      1 ;     HE - NH1
5171   5176      1 ;     HE - NH2
5173   5177      1 ;    NH1 - HH21
5173   5178      1 ;    NH1 - HH22
5174   5176      1 ;   HH11 - NH2
5175   5176      1 ;   HH12 - NH2
5179   5184      1 ;      C - HA
```

```
5179   5185       1 ;       C - CB
5179   5193       1 ;       C - C
5180   5182       1 ;       O - H
5180   5183       1 ;       O - CA
5181   5186       1 ;       N - HB2
5181   5187       1 ;       N - HB3
5181   5188       1 ;       N - CG
5181   5194       1 ;       N - O
5181   5195       1 ;       N - N
5182   5184       1 ;       H - HA
5182   5185       1 ;       H - CB
5182   5193       1 ;       H - C
5183   5189       1 ;      CA - OD1
5183   5190       1 ;      CA - ND2
5183   5196       1 ;      CA - H
5183   5197       1 ;      CA - CA
5184   5186       1 ;      HA - HB2
5184   5187       1 ;      HA - HB3
5184   5188       1 ;      HA - CG
5184   5194       1 ;      HA - O
5184   5195       1 ;      HA - N
5185   5191       1 ;      CB - HD21
5185   5192       1 ;      CB - HD22
5185   5194       1 ;      CB - O
5185   5195       1 ;      CB - N
5186   5189       1 ;     HB2 - OD1
5186   5190       1 ;     HB2 - ND2
5186   5193       1 ;     HB2 - C
5187   5189       1 ;     HB3 - OD1
5187   5190       1 ;     HB3 - ND2
5187   5193       1 ;     HB3 - C
5188   5193       1 ;      CG - C
5189   5191       1 ;     OD1 - HD21
5189   5192       1 ;     OD1 - HD22
5193   5198       1 ;       C - HA
5193   5199       1 ;       C - CB
5193   5212       1 ;       C - C
5194   5196       1 ;       O - H
5194   5197       1 ;       O - CA
5195   5200       1 ;       N - HB
5195   5201       1 ;       N - CG2
5195   5205       1 ;       N - CG1
5195   5213       1 ;       N - O
5195   5214       1 ;       N - N
5196   5198       1 ;       H - HA
5196   5199       1 ;       H - CB
5196   5212       1 ;       H - C
5197   5202       1 ;      CA - HG21
5197   5203       1 ;      CA - HG22
5197   5204       1 ;      CA - HG23
5197   5206       1 ;      CA - HG12
5197   5207       1 ;      CA - HG13
```

```
  5197   5208       1 ;      CA - CD1
  5197   5215       1 ;      CA - H
  5197   5216       1 ;      CA - CA
  5198   5200       1 ;      HA - HB
  5198   5201       1 ;      HA - CG2
  5198   5205       1 ;      HA - CG1
  5198   5213       1 ;      HA - O
  5198   5214       1 ;      HA - N
  5199   5209       1 ;      CB - HD11
  5199   5210       1 ;      CB - HD12
  5199   5211       1 ;      CB - HD13
  5199   5213       1 ;      CB - O
  5199   5214       1 ;      CB - N
  5200   5202       1 ;      HB - HG21
  5200   5203       1 ;      HB - HG22
  5200   5204       1 ;      HB - HG23
  5200   5206       1 ;      HB - HG12
  5200   5207       1 ;      HB - HG13
  5200   5208       1 ;      HB - CD1
  5200   5212       1 ;      HB - C
  5201   5206       1 ;     CG2 - HG12
  5201   5207       1 ;     CG2 - HG13
  5201   5208       1 ;     CG2 - CD1
  5201   5212       1 ;     CG2 - C
  5202   5205       1 ;    HG21 - CG1
  5203   5205       1 ;    HG22 - CG1
  5204   5205       1 ;    HG23 - CG1
  5205   5212       1 ;     CG1 - C
  5206   5209       1 ;    HG12 - HD11
  5206   5210       1 ;    HG12 - HD12
  5206   5211       1 ;    HG12 - HD13
  5207   5209       1 ;    HG13 - HD11
  5207   5210       1 ;    HG13 - HD12
  5207   5211       1 ;    HG13 - HD13
  5212   5217       1 ;       C - HA
  5212   5218       1 ;       C - CB
  5212   5232       1 ;       C - C
  5213   5215       1 ;       O - H
  5213   5216       1 ;       O - CA
  5214   5219       1 ;       N - HB2
  5214   5220       1 ;       N - HB3
  5214   5221       1 ;       N - CG
  5214   5233       1 ;       N - O
  5214   5234       1 ;       N - N
  5215   5217       1 ;       H - HA
  5215   5218       1 ;       H - CB
  5215   5232       1 ;       H - C
  5216   5222       1 ;      CA - CD1
  5216   5230       1 ;      CA - CD2
  5216   5235       1 ;      CA - H
  5216   5236       1 ;      CA - CA
  5217   5219       1 ;      HA - HB2
```

| | | | | | |
|---|---|---|---|---|---|
| 5217 | 5220 | 1 ; | HA | - | HB3 |
| 5217 | 5221 | 1 ; | HA | - | CG |
| 5217 | 5233 | 1 ; | HA | - | O |
| 5217 | 5234 | 1 ; | HA | - | N |
| 5218 | 5223 | 1 ; | CB | - | HD1 |
| 5218 | 5224 | 1 ; | CB | - | CE1 |
| 5218 | 5228 | 1 ; | CB | - | CE2 |
| 5218 | 5231 | 1 ; | CB | - | HD2 |
| 5218 | 5233 | 1 ; | CB | - | O |
| 5218 | 5234 | 1 ; | CB | - | N |
| 5219 | 5222 | 1 ; | HB2 | - | CD1 |
| 5219 | 5230 | 1 ; | HB2 | - | CD2 |
| 5219 | 5232 | 1 ; | HB2 | - | C |
| 5220 | 5222 | 1 ; | HB3 | - | CD1 |
| 5220 | 5230 | 1 ; | HB3 | - | CD2 |
| 5220 | 5232 | 1 ; | HB3 | - | C |
| 5221 | 5225 | 1 ; | CG | - | HE1 |
| 5221 | 5226 | 1 ; | CG | - | CZ |
| 5221 | 5229 | 1 ; | CG | - | HE2 |
| 5221 | 5232 | 1 ; | CG | - | C |
| 5222 | 5227 | 1 ; | CD1 | - | HZ |
| 5222 | 5228 | 1 ; | CD1 | - | CE2 |
| 5222 | 5231 | 1 ; | CD1 | - | HD2 |
| 5223 | 5225 | 1 ; | HD1 | - | HE1 |
| 5223 | 5226 | 1 ; | HD1 | - | CZ |
| 5223 | 5230 | 1 ; | HD1 | - | CD2 |
| 5224 | 5229 | 1 ; | CE1 | - | HE2 |
| 5224 | 5230 | 1 ; | CE1 | - | CD2 |
| 5225 | 5227 | 1 ; | HE1 | - | HZ |
| 5225 | 5228 | 1 ; | HE1 | - | CE2 |
| 5226 | 5231 | 1 ; | CZ | - | HD2 |
| 5227 | 5229 | 1 ; | HZ | - | HE2 |
| 5227 | 5230 | 1 ; | HZ | - | CD2 |
| 5229 | 5231 | 1 ; | HE2 | - | HD2 |
| 5232 | 5237 | 1 ; | C | - | HA |
| 5232 | 5238 | 1 ; | C | - | CB |
| 5232 | 5251 | 1 ; | C | - | C |
| 5233 | 5235 | 1 ; | O | - | H |
| 5233 | 5236 | 1 ; | O | - | CA |
| 5234 | 5239 | 1 ; | N | - | HB |
| 5234 | 5240 | 1 ; | N | - | CG2 |
| 5234 | 5244 | 1 ; | N | - | CG1 |
| 5234 | 5252 | 1 ; | N | - | O |
| 5234 | 5253 | 1 ; | N | - | N |
| 5235 | 5237 | 1 ; | H | - | HA |
| 5235 | 5238 | 1 ; | H | - | CB |
| 5235 | 5251 | 1 ; | H | - | C |
| 5236 | 5241 | 1 ; | CA | - | HG21 |
| 5236 | 5242 | 1 ; | CA | - | HG22 |
| 5236 | 5243 | 1 ; | CA | - | HG23 |
| 5236 | 5245 | 1 ; | CA | - | HG12 |
| 5236 | 5246 | 1 ; | CA | - | HG13 |

```
5236   5247       1 ;      CA - CD1
5236   5254       1 ;      CA - H
5236   5255       1 ;      CA - CA
5237   5239       1 ;      HA - HB
5237   5240       1 ;      HA - CG2
5237   5244       1 ;      HA - CG1
5237   5252       1 ;      HA - O
5237   5253       1 ;      HA - N
5238   5248       1 ;      CB - HD11
5238   5249       1 ;      CB - HD12
5238   5250       1 ;      CB - HD13
5238   5252       1 ;      CB - O
5238   5253       1 ;      CB - N
5239   5241       1 ;      HB - HG21
5239   5242       1 ;      HB - HG22
5239   5243       1 ;      HB - HG23
5239   5245       1 ;      HB - HG12
5239   5246       1 ;      HB - HG13
5239   5247       1 ;      HB - CD1
5239   5251       1 ;      HB - C
5240   5245       1 ;     CG2 - HG12
5240   5246       1 ;     CG2 - HG13
5240   5247       1 ;     CG2 - CD1
5240   5251       1 ;     CG2 - C
5241   5244       1 ;    HG21 - CG1
5242   5244       1 ;    HG22 - CG1
5243   5244       1 ;    HG23 - CG1
5244   5251       1 ;     CG1 - C
5245   5248       1 ;    HG12 - HD11
5245   5249       1 ;    HG12 - HD12
5245   5250       1 ;    HG12 - HD13
5246   5248       1 ;    HG13 - HD11
5246   5249       1 ;    HG13 - HD12
5246   5250       1 ;    HG13 - HD13
5251   5256       1 ;       C - HA
5251   5257       1 ;       C - CB
5251   5261       1 ;       C - C
5252   5254       1 ;       O - H
5252   5255       1 ;       O - CA
5253   5258       1 ;       N - HB1
5253   5259       1 ;       N - HB2
5253   5260       1 ;       N - HB3
5253   5262       1 ;       N - O
5253   5263       1 ;       N - N
5254   5256       1 ;       H - HA
5254   5257       1 ;       H - CB
5254   5261       1 ;       H - C
5255   5264       1 ;      CA - H
5255   5265       1 ;      CA - CA
5256   5258       1 ;      HA - HB1
5256   5259       1 ;      HA - HB2
5256   5260       1 ;      HA - HB3
```

```
5256   5262       1 ;      HA - O
5256   5263       1 ;      HA - N
5257   5262       1 ;      CB - O
5257   5263       1 ;      CB - N
5258   5261       1 ;     HB1 - C
5259   5261       1 ;     HB2 - C
5260   5261       1 ;     HB3 - C
5261   5266       1 ;       C - HA
5261   5267       1 ;       C - CB
5261   5280       1 ;       C - C
5262   5264       1 ;       O - H
5262   5265       1 ;       O - CA
5263   5268       1 ;       N - HB2
5263   5269       1 ;       N - HB3
5263   5270       1 ;       N - CG
5263   5281       1 ;       N - O
5263   5282       1 ;       N - N
5264   5266       1 ;       H - HA
5264   5267       1 ;       H - CB
5264   5280       1 ;       H - C
5265   5271       1 ;      CA - HG
5265   5272       1 ;      CA - CD1
5265   5276       1 ;      CA - CD2
5265   5283       1 ;      CA - H
5265   5284       1 ;      CA - CA
5266   5268       1 ;      HA - HB2
5266   5269       1 ;      HA - HB3
5266   5270       1 ;      HA - CG
5266   5281       1 ;      HA - O
5266   5282       1 ;      HA - N
5267   5273       1 ;      CB - HD11
5267   5274       1 ;      CB - HD12
5267   5275       1 ;      CB - HD13
5267   5277       1 ;      CB - HD21
5267   5278       1 ;      CB - HD22
5267   5279       1 ;      CB - HD23
5267   5281       1 ;      CB - O
5267   5282       1 ;      CB - N
5268   5271       1 ;     HB2 - HG
5268   5272       1 ;     HB2 - CD1
5268   5276       1 ;     HB2 - CD2
5268   5280       1 ;     HB2 - C
5269   5271       1 ;     HB3 - HG
5269   5272       1 ;     HB3 - CD1
5269   5276       1 ;     HB3 - CD2
5269   5280       1 ;     HB3 - C
5270   5280       1 ;      CG - C
5271   5273       1 ;      HG - HD11
5271   5274       1 ;      HG - HD12
5271   5275       1 ;      HG - HD13
5271   5277       1 ;      HG - HD21
5271   5278       1 ;      HG - HD22
```

| | | | | | |
|---|---|---|---|---|---|
| 5271 | 5279 | 1 ; | HG | - | HD23 |
| 5272 | 5277 | 1 ; | CD1 | - | HD21 |
| 5272 | 5278 | 1 ; | CD1 | - | HD22 |
| 5272 | 5279 | 1 ; | CD1 | - | HD23 |
| 5273 | 5276 | 1 ; | HD11 | - | CD2 |
| 5274 | 5276 | 1 ; | HD12 | - | CD2 |
| 5275 | 5276 | 1 ; | HD13 | - | CD2 |
| 5280 | 5285 | 1 ; | C | - | HA |
| 5280 | 5286 | 1 ; | C | - | CB |
| 5280 | 5292 | 1 ; | C | - | C |
| 5281 | 5283 | 1 ; | O | - | H |
| 5281 | 5284 | 1 ; | O | - | CA |
| 5282 | 5287 | 1 ; | N | - | HB2 |
| 5282 | 5288 | 1 ; | N | - | HB3 |
| 5282 | 5289 | 1 ; | N | - | CG |
| 5282 | 5293 | 1 ; | N | - | O |
| 5282 | 5294 | 1 ; | N | - | N |
| 5283 | 5285 | 1 ; | H | - | HA |
| 5283 | 5286 | 1 ; | H | - | CB |
| 5283 | 5292 | 1 ; | H | - | C |
| 5284 | 5290 | 1 ; | CA | - | OD1 |
| 5284 | 5291 | 1 ; | CA | - | OD2 |
| 5284 | 5295 | 1 ; | CA | - | H |
| 5284 | 5296 | 1 ; | CA | - | CA |
| 5285 | 5287 | 1 ; | HA | - | HB2 |
| 5285 | 5288 | 1 ; | HA | - | HB3 |
| 5285 | 5289 | 1 ; | HA | - | CG |
| 5285 | 5293 | 1 ; | HA | - | O |
| 5285 | 5294 | 1 ; | HA | - | N |
| 5286 | 5293 | 1 ; | CB | - | O |
| 5286 | 5294 | 1 ; | CB | - | N |
| 5287 | 5290 | 1 ; | HB2 | - | OD1 |
| 5287 | 5291 | 1 ; | HB2 | - | OD2 |
| 5287 | 5292 | 1 ; | HB2 | - | C |
| 5288 | 5290 | 1 ; | HB3 | - | OD1 |
| 5288 | 5291 | 1 ; | HB3 | - | OD2 |
| 5288 | 5292 | 1 ; | HB3 | - | C |
| 5289 | 5292 | 1 ; | CG | - | C |
| 5292 | 5297 | 1 ; | C | - | HA |
| 5292 | 5298 | 1 ; | C | - | CB |
| 5292 | 5308 | 1 ; | C | - | C |
| 5293 | 5295 | 1 ; | O | - | H |
| 5293 | 5296 | 1 ; | O | - | CA |
| 5294 | 5299 | 1 ; | N | - | HB |
| 5294 | 5300 | 1 ; | N | - | CG1 |
| 5294 | 5304 | 1 ; | N | - | CG2 |
| 5294 | 5309 | 1 ; | N | - | O |
| 5294 | 5310 | 1 ; | N | - | N |
| 5295 | 5297 | 1 ; | H | - | HA |
| 5295 | 5298 | 1 ; | H | - | CB |
| 5295 | 5308 | 1 ; | H | - | C |
| 5296 | 5301 | 1 ; | CA | - | HG11 |

| | | | | | |
|---|---|---|---|---|---|
| 5296 | 5302 | 1 ; | CA | - | HG12 |
| 5296 | 5303 | 1 ; | CA | - | HG13 |
| 5296 | 5305 | 1 ; | CA | - | HG21 |
| 5296 | 5306 | 1 ; | CA | - | HG22 |
| 5296 | 5307 | 1 ; | CA | - | HG23 |
| 5296 | 5311 | 1 ; | CA | - | H |
| 5296 | 5312 | 1 ; | CA | - | CA |
| 5297 | 5299 | 1 ; | HA | - | HB |
| 5297 | 5300 | 1 ; | HA | - | CG1 |
| 5297 | 5304 | 1 ; | HA | - | CG2 |
| 5297 | 5309 | 1 ; | HA | - | O |
| 5297 | 5310 | 1 ; | HA | - | N |
| 5298 | 5309 | 1 ; | CB | - | O |
| 5298 | 5310 | 1 ; | CB | - | N |
| 5299 | 5301 | 1 ; | HB | - | HG11 |
| 5299 | 5302 | 1 ; | HB | - | HG12 |
| 5299 | 5303 | 1 ; | HB | - | HG13 |
| 5299 | 5305 | 1 ; | HB | - | HG21 |
| 5299 | 5306 | 1 ; | HB | - | HG22 |
| 5299 | 5307 | 1 ; | HB | - | HG23 |
| 5299 | 5308 | 1 ; | HB | - | C |
| 5300 | 5305 | 1 ; | CG1 | - | HG21 |
| 5300 | 5306 | 1 ; | CG1 | - | HG22 |
| 5300 | 5307 | 1 ; | CG1 | - | HG23 |
| 5300 | 5308 | 1 ; | CG1 | - | C |
| 5301 | 5304 | 1 ; | HG11 | - | CG2 |
| 5302 | 5304 | 1 ; | HG12 | - | CG2 |
| 5303 | 5304 | 1 ; | HG13 | - | CG2 |
| 5304 | 5308 | 1 ; | CG2 | - | C |
| 5308 | 5313 | 1 ; | C | - | HA |
| 5308 | 5314 | 1 ; | C | - | CB |
| 5308 | 5320 | 1 ; | C | - | C |
| 5309 | 5311 | 1 ; | O | - | H |
| 5309 | 5312 | 1 ; | O | - | CA |
| 5310 | 5315 | 1 ; | N | - | HB2 |
| 5310 | 5316 | 1 ; | N | - | HB3 |
| 5310 | 5317 | 1 ; | N | - | CG |
| 5310 | 5321 | 1 ; | N | - | O |
| 5310 | 5322 | 1 ; | N | - | N |
| 5311 | 5313 | 1 ; | H | - | HA |
| 5311 | 5314 | 1 ; | H | - | CB |
| 5311 | 5320 | 1 ; | H | - | C |
| 5312 | 5318 | 1 ; | CA | - | OD1 |
| 5312 | 5319 | 1 ; | CA | - | OD2 |
| 5312 | 5323 | 1 ; | CA | - | H |
| 5312 | 5324 | 1 ; | CA | - | CA |
| 5313 | 5315 | 1 ; | HA | - | HB2 |
| 5313 | 5316 | 1 ; | HA | - | HB3 |
| 5313 | 5317 | 1 ; | HA | - | CG |
| 5313 | 5321 | 1 ; | HA | - | O |
| 5313 | 5322 | 1 ; | HA | - | N |
| 5314 | 5321 | 1 ; | CB | - | O |

```
5314   5322       1 ;      CB - N
5315   5318       1 ;     HB2 - OD1
5315   5319       1 ;     HB2 - OD2
5315   5320       1 ;     HB2 - C
5316   5318       1 ;     HB3 - OD1
5316   5319       1 ;     HB3 - OD2
5316   5320       1 ;     HB3 - C
5317   5320       1 ;      CG - C
5320   5325       1 ;       C - HA
5320   5326       1 ;       C - CB
5320   5334       1 ;       C - C
5321   5323       1 ;       O - H
5321   5324       1 ;       O - CA
5322   5327       1 ;       N - HB2
5322   5328       1 ;       N - HB3
5322   5329       1 ;       N - CG
5322   5335       1 ;       N - O
5322   5336       1 ;       N - N
5323   5325       1 ;       H - HA
5323   5326       1 ;       H - CB
5323   5334       1 ;       H - C
5324   5330       1 ;      CA - OD1
5324   5331       1 ;      CA - ND2
5324   5337       1 ;      CA - H
5324   5338       1 ;      CA - CA
5325   5327       1 ;      HA - HB2
5325   5328       1 ;      HA - HB3
5325   5329       1 ;      HA - CG
5325   5335       1 ;      HA - O
5325   5336       1 ;      HA - N
5326   5332       1 ;      CB - HD21
5326   5333       1 ;      CB - HD22
5326   5335       1 ;      CB - O
5326   5336       1 ;      CB - N
5327   5330       1 ;     HB2 - OD1
5327   5331       1 ;     HB2 - ND2
5327   5334       1 ;     HB2 - C
5328   5330       1 ;     HB3 - OD1
5328   5331       1 ;     HB3 - ND2
5328   5334       1 ;     HB3 - C
5329   5334       1 ;      CG - C
5330   5332       1 ;     OD1 - HD21
5330   5333       1 ;     OD1 - HD22
5334   5339       1 ;       C - HA
5334   5340       1 ;       C - CB
5334   5345       1 ;       C - C
5335   5337       1 ;       O - H
5335   5338       1 ;       O - CA
5336   5341       1 ;       N - HB2
5336   5342       1 ;       N - HB3
5336   5343       1 ;       N - OG
5336   5346       1 ;       N - O
```

```
5336   5347       1 ;       N - N
5337   5339       1 ;       H - HA
5337   5340       1 ;       H - CB
5337   5345       1 ;       H - C
5338   5344       1 ;      CA - HG
5338   5348       1 ;      CA - H
5338   5349       1 ;      CA - CA
5339   5341       1 ;      HA - HB2
5339   5342       1 ;      HA - HB3
5339   5343       1 ;      HA - OG
5339   5346       1 ;      HA - O
5339   5347       1 ;      HA - N
5340   5346       1 ;      CB - O
5340   5347       1 ;      CB - N
5341   5344       1 ;     HB2 - HG
5341   5345       1 ;     HB2 - C
5342   5344       1 ;     HB3 - HG
5342   5345       1 ;     HB3 - C
5343   5345       1 ;      OG - C
5345   5350       1 ;       C - HA2
5345   5351       1 ;       C - HA3
5345   5352       1 ;       C - C
5346   5348       1 ;       O - H
5346   5349       1 ;       O - CA
5347   5353       1 ;       N - O
5347   5354       1 ;       N - N
5348   5350       1 ;       H - HA2
5348   5351       1 ;       H - HA3
5348   5352       1 ;       H - C
5349   5355       1 ;      CA - H
5349   5356       1 ;      CA - CA
5350   5353       1 ;     HA2 - O
5350   5354       1 ;     HA2 - N
5351   5353       1 ;     HA3 - O
5351   5354       1 ;     HA3 - N
5352   5357       1 ;       C - HA
5352   5358       1 ;       C - CB
5352   5366       1 ;       C - C
5353   5355       1 ;       O - H
5353   5356       1 ;       O - CA
5354   5359       1 ;       N - HB
5354   5360       1 ;       N - CG2
5354   5364       1 ;       N - OG1
5354   5367       1 ;       N - O
5354   5368       1 ;       N - N
5355   5357       1 ;       H - HA
5355   5358       1 ;       H - CB
5355   5366       1 ;       H - C
5356   5361       1 ;      CA - HG21
5356   5362       1 ;      CA - HG22
5356   5363       1 ;      CA - HG23
5356   5365       1 ;      CA - HG1
```

```
5356   5369      1 ;     CA - H
5356   5370      1 ;     CA - CA
5357   5359      1 ;     HA - HB
5357   5360      1 ;     HA - CG2
5357   5364      1 ;     HA - OG1
5357   5367      1 ;     HA - O
5357   5368      1 ;     HA - N
5358   5367      1 ;     CB - O
5358   5368      1 ;     CB - N
5359   5361      1 ;     HB - HG21
5359   5362      1 ;     HB - HG22
5359   5363      1 ;     HB - HG23
5359   5365      1 ;     HB - HG1
5359   5366      1 ;     HB - C
5360   5365      1 ;    CG2 - HG1
5360   5366      1 ;    CG2 - C
5361   5364      1 ;   HG21 - OG1
5362   5364      1 ;   HG22 - OG1
5363   5364      1 ;   HG23 - OG1
5364   5366      1 ;    OG1 - C
5366   5371      1 ;      C - HA
5366   5372      1 ;      C - CB
5366   5385      1 ;      C - C
5367   5369      1 ;      O - H
5367   5370      1 ;      O - CA
5368   5373      1 ;      N - HB2
5368   5374      1 ;      N - HB3
5368   5375      1 ;      N - CG
5368   5386      1 ;      N - O
5368   5387      1 ;      N - N
5369   5371      1 ;      H - HA
5369   5372      1 ;      H - CB
5369   5385      1 ;      H - C
5370   5376      1 ;     CA - HG
5370   5377      1 ;     CA - CD1
5370   5381      1 ;     CA - CD2
5370   5388      1 ;     CA - H
5370   5389      1 ;     CA - CA
5371   5373      1 ;     HA - HB2
5371   5374      1 ;     HA - HB3
5371   5375      1 ;     HA - CG
5371   5386      1 ;     HA - O
5371   5387      1 ;     HA - N
5372   5378      1 ;     CB - HD11
5372   5379      1 ;     CB - HD12
5372   5380      1 ;     CB - HD13
5372   5382      1 ;     CB - HD21
5372   5383      1 ;     CB - HD22
5372   5384      1 ;     CB - HD23
5372   5386      1 ;     CB - O
5372   5387      1 ;     CB - N
5373   5376      1 ;    HB2 - HG
```

| | | | |
|---|---|---|---|
| 5373 | 5377 | 1 ; | HB2 - CD1 |
| 5373 | 5381 | 1 ; | HB2 - CD2 |
| 5373 | 5385 | 1 ; | HB2 - C |
| 5374 | 5376 | 1 ; | HB3 - HG |
| 5374 | 5377 | 1 ; | HB3 - CD1 |
| 5374 | 5381 | 1 ; | HB3 - CD2 |
| 5374 | 5385 | 1 ; | HB3 - C |
| 5375 | 5385 | 1 ; | CG - C |
| 5376 | 5378 | 1 ; | HG - HD11 |
| 5376 | 5379 | 1 ; | HG - HD12 |
| 5376 | 5380 | 1 ; | HG - HD13 |
| 5376 | 5382 | 1 ; | HG - HD21 |
| 5376 | 5383 | 1 ; | HG - HD22 |
| 5376 | 5384 | 1 ; | HG - HD23 |
| 5377 | 5382 | 1 ; | CD1 - HD21 |
| 5377 | 5383 | 1 ; | CD1 - HD22 |
| 5377 | 5384 | 1 ; | CD1 - HD23 |
| 5378 | 5381 | 1 ; | HD11 - CD2 |
| 5379 | 5381 | 1 ; | HD12 - CD2 |
| 5380 | 5381 | 1 ; | HD13 - CD2 |
| 5385 | 5390 | 1 ; | C - HA |
| 5385 | 5391 | 1 ; | C - CB |
| 5385 | 5396 | 1 ; | C - C |
| 5386 | 5388 | 1 ; | O - H |
| 5386 | 5389 | 1 ; | O - CA |
| 5387 | 5392 | 1 ; | N - HB2 |
| 5387 | 5393 | 1 ; | N - HB3 |
| 5387 | 5394 | 1 ; | N - OG |
| 5387 | 5397 | 1 ; | N - O |
| 5387 | 5398 | 1 ; | N - N |
| 5388 | 5390 | 1 ; | H - HA |
| 5388 | 5391 | 1 ; | H - CB |
| 5388 | 5396 | 1 ; | H - C |
| 5389 | 5395 | 1 ; | CA - HG |
| 5389 | 5399 | 1 ; | CA - H |
| 5389 | 5400 | 1 ; | CA - CA |
| 5390 | 5392 | 1 ; | HA - HB2 |
| 5390 | 5393 | 1 ; | HA - HB3 |
| 5390 | 5394 | 1 ; | HA - OG |
| 5390 | 5397 | 1 ; | HA - O |
| 5390 | 5398 | 1 ; | HA - N |
| 5391 | 5397 | 1 ; | CB - O |
| 5391 | 5398 | 1 ; | CB - N |
| 5392 | 5395 | 1 ; | HB2 - HG |
| 5392 | 5396 | 1 ; | HB2 - C |
| 5393 | 5395 | 1 ; | HB3 - HG |
| 5393 | 5396 | 1 ; | HB3 - C |
| 5394 | 5396 | 1 ; | OG - C |
| 5396 | 5401 | 1 ; | C - HA |
| 5396 | 5402 | 1 ; | C - CB |
| 5396 | 5407 | 1 ; | C - C |
| 5397 | 5399 | 1 ; | O - H |

| | | | |
|---|---|---|---|
| 5397 | 5400 | 1 ; | O - CA |
| 5398 | 5403 | 1 ; | N - HB2 |
| 5398 | 5404 | 1 ; | N - HB3 |
| 5398 | 5405 | 1 ; | N - OG |
| 5398 | 5408 | 1 ; | N - O |
| 5398 | 5409 | 1 ; | N - N |
| 5399 | 5401 | 1 ; | H - HA |
| 5399 | 5402 | 1 ; | H - CB |
| 5399 | 5407 | 1 ; | H - C |
| 5400 | 5406 | 1 ; | CA - HG |
| 5400 | 5410 | 1 ; | CA - H |
| 5400 | 5411 | 1 ; | CA - CA |
| 5401 | 5403 | 1 ; | HA - HB2 |
| 5401 | 5404 | 1 ; | HA - HB3 |
| 5401 | 5405 | 1 ; | HA - OG |
| 5401 | 5408 | 1 ; | HA - O |
| 5401 | 5409 | 1 ; | HA - N |
| 5402 | 5408 | 1 ; | CB - O |
| 5402 | 5409 | 1 ; | CB - N |
| 5403 | 5406 | 1 ; | HB2 - HG |
| 5403 | 5407 | 1 ; | HB2 - C |
| 5404 | 5406 | 1 ; | HB3 - HG |
| 5404 | 5407 | 1 ; | HB3 - C |
| 5405 | 5407 | 1 ; | OG - C |
| 5407 | 5412 | 1 ; | C - HA |
| 5407 | 5413 | 1 ; | C - CB |
| 5407 | 5424 | 1 ; | C - C |
| 5408 | 5410 | 1 ; | O - H |
| 5408 | 5411 | 1 ; | O - CA |
| 5409 | 5414 | 1 ; | N - HB2 |
| 5409 | 5415 | 1 ; | N - HB3 |
| 5409 | 5416 | 1 ; | N - CG |
| 5409 | 5425 | 1 ; | N - O |
| 5409 | 5426 | 1 ; | N - N |
| 5410 | 5412 | 1 ; | H - HA |
| 5410 | 5413 | 1 ; | H - CB |
| 5410 | 5424 | 1 ; | H - C |
| 5411 | 5417 | 1 ; | CA - HG2 |
| 5411 | 5418 | 1 ; | CA - HG3 |
| 5411 | 5419 | 1 ; | CA - CD |
| 5411 | 5427 | 1 ; | CA - H |
| 5411 | 5428 | 1 ; | CA - CA |
| 5412 | 5414 | 1 ; | HA - HB2 |
| 5412 | 5415 | 1 ; | HA - HB3 |
| 5412 | 5416 | 1 ; | HA - CG |
| 5412 | 5425 | 1 ; | HA - O |
| 5412 | 5426 | 1 ; | HA - N |
| 5413 | 5420 | 1 ; | CB - OE1 |
| 5413 | 5421 | 1 ; | CB - NE2 |
| 5413 | 5425 | 1 ; | CB - O |
| 5413 | 5426 | 1 ; | CB - N |
| 5414 | 5417 | 1 ; | HB2 - HG2 |

```
5414   5418       1 ;    HB2 - HG3
5414   5419       1 ;    HB2 - CD
5414   5424       1 ;    HB2 - C
5415   5417       1 ;    HB3 - HG2
5415   5418       1 ;    HB3 - HG3
5415   5419       1 ;    HB3 - CD
5415   5424       1 ;    HB3 - C
5416   5422       1 ;     CG - HE21
5416   5423       1 ;     CG - HE22
5416   5424       1 ;     CG - C
5417   5420       1 ;    HG2 - OE1
5417   5421       1 ;    HG2 - NE2
5418   5420       1 ;    HG3 - OE1
5418   5421       1 ;    HG3 - NE2
5420   5422       1 ;    OE1 - HE21
5420   5423       1 ;    OE1 - HE22
5424   5429       1 ;      C - HA
5424   5430       1 ;      C - CB
5424   5440       1 ;      C - C
5425   5427       1 ;      O - H
5425   5428       1 ;      O - CA
5426   5431       1 ;      N - HB2
5426   5432       1 ;      N - HB3
5426   5433       1 ;      N - CG
5426   5441       1 ;      N - O
5426   5442       1 ;      N - N
5427   5429       1 ;      H - HA
5427   5430       1 ;      H - CB
5427   5440       1 ;      H - C
5428   5434       1 ;     CA - HG2
5428   5435       1 ;     CA - HG3
5428   5436       1 ;     CA - CD
5428   5443       1 ;     CA - H
5428   5444       1 ;     CA - CA
5429   5431       1 ;     HA - HB2
5429   5432       1 ;     HA - HB3
5429   5433       1 ;     HA - CG
5429   5441       1 ;     HA - O
5429   5442       1 ;     HA - N
5430   5437       1 ;     CB - OE1
5430   5438       1 ;     CB - OE2
5430   5441       1 ;     CB - O
5430   5442       1 ;     CB - N
5431   5434       1 ;    HB2 - HG2
5431   5435       1 ;    HB2 - HG3
5431   5436       1 ;    HB2 - CD
5431   5440       1 ;    HB2 - C
5432   5434       1 ;    HB3 - HG2
5432   5435       1 ;    HB3 - HG3
5432   5436       1 ;    HB3 - CD
5432   5440       1 ;    HB3 - C
5433   5439       1 ;     CG - HE2
```

| | | | | | |
|---|---|---|---|---|---|
| 5433 | 5440 | 1 ; | CG | - | C |
| 5434 | 5437 | 1 ; | HG2 | - | OE1 |
| 5434 | 5438 | 1 ; | HG2 | - | OE2 |
| 5435 | 5437 | 1 ; | HG3 | - | OE1 |
| 5435 | 5438 | 1 ; | HG3 | - | OE2 |
| 5437 | 5439 | 1 ; | OE1 | - | HE2 |
| 5440 | 5445 | 1 ; | C | - | HA |
| 5440 | 5446 | 1 ; | C | - | CB |
| 5440 | 5459 | 1 ; | C | - | C |
| 5441 | 5443 | 1 ; | O | - | H |
| 5441 | 5444 | 1 ; | O | - | CA |
| 5442 | 5447 | 1 ; | N | - | HB |
| 5442 | 5448 | 1 ; | N | - | CG2 |
| 5442 | 5452 | 1 ; | N | - | CG1 |
| 5442 | 5460 | 1 ; | N | - | O |
| 5442 | 5461 | 1 ; | N | - | N |
| 5443 | 5445 | 1 ; | H | - | HA |
| 5443 | 5446 | 1 ; | H | - | CB |
| 5443 | 5459 | 1 ; | H | - | C |
| 5444 | 5449 | 1 ; | CA | - | HG21 |
| 5444 | 5450 | 1 ; | CA | - | HG22 |
| 5444 | 5451 | 1 ; | CA | - | HG23 |
| 5444 | 5453 | 1 ; | CA | - | HG12 |
| 5444 | 5454 | 1 ; | CA | - | HG13 |
| 5444 | 5455 | 1 ; | CA | - | CD1 |
| 5444 | 5462 | 1 ; | CA | - | H |
| 5444 | 5463 | 1 ; | CA | - | CA |
| 5445 | 5447 | 1 ; | HA | - | HB |
| 5445 | 5448 | 1 ; | HA | - | CG2 |
| 5445 | 5452 | 1 ; | HA | - | CG1 |
| 5445 | 5460 | 1 ; | HA | - | O |
| 5445 | 5461 | 1 ; | HA | - | N |
| 5446 | 5456 | 1 ; | CB | - | HD11 |
| 5446 | 5457 | 1 ; | CB | - | HD12 |
| 5446 | 5458 | 1 ; | CB | - | HD13 |
| 5446 | 5460 | 1 ; | CB | - | O |
| 5446 | 5461 | 1 ; | CB | - | N |
| 5447 | 5449 | 1 ; | HB | - | HG21 |
| 5447 | 5450 | 1 ; | HB | - | HG22 |
| 5447 | 5451 | 1 ; | HB | - | HG23 |
| 5447 | 5453 | 1 ; | HB | - | HG12 |
| 5447 | 5454 | 1 ; | HB | - | HG13 |
| 5447 | 5455 | 1 ; | HB | - | CD1 |
| 5447 | 5459 | 1 ; | HB | - | C |
| 5448 | 5453 | 1 ; | CG2 | - | HG12 |
| 5448 | 5454 | 1 ; | CG2 | - | HG13 |
| 5448 | 5455 | 1 ; | CG2 | - | CD1 |
| 5448 | 5459 | 1 ; | CG2 | - | C |
| 5449 | 5452 | 1 ; | HG21 | - | CG1 |
| 5450 | 5452 | 1 ; | HG22 | - | CG1 |
| 5451 | 5452 | 1 ; | HG23 | - | CG1 |
| 5452 | 5459 | 1 ; | CG1 | - | C |

```
5453   5456       1 ;   HG12 - HD11
5453   5457       1 ;   HG12 - HD12
5453   5458       1 ;   HG12 - HD13
5454   5456       1 ;   HG13 - HD11
5454   5457       1 ;   HG13 - HD12
5454   5458       1 ;   HG13 - HD13
5459   5464       1 ;      C - HA
5459   5465       1 ;      C - CB
5459   5478       1 ;      C - C
5460   5462       1 ;      O - H
5460   5463       1 ;      O - CA
5461   5466       1 ;      N - HB2
5461   5467       1 ;      N - HB3
5461   5468       1 ;      N - CG
5461   5479       1 ;      N - O
5461   5480       1 ;      N - N
5462   5464       1 ;      H - HA
5462   5465       1 ;      H - CB
5462   5478       1 ;      H - C
5463   5469       1 ;     CA - HG
5463   5470       1 ;     CA - CD1
5463   5474       1 ;     CA - CD2
5463   5481       1 ;     CA - H
5463   5482       1 ;     CA - CA
5464   5466       1 ;     HA - HB2
5464   5467       1 ;     HA - HB3
5464   5468       1 ;     HA - CG
5464   5479       1 ;     HA - O
5464   5480       1 ;     HA - N
5465   5471       1 ;     CB - HD11
5465   5472       1 ;     CB - HD12
5465   5473       1 ;     CB - HD13
5465   5475       1 ;     CB - HD21
5465   5476       1 ;     CB - HD22
5465   5477       1 ;     CB - HD23
5465   5479       1 ;     CB - O
5465   5480       1 ;     CB - N
5466   5469       1 ;    HB2 - HG
5466   5470       1 ;    HB2 - CD1
5466   5474       1 ;    HB2 - CD2
5466   5478       1 ;    HB2 - C
5467   5469       1 ;    HB3 - HG
5467   5470       1 ;    HB3 - CD1
5467   5474       1 ;    HB3 - CD2
5467   5478       1 ;    HB3 - C
5468   5478       1 ;     CG - C
5469   5471       1 ;     HG - HD11
5469   5472       1 ;     HG - HD12
5469   5473       1 ;     HG - HD13
5469   5475       1 ;     HG - HD21
5469   5476       1 ;     HG - HD22
5469   5477       1 ;     HG - HD23
```

```
5470   5475      1 ;    CD1 - HD21
5470   5476      1 ;    CD1 - HD22
5470   5477      1 ;    CD1 - HD23
5471   5474      1 ;   HD11 - CD2
5472   5474      1 ;   HD12 - CD2
5473   5474      1 ;   HD13 - CD2
5478   5483      1 ;      C - HA
5478   5484      1 ;      C - CB
5478   5490      1 ;      C - C
5479   5481      1 ;      O - H
5479   5482      1 ;      O - CA
5480   5485      1 ;      N - HB2
5480   5486      1 ;      N - HB3
5480   5487      1 ;      N - CG
5480   5491      1 ;      N - O
5480   5492      1 ;      N - N
5481   5483      1 ;      H - HA
5481   5484      1 ;      H - CB
5481   5490      1 ;      H - C
5482   5488      1 ;     CA - OD1
5482   5489      1 ;     CA - OD2
5482   5493      1 ;     CA - H
5482   5494      1 ;     CA - CA
5483   5485      1 ;     HA - HB2
5483   5486      1 ;     HA - HB3
5483   5487      1 ;     HA - CG
5483   5491      1 ;     HA - O
5483   5492      1 ;     HA - N
5484   5491      1 ;     CB - O
5484   5492      1 ;     CB - N
5485   5488      1 ;    HB2 - OD1
5485   5489      1 ;    HB2 - OD2
5485   5490      1 ;    HB2 - C
5486   5488      1 ;    HB3 - OD1
5486   5489      1 ;    HB3 - OD2
5486   5490      1 ;    HB3 - C
5487   5490      1 ;     CG - C
5490   5495      1 ;      C - HA2
5490   5496      1 ;      C - HA3
5490   5497      1 ;      C - C
5491   5493      1 ;      O - H
5491   5494      1 ;      O - CA
5492   5498      1 ;      N - O
5492   5499      1 ;      N - N
5493   5495      1 ;      H - HA2
5493   5496      1 ;      H - HA3
5493   5497      1 ;      H - C
5494   5500      1 ;     CA - H
5494   5501      1 ;     CA - CA
5495   5498      1 ;    HA2 - O
5495   5499      1 ;    HA2 - N
5496   5498      1 ;    HA3 - O
```

```
5496   5499       1 ;    HA3 - N
5497   5502       1 ;      C - HA
5497   5503       1 ;      C - CB
5497   5516       1 ;      C - C
5498   5500       1 ;      O - H
5498   5501       1 ;      O - CA
5499   5504       1 ;      N - HB2
5499   5505       1 ;      N - HB3
5499   5506       1 ;      N - CG
5499   5517       1 ;      N - O
5499   5518       1 ;      N - N
5500   5502       1 ;      H - HA
5500   5503       1 ;      H - CB
5500   5516       1 ;      H - C
5501   5507       1 ;     CA - HG
5501   5508       1 ;     CA - CD1
5501   5512       1 ;     CA - CD2
5501   5519       1 ;     CA - H
5501   5520       1 ;     CA - CA
5502   5504       1 ;     HA - HB2
5502   5505       1 ;     HA - HB3
5502   5506       1 ;     HA - CG
5502   5517       1 ;     HA - O
5502   5518       1 ;     HA - N
5503   5509       1 ;     CB - HD11
5503   5510       1 ;     CB - HD12
5503   5511       1 ;     CB - HD13
5503   5513       1 ;     CB - HD21
5503   5514       1 ;     CB - HD22
5503   5515       1 ;     CB - HD23
5503   5517       1 ;     CB - O
5503   5518       1 ;     CB - N
5504   5507       1 ;    HB2 - HG
5504   5508       1 ;    HB2 - CD1
5504   5512       1 ;    HB2 - CD2
5504   5516       1 ;    HB2 - C
5505   5507       1 ;    HB3 - HG
5505   5508       1 ;    HB3 - CD1
5505   5512       1 ;    HB3 - CD2
5505   5516       1 ;    HB3 - C
5506   5516       1 ;     CG - C
5507   5509       1 ;     HG - HD11
5507   5510       1 ;     HG - HD12
5507   5511       1 ;     HG - HD13
5507   5513       1 ;     HG - HD21
5507   5514       1 ;     HG - HD22
5507   5515       1 ;     HG - HD23
5508   5513       1 ;    CD1 - HD21
5508   5514       1 ;    CD1 - HD22
5508   5515       1 ;    CD1 - HD23
5509   5512       1 ;   HD11 - CD2
5510   5512       1 ;   HD12 - CD2
```

```
5511   5512       1 ;   HD13 - CD2
5516   5521       1 ;      C - HA
5516   5522       1 ;      C - CB
5516   5538       1 ;      C - C
5517   5519       1 ;      O - H
5517   5520       1 ;      O - CA
5518   5523       1 ;      N - HB2
5518   5524       1 ;      N - HB3
5518   5525       1 ;      N - CG
5518   5539       1 ;      N - O
5518   5540       1 ;      N - N
5519   5521       1 ;      H - HA
5519   5522       1 ;      H - CB
5519   5538       1 ;      H - C
5520   5526       1 ;     CA - HG2
5520   5527       1 ;     CA - HG3
5520   5528       1 ;     CA - CD
5520   5541       1 ;     CA - H
5520   5542       1 ;     CA - CA
5521   5523       1 ;     HA - HB2
5521   5524       1 ;     HA - HB3
5521   5525       1 ;     HA - CG
5521   5539       1 ;     HA - O
5521   5540       1 ;     HA - N
5522   5529       1 ;     CB - HD2
5522   5530       1 ;     CB - HD3
5522   5531       1 ;     CB - CE
5522   5539       1 ;     CB - O
5522   5540       1 ;     CB - N
5523   5526       1 ;    HB2 - HG2
5523   5527       1 ;    HB2 - HG3
5523   5528       1 ;    HB2 - CD
5523   5538       1 ;    HB2 - C
5524   5526       1 ;    HB3 - HG2
5524   5527       1 ;    HB3 - HG3
5524   5528       1 ;    HB3 - CD
5524   5538       1 ;    HB3 - C
5525   5532       1 ;     CG - HE2
5525   5533       1 ;     CG - HE3
5525   5534       1 ;     CG - NZ
5525   5538       1 ;     CG - C
5526   5529       1 ;    HG2 - HD2
5526   5530       1 ;    HG2 - HD3
5526   5531       1 ;    HG2 - CE
5527   5529       1 ;    HG3 - HD2
5527   5530       1 ;    HG3 - HD3
5527   5531       1 ;    HG3 - CE
5528   5535       1 ;     CD - HZ1
5528   5536       1 ;     CD - HZ2
5528   5537       1 ;     CD - HZ3
5529   5532       1 ;    HD2 - HE2
5529   5533       1 ;    HD2 - HE3
```

| | | | |
|---|---|---|---|
| 5529 | 5534 | 1 ; | HD2 - NZ |
| 5530 | 5532 | 1 ; | HD3 - HE2 |
| 5530 | 5533 | 1 ; | HD3 - HE3 |
| 5530 | 5534 | 1 ; | HD3 - NZ |
| 5532 | 5535 | 1 ; | HE2 - HZ1 |
| 5532 | 5536 | 1 ; | HE2 - HZ2 |
| 5532 | 5537 | 1 ; | HE2 - HZ3 |
| 5533 | 5535 | 1 ; | HE3 - HZ1 |
| 5533 | 5536 | 1 ; | HE3 - HZ2 |
| 5533 | 5537 | 1 ; | HE3 - HZ3 |
| 5538 | 5543 | 1 ; | C - HA |
| 5538 | 5544 | 1 ; | C - CB |
| 5538 | 5560 | 1 ; | C - C |
| 5539 | 5541 | 1 ; | O - H |
| 5539 | 5542 | 1 ; | O - CA |
| 5540 | 5545 | 1 ; | N - HB2 |
| 5540 | 5546 | 1 ; | N - HB3 |
| 5540 | 5547 | 1 ; | N - CG |
| 5540 | 5561 | 1 ; | N - O |
| 5540 | 5562 | 1 ; | N - N |
| 5541 | 5543 | 1 ; | H - HA |
| 5541 | 5544 | 1 ; | H - CB |
| 5541 | 5560 | 1 ; | H - C |
| 5542 | 5548 | 1 ; | CA - HG2 |
| 5542 | 5549 | 1 ; | CA - HG3 |
| 5542 | 5550 | 1 ; | CA - CD |
| 5542 | 5563 | 1 ; | CA - H |
| 5542 | 5564 | 1 ; | CA - CA |
| 5543 | 5545 | 1 ; | HA - HB2 |
| 5543 | 5546 | 1 ; | HA - HB3 |
| 5543 | 5547 | 1 ; | HA - CG |
| 5543 | 5561 | 1 ; | HA - O |
| 5543 | 5562 | 1 ; | HA - N |
| 5544 | 5551 | 1 ; | CB - HD2 |
| 5544 | 5552 | 1 ; | CB - HD3 |
| 5544 | 5553 | 1 ; | CB - CE |
| 5544 | 5561 | 1 ; | CB - O |
| 5544 | 5562 | 1 ; | CB - N |
| 5545 | 5548 | 1 ; | HB2 - HG2 |
| 5545 | 5549 | 1 ; | HB2 - HG3 |
| 5545 | 5550 | 1 ; | HB2 - CD |
| 5545 | 5560 | 1 ; | HB2 - C |
| 5546 | 5548 | 1 ; | HB3 - HG2 |
| 5546 | 5549 | 1 ; | HB3 - HG3 |
| 5546 | 5550 | 1 ; | HB3 - CD |
| 5546 | 5560 | 1 ; | HB3 - C |
| 5547 | 5554 | 1 ; | CG - HE2 |
| 5547 | 5555 | 1 ; | CG - HE3 |
| 5547 | 5556 | 1 ; | CG - NZ |
| 5547 | 5560 | 1 ; | CG - C |
| 5548 | 5551 | 1 ; | HG2 - HD2 |
| 5548 | 5552 | 1 ; | HG2 - HD3 |

| | | | |
|---|---|---|---|
| 5548 | 5553 | 1 ; | HG2 - CE |
| 5549 | 5551 | 1 ; | HG3 - HD2 |
| 5549 | 5552 | 1 ; | HG3 - HD3 |
| 5549 | 5553 | 1 ; | HG3 - CE |
| 5550 | 5557 | 1 ; | CD - HZ1 |
| 5550 | 5558 | 1 ; | CD - HZ2 |
| 5550 | 5559 | 1 ; | CD - HZ3 |
| 5551 | 5554 | 1 ; | HD2 - HE2 |
| 5551 | 5555 | 1 ; | HD2 - HE3 |
| 5551 | 5556 | 1 ; | HD2 - NZ |
| 5552 | 5554 | 1 ; | HD3 - HE2 |
| 5552 | 5555 | 1 ; | HD3 - HE3 |
| 5552 | 5556 | 1 ; | HD3 - NZ |
| 5554 | 5557 | 1 ; | HE2 - HZ1 |
| 5554 | 5558 | 1 ; | HE2 - HZ2 |
| 5554 | 5559 | 1 ; | HE2 - HZ3 |
| 5555 | 5557 | 1 ; | HE3 - HZ1 |
| 5555 | 5558 | 1 ; | HE3 - HZ2 |
| 5555 | 5559 | 1 ; | HE3 - HZ3 |
| 5560 | 5565 | 1 ; | C - HA |
| 5560 | 5566 | 1 ; | C - CB |
| 5560 | 5579 | 1 ; | C - C |
| 5561 | 5563 | 1 ; | O - H |
| 5561 | 5564 | 1 ; | O - CA |
| 5562 | 5567 | 1 ; | N - HB |
| 5562 | 5568 | 1 ; | N - CG2 |
| 5562 | 5572 | 1 ; | N - CG1 |
| 5562 | 5580 | 1 ; | N - O |
| 5562 | 5581 | 1 ; | N - N |
| 5563 | 5565 | 1 ; | H - HA |
| 5563 | 5566 | 1 ; | H - CB |
| 5563 | 5579 | 1 ; | H - C |
| 5564 | 5569 | 1 ; | CA - HG21 |
| 5564 | 5570 | 1 ; | CA - HG22 |
| 5564 | 5571 | 1 ; | CA - HG23 |
| 5564 | 5573 | 1 ; | CA - HG12 |
| 5564 | 5574 | 1 ; | CA - HG13 |
| 5564 | 5575 | 1 ; | CA - CD1 |
| 5564 | 5582 | 1 ; | CA - H |
| 5564 | 5583 | 1 ; | CA - CA |
| 5565 | 5567 | 1 ; | HA - HB |
| 5565 | 5568 | 1 ; | HA - CG2 |
| 5565 | 5572 | 1 ; | HA - CG1 |
| 5565 | 5580 | 1 ; | HA - O |
| 5565 | 5581 | 1 ; | HA - N |
| 5566 | 5576 | 1 ; | CB - HD11 |
| 5566 | 5577 | 1 ; | CB - HD12 |
| 5566 | 5578 | 1 ; | CB - HD13 |
| 5566 | 5580 | 1 ; | CB - O |
| 5566 | 5581 | 1 ; | CB - N |
| 5567 | 5569 | 1 ; | HB - HG21 |
| 5567 | 5570 | 1 ; | HB - HG22 |

```
5567   5571      1 ;     HB - HG23
5567   5573      1 ;     HB - HG12
5567   5574      1 ;     HB - HG13
5567   5575      1 ;     HB - CD1
5567   5579      1 ;     HB - C
5568   5573      1 ;    CG2 - HG12
5568   5574      1 ;    CG2 - HG13
5568   5575      1 ;    CG2 - CD1
5568   5579      1 ;    CG2 - C
5569   5572      1 ;   HG21 - CG1
5570   5572      1 ;   HG22 - CG1
5571   5572      1 ;   HG23 - CG1
5572   5579      1 ;    CG1 - C
5573   5576      1 ;   HG12 - HD11
5573   5577      1 ;   HG12 - HD12
5573   5578      1 ;   HG12 - HD13
5574   5576      1 ;   HG13 - HD11
5574   5577      1 ;   HG13 - HD12
5574   5578      1 ;   HG13 - HD13
5579   5584      1 ;      C - HA2
5579   5585      1 ;      C - HA3
5579   5586      1 ;      C - C
5580   5582      1 ;      O - H
5580   5583      1 ;      O - CA
5581   5587      1 ;      N - O
5581   5588      1 ;      N - N
5582   5584      1 ;      H - HA2
5582   5585      1 ;      H - HA3
5582   5586      1 ;      H - C
5583   5589      1 ;     CA - H
5583   5590      1 ;     CA - CA
5584   5587      1 ;    HA2 - O
5584   5588      1 ;    HA2 - N
5585   5587      1 ;    HA3 - O
5585   5588      1 ;    HA3 - N
5586   5591      1 ;      C - HA
5586   5592      1 ;      C - CB
5586   5607      1 ;      C - C
5587   5589      1 ;      O - H
5587   5590      1 ;      O - CA
5588   5593      1 ;      N - HB2
5588   5594      1 ;      N - HB3
5588   5595      1 ;      N - CG
5588   5608      1 ;      N - O
5588   5609      1 ;      N - N
5589   5591      1 ;      H - HA
5589   5592      1 ;      H - CB
5589   5607      1 ;      H - C
5590   5596      1 ;     CA - CD1
5590   5605      1 ;     CA - CD2
5590   5610      1 ;     CA - H
5590   5611      1 ;     CA - CA
```

| | | | | | |
|---|---|---|---|---|---|
| 5591 | 5593 | 1 ; | HA | - | HB2 |
| 5591 | 5594 | 1 ; | HA | - | HB3 |
| 5591 | 5595 | 1 ; | HA | - | CG |
| 5591 | 5608 | 1 ; | HA | - | O |
| 5591 | 5609 | 1 ; | HA | - | N |
| 5592 | 5597 | 1 ; | CB | - | HD1 |
| 5592 | 5598 | 1 ; | CB | - | CE1 |
| 5592 | 5603 | 1 ; | CB | - | CE2 |
| 5592 | 5606 | 1 ; | CB | - | HD2 |
| 5592 | 5608 | 1 ; | CB | - | O |
| 5592 | 5609 | 1 ; | CB | - | N |
| 5593 | 5596 | 1 ; | HB2 | - | CD1 |
| 5593 | 5605 | 1 ; | HB2 | - | CD2 |
| 5593 | 5607 | 1 ; | HB2 | - | C |
| 5594 | 5596 | 1 ; | HB3 | - | CD1 |
| 5594 | 5605 | 1 ; | HB3 | - | CD2 |
| 5594 | 5607 | 1 ; | HB3 | - | C |
| 5595 | 5599 | 1 ; | CG | - | HE1 |
| 5595 | 5600 | 1 ; | CG | - | CZ |
| 5595 | 5604 | 1 ; | CG | - | HE2 |
| 5595 | 5607 | 1 ; | CG | - | C |
| 5596 | 5601 | 1 ; | CD1 | - | OH |
| 5596 | 5603 | 1 ; | CD1 | - | CE2 |
| 5596 | 5606 | 1 ; | CD1 | - | HD2 |
| 5597 | 5599 | 1 ; | HD1 | - | HE1 |
| 5597 | 5600 | 1 ; | HD1 | - | CZ |
| 5597 | 5605 | 1 ; | HD1 | - | CD2 |
| 5598 | 5602 | 1 ; | CE1 | - | HH |
| 5598 | 5604 | 1 ; | CE1 | - | HE2 |
| 5598 | 5605 | 1 ; | CE1 | - | CD2 |
| 5599 | 5601 | 1 ; | HE1 | - | OH |
| 5599 | 5603 | 1 ; | HE1 | - | CE2 |
| 5600 | 5606 | 1 ; | CZ | - | HD2 |
| 5601 | 5604 | 1 ; | OH | - | HE2 |
| 5601 | 5605 | 1 ; | OH | - | CD2 |
| 5602 | 5603 | 1 ; | HH | - | CE2 |
| 5604 | 5606 | 1 ; | HE2 | - | HD2 |
| 5607 | 5612 | 1 ; | C | - | HA |
| 5607 | 5613 | 1 ; | C | - | CB |
| 5607 | 5624 | 1 ; | C | - | C |
| 5608 | 5610 | 1 ; | O | - | H |
| 5608 | 5611 | 1 ; | O | - | CA |
| 5609 | 5614 | 1 ; | N | - | HB2 |
| 5609 | 5615 | 1 ; | N | - | HB3 |
| 5609 | 5616 | 1 ; | N | - | CG |
| 5609 | 5625 | 1 ; | N | - | O |
| 5609 | 5626 | 1 ; | N | - | N |
| 5610 | 5612 | 1 ; | H | - | HA |
| 5610 | 5613 | 1 ; | H | - | CB |
| 5610 | 5624 | 1 ; | H | - | C |
| 5611 | 5617 | 1 ; | CA | - | HG2 |
| 5611 | 5618 | 1 ; | CA | - | HG3 |

```
5611   5619       1 ;      CA - CD
5611   5627       1 ;      CA - H
5611   5628       1 ;      CA - CA
5612   5614       1 ;      HA - HB2
5612   5615       1 ;      HA - HB3
5612   5616       1 ;      HA - CG
5612   5625       1 ;      HA - O
5612   5626       1 ;      HA - N
5613   5620       1 ;      CB - OE1
5613   5621       1 ;      CB - NE2
5613   5625       1 ;      CB - O
5613   5626       1 ;      CB - N
5614   5617       1 ;     HB2 - HG2
5614   5618       1 ;     HB2 - HG3
5614   5619       1 ;     HB2 - CD
5614   5624       1 ;     HB2 - C
5615   5617       1 ;     HB3 - HG2
5615   5618       1 ;     HB3 - HG3
5615   5619       1 ;     HB3 - CD
5615   5624       1 ;     HB3 - C
5616   5622       1 ;      CG - HE21
5616   5623       1 ;      CG - HE22
5616   5624       1 ;      CG - C
5617   5620       1 ;     HG2 - OE1
5617   5621       1 ;     HG2 - NE2
5618   5620       1 ;     HG3 - OE1
5618   5621       1 ;     HG3 - NE2
5620   5622       1 ;     OE1 - HE21
5620   5623       1 ;     OE1 - HE22
5624   5629       1 ;       C - HA
5624   5630       1 ;       C - CB
5624   5646       1 ;       C - C
5625   5627       1 ;       O - H
5625   5628       1 ;       O - CA
5626   5631       1 ;       N - HB2
5626   5632       1 ;       N - HB3
5626   5633       1 ;       N - CG
5626   5647       1 ;       N - O
5626   5648       1 ;       N - N
5627   5629       1 ;       H - HA
5627   5630       1 ;       H - CB
5627   5646       1 ;       H - C
5628   5634       1 ;      CA - HG2
5628   5635       1 ;      CA - HG3
5628   5636       1 ;      CA - CD
5628   5649       1 ;      CA - H
5628   5650       1 ;      CA - CA
5629   5631       1 ;      HA - HB2
5629   5632       1 ;      HA - HB3
5629   5633       1 ;      HA - CG
5629   5647       1 ;      HA - O
5629   5648       1 ;      HA - N
```

```
5630   5637      1 ;     CB - HD2
5630   5638      1 ;     CB - HD3
5630   5639      1 ;     CB - CE
5630   5647      1 ;     CB - O
5630   5648      1 ;     CB - N
5631   5634      1 ;    HB2 - HG2
5631   5635      1 ;    HB2 - HG3
5631   5636      1 ;    HB2 - CD
5631   5646      1 ;    HB2 - C
5632   5634      1 ;    HB3 - HG2
5632   5635      1 ;    HB3 - HG3
5632   5636      1 ;    HB3 - CD
5632   5646      1 ;    HB3 - C
5633   5640      1 ;     CG - HE2
5633   5641      1 ;     CG - HE3
5633   5642      1 ;     CG - NZ
5633   5646      1 ;     CG - C
5634   5637      1 ;    HG2 - HD2
5634   5638      1 ;    HG2 - HD3
5634   5639      1 ;    HG2 - CE
5635   5637      1 ;    HG3 - HD2
5635   5638      1 ;    HG3 - HD3
5635   5639      1 ;    HG3 - CE
5636   5643      1 ;     CD - HZ1
5636   5644      1 ;     CD - HZ2
5636   5645      1 ;     CD - HZ3
5637   5640      1 ;    HD2 - HE2
5637   5641      1 ;    HD2 - HE3
5637   5642      1 ;    HD2 - NZ
5638   5640      1 ;    HD3 - HE2
5638   5641      1 ;    HD3 - HE3
5638   5642      1 ;    HD3 - NZ
5640   5643      1 ;    HE2 - HZ1
5640   5644      1 ;    HE2 - HZ2
5640   5645      1 ;    HE2 - HZ3
5641   5643      1 ;    HE3 - HZ1
5641   5644      1 ;    HE3 - HZ2
5641   5645      1 ;    HE3 - HZ3
5646   5651      1 ;      C - HA
5646   5652      1 ;      C - CB
5646   5665      1 ;      C - C
5647   5649      1 ;      O - H
5647   5650      1 ;      O - CA
5648   5653      1 ;      N - HB
5648   5654      1 ;      N - CG2
5648   5658      1 ;      N - CG1
5648   5666      1 ;      N - O
5648   5667      1 ;      N - N
5649   5651      1 ;      H - HA
5649   5652      1 ;      H - CB
5649   5665      1 ;      H - C
5650   5655      1 ;     CA - HG21
```

```
5650   5656       1 ;      CA - HG22
5650   5657       1 ;      CA - HG23
5650   5659       1 ;      CA - HG12
5650   5660       1 ;      CA - HG13
5650   5661       1 ;      CA - CD1
5650   5668       1 ;      CA - CD
5650   5677       1 ;      CA - CA
5651   5653       1 ;      HA - HB
5651   5654       1 ;      HA - CG2
5651   5658       1 ;      HA - CG1
5651   5666       1 ;      HA - O
5651   5667       1 ;      HA - N
5652   5662       1 ;      CB - HD11
5652   5663       1 ;      CB - HD12
5652   5664       1 ;      CB - HD13
5652   5666       1 ;      CB - O
5652   5667       1 ;      CB - N
5653   5655       1 ;      HB - HG21
5653   5656       1 ;      HB - HG22
5653   5657       1 ;      HB - HG23
5653   5659       1 ;      HB - HG12
5653   5660       1 ;      HB - HG13
5653   5661       1 ;      HB - CD1
5653   5665       1 ;      HB - C
5654   5659       1 ;     CG2 - HG12
5654   5660       1 ;     CG2 - HG13
5654   5661       1 ;     CG2 - CD1
5654   5665       1 ;     CG2 - C
5655   5658       1 ;    HG21 - CG1
5656   5658       1 ;    HG22 - CG1
5657   5658       1 ;    HG23 - CG1
5658   5665       1 ;     CG1 - C
5659   5662       1 ;    HG12 - HD11
5659   5663       1 ;    HG12 - HD12
5659   5664       1 ;    HG12 - HD13
5660   5662       1 ;    HG13 - HD11
5660   5663       1 ;    HG13 - HD12
5660   5664       1 ;    HG13 - HD13
5665   5669       1 ;       C - HD2
5665   5670       1 ;       C - HD3
5665   5671       1 ;       C - CG
5665   5674       1 ;       C - CB
5665   5678       1 ;       C - HA
5665   5679       1 ;       C - C
5666   5668       1 ;       O - CD
5666   5677       1 ;       O - CA
5667   5672       1 ;       N - HG2
5667   5673       1 ;       N - HG3
5667   5675       1 ;       N - HB2
5667   5676       1 ;       N - HB3
5667   5680       1 ;       N - O
5667   5681       1 ;       N - N
```

```
5668   5675       1 ;      CD - HB2
5668   5676       1 ;      CD - HB3
5668   5678       1 ;      CD - HA
5668   5679       1 ;      CD - C
5669   5672       1 ;     HD2 - HG2
5669   5673       1 ;     HD2 - HG3
5669   5674       1 ;     HD2 - CB
5669   5677       1 ;     HD2 - CA
5670   5672       1 ;     HD3 - HG2
5670   5673       1 ;     HD3 - HG3
5670   5674       1 ;     HD3 - CB
5670   5677       1 ;     HD3 - CA
5671   5678       1 ;      CG - HA
5671   5679       1 ;      CG - C
5672   5675       1 ;     HG2 - HB2
5672   5676       1 ;     HG2 - HB3
5672   5677       1 ;     HG2 - CA
5673   5675       1 ;     HG3 - HB2
5673   5676       1 ;     HG3 - HB3
5673   5677       1 ;     HG3 - CA
5674   5680       1 ;      CB - O
5674   5681       1 ;      CB - N
5675   5678       1 ;     HB2 - HA
5675   5679       1 ;     HB2 - C
5676   5678       1 ;     HB3 - HA
5676   5679       1 ;     HB3 - C
5677   5682       1 ;      CA - CD
5677   5691       1 ;      CA - CA
5678   5680       1 ;      HA - O
5678   5681       1 ;      HA - N
5679   5683       1 ;       C - HD2
5679   5684       1 ;       C - HD3
5679   5685       1 ;       C - CG
5679   5688       1 ;       C - CB
5679   5692       1 ;       C - HA
5679   5693       1 ;       C - C
5680   5682       1 ;       O - CD
5680   5691       1 ;       O - CA
5681   5686       1 ;       N - HG2
5681   5687       1 ;       N - HG3
5681   5689       1 ;       N - HB2
5681   5690       1 ;       N - HB3
5681   5694       1 ;       N - O
5681   5695       1 ;       N - N
5682   5689       1 ;      CD - HB2
5682   5690       1 ;      CD - HB3
5682   5692       1 ;      CD - HA
5682   5693       1 ;      CD - C
5683   5686       1 ;     HD2 - HG2
5683   5687       1 ;     HD2 - HG3
5683   5688       1 ;     HD2 - CB
5683   5691       1 ;     HD2 - CA
```

| | | | |
|---|---|---|---|
| 5684 | 5686 | 1 ; | HD3 - HG2 |
| 5684 | 5687 | 1 ; | HD3 - HG3 |
| 5684 | 5688 | 1 ; | HD3 - CB |
| 5684 | 5691 | 1 ; | HD3 - CA |
| 5685 | 5692 | 1 ; | CG - HA |
| 5685 | 5693 | 1 ; | CG - C |
| 5686 | 5689 | 1 ; | HG2 - HB2 |
| 5686 | 5690 | 1 ; | HG2 - HB3 |
| 5686 | 5691 | 1 ; | HG2 - CA |
| 5687 | 5689 | 1 ; | HG3 - HB2 |
| 5687 | 5690 | 1 ; | HG3 - HB3 |
| 5687 | 5691 | 1 ; | HG3 - CA |
| 5688 | 5694 | 1 ; | CB - O |
| 5688 | 5695 | 1 ; | CB - N |
| 5689 | 5692 | 1 ; | HB2 - HA |
| 5689 | 5693 | 1 ; | HB2 - C |
| 5690 | 5692 | 1 ; | HB3 - HA |
| 5690 | 5693 | 1 ; | HB3 - C |
| 5691 | 5696 | 1 ; | CA - H |
| 5691 | 5697 | 1 ; | CA - CA |
| 5692 | 5694 | 1 ; | HA - O |
| 5692 | 5695 | 1 ; | HA - N |
| 5693 | 5698 | 1 ; | C - HA |
| 5693 | 5699 | 1 ; | C - CB |
| 5693 | 5705 | 1 ; | C - C |
| 5694 | 5696 | 1 ; | O - H |
| 5694 | 5697 | 1 ; | O - CA |
| 5695 | 5700 | 1 ; | N - HB2 |
| 5695 | 5701 | 1 ; | N - HB3 |
| 5695 | 5702 | 1 ; | N - CG |
| 5695 | 5706 | 1 ; | N - O |
| 5695 | 5707 | 1 ; | N - N |
| 5696 | 5698 | 1 ; | H - HA |
| 5696 | 5699 | 1 ; | H - CB |
| 5696 | 5705 | 1 ; | H - C |
| 5697 | 5703 | 1 ; | CA - OD1 |
| 5697 | 5704 | 1 ; | CA - OD2 |
| 5697 | 5708 | 1 ; | CA - H |
| 5697 | 5709 | 1 ; | CA - CA |
| 5698 | 5700 | 1 ; | HA - HB2 |
| 5698 | 5701 | 1 ; | HA - HB3 |
| 5698 | 5702 | 1 ; | HA - CG |
| 5698 | 5706 | 1 ; | HA - O |
| 5698 | 5707 | 1 ; | HA - N |
| 5699 | 5706 | 1 ; | CB - O |
| 5699 | 5707 | 1 ; | CB - N |
| 5700 | 5703 | 1 ; | HB2 - OD1 |
| 5700 | 5704 | 1 ; | HB2 - OD2 |
| 5700 | 5705 | 1 ; | HB2 - C |
| 5701 | 5703 | 1 ; | HB3 - OD1 |
| 5701 | 5704 | 1 ; | HB3 - OD2 |
| 5701 | 5705 | 1 ; | HB3 - C |

```
5702   5705       1 ;      CG - C
5705   5710       1 ;       C - HA
5705   5711       1 ;       C - CB
5705   5724       1 ;       C - C
5706   5708       1 ;       O - H
5706   5709       1 ;       O - CA
5707   5712       1 ;       N - HB
5707   5713       1 ;       N - CG2
5707   5717       1 ;       N - CG1
5707   5725       1 ;       N - O
5707   5726       1 ;       N - N
5708   5710       1 ;       H - HA
5708   5711       1 ;       H - CB
5708   5724       1 ;       H - C
5709   5714       1 ;      CA - HG21
5709   5715       1 ;      CA - HG22
5709   5716       1 ;      CA - HG23
5709   5718       1 ;      CA - HG12
5709   5719       1 ;      CA - HG13
5709   5720       1 ;      CA - CD1
5709   5727       1 ;      CA - H
5709   5728       1 ;      CA - CA
5710   5712       1 ;      HA - HB
5710   5713       1 ;      HA - CG2
5710   5717       1 ;      HA - CG1
5710   5725       1 ;      HA - O
5710   5726       1 ;      HA - N
5711   5721       1 ;      CB - HD11
5711   5722       1 ;      CB - HD12
5711   5723       1 ;      CB - HD13
5711   5725       1 ;      CB - O
5711   5726       1 ;      CB - N
5712   5714       1 ;      HB - HG21
5712   5715       1 ;      HB - HG22
5712   5716       1 ;      HB - HG23
5712   5718       1 ;      HB - HG12
5712   5719       1 ;      HB - HG13
5712   5720       1 ;      HB - CD1
5712   5724       1 ;      HB - C
5713   5718       1 ;     CG2 - HG12
5713   5719       1 ;     CG2 - HG13
5713   5720       1 ;     CG2 - CD1
5713   5724       1 ;     CG2 - C
5714   5717       1 ;    HG21 - CG1
5715   5717       1 ;    HG22 - CG1
5716   5717       1 ;    HG23 - CG1
5717   5724       1 ;     CG1 - C
5718   5721       1 ;    HG12 - HD11
5718   5722       1 ;    HG12 - HD12
5718   5723       1 ;    HG12 - HD13
5719   5721       1 ;    HG13 - HD11
5719   5722       1 ;    HG13 - HD12
```

| | | | | | |
|---|---|---|---|---|---|
| 5719 | 5723 | 1 ; | HG13 | - | HD13 |
| 5724 | 5729 | 1 ; | C | - | HA |
| 5724 | 5730 | 1 ; | C | - | CB |
| 5724 | 5741 | 1 ; | C | - | C |
| 5725 | 5727 | 1 ; | O | - | H |
| 5725 | 5728 | 1 ; | O | - | CA |
| 5726 | 5731 | 1 ; | N | - | HB2 |
| 5726 | 5732 | 1 ; | N | - | HB3 |
| 5726 | 5733 | 1 ; | N | - | CG |
| 5726 | 5742 | 1 ; | N | - | O |
| 5726 | 5743 | 1 ; | N | - | N |
| 5727 | 5729 | 1 ; | H | - | HA |
| 5727 | 5730 | 1 ; | H | - | CB |
| 5727 | 5741 | 1 ; | H | - | C |
| 5728 | 5734 | 1 ; | CA | - | ND1 |
| 5728 | 5739 | 1 ; | CA | - | CD2 |
| 5728 | 5744 | 1 ; | CA | - | H |
| 5728 | 5745 | 1 ; | CA | - | CA |
| 5729 | 5731 | 1 ; | HA | - | HB2 |
| 5729 | 5732 | 1 ; | HA | - | HB3 |
| 5729 | 5733 | 1 ; | HA | - | CG |
| 5729 | 5742 | 1 ; | HA | - | O |
| 5729 | 5743 | 1 ; | HA | - | N |
| 5730 | 5735 | 1 ; | CB | - | CE1 |
| 5730 | 5737 | 1 ; | CB | - | NE2 |
| 5730 | 5740 | 1 ; | CB | - | HD2 |
| 5730 | 5742 | 1 ; | CB | - | O |
| 5730 | 5743 | 1 ; | CB | - | N |
| 5731 | 5734 | 1 ; | HB2 | - | ND1 |
| 5731 | 5739 | 1 ; | HB2 | - | CD2 |
| 5731 | 5741 | 1 ; | HB2 | - | C |
| 5732 | 5734 | 1 ; | HB3 | - | ND1 |
| 5732 | 5739 | 1 ; | HB3 | - | CD2 |
| 5732 | 5741 | 1 ; | HB3 | - | C |
| 5733 | 5736 | 1 ; | CG | - | HE1 |
| 5733 | 5738 | 1 ; | CG | - | HE2 |
| 5733 | 5741 | 1 ; | CG | - | C |
| 5734 | 5738 | 1 ; | ND1 | - | HE2 |
| 5734 | 5740 | 1 ; | ND1 | - | HD2 |
| 5735 | 5740 | 1 ; | CE1 | - | HD2 |
| 5736 | 5738 | 1 ; | HE1 | - | HE2 |
| 5736 | 5739 | 1 ; | HE1 | - | CD2 |
| 5738 | 5740 | 1 ; | HE2 | - | HD2 |
| 5741 | 5746 | 1 ; | C | - | HA |
| 5741 | 5747 | 1 ; | C | - | CB |
| 5741 | 5758 | 1 ; | C | - | C |
| 5742 | 5744 | 1 ; | O | - | H |
| 5742 | 5745 | 1 ; | O | - | CA |
| 5743 | 5748 | 1 ; | N | - | HB2 |
| 5743 | 5749 | 1 ; | N | - | HB3 |
| 5743 | 5750 | 1 ; | N | - | CG |
| 5743 | 5759 | 1 ; | N | - | O |

```
5743   5760       1 ;       N - N
5744   5746       1 ;       H - HA
5744   5747       1 ;       H - CB
5744   5758       1 ;       H - C
5745   5751       1 ;      CA - HG2
5745   5752       1 ;      CA - HG3
5745   5753       1 ;      CA - CD
5745   5761       1 ;      CA - H
5745   5762       1 ;      CA - CA
5746   5748       1 ;      HA - HB2
5746   5749       1 ;      HA - HB3
5746   5750       1 ;      HA - CG
5746   5759       1 ;      HA - O
5746   5760       1 ;      HA - N
5747   5754       1 ;      CB - OE1
5747   5755       1 ;      CB - NE2
5747   5759       1 ;      CB - O
5747   5760       1 ;      CB - N
5748   5751       1 ;     HB2 - HG2
5748   5752       1 ;     HB2 - HG3
5748   5753       1 ;     HB2 - CD
5748   5758       1 ;     HB2 - C
5749   5751       1 ;     HB3 - HG2
5749   5752       1 ;     HB3 - HG3
5749   5753       1 ;     HB3 - CD
5749   5758       1 ;     HB3 - C
5750   5756       1 ;      CG - HE21
5750   5757       1 ;      CG - HE22
5750   5758       1 ;      CG - C
5751   5754       1 ;     HG2 - OE1
5751   5755       1 ;     HG2 - NE2
5752   5754       1 ;     HG3 - OE1
5752   5755       1 ;     HG3 - NE2
5754   5756       1 ;     OE1 - HE21
5754   5757       1 ;     OE1 - HE22
5758   5763       1 ;       C - HA
5758   5764       1 ;       C - CB
5758   5774       1 ;       C - C
5759   5761       1 ;       O - H
5759   5762       1 ;       O - CA
5760   5765       1 ;       N - HB
5760   5766       1 ;       N - CG1
5760   5770       1 ;       N - CG2
5760   5775       1 ;       N - O
5760   5776       1 ;       N - N
5761   5763       1 ;       H - HA
5761   5764       1 ;       H - CB
5761   5774       1 ;       H - C
5762   5767       1 ;      CA - HG11
5762   5768       1 ;      CA - HG12
5762   5769       1 ;      CA - HG13
5762   5771       1 ;      CA - HG21
```

| | | | |
|---|---|---|---|
| 5762 | 5772 | 1 ; | CA - HG22 |
| 5762 | 5773 | 1 ; | CA - HG23 |
| 5762 | 5777 | 1 ; | CA - H |
| 5762 | 5778 | 1 ; | CA - CA |
| 5763 | 5765 | 1 ; | HA - HB |
| 5763 | 5766 | 1 ; | HA - CG1 |
| 5763 | 5770 | 1 ; | HA - CG2 |
| 5763 | 5775 | 1 ; | HA - O |
| 5763 | 5776 | 1 ; | HA - N |
| 5764 | 5775 | 1 ; | CB - O |
| 5764 | 5776 | 1 ; | CB - N |
| 5765 | 5767 | 1 ; | HB - HG11 |
| 5765 | 5768 | 1 ; | HB - HG12 |
| 5765 | 5769 | 1 ; | HB - HG13 |
| 5765 | 5771 | 1 ; | HB - HG21 |
| 5765 | 5772 | 1 ; | HB - HG22 |
| 5765 | 5773 | 1 ; | HB - HG23 |
| 5765 | 5774 | 1 ; | HB - C |
| 5766 | 5771 | 1 ; | CG1 - HG21 |
| 5766 | 5772 | 1 ; | CG1 - HG22 |
| 5766 | 5773 | 1 ; | CG1 - HG23 |
| 5766 | 5774 | 1 ; | CG1 - C |
| 5767 | 5770 | 1 ; | HG11 - CG2 |
| 5768 | 5770 | 1 ; | HG12 - CG2 |
| 5769 | 5770 | 1 ; | HG13 - CG2 |
| 5770 | 5774 | 1 ; | CG2 - C |
| 5774 | 5779 | 1 ; | C - HA |
| 5774 | 5780 | 1 ; | C - CB |
| 5774 | 5793 | 1 ; | C - C |
| 5775 | 5777 | 1 ; | O - H |
| 5775 | 5778 | 1 ; | O - CA |
| 5776 | 5781 | 1 ; | N - HB2 |
| 5776 | 5782 | 1 ; | N - HB3 |
| 5776 | 5783 | 1 ; | N - CG |
| 5776 | 5794 | 1 ; | N - O |
| 5776 | 5795 | 1 ; | N - N |
| 5777 | 5779 | 1 ; | H - HA |
| 5777 | 5780 | 1 ; | H - CB |
| 5777 | 5793 | 1 ; | H - C |
| 5778 | 5784 | 1 ; | CA - HG |
| 5778 | 5785 | 1 ; | CA - CD1 |
| 5778 | 5789 | 1 ; | CA - CD2 |
| 5778 | 5796 | 1 ; | CA - H |
| 5778 | 5797 | 1 ; | CA - CA |
| 5779 | 5781 | 1 ; | HA - HB2 |
| 5779 | 5782 | 1 ; | HA - HB3 |
| 5779 | 5783 | 1 ; | HA - CG |
| 5779 | 5794 | 1 ; | HA - O |
| 5779 | 5795 | 1 ; | HA - N |
| 5780 | 5786 | 1 ; | CB - HD11 |
| 5780 | 5787 | 1 ; | CB - HD12 |
| 5780 | 5788 | 1 ; | CB - HD13 |

| | | | | |
|---|---|---|---|---|
| 5780 | 5790 | 1 ; | CB - | HD21 |
| 5780 | 5791 | 1 ; | CB - | HD22 |
| 5780 | 5792 | 1 ; | CB - | HD23 |
| 5780 | 5794 | 1 ; | CB - | O |
| 5780 | 5795 | 1 ; | CB - | N |
| 5781 | 5784 | 1 ; | HB2 - | HG |
| 5781 | 5785 | 1 ; | HB2 - | CD1 |
| 5781 | 5789 | 1 ; | HB2 - | CD2 |
| 5781 | 5793 | 1 ; | HB2 - | C |
| 5782 | 5784 | 1 ; | HB3 - | HG |
| 5782 | 5785 | 1 ; | HB3 - | CD1 |
| 5782 | 5789 | 1 ; | HB3 - | CD2 |
| 5782 | 5793 | 1 ; | HB3 - | C |
| 5783 | 5793 | 1 ; | CG - | C |
| 5784 | 5786 | 1 ; | HG - | HD11 |
| 5784 | 5787 | 1 ; | HG - | HD12 |
| 5784 | 5788 | 1 ; | HG - | HD13 |
| 5784 | 5790 | 1 ; | HG - | HD21 |
| 5784 | 5791 | 1 ; | HG - | HD22 |
| 5784 | 5792 | 1 ; | HG - | HD23 |
| 5785 | 5790 | 1 ; | CD1 - | HD21 |
| 5785 | 5791 | 1 ; | CD1 - | HD22 |
| 5785 | 5792 | 1 ; | CD1 - | HD23 |
| 5786 | 5789 | 1 ; | HD11 - | CD2 |
| 5787 | 5789 | 1 ; | HD12 - | CD2 |
| 5788 | 5789 | 1 ; | HD13 - | CD2 |
| 5793 | 5798 | 1 ; | C - | HA |
| 5793 | 5799 | 1 ; | C - | CB |
| 5793 | 5817 | 1 ; | C - | C |
| 5794 | 5796 | 1 ; | O - | H |
| 5794 | 5797 | 1 ; | O - | CA |
| 5795 | 5800 | 1 ; | N - | HB2 |
| 5795 | 5801 | 1 ; | N - | HB3 |
| 5795 | 5802 | 1 ; | N - | CG |
| 5795 | 5818 | 1 ; | N - | O |
| 5795 | 5819 | 1 ; | N - | N |
| 5796 | 5798 | 1 ; | H - | HA |
| 5796 | 5799 | 1 ; | H - | CB |
| 5796 | 5817 | 1 ; | H - | C |
| 5797 | 5803 | 1 ; | CA - | HG2 |
| 5797 | 5804 | 1 ; | CA - | HG3 |
| 5797 | 5805 | 1 ; | CA - | CD |
| 5797 | 5820 | 1 ; | CA - | H |
| 5797 | 5821 | 1 ; | CA - | CA |
| 5798 | 5800 | 1 ; | HA - | HB2 |
| 5798 | 5801 | 1 ; | HA - | HB3 |
| 5798 | 5802 | 1 ; | HA - | CG |
| 5798 | 5818 | 1 ; | HA - | O |
| 5798 | 5819 | 1 ; | HA - | N |
| 5799 | 5806 | 1 ; | CB - | HD2 |
| 5799 | 5807 | 1 ; | CB - | HD3 |
| 5799 | 5808 | 1 ; | CB - | NE |

```
5799   5818       1 ;      CB - O
5799   5819       1 ;      CB - N
5800   5803       1 ;     HB2 - HG2
5800   5804       1 ;     HB2 - HG3
5800   5805       1 ;     HB2 - CD
5800   5817       1 ;     HB2 - C
5801   5803       1 ;     HB3 - HG2
5801   5804       1 ;     HB3 - HG3
5801   5805       1 ;     HB3 - CD
5801   5817       1 ;     HB3 - C
5802   5809       1 ;      CG - HE
5802   5810       1 ;      CG - CZ
5802   5817       1 ;      CG - C
5803   5806       1 ;     HG2 - HD2
5803   5807       1 ;     HG2 - HD3
5803   5808       1 ;     HG2 - NE
5804   5806       1 ;     HG3 - HD2
5804   5807       1 ;     HG3 - HD3
5804   5808       1 ;     HG3 - NE
5805   5811       1 ;      CD - NH1
5805   5814       1 ;      CD - NH2
5806   5809       1 ;     HD2 - HE
5806   5810       1 ;     HD2 - CZ
5807   5809       1 ;     HD3 - HE
5807   5810       1 ;     HD3 - CZ
5808   5812       1 ;      NE - HH11
5808   5813       1 ;      NE - HH12
5808   5815       1 ;      NE - HH21
5808   5816       1 ;      NE - HH22
5809   5811       1 ;      HE - NH1
5809   5814       1 ;      HE - NH2
5811   5815       1 ;     NH1 - HH21
5811   5816       1 ;     NH1 - HH22
5812   5814       1 ;    HH11 - NH2
5813   5814       1 ;    HH12 - NH2
5817   5822       1 ;       C - HA
5817   5823       1 ;       C - CB
5817   5829       1 ;       C - C
5818   5820       1 ;       O - H
5818   5821       1 ;       O - CA
5819   5824       1 ;       N - HB2
5819   5825       1 ;       N - HB3
5819   5826       1 ;       N - CG
5819   5830       1 ;       N - O
5819   5831       1 ;       N - N
5820   5822       1 ;       H - HA
5820   5823       1 ;       H - CB
5820   5829       1 ;       H - C
5821   5827       1 ;      CA - OD1
5821   5828       1 ;      CA - OD2
5821   5832       1 ;      CA - H
5821   5833       1 ;      CA - CA
```

```
5822   5824      1 ;      HA - HB2
5822   5825      1 ;      HA - HB3
5822   5826      1 ;      HA - CG
5822   5830      1 ;      HA - O
5822   5831      1 ;      HA - N
5823   5830      1 ;      CB - O
5823   5831      1 ;      CB - N
5824   5827      1 ;     HB2 - OD1
5824   5828      1 ;     HB2 - OD2
5824   5829      1 ;     HB2 - C
5825   5827      1 ;     HB3 - OD1
5825   5828      1 ;     HB3 - OD2
5825   5829      1 ;     HB3 - C
5826   5829      1 ;      CG - C
5829   5834      1 ;       C - HA
5829   5835      1 ;       C - CB
5829   5848      1 ;       C - C
5830   5832      1 ;       O - H
5830   5833      1 ;       O - CA
5831   5836      1 ;       N - HB
5831   5837      1 ;       N - CG2
5831   5841      1 ;       N - CG1
5831   5849      1 ;       N - O
5831   5850      1 ;       N - N
5832   5834      1 ;       H - HA
5832   5835      1 ;       H - CB
5832   5848      1 ;       H - C
5833   5838      1 ;      CA - HG21
5833   5839      1 ;      CA - HG22
5833   5840      1 ;      CA - HG23
5833   5842      1 ;      CA - HG12
5833   5843      1 ;      CA - HG13
5833   5844      1 ;      CA - CD1
5833   5851      1 ;      CA - H
5833   5852      1 ;      CA - CA
5834   5836      1 ;      HA - HB
5834   5837      1 ;      HA - CG2
5834   5841      1 ;      HA - CG1
5834   5849      1 ;      HA - O
5834   5850      1 ;      HA - N
5835   5845      1 ;      CB - HD11
5835   5846      1 ;      CB - HD12
5835   5847      1 ;      CB - HD13
5835   5849      1 ;      CB - O
5835   5850      1 ;      CB - N
5836   5838      1 ;      HB - HG21
5836   5839      1 ;      HB - HG22
5836   5840      1 ;      HB - HG23
5836   5842      1 ;      HB - HG12
5836   5843      1 ;      HB - HG13
5836   5844      1 ;      HB - CD1
5836   5848      1 ;      HB - C
```

```
 5837   5842       1 ;     CG2 - HG12
 5837   5843       1 ;     CG2 - HG13
 5837   5844       1 ;     CG2 - CD1
 5837   5848       1 ;     CG2 - C
 5838   5841       1 ;    HG21 - CG1
 5839   5841       1 ;    HG22 - CG1
 5840   5841       1 ;    HG23 - CG1
 5841   5848       1 ;     CG1 - C
 5842   5845       1 ;    HG12 - HD11
 5842   5846       1 ;    HG12 - HD12
 5842   5847       1 ;    HG12 - HD13
 5843   5845       1 ;    HG13 - HD11
 5843   5846       1 ;    HG13 - HD12
 5843   5847       1 ;    HG13 - HD13
 5848   5853       1 ;       C - HA
 5848   5854       1 ;       C - CB
 5848   5860       1 ;       C - C
 5849   5851       1 ;       O - H
 5849   5852       1 ;       O - CA
 5850   5855       1 ;       N - HB2
 5850   5856       1 ;       N - HB3
 5850   5857       1 ;       N - CG
 5850   5861       1 ;       N - O
 5850   5862       1 ;       N - N
 5851   5853       1 ;       H - HA
 5851   5854       1 ;       H - CB
 5851   5860       1 ;       H - C
 5852   5858       1 ;      CA - OD1
 5852   5859       1 ;      CA - OD2
 5852   5863       1 ;      CA - H
 5852   5864       1 ;      CA - CA
 5853   5855       1 ;      HA - HB2
 5853   5856       1 ;      HA - HB3
 5853   5857       1 ;      HA - CG
 5853   5861       1 ;      HA - O
 5853   5862       1 ;      HA - N
 5854   5861       1 ;      CB - O
 5854   5862       1 ;      CB - N
 5855   5858       1 ;     HB2 - OD1
 5855   5859       1 ;     HB2 - OD2
 5855   5860       1 ;     HB2 - C
 5856   5858       1 ;     HB3 - OD1
 5856   5859       1 ;     HB3 - OD2
 5856   5860       1 ;     HB3 - C
 5857   5860       1 ;      CG - C
 5860   5865       1 ;       C - HA
 5860   5866       1 ;       C - CB
 5860   5871       1 ;       C - C
 5861   5863       1 ;       O - H
 5861   5864       1 ;       O - CA
 5862   5867       1 ;       N - HB2
 5862   5868       1 ;       N - HB3
```

```
5862   5869       1 ;       N - OG
5862   5872       1 ;       N - O
5862   5873       1 ;       N - N
5863   5865       1 ;       H - HA
5863   5866       1 ;       H - CB
5863   5871       1 ;       H - C
5864   5870       1 ;      CA - HG
5864   5874       1 ;      CA - H
5864   5875       1 ;      CA - CA
5865   5867       1 ;      HA - HB2
5865   5868       1 ;      HA - HB3
5865   5869       1 ;      HA - OG
5865   5872       1 ;      HA - O
5865   5873       1 ;      HA - N
5866   5872       1 ;      CB - O
5866   5873       1 ;      CB - N
5867   5870       1 ;     HB2 - HG
5867   5871       1 ;     HB2 - C
5868   5870       1 ;     HB3 - HG
5868   5871       1 ;     HB3 - C
5869   5871       1 ;      OG - C
5871   5876       1 ;       C - HA
5871   5877       1 ;       C - CB
5871   5885       1 ;       C - C
5872   5874       1 ;       O - H
5872   5875       1 ;       O - CA
5873   5878       1 ;       N - HB2
5873   5879       1 ;       N - HB3
5873   5880       1 ;       N - CG
5873   5886       1 ;       N - O
5873   5887       1 ;       N - N
5874   5876       1 ;       H - HA
5874   5877       1 ;       H - CB
5874   5885       1 ;       H - C
5875   5881       1 ;      CA - OD1
5875   5882       1 ;      CA - ND2
5875   5888       1 ;      CA - H
5875   5889       1 ;      CA - CA
5876   5878       1 ;      HA - HB2
5876   5879       1 ;      HA - HB3
5876   5880       1 ;      HA - CG
5876   5886       1 ;      HA - O
5876   5887       1 ;      HA - N
5877   5883       1 ;      CB - HD21
5877   5884       1 ;      CB - HD22
5877   5886       1 ;      CB - O
5877   5887       1 ;      CB - N
5878   5881       1 ;     HB2 - OD1
5878   5882       1 ;     HB2 - ND2
5878   5885       1 ;     HB2 - C
5879   5881       1 ;     HB3 - OD1
5879   5882       1 ;     HB3 - ND2
```

```
5879   5885      1 ;    HB3 - C
5880   5885      1 ;     CG - C
5881   5883      1 ;    OD1 - HD21
5881   5884      1 ;    OD1 - HD22
5885   5890      1 ;      C - HA
5885   5891      1 ;      C - CB
5885   5895      1 ;      C - C
5886   5888      1 ;      O - H
5886   5889      1 ;      O - CA
5887   5892      1 ;      N - HB1
5887   5893      1 ;      N - HB2
5887   5894      1 ;      N - HB3
5887   5896      1 ;      N - O
5887   5897      1 ;      N - N
5888   5890      1 ;      H - HA
5888   5891      1 ;      H - CB
5888   5895      1 ;      H - C
5889   5898      1 ;     CA - H
5889   5899      1 ;     CA - CA
5890   5892      1 ;     HA - HB1
5890   5893      1 ;     HA - HB2
5890   5894      1 ;     HA - HB3
5890   5896      1 ;     HA - O
5890   5897      1 ;     HA - N
5891   5896      1 ;     CB - O
5891   5897      1 ;     CB - N
5892   5895      1 ;    HB1 - C
5893   5895      1 ;    HB2 - C
5894   5895      1 ;    HB3 - C
5895   5900      1 ;      C - HA
5895   5901      1 ;      C - CB
5895   5906      1 ;      C - C
5896   5898      1 ;      O - H
5896   5899      1 ;      O - CA
5897   5902      1 ;      N - HB2
5897   5903      1 ;      N - HB3
5897   5904      1 ;      N - OG
5897   5907      1 ;      N - O
5897   5908      1 ;      N - N
5898   5900      1 ;      H - HA
5898   5901      1 ;      H - CB
5898   5906      1 ;      H - C
5899   5905      1 ;     CA - HG
5899   5909      1 ;     CA - H
5899   5910      1 ;     CA - CA
5900   5902      1 ;     HA - HB2
5900   5903      1 ;     HA - HB3
5900   5904      1 ;     HA - OG
5900   5907      1 ;     HA - O
5900   5908      1 ;     HA - N
5901   5907      1 ;     CB - O
5901   5908      1 ;     CB - N
```

| | | | |
|---|---|---|---|
| 5902 | 5905 | 1 ; | HB2 - HG |
| 5902 | 5906 | 1 ; | HB2 - C |
| 5903 | 5905 | 1 ; | HB3 - HG |
| 5903 | 5906 | 1 ; | HB3 - C |
| 5904 | 5906 | 1 ; | OG - C |
| 5906 | 5911 | 1 ; | C - HA2 |
| 5906 | 5912 | 1 ; | C - HA3 |
| 5906 | 5913 | 1 ; | C - C |
| 5907 | 5909 | 1 ; | O - H |
| 5907 | 5910 | 1 ; | O - CA |
| 5908 | 5914 | 1 ; | N - O |
| 5908 | 5915 | 1 ; | N - N |
| 5909 | 5911 | 1 ; | H - HA2 |
| 5909 | 5912 | 1 ; | H - HA3 |
| 5909 | 5913 | 1 ; | H - C |
| 5910 | 5916 | 1 ; | CA - H |
| 5910 | 5917 | 1 ; | CA - CA |
| 5911 | 5914 | 1 ; | HA2 - O |
| 5911 | 5915 | 1 ; | HA2 - N |
| 5912 | 5914 | 1 ; | HA3 - O |
| 5912 | 5915 | 1 ; | HA3 - N |
| 5913 | 5918 | 1 ; | C - HA |
| 5913 | 5919 | 1 ; | C - CB |
| 5913 | 5930 | 1 ; | C - C |
| 5914 | 5916 | 1 ; | O - H |
| 5914 | 5917 | 1 ; | O - CA |
| 5915 | 5920 | 1 ; | N - HB2 |
| 5915 | 5921 | 1 ; | N - HB3 |
| 5915 | 5922 | 1 ; | N - CG |
| 5915 | 5931 | 1 ; | N - O |
| 5915 | 5932 | 1 ; | N - N |
| 5916 | 5918 | 1 ; | H - HA |
| 5916 | 5919 | 1 ; | H - CB |
| 5916 | 5930 | 1 ; | H - C |
| 5917 | 5923 | 1 ; | CA - HG2 |
| 5917 | 5924 | 1 ; | CA - HG3 |
| 5917 | 5925 | 1 ; | CA - CD |
| 5917 | 5933 | 1 ; | CA - H |
| 5917 | 5934 | 1 ; | CA - CA |
| 5918 | 5920 | 1 ; | HA - HB2 |
| 5918 | 5921 | 1 ; | HA - HB3 |
| 5918 | 5922 | 1 ; | HA - CG |
| 5918 | 5931 | 1 ; | HA - O |
| 5918 | 5932 | 1 ; | HA - N |
| 5919 | 5926 | 1 ; | CB - OE1 |
| 5919 | 5927 | 1 ; | CB - NE2 |
| 5919 | 5931 | 1 ; | CB - O |
| 5919 | 5932 | 1 ; | CB - N |
| 5920 | 5923 | 1 ; | HB2 - HG2 |
| 5920 | 5924 | 1 ; | HB2 - HG3 |
| 5920 | 5925 | 1 ; | HB2 - CD |
| 5920 | 5930 | 1 ; | HB2 - C |

```
5921   5923      1 ;    HB3 - HG2
5921   5924      1 ;    HB3 - HG3
5921   5925      1 ;    HB3 - CD
5921   5930      1 ;    HB3 - C
5922   5928      1 ;     CG - HE21
5922   5929      1 ;     CG - HE22
5922   5930      1 ;     CG - C
5923   5926      1 ;    HG2 - OE1
5923   5927      1 ;    HG2 - NE2
5924   5926      1 ;    HG3 - OE1
5924   5927      1 ;    HG3 - NE2
5926   5928      1 ;    OE1 - HE21
5926   5929      1 ;    OE1 - HE22
5930   5935      1 ;      C - HA
5930   5936      1 ;      C - CB
5930   5949      1 ;      C - C
5931   5933      1 ;      O - H
5931   5934      1 ;      O - CA
5932   5937      1 ;      N - HB
5932   5938      1 ;      N - CG2
5932   5942      1 ;      N - CG1
5932   5950      1 ;      N - O
5932   5951      1 ;      N - N
5933   5935      1 ;      H - HA
5933   5936      1 ;      H - CB
5933   5949      1 ;      H - C
5934   5939      1 ;     CA - HG21
5934   5940      1 ;     CA - HG22
5934   5941      1 ;     CA - HG23
5934   5943      1 ;     CA - HG12
5934   5944      1 ;     CA - HG13
5934   5945      1 ;     CA - CD1
5934   5952      1 ;     CA - H
5934   5953      1 ;     CA - CA
5935   5937      1 ;     HA - HB
5935   5938      1 ;     HA - CG2
5935   5942      1 ;     HA - CG1
5935   5950      1 ;     HA - O
5935   5951      1 ;     HA - N
5936   5946      1 ;     CB - HD11
5936   5947      1 ;     CB - HD12
5936   5948      1 ;     CB - HD13
5936   5950      1 ;     CB - O
5936   5951      1 ;     CB - N
5937   5939      1 ;     HB - HG21
5937   5940      1 ;     HB - HG22
5937   5941      1 ;     HB - HG23
5937   5943      1 ;     HB - HG12
5937   5944      1 ;     HB - HG13
5937   5945      1 ;     HB - CD1
5937   5949      1 ;     HB - C
5938   5943      1 ;    CG2 - HG12
```

```
5938   5944       1 ;    CG2 - HG13
5938   5945       1 ;    CG2 - CD1
5938   5949       1 ;    CG2 - C
5939   5942       1 ;   HG21 - CG1
5940   5942       1 ;   HG22 - CG1
5941   5942       1 ;   HG23 - CG1
5942   5949       1 ;    CG1 - C
5943   5946       1 ;   HG12 - HD11
5943   5947       1 ;   HG12 - HD12
5943   5948       1 ;   HG12 - HD13
5944   5946       1 ;   HG13 - HD11
5944   5947       1 ;   HG13 - HD12
5944   5948       1 ;   HG13 - HD13
5949   5954       1 ;      C - HA
5949   5955       1 ;      C - CB
5949   5966       1 ;      C - C
5950   5952       1 ;      O - H
5950   5953       1 ;      O - CA
5951   5956       1 ;      N - HB2
5951   5957       1 ;      N - HB3
5951   5958       1 ;      N - CG
5951   5967       1 ;      N - O
5951   5968       1 ;      N - N
5952   5954       1 ;      H - HA
5952   5955       1 ;      H - CB
5952   5966       1 ;      H - C
5953   5959       1 ;     CA - ND1
5953   5964       1 ;     CA - CD2
5953   5969       1 ;     CA - H
5953   5970       1 ;     CA - CA
5954   5956       1 ;     HA - HB2
5954   5957       1 ;     HA - HB3
5954   5958       1 ;     HA - CG
5954   5967       1 ;     HA - O
5954   5968       1 ;     HA - N
5955   5960       1 ;     CB - CE1
5955   5962       1 ;     CB - NE2
5955   5965       1 ;     CB - HD2
5955   5967       1 ;     CB - O
5955   5968       1 ;     CB - N
5956   5959       1 ;    HB2 - ND1
5956   5964       1 ;    HB2 - CD2
5956   5966       1 ;    HB2 - C
5957   5959       1 ;    HB3 - ND1
5957   5964       1 ;    HB3 - CD2
5957   5966       1 ;    HB3 - C
5958   5961       1 ;     CG - HE1
5958   5963       1 ;     CG - HE2
5958   5966       1 ;     CG - C
5959   5963       1 ;    ND1 - HE2
5959   5965       1 ;    ND1 - HD2
5960   5965       1 ;    CE1 - HD2
```

```
5961   5963      1 ;    HE1 - HE2
5961   5964      1 ;    HE1 - CD2
5963   5965      1 ;    HE2 - HD2
5966   5971      1 ;      C - HA
5966   5972      1 ;      C - CB
5966   5987      1 ;      C - C
5967   5969      1 ;      O - H
5967   5970      1 ;      O - CA
5968   5973      1 ;      N - HB2
5968   5974      1 ;      N - HB3
5968   5975      1 ;      N - CG
5968   5988      1 ;      N - O
5968   5989      1 ;      N - N
5969   5971      1 ;      H - HA
5969   5972      1 ;      H - CB
5969   5987      1 ;      H - C
5970   5976      1 ;     CA - CD1
5970   5985      1 ;     CA - CD2
5970   5990      1 ;     CA - H
5970   5991      1 ;     CA - CA
5971   5973      1 ;     HA - HB2
5971   5974      1 ;     HA - HB3
5971   5975      1 ;     HA - CG
5971   5988      1 ;     HA - O
5971   5989      1 ;     HA - N
5972   5977      1 ;     CB - HD1
5972   5978      1 ;     CB - CE1
5972   5983      1 ;     CB - CE2
5972   5986      1 ;     CB - HD2
5972   5988      1 ;     CB - O
5972   5989      1 ;     CB - N
5973   5976      1 ;    HB2 - CD1
5973   5985      1 ;    HB2 - CD2
5973   5987      1 ;    HB2 - C
5974   5976      1 ;    HB3 - CD1
5974   5985      1 ;    HB3 - CD2
5974   5987      1 ;    HB3 - C
5975   5979      1 ;     CG - HE1
5975   5980      1 ;     CG - CZ
5975   5984      1 ;     CG - HE2
5975   5987      1 ;     CG - C
5976   5981      1 ;    CD1 - OH
5976   5983      1 ;    CD1 - CE2
5976   5986      1 ;    CD1 - HD2
5977   5979      1 ;    HD1 - HE1
5977   5980      1 ;    HD1 - CZ
5977   5985      1 ;    HD1 - CD2
5978   5982      1 ;    CE1 - HH
5978   5984      1 ;    CE1 - HE2
5978   5985      1 ;    CE1 - CD2
5979   5981      1 ;    HE1 - OH
5979   5983      1 ;    HE1 - CE2
```

```
 5980   5986       1 ;      CZ - HD2
 5981   5984       1 ;      OH - HE2
 5981   5985       1 ;      OH - CD2
 5982   5983       1 ;      HH - CE2
 5984   5986       1 ;     HE2 - HD2
 5987   5992       1 ;       C - HA
 5987   5993       1 ;       C - CB
 5987   6001       1 ;       C - C
 5988   5990       1 ;       O - H
 5988   5991       1 ;       O - CA
 5989   5994       1 ;       N - HB
 5989   5995       1 ;       N - CG2
 5989   5999       1 ;       N - OG1
 5989   6002       1 ;       N - O
 5989   6003       1 ;       N - N
 5990   5992       1 ;       H - HA
 5990   5993       1 ;       H - CB
 5990   6001       1 ;       H - C
 5991   5996       1 ;      CA - HG21
 5991   5997       1 ;      CA - HG22
 5991   5998       1 ;      CA - HG23
 5991   6000       1 ;      CA - HG1
 5991   6004       1 ;      CA - H
 5991   6005       1 ;      CA - CA
 5992   5994       1 ;      HA - HB
 5992   5995       1 ;      HA - CG2
 5992   5999       1 ;      HA - OG1
 5992   6002       1 ;      HA - O
 5992   6003       1 ;      HA - N
 5993   6002       1 ;      CB - O
 5993   6003       1 ;      CB - N
 5994   5996       1 ;      HB - HG21
 5994   5997       1 ;      HB - HG22
 5994   5998       1 ;      HB - HG23
 5994   6000       1 ;      HB - HG1
 5994   6001       1 ;      HB - C
 5995   6000       1 ;     CG2 - HG1
 5995   6001       1 ;     CG2 - C
 5996   5999       1 ;    HG21 - OG1
 5997   5999       1 ;    HG22 - OG1
 5998   5999       1 ;    HG23 - OG1
 5999   6001       1 ;     OG1 - C
 6001   6006       1 ;       C - HA
 6001   6007       1 ;       C - CB
 6001   6013       1 ;       C - C
 6002   6004       1 ;       O - H
 6002   6005       1 ;       O - CA
 6003   6008       1 ;       N - HB2
 6003   6009       1 ;       N - HB3
 6003   6010       1 ;       N - CG
 6003   6014       1 ;       N - O
 6003   6015       1 ;       N - N
```

```
6004   6006       1 ;      H - HA
6004   6007       1 ;      H - CB
6004   6013       1 ;      H - C
6005   6011       1 ;     CA - OD1
6005   6012       1 ;     CA - OD2
6005   6016       1 ;     CA - H
6005   6017       1 ;     CA - CA
6006   6008       1 ;     HA - HB2
6006   6009       1 ;     HA - HB3
6006   6010       1 ;     HA - CG
6006   6014       1 ;     HA - O
6006   6015       1 ;     HA - N
6007   6014       1 ;     CB - O
6007   6015       1 ;     CB - N
6008   6011       1 ;    HB2 - OD1
6008   6012       1 ;    HB2 - OD2
6008   6013       1 ;    HB2 - C
6009   6011       1 ;    HB3 - OD1
6009   6012       1 ;    HB3 - OD2
6009   6013       1 ;    HB3 - C
6010   6013       1 ;     CG - C
6013   6018       1 ;      C - HA
6013   6019       1 ;      C - CB
6013   6033       1 ;      C - C
6014   6016       1 ;      O - H
6014   6017       1 ;      O - CA
6015   6020       1 ;      N - HB2
6015   6021       1 ;      N - HB3
6015   6022       1 ;      N - CG
6015   6034       1 ;      N - O
6015   6035       1 ;      N - N
6016   6018       1 ;      H - HA
6016   6019       1 ;      H - CB
6016   6033       1 ;      H - C
6017   6023       1 ;     CA - CD1
6017   6031       1 ;     CA - CD2
6017   6036       1 ;     CA - H
6017   6037       1 ;     CA - CA
6018   6020       1 ;     HA - HB2
6018   6021       1 ;     HA - HB3
6018   6022       1 ;     HA - CG
6018   6034       1 ;     HA - O
6018   6035       1 ;     HA - N
6019   6024       1 ;     CB - HD1
6019   6025       1 ;     CB - CE1
6019   6029       1 ;     CB - CE2
6019   6032       1 ;     CB - HD2
6019   6034       1 ;     CB - O
6019   6035       1 ;     CB - N
6020   6023       1 ;    HB2 - CD1
6020   6031       1 ;    HB2 - CD2
6020   6033       1 ;    HB2 - C
```

```
6021   6023      1 ;    HB3 - CD1
6021   6031      1 ;    HB3 - CD2
6021   6033      1 ;    HB3 - C
6022   6026      1 ;     CG - HE1
6022   6027      1 ;     CG - CZ
6022   6030      1 ;     CG - HE2
6022   6033      1 ;     CG - C
6023   6028      1 ;    CD1 - HZ
6023   6029      1 ;    CD1 - CE2
6023   6032      1 ;    CD1 - HD2
6024   6026      1 ;    HD1 - HE1
6024   6027      1 ;    HD1 - CZ
6024   6031      1 ;    HD1 - CD2
6025   6030      1 ;    CE1 - HE2
6025   6031      1 ;    CE1 - CD2
6026   6028      1 ;    HE1 - HZ
6026   6029      1 ;    HE1 - CE2
6027   6032      1 ;     CZ - HD2
6028   6030      1 ;     HZ - HE2
6028   6031      1 ;     HZ - CD2
6030   6032      1 ;    HE2 - HD2
6033   6038      1 ;      C - HA
6033   6039      1 ;      C - CB
6033   6052      1 ;      C - C
6034   6036      1 ;      O - H
6034   6037      1 ;      O - CA
6035   6040      1 ;      N - HB2
6035   6041      1 ;      N - HB3
6035   6042      1 ;      N - CG
6035   6053      1 ;      N - O
6035   6054      1 ;      N - N
6036   6038      1 ;      H - HA
6036   6039      1 ;      H - CB
6036   6052      1 ;      H - C
6037   6043      1 ;     CA - HG
6037   6044      1 ;     CA - CD1
6037   6048      1 ;     CA - CD2
6037   6055      1 ;     CA - H
6037   6056      1 ;     CA - CA
6038   6040      1 ;     HA - HB2
6038   6041      1 ;     HA - HB3
6038   6042      1 ;     HA - CG
6038   6053      1 ;     HA - O
6038   6054      1 ;     HA - N
6039   6045      1 ;     CB - HD11
6039   6046      1 ;     CB - HD12
6039   6047      1 ;     CB - HD13
6039   6049      1 ;     CB - HD21
6039   6050      1 ;     CB - HD22
6039   6051      1 ;     CB - HD23
6039   6053      1 ;     CB - O
6039   6054      1 ;     CB - N
```

```
6040   6043      1 ;    HB2 - HG
6040   6044      1 ;    HB2 - CD1
6040   6048      1 ;    HB2 - CD2
6040   6052      1 ;    HB2 - C
6041   6043      1 ;    HB3 - HG
6041   6044      1 ;    HB3 - CD1
6041   6048      1 ;    HB3 - CD2
6041   6052      1 ;    HB3 - C
6042   6052      1 ;     CG - C
6043   6045      1 ;     HG - HD11
6043   6046      1 ;     HG - HD12
6043   6047      1 ;     HG - HD13
6043   6049      1 ;     HG - HD21
6043   6050      1 ;     HG - HD22
6043   6051      1 ;     HG - HD23
6044   6049      1 ;    CD1 - HD21
6044   6050      1 ;    CD1 - HD22
6044   6051      1 ;    CD1 - HD23
6045   6048      1 ;   HD11 - CD2
6046   6048      1 ;   HD12 - CD2
6047   6048      1 ;   HD13 - CD2
6052   6057      1 ;      C - HA
6052   6058      1 ;      C - CB
6052   6062      1 ;      C - C
6053   6055      1 ;      O - H
6053   6056      1 ;      O - CA
6054   6059      1 ;      N - HB1
6054   6060      1 ;      N - HB2
6054   6061      1 ;      N - HB3
6054   6063      1 ;      N - O
6054   6064      1 ;      N - N
6055   6057      1 ;      H - HA
6055   6058      1 ;      H - CB
6055   6062      1 ;      H - C
6056   6065      1 ;     CA - H
6056   6066      1 ;     CA - CA
6057   6059      1 ;     HA - HB1
6057   6060      1 ;     HA - HB2
6057   6061      1 ;     HA - HB3
6057   6063      1 ;     HA - O
6057   6064      1 ;     HA - N
6058   6063      1 ;     CB - O
6058   6064      1 ;     CB - N
6059   6062      1 ;    HB1 - C
6060   6062      1 ;    HB2 - C
6061   6062      1 ;    HB3 - C
6062   6067      1 ;      C - HA
6062   6068      1 ;      C - CB
6062   6072      1 ;      C - C
6063   6065      1 ;      O - H
6063   6066      1 ;      O - CA
6064   6069      1 ;      N - HB1
```

```
6064   6070      1 ;      N - HB2
6064   6071      1 ;      N - HB3
6064   6073      1 ;      N - O
6064   6074      1 ;      N - N
6065   6067      1 ;      H - HA
6065   6068      1 ;      H - CB
6065   6072      1 ;      H - C
6066   6075      1 ;     CA - H
6066   6076      1 ;     CA - CA
6067   6069      1 ;     HA - HB1
6067   6070      1 ;     HA - HB2
6067   6071      1 ;     HA - HB3
6067   6073      1 ;     HA - O
6067   6074      1 ;     HA - N
6068   6073      1 ;     CB - O
6068   6074      1 ;     CB - N
6069   6072      1 ;    HB1 - C
6070   6072      1 ;    HB2 - C
6071   6072      1 ;    HB3 - C
6072   6077      1 ;      C - HA
6072   6078      1 ;      C - CB
6072   6086      1 ;      C - C
6073   6075      1 ;      O - H
6073   6076      1 ;      O - CA
6074   6079      1 ;      N - HB
6074   6080      1 ;      N - CG2
6074   6084      1 ;      N - OG1
6074   6087      1 ;      N - O
6074   6088      1 ;      N - N
6075   6077      1 ;      H - HA
6075   6078      1 ;      H - CB
6075   6086      1 ;      H - C
6076   6081      1 ;     CA - HG21
6076   6082      1 ;     CA - HG22
6076   6083      1 ;     CA - HG23
6076   6085      1 ;     CA - HG1
6076   6089      1 ;     CA - H
6076   6090      1 ;     CA - CA
6077   6079      1 ;     HA - HB
6077   6080      1 ;     HA - CG2
6077   6084      1 ;     HA - OG1
6077   6087      1 ;     HA - O
6077   6088      1 ;     HA - N
6078   6087      1 ;     CB - O
6078   6088      1 ;     CB - N
6079   6081      1 ;     HB - HG21
6079   6082      1 ;     HB - HG22
6079   6083      1 ;     HB - HG23
6079   6085      1 ;     HB - HG1
6079   6086      1 ;     HB - C
6080   6085      1 ;    CG2 - HG1
6080   6086      1 ;    CG2 - C
```

```
6081   6084       1 ;   HG21 - OG1
6082   6084       1 ;   HG22 - OG1
6083   6084       1 ;   HG23 - OG1
6084   6086       1 ;    OG1 - C
6086   6091       1 ;      C - HA
6086   6092       1 ;      C - CB
6086   6105       1 ;      C - C
6087   6089       1 ;      O - H
6087   6090       1 ;      O - CA
6088   6093       1 ;      N - HB
6088   6094       1 ;      N - CG2
6088   6098       1 ;      N - CG1
6088   6106       1 ;      N - O
6088   6107       1 ;      N - N
6089   6091       1 ;      H - HA
6089   6092       1 ;      H - CB
6089   6105       1 ;      H - C
6090   6095       1 ;     CA - HG21
6090   6096       1 ;     CA - HG22
6090   6097       1 ;     CA - HG23
6090   6099       1 ;     CA - HG12
6090   6100       1 ;     CA - HG13
6090   6101       1 ;     CA - CD1
6090   6108       1 ;     CA - H
6090   6109       1 ;     CA - CA
6091   6093       1 ;     HA - HB
6091   6094       1 ;     HA - CG2
6091   6098       1 ;     HA - CG1
6091   6106       1 ;     HA - O
6091   6107       1 ;     HA - N
6092   6102       1 ;     CB - HD11
6092   6103       1 ;     CB - HD12
6092   6104       1 ;     CB - HD13
6092   6106       1 ;     CB - O
6092   6107       1 ;     CB - N
6093   6095       1 ;     HB - HG21
6093   6096       1 ;     HB - HG22
6093   6097       1 ;     HB - HG23
6093   6099       1 ;     HB - HG12
6093   6100       1 ;     HB - HG13
6093   6101       1 ;     HB - CD1
6093   6105       1 ;     HB - C
6094   6099       1 ;    CG2 - HG12
6094   6100       1 ;    CG2 - HG13
6094   6101       1 ;    CG2 - CD1
6094   6105       1 ;    CG2 - C
6095   6098       1 ;   HG21 - CG1
6096   6098       1 ;   HG22 - CG1
6097   6098       1 ;   HG23 - CG1
6098   6105       1 ;    CG1 - C
6099   6102       1 ;   HG12 - HD11
6099   6103       1 ;   HG12 - HD12
```

```
6099   6104       1 ;   HG12 - HD13
6100   6102       1 ;   HG13 - HD11
6100   6103       1 ;   HG13 - HD12
6100   6104       1 ;   HG13 - HD13
6105   6110       1 ;      C - HA
6105   6111       1 ;      C - CB
6105   6117       1 ;      C - C
6106   6108       1 ;      O - H
6106   6109       1 ;      O - CA
6107   6112       1 ;      N - HB2
6107   6113       1 ;      N - HB3
6107   6114       1 ;      N - CG
6107   6118       1 ;      N - O
6107   6119       1 ;      N - N
6108   6110       1 ;      H - HA
6108   6111       1 ;      H - CB
6108   6117       1 ;      H - C
6109   6115       1 ;     CA - OD1
6109   6116       1 ;     CA - OD2
6109   6120       1 ;     CA - H
6109   6121       1 ;     CA - CA
6110   6112       1 ;     HA - HB2
6110   6113       1 ;     HA - HB3
6110   6114       1 ;     HA - CG
6110   6118       1 ;     HA - O
6110   6119       1 ;     HA - N
6111   6118       1 ;     CB - O
6111   6119       1 ;     CB - N
6112   6115       1 ;    HB2 - OD1
6112   6116       1 ;    HB2 - OD2
6112   6117       1 ;    HB2 - C
6113   6115       1 ;    HB3 - OD1
6113   6116       1 ;    HB3 - OD2
6113   6117       1 ;    HB3 - C
6114   6117       1 ;     CG - C
6117   6122       1 ;      C - HA
6117   6123       1 ;      C - CB
6117   6139       1 ;      C - C
6118   6120       1 ;      O - H
6118   6121       1 ;      O - CA
6119   6124       1 ;      N - HB2
6119   6125       1 ;      N - HB3
6119   6126       1 ;      N - CG
6119   6140       1 ;      N - O
6119   6141       1 ;      N - N
6120   6122       1 ;      H - HA
6120   6123       1 ;      H - CB
6120   6139       1 ;      H - C
6121   6127       1 ;     CA - HG2
6121   6128       1 ;     CA - HG3
6121   6129       1 ;     CA - CD
6121   6142       1 ;     CA - H
```

```
6121   6143       1 ;      CA - CA
6122   6124       1 ;      HA - HB2
6122   6125       1 ;      HA - HB3
6122   6126       1 ;      HA - CG
6122   6140       1 ;      HA - O
6122   6141       1 ;      HA - N
6123   6130       1 ;      CB - HD2
6123   6131       1 ;      CB - HD3
6123   6132       1 ;      CB - CE
6123   6140       1 ;      CB - O
6123   6141       1 ;      CB - N
6124   6127       1 ;     HB2 - HG2
6124   6128       1 ;     HB2 - HG3
6124   6129       1 ;     HB2 - CD
6124   6139       1 ;     HB2 - C
6125   6127       1 ;     HB3 - HG2
6125   6128       1 ;     HB3 - HG3
6125   6129       1 ;     HB3 - CD
6125   6139       1 ;     HB3 - C
6126   6133       1 ;      CG - HE2
6126   6134       1 ;      CG - HE3
6126   6135       1 ;      CG - NZ
6126   6139       1 ;      CG - C
6127   6130       1 ;     HG2 - HD2
6127   6131       1 ;     HG2 - HD3
6127   6132       1 ;     HG2 - CE
6128   6130       1 ;     HG3 - HD2
6128   6131       1 ;     HG3 - HD3
6128   6132       1 ;     HG3 - CE
6129   6136       1 ;      CD - HZ1
6129   6137       1 ;      CD - HZ2
6129   6138       1 ;      CD - HZ3
6130   6133       1 ;     HD2 - HE2
6130   6134       1 ;     HD2 - HE3
6130   6135       1 ;     HD2 - NZ
6131   6133       1 ;     HD3 - HE2
6131   6134       1 ;     HD3 - HE3
6131   6135       1 ;     HD3 - NZ
6133   6136       1 ;     HE2 - HZ1
6133   6137       1 ;     HE2 - HZ2
6133   6138       1 ;     HE2 - HZ3
6134   6136       1 ;     HE3 - HZ1
6134   6137       1 ;     HE3 - HZ2
6134   6138       1 ;     HE3 - HZ3
6139   6144       1 ;       C - HA
6139   6145       1 ;       C - CB
6139   6156       1 ;       C - C
6140   6142       1 ;       O - H
6140   6143       1 ;       O - CA
6141   6146       1 ;       N - HB2
6141   6147       1 ;       N - HB3
6141   6148       1 ;       N - CG
```

| | | | |
|---|---|---|---|
| 6141 | 6157 | 1 ; | N - O |
| 6141 | 6158 | 1 ; | N - N |
| 6142 | 6144 | 1 ; | H - HA |
| 6142 | 6145 | 1 ; | H - CB |
| 6142 | 6156 | 1 ; | H - C |
| 6143 | 6149 | 1 ; | CA - HG2 |
| 6143 | 6150 | 1 ; | CA - HG3 |
| 6143 | 6151 | 1 ; | CA - CD |
| 6143 | 6159 | 1 ; | CA - H |
| 6143 | 6160 | 1 ; | CA - CA |
| 6144 | 6146 | 1 ; | HA - HB2 |
| 6144 | 6147 | 1 ; | HA - HB3 |
| 6144 | 6148 | 1 ; | HA - CG |
| 6144 | 6157 | 1 ; | HA - O |
| 6144 | 6158 | 1 ; | HA - N |
| 6145 | 6152 | 1 ; | CB - OE1 |
| 6145 | 6153 | 1 ; | CB - NE2 |
| 6145 | 6157 | 1 ; | CB - O |
| 6145 | 6158 | 1 ; | CB - N |
| 6146 | 6149 | 1 ; | HB2 - HG2 |
| 6146 | 6150 | 1 ; | HB2 - HG3 |
| 6146 | 6151 | 1 ; | HB2 - CD |
| 6146 | 6156 | 1 ; | HB2 - C |
| 6147 | 6149 | 1 ; | HB3 - HG2 |
| 6147 | 6150 | 1 ; | HB3 - HG3 |
| 6147 | 6151 | 1 ; | HB3 - CD |
| 6147 | 6156 | 1 ; | HB3 - C |
| 6148 | 6154 | 1 ; | CG - HE21 |
| 6148 | 6155 | 1 ; | CG - HE22 |
| 6148 | 6156 | 1 ; | CG - C |
| 6149 | 6152 | 1 ; | HG2 - OE1 |
| 6149 | 6153 | 1 ; | HG2 - NE2 |
| 6150 | 6152 | 1 ; | HG3 - OE1 |
| 6150 | 6153 | 1 ; | HG3 - NE2 |
| 6152 | 6154 | 1 ; | OE1 - HE21 |
| 6152 | 6155 | 1 ; | OE1 - HE22 |
| 6156 | 6161 | 1 ; | C - HA |
| 6156 | 6162 | 1 ; | C - CB |
| 6156 | 6170 | 1 ; | C - C |
| 6157 | 6159 | 1 ; | O - H |
| 6157 | 6160 | 1 ; | O - CA |
| 6158 | 6163 | 1 ; | N - HB |
| 6158 | 6164 | 1 ; | N - CG2 |
| 6158 | 6168 | 1 ; | N - OG1 |
| 6158 | 6171 | 1 ; | N - O |
| 6158 | 6172 | 1 ; | N - N |
| 6159 | 6161 | 1 ; | H - HA |
| 6159 | 6162 | 1 ; | H - CB |
| 6159 | 6170 | 1 ; | H - C |
| 6160 | 6165 | 1 ; | CA - HG21 |
| 6160 | 6166 | 1 ; | CA - HG22 |
| 6160 | 6167 | 1 ; | CA - HG23 |

```
6160   6169      1 ;     CA - HG1
6160   6173      1 ;     CA - H
6160   6174      1 ;     CA - CA
6161   6163      1 ;     HA - HB
6161   6164      1 ;     HA - CG2
6161   6168      1 ;     HA - OG1
6161   6171      1 ;     HA - O
6161   6172      1 ;     HA - N
6162   6171      1 ;     CB - O
6162   6172      1 ;     CB - N
6163   6165      1 ;     HB - HG21
6163   6166      1 ;     HB - HG22
6163   6167      1 ;     HB - HG23
6163   6169      1 ;     HB - HG1
6163   6170      1 ;     HB - C
6164   6169      1 ;    CG2 - HG1
6164   6170      1 ;    CG2 - C
6165   6168      1 ;   HG21 - OG1
6166   6168      1 ;   HG22 - OG1
6167   6168      1 ;   HG23 - OG1
6168   6170      1 ;    OG1 - C
6170   6175      1 ;      C - HA
6170   6176      1 ;      C - CB
6170   6191      1 ;      C - C
6171   6173      1 ;      O - H
6171   6174      1 ;      O - CA
6172   6177      1 ;      N - HB2
6172   6178      1 ;      N - HB3
6172   6179      1 ;      N - CG
6172   6192      1 ;      N - O
6172   6193      1 ;      N - N
6173   6175      1 ;      H - HA
6173   6176      1 ;      H - CB
6173   6191      1 ;      H - C
6174   6180      1 ;     CA - CD1
6174   6189      1 ;     CA - CD2
6174   6194      1 ;     CA - H
6174   6195      1 ;     CA - CA
6175   6177      1 ;     HA - HB2
6175   6178      1 ;     HA - HB3
6175   6179      1 ;     HA - CG
6175   6192      1 ;     HA - O
6175   6193      1 ;     HA - N
6176   6181      1 ;     CB - HD1
6176   6182      1 ;     CB - CE1
6176   6187      1 ;     CB - CE2
6176   6190      1 ;     CB - HD2
6176   6192      1 ;     CB - O
6176   6193      1 ;     CB - N
6177   6180      1 ;    HB2 - CD1
6177   6189      1 ;    HB2 - CD2
6177   6191      1 ;    HB2 - C
```

```
6178   6180      1 ;    HB3 - CD1
6178   6189      1 ;    HB3 - CD2
6178   6191      1 ;    HB3 - C
6179   6183      1 ;     CG - HE1
6179   6184      1 ;     CG - CZ
6179   6188      1 ;     CG - HE2
6179   6191      1 ;     CG - C
6180   6185      1 ;    CD1 - OH
6180   6187      1 ;    CD1 - CE2
6180   6190      1 ;    CD1 - HD2
6181   6183      1 ;    HD1 - HE1
6181   6184      1 ;    HD1 - CZ
6181   6189      1 ;    HD1 - CD2
6182   6186      1 ;    CE1 - HH
6182   6188      1 ;    CE1 - HE2
6182   6189      1 ;    CE1 - CD2
6183   6185      1 ;    HE1 - OH
6183   6187      1 ;    HE1 - CE2
6184   6190      1 ;     CZ - HD2
6185   6188      1 ;     OH - HE2
6185   6189      1 ;     OH - CD2
6186   6187      1 ;     HH - CE2
6188   6190      1 ;    HE2 - HD2
6191   6196      1 ;      C - HA
6191   6197      1 ;      C - CB
6191   6210      1 ;      C - C
6192   6194      1 ;      O - H
6192   6195      1 ;      O - CA
6193   6198      1 ;      N - HB2
6193   6199      1 ;      N - HB3
6193   6200      1 ;      N - CG
6193   6211      1 ;      N - O
6193   6212      1 ;      N - N
6194   6196      1 ;      H - HA
6194   6197      1 ;      H - CB
6194   6210      1 ;      H - C
6195   6201      1 ;     CA - HG
6195   6202      1 ;     CA - CD1
6195   6206      1 ;     CA - CD2
6195   6213      1 ;     CA - H
6195   6214      1 ;     CA - CA
6196   6198      1 ;     HA - HB2
6196   6199      1 ;     HA - HB3
6196   6200      1 ;     HA - CG
6196   6211      1 ;     HA - O
6196   6212      1 ;     HA - N
6197   6203      1 ;     CB - HD11
6197   6204      1 ;     CB - HD12
6197   6205      1 ;     CB - HD13
6197   6207      1 ;     CB - HD21
6197   6208      1 ;     CB - HD22
6197   6209      1 ;     CB - HD23
```

| | | | | | |
|---|---|---|---|---|---|
| 6197 | 6211 | 1 ; | CB | - | O |
| 6197 | 6212 | 1 ; | CB | - | N |
| 6198 | 6201 | 1 ; | HB2 | - | HG |
| 6198 | 6202 | 1 ; | HB2 | - | CD1 |
| 6198 | 6206 | 1 ; | HB2 | - | CD2 |
| 6198 | 6210 | 1 ; | HB2 | - | C |
| 6199 | 6201 | 1 ; | HB3 | - | HG |
| 6199 | 6202 | 1 ; | HB3 | - | CD1 |
| 6199 | 6206 | 1 ; | HB3 | - | CD2 |
| 6199 | 6210 | 1 ; | HB3 | - | C |
| 6200 | 6210 | 1 ; | CG | - | C |
| 6201 | 6203 | 1 ; | HG | - | HD11 |
| 6201 | 6204 | 1 ; | HG | - | HD12 |
| 6201 | 6205 | 1 ; | HG | - | HD13 |
| 6201 | 6207 | 1 ; | HG | - | HD21 |
| 6201 | 6208 | 1 ; | HG | - | HD22 |
| 6201 | 6209 | 1 ; | HG | - | HD23 |
| 6202 | 6207 | 1 ; | CD1 | - | HD21 |
| 6202 | 6208 | 1 ; | CD1 | - | HD22 |
| 6202 | 6209 | 1 ; | CD1 | - | HD23 |
| 6203 | 6206 | 1 ; | HD11 | - | CD2 |
| 6204 | 6206 | 1 ; | HD12 | - | CD2 |
| 6205 | 6206 | 1 ; | HD13 | - | CD2 |
| 6210 | 6215 | 1 ; | C | - | HA |
| 6210 | 6216 | 1 ; | C | - | CB |
| 6210 | 6232 | 1 ; | C | - | C |
| 6211 | 6213 | 1 ; | O | - | H |
| 6211 | 6214 | 1 ; | O | - | CA |
| 6212 | 6217 | 1 ; | N | - | HB2 |
| 6212 | 6218 | 1 ; | N | - | HB3 |
| 6212 | 6219 | 1 ; | N | - | CG |
| 6212 | 6233 | 1 ; | N | - | O |
| 6212 | 6234 | 1 ; | N | - | N |
| 6213 | 6215 | 1 ; | H | - | HA |
| 6213 | 6216 | 1 ; | H | - | CB |
| 6213 | 6232 | 1 ; | H | - | C |
| 6214 | 6220 | 1 ; | CA | - | HG2 |
| 6214 | 6221 | 1 ; | CA | - | HG3 |
| 6214 | 6222 | 1 ; | CA | - | CD |
| 6214 | 6235 | 1 ; | CA | - | H |
| 6214 | 6236 | 1 ; | CA | - | CA |
| 6215 | 6217 | 1 ; | HA | - | HB2 |
| 6215 | 6218 | 1 ; | HA | - | HB3 |
| 6215 | 6219 | 1 ; | HA | - | CG |
| 6215 | 6233 | 1 ; | HA | - | O |
| 6215 | 6234 | 1 ; | HA | - | N |
| 6216 | 6223 | 1 ; | CB | - | HD2 |
| 6216 | 6224 | 1 ; | CB | - | HD3 |
| 6216 | 6225 | 1 ; | CB | - | CE |
| 6216 | 6233 | 1 ; | CB | - | O |
| 6216 | 6234 | 1 ; | CB | - | N |
| 6217 | 6220 | 1 ; | HB2 | - | HG2 |

```
6217   6221       1 ;     HB2 - HG3
6217   6222       1 ;     HB2 - CD
6217   6232       1 ;     HB2 - C
6218   6220       1 ;     HB3 - HG2
6218   6221       1 ;     HB3 - HG3
6218   6222       1 ;     HB3 - CD
6218   6232       1 ;     HB3 - C
6219   6226       1 ;      CG - HE2
6219   6227       1 ;      CG - HE3
6219   6228       1 ;      CG - NZ
6219   6232       1 ;      CG - C
6220   6223       1 ;     HG2 - HD2
6220   6224       1 ;     HG2 - HD3
6220   6225       1 ;     HG2 - CE
6221   6223       1 ;     HG3 - HD2
6221   6224       1 ;     HG3 - HD3
6221   6225       1 ;     HG3 - CE
6222   6229       1 ;      CD - HZ1
6222   6230       1 ;      CD - HZ2
6222   6231       1 ;      CD - HZ3
6223   6226       1 ;     HD2 - HE2
6223   6227       1 ;     HD2 - HE3
6223   6228       1 ;     HD2 - NZ
6224   6226       1 ;     HD3 - HE2
6224   6227       1 ;     HD3 - HE3
6224   6228       1 ;     HD3 - NZ
6226   6229       1 ;     HE2 - HZ1
6226   6230       1 ;     HE2 - HZ2
6226   6231       1 ;     HE2 - HZ3
6227   6229       1 ;     HE3 - HZ1
6227   6230       1 ;     HE3 - HZ2
6227   6231       1 ;     HE3 - HZ3
6232   6237       1 ;       C - HA
6232   6238       1 ;       C - CB
6232   6254       1 ;       C - C
6233   6235       1 ;       O - H
6233   6236       1 ;       O - CA
6234   6239       1 ;       N - HB2
6234   6240       1 ;       N - HB3
6234   6241       1 ;       N - CG
6234   6255       1 ;       N - O
6234   6256       1 ;       N - N
6235   6237       1 ;       H - HA
6235   6238       1 ;       H - CB
6235   6254       1 ;       H - C
6236   6242       1 ;      CA - HG2
6236   6243       1 ;      CA - HG3
6236   6244       1 ;      CA - CD
6236   6257       1 ;      CA - H
6236   6258       1 ;      CA - CA
6237   6239       1 ;      HA - HB2
6237   6240       1 ;      HA - HB3
```

| | | | |
|---|---|---|---|
| 6237 | 6241 | 1 ; | HA - CG |
| 6237 | 6255 | 1 ; | HA - O |
| 6237 | 6256 | 1 ; | HA - N |
| 6238 | 6245 | 1 ; | CB - HD2 |
| 6238 | 6246 | 1 ; | CB - HD3 |
| 6238 | 6247 | 1 ; | CB - CE |
| 6238 | 6255 | 1 ; | CB - O |
| 6238 | 6256 | 1 ; | CB - N |
| 6239 | 6242 | 1 ; | HB2 - HG2 |
| 6239 | 6243 | 1 ; | HB2 - HG3 |
| 6239 | 6244 | 1 ; | HB2 - CD |
| 6239 | 6254 | 1 ; | HB2 - C |
| 6240 | 6242 | 1 ; | HB3 - HG2 |
| 6240 | 6243 | 1 ; | HB3 - HG3 |
| 6240 | 6244 | 1 ; | HB3 - CD |
| 6240 | 6254 | 1 ; | HB3 - C |
| 6241 | 6248 | 1 ; | CG - HE2 |
| 6241 | 6249 | 1 ; | CG - HE3 |
| 6241 | 6250 | 1 ; | CG - NZ |
| 6241 | 6254 | 1 ; | CG - C |
| 6242 | 6245 | 1 ; | HG2 - HD2 |
| 6242 | 6246 | 1 ; | HG2 - HD3 |
| 6242 | 6247 | 1 ; | HG2 - CE |
| 6243 | 6245 | 1 ; | HG3 - HD2 |
| 6243 | 6246 | 1 ; | HG3 - HD3 |
| 6243 | 6247 | 1 ; | HG3 - CE |
| 6244 | 6251 | 1 ; | CD - HZ1 |
| 6244 | 6252 | 1 ; | CD - HZ2 |
| 6244 | 6253 | 1 ; | CD - HZ3 |
| 6245 | 6248 | 1 ; | HD2 - HE2 |
| 6245 | 6249 | 1 ; | HD2 - HE3 |
| 6245 | 6250 | 1 ; | HD2 - NZ |
| 6246 | 6248 | 1 ; | HD3 - HE2 |
| 6246 | 6249 | 1 ; | HD3 - HE3 |
| 6246 | 6250 | 1 ; | HD3 - NZ |
| 6248 | 6251 | 1 ; | HE2 - HZ1 |
| 6248 | 6252 | 1 ; | HE2 - HZ2 |
| 6248 | 6253 | 1 ; | HE2 - HZ3 |
| 6249 | 6251 | 1 ; | HE3 - HZ1 |
| 6249 | 6252 | 1 ; | HE3 - HZ2 |
| 6249 | 6253 | 1 ; | HE3 - HZ3 |
| 6254 | 6259 | 1 ; | C - HA |
| 6254 | 6260 | 1 ; | C - CB |
| 6254 | 6269 | 1 ; | C - C |
| 6255 | 6257 | 1 ; | O - H |
| 6255 | 6258 | 1 ; | O - CA |
| 6256 | 6261 | 1 ; | N - HB2 |
| 6256 | 6262 | 1 ; | N - HB3 |
| 6256 | 6263 | 1 ; | N - CG |
| 6256 | 6270 | 1 ; | N - O |
| 6256 | 6271 | 1 ; | N - N |
| 6257 | 6259 | 1 ; | H - HA |

```
6257   6260       1 ;       H - CB
6257   6269       1 ;       H - C
6258   6264       1 ;      CA - HG2
6258   6265       1 ;      CA - HG3
6258   6266       1 ;      CA - CD
6258   6272       1 ;      CA - H
6258   6273       1 ;      CA - CA
6259   6261       1 ;      HA - HB2
6259   6262       1 ;      HA - HB3
6259   6263       1 ;      HA - CG
6259   6270       1 ;      HA - O
6259   6271       1 ;      HA - N
6260   6267       1 ;      CB - OE1
6260   6268       1 ;      CB - OE2
6260   6270       1 ;      CB - O
6260   6271       1 ;      CB - N
6261   6264       1 ;     HB2 - HG2
6261   6265       1 ;     HB2 - HG3
6261   6266       1 ;     HB2 - CD
6261   6269       1 ;     HB2 - C
6262   6264       1 ;     HB3 - HG2
6262   6265       1 ;     HB3 - HG3
6262   6266       1 ;     HB3 - CD
6262   6269       1 ;     HB3 - C
6263   6269       1 ;      CG - C
6264   6267       1 ;     HG2 - OE1
6264   6268       1 ;     HG2 - OE2
6265   6267       1 ;     HG3 - OE1
6265   6268       1 ;     HG3 - OE2
6269   6274       1 ;       C - HA
6269   6275       1 ;       C - CB
6269   6285       1 ;       C - C
6270   6272       1 ;       O - H
6270   6273       1 ;       O - CA
6271   6276       1 ;       N - HB
6271   6277       1 ;       N - CG1
6271   6281       1 ;       N - CG2
6271   6286       1 ;       N - O
6271   6287       1 ;       N - N
6272   6274       1 ;       H - HA
6272   6275       1 ;       H - CB
6272   6285       1 ;       H - C
6273   6278       1 ;      CA - HG11
6273   6279       1 ;      CA - HG12
6273   6280       1 ;      CA - HG13
6273   6282       1 ;      CA - HG21
6273   6283       1 ;      CA - HG22
6273   6284       1 ;      CA - HG23
6273   6288       1 ;      CA - H
6273   6289       1 ;      CA - CA
6274   6276       1 ;      HA - HB
6274   6277       1 ;      HA - CG1
```

```
6274   6281       1 ;      HA - CG2
6274   6286       1 ;      HA - O
6274   6287       1 ;      HA - N
6275   6286       1 ;      CB - O
6275   6287       1 ;      CB - N
6276   6278       1 ;      HB - HG11
6276   6279       1 ;      HB - HG12
6276   6280       1 ;      HB - HG13
6276   6282       1 ;      HB - HG21
6276   6283       1 ;      HB - HG22
6276   6284       1 ;      HB - HG23
6276   6285       1 ;      HB - C
6277   6282       1 ;     CG1 - HG21
6277   6283       1 ;     CG1 - HG22
6277   6284       1 ;     CG1 - HG23
6277   6285       1 ;     CG1 - C
6278   6281       1 ;    HG11 - CG2
6279   6281       1 ;    HG12 - CG2
6280   6281       1 ;    HG13 - CG2
6281   6285       1 ;     CG2 - C
6285   6290       1 ;       C - HA
6285   6291       1 ;       C - CB
6285   6296       1 ;       C - C
6286   6288       1 ;       O - H
6286   6289       1 ;       O - CA
6287   6292       1 ;       N - HB2
6287   6293       1 ;       N - HB3
6287   6294       1 ;       N - SG
6287   6297       1 ;       N - O
6287   6298       1 ;       N - N
6288   6290       1 ;       H - HA
6288   6291       1 ;       H - CB
6288   6296       1 ;       H - C
6289   6295       1 ;      CA - HG
6289   6299       1 ;      CA - H
6289   6300       1 ;      CA - CA
6290   6292       1 ;      HA - HB2
6290   6293       1 ;      HA - HB3
6290   6294       1 ;      HA - SG
6290   6297       1 ;      HA - O
6290   6298       1 ;      HA - N
6291   6297       1 ;      CB - O
6291   6298       1 ;      CB - N
6292   6295       1 ;     HB2 - HG
6292   6296       1 ;     HB2 - C
6293   6295       1 ;     HB3 - HG
6293   6296       1 ;     HB3 - C
6294   6296       1 ;      SG - C
6296   6301       1 ;       C - HA
6296   6302       1 ;       C - CB
6296   6315       1 ;       C - C
6297   6299       1 ;       O - H
```

| | | | | |
|---|---|---|---|---|
| 6297 | 6300 | 1 ; | O - | CA |
| 6298 | 6303 | 1 ; | N - | HB2 |
| 6298 | 6304 | 1 ; | N - | HB3 |
| 6298 | 6305 | 1 ; | N - | CG |
| 6298 | 6316 | 1 ; | N - | O |
| 6298 | 6317 | 1 ; | N - | N |
| 6299 | 6301 | 1 ; | H - | HA |
| 6299 | 6302 | 1 ; | H - | CB |
| 6299 | 6315 | 1 ; | H - | C |
| 6300 | 6306 | 1 ; | CA - | HG |
| 6300 | 6307 | 1 ; | CA - | CD1 |
| 6300 | 6311 | 1 ; | CA - | CD2 |
| 6300 | 6318 | 1 ; | CA - | H |
| 6300 | 6319 | 1 ; | CA - | CA |
| 6301 | 6303 | 1 ; | HA - | HB2 |
| 6301 | 6304 | 1 ; | HA - | HB3 |
| 6301 | 6305 | 1 ; | HA - | CG |
| 6301 | 6316 | 1 ; | HA - | O |
| 6301 | 6317 | 1 ; | HA - | N |
| 6302 | 6308 | 1 ; | CB - | HD11 |
| 6302 | 6309 | 1 ; | CB - | HD12 |
| 6302 | 6310 | 1 ; | CB - | HD13 |
| 6302 | 6312 | 1 ; | CB - | HD21 |
| 6302 | 6313 | 1 ; | CB - | HD22 |
| 6302 | 6314 | 1 ; | CB - | HD23 |
| 6302 | 6316 | 1 ; | CB - | O |
| 6302 | 6317 | 1 ; | CB - | N |
| 6303 | 6306 | 1 ; | HB2 - | HG |
| 6303 | 6307 | 1 ; | HB2 - | CD1 |
| 6303 | 6311 | 1 ; | HB2 - | CD2 |
| 6303 | 6315 | 1 ; | HB2 - | C |
| 6304 | 6306 | 1 ; | HB3 - | HG |
| 6304 | 6307 | 1 ; | HB3 - | CD1 |
| 6304 | 6311 | 1 ; | HB3 - | CD2 |
| 6304 | 6315 | 1 ; | HB3 - | C |
| 6305 | 6315 | 1 ; | CG - | C |
| 6306 | 6308 | 1 ; | HG - | HD11 |
| 6306 | 6309 | 1 ; | HG - | HD12 |
| 6306 | 6310 | 1 ; | HG - | HD13 |
| 6306 | 6312 | 1 ; | HG - | HD21 |
| 6306 | 6313 | 1 ; | HG - | HD22 |
| 6306 | 6314 | 1 ; | HG - | HD23 |
| 6307 | 6312 | 1 ; | CD1 - | HD21 |
| 6307 | 6313 | 1 ; | CD1 - | HD22 |
| 6307 | 6314 | 1 ; | CD1 - | HD23 |
| 6308 | 6311 | 1 ; | HD11 - | CD2 |
| 6309 | 6311 | 1 ; | HD12 - | CD2 |
| 6310 | 6311 | 1 ; | HD13 - | CD2 |
| 6315 | 6320 | 1 ; | C - | HA |
| 6315 | 6321 | 1 ; | C - | CB |
| 6315 | 6334 | 1 ; | C - | C |
| 6316 | 6318 | 1 ; | O - | H |

```
6316   6319      1 ;      O - CA
6317   6322      1 ;      N - HB
6317   6323      1 ;      N - CG2
6317   6327      1 ;      N - CG1
6317   6335      1 ;      N - O
6317   6336      1 ;      N - N
6318   6320      1 ;      H - HA
6318   6321      1 ;      H - CB
6318   6334      1 ;      H - C
6319   6324      1 ;     CA - HG21
6319   6325      1 ;     CA - HG22
6319   6326      1 ;     CA - HG23
6319   6328      1 ;     CA - HG12
6319   6329      1 ;     CA - HG13
6319   6330      1 ;     CA - CD1
6319   6337      1 ;     CA - CD
6319   6346      1 ;     CA - CA
6320   6322      1 ;     HA - HB
6320   6323      1 ;     HA - CG2
6320   6327      1 ;     HA - CG1
6320   6335      1 ;     HA - O
6320   6336      1 ;     HA - N
6321   6331      1 ;     CB - HD11
6321   6332      1 ;     CB - HD12
6321   6333      1 ;     CB - HD13
6321   6335      1 ;     CB - O
6321   6336      1 ;     CB - N
6322   6324      1 ;     HB - HG21
6322   6325      1 ;     HB - HG22
6322   6326      1 ;     HB - HG23
6322   6328      1 ;     HB - HG12
6322   6329      1 ;     HB - HG13
6322   6330      1 ;     HB - CD1
6322   6334      1 ;     HB - C
6323   6328      1 ;    CG2 - HG12
6323   6329      1 ;    CG2 - HG13
6323   6330      1 ;    CG2 - CD1
6323   6334      1 ;    CG2 - C
6324   6327      1 ;   HG21 - CG1
6325   6327      1 ;   HG22 - CG1
6326   6327      1 ;   HG23 - CG1
6327   6334      1 ;    CG1 - C
6328   6331      1 ;   HG12 - HD11
6328   6332      1 ;   HG12 - HD12
6328   6333      1 ;   HG12 - HD13
6329   6331      1 ;   HG13 - HD11
6329   6332      1 ;   HG13 - HD12
6329   6333      1 ;   HG13 - HD13
6334   6338      1 ;      C - HD2
6334   6339      1 ;      C - HD3
6334   6340      1 ;      C - CG
6334   6343      1 ;      C - CB
```

| | | | |
|---|---|---|---|
| 6334 | 6347 | 1 ; | C - HA |
| 6334 | 6348 | 1 ; | C - C |
| 6335 | 6337 | 1 ; | O - CD |
| 6335 | 6346 | 1 ; | O - CA |
| 6336 | 6341 | 1 ; | N - HG2 |
| 6336 | 6342 | 1 ; | N - HG3 |
| 6336 | 6344 | 1 ; | N - HB2 |
| 6336 | 6345 | 1 ; | N - HB3 |
| 6336 | 6349 | 1 ; | N - O |
| 6336 | 6350 | 1 ; | N - N |
| 6337 | 6344 | 1 ; | CD - HB2 |
| 6337 | 6345 | 1 ; | CD - HB3 |
| 6337 | 6347 | 1 ; | CD - HA |
| 6337 | 6348 | 1 ; | CD - C |
| 6338 | 6341 | 1 ; | HD2 - HG2 |
| 6338 | 6342 | 1 ; | HD2 - HG3 |
| 6338 | 6343 | 1 ; | HD2 - CB |
| 6338 | 6346 | 1 ; | HD2 - CA |
| 6339 | 6341 | 1 ; | HD3 - HG2 |
| 6339 | 6342 | 1 ; | HD3 - HG3 |
| 6339 | 6343 | 1 ; | HD3 - CB |
| 6339 | 6346 | 1 ; | HD3 - CA |
| 6340 | 6347 | 1 ; | CG - HA |
| 6340 | 6348 | 1 ; | CG - C |
| 6341 | 6344 | 1 ; | HG2 - HB2 |
| 6341 | 6345 | 1 ; | HG2 - HB3 |
| 6341 | 6346 | 1 ; | HG2 - CA |
| 6342 | 6344 | 1 ; | HG3 - HB2 |
| 6342 | 6345 | 1 ; | HG3 - HB3 |
| 6342 | 6346 | 1 ; | HG3 - CA |
| 6343 | 6349 | 1 ; | CB - O |
| 6343 | 6350 | 1 ; | CB - N |
| 6344 | 6347 | 1 ; | HB2 - HA |
| 6344 | 6348 | 1 ; | HB2 - C |
| 6345 | 6347 | 1 ; | HB3 - HA |
| 6345 | 6348 | 1 ; | HB3 - C |
| 6346 | 6351 | 1 ; | CA - H |
| 6346 | 6352 | 1 ; | CA - CA |
| 6347 | 6349 | 1 ; | HA - O |
| 6347 | 6350 | 1 ; | HA - N |
| 6348 | 6353 | 1 ; | C - HA |
| 6348 | 6354 | 1 ; | C - CB |
| 6348 | 6368 | 1 ; | C - C |
| 6349 | 6351 | 1 ; | O - H |
| 6349 | 6352 | 1 ; | O - CA |
| 6350 | 6355 | 1 ; | N - HB2 |
| 6350 | 6356 | 1 ; | N - HB3 |
| 6350 | 6357 | 1 ; | N - CG |
| 6350 | 6369 | 1 ; | N - O |
| 6350 | 6370 | 1 ; | N - N |
| 6351 | 6353 | 1 ; | H - HA |
| 6351 | 6354 | 1 ; | H - CB |

```
6351   6368       1 ;      H - C
6352   6358       1 ;     CA - CD1
6352   6366       1 ;     CA - CD2
6352   6371       1 ;     CA - H
6352   6372       1 ;     CA - CA
6353   6355       1 ;     HA - HB2
6353   6356       1 ;     HA - HB3
6353   6357       1 ;     HA - CG
6353   6369       1 ;     HA - O
6353   6370       1 ;     HA - N
6354   6359       1 ;     CB - HD1
6354   6360       1 ;     CB - CE1
6354   6364       1 ;     CB - CE2
6354   6367       1 ;     CB - HD2
6354   6369       1 ;     CB - O
6354   6370       1 ;     CB - N
6355   6358       1 ;    HB2 - CD1
6355   6366       1 ;    HB2 - CD2
6355   6368       1 ;    HB2 - C
6356   6358       1 ;    HB3 - CD1
6356   6366       1 ;    HB3 - CD2
6356   6368       1 ;    HB3 - C
6357   6361       1 ;     CG - HE1
6357   6362       1 ;     CG - CZ
6357   6365       1 ;     CG - HE2
6357   6368       1 ;     CG - C
6358   6363       1 ;    CD1 - HZ
6358   6364       1 ;    CD1 - CE2
6358   6367       1 ;    CD1 - HD2
6359   6361       1 ;    HD1 - HE1
6359   6362       1 ;    HD1 - CZ
6359   6366       1 ;    HD1 - CD2
6360   6365       1 ;    CE1 - HE2
6360   6366       1 ;    CE1 - CD2
6361   6363       1 ;    HE1 - HZ
6361   6364       1 ;    HE1 - CE2
6362   6367       1 ;     CZ - HD2
6363   6365       1 ;     HZ - HE2
6363   6366       1 ;     HZ - CD2
6365   6367       1 ;    HE2 - HD2
6368   6373       1 ;      C - HA
6368   6374       1 ;      C - CB
6368   6390       1 ;      C - C
6369   6371       1 ;      O - H
6369   6372       1 ;      O - CA
6370   6375       1 ;      N - HB2
6370   6376       1 ;      N - HB3
6370   6377       1 ;      N - CG
6370   6391       1 ;      N - O
6370   6392       1 ;      N - N
6371   6373       1 ;      H - HA
6371   6374       1 ;      H - CB
```

```
6371   6390       1 ;       H - C
6372   6378       1 ;      CA - HG2
6372   6379       1 ;      CA - HG3
6372   6380       1 ;      CA - CD
6372   6393       1 ;      CA - H
6372   6394       1 ;      CA - CA
6373   6375       1 ;      HA - HB2
6373   6376       1 ;      HA - HB3
6373   6377       1 ;      HA - CG
6373   6391       1 ;      HA - O
6373   6392       1 ;      HA - N
6374   6381       1 ;      CB - HD2
6374   6382       1 ;      CB - HD3
6374   6383       1 ;      CB - CE
6374   6391       1 ;      CB - O
6374   6392       1 ;      CB - N
6375   6378       1 ;     HB2 - HG2
6375   6379       1 ;     HB2 - HG3
6375   6380       1 ;     HB2 - CD
6375   6390       1 ;     HB2 - C
6376   6378       1 ;     HB3 - HG2
6376   6379       1 ;     HB3 - HG3
6376   6380       1 ;     HB3 - CD
6376   6390       1 ;     HB3 - C
6377   6384       1 ;      CG - HE2
6377   6385       1 ;      CG - HE3
6377   6386       1 ;      CG - NZ
6377   6390       1 ;      CG - C
6378   6381       1 ;     HG2 - HD2
6378   6382       1 ;     HG2 - HD3
6378   6383       1 ;     HG2 - CE
6379   6381       1 ;     HG3 - HD2
6379   6382       1 ;     HG3 - HD3
6379   6383       1 ;     HG3 - CE
6380   6387       1 ;      CD - HZ1
6380   6388       1 ;      CD - HZ2
6380   6389       1 ;      CD - HZ3
6381   6384       1 ;     HD2 - HE2
6381   6385       1 ;     HD2 - HE3
6381   6386       1 ;     HD2 - NZ
6382   6384       1 ;     HD3 - HE2
6382   6385       1 ;     HD3 - HE3
6382   6386       1 ;     HD3 - NZ
6384   6387       1 ;     HE2 - HZ1
6384   6388       1 ;     HE2 - HZ2
6384   6389       1 ;     HE2 - HZ3
6385   6387       1 ;     HE3 - HZ1
6385   6388       1 ;     HE3 - HZ2
6385   6389       1 ;     HE3 - HZ3
6390   6395       1 ;       C - HA
6390   6396       1 ;       C - CB
6390   6410       1 ;       C - C
```

| | | | |
|---|---|---|---|
| 6391 | 6393 | 1 ; | O - H |
| 6391 | 6394 | 1 ; | O - CA |
| 6392 | 6397 | 1 ; | N - HB2 |
| 6392 | 6398 | 1 ; | N - HB3 |
| 6392 | 6399 | 1 ; | N - CG |
| 6392 | 6411 | 1 ; | N - O |
| 6392 | 6412 | 1 ; | N - N |
| 6393 | 6395 | 1 ; | H - HA |
| 6393 | 6396 | 1 ; | H - CB |
| 6393 | 6410 | 1 ; | H - C |
| 6394 | 6400 | 1 ; | CA - CD1 |
| 6394 | 6408 | 1 ; | CA - CD2 |
| 6394 | 6413 | 1 ; | CA - H |
| 6394 | 6414 | 1 ; | CA - CA |
| 6395 | 6397 | 1 ; | HA - HB2 |
| 6395 | 6398 | 1 ; | HA - HB3 |
| 6395 | 6399 | 1 ; | HA - CG |
| 6395 | 6411 | 1 ; | HA - O |
| 6395 | 6412 | 1 ; | HA - N |
| 6396 | 6401 | 1 ; | CB - HD1 |
| 6396 | 6402 | 1 ; | CB - CE1 |
| 6396 | 6406 | 1 ; | CB - CE2 |
| 6396 | 6409 | 1 ; | CB - HD2 |
| 6396 | 6411 | 1 ; | CB - O |
| 6396 | 6412 | 1 ; | CB - N |
| 6397 | 6400 | 1 ; | HB2 - CD1 |
| 6397 | 6408 | 1 ; | HB2 - CD2 |
| 6397 | 6410 | 1 ; | HB2 - C |
| 6398 | 6400 | 1 ; | HB3 - CD1 |
| 6398 | 6408 | 1 ; | HB3 - CD2 |
| 6398 | 6410 | 1 ; | HB3 - C |
| 6399 | 6403 | 1 ; | CG - HE1 |
| 6399 | 6404 | 1 ; | CG - CZ |
| 6399 | 6407 | 1 ; | CG - HE2 |
| 6399 | 6410 | 1 ; | CG - C |
| 6400 | 6405 | 1 ; | CD1 - HZ |
| 6400 | 6406 | 1 ; | CD1 - CE2 |
| 6400 | 6409 | 1 ; | CD1 - HD2 |
| 6401 | 6403 | 1 ; | HD1 - HE1 |
| 6401 | 6404 | 1 ; | HD1 - CZ |
| 6401 | 6408 | 1 ; | HD1 - CD2 |
| 6402 | 6407 | 1 ; | CE1 - HE2 |
| 6402 | 6408 | 1 ; | CE1 - CD2 |
| 6403 | 6405 | 1 ; | HE1 - HZ |
| 6403 | 6406 | 1 ; | HE1 - CE2 |
| 6404 | 6409 | 1 ; | CZ - HD2 |
| 6405 | 6407 | 1 ; | HZ - HE2 |
| 6405 | 6408 | 1 ; | HZ - CD2 |
| 6407 | 6409 | 1 ; | HE2 - HD2 |
| 6410 | 6415 | 1 ; | C - HA |
| 6410 | 6416 | 1 ; | C - CB |
| 6410 | 6430 | 1 ; | C - C |

```
6411   6413     1 ;     O - H
6411   6414     1 ;     O - CA
6412   6417     1 ;     N - HB2
6412   6418     1 ;     N - HB3
6412   6419     1 ;     N - CG
6412   6431     1 ;     N - O
6412   6432     1 ;     N - N
6413   6415     1 ;     H - HA
6413   6416     1 ;     H - CB
6413   6430     1 ;     H - C
6414   6420     1 ;    CA - CD1
6414   6428     1 ;    CA - CD2
6414   6433     1 ;    CA - H
6414   6434     1 ;    CA - CA
6415   6417     1 ;    HA - HB2
6415   6418     1 ;    HA - HB3
6415   6419     1 ;    HA - CG
6415   6431     1 ;    HA - O
6415   6432     1 ;    HA - N
6416   6421     1 ;    CB - HD1
6416   6422     1 ;    CB - CE1
6416   6426     1 ;    CB - CE2
6416   6429     1 ;    CB - HD2
6416   6431     1 ;    CB - O
6416   6432     1 ;    CB - N
6417   6420     1 ;   HB2 - CD1
6417   6428     1 ;   HB2 - CD2
6417   6430     1 ;   HB2 - C
6418   6420     1 ;   HB3 - CD1
6418   6428     1 ;   HB3 - CD2
6418   6430     1 ;   HB3 - C
6419   6423     1 ;    CG - HE1
6419   6424     1 ;    CG - CZ
6419   6427     1 ;    CG - HE2
6419   6430     1 ;    CG - C
6420   6425     1 ;   CD1 - HZ
6420   6426     1 ;   CD1 - CE2
6420   6429     1 ;   CD1 - HD2
6421   6423     1 ;   HD1 - HE1
6421   6424     1 ;   HD1 - CZ
6421   6428     1 ;   HD1 - CD2
6422   6427     1 ;   CE1 - HE2
6422   6428     1 ;   CE1 - CD2
6423   6425     1 ;   HE1 - HZ
6423   6426     1 ;   HE1 - CE2
6424   6429     1 ;    CZ - HD2
6425   6427     1 ;    HZ - HE2
6425   6428     1 ;    HZ - CD2
6427   6429     1 ;   HE2 - HD2
6430   6435     1 ;     C - HA
6430   6436     1 ;     C - CB
6430   6442     1 ;     C - C
```

```
6431   6433       1 ;       O - H
6431   6434       1 ;       O - CA
6432   6437       1 ;       N - HB2
6432   6438       1 ;       N - HB3
6432   6439       1 ;       N - CG
6432   6443       1 ;       N - O
6432   6444       1 ;       N - N
6433   6435       1 ;       H - HA
6433   6436       1 ;       H - CB
6433   6442       1 ;       H - C
6434   6440       1 ;      CA - OD1
6434   6441       1 ;      CA - OD2
6434   6445       1 ;      CA - H
6434   6446       1 ;      CA - CA
6435   6437       1 ;      HA - HB2
6435   6438       1 ;      HA - HB3
6435   6439       1 ;      HA - CG
6435   6443       1 ;      HA - O
6435   6444       1 ;      HA - N
6436   6443       1 ;      CB - O
6436   6444       1 ;      CB - N
6437   6440       1 ;     HB2 - OD1
6437   6441       1 ;     HB2 - OD2
6437   6442       1 ;     HB2 - C
6438   6440       1 ;     HB3 - OD1
6438   6441       1 ;     HB3 - OD2
6438   6442       1 ;     HB3 - C
6439   6442       1 ;      CG - C
6442   6447       1 ;       C - HA
6442   6448       1 ;       C - CB
6442   6461       1 ;       C - C
6443   6445       1 ;       O - H
6443   6446       1 ;       O - CA
6444   6449       1 ;       N - HB
6444   6450       1 ;       N - CG2
6444   6454       1 ;       N - CG1
6444   6462       1 ;       N - O
6444   6463       1 ;       N - N
6445   6447       1 ;       H - HA
6445   6448       1 ;       H - CB
6445   6461       1 ;       H - C
6446   6451       1 ;      CA - HG21
6446   6452       1 ;      CA - HG22
6446   6453       1 ;      CA - HG23
6446   6455       1 ;      CA - HG12
6446   6456       1 ;      CA - HG13
6446   6457       1 ;      CA - CD1
6446   6464       1 ;      CA - H
6446   6465       1 ;      CA - CA
6447   6449       1 ;      HA - HB
6447   6450       1 ;      HA - CG2
6447   6454       1 ;      HA - CG1
```

```
6447   6462       1 ;      HA - O
6447   6463       1 ;      HA - N
6448   6458       1 ;      CB - HD11
6448   6459       1 ;      CB - HD12
6448   6460       1 ;      CB - HD13
6448   6462       1 ;      CB - O
6448   6463       1 ;      CB - N
6449   6451       1 ;      HB - HG21
6449   6452       1 ;      HB - HG22
6449   6453       1 ;      HB - HG23
6449   6455       1 ;      HB - HG12
6449   6456       1 ;      HB - HG13
6449   6457       1 ;      HB - CD1
6449   6461       1 ;      HB - C
6450   6455       1 ;     CG2 - HG12
6450   6456       1 ;     CG2 - HG13
6450   6457       1 ;     CG2 - CD1
6450   6461       1 ;     CG2 - C
6451   6454       1 ;    HG21 - CG1
6452   6454       1 ;    HG22 - CG1
6453   6454       1 ;    HG23 - CG1
6454   6461       1 ;     CG1 - C
6455   6458       1 ;    HG12 - HD11
6455   6459       1 ;    HG12 - HD12
6455   6460       1 ;    HG12 - HD13
6456   6458       1 ;    HG13 - HD11
6456   6459       1 ;    HG13 - HD12
6456   6460       1 ;    HG13 - HD13
6461   6466       1 ;       C - HA
6461   6467       1 ;       C - CB
6461   6473       1 ;       C - C
6462   6464       1 ;       O - H
6462   6465       1 ;       O - CA
6463   6468       1 ;       N - HB2
6463   6469       1 ;       N - HB3
6463   6470       1 ;       N - CG
6463   6474       1 ;       N - O
6463   6475       1 ;       N - N
6464   6466       1 ;       H - HA
6464   6467       1 ;       H - CB
6464   6473       1 ;       H - C
6465   6471       1 ;      CA - OD1
6465   6472       1 ;      CA - OD2
6465   6476       1 ;      CA - H
6465   6477       1 ;      CA - CA
6466   6468       1 ;      HA - HB2
6466   6469       1 ;      HA - HB3
6466   6470       1 ;      HA - CG
6466   6474       1 ;      HA - O
6466   6475       1 ;      HA - N
6467   6474       1 ;      CB - O
6467   6475       1 ;      CB - N
```

```
6468   6471      1 ;    HB2 - OD1
6468   6472      1 ;    HB2 - OD2
6468   6473      1 ;    HB2 - C
6469   6471      1 ;    HB3 - OD1
6469   6472      1 ;    HB3 - OD2
6469   6473      1 ;    HB3 - C
6470   6473      1 ;     CG - C
6473   6478      1 ;      C - HA2
6473   6479      1 ;      C - HA3
6473   6480      1 ;      C - C
6474   6476      1 ;      O - H
6474   6477      1 ;      O - CA
6475   6481      1 ;      N - O
6475   6482      1 ;      N - N
6476   6478      1 ;      H - HA2
6476   6479      1 ;      H - HA3
6476   6480      1 ;      H - C
6477   6483      1 ;     CA - H
6477   6484      1 ;     CA - CA
6478   6481      1 ;    HA2 - O
6478   6482      1 ;    HA2 - N
6479   6481      1 ;    HA3 - O
6479   6482      1 ;    HA3 - N
6480   6485      1 ;      C - HA
6480   6486      1 ;      C - CB
6480   6494      1 ;      C - C
6481   6483      1 ;      O - H
6481   6484      1 ;      O - CA
6482   6487      1 ;      N - HB2
6482   6488      1 ;      N - HB3
6482   6489      1 ;      N - CG
6482   6495      1 ;      N - O
6482   6496      1 ;      N - N
6483   6485      1 ;      H - HA
6483   6486      1 ;      H - CB
6483   6494      1 ;      H - C
6484   6490      1 ;     CA - OD1
6484   6491      1 ;     CA - ND2
6484   6497      1 ;     CA - H
6484   6498      1 ;     CA - CA
6485   6487      1 ;     HA - HB2
6485   6488      1 ;     HA - HB3
6485   6489      1 ;     HA - CG
6485   6495      1 ;     HA - O
6485   6496      1 ;     HA - N
6486   6492      1 ;     CB - HD21
6486   6493      1 ;     CB - HD22
6486   6495      1 ;     CB - O
6486   6496      1 ;     CB - N
6487   6490      1 ;    HB2 - OD1
6487   6491      1 ;    HB2 - ND2
6487   6494      1 ;    HB2 - C
```

| | | | | | |
|---|---|---|---|---|---|
| 6488 | 6490 | 1 ; | HB3 | - | OD1 |
| 6488 | 6491 | 1 ; | HB3 | - | ND2 |
| 6488 | 6494 | 1 ; | HB3 | - | C |
| 6489 | 6494 | 1 ; | CG | - | C |
| 6490 | 6492 | 1 ; | OD1 | - | HD21 |
| 6490 | 6493 | 1 ; | OD1 | - | HD22 |
| 6494 | 6499 | 1 ; | C | - | HA2 |
| 6494 | 6500 | 1 ; | C | - | HA3 |
| 6494 | 6501 | 1 ; | C | - | C |
| 6495 | 6497 | 1 ; | O | - | H |
| 6495 | 6498 | 1 ; | O | - | CA |
| 6496 | 6502 | 1 ; | N | - | O |
| 6496 | 6503 | 1 ; | N | - | N |
| 6497 | 6499 | 1 ; | H | - | HA2 |
| 6497 | 6500 | 1 ; | H | - | HA3 |
| 6497 | 6501 | 1 ; | H | - | C |
| 6498 | 6504 | 1 ; | CA | - | H |
| 6498 | 6505 | 1 ; | CA | - | CA |
| 6499 | 6502 | 1 ; | HA2 | - | O |
| 6499 | 6503 | 1 ; | HA2 | - | N |
| 6500 | 6502 | 1 ; | HA3 | - | O |
| 6500 | 6503 | 1 ; | HA3 | - | N |
| 6501 | 6506 | 1 ; | C | - | HA |
| 6501 | 6507 | 1 ; | C | - | CB |
| 6501 | 6523 | 1 ; | C | - | C |
| 6502 | 6504 | 1 ; | O | - | H |
| 6502 | 6505 | 1 ; | O | - | CA |
| 6503 | 6508 | 1 ; | N | - | HB2 |
| 6503 | 6509 | 1 ; | N | - | HB3 |
| 6503 | 6510 | 1 ; | N | - | CG |
| 6503 | 6524 | 1 ; | N | - | O |
| 6503 | 6525 | 1 ; | N | - | N |
| 6504 | 6506 | 1 ; | H | - | HA |
| 6504 | 6507 | 1 ; | H | - | CB |
| 6504 | 6523 | 1 ; | H | - | C |
| 6505 | 6511 | 1 ; | CA | - | HG2 |
| 6505 | 6512 | 1 ; | CA | - | HG3 |
| 6505 | 6513 | 1 ; | CA | - | CD |
| 6505 | 6526 | 1 ; | CA | - | H |
| 6505 | 6527 | 1 ; | CA | - | CA |
| 6506 | 6508 | 1 ; | HA | - | HB2 |
| 6506 | 6509 | 1 ; | HA | - | HB3 |
| 6506 | 6510 | 1 ; | HA | - | CG |
| 6506 | 6524 | 1 ; | HA | - | O |
| 6506 | 6525 | 1 ; | HA | - | N |
| 6507 | 6514 | 1 ; | CB | - | HD2 |
| 6507 | 6515 | 1 ; | CB | - | HD3 |
| 6507 | 6516 | 1 ; | CB | - | CE |
| 6507 | 6524 | 1 ; | CB | - | O |
| 6507 | 6525 | 1 ; | CB | - | N |
| 6508 | 6511 | 1 ; | HB2 | - | HG2 |
| 6508 | 6512 | 1 ; | HB2 | - | HG3 |

| | | | | | |
|---|---|---|---|---|---|
| 6508 | 6513 | 1 ; | HB2 | - | CD |
| 6508 | 6523 | 1 ; | HB2 | - | C |
| 6509 | 6511 | 1 ; | HB3 | - | HG2 |
| 6509 | 6512 | 1 ; | HB3 | - | HG3 |
| 6509 | 6513 | 1 ; | HB3 | - | CD |
| 6509 | 6523 | 1 ; | HB3 | - | C |
| 6510 | 6517 | 1 ; | CG | - | HE2 |
| 6510 | 6518 | 1 ; | CG | - | HE3 |
| 6510 | 6519 | 1 ; | CG | - | NZ |
| 6510 | 6523 | 1 ; | CG | - | C |
| 6511 | 6514 | 1 ; | HG2 | - | HD2 |
| 6511 | 6515 | 1 ; | HG2 | - | HD3 |
| 6511 | 6516 | 1 ; | HG2 | - | CE |
| 6512 | 6514 | 1 ; | HG3 | - | HD2 |
| 6512 | 6515 | 1 ; | HG3 | - | HD3 |
| 6512 | 6516 | 1 ; | HG3 | - | CE |
| 6513 | 6520 | 1 ; | CD | - | HZ1 |
| 6513 | 6521 | 1 ; | CD | - | HZ2 |
| 6513 | 6522 | 1 ; | CD | - | HZ3 |
| 6514 | 6517 | 1 ; | HD2 | - | HE2 |
| 6514 | 6518 | 1 ; | HD2 | - | HE3 |
| 6514 | 6519 | 1 ; | HD2 | - | NZ |
| 6515 | 6517 | 1 ; | HD3 | - | HE2 |
| 6515 | 6518 | 1 ; | HD3 | - | HE3 |
| 6515 | 6519 | 1 ; | HD3 | - | NZ |
| 6517 | 6520 | 1 ; | HE2 | - | HZ1 |
| 6517 | 6521 | 1 ; | HE2 | - | HZ2 |
| 6517 | 6522 | 1 ; | HE2 | - | HZ3 |
| 6518 | 6520 | 1 ; | HE3 | - | HZ1 |
| 6518 | 6521 | 1 ; | HE3 | - | HZ2 |
| 6518 | 6522 | 1 ; | HE3 | - | HZ3 |
| 6523 | 6528 | 1 ; | C | - | HA |
| 6523 | 6529 | 1 ; | C | - | CB |
| 6523 | 6542 | 1 ; | C | - | C |
| 6524 | 6526 | 1 ; | O | - | H |
| 6524 | 6527 | 1 ; | O | - | CA |
| 6525 | 6530 | 1 ; | N | - | HB |
| 6525 | 6531 | 1 ; | N | - | CG2 |
| 6525 | 6535 | 1 ; | N | - | CG1 |
| 6525 | 6543 | 1 ; | N | - | O |
| 6525 | 6544 | 1 ; | N | - | N |
| 6526 | 6528 | 1 ; | H | - | HA |
| 6526 | 6529 | 1 ; | H | - | CB |
| 6526 | 6542 | 1 ; | H | - | C |
| 6527 | 6532 | 1 ; | CA | - | HG21 |
| 6527 | 6533 | 1 ; | CA | - | HG22 |
| 6527 | 6534 | 1 ; | CA | - | HG23 |
| 6527 | 6536 | 1 ; | CA | - | HG12 |
| 6527 | 6537 | 1 ; | CA | - | HG13 |
| 6527 | 6538 | 1 ; | CA | - | CD1 |
| 6527 | 6545 | 1 ; | CA | - | H |
| 6527 | 6546 | 1 ; | CA | - | CA |

```
6528   6530      1 ;      HA - HB
6528   6531      1 ;      HA - CG2
6528   6535      1 ;      HA - CG1
6528   6543      1 ;      HA - O
6528   6544      1 ;      HA - N
6529   6539      1 ;      CB - HD11
6529   6540      1 ;      CB - HD12
6529   6541      1 ;      CB - HD13
6529   6543      1 ;      CB - O
6529   6544      1 ;      CB - N
6530   6532      1 ;      HB - HG21
6530   6533      1 ;      HB - HG22
6530   6534      1 ;      HB - HG23
6530   6536      1 ;      HB - HG12
6530   6537      1 ;      HB - HG13
6530   6538      1 ;      HB - CD1
6530   6542      1 ;      HB - C
6531   6536      1 ;     CG2 - HG12
6531   6537      1 ;     CG2 - HG13
6531   6538      1 ;     CG2 - CD1
6531   6542      1 ;     CG2 - C
6532   6535      1 ;    HG21 - CG1
6533   6535      1 ;    HG22 - CG1
6534   6535      1 ;    HG23 - CG1
6535   6542      1 ;     CG1 - C
6536   6539      1 ;    HG12 - HD11
6536   6540      1 ;    HG12 - HD12
6536   6541      1 ;    HG12 - HD13
6537   6539      1 ;    HG13 - HD11
6537   6540      1 ;    HG13 - HD12
6537   6541      1 ;    HG13 - HD13
6542   6547      1 ;       C - HA
6542   6548      1 ;       C - CB
6542   6553      1 ;       C - C
6543   6545      1 ;       O - H
6543   6546      1 ;       O - CA
6544   6549      1 ;       N - HB2
6544   6550      1 ;       N - HB3
6544   6551      1 ;       N - OG
6544   6554      1 ;       N - O
6544   6555      1 ;       N - N
6545   6547      1 ;       H - HA
6545   6548      1 ;       H - CB
6545   6553      1 ;       H - C
6546   6552      1 ;      CA - HG
6546   6556      1 ;      CA - H
6546   6557      1 ;      CA - CA
6547   6549      1 ;      HA - HB2
6547   6550      1 ;      HA - HB3
6547   6551      1 ;      HA - OG
6547   6554      1 ;      HA - O
6547   6555      1 ;      HA - N
```

```
6548   6554       1 ;      CB - O
6548   6555       1 ;      CB - N
6549   6552       1 ;     HB2 - HG
6549   6553       1 ;     HB2 - C
6550   6552       1 ;     HB3 - HG
6550   6553       1 ;     HB3 - C
6551   6553       1 ;      OG - C
6553   6558       1 ;       C - HA
6553   6559       1 ;       C - CB
6553   6569       1 ;       C - C
6554   6556       1 ;       O - H
6554   6557       1 ;       O - CA
6555   6560       1 ;       N - HB
6555   6561       1 ;       N - CG1
6555   6565       1 ;       N - CG2
6555   6570       1 ;       N - O
6555   6571       1 ;       N - N
6556   6558       1 ;       H - HA
6556   6559       1 ;       H - CB
6556   6569       1 ;       H - C
6557   6562       1 ;      CA - HG11
6557   6563       1 ;      CA - HG12
6557   6564       1 ;      CA - HG13
6557   6566       1 ;      CA - HG21
6557   6567       1 ;      CA - HG22
6557   6568       1 ;      CA - HG23
6557   6572       1 ;      CA - H
6557   6573       1 ;      CA - CA
6558   6560       1 ;      HA - HB
6558   6561       1 ;      HA - CG1
6558   6565       1 ;      HA - CG2
6558   6570       1 ;      HA - O
6558   6571       1 ;      HA - N
6559   6570       1 ;      CB - O
6559   6571       1 ;      CB - N
6560   6562       1 ;      HB - HG11
6560   6563       1 ;      HB - HG12
6560   6564       1 ;      HB - HG13
6560   6566       1 ;      HB - HG21
6560   6567       1 ;      HB - HG22
6560   6568       1 ;      HB - HG23
6560   6569       1 ;      HB - C
6561   6566       1 ;     CG1 - HG21
6561   6567       1 ;     CG1 - HG22
6561   6568       1 ;     CG1 - HG23
6561   6569       1 ;     CG1 - C
6562   6565       1 ;    HG11 - CG2
6563   6565       1 ;    HG12 - CG2
6564   6565       1 ;    HG13 - CG2
6565   6569       1 ;     CG2 - C
6569   6574       1 ;       C - HA
6569   6575       1 ;       C - CB
```

| | | | | |
|---|---|---|---|---|
| 6569 | 6584 | 1 ; | C - | C |
| 6570 | 6572 | 1 ; | O - | H |
| 6570 | 6573 | 1 ; | O - | CA |
| 6571 | 6576 | 1 ; | N - | HB2 |
| 6571 | 6577 | 1 ; | N - | HB3 |
| 6571 | 6578 | 1 ; | N - | CG |
| 6571 | 6585 | 1 ; | N - | O |
| 6571 | 6586 | 1 ; | N - | N |
| 6572 | 6574 | 1 ; | H - | HA |
| 6572 | 6575 | 1 ; | H - | CB |
| 6572 | 6584 | 1 ; | H - | C |
| 6573 | 6579 | 1 ; | CA - | HG2 |
| 6573 | 6580 | 1 ; | CA - | HG3 |
| 6573 | 6581 | 1 ; | CA - | CD |
| 6573 | 6587 | 1 ; | CA - | H |
| 6573 | 6588 | 1 ; | CA - | CA |
| 6574 | 6576 | 1 ; | HA - | HB2 |
| 6574 | 6577 | 1 ; | HA - | HB3 |
| 6574 | 6578 | 1 ; | HA - | CG |
| 6574 | 6585 | 1 ; | HA - | O |
| 6574 | 6586 | 1 ; | HA - | N |
| 6575 | 6582 | 1 ; | CB - | OE1 |
| 6575 | 6583 | 1 ; | CB - | OE2 |
| 6575 | 6585 | 1 ; | CB - | O |
| 6575 | 6586 | 1 ; | CB - | N |
| 6576 | 6579 | 1 ; | HB2 - | HG2 |
| 6576 | 6580 | 1 ; | HB2 - | HG3 |
| 6576 | 6581 | 1 ; | HB2 - | CD |
| 6576 | 6584 | 1 ; | HB2 - | C |
| 6577 | 6579 | 1 ; | HB3 - | HG2 |
| 6577 | 6580 | 1 ; | HB3 - | HG3 |
| 6577 | 6581 | 1 ; | HB3 - | CD |
| 6577 | 6584 | 1 ; | HB3 - | C |
| 6578 | 6584 | 1 ; | CG - | C |
| 6579 | 6582 | 1 ; | HG2 - | OE1 |
| 6579 | 6583 | 1 ; | HG2 - | OE2 |
| 6580 | 6582 | 1 ; | HG3 - | OE1 |
| 6580 | 6583 | 1 ; | HG3 - | OE2 |
| 6584 | 6589 | 1 ; | C - | HA |
| 6584 | 6590 | 1 ; | C - | CB |
| 6584 | 6600 | 1 ; | C - | C |
| 6585 | 6587 | 1 ; | O - | H |
| 6585 | 6588 | 1 ; | O - | CA |
| 6586 | 6591 | 1 ; | N - | HB2 |
| 6586 | 6592 | 1 ; | N - | HB3 |
| 6586 | 6593 | 1 ; | N - | CG |
| 6586 | 6601 | 1 ; | N - | O |
| 6586 | 6602 | 1 ; | N - | N |
| 6587 | 6589 | 1 ; | H - | HA |
| 6587 | 6590 | 1 ; | H - | CB |
| 6587 | 6600 | 1 ; | H - | C |
| 6588 | 6594 | 1 ; | CA - | HG2 |

```
6588   6595       1 ;      CA - HG3
6588   6596       1 ;      CA - CD
6588   6603       1 ;      CA - H
6588   6604       1 ;      CA - CA
6589   6591       1 ;      HA - HB2
6589   6592       1 ;      HA - HB3
6589   6593       1 ;      HA - CG
6589   6601       1 ;      HA - O
6589   6602       1 ;      HA - N
6590   6597       1 ;      CB - OE1
6590   6598       1 ;      CB - OE2
6590   6601       1 ;      CB - O
6590   6602       1 ;      CB - N
6591   6594       1 ;     HB2 - HG2
6591   6595       1 ;     HB2 - HG3
6591   6596       1 ;     HB2 - CD
6591   6600       1 ;     HB2 - C
6592   6594       1 ;     HB3 - HG2
6592   6595       1 ;     HB3 - HG3
6592   6596       1 ;     HB3 - CD
6592   6600       1 ;     HB3 - C
6593   6599       1 ;      CG - HE2
6593   6600       1 ;      CG - C
6594   6597       1 ;     HG2 - OE1
6594   6598       1 ;     HG2 - OE2
6595   6597       1 ;     HG3 - OE1
6595   6598       1 ;     HG3 - OE2
6597   6599       1 ;     OE1 - HE2
6600   6605       1 ;       C - HA
6600   6606       1 ;       C - CB
6600   6619       1 ;       C - C
6601   6603       1 ;       O - H
6601   6604       1 ;       O - CA
6602   6607       1 ;       N - HB2
6602   6608       1 ;       N - HB3
6602   6609       1 ;       N - CG
6602   6620       1 ;       N - O
6602   6621       1 ;       N - N
6603   6605       1 ;       H - HA
6603   6606       1 ;       H - CB
6603   6619       1 ;       H - C
6604   6610       1 ;      CA - HG
6604   6611       1 ;      CA - CD1
6604   6615       1 ;      CA - CD2
6604   6622       1 ;      CA - H
6604   6623       1 ;      CA - CA
6605   6607       1 ;      HA - HB2
6605   6608       1 ;      HA - HB3
6605   6609       1 ;      HA - CG
6605   6620       1 ;      HA - O
6605   6621       1 ;      HA - N
6606   6612       1 ;      CB - HD11
```

```
6606   6613      1 ;     CB - HD12
6606   6614      1 ;     CB - HD13
6606   6616      1 ;     CB - HD21
6606   6617      1 ;     CB - HD22
6606   6618      1 ;     CB - HD23
6606   6620      1 ;     CB - O
6606   6621      1 ;     CB - N
6607   6610      1 ;    HB2 - HG
6607   6611      1 ;    HB2 - CD1
6607   6615      1 ;    HB2 - CD2
6607   6619      1 ;    HB2 - C
6608   6610      1 ;    HB3 - HG
6608   6611      1 ;    HB3 - CD1
6608   6615      1 ;    HB3 - CD2
6608   6619      1 ;    HB3 - C
6609   6619      1 ;     CG - C
6610   6612      1 ;     HG - HD11
6610   6613      1 ;     HG - HD12
6610   6614      1 ;     HG - HD13
6610   6616      1 ;     HG - HD21
6610   6617      1 ;     HG - HD22
6610   6618      1 ;     HG - HD23
6611   6616      1 ;    CD1 - HD21
6611   6617      1 ;    CD1 - HD22
6611   6618      1 ;    CD1 - HD23
6612   6615      1 ;   HD11 - CD2
6613   6615      1 ;   HD12 - CD2
6614   6615      1 ;   HD13 - CD2
6619   6624      1 ;      C - HA
6619   6625      1 ;      C - CB
6619   6641      1 ;      C - C
6620   6622      1 ;      O - H
6620   6623      1 ;      O - CA
6621   6626      1 ;      N - HB2
6621   6627      1 ;      N - HB3
6621   6628      1 ;      N - CG
6621   6642      1 ;      N - O
6621   6643      1 ;      N - N
6622   6624      1 ;      H - HA
6622   6625      1 ;      H - CB
6622   6641      1 ;      H - C
6623   6629      1 ;     CA - HG2
6623   6630      1 ;     CA - HG3
6623   6631      1 ;     CA - CD
6623   6644      1 ;     CA - H
6623   6645      1 ;     CA - CA
6624   6626      1 ;     HA - HB2
6624   6627      1 ;     HA - HB3
6624   6628      1 ;     HA - CG
6624   6642      1 ;     HA - O
6624   6643      1 ;     HA - N
6625   6632      1 ;     CB - HD2
```

```
6625   6633       1 ;      CB - HD3
6625   6634       1 ;      CB - CE
6625   6642       1 ;      CB - O
6625   6643       1 ;      CB - N
6626   6629       1 ;     HB2 - HG2
6626   6630       1 ;     HB2 - HG3
6626   6631       1 ;     HB2 - CD
6626   6641       1 ;     HB2 - C
6627   6629       1 ;     HB3 - HG2
6627   6630       1 ;     HB3 - HG3
6627   6631       1 ;     HB3 - CD
6627   6641       1 ;     HB3 - C
6628   6635       1 ;      CG - HE2
6628   6636       1 ;      CG - HE3
6628   6637       1 ;      CG - NZ
6628   6641       1 ;      CG - C
6629   6632       1 ;     HG2 - HD2
6629   6633       1 ;     HG2 - HD3
6629   6634       1 ;     HG2 - CE
6630   6632       1 ;     HG3 - HD2
6630   6633       1 ;     HG3 - HD3
6630   6634       1 ;     HG3 - CE
6631   6638       1 ;      CD - HZ1
6631   6639       1 ;      CD - HZ2
6631   6640       1 ;      CD - HZ3
6632   6635       1 ;     HD2 - HE2
6632   6636       1 ;     HD2 - HE3
6632   6637       1 ;     HD2 - NZ
6633   6635       1 ;     HD3 - HE2
6633   6636       1 ;     HD3 - HE3
6633   6637       1 ;     HD3 - NZ
6635   6638       1 ;     HE2 - HZ1
6635   6639       1 ;     HE2 - HZ2
6635   6640       1 ;     HE2 - HZ3
6636   6638       1 ;     HE3 - HZ1
6636   6639       1 ;     HE3 - HZ2
6636   6640       1 ;     HE3 - HZ3
6641   6646       1 ;       C - HA
6641   6647       1 ;       C - CB
6641   6665       1 ;       C - C
6642   6644       1 ;       O - H
6642   6645       1 ;       O - CA
6643   6648       1 ;       N - HB2
6643   6649       1 ;       N - HB3
6643   6650       1 ;       N - CG
6643   6666       1 ;       N - O
6643   6667       1 ;       N - N
6644   6646       1 ;       H - HA
6644   6647       1 ;       H - CB
6644   6665       1 ;       H - C
6645   6651       1 ;      CA - HG2
6645   6652       1 ;      CA - HG3
```

```
6645   6653      1 ;      CA - CD
6645   6668      1 ;      CA - H
6645   6669      1 ;      CA - CA
6646   6648      1 ;      HA - HB2
6646   6649      1 ;      HA - HB3
6646   6650      1 ;      HA - CG
6646   6666      1 ;      HA - O
6646   6667      1 ;      HA - N
6647   6654      1 ;      CB - HD2
6647   6655      1 ;      CB - HD3
6647   6656      1 ;      CB - NE
6647   6666      1 ;      CB - O
6647   6667      1 ;      CB - N
6648   6651      1 ;     HB2 - HG2
6648   6652      1 ;     HB2 - HG3
6648   6653      1 ;     HB2 - CD
6648   6665      1 ;     HB2 - C
6649   6651      1 ;     HB3 - HG2
6649   6652      1 ;     HB3 - HG3
6649   6653      1 ;     HB3 - CD
6649   6665      1 ;     HB3 - C
6650   6657      1 ;      CG - HE
6650   6658      1 ;      CG - CZ
6650   6665      1 ;      CG - C
6651   6654      1 ;     HG2 - HD2
6651   6655      1 ;     HG2 - HD3
6651   6656      1 ;     HG2 - NE
6652   6654      1 ;     HG3 - HD2
6652   6655      1 ;     HG3 - HD3
6652   6656      1 ;     HG3 - NE
6653   6659      1 ;      CD - NH1
6653   6662      1 ;      CD - NH2
6654   6657      1 ;     HD2 - HE
6654   6658      1 ;     HD2 - CZ
6655   6657      1 ;     HD3 - HE
6655   6658      1 ;     HD3 - CZ
6656   6660      1 ;      NE - HH11
6656   6661      1 ;      NE - HH12
6656   6663      1 ;      NE - HH21
6656   6664      1 ;      NE - HH22
6657   6659      1 ;      HE - NH1
6657   6662      1 ;      HE - NH2
6659   6663      1 ;     NH1 - HH21
6659   6664      1 ;     NH1 - HH22
6660   6662      1 ;    HH11 - NH2
6661   6662      1 ;    HH12 - NH2
6665   6670      1 ;       C - HA
6665   6671      1 ;       C - CB
6665   6684      1 ;       C - C
6666   6668      1 ;       O - H
6666   6669      1 ;       O - CA
6667   6672      1 ;       N - HB
```

```
 6667   6673       1 ;       N - CG2
 6667   6677       1 ;       N - CG1
 6667   6685       1 ;       N - O
 6667   6686       1 ;       N - N
 6668   6670       1 ;       H - HA
 6668   6671       1 ;       H - CB
 6668   6684       1 ;       H - C
 6669   6674       1 ;      CA - HG21
 6669   6675       1 ;      CA - HG22
 6669   6676       1 ;      CA - HG23
 6669   6678       1 ;      CA - HG12
 6669   6679       1 ;      CA - HG13
 6669   6680       1 ;      CA - CD1
 6669   6687       1 ;      CA - H
 6669   6688       1 ;      CA - CA
 6670   6672       1 ;      HA - HB
 6670   6673       1 ;      HA - CG2
 6670   6677       1 ;      HA - CG1
 6670   6685       1 ;      HA - O
 6670   6686       1 ;      HA - N
 6671   6681       1 ;      CB - HD11
 6671   6682       1 ;      CB - HD12
 6671   6683       1 ;      CB - HD13
 6671   6685       1 ;      CB - O
 6671   6686       1 ;      CB - N
 6672   6674       1 ;      HB - HG21
 6672   6675       1 ;      HB - HG22
 6672   6676       1 ;      HB - HG23
 6672   6678       1 ;      HB - HG12
 6672   6679       1 ;      HB - HG13
 6672   6680       1 ;      HB - CD1
 6672   6684       1 ;      HB - C
 6673   6678       1 ;     CG2 - HG12
 6673   6679       1 ;     CG2 - HG13
 6673   6680       1 ;     CG2 - CD1
 6673   6684       1 ;     CG2 - C
 6674   6677       1 ;    HG21 - CG1
 6675   6677       1 ;    HG22 - CG1
 6676   6677       1 ;    HG23 - CG1
 6677   6684       1 ;     CG1 - C
 6678   6681       1 ;    HG12 - HD11
 6678   6682       1 ;    HG12 - HD12
 6678   6683       1 ;    HG12 - HD13
 6679   6681       1 ;    HG13 - HD11
 6679   6682       1 ;    HG13 - HD12
 6679   6683       1 ;    HG13 - HD13
 6684   6689       1 ;       C - HA
 6684   6690       1 ;       C - CB
 6684   6704       1 ;       C - C
 6685   6687       1 ;       O - H
 6685   6688       1 ;       O - CA
 6686   6691       1 ;       N - HB2
```

```
6686   6692      1 ;      N - HB3
6686   6693      1 ;      N - CG
6686   6705      1 ;      N - O
6686   6706      1 ;      N - N
6687   6689      1 ;      H - HA
6687   6690      1 ;      H - CB
6687   6704      1 ;      H - C
6688   6694      1 ;     CA - CD1
6688   6702      1 ;     CA - CD2
6688   6707      1 ;     CA - H
6688   6708      1 ;     CA - CA
6689   6691      1 ;     HA - HB2
6689   6692      1 ;     HA - HB3
6689   6693      1 ;     HA - CG
6689   6705      1 ;     HA - O
6689   6706      1 ;     HA - N
6690   6695      1 ;     CB - HD1
6690   6696      1 ;     CB - CE1
6690   6700      1 ;     CB - CE2
6690   6703      1 ;     CB - HD2
6690   6705      1 ;     CB - O
6690   6706      1 ;     CB - N
6691   6694      1 ;    HB2 - CD1
6691   6702      1 ;    HB2 - CD2
6691   6704      1 ;    HB2 - C
6692   6694      1 ;    HB3 - CD1
6692   6702      1 ;    HB3 - CD2
6692   6704      1 ;    HB3 - C
6693   6697      1 ;     CG - HE1
6693   6698      1 ;     CG - CZ
6693   6701      1 ;     CG - HE2
6693   6704      1 ;     CG - C
6694   6699      1 ;    CD1 - HZ
6694   6700      1 ;    CD1 - CE2
6694   6703      1 ;    CD1 - HD2
6695   6697      1 ;    HD1 - HE1
6695   6698      1 ;    HD1 - CZ
6695   6702      1 ;    HD1 - CD2
6696   6701      1 ;    CE1 - HE2
6696   6702      1 ;    CE1 - CD2
6697   6699      1 ;    HE1 - HZ
6697   6700      1 ;    HE1 - CE2
6698   6703      1 ;     CZ - HD2
6699   6701      1 ;     HZ - HE2
6699   6702      1 ;     HZ - CD2
6701   6703      1 ;    HE2 - HD2
6704   6709      1 ;      C - HA2
6704   6710      1 ;      C - HA3
6704   6711      1 ;      C - C
6705   6707      1 ;      O - H
6705   6708      1 ;      O - CA
6706   6712      1 ;      N - O
```

| | | | | | |
|---|---|---|---|---|---|
| 6706 | 6713 | 1 ; | N | - | N |
| 6707 | 6709 | 1 ; | H | - | HA2 |
| 6707 | 6710 | 1 ; | H | - | HA3 |
| 6707 | 6711 | 1 ; | H | - | C |
| 6708 | 6714 | 1 ; | CA | - | H |
| 6708 | 6715 | 1 ; | CA | - | CA |
| 6709 | 6712 | 1 ; | HA2 | - | O |
| 6709 | 6713 | 1 ; | HA2 | - | N |
| 6710 | 6712 | 1 ; | HA3 | - | O |
| 6710 | 6713 | 1 ; | HA3 | - | N |
| 6711 | 6716 | 1 ; | C | - | HA |
| 6711 | 6717 | 1 ; | C | - | CB |
| 6711 | 6735 | 1 ; | C | - | C |
| 6712 | 6714 | 1 ; | O | - | H |
| 6712 | 6715 | 1 ; | O | - | CA |
| 6713 | 6718 | 1 ; | N | - | HB2 |
| 6713 | 6719 | 1 ; | N | - | HB3 |
| 6713 | 6720 | 1 ; | N | - | CG |
| 6713 | 6736 | 1 ; | N | - | O |
| 6713 | 6737 | 1 ; | N | - | N |
| 6714 | 6716 | 1 ; | H | - | HA |
| 6714 | 6717 | 1 ; | H | - | CB |
| 6714 | 6735 | 1 ; | H | - | C |
| 6715 | 6721 | 1 ; | CA | - | HG2 |
| 6715 | 6722 | 1 ; | CA | - | HG3 |
| 6715 | 6723 | 1 ; | CA | - | CD |
| 6715 | 6738 | 1 ; | CA | - | H |
| 6715 | 6739 | 1 ; | CA | - | CA |
| 6716 | 6718 | 1 ; | HA | - | HB2 |
| 6716 | 6719 | 1 ; | HA | - | HB3 |
| 6716 | 6720 | 1 ; | HA | - | CG |
| 6716 | 6736 | 1 ; | HA | - | O |
| 6716 | 6737 | 1 ; | HA | - | N |
| 6717 | 6724 | 1 ; | CB | - | HD2 |
| 6717 | 6725 | 1 ; | CB | - | HD3 |
| 6717 | 6726 | 1 ; | CB | - | NE |
| 6717 | 6736 | 1 ; | CB | - | O |
| 6717 | 6737 | 1 ; | CB | - | N |
| 6718 | 6721 | 1 ; | HB2 | - | HG2 |
| 6718 | 6722 | 1 ; | HB2 | - | HG3 |
| 6718 | 6723 | 1 ; | HB2 | - | CD |
| 6718 | 6735 | 1 ; | HB2 | - | C |
| 6719 | 6721 | 1 ; | HB3 | - | HG2 |
| 6719 | 6722 | 1 ; | HB3 | - | HG3 |
| 6719 | 6723 | 1 ; | HB3 | - | CD |
| 6719 | 6735 | 1 ; | HB3 | - | C |
| 6720 | 6727 | 1 ; | CG | - | HE |
| 6720 | 6728 | 1 ; | CG | - | CZ |
| 6720 | 6735 | 1 ; | CG | - | C |
| 6721 | 6724 | 1 ; | HG2 | - | HD2 |
| 6721 | 6725 | 1 ; | HG2 | - | HD3 |
| 6721 | 6726 | 1 ; | HG2 | - | NE |

```
6722   6724       1 ;    HG3 - HD2
6722   6725       1 ;    HG3 - HD3
6722   6726       1 ;    HG3 - NE
6723   6729       1 ;     CD - NH1
6723   6732       1 ;     CD - NH2
6724   6727       1 ;    HD2 - HE
6724   6728       1 ;    HD2 - CZ
6725   6727       1 ;    HD3 - HE
6725   6728       1 ;    HD3 - CZ
6726   6730       1 ;     NE - HH11
6726   6731       1 ;     NE - HH12
6726   6733       1 ;     NE - HH21
6726   6734       1 ;     NE - HH22
6727   6729       1 ;     HE - NH1
6727   6732       1 ;     HE - NH2
6729   6733       1 ;    NH1 - HH21
6729   6734       1 ;    NH1 - HH22
6730   6732       1 ;   HH11 - NH2
6731   6732       1 ;   HH12 - NH2
6735   6740       1 ;      C - HA
6735   6741       1 ;      C - CB
6735   6747       1 ;      C - C
6736   6738       1 ;      O - H
6736   6739       1 ;      O - CA
6737   6742       1 ;      N - HB2
6737   6743       1 ;      N - HB3
6737   6744       1 ;      N - CG
6737   6748       1 ;      N - O
6737   6749       1 ;      N - N
6738   6740       1 ;      H - HA
6738   6741       1 ;      H - CB
6738   6747       1 ;      H - C
6739   6745       1 ;     CA - OD1
6739   6746       1 ;     CA - OD2
6739   6750       1 ;     CA - H
6739   6751       1 ;     CA - CA
6740   6742       1 ;     HA - HB2
6740   6743       1 ;     HA - HB3
6740   6744       1 ;     HA - CG
6740   6748       1 ;     HA - O
6740   6749       1 ;     HA - N
6741   6748       1 ;     CB - O
6741   6749       1 ;     CB - N
6742   6745       1 ;    HB2 - OD1
6742   6746       1 ;    HB2 - OD2
6742   6747       1 ;    HB2 - C
6743   6745       1 ;    HB3 - OD1
6743   6746       1 ;    HB3 - OD2
6743   6747       1 ;    HB3 - C
6744   6747       1 ;     CG - C
6747   6752       1 ;      C - HA
6747   6753       1 ;      C - CB
```

| | | | | | |
|---|---|---|---|---|---|
| 6747 | 6759 | 1 ; | C | - | C |
| 6748 | 6750 | 1 ; | O | - | H |
| 6748 | 6751 | 1 ; | O | - | CA |
| 6749 | 6754 | 1 ; | N | - | HB2 |
| 6749 | 6755 | 1 ; | N | - | HB3 |
| 6749 | 6756 | 1 ; | N | - | CG |
| 6749 | 6760 | 1 ; | N | - | O |
| 6749 | 6761 | 1 ; | N | - | N |
| 6750 | 6752 | 1 ; | H | - | HA |
| 6750 | 6753 | 1 ; | H | - | CB |
| 6750 | 6759 | 1 ; | H | - | C |
| 6751 | 6757 | 1 ; | CA | - | OD1 |
| 6751 | 6758 | 1 ; | CA | - | OD2 |
| 6751 | 6762 | 1 ; | CA | - | H |
| 6751 | 6763 | 1 ; | CA | - | CA |
| 6752 | 6754 | 1 ; | HA | - | HB2 |
| 6752 | 6755 | 1 ; | HA | - | HB3 |
| 6752 | 6756 | 1 ; | HA | - | CG |
| 6752 | 6760 | 1 ; | HA | - | O |
| 6752 | 6761 | 1 ; | HA | - | N |
| 6753 | 6760 | 1 ; | CB | - | O |
| 6753 | 6761 | 1 ; | CB | - | N |
| 6754 | 6757 | 1 ; | HB2 | - | OD1 |
| 6754 | 6758 | 1 ; | HB2 | - | OD2 |
| 6754 | 6759 | 1 ; | HB2 | - | C |
| 6755 | 6757 | 1 ; | HB3 | - | OD1 |
| 6755 | 6758 | 1 ; | HB3 | - | OD2 |
| 6755 | 6759 | 1 ; | HB3 | - | C |
| 6756 | 6759 | 1 ; | CG | - | C |
| 6759 | 6764 | 1 ; | C | - | HA |
| 6759 | 6765 | 1 ; | C | - | CB |
| 6759 | 6778 | 1 ; | C | - | C |
| 6760 | 6762 | 1 ; | O | - | H |
| 6760 | 6763 | 1 ; | O | - | CA |
| 6761 | 6766 | 1 ; | N | - | HB |
| 6761 | 6767 | 1 ; | N | - | CG2 |
| 6761 | 6771 | 1 ; | N | - | CG1 |
| 6761 | 6779 | 1 ; | N | - | O |
| 6761 | 6780 | 1 ; | N | - | N |
| 6762 | 6764 | 1 ; | H | - | HA |
| 6762 | 6765 | 1 ; | H | - | CB |
| 6762 | 6778 | 1 ; | H | - | C |
| 6763 | 6768 | 1 ; | CA | - | HG21 |
| 6763 | 6769 | 1 ; | CA | - | HG22 |
| 6763 | 6770 | 1 ; | CA | - | HG23 |
| 6763 | 6772 | 1 ; | CA | - | HG12 |
| 6763 | 6773 | 1 ; | CA | - | HG13 |
| 6763 | 6774 | 1 ; | CA | - | CD1 |
| 6763 | 6781 | 1 ; | CA | - | H |
| 6763 | 6782 | 1 ; | CA | - | CA |
| 6764 | 6766 | 1 ; | HA | - | HB |
| 6764 | 6767 | 1 ; | HA | - | CG2 |

```
6764   6771      1 ;     HA - CG1
6764   6779      1 ;     HA - O
6764   6780      1 ;     HA - N
6765   6775      1 ;     CB - HD11
6765   6776      1 ;     CB - HD12
6765   6777      1 ;     CB - HD13
6765   6779      1 ;     CB - O
6765   6780      1 ;     CB - N
6766   6768      1 ;     HB - HG21
6766   6769      1 ;     HB - HG22
6766   6770      1 ;     HB - HG23
6766   6772      1 ;     HB - HG12
6766   6773      1 ;     HB - HG13
6766   6774      1 ;     HB - CD1
6766   6778      1 ;     HB - C
6767   6772      1 ;    CG2 - HG12
6767   6773      1 ;    CG2 - HG13
6767   6774      1 ;    CG2 - CD1
6767   6778      1 ;    CG2 - C
6768   6771      1 ;   HG21 - CG1
6769   6771      1 ;   HG22 - CG1
6770   6771      1 ;   HG23 - CG1
6771   6778      1 ;    CG1 - C
6772   6775      1 ;   HG12 - HD11
6772   6776      1 ;   HG12 - HD12
6772   6777      1 ;   HG12 - HD13
6773   6775      1 ;   HG13 - HD11
6773   6776      1 ;   HG13 - HD12
6773   6777      1 ;   HG13 - HD13
6778   6783      1 ;      C - HA
6778   6784      1 ;      C - CB
6778   6793      1 ;      C - C
6779   6781      1 ;      O - H
6779   6782      1 ;      O - CA
6780   6785      1 ;      N - HB2
6780   6786      1 ;      N - HB3
6780   6787      1 ;      N - CG
6780   6794      1 ;      N - O
6780   6795      1 ;      N - N
6781   6783      1 ;      H - HA
6781   6784      1 ;      H - CB
6781   6793      1 ;      H - C
6782   6788      1 ;     CA - HG2
6782   6789      1 ;     CA - HG3
6782   6790      1 ;     CA - CD
6782   6796      1 ;     CA - H
6782   6797      1 ;     CA - CA
6783   6785      1 ;     HA - HB2
6783   6786      1 ;     HA - HB3
6783   6787      1 ;     HA - CG
6783   6794      1 ;     HA - O
6783   6795      1 ;     HA - N
```

```
6784   6791       1 ;      CB - OE1
6784   6792       1 ;      CB - OE2
6784   6794       1 ;      CB - O
6784   6795       1 ;      CB - N
6785   6788       1 ;     HB2 - HG2
6785   6789       1 ;     HB2 - HG3
6785   6790       1 ;     HB2 - CD
6785   6793       1 ;     HB2 - C
6786   6788       1 ;     HB3 - HG2
6786   6789       1 ;     HB3 - HG3
6786   6790       1 ;     HB3 - CD
6786   6793       1 ;     HB3 - C
6787   6793       1 ;      CG - C
6788   6791       1 ;     HG2 - OE1
6788   6792       1 ;     HG2 - OE2
6789   6791       1 ;     HG3 - OE1
6789   6792       1 ;     HG3 - OE2
6793   6798       1 ;       C - HA
6793   6799       1 ;       C - CB
6793   6807       1 ;       C - C
6794   6796       1 ;       O - H
6794   6797       1 ;       O - CA
6795   6800       1 ;       N - HB2
6795   6801       1 ;       N - HB3
6795   6802       1 ;       N - CG
6795   6808       1 ;       N - O
6795   6809       1 ;       N - N
6796   6798       1 ;       H - HA
6796   6799       1 ;       H - CB
6796   6807       1 ;       H - C
6797   6803       1 ;      CA - OD1
6797   6804       1 ;      CA - ND2
6797   6810       1 ;      CA - CD
6797   6819       1 ;      CA - CA
6798   6800       1 ;      HA - HB2
6798   6801       1 ;      HA - HB3
6798   6802       1 ;      HA - CG
6798   6808       1 ;      HA - O
6798   6809       1 ;      HA - N
6799   6805       1 ;      CB - HD21
6799   6806       1 ;      CB - HD22
6799   6808       1 ;      CB - O
6799   6809       1 ;      CB - N
6800   6803       1 ;     HB2 - OD1
6800   6804       1 ;     HB2 - ND2
6800   6807       1 ;     HB2 - C
6801   6803       1 ;     HB3 - OD1
6801   6804       1 ;     HB3 - ND2
6801   6807       1 ;     HB3 - C
6802   6807       1 ;      CG - C
6803   6805       1 ;     OD1 - HD21
6803   6806       1 ;     OD1 - HD22
```

```
6807   6811       1 ;       C - HD2
6807   6812       1 ;       C - HD3
6807   6813       1 ;       C - CG
6807   6816       1 ;       C - CB
6807   6820       1 ;       C - HA
6807   6821       1 ;       C - C
6808   6810       1 ;       O - CD
6808   6819       1 ;       O - CA
6809   6814       1 ;       N - HG2
6809   6815       1 ;       N - HG3
6809   6817       1 ;       N - HB2
6809   6818       1 ;       N - HB3
6809   6822       1 ;       N - O
6809   6823       1 ;       N - N
6810   6817       1 ;      CD - HB2
6810   6818       1 ;      CD - HB3
6810   6820       1 ;      CD - HA
6810   6821       1 ;      CD - C
6811   6814       1 ;     HD2 - HG2
6811   6815       1 ;     HD2 - HG3
6811   6816       1 ;     HD2 - CB
6811   6819       1 ;     HD2 - CA
6812   6814       1 ;     HD3 - HG2
6812   6815       1 ;     HD3 - HG3
6812   6816       1 ;     HD3 - CB
6812   6819       1 ;     HD3 - CA
6813   6820       1 ;      CG - HA
6813   6821       1 ;      CG - C
6814   6817       1 ;     HG2 - HB2
6814   6818       1 ;     HG2 - HB3
6814   6819       1 ;     HG2 - CA
6815   6817       1 ;     HG3 - HB2
6815   6818       1 ;     HG3 - HB3
6815   6819       1 ;     HG3 - CA
6816   6822       1 ;      CB - O
6816   6823       1 ;      CB - N
6817   6820       1 ;     HB2 - HA
6817   6821       1 ;     HB2 - C
6818   6820       1 ;     HB3 - HA
6818   6821       1 ;     HB3 - C
6819   6824       1 ;      CA - H
6819   6825       1 ;      CA - CA
6820   6822       1 ;      HA - O
6820   6823       1 ;      HA - N
6821   6826       1 ;       C - HA
6821   6827       1 ;       C - CB
6821   6840       1 ;       C - C
6822   6824       1 ;       O - H
6822   6825       1 ;       O - CA
6823   6828       1 ;       N - HB2
6823   6829       1 ;       N - HB3
6823   6830       1 ;       N - CG
```

```
6823   6841       1 ;       N - O
6823   6842       1 ;       N - N
6824   6826       1 ;       H - HA
6824   6827       1 ;       H - CB
6824   6840       1 ;       H - C
6825   6831       1 ;      CA - HG
6825   6832       1 ;      CA - CD1
6825   6836       1 ;      CA - CD2
6825   6843       1 ;      CA - H
6825   6844       1 ;      CA - CA
6826   6828       1 ;      HA - HB2
6826   6829       1 ;      HA - HB3
6826   6830       1 ;      HA - CG
6826   6841       1 ;      HA - O
6826   6842       1 ;      HA - N
6827   6833       1 ;      CB - HD11
6827   6834       1 ;      CB - HD12
6827   6835       1 ;      CB - HD13
6827   6837       1 ;      CB - HD21
6827   6838       1 ;      CB - HD22
6827   6839       1 ;      CB - HD23
6827   6841       1 ;      CB - O
6827   6842       1 ;      CB - N
6828   6831       1 ;     HB2 - HG
6828   6832       1 ;     HB2 - CD1
6828   6836       1 ;     HB2 - CD2
6828   6840       1 ;     HB2 - C
6829   6831       1 ;     HB3 - HG
6829   6832       1 ;     HB3 - CD1
6829   6836       1 ;     HB3 - CD2
6829   6840       1 ;     HB3 - C
6830   6840       1 ;      CG - C
6831   6833       1 ;      HG - HD11
6831   6834       1 ;      HG - HD12
6831   6835       1 ;      HG - HD13
6831   6837       1 ;      HG - HD21
6831   6838       1 ;      HG - HD22
6831   6839       1 ;      HG - HD23
6832   6837       1 ;     CD1 - HD21
6832   6838       1 ;     CD1 - HD22
6832   6839       1 ;     CD1 - HD23
6833   6836       1 ;    HD11 - CD2
6834   6836       1 ;    HD12 - CD2
6835   6836       1 ;    HD13 - CD2
6840   6845       1 ;       C - HA
6840   6846       1 ;       C - CB
6840   6859       1 ;       C - C
6841   6843       1 ;       O - H
6841   6844       1 ;       O - CA
6842   6847       1 ;       N - HB
6842   6848       1 ;       N - CG2
6842   6852       1 ;       N - CG1
```

```
6842   6860      1 ;      N - O
6842   6861      1 ;      N - N
6843   6845      1 ;      H - HA
6843   6846      1 ;      H - CB
6843   6859      1 ;      H - C
6844   6849      1 ;     CA - HG21
6844   6850      1 ;     CA - HG22
6844   6851      1 ;     CA - HG23
6844   6853      1 ;     CA - HG12
6844   6854      1 ;     CA - HG13
6844   6855      1 ;     CA - CD1
6844   6862      1 ;     CA - H
6844   6863      1 ;     CA - CA
6845   6847      1 ;     HA - HB
6845   6848      1 ;     HA - CG2
6845   6852      1 ;     HA - CG1
6845   6860      1 ;     HA - O
6845   6861      1 ;     HA - N
6846   6856      1 ;     CB - HD11
6846   6857      1 ;     CB - HD12
6846   6858      1 ;     CB - HD13
6846   6860      1 ;     CB - O
6846   6861      1 ;     CB - N
6847   6849      1 ;     HB - HG21
6847   6850      1 ;     HB - HG22
6847   6851      1 ;     HB - HG23
6847   6853      1 ;     HB - HG12
6847   6854      1 ;     HB - HG13
6847   6855      1 ;     HB - CD1
6847   6859      1 ;     HB - C
6848   6853      1 ;    CG2 - HG12
6848   6854      1 ;    CG2 - HG13
6848   6855      1 ;    CG2 - CD1
6848   6859      1 ;    CG2 - C
6849   6852      1 ;   HG21 - CG1
6850   6852      1 ;   HG22 - CG1
6851   6852      1 ;   HG23 - CG1
6852   6859      1 ;    CG1 - C
6853   6856      1 ;   HG12 - HD11
6853   6857      1 ;   HG12 - HD12
6853   6858      1 ;   HG12 - HD13
6854   6856      1 ;   HG13 - HD11
6854   6857      1 ;   HG13 - HD12
6854   6858      1 ;   HG13 - HD13
6859   6864      1 ;      C - HA
6859   6865      1 ;      C - CB
6859   6871      1 ;      C - C
6860   6862      1 ;      O - H
6860   6863      1 ;      O - CA
6861   6866      1 ;      N - HB2
6861   6867      1 ;      N - HB3
6861   6868      1 ;      N - CG
```

```
6861   6872       1 ;       N - O
6861   6873       1 ;       N - N
6862   6864       1 ;       H - HA
6862   6865       1 ;       H - CB
6862   6871       1 ;       H - C
6863   6869       1 ;      CA - OD1
6863   6870       1 ;      CA - OD2
6863   6874       1 ;      CA - H
6863   6875       1 ;      CA - CA
6864   6866       1 ;      HA - HB2
6864   6867       1 ;      HA - HB3
6864   6868       1 ;      HA - CG
6864   6872       1 ;      HA - O
6864   6873       1 ;      HA - N
6865   6872       1 ;      CB - O
6865   6873       1 ;      CB - N
6866   6869       1 ;     HB2 - OD1
6866   6870       1 ;     HB2 - OD2
6866   6871       1 ;     HB2 - C
6867   6869       1 ;     HB3 - OD1
6867   6870       1 ;     HB3 - OD2
6867   6871       1 ;     HB3 - C
6868   6871       1 ;      CG - C
6871   6876       1 ;       C - HA
6871   6877       1 ;       C - CB
6871   6893       1 ;       C - C
6872   6874       1 ;       O - H
6872   6875       1 ;       O - CA
6873   6878       1 ;       N - HB2
6873   6879       1 ;       N - HB3
6873   6880       1 ;       N - CG
6873   6894       1 ;       N - O
6873   6895       1 ;       N - N
6874   6876       1 ;       H - HA
6874   6877       1 ;       H - CB
6874   6893       1 ;       H - C
6875   6881       1 ;      CA - HG2
6875   6882       1 ;      CA - HG3
6875   6883       1 ;      CA - CD
6875   6896       1 ;      CA - H
6875   6897       1 ;      CA - CA
6876   6878       1 ;      HA - HB2
6876   6879       1 ;      HA - HB3
6876   6880       1 ;      HA - CG
6876   6894       1 ;      HA - O
6876   6895       1 ;      HA - N
6877   6884       1 ;      CB - HD2
6877   6885       1 ;      CB - HD3
6877   6886       1 ;      CB - CE
6877   6894       1 ;      CB - O
6877   6895       1 ;      CB - N
6878   6881       1 ;     HB2 - HG2
```

| | | | | |
|---|---|---|---|---|
| 6878 | 6882 | 1 ; | HB2 - | HG3 |
| 6878 | 6883 | 1 ; | HB2 - | CD |
| 6878 | 6893 | 1 ; | HB2 - | C |
| 6879 | 6881 | 1 ; | HB3 - | HG2 |
| 6879 | 6882 | 1 ; | HB3 - | HG3 |
| 6879 | 6883 | 1 ; | HB3 - | CD |
| 6879 | 6893 | 1 ; | HB3 - | C |
| 6880 | 6887 | 1 ; | CG - | HE2 |
| 6880 | 6888 | 1 ; | CG - | HE3 |
| 6880 | 6889 | 1 ; | CG - | NZ |
| 6880 | 6893 | 1 ; | CG - | C |
| 6881 | 6884 | 1 ; | HG2 - | HD2 |
| 6881 | 6885 | 1 ; | HG2 - | HD3 |
| 6881 | 6886 | 1 ; | HG2 - | CE |
| 6882 | 6884 | 1 ; | HG3 - | HD2 |
| 6882 | 6885 | 1 ; | HG3 - | HD3 |
| 6882 | 6886 | 1 ; | HG3 - | CE |
| 6883 | 6890 | 1 ; | CD - | HZ1 |
| 6883 | 6891 | 1 ; | CD - | HZ2 |
| 6883 | 6892 | 1 ; | CD - | HZ3 |
| 6884 | 6887 | 1 ; | HD2 - | HE2 |
| 6884 | 6888 | 1 ; | HD2 - | HE3 |
| 6884 | 6889 | 1 ; | HD2 - | NZ |
| 6885 | 6887 | 1 ; | HD3 - | HE2 |
| 6885 | 6888 | 1 ; | HD3 - | HE3 |
| 6885 | 6889 | 1 ; | HD3 - | NZ |
| 6887 | 6890 | 1 ; | HE2 - | HZ1 |
| 6887 | 6891 | 1 ; | HE2 - | HZ2 |
| 6887 | 6892 | 1 ; | HE2 - | HZ3 |
| 6888 | 6890 | 1 ; | HE3 - | HZ1 |
| 6888 | 6891 | 1 ; | HE3 - | HZ2 |
| 6888 | 6892 | 1 ; | HE3 - | HZ3 |
| 6893 | 6898 | 1 ; | C - | HA |
| 6893 | 6899 | 1 ; | C - | CB |
| 6893 | 6903 | 1 ; | C - | C |
| 6894 | 6896 | 1 ; | O - | H |
| 6894 | 6897 | 1 ; | O - | CA |
| 6895 | 6900 | 1 ; | N - | HB1 |
| 6895 | 6901 | 1 ; | N - | HB2 |
| 6895 | 6902 | 1 ; | N - | HB3 |
| 6895 | 6904 | 1 ; | N - | O |
| 6895 | 6905 | 1 ; | N - | N |
| 6896 | 6898 | 1 ; | H - | HA |
| 6896 | 6899 | 1 ; | H - | CB |
| 6896 | 6903 | 1 ; | H - | C |
| 6897 | 6906 | 1 ; | CA - | H |
| 6897 | 6907 | 1 ; | CA - | CA |
| 6898 | 6900 | 1 ; | HA - | HB1 |
| 6898 | 6901 | 1 ; | HA - | HB2 |
| 6898 | 6902 | 1 ; | HA - | HB3 |
| 6898 | 6904 | 1 ; | HA - | O |
| 6898 | 6905 | 1 ; | HA - | N |

| | | | | | |
|---|---|---|---|---|---|
| 6899 | 6904 | 1 ; | CB | - | O |
| 6899 | 6905 | 1 ; | CB | - | N |
| 6900 | 6903 | 1 ; | HB1 | - | C |
| 6901 | 6903 | 1 ; | HB2 | - | C |
| 6902 | 6903 | 1 ; | HB3 | - | C |
| 6903 | 6908 | 1 ; | C | - | HA |
| 6903 | 6909 | 1 ; | C | - | CB |
| 6903 | 6922 | 1 ; | C | - | C |
| 6904 | 6906 | 1 ; | O | - | H |
| 6904 | 6907 | 1 ; | O | - | CA |
| 6905 | 6910 | 1 ; | N | - | HB |
| 6905 | 6911 | 1 ; | N | - | CG2 |
| 6905 | 6915 | 1 ; | N | - | CG1 |
| 6905 | 6923 | 1 ; | N | - | O |
| 6905 | 6924 | 1 ; | N | - | N |
| 6906 | 6908 | 1 ; | H | - | HA |
| 6906 | 6909 | 1 ; | H | - | CB |
| 6906 | 6922 | 1 ; | H | - | C |
| 6907 | 6912 | 1 ; | CA | - | HG21 |
| 6907 | 6913 | 1 ; | CA | - | HG22 |
| 6907 | 6914 | 1 ; | CA | - | HG23 |
| 6907 | 6916 | 1 ; | CA | - | HG12 |
| 6907 | 6917 | 1 ; | CA | - | HG13 |
| 6907 | 6918 | 1 ; | CA | - | CD1 |
| 6907 | 6925 | 1 ; | CA | - | H |
| 6907 | 6926 | 1 ; | CA | - | CA |
| 6908 | 6910 | 1 ; | HA | - | HB |
| 6908 | 6911 | 1 ; | HA | - | CG2 |
| 6908 | 6915 | 1 ; | HA | - | CG1 |
| 6908 | 6923 | 1 ; | HA | - | O |
| 6908 | 6924 | 1 ; | HA | - | N |
| 6909 | 6919 | 1 ; | CB | - | HD11 |
| 6909 | 6920 | 1 ; | CB | - | HD12 |
| 6909 | 6921 | 1 ; | CB | - | HD13 |
| 6909 | 6923 | 1 ; | CB | - | O |
| 6909 | 6924 | 1 ; | CB | - | N |
| 6910 | 6912 | 1 ; | HB | - | HG21 |
| 6910 | 6913 | 1 ; | HB | - | HG22 |
| 6910 | 6914 | 1 ; | HB | - | HG23 |
| 6910 | 6916 | 1 ; | HB | - | HG12 |
| 6910 | 6917 | 1 ; | HB | - | HG13 |
| 6910 | 6918 | 1 ; | HB | - | CD1 |
| 6910 | 6922 | 1 ; | HB | - | C |
| 6911 | 6916 | 1 ; | CG2 | - | HG12 |
| 6911 | 6917 | 1 ; | CG2 | - | HG13 |
| 6911 | 6918 | 1 ; | CG2 | - | CD1 |
| 6911 | 6922 | 1 ; | CG2 | - | C |
| 6912 | 6915 | 1 ; | HG21 | - | CG1 |
| 6913 | 6915 | 1 ; | HG22 | - | CG1 |
| 6914 | 6915 | 1 ; | HG23 | - | CG1 |
| 6915 | 6922 | 1 ; | CG1 | - | C |
| 6916 | 6919 | 1 ; | HG12 | - | HD11 |

```
6916   6920      1 ;   HG12 - HD12
6916   6921      1 ;   HG12 - HD13
6917   6919      1 ;   HG13 - HD11
6917   6920      1 ;   HG13 - HD12
6917   6921      1 ;   HG13 - HD13
6922   6927      1 ;      C - HA
6922   6928      1 ;      C - CB
6922   6934      1 ;      C - C
6923   6925      1 ;      O - H
6923   6926      1 ;      O - CA
6924   6929      1 ;      N - HB2
6924   6930      1 ;      N - HB3
6924   6931      1 ;      N - CG
6924   6935      1 ;      N - O
6924   6936      1 ;      N - N
6925   6927      1 ;      H - HA
6925   6928      1 ;      H - CB
6925   6934      1 ;      H - C
6926   6932      1 ;     CA - OD1
6926   6933      1 ;     CA - OD2
6926   6937      1 ;     CA - H
6926   6938      1 ;     CA - CA
6927   6929      1 ;     HA - HB2
6927   6930      1 ;     HA - HB3
6927   6931      1 ;     HA - CG
6927   6935      1 ;     HA - O
6927   6936      1 ;     HA - N
6928   6935      1 ;     CB - O
6928   6936      1 ;     CB - N
6929   6932      1 ;    HB2 - OD1
6929   6933      1 ;    HB2 - OD2
6929   6934      1 ;    HB2 - C
6930   6932      1 ;    HB3 - OD1
6930   6933      1 ;    HB3 - OD2
6930   6934      1 ;    HB3 - C
6931   6934      1 ;     CG - C
6934   6939      1 ;      C - HA
6934   6940      1 ;      C - CB
6934   6945      1 ;      C - C
6935   6937      1 ;      O - H
6935   6938      1 ;      O - CA
6936   6941      1 ;      N - HB2
6936   6942      1 ;      N - HB3
6936   6943      1 ;      N - OG
6936   6946      1 ;      N - O
6936   6947      1 ;      N - N
6937   6939      1 ;      H - HA
6937   6940      1 ;      H - CB
6937   6945      1 ;      H - C
6938   6944      1 ;     CA - HG
6938   6948      1 ;     CA - H
6938   6949      1 ;     CA - CA
```

| | | | | |
|---|---|---|---|---|
| 6939 | 6941 | 1 | ; | HA - HB2 |
| 6939 | 6942 | 1 | ; | HA - HB3 |
| 6939 | 6943 | 1 | ; | HA - OG |
| 6939 | 6946 | 1 | ; | HA - O |
| 6939 | 6947 | 1 | ; | HA - N |
| 6940 | 6946 | 1 | ; | CB - O |
| 6940 | 6947 | 1 | ; | CB - N |
| 6941 | 6944 | 1 | ; | HB2 - HG |
| 6941 | 6945 | 1 | ; | HB2 - C |
| 6942 | 6944 | 1 | ; | HB3 - HG |
| 6942 | 6945 | 1 | ; | HB3 - C |
| 6943 | 6945 | 1 | ; | OG - C |
| 6945 | 6950 | 1 | ; | C - HA |
| 6945 | 6951 | 1 | ; | C - CB |
| 6945 | 6964 | 1 | ; | C - C |
| 6946 | 6948 | 1 | ; | O - H |
| 6946 | 6949 | 1 | ; | O - CA |
| 6947 | 6952 | 1 | ; | N - HB2 |
| 6947 | 6953 | 1 | ; | N - HB3 |
| 6947 | 6954 | 1 | ; | N - CG |
| 6947 | 6965 | 1 | ; | N - O |
| 6947 | 6966 | 1 | ; | N - N |
| 6948 | 6950 | 1 | ; | H - HA |
| 6948 | 6951 | 1 | ; | H - CB |
| 6948 | 6964 | 1 | ; | H - C |
| 6949 | 6955 | 1 | ; | CA - HG |
| 6949 | 6956 | 1 | ; | CA - CD1 |
| 6949 | 6960 | 1 | ; | CA - CD2 |
| 6949 | 6967 | 1 | ; | CA - H |
| 6949 | 6968 | 1 | ; | CA - CA |
| 6950 | 6952 | 1 | ; | HA - HB2 |
| 6950 | 6953 | 1 | ; | HA - HB3 |
| 6950 | 6954 | 1 | ; | HA - CG |
| 6950 | 6965 | 1 | ; | HA - O |
| 6950 | 6966 | 1 | ; | HA - N |
| 6951 | 6957 | 1 | ; | CB - HD11 |
| 6951 | 6958 | 1 | ; | CB - HD12 |
| 6951 | 6959 | 1 | ; | CB - HD13 |
| 6951 | 6961 | 1 | ; | CB - HD21 |
| 6951 | 6962 | 1 | ; | CB - HD22 |
| 6951 | 6963 | 1 | ; | CB - HD23 |
| 6951 | 6965 | 1 | ; | CB - O |
| 6951 | 6966 | 1 | ; | CB - N |
| 6952 | 6955 | 1 | ; | HB2 - HG |
| 6952 | 6956 | 1 | ; | HB2 - CD1 |
| 6952 | 6960 | 1 | ; | HB2 - CD2 |
| 6952 | 6964 | 1 | ; | HB2 - C |
| 6953 | 6955 | 1 | ; | HB3 - HG |
| 6953 | 6956 | 1 | ; | HB3 - CD1 |
| 6953 | 6960 | 1 | ; | HB3 - CD2 |
| 6953 | 6964 | 1 | ; | HB3 - C |
| 6954 | 6964 | 1 | ; | CG - C |

```
6955   6957       1 ;      HG - HD11
6955   6958       1 ;      HG - HD12
6955   6959       1 ;      HG - HD13
6955   6961       1 ;      HG - HD21
6955   6962       1 ;      HG - HD22
6955   6963       1 ;      HG - HD23
6956   6961       1 ;     CD1 - HD21
6956   6962       1 ;     CD1 - HD22
6956   6963       1 ;     CD1 - HD23
6957   6960       1 ;    HD11 - CD2
6958   6960       1 ;    HD12 - CD2
6959   6960       1 ;    HD13 - CD2
6964   6969       1 ;       C - HA
6964   6970       1 ;       C - CB
6964   6983       1 ;       C - C
6965   6967       1 ;       O - H
6965   6968       1 ;       O - CA
6966   6971       1 ;       N - HB2
6966   6972       1 ;       N - HB3
6966   6973       1 ;       N - CG
6966   6984       1 ;       N - O
6966   6985       1 ;       N - N
6967   6969       1 ;       H - HA
6967   6970       1 ;       H - CB
6967   6983       1 ;       H - C
6968   6974       1 ;      CA - HG
6968   6975       1 ;      CA - CD1
6968   6979       1 ;      CA - CD2
6968   6986       1 ;      CA - H
6968   6987       1 ;      CA - CA
6969   6971       1 ;      HA - HB2
6969   6972       1 ;      HA - HB3
6969   6973       1 ;      HA - CG
6969   6984       1 ;      HA - O
6969   6985       1 ;      HA - N
6970   6976       1 ;      CB - HD11
6970   6977       1 ;      CB - HD12
6970   6978       1 ;      CB - HD13
6970   6980       1 ;      CB - HD21
6970   6981       1 ;      CB - HD22
6970   6982       1 ;      CB - HD23
6970   6984       1 ;      CB - O
6970   6985       1 ;      CB - N
6971   6974       1 ;     HB2 - HG
6971   6975       1 ;     HB2 - CD1
6971   6979       1 ;     HB2 - CD2
6971   6983       1 ;     HB2 - C
6972   6974       1 ;     HB3 - HG
6972   6975       1 ;     HB3 - CD1
6972   6979       1 ;     HB3 - CD2
6972   6983       1 ;     HB3 - C
6973   6983       1 ;      CG - C
```

```
6974   6976      1 ;     HG - HD11
6974   6977      1 ;     HG - HD12
6974   6978      1 ;     HG - HD13
6974   6980      1 ;     HG - HD21
6974   6981      1 ;     HG - HD22
6974   6982      1 ;     HG - HD23
6975   6980      1 ;    CD1 - HD21
6975   6981      1 ;    CD1 - HD22
6975   6982      1 ;    CD1 - HD23
6976   6979      1 ;   HD11 - CD2
6977   6979      1 ;   HD12 - CD2
6978   6979      1 ;   HD13 - CD2
6983   6988      1 ;      C - HA
6983   6989      1 ;      C - CB
6983   7000      1 ;      C - C
6984   6986      1 ;      O - H
6984   6987      1 ;      O - CA
6985   6990      1 ;      N - HB2
6985   6991      1 ;      N - HB3
6985   6992      1 ;      N - CG
6985   7001      1 ;      N - O
6985   7002      1 ;      N - N
6986   6988      1 ;      H - HA
6986   6989      1 ;      H - CB
6986   7000      1 ;      H - C
6987   6993      1 ;     CA - HG2
6987   6994      1 ;     CA - HG3
6987   6995      1 ;     CA - CD
6987   7003      1 ;     CA - H
6987   7004      1 ;     CA - CA
6988   6990      1 ;     HA - HB2
6988   6991      1 ;     HA - HB3
6988   6992      1 ;     HA - CG
6988   7001      1 ;     HA - O
6988   7002      1 ;     HA - N
6989   6996      1 ;     CB - OE1
6989   6997      1 ;     CB - NE2
6989   7001      1 ;     CB - O
6989   7002      1 ;     CB - N
6990   6993      1 ;    HB2 - HG2
6990   6994      1 ;    HB2 - HG3
6990   6995      1 ;    HB2 - CD
6990   7000      1 ;    HB2 - C
6991   6993      1 ;    HB3 - HG2
6991   6994      1 ;    HB3 - HG3
6991   6995      1 ;    HB3 - CD
6991   7000      1 ;    HB3 - C
6992   6998      1 ;     CG - HE21
6992   6999      1 ;     CG - HE22
6992   7000      1 ;     CG - C
6993   6996      1 ;    HG2 - OE1
6993   6997      1 ;    HG2 - NE2
```

```
6994   6996       1 ;    HG3 - OE1
6994   6997       1 ;    HG3 - NE2
6996   6998       1 ;    OE1 - HE21
6996   6999       1 ;    OE1 - HE22
7000   7005       1 ;      C - HA
7000   7006       1 ;      C - CB
7000   7015       1 ;      C - C
7001   7003       1 ;      O - H
7001   7004       1 ;      O - CA
7002   7007       1 ;      N - HB2
7002   7008       1 ;      N - HB3
7002   7009       1 ;      N - CG
7002   7016       1 ;      N - O
7002   7017       1 ;      N - N
7003   7005       1 ;      H - HA
7003   7006       1 ;      H - CB
7003   7015       1 ;      H - C
7004   7010       1 ;     CA - HG2
7004   7011       1 ;     CA - HG3
7004   7012       1 ;     CA - CD
7004   7018       1 ;     CA - H
7004   7019       1 ;     CA - CA
7005   7007       1 ;     HA - HB2
7005   7008       1 ;     HA - HB3
7005   7009       1 ;     HA - CG
7005   7016       1 ;     HA - O
7005   7017       1 ;     HA - N
7006   7013       1 ;     CB - OE1
7006   7014       1 ;     CB - OE2
7006   7016       1 ;     CB - O
7006   7017       1 ;     CB - N
7007   7010       1 ;    HB2 - HG2
7007   7011       1 ;    HB2 - HG3
7007   7012       1 ;    HB2 - CD
7007   7015       1 ;    HB2 - C
7008   7010       1 ;    HB3 - HG2
7008   7011       1 ;    HB3 - HG3
7008   7012       1 ;    HB3 - CD
7008   7015       1 ;    HB3 - C
7009   7015       1 ;     CG - C
7010   7013       1 ;    HG2 - OE1
7010   7014       1 ;    HG2 - OE2
7011   7013       1 ;    HG3 - OE1
7011   7014       1 ;    HG3 - OE2
7015   7020       1 ;      C - HA
7015   7021       1 ;      C - CB
7015   7031       1 ;      C - C
7016   7018       1 ;      O - H
7016   7019       1 ;      O - CA
7017   7022       1 ;      N - HB
7017   7023       1 ;      N - CG1
7017   7027       1 ;      N - CG2
```

| | | | | | |
|---|---|---|---|---|---|
| 7017 | 7032 | 1 ; | N | - | O |
| 7017 | 7033 | 1 ; | N | - | N |
| 7018 | 7020 | 1 ; | H | - | HA |
| 7018 | 7021 | 1 ; | H | - | CB |
| 7018 | 7031 | 1 ; | H | - | C |
| 7019 | 7024 | 1 ; | CA | - | HG11 |
| 7019 | 7025 | 1 ; | CA | - | HG12 |
| 7019 | 7026 | 1 ; | CA | - | HG13 |
| 7019 | 7028 | 1 ; | CA | - | HG21 |
| 7019 | 7029 | 1 ; | CA | - | HG22 |
| 7019 | 7030 | 1 ; | CA | - | HG23 |
| 7019 | 7034 | 1 ; | CA | - | H |
| 7019 | 7035 | 1 ; | CA | - | CA |
| 7020 | 7022 | 1 ; | HA | - | HB |
| 7020 | 7023 | 1 ; | HA | - | CG1 |
| 7020 | 7027 | 1 ; | HA | - | CG2 |
| 7020 | 7032 | 1 ; | HA | - | O |
| 7020 | 7033 | 1 ; | HA | - | N |
| 7021 | 7032 | 1 ; | CB | - | O |
| 7021 | 7033 | 1 ; | CB | - | N |
| 7022 | 7024 | 1 ; | HB | - | HG11 |
| 7022 | 7025 | 1 ; | HB | - | HG12 |
| 7022 | 7026 | 1 ; | HB | - | HG13 |
| 7022 | 7028 | 1 ; | HB | - | HG21 |
| 7022 | 7029 | 1 ; | HB | - | HG22 |
| 7022 | 7030 | 1 ; | HB | - | HG23 |
| 7022 | 7031 | 1 ; | HB | - | C |
| 7023 | 7028 | 1 ; | CG1 | - | HG21 |
| 7023 | 7029 | 1 ; | CG1 | - | HG22 |
| 7023 | 7030 | 1 ; | CG1 | - | HG23 |
| 7023 | 7031 | 1 ; | CG1 | - | C |
| 7024 | 7027 | 1 ; | HG11 | - | CG2 |
| 7025 | 7027 | 1 ; | HG12 | - | CG2 |
| 7026 | 7027 | 1 ; | HG13 | - | CG2 |
| 7027 | 7031 | 1 ; | CG2 | - | C |
| 7031 | 7036 | 1 ; | C | - | HA |
| 7031 | 7037 | 1 ; | C | - | CB |
| 7031 | 7043 | 1 ; | C | - | C |
| 7032 | 7034 | 1 ; | O | - | H |
| 7032 | 7035 | 1 ; | O | - | CA |
| 7033 | 7038 | 1 ; | N | - | HB2 |
| 7033 | 7039 | 1 ; | N | - | HB3 |
| 7033 | 7040 | 1 ; | N | - | CG |
| 7033 | 7044 | 1 ; | N | - | O |
| 7033 | 7045 | 1 ; | N | - | N |
| 7034 | 7036 | 1 ; | H | - | HA |
| 7034 | 7037 | 1 ; | H | - | CB |
| 7034 | 7043 | 1 ; | H | - | C |
| 7035 | 7041 | 1 ; | CA | - | OD1 |
| 7035 | 7042 | 1 ; | CA | - | OD2 |
| 7035 | 7046 | 1 ; | CA | - | H |
| 7035 | 7047 | 1 ; | CA | - | CA |

```
7036   7038      1 ;     HA - HB2
7036   7039      1 ;     HA - HB3
7036   7040      1 ;     HA - CG
7036   7044      1 ;     HA - O
7036   7045      1 ;     HA - N
7037   7044      1 ;     CB - O
7037   7045      1 ;     CB - N
7038   7041      1 ;    HB2 - OD1
7038   7042      1 ;    HB2 - OD2
7038   7043      1 ;    HB2 - C
7039   7041      1 ;    HB3 - OD1
7039   7042      1 ;    HB3 - OD2
7039   7043      1 ;    HB3 - C
7040   7043      1 ;     CG - C
7043   7048      1 ;      C - HA
7043   7049      1 ;      C - CB
7043   7062      1 ;      C - C
7044   7046      1 ;      O - H
7044   7047      1 ;      O - CA
7045   7050      1 ;      N - HB2
7045   7051      1 ;      N - HB3
7045   7052      1 ;      N - CG
7045   7063      1 ;      N - O
7045   7064      1 ;      N - N
7046   7048      1 ;      H - HA
7046   7049      1 ;      H - CB
7046   7062      1 ;      H - C
7047   7053      1 ;     CA - HG
7047   7054      1 ;     CA - CD1
7047   7058      1 ;     CA - CD2
7047   7065      1 ;     CA - H
7047   7066      1 ;     CA - CA
7048   7050      1 ;     HA - HB2
7048   7051      1 ;     HA - HB3
7048   7052      1 ;     HA - CG
7048   7063      1 ;     HA - O
7048   7064      1 ;     HA - N
7049   7055      1 ;     CB - HD11
7049   7056      1 ;     CB - HD12
7049   7057      1 ;     CB - HD13
7049   7059      1 ;     CB - HD21
7049   7060      1 ;     CB - HD22
7049   7061      1 ;     CB - HD23
7049   7063      1 ;     CB - O
7049   7064      1 ;     CB - N
7050   7053      1 ;    HB2 - HG
7050   7054      1 ;    HB2 - CD1
7050   7058      1 ;    HB2 - CD2
7050   7062      1 ;    HB2 - C
7051   7053      1 ;    HB3 - HG
7051   7054      1 ;    HB3 - CD1
7051   7058      1 ;    HB3 - CD2
```

```
7051   7062       1 ;     HB3 - C
7052   7062       1 ;      CG - C
7053   7055       1 ;      HG - HD11
7053   7056       1 ;      HG - HD12
7053   7057       1 ;      HG - HD13
7053   7059       1 ;      HG - HD21
7053   7060       1 ;      HG - HD22
7053   7061       1 ;      HG - HD23
7054   7059       1 ;     CD1 - HD21
7054   7060       1 ;     CD1 - HD22
7054   7061       1 ;     CD1 - HD23
7055   7058       1 ;    HD11 - CD2
7056   7058       1 ;    HD12 - CD2
7057   7058       1 ;    HD13 - CD2
7062   7067       1 ;       C - HA
7062   7068       1 ;       C - CB
7062   7076       1 ;       C - C
7063   7065       1 ;       O - H
7063   7066       1 ;       O - CA
7064   7069       1 ;       N - HB2
7064   7070       1 ;       N - HB3
7064   7071       1 ;       N - CG
7064   7077       1 ;       N - O
7064   7078       1 ;       N - N
7065   7067       1 ;       H - HA
7065   7068       1 ;       H - CB
7065   7076       1 ;       H - C
7066   7072       1 ;      CA - OD1
7066   7073       1 ;      CA - ND2
7066   7079       1 ;      CA - H
7066   7080       1 ;      CA - CA
7067   7069       1 ;      HA - HB2
7067   7070       1 ;      HA - HB3
7067   7071       1 ;      HA - CG
7067   7077       1 ;      HA - O
7067   7078       1 ;      HA - N
7068   7074       1 ;      CB - HD21
7068   7075       1 ;      CB - HD22
7068   7077       1 ;      CB - O
7068   7078       1 ;      CB - N
7069   7072       1 ;     HB2 - OD1
7069   7073       1 ;     HB2 - ND2
7069   7076       1 ;     HB2 - C
7070   7072       1 ;     HB3 - OD1
7070   7073       1 ;     HB3 - ND2
7070   7076       1 ;     HB3 - C
7071   7076       1 ;      CG - C
7072   7074       1 ;     OD1 - HD21
7072   7075       1 ;     OD1 - HD22
7076   7081       1 ;       C - HA2
7076   7082       1 ;       C - HA3
7076   7083       1 ;       C - C
```

| | | | |
|---|---|---|---|
| 7077 | 7079 | 1 ; | O - H |
| 7077 | 7080 | 1 ; | O - CA |
| 7078 | 7084 | 1 ; | N - O |
| 7078 | 7085 | 1 ; | N - N |
| 7079 | 7081 | 1 ; | H - HA2 |
| 7079 | 7082 | 1 ; | H - HA3 |
| 7079 | 7083 | 1 ; | H - C |
| 7080 | 7086 | 1 ; | CA - H |
| 7080 | 7087 | 1 ; | CA - CA |
| 7081 | 7084 | 1 ; | HA2 - O |
| 7081 | 7085 | 1 ; | HA2 - N |
| 7082 | 7084 | 1 ; | HA3 - O |
| 7082 | 7085 | 1 ; | HA3 - N |
| 7083 | 7088 | 1 ; | C - HA |
| 7083 | 7089 | 1 ; | C - CB |
| 7083 | 7095 | 1 ; | C - C |
| 7084 | 7086 | 1 ; | O - H |
| 7084 | 7087 | 1 ; | O - CA |
| 7085 | 7090 | 1 ; | N - HB2 |
| 7085 | 7091 | 1 ; | N - HB3 |
| 7085 | 7092 | 1 ; | N - CG |
| 7085 | 7096 | 1 ; | N - O |
| 7085 | 7097 | 1 ; | N - N |
| 7086 | 7088 | 1 ; | H - HA |
| 7086 | 7089 | 1 ; | H - CB |
| 7086 | 7095 | 1 ; | H - C |
| 7087 | 7093 | 1 ; | CA - OD1 |
| 7087 | 7094 | 1 ; | CA - OD2 |
| 7087 | 7098 | 1 ; | CA - H |
| 7087 | 7099 | 1 ; | CA - CA |
| 7088 | 7090 | 1 ; | HA - HB2 |
| 7088 | 7091 | 1 ; | HA - HB3 |
| 7088 | 7092 | 1 ; | HA - CG |
| 7088 | 7096 | 1 ; | HA - O |
| 7088 | 7097 | 1 ; | HA - N |
| 7089 | 7096 | 1 ; | CB - O |
| 7089 | 7097 | 1 ; | CB - N |
| 7090 | 7093 | 1 ; | HB2 - OD1 |
| 7090 | 7094 | 1 ; | HB2 - OD2 |
| 7090 | 7095 | 1 ; | HB2 - C |
| 7091 | 7093 | 1 ; | HB3 - OD1 |
| 7091 | 7094 | 1 ; | HB3 - OD2 |
| 7091 | 7095 | 1 ; | HB3 - C |
| 7092 | 7095 | 1 ; | CG - C |
| 7095 | 7100 | 1 ; | C - HA2 |
| 7095 | 7101 | 1 ; | C - HA3 |
| 7095 | 7102 | 1 ; | C - C |
| 7096 | 7098 | 1 ; | O - H |
| 7096 | 7099 | 1 ; | O - CA |
| 7097 | 7103 | 1 ; | N - O |
| 7097 | 7104 | 1 ; | N - N |
| 7098 | 7100 | 1 ; | H - HA2 |

```
7098   7101      1 ;      H - HA3
7098   7102      1 ;      H - C
7099   7105      1 ;     CA - H
7099   7106      1 ;     CA - CA
7100   7103      1 ;    HA2 - O
7100   7104      1 ;    HA2 - N
7101   7103      1 ;    HA3 - O
7101   7104      1 ;    HA3 - N
7102   7107      1 ;      C - HA
7102   7108      1 ;      C - CB
7102   7117      1 ;      C - C
7103   7105      1 ;      O - H
7103   7106      1 ;      O - CA
7104   7109      1 ;      N - HB2
7104   7110      1 ;      N - HB3
7104   7111      1 ;      N - CG
7104   7118      1 ;      N - O
7104   7119      1 ;      N - N
7105   7107      1 ;      H - HA
7105   7108      1 ;      H - CB
7105   7117      1 ;      H - C
7106   7112      1 ;     CA - HG2
7106   7113      1 ;     CA - HG3
7106   7114      1 ;     CA - CD
7106   7120      1 ;     CA - H
7106   7121      1 ;     CA - CA
7107   7109      1 ;     HA - HB2
7107   7110      1 ;     HA - HB3
7107   7111      1 ;     HA - CG
7107   7118      1 ;     HA - O
7107   7119      1 ;     HA - N
7108   7115      1 ;     CB - OE1
7108   7116      1 ;     CB - OE2
7108   7118      1 ;     CB - O
7108   7119      1 ;     CB - N
7109   7112      1 ;    HB2 - HG2
7109   7113      1 ;    HB2 - HG3
7109   7114      1 ;    HB2 - CD
7109   7117      1 ;    HB2 - C
7110   7112      1 ;    HB3 - HG2
7110   7113      1 ;    HB3 - HG3
7110   7114      1 ;    HB3 - CD
7110   7117      1 ;    HB3 - C
7111   7117      1 ;     CG - C
7112   7115      1 ;    HG2 - OE1
7112   7116      1 ;    HG2 - OE2
7113   7115      1 ;    HG3 - OE1
7113   7116      1 ;    HG3 - OE2
7117   7122      1 ;      C - HA
7117   7123      1 ;      C - CB
7117   7136      1 ;      C - C
7118   7120      1 ;      O - H
```

| | | | | |
|---|---|---|---|---|
| 7118 | 7121 | 1 ; | O - | CA |
| 7119 | 7124 | 1 ; | N - | HB |
| 7119 | 7125 | 1 ; | N - | CG2 |
| 7119 | 7129 | 1 ; | N - | CG1 |
| 7119 | 7137 | 1 ; | N - | O |
| 7119 | 7138 | 1 ; | N - | N |
| 7120 | 7122 | 1 ; | H - | HA |
| 7120 | 7123 | 1 ; | H - | CB |
| 7120 | 7136 | 1 ; | H - | C |
| 7121 | 7126 | 1 ; | CA - | HG21 |
| 7121 | 7127 | 1 ; | CA - | HG22 |
| 7121 | 7128 | 1 ; | CA - | HG23 |
| 7121 | 7130 | 1 ; | CA - | HG12 |
| 7121 | 7131 | 1 ; | CA - | HG13 |
| 7121 | 7132 | 1 ; | CA - | CD1 |
| 7121 | 7139 | 1 ; | CA - | H |
| 7121 | 7140 | 1 ; | CA - | CA |
| 7122 | 7124 | 1 ; | HA - | HB |
| 7122 | 7125 | 1 ; | HA - | CG2 |
| 7122 | 7129 | 1 ; | HA - | CG1 |
| 7122 | 7137 | 1 ; | HA - | O |
| 7122 | 7138 | 1 ; | HA - | N |
| 7123 | 7133 | 1 ; | CB - | HD11 |
| 7123 | 7134 | 1 ; | CB - | HD12 |
| 7123 | 7135 | 1 ; | CB - | HD13 |
| 7123 | 7137 | 1 ; | CB - | O |
| 7123 | 7138 | 1 ; | CB - | N |
| 7124 | 7126 | 1 ; | HB - | HG21 |
| 7124 | 7127 | 1 ; | HB - | HG22 |
| 7124 | 7128 | 1 ; | HB - | HG23 |
| 7124 | 7130 | 1 ; | HB - | HG12 |
| 7124 | 7131 | 1 ; | HB - | HG13 |
| 7124 | 7132 | 1 ; | HB - | CD1 |
| 7124 | 7136 | 1 ; | HB - | C |
| 7125 | 7130 | 1 ; | CG2 - | HG12 |
| 7125 | 7131 | 1 ; | CG2 - | HG13 |
| 7125 | 7132 | 1 ; | CG2 - | CD1 |
| 7125 | 7136 | 1 ; | CG2 - | C |
| 7126 | 7129 | 1 ; | HG21 - | CG1 |
| 7127 | 7129 | 1 ; | HG22 - | CG1 |
| 7128 | 7129 | 1 ; | HG23 - | CG1 |
| 7129 | 7136 | 1 ; | CG1 - | C |
| 7130 | 7133 | 1 ; | HG12 - | HD11 |
| 7130 | 7134 | 1 ; | HG12 - | HD12 |
| 7130 | 7135 | 1 ; | HG12 - | HD13 |
| 7131 | 7133 | 1 ; | HG13 - | HD11 |
| 7131 | 7134 | 1 ; | HG13 - | HD12 |
| 7131 | 7135 | 1 ; | HG13 - | HD13 |
| 7136 | 7141 | 1 ; | C - | HA |
| 7136 | 7142 | 1 ; | C - | CB |
| 7136 | 7148 | 1 ; | C - | C |
| 7137 | 7139 | 1 ; | O - | H |

```
 7137   7140       1 ;       O - CA
 7138   7143       1 ;       N - HB2
 7138   7144       1 ;       N - HB3
 7138   7145       1 ;       N - CG
 7138   7149       1 ;       N - O
 7138   7150       1 ;       N - N
 7139   7141       1 ;       H - HA
 7139   7142       1 ;       H - CB
 7139   7148       1 ;       H - C
 7140   7146       1 ;      CA - OD1
 7140   7147       1 ;      CA - OD2
 7140   7151       1 ;      CA - H
 7140   7152       1 ;      CA - CA
 7141   7143       1 ;      HA - HB2
 7141   7144       1 ;      HA - HB3
 7141   7145       1 ;      HA - CG
 7141   7149       1 ;      HA - O
 7141   7150       1 ;      HA - N
 7142   7149       1 ;      CB - O
 7142   7150       1 ;      CB - N
 7143   7146       1 ;     HB2 - OD1
 7143   7147       1 ;     HB2 - OD2
 7143   7148       1 ;     HB2 - C
 7144   7146       1 ;     HB3 - OD1
 7144   7147       1 ;     HB3 - OD2
 7144   7148       1 ;     HB3 - C
 7145   7148       1 ;      CG - C
 7148   7153       1 ;       C - HA
 7148   7154       1 ;       C - CB
 7148   7168       1 ;       C - C
 7149   7151       1 ;       O - H
 7149   7152       1 ;       O - CA
 7150   7155       1 ;       N - HB2
 7150   7156       1 ;       N - HB3
 7150   7157       1 ;       N - CG
 7150   7169       1 ;       N - O
 7150   7170       1 ;       N - N
 7151   7153       1 ;       H - HA
 7151   7154       1 ;       H - CB
 7151   7168       1 ;       H - C
 7152   7158       1 ;      CA - CD1
 7152   7166       1 ;      CA - CD2
 7152   7171       1 ;      CA - H
 7152   7172       1 ;      CA - CA
 7153   7155       1 ;      HA - HB2
 7153   7156       1 ;      HA - HB3
 7153   7157       1 ;      HA - CG
 7153   7169       1 ;      HA - O
 7153   7170       1 ;      HA - N
 7154   7159       1 ;      CB - HD1
 7154   7160       1 ;      CB - CE1
 7154   7164       1 ;      CB - CE2
```

| | | | |
|---|---|---|---|
| 7154 | 7167 | 1 ; | CB - HD2 |
| 7154 | 7169 | 1 ; | CB - O |
| 7154 | 7170 | 1 ; | CB - N |
| 7155 | 7158 | 1 ; | HB2 - CD1 |
| 7155 | 7166 | 1 ; | HB2 - CD2 |
| 7155 | 7168 | 1 ; | HB2 - C |
| 7156 | 7158 | 1 ; | HB3 - CD1 |
| 7156 | 7166 | 1 ; | HB3 - CD2 |
| 7156 | 7168 | 1 ; | HB3 - C |
| 7157 | 7161 | 1 ; | CG - HE1 |
| 7157 | 7162 | 1 ; | CG - CZ |
| 7157 | 7165 | 1 ; | CG - HE2 |
| 7157 | 7168 | 1 ; | CG - C |
| 7158 | 7163 | 1 ; | CD1 - HZ |
| 7158 | 7164 | 1 ; | CD1 - CE2 |
| 7158 | 7167 | 1 ; | CD1 - HD2 |
| 7159 | 7161 | 1 ; | HD1 - HE1 |
| 7159 | 7162 | 1 ; | HD1 - CZ |
| 7159 | 7166 | 1 ; | HD1 - CD2 |
| 7160 | 7165 | 1 ; | CE1 - HE2 |
| 7160 | 7166 | 1 ; | CE1 - CD2 |
| 7161 | 7163 | 1 ; | HE1 - HZ |
| 7161 | 7164 | 1 ; | HE1 - CE2 |
| 7162 | 7167 | 1 ; | CZ - HD2 |
| 7163 | 7165 | 1 ; | HZ - HE2 |
| 7163 | 7166 | 1 ; | HZ - CD2 |
| 7165 | 7167 | 1 ; | HE2 - HD2 |
| 7168 | 7173 | 1 ; | C - HA |
| 7168 | 7174 | 1 ; | C - CB |
| 7168 | 7186 | 1 ; | C - C |
| 7169 | 7171 | 1 ; | O - H |
| 7169 | 7172 | 1 ; | O - CA |
| 7170 | 7175 | 1 ; | N - HB2 |
| 7170 | 7176 | 1 ; | N - HB3 |
| 7170 | 7177 | 1 ; | N - CG |
| 7170 | 7187 | 1 ; | N - O |
| 7170 | 7188 | 1 ; | N - N |
| 7171 | 7173 | 1 ; | H - HA |
| 7171 | 7174 | 1 ; | H - CB |
| 7171 | 7186 | 1 ; | H - C |
| 7172 | 7178 | 1 ; | CA - ND1 |
| 7172 | 7184 | 1 ; | CA - CD2 |
| 7172 | 7189 | 1 ; | CA - H |
| 7172 | 7190 | 1 ; | CA - CA |
| 7173 | 7175 | 1 ; | HA - HB2 |
| 7173 | 7176 | 1 ; | HA - HB3 |
| 7173 | 7177 | 1 ; | HA - CG |
| 7173 | 7187 | 1 ; | HA - O |
| 7173 | 7188 | 1 ; | HA - N |
| 7174 | 7179 | 1 ; | CB - HD1 |
| 7174 | 7180 | 1 ; | CB - CE1 |
| 7174 | 7182 | 1 ; | CB - NE2 |

| | | | |
|---|---|---|---|
| 7174 | 7185 | 1 ; | CB - HD2 |
| 7174 | 7187 | 1 ; | CB - O |
| 7174 | 7188 | 1 ; | CB - N |
| 7175 | 7178 | 1 ; | HB2 - ND1 |
| 7175 | 7184 | 1 ; | HB2 - CD2 |
| 7175 | 7186 | 1 ; | HB2 - C |
| 7176 | 7178 | 1 ; | HB3 - ND1 |
| 7176 | 7184 | 1 ; | HB3 - CD2 |
| 7176 | 7186 | 1 ; | HB3 - C |
| 7177 | 7181 | 1 ; | CG - HE1 |
| 7177 | 7183 | 1 ; | CG - HE2 |
| 7177 | 7186 | 1 ; | CG - C |
| 7178 | 7183 | 1 ; | ND1 - HE2 |
| 7178 | 7185 | 1 ; | ND1 - HD2 |
| 7179 | 7181 | 1 ; | HD1 - HE1 |
| 7179 | 7182 | 1 ; | HD1 - NE2 |
| 7179 | 7184 | 1 ; | HD1 - CD2 |
| 7180 | 7185 | 1 ; | CE1 - HD2 |
| 7181 | 7183 | 1 ; | HE1 - HE2 |
| 7181 | 7184 | 1 ; | HE1 - CD2 |
| 7183 | 7185 | 1 ; | HE2 - HD2 |
| 7186 | 7191 | 1 ; | C - HA |
| 7186 | 7192 | 1 ; | C - CB |
| 7186 | 7202 | 1 ; | C - C |
| 7187 | 7189 | 1 ; | O - H |
| 7187 | 7190 | 1 ; | O - CA |
| 7188 | 7193 | 1 ; | N - HB2 |
| 7188 | 7194 | 1 ; | N - HB3 |
| 7188 | 7195 | 1 ; | N - CG |
| 7188 | 7203 | 1 ; | N - O |
| 7188 | 7204 | 1 ; | N - N |
| 7189 | 7191 | 1 ; | H - HA |
| 7189 | 7192 | 1 ; | H - CB |
| 7189 | 7202 | 1 ; | H - C |
| 7190 | 7196 | 1 ; | CA - HG2 |
| 7190 | 7197 | 1 ; | CA - HG3 |
| 7190 | 7198 | 1 ; | CA - CD |
| 7190 | 7205 | 1 ; | CA - H |
| 7190 | 7206 | 1 ; | CA - CA |
| 7191 | 7193 | 1 ; | HA - HB2 |
| 7191 | 7194 | 1 ; | HA - HB3 |
| 7191 | 7195 | 1 ; | HA - CG |
| 7191 | 7203 | 1 ; | HA - O |
| 7191 | 7204 | 1 ; | HA - N |
| 7192 | 7199 | 1 ; | CB - OE1 |
| 7192 | 7200 | 1 ; | CB - OE2 |
| 7192 | 7203 | 1 ; | CB - O |
| 7192 | 7204 | 1 ; | CB - N |
| 7193 | 7196 | 1 ; | HB2 - HG2 |
| 7193 | 7197 | 1 ; | HB2 - HG3 |
| 7193 | 7198 | 1 ; | HB2 - CD |
| 7193 | 7202 | 1 ; | HB2 - C |

```
 7194   7196       1 ;    HB3 - HG2
 7194   7197       1 ;    HB3 - HG3
 7194   7198       1 ;    HB3 - CD
 7194   7202       1 ;    HB3 - C
 7195   7201       1 ;     CG - HE2
 7195   7202       1 ;     CG - C
 7196   7199       1 ;    HG2 - OE1
 7196   7200       1 ;    HG2 - OE2
 7197   7199       1 ;    HG3 - OE1
 7197   7200       1 ;    HG3 - OE2
 7199   7201       1 ;    OE1 - HE2
 7202   7207       1 ;      C - HA
 7202   7208       1 ;      C - CB
 7202   7222       1 ;      C - C
 7203   7205       1 ;      O - H
 7203   7206       1 ;      O - CA
 7204   7209       1 ;      N - HB2
 7204   7210       1 ;      N - HB3
 7204   7211       1 ;      N - CG
 7204   7223       1 ;      N - O
 7204   7224       1 ;      N - N
 7205   7207       1 ;      H - HA
 7205   7208       1 ;      H - CB
 7205   7222       1 ;      H - C
 7206   7212       1 ;     CA - CD1
 7206   7220       1 ;     CA - CD2
 7206   7225       1 ;     CA - H
 7206   7226       1 ;     CA - CA
 7207   7209       1 ;     HA - HB2
 7207   7210       1 ;     HA - HB3
 7207   7211       1 ;     HA - CG
 7207   7223       1 ;     HA - O
 7207   7224       1 ;     HA - N
 7208   7213       1 ;     CB - HD1
 7208   7214       1 ;     CB - CE1
 7208   7218       1 ;     CB - CE2
 7208   7221       1 ;     CB - HD2
 7208   7223       1 ;     CB - O
 7208   7224       1 ;     CB - N
 7209   7212       1 ;    HB2 - CD1
 7209   7220       1 ;    HB2 - CD2
 7209   7222       1 ;    HB2 - C
 7210   7212       1 ;    HB3 - CD1
 7210   7220       1 ;    HB3 - CD2
 7210   7222       1 ;    HB3 - C
 7211   7215       1 ;     CG - HE1
 7211   7216       1 ;     CG - CZ
 7211   7219       1 ;     CG - HE2
 7211   7222       1 ;     CG - C
 7212   7217       1 ;    CD1 - HZ
 7212   7218       1 ;    CD1 - CE2
 7212   7221       1 ;    CD1 - HD2
```

```
  7213   7215       1 ;     HD1 - HE1
  7213   7216       1 ;     HD1 - CZ
  7213   7220       1 ;     HD1 - CD2
  7214   7219       1 ;     CE1 - HE2
  7214   7220       1 ;     CE1 - CD2
  7215   7217       1 ;     HE1 - HZ
  7215   7218       1 ;     HE1 - CE2
  7216   7221       1 ;      CZ - HD2
  7217   7219       1 ;      HZ - HE2
  7217   7220       1 ;      HZ - CD2
  7219   7221       1 ;     HE2 - HD2
  7222   7227       1 ;       C - HA
  7222   7228       1 ;       C - CB
  7222   7239       1 ;       C - C
  7223   7225       1 ;       O - H
  7223   7226       1 ;       O - CA
  7224   7229       1 ;       N - HB2
  7224   7230       1 ;       N - HB3
  7224   7231       1 ;       N - CG
  7224   7240       1 ;       N - O
  7224   7241       1 ;       N - N
  7225   7227       1 ;       H - HA
  7225   7228       1 ;       H - CB
  7225   7239       1 ;       H - C
  7226   7232       1 ;      CA - HG2
  7226   7233       1 ;      CA - HG3
  7226   7234       1 ;      CA - SD
  7226   7242       1 ;      CA - H
  7226   7243       1 ;      CA - CA
  7227   7229       1 ;      HA - HB2
  7227   7230       1 ;      HA - HB3
  7227   7231       1 ;      HA - CG
  7227   7240       1 ;      HA - O
  7227   7241       1 ;      HA - N
  7228   7235       1 ;      CB - CE
  7228   7240       1 ;      CB - O
  7228   7241       1 ;      CB - N
  7229   7232       1 ;     HB2 - HG2
  7229   7233       1 ;     HB2 - HG3
  7229   7234       1 ;     HB2 - SD
  7229   7239       1 ;     HB2 - C
  7230   7232       1 ;     HB3 - HG2
  7230   7233       1 ;     HB3 - HG3
  7230   7234       1 ;     HB3 - SD
  7230   7239       1 ;     HB3 - C
  7231   7236       1 ;      CG - HE1
  7231   7237       1 ;      CG - HE2
  7231   7238       1 ;      CG - HE3
  7231   7239       1 ;      CG - C
  7232   7235       1 ;     HG2 - CE
  7233   7235       1 ;     HG3 - CE
  7239   7244       1 ;       C - HA
```

```
7239   7245      1 ;      C - CB
7239   7258      1 ;      C - C
7240   7242      1 ;      O - H
7240   7243      1 ;      O - CA
7241   7246      1 ;      N - HB2
7241   7247      1 ;      N - HB3
7241   7248      1 ;      N - CG
7241   7259      1 ;      N - O
7241   7260      1 ;      N - N
7242   7244      1 ;      H - HA
7242   7245      1 ;      H - CB
7242   7258      1 ;      H - C
7243   7249      1 ;     CA - HG
7243   7250      1 ;     CA - CD1
7243   7254      1 ;     CA - CD2
7243   7261      1 ;     CA - H
7243   7262      1 ;     CA - CA
7244   7246      1 ;     HA - HB2
7244   7247      1 ;     HA - HB3
7244   7248      1 ;     HA - CG
7244   7259      1 ;     HA - O
7244   7260      1 ;     HA - N
7245   7251      1 ;     CB - HD11
7245   7252      1 ;     CB - HD12
7245   7253      1 ;     CB - HD13
7245   7255      1 ;     CB - HD21
7245   7256      1 ;     CB - HD22
7245   7257      1 ;     CB - HD23
7245   7259      1 ;     CB - O
7245   7260      1 ;     CB - N
7246   7249      1 ;    HB2 - HG
7246   7250      1 ;    HB2 - CD1
7246   7254      1 ;    HB2 - CD2
7246   7258      1 ;    HB2 - C
7247   7249      1 ;    HB3 - HG
7247   7250      1 ;    HB3 - CD1
7247   7254      1 ;    HB3 - CD2
7247   7258      1 ;    HB3 - C
7248   7258      1 ;     CG - C
7249   7251      1 ;     HG - HD11
7249   7252      1 ;     HG - HD12
7249   7253      1 ;     HG - HD13
7249   7255      1 ;     HG - HD21
7249   7256      1 ;     HG - HD22
7249   7257      1 ;     HG - HD23
7250   7255      1 ;    CD1 - HD21
7250   7256      1 ;    CD1 - HD22
7250   7257      1 ;    CD1 - HD23
7251   7254      1 ;   HD11 - CD2
7252   7254      1 ;   HD12 - CD2
7253   7254      1 ;   HD13 - CD2
7258   7263      1 ;      C - HA
```

```
7258   7264       1 ;       C - CB
7258   7275       1 ;       C - C
7259   7261       1 ;       O - H
7259   7262       1 ;       O - CA
7260   7265       1 ;       N - HB2
7260   7266       1 ;       N - HB3
7260   7267       1 ;       N - CG
7260   7276       1 ;       N - O
7260   7277       1 ;       N - N
7261   7263       1 ;       H - HA
7261   7264       1 ;       H - CB
7261   7275       1 ;       H - C
7262   7268       1 ;      CA - HG2
7262   7269       1 ;      CA - HG3
7262   7270       1 ;      CA - SD
7262   7278       1 ;      CA - H
7262   7279       1 ;      CA - CA
7263   7265       1 ;      HA - HB2
7263   7266       1 ;      HA - HB3
7263   7267       1 ;      HA - CG
7263   7276       1 ;      HA - O
7263   7277       1 ;      HA - N
7264   7271       1 ;      CB - CE
7264   7276       1 ;      CB - O
7264   7277       1 ;      CB - N
7265   7268       1 ;     HB2 - HG2
7265   7269       1 ;     HB2 - HG3
7265   7270       1 ;     HB2 - SD
7265   7275       1 ;     HB2 - C
7266   7268       1 ;     HB3 - HG2
7266   7269       1 ;     HB3 - HG3
7266   7270       1 ;     HB3 - SD
7266   7275       1 ;     HB3 - C
7267   7272       1 ;      CG - HE1
7267   7273       1 ;      CG - HE2
7267   7274       1 ;      CG - HE3
7267   7275       1 ;      CG - C
7268   7271       1 ;     HG2 - CE
7269   7271       1 ;     HG3 - CE
7275   7280       1 ;       C - HA
7275   7281       1 ;       C - CB
7275   7292       1 ;       C - C
7276   7278       1 ;       O - H
7276   7279       1 ;       O - CA
7277   7282       1 ;       N - HB2
7277   7283       1 ;       N - HB3
7277   7284       1 ;       N - CG
7277   7293       1 ;       N - O
7277   7294       1 ;       N - N
7278   7280       1 ;       H - HA
7278   7281       1 ;       H - CB
7278   7292       1 ;       H - C
```

```
7279   7285      1 ;     CA - HG2
7279   7286      1 ;     CA - HG3
7279   7287      1 ;     CA - SD
7279   7295      1 ;     CA - H
7279   7296      1 ;     CA - CA
7280   7282      1 ;     HA - HB2
7280   7283      1 ;     HA - HB3
7280   7284      1 ;     HA - CG
7280   7293      1 ;     HA - O
7280   7294      1 ;     HA - N
7281   7288      1 ;     CB - CE
7281   7293      1 ;     CB - O
7281   7294      1 ;     CB - N
7282   7285      1 ;    HB2 - HG2
7282   7286      1 ;    HB2 - HG3
7282   7287      1 ;    HB2 - SD
7282   7292      1 ;    HB2 - C
7283   7285      1 ;    HB3 - HG2
7283   7286      1 ;    HB3 - HG3
7283   7287      1 ;    HB3 - SD
7283   7292      1 ;    HB3 - C
7284   7289      1 ;     CG - HE1
7284   7290      1 ;     CG - HE2
7284   7291      1 ;     CG - HE3
7284   7292      1 ;     CG - C
7285   7288      1 ;    HG2 - CE
7286   7288      1 ;    HG3 - CE
7292   7297      1 ;      C - HA
7292   7298      1 ;      C - CB
7292   7303      1 ;      C - C
7293   7295      1 ;      O - H
7293   7296      1 ;      O - CA
7294   7299      1 ;      N - HB2
7294   7300      1 ;      N - HB3
7294   7301      1 ;      N - OG
7294   7304      1 ;      N - O
7294   7305      1 ;      N - N
7295   7297      1 ;      H - HA
7295   7298      1 ;      H - CB
7295   7303      1 ;      H - C
7296   7302      1 ;     CA - HG
7296   7306      1 ;     CA - H
7296   7307      1 ;     CA - CA
7297   7299      1 ;     HA - HB2
7297   7300      1 ;     HA - HB3
7297   7301      1 ;     HA - OG
7297   7304      1 ;     HA - O
7297   7305      1 ;     HA - N
7298   7304      1 ;     CB - O
7298   7305      1 ;     CB - N
7299   7302      1 ;    HB2 - HG
7299   7303      1 ;    HB2 - C
```

```
7300   7302       1 ;     HB3 - HG
7300   7303       1 ;     HB3 - C
7301   7303       1 ;      OG - C
7303   7308       1 ;       C - HA
7303   7309       1 ;       C - CB
7303   7325       1 ;       C - C
7304   7306       1 ;       O - H
7304   7307       1 ;       O - CA
7305   7310       1 ;       N - HB2
7305   7311       1 ;       N - HB3
7305   7312       1 ;       N - CG
7305   7326       1 ;       N - O
7305   7327       1 ;       N - N
7306   7308       1 ;       H - HA
7306   7309       1 ;       H - CB
7306   7325       1 ;       H - C
7307   7313       1 ;      CA - HG2
7307   7314       1 ;      CA - HG3
7307   7315       1 ;      CA - CD
7307   7328       1 ;      CA - H
7307   7329       1 ;      CA - CA
7308   7310       1 ;      HA - HB2
7308   7311       1 ;      HA - HB3
7308   7312       1 ;      HA - CG
7308   7326       1 ;      HA - O
7308   7327       1 ;      HA - N
7309   7316       1 ;      CB - HD2
7309   7317       1 ;      CB - HD3
7309   7318       1 ;      CB - CE
7309   7326       1 ;      CB - O
7309   7327       1 ;      CB - N
7310   7313       1 ;     HB2 - HG2
7310   7314       1 ;     HB2 - HG3
7310   7315       1 ;     HB2 - CD
7310   7325       1 ;     HB2 - C
7311   7313       1 ;     HB3 - HG2
7311   7314       1 ;     HB3 - HG3
7311   7315       1 ;     HB3 - CD
7311   7325       1 ;     HB3 - C
7312   7319       1 ;      CG - HE2
7312   7320       1 ;      CG - HE3
7312   7321       1 ;      CG - NZ
7312   7325       1 ;      CG - C
7313   7316       1 ;     HG2 - HD2
7313   7317       1 ;     HG2 - HD3
7313   7318       1 ;     HG2 - CE
7314   7316       1 ;     HG3 - HD2
7314   7317       1 ;     HG3 - HD3
7314   7318       1 ;     HG3 - CE
7315   7322       1 ;      CD - HZ1
7315   7323       1 ;      CD - HZ2
7315   7324       1 ;      CD - HZ3
```

| | | | |
|---|---|---|---|
| 7316 | 7319 | 1 ; | HD2 - HE2 |
| 7316 | 7320 | 1 ; | HD2 - HE3 |
| 7316 | 7321 | 1 ; | HD2 - NZ |
| 7317 | 7319 | 1 ; | HD3 - HE2 |
| 7317 | 7320 | 1 ; | HD3 - HE3 |
| 7317 | 7321 | 1 ; | HD3 - NZ |
| 7319 | 7322 | 1 ; | HE2 - HZ1 |
| 7319 | 7323 | 1 ; | HE2 - HZ2 |
| 7319 | 7324 | 1 ; | HE2 - HZ3 |
| 7320 | 7322 | 1 ; | HE3 - HZ1 |
| 7320 | 7323 | 1 ; | HE3 - HZ2 |
| 7320 | 7324 | 1 ; | HE3 - HZ3 |
| 7325 | 7330 | 1 ; | C - HA |
| 7325 | 7331 | 1 ; | C - CB |
| 7325 | 7347 | 1 ; | C - C |
| 7326 | 7328 | 1 ; | O - H |
| 7326 | 7329 | 1 ; | O - CA |
| 7327 | 7332 | 1 ; | N - HB2 |
| 7327 | 7333 | 1 ; | N - HB3 |
| 7327 | 7334 | 1 ; | N - CG |
| 7327 | 7348 | 1 ; | N - O |
| 7327 | 7349 | 1 ; | N - N |
| 7328 | 7330 | 1 ; | H - HA |
| 7328 | 7331 | 1 ; | H - CB |
| 7328 | 7347 | 1 ; | H - C |
| 7329 | 7335 | 1 ; | CA - HG2 |
| 7329 | 7336 | 1 ; | CA - HG3 |
| 7329 | 7337 | 1 ; | CA - CD |
| 7329 | 7350 | 1 ; | CA - H |
| 7329 | 7351 | 1 ; | CA - CA |
| 7330 | 7332 | 1 ; | HA - HB2 |
| 7330 | 7333 | 1 ; | HA - HB3 |
| 7330 | 7334 | 1 ; | HA - CG |
| 7330 | 7348 | 1 ; | HA - O |
| 7330 | 7349 | 1 ; | HA - N |
| 7331 | 7338 | 1 ; | CB - HD2 |
| 7331 | 7339 | 1 ; | CB - HD3 |
| 7331 | 7340 | 1 ; | CB - CE |
| 7331 | 7348 | 1 ; | CB - O |
| 7331 | 7349 | 1 ; | CB - N |
| 7332 | 7335 | 1 ; | HB2 - HG2 |
| 7332 | 7336 | 1 ; | HB2 - HG3 |
| 7332 | 7337 | 1 ; | HB2 - CD |
| 7332 | 7347 | 1 ; | HB2 - C |
| 7333 | 7335 | 1 ; | HB3 - HG2 |
| 7333 | 7336 | 1 ; | HB3 - HG3 |
| 7333 | 7337 | 1 ; | HB3 - CD |
| 7333 | 7347 | 1 ; | HB3 - C |
| 7334 | 7341 | 1 ; | CG - HE2 |
| 7334 | 7342 | 1 ; | CG - HE3 |
| 7334 | 7343 | 1 ; | CG - NZ |
| 7334 | 7347 | 1 ; | CG - C |

| | | | |
|---|---|---|---|
| 7335 | 7338 | 1 ; | HG2 - HD2 |
| 7335 | 7339 | 1 ; | HG2 - HD3 |
| 7335 | 7340 | 1 ; | HG2 - CE |
| 7336 | 7338 | 1 ; | HG3 - HD2 |
| 7336 | 7339 | 1 ; | HG3 - HD3 |
| 7336 | 7340 | 1 ; | HG3 - CE |
| 7337 | 7344 | 1 ; | CD - HZ1 |
| 7337 | 7345 | 1 ; | CD - HZ2 |
| 7337 | 7346 | 1 ; | CD - HZ3 |
| 7338 | 7341 | 1 ; | HD2 - HE2 |
| 7338 | 7342 | 1 ; | HD2 - HE3 |
| 7338 | 7343 | 1 ; | HD2 - NZ |
| 7339 | 7341 | 1 ; | HD3 - HE2 |
| 7339 | 7342 | 1 ; | HD3 - HE3 |
| 7339 | 7343 | 1 ; | HD3 - NZ |
| 7341 | 7344 | 1 ; | HE2 - HZ1 |
| 7341 | 7345 | 1 ; | HE2 - HZ2 |
| 7341 | 7346 | 1 ; | HE2 - HZ3 |
| 7342 | 7344 | 1 ; | HE3 - HZ1 |
| 7342 | 7345 | 1 ; | HE3 - HZ2 |
| 7342 | 7346 | 1 ; | HE3 - HZ3 |
| 7347 | 7352 | 1 ; | C - HA |
| 7347 | 7353 | 1 ; | C - CB |
| 7347 | 7369 | 1 ; | C - C |
| 7348 | 7350 | 1 ; | O - H |
| 7348 | 7351 | 1 ; | O - CA |
| 7349 | 7354 | 1 ; | N - HB2 |
| 7349 | 7355 | 1 ; | N - HB3 |
| 7349 | 7356 | 1 ; | N - CG |
| 7349 | 7370 | 1 ; | N - O |
| 7349 | 7371 | 1 ; | N - OXT |
| 7350 | 7352 | 1 ; | H - HA |
| 7350 | 7353 | 1 ; | H - CB |
| 7350 | 7369 | 1 ; | H - C |
| 7351 | 7357 | 1 ; | CA - HG2 |
| 7351 | 7358 | 1 ; | CA - HG3 |
| 7351 | 7359 | 1 ; | CA - CD |
| 7352 | 7354 | 1 ; | HA - HB2 |
| 7352 | 7355 | 1 ; | HA - HB3 |
| 7352 | 7356 | 1 ; | HA - CG |
| 7352 | 7370 | 1 ; | HA - O |
| 7352 | 7371 | 1 ; | HA - OXT |
| 7353 | 7360 | 1 ; | CB - HD2 |
| 7353 | 7361 | 1 ; | CB - HD3 |
| 7353 | 7362 | 1 ; | CB - CE |
| 7353 | 7370 | 1 ; | CB - O |
| 7353 | 7371 | 1 ; | CB - OXT |
| 7354 | 7357 | 1 ; | HB2 - HG2 |
| 7354 | 7358 | 1 ; | HB2 - HG3 |
| 7354 | 7359 | 1 ; | HB2 - CD |
| 7354 | 7369 | 1 ; | HB2 - C |
| 7355 | 7357 | 1 ; | HB3 - HG2 |

| | | | |
|---|---|---|---|
| 7355 | 7358 | 1 ; | HB3 - HG3 |
| 7355 | 7359 | 1 ; | HB3 - CD |
| 7355 | 7369 | 1 ; | HB3 - C |
| 7356 | 7363 | 1 ; | CG - HE2 |
| 7356 | 7364 | 1 ; | CG - HE3 |
| 7356 | 7365 | 1 ; | CG - NZ |
| 7356 | 7369 | 1 ; | CG - C |
| 7357 | 7360 | 1 ; | HG2 - HD2 |
| 7357 | 7361 | 1 ; | HG2 - HD3 |
| 7357 | 7362 | 1 ; | HG2 - CE |
| 7358 | 7360 | 1 ; | HG3 - HD2 |
| 7358 | 7361 | 1 ; | HG3 - HD3 |
| 7358 | 7362 | 1 ; | HG3 - CE |
| 7359 | 7366 | 1 ; | CD - HZ1 |
| 7359 | 7367 | 1 ; | CD - HZ2 |
| 7359 | 7368 | 1 ; | CD - HZ3 |
| 7360 | 7363 | 1 ; | HD2 - HE2 |
| 7360 | 7364 | 1 ; | HD2 - HE3 |
| 7360 | 7365 | 1 ; | HD2 - NZ |
| 7361 | 7363 | 1 ; | HD3 - HE2 |
| 7361 | 7364 | 1 ; | HD3 - HE3 |
| 7361 | 7365 | 1 ; | HD3 - NZ |
| 7363 | 7366 | 1 ; | HE2 - HZ1 |
| 7363 | 7367 | 1 ; | HE2 - HZ2 |
| 7363 | 7368 | 1 ; | HE2 - HZ3 |
| 7364 | 7366 | 1 ; | HE3 - HZ1 |
| 7364 | 7367 | 1 ; | HE3 - HZ2 |
| 7364 | 7368 | 1 ; | HE3 - HZ3 |
| 7372 | 7375 | 1 ; | C1 - C4 |
| 7372 | 7379 | 1 ; | C1 - C8 |
| 7372 | 7414 | 1 ; | C1 - H43 |
| 7373 | 7376 | 1 ; | C2 - C5 |
| 7373 | 7387 | 1 ; | C2 - N16 |
| 7373 | 7415 | 1 ; | C2 - H44 |
| 7373 | 7430 | 1 ; | C2 - H59 |
| 7374 | 7377 | 1 ; | C3 - C6 |
| 7374 | 7378 | 1 ; | C3 - C7 |
| 7374 | 7379 | 1 ; | C3 - C8 |
| 7374 | 7388 | 1 ; | C3 - O17 |
| 7374 | 7436 | 1 ; | C3 - RU |
| 7375 | 7380 | 1 ; | C4 - C9 |
| 7375 | 7386 | 1 ; | C4 - N15 |
| 7375 | 7389 | 1 ; | C4 - C18 |
| 7375 | 7390 | 1 ; | C4 - C19 |
| 7375 | 7391 | 1 ; | C4 - C20 |
| 7375 | 7392 | 1 ; | C4 - C21 |
| 7375 | 7393 | 1 ; | C4 - C22 |
| 7375 | 7394 | 1 ; | C4 - C23 |
| 7375 | 7405 | 1 ; | C4 - C34 |
| 7375 | 7413 | 1 ; | C4 - H42 |
| 7375 | 7415 | 1 ; | C4 - H44 |
| 7376 | 7380 | 1 ; | C5 - C9 |

```
7376   7388       1 ;      C5 - O17
7376   7406       1 ;      C5 - C35
7376   7412       1 ;      C5 - C41
7376   7414       1 ;      C5 - H43
7376   7436       1 ;      C5 - RU
7377   7378       1 ;      C6 - C7
7377   7387       1 ;      C6 - N16
7377   7405       1 ;      C6 - C34
7377   7413       1 ;      C6 - H42
7377   7430       1 ;      C6 - H59
7378   7382       1 ;      C7 - C11
7378   7384       1 ;      C7 - C13
7378   7389       1 ;      C7 - C18
7378   7390       1 ;      C7 - C19
7378   7391       1 ;      C7 - C20
7378   7392       1 ;      C7 - C21
7378   7393       1 ;      C7 - C22
7378   7394       1 ;      C7 - C23
7378   7406       1 ;      C7 - C35
7378   7412       1 ;      C7 - C41
7379   7381       1 ;      C8 - C10
7379   7386       1 ;      C8 - N15
7379   7407       1 ;      C8 - N36
7379   7408       1 ;      C8 - O37
7379   7410       1 ;      C8 - C39
7379   7415       1 ;      C8 - H44
7379   7436       1 ;      C8 - RU
7380   7383       1 ;      C9 - C12
7380   7389       1 ;      C9 - C18
7380   7390       1 ;      C9 - C19
7380   7391       1 ;      C9 - C20
7380   7392       1 ;      C9 - C21
7380   7393       1 ;      C9 - C22
7380   7394       1 ;      C9 - C23
7380   7405       1 ;      C9 - C34
7380   7410       1 ;      C9 - C39
7380   7416       1 ;      C9 - H45
7380   7417       1 ;      C9 - H46
7381   7382       1 ;     C10 - C11
7381   7385       1 ;     C10 - F14
7381   7387       1 ;     C10 - N16
7381   7406       1 ;     C10 - C35
7381   7407       1 ;     C10 - N36
7381   7411       1 ;     C10 - O40
7381   7436       1 ;     C10 - RU
7382   7387       1 ;     C11 - N16
7382   7389       1 ;     C11 - C18
7382   7390       1 ;     C11 - C19
7382   7391       1 ;     C11 - C20
7382   7392       1 ;     C11 - C21
7382   7393       1 ;     C11 - C22
7382   7394       1 ;     C11 - C23
```

| | | | | |
|---|---|---|---|---|
| 7382 | 7417 | 1 | ; | C11 - H46 |
| 7383 | 7412 | 1 | ; | C12 - C41 |
| 7383 | 7436 | 1 | ; | C12 - RU |
| 7384 | 7386 | 1 | ; | C13 - N15 |
| 7384 | 7405 | 1 | ; | C13 - C34 |
| 7384 | 7410 | 1 | ; | C13 - C39 |
| 7384 | 7416 | 1 | ; | C13 - H45 |
| 7385 | 7386 | 1 | ; | F14 - N15 |
| 7385 | 7416 | 1 | ; | F14 - H45 |
| 7385 | 7417 | 1 | ; | F14 - H46 |
| 7386 | 7395 | 1 | ; | N15 - O24 |
| 7386 | 7396 | 1 | ; | N15 - C25 |
| 7386 | 7412 | 1 | ; | N15 - C41 |
| 7386 | 7418 | 1 | ; | N15 - H47 |
| 7386 | 7419 | 1 | ; | N15 - H48 |
| 7386 | 7420 | 1 | ; | N15 - H49 |
| 7386 | 7435 | 1 | ; | N15 - H64 |
| 7387 | 7395 | 1 | ; | N16 - O24 |
| 7387 | 7396 | 1 | ; | N16 - C25 |
| 7387 | 7405 | 1 | ; | N16 - C34 |
| 7387 | 7414 | 1 | ; | N16 - H43 |
| 7387 | 7418 | 1 | ; | N16 - H47 |
| 7387 | 7419 | 1 | ; | N16 - H48 |
| 7387 | 7420 | 1 | ; | N16 - H49 |
| 7387 | 7435 | 1 | ; | N16 - H64 |
| 7388 | 7413 | 1 | ; | O17 - H42 |
| 7388 | 7415 | 1 | ; | O17 - H44 |
| 7389 | 7395 | 1 | ; | C18 - O24 |
| 7389 | 7397 | 1 | ; | C18 - N26 |
| 7389 | 7398 | 1 | ; | C18 - O27 |
| 7389 | 7419 | 1 | ; | C18 - H48 |
| 7389 | 7435 | 1 | ; | C18 - H64 |
| 7390 | 7395 | 1 | ; | C19 - O24 |
| 7390 | 7399 | 1 | ; | C19 - C28 |
| 7390 | 7419 | 1 | ; | C19 - H48 |
| 7390 | 7420 | 1 | ; | C19 - H49 |
| 7390 | 7434 | 1 | ; | C19 - H63 |
| 7391 | 7395 | 1 | ; | C20 - O24 |
| 7391 | 7397 | 1 | ; | C20 - N26 |
| 7391 | 7398 | 1 | ; | C20 - O27 |
| 7391 | 7418 | 1 | ; | C20 - H47 |
| 7391 | 7420 | 1 | ; | C20 - H49 |
| 7392 | 7395 | 1 | ; | C21 - O24 |
| 7392 | 7396 | 1 | ; | C21 - C25 |
| 7392 | 7418 | 1 | ; | C21 - H47 |
| 7393 | 7395 | 1 | ; | C22 - O24 |
| 7393 | 7396 | 1 | ; | C22 - C25 |
| 7393 | 7435 | 1 | ; | C22 - H64 |
| 7394 | 7396 | 1 | ; | C23 - C25 |
| 7394 | 7418 | 1 | ; | C23 - H47 |
| 7394 | 7419 | 1 | ; | C23 - H48 |
| 7394 | 7420 | 1 | ; | C23 - H49 |

| | | | |
|---|---|---|---|
| 7394 | 7435 | 1 ; | C23 - H64 |
| 7396 | 7400 | 1 ; | C25 - C29 |
| 7396 | 7401 | 1 ; | C25 - C30 |
| 7396 | 7403 | 1 ; | C25 - C32 |
| 7396 | 7418 | 1 ; | C25 - H47 |
| 7396 | 7435 | 1 ; | C25 - H64 |
| 7397 | 7402 | 1 ; | N26 - O31 |
| 7397 | 7404 | 1 ; | N26 - O33 |
| 7397 | 7421 | 1 ; | N26 - H50 |
| 7397 | 7422 | 1 ; | N26 - H51 |
| 7397 | 7423 | 1 ; | N26 - H52 |
| 7397 | 7424 | 1 ; | N26 - H53 |
| 7397 | 7431 | 1 ; | N26 - H60 |
| 7397 | 7432 | 1 ; | N26 - H61 |
| 7397 | 7433 | 1 ; | N26 - H62 |
| 7397 | 7436 | 1 ; | N26 - RU |
| 7398 | 7399 | 1 ; | O27 - C28 |
| 7398 | 7434 | 1 ; | O27 - H63 |
| 7398 | 7436 | 1 ; | O27 - RU |
| 7399 | 7425 | 1 ; | C28 - H54 |
| 7399 | 7429 | 1 ; | C28 - H58 |
| 7400 | 7402 | 1 ; | C29 - O31 |
| 7400 | 7404 | 1 ; | C29 - O33 |
| 7400 | 7421 | 1 ; | C29 - H50 |
| 7400 | 7422 | 1 ; | C29 - H51 |
| 7400 | 7423 | 1 ; | C29 - H52 |
| 7400 | 7424 | 1 ; | C29 - H53 |
| 7400 | 7434 | 1 ; | C29 - H63 |
| 7401 | 7404 | 1 ; | C30 - O33 |
| 7401 | 7423 | 1 ; | C30 - H52 |
| 7401 | 7424 | 1 ; | C30 - H53 |
| 7401 | 7431 | 1 ; | C30 - H60 |
| 7401 | 7432 | 1 ; | C30 - H61 |
| 7401 | 7433 | 1 ; | C30 - H62 |
| 7401 | 7434 | 1 ; | C30 - H63 |
| 7402 | 7403 | 1 ; | O31 - C32 |
| 7403 | 7421 | 1 ; | C32 - H50 |
| 7403 | 7422 | 1 ; | C32 - H51 |
| 7403 | 7431 | 1 ; | C32 - H60 |
| 7403 | 7432 | 1 ; | C32 - H61 |
| 7403 | 7433 | 1 ; | C32 - H62 |
| 7403 | 7434 | 1 ; | C32 - H63 |
| 7405 | 7409 | 1 ; | C34 - C38 |
| 7405 | 7411 | 1 ; | C34 - O40 |
| 7406 | 7411 | 1 ; | C35 - O40 |
| 7406 | 7426 | 1 ; | C35 - H55 |
| 7406 | 7427 | 1 ; | C35 - H56 |
| 7406 | 7428 | 1 ; | C35 - H57 |
| 7408 | 7409 | 1 ; | O37 - C38 |
| 7408 | 7410 | 1 ; | O37 - C39 |
| 7408 | 7412 | 1 ; | O37 - C41 |
| 7409 | 7411 | 1 ; | C38 - O40 |

```
  7409   7412       1 ;    C38 - C41
  7410   7426       1 ;    C39 - H55
  7410   7427       1 ;    C39 - H56
  7410   7428       1 ;    C39 - H57
  7412   7417       1 ;    C41 - H46
  7413   7414       1 ;    H42 - H43
  7416   7436       1 ;    H45 - RU
  7418   7420       1 ;    H47 - H49
  7419   7420       1 ;    H48 - H49
  7419   7435       1 ;    H48 - H64
  7421   7425       1 ;    H50 - H54
  7422   7425       1 ;    H51 - H54
  7423   7429       1 ;    H52 - H58
  7424   7429       1 ;    H53 - H58

[ angles ]
;   ai     aj     ak    funct   theta         cth
     1      5      6      1    1.0950e+02    4.1840e+02 ;      N - CA
- HA
     1      5      7      1    1.1120e+02    6.6944e+02 ;      N - CA
- CB
     1      5     20      1    1.1120e+02    6.6944e+02 ;      N - CA
- C
     2      1      3      1    1.0950e+02    2.9288e+02 ;     H1 - N
- H2
     2      1      4      1    1.0950e+02    2.9288e+02 ;     H1 - N
- H3
     2      1      5      1    1.0950e+02    4.1840e+02 ;     H1 - N
- CA
     3      1      4      1    1.0950e+02    2.9288e+02 ;     H2 - N
- H3
     3      1      5      1    1.0950e+02    4.1840e+02 ;     H2 - N
- CA
     4      1      5      1    1.0950e+02    4.1840e+02 ;     H3 - N
- CA
     5      7      8      1    1.0950e+02    4.1840e+02 ;     CA - CB
- HB2
     5      7      9      1    1.0950e+02    4.1840e+02 ;     CA - CB
- HB3
     5      7     10      1    1.0950e+02    3.3472e+02 ;     CA - CB
- CG
     5     20     21      1    1.2040e+02    6.6944e+02 ;     CA - C
- O
     5     20     22      1    1.1660e+02    5.8576e+02 ;     CA - C
- N
     6      5      7      1    1.0950e+02    4.1840e+02 ;     HA - CA
- CB
     6      5     20      1    1.0950e+02    4.1840e+02 ;     HA - CA
- C
     7      5     20      1    1.1110e+02    5.2718e+02 ;     CB - CA
- C
```

```
      62      69      71       1    1.1660e+02    5.8576e+02 ;     CA - C
  - N
      63      62      64       1    1.0950e+02    4.1840e+02 ;     HA - CA
  - CB
      63      62      69       1    1.0950e+02    4.1840e+02 ;     HA - CA
  - C
      64      62      69       1    1.1110e+02    5.2718e+02 ;     CB - CA
  - C
      64      67      68       1    9.6000e+01    3.5982e+02 ;     CB - SG
  - HG
      65      64      66       1    1.0950e+02    2.9288e+02 ;    HB2 - CB
  - HB3
      65      64      67       1    1.0950e+02    4.1840e+02 ;    HB2 - CB
  - SG
      66      64      67       1    1.0950e+02    4.1840e+02 ;    HB3 - CB
  - SG
      69      71      72       1    1.2000e+02    4.1840e+02 ;      C - N
  - H
      69      71      73       1    1.2190e+02    4.1840e+02 ;      C - N
  - CA
      70      69      71       1    1.2290e+02    6.6944e+02 ;      O - C
  - N
      71      73      74       1    1.0950e+02    4.1840e+02 ;      N - CA
  - HA
      71      73      75       1    1.0970e+02    6.6944e+02 ;      N - CA
  - CB
      71      73      87       1    1.1010e+02    5.2718e+02 ;      N - CA
  - C
      72      71      73       1    1.1804e+02    4.1840e+02 ;      H - N
  - CA
      73      75      76       1    1.0950e+02    4.1840e+02 ;     CA - CB
  - HB2
      73      75      77       1    1.0950e+02    4.1840e+02 ;     CA - CB
  - HB3
      73      75      78       1    1.1310e+02    5.2718e+02 ;     CA - CB
  - CG
      73      87      88       1    1.2040e+02    6.6944e+02 ;     CA - C
  - O
      73      87      89       1    1.1660e+02    5.8576e+02 ;     CA - C
  - N
      74      73      75       1    1.0950e+02    4.1840e+02 ;     HA - CA
  - CB
      74      73      87       1    1.0950e+02    4.1840e+02 ;     HA - CA
  - C
      75      73      87       1    1.1110e+02    5.2718e+02 ;     CB - CA
  - C
      75      78      79       1    1.2000e+02    5.8576e+02 ;     CB - CG
  - ND1
      75      78      85       1    1.2000e+02    5.8576e+02 ;     CB - CG
  - CD2
      76      75      77       1    1.0950e+02    2.9288e+02 ;    HB2 - CB
  - HB3
```

```
     76      75      78       1    1.0950e+02    4.1840e+02 ;    HB2 - CB
- CG
     77      75      78       1    1.0950e+02    4.1840e+02 ;    HB3 - CB
- CG
     78      79      80       1    1.2000e+02    4.1840e+02 ;     CG - ND1
- HD1
     78      79      81       1    1.2000e+02    5.8576e+02 ;     CG - ND1
- CE1
     78      85      83       1    1.2000e+02    5.8576e+02 ;     CG - CD2
- NE2
     78      85      86       1    1.2000e+02    4.1840e+02 ;     CG - CD2
- HD2
     79      78      85       1    1.2000e+02    5.8576e+02 ;    ND1 - CG
- CD2
     79      81      82       1    1.2000e+02    4.1840e+02 ;    ND1 - CE1
- HE1
     79      81      83       1    1.2000e+02    5.8576e+02 ;    ND1 - CE1
- NE2
     80      79      81       1    1.2000e+02    4.1840e+02 ;    HD1 - ND1
- CE1
     81      83      84       1    1.2000e+02    4.1840e+02 ;    CE1 - NE2
- HE2
     81      83      85       1    1.2000e+02    5.8576e+02 ;    CE1 - NE2
- CD2
     82      81      83       1    1.2000e+02    4.1840e+02 ;    HE1 - CE1
- NE2
     83      85      86       1    1.2000e+02    4.1840e+02 ;    NE2 - CD2
- HD2
     84      83      85       1    1.2000e+02    4.1840e+02 ;    HE2 - NE2
- CD2
     87      89      90       1    1.2000e+02    4.1840e+02 ;      C - N
- H
     87      89      91       1    1.2190e+02    4.1840e+02 ;      C - N
- CA
     88      87      89       1    1.2290e+02    6.6944e+02 ;      O - C
- N
     89      91      92       1    1.0950e+02    4.1840e+02 ;      N - CA
- HA
     89      91      93       1    1.0970e+02    6.6944e+02 ;      N - CA
- CB
     89      91      98       1    1.1010e+02    5.2718e+02 ;      N - CA
- C
     90      89      91       1    1.1804e+02    4.1840e+02 ;      H - N
- CA
     91      93      94       1    1.0950e+02    4.1840e+02 ;     CA - CB
- HB2
     91      93      95       1    1.0950e+02    4.1840e+02 ;     CA - CB
- HB3
     91      93      96       1    1.0950e+02    4.1840e+02 ;     CA - CB
- OG
     91      98      99       1    1.2040e+02    6.6944e+02 ;     CA - C
- O
```

```
    91     98    100      1    1.1660e+02    5.8576e+02 ;     CA - C
- N
    92     91     93      1    1.0950e+02    4.1840e+02 ;     HA - CA
- CB
    92     91     98      1    1.0950e+02    4.1840e+02 ;     HA - CA
- C
    93     91     98      1    1.1110e+02    5.2718e+02 ;     CB - CA
- C
    93     96     97      1    1.0850e+02    4.6024e+02 ;     CB - OG
- HG
    94     93     95      1    1.0950e+02    2.9288e+02 ;    HB2 - CB
- HB3
    94     93     96      1    1.0950e+02    4.1840e+02 ;    HB2 - CB
- OG
    95     93     96      1    1.0950e+02    4.1840e+02 ;    HB3 - CB
- OG
    98    100    101      1    1.2000e+02    4.1840e+02 ;      C - N
- H
    98    100    102      1    1.2190e+02    4.1840e+02 ;      C - N
- CA
    99     98    100      1    1.2290e+02    6.6944e+02 ;      O - C
- N
   100    102    103      1    1.0950e+02    4.1840e+02 ;      N - CA
- HA2
   100    102    104      1    1.0950e+02    4.1840e+02 ;      N - CA
- HA3
   100    102    105      1    1.1010e+02    5.2718e+02 ;      N - CA
- C
   101    100    102      1    1.1804e+02    4.1840e+02 ;      H - N
- CA
   102    105    106      1    1.2040e+02    6.6944e+02 ;     CA - C
- O
   102    105    107      1    1.1660e+02    5.8576e+02 ;     CA - C
- N
   103    102    104      1    1.0950e+02    2.9288e+02 ;    HA2 - CA
- HA3
   103    102    105      1    1.0950e+02    4.1840e+02 ;    HA2 - CA
- C
   104    102    105      1    1.0950e+02    4.1840e+02 ;    HA3 - CA
- C
   105    107    108      1    1.2000e+02    4.1840e+02 ;      C - N
- H
   105    107    109      1    1.2190e+02    4.1840e+02 ;      C - N
- CA
   106    105    107      1    1.2290e+02    6.6944e+02 ;      O - C
- N
   107    109    110      1    1.0950e+02    4.1840e+02 ;      N - CA
- HA
   107    109    111      1    1.0970e+02    6.6944e+02 ;      N - CA
- CB
   107    109    127      1    1.1010e+02    5.2718e+02 ;      N - CA
- C
```

```
   108    107    109      1    1.1804e+02    4.1840e+02 ;      H - N
- CA
   109    111    112      1    1.0950e+02    4.1840e+02 ;     CA - CB
- HB2
   109    111    113      1    1.0950e+02    4.1840e+02 ;     CA - CB
- HB3
   109    111    114      1    1.0950e+02    3.3472e+02 ;     CA - CB
- CG
   109    127    128      1    1.2040e+02    6.6944e+02 ;     CA - C
- O
   109    127    129      1    1.1660e+02    5.8576e+02 ;     CA - C
- N
   110    109    111      1    1.0950e+02    4.1840e+02 ;     HA - CA
- CB
   110    109    127      1    1.0950e+02    4.1840e+02 ;     HA - CA
- C
   111    109    127      1    1.1110e+02    5.2718e+02 ;     CB - CA
- C
   111    114    115      1    1.0950e+02    4.1840e+02 ;     CB - CG
- HG2
   111    114    116      1    1.0950e+02    4.1840e+02 ;     CB - CG
- HG3
   111    114    117      1    1.0950e+02    3.3472e+02 ;     CB - CG
- CD
   112    111    113      1    1.0950e+02    2.9288e+02 ;    HB2 - CB
- HB3
   112    111    114      1    1.0950e+02    4.1840e+02 ;    HB2 - CB
- CG
   113    111    114      1    1.0950e+02    4.1840e+02 ;    HB3 - CB
- CG
   114    117    118      1    1.0950e+02    4.1840e+02 ;     CG - CD
- HD2
   114    117    119      1    1.0950e+02    4.1840e+02 ;     CG - CD
- HD3
   114    117    120      1    1.0950e+02    3.3472e+02 ;     CG - CD
- CE
   115    114    116      1    1.0950e+02    2.9288e+02 ;    HG2 - CG
- HG3
   115    114    117      1    1.0950e+02    4.1840e+02 ;    HG2 - CG
- CD
   116    114    117      1    1.0950e+02    4.1840e+02 ;    HG3 - CG
- CD
   117    120    121      1    1.0950e+02    4.1840e+02 ;     CD - CE
- HE2
   117    120    122      1    1.0950e+02    4.1840e+02 ;     CD - CE
- HE3
   117    120    123      1    1.1120e+02    6.6944e+02 ;     CD - CE
- NZ
   118    117    119      1    1.0950e+02    2.9288e+02 ;    HD2 - CD
- HD3
   118    117    120      1    1.0950e+02    4.1840e+02 ;    HD2 - CD
- CE
```

```
   119    117    120      1    1.0950e+02    4.1840e+02 ;    HD3 - CD
- CE
   120    123    124      1    1.0950e+02    4.1840e+02 ;     CE - NZ
- HZ1
   120    123    125      1    1.0950e+02    4.1840e+02 ;     CE - NZ
- HZ2
   120    123    126      1    1.0950e+02    4.1840e+02 ;     CE - NZ
- HZ3
   121    120    122      1    1.0950e+02    2.9288e+02 ;    HE2 - CE
- HE3
   121    120    123      1    1.0950e+02    4.1840e+02 ;    HE2 - CE
- NZ
   122    120    123      1    1.0950e+02    4.1840e+02 ;    HE3 - CE
- NZ
   124    123    125      1    1.0950e+02    2.9288e+02 ;    HZ1 - NZ
- HZ2
   124    123    126      1    1.0950e+02    2.9288e+02 ;    HZ1 - NZ
- HZ3
   125    123    126      1    1.0950e+02    2.9288e+02 ;    HZ2 - NZ
- HZ3
   127    129    130      1    1.2000e+02    4.1840e+02 ;      C - N
- H
   127    129    131      1    1.2190e+02    4.1840e+02 ;      C - N
- CA
   128    127    129      1    1.2290e+02    6.6944e+02 ;      O - C
- N
   129    131    132      1    1.0950e+02    4.1840e+02 ;      N - CA
- HA
   129    131    133      1    1.0970e+02    6.6944e+02 ;      N - CA
- CB
   129    131    146      1    1.1010e+02    5.2718e+02 ;      N - CA
- C
   130    129    131      1    1.1804e+02    4.1840e+02 ;      H - N
- CA
   131    133    134      1    1.0950e+02    4.1840e+02 ;     CA - CB
- HB2
   131    133    135      1    1.0950e+02    4.1840e+02 ;     CA - CB
- HB3
   131    133    136      1    1.0950e+02    3.3472e+02 ;     CA - CB
- CG
   131    146    147      1    1.2040e+02    6.6944e+02 ;     CA - C
- O
   131    146    148      1    1.1660e+02    5.8576e+02 ;     CA - C
- N
   132    131    133      1    1.0950e+02    4.1840e+02 ;     HA - CA
- CB
   132    131    146      1    1.0950e+02    4.1840e+02 ;     HA - CA
- C
   133    131    146      1    1.1110e+02    5.2718e+02 ;     CB - CA
- C
   133    136    137      1    1.0950e+02    4.1840e+02 ;     CB - CG
- HG
```

133 136 138 1 1.0950e+02 3.3472e+02 ; CB - CG
- CD1
133 136 142 1 1.0950e+02 3.3472e+02 ; CB - CG
- CD2
134 133 135 1 1.0950e+02 2.9288e+02 ; HB2 - CB
- HB3
134 133 136 1 1.0950e+02 4.1840e+02 ; HB2 - CB
- CG
135 133 136 1 1.0950e+02 4.1840e+02 ; HB3 - CB
- CG
136 138 139 1 1.0950e+02 4.1840e+02 ; CG - CD1
- HD11
136 138 140 1 1.0950e+02 4.1840e+02 ; CG - CD1
- HD12
136 138 141 1 1.0950e+02 4.1840e+02 ; CG - CD1
- HD13
136 142 143 1 1.0950e+02 4.1840e+02 ; CG - CD2
- HD21
136 142 144 1 1.0950e+02 4.1840e+02 ; CG - CD2
- HD22
136 142 145 1 1.0950e+02 4.1840e+02 ; CG - CD2
- HD23
137 136 138 1 1.0950e+02 4.1840e+02 ; HG - CG
- CD1
137 136 142 1 1.0950e+02 4.1840e+02 ; HG - CG
- CD2
138 136 142 1 1.0950e+02 3.3472e+02 ; CD1 - CG
- CD2
139 138 140 1 1.0950e+02 2.9288e+02 ; HD11 - CD1
- HD12
139 138 141 1 1.0950e+02 2.9288e+02 ; HD11 - CD1
- HD13
140 138 141 1 1.0950e+02 2.9288e+02 ; HD12 - CD1
- HD13
143 142 144 1 1.0950e+02 2.9288e+02 ; HD21 - CD2
- HD22
143 142 145 1 1.0950e+02 2.9288e+02 ; HD21 - CD2
- HD23
144 142 145 1 1.0950e+02 2.9288e+02 ; HD22 - CD2
- HD23
146 148 149 1 1.2000e+02 4.1840e+02 ; C - N
- H
146 148 150 1 1.2190e+02 4.1840e+02 ; C - N
- CA
147 146 148 1 1.2290e+02 6.6944e+02 ; O - C
- N
148 150 151 1 1.0950e+02 4.1840e+02 ; N - CA
- HA
148 150 152 1 1.0970e+02 6.6944e+02 ; N - CA
- CB
148 150 161 1 1.1010e+02 5.2718e+02 ; N - CA
- C

```
   149    148    150     1    1.1804e+02    4.1840e+02 ;      H - N
- CA
   150    152    153     1    1.0950e+02    4.1840e+02 ;     CA - CB
- HB2
   150    152    154     1    1.0950e+02    4.1840e+02 ;     CA - CB
- HB3
   150    152    155     1    1.0950e+02    3.3472e+02 ;     CA - CB
- CG
   150    161    162     1    1.2040e+02    6.6944e+02 ;     CA - C
- O
   150    161    163     1    1.1660e+02    5.8576e+02 ;     CA - C
- N
   151    150    152     1    1.0950e+02    4.1840e+02 ;     HA - CA
- CB
   151    150    161     1    1.0950e+02    4.1840e+02 ;     HA - CA
- C
   152    150    161     1    1.1110e+02    5.2718e+02 ;     CB - CA
- C
   152    155    156     1    1.0950e+02    4.1840e+02 ;     CB - CG
- HG2
   152    155    157     1    1.0950e+02    4.1840e+02 ;     CB - CG
- HG3
   152    155    158     1    1.1110e+02    5.2718e+02 ;     CB - CG
- CD
   153    152    154     1    1.0950e+02    2.9288e+02 ;    HB2 - CB
- HB3
   153    152    155     1    1.0950e+02    4.1840e+02 ;    HB2 - CB
- CG
   154    152    155     1    1.0950e+02    4.1840e+02 ;    HB3 - CB
- CG
   155    158    159     1    1.1700e+02    5.8576e+02 ;     CG - CD
- OE1
   155    158    160     1    1.1700e+02    5.8576e+02 ;     CG - CD
- OE2
   156    155    157     1    1.0950e+02    2.9288e+02 ;    HG2 - CG
- HG3
   156    155    158     1    1.0950e+02    4.1840e+02 ;    HG2 - CG
- CD
   157    155    158     1    1.0950e+02    4.1840e+02 ;    HG3 - CG
- CD
   159    158    160     1    1.2600e+02    6.6944e+02 ;    OE1 - CD
- OE2
   161    163    164     1    1.2000e+02    4.1840e+02 ;      C - N
- H
   161    163    165     1    1.2190e+02    4.1840e+02 ;      C - N
- CA
   162    161    163     1    1.2290e+02    6.6944e+02 ;      O - C
- N
   163    165    166     1    1.0950e+02    4.1840e+02 ;      N - CA
- HA
   163    165    167     1    1.0970e+02    6.6944e+02 ;      N - CA
- CB
```

163 165 173 1 1.1010e+02 5.2718e+02 ; N - CA
- C
164 163 165 1 1.1804e+02 4.1840e+02 ; H - N
- CA
165 167 168 1 1.0950e+02 4.1840e+02 ; CA - CB
- HB2
165 167 169 1 1.0950e+02 4.1840e+02 ; CA - CB
- HB3
165 167 170 1 1.1110e+02 5.2718e+02 ; CA - CB
- CG
165 173 174 1 1.2040e+02 6.6944e+02 ; CA - C
- O
165 173 175 1 1.1660e+02 5.8576e+02 ; CA - C
- N
166 165 167 1 1.0950e+02 4.1840e+02 ; HA - CA
- CB
166 165 173 1 1.0950e+02 4.1840e+02 ; HA - CA
- C
167 165 173 1 1.1110e+02 5.2718e+02 ; CB - CA
- C
167 170 171 1 1.1700e+02 5.8576e+02 ; CB - CG
- OD1
167 170 172 1 1.1700e+02 5.8576e+02 ; CB - CG
- OD2
168 167 169 1 1.0950e+02 2.9288e+02 ; HB2 - CB
- HB3
168 167 170 1 1.0950e+02 4.1840e+02 ; HB2 - CB
- CG
169 167 170 1 1.0950e+02 4.1840e+02 ; HB3 - CB
- CG
171 170 172 1 1.2600e+02 6.6944e+02 ; OD1 - CG
- OD2
173 175 176 1 1.2000e+02 4.1840e+02 ; C - N
- H
173 175 177 1 1.2190e+02 4.1840e+02 ; C - N
- CA
174 173 175 1 1.2290e+02 6.6944e+02 ; O - C
- N
175 177 178 1 1.0950e+02 4.1840e+02 ; N - CA
- HA
175 177 179 1 1.0970e+02 6.6944e+02 ; N - CA
- CB
175 177 195 1 1.1010e+02 5.2718e+02 ; N - CA
- C
176 175 177 1 1.1804e+02 4.1840e+02 ; H - N
- CA
177 179 180 1 1.0950e+02 4.1840e+02 ; CA - CB
- HB2
177 179 181 1 1.0950e+02 4.1840e+02 ; CA - CB
- HB3
177 179 182 1 1.0950e+02 3.3472e+02 ; CA - CB
- CG

```
   177    195    196      1    1.2040e+02    6.6944e+02 ;     CA - C
- O
   177    195    197      1    1.1660e+02    5.8576e+02 ;     CA - C
- N
   178    177    179      1    1.0950e+02    4.1840e+02 ;     HA - CA
- CB
   178    177    195      1    1.0950e+02    4.1840e+02 ;     HA - CA
- C
   179    177    195      1    1.1110e+02    5.2718e+02 ;     CB - CA
- C
   179    182    183      1    1.0950e+02    4.1840e+02 ;     CB - CG
- HG2
   179    182    184      1    1.0950e+02    4.1840e+02 ;     CB - CG
- HG3
   179    182    185      1    1.0950e+02    3.3472e+02 ;     CB - CG
- CD
   180    179    181      1    1.0950e+02    2.9288e+02 ;    HB2 - CB
- HB3
   180    179    182      1    1.0950e+02    4.1840e+02 ;    HB2 - CB
- CG
   181    179    182      1    1.0950e+02    4.1840e+02 ;    HB3 - CB
- CG
   182    185    186      1    1.0950e+02    4.1840e+02 ;     CG - CD
- HD2
   182    185    187      1    1.0950e+02    4.1840e+02 ;     CG - CD
- HD3
   182    185    188      1    1.0950e+02    3.3472e+02 ;     CG - CD
- CE
   183    182    184      1    1.0950e+02    2.9288e+02 ;    HG2 - CG
- HG3
   183    182    185      1    1.0950e+02    4.1840e+02 ;    HG2 - CG
- CD
   184    182    185      1    1.0950e+02    4.1840e+02 ;    HG3 - CG
- CD
   185    188    189      1    1.0950e+02    4.1840e+02 ;     CD - CE
- HE2
   185    188    190      1    1.0950e+02    4.1840e+02 ;     CD - CE
- HE3
   185    188    191      1    1.1120e+02    6.6944e+02 ;     CD - CE
- NZ
   186    185    187      1    1.0950e+02    2.9288e+02 ;    HD2 - CD
- HD3
   186    185    188      1    1.0950e+02    4.1840e+02 ;    HD2 - CD
- CE
   187    185    188      1    1.0950e+02    4.1840e+02 ;    HD3 - CD
- CE
   188    191    192      1    1.0950e+02    4.1840e+02 ;     CE - NZ
- HZ1
   188    191    193      1    1.0950e+02    4.1840e+02 ;     CE - NZ
- HZ2
   188    191    194      1    1.0950e+02    4.1840e+02 ;     CE - NZ
- HZ3
```

```
   189    188    190      1    1.0950e+02    2.9288e+02 ;    HE2 - CE - HE3
   189    188    191      1    1.0950e+02    4.1840e+02 ;    HE2 - CE - NZ
   190    188    191      1    1.0950e+02    4.1840e+02 ;    HE3 - CE - NZ
   192    191    193      1    1.0950e+02    2.9288e+02 ;    HZ1 - NZ - HZ2
   192    191    194      1    1.0950e+02    2.9288e+02 ;    HZ1 - NZ - HZ3
   193    191    194      1    1.0950e+02    2.9288e+02 ;    HZ2 - NZ - HZ3
   195    197    198      1    1.2000e+02    4.1840e+02 ;      C - N - H
   195    197    199      1    1.2190e+02    4.1840e+02 ;      C - N - CA
   196    195    197      1    1.2290e+02    6.6944e+02 ;      O - C - N
   197    199    200      1    1.0950e+02    4.1840e+02 ;      N - CA - HA
   197    199    201      1    1.0970e+02    6.6944e+02 ;      N - CA - CB
   197    199    216      1    1.1010e+02    5.2718e+02 ;      N - CA - C
   198    197    199      1    1.1804e+02    4.1840e+02 ;      H - N - CA
   199    201    202      1    1.0950e+02    4.1840e+02 ;     CA - CB - HB2
   199    201    203      1    1.0950e+02    4.1840e+02 ;     CA - CB - HB3
   199    201    204      1    1.1400e+02    5.2718e+02 ;     CA - CB - CG
   199    216    217      1    1.2040e+02    6.6944e+02 ;     CA - C - O
   199    216    218      1    1.1660e+02    5.8576e+02 ;     CA - C - N
   200    199    201      1    1.0950e+02    4.1840e+02 ;     HA - CA - CB
   200    199    216      1    1.0950e+02    4.1840e+02 ;     HA - CA - C
   201    199    216      1    1.1110e+02    5.2718e+02 ;     CB - CA - C
   201    204    205      1    1.2000e+02    5.8576e+02 ;     CB - CG - CD1
   201    204    214      1    1.2000e+02    5.8576e+02 ;     CB - CG - CD2
   202    201    203      1    1.0950e+02    2.9288e+02 ;    HB2 - CB - HB3
   202    201    204      1    1.0950e+02    4.1840e+02 ;    HB2 - CB - CG
   203    201    204      1    1.0950e+02    4.1840e+02 ;    HB3 - CB - CG
```

204 205 206 1 1.2000e+02 4.1840e+02 ; CG - CD1
- HD1
204 205 207 1 1.2000e+02 5.2718e+02 ; CG - CD1
- CE1
204 214 212 1 1.2000e+02 5.2718e+02 ; CG - CD2
- CE2
204 214 215 1 1.2000e+02 4.1840e+02 ; CG - CD2
- HD2
205 204 214 1 1.2000e+02 5.2718e+02 ; CD1 - CG
- CD2
205 207 208 1 1.2000e+02 4.1840e+02 ; CD1 - CE1
- HE1
205 207 209 1 1.2000e+02 5.2718e+02 ; CD1 - CE1
- CZ
206 205 207 1 1.2000e+02 4.1840e+02 ; HD1 - CD1
- CE1
207 209 210 1 1.2000e+02 5.8576e+02 ; CE1 - CZ
- OH
207 209 212 1 1.2000e+02 5.2718e+02 ; CE1 - CZ
- CE2
208 207 209 1 1.2000e+02 4.1840e+02 ; HE1 - CE1
- CZ
209 210 211 1 1.1300e+02 4.1840e+02 ; CZ - OH
- HH
209 212 213 1 1.2000e+02 4.1840e+02 ; CZ - CE2
- HE2
209 212 214 1 1.2000e+02 5.2718e+02 ; CZ - CE2
- CD2
210 209 212 1 1.2000e+02 5.8576e+02 ; OH - CZ
- CE2
212 214 215 1 1.2000e+02 4.1840e+02 ; CE2 - CD2
- HD2
213 212 214 1 1.2000e+02 4.1840e+02 ; HE2 - CE2
- CD2
216 218 219 1 1.2000e+02 4.1840e+02 ; C - N
- H
216 218 220 1 1.2190e+02 4.1840e+02 ; C - N
- CA
217 216 218 1 1.2290e+02 6.6944e+02 ; O - C
- N
218 220 221 1 1.0950e+02 4.1840e+02 ; N - CA
- HA
218 220 222 1 1.0970e+02 6.6944e+02 ; N - CA
- CB
218 220 235 1 1.1010e+02 5.2718e+02 ; N - CA
- C
219 218 220 1 1.1804e+02 4.1840e+02 ; H - N
- CA
220 222 223 1 1.0950e+02 4.1840e+02 ; CA - CB
- HB
220 222 224 1 1.0950e+02 3.3472e+02 ; CA - CB
- CG2

220 222 228 1 1.0950e+02 3.3472e+02 ; CA - CB
- CG1
220 235 236 1 1.2040e+02 6.6944e+02 ; CA - C
- O
220 235 237 1 1.1660e+02 5.8576e+02 ; CA - C
- N
221 220 222 1 1.0950e+02 4.1840e+02 ; HA - CA
- CB
221 220 235 1 1.0950e+02 4.1840e+02 ; HA - CA
- C
222 220 235 1 1.1110e+02 5.2718e+02 ; CB - CA
- C
222 224 225 1 1.0950e+02 4.1840e+02 ; CB - CG2
- HG21
222 224 226 1 1.0950e+02 4.1840e+02 ; CB - CG2
- HG22
222 224 227 1 1.0950e+02 4.1840e+02 ; CB - CG2
- HG23
222 228 229 1 1.0950e+02 4.1840e+02 ; CB - CG1
- HG12
222 228 230 1 1.0950e+02 4.1840e+02 ; CB - CG1
- HG13
222 228 231 1 1.0950e+02 3.3472e+02 ; CB - CG1
- CD1
223 222 224 1 1.0950e+02 4.1840e+02 ; HB - CB
- CG2
223 222 228 1 1.0950e+02 4.1840e+02 ; HB - CB
- CG1
224 222 228 1 1.0950e+02 3.3472e+02 ; CG2 - CB
- CG1
225 224 226 1 1.0950e+02 2.9288e+02 ; HG21 - CG2
- HG22
225 224 227 1 1.0950e+02 2.9288e+02 ; HG21 - CG2
- HG23
226 224 227 1 1.0950e+02 2.9288e+02 ; HG22 - CG2
- HG23
228 231 232 1 1.0950e+02 4.1840e+02 ; CG1 - CD1
- HD11
228 231 233 1 1.0950e+02 4.1840e+02 ; CG1 - CD1
- HD12
228 231 234 1 1.0950e+02 4.1840e+02 ; CG1 - CD1
- HD13
229 228 230 1 1.0950e+02 2.9288e+02 ; HG12 - CG1
- HG13
229 228 231 1 1.0950e+02 4.1840e+02 ; HG12 - CG1
- CD1
230 228 231 1 1.0950e+02 4.1840e+02 ; HG13 - CG1
- CD1
232 231 233 1 1.0950e+02 2.9288e+02 ; HD11 - CD1
- HD12
232 231 234 1 1.0950e+02 2.9288e+02 ; HD11 - CD1
- HD13

```
   233   231   234     1   1.0950e+02   2.9288e+02 ;   HD12 - CD1
- HD13
   235   237   238     1   1.2000e+02   4.1840e+02 ;      C - N
- H
   235   237   239     1   1.2190e+02   4.1840e+02 ;      C - N
- CA
   236   235   237     1   1.2290e+02   6.6944e+02 ;      O - C
- N
   237   239   240     1   1.0950e+02   4.1840e+02 ;      N - CA
- HA
   237   239   241     1   1.0970e+02   6.6944e+02 ;      N - CA
- CB
   237   239   254     1   1.1010e+02   5.2718e+02 ;      N - CA
- C
   238   237   239     1   1.1804e+02   4.1840e+02 ;      H - N
- CA
   239   241   242     1   1.0950e+02   4.1840e+02 ;     CA - CB
- HB
   239   241   243     1   1.0950e+02   3.3472e+02 ;     CA - CB
- CG2
   239   241   247     1   1.0950e+02   3.3472e+02 ;     CA - CB
- CG1
   239   254   255     1   1.2040e+02   6.6944e+02 ;     CA - C
- O
   239   254   256     1   1.1660e+02   5.8576e+02 ;     CA - C
- N
   240   239   241     1   1.0950e+02   4.1840e+02 ;     HA - CA
- CB
   240   239   254     1   1.0950e+02   4.1840e+02 ;     HA - CA
- C
   241   239   254     1   1.1110e+02   5.2718e+02 ;     CB - CA
- C
   241   243   244     1   1.0950e+02   4.1840e+02 ;     CB - CG2
- HG21
   241   243   245     1   1.0950e+02   4.1840e+02 ;     CB - CG2
- HG22
   241   243   246     1   1.0950e+02   4.1840e+02 ;     CB - CG2
- HG23
   241   247   248     1   1.0950e+02   4.1840e+02 ;     CB - CG1
- HG12
   241   247   249     1   1.0950e+02   4.1840e+02 ;     CB - CG1
- HG13
   241   247   250     1   1.0950e+02   3.3472e+02 ;     CB - CG1
- CD1
   242   241   243     1   1.0950e+02   4.1840e+02 ;     HB - CB
- CG2
   242   241   247     1   1.0950e+02   4.1840e+02 ;     HB - CB
- CG1
   243   241   247     1   1.0950e+02   3.3472e+02 ;    CG2 - CB
- CG1
   244   243   245     1   1.0950e+02   2.9288e+02 ;   HG21 - CG2
- HG22
```

```
   244    243    246      1    1.0950e+02    2.9288e+02 ;   HG21 - CG2
- HG23
   245    243    246      1    1.0950e+02    2.9288e+02 ;   HG22 - CG2
- HG23
   247    250    251      1    1.0950e+02    4.1840e+02 ;    CG1 - CD1
- HD11
   247    250    252      1    1.0950e+02    4.1840e+02 ;    CG1 - CD1
- HD12
   247    250    253      1    1.0950e+02    4.1840e+02 ;    CG1 - CD1
- HD13
   248    247    249      1    1.0950e+02    2.9288e+02 ;   HG12 - CG1
- HG13
   248    247    250      1    1.0950e+02    4.1840e+02 ;   HG12 - CG1
- CD1
   249    247    250      1    1.0950e+02    4.1840e+02 ;   HG13 - CG1
- CD1
   251    250    252      1    1.0950e+02    2.9288e+02 ;   HD11 - CD1
- HD12
   251    250    253      1    1.0950e+02    2.9288e+02 ;   HD11 - CD1
- HD13
   252    250    253      1    1.0950e+02    2.9288e+02 ;   HD12 - CD1
- HD13
   254    256    257      1    1.2000e+02    4.1840e+02 ;      C - N
- H
   254    256    258      1    1.2190e+02    4.1840e+02 ;      C - N
- CA
   255    254    256      1    1.2290e+02    6.6944e+02 ;      O - C
- N
   256    258    259      1    1.0950e+02    4.1840e+02 ;      N - CA
- HA
   256    258    260      1    1.0970e+02    6.6944e+02 ;      N - CA
- CB
   256    258    266      1    1.1010e+02    5.2718e+02 ;      N - CA
- C
   257    256    258      1    1.1804e+02    4.1840e+02 ;      H - N
- CA
   258    260    261      1    1.0950e+02    4.1840e+02 ;     CA - CB
- HB2
   258    260    262      1    1.0950e+02    4.1840e+02 ;     CA - CB
- HB3
   258    260    263      1    1.1110e+02    5.2718e+02 ;     CA - CB
- CG
   258    266    267      1    1.2040e+02    6.6944e+02 ;     CA - C
- O
   258    266    268      1    1.1660e+02    5.8576e+02 ;     CA - C
- N
   259    258    260      1    1.0950e+02    4.1840e+02 ;     HA - CA
- CB
   259    258    266      1    1.0950e+02    4.1840e+02 ;     HA - CA
- C
   260    258    266      1    1.1110e+02    5.2718e+02 ;     CB - CA
- C
```

| | | | | | | | |
|---|---|---|---|---|---|---|---|
| 260 | 263 | 264 | 1 | 1.1700e+02 | 5.8576e+02 | ; | CB - CG - OD1 |
| 260 | 263 | 265 | 1 | 1.1700e+02 | 5.8576e+02 | ; | CB - CG - OD2 |
| 261 | 260 | 262 | 1 | 1.0950e+02 | 2.9288e+02 | ; | HB2 - CB - HB3 |
| 261 | 260 | 263 | 1 | 1.0950e+02 | 4.1840e+02 | ; | HB2 - CB - CG |
| 262 | 260 | 263 | 1 | 1.0950e+02 | 4.1840e+02 | ; | HB3 - CB - CG |
| 264 | 263 | 265 | 1 | 1.2600e+02 | 6.6944e+02 | ; | OD1 - CG - OD2 |
| 266 | 268 | 269 | 1 | 1.2000e+02 | 4.1840e+02 | ; | C - N - H |
| 266 | 268 | 270 | 1 | 1.2190e+02 | 4.1840e+02 | ; | C - N - CA |
| 267 | 266 | 268 | 1 | 1.2290e+02 | 6.6944e+02 | ; | O - C - N |
| 268 | 270 | 271 | 1 | 1.0950e+02 | 4.1840e+02 | ; | N - CA - HA |
| 268 | 270 | 272 | 1 | 1.0970e+02 | 6.6944e+02 | ; | N - CA - CB |
| 268 | 270 | 281 | 1 | 1.1010e+02 | 5.2718e+02 | ; | N - CA - C |
| 269 | 268 | 270 | 1 | 1.1804e+02 | 4.1840e+02 | ; | H - N - CA |
| 270 | 272 | 273 | 1 | 1.0950e+02 | 4.1840e+02 | ; | CA - CB - HB2 |
| 270 | 272 | 274 | 1 | 1.0950e+02 | 4.1840e+02 | ; | CA - CB - HB3 |
| 270 | 272 | 275 | 1 | 1.0950e+02 | 3.3472e+02 | ; | CA - CB - CG |
| 270 | 281 | 282 | 1 | 1.2040e+02 | 6.6944e+02 | ; | CA - C - O |
| 270 | 281 | 283 | 1 | 1.1660e+02 | 5.8576e+02 | ; | CA - C - N |
| 271 | 270 | 272 | 1 | 1.0950e+02 | 4.1840e+02 | ; | HA - CA - CB |
| 271 | 270 | 281 | 1 | 1.0950e+02 | 4.1840e+02 | ; | HA - CA - C |
| 272 | 270 | 281 | 1 | 1.1110e+02 | 5.2718e+02 | ; | CB - CA - C |
| 272 | 275 | 276 | 1 | 1.0950e+02 | 4.1840e+02 | ; | CB - CG - HG2 |
| 272 | 275 | 277 | 1 | 1.0950e+02 | 4.1840e+02 | ; | CB - CG - HG3 |
| 272 | 275 | 278 | 1 | 1.1110e+02 | 5.2718e+02 | ; | CB - CG - CD |
| 273 | 272 | 274 | 1 | 1.0950e+02 | 2.9288e+02 | ; | HB2 - CB - HB3 |
| 273 | 272 | 275 | 1 | 1.0950e+02 | 4.1840e+02 | ; | HB2 - CB - CG |

```
   274    272    275      1    1.0950e+02    4.1840e+02 ;    HB3 - CB
- CG
   275    278    279      1    1.1700e+02    5.8576e+02 ;     CG - CD
- OE1
   275    278    280      1    1.1700e+02    5.8576e+02 ;     CG - CD
- OE2
   276    275    277      1    1.0950e+02    2.9288e+02 ;    HG2 - CG
- HG3
   276    275    278      1    1.0950e+02    4.1840e+02 ;    HG2 - CG
- CD
   277    275    278      1    1.0950e+02    4.1840e+02 ;    HG3 - CG
- CD
   279    278    280      1    1.2600e+02    6.6944e+02 ;    OE1 - CD
- OE2
   281    283    284      1    1.2000e+02    4.1840e+02 ;      C - N
- H
   281    283    285      1    1.2190e+02    4.1840e+02 ;      C - N
- CA
   282    281    283      1    1.2290e+02    6.6944e+02 ;      O - C
- N
   283    285    286      1    1.0950e+02    4.1840e+02 ;      N - CA
- HA
   283    285    287      1    1.0970e+02    6.6944e+02 ;      N - CA
- CB
   283    285    303      1    1.1010e+02    5.2718e+02 ;      N - CA
- C
   284    283    285      1    1.1804e+02    4.1840e+02 ;      H - N
- CA
   285    287    288      1    1.0950e+02    4.1840e+02 ;     CA - CB
- HB2
   285    287    289      1    1.0950e+02    4.1840e+02 ;     CA - CB
- HB3
   285    287    290      1    1.0950e+02    3.3472e+02 ;     CA - CB
- CG
   285    303    304      1    1.2040e+02    6.6944e+02 ;     CA - C
- O
   285    303    305      1    1.1660e+02    5.8576e+02 ;     CA - C
- N
   286    285    287      1    1.0950e+02    4.1840e+02 ;     HA - CA
- CB
   286    285    303      1    1.0950e+02    4.1840e+02 ;     HA - CA
- C
   287    285    303      1    1.1110e+02    5.2718e+02 ;     CB - CA
- C
   287    290    291      1    1.0950e+02    4.1840e+02 ;     CB - CG
- HG2
   287    290    292      1    1.0950e+02    4.1840e+02 ;     CB - CG
- HG3
   287    290    293      1    1.0950e+02    3.3472e+02 ;     CB - CG
- CD
   288    287    289      1    1.0950e+02    2.9288e+02 ;    HB2 - CB
- HB3
```

```
   288    287    290      1    1.0950e+02    4.1840e+02 ;    HB2 - CB
 - CG
   289    287    290      1    1.0950e+02    4.1840e+02 ;    HB3 - CB
 - CG
   290    293    294      1    1.0950e+02    4.1840e+02 ;     CG - CD
 - HD2
   290    293    295      1    1.0950e+02    4.1840e+02 ;     CG - CD
 - HD3
   290    293    296      1    1.0950e+02    3.3472e+02 ;     CG - CD
 - CE
   291    290    292      1    1.0950e+02    2.9288e+02 ;    HG2 - CG
 - HG3
   291    290    293      1    1.0950e+02    4.1840e+02 ;    HG2 - CG
 - CD
   292    290    293      1    1.0950e+02    4.1840e+02 ;    HG3 - CG
 - CD
   293    296    297      1    1.0950e+02    4.1840e+02 ;     CD - CE
 - HE2
   293    296    298      1    1.0950e+02    4.1840e+02 ;     CD - CE
 - HE3
   293    296    299      1    1.1120e+02    6.6944e+02 ;     CD - CE
 - NZ
   294    293    295      1    1.0950e+02    2.9288e+02 ;    HD2 - CD
 - HD3
   294    293    296      1    1.0950e+02    4.1840e+02 ;    HD2 - CD
 - CE
   295    293    296      1    1.0950e+02    4.1840e+02 ;    HD3 - CD
 - CE
   296    299    300      1    1.0950e+02    4.1840e+02 ;     CE - NZ
 - HZ1
   296    299    301      1    1.0950e+02    4.1840e+02 ;     CE - NZ
 - HZ2
   296    299    302      1    1.0950e+02    4.1840e+02 ;     CE - NZ
 - HZ3
   297    296    298      1    1.0950e+02    2.9288e+02 ;    HE2 - CE
 - HE3
   297    296    299      1    1.0950e+02    4.1840e+02 ;    HE2 - CE
 - NZ
   298    296    299      1    1.0950e+02    4.1840e+02 ;    HE3 - CE
 - NZ
   300    299    301      1    1.0950e+02    2.9288e+02 ;    HZ1 - NZ
 - HZ2
   300    299    302      1    1.0950e+02    2.9288e+02 ;    HZ1 - NZ
 - HZ3
   301    299    302      1    1.0950e+02    2.9288e+02 ;    HZ2 - NZ
 - HZ3
   303    305    306      1    1.2000e+02    4.1840e+02 ;      C - N
 - H
   303    305    307      1    1.2190e+02    4.1840e+02 ;      C - N
 - CA
   304    303    305      1    1.2290e+02    6.6944e+02 ;      O - C
 - N
```

```
   305    307    308      1    1.0950e+02    4.1840e+02 ;      N - CA
- HA
   305    307    309      1    1.0970e+02    6.6944e+02 ;      N - CA
- CB
   305    307    322      1    1.1010e+02    5.2718e+02 ;      N - CA
- C
   306    305    307      1    1.1804e+02    4.1840e+02 ;      H - N
- CA
   307    309    310      1    1.0950e+02    4.1840e+02 ;     CA - CB
- HB2
   307    309    311      1    1.0950e+02    4.1840e+02 ;     CA - CB
- HB3
   307    309    312      1    1.0950e+02    3.3472e+02 ;     CA - CB
- CG
   307    322    323      1    1.2040e+02    6.6944e+02 ;     CA - C
- O
   307    322    324      1    1.1660e+02    5.8576e+02 ;     CA - C
- N
   308    307    309      1    1.0950e+02    4.1840e+02 ;     HA - CA
- CB
   308    307    322      1    1.0950e+02    4.1840e+02 ;     HA - CA
- C
   309    307    322      1    1.1110e+02    5.2718e+02 ;     CB - CA
- C
   309    312    313      1    1.0950e+02    4.1840e+02 ;     CB - CG
- HG
   309    312    314      1    1.0950e+02    3.3472e+02 ;     CB - CG
- CD1
   309    312    318      1    1.0950e+02    3.3472e+02 ;     CB - CG
- CD2
   310    309    311      1    1.0950e+02    2.9288e+02 ;    HB2 - CB
- HB3
   310    309    312      1    1.0950e+02    4.1840e+02 ;    HB2 - CB
- CG
   311    309    312      1    1.0950e+02    4.1840e+02 ;    HB3 - CB
- CG
   312    314    315      1    1.0950e+02    4.1840e+02 ;     CG - CD1
- HD11
   312    314    316      1    1.0950e+02    4.1840e+02 ;     CG - CD1
- HD12
   312    314    317      1    1.0950e+02    4.1840e+02 ;     CG - CD1
- HD13
   312    318    319      1    1.0950e+02    4.1840e+02 ;     CG - CD2
- HD21
   312    318    320      1    1.0950e+02    4.1840e+02 ;     CG - CD2
- HD22
   312    318    321      1    1.0950e+02    4.1840e+02 ;     CG - CD2
- HD23
   313    312    314      1    1.0950e+02    4.1840e+02 ;     HG - CG
- CD1
   313    312    318      1    1.0950e+02    4.1840e+02 ;     HG - CG
- CD2
```

```
   314    312    318      1    1.0950e+02    3.3472e+02 ;    CD1 - CG
- CD2
   315    314    316      1    1.0950e+02    2.9288e+02 ;   HD11 - CD1
- HD12
   315    314    317      1    1.0950e+02    2.9288e+02 ;   HD11 - CD1
- HD13
   316    314    317      1    1.0950e+02    2.9288e+02 ;   HD12 - CD1
- HD13
   319    318    320      1    1.0950e+02    2.9288e+02 ;   HD21 - CD2
- HD22
   319    318    321      1    1.0950e+02    2.9288e+02 ;   HD21 - CD2
- HD23
   320    318    321      1    1.0950e+02    2.9288e+02 ;   HD22 - CD2
- HD23
   322    324    325      1    1.2000e+02    4.1840e+02 ;      C - N
- H
   322    324    326      1    1.2190e+02    4.1840e+02 ;      C - N
- CA
   323    322    324      1    1.2290e+02    6.6944e+02 ;      O - C
- N
   324    326    327      1    1.0950e+02    4.1840e+02 ;      N - CA
- HA2
   324    326    328      1    1.0950e+02    4.1840e+02 ;      N - CA
- HA3
   324    326    329      1    1.1010e+02    5.2718e+02 ;      N - CA
- C
   325    324    326      1    1.1804e+02    4.1840e+02 ;      H - N
- CA
   326    329    330      1    1.2040e+02    6.6944e+02 ;     CA - C
- O
   326    329    331      1    1.1660e+02    5.8576e+02 ;     CA - C
- N
   327    326    328      1    1.0950e+02    2.9288e+02 ;    HA2 - CA
- HA3
   327    326    329      1    1.0950e+02    4.1840e+02 ;    HA2 - CA
- C
   328    326    329      1    1.0950e+02    4.1840e+02 ;    HA3 - CA
- C
   329    331    332      1    1.2000e+02    4.1840e+02 ;      C - N
- H
   329    331    333      1    1.2190e+02    4.1840e+02 ;      C - N
- CA
   330    329    331      1    1.2290e+02    6.6944e+02 ;      O - C
- N
   331    333    334      1    1.0950e+02    4.1840e+02 ;      N - CA
- HA
   331    333    335      1    1.0970e+02    6.6944e+02 ;      N - CA
- CB
   331    333    346      1    1.1010e+02    5.2718e+02 ;      N - CA
- C
   332    331    333      1    1.1804e+02    4.1840e+02 ;      H - N
- CA
```

```
   333   335   336     1    1.0950e+02   4.1840e+02 ;     CA - CB
- HB2
   333   335   337     1    1.0950e+02   4.1840e+02 ;     CA - CB
- HB3
   333   335   338     1    1.0950e+02   3.3472e+02 ;     CA - CB
- CG
   333   346   347     1    1.2040e+02   6.6944e+02 ;     CA - C
- O
   333   346   348     1    1.1660e+02   5.8576e+02 ;     CA - C
- N
   334   333   335     1    1.0950e+02   4.1840e+02 ;     HA - CA
- CB
   334   333   346     1    1.0950e+02   4.1840e+02 ;     HA - CA
- C
   335   333   346     1    1.1110e+02   5.2718e+02 ;     CB - CA
- C
   335   338   339     1    1.0950e+02   4.1840e+02 ;     CB - CG
- HG2
   335   338   340     1    1.0950e+02   4.1840e+02 ;     CB - CG
- HG3
   335   338   341     1    1.1110e+02   5.2718e+02 ;     CB - CG
- CD
   336   335   337     1    1.0950e+02   2.9288e+02 ;    HB2 - CB
- HB3
   336   335   338     1    1.0950e+02   4.1840e+02 ;    HB2 - CB
- CG
   337   335   338     1    1.0950e+02   4.1840e+02 ;    HB3 - CB
- CG
   338   341   342     1    1.2040e+02   6.6944e+02 ;     CG - CD
- OE1
   338   341   343     1    1.1660e+02   5.8576e+02 ;     CG - CD
- NE2
   339   338   340     1    1.0950e+02   2.9288e+02 ;    HG2 - CG
- HG3
   339   338   341     1    1.0950e+02   4.1840e+02 ;    HG2 - CG
- CD
   340   338   341     1    1.0950e+02   4.1840e+02 ;    HG3 - CG
- CD
   341   343   344     1    1.2000e+02   4.1840e+02 ;     CD - NE2
- HE21
   341   343   345     1    1.2000e+02   4.1840e+02 ;     CD - NE2
- HE22
   342   341   343     1    1.2290e+02   6.6944e+02 ;    OE1 - CD
- NE2
   344   343   345     1    1.2000e+02   2.9288e+02 ;   HE21 - NE2
- HE22
   346   348   349     1    1.2000e+02   4.1840e+02 ;      C - N
- H
   346   348   350     1    1.2190e+02   4.1840e+02 ;      C - N
- CA
   347   346   348     1    1.2290e+02   6.6944e+02 ;      O - C
- N
```

```
   348    350    351      1    1.0950e+02    4.1840e+02 ;      N - CA
- HA2
   348    350    352      1    1.0950e+02    4.1840e+02 ;      N - CA
- HA3
   348    350    353      1    1.1010e+02    5.2718e+02 ;      N - CA
- C
   349    348    350      1    1.1804e+02    4.1840e+02 ;      H - N
- CA
   350    353    354      1    1.2040e+02    6.6944e+02 ;     CA - C
- O
   350    353    355      1    1.1660e+02    5.8576e+02 ;     CA - C
- N
   351    350    352      1    1.0950e+02    2.9288e+02 ;    HA2 - CA
- HA3
   351    350    353      1    1.0950e+02    4.1840e+02 ;    HA2 - CA
- C
   352    350    353      1    1.0950e+02    4.1840e+02 ;    HA3 - CA
- C
   353    355    356      1    1.2000e+02    4.1840e+02 ;      C - N
- H
   353    355    357      1    1.2190e+02    4.1840e+02 ;      C - N
- CA
   354    353    355      1    1.2290e+02    6.6944e+02 ;      O - C
- N
   355    357    358      1    1.0950e+02    4.1840e+02 ;      N - CA
- HA
   355    357    359      1    1.0970e+02    6.6944e+02 ;      N - CA
- CB
   355    357    367      1    1.1010e+02    5.2718e+02 ;      N - CA
- C
   356    355    357      1    1.1804e+02    4.1840e+02 ;      H - N
- CA
   357    359    360      1    1.0950e+02    4.1840e+02 ;     CA - CB
- HB
   357    359    361      1    1.0950e+02    3.3472e+02 ;     CA - CB
- CG2
   357    359    365      1    1.0950e+02    4.1840e+02 ;     CA - CB
- OG1
   357    367    368      1    1.2040e+02    6.6944e+02 ;     CA - C
- O
   357    367    369      1    1.1660e+02    5.8576e+02 ;     CA - C
- N
   358    357    359      1    1.0950e+02    4.1840e+02 ;     HA - CA
- CB
   358    357    367      1    1.0950e+02    4.1840e+02 ;     HA - CA
- C
   359    357    367      1    1.1110e+02    5.2718e+02 ;     CB - CA
- C
   359    361    362      1    1.0950e+02    4.1840e+02 ;     CB - CG2
- HG21
   359    361    363      1    1.0950e+02    4.1840e+02 ;     CB - CG2
- HG22
```

```
   359    361    364      1    1.0950e+02    4.1840e+02 ;     CB - CG2
- HG23
   359    365    366      1    1.0850e+02    4.6024e+02 ;     CB - OG1
- HG1
   360    359    361      1    1.0950e+02    4.1840e+02 ;     HB - CB
- CG2
   360    359    365      1    1.0950e+02    4.1840e+02 ;     HB - CB
- OG1
   361    359    365      1    1.0950e+02    4.1840e+02 ;    CG2 - CB
- OG1
   362    361    363      1    1.0950e+02    2.9288e+02 ;   HG21 - CG2
- HG22
   362    361    364      1    1.0950e+02    2.9288e+02 ;   HG21 - CG2
- HG23
   363    361    364      1    1.0950e+02    2.9288e+02 ;   HG22 - CG2
- HG23
   367    369    370      1    1.2000e+02    4.1840e+02 ;      C - N
- H
   367    369    371      1    1.2190e+02    4.1840e+02 ;      C - N
- CA
   368    367    369      1    1.2290e+02    6.6944e+02 ;      O - C
- N
   369    371    372      1    1.0950e+02    4.1840e+02 ;      N - CA
- HA
   369    371    373      1    1.0970e+02    6.6944e+02 ;      N - CA
- CB
   369    371    388      1    1.1010e+02    5.2718e+02 ;      N - CA
- C
   370    369    371      1    1.1804e+02    4.1840e+02 ;      H - N
- CA
   371    373    374      1    1.0950e+02    4.1840e+02 ;     CA - CB
- HB2
   371    373    375      1    1.0950e+02    4.1840e+02 ;     CA - CB
- HB3
   371    373    376      1    1.1400e+02    5.2718e+02 ;     CA - CB
- CG
   371    388    389      1    1.2040e+02    6.6944e+02 ;     CA - C
- O
   371    388    390      1    1.1660e+02    5.8576e+02 ;     CA - C
- N
   372    371    373      1    1.0950e+02    4.1840e+02 ;     HA - CA
- CB
   372    371    388      1    1.0950e+02    4.1840e+02 ;     HA - CA
- C
   373    371    388      1    1.1110e+02    5.2718e+02 ;     CB - CA
- C
   373    376    377      1    1.2000e+02    5.8576e+02 ;     CB - CG
- CD1
   373    376    386      1    1.2000e+02    5.8576e+02 ;     CB - CG
- CD2
   374    373    375      1    1.0950e+02    2.9288e+02 ;    HB2 - CB
- HB3
```

```
   374    373    376       1    1.0950e+02    4.1840e+02 ;    HB2 - CB
- CG
   375    373    376       1    1.0950e+02    4.1840e+02 ;    HB3 - CB
- CG
   376    377    378       1    1.2000e+02    4.1840e+02 ;     CG - CD1
- HD1
   376    377    379       1    1.2000e+02    5.2718e+02 ;     CG - CD1
- CE1
   376    386    384       1    1.2000e+02    5.2718e+02 ;     CG - CD2
- CE2
   376    386    387       1    1.2000e+02    4.1840e+02 ;     CG - CD2
- HD2
   377    376    386       1    1.2000e+02    5.2718e+02 ;    CD1 - CG
- CD2
   377    379    380       1    1.2000e+02    4.1840e+02 ;    CD1 - CE1
- HE1
   377    379    381       1    1.2000e+02    5.2718e+02 ;    CD1 - CE1
- CZ
   378    377    379       1    1.2000e+02    4.1840e+02 ;    HD1 - CD1
- CE1
   379    381    382       1    1.2000e+02    5.8576e+02 ;    CE1 - CZ
- OH
   379    381    384       1    1.2000e+02    5.2718e+02 ;    CE1 - CZ
- CE2
   380    379    381       1    1.2000e+02    4.1840e+02 ;    HE1 - CE1
- CZ
   381    382    383       1    1.1300e+02    4.1840e+02 ;     CZ - OH
- HH
   381    384    385       1    1.2000e+02    4.1840e+02 ;     CZ - CE2
- HE2
   381    384    386       1    1.2000e+02    5.2718e+02 ;     CZ - CE2
- CD2
   382    381    384       1    1.2000e+02    5.8576e+02 ;     OH - CZ
- CE2
   384    386    387       1    1.2000e+02    4.1840e+02 ;    CE2 - CD2
- HD2
   385    384    386       1    1.2000e+02    4.1840e+02 ;    HE2 - CE2
- CD2
   388    390    391       1    1.2000e+02    4.1840e+02 ;      C - N
- H
   388    390    392       1    1.2190e+02    4.1840e+02 ;      C - N
- CA
   389    388    390       1    1.2290e+02    6.6944e+02 ;      O - C
- N
   390    392    393       1    1.0950e+02    4.1840e+02 ;      N - CA
- HA2
   390    392    394       1    1.0950e+02    4.1840e+02 ;      N - CA
- HA3
   390    392    395       1    1.1010e+02    5.2718e+02 ;      N - CA
- C
   391    390    392       1    1.1804e+02    4.1840e+02 ;      H - N
- CA
```

```
   392    395    396       1    1.2040e+02    6.6944e+02 ;     CA - C
- O
   392    395    397       1    1.1660e+02    5.8576e+02 ;     CA - C
- N
   393    392    394       1    1.0950e+02    2.9288e+02 ;    HA2 - CA
- HA3
   393    392    395       1    1.0950e+02    4.1840e+02 ;    HA2 - CA
- C
   394    392    395       1    1.0950e+02    4.1840e+02 ;    HA3 - CA
- C
   395    397    398       1    1.2000e+02    4.1840e+02 ;      C - N
- H
   395    397    399       1    1.2190e+02    4.1840e+02 ;      C - N
- CA
   396    395    397       1    1.2290e+02    6.6944e+02 ;      O - C
- N
   397    399    400       1    1.0950e+02    4.1840e+02 ;      N - CA
- HA
   397    399    401       1    1.0970e+02    6.6944e+02 ;      N - CA
- CB
   397    399    406       1    1.1010e+02    5.2718e+02 ;      N - CA
- C
   398    397    399       1    1.1804e+02    4.1840e+02 ;      H - N
- CA
   399    401    402       1    1.0950e+02    4.1840e+02 ;     CA - CB
- HB2
   399    401    403       1    1.0950e+02    4.1840e+02 ;     CA - CB
- HB3
   399    401    404       1    1.0860e+02    4.1840e+02 ;     CA - CB
- SG
   399    406    407       1    1.2040e+02    6.6944e+02 ;     CA - C
- O
   399    406    408       1    1.1660e+02    5.8576e+02 ;     CA - C
- N
   400    399    401       1    1.0950e+02    4.1840e+02 ;     HA - CA
- CB
   400    399    406       1    1.0950e+02    4.1840e+02 ;     HA - CA
- C
   401    399    406       1    1.1110e+02    5.2718e+02 ;     CB - CA
- C
   401    404    405       1    9.6000e+01    3.5982e+02 ;     CB - SG
- HG
   402    401    403       1    1.0950e+02    2.9288e+02 ;    HB2 - CB
- HB3
   402    401    404       1    1.0950e+02    4.1840e+02 ;    HB2 - CB
- SG
   403    401    404       1    1.0950e+02    4.1840e+02 ;    HB3 - CB
- SG
   406    408    409       1    1.2000e+02    4.1840e+02 ;      C - N
- H
   406    408    410       1    1.2190e+02    4.1840e+02 ;      C - N
- CA
```

```
   407    406    408      1    1.2290e+02    6.6944e+02 ;      O - C
 - N
   408    410    411      1    1.0950e+02    4.1840e+02 ;      N - CA
 - HA
   408    410    412      1    1.0970e+02    6.6944e+02 ;      N - CA
 - CB
   408    410    422      1    1.1010e+02    5.2718e+02 ;      N - CA
 - C
   409    408    410      1    1.1804e+02    4.1840e+02 ;      H - N
 - CA
   410    412    413      1    1.0950e+02    4.1840e+02 ;     CA - CB
 - HB
   410    412    414      1    1.0950e+02    3.3472e+02 ;     CA - CB
 - CG1
   410    412    418      1    1.0950e+02    3.3472e+02 ;     CA - CB
 - CG2
   410    422    423      1    1.2040e+02    6.6944e+02 ;     CA - C
 - O
   410    422    424      1    1.1660e+02    5.8576e+02 ;     CA - C
 - N
   411    410    412      1    1.0950e+02    4.1840e+02 ;     HA - CA
 - CB
   411    410    422      1    1.0950e+02    4.1840e+02 ;     HA - CA
 - C
   412    410    422      1    1.1110e+02    5.2718e+02 ;     CB - CA
 - C
   412    414    415      1    1.0950e+02    4.1840e+02 ;     CB - CG1
 - HG11
   412    414    416      1    1.0950e+02    4.1840e+02 ;     CB - CG1
 - HG12
   412    414    417      1    1.0950e+02    4.1840e+02 ;     CB - CG1
 - HG13
   412    418    419      1    1.0950e+02    4.1840e+02 ;     CB - CG2
 - HG21
   412    418    420      1    1.0950e+02    4.1840e+02 ;     CB - CG2
 - HG22
   412    418    421      1    1.0950e+02    4.1840e+02 ;     CB - CG2
 - HG23
   413    412    414      1    1.0950e+02    4.1840e+02 ;     HB - CB
 - CG1
   413    412    418      1    1.0950e+02    4.1840e+02 ;     HB - CB
 - CG2
   414    412    418      1    1.0950e+02    3.3472e+02 ;    CG1 - CB
 - CG2
   415    414    416      1    1.0950e+02    2.9288e+02 ;   HG11 - CG1
 - HG12
   415    414    417      1    1.0950e+02    2.9288e+02 ;   HG11 - CG1
 - HG13
   416    414    417      1    1.0950e+02    2.9288e+02 ;   HG12 - CG1
 - HG13
   419    418    420      1    1.0950e+02    2.9288e+02 ;   HG21 - CG2
 - HG22
```

419 418 421 1 1.0950e+02 2.9288e+02 ; HG21 - CG2
- HG23
420 418 421 1 1.0950e+02 2.9288e+02 ; HG22 - CG2
- HG23
422 424 425 1 1.2000e+02 4.1840e+02 ; C - N
- H
422 424 426 1 1.2190e+02 4.1840e+02 ; C - N
- CA
423 422 424 1 1.2290e+02 6.6944e+02 ; O - C
- N
424 426 427 1 1.0950e+02 4.1840e+02 ; N - CA
- HA
424 426 428 1 1.0970e+02 6.6944e+02 ; N - CA
- CB
424 426 443 1 1.1010e+02 5.2718e+02 ; N - CA
- C
425 424 426 1 1.1804e+02 4.1840e+02 ; H - N
- CA
426 428 429 1 1.0950e+02 4.1840e+02 ; CA - CB
- HB2
426 428 430 1 1.0950e+02 4.1840e+02 ; CA - CB
- HB3
426 428 431 1 1.1400e+02 5.2718e+02 ; CA - CB
- CG
426 443 444 1 1.2040e+02 6.6944e+02 ; CA - C
- O
426 443 445 1 1.1660e+02 5.8576e+02 ; CA - C
- N
427 426 428 1 1.0950e+02 4.1840e+02 ; HA - CA
- CB
427 426 443 1 1.0950e+02 4.1840e+02 ; HA - CA
- C
428 426 443 1 1.1110e+02 5.2718e+02 ; CB - CA
- C
428 431 432 1 1.2000e+02 5.8576e+02 ; CB - CG
- CD1
428 431 441 1 1.2000e+02 5.8576e+02 ; CB - CG
- CD2
429 428 430 1 1.0950e+02 2.9288e+02 ; HB2 - CB
- HB3
429 428 431 1 1.0950e+02 4.1840e+02 ; HB2 - CB
- CG
430 428 431 1 1.0950e+02 4.1840e+02 ; HB3 - CB
- CG
431 432 433 1 1.2000e+02 4.1840e+02 ; CG - CD1
- HD1
431 432 434 1 1.2000e+02 5.2718e+02 ; CG - CD1
- CE1
431 441 439 1 1.2000e+02 5.2718e+02 ; CG - CD2
- CE2
431 441 442 1 1.2000e+02 4.1840e+02 ; CG - CD2
- HD2

```
   432    431    441      1    1.2000e+02    5.2718e+02 ;    CD1 - CG
- CD2
   432    434    435      1    1.2000e+02    4.1840e+02 ;    CD1 - CE1
- HE1
   432    434    436      1    1.2000e+02    5.2718e+02 ;    CD1 - CE1
- CZ
   433    432    434      1    1.2000e+02    4.1840e+02 ;    HD1 - CD1
- CE1
   434    436    437      1    1.2000e+02    5.8576e+02 ;    CE1 - CZ
- OH
   434    436    439      1    1.2000e+02    5.2718e+02 ;    CE1 - CZ
- CE2
   435    434    436      1    1.2000e+02    4.1840e+02 ;    HE1 - CE1
- CZ
   436    437    438      1    1.1300e+02    4.1840e+02 ;     CZ - OH
- HH
   436    439    440      1    1.2000e+02    4.1840e+02 ;     CZ - CE2
- HE2
   436    439    441      1    1.2000e+02    5.2718e+02 ;     CZ - CE2
- CD2
   437    436    439      1    1.2000e+02    5.8576e+02 ;     OH - CZ
- CE2
   439    441    442      1    1.2000e+02    4.1840e+02 ;    CE2 - CD2
- HD2
   440    439    441      1    1.2000e+02    4.1840e+02 ;    HE2 - CE2
- CD2
   443    445    446      1    1.2000e+02    4.1840e+02 ;      C - N
- H
   443    445    447      1    1.2190e+02    4.1840e+02 ;      C - N
- CA
   444    443    445      1    1.2290e+02    6.6944e+02 ;      O - C
- N
   445    447    448      1    1.0950e+02    4.1840e+02 ;      N - CA
- HA
   445    447    449      1    1.0970e+02    6.6944e+02 ;      N - CA
- CB
   445    447    465      1    1.1010e+02    5.2718e+02 ;      N - CA
- C
   446    445    447      1    1.1804e+02    4.1840e+02 ;      H - N
- CA
   447    449    450      1    1.0950e+02    4.1840e+02 ;     CA - CB
- HB2
   447    449    451      1    1.0950e+02    4.1840e+02 ;     CA - CB
- HB3
   447    449    452      1    1.0950e+02    3.3472e+02 ;     CA - CB
- CG
   447    465    466      1    1.2040e+02    6.6944e+02 ;     CA - C
- O
   447    465    467      1    1.1660e+02    5.8576e+02 ;     CA - C
- N
   448    447    449      1    1.0950e+02    4.1840e+02 ;     HA - CA
- CB
```

```
   448    447    465      1   1.0950e+02   4.1840e+02 ;     HA - CA
 - C
   449    447    465      1   1.1110e+02   5.2718e+02 ;     CB - CA
 - C
   449    452    453      1   1.0950e+02   4.1840e+02 ;     CB - CG
 - HG2
   449    452    454      1   1.0950e+02   4.1840e+02 ;     CB - CG
 - HG3
   449    452    455      1   1.0950e+02   3.3472e+02 ;     CB - CG
 - CD
   450    449    451      1   1.0950e+02   2.9288e+02 ;    HB2 - CB
 - HB3
   450    449    452      1   1.0950e+02   4.1840e+02 ;    HB2 - CB
 - CG
   451    449    452      1   1.0950e+02   4.1840e+02 ;    HB3 - CB
 - CG
   452    455    456      1   1.0950e+02   4.1840e+02 ;     CG - CD
 - HD2
   452    455    457      1   1.0950e+02   4.1840e+02 ;     CG - CD
 - HD3
   452    455    458      1   1.0950e+02   3.3472e+02 ;     CG - CD
 - CE
   453    452    454      1   1.0950e+02   2.9288e+02 ;    HG2 - CG
 - HG3
   453    452    455      1   1.0950e+02   4.1840e+02 ;    HG2 - CG
 - CD
   454    452    455      1   1.0950e+02   4.1840e+02 ;    HG3 - CG
 - CD
   455    458    459      1   1.0950e+02   4.1840e+02 ;     CD - CE
 - HE2
   455    458    460      1   1.0950e+02   4.1840e+02 ;     CD - CE
 - HE3
   455    458    461      1   1.1120e+02   6.6944e+02 ;     CD - CE
 - NZ
   456    455    457      1   1.0950e+02   2.9288e+02 ;    HD2 - CD
 - HD3
   456    455    458      1   1.0950e+02   4.1840e+02 ;    HD2 - CD
 - CE
   457    455    458      1   1.0950e+02   4.1840e+02 ;    HD3 - CD
 - CE
   458    461    462      1   1.0950e+02   4.1840e+02 ;     CE - NZ
 - HZ1
   458    461    463      1   1.0950e+02   4.1840e+02 ;     CE - NZ
 - HZ2
   458    461    464      1   1.0950e+02   4.1840e+02 ;     CE - NZ
 - HZ3
   459    458    460      1   1.0950e+02   2.9288e+02 ;    HE2 - CE
 - HE3
   459    458    461      1   1.0950e+02   4.1840e+02 ;    HE2 - CE
 - NZ
   460    458    461      1   1.0950e+02   4.1840e+02 ;    HE3 - CE
 - NZ
```

```
   462    461    463      1    1.0950e+02    2.9288e+02 ;    HZ1 - NZ
- HZ2
   462    461    464      1    1.0950e+02    2.9288e+02 ;    HZ1 - NZ
- HZ3
   463    461    464      1    1.0950e+02    2.9288e+02 ;    HZ2 - NZ
- HZ3
   465    467    468      1    1.2000e+02    4.1840e+02 ;      C - N
- H
   465    467    469      1    1.2190e+02    4.1840e+02 ;      C - N
- CA
   466    465    467      1    1.2290e+02    6.6944e+02 ;      O - C
- N
   467    469    470      1    1.0950e+02    4.1840e+02 ;      N - CA
- HA2
   467    469    471      1    1.0950e+02    4.1840e+02 ;      N - CA
- HA3
   467    469    472      1    1.1010e+02    5.2718e+02 ;      N - CA
- C
   468    467    469      1    1.1804e+02    4.1840e+02 ;      H - N
- CA
   469    472    473      1    1.2040e+02    6.6944e+02 ;     CA - C
- O
   469    472    474      1    1.1660e+02    5.8576e+02 ;     CA - C
- N
   470    469    471      1    1.0950e+02    2.9288e+02 ;    HA2 - CA
- HA3
   470    469    472      1    1.0950e+02    4.1840e+02 ;    HA2 - CA
- C
   471    469    472      1    1.0950e+02    4.1840e+02 ;    HA3 - CA
- C
   472    474    475      1    1.2000e+02    4.1840e+02 ;      C - N
- H
   472    474    476      1    1.2190e+02    4.1840e+02 ;      C - N
- CA
   473    472    474      1    1.2290e+02    6.6944e+02 ;      O - C
- N
   474    476    477      1    1.0950e+02    4.1840e+02 ;      N - CA
- HA
   474    476    478      1    1.0970e+02    6.6944e+02 ;      N - CA
- CB
   474    476    491      1    1.1010e+02    5.2718e+02 ;      N - CA
- C
   475    474    476      1    1.1804e+02    4.1840e+02 ;      H - N
- CA
   476    478    479      1    1.0950e+02    4.1840e+02 ;     CA - CB
- HB
   476    478    480      1    1.0950e+02    3.3472e+02 ;     CA - CB
- CG2
   476    478    484      1    1.0950e+02    3.3472e+02 ;     CA - CB
- CG1
   476    491    492      1    1.2040e+02    6.6944e+02 ;     CA - C
- O
```

```
   476    491    493      1    1.1660e+02    5.8576e+02 ;     CA - C
- N
   477    476    478      1    1.0950e+02    4.1840e+02 ;     HA - CA
- CB
   477    476    491      1    1.0950e+02    4.1840e+02 ;     HA - CA
- C
   478    476    491      1    1.1110e+02    5.2718e+02 ;     CB - CA
- C
   478    480    481      1    1.0950e+02    4.1840e+02 ;     CB - CG2
- HG21
   478    480    482      1    1.0950e+02    4.1840e+02 ;     CB - CG2
- HG22
   478    480    483      1    1.0950e+02    4.1840e+02 ;     CB - CG2
- HG23
   478    484    485      1    1.0950e+02    4.1840e+02 ;     CB - CG1
- HG12
   478    484    486      1    1.0950e+02    4.1840e+02 ;     CB - CG1
- HG13
   478    484    487      1    1.0950e+02    3.3472e+02 ;     CB - CG1
- CD1
   479    478    480      1    1.0950e+02    4.1840e+02 ;     HB - CB
- CG2
   479    478    484      1    1.0950e+02    4.1840e+02 ;     HB - CB
- CG1
   480    478    484      1    1.0950e+02    3.3472e+02 ;    CG2 - CB
- CG1
   481    480    482      1    1.0950e+02    2.9288e+02 ;   HG21 - CG2
- HG22
   481    480    483      1    1.0950e+02    2.9288e+02 ;   HG21 - CG2
- HG23
   482    480    483      1    1.0950e+02    2.9288e+02 ;   HG22 - CG2
- HG23
   484    487    488      1    1.0950e+02    4.1840e+02 ;    CG1 - CD1
- HD11
   484    487    489      1    1.0950e+02    4.1840e+02 ;    CG1 - CD1
- HD12
   484    487    490      1    1.0950e+02    4.1840e+02 ;    CG1 - CD1
- HD13
   485    484    486      1    1.0950e+02    2.9288e+02 ;   HG12 - CG1
- HG13
   485    484    487      1    1.0950e+02    4.1840e+02 ;   HG12 - CG1
- CD1
   486    484    487      1    1.0950e+02    4.1840e+02 ;   HG13 - CG1
- CD1
   488    487    489      1    1.0950e+02    2.9288e+02 ;   HD11 - CD1
- HD12
   488    487    490      1    1.0950e+02    2.9288e+02 ;   HD11 - CD1
- HD13
   489    487    490      1    1.0950e+02    2.9288e+02 ;   HD12 - CD1
- HD13
   491    493    494      1    1.2000e+02    4.1840e+02 ;      C - N
- H
```

```
  491   493   495     1   1.2190e+02   4.1840e+02 ;      C - N - CA
  492   491   493     1   1.2290e+02   6.6944e+02 ;      O - C - N
  493   495   496     1   1.0950e+02   4.1840e+02 ;      N - CA - HA
  493   495   497     1   1.0970e+02   6.6944e+02 ;      N - CA - CB
  493   495   503     1   1.1010e+02   5.2718e+02 ;      N - CA - C
  494   493   495     1   1.1804e+02   4.1840e+02 ;      H - N - CA
  495   497   498     1   1.0950e+02   4.1840e+02 ;     CA - CB - HB2
  495   497   499     1   1.0950e+02   4.1840e+02 ;     CA - CB - HB3
  495   497   500     1   1.1110e+02   5.2718e+02 ;     CA - CB - CG
  495   503   504     1   1.2040e+02   6.6944e+02 ;     CA - C - O
  495   503   505     1   1.1660e+02   5.8576e+02 ;     CA - C - N
  496   495   497     1   1.0950e+02   4.1840e+02 ;     HA - CA - CB
  496   495   503     1   1.0950e+02   4.1840e+02 ;     HA - CA - C
  497   495   503     1   1.1110e+02   5.2718e+02 ;     CB - CA - C
  497   500   501     1   1.1700e+02   5.8576e+02 ;     CB - CG - OD1
  497   500   502     1   1.1700e+02   5.8576e+02 ;     CB - CG - OD2
  498   497   499     1   1.0950e+02   2.9288e+02 ;    HB2 - CB - HB3
  498   497   500     1   1.0950e+02   4.1840e+02 ;    HB2 - CB - CG
  499   497   500     1   1.0950e+02   4.1840e+02 ;    HB3 - CB - CG
  501   500   502     1   1.2600e+02   6.6944e+02 ;    OD1 - CG - OD2
  503   505   506     1   1.2000e+02   4.1840e+02 ;      C - N - H
  503   505   507     1   1.2190e+02   4.1840e+02 ;      C - N - CA
  504   503   505     1   1.2290e+02   6.6944e+02 ;      O - C - N
  505   507   508     1   1.0950e+02   4.1840e+02 ;      N - CA - HA
  505   507   509     1   1.0970e+02   6.6944e+02 ;      N - CA - CB
  505   507   525     1   1.1010e+02   5.2718e+02 ;      N - CA - C
```

```
   506    505    507      1    1.1804e+02    4.1840e+02 ;      H - N
- CA
   507    509    510      1    1.0950e+02    4.1840e+02 ;     CA - CB
- HB2
   507    509    511      1    1.0950e+02    4.1840e+02 ;     CA - CB
- HB3
   507    509    512      1    1.0950e+02    3.3472e+02 ;     CA - CB
- CG
   507    525    526      1    1.2040e+02    6.6944e+02 ;     CA - C
- O
   507    525    527      1    1.1660e+02    5.8576e+02 ;     CA - C
- N
   508    507    509      1    1.0950e+02    4.1840e+02 ;     HA - CA
- CB
   508    507    525      1    1.0950e+02    4.1840e+02 ;     HA - CA
- C
   509    507    525      1    1.1110e+02    5.2718e+02 ;     CB - CA
- C
   509    512    513      1    1.0950e+02    4.1840e+02 ;     CB - CG
- HG2
   509    512    514      1    1.0950e+02    4.1840e+02 ;     CB - CG
- HG3
   509    512    515      1    1.0950e+02    3.3472e+02 ;     CB - CG
- CD
   510    509    511      1    1.0950e+02    2.9288e+02 ;    HB2 - CB
- HB3
   510    509    512      1    1.0950e+02    4.1840e+02 ;    HB2 - CB
- CG
   511    509    512      1    1.0950e+02    4.1840e+02 ;    HB3 - CB
- CG
   512    515    516      1    1.0950e+02    4.1840e+02 ;     CG - CD
- HD2
   512    515    517      1    1.0950e+02    4.1840e+02 ;     CG - CD
- HD3
   512    515    518      1    1.0950e+02    3.3472e+02 ;     CG - CD
- CE
   513    512    514      1    1.0950e+02    2.9288e+02 ;    HG2 - CG
- HG3
   513    512    515      1    1.0950e+02    4.1840e+02 ;    HG2 - CG
- CD
   514    512    515      1    1.0950e+02    4.1840e+02 ;    HG3 - CG
- CD
   515    518    519      1    1.0950e+02    4.1840e+02 ;     CD - CE
- HE2
   515    518    520      1    1.0950e+02    4.1840e+02 ;     CD - CE
- HE3
   515    518    521      1    1.1120e+02    6.6944e+02 ;     CD - CE
- NZ
   516    515    517      1    1.0950e+02    2.9288e+02 ;    HD2 - CD
- HD3
   516    515    518      1    1.0950e+02    4.1840e+02 ;    HD2 - CD
- CE
```

517 515 518 1 1.0950e+02 4.1840e+02 ; HD3 - CD
- CE
518 521 522 1 1.0950e+02 4.1840e+02 ; CE - NZ
- HZ1
518 521 523 1 1.0950e+02 4.1840e+02 ; CE - NZ
- HZ2
518 521 524 1 1.0950e+02 4.1840e+02 ; CE - NZ
- HZ3
519 518 520 1 1.0950e+02 2.9288e+02 ; HE2 - CE
- HE3
519 518 521 1 1.0950e+02 4.1840e+02 ; HE2 - CE
- NZ
520 518 521 1 1.0950e+02 4.1840e+02 ; HE3 - CE
- NZ
522 521 523 1 1.0950e+02 2.9288e+02 ; HZ1 - NZ
- HZ2
522 521 524 1 1.0950e+02 2.9288e+02 ; HZ1 - NZ
- HZ3
523 521 524 1 1.0950e+02 2.9288e+02 ; HZ2 - NZ
- HZ3
525 527 528 1 1.2000e+02 4.1840e+02 ; C - N
- H
525 527 529 1 1.2190e+02 4.1840e+02 ; C - N
- CA
526 525 527 1 1.2290e+02 6.6944e+02 ; O - C
- N
527 529 530 1 1.0950e+02 4.1840e+02 ; N - CA
- HA
527 529 531 1 1.0970e+02 6.6944e+02 ; N - CA
- CB
527 529 541 1 1.1010e+02 5.2718e+02 ; N - CA
- C
528 527 529 1 1.1804e+02 4.1840e+02 ; H - N
- CA
529 531 532 1 1.0950e+02 4.1840e+02 ; CA - CB
- HB
529 531 533 1 1.0950e+02 3.3472e+02 ; CA - CB
- CG1
529 531 537 1 1.0950e+02 3.3472e+02 ; CA - CB
- CG2
529 541 542 1 1.2040e+02 6.6944e+02 ; CA - C
- O
529 541 543 1 1.1660e+02 5.8576e+02 ; CA - C
- N
530 529 531 1 1.0950e+02 4.1840e+02 ; HA - CA
- CB
530 529 541 1 1.0950e+02 4.1840e+02 ; HA - CA
- C
531 529 541 1 1.1110e+02 5.2718e+02 ; CB - CA
- C
531 533 534 1 1.0950e+02 4.1840e+02 ; CB - CG1
- HG11

```
   531   533   535     1    1.0950e+02    4.1840e+02 ;     CB - CG1
- HG12
   531   533   536     1    1.0950e+02    4.1840e+02 ;     CB - CG1
- HG13
   531   537   538     1    1.0950e+02    4.1840e+02 ;     CB - CG2
- HG21
   531   537   539     1    1.0950e+02    4.1840e+02 ;     CB - CG2
- HG22
   531   537   540     1    1.0950e+02    4.1840e+02 ;     CB - CG2
- HG23
   532   531   533     1    1.0950e+02    4.1840e+02 ;     HB - CB
- CG1
   532   531   537     1    1.0950e+02    4.1840e+02 ;     HB - CB
- CG2
   533   531   537     1    1.0950e+02    3.3472e+02 ;    CG1 - CB
- CG2
   534   533   535     1    1.0950e+02    2.9288e+02 ;   HG11 - CG1
- HG12
   534   533   536     1    1.0950e+02    2.9288e+02 ;   HG11 - CG1
- HG13
   535   533   536     1    1.0950e+02    2.9288e+02 ;   HG12 - CG1
- HG13
   538   537   539     1    1.0950e+02    2.9288e+02 ;   HG21 - CG2
- HG22
   538   537   540     1    1.0950e+02    2.9288e+02 ;   HG21 - CG2
- HG23
   539   537   540     1    1.0950e+02    2.9288e+02 ;   HG22 - CG2
- HG23
   541   543   544     1    1.2000e+02    4.1840e+02 ;      C - N
- H
   541   543   545     1    1.2190e+02    4.1840e+02 ;      C - N
- CA
   542   541   543     1    1.2290e+02    6.6944e+02 ;      O - C
- N
   543   545   546     1    1.0950e+02    4.1840e+02 ;      N - CA
- HA
   543   545   547     1    1.0970e+02    6.6944e+02 ;      N - CA
- CB
   543   545   555     1    1.1010e+02    5.2718e+02 ;      N - CA
- C
   544   543   545     1    1.1804e+02    4.1840e+02 ;      H - N
- CA
   545   547   548     1    1.0950e+02    4.1840e+02 ;     CA - CB
- HB
   545   547   549     1    1.0950e+02    3.3472e+02 ;     CA - CB
- CG2
   545   547   553     1    1.0950e+02    4.1840e+02 ;     CA - CB
- OG1
   545   555   556     1    1.2040e+02    6.6944e+02 ;     CA - C
- O
   545   555   557     1    1.1660e+02    5.8576e+02 ;     CA - C
- N
```

546 545 547 1 1.0950e+02 4.1840e+02 ; HA - CA
- CB
546 545 555 1 1.0950e+02 4.1840e+02 ; HA - CA
- C
547 545 555 1 1.1110e+02 5.2718e+02 ; CB - CA
- C
547 549 550 1 1.0950e+02 4.1840e+02 ; CB - CG2
- HG21
547 549 551 1 1.0950e+02 4.1840e+02 ; CB - CG2
- HG22
547 549 552 1 1.0950e+02 4.1840e+02 ; CB - CG2
- HG23
547 553 554 1 1.0850e+02 4.6024e+02 ; CB - OG1
- HG1
548 547 549 1 1.0950e+02 4.1840e+02 ; HB - CB
- CG2
548 547 553 1 1.0950e+02 4.1840e+02 ; HB - CB
- OG1
549 547 553 1 1.0950e+02 4.1840e+02 ; CG2 - CB
- OG1
550 549 551 1 1.0950e+02 2.9288e+02 ; HG21 - CG2
- HG22
550 549 552 1 1.0950e+02 2.9288e+02 ; HG21 - CG2
- HG23
551 549 552 1 1.0950e+02 2.9288e+02 ; HG22 - CG2
- HG23
555 557 558 1 1.2000e+02 4.1840e+02 ; C - N
- H
555 557 559 1 1.2190e+02 4.1840e+02 ; C - N
- CA
556 555 557 1 1.2290e+02 6.6944e+02 ; O - C
- N
557 559 560 1 1.0950e+02 4.1840e+02 ; N - CA
- HA
557 559 561 1 1.0970e+02 6.6944e+02 ; N - CA
- CB
557 559 569 1 1.1010e+02 5.2718e+02 ; N - CA
- C
558 557 559 1 1.1804e+02 4.1840e+02 ; H - N
- CA
559 561 562 1 1.0950e+02 4.1840e+02 ; CA - CB
- HB2
559 561 563 1 1.0950e+02 4.1840e+02 ; CA - CB
- HB3
559 561 564 1 1.1110e+02 5.2718e+02 ; CA - CB
- CG
559 569 570 1 1.2040e+02 6.6944e+02 ; CA - C
- O
559 569 571 1 1.1660e+02 5.8576e+02 ; CA - C
- N
560 559 561 1 1.0950e+02 4.1840e+02 ; HA - CA
- CB

```
   560    559    569      1    1.0950e+02    4.1840e+02 ;     HA - CA
- C
   561    559    569      1    1.1110e+02    5.2718e+02 ;     CB - CA
- C
   561    564    565      1    1.2040e+02    6.6944e+02 ;     CB - CG
- OD1
   561    564    566      1    1.1660e+02    5.8576e+02 ;     CB - CG
- ND2
   562    561    563      1    1.0950e+02    2.9288e+02 ;    HB2 - CB
- HB3
   562    561    564      1    1.0950e+02    4.1840e+02 ;    HB2 - CB
- CG
   563    561    564      1    1.0950e+02    4.1840e+02 ;    HB3 - CB
- CG
   564    566    567      1    1.2000e+02    4.1840e+02 ;     CG - ND2
- HD21
   564    566    568      1    1.2000e+02    4.1840e+02 ;     CG - ND2
- HD22
   565    564    566      1    1.2290e+02    6.6944e+02 ;    OD1 - CG
- ND2
   567    566    568      1    1.2000e+02    2.9288e+02 ;   HD21 - ND2
- HD22
   569    571    572      1    1.2000e+02    4.1840e+02 ;      C - N
- H
   569    571    573      1    1.2190e+02    4.1840e+02 ;      C - N
- CA
   570    569    571      1    1.2290e+02    6.6944e+02 ;      O - C
- N
   571    573    574      1    1.0950e+02    4.1840e+02 ;      N - CA
- HA
   571    573    575      1    1.0970e+02    6.6944e+02 ;      N - CA
- CB
   571    573    586      1    1.1010e+02    5.2718e+02 ;      N - CA
- C
   572    571    573      1    1.1804e+02    4.1840e+02 ;      H - N
- CA
   573    575    576      1    1.0950e+02    4.1840e+02 ;     CA - CB
- HB2
   573    575    577      1    1.0950e+02    4.1840e+02 ;     CA - CB
- HB3
   573    575    578      1    1.0950e+02    3.3472e+02 ;     CA - CB
- CG
   573    586    587      1    1.2040e+02    6.6944e+02 ;     CA - C
- O
   573    586    588      1    1.1660e+02    5.8576e+02 ;     CA - C
- N
   574    573    575      1    1.0950e+02    4.1840e+02 ;     HA - CA
- CB
   574    573    586      1    1.0950e+02    4.1840e+02 ;     HA - CA
- C
   575    573    586      1    1.1110e+02    5.2718e+02 ;     CB - CA
- C
```

575 578 579 1 1.0950e+02 4.1840e+02 ; CB - CG
- HG2
575 578 580 1 1.0950e+02 4.1840e+02 ; CB - CG
- HG3
575 578 581 1 1.1110e+02 5.2718e+02 ; CB - CG
- CD
576 575 577 1 1.0950e+02 2.9288e+02 ; HB2 - CB
- HB3
576 575 578 1 1.0950e+02 4.1840e+02 ; HB2 - CB
- CG
577 575 578 1 1.0950e+02 4.1840e+02 ; HB3 - CB
- CG
578 581 582 1 1.2040e+02 6.6944e+02 ; CG - CD
- OE1
578 581 583 1 1.1660e+02 5.8576e+02 ; CG - CD
- NE2
579 578 580 1 1.0950e+02 2.9288e+02 ; HG2 - CG
- HG3
579 578 581 1 1.0950e+02 4.1840e+02 ; HG2 - CG
- CD
580 578 581 1 1.0950e+02 4.1840e+02 ; HG3 - CG
- CD
581 583 584 1 1.2000e+02 4.1840e+02 ; CD - NE2
- HE21
581 583 585 1 1.2000e+02 4.1840e+02 ; CD - NE2
- HE22
582 581 583 1 1.2290e+02 6.6944e+02 ; OE1 - CD
- NE2
584 583 585 1 1.2000e+02 2.9288e+02 ; HE21 - NE2
- HE22
586 588 589 1 1.2000e+02 4.1840e+02 ; C - N
- H
586 588 590 1 1.2190e+02 4.1840e+02 ; C - N
- CA
587 586 588 1 1.2290e+02 6.6944e+02 ; O - C
- N
588 590 591 1 1.0950e+02 4.1840e+02 ; N - CA
- HA
588 590 592 1 1.0970e+02 6.6944e+02 ; N - CA
- CB
588 590 605 1 1.1010e+02 5.2718e+02 ; N - CA
- C
589 588 590 1 1.1804e+02 4.1840e+02 ; H - N
- CA
590 592 593 1 1.0950e+02 4.1840e+02 ; CA - CB
- HB2
590 592 594 1 1.0950e+02 4.1840e+02 ; CA - CB
- HB3
590 592 595 1 1.0950e+02 3.3472e+02 ; CA - CB
- CG
590 605 606 1 1.2040e+02 6.6944e+02 ; CA - C
- O

```
   590    605    607      1    1.1660e+02    5.8576e+02 ;     CA - C
- N
   591    590    592      1    1.0950e+02    4.1840e+02 ;     HA - CA
- CB
   591    590    605      1    1.0950e+02    4.1840e+02 ;     HA - CA
- C
   592    590    605      1    1.1110e+02    5.2718e+02 ;     CB - CA
- C
   592    595    596      1    1.0950e+02    4.1840e+02 ;     CB - CG
- HG
   592    595    597      1    1.0950e+02    3.3472e+02 ;     CB - CG
- CD1
   592    595    601      1    1.0950e+02    3.3472e+02 ;     CB - CG
- CD2
   593    592    594      1    1.0950e+02    2.9288e+02 ;    HB2 - CB
- HB3
   593    592    595      1    1.0950e+02    4.1840e+02 ;    HB2 - CB
- CG
   594    592    595      1    1.0950e+02    4.1840e+02 ;    HB3 - CB
- CG
   595    597    598      1    1.0950e+02    4.1840e+02 ;     CG - CD1
- HD11
   595    597    599      1    1.0950e+02    4.1840e+02 ;     CG - CD1
- HD12
   595    597    600      1    1.0950e+02    4.1840e+02 ;     CG - CD1
- HD13
   595    601    602      1    1.0950e+02    4.1840e+02 ;     CG - CD2
- HD21
   595    601    603      1    1.0950e+02    4.1840e+02 ;     CG - CD2
- HD22
   595    601    604      1    1.0950e+02    4.1840e+02 ;     CG - CD2
- HD23
   596    595    597      1    1.0950e+02    4.1840e+02 ;     HG - CG
- CD1
   596    595    601      1    1.0950e+02    4.1840e+02 ;     HG - CG
- CD2
   597    595    601      1    1.0950e+02    3.3472e+02 ;    CD1 - CG
- CD2
   598    597    599      1    1.0950e+02    2.9288e+02 ;   HD11 - CD1
- HD12
   598    597    600      1    1.0950e+02    2.9288e+02 ;   HD11 - CD1
- HD13
   599    597    600      1    1.0950e+02    2.9288e+02 ;   HD12 - CD1
- HD13
   602    601    603      1    1.0950e+02    2.9288e+02 ;   HD21 - CD2
- HD22
   602    601    604      1    1.0950e+02    2.9288e+02 ;   HD21 - CD2
- HD23
   603    601    604      1    1.0950e+02    2.9288e+02 ;   HD22 - CD2
- HD23
   605    607    608      1    1.2000e+02    4.1840e+02 ;      C - N
- H
```

```
   605    607    609      1    1.2190e+02    4.1840e+02 ;      C - N
 - CA
   606    605    607      1    1.2290e+02    6.6944e+02 ;      O - C
 - N
   607    609    610      1    1.0950e+02    4.1840e+02 ;      N - CA
 - HA
   607    609    611      1    1.0970e+02    6.6944e+02 ;      N - CA
 - CB
   607    609    626      1    1.1010e+02    5.2718e+02 ;      N - CA
 - C
   608    607    609      1    1.1804e+02    4.1840e+02 ;      H - N
 - CA
   609    611    612      1    1.0950e+02    4.1840e+02 ;     CA - CB
 - HB2
   609    611    613      1    1.0950e+02    4.1840e+02 ;     CA - CB
 - HB3
   609    611    614      1    1.1400e+02    5.2718e+02 ;     CA - CB
 - CG
   609    626    627      1    1.2040e+02    6.6944e+02 ;     CA - C
 - O
   609    626    628      1    1.1660e+02    5.8576e+02 ;     CA - C
 - N
   610    609    611      1    1.0950e+02    4.1840e+02 ;     HA - CA
 - CB
   610    609    626      1    1.0950e+02    4.1840e+02 ;     HA - CA
 - C
   611    609    626      1    1.1110e+02    5.2718e+02 ;     CB - CA
 - C
   611    614    615      1    1.2000e+02    5.8576e+02 ;     CB - CG
 - CD1
   611    614    624      1    1.2000e+02    5.8576e+02 ;     CB - CG
 - CD2
   612    611    613      1    1.0950e+02    2.9288e+02 ;    HB2 - CB
 - HB3
   612    611    614      1    1.0950e+02    4.1840e+02 ;    HB2 - CB
 - CG
   613    611    614      1    1.0950e+02    4.1840e+02 ;    HB3 - CB
 - CG
   614    615    616      1    1.2000e+02    4.1840e+02 ;     CG - CD1
 - HD1
   614    615    617      1    1.2000e+02    5.2718e+02 ;     CG - CD1
 - CE1
   614    624    622      1    1.2000e+02    5.2718e+02 ;     CG - CD2
 - CE2
   614    624    625      1    1.2000e+02    4.1840e+02 ;     CG - CD2
 - HD2
   615    614    624      1    1.2000e+02    5.2718e+02 ;    CD1 - CG
 - CD2
   615    617    618      1    1.2000e+02    4.1840e+02 ;    CD1 - CE1
 - HE1
   615    617    619      1    1.2000e+02    5.2718e+02 ;    CD1 - CE1
 - CZ
```

616 615 617 1 1.2000e+02 4.1840e+02 ; HD1 - CD1
- CE1
617 619 620 1 1.2000e+02 5.8576e+02 ; CE1 - CZ
- OH
617 619 622 1 1.2000e+02 5.2718e+02 ; CE1 - CZ
- CE2
618 617 619 1 1.2000e+02 4.1840e+02 ; HE1 - CE1
- CZ
619 620 621 1 1.1300e+02 4.1840e+02 ; CZ - OH
- HH
619 622 623 1 1.2000e+02 4.1840e+02 ; CZ - CE2
- HE2
619 622 624 1 1.2000e+02 5.2718e+02 ; CZ - CE2
- CD2
620 619 622 1 1.2000e+02 5.8576e+02 ; OH - CZ
- CE2
622 624 625 1 1.2000e+02 4.1840e+02 ; CE2 - CD2
- HD2
623 622 624 1 1.2000e+02 4.1840e+02 ; HE2 - CE2
- CD2
626 628 629 1 1.2000e+02 4.1840e+02 ; C - N
- H
626 628 630 1 1.2190e+02 4.1840e+02 ; C - N
- CA
627 626 628 1 1.2290e+02 6.6944e+02 ; O - C
- N
628 630 631 1 1.0950e+02 4.1840e+02 ; N - CA
- HA
628 630 632 1 1.0970e+02 6.6944e+02 ; N - CA
- CB
628 630 636 1 1.1010e+02 5.2718e+02 ; N - CA
- C
629 628 630 1 1.1804e+02 4.1840e+02 ; H - N
- CA
630 632 633 1 1.0950e+02 4.1840e+02 ; CA - CB
- HB1
630 632 634 1 1.0950e+02 4.1840e+02 ; CA - CB
- HB2
630 632 635 1 1.0950e+02 4.1840e+02 ; CA - CB
- HB3
630 636 637 1 1.2040e+02 6.6944e+02 ; CA - C
- O
630 636 638 1 1.1660e+02 5.8576e+02 ; CA - C
- N
631 630 632 1 1.0950e+02 4.1840e+02 ; HA - CA
- CB
631 630 636 1 1.0950e+02 4.1840e+02 ; HA - CA
- C
632 630 636 1 1.1110e+02 5.2718e+02 ; CB - CA
- C
633 632 634 1 1.0950e+02 2.9288e+02 ; HB1 - CB
- HB2

```
   633    632    635      1    1.0950e+02    2.9288e+02 ;    HB1 - CB
- HB3
   634    632    635      1    1.0950e+02    2.9288e+02 ;    HB2 - CB
- HB3
   636    638    639      1    1.2000e+02    4.1840e+02 ;      C - N
- H
   636    638    640      1    1.2190e+02    4.1840e+02 ;      C - N
- CA
   637    636    638      1    1.2290e+02    6.6944e+02 ;      O - C
- N
   638    640    641      1    1.0950e+02    4.1840e+02 ;      N - CA
- HA
   638    640    642      1    1.0970e+02    6.6944e+02 ;      N - CA
- CB
   638    640    655      1    1.1010e+02    5.2718e+02 ;      N - CA
- C
   639    638    640      1    1.1804e+02    4.1840e+02 ;      H - N
- CA
   640    642    643      1    1.0950e+02    4.1840e+02 ;     CA - CB
- HB
   640    642    644      1    1.0950e+02    3.3472e+02 ;     CA - CB
- CG2
   640    642    648      1    1.0950e+02    3.3472e+02 ;     CA - CB
- CG1
   640    655    656      1    1.2040e+02    6.6944e+02 ;     CA - C
- O
   640    655    657      1    1.1660e+02    5.8576e+02 ;     CA - C
- N
   641    640    642      1    1.0950e+02    4.1840e+02 ;     HA - CA
- CB
   641    640    655      1    1.0950e+02    4.1840e+02 ;     HA - CA
- C
   642    640    655      1    1.1110e+02    5.2718e+02 ;     CB - CA
- C
   642    644    645      1    1.0950e+02    4.1840e+02 ;     CB - CG2
- HG21
   642    644    646      1    1.0950e+02    4.1840e+02 ;     CB - CG2
- HG22
   642    644    647      1    1.0950e+02    4.1840e+02 ;     CB - CG2
- HG23
   642    648    649      1    1.0950e+02    4.1840e+02 ;     CB - CG1
- HG12
   642    648    650      1    1.0950e+02    4.1840e+02 ;     CB - CG1
- HG13
   642    648    651      1    1.0950e+02    3.3472e+02 ;     CB - CG1
- CD1
   643    642    644      1    1.0950e+02    4.1840e+02 ;     HB - CB
- CG2
   643    642    648      1    1.0950e+02    4.1840e+02 ;     HB - CB
- CG1
   644    642    648      1    1.0950e+02    3.3472e+02 ;    CG2 - CB
- CG1
```

645 644 646 1 1.0950e+02 2.9288e+02 ; HG21 - CG2
- HG22
645 644 647 1 1.0950e+02 2.9288e+02 ; HG21 - CG2
- HG23
646 644 647 1 1.0950e+02 2.9288e+02 ; HG22 - CG2
- HG23
648 651 652 1 1.0950e+02 4.1840e+02 ; CG1 - CD1
- HD11
648 651 653 1 1.0950e+02 4.1840e+02 ; CG1 - CD1
- HD12
648 651 654 1 1.0950e+02 4.1840e+02 ; CG1 - CD1
- HD13
649 648 650 1 1.0950e+02 2.9288e+02 ; HG12 - CG1
- HG13
649 648 651 1 1.0950e+02 4.1840e+02 ; HG12 - CG1
- CD1
650 648 651 1 1.0950e+02 4.1840e+02 ; HG13 - CG1
- CD1
652 651 653 1 1.0950e+02 2.9288e+02 ; HD11 - CD1
- HD12
652 651 654 1 1.0950e+02 2.9288e+02 ; HD11 - CD1
- HD13
653 651 654 1 1.0950e+02 2.9288e+02 ; HD12 - CD1
- HD13
655 657 658 1 1.2000e+02 4.1840e+02 ; C - N
- H
655 657 659 1 1.2190e+02 4.1840e+02 ; C - N
- CA
656 655 657 1 1.2290e+02 6.6944e+02 ; O - C
- N
657 659 660 1 1.0950e+02 4.1840e+02 ; N - CA
- HA
657 659 661 1 1.0970e+02 6.6944e+02 ; N - CA
- CB
657 659 677 1 1.1010e+02 5.2718e+02 ; N - CA
- C
658 657 659 1 1.1804e+02 4.1840e+02 ; H - N
- CA
659 661 662 1 1.0950e+02 4.1840e+02 ; CA - CB
- HB2
659 661 663 1 1.0950e+02 4.1840e+02 ; CA - CB
- HB3
659 661 664 1 1.0950e+02 3.3472e+02 ; CA - CB
- CG
659 677 678 1 1.2040e+02 6.6944e+02 ; CA - C
- O
659 677 679 1 1.1660e+02 5.8576e+02 ; CA - C
- N
660 659 661 1 1.0950e+02 4.1840e+02 ; HA - CA
- CB
660 659 677 1 1.0950e+02 4.1840e+02 ; HA - CA
- C

```
  661    659    677      1    1.1110e+02    5.2718e+02 ;     CB - CA
- C
  661    664    665      1    1.0950e+02    4.1840e+02 ;     CB - CG
- HG2
  661    664    666      1    1.0950e+02    4.1840e+02 ;     CB - CG
- HG3
  661    664    667      1    1.0950e+02    3.3472e+02 ;     CB - CG
- CD
  662    661    663      1    1.0950e+02    2.9288e+02 ;    HB2 - CB
- HB3
  662    661    664      1    1.0950e+02    4.1840e+02 ;    HB2 - CB
- CG
  663    661    664      1    1.0950e+02    4.1840e+02 ;    HB3 - CB
- CG
  664    667    668      1    1.0950e+02    4.1840e+02 ;     CG - CD
- HD2
  664    667    669      1    1.0950e+02    4.1840e+02 ;     CG - CD
- HD3
  664    667    670      1    1.0950e+02    3.3472e+02 ;     CG - CD
- CE
  665    664    666      1    1.0950e+02    2.9288e+02 ;    HG2 - CG
- HG3
  665    664    667      1    1.0950e+02    4.1840e+02 ;    HG2 - CG
- CD
  666    664    667      1    1.0950e+02    4.1840e+02 ;    HG3 - CG
- CD
  667    670    671      1    1.0950e+02    4.1840e+02 ;     CD - CE
- HE2
  667    670    672      1    1.0950e+02    4.1840e+02 ;     CD - CE
- HE3
  667    670    673      1    1.1120e+02    6.6944e+02 ;     CD - CE
- NZ
  668    667    669      1    1.0950e+02    2.9288e+02 ;    HD2 - CD
- HD3
  668    667    670      1    1.0950e+02    4.1840e+02 ;    HD2 - CD
- CE
  669    667    670      1    1.0950e+02    4.1840e+02 ;    HD3 - CD
- CE
  670    673    674      1    1.0950e+02    4.1840e+02 ;     CE - NZ
- HZ1
  670    673    675      1    1.0950e+02    4.1840e+02 ;     CE - NZ
- HZ2
  670    673    676      1    1.0950e+02    4.1840e+02 ;     CE - NZ
- HZ3
  671    670    672      1    1.0950e+02    2.9288e+02 ;    HE2 - CE
- HE3
  671    670    673      1    1.0950e+02    4.1840e+02 ;    HE2 - CE
- NZ
  672    670    673      1    1.0950e+02    4.1840e+02 ;    HE3 - CE
- NZ
  674    673    675      1    1.0950e+02    2.9288e+02 ;    HZ1 - NZ
- HZ2
```

674 673 676 1 1.0950e+02 2.9288e+02 ; HZ1 - NZ
- HZ3
675 673 676 1 1.0950e+02 2.9288e+02 ; HZ2 - NZ
- HZ3
677 679 680 1 1.2000e+02 4.1840e+02 ; C - N
- H
677 679 681 1 1.2190e+02 4.1840e+02 ; C - N
- CA
678 677 679 1 1.2290e+02 6.6944e+02 ; O - C
- N
679 681 682 1 1.0950e+02 4.1840e+02 ; N - CA
- HA
679 681 683 1 1.0970e+02 6.6944e+02 ; N - CA
- CB
679 681 692 1 1.1010e+02 5.2718e+02 ; N - CA
- C
680 679 681 1 1.1804e+02 4.1840e+02 ; H - N
- CA
681 683 684 1 1.0950e+02 4.1840e+02 ; CA - CB
- HB2
681 683 685 1 1.0950e+02 4.1840e+02 ; CA - CB
- HB3
681 683 686 1 1.0950e+02 3.3472e+02 ; CA - CB
- CG
681 692 693 1 1.2040e+02 6.6944e+02 ; CA - C
- O
681 692 694 1 1.1660e+02 5.8576e+02 ; CA - C
- N
682 681 683 1 1.0950e+02 4.1840e+02 ; HA - CA
- CB
682 681 692 1 1.0950e+02 4.1840e+02 ; HA - CA
- C
683 681 692 1 1.1110e+02 5.2718e+02 ; CB - CA
- C
683 686 687 1 1.0950e+02 4.1840e+02 ; CB - CG
- HG2
683 686 688 1 1.0950e+02 4.1840e+02 ; CB - CG
- HG3
683 686 689 1 1.1110e+02 5.2718e+02 ; CB - CG
- CD
684 683 685 1 1.0950e+02 2.9288e+02 ; HB2 - CB
- HB3
684 683 686 1 1.0950e+02 4.1840e+02 ; HB2 - CB
- CG
685 683 686 1 1.0950e+02 4.1840e+02 ; HB3 - CB
- CG
686 689 690 1 1.1700e+02 5.8576e+02 ; CG - CD
- OE1
686 689 691 1 1.1700e+02 5.8576e+02 ; CG - CD
- OE2
687 686 688 1 1.0950e+02 2.9288e+02 ; HG2 - CG
- HG3

```
   687    686    689      1    1.0950e+02    4.1840e+02 ;    HG2 - CG
- CD
   688    686    689      1    1.0950e+02    4.1840e+02 ;    HG3 - CG
- CD
   690    689    691      1    1.2600e+02    6.6944e+02 ;    OE1 - CD
- OE2
   692    694    695      1    1.2000e+02    4.1840e+02 ;      C - N
- H
   692    694    696      1    1.2190e+02    4.1840e+02 ;      C - N
- CA
   693    692    694      1    1.2290e+02    6.6944e+02 ;      O - C
- N
   694    696    697      1    1.0950e+02    4.1840e+02 ;      N - CA
- HA
   694    696    698      1    1.0970e+02    6.6944e+02 ;      N - CA
- CB
   694    696    707      1    1.1010e+02    5.2718e+02 ;      N - CA
- C
   695    694    696      1    1.1804e+02    4.1840e+02 ;      H - N
- CA
   696    698    699      1    1.0950e+02    4.1840e+02 ;     CA - CB
- HB2
   696    698    700      1    1.0950e+02    4.1840e+02 ;     CA - CB
- HB3
   696    698    701      1    1.0950e+02    3.3472e+02 ;     CA - CB
- CG
   696    707    708      1    1.2040e+02    6.6944e+02 ;     CA - C
- O
   696    707    709      1    1.1660e+02    5.8576e+02 ;     CA - C
- N
   697    696    698      1    1.0950e+02    4.1840e+02 ;     HA - CA
- CB
   697    696    707      1    1.0950e+02    4.1840e+02 ;     HA - CA
- C
   698    696    707      1    1.1110e+02    5.2718e+02 ;     CB - CA
- C
   698    701    702      1    1.0950e+02    4.1840e+02 ;     CB - CG
- HG2
   698    701    703      1    1.0950e+02    4.1840e+02 ;     CB - CG
- HG3
   698    701    704      1    1.1110e+02    5.2718e+02 ;     CB - CG
- CD
   699    698    700      1    1.0950e+02    2.9288e+02 ;    HB2 - CB
- HB3
   699    698    701      1    1.0950e+02    4.1840e+02 ;    HB2 - CB
- CG
   700    698    701      1    1.0950e+02    4.1840e+02 ;    HB3 - CB
- CG
   701    704    705      1    1.1700e+02    5.8576e+02 ;     CG - CD
- OE1
   701    704    706      1    1.1700e+02    5.8576e+02 ;     CG - CD
- OE2
```

```
  2054   2053   2056      1    1.0950e+02    2.9288e+02 ;   HD11 - CD1
- HD13
  2055   2053   2056      1    1.0950e+02    2.9288e+02 ;   HD12 - CD1
- HD13
  2058   2057   2059      1    1.0950e+02    2.9288e+02 ;   HD21 - CD2
- HD22
  2058   2057   2060      1    1.0950e+02    2.9288e+02 ;   HD21 - CD2
- HD23
  2059   2057   2060      1    1.0950e+02    2.9288e+02 ;   HD22 - CD2
- HD23
  2061   2063   2064      1    1.2000e+02    4.1840e+02 ;      C - N
- H
  2061   2063   2065      1    1.2190e+02    4.1840e+02 ;      C - N
- CA
  2062   2061   2063      1    1.2290e+02    6.6944e+02 ;      O - C
- N
  2063   2065   2066      1    1.0950e+02    4.1840e+02 ;      N - CA
- HA
  2063   2065   2067      1    1.0970e+02    6.6944e+02 ;      N - CA
- CB
  2063   2065   2078      1    1.1010e+02    5.2718e+02 ;      N - CA
- C
  2064   2063   2065      1    1.1804e+02    4.1840e+02 ;      H - N
- CA
  2065   2067   2068      1    1.0950e+02    4.1840e+02 ;     CA - CB
- HB2
  2065   2067   2069      1    1.0950e+02    4.1840e+02 ;     CA - CB
- HB3
  2065   2067   2070      1    1.1310e+02    5.2718e+02 ;     CA - CB
- CG
  2065   2078   2079      1    1.2040e+02    6.6944e+02 ;     CA - C
- O
  2065   2078   2080      1    1.1660e+02    5.8576e+02 ;     CA - C
- N
  2066   2065   2067      1    1.0950e+02    4.1840e+02 ;     HA - CA
- CB
  2066   2065   2078      1    1.0950e+02    4.1840e+02 ;     HA - CA
- C
  2067   2065   2078      1    1.1110e+02    5.2718e+02 ;     CB - CA
- C
  2067   2070   2071      1    1.2000e+02    5.8576e+02 ;     CB - CG
- ND1
  2067   2070   2076      1    1.2000e+02    5.8576e+02 ;     CB - CG
- CD2
  2068   2067   2069      1    1.0950e+02    2.9288e+02 ;    HB2 - CB
- HB3
  2068   2067   2070      1    1.0950e+02    4.1840e+02 ;    HB2 - CB
- CG
  2069   2067   2070      1    1.0950e+02    4.1840e+02 ;    HB3 - CB
- CG
  2070   2071   2072      1    1.1700e+02    5.8576e+02 ;     CG - ND1
- CE1
```

```
  2070   2076   2074      1    1.2000e+02    5.8576e+02 ;     CG - CD2
- NE2
  2070   2076   2077      1    1.2000e+02    4.1840e+02 ;     CG - CD2
- HD2
  2071   2070   2076      1    1.2000e+02    5.8576e+02 ;    ND1 - CG
- CD2
  2071   2072   2073      1    1.2000e+02    4.1840e+02 ;    ND1 - CE1
- HE1
  2071   2072   2074      1    1.2000e+02    5.8576e+02 ;    ND1 - CE1
- NE2
  2072   2074   2075      1    1.2000e+02    4.1840e+02 ;    CE1 - NE2
- HE2
  2072   2074   2076      1    1.2000e+02    5.8576e+02 ;    CE1 - NE2
- CD2
  2073   2072   2074      1    1.2000e+02    4.1840e+02 ;    HE1 - CE1
- NE2
  2074   2076   2077      1    1.2000e+02    4.1840e+02 ;    NE2 - CD2
- HD2
  2075   2074   2076      1    1.2000e+02    4.1840e+02 ;    HE2 - NE2
- CD2
  2078   2080   2081      1    1.2000e+02    4.1840e+02 ;      C - N
- H
  2078   2080   2082      1    1.2190e+02    4.1840e+02 ;      C - N
- CA
  2079   2078   2080      1    1.2290e+02    6.6944e+02 ;      O - C
- N
  2080   2082   2083      1    1.0950e+02    4.1840e+02 ;      N - CA
- HA
  2080   2082   2084      1    1.0970e+02    6.6944e+02 ;      N - CA
- CB
  2080   2082   2089      1    1.1010e+02    5.2718e+02 ;      N - CA
- C
  2081   2080   2082      1    1.1804e+02    4.1840e+02 ;      H - N
- CA
  2082   2084   2085      1    1.0950e+02    4.1840e+02 ;     CA - CB
- HB2
  2082   2084   2086      1    1.0950e+02    4.1840e+02 ;     CA - CB
- HB3
  2082   2084   2087      1    1.0950e+02    4.1840e+02 ;     CA - CB
- OG
  2082   2089   2090      1    1.2040e+02    6.6944e+02 ;     CA - C
- O
  2082   2089   2091      1    1.1660e+02    5.8576e+02 ;     CA - C
- N
  2083   2082   2084      1    1.0950e+02    4.1840e+02 ;     HA - CA
- CB
  2083   2082   2089      1    1.0950e+02    4.1840e+02 ;     HA - CA
- C
  2084   2082   2089      1    1.1110e+02    5.2718e+02 ;     CB - CA
- C
  2084   2087   2088      1    1.0850e+02    4.6024e+02 ;     CB - OG
- HG
```

2085 2084 2086 1 1.0950e+02 2.9288e+02 ; HB2 - CB
- HB3
2085 2084 2087 1 1.0950e+02 4.1840e+02 ; HB2 - CB
- OG
2086 2084 2087 1 1.0950e+02 4.1840e+02 ; HB3 - CB
- OG
2089 2091 2092 1 1.2000e+02 4.1840e+02 ; C - N
- H
2089 2091 2093 1 1.2190e+02 4.1840e+02 ; C - N
- CA
2090 2089 2091 1 1.2290e+02 6.6944e+02 ; O - C
- N
2091 2093 2094 1 1.0950e+02 4.1840e+02 ; N - CA
- HA
2091 2093 2095 1 1.0970e+02 6.6944e+02 ; N - CA
- CB
2091 2093 2108 1 1.1010e+02 5.2718e+02 ; N - CA
- C
2092 2091 2093 1 1.1804e+02 4.1840e+02 ; H - N
- CA
2093 2095 2096 1 1.0950e+02 4.1840e+02 ; CA - CB
- HB2
2093 2095 2097 1 1.0950e+02 4.1840e+02 ; CA - CB
- HB3
2093 2095 2098 1 1.0950e+02 3.3472e+02 ; CA - CB
- CG
2093 2108 2109 1 1.2040e+02 6.6944e+02 ; CA - C
- O
2093 2108 2110 1 1.1660e+02 5.8576e+02 ; CA - C
- N
2094 2093 2095 1 1.0950e+02 4.1840e+02 ; HA - CA
- CB
2094 2093 2108 1 1.0950e+02 4.1840e+02 ; HA - CA
- C
2095 2093 2108 1 1.1110e+02 5.2718e+02 ; CB - CA
- C
2095 2098 2099 1 1.0950e+02 4.1840e+02 ; CB - CG
- HG
2095 2098 2100 1 1.0950e+02 3.3472e+02 ; CB - CG
- CD1
2095 2098 2104 1 1.0950e+02 3.3472e+02 ; CB - CG
- CD2
2096 2095 2097 1 1.0950e+02 2.9288e+02 ; HB2 - CB
- HB3
2096 2095 2098 1 1.0950e+02 4.1840e+02 ; HB2 - CB
- CG
2097 2095 2098 1 1.0950e+02 4.1840e+02 ; HB3 - CB
- CG
2098 2100 2101 1 1.0950e+02 4.1840e+02 ; CG - CD1
- HD11
2098 2100 2102 1 1.0950e+02 4.1840e+02 ; CG - CD1
- HD12

2098 2100 2103 1 1.0950e+02 4.1840e+02 ; CG - CD1
- HD13
2098 2104 2105 1 1.0950e+02 4.1840e+02 ; CG - CD2
- HD21
2098 2104 2106 1 1.0950e+02 4.1840e+02 ; CG - CD2
- HD22
2098 2104 2107 1 1.0950e+02 4.1840e+02 ; CG - CD2
- HD23
2099 2098 2100 1 1.0950e+02 4.1840e+02 ; HG - CG
- CD1
2099 2098 2104 1 1.0950e+02 4.1840e+02 ; HG - CG
- CD2
2100 2098 2104 1 1.0950e+02 3.3472e+02 ; CD1 - CG
- CD2
2101 2100 2102 1 1.0950e+02 2.9288e+02 ; HD11 - CD1
- HD12
2101 2100 2103 1 1.0950e+02 2.9288e+02 ; HD11 - CD1
- HD13
2102 2100 2103 1 1.0950e+02 2.9288e+02 ; HD12 - CD1
- HD13
2105 2104 2106 1 1.0950e+02 2.9288e+02 ; HD21 - CD2
- HD22
2105 2104 2107 1 1.0950e+02 2.9288e+02 ; HD21 - CD2
- HD23
2106 2104 2107 1 1.0950e+02 2.9288e+02 ; HD22 - CD2
- HD23
2108 2110 2111 1 1.2000e+02 4.1840e+02 ; C - N
- H
2108 2110 2112 1 1.2190e+02 4.1840e+02 ; C - N
- CA
2109 2108 2110 1 1.2290e+02 6.6944e+02 ; O - C
- N
2110 2112 2113 1 1.0950e+02 4.1840e+02 ; N - CA
- HA
2110 2112 2114 1 1.0970e+02 6.6944e+02 ; N - CA
- CB
2110 2112 2122 1 1.1010e+02 5.2718e+02 ; N - CA
- C
2111 2110 2112 1 1.1804e+02 4.1840e+02 ; H - N
- CA
2112 2114 2115 1 1.0950e+02 4.1840e+02 ; CA - CB
- HB2
2112 2114 2116 1 1.0950e+02 4.1840e+02 ; CA - CB
- HB3
2112 2114 2117 1 1.1110e+02 5.2718e+02 ; CA - CB
- CG
2112 2122 2123 1 1.2040e+02 6.6944e+02 ; CA - C
- O
2112 2122 2124 1 1.1660e+02 5.8576e+02 ; CA - C
- N
2113 2112 2114 1 1.0950e+02 4.1840e+02 ; HA - CA
- CB

```
  2113   2112   2122       1    1.0950e+02    4.1840e+02 ;     HA - CA
- C
  2114   2112   2122       1    1.1110e+02    5.2718e+02 ;     CB - CA
- C
  2114   2117   2118       1    1.2040e+02    6.6944e+02 ;     CB - CG
- OD1
  2114   2117   2119       1    1.1660e+02    5.8576e+02 ;     CB - CG
- ND2
  2115   2114   2116       1    1.0950e+02    2.9288e+02 ;    HB2 - CB
- HB3
  2115   2114   2117       1    1.0950e+02    4.1840e+02 ;    HB2 - CB
- CG
  2116   2114   2117       1    1.0950e+02    4.1840e+02 ;    HB3 - CB
- CG
  2117   2119   2120       1    1.2000e+02    4.1840e+02 ;     CG - ND2
- HD21
  2117   2119   2121       1    1.2000e+02    4.1840e+02 ;     CG - ND2
- HD22
  2118   2117   2119       1    1.2290e+02    6.6944e+02 ;    OD1 - CG
- ND2
  2120   2119   2121       1    1.2000e+02    2.9288e+02 ;   HD21 - ND2
- HD22
  2122   2124   2125       1    1.2000e+02    4.1840e+02 ;      C - N
- H
  2122   2124   2126       1    1.2190e+02    4.1840e+02 ;      C - N
- CA
  2123   2122   2124       1    1.2290e+02    6.6944e+02 ;      O - C
- N
  2124   2126   2127       1    1.0950e+02    4.1840e+02 ;      N - CA
- HA
  2124   2126   2128       1    1.0970e+02    6.6944e+02 ;      N - CA
- CB
  2124   2126   2141       1    1.1010e+02    5.2718e+02 ;      N - CA
- C
  2125   2124   2126       1    1.1804e+02    4.1840e+02 ;      H - N
- CA
  2126   2128   2129       1    1.0950e+02    4.1840e+02 ;     CA - CB
- HB
  2126   2128   2130       1    1.0950e+02    3.3472e+02 ;     CA - CB
- CG2
  2126   2128   2134       1    1.0950e+02    3.3472e+02 ;     CA - CB
- CG1
  2126   2141   2142       1    1.2040e+02    6.6944e+02 ;     CA - C
- O
  2126   2141   2143       1    1.1660e+02    5.8576e+02 ;     CA - C
- N
  2127   2126   2128       1    1.0950e+02    4.1840e+02 ;     HA - CA
- CB
  2127   2126   2141       1    1.0950e+02    4.1840e+02 ;     HA - CA
- C
  2128   2126   2141       1    1.1110e+02    5.2718e+02 ;     CB - CA
- C
```

2128 2130 2131 1 1.0950e+02 4.1840e+02 ; CB - CG2
- HG21
2128 2130 2132 1 1.0950e+02 4.1840e+02 ; CB - CG2
- HG22
2128 2130 2133 1 1.0950e+02 4.1840e+02 ; CB - CG2
- HG23
2128 2134 2135 1 1.0950e+02 4.1840e+02 ; CB - CG1
- HG12
2128 2134 2136 1 1.0950e+02 4.1840e+02 ; CB - CG1
- HG13
2128 2134 2137 1 1.0950e+02 3.3472e+02 ; CB - CG1
- CD1
2129 2128 2130 1 1.0950e+02 4.1840e+02 ; HB - CB
- CG2
2129 2128 2134 1 1.0950e+02 4.1840e+02 ; HB - CB
- CG1
2130 2128 2134 1 1.0950e+02 3.3472e+02 ; CG2 - CB
- CG1
2131 2130 2132 1 1.0950e+02 2.9288e+02 ; HG21 - CG2
- HG22
2131 2130 2133 1 1.0950e+02 2.9288e+02 ; HG21 - CG2
- HG23
2132 2130 2133 1 1.0950e+02 2.9288e+02 ; HG22 - CG2
- HG23
2134 2137 2138 1 1.0950e+02 4.1840e+02 ; CG1 - CD1
- HD11
2134 2137 2139 1 1.0950e+02 4.1840e+02 ; CG1 - CD1
- HD12
2134 2137 2140 1 1.0950e+02 4.1840e+02 ; CG1 - CD1
- HD13
2135 2134 2136 1 1.0950e+02 2.9288e+02 ; HG12 - CG1
- HG13
2135 2134 2137 1 1.0950e+02 4.1840e+02 ; HG12 - CG1
- CD1
2136 2134 2137 1 1.0950e+02 4.1840e+02 ; HG13 - CG1
- CD1
2138 2137 2139 1 1.0950e+02 2.9288e+02 ; HD11 - CD1
- HD12
2138 2137 2140 1 1.0950e+02 2.9288e+02 ; HD11 - CD1
- HD13
2139 2137 2140 1 1.0950e+02 2.9288e+02 ; HD12 - CD1
- HD13
2141 2143 2144 1 1.2000e+02 4.1840e+02 ; C - N
- H
2141 2143 2145 1 1.2190e+02 4.1840e+02 ; C - N
- CA
2142 2141 2143 1 1.2290e+02 6.6944e+02 ; O - C
- N
2143 2145 2146 1 1.0950e+02 4.1840e+02 ; N - CA
- HA
2143 2145 2147 1 1.0970e+02 6.6944e+02 ; N - CA
- CB

2143 2145 2157 1 1.1010e+02 5.2718e+02 ; N - CA
- C
2144 2143 2145 1 1.1804e+02 4.1840e+02 ; H - N
- CA
2145 2147 2148 1 1.0950e+02 4.1840e+02 ; CA - CB
- HB
2145 2147 2149 1 1.0950e+02 3.3472e+02 ; CA - CB
- CG1
2145 2147 2153 1 1.0950e+02 3.3472e+02 ; CA - CB
- CG2
2145 2157 2158 1 1.2040e+02 6.6944e+02 ; CA - C
- O
2145 2157 2159 1 1.1660e+02 5.8576e+02 ; CA - C
- N
2146 2145 2147 1 1.0950e+02 4.1840e+02 ; HA - CA
- CB
2146 2145 2157 1 1.0950e+02 4.1840e+02 ; HA - CA
- C
2147 2145 2157 1 1.1110e+02 5.2718e+02 ; CB - CA
- C
2147 2149 2150 1 1.0950e+02 4.1840e+02 ; CB - CG1
- HG11
2147 2149 2151 1 1.0950e+02 4.1840e+02 ; CB - CG1
- HG12
2147 2149 2152 1 1.0950e+02 4.1840e+02 ; CB - CG1
- HG13
2147 2153 2154 1 1.0950e+02 4.1840e+02 ; CB - CG2
- HG21
2147 2153 2155 1 1.0950e+02 4.1840e+02 ; CB - CG2
- HG22
2147 2153 2156 1 1.0950e+02 4.1840e+02 ; CB - CG2
- HG23
2148 2147 2149 1 1.0950e+02 4.1840e+02 ; HB - CB
- CG1
2148 2147 2153 1 1.0950e+02 4.1840e+02 ; HB - CB
- CG2
2149 2147 2153 1 1.0950e+02 3.3472e+02 ; CG1 - CB
- CG2
2150 2149 2151 1 1.0950e+02 2.9288e+02 ; HG11 - CG1
- HG12
2150 2149 2152 1 1.0950e+02 2.9288e+02 ; HG11 - CG1
- HG13
2151 2149 2152 1 1.0950e+02 2.9288e+02 ; HG12 - CG1
- HG13
2154 2153 2155 1 1.0950e+02 2.9288e+02 ; HG21 - CG2
- HG22
2154 2153 2156 1 1.0950e+02 2.9288e+02 ; HG21 - CG2
- HG23
2155 2153 2156 1 1.0950e+02 2.9288e+02 ; HG22 - CG2
- HG23
2157 2159 2160 1 1.2000e+02 4.1840e+02 ; C - N
- H

2157 2159 2161 1 1.2190e+02 4.1840e+02 ; C - N
- CA
2158 2157 2159 1 1.2290e+02 6.6944e+02 ; O - C
- N
2159 2161 2162 1 1.0950e+02 4.1840e+02 ; N - CA
- HA
2159 2161 2163 1 1.0970e+02 6.6944e+02 ; N - CA
- CB
2159 2161 2174 1 1.1010e+02 5.2718e+02 ; N - CA
- C
2160 2159 2161 1 1.1804e+02 4.1840e+02 ; H - N
- CA
2161 2163 2164 1 1.0950e+02 4.1840e+02 ; CA - CB
- HB2
2161 2163 2165 1 1.0950e+02 4.1840e+02 ; CA - CB
- HB3
2161 2163 2166 1 1.1310e+02 5.2718e+02 ; CA - CB
- CG
2161 2174 2175 1 1.2040e+02 6.6944e+02 ; CA - C
- O
2161 2174 2176 1 1.1660e+02 5.8576e+02 ; CA - C
- N
2162 2161 2163 1 1.0950e+02 4.1840e+02 ; HA - CA
- CB
2162 2161 2174 1 1.0950e+02 4.1840e+02 ; HA - CA
- C
2163 2161 2174 1 1.1110e+02 5.2718e+02 ; CB - CA
- C
2163 2166 2167 1 1.2000e+02 5.8576e+02 ; CB - CG
- ND1
2163 2166 2172 1 1.2000e+02 5.8576e+02 ; CB - CG
- CD2
2164 2163 2165 1 1.0950e+02 2.9288e+02 ; HB2 - CB
- HB3
2164 2163 2166 1 1.0950e+02 4.1840e+02 ; HB2 - CB
- CG
2165 2163 2166 1 1.0950e+02 4.1840e+02 ; HB3 - CB
- CG
2166 2167 2168 1 1.1700e+02 5.8576e+02 ; CG - ND1
- CE1
2166 2172 2170 1 1.2000e+02 5.8576e+02 ; CG - CD2
- NE2
2166 2172 2173 1 1.2000e+02 4.1840e+02 ; CG - CD2
- HD2
2167 2166 2172 1 1.2000e+02 5.8576e+02 ; ND1 - CG
- CD2
2167 2168 2169 1 1.2000e+02 4.1840e+02 ; ND1 - CE1
- HE1
2167 2168 2170 1 1.2000e+02 5.8576e+02 ; ND1 - CE1
- NE2
2168 2170 2171 1 1.2000e+02 4.1840e+02 ; CE1 - NE2
- HE2

| | | | | | | | |
|---|---|---|---|---|---|---|---|
| 2168 | 2170 | 2172 | 1 | 1.2000e+02 | 5.8576e+02 | ; | CE1 - NE2 - CD2 |
| 2169 | 2168 | 2170 | 1 | 1.2000e+02 | 4.1840e+02 | ; | HE1 - CE1 - NE2 |
| 2170 | 2172 | 2173 | 1 | 1.2000e+02 | 4.1840e+02 | ; | NE2 - CD2 - HD2 |
| 2171 | 2170 | 2172 | 1 | 1.2000e+02 | 4.1840e+02 | ; | HE2 - NE2 - CD2 |
| 2174 | 2176 | 2177 | 1 | 1.2000e+02 | 4.1840e+02 | ; | C - N - H |
| 2174 | 2176 | 2178 | 1 | 1.2190e+02 | 4.1840e+02 | ; | C - N - CA |
| 2175 | 2174 | 2176 | 1 | 1.2290e+02 | 6.6944e+02 | ; | O - C - N |
| 2176 | 2178 | 2179 | 1 | 1.0950e+02 | 4.1840e+02 | ; | N - CA - HA |
| 2176 | 2178 | 2180 | 1 | 1.0970e+02 | 6.6944e+02 | ; | N - CA - CB |
| 2176 | 2178 | 2198 | 1 | 1.1010e+02 | 5.2718e+02 | ; | N - CA - C |
| 2177 | 2176 | 2178 | 1 | 1.1804e+02 | 4.1840e+02 | ; | H - N - CA |
| 2178 | 2180 | 2181 | 1 | 1.0950e+02 | 4.1840e+02 | ; | CA - CB - HB2 |
| 2178 | 2180 | 2182 | 1 | 1.0950e+02 | 4.1840e+02 | ; | CA - CB - HB3 |
| 2178 | 2180 | 2183 | 1 | 1.0950e+02 | 3.3472e+02 | ; | CA - CB - CG |
| 2178 | 2198 | 2199 | 1 | 1.2040e+02 | 6.6944e+02 | ; | CA - C - O |
| 2178 | 2198 | 2200 | 1 | 1.1660e+02 | 5.8576e+02 | ; | CA - C - N |
| 2179 | 2178 | 2180 | 1 | 1.0950e+02 | 4.1840e+02 | ; | HA - CA - CB |
| 2179 | 2178 | 2198 | 1 | 1.0950e+02 | 4.1840e+02 | ; | HA - CA - C |
| 2180 | 2178 | 2198 | 1 | 1.1110e+02 | 5.2718e+02 | ; | CB - CA - C |
| 2180 | 2183 | 2184 | 1 | 1.0950e+02 | 4.1840e+02 | ; | CB - CG - HG2 |
| 2180 | 2183 | 2185 | 1 | 1.0950e+02 | 4.1840e+02 | ; | CB - CG - HG3 |
| 2180 | 2183 | 2186 | 1 | 1.0950e+02 | 3.3472e+02 | ; | CB - CG - CD |
| 2181 | 2180 | 2182 | 1 | 1.0950e+02 | 2.9288e+02 | ; | HB2 - CB - HB3 |
| 2181 | 2180 | 2183 | 1 | 1.0950e+02 | 4.1840e+02 | ; | HB2 - CB - CG |
| 2182 | 2180 | 2183 | 1 | 1.0950e+02 | 4.1840e+02 | ; | HB3 - CB - CG |
| 2183 | 2186 | 2187 | 1 | 1.0950e+02 | 4.1840e+02 | ; | CG - CD - HD2 |

```
  2183   2186   2188       1    1.0950e+02    4.1840e+02 ;     CG - CD
- HD3
  2183   2186   2189       1    1.1120e+02    6.6944e+02 ;     CG - CD
- NE
  2184   2183   2185       1    1.0950e+02    2.9288e+02 ;    HG2 - CG
- HG3
  2184   2183   2186       1    1.0950e+02    4.1840e+02 ;    HG2 - CG
- CD
  2185   2183   2186       1    1.0950e+02    4.1840e+02 ;    HG3 - CG
- CD
  2186   2189   2190       1    1.1840e+02    4.1840e+02 ;     CD - NE
- HE
  2186   2189   2191       1    1.2320e+02    4.1840e+02 ;     CD - NE
- CZ
  2187   2186   2188       1    1.0950e+02    2.9288e+02 ;    HD2 - CD
- HD3
  2187   2186   2189       1    1.0950e+02    4.1840e+02 ;    HD2 - CD
- NE
  2188   2186   2189       1    1.0950e+02    4.1840e+02 ;    HD3 - CD
- NE
  2189   2191   2192       1    1.2000e+02    5.8576e+02 ;     NE - CZ
- NH1
  2189   2191   2195       1    1.2000e+02    5.8576e+02 ;     NE - CZ
- NH2
  2190   2189   2191       1    1.2000e+02    4.1840e+02 ;     HE - NE
- CZ
  2191   2192   2193       1    1.2000e+02    4.1840e+02 ;     CZ - NH1
- HH11
  2191   2192   2194       1    1.2000e+02    4.1840e+02 ;     CZ - NH1
- HH12
  2191   2195   2196       1    1.2000e+02    4.1840e+02 ;     CZ - NH2
- HH21
  2191   2195   2197       1    1.2000e+02    4.1840e+02 ;     CZ - NH2
- HH22
  2192   2191   2195       1    1.2000e+02    5.8576e+02 ;    NH1 - CZ
- NH2
  2193   2192   2194       1    1.2000e+02    2.9288e+02 ;   HH11 - NH1
- HH12
  2196   2195   2197       1    1.2000e+02    2.9288e+02 ;   HH21 - NH2
- HH22
  2198   2200   2201       1    1.2000e+02    4.1840e+02 ;      C - N
- H
  2198   2200   2202       1    1.2190e+02    4.1840e+02 ;      C - N
- CA
  2199   2198   2200       1    1.2290e+02    6.6944e+02 ;      O - C
- N
  2200   2202   2203       1    1.0950e+02    4.1840e+02 ;      N - CA
- HA
  2200   2202   2204       1    1.0970e+02    6.6944e+02 ;      N - CA
- CB
  2200   2202   2210       1    1.1010e+02    5.2718e+02 ;      N - CA
- C
```

```
  2201   2200   2202       1    1.1804e+02    4.1840e+02 ;      H - N
- CA
  2202   2204   2205       1    1.0950e+02    4.1840e+02 ;     CA - CB
- HB2
  2202   2204   2206       1    1.0950e+02    4.1840e+02 ;     CA - CB
- HB3
  2202   2204   2207       1    1.1110e+02    5.2718e+02 ;     CA - CB
- CG
  2202   2210   2211       1    1.2040e+02    6.6944e+02 ;     CA - C
- O
  2202   2210   2212       1    1.1660e+02    5.8576e+02 ;     CA - C
- N
  2203   2202   2204       1    1.0950e+02    4.1840e+02 ;     HA - CA
- CB
  2203   2202   2210       1    1.0950e+02    4.1840e+02 ;     HA - CA
- C
  2204   2202   2210       1    1.1110e+02    5.2718e+02 ;     CB - CA
- C
  2204   2207   2208       1    1.1700e+02    5.8576e+02 ;     CB - CG
- OD1
  2204   2207   2209       1    1.1700e+02    5.8576e+02 ;     CB - CG
- OD2
  2205   2204   2206       1    1.0950e+02    2.9288e+02 ;    HB2 - CB
- HB3
  2205   2204   2207       1    1.0950e+02    4.1840e+02 ;    HB2 - CB
- CG
  2206   2204   2207       1    1.0950e+02    4.1840e+02 ;    HB3 - CB
- CG
  2208   2207   2209       1    1.2600e+02    6.6944e+02 ;    OD1 - CG
- OD2
  2210   2212   2213       1    1.2000e+02    4.1840e+02 ;      C - N
- H
  2210   2212   2214       1    1.2190e+02    4.1840e+02 ;      C - N
- CA
  2211   2210   2212       1    1.2290e+02    6.6944e+02 ;      O - C
- N
  2212   2214   2215       1    1.0950e+02    4.1840e+02 ;      N - CA
- HA
  2212   2214   2216       1    1.0970e+02    6.6944e+02 ;      N - CA
- CB
  2212   2214   2229       1    1.1010e+02    5.2718e+02 ;      N - CA
- C
  2213   2212   2214       1    1.1804e+02    4.1840e+02 ;      H - N
- CA
  2214   2216   2217       1    1.0950e+02    4.1840e+02 ;     CA - CB
- HB2
  2214   2216   2218       1    1.0950e+02    4.1840e+02 ;     CA - CB
- HB3
  2214   2216   2219       1    1.0950e+02    3.3472e+02 ;     CA - CB
- CG
  2214   2229   2230       1    1.2040e+02    6.6944e+02 ;     CA - C
- O
```

```
  2214   2229   2231      1    1.1660e+02    5.8576e+02 ;     CA - C
- N
  2215   2214   2216      1    1.0950e+02    4.1840e+02 ;     HA - CA
- CB
  2215   2214   2229      1    1.0950e+02    4.1840e+02 ;     HA - CA
- C
  2216   2214   2229      1    1.1110e+02    5.2718e+02 ;     CB - CA
- C
  2216   2219   2220      1    1.0950e+02    4.1840e+02 ;     CB - CG
- HG
  2216   2219   2221      1    1.0950e+02    3.3472e+02 ;     CB - CG
- CD1
  2216   2219   2225      1    1.0950e+02    3.3472e+02 ;     CB - CG
- CD2
  2217   2216   2218      1    1.0950e+02    2.9288e+02 ;    HB2 - CB
- HB3
  2217   2216   2219      1    1.0950e+02    4.1840e+02 ;    HB2 - CB
- CG
  2218   2216   2219      1    1.0950e+02    4.1840e+02 ;    HB3 - CB
- CG
  2219   2221   2222      1    1.0950e+02    4.1840e+02 ;     CG - CD1
- HD11
  2219   2221   2223      1    1.0950e+02    4.1840e+02 ;     CG - CD1
- HD12
  2219   2221   2224      1    1.0950e+02    4.1840e+02 ;     CG - CD1
- HD13
  2219   2225   2226      1    1.0950e+02    4.1840e+02 ;     CG - CD2
- HD21
  2219   2225   2227      1    1.0950e+02    4.1840e+02 ;     CG - CD2
- HD22
  2219   2225   2228      1    1.0950e+02    4.1840e+02 ;     CG - CD2
- HD23
  2220   2219   2221      1    1.0950e+02    4.1840e+02 ;     HG - CG
- CD1
  2220   2219   2225      1    1.0950e+02    4.1840e+02 ;     HG - CG
- CD2
  2221   2219   2225      1    1.0950e+02    3.3472e+02 ;    CD1 - CG
- CD2
  2222   2221   2223      1    1.0950e+02    2.9288e+02 ;   HD11 - CD1
- HD12
  2222   2221   2224      1    1.0950e+02    2.9288e+02 ;   HD11 - CD1
- HD13
  2223   2221   2224      1    1.0950e+02    2.9288e+02 ;   HD12 - CD1
- HD13
  2226   2225   2227      1    1.0950e+02    2.9288e+02 ;   HD21 - CD2
- HD22
  2226   2225   2228      1    1.0950e+02    2.9288e+02 ;   HD21 - CD2
- HD23
  2227   2225   2228      1    1.0950e+02    2.9288e+02 ;   HD22 - CD2
- HD23
  2229   2231   2232      1    1.2000e+02    4.1840e+02 ;      C - N
- H
```

2229 2231 2233 1 1.2190e+02 4.1840e+02 ; C - N
- CA
2230 2229 2231 1 1.2290e+02 6.6944e+02 ; O - C
- N
2231 2233 2234 1 1.0950e+02 4.1840e+02 ; N - CA
- HA
2231 2233 2235 1 1.0970e+02 6.6944e+02 ; N - CA
- CB
2231 2233 2251 1 1.1010e+02 5.2718e+02 ; N - CA
- C
2232 2231 2233 1 1.1804e+02 4.1840e+02 ; H - N
- CA
2233 2235 2236 1 1.0950e+02 4.1840e+02 ; CA - CB
- HB2
2233 2235 2237 1 1.0950e+02 4.1840e+02 ; CA - CB
- HB3
2233 2235 2238 1 1.0950e+02 3.3472e+02 ; CA - CB
- CG
2233 2251 2252 1 1.2040e+02 6.6944e+02 ; CA - C
- O
2233 2251 2253 1 1.1660e+02 5.8576e+02 ; CA - C
- N
2234 2233 2235 1 1.0950e+02 4.1840e+02 ; HA - CA
- CB
2234 2233 2251 1 1.0950e+02 4.1840e+02 ; HA - CA
- C
2235 2233 2251 1 1.1110e+02 5.2718e+02 ; CB - CA
- C
2235 2238 2239 1 1.0950e+02 4.1840e+02 ; CB - CG
- HG2
2235 2238 2240 1 1.0950e+02 4.1840e+02 ; CB - CG
- HG3
2235 2238 2241 1 1.0950e+02 3.3472e+02 ; CB - CG
- CD
2236 2235 2237 1 1.0950e+02 2.9288e+02 ; HB2 - CB
- HB3
2236 2235 2238 1 1.0950e+02 4.1840e+02 ; HB2 - CB
- CG
2237 2235 2238 1 1.0950e+02 4.1840e+02 ; HB3 - CB
- CG
2238 2241 2242 1 1.0950e+02 4.1840e+02 ; CG - CD
- HD2
2238 2241 2243 1 1.0950e+02 4.1840e+02 ; CG - CD
- HD3
2238 2241 2244 1 1.0950e+02 3.3472e+02 ; CG - CD
- CE
2239 2238 2240 1 1.0950e+02 2.9288e+02 ; HG2 - CG
- HG3
2239 2238 2241 1 1.0950e+02 4.1840e+02 ; HG2 - CG
- CD
2240 2238 2241 1 1.0950e+02 4.1840e+02 ; HG3 - CG
- CD

2241 2244 2245 1 1.0950e+02 4.1840e+02 ; CD - CE
- HE2
2241 2244 2246 1 1.0950e+02 4.1840e+02 ; CD - CE
- HE3
2241 2244 2247 1 1.1120e+02 6.6944e+02 ; CD - CE
- NZ
2242 2241 2243 1 1.0950e+02 2.9288e+02 ; HD2 - CD
- HD3
2242 2241 2244 1 1.0950e+02 4.1840e+02 ; HD2 - CD
- CE
2243 2241 2244 1 1.0950e+02 4.1840e+02 ; HD3 - CD
- CE
2244 2247 2248 1 1.0950e+02 4.1840e+02 ; CE - NZ
- HZ1
2244 2247 2249 1 1.0950e+02 4.1840e+02 ; CE - NZ
- HZ2
2244 2247 2250 1 1.0950e+02 4.1840e+02 ; CE - NZ
- HZ3
2245 2244 2246 1 1.0950e+02 2.9288e+02 ; HE2 - CE
- HE3
2245 2244 2247 1 1.0950e+02 4.1840e+02 ; HE2 - CE
- NZ
2246 2244 2247 1 1.0950e+02 4.1840e+02 ; HE3 - CE
- NZ
2248 2247 2249 1 1.0950e+02 2.9288e+02 ; HZ1 - NZ
- HZ2
2248 2247 2250 1 1.0950e+02 2.9288e+02 ; HZ1 - NZ
- HZ3
2249 2247 2250 1 1.0950e+02 2.9288e+02 ; HZ2 - NZ
- HZ3
2251 2253 2254 1 1.2190e+02 4.1840e+02 ; C - N
- CD
2251 2253 2263 1 1.2190e+02 4.1840e+02 ; C - N
- CA
2252 2251 2253 1 1.2290e+02 6.6944e+02 ; O - C
- N
2253 2254 2255 1 1.0950e+02 4.1840e+02 ; N - CD
- HD2
2253 2254 2256 1 1.0950e+02 4.1840e+02 ; N - CD
- HD3
2253 2254 2257 1 1.0970e+02 6.6944e+02 ; N - CD
- CG
2253 2263 2260 1 1.0970e+02 6.6944e+02 ; N - CA
- CB
2253 2263 2264 1 1.0950e+02 4.1840e+02 ; N - CA
- HA
2253 2263 2265 1 1.1010e+02 5.2718e+02 ; N - CA
- C
2254 2253 2263 1 1.1800e+02 4.1840e+02 ; CD - N
- CA
2254 2257 2258 1 1.0950e+02 4.1840e+02 ; CD - CG
- HG2

2254 2257 2259 1 1.0950e+02 4.1840e+02 ; CD - CG
- HG3
2254 2257 2260 1 1.0950e+02 3.3472e+02 ; CD - CG
- CB
2255 2254 2256 1 1.0950e+02 2.9288e+02 ; HD2 - CD
- HD3
2255 2254 2257 1 1.0950e+02 4.1840e+02 ; HD2 - CD
- CG
2256 2254 2257 1 1.0950e+02 4.1840e+02 ; HD3 - CD
- CG
2257 2260 2261 1 1.0950e+02 4.1840e+02 ; CG - CB
- HB2
2257 2260 2262 1 1.0950e+02 4.1840e+02 ; CG - CB
- HB3
2257 2260 2263 1 1.0950e+02 3.3472e+02 ; CG - CB
- CA
2258 2257 2259 1 1.0950e+02 2.9288e+02 ; HG2 - CG
- HG3
2258 2257 2260 1 1.0950e+02 4.1840e+02 ; HG2 - CG
- CB
2259 2257 2260 1 1.0950e+02 4.1840e+02 ; HG3 - CG
- CB
2260 2263 2264 1 1.0950e+02 4.1840e+02 ; CB - CA
- HA
2260 2263 2265 1 1.1110e+02 5.2718e+02 ; CB - CA
- C
2261 2260 2262 1 1.0950e+02 2.9288e+02 ; HB2 - CB
- HB3
2261 2260 2263 1 1.0950e+02 4.1840e+02 ; HB2 - CB
- CA
2262 2260 2263 1 1.0950e+02 4.1840e+02 ; HB3 - CB
- CA
2263 2265 2266 1 1.2040e+02 6.6944e+02 ; CA - C
- O
2263 2265 2267 1 1.1660e+02 5.8576e+02 ; CA - C
- N
2264 2263 2265 1 1.0950e+02 4.1840e+02 ; HA - CA
- C
2265 2267 2268 1 1.2000e+02 4.1840e+02 ; C - N
- H
2265 2267 2269 1 1.2190e+02 4.1840e+02 ; C - N
- CA
2266 2265 2267 1 1.2290e+02 6.6944e+02 ; O - C
- N
2267 2269 2270 1 1.0950e+02 4.1840e+02 ; N - CA
- HA
2267 2269 2271 1 1.0970e+02 6.6944e+02 ; N - CA
- CB
2267 2269 2280 1 1.1010e+02 5.2718e+02 ; N - CA
- C
2268 2267 2269 1 1.1804e+02 4.1840e+02 ; H - N
- CA

```
  2269   2271   2272       1    1.0950e+02    4.1840e+02 ;     CA - CB
- HB2
  2269   2271   2273       1    1.0950e+02    4.1840e+02 ;     CA - CB
- HB3
  2269   2271   2274       1    1.0950e+02    3.3472e+02 ;     CA - CB
- CG
  2269   2280   2281       1    1.2040e+02    6.6944e+02 ;     CA - C
- O
  2269   2280   2282       1    1.1660e+02    5.8576e+02 ;     CA - C
- N
  2270   2269   2271       1    1.0950e+02    4.1840e+02 ;     HA - CA
- CB
  2270   2269   2280       1    1.0950e+02    4.1840e+02 ;     HA - CA
- C
  2271   2269   2280       1    1.1110e+02    5.2718e+02 ;     CB - CA
- C
  2271   2274   2275       1    1.0950e+02    4.1840e+02 ;     CB - CG
- HG2
  2271   2274   2276       1    1.0950e+02    4.1840e+02 ;     CB - CG
- HG3
  2271   2274   2277       1    1.1110e+02    5.2718e+02 ;     CB - CG
- CD
  2272   2271   2273       1    1.0950e+02    2.9288e+02 ;    HB2 - CB
- HB3
  2272   2271   2274       1    1.0950e+02    4.1840e+02 ;    HB2 - CB
- CG
  2273   2271   2274       1    1.0950e+02    4.1840e+02 ;    HB3 - CB
- CG
  2274   2277   2278       1    1.1700e+02    5.8576e+02 ;     CG - CD
- OE1
  2274   2277   2279       1    1.1700e+02    5.8576e+02 ;     CG - CD
- OE2
  2275   2274   2276       1    1.0950e+02    2.9288e+02 ;    HG2 - CG
- HG3
  2275   2274   2277       1    1.0950e+02    4.1840e+02 ;    HG2 - CG
- CD
  2276   2274   2277       1    1.0950e+02    4.1840e+02 ;    HG3 - CG
- CD
  2278   2277   2279       1    1.2600e+02    6.6944e+02 ;    OE1 - CD
- OE2
  2280   2282   2283       1    1.2000e+02    4.1840e+02 ;      C - N
- H
  2280   2282   2284       1    1.2190e+02    4.1840e+02 ;      C - N
- CA
  2281   2280   2282       1    1.2290e+02    6.6944e+02 ;      O - C
- N
  2282   2284   2285       1    1.0950e+02    4.1840e+02 ;      N - CA
- HA
  2282   2284   2286       1    1.0970e+02    6.6944e+02 ;      N - CA
- CB
  2282   2284   2294       1    1.1010e+02    5.2718e+02 ;      N - CA
- C
```

2283 2282 2284 1 1.1804e+02 4.1840e+02 ; H - N
- CA
2284 2286 2287 1 1.0950e+02 4.1840e+02 ; CA - CB
- HB2
2284 2286 2288 1 1.0950e+02 4.1840e+02 ; CA - CB
- HB3
2284 2286 2289 1 1.1110e+02 5.2718e+02 ; CA - CB
- CG
2284 2294 2295 1 1.2040e+02 6.6944e+02 ; CA - C
- O
2284 2294 2296 1 1.1660e+02 5.8576e+02 ; CA - C
- N
2285 2284 2286 1 1.0950e+02 4.1840e+02 ; HA - CA
- CB
2285 2284 2294 1 1.0950e+02 4.1840e+02 ; HA - CA
- C
2286 2284 2294 1 1.1110e+02 5.2718e+02 ; CB - CA
- C
2286 2289 2290 1 1.2040e+02 6.6944e+02 ; CB - CG
- OD1
2286 2289 2291 1 1.1660e+02 5.8576e+02 ; CB - CG
- ND2
2287 2286 2288 1 1.0950e+02 2.9288e+02 ; HB2 - CB
- HB3
2287 2286 2289 1 1.0950e+02 4.1840e+02 ; HB2 - CB
- CG
2288 2286 2289 1 1.0950e+02 4.1840e+02 ; HB3 - CB
- CG
2289 2291 2292 1 1.2000e+02 4.1840e+02 ; CG - ND2
- HD21
2289 2291 2293 1 1.2000e+02 4.1840e+02 ; CG - ND2
- HD22
2290 2289 2291 1 1.2290e+02 6.6944e+02 ; OD1 - CG
- ND2
2292 2291 2293 1 1.2000e+02 2.9288e+02 ; HD21 - ND2
- HD22
2294 2296 2297 1 1.2000e+02 4.1840e+02 ; C - N
- H
2294 2296 2298 1 1.2190e+02 4.1840e+02 ; C - N
- CA
2295 2294 2296 1 1.2290e+02 6.6944e+02 ; O - C
- N
2296 2298 2299 1 1.0950e+02 4.1840e+02 ; N - CA
- HA
2296 2298 2300 1 1.0970e+02 6.6944e+02 ; N - CA
- CB
2296 2298 2314 1 1.1010e+02 5.2718e+02 ; N - CA
- C
2297 2296 2298 1 1.1804e+02 4.1840e+02 ; H - N
- CA
2298 2300 2301 1 1.0950e+02 4.1840e+02 ; CA - CB
- HB2

```
  2298   2300   2302       1    1.0950e+02    4.1840e+02 ;     CA - CB
- HB3
  2298   2300   2303       1    1.1400e+02    5.2718e+02 ;     CA - CB
- CG
  2298   2314   2315       1    1.2040e+02    6.6944e+02 ;     CA - C
- O
  2298   2314   2316       1    1.1660e+02    5.8576e+02 ;     CA - C
- N
  2299   2298   2300       1    1.0950e+02    4.1840e+02 ;     HA - CA
- CB
  2299   2298   2314       1    1.0950e+02    4.1840e+02 ;     HA - CA
- C
  2300   2298   2314       1    1.1110e+02    5.2718e+02 ;     CB - CA
- C
  2300   2303   2304       1    1.2000e+02    5.8576e+02 ;     CB - CG
- CD1
  2300   2303   2312       1    1.2000e+02    5.8576e+02 ;     CB - CG
- CD2
  2301   2300   2302       1    1.0950e+02    2.9288e+02 ;    HB2 - CB
- HB3
  2301   2300   2303       1    1.0950e+02    4.1840e+02 ;    HB2 - CB
- CG
  2302   2300   2303       1    1.0950e+02    4.1840e+02 ;    HB3 - CB
- CG
  2303   2304   2305       1    1.2000e+02    4.1840e+02 ;     CG - CD1
- HD1
  2303   2304   2306       1    1.2000e+02    5.2718e+02 ;     CG - CD1
- CE1
  2303   2312   2310       1    1.2000e+02    5.2718e+02 ;     CG - CD2
- CE2
  2303   2312   2313       1    1.2000e+02    4.1840e+02 ;     CG - CD2
- HD2
  2304   2303   2312       1    1.2000e+02    5.2718e+02 ;    CD1 - CG
- CD2
  2304   2306   2307       1    1.2000e+02    4.1840e+02 ;    CD1 - CE1
- HE1
  2304   2306   2308       1    1.2000e+02    5.2718e+02 ;    CD1 - CE1
- CZ
  2305   2304   2306       1    1.2000e+02    4.1840e+02 ;    HD1 - CD1
- CE1
  2306   2308   2309       1    1.2000e+02    4.1840e+02 ;    CE1 - CZ
- HZ
  2306   2308   2310       1    1.2000e+02    5.2718e+02 ;    CE1 - CZ
- CE2
  2307   2306   2308       1    1.2000e+02    4.1840e+02 ;    HE1 - CE1
- CZ
  2308   2310   2311       1    1.2000e+02    4.1840e+02 ;     CZ - CE2
- HE2
  2308   2310   2312       1    1.2000e+02    5.2718e+02 ;     CZ - CE2
- CD2
  2309   2308   2310       1    1.2000e+02    4.1840e+02 ;     HZ - CZ
- CE2
```

2310 2312 2313 1 1.2000e+02 4.1840e+02 ; CE2 - CD2
- HD2
2311 2310 2312 1 1.2000e+02 4.1840e+02 ; HE2 - CE2
- CD2
2314 2316 2317 1 1.2000e+02 4.1840e+02 ; C - N
- H
2314 2316 2318 1 1.2190e+02 4.1840e+02 ; C - N
- CA
2315 2314 2316 1 1.2290e+02 6.6944e+02 ; O - C
- N
2316 2318 2319 1 1.0950e+02 4.1840e+02 ; N - CA
- HA
2316 2318 2320 1 1.0970e+02 6.6944e+02 ; N - CA
- CB
2316 2318 2333 1 1.1010e+02 5.2718e+02 ; N - CA
- C
2317 2316 2318 1 1.1804e+02 4.1840e+02 ; H - N
- CA
2318 2320 2321 1 1.0950e+02 4.1840e+02 ; CA - CB
- HB2
2318 2320 2322 1 1.0950e+02 4.1840e+02 ; CA - CB
- HB3
2318 2320 2323 1 1.0950e+02 3.3472e+02 ; CA - CB
- CG
2318 2333 2334 1 1.2040e+02 6.6944e+02 ; CA - C
- O
2318 2333 2335 1 1.1660e+02 5.8576e+02 ; CA - C
- N
2319 2318 2320 1 1.0950e+02 4.1840e+02 ; HA - CA
- CB
2319 2318 2333 1 1.0950e+02 4.1840e+02 ; HA - CA
- C
2320 2318 2333 1 1.1110e+02 5.2718e+02 ; CB - CA
- C
2320 2323 2324 1 1.0950e+02 4.1840e+02 ; CB - CG
- HG
2320 2323 2325 1 1.0950e+02 3.3472e+02 ; CB - CG
- CD1
2320 2323 2329 1 1.0950e+02 3.3472e+02 ; CB - CG
- CD2
2321 2320 2322 1 1.0950e+02 2.9288e+02 ; HB2 - CB
- HB3
2321 2320 2323 1 1.0950e+02 4.1840e+02 ; HB2 - CB
- CG
2322 2320 2323 1 1.0950e+02 4.1840e+02 ; HB3 - CB
- CG
2323 2325 2326 1 1.0950e+02 4.1840e+02 ; CG - CD1
- HD11
2323 2325 2327 1 1.0950e+02 4.1840e+02 ; CG - CD1
- HD12
2323 2325 2328 1 1.0950e+02 4.1840e+02 ; CG - CD1
- HD13

```
  2323   2329   2330      1    1.0950e+02    4.1840e+02 ;     CG - CD2
- HD21
  2323   2329   2331      1    1.0950e+02    4.1840e+02 ;     CG - CD2
- HD22
  2323   2329   2332      1    1.0950e+02    4.1840e+02 ;     CG - CD2
- HD23
  2324   2323   2325      1    1.0950e+02    4.1840e+02 ;     HG - CG
- CD1
  2324   2323   2329      1    1.0950e+02    4.1840e+02 ;     HG - CG
- CD2
  2325   2323   2329      1    1.0950e+02    3.3472e+02 ;    CD1 - CG
- CD2
  2326   2325   2327      1    1.0950e+02    2.9288e+02 ;   HD11 - CD1
- HD12
  2326   2325   2328      1    1.0950e+02    2.9288e+02 ;   HD11 - CD1
- HD13
  2327   2325   2328      1    1.0950e+02    2.9288e+02 ;   HD12 - CD1
- HD13
  2330   2329   2331      1    1.0950e+02    2.9288e+02 ;   HD21 - CD2
- HD22
  2330   2329   2332      1    1.0950e+02    2.9288e+02 ;   HD21 - CD2
- HD23
  2331   2329   2332      1    1.0950e+02    2.9288e+02 ;   HD22 - CD2
- HD23
  2333   2335   2336      1    1.2000e+02    4.1840e+02 ;      C - N
- H
  2333   2335   2337      1    1.2190e+02    4.1840e+02 ;      C - N
- CA
  2334   2333   2335      1    1.2290e+02    6.6944e+02 ;      O - C
- N
  2335   2337   2338      1    1.0950e+02    4.1840e+02 ;      N - CA
- HA
  2335   2337   2339      1    1.0970e+02    6.6944e+02 ;      N - CA
- CB
  2335   2337   2353      1    1.1010e+02    5.2718e+02 ;      N - CA
- C
  2336   2335   2337      1    1.1804e+02    4.1840e+02 ;      H - N
- CA
  2337   2339   2340      1    1.0950e+02    4.1840e+02 ;     CA - CB
- HB2
  2337   2339   2341      1    1.0950e+02    4.1840e+02 ;     CA - CB
- HB3
  2337   2339   2342      1    1.1400e+02    5.2718e+02 ;     CA - CB
- CG
  2337   2353   2354      1    1.2040e+02    6.6944e+02 ;     CA - C
- O
  2337   2353   2355      1    1.1660e+02    5.8576e+02 ;     CA - C
- N
  2338   2337   2339      1    1.0950e+02    4.1840e+02 ;     HA - CA
- CB
  2338   2337   2353      1    1.0950e+02    4.1840e+02 ;     HA - CA
- C
```

2339 2337 2353 1 1.1110e+02 5.2718e+02 ; CB - CA
- C
2339 2342 2343 1 1.2000e+02 5.8576e+02 ; CB - CG
- CD1
2339 2342 2351 1 1.2000e+02 5.8576e+02 ; CB - CG
- CD2
2340 2339 2341 1 1.0950e+02 2.9288e+02 ; HB2 - CB
- HB3
2340 2339 2342 1 1.0950e+02 4.1840e+02 ; HB2 - CB
- CG
2341 2339 2342 1 1.0950e+02 4.1840e+02 ; HB3 - CB
- CG
2342 2343 2344 1 1.2000e+02 4.1840e+02 ; CG - CD1
- HD1
2342 2343 2345 1 1.2000e+02 5.2718e+02 ; CG - CD1
- CE1
2342 2351 2349 1 1.2000e+02 5.2718e+02 ; CG - CD2
- CE2
2342 2351 2352 1 1.2000e+02 4.1840e+02 ; CG - CD2
- HD2
2343 2342 2351 1 1.2000e+02 5.2718e+02 ; CD1 - CG
- CD2
2343 2345 2346 1 1.2000e+02 4.1840e+02 ; CD1 - CE1
- HE1
2343 2345 2347 1 1.2000e+02 5.2718e+02 ; CD1 - CE1
- CZ
2344 2343 2345 1 1.2000e+02 4.1840e+02 ; HD1 - CD1
- CE1
2345 2347 2348 1 1.2000e+02 4.1840e+02 ; CE1 - CZ
- HZ
2345 2347 2349 1 1.2000e+02 5.2718e+02 ; CE1 - CZ
- CE2
2346 2345 2347 1 1.2000e+02 4.1840e+02 ; HE1 - CE1
- CZ
2347 2349 2350 1 1.2000e+02 4.1840e+02 ; CZ - CE2
- HE2
2347 2349 2351 1 1.2000e+02 5.2718e+02 ; CZ - CE2
- CD2
2348 2347 2349 1 1.2000e+02 4.1840e+02 ; HZ - CZ
- CE2
2349 2351 2352 1 1.2000e+02 4.1840e+02 ; CE2 - CD2
- HD2
2350 2349 2351 1 1.2000e+02 4.1840e+02 ; HE2 - CE2
- CD2
2353 2355 2356 1 1.2000e+02 4.1840e+02 ; C - N
- H
2353 2355 2357 1 1.2190e+02 4.1840e+02 ; C - N
- CA
2354 2353 2355 1 1.2290e+02 6.6944e+02 ; O - C
- N
2355 2357 2358 1 1.0950e+02 4.1840e+02 ; N - CA
- HA

```
  2355   2357   2359       1    1.0970e+02    6.6944e+02 ;       N - CA
- CB
  2355   2357   2370       1    1.1010e+02    5.2718e+02 ;       N - CA
- C
  2356   2355   2357       1    1.1804e+02    4.1840e+02 ;       H - N
- CA
  2357   2359   2360       1    1.0950e+02    4.1840e+02 ;      CA - CB
- HB2
  2357   2359   2361       1    1.0950e+02    4.1840e+02 ;      CA - CB
- HB3
  2357   2359   2362       1    1.0950e+02    3.3472e+02 ;      CA - CB
- CG
  2357   2370   2371       1    1.2040e+02    6.6944e+02 ;      CA - C
- O
  2357   2370   2372       1    1.1660e+02    5.8576e+02 ;      CA - C
- N
  2358   2357   2359       1    1.0950e+02    4.1840e+02 ;      HA - CA
- CB
  2358   2357   2370       1    1.0950e+02    4.1840e+02 ;      HA - CA
- C
  2359   2357   2370       1    1.1110e+02    5.2718e+02 ;      CB - CA
- C
  2359   2362   2363       1    1.0950e+02    4.1840e+02 ;      CB - CG
- HG2
  2359   2362   2364       1    1.0950e+02    4.1840e+02 ;      CB - CG
- HG3
  2359   2362   2365       1    1.1110e+02    5.2718e+02 ;      CB - CG
- CD
  2360   2359   2361       1    1.0950e+02    2.9288e+02 ;     HB2 - CB
- HB3
  2360   2359   2362       1    1.0950e+02    4.1840e+02 ;     HB2 - CB
- CG
  2361   2359   2362       1    1.0950e+02    4.1840e+02 ;     HB3 - CB
- CG
  2362   2365   2366       1    1.2040e+02    6.6944e+02 ;      CG - CD
- OE1
  2362   2365   2367       1    1.1660e+02    5.8576e+02 ;      CG - CD
- NE2
  2363   2362   2364       1    1.0950e+02    2.9288e+02 ;     HG2 - CG
- HG3
  2363   2362   2365       1    1.0950e+02    4.1840e+02 ;     HG2 - CG
- CD
  2364   2362   2365       1    1.0950e+02    4.1840e+02 ;     HG3 - CG
- CD
  2365   2367   2368       1    1.2000e+02    4.1840e+02 ;      CD - NE2
- HE21
  2365   2367   2369       1    1.2000e+02    4.1840e+02 ;      CD - NE2
- HE22
  2366   2365   2367       1    1.2290e+02    6.6944e+02 ;     OE1 - CD
- NE2
  2368   2367   2369       1    1.2000e+02    2.9288e+02 ;    HE21 - NE2
- HE22
```

```
  2370   2372   2373       1    1.2000e+02    4.1840e+02 ;      C - N
- H
  2370   2372   2374       1    1.2190e+02    4.1840e+02 ;      C - N
- CA
  2371   2370   2372       1    1.2290e+02    6.6944e+02 ;      O - C
- N
  2372   2374   2375       1    1.0950e+02    4.1840e+02 ;      N - CA
- HA
  2372   2374   2376       1    1.0970e+02    6.6944e+02 ;      N - CA
- CB
  2372   2374   2381       1    1.1010e+02    5.2718e+02 ;      N - CA
- C
  2373   2372   2374       1    1.1804e+02    4.1840e+02 ;      H - N
- CA
  2374   2376   2377       1    1.0950e+02    4.1840e+02 ;     CA - CB
- HB2
  2374   2376   2378       1    1.0950e+02    4.1840e+02 ;     CA - CB
- HB3
  2374   2376   2379       1    1.0950e+02    4.1840e+02 ;     CA - CB
- OG
  2374   2381   2382       1    1.2040e+02    6.6944e+02 ;     CA - C
- O
  2374   2381   2383       1    1.1660e+02    5.8576e+02 ;     CA - C
- N
  2375   2374   2376       1    1.0950e+02    4.1840e+02 ;     HA - CA
- CB
  2375   2374   2381       1    1.0950e+02    4.1840e+02 ;     HA - CA
- C
  2376   2374   2381       1    1.1110e+02    5.2718e+02 ;     CB - CA
- C
  2376   2379   2380       1    1.0850e+02    4.6024e+02 ;     CB - OG
- HG
  2377   2376   2378       1    1.0950e+02    2.9288e+02 ;    HB2 - CB
- HB3
  2377   2376   2379       1    1.0950e+02    4.1840e+02 ;    HB2 - CB
- OG
  2378   2376   2379       1    1.0950e+02    4.1840e+02 ;    HB3 - CB
- OG
  2381   2383   2384       1    1.2000e+02    4.1840e+02 ;      C - N
- H
  2381   2383   2385       1    1.2190e+02    4.1840e+02 ;      C - N
- CA
  2382   2381   2383       1    1.2290e+02    6.6944e+02 ;      O - C
- N
  2383   2385   2386       1    1.0950e+02    4.1840e+02 ;      N - CA
- HA
  2383   2385   2387       1    1.0970e+02    6.6944e+02 ;      N - CA
- CB
  2383   2385   2396       1    1.1010e+02    5.2718e+02 ;      N - CA
- C
  2384   2383   2385       1    1.1804e+02    4.1840e+02 ;      H - N
- CA
```

2385 2387 2388 1 1.0950e+02 4.1840e+02 ; CA - CB
- HB2
2385 2387 2389 1 1.0950e+02 4.1840e+02 ; CA - CB
- HB3
2385 2387 2390 1 1.0950e+02 3.3472e+02 ; CA - CB
- CG
2385 2396 2397 1 1.2040e+02 6.6944e+02 ; CA - C
- O
2385 2396 2398 1 1.1660e+02 5.8576e+02 ; CA - C
- N
2386 2385 2387 1 1.0950e+02 4.1840e+02 ; HA - CA
- CB
2386 2385 2396 1 1.0950e+02 4.1840e+02 ; HA - CA
- C
2387 2385 2396 1 1.1110e+02 5.2718e+02 ; CB - CA
- C
2387 2390 2391 1 1.0950e+02 4.1840e+02 ; CB - CG
- HG2
2387 2390 2392 1 1.0950e+02 4.1840e+02 ; CB - CG
- HG3
2387 2390 2393 1 1.1110e+02 5.2718e+02 ; CB - CG
- CD
2388 2387 2389 1 1.0950e+02 2.9288e+02 ; HB2 - CB
- HB3
2388 2387 2390 1 1.0950e+02 4.1840e+02 ; HB2 - CB
- CG
2389 2387 2390 1 1.0950e+02 4.1840e+02 ; HB3 - CB
- CG
2390 2393 2394 1 1.1700e+02 5.8576e+02 ; CG - CD
- OE1
2390 2393 2395 1 1.1700e+02 5.8576e+02 ; CG - CD
- OE2
2391 2390 2392 1 1.0950e+02 2.9288e+02 ; HG2 - CG
- HG3
2391 2390 2393 1 1.0950e+02 4.1840e+02 ; HG2 - CG
- CD
2392 2390 2393 1 1.0950e+02 4.1840e+02 ; HG3 - CG
- CD
2394 2393 2395 1 1.2600e+02 6.6944e+02 ; OE1 - CD
- OE2
2396 2398 2399 1 1.2000e+02 4.1840e+02 ; C - N
- H
2396 2398 2400 1 1.2190e+02 4.1840e+02 ; C - N
- CA
2397 2396 2398 1 1.2290e+02 6.6944e+02 ; O - C
- N
2398 2400 2401 1 1.0950e+02 4.1840e+02 ; N - CA
- HA
2398 2400 2402 1 1.0970e+02 6.6944e+02 ; N - CA
- CB
2398 2400 2410 1 1.1010e+02 5.2718e+02 ; N - CA
- C

2399 2398 2400 1 1.1804e+02 4.1840e+02 ; H - N
- CA
2400 2402 2403 1 1.0950e+02 4.1840e+02 ; CA - CB
- HB2
2400 2402 2404 1 1.0950e+02 4.1840e+02 ; CA - CB
- HB3
2400 2402 2405 1 1.1110e+02 5.2718e+02 ; CA - CB
- CG
2400 2410 2411 1 1.2040e+02 6.6944e+02 ; CA - C
- O
2400 2410 2412 1 1.1660e+02 5.8576e+02 ; CA - C
- N
2401 2400 2402 1 1.0950e+02 4.1840e+02 ; HA - CA
- CB
2401 2400 2410 1 1.0950e+02 4.1840e+02 ; HA - CA
- C
2402 2400 2410 1 1.1110e+02 5.2718e+02 ; CB - CA
- C
2402 2405 2406 1 1.2040e+02 6.6944e+02 ; CB - CG
- OD1
2402 2405 2407 1 1.1660e+02 5.8576e+02 ; CB - CG
- ND2
2403 2402 2404 1 1.0950e+02 2.9288e+02 ; HB2 - CB
- HB3
2403 2402 2405 1 1.0950e+02 4.1840e+02 ; HB2 - CB
- CG
2404 2402 2405 1 1.0950e+02 4.1840e+02 ; HB3 - CB
- CG
2405 2407 2408 1 1.2000e+02 4.1840e+02 ; CG - ND2
- HD21
2405 2407 2409 1 1.2000e+02 4.1840e+02 ; CG - ND2
- HD22
2406 2405 2407 1 1.2290e+02 6.6944e+02 ; OD1 - CG
- ND2
2408 2407 2409 1 1.2000e+02 2.9288e+02 ; HD21 - ND2
- HD22
2410 2412 2413 1 1.2000e+02 4.1840e+02 ; C - N
- H
2410 2412 2414 1 1.2190e+02 4.1840e+02 ; C - N
- CA
2411 2410 2412 1 1.2290e+02 6.6944e+02 ; O - C
- N
2412 2414 2415 1 1.0950e+02 4.1840e+02 ; N - CA
- HA
2412 2414 2416 1 1.0970e+02 6.6944e+02 ; N - CA
- CB
2412 2414 2432 1 1.1010e+02 5.2718e+02 ; N - CA
- C
2413 2412 2414 1 1.1804e+02 4.1840e+02 ; H - N
- CA
2414 2416 2417 1 1.0950e+02 4.1840e+02 ; CA - CB
- HB2

2414 2416 2418 1 1.0950e+02 4.1840e+02 ; CA - CB
- HB3
2414 2416 2419 1 1.0950e+02 3.3472e+02 ; CA - CB
- CG
2414 2432 2433 1 1.2040e+02 6.6944e+02 ; CA - C
- O
2414 2432 2434 1 1.1660e+02 5.8576e+02 ; CA - C
- N
2415 2414 2416 1 1.0950e+02 4.1840e+02 ; HA - CA
- CB
2415 2414 2432 1 1.0950e+02 4.1840e+02 ; HA - CA
- C
2416 2414 2432 1 1.1110e+02 5.2718e+02 ; CB - CA
- C
2416 2419 2420 1 1.0950e+02 4.1840e+02 ; CB - CG
- HG2
2416 2419 2421 1 1.0950e+02 4.1840e+02 ; CB - CG
- HG3
2416 2419 2422 1 1.0950e+02 3.3472e+02 ; CB - CG
- CD
2417 2416 2418 1 1.0950e+02 2.9288e+02 ; HB2 - CB
- HB3
2417 2416 2419 1 1.0950e+02 4.1840e+02 ; HB2 - CB
- CG
2418 2416 2419 1 1.0950e+02 4.1840e+02 ; HB3 - CB
- CG
2419 2422 2423 1 1.0950e+02 4.1840e+02 ; CG - CD
- HD2
2419 2422 2424 1 1.0950e+02 4.1840e+02 ; CG - CD
- HD3
2419 2422 2425 1 1.0950e+02 3.3472e+02 ; CG - CD
- CE
2420 2419 2421 1 1.0950e+02 2.9288e+02 ; HG2 - CG
- HG3
2420 2419 2422 1 1.0950e+02 4.1840e+02 ; HG2 - CG
- CD
2421 2419 2422 1 1.0950e+02 4.1840e+02 ; HG3 - CG
- CD
2422 2425 2426 1 1.0950e+02 4.1840e+02 ; CD - CE
- HE2
2422 2425 2427 1 1.0950e+02 4.1840e+02 ; CD - CE
- HE3
2422 2425 2428 1 1.1120e+02 6.6944e+02 ; CD - CE
- NZ
2423 2422 2424 1 1.0950e+02 2.9288e+02 ; HD2 - CD
- HD3
2423 2422 2425 1 1.0950e+02 4.1840e+02 ; HD2 - CD
- CE
2424 2422 2425 1 1.0950e+02 4.1840e+02 ; HD3 - CD
- CE
2425 2428 2429 1 1.0950e+02 4.1840e+02 ; CE - NZ
- HZ1

2425 2428 2430 1 1.0950e+02 4.1840e+02 ; CE - NZ
- HZ2
2425 2428 2431 1 1.0950e+02 4.1840e+02 ; CE - NZ
- HZ3
2426 2425 2427 1 1.0950e+02 2.9288e+02 ; HE2 - CE
- HE3
2426 2425 2428 1 1.0950e+02 4.1840e+02 ; HE2 - CE
- NZ
2427 2425 2428 1 1.0950e+02 4.1840e+02 ; HE3 - CE
- NZ
2429 2428 2430 1 1.0950e+02 2.9288e+02 ; HZ1 - NZ
- HZ2
2429 2428 2431 1 1.0950e+02 2.9288e+02 ; HZ1 - NZ
- HZ3
2430 2428 2431 1 1.0950e+02 2.9288e+02 ; HZ2 - NZ
- HZ3
2432 2434 2435 1 1.2000e+02 4.1840e+02 ; C - N
- H
2432 2434 2436 1 1.2190e+02 4.1840e+02 ; C - N
- CA
2433 2432 2434 1 1.2290e+02 6.6944e+02 ; O - C
- N
2434 2436 2437 1 1.0950e+02 4.1840e+02 ; N - CA
- HA
2434 2436 2438 1 1.0970e+02 6.6944e+02 ; N - CA
- CB
2434 2436 2444 1 1.1010e+02 5.2718e+02 ; N - CA
- C
2435 2434 2436 1 1.1804e+02 4.1840e+02 ; H - N
- CA
2436 2438 2439 1 1.0950e+02 4.1840e+02 ; CA - CB
- HB2
2436 2438 2440 1 1.0950e+02 4.1840e+02 ; CA - CB
- HB3
2436 2438 2441 1 1.1110e+02 5.2718e+02 ; CA - CB
- CG
2436 2444 2445 1 1.2040e+02 6.6944e+02 ; CA - C
- O
2436 2444 2446 1 1.1660e+02 5.8576e+02 ; CA - C
- N
2437 2436 2438 1 1.0950e+02 4.1840e+02 ; HA - CA
- CB
2437 2436 2444 1 1.0950e+02 4.1840e+02 ; HA - CA
- C
2438 2436 2444 1 1.1110e+02 5.2718e+02 ; CB - CA
- C
2438 2441 2442 1 1.1700e+02 5.8576e+02 ; CB - CG
- OD1
2438 2441 2443 1 1.1700e+02 5.8576e+02 ; CB - CG
- OD2
2439 2438 2440 1 1.0950e+02 2.9288e+02 ; HB2 - CB
- HB3

```
  2439   2438   2441      1    1.0950e+02    4.1840e+02 ;    HB2 - CB
- CG
  2440   2438   2441      1    1.0950e+02    4.1840e+02 ;    HB3 - CB
- CG
  2442   2441   2443      1    1.2600e+02    6.6944e+02 ;    OD1 - CG
- OD2
  2444   2446   2447      1    1.2000e+02    4.1840e+02 ;      C - N
- H
  2444   2446   2448      1    1.2190e+02    4.1840e+02 ;      C - N
- CA
  2445   2444   2446      1    1.2290e+02    6.6944e+02 ;      O - C
- N
  2446   2448   2449      1    1.0950e+02    4.1840e+02 ;      N - CA
- HA
  2446   2448   2450      1    1.0970e+02    6.6944e+02 ;      N - CA
- CB
  2446   2448   2455      1    1.1010e+02    5.2718e+02 ;      N - CA
- C
  2447   2446   2448      1    1.1804e+02    4.1840e+02 ;      H - N
- CA
  2448   2450   2451      1    1.0950e+02    4.1840e+02 ;     CA - CB
- HB2
  2448   2450   2452      1    1.0950e+02    4.1840e+02 ;     CA - CB
- HB3
  2448   2450   2453      1    1.0950e+02    4.1840e+02 ;     CA - CB
- OG
  2448   2455   2456      1    1.2040e+02    6.6944e+02 ;     CA - C
- O
  2448   2455   2457      1    1.1660e+02    5.8576e+02 ;     CA - C
- N
  2449   2448   2450      1    1.0950e+02    4.1840e+02 ;     HA - CA
- CB
  2449   2448   2455      1    1.0950e+02    4.1840e+02 ;     HA - CA
- C
  2450   2448   2455      1    1.1110e+02    5.2718e+02 ;     CB - CA
- C
  2450   2453   2454      1    1.0850e+02    4.6024e+02 ;     CB - OG
- HG
  2451   2450   2452      1    1.0950e+02    2.9288e+02 ;    HB2 - CB
- HB3
  2451   2450   2453      1    1.0950e+02    4.1840e+02 ;    HB2 - CB
- OG
  2452   2450   2453      1    1.0950e+02    4.1840e+02 ;    HB3 - CB
- OG
  2455   2457   2458      1    1.2000e+02    4.1840e+02 ;      C - N
- H
  2455   2457   2459      1    1.2190e+02    4.1840e+02 ;      C - N
- CA
  2456   2455   2457      1    1.2290e+02    6.6944e+02 ;      O - C
- N
  2457   2459   2460      1    1.0950e+02    4.1840e+02 ;      N - CA
- HA
```

2457 2459 2461 1 1.0970e+02 6.6944e+02 ; N - CA
- CB
2457 2459 2474 1 1.1010e+02 5.2718e+02 ; N - CA
- C
2458 2457 2459 1 1.1804e+02 4.1840e+02 ; H - N
- CA
2459 2461 2462 1 1.0950e+02 4.1840e+02 ; CA - CB
- HB2
2459 2461 2463 1 1.0950e+02 4.1840e+02 ; CA - CB
- HB3
2459 2461 2464 1 1.0950e+02 3.3472e+02 ; CA - CB
- CG
2459 2474 2475 1 1.2040e+02 6.6944e+02 ; CA - C
- O
2459 2474 2476 1 1.1660e+02 5.8576e+02 ; CA - C
- N
2460 2459 2461 1 1.0950e+02 4.1840e+02 ; HA - CA
- CB
2460 2459 2474 1 1.0950e+02 4.1840e+02 ; HA - CA
- C
2461 2459 2474 1 1.1110e+02 5.2718e+02 ; CB - CA
- C
2461 2464 2465 1 1.0950e+02 4.1840e+02 ; CB - CG
- HG
2461 2464 2466 1 1.0950e+02 3.3472e+02 ; CB - CG
- CD1
2461 2464 2470 1 1.0950e+02 3.3472e+02 ; CB - CG
- CD2
2462 2461 2463 1 1.0950e+02 2.9288e+02 ; HB2 - CB
- HB3
2462 2461 2464 1 1.0950e+02 4.1840e+02 ; HB2 - CB
- CG
2463 2461 2464 1 1.0950e+02 4.1840e+02 ; HB3 - CB
- CG
2464 2466 2467 1 1.0950e+02 4.1840e+02 ; CG - CD1
- HD11
2464 2466 2468 1 1.0950e+02 4.1840e+02 ; CG - CD1
- HD12
2464 2466 2469 1 1.0950e+02 4.1840e+02 ; CG - CD1
- HD13
2464 2470 2471 1 1.0950e+02 4.1840e+02 ; CG - CD2
- HD21
2464 2470 2472 1 1.0950e+02 4.1840e+02 ; CG - CD2
- HD22
2464 2470 2473 1 1.0950e+02 4.1840e+02 ; CG - CD2
- HD23
2465 2464 2466 1 1.0950e+02 4.1840e+02 ; HG - CG
- CD1
2465 2464 2470 1 1.0950e+02 4.1840e+02 ; HG - CG
- CD2
2466 2464 2470 1 1.0950e+02 3.3472e+02 ; CD1 - CG
- CD2

2467 2466 2468 1 1.0950e+02 2.9288e+02 ; HD11 - CD1 - HD12
2467 2466 2469 1 1.0950e+02 2.9288e+02 ; HD11 - CD1 - HD13
2468 2466 2469 1 1.0950e+02 2.9288e+02 ; HD12 - CD1 - HD13
2471 2470 2472 1 1.0950e+02 2.9288e+02 ; HD21 - CD2 - HD22
2471 2470 2473 1 1.0950e+02 2.9288e+02 ; HD21 - CD2 - HD23
2472 2470 2473 1 1.0950e+02 2.9288e+02 ; HD22 - CD2 - HD23
2474 2476 2477 1 1.2000e+02 4.1840e+02 ; C - N - H
2474 2476 2478 1 1.2190e+02 4.1840e+02 ; C - N - CA
2475 2474 2476 1 1.2290e+02 6.6944e+02 ; O - C - N
2476 2478 2479 1 1.0950e+02 4.1840e+02 ; N - CA - HA
2476 2478 2480 1 1.0970e+02 6.6944e+02 ; N - CA - CB
2476 2478 2493 1 1.1010e+02 5.2718e+02 ; N - CA - C
2477 2476 2478 1 1.1804e+02 4.1840e+02 ; H - N - CA
2478 2480 2481 1 1.0950e+02 4.1840e+02 ; CA - CB - HB2
2478 2480 2482 1 1.0950e+02 4.1840e+02 ; CA - CB - HB3
2478 2480 2483 1 1.0950e+02 3.3472e+02 ; CA - CB - CG
2478 2493 2494 1 1.2040e+02 6.6944e+02 ; CA - C - O
2478 2493 2495 1 1.1660e+02 5.8576e+02 ; CA - C - N
2479 2478 2480 1 1.0950e+02 4.1840e+02 ; HA - CA - CB
2479 2478 2493 1 1.0950e+02 4.1840e+02 ; HA - CA - C
2480 2478 2493 1 1.1110e+02 5.2718e+02 ; CB - CA - C
2480 2483 2484 1 1.0950e+02 4.1840e+02 ; CB - CG - HG
2480 2483 2485 1 1.0950e+02 3.3472e+02 ; CB - CG - CD1
2480 2483 2489 1 1.0950e+02 3.3472e+02 ; CB - CG - CD2
2481 2480 2482 1 1.0950e+02 2.9288e+02 ; HB2 - CB - HB3
2481 2480 2483 1 1.0950e+02 4.1840e+02 ; HB2 - CB - CG

```
  2482   2480   2483      1    1.0950e+02    4.1840e+02 ;    HB3 - CB
- CG
  2483   2485   2486      1    1.0950e+02    4.1840e+02 ;     CG - CD1
- HD11
  2483   2485   2487      1    1.0950e+02    4.1840e+02 ;     CG - CD1
- HD12
  2483   2485   2488      1    1.0950e+02    4.1840e+02 ;     CG - CD1
- HD13
  2483   2489   2490      1    1.0950e+02    4.1840e+02 ;     CG - CD2
- HD21
  2483   2489   2491      1    1.0950e+02    4.1840e+02 ;     CG - CD2
- HD22
  2483   2489   2492      1    1.0950e+02    4.1840e+02 ;     CG - CD2
- HD23
  2484   2483   2485      1    1.0950e+02    4.1840e+02 ;     HG - CG
- CD1
  2484   2483   2489      1    1.0950e+02    4.1840e+02 ;     HG - CG
- CD2
  2485   2483   2489      1    1.0950e+02    3.3472e+02 ;    CD1 - CG
- CD2
  2486   2485   2487      1    1.0950e+02    2.9288e+02 ;   HD11 - CD1
- HD12
  2486   2485   2488      1    1.0950e+02    2.9288e+02 ;   HD11 - CD1
- HD13
  2487   2485   2488      1    1.0950e+02    2.9288e+02 ;   HD12 - CD1
- HD13
  2490   2489   2491      1    1.0950e+02    2.9288e+02 ;   HD21 - CD2
- HD22
  2490   2489   2492      1    1.0950e+02    2.9288e+02 ;   HD21 - CD2
- HD23
  2491   2489   2492      1    1.0950e+02    2.9288e+02 ;   HD22 - CD2
- HD23
  2493   2495   2496      1    1.2000e+02    4.1840e+02 ;      C - N
- H
  2493   2495   2497      1    1.2190e+02    4.1840e+02 ;      C - N
- CA
  2494   2493   2495      1    1.2290e+02    6.6944e+02 ;      O - C
- N
  2495   2497   2498      1    1.0950e+02    4.1840e+02 ;      N - CA
- HA
  2495   2497   2499      1    1.0970e+02    6.6944e+02 ;      N - CA
- CB
  2495   2497   2515      1    1.1010e+02    5.2718e+02 ;      N - CA
- C
  2496   2495   2497      1    1.1804e+02    4.1840e+02 ;      H - N
- CA
  2497   2499   2500      1    1.0950e+02    4.1840e+02 ;     CA - CB
- HB2
  2497   2499   2501      1    1.0950e+02    4.1840e+02 ;     CA - CB
- HB3
  2497   2499   2502      1    1.0950e+02    3.3472e+02 ;     CA - CB
- CG
```

2497 2515 2516 1 1.2040e+02 6.6944e+02 ; CA - C
- O
2497 2515 2517 1 1.1660e+02 5.8576e+02 ; CA - C
- N
2498 2497 2499 1 1.0950e+02 4.1840e+02 ; HA - CA
- CB
2498 2497 2515 1 1.0950e+02 4.1840e+02 ; HA - CA
- C
2499 2497 2515 1 1.1110e+02 5.2718e+02 ; CB - CA
- C
2499 2502 2503 1 1.0950e+02 4.1840e+02 ; CB - CG
- HG2
2499 2502 2504 1 1.0950e+02 4.1840e+02 ; CB - CG
- HG3
2499 2502 2505 1 1.0950e+02 3.3472e+02 ; CB - CG
- CD
2500 2499 2501 1 1.0950e+02 2.9288e+02 ; HB2 - CB
- HB3
2500 2499 2502 1 1.0950e+02 4.1840e+02 ; HB2 - CB
- CG
2501 2499 2502 1 1.0950e+02 4.1840e+02 ; HB3 - CB
- CG
2502 2505 2506 1 1.0950e+02 4.1840e+02 ; CG - CD
- HD2
2502 2505 2507 1 1.0950e+02 4.1840e+02 ; CG - CD
- HD3
2502 2505 2508 1 1.0950e+02 3.3472e+02 ; CG - CD
- CE
2503 2502 2504 1 1.0950e+02 2.9288e+02 ; HG2 - CG
- HG3
2503 2502 2505 1 1.0950e+02 4.1840e+02 ; HG2 - CG
- CD
2504 2502 2505 1 1.0950e+02 4.1840e+02 ; HG3 - CG
- CD
2505 2508 2509 1 1.0950e+02 4.1840e+02 ; CD - CE
- HE2
2505 2508 2510 1 1.0950e+02 4.1840e+02 ; CD - CE
- HE3
2505 2508 2511 1 1.1120e+02 6.6944e+02 ; CD - CE
- NZ
2506 2505 2507 1 1.0950e+02 2.9288e+02 ; HD2 - CD
- HD3
2506 2505 2508 1 1.0950e+02 4.1840e+02 ; HD2 - CD
- CE
2507 2505 2508 1 1.0950e+02 4.1840e+02 ; HD3 - CD
- CE
2508 2511 2512 1 1.0950e+02 4.1840e+02 ; CE - NZ
- HZ1
2508 2511 2513 1 1.0950e+02 4.1840e+02 ; CE - NZ
- HZ2
2508 2511 2514 1 1.0950e+02 4.1840e+02 ; CE - NZ
- HZ3

2509 2508 2510 1 1.0950e+02 2.9288e+02 ; HE2 - CE
- HE3
2509 2508 2511 1 1.0950e+02 4.1840e+02 ; HE2 - CE
- NZ
2510 2508 2511 1 1.0950e+02 4.1840e+02 ; HE3 - CE
- NZ
2512 2511 2513 1 1.0950e+02 2.9288e+02 ; HZ1 - NZ
- HZ2
2512 2511 2514 1 1.0950e+02 2.9288e+02 ; HZ1 - NZ
- HZ3
2513 2511 2514 1 1.0950e+02 2.9288e+02 ; HZ2 - NZ
- HZ3
2515 2517 2518 1 1.2000e+02 4.1840e+02 ; C - N
- H
2515 2517 2519 1 1.2190e+02 4.1840e+02 ; C - N
- CA
2516 2515 2517 1 1.2290e+02 6.6944e+02 ; O - C
- N
2517 2519 2520 1 1.0950e+02 4.1840e+02 ; N - CA
- HA
2517 2519 2521 1 1.0970e+02 6.6944e+02 ; N - CA
- CB
2517 2519 2534 1 1.1010e+02 5.2718e+02 ; N - CA
- C
2518 2517 2519 1 1.1804e+02 4.1840e+02 ; H - N
- CA
2519 2521 2522 1 1.0950e+02 4.1840e+02 ; CA - CB
- HB
2519 2521 2523 1 1.0950e+02 3.3472e+02 ; CA - CB
- CG2
2519 2521 2527 1 1.0950e+02 3.3472e+02 ; CA - CB
- CG1
2519 2534 2535 1 1.2040e+02 6.6944e+02 ; CA - C
- O
2519 2534 2536 1 1.1660e+02 5.8576e+02 ; CA - C
- N
2520 2519 2521 1 1.0950e+02 4.1840e+02 ; HA - CA
- CB
2520 2519 2534 1 1.0950e+02 4.1840e+02 ; HA - CA
- C
2521 2519 2534 1 1.1110e+02 5.2718e+02 ; CB - CA
- C
2521 2523 2524 1 1.0950e+02 4.1840e+02 ; CB - CG2
- HG21
2521 2523 2525 1 1.0950e+02 4.1840e+02 ; CB - CG2
- HG22
2521 2523 2526 1 1.0950e+02 4.1840e+02 ; CB - CG2
- HG23
2521 2527 2528 1 1.0950e+02 4.1840e+02 ; CB - CG1
- HG12
2521 2527 2529 1 1.0950e+02 4.1840e+02 ; CB - CG1
- HG13

2521 2527 2530 1 1.0950e+02 3.3472e+02 ; CB - CG1
- CD1
2522 2521 2523 1 1.0950e+02 4.1840e+02 ; HB - CB
- CG2
2522 2521 2527 1 1.0950e+02 4.1840e+02 ; HB - CB
- CG1
2523 2521 2527 1 1.0950e+02 3.3472e+02 ; CG2 - CB
- CG1
2524 2523 2525 1 1.0950e+02 2.9288e+02 ; HG21 - CG2
- HG22
2524 2523 2526 1 1.0950e+02 2.9288e+02 ; HG21 - CG2
- HG23
2525 2523 2526 1 1.0950e+02 2.9288e+02 ; HG22 - CG2
- HG23
2527 2530 2531 1 1.0950e+02 4.1840e+02 ; CG1 - CD1
- HD11
2527 2530 2532 1 1.0950e+02 4.1840e+02 ; CG1 - CD1
- HD12
2527 2530 2533 1 1.0950e+02 4.1840e+02 ; CG1 - CD1
- HD13
2528 2527 2529 1 1.0950e+02 2.9288e+02 ; HG12 - CG1
- HG13
2528 2527 2530 1 1.0950e+02 4.1840e+02 ; HG12 - CG1
- CD1
2529 2527 2530 1 1.0950e+02 4.1840e+02 ; HG13 - CG1
- CD1
2531 2530 2532 1 1.0950e+02 2.9288e+02 ; HD11 - CD1
- HD12
2531 2530 2533 1 1.0950e+02 2.9288e+02 ; HD11 - CD1
- HD13
2532 2530 2533 1 1.0950e+02 2.9288e+02 ; HD12 - CD1
- HD13
2534 2536 2537 1 1.2000e+02 4.1840e+02 ; C - N
- H
2534 2536 2538 1 1.2190e+02 4.1840e+02 ; C - N
- CA
2535 2534 2536 1 1.2290e+02 6.6944e+02 ; O - C
- N
2536 2538 2539 1 1.0950e+02 4.1840e+02 ; N - CA
- HA
2536 2538 2540 1 1.0970e+02 6.6944e+02 ; N - CA
- CB
2536 2538 2553 1 1.1010e+02 5.2718e+02 ; N - CA
- C
2537 2536 2538 1 1.1804e+02 4.1840e+02 ; H - N
- CA
2538 2540 2541 1 1.0950e+02 4.1840e+02 ; CA - CB
- HB
2538 2540 2542 1 1.0950e+02 3.3472e+02 ; CA - CB
- CG2
2538 2540 2546 1 1.0950e+02 3.3472e+02 ; CA - CB
- CG1

2538 2553 2554 1 1.2040e+02 6.6944e+02 ; CA - C
- O
2538 2553 2555 1 1.1660e+02 5.8576e+02 ; CA - C
- N
2539 2538 2540 1 1.0950e+02 4.1840e+02 ; HA - CA
- CB
2539 2538 2553 1 1.0950e+02 4.1840e+02 ; HA - CA
- C
2540 2538 2553 1 1.1110e+02 5.2718e+02 ; CB - CA
- C
2540 2542 2543 1 1.0950e+02 4.1840e+02 ; CB - CG2
- HG21
2540 2542 2544 1 1.0950e+02 4.1840e+02 ; CB - CG2
- HG22
2540 2542 2545 1 1.0950e+02 4.1840e+02 ; CB - CG2
- HG23
2540 2546 2547 1 1.0950e+02 4.1840e+02 ; CB - CG1
- HG12
2540 2546 2548 1 1.0950e+02 4.1840e+02 ; CB - CG1
- HG13
2540 2546 2549 1 1.0950e+02 3.3472e+02 ; CB - CG1
- CD1
2541 2540 2542 1 1.0950e+02 4.1840e+02 ; HB - CB
- CG2
2541 2540 2546 1 1.0950e+02 4.1840e+02 ; HB - CB
- CG1
2542 2540 2546 1 1.0950e+02 3.3472e+02 ; CG2 - CB
- CG1
2543 2542 2544 1 1.0950e+02 2.9288e+02 ; HG21 - CG2
- HG22
2543 2542 2545 1 1.0950e+02 2.9288e+02 ; HG21 - CG2
- HG23
2544 2542 2545 1 1.0950e+02 2.9288e+02 ; HG22 - CG2
- HG23
2546 2549 2550 1 1.0950e+02 4.1840e+02 ; CG1 - CD1
- HD11
2546 2549 2551 1 1.0950e+02 4.1840e+02 ; CG1 - CD1
- HD12
2546 2549 2552 1 1.0950e+02 4.1840e+02 ; CG1 - CD1
- HD13
2547 2546 2548 1 1.0950e+02 2.9288e+02 ; HG12 - CG1
- HG13
2547 2546 2549 1 1.0950e+02 4.1840e+02 ; HG12 - CG1
- CD1
2548 2546 2549 1 1.0950e+02 4.1840e+02 ; HG13 - CG1
- CD1
2550 2549 2551 1 1.0950e+02 2.9288e+02 ; HD11 - CD1
- HD12
2550 2549 2552 1 1.0950e+02 2.9288e+02 ; HD11 - CD1
- HD13
2551 2549 2552 1 1.0950e+02 2.9288e+02 ; HD12 - CD1
- HD13

2553 2555 2556 1 1.2000e+02 4.1840e+02 ; C - N
- H
2553 2555 2557 1 1.2190e+02 4.1840e+02 ; C - N
- CA
2554 2553 2555 1 1.2290e+02 6.6944e+02 ; O - C
- N
2555 2557 2558 1 1.0950e+02 4.1840e+02 ; N - CA
- HA
2555 2557 2559 1 1.0970e+02 6.6944e+02 ; N - CA
- CB
2555 2557 2565 1 1.1010e+02 5.2718e+02 ; N - CA
- C
2556 2555 2557 1 1.1804e+02 4.1840e+02 ; H - N
- CA
2557 2559 2560 1 1.0950e+02 4.1840e+02 ; CA - CB
- HB2
2557 2559 2561 1 1.0950e+02 4.1840e+02 ; CA - CB
- HB3
2557 2559 2562 1 1.1110e+02 5.2718e+02 ; CA - CB
- CG
2557 2565 2566 1 1.2040e+02 6.6944e+02 ; CA - C
- O
2557 2565 2567 1 1.1660e+02 5.8576e+02 ; CA - C
- N
2558 2557 2559 1 1.0950e+02 4.1840e+02 ; HA - CA
- CB
2558 2557 2565 1 1.0950e+02 4.1840e+02 ; HA - CA
- C
2559 2557 2565 1 1.1110e+02 5.2718e+02 ; CB - CA
- C
2559 2562 2563 1 1.1700e+02 5.8576e+02 ; CB - CG
- OD1
2559 2562 2564 1 1.1700e+02 5.8576e+02 ; CB - CG
- OD2
2560 2559 2561 1 1.0950e+02 2.9288e+02 ; HB2 - CB
- HB3
2560 2559 2562 1 1.0950e+02 4.1840e+02 ; HB2 - CB
- CG
2561 2559 2562 1 1.0950e+02 4.1840e+02 ; HB3 - CB
- CG
2563 2562 2564 1 1.2600e+02 6.6944e+02 ; OD1 - CG
- OD2
2565 2567 2568 1 1.2000e+02 4.1840e+02 ; C - N
- H
2565 2567 2569 1 1.2190e+02 4.1840e+02 ; C - N
- CA
2566 2565 2567 1 1.2290e+02 6.6944e+02 ; O - C
- N
2567 2569 2570 1 1.0950e+02 4.1840e+02 ; N - CA
- HA
2567 2569 2571 1 1.0970e+02 6.6944e+02 ; N - CA
- CB

```
  2567   2569   2585       1    1.1010e+02    5.2718e+02 ;      N - CA
- C
  2568   2567   2569       1    1.1804e+02    4.1840e+02 ;      H - N
- CA
  2569   2571   2572       1    1.0950e+02    4.1840e+02 ;     CA - CB
- HB2
  2569   2571   2573       1    1.0950e+02    4.1840e+02 ;     CA - CB
- HB3
  2569   2571   2574       1    1.1400e+02    5.2718e+02 ;     CA - CB
- CG
  2569   2585   2586       1    1.2040e+02    6.6944e+02 ;     CA - C
- O
  2569   2585   2587       1    1.1660e+02    5.8576e+02 ;     CA - C
- N
  2570   2569   2571       1    1.0950e+02    4.1840e+02 ;     HA - CA
- CB
  2570   2569   2585       1    1.0950e+02    4.1840e+02 ;     HA - CA
- C
  2571   2569   2585       1    1.1110e+02    5.2718e+02 ;     CB - CA
- C
  2571   2574   2575       1    1.2000e+02    5.8576e+02 ;     CB - CG
- CD1
  2571   2574   2583       1    1.2000e+02    5.8576e+02 ;     CB - CG
- CD2
  2572   2571   2573       1    1.0950e+02    2.9288e+02 ;    HB2 - CB
- HB3
  2572   2571   2574       1    1.0950e+02    4.1840e+02 ;    HB2 - CB
- CG
  2573   2571   2574       1    1.0950e+02    4.1840e+02 ;    HB3 - CB
- CG
  2574   2575   2576       1    1.2000e+02    4.1840e+02 ;     CG - CD1
- HD1
  2574   2575   2577       1    1.2000e+02    5.2718e+02 ;     CG - CD1
- CE1
  2574   2583   2581       1    1.2000e+02    5.2718e+02 ;     CG - CD2
- CE2
  2574   2583   2584       1    1.2000e+02    4.1840e+02 ;     CG - CD2
- HD2
  2575   2574   2583       1    1.2000e+02    5.2718e+02 ;    CD1 - CG
- CD2
  2575   2577   2578       1    1.2000e+02    4.1840e+02 ;    CD1 - CE1
- HE1
  2575   2577   2579       1    1.2000e+02    5.2718e+02 ;    CD1 - CE1
- CZ
  2576   2575   2577       1    1.2000e+02    4.1840e+02 ;    HD1 - CD1
- CE1
  2577   2579   2580       1    1.2000e+02    4.1840e+02 ;    CE1 - CZ
- HZ
  2577   2579   2581       1    1.2000e+02    5.2718e+02 ;    CE1 - CZ
- CE2
  2578   2577   2579       1    1.2000e+02    4.1840e+02 ;    HE1 - CE1
- CZ
```

```
   2579   2581   2582       1    1.2000e+02    4.1840e+02 ;     CZ - CE2
 - HE2
   2579   2581   2583       1    1.2000e+02    5.2718e+02 ;     CZ - CE2
 - CD2
   2580   2579   2581       1    1.2000e+02    4.1840e+02 ;     HZ - CZ
 - CE2
   2581   2583   2584       1    1.2000e+02    4.1840e+02 ;    CE2 - CD2
 - HD2
   2582   2581   2583       1    1.2000e+02    4.1840e+02 ;    HE2 - CE2
 - CD2
   2585   2587   2588       1    1.2000e+02    4.1840e+02 ;      C - N
 - H
   2585   2587   2589       1    1.2190e+02    4.1840e+02 ;      C - N
 - CA
   2586   2585   2587       1    1.2290e+02    6.6944e+02 ;      O - C
 - N
   2587   2589   2590       1    1.0950e+02    4.1840e+02 ;      N - CA
 - HA2
   2587   2589   2591       1    1.0950e+02    4.1840e+02 ;      N - CA
 - HA3
   2587   2589   2592       1    1.1010e+02    5.2718e+02 ;      N - CA
 - C
   2588   2587   2589       1    1.1804e+02    4.1840e+02 ;      H - N
 - CA
   2589   2592   2593       1    1.2040e+02    6.6944e+02 ;     CA - C
 - O
   2589   2592   2594       1    1.1660e+02    5.8576e+02 ;     CA - C
 - N
   2590   2589   2591       1    1.0950e+02    2.9288e+02 ;    HA2 - CA
 - HA3
   2590   2589   2592       1    1.0950e+02    4.1840e+02 ;    HA2 - CA
 - C
   2591   2589   2592       1    1.0950e+02    4.1840e+02 ;    HA3 - CA
 - C
   2592   2594   2595       1    1.2000e+02    4.1840e+02 ;      C - N
 - H
   2592   2594   2596       1    1.2190e+02    4.1840e+02 ;      C - N
 - CA
   2593   2592   2594       1    1.2290e+02    6.6944e+02 ;      O - C
 - N
   2594   2596   2597       1    1.0950e+02    4.1840e+02 ;      N - CA
 - HA
   2594   2596   2598       1    1.0970e+02    6.6944e+02 ;      N - CA
 - CB
   2594   2596   2611       1    1.1010e+02    5.2718e+02 ;      N - CA
 - C
   2595   2594   2596       1    1.1804e+02    4.1840e+02 ;      H - N
 - CA
   2596   2598   2599       1    1.0950e+02    4.1840e+02 ;     CA - CB
 - HB2
   2596   2598   2600       1    1.0950e+02    4.1840e+02 ;     CA - CB
 - HB3
```

2596 2598 2601 1 1.0950e+02 3.3472e+02 ; CA - CB
- CG
2596 2611 2612 1 1.2040e+02 6.6944e+02 ; CA - C
- O
2596 2611 2613 1 1.1660e+02 5.8576e+02 ; CA - C
- N
2597 2596 2598 1 1.0950e+02 4.1840e+02 ; HA - CA
- CB
2597 2596 2611 1 1.0950e+02 4.1840e+02 ; HA - CA
- C
2598 2596 2611 1 1.1110e+02 5.2718e+02 ; CB - CA
- C
2598 2601 2602 1 1.0950e+02 4.1840e+02 ; CB - CG
- HG
2598 2601 2603 1 1.0950e+02 3.3472e+02 ; CB - CG
- CD1
2598 2601 2607 1 1.0950e+02 3.3472e+02 ; CB - CG
- CD2
2599 2598 2600 1 1.0950e+02 2.9288e+02 ; HB2 - CB
- HB3
2599 2598 2601 1 1.0950e+02 4.1840e+02 ; HB2 - CB
- CG
2600 2598 2601 1 1.0950e+02 4.1840e+02 ; HB3 - CB
- CG
2601 2603 2604 1 1.0950e+02 4.1840e+02 ; CG - CD1
- HD11
2601 2603 2605 1 1.0950e+02 4.1840e+02 ; CG - CD1
- HD12
2601 2603 2606 1 1.0950e+02 4.1840e+02 ; CG - CD1
- HD13
2601 2607 2608 1 1.0950e+02 4.1840e+02 ; CG - CD2
- HD21
2601 2607 2609 1 1.0950e+02 4.1840e+02 ; CG - CD2
- HD22
2601 2607 2610 1 1.0950e+02 4.1840e+02 ; CG - CD2
- HD23
2602 2601 2603 1 1.0950e+02 4.1840e+02 ; HG - CG
- CD1
2602 2601 2607 1 1.0950e+02 4.1840e+02 ; HG - CG
- CD2
2603 2601 2607 1 1.0950e+02 3.3472e+02 ; CD1 - CG
- CD2
2604 2603 2605 1 1.0950e+02 2.9288e+02 ; HD11 - CD1
- HD12
2604 2603 2606 1 1.0950e+02 2.9288e+02 ; HD11 - CD1
- HD13
2605 2603 2606 1 1.0950e+02 2.9288e+02 ; HD12 - CD1
- HD13
2608 2607 2609 1 1.0950e+02 2.9288e+02 ; HD21 - CD2
- HD22
2608 2607 2610 1 1.0950e+02 2.9288e+02 ; HD21 - CD2
- HD23

2609 2607 2610 1 1.0950e+02 2.9288e+02 ; HD22 - CD2
- HD23
2611 2613 2614 1 1.2000e+02 4.1840e+02 ; C - N
- H
2611 2613 2615 1 1.2190e+02 4.1840e+02 ; C - N
- CA
2612 2611 2613 1 1.2290e+02 6.6944e+02 ; O - C
- N
2613 2615 2616 1 1.0950e+02 4.1840e+02 ; N - CA
- HA
2613 2615 2617 1 1.0970e+02 6.6944e+02 ; N - CA
- CB
2613 2615 2622 1 1.1010e+02 5.2718e+02 ; N - CA
- C
2614 2613 2615 1 1.1804e+02 4.1840e+02 ; H - N
- CA
2615 2617 2618 1 1.0950e+02 4.1840e+02 ; CA - CB
- HB2
2615 2617 2619 1 1.0950e+02 4.1840e+02 ; CA - CB
- HB3
2615 2617 2620 1 1.0950e+02 4.1840e+02 ; CA - CB
- OG
2615 2622 2623 1 1.2040e+02 6.6944e+02 ; CA - C
- O
2615 2622 2624 1 1.1660e+02 5.8576e+02 ; CA - C
- N
2616 2615 2617 1 1.0950e+02 4.1840e+02 ; HA - CA
- CB
2616 2615 2622 1 1.0950e+02 4.1840e+02 ; HA - CA
- C
2617 2615 2622 1 1.1110e+02 5.2718e+02 ; CB - CA
- C
2617 2620 2621 1 1.0850e+02 4.6024e+02 ; CB - OG
- HG
2618 2617 2619 1 1.0950e+02 2.9288e+02 ; HB2 - CB
- HB3
2618 2617 2620 1 1.0950e+02 4.1840e+02 ; HB2 - CB
- OG
2619 2617 2620 1 1.0950e+02 4.1840e+02 ; HB3 - CB
- OG
2622 2624 2625 1 1.2000e+02 4.1840e+02 ; C - N
- H
2622 2624 2626 1 1.2190e+02 4.1840e+02 ; C - N
- CA
2623 2622 2624 1 1.2290e+02 6.6944e+02 ; O - C
- N
2624 2626 2627 1 1.0950e+02 4.1840e+02 ; N - CA
- HA
2624 2626 2628 1 1.0970e+02 6.6944e+02 ; N - CA
- CB
2624 2626 2644 1 1.1010e+02 5.2718e+02 ; N - CA
- C

```
  2625   2624   2626       1    1.1804e+02    4.1840e+02 ;      H - N
- CA
  2626   2628   2629       1    1.0950e+02    4.1840e+02 ;     CA - CB
- HB2
  2626   2628   2630       1    1.0950e+02    4.1840e+02 ;     CA - CB
- HB3
  2626   2628   2631       1    1.0950e+02    3.3472e+02 ;     CA - CB
- CG
  2626   2644   2645       1    1.2040e+02    6.6944e+02 ;     CA - C
- O
  2626   2644   2646       1    1.1660e+02    5.8576e+02 ;     CA - C
- N
  2627   2626   2628       1    1.0950e+02    4.1840e+02 ;     HA - CA
- CB
  2627   2626   2644       1    1.0950e+02    4.1840e+02 ;     HA - CA
- C
  2628   2626   2644       1    1.1110e+02    5.2718e+02 ;     CB - CA
- C
  2628   2631   2632       1    1.0950e+02    4.1840e+02 ;     CB - CG
- HG2
  2628   2631   2633       1    1.0950e+02    4.1840e+02 ;     CB - CG
- HG3
  2628   2631   2634       1    1.0950e+02    3.3472e+02 ;     CB - CG
- CD
  2629   2628   2630       1    1.0950e+02    2.9288e+02 ;    HB2 - CB
- HB3
  2629   2628   2631       1    1.0950e+02    4.1840e+02 ;    HB2 - CB
- CG
  2630   2628   2631       1    1.0950e+02    4.1840e+02 ;    HB3 - CB
- CG
  2631   2634   2635       1    1.0950e+02    4.1840e+02 ;     CG - CD
- HD2
  2631   2634   2636       1    1.0950e+02    4.1840e+02 ;     CG - CD
- HD3
  2631   2634   2637       1    1.0950e+02    3.3472e+02 ;     CG - CD
- CE
  2632   2631   2633       1    1.0950e+02    2.9288e+02 ;    HG2 - CG
- HG3
  2632   2631   2634       1    1.0950e+02    4.1840e+02 ;    HG2 - CG
- CD
  2633   2631   2634       1    1.0950e+02    4.1840e+02 ;    HG3 - CG
- CD
  2634   2637   2638       1    1.0950e+02    4.1840e+02 ;     CD - CE
- HE2
  2634   2637   2639       1    1.0950e+02    4.1840e+02 ;     CD - CE
- HE3
  2634   2637   2640       1    1.1120e+02    6.6944e+02 ;     CD - CE
- NZ
  2635   2634   2636       1    1.0950e+02    2.9288e+02 ;    HD2 - CD
- HD3
  2635   2634   2637       1    1.0950e+02    4.1840e+02 ;    HD2 - CD
- CE
```

```
  2636   2634   2637       1    1.0950e+02    4.1840e+02 ;    HD3 - CD
- CE
  2637   2640   2641       1    1.0950e+02    4.1840e+02 ;     CE - NZ
- HZ1
  2637   2640   2642       1    1.0950e+02    4.1840e+02 ;     CE - NZ
- HZ2
  2637   2640   2643       1    1.0950e+02    4.1840e+02 ;     CE - NZ
- HZ3
  2638   2637   2639       1    1.0950e+02    2.9288e+02 ;    HE2 - CE
- HE3
  2638   2637   2640       1    1.0950e+02    4.1840e+02 ;    HE2 - CE
- NZ
  2639   2637   2640       1    1.0950e+02    4.1840e+02 ;    HE3 - CE
- NZ
  2641   2640   2642       1    1.0950e+02    2.9288e+02 ;    HZ1 - NZ
- HZ2
  2641   2640   2643       1    1.0950e+02    2.9288e+02 ;    HZ1 - NZ
- HZ3
  2642   2640   2643       1    1.0950e+02    2.9288e+02 ;    HZ2 - NZ
- HZ3
  2644   2646   2647       1    1.2000e+02    4.1840e+02 ;      C - N
- H
  2644   2646   2648       1    1.2190e+02    4.1840e+02 ;      C - N
- CA
  2645   2644   2646       1    1.2290e+02    6.6944e+02 ;      O - C
- N
  2646   2648   2649       1    1.0950e+02    4.1840e+02 ;      N - CA
- HA
  2646   2648   2650       1    1.0970e+02    6.6944e+02 ;      N - CA
- CB
  2646   2648   2658       1    1.1010e+02    5.2718e+02 ;      N - CA
- C
  2647   2646   2648       1    1.1804e+02    4.1840e+02 ;      H - N
- CA
  2648   2650   2651       1    1.0950e+02    4.1840e+02 ;     CA - CB
- HB2
  2648   2650   2652       1    1.0950e+02    4.1840e+02 ;     CA - CB
- HB3
  2648   2650   2653       1    1.1110e+02    5.2718e+02 ;     CA - CB
- CG
  2648   2658   2659       1    1.2040e+02    6.6944e+02 ;     CA - C
- O
  2648   2658   2660       1    1.1660e+02    5.8576e+02 ;     CA - C
- N
  2649   2648   2650       1    1.0950e+02    4.1840e+02 ;     HA - CA
- CB
  2649   2648   2658       1    1.0950e+02    4.1840e+02 ;     HA - CA
- C
  2650   2648   2658       1    1.1110e+02    5.2718e+02 ;     CB - CA
- C
  2650   2653   2654       1    1.2040e+02    6.6944e+02 ;     CB - CG
- OD1
```

2650 2653 2655 1 1.1660e+02 5.8576e+02 ; CB - CG
- ND2
2651 2650 2652 1 1.0950e+02 2.9288e+02 ; HB2 - CB
- HB3
2651 2650 2653 1 1.0950e+02 4.1840e+02 ; HB2 - CB
- CG
2652 2650 2653 1 1.0950e+02 4.1840e+02 ; HB3 - CB
- CG
2653 2655 2656 1 1.2000e+02 4.1840e+02 ; CG - ND2
- HD21
2653 2655 2657 1 1.2000e+02 4.1840e+02 ; CG - ND2
- HD22
2654 2653 2655 1 1.2290e+02 6.6944e+02 ; OD1 - CG
- ND2
2656 2655 2657 1 1.2000e+02 2.9288e+02 ; HD21 - ND2
- HD22
2658 2660 2661 1 1.2000e+02 4.1840e+02 ; C - N
- H
2658 2660 2662 1 1.2190e+02 4.1840e+02 ; C - N
- CA
2659 2658 2660 1 1.2290e+02 6.6944e+02 ; O - C
- N
2660 2662 2663 1 1.0950e+02 4.1840e+02 ; N - CA
- HA
2660 2662 2664 1 1.0970e+02 6.6944e+02 ; N - CA
- CB
2660 2662 2677 1 1.1010e+02 5.2718e+02 ; N - CA
- C
2661 2660 2662 1 1.1804e+02 4.1840e+02 ; H - N
- CA
2662 2664 2665 1 1.0950e+02 4.1840e+02 ; CA - CB
- HB2
2662 2664 2666 1 1.0950e+02 4.1840e+02 ; CA - CB
- HB3
2662 2664 2667 1 1.0950e+02 3.3472e+02 ; CA - CB
- CG
2662 2677 2678 1 1.2040e+02 6.6944e+02 ; CA - C
- O
2662 2677 2679 1 1.1660e+02 5.8576e+02 ; CA - C
- N
2663 2662 2664 1 1.0950e+02 4.1840e+02 ; HA - CA
- CB
2663 2662 2677 1 1.0950e+02 4.1840e+02 ; HA - CA
- C
2664 2662 2677 1 1.1110e+02 5.2718e+02 ; CB - CA
- C
2664 2667 2668 1 1.0950e+02 4.1840e+02 ; CB - CG
- HG
2664 2667 2669 1 1.0950e+02 3.3472e+02 ; CB - CG
- CD1
2664 2667 2673 1 1.0950e+02 3.3472e+02 ; CB - CG
- CD2

| | | | | | | | |
|---|---|---|---|---|---|---|---|
| 2665 | 2664 | 2666 | 1 | 1.0950e+02 | 2.9288e+02 | ; | HB2 - CB - HB3 |
| 2665 | 2664 | 2667 | 1 | 1.0950e+02 | 4.1840e+02 | ; | HB2 - CB - CG |
| 2666 | 2664 | 2667 | 1 | 1.0950e+02 | 4.1840e+02 | ; | HB3 - CB - CG |
| 2667 | 2669 | 2670 | 1 | 1.0950e+02 | 4.1840e+02 | ; | CG - CD1 - HD11 |
| 2667 | 2669 | 2671 | 1 | 1.0950e+02 | 4.1840e+02 | ; | CG - CD1 - HD12 |
| 2667 | 2669 | 2672 | 1 | 1.0950e+02 | 4.1840e+02 | ; | CG - CD1 - HD13 |
| 2667 | 2673 | 2674 | 1 | 1.0950e+02 | 4.1840e+02 | ; | CG - CD2 - HD21 |
| 2667 | 2673 | 2675 | 1 | 1.0950e+02 | 4.1840e+02 | ; | CG - CD2 - HD22 |
| 2667 | 2673 | 2676 | 1 | 1.0950e+02 | 4.1840e+02 | ; | CG - CD2 - HD23 |
| 2668 | 2667 | 2669 | 1 | 1.0950e+02 | 4.1840e+02 | ; | HG - CG - CD1 |
| 2668 | 2667 | 2673 | 1 | 1.0950e+02 | 4.1840e+02 | ; | HG - CG - CD2 |
| 2669 | 2667 | 2673 | 1 | 1.0950e+02 | 3.3472e+02 | ; | CD1 - CG - CD2 |
| 2670 | 2669 | 2671 | 1 | 1.0950e+02 | 2.9288e+02 | ; | HD11 - CD1 - HD12 |
| 2670 | 2669 | 2672 | 1 | 1.0950e+02 | 2.9288e+02 | ; | HD11 - CD1 - HD13 |
| 2671 | 2669 | 2672 | 1 | 1.0950e+02 | 2.9288e+02 | ; | HD12 - CD1 - HD13 |
| 2674 | 2673 | 2675 | 1 | 1.0950e+02 | 2.9288e+02 | ; | HD21 - CD2 - HD22 |
| 2674 | 2673 | 2676 | 1 | 1.0950e+02 | 2.9288e+02 | ; | HD21 - CD2 - HD23 |
| 2675 | 2673 | 2676 | 1 | 1.0950e+02 | 2.9288e+02 | ; | HD22 - CD2 - HD23 |
| 2677 | 2679 | 2680 | 1 | 1.2000e+02 | 4.1840e+02 | ; | C - N - H |
| 2677 | 2679 | 2681 | 1 | 1.2190e+02 | 4.1840e+02 | ; | C - N - CA |
| 2678 | 2677 | 2679 | 1 | 1.2290e+02 | 6.6944e+02 | ; | O - C - N |
| 2679 | 2681 | 2682 | 1 | 1.0950e+02 | 4.1840e+02 | ; | N - CA - HA2 |
| 2679 | 2681 | 2683 | 1 | 1.0950e+02 | 4.1840e+02 | ; | N - CA - HA3 |
| 2679 | 2681 | 2684 | 1 | 1.1010e+02 | 5.2718e+02 | ; | N - CA - C |
| 2680 | 2679 | 2681 | 1 | 1.1804e+02 | 4.1840e+02 | ; | H - N - CA |
| 2681 | 2684 | 2685 | 1 | 1.2040e+02 | 6.6944e+02 | ; | CA - C - O |

2681 2684 2686 1 1.1660e+02 5.8576e+02 ; CA - C
- N
2682 2681 2683 1 1.0950e+02 2.9288e+02 ; HA2 - CA
- HA3
2682 2681 2684 1 1.0950e+02 4.1840e+02 ; HA2 - CA
- C
2683 2681 2684 1 1.0950e+02 4.1840e+02 ; HA3 - CA
- C
2684 2686 2687 1 1.2000e+02 4.1840e+02 ; C - N
- H
2684 2686 2688 1 1.2190e+02 4.1840e+02 ; C - N
- CA
2685 2684 2686 1 1.2290e+02 6.6944e+02 ; O - C
- N
2686 2688 2689 1 1.0950e+02 4.1840e+02 ; N - CA
- HA
2686 2688 2690 1 1.0970e+02 6.6944e+02 ; N - CA
- CB
2686 2688 2698 1 1.1010e+02 5.2718e+02 ; N - CA
- C
2687 2686 2688 1 1.1804e+02 4.1840e+02 ; H - N
- CA
2688 2690 2691 1 1.0950e+02 4.1840e+02 ; CA - CB
- HB
2688 2690 2692 1 1.0950e+02 3.3472e+02 ; CA - CB
- CG2
2688 2690 2696 1 1.0950e+02 4.1840e+02 ; CA - CB
- OG1
2688 2698 2699 1 1.2040e+02 6.6944e+02 ; CA - C
- O
2688 2698 2700 1 1.1660e+02 5.8576e+02 ; CA - C
- N
2689 2688 2690 1 1.0950e+02 4.1840e+02 ; HA - CA
- CB
2689 2688 2698 1 1.0950e+02 4.1840e+02 ; HA - CA
- C
2690 2688 2698 1 1.1110e+02 5.2718e+02 ; CB - CA
- C
2690 2692 2693 1 1.0950e+02 4.1840e+02 ; CB - CG2
- HG21
2690 2692 2694 1 1.0950e+02 4.1840e+02 ; CB - CG2
- HG22
2690 2692 2695 1 1.0950e+02 4.1840e+02 ; CB - CG2
- HG23
2690 2696 2697 1 1.0850e+02 4.6024e+02 ; CB - OG1
- HG1
2691 2690 2692 1 1.0950e+02 4.1840e+02 ; HB - CB
- CG2
2691 2690 2696 1 1.0950e+02 4.1840e+02 ; HB - CB
- OG1
2692 2690 2696 1 1.0950e+02 4.1840e+02 ; CG2 - CB
- OG1

```
  2693   2692   2694      1    1.0950e+02    2.9288e+02 ;   HG21 - CG2
- HG22
  2693   2692   2695      1    1.0950e+02    2.9288e+02 ;   HG21 - CG2
- HG23
  2694   2692   2695      1    1.0950e+02    2.9288e+02 ;   HG22 - CG2
- HG23
  2698   2700   2701      1    1.2000e+02    4.1840e+02 ;      C - N
- H
  2698   2700   2702      1    1.2190e+02    4.1840e+02 ;      C - N
- CA
  2699   2698   2700      1    1.2290e+02    6.6944e+02 ;      O - C
- N
  2700   2702   2703      1    1.0950e+02    4.1840e+02 ;      N - CA
- HA2
  2700   2702   2704      1    1.0950e+02    4.1840e+02 ;      N - CA
- HA3
  2700   2702   2705      1    1.1010e+02    5.2718e+02 ;      N - CA
- C
  2701   2700   2702      1    1.1804e+02    4.1840e+02 ;      H - N
- CA
  2702   2705   2706      1    1.2040e+02    6.6944e+02 ;     CA - C
- O
  2702   2705   2707      1    1.1660e+02    5.8576e+02 ;     CA - C
- N
  2703   2702   2704      1    1.0950e+02    2.9288e+02 ;    HA2 - CA
- HA3
  2703   2702   2705      1    1.0950e+02    4.1840e+02 ;    HA2 - CA
- C
  2704   2702   2705      1    1.0950e+02    4.1840e+02 ;    HA3 - CA
- C
  2705   2707   2708      1    1.2000e+02    4.1840e+02 ;      C - N
- H
  2705   2707   2709      1    1.2190e+02    4.1840e+02 ;      C - N
- CA
  2706   2705   2707      1    1.2290e+02    6.6944e+02 ;      O - C
- N
  2707   2709   2710      1    1.0950e+02    4.1840e+02 ;      N - CA
- HA
  2707   2709   2711      1    1.0970e+02    6.6944e+02 ;      N - CA
- CB
  2707   2709   2720      1    1.1010e+02    5.2718e+02 ;      N - CA
- C
  2708   2707   2709      1    1.1804e+02    4.1840e+02 ;      H - N
- CA
  2709   2711   2712      1    1.0950e+02    4.1840e+02 ;     CA - CB
- HB2
  2709   2711   2713      1    1.0950e+02    4.1840e+02 ;     CA - CB
- HB3
  2709   2711   2714      1    1.0950e+02    3.3472e+02 ;     CA - CB
- CG
  2709   2720   2721      1    1.2040e+02    6.6944e+02 ;     CA - C
- O
```

2709 2720 2722 1 1.1660e+02 5.8576e+02 ; CA - C
- N
2710 2709 2711 1 1.0950e+02 4.1840e+02 ; HA - CA
- CB
2710 2709 2720 1 1.0950e+02 4.1840e+02 ; HA - CA
- C
2711 2709 2720 1 1.1110e+02 5.2718e+02 ; CB - CA
- C
2711 2714 2715 1 1.0950e+02 4.1840e+02 ; CB - CG
- HG2
2711 2714 2716 1 1.0950e+02 4.1840e+02 ; CB - CG
- HG3
2711 2714 2717 1 1.1110e+02 5.2718e+02 ; CB - CG
- CD
2712 2711 2713 1 1.0950e+02 2.9288e+02 ; HB2 - CB
- HB3
2712 2711 2714 1 1.0950e+02 4.1840e+02 ; HB2 - CB
- CG
2713 2711 2714 1 1.0950e+02 4.1840e+02 ; HB3 - CB
- CG
2714 2717 2718 1 1.1700e+02 5.8576e+02 ; CG - CD
- OE1
2714 2717 2719 1 1.1700e+02 5.8576e+02 ; CG - CD
- OE2
2715 2714 2716 1 1.0950e+02 2.9288e+02 ; HG2 - CG
- HG3
2715 2714 2717 1 1.0950e+02 4.1840e+02 ; HG2 - CG
- CD
2716 2714 2717 1 1.0950e+02 4.1840e+02 ; HG3 - CG
- CD
2718 2717 2719 1 1.2600e+02 6.6944e+02 ; OE1 - CD
- OE2
2720 2722 2723 1 1.2000e+02 4.1840e+02 ; C - N
- H
2720 2722 2724 1 1.2190e+02 4.1840e+02 ; C - N
- CA
2721 2720 2722 1 1.2290e+02 6.6944e+02 ; O - C
- N
2722 2724 2725 1 1.0950e+02 4.1840e+02 ; N - CA
- HA
2722 2724 2726 1 1.0970e+02 6.6944e+02 ; N - CA
- CB
2722 2724 2740 1 1.1010e+02 5.2718e+02 ; N - CA
- C
2723 2722 2724 1 1.1804e+02 4.1840e+02 ; H - N
- CA
2724 2726 2727 1 1.0950e+02 4.1840e+02 ; CA - CB
- HB2
2724 2726 2728 1 1.0950e+02 4.1840e+02 ; CA - CB
- HB3
2724 2726 2729 1 1.1400e+02 5.2718e+02 ; CA - CB
- CG

```
  2724   2740   2741       1    1.2040e+02    6.6944e+02 ;     CA - C
- O
  2724   2740   2742       1    1.1660e+02    5.8576e+02 ;     CA - C
- N
  2725   2724   2726       1    1.0950e+02    4.1840e+02 ;     HA - CA
- CB
  2725   2724   2740       1    1.0950e+02    4.1840e+02 ;     HA - CA
- C
  2726   2724   2740       1    1.1110e+02    5.2718e+02 ;     CB - CA
- C
  2726   2729   2730       1    1.2000e+02    5.8576e+02 ;     CB - CG
- CD1
  2726   2729   2738       1    1.2000e+02    5.8576e+02 ;     CB - CG
- CD2
  2727   2726   2728       1    1.0950e+02    2.9288e+02 ;    HB2 - CB
- HB3
  2727   2726   2729       1    1.0950e+02    4.1840e+02 ;    HB2 - CB
- CG
  2728   2726   2729       1    1.0950e+02    4.1840e+02 ;    HB3 - CB
- CG
  2729   2730   2731       1    1.2000e+02    4.1840e+02 ;     CG - CD1
- HD1
  2729   2730   2732       1    1.2000e+02    5.2718e+02 ;     CG - CD1
- CE1
  2729   2738   2736       1    1.2000e+02    5.2718e+02 ;     CG - CD2
- CE2
  2729   2738   2739       1    1.2000e+02    4.1840e+02 ;     CG - CD2
- HD2
  2730   2729   2738       1    1.2000e+02    5.2718e+02 ;    CD1 - CG
- CD2
  2730   2732   2733       1    1.2000e+02    4.1840e+02 ;    CD1 - CE1
- HE1
  2730   2732   2734       1    1.2000e+02    5.2718e+02 ;    CD1 - CE1
- CZ
  2731   2730   2732       1    1.2000e+02    4.1840e+02 ;    HD1 - CD1
- CE1
  2732   2734   2735       1    1.2000e+02    4.1840e+02 ;    CE1 - CZ
- HZ
  2732   2734   2736       1    1.2000e+02    5.2718e+02 ;    CE1 - CZ
- CE2
  2733   2732   2734       1    1.2000e+02    4.1840e+02 ;    HE1 - CE1
- CZ
  2734   2736   2737       1    1.2000e+02    4.1840e+02 ;     CZ - CE2
- HE2
  2734   2736   2738       1    1.2000e+02    5.2718e+02 ;     CZ - CE2
- CD2
  2735   2734   2736       1    1.2000e+02    4.1840e+02 ;     HZ - CZ
- CE2
  2736   2738   2739       1    1.2000e+02    4.1840e+02 ;    CE2 - CD2
- HD2
  2737   2736   2738       1    1.2000e+02    4.1840e+02 ;    HE2 - CE2
- CD2
```

```
  2740   2742   2743       1    1.2000e+02    4.1840e+02 ;      C - N
- H
  2740   2742   2744       1    1.2190e+02    4.1840e+02 ;      C - N
- CA
  2741   2740   2742       1    1.2290e+02    6.6944e+02 ;      O - C
- N
  2742   2744   2745       1    1.0950e+02    4.1840e+02 ;      N - CA
- HA
  2742   2744   2746       1    1.0970e+02    6.6944e+02 ;      N - CA
- CB
  2742   2744   2754       1    1.1010e+02    5.2718e+02 ;      N - CA
- C
  2743   2742   2744       1    1.1804e+02    4.1840e+02 ;      H - N
- CA
  2744   2746   2747       1    1.0950e+02    4.1840e+02 ;     CA - CB
- HB
  2744   2746   2748       1    1.0950e+02    3.3472e+02 ;     CA - CB
- CG2
  2744   2746   2752       1    1.0950e+02    4.1840e+02 ;     CA - CB
- OG1
  2744   2754   2755       1    1.2040e+02    6.6944e+02 ;     CA - C
- O
  2744   2754   2756       1    1.1660e+02    5.8576e+02 ;     CA - C
- N
  2745   2744   2746       1    1.0950e+02    4.1840e+02 ;     HA - CA
- CB
  2745   2744   2754       1    1.0950e+02    4.1840e+02 ;     HA - CA
- C
  2746   2744   2754       1    1.1110e+02    5.2718e+02 ;     CB - CA
- C
  2746   2748   2749       1    1.0950e+02    4.1840e+02 ;     CB - CG2
- HG21
  2746   2748   2750       1    1.0950e+02    4.1840e+02 ;     CB - CG2
- HG22
  2746   2748   2751       1    1.0950e+02    4.1840e+02 ;     CB - CG2
- HG23
  2746   2752   2753       1    1.0850e+02    4.6024e+02 ;     CB - OG1
- HG1
  2747   2746   2748       1    1.0950e+02    4.1840e+02 ;     HB - CB
- CG2
  2747   2746   2752       1    1.0950e+02    4.1840e+02 ;     HB - CB
- OG1
  2748   2746   2752       1    1.0950e+02    4.1840e+02 ;    CG2 - CB
- OG1
  2749   2748   2750       1    1.0950e+02    2.9288e+02 ;   HG21 - CG2
- HG22
  2749   2748   2751       1    1.0950e+02    2.9288e+02 ;   HG21 - CG2
- HG23
  2750   2748   2751       1    1.0950e+02    2.9288e+02 ;   HG22 - CG2
- HG23
  2754   2756   2757       1    1.2000e+02    4.1840e+02 ;      C - N
- H
```

```
  2754   2756   2758       1    1.2190e+02    4.1840e+02 ;      C - N
- CA
  2755   2754   2756       1    1.2290e+02    6.6944e+02 ;      O - C
- N
  2756   2758   2759       1    1.0950e+02    4.1840e+02 ;      N - CA
- HA
  2756   2758   2760       1    1.0970e+02    6.6944e+02 ;      N - CA
- CB
  2756   2758   2768       1    1.1010e+02    5.2718e+02 ;      N - CA
- C
  2757   2756   2758       1    1.1804e+02    4.1840e+02 ;      H - N
- CA
  2758   2760   2761       1    1.0950e+02    4.1840e+02 ;     CA - CB
- HB
  2758   2760   2762       1    1.0950e+02    3.3472e+02 ;     CA - CB
- CG2
  2758   2760   2766       1    1.0950e+02    4.1840e+02 ;     CA - CB
- OG1
  2758   2768   2769       1    1.2040e+02    6.6944e+02 ;     CA - C
- O
  2758   2768   2770       1    1.1660e+02    5.8576e+02 ;     CA - C
- N
  2759   2758   2760       1    1.0950e+02    4.1840e+02 ;     HA - CA
- CB
  2759   2758   2768       1    1.0950e+02    4.1840e+02 ;     HA - CA
- C
  2760   2758   2768       1    1.1110e+02    5.2718e+02 ;     CB - CA
- C
  2760   2762   2763       1    1.0950e+02    4.1840e+02 ;     CB - CG2
- HG21
  2760   2762   2764       1    1.0950e+02    4.1840e+02 ;     CB - CG2
- HG22
  2760   2762   2765       1    1.0950e+02    4.1840e+02 ;     CB - CG2
- HG23
  2760   2766   2767       1    1.0850e+02    4.6024e+02 ;     CB - OG1
- HG1
  2761   2760   2762       1    1.0950e+02    4.1840e+02 ;     HB - CB
- CG2
  2761   2760   2766       1    1.0950e+02    4.1840e+02 ;     HB - CB
- OG1
  2762   2760   2766       1    1.0950e+02    4.1840e+02 ;    CG2 - CB
- OG1
  2763   2762   2764       1    1.0950e+02    2.9288e+02 ;   HG21 - CG2
- HG22
  2763   2762   2765       1    1.0950e+02    2.9288e+02 ;   HG21 - CG2
- HG23
  2764   2762   2765       1    1.0950e+02    2.9288e+02 ;   HG22 - CG2
- HG23
  2768   2770   2771       1    1.2000e+02    4.1840e+02 ;      C - N
- H
  2768   2770   2772       1    1.2190e+02    4.1840e+02 ;      C - N
- CA
```

2769 2768 2770 1 1.2290e+02 6.6944e+02 ; O - C
- N
2770 2772 2773 1 1.0950e+02 4.1840e+02 ; N - CA
- HA
2770 2772 2774 1 1.0970e+02 6.6944e+02 ; N - CA
- CB
2770 2772 2782 1 1.1010e+02 5.2718e+02 ; N - CA
- C
2771 2770 2772 1 1.1804e+02 4.1840e+02 ; H - N
- CA
2772 2774 2775 1 1.0950e+02 4.1840e+02 ; CA - CB
- HB
2772 2774 2776 1 1.0950e+02 3.3472e+02 ; CA - CB
- CG2
2772 2774 2780 1 1.0950e+02 4.1840e+02 ; CA - CB
- OG1
2772 2782 2783 1 1.2040e+02 6.6944e+02 ; CA - C
- O
2772 2782 2784 1 1.1660e+02 5.8576e+02 ; CA - C
- N
2773 2772 2774 1 1.0950e+02 4.1840e+02 ; HA - CA
- CB
2773 2772 2782 1 1.0950e+02 4.1840e+02 ; HA - CA
- C
2774 2772 2782 1 1.1110e+02 5.2718e+02 ; CB - CA
- C
2774 2776 2777 1 1.0950e+02 4.1840e+02 ; CB - CG2
- HG21
2774 2776 2778 1 1.0950e+02 4.1840e+02 ; CB - CG2
- HG22
2774 2776 2779 1 1.0950e+02 4.1840e+02 ; CB - CG2
- HG23
2774 2780 2781 1 1.0850e+02 4.6024e+02 ; CB - OG1
- HG1
2775 2774 2776 1 1.0950e+02 4.1840e+02 ; HB - CB
- CG2
2775 2774 2780 1 1.0950e+02 4.1840e+02 ; HB - CB
- OG1
2776 2774 2780 1 1.0950e+02 4.1840e+02 ; CG2 - CB
- OG1
2777 2776 2778 1 1.0950e+02 2.9288e+02 ; HG21 - CG2
- HG22
2777 2776 2779 1 1.0950e+02 2.9288e+02 ; HG21 - CG2
- HG23
2778 2776 2779 1 1.0950e+02 2.9288e+02 ; HG22 - CG2
- HG23
2782 2784 2785 1 1.2000e+02 4.1840e+02 ; C - N
- H
2782 2784 2786 1 1.2190e+02 4.1840e+02 ; C - N
- CA
2783 2782 2784 1 1.2290e+02 6.6944e+02 ; O - C
- N

2784 2786 2787 1 1.0950e+02 4.1840e+02 ; N - CA
- HA
2784 2786 2788 1 1.0970e+02 6.6944e+02 ; N - CA
- CB
2784 2786 2804 1 1.1010e+02 5.2718e+02 ; N - CA
- C
2785 2784 2786 1 1.1804e+02 4.1840e+02 ; H - N
- CA
2786 2788 2789 1 1.0950e+02 4.1840e+02 ; CA - CB
- HB2
2786 2788 2790 1 1.0950e+02 4.1840e+02 ; CA - CB
- HB3
2786 2788 2791 1 1.0950e+02 3.3472e+02 ; CA - CB
- CG
2786 2804 2805 1 1.2040e+02 6.6944e+02 ; CA - C
- O
2786 2804 2806 1 1.1660e+02 5.8576e+02 ; CA - C
- N
2787 2786 2788 1 1.0950e+02 4.1840e+02 ; HA - CA
- CB
2787 2786 2804 1 1.0950e+02 4.1840e+02 ; HA - CA
- C
2788 2786 2804 1 1.1110e+02 5.2718e+02 ; CB - CA
- C
2788 2791 2792 1 1.0950e+02 4.1840e+02 ; CB - CG
- HG2
2788 2791 2793 1 1.0950e+02 4.1840e+02 ; CB - CG
- HG3
2788 2791 2794 1 1.0950e+02 3.3472e+02 ; CB - CG
- CD
2789 2788 2790 1 1.0950e+02 2.9288e+02 ; HB2 - CB
- HB3
2789 2788 2791 1 1.0950e+02 4.1840e+02 ; HB2 - CB
- CG
2790 2788 2791 1 1.0950e+02 4.1840e+02 ; HB3 - CB
- CG
2791 2794 2795 1 1.0950e+02 4.1840e+02 ; CG - CD
- HD2
2791 2794 2796 1 1.0950e+02 4.1840e+02 ; CG - CD
- HD3
2791 2794 2797 1 1.0950e+02 3.3472e+02 ; CG - CD
- CE
2792 2791 2793 1 1.0950e+02 2.9288e+02 ; HG2 - CG
- HG3
2792 2791 2794 1 1.0950e+02 4.1840e+02 ; HG2 - CG
- CD
2793 2791 2794 1 1.0950e+02 4.1840e+02 ; HG3 - CG
- CD
2794 2797 2798 1 1.0950e+02 4.1840e+02 ; CD - CE
- HE2
2794 2797 2799 1 1.0950e+02 4.1840e+02 ; CD - CE
- HE3

| | | | | | | | |
|---|---|---|---|---|---|---|---|
| 2794 | 2797 | 2800 | 1 | 1.1120e+02 | 6.6944e+02 | ; | CD - CE - NZ |
| 2795 | 2794 | 2796 | 1 | 1.0950e+02 | 2.9288e+02 | ; | HD2 - CD - HD3 |
| 2795 | 2794 | 2797 | 1 | 1.0950e+02 | 4.1840e+02 | ; | HD2 - CD - CE |
| 2796 | 2794 | 2797 | 1 | 1.0950e+02 | 4.1840e+02 | ; | HD3 - CD - CE |
| 2797 | 2800 | 2801 | 1 | 1.0950e+02 | 4.1840e+02 | ; | CE - NZ - HZ1 |
| 2797 | 2800 | 2802 | 1 | 1.0950e+02 | 4.1840e+02 | ; | CE - NZ - HZ2 |
| 2797 | 2800 | 2803 | 1 | 1.0950e+02 | 4.1840e+02 | ; | CE - NZ - HZ3 |
| 2798 | 2797 | 2799 | 1 | 1.0950e+02 | 2.9288e+02 | ; | HE2 - CE - HE3 |
| 2798 | 2797 | 2800 | 1 | 1.0950e+02 | 4.1840e+02 | ; | HE2 - CE - NZ |
| 2799 | 2797 | 2800 | 1 | 1.0950e+02 | 4.1840e+02 | ; | HE3 - CE - NZ |
| 2801 | 2800 | 2802 | 1 | 1.0950e+02 | 2.9288e+02 | ; | HZ1 - NZ - HZ2 |
| 2801 | 2800 | 2803 | 1 | 1.0950e+02 | 2.9288e+02 | ; | HZ1 - NZ - HZ3 |
| 2802 | 2800 | 2803 | 1 | 1.0950e+02 | 2.9288e+02 | ; | HZ2 - NZ - HZ3 |
| 2804 | 2806 | 2807 | 1 | 1.2000e+02 | 4.1840e+02 | ; | C - N - H |
| 2804 | 2806 | 2808 | 1 | 1.2190e+02 | 4.1840e+02 | ; | C - N - CA |
| 2805 | 2804 | 2806 | 1 | 1.2290e+02 | 6.6944e+02 | ; | O - C - N |
| 2806 | 2808 | 2809 | 1 | 1.0950e+02 | 4.1840e+02 | ; | N - CA - HA |
| 2806 | 2808 | 2810 | 1 | 1.0970e+02 | 6.6944e+02 | ; | N - CA - CB |
| 2806 | 2808 | 2814 | 1 | 1.1010e+02 | 5.2718e+02 | ; | N - CA - C |
| 2807 | 2806 | 2808 | 1 | 1.1804e+02 | 4.1840e+02 | ; | H - N - CA |
| 2808 | 2810 | 2811 | 1 | 1.0950e+02 | 4.1840e+02 | ; | CA - CB - HB1 |
| 2808 | 2810 | 2812 | 1 | 1.0950e+02 | 4.1840e+02 | ; | CA - CB - HB2 |
| 2808 | 2810 | 2813 | 1 | 1.0950e+02 | 4.1840e+02 | ; | CA - CB - HB3 |
| 2808 | 2814 | 2815 | 1 | 1.2040e+02 | 6.6944e+02 | ; | CA - C - O |
| 2808 | 2814 | 2816 | 1 | 1.1660e+02 | 5.8576e+02 | ; | CA - C - N |
| 2809 | 2808 | 2810 | 1 | 1.0950e+02 | 4.1840e+02 | ; | HA - CA - CB |

| | | | | | | | |
|---|---|---|---|---|---|---|---|
| 2809 | 2808 | 2814 | 1 | 1.0950e+02 | 4.1840e+02 | ; | HA - CA - C |
| 2810 | 2808 | 2814 | 1 | 1.1110e+02 | 5.2718e+02 | ; | CB - CA - C |
| 2811 | 2810 | 2812 | 1 | 1.0950e+02 | 2.9288e+02 | ; | HB1 - CB - HB2 |
| 2811 | 2810 | 2813 | 1 | 1.0950e+02 | 2.9288e+02 | ; | HB1 - CB - HB3 |
| 2812 | 2810 | 2813 | 1 | 1.0950e+02 | 2.9288e+02 | ; | HB2 - CB - HB3 |
| 2814 | 2816 | 2817 | 1 | 1.2000e+02 | 4.1840e+02 | ; | C - N - H |
| 2814 | 2816 | 2818 | 1 | 1.2190e+02 | 4.1840e+02 | ; | C - N - CA |
| 2815 | 2814 | 2816 | 1 | 1.2290e+02 | 6.6944e+02 | ; | O - C - N |
| 2816 | 2818 | 2819 | 1 | 1.0950e+02 | 4.1840e+02 | ; | N - CA - HA2 |
| 2816 | 2818 | 2820 | 1 | 1.0950e+02 | 4.1840e+02 | ; | N - CA - HA3 |
| 2816 | 2818 | 2821 | 1 | 1.1010e+02 | 5.2718e+02 | ; | N - CA - C |
| 2817 | 2816 | 2818 | 1 | 1.1804e+02 | 4.1840e+02 | ; | H - N - CA |
| 2818 | 2821 | 2822 | 1 | 1.2040e+02 | 6.6944e+02 | ; | CA - C - O |
| 2818 | 2821 | 2823 | 1 | 1.1660e+02 | 5.8576e+02 | ; | CA - C - N |
| 2819 | 2818 | 2820 | 1 | 1.0950e+02 | 2.9288e+02 | ; | HA2 - CA - HA3 |
| 2819 | 2818 | 2821 | 1 | 1.0950e+02 | 4.1840e+02 | ; | HA2 - CA - C |
| 2820 | 2818 | 2821 | 1 | 1.0950e+02 | 4.1840e+02 | ; | HA3 - CA - C |
| 2821 | 2823 | 2824 | 1 | 1.2000e+02 | 4.1840e+02 | ; | C - N - H |
| 2821 | 2823 | 2825 | 1 | 1.2190e+02 | 4.1840e+02 | ; | C - N - CA |
| 2822 | 2821 | 2823 | 1 | 1.2290e+02 | 6.6944e+02 | ; | O - C - N |
| 2823 | 2825 | 2826 | 1 | 1.0950e+02 | 4.1840e+02 | ; | N - CA - HA |
| 2823 | 2825 | 2827 | 1 | 1.0970e+02 | 6.6944e+02 | ; | N - CA - CB |
| 2823 | 2825 | 2835 | 1 | 1.1010e+02 | 5.2718e+02 | ; | N - CA - C |
| 2824 | 2823 | 2825 | 1 | 1.1804e+02 | 4.1840e+02 | ; | H - N - CA |
| 2825 | 2827 | 2828 | 1 | 1.0950e+02 | 4.1840e+02 | ; | CA - CB - HB |
| 2825 | 2827 | 2829 | 1 | 1.0950e+02 | 3.3472e+02 | ; | CA - CB - CG2 |

```
  2825   2827   2833      1    1.0950e+02    4.1840e+02 ;     CA - CB
- OG1
  2825   2835   2836      1    1.2040e+02    6.6944e+02 ;     CA - C
- O
  2825   2835   2837      1    1.1660e+02    5.8576e+02 ;     CA - C
- N
  2826   2825   2827      1    1.0950e+02    4.1840e+02 ;     HA - CA
- CB
  2826   2825   2835      1    1.0950e+02    4.1840e+02 ;     HA - CA
- C
  2827   2825   2835      1    1.1110e+02    5.2718e+02 ;     CB - CA
- C
  2827   2829   2830      1    1.0950e+02    4.1840e+02 ;     CB - CG2
- HG21
  2827   2829   2831      1    1.0950e+02    4.1840e+02 ;     CB - CG2
- HG22
  2827   2829   2832      1    1.0950e+02    4.1840e+02 ;     CB - CG2
- HG23
  2827   2833   2834      1    1.0850e+02    4.6024e+02 ;     CB - OG1
- HG1
  2828   2827   2829      1    1.0950e+02    4.1840e+02 ;     HB - CB
- CG2
  2828   2827   2833      1    1.0950e+02    4.1840e+02 ;     HB - CB
- OG1
  2829   2827   2833      1    1.0950e+02    4.1840e+02 ;    CG2 - CB
- OG1
  2830   2829   2831      1    1.0950e+02    2.9288e+02 ;   HG21 - CG2
- HG22
  2830   2829   2832      1    1.0950e+02    2.9288e+02 ;   HG21 - CG2
- HG23
  2831   2829   2832      1    1.0950e+02    2.9288e+02 ;   HG22 - CG2
- HG23
  2835   2837   2838      1    1.2190e+02    4.1840e+02 ;      C - N
- CD
  2835   2837   2847      1    1.2190e+02    4.1840e+02 ;      C - N
- CA
  2836   2835   2837      1    1.2290e+02    6.6944e+02 ;      O - C
- N
  2837   2838   2839      1    1.0950e+02    4.1840e+02 ;      N - CD
- HD2
  2837   2838   2840      1    1.0950e+02    4.1840e+02 ;      N - CD
- HD3
  2837   2838   2841      1    1.0970e+02    6.6944e+02 ;      N - CD
- CG
  2837   2847   2844      1    1.0970e+02    6.6944e+02 ;      N - CA
- CB
  2837   2847   2848      1    1.0950e+02    4.1840e+02 ;      N - CA
- HA
  2837   2847   2849      1    1.1010e+02    5.2718e+02 ;      N - CA
- C
  2838   2837   2847      1    1.1800e+02    4.1840e+02 ;     CD - N
- CA
```

| | | | | | | | |
|---|---|---|---|---|---|---|---|
| 2838 | 2841 | 2842 | 1 | 1.0950e+02 | 4.1840e+02 | ; | CD - CG - HG2 |
| 2838 | 2841 | 2843 | 1 | 1.0950e+02 | 4.1840e+02 | ; | CD - CG - HG3 |
| 2838 | 2841 | 2844 | 1 | 1.0950e+02 | 3.3472e+02 | ; | CD - CG - CB |
| 2839 | 2838 | 2840 | 1 | 1.0950e+02 | 2.9288e+02 | ; | HD2 - CD - HD3 |
| 2839 | 2838 | 2841 | 1 | 1.0950e+02 | 4.1840e+02 | ; | HD2 - CD - CG |
| 2840 | 2838 | 2841 | 1 | 1.0950e+02 | 4.1840e+02 | ; | HD3 - CD - CG |
| 2841 | 2844 | 2845 | 1 | 1.0950e+02 | 4.1840e+02 | ; | CG - CB - HB2 |
| 2841 | 2844 | 2846 | 1 | 1.0950e+02 | 4.1840e+02 | ; | CG - CB - HB3 |
| 2841 | 2844 | 2847 | 1 | 1.0950e+02 | 3.3472e+02 | ; | CG - CB - CA |
| 2842 | 2841 | 2843 | 1 | 1.0950e+02 | 2.9288e+02 | ; | HG2 - CG - HG3 |
| 2842 | 2841 | 2844 | 1 | 1.0950e+02 | 4.1840e+02 | ; | HG2 - CG - CB |
| 2843 | 2841 | 2844 | 1 | 1.0950e+02 | 4.1840e+02 | ; | HG3 - CG - CB |
| 2844 | 2847 | 2848 | 1 | 1.0950e+02 | 4.1840e+02 | ; | CB - CA - HA |
| 2844 | 2847 | 2849 | 1 | 1.1110e+02 | 5.2718e+02 | ; | CB - CA - C |
| 2845 | 2844 | 2846 | 1 | 1.0950e+02 | 2.9288e+02 | ; | HB2 - CB - HB3 |
| 2845 | 2844 | 2847 | 1 | 1.0950e+02 | 4.1840e+02 | ; | HB2 - CB - CA |
| 2846 | 2844 | 2847 | 1 | 1.0950e+02 | 4.1840e+02 | ; | HB3 - CB - CA |
| 2847 | 2849 | 2850 | 1 | 1.2040e+02 | 6.6944e+02 | ; | CA - C - O |
| 2847 | 2849 | 2851 | 1 | 1.1660e+02 | 5.8576e+02 | ; | CA - C - N |
| 2848 | 2847 | 2849 | 1 | 1.0950e+02 | 4.1840e+02 | ; | HA - CA - C |
| 2849 | 2851 | 2852 | 1 | 1.2000e+02 | 4.1840e+02 | ; | C - N - H |
| 2849 | 2851 | 2853 | 1 | 1.2190e+02 | 4.1840e+02 | ; | C - N - CA |
| 2850 | 2849 | 2851 | 1 | 1.2290e+02 | 6.6944e+02 | ; | O - C - N |
| 2851 | 2853 | 2854 | 1 | 1.0950e+02 | 4.1840e+02 | ; | N - CA - HA |
| 2851 | 2853 | 2855 | 1 | 1.0970e+02 | 6.6944e+02 | ; | N - CA - CB |
| 2851 | 2853 | 2870 | 1 | 1.1010e+02 | 5.2718e+02 | ; | N - CA - C |

2852 2851 2853 1 1.1804e+02 4.1840e+02 ; H - N
- CA
2853 2855 2856 1 1.0950e+02 4.1840e+02 ; CA - CB
- HB2
2853 2855 2857 1 1.0950e+02 4.1840e+02 ; CA - CB
- HB3
2853 2855 2858 1 1.1400e+02 5.2718e+02 ; CA - CB
- CG
2853 2870 2871 1 1.2040e+02 6.6944e+02 ; CA - C
- O
2853 2870 2872 1 1.1660e+02 5.8576e+02 ; CA - C
- N
2854 2853 2855 1 1.0950e+02 4.1840e+02 ; HA - CA
- CB
2854 2853 2870 1 1.0950e+02 4.1840e+02 ; HA - CA
- C
2855 2853 2870 1 1.1110e+02 5.2718e+02 ; CB - CA
- C
2855 2858 2859 1 1.2000e+02 5.8576e+02 ; CB - CG
- CD1
2855 2858 2868 1 1.2000e+02 5.8576e+02 ; CB - CG
- CD2
2856 2855 2857 1 1.0950e+02 2.9288e+02 ; HB2 - CB
- HB3
2856 2855 2858 1 1.0950e+02 4.1840e+02 ; HB2 - CB
- CG
2857 2855 2858 1 1.0950e+02 4.1840e+02 ; HB3 - CB
- CG
2858 2859 2860 1 1.2000e+02 4.1840e+02 ; CG - CD1
- HD1
2858 2859 2861 1 1.2000e+02 5.2718e+02 ; CG - CD1
- CE1
2858 2868 2866 1 1.2000e+02 5.2718e+02 ; CG - CD2
- CE2
2858 2868 2869 1 1.2000e+02 4.1840e+02 ; CG - CD2
- HD2
2859 2858 2868 1 1.2000e+02 5.2718e+02 ; CD1 - CG
- CD2
2859 2861 2862 1 1.2000e+02 4.1840e+02 ; CD1 - CE1
- HE1
2859 2861 2863 1 1.2000e+02 5.2718e+02 ; CD1 - CE1
- CZ
2860 2859 2861 1 1.2000e+02 4.1840e+02 ; HD1 - CD1
- CE1
2861 2863 2864 1 1.2000e+02 5.8576e+02 ; CE1 - CZ
- OH
2861 2863 2866 1 1.2000e+02 5.2718e+02 ; CE1 - CZ
- CE2
2862 2861 2863 1 1.2000e+02 4.1840e+02 ; HE1 - CE1
- CZ
2863 2864 2865 1 1.1300e+02 4.1840e+02 ; CZ - OH
- HH

```
  2863   2866   2867      1    1.2000e+02    4.1840e+02 ;     CZ - CE2
- HE2
  2863   2866   2868      1    1.2000e+02    5.2718e+02 ;     CZ - CE2
- CD2
  2864   2863   2866      1    1.2000e+02    5.8576e+02 ;     OH - CZ
- CE2
  2866   2868   2869      1    1.2000e+02    4.1840e+02 ;    CE2 - CD2
- HD2
  2867   2866   2868      1    1.2000e+02    4.1840e+02 ;    HE2 - CE2
- CD2
  2870   2872   2873      1    1.2000e+02    4.1840e+02 ;      C - N
- H
  2870   2872   2874      1    1.2190e+02    4.1840e+02 ;      C - N
- CA
  2871   2870   2872      1    1.2290e+02    6.6944e+02 ;      O - C
- N
  2872   2874   2875      1    1.0950e+02    4.1840e+02 ;      N - CA
- HA
  2872   2874   2876      1    1.0970e+02    6.6944e+02 ;      N - CA
- CB
  2872   2874   2891      1    1.1010e+02    5.2718e+02 ;      N - CA
- C
  2873   2872   2874      1    1.1804e+02    4.1840e+02 ;      H - N
- CA
  2874   2876   2877      1    1.0950e+02    4.1840e+02 ;     CA - CB
- HB2
  2874   2876   2878      1    1.0950e+02    4.1840e+02 ;     CA - CB
- HB3
  2874   2876   2879      1    1.1400e+02    5.2718e+02 ;     CA - CB
- CG
  2874   2891   2892      1    1.2040e+02    6.6944e+02 ;     CA - C
- O
  2874   2891   2893      1    1.1660e+02    5.8576e+02 ;     CA - C
- N
  2875   2874   2876      1    1.0950e+02    4.1840e+02 ;     HA - CA
- CB
  2875   2874   2891      1    1.0950e+02    4.1840e+02 ;     HA - CA
- C
  2876   2874   2891      1    1.1110e+02    5.2718e+02 ;     CB - CA
- C
  2876   2879   2880      1    1.2000e+02    5.8576e+02 ;     CB - CG
- CD1
  2876   2879   2889      1    1.2000e+02    5.8576e+02 ;     CB - CG
- CD2
  2877   2876   2878      1    1.0950e+02    2.9288e+02 ;    HB2 - CB
- HB3
  2877   2876   2879      1    1.0950e+02    4.1840e+02 ;    HB2 - CB
- CG
  2878   2876   2879      1    1.0950e+02    4.1840e+02 ;    HB3 - CB
- CG
  2879   2880   2881      1    1.2000e+02    4.1840e+02 ;     CG - CD1
- HD1
```

2879 2880 2882 1 1.2000e+02 5.2718e+02 ; CG - CD1
- CE1
2879 2889 2887 1 1.2000e+02 5.2718e+02 ; CG - CD2
- CE2
2879 2889 2890 1 1.2000e+02 4.1840e+02 ; CG - CD2
- HD2
2880 2879 2889 1 1.2000e+02 5.2718e+02 ; CD1 - CG
- CD2
2880 2882 2883 1 1.2000e+02 4.1840e+02 ; CD1 - CE1
- HE1
2880 2882 2884 1 1.2000e+02 5.2718e+02 ; CD1 - CE1
- CZ
2881 2880 2882 1 1.2000e+02 4.1840e+02 ; HD1 - CD1
- CE1
2882 2884 2885 1 1.2000e+02 5.8576e+02 ; CE1 - CZ
- OH
2882 2884 2887 1 1.2000e+02 5.2718e+02 ; CE1 - CZ
- CE2
2883 2882 2884 1 1.2000e+02 4.1840e+02 ; HE1 - CE1
- CZ
2884 2885 2886 1 1.1300e+02 4.1840e+02 ; CZ - OH
- HH
2884 2887 2888 1 1.2000e+02 4.1840e+02 ; CZ - CE2
- HE2
2884 2887 2889 1 1.2000e+02 5.2718e+02 ; CZ - CE2
- CD2
2885 2884 2887 1 1.2000e+02 5.8576e+02 ; OH - CZ
- CE2
2887 2889 2890 1 1.2000e+02 4.1840e+02 ; CE2 - CD2
- HD2
2888 2887 2889 1 1.2000e+02 4.1840e+02 ; HE2 - CE2
- CD2
2891 2893 2894 1 1.2000e+02 4.1840e+02 ; C - N
- H
2891 2893 2895 1 1.2190e+02 4.1840e+02 ; C - N
- CA
2892 2891 2893 1 1.2290e+02 6.6944e+02 ; O - C
- N
2893 2895 2896 1 1.0950e+02 4.1840e+02 ; N - CA
- HA
2893 2895 2897 1 1.0970e+02 6.6944e+02 ; N - CA
- CB
2893 2895 2907 1 1.1010e+02 5.2718e+02 ; N - CA
- C
2894 2893 2895 1 1.1804e+02 4.1840e+02 ; H - N
- CA
2895 2897 2898 1 1.0950e+02 4.1840e+02 ; CA - CB
- HB
2895 2897 2899 1 1.0950e+02 3.3472e+02 ; CA - CB
- CG1
2895 2897 2903 1 1.0950e+02 3.3472e+02 ; CA - CB
- CG2

```
  2895   2907   2908       1    1.2040e+02    6.6944e+02 ;     CA - C
- O
  2895   2907   2909       1    1.1660e+02    5.8576e+02 ;     CA - C
- N
  2896   2895   2897       1    1.0950e+02    4.1840e+02 ;     HA - CA
- CB
  2896   2895   2907       1    1.0950e+02    4.1840e+02 ;     HA - CA
- C
  2897   2895   2907       1    1.1110e+02    5.2718e+02 ;     CB - CA
- C
  2897   2899   2900       1    1.0950e+02    4.1840e+02 ;     CB - CG1
- HG11
  2897   2899   2901       1    1.0950e+02    4.1840e+02 ;     CB - CG1
- HG12
  2897   2899   2902       1    1.0950e+02    4.1840e+02 ;     CB - CG1
- HG13
  2897   2903   2904       1    1.0950e+02    4.1840e+02 ;     CB - CG2
- HG21
  2897   2903   2905       1    1.0950e+02    4.1840e+02 ;     CB - CG2
- HG22
  2897   2903   2906       1    1.0950e+02    4.1840e+02 ;     CB - CG2
- HG23
  2898   2897   2899       1    1.0950e+02    4.1840e+02 ;     HB - CB
- CG1
  2898   2897   2903       1    1.0950e+02    4.1840e+02 ;     HB - CB
- CG2
  2899   2897   2903       1    1.0950e+02    3.3472e+02 ;    CG1 - CB
- CG2
  2900   2899   2901       1    1.0950e+02    2.9288e+02 ;   HG11 - CG1
- HG12
  2900   2899   2902       1    1.0950e+02    2.9288e+02 ;   HG11 - CG1
- HG13
  2901   2899   2902       1    1.0950e+02    2.9288e+02 ;   HG12 - CG1
- HG13
  2904   2903   2905       1    1.0950e+02    2.9288e+02 ;   HG21 - CG2
- HG22
  2904   2903   2906       1    1.0950e+02    2.9288e+02 ;   HG21 - CG2
- HG23
  2905   2903   2906       1    1.0950e+02    2.9288e+02 ;   HG22 - CG2
- HG23
  2907   2909   2910       1    1.2000e+02    4.1840e+02 ;      C - N
- H
  2907   2909   2911       1    1.2190e+02    4.1840e+02 ;      C - N
- CA
  2908   2907   2909       1    1.2290e+02    6.6944e+02 ;      O - C
- N
  2909   2911   2912       1    1.0950e+02    4.1840e+02 ;      N - CA
- HA
  2909   2911   2913       1    1.0970e+02    6.6944e+02 ;      N - CA
- CB
  2909   2911   2917       1    1.1010e+02    5.2718e+02 ;      N - CA
- C
```

```
  2910   2909   2911      1    1.1804e+02    4.1840e+02 ;      H - N
- CA
  2911   2913   2914      1    1.0950e+02    4.1840e+02 ;     CA - CB
- HB1
  2911   2913   2915      1    1.0950e+02    4.1840e+02 ;     CA - CB
- HB2
  2911   2913   2916      1    1.0950e+02    4.1840e+02 ;     CA - CB
- HB3
  2911   2917   2918      1    1.2040e+02    6.6944e+02 ;     CA - C
- O
  2911   2917   2919      1    1.1660e+02    5.8576e+02 ;     CA - C
- N
  2912   2911   2913      1    1.0950e+02    4.1840e+02 ;     HA - CA
- CB
  2912   2911   2917      1    1.0950e+02    4.1840e+02 ;     HA - CA
- C
  2913   2911   2917      1    1.1110e+02    5.2718e+02 ;     CB - CA
- C
  2914   2913   2915      1    1.0950e+02    2.9288e+02 ;    HB1 - CB
- HB2
  2914   2913   2916      1    1.0950e+02    2.9288e+02 ;    HB1 - CB
- HB3
  2915   2913   2916      1    1.0950e+02    2.9288e+02 ;    HB2 - CB
- HB3
  2917   2919   2920      1    1.2190e+02    4.1840e+02 ;      C - N
- CD
  2917   2919   2929      1    1.2190e+02    4.1840e+02 ;      C - N
- CA
  2918   2917   2919      1    1.2290e+02    6.6944e+02 ;      O - C
- N
  2919   2920   2921      1    1.0950e+02    4.1840e+02 ;      N - CD
- HD2
  2919   2920   2922      1    1.0950e+02    4.1840e+02 ;      N - CD
- HD3
  2919   2920   2923      1    1.0970e+02    6.6944e+02 ;      N - CD
- CG
  2919   2929   2926      1    1.0970e+02    6.6944e+02 ;      N - CA
- CB
  2919   2929   2930      1    1.0950e+02    4.1840e+02 ;      N - CA
- HA
  2919   2929   2931      1    1.1010e+02    5.2718e+02 ;      N - CA
- C
  2920   2919   2929      1    1.1800e+02    4.1840e+02 ;     CD - N
- CA
  2920   2923   2924      1    1.0950e+02    4.1840e+02 ;     CD - CG
- HG2
  2920   2923   2925      1    1.0950e+02    4.1840e+02 ;     CD - CG
- HG3
  2920   2923   2926      1    1.0950e+02    3.3472e+02 ;     CD - CG
- CB
  2921   2920   2922      1    1.0950e+02    2.9288e+02 ;    HD2 - CD
- HD3
```

```
  2921   2920   2923       1    1.0950e+02    4.1840e+02 ;    HD2 - CD
- CG
  2922   2920   2923       1    1.0950e+02    4.1840e+02 ;    HD3 - CD
- CG
  2923   2926   2927       1    1.0950e+02    4.1840e+02 ;     CG - CB
- HB2
  2923   2926   2928       1    1.0950e+02    4.1840e+02 ;     CG - CB
- HB3
  2923   2926   2929       1    1.0950e+02    3.3472e+02 ;     CG - CB
- CA
  2924   2923   2925       1    1.0950e+02    2.9288e+02 ;    HG2 - CG
- HG3
  2924   2923   2926       1    1.0950e+02    4.1840e+02 ;    HG2 - CG
- CB
  2925   2923   2926       1    1.0950e+02    4.1840e+02 ;    HG3 - CG
- CB
  2926   2929   2930       1    1.0950e+02    4.1840e+02 ;     CB - CA
- HA
  2926   2929   2931       1    1.1110e+02    5.2718e+02 ;     CB - CA
- C
  2927   2926   2928       1    1.0950e+02    2.9288e+02 ;    HB2 - CB
- HB3
  2927   2926   2929       1    1.0950e+02    4.1840e+02 ;    HB2 - CB
- CA
  2928   2926   2929       1    1.0950e+02    4.1840e+02 ;    HB3 - CB
- CA
  2929   2931   2932       1    1.2040e+02    6.6944e+02 ;     CA - C
- O
  2929   2931   2933       1    1.1660e+02    5.8576e+02 ;     CA - C
- N
  2930   2929   2931       1    1.0950e+02    4.1840e+02 ;     HA - CA
- C
  2931   2933   2934       1    1.2000e+02    4.1840e+02 ;      C - N
- H
  2931   2933   2935       1    1.2190e+02    4.1840e+02 ;      C - N
- CA
  2932   2931   2933       1    1.2290e+02    6.6944e+02 ;      O - C
- N
  2933   2935   2936       1    1.0950e+02    4.1840e+02 ;      N - CA
- HA
  2933   2935   2937       1    1.0970e+02    6.6944e+02 ;      N - CA
- CB
  2933   2935   2948       1    1.1010e+02    5.2718e+02 ;      N - CA
- C
  2934   2933   2935       1    1.1804e+02    4.1840e+02 ;      H - N
- CA
  2935   2937   2938       1    1.0950e+02    4.1840e+02 ;     CA - CB
- HB2
  2935   2937   2939       1    1.0950e+02    4.1840e+02 ;     CA - CB
- HB3
  2935   2937   2940       1    1.0950e+02    3.3472e+02 ;     CA - CB
- CG
```

```
   2935   2948   2949      1    1.2040e+02    6.6944e+02 ;     CA - C
 - O
   2935   2948   2950      1    1.1660e+02    5.8576e+02 ;     CA - C
 - N
   2936   2935   2937      1    1.0950e+02    4.1840e+02 ;     HA - CA
 - CB
   2936   2935   2948      1    1.0950e+02    4.1840e+02 ;     HA - CA
 - C
   2937   2935   2948      1    1.1110e+02    5.2718e+02 ;     CB - CA
 - C
   2937   2940   2941      1    1.0950e+02    4.1840e+02 ;     CB - CG
 - HG2
   2937   2940   2942      1    1.0950e+02    4.1840e+02 ;     CB - CG
 - HG3
   2937   2940   2943      1    1.1110e+02    5.2718e+02 ;     CB - CG
 - CD
   2938   2937   2939      1    1.0950e+02    2.9288e+02 ;    HB2 - CB
 - HB3
   2938   2937   2940      1    1.0950e+02    4.1840e+02 ;    HB2 - CB
 - CG
   2939   2937   2940      1    1.0950e+02    4.1840e+02 ;    HB3 - CB
 - CG
   2940   2943   2944      1    1.2040e+02    6.6944e+02 ;     CG - CD
 - OE1
   2940   2943   2945      1    1.1660e+02    5.8576e+02 ;     CG - CD
 - NE2
   2941   2940   2942      1    1.0950e+02    2.9288e+02 ;    HG2 - CG
 - HG3
   2941   2940   2943      1    1.0950e+02    4.1840e+02 ;    HG2 - CG
 - CD
   2942   2940   2943      1    1.0950e+02    4.1840e+02 ;    HG3 - CG
 - CD
   2943   2945   2946      1    1.2000e+02    4.1840e+02 ;     CD - NE2
 - HE21
   2943   2945   2947      1    1.2000e+02    4.1840e+02 ;     CD - NE2
 - HE22
   2944   2943   2945      1    1.2290e+02    6.6944e+02 ;    OE1 - CD
 - NE2
   2946   2945   2947      1    1.2000e+02    2.9288e+02 ;   HE21 - NE2
 - HE22
   2948   2950   2951      1    1.2000e+02    4.1840e+02 ;      C - N
 - H
   2948   2950   2952      1    1.2190e+02    4.1840e+02 ;      C - N
 - CA
   2949   2948   2950      1    1.2290e+02    6.6944e+02 ;      O - C
 - N
   2950   2952   2953      1    1.0950e+02    4.1840e+02 ;      N - CA
 - HA
   2950   2952   2954      1    1.0970e+02    6.6944e+02 ;      N - CA
 - CB
   2950   2952   2964      1    1.1010e+02    5.2718e+02 ;      N - CA
 - C
```

```
  2951   2950   2952      1    1.1804e+02    4.1840e+02 ;      H - N
- CA
  2952   2954   2955      1    1.0950e+02    4.1840e+02 ;     CA - CB
- HB
  2952   2954   2956      1    1.0950e+02    3.3472e+02 ;     CA - CB
- CG1
  2952   2954   2960      1    1.0950e+02    3.3472e+02 ;     CA - CB
- CG2
  2952   2964   2965      1    1.2040e+02    6.6944e+02 ;     CA - C
- O
  2952   2964   2966      1    1.1660e+02    5.8576e+02 ;     CA - C
- N
  2953   2952   2954      1    1.0950e+02    4.1840e+02 ;     HA - CA
- CB
  2953   2952   2964      1    1.0950e+02    4.1840e+02 ;     HA - CA
- C
  2954   2952   2964      1    1.1110e+02    5.2718e+02 ;     CB - CA
- C
  2954   2956   2957      1    1.0950e+02    4.1840e+02 ;     CB - CG1
- HG11
  2954   2956   2958      1    1.0950e+02    4.1840e+02 ;     CB - CG1
- HG12
  2954   2956   2959      1    1.0950e+02    4.1840e+02 ;     CB - CG1
- HG13
  2954   2960   2961      1    1.0950e+02    4.1840e+02 ;     CB - CG2
- HG21
  2954   2960   2962      1    1.0950e+02    4.1840e+02 ;     CB - CG2
- HG22
  2954   2960   2963      1    1.0950e+02    4.1840e+02 ;     CB - CG2
- HG23
  2955   2954   2956      1    1.0950e+02    4.1840e+02 ;     HB - CB
- CG1
  2955   2954   2960      1    1.0950e+02    4.1840e+02 ;     HB - CB
- CG2
  2956   2954   2960      1    1.0950e+02    3.3472e+02 ;    CG1 - CB
- CG2
  2957   2956   2958      1    1.0950e+02    2.9288e+02 ;   HG11 - CG1
- HG12
  2957   2956   2959      1    1.0950e+02    2.9288e+02 ;   HG11 - CG1
- HG13
  2958   2956   2959      1    1.0950e+02    2.9288e+02 ;   HG12 - CG1
- HG13
  2961   2960   2962      1    1.0950e+02    2.9288e+02 ;   HG21 - CG2
- HG22
  2961   2960   2963      1    1.0950e+02    2.9288e+02 ;   HG21 - CG2
- HG23
  2962   2960   2963      1    1.0950e+02    2.9288e+02 ;   HG22 - CG2
- HG23
  2964   2966   2967      1    1.2000e+02    4.1840e+02 ;      C - N
- H
  2964   2966   2968      1    1.2190e+02    4.1840e+02 ;      C - N
- CA
```

```
 2965  2964  2966    1   1.2290e+02   6.6944e+02 ;      O - C
- N
 2966  2968  2969    1   1.0950e+02   4.1840e+02 ;      N - CA
- HA
 2966  2968  2970    1   1.0970e+02   6.6944e+02 ;      N - CA
- CB
 2966  2968  2983    1   1.1010e+02   5.2718e+02 ;      N - CA
- C
 2967  2966  2968    1   1.1804e+02   4.1840e+02 ;      H - N
- CA
 2968  2970  2971    1   1.0950e+02   4.1840e+02 ;     CA - CB
- HB2
 2968  2970  2972    1   1.0950e+02   4.1840e+02 ;     CA - CB
- HB3
 2968  2970  2973    1   1.0950e+02   3.3472e+02 ;     CA - CB
- CG
 2968  2983  2984    1   1.2040e+02   6.6944e+02 ;     CA - C
- O
 2968  2983  2985    1   1.1660e+02   5.8576e+02 ;     CA - C
- N
 2969  2968  2970    1   1.0950e+02   4.1840e+02 ;     HA - CA
- CB
 2969  2968  2983    1   1.0950e+02   4.1840e+02 ;     HA - CA
- C
 2970  2968  2983    1   1.1110e+02   5.2718e+02 ;     CB - CA
- C
 2970  2973  2974    1   1.0950e+02   4.1840e+02 ;     CB - CG
- HG
 2970  2973  2975    1   1.0950e+02   3.3472e+02 ;     CB - CG
- CD1
 2970  2973  2979    1   1.0950e+02   3.3472e+02 ;     CB - CG
- CD2
 2971  2970  2972    1   1.0950e+02   2.9288e+02 ;    HB2 - CB
- HB3
 2971  2970  2973    1   1.0950e+02   4.1840e+02 ;    HB2 - CB
- CG
 2972  2970  2973    1   1.0950e+02   4.1840e+02 ;    HB3 - CB
- CG
 2973  2975  2976    1   1.0950e+02   4.1840e+02 ;     CG - CD1
- HD11
 2973  2975  2977    1   1.0950e+02   4.1840e+02 ;     CG - CD1
- HD12
 2973  2975  2978    1   1.0950e+02   4.1840e+02 ;     CG - CD1
- HD13
 2973  2979  2980    1   1.0950e+02   4.1840e+02 ;     CG - CD2
- HD21
 2973  2979  2981    1   1.0950e+02   4.1840e+02 ;     CG - CD2
- HD22
 2973  2979  2982    1   1.0950e+02   4.1840e+02 ;     CG - CD2
- HD23
 2974  2973  2975    1   1.0950e+02   4.1840e+02 ;     HG - CG
- CD1
```

```
  2974   2973   2979      1    1.0950e+02    4.1840e+02 ;     HG - CG
- CD2
  2975   2973   2979      1    1.0950e+02    3.3472e+02 ;    CD1 - CG
- CD2
  2976   2975   2977      1    1.0950e+02    2.9288e+02 ;   HD11 - CD1
- HD12
  2976   2975   2978      1    1.0950e+02    2.9288e+02 ;   HD11 - CD1
- HD13
  2977   2975   2978      1    1.0950e+02    2.9288e+02 ;   HD12 - CD1
- HD13
  2980   2979   2981      1    1.0950e+02    2.9288e+02 ;   HD21 - CD2
- HD22
  2980   2979   2982      1    1.0950e+02    2.9288e+02 ;   HD21 - CD2
- HD23
  2981   2979   2982      1    1.0950e+02    2.9288e+02 ;   HD22 - CD2
- HD23
  2983   2985   2986      1    1.2000e+02    4.1840e+02 ;      C - N
- H
  2983   2985   2987      1    1.2190e+02    4.1840e+02 ;      C - N
- CA
  2984   2983   2985      1    1.2290e+02    6.6944e+02 ;      O - C
- N
  2985   2987   2988      1    1.0950e+02    4.1840e+02 ;      N - CA
- HA
  2985   2987   2989      1    1.0970e+02    6.6944e+02 ;      N - CA
- CB
  2985   2987   2995      1    1.1010e+02    5.2718e+02 ;      N - CA
- C
  2986   2985   2987      1    1.1804e+02    4.1840e+02 ;      H - N
- CA
  2987   2989   2990      1    1.0950e+02    4.1840e+02 ;     CA - CB
- HB2
  2987   2989   2991      1    1.0950e+02    4.1840e+02 ;     CA - CB
- HB3
  2987   2989   2992      1    1.1110e+02    5.2718e+02 ;     CA - CB
- CG
  2987   2995   2996      1    1.2040e+02    6.6944e+02 ;     CA - C
- O
  2987   2995   2997      1    1.1660e+02    5.8576e+02 ;     CA - C
- N
  2988   2987   2989      1    1.0950e+02    4.1840e+02 ;     HA - CA
- CB
  2988   2987   2995      1    1.0950e+02    4.1840e+02 ;     HA - CA
- C
  2989   2987   2995      1    1.1110e+02    5.2718e+02 ;     CB - CA
- C
  2989   2992   2993      1    1.1700e+02    5.8576e+02 ;     CB - CG
- OD1
  2989   2992   2994      1    1.1700e+02    5.8576e+02 ;     CB - CG
- OD2
  2990   2989   2991      1    1.0950e+02    2.9288e+02 ;    HB2 - CB
- HB3
```

```
  2990   2989   2992       1    1.0950e+02    4.1840e+02 ;    HB2 - CB
- CG
  2991   2989   2992       1    1.0950e+02    4.1840e+02 ;    HB3 - CB
- CG
  2993   2992   2994       1    1.2600e+02    6.6944e+02 ;    OD1 - CG
- OD2
  2995   2997   2998       1    1.2000e+02    4.1840e+02 ;      C - N
- H
  2995   2997   2999       1    1.2190e+02    4.1840e+02 ;      C - N
- CA
  2996   2995   2997       1    1.2290e+02    6.6944e+02 ;      O - C
- N
  2997   2999   3000       1    1.0950e+02    4.1840e+02 ;      N - CA
- HA2
  2997   2999   3001       1    1.0950e+02    4.1840e+02 ;      N - CA
- HA3
  2997   2999   3002       1    1.1010e+02    5.2718e+02 ;      N - CA
- C
  2998   2997   2999       1    1.1804e+02    4.1840e+02 ;      H - N
- CA
  2999   3002   3003       1    1.2040e+02    6.6944e+02 ;     CA - C
- O
  2999   3002   3004       1    1.1660e+02    5.8576e+02 ;     CA - C
- N
  3000   2999   3001       1    1.0950e+02    2.9288e+02 ;    HA2 - CA
- HA3
  3000   2999   3002       1    1.0950e+02    4.1840e+02 ;    HA2 - CA
- C
  3001   2999   3002       1    1.0950e+02    4.1840e+02 ;    HA3 - CA
- C
  3002   3004   3005       1    1.2000e+02    4.1840e+02 ;      C - N
- H
  3002   3004   3006       1    1.2190e+02    4.1840e+02 ;      C - N
- CA
  3003   3002   3004       1    1.2290e+02    6.6944e+02 ;      O - C
- N
  3004   3006   3007       1    1.0950e+02    4.1840e+02 ;      N - CA
- HA
  3004   3006   3008       1    1.0970e+02    6.6944e+02 ;      N - CA
- CB
  3004   3006   3024       1    1.1010e+02    5.2718e+02 ;      N - CA
- C
  3005   3004   3006       1    1.1804e+02    4.1840e+02 ;      H - N
- CA
  3006   3008   3009       1    1.0950e+02    4.1840e+02 ;     CA - CB
- HB2
  3006   3008   3010       1    1.0950e+02    4.1840e+02 ;     CA - CB
- HB3
  3006   3008   3011       1    1.0950e+02    3.3472e+02 ;     CA - CB
- CG
  3006   3024   3025       1    1.2040e+02    6.6944e+02 ;     CA - C
- O
```

3006 3024 3026 1 1.1660e+02 5.8576e+02 ; CA - C
- N
3007 3006 3008 1 1.0950e+02 4.1840e+02 ; HA - CA
- CB
3007 3006 3024 1 1.0950e+02 4.1840e+02 ; HA - CA
- C
3008 3006 3024 1 1.1110e+02 5.2718e+02 ; CB - CA
- C
3008 3011 3012 1 1.0950e+02 4.1840e+02 ; CB - CG
- HG2
3008 3011 3013 1 1.0950e+02 4.1840e+02 ; CB - CG
- HG3
3008 3011 3014 1 1.0950e+02 3.3472e+02 ; CB - CG
- CD
3009 3008 3010 1 1.0950e+02 2.9288e+02 ; HB2 - CB
- HB3
3009 3008 3011 1 1.0950e+02 4.1840e+02 ; HB2 - CB
- CG
3010 3008 3011 1 1.0950e+02 4.1840e+02 ; HB3 - CB
- CG
3011 3014 3015 1 1.0950e+02 4.1840e+02 ; CG - CD
- HD2
3011 3014 3016 1 1.0950e+02 4.1840e+02 ; CG - CD
- HD3
3011 3014 3017 1 1.0950e+02 3.3472e+02 ; CG - CD
- CE
3012 3011 3013 1 1.0950e+02 2.9288e+02 ; HG2 - CG
- HG3
3012 3011 3014 1 1.0950e+02 4.1840e+02 ; HG2 - CG
- CD
3013 3011 3014 1 1.0950e+02 4.1840e+02 ; HG3 - CG
- CD
3014 3017 3018 1 1.0950e+02 4.1840e+02 ; CD - CE
- HE2
3014 3017 3019 1 1.0950e+02 4.1840e+02 ; CD - CE
- HE3
3014 3017 3020 1 1.1120e+02 6.6944e+02 ; CD - CE
- NZ
3015 3014 3016 1 1.0950e+02 2.9288e+02 ; HD2 - CD
- HD3
3015 3014 3017 1 1.0950e+02 4.1840e+02 ; HD2 - CD
- CE
3016 3014 3017 1 1.0950e+02 4.1840e+02 ; HD3 - CD
- CE
3017 3020 3021 1 1.0950e+02 4.1840e+02 ; CE - NZ
- HZ1
3017 3020 3022 1 1.0950e+02 4.1840e+02 ; CE - NZ
- HZ2
3017 3020 3023 1 1.0950e+02 4.1840e+02 ; CE - NZ
- HZ3
3018 3017 3019 1 1.0950e+02 2.9288e+02 ; HE2 - CE
- HE3

```
  3018   3017   3020      1    1.0950e+02    4.1840e+02 ;    HE2 - CE
- NZ
  3019   3017   3020      1    1.0950e+02    4.1840e+02 ;    HE3 - CE
- NZ
  3021   3020   3022      1    1.0950e+02    2.9288e+02 ;    HZ1 - NZ
- HZ2
  3021   3020   3023      1    1.0950e+02    2.9288e+02 ;    HZ1 - NZ
- HZ3
  3022   3020   3023      1    1.0950e+02    2.9288e+02 ;    HZ2 - NZ
- HZ3
  3024   3026   3027      1    1.2000e+02    4.1840e+02 ;      C - N
- H
  3024   3026   3028      1    1.2190e+02    4.1840e+02 ;      C - N
- CA
  3025   3024   3026      1    1.2290e+02    6.6944e+02 ;      O - C
- N
  3026   3028   3029      1    1.0950e+02    4.1840e+02 ;      N - CA
- HA
  3026   3028   3030      1    1.0970e+02    6.6944e+02 ;      N - CA
- CB
  3026   3028   3045      1    1.1010e+02    5.2718e+02 ;      N - CA
- C
  3027   3026   3028      1    1.1804e+02    4.1840e+02 ;      H - N
- CA
  3028   3030   3031      1    1.0950e+02    4.1840e+02 ;     CA - CB
- HB2
  3028   3030   3032      1    1.0950e+02    4.1840e+02 ;     CA - CB
- HB3
  3028   3030   3033      1    1.1400e+02    5.2718e+02 ;     CA - CB
- CG
  3028   3045   3046      1    1.2040e+02    6.6944e+02 ;     CA - C
- O
  3028   3045   3047      1    1.1660e+02    5.8576e+02 ;     CA - C
- N
  3029   3028   3030      1    1.0950e+02    4.1840e+02 ;     HA - CA
- CB
  3029   3028   3045      1    1.0950e+02    4.1840e+02 ;     HA - CA
- C
  3030   3028   3045      1    1.1110e+02    5.2718e+02 ;     CB - CA
- C
  3030   3033   3034      1    1.2000e+02    5.8576e+02 ;     CB - CG
- CD1
  3030   3033   3043      1    1.2000e+02    5.8576e+02 ;     CB - CG
- CD2
  3031   3030   3032      1    1.0950e+02    2.9288e+02 ;    HB2 - CB
- HB3
  3031   3030   3033      1    1.0950e+02    4.1840e+02 ;    HB2 - CB
- CG
  3032   3030   3033      1    1.0950e+02    4.1840e+02 ;    HB3 - CB
- CG
  3033   3034   3035      1    1.2000e+02    4.1840e+02 ;     CG - CD1
- HD1
```

```
  3033   3034   3036       1    1.2000e+02    5.2718e+02 ;     CG - CD1
- CE1
  3033   3043   3041       1    1.2000e+02    5.2718e+02 ;     CG - CD2
- CE2
  3033   3043   3044       1    1.2000e+02    4.1840e+02 ;     CG - CD2
- HD2
  3034   3033   3043       1    1.2000e+02    5.2718e+02 ;    CD1 - CG
- CD2
  3034   3036   3037       1    1.2000e+02    4.1840e+02 ;    CD1 - CE1
- HE1
  3034   3036   3038       1    1.2000e+02    5.2718e+02 ;    CD1 - CE1
- CZ
  3035   3034   3036       1    1.2000e+02    4.1840e+02 ;    HD1 - CD1
- CE1
  3036   3038   3039       1    1.2000e+02    5.8576e+02 ;    CE1 - CZ
- OH
  3036   3038   3041       1    1.2000e+02    5.2718e+02 ;    CE1 - CZ
- CE2
  3037   3036   3038       1    1.2000e+02    4.1840e+02 ;    HE1 - CE1
- CZ
  3038   3039   3040       1    1.1300e+02    4.1840e+02 ;     CZ - OH
- HH
  3038   3041   3042       1    1.2000e+02    4.1840e+02 ;     CZ - CE2
- HE2
  3038   3041   3043       1    1.2000e+02    5.2718e+02 ;     CZ - CE2
- CD2
  3039   3038   3041       1    1.2000e+02    5.8576e+02 ;     OH - CZ
- CE2
  3041   3043   3044       1    1.2000e+02    4.1840e+02 ;    CE2 - CD2
- HD2
  3042   3041   3043       1    1.2000e+02    4.1840e+02 ;    HE2 - CE2
- CD2
  3045   3047   3048       1    1.2000e+02    4.1840e+02 ;      C - N
- H
  3045   3047   3049       1    1.2190e+02    4.1840e+02 ;      C - N
- CA
  3046   3045   3047       1    1.2290e+02    6.6944e+02 ;      O - C
- N
  3047   3049   3050       1    1.0950e+02    4.1840e+02 ;      N - CA
- HA
  3047   3049   3051       1    1.0970e+02    6.6944e+02 ;      N - CA
- CB
  3047   3049   3057       1    1.1010e+02    5.2718e+02 ;      N - CA
- C
  3048   3047   3049       1    1.1804e+02    4.1840e+02 ;      H - N
- CA
  3049   3051   3052       1    1.0950e+02    4.1840e+02 ;     CA - CB
- HB2
  3049   3051   3053       1    1.0950e+02    4.1840e+02 ;     CA - CB
- HB3
  3049   3051   3054       1    1.1110e+02    5.2718e+02 ;     CA - CB
- CG
```

3049 3057 3058 1 1.2040e+02 6.6944e+02 ; CA - C
- O
3049 3057 3059 1 1.1660e+02 5.8576e+02 ; CA - C
- N
3050 3049 3051 1 1.0950e+02 4.1840e+02 ; HA - CA
- CB
3050 3049 3057 1 1.0950e+02 4.1840e+02 ; HA - CA
- C
3051 3049 3057 1 1.1110e+02 5.2718e+02 ; CB - CA
- C
3051 3054 3055 1 1.1700e+02 5.8576e+02 ; CB - CG
- OD1
3051 3054 3056 1 1.1700e+02 5.8576e+02 ; CB - CG
- OD2
3052 3051 3053 1 1.0950e+02 2.9288e+02 ; HB2 - CB
- HB3
3052 3051 3054 1 1.0950e+02 4.1840e+02 ; HB2 - CB
- CG
3053 3051 3054 1 1.0950e+02 4.1840e+02 ; HB3 - CB
- CG
3055 3054 3056 1 1.2600e+02 6.6944e+02 ; OD1 - CG
- OD2
3057 3059 3060 1 1.2000e+02 4.1840e+02 ; C - N
- H
3057 3059 3061 1 1.2190e+02 4.1840e+02 ; C - N
- CA
3058 3057 3059 1 1.2290e+02 6.6944e+02 ; O - C
- N
3059 3061 3062 1 1.0950e+02 4.1840e+02 ; N - CA
- HA
3059 3061 3063 1 1.0970e+02 6.6944e+02 ; N - CA
- CB
3059 3061 3079 1 1.1010e+02 5.2718e+02 ; N - CA
- C
3060 3059 3061 1 1.1804e+02 4.1840e+02 ; H - N
- CA
3061 3063 3064 1 1.0950e+02 4.1840e+02 ; CA - CB
- HB2
3061 3063 3065 1 1.0950e+02 4.1840e+02 ; CA - CB
- HB3
3061 3063 3066 1 1.0950e+02 3.3472e+02 ; CA - CB
- CG
3061 3079 3080 1 1.2040e+02 6.6944e+02 ; CA - C
- O
3061 3079 3081 1 1.1660e+02 5.8576e+02 ; CA - C
- N
3062 3061 3063 1 1.0950e+02 4.1840e+02 ; HA - CA
- CB
3062 3061 3079 1 1.0950e+02 4.1840e+02 ; HA - CA
- C
3063 3061 3079 1 1.1110e+02 5.2718e+02 ; CB - CA
- C

```
  3063   3066   3067      1    1.0950e+02    4.1840e+02 ;     CB - CG
- HG2
  3063   3066   3068      1    1.0950e+02    4.1840e+02 ;     CB - CG
- HG3
  3063   3066   3069      1    1.0950e+02    3.3472e+02 ;     CB - CG
- CD
  3064   3063   3065      1    1.0950e+02    2.9288e+02 ;    HB2 - CB
- HB3
  3064   3063   3066      1    1.0950e+02    4.1840e+02 ;    HB2 - CB
- CG
  3065   3063   3066      1    1.0950e+02    4.1840e+02 ;    HB3 - CB
- CG
  3066   3069   3070      1    1.0950e+02    4.1840e+02 ;     CG - CD
- HD2
  3066   3069   3071      1    1.0950e+02    4.1840e+02 ;     CG - CD
- HD3
  3066   3069   3072      1    1.0950e+02    3.3472e+02 ;     CG - CD
- CE
  3067   3066   3068      1    1.0950e+02    2.9288e+02 ;    HG2 - CG
- HG3
  3067   3066   3069      1    1.0950e+02    4.1840e+02 ;    HG2 - CG
- CD
  3068   3066   3069      1    1.0950e+02    4.1840e+02 ;    HG3 - CG
- CD
  3069   3072   3073      1    1.0950e+02    4.1840e+02 ;     CD - CE
- HE2
  3069   3072   3074      1    1.0950e+02    4.1840e+02 ;     CD - CE
- HE3
  3069   3072   3075      1    1.1120e+02    6.6944e+02 ;     CD - CE
- NZ
  3070   3069   3071      1    1.0950e+02    2.9288e+02 ;    HD2 - CD
- HD3
  3070   3069   3072      1    1.0950e+02    4.1840e+02 ;    HD2 - CD
- CE
  3071   3069   3072      1    1.0950e+02    4.1840e+02 ;    HD3 - CD
- CE
  3072   3075   3076      1    1.0950e+02    4.1840e+02 ;     CE - NZ
- HZ1
  3072   3075   3077      1    1.0950e+02    4.1840e+02 ;     CE - NZ
- HZ2
  3072   3075   3078      1    1.0950e+02    4.1840e+02 ;     CE - NZ
- HZ3
  3073   3072   3074      1    1.0950e+02    2.9288e+02 ;    HE2 - CE
- HE3
  3073   3072   3075      1    1.0950e+02    4.1840e+02 ;    HE2 - CE
- NZ
  3074   3072   3075      1    1.0950e+02    4.1840e+02 ;    HE3 - CE
- NZ
  3076   3075   3077      1    1.0950e+02    2.9288e+02 ;    HZ1 - NZ
- HZ2
  3076   3075   3078      1    1.0950e+02    2.9288e+02 ;    HZ1 - NZ
- HZ3
```

3077 3075 3078 1 1.0950e+02 2.9288e+02 ; HZ2 - NZ
- HZ3
3079 3081 3082 1 1.2000e+02 4.1840e+02 ; C - N
- H
3079 3081 3083 1 1.2190e+02 4.1840e+02 ; C - N
- CA
3080 3079 3081 1 1.2290e+02 6.6944e+02 ; O - C
- N
3081 3083 3084 1 1.0950e+02 4.1840e+02 ; N - CA
- HA
3081 3083 3085 1 1.0970e+02 6.6944e+02 ; N - CA
- CB
3081 3083 3100 1 1.1010e+02 5.2718e+02 ; N - CA
- C
3082 3081 3083 1 1.1804e+02 4.1840e+02 ; H - N
- CA
3083 3085 3086 1 1.0950e+02 4.1840e+02 ; CA - CB
- HB2
3083 3085 3087 1 1.0950e+02 4.1840e+02 ; CA - CB
- HB3
3083 3085 3088 1 1.0950e+02 3.3472e+02 ; CA - CB
- CG
3083 3100 3101 1 1.2040e+02 6.6944e+02 ; CA - C
- O
3083 3100 3102 1 1.1660e+02 5.8576e+02 ; CA - C
- N
3084 3083 3085 1 1.0950e+02 4.1840e+02 ; HA - CA
- CB
3084 3083 3100 1 1.0950e+02 4.1840e+02 ; HA - CA
- C
3085 3083 3100 1 1.1110e+02 5.2718e+02 ; CB - CA
- C
3085 3088 3089 1 1.0950e+02 4.1840e+02 ; CB - CG
- HG2
3085 3088 3090 1 1.0950e+02 4.1840e+02 ; CB - CG
- HG3
3085 3088 3091 1 1.0950e+02 3.3472e+02 ; CB - CG
- CD
3086 3085 3087 1 1.0950e+02 2.9288e+02 ; HB2 - CB
- HB3
3086 3085 3088 1 1.0950e+02 4.1840e+02 ; HB2 - CB
- CG
3087 3085 3088 1 1.0950e+02 4.1840e+02 ; HB3 - CB
- CG
3088 3091 3092 1 1.0950e+02 4.1840e+02 ; CG - CD
- HD2
3088 3091 3093 1 1.0950e+02 4.1840e+02 ; CG - CD
- HD3
3088 3091 3094 1 1.0950e+02 3.3472e+02 ; CG - CD
- CE
3089 3088 3090 1 1.0950e+02 2.9288e+02 ; HG2 - CG
- HG3

```
  3089   3088   3091       1    1.0950e+02    4.1840e+02 ;    HG2 - CG
- CD
  3090   3088   3091       1    1.0950e+02    4.1840e+02 ;    HG3 - CG
- CD
  3091   3094   3095       1    1.0950e+02    4.1840e+02 ;     CD - CE
- HE2
  3091   3094   3096       1    1.0950e+02    4.1840e+02 ;     CD - CE
- HE3
  3091   3094   3097       1    1.1120e+02    6.6944e+02 ;     CD - CE
- NZ
  3092   3091   3093       1    1.0950e+02    2.9288e+02 ;    HD2 - CD
- HD3
  3092   3091   3094       1    1.0950e+02    4.1840e+02 ;    HD2 - CD
- CE
  3093   3091   3094       1    1.0950e+02    4.1840e+02 ;    HD3 - CD
- CE
  3094   3097   3098       1    1.0950e+02    4.1840e+02 ;     CE - NZ
- HZ2
  3094   3097   3099       1    1.0950e+02    4.1840e+02 ;     CE - NZ
- HZ3
  3095   3094   3096       1    1.0950e+02    2.9288e+02 ;    HE2 - CE
- HE3
  3095   3094   3097       1    1.0950e+02    4.1840e+02 ;    HE2 - CE
- NZ
  3096   3094   3097       1    1.0950e+02    4.1840e+02 ;    HE3 - CE
- NZ
  3098   3097   3099       1    1.0950e+02    2.9288e+02 ;    HZ2 - NZ
- HZ3
  3100   3102   3103       1    1.2000e+02    4.1840e+02 ;      C - N
- H
  3100   3102   3104       1    1.2190e+02    4.1840e+02 ;      C - N
- CA
  3101   3100   3102       1    1.2290e+02    6.6944e+02 ;      O - C
- N
  3102   3104   3105       1    1.0950e+02    4.1840e+02 ;      N - CA
- HA
  3102   3104   3106       1    1.0970e+02    6.6944e+02 ;      N - CA
- CB
  3102   3104   3111       1    1.1010e+02    5.2718e+02 ;      N - CA
- C
  3103   3102   3104       1    1.1804e+02    4.1840e+02 ;      H - N
- CA
  3104   3106   3107       1    1.0950e+02    4.1840e+02 ;     CA - CB
- HB2
  3104   3106   3108       1    1.0950e+02    4.1840e+02 ;     CA - CB
- HB3
  3104   3106   3109       1    1.0860e+02    4.1840e+02 ;     CA - CB
- SG
  3104   3111   3112       1    1.2040e+02    6.6944e+02 ;     CA - C
- O
  3104   3111   3113       1    1.1660e+02    5.8576e+02 ;     CA - C
- N
```

3105 3104 3106 1 1.0950e+02 4.1840e+02 ; HA - CA
- CB
3105 3104 3111 1 1.0950e+02 4.1840e+02 ; HA - CA
- C
3106 3104 3111 1 1.1110e+02 5.2718e+02 ; CB - CA
- C
3106 3109 3110 1 9.6000e+01 3.5982e+02 ; CB - SG
- HG
3107 3106 3108 1 1.0950e+02 2.9288e+02 ; HB2 - CB
- HB3
3107 3106 3109 1 1.0950e+02 4.1840e+02 ; HB2 - CB
- SG
3108 3106 3109 1 1.0950e+02 4.1840e+02 ; HB3 - CB
- SG
3111 3113 3114 1 1.2000e+02 4.1840e+02 ; C - N
- H
3111 3113 3115 1 1.2190e+02 4.1840e+02 ; C - N
- CA
3112 3111 3113 1 1.2290e+02 6.6944e+02 ; O - C
- N
3113 3115 3116 1 1.0950e+02 4.1840e+02 ; N - CA
- HA
3113 3115 3117 1 1.0970e+02 6.6944e+02 ; N - CA
- CB
3113 3115 3123 1 1.1010e+02 5.2718e+02 ; N - CA
- C
3114 3113 3115 1 1.1804e+02 4.1840e+02 ; H - N
- CA
3115 3117 3118 1 1.0950e+02 4.1840e+02 ; CA - CB
- HB2
3115 3117 3119 1 1.0950e+02 4.1840e+02 ; CA - CB
- HB3
3115 3117 3120 1 1.1110e+02 5.2718e+02 ; CA - CB
- CG
3115 3123 3124 1 1.2040e+02 6.6944e+02 ; CA - C
- O
3115 3123 3125 1 1.1660e+02 5.8576e+02 ; CA - C
- N
3116 3115 3117 1 1.0950e+02 4.1840e+02 ; HA - CA
- CB
3116 3115 3123 1 1.0950e+02 4.1840e+02 ; HA - CA
- C
3117 3115 3123 1 1.1110e+02 5.2718e+02 ; CB - CA
- C
3117 3120 3121 1 1.1700e+02 5.8576e+02 ; CB - CG
- OD1
3117 3120 3122 1 1.1700e+02 5.8576e+02 ; CB - CG
- OD2
3118 3117 3119 1 1.0950e+02 2.9288e+02 ; HB2 - CB
- HB3
3118 3117 3120 1 1.0950e+02 4.1840e+02 ; HB2 - CB
- CG

```
  3119   3117   3120        1    1.0950e+02    4.1840e+02 ;    HB3 - CB
- CG
  3121   3120   3122        1    1.2600e+02    6.6944e+02 ;    OD1 - CG
- OD2
  3123   3125   3126        1    1.2000e+02    4.1840e+02 ;      C - N
- H
  3123   3125   3127        1    1.2190e+02    4.1840e+02 ;      C - N
- CA
  3124   3123   3125        1    1.2290e+02    6.6944e+02 ;      O - C
- N
  3125   3127   3128        1    1.0950e+02    4.1840e+02 ;      N - CA
- HA
  3125   3127   3129        1    1.0970e+02    6.6944e+02 ;      N - CA
- CB
  3125   3127   3142        1    1.1010e+02    5.2718e+02 ;      N - CA
- C
  3126   3125   3127        1    1.1804e+02    4.1840e+02 ;      H - N
- CA
  3127   3129   3130        1    1.0950e+02    4.1840e+02 ;     CA - CB
- HB
  3127   3129   3131        1    1.0950e+02    3.3472e+02 ;     CA - CB
- CG2
  3127   3129   3135        1    1.0950e+02    3.3472e+02 ;     CA - CB
- CG1
  3127   3142   3143        1    1.2040e+02    6.6944e+02 ;     CA - C
- O
  3127   3142   3144        1    1.1660e+02    5.8576e+02 ;     CA - C
- N
  3128   3127   3129        1    1.0950e+02    4.1840e+02 ;     HA - CA
- CB
  3128   3127   3142        1    1.0950e+02    4.1840e+02 ;     HA - CA
- C
  3129   3127   3142        1    1.1110e+02    5.2718e+02 ;     CB - CA
- C
  3129   3131   3132        1    1.0950e+02    4.1840e+02 ;     CB - CG2
- HG21
  3129   3131   3133        1    1.0950e+02    4.1840e+02 ;     CB - CG2
- HG22
  3129   3131   3134        1    1.0950e+02    4.1840e+02 ;     CB - CG2
- HG23
  3129   3135   3136        1    1.0950e+02    4.1840e+02 ;     CB - CG1
- HG12
  3129   3135   3137        1    1.0950e+02    4.1840e+02 ;     CB - CG1
- HG13
  3129   3135   3138        1    1.0950e+02    3.3472e+02 ;     CB - CG1
- CD1
  3130   3129   3131        1    1.0950e+02    4.1840e+02 ;     HB - CB
- CG2
  3130   3129   3135        1    1.0950e+02    4.1840e+02 ;     HB - CB
- CG1
  3131   3129   3135        1    1.0950e+02    3.3472e+02 ;    CG2 - CB
- CG1
```

3132 3131 3133 1 1.0950e+02 2.9288e+02 ; HG21 - CG2
- HG22
3132 3131 3134 1 1.0950e+02 2.9288e+02 ; HG21 - CG2
- HG23
3133 3131 3134 1 1.0950e+02 2.9288e+02 ; HG22 - CG2
- HG23
3135 3138 3139 1 1.0950e+02 4.1840e+02 ; CG1 - CD1
- HD11
3135 3138 3140 1 1.0950e+02 4.1840e+02 ; CG1 - CD1
- HD12
3135 3138 3141 1 1.0950e+02 4.1840e+02 ; CG1 - CD1
- HD13
3136 3135 3137 1 1.0950e+02 2.9288e+02 ; HG12 - CG1
- HG13
3136 3135 3138 1 1.0950e+02 4.1840e+02 ; HG12 - CG1
- CD1
3137 3135 3138 1 1.0950e+02 4.1840e+02 ; HG13 - CG1
- CD1
3139 3138 3140 1 1.0950e+02 2.9288e+02 ; HD11 - CD1
- HD12
3139 3138 3141 1 1.0950e+02 2.9288e+02 ; HD11 - CD1
- HD13
3140 3138 3141 1 1.0950e+02 2.9288e+02 ; HD12 - CD1
- HD13
3142 3144 3145 1 1.2000e+02 4.1840e+02 ; C - N
- H
3142 3144 3146 1 1.2190e+02 4.1840e+02 ; C - N
- CA
3143 3142 3144 1 1.2290e+02 6.6944e+02 ; O - C
- N
3144 3146 3147 1 1.0950e+02 4.1840e+02 ; N - CA
- HA
3144 3146 3148 1 1.0970e+02 6.6944e+02 ; N - CA
- CB
3144 3146 3166 1 1.1010e+02 5.2718e+02 ; N - CA
- C
3145 3144 3146 1 1.1804e+02 4.1840e+02 ; H - N
- CA
3146 3148 3149 1 1.0950e+02 4.1840e+02 ; CA - CB
- HB2
3146 3148 3150 1 1.0950e+02 4.1840e+02 ; CA - CB
- HB3
3146 3148 3151 1 1.1560e+02 5.2718e+02 ; CA - CB
- CG
3146 3166 3167 1 1.2040e+02 6.6944e+02 ; CA - C
- O
3146 3166 3168 1 1.1660e+02 5.8576e+02 ; CA - C
- N
3147 3146 3148 1 1.0950e+02 4.1840e+02 ; HA - CA
- CB
3147 3146 3166 1 1.0950e+02 4.1840e+02 ; HA - CA
- C

3148 3146 3166 1 1.1110e+02 5.2718e+02 ; CB - CA
- C
3148 3151 3152 1 1.2500e+02 5.8576e+02 ; CB - CG
- CD1
3148 3151 3165 1 1.2860e+02 5.8576e+02 ; CB - CG
- CD2
3149 3148 3150 1 1.0950e+02 2.9288e+02 ; HB2 - CB
- HB3
3149 3148 3151 1 1.0950e+02 4.1840e+02 ; HB2 - CB
- CG
3150 3148 3151 1 1.0950e+02 4.1840e+02 ; HB3 - CB
- CG
3151 3152 3153 1 1.2000e+02 4.1840e+02 ; CG - CD1
- HD1
3151 3152 3154 1 1.0870e+02 5.8576e+02 ; CG - CD1
- NE1
3151 3165 3156 1 1.0880e+02 5.2718e+02 ; CG - CD2
- CE2
3151 3165 3163 1 1.3490e+02 5.2718e+02 ; CG - CD2
- CE3
3152 3151 3165 1 1.0640e+02 5.2718e+02 ; CD1 - CG
- CD2
3152 3154 3155 1 1.2000e+02 4.1840e+02 ; CD1 - NE1
- HE1
3152 3154 3156 1 1.1160e+02 5.8576e+02 ; CD1 - NE1
- CE2
3153 3152 3154 1 1.2000e+02 4.1840e+02 ; HD1 - CD1
- NE1
3154 3156 3157 1 1.3280e+02 5.8576e+02 ; NE1 - CE2
- CZ2
3154 3156 3165 1 1.0440e+02 5.8576e+02 ; NE1 - CE2
- CD2
3155 3154 3156 1 1.2310e+02 4.1840e+02 ; HE1 - NE1
- CE2
3156 3157 3158 1 1.2000e+02 4.1840e+02 ; CE2 - CZ2
- HZ2
3156 3157 3159 1 1.2000e+02 5.2718e+02 ; CE2 - CZ2
- CH2
3156 3165 3163 1 1.1620e+02 5.2718e+02 ; CE2 - CD2
- CE3
3157 3156 3165 1 1.2270e+02 5.2718e+02 ; CZ2 - CE2
- CD2
3157 3159 3160 1 1.2000e+02 4.1840e+02 ; CZ2 - CH2
- HH2
3157 3159 3161 1 1.2000e+02 5.2718e+02 ; CZ2 - CH2
- CZ3
3158 3157 3159 1 1.2000e+02 4.1840e+02 ; HZ2 - CZ2
- CH2
3159 3161 3162 1 1.2000e+02 4.1840e+02 ; CH2 - CZ3
- HZ3
3159 3161 3163 1 1.2000e+02 5.2718e+02 ; CH2 - CZ3
- CE3

3160 3159 3161 1 1.2000e+02 4.1840e+02 ; HH2 - CH2 - CZ3
3161 3163 3164 1 1.2000e+02 4.1840e+02 ; CZ3 - CE3 - HE3
3161 3163 3165 1 1.2000e+02 5.2718e+02 ; CZ3 - CE3 - CD2
3162 3161 3163 1 1.2000e+02 4.1840e+02 ; HZ3 - CZ3 - CE3
3164 3163 3165 1 1.2000e+02 4.1840e+02 ; HE3 - CE3 - CD2
3166 3168 3169 1 1.2000e+02 4.1840e+02 ; C - N - H
3166 3168 3170 1 1.2190e+02 4.1840e+02 ; C - N - CA
3167 3166 3168 1 1.2290e+02 6.6944e+02 ; O - C - N
3168 3170 3171 1 1.0950e+02 4.1840e+02 ; N - CA - HA
3168 3170 3172 1 1.0970e+02 6.6944e+02 ; N - CA - CB
3168 3170 3177 1 1.1010e+02 5.2718e+02 ; N - CA - C
3169 3168 3170 1 1.1804e+02 4.1840e+02 ; H - N - CA
3170 3172 3173 1 1.0950e+02 4.1840e+02 ; CA - CB - HB2
3170 3172 3174 1 1.0950e+02 4.1840e+02 ; CA - CB - HB3
3170 3172 3175 1 1.0950e+02 4.1840e+02 ; CA - CB - OG
3170 3177 3178 1 1.2040e+02 6.6944e+02 ; CA - C - O
3170 3177 3179 1 1.1660e+02 5.8576e+02 ; CA - C - N
3171 3170 3172 1 1.0950e+02 4.1840e+02 ; HA - CA - CB
3171 3170 3177 1 1.0950e+02 4.1840e+02 ; HA - CA - C
3172 3170 3177 1 1.1110e+02 5.2718e+02 ; CB - CA - C
3172 3175 3176 1 1.0850e+02 4.6024e+02 ; CB - OG - HG
3173 3172 3174 1 1.0950e+02 2.9288e+02 ; HB2 - CB - HB3
3173 3172 3175 1 1.0950e+02 4.1840e+02 ; HB2 - CB - OG
3174 3172 3175 1 1.0950e+02 4.1840e+02 ; HB3 - CB - OG
3177 3179 3180 1 1.2000e+02 4.1840e+02 ; C - N - H
3177 3179 3181 1 1.2190e+02 4.1840e+02 ; C - N - CA

3178 3177 3179 1 1.2290e+02 6.6944e+02 ; O - C
- N
3179 3181 3182 1 1.0950e+02 4.1840e+02 ; N - CA
- HA
3179 3181 3183 1 1.0970e+02 6.6944e+02 ; N - CA
- CB
3179 3181 3188 1 1.1010e+02 5.2718e+02 ; N - CA
- C
3180 3179 3181 1 1.1804e+02 4.1840e+02 ; H - N
- CA
3181 3183 3184 1 1.0950e+02 4.1840e+02 ; CA - CB
- HB2
3181 3183 3185 1 1.0950e+02 4.1840e+02 ; CA - CB
- HB3
3181 3183 3186 1 1.0950e+02 4.1840e+02 ; CA - CB
- OG
3181 3188 3189 1 1.2040e+02 6.6944e+02 ; CA - C
- O
3181 3188 3190 1 1.1660e+02 5.8576e+02 ; CA - C
- N
3182 3181 3183 1 1.0950e+02 4.1840e+02 ; HA - CA
- CB
3182 3181 3188 1 1.0950e+02 4.1840e+02 ; HA - CA
- C
3183 3181 3188 1 1.1110e+02 5.2718e+02 ; CB - CA
- C
3183 3186 3187 1 1.0850e+02 4.6024e+02 ; CB - OG
- HG
3184 3183 3185 1 1.0950e+02 2.9288e+02 ; HB2 - CB
- HB3
3184 3183 3186 1 1.0950e+02 4.1840e+02 ; HB2 - CB
- OG
3185 3183 3186 1 1.0950e+02 4.1840e+02 ; HB3 - CB
- OG
3188 3190 3191 1 1.2000e+02 4.1840e+02 ; C - N
- H
3188 3190 3192 1 1.2190e+02 4.1840e+02 ; C - N
- CA
3189 3188 3190 1 1.2290e+02 6.6944e+02 ; O - C
- N
3190 3192 3193 1 1.0950e+02 4.1840e+02 ; N - CA
- HA2
3190 3192 3194 1 1.0950e+02 4.1840e+02 ; N - CA
- HA3
3190 3192 3195 1 1.1010e+02 5.2718e+02 ; N - CA
- C
3191 3190 3192 1 1.1804e+02 4.1840e+02 ; H - N
- CA
3192 3195 3196 1 1.2040e+02 6.6944e+02 ; CA - C
- O
3192 3195 3197 1 1.1660e+02 5.8576e+02 ; CA - C
- N

3193 3192 3194 1 1.0950e+02 2.9288e+02 ; HA2 - CA
- HA3
3193 3192 3195 1 1.0950e+02 4.1840e+02 ; HA2 - CA
- C
3194 3192 3195 1 1.0950e+02 4.1840e+02 ; HA3 - CA
- C
3195 3197 3198 1 1.2000e+02 4.1840e+02 ; C - N
- H
3195 3197 3199 1 1.2190e+02 4.1840e+02 ; C - N
- CA
3196 3195 3197 1 1.2290e+02 6.6944e+02 ; O - C
- N
3197 3199 3200 1 1.0950e+02 4.1840e+02 ; N - CA
- HA
3197 3199 3201 1 1.0970e+02 6.6944e+02 ; N - CA
- CB
3197 3199 3211 1 1.1010e+02 5.2718e+02 ; N - CA
- C
3198 3197 3199 1 1.1804e+02 4.1840e+02 ; H - N
- CA
3199 3201 3202 1 1.0950e+02 4.1840e+02 ; CA - CB
- HB
3199 3201 3203 1 1.0950e+02 3.3472e+02 ; CA - CB
- CG1
3199 3201 3207 1 1.0950e+02 3.3472e+02 ; CA - CB
- CG2
3199 3211 3212 1 1.2040e+02 6.6944e+02 ; CA - C
- O
3199 3211 3213 1 1.1660e+02 5.8576e+02 ; CA - C
- N
3200 3199 3201 1 1.0950e+02 4.1840e+02 ; HA - CA
- CB
3200 3199 3211 1 1.0950e+02 4.1840e+02 ; HA - CA
- C
3201 3199 3211 1 1.1110e+02 5.2718e+02 ; CB - CA
- C
3201 3203 3204 1 1.0950e+02 4.1840e+02 ; CB - CG1
- HG11
3201 3203 3205 1 1.0950e+02 4.1840e+02 ; CB - CG1
- HG12
3201 3203 3206 1 1.0950e+02 4.1840e+02 ; CB - CG1
- HG13
3201 3207 3208 1 1.0950e+02 4.1840e+02 ; CB - CG2
- HG21
3201 3207 3209 1 1.0950e+02 4.1840e+02 ; CB - CG2
- HG22
3201 3207 3210 1 1.0950e+02 4.1840e+02 ; CB - CG2
- HG23
3202 3201 3203 1 1.0950e+02 4.1840e+02 ; HB - CB
- CG1
3202 3201 3207 1 1.0950e+02 4.1840e+02 ; HB - CB
- CG2

| | | | | | | | |
|---|---|---|---|---|---|---|---|
| 3203 | 3201 | 3207 | 1 | 1.0950e+02 | 3.3472e+02 | ; | CG1 - CB - CG2 |
| 3204 | 3203 | 3205 | 1 | 1.0950e+02 | 2.9288e+02 | ; | HG11 - CG1 - HG12 |
| 3204 | 3203 | 3206 | 1 | 1.0950e+02 | 2.9288e+02 | ; | HG11 - CG1 - HG13 |
| 3205 | 3203 | 3206 | 1 | 1.0950e+02 | 2.9288e+02 | ; | HG12 - CG1 - HG13 |
| 3208 | 3207 | 3209 | 1 | 1.0950e+02 | 2.9288e+02 | ; | HG21 - CG2 - HG22 |
| 3208 | 3207 | 3210 | 1 | 1.0950e+02 | 2.9288e+02 | ; | HG21 - CG2 - HG23 |
| 3209 | 3207 | 3210 | 1 | 1.0950e+02 | 2.9288e+02 | ; | HG22 - CG2 - HG23 |
| 3211 | 3213 | 3214 | 1 | 1.2000e+02 | 4.1840e+02 | ; | C - N - H |
| 3211 | 3213 | 3215 | 1 | 1.2190e+02 | 4.1840e+02 | ; | C - N - CA |
| 3212 | 3211 | 3213 | 1 | 1.2290e+02 | 6.6944e+02 | ; | O - C - N |
| 3213 | 3215 | 3216 | 1 | 1.0950e+02 | 4.1840e+02 | ; | N - CA - HA |
| 3213 | 3215 | 3217 | 1 | 1.0970e+02 | 6.6944e+02 | ; | N - CA - CB |
| 3213 | 3215 | 3230 | 1 | 1.1010e+02 | 5.2718e+02 | ; | N - CA - C |
| 3214 | 3213 | 3215 | 1 | 1.1804e+02 | 4.1840e+02 | ; | H - N - CA |
| 3215 | 3217 | 3218 | 1 | 1.0950e+02 | 4.1840e+02 | ; | CA - CB - HB |
| 3215 | 3217 | 3219 | 1 | 1.0950e+02 | 3.3472e+02 | ; | CA - CB - CG2 |
| 3215 | 3217 | 3223 | 1 | 1.0950e+02 | 3.3472e+02 | ; | CA - CB - CG1 |
| 3215 | 3230 | 3231 | 1 | 1.2040e+02 | 6.6944e+02 | ; | CA - C - O |
| 3215 | 3230 | 3232 | 1 | 1.1660e+02 | 5.8576e+02 | ; | CA - C - N |
| 3216 | 3215 | 3217 | 1 | 1.0950e+02 | 4.1840e+02 | ; | HA - CA - CB |
| 3216 | 3215 | 3230 | 1 | 1.0950e+02 | 4.1840e+02 | ; | HA - CA - C |
| 3217 | 3215 | 3230 | 1 | 1.1110e+02 | 5.2718e+02 | ; | CB - CA - C |
| 3217 | 3219 | 3220 | 1 | 1.0950e+02 | 4.1840e+02 | ; | CB - CG2 - HG21 |
| 3217 | 3219 | 3221 | 1 | 1.0950e+02 | 4.1840e+02 | ; | CB - CG2 - HG22 |
| 3217 | 3219 | 3222 | 1 | 1.0950e+02 | 4.1840e+02 | ; | CB - CG2 - HG23 |
| 3217 | 3223 | 3224 | 1 | 1.0950e+02 | 4.1840e+02 | ; | CB - CG1 - HG12 |

3217 3223 3225 1 1.0950e+02 4.1840e+02 ; CB - CG1
- HG13
3217 3223 3226 1 1.0950e+02 3.3472e+02 ; CB - CG1
- CD1
3218 3217 3219 1 1.0950e+02 4.1840e+02 ; HB - CB
- CG2
3218 3217 3223 1 1.0950e+02 4.1840e+02 ; HB - CB
- CG1
3219 3217 3223 1 1.0950e+02 3.3472e+02 ; CG2 - CB
- CG1
3220 3219 3221 1 1.0950e+02 2.9288e+02 ; HG21 - CG2
- HG22
3220 3219 3222 1 1.0950e+02 2.9288e+02 ; HG21 - CG2
- HG23
3221 3219 3222 1 1.0950e+02 2.9288e+02 ; HG22 - CG2
- HG23
3223 3226 3227 1 1.0950e+02 4.1840e+02 ; CG1 - CD1
- HD11
3223 3226 3228 1 1.0950e+02 4.1840e+02 ; CG1 - CD1
- HD12
3223 3226 3229 1 1.0950e+02 4.1840e+02 ; CG1 - CD1
- HD13
3224 3223 3225 1 1.0950e+02 2.9288e+02 ; HG12 - CG1
- HG13
3224 3223 3226 1 1.0950e+02 4.1840e+02 ; HG12 - CG1
- CD1
3225 3223 3226 1 1.0950e+02 4.1840e+02 ; HG13 - CG1
- CD1
3227 3226 3228 1 1.0950e+02 2.9288e+02 ; HD11 - CD1
- HD12
3227 3226 3229 1 1.0950e+02 2.9288e+02 ; HD11 - CD1
- HD13
3228 3226 3229 1 1.0950e+02 2.9288e+02 ; HD12 - CD1
- HD13
3230 3232 3233 1 1.2000e+02 4.1840e+02 ; C - N
- H
3230 3232 3234 1 1.2190e+02 4.1840e+02 ; C - N
- CA
3231 3230 3232 1 1.2290e+02 6.6944e+02 ; O - C
- N
3232 3234 3235 1 1.0950e+02 4.1840e+02 ; N - CA
- HA
3232 3234 3236 1 1.0970e+02 6.6944e+02 ; N - CA
- CB
3232 3234 3247 1 1.1010e+02 5.2718e+02 ; N - CA
- C
3233 3232 3234 1 1.1804e+02 4.1840e+02 ; H - N
- CA
3234 3236 3237 1 1.0950e+02 4.1840e+02 ; CA - CB
- HB2
3234 3236 3238 1 1.0950e+02 4.1840e+02 ; CA - CB
- HB3

3234 3236 3239 1 1.0950e+02 3.3472e+02 ; CA - CB
- CG
3234 3247 3248 1 1.2040e+02 6.6944e+02 ; CA - C
- O
3234 3247 3249 1 1.1660e+02 5.8576e+02 ; CA - C
- N
3235 3234 3236 1 1.0950e+02 4.1840e+02 ; HA - CA
- CB
3235 3234 3247 1 1.0950e+02 4.1840e+02 ; HA - CA
- C
3236 3234 3247 1 1.1110e+02 5.2718e+02 ; CB - CA
- C
3236 3239 3240 1 1.0950e+02 4.1840e+02 ; CB - CG
- HG2
3236 3239 3241 1 1.0950e+02 4.1840e+02 ; CB - CG
- HG3
3236 3239 3242 1 1.1470e+02 4.1840e+02 ; CB - CG
- SD
3237 3236 3238 1 1.0950e+02 2.9288e+02 ; HB2 - CB
- HB3
3237 3236 3239 1 1.0950e+02 4.1840e+02 ; HB2 - CB
- CG
3238 3236 3239 1 1.0950e+02 4.1840e+02 ; HB3 - CB
- CG
3239 3242 3243 1 9.8900e+01 5.1882e+02 ; CG - SD
- CE
3240 3239 3241 1 1.0950e+02 2.9288e+02 ; HG2 - CG
- HG3
3240 3239 3242 1 1.0950e+02 4.1840e+02 ; HG2 - CG
- SD
3241 3239 3242 1 1.0950e+02 4.1840e+02 ; HG3 - CG
- SD
3242 3243 3244 1 1.0950e+02 4.1840e+02 ; SD - CE
- HE1
3242 3243 3245 1 1.0950e+02 4.1840e+02 ; SD - CE
- HE2
3242 3243 3246 1 1.0950e+02 4.1840e+02 ; SD - CE
- HE3
3244 3243 3245 1 1.0950e+02 2.9288e+02 ; HE1 - CE
- HE2
3244 3243 3246 1 1.0950e+02 2.9288e+02 ; HE1 - CE
- HE3
3245 3243 3246 1 1.0950e+02 2.9288e+02 ; HE2 - CE
- HE3
3247 3249 3250 1 1.2000e+02 4.1840e+02 ; C - N
- H
3247 3249 3251 1 1.2190e+02 4.1840e+02 ; C - N
- CA
3248 3247 3249 1 1.2290e+02 6.6944e+02 ; O - C
- N
3249 3251 3252 1 1.0950e+02 4.1840e+02 ; N - CA
- HA

3249 3251 3253 1 1.0970e+02 6.6944e+02 ; N - CA
- CB
3249 3251 3268 1 1.1010e+02 5.2718e+02 ; N - CA
- C
3250 3249 3251 1 1.1804e+02 4.1840e+02 ; H - N
- CA
3251 3253 3254 1 1.0950e+02 4.1840e+02 ; CA - CB
- HB2
3251 3253 3255 1 1.0950e+02 4.1840e+02 ; CA - CB
- HB3
3251 3253 3256 1 1.1400e+02 5.2718e+02 ; CA - CB
- CG
3251 3268 3269 1 1.2040e+02 6.6944e+02 ; CA - C
- O
3251 3268 3270 1 1.1660e+02 5.8576e+02 ; CA - C
- N
3252 3251 3253 1 1.0950e+02 4.1840e+02 ; HA - CA
- CB
3252 3251 3268 1 1.0950e+02 4.1840e+02 ; HA - CA
- C
3253 3251 3268 1 1.1110e+02 5.2718e+02 ; CB - CA
- C
3253 3256 3257 1 1.2000e+02 5.8576e+02 ; CB - CG
- CD1
3253 3256 3266 1 1.2000e+02 5.8576e+02 ; CB - CG
- CD2
3254 3253 3255 1 1.0950e+02 2.9288e+02 ; HB2 - CB
- HB3
3254 3253 3256 1 1.0950e+02 4.1840e+02 ; HB2 - CB
- CG
3255 3253 3256 1 1.0950e+02 4.1840e+02 ; HB3 - CB
- CG
3256 3257 3258 1 1.2000e+02 4.1840e+02 ; CG - CD1
- HD1
3256 3257 3259 1 1.2000e+02 5.2718e+02 ; CG - CD1
- CE1
3256 3266 3264 1 1.2000e+02 5.2718e+02 ; CG - CD2
- CE2
3256 3266 3267 1 1.2000e+02 4.1840e+02 ; CG - CD2
- HD2
3257 3256 3266 1 1.2000e+02 5.2718e+02 ; CD1 - CG
- CD2
3257 3259 3260 1 1.2000e+02 4.1840e+02 ; CD1 - CE1
- HE1
3257 3259 3261 1 1.2000e+02 5.2718e+02 ; CD1 - CE1
- CZ
3258 3257 3259 1 1.2000e+02 4.1840e+02 ; HD1 - CD1
- CE1
3259 3261 3262 1 1.2000e+02 5.8576e+02 ; CE1 - CZ
- OH
3259 3261 3264 1 1.2000e+02 5.2718e+02 ; CE1 - CZ
- CE2

3260 3259 3261 1 1.2000e+02 4.1840e+02 ; HE1 - CE1
- CZ
3261 3262 3263 1 1.1300e+02 4.1840e+02 ; CZ - OH
- HH
3261 3264 3265 1 1.2000e+02 4.1840e+02 ; CZ - CE2
- HE2
3261 3264 3266 1 1.2000e+02 5.2718e+02 ; CZ - CE2
- CD2
3262 3261 3264 1 1.2000e+02 5.8576e+02 ; OH - CZ
- CE2
3264 3266 3267 1 1.2000e+02 4.1840e+02 ; CE2 - CD2
- HD2
3265 3264 3266 1 1.2000e+02 4.1840e+02 ; HE2 - CE2
- CD2
3268 3270 3271 1 1.2000e+02 4.1840e+02 ; C - N
- H
3268 3270 3272 1 1.2190e+02 4.1840e+02 ; C - N
- CA
3269 3268 3270 1 1.2290e+02 6.6944e+02 ; O - C
- N
3270 3272 3273 1 1.0950e+02 4.1840e+02 ; N - CA
- HA
3270 3272 3274 1 1.0970e+02 6.6944e+02 ; N - CA
- CB
3270 3272 3282 1 1.1010e+02 5.2718e+02 ; N - CA
- C
3271 3270 3272 1 1.1804e+02 4.1840e+02 ; H - N
- CA
3272 3274 3275 1 1.0950e+02 4.1840e+02 ; CA - CB
- HB
3272 3274 3276 1 1.0950e+02 3.3472e+02 ; CA - CB
- CG2
3272 3274 3280 1 1.0950e+02 4.1840e+02 ; CA - CB
- OG1
3272 3282 3283 1 1.2040e+02 6.6944e+02 ; CA - C
- O
3272 3282 3284 1 1.1660e+02 5.8576e+02 ; CA - C
- N
3273 3272 3274 1 1.0950e+02 4.1840e+02 ; HA - CA
- CB
3273 3272 3282 1 1.0950e+02 4.1840e+02 ; HA - CA
- C
3274 3272 3282 1 1.1110e+02 5.2718e+02 ; CB - CA
- C
3274 3276 3277 1 1.0950e+02 4.1840e+02 ; CB - CG2
- HG21
3274 3276 3278 1 1.0950e+02 4.1840e+02 ; CB - CG2
- HG22
3274 3276 3279 1 1.0950e+02 4.1840e+02 ; CB - CG2
- HG23
3274 3280 3281 1 1.0850e+02 4.6024e+02 ; CB - OG1
- HG1

```
  3275   3274   3276      1    1.0950e+02    4.1840e+02 ;     HB - CB
- CG2
  3275   3274   3280      1    1.0950e+02    4.1840e+02 ;     HB - CB
- OG1
  3276   3274   3280      1    1.0950e+02    4.1840e+02 ;    CG2 - CB
- OG1
  3277   3276   3278      1    1.0950e+02    2.9288e+02 ;   HG21 - CG2
- HG22
  3277   3276   3279      1    1.0950e+02    2.9288e+02 ;   HG21 - CG2
- HG23
  3278   3276   3279      1    1.0950e+02    2.9288e+02 ;   HG22 - CG2
- HG23
  3282   3284   3285      1    1.2000e+02    4.1840e+02 ;      C - N
- H
  3282   3284   3286      1    1.2190e+02    4.1840e+02 ;      C - N
- CA
  3283   3282   3284      1    1.2290e+02    6.6944e+02 ;      O - C
- N
  3284   3286   3287      1    1.0950e+02    4.1840e+02 ;      N - CA
- HA
  3284   3286   3288      1    1.0970e+02    6.6944e+02 ;      N - CA
- CB
  3284   3286   3301      1    1.1010e+02    5.2718e+02 ;      N - CA
- C
  3285   3284   3286      1    1.1804e+02    4.1840e+02 ;      H - N
- CA
  3286   3288   3289      1    1.0950e+02    4.1840e+02 ;     CA - CB
- HB2
  3286   3288   3290      1    1.0950e+02    4.1840e+02 ;     CA - CB
- HB3
  3286   3288   3291      1    1.0950e+02    3.3472e+02 ;     CA - CB
- CG
  3286   3301   3302      1    1.2040e+02    6.6944e+02 ;     CA - C
- O
  3286   3301   3303      1    1.1660e+02    5.8576e+02 ;     CA - C
- N
  3287   3286   3288      1    1.0950e+02    4.1840e+02 ;     HA - CA
- CB
  3287   3286   3301      1    1.0950e+02    4.1840e+02 ;     HA - CA
- C
  3288   3286   3301      1    1.1110e+02    5.2718e+02 ;     CB - CA
- C
  3288   3291   3292      1    1.0950e+02    4.1840e+02 ;     CB - CG
- HG
  3288   3291   3293      1    1.0950e+02    3.3472e+02 ;     CB - CG
- CD1
  3288   3291   3297      1    1.0950e+02    3.3472e+02 ;     CB - CG
- CD2
  3289   3288   3290      1    1.0950e+02    2.9288e+02 ;    HB2 - CB
- HB3
  3289   3288   3291      1    1.0950e+02    4.1840e+02 ;    HB2 - CB
- CG
```

3290 3288 3291 1 1.0950e+02 4.1840e+02 ; HB3 - CB
- CG
3291 3293 3294 1 1.0950e+02 4.1840e+02 ; CG - CD1
- HD11
3291 3293 3295 1 1.0950e+02 4.1840e+02 ; CG - CD1
- HD12
3291 3293 3296 1 1.0950e+02 4.1840e+02 ; CG - CD1
- HD13
3291 3297 3298 1 1.0950e+02 4.1840e+02 ; CG - CD2
- HD21
3291 3297 3299 1 1.0950e+02 4.1840e+02 ; CG - CD2
- HD22
3291 3297 3300 1 1.0950e+02 4.1840e+02 ; CG - CD2
- HD23
3292 3291 3293 1 1.0950e+02 4.1840e+02 ; HG - CG
- CD1
3292 3291 3297 1 1.0950e+02 4.1840e+02 ; HG - CG
- CD2
3293 3291 3297 1 1.0950e+02 3.3472e+02 ; CD1 - CG
- CD2
3294 3293 3295 1 1.0950e+02 2.9288e+02 ; HD11 - CD1
- HD12
3294 3293 3296 1 1.0950e+02 2.9288e+02 ; HD11 - CD1
- HD13
3295 3293 3296 1 1.0950e+02 2.9288e+02 ; HD12 - CD1
- HD13
3298 3297 3299 1 1.0950e+02 2.9288e+02 ; HD21 - CD2
- HD22
3298 3297 3300 1 1.0950e+02 2.9288e+02 ; HD21 - CD2
- HD23
3299 3297 3300 1 1.0950e+02 2.9288e+02 ; HD22 - CD2
- HD23
3301 3303 3304 1 1.2000e+02 4.1840e+02 ; C - N
- H
3301 3303 3305 1 1.2190e+02 4.1840e+02 ; C - N
- CA
3302 3301 3303 1 1.2290e+02 6.6944e+02 ; O - C
- N
3303 3305 3306 1 1.0950e+02 4.1840e+02 ; N - CA
- HA
3303 3305 3307 1 1.0970e+02 6.6944e+02 ; N - CA
- CB
3303 3305 3320 1 1.1010e+02 5.2718e+02 ; N - CA
- C
3304 3303 3305 1 1.1804e+02 4.1840e+02 ; H - N
- CA
3305 3307 3308 1 1.0950e+02 4.1840e+02 ; CA - CB
- HB2
3305 3307 3309 1 1.0950e+02 4.1840e+02 ; CA - CB
- HB3
3305 3307 3310 1 1.0950e+02 3.3472e+02 ; CA - CB
- CG

```
  3305   3320   3321       1    1.2040e+02    6.6944e+02 ;     CA - C
- O
  3305   3320   3322       1    1.1660e+02    5.8576e+02 ;     CA - C
- N
  3306   3305   3307       1    1.0950e+02    4.1840e+02 ;     HA - CA
- CB
  3306   3305   3320       1    1.0950e+02    4.1840e+02 ;     HA - CA
- C
  3307   3305   3320       1    1.1110e+02    5.2718e+02 ;     CB - CA
- C
  3307   3310   3311       1    1.0950e+02    4.1840e+02 ;     CB - CG
- HG
  3307   3310   3312       1    1.0950e+02    3.3472e+02 ;     CB - CG
- CD1
  3307   3310   3316       1    1.0950e+02    3.3472e+02 ;     CB - CG
- CD2
  3308   3307   3309       1    1.0950e+02    2.9288e+02 ;    HB2 - CB
- HB3
  3308   3307   3310       1    1.0950e+02    4.1840e+02 ;    HB2 - CB
- CG
  3309   3307   3310       1    1.0950e+02    4.1840e+02 ;    HB3 - CB
- CG
  3310   3312   3313       1    1.0950e+02    4.1840e+02 ;     CG - CD1
- HD11
  3310   3312   3314       1    1.0950e+02    4.1840e+02 ;     CG - CD1
- HD12
  3310   3312   3315       1    1.0950e+02    4.1840e+02 ;     CG - CD1
- HD13
  3310   3316   3317       1    1.0950e+02    4.1840e+02 ;     CG - CD2
- HD21
  3310   3316   3318       1    1.0950e+02    4.1840e+02 ;     CG - CD2
- HD22
  3310   3316   3319       1    1.0950e+02    4.1840e+02 ;     CG - CD2
- HD23
  3311   3310   3312       1    1.0950e+02    4.1840e+02 ;     HG - CG
- CD1
  3311   3310   3316       1    1.0950e+02    4.1840e+02 ;     HG - CG
- CD2
  3312   3310   3316       1    1.0950e+02    3.3472e+02 ;    CD1 - CG
- CD2
  3313   3312   3314       1    1.0950e+02    2.9288e+02 ;   HD11 - CD1
- HD12
  3313   3312   3315       1    1.0950e+02    2.9288e+02 ;   HD11 - CD1
- HD13
  3314   3312   3315       1    1.0950e+02    2.9288e+02 ;   HD12 - CD1
- HD13
  3317   3316   3318       1    1.0950e+02    2.9288e+02 ;   HD21 - CD2
- HD22
  3317   3316   3319       1    1.0950e+02    2.9288e+02 ;   HD21 - CD2
- HD23
  3318   3316   3319       1    1.0950e+02    2.9288e+02 ;   HD22 - CD2
- HD23
```

3320 3322 3323 1 1.2000e+02 4.1840e+02 ; C - N
- H
3320 3322 3324 1 1.2190e+02 4.1840e+02 ; C - N
- CA
3321 3320 3322 1 1.2290e+02 6.6944e+02 ; O - C
- N
3322 3324 3325 1 1.0950e+02 4.1840e+02 ; N - CA
- HA
3322 3324 3326 1 1.0970e+02 6.6944e+02 ; N - CA
- CB
3322 3324 3331 1 1.1010e+02 5.2718e+02 ; N - CA
- C
3323 3322 3324 1 1.1804e+02 4.1840e+02 ; H - N
- CA
3324 3326 3327 1 1.0950e+02 4.1840e+02 ; CA - CB
- HB2
3324 3326 3328 1 1.0950e+02 4.1840e+02 ; CA - CB
- HB3
3324 3326 3329 1 1.0860e+02 4.1840e+02 ; CA - CB
- SG
3324 3331 3332 1 1.2040e+02 6.6944e+02 ; CA - C
- O
3324 3331 3333 1 1.1660e+02 5.8576e+02 ; CA - C
- N
3325 3324 3326 1 1.0950e+02 4.1840e+02 ; HA - CA
- CB
3325 3324 3331 1 1.0950e+02 4.1840e+02 ; HA - CA
- C
3326 3324 3331 1 1.1110e+02 5.2718e+02 ; CB - CA
- C
3326 3329 3330 1 9.6000e+01 3.5982e+02 ; CB - SG
- HG
3327 3326 3328 1 1.0950e+02 2.9288e+02 ; HB2 - CB
- HB3
3327 3326 3329 1 1.0950e+02 4.1840e+02 ; HB2 - CB
- SG
3328 3326 3329 1 1.0950e+02 4.1840e+02 ; HB3 - CB
- SG
3331 3333 3334 1 1.2000e+02 4.1840e+02 ; C - N
- H
3331 3333 3335 1 1.2190e+02 4.1840e+02 ; C - N
- CA
3332 3331 3333 1 1.2290e+02 6.6944e+02 ; O - C
- N
3333 3335 3336 1 1.0950e+02 4.1840e+02 ; N - CA
- HA2
3333 3335 3337 1 1.0950e+02 4.1840e+02 ; N - CA
- HA3
3333 3335 3338 1 1.1010e+02 5.2718e+02 ; N - CA
- C
3334 3333 3335 1 1.1804e+02 4.1840e+02 ; H - N
- CA

3335 3338 3339 1 1.2040e+02 6.6944e+02 ; CA - C
- O
3335 3338 3340 1 1.1660e+02 5.8576e+02 ; CA - C
- N
3336 3335 3337 1 1.0950e+02 2.9288e+02 ; HA2 - CA
- HA3
3336 3335 3338 1 1.0950e+02 4.1840e+02 ; HA2 - CA
- C
3337 3335 3338 1 1.0950e+02 4.1840e+02 ; HA3 - CA
- C
3338 3340 3341 1 1.2000e+02 4.1840e+02 ; C - N
- H
3338 3340 3342 1 1.2190e+02 4.1840e+02 ; C - N
- CA
3339 3338 3340 1 1.2290e+02 6.6944e+02 ; O - C
- N
3340 3342 3343 1 1.0950e+02 4.1840e+02 ; N - CA
- HA
3340 3342 3344 1 1.0970e+02 6.6944e+02 ; N - CA
- CB
3340 3342 3359 1 1.1010e+02 5.2718e+02 ; N - CA
- C
3341 3340 3342 1 1.1804e+02 4.1840e+02 ; H - N
- CA
3342 3344 3345 1 1.0950e+02 4.1840e+02 ; CA - CB
- HB2
3342 3344 3346 1 1.0950e+02 4.1840e+02 ; CA - CB
- HB3
3342 3344 3347 1 1.1400e+02 5.2718e+02 ; CA - CB
- CG
3342 3359 3360 1 1.2040e+02 6.6944e+02 ; CA - C
- O
3342 3359 3361 1 1.1660e+02 5.8576e+02 ; CA - C
- N
3343 3342 3344 1 1.0950e+02 4.1840e+02 ; HA - CA
- CB
3343 3342 3359 1 1.0950e+02 4.1840e+02 ; HA - CA
- C
3344 3342 3359 1 1.1110e+02 5.2718e+02 ; CB - CA
- C
3344 3347 3348 1 1.2000e+02 5.8576e+02 ; CB - CG
- CD1
3344 3347 3357 1 1.2000e+02 5.8576e+02 ; CB - CG
- CD2
3345 3344 3346 1 1.0950e+02 2.9288e+02 ; HB2 - CB
- HB3
3345 3344 3347 1 1.0950e+02 4.1840e+02 ; HB2 - CB
- CG
3346 3344 3347 1 1.0950e+02 4.1840e+02 ; HB3 - CB
- CG
3347 3348 3349 1 1.2000e+02 4.1840e+02 ; CG - CD1
- HD1

3347   3348   3350      1    1.2000e+02    5.2718e+02 ;     CG - CD1
- CE1
3347   3357   3355      1    1.2000e+02    5.2718e+02 ;     CG - CD2
- CE2
3347   3357   3358      1    1.2000e+02    4.1840e+02 ;     CG - CD2
- HD2
3348   3347   3357      1    1.2000e+02    5.2718e+02 ;    CD1 - CG
- CD2
3348   3350   3351      1    1.2000e+02    4.1840e+02 ;    CD1 - CE1
- HE1
3348   3350   3352      1    1.2000e+02    5.2718e+02 ;    CD1 - CE1
- CZ
3349   3348   3350      1    1.2000e+02    4.1840e+02 ;    HD1 - CD1
- CE1
3350   3352   3353      1    1.2000e+02    5.8576e+02 ;    CE1 - CZ
- OH
3350   3352   3355      1    1.2000e+02    5.2718e+02 ;    CE1 - CZ
- CE2
3351   3350   3352      1    1.2000e+02    4.1840e+02 ;    HE1 - CE1
- CZ
3352   3353   3354      1    1.1300e+02    4.1840e+02 ;     CZ - OH
- HH
3352   3355   3356      1    1.2000e+02    4.1840e+02 ;     CZ - CE2
- HE2
3352   3355   3357      1    1.2000e+02    5.2718e+02 ;     CZ - CE2
- CD2
3353   3352   3355      1    1.2000e+02    5.8576e+02 ;     OH - CZ
- CE2
3355   3357   3358      1    1.2000e+02    4.1840e+02 ;    CE2 - CD2
- HD2
3356   3355   3357      1    1.2000e+02    4.1840e+02 ;    HE2 - CE2
- CD2
3359   3361   3362      1    1.2190e+02    4.1840e+02 ;      C - N
- CD
3359   3361   3371      1    1.2190e+02    4.1840e+02 ;      C - N
- CA
3360   3359   3361      1    1.2290e+02    6.6944e+02 ;      O - C
- N
3361   3362   3363      1    1.0950e+02    4.1840e+02 ;      N - CD
- HD2
3361   3362   3364      1    1.0950e+02    4.1840e+02 ;      N - CD
- HD3
3361   3362   3365      1    1.0970e+02    6.6944e+02 ;      N - CD
- CG
3361   3371   3368      1    1.0970e+02    6.6944e+02 ;      N - CA
- CB
3361   3371   3372      1    1.0950e+02    4.1840e+02 ;      N - CA
- HA
3361   3371   3373      1    1.1010e+02    5.2718e+02 ;      N - CA
- C
3362   3361   3371      1    1.1800e+02    4.1840e+02 ;     CD - N
- CA

3362 3365 3366 1 1.0950e+02 4.1840e+02 ; CD - CG
- HG2
3362 3365 3367 1 1.0950e+02 4.1840e+02 ; CD - CG
- HG3
3362 3365 3368 1 1.0950e+02 3.3472e+02 ; CD - CG
- CB
3363 3362 3364 1 1.0950e+02 2.9288e+02 ; HD2 - CD
- HD3
3363 3362 3365 1 1.0950e+02 4.1840e+02 ; HD2 - CD
- CG
3364 3362 3365 1 1.0950e+02 4.1840e+02 ; HD3 - CD
- CG
3365 3368 3369 1 1.0950e+02 4.1840e+02 ; CG - CB
- HB2
3365 3368 3370 1 1.0950e+02 4.1840e+02 ; CG - CB
- HB3
3365 3368 3371 1 1.0950e+02 3.3472e+02 ; CG - CB
- CA
3366 3365 3367 1 1.0950e+02 2.9288e+02 ; HG2 - CG
- HG3
3366 3365 3368 1 1.0950e+02 4.1840e+02 ; HG2 - CG
- CB
3367 3365 3368 1 1.0950e+02 4.1840e+02 ; HG3 - CG
- CB
3368 3371 3372 1 1.0950e+02 4.1840e+02 ; CB - CA
- HA
3368 3371 3373 1 1.1110e+02 5.2718e+02 ; CB - CA
- C
3369 3368 3370 1 1.0950e+02 2.9288e+02 ; HB2 - CB
- HB3
3369 3368 3371 1 1.0950e+02 4.1840e+02 ; HB2 - CB
- CA
3370 3368 3371 1 1.0950e+02 4.1840e+02 ; HB3 - CB
- CA
3371 3373 3374 1 1.2040e+02 6.6944e+02 ; CA - C
- O
3371 3373 3375 1 1.1660e+02 5.8576e+02 ; CA - C
- N
3372 3371 3373 1 1.0950e+02 4.1840e+02 ; HA - CA
- C
3373 3375 3376 1 1.2190e+02 4.1840e+02 ; C - N
- CD
3373 3375 3385 1 1.2190e+02 4.1840e+02 ; C - N
- CA
3374 3373 3375 1 1.2290e+02 6.6944e+02 ; O - C
- N
3375 3376 3377 1 1.0950e+02 4.1840e+02 ; N - CD
- HD2
3375 3376 3378 1 1.0950e+02 4.1840e+02 ; N - CD
- HD3
3375 3376 3379 1 1.0970e+02 6.6944e+02 ; N - CD
- CG

3375 3385 3382 1 1.0970e+02 6.6944e+02 ; N - CA
- CB
3375 3385 3386 1 1.0950e+02 4.1840e+02 ; N - CA
- HA
3375 3385 3387 1 1.1010e+02 5.2718e+02 ; N - CA
- C
3376 3375 3385 1 1.1800e+02 4.1840e+02 ; CD - N
- CA
3376 3379 3380 1 1.0950e+02 4.1840e+02 ; CD - CG
- HG2
3376 3379 3381 1 1.0950e+02 4.1840e+02 ; CD - CG
- HG3
3376 3379 3382 1 1.0950e+02 3.3472e+02 ; CD - CG
- CB
3377 3376 3378 1 1.0950e+02 2.9288e+02 ; HD2 - CD
- HD3
3377 3376 3379 1 1.0950e+02 4.1840e+02 ; HD2 - CD
- CG
3378 3376 3379 1 1.0950e+02 4.1840e+02 ; HD3 - CD
- CG
3379 3382 3383 1 1.0950e+02 4.1840e+02 ; CG - CB
- HB2
3379 3382 3384 1 1.0950e+02 4.1840e+02 ; CG - CB
- HB3
3379 3382 3385 1 1.0950e+02 3.3472e+02 ; CG - CB
- CA
3380 3379 3381 1 1.0950e+02 2.9288e+02 ; HG2 - CG
- HG3
3380 3379 3382 1 1.0950e+02 4.1840e+02 ; HG2 - CG
- CB
3381 3379 3382 1 1.0950e+02 4.1840e+02 ; HG3 - CG
- CB
3382 3385 3386 1 1.0950e+02 4.1840e+02 ; CB - CA
- HA
3382 3385 3387 1 1.1110e+02 5.2718e+02 ; CB - CA
- C
3383 3382 3384 1 1.0950e+02 2.9288e+02 ; HB2 - CB
- HB3
3383 3382 3385 1 1.0950e+02 4.1840e+02 ; HB2 - CB
- CA
3384 3382 3385 1 1.0950e+02 4.1840e+02 ; HB3 - CB
- CA
3385 3387 3388 1 1.2040e+02 6.6944e+02 ; CA - C
- O
3385 3387 3389 1 1.1660e+02 5.8576e+02 ; CA - C
- N
3386 3385 3387 1 1.0950e+02 4.1840e+02 ; HA - CA
- C
3387 3389 3390 1 1.2000e+02 4.1840e+02 ; C - N
- H
3387 3389 3391 1 1.2190e+02 4.1840e+02 ; C - N
- CA

```
  3388   3387   3389       1    1.2290e+02    6.6944e+02 ;      O - C
- N
  3389   3391   3392       1    1.0950e+02    4.1840e+02 ;      N - CA
- HA
  3389   3391   3393       1    1.0970e+02    6.6944e+02 ;      N - CA
- CB
  3389   3391   3407       1    1.1010e+02    5.2718e+02 ;      N - CA
- C
  3390   3389   3391       1    1.1804e+02    4.1840e+02 ;      H - N
- CA
  3391   3393   3394       1    1.0950e+02    4.1840e+02 ;     CA - CB
- HB2
  3391   3393   3395       1    1.0950e+02    4.1840e+02 ;     CA - CB
- HB3
  3391   3393   3396       1    1.1400e+02    5.2718e+02 ;     CA - CB
- CG
  3391   3407   3408       1    1.2040e+02    6.6944e+02 ;     CA - C
- O
  3391   3407   3409       1    1.1660e+02    5.8576e+02 ;     CA - C
- N
  3392   3391   3393       1    1.0950e+02    4.1840e+02 ;     HA - CA
- CB
  3392   3391   3407       1    1.0950e+02    4.1840e+02 ;     HA - CA
- C
  3393   3391   3407       1    1.1110e+02    5.2718e+02 ;     CB - CA
- C
  3393   3396   3397       1    1.2000e+02    5.8576e+02 ;     CB - CG
- CD1
  3393   3396   3405       1    1.2000e+02    5.8576e+02 ;     CB - CG
- CD2
  3394   3393   3395       1    1.0950e+02    2.9288e+02 ;    HB2 - CB
- HB3
  3394   3393   3396       1    1.0950e+02    4.1840e+02 ;    HB2 - CB
- CG
  3395   3393   3396       1    1.0950e+02    4.1840e+02 ;    HB3 - CB
- CG
  3396   3397   3398       1    1.2000e+02    4.1840e+02 ;     CG - CD1
- HD1
  3396   3397   3399       1    1.2000e+02    5.2718e+02 ;     CG - CD1
- CE1
  3396   3405   3403       1    1.2000e+02    5.2718e+02 ;     CG - CD2
- CE2
  3396   3405   3406       1    1.2000e+02    4.1840e+02 ;     CG - CD2
- HD2
  3397   3396   3405       1    1.2000e+02    5.2718e+02 ;    CD1 - CG
- CD2
  3397   3399   3400       1    1.2000e+02    4.1840e+02 ;    CD1 - CE1
- HE1
  3397   3399   3401       1    1.2000e+02    5.2718e+02 ;    CD1 - CE1
- CZ
  3398   3397   3399       1    1.2000e+02    4.1840e+02 ;    HD1 - CD1
- CE1
```

```
  3399   3401   3402       1    1.2000e+02    4.1840e+02 ;    CE1 - CZ
- HZ
  3399   3401   3403       1    1.2000e+02    5.2718e+02 ;    CE1 - CZ
- CE2
  3400   3399   3401       1    1.2000e+02    4.1840e+02 ;    HE1 - CE1
- CZ
  3401   3403   3404       1    1.2000e+02    4.1840e+02 ;     CZ - CE2
- HE2
  3401   3403   3405       1    1.2000e+02    5.2718e+02 ;     CZ - CE2
- CD2
  3402   3401   3403       1    1.2000e+02    4.1840e+02 ;     HZ - CZ
- CE2
  3403   3405   3406       1    1.2000e+02    4.1840e+02 ;    CE2 - CD2
- HD2
  3404   3403   3405       1    1.2000e+02    4.1840e+02 ;    HE2 - CE2
- CD2
  3407   3409   3410       1    1.2000e+02    4.1840e+02 ;      C - N
- H
  3407   3409   3411       1    1.2190e+02    4.1840e+02 ;      C - N
- CA
  3408   3407   3409       1    1.2290e+02    6.6944e+02 ;      O - C
- N
  3409   3411   3412       1    1.0950e+02    4.1840e+02 ;      N - CA
- HA
  3409   3411   3413       1    1.0970e+02    6.6944e+02 ;      N - CA
- CB
  3409   3411   3428       1    1.1010e+02    5.2718e+02 ;      N - CA
- C
  3410   3409   3411       1    1.1804e+02    4.1840e+02 ;      H - N
- CA
  3411   3413   3414       1    1.0950e+02    4.1840e+02 ;     CA - CB
- HB2
  3411   3413   3415       1    1.0950e+02    4.1840e+02 ;     CA - CB
- HB3
  3411   3413   3416       1    1.1400e+02    5.2718e+02 ;     CA - CB
- CG
  3411   3428   3429       1    1.2040e+02    6.6944e+02 ;     CA - C
- O
  3411   3428   3430       1    1.1660e+02    5.8576e+02 ;     CA - C
- N
  3412   3411   3413       1    1.0950e+02    4.1840e+02 ;     HA - CA
- CB
  3412   3411   3428       1    1.0950e+02    4.1840e+02 ;     HA - CA
- C
  3413   3411   3428       1    1.1110e+02    5.2718e+02 ;     CB - CA
- C
  3413   3416   3417       1    1.2000e+02    5.8576e+02 ;     CB - CG
- CD1
  3413   3416   3426       1    1.2000e+02    5.8576e+02 ;     CB - CG
- CD2
  3414   3413   3415       1    1.0950e+02    2.9288e+02 ;    HB2 - CB
- HB3
```

3414 3413 3416 1 1.0950e+02 4.1840e+02 ; HB2 - CB
- CG
3415 3413 3416 1 1.0950e+02 4.1840e+02 ; HB3 - CB
- CG
3416 3417 3418 1 1.2000e+02 4.1840e+02 ; CG - CD1
- HD1
3416 3417 3419 1 1.2000e+02 5.2718e+02 ; CG - CD1
- CE1
3416 3426 3424 1 1.2000e+02 5.2718e+02 ; CG - CD2
- CE2
3416 3426 3427 1 1.2000e+02 4.1840e+02 ; CG - CD2
- HD2
3417 3416 3426 1 1.2000e+02 5.2718e+02 ; CD1 - CG
- CD2
3417 3419 3420 1 1.2000e+02 4.1840e+02 ; CD1 - CE1
- HE1
3417 3419 3421 1 1.2000e+02 5.2718e+02 ; CD1 - CE1
- CZ
3418 3417 3419 1 1.2000e+02 4.1840e+02 ; HD1 - CD1
- CE1
3419 3421 3422 1 1.2000e+02 5.8576e+02 ; CE1 - CZ
- OH
3419 3421 3424 1 1.2000e+02 5.2718e+02 ; CE1 - CZ
- CE2
3420 3419 3421 1 1.2000e+02 4.1840e+02 ; HE1 - CE1
- CZ
3421 3422 3423 1 1.1300e+02 4.1840e+02 ; CZ - OH
- HH
3421 3424 3425 1 1.2000e+02 4.1840e+02 ; CZ - CE2
- HE2
3421 3424 3426 1 1.2000e+02 5.2718e+02 ; CZ - CE2
- CD2
3422 3421 3424 1 1.2000e+02 5.8576e+02 ; OH - CZ
- CE2
3424 3426 3427 1 1.2000e+02 4.1840e+02 ; CE2 - CD2
- HD2
3425 3424 3426 1 1.2000e+02 4.1840e+02 ; HE2 - CE2
- CD2
3428 3430 3431 1 1.2000e+02 4.1840e+02 ; C - N
- H
3428 3430 3432 1 1.2190e+02 4.1840e+02 ; C - N
- CA
3429 3428 3430 1 1.2290e+02 6.6944e+02 ; O - C
- N
3430 3432 3433 1 1.0950e+02 4.1840e+02 ; N - CA
- HA2
3430 3432 3434 1 1.0950e+02 4.1840e+02 ; N - CA
- HA3
3430 3432 3435 1 1.1010e+02 5.2718e+02 ; N - CA
- C
3431 3430 3432 1 1.1804e+02 4.1840e+02 ; H - N
- CA

```
  3432   3435   3436      1    1.2040e+02    6.6944e+02 ;     CA - C
- O
  3432   3435   3437      1    1.1660e+02    5.8576e+02 ;     CA - C
- N
  3433   3432   3434      1    1.0950e+02    2.9288e+02 ;    HA2 - CA
- HA3
  3433   3432   3435      1    1.0950e+02    4.1840e+02 ;    HA2 - CA
- C
  3434   3432   3435      1    1.0950e+02    4.1840e+02 ;    HA3 - CA
- C
  3435   3437   3438      1    1.2000e+02    4.1840e+02 ;      C - N
- H
  3435   3437   3439      1    1.2190e+02    4.1840e+02 ;      C - N
- CA
  3436   3435   3437      1    1.2290e+02    6.6944e+02 ;      O - C
- N
  3437   3439   3440      1    1.0950e+02    4.1840e+02 ;      N - CA
- HA
  3437   3439   3441      1    1.0970e+02    6.6944e+02 ;      N - CA
- CB
  3437   3439   3447      1    1.1010e+02    5.2718e+02 ;      N - CA
- C
  3438   3437   3439      1    1.1804e+02    4.1840e+02 ;      H - N
- CA
  3439   3441   3442      1    1.0950e+02    4.1840e+02 ;     CA - CB
- HB2
  3439   3441   3443      1    1.0950e+02    4.1840e+02 ;     CA - CB
- HB3
  3439   3441   3444      1    1.1110e+02    5.2718e+02 ;     CA - CB
- CG
  3439   3447   3448      1    1.2040e+02    6.6944e+02 ;     CA - C
- O
  3439   3447   3449      1    1.1660e+02    5.8576e+02 ;     CA - C
- N
  3440   3439   3441      1    1.0950e+02    4.1840e+02 ;     HA - CA
- CB
  3440   3439   3447      1    1.0950e+02    4.1840e+02 ;     HA - CA
- C
  3441   3439   3447      1    1.1110e+02    5.2718e+02 ;     CB - CA
- C
  3441   3444   3445      1    1.1700e+02    5.8576e+02 ;     CB - CG
- OD1
  3441   3444   3446      1    1.1700e+02    5.8576e+02 ;     CB - CG
- OD2
  3442   3441   3443      1    1.0950e+02    2.9288e+02 ;    HB2 - CB
- HB3
  3442   3441   3444      1    1.0950e+02    4.1840e+02 ;    HB2 - CB
- CG
  3443   3441   3444      1    1.0950e+02    4.1840e+02 ;    HB3 - CB
- CG
  3445   3444   3446      1    1.2600e+02    6.6944e+02 ;    OD1 - CG
- OD2
```

```
  3447   3449   3450       1    1.2000e+02    4.1840e+02 ;      C - N
- H
  3447   3449   3451       1    1.2190e+02    4.1840e+02 ;      C - N
- CA
  3448   3447   3449       1    1.2290e+02    6.6944e+02 ;      O - C
- N
  3449   3451   3452       1    1.0950e+02    4.1840e+02 ;      N - CA
- HA
  3449   3451   3453       1    1.0970e+02    6.6944e+02 ;      N - CA
- CB
  3449   3451   3461       1    1.1010e+02    5.2718e+02 ;      N - CA
- C
  3450   3449   3451       1    1.1804e+02    4.1840e+02 ;      H - N
- CA
  3451   3453   3454       1    1.0950e+02    4.1840e+02 ;     CA - CB
- HB
  3451   3453   3455       1    1.0950e+02    3.3472e+02 ;     CA - CB
- CG2
  3451   3453   3459       1    1.0950e+02    4.1840e+02 ;     CA - CB
- OG1
  3451   3461   3462       1    1.2040e+02    6.6944e+02 ;     CA - C
- O
  3451   3461   3463       1    1.1660e+02    5.8576e+02 ;     CA - C
- N
  3452   3451   3453       1    1.0950e+02    4.1840e+02 ;     HA - CA
- CB
  3452   3451   3461       1    1.0950e+02    4.1840e+02 ;     HA - CA
- C
  3453   3451   3461       1    1.1110e+02    5.2718e+02 ;     CB - CA
- C
  3453   3455   3456       1    1.0950e+02    4.1840e+02 ;     CB - CG2
- HG21
  3453   3455   3457       1    1.0950e+02    4.1840e+02 ;     CB - CG2
- HG22
  3453   3455   3458       1    1.0950e+02    4.1840e+02 ;     CB - CG2
- HG23
  3453   3459   3460       1    1.0850e+02    4.6024e+02 ;     CB - OG1
- HG1
  3454   3453   3455       1    1.0950e+02    4.1840e+02 ;     HB - CB
- CG2
  3454   3453   3459       1    1.0950e+02    4.1840e+02 ;     HB - CB
- OG1
  3455   3453   3459       1    1.0950e+02    4.1840e+02 ;    CG2 - CB
- OG1
  3456   3455   3457       1    1.0950e+02    2.9288e+02 ;   HG21 - CG2
- HG22
  3456   3455   3458       1    1.0950e+02    2.9288e+02 ;   HG21 - CG2
- HG23
  3457   3455   3458       1    1.0950e+02    2.9288e+02 ;   HG22 - CG2
- HG23
  3461   3463   3464       1    1.2000e+02    4.1840e+02 ;      C - N
- H
```

```
  3461   3463   3465      1    1.2190e+02    4.1840e+02 ;       C - N
- CA
  3462   3461   3463      1    1.2290e+02    6.6944e+02 ;       O - C
- N
  3463   3465   3466      1    1.0950e+02    4.1840e+02 ;       N - CA
- HA
  3463   3465   3467      1    1.0970e+02    6.6944e+02 ;       N - CA
- CB
  3463   3465   3473      1    1.1010e+02    5.2718e+02 ;       N - CA
- C
  3464   3463   3465      1    1.1804e+02    4.1840e+02 ;       H - N
- CA
  3465   3467   3468      1    1.0950e+02    4.1840e+02 ;      CA - CB
- HB2
  3465   3467   3469      1    1.0950e+02    4.1840e+02 ;      CA - CB
- HB3
  3465   3467   3470      1    1.1110e+02    5.2718e+02 ;      CA - CB
- CG
  3465   3473   3474      1    1.2040e+02    6.6944e+02 ;      CA - C
- O
  3465   3473   3475      1    1.1660e+02    5.8576e+02 ;      CA - C
- N
  3466   3465   3467      1    1.0950e+02    4.1840e+02 ;      HA - CA
- CB
  3466   3465   3473      1    1.0950e+02    4.1840e+02 ;      HA - CA
- C
  3467   3465   3473      1    1.1110e+02    5.2718e+02 ;      CB - CA
- C
  3467   3470   3471      1    1.1700e+02    5.8576e+02 ;      CB - CG
- OD1
  3467   3470   3472      1    1.1700e+02    5.8576e+02 ;      CB - CG
- OD2
  3468   3467   3469      1    1.0950e+02    2.9288e+02 ;     HB2 - CB
- HB3
  3468   3467   3470      1    1.0950e+02    4.1840e+02 ;     HB2 - CB
- CG
  3469   3467   3470      1    1.0950e+02    4.1840e+02 ;     HB3 - CB
- CG
  3471   3470   3472      1    1.2600e+02    6.6944e+02 ;     OD1 - CG
- OD2
  3473   3475   3476      1    1.2000e+02    4.1840e+02 ;       C - N
- H
  3473   3475   3477      1    1.2190e+02    4.1840e+02 ;       C - N
- CA
  3474   3473   3475      1    1.2290e+02    6.6944e+02 ;       O - C
- N
  3475   3477   3478      1    1.0950e+02    4.1840e+02 ;       N - CA
- HA
  3475   3477   3479      1    1.0970e+02    6.6944e+02 ;       N - CA
- CB
  3475   3477   3487      1    1.1010e+02    5.2718e+02 ;       N - CA
- C
```

3476 3475 3477 1 1.1804e+02 4.1840e+02 ; H - N
- CA
3477 3479 3480 1 1.0950e+02 4.1840e+02 ; CA - CB
- HB2
3477 3479 3481 1 1.0950e+02 4.1840e+02 ; CA - CB
- HB3
3477 3479 3482 1 1.1110e+02 5.2718e+02 ; CA - CB
- CG
3477 3487 3488 1 1.2040e+02 6.6944e+02 ; CA - C
- O
3477 3487 3489 1 1.1660e+02 5.8576e+02 ; CA - C
- N
3478 3477 3479 1 1.0950e+02 4.1840e+02 ; HA - CA
- CB
3478 3477 3487 1 1.0950e+02 4.1840e+02 ; HA - CA
- C
3479 3477 3487 1 1.1110e+02 5.2718e+02 ; CB - CA
- C
3479 3482 3483 1 1.2040e+02 6.6944e+02 ; CB - CG
- OD1
3479 3482 3484 1 1.1660e+02 5.8576e+02 ; CB - CG
- ND2
3480 3479 3481 1 1.0950e+02 2.9288e+02 ; HB2 - CB
- HB3
3480 3479 3482 1 1.0950e+02 4.1840e+02 ; HB2 - CB
- CG
3481 3479 3482 1 1.0950e+02 4.1840e+02 ; HB3 - CB
- CG
3482 3484 3485 1 1.2000e+02 4.1840e+02 ; CG - ND2
- HD21
3482 3484 3486 1 1.2000e+02 4.1840e+02 ; CG - ND2
- HD22
3483 3482 3484 1 1.2290e+02 6.6944e+02 ; OD1 - CG
- ND2
3485 3484 3486 1 1.2000e+02 2.9288e+02 ; HD21 - ND2
- HD22
3487 3489 3490 1 1.2000e+02 4.1840e+02 ; C - N
- H
3487 3489 3491 1 1.2190e+02 4.1840e+02 ; C - N
- CA
3488 3487 3489 1 1.2290e+02 6.6944e+02 ; O - C
- N
3489 3491 3492 1 1.0950e+02 4.1840e+02 ; N - CA
- HA
3489 3491 3493 1 1.0970e+02 6.6944e+02 ; N - CA
- CB
3489 3491 3502 1 1.1010e+02 5.2718e+02 ; N - CA
- C
3490 3489 3491 1 1.1804e+02 4.1840e+02 ; H - N
- CA
3491 3493 3494 1 1.0950e+02 4.1840e+02 ; CA - CB
- HB2

```
  3491   3493   3495       1    1.0950e+02    4.1840e+02 ;     CA - CB
- HB3
  3491   3493   3496       1    1.0950e+02    3.3472e+02 ;     CA - CB
- CG
  3491   3502   3503       1    1.2040e+02    6.6944e+02 ;     CA - C
- O
  3491   3502   3504       1    1.1660e+02    5.8576e+02 ;     CA - C
- N
  3492   3491   3493       1    1.0950e+02    4.1840e+02 ;     HA - CA
- CB
  3492   3491   3502       1    1.0950e+02    4.1840e+02 ;     HA - CA
- C
  3493   3491   3502       1    1.1110e+02    5.2718e+02 ;     CB - CA
- C
  3493   3496   3497       1    1.0950e+02    4.1840e+02 ;     CB - CG
- HG2
  3493   3496   3498       1    1.0950e+02    4.1840e+02 ;     CB - CG
- HG3
  3493   3496   3499       1    1.1110e+02    5.2718e+02 ;     CB - CG
- CD
  3494   3493   3495       1    1.0950e+02    2.9288e+02 ;    HB2 - CB
- HB3
  3494   3493   3496       1    1.0950e+02    4.1840e+02 ;    HB2 - CB
- CG
  3495   3493   3496       1    1.0950e+02    4.1840e+02 ;    HB3 - CB
- CG
  3496   3499   3500       1    1.1700e+02    5.8576e+02 ;     CG - CD
- OE1
  3496   3499   3501       1    1.1700e+02    5.8576e+02 ;     CG - CD
- OE2
  3497   3496   3498       1    1.0950e+02    2.9288e+02 ;    HG2 - CG
- HG3
  3497   3496   3499       1    1.0950e+02    4.1840e+02 ;    HG2 - CG
- CD
  3498   3496   3499       1    1.0950e+02    4.1840e+02 ;    HG3 - CG
- CD
  3500   3499   3501       1    1.2600e+02    6.6944e+02 ;    OE1 - CD
- OE2
  3502   3504   3505       1    1.2000e+02    4.1840e+02 ;      C - N
- H
  3502   3504   3506       1    1.2190e+02    4.1840e+02 ;      C - N
- CA
  3503   3502   3504       1    1.2290e+02    6.6944e+02 ;      O - C
- N
  3504   3506   3507       1    1.0950e+02    4.1840e+02 ;      N - CA
- HA
  3504   3506   3508       1    1.0970e+02    6.6944e+02 ;      N - CA
- CB
  3504   3506   3518       1    1.1010e+02    5.2718e+02 ;      N - CA
- C
  3505   3504   3506       1    1.1804e+02    4.1840e+02 ;      H - N
- CA
```

3506 3508 3509 1 1.0950e+02 4.1840e+02 ; CA - CB
- HB
3506 3508 3510 1 1.0950e+02 3.3472e+02 ; CA - CB
- CG1
3506 3508 3514 1 1.0950e+02 3.3472e+02 ; CA - CB
- CG2
3506 3518 3519 1 1.2040e+02 6.6944e+02 ; CA - C
- O
3506 3518 3520 1 1.1660e+02 5.8576e+02 ; CA - C
- N
3507 3506 3508 1 1.0950e+02 4.1840e+02 ; HA - CA
- CB
3507 3506 3518 1 1.0950e+02 4.1840e+02 ; HA - CA
- C
3508 3506 3518 1 1.1110e+02 5.2718e+02 ; CB - CA
- C
3508 3510 3511 1 1.0950e+02 4.1840e+02 ; CB - CG1
- HG11
3508 3510 3512 1 1.0950e+02 4.1840e+02 ; CB - CG1
- HG12
3508 3510 3513 1 1.0950e+02 4.1840e+02 ; CB - CG1
- HG13
3508 3514 3515 1 1.0950e+02 4.1840e+02 ; CB - CG2
- HG21
3508 3514 3516 1 1.0950e+02 4.1840e+02 ; CB - CG2
- HG22
3508 3514 3517 1 1.0950e+02 4.1840e+02 ; CB - CG2
- HG23
3509 3508 3510 1 1.0950e+02 4.1840e+02 ; HB - CB
- CG1
3509 3508 3514 1 1.0950e+02 4.1840e+02 ; HB - CB
- CG2
3510 3508 3514 1 1.0950e+02 3.3472e+02 ; CG1 - CB
- CG2
3511 3510 3512 1 1.0950e+02 2.9288e+02 ; HG11 - CG1
- HG12
3511 3510 3513 1 1.0950e+02 2.9288e+02 ; HG11 - CG1
- HG13
3512 3510 3513 1 1.0950e+02 2.9288e+02 ; HG12 - CG1
- HG13
3515 3514 3516 1 1.0950e+02 2.9288e+02 ; HG21 - CG2
- HG22
3515 3514 3517 1 1.0950e+02 2.9288e+02 ; HG21 - CG2
- HG23
3516 3514 3517 1 1.0950e+02 2.9288e+02 ; HG22 - CG2
- HG23
3518 3520 3521 1 1.2000e+02 4.1840e+02 ; C - N
- H
3518 3520 3522 1 1.2190e+02 4.1840e+02 ; C - N
- CA
3519 3518 3520 1 1.2290e+02 6.6944e+02 ; O - C
- N

3520 3522 3523 1 1.0950e+02 4.1840e+02 ; N - CA
- HA
3520 3522 3524 1 1.0970e+02 6.6944e+02 ; N - CA
- CB
3520 3522 3537 1 1.1010e+02 5.2718e+02 ; N - CA
- C
3521 3520 3522 1 1.1804e+02 4.1840e+02 ; H - N
- CA
3522 3524 3525 1 1.0950e+02 4.1840e+02 ; CA - CB
- HB2
3522 3524 3526 1 1.0950e+02 4.1840e+02 ; CA - CB
- HB3
3522 3524 3527 1 1.0950e+02 3.3472e+02 ; CA - CB
- CG
3522 3537 3538 1 1.2040e+02 6.6944e+02 ; CA - C
- O
3522 3537 3539 1 1.1660e+02 5.8576e+02 ; CA - C
- N
3523 3522 3524 1 1.0950e+02 4.1840e+02 ; HA - CA
- CB
3523 3522 3537 1 1.0950e+02 4.1840e+02 ; HA - CA
- C
3524 3522 3537 1 1.1110e+02 5.2718e+02 ; CB - CA
- C
3524 3527 3528 1 1.0950e+02 4.1840e+02 ; CB - CG
- HG
3524 3527 3529 1 1.0950e+02 3.3472e+02 ; CB - CG
- CD1
3524 3527 3533 1 1.0950e+02 3.3472e+02 ; CB - CG
- CD2
3525 3524 3526 1 1.0950e+02 2.9288e+02 ; HB2 - CB
- HB3
3525 3524 3527 1 1.0950e+02 4.1840e+02 ; HB2 - CB
- CG
3526 3524 3527 1 1.0950e+02 4.1840e+02 ; HB3 - CB
- CG
3527 3529 3530 1 1.0950e+02 4.1840e+02 ; CG - CD1
- HD11
3527 3529 3531 1 1.0950e+02 4.1840e+02 ; CG - CD1
- HD12
3527 3529 3532 1 1.0950e+02 4.1840e+02 ; CG - CD1
- HD13
3527 3533 3534 1 1.0950e+02 4.1840e+02 ; CG - CD2
- HD21
3527 3533 3535 1 1.0950e+02 4.1840e+02 ; CG - CD2
- HD22
3527 3533 3536 1 1.0950e+02 4.1840e+02 ; CG - CD2
- HD23
3528 3527 3529 1 1.0950e+02 4.1840e+02 ; HG - CG
- CD1
3528 3527 3533 1 1.0950e+02 4.1840e+02 ; HG - CG
- CD2

3529 3527 3533 1 1.0950e+02 3.3472e+02 ; CD1 - CG
- CD2
3530 3529 3531 1 1.0950e+02 2.9288e+02 ; HD11 - CD1
- HD12
3530 3529 3532 1 1.0950e+02 2.9288e+02 ; HD11 - CD1
- HD13
3531 3529 3532 1 1.0950e+02 2.9288e+02 ; HD12 - CD1
- HD13
3534 3533 3535 1 1.0950e+02 2.9288e+02 ; HD21 - CD2
- HD22
3534 3533 3536 1 1.0950e+02 2.9288e+02 ; HD21 - CD2
- HD23
3535 3533 3536 1 1.0950e+02 2.9288e+02 ; HD22 - CD2
- HD23
3537 3539 3540 1 1.2000e+02 4.1840e+02 ; C - N
- H
3537 3539 3541 1 1.2190e+02 4.1840e+02 ; C - N
- CA
3538 3537 3539 1 1.2290e+02 6.6944e+02 ; O - C
- N
3539 3541 3542 1 1.0950e+02 4.1840e+02 ; N - CA
- HA
3539 3541 3543 1 1.0970e+02 6.6944e+02 ; N - CA
- CB
3539 3541 3559 1 1.1010e+02 5.2718e+02 ; N - CA
- C
3540 3539 3541 1 1.1804e+02 4.1840e+02 ; H - N
- CA
3541 3543 3544 1 1.0950e+02 4.1840e+02 ; CA - CB
- HB2
3541 3543 3545 1 1.0950e+02 4.1840e+02 ; CA - CB
- HB3
3541 3543 3546 1 1.0950e+02 3.3472e+02 ; CA - CB
- CG
3541 3559 3560 1 1.2040e+02 6.6944e+02 ; CA - C
- O
3541 3559 3561 1 1.1660e+02 5.8576e+02 ; CA - C
- N
3542 3541 3543 1 1.0950e+02 4.1840e+02 ; HA - CA
- CB
3542 3541 3559 1 1.0950e+02 4.1840e+02 ; HA - CA
- C
3543 3541 3559 1 1.1110e+02 5.2718e+02 ; CB - CA
- C
3543 3546 3547 1 1.0950e+02 4.1840e+02 ; CB - CG
- HG2
3543 3546 3548 1 1.0950e+02 4.1840e+02 ; CB - CG
- HG3
3543 3546 3549 1 1.0950e+02 3.3472e+02 ; CB - CG
- CD
3544 3543 3545 1 1.0950e+02 2.9288e+02 ; HB2 - CB
- HB3

```
  3544   3543   3546       1    1.0950e+02    4.1840e+02 ;    HB2 - CB
- CG
  3545   3543   3546       1    1.0950e+02    4.1840e+02 ;    HB3 - CB
- CG
  3546   3549   3550       1    1.0950e+02    4.1840e+02 ;     CG - CD
- HD2
  3546   3549   3551       1    1.0950e+02    4.1840e+02 ;     CG - CD
- HD3
  3546   3549   3552       1    1.0950e+02    3.3472e+02 ;     CG - CD
- CE
  3547   3546   3548       1    1.0950e+02    2.9288e+02 ;    HG2 - CG
- HG3
  3547   3546   3549       1    1.0950e+02    4.1840e+02 ;    HG2 - CG
- CD
  3548   3546   3549       1    1.0950e+02    4.1840e+02 ;    HG3 - CG
- CD
  3549   3552   3553       1    1.0950e+02    4.1840e+02 ;     CD - CE
- HE2
  3549   3552   3554       1    1.0950e+02    4.1840e+02 ;     CD - CE
- HE3
  3549   3552   3555       1    1.1120e+02    6.6944e+02 ;     CD - CE
- NZ
  3550   3549   3551       1    1.0950e+02    2.9288e+02 ;    HD2 - CD
- HD3
  3550   3549   3552       1    1.0950e+02    4.1840e+02 ;    HD2 - CD
- CE
  3551   3549   3552       1    1.0950e+02    4.1840e+02 ;    HD3 - CD
- CE
  3552   3555   3556       1    1.0950e+02    4.1840e+02 ;     CE - NZ
- HZ1
  3552   3555   3557       1    1.0950e+02    4.1840e+02 ;     CE - NZ
- HZ2
  3552   3555   3558       1    1.0950e+02    4.1840e+02 ;     CE - NZ
- HZ3
  3553   3552   3554       1    1.0950e+02    2.9288e+02 ;    HE2 - CE
- HE3
  3553   3552   3555       1    1.0950e+02    4.1840e+02 ;    HE2 - CE
- NZ
  3554   3552   3555       1    1.0950e+02    4.1840e+02 ;    HE3 - CE
- NZ
  3556   3555   3557       1    1.0950e+02    2.9288e+02 ;    HZ1 - NZ
- HZ2
  3556   3555   3558       1    1.0950e+02    2.9288e+02 ;    HZ1 - NZ
- HZ3
  3557   3555   3558       1    1.0950e+02    2.9288e+02 ;    HZ2 - NZ
- HZ3
  3559   3561   3562       1    1.2000e+02    4.1840e+02 ;      C - N
- H
  3559   3561   3563       1    1.2190e+02    4.1840e+02 ;      C - N
- CA
  3560   3559   3561       1    1.2290e+02    6.6944e+02 ;      O - C
- N
```

3561 3563 3564 1 1.0950e+02 4.1840e+02 ; N - CA - HA
3561 3563 3565 1 1.0970e+02 6.6944e+02 ; N - CA - CB
3561 3563 3581 1 1.1010e+02 5.2718e+02 ; N - CA - C
3562 3561 3563 1 1.1804e+02 4.1840e+02 ; H - N - CA
3563 3565 3566 1 1.0950e+02 4.1840e+02 ; CA - CB - HB2
3563 3565 3567 1 1.0950e+02 4.1840e+02 ; CA - CB - HB3
3563 3565 3568 1 1.0950e+02 3.3472e+02 ; CA - CB - CG
3563 3581 3582 1 1.2040e+02 6.6944e+02 ; CA - C - O
3563 3581 3583 1 1.1660e+02 5.8576e+02 ; CA - C - N
3564 3563 3565 1 1.0950e+02 4.1840e+02 ; HA - CA - CB
3564 3563 3581 1 1.0950e+02 4.1840e+02 ; HA - CA - C
3565 3563 3581 1 1.1110e+02 5.2718e+02 ; CB - CA - C
3565 3568 3569 1 1.0950e+02 4.1840e+02 ; CB - CG - HG2
3565 3568 3570 1 1.0950e+02 4.1840e+02 ; CB - CG - HG3
3565 3568 3571 1 1.0950e+02 3.3472e+02 ; CB - CG - CD
3566 3565 3567 1 1.0950e+02 2.9288e+02 ; HB2 - CB - HB3
3566 3565 3568 1 1.0950e+02 4.1840e+02 ; HB2 - CB - CG
3567 3565 3568 1 1.0950e+02 4.1840e+02 ; HB3 - CB - CG
3568 3571 3572 1 1.0950e+02 4.1840e+02 ; CG - CD - HD2
3568 3571 3573 1 1.0950e+02 4.1840e+02 ; CG - CD - HD3
3568 3571 3574 1 1.0950e+02 3.3472e+02 ; CG - CD - CE
3569 3568 3570 1 1.0950e+02 2.9288e+02 ; HG2 - CG - HG3
3569 3568 3571 1 1.0950e+02 4.1840e+02 ; HG2 - CG - CD
3570 3568 3571 1 1.0950e+02 4.1840e+02 ; HG3 - CG - CD
3571 3574 3575 1 1.0950e+02 4.1840e+02 ; CD - CE - HE2
3571 3574 3576 1 1.0950e+02 4.1840e+02 ; CD - CE - HE3

3571 3574 3577 1 1.1120e+02 6.6944e+02 ; CD - CE
- NZ
3572 3571 3573 1 1.0950e+02 2.9288e+02 ; HD2 - CD
- HD3
3572 3571 3574 1 1.0950e+02 4.1840e+02 ; HD2 - CD
- CE
3573 3571 3574 1 1.0950e+02 4.1840e+02 ; HD3 - CD
- CE
3574 3577 3578 1 1.0950e+02 4.1840e+02 ; CE - NZ
- HZ1
3574 3577 3579 1 1.0950e+02 4.1840e+02 ; CE - NZ
- HZ2
3574 3577 3580 1 1.0950e+02 4.1840e+02 ; CE - NZ
- HZ3
3575 3574 3576 1 1.0950e+02 2.9288e+02 ; HE2 - CE
- HE3
3575 3574 3577 1 1.0950e+02 4.1840e+02 ; HE2 - CE
- NZ
3576 3574 3577 1 1.0950e+02 4.1840e+02 ; HE3 - CE
- NZ
3578 3577 3579 1 1.0950e+02 2.9288e+02 ; HZ1 - NZ
- HZ2
3578 3577 3580 1 1.0950e+02 2.9288e+02 ; HZ1 - NZ
- HZ3
3579 3577 3580 1 1.0950e+02 2.9288e+02 ; HZ2 - NZ
- HZ3
3581 3583 3584 1 1.2000e+02 4.1840e+02 ; C - N
- H
3581 3583 3585 1 1.2190e+02 4.1840e+02 ; C - N
- CA
3582 3581 3583 1 1.2290e+02 6.6944e+02 ; O - C
- N
3583 3585 3586 1 1.0950e+02 4.1840e+02 ; N - CA
- HA
3583 3585 3587 1 1.0970e+02 6.6944e+02 ; N - CA
- CB
3583 3585 3597 1 1.1010e+02 5.2718e+02 ; N - CA
- C
3584 3583 3585 1 1.1804e+02 4.1840e+02 ; H - N
- CA
3585 3587 3588 1 1.0950e+02 4.1840e+02 ; CA - CB
- HB
3585 3587 3589 1 1.0950e+02 3.3472e+02 ; CA - CB
- CG1
3585 3587 3593 1 1.0950e+02 3.3472e+02 ; CA - CB
- CG2
3585 3597 3598 1 1.2040e+02 6.6944e+02 ; CA - C
- O
3585 3597 3599 1 1.1660e+02 5.8576e+02 ; CA - C
- N
3586 3585 3587 1 1.0950e+02 4.1840e+02 ; HA - CA
- CB

3586 3585 3597 1 1.0950e+02 4.1840e+02 ; HA - CA
- C
3587 3585 3597 1 1.1110e+02 5.2718e+02 ; CB - CA
- C
3587 3589 3590 1 1.0950e+02 4.1840e+02 ; CB - CG1
- HG11
3587 3589 3591 1 1.0950e+02 4.1840e+02 ; CB - CG1
- HG12
3587 3589 3592 1 1.0950e+02 4.1840e+02 ; CB - CG1
- HG13
3587 3593 3594 1 1.0950e+02 4.1840e+02 ; CB - CG2
- HG21
3587 3593 3595 1 1.0950e+02 4.1840e+02 ; CB - CG2
- HG22
3587 3593 3596 1 1.0950e+02 4.1840e+02 ; CB - CG2
- HG23
3588 3587 3589 1 1.0950e+02 4.1840e+02 ; HB - CB
- CG1
3588 3587 3593 1 1.0950e+02 4.1840e+02 ; HB - CB
- CG2
3589 3587 3593 1 1.0950e+02 3.3472e+02 ; CG1 - CB
- CG2
3590 3589 3591 1 1.0950e+02 2.9288e+02 ; HG11 - CG1
- HG12
3590 3589 3592 1 1.0950e+02 2.9288e+02 ; HG11 - CG1
- HG13
3591 3589 3592 1 1.0950e+02 2.9288e+02 ; HG12 - CG1
- HG13
3594 3593 3595 1 1.0950e+02 2.9288e+02 ; HG21 - CG2
- HG22
3594 3593 3596 1 1.0950e+02 2.9288e+02 ; HG21 - CG2
- HG23
3595 3593 3596 1 1.0950e+02 2.9288e+02 ; HG22 - CG2
- HG23
3597 3599 3600 1 1.2000e+02 4.1840e+02 ; C - N
- H
3597 3599 3601 1 1.2190e+02 4.1840e+02 ; C - N
- CA
3598 3597 3599 1 1.2290e+02 6.6944e+02 ; O - C
- N
3599 3601 3602 1 1.0950e+02 4.1840e+02 ; N - CA
- HA
3599 3601 3603 1 1.0970e+02 6.6944e+02 ; N - CA
- CB
3599 3601 3619 1 1.1010e+02 5.2718e+02 ; N - CA
- C
3600 3599 3601 1 1.1804e+02 4.1840e+02 ; H - N
- CA
3601 3603 3604 1 1.0950e+02 4.1840e+02 ; CA - CB
- HB2
3601 3603 3605 1 1.0950e+02 4.1840e+02 ; CA - CB
- HB3

```
  3601   3603   3606       1    1.0950e+02    3.3472e+02 ;     CA - CB
- CG
  3601   3619   3620       1    1.2040e+02    6.6944e+02 ;     CA - C
- O
  3601   3619   3621       1    1.1660e+02    5.8576e+02 ;     CA - C
- N
  3602   3601   3603       1    1.0950e+02    4.1840e+02 ;     HA - CA
- CB
  3602   3601   3619       1    1.0950e+02    4.1840e+02 ;     HA - CA
- C
  3603   3601   3619       1    1.1110e+02    5.2718e+02 ;     CB - CA
- C
  3603   3606   3607       1    1.0950e+02    4.1840e+02 ;     CB - CG
- HG2
  3603   3606   3608       1    1.0950e+02    4.1840e+02 ;     CB - CG
- HG3
  3603   3606   3609       1    1.0950e+02    3.3472e+02 ;     CB - CG
- CD
  3604   3603   3605       1    1.0950e+02    2.9288e+02 ;    HB2 - CB
- HB3
  3604   3603   3606       1    1.0950e+02    4.1840e+02 ;    HB2 - CB
- CG
  3605   3603   3606       1    1.0950e+02    4.1840e+02 ;    HB3 - CB
- CG
  3606   3609   3610       1    1.0950e+02    4.1840e+02 ;     CG - CD
- HD2
  3606   3609   3611       1    1.0950e+02    4.1840e+02 ;     CG - CD
- HD3
  3606   3609   3612       1    1.0950e+02    3.3472e+02 ;     CG - CD
- CE
  3607   3606   3608       1    1.0950e+02    2.9288e+02 ;    HG2 - CG
- HG3
  3607   3606   3609       1    1.0950e+02    4.1840e+02 ;    HG2 - CG
- CD
  3608   3606   3609       1    1.0950e+02    4.1840e+02 ;    HG3 - CG
- CD
  3609   3612   3613       1    1.0950e+02    4.1840e+02 ;     CD - CE
- HE2
  3609   3612   3614       1    1.0950e+02    4.1840e+02 ;     CD - CE
- HE3
  3609   3612   3615       1    1.1120e+02    6.6944e+02 ;     CD - CE
- NZ
  3610   3609   3611       1    1.0950e+02    2.9288e+02 ;    HD2 - CD
- HD3
  3610   3609   3612       1    1.0950e+02    4.1840e+02 ;    HD2 - CD
- CE
  3611   3609   3612       1    1.0950e+02    4.1840e+02 ;    HD3 - CD
- CE
  3612   3615   3616       1    1.0950e+02    4.1840e+02 ;     CE - NZ
- HZ1
  3612   3615   3617       1    1.0950e+02    4.1840e+02 ;     CE - NZ
- HZ2
```

3612 3615 3618 1 1.0950e+02 4.1840e+02 ; CE - NZ
- HZ3
3613 3612 3614 1 1.0950e+02 2.9288e+02 ; HE2 - CE
- HE3
3613 3612 3615 1 1.0950e+02 4.1840e+02 ; HE2 - CE
- NZ
3614 3612 3615 1 1.0950e+02 4.1840e+02 ; HE3 - CE
- NZ
3616 3615 3617 1 1.0950e+02 2.9288e+02 ; HZ1 - NZ
- HZ2
3616 3615 3618 1 1.0950e+02 2.9288e+02 ; HZ1 - NZ
- HZ3
3617 3615 3618 1 1.0950e+02 2.9288e+02 ; HZ2 - NZ
- HZ3
3619 3621 3622 1 1.2000e+02 4.1840e+02 ; C - N
- H
3619 3621 3623 1 1.2190e+02 4.1840e+02 ; C - N
- CA
3620 3619 3621 1 1.2290e+02 6.6944e+02 ; O - C
- N
3621 3623 3624 1 1.0950e+02 4.1840e+02 ; N - CA
- HA
3621 3623 3625 1 1.0970e+02 6.6944e+02 ; N - CA
- CB
3621 3623 3641 1 1.1010e+02 5.2718e+02 ; N - CA
- C
3622 3621 3623 1 1.1804e+02 4.1840e+02 ; H - N
- CA
3623 3625 3626 1 1.0950e+02 4.1840e+02 ; CA - CB
- HB2
3623 3625 3627 1 1.0950e+02 4.1840e+02 ; CA - CB
- HB3
3623 3625 3628 1 1.0950e+02 3.3472e+02 ; CA - CB
- CG
3623 3641 3642 1 1.2040e+02 6.6944e+02 ; CA - C
- O
3623 3641 3643 1 1.1660e+02 5.8576e+02 ; CA - C
- N
3624 3623 3625 1 1.0950e+02 4.1840e+02 ; HA - CA
- CB
3624 3623 3641 1 1.0950e+02 4.1840e+02 ; HA - CA
- C
3625 3623 3641 1 1.1110e+02 5.2718e+02 ; CB - CA
- C
3625 3628 3629 1 1.0950e+02 4.1840e+02 ; CB - CG
- HG2
3625 3628 3630 1 1.0950e+02 4.1840e+02 ; CB - CG
- HG3
3625 3628 3631 1 1.0950e+02 3.3472e+02 ; CB - CG
- CD
3626 3625 3627 1 1.0950e+02 2.9288e+02 ; HB2 - CB
- HB3

3626 3625 3628 1 1.0950e+02 4.1840e+02 ; HB2 - CB
- CG
3627 3625 3628 1 1.0950e+02 4.1840e+02 ; HB3 - CB
- CG
3628 3631 3632 1 1.0950e+02 4.1840e+02 ; CG - CD
- HD2
3628 3631 3633 1 1.0950e+02 4.1840e+02 ; CG - CD
- HD3
3628 3631 3634 1 1.0950e+02 3.3472e+02 ; CG - CD
- CE
3629 3628 3630 1 1.0950e+02 2.9288e+02 ; HG2 - CG
- HG3
3629 3628 3631 1 1.0950e+02 4.1840e+02 ; HG2 - CG
- CD
3630 3628 3631 1 1.0950e+02 4.1840e+02 ; HG3 - CG
- CD
3631 3634 3635 1 1.0950e+02 4.1840e+02 ; CD - CE
- HE2
3631 3634 3636 1 1.0950e+02 4.1840e+02 ; CD - CE
- HE3
3631 3634 3637 1 1.1120e+02 6.6944e+02 ; CD - CE
- NZ
3632 3631 3633 1 1.0950e+02 2.9288e+02 ; HD2 - CD
- HD3
3632 3631 3634 1 1.0950e+02 4.1840e+02 ; HD2 - CD
- CE
3633 3631 3634 1 1.0950e+02 4.1840e+02 ; HD3 - CD
- CE
3634 3637 3638 1 1.0950e+02 4.1840e+02 ; CE - NZ
- HZ1
3634 3637 3639 1 1.0950e+02 4.1840e+02 ; CE - NZ
- HZ2
3634 3637 3640 1 1.0950e+02 4.1840e+02 ; CE - NZ
- HZ3
3635 3634 3636 1 1.0950e+02 2.9288e+02 ; HE2 - CE
- HE3
3635 3634 3637 1 1.0950e+02 4.1840e+02 ; HE2 - CE
- NZ
3636 3634 3637 1 1.0950e+02 4.1840e+02 ; HE3 - CE
- NZ
3638 3637 3639 1 1.0950e+02 2.9288e+02 ; HZ1 - NZ
- HZ2
3638 3637 3640 1 1.0950e+02 2.9288e+02 ; HZ1 - NZ
- HZ3
3639 3637 3640 1 1.0950e+02 2.9288e+02 ; HZ2 - NZ
- HZ3
3641 3643 3644 1 1.2000e+02 4.1840e+02 ; C - N
- H
3641 3643 3645 1 1.2190e+02 4.1840e+02 ; C - N
- CA
3642 3641 3643 1 1.2290e+02 6.6944e+02 ; O - C
- N

```
  3643   3645   3646      1    1.0950e+02    4.1840e+02 ;      N - CA
- HA2
  3643   3645   3647      1    1.0950e+02    4.1840e+02 ;      N - CA
- HA3
  3643   3645   3648      1    1.1010e+02    5.2718e+02 ;      N - CA
- C
  3644   3643   3645      1    1.1804e+02    4.1840e+02 ;      H - N
- CA
  3645   3648   3649      1    1.2040e+02    6.6944e+02 ;     CA - C
- O
  3645   3648   3650      1    1.1660e+02    5.8576e+02 ;     CA - C
- N
  3646   3645   3647      1    1.0950e+02    2.9288e+02 ;    HA2 - CA
- HA3
  3646   3645   3648      1    1.0950e+02    4.1840e+02 ;    HA2 - CA
- C
  3647   3645   3648      1    1.0950e+02    4.1840e+02 ;    HA3 - CA
- C
  3648   3650   3651      1    1.2000e+02    4.1840e+02 ;      C - N
- H
  3648   3650   3652      1    1.2190e+02    4.1840e+02 ;      C - N
- CA
  3649   3648   3650      1    1.2290e+02    6.6944e+02 ;      O - C
- N
  3650   3652   3653      1    1.0950e+02    4.1840e+02 ;      N - CA
- HA
  3650   3652   3654      1    1.0970e+02    6.6944e+02 ;      N - CA
- CB
  3650   3652   3663      1    1.1010e+02    5.2718e+02 ;      N - CA
- C
  3651   3650   3652      1    1.1804e+02    4.1840e+02 ;      H - N
- CA
  3652   3654   3655      1    1.0950e+02    4.1840e+02 ;     CA - CB
- HB2
  3652   3654   3656      1    1.0950e+02    4.1840e+02 ;     CA - CB
- HB3
  3652   3654   3657      1    1.0950e+02    3.3472e+02 ;     CA - CB
- CG
  3652   3663   3664      1    1.2040e+02    6.6944e+02 ;     CA - C
- O
  3652   3663   3665      1    1.1660e+02    5.8576e+02 ;     CA - C
- N
  3653   3652   3654      1    1.0950e+02    4.1840e+02 ;     HA - CA
- CB
  3653   3652   3663      1    1.0950e+02    4.1840e+02 ;     HA - CA
- C
  3654   3652   3663      1    1.1110e+02    5.2718e+02 ;     CB - CA
- C
  3654   3657   3658      1    1.0950e+02    4.1840e+02 ;     CB - CG
- HG2
  3654   3657   3659      1    1.0950e+02    4.1840e+02 ;     CB - CG
- HG3
```

3654 3657 3660 1 1.1110e+02 5.2718e+02 ; CB - CG
- CD
3655 3654 3656 1 1.0950e+02 2.9288e+02 ; HB2 - CB
- HB3
3655 3654 3657 1 1.0950e+02 4.1840e+02 ; HB2 - CB
- CG
3656 3654 3657 1 1.0950e+02 4.1840e+02 ; HB3 - CB
- CG
3657 3660 3661 1 1.1700e+02 5.8576e+02 ; CG - CD
- OE1
3657 3660 3662 1 1.1700e+02 5.8576e+02 ; CG - CD
- OE2
3658 3657 3659 1 1.0950e+02 2.9288e+02 ; HG2 - CG
- HG3
3658 3657 3660 1 1.0950e+02 4.1840e+02 ; HG2 - CG
- CD
3659 3657 3660 1 1.0950e+02 4.1840e+02 ; HG3 - CG
- CD
3661 3660 3662 1 1.2600e+02 6.6944e+02 ; OE1 - CD
- OE2
3663 3665 3666 1 1.2000e+02 4.1840e+02 ; C - N
- H
3663 3665 3667 1 1.2190e+02 4.1840e+02 ; C - N
- CA
3664 3663 3665 1 1.2290e+02 6.6944e+02 ; O - C
- N
3665 3667 3668 1 1.0950e+02 4.1840e+02 ; N - CA
- HA
3665 3667 3669 1 1.0970e+02 6.6944e+02 ; N - CA
- CB
3665 3667 3683 1 1.1010e+02 5.2718e+02 ; N - CA
- C
3666 3665 3667 1 1.1804e+02 4.1840e+02 ; H - N
- CA
3667 3669 3670 1 1.0950e+02 4.1840e+02 ; CA - CB
- HB2
3667 3669 3671 1 1.0950e+02 4.1840e+02 ; CA - CB
- HB3
3667 3669 3672 1 1.1400e+02 5.2718e+02 ; CA - CB
- CG
3667 3683 3684 1 1.2040e+02 6.6944e+02 ; CA - C
- O
3667 3683 3685 1 1.1660e+02 5.8576e+02 ; CA - C
- N
3668 3667 3669 1 1.0950e+02 4.1840e+02 ; HA - CA
- CB
3668 3667 3683 1 1.0950e+02 4.1840e+02 ; HA - CA
- C
3669 3667 3683 1 1.1110e+02 5.2718e+02 ; CB - CA
- C
3669 3672 3673 1 1.2000e+02 5.8576e+02 ; CB - CG
- CD1

```
  3669   3672   3681       1    1.2000e+02    5.8576e+02 ;     CB - CG
- CD2
  3670   3669   3671       1    1.0950e+02    2.9288e+02 ;    HB2 - CB
- HB3
  3670   3669   3672       1    1.0950e+02    4.1840e+02 ;    HB2 - CB
- CG
  3671   3669   3672       1    1.0950e+02    4.1840e+02 ;    HB3 - CB
- CG
  3672   3673   3674       1    1.2000e+02    4.1840e+02 ;     CG - CD1
- HD1
  3672   3673   3675       1    1.2000e+02    5.2718e+02 ;     CG - CD1
- CE1
  3672   3681   3679       1    1.2000e+02    5.2718e+02 ;     CG - CD2
- CE2
  3672   3681   3682       1    1.2000e+02    4.1840e+02 ;     CG - CD2
- HD2
  3673   3672   3681       1    1.2000e+02    5.2718e+02 ;    CD1 - CG
- CD2
  3673   3675   3676       1    1.2000e+02    4.1840e+02 ;    CD1 - CE1
- HE1
  3673   3675   3677       1    1.2000e+02    5.2718e+02 ;    CD1 - CE1
- CZ
  3674   3673   3675       1    1.2000e+02    4.1840e+02 ;    HD1 - CD1
- CE1
  3675   3677   3678       1    1.2000e+02    4.1840e+02 ;    CE1 - CZ
- HZ
  3675   3677   3679       1    1.2000e+02    5.2718e+02 ;    CE1 - CZ
- CE2
  3676   3675   3677       1    1.2000e+02    4.1840e+02 ;    HE1 - CE1
- CZ
  3677   3679   3680       1    1.2000e+02    4.1840e+02 ;     CZ - CE2
- HE2
  3677   3679   3681       1    1.2000e+02    5.2718e+02 ;     CZ - CE2
- CD2
  3678   3677   3679       1    1.2000e+02    4.1840e+02 ;     HZ - CZ
- CE2
  3679   3681   3682       1    1.2000e+02    4.1840e+02 ;    CE2 - CD2
- HD2
  3680   3679   3681       1    1.2000e+02    4.1840e+02 ;    HE2 - CE2
- CD2
  3683   3685   3686       1    1.2000e+02    4.1840e+02 ;      C - N
- H
  3683   3685   3687       1    1.2190e+02    4.1840e+02 ;      C - N
- CA
  3684   3683   3685       1    1.2290e+02    6.6944e+02 ;      O - C
- N
  3685   3687   3688       1    1.0950e+02    4.1840e+02 ;      N - CA
- HA
  3685   3687   3689       1    1.0970e+02    6.6944e+02 ;      N - CA
- CB
  3685   3687   3694       1    1.1010e+02    5.2718e+02 ;      N - CA
- C
```

3686 3685 3687 1 1.1804e+02 4.1840e+02 ; H - N
- CA
3687 3689 3690 1 1.0950e+02 4.1840e+02 ; CA - CB
- HB2
3687 3689 3691 1 1.0950e+02 4.1840e+02 ; CA - CB
- HB3
3687 3689 3692 1 1.0860e+02 4.1840e+02 ; CA - CB
- SG
3687 3694 3695 1 1.2040e+02 6.6944e+02 ; CA - C
- O
3687 3694 3696 1 1.1660e+02 5.8576e+02 ; CA - C
- N
3688 3687 3689 1 1.0950e+02 4.1840e+02 ; HA - CA
- CB
3688 3687 3694 1 1.0950e+02 4.1840e+02 ; HA - CA
- C
3689 3687 3694 1 1.1110e+02 5.2718e+02 ; CB - CA
- C
3689 3692 3693 1 9.6000e+01 3.5982e+02 ; CB - SG
- HG
3690 3689 3691 1 1.0950e+02 2.9288e+02 ; HB2 - CB
- HB3
3690 3689 3692 1 1.0950e+02 4.1840e+02 ; HB2 - CB
- SG
3691 3689 3692 1 1.0950e+02 4.1840e+02 ; HB3 - CB
- SG
3694 3696 3697 1 1.2000e+02 4.1840e+02 ; C - N
- H
3694 3696 3698 1 1.2190e+02 4.1840e+02 ; C - N
- CA
3695 3694 3696 1 1.2290e+02 6.6944e+02 ; O - C
- N
3696 3698 3699 1 1.0950e+02 4.1840e+02 ; N - CA
- HA
3696 3698 3700 1 1.0970e+02 6.6944e+02 ; N - CA
- CB
3696 3698 3714 1 1.1010e+02 5.2718e+02 ; N - CA
- C
3697 3696 3698 1 1.1804e+02 4.1840e+02 ; H - N
- CA
3698 3700 3701 1 1.0950e+02 4.1840e+02 ; CA - CB
- HB2
3698 3700 3702 1 1.0950e+02 4.1840e+02 ; CA - CB
- HB3
3698 3700 3703 1 1.1400e+02 5.2718e+02 ; CA - CB
- CG
3698 3714 3715 1 1.2040e+02 6.6944e+02 ; CA - C
- O
3698 3714 3716 1 1.1660e+02 5.8576e+02 ; CA - C
- N
3699 3698 3700 1 1.0950e+02 4.1840e+02 ; HA - CA
- CB

| | | | | | | | |
|---|---|---|---|---|---|---|---|
| 3699 | 3698 | 3714 | 1 | 1.0950e+02 | 4.1840e+02 | ; | HA - CA - C |
| 3700 | 3698 | 3714 | 1 | 1.1110e+02 | 5.2718e+02 | ; | CB - CA - C |
| 3700 | 3703 | 3704 | 1 | 1.2000e+02 | 5.8576e+02 | ; | CB - CG - CD1 |
| 3700 | 3703 | 3712 | 1 | 1.2000e+02 | 5.8576e+02 | ; | CB - CG - CD2 |
| 3701 | 3700 | 3702 | 1 | 1.0950e+02 | 2.9288e+02 | ; | HB2 - CB - HB3 |
| 3701 | 3700 | 3703 | 1 | 1.0950e+02 | 4.1840e+02 | ; | HB2 - CB - CG |
| 3702 | 3700 | 3703 | 1 | 1.0950e+02 | 4.1840e+02 | ; | HB3 - CB - CG |
| 3703 | 3704 | 3705 | 1 | 1.2000e+02 | 4.1840e+02 | ; | CG - CD1 - HD1 |
| 3703 | 3704 | 3706 | 1 | 1.2000e+02 | 5.2718e+02 | ; | CG - CD1 - CE1 |
| 3703 | 3712 | 3710 | 1 | 1.2000e+02 | 5.2718e+02 | ; | CG - CD2 - CE2 |
| 3703 | 3712 | 3713 | 1 | 1.2000e+02 | 4.1840e+02 | ; | CG - CD2 - HD2 |
| 3704 | 3703 | 3712 | 1 | 1.2000e+02 | 5.2718e+02 | ; | CD1 - CG - CD2 |
| 3704 | 3706 | 3707 | 1 | 1.2000e+02 | 4.1840e+02 | ; | CD1 - CE1 - HE1 |
| 3704 | 3706 | 3708 | 1 | 1.2000e+02 | 5.2718e+02 | ; | CD1 - CE1 - CZ |
| 3705 | 3704 | 3706 | 1 | 1.2000e+02 | 4.1840e+02 | ; | HD1 - CD1 - CE1 |
| 3706 | 3708 | 3709 | 1 | 1.2000e+02 | 4.1840e+02 | ; | CE1 - CZ - HZ |
| 3706 | 3708 | 3710 | 1 | 1.2000e+02 | 5.2718e+02 | ; | CE1 - CZ - CE2 |
| 3707 | 3706 | 3708 | 1 | 1.2000e+02 | 4.1840e+02 | ; | HE1 - CE1 - CZ |
| 3708 | 3710 | 3711 | 1 | 1.2000e+02 | 4.1840e+02 | ; | CZ - CE2 - HE2 |
| 3708 | 3710 | 3712 | 1 | 1.2000e+02 | 5.2718e+02 | ; | CZ - CE2 - CD2 |
| 3709 | 3708 | 3710 | 1 | 1.2000e+02 | 4.1840e+02 | ; | HZ - CZ - CE2 |
| 3710 | 3712 | 3713 | 1 | 1.2000e+02 | 4.1840e+02 | ; | CE2 - CD2 - HD2 |
| 3711 | 3710 | 3712 | 1 | 1.2000e+02 | 4.1840e+02 | ; | HE2 - CE2 - CD2 |
| 3714 | 3716 | 3717 | 1 | 1.2000e+02 | 4.1840e+02 | ; | C - N - H |
| 3714 | 3716 | 3718 | 1 | 1.2190e+02 | 4.1840e+02 | ; | C - N - CA |
| 3715 | 3714 | 3716 | 1 | 1.2290e+02 | 6.6944e+02 | ; | O - C - N |

3716 3718 3719 1 1.0950e+02 4.1840e+02 ; N - CA
- HA
3716 3718 3720 1 1.0970e+02 6.6944e+02 ; N - CA
- CB
3716 3718 3735 1 1.1010e+02 5.2718e+02 ; N - CA
- C
3717 3716 3718 1 1.1804e+02 4.1840e+02 ; H - N
- CA
3718 3720 3721 1 1.0950e+02 4.1840e+02 ; CA - CB
- HB2
3718 3720 3722 1 1.0950e+02 4.1840e+02 ; CA - CB
- HB3
3718 3720 3723 1 1.1400e+02 5.2718e+02 ; CA - CB
- CG
3718 3735 3736 1 1.2040e+02 6.6944e+02 ; CA - C
- O
3718 3735 3737 1 1.1660e+02 5.8576e+02 ; CA - C
- N
3719 3718 3720 1 1.0950e+02 4.1840e+02 ; HA - CA
- CB
3719 3718 3735 1 1.0950e+02 4.1840e+02 ; HA - CA
- C
3720 3718 3735 1 1.1110e+02 5.2718e+02 ; CB - CA
- C
3720 3723 3724 1 1.2000e+02 5.8576e+02 ; CB - CG
- CD1
3720 3723 3733 1 1.2000e+02 5.8576e+02 ; CB - CG
- CD2
3721 3720 3722 1 1.0950e+02 2.9288e+02 ; HB2 - CB
- HB3
3721 3720 3723 1 1.0950e+02 4.1840e+02 ; HB2 - CB
- CG
3722 3720 3723 1 1.0950e+02 4.1840e+02 ; HB3 - CB
- CG
3723 3724 3725 1 1.2000e+02 4.1840e+02 ; CG - CD1
- HD1
3723 3724 3726 1 1.2000e+02 5.2718e+02 ; CG - CD1
- CE1
3723 3733 3731 1 1.2000e+02 5.2718e+02 ; CG - CD2
- CE2
3723 3733 3734 1 1.2000e+02 4.1840e+02 ; CG - CD2
- HD2
3724 3723 3733 1 1.2000e+02 5.2718e+02 ; CD1 - CG
- CD2
3724 3726 3727 1 1.2000e+02 4.1840e+02 ; CD1 - CE1
- HE1
3724 3726 3728 1 1.2000e+02 5.2718e+02 ; CD1 - CE1
- CZ
3725 3724 3726 1 1.2000e+02 4.1840e+02 ; HD1 - CD1
- CE1
3726 3728 3729 1 1.2000e+02 5.8576e+02 ; CE1 - CZ
- OH

3726 3728 3731 1 1.2000e+02 5.2718e+02 ; CE1 - CZ
- CE2
3727 3726 3728 1 1.2000e+02 4.1840e+02 ; HE1 - CE1
- CZ
3728 3729 3730 1 1.1300e+02 4.1840e+02 ; CZ - OH
- HH
3728 3731 3732 1 1.2000e+02 4.1840e+02 ; CZ - CE2
- HE2
3728 3731 3733 1 1.2000e+02 5.2718e+02 ; CZ - CE2
- CD2
3729 3728 3731 1 1.2000e+02 5.8576e+02 ; OH - CZ
- CE2
3731 3733 3734 1 1.2000e+02 4.1840e+02 ; CE2 - CD2
- HD2
3732 3731 3733 1 1.2000e+02 4.1840e+02 ; HE2 - CE2
- CD2
3735 3737 3738 1 1.2000e+02 4.1840e+02 ; C - N
- H
3735 3737 3739 1 1.2190e+02 4.1840e+02 ; C - N
- CA
3736 3735 3737 1 1.2290e+02 6.6944e+02 ; O - C
- N
3737 3739 3740 1 1.0950e+02 4.1840e+02 ; N - CA
- HA
3737 3739 3741 1 1.0970e+02 6.6944e+02 ; N - CA
- CB
3737 3739 3750 1 1.1010e+02 5.2718e+02 ; N - CA
- C
3738 3737 3739 1 1.1804e+02 4.1840e+02 ; H - N
- CA
3739 3741 3742 1 1.0950e+02 4.1840e+02 ; CA - CB
- HB2
3739 3741 3743 1 1.0950e+02 4.1840e+02 ; CA - CB
- HB3
3739 3741 3744 1 1.0950e+02 3.3472e+02 ; CA - CB
- CG
3739 3750 3751 1 1.2040e+02 6.6944e+02 ; CA - C
- O
3739 3750 3752 1 1.1660e+02 5.8576e+02 ; CA - C
- N
3740 3739 3741 1 1.0950e+02 4.1840e+02 ; HA - CA
- CB
3740 3739 3750 1 1.0950e+02 4.1840e+02 ; HA - CA
- C
3741 3739 3750 1 1.1110e+02 5.2718e+02 ; CB - CA
- C
3741 3744 3745 1 1.0950e+02 4.1840e+02 ; CB - CG
- HG2
3741 3744 3746 1 1.0950e+02 4.1840e+02 ; CB - CG
- HG3
3741 3744 3747 1 1.1110e+02 5.2718e+02 ; CB - CG
- CD

3742 3741 3743 1 1.0950e+02 2.9288e+02 ; HB2 - CB
- HB3
3742 3741 3744 1 1.0950e+02 4.1840e+02 ; HB2 - CB
- CG
3743 3741 3744 1 1.0950e+02 4.1840e+02 ; HB3 - CB
- CG
3744 3747 3748 1 1.1700e+02 5.8576e+02 ; CG - CD
- OE1
3744 3747 3749 1 1.1700e+02 5.8576e+02 ; CG - CD
- OE2
3745 3744 3746 1 1.0950e+02 2.9288e+02 ; HG2 - CG
- HG3
3745 3744 3747 1 1.0950e+02 4.1840e+02 ; HG2 - CG
- CD
3746 3744 3747 1 1.0950e+02 4.1840e+02 ; HG3 - CG
- CD
3748 3747 3749 1 1.2600e+02 6.6944e+02 ; OE1 - CD
- OE2
3750 3752 3753 1 1.2000e+02 4.1840e+02 ; C - N
- H
3750 3752 3754 1 1.2190e+02 4.1840e+02 ; C - N
- CA
3751 3750 3752 1 1.2290e+02 6.6944e+02 ; O - C
- N
3752 3754 3755 1 1.0950e+02 4.1840e+02 ; N - CA
- HA
3752 3754 3756 1 1.0970e+02 6.6944e+02 ; N - CA
- CB
3752 3754 3764 1 1.1010e+02 5.2718e+02 ; N - CA
- C
3753 3752 3754 1 1.1804e+02 4.1840e+02 ; H - N
- CA
3754 3756 3757 1 1.0950e+02 4.1840e+02 ; CA - CB
- HB2
3754 3756 3758 1 1.0950e+02 4.1840e+02 ; CA - CB
- HB3
3754 3756 3759 1 1.1110e+02 5.2718e+02 ; CA - CB
- CG
3754 3764 3765 1 1.2040e+02 6.6944e+02 ; CA - C
- O
3754 3764 3766 1 1.1660e+02 5.8576e+02 ; CA - C
- N
3755 3754 3756 1 1.0950e+02 4.1840e+02 ; HA - CA
- CB
3755 3754 3764 1 1.0950e+02 4.1840e+02 ; HA - CA
- C
3756 3754 3764 1 1.1110e+02 5.2718e+02 ; CB - CA
- C
3756 3759 3760 1 1.2040e+02 6.6944e+02 ; CB - CG
- OD1
3756 3759 3761 1 1.1660e+02 5.8576e+02 ; CB - CG
- ND2

| | | | | | | | |
|---|---|---|---|---|---|---|---|
| 3757 | 3756 | 3758 | 1 | 1.0950e+02 | 2.9288e+02 | ; | HB2 - CB - HB3 |
| 3757 | 3756 | 3759 | 1 | 1.0950e+02 | 4.1840e+02 | ; | HB2 - CB - CG |
| 3758 | 3756 | 3759 | 1 | 1.0950e+02 | 4.1840e+02 | ; | HB3 - CB - CG |
| 3759 | 3761 | 3762 | 1 | 1.2000e+02 | 4.1840e+02 | ; | CG - ND2 - HD21 |
| 3759 | 3761 | 3763 | 1 | 1.2000e+02 | 4.1840e+02 | ; | CG - ND2 - HD22 |
| 3760 | 3759 | 3761 | 1 | 1.2290e+02 | 6.6944e+02 | ; | OD1 - CG - ND2 |
| 3762 | 3761 | 3763 | 1 | 1.2000e+02 | 2.9288e+02 | ; | HD21 - ND2 - HD22 |
| 3764 | 3766 | 3767 | 1 | 1.2000e+02 | 4.1840e+02 | ; | C - N - H |
| 3764 | 3766 | 3768 | 1 | 1.2190e+02 | 4.1840e+02 | ; | C - N - CA |
| 3765 | 3764 | 3766 | 1 | 1.2290e+02 | 6.6944e+02 | ; | O - C - N |
| 3766 | 3768 | 3769 | 1 | 1.0950e+02 | 4.1840e+02 | ; | N - CA - HA |
| 3766 | 3768 | 3770 | 1 | 1.0970e+02 | 6.6944e+02 | ; | N - CA - CB |
| 3766 | 3768 | 3776 | 1 | 1.1010e+02 | 5.2718e+02 | ; | N - CA - C |
| 3767 | 3766 | 3768 | 1 | 1.1804e+02 | 4.1840e+02 | ; | H - N - CA |
| 3768 | 3770 | 3771 | 1 | 1.0950e+02 | 4.1840e+02 | ; | CA - CB - HB2 |
| 3768 | 3770 | 3772 | 1 | 1.0950e+02 | 4.1840e+02 | ; | CA - CB - HB3 |
| 3768 | 3770 | 3773 | 1 | 1.1110e+02 | 5.2718e+02 | ; | CA - CB - CG |
| 3768 | 3776 | 3777 | 1 | 1.2040e+02 | 6.6944e+02 | ; | CA - C - O |
| 3768 | 3776 | 3778 | 1 | 1.1660e+02 | 5.8576e+02 | ; | CA - C - N |
| 3769 | 3768 | 3770 | 1 | 1.0950e+02 | 4.1840e+02 | ; | HA - CA - CB |
| 3769 | 3768 | 3776 | 1 | 1.0950e+02 | 4.1840e+02 | ; | HA - CA - C |
| 3770 | 3768 | 3776 | 1 | 1.1110e+02 | 5.2718e+02 | ; | CB - CA - C |
| 3770 | 3773 | 3774 | 1 | 1.1700e+02 | 5.8576e+02 | ; | CB - CG - OD1 |
| 3770 | 3773 | 3775 | 1 | 1.1700e+02 | 5.8576e+02 | ; | CB - CG - OD2 |
| 3771 | 3770 | 3772 | 1 | 1.0950e+02 | 2.9288e+02 | ; | HB2 - CB - HB3 |
| 3771 | 3770 | 3773 | 1 | 1.0950e+02 | 4.1840e+02 | ; | HB2 - CB - CG |

3772 3770 3773 1 1.0950e+02 4.1840e+02 ; HB3 - CB
- CG
3774 3773 3775 1 1.2600e+02 6.6944e+02 ; OD1 - CG
- OD2
3776 3778 3779 1 1.2000e+02 4.1840e+02 ; C - N
- H
3776 3778 3780 1 1.2190e+02 4.1840e+02 ; C - N
- CA
3777 3776 3778 1 1.2290e+02 6.6944e+02 ; O - C
- N
3778 3780 3781 1 1.0950e+02 4.1840e+02 ; N - CA
- HA
3778 3780 3782 1 1.0970e+02 6.6944e+02 ; N - CA
- CB
3778 3780 3800 1 1.1010e+02 5.2718e+02 ; N - CA
- C
3779 3778 3780 1 1.1804e+02 4.1840e+02 ; H - N
- CA
3780 3782 3783 1 1.0950e+02 4.1840e+02 ; CA - CB
- HB2
3780 3782 3784 1 1.0950e+02 4.1840e+02 ; CA - CB
- HB3
3780 3782 3785 1 1.1560e+02 5.2718e+02 ; CA - CB
- CG
3780 3800 3801 1 1.2040e+02 6.6944e+02 ; CA - C
- O
3780 3800 3802 1 1.1660e+02 5.8576e+02 ; CA - C
- N
3781 3780 3782 1 1.0950e+02 4.1840e+02 ; HA - CA
- CB
3781 3780 3800 1 1.0950e+02 4.1840e+02 ; HA - CA
- C
3782 3780 3800 1 1.1110e+02 5.2718e+02 ; CB - CA
- C
3782 3785 3786 1 1.2500e+02 5.8576e+02 ; CB - CG
- CD1
3782 3785 3799 1 1.2860e+02 5.8576e+02 ; CB - CG
- CD2
3783 3782 3784 1 1.0950e+02 2.9288e+02 ; HB2 - CB
- HB3
3783 3782 3785 1 1.0950e+02 4.1840e+02 ; HB2 - CB
- CG
3784 3782 3785 1 1.0950e+02 4.1840e+02 ; HB3 - CB
- CG
3785 3786 3787 1 1.2000e+02 4.1840e+02 ; CG - CD1
- HD1
3785 3786 3788 1 1.0870e+02 5.8576e+02 ; CG - CD1
- NE1
3785 3799 3790 1 1.0880e+02 5.2718e+02 ; CG - CD2
- CE2
3785 3799 3797 1 1.3490e+02 5.2718e+02 ; CG - CD2
- CE3

```
  3786   3785   3799      1    1.0640e+02    5.2718e+02 ;    CD1 - CG
- CD2
  3786   3788   3789      1    1.2000e+02    4.1840e+02 ;    CD1 - NE1
- HE1
  3786   3788   3790      1    1.1160e+02    5.8576e+02 ;    CD1 - NE1
- CE2
  3787   3786   3788      1    1.2000e+02    4.1840e+02 ;    HD1 - CD1
- NE1
  3788   3790   3791      1    1.3280e+02    5.8576e+02 ;    NE1 - CE2
- CZ2
  3788   3790   3799      1    1.0440e+02    5.8576e+02 ;    NE1 - CE2
- CD2
  3789   3788   3790      1    1.2310e+02    4.1840e+02 ;    HE1 - NE1
- CE2
  3790   3791   3792      1    1.2000e+02    4.1840e+02 ;    CE2 - CZ2
- HZ2
  3790   3791   3793      1    1.2000e+02    5.2718e+02 ;    CE2 - CZ2
- CH2
  3790   3799   3797      1    1.1620e+02    5.2718e+02 ;    CE2 - CD2
- CE3
  3791   3790   3799      1    1.2270e+02    5.2718e+02 ;    CZ2 - CE2
- CD2
  3791   3793   3794      1    1.2000e+02    4.1840e+02 ;    CZ2 - CH2
- HH2
  3791   3793   3795      1    1.2000e+02    5.2718e+02 ;    CZ2 - CH2
- CZ3
  3792   3791   3793      1    1.2000e+02    4.1840e+02 ;    HZ2 - CZ2
- CH2
  3793   3795   3796      1    1.2000e+02    4.1840e+02 ;    CH2 - CZ3
- HZ3
  3793   3795   3797      1    1.2000e+02    5.2718e+02 ;    CH2 - CZ3
- CE3
  3794   3793   3795      1    1.2000e+02    4.1840e+02 ;    HH2 - CH2
- CZ3
  3795   3797   3798      1    1.2000e+02    4.1840e+02 ;    CZ3 - CE3
- HE3
  3795   3797   3799      1    1.2000e+02    5.2718e+02 ;    CZ3 - CE3
- CD2
  3796   3795   3797      1    1.2000e+02    4.1840e+02 ;    HZ3 - CZ3
- CE3
  3798   3797   3799      1    1.2000e+02    4.1840e+02 ;    HE3 - CE3
- CD2
  3800   3802   3803      1    1.2000e+02    4.1840e+02 ;      C - N
- H
  3800   3802   3804      1    1.2190e+02    4.1840e+02 ;      C - N
- CA
  3801   3800   3802      1    1.2290e+02    6.6944e+02 ;      O - C
- N
  3802   3804   3805      1    1.0950e+02    4.1840e+02 ;      N - CA
- HA2
  3802   3804   3806      1    1.0950e+02    4.1840e+02 ;      N - CA
- HA3
```

```
  3802   3804   3807       1    1.1010e+02    5.2718e+02 ;      N - CA
- C
  3803   3802   3804       1    1.1804e+02    4.1840e+02 ;      H - N
- CA
  3804   3807   3808       1    1.2040e+02    6.6944e+02 ;     CA - C
- O
  3804   3807   3809       1    1.1660e+02    5.8576e+02 ;     CA - C
- N
  3805   3804   3806       1    1.0950e+02    2.9288e+02 ;    HA2 - CA
- HA3
  3805   3804   3807       1    1.0950e+02    4.1840e+02 ;    HA2 - CA
- C
  3806   3804   3807       1    1.0950e+02    4.1840e+02 ;    HA3 - CA
- C
  3807   3809   3810       1    1.2000e+02    4.1840e+02 ;      C - N
- H
  3807   3809   3811       1    1.2190e+02    4.1840e+02 ;      C - N
- CA
  3808   3807   3809       1    1.2290e+02    6.6944e+02 ;      O - C
- N
  3809   3811   3812       1    1.0950e+02    4.1840e+02 ;      N - CA
- HA
  3809   3811   3813       1    1.0970e+02    6.6944e+02 ;      N - CA
- CB
  3809   3811   3818       1    1.1010e+02    5.2718e+02 ;      N - CA
- C
  3810   3809   3811       1    1.1804e+02    4.1840e+02 ;      H - N
- CA
  3811   3813   3814       1    1.0950e+02    4.1840e+02 ;     CA - CB
- HB2
  3811   3813   3815       1    1.0950e+02    4.1840e+02 ;     CA - CB
- HB3
  3811   3813   3816       1    1.0950e+02    4.1840e+02 ;     CA - CB
- OG
  3811   3818   3819       1    1.2040e+02    6.6944e+02 ;     CA - C
- O
  3811   3818   3820       1    1.1660e+02    5.8576e+02 ;     CA - C
- N
  3812   3811   3813       1    1.0950e+02    4.1840e+02 ;     HA - CA
- CB
  3812   3811   3818       1    1.0950e+02    4.1840e+02 ;     HA - CA
- C
  3813   3811   3818       1    1.1110e+02    5.2718e+02 ;     CB - CA
- C
  3813   3816   3817       1    1.0850e+02    4.6024e+02 ;     CB - OG
- HG
  3814   3813   3815       1    1.0950e+02    2.9288e+02 ;    HB2 - CB
- HB3
  3814   3813   3816       1    1.0950e+02    4.1840e+02 ;    HB2 - CB
- OG
  3815   3813   3816       1    1.0950e+02    4.1840e+02 ;    HB3 - CB
- OG
```

3818 3820 3821 1 1.2000e+02 4.1840e+02 ; C - N
- H
3818 3820 3822 1 1.2190e+02 4.1840e+02 ; C - N
- CA
3819 3818 3820 1 1.2290e+02 6.6944e+02 ; O - C
- N
3820 3822 3823 1 1.0950e+02 4.1840e+02 ; N - CA
- HA
3820 3822 3824 1 1.0970e+02 6.6944e+02 ; N - CA
- CB
3820 3822 3837 1 1.1010e+02 5.2718e+02 ; N - CA
- C
3821 3820 3822 1 1.1804e+02 4.1840e+02 ; H - N
- CA
3822 3824 3825 1 1.0950e+02 4.1840e+02 ; CA - CB
- HB
3822 3824 3826 1 1.0950e+02 3.3472e+02 ; CA - CB
- CG2
3822 3824 3830 1 1.0950e+02 3.3472e+02 ; CA - CB
- CG1
3822 3837 3838 1 1.2040e+02 6.6944e+02 ; CA - C
- O
3822 3837 3839 1 1.1660e+02 5.8576e+02 ; CA - C
- N
3823 3822 3824 1 1.0950e+02 4.1840e+02 ; HA - CA
- CB
3823 3822 3837 1 1.0950e+02 4.1840e+02 ; HA - CA
- C
3824 3822 3837 1 1.1110e+02 5.2718e+02 ; CB - CA
- C
3824 3826 3827 1 1.0950e+02 4.1840e+02 ; CB - CG2
- HG21
3824 3826 3828 1 1.0950e+02 4.1840e+02 ; CB - CG2
- HG22
3824 3826 3829 1 1.0950e+02 4.1840e+02 ; CB - CG2
- HG23
3824 3830 3831 1 1.0950e+02 4.1840e+02 ; CB - CG1
- HG12
3824 3830 3832 1 1.0950e+02 4.1840e+02 ; CB - CG1
- HG13
3824 3830 3833 1 1.0950e+02 3.3472e+02 ; CB - CG1
- CD1
3825 3824 3826 1 1.0950e+02 4.1840e+02 ; HB - CB
- CG2
3825 3824 3830 1 1.0950e+02 4.1840e+02 ; HB - CB
- CG1
3826 3824 3830 1 1.0950e+02 3.3472e+02 ; CG2 - CB
- CG1
3827 3826 3828 1 1.0950e+02 2.9288e+02 ; HG21 - CG2
- HG22
3827 3826 3829 1 1.0950e+02 2.9288e+02 ; HG21 - CG2
- HG23

3828 3826 3829 1 1.0950e+02 2.9288e+02 ; HG22 - CG2
- HG23
3830 3833 3834 1 1.0950e+02 4.1840e+02 ; CG1 - CD1
- HD11
3830 3833 3835 1 1.0950e+02 4.1840e+02 ; CG1 - CD1
- HD12
3830 3833 3836 1 1.0950e+02 4.1840e+02 ; CG1 - CD1
- HD13
3831 3830 3832 1 1.0950e+02 2.9288e+02 ; HG12 - CG1
- HG13
3831 3830 3833 1 1.0950e+02 4.1840e+02 ; HG12 - CG1
- CD1
3832 3830 3833 1 1.0950e+02 4.1840e+02 ; HG13 - CG1
- CD1
3834 3833 3835 1 1.0950e+02 2.9288e+02 ; HD11 - CD1
- HD12
3834 3833 3836 1 1.0950e+02 2.9288e+02 ; HD11 - CD1
- HD13
3835 3833 3836 1 1.0950e+02 2.9288e+02 ; HD12 - CD1
- HD13
3837 3839 3840 1 1.2000e+02 4.1840e+02 ; C - N
- H
3837 3839 3841 1 1.2190e+02 4.1840e+02 ; C - N
- CA
3838 3837 3839 1 1.2290e+02 6.6944e+02 ; O - C
- N
3839 3841 3842 1 1.0950e+02 4.1840e+02 ; N - CA
- HA
3839 3841 3843 1 1.0970e+02 6.6944e+02 ; N - CA
- CB
3839 3841 3848 1 1.1010e+02 5.2718e+02 ; N - CA
- C
3840 3839 3841 1 1.1804e+02 4.1840e+02 ; H - N
- CA
3841 3843 3844 1 1.0950e+02 4.1840e+02 ; CA - CB
- HB2
3841 3843 3845 1 1.0950e+02 4.1840e+02 ; CA - CB
- HB3
3841 3843 3846 1 1.0950e+02 4.1840e+02 ; CA - CB
- OG
3841 3848 3849 1 1.2040e+02 6.6944e+02 ; CA - C
- O
3841 3848 3850 1 1.1660e+02 5.8576e+02 ; CA - C
- N
3842 3841 3843 1 1.0950e+02 4.1840e+02 ; HA - CA
- CB
3842 3841 3848 1 1.0950e+02 4.1840e+02 ; HA - CA
- C
3843 3841 3848 1 1.1110e+02 5.2718e+02 ; CB - CA
- C
3843 3846 3847 1 1.0850e+02 4.6024e+02 ; CB - OG
- HG

| | | | | | | | |
|---|---|---|---|---|---|---|---|
| 3844 | 3843 | 3845 | 1 | 1.0950e+02 | 2.9288e+02 | ; | HB2 - CB - HB3 |
| 3844 | 3843 | 3846 | 1 | 1.0950e+02 | 4.1840e+02 | ; | HB2 - CB - OG |
| 3845 | 3843 | 3846 | 1 | 1.0950e+02 | 4.1840e+02 | ; | HB3 - CB - OG |
| 3848 | 3850 | 3851 | 1 | 1.2000e+02 | 4.1840e+02 | ; | C - N - H |
| 3848 | 3850 | 3852 | 1 | 1.2190e+02 | 4.1840e+02 | ; | C - N - CA |
| 3849 | 3848 | 3850 | 1 | 1.2290e+02 | 6.6944e+02 | ; | O - C - N |
| 3850 | 3852 | 3853 | 1 | 1.0950e+02 | 4.1840e+02 | ; | N - CA - HA |
| 3850 | 3852 | 3854 | 1 | 1.0970e+02 | 6.6944e+02 | ; | N - CA - CB |
| 3850 | 3852 | 3859 | 1 | 1.1010e+02 | 5.2718e+02 | ; | N - CA - C |
| 3851 | 3850 | 3852 | 1 | 1.1804e+02 | 4.1840e+02 | ; | H - N - CA |
| 3852 | 3854 | 3855 | 1 | 1.0950e+02 | 4.1840e+02 | ; | CA - CB - HB2 |
| 3852 | 3854 | 3856 | 1 | 1.0950e+02 | 4.1840e+02 | ; | CA - CB - HB3 |
| 3852 | 3854 | 3857 | 1 | 1.0950e+02 | 4.1840e+02 | ; | CA - CB - OG |
| 3852 | 3859 | 3860 | 1 | 1.2040e+02 | 6.6944e+02 | ; | CA - C - O |
| 3852 | 3859 | 3861 | 1 | 1.1660e+02 | 5.8576e+02 | ; | CA - C - N |
| 3853 | 3852 | 3854 | 1 | 1.0950e+02 | 4.1840e+02 | ; | HA - CA - CB |
| 3853 | 3852 | 3859 | 1 | 1.0950e+02 | 4.1840e+02 | ; | HA - CA - C |
| 3854 | 3852 | 3859 | 1 | 1.1110e+02 | 5.2718e+02 | ; | CB - CA - C |
| 3854 | 3857 | 3858 | 1 | 1.0850e+02 | 4.6024e+02 | ; | CB - OG - HG |
| 3855 | 3854 | 3856 | 1 | 1.0950e+02 | 2.9288e+02 | ; | HB2 - CB - HB3 |
| 3855 | 3854 | 3857 | 1 | 1.0950e+02 | 4.1840e+02 | ; | HB2 - CB - OG |
| 3856 | 3854 | 3857 | 1 | 1.0950e+02 | 4.1840e+02 | ; | HB3 - CB - OG |
| 3859 | 3861 | 3862 | 1 | 1.2000e+02 | 4.1840e+02 | ; | C - N - H |
| 3859 | 3861 | 3863 | 1 | 1.2190e+02 | 4.1840e+02 | ; | C - N - CA |
| 3860 | 3859 | 3861 | 1 | 1.2290e+02 | 6.6944e+02 | ; | O - C - N |
| 3861 | 3863 | 3864 | 1 | 1.0950e+02 | 4.1840e+02 | ; | N - CA - HA |

3861 3863 3865 1 1.0970e+02 6.6944e+02 ; N - CA
- CB
3861 3863 3871 1 1.1010e+02 5.2718e+02 ; N - CA
- C
3862 3861 3863 1 1.1804e+02 4.1840e+02 ; H - N
- CA
3863 3865 3866 1 1.0950e+02 4.1840e+02 ; CA - CB
- HB2
3863 3865 3867 1 1.0950e+02 4.1840e+02 ; CA - CB
- HB3
3863 3865 3868 1 1.1110e+02 5.2718e+02 ; CA - CB
- CG
3863 3871 3872 1 1.2040e+02 6.6944e+02 ; CA - C
- O
3863 3871 3873 1 1.1660e+02 5.8576e+02 ; CA - C
- N
3864 3863 3865 1 1.0950e+02 4.1840e+02 ; HA - CA
- CB
3864 3863 3871 1 1.0950e+02 4.1840e+02 ; HA - CA
- C
3865 3863 3871 1 1.1110e+02 5.2718e+02 ; CB - CA
- C
3865 3868 3869 1 1.1700e+02 5.8576e+02 ; CB - CG
- OD1
3865 3868 3870 1 1.1700e+02 5.8576e+02 ; CB - CG
- OD2
3866 3865 3867 1 1.0950e+02 2.9288e+02 ; HB2 - CB
- HB3
3866 3865 3868 1 1.0950e+02 4.1840e+02 ; HB2 - CB
- CG
3867 3865 3868 1 1.0950e+02 4.1840e+02 ; HB3 - CB
- CG
3869 3868 3870 1 1.2600e+02 6.6944e+02 ; OD1 - CG
- OD2
3871 3873 3874 1 1.2000e+02 4.1840e+02 ; C - N
- H
3871 3873 3875 1 1.2190e+02 4.1840e+02 ; C - N
- CA
3872 3871 3873 1 1.2290e+02 6.6944e+02 ; O - C
- N
3873 3875 3876 1 1.0950e+02 4.1840e+02 ; N - CA
- HA
3873 3875 3877 1 1.0970e+02 6.6944e+02 ; N - CA
- CB
3873 3875 3881 1 1.1010e+02 5.2718e+02 ; N - CA
- C
3874 3873 3875 1 1.1804e+02 4.1840e+02 ; H - N
- CA
3875 3877 3878 1 1.0950e+02 4.1840e+02 ; CA - CB
- HB1
3875 3877 3879 1 1.0950e+02 4.1840e+02 ; CA - CB
- HB2

3875 3877 3880 1 1.0950e+02 4.1840e+02 ; CA - CB
- HB3
3875 3881 3882 1 1.2040e+02 6.6944e+02 ; CA - C
- O
3875 3881 3883 1 1.1660e+02 5.8576e+02 ; CA - C
- N
3876 3875 3877 1 1.0950e+02 4.1840e+02 ; HA - CA
- CB
3876 3875 3881 1 1.0950e+02 4.1840e+02 ; HA - CA
- C
3877 3875 3881 1 1.1110e+02 5.2718e+02 ; CB - CA
- C
3878 3877 3879 1 1.0950e+02 2.9288e+02 ; HB1 - CB
- HB2
3878 3877 3880 1 1.0950e+02 2.9288e+02 ; HB1 - CB
- HB3
3879 3877 3880 1 1.0950e+02 2.9288e+02 ; HB2 - CB
- HB3
3881 3883 3884 1 1.2000e+02 4.1840e+02 ; C - N
- H
3881 3883 3885 1 1.2190e+02 4.1840e+02 ; C - N
- CA
3882 3881 3883 1 1.2290e+02 6.6944e+02 ; O - C
- N
3883 3885 3886 1 1.0950e+02 4.1840e+02 ; N - CA
- HA
3883 3885 3887 1 1.0970e+02 6.6944e+02 ; N - CA
- CB
3883 3885 3903 1 1.1010e+02 5.2718e+02 ; N - CA
- C
3884 3883 3885 1 1.1804e+02 4.1840e+02 ; H - N
- CA
3885 3887 3888 1 1.0950e+02 4.1840e+02 ; CA - CB
- HB2
3885 3887 3889 1 1.0950e+02 4.1840e+02 ; CA - CB
- HB3
3885 3887 3890 1 1.0950e+02 3.3472e+02 ; CA - CB
- CG
3885 3903 3904 1 1.2040e+02 6.6944e+02 ; CA - C
- O
3885 3903 3905 1 1.1660e+02 5.8576e+02 ; CA - C
- N
3886 3885 3887 1 1.0950e+02 4.1840e+02 ; HA - CA
- CB
3886 3885 3903 1 1.0950e+02 4.1840e+02 ; HA - CA
- C
3887 3885 3903 1 1.1110e+02 5.2718e+02 ; CB - CA
- C
3887 3890 3891 1 1.0950e+02 4.1840e+02 ; CB - CG
- HG2
3887 3890 3892 1 1.0950e+02 4.1840e+02 ; CB - CG
- HG3

```
  3887   3890   3893      1    1.0950e+02    3.3472e+02 ;     CB - CG
- CD
  3888   3887   3889      1    1.0950e+02    2.9288e+02 ;    HB2 - CB
- HB3
  3888   3887   3890      1    1.0950e+02    4.1840e+02 ;    HB2 - CB
- CG
  3889   3887   3890      1    1.0950e+02    4.1840e+02 ;    HB3 - CB
- CG
  3890   3893   3894      1    1.0950e+02    4.1840e+02 ;     CG - CD
- HD2
  3890   3893   3895      1    1.0950e+02    4.1840e+02 ;     CG - CD
- HD3
  3890   3893   3896      1    1.0950e+02    3.3472e+02 ;     CG - CD
- CE
  3891   3890   3892      1    1.0950e+02    2.9288e+02 ;    HG2 - CG
- HG3
  3891   3890   3893      1    1.0950e+02    4.1840e+02 ;    HG2 - CG
- CD
  3892   3890   3893      1    1.0950e+02    4.1840e+02 ;    HG3 - CG
- CD
  3893   3896   3897      1    1.0950e+02    4.1840e+02 ;     CD - CE
- HE2
  3893   3896   3898      1    1.0950e+02    4.1840e+02 ;     CD - CE
- HE3
  3893   3896   3899      1    1.1120e+02    6.6944e+02 ;     CD - CE
- NZ
  3894   3893   3895      1    1.0950e+02    2.9288e+02 ;    HD2 - CD
- HD3
  3894   3893   3896      1    1.0950e+02    4.1840e+02 ;    HD2 - CD
- CE
  3895   3893   3896      1    1.0950e+02    4.1840e+02 ;    HD3 - CD
- CE
  3896   3899   3900      1    1.0950e+02    4.1840e+02 ;     CE - NZ
- HZ1
  3896   3899   3901      1    1.0950e+02    4.1840e+02 ;     CE - NZ
- HZ2
  3896   3899   3902      1    1.0950e+02    4.1840e+02 ;     CE - NZ
- HZ3
  3897   3896   3898      1    1.0950e+02    2.9288e+02 ;    HE2 - CE
- HE3
  3897   3896   3899      1    1.0950e+02    4.1840e+02 ;    HE2 - CE
- NZ
  3898   3896   3899      1    1.0950e+02    4.1840e+02 ;    HE3 - CE
- NZ
  3900   3899   3901      1    1.0950e+02    2.9288e+02 ;    HZ1 - NZ
- HZ2
  3900   3899   3902      1    1.0950e+02    2.9288e+02 ;    HZ1 - NZ
- HZ3
  3901   3899   3902      1    1.0950e+02    2.9288e+02 ;    HZ2 - NZ
- HZ3
  3903   3905   3906      1    1.2000e+02    4.1840e+02 ;      C - N
- H
```

```
  3903   3905   3907       1    1.2190e+02    4.1840e+02 ;      C - N
- CA
  3904   3903   3905       1    1.2290e+02    6.6944e+02 ;      O - C
- N
  3905   3907   3908       1    1.0950e+02    4.1840e+02 ;      N - CA
- HA
  3905   3907   3909       1    1.0970e+02    6.6944e+02 ;      N - CA
- CB
  3905   3907   3917       1    1.1010e+02    5.2718e+02 ;      N - CA
- C
  3906   3905   3907       1    1.1804e+02    4.1840e+02 ;      H - N
- CA
  3907   3909   3910       1    1.0950e+02    4.1840e+02 ;     CA - CB
- HB2
  3907   3909   3911       1    1.0950e+02    4.1840e+02 ;     CA - CB
- HB3
  3907   3909   3912       1    1.1110e+02    5.2718e+02 ;     CA - CB
- CG
  3907   3917   3918       1    1.2040e+02    6.6944e+02 ;     CA - C
- O
  3907   3917   3919       1    1.1660e+02    5.8576e+02 ;     CA - C
- N
  3908   3907   3909       1    1.0950e+02    4.1840e+02 ;     HA - CA
- CB
  3908   3907   3917       1    1.0950e+02    4.1840e+02 ;     HA - CA
- C
  3909   3907   3917       1    1.1110e+02    5.2718e+02 ;     CB - CA
- C
  3909   3912   3913       1    1.2040e+02    6.6944e+02 ;     CB - CG
- OD1
  3909   3912   3914       1    1.1660e+02    5.8576e+02 ;     CB - CG
- ND2
  3910   3909   3911       1    1.0950e+02    2.9288e+02 ;    HB2 - CB
- HB3
  3910   3909   3912       1    1.0950e+02    4.1840e+02 ;    HB2 - CB
- CG
  3911   3909   3912       1    1.0950e+02    4.1840e+02 ;    HB3 - CB
- CG
  3912   3914   3915       1    1.2000e+02    4.1840e+02 ;     CG - ND2
- HD21
  3912   3914   3916       1    1.2000e+02    4.1840e+02 ;     CG - ND2
- HD22
  3913   3912   3914       1    1.2290e+02    6.6944e+02 ;    OD1 - CG
- ND2
  3915   3914   3916       1    1.2000e+02    2.9288e+02 ;   HD21 - ND2
- HD22
  3917   3919   3920       1    1.2000e+02    4.1840e+02 ;      C - N
- H
  3917   3919   3921       1    1.2190e+02    4.1840e+02 ;      C - N
- CA
  3918   3917   3919       1    1.2290e+02    6.6944e+02 ;      O - C
- N
```

```
  3919   3921   3922       1    1.0950e+02    4.1840e+02 ;      N - CA
- HA
  3919   3921   3923       1    1.0970e+02    6.6944e+02 ;      N - CA
- CB
  3919   3921   3936       1    1.1010e+02    5.2718e+02 ;      N - CA
- C
  3920   3919   3921       1    1.1804e+02    4.1840e+02 ;      H - N
- CA
  3921   3923   3924       1    1.0950e+02    4.1840e+02 ;     CA - CB
- HB2
  3921   3923   3925       1    1.0950e+02    4.1840e+02 ;     CA - CB
- HB3
  3921   3923   3926       1    1.0950e+02    3.3472e+02 ;     CA - CB
- CG
  3921   3936   3937       1    1.2040e+02    6.6944e+02 ;     CA - C
- O
  3921   3936   3938       1    1.1660e+02    5.8576e+02 ;     CA - C
- N
  3922   3921   3923       1    1.0950e+02    4.1840e+02 ;     HA - CA
- CB
  3922   3921   3936       1    1.0950e+02    4.1840e+02 ;     HA - CA
- C
  3923   3921   3936       1    1.1110e+02    5.2718e+02 ;     CB - CA
- C
  3923   3926   3927       1    1.0950e+02    4.1840e+02 ;     CB - CG
- HG
  3923   3926   3928       1    1.0950e+02    3.3472e+02 ;     CB - CG
- CD1
  3923   3926   3932       1    1.0950e+02    3.3472e+02 ;     CB - CG
- CD2
  3924   3923   3925       1    1.0950e+02    2.9288e+02 ;    HB2 - CB
- HB3
  3924   3923   3926       1    1.0950e+02    4.1840e+02 ;    HB2 - CB
- CG
  3925   3923   3926       1    1.0950e+02    4.1840e+02 ;    HB3 - CB
- CG
  3926   3928   3929       1    1.0950e+02    4.1840e+02 ;     CG - CD1
- HD11
  3926   3928   3930       1    1.0950e+02    4.1840e+02 ;     CG - CD1
- HD12
  3926   3928   3931       1    1.0950e+02    4.1840e+02 ;     CG - CD1
- HD13
  3926   3932   3933       1    1.0950e+02    4.1840e+02 ;     CG - CD2
- HD21
  3926   3932   3934       1    1.0950e+02    4.1840e+02 ;     CG - CD2
- HD22
  3926   3932   3935       1    1.0950e+02    4.1840e+02 ;     CG - CD2
- HD23
  3927   3926   3928       1    1.0950e+02    4.1840e+02 ;     HG - CG
- CD1
  3927   3926   3932       1    1.0950e+02    4.1840e+02 ;     HG - CG
- CD2
```

3928 3926 3932 1 1.0950e+02 3.3472e+02 ; CD1 - CG
- CD2
3929 3928 3930 1 1.0950e+02 2.9288e+02 ; HD11 - CD1
- HD12
3929 3928 3931 1 1.0950e+02 2.9288e+02 ; HD11 - CD1
- HD13
3930 3928 3931 1 1.0950e+02 2.9288e+02 ; HD12 - CD1
- HD13
3933 3932 3934 1 1.0950e+02 2.9288e+02 ; HD21 - CD2
- HD22
3933 3932 3935 1 1.0950e+02 2.9288e+02 ; HD21 - CD2
- HD23
3934 3932 3935 1 1.0950e+02 2.9288e+02 ; HD22 - CD2
- HD23
3936 3938 3939 1 1.2000e+02 4.1840e+02 ; C - N
- H
3936 3938 3940 1 1.2190e+02 4.1840e+02 ; C - N
- CA
3937 3936 3938 1 1.2290e+02 6.6944e+02 ; O - C
- N
3938 3940 3941 1 1.0950e+02 4.1840e+02 ; N - CA
- HA
3938 3940 3942 1 1.0970e+02 6.6944e+02 ; N - CA
- CB
3938 3940 3955 1 1.1010e+02 5.2718e+02 ; N - CA
- C
3939 3938 3940 1 1.1804e+02 4.1840e+02 ; H - N
- CA
3940 3942 3943 1 1.0950e+02 4.1840e+02 ; CA - CB
- HB
3940 3942 3944 1 1.0950e+02 3.3472e+02 ; CA - CB
- CG2
3940 3942 3948 1 1.0950e+02 3.3472e+02 ; CA - CB
- CG1
3940 3955 3956 1 1.2040e+02 6.6944e+02 ; CA - C
- O
3940 3955 3957 1 1.1660e+02 5.8576e+02 ; CA - C
- N
3941 3940 3942 1 1.0950e+02 4.1840e+02 ; HA - CA
- CB
3941 3940 3955 1 1.0950e+02 4.1840e+02 ; HA - CA
- C
3942 3940 3955 1 1.1110e+02 5.2718e+02 ; CB - CA
- C
3942 3944 3945 1 1.0950e+02 4.1840e+02 ; CB - CG2
- HG21
3942 3944 3946 1 1.0950e+02 4.1840e+02 ; CB - CG2
- HG22
3942 3944 3947 1 1.0950e+02 4.1840e+02 ; CB - CG2
- HG23
3942 3948 3949 1 1.0950e+02 4.1840e+02 ; CB - CG1
- HG12

3942 3948 3950 1 1.0950e+02 4.1840e+02 ; CB - CG1
- HG13
3942 3948 3951 1 1.0950e+02 3.3472e+02 ; CB - CG1
- CD1
3943 3942 3944 1 1.0950e+02 4.1840e+02 ; HB - CB
- CG2
3943 3942 3948 1 1.0950e+02 4.1840e+02 ; HB - CB
- CG1
3944 3942 3948 1 1.0950e+02 3.3472e+02 ; CG2 - CB
- CG1
3945 3944 3946 1 1.0950e+02 2.9288e+02 ; HG21 - CG2
- HG22
3945 3944 3947 1 1.0950e+02 2.9288e+02 ; HG21 - CG2
- HG23
3946 3944 3947 1 1.0950e+02 2.9288e+02 ; HG22 - CG2
- HG23
3948 3951 3952 1 1.0950e+02 4.1840e+02 ; CG1 - CD1
- HD11
3948 3951 3953 1 1.0950e+02 4.1840e+02 ; CG1 - CD1
- HD12
3948 3951 3954 1 1.0950e+02 4.1840e+02 ; CG1 - CD1
- HD13
3949 3948 3950 1 1.0950e+02 2.9288e+02 ; HG12 - CG1
- HG13
3949 3948 3951 1 1.0950e+02 4.1840e+02 ; HG12 - CG1
- CD1
3950 3948 3951 1 1.0950e+02 4.1840e+02 ; HG13 - CG1
- CD1
3952 3951 3953 1 1.0950e+02 2.9288e+02 ; HD11 - CD1
- HD12
3952 3951 3954 1 1.0950e+02 2.9288e+02 ; HD11 - CD1
- HD13
3953 3951 3954 1 1.0950e+02 2.9288e+02 ; HD12 - CD1
- HD13
3955 3957 3958 1 1.2000e+02 4.1840e+02 ; C - N
- H
3955 3957 3959 1 1.2190e+02 4.1840e+02 ; C - N
- CA
3956 3955 3957 1 1.2290e+02 6.6944e+02 ; O - C
- N
3957 3959 3960 1 1.0950e+02 4.1840e+02 ; N - CA
- HA
3957 3959 3961 1 1.0970e+02 6.6944e+02 ; N - CA
- CB
3957 3959 3969 1 1.1010e+02 5.2718e+02 ; N - CA
- C
3958 3957 3959 1 1.1804e+02 4.1840e+02 ; H - N
- CA
3959 3961 3962 1 1.0950e+02 4.1840e+02 ; CA - CB
- HB
3959 3961 3963 1 1.0950e+02 3.3472e+02 ; CA - CB
- CG2

3959 3961 3967 1 1.0950e+02 4.1840e+02 ; CA - CB
- OG1
3959 3969 3970 1 1.2040e+02 6.6944e+02 ; CA - C
- O
3959 3969 3971 1 1.1660e+02 5.8576e+02 ; CA - C
- N
3960 3959 3961 1 1.0950e+02 4.1840e+02 ; HA - CA
- CB
3960 3959 3969 1 1.0950e+02 4.1840e+02 ; HA - CA
- C
3961 3959 3969 1 1.1110e+02 5.2718e+02 ; CB - CA
- C
3961 3963 3964 1 1.0950e+02 4.1840e+02 ; CB - CG2
- HG21
3961 3963 3965 1 1.0950e+02 4.1840e+02 ; CB - CG2
- HG22
3961 3963 3966 1 1.0950e+02 4.1840e+02 ; CB - CG2
- HG23
3961 3967 3968 1 1.0850e+02 4.6024e+02 ; CB - OG1
- HG1
3962 3961 3963 1 1.0950e+02 4.1840e+02 ; HB - CB
- CG2
3962 3961 3967 1 1.0950e+02 4.1840e+02 ; HB - CB
- OG1
3963 3961 3967 1 1.0950e+02 4.1840e+02 ; CG2 - CB
- OG1
3964 3963 3965 1 1.0950e+02 2.9288e+02 ; HG21 - CG2
- HG22
3964 3963 3966 1 1.0950e+02 2.9288e+02 ; HG21 - CG2
- HG23
3965 3963 3966 1 1.0950e+02 2.9288e+02 ; HG22 - CG2
- HG23
3969 3971 3972 1 1.2000e+02 4.1840e+02 ; C - N
- H
3969 3971 3973 1 1.2190e+02 4.1840e+02 ; C - N
- CA
3970 3969 3971 1 1.2290e+02 6.6944e+02 ; O - C
- N
3971 3973 3974 1 1.0950e+02 4.1840e+02 ; N - CA
- HA
3971 3973 3975 1 1.0970e+02 6.6944e+02 ; N - CA
- CB
3971 3973 3991 1 1.1010e+02 5.2718e+02 ; N - CA
- C
3972 3971 3973 1 1.1804e+02 4.1840e+02 ; H - N
- CA
3973 3975 3976 1 1.0950e+02 4.1840e+02 ; CA - CB
- HB2
3973 3975 3977 1 1.0950e+02 4.1840e+02 ; CA - CB
- HB3
3973 3975 3978 1 1.0950e+02 3.3472e+02 ; CA - CB
- CG

```
  3973   3991   3992       1    1.2040e+02    6.6944e+02 ;     CA - C
- O
  3973   3991   3993       1    1.1660e+02    5.8576e+02 ;     CA - C
- N
  3974   3973   3975       1    1.0950e+02    4.1840e+02 ;     HA - CA
- CB
  3974   3973   3991       1    1.0950e+02    4.1840e+02 ;     HA - CA
- C
  3975   3973   3991       1    1.1110e+02    5.2718e+02 ;     CB - CA
- C
  3975   3978   3979       1    1.0950e+02    4.1840e+02 ;     CB - CG
- HG2
  3975   3978   3980       1    1.0950e+02    4.1840e+02 ;     CB - CG
- HG3
  3975   3978   3981       1    1.0950e+02    3.3472e+02 ;     CB - CG
- CD
  3976   3975   3977       1    1.0950e+02    2.9288e+02 ;    HB2 - CB
- HB3
  3976   3975   3978       1    1.0950e+02    4.1840e+02 ;    HB2 - CB
- CG
  3977   3975   3978       1    1.0950e+02    4.1840e+02 ;    HB3 - CB
- CG
  3978   3981   3982       1    1.0950e+02    4.1840e+02 ;     CG - CD
- HD2
  3978   3981   3983       1    1.0950e+02    4.1840e+02 ;     CG - CD
- HD3
  3978   3981   3984       1    1.0950e+02    3.3472e+02 ;     CG - CD
- CE
  3979   3978   3980       1    1.0950e+02    2.9288e+02 ;    HG2 - CG
- HG3
  3979   3978   3981       1    1.0950e+02    4.1840e+02 ;    HG2 - CG
- CD
  3980   3978   3981       1    1.0950e+02    4.1840e+02 ;    HG3 - CG
- CD
  3981   3984   3985       1    1.0950e+02    4.1840e+02 ;     CD - CE
- HE2
  3981   3984   3986       1    1.0950e+02    4.1840e+02 ;     CD - CE
- HE3
  3981   3984   3987       1    1.1120e+02    6.6944e+02 ;     CD - CE
- NZ
  3982   3981   3983       1    1.0950e+02    2.9288e+02 ;    HD2 - CD
- HD3
  3982   3981   3984       1    1.0950e+02    4.1840e+02 ;    HD2 - CD
- CE
  3983   3981   3984       1    1.0950e+02    4.1840e+02 ;    HD3 - CD
- CE
  3984   3987   3988       1    1.0950e+02    4.1840e+02 ;     CE - NZ
- HZ1
  3984   3987   3989       1    1.0950e+02    4.1840e+02 ;     CE - NZ
- HZ2
  3984   3987   3990       1    1.0950e+02    4.1840e+02 ;     CE - NZ
- HZ3
```

3985 3984 3986 1 1.0950e+02 2.9288e+02 ; HE2 - CE
- HE3
3985 3984 3987 1 1.0950e+02 4.1840e+02 ; HE2 - CE
- NZ
3986 3984 3987 1 1.0950e+02 4.1840e+02 ; HE3 - CE
- NZ
3988 3987 3989 1 1.0950e+02 2.9288e+02 ; HZ1 - NZ
- HZ2
3988 3987 3990 1 1.0950e+02 2.9288e+02 ; HZ1 - NZ
- HZ3
3989 3987 3990 1 1.0950e+02 2.9288e+02 ; HZ2 - NZ
- HZ3
3991 3993 3994 1 1.2000e+02 4.1840e+02 ; C - N
- H
3991 3993 3995 1 1.2190e+02 4.1840e+02 ; C - N
- CA
3992 3991 3993 1 1.2290e+02 6.6944e+02 ; O - C
- N
3993 3995 3996 1 1.0950e+02 4.1840e+02 ; N - CA
- HA
3993 3995 3997 1 1.0970e+02 6.6944e+02 ; N - CA
- CB
3993 3995 4010 1 1.1010e+02 5.2718e+02 ; N - CA
- C
3994 3993 3995 1 1.1804e+02 4.1840e+02 ; H - N
- CA
3995 3997 3998 1 1.0950e+02 4.1840e+02 ; CA - CB
- HB2
3995 3997 3999 1 1.0950e+02 4.1840e+02 ; CA - CB
- HB3
3995 3997 4000 1 1.0950e+02 3.3472e+02 ; CA - CB
- CG
3995 4010 4011 1 1.2040e+02 6.6944e+02 ; CA - C
- O
3995 4010 4012 1 1.1660e+02 5.8576e+02 ; CA - C
- N
3996 3995 3997 1 1.0950e+02 4.1840e+02 ; HA - CA
- CB
3996 3995 4010 1 1.0950e+02 4.1840e+02 ; HA - CA
- C
3997 3995 4010 1 1.1110e+02 5.2718e+02 ; CB - CA
- C
3997 4000 4001 1 1.0950e+02 4.1840e+02 ; CB - CG
- HG
3997 4000 4002 1 1.0950e+02 3.3472e+02 ; CB - CG
- CD1
3997 4000 4006 1 1.0950e+02 3.3472e+02 ; CB - CG
- CD2
3998 3997 3999 1 1.0950e+02 2.9288e+02 ; HB2 - CB
- HB3
3998 3997 4000 1 1.0950e+02 4.1840e+02 ; HB2 - CB
- CG

```
  3999   3997   4000      1    1.0950e+02    4.1840e+02 ;    HB3 - CB
- CG
  4000   4002   4003      1    1.0950e+02    4.1840e+02 ;     CG - CD1
- HD11
  4000   4002   4004      1    1.0950e+02    4.1840e+02 ;     CG - CD1
- HD12
  4000   4002   4005      1    1.0950e+02    4.1840e+02 ;     CG - CD1
- HD13
  4000   4006   4007      1    1.0950e+02    4.1840e+02 ;     CG - CD2
- HD21
  4000   4006   4008      1    1.0950e+02    4.1840e+02 ;     CG - CD2
- HD22
  4000   4006   4009      1    1.0950e+02    4.1840e+02 ;     CG - CD2
- HD23
  4001   4000   4002      1    1.0950e+02    4.1840e+02 ;     HG - CG
- CD1
  4001   4000   4006      1    1.0950e+02    4.1840e+02 ;     HG - CG
- CD2
  4002   4000   4006      1    1.0950e+02    3.3472e+02 ;    CD1 - CG
- CD2
  4003   4002   4004      1    1.0950e+02    2.9288e+02 ;   HD11 - CD1
- HD12
  4003   4002   4005      1    1.0950e+02    2.9288e+02 ;   HD11 - CD1
- HD13
  4004   4002   4005      1    1.0950e+02    2.9288e+02 ;   HD12 - CD1
- HD13
  4007   4006   4008      1    1.0950e+02    2.9288e+02 ;   HD21 - CD2
- HD22
  4007   4006   4009      1    1.0950e+02    2.9288e+02 ;   HD21 - CD2
- HD23
  4008   4006   4009      1    1.0950e+02    2.9288e+02 ;   HD22 - CD2
- HD23
  4010   4012   4013      1    1.2000e+02    4.1840e+02 ;      C - N
- H
  4010   4012   4014      1    1.2190e+02    4.1840e+02 ;      C - N
- CA
  4011   4010   4012      1    1.2290e+02    6.6944e+02 ;      O - C
- N
  4012   4014   4015      1    1.0950e+02    4.1840e+02 ;      N - CA
- HA
  4012   4014   4016      1    1.0970e+02    6.6944e+02 ;      N - CA
- CB
  4012   4014   4029      1    1.1010e+02    5.2718e+02 ;      N - CA
- C
  4013   4012   4014      1    1.1804e+02    4.1840e+02 ;      H - N
- CA
  4014   4016   4017      1    1.0950e+02    4.1840e+02 ;     CA - CB
- HB2
  4014   4016   4018      1    1.0950e+02    4.1840e+02 ;     CA - CB
- HB3
  4014   4016   4019      1    1.0950e+02    3.3472e+02 ;     CA - CB
- CG
```

4014 4029 4030 1 1.2040e+02 6.6944e+02 ; CA - C
- O
4014 4029 4031 1 1.1660e+02 5.8576e+02 ; CA - C
- N
4015 4014 4016 1 1.0950e+02 4.1840e+02 ; HA - CA
- CB
4015 4014 4029 1 1.0950e+02 4.1840e+02 ; HA - CA
- C
4016 4014 4029 1 1.1110e+02 5.2718e+02 ; CB - CA
- C
4016 4019 4020 1 1.0950e+02 4.1840e+02 ; CB - CG
- HG
4016 4019 4021 1 1.0950e+02 3.3472e+02 ; CB - CG
- CD1
4016 4019 4025 1 1.0950e+02 3.3472e+02 ; CB - CG
- CD2
4017 4016 4018 1 1.0950e+02 2.9288e+02 ; HB2 - CB
- HB3
4017 4016 4019 1 1.0950e+02 4.1840e+02 ; HB2 - CB
- CG
4018 4016 4019 1 1.0950e+02 4.1840e+02 ; HB3 - CB
- CG
4019 4021 4022 1 1.0950e+02 4.1840e+02 ; CG - CD1
- HD11
4019 4021 4023 1 1.0950e+02 4.1840e+02 ; CG - CD1
- HD12
4019 4021 4024 1 1.0950e+02 4.1840e+02 ; CG - CD1
- HD13
4019 4025 4026 1 1.0950e+02 4.1840e+02 ; CG - CD2
- HD21
4019 4025 4027 1 1.0950e+02 4.1840e+02 ; CG - CD2
- HD22
4019 4025 4028 1 1.0950e+02 4.1840e+02 ; CG - CD2
- HD23
4020 4019 4021 1 1.0950e+02 4.1840e+02 ; HG - CG
- CD1
4020 4019 4025 1 1.0950e+02 4.1840e+02 ; HG - CG
- CD2
4021 4019 4025 1 1.0950e+02 3.3472e+02 ; CD1 - CG
- CD2
4022 4021 4023 1 1.0950e+02 2.9288e+02 ; HD11 - CD1
- HD12
4022 4021 4024 1 1.0950e+02 2.9288e+02 ; HD11 - CD1
- HD13
4023 4021 4024 1 1.0950e+02 2.9288e+02 ; HD12 - CD1
- HD13
4026 4025 4027 1 1.0950e+02 2.9288e+02 ; HD21 - CD2
- HD22
4026 4025 4028 1 1.0950e+02 2.9288e+02 ; HD21 - CD2
- HD23
4027 4025 4028 1 1.0950e+02 2.9288e+02 ; HD22 - CD2
- HD23

4029 4031 4032 1 1.2000e+02 4.1840e+02 ; C - N
- H
4029 4031 4033 1 1.2190e+02 4.1840e+02 ; C - N
- CA
4030 4029 4031 1 1.2290e+02 6.6944e+02 ; O - C
- N
4031 4033 4034 1 1.0950e+02 4.1840e+02 ; N - CA
- HA
4031 4033 4035 1 1.0970e+02 6.6944e+02 ; N - CA
- CB
4031 4033 4043 1 1.1010e+02 5.2718e+02 ; N - CA
- C
4032 4031 4033 1 1.1804e+02 4.1840e+02 ; H - N
- CA
4033 4035 4036 1 1.0950e+02 4.1840e+02 ; CA - CB
- HB
4033 4035 4037 1 1.0950e+02 3.3472e+02 ; CA - CB
- CG2
4033 4035 4041 1 1.0950e+02 4.1840e+02 ; CA - CB
- OG1
4033 4043 4044 1 1.2040e+02 6.6944e+02 ; CA - C
- O
4033 4043 4045 1 1.1660e+02 5.8576e+02 ; CA - C
- N
4034 4033 4035 1 1.0950e+02 4.1840e+02 ; HA - CA
- CB
4034 4033 4043 1 1.0950e+02 4.1840e+02 ; HA - CA
- C
4035 4033 4043 1 1.1110e+02 5.2718e+02 ; CB - CA
- C
4035 4037 4038 1 1.0950e+02 4.1840e+02 ; CB - CG2
- HG21
4035 4037 4039 1 1.0950e+02 4.1840e+02 ; CB - CG2
- HG22
4035 4037 4040 1 1.0950e+02 4.1840e+02 ; CB - CG2
- HG23
4035 4041 4042 1 1.0850e+02 4.6024e+02 ; CB - OG1
- HG1
4036 4035 4037 1 1.0950e+02 4.1840e+02 ; HB - CB
- CG2
4036 4035 4041 1 1.0950e+02 4.1840e+02 ; HB - CB
- OG1
4037 4035 4041 1 1.0950e+02 4.1840e+02 ; CG2 - CB
- OG1
4038 4037 4039 1 1.0950e+02 2.9288e+02 ; HG21 - CG2
- HG22
4038 4037 4040 1 1.0950e+02 2.9288e+02 ; HG21 - CG2
- HG23
4039 4037 4040 1 1.0950e+02 2.9288e+02 ; HG22 - CG2
- HG23
4043 4045 4046 1 1.2000e+02 4.1840e+02 ; C - N
- H

4043 4045 4047 1 1.2190e+02 4.1840e+02 ; C - N
- CA
4044 4043 4045 1 1.2290e+02 6.6944e+02 ; O - C
- N
4045 4047 4048 1 1.0950e+02 4.1840e+02 ; N - CA
- HA
4045 4047 4049 1 1.0970e+02 6.6944e+02 ; N - CA
- CB
4045 4047 4064 1 1.1010e+02 5.2718e+02 ; N - CA
- C
4046 4045 4047 1 1.1804e+02 4.1840e+02 ; H - N
- CA
4047 4049 4050 1 1.0950e+02 4.1840e+02 ; CA - CB
- HB2
4047 4049 4051 1 1.0950e+02 4.1840e+02 ; CA - CB
- HB3
4047 4049 4052 1 1.1400e+02 5.2718e+02 ; CA - CB
- CG
4047 4064 4065 1 1.2040e+02 6.6944e+02 ; CA - C
- O
4047 4064 4066 1 1.1660e+02 5.8576e+02 ; CA - C
- N
4048 4047 4049 1 1.0950e+02 4.1840e+02 ; HA - CA
- CB
4048 4047 4064 1 1.0950e+02 4.1840e+02 ; HA - CA
- C
4049 4047 4064 1 1.1110e+02 5.2718e+02 ; CB - CA
- C
4049 4052 4053 1 1.2000e+02 5.8576e+02 ; CB - CG
- CD1
4049 4052 4062 1 1.2000e+02 5.8576e+02 ; CB - CG
- CD2
4050 4049 4051 1 1.0950e+02 2.9288e+02 ; HB2 - CB
- HB3
4050 4049 4052 1 1.0950e+02 4.1840e+02 ; HB2 - CB
- CG
4051 4049 4052 1 1.0950e+02 4.1840e+02 ; HB3 - CB
- CG
4052 4053 4054 1 1.2000e+02 4.1840e+02 ; CG - CD1
- HD1
4052 4053 4055 1 1.2000e+02 5.2718e+02 ; CG - CD1
- CE1
4052 4062 4060 1 1.2000e+02 5.2718e+02 ; CG - CD2
- CE2
4052 4062 4063 1 1.2000e+02 4.1840e+02 ; CG - CD2
- HD2
4053 4052 4062 1 1.2000e+02 5.2718e+02 ; CD1 - CG
- CD2
4053 4055 4056 1 1.2000e+02 4.1840e+02 ; CD1 - CE1
- HE1
4053 4055 4057 1 1.2000e+02 5.2718e+02 ; CD1 - CE1
- CZ

```
  4054   4053   4055       1    1.2000e+02    4.1840e+02 ;    HD1 - CD1
- CE1
  4055   4057   4058       1    1.2000e+02    5.8576e+02 ;    CE1 - CZ
- OH
  4055   4057   4060       1    1.2000e+02    5.2718e+02 ;    CE1 - CZ
- CE2
  4056   4055   4057       1    1.2000e+02    4.1840e+02 ;    HE1 - CE1
- CZ
  4057   4058   4059       1    1.1300e+02    4.1840e+02 ;     CZ - OH
- HH
  4057   4060   4061       1    1.2000e+02    4.1840e+02 ;     CZ - CE2
- HE2
  4057   4060   4062       1    1.2000e+02    5.2718e+02 ;     CZ - CE2
- CD2
  4058   4057   4060       1    1.2000e+02    5.8576e+02 ;     OH - CZ
- CE2
  4060   4062   4063       1    1.2000e+02    4.1840e+02 ;    CE2 - CD2
- HD2
  4061   4060   4062       1    1.2000e+02    4.1840e+02 ;    HE2 - CE2
- CD2
  4064   4066   4067       1    1.2000e+02    4.1840e+02 ;      C - N
- H
  4064   4066   4068       1    1.2190e+02    4.1840e+02 ;      C - N
- CA
  4065   4064   4066       1    1.2290e+02    6.6944e+02 ;      O - C
- N
  4066   4068   4069       1    1.0950e+02    4.1840e+02 ;      N - CA
- HA
  4066   4068   4070       1    1.0970e+02    6.6944e+02 ;      N - CA
- CB
  4066   4068   4078       1    1.1010e+02    5.2718e+02 ;      N - CA
- C
  4067   4066   4068       1    1.1804e+02    4.1840e+02 ;      H - N
- CA
  4068   4070   4071       1    1.0950e+02    4.1840e+02 ;     CA - CB
- HB2
  4068   4070   4072       1    1.0950e+02    4.1840e+02 ;     CA - CB
- HB3
  4068   4070   4073       1    1.1110e+02    5.2718e+02 ;     CA - CB
- CG
  4068   4078   4079       1    1.2040e+02    6.6944e+02 ;     CA - C
- O
  4068   4078   4080       1    1.1660e+02    5.8576e+02 ;     CA - C
- N
  4069   4068   4070       1    1.0950e+02    4.1840e+02 ;     HA - CA
- CB
  4069   4068   4078       1    1.0950e+02    4.1840e+02 ;     HA - CA
- C
  4070   4068   4078       1    1.1110e+02    5.2718e+02 ;     CB - CA
- C
  4070   4073   4074       1    1.2040e+02    6.6944e+02 ;     CB - CG
- OD1
```

4070 4073 4075 1 1.1660e+02 5.8576e+02 ; CB - CG
- ND2
4071 4070 4072 1 1.0950e+02 2.9288e+02 ; HB2 - CB
- HB3
4071 4070 4073 1 1.0950e+02 4.1840e+02 ; HB2 - CB
- CG
4072 4070 4073 1 1.0950e+02 4.1840e+02 ; HB3 - CB
- CG
4073 4075 4076 1 1.2000e+02 4.1840e+02 ; CG - ND2
- HD21
4073 4075 4077 1 1.2000e+02 4.1840e+02 ; CG - ND2
- HD22
4074 4073 4075 1 1.2290e+02 6.6944e+02 ; OD1 - CG
- ND2
4076 4075 4077 1 1.2000e+02 2.9288e+02 ; HD21 - ND2
- HD22
4078 4080 4081 1 1.2190e+02 4.1840e+02 ; C - N
- CD
4078 4080 4090 1 1.2190e+02 4.1840e+02 ; C - N
- CA
4079 4078 4080 1 1.2290e+02 6.6944e+02 ; O - C
- N
4080 4081 4082 1 1.0950e+02 4.1840e+02 ; N - CD
- HD2
4080 4081 4083 1 1.0950e+02 4.1840e+02 ; N - CD
- HD3
4080 4081 4084 1 1.0970e+02 6.6944e+02 ; N - CD
- CG
4080 4090 4087 1 1.0970e+02 6.6944e+02 ; N - CA
- CB
4080 4090 4091 1 1.0950e+02 4.1840e+02 ; N - CA
- HA
4080 4090 4092 1 1.1010e+02 5.2718e+02 ; N - CA
- C
4081 4080 4090 1 1.1800e+02 4.1840e+02 ; CD - N
- CA
4081 4084 4085 1 1.0950e+02 4.1840e+02 ; CD - CG
- HG2
4081 4084 4086 1 1.0950e+02 4.1840e+02 ; CD - CG
- HG3
4081 4084 4087 1 1.0950e+02 3.3472e+02 ; CD - CG
- CB
4082 4081 4083 1 1.0950e+02 2.9288e+02 ; HD2 - CD
- HD3
4082 4081 4084 1 1.0950e+02 4.1840e+02 ; HD2 - CD
- CG
4083 4081 4084 1 1.0950e+02 4.1840e+02 ; HD3 - CD
- CG
4084 4087 4088 1 1.0950e+02 4.1840e+02 ; CG - CB
- HB2
4084 4087 4089 1 1.0950e+02 4.1840e+02 ; CG - CB
- HB3

4084 4087 4090 1 1.0950e+02 3.3472e+02 ; CG - CB
- CA
4085 4084 4086 1 1.0950e+02 2.9288e+02 ; HG2 - CG
- HG3
4085 4084 4087 1 1.0950e+02 4.1840e+02 ; HG2 - CG
- CB
4086 4084 4087 1 1.0950e+02 4.1840e+02 ; HG3 - CG
- CB
4087 4090 4091 1 1.0950e+02 4.1840e+02 ; CB - CA
- HA
4087 4090 4092 1 1.1110e+02 5.2718e+02 ; CB - CA
- C
4088 4087 4089 1 1.0950e+02 2.9288e+02 ; HB2 - CB
- HB3
4088 4087 4090 1 1.0950e+02 4.1840e+02 ; HB2 - CB
- CA
4089 4087 4090 1 1.0950e+02 4.1840e+02 ; HB3 - CB
- CA
4090 4092 4093 1 1.2040e+02 6.6944e+02 ; CA - C
- O
4090 4092 4094 1 1.1660e+02 5.8576e+02 ; CA - C
- N
4091 4090 4092 1 1.0950e+02 4.1840e+02 ; HA - CA
- C
4092 4094 4095 1 1.2000e+02 4.1840e+02 ; C - N
- H
4092 4094 4096 1 1.2190e+02 4.1840e+02 ; C - N
- CA
4093 4092 4094 1 1.2290e+02 6.6944e+02 ; O - C
- N
4094 4096 4097 1 1.0950e+02 4.1840e+02 ; N - CA
- HA
4094 4096 4098 1 1.0970e+02 6.6944e+02 ; N - CA
- CB
4094 4096 4106 1 1.1010e+02 5.2718e+02 ; N - CA
- C
4095 4094 4096 1 1.1804e+02 4.1840e+02 ; H - N
- CA
4096 4098 4099 1 1.0950e+02 4.1840e+02 ; CA - CB
- HB2
4096 4098 4100 1 1.0950e+02 4.1840e+02 ; CA - CB
- HB3
4096 4098 4101 1 1.1110e+02 5.2718e+02 ; CA - CB
- CG
4096 4106 4107 1 1.2040e+02 6.6944e+02 ; CA - C
- O
4096 4106 4108 1 1.1660e+02 5.8576e+02 ; CA - C
- N
4097 4096 4098 1 1.0950e+02 4.1840e+02 ; HA - CA
- CB
4097 4096 4106 1 1.0950e+02 4.1840e+02 ; HA - CA
- C

```
  4098   4096   4106       1    1.1110e+02    5.2718e+02 ;      CB - CA
- C
  4098   4101   4102       1    1.2040e+02    6.6944e+02 ;      CB - CG
- OD1
  4098   4101   4103       1    1.1660e+02    5.8576e+02 ;      CB - CG
- ND2
  4099   4098   4100       1    1.0950e+02    2.9288e+02 ;     HB2 - CB
- HB3
  4099   4098   4101       1    1.0950e+02    4.1840e+02 ;     HB2 - CB
- CG
  4100   4098   4101       1    1.0950e+02    4.1840e+02 ;     HB3 - CB
- CG
  4101   4103   4104       1    1.2000e+02    4.1840e+02 ;      CG - ND2
- HD21
  4101   4103   4105       1    1.2000e+02    4.1840e+02 ;      CG - ND2
- HD22
  4102   4101   4103       1    1.2290e+02    6.6944e+02 ;     OD1 - CG
- ND2
  4104   4103   4105       1    1.2000e+02    2.9288e+02 ;    HD21 - ND2
- HD22
  4106   4108   4109       1    1.2000e+02    4.1840e+02 ;       C - N
- H
  4106   4108   4110       1    1.2190e+02    4.1840e+02 ;       C - N
- CA
  4107   4106   4108       1    1.2290e+02    6.6944e+02 ;       O - C
- N
  4108   4110   4111       1    1.0950e+02    4.1840e+02 ;       N - CA
- HA
  4108   4110   4112       1    1.0970e+02    6.6944e+02 ;       N - CA
- CB
  4108   4110   4121       1    1.1010e+02    5.2718e+02 ;       N - CA
- C
  4109   4108   4110       1    1.1804e+02    4.1840e+02 ;       H - N
- CA
  4110   4112   4113       1    1.0950e+02    4.1840e+02 ;      CA - CB
- HB2
  4110   4112   4114       1    1.0950e+02    4.1840e+02 ;      CA - CB
- HB3
  4110   4112   4115       1    1.0950e+02    3.3472e+02 ;      CA - CB
- CG
  4110   4121   4122       1    1.2040e+02    6.6944e+02 ;      CA - C
- O
  4110   4121   4123       1    1.1660e+02    5.8576e+02 ;      CA - C
- N
  4111   4110   4112       1    1.0950e+02    4.1840e+02 ;      HA - CA
- CB
  4111   4110   4121       1    1.0950e+02    4.1840e+02 ;      HA - CA
- C
  4112   4110   4121       1    1.1110e+02    5.2718e+02 ;      CB - CA
- C
  4112   4115   4116       1    1.0950e+02    4.1840e+02 ;      CB - CG
- HG2
```

```
  4112   4115   4117      1    1.0950e+02    4.1840e+02 ;     CB - CG
- HG3
  4112   4115   4118      1    1.1110e+02    5.2718e+02 ;     CB - CG
- CD
  4113   4112   4114      1    1.0950e+02    2.9288e+02 ;    HB2 - CB
- HB3
  4113   4112   4115      1    1.0950e+02    4.1840e+02 ;    HB2 - CB
- CG
  4114   4112   4115      1    1.0950e+02    4.1840e+02 ;    HB3 - CB
- CG
  4115   4118   4119      1    1.1700e+02    5.8576e+02 ;     CG - CD
- OE1
  4115   4118   4120      1    1.1700e+02    5.8576e+02 ;     CG - CD
- OE2
  4116   4115   4117      1    1.0950e+02    2.9288e+02 ;    HG2 - CG
- HG3
  4116   4115   4118      1    1.0950e+02    4.1840e+02 ;    HG2 - CG
- CD
  4117   4115   4118      1    1.0950e+02    4.1840e+02 ;    HG3 - CG
- CD
  4119   4118   4120      1    1.2600e+02    6.6944e+02 ;    OE1 - CD
- OE2
  4121   4123   4124      1    1.2000e+02    4.1840e+02 ;      C - N
- H
  4121   4123   4125      1    1.2190e+02    4.1840e+02 ;      C - N
- CA
  4122   4121   4123      1    1.2290e+02    6.6944e+02 ;      O - C
- N
  4123   4125   4126      1    1.0950e+02    4.1840e+02 ;      N - CA
- HA
  4123   4125   4127      1    1.0970e+02    6.6944e+02 ;      N - CA
- CB
  4123   4125   4145      1    1.1010e+02    5.2718e+02 ;      N - CA
- C
  4124   4123   4125      1    1.1804e+02    4.1840e+02 ;      H - N
- CA
  4125   4127   4128      1    1.0950e+02    4.1840e+02 ;     CA - CB
- HB2
  4125   4127   4129      1    1.0950e+02    4.1840e+02 ;     CA - CB
- HB3
  4125   4127   4130      1    1.0950e+02    3.3472e+02 ;     CA - CB
- CG
  4125   4145   4146      1    1.2040e+02    6.6944e+02 ;     CA - C
- O
  4125   4145   4147      1    1.1660e+02    5.8576e+02 ;     CA - C
- N
  4126   4125   4127      1    1.0950e+02    4.1840e+02 ;     HA - CA
- CB
  4126   4125   4145      1    1.0950e+02    4.1840e+02 ;     HA - CA
- C
  4127   4125   4145      1    1.1110e+02    5.2718e+02 ;     CB - CA
- C
```

```
  4127   4130   4131      1    1.0950e+02    4.1840e+02 ;     CB - CG
- HG2
  4127   4130   4132      1    1.0950e+02    4.1840e+02 ;     CB - CG
- HG3
  4127   4130   4133      1    1.0950e+02    3.3472e+02 ;     CB - CG
- CD
  4128   4127   4129      1    1.0950e+02    2.9288e+02 ;    HB2 - CB
- HB3
  4128   4127   4130      1    1.0950e+02    4.1840e+02 ;    HB2 - CB
- CG
  4129   4127   4130      1    1.0950e+02    4.1840e+02 ;    HB3 - CB
- CG
  4130   4133   4134      1    1.0950e+02    4.1840e+02 ;     CG - CD
- HD2
  4130   4133   4135      1    1.0950e+02    4.1840e+02 ;     CG - CD
- HD3
  4130   4133   4136      1    1.1120e+02    6.6944e+02 ;     CG - CD
- NE
  4131   4130   4132      1    1.0950e+02    2.9288e+02 ;    HG2 - CG
- HG3
  4131   4130   4133      1    1.0950e+02    4.1840e+02 ;    HG2 - CG
- CD
  4132   4130   4133      1    1.0950e+02    4.1840e+02 ;    HG3 - CG
- CD
  4133   4136   4137      1    1.1840e+02    4.1840e+02 ;     CD - NE
- HE
  4133   4136   4138      1    1.2320e+02    4.1840e+02 ;     CD - NE
- CZ
  4134   4133   4135      1    1.0950e+02    2.9288e+02 ;    HD2 - CD
- HD3
  4134   4133   4136      1    1.0950e+02    4.1840e+02 ;    HD2 - CD
- NE
  4135   4133   4136      1    1.0950e+02    4.1840e+02 ;    HD3 - CD
- NE
  4136   4138   4139      1    1.2000e+02    5.8576e+02 ;     NE - CZ
- NH1
  4136   4138   4142      1    1.2000e+02    5.8576e+02 ;     NE - CZ
- NH2
  4137   4136   4138      1    1.2000e+02    4.1840e+02 ;     HE - NE
- CZ
  4138   4139   4140      1    1.2000e+02    4.1840e+02 ;     CZ - NH1
- HH11
  4138   4139   4141      1    1.2000e+02    4.1840e+02 ;     CZ - NH1
- HH12
  4138   4142   4143      1    1.2000e+02    4.1840e+02 ;     CZ - NH2
- HH21
  4138   4142   4144      1    1.2000e+02    4.1840e+02 ;     CZ - NH2
- HH22
  4139   4138   4142      1    1.2000e+02    5.8576e+02 ;    NH1 - CZ
- NH2
  4140   4139   4141      1    1.2000e+02    2.9288e+02 ;   HH11 - NH1
- HH12
```

```
  4143   4142   4144       1    1.2000e+02    2.9288e+02 ;   HH21 - NH2
- HH22
  4145   4147   4148       1    1.2000e+02    4.1840e+02 ;      C - N
- H
  4145   4147   4149       1    1.2190e+02    4.1840e+02 ;      C - N
- CA
  4146   4145   4147       1    1.2290e+02    6.6944e+02 ;      O - C
- N
  4147   4149   4150       1    1.0950e+02    4.1840e+02 ;      N - CA
- HA
  4147   4149   4151       1    1.0970e+02    6.6944e+02 ;      N - CA
- CB
  4147   4149   4156       1    1.1010e+02    5.2718e+02 ;      N - CA
- C
  4148   4147   4149       1    1.1804e+02    4.1840e+02 ;      H - N
- CA
  4149   4151   4152       1    1.0950e+02    4.1840e+02 ;     CA - CB
- HB2
  4149   4151   4153       1    1.0950e+02    4.1840e+02 ;     CA - CB
- HB3
  4149   4151   4154       1    1.0860e+02    4.1840e+02 ;     CA - CB
- SG
  4149   4156   4157       1    1.2040e+02    6.6944e+02 ;     CA - C
- O
  4149   4156   4158       1    1.1660e+02    5.8576e+02 ;     CA - C
- N
  4150   4149   4151       1    1.0950e+02    4.1840e+02 ;     HA - CA
- CB
  4150   4149   4156       1    1.0950e+02    4.1840e+02 ;     HA - CA
- C
  4151   4149   4156       1    1.1110e+02    5.2718e+02 ;     CB - CA
- C
  4151   4154   4155       1    9.6000e+01    3.5982e+02 ;     CB - SG
- HG
  4152   4151   4153       1    1.0950e+02    2.9288e+02 ;    HB2 - CB
- HB3
  4152   4151   4154       1    1.0950e+02    4.1840e+02 ;    HB2 - CB
- SG
  4153   4151   4154       1    1.0950e+02    4.1840e+02 ;    HB3 - CB
- SG
  4156   4158   4159       1    1.2000e+02    4.1840e+02 ;      C - N
- H
  4156   4158   4160       1    1.2190e+02    4.1840e+02 ;      C - N
- CA
  4157   4156   4158       1    1.2290e+02    6.6944e+02 ;      O - C
- N
  4158   4160   4161       1    1.0950e+02    4.1840e+02 ;      N - CA
- HA
  4158   4160   4162       1    1.0970e+02    6.6944e+02 ;      N - CA
- CB
  4158   4160   4170       1    1.1010e+02    5.2718e+02 ;      N - CA
- C
```

```
   4159   4158   4160       1    1.1804e+02    4.1840e+02 ;      H - N
 - CA
   4160   4162   4163       1    1.0950e+02    4.1840e+02 ;     CA - CB
 - HB
   4160   4162   4164       1    1.0950e+02    3.3472e+02 ;     CA - CB
 - CG2
   4160   4162   4168       1    1.0950e+02    4.1840e+02 ;     CA - CB
 - OG1
   4160   4170   4171       1    1.2040e+02    6.6944e+02 ;     CA - C
 - O
   4160   4170   4172       1    1.1660e+02    5.8576e+02 ;     CA - C
 - N
   4161   4160   4162       1    1.0950e+02    4.1840e+02 ;     HA - CA
 - CB
   4161   4160   4170       1    1.0950e+02    4.1840e+02 ;     HA - CA
 - C
   4162   4160   4170       1    1.1110e+02    5.2718e+02 ;     CB - CA
 - C
   4162   4164   4165       1    1.0950e+02    4.1840e+02 ;     CB - CG2
 - HG21
   4162   4164   4166       1    1.0950e+02    4.1840e+02 ;     CB - CG2
 - HG22
   4162   4164   4167       1    1.0950e+02    4.1840e+02 ;     CB - CG2
 - HG23
   4162   4168   4169       1    1.0850e+02    4.6024e+02 ;     CB - OG1
 - HG1
   4163   4162   4164       1    1.0950e+02    4.1840e+02 ;     HB - CB
 - CG2
   4163   4162   4168       1    1.0950e+02    4.1840e+02 ;     HB - CB
 - OG1
   4164   4162   4168       1    1.0950e+02    4.1840e+02 ;    CG2 - CB
 - OG1
   4165   4164   4166       1    1.0950e+02    2.9288e+02 ;   HG21 - CG2
 - HG22
   4165   4164   4167       1    1.0950e+02    2.9288e+02 ;   HG21 - CG2
 - HG23
   4166   4164   4167       1    1.0950e+02    2.9288e+02 ;   HG22 - CG2
 - HG23
   4170   4172   4173       1    1.2000e+02    4.1840e+02 ;      C - N
 - H
   4170   4172   4174       1    1.2190e+02    4.1840e+02 ;      C - N
 - CA
   4171   4170   4172       1    1.2290e+02    6.6944e+02 ;      O - C
 - N
   4172   4174   4175       1    1.0950e+02    4.1840e+02 ;      N - CA
 - HA
   4172   4174   4176       1    1.0970e+02    6.6944e+02 ;      N - CA
 - CB
   4172   4174   4189       1    1.1010e+02    5.2718e+02 ;      N - CA
 - C
   4173   4172   4174       1    1.1804e+02    4.1840e+02 ;      H - N
 - CA
```

4174 4176 4177 1 1.0950e+02 4.1840e+02 ; CA - CB - HB
4174 4176 4178 1 1.0950e+02 3.3472e+02 ; CA - CB - CG2
4174 4176 4182 1 1.0950e+02 3.3472e+02 ; CA - CB - CG1
4174 4189 4190 1 1.2040e+02 6.6944e+02 ; CA - C - O
4174 4189 4191 1 1.1660e+02 5.8576e+02 ; CA - C - N
4175 4174 4176 1 1.0950e+02 4.1840e+02 ; HA - CA - CB
4175 4174 4189 1 1.0950e+02 4.1840e+02 ; HA - CA - C
4176 4174 4189 1 1.1110e+02 5.2718e+02 ; CB - CA - C
4176 4178 4179 1 1.0950e+02 4.1840e+02 ; CB - CG2 - HG21
4176 4178 4180 1 1.0950e+02 4.1840e+02 ; CB - CG2 - HG22
4176 4178 4181 1 1.0950e+02 4.1840e+02 ; CB - CG2 - HG23
4176 4182 4183 1 1.0950e+02 4.1840e+02 ; CB - CG1 - HG12
4176 4182 4184 1 1.0950e+02 4.1840e+02 ; CB - CG1 - HG13
4176 4182 4185 1 1.0950e+02 3.3472e+02 ; CB - CG1 - CD1
4177 4176 4178 1 1.0950e+02 4.1840e+02 ; HB - CB - CG2
4177 4176 4182 1 1.0950e+02 4.1840e+02 ; HB - CB - CG1
4178 4176 4182 1 1.0950e+02 3.3472e+02 ; CG2 - CB - CG1
4179 4178 4180 1 1.0950e+02 2.9288e+02 ; HG21 - CG2 - HG22
4179 4178 4181 1 1.0950e+02 2.9288e+02 ; HG21 - CG2 - HG23
4180 4178 4181 1 1.0950e+02 2.9288e+02 ; HG22 - CG2 - HG23
4182 4185 4186 1 1.0950e+02 4.1840e+02 ; CG1 - CD1 - HD11
4182 4185 4187 1 1.0950e+02 4.1840e+02 ; CG1 - CD1 - HD12
4182 4185 4188 1 1.0950e+02 4.1840e+02 ; CG1 - CD1 - HD13
4183 4182 4184 1 1.0950e+02 2.9288e+02 ; HG12 - CG1 - HG13
4183 4182 4185 1 1.0950e+02 4.1840e+02 ; HG12 - CG1 - CD1
4184 4182 4185 1 1.0950e+02 4.1840e+02 ; HG13 - CG1 - CD1

4186 4185 4187 1 1.0950e+02 2.9288e+02 ; HD11 - CD1 - HD12
4186 4185 4188 1 1.0950e+02 2.9288e+02 ; HD11 - CD1 - HD13
4187 4185 4188 1 1.0950e+02 2.9288e+02 ; HD12 - CD1 - HD13
4189 4191 4192 1 1.2000e+02 4.1840e+02 ; C - N - H
4189 4191 4193 1 1.2190e+02 4.1840e+02 ; C - N - CA
4190 4189 4191 1 1.2290e+02 6.6944e+02 ; O - C - N
4191 4193 4194 1 1.0950e+02 4.1840e+02 ; N - CA - HA
4191 4193 4195 1 1.0970e+02 6.6944e+02 ; N - CA - CB
4191 4193 4204 1 1.1010e+02 5.2718e+02 ; N - CA - C
4192 4191 4193 1 1.1804e+02 4.1840e+02 ; H - N - CA
4193 4195 4196 1 1.0950e+02 4.1840e+02 ; CA - CB - HB2
4193 4195 4197 1 1.0950e+02 4.1840e+02 ; CA - CB - HB3
4193 4195 4198 1 1.0950e+02 3.3472e+02 ; CA - CB - CG
4193 4204 4205 1 1.2040e+02 6.6944e+02 ; CA - C - O
4193 4204 4206 1 1.1660e+02 5.8576e+02 ; CA - C - N
4194 4193 4195 1 1.0950e+02 4.1840e+02 ; HA - CA - CB
4194 4193 4204 1 1.0950e+02 4.1840e+02 ; HA - CA - C
4195 4193 4204 1 1.1110e+02 5.2718e+02 ; CB - CA - C
4195 4198 4199 1 1.0950e+02 4.1840e+02 ; CB - CG - HG2
4195 4198 4200 1 1.0950e+02 4.1840e+02 ; CB - CG - HG3
4195 4198 4201 1 1.1110e+02 5.2718e+02 ; CB - CG - CD
4196 4195 4197 1 1.0950e+02 2.9288e+02 ; HB2 - CB - HB3
4196 4195 4198 1 1.0950e+02 4.1840e+02 ; HB2 - CB - CG
4197 4195 4198 1 1.0950e+02 4.1840e+02 ; HB3 - CB - CG
4198 4201 4202 1 1.1700e+02 5.8576e+02 ; CG - CD - OE1
4198 4201 4203 1 1.1700e+02 5.8576e+02 ; CG - CD - OE2

4199 4198 4200 1 1.0950e+02 2.9288e+02 ; HG2 - CG
- HG3
4199 4198 4201 1 1.0950e+02 4.1840e+02 ; HG2 - CG
- CD
4200 4198 4201 1 1.0950e+02 4.1840e+02 ; HG3 - CG
- CD
4202 4201 4203 1 1.2600e+02 6.6944e+02 ; OE1 - CD
- OE2
4204 4206 4207 1 1.2000e+02 4.1840e+02 ; C - N
- H
4204 4206 4208 1 1.2190e+02 4.1840e+02 ; C - N
- CA
4205 4204 4206 1 1.2290e+02 6.6944e+02 ; O - C
- N
4206 4208 4209 1 1.0950e+02 4.1840e+02 ; N - CA
- HA
4206 4208 4210 1 1.0970e+02 6.6944e+02 ; N - CA
- CB
4206 4208 4219 1 1.1010e+02 5.2718e+02 ; N - CA
- C
4207 4206 4208 1 1.1804e+02 4.1840e+02 ; H - N
- CA
4208 4210 4211 1 1.0950e+02 4.1840e+02 ; CA - CB
- HB2
4208 4210 4212 1 1.0950e+02 4.1840e+02 ; CA - CB
- HB3
4208 4210 4213 1 1.0950e+02 3.3472e+02 ; CA - CB
- CG
4208 4219 4220 1 1.2040e+02 6.6944e+02 ; CA - C
- O
4208 4219 4221 1 1.1660e+02 5.8576e+02 ; CA - C
- N
4209 4208 4210 1 1.0950e+02 4.1840e+02 ; HA - CA
- CB
4209 4208 4219 1 1.0950e+02 4.1840e+02 ; HA - CA
- C
4210 4208 4219 1 1.1110e+02 5.2718e+02 ; CB - CA
- C
4210 4213 4214 1 1.0950e+02 4.1840e+02 ; CB - CG
- HG2
4210 4213 4215 1 1.0950e+02 4.1840e+02 ; CB - CG
- HG3
4210 4213 4216 1 1.1110e+02 5.2718e+02 ; CB - CG
- CD
4211 4210 4212 1 1.0950e+02 2.9288e+02 ; HB2 - CB
- HB3
4211 4210 4213 1 1.0950e+02 4.1840e+02 ; HB2 - CB
- CG
4212 4210 4213 1 1.0950e+02 4.1840e+02 ; HB3 - CB
- CG
4213 4216 4217 1 1.1700e+02 5.8576e+02 ; CG - CD
- OE1

4213 4216 4218 1 1.1700e+02 5.8576e+02 ; CG - CD
- OE2
4214 4213 4215 1 1.0950e+02 2.9288e+02 ; HG2 - CG
- HG3
4214 4213 4216 1 1.0950e+02 4.1840e+02 ; HG2 - CG
- CD
4215 4213 4216 1 1.0950e+02 4.1840e+02 ; HG3 - CG
- CD
4217 4216 4218 1 1.2600e+02 6.6944e+02 ; OE1 - CD
- OE2
4219 4221 4222 1 1.2000e+02 4.1840e+02 ; C - N
- H
4219 4221 4223 1 1.2190e+02 4.1840e+02 ; C - N
- CA
4220 4219 4221 1 1.2290e+02 6.6944e+02 ; O - C
- N
4221 4223 4224 1 1.0950e+02 4.1840e+02 ; N - CA
- HA
4221 4223 4225 1 1.0970e+02 6.6944e+02 ; N - CA
- CB
4221 4223 4229 1 1.1010e+02 5.2718e+02 ; N - CA
- C
4222 4221 4223 1 1.1804e+02 4.1840e+02 ; H - N
- CA
4223 4225 4226 1 1.0950e+02 4.1840e+02 ; CA - CB
- HB1
4223 4225 4227 1 1.0950e+02 4.1840e+02 ; CA - CB
- HB2
4223 4225 4228 1 1.0950e+02 4.1840e+02 ; CA - CB
- HB3
4223 4229 4230 1 1.2040e+02 6.6944e+02 ; CA - C
- O
4223 4229 4231 1 1.1660e+02 5.8576e+02 ; CA - C
- N
4224 4223 4225 1 1.0950e+02 4.1840e+02 ; HA - CA
- CB
4224 4223 4229 1 1.0950e+02 4.1840e+02 ; HA - CA
- C
4225 4223 4229 1 1.1110e+02 5.2718e+02 ; CB - CA
- C
4226 4225 4227 1 1.0950e+02 2.9288e+02 ; HB1 - CB
- HB2
4226 4225 4228 1 1.0950e+02 2.9288e+02 ; HB1 - CB
- HB3
4227 4225 4228 1 1.0950e+02 2.9288e+02 ; HB2 - CB
- HB3
4229 4231 4232 1 1.2000e+02 4.1840e+02 ; C - N
- H
4229 4231 4233 1 1.2190e+02 4.1840e+02 ; C - N
- CA
4230 4229 4231 1 1.2290e+02 6.6944e+02 ; O - C
- N

4231 4233 4234 1 1.0950e+02 4.1840e+02 ; N - CA
- HA
4231 4233 4235 1 1.0970e+02 6.6944e+02 ; N - CA
- CB
4231 4233 4248 1 1.1010e+02 5.2718e+02 ; N - CA
- C
4232 4231 4233 1 1.1804e+02 4.1840e+02 ; H - N
- CA
4233 4235 4236 1 1.0950e+02 4.1840e+02 ; CA - CB
- HB2
4233 4235 4237 1 1.0950e+02 4.1840e+02 ; CA - CB
- HB3
4233 4235 4238 1 1.0950e+02 3.3472e+02 ; CA - CB
- CG
4233 4248 4249 1 1.2040e+02 6.6944e+02 ; CA - C
- O
4233 4248 4250 1 1.1660e+02 5.8576e+02 ; CA - C
- N
4234 4233 4235 1 1.0950e+02 4.1840e+02 ; HA - CA
- CB
4234 4233 4248 1 1.0950e+02 4.1840e+02 ; HA - CA
- C
4235 4233 4248 1 1.1110e+02 5.2718e+02 ; CB - CA
- C
4235 4238 4239 1 1.0950e+02 4.1840e+02 ; CB - CG
- HG
4235 4238 4240 1 1.0950e+02 3.3472e+02 ; CB - CG
- CD1
4235 4238 4244 1 1.0950e+02 3.3472e+02 ; CB - CG
- CD2
4236 4235 4237 1 1.0950e+02 2.9288e+02 ; HB2 - CB
- HB3
4236 4235 4238 1 1.0950e+02 4.1840e+02 ; HB2 - CB
- CG
4237 4235 4238 1 1.0950e+02 4.1840e+02 ; HB3 - CB
- CG
4238 4240 4241 1 1.0950e+02 4.1840e+02 ; CG - CD1
- HD11
4238 4240 4242 1 1.0950e+02 4.1840e+02 ; CG - CD1
- HD12
4238 4240 4243 1 1.0950e+02 4.1840e+02 ; CG - CD1
- HD13
4238 4244 4245 1 1.0950e+02 4.1840e+02 ; CG - CD2
- HD21
4238 4244 4246 1 1.0950e+02 4.1840e+02 ; CG - CD2
- HD22
4238 4244 4247 1 1.0950e+02 4.1840e+02 ; CG - CD2
- HD23
4239 4238 4240 1 1.0950e+02 4.1840e+02 ; HG - CG
- CD1
4239 4238 4244 1 1.0950e+02 4.1840e+02 ; HG - CG
- CD2

| | | | | | | | |
|---|---|---|---|---|---|---|---|
| 4240 | 4238 | 4244 | 1 | 1.0950e+02 | 3.3472e+02 | ; | CD1 - CG - CD2 |
| 4241 | 4240 | 4242 | 1 | 1.0950e+02 | 2.9288e+02 | ; | HD11 - CD1 - HD12 |
| 4241 | 4240 | 4243 | 1 | 1.0950e+02 | 2.9288e+02 | ; | HD11 - CD1 - HD13 |
| 4242 | 4240 | 4243 | 1 | 1.0950e+02 | 2.9288e+02 | ; | HD12 - CD1 - HD13 |
| 4245 | 4244 | 4246 | 1 | 1.0950e+02 | 2.9288e+02 | ; | HD21 - CD2 - HD22 |
| 4245 | 4244 | 4247 | 1 | 1.0950e+02 | 2.9288e+02 | ; | HD21 - CD2 - HD23 |
| 4246 | 4244 | 4247 | 1 | 1.0950e+02 | 2.9288e+02 | ; | HD22 - CD2 - HD23 |
| 4248 | 4250 | 4251 | 1 | 1.2000e+02 | 4.1840e+02 | ; | C - N - H |
| 4248 | 4250 | 4252 | 1 | 1.2190e+02 | 4.1840e+02 | ; | C - N - CA |
| 4249 | 4248 | 4250 | 1 | 1.2290e+02 | 6.6944e+02 | ; | O - C - N |
| 4250 | 4252 | 4253 | 1 | 1.0950e+02 | 4.1840e+02 | ; | N - CA - HA |
| 4250 | 4252 | 4254 | 1 | 1.0970e+02 | 6.6944e+02 | ; | N - CA - CB |
| 4250 | 4252 | 4262 | 1 | 1.1010e+02 | 5.2718e+02 | ; | N - CA - C |
| 4251 | 4250 | 4252 | 1 | 1.1804e+02 | 4.1840e+02 | ; | H - N - CA |
| 4252 | 4254 | 4255 | 1 | 1.0950e+02 | 4.1840e+02 | ; | CA - CB - HB2 |
| 4252 | 4254 | 4256 | 1 | 1.0950e+02 | 4.1840e+02 | ; | CA - CB - HB3 |
| 4252 | 4254 | 4257 | 1 | 1.1110e+02 | 5.2718e+02 | ; | CA - CB - CG |
| 4252 | 4262 | 4263 | 1 | 1.2040e+02 | 6.6944e+02 | ; | CA - C - O |
| 4252 | 4262 | 4264 | 1 | 1.1660e+02 | 5.8576e+02 | ; | CA - C - N |
| 4253 | 4252 | 4254 | 1 | 1.0950e+02 | 4.1840e+02 | ; | HA - CA - CB |
| 4253 | 4252 | 4262 | 1 | 1.0950e+02 | 4.1840e+02 | ; | HA - CA - C |
| 4254 | 4252 | 4262 | 1 | 1.1110e+02 | 5.2718e+02 | ; | CB - CA - C |
| 4254 | 4257 | 4258 | 1 | 1.2040e+02 | 6.6944e+02 | ; | CB - CG - OD1 |
| 4254 | 4257 | 4259 | 1 | 1.1660e+02 | 5.8576e+02 | ; | CB - CG - ND2 |
| 4255 | 4254 | 4256 | 1 | 1.0950e+02 | 2.9288e+02 | ; | HB2 - CB - HB3 |
| 4255 | 4254 | 4257 | 1 | 1.0950e+02 | 4.1840e+02 | ; | HB2 - CB - CG |

| | | | | | | | |
|---|---|---|---|---|---|---|---|
| 4256 | 4254 | 4257 | 1 | 1.0950e+02 | 4.1840e+02 | ; | HB3 - CB - CG |
| 4257 | 4259 | 4260 | 1 | 1.2000e+02 | 4.1840e+02 | ; | CG - ND2 - HD21 |
| 4257 | 4259 | 4261 | 1 | 1.2000e+02 | 4.1840e+02 | ; | CG - ND2 - HD22 |
| 4258 | 4257 | 4259 | 1 | 1.2290e+02 | 6.6944e+02 | ; | OD1 - CG - ND2 |
| 4260 | 4259 | 4261 | 1 | 1.2000e+02 | 2.9288e+02 | ; | HD21 - ND2 - HD22 |
| 4262 | 4264 | 4265 | 1 | 1.2000e+02 | 4.1840e+02 | ; | C - N - H |
| 4262 | 4264 | 4266 | 1 | 1.2190e+02 | 4.1840e+02 | ; | C - N - CA |
| 4263 | 4262 | 4264 | 1 | 1.2290e+02 | 6.6944e+02 | ; | O - C - N |
| 4264 | 4266 | 4267 | 1 | 1.0950e+02 | 4.1840e+02 | ; | N - CA - HA |
| 4264 | 4266 | 4268 | 1 | 1.0970e+02 | 6.6944e+02 | ; | N - CA - CB |
| 4264 | 4266 | 4280 | 1 | 1.1010e+02 | 5.2718e+02 | ; | N - CA - C |
| 4265 | 4264 | 4266 | 1 | 1.1804e+02 | 4.1840e+02 | ; | H - N - CA |
| 4266 | 4268 | 4269 | 1 | 1.0950e+02 | 4.1840e+02 | ; | CA - CB - HB2 |
| 4266 | 4268 | 4270 | 1 | 1.0950e+02 | 4.1840e+02 | ; | CA - CB - HB3 |
| 4266 | 4268 | 4271 | 1 | 1.1310e+02 | 5.2718e+02 | ; | CA - CB - CG |
| 4266 | 4280 | 4281 | 1 | 1.2040e+02 | 6.6944e+02 | ; | CA - C - O |
| 4266 | 4280 | 4282 | 1 | 1.1660e+02 | 5.8576e+02 | ; | CA - C - N |
| 4267 | 4266 | 4268 | 1 | 1.0950e+02 | 4.1840e+02 | ; | HA - CA - CB |
| 4267 | 4266 | 4280 | 1 | 1.0950e+02 | 4.1840e+02 | ; | HA - CA - C |
| 4268 | 4266 | 4280 | 1 | 1.1110e+02 | 5.2718e+02 | ; | CB - CA - C |
| 4268 | 4271 | 4272 | 1 | 1.2000e+02 | 5.8576e+02 | ; | CB - CG - ND1 |
| 4268 | 4271 | 4278 | 1 | 1.2000e+02 | 5.8576e+02 | ; | CB - CG - CD2 |
| 4269 | 4268 | 4270 | 1 | 1.0950e+02 | 2.9288e+02 | ; | HB2 - CB - HB3 |
| 4269 | 4268 | 4271 | 1 | 1.0950e+02 | 4.1840e+02 | ; | HB2 - CB - CG |
| 4270 | 4268 | 4271 | 1 | 1.0950e+02 | 4.1840e+02 | ; | HB3 - CB - CG |
| 4271 | 4272 | 4273 | 1 | 1.2000e+02 | 4.1840e+02 | ; | CG - ND1 - HD1 |

| | | | | | | | |
|---|---|---|---|---|---|---|---|
| 4271 | 4272 | 4274 | 1 | 1.2000e+02 | 5.8576e+02 | ; | CG - ND1 - CE1 |
| 4271 | 4278 | 4276 | 1 | 1.2000e+02 | 5.8576e+02 | ; | CG - CD2 - NE2 |
| 4271 | 4278 | 4279 | 1 | 1.2000e+02 | 4.1840e+02 | ; | CG - CD2 - HD2 |
| 4272 | 4271 | 4278 | 1 | 1.2000e+02 | 5.8576e+02 | ; | ND1 - CG - CD2 |
| 4272 | 4274 | 4275 | 1 | 1.2000e+02 | 4.1840e+02 | ; | ND1 - CE1 - HE1 |
| 4272 | 4274 | 4276 | 1 | 1.2000e+02 | 5.8576e+02 | ; | ND1 - CE1 - NE2 |
| 4273 | 4272 | 4274 | 1 | 1.2000e+02 | 4.1840e+02 | ; | HD1 - ND1 - CE1 |
| 4274 | 4276 | 4277 | 1 | 1.2000e+02 | 4.1840e+02 | ; | CE1 - NE2 - HE2 |
| 4274 | 4276 | 4278 | 1 | 1.2000e+02 | 5.8576e+02 | ; | CE1 - NE2 - CD2 |
| 4275 | 4274 | 4276 | 1 | 1.2000e+02 | 4.1840e+02 | ; | HE1 - CE1 - NE2 |
| 4276 | 4278 | 4279 | 1 | 1.2000e+02 | 4.1840e+02 | ; | NE2 - CD2 - HD2 |
| 4277 | 4276 | 4278 | 1 | 1.2000e+02 | 4.1840e+02 | ; | HE2 - NE2 - CD2 |
| 4280 | 4282 | 4283 | 1 | 1.2190e+02 | 4.1840e+02 | ; | C - N - CD |
| 4280 | 4282 | 4292 | 1 | 1.2190e+02 | 4.1840e+02 | ; | C - N - CA |
| 4281 | 4280 | 4282 | 1 | 1.2290e+02 | 6.6944e+02 | ; | O - C - N |
| 4282 | 4283 | 4284 | 1 | 1.0950e+02 | 4.1840e+02 | ; | N - CD - HD2 |
| 4282 | 4283 | 4285 | 1 | 1.0950e+02 | 4.1840e+02 | ; | N - CD - HD3 |
| 4282 | 4283 | 4286 | 1 | 1.0970e+02 | 6.6944e+02 | ; | N - CD - CG |
| 4282 | 4292 | 4289 | 1 | 1.0970e+02 | 6.6944e+02 | ; | N - CA - CB |
| 4282 | 4292 | 4293 | 1 | 1.0950e+02 | 4.1840e+02 | ; | N - CA - HA |
| 4282 | 4292 | 4294 | 1 | 1.1010e+02 | 5.2718e+02 | ; | N - CA - C |
| 4283 | 4282 | 4292 | 1 | 1.1800e+02 | 4.1840e+02 | ; | CD - N - CA |
| 4283 | 4286 | 4287 | 1 | 1.0950e+02 | 4.1840e+02 | ; | CD - CG - HG2 |
| 4283 | 4286 | 4288 | 1 | 1.0950e+02 | 4.1840e+02 | ; | CD - CG - HG3 |
| 4283 | 4286 | 4289 | 1 | 1.0950e+02 | 3.3472e+02 | ; | CD - CG - CB |
| 4284 | 4283 | 4285 | 1 | 1.0950e+02 | 2.9288e+02 | ; | HD2 - CD - HD3 |

```
  4284   4283   4286      1    1.0950e+02    4.1840e+02 ;    HD2 - CD 
- CG
  4285   4283   4286      1    1.0950e+02    4.1840e+02 ;    HD3 - CD 
- CG
  4286   4289   4290      1    1.0950e+02    4.1840e+02 ;     CG - CB 
- HB2
  4286   4289   4291      1    1.0950e+02    4.1840e+02 ;     CG - CB 
- HB3
  4286   4289   4292      1    1.0950e+02    3.3472e+02 ;     CG - CB 
- CA
  4287   4286   4288      1    1.0950e+02    2.9288e+02 ;    HG2 - CG 
- HG3
  4287   4286   4289      1    1.0950e+02    4.1840e+02 ;    HG2 - CG 
- CB
  4288   4286   4289      1    1.0950e+02    4.1840e+02 ;    HG3 - CG 
- CB
  4289   4292   4293      1    1.0950e+02    4.1840e+02 ;     CB - CA 
- HA
  4289   4292   4294      1    1.1110e+02    5.2718e+02 ;     CB - CA 
- C
  4290   4289   4291      1    1.0950e+02    2.9288e+02 ;    HB2 - CB 
- HB3
  4290   4289   4292      1    1.0950e+02    4.1840e+02 ;    HB2 - CB 
- CA
  4291   4289   4292      1    1.0950e+02    4.1840e+02 ;    HB3 - CB 
- CA
  4292   4294   4295      1    1.2040e+02    6.6944e+02 ;     CA - C 
- O
  4292   4294   4296      1    1.1660e+02    5.8576e+02 ;     CA - C 
- N
  4293   4292   4294      1    1.0950e+02    4.1840e+02 ;     HA - CA 
- C
  4294   4296   4297      1    1.2000e+02    4.1840e+02 ;      C - N 
- H
  4294   4296   4298      1    1.2190e+02    4.1840e+02 ;      C - N 
- CA
  4295   4294   4296      1    1.2290e+02    6.6944e+02 ;      O - C 
- N
  4296   4298   4299      1    1.0950e+02    4.1840e+02 ;      N - CA 
- HA
  4296   4298   4300      1    1.0970e+02    6.6944e+02 ;      N - CA 
- CB
  4296   4298   4318      1    1.1010e+02    5.2718e+02 ;      N - CA 
- C
  4297   4296   4298      1    1.1804e+02    4.1840e+02 ;      H - N 
- CA
  4298   4300   4301      1    1.0950e+02    4.1840e+02 ;     CA - CB 
- HB2
  4298   4300   4302      1    1.0950e+02    4.1840e+02 ;     CA - CB 
- HB3
  4298   4300   4303      1    1.1560e+02    5.2718e+02 ;     CA - CB 
- CG
```

```
  4298   4318   4319      1    1.2040e+02    6.6944e+02 ;     CA - C
- O
  4298   4318   4320      1    1.1660e+02    5.8576e+02 ;     CA - C
- N
  4299   4298   4300      1    1.0950e+02    4.1840e+02 ;     HA - CA
- CB
  4299   4298   4318      1    1.0950e+02    4.1840e+02 ;     HA - CA
- C
  4300   4298   4318      1    1.1110e+02    5.2718e+02 ;     CB - CA
- C
  4300   4303   4304      1    1.2500e+02    5.8576e+02 ;     CB - CG
- CD1
  4300   4303   4317      1    1.2860e+02    5.8576e+02 ;     CB - CG
- CD2
  4301   4300   4302      1    1.0950e+02    2.9288e+02 ;    HB2 - CB
- HB3
  4301   4300   4303      1    1.0950e+02    4.1840e+02 ;    HB2 - CB
- CG
  4302   4300   4303      1    1.0950e+02    4.1840e+02 ;    HB3 - CB
- CG
  4303   4304   4305      1    1.2000e+02    4.1840e+02 ;     CG - CD1
- HD1
  4303   4304   4306      1    1.0870e+02    5.8576e+02 ;     CG - CD1
- NE1
  4303   4317   4308      1    1.0880e+02    5.2718e+02 ;     CG - CD2
- CE2
  4303   4317   4315      1    1.3490e+02    5.2718e+02 ;     CG - CD2
- CE3
  4304   4303   4317      1    1.0640e+02    5.2718e+02 ;    CD1 - CG
- CD2
  4304   4306   4307      1    1.2000e+02    4.1840e+02 ;    CD1 - NE1
- HE1
  4304   4306   4308      1    1.1160e+02    5.8576e+02 ;    CD1 - NE1
- CE2
  4305   4304   4306      1    1.2000e+02    4.1840e+02 ;    HD1 - CD1
- NE1
  4306   4308   4309      1    1.3280e+02    5.8576e+02 ;    NE1 - CE2
- CZ2
  4306   4308   4317      1    1.0440e+02    5.8576e+02 ;    NE1 - CE2
- CD2
  4307   4306   4308      1    1.2310e+02    4.1840e+02 ;    HE1 - NE1
- CE2
  4308   4309   4310      1    1.2000e+02    4.1840e+02 ;    CE2 - CZ2
- HZ2
  4308   4309   4311      1    1.2000e+02    5.2718e+02 ;    CE2 - CZ2
- CH2
  4308   4317   4315      1    1.1620e+02    5.2718e+02 ;    CE2 - CD2
- CE3
  4309   4308   4317      1    1.2270e+02    5.2718e+02 ;    CZ2 - CE2
- CD2
  4309   4311   4312      1    1.2000e+02    4.1840e+02 ;    CZ2 - CH2
- HH2
```

4309 4311 4313 1 1.2000e+02 5.2718e+02 ; CZ2 - CH2
- CZ3
4310 4309 4311 1 1.2000e+02 4.1840e+02 ; HZ2 - CZ2
- CH2
4311 4313 4314 1 1.2000e+02 4.1840e+02 ; CH2 - CZ3
- HZ3
4311 4313 4315 1 1.2000e+02 5.2718e+02 ; CH2 - CZ3
- CE3
4312 4311 4313 1 1.2000e+02 4.1840e+02 ; HH2 - CH2
- CZ3
4313 4315 4316 1 1.2000e+02 4.1840e+02 ; CZ3 - CE3
- HE3
4313 4315 4317 1 1.2000e+02 5.2718e+02 ; CZ3 - CE3
- CD2
4314 4313 4315 1 1.2000e+02 4.1840e+02 ; HZ3 - CZ3
- CE3
4316 4315 4317 1 1.2000e+02 4.1840e+02 ; HE3 - CE3
- CD2
4318 4320 4321 1 1.2000e+02 4.1840e+02 ; C - N
- H
4318 4320 4322 1 1.2190e+02 4.1840e+02 ; C - N
- CA
4319 4318 4320 1 1.2290e+02 6.6944e+02 ; O - C
- N
4320 4322 4323 1 1.0950e+02 4.1840e+02 ; N - CA
- HA
4320 4322 4324 1 1.0970e+02 6.6944e+02 ; N - CA
- CB
4320 4322 4337 1 1.1010e+02 5.2718e+02 ; N - CA
- C
4321 4320 4322 1 1.1804e+02 4.1840e+02 ; H - N
- CA
4322 4324 4325 1 1.0950e+02 4.1840e+02 ; CA - CB
- HB
4322 4324 4326 1 1.0950e+02 3.3472e+02 ; CA - CB
- CG2
4322 4324 4330 1 1.0950e+02 3.3472e+02 ; CA - CB
- CG1
4322 4337 4338 1 1.2040e+02 6.6944e+02 ; CA - C
- O
4322 4337 4339 1 1.1660e+02 5.8576e+02 ; CA - C
- N
4323 4322 4324 1 1.0950e+02 4.1840e+02 ; HA - CA
- CB
4323 4322 4337 1 1.0950e+02 4.1840e+02 ; HA - CA
- C
4324 4322 4337 1 1.1110e+02 5.2718e+02 ; CB - CA
- C
4324 4326 4327 1 1.0950e+02 4.1840e+02 ; CB - CG2
- HG21
4324 4326 4328 1 1.0950e+02 4.1840e+02 ; CB - CG2
- HG22

4324 4326 4329 1 1.0950e+02 4.1840e+02 ; CB - CG2
- HG23
4324 4330 4331 1 1.0950e+02 4.1840e+02 ; CB - CG1
- HG12
4324 4330 4332 1 1.0950e+02 4.1840e+02 ; CB - CG1
- HG13
4324 4330 4333 1 1.0950e+02 3.3472e+02 ; CB - CG1
- CD1
4325 4324 4326 1 1.0950e+02 4.1840e+02 ; HB - CB
- CG2
4325 4324 4330 1 1.0950e+02 4.1840e+02 ; HB - CB
- CG1
4326 4324 4330 1 1.0950e+02 3.3472e+02 ; CG2 - CB
- CG1
4327 4326 4328 1 1.0950e+02 2.9288e+02 ; HG21 - CG2
- HG22
4327 4326 4329 1 1.0950e+02 2.9288e+02 ; HG21 - CG2
- HG23
4328 4326 4329 1 1.0950e+02 2.9288e+02 ; HG22 - CG2
- HG23
4330 4333 4334 1 1.0950e+02 4.1840e+02 ; CG1 - CD1
- HD11
4330 4333 4335 1 1.0950e+02 4.1840e+02 ; CG1 - CD1
- HD12
4330 4333 4336 1 1.0950e+02 4.1840e+02 ; CG1 - CD1
- HD13
4331 4330 4332 1 1.0950e+02 2.9288e+02 ; HG12 - CG1
- HG13
4331 4330 4333 1 1.0950e+02 4.1840e+02 ; HG12 - CG1
- CD1
4332 4330 4333 1 1.0950e+02 4.1840e+02 ; HG13 - CG1
- CD1
4334 4333 4335 1 1.0950e+02 2.9288e+02 ; HD11 - CD1
- HD12
4334 4333 4336 1 1.0950e+02 2.9288e+02 ; HD11 - CD1
- HD13
4335 4333 4336 1 1.0950e+02 2.9288e+02 ; HD12 - CD1
- HD13
4337 4339 4340 1 1.2000e+02 4.1840e+02 ; C - N
- H
4337 4339 4341 1 1.2190e+02 4.1840e+02 ; C - N
- CA
4338 4337 4339 1 1.2290e+02 6.6944e+02 ; O - C
- N
4339 4341 4342 1 1.0950e+02 4.1840e+02 ; N - CA
- HA
4339 4341 4343 1 1.0970e+02 6.6944e+02 ; N - CA
- CB
4339 4341 4351 1 1.1010e+02 5.2718e+02 ; N - CA
- C
4340 4339 4341 1 1.1804e+02 4.1840e+02 ; H - N
- CA

4341 4343 4344 1 1.0950e+02 4.1840e+02 ; CA - CB
- HB
4341 4343 4345 1 1.0950e+02 3.3472e+02 ; CA - CB
- CG2
4341 4343 4349 1 1.0950e+02 4.1840e+02 ; CA - CB
- OG1
4341 4351 4352 1 1.2040e+02 6.6944e+02 ; CA - C
- O
4341 4351 4353 1 1.1660e+02 5.8576e+02 ; CA - C
- N
4342 4341 4343 1 1.0950e+02 4.1840e+02 ; HA - CA
- CB
4342 4341 4351 1 1.0950e+02 4.1840e+02 ; HA - CA
- C
4343 4341 4351 1 1.1110e+02 5.2718e+02 ; CB - CA
- C
4343 4345 4346 1 1.0950e+02 4.1840e+02 ; CB - CG2
- HG21
4343 4345 4347 1 1.0950e+02 4.1840e+02 ; CB - CG2
- HG22
4343 4345 4348 1 1.0950e+02 4.1840e+02 ; CB - CG2
- HG23
4343 4349 4350 1 1.0850e+02 4.6024e+02 ; CB - OG1
- HG1
4344 4343 4345 1 1.0950e+02 4.1840e+02 ; HB - CB
- CG2
4344 4343 4349 1 1.0950e+02 4.1840e+02 ; HB - CB
- OG1
4345 4343 4349 1 1.0950e+02 4.1840e+02 ; CG2 - CB
- OG1
4346 4345 4347 1 1.0950e+02 2.9288e+02 ; HG21 - CG2
- HG22
4346 4345 4348 1 1.0950e+02 2.9288e+02 ; HG21 - CG2
- HG23
4347 4345 4348 1 1.0950e+02 2.9288e+02 ; HG22 - CG2
- HG23
4351 4353 4354 1 1.2000e+02 4.1840e+02 ; C - N
- H
4351 4353 4355 1 1.2190e+02 4.1840e+02 ; C - N
- CA
4352 4351 4353 1 1.2290e+02 6.6944e+02 ; O - C
- N
4353 4355 4356 1 1.0950e+02 4.1840e+02 ; N - CA
- HA
4353 4355 4357 1 1.0970e+02 6.6944e+02 ; N - CA
- CB
4353 4355 4368 1 1.1010e+02 5.2718e+02 ; N - CA
- C
4354 4353 4355 1 1.1804e+02 4.1840e+02 ; H - N
- CA
4355 4357 4358 1 1.0950e+02 4.1840e+02 ; CA - CB
- HB2

```
  4355   4357   4359      1    1.0950e+02    4.1840e+02 ;     CA - CB
- HB3
  4355   4357   4360      1    1.0950e+02    3.3472e+02 ;     CA - CB
- CG
  4355   4368   4369      1    1.2040e+02    6.6944e+02 ;     CA - C
- O
  4355   4368   4370      1    1.1660e+02    5.8576e+02 ;     CA - C
- N
  4356   4355   4357      1    1.0950e+02    4.1840e+02 ;     HA - CA
- CB
  4356   4355   4368      1    1.0950e+02    4.1840e+02 ;     HA - CA
- C
  4357   4355   4368      1    1.1110e+02    5.2718e+02 ;     CB - CA
- C
  4357   4360   4361      1    1.0950e+02    4.1840e+02 ;     CB - CG
- HG2
  4357   4360   4362      1    1.0950e+02    4.1840e+02 ;     CB - CG
- HG3
  4357   4360   4363      1    1.1110e+02    5.2718e+02 ;     CB - CG
- CD
  4358   4357   4359      1    1.0950e+02    2.9288e+02 ;    HB2 - CB
- HB3
  4358   4357   4360      1    1.0950e+02    4.1840e+02 ;    HB2 - CB
- CG
  4359   4357   4360      1    1.0950e+02    4.1840e+02 ;    HB3 - CB
- CG
  4360   4363   4364      1    1.2040e+02    6.6944e+02 ;     CG - CD
- OE1
  4360   4363   4365      1    1.1660e+02    5.8576e+02 ;     CG - CD
- NE2
  4361   4360   4362      1    1.0950e+02    2.9288e+02 ;    HG2 - CG
- HG3
  4361   4360   4363      1    1.0950e+02    4.1840e+02 ;    HG2 - CG
- CD
  4362   4360   4363      1    1.0950e+02    4.1840e+02 ;    HG3 - CG
- CD
  4363   4365   4366      1    1.2000e+02    4.1840e+02 ;     CD - NE2
- HE21
  4363   4365   4367      1    1.2000e+02    4.1840e+02 ;     CD - NE2
- HE22
  4364   4363   4365      1    1.2290e+02    6.6944e+02 ;    OE1 - CD
- NE2
  4366   4365   4367      1    1.2000e+02    2.9288e+02 ;   HE21 - NE2
- HE22
  4368   4370   4371      1    1.2000e+02    4.1840e+02 ;      C - N
- H
  4368   4370   4372      1    1.2190e+02    4.1840e+02 ;      C - N
- CA
  4369   4368   4370      1    1.2290e+02    6.6944e+02 ;      O - C
- N
  4370   4372   4373      1    1.0950e+02    4.1840e+02 ;      N - CA
- HA
```

4370 4372 4374 1 1.0970e+02 6.6944e+02 ; N - CA
- CB
4370 4372 4385 1 1.1010e+02 5.2718e+02 ; N - CA
- C
4371 4370 4372 1 1.1804e+02 4.1840e+02 ; H - N
- CA
4372 4374 4375 1 1.0950e+02 4.1840e+02 ; CA - CB
- HB2
4372 4374 4376 1 1.0950e+02 4.1840e+02 ; CA - CB
- HB3
4372 4374 4377 1 1.0950e+02 3.3472e+02 ; CA - CB
- CG
4372 4385 4386 1 1.2040e+02 6.6944e+02 ; CA - C
- O
4372 4385 4387 1 1.1660e+02 5.8576e+02 ; CA - C
- N
4373 4372 4374 1 1.0950e+02 4.1840e+02 ; HA - CA
- CB
4373 4372 4385 1 1.0950e+02 4.1840e+02 ; HA - CA
- C
4374 4372 4385 1 1.1110e+02 5.2718e+02 ; CB - CA
- C
4374 4377 4378 1 1.0950e+02 4.1840e+02 ; CB - CG
- HG2
4374 4377 4379 1 1.0950e+02 4.1840e+02 ; CB - CG
- HG3
4374 4377 4380 1 1.1470e+02 4.1840e+02 ; CB - CG
- SD
4375 4374 4376 1 1.0950e+02 2.9288e+02 ; HB2 - CB
- HB3
4375 4374 4377 1 1.0950e+02 4.1840e+02 ; HB2 - CB
- CG
4376 4374 4377 1 1.0950e+02 4.1840e+02 ; HB3 - CB
- CG
4377 4380 4381 1 9.8900e+01 5.1882e+02 ; CG - SD
- CE
4378 4377 4379 1 1.0950e+02 2.9288e+02 ; HG2 - CG
- HG3
4378 4377 4380 1 1.0950e+02 4.1840e+02 ; HG2 - CG
- SD
4379 4377 4380 1 1.0950e+02 4.1840e+02 ; HG3 - CG
- SD
4380 4381 4382 1 1.0950e+02 4.1840e+02 ; SD - CE
- HE1
4380 4381 4383 1 1.0950e+02 4.1840e+02 ; SD - CE
- HE2
4380 4381 4384 1 1.0950e+02 4.1840e+02 ; SD - CE
- HE3
4382 4381 4383 1 1.0950e+02 2.9288e+02 ; HE1 - CE
- HE2
4382 4381 4384 1 1.0950e+02 2.9288e+02 ; HE1 - CE
- HE3

```
  4383   4381   4384       1    1.0950e+02    2.9288e+02 ;    HE2 - CE
- HE3
  4385   4387   4388       1    1.2000e+02    4.1840e+02 ;      C - N
- H
  4385   4387   4389       1    1.2190e+02    4.1840e+02 ;      C - N
- CA
  4386   4385   4387       1    1.2290e+02    6.6944e+02 ;      O - C
- N
  4387   4389   4390       1    1.0950e+02    4.1840e+02 ;      N - CA
- HA
  4387   4389   4391       1    1.0970e+02    6.6944e+02 ;      N - CA
- CB
  4387   4389   4399       1    1.1010e+02    5.2718e+02 ;      N - CA
- C
  4388   4387   4389       1    1.1804e+02    4.1840e+02 ;      H - N
- CA
  4389   4391   4392       1    1.0950e+02    4.1840e+02 ;     CA - CB
- HB
  4389   4391   4393       1    1.0950e+02    3.3472e+02 ;     CA - CB
- CG2
  4389   4391   4397       1    1.0950e+02    4.1840e+02 ;     CA - CB
- OG1
  4389   4399   4400       1    1.2040e+02    6.6944e+02 ;     CA - C
- O
  4389   4399   4401       1    1.1660e+02    5.8576e+02 ;     CA - C
- N
  4390   4389   4391       1    1.0950e+02    4.1840e+02 ;     HA - CA
- CB
  4390   4389   4399       1    1.0950e+02    4.1840e+02 ;     HA - CA
- C
  4391   4389   4399       1    1.1110e+02    5.2718e+02 ;     CB - CA
- C
  4391   4393   4394       1    1.0950e+02    4.1840e+02 ;     CB - CG2
- HG21
  4391   4393   4395       1    1.0950e+02    4.1840e+02 ;     CB - CG2
- HG22
  4391   4393   4396       1    1.0950e+02    4.1840e+02 ;     CB - CG2
- HG23
  4391   4397   4398       1    1.0850e+02    4.6024e+02 ;     CB - OG1
- HG1
  4392   4391   4393       1    1.0950e+02    4.1840e+02 ;     HB - CB
- CG2
  4392   4391   4397       1    1.0950e+02    4.1840e+02 ;     HB - CB
- OG1
  4393   4391   4397       1    1.0950e+02    4.1840e+02 ;    CG2 - CB
- OG1
  4394   4393   4395       1    1.0950e+02    2.9288e+02 ;   HG21 - CG2
- HG22
  4394   4393   4396       1    1.0950e+02    2.9288e+02 ;   HG21 - CG2
- HG23
  4395   4393   4396       1    1.0950e+02    2.9288e+02 ;   HG22 - CG2
- HG23
```

```
  4399   4401   4402       1    1.2000e+02    4.1840e+02 ;      C - N
- H
  4399   4401   4403       1    1.2190e+02    4.1840e+02 ;      C - N
- CA
  4400   4399   4401       1    1.2290e+02    6.6944e+02 ;      O - C
- N
  4401   4403   4404       1    1.0950e+02    4.1840e+02 ;      N - CA
- HA
  4401   4403   4405       1    1.0970e+02    6.6944e+02 ;      N - CA
- CB
  4401   4403   4421       1    1.1010e+02    5.2718e+02 ;      N - CA
- C
  4402   4401   4403       1    1.1804e+02    4.1840e+02 ;      H - N
- CA
  4403   4405   4406       1    1.0950e+02    4.1840e+02 ;     CA - CB
- HB2
  4403   4405   4407       1    1.0950e+02    4.1840e+02 ;     CA - CB
- HB3
  4403   4405   4408       1    1.0950e+02    3.3472e+02 ;     CA - CB
- CG
  4403   4421   4422       1    1.2040e+02    6.6944e+02 ;     CA - C
- O
  4403   4421   4423       1    1.1660e+02    5.8576e+02 ;     CA - C
- N
  4404   4403   4405       1    1.0950e+02    4.1840e+02 ;     HA - CA
- CB
  4404   4403   4421       1    1.0950e+02    4.1840e+02 ;     HA - CA
- C
  4405   4403   4421       1    1.1110e+02    5.2718e+02 ;     CB - CA
- C
  4405   4408   4409       1    1.0950e+02    4.1840e+02 ;     CB - CG
- HG2
  4405   4408   4410       1    1.0950e+02    4.1840e+02 ;     CB - CG
- HG3
  4405   4408   4411       1    1.0950e+02    3.3472e+02 ;     CB - CG
- CD
  4406   4405   4407       1    1.0950e+02    2.9288e+02 ;    HB2 - CB
- HB3
  4406   4405   4408       1    1.0950e+02    4.1840e+02 ;    HB2 - CB
- CG
  4407   4405   4408       1    1.0950e+02    4.1840e+02 ;    HB3 - CB
- CG
  4408   4411   4412       1    1.0950e+02    4.1840e+02 ;     CG - CD
- HD2
  4408   4411   4413       1    1.0950e+02    4.1840e+02 ;     CG - CD
- HD3
  4408   4411   4414       1    1.0950e+02    3.3472e+02 ;     CG - CD
- CE
  4409   4408   4410       1    1.0950e+02    2.9288e+02 ;    HG2 - CG
- HG3
  4409   4408   4411       1    1.0950e+02    4.1840e+02 ;    HG2 - CG
- CD
```

```
  4410   4408   4411       1    1.0950e+02    4.1840e+02 ;    HG3 - CG
- CD
  4411   4414   4415       1    1.0950e+02    4.1840e+02 ;     CD - CE
- HE2
  4411   4414   4416       1    1.0950e+02    4.1840e+02 ;     CD - CE
- HE3
  4411   4414   4417       1    1.1120e+02    6.6944e+02 ;     CD - CE
- NZ
  4412   4411   4413       1    1.0950e+02    2.9288e+02 ;    HD2 - CD
- HD3
  4412   4411   4414       1    1.0950e+02    4.1840e+02 ;    HD2 - CD
- CE
  4413   4411   4414       1    1.0950e+02    4.1840e+02 ;    HD3 - CD
- CE
  4414   4417   4418       1    1.0950e+02    4.1840e+02 ;     CE - NZ
- HZ1
  4414   4417   4419       1    1.0950e+02    4.1840e+02 ;     CE - NZ
- HZ2
  4414   4417   4420       1    1.0950e+02    4.1840e+02 ;     CE - NZ
- HZ3
  4415   4414   4416       1    1.0950e+02    2.9288e+02 ;    HE2 - CE
- HE3
  4415   4414   4417       1    1.0950e+02    4.1840e+02 ;    HE2 - CE
- NZ
  4416   4414   4417       1    1.0950e+02    4.1840e+02 ;    HE3 - CE
- NZ
  4418   4417   4419       1    1.0950e+02    2.9288e+02 ;    HZ1 - NZ
- HZ2
  4418   4417   4420       1    1.0950e+02    2.9288e+02 ;    HZ1 - NZ
- HZ3
  4419   4417   4420       1    1.0950e+02    2.9288e+02 ;    HZ2 - NZ
- HZ3
  4421   4423   4424       1    1.2000e+02    4.1840e+02 ;      C - N
- H
  4421   4423   4425       1    1.2190e+02    4.1840e+02 ;      C - N
- CA
  4422   4421   4423       1    1.2290e+02    6.6944e+02 ;      O - C
- N
  4423   4425   4426       1    1.0950e+02    4.1840e+02 ;      N - CA
- HA
  4423   4425   4427       1    1.0970e+02    6.6944e+02 ;      N - CA
- CB
  4423   4425   4432       1    1.1010e+02    5.2718e+02 ;      N - CA
- C
  4424   4423   4425       1    1.1804e+02    4.1840e+02 ;      H - N
- CA
  4425   4427   4428       1    1.0950e+02    4.1840e+02 ;     CA - CB
- HB2
  4425   4427   4429       1    1.0950e+02    4.1840e+02 ;     CA - CB
- HB3
  4425   4427   4430       1    1.0950e+02    4.1840e+02 ;     CA - CB
- OG
```

4425 4432 4433 1 1.2040e+02 6.6944e+02 ; CA - C
- O
4425 4432 4434 1 1.1660e+02 5.8576e+02 ; CA - C
- N
4426 4425 4427 1 1.0950e+02 4.1840e+02 ; HA - CA
- CB
4426 4425 4432 1 1.0950e+02 4.1840e+02 ; HA - CA
- C
4427 4425 4432 1 1.1110e+02 5.2718e+02 ; CB - CA
- C
4427 4430 4431 1 1.0850e+02 4.6024e+02 ; CB - OG
- HG
4428 4427 4429 1 1.0950e+02 2.9288e+02 ; HB2 - CB
- HB3
4428 4427 4430 1 1.0950e+02 4.1840e+02 ; HB2 - CB
- OG
4429 4427 4430 1 1.0950e+02 4.1840e+02 ; HB3 - CB
- OG
4432 4434 4435 1 1.2000e+02 4.1840e+02 ; C - N
- H
4432 4434 4436 1 1.2190e+02 4.1840e+02 ; C - N
- CA
4433 4432 4434 1 1.2290e+02 6.6944e+02 ; O - C
- N
4434 4436 4437 1 1.0950e+02 4.1840e+02 ; N - CA
- HA
4434 4436 4438 1 1.0970e+02 6.6944e+02 ; N - CA
- CB
4434 4436 4449 1 1.1010e+02 5.2718e+02 ; N - CA
- C
4435 4434 4436 1 1.1804e+02 4.1840e+02 ; H - N
- CA
4436 4438 4439 1 1.0950e+02 4.1840e+02 ; CA - CB
- HB2
4436 4438 4440 1 1.0950e+02 4.1840e+02 ; CA - CB
- HB3
4436 4438 4441 1 1.1310e+02 5.2718e+02 ; CA - CB
- CG
4436 4449 4450 1 1.2040e+02 6.6944e+02 ; CA - C
- O
4436 4449 4451 1 1.1660e+02 5.8576e+02 ; CA - C
- N
4437 4436 4438 1 1.0950e+02 4.1840e+02 ; HA - CA
- CB
4437 4436 4449 1 1.0950e+02 4.1840e+02 ; HA - CA
- C
4438 4436 4449 1 1.1110e+02 5.2718e+02 ; CB - CA
- C
4438 4441 4442 1 1.2000e+02 5.8576e+02 ; CB - CG
- ND1
4438 4441 4447 1 1.2000e+02 5.8576e+02 ; CB - CG
- CD2

4439 4438 4440 1 1.0950e+02 2.9288e+02 ; HB2 - CB
- HB3
4439 4438 4441 1 1.0950e+02 4.1840e+02 ; HB2 - CB
- CG
4440 4438 4441 1 1.0950e+02 4.1840e+02 ; HB3 - CB
- CG
4441 4442 4443 1 1.1700e+02 5.8576e+02 ; CG - ND1
- CE1
4441 4447 4445 1 1.2000e+02 5.8576e+02 ; CG - CD2
- NE2
4441 4447 4448 1 1.2000e+02 4.1840e+02 ; CG - CD2
- HD2
4442 4441 4447 1 1.2000e+02 5.8576e+02 ; ND1 - CG
- CD2
4442 4443 4444 1 1.2000e+02 4.1840e+02 ; ND1 - CE1
- HE1
4442 4443 4445 1 1.2000e+02 5.8576e+02 ; ND1 - CE1
- NE2
4443 4445 4446 1 1.2000e+02 4.1840e+02 ; CE1 - NE2
- HE2
4443 4445 4447 1 1.2000e+02 5.8576e+02 ; CE1 - NE2
- CD2
4444 4443 4445 1 1.2000e+02 4.1840e+02 ; HE1 - CE1
- NE2
4445 4447 4448 1 1.2000e+02 4.1840e+02 ; NE2 - CD2
- HD2
4446 4445 4447 1 1.2000e+02 4.1840e+02 ; HE2 - NE2
- CD2
4449 4451 4452 1 1.2000e+02 4.1840e+02 ; C - N
- H
4449 4451 4453 1 1.2190e+02 4.1840e+02 ; C - N
- CA
4450 4449 4451 1 1.2290e+02 6.6944e+02 ; O - C
- N
4451 4453 4454 1 1.0950e+02 4.1840e+02 ; N - CA
- HA
4451 4453 4455 1 1.0970e+02 6.6944e+02 ; N - CA
- CB
4451 4453 4464 1 1.1010e+02 5.2718e+02 ; N - CA
- C
4452 4451 4453 1 1.1804e+02 4.1840e+02 ; H - N
- CA
4453 4455 4456 1 1.0950e+02 4.1840e+02 ; CA - CB
- HB2
4453 4455 4457 1 1.0950e+02 4.1840e+02 ; CA - CB
- HB3
4453 4455 4458 1 1.0950e+02 3.3472e+02 ; CA - CB
- CG
4453 4464 4465 1 1.2040e+02 6.6944e+02 ; CA - C
- O
4453 4464 4466 1 1.1660e+02 5.8576e+02 ; CA - C
- N

4454 4453 4455 1 1.0950e+02 4.1840e+02 ; HA - CA - CB
4454 4453 4464 1 1.0950e+02 4.1840e+02 ; HA - CA - C
4455 4453 4464 1 1.1110e+02 5.2718e+02 ; CB - CA - C
4455 4458 4459 1 1.0950e+02 4.1840e+02 ; CB - CG - HG2
4455 4458 4460 1 1.0950e+02 4.1840e+02 ; CB - CG - HG3
4455 4458 4461 1 1.1110e+02 5.2718e+02 ; CB - CG - CD
4456 4455 4457 1 1.0950e+02 2.9288e+02 ; HB2 - CB - HB3
4456 4455 4458 1 1.0950e+02 4.1840e+02 ; HB2 - CB - CG
4457 4455 4458 1 1.0950e+02 4.1840e+02 ; HB3 - CB - CG
4458 4461 4462 1 1.1700e+02 5.8576e+02 ; CG - CD - OE1
4458 4461 4463 1 1.1700e+02 5.8576e+02 ; CG - CD - OE2
4459 4458 4460 1 1.0950e+02 2.9288e+02 ; HG2 - CG - HG3
4459 4458 4461 1 1.0950e+02 4.1840e+02 ; HG2 - CG - CD
4460 4458 4461 1 1.0950e+02 4.1840e+02 ; HG3 - CG - CD
4462 4461 4463 1 1.2600e+02 6.6944e+02 ; OE1 - CD - OE2
4464 4466 4467 1 1.2000e+02 4.1840e+02 ; C - N - H
4464 4466 4468 1 1.2190e+02 4.1840e+02 ; C - N - CA
4465 4464 4466 1 1.2290e+02 6.6944e+02 ; O - C - N
4466 4468 4469 1 1.0950e+02 4.1840e+02 ; N - CA - HA
4466 4468 4470 1 1.0970e+02 6.6944e+02 ; N - CA - CB
4466 4468 4481 1 1.1010e+02 5.2718e+02 ; N - CA - C
4467 4466 4468 1 1.1804e+02 4.1840e+02 ; H - N - CA
4468 4470 4471 1 1.0950e+02 4.1840e+02 ; CA - CB - HB2
4468 4470 4472 1 1.0950e+02 4.1840e+02 ; CA - CB - HB3
4468 4470 4473 1 1.1310e+02 5.2718e+02 ; CA - CB - CG
4468 4481 4482 1 1.2040e+02 6.6944e+02 ; CA - C - O

4468 4481 4483 1 1.1660e+02 5.8576e+02 ; CA - C
- N
4469 4468 4470 1 1.0950e+02 4.1840e+02 ; HA - CA
- CB
4469 4468 4481 1 1.0950e+02 4.1840e+02 ; HA - CA
- C
4470 4468 4481 1 1.1110e+02 5.2718e+02 ; CB - CA
- C
4470 4473 4474 1 1.2000e+02 5.8576e+02 ; CB - CG
- ND1
4470 4473 4479 1 1.2000e+02 5.8576e+02 ; CB - CG
- CD2
4471 4470 4472 1 1.0950e+02 2.9288e+02 ; HB2 - CB
- HB3
4471 4470 4473 1 1.0950e+02 4.1840e+02 ; HB2 - CB
- CG
4472 4470 4473 1 1.0950e+02 4.1840e+02 ; HB3 - CB
- CG
4473 4474 4475 1 1.1700e+02 5.8576e+02 ; CG - ND1
- CE1
4473 4479 4477 1 1.2000e+02 5.8576e+02 ; CG - CD2
- NE2
4473 4479 4480 1 1.2000e+02 4.1840e+02 ; CG - CD2
- HD2
4474 4473 4479 1 1.2000e+02 5.8576e+02 ; ND1 - CG
- CD2
4474 4475 4476 1 1.2000e+02 4.1840e+02 ; ND1 - CE1
- HE1
4474 4475 4477 1 1.2000e+02 5.8576e+02 ; ND1 - CE1
- NE2
4475 4477 4478 1 1.2000e+02 4.1840e+02 ; CE1 - NE2
- HE2
4475 4477 4479 1 1.2000e+02 5.8576e+02 ; CE1 - NE2
- CD2
4476 4475 4477 1 1.2000e+02 4.1840e+02 ; HE1 - CE1
- NE2
4477 4479 4480 1 1.2000e+02 4.1840e+02 ; NE2 - CD2
- HD2
4478 4477 4479 1 1.2000e+02 4.1840e+02 ; HE2 - NE2
- CD2
4481 4483 4484 1 1.2000e+02 4.1840e+02 ; C - N
- H
4481 4483 4485 1 1.2190e+02 4.1840e+02 ; C - N
- CA
4482 4481 4483 1 1.2290e+02 6.6944e+02 ; O - C
- N
4483 4485 4486 1 1.0950e+02 4.1840e+02 ; N - CA
- HA
4483 4485 4487 1 1.0970e+02 6.6944e+02 ; N - CA
- CB
4483 4485 4497 1 1.1010e+02 5.2718e+02 ; N - CA
- C

```
  4484   4483   4485      1    1.1804e+02    4.1840e+02 ;      H - N
- CA
  4485   4487   4488      1    1.0950e+02    4.1840e+02 ;     CA - CB
- HB
  4485   4487   4489      1    1.0950e+02    3.3472e+02 ;     CA - CB
- CG1
  4485   4487   4493      1    1.0950e+02    3.3472e+02 ;     CA - CB
- CG2
  4485   4497   4498      1    1.2040e+02    6.6944e+02 ;     CA - C
- O
  4485   4497   4499      1    1.1660e+02    5.8576e+02 ;     CA - C
- N
  4486   4485   4487      1    1.0950e+02    4.1840e+02 ;     HA - CA
- CB
  4486   4485   4497      1    1.0950e+02    4.1840e+02 ;     HA - CA
- C
  4487   4485   4497      1    1.1110e+02    5.2718e+02 ;     CB - CA
- C
  4487   4489   4490      1    1.0950e+02    4.1840e+02 ;     CB - CG1
- HG11
  4487   4489   4491      1    1.0950e+02    4.1840e+02 ;     CB - CG1
- HG12
  4487   4489   4492      1    1.0950e+02    4.1840e+02 ;     CB - CG1
- HG13
  4487   4493   4494      1    1.0950e+02    4.1840e+02 ;     CB - CG2
- HG21
  4487   4493   4495      1    1.0950e+02    4.1840e+02 ;     CB - CG2
- HG22
  4487   4493   4496      1    1.0950e+02    4.1840e+02 ;     CB - CG2
- HG23
  4488   4487   4489      1    1.0950e+02    4.1840e+02 ;     HB - CB
- CG1
  4488   4487   4493      1    1.0950e+02    4.1840e+02 ;     HB - CB
- CG2
  4489   4487   4493      1    1.0950e+02    3.3472e+02 ;    CG1 - CB
- CG2
  4490   4489   4491      1    1.0950e+02    2.9288e+02 ;   HG11 - CG1
- HG12
  4490   4489   4492      1    1.0950e+02    2.9288e+02 ;   HG11 - CG1
- HG13
  4491   4489   4492      1    1.0950e+02    2.9288e+02 ;   HG12 - CG1
- HG13
  4494   4493   4495      1    1.0950e+02    2.9288e+02 ;   HG21 - CG2
- HG22
  4494   4493   4496      1    1.0950e+02    2.9288e+02 ;   HG21 - CG2
- HG23
  4495   4493   4496      1    1.0950e+02    2.9288e+02 ;   HG22 - CG2
- HG23
  4497   4499   4500      1    1.2000e+02    4.1840e+02 ;      C - N
- H
  4497   4499   4501      1    1.2190e+02    4.1840e+02 ;      C - N
- CA
```

```
  4498   4497   4499       1    1.2290e+02    6.6944e+02 ;      O - C
- N
  4499   4501   4502       1    1.0950e+02    4.1840e+02 ;      N - CA
- HA
  4499   4501   4503       1    1.0970e+02    6.6944e+02 ;      N - CA
- CB
  4499   4501   4512       1    1.1010e+02    5.2718e+02 ;      N - CA
- C
  4500   4499   4501       1    1.1804e+02    4.1840e+02 ;      H - N
- CA
  4501   4503   4504       1    1.0950e+02    4.1840e+02 ;     CA - CB
- HB2
  4501   4503   4505       1    1.0950e+02    4.1840e+02 ;     CA - CB
- HB3
  4501   4503   4506       1    1.0950e+02    3.3472e+02 ;     CA - CB
- CG
  4501   4512   4513       1    1.2040e+02    6.6944e+02 ;     CA - C
- O
  4501   4512   4514       1    1.1660e+02    5.8576e+02 ;     CA - C
- N
  4502   4501   4503       1    1.0950e+02    4.1840e+02 ;     HA - CA
- CB
  4502   4501   4512       1    1.0950e+02    4.1840e+02 ;     HA - CA
- C
  4503   4501   4512       1    1.1110e+02    5.2718e+02 ;     CB - CA
- C
  4503   4506   4507       1    1.0950e+02    4.1840e+02 ;     CB - CG
- HG2
  4503   4506   4508       1    1.0950e+02    4.1840e+02 ;     CB - CG
- HG3
  4503   4506   4509       1    1.1110e+02    5.2718e+02 ;     CB - CG
- CD
  4504   4503   4505       1    1.0950e+02    2.9288e+02 ;    HB2 - CB
- HB3
  4504   4503   4506       1    1.0950e+02    4.1840e+02 ;    HB2 - CB
- CG
  4505   4503   4506       1    1.0950e+02    4.1840e+02 ;    HB3 - CB
- CG
  4506   4509   4510       1    1.1700e+02    5.8576e+02 ;     CG - CD
- OE1
  4506   4509   4511       1    1.1700e+02    5.8576e+02 ;     CG - CD
- OE2
  4507   4506   4508       1    1.0950e+02    2.9288e+02 ;    HG2 - CG
- HG3
  4507   4506   4509       1    1.0950e+02    4.1840e+02 ;    HG2 - CG
- CD
  4508   4506   4509       1    1.0950e+02    4.1840e+02 ;    HG3 - CG
- CD
  4510   4509   4511       1    1.2600e+02    6.6944e+02 ;    OE1 - CD
- OE2
  4512   4514   4515       1    1.2000e+02    4.1840e+02 ;      C - N
- H
```

| | | | | | | | | |
|---|---|---|---|---|---|---|---|---|
| 4512 | 4514 | 4516 | 1 | 1.2190e+02 | 4.1840e+02 ; | C | - N | - CA |
| 4513 | 4512 | 4514 | 1 | 1.2290e+02 | 6.6944e+02 ; | O | - C | - N |
| 4514 | 4516 | 4517 | 1 | 1.0950e+02 | 4.1840e+02 ; | N | - CA | - HA |
| 4514 | 4516 | 4518 | 1 | 1.0970e+02 | 6.6944e+02 ; | N | - CA | - CB |
| 4514 | 4516 | 4531 | 1 | 1.1010e+02 | 5.2718e+02 ; | N | - CA | - C |
| 4515 | 4514 | 4516 | 1 | 1.1804e+02 | 4.1840e+02 ; | H | - N | - CA |
| 4516 | 4518 | 4519 | 1 | 1.0950e+02 | 4.1840e+02 ; | CA | - CB | - HB2 |
| 4516 | 4518 | 4520 | 1 | 1.0950e+02 | 4.1840e+02 ; | CA | - CB | - HB3 |
| 4516 | 4518 | 4521 | 1 | 1.0950e+02 | 3.3472e+02 ; | CA | - CB | - CG |
| 4516 | 4531 | 4532 | 1 | 1.2040e+02 | 6.6944e+02 ; | CA | - C | - O |
| 4516 | 4531 | 4533 | 1 | 1.1660e+02 | 5.8576e+02 ; | CA | - C | - N |
| 4517 | 4516 | 4518 | 1 | 1.0950e+02 | 4.1840e+02 ; | HA | - CA | - CB |
| 4517 | 4516 | 4531 | 1 | 1.0950e+02 | 4.1840e+02 ; | HA | - CA | - C |
| 4518 | 4516 | 4531 | 1 | 1.1110e+02 | 5.2718e+02 ; | CB | - CA | - C |
| 4518 | 4521 | 4522 | 1 | 1.0950e+02 | 4.1840e+02 ; | CB | - CG | - HG |
| 4518 | 4521 | 4523 | 1 | 1.0950e+02 | 3.3472e+02 ; | CB | - CG | - CD1 |
| 4518 | 4521 | 4527 | 1 | 1.0950e+02 | 3.3472e+02 ; | CB | - CG | - CD2 |
| 4519 | 4518 | 4520 | 1 | 1.0950e+02 | 2.9288e+02 ; | HB2 | - CB | - HB3 |
| 4519 | 4518 | 4521 | 1 | 1.0950e+02 | 4.1840e+02 ; | HB2 | - CB | - CG |
| 4520 | 4518 | 4521 | 1 | 1.0950e+02 | 4.1840e+02 ; | HB3 | - CB | - CG |
| 4521 | 4523 | 4524 | 1 | 1.0950e+02 | 4.1840e+02 ; | CG | - CD1 | - HD11 |
| 4521 | 4523 | 4525 | 1 | 1.0950e+02 | 4.1840e+02 ; | CG | - CD1 | - HD12 |
| 4521 | 4523 | 4526 | 1 | 1.0950e+02 | 4.1840e+02 ; | CG | - CD1 | - HD13 |
| 4521 | 4527 | 4528 | 1 | 1.0950e+02 | 4.1840e+02 ; | CG | - CD2 | - HD21 |
| 4521 | 4527 | 4529 | 1 | 1.0950e+02 | 4.1840e+02 ; | CG | - CD2 | - HD22 |
| 4521 | 4527 | 4530 | 1 | 1.0950e+02 | 4.1840e+02 ; | CG | - CD2 | - HD23 |

```
  4522   4521   4523       1    1.0950e+02    4.1840e+02 ;     HG - CG - CD1
  4522   4521   4527       1    1.0950e+02    4.1840e+02 ;     HG - CG - CD2
  4523   4521   4527       1    1.0950e+02    3.3472e+02 ;    CD1 - CG - CD2
  4524   4523   4525       1    1.0950e+02    2.9288e+02 ;   HD11 - CD1 - HD12
  4524   4523   4526       1    1.0950e+02    2.9288e+02 ;   HD11 - CD1 - HD13
  4525   4523   4526       1    1.0950e+02    2.9288e+02 ;   HD12 - CD1 - HD13
  4528   4527   4529       1    1.0950e+02    2.9288e+02 ;   HD21 - CD2 - HD22
  4528   4527   4530       1    1.0950e+02    2.9288e+02 ;   HD21 - CD2 - HD23
  4529   4527   4530       1    1.0950e+02    2.9288e+02 ;   HD22 - CD2 - HD23
  4531   4533   4534       1    1.2000e+02    4.1840e+02 ;      C - N - H
  4531   4533   4535       1    1.2190e+02    4.1840e+02 ;      C - N - CA
  4532   4531   4533       1    1.2290e+02    6.6944e+02 ;      O - C - N
  4533   4535   4536       1    1.0950e+02    4.1840e+02 ;      N - CA - HA
  4533   4535   4537       1    1.0970e+02    6.6944e+02 ;      N - CA - CB
  4533   4535   4542       1    1.1010e+02    5.2718e+02 ;      N - CA - C
  4534   4533   4535       1    1.1804e+02    4.1840e+02 ;      H - N - CA
  4535   4537   4538       1    1.0950e+02    4.1840e+02 ;     CA - CB - HB2
  4535   4537   4539       1    1.0950e+02    4.1840e+02 ;     CA - CB - HB3
  4535   4537   4540       1    1.0950e+02    4.1840e+02 ;     CA - CB - OG
  4535   4542   4543       1    1.2040e+02    6.6944e+02 ;     CA - C - O
  4535   4542   4544       1    1.1660e+02    5.8576e+02 ;     CA - C - N
  4536   4535   4537       1    1.0950e+02    4.1840e+02 ;     HA - CA - CB
  4536   4535   4542       1    1.0950e+02    4.1840e+02 ;     HA - CA - C
  4537   4535   4542       1    1.1110e+02    5.2718e+02 ;     CB - CA - C
  4537   4540   4541       1    1.0850e+02    4.6024e+02 ;     CB - OG - HG
  4538   4537   4539       1    1.0950e+02    2.9288e+02 ;    HB2 - CB - HB3
```

| | | | | | | |
|---|---|---|---|---|---|---|
| 4538 | 4537 | 4540 | 1 | 1.0950e+02 | 4.1840e+02 ; | HB2 - CB - OG |
| 4539 | 4537 | 4540 | 1 | 1.0950e+02 | 4.1840e+02 ; | HB3 - CB - OG |
| 4542 | 4544 | 4545 | 1 | 1.2000e+02 | 4.1840e+02 ; | C - N - H |
| 4542 | 4544 | 4546 | 1 | 1.2190e+02 | 4.1840e+02 ; | C - N - CA |
| 4543 | 4542 | 4544 | 1 | 1.2290e+02 | 6.6944e+02 ; | O - C - N |
| 4544 | 4546 | 4547 | 1 | 1.0950e+02 | 4.1840e+02 ; | N - CA - HA |
| 4544 | 4546 | 4548 | 1 | 1.0970e+02 | 6.6944e+02 ; | N - CA - CB |
| 4544 | 4546 | 4553 | 1 | 1.1010e+02 | 5.2718e+02 ; | N - CA - C |
| 4545 | 4544 | 4546 | 1 | 1.1804e+02 | 4.1840e+02 ; | H - N - CA |
| 4546 | 4548 | 4549 | 1 | 1.0950e+02 | 4.1840e+02 ; | CA - CB - HB2 |
| 4546 | 4548 | 4550 | 1 | 1.0950e+02 | 4.1840e+02 ; | CA - CB - HB3 |
| 4546 | 4548 | 4551 | 1 | 1.0950e+02 | 4.1840e+02 ; | CA - CB - OG |
| 4546 | 4553 | 4554 | 1 | 1.2040e+02 | 6.6944e+02 ; | CA - C - O |
| 4546 | 4553 | 4555 | 1 | 1.1660e+02 | 5.8576e+02 ; | CA - C - N |
| 4547 | 4546 | 4548 | 1 | 1.0950e+02 | 4.1840e+02 ; | HA - CA - CB |
| 4547 | 4546 | 4553 | 1 | 1.0950e+02 | 4.1840e+02 ; | HA - CA - C |
| 4548 | 4546 | 4553 | 1 | 1.1110e+02 | 5.2718e+02 ; | CB - CA - C |
| 4548 | 4551 | 4552 | 1 | 1.0850e+02 | 4.6024e+02 ; | CB - OG - HG |
| 4549 | 4548 | 4550 | 1 | 1.0950e+02 | 2.9288e+02 ; | HB2 - CB - HB3 |
| 4549 | 4548 | 4551 | 1 | 1.0950e+02 | 4.1840e+02 ; | HB2 - CB - OG |
| 4550 | 4548 | 4551 | 1 | 1.0950e+02 | 4.1840e+02 ; | HB3 - CB - OG |
| 4553 | 4555 | 4556 | 1 | 1.2000e+02 | 4.1840e+02 ; | C - N - H |
| 4553 | 4555 | 4557 | 1 | 1.2190e+02 | 4.1840e+02 ; | C - N - CA |
| 4554 | 4553 | 4555 | 1 | 1.2290e+02 | 6.6944e+02 ; | O - C - N |
| 4555 | 4557 | 4558 | 1 | 1.0950e+02 | 4.1840e+02 ; | N - CA - HA |
| 4555 | 4557 | 4559 | 1 | 1.0970e+02 | 6.6944e+02 ; | N - CA - CB |

```
  4555   4557   4567       1    1.1010e+02    5.2718e+02 ;      N - CA
- C
  4556   4555   4557       1    1.1804e+02    4.1840e+02 ;      H - N
- CA
  4557   4559   4560       1    1.0950e+02    4.1840e+02 ;     CA - CB
- HB
  4557   4559   4561       1    1.0950e+02    3.3472e+02 ;     CA - CB
- CG2
  4557   4559   4565       1    1.0950e+02    4.1840e+02 ;     CA - CB
- OG1
  4557   4567   4568       1    1.2040e+02    6.6944e+02 ;     CA - C
- O
  4557   4567   4569       1    1.1660e+02    5.8576e+02 ;     CA - C
- N
  4558   4557   4559       1    1.0950e+02    4.1840e+02 ;     HA - CA
- CB
  4558   4557   4567       1    1.0950e+02    4.1840e+02 ;     HA - CA
- C
  4559   4557   4567       1    1.1110e+02    5.2718e+02 ;     CB - CA
- C
  4559   4561   4562       1    1.0950e+02    4.1840e+02 ;     CB - CG2
- HG21
  4559   4561   4563       1    1.0950e+02    4.1840e+02 ;     CB - CG2
- HG22
  4559   4561   4564       1    1.0950e+02    4.1840e+02 ;     CB - CG2
- HG23
  4559   4565   4566       1    1.0850e+02    4.6024e+02 ;     CB - OG1
- HG1
  4560   4559   4561       1    1.0950e+02    4.1840e+02 ;     HB - CB
- CG2
  4560   4559   4565       1    1.0950e+02    4.1840e+02 ;     HB - CB
- OG1
  4561   4559   4565       1    1.0950e+02    4.1840e+02 ;    CG2 - CB
- OG1
  4562   4561   4563       1    1.0950e+02    2.9288e+02 ;   HG21 - CG2
- HG22
  4562   4561   4564       1    1.0950e+02    2.9288e+02 ;   HG21 - CG2
- HG23
  4563   4561   4564       1    1.0950e+02    2.9288e+02 ;   HG22 - CG2
- HG23
  4567   4569   4570       1    1.2000e+02    4.1840e+02 ;      C - N
- H
  4567   4569   4571       1    1.2190e+02    4.1840e+02 ;      C - N
- CA
  4568   4567   4569       1    1.2290e+02    6.6944e+02 ;      O - C
- N
  4569   4571   4572       1    1.0950e+02    4.1840e+02 ;      N - CA
- HA
  4569   4571   4573       1    1.0970e+02    6.6944e+02 ;      N - CA
- CB
  4569   4571   4586       1    1.1010e+02    5.2718e+02 ;      N - CA
- C
```

```
  4570   4569   4571       1    1.1804e+02    4.1840e+02 ;      H - N
- CA
  4571   4573   4574       1    1.0950e+02    4.1840e+02 ;     CA - CB
- HB2
  4571   4573   4575       1    1.0950e+02    4.1840e+02 ;     CA - CB
- HB3
  4571   4573   4576       1    1.0950e+02    3.3472e+02 ;     CA - CB
- CG
  4571   4586   4587       1    1.2040e+02    6.6944e+02 ;     CA - C
- O
  4571   4586   4588       1    1.1660e+02    5.8576e+02 ;     CA - C
- N
  4572   4571   4573       1    1.0950e+02    4.1840e+02 ;     HA - CA
- CB
  4572   4571   4586       1    1.0950e+02    4.1840e+02 ;     HA - CA
- C
  4573   4571   4586       1    1.1110e+02    5.2718e+02 ;     CB - CA
- C
  4573   4576   4577       1    1.0950e+02    4.1840e+02 ;     CB - CG
- HG
  4573   4576   4578       1    1.0950e+02    3.3472e+02 ;     CB - CG
- CD1
  4573   4576   4582       1    1.0950e+02    3.3472e+02 ;     CB - CG
- CD2
  4574   4573   4575       1    1.0950e+02    2.9288e+02 ;    HB2 - CB
- HB3
  4574   4573   4576       1    1.0950e+02    4.1840e+02 ;    HB2 - CB
- CG
  4575   4573   4576       1    1.0950e+02    4.1840e+02 ;    HB3 - CB
- CG
  4576   4578   4579       1    1.0950e+02    4.1840e+02 ;     CG - CD1
- HD11
  4576   4578   4580       1    1.0950e+02    4.1840e+02 ;     CG - CD1
- HD12
  4576   4578   4581       1    1.0950e+02    4.1840e+02 ;     CG - CD1
- HD13
  4576   4582   4583       1    1.0950e+02    4.1840e+02 ;     CG - CD2
- HD21
  4576   4582   4584       1    1.0950e+02    4.1840e+02 ;     CG - CD2
- HD22
  4576   4582   4585       1    1.0950e+02    4.1840e+02 ;     CG - CD2
- HD23
  4577   4576   4578       1    1.0950e+02    4.1840e+02 ;     HG - CG
- CD1
  4577   4576   4582       1    1.0950e+02    4.1840e+02 ;     HG - CG
- CD2
  4578   4576   4582       1    1.0950e+02    3.3472e+02 ;    CD1 - CG
- CD2
  4579   4578   4580       1    1.0950e+02    2.9288e+02 ;   HD11 - CD1
- HD12
  4579   4578   4581       1    1.0950e+02    2.9288e+02 ;   HD11 - CD1
- HD13
```

```
  4580   4578   4581      1    1.0950e+02    2.9288e+02 ;   HD12 - CD1
- HD13
  4583   4582   4584      1    1.0950e+02    2.9288e+02 ;   HD21 - CD2
- HD22
  4583   4582   4585      1    1.0950e+02    2.9288e+02 ;   HD21 - CD2
- HD23
  4584   4582   4585      1    1.0950e+02    2.9288e+02 ;   HD22 - CD2
- HD23
  4586   4588   4589      1    1.2000e+02    4.1840e+02 ;      C - N
- H
  4586   4588   4590      1    1.2190e+02    4.1840e+02 ;      C - N
- CA
  4587   4586   4588      1    1.2290e+02    6.6944e+02 ;      O - C
- N
  4588   4590   4591      1    1.0950e+02    4.1840e+02 ;      N - CA
- HA
  4588   4590   4592      1    1.0970e+02    6.6944e+02 ;      N - CA
- CB
  4588   4590   4605      1    1.1010e+02    5.2718e+02 ;      N - CA
- C
  4589   4588   4590      1    1.1804e+02    4.1840e+02 ;      H - N
- CA
  4590   4592   4593      1    1.0950e+02    4.1840e+02 ;     CA - CB
- HB2
  4590   4592   4594      1    1.0950e+02    4.1840e+02 ;     CA - CB
- HB3
  4590   4592   4595      1    1.0950e+02    3.3472e+02 ;     CA - CB
- CG
  4590   4605   4606      1    1.2040e+02    6.6944e+02 ;     CA - C
- O
  4590   4605   4607      1    1.1660e+02    5.8576e+02 ;     CA - C
- N
  4591   4590   4592      1    1.0950e+02    4.1840e+02 ;     HA - CA
- CB
  4591   4590   4605      1    1.0950e+02    4.1840e+02 ;     HA - CA
- C
  4592   4590   4605      1    1.1110e+02    5.2718e+02 ;     CB - CA
- C
  4592   4595   4596      1    1.0950e+02    4.1840e+02 ;     CB - CG
- HG
  4592   4595   4597      1    1.0950e+02    3.3472e+02 ;     CB - CG
- CD1
  4592   4595   4601      1    1.0950e+02    3.3472e+02 ;     CB - CG
- CD2
  4593   4592   4594      1    1.0950e+02    2.9288e+02 ;    HB2 - CB
- HB3
  4593   4592   4595      1    1.0950e+02    4.1840e+02 ;    HB2 - CB
- CG
  4594   4592   4595      1    1.0950e+02    4.1840e+02 ;    HB3 - CB
- CG
  4595   4597   4598      1    1.0950e+02    4.1840e+02 ;     CG - CD1
- HD11
```

```
 4595  4597  4599     1    1.0950e+02    4.1840e+02 ;     CG - CD1
- HD12
 4595  4597  4600     1    1.0950e+02    4.1840e+02 ;     CG - CD1
- HD13
 4595  4601  4602     1    1.0950e+02    4.1840e+02 ;     CG - CD2
- HD21
 4595  4601  4603     1    1.0950e+02    4.1840e+02 ;     CG - CD2
- HD22
 4595  4601  4604     1    1.0950e+02    4.1840e+02 ;     CG - CD2
- HD23
 4596  4595  4597     1    1.0950e+02    4.1840e+02 ;     HG - CG
- CD1
 4596  4595  4601     1    1.0950e+02    4.1840e+02 ;     HG - CG
- CD2
 4597  4595  4601     1    1.0950e+02    3.3472e+02 ;    CD1 - CG
- CD2
 4598  4597  4599     1    1.0950e+02    2.9288e+02 ;   HD11 - CD1
- HD12
 4598  4597  4600     1    1.0950e+02    2.9288e+02 ;   HD11 - CD1
- HD13
 4599  4597  4600     1    1.0950e+02    2.9288e+02 ;   HD12 - CD1
- HD13
 4602  4601  4603     1    1.0950e+02    2.9288e+02 ;   HD21 - CD2
- HD22
 4602  4601  4604     1    1.0950e+02    2.9288e+02 ;   HD21 - CD2
- HD23
 4603  4601  4604     1    1.0950e+02    2.9288e+02 ;   HD22 - CD2
- HD23
 4605  4607  4608     1    1.2000e+02    4.1840e+02 ;      C - N
- H
 4605  4607  4609     1    1.2190e+02    4.1840e+02 ;      C - N
- CA
 4606  4605  4607     1    1.2290e+02    6.6944e+02 ;      O - C
- N
 4607  4609  4610     1    1.0950e+02    4.1840e+02 ;      N - CA
- HA
 4607  4609  4611     1    1.0970e+02    6.6944e+02 ;      N - CA
- CB
 4607  4609  4627     1    1.1010e+02    5.2718e+02 ;      N - CA
- C
 4608  4607  4609     1    1.1804e+02    4.1840e+02 ;      H - N
- CA
 4609  4611  4612     1    1.0950e+02    4.1840e+02 ;     CA - CB
- HB2
 4609  4611  4613     1    1.0950e+02    4.1840e+02 ;     CA - CB
- HB3
 4609  4611  4614     1    1.0950e+02    3.3472e+02 ;     CA - CB
- CG
 4609  4627  4628     1    1.2040e+02    6.6944e+02 ;     CA - C
- O
 4609  4627  4629     1    1.1660e+02    5.8576e+02 ;     CA - C
- N
```

4610 4609 4611 1 1.0950e+02 4.1840e+02 ; HA - CA - CB
4610 4609 4627 1 1.0950e+02 4.1840e+02 ; HA - CA - C
4611 4609 4627 1 1.1110e+02 5.2718e+02 ; CB - CA - C
4611 4614 4615 1 1.0950e+02 4.1840e+02 ; CB - CG - HG2
4611 4614 4616 1 1.0950e+02 4.1840e+02 ; CB - CG - HG3
4611 4614 4617 1 1.0950e+02 3.3472e+02 ; CB - CG - CD
4612 4611 4613 1 1.0950e+02 2.9288e+02 ; HB2 - CB - HB3
4612 4611 4614 1 1.0950e+02 4.1840e+02 ; HB2 - CB - CG
4613 4611 4614 1 1.0950e+02 4.1840e+02 ; HB3 - CB - CG
4614 4617 4618 1 1.0950e+02 4.1840e+02 ; CG - CD - HD2
4614 4617 4619 1 1.0950e+02 4.1840e+02 ; CG - CD - HD3
4614 4617 4620 1 1.0950e+02 3.3472e+02 ; CG - CD - CE
4615 4614 4616 1 1.0950e+02 2.9288e+02 ; HG2 - CG - HG3
4615 4614 4617 1 1.0950e+02 4.1840e+02 ; HG2 - CG - CD
4616 4614 4617 1 1.0950e+02 4.1840e+02 ; HG3 - CG - CD
4617 4620 4621 1 1.0950e+02 4.1840e+02 ; CD - CE - HE2
4617 4620 4622 1 1.0950e+02 4.1840e+02 ; CD - CE - HE3
4617 4620 4623 1 1.1120e+02 6.6944e+02 ; CD - CE - NZ
4618 4617 4619 1 1.0950e+02 2.9288e+02 ; HD2 - CD - HD3
4618 4617 4620 1 1.0950e+02 4.1840e+02 ; HD2 - CD - CE
4619 4617 4620 1 1.0950e+02 4.1840e+02 ; HD3 - CD - CE
4620 4623 4624 1 1.0950e+02 4.1840e+02 ; CE - NZ - HZ1
4620 4623 4625 1 1.0950e+02 4.1840e+02 ; CE - NZ - HZ2
4620 4623 4626 1 1.0950e+02 4.1840e+02 ; CE - NZ - HZ3
4621 4620 4622 1 1.0950e+02 2.9288e+02 ; HE2 - CE - HE3
4621 4620 4623 1 1.0950e+02 4.1840e+02 ; HE2 - CE - NZ

```
  4622   4620   4623       1    1.0950e+02    4.1840e+02 ;    HE3 - CE
- NZ
  4624   4623   4625       1    1.0950e+02    2.9288e+02 ;    HZ1 - NZ
- HZ2
  4624   4623   4626       1    1.0950e+02    2.9288e+02 ;    HZ1 - NZ
- HZ3
  4625   4623   4626       1    1.0950e+02    2.9288e+02 ;    HZ2 - NZ
- HZ3
  4627   4629   4630       1    1.2000e+02    4.1840e+02 ;      C - N
- H
  4627   4629   4631       1    1.2190e+02    4.1840e+02 ;      C - N
- CA
  4628   4627   4629       1    1.2290e+02    6.6944e+02 ;      O - C
- N
  4629   4631   4632       1    1.0950e+02    4.1840e+02 ;      N - CA
- HA
  4629   4631   4633       1    1.0970e+02    6.6944e+02 ;      N - CA
- CB
  4629   4631   4641       1    1.1010e+02    5.2718e+02 ;      N - CA
- C
  4630   4629   4631       1    1.1804e+02    4.1840e+02 ;      H - N
- CA
  4631   4633   4634       1    1.0950e+02    4.1840e+02 ;     CA - CB
- HB2
  4631   4633   4635       1    1.0950e+02    4.1840e+02 ;     CA - CB
- HB3
  4631   4633   4636       1    1.1110e+02    5.2718e+02 ;     CA - CB
- CG
  4631   4641   4642       1    1.2040e+02    6.6944e+02 ;     CA - C
- O
  4631   4641   4643       1    1.1660e+02    5.8576e+02 ;     CA - C
- N
  4632   4631   4633       1    1.0950e+02    4.1840e+02 ;     HA - CA
- CB
  4632   4631   4641       1    1.0950e+02    4.1840e+02 ;     HA - CA
- C
  4633   4631   4641       1    1.1110e+02    5.2718e+02 ;     CB - CA
- C
  4633   4636   4637       1    1.2040e+02    6.6944e+02 ;     CB - CG
- OD1
  4633   4636   4638       1    1.1660e+02    5.8576e+02 ;     CB - CG
- ND2
  4634   4633   4635       1    1.0950e+02    2.9288e+02 ;    HB2 - CB
- HB3
  4634   4633   4636       1    1.0950e+02    4.1840e+02 ;    HB2 - CB
- CG
  4635   4633   4636       1    1.0950e+02    4.1840e+02 ;    HB3 - CB
- CG
  4636   4638   4639       1    1.2000e+02    4.1840e+02 ;     CG - ND2
- HD21
  4636   4638   4640       1    1.2000e+02    4.1840e+02 ;     CG - ND2
- HD22
```

4637 4636 4638 1 1.2290e+02 6.6944e+02 ; OD1 - CG
- ND2
4639 4638 4640 1 1.2000e+02 2.9288e+02 ; HD21 - ND2
- HD22
4641 4643 4644 1 1.2000e+02 4.1840e+02 ; C - N
- H
4641 4643 4645 1 1.2190e+02 4.1840e+02 ; C - N
- CA
4642 4641 4643 1 1.2290e+02 6.6944e+02 ; O - C
- N
4643 4645 4646 1 1.0950e+02 4.1840e+02 ; N - CA
- HA
4643 4645 4647 1 1.0970e+02 6.6944e+02 ; N - CA
- CB
4643 4645 4660 1 1.1010e+02 5.2718e+02 ; N - CA
- C
4644 4643 4645 1 1.1804e+02 4.1840e+02 ; H - N
- CA
4645 4647 4648 1 1.0950e+02 4.1840e+02 ; CA - CB
- HB2
4645 4647 4649 1 1.0950e+02 4.1840e+02 ; CA - CB
- HB3
4645 4647 4650 1 1.0950e+02 3.3472e+02 ; CA - CB
- CG
4645 4660 4661 1 1.2040e+02 6.6944e+02 ; CA - C
- O
4645 4660 4662 1 1.1660e+02 5.8576e+02 ; CA - C
- N
4646 4645 4647 1 1.0950e+02 4.1840e+02 ; HA - CA
- CB
4646 4645 4660 1 1.0950e+02 4.1840e+02 ; HA - CA
- C
4647 4645 4660 1 1.1110e+02 5.2718e+02 ; CB - CA
- C
4647 4650 4651 1 1.0950e+02 4.1840e+02 ; CB - CG
- HG
4647 4650 4652 1 1.0950e+02 3.3472e+02 ; CB - CG
- CD1
4647 4650 4656 1 1.0950e+02 3.3472e+02 ; CB - CG
- CD2
4648 4647 4649 1 1.0950e+02 2.9288e+02 ; HB2 - CB
- HB3
4648 4647 4650 1 1.0950e+02 4.1840e+02 ; HB2 - CB
- CG
4649 4647 4650 1 1.0950e+02 4.1840e+02 ; HB3 - CB
- CG
4650 4652 4653 1 1.0950e+02 4.1840e+02 ; CG - CD1
- HD11
4650 4652 4654 1 1.0950e+02 4.1840e+02 ; CG - CD1
- HD12
4650 4652 4655 1 1.0950e+02 4.1840e+02 ; CG - CD1
- HD13

```
  4650   4656   4657       1    1.0950e+02    4.1840e+02 ;     CG - CD2
- HD21
  4650   4656   4658       1    1.0950e+02    4.1840e+02 ;     CG - CD2
- HD22
  4650   4656   4659       1    1.0950e+02    4.1840e+02 ;     CG - CD2
- HD23
  4651   4650   4652       1    1.0950e+02    4.1840e+02 ;     HG - CG
- CD1
  4651   4650   4656       1    1.0950e+02    4.1840e+02 ;     HG - CG
- CD2
  4652   4650   4656       1    1.0950e+02    3.3472e+02 ;    CD1 - CG
- CD2
  4653   4652   4654       1    1.0950e+02    2.9288e+02 ;   HD11 - CD1
- HD12
  4653   4652   4655       1    1.0950e+02    2.9288e+02 ;   HD11 - CD1
- HD13
  4654   4652   4655       1    1.0950e+02    2.9288e+02 ;   HD12 - CD1
- HD13
  4657   4656   4658       1    1.0950e+02    2.9288e+02 ;   HD21 - CD2
- HD22
  4657   4656   4659       1    1.0950e+02    2.9288e+02 ;   HD21 - CD2
- HD23
  4658   4656   4659       1    1.0950e+02    2.9288e+02 ;   HD22 - CD2
- HD23
  4660   4662   4663       1    1.2000e+02    4.1840e+02 ;      C - N
- H
  4660   4662   4664       1    1.2190e+02    4.1840e+02 ;      C - N
- CA
  4661   4660   4662       1    1.2290e+02    6.6944e+02 ;      O - C
- N
  4662   4664   4665       1    1.0950e+02    4.1840e+02 ;      N - CA
- HA
  4662   4664   4666       1    1.0970e+02    6.6944e+02 ;      N - CA
- CB
  4662   4664   4682       1    1.1010e+02    5.2718e+02 ;      N - CA
- C
  4663   4662   4664       1    1.1804e+02    4.1840e+02 ;      H - N
- CA
  4664   4666   4667       1    1.0950e+02    4.1840e+02 ;     CA - CB
- HB2
  4664   4666   4668       1    1.0950e+02    4.1840e+02 ;     CA - CB
- HB3
  4664   4666   4669       1    1.0950e+02    3.3472e+02 ;     CA - CB
- CG
  4664   4682   4683       1    1.2040e+02    6.6944e+02 ;     CA - C
- O
  4664   4682   4684       1    1.1660e+02    5.8576e+02 ;     CA - C
- N
  4665   4664   4666       1    1.0950e+02    4.1840e+02 ;     HA - CA
- CB
  4665   4664   4682       1    1.0950e+02    4.1840e+02 ;     HA - CA
- C
```

4666 4664 4682 1 1.1110e+02 5.2718e+02 ; CB - CA
- C
4666 4669 4670 1 1.0950e+02 4.1840e+02 ; CB - CG
- HG2
4666 4669 4671 1 1.0950e+02 4.1840e+02 ; CB - CG
- HG3
4666 4669 4672 1 1.0950e+02 3.3472e+02 ; CB - CG
- CD
4667 4666 4668 1 1.0950e+02 2.9288e+02 ; HB2 - CB
- HB3
4667 4666 4669 1 1.0950e+02 4.1840e+02 ; HB2 - CB
- CG
4668 4666 4669 1 1.0950e+02 4.1840e+02 ; HB3 - CB
- CG
4669 4672 4673 1 1.0950e+02 4.1840e+02 ; CG - CD
- HD2
4669 4672 4674 1 1.0950e+02 4.1840e+02 ; CG - CD
- HD3
4669 4672 4675 1 1.0950e+02 3.3472e+02 ; CG - CD
- CE
4670 4669 4671 1 1.0950e+02 2.9288e+02 ; HG2 - CG
- HG3
4670 4669 4672 1 1.0950e+02 4.1840e+02 ; HG2 - CG
- CD
4671 4669 4672 1 1.0950e+02 4.1840e+02 ; HG3 - CG
- CD
4672 4675 4676 1 1.0950e+02 4.1840e+02 ; CD - CE
- HE2
4672 4675 4677 1 1.0950e+02 4.1840e+02 ; CD - CE
- HE3
4672 4675 4678 1 1.1120e+02 6.6944e+02 ; CD - CE
- NZ
4673 4672 4674 1 1.0950e+02 2.9288e+02 ; HD2 - CD
- HD3
4673 4672 4675 1 1.0950e+02 4.1840e+02 ; HD2 - CD
- CE
4674 4672 4675 1 1.0950e+02 4.1840e+02 ; HD3 - CD
- CE
4675 4678 4679 1 1.0950e+02 4.1840e+02 ; CE - NZ
- HZ1
4675 4678 4680 1 1.0950e+02 4.1840e+02 ; CE - NZ
- HZ2
4675 4678 4681 1 1.0950e+02 4.1840e+02 ; CE - NZ
- HZ3
4676 4675 4677 1 1.0950e+02 2.9288e+02 ; HE2 - CE
- HE3
4676 4675 4678 1 1.0950e+02 4.1840e+02 ; HE2 - CE
- NZ
4677 4675 4678 1 1.0950e+02 4.1840e+02 ; HE3 - CE
- NZ
4679 4678 4680 1 1.0950e+02 2.9288e+02 ; HZ1 - NZ
- HZ2

```
  4679   4678   4681       1    1.0950e+02    2.9288e+02 ;    HZ1 - NZ
- HZ3
  4680   4678   4681       1    1.0950e+02    2.9288e+02 ;    HZ2 - NZ
- HZ3
  4682   4684   4685       1    1.2000e+02    4.1840e+02 ;      C - N
- H
  4682   4684   4686       1    1.2190e+02    4.1840e+02 ;      C - N
- CA
  4683   4682   4684       1    1.2290e+02    6.6944e+02 ;      O - C
- N
  4684   4686   4687       1    1.0950e+02    4.1840e+02 ;      N - CA
- HA
  4684   4686   4688       1    1.0970e+02    6.6944e+02 ;      N - CA
- CB
  4684   4686   4696       1    1.1010e+02    5.2718e+02 ;      N - CA
- C
  4685   4684   4686       1    1.1804e+02    4.1840e+02 ;      H - N
- CA
  4686   4688   4689       1    1.0950e+02    4.1840e+02 ;     CA - CB
- HB2
  4686   4688   4690       1    1.0950e+02    4.1840e+02 ;     CA - CB
- HB3
  4686   4688   4691       1    1.1110e+02    5.2718e+02 ;     CA - CB
- CG
  4686   4696   4697       1    1.2040e+02    6.6944e+02 ;     CA - C
- O
  4686   4696   4698       1    1.1660e+02    5.8576e+02 ;     CA - C
- N
  4687   4686   4688       1    1.0950e+02    4.1840e+02 ;     HA - CA
- CB
  4687   4686   4696       1    1.0950e+02    4.1840e+02 ;     HA - CA
- C
  4688   4686   4696       1    1.1110e+02    5.2718e+02 ;     CB - CA
- C
  4688   4691   4692       1    1.2040e+02    6.6944e+02 ;     CB - CG
- OD1
  4688   4691   4693       1    1.1660e+02    5.8576e+02 ;     CB - CG
- ND2
  4689   4688   4690       1    1.0950e+02    2.9288e+02 ;    HB2 - CB
- HB3
  4689   4688   4691       1    1.0950e+02    4.1840e+02 ;    HB2 - CB
- CG
  4690   4688   4691       1    1.0950e+02    4.1840e+02 ;    HB3 - CB
- CG
  4691   4693   4694       1    1.2000e+02    4.1840e+02 ;     CG - ND2
- HD21
  4691   4693   4695       1    1.2000e+02    4.1840e+02 ;     CG - ND2
- HD22
  4692   4691   4693       1    1.2290e+02    6.6944e+02 ;    OD1 - CG
- ND2
  4694   4693   4695       1    1.2000e+02    2.9288e+02 ;   HD21 - ND2
- HD22
```

4696 4698 4699 1 1.2000e+02 4.1840e+02 ; C - N
- H
4696 4698 4700 1 1.2190e+02 4.1840e+02 ; C - N
- CA
4697 4696 4698 1 1.2290e+02 6.6944e+02 ; O - C
- N
4698 4700 4701 1 1.0950e+02 4.1840e+02 ; N - CA
- HA
4698 4700 4702 1 1.0970e+02 6.6944e+02 ; N - CA
- CB
4698 4700 4716 1 1.1010e+02 5.2718e+02 ; N - CA
- C
4699 4698 4700 1 1.1804e+02 4.1840e+02 ; H - N
- CA
4700 4702 4703 1 1.0950e+02 4.1840e+02 ; CA - CB
- HB2
4700 4702 4704 1 1.0950e+02 4.1840e+02 ; CA - CB
- HB3
4700 4702 4705 1 1.1400e+02 5.2718e+02 ; CA - CB
- CG
4700 4716 4717 1 1.2040e+02 6.6944e+02 ; CA - C
- O
4700 4716 4718 1 1.1660e+02 5.8576e+02 ; CA - C
- N
4701 4700 4702 1 1.0950e+02 4.1840e+02 ; HA - CA
- CB
4701 4700 4716 1 1.0950e+02 4.1840e+02 ; HA - CA
- C
4702 4700 4716 1 1.1110e+02 5.2718e+02 ; CB - CA
- C
4702 4705 4706 1 1.2000e+02 5.8576e+02 ; CB - CG
- CD1
4702 4705 4714 1 1.2000e+02 5.8576e+02 ; CB - CG
- CD2
4703 4702 4704 1 1.0950e+02 2.9288e+02 ; HB2 - CB
- HB3
4703 4702 4705 1 1.0950e+02 4.1840e+02 ; HB2 - CB
- CG
4704 4702 4705 1 1.0950e+02 4.1840e+02 ; HB3 - CB
- CG
4705 4706 4707 1 1.2000e+02 4.1840e+02 ; CG - CD1
- HD1
4705 4706 4708 1 1.2000e+02 5.2718e+02 ; CG - CD1
- CE1
4705 4714 4712 1 1.2000e+02 5.2718e+02 ; CG - CD2
- CE2
4705 4714 4715 1 1.2000e+02 4.1840e+02 ; CG - CD2
- HD2
4706 4705 4714 1 1.2000e+02 5.2718e+02 ; CD1 - CG
- CD2
4706 4708 4709 1 1.2000e+02 4.1840e+02 ; CD1 - CE1
- HE1

4706 4708 4710 1 1.2000e+02 5.2718e+02 ; CD1 - CE1
- CZ
4707 4706 4708 1 1.2000e+02 4.1840e+02 ; HD1 - CD1
- CE1
4708 4710 4711 1 1.2000e+02 4.1840e+02 ; CE1 - CZ
- HZ
4708 4710 4712 1 1.2000e+02 5.2718e+02 ; CE1 - CZ
- CE2
4709 4708 4710 1 1.2000e+02 4.1840e+02 ; HE1 - CE1
- CZ
4710 4712 4713 1 1.2000e+02 4.1840e+02 ; CZ - CE2
- HE2
4710 4712 4714 1 1.2000e+02 5.2718e+02 ; CZ - CE2
- CD2
4711 4710 4712 1 1.2000e+02 4.1840e+02 ; HZ - CZ
- CE2
4712 4714 4715 1 1.2000e+02 4.1840e+02 ; CE2 - CD2
- HD2
4713 4712 4714 1 1.2000e+02 4.1840e+02 ; HE2 - CE2
- CD2
4716 4718 4719 1 1.2000e+02 4.1840e+02 ; C - N
- H
4716 4718 4720 1 1.2190e+02 4.1840e+02 ; C - N
- CA
4717 4716 4718 1 1.2290e+02 6.6944e+02 ; O - C
- N
4718 4720 4721 1 1.0950e+02 4.1840e+02 ; N - CA
- HA
4718 4720 4722 1 1.0970e+02 6.6944e+02 ; N - CA
- CB
4718 4720 4738 1 1.1010e+02 5.2718e+02 ; N - CA
- C
4719 4718 4720 1 1.1804e+02 4.1840e+02 ; H - N
- CA
4720 4722 4723 1 1.0950e+02 4.1840e+02 ; CA - CB
- HB2
4720 4722 4724 1 1.0950e+02 4.1840e+02 ; CA - CB
- HB3
4720 4722 4725 1 1.0950e+02 3.3472e+02 ; CA - CB
- CG
4720 4738 4739 1 1.2040e+02 6.6944e+02 ; CA - C
- O
4720 4738 4740 1 1.1660e+02 5.8576e+02 ; CA - C
- N
4721 4720 4722 1 1.0950e+02 4.1840e+02 ; HA - CA
- CB
4721 4720 4738 1 1.0950e+02 4.1840e+02 ; HA - CA
- C
4722 4720 4738 1 1.1110e+02 5.2718e+02 ; CB - CA
- C
4722 4725 4726 1 1.0950e+02 4.1840e+02 ; CB - CG
- HG2

4722 4725 4727 1 1.0950e+02 4.1840e+02 ; CB - CG
- HG3
4722 4725 4728 1 1.0950e+02 3.3472e+02 ; CB - CG
- CD
4723 4722 4724 1 1.0950e+02 2.9288e+02 ; HB2 - CB
- HB3
4723 4722 4725 1 1.0950e+02 4.1840e+02 ; HB2 - CB
- CG
4724 4722 4725 1 1.0950e+02 4.1840e+02 ; HB3 - CB
- CG
4725 4728 4729 1 1.0950e+02 4.1840e+02 ; CG - CD
- HD2
4725 4728 4730 1 1.0950e+02 4.1840e+02 ; CG - CD
- HD3
4725 4728 4731 1 1.0950e+02 3.3472e+02 ; CG - CD
- CE
4726 4725 4727 1 1.0950e+02 2.9288e+02 ; HG2 - CG
- HG3
4726 4725 4728 1 1.0950e+02 4.1840e+02 ; HG2 - CG
- CD
4727 4725 4728 1 1.0950e+02 4.1840e+02 ; HG3 - CG
- CD
4728 4731 4732 1 1.0950e+02 4.1840e+02 ; CD - CE
- HE2
4728 4731 4733 1 1.0950e+02 4.1840e+02 ; CD - CE
- HE3
4728 4731 4734 1 1.1120e+02 6.6944e+02 ; CD - CE
- NZ
4729 4728 4730 1 1.0950e+02 2.9288e+02 ; HD2 - CD
- HD3
4729 4728 4731 1 1.0950e+02 4.1840e+02 ; HD2 - CD
- CE
4730 4728 4731 1 1.0950e+02 4.1840e+02 ; HD3 - CD
- CE
4731 4734 4735 1 1.0950e+02 4.1840e+02 ; CE - NZ
- HZ1
4731 4734 4736 1 1.0950e+02 4.1840e+02 ; CE - NZ
- HZ2
4731 4734 4737 1 1.0950e+02 4.1840e+02 ; CE - NZ
- HZ3
4732 4731 4733 1 1.0950e+02 2.9288e+02 ; HE2 - CE
- HE3
4732 4731 4734 1 1.0950e+02 4.1840e+02 ; HE2 - CE
- NZ
4733 4731 4734 1 1.0950e+02 4.1840e+02 ; HE3 - CE
- NZ
4735 4734 4736 1 1.0950e+02 2.9288e+02 ; HZ1 - NZ
- HZ2
4735 4734 4737 1 1.0950e+02 2.9288e+02 ; HZ1 - NZ
- HZ3
4736 4734 4737 1 1.0950e+02 2.9288e+02 ; HZ2 - NZ
- HZ3

4738 4740 4741 1 1.2000e+02 4.1840e+02 ; C - N
- H
4738 4740 4742 1 1.2190e+02 4.1840e+02 ; C - N
- CA
4739 4738 4740 1 1.2290e+02 6.6944e+02 ; O - C
- N
4740 4742 4743 1 1.0950e+02 4.1840e+02 ; N - CA
- HA
4740 4742 4744 1 1.0970e+02 6.6944e+02 ; N - CA
- CB
4740 4742 4760 1 1.1010e+02 5.2718e+02 ; N - CA
- C
4741 4740 4742 1 1.1804e+02 4.1840e+02 ; H - N
- CA
4742 4744 4745 1 1.0950e+02 4.1840e+02 ; CA - CB
- HB2
4742 4744 4746 1 1.0950e+02 4.1840e+02 ; CA - CB
- HB3
4742 4744 4747 1 1.0950e+02 3.3472e+02 ; CA - CB
- CG
4742 4760 4761 1 1.2040e+02 6.6944e+02 ; CA - C
- O
4742 4760 4762 1 1.1660e+02 5.8576e+02 ; CA - C
- N
4743 4742 4744 1 1.0950e+02 4.1840e+02 ; HA - CA
- CB
4743 4742 4760 1 1.0950e+02 4.1840e+02 ; HA - CA
- C
4744 4742 4760 1 1.1110e+02 5.2718e+02 ; CB - CA
- C
4744 4747 4748 1 1.0950e+02 4.1840e+02 ; CB - CG
- HG2
4744 4747 4749 1 1.0950e+02 4.1840e+02 ; CB - CG
- HG3
4744 4747 4750 1 1.0950e+02 3.3472e+02 ; CB - CG
- CD
4745 4744 4746 1 1.0950e+02 2.9288e+02 ; HB2 - CB
- HB3
4745 4744 4747 1 1.0950e+02 4.1840e+02 ; HB2 - CB
- CG
4746 4744 4747 1 1.0950e+02 4.1840e+02 ; HB3 - CB
- CG
4747 4750 4751 1 1.0950e+02 4.1840e+02 ; CG - CD
- HD2
4747 4750 4752 1 1.0950e+02 4.1840e+02 ; CG - CD
- HD3
4747 4750 4753 1 1.0950e+02 3.3472e+02 ; CG - CD
- CE
4748 4747 4749 1 1.0950e+02 2.9288e+02 ; HG2 - CG
- HG3
4748 4747 4750 1 1.0950e+02 4.1840e+02 ; HG2 - CG
- CD

4749 4747 4750 1 1.0950e+02 4.1840e+02 ; HG3 - CG
- CD
4750 4753 4754 1 1.0950e+02 4.1840e+02 ; CD - CE
- HE2
4750 4753 4755 1 1.0950e+02 4.1840e+02 ; CD - CE
- HE3
4750 4753 4756 1 1.1120e+02 6.6944e+02 ; CD - CE
- NZ
4751 4750 4752 1 1.0950e+02 2.9288e+02 ; HD2 - CD
- HD3
4751 4750 4753 1 1.0950e+02 4.1840e+02 ; HD2 - CD
- CE
4752 4750 4753 1 1.0950e+02 4.1840e+02 ; HD3 - CD
- CE
4753 4756 4757 1 1.0950e+02 4.1840e+02 ; CE - NZ
- HZ1
4753 4756 4758 1 1.0950e+02 4.1840e+02 ; CE - NZ
- HZ2
4753 4756 4759 1 1.0950e+02 4.1840e+02 ; CE - NZ
- HZ3
4754 4753 4755 1 1.0950e+02 2.9288e+02 ; HE2 - CE
- HE3
4754 4753 4756 1 1.0950e+02 4.1840e+02 ; HE2 - CE
- NZ
4755 4753 4756 1 1.0950e+02 4.1840e+02 ; HE3 - CE
- NZ
4757 4756 4758 1 1.0950e+02 2.9288e+02 ; HZ1 - NZ
- HZ2
4757 4756 4759 1 1.0950e+02 2.9288e+02 ; HZ1 - NZ
- HZ3
4758 4756 4759 1 1.0950e+02 2.9288e+02 ; HZ2 - NZ
- HZ3
4760 4762 4763 1 1.2000e+02 4.1840e+02 ; C - N
- H
4760 4762 4764 1 1.2190e+02 4.1840e+02 ; C - N
- CA
4761 4760 4762 1 1.2290e+02 6.6944e+02 ; O - C
- N
4762 4764 4765 1 1.0950e+02 4.1840e+02 ; N - CA
- HA
4762 4764 4766 1 1.0970e+02 6.6944e+02 ; N - CA
- CB
4762 4764 4775 1 1.1010e+02 5.2718e+02 ; N - CA
- C
4763 4762 4764 1 1.1804e+02 4.1840e+02 ; H - N
- CA
4764 4766 4767 1 1.0950e+02 4.1840e+02 ; CA - CB
- HB2
4764 4766 4768 1 1.0950e+02 4.1840e+02 ; CA - CB
- HB3
4764 4766 4769 1 1.0950e+02 3.3472e+02 ; CA - CB
- CG

4764 4775 4776 1 1.2040e+02 6.6944e+02 ; CA - C
- O
4764 4775 4777 1 1.1660e+02 5.8576e+02 ; CA - C
- N
4765 4764 4766 1 1.0950e+02 4.1840e+02 ; HA - CA
- CB
4765 4764 4775 1 1.0950e+02 4.1840e+02 ; HA - CA
- C
4766 4764 4775 1 1.1110e+02 5.2718e+02 ; CB - CA
- C
4766 4769 4770 1 1.0950e+02 4.1840e+02 ; CB - CG
- HG2
4766 4769 4771 1 1.0950e+02 4.1840e+02 ; CB - CG
- HG3
4766 4769 4772 1 1.1110e+02 5.2718e+02 ; CB - CG
- CD
4767 4766 4768 1 1.0950e+02 2.9288e+02 ; HB2 - CB
- HB3
4767 4766 4769 1 1.0950e+02 4.1840e+02 ; HB2 - CB
- CG
4768 4766 4769 1 1.0950e+02 4.1840e+02 ; HB3 - CB
- CG
4769 4772 4773 1 1.1700e+02 5.8576e+02 ; CG - CD
- OE1
4769 4772 4774 1 1.1700e+02 5.8576e+02 ; CG - CD
- OE2
4770 4769 4771 1 1.0950e+02 2.9288e+02 ; HG2 - CG
- HG3
4770 4769 4772 1 1.0950e+02 4.1840e+02 ; HG2 - CG
- CD
4771 4769 4772 1 1.0950e+02 4.1840e+02 ; HG3 - CG
- CD
4773 4772 4774 1 1.2600e+02 6.6944e+02 ; OE1 - CD
- OE2
4775 4777 4778 1 1.2000e+02 4.1840e+02 ; C - N
- H
4775 4777 4779 1 1.2190e+02 4.1840e+02 ; C - N
- CA
4776 4775 4777 1 1.2290e+02 6.6944e+02 ; O - C
- N
4777 4779 4780 1 1.0950e+02 4.1840e+02 ; N - CA
- HA
4777 4779 4781 1 1.0970e+02 6.6944e+02 ; N - CA
- CB
4777 4779 4789 1 1.1010e+02 5.2718e+02 ; N - CA
- C
4778 4777 4779 1 1.1804e+02 4.1840e+02 ; H - N
- CA
4779 4781 4782 1 1.0950e+02 4.1840e+02 ; CA - CB
- HB2
4779 4781 4783 1 1.0950e+02 4.1840e+02 ; CA - CB
- HB3

```
  4779   4781   4784      1    1.1110e+02    5.2718e+02 ;     CA - CB
- CG
  4779   4789   4790      1    1.2040e+02    6.6944e+02 ;     CA - C
- O
  4779   4789   4791      1    1.1660e+02    5.8576e+02 ;     CA - C
- N
  4780   4779   4781      1    1.0950e+02    4.1840e+02 ;     HA - CA
- CB
  4780   4779   4789      1    1.0950e+02    4.1840e+02 ;     HA - CA
- C
  4781   4779   4789      1    1.1110e+02    5.2718e+02 ;     CB - CA
- C
  4781   4784   4785      1    1.2040e+02    6.6944e+02 ;     CB - CG
- OD1
  4781   4784   4786      1    1.1660e+02    5.8576e+02 ;     CB - CG
- ND2
  4782   4781   4783      1    1.0950e+02    2.9288e+02 ;    HB2 - CB
- HB3
  4782   4781   4784      1    1.0950e+02    4.1840e+02 ;    HB2 - CB
- CG
  4783   4781   4784      1    1.0950e+02    4.1840e+02 ;    HB3 - CB
- CG
  4784   4786   4787      1    1.2000e+02    4.1840e+02 ;     CG - ND2
- HD21
  4784   4786   4788      1    1.2000e+02    4.1840e+02 ;     CG - ND2
- HD22
  4785   4784   4786      1    1.2290e+02    6.6944e+02 ;    OD1 - CG
- ND2
  4787   4786   4788      1    1.2000e+02    2.9288e+02 ;   HD21 - ND2
- HD22
  4789   4791   4792      1    1.2000e+02    4.1840e+02 ;      C - N
- H
  4789   4791   4793      1    1.2190e+02    4.1840e+02 ;      C - N
- CA
  4790   4789   4791      1    1.2290e+02    6.6944e+02 ;      O - C
- N
  4791   4793   4794      1    1.0950e+02    4.1840e+02 ;      N - CA
- HA
  4791   4793   4795      1    1.0970e+02    6.6944e+02 ;      N - CA
- CB
  4791   4793   4804      1    1.1010e+02    5.2718e+02 ;      N - CA
- C
  4792   4791   4793      1    1.1804e+02    4.1840e+02 ;      H - N
- CA
  4793   4795   4796      1    1.0950e+02    4.1840e+02 ;     CA - CB
- HB2
  4793   4795   4797      1    1.0950e+02    4.1840e+02 ;     CA - CB
- HB3
  4793   4795   4798      1    1.0950e+02    3.3472e+02 ;     CA - CB
- CG
  4793   4804   4805      1    1.2040e+02    6.6944e+02 ;     CA - C
- O
```

```
  4793   4804   4806      1    1.1660e+02    5.8576e+02 ;     CA - C
- N
  4794   4793   4795      1    1.0950e+02    4.1840e+02 ;     HA - CA
- CB
  4794   4793   4804      1    1.0950e+02    4.1840e+02 ;     HA - CA
- C
  4795   4793   4804      1    1.1110e+02    5.2718e+02 ;     CB - CA
- C
  4795   4798   4799      1    1.0950e+02    4.1840e+02 ;     CB - CG
- HG2
  4795   4798   4800      1    1.0950e+02    4.1840e+02 ;     CB - CG
- HG3
  4795   4798   4801      1    1.1110e+02    5.2718e+02 ;     CB - CG
- CD
  4796   4795   4797      1    1.0950e+02    2.9288e+02 ;    HB2 - CB
- HB3
  4796   4795   4798      1    1.0950e+02    4.1840e+02 ;    HB2 - CB
- CG
  4797   4795   4798      1    1.0950e+02    4.1840e+02 ;    HB3 - CB
- CG
  4798   4801   4802      1    1.1700e+02    5.8576e+02 ;     CG - CD
- OE1
  4798   4801   4803      1    1.1700e+02    5.8576e+02 ;     CG - CD
- OE2
  4799   4798   4800      1    1.0950e+02    2.9288e+02 ;    HG2 - CG
- HG3
  4799   4798   4801      1    1.0950e+02    4.1840e+02 ;    HG2 - CG
- CD
  4800   4798   4801      1    1.0950e+02    4.1840e+02 ;    HG3 - CG
- CD
  4802   4801   4803      1    1.2600e+02    6.6944e+02 ;    OE1 - CD
- OE2
  4804   4806   4807      1    1.2000e+02    4.1840e+02 ;      C - N
- H
  4804   4806   4808      1    1.2190e+02    4.1840e+02 ;      C - N
- CA
  4805   4804   4806      1    1.2290e+02    6.6944e+02 ;      O - C
- N
  4806   4808   4809      1    1.0950e+02    4.1840e+02 ;      N - CA
- HA
  4806   4808   4810      1    1.0970e+02    6.6944e+02 ;      N - CA
- CB
  4806   4808   4823      1    1.1010e+02    5.2718e+02 ;      N - CA
- C
  4807   4806   4808      1    1.1804e+02    4.1840e+02 ;      H - N
- CA
  4808   4810   4811      1    1.0950e+02    4.1840e+02 ;     CA - CB
- HB2
  4808   4810   4812      1    1.0950e+02    4.1840e+02 ;     CA - CB
- HB3
  4808   4810   4813      1    1.0950e+02    3.3472e+02 ;     CA - CB
- CG
```

```
  4808   4823   4824      1    1.2040e+02    6.6944e+02 ;     CA - C
- O
  4808   4823   4825      1    1.1660e+02    5.8576e+02 ;     CA - C
- N
  4809   4808   4810      1    1.0950e+02    4.1840e+02 ;     HA - CA
- CB
  4809   4808   4823      1    1.0950e+02    4.1840e+02 ;     HA - CA
- C
  4810   4808   4823      1    1.1110e+02    5.2718e+02 ;     CB - CA
- C
  4810   4813   4814      1    1.0950e+02    4.1840e+02 ;     CB - CG
- HG
  4810   4813   4815      1    1.0950e+02    3.3472e+02 ;     CB - CG
- CD1
  4810   4813   4819      1    1.0950e+02    3.3472e+02 ;     CB - CG
- CD2
  4811   4810   4812      1    1.0950e+02    2.9288e+02 ;    HB2 - CB
- HB3
  4811   4810   4813      1    1.0950e+02    4.1840e+02 ;    HB2 - CB
- CG
  4812   4810   4813      1    1.0950e+02    4.1840e+02 ;    HB3 - CB
- CG
  4813   4815   4816      1    1.0950e+02    4.1840e+02 ;     CG - CD1
- HD11
  4813   4815   4817      1    1.0950e+02    4.1840e+02 ;     CG - CD1
- HD12
  4813   4815   4818      1    1.0950e+02    4.1840e+02 ;     CG - CD1
- HD13
  4813   4819   4820      1    1.0950e+02    4.1840e+02 ;     CG - CD2
- HD21
  4813   4819   4821      1    1.0950e+02    4.1840e+02 ;     CG - CD2
- HD22
  4813   4819   4822      1    1.0950e+02    4.1840e+02 ;     CG - CD2
- HD23
  4814   4813   4815      1    1.0950e+02    4.1840e+02 ;     HG - CG
- CD1
  4814   4813   4819      1    1.0950e+02    4.1840e+02 ;     HG - CG
- CD2
  4815   4813   4819      1    1.0950e+02    3.3472e+02 ;    CD1 - CG
- CD2
  4816   4815   4817      1    1.0950e+02    2.9288e+02 ;   HD11 - CD1
- HD12
  4816   4815   4818      1    1.0950e+02    2.9288e+02 ;   HD11 - CD1
- HD13
  4817   4815   4818      1    1.0950e+02    2.9288e+02 ;   HD12 - CD1
- HD13
  4820   4819   4821      1    1.0950e+02    2.9288e+02 ;   HD21 - CD2
- HD22
  4820   4819   4822      1    1.0950e+02    2.9288e+02 ;   HD21 - CD2
- HD23
  4821   4819   4822      1    1.0950e+02    2.9288e+02 ;   HD22 - CD2
- HD23
```

4823 4825 4826 1 1.2000e+02 4.1840e+02 ; C - N
- H
4823 4825 4827 1 1.2190e+02 4.1840e+02 ; C - N
- CA
4824 4823 4825 1 1.2290e+02 6.6944e+02 ; O - C
- N
4825 4827 4828 1 1.0950e+02 4.1840e+02 ; N - CA
- HA
4825 4827 4829 1 1.0970e+02 6.6944e+02 ; N - CA
- CB
4825 4827 4845 1 1.1010e+02 5.2718e+02 ; N - CA
- C
4826 4825 4827 1 1.1804e+02 4.1840e+02 ; H - N
- CA
4827 4829 4830 1 1.0950e+02 4.1840e+02 ; CA - CB
- HB2
4827 4829 4831 1 1.0950e+02 4.1840e+02 ; CA - CB
- HB3
4827 4829 4832 1 1.0950e+02 3.3472e+02 ; CA - CB
- CG
4827 4845 4846 1 1.2040e+02 6.6944e+02 ; CA - C
- O
4827 4845 4847 1 1.1660e+02 5.8576e+02 ; CA - C
- N
4828 4827 4829 1 1.0950e+02 4.1840e+02 ; HA - CA
- CB
4828 4827 4845 1 1.0950e+02 4.1840e+02 ; HA - CA
- C
4829 4827 4845 1 1.1110e+02 5.2718e+02 ; CB - CA
- C
4829 4832 4833 1 1.0950e+02 4.1840e+02 ; CB - CG
- HG2
4829 4832 4834 1 1.0950e+02 4.1840e+02 ; CB - CG
- HG3
4829 4832 4835 1 1.0950e+02 3.3472e+02 ; CB - CG
- CD
4830 4829 4831 1 1.0950e+02 2.9288e+02 ; HB2 - CB
- HB3
4830 4829 4832 1 1.0950e+02 4.1840e+02 ; HB2 - CB
- CG
4831 4829 4832 1 1.0950e+02 4.1840e+02 ; HB3 - CB
- CG
4832 4835 4836 1 1.0950e+02 4.1840e+02 ; CG - CD
- HD2
4832 4835 4837 1 1.0950e+02 4.1840e+02 ; CG - CD
- HD3
4832 4835 4838 1 1.0950e+02 3.3472e+02 ; CG - CD
- CE
4833 4832 4834 1 1.0950e+02 2.9288e+02 ; HG2 - CG
- HG3
4833 4832 4835 1 1.0950e+02 4.1840e+02 ; HG2 - CG
- CD

```
  4834   4832   4835      1    1.0950e+02    4.1840e+02 ;    HG3 - CG
- CD
  4835   4838   4839      1    1.0950e+02    4.1840e+02 ;     CD - CE
- HE2
  4835   4838   4840      1    1.0950e+02    4.1840e+02 ;     CD - CE
- HE3
  4835   4838   4841      1    1.1120e+02    6.6944e+02 ;     CD - CE
- NZ
  4836   4835   4837      1    1.0950e+02    2.9288e+02 ;    HD2 - CD
- HD3
  4836   4835   4838      1    1.0950e+02    4.1840e+02 ;    HD2 - CD
- CE
  4837   4835   4838      1    1.0950e+02    4.1840e+02 ;    HD3 - CD
- CE
  4838   4841   4842      1    1.0950e+02    4.1840e+02 ;     CE - NZ
- HZ1
  4838   4841   4843      1    1.0950e+02    4.1840e+02 ;     CE - NZ
- HZ2
  4838   4841   4844      1    1.0950e+02    4.1840e+02 ;     CE - NZ
- HZ3
  4839   4838   4840      1    1.0950e+02    2.9288e+02 ;    HE2 - CE
- HE3
  4839   4838   4841      1    1.0950e+02    4.1840e+02 ;    HE2 - CE
- NZ
  4840   4838   4841      1    1.0950e+02    4.1840e+02 ;    HE3 - CE
- NZ
  4842   4841   4843      1    1.0950e+02    2.9288e+02 ;    HZ1 - NZ
- HZ2
  4842   4841   4844      1    1.0950e+02    2.9288e+02 ;    HZ1 - NZ
- HZ3
  4843   4841   4844      1    1.0950e+02    2.9288e+02 ;    HZ2 - NZ
- HZ3
  4845   4847   4848      1    1.2000e+02    4.1840e+02 ;      C - N
- H
  4845   4847   4849      1    1.2190e+02    4.1840e+02 ;      C - N
- CA
  4846   4845   4847      1    1.2290e+02    6.6944e+02 ;      O - C
- N
  4847   4849   4850      1    1.0950e+02    4.1840e+02 ;      N - CA
- HA
  4847   4849   4851      1    1.0970e+02    6.6944e+02 ;      N - CA
- CB
  4847   4849   4867      1    1.1010e+02    5.2718e+02 ;      N - CA
- C
  4848   4847   4849      1    1.1804e+02    4.1840e+02 ;      H - N
- CA
  4849   4851   4852      1    1.0950e+02    4.1840e+02 ;     CA - CB
- HB2
  4849   4851   4853      1    1.0950e+02    4.1840e+02 ;     CA - CB
- HB3
  4849   4851   4854      1    1.0950e+02    3.3472e+02 ;     CA - CB
- CG
```

4849 4867 4868 1 1.2040e+02 6.6944e+02 ; CA - C
- O
4849 4867 4869 1 1.1660e+02 5.8576e+02 ; CA - C
- N
4850 4849 4851 1 1.0950e+02 4.1840e+02 ; HA - CA
- CB
4850 4849 4867 1 1.0950e+02 4.1840e+02 ; HA - CA
- C
4851 4849 4867 1 1.1110e+02 5.2718e+02 ; CB - CA
- C
4851 4854 4855 1 1.0950e+02 4.1840e+02 ; CB - CG
- HG2
4851 4854 4856 1 1.0950e+02 4.1840e+02 ; CB - CG
- HG3
4851 4854 4857 1 1.0950e+02 3.3472e+02 ; CB - CG
- CD
4852 4851 4853 1 1.0950e+02 2.9288e+02 ; HB2 - CB
- HB3
4852 4851 4854 1 1.0950e+02 4.1840e+02 ; HB2 - CB
- CG
4853 4851 4854 1 1.0950e+02 4.1840e+02 ; HB3 - CB
- CG
4854 4857 4858 1 1.0950e+02 4.1840e+02 ; CG - CD
- HD2
4854 4857 4859 1 1.0950e+02 4.1840e+02 ; CG - CD
- HD3
4854 4857 4860 1 1.0950e+02 3.3472e+02 ; CG - CD
- CE
4855 4854 4856 1 1.0950e+02 2.9288e+02 ; HG2 - CG
- HG3
4855 4854 4857 1 1.0950e+02 4.1840e+02 ; HG2 - CG
- CD
4856 4854 4857 1 1.0950e+02 4.1840e+02 ; HG3 - CG
- CD
4857 4860 4861 1 1.0950e+02 4.1840e+02 ; CD - CE
- HE2
4857 4860 4862 1 1.0950e+02 4.1840e+02 ; CD - CE
- HE3
4857 4860 4863 1 1.1120e+02 6.6944e+02 ; CD - CE
- NZ
4858 4857 4859 1 1.0950e+02 2.9288e+02 ; HD2 - CD
- HD3
4858 4857 4860 1 1.0950e+02 4.1840e+02 ; HD2 - CD
- CE
4859 4857 4860 1 1.0950e+02 4.1840e+02 ; HD3 - CD
- CE
4860 4863 4864 1 1.0950e+02 4.1840e+02 ; CE - NZ
- HZ1
4860 4863 4865 1 1.0950e+02 4.1840e+02 ; CE - NZ
- HZ2
4860 4863 4866 1 1.0950e+02 4.1840e+02 ; CE - NZ
- HZ3

4861 4860 4862 1 1.0950e+02 2.9288e+02 ; HE2 - CE
- HE3
4861 4860 4863 1 1.0950e+02 4.1840e+02 ; HE2 - CE
- NZ
4862 4860 4863 1 1.0950e+02 4.1840e+02 ; HE3 - CE
- NZ
4864 4863 4865 1 1.0950e+02 2.9288e+02 ; HZ1 - NZ
- HZ2
4864 4863 4866 1 1.0950e+02 2.9288e+02 ; HZ1 - NZ
- HZ3
4865 4863 4866 1 1.0950e+02 2.9288e+02 ; HZ2 - NZ
- HZ3
4867 4869 4870 1 1.2000e+02 4.1840e+02 ; C - N
- H
4867 4869 4871 1 1.2190e+02 4.1840e+02 ; C - N
- CA
4868 4867 4869 1 1.2290e+02 6.6944e+02 ; O - C
- N
4869 4871 4872 1 1.0950e+02 4.1840e+02 ; N - CA
- HA
4869 4871 4873 1 1.0970e+02 6.6944e+02 ; N - CA
- CB
4869 4871 4886 1 1.1010e+02 5.2718e+02 ; N - CA
- C
4870 4869 4871 1 1.1804e+02 4.1840e+02 ; H - N
- CA
4871 4873 4874 1 1.0950e+02 4.1840e+02 ; CA - CB
- HB
4871 4873 4875 1 1.0950e+02 3.3472e+02 ; CA - CB
- CG2
4871 4873 4879 1 1.0950e+02 3.3472e+02 ; CA - CB
- CG1
4871 4886 4887 1 1.2040e+02 6.6944e+02 ; CA - C
- O
4871 4886 4888 1 1.1660e+02 5.8576e+02 ; CA - C
- N
4872 4871 4873 1 1.0950e+02 4.1840e+02 ; HA - CA
- CB
4872 4871 4886 1 1.0950e+02 4.1840e+02 ; HA - CA
- C
4873 4871 4886 1 1.1110e+02 5.2718e+02 ; CB - CA
- C
4873 4875 4876 1 1.0950e+02 4.1840e+02 ; CB - CG2
- HG21
4873 4875 4877 1 1.0950e+02 4.1840e+02 ; CB - CG2
- HG22
4873 4875 4878 1 1.0950e+02 4.1840e+02 ; CB - CG2
- HG23
4873 4879 4880 1 1.0950e+02 4.1840e+02 ; CB - CG1
- HG12
4873 4879 4881 1 1.0950e+02 4.1840e+02 ; CB - CG1
- HG13

4873 4879 4882 1 1.0950e+02 3.3472e+02 ; CB - CG1
- CD1
4874 4873 4875 1 1.0950e+02 4.1840e+02 ; HB - CB
- CG2
4874 4873 4879 1 1.0950e+02 4.1840e+02 ; HB - CB
- CG1
4875 4873 4879 1 1.0950e+02 3.3472e+02 ; CG2 - CB
- CG1
4876 4875 4877 1 1.0950e+02 2.9288e+02 ; HG21 - CG2
- HG22
4876 4875 4878 1 1.0950e+02 2.9288e+02 ; HG21 - CG2
- HG23
4877 4875 4878 1 1.0950e+02 2.9288e+02 ; HG22 - CG2
- HG23
4879 4882 4883 1 1.0950e+02 4.1840e+02 ; CG1 - CD1
- HD11
4879 4882 4884 1 1.0950e+02 4.1840e+02 ; CG1 - CD1
- HD12
4879 4882 4885 1 1.0950e+02 4.1840e+02 ; CG1 - CD1
- HD13
4880 4879 4881 1 1.0950e+02 2.9288e+02 ; HG12 - CG1
- HG13
4880 4879 4882 1 1.0950e+02 4.1840e+02 ; HG12 - CG1
- CD1
4881 4879 4882 1 1.0950e+02 4.1840e+02 ; HG13 - CG1
- CD1
4883 4882 4884 1 1.0950e+02 2.9288e+02 ; HD11 - CD1
- HD12
4883 4882 4885 1 1.0950e+02 2.9288e+02 ; HD11 - CD1
- HD13
4884 4882 4885 1 1.0950e+02 2.9288e+02 ; HD12 - CD1
- HD13
4886 4888 4889 1 1.2000e+02 4.1840e+02 ; C - N
- H
4886 4888 4890 1 1.2190e+02 4.1840e+02 ; C - N
- CA
4887 4886 4888 1 1.2290e+02 6.6944e+02 ; O - C
- N
4888 4890 4891 1 1.0950e+02 4.1840e+02 ; N - CA
- HA
4888 4890 4892 1 1.0970e+02 6.6944e+02 ; N - CA
- CB
4888 4890 4896 1 1.1010e+02 5.2718e+02 ; N - CA
- C
4889 4888 4890 1 1.1804e+02 4.1840e+02 ; H - N
- CA
4890 4892 4893 1 1.0950e+02 4.1840e+02 ; CA - CB
- HB1
4890 4892 4894 1 1.0950e+02 4.1840e+02 ; CA - CB
- HB2
4890 4892 4895 1 1.0950e+02 4.1840e+02 ; CA - CB
- HB3

4890 4896 4897 1 1.2040e+02 6.6944e+02 ; CA - C
- O
4890 4896 4898 1 1.1660e+02 5.8576e+02 ; CA - C
- N
4891 4890 4892 1 1.0950e+02 4.1840e+02 ; HA - CA
- CB
4891 4890 4896 1 1.0950e+02 4.1840e+02 ; HA - CA
- C
4892 4890 4896 1 1.1110e+02 5.2718e+02 ; CB - CA
- C
4893 4892 4894 1 1.0950e+02 2.9288e+02 ; HB1 - CB
- HB2
4893 4892 4895 1 1.0950e+02 2.9288e+02 ; HB1 - CB
- HB3
4894 4892 4895 1 1.0950e+02 2.9288e+02 ; HB2 - CB
- HB3
4896 4898 4899 1 1.2000e+02 4.1840e+02 ; C - N
- H
4896 4898 4900 1 1.2190e+02 4.1840e+02 ; C - N
- CA
4897 4896 4898 1 1.2290e+02 6.6944e+02 ; O - C
- N
4898 4900 4901 1 1.0950e+02 4.1840e+02 ; N - CA
- HA
4898 4900 4902 1 1.0970e+02 6.6944e+02 ; N - CA
- CB
4898 4900 4915 1 1.1010e+02 5.2718e+02 ; N - CA
- C
4899 4898 4900 1 1.1804e+02 4.1840e+02 ; H - N
- CA
4900 4902 4903 1 1.0950e+02 4.1840e+02 ; CA - CB
- HB2
4900 4902 4904 1 1.0950e+02 4.1840e+02 ; CA - CB
- HB3
4900 4902 4905 1 1.0950e+02 3.3472e+02 ; CA - CB
- CG
4900 4915 4916 1 1.2040e+02 6.6944e+02 ; CA - C
- O
4900 4915 4917 1 1.1660e+02 5.8576e+02 ; CA - C
- N
4901 4900 4902 1 1.0950e+02 4.1840e+02 ; HA - CA
- CB
4901 4900 4915 1 1.0950e+02 4.1840e+02 ; HA - CA
- C
4902 4900 4915 1 1.1110e+02 5.2718e+02 ; CB - CA
- C
4902 4905 4906 1 1.0950e+02 4.1840e+02 ; CB - CG
- HG
4902 4905 4907 1 1.0950e+02 3.3472e+02 ; CB - CG
- CD1
4902 4905 4911 1 1.0950e+02 3.3472e+02 ; CB - CG
- CD2

```
  4903   4902   4904      1    1.0950e+02    2.9288e+02 ;    HB2 - CB
- HB3
  4903   4902   4905      1    1.0950e+02    4.1840e+02 ;    HB2 - CB
- CG
  4904   4902   4905      1    1.0950e+02    4.1840e+02 ;    HB3 - CB
- CG
  4905   4907   4908      1    1.0950e+02    4.1840e+02 ;     CG - CD1
- HD11
  4905   4907   4909      1    1.0950e+02    4.1840e+02 ;     CG - CD1
- HD12
  4905   4907   4910      1    1.0950e+02    4.1840e+02 ;     CG - CD1
- HD13
  4905   4911   4912      1    1.0950e+02    4.1840e+02 ;     CG - CD2
- HD21
  4905   4911   4913      1    1.0950e+02    4.1840e+02 ;     CG - CD2
- HD22
  4905   4911   4914      1    1.0950e+02    4.1840e+02 ;     CG - CD2
- HD23
  4906   4905   4907      1    1.0950e+02    4.1840e+02 ;     HG - CG
- CD1
  4906   4905   4911      1    1.0950e+02    4.1840e+02 ;     HG - CG
- CD2
  4907   4905   4911      1    1.0950e+02    3.3472e+02 ;    CD1 - CG
- CD2
  4908   4907   4909      1    1.0950e+02    2.9288e+02 ;   HD11 - CD1
- HD12
  4908   4907   4910      1    1.0950e+02    2.9288e+02 ;   HD11 - CD1
- HD13
  4909   4907   4910      1    1.0950e+02    2.9288e+02 ;   HD12 - CD1
- HD13
  4912   4911   4913      1    1.0950e+02    2.9288e+02 ;   HD21 - CD2
- HD22
  4912   4911   4914      1    1.0950e+02    2.9288e+02 ;   HD21 - CD2
- HD23
  4913   4911   4914      1    1.0950e+02    2.9288e+02 ;   HD22 - CD2
- HD23
  4915   4917   4918      1    1.2000e+02    4.1840e+02 ;      C - N
- H
  4915   4917   4919      1    1.2190e+02    4.1840e+02 ;      C - N
- CA
  4916   4915   4917      1    1.2290e+02    6.6944e+02 ;      O - C
- N
  4917   4919   4920      1    1.0950e+02    4.1840e+02 ;      N - CA
- HA
  4917   4919   4921      1    1.0970e+02    6.6944e+02 ;      N - CA
- CB
  4917   4919   4929      1    1.1010e+02    5.2718e+02 ;      N - CA
- C
  4918   4917   4919      1    1.1804e+02    4.1840e+02 ;      H - N
- CA
  4919   4921   4922      1    1.0950e+02    4.1840e+02 ;     CA - CB
- HB
```

```
  4919   4921   4923       1    1.0950e+02    3.3472e+02 ;     CA - CB
- CG2
  4919   4921   4927       1    1.0950e+02    4.1840e+02 ;     CA - CB
- OG1
  4919   4929   4930       1    1.2040e+02    6.6944e+02 ;     CA - C
- O
  4919   4929   4931       1    1.1660e+02    5.8576e+02 ;     CA - C
- N
  4920   4919   4921       1    1.0950e+02    4.1840e+02 ;     HA - CA
- CB
  4920   4919   4929       1    1.0950e+02    4.1840e+02 ;     HA - CA
- C
  4921   4919   4929       1    1.1110e+02    5.2718e+02 ;     CB - CA
- C
  4921   4923   4924       1    1.0950e+02    4.1840e+02 ;     CB - CG2
- HG21
  4921   4923   4925       1    1.0950e+02    4.1840e+02 ;     CB - CG2
- HG22
  4921   4923   4926       1    1.0950e+02    4.1840e+02 ;     CB - CG2
- HG23
  4921   4927   4928       1    1.0850e+02    4.6024e+02 ;     CB - OG1
- HG1
  4922   4921   4923       1    1.0950e+02    4.1840e+02 ;     HB - CB
- CG2
  4922   4921   4927       1    1.0950e+02    4.1840e+02 ;     HB - CB
- OG1
  4923   4921   4927       1    1.0950e+02    4.1840e+02 ;    CG2 - CB
- OG1
  4924   4923   4925       1    1.0950e+02    2.9288e+02 ;   HG21 - CG2
- HG22
  4924   4923   4926       1    1.0950e+02    2.9288e+02 ;   HG21 - CG2
- HG23
  4925   4923   4926       1    1.0950e+02    2.9288e+02 ;   HG22 - CG2
- HG23
  4929   4931   4932       1    1.2000e+02    4.1840e+02 ;      C - N
- H
  4929   4931   4933       1    1.2190e+02    4.1840e+02 ;      C - N
- CA
  4930   4929   4931       1    1.2290e+02    6.6944e+02 ;      O - C
- N
  4931   4933   4934       1    1.0950e+02    4.1840e+02 ;      N - CA
- HA
  4931   4933   4935       1    1.0970e+02    6.6944e+02 ;      N - CA
- CB
  4931   4933   4948       1    1.1010e+02    5.2718e+02 ;      N - CA
- C
  4932   4931   4933       1    1.1804e+02    4.1840e+02 ;      H - N
- CA
  4933   4935   4936       1    1.0950e+02    4.1840e+02 ;     CA - CB
- HB
  4933   4935   4937       1    1.0950e+02    3.3472e+02 ;     CA - CB
- CG2
```

4933 4935 4941 1 1.0950e+02 3.3472e+02 ; CA - CB
- CG1
4933 4948 4949 1 1.2040e+02 6.6944e+02 ; CA - C
- O
4933 4948 4950 1 1.1660e+02 5.8576e+02 ; CA - C
- N
4934 4933 4935 1 1.0950e+02 4.1840e+02 ; HA - CA
- CB
4934 4933 4948 1 1.0950e+02 4.1840e+02 ; HA - CA
- C
4935 4933 4948 1 1.1110e+02 5.2718e+02 ; CB - CA
- C
4935 4937 4938 1 1.0950e+02 4.1840e+02 ; CB - CG2
- HG21
4935 4937 4939 1 1.0950e+02 4.1840e+02 ; CB - CG2
- HG22
4935 4937 4940 1 1.0950e+02 4.1840e+02 ; CB - CG2
- HG23
4935 4941 4942 1 1.0950e+02 4.1840e+02 ; CB - CG1
- HG12
4935 4941 4943 1 1.0950e+02 4.1840e+02 ; CB - CG1
- HG13
4935 4941 4944 1 1.0950e+02 3.3472e+02 ; CB - CG1
- CD1
4936 4935 4937 1 1.0950e+02 4.1840e+02 ; HB - CB
- CG2
4936 4935 4941 1 1.0950e+02 4.1840e+02 ; HB - CB
- CG1
4937 4935 4941 1 1.0950e+02 3.3472e+02 ; CG2 - CB
- CG1
4938 4937 4939 1 1.0950e+02 2.9288e+02 ; HG21 - CG2
- HG22
4938 4937 4940 1 1.0950e+02 2.9288e+02 ; HG21 - CG2
- HG23
4939 4937 4940 1 1.0950e+02 2.9288e+02 ; HG22 - CG2
- HG23
4941 4944 4945 1 1.0950e+02 4.1840e+02 ; CG1 - CD1
- HD11
4941 4944 4946 1 1.0950e+02 4.1840e+02 ; CG1 - CD1
- HD12
4941 4944 4947 1 1.0950e+02 4.1840e+02 ; CG1 - CD1
- HD13
4942 4941 4943 1 1.0950e+02 2.9288e+02 ; HG12 - CG1
- HG13
4942 4941 4944 1 1.0950e+02 4.1840e+02 ; HG12 - CG1
- CD1
4943 4941 4944 1 1.0950e+02 4.1840e+02 ; HG13 - CG1
- CD1
4945 4944 4946 1 1.0950e+02 2.9288e+02 ; HD11 - CD1
- HD12
4945 4944 4947 1 1.0950e+02 2.9288e+02 ; HD11 - CD1
- HD13

```
  4946   4944   4947       1    1.0950e+02    2.9288e+02 ;   HD12 - CD1
- HD13
  4948   4950   4951       1    1.2000e+02    4.1840e+02 ;      C - N
- H
  4948   4950   4952       1    1.2190e+02    4.1840e+02 ;      C - N
- CA
  4949   4948   4950       1    1.2290e+02    6.6944e+02 ;      O - C
- N
  4950   4952   4953       1    1.0950e+02    4.1840e+02 ;      N - CA
- HA
  4950   4952   4954       1    1.0970e+02    6.6944e+02 ;      N - CA
- CB
  4950   4952   4967       1    1.1010e+02    5.2718e+02 ;      N - CA
- C
  4951   4950   4952       1    1.1804e+02    4.1840e+02 ;      H - N
- CA
  4952   4954   4955       1    1.0950e+02    4.1840e+02 ;     CA - CB
- HB
  4952   4954   4956       1    1.0950e+02    3.3472e+02 ;     CA - CB
- CG2
  4952   4954   4960       1    1.0950e+02    3.3472e+02 ;     CA - CB
- CG1
  4952   4967   4968       1    1.2040e+02    6.6944e+02 ;     CA - C
- O
  4952   4967   4969       1    1.1660e+02    5.8576e+02 ;     CA - C
- N
  4953   4952   4954       1    1.0950e+02    4.1840e+02 ;     HA - CA
- CB
  4953   4952   4967       1    1.0950e+02    4.1840e+02 ;     HA - CA
- C
  4954   4952   4967       1    1.1110e+02    5.2718e+02 ;     CB - CA
- C
  4954   4956   4957       1    1.0950e+02    4.1840e+02 ;     CB - CG2
- HG21
  4954   4956   4958       1    1.0950e+02    4.1840e+02 ;     CB - CG2
- HG22
  4954   4956   4959       1    1.0950e+02    4.1840e+02 ;     CB - CG2
- HG23
  4954   4960   4961       1    1.0950e+02    4.1840e+02 ;     CB - CG1
- HG12
  4954   4960   4962       1    1.0950e+02    4.1840e+02 ;     CB - CG1
- HG13
  4954   4960   4963       1    1.0950e+02    3.3472e+02 ;     CB - CG1
- CD1
  4955   4954   4956       1    1.0950e+02    4.1840e+02 ;     HB - CB
- CG2
  4955   4954   4960       1    1.0950e+02    4.1840e+02 ;     HB - CB
- CG1
  4956   4954   4960       1    1.0950e+02    3.3472e+02 ;    CG2 - CB
- CG1
  4957   4956   4958       1    1.0950e+02    2.9288e+02 ;   HG21 - CG2
- HG22
```

```
  4957   4956   4959      1    1.0950e+02    2.9288e+02 ;   HG21 - CG2
- HG23
  4958   4956   4959      1    1.0950e+02    2.9288e+02 ;   HG22 - CG2
- HG23
  4960   4963   4964      1    1.0950e+02    4.1840e+02 ;    CG1 - CD1
- HD11
  4960   4963   4965      1    1.0950e+02    4.1840e+02 ;    CG1 - CD1
- HD12
  4960   4963   4966      1    1.0950e+02    4.1840e+02 ;    CG1 - CD1
- HD13
  4961   4960   4962      1    1.0950e+02    2.9288e+02 ;   HG12 - CG1
- HG13
  4961   4960   4963      1    1.0950e+02    4.1840e+02 ;   HG12 - CG1
- CD1
  4962   4960   4963      1    1.0950e+02    4.1840e+02 ;   HG13 - CG1
- CD1
  4964   4963   4965      1    1.0950e+02    2.9288e+02 ;   HD11 - CD1
- HD12
  4964   4963   4966      1    1.0950e+02    2.9288e+02 ;   HD11 - CD1
- HD13
  4965   4963   4966      1    1.0950e+02    2.9288e+02 ;   HD12 - CD1
- HD13
  4967   4969   4970      1    1.2000e+02    4.1840e+02 ;      C - N
- H
  4967   4969   4971      1    1.2190e+02    4.1840e+02 ;      C - N
- CA
  4968   4967   4969      1    1.2290e+02    6.6944e+02 ;      O - C
- N
  4969   4971   4972      1    1.0950e+02    4.1840e+02 ;      N - CA
- HA
  4969   4971   4973      1    1.0970e+02    6.6944e+02 ;      N - CA
- CB
  4969   4971   4977      1    1.1010e+02    5.2718e+02 ;      N - CA
- C
  4970   4969   4971      1    1.1804e+02    4.1840e+02 ;      H - N
- CA
  4971   4973   4974      1    1.0950e+02    4.1840e+02 ;     CA - CB
- HB1
  4971   4973   4975      1    1.0950e+02    4.1840e+02 ;     CA - CB
- HB2
  4971   4973   4976      1    1.0950e+02    4.1840e+02 ;     CA - CB
- HB3
  4971   4977   4978      1    1.2040e+02    6.6944e+02 ;     CA - C
- O
  4971   4977   4979      1    1.1660e+02    5.8576e+02 ;     CA - C
- N
  4972   4971   4973      1    1.0950e+02    4.1840e+02 ;     HA - CA
- CB
  4972   4971   4977      1    1.0950e+02    4.1840e+02 ;     HA - CA
- C
  4973   4971   4977      1    1.1110e+02    5.2718e+02 ;     CB - CA
- C
```

4974 4973 4975 1 1.0950e+02 2.9288e+02 ; HB1 - CB - HB2
4974 4973 4976 1 1.0950e+02 2.9288e+02 ; HB1 - CB - HB3
4975 4973 4976 1 1.0950e+02 2.9288e+02 ; HB2 - CB - HB3
4977 4979 4980 1 1.2000e+02 4.1840e+02 ; C - N - H
4977 4979 4981 1 1.2190e+02 4.1840e+02 ; C - N - CA
4978 4977 4979 1 1.2290e+02 6.6944e+02 ; O - C - N
4979 4981 4982 1 1.0950e+02 4.1840e+02 ; N - CA - HA
4979 4981 4983 1 1.0970e+02 6.6944e+02 ; N - CA - CB
4979 4981 4999 1 1.1010e+02 5.2718e+02 ; N - CA - C
4980 4979 4981 1 1.1804e+02 4.1840e+02 ; H - N - CA
4981 4983 4984 1 1.0950e+02 4.1840e+02 ; CA - CB - HB2
4981 4983 4985 1 1.0950e+02 4.1840e+02 ; CA - CB - HB3
4981 4983 4986 1 1.0950e+02 3.3472e+02 ; CA - CB - CG
4981 4999 5000 1 1.2040e+02 6.6944e+02 ; CA - C - O
4981 4999 5001 1 1.1660e+02 5.8576e+02 ; CA - C - N
4982 4981 4983 1 1.0950e+02 4.1840e+02 ; HA - CA - CB
4982 4981 4999 1 1.0950e+02 4.1840e+02 ; HA - CA - C
4983 4981 4999 1 1.1110e+02 5.2718e+02 ; CB - CA - C
4983 4986 4987 1 1.0950e+02 4.1840e+02 ; CB - CG - HG2
4983 4986 4988 1 1.0950e+02 4.1840e+02 ; CB - CG - HG3
4983 4986 4989 1 1.0950e+02 3.3472e+02 ; CB - CG - CD
4984 4983 4985 1 1.0950e+02 2.9288e+02 ; HB2 - CB - HB3
4984 4983 4986 1 1.0950e+02 4.1840e+02 ; HB2 - CB - CG
4985 4983 4986 1 1.0950e+02 4.1840e+02 ; HB3 - CB - CG
4986 4989 4990 1 1.0950e+02 4.1840e+02 ; CG - CD - HD2
4986 4989 4991 1 1.0950e+02 4.1840e+02 ; CG - CD - HD3

4986 4989 4992 1 1.0950e+02 3.3472e+02 ; CG - CD - CE
4987 4986 4988 1 1.0950e+02 2.9288e+02 ; HG2 - CG - HG3
4987 4986 4989 1 1.0950e+02 4.1840e+02 ; HG2 - CG - CD
4988 4986 4989 1 1.0950e+02 4.1840e+02 ; HG3 - CG - CD
4989 4992 4993 1 1.0950e+02 4.1840e+02 ; CD - CE - HE2
4989 4992 4994 1 1.0950e+02 4.1840e+02 ; CD - CE - HE3
4989 4992 4995 1 1.1120e+02 6.6944e+02 ; CD - CE - NZ
4990 4989 4991 1 1.0950e+02 2.9288e+02 ; HD2 - CD - HD3
4990 4989 4992 1 1.0950e+02 4.1840e+02 ; HD2 - CD - CE
4991 4989 4992 1 1.0950e+02 4.1840e+02 ; HD3 - CD - CE
4992 4995 4996 1 1.0950e+02 4.1840e+02 ; CE - NZ - HZ1
4992 4995 4997 1 1.0950e+02 4.1840e+02 ; CE - NZ - HZ2
4992 4995 4998 1 1.0950e+02 4.1840e+02 ; CE - NZ - HZ3
4993 4992 4994 1 1.0950e+02 2.9288e+02 ; HE2 - CE - HE3
4993 4992 4995 1 1.0950e+02 4.1840e+02 ; HE2 - CE - NZ
4994 4992 4995 1 1.0950e+02 4.1840e+02 ; HE3 - CE - NZ
4996 4995 4997 1 1.0950e+02 2.9288e+02 ; HZ1 - NZ - HZ2
4996 4995 4998 1 1.0950e+02 2.9288e+02 ; HZ1 - NZ - HZ3
4997 4995 4998 1 1.0950e+02 2.9288e+02 ; HZ2 - NZ - HZ3
4999 5001 5002 1 1.2000e+02 4.1840e+02 ; C - N - H
4999 5001 5003 1 1.2190e+02 4.1840e+02 ; C - N - CA
5000 4999 5001 1 1.2290e+02 6.6944e+02 ; O - C - N
5001 5003 5004 1 1.0950e+02 4.1840e+02 ; N - CA - HA
5001 5003 5005 1 1.0970e+02 6.6944e+02 ; N - CA - CB
5001 5003 5016 1 1.1010e+02 5.2718e+02 ; N - CA - C
5002 5001 5003 1 1.1804e+02 4.1840e+02 ; H - N - CA

5003 5005 5006 1 1.0950e+02 4.1840e+02 ; CA - CB
- HB2
5003 5005 5007 1 1.0950e+02 4.1840e+02 ; CA - CB
- HB3
5003 5005 5008 1 1.1310e+02 5.2718e+02 ; CA - CB
- CG
5003 5016 5017 1 1.2040e+02 6.6944e+02 ; CA - C
- O
5003 5016 5018 1 1.1660e+02 5.8576e+02 ; CA - C
- N
5004 5003 5005 1 1.0950e+02 4.1840e+02 ; HA - CA
- CB
5004 5003 5016 1 1.0950e+02 4.1840e+02 ; HA - CA
- C
5005 5003 5016 1 1.1110e+02 5.2718e+02 ; CB - CA
- C
5005 5008 5009 1 1.2000e+02 5.8576e+02 ; CB - CG
- ND1
5005 5008 5014 1 1.2000e+02 5.8576e+02 ; CB - CG
- CD2
5006 5005 5007 1 1.0950e+02 2.9288e+02 ; HB2 - CB
- HB3
5006 5005 5008 1 1.0950e+02 4.1840e+02 ; HB2 - CB
- CG
5007 5005 5008 1 1.0950e+02 4.1840e+02 ; HB3 - CB
- CG
5008 5009 5010 1 1.1700e+02 5.8576e+02 ; CG - ND1
- CE1
5008 5014 5012 1 1.2000e+02 5.8576e+02 ; CG - CD2
- NE2
5008 5014 5015 1 1.2000e+02 4.1840e+02 ; CG - CD2
- HD2
5009 5008 5014 1 1.2000e+02 5.8576e+02 ; ND1 - CG
- CD2
5009 5010 5011 1 1.2000e+02 4.1840e+02 ; ND1 - CE1
- HE1
5009 5010 5012 1 1.2000e+02 5.8576e+02 ; ND1 - CE1
- NE2
5010 5012 5013 1 1.2000e+02 4.1840e+02 ; CE1 - NE2
- HE2
5010 5012 5014 1 1.2000e+02 5.8576e+02 ; CE1 - NE2
- CD2
5011 5010 5012 1 1.2000e+02 4.1840e+02 ; HE1 - CE1
- NE2
5012 5014 5015 1 1.2000e+02 4.1840e+02 ; NE2 - CD2
- HD2
5013 5012 5014 1 1.2000e+02 4.1840e+02 ; HE2 - NE2
- CD2
5016 5018 5019 1 1.2000e+02 4.1840e+02 ; C - N
- H
5016 5018 5020 1 1.2190e+02 4.1840e+02 ; C - N
- CA

```
  5017   5016   5018      1    1.2290e+02    6.6944e+02 ;      O - C
- N
  5018   5020   5021      1    1.0950e+02    4.1840e+02 ;      N - CA
- HA
  5018   5020   5022      1    1.0970e+02    6.6944e+02 ;      N - CA
- CB
  5018   5020   5035      1    1.1010e+02    5.2718e+02 ;      N - CA
- C
  5019   5018   5020      1    1.1804e+02    4.1840e+02 ;      H - N
- CA
  5020   5022   5023      1    1.0950e+02    4.1840e+02 ;     CA - CB
- HB2
  5020   5022   5024      1    1.0950e+02    4.1840e+02 ;     CA - CB
- HB3
  5020   5022   5025      1    1.0950e+02    3.3472e+02 ;     CA - CB
- CG
  5020   5035   5036      1    1.2040e+02    6.6944e+02 ;     CA - C
- O
  5020   5035   5037      1    1.1660e+02    5.8576e+02 ;     CA - C
- N
  5021   5020   5022      1    1.0950e+02    4.1840e+02 ;     HA - CA
- CB
  5021   5020   5035      1    1.0950e+02    4.1840e+02 ;     HA - CA
- C
  5022   5020   5035      1    1.1110e+02    5.2718e+02 ;     CB - CA
- C
  5022   5025   5026      1    1.0950e+02    4.1840e+02 ;     CB - CG
- HG
  5022   5025   5027      1    1.0950e+02    3.3472e+02 ;     CB - CG
- CD1
  5022   5025   5031      1    1.0950e+02    3.3472e+02 ;     CB - CG
- CD2
  5023   5022   5024      1    1.0950e+02    2.9288e+02 ;    HB2 - CB
- HB3
  5023   5022   5025      1    1.0950e+02    4.1840e+02 ;    HB2 - CB
- CG
  5024   5022   5025      1    1.0950e+02    4.1840e+02 ;    HB3 - CB
- CG
  5025   5027   5028      1    1.0950e+02    4.1840e+02 ;     CG - CD1
- HD11
  5025   5027   5029      1    1.0950e+02    4.1840e+02 ;     CG - CD1
- HD12
  5025   5027   5030      1    1.0950e+02    4.1840e+02 ;     CG - CD1
- HD13
  5025   5031   5032      1    1.0950e+02    4.1840e+02 ;     CG - CD2
- HD21
  5025   5031   5033      1    1.0950e+02    4.1840e+02 ;     CG - CD2
- HD22
  5025   5031   5034      1    1.0950e+02    4.1840e+02 ;     CG - CD2
- HD23
  5026   5025   5027      1    1.0950e+02    4.1840e+02 ;     HG - CG
- CD1
```

5026 5025 5031 1 1.0950e+02 4.1840e+02 ; HG - CG
- CD2
5027 5025 5031 1 1.0950e+02 3.3472e+02 ; CD1 - CG
- CD2
5028 5027 5029 1 1.0950e+02 2.9288e+02 ; HD11 - CD1
- HD12
5028 5027 5030 1 1.0950e+02 2.9288e+02 ; HD11 - CD1
- HD13
5029 5027 5030 1 1.0950e+02 2.9288e+02 ; HD12 - CD1
- HD13
5032 5031 5033 1 1.0950e+02 2.9288e+02 ; HD21 - CD2
- HD22
5032 5031 5034 1 1.0950e+02 2.9288e+02 ; HD21 - CD2
- HD23
5033 5031 5034 1 1.0950e+02 2.9288e+02 ; HD22 - CD2
- HD23
5035 5037 5038 1 1.2000e+02 4.1840e+02 ; C - N
- H
5035 5037 5039 1 1.2190e+02 4.1840e+02 ; C - N
- CA
5036 5035 5037 1 1.2290e+02 6.6944e+02 ; O - C
- N
5037 5039 5040 1 1.0950e+02 4.1840e+02 ; N - CA
- HA
5037 5039 5041 1 1.0970e+02 6.6944e+02 ; N - CA
- CB
5037 5039 5046 1 1.1010e+02 5.2718e+02 ; N - CA
- C
5038 5037 5039 1 1.1804e+02 4.1840e+02 ; H - N
- CA
5039 5041 5042 1 1.0950e+02 4.1840e+02 ; CA - CB
- HB2
5039 5041 5043 1 1.0950e+02 4.1840e+02 ; CA - CB
- HB3
5039 5041 5044 1 1.0860e+02 4.1840e+02 ; CA - CB
- SG
5039 5046 5047 1 1.2040e+02 6.6944e+02 ; CA - C
- O
5039 5046 5048 1 1.1660e+02 5.8576e+02 ; CA - C
- N
5040 5039 5041 1 1.0950e+02 4.1840e+02 ; HA - CA
- CB
5040 5039 5046 1 1.0950e+02 4.1840e+02 ; HA - CA
- C
5041 5039 5046 1 1.1110e+02 5.2718e+02 ; CB - CA
- C
5041 5044 5045 1 9.6000e+01 3.5982e+02 ; CB - SG
- HG
5042 5041 5043 1 1.0950e+02 2.9288e+02 ; HB2 - CB
- HB3
5042 5041 5044 1 1.0950e+02 4.1840e+02 ; HB2 - CB
- SG

5043 5041 5044 1 1.0950e+02 4.1840e+02 ; HB3 - CB
- SG
5046 5048 5049 1 1.2000e+02 4.1840e+02 ; C - N
- H
5046 5048 5050 1 1.2190e+02 4.1840e+02 ; C - N
- CA
5047 5046 5048 1 1.2290e+02 6.6944e+02 ; O - C
- N
5048 5050 5051 1 1.0950e+02 4.1840e+02 ; N - CA
- HA
5048 5050 5052 1 1.0970e+02 6.6944e+02 ; N - CA
- CB
5048 5050 5058 1 1.1010e+02 5.2718e+02 ; N - CA
- C
5049 5048 5050 1 1.1804e+02 4.1840e+02 ; H - N
- CA
5050 5052 5053 1 1.0950e+02 4.1840e+02 ; CA - CB
- HB2
5050 5052 5054 1 1.0950e+02 4.1840e+02 ; CA - CB
- HB3
5050 5052 5055 1 1.1110e+02 5.2718e+02 ; CA - CB
- CG
5050 5058 5059 1 1.2040e+02 6.6944e+02 ; CA - C
- O
5050 5058 5060 1 1.1660e+02 5.8576e+02 ; CA - C
- N
5051 5050 5052 1 1.0950e+02 4.1840e+02 ; HA - CA
- CB
5051 5050 5058 1 1.0950e+02 4.1840e+02 ; HA - CA
- C
5052 5050 5058 1 1.1110e+02 5.2718e+02 ; CB - CA
- C
5052 5055 5056 1 1.1700e+02 5.8576e+02 ; CB - CG
- OD1
5052 5055 5057 1 1.1700e+02 5.8576e+02 ; CB - CG
- OD2
5053 5052 5054 1 1.0950e+02 2.9288e+02 ; HB2 - CB
- HB3
5053 5052 5055 1 1.0950e+02 4.1840e+02 ; HB2 - CB
- CG
5054 5052 5055 1 1.0950e+02 4.1840e+02 ; HB3 - CB
- CG
5056 5055 5057 1 1.2600e+02 6.6944e+02 ; OD1 - CG
- OD2
5058 5060 5061 1 1.2000e+02 4.1840e+02 ; C - N
- H
5058 5060 5062 1 1.2190e+02 4.1840e+02 ; C - N
- CA
5059 5058 5060 1 1.2290e+02 6.6944e+02 ; O - C
- N
5060 5062 5063 1 1.0950e+02 4.1840e+02 ; N - CA
- HA

5060 5062 5064 1 1.0970e+02 6.6944e+02 ; N - CA
- CB
5060 5062 5074 1 1.1010e+02 5.2718e+02 ; N - CA
- C
5061 5060 5062 1 1.1804e+02 4.1840e+02 ; H - N
- CA
5062 5064 5065 1 1.0950e+02 4.1840e+02 ; CA - CB
- HB
5062 5064 5066 1 1.0950e+02 3.3472e+02 ; CA - CB
- CG1
5062 5064 5070 1 1.0950e+02 3.3472e+02 ; CA - CB
- CG2
5062 5074 5075 1 1.2040e+02 6.6944e+02 ; CA - C
- O
5062 5074 5076 1 1.1660e+02 5.8576e+02 ; CA - C
- N
5063 5062 5064 1 1.0950e+02 4.1840e+02 ; HA - CA
- CB
5063 5062 5074 1 1.0950e+02 4.1840e+02 ; HA - CA
- C
5064 5062 5074 1 1.1110e+02 5.2718e+02 ; CB - CA
- C
5064 5066 5067 1 1.0950e+02 4.1840e+02 ; CB - CG1
- HG11
5064 5066 5068 1 1.0950e+02 4.1840e+02 ; CB - CG1
- HG12
5064 5066 5069 1 1.0950e+02 4.1840e+02 ; CB - CG1
- HG13
5064 5070 5071 1 1.0950e+02 4.1840e+02 ; CB - CG2
- HG21
5064 5070 5072 1 1.0950e+02 4.1840e+02 ; CB - CG2
- HG22
5064 5070 5073 1 1.0950e+02 4.1840e+02 ; CB - CG2
- HG23
5065 5064 5066 1 1.0950e+02 4.1840e+02 ; HB - CB
- CG1
5065 5064 5070 1 1.0950e+02 4.1840e+02 ; HB - CB
- CG2
5066 5064 5070 1 1.0950e+02 3.3472e+02 ; CG1 - CB
- CG2
5067 5066 5068 1 1.0950e+02 2.9288e+02 ; HG11 - CG1
- HG12
5067 5066 5069 1 1.0950e+02 2.9288e+02 ; HG11 - CG1
- HG13
5068 5066 5069 1 1.0950e+02 2.9288e+02 ; HG12 - CG1
- HG13
5071 5070 5072 1 1.0950e+02 2.9288e+02 ; HG21 - CG2
- HG22
5071 5070 5073 1 1.0950e+02 2.9288e+02 ; HG21 - CG2
- HG23
5072 5070 5073 1 1.0950e+02 2.9288e+02 ; HG22 - CG2
- HG23

5074 5076 5077 1 1.2000e+02 4.1840e+02 ; C - N
- H
5074 5076 5078 1 1.2190e+02 4.1840e+02 ; C - N
- CA
5075 5074 5076 1 1.2290e+02 6.6944e+02 ; O - C
- N
5076 5078 5079 1 1.0950e+02 4.1840e+02 ; N - CA
- HA
5076 5078 5080 1 1.0970e+02 6.6944e+02 ; N - CA
- CB
5076 5078 5089 1 1.1010e+02 5.2718e+02 ; N - CA
- C
5077 5076 5078 1 1.1804e+02 4.1840e+02 ; H - N
- CA
5078 5080 5081 1 1.0950e+02 4.1840e+02 ; CA - CB
- HB2
5078 5080 5082 1 1.0950e+02 4.1840e+02 ; CA - CB
- HB3
5078 5080 5083 1 1.0950e+02 3.3472e+02 ; CA - CB
- CG
5078 5089 5090 1 1.2040e+02 6.6944e+02 ; CA - C
- O
5078 5089 5091 1 1.1660e+02 5.8576e+02 ; CA - C
- N
5079 5078 5080 1 1.0950e+02 4.1840e+02 ; HA - CA
- CB
5079 5078 5089 1 1.0950e+02 4.1840e+02 ; HA - CA
- C
5080 5078 5089 1 1.1110e+02 5.2718e+02 ; CB - CA
- C
5080 5083 5084 1 1.0950e+02 4.1840e+02 ; CB - CG
- HG2
5080 5083 5085 1 1.0950e+02 4.1840e+02 ; CB - CG
- HG3
5080 5083 5086 1 1.1110e+02 5.2718e+02 ; CB - CG
- CD
5081 5080 5082 1 1.0950e+02 2.9288e+02 ; HB2 - CB
- HB3
5081 5080 5083 1 1.0950e+02 4.1840e+02 ; HB2 - CB
- CG
5082 5080 5083 1 1.0950e+02 4.1840e+02 ; HB3 - CB
- CG
5083 5086 5087 1 1.1700e+02 5.8576e+02 ; CG - CD
- OE1
5083 5086 5088 1 1.1700e+02 5.8576e+02 ; CG - CD
- OE2
5084 5083 5085 1 1.0950e+02 2.9288e+02 ; HG2 - CG
- HG3
5084 5083 5086 1 1.0950e+02 4.1840e+02 ; HG2 - CG
- CD
5085 5083 5086 1 1.0950e+02 4.1840e+02 ; HG3 - CG
- CD

5087 5086 5088 1 1.2600e+02 6.6944e+02 ; OE1 - CD
- OE2
5089 5091 5092 1 1.2000e+02 4.1840e+02 ; C - N
- H
5089 5091 5093 1 1.2190e+02 4.1840e+02 ; C - N
- CA
5090 5089 5091 1 1.2290e+02 6.6944e+02 ; O - C
- N
5091 5093 5094 1 1.0950e+02 4.1840e+02 ; N - CA
- HA
5091 5093 5095 1 1.0970e+02 6.6944e+02 ; N - CA
- CB
5091 5093 5108 1 1.1010e+02 5.2718e+02 ; N - CA
- C
5092 5091 5093 1 1.1804e+02 4.1840e+02 ; H - N
- CA
5093 5095 5096 1 1.0950e+02 4.1840e+02 ; CA - CB
- HB
5093 5095 5097 1 1.0950e+02 3.3472e+02 ; CA - CB
- CG2
5093 5095 5101 1 1.0950e+02 3.3472e+02 ; CA - CB
- CG1
5093 5108 5109 1 1.2040e+02 6.6944e+02 ; CA - C
- O
5093 5108 5110 1 1.1660e+02 5.8576e+02 ; CA - C
- N
5094 5093 5095 1 1.0950e+02 4.1840e+02 ; HA - CA
- CB
5094 5093 5108 1 1.0950e+02 4.1840e+02 ; HA - CA
- C
5095 5093 5108 1 1.1110e+02 5.2718e+02 ; CB - CA
- C
5095 5097 5098 1 1.0950e+02 4.1840e+02 ; CB - CG2
- HG21
5095 5097 5099 1 1.0950e+02 4.1840e+02 ; CB - CG2
- HG22
5095 5097 5100 1 1.0950e+02 4.1840e+02 ; CB - CG2
- HG23
5095 5101 5102 1 1.0950e+02 4.1840e+02 ; CB - CG1
- HG12
5095 5101 5103 1 1.0950e+02 4.1840e+02 ; CB - CG1
- HG13
5095 5101 5104 1 1.0950e+02 3.3472e+02 ; CB - CG1
- CD1
5096 5095 5097 1 1.0950e+02 4.1840e+02 ; HB - CB
- CG2
5096 5095 5101 1 1.0950e+02 4.1840e+02 ; HB - CB
- CG1
5097 5095 5101 1 1.0950e+02 3.3472e+02 ; CG2 - CB
- CG1
5098 5097 5099 1 1.0950e+02 2.9288e+02 ; HG21 - CG2
- HG22

```
  5098   5097   5100      1    1.0950e+02    2.9288e+02 ;   HG21 - CG2
- HG23
  5099   5097   5100      1    1.0950e+02    2.9288e+02 ;   HG22 - CG2
- HG23
  5101   5104   5105      1    1.0950e+02    4.1840e+02 ;    CG1 - CD1
- HD11
  5101   5104   5106      1    1.0950e+02    4.1840e+02 ;    CG1 - CD1
- HD12
  5101   5104   5107      1    1.0950e+02    4.1840e+02 ;    CG1 - CD1
- HD13
  5102   5101   5103      1    1.0950e+02    2.9288e+02 ;   HG12 - CG1
- HG13
  5102   5101   5104      1    1.0950e+02    4.1840e+02 ;   HG12 - CG1
- CD1
  5103   5101   5104      1    1.0950e+02    4.1840e+02 ;   HG13 - CG1
- CD1
  5105   5104   5106      1    1.0950e+02    2.9288e+02 ;   HD11 - CD1
- HD12
  5105   5104   5107      1    1.0950e+02    2.9288e+02 ;   HD11 - CD1
- HD13
  5106   5104   5107      1    1.0950e+02    2.9288e+02 ;   HD12 - CD1
- HD13
  5108   5110   5111      1    1.2000e+02    4.1840e+02 ;      C - N
- H
  5108   5110   5112      1    1.2190e+02    4.1840e+02 ;      C - N
- CA
  5109   5108   5110      1    1.2290e+02    6.6944e+02 ;      O - C
- N
  5110   5112   5113      1    1.0950e+02    4.1840e+02 ;      N - CA
- HA
  5110   5112   5114      1    1.0970e+02    6.6944e+02 ;      N - CA
- CB
  5110   5112   5122      1    1.1010e+02    5.2718e+02 ;      N - CA
- C
  5111   5110   5112      1    1.1804e+02    4.1840e+02 ;      H - N
- CA
  5112   5114   5115      1    1.0950e+02    4.1840e+02 ;     CA - CB
- HB2
  5112   5114   5116      1    1.0950e+02    4.1840e+02 ;     CA - CB
- HB3
  5112   5114   5117      1    1.1110e+02    5.2718e+02 ;     CA - CB
- CG
  5112   5122   5123      1    1.2040e+02    6.6944e+02 ;     CA - C
- O
  5112   5122   5124      1    1.1660e+02    5.8576e+02 ;     CA - C
- N
  5113   5112   5114      1    1.0950e+02    4.1840e+02 ;     HA - CA
- CB
  5113   5112   5122      1    1.0950e+02    4.1840e+02 ;     HA - CA
- C
  5114   5112   5122      1    1.1110e+02    5.2718e+02 ;     CB - CA
- C
```

```
  5114   5117   5118      1    1.2040e+02    6.6944e+02 ;     CB - CG
- OD1
  5114   5117   5119      1    1.1660e+02    5.8576e+02 ;     CB - CG
- ND2
  5115   5114   5116      1    1.0950e+02    2.9288e+02 ;    HB2 - CB
- HB3
  5115   5114   5117      1    1.0950e+02    4.1840e+02 ;    HB2 - CB
- CG
  5116   5114   5117      1    1.0950e+02    4.1840e+02 ;    HB3 - CB
- CG
  5117   5119   5120      1    1.2000e+02    4.1840e+02 ;     CG - ND2
- HD21
  5117   5119   5121      1    1.2000e+02    4.1840e+02 ;     CG - ND2
- HD22
  5118   5117   5119      1    1.2290e+02    6.6944e+02 ;    OD1 - CG
- ND2
  5120   5119   5121      1    1.2000e+02    2.9288e+02 ;   HD21 - ND2
- HD22
  5122   5124   5125      1    1.2000e+02    4.1840e+02 ;      C - N
- H
  5122   5124   5126      1    1.2190e+02    4.1840e+02 ;      C - N
- CA
  5123   5122   5124      1    1.2290e+02    6.6944e+02 ;      O - C
- N
  5124   5126   5127      1    1.0950e+02    4.1840e+02 ;      N - CA
- HA
  5124   5126   5128      1    1.0970e+02    6.6944e+02 ;      N - CA
- CB
  5124   5126   5136      1    1.1010e+02    5.2718e+02 ;      N - CA
- C
  5125   5124   5126      1    1.1804e+02    4.1840e+02 ;      H - N
- CA
  5126   5128   5129      1    1.0950e+02    4.1840e+02 ;     CA - CB
- HB2
  5126   5128   5130      1    1.0950e+02    4.1840e+02 ;     CA - CB
- HB3
  5126   5128   5131      1    1.1110e+02    5.2718e+02 ;     CA - CB
- CG
  5126   5136   5137      1    1.2040e+02    6.6944e+02 ;     CA - C
- O
  5126   5136   5138      1    1.1660e+02    5.8576e+02 ;     CA - C
- N
  5127   5126   5128      1    1.0950e+02    4.1840e+02 ;     HA - CA
- CB
  5127   5126   5136      1    1.0950e+02    4.1840e+02 ;     HA - CA
- C
  5128   5126   5136      1    1.1110e+02    5.2718e+02 ;     CB - CA
- C
  5128   5131   5132      1    1.2040e+02    6.6944e+02 ;     CB - CG
- OD1
  5128   5131   5133      1    1.1660e+02    5.8576e+02 ;     CB - CG
- ND2
```

```
  5129   5128   5130      1    1.0950e+02    2.9288e+02 ;    HB2 - CB
- HB3
  5129   5128   5131      1    1.0950e+02    4.1840e+02 ;    HB2 - CB
- CG
  5130   5128   5131      1    1.0950e+02    4.1840e+02 ;    HB3 - CB
- CG
  5131   5133   5134      1    1.2000e+02    4.1840e+02 ;     CG - ND2
- HD21
  5131   5133   5135      1    1.2000e+02    4.1840e+02 ;     CG - ND2
- HD22
  5132   5131   5133      1    1.2290e+02    6.6944e+02 ;    OD1 - CG
- ND2
  5134   5133   5135      1    1.2000e+02    2.9288e+02 ;   HD21 - ND2
- HD22
  5136   5138   5139      1    1.2000e+02    4.1840e+02 ;      C - N
- H
  5136   5138   5140      1    1.2190e+02    4.1840e+02 ;      C - N
- CA
  5137   5136   5138      1    1.2290e+02    6.6944e+02 ;      O - C
- N
  5138   5140   5141      1    1.0950e+02    4.1840e+02 ;      N - CA
- HA
  5138   5140   5142      1    1.0970e+02    6.6944e+02 ;      N - CA
- CB
  5138   5140   5155      1    1.1010e+02    5.2718e+02 ;      N - CA
- C
  5139   5138   5140      1    1.1804e+02    4.1840e+02 ;      H - N
- CA
  5140   5142   5143      1    1.0950e+02    4.1840e+02 ;     CA - CB
- HB2
  5140   5142   5144      1    1.0950e+02    4.1840e+02 ;     CA - CB
- HB3
  5140   5142   5145      1    1.0950e+02    3.3472e+02 ;     CA - CB
- CG
  5140   5155   5156      1    1.2040e+02    6.6944e+02 ;     CA - C
- O
  5140   5155   5157      1    1.1660e+02    5.8576e+02 ;     CA - C
- N
  5141   5140   5142      1    1.0950e+02    4.1840e+02 ;     HA - CA
- CB
  5141   5140   5155      1    1.0950e+02    4.1840e+02 ;     HA - CA
- C
  5142   5140   5155      1    1.1110e+02    5.2718e+02 ;     CB - CA
- C
  5142   5145   5146      1    1.0950e+02    4.1840e+02 ;     CB - CG
- HG
  5142   5145   5147      1    1.0950e+02    3.3472e+02 ;     CB - CG
- CD1
  5142   5145   5151      1    1.0950e+02    3.3472e+02 ;     CB - CG
- CD2
  5143   5142   5144      1    1.0950e+02    2.9288e+02 ;    HB2 - CB
- HB3
```

5143 5142 5145 1 1.0950e+02 4.1840e+02 ; HB2 - CB
- CG
5144 5142 5145 1 1.0950e+02 4.1840e+02 ; HB3 - CB
- CG
5145 5147 5148 1 1.0950e+02 4.1840e+02 ; CG - CD1
- HD11
5145 5147 5149 1 1.0950e+02 4.1840e+02 ; CG - CD1
- HD12
5145 5147 5150 1 1.0950e+02 4.1840e+02 ; CG - CD1
- HD13
5145 5151 5152 1 1.0950e+02 4.1840e+02 ; CG - CD2
- HD21
5145 5151 5153 1 1.0950e+02 4.1840e+02 ; CG - CD2
- HD22
5145 5151 5154 1 1.0950e+02 4.1840e+02 ; CG - CD2
- HD23
5146 5145 5147 1 1.0950e+02 4.1840e+02 ; HG - CG
- CD1
5146 5145 5151 1 1.0950e+02 4.1840e+02 ; HG - CG
- CD2
5147 5145 5151 1 1.0950e+02 3.3472e+02 ; CD1 - CG
- CD2
5148 5147 5149 1 1.0950e+02 2.9288e+02 ; HD11 - CD1
- HD12
5148 5147 5150 1 1.0950e+02 2.9288e+02 ; HD11 - CD1
- HD13
5149 5147 5150 1 1.0950e+02 2.9288e+02 ; HD12 - CD1
- HD13
5152 5151 5153 1 1.0950e+02 2.9288e+02 ; HD21 - CD2
- HD22
5152 5151 5154 1 1.0950e+02 2.9288e+02 ; HD21 - CD2
- HD23
5153 5151 5154 1 1.0950e+02 2.9288e+02 ; HD22 - CD2
- HD23
5155 5157 5158 1 1.2000e+02 4.1840e+02 ; C - N
- H
5155 5157 5159 1 1.2190e+02 4.1840e+02 ; C - N
- CA
5156 5155 5157 1 1.2290e+02 6.6944e+02 ; O - C
- N
5157 5159 5160 1 1.0950e+02 4.1840e+02 ; N - CA
- HA
5157 5159 5161 1 1.0970e+02 6.6944e+02 ; N - CA
- CB
5157 5159 5179 1 1.1010e+02 5.2718e+02 ; N - CA
- C
5158 5157 5159 1 1.1804e+02 4.1840e+02 ; H - N
- CA
5159 5161 5162 1 1.0950e+02 4.1840e+02 ; CA - CB
- HB2
5159 5161 5163 1 1.0950e+02 4.1840e+02 ; CA - CB
- HB3

5159 5161 5164 1 1.0950e+02 3.3472e+02 ; CA - CB
- CG
5159 5179 5180 1 1.2040e+02 6.6944e+02 ; CA - C
- O
5159 5179 5181 1 1.1660e+02 5.8576e+02 ; CA - C
- N
5160 5159 5161 1 1.0950e+02 4.1840e+02 ; HA - CA
- CB
5160 5159 5179 1 1.0950e+02 4.1840e+02 ; HA - CA
- C
5161 5159 5179 1 1.1110e+02 5.2718e+02 ; CB - CA
- C
5161 5164 5165 1 1.0950e+02 4.1840e+02 ; CB - CG
- HG2
5161 5164 5166 1 1.0950e+02 4.1840e+02 ; CB - CG
- HG3
5161 5164 5167 1 1.0950e+02 3.3472e+02 ; CB - CG
- CD
5162 5161 5163 1 1.0950e+02 2.9288e+02 ; HB2 - CB
- HB3
5162 5161 5164 1 1.0950e+02 4.1840e+02 ; HB2 - CB
- CG
5163 5161 5164 1 1.0950e+02 4.1840e+02 ; HB3 - CB
- CG
5164 5167 5168 1 1.0950e+02 4.1840e+02 ; CG - CD
- HD2
5164 5167 5169 1 1.0950e+02 4.1840e+02 ; CG - CD
- HD3
5164 5167 5170 1 1.1120e+02 6.6944e+02 ; CG - CD
- NE
5165 5164 5166 1 1.0950e+02 2.9288e+02 ; HG2 - CG
- HG3
5165 5164 5167 1 1.0950e+02 4.1840e+02 ; HG2 - CG
- CD
5166 5164 5167 1 1.0950e+02 4.1840e+02 ; HG3 - CG
- CD
5167 5170 5171 1 1.1840e+02 4.1840e+02 ; CD - NE
- HE
5167 5170 5172 1 1.2320e+02 4.1840e+02 ; CD - NE
- CZ
5168 5167 5169 1 1.0950e+02 2.9288e+02 ; HD2 - CD
- HD3
5168 5167 5170 1 1.0950e+02 4.1840e+02 ; HD2 - CD
- NE
5169 5167 5170 1 1.0950e+02 4.1840e+02 ; HD3 - CD
- NE
5170 5172 5173 1 1.2000e+02 5.8576e+02 ; NE - CZ
- NH1
5170 5172 5176 1 1.2000e+02 5.8576e+02 ; NE - CZ
- NH2
5171 5170 5172 1 1.2000e+02 4.1840e+02 ; HE - NE
- CZ

```
  5172   5173   5174      1    1.2000e+02    4.1840e+02 ;     CZ - NH1 - HH11
  5172   5173   5175      1    1.2000e+02    4.1840e+02 ;     CZ - NH1 - HH12
  5172   5176   5177      1    1.2000e+02    4.1840e+02 ;     CZ - NH2 - HH21
  5172   5176   5178      1    1.2000e+02    4.1840e+02 ;     CZ - NH2 - HH22
  5173   5172   5176      1    1.2000e+02    5.8576e+02 ;    NH1 - CZ - NH2
  5174   5173   5175      1    1.2000e+02    2.9288e+02 ;   HH11 - NH1 - HH12
  5177   5176   5178      1    1.2000e+02    2.9288e+02 ;   HH21 - NH2 - HH22
  5179   5181   5182      1    1.2000e+02    4.1840e+02 ;      C - N - H
  5179   5181   5183      1    1.2190e+02    4.1840e+02 ;      C - N - CA
  5180   5179   5181      1    1.2290e+02    6.6944e+02 ;      O - C - N
  5181   5183   5184      1    1.0950e+02    4.1840e+02 ;      N - CA - HA
  5181   5183   5185      1    1.0970e+02    6.6944e+02 ;      N - CA - CB
  5181   5183   5193      1    1.1010e+02    5.2718e+02 ;      N - CA - C
  5182   5181   5183      1    1.1804e+02    4.1840e+02 ;      H - N - CA
  5183   5185   5186      1    1.0950e+02    4.1840e+02 ;     CA - CB - HB2
  5183   5185   5187      1    1.0950e+02    4.1840e+02 ;     CA - CB - HB3
  5183   5185   5188      1    1.1110e+02    5.2718e+02 ;     CA - CB - CG
  5183   5193   5194      1    1.2040e+02    6.6944e+02 ;     CA - C - O
  5183   5193   5195      1    1.1660e+02    5.8576e+02 ;     CA - C - N
  5184   5183   5185      1    1.0950e+02    4.1840e+02 ;     HA - CA - CB
  5184   5183   5193      1    1.0950e+02    4.1840e+02 ;     HA - CA - C
  5185   5183   5193      1    1.1110e+02    5.2718e+02 ;     CB - CA - C
  5185   5188   5189      1    1.2040e+02    6.6944e+02 ;     CB - CG - OD1
  5185   5188   5190      1    1.1660e+02    5.8576e+02 ;     CB - CG - ND2
  5186   5185   5187      1    1.0950e+02    2.9288e+02 ;    HB2 - CB - HB3
  5186   5185   5188      1    1.0950e+02    4.1840e+02 ;    HB2 - CB - CG
```

5187 5185 5188 1 1.0950e+02 4.1840e+02 ; HB3 - CB
- CG
5188 5190 5191 1 1.2000e+02 4.1840e+02 ; CG - ND2
- HD21
5188 5190 5192 1 1.2000e+02 4.1840e+02 ; CG - ND2
- HD22
5189 5188 5190 1 1.2290e+02 6.6944e+02 ; OD1 - CG
- ND2
5191 5190 5192 1 1.2000e+02 2.9288e+02 ; HD21 - ND2
- HD22
5193 5195 5196 1 1.2000e+02 4.1840e+02 ; C - N
- H
5193 5195 5197 1 1.2190e+02 4.1840e+02 ; C - N
- CA
5194 5193 5195 1 1.2290e+02 6.6944e+02 ; O - C
- N
5195 5197 5198 1 1.0950e+02 4.1840e+02 ; N - CA
- HA
5195 5197 5199 1 1.0970e+02 6.6944e+02 ; N - CA
- CB
5195 5197 5212 1 1.1010e+02 5.2718e+02 ; N - CA
- C
5196 5195 5197 1 1.1804e+02 4.1840e+02 ; H - N
- CA
5197 5199 5200 1 1.0950e+02 4.1840e+02 ; CA - CB
- HB
5197 5199 5201 1 1.0950e+02 3.3472e+02 ; CA - CB
- CG2
5197 5199 5205 1 1.0950e+02 3.3472e+02 ; CA - CB
- CG1
5197 5212 5213 1 1.2040e+02 6.6944e+02 ; CA - C
- O
5197 5212 5214 1 1.1660e+02 5.8576e+02 ; CA - C
- N
5198 5197 5199 1 1.0950e+02 4.1840e+02 ; HA - CA
- CB
5198 5197 5212 1 1.0950e+02 4.1840e+02 ; HA - CA
- C
5199 5197 5212 1 1.1110e+02 5.2718e+02 ; CB - CA
- C
5199 5201 5202 1 1.0950e+02 4.1840e+02 ; CB - CG2
- HG21
5199 5201 5203 1 1.0950e+02 4.1840e+02 ; CB - CG2
- HG22
5199 5201 5204 1 1.0950e+02 4.1840e+02 ; CB - CG2
- HG23
5199 5205 5206 1 1.0950e+02 4.1840e+02 ; CB - CG1
- HG12
5199 5205 5207 1 1.0950e+02 4.1840e+02 ; CB - CG1
- HG13
5199 5205 5208 1 1.0950e+02 3.3472e+02 ; CB - CG1
- CD1

```
  5200   5199   5201      1    1.0950e+02    4.1840e+02 ;     HB - CB
- CG2
  5200   5199   5205      1    1.0950e+02    4.1840e+02 ;     HB - CB
- CG1
  5201   5199   5205      1    1.0950e+02    3.3472e+02 ;    CG2 - CB
- CG1
  5202   5201   5203      1    1.0950e+02    2.9288e+02 ;   HG21 - CG2
- HG22
  5202   5201   5204      1    1.0950e+02    2.9288e+02 ;   HG21 - CG2
- HG23
  5203   5201   5204      1    1.0950e+02    2.9288e+02 ;   HG22 - CG2
- HG23
  5205   5208   5209      1    1.0950e+02    4.1840e+02 ;    CG1 - CD1
- HD11
  5205   5208   5210      1    1.0950e+02    4.1840e+02 ;    CG1 - CD1
- HD12
  5205   5208   5211      1    1.0950e+02    4.1840e+02 ;    CG1 - CD1
- HD13
  5206   5205   5207      1    1.0950e+02    2.9288e+02 ;   HG12 - CG1
- HG13
  5206   5205   5208      1    1.0950e+02    4.1840e+02 ;   HG12 - CG1
- CD1
  5207   5205   5208      1    1.0950e+02    4.1840e+02 ;   HG13 - CG1
- CD1
  5209   5208   5210      1    1.0950e+02    2.9288e+02 ;   HD11 - CD1
- HD12
  5209   5208   5211      1    1.0950e+02    2.9288e+02 ;   HD11 - CD1
- HD13
  5210   5208   5211      1    1.0950e+02    2.9288e+02 ;   HD12 - CD1
- HD13
  5212   5214   5215      1    1.2000e+02    4.1840e+02 ;      C - N
- H
  5212   5214   5216      1    1.2190e+02    4.1840e+02 ;      C - N
- CA
  5213   5212   5214      1    1.2290e+02    6.6944e+02 ;      O - C
- N
  5214   5216   5217      1    1.0950e+02    4.1840e+02 ;      N - CA
- HA
  5214   5216   5218      1    1.0970e+02    6.6944e+02 ;      N - CA
- CB
  5214   5216   5232      1    1.1010e+02    5.2718e+02 ;      N - CA
- C
  5215   5214   5216      1    1.1804e+02    4.1840e+02 ;      H - N
- CA
  5216   5218   5219      1    1.0950e+02    4.1840e+02 ;     CA - CB
- HB2
  5216   5218   5220      1    1.0950e+02    4.1840e+02 ;     CA - CB
- HB3
  5216   5218   5221      1    1.1400e+02    5.2718e+02 ;     CA - CB
- CG
  5216   5232   5233      1    1.2040e+02    6.6944e+02 ;     CA - C
- O
```

| | | | | | | |
|---|---|---|---|---|---|---|
| 5216 | 5232 | 5234 | 1 | 1.1660e+02 | 5.8576e+02 ; | CA - C - N |
| 5217 | 5216 | 5218 | 1 | 1.0950e+02 | 4.1840e+02 ; | HA - CA - CB |
| 5217 | 5216 | 5232 | 1 | 1.0950e+02 | 4.1840e+02 ; | HA - CA - C |
| 5218 | 5216 | 5232 | 1 | 1.1110e+02 | 5.2718e+02 ; | CB - CA - C |
| 5218 | 5221 | 5222 | 1 | 1.2000e+02 | 5.8576e+02 ; | CB - CG - CD1 |
| 5218 | 5221 | 5230 | 1 | 1.2000e+02 | 5.8576e+02 ; | CB - CG - CD2 |
| 5219 | 5218 | 5220 | 1 | 1.0950e+02 | 2.9288e+02 ; | HB2 - CB - HB3 |
| 5219 | 5218 | 5221 | 1 | 1.0950e+02 | 4.1840e+02 ; | HB2 - CB - CG |
| 5220 | 5218 | 5221 | 1 | 1.0950e+02 | 4.1840e+02 ; | HB3 - CB - CG |
| 5221 | 5222 | 5223 | 1 | 1.2000e+02 | 4.1840e+02 ; | CG - CD1 - HD1 |
| 5221 | 5222 | 5224 | 1 | 1.2000e+02 | 5.2718e+02 ; | CG - CD1 - CE1 |
| 5221 | 5230 | 5228 | 1 | 1.2000e+02 | 5.2718e+02 ; | CG - CD2 - CE2 |
| 5221 | 5230 | 5231 | 1 | 1.2000e+02 | 4.1840e+02 ; | CG - CD2 - HD2 |
| 5222 | 5221 | 5230 | 1 | 1.2000e+02 | 5.2718e+02 ; | CD1 - CG - CD2 |
| 5222 | 5224 | 5225 | 1 | 1.2000e+02 | 4.1840e+02 ; | CD1 - CE1 - HE1 |
| 5222 | 5224 | 5226 | 1 | 1.2000e+02 | 5.2718e+02 ; | CD1 - CE1 - CZ |
| 5223 | 5222 | 5224 | 1 | 1.2000e+02 | 4.1840e+02 ; | HD1 - CD1 - CE1 |
| 5224 | 5226 | 5227 | 1 | 1.2000e+02 | 4.1840e+02 ; | CE1 - CZ - HZ |
| 5224 | 5226 | 5228 | 1 | 1.2000e+02 | 5.2718e+02 ; | CE1 - CZ - CE2 |
| 5225 | 5224 | 5226 | 1 | 1.2000e+02 | 4.1840e+02 ; | HE1 - CE1 - CZ |
| 5226 | 5228 | 5229 | 1 | 1.2000e+02 | 4.1840e+02 ; | CZ - CE2 - HE2 |
| 5226 | 5228 | 5230 | 1 | 1.2000e+02 | 5.2718e+02 ; | CZ - CE2 - CD2 |
| 5227 | 5226 | 5228 | 1 | 1.2000e+02 | 4.1840e+02 ; | HZ - CZ - CE2 |
| 5228 | 5230 | 5231 | 1 | 1.2000e+02 | 4.1840e+02 ; | CE2 - CD2 - HD2 |
| 5229 | 5228 | 5230 | 1 | 1.2000e+02 | 4.1840e+02 ; | HE2 - CE2 - CD2 |
| 5232 | 5234 | 5235 | 1 | 1.2000e+02 | 4.1840e+02 ; | C - N - H |

```
  5232   5234   5236      1    1.2190e+02    4.1840e+02 ;      C - N
- CA
  5233   5232   5234      1    1.2290e+02    6.6944e+02 ;      O - C
- N
  5234   5236   5237      1    1.0950e+02    4.1840e+02 ;      N - CA
- HA
  5234   5236   5238      1    1.0970e+02    6.6944e+02 ;      N - CA
- CB
  5234   5236   5251      1    1.1010e+02    5.2718e+02 ;      N - CA
- C
  5235   5234   5236      1    1.1804e+02    4.1840e+02 ;      H - N
- CA
  5236   5238   5239      1    1.0950e+02    4.1840e+02 ;     CA - CB
- HB
  5236   5238   5240      1    1.0950e+02    3.3472e+02 ;     CA - CB
- CG2
  5236   5238   5244      1    1.0950e+02    3.3472e+02 ;     CA - CB
- CG1
  5236   5251   5252      1    1.2040e+02    6.6944e+02 ;     CA - C
- O
  5236   5251   5253      1    1.1660e+02    5.8576e+02 ;     CA - C
- N
  5237   5236   5238      1    1.0950e+02    4.1840e+02 ;     HA - CA
- CB
  5237   5236   5251      1    1.0950e+02    4.1840e+02 ;     HA - CA
- C
  5238   5236   5251      1    1.1110e+02    5.2718e+02 ;     CB - CA
- C
  5238   5240   5241      1    1.0950e+02    4.1840e+02 ;     CB - CG2
- HG21
  5238   5240   5242      1    1.0950e+02    4.1840e+02 ;     CB - CG2
- HG22
  5238   5240   5243      1    1.0950e+02    4.1840e+02 ;     CB - CG2
- HG23
  5238   5244   5245      1    1.0950e+02    4.1840e+02 ;     CB - CG1
- HG12
  5238   5244   5246      1    1.0950e+02    4.1840e+02 ;     CB - CG1
- HG13
  5238   5244   5247      1    1.0950e+02    3.3472e+02 ;     CB - CG1
- CD1
  5239   5238   5240      1    1.0950e+02    4.1840e+02 ;     HB - CB
- CG2
  5239   5238   5244      1    1.0950e+02    4.1840e+02 ;     HB - CB
- CG1
  5240   5238   5244      1    1.0950e+02    3.3472e+02 ;    CG2 - CB
- CG1
  5241   5240   5242      1    1.0950e+02    2.9288e+02 ;   HG21 - CG2
- HG22
  5241   5240   5243      1    1.0950e+02    2.9288e+02 ;   HG21 - CG2
- HG23
  5242   5240   5243      1    1.0950e+02    2.9288e+02 ;   HG22 - CG2
- HG23
```

5244 5247 5248 1 1.0950e+02 4.1840e+02 ; CG1 - CD1
- HD11
5244 5247 5249 1 1.0950e+02 4.1840e+02 ; CG1 - CD1
- HD12
5244 5247 5250 1 1.0950e+02 4.1840e+02 ; CG1 - CD1
- HD13
5245 5244 5246 1 1.0950e+02 2.9288e+02 ; HG12 - CG1
- HG13
5245 5244 5247 1 1.0950e+02 4.1840e+02 ; HG12 - CG1
- CD1
5246 5244 5247 1 1.0950e+02 4.1840e+02 ; HG13 - CG1
- CD1
5248 5247 5249 1 1.0950e+02 2.9288e+02 ; HD11 - CD1
- HD12
5248 5247 5250 1 1.0950e+02 2.9288e+02 ; HD11 - CD1
- HD13
5249 5247 5250 1 1.0950e+02 2.9288e+02 ; HD12 - CD1
- HD13
5251 5253 5254 1 1.2000e+02 4.1840e+02 ; C - N
- H
5251 5253 5255 1 1.2190e+02 4.1840e+02 ; C - N
- CA
5252 5251 5253 1 1.2290e+02 6.6944e+02 ; O - C
- N
5253 5255 5256 1 1.0950e+02 4.1840e+02 ; N - CA
- HA
5253 5255 5257 1 1.0970e+02 6.6944e+02 ; N - CA
- CB
5253 5255 5261 1 1.1010e+02 5.2718e+02 ; N - CA
- C
5254 5253 5255 1 1.1804e+02 4.1840e+02 ; H - N
- CA
5255 5257 5258 1 1.0950e+02 4.1840e+02 ; CA - CB
- HB1
5255 5257 5259 1 1.0950e+02 4.1840e+02 ; CA - CB
- HB2
5255 5257 5260 1 1.0950e+02 4.1840e+02 ; CA - CB
- HB3
5255 5261 5262 1 1.2040e+02 6.6944e+02 ; CA - C
- O
5255 5261 5263 1 1.1660e+02 5.8576e+02 ; CA - C
- N
5256 5255 5257 1 1.0950e+02 4.1840e+02 ; HA - CA
- CB
5256 5255 5261 1 1.0950e+02 4.1840e+02 ; HA - CA
- C
5257 5255 5261 1 1.1110e+02 5.2718e+02 ; CB - CA
- C
5258 5257 5259 1 1.0950e+02 2.9288e+02 ; HB1 - CB
- HB2
5258 5257 5260 1 1.0950e+02 2.9288e+02 ; HB1 - CB
- HB3

```
  5259   5257   5260      1    1.0950e+02    2.9288e+02 ;    HB2 - CB
- HB3
  5261   5263   5264      1    1.2000e+02    4.1840e+02 ;      C - N
- H
  5261   5263   5265      1    1.2190e+02    4.1840e+02 ;      C - N
- CA
  5262   5261   5263      1    1.2290e+02    6.6944e+02 ;      O - C
- N
  5263   5265   5266      1    1.0950e+02    4.1840e+02 ;      N - CA
- HA
  5263   5265   5267      1    1.0970e+02    6.6944e+02 ;      N - CA
- CB
  5263   5265   5280      1    1.1010e+02    5.2718e+02 ;      N - CA
- C
  5264   5263   5265      1    1.1804e+02    4.1840e+02 ;      H - N
- CA
  5265   5267   5268      1    1.0950e+02    4.1840e+02 ;     CA - CB
- HB2
  5265   5267   5269      1    1.0950e+02    4.1840e+02 ;     CA - CB
- HB3
  5265   5267   5270      1    1.0950e+02    3.3472e+02 ;     CA - CB
- CG
  5265   5280   5281      1    1.2040e+02    6.6944e+02 ;     CA - C
- O
  5265   5280   5282      1    1.1660e+02    5.8576e+02 ;     CA - C
- N
  5266   5265   5267      1    1.0950e+02    4.1840e+02 ;     HA - CA
- CB
  5266   5265   5280      1    1.0950e+02    4.1840e+02 ;     HA - CA
- C
  5267   5265   5280      1    1.1110e+02    5.2718e+02 ;     CB - CA
- C
  5267   5270   5271      1    1.0950e+02    4.1840e+02 ;     CB - CG
- HG
  5267   5270   5272      1    1.0950e+02    3.3472e+02 ;     CB - CG
- CD1
  5267   5270   5276      1    1.0950e+02    3.3472e+02 ;     CB - CG
- CD2
  5268   5267   5269      1    1.0950e+02    2.9288e+02 ;    HB2 - CB
- HB3
  5268   5267   5270      1    1.0950e+02    4.1840e+02 ;    HB2 - CB
- CG
  5269   5267   5270      1    1.0950e+02    4.1840e+02 ;    HB3 - CB
- CG
  5270   5272   5273      1    1.0950e+02    4.1840e+02 ;     CG - CD1
- HD11
  5270   5272   5274      1    1.0950e+02    4.1840e+02 ;     CG - CD1
- HD12
  5270   5272   5275      1    1.0950e+02    4.1840e+02 ;     CG - CD1
- HD13
  5270   5276   5277      1    1.0950e+02    4.1840e+02 ;     CG - CD2
- HD21
```

5270 5276 5278 1 1.0950e+02 4.1840e+02 ; CG - CD2
- HD22
5270 5276 5279 1 1.0950e+02 4.1840e+02 ; CG - CD2
- HD23
5271 5270 5272 1 1.0950e+02 4.1840e+02 ; HG - CG
- CD1
5271 5270 5276 1 1.0950e+02 4.1840e+02 ; HG - CG
- CD2
5272 5270 5276 1 1.0950e+02 3.3472e+02 ; CD1 - CG
- CD2
5273 5272 5274 1 1.0950e+02 2.9288e+02 ; HD11 - CD1
- HD12
5273 5272 5275 1 1.0950e+02 2.9288e+02 ; HD11 - CD1
- HD13
5274 5272 5275 1 1.0950e+02 2.9288e+02 ; HD12 - CD1
- HD13
5277 5276 5278 1 1.0950e+02 2.9288e+02 ; HD21 - CD2
- HD22
5277 5276 5279 1 1.0950e+02 2.9288e+02 ; HD21 - CD2
- HD23
5278 5276 5279 1 1.0950e+02 2.9288e+02 ; HD22 - CD2
- HD23
5280 5282 5283 1 1.2000e+02 4.1840e+02 ; C - N
- H
5280 5282 5284 1 1.2190e+02 4.1840e+02 ; C - N
- CA
5281 5280 5282 1 1.2290e+02 6.6944e+02 ; O - C
- N
5282 5284 5285 1 1.0950e+02 4.1840e+02 ; N - CA
- HA
5282 5284 5286 1 1.0970e+02 6.6944e+02 ; N - CA
- CB
5282 5284 5292 1 1.1010e+02 5.2718e+02 ; N - CA
- C
5283 5282 5284 1 1.1804e+02 4.1840e+02 ; H - N
- CA
5284 5286 5287 1 1.0950e+02 4.1840e+02 ; CA - CB
- HB2
5284 5286 5288 1 1.0950e+02 4.1840e+02 ; CA - CB
- HB3
5284 5286 5289 1 1.1110e+02 5.2718e+02 ; CA - CB
- CG
5284 5292 5293 1 1.2040e+02 6.6944e+02 ; CA - C
- O
5284 5292 5294 1 1.1660e+02 5.8576e+02 ; CA - C
- N
5285 5284 5286 1 1.0950e+02 4.1840e+02 ; HA - CA
- CB
5285 5284 5292 1 1.0950e+02 4.1840e+02 ; HA - CA
- C
5286 5284 5292 1 1.1110e+02 5.2718e+02 ; CB - CA
- C

| | | | | | | | | | | | |
|---|---|---|---|---|---|---|---|---|---|---|---|
| 5286 | 5289 | 5290 | 1 | 1.1700e+02 | 5.8576e+02 | ; | CB | - | CG | - | OD1 |
| 5286 | 5289 | 5291 | 1 | 1.1700e+02 | 5.8576e+02 | ; | CB | - | CG | - | OD2 |
| 5287 | 5286 | 5288 | 1 | 1.0950e+02 | 2.9288e+02 | ; | HB2 | - | CB | - | HB3 |
| 5287 | 5286 | 5289 | 1 | 1.0950e+02 | 4.1840e+02 | ; | HB2 | - | CB | - | CG |
| 5288 | 5286 | 5289 | 1 | 1.0950e+02 | 4.1840e+02 | ; | HB3 | - | CB | - | CG |
| 5290 | 5289 | 5291 | 1 | 1.2600e+02 | 6.6944e+02 | ; | OD1 | - | CG | - | OD2 |
| 5292 | 5294 | 5295 | 1 | 1.2000e+02 | 4.1840e+02 | ; | C | - | N | - | H |
| 5292 | 5294 | 5296 | 1 | 1.2190e+02 | 4.1840e+02 | ; | C | - | N | - | CA |
| 5293 | 5292 | 5294 | 1 | 1.2290e+02 | 6.6944e+02 | ; | O | - | C | - | N |
| 5294 | 5296 | 5297 | 1 | 1.0950e+02 | 4.1840e+02 | ; | N | - | CA | - | HA |
| 5294 | 5296 | 5298 | 1 | 1.0970e+02 | 6.6944e+02 | ; | N | - | CA | - | CB |
| 5294 | 5296 | 5308 | 1 | 1.1010e+02 | 5.2718e+02 | ; | N | - | CA | - | C |
| 5295 | 5294 | 5296 | 1 | 1.1804e+02 | 4.1840e+02 | ; | H | - | N | - | CA |
| 5296 | 5298 | 5299 | 1 | 1.0950e+02 | 4.1840e+02 | ; | CA | - | CB | - | HB |
| 5296 | 5298 | 5300 | 1 | 1.0950e+02 | 3.3472e+02 | ; | CA | - | CB | - | CG1 |
| 5296 | 5298 | 5304 | 1 | 1.0950e+02 | 3.3472e+02 | ; | CA | - | CB | - | CG2 |
| 5296 | 5308 | 5309 | 1 | 1.2040e+02 | 6.6944e+02 | ; | CA | - | C | - | O |
| 5296 | 5308 | 5310 | 1 | 1.1660e+02 | 5.8576e+02 | ; | CA | - | C | - | N |
| 5297 | 5296 | 5298 | 1 | 1.0950e+02 | 4.1840e+02 | ; | HA | - | CA | - | CB |
| 5297 | 5296 | 5308 | 1 | 1.0950e+02 | 4.1840e+02 | ; | HA | - | CA | - | C |
| 5298 | 5296 | 5308 | 1 | 1.1110e+02 | 5.2718e+02 | ; | CB | - | CA | - | C |
| 5298 | 5300 | 5301 | 1 | 1.0950e+02 | 4.1840e+02 | ; | CB | - | CG1 | - | HG11 |
| 5298 | 5300 | 5302 | 1 | 1.0950e+02 | 4.1840e+02 | ; | CB | - | CG1 | - | HG12 |
| 5298 | 5300 | 5303 | 1 | 1.0950e+02 | 4.1840e+02 | ; | CB | - | CG1 | - | HG13 |
| 5298 | 5304 | 5305 | 1 | 1.0950e+02 | 4.1840e+02 | ; | CB | - | CG2 | - | HG21 |
| 5298 | 5304 | 5306 | 1 | 1.0950e+02 | 4.1840e+02 | ; | CB | - | CG2 | - | HG22 |

5298 5304 5307 1 1.0950e+02 4.1840e+02 ; CB - CG2
- HG23
5299 5298 5300 1 1.0950e+02 4.1840e+02 ; HB - CB
- CG1
5299 5298 5304 1 1.0950e+02 4.1840e+02 ; HB - CB
- CG2
5300 5298 5304 1 1.0950e+02 3.3472e+02 ; CG1 - CB
- CG2
5301 5300 5302 1 1.0950e+02 2.9288e+02 ; HG11 - CG1
- HG12
5301 5300 5303 1 1.0950e+02 2.9288e+02 ; HG11 - CG1
- HG13
5302 5300 5303 1 1.0950e+02 2.9288e+02 ; HG12 - CG1
- HG13
5305 5304 5306 1 1.0950e+02 2.9288e+02 ; HG21 - CG2
- HG22
5305 5304 5307 1 1.0950e+02 2.9288e+02 ; HG21 - CG2
- HG23
5306 5304 5307 1 1.0950e+02 2.9288e+02 ; HG22 - CG2
- HG23
5308 5310 5311 1 1.2000e+02 4.1840e+02 ; C - N
- H
5308 5310 5312 1 1.2190e+02 4.1840e+02 ; C - N
- CA
5309 5308 5310 1 1.2290e+02 6.6944e+02 ; O - C
- N
5310 5312 5313 1 1.0950e+02 4.1840e+02 ; N - CA
- HA
5310 5312 5314 1 1.0970e+02 6.6944e+02 ; N - CA
- CB
5310 5312 5320 1 1.1010e+02 5.2718e+02 ; N - CA
- C
5311 5310 5312 1 1.1804e+02 4.1840e+02 ; H - N
- CA
5312 5314 5315 1 1.0950e+02 4.1840e+02 ; CA - CB
- HB2
5312 5314 5316 1 1.0950e+02 4.1840e+02 ; CA - CB
- HB3
5312 5314 5317 1 1.1110e+02 5.2718e+02 ; CA - CB
- CG
5312 5320 5321 1 1.2040e+02 6.6944e+02 ; CA - C
- O
5312 5320 5322 1 1.1660e+02 5.8576e+02 ; CA - C
- N
5313 5312 5314 1 1.0950e+02 4.1840e+02 ; HA - CA
- CB
5313 5312 5320 1 1.0950e+02 4.1840e+02 ; HA - CA
- C
5314 5312 5320 1 1.1110e+02 5.2718e+02 ; CB - CA
- C
5314 5317 5318 1 1.1700e+02 5.8576e+02 ; CB - CG
- OD1

5314 5317 5319 1 1.1700e+02 5.8576e+02 ; CB - CG
- OD2
5315 5314 5316 1 1.0950e+02 2.9288e+02 ; HB2 - CB
- HB3
5315 5314 5317 1 1.0950e+02 4.1840e+02 ; HB2 - CB
- CG
5316 5314 5317 1 1.0950e+02 4.1840e+02 ; HB3 - CB
- CG
5318 5317 5319 1 1.2600e+02 6.6944e+02 ; OD1 - CG
- OD2
5320 5322 5323 1 1.2000e+02 4.1840e+02 ; C - N
- H
5320 5322 5324 1 1.2190e+02 4.1840e+02 ; C - N
- CA
5321 5320 5322 1 1.2290e+02 6.6944e+02 ; O - C
- N
5322 5324 5325 1 1.0950e+02 4.1840e+02 ; N - CA
- HA
5322 5324 5326 1 1.0970e+02 6.6944e+02 ; N - CA
- CB
5322 5324 5334 1 1.1010e+02 5.2718e+02 ; N - CA
- C
5323 5322 5324 1 1.1804e+02 4.1840e+02 ; H - N
- CA
5324 5326 5327 1 1.0950e+02 4.1840e+02 ; CA - CB
- HB2
5324 5326 5328 1 1.0950e+02 4.1840e+02 ; CA - CB
- HB3
5324 5326 5329 1 1.1110e+02 5.2718e+02 ; CA - CB
- CG
5324 5334 5335 1 1.2040e+02 6.6944e+02 ; CA - C
- O
5324 5334 5336 1 1.1660e+02 5.8576e+02 ; CA - C
- N
5325 5324 5326 1 1.0950e+02 4.1840e+02 ; HA - CA
- CB
5325 5324 5334 1 1.0950e+02 4.1840e+02 ; HA - CA
- C
5326 5324 5334 1 1.1110e+02 5.2718e+02 ; CB - CA
- C
5326 5329 5330 1 1.2040e+02 6.6944e+02 ; CB - CG
- OD1
5326 5329 5331 1 1.1660e+02 5.8576e+02 ; CB - CG
- ND2
5327 5326 5328 1 1.0950e+02 2.9288e+02 ; HB2 - CB
- HB3
5327 5326 5329 1 1.0950e+02 4.1840e+02 ; HB2 - CB
- CG
5328 5326 5329 1 1.0950e+02 4.1840e+02 ; HB3 - CB
- CG
5329 5331 5332 1 1.2000e+02 4.1840e+02 ; CG - ND2
- HD21

5329 5331 5333 1 1.2000e+02 4.1840e+02 ; CG - ND2
- HD22
5330 5329 5331 1 1.2290e+02 6.6944e+02 ; OD1 - CG
- ND2
5332 5331 5333 1 1.2000e+02 2.9288e+02 ; HD21 - ND2
- HD22
5334 5336 5337 1 1.2000e+02 4.1840e+02 ; C - N
- H
5334 5336 5338 1 1.2190e+02 4.1840e+02 ; C - N
- CA
5335 5334 5336 1 1.2290e+02 6.6944e+02 ; O - C
- N
5336 5338 5339 1 1.0950e+02 4.1840e+02 ; N - CA
- HA
5336 5338 5340 1 1.0970e+02 6.6944e+02 ; N - CA
- CB
5336 5338 5345 1 1.1010e+02 5.2718e+02 ; N - CA
- C
5337 5336 5338 1 1.1804e+02 4.1840e+02 ; H - N
- CA
5338 5340 5341 1 1.0950e+02 4.1840e+02 ; CA - CB
- HB2
5338 5340 5342 1 1.0950e+02 4.1840e+02 ; CA - CB
- HB3
5338 5340 5343 1 1.0950e+02 4.1840e+02 ; CA - CB
- OG
5338 5345 5346 1 1.2040e+02 6.6944e+02 ; CA - C
- O
5338 5345 5347 1 1.1660e+02 5.8576e+02 ; CA - C
- N
5339 5338 5340 1 1.0950e+02 4.1840e+02 ; HA - CA
- CB
5339 5338 5345 1 1.0950e+02 4.1840e+02 ; HA - CA
- C
5340 5338 5345 1 1.1110e+02 5.2718e+02 ; CB - CA
- C
5340 5343 5344 1 1.0850e+02 4.6024e+02 ; CB - OG
- HG
5341 5340 5342 1 1.0950e+02 2.9288e+02 ; HB2 - CB
- HB3
5341 5340 5343 1 1.0950e+02 4.1840e+02 ; HB2 - CB
- OG
5342 5340 5343 1 1.0950e+02 4.1840e+02 ; HB3 - CB
- OG
5345 5347 5348 1 1.2000e+02 4.1840e+02 ; C - N
- H
5345 5347 5349 1 1.2190e+02 4.1840e+02 ; C - N
- CA
5346 5345 5347 1 1.2290e+02 6.6944e+02 ; O - C
- N
5347 5349 5350 1 1.0950e+02 4.1840e+02 ; N - CA
- HA2

5347 5349 5351 1 1.0950e+02 4.1840e+02 ; N - CA
- HA3
5347 5349 5352 1 1.1010e+02 5.2718e+02 ; N - CA
- C
5348 5347 5349 1 1.1804e+02 4.1840e+02 ; H - N
- CA
5349 5352 5353 1 1.2040e+02 6.6944e+02 ; CA - C
- O
5349 5352 5354 1 1.1660e+02 5.8576e+02 ; CA - C
- N
5350 5349 5351 1 1.0950e+02 2.9288e+02 ; HA2 - CA
- HA3
5350 5349 5352 1 1.0950e+02 4.1840e+02 ; HA2 - CA
- C
5351 5349 5352 1 1.0950e+02 4.1840e+02 ; HA3 - CA
- C
5352 5354 5355 1 1.2000e+02 4.1840e+02 ; C - N
- H
5352 5354 5356 1 1.2190e+02 4.1840e+02 ; C - N
- CA
5353 5352 5354 1 1.2290e+02 6.6944e+02 ; O - C
- N
5354 5356 5357 1 1.0950e+02 4.1840e+02 ; N - CA
- HA
5354 5356 5358 1 1.0970e+02 6.6944e+02 ; N - CA
- CB
5354 5356 5366 1 1.1010e+02 5.2718e+02 ; N - CA
- C
5355 5354 5356 1 1.1804e+02 4.1840e+02 ; H - N
- CA
5356 5358 5359 1 1.0950e+02 4.1840e+02 ; CA - CB
- HB
5356 5358 5360 1 1.0950e+02 3.3472e+02 ; CA - CB
- CG2
5356 5358 5364 1 1.0950e+02 4.1840e+02 ; CA - CB
- OG1
5356 5366 5367 1 1.2040e+02 6.6944e+02 ; CA - C
- O
5356 5366 5368 1 1.1660e+02 5.8576e+02 ; CA - C
- N
5357 5356 5358 1 1.0950e+02 4.1840e+02 ; HA - CA
- CB
5357 5356 5366 1 1.0950e+02 4.1840e+02 ; HA - CA
- C
5358 5356 5366 1 1.1110e+02 5.2718e+02 ; CB - CA
- C
5358 5360 5361 1 1.0950e+02 4.1840e+02 ; CB - CG2
- HG21
5358 5360 5362 1 1.0950e+02 4.1840e+02 ; CB - CG2
- HG22
5358 5360 5363 1 1.0950e+02 4.1840e+02 ; CB - CG2
- HG23

5358 5364 5365 1 1.0850e+02 4.6024e+02 ; CB - OG1
- HG1
5359 5358 5360 1 1.0950e+02 4.1840e+02 ; HB - CB
- CG2
5359 5358 5364 1 1.0950e+02 4.1840e+02 ; HB - CB
- OG1
5360 5358 5364 1 1.0950e+02 4.1840e+02 ; CG2 - CB
- OG1
5361 5360 5362 1 1.0950e+02 2.9288e+02 ; HG21 - CG2
- HG22
5361 5360 5363 1 1.0950e+02 2.9288e+02 ; HG21 - CG2
- HG23
5362 5360 5363 1 1.0950e+02 2.9288e+02 ; HG22 - CG2
- HG23
5366 5368 5369 1 1.2000e+02 4.1840e+02 ; C - N
- H
5366 5368 5370 1 1.2190e+02 4.1840e+02 ; C - N
- CA
5367 5366 5368 1 1.2290e+02 6.6944e+02 ; O - C
- N
5368 5370 5371 1 1.0950e+02 4.1840e+02 ; N - CA
- HA
5368 5370 5372 1 1.0970e+02 6.6944e+02 ; N - CA
- CB
5368 5370 5385 1 1.1010e+02 5.2718e+02 ; N - CA
- C
5369 5368 5370 1 1.1804e+02 4.1840e+02 ; H - N
- CA
5370 5372 5373 1 1.0950e+02 4.1840e+02 ; CA - CB
- HB2
5370 5372 5374 1 1.0950e+02 4.1840e+02 ; CA - CB
- HB3
5370 5372 5375 1 1.0950e+02 3.3472e+02 ; CA - CB
- CG
5370 5385 5386 1 1.2040e+02 6.6944e+02 ; CA - C
- O
5370 5385 5387 1 1.1660e+02 5.8576e+02 ; CA - C
- N
5371 5370 5372 1 1.0950e+02 4.1840e+02 ; HA - CA
- CB
5371 5370 5385 1 1.0950e+02 4.1840e+02 ; HA - CA
- C
5372 5370 5385 1 1.1110e+02 5.2718e+02 ; CB - CA
- C
5372 5375 5376 1 1.0950e+02 4.1840e+02 ; CB - CG
- HG
5372 5375 5377 1 1.0950e+02 3.3472e+02 ; CB - CG
- CD1
5372 5375 5381 1 1.0950e+02 3.3472e+02 ; CB - CG
- CD2
5373 5372 5374 1 1.0950e+02 2.9288e+02 ; HB2 - CB
- HB3

```
  5373   5372   5375      1    1.0950e+02    4.1840e+02 ;    HB2 - CB
- CG
  5374   5372   5375      1    1.0950e+02    4.1840e+02 ;    HB3 - CB
- CG
  5375   5377   5378      1    1.0950e+02    4.1840e+02 ;     CG - CD1
- HD11
  5375   5377   5379      1    1.0950e+02    4.1840e+02 ;     CG - CD1
- HD12
  5375   5377   5380      1    1.0950e+02    4.1840e+02 ;     CG - CD1
- HD13
  5375   5381   5382      1    1.0950e+02    4.1840e+02 ;     CG - CD2
- HD21
  5375   5381   5383      1    1.0950e+02    4.1840e+02 ;     CG - CD2
- HD22
  5375   5381   5384      1    1.0950e+02    4.1840e+02 ;     CG - CD2
- HD23
  5376   5375   5377      1    1.0950e+02    4.1840e+02 ;     HG - CG
- CD1
  5376   5375   5381      1    1.0950e+02    4.1840e+02 ;     HG - CG
- CD2
  5377   5375   5381      1    1.0950e+02    3.3472e+02 ;    CD1 - CG
- CD2
  5378   5377   5379      1    1.0950e+02    2.9288e+02 ;   HD11 - CD1
- HD12
  5378   5377   5380      1    1.0950e+02    2.9288e+02 ;   HD11 - CD1
- HD13
  5379   5377   5380      1    1.0950e+02    2.9288e+02 ;   HD12 - CD1
- HD13
  5382   5381   5383      1    1.0950e+02    2.9288e+02 ;   HD21 - CD2
- HD22
  5382   5381   5384      1    1.0950e+02    2.9288e+02 ;   HD21 - CD2
- HD23
  5383   5381   5384      1    1.0950e+02    2.9288e+02 ;   HD22 - CD2
- HD23
  5385   5387   5388      1    1.2000e+02    4.1840e+02 ;      C - N
- H
  5385   5387   5389      1    1.2190e+02    4.1840e+02 ;      C - N
- CA
  5386   5385   5387      1    1.2290e+02    6.6944e+02 ;      O - C
- N
  5387   5389   5390      1    1.0950e+02    4.1840e+02 ;      N - CA
- HA
  5387   5389   5391      1    1.0970e+02    6.6944e+02 ;      N - CA
- CB
  5387   5389   5396      1    1.1010e+02    5.2718e+02 ;      N - CA
- C
  5388   5387   5389      1    1.1804e+02    4.1840e+02 ;      H - N
- CA
  5389   5391   5392      1    1.0950e+02    4.1840e+02 ;     CA - CB
- HB2
  5389   5391   5393      1    1.0950e+02    4.1840e+02 ;     CA - CB
- HB3
```

5389 5391 5394 1 1.0950e+02 4.1840e+02 ; CA - CB
- OG
5389 5396 5397 1 1.2040e+02 6.6944e+02 ; CA - C
- O
5389 5396 5398 1 1.1660e+02 5.8576e+02 ; CA - C
- N
5390 5389 5391 1 1.0950e+02 4.1840e+02 ; HA - CA
- CB
5390 5389 5396 1 1.0950e+02 4.1840e+02 ; HA - CA
- C
5391 5389 5396 1 1.1110e+02 5.2718e+02 ; CB - CA
- C
5391 5394 5395 1 1.0850e+02 4.6024e+02 ; CB - OG
- HG
5392 5391 5393 1 1.0950e+02 2.9288e+02 ; HB2 - CB
- HB3
5392 5391 5394 1 1.0950e+02 4.1840e+02 ; HB2 - CB
- OG
5393 5391 5394 1 1.0950e+02 4.1840e+02 ; HB3 - CB
- OG
5396 5398 5399 1 1.2000e+02 4.1840e+02 ; C - N
- H
5396 5398 5400 1 1.2190e+02 4.1840e+02 ; C - N
- CA
5397 5396 5398 1 1.2290e+02 6.6944e+02 ; O - C
- N
5398 5400 5401 1 1.0950e+02 4.1840e+02 ; N - CA
- HA
5398 5400 5402 1 1.0970e+02 6.6944e+02 ; N - CA
- CB
5398 5400 5407 1 1.1010e+02 5.2718e+02 ; N - CA
- C
5399 5398 5400 1 1.1804e+02 4.1840e+02 ; H - N
- CA
5400 5402 5403 1 1.0950e+02 4.1840e+02 ; CA - CB
- HB2
5400 5402 5404 1 1.0950e+02 4.1840e+02 ; CA - CB
- HB3
5400 5402 5405 1 1.0950e+02 4.1840e+02 ; CA - CB
- OG
5400 5407 5408 1 1.2040e+02 6.6944e+02 ; CA - C
- O
5400 5407 5409 1 1.1660e+02 5.8576e+02 ; CA - C
- N
5401 5400 5402 1 1.0950e+02 4.1840e+02 ; HA - CA
- CB
5401 5400 5407 1 1.0950e+02 4.1840e+02 ; HA - CA
- C
5402 5400 5407 1 1.1110e+02 5.2718e+02 ; CB - CA
- C
5402 5405 5406 1 1.0850e+02 4.6024e+02 ; CB - OG
- HG

```
  5403   5402   5404       1    1.0950e+02    2.9288e+02 ;    HB2 - CB
- HB3
  5403   5402   5405       1    1.0950e+02    4.1840e+02 ;    HB2 - CB
- OG
  5404   5402   5405       1    1.0950e+02    4.1840e+02 ;    HB3 - CB
- OG
  5407   5409   5410       1    1.2000e+02    4.1840e+02 ;      C - N
- H
  5407   5409   5411       1    1.2190e+02    4.1840e+02 ;      C - N
- CA
  5408   5407   5409       1    1.2290e+02    6.6944e+02 ;      O - C
- N
  5409   5411   5412       1    1.0950e+02    4.1840e+02 ;      N - CA
- HA
  5409   5411   5413       1    1.0970e+02    6.6944e+02 ;      N - CA
- CB
  5409   5411   5424       1    1.1010e+02    5.2718e+02 ;      N - CA
- C
  5410   5409   5411       1    1.1804e+02    4.1840e+02 ;      H - N
- CA
  5411   5413   5414       1    1.0950e+02    4.1840e+02 ;     CA - CB
- HB2
  5411   5413   5415       1    1.0950e+02    4.1840e+02 ;     CA - CB
- HB3
  5411   5413   5416       1    1.0950e+02    3.3472e+02 ;     CA - CB
- CG
  5411   5424   5425       1    1.2040e+02    6.6944e+02 ;     CA - C
- O
  5411   5424   5426       1    1.1660e+02    5.8576e+02 ;     CA - C
- N
  5412   5411   5413       1    1.0950e+02    4.1840e+02 ;     HA - CA
- CB
  5412   5411   5424       1    1.0950e+02    4.1840e+02 ;     HA - CA
- C
  5413   5411   5424       1    1.1110e+02    5.2718e+02 ;     CB - CA
- C
  5413   5416   5417       1    1.0950e+02    4.1840e+02 ;     CB - CG
- HG2
  5413   5416   5418       1    1.0950e+02    4.1840e+02 ;     CB - CG
- HG3
  5413   5416   5419       1    1.1110e+02    5.2718e+02 ;     CB - CG
- CD
  5414   5413   5415       1    1.0950e+02    2.9288e+02 ;    HB2 - CB
- HB3
  5414   5413   5416       1    1.0950e+02    4.1840e+02 ;    HB2 - CB
- CG
  5415   5413   5416       1    1.0950e+02    4.1840e+02 ;    HB3 - CB
- CG
  5416   5419   5420       1    1.2040e+02    6.6944e+02 ;     CG - CD
- OE1
  5416   5419   5421       1    1.1660e+02    5.8576e+02 ;     CG - CD
- NE2
```

5417 5416 5418 1 1.0950e+02 2.9288e+02 ; HG2 - CG
- HG3
5417 5416 5419 1 1.0950e+02 4.1840e+02 ; HG2 - CG
- CD
5418 5416 5419 1 1.0950e+02 4.1840e+02 ; HG3 - CG
- CD
5419 5421 5422 1 1.2000e+02 4.1840e+02 ; CD - NE2
- HE21
5419 5421 5423 1 1.2000e+02 4.1840e+02 ; CD - NE2
- HE22
5420 5419 5421 1 1.2290e+02 6.6944e+02 ; OE1 - CD
- NE2
5422 5421 5423 1 1.2000e+02 2.9288e+02 ; HE21 - NE2
- HE22
5424 5426 5427 1 1.2000e+02 4.1840e+02 ; C - N
- H
5424 5426 5428 1 1.2190e+02 4.1840e+02 ; C - N
- CA
5425 5424 5426 1 1.2290e+02 6.6944e+02 ; O - C
- N
5426 5428 5429 1 1.0950e+02 4.1840e+02 ; N - CA
- HA
5426 5428 5430 1 1.0970e+02 6.6944e+02 ; N - CA
- CB
5426 5428 5440 1 1.1010e+02 5.2718e+02 ; N - CA
- C
5427 5426 5428 1 1.1804e+02 4.1840e+02 ; H - N
- CA
5428 5430 5431 1 1.0950e+02 4.1840e+02 ; CA - CB
- HB2
5428 5430 5432 1 1.0950e+02 4.1840e+02 ; CA - CB
- HB3
5428 5430 5433 1 1.0950e+02 3.3472e+02 ; CA - CB
- CG
5428 5440 5441 1 1.2040e+02 6.6944e+02 ; CA - C
- O
5428 5440 5442 1 1.1660e+02 5.8576e+02 ; CA - C
- N
5429 5428 5430 1 1.0950e+02 4.1840e+02 ; HA - CA
- CB
5429 5428 5440 1 1.0950e+02 4.1840e+02 ; HA - CA
- C
5430 5428 5440 1 1.1110e+02 5.2718e+02 ; CB - CA
- C
5430 5433 5434 1 1.0950e+02 4.1840e+02 ; CB - CG
- HG2
5430 5433 5435 1 1.0950e+02 4.1840e+02 ; CB - CG
- HG3
5430 5433 5436 1 1.1110e+02 5.2718e+02 ; CB - CG
- CD
5431 5430 5432 1 1.0950e+02 2.9288e+02 ; HB2 - CB
- HB3

5431 5430 5433 1 1.0950e+02 4.1840e+02 ; HB2 - CB
- CG
5432 5430 5433 1 1.0950e+02 4.1840e+02 ; HB3 - CB
- CG
5433 5436 5437 1 1.2040e+02 6.6944e+02 ; CG - CD
- OE1
5433 5436 5438 1 1.1000e+02 6.6944e+02 ; CG - CD
- OE2
5434 5433 5435 1 1.0950e+02 2.9288e+02 ; HG2 - CG
- HG3
5434 5433 5436 1 1.0950e+02 4.1840e+02 ; HG2 - CG
- CD
5435 5433 5436 1 1.0950e+02 4.1840e+02 ; HG3 - CG
- CD
5436 5438 5439 1 1.1300e+02 4.1840e+02 ; CD - OE2
- HE2
5437 5436 5438 1 1.2000e+02 6.6944e+02 ; OE1 - CD
- OE2
5440 5442 5443 1 1.2000e+02 4.1840e+02 ; C - N
- H
5440 5442 5444 1 1.2190e+02 4.1840e+02 ; C - N
- CA
5441 5440 5442 1 1.2290e+02 6.6944e+02 ; O - C
- N
5442 5444 5445 1 1.0950e+02 4.1840e+02 ; N - CA
- HA
5442 5444 5446 1 1.0970e+02 6.6944e+02 ; N - CA
- CB
5442 5444 5459 1 1.1010e+02 5.2718e+02 ; N - CA
- C
5443 5442 5444 1 1.1804e+02 4.1840e+02 ; H - N
- CA
5444 5446 5447 1 1.0950e+02 4.1840e+02 ; CA - CB
- HB
5444 5446 5448 1 1.0950e+02 3.3472e+02 ; CA - CB
- CG2
5444 5446 5452 1 1.0950e+02 3.3472e+02 ; CA - CB
- CG1
5444 5459 5460 1 1.2040e+02 6.6944e+02 ; CA - C
- O
5444 5459 5461 1 1.1660e+02 5.8576e+02 ; CA - C
- N
5445 5444 5446 1 1.0950e+02 4.1840e+02 ; HA - CA
- CB
5445 5444 5459 1 1.0950e+02 4.1840e+02 ; HA - CA
- C
5446 5444 5459 1 1.1110e+02 5.2718e+02 ; CB - CA
- C
5446 5448 5449 1 1.0950e+02 4.1840e+02 ; CB - CG2
- HG21
5446 5448 5450 1 1.0950e+02 4.1840e+02 ; CB - CG2
- HG22

5446 5448 5451 1 1.0950e+02 4.1840e+02 ; CB - CG2
- HG23
5446 5452 5453 1 1.0950e+02 4.1840e+02 ; CB - CG1
- HG12
5446 5452 5454 1 1.0950e+02 4.1840e+02 ; CB - CG1
- HG13
5446 5452 5455 1 1.0950e+02 3.3472e+02 ; CB - CG1
- CD1
5447 5446 5448 1 1.0950e+02 4.1840e+02 ; HB - CB
- CG2
5447 5446 5452 1 1.0950e+02 4.1840e+02 ; HB - CB
- CG1
5448 5446 5452 1 1.0950e+02 3.3472e+02 ; CG2 - CB
- CG1
5449 5448 5450 1 1.0950e+02 2.9288e+02 ; HG21 - CG2
- HG22
5449 5448 5451 1 1.0950e+02 2.9288e+02 ; HG21 - CG2
- HG23
5450 5448 5451 1 1.0950e+02 2.9288e+02 ; HG22 - CG2
- HG23
5452 5455 5456 1 1.0950e+02 4.1840e+02 ; CG1 - CD1
- HD11
5452 5455 5457 1 1.0950e+02 4.1840e+02 ; CG1 - CD1
- HD12
5452 5455 5458 1 1.0950e+02 4.1840e+02 ; CG1 - CD1
- HD13
5453 5452 5454 1 1.0950e+02 2.9288e+02 ; HG12 - CG1
- HG13
5453 5452 5455 1 1.0950e+02 4.1840e+02 ; HG12 - CG1
- CD1
5454 5452 5455 1 1.0950e+02 4.1840e+02 ; HG13 - CG1
- CD1
5456 5455 5457 1 1.0950e+02 2.9288e+02 ; HD11 - CD1
- HD12
5456 5455 5458 1 1.0950e+02 2.9288e+02 ; HD11 - CD1
- HD13
5457 5455 5458 1 1.0950e+02 2.9288e+02 ; HD12 - CD1
- HD13
5459 5461 5462 1 1.2000e+02 4.1840e+02 ; C - N
- H
5459 5461 5463 1 1.2190e+02 4.1840e+02 ; C - N
- CA
5460 5459 5461 1 1.2290e+02 6.6944e+02 ; O - C
- N
5461 5463 5464 1 1.0950e+02 4.1840e+02 ; N - CA
- HA
5461 5463 5465 1 1.0970e+02 6.6944e+02 ; N - CA
- CB
5461 5463 5478 1 1.1010e+02 5.2718e+02 ; N - CA
- C
5462 5461 5463 1 1.1804e+02 4.1840e+02 ; H - N
- CA

5463 5465 5466 1 1.0950e+02 4.1840e+02 ; CA - CB - HB2
5463 5465 5467 1 1.0950e+02 4.1840e+02 ; CA - CB - HB3
5463 5465 5468 1 1.0950e+02 3.3472e+02 ; CA - CB - CG
5463 5478 5479 1 1.2040e+02 6.6944e+02 ; CA - C - O
5463 5478 5480 1 1.1660e+02 5.8576e+02 ; CA - C - N
5464 5463 5465 1 1.0950e+02 4.1840e+02 ; HA - CA - CB
5464 5463 5478 1 1.0950e+02 4.1840e+02 ; HA - CA - C
5465 5463 5478 1 1.1110e+02 5.2718e+02 ; CB - CA - C
5465 5468 5469 1 1.0950e+02 4.1840e+02 ; CB - CG - HG
5465 5468 5470 1 1.0950e+02 3.3472e+02 ; CB - CG - CD1
5465 5468 5474 1 1.0950e+02 3.3472e+02 ; CB - CG - CD2
5466 5465 5467 1 1.0950e+02 2.9288e+02 ; HB2 - CB - HB3
5466 5465 5468 1 1.0950e+02 4.1840e+02 ; HB2 - CB - CG
5467 5465 5468 1 1.0950e+02 4.1840e+02 ; HB3 - CB - CG
5468 5470 5471 1 1.0950e+02 4.1840e+02 ; CG - CD1 - HD11
5468 5470 5472 1 1.0950e+02 4.1840e+02 ; CG - CD1 - HD12
5468 5470 5473 1 1.0950e+02 4.1840e+02 ; CG - CD1 - HD13
5468 5474 5475 1 1.0950e+02 4.1840e+02 ; CG - CD2 - HD21
5468 5474 5476 1 1.0950e+02 4.1840e+02 ; CG - CD2 - HD22
5468 5474 5477 1 1.0950e+02 4.1840e+02 ; CG - CD2 - HD23
5469 5468 5470 1 1.0950e+02 4.1840e+02 ; HG - CG - CD1
5469 5468 5474 1 1.0950e+02 4.1840e+02 ; HG - CG - CD2
5470 5468 5474 1 1.0950e+02 3.3472e+02 ; CD1 - CG - CD2
5471 5470 5472 1 1.0950e+02 2.9288e+02 ; HD11 - CD1 - HD12
5471 5470 5473 1 1.0950e+02 2.9288e+02 ; HD11 - CD1 - HD13
5472 5470 5473 1 1.0950e+02 2.9288e+02 ; HD12 - CD1 - HD13

5475 5474 5476 1 1.0950e+02 2.9288e+02 ; HD21 - CD2
- HD22
5475 5474 5477 1 1.0950e+02 2.9288e+02 ; HD21 - CD2
- HD23
5476 5474 5477 1 1.0950e+02 2.9288e+02 ; HD22 - CD2
- HD23
5478 5480 5481 1 1.2000e+02 4.1840e+02 ; C - N
- H
5478 5480 5482 1 1.2190e+02 4.1840e+02 ; C - N
- CA
5479 5478 5480 1 1.2290e+02 6.6944e+02 ; O - C
- N
5480 5482 5483 1 1.0950e+02 4.1840e+02 ; N - CA
- HA
5480 5482 5484 1 1.0970e+02 6.6944e+02 ; N - CA
- CB
5480 5482 5490 1 1.1010e+02 5.2718e+02 ; N - CA
- C
5481 5480 5482 1 1.1804e+02 4.1840e+02 ; H - N
- CA
5482 5484 5485 1 1.0950e+02 4.1840e+02 ; CA - CB
- HB2
5482 5484 5486 1 1.0950e+02 4.1840e+02 ; CA - CB
- HB3
5482 5484 5487 1 1.1110e+02 5.2718e+02 ; CA - CB
- CG
5482 5490 5491 1 1.2040e+02 6.6944e+02 ; CA - C
- O
5482 5490 5492 1 1.1660e+02 5.8576e+02 ; CA - C
- N
5483 5482 5484 1 1.0950e+02 4.1840e+02 ; HA - CA
- CB
5483 5482 5490 1 1.0950e+02 4.1840e+02 ; HA - CA
- C
5484 5482 5490 1 1.1110e+02 5.2718e+02 ; CB - CA
- C
5484 5487 5488 1 1.1700e+02 5.8576e+02 ; CB - CG
- OD1
5484 5487 5489 1 1.1700e+02 5.8576e+02 ; CB - CG
- OD2
5485 5484 5486 1 1.0950e+02 2.9288e+02 ; HB2 - CB
- HB3
5485 5484 5487 1 1.0950e+02 4.1840e+02 ; HB2 - CB
- CG
5486 5484 5487 1 1.0950e+02 4.1840e+02 ; HB3 - CB
- CG
5488 5487 5489 1 1.2600e+02 6.6944e+02 ; OD1 - CG
- OD2
5490 5492 5493 1 1.2000e+02 4.1840e+02 ; C - N
- H
5490 5492 5494 1 1.2190e+02 4.1840e+02 ; C - N
- CA

5491 5490 5492 1 1.2290e+02 6.6944e+02 ; O - C
- N
5492 5494 5495 1 1.0950e+02 4.1840e+02 ; N - CA
- HA2
5492 5494 5496 1 1.0950e+02 4.1840e+02 ; N - CA
- HA3
5492 5494 5497 1 1.1010e+02 5.2718e+02 ; N - CA
- C
5493 5492 5494 1 1.1804e+02 4.1840e+02 ; H - N
- CA
5494 5497 5498 1 1.2040e+02 6.6944e+02 ; CA - C
- O
5494 5497 5499 1 1.1660e+02 5.8576e+02 ; CA - C
- N
5495 5494 5496 1 1.0950e+02 2.9288e+02 ; HA2 - CA
- HA3
5495 5494 5497 1 1.0950e+02 4.1840e+02 ; HA2 - CA
- C
5496 5494 5497 1 1.0950e+02 4.1840e+02 ; HA3 - CA
- C
5497 5499 5500 1 1.2000e+02 4.1840e+02 ; C - N
- H
5497 5499 5501 1 1.2190e+02 4.1840e+02 ; C - N
- CA
5498 5497 5499 1 1.2290e+02 6.6944e+02 ; O - C
- N
5499 5501 5502 1 1.0950e+02 4.1840e+02 ; N - CA
- HA
5499 5501 5503 1 1.0970e+02 6.6944e+02 ; N - CA
- CB
5499 5501 5516 1 1.1010e+02 5.2718e+02 ; N - CA
- C
5500 5499 5501 1 1.1804e+02 4.1840e+02 ; H - N
- CA
5501 5503 5504 1 1.0950e+02 4.1840e+02 ; CA - CB
- HB2
5501 5503 5505 1 1.0950e+02 4.1840e+02 ; CA - CB
- HB3
5501 5503 5506 1 1.0950e+02 3.3472e+02 ; CA - CB
- CG
5501 5516 5517 1 1.2040e+02 6.6944e+02 ; CA - C
- O
5501 5516 5518 1 1.1660e+02 5.8576e+02 ; CA - C
- N
5502 5501 5503 1 1.0950e+02 4.1840e+02 ; HA - CA
- CB
5502 5501 5516 1 1.0950e+02 4.1840e+02 ; HA - CA
- C
5503 5501 5516 1 1.1110e+02 5.2718e+02 ; CB - CA
- C
5503 5506 5507 1 1.0950e+02 4.1840e+02 ; CB - CG
- HG

5503 5506 5508 1 1.0950e+02 3.3472e+02 ; CB - CG
- CD1
5503 5506 5512 1 1.0950e+02 3.3472e+02 ; CB - CG
- CD2
5504 5503 5505 1 1.0950e+02 2.9288e+02 ; HB2 - CB
- HB3
5504 5503 5506 1 1.0950e+02 4.1840e+02 ; HB2 - CB
- CG
5505 5503 5506 1 1.0950e+02 4.1840e+02 ; HB3 - CB
- CG
5506 5508 5509 1 1.0950e+02 4.1840e+02 ; CG - CD1
- HD11
5506 5508 5510 1 1.0950e+02 4.1840e+02 ; CG - CD1
- HD12
5506 5508 5511 1 1.0950e+02 4.1840e+02 ; CG - CD1
- HD13
5506 5512 5513 1 1.0950e+02 4.1840e+02 ; CG - CD2
- HD21
5506 5512 5514 1 1.0950e+02 4.1840e+02 ; CG - CD2
- HD22
5506 5512 5515 1 1.0950e+02 4.1840e+02 ; CG - CD2
- HD23
5507 5506 5508 1 1.0950e+02 4.1840e+02 ; HG - CG
- CD1
5507 5506 5512 1 1.0950e+02 4.1840e+02 ; HG - CG
- CD2
5508 5506 5512 1 1.0950e+02 3.3472e+02 ; CD1 - CG
- CD2
5509 5508 5510 1 1.0950e+02 2.9288e+02 ; HD11 - CD1
- HD12
5509 5508 5511 1 1.0950e+02 2.9288e+02 ; HD11 - CD1
- HD13
5510 5508 5511 1 1.0950e+02 2.9288e+02 ; HD12 - CD1
- HD13
5513 5512 5514 1 1.0950e+02 2.9288e+02 ; HD21 - CD2
- HD22
5513 5512 5515 1 1.0950e+02 2.9288e+02 ; HD21 - CD2
- HD23
5514 5512 5515 1 1.0950e+02 2.9288e+02 ; HD22 - CD2
- HD23
5516 5518 5519 1 1.2000e+02 4.1840e+02 ; C - N
- H
5516 5518 5520 1 1.2190e+02 4.1840e+02 ; C - N
- CA
5517 5516 5518 1 1.2290e+02 6.6944e+02 ; O - C
- N
5518 5520 5521 1 1.0950e+02 4.1840e+02 ; N - CA
- HA
5518 5520 5522 1 1.0970e+02 6.6944e+02 ; N - CA
- CB
5518 5520 5538 1 1.1010e+02 5.2718e+02 ; N - CA
- C

5519 5518 5520 1 1.1804e+02 4.1840e+02 ; H - N
- CA
5520 5522 5523 1 1.0950e+02 4.1840e+02 ; CA - CB
- HB2
5520 5522 5524 1 1.0950e+02 4.1840e+02 ; CA - CB
- HB3
5520 5522 5525 1 1.0950e+02 3.3472e+02 ; CA - CB
- CG
5520 5538 5539 1 1.2040e+02 6.6944e+02 ; CA - C
- O
5520 5538 5540 1 1.1660e+02 5.8576e+02 ; CA - C
- N
5521 5520 5522 1 1.0950e+02 4.1840e+02 ; HA - CA
- CB
5521 5520 5538 1 1.0950e+02 4.1840e+02 ; HA - CA
- C
5522 5520 5538 1 1.1110e+02 5.2718e+02 ; CB - CA
- C
5522 5525 5526 1 1.0950e+02 4.1840e+02 ; CB - CG
- HG2
5522 5525 5527 1 1.0950e+02 4.1840e+02 ; CB - CG
- HG3
5522 5525 5528 1 1.0950e+02 3.3472e+02 ; CB - CG
- CD
5523 5522 5524 1 1.0950e+02 2.9288e+02 ; HB2 - CB
- HB3
5523 5522 5525 1 1.0950e+02 4.1840e+02 ; HB2 - CB
- CG
5524 5522 5525 1 1.0950e+02 4.1840e+02 ; HB3 - CB
- CG
5525 5528 5529 1 1.0950e+02 4.1840e+02 ; CG - CD
- HD2
5525 5528 5530 1 1.0950e+02 4.1840e+02 ; CG - CD
- HD3
5525 5528 5531 1 1.0950e+02 3.3472e+02 ; CG - CD
- CE
5526 5525 5527 1 1.0950e+02 2.9288e+02 ; HG2 - CG
- HG3
5526 5525 5528 1 1.0950e+02 4.1840e+02 ; HG2 - CG
- CD
5527 5525 5528 1 1.0950e+02 4.1840e+02 ; HG3 - CG
- CD
5528 5531 5532 1 1.0950e+02 4.1840e+02 ; CD - CE
- HE2
5528 5531 5533 1 1.0950e+02 4.1840e+02 ; CD - CE
- HE3
5528 5531 5534 1 1.1120e+02 6.6944e+02 ; CD - CE
- NZ
5529 5528 5530 1 1.0950e+02 2.9288e+02 ; HD2 - CD
- HD3
5529 5528 5531 1 1.0950e+02 4.1840e+02 ; HD2 - CD
- CE

| | | | | | | | |
|---|---|---|---|---|---|---|---|
| 5530 | 5528 | 5531 | 1 | 1.0950e+02 | 4.1840e+02 | ; | HD3 - CD - CE |
| 5531 | 5534 | 5535 | 1 | 1.0950e+02 | 4.1840e+02 | ; | CE - NZ - HZ1 |
| 5531 | 5534 | 5536 | 1 | 1.0950e+02 | 4.1840e+02 | ; | CE - NZ - HZ2 |
| 5531 | 5534 | 5537 | 1 | 1.0950e+02 | 4.1840e+02 | ; | CE - NZ - HZ3 |
| 5532 | 5531 | 5533 | 1 | 1.0950e+02 | 2.9288e+02 | ; | HE2 - CE - HE3 |
| 5532 | 5531 | 5534 | 1 | 1.0950e+02 | 4.1840e+02 | ; | HE2 - CE - NZ |
| 5533 | 5531 | 5534 | 1 | 1.0950e+02 | 4.1840e+02 | ; | HE3 - CE - NZ |
| 5535 | 5534 | 5536 | 1 | 1.0950e+02 | 2.9288e+02 | ; | HZ1 - NZ - HZ2 |
| 5535 | 5534 | 5537 | 1 | 1.0950e+02 | 2.9288e+02 | ; | HZ1 - NZ - HZ3 |
| 5536 | 5534 | 5537 | 1 | 1.0950e+02 | 2.9288e+02 | ; | HZ2 - NZ - HZ3 |
| 5538 | 5540 | 5541 | 1 | 1.2000e+02 | 4.1840e+02 | ; | C - N - H |
| 5538 | 5540 | 5542 | 1 | 1.2190e+02 | 4.1840e+02 | ; | C - N - CA |
| 5539 | 5538 | 5540 | 1 | 1.2290e+02 | 6.6944e+02 | ; | O - C - N |
| 5540 | 5542 | 5543 | 1 | 1.0950e+02 | 4.1840e+02 | ; | N - CA - HA |
| 5540 | 5542 | 5544 | 1 | 1.0970e+02 | 6.6944e+02 | ; | N - CA - CB |
| 5540 | 5542 | 5560 | 1 | 1.1010e+02 | 5.2718e+02 | ; | N - CA - C |
| 5541 | 5540 | 5542 | 1 | 1.1804e+02 | 4.1840e+02 | ; | H - N - CA |
| 5542 | 5544 | 5545 | 1 | 1.0950e+02 | 4.1840e+02 | ; | CA - CB - HB2 |
| 5542 | 5544 | 5546 | 1 | 1.0950e+02 | 4.1840e+02 | ; | CA - CB - HB3 |
| 5542 | 5544 | 5547 | 1 | 1.0950e+02 | 3.3472e+02 | ; | CA - CB - CG |
| 5542 | 5560 | 5561 | 1 | 1.2040e+02 | 6.6944e+02 | ; | CA - C - O |
| 5542 | 5560 | 5562 | 1 | 1.1660e+02 | 5.8576e+02 | ; | CA - C - N |
| 5543 | 5542 | 5544 | 1 | 1.0950e+02 | 4.1840e+02 | ; | HA - CA - CB |
| 5543 | 5542 | 5560 | 1 | 1.0950e+02 | 4.1840e+02 | ; | HA - CA - C |
| 5544 | 5542 | 5560 | 1 | 1.1110e+02 | 5.2718e+02 | ; | CB - CA - C |
| 5544 | 5547 | 5548 | 1 | 1.0950e+02 | 4.1840e+02 | ; | CB - CG - HG2 |

5544 5547 5549 1 1.0950e+02 4.1840e+02 ; CB - CG
- HG3
5544 5547 5550 1 1.0950e+02 3.3472e+02 ; CB - CG
- CD
5545 5544 5546 1 1.0950e+02 2.9288e+02 ; HB2 - CB
- HB3
5545 5544 5547 1 1.0950e+02 4.1840e+02 ; HB2 - CB
- CG
5546 5544 5547 1 1.0950e+02 4.1840e+02 ; HB3 - CB
- CG
5547 5550 5551 1 1.0950e+02 4.1840e+02 ; CG - CD
- HD2
5547 5550 5552 1 1.0950e+02 4.1840e+02 ; CG - CD
- HD3
5547 5550 5553 1 1.0950e+02 3.3472e+02 ; CG - CD
- CE
5548 5547 5549 1 1.0950e+02 2.9288e+02 ; HG2 - CG
- HG3
5548 5547 5550 1 1.0950e+02 4.1840e+02 ; HG2 - CG
- CD
5549 5547 5550 1 1.0950e+02 4.1840e+02 ; HG3 - CG
- CD
5550 5553 5554 1 1.0950e+02 4.1840e+02 ; CD - CE
- HE2
5550 5553 5555 1 1.0950e+02 4.1840e+02 ; CD - CE
- HE3
5550 5553 5556 1 1.1120e+02 6.6944e+02 ; CD - CE
- NZ
5551 5550 5552 1 1.0950e+02 2.9288e+02 ; HD2 - CD
- HD3
5551 5550 5553 1 1.0950e+02 4.1840e+02 ; HD2 - CD
- CE
5552 5550 5553 1 1.0950e+02 4.1840e+02 ; HD3 - CD
- CE
5553 5556 5557 1 1.0950e+02 4.1840e+02 ; CE - NZ
- HZ1
5553 5556 5558 1 1.0950e+02 4.1840e+02 ; CE - NZ
- HZ2
5553 5556 5559 1 1.0950e+02 4.1840e+02 ; CE - NZ
- HZ3
5554 5553 5555 1 1.0950e+02 2.9288e+02 ; HE2 - CE
- HE3
5554 5553 5556 1 1.0950e+02 4.1840e+02 ; HE2 - CE
- NZ
5555 5553 5556 1 1.0950e+02 4.1840e+02 ; HE3 - CE
- NZ
5557 5556 5558 1 1.0950e+02 2.9288e+02 ; HZ1 - NZ
- HZ2
5557 5556 5559 1 1.0950e+02 2.9288e+02 ; HZ1 - NZ
- HZ3
5558 5556 5559 1 1.0950e+02 2.9288e+02 ; HZ2 - NZ
- HZ3

5560 5562 5563 1 1.2000e+02 4.1840e+02 ; C - N
- H
5560 5562 5564 1 1.2190e+02 4.1840e+02 ; C - N
- CA
5561 5560 5562 1 1.2290e+02 6.6944e+02 ; O - C
- N
5562 5564 5565 1 1.0950e+02 4.1840e+02 ; N - CA
- HA
5562 5564 5566 1 1.0970e+02 6.6944e+02 ; N - CA
- CB
5562 5564 5579 1 1.1010e+02 5.2718e+02 ; N - CA
- C
5563 5562 5564 1 1.1804e+02 4.1840e+02 ; H - N
- CA
5564 5566 5567 1 1.0950e+02 4.1840e+02 ; CA - CB
- HB
5564 5566 5568 1 1.0950e+02 3.3472e+02 ; CA - CB
- CG2
5564 5566 5572 1 1.0950e+02 3.3472e+02 ; CA - CB
- CG1
5564 5579 5580 1 1.2040e+02 6.6944e+02 ; CA - C
- O
5564 5579 5581 1 1.1660e+02 5.8576e+02 ; CA - C
- N
5565 5564 5566 1 1.0950e+02 4.1840e+02 ; HA - CA
- CB
5565 5564 5579 1 1.0950e+02 4.1840e+02 ; HA - CA
- C
5566 5564 5579 1 1.1110e+02 5.2718e+02 ; CB - CA
- C
5566 5568 5569 1 1.0950e+02 4.1840e+02 ; CB - CG2
- HG21
5566 5568 5570 1 1.0950e+02 4.1840e+02 ; CB - CG2
- HG22
5566 5568 5571 1 1.0950e+02 4.1840e+02 ; CB - CG2
- HG23
5566 5572 5573 1 1.0950e+02 4.1840e+02 ; CB - CG1
- HG12
5566 5572 5574 1 1.0950e+02 4.1840e+02 ; CB - CG1
- HG13
5566 5572 5575 1 1.0950e+02 3.3472e+02 ; CB - CG1
- CD1
5567 5566 5568 1 1.0950e+02 4.1840e+02 ; HB - CB
- CG2
5567 5566 5572 1 1.0950e+02 4.1840e+02 ; HB - CB
- CG1
5568 5566 5572 1 1.0950e+02 3.3472e+02 ; CG2 - CB
- CG1
5569 5568 5570 1 1.0950e+02 2.9288e+02 ; HG21 - CG2
- HG22
5569 5568 5571 1 1.0950e+02 2.9288e+02 ; HG21 - CG2
- HG23

5570 5568 5571 1 1.0950e+02 2.9288e+02 ; HG22 - CG2
- HG23
5572 5575 5576 1 1.0950e+02 4.1840e+02 ; CG1 - CD1
- HD11
5572 5575 5577 1 1.0950e+02 4.1840e+02 ; CG1 - CD1
- HD12
5572 5575 5578 1 1.0950e+02 4.1840e+02 ; CG1 - CD1
- HD13
5573 5572 5574 1 1.0950e+02 2.9288e+02 ; HG12 - CG1
- HG13
5573 5572 5575 1 1.0950e+02 4.1840e+02 ; HG12 - CG1
- CD1
5574 5572 5575 1 1.0950e+02 4.1840e+02 ; HG13 - CG1
- CD1
5576 5575 5577 1 1.0950e+02 2.9288e+02 ; HD11 - CD1
- HD12
5576 5575 5578 1 1.0950e+02 2.9288e+02 ; HD11 - CD1
- HD13
5577 5575 5578 1 1.0950e+02 2.9288e+02 ; HD12 - CD1
- HD13
5579 5581 5582 1 1.2000e+02 4.1840e+02 ; C - N
- H
5579 5581 5583 1 1.2190e+02 4.1840e+02 ; C - N
- CA
5580 5579 5581 1 1.2290e+02 6.6944e+02 ; O - C
- N
5581 5583 5584 1 1.0950e+02 4.1840e+02 ; N - CA
- HA2
5581 5583 5585 1 1.0950e+02 4.1840e+02 ; N - CA
- HA3
5581 5583 5586 1 1.1010e+02 5.2718e+02 ; N - CA
- C
5582 5581 5583 1 1.1804e+02 4.1840e+02 ; H - N
- CA
5583 5586 5587 1 1.2040e+02 6.6944e+02 ; CA - C
- O
5583 5586 5588 1 1.1660e+02 5.8576e+02 ; CA - C
- N
5584 5583 5585 1 1.0950e+02 2.9288e+02 ; HA2 - CA
- HA3
5584 5583 5586 1 1.0950e+02 4.1840e+02 ; HA2 - CA
- C
5585 5583 5586 1 1.0950e+02 4.1840e+02 ; HA3 - CA
- C
5586 5588 5589 1 1.2000e+02 4.1840e+02 ; C - N
- H
5586 5588 5590 1 1.2190e+02 4.1840e+02 ; C - N
- CA
5587 5586 5588 1 1.2290e+02 6.6944e+02 ; O - C
- N
5588 5590 5591 1 1.0950e+02 4.1840e+02 ; N - CA
- HA

```
  5588   5590   5592      1    1.0970e+02    6.6944e+02 ;      N - CA
- CB
  5588   5590   5607      1    1.1010e+02    5.2718e+02 ;      N - CA
- C
  5589   5588   5590      1    1.1804e+02    4.1840e+02 ;      H - N
- CA
  5590   5592   5593      1    1.0950e+02    4.1840e+02 ;     CA - CB
- HB2
  5590   5592   5594      1    1.0950e+02    4.1840e+02 ;     CA - CB
- HB3
  5590   5592   5595      1    1.1400e+02    5.2718e+02 ;     CA - CB
- CG
  5590   5607   5608      1    1.2040e+02    6.6944e+02 ;     CA - C
- O
  5590   5607   5609      1    1.1660e+02    5.8576e+02 ;     CA - C
- N
  5591   5590   5592      1    1.0950e+02    4.1840e+02 ;     HA - CA
- CB
  5591   5590   5607      1    1.0950e+02    4.1840e+02 ;     HA - CA
- C
  5592   5590   5607      1    1.1110e+02    5.2718e+02 ;     CB - CA
- C
  5592   5595   5596      1    1.2000e+02    5.8576e+02 ;     CB - CG
- CD1
  5592   5595   5605      1    1.2000e+02    5.8576e+02 ;     CB - CG
- CD2
  5593   5592   5594      1    1.0950e+02    2.9288e+02 ;    HB2 - CB
- HB3
  5593   5592   5595      1    1.0950e+02    4.1840e+02 ;    HB2 - CB
- CG
  5594   5592   5595      1    1.0950e+02    4.1840e+02 ;    HB3 - CB
- CG
  5595   5596   5597      1    1.2000e+02    4.1840e+02 ;     CG - CD1
- HD1
  5595   5596   5598      1    1.2000e+02    5.2718e+02 ;     CG - CD1
- CE1
  5595   5605   5603      1    1.2000e+02    5.2718e+02 ;     CG - CD2
- CE2
  5595   5605   5606      1    1.2000e+02    4.1840e+02 ;     CG - CD2
- HD2
  5596   5595   5605      1    1.2000e+02    5.2718e+02 ;    CD1 - CG
- CD2
  5596   5598   5599      1    1.2000e+02    4.1840e+02 ;    CD1 - CE1
- HE1
  5596   5598   5600      1    1.2000e+02    5.2718e+02 ;    CD1 - CE1
- CZ
  5597   5596   5598      1    1.2000e+02    4.1840e+02 ;    HD1 - CD1
- CE1
  5598   5600   5601      1    1.2000e+02    5.8576e+02 ;    CE1 - CZ
- OH
  5598   5600   5603      1    1.2000e+02    5.2718e+02 ;    CE1 - CZ
- CE2
```

5599 5598 5600 1 1.2000e+02 4.1840e+02 ; HE1 - CE1
- CZ
5600 5601 5602 1 1.1300e+02 4.1840e+02 ; CZ - OH
- HH
5600 5603 5604 1 1.2000e+02 4.1840e+02 ; CZ - CE2
- HE2
5600 5603 5605 1 1.2000e+02 5.2718e+02 ; CZ - CE2
- CD2
5601 5600 5603 1 1.2000e+02 5.8576e+02 ; OH - CZ
- CE2
5603 5605 5606 1 1.2000e+02 4.1840e+02 ; CE2 - CD2
- HD2
5604 5603 5605 1 1.2000e+02 4.1840e+02 ; HE2 - CE2
- CD2
5607 5609 5610 1 1.2000e+02 4.1840e+02 ; C - N
- H
5607 5609 5611 1 1.2190e+02 4.1840e+02 ; C - N
- CA
5608 5607 5609 1 1.2290e+02 6.6944e+02 ; O - C
- N
5609 5611 5612 1 1.0950e+02 4.1840e+02 ; N - CA
- HA
5609 5611 5613 1 1.0970e+02 6.6944e+02 ; N - CA
- CB
5609 5611 5624 1 1.1010e+02 5.2718e+02 ; N - CA
- C
5610 5609 5611 1 1.1804e+02 4.1840e+02 ; H - N
- CA
5611 5613 5614 1 1.0950e+02 4.1840e+02 ; CA - CB
- HB2
5611 5613 5615 1 1.0950e+02 4.1840e+02 ; CA - CB
- HB3
5611 5613 5616 1 1.0950e+02 3.3472e+02 ; CA - CB
- CG
5611 5624 5625 1 1.2040e+02 6.6944e+02 ; CA - C
- O
5611 5624 5626 1 1.1660e+02 5.8576e+02 ; CA - C
- N
5612 5611 5613 1 1.0950e+02 4.1840e+02 ; HA - CA
- CB
5612 5611 5624 1 1.0950e+02 4.1840e+02 ; HA - CA
- C
5613 5611 5624 1 1.1110e+02 5.2718e+02 ; CB - CA
- C
5613 5616 5617 1 1.0950e+02 4.1840e+02 ; CB - CG
- HG2
5613 5616 5618 1 1.0950e+02 4.1840e+02 ; CB - CG
- HG3
5613 5616 5619 1 1.1110e+02 5.2718e+02 ; CB - CG
- CD
5614 5613 5615 1 1.0950e+02 2.9288e+02 ; HB2 - CB
- HB3

5614 5613 5616 1 1.0950e+02 4.1840e+02 ; HB2 - CB
- CG
5615 5613 5616 1 1.0950e+02 4.1840e+02 ; HB3 - CB
- CG
5616 5619 5620 1 1.2040e+02 6.6944e+02 ; CG - CD
- OE1
5616 5619 5621 1 1.1660e+02 5.8576e+02 ; CG - CD
- NE2
5617 5616 5618 1 1.0950e+02 2.9288e+02 ; HG2 - CG
- HG3
5617 5616 5619 1 1.0950e+02 4.1840e+02 ; HG2 - CG
- CD
5618 5616 5619 1 1.0950e+02 4.1840e+02 ; HG3 - CG
- CD
5619 5621 5622 1 1.2000e+02 4.1840e+02 ; CD - NE2
- HE21
5619 5621 5623 1 1.2000e+02 4.1840e+02 ; CD - NE2
- HE22
5620 5619 5621 1 1.2290e+02 6.6944e+02 ; OE1 - CD
- NE2
5622 5621 5623 1 1.2000e+02 2.9288e+02 ; HE21 - NE2
- HE22
5624 5626 5627 1 1.2000e+02 4.1840e+02 ; C - N
- H
5624 5626 5628 1 1.2190e+02 4.1840e+02 ; C - N
- CA
5625 5624 5626 1 1.2290e+02 6.6944e+02 ; O - C
- N
5626 5628 5629 1 1.0950e+02 4.1840e+02 ; N - CA
- HA
5626 5628 5630 1 1.0970e+02 6.6944e+02 ; N - CA
- CB
5626 5628 5646 1 1.1010e+02 5.2718e+02 ; N - CA
- C
5627 5626 5628 1 1.1804e+02 4.1840e+02 ; H - N
- CA
5628 5630 5631 1 1.0950e+02 4.1840e+02 ; CA - CB
- HB2
5628 5630 5632 1 1.0950e+02 4.1840e+02 ; CA - CB
- HB3
5628 5630 5633 1 1.0950e+02 3.3472e+02 ; CA - CB
- CG
5628 5646 5647 1 1.2040e+02 6.6944e+02 ; CA - C
- O
5628 5646 5648 1 1.1660e+02 5.8576e+02 ; CA - C
- N
5629 5628 5630 1 1.0950e+02 4.1840e+02 ; HA - CA
- CB
5629 5628 5646 1 1.0950e+02 4.1840e+02 ; HA - CA
- C
5630 5628 5646 1 1.1110e+02 5.2718e+02 ; CB - CA
- C

```
  5630   5633   5634       1    1.0950e+02    4.1840e+02 ;     CB - CG
- HG2
  5630   5633   5635       1    1.0950e+02    4.1840e+02 ;     CB - CG
- HG3
  5630   5633   5636       1    1.0950e+02    3.3472e+02 ;     CB - CG
- CD
  5631   5630   5632       1    1.0950e+02    2.9288e+02 ;    HB2 - CB
- HB3
  5631   5630   5633       1    1.0950e+02    4.1840e+02 ;    HB2 - CB
- CG
  5632   5630   5633       1    1.0950e+02    4.1840e+02 ;    HB3 - CB
- CG
  5633   5636   5637       1    1.0950e+02    4.1840e+02 ;     CG - CD
- HD2
  5633   5636   5638       1    1.0950e+02    4.1840e+02 ;     CG - CD
- HD3
  5633   5636   5639       1    1.0950e+02    3.3472e+02 ;     CG - CD
- CE
  5634   5633   5635       1    1.0950e+02    2.9288e+02 ;    HG2 - CG
- HG3
  5634   5633   5636       1    1.0950e+02    4.1840e+02 ;    HG2 - CG
- CD
  5635   5633   5636       1    1.0950e+02    4.1840e+02 ;    HG3 - CG
- CD
  5636   5639   5640       1    1.0950e+02    4.1840e+02 ;     CD - CE
- HE2
  5636   5639   5641       1    1.0950e+02    4.1840e+02 ;     CD - CE
- HE3
  5636   5639   5642       1    1.1120e+02    6.6944e+02 ;     CD - CE
- NZ
  5637   5636   5638       1    1.0950e+02    2.9288e+02 ;    HD2 - CD
- HD3
  5637   5636   5639       1    1.0950e+02    4.1840e+02 ;    HD2 - CD
- CE
  5638   5636   5639       1    1.0950e+02    4.1840e+02 ;    HD3 - CD
- CE
  5639   5642   5643       1    1.0950e+02    4.1840e+02 ;     CE - NZ
- HZ1
  5639   5642   5644       1    1.0950e+02    4.1840e+02 ;     CE - NZ
- HZ2
  5639   5642   5645       1    1.0950e+02    4.1840e+02 ;     CE - NZ
- HZ3
  5640   5639   5641       1    1.0950e+02    2.9288e+02 ;    HE2 - CE
- HE3
  5640   5639   5642       1    1.0950e+02    4.1840e+02 ;    HE2 - CE
- NZ
  5641   5639   5642       1    1.0950e+02    4.1840e+02 ;    HE3 - CE
- NZ
  5643   5642   5644       1    1.0950e+02    2.9288e+02 ;    HZ1 - NZ
- HZ2
  5643   5642   5645       1    1.0950e+02    2.9288e+02 ;    HZ1 - NZ
- HZ3
```

5644 5642 5645 1 1.0950e+02 2.9288e+02 ; HZ2 - NZ
- HZ3
5646 5648 5649 1 1.2000e+02 4.1840e+02 ; C - N
- H
5646 5648 5650 1 1.2190e+02 4.1840e+02 ; C - N
- CA
5647 5646 5648 1 1.2290e+02 6.6944e+02 ; O - C
- N
5648 5650 5651 1 1.0950e+02 4.1840e+02 ; N - CA
- HA
5648 5650 5652 1 1.0970e+02 6.6944e+02 ; N - CA
- CB
5648 5650 5665 1 1.1010e+02 5.2718e+02 ; N - CA
- C
5649 5648 5650 1 1.1804e+02 4.1840e+02 ; H - N
- CA
5650 5652 5653 1 1.0950e+02 4.1840e+02 ; CA - CB
- HB
5650 5652 5654 1 1.0950e+02 3.3472e+02 ; CA - CB
- CG2
5650 5652 5658 1 1.0950e+02 3.3472e+02 ; CA - CB
- CG1
5650 5665 5666 1 1.2040e+02 6.6944e+02 ; CA - C
- O
5650 5665 5667 1 1.1660e+02 5.8576e+02 ; CA - C
- N
5651 5650 5652 1 1.0950e+02 4.1840e+02 ; HA - CA
- CB
5651 5650 5665 1 1.0950e+02 4.1840e+02 ; HA - CA
- C
5652 5650 5665 1 1.1110e+02 5.2718e+02 ; CB - CA
- C
5652 5654 5655 1 1.0950e+02 4.1840e+02 ; CB - CG2
- HG21
5652 5654 5656 1 1.0950e+02 4.1840e+02 ; CB - CG2
- HG22
5652 5654 5657 1 1.0950e+02 4.1840e+02 ; CB - CG2
- HG23
5652 5658 5659 1 1.0950e+02 4.1840e+02 ; CB - CG1
- HG12
5652 5658 5660 1 1.0950e+02 4.1840e+02 ; CB - CG1
- HG13
5652 5658 5661 1 1.0950e+02 3.3472e+02 ; CB - CG1
- CD1
5653 5652 5654 1 1.0950e+02 4.1840e+02 ; HB - CB
- CG2
5653 5652 5658 1 1.0950e+02 4.1840e+02 ; HB - CB
- CG1
5654 5652 5658 1 1.0950e+02 3.3472e+02 ; CG2 - CB
- CG1
5655 5654 5656 1 1.0950e+02 2.9288e+02 ; HG21 - CG2
- HG22

5655 5654 5657 1 1.0950e+02 2.9288e+02 ; HG21 - CG2
- HG23
5656 5654 5657 1 1.0950e+02 2.9288e+02 ; HG22 - CG2
- HG23
5658 5661 5662 1 1.0950e+02 4.1840e+02 ; CG1 - CD1
- HD11
5658 5661 5663 1 1.0950e+02 4.1840e+02 ; CG1 - CD1
- HD12
5658 5661 5664 1 1.0950e+02 4.1840e+02 ; CG1 - CD1
- HD13
5659 5658 5660 1 1.0950e+02 2.9288e+02 ; HG12 - CG1
- HG13
5659 5658 5661 1 1.0950e+02 4.1840e+02 ; HG12 - CG1
- CD1
5660 5658 5661 1 1.0950e+02 4.1840e+02 ; HG13 - CG1
- CD1
5662 5661 5663 1 1.0950e+02 2.9288e+02 ; HD11 - CD1
- HD12
5662 5661 5664 1 1.0950e+02 2.9288e+02 ; HD11 - CD1
- HD13
5663 5661 5664 1 1.0950e+02 2.9288e+02 ; HD12 - CD1
- HD13
5665 5667 5668 1 1.2190e+02 4.1840e+02 ; C - N
- CD
5665 5667 5677 1 1.2190e+02 4.1840e+02 ; C - N
- CA
5666 5665 5667 1 1.2290e+02 6.6944e+02 ; O - C
- N
5667 5668 5669 1 1.0950e+02 4.1840e+02 ; N - CD
- HD2
5667 5668 5670 1 1.0950e+02 4.1840e+02 ; N - CD
- HD3
5667 5668 5671 1 1.0970e+02 6.6944e+02 ; N - CD
- CG
5667 5677 5674 1 1.0970e+02 6.6944e+02 ; N - CA
- CB
5667 5677 5678 1 1.0950e+02 4.1840e+02 ; N - CA
- HA
5667 5677 5679 1 1.1010e+02 5.2718e+02 ; N - CA
- C
5668 5667 5677 1 1.1800e+02 4.1840e+02 ; CD - N
- CA
5668 5671 5672 1 1.0950e+02 4.1840e+02 ; CD - CG
- HG2
5668 5671 5673 1 1.0950e+02 4.1840e+02 ; CD - CG
- HG3
5668 5671 5674 1 1.0950e+02 3.3472e+02 ; CD - CG
- CB
5669 5668 5670 1 1.0950e+02 2.9288e+02 ; HD2 - CD
- HD3
5669 5668 5671 1 1.0950e+02 4.1840e+02 ; HD2 - CD
- CG

5670 5668 5671 1 1.0950e+02 4.1840e+02 ; HD3 - CD
- CG
5671 5674 5675 1 1.0950e+02 4.1840e+02 ; CG - CB
- HB2
5671 5674 5676 1 1.0950e+02 4.1840e+02 ; CG - CB
- HB3
5671 5674 5677 1 1.0950e+02 3.3472e+02 ; CG - CB
- CA
5672 5671 5673 1 1.0950e+02 2.9288e+02 ; HG2 - CG
- HG3
5672 5671 5674 1 1.0950e+02 4.1840e+02 ; HG2 - CG
- CB
5673 5671 5674 1 1.0950e+02 4.1840e+02 ; HG3 - CG
- CB
5674 5677 5678 1 1.0950e+02 4.1840e+02 ; CB - CA
- HA
5674 5677 5679 1 1.1110e+02 5.2718e+02 ; CB - CA
- C
5675 5674 5676 1 1.0950e+02 2.9288e+02 ; HB2 - CB
- HB3
5675 5674 5677 1 1.0950e+02 4.1840e+02 ; HB2 - CB
- CA
5676 5674 5677 1 1.0950e+02 4.1840e+02 ; HB3 - CB
- CA
5677 5679 5680 1 1.2040e+02 6.6944e+02 ; CA - C
- O
5677 5679 5681 1 1.1660e+02 5.8576e+02 ; CA - C
- N
5678 5677 5679 1 1.0950e+02 4.1840e+02 ; HA - CA
- C
5679 5681 5682 1 1.2190e+02 4.1840e+02 ; C - N
- CD
5679 5681 5691 1 1.2190e+02 4.1840e+02 ; C - N
- CA
5680 5679 5681 1 1.2290e+02 6.6944e+02 ; O - C
- N
5681 5682 5683 1 1.0950e+02 4.1840e+02 ; N - CD
- HD2
5681 5682 5684 1 1.0950e+02 4.1840e+02 ; N - CD
- HD3
5681 5682 5685 1 1.0970e+02 6.6944e+02 ; N - CD
- CG
5681 5691 5688 1 1.0970e+02 6.6944e+02 ; N - CA
- CB
5681 5691 5692 1 1.0950e+02 4.1840e+02 ; N - CA
- HA
5681 5691 5693 1 1.1010e+02 5.2718e+02 ; N - CA
- C
5682 5681 5691 1 1.1800e+02 4.1840e+02 ; CD - N
- CA
5682 5685 5686 1 1.0950e+02 4.1840e+02 ; CD - CG
- HG2

| | | | | | | | |
|---|---|---|---|---|---|---|---|
| 5682 | 5685 | 5687 | 1 | 1.0950e+02 | 4.1840e+02 | ; | CD - CG - HG3 |
| 5682 | 5685 | 5688 | 1 | 1.0950e+02 | 3.3472e+02 | ; | CD - CG - CB |
| 5683 | 5682 | 5684 | 1 | 1.0950e+02 | 2.9288e+02 | ; | HD2 - CD - HD3 |
| 5683 | 5682 | 5685 | 1 | 1.0950e+02 | 4.1840e+02 | ; | HD2 - CD - CG |
| 5684 | 5682 | 5685 | 1 | 1.0950e+02 | 4.1840e+02 | ; | HD3 - CD - CG |
| 5685 | 5688 | 5689 | 1 | 1.0950e+02 | 4.1840e+02 | ; | CG - CB - HB2 |
| 5685 | 5688 | 5690 | 1 | 1.0950e+02 | 4.1840e+02 | ; | CG - CB - HB3 |
| 5685 | 5688 | 5691 | 1 | 1.0950e+02 | 3.3472e+02 | ; | CG - CB - CA |
| 5686 | 5685 | 5687 | 1 | 1.0950e+02 | 2.9288e+02 | ; | HG2 - CG - HG3 |
| 5686 | 5685 | 5688 | 1 | 1.0950e+02 | 4.1840e+02 | ; | HG2 - CG - CB |
| 5687 | 5685 | 5688 | 1 | 1.0950e+02 | 4.1840e+02 | ; | HG3 - CG - CB |
| 5688 | 5691 | 5692 | 1 | 1.0950e+02 | 4.1840e+02 | ; | CB - CA - HA |
| 5688 | 5691 | 5693 | 1 | 1.1110e+02 | 5.2718e+02 | ; | CB - CA - C |
| 5689 | 5688 | 5690 | 1 | 1.0950e+02 | 2.9288e+02 | ; | HB2 - CB - HB3 |
| 5689 | 5688 | 5691 | 1 | 1.0950e+02 | 4.1840e+02 | ; | HB2 - CB - CA |
| 5690 | 5688 | 5691 | 1 | 1.0950e+02 | 4.1840e+02 | ; | HB3 - CB - CA |
| 5691 | 5693 | 5694 | 1 | 1.2040e+02 | 6.6944e+02 | ; | CA - C - O |
| 5691 | 5693 | 5695 | 1 | 1.1660e+02 | 5.8576e+02 | ; | CA - C - N |
| 5692 | 5691 | 5693 | 1 | 1.0950e+02 | 4.1840e+02 | ; | HA - CA - C |
| 5693 | 5695 | 5696 | 1 | 1.2000e+02 | 4.1840e+02 | ; | C - N - H |
| 5693 | 5695 | 5697 | 1 | 1.2190e+02 | 4.1840e+02 | ; | C - N - CA |
| 5694 | 5693 | 5695 | 1 | 1.2290e+02 | 6.6944e+02 | ; | O - C - N |
| 5695 | 5697 | 5698 | 1 | 1.0950e+02 | 4.1840e+02 | ; | N - CA - HA |
| 5695 | 5697 | 5699 | 1 | 1.0970e+02 | 6.6944e+02 | ; | N - CA - CB |
| 5695 | 5697 | 5705 | 1 | 1.1010e+02 | 5.2718e+02 | ; | N - CA - C |
| 5696 | 5695 | 5697 | 1 | 1.1804e+02 | 4.1840e+02 | ; | H - N - CA |

| | | | | | | |
|---|---|---|---|---|---|---|
| 5697 | 5699 | 5700 | 1 | 1.0950e+02 | 4.1840e+02 | ; CA - CB - HB2 |
| 5697 | 5699 | 5701 | 1 | 1.0950e+02 | 4.1840e+02 | ; CA - CB - HB3 |
| 5697 | 5699 | 5702 | 1 | 1.1110e+02 | 5.2718e+02 | ; CA - CB - CG |
| 5697 | 5705 | 5706 | 1 | 1.2040e+02 | 6.6944e+02 | ; CA - C - O |
| 5697 | 5705 | 5707 | 1 | 1.1660e+02 | 5.8576e+02 | ; CA - C - N |
| 5698 | 5697 | 5699 | 1 | 1.0950e+02 | 4.1840e+02 | ; HA - CA - CB |
| 5698 | 5697 | 5705 | 1 | 1.0950e+02 | 4.1840e+02 | ; HA - CA - C |
| 5699 | 5697 | 5705 | 1 | 1.1110e+02 | 5.2718e+02 | ; CB - CA - C |
| 5699 | 5702 | 5703 | 1 | 1.1700e+02 | 5.8576e+02 | ; CB - CG - OD1 |
| 5699 | 5702 | 5704 | 1 | 1.1700e+02 | 5.8576e+02 | ; CB - CG - OD2 |
| 5700 | 5699 | 5701 | 1 | 1.0950e+02 | 2.9288e+02 | ; HB2 - CB - HB3 |
| 5700 | 5699 | 5702 | 1 | 1.0950e+02 | 4.1840e+02 | ; HB2 - CB - CG |
| 5701 | 5699 | 5702 | 1 | 1.0950e+02 | 4.1840e+02 | ; HB3 - CB - CG |
| 5703 | 5702 | 5704 | 1 | 1.2600e+02 | 6.6944e+02 | ; OD1 - CG - OD2 |
| 5705 | 5707 | 5708 | 1 | 1.2000e+02 | 4.1840e+02 | ; C - N - H |
| 5705 | 5707 | 5709 | 1 | 1.2190e+02 | 4.1840e+02 | ; C - N - CA |
| 5706 | 5705 | 5707 | 1 | 1.2290e+02 | 6.6944e+02 | ; O - C - N |
| 5707 | 5709 | 5710 | 1 | 1.0950e+02 | 4.1840e+02 | ; N - CA - HA |
| 5707 | 5709 | 5711 | 1 | 1.0970e+02 | 6.6944e+02 | ; N - CA - CB |
| 5707 | 5709 | 5724 | 1 | 1.1010e+02 | 5.2718e+02 | ; N - CA - C |
| 5708 | 5707 | 5709 | 1 | 1.1804e+02 | 4.1840e+02 | ; H - N - CA |
| 5709 | 5711 | 5712 | 1 | 1.0950e+02 | 4.1840e+02 | ; CA - CB - HB |
| 5709 | 5711 | 5713 | 1 | 1.0950e+02 | 3.3472e+02 | ; CA - CB - CG2 |
| 5709 | 5711 | 5717 | 1 | 1.0950e+02 | 3.3472e+02 | ; CA - CB - CG1 |
| 5709 | 5724 | 5725 | 1 | 1.2040e+02 | 6.6944e+02 | ; CA - C - O |
| 5709 | 5724 | 5726 | 1 | 1.1660e+02 | 5.8576e+02 | ; CA - C - N |

```
  5710   5709   5711      1    1.0950e+02    4.1840e+02 ;     HA - CA
- CB
  5710   5709   5724      1    1.0950e+02    4.1840e+02 ;     HA - CA
- C
  5711   5709   5724      1    1.1110e+02    5.2718e+02 ;     CB - CA
- C
  5711   5713   5714      1    1.0950e+02    4.1840e+02 ;     CB - CG2
- HG21
  5711   5713   5715      1    1.0950e+02    4.1840e+02 ;     CB - CG2
- HG22
  5711   5713   5716      1    1.0950e+02    4.1840e+02 ;     CB - CG2
- HG23
  5711   5717   5718      1    1.0950e+02    4.1840e+02 ;     CB - CG1
- HG12
  5711   5717   5719      1    1.0950e+02    4.1840e+02 ;     CB - CG1
- HG13
  5711   5717   5720      1    1.0950e+02    3.3472e+02 ;     CB - CG1
- CD1
  5712   5711   5713      1    1.0950e+02    4.1840e+02 ;     HB - CB
- CG2
  5712   5711   5717      1    1.0950e+02    4.1840e+02 ;     HB - CB
- CG1
  5713   5711   5717      1    1.0950e+02    3.3472e+02 ;    CG2 - CB
- CG1
  5714   5713   5715      1    1.0950e+02    2.9288e+02 ;   HG21 - CG2
- HG22
  5714   5713   5716      1    1.0950e+02    2.9288e+02 ;   HG21 - CG2
- HG23
  5715   5713   5716      1    1.0950e+02    2.9288e+02 ;   HG22 - CG2
- HG23
  5717   5720   5721      1    1.0950e+02    4.1840e+02 ;    CG1 - CD1
- HD11
  5717   5720   5722      1    1.0950e+02    4.1840e+02 ;    CG1 - CD1
- HD12
  5717   5720   5723      1    1.0950e+02    4.1840e+02 ;    CG1 - CD1
- HD13
  5718   5717   5719      1    1.0950e+02    2.9288e+02 ;   HG12 - CG1
- HG13
  5718   5717   5720      1    1.0950e+02    4.1840e+02 ;   HG12 - CG1
- CD1
  5719   5717   5720      1    1.0950e+02    4.1840e+02 ;   HG13 - CG1
- CD1
  5721   5720   5722      1    1.0950e+02    2.9288e+02 ;   HD11 - CD1
- HD12
  5721   5720   5723      1    1.0950e+02    2.9288e+02 ;   HD11 - CD1
- HD13
  5722   5720   5723      1    1.0950e+02    2.9288e+02 ;   HD12 - CD1
- HD13
  5724   5726   5727      1    1.2000e+02    4.1840e+02 ;      C - N
- H
  5724   5726   5728      1    1.2190e+02    4.1840e+02 ;      C - N
- CA
```

```
  5725   5724   5726      1    1.2290e+02    6.6944e+02 ;      O - C
- N
  5726   5728   5729      1    1.0950e+02    4.1840e+02 ;      N - CA
- HA
  5726   5728   5730      1    1.0970e+02    6.6944e+02 ;      N - CA
- CB
  5726   5728   5741      1    1.1010e+02    5.2718e+02 ;      N - CA
- C
  5727   5726   5728      1    1.1804e+02    4.1840e+02 ;      H - N
- CA
  5728   5730   5731      1    1.0950e+02    4.1840e+02 ;     CA - CB
- HB2
  5728   5730   5732      1    1.0950e+02    4.1840e+02 ;     CA - CB
- HB3
  5728   5730   5733      1    1.1310e+02    5.2718e+02 ;     CA - CB
- CG
  5728   5741   5742      1    1.2040e+02    6.6944e+02 ;     CA - C
- O
  5728   5741   5743      1    1.1660e+02    5.8576e+02 ;     CA - C
- N
  5729   5728   5730      1    1.0950e+02    4.1840e+02 ;     HA - CA
- CB
  5729   5728   5741      1    1.0950e+02    4.1840e+02 ;     HA - CA
- C
  5730   5728   5741      1    1.1110e+02    5.2718e+02 ;     CB - CA
- C
  5730   5733   5734      1    1.2000e+02    5.8576e+02 ;     CB - CG
- ND1
  5730   5733   5739      1    1.2000e+02    5.8576e+02 ;     CB - CG
- CD2
  5731   5730   5732      1    1.0950e+02    2.9288e+02 ;    HB2 - CB
- HB3
  5731   5730   5733      1    1.0950e+02    4.1840e+02 ;    HB2 - CB
- CG
  5732   5730   5733      1    1.0950e+02    4.1840e+02 ;    HB3 - CB
- CG
  5733   5734   5735      1    1.1700e+02    5.8576e+02 ;     CG - ND1
- CE1
  5733   5739   5737      1    1.2000e+02    5.8576e+02 ;     CG - CD2
- NE2
  5733   5739   5740      1    1.2000e+02    4.1840e+02 ;     CG - CD2
- HD2
  5734   5733   5739      1    1.2000e+02    5.8576e+02 ;    ND1 - CG
- CD2
  5734   5735   5736      1    1.2000e+02    4.1840e+02 ;    ND1 - CE1
- HE1
  5734   5735   5737      1    1.2000e+02    5.8576e+02 ;    ND1 - CE1
- NE2
  5735   5737   5738      1    1.2000e+02    4.1840e+02 ;    CE1 - NE2
- HE2
  5735   5737   5739      1    1.2000e+02    5.8576e+02 ;    CE1 - NE2
- CD2
```

5736 5735 5737 1 1.2000e+02 4.1840e+02 ; HE1 - CE1
- NE2
5737 5739 5740 1 1.2000e+02 4.1840e+02 ; NE2 - CD2
- HD2
5738 5737 5739 1 1.2000e+02 4.1840e+02 ; HE2 - NE2
- CD2
5741 5743 5744 1 1.2000e+02 4.1840e+02 ; C - N
- H
5741 5743 5745 1 1.2190e+02 4.1840e+02 ; C - N
- CA
5742 5741 5743 1 1.2290e+02 6.6944e+02 ; O - C
- N
5743 5745 5746 1 1.0950e+02 4.1840e+02 ; N - CA
- HA
5743 5745 5747 1 1.0970e+02 6.6944e+02 ; N - CA
- CB
5743 5745 5758 1 1.1010e+02 5.2718e+02 ; N - CA
- C
5744 5743 5745 1 1.1804e+02 4.1840e+02 ; H - N
- CA
5745 5747 5748 1 1.0950e+02 4.1840e+02 ; CA - CB
- HB2
5745 5747 5749 1 1.0950e+02 4.1840e+02 ; CA - CB
- HB3
5745 5747 5750 1 1.0950e+02 3.3472e+02 ; CA - CB
- CG
5745 5758 5759 1 1.2040e+02 6.6944e+02 ; CA - C
- O
5745 5758 5760 1 1.1660e+02 5.8576e+02 ; CA - C
- N
5746 5745 5747 1 1.0950e+02 4.1840e+02 ; HA - CA
- CB
5746 5745 5758 1 1.0950e+02 4.1840e+02 ; HA - CA
- C
5747 5745 5758 1 1.1110e+02 5.2718e+02 ; CB - CA
- C
5747 5750 5751 1 1.0950e+02 4.1840e+02 ; CB - CG
- HG2
5747 5750 5752 1 1.0950e+02 4.1840e+02 ; CB - CG
- HG3
5747 5750 5753 1 1.1110e+02 5.2718e+02 ; CB - CG
- CD
5748 5747 5749 1 1.0950e+02 2.9288e+02 ; HB2 - CB
- HB3
5748 5747 5750 1 1.0950e+02 4.1840e+02 ; HB2 - CB
- CG
5749 5747 5750 1 1.0950e+02 4.1840e+02 ; HB3 - CB
- CG
5750 5753 5754 1 1.2040e+02 6.6944e+02 ; CG - CD
- OE1
5750 5753 5755 1 1.1660e+02 5.8576e+02 ; CG - CD
- NE2

5751 5750 5752 1 1.0950e+02 2.9288e+02 ; HG2 - CG
- HG3
5751 5750 5753 1 1.0950e+02 4.1840e+02 ; HG2 - CG
- CD
5752 5750 5753 1 1.0950e+02 4.1840e+02 ; HG3 - CG
- CD
5753 5755 5756 1 1.2000e+02 4.1840e+02 ; CD - NE2
- HE21
5753 5755 5757 1 1.2000e+02 4.1840e+02 ; CD - NE2
- HE22
5754 5753 5755 1 1.2290e+02 6.6944e+02 ; OE1 - CD
- NE2
5756 5755 5757 1 1.2000e+02 2.9288e+02 ; HE21 - NE2
- HE22
5758 5760 5761 1 1.2000e+02 4.1840e+02 ; C - N
- H
5758 5760 5762 1 1.2190e+02 4.1840e+02 ; C - N
- CA
5759 5758 5760 1 1.2290e+02 6.6944e+02 ; O - C
- N
5760 5762 5763 1 1.0950e+02 4.1840e+02 ; N - CA
- HA
5760 5762 5764 1 1.0970e+02 6.6944e+02 ; N - CA
- CB
5760 5762 5774 1 1.1010e+02 5.2718e+02 ; N - CA
- C
5761 5760 5762 1 1.1804e+02 4.1840e+02 ; H - N
- CA
5762 5764 5765 1 1.0950e+02 4.1840e+02 ; CA - CB
- HB
5762 5764 5766 1 1.0950e+02 3.3472e+02 ; CA - CB
- CG1
5762 5764 5770 1 1.0950e+02 3.3472e+02 ; CA - CB
- CG2
5762 5774 5775 1 1.2040e+02 6.6944e+02 ; CA - C
- O
5762 5774 5776 1 1.1660e+02 5.8576e+02 ; CA - C
- N
5763 5762 5764 1 1.0950e+02 4.1840e+02 ; HA - CA
- CB
5763 5762 5774 1 1.0950e+02 4.1840e+02 ; HA - CA
- C
5764 5762 5774 1 1.1110e+02 5.2718e+02 ; CB - CA
- C
5764 5766 5767 1 1.0950e+02 4.1840e+02 ; CB - CG1
- HG11
5764 5766 5768 1 1.0950e+02 4.1840e+02 ; CB - CG1
- HG12
5764 5766 5769 1 1.0950e+02 4.1840e+02 ; CB - CG1
- HG13
5764 5770 5771 1 1.0950e+02 4.1840e+02 ; CB - CG2
- HG21

5764 5770 5772 1 1.0950e+02 4.1840e+02 ; CB - CG2
- HG22
5764 5770 5773 1 1.0950e+02 4.1840e+02 ; CB - CG2
- HG23
5765 5764 5766 1 1.0950e+02 4.1840e+02 ; HB - CB
- CG1
5765 5764 5770 1 1.0950e+02 4.1840e+02 ; HB - CB
- CG2
5766 5764 5770 1 1.0950e+02 3.3472e+02 ; CG1 - CB
- CG2
5767 5766 5768 1 1.0950e+02 2.9288e+02 ; HG11 - CG1
- HG12
5767 5766 5769 1 1.0950e+02 2.9288e+02 ; HG11 - CG1
- HG13
5768 5766 5769 1 1.0950e+02 2.9288e+02 ; HG12 - CG1
- HG13
5771 5770 5772 1 1.0950e+02 2.9288e+02 ; HG21 - CG2
- HG22
5771 5770 5773 1 1.0950e+02 2.9288e+02 ; HG21 - CG2
- HG23
5772 5770 5773 1 1.0950e+02 2.9288e+02 ; HG22 - CG2
- HG23
5774 5776 5777 1 1.2000e+02 4.1840e+02 ; C - N
- H
5774 5776 5778 1 1.2190e+02 4.1840e+02 ; C - N
- CA
5775 5774 5776 1 1.2290e+02 6.6944e+02 ; O - C
- N
5776 5778 5779 1 1.0950e+02 4.1840e+02 ; N - CA
- HA
5776 5778 5780 1 1.0970e+02 6.6944e+02 ; N - CA
- CB
5776 5778 5793 1 1.1010e+02 5.2718e+02 ; N - CA
- C
5777 5776 5778 1 1.1804e+02 4.1840e+02 ; H - N
- CA
5778 5780 5781 1 1.0950e+02 4.1840e+02 ; CA - CB
- HB2
5778 5780 5782 1 1.0950e+02 4.1840e+02 ; CA - CB
- HB3
5778 5780 5783 1 1.0950e+02 3.3472e+02 ; CA - CB
- CG
5778 5793 5794 1 1.2040e+02 6.6944e+02 ; CA - C
- O
5778 5793 5795 1 1.1660e+02 5.8576e+02 ; CA - C
- N
5779 5778 5780 1 1.0950e+02 4.1840e+02 ; HA - CA
- CB
5779 5778 5793 1 1.0950e+02 4.1840e+02 ; HA - CA
- C
5780 5778 5793 1 1.1110e+02 5.2718e+02 ; CB - CA
- C

```
  5780   5783   5784      1    1.0950e+02    4.1840e+02 ;     CB - CG
- HG
  5780   5783   5785      1    1.0950e+02    3.3472e+02 ;     CB - CG
- CD1
  5780   5783   5789      1    1.0950e+02    3.3472e+02 ;     CB - CG
- CD2
  5781   5780   5782      1    1.0950e+02    2.9288e+02 ;    HB2 - CB
- HB3
  5781   5780   5783      1    1.0950e+02    4.1840e+02 ;    HB2 - CB
- CG
  5782   5780   5783      1    1.0950e+02    4.1840e+02 ;    HB3 - CB
- CG
  5783   5785   5786      1    1.0950e+02    4.1840e+02 ;     CG - CD1
- HD11
  5783   5785   5787      1    1.0950e+02    4.1840e+02 ;     CG - CD1
- HD12
  5783   5785   5788      1    1.0950e+02    4.1840e+02 ;     CG - CD1
- HD13
  5783   5789   5790      1    1.0950e+02    4.1840e+02 ;     CG - CD2
- HD21
  5783   5789   5791      1    1.0950e+02    4.1840e+02 ;     CG - CD2
- HD22
  5783   5789   5792      1    1.0950e+02    4.1840e+02 ;     CG - CD2
- HD23
  5784   5783   5785      1    1.0950e+02    4.1840e+02 ;     HG - CG
- CD1
  5784   5783   5789      1    1.0950e+02    4.1840e+02 ;     HG - CG
- CD2
  5785   5783   5789      1    1.0950e+02    3.3472e+02 ;    CD1 - CG
- CD2
  5786   5785   5787      1    1.0950e+02    2.9288e+02 ;   HD11 - CD1
- HD12
  5786   5785   5788      1    1.0950e+02    2.9288e+02 ;   HD11 - CD1
- HD13
  5787   5785   5788      1    1.0950e+02    2.9288e+02 ;   HD12 - CD1
- HD13
  5790   5789   5791      1    1.0950e+02    2.9288e+02 ;   HD21 - CD2
- HD22
  5790   5789   5792      1    1.0950e+02    2.9288e+02 ;   HD21 - CD2
- HD23
  5791   5789   5792      1    1.0950e+02    2.9288e+02 ;   HD22 - CD2
- HD23
  5793   5795   5796      1    1.2000e+02    4.1840e+02 ;      C - N
- H
  5793   5795   5797      1    1.2190e+02    4.1840e+02 ;      C - N
- CA
  5794   5793   5795      1    1.2290e+02    6.6944e+02 ;      O - C
- N
  5795   5797   5798      1    1.0950e+02    4.1840e+02 ;      N - CA
- HA
  5795   5797   5799      1    1.0970e+02    6.6944e+02 ;      N - CA
- CB
```

5795 5797 5817 1 1.1010e+02 5.2718e+02 ; N - CA
- C
5796 5795 5797 1 1.1804e+02 4.1840e+02 ; H - N
- CA
5797 5799 5800 1 1.0950e+02 4.1840e+02 ; CA - CB
- HB2
5797 5799 5801 1 1.0950e+02 4.1840e+02 ; CA - CB
- HB3
5797 5799 5802 1 1.0950e+02 3.3472e+02 ; CA - CB
- CG
5797 5817 5818 1 1.2040e+02 6.6944e+02 ; CA - C
- O
5797 5817 5819 1 1.1660e+02 5.8576e+02 ; CA - C
- N
5798 5797 5799 1 1.0950e+02 4.1840e+02 ; HA - CA
- CB
5798 5797 5817 1 1.0950e+02 4.1840e+02 ; HA - CA
- C
5799 5797 5817 1 1.1110e+02 5.2718e+02 ; CB - CA
- C
5799 5802 5803 1 1.0950e+02 4.1840e+02 ; CB - CG
- HG2
5799 5802 5804 1 1.0950e+02 4.1840e+02 ; CB - CG
- HG3
5799 5802 5805 1 1.0950e+02 3.3472e+02 ; CB - CG
- CD
5800 5799 5801 1 1.0950e+02 2.9288e+02 ; HB2 - CB
- HB3
5800 5799 5802 1 1.0950e+02 4.1840e+02 ; HB2 - CB
- CG
5801 5799 5802 1 1.0950e+02 4.1840e+02 ; HB3 - CB
- CG
5802 5805 5806 1 1.0950e+02 4.1840e+02 ; CG - CD
- HD2
5802 5805 5807 1 1.0950e+02 4.1840e+02 ; CG - CD
- HD3
5802 5805 5808 1 1.1120e+02 6.6944e+02 ; CG - CD
- NE
5803 5802 5804 1 1.0950e+02 2.9288e+02 ; HG2 - CG
- HG3
5803 5802 5805 1 1.0950e+02 4.1840e+02 ; HG2 - CG
- CD
5804 5802 5805 1 1.0950e+02 4.1840e+02 ; HG3 - CG
- CD
5805 5808 5809 1 1.1840e+02 4.1840e+02 ; CD - NE
- HE
5805 5808 5810 1 1.2320e+02 4.1840e+02 ; CD - NE
- CZ
5806 5805 5807 1 1.0950e+02 2.9288e+02 ; HD2 - CD
- HD3
5806 5805 5808 1 1.0950e+02 4.1840e+02 ; HD2 - CD
- NE

5807 5805 5808 1 1.0950e+02 4.1840e+02 ; HD3 - CD
- NE
5808 5810 5811 1 1.2000e+02 5.8576e+02 ; NE - CZ
- NH1
5808 5810 5814 1 1.2000e+02 5.8576e+02 ; NE - CZ
- NH2
5809 5808 5810 1 1.2000e+02 4.1840e+02 ; HE - NE
- CZ
5810 5811 5812 1 1.2000e+02 4.1840e+02 ; CZ - NH1
- HH11
5810 5811 5813 1 1.2000e+02 4.1840e+02 ; CZ - NH1
- HH12
5810 5814 5815 1 1.2000e+02 4.1840e+02 ; CZ - NH2
- HH21
5810 5814 5816 1 1.2000e+02 4.1840e+02 ; CZ - NH2
- HH22
5811 5810 5814 1 1.2000e+02 5.8576e+02 ; NH1 - CZ
- NH2
5812 5811 5813 1 1.2000e+02 2.9288e+02 ; HH11 - NH1
- HH12
5815 5814 5816 1 1.2000e+02 2.9288e+02 ; HH21 - NH2
- HH22
5817 5819 5820 1 1.2000e+02 4.1840e+02 ; C - N
- H
5817 5819 5821 1 1.2190e+02 4.1840e+02 ; C - N
- CA
5818 5817 5819 1 1.2290e+02 6.6944e+02 ; O - C
- N
5819 5821 5822 1 1.0950e+02 4.1840e+02 ; N - CA
- HA
5819 5821 5823 1 1.0970e+02 6.6944e+02 ; N - CA
- CB
5819 5821 5829 1 1.1010e+02 5.2718e+02 ; N - CA
- C
5820 5819 5821 1 1.1804e+02 4.1840e+02 ; H - N
- CA
5821 5823 5824 1 1.0950e+02 4.1840e+02 ; CA - CB
- HB2
5821 5823 5825 1 1.0950e+02 4.1840e+02 ; CA - CB
- HB3
5821 5823 5826 1 1.1110e+02 5.2718e+02 ; CA - CB
- CG
5821 5829 5830 1 1.2040e+02 6.6944e+02 ; CA - C
- O
5821 5829 5831 1 1.1660e+02 5.8576e+02 ; CA - C
- N
5822 5821 5823 1 1.0950e+02 4.1840e+02 ; HA - CA
- CB
5822 5821 5829 1 1.0950e+02 4.1840e+02 ; HA - CA
- C
5823 5821 5829 1 1.1110e+02 5.2718e+02 ; CB - CA
- C

```
  5823   5826   5827      1    1.1700e+02    5.8576e+02 ;     CB - CG
- OD1
  5823   5826   5828      1    1.1700e+02    5.8576e+02 ;     CB - CG
- OD2
  5824   5823   5825      1    1.0950e+02    2.9288e+02 ;    HB2 - CB
- HB3
  5824   5823   5826      1    1.0950e+02    4.1840e+02 ;    HB2 - CB
- CG
  5825   5823   5826      1    1.0950e+02    4.1840e+02 ;    HB3 - CB
- CG
  5827   5826   5828      1    1.2600e+02    6.6944e+02 ;    OD1 - CG
- OD2
  5829   5831   5832      1    1.2000e+02    4.1840e+02 ;      C - N
- H
  5829   5831   5833      1    1.2190e+02    4.1840e+02 ;      C - N
- CA
  5830   5829   5831      1    1.2290e+02    6.6944e+02 ;      O - C
- N
  5831   5833   5834      1    1.0950e+02    4.1840e+02 ;      N - CA
- HA
  5831   5833   5835      1    1.0970e+02    6.6944e+02 ;      N - CA
- CB
  5831   5833   5848      1    1.1010e+02    5.2718e+02 ;      N - CA
- C
  5832   5831   5833      1    1.1804e+02    4.1840e+02 ;      H - N
- CA
  5833   5835   5836      1    1.0950e+02    4.1840e+02 ;     CA - CB
- HB
  5833   5835   5837      1    1.0950e+02    3.3472e+02 ;     CA - CB
- CG2
  5833   5835   5841      1    1.0950e+02    3.3472e+02 ;     CA - CB
- CG1
  5833   5848   5849      1    1.2040e+02    6.6944e+02 ;     CA - C
- O
  5833   5848   5850      1    1.1660e+02    5.8576e+02 ;     CA - C
- N
  5834   5833   5835      1    1.0950e+02    4.1840e+02 ;     HA - CA
- CB
  5834   5833   5848      1    1.0950e+02    4.1840e+02 ;     HA - CA
- C
  5835   5833   5848      1    1.1110e+02    5.2718e+02 ;     CB - CA
- C
  5835   5837   5838      1    1.0950e+02    4.1840e+02 ;     CB - CG2
- HG21
  5835   5837   5839      1    1.0950e+02    4.1840e+02 ;     CB - CG2
- HG22
  5835   5837   5840      1    1.0950e+02    4.1840e+02 ;     CB - CG2
- HG23
  5835   5841   5842      1    1.0950e+02    4.1840e+02 ;     CB - CG1
- HG12
  5835   5841   5843      1    1.0950e+02    4.1840e+02 ;     CB - CG1
- HG13
```

5835 5841 5844 1 1.0950e+02 3.3472e+02 ; CB - CG1
- CD1
5836 5835 5837 1 1.0950e+02 4.1840e+02 ; HB - CB
- CG2
5836 5835 5841 1 1.0950e+02 4.1840e+02 ; HB - CB
- CG1
5837 5835 5841 1 1.0950e+02 3.3472e+02 ; CG2 - CB
- CG1
5838 5837 5839 1 1.0950e+02 2.9288e+02 ; HG21 - CG2
- HG22
5838 5837 5840 1 1.0950e+02 2.9288e+02 ; HG21 - CG2
- HG23
5839 5837 5840 1 1.0950e+02 2.9288e+02 ; HG22 - CG2
- HG23
5841 5844 5845 1 1.0950e+02 4.1840e+02 ; CG1 - CD1
- HD11
5841 5844 5846 1 1.0950e+02 4.1840e+02 ; CG1 - CD1
- HD12
5841 5844 5847 1 1.0950e+02 4.1840e+02 ; CG1 - CD1
- HD13
5842 5841 5843 1 1.0950e+02 2.9288e+02 ; HG12 - CG1
- HG13
5842 5841 5844 1 1.0950e+02 4.1840e+02 ; HG12 - CG1
- CD1
5843 5841 5844 1 1.0950e+02 4.1840e+02 ; HG13 - CG1
- CD1
5845 5844 5846 1 1.0950e+02 2.9288e+02 ; HD11 - CD1
- HD12
5845 5844 5847 1 1.0950e+02 2.9288e+02 ; HD11 - CD1
- HD13
5846 5844 5847 1 1.0950e+02 2.9288e+02 ; HD12 - CD1
- HD13
5848 5850 5851 1 1.2000e+02 4.1840e+02 ; C - N
- H
5848 5850 5852 1 1.2190e+02 4.1840e+02 ; C - N
- CA
5849 5848 5850 1 1.2290e+02 6.6944e+02 ; O - C
- N
5850 5852 5853 1 1.0950e+02 4.1840e+02 ; N - CA
- HA
5850 5852 5854 1 1.0970e+02 6.6944e+02 ; N - CA
- CB
5850 5852 5860 1 1.1010e+02 5.2718e+02 ; N - CA
- C
5851 5850 5852 1 1.1804e+02 4.1840e+02 ; H - N
- CA
5852 5854 5855 1 1.0950e+02 4.1840e+02 ; CA - CB
- HB2
5852 5854 5856 1 1.0950e+02 4.1840e+02 ; CA - CB
- HB3
5852 5854 5857 1 1.1110e+02 5.2718e+02 ; CA - CB
- CG

5852 5860 5861 1 1.2040e+02 6.6944e+02 ; CA - C
- O
5852 5860 5862 1 1.1660e+02 5.8576e+02 ; CA - C
- N
5853 5852 5854 1 1.0950e+02 4.1840e+02 ; HA - CA
- CB
5853 5852 5860 1 1.0950e+02 4.1840e+02 ; HA - CA
- C
5854 5852 5860 1 1.1110e+02 5.2718e+02 ; CB - CA
- C
5854 5857 5858 1 1.1700e+02 5.8576e+02 ; CB - CG
- OD1
5854 5857 5859 1 1.1700e+02 5.8576e+02 ; CB - CG
- OD2
5855 5854 5856 1 1.0950e+02 2.9288e+02 ; HB2 - CB
- HB3
5855 5854 5857 1 1.0950e+02 4.1840e+02 ; HB2 - CB
- CG
5856 5854 5857 1 1.0950e+02 4.1840e+02 ; HB3 - CB
- CG
5858 5857 5859 1 1.2600e+02 6.6944e+02 ; OD1 - CG
- OD2
5860 5862 5863 1 1.2000e+02 4.1840e+02 ; C - N
- H
5860 5862 5864 1 1.2190e+02 4.1840e+02 ; C - N
- CA
5861 5860 5862 1 1.2290e+02 6.6944e+02 ; O - C
- N
5862 5864 5865 1 1.0950e+02 4.1840e+02 ; N - CA
- HA
5862 5864 5866 1 1.0970e+02 6.6944e+02 ; N - CA
- CB
5862 5864 5871 1 1.1010e+02 5.2718e+02 ; N - CA
- C
5863 5862 5864 1 1.1804e+02 4.1840e+02 ; H - N
- CA
5864 5866 5867 1 1.0950e+02 4.1840e+02 ; CA - CB
- HB2
5864 5866 5868 1 1.0950e+02 4.1840e+02 ; CA - CB
- HB3
5864 5866 5869 1 1.0950e+02 4.1840e+02 ; CA - CB
- OG
5864 5871 5872 1 1.2040e+02 6.6944e+02 ; CA - C
- O
5864 5871 5873 1 1.1660e+02 5.8576e+02 ; CA - C
- N
5865 5864 5866 1 1.0950e+02 4.1840e+02 ; HA - CA
- CB
5865 5864 5871 1 1.0950e+02 4.1840e+02 ; HA - CA
- C
5866 5864 5871 1 1.1110e+02 5.2718e+02 ; CB - CA
- C

5866 5869 5870 1 1.0850e+02 4.6024e+02 ; CB - OG
- HG
5867 5866 5868 1 1.0950e+02 2.9288e+02 ; HB2 - CB
- HB3
5867 5866 5869 1 1.0950e+02 4.1840e+02 ; HB2 - CB
- OG
5868 5866 5869 1 1.0950e+02 4.1840e+02 ; HB3 - CB
- OG
5871 5873 5874 1 1.2000e+02 4.1840e+02 ; C - N
- H
5871 5873 5875 1 1.2190e+02 4.1840e+02 ; C - N
- CA
5872 5871 5873 1 1.2290e+02 6.6944e+02 ; O - C
- N
5873 5875 5876 1 1.0950e+02 4.1840e+02 ; N - CA
- HA
5873 5875 5877 1 1.0970e+02 6.6944e+02 ; N - CA
- CB
5873 5875 5885 1 1.1010e+02 5.2718e+02 ; N - CA
- C
5874 5873 5875 1 1.1804e+02 4.1840e+02 ; H - N
- CA
5875 5877 5878 1 1.0950e+02 4.1840e+02 ; CA - CB
- HB2
5875 5877 5879 1 1.0950e+02 4.1840e+02 ; CA - CB
- HB3
5875 5877 5880 1 1.1110e+02 5.2718e+02 ; CA - CB
- CG
5875 5885 5886 1 1.2040e+02 6.6944e+02 ; CA - C
- O
5875 5885 5887 1 1.1660e+02 5.8576e+02 ; CA - C
- N
5876 5875 5877 1 1.0950e+02 4.1840e+02 ; HA - CA
- CB
5876 5875 5885 1 1.0950e+02 4.1840e+02 ; HA - CA
- C
5877 5875 5885 1 1.1110e+02 5.2718e+02 ; CB - CA
- C
5877 5880 5881 1 1.2040e+02 6.6944e+02 ; CB - CG
- OD1
5877 5880 5882 1 1.1660e+02 5.8576e+02 ; CB - CG
- ND2
5878 5877 5879 1 1.0950e+02 2.9288e+02 ; HB2 - CB
- HB3
5878 5877 5880 1 1.0950e+02 4.1840e+02 ; HB2 - CB
- CG
5879 5877 5880 1 1.0950e+02 4.1840e+02 ; HB3 - CB
- CG
5880 5882 5883 1 1.2000e+02 4.1840e+02 ; CG - ND2
- HD21
5880 5882 5884 1 1.2000e+02 4.1840e+02 ; CG - ND2
- HD22

```
  5881   5880   5882       1    1.2290e+02    6.6944e+02 ;    OD1 - CG
- ND2
  5883   5882   5884       1    1.2000e+02    2.9288e+02 ;   HD21 - ND2
- HD22
  5885   5887   5888       1    1.2000e+02    4.1840e+02 ;      C - N
- H
  5885   5887   5889       1    1.2190e+02    4.1840e+02 ;      C - N
- CA
  5886   5885   5887       1    1.2290e+02    6.6944e+02 ;      O - C
- N
  5887   5889   5890       1    1.0950e+02    4.1840e+02 ;      N - CA
- HA
  5887   5889   5891       1    1.0970e+02    6.6944e+02 ;      N - CA
- CB
  5887   5889   5895       1    1.1010e+02    5.2718e+02 ;      N - CA
- C
  5888   5887   5889       1    1.1804e+02    4.1840e+02 ;      H - N
- CA
  5889   5891   5892       1    1.0950e+02    4.1840e+02 ;     CA - CB
- HB1
  5889   5891   5893       1    1.0950e+02    4.1840e+02 ;     CA - CB
- HB2
  5889   5891   5894       1    1.0950e+02    4.1840e+02 ;     CA - CB
- HB3
  5889   5895   5896       1    1.2040e+02    6.6944e+02 ;     CA - C
- O
  5889   5895   5897       1    1.1660e+02    5.8576e+02 ;     CA - C
- N
  5890   5889   5891       1    1.0950e+02    4.1840e+02 ;     HA - CA
- CB
  5890   5889   5895       1    1.0950e+02    4.1840e+02 ;     HA - CA
- C
  5891   5889   5895       1    1.1110e+02    5.2718e+02 ;     CB - CA
- C
  5892   5891   5893       1    1.0950e+02    2.9288e+02 ;    HB1 - CB
- HB2
  5892   5891   5894       1    1.0950e+02    2.9288e+02 ;    HB1 - CB
- HB3
  5893   5891   5894       1    1.0950e+02    2.9288e+02 ;    HB2 - CB
- HB3
  5895   5897   5898       1    1.2000e+02    4.1840e+02 ;      C - N
- H
  5895   5897   5899       1    1.2190e+02    4.1840e+02 ;      C - N
- CA
  5896   5895   5897       1    1.2290e+02    6.6944e+02 ;      O - C
- N
  5897   5899   5900       1    1.0950e+02    4.1840e+02 ;      N - CA
- HA
  5897   5899   5901       1    1.0970e+02    6.6944e+02 ;      N - CA
- CB
  5897   5899   5906       1    1.1010e+02    5.2718e+02 ;      N - CA
- C
```

5898 5897 5899 1 1.1804e+02 4.1840e+02 ; H - N
- CA
5899 5901 5902 1 1.0950e+02 4.1840e+02 ; CA - CB
- HB2
5899 5901 5903 1 1.0950e+02 4.1840e+02 ; CA - CB
- HB3
5899 5901 5904 1 1.0950e+02 4.1840e+02 ; CA - CB
- OG
5899 5906 5907 1 1.2040e+02 6.6944e+02 ; CA - C
- O
5899 5906 5908 1 1.1660e+02 5.8576e+02 ; CA - C
- N
5900 5899 5901 1 1.0950e+02 4.1840e+02 ; HA - CA
- CB
5900 5899 5906 1 1.0950e+02 4.1840e+02 ; HA - CA
- C
5901 5899 5906 1 1.1110e+02 5.2718e+02 ; CB - CA
- C
5901 5904 5905 1 1.0850e+02 4.6024e+02 ; CB - OG
- HG
5902 5901 5903 1 1.0950e+02 2.9288e+02 ; HB2 - CB
- HB3
5902 5901 5904 1 1.0950e+02 4.1840e+02 ; HB2 - CB
- OG
5903 5901 5904 1 1.0950e+02 4.1840e+02 ; HB3 - CB
- OG
5906 5908 5909 1 1.2000e+02 4.1840e+02 ; C - N
- H
5906 5908 5910 1 1.2190e+02 4.1840e+02 ; C - N
- CA
5907 5906 5908 1 1.2290e+02 6.6944e+02 ; O - C
- N
5908 5910 5911 1 1.0950e+02 4.1840e+02 ; N - CA
- HA2
5908 5910 5912 1 1.0950e+02 4.1840e+02 ; N - CA
- HA3
5908 5910 5913 1 1.1010e+02 5.2718e+02 ; N - CA
- C
5909 5908 5910 1 1.1804e+02 4.1840e+02 ; H - N
- CA
5910 5913 5914 1 1.2040e+02 6.6944e+02 ; CA - C
- O
5910 5913 5915 1 1.1660e+02 5.8576e+02 ; CA - C
- N
5911 5910 5912 1 1.0950e+02 2.9288e+02 ; HA2 - CA
- HA3
5911 5910 5913 1 1.0950e+02 4.1840e+02 ; HA2 - CA
- C
5912 5910 5913 1 1.0950e+02 4.1840e+02 ; HA3 - CA
- C
5913 5915 5916 1 1.2000e+02 4.1840e+02 ; C - N
- H

5913 5915 5917 1 1.2190e+02 4.1840e+02 ; C - N
- CA
5914 5913 5915 1 1.2290e+02 6.6944e+02 ; O - C
- N
5915 5917 5918 1 1.0950e+02 4.1840e+02 ; N - CA
- HA
5915 5917 5919 1 1.0970e+02 6.6944e+02 ; N - CA
- CB
5915 5917 5930 1 1.1010e+02 5.2718e+02 ; N - CA
- C
5916 5915 5917 1 1.1804e+02 4.1840e+02 ; H - N
- CA
5917 5919 5920 1 1.0950e+02 4.1840e+02 ; CA - CB
- HB2
5917 5919 5921 1 1.0950e+02 4.1840e+02 ; CA - CB
- HB3
5917 5919 5922 1 1.0950e+02 3.3472e+02 ; CA - CB
- CG
5917 5930 5931 1 1.2040e+02 6.6944e+02 ; CA - C
- O
5917 5930 5932 1 1.1660e+02 5.8576e+02 ; CA - C
- N
5918 5917 5919 1 1.0950e+02 4.1840e+02 ; HA - CA
- CB
5918 5917 5930 1 1.0950e+02 4.1840e+02 ; HA - CA
- C
5919 5917 5930 1 1.1110e+02 5.2718e+02 ; CB - CA
- C
5919 5922 5923 1 1.0950e+02 4.1840e+02 ; CB - CG
- HG2
5919 5922 5924 1 1.0950e+02 4.1840e+02 ; CB - CG
- HG3
5919 5922 5925 1 1.1110e+02 5.2718e+02 ; CB - CG
- CD
5920 5919 5921 1 1.0950e+02 2.9288e+02 ; HB2 - CB
- HB3
5920 5919 5922 1 1.0950e+02 4.1840e+02 ; HB2 - CB
- CG
5921 5919 5922 1 1.0950e+02 4.1840e+02 ; HB3 - CB
- CG
5922 5925 5926 1 1.2040e+02 6.6944e+02 ; CG - CD
- OE1
5922 5925 5927 1 1.1660e+02 5.8576e+02 ; CG - CD
- NE2
5923 5922 5924 1 1.0950e+02 2.9288e+02 ; HG2 - CG
- HG3
5923 5922 5925 1 1.0950e+02 4.1840e+02 ; HG2 - CG
- CD
5924 5922 5925 1 1.0950e+02 4.1840e+02 ; HG3 - CG
- CD
5925 5927 5928 1 1.2000e+02 4.1840e+02 ; CD - NE2
- HE21

```
  5925   5927   5929      1    1.2000e+02    4.1840e+02 ;     CD - NE2
- HE22
  5926   5925   5927      1    1.2290e+02    6.6944e+02 ;    OE1 - CD
- NE2
  5928   5927   5929      1    1.2000e+02    2.9288e+02 ;   HE21 - NE2
- HE22
  5930   5932   5933      1    1.2000e+02    4.1840e+02 ;      C - N
- H
  5930   5932   5934      1    1.2190e+02    4.1840e+02 ;      C - N
- CA
  5931   5930   5932      1    1.2290e+02    6.6944e+02 ;      O - C
- N
  5932   5934   5935      1    1.0950e+02    4.1840e+02 ;      N - CA
- HA
  5932   5934   5936      1    1.0970e+02    6.6944e+02 ;      N - CA
- CB
  5932   5934   5949      1    1.1010e+02    5.2718e+02 ;      N - CA
- C
  5933   5932   5934      1    1.1804e+02    4.1840e+02 ;      H - N
- CA
  5934   5936   5937      1    1.0950e+02    4.1840e+02 ;     CA - CB
- HB
  5934   5936   5938      1    1.0950e+02    3.3472e+02 ;     CA - CB
- CG2
  5934   5936   5942      1    1.0950e+02    3.3472e+02 ;     CA - CB
- CG1
  5934   5949   5950      1    1.2040e+02    6.6944e+02 ;     CA - C
- O
  5934   5949   5951      1    1.1660e+02    5.8576e+02 ;     CA - C
- N
  5935   5934   5936      1    1.0950e+02    4.1840e+02 ;     HA - CA
- CB
  5935   5934   5949      1    1.0950e+02    4.1840e+02 ;     HA - CA
- C
  5936   5934   5949      1    1.1110e+02    5.2718e+02 ;     CB - CA
- C
  5936   5938   5939      1    1.0950e+02    4.1840e+02 ;     CB - CG2
- HG21
  5936   5938   5940      1    1.0950e+02    4.1840e+02 ;     CB - CG2
- HG22
  5936   5938   5941      1    1.0950e+02    4.1840e+02 ;     CB - CG2
- HG23
  5936   5942   5943      1    1.0950e+02    4.1840e+02 ;     CB - CG1
- HG12
  5936   5942   5944      1    1.0950e+02    4.1840e+02 ;     CB - CG1
- HG13
  5936   5942   5945      1    1.0950e+02    3.3472e+02 ;     CB - CG1
- CD1
  5937   5936   5938      1    1.0950e+02    4.1840e+02 ;     HB - CB
- CG2
  5937   5936   5942      1    1.0950e+02    4.1840e+02 ;     HB - CB
- CG1
```

5938 5936 5942 1 1.0950e+02 3.3472e+02 ; CG2 - CB
- CG1
5939 5938 5940 1 1.0950e+02 2.9288e+02 ; HG21 - CG2
- HG22
5939 5938 5941 1 1.0950e+02 2.9288e+02 ; HG21 - CG2
- HG23
5940 5938 5941 1 1.0950e+02 2.9288e+02 ; HG22 - CG2
- HG23
5942 5945 5946 1 1.0950e+02 4.1840e+02 ; CG1 - CD1
- HD11
5942 5945 5947 1 1.0950e+02 4.1840e+02 ; CG1 - CD1
- HD12
5942 5945 5948 1 1.0950e+02 4.1840e+02 ; CG1 - CD1
- HD13
5943 5942 5944 1 1.0950e+02 2.9288e+02 ; HG12 - CG1
- HG13
5943 5942 5945 1 1.0950e+02 4.1840e+02 ; HG12 - CG1
- CD1
5944 5942 5945 1 1.0950e+02 4.1840e+02 ; HG13 - CG1
- CD1
5946 5945 5947 1 1.0950e+02 2.9288e+02 ; HD11 - CD1
- HD12
5946 5945 5948 1 1.0950e+02 2.9288e+02 ; HD11 - CD1
- HD13
5947 5945 5948 1 1.0950e+02 2.9288e+02 ; HD12 - CD1
- HD13
5949 5951 5952 1 1.2000e+02 4.1840e+02 ; C - N
- H
5949 5951 5953 1 1.2190e+02 4.1840e+02 ; C - N
- CA
5950 5949 5951 1 1.2290e+02 6.6944e+02 ; O - C
- N
5951 5953 5954 1 1.0950e+02 4.1840e+02 ; N - CA
- HA
5951 5953 5955 1 1.0970e+02 6.6944e+02 ; N - CA
- CB
5951 5953 5966 1 1.1010e+02 5.2718e+02 ; N - CA
- C
5952 5951 5953 1 1.1804e+02 4.1840e+02 ; H - N
- CA
5953 5955 5956 1 1.0950e+02 4.1840e+02 ; CA - CB
- HB2
5953 5955 5957 1 1.0950e+02 4.1840e+02 ; CA - CB
- HB3
5953 5955 5958 1 1.1310e+02 5.2718e+02 ; CA - CB
- CG
5953 5966 5967 1 1.2040e+02 6.6944e+02 ; CA - C
- O
5953 5966 5968 1 1.1660e+02 5.8576e+02 ; CA - C
- N
5954 5953 5955 1 1.0950e+02 4.1840e+02 ; HA - CA
- CB

```
  5954   5953   5966      1    1.0950e+02    4.1840e+02 ;     HA - CA
- C
  5955   5953   5966      1    1.1110e+02    5.2718e+02 ;     CB - CA
- C
  5955   5958   5959      1    1.2000e+02    5.8576e+02 ;     CB - CG
- ND1
  5955   5958   5964      1    1.2000e+02    5.8576e+02 ;     CB - CG
- CD2
  5956   5955   5957      1    1.0950e+02    2.9288e+02 ;    HB2 - CB
- HB3
  5956   5955   5958      1    1.0950e+02    4.1840e+02 ;    HB2 - CB
- CG
  5957   5955   5958      1    1.0950e+02    4.1840e+02 ;    HB3 - CB
- CG
  5958   5959   5960      1    1.1700e+02    5.8576e+02 ;     CG - ND1
- CE1
  5958   5964   5962      1    1.2000e+02    5.8576e+02 ;     CG - CD2
- NE2
  5958   5964   5965      1    1.2000e+02    4.1840e+02 ;     CG - CD2
- HD2
  5959   5958   5964      1    1.2000e+02    5.8576e+02 ;    ND1 - CG
- CD2
  5959   5960   5961      1    1.2000e+02    4.1840e+02 ;    ND1 - CE1
- HE1
  5959   5960   5962      1    1.2000e+02    5.8576e+02 ;    ND1 - CE1
- NE2
  5960   5962   5963      1    1.2000e+02    4.1840e+02 ;    CE1 - NE2
- HE2
  5960   5962   5964      1    1.2000e+02    5.8576e+02 ;    CE1 - NE2
- CD2
  5961   5960   5962      1    1.2000e+02    4.1840e+02 ;    HE1 - CE1
- NE2
  5962   5964   5965      1    1.2000e+02    4.1840e+02 ;    NE2 - CD2
- HD2
  5963   5962   5964      1    1.2000e+02    4.1840e+02 ;    HE2 - NE2
- CD2
  5966   5968   5969      1    1.2000e+02    4.1840e+02 ;      C - N
- H
  5966   5968   5970      1    1.2190e+02    4.1840e+02 ;      C - N
- CA
  5967   5966   5968      1    1.2290e+02    6.6944e+02 ;      O - C
- N
  5968   5970   5971      1    1.0950e+02    4.1840e+02 ;      N - CA
- HA
  5968   5970   5972      1    1.0970e+02    6.6944e+02 ;      N - CA
- CB
  5968   5970   5987      1    1.1010e+02    5.2718e+02 ;      N - CA
- C
  5969   5968   5970      1    1.1804e+02    4.1840e+02 ;      H - N
- CA
  5970   5972   5973      1    1.0950e+02    4.1840e+02 ;     CA - CB
- HB2
```

5970 5972 5974 1 1.0950e+02 4.1840e+02 ; CA - CB
- HB3
5970 5972 5975 1 1.1400e+02 5.2718e+02 ; CA - CB
- CG
5970 5987 5988 1 1.2040e+02 6.6944e+02 ; CA - C
- O
5970 5987 5989 1 1.1660e+02 5.8576e+02 ; CA - C
- N
5971 5970 5972 1 1.0950e+02 4.1840e+02 ; HA - CA
- CB
5971 5970 5987 1 1.0950e+02 4.1840e+02 ; HA - CA
- C
5972 5970 5987 1 1.1110e+02 5.2718e+02 ; CB - CA
- C
5972 5975 5976 1 1.2000e+02 5.8576e+02 ; CB - CG
- CD1
5972 5975 5985 1 1.2000e+02 5.8576e+02 ; CB - CG
- CD2
5973 5972 5974 1 1.0950e+02 2.9288e+02 ; HB2 - CB
- HB3
5973 5972 5975 1 1.0950e+02 4.1840e+02 ; HB2 - CB
- CG
5974 5972 5975 1 1.0950e+02 4.1840e+02 ; HB3 - CB
- CG
5975 5976 5977 1 1.2000e+02 4.1840e+02 ; CG - CD1
- HD1
5975 5976 5978 1 1.2000e+02 5.2718e+02 ; CG - CD1
- CE1
5975 5985 5983 1 1.2000e+02 5.2718e+02 ; CG - CD2
- CE2
5975 5985 5986 1 1.2000e+02 4.1840e+02 ; CG - CD2
- HD2
5976 5975 5985 1 1.2000e+02 5.2718e+02 ; CD1 - CG
- CD2
5976 5978 5979 1 1.2000e+02 4.1840e+02 ; CD1 - CE1
- HE1
5976 5978 5980 1 1.2000e+02 5.2718e+02 ; CD1 - CE1
- CZ
5977 5976 5978 1 1.2000e+02 4.1840e+02 ; HD1 - CD1
- CE1
5978 5980 5981 1 1.2000e+02 5.8576e+02 ; CE1 - CZ
- OH
5978 5980 5983 1 1.2000e+02 5.2718e+02 ; CE1 - CZ
- CE2
5979 5978 5980 1 1.2000e+02 4.1840e+02 ; HE1 - CE1
- CZ
5980 5981 5982 1 1.1300e+02 4.1840e+02 ; CZ - OH
- HH
5980 5983 5984 1 1.2000e+02 4.1840e+02 ; CZ - CE2
- HE2
5980 5983 5985 1 1.2000e+02 5.2718e+02 ; CZ - CE2
- CD2

5981 5980 5983 1 1.2000e+02 5.8576e+02 ; OH - CZ
- CE2
5983 5985 5986 1 1.2000e+02 4.1840e+02 ; CE2 - CD2
- HD2
5984 5983 5985 1 1.2000e+02 4.1840e+02 ; HE2 - CE2
- CD2
5987 5989 5990 1 1.2000e+02 4.1840e+02 ; C - N
- H
5987 5989 5991 1 1.2190e+02 4.1840e+02 ; C - N
- CA
5988 5987 5989 1 1.2290e+02 6.6944e+02 ; O - C
- N
5989 5991 5992 1 1.0950e+02 4.1840e+02 ; N - CA
- HA
5989 5991 5993 1 1.0970e+02 6.6944e+02 ; N - CA
- CB
5989 5991 6001 1 1.1010e+02 5.2718e+02 ; N - CA
- C
5990 5989 5991 1 1.1804e+02 4.1840e+02 ; H - N
- CA
5991 5993 5994 1 1.0950e+02 4.1840e+02 ; CA - CB
- HB
5991 5993 5995 1 1.0950e+02 3.3472e+02 ; CA - CB
- CG2
5991 5993 5999 1 1.0950e+02 4.1840e+02 ; CA - CB
- OG1
5991 6001 6002 1 1.2040e+02 6.6944e+02 ; CA - C
- O
5991 6001 6003 1 1.1660e+02 5.8576e+02 ; CA - C
- N
5992 5991 5993 1 1.0950e+02 4.1840e+02 ; HA - CA
- CB
5992 5991 6001 1 1.0950e+02 4.1840e+02 ; HA - CA
- C
5993 5991 6001 1 1.1110e+02 5.2718e+02 ; CB - CA
- C
5993 5995 5996 1 1.0950e+02 4.1840e+02 ; CB - CG2
- HG21
5993 5995 5997 1 1.0950e+02 4.1840e+02 ; CB - CG2
- HG22
5993 5995 5998 1 1.0950e+02 4.1840e+02 ; CB - CG2
- HG23
5993 5999 6000 1 1.0850e+02 4.6024e+02 ; CB - OG1
- HG1
5994 5993 5995 1 1.0950e+02 4.1840e+02 ; HB - CB
- CG2
5994 5993 5999 1 1.0950e+02 4.1840e+02 ; HB - CB
- OG1
5995 5993 5999 1 1.0950e+02 4.1840e+02 ; CG2 - CB
- OG1
5996 5995 5997 1 1.0950e+02 2.9288e+02 ; HG21 - CG2
- HG22

5996 5995 5998 1 1.0950e+02 2.9288e+02 ; HG21 - CG2
- HG23
5997 5995 5998 1 1.0950e+02 2.9288e+02 ; HG22 - CG2
- HG23
6001 6003 6004 1 1.2000e+02 4.1840e+02 ; C - N
- H
6001 6003 6005 1 1.2190e+02 4.1840e+02 ; C - N
- CA
6002 6001 6003 1 1.2290e+02 6.6944e+02 ; O - C
- N
6003 6005 6006 1 1.0950e+02 4.1840e+02 ; N - CA
- HA
6003 6005 6007 1 1.0970e+02 6.6944e+02 ; N - CA
- CB
6003 6005 6013 1 1.1010e+02 5.2718e+02 ; N - CA
- C
6004 6003 6005 1 1.1804e+02 4.1840e+02 ; H - N
- CA
6005 6007 6008 1 1.0950e+02 4.1840e+02 ; CA - CB
- HB2
6005 6007 6009 1 1.0950e+02 4.1840e+02 ; CA - CB
- HB3
6005 6007 6010 1 1.1110e+02 5.2718e+02 ; CA - CB
- CG
6005 6013 6014 1 1.2040e+02 6.6944e+02 ; CA - C
- O
6005 6013 6015 1 1.1660e+02 5.8576e+02 ; CA - C
- N
6006 6005 6007 1 1.0950e+02 4.1840e+02 ; HA - CA
- CB
6006 6005 6013 1 1.0950e+02 4.1840e+02 ; HA - CA
- C
6007 6005 6013 1 1.1110e+02 5.2718e+02 ; CB - CA
- C
6007 6010 6011 1 1.1700e+02 5.8576e+02 ; CB - CG
- OD1
6007 6010 6012 1 1.1700e+02 5.8576e+02 ; CB - CG
- OD2
6008 6007 6009 1 1.0950e+02 2.9288e+02 ; HB2 - CB
- HB3
6008 6007 6010 1 1.0950e+02 4.1840e+02 ; HB2 - CB
- CG
6009 6007 6010 1 1.0950e+02 4.1840e+02 ; HB3 - CB
- CG
6011 6010 6012 1 1.2600e+02 6.6944e+02 ; OD1 - CG
- OD2
6013 6015 6016 1 1.2000e+02 4.1840e+02 ; C - N
- H
6013 6015 6017 1 1.2190e+02 4.1840e+02 ; C - N
- CA
6014 6013 6015 1 1.2290e+02 6.6944e+02 ; O - C
- N

```
  6015   6017   6018       1    1.0950e+02    4.1840e+02 ;      N - CA
- HA
  6015   6017   6019       1    1.0970e+02    6.6944e+02 ;      N - CA
- CB
  6015   6017   6033       1    1.1010e+02    5.2718e+02 ;      N - CA
- C
  6016   6015   6017       1    1.1804e+02    4.1840e+02 ;      H - N
- CA
  6017   6019   6020       1    1.0950e+02    4.1840e+02 ;     CA - CB
- HB2
  6017   6019   6021       1    1.0950e+02    4.1840e+02 ;     CA - CB
- HB3
  6017   6019   6022       1    1.1400e+02    5.2718e+02 ;     CA - CB
- CG
  6017   6033   6034       1    1.2040e+02    6.6944e+02 ;     CA - C
- O
  6017   6033   6035       1    1.1660e+02    5.8576e+02 ;     CA - C
- N
  6018   6017   6019       1    1.0950e+02    4.1840e+02 ;     HA - CA
- CB
  6018   6017   6033       1    1.0950e+02    4.1840e+02 ;     HA - CA
- C
  6019   6017   6033       1    1.1110e+02    5.2718e+02 ;     CB - CA
- C
  6019   6022   6023       1    1.2000e+02    5.8576e+02 ;     CB - CG
- CD1
  6019   6022   6031       1    1.2000e+02    5.8576e+02 ;     CB - CG
- CD2
  6020   6019   6021       1    1.0950e+02    2.9288e+02 ;    HB2 - CB
- HB3
  6020   6019   6022       1    1.0950e+02    4.1840e+02 ;    HB2 - CB
- CG
  6021   6019   6022       1    1.0950e+02    4.1840e+02 ;    HB3 - CB
- CG
  6022   6023   6024       1    1.2000e+02    4.1840e+02 ;     CG - CD1
- HD1
  6022   6023   6025       1    1.2000e+02    5.2718e+02 ;     CG - CD1
- CE1
  6022   6031   6029       1    1.2000e+02    5.2718e+02 ;     CG - CD2
- CE2
  6022   6031   6032       1    1.2000e+02    4.1840e+02 ;     CG - CD2
- HD2
  6023   6022   6031       1    1.2000e+02    5.2718e+02 ;    CD1 - CG
- CD2
  6023   6025   6026       1    1.2000e+02    4.1840e+02 ;    CD1 - CE1
- HE1
  6023   6025   6027       1    1.2000e+02    5.2718e+02 ;    CD1 - CE1
- CZ
  6024   6023   6025       1    1.2000e+02    4.1840e+02 ;    HD1 - CD1
- CE1
  6025   6027   6028       1    1.2000e+02    4.1840e+02 ;    CE1 - CZ
- HZ
```

```
  6025   6027   6029      1    1.2000e+02    5.2718e+02 ;    CE1 - CZ
- CE2
  6026   6025   6027      1    1.2000e+02    4.1840e+02 ;    HE1 - CE1
- CZ
  6027   6029   6030      1    1.2000e+02    4.1840e+02 ;     CZ - CE2
- HE2
  6027   6029   6031      1    1.2000e+02    5.2718e+02 ;     CZ - CE2
- CD2
  6028   6027   6029      1    1.2000e+02    4.1840e+02 ;     HZ - CZ
- CE2
  6029   6031   6032      1    1.2000e+02    4.1840e+02 ;    CE2 - CD2
- HD2
  6030   6029   6031      1    1.2000e+02    4.1840e+02 ;    HE2 - CE2
- CD2
  6033   6035   6036      1    1.2000e+02    4.1840e+02 ;      C - N
- H
  6033   6035   6037      1    1.2190e+02    4.1840e+02 ;      C - N
- CA
  6034   6033   6035      1    1.2290e+02    6.6944e+02 ;      O - C
- N
  6035   6037   6038      1    1.0950e+02    4.1840e+02 ;      N - CA
- HA
  6035   6037   6039      1    1.0970e+02    6.6944e+02 ;      N - CA
- CB
  6035   6037   6052      1    1.1010e+02    5.2718e+02 ;      N - CA
- C
  6036   6035   6037      1    1.1804e+02    4.1840e+02 ;      H - N
- CA
  6037   6039   6040      1    1.0950e+02    4.1840e+02 ;     CA - CB
- HB2
  6037   6039   6041      1    1.0950e+02    4.1840e+02 ;     CA - CB
- HB3
  6037   6039   6042      1    1.0950e+02    3.3472e+02 ;     CA - CB
- CG
  6037   6052   6053      1    1.2040e+02    6.6944e+02 ;     CA - C
- O
  6037   6052   6054      1    1.1660e+02    5.8576e+02 ;     CA - C
- N
  6038   6037   6039      1    1.0950e+02    4.1840e+02 ;     HA - CA
- CB
  6038   6037   6052      1    1.0950e+02    4.1840e+02 ;     HA - CA
- C
  6039   6037   6052      1    1.1110e+02    5.2718e+02 ;     CB - CA
- C
  6039   6042   6043      1    1.0950e+02    4.1840e+02 ;     CB - CG
- HG
  6039   6042   6044      1    1.0950e+02    3.3472e+02 ;     CB - CG
- CD1
  6039   6042   6048      1    1.0950e+02    3.3472e+02 ;     CB - CG
- CD2
  6040   6039   6041      1    1.0950e+02    2.9288e+02 ;    HB2 - CB
- HB3
```

```
  6040   6039   6042      1    1.0950e+02    4.1840e+02 ;    HB2 - CB
- CG
  6041   6039   6042      1    1.0950e+02    4.1840e+02 ;    HB3 - CB
- CG
  6042   6044   6045      1    1.0950e+02    4.1840e+02 ;     CG - CD1
- HD11
  6042   6044   6046      1    1.0950e+02    4.1840e+02 ;     CG - CD1
- HD12
  6042   6044   6047      1    1.0950e+02    4.1840e+02 ;     CG - CD1
- HD13
  6042   6048   6049      1    1.0950e+02    4.1840e+02 ;     CG - CD2
- HD21
  6042   6048   6050      1    1.0950e+02    4.1840e+02 ;     CG - CD2
- HD22
  6042   6048   6051      1    1.0950e+02    4.1840e+02 ;     CG - CD2
- HD23
  6043   6042   6044      1    1.0950e+02    4.1840e+02 ;     HG - CG
- CD1
  6043   6042   6048      1    1.0950e+02    4.1840e+02 ;     HG - CG
- CD2
  6044   6042   6048      1    1.0950e+02    3.3472e+02 ;    CD1 - CG
- CD2
  6045   6044   6046      1    1.0950e+02    2.9288e+02 ;   HD11 - CD1
- HD12
  6045   6044   6047      1    1.0950e+02    2.9288e+02 ;   HD11 - CD1
- HD13
  6046   6044   6047      1    1.0950e+02    2.9288e+02 ;   HD12 - CD1
- HD13
  6049   6048   6050      1    1.0950e+02    2.9288e+02 ;   HD21 - CD2
- HD22
  6049   6048   6051      1    1.0950e+02    2.9288e+02 ;   HD21 - CD2
- HD23
  6050   6048   6051      1    1.0950e+02    2.9288e+02 ;   HD22 - CD2
- HD23
  6052   6054   6055      1    1.2000e+02    4.1840e+02 ;      C - N
- H
  6052   6054   6056      1    1.2190e+02    4.1840e+02 ;      C - N
- CA
  6053   6052   6054      1    1.2290e+02    6.6944e+02 ;      O - C
- N
  6054   6056   6057      1    1.0950e+02    4.1840e+02 ;      N - CA
- HA
  6054   6056   6058      1    1.0970e+02    6.6944e+02 ;      N - CA
- CB
  6054   6056   6062      1    1.1010e+02    5.2718e+02 ;      N - CA
- C
  6055   6054   6056      1    1.1804e+02    4.1840e+02 ;      H - N
- CA
  6056   6058   6059      1    1.0950e+02    4.1840e+02 ;     CA - CB
- HB1
  6056   6058   6060      1    1.0950e+02    4.1840e+02 ;     CA - CB
- HB2
```

6056 6058 6061 1 1.0950e+02 4.1840e+02 ; CA - CB
- HB3
6056 6062 6063 1 1.2040e+02 6.6944e+02 ; CA - C
- O
6056 6062 6064 1 1.1660e+02 5.8576e+02 ; CA - C
- N
6057 6056 6058 1 1.0950e+02 4.1840e+02 ; HA - CA
- CB
6057 6056 6062 1 1.0950e+02 4.1840e+02 ; HA - CA
- C
6058 6056 6062 1 1.1110e+02 5.2718e+02 ; CB - CA
- C
6059 6058 6060 1 1.0950e+02 2.9288e+02 ; HB1 - CB
- HB2
6059 6058 6061 1 1.0950e+02 2.9288e+02 ; HB1 - CB
- HB3
6060 6058 6061 1 1.0950e+02 2.9288e+02 ; HB2 - CB
- HB3
6062 6064 6065 1 1.2000e+02 4.1840e+02 ; C - N
- H
6062 6064 6066 1 1.2190e+02 4.1840e+02 ; C - N
- CA
6063 6062 6064 1 1.2290e+02 6.6944e+02 ; O - C
- N
6064 6066 6067 1 1.0950e+02 4.1840e+02 ; N - CA
- HA
6064 6066 6068 1 1.0970e+02 6.6944e+02 ; N - CA
- CB
6064 6066 6072 1 1.1010e+02 5.2718e+02 ; N - CA
- C
6065 6064 6066 1 1.1804e+02 4.1840e+02 ; H - N
- CA
6066 6068 6069 1 1.0950e+02 4.1840e+02 ; CA - CB
- HB1
6066 6068 6070 1 1.0950e+02 4.1840e+02 ; CA - CB
- HB2
6066 6068 6071 1 1.0950e+02 4.1840e+02 ; CA - CB
- HB3
6066 6072 6073 1 1.2040e+02 6.6944e+02 ; CA - C
- O
6066 6072 6074 1 1.1660e+02 5.8576e+02 ; CA - C
- N
6067 6066 6068 1 1.0950e+02 4.1840e+02 ; HA - CA
- CB
6067 6066 6072 1 1.0950e+02 4.1840e+02 ; HA - CA
- C
6068 6066 6072 1 1.1110e+02 5.2718e+02 ; CB - CA
- C
6069 6068 6070 1 1.0950e+02 2.9288e+02 ; HB1 - CB
- HB2
6069 6068 6071 1 1.0950e+02 2.9288e+02 ; HB1 - CB
- HB3

```
  6070   6068   6071       1    1.0950e+02    2.9288e+02 ;    HB2 - CB
- HB3
  6072   6074   6075       1    1.2000e+02    4.1840e+02 ;      C - N
- H
  6072   6074   6076       1    1.2190e+02    4.1840e+02 ;      C - N
- CA
  6073   6072   6074       1    1.2290e+02    6.6944e+02 ;      O - C
- N
  6074   6076   6077       1    1.0950e+02    4.1840e+02 ;      N - CA
- HA
  6074   6076   6078       1    1.0970e+02    6.6944e+02 ;      N - CA
- CB
  6074   6076   6086       1    1.1010e+02    5.2718e+02 ;      N - CA
- C
  6075   6074   6076       1    1.1804e+02    4.1840e+02 ;      H - N
- CA
  6076   6078   6079       1    1.0950e+02    4.1840e+02 ;     CA - CB
- HB
  6076   6078   6080       1    1.0950e+02    3.3472e+02 ;     CA - CB
- CG2
  6076   6078   6084       1    1.0950e+02    4.1840e+02 ;     CA - CB
- OG1
  6076   6086   6087       1    1.2040e+02    6.6944e+02 ;     CA - C
- O
  6076   6086   6088       1    1.1660e+02    5.8576e+02 ;     CA - C
- N
  6077   6076   6078       1    1.0950e+02    4.1840e+02 ;     HA - CA
- CB
  6077   6076   6086       1    1.0950e+02    4.1840e+02 ;     HA - CA
- C
  6078   6076   6086       1    1.1110e+02    5.2718e+02 ;     CB - CA
- C
  6078   6080   6081       1    1.0950e+02    4.1840e+02 ;     CB - CG2
- HG21
  6078   6080   6082       1    1.0950e+02    4.1840e+02 ;     CB - CG2
- HG22
  6078   6080   6083       1    1.0950e+02    4.1840e+02 ;     CB - CG2
- HG23
  6078   6084   6085       1    1.0850e+02    4.6024e+02 ;     CB - OG1
- HG1
  6079   6078   6080       1    1.0950e+02    4.1840e+02 ;     HB - CB
- CG2
  6079   6078   6084       1    1.0950e+02    4.1840e+02 ;     HB - CB
- OG1
  6080   6078   6084       1    1.0950e+02    4.1840e+02 ;    CG2 - CB
- OG1
  6081   6080   6082       1    1.0950e+02    2.9288e+02 ;   HG21 - CG2
- HG22
  6081   6080   6083       1    1.0950e+02    2.9288e+02 ;   HG21 - CG2
- HG23
  6082   6080   6083       1    1.0950e+02    2.9288e+02 ;   HG22 - CG2
- HG23
```

```
  6086   6088   6089       1    1.2000e+02    4.1840e+02 ;      C - N
- H
  6086   6088   6090       1    1.2190e+02    4.1840e+02 ;      C - N
- CA
  6087   6086   6088       1    1.2290e+02    6.6944e+02 ;      O - C
- N
  6088   6090   6091       1    1.0950e+02    4.1840e+02 ;      N - CA
- HA
  6088   6090   6092       1    1.0970e+02    6.6944e+02 ;      N - CA
- CB
  6088   6090   6105       1    1.1010e+02    5.2718e+02 ;      N - CA
- C
  6089   6088   6090       1    1.1804e+02    4.1840e+02 ;      H - N
- CA
  6090   6092   6093       1    1.0950e+02    4.1840e+02 ;     CA - CB
- HB
  6090   6092   6094       1    1.0950e+02    3.3472e+02 ;     CA - CB
- CG2
  6090   6092   6098       1    1.0950e+02    3.3472e+02 ;     CA - CB
- CG1
  6090   6105   6106       1    1.2040e+02    6.6944e+02 ;     CA - C
- O
  6090   6105   6107       1    1.1660e+02    5.8576e+02 ;     CA - C
- N
  6091   6090   6092       1    1.0950e+02    4.1840e+02 ;     HA - CA
- CB
  6091   6090   6105       1    1.0950e+02    4.1840e+02 ;     HA - CA
- C
  6092   6090   6105       1    1.1110e+02    5.2718e+02 ;     CB - CA
- C
  6092   6094   6095       1    1.0950e+02    4.1840e+02 ;     CB - CG2
- HG21
  6092   6094   6096       1    1.0950e+02    4.1840e+02 ;     CB - CG2
- HG22
  6092   6094   6097       1    1.0950e+02    4.1840e+02 ;     CB - CG2
- HG23
  6092   6098   6099       1    1.0950e+02    4.1840e+02 ;     CB - CG1
- HG12
  6092   6098   6100       1    1.0950e+02    4.1840e+02 ;     CB - CG1
- HG13
  6092   6098   6101       1    1.0950e+02    3.3472e+02 ;     CB - CG1
- CD1
  6093   6092   6094       1    1.0950e+02    4.1840e+02 ;     HB - CB
- CG2
  6093   6092   6098       1    1.0950e+02    4.1840e+02 ;     HB - CB
- CG1
  6094   6092   6098       1    1.0950e+02    3.3472e+02 ;    CG2 - CB
- CG1
  6095   6094   6096       1    1.0950e+02    2.9288e+02 ;   HG21 - CG2
- HG22
  6095   6094   6097       1    1.0950e+02    2.9288e+02 ;   HG21 - CG2
- HG23
```

```
  6096   6094   6097       1    1.0950e+02    2.9288e+02 ;   HG22 - CG2
- HG23
  6098   6101   6102       1    1.0950e+02    4.1840e+02 ;    CG1 - CD1
- HD11
  6098   6101   6103       1    1.0950e+02    4.1840e+02 ;    CG1 - CD1
- HD12
  6098   6101   6104       1    1.0950e+02    4.1840e+02 ;    CG1 - CD1
- HD13
  6099   6098   6100       1    1.0950e+02    2.9288e+02 ;   HG12 - CG1
- HG13
  6099   6098   6101       1    1.0950e+02    4.1840e+02 ;   HG12 - CG1
- CD1
  6100   6098   6101       1    1.0950e+02    4.1840e+02 ;   HG13 - CG1
- CD1
  6102   6101   6103       1    1.0950e+02    2.9288e+02 ;   HD11 - CD1
- HD12
  6102   6101   6104       1    1.0950e+02    2.9288e+02 ;   HD11 - CD1
- HD13
  6103   6101   6104       1    1.0950e+02    2.9288e+02 ;   HD12 - CD1
- HD13
  6105   6107   6108       1    1.2000e+02    4.1840e+02 ;      C - N
- H
  6105   6107   6109       1    1.2190e+02    4.1840e+02 ;      C - N
- CA
  6106   6105   6107       1    1.2290e+02    6.6944e+02 ;      O - C
- N
  6107   6109   6110       1    1.0950e+02    4.1840e+02 ;      N - CA
- HA
  6107   6109   6111       1    1.0970e+02    6.6944e+02 ;      N - CA
- CB
  6107   6109   6117       1    1.1010e+02    5.2718e+02 ;      N - CA
- C
  6108   6107   6109       1    1.1804e+02    4.1840e+02 ;      H - N
- CA
  6109   6111   6112       1    1.0950e+02    4.1840e+02 ;     CA - CB
- HB2
  6109   6111   6113       1    1.0950e+02    4.1840e+02 ;     CA - CB
- HB3
  6109   6111   6114       1    1.1110e+02    5.2718e+02 ;     CA - CB
- CG
  6109   6117   6118       1    1.2040e+02    6.6944e+02 ;     CA - C
- O
  6109   6117   6119       1    1.1660e+02    5.8576e+02 ;     CA - C
- N
  6110   6109   6111       1    1.0950e+02    4.1840e+02 ;     HA - CA
- CB
  6110   6109   6117       1    1.0950e+02    4.1840e+02 ;     HA - CA
- C
  6111   6109   6117       1    1.1110e+02    5.2718e+02 ;     CB - CA
- C
  6111   6114   6115       1    1.1700e+02    5.8576e+02 ;     CB - CG
- OD1
```

6111 6114 6116 1 1.1700e+02 5.8576e+02 ; CB - CG
- OD2
6112 6111 6113 1 1.0950e+02 2.9288e+02 ; HB2 - CB
- HB3
6112 6111 6114 1 1.0950e+02 4.1840e+02 ; HB2 - CB
- CG
6113 6111 6114 1 1.0950e+02 4.1840e+02 ; HB3 - CB
- CG
6115 6114 6116 1 1.2600e+02 6.6944e+02 ; OD1 - CG
- OD2
6117 6119 6120 1 1.2000e+02 4.1840e+02 ; C - N
- H
6117 6119 6121 1 1.2190e+02 4.1840e+02 ; C - N
- CA
6118 6117 6119 1 1.2290e+02 6.6944e+02 ; O - C
- N
6119 6121 6122 1 1.0950e+02 4.1840e+02 ; N - CA
- HA
6119 6121 6123 1 1.0970e+02 6.6944e+02 ; N - CA
- CB
6119 6121 6139 1 1.1010e+02 5.2718e+02 ; N - CA
- C
6120 6119 6121 1 1.1804e+02 4.1840e+02 ; H - N
- CA
6121 6123 6124 1 1.0950e+02 4.1840e+02 ; CA - CB
- HB2
6121 6123 6125 1 1.0950e+02 4.1840e+02 ; CA - CB
- HB3
6121 6123 6126 1 1.0950e+02 3.3472e+02 ; CA - CB
- CG
6121 6139 6140 1 1.2040e+02 6.6944e+02 ; CA - C
- O
6121 6139 6141 1 1.1660e+02 5.8576e+02 ; CA - C
- N
6122 6121 6123 1 1.0950e+02 4.1840e+02 ; HA - CA
- CB
6122 6121 6139 1 1.0950e+02 4.1840e+02 ; HA - CA
- C
6123 6121 6139 1 1.1110e+02 5.2718e+02 ; CB - CA
- C
6123 6126 6127 1 1.0950e+02 4.1840e+02 ; CB - CG
- HG2
6123 6126 6128 1 1.0950e+02 4.1840e+02 ; CB - CG
- HG3
6123 6126 6129 1 1.0950e+02 3.3472e+02 ; CB - CG
- CD
6124 6123 6125 1 1.0950e+02 2.9288e+02 ; HB2 - CB
- HB3
6124 6123 6126 1 1.0950e+02 4.1840e+02 ; HB2 - CB
- CG
6125 6123 6126 1 1.0950e+02 4.1840e+02 ; HB3 - CB
- CG

```
  6126   6129   6130      1    1.0950e+02    4.1840e+02 ;     CG - CD
- HD2
  6126   6129   6131      1    1.0950e+02    4.1840e+02 ;     CG - CD
- HD3
  6126   6129   6132      1    1.0950e+02    3.3472e+02 ;     CG - CD
- CE
  6127   6126   6128      1    1.0950e+02    2.9288e+02 ;    HG2 - CG
- HG3
  6127   6126   6129      1    1.0950e+02    4.1840e+02 ;    HG2 - CG
- CD
  6128   6126   6129      1    1.0950e+02    4.1840e+02 ;    HG3 - CG
- CD
  6129   6132   6133      1    1.0950e+02    4.1840e+02 ;     CD - CE
- HE2
  6129   6132   6134      1    1.0950e+02    4.1840e+02 ;     CD - CE
- HE3
  6129   6132   6135      1    1.1120e+02    6.6944e+02 ;     CD - CE
- NZ
  6130   6129   6131      1    1.0950e+02    2.9288e+02 ;    HD2 - CD
- HD3
  6130   6129   6132      1    1.0950e+02    4.1840e+02 ;    HD2 - CD
- CE
  6131   6129   6132      1    1.0950e+02    4.1840e+02 ;    HD3 - CD
- CE
  6132   6135   6136      1    1.0950e+02    4.1840e+02 ;     CE - NZ
- HZ1
  6132   6135   6137      1    1.0950e+02    4.1840e+02 ;     CE - NZ
- HZ2
  6132   6135   6138      1    1.0950e+02    4.1840e+02 ;     CE - NZ
- HZ3
  6133   6132   6134      1    1.0950e+02    2.9288e+02 ;    HE2 - CE
- HE3
  6133   6132   6135      1    1.0950e+02    4.1840e+02 ;    HE2 - CE
- NZ
  6134   6132   6135      1    1.0950e+02    4.1840e+02 ;    HE3 - CE
- NZ
  6136   6135   6137      1    1.0950e+02    2.9288e+02 ;    HZ1 - NZ
- HZ2
  6136   6135   6138      1    1.0950e+02    2.9288e+02 ;    HZ1 - NZ
- HZ3
  6137   6135   6138      1    1.0950e+02    2.9288e+02 ;    HZ2 - NZ
- HZ3
  6139   6141   6142      1    1.2000e+02    4.1840e+02 ;      C - N
- H
  6139   6141   6143      1    1.2190e+02    4.1840e+02 ;      C - N
- CA
  6140   6139   6141      1    1.2290e+02    6.6944e+02 ;      O - C
- N
  6141   6143   6144      1    1.0950e+02    4.1840e+02 ;      N - CA
- HA
  6141   6143   6145      1    1.0970e+02    6.6944e+02 ;      N - CA
- CB
```

```
  6141   6143   6156       1    1.1010e+02    5.2718e+02 ;      N - CA
- C
  6142   6141   6143       1    1.1804e+02    4.1840e+02 ;      H - N
- CA
  6143   6145   6146       1    1.0950e+02    4.1840e+02 ;     CA - CB
- HB2
  6143   6145   6147       1    1.0950e+02    4.1840e+02 ;     CA - CB
- HB3
  6143   6145   6148       1    1.0950e+02    3.3472e+02 ;     CA - CB
- CG
  6143   6156   6157       1    1.2040e+02    6.6944e+02 ;     CA - C
- O
  6143   6156   6158       1    1.1660e+02    5.8576e+02 ;     CA - C
- N
  6144   6143   6145       1    1.0950e+02    4.1840e+02 ;     HA - CA
- CB
  6144   6143   6156       1    1.0950e+02    4.1840e+02 ;     HA - CA
- C
  6145   6143   6156       1    1.1110e+02    5.2718e+02 ;     CB - CA
- C
  6145   6148   6149       1    1.0950e+02    4.1840e+02 ;     CB - CG
- HG2
  6145   6148   6150       1    1.0950e+02    4.1840e+02 ;     CB - CG
- HG3
  6145   6148   6151       1    1.1110e+02    5.2718e+02 ;     CB - CG
- CD
  6146   6145   6147       1    1.0950e+02    2.9288e+02 ;    HB2 - CB
- HB3
  6146   6145   6148       1    1.0950e+02    4.1840e+02 ;    HB2 - CB
- CG
  6147   6145   6148       1    1.0950e+02    4.1840e+02 ;    HB3 - CB
- CG
  6148   6151   6152       1    1.2040e+02    6.6944e+02 ;     CG - CD
- OE1
  6148   6151   6153       1    1.1660e+02    5.8576e+02 ;     CG - CD
- NE2
  6149   6148   6150       1    1.0950e+02    2.9288e+02 ;    HG2 - CG
- HG3
  6149   6148   6151       1    1.0950e+02    4.1840e+02 ;    HG2 - CG
- CD
  6150   6148   6151       1    1.0950e+02    4.1840e+02 ;    HG3 - CG
- CD
  6151   6153   6154       1    1.2000e+02    4.1840e+02 ;     CD - NE2
- HE21
  6151   6153   6155       1    1.2000e+02    4.1840e+02 ;     CD - NE2
- HE22
  6152   6151   6153       1    1.2290e+02    6.6944e+02 ;    OE1 - CD
- NE2
  6154   6153   6155       1    1.2000e+02    2.9288e+02 ;   HE21 - NE2
- HE22
  6156   6158   6159       1    1.2000e+02    4.1840e+02 ;      C - N
- H
```

6156 6158 6160 1 1.2190e+02 4.1840e+02 ; C - N
- CA
6157 6156 6158 1 1.2290e+02 6.6944e+02 ; O - C
- N
6158 6160 6161 1 1.0950e+02 4.1840e+02 ; N - CA
- HA
6158 6160 6162 1 1.0970e+02 6.6944e+02 ; N - CA
- CB
6158 6160 6170 1 1.1010e+02 5.2718e+02 ; N - CA
- C
6159 6158 6160 1 1.1804e+02 4.1840e+02 ; H - N
- CA
6160 6162 6163 1 1.0950e+02 4.1840e+02 ; CA - CB
- HB
6160 6162 6164 1 1.0950e+02 3.3472e+02 ; CA - CB
- CG2
6160 6162 6168 1 1.0950e+02 4.1840e+02 ; CA - CB
- OG1
6160 6170 6171 1 1.2040e+02 6.6944e+02 ; CA - C
- O
6160 6170 6172 1 1.1660e+02 5.8576e+02 ; CA - C
- N
6161 6160 6162 1 1.0950e+02 4.1840e+02 ; HA - CA
- CB
6161 6160 6170 1 1.0950e+02 4.1840e+02 ; HA - CA
- C
6162 6160 6170 1 1.1110e+02 5.2718e+02 ; CB - CA
- C
6162 6164 6165 1 1.0950e+02 4.1840e+02 ; CB - CG2
- HG21
6162 6164 6166 1 1.0950e+02 4.1840e+02 ; CB - CG2
- HG22
6162 6164 6167 1 1.0950e+02 4.1840e+02 ; CB - CG2
- HG23
6162 6168 6169 1 1.0850e+02 4.6024e+02 ; CB - OG1
- HG1
6163 6162 6164 1 1.0950e+02 4.1840e+02 ; HB - CB
- CG2
6163 6162 6168 1 1.0950e+02 4.1840e+02 ; HB - CB
- OG1
6164 6162 6168 1 1.0950e+02 4.1840e+02 ; CG2 - CB
- OG1
6165 6164 6166 1 1.0950e+02 2.9288e+02 ; HG21 - CG2
- HG22
6165 6164 6167 1 1.0950e+02 2.9288e+02 ; HG21 - CG2
- HG23
6166 6164 6167 1 1.0950e+02 2.9288e+02 ; HG22 - CG2
- HG23
6170 6172 6173 1 1.2000e+02 4.1840e+02 ; C - N
- H
6170 6172 6174 1 1.2190e+02 4.1840e+02 ; C - N
- CA

```
  6171   6170   6172      1    1.2290e+02    6.6944e+02 ;      O - C
- N
  6172   6174   6175      1    1.0950e+02    4.1840e+02 ;      N - CA
- HA
  6172   6174   6176      1    1.0970e+02    6.6944e+02 ;      N - CA
- CB
  6172   6174   6191      1    1.1010e+02    5.2718e+02 ;      N - CA
- C
  6173   6172   6174      1    1.1804e+02    4.1840e+02 ;      H - N
- CA
  6174   6176   6177      1    1.0950e+02    4.1840e+02 ;     CA - CB
- HB2
  6174   6176   6178      1    1.0950e+02    4.1840e+02 ;     CA - CB
- HB3
  6174   6176   6179      1    1.1400e+02    5.2718e+02 ;     CA - CB
- CG
  6174   6191   6192      1    1.2040e+02    6.6944e+02 ;     CA - C
- O
  6174   6191   6193      1    1.1660e+02    5.8576e+02 ;     CA - C
- N
  6175   6174   6176      1    1.0950e+02    4.1840e+02 ;     HA - CA
- CB
  6175   6174   6191      1    1.0950e+02    4.1840e+02 ;     HA - CA
- C
  6176   6174   6191      1    1.1110e+02    5.2718e+02 ;     CB - CA
- C
  6176   6179   6180      1    1.2000e+02    5.8576e+02 ;     CB - CG
- CD1
  6176   6179   6189      1    1.2000e+02    5.8576e+02 ;     CB - CG
- CD2
  6177   6176   6178      1    1.0950e+02    2.9288e+02 ;    HB2 - CB
- HB3
  6177   6176   6179      1    1.0950e+02    4.1840e+02 ;    HB2 - CB
- CG
  6178   6176   6179      1    1.0950e+02    4.1840e+02 ;    HB3 - CB
- CG
  6179   6180   6181      1    1.2000e+02    4.1840e+02 ;     CG - CD1
- HD1
  6179   6180   6182      1    1.2000e+02    5.2718e+02 ;     CG - CD1
- CE1
  6179   6189   6187      1    1.2000e+02    5.2718e+02 ;     CG - CD2
- CE2
  6179   6189   6190      1    1.2000e+02    4.1840e+02 ;     CG - CD2
- HD2
  6180   6179   6189      1    1.2000e+02    5.2718e+02 ;    CD1 - CG
- CD2
  6180   6182   6183      1    1.2000e+02    4.1840e+02 ;    CD1 - CE1
- HE1
  6180   6182   6184      1    1.2000e+02    5.2718e+02 ;    CD1 - CE1
- CZ
  6181   6180   6182      1    1.2000e+02    4.1840e+02 ;    HD1 - CD1
- CE1
```

6182 6184 6185 1 1.2000e+02 5.8576e+02 ; CE1 - CZ
- OH
6182 6184 6187 1 1.2000e+02 5.2718e+02 ; CE1 - CZ
- CE2
6183 6182 6184 1 1.2000e+02 4.1840e+02 ; HE1 - CE1
- CZ
6184 6185 6186 1 1.1300e+02 4.1840e+02 ; CZ - OH
- HH
6184 6187 6188 1 1.2000e+02 4.1840e+02 ; CZ - CE2
- HE2
6184 6187 6189 1 1.2000e+02 5.2718e+02 ; CZ - CE2
- CD2
6185 6184 6187 1 1.2000e+02 5.8576e+02 ; OH - CZ
- CE2
6187 6189 6190 1 1.2000e+02 4.1840e+02 ; CE2 - CD2
- HD2
6188 6187 6189 1 1.2000e+02 4.1840e+02 ; HE2 - CE2
- CD2
6191 6193 6194 1 1.2000e+02 4.1840e+02 ; C - N
- H
6191 6193 6195 1 1.2190e+02 4.1840e+02 ; C - N
- CA
6192 6191 6193 1 1.2290e+02 6.6944e+02 ; O - C
- N
6193 6195 6196 1 1.0950e+02 4.1840e+02 ; N - CA
- HA
6193 6195 6197 1 1.0970e+02 6.6944e+02 ; N - CA
- CB
6193 6195 6210 1 1.1010e+02 5.2718e+02 ; N - CA
- C
6194 6193 6195 1 1.1804e+02 4.1840e+02 ; H - N
- CA
6195 6197 6198 1 1.0950e+02 4.1840e+02 ; CA - CB
- HB2
6195 6197 6199 1 1.0950e+02 4.1840e+02 ; CA - CB
- HB3
6195 6197 6200 1 1.0950e+02 3.3472e+02 ; CA - CB
- CG
6195 6210 6211 1 1.2040e+02 6.6944e+02 ; CA - C
- O
6195 6210 6212 1 1.1660e+02 5.8576e+02 ; CA - C
- N
6196 6195 6197 1 1.0950e+02 4.1840e+02 ; HA - CA
- CB
6196 6195 6210 1 1.0950e+02 4.1840e+02 ; HA - CA
- C
6197 6195 6210 1 1.1110e+02 5.2718e+02 ; CB - CA
- C
6197 6200 6201 1 1.0950e+02 4.1840e+02 ; CB - CG
- HG
6197 6200 6202 1 1.0950e+02 3.3472e+02 ; CB - CG
- CD1

```
  6197   6200   6206      1    1.0950e+02    3.3472e+02 ;     CB - CG
- CD2
  6198   6197   6199      1    1.0950e+02    2.9288e+02 ;    HB2 - CB
- HB3
  6198   6197   6200      1    1.0950e+02    4.1840e+02 ;    HB2 - CB
- CG
  6199   6197   6200      1    1.0950e+02    4.1840e+02 ;    HB3 - CB
- CG
  6200   6202   6203      1    1.0950e+02    4.1840e+02 ;     CG - CD1
- HD11
  6200   6202   6204      1    1.0950e+02    4.1840e+02 ;     CG - CD1
- HD12
  6200   6202   6205      1    1.0950e+02    4.1840e+02 ;     CG - CD1
- HD13
  6200   6206   6207      1    1.0950e+02    4.1840e+02 ;     CG - CD2
- HD21
  6200   6206   6208      1    1.0950e+02    4.1840e+02 ;     CG - CD2
- HD22
  6200   6206   6209      1    1.0950e+02    4.1840e+02 ;     CG - CD2
- HD23
  6201   6200   6202      1    1.0950e+02    4.1840e+02 ;     HG - CG
- CD1
  6201   6200   6206      1    1.0950e+02    4.1840e+02 ;     HG - CG
- CD2
  6202   6200   6206      1    1.0950e+02    3.3472e+02 ;    CD1 - CG
- CD2
  6203   6202   6204      1    1.0950e+02    2.9288e+02 ;   HD11 - CD1
- HD12
  6203   6202   6205      1    1.0950e+02    2.9288e+02 ;   HD11 - CD1
- HD13
  6204   6202   6205      1    1.0950e+02    2.9288e+02 ;   HD12 - CD1
- HD13
  6207   6206   6208      1    1.0950e+02    2.9288e+02 ;   HD21 - CD2
- HD22
  6207   6206   6209      1    1.0950e+02    2.9288e+02 ;   HD21 - CD2
- HD23
  6208   6206   6209      1    1.0950e+02    2.9288e+02 ;   HD22 - CD2
- HD23
  6210   6212   6213      1    1.2000e+02    4.1840e+02 ;      C - N
- H
  6210   6212   6214      1    1.2190e+02    4.1840e+02 ;      C - N
- CA
  6211   6210   6212      1    1.2290e+02    6.6944e+02 ;      O - C
- N
  6212   6214   6215      1    1.0950e+02    4.1840e+02 ;      N - CA
- HA
  6212   6214   6216      1    1.0970e+02    6.6944e+02 ;      N - CA
- CB
  6212   6214   6232      1    1.1010e+02    5.2718e+02 ;      N - CA
- C
  6213   6212   6214      1    1.1804e+02    4.1840e+02 ;      H - N
- CA
```

6214 6216 6217 1 1.0950e+02 4.1840e+02 ; CA - CB
- HB2
6214 6216 6218 1 1.0950e+02 4.1840e+02 ; CA - CB
- HB3
6214 6216 6219 1 1.0950e+02 3.3472e+02 ; CA - CB
- CG
6214 6232 6233 1 1.2040e+02 6.6944e+02 ; CA - C
- O
6214 6232 6234 1 1.1660e+02 5.8576e+02 ; CA - C
- N
6215 6214 6216 1 1.0950e+02 4.1840e+02 ; HA - CA
- CB
6215 6214 6232 1 1.0950e+02 4.1840e+02 ; HA - CA
- C
6216 6214 6232 1 1.1110e+02 5.2718e+02 ; CB - CA
- C
6216 6219 6220 1 1.0950e+02 4.1840e+02 ; CB - CG
- HG2
6216 6219 6221 1 1.0950e+02 4.1840e+02 ; CB - CG
- HG3
6216 6219 6222 1 1.0950e+02 3.3472e+02 ; CB - CG
- CD
6217 6216 6218 1 1.0950e+02 2.9288e+02 ; HB2 - CB
- HB3
6217 6216 6219 1 1.0950e+02 4.1840e+02 ; HB2 - CB
- CG
6218 6216 6219 1 1.0950e+02 4.1840e+02 ; HB3 - CB
- CG
6219 6222 6223 1 1.0950e+02 4.1840e+02 ; CG - CD
- HD2
6219 6222 6224 1 1.0950e+02 4.1840e+02 ; CG - CD
- HD3
6219 6222 6225 1 1.0950e+02 3.3472e+02 ; CG - CD
- CE
6220 6219 6221 1 1.0950e+02 2.9288e+02 ; HG2 - CG
- HG3
6220 6219 6222 1 1.0950e+02 4.1840e+02 ; HG2 - CG
- CD
6221 6219 6222 1 1.0950e+02 4.1840e+02 ; HG3 - CG
- CD
6222 6225 6226 1 1.0950e+02 4.1840e+02 ; CD - CE
- HE2
6222 6225 6227 1 1.0950e+02 4.1840e+02 ; CD - CE
- HE3
6222 6225 6228 1 1.1120e+02 6.6944e+02 ; CD - CE
- NZ
6223 6222 6224 1 1.0950e+02 2.9288e+02 ; HD2 - CD
- HD3
6223 6222 6225 1 1.0950e+02 4.1840e+02 ; HD2 - CD
- CE
6224 6222 6225 1 1.0950e+02 4.1840e+02 ; HD3 - CD
- CE

```
  6225   6228   6229       1    1.0950e+02    4.1840e+02 ;     CE - NZ
- HZ1
  6225   6228   6230       1    1.0950e+02    4.1840e+02 ;     CE - NZ
- HZ2
  6225   6228   6231       1    1.0950e+02    4.1840e+02 ;     CE - NZ
- HZ3
  6226   6225   6227       1    1.0950e+02    2.9288e+02 ;    HE2 - CE
- HE3
  6226   6225   6228       1    1.0950e+02    4.1840e+02 ;    HE2 - CE
- NZ
  6227   6225   6228       1    1.0950e+02    4.1840e+02 ;    HE3 - CE
- NZ
  6229   6228   6230       1    1.0950e+02    2.9288e+02 ;    HZ1 - NZ
- HZ2
  6229   6228   6231       1    1.0950e+02    2.9288e+02 ;    HZ1 - NZ
- HZ3
  6230   6228   6231       1    1.0950e+02    2.9288e+02 ;    HZ2 - NZ
- HZ3
  6232   6234   6235       1    1.2000e+02    4.1840e+02 ;      C - N
- H
  6232   6234   6236       1    1.2190e+02    4.1840e+02 ;      C - N
- CA
  6233   6232   6234       1    1.2290e+02    6.6944e+02 ;      O - C
- N
  6234   6236   6237       1    1.0950e+02    4.1840e+02 ;      N - CA
- HA
  6234   6236   6238       1    1.0970e+02    6.6944e+02 ;      N - CA
- CB
  6234   6236   6254       1    1.1010e+02    5.2718e+02 ;      N - CA
- C
  6235   6234   6236       1    1.1804e+02    4.1840e+02 ;      H - N
- CA
  6236   6238   6239       1    1.0950e+02    4.1840e+02 ;     CA - CB
- HB2
  6236   6238   6240       1    1.0950e+02    4.1840e+02 ;     CA - CB
- HB3
  6236   6238   6241       1    1.0950e+02    3.3472e+02 ;     CA - CB
- CG
  6236   6254   6255       1    1.2040e+02    6.6944e+02 ;     CA - C
- O
  6236   6254   6256       1    1.1660e+02    5.8576e+02 ;     CA - C
- N
  6237   6236   6238       1    1.0950e+02    4.1840e+02 ;     HA - CA
- CB
  6237   6236   6254       1    1.0950e+02    4.1840e+02 ;     HA - CA
- C
  6238   6236   6254       1    1.1110e+02    5.2718e+02 ;     CB - CA
- C
  6238   6241   6242       1    1.0950e+02    4.1840e+02 ;     CB - CG
- HG2
  6238   6241   6243       1    1.0950e+02    4.1840e+02 ;     CB - CG
- HG3
```

6238 6241 6244 1 1.0950e+02 3.3472e+02 ; CB - CG
- CD
6239 6238 6240 1 1.0950e+02 2.9288e+02 ; HB2 - CB
- HB3
6239 6238 6241 1 1.0950e+02 4.1840e+02 ; HB2 - CB
- CG
6240 6238 6241 1 1.0950e+02 4.1840e+02 ; HB3 - CB
- CG
6241 6244 6245 1 1.0950e+02 4.1840e+02 ; CG - CD
- HD2
6241 6244 6246 1 1.0950e+02 4.1840e+02 ; CG - CD
- HD3
6241 6244 6247 1 1.0950e+02 3.3472e+02 ; CG - CD
- CE
6242 6241 6243 1 1.0950e+02 2.9288e+02 ; HG2 - CG
- HG3
6242 6241 6244 1 1.0950e+02 4.1840e+02 ; HG2 - CG
- CD
6243 6241 6244 1 1.0950e+02 4.1840e+02 ; HG3 - CG
- CD
6244 6247 6248 1 1.0950e+02 4.1840e+02 ; CD - CE
- HE2
6244 6247 6249 1 1.0950e+02 4.1840e+02 ; CD - CE
- HE3
6244 6247 6250 1 1.1120e+02 6.6944e+02 ; CD - CE
- NZ
6245 6244 6246 1 1.0950e+02 2.9288e+02 ; HD2 - CD
- HD3
6245 6244 6247 1 1.0950e+02 4.1840e+02 ; HD2 - CD
- CE
6246 6244 6247 1 1.0950e+02 4.1840e+02 ; HD3 - CD
- CE
6247 6250 6251 1 1.0950e+02 4.1840e+02 ; CE - NZ
- HZ1
6247 6250 6252 1 1.0950e+02 4.1840e+02 ; CE - NZ
- HZ2
6247 6250 6253 1 1.0950e+02 4.1840e+02 ; CE - NZ
- HZ3
6248 6247 6249 1 1.0950e+02 2.9288e+02 ; HE2 - CE
- HE3
6248 6247 6250 1 1.0950e+02 4.1840e+02 ; HE2 - CE
- NZ
6249 6247 6250 1 1.0950e+02 4.1840e+02 ; HE3 - CE
- NZ
6251 6250 6252 1 1.0950e+02 2.9288e+02 ; HZ1 - NZ
- HZ2
6251 6250 6253 1 1.0950e+02 2.9288e+02 ; HZ1 - NZ
- HZ3
6252 6250 6253 1 1.0950e+02 2.9288e+02 ; HZ2 - NZ
- HZ3
6254 6256 6257 1 1.2000e+02 4.1840e+02 ; C - N
- H

6254 6256 6258 1 1.2190e+02 4.1840e+02 ; C - N
- CA
6255 6254 6256 1 1.2290e+02 6.6944e+02 ; O - C
- N
6256 6258 6259 1 1.0950e+02 4.1840e+02 ; N - CA
- HA
6256 6258 6260 1 1.0970e+02 6.6944e+02 ; N - CA
- CB
6256 6258 6269 1 1.1010e+02 5.2718e+02 ; N - CA
- C
6257 6256 6258 1 1.1804e+02 4.1840e+02 ; H - N
- CA
6258 6260 6261 1 1.0950e+02 4.1840e+02 ; CA - CB
- HB2
6258 6260 6262 1 1.0950e+02 4.1840e+02 ; CA - CB
- HB3
6258 6260 6263 1 1.0950e+02 3.3472e+02 ; CA - CB
- CG
6258 6269 6270 1 1.2040e+02 6.6944e+02 ; CA - C
- O
6258 6269 6271 1 1.1660e+02 5.8576e+02 ; CA - C
- N
6259 6258 6260 1 1.0950e+02 4.1840e+02 ; HA - CA
- CB
6259 6258 6269 1 1.0950e+02 4.1840e+02 ; HA - CA
- C
6260 6258 6269 1 1.1110e+02 5.2718e+02 ; CB - CA
- C
6260 6263 6264 1 1.0950e+02 4.1840e+02 ; CB - CG
- HG2
6260 6263 6265 1 1.0950e+02 4.1840e+02 ; CB - CG
- HG3
6260 6263 6266 1 1.1110e+02 5.2718e+02 ; CB - CG
- CD
6261 6260 6262 1 1.0950e+02 2.9288e+02 ; HB2 - CB
- HB3
6261 6260 6263 1 1.0950e+02 4.1840e+02 ; HB2 - CB
- CG
6262 6260 6263 1 1.0950e+02 4.1840e+02 ; HB3 - CB
- CG
6263 6266 6267 1 1.1700e+02 5.8576e+02 ; CG - CD
- OE1
6263 6266 6268 1 1.1700e+02 5.8576e+02 ; CG - CD
- OE2
6264 6263 6265 1 1.0950e+02 2.9288e+02 ; HG2 - CG
- HG3
6264 6263 6266 1 1.0950e+02 4.1840e+02 ; HG2 - CG
- CD
6265 6263 6266 1 1.0950e+02 4.1840e+02 ; HG3 - CG
- CD
6267 6266 6268 1 1.2600e+02 6.6944e+02 ; OE1 - CD
- OE2

6269 6271 6272 1 1.2000e+02 4.1840e+02 ; C - N
- H
6269 6271 6273 1 1.2190e+02 4.1840e+02 ; C - N
- CA
6270 6269 6271 1 1.2290e+02 6.6944e+02 ; O - C
- N
6271 6273 6274 1 1.0950e+02 4.1840e+02 ; N - CA
- HA
6271 6273 6275 1 1.0970e+02 6.6944e+02 ; N - CA
- CB
6271 6273 6285 1 1.1010e+02 5.2718e+02 ; N - CA
- C
6272 6271 6273 1 1.1804e+02 4.1840e+02 ; H - N
- CA
6273 6275 6276 1 1.0950e+02 4.1840e+02 ; CA - CB
- HB
6273 6275 6277 1 1.0950e+02 3.3472e+02 ; CA - CB
- CG1
6273 6275 6281 1 1.0950e+02 3.3472e+02 ; CA - CB
- CG2
6273 6285 6286 1 1.2040e+02 6.6944e+02 ; CA - C
- O
6273 6285 6287 1 1.1660e+02 5.8576e+02 ; CA - C
- N
6274 6273 6275 1 1.0950e+02 4.1840e+02 ; HA - CA
- CB
6274 6273 6285 1 1.0950e+02 4.1840e+02 ; HA - CA
- C
6275 6273 6285 1 1.1110e+02 5.2718e+02 ; CB - CA
- C
6275 6277 6278 1 1.0950e+02 4.1840e+02 ; CB - CG1
- HG11
6275 6277 6279 1 1.0950e+02 4.1840e+02 ; CB - CG1
- HG12
6275 6277 6280 1 1.0950e+02 4.1840e+02 ; CB - CG1
- HG13
6275 6281 6282 1 1.0950e+02 4.1840e+02 ; CB - CG2
- HG21
6275 6281 6283 1 1.0950e+02 4.1840e+02 ; CB - CG2
- HG22
6275 6281 6284 1 1.0950e+02 4.1840e+02 ; CB - CG2
- HG23
6276 6275 6277 1 1.0950e+02 4.1840e+02 ; HB - CB
- CG1
6276 6275 6281 1 1.0950e+02 4.1840e+02 ; HB - CB
- CG2
6277 6275 6281 1 1.0950e+02 3.3472e+02 ; CG1 - CB
- CG2
6278 6277 6279 1 1.0950e+02 2.9288e+02 ; HG11 - CG1
- HG12
6278 6277 6280 1 1.0950e+02 2.9288e+02 ; HG11 - CG1
- HG13

6279 6277 6280 1 1.0950e+02 2.9288e+02 ; HG12 - CG1
- HG13
6282 6281 6283 1 1.0950e+02 2.9288e+02 ; HG21 - CG2
- HG22
6282 6281 6284 1 1.0950e+02 2.9288e+02 ; HG21 - CG2
- HG23
6283 6281 6284 1 1.0950e+02 2.9288e+02 ; HG22 - CG2
- HG23
6285 6287 6288 1 1.2000e+02 4.1840e+02 ; C - N
- H
6285 6287 6289 1 1.2190e+02 4.1840e+02 ; C - N
- CA
6286 6285 6287 1 1.2290e+02 6.6944e+02 ; O - C
- N
6287 6289 6290 1 1.0950e+02 4.1840e+02 ; N - CA
- HA
6287 6289 6291 1 1.0970e+02 6.6944e+02 ; N - CA
- CB
6287 6289 6296 1 1.1010e+02 5.2718e+02 ; N - CA
- C
6288 6287 6289 1 1.1804e+02 4.1840e+02 ; H - N
- CA
6289 6291 6292 1 1.0950e+02 4.1840e+02 ; CA - CB
- HB2
6289 6291 6293 1 1.0950e+02 4.1840e+02 ; CA - CB
- HB3
6289 6291 6294 1 1.0860e+02 4.1840e+02 ; CA - CB
- SG
6289 6296 6297 1 1.2040e+02 6.6944e+02 ; CA - C
- O
6289 6296 6298 1 1.1660e+02 5.8576e+02 ; CA - C
- N
6290 6289 6291 1 1.0950e+02 4.1840e+02 ; HA - CA
- CB
6290 6289 6296 1 1.0950e+02 4.1840e+02 ; HA - CA
- C
6291 6289 6296 1 1.1110e+02 5.2718e+02 ; CB - CA
- C
6291 6294 6295 1 9.6000e+01 3.5982e+02 ; CB - SG
- HG
6292 6291 6293 1 1.0950e+02 2.9288e+02 ; HB2 - CB
- HB3
6292 6291 6294 1 1.0950e+02 4.1840e+02 ; HB2 - CB
- SG
6293 6291 6294 1 1.0950e+02 4.1840e+02 ; HB3 - CB
- SG
6296 6298 6299 1 1.2000e+02 4.1840e+02 ; C - N
- H
6296 6298 6300 1 1.2190e+02 4.1840e+02 ; C - N
- CA
6297 6296 6298 1 1.2290e+02 6.6944e+02 ; O - C
- N

```
  6298   6300   6301      1    1.0950e+02    4.1840e+02 ;      N - CA
- HA
  6298   6300   6302      1    1.0970e+02    6.6944e+02 ;      N - CA
- CB
  6298   6300   6315      1    1.1010e+02    5.2718e+02 ;      N - CA
- C
  6299   6298   6300      1    1.1804e+02    4.1840e+02 ;      H - N
- CA
  6300   6302   6303      1    1.0950e+02    4.1840e+02 ;     CA - CB
- HB2
  6300   6302   6304      1    1.0950e+02    4.1840e+02 ;     CA - CB
- HB3
  6300   6302   6305      1    1.0950e+02    3.3472e+02 ;     CA - CB
- CG
  6300   6315   6316      1    1.2040e+02    6.6944e+02 ;     CA - C
- O
  6300   6315   6317      1    1.1660e+02    5.8576e+02 ;     CA - C
- N
  6301   6300   6302      1    1.0950e+02    4.1840e+02 ;     HA - CA
- CB
  6301   6300   6315      1    1.0950e+02    4.1840e+02 ;     HA - CA
- C
  6302   6300   6315      1    1.1110e+02    5.2718e+02 ;     CB - CA
- C
  6302   6305   6306      1    1.0950e+02    4.1840e+02 ;     CB - CG
- HG
  6302   6305   6307      1    1.0950e+02    3.3472e+02 ;     CB - CG
- CD1
  6302   6305   6311      1    1.0950e+02    3.3472e+02 ;     CB - CG
- CD2
  6303   6302   6304      1    1.0950e+02    2.9288e+02 ;    HB2 - CB
- HB3
  6303   6302   6305      1    1.0950e+02    4.1840e+02 ;    HB2 - CB
- CG
  6304   6302   6305      1    1.0950e+02    4.1840e+02 ;    HB3 - CB
- CG
  6305   6307   6308      1    1.0950e+02    4.1840e+02 ;     CG - CD1
- HD11
  6305   6307   6309      1    1.0950e+02    4.1840e+02 ;     CG - CD1
- HD12
  6305   6307   6310      1    1.0950e+02    4.1840e+02 ;     CG - CD1
- HD13
  6305   6311   6312      1    1.0950e+02    4.1840e+02 ;     CG - CD2
- HD21
  6305   6311   6313      1    1.0950e+02    4.1840e+02 ;     CG - CD2
- HD22
  6305   6311   6314      1    1.0950e+02    4.1840e+02 ;     CG - CD2
- HD23
  6306   6305   6307      1    1.0950e+02    4.1840e+02 ;     HG - CG
- CD1
  6306   6305   6311      1    1.0950e+02    4.1840e+02 ;     HG - CG
- CD2
```

6307 6305 6311 1 1.0950e+02 3.3472e+02 ; CD1 - CG
- CD2
6308 6307 6309 1 1.0950e+02 2.9288e+02 ; HD11 - CD1
- HD12
6308 6307 6310 1 1.0950e+02 2.9288e+02 ; HD11 - CD1
- HD13
6309 6307 6310 1 1.0950e+02 2.9288e+02 ; HD12 - CD1
- HD13
6312 6311 6313 1 1.0950e+02 2.9288e+02 ; HD21 - CD2
- HD22
6312 6311 6314 1 1.0950e+02 2.9288e+02 ; HD21 - CD2
- HD23
6313 6311 6314 1 1.0950e+02 2.9288e+02 ; HD22 - CD2
- HD23
6315 6317 6318 1 1.2000e+02 4.1840e+02 ; C - N
- H
6315 6317 6319 1 1.2190e+02 4.1840e+02 ; C - N
- CA
6316 6315 6317 1 1.2290e+02 6.6944e+02 ; O - C
- N
6317 6319 6320 1 1.0950e+02 4.1840e+02 ; N - CA
- HA
6317 6319 6321 1 1.0970e+02 6.6944e+02 ; N - CA
- CB
6317 6319 6334 1 1.1010e+02 5.2718e+02 ; N - CA
- C
6318 6317 6319 1 1.1804e+02 4.1840e+02 ; H - N
- CA
6319 6321 6322 1 1.0950e+02 4.1840e+02 ; CA - CB
- HB
6319 6321 6323 1 1.0950e+02 3.3472e+02 ; CA - CB
- CG2
6319 6321 6327 1 1.0950e+02 3.3472e+02 ; CA - CB
- CG1
6319 6334 6335 1 1.2040e+02 6.6944e+02 ; CA - C
- O
6319 6334 6336 1 1.1660e+02 5.8576e+02 ; CA - C
- N
6320 6319 6321 1 1.0950e+02 4.1840e+02 ; HA - CA
- CB
6320 6319 6334 1 1.0950e+02 4.1840e+02 ; HA - CA
- C
6321 6319 6334 1 1.1110e+02 5.2718e+02 ; CB - CA
- C
6321 6323 6324 1 1.0950e+02 4.1840e+02 ; CB - CG2
- HG21
6321 6323 6325 1 1.0950e+02 4.1840e+02 ; CB - CG2
- HG22
6321 6323 6326 1 1.0950e+02 4.1840e+02 ; CB - CG2
- HG23
6321 6327 6328 1 1.0950e+02 4.1840e+02 ; CB - CG1
- HG12

```
  6321   6327   6329       1    1.0950e+02    4.1840e+02 ;     CB - CG1
- HG13
  6321   6327   6330       1    1.0950e+02    3.3472e+02 ;     CB - CG1
- CD1
  6322   6321   6323       1    1.0950e+02    4.1840e+02 ;     HB - CB
- CG2
  6322   6321   6327       1    1.0950e+02    4.1840e+02 ;     HB - CB
- CG1
  6323   6321   6327       1    1.0950e+02    3.3472e+02 ;    CG2 - CB
- CG1
  6324   6323   6325       1    1.0950e+02    2.9288e+02 ;   HG21 - CG2
- HG22
  6324   6323   6326       1    1.0950e+02    2.9288e+02 ;   HG21 - CG2
- HG23
  6325   6323   6326       1    1.0950e+02    2.9288e+02 ;   HG22 - CG2
- HG23
  6327   6330   6331       1    1.0950e+02    4.1840e+02 ;    CG1 - CD1
- HD11
  6327   6330   6332       1    1.0950e+02    4.1840e+02 ;    CG1 - CD1
- HD12
  6327   6330   6333       1    1.0950e+02    4.1840e+02 ;    CG1 - CD1
- HD13
  6328   6327   6329       1    1.0950e+02    2.9288e+02 ;   HG12 - CG1
- HG13
  6328   6327   6330       1    1.0950e+02    4.1840e+02 ;   HG12 - CG1
- CD1
  6329   6327   6330       1    1.0950e+02    4.1840e+02 ;   HG13 - CG1
- CD1
  6331   6330   6332       1    1.0950e+02    2.9288e+02 ;   HD11 - CD1
- HD12
  6331   6330   6333       1    1.0950e+02    2.9288e+02 ;   HD11 - CD1
- HD13
  6332   6330   6333       1    1.0950e+02    2.9288e+02 ;   HD12 - CD1
- HD13
  6334   6336   6337       1    1.2190e+02    4.1840e+02 ;      C - N
- CD
  6334   6336   6346       1    1.2190e+02    4.1840e+02 ;      C - N
- CA
  6335   6334   6336       1    1.2290e+02    6.6944e+02 ;      O - C
- N
  6336   6337   6338       1    1.0950e+02    4.1840e+02 ;      N - CD
- HD2
  6336   6337   6339       1    1.0950e+02    4.1840e+02 ;      N - CD
- HD3
  6336   6337   6340       1    1.0970e+02    6.6944e+02 ;      N - CD
- CG
  6336   6346   6343       1    1.0970e+02    6.6944e+02 ;      N - CA
- CB
  6336   6346   6347       1    1.0950e+02    4.1840e+02 ;      N - CA
- HA
  6336   6346   6348       1    1.1010e+02    5.2718e+02 ;      N - CA
- C
```

| | | | | | | | |
|---|---|---|---|---|---|---|---|
| 6337 | 6336 | 6346 | 1 | 1.1800e+02 | 4.1840e+02 | ; | CD - N - CA |
| 6337 | 6340 | 6341 | 1 | 1.0950e+02 | 4.1840e+02 | ; | CD - CG - HG2 |
| 6337 | 6340 | 6342 | 1 | 1.0950e+02 | 4.1840e+02 | ; | CD - CG - HG3 |
| 6337 | 6340 | 6343 | 1 | 1.0950e+02 | 3.3472e+02 | ; | CD - CG - CB |
| 6338 | 6337 | 6339 | 1 | 1.0950e+02 | 2.9288e+02 | ; | HD2 - CD - HD3 |
| 6338 | 6337 | 6340 | 1 | 1.0950e+02 | 4.1840e+02 | ; | HD2 - CD - CG |
| 6339 | 6337 | 6340 | 1 | 1.0950e+02 | 4.1840e+02 | ; | HD3 - CD - CG |
| 6340 | 6343 | 6344 | 1 | 1.0950e+02 | 4.1840e+02 | ; | CG - CB - HB2 |
| 6340 | 6343 | 6345 | 1 | 1.0950e+02 | 4.1840e+02 | ; | CG - CB - HB3 |
| 6340 | 6343 | 6346 | 1 | 1.0950e+02 | 3.3472e+02 | ; | CG - CB - CA |
| 6341 | 6340 | 6342 | 1 | 1.0950e+02 | 2.9288e+02 | ; | HG2 - CG - HG3 |
| 6341 | 6340 | 6343 | 1 | 1.0950e+02 | 4.1840e+02 | ; | HG2 - CG - CB |
| 6342 | 6340 | 6343 | 1 | 1.0950e+02 | 4.1840e+02 | ; | HG3 - CG - CB |
| 6343 | 6346 | 6347 | 1 | 1.0950e+02 | 4.1840e+02 | ; | CB - CA - HA |
| 6343 | 6346 | 6348 | 1 | 1.1110e+02 | 5.2718e+02 | ; | CB - CA - C |
| 6344 | 6343 | 6345 | 1 | 1.0950e+02 | 2.9288e+02 | ; | HB2 - CB - HB3 |
| 6344 | 6343 | 6346 | 1 | 1.0950e+02 | 4.1840e+02 | ; | HB2 - CB - CA |
| 6345 | 6343 | 6346 | 1 | 1.0950e+02 | 4.1840e+02 | ; | HB3 - CB - CA |
| 6346 | 6348 | 6349 | 1 | 1.2040e+02 | 6.6944e+02 | ; | CA - C - O |
| 6346 | 6348 | 6350 | 1 | 1.1660e+02 | 5.8576e+02 | ; | CA - C - N |
| 6347 | 6346 | 6348 | 1 | 1.0950e+02 | 4.1840e+02 | ; | HA - CA - C |
| 6348 | 6350 | 6351 | 1 | 1.2000e+02 | 4.1840e+02 | ; | C - N - H |
| 6348 | 6350 | 6352 | 1 | 1.2190e+02 | 4.1840e+02 | ; | C - N - CA |
| 6349 | 6348 | 6350 | 1 | 1.2290e+02 | 6.6944e+02 | ; | O - C - N |
| 6350 | 6352 | 6353 | 1 | 1.0950e+02 | 4.1840e+02 | ; | N - CA - HA |
| 6350 | 6352 | 6354 | 1 | 1.0970e+02 | 6.6944e+02 | ; | N - CA - CB |

| | | | | | | | |
|---|---|---|---|---|---|---|---|
| 6350 | 6352 | 6368 | 1 | 1.1010e+02 | 5.2718e+02 | ; | N - CA - C |
| 6351 | 6350 | 6352 | 1 | 1.1804e+02 | 4.1840e+02 | ; | H - N - CA |
| 6352 | 6354 | 6355 | 1 | 1.0950e+02 | 4.1840e+02 | ; | CA - CB - HB2 |
| 6352 | 6354 | 6356 | 1 | 1.0950e+02 | 4.1840e+02 | ; | CA - CB - HB3 |
| 6352 | 6354 | 6357 | 1 | 1.1400e+02 | 5.2718e+02 | ; | CA - CB - CG |
| 6352 | 6368 | 6369 | 1 | 1.2040e+02 | 6.6944e+02 | ; | CA - C - O |
| 6352 | 6368 | 6370 | 1 | 1.1660e+02 | 5.8576e+02 | ; | CA - C - N |
| 6353 | 6352 | 6354 | 1 | 1.0950e+02 | 4.1840e+02 | ; | HA - CA - CB |
| 6353 | 6352 | 6368 | 1 | 1.0950e+02 | 4.1840e+02 | ; | HA - CA - C |
| 6354 | 6352 | 6368 | 1 | 1.1110e+02 | 5.2718e+02 | ; | CB - CA - C |
| 6354 | 6357 | 6358 | 1 | 1.2000e+02 | 5.8576e+02 | ; | CB - CG - CD1 |
| 6354 | 6357 | 6366 | 1 | 1.2000e+02 | 5.8576e+02 | ; | CB - CG - CD2 |
| 6355 | 6354 | 6356 | 1 | 1.0950e+02 | 2.9288e+02 | ; | HB2 - CB - HB3 |
| 6355 | 6354 | 6357 | 1 | 1.0950e+02 | 4.1840e+02 | ; | HB2 - CB - CG |
| 6356 | 6354 | 6357 | 1 | 1.0950e+02 | 4.1840e+02 | ; | HB3 - CB - CG |
| 6357 | 6358 | 6359 | 1 | 1.2000e+02 | 4.1840e+02 | ; | CG - CD1 - HD1 |
| 6357 | 6358 | 6360 | 1 | 1.2000e+02 | 5.2718e+02 | ; | CG - CD1 - CE1 |
| 6357 | 6366 | 6364 | 1 | 1.2000e+02 | 5.2718e+02 | ; | CG - CD2 - CE2 |
| 6357 | 6366 | 6367 | 1 | 1.2000e+02 | 4.1840e+02 | ; | CG - CD2 - HD2 |
| 6358 | 6357 | 6366 | 1 | 1.2000e+02 | 5.2718e+02 | ; | CD1 - CG - CD2 |
| 6358 | 6360 | 6361 | 1 | 1.2000e+02 | 4.1840e+02 | ; | CD1 - CE1 - HE1 |
| 6358 | 6360 | 6362 | 1 | 1.2000e+02 | 5.2718e+02 | ; | CD1 - CE1 - CZ |
| 6359 | 6358 | 6360 | 1 | 1.2000e+02 | 4.1840e+02 | ; | HD1 - CD1 - CE1 |
| 6360 | 6362 | 6363 | 1 | 1.2000e+02 | 4.1840e+02 | ; | CE1 - CZ - HZ |
| 6360 | 6362 | 6364 | 1 | 1.2000e+02 | 5.2718e+02 | ; | CE1 - CZ - CE2 |
| 6361 | 6360 | 6362 | 1 | 1.2000e+02 | 4.1840e+02 | ; | HE1 - CE1 - CZ |

6362 6364 6365 1 1.2000e+02 4.1840e+02 ; CZ - CE2
- HE2
6362 6364 6366 1 1.2000e+02 5.2718e+02 ; CZ - CE2
- CD2
6363 6362 6364 1 1.2000e+02 4.1840e+02 ; HZ - CZ
- CE2
6364 6366 6367 1 1.2000e+02 4.1840e+02 ; CE2 - CD2
- HD2
6365 6364 6366 1 1.2000e+02 4.1840e+02 ; HE2 - CE2
- CD2
6368 6370 6371 1 1.2000e+02 4.1840e+02 ; C - N
- H
6368 6370 6372 1 1.2190e+02 4.1840e+02 ; C - N
- CA
6369 6368 6370 1 1.2290e+02 6.6944e+02 ; O - C
- N
6370 6372 6373 1 1.0950e+02 4.1840e+02 ; N - CA
- HA
6370 6372 6374 1 1.0970e+02 6.6944e+02 ; N - CA
- CB
6370 6372 6390 1 1.1010e+02 5.2718e+02 ; N - CA
- C
6371 6370 6372 1 1.1804e+02 4.1840e+02 ; H - N
- CA
6372 6374 6375 1 1.0950e+02 4.1840e+02 ; CA - CB
- HB2
6372 6374 6376 1 1.0950e+02 4.1840e+02 ; CA - CB
- HB3
6372 6374 6377 1 1.0950e+02 3.3472e+02 ; CA - CB
- CG
6372 6390 6391 1 1.2040e+02 6.6944e+02 ; CA - C
- O
6372 6390 6392 1 1.1660e+02 5.8576e+02 ; CA - C
- N
6373 6372 6374 1 1.0950e+02 4.1840e+02 ; HA - CA
- CB
6373 6372 6390 1 1.0950e+02 4.1840e+02 ; HA - CA
- C
6374 6372 6390 1 1.1110e+02 5.2718e+02 ; CB - CA
- C
6374 6377 6378 1 1.0950e+02 4.1840e+02 ; CB - CG
- HG2
6374 6377 6379 1 1.0950e+02 4.1840e+02 ; CB - CG
- HG3
6374 6377 6380 1 1.0950e+02 3.3472e+02 ; CB - CG
- CD
6375 6374 6376 1 1.0950e+02 2.9288e+02 ; HB2 - CB
- HB3
6375 6374 6377 1 1.0950e+02 4.1840e+02 ; HB2 - CB
- CG
6376 6374 6377 1 1.0950e+02 4.1840e+02 ; HB3 - CB
- CG

6377 6380 6381 1 1.0950e+02 4.1840e+02 ; CG - CD
- HD2
6377 6380 6382 1 1.0950e+02 4.1840e+02 ; CG - CD
- HD3
6377 6380 6383 1 1.0950e+02 3.3472e+02 ; CG - CD
- CE
6378 6377 6379 1 1.0950e+02 2.9288e+02 ; HG2 - CG
- HG3
6378 6377 6380 1 1.0950e+02 4.1840e+02 ; HG2 - CG
- CD
6379 6377 6380 1 1.0950e+02 4.1840e+02 ; HG3 - CG
- CD
6380 6383 6384 1 1.0950e+02 4.1840e+02 ; CD - CE
- HE2
6380 6383 6385 1 1.0950e+02 4.1840e+02 ; CD - CE
- HE3
6380 6383 6386 1 1.1120e+02 6.6944e+02 ; CD - CE
- NZ
6381 6380 6382 1 1.0950e+02 2.9288e+02 ; HD2 - CD
- HD3
6381 6380 6383 1 1.0950e+02 4.1840e+02 ; HD2 - CD
- CE
6382 6380 6383 1 1.0950e+02 4.1840e+02 ; HD3 - CD
- CE
6383 6386 6387 1 1.0950e+02 4.1840e+02 ; CE - NZ
- HZ1
6383 6386 6388 1 1.0950e+02 4.1840e+02 ; CE - NZ
- HZ2
6383 6386 6389 1 1.0950e+02 4.1840e+02 ; CE - NZ
- HZ3
6384 6383 6385 1 1.0950e+02 2.9288e+02 ; HE2 - CE
- HE3
6384 6383 6386 1 1.0950e+02 4.1840e+02 ; HE2 - CE
- NZ
6385 6383 6386 1 1.0950e+02 4.1840e+02 ; HE3 - CE
- NZ
6387 6386 6388 1 1.0950e+02 2.9288e+02 ; HZ1 - NZ
- HZ2
6387 6386 6389 1 1.0950e+02 2.9288e+02 ; HZ1 - NZ
- HZ3
6388 6386 6389 1 1.0950e+02 2.9288e+02 ; HZ2 - NZ
- HZ3
6390 6392 6393 1 1.2000e+02 4.1840e+02 ; C - N
- H
6390 6392 6394 1 1.2190e+02 4.1840e+02 ; C - N
- CA
6391 6390 6392 1 1.2290e+02 6.6944e+02 ; O - C
- N
6392 6394 6395 1 1.0950e+02 4.1840e+02 ; N - CA
- HA
6392 6394 6396 1 1.0970e+02 6.6944e+02 ; N - CA
- CB

6392 6394 6410 1 1.1010e+02 5.2718e+02 ; N - CA
- C
6393 6392 6394 1 1.1804e+02 4.1840e+02 ; H - N
- CA
6394 6396 6397 1 1.0950e+02 4.1840e+02 ; CA - CB
- HB2
6394 6396 6398 1 1.0950e+02 4.1840e+02 ; CA - CB
- HB3
6394 6396 6399 1 1.1400e+02 5.2718e+02 ; CA - CB
- CG
6394 6410 6411 1 1.2040e+02 6.6944e+02 ; CA - C
- O
6394 6410 6412 1 1.1660e+02 5.8576e+02 ; CA - C
- N
6395 6394 6396 1 1.0950e+02 4.1840e+02 ; HA - CA
- CB
6395 6394 6410 1 1.0950e+02 4.1840e+02 ; HA - CA
- C
6396 6394 6410 1 1.1110e+02 5.2718e+02 ; CB - CA
- C
6396 6399 6400 1 1.2000e+02 5.8576e+02 ; CB - CG
- CD1
6396 6399 6408 1 1.2000e+02 5.8576e+02 ; CB - CG
- CD2
6397 6396 6398 1 1.0950e+02 2.9288e+02 ; HB2 - CB
- HB3
6397 6396 6399 1 1.0950e+02 4.1840e+02 ; HB2 - CB
- CG
6398 6396 6399 1 1.0950e+02 4.1840e+02 ; HB3 - CB
- CG
6399 6400 6401 1 1.2000e+02 4.1840e+02 ; CG - CD1
- HD1
6399 6400 6402 1 1.2000e+02 5.2718e+02 ; CG - CD1
- CE1
6399 6408 6406 1 1.2000e+02 5.2718e+02 ; CG - CD2
- CE2
6399 6408 6409 1 1.2000e+02 4.1840e+02 ; CG - CD2
- HD2
6400 6399 6408 1 1.2000e+02 5.2718e+02 ; CD1 - CG
- CD2
6400 6402 6403 1 1.2000e+02 4.1840e+02 ; CD1 - CE1
- HE1
6400 6402 6404 1 1.2000e+02 5.2718e+02 ; CD1 - CE1
- CZ
6401 6400 6402 1 1.2000e+02 4.1840e+02 ; HD1 - CD1
- CE1
6402 6404 6405 1 1.2000e+02 4.1840e+02 ; CE1 - CZ
- HZ
6402 6404 6406 1 1.2000e+02 5.2718e+02 ; CE1 - CZ
- CE2
6403 6402 6404 1 1.2000e+02 4.1840e+02 ; HE1 - CE1
- CZ

```
  6404   6406   6407       1    1.2000e+02    4.1840e+02 ;     CZ - CE2
- HE2
  6404   6406   6408       1    1.2000e+02    5.2718e+02 ;     CZ - CE2
- CD2
  6405   6404   6406       1    1.2000e+02    4.1840e+02 ;     HZ - CZ
- CE2
  6406   6408   6409       1    1.2000e+02    4.1840e+02 ;    CE2 - CD2
- HD2
  6407   6406   6408       1    1.2000e+02    4.1840e+02 ;    HE2 - CE2
- CD2
  6410   6412   6413       1    1.2000e+02    4.1840e+02 ;      C - N
- H
  6410   6412   6414       1    1.2190e+02    4.1840e+02 ;      C - N
- CA
  6411   6410   6412       1    1.2290e+02    6.6944e+02 ;      O - C
- N
  6412   6414   6415       1    1.0950e+02    4.1840e+02 ;      N - CA
- HA
  6412   6414   6416       1    1.0970e+02    6.6944e+02 ;      N - CA
- CB
  6412   6414   6430       1    1.1010e+02    5.2718e+02 ;      N - CA
- C
  6413   6412   6414       1    1.1804e+02    4.1840e+02 ;      H - N
- CA
  6414   6416   6417       1    1.0950e+02    4.1840e+02 ;     CA - CB
- HB2
  6414   6416   6418       1    1.0950e+02    4.1840e+02 ;     CA - CB
- HB3
  6414   6416   6419       1    1.1400e+02    5.2718e+02 ;     CA - CB
- CG
  6414   6430   6431       1    1.2040e+02    6.6944e+02 ;     CA - C
- O
  6414   6430   6432       1    1.1660e+02    5.8576e+02 ;     CA - C
- N
  6415   6414   6416       1    1.0950e+02    4.1840e+02 ;     HA - CA
- CB
  6415   6414   6430       1    1.0950e+02    4.1840e+02 ;     HA - CA
- C
  6416   6414   6430       1    1.1110e+02    5.2718e+02 ;     CB - CA
- C
  6416   6419   6420       1    1.2000e+02    5.8576e+02 ;     CB - CG
- CD1
  6416   6419   6428       1    1.2000e+02    5.8576e+02 ;     CB - CG
- CD2
  6417   6416   6418       1    1.0950e+02    2.9288e+02 ;    HB2 - CB
- HB3
  6417   6416   6419       1    1.0950e+02    4.1840e+02 ;    HB2 - CB
- CG
  6418   6416   6419       1    1.0950e+02    4.1840e+02 ;    HB3 - CB
- CG
  6419   6420   6421       1    1.2000e+02    4.1840e+02 ;     CG - CD1
- HD1
```

```
  6419   6420   6422       1    1.2000e+02    5.2718e+02 ;     CG - CD1
- CE1
  6419   6428   6426       1    1.2000e+02    5.2718e+02 ;     CG - CD2
- CE2
  6419   6428   6429       1    1.2000e+02    4.1840e+02 ;     CG - CD2
- HD2
  6420   6419   6428       1    1.2000e+02    5.2718e+02 ;    CD1 - CG
- CD2
  6420   6422   6423       1    1.2000e+02    4.1840e+02 ;    CD1 - CE1
- HE1
  6420   6422   6424       1    1.2000e+02    5.2718e+02 ;    CD1 - CE1
- CZ
  6421   6420   6422       1    1.2000e+02    4.1840e+02 ;    HD1 - CD1
- CE1
  6422   6424   6425       1    1.2000e+02    4.1840e+02 ;    CE1 - CZ
- HZ
  6422   6424   6426       1    1.2000e+02    5.2718e+02 ;    CE1 - CZ
- CE2
  6423   6422   6424       1    1.2000e+02    4.1840e+02 ;    HE1 - CE1
- CZ
  6424   6426   6427       1    1.2000e+02    4.1840e+02 ;     CZ - CE2
- HE2
  6424   6426   6428       1    1.2000e+02    5.2718e+02 ;     CZ - CE2
- CD2
  6425   6424   6426       1    1.2000e+02    4.1840e+02 ;     HZ - CZ
- CE2
  6426   6428   6429       1    1.2000e+02    4.1840e+02 ;    CE2 - CD2
- HD2
  6427   6426   6428       1    1.2000e+02    4.1840e+02 ;    HE2 - CE2
- CD2
  6430   6432   6433       1    1.2000e+02    4.1840e+02 ;      C - N
- H
  6430   6432   6434       1    1.2190e+02    4.1840e+02 ;      C - N
- CA
  6431   6430   6432       1    1.2290e+02    6.6944e+02 ;      O - C
- N
  6432   6434   6435       1    1.0950e+02    4.1840e+02 ;      N - CA
- HA
  6432   6434   6436       1    1.0970e+02    6.6944e+02 ;      N - CA
- CB
  6432   6434   6442       1    1.1010e+02    5.2718e+02 ;      N - CA
- C
  6433   6432   6434       1    1.1804e+02    4.1840e+02 ;      H - N
- CA
  6434   6436   6437       1    1.0950e+02    4.1840e+02 ;     CA - CB
- HB2
  6434   6436   6438       1    1.0950e+02    4.1840e+02 ;     CA - CB
- HB3
  6434   6436   6439       1    1.1110e+02    5.2718e+02 ;     CA - CB
- CG
  6434   6442   6443       1    1.2040e+02    6.6944e+02 ;     CA - C
- O
```

6434 6442 6444 1 1.1660e+02 5.8576e+02 ; CA - C
- N
6435 6434 6436 1 1.0950e+02 4.1840e+02 ; HA - CA
- CB
6435 6434 6442 1 1.0950e+02 4.1840e+02 ; HA - CA
- C
6436 6434 6442 1 1.1110e+02 5.2718e+02 ; CB - CA
- C
6436 6439 6440 1 1.1700e+02 5.8576e+02 ; CB - CG
- OD1
6436 6439 6441 1 1.1700e+02 5.8576e+02 ; CB - CG
- OD2
6437 6436 6438 1 1.0950e+02 2.9288e+02 ; HB2 - CB
- HB3
6437 6436 6439 1 1.0950e+02 4.1840e+02 ; HB2 - CB
- CG
6438 6436 6439 1 1.0950e+02 4.1840e+02 ; HB3 - CB
- CG
6440 6439 6441 1 1.2600e+02 6.6944e+02 ; OD1 - CG
- OD2
6442 6444 6445 1 1.2000e+02 4.1840e+02 ; C - N
- H
6442 6444 6446 1 1.2190e+02 4.1840e+02 ; C - N
- CA
6443 6442 6444 1 1.2290e+02 6.6944e+02 ; O - C
- N
6444 6446 6447 1 1.0950e+02 4.1840e+02 ; N - CA
- HA
6444 6446 6448 1 1.0970e+02 6.6944e+02 ; N - CA
- CB
6444 6446 6461 1 1.1010e+02 5.2718e+02 ; N - CA
- C
6445 6444 6446 1 1.1804e+02 4.1840e+02 ; H - N
- CA
6446 6448 6449 1 1.0950e+02 4.1840e+02 ; CA - CB
- HB
6446 6448 6450 1 1.0950e+02 3.3472e+02 ; CA - CB
- CG2
6446 6448 6454 1 1.0950e+02 3.3472e+02 ; CA - CB
- CG1
6446 6461 6462 1 1.2040e+02 6.6944e+02 ; CA - C
- O
6446 6461 6463 1 1.1660e+02 5.8576e+02 ; CA - C
- N
6447 6446 6448 1 1.0950e+02 4.1840e+02 ; HA - CA
- CB
6447 6446 6461 1 1.0950e+02 4.1840e+02 ; HA - CA
- C
6448 6446 6461 1 1.1110e+02 5.2718e+02 ; CB - CA
- C
6448 6450 6451 1 1.0950e+02 4.1840e+02 ; CB - CG2
- HG21

```
  6448   6450   6452      1    1.0950e+02    4.1840e+02 ;     CB - CG2
- HG22
  6448   6450   6453      1    1.0950e+02    4.1840e+02 ;     CB - CG2
- HG23
  6448   6454   6455      1    1.0950e+02    4.1840e+02 ;     CB - CG1
- HG12
  6448   6454   6456      1    1.0950e+02    4.1840e+02 ;     CB - CG1
- HG13
  6448   6454   6457      1    1.0950e+02    3.3472e+02 ;     CB - CG1
- CD1
  6449   6448   6450      1    1.0950e+02    4.1840e+02 ;     HB - CB
- CG2
  6449   6448   6454      1    1.0950e+02    4.1840e+02 ;     HB - CB
- CG1
  6450   6448   6454      1    1.0950e+02    3.3472e+02 ;    CG2 - CB
- CG1
  6451   6450   6452      1    1.0950e+02    2.9288e+02 ;   HG21 - CG2
- HG22
  6451   6450   6453      1    1.0950e+02    2.9288e+02 ;   HG21 - CG2
- HG23
  6452   6450   6453      1    1.0950e+02    2.9288e+02 ;   HG22 - CG2
- HG23
  6454   6457   6458      1    1.0950e+02    4.1840e+02 ;    CG1 - CD1
- HD11
  6454   6457   6459      1    1.0950e+02    4.1840e+02 ;    CG1 - CD1
- HD12
  6454   6457   6460      1    1.0950e+02    4.1840e+02 ;    CG1 - CD1
- HD13
  6455   6454   6456      1    1.0950e+02    2.9288e+02 ;   HG12 - CG1
- HG13
  6455   6454   6457      1    1.0950e+02    4.1840e+02 ;   HG12 - CG1
- CD1
  6456   6454   6457      1    1.0950e+02    4.1840e+02 ;   HG13 - CG1
- CD1
  6458   6457   6459      1    1.0950e+02    2.9288e+02 ;   HD11 - CD1
- HD12
  6458   6457   6460      1    1.0950e+02    2.9288e+02 ;   HD11 - CD1
- HD13
  6459   6457   6460      1    1.0950e+02    2.9288e+02 ;   HD12 - CD1
- HD13
  6461   6463   6464      1    1.2000e+02    4.1840e+02 ;      C - N
- H
  6461   6463   6465      1    1.2190e+02    4.1840e+02 ;      C - N
- CA
  6462   6461   6463      1    1.2290e+02    6.6944e+02 ;      O - C
- N
  6463   6465   6466      1    1.0950e+02    4.1840e+02 ;      N - CA
- HA
  6463   6465   6467      1    1.0970e+02    6.6944e+02 ;      N - CA
- CB
  6463   6465   6473      1    1.1010e+02    5.2718e+02 ;      N - CA
- C
```

```
  6464   6463   6465      1    1.1804e+02    4.1840e+02 ;      H - N
- CA
  6465   6467   6468      1    1.0950e+02    4.1840e+02 ;     CA - CB
- HB2
  6465   6467   6469      1    1.0950e+02    4.1840e+02 ;     CA - CB
- HB3
  6465   6467   6470      1    1.1110e+02    5.2718e+02 ;     CA - CB
- CG
  6465   6473   6474      1    1.2040e+02    6.6944e+02 ;     CA - C
- O
  6465   6473   6475      1    1.1660e+02    5.8576e+02 ;     CA - C
- N
  6466   6465   6467      1    1.0950e+02    4.1840e+02 ;     HA - CA
- CB
  6466   6465   6473      1    1.0950e+02    4.1840e+02 ;     HA - CA
- C
  6467   6465   6473      1    1.1110e+02    5.2718e+02 ;     CB - CA
- C
  6467   6470   6471      1    1.1700e+02    5.8576e+02 ;     CB - CG
- OD1
  6467   6470   6472      1    1.1700e+02    5.8576e+02 ;     CB - CG
- OD2
  6468   6467   6469      1    1.0950e+02    2.9288e+02 ;    HB2 - CB
- HB3
  6468   6467   6470      1    1.0950e+02    4.1840e+02 ;    HB2 - CB
- CG
  6469   6467   6470      1    1.0950e+02    4.1840e+02 ;    HB3 - CB
- CG
  6471   6470   6472      1    1.2600e+02    6.6944e+02 ;    OD1 - CG
- OD2
  6473   6475   6476      1    1.2000e+02    4.1840e+02 ;      C - N
- H
  6473   6475   6477      1    1.2190e+02    4.1840e+02 ;      C - N
- CA
  6474   6473   6475      1    1.2290e+02    6.6944e+02 ;      O - C
- N
  6475   6477   6478      1    1.0950e+02    4.1840e+02 ;      N - CA
- HA2
  6475   6477   6479      1    1.0950e+02    4.1840e+02 ;      N - CA
- HA3
  6475   6477   6480      1    1.1010e+02    5.2718e+02 ;      N - CA
- C
  6476   6475   6477      1    1.1804e+02    4.1840e+02 ;      H - N
- CA
  6477   6480   6481      1    1.2040e+02    6.6944e+02 ;     CA - C
- O
  6477   6480   6482      1    1.1660e+02    5.8576e+02 ;     CA - C
- N
  6478   6477   6479      1    1.0950e+02    2.9288e+02 ;    HA2 - CA
- HA3
  6478   6477   6480      1    1.0950e+02    4.1840e+02 ;    HA2 - CA
- C
```

```
  6479   6477   6480       1    1.0950e+02    4.1840e+02 ;    HA3 - CA
- C
  6480   6482   6483       1    1.2000e+02    4.1840e+02 ;      C - N
- H
  6480   6482   6484       1    1.2190e+02    4.1840e+02 ;      C - N
- CA
  6481   6480   6482       1    1.2290e+02    6.6944e+02 ;      O - C
- N
  6482   6484   6485       1    1.0950e+02    4.1840e+02 ;      N - CA
- HA
  6482   6484   6486       1    1.0970e+02    6.6944e+02 ;      N - CA
- CB
  6482   6484   6494       1    1.1010e+02    5.2718e+02 ;      N - CA
- C
  6483   6482   6484       1    1.1804e+02    4.1840e+02 ;      H - N
- CA
  6484   6486   6487       1    1.0950e+02    4.1840e+02 ;     CA - CB
- HB2
  6484   6486   6488       1    1.0950e+02    4.1840e+02 ;     CA - CB
- HB3
  6484   6486   6489       1    1.1110e+02    5.2718e+02 ;     CA - CB
- CG
  6484   6494   6495       1    1.2040e+02    6.6944e+02 ;     CA - C
- O
  6484   6494   6496       1    1.1660e+02    5.8576e+02 ;     CA - C
- N
  6485   6484   6486       1    1.0950e+02    4.1840e+02 ;     HA - CA
- CB
  6485   6484   6494       1    1.0950e+02    4.1840e+02 ;     HA - CA
- C
  6486   6484   6494       1    1.1110e+02    5.2718e+02 ;     CB - CA
- C
  6486   6489   6490       1    1.2040e+02    6.6944e+02 ;     CB - CG
- OD1
  6486   6489   6491       1    1.1660e+02    5.8576e+02 ;     CB - CG
- ND2
  6487   6486   6488       1    1.0950e+02    2.9288e+02 ;    HB2 - CB
- HB3
  6487   6486   6489       1    1.0950e+02    4.1840e+02 ;    HB2 - CB
- CG
  6488   6486   6489       1    1.0950e+02    4.1840e+02 ;    HB3 - CB
- CG
  6489   6491   6492       1    1.2000e+02    4.1840e+02 ;     CG - ND2
- HD21
  6489   6491   6493       1    1.2000e+02    4.1840e+02 ;     CG - ND2
- HD22
  6490   6489   6491       1    1.2290e+02    6.6944e+02 ;    OD1 - CG
- ND2
  6492   6491   6493       1    1.2000e+02    2.9288e+02 ;   HD21 - ND2
- HD22
  6494   6496   6497       1    1.2000e+02    4.1840e+02 ;      C - N
- H
```

6494 6496 6498 1 1.2190e+02 4.1840e+02 ; C - N
- CA
6495 6494 6496 1 1.2290e+02 6.6944e+02 ; O - C
- N
6496 6498 6499 1 1.0950e+02 4.1840e+02 ; N - CA
- HA2
6496 6498 6500 1 1.0950e+02 4.1840e+02 ; N - CA
- HA3
6496 6498 6501 1 1.1010e+02 5.2718e+02 ; N - CA
- C
6497 6496 6498 1 1.1804e+02 4.1840e+02 ; H - N
- CA
6498 6501 6502 1 1.2040e+02 6.6944e+02 ; CA - C
- O
6498 6501 6503 1 1.1660e+02 5.8576e+02 ; CA - C
- N
6499 6498 6500 1 1.0950e+02 2.9288e+02 ; HA2 - CA
- HA3
6499 6498 6501 1 1.0950e+02 4.1840e+02 ; HA2 - CA
- C
6500 6498 6501 1 1.0950e+02 4.1840e+02 ; HA3 - CA
- C
6501 6503 6504 1 1.2000e+02 4.1840e+02 ; C - N
- H
6501 6503 6505 1 1.2190e+02 4.1840e+02 ; C - N
- CA
6502 6501 6503 1 1.2290e+02 6.6944e+02 ; O - C
- N
6503 6505 6506 1 1.0950e+02 4.1840e+02 ; N - CA
- HA
6503 6505 6507 1 1.0970e+02 6.6944e+02 ; N - CA
- CB
6503 6505 6523 1 1.1010e+02 5.2718e+02 ; N - CA
- C
6504 6503 6505 1 1.1804e+02 4.1840e+02 ; H - N
- CA
6505 6507 6508 1 1.0950e+02 4.1840e+02 ; CA - CB
- HB2
6505 6507 6509 1 1.0950e+02 4.1840e+02 ; CA - CB
- HB3
6505 6507 6510 1 1.0950e+02 3.3472e+02 ; CA - CB
- CG
6505 6523 6524 1 1.2040e+02 6.6944e+02 ; CA - C
- O
6505 6523 6525 1 1.1660e+02 5.8576e+02 ; CA - C
- N
6506 6505 6507 1 1.0950e+02 4.1840e+02 ; HA - CA
- CB
6506 6505 6523 1 1.0950e+02 4.1840e+02 ; HA - CA
- C
6507 6505 6523 1 1.1110e+02 5.2718e+02 ; CB - CA
- C

6507 6510 6511 1 1.0950e+02 4.1840e+02 ; CB - CG
- HG2
6507 6510 6512 1 1.0950e+02 4.1840e+02 ; CB - CG
- HG3
6507 6510 6513 1 1.0950e+02 3.3472e+02 ; CB - CG
- CD
6508 6507 6509 1 1.0950e+02 2.9288e+02 ; HB2 - CB
- HB3
6508 6507 6510 1 1.0950e+02 4.1840e+02 ; HB2 - CB
- CG
6509 6507 6510 1 1.0950e+02 4.1840e+02 ; HB3 - CB
- CG
6510 6513 6514 1 1.0950e+02 4.1840e+02 ; CG - CD
- HD2
6510 6513 6515 1 1.0950e+02 4.1840e+02 ; CG - CD
- HD3
6510 6513 6516 1 1.0950e+02 3.3472e+02 ; CG - CD
- CE
6511 6510 6512 1 1.0950e+02 2.9288e+02 ; HG2 - CG
- HG3
6511 6510 6513 1 1.0950e+02 4.1840e+02 ; HG2 - CG
- CD
6512 6510 6513 1 1.0950e+02 4.1840e+02 ; HG3 - CG
- CD
6513 6516 6517 1 1.0950e+02 4.1840e+02 ; CD - CE
- HE2
6513 6516 6518 1 1.0950e+02 4.1840e+02 ; CD - CE
- HE3
6513 6516 6519 1 1.1120e+02 6.6944e+02 ; CD - CE
- NZ
6514 6513 6515 1 1.0950e+02 2.9288e+02 ; HD2 - CD
- HD3
6514 6513 6516 1 1.0950e+02 4.1840e+02 ; HD2 - CD
- CE
6515 6513 6516 1 1.0950e+02 4.1840e+02 ; HD3 - CD
- CE
6516 6519 6520 1 1.0950e+02 4.1840e+02 ; CE - NZ
- HZ1
6516 6519 6521 1 1.0950e+02 4.1840e+02 ; CE - NZ
- HZ2
6516 6519 6522 1 1.0950e+02 4.1840e+02 ; CE - NZ
- HZ3
6517 6516 6518 1 1.0950e+02 2.9288e+02 ; HE2 - CE
- HE3
6517 6516 6519 1 1.0950e+02 4.1840e+02 ; HE2 - CE
- NZ
6518 6516 6519 1 1.0950e+02 4.1840e+02 ; HE3 - CE
- NZ
6520 6519 6521 1 1.0950e+02 2.9288e+02 ; HZ1 - NZ
- HZ2
6520 6519 6522 1 1.0950e+02 2.9288e+02 ; HZ1 - NZ
- HZ3

```
  6521   6519   6522       1    1.0950e+02    2.9288e+02 ;    HZ2 - NZ
- HZ3
  6523   6525   6526       1    1.2000e+02    4.1840e+02 ;      C - N
- H
  6523   6525   6527       1    1.2190e+02    4.1840e+02 ;      C - N
- CA
  6524   6523   6525       1    1.2290e+02    6.6944e+02 ;      O - C
- N
  6525   6527   6528       1    1.0950e+02    4.1840e+02 ;      N - CA
- HA
  6525   6527   6529       1    1.0970e+02    6.6944e+02 ;      N - CA
- CB
  6525   6527   6542       1    1.1010e+02    5.2718e+02 ;      N - CA
- C
  6526   6525   6527       1    1.1804e+02    4.1840e+02 ;      H - N
- CA
  6527   6529   6530       1    1.0950e+02    4.1840e+02 ;     CA - CB
- HB
  6527   6529   6531       1    1.0950e+02    3.3472e+02 ;     CA - CB
- CG2
  6527   6529   6535       1    1.0950e+02    3.3472e+02 ;     CA - CB
- CG1
  6527   6542   6543       1    1.2040e+02    6.6944e+02 ;     CA - C
- O
  6527   6542   6544       1    1.1660e+02    5.8576e+02 ;     CA - C
- N
  6528   6527   6529       1    1.0950e+02    4.1840e+02 ;     HA - CA
- CB
  6528   6527   6542       1    1.0950e+02    4.1840e+02 ;     HA - CA
- C
  6529   6527   6542       1    1.1110e+02    5.2718e+02 ;     CB - CA
- C
  6529   6531   6532       1    1.0950e+02    4.1840e+02 ;     CB - CG2
- HG21
  6529   6531   6533       1    1.0950e+02    4.1840e+02 ;     CB - CG2
- HG22
  6529   6531   6534       1    1.0950e+02    4.1840e+02 ;     CB - CG2
- HG23
  6529   6535   6536       1    1.0950e+02    4.1840e+02 ;     CB - CG1
- HG12
  6529   6535   6537       1    1.0950e+02    4.1840e+02 ;     CB - CG1
- HG13
  6529   6535   6538       1    1.0950e+02    3.3472e+02 ;     CB - CG1
- CD1
  6530   6529   6531       1    1.0950e+02    4.1840e+02 ;     HB - CB
- CG2
  6530   6529   6535       1    1.0950e+02    4.1840e+02 ;     HB - CB
- CG1
  6531   6529   6535       1    1.0950e+02    3.3472e+02 ;    CG2 - CB
- CG1
  6532   6531   6533       1    1.0950e+02    2.9288e+02 ;   HG21 - CG2
- HG22
```

```
  6532   6531   6534       1    1.0950e+02    2.9288e+02 ;   HG21 - CG2
- HG23
  6533   6531   6534       1    1.0950e+02    2.9288e+02 ;   HG22 - CG2
- HG23
  6535   6538   6539       1    1.0950e+02    4.1840e+02 ;    CG1 - CD1
- HD11
  6535   6538   6540       1    1.0950e+02    4.1840e+02 ;    CG1 - CD1
- HD12
  6535   6538   6541       1    1.0950e+02    4.1840e+02 ;    CG1 - CD1
- HD13
  6536   6535   6537       1    1.0950e+02    2.9288e+02 ;   HG12 - CG1
- HG13
  6536   6535   6538       1    1.0950e+02    4.1840e+02 ;   HG12 - CG1
- CD1
  6537   6535   6538       1    1.0950e+02    4.1840e+02 ;   HG13 - CG1
- CD1
  6539   6538   6540       1    1.0950e+02    2.9288e+02 ;   HD11 - CD1
- HD12
  6539   6538   6541       1    1.0950e+02    2.9288e+02 ;   HD11 - CD1
- HD13
  6540   6538   6541       1    1.0950e+02    2.9288e+02 ;   HD12 - CD1
- HD13
  6542   6544   6545       1    1.2000e+02    4.1840e+02 ;      C - N
- H
  6542   6544   6546       1    1.2190e+02    4.1840e+02 ;      C - N
- CA
  6543   6542   6544       1    1.2290e+02    6.6944e+02 ;      O - C
- N
  6544   6546   6547       1    1.0950e+02    4.1840e+02 ;      N - CA
- HA
  6544   6546   6548       1    1.0970e+02    6.6944e+02 ;      N - CA
- CB
  6544   6546   6553       1    1.1010e+02    5.2718e+02 ;      N - CA
- C
  6545   6544   6546       1    1.1804e+02    4.1840e+02 ;      H - N
- CA
  6546   6548   6549       1    1.0950e+02    4.1840e+02 ;     CA - CB
- HB2
  6546   6548   6550       1    1.0950e+02    4.1840e+02 ;     CA - CB
- HB3
  6546   6548   6551       1    1.0950e+02    4.1840e+02 ;     CA - CB
- OG
  6546   6553   6554       1    1.2040e+02    6.6944e+02 ;     CA - C
- O
  6546   6553   6555       1    1.1660e+02    5.8576e+02 ;     CA - C
- N
  6547   6546   6548       1    1.0950e+02    4.1840e+02 ;     HA - CA
- CB
  6547   6546   6553       1    1.0950e+02    4.1840e+02 ;     HA - CA
- C
  6548   6546   6553       1    1.1110e+02    5.2718e+02 ;     CB - CA
- C
```

6548 6551 6552 1 1.0850e+02 4.6024e+02 ; CB - OG
- HG
6549 6548 6550 1 1.0950e+02 2.9288e+02 ; HB2 - CB
- HB3
6549 6548 6551 1 1.0950e+02 4.1840e+02 ; HB2 - CB
- OG
6550 6548 6551 1 1.0950e+02 4.1840e+02 ; HB3 - CB
- OG
6553 6555 6556 1 1.2000e+02 4.1840e+02 ; C - N
- H
6553 6555 6557 1 1.2190e+02 4.1840e+02 ; C - N
- CA
6554 6553 6555 1 1.2290e+02 6.6944e+02 ; O - C
- N
6555 6557 6558 1 1.0950e+02 4.1840e+02 ; N - CA
- HA
6555 6557 6559 1 1.0970e+02 6.6944e+02 ; N - CA
- CB
6555 6557 6569 1 1.1010e+02 5.2718e+02 ; N - CA
- C
6556 6555 6557 1 1.1804e+02 4.1840e+02 ; H - N
- CA
6557 6559 6560 1 1.0950e+02 4.1840e+02 ; CA - CB
- HB
6557 6559 6561 1 1.0950e+02 3.3472e+02 ; CA - CB
- CG1
6557 6559 6565 1 1.0950e+02 3.3472e+02 ; CA - CB
- CG2
6557 6569 6570 1 1.2040e+02 6.6944e+02 ; CA - C
- O
6557 6569 6571 1 1.1660e+02 5.8576e+02 ; CA - C
- N
6558 6557 6559 1 1.0950e+02 4.1840e+02 ; HA - CA
- CB
6558 6557 6569 1 1.0950e+02 4.1840e+02 ; HA - CA
- C
6559 6557 6569 1 1.1110e+02 5.2718e+02 ; CB - CA
- C
6559 6561 6562 1 1.0950e+02 4.1840e+02 ; CB - CG1
- HG11
6559 6561 6563 1 1.0950e+02 4.1840e+02 ; CB - CG1
- HG12
6559 6561 6564 1 1.0950e+02 4.1840e+02 ; CB - CG1
- HG13
6559 6565 6566 1 1.0950e+02 4.1840e+02 ; CB - CG2
- HG21
6559 6565 6567 1 1.0950e+02 4.1840e+02 ; CB - CG2
- HG22
6559 6565 6568 1 1.0950e+02 4.1840e+02 ; CB - CG2
- HG23
6560 6559 6561 1 1.0950e+02 4.1840e+02 ; HB - CB
- CG1

```
  6560   6559   6565      1    1.0950e+02    4.1840e+02 ;     HB - CB
- CG2
  6561   6559   6565      1    1.0950e+02    3.3472e+02 ;    CG1 - CB
- CG2
  6562   6561   6563      1    1.0950e+02    2.9288e+02 ;   HG11 - CG1
- HG12
  6562   6561   6564      1    1.0950e+02    2.9288e+02 ;   HG11 - CG1
- HG13
  6563   6561   6564      1    1.0950e+02    2.9288e+02 ;   HG12 - CG1
- HG13
  6566   6565   6567      1    1.0950e+02    2.9288e+02 ;   HG21 - CG2
- HG22
  6566   6565   6568      1    1.0950e+02    2.9288e+02 ;   HG21 - CG2
- HG23
  6567   6565   6568      1    1.0950e+02    2.9288e+02 ;   HG22 - CG2
- HG23
  6569   6571   6572      1    1.2000e+02    4.1840e+02 ;      C - N
- H
  6569   6571   6573      1    1.2190e+02    4.1840e+02 ;      C - N
- CA
  6570   6569   6571      1    1.2290e+02    6.6944e+02 ;      O - C
- N
  6571   6573   6574      1    1.0950e+02    4.1840e+02 ;      N - CA
- HA
  6571   6573   6575      1    1.0970e+02    6.6944e+02 ;      N - CA
- CB
  6571   6573   6584      1    1.1010e+02    5.2718e+02 ;      N - CA
- C
  6572   6571   6573      1    1.1804e+02    4.1840e+02 ;      H - N
- CA
  6573   6575   6576      1    1.0950e+02    4.1840e+02 ;     CA - CB
- HB2
  6573   6575   6577      1    1.0950e+02    4.1840e+02 ;     CA - CB
- HB3
  6573   6575   6578      1    1.0950e+02    3.3472e+02 ;     CA - CB
- CG
  6573   6584   6585      1    1.2040e+02    6.6944e+02 ;     CA - C
- O
  6573   6584   6586      1    1.1660e+02    5.8576e+02 ;     CA - C
- N
  6574   6573   6575      1    1.0950e+02    4.1840e+02 ;     HA - CA
- CB
  6574   6573   6584      1    1.0950e+02    4.1840e+02 ;     HA - CA
- C
  6575   6573   6584      1    1.1110e+02    5.2718e+02 ;     CB - CA
- C
  6575   6578   6579      1    1.0950e+02    4.1840e+02 ;     CB - CG
- HG2
  6575   6578   6580      1    1.0950e+02    4.1840e+02 ;     CB - CG
- HG3
  6575   6578   6581      1    1.1110e+02    5.2718e+02 ;     CB - CG
- CD
```

```
  6576   6575   6577       1    1.0950e+02    2.9288e+02 ;    HB2 - CB
- HB3
  6576   6575   6578       1    1.0950e+02    4.1840e+02 ;    HB2 - CB
- CG
  6577   6575   6578       1    1.0950e+02    4.1840e+02 ;    HB3 - CB
- CG
  6578   6581   6582       1    1.1700e+02    5.8576e+02 ;     CG - CD
- OE1
  6578   6581   6583       1    1.1700e+02    5.8576e+02 ;     CG - CD
- OE2
  6579   6578   6580       1    1.0950e+02    2.9288e+02 ;    HG2 - CG
- HG3
  6579   6578   6581       1    1.0950e+02    4.1840e+02 ;    HG2 - CG
- CD
  6580   6578   6581       1    1.0950e+02    4.1840e+02 ;    HG3 - CG
- CD
  6582   6581   6583       1    1.2600e+02    6.6944e+02 ;    OE1 - CD
- OE2
  6584   6586   6587       1    1.2000e+02    4.1840e+02 ;      C - N
- H
  6584   6586   6588       1    1.2190e+02    4.1840e+02 ;      C - N
- CA
  6585   6584   6586       1    1.2290e+02    6.6944e+02 ;      O - C
- N
  6586   6588   6589       1    1.0950e+02    4.1840e+02 ;      N - CA
- HA
  6586   6588   6590       1    1.0970e+02    6.6944e+02 ;      N - CA
- CB
  6586   6588   6600       1    1.1010e+02    5.2718e+02 ;      N - CA
- C
  6587   6586   6588       1    1.1804e+02    4.1840e+02 ;      H - N
- CA
  6588   6590   6591       1    1.0950e+02    4.1840e+02 ;     CA - CB
- HB2
  6588   6590   6592       1    1.0950e+02    4.1840e+02 ;     CA - CB
- HB3
  6588   6590   6593       1    1.0950e+02    3.3472e+02 ;     CA - CB
- CG
  6588   6600   6601       1    1.2040e+02    6.6944e+02 ;     CA - C
- O
  6588   6600   6602       1    1.1660e+02    5.8576e+02 ;     CA - C
- N
  6589   6588   6590       1    1.0950e+02    4.1840e+02 ;     HA - CA
- CB
  6589   6588   6600       1    1.0950e+02    4.1840e+02 ;     HA - CA
- C
  6590   6588   6600       1    1.1110e+02    5.2718e+02 ;     CB - CA
- C
  6590   6593   6594       1    1.0950e+02    4.1840e+02 ;     CB - CG
- HG2
  6590   6593   6595       1    1.0950e+02    4.1840e+02 ;     CB - CG
- HG3
```

```
  6590   6593   6596       1    1.1110e+02    5.2718e+02 ;     CB - CG
- CD
  6591   6590   6592       1    1.0950e+02    2.9288e+02 ;    HB2 - CB
- HB3
  6591   6590   6593       1    1.0950e+02    4.1840e+02 ;    HB2 - CB
- CG
  6592   6590   6593       1    1.0950e+02    4.1840e+02 ;    HB3 - CB
- CG
  6593   6596   6597       1    1.2040e+02    6.6944e+02 ;     CG - CD
- OE1
  6593   6596   6598       1    1.1000e+02    6.6944e+02 ;     CG - CD
- OE2
  6594   6593   6595       1    1.0950e+02    2.9288e+02 ;    HG2 - CG
- HG3
  6594   6593   6596       1    1.0950e+02    4.1840e+02 ;    HG2 - CG
- CD
  6595   6593   6596       1    1.0950e+02    4.1840e+02 ;    HG3 - CG
- CD
  6596   6598   6599       1    1.1300e+02    4.1840e+02 ;     CD - OE2
- HE2
  6597   6596   6598       1    1.2000e+02    6.6944e+02 ;    OE1 - CD
- OE2
  6600   6602   6603       1    1.2000e+02    4.1840e+02 ;      C - N
- H
  6600   6602   6604       1    1.2190e+02    4.1840e+02 ;      C - N
- CA
  6601   6600   6602       1    1.2290e+02    6.6944e+02 ;      O - C
- N
  6602   6604   6605       1    1.0950e+02    4.1840e+02 ;      N - CA
- HA
  6602   6604   6606       1    1.0970e+02    6.6944e+02 ;      N - CA
- CB
  6602   6604   6619       1    1.1010e+02    5.2718e+02 ;      N - CA
- C
  6603   6602   6604       1    1.1804e+02    4.1840e+02 ;      H - N
- CA
  6604   6606   6607       1    1.0950e+02    4.1840e+02 ;     CA - CB
- HB2
  6604   6606   6608       1    1.0950e+02    4.1840e+02 ;     CA - CB
- HB3
  6604   6606   6609       1    1.0950e+02    3.3472e+02 ;     CA - CB
- CG
  6604   6619   6620       1    1.2040e+02    6.6944e+02 ;     CA - C
- O
  6604   6619   6621       1    1.1660e+02    5.8576e+02 ;     CA - C
- N
  6605   6604   6606       1    1.0950e+02    4.1840e+02 ;     HA - CA
- CB
  6605   6604   6619       1    1.0950e+02    4.1840e+02 ;     HA - CA
- C
  6606   6604   6619       1    1.1110e+02    5.2718e+02 ;     CB - CA
- C
```

6606 6609 6610 1 1.0950e+02 4.1840e+02 ; CB - CG
- HG
6606 6609 6611 1 1.0950e+02 3.3472e+02 ; CB - CG
- CD1
6606 6609 6615 1 1.0950e+02 3.3472e+02 ; CB - CG
- CD2
6607 6606 6608 1 1.0950e+02 2.9288e+02 ; HB2 - CB
- HB3
6607 6606 6609 1 1.0950e+02 4.1840e+02 ; HB2 - CB
- CG
6608 6606 6609 1 1.0950e+02 4.1840e+02 ; HB3 - CB
- CG
6609 6611 6612 1 1.0950e+02 4.1840e+02 ; CG - CD1
- HD11
6609 6611 6613 1 1.0950e+02 4.1840e+02 ; CG - CD1
- HD12
6609 6611 6614 1 1.0950e+02 4.1840e+02 ; CG - CD1
- HD13
6609 6615 6616 1 1.0950e+02 4.1840e+02 ; CG - CD2
- HD21
6609 6615 6617 1 1.0950e+02 4.1840e+02 ; CG - CD2
- HD22
6609 6615 6618 1 1.0950e+02 4.1840e+02 ; CG - CD2
- HD23
6610 6609 6611 1 1.0950e+02 4.1840e+02 ; HG - CG
- CD1
6610 6609 6615 1 1.0950e+02 4.1840e+02 ; HG - CG
- CD2
6611 6609 6615 1 1.0950e+02 3.3472e+02 ; CD1 - CG
- CD2
6612 6611 6613 1 1.0950e+02 2.9288e+02 ; HD11 - CD1
- HD12
6612 6611 6614 1 1.0950e+02 2.9288e+02 ; HD11 - CD1
- HD13
6613 6611 6614 1 1.0950e+02 2.9288e+02 ; HD12 - CD1
- HD13
6616 6615 6617 1 1.0950e+02 2.9288e+02 ; HD21 - CD2
- HD22
6616 6615 6618 1 1.0950e+02 2.9288e+02 ; HD21 - CD2
- HD23
6617 6615 6618 1 1.0950e+02 2.9288e+02 ; HD22 - CD2
- HD23
6619 6621 6622 1 1.2000e+02 4.1840e+02 ; C - N
- H
6619 6621 6623 1 1.2190e+02 4.1840e+02 ; C - N
- CA
6620 6619 6621 1 1.2290e+02 6.6944e+02 ; O - C
- N
6621 6623 6624 1 1.0950e+02 4.1840e+02 ; N - CA
- HA
6621 6623 6625 1 1.0970e+02 6.6944e+02 ; N - CA
- CB

6621 6623 6641 1 1.1010e+02 5.2718e+02 ; N - CA
- C
6622 6621 6623 1 1.1804e+02 4.1840e+02 ; H - N
- CA
6623 6625 6626 1 1.0950e+02 4.1840e+02 ; CA - CB
- HB2
6623 6625 6627 1 1.0950e+02 4.1840e+02 ; CA - CB
- HB3
6623 6625 6628 1 1.0950e+02 3.3472e+02 ; CA - CB
- CG
6623 6641 6642 1 1.2040e+02 6.6944e+02 ; CA - C
- O
6623 6641 6643 1 1.1660e+02 5.8576e+02 ; CA - C
- N
6624 6623 6625 1 1.0950e+02 4.1840e+02 ; HA - CA
- CB
6624 6623 6641 1 1.0950e+02 4.1840e+02 ; HA - CA
- C
6625 6623 6641 1 1.1110e+02 5.2718e+02 ; CB - CA
- C
6625 6628 6629 1 1.0950e+02 4.1840e+02 ; CB - CG
- HG2
6625 6628 6630 1 1.0950e+02 4.1840e+02 ; CB - CG
- HG3
6625 6628 6631 1 1.0950e+02 3.3472e+02 ; CB - CG
- CD
6626 6625 6627 1 1.0950e+02 2.9288e+02 ; HB2 - CB
- HB3
6626 6625 6628 1 1.0950e+02 4.1840e+02 ; HB2 - CB
- CG
6627 6625 6628 1 1.0950e+02 4.1840e+02 ; HB3 - CB
- CG
6628 6631 6632 1 1.0950e+02 4.1840e+02 ; CG - CD
- HD2
6628 6631 6633 1 1.0950e+02 4.1840e+02 ; CG - CD
- HD3
6628 6631 6634 1 1.0950e+02 3.3472e+02 ; CG - CD
- CE
6629 6628 6630 1 1.0950e+02 2.9288e+02 ; HG2 - CG
- HG3
6629 6628 6631 1 1.0950e+02 4.1840e+02 ; HG2 - CG
- CD
6630 6628 6631 1 1.0950e+02 4.1840e+02 ; HG3 - CG
- CD
6631 6634 6635 1 1.0950e+02 4.1840e+02 ; CD - CE
- HE2
6631 6634 6636 1 1.0950e+02 4.1840e+02 ; CD - CE
- HE3
6631 6634 6637 1 1.1120e+02 6.6944e+02 ; CD - CE
- NZ
6632 6631 6633 1 1.0950e+02 2.9288e+02 ; HD2 - CD
- HD3

```
  6632   6631   6634       1    1.0950e+02    4.1840e+02 ;    HD2 - CD
- CE
  6633   6631   6634       1    1.0950e+02    4.1840e+02 ;    HD3 - CD
- CE
  6634   6637   6638       1    1.0950e+02    4.1840e+02 ;     CE - NZ
- HZ1
  6634   6637   6639       1    1.0950e+02    4.1840e+02 ;     CE - NZ
- HZ2
  6634   6637   6640       1    1.0950e+02    4.1840e+02 ;     CE - NZ
- HZ3
  6635   6634   6636       1    1.0950e+02    2.9288e+02 ;    HE2 - CE
- HE3
  6635   6634   6637       1    1.0950e+02    4.1840e+02 ;    HE2 - CE
- NZ
  6636   6634   6637       1    1.0950e+02    4.1840e+02 ;    HE3 - CE
- NZ
  6638   6637   6639       1    1.0950e+02    2.9288e+02 ;    HZ1 - NZ
- HZ2
  6638   6637   6640       1    1.0950e+02    2.9288e+02 ;    HZ1 - NZ
- HZ3
  6639   6637   6640       1    1.0950e+02    2.9288e+02 ;    HZ2 - NZ
- HZ3
  6641   6643   6644       1    1.2000e+02    4.1840e+02 ;      C - N
- H
  6641   6643   6645       1    1.2190e+02    4.1840e+02 ;      C - N
- CA
  6642   6641   6643       1    1.2290e+02    6.6944e+02 ;      O - C
- N
  6643   6645   6646       1    1.0950e+02    4.1840e+02 ;      N - CA
- HA
  6643   6645   6647       1    1.0970e+02    6.6944e+02 ;      N - CA
- CB
  6643   6645   6665       1    1.1010e+02    5.2718e+02 ;      N - CA
- C
  6644   6643   6645       1    1.1804e+02    4.1840e+02 ;      H - N
- CA
  6645   6647   6648       1    1.0950e+02    4.1840e+02 ;     CA - CB
- HB2
  6645   6647   6649       1    1.0950e+02    4.1840e+02 ;     CA - CB
- HB3
  6645   6647   6650       1    1.0950e+02    3.3472e+02 ;     CA - CB
- CG
  6645   6665   6666       1    1.2040e+02    6.6944e+02 ;     CA - C
- O
  6645   6665   6667       1    1.1660e+02    5.8576e+02 ;     CA - C
- N
  6646   6645   6647       1    1.0950e+02    4.1840e+02 ;     HA - CA
- CB
  6646   6645   6665       1    1.0950e+02    4.1840e+02 ;     HA - CA
- C
  6647   6645   6665       1    1.1110e+02    5.2718e+02 ;     CB - CA
- C
```

6647 6650 6651 1 1.0950e+02 4.1840e+02 ; CB - CG
- HG2
6647 6650 6652 1 1.0950e+02 4.1840e+02 ; CB - CG
- HG3
6647 6650 6653 1 1.0950e+02 3.3472e+02 ; CB - CG
- CD
6648 6647 6649 1 1.0950e+02 2.9288e+02 ; HB2 - CB
- HB3
6648 6647 6650 1 1.0950e+02 4.1840e+02 ; HB2 - CB
- CG
6649 6647 6650 1 1.0950e+02 4.1840e+02 ; HB3 - CB
- CG
6650 6653 6654 1 1.0950e+02 4.1840e+02 ; CG - CD
- HD2
6650 6653 6655 1 1.0950e+02 4.1840e+02 ; CG - CD
- HD3
6650 6653 6656 1 1.1120e+02 6.6944e+02 ; CG - CD
- NE
6651 6650 6652 1 1.0950e+02 2.9288e+02 ; HG2 - CG
- HG3
6651 6650 6653 1 1.0950e+02 4.1840e+02 ; HG2 - CG
- CD
6652 6650 6653 1 1.0950e+02 4.1840e+02 ; HG3 - CG
- CD
6653 6656 6657 1 1.1840e+02 4.1840e+02 ; CD - NE
- HE
6653 6656 6658 1 1.2320e+02 4.1840e+02 ; CD - NE
- CZ
6654 6653 6655 1 1.0950e+02 2.9288e+02 ; HD2 - CD
- HD3
6654 6653 6656 1 1.0950e+02 4.1840e+02 ; HD2 - CD
- NE
6655 6653 6656 1 1.0950e+02 4.1840e+02 ; HD3 - CD
- NE
6656 6658 6659 1 1.2000e+02 5.8576e+02 ; NE - CZ
- NH1
6656 6658 6662 1 1.2000e+02 5.8576e+02 ; NE - CZ
- NH2
6657 6656 6658 1 1.2000e+02 4.1840e+02 ; HE - NE
- CZ
6658 6659 6660 1 1.2000e+02 4.1840e+02 ; CZ - NH1
- HH11
6658 6659 6661 1 1.2000e+02 4.1840e+02 ; CZ - NH1
- HH12
6658 6662 6663 1 1.2000e+02 4.1840e+02 ; CZ - NH2
- HH21
6658 6662 6664 1 1.2000e+02 4.1840e+02 ; CZ - NH2
- HH22
6659 6658 6662 1 1.2000e+02 5.8576e+02 ; NH1 - CZ
- NH2
6660 6659 6661 1 1.2000e+02 2.9288e+02 ; HH11 - NH1
- HH12

```
  6663   6662   6664       1    1.2000e+02    2.9288e+02 ;   HH21 - NH2
- HH22
  6665   6667   6668       1    1.2000e+02    4.1840e+02 ;      C - N
- H
  6665   6667   6669       1    1.2190e+02    4.1840e+02 ;      C - N
- CA
  6666   6665   6667       1    1.2290e+02    6.6944e+02 ;      O - C
- N
  6667   6669   6670       1    1.0950e+02    4.1840e+02 ;      N - CA
- HA
  6667   6669   6671       1    1.0970e+02    6.6944e+02 ;      N - CA
- CB
  6667   6669   6684       1    1.1010e+02    5.2718e+02 ;      N - CA
- C
  6668   6667   6669       1    1.1804e+02    4.1840e+02 ;      H - N
- CA
  6669   6671   6672       1    1.0950e+02    4.1840e+02 ;     CA - CB
- HB
  6669   6671   6673       1    1.0950e+02    3.3472e+02 ;     CA - CB
- CG2
  6669   6671   6677       1    1.0950e+02    3.3472e+02 ;     CA - CB
- CG1
  6669   6684   6685       1    1.2040e+02    6.6944e+02 ;     CA - C
- O
  6669   6684   6686       1    1.1660e+02    5.8576e+02 ;     CA - C
- N
  6670   6669   6671       1    1.0950e+02    4.1840e+02 ;     HA - CA
- CB
  6670   6669   6684       1    1.0950e+02    4.1840e+02 ;     HA - CA
- C
  6671   6669   6684       1    1.1110e+02    5.2718e+02 ;     CB - CA
- C
  6671   6673   6674       1    1.0950e+02    4.1840e+02 ;     CB - CG2
- HG21
  6671   6673   6675       1    1.0950e+02    4.1840e+02 ;     CB - CG2
- HG22
  6671   6673   6676       1    1.0950e+02    4.1840e+02 ;     CB - CG2
- HG23
  6671   6677   6678       1    1.0950e+02    4.1840e+02 ;     CB - CG1
- HG12
  6671   6677   6679       1    1.0950e+02    4.1840e+02 ;     CB - CG1
- HG13
  6671   6677   6680       1    1.0950e+02    3.3472e+02 ;     CB - CG1
- CD1
  6672   6671   6673       1    1.0950e+02    4.1840e+02 ;     HB - CB
- CG2
  6672   6671   6677       1    1.0950e+02    4.1840e+02 ;     HB - CB
- CG1
  6673   6671   6677       1    1.0950e+02    3.3472e+02 ;    CG2 - CB
- CG1
  6674   6673   6675       1    1.0950e+02    2.9288e+02 ;   HG21 - CG2
- HG22
```

```
   6674   6673   6676      1    1.0950e+02    2.9288e+02 ;   HG21 - CG2
 - HG23
   6675   6673   6676      1    1.0950e+02    2.9288e+02 ;   HG22 - CG2
 - HG23
   6677   6680   6681      1    1.0950e+02    4.1840e+02 ;    CG1 - CD1
 - HD11
   6677   6680   6682      1    1.0950e+02    4.1840e+02 ;    CG1 - CD1
 - HD12
   6677   6680   6683      1    1.0950e+02    4.1840e+02 ;    CG1 - CD1
 - HD13
   6678   6677   6679      1    1.0950e+02    2.9288e+02 ;   HG12 - CG1
 - HG13
   6678   6677   6680      1    1.0950e+02    4.1840e+02 ;   HG12 - CG1
 - CD1
   6679   6677   6680      1    1.0950e+02    4.1840e+02 ;   HG13 - CG1
 - CD1
   6681   6680   6682      1    1.0950e+02    2.9288e+02 ;   HD11 - CD1
 - HD12
   6681   6680   6683      1    1.0950e+02    2.9288e+02 ;   HD11 - CD1
 - HD13
   6682   6680   6683      1    1.0950e+02    2.9288e+02 ;   HD12 - CD1
 - HD13
   6684   6686   6687      1    1.2000e+02    4.1840e+02 ;      C - N
 - H
   6684   6686   6688      1    1.2190e+02    4.1840e+02 ;      C - N
 - CA
   6685   6684   6686      1    1.2290e+02    6.6944e+02 ;      O - C
 - N
   6686   6688   6689      1    1.0950e+02    4.1840e+02 ;      N - CA
 - HA
   6686   6688   6690      1    1.0970e+02    6.6944e+02 ;      N - CA
 - CB
   6686   6688   6704      1    1.1010e+02    5.2718e+02 ;      N - CA
 - C
   6687   6686   6688      1    1.1804e+02    4.1840e+02 ;      H - N
 - CA
   6688   6690   6691      1    1.0950e+02    4.1840e+02 ;     CA - CB
 - HB2
   6688   6690   6692      1    1.0950e+02    4.1840e+02 ;     CA - CB
 - HB3
   6688   6690   6693      1    1.1400e+02    5.2718e+02 ;     CA - CB
 - CG
   6688   6704   6705      1    1.2040e+02    6.6944e+02 ;     CA - C
 - O
   6688   6704   6706      1    1.1660e+02    5.8576e+02 ;     CA - C
 - N
   6689   6688   6690      1    1.0950e+02    4.1840e+02 ;     HA - CA
 - CB
   6689   6688   6704      1    1.0950e+02    4.1840e+02 ;     HA - CA
 - C
   6690   6688   6704      1    1.1110e+02    5.2718e+02 ;     CB - CA
 - C
```

| | | | | | | | |
|---|---|---|---|---|---|---|---|
| 6690 | 6693 | 6694 | 1 | 1.2000e+02 | 5.8576e+02 | ; | CB - CG - CD1 |
| 6690 | 6693 | 6702 | 1 | 1.2000e+02 | 5.8576e+02 | ; | CB - CG - CD2 |
| 6691 | 6690 | 6692 | 1 | 1.0950e+02 | 2.9288e+02 | ; | HB2 - CB - HB3 |
| 6691 | 6690 | 6693 | 1 | 1.0950e+02 | 4.1840e+02 | ; | HB2 - CB - CG |
| 6692 | 6690 | 6693 | 1 | 1.0950e+02 | 4.1840e+02 | ; | HB3 - CB - CG |
| 6693 | 6694 | 6695 | 1 | 1.2000e+02 | 4.1840e+02 | ; | CG - CD1 - HD1 |
| 6693 | 6694 | 6696 | 1 | 1.2000e+02 | 5.2718e+02 | ; | CG - CD1 - CE1 |
| 6693 | 6702 | 6700 | 1 | 1.2000e+02 | 5.2718e+02 | ; | CG - CD2 - CE2 |
| 6693 | 6702 | 6703 | 1 | 1.2000e+02 | 4.1840e+02 | ; | CG - CD2 - HD2 |
| 6694 | 6693 | 6702 | 1 | 1.2000e+02 | 5.2718e+02 | ; | CD1 - CG - CD2 |
| 6694 | 6696 | 6697 | 1 | 1.2000e+02 | 4.1840e+02 | ; | CD1 - CE1 - HE1 |
| 6694 | 6696 | 6698 | 1 | 1.2000e+02 | 5.2718e+02 | ; | CD1 - CE1 - CZ |
| 6695 | 6694 | 6696 | 1 | 1.2000e+02 | 4.1840e+02 | ; | HD1 - CD1 - CE1 |
| 6696 | 6698 | 6699 | 1 | 1.2000e+02 | 4.1840e+02 | ; | CE1 - CZ - HZ |
| 6696 | 6698 | 6700 | 1 | 1.2000e+02 | 5.2718e+02 | ; | CE1 - CZ - CE2 |
| 6697 | 6696 | 6698 | 1 | 1.2000e+02 | 4.1840e+02 | ; | HE1 - CE1 - CZ |
| 6698 | 6700 | 6701 | 1 | 1.2000e+02 | 4.1840e+02 | ; | CZ - CE2 - HE2 |
| 6698 | 6700 | 6702 | 1 | 1.2000e+02 | 5.2718e+02 | ; | CZ - CE2 - CD2 |
| 6699 | 6698 | 6700 | 1 | 1.2000e+02 | 4.1840e+02 | ; | HZ - CZ - CE2 |
| 6700 | 6702 | 6703 | 1 | 1.2000e+02 | 4.1840e+02 | ; | CE2 - CD2 - HD2 |
| 6701 | 6700 | 6702 | 1 | 1.2000e+02 | 4.1840e+02 | ; | HE2 - CE2 - CD2 |
| 6704 | 6706 | 6707 | 1 | 1.2000e+02 | 4.1840e+02 | ; | C - N - H |
| 6704 | 6706 | 6708 | 1 | 1.2190e+02 | 4.1840e+02 | ; | C - N - CA |
| 6705 | 6704 | 6706 | 1 | 1.2290e+02 | 6.6944e+02 | ; | O - C - N |
| 6706 | 6708 | 6709 | 1 | 1.0950e+02 | 4.1840e+02 | ; | N - CA - HA2 |
| 6706 | 6708 | 6710 | 1 | 1.0950e+02 | 4.1840e+02 | ; | N - CA - HA3 |

| | | | | | | |
|---|---|---|---|---|---|---|
| 6706 | 6708 | 6711 | 1 | 1.1010e+02 | 5.2718e+02 ; | N - CA - C |
| 6707 | 6706 | 6708 | 1 | 1.1804e+02 | 4.1840e+02 ; | H - N - CA |
| 6708 | 6711 | 6712 | 1 | 1.2040e+02 | 6.6944e+02 ; | CA - C - O |
| 6708 | 6711 | 6713 | 1 | 1.1660e+02 | 5.8576e+02 ; | CA - C - N |
| 6709 | 6708 | 6710 | 1 | 1.0950e+02 | 2.9288e+02 ; | HA2 - CA - HA3 |
| 6709 | 6708 | 6711 | 1 | 1.0950e+02 | 4.1840e+02 ; | HA2 - CA - C |
| 6710 | 6708 | 6711 | 1 | 1.0950e+02 | 4.1840e+02 ; | HA3 - CA - C |
| 6711 | 6713 | 6714 | 1 | 1.2000e+02 | 4.1840e+02 ; | C - N - H |
| 6711 | 6713 | 6715 | 1 | 1.2190e+02 | 4.1840e+02 ; | C - N - CA |
| 6712 | 6711 | 6713 | 1 | 1.2290e+02 | 6.6944e+02 ; | O - C - N |
| 6713 | 6715 | 6716 | 1 | 1.0950e+02 | 4.1840e+02 ; | N - CA - HA |
| 6713 | 6715 | 6717 | 1 | 1.0970e+02 | 6.6944e+02 ; | N - CA - CB |
| 6713 | 6715 | 6735 | 1 | 1.1010e+02 | 5.2718e+02 ; | N - CA - C |
| 6714 | 6713 | 6715 | 1 | 1.1804e+02 | 4.1840e+02 ; | H - N - CA |
| 6715 | 6717 | 6718 | 1 | 1.0950e+02 | 4.1840e+02 ; | CA - CB - HB2 |
| 6715 | 6717 | 6719 | 1 | 1.0950e+02 | 4.1840e+02 ; | CA - CB - HB3 |
| 6715 | 6717 | 6720 | 1 | 1.0950e+02 | 3.3472e+02 ; | CA - CB - CG |
| 6715 | 6735 | 6736 | 1 | 1.2040e+02 | 6.6944e+02 ; | CA - C - O |
| 6715 | 6735 | 6737 | 1 | 1.1660e+02 | 5.8576e+02 ; | CA - C - N |
| 6716 | 6715 | 6717 | 1 | 1.0950e+02 | 4.1840e+02 ; | HA - CA - CB |
| 6716 | 6715 | 6735 | 1 | 1.0950e+02 | 4.1840e+02 ; | HA - CA - C |
| 6717 | 6715 | 6735 | 1 | 1.1110e+02 | 5.2718e+02 ; | CB - CA - C |
| 6717 | 6720 | 6721 | 1 | 1.0950e+02 | 4.1840e+02 ; | CB - CG - HG2 |
| 6717 | 6720 | 6722 | 1 | 1.0950e+02 | 4.1840e+02 ; | CB - CG - HG3 |
| 6717 | 6720 | 6723 | 1 | 1.0950e+02 | 3.3472e+02 ; | CB - CG - CD |
| 6718 | 6717 | 6719 | 1 | 1.0950e+02 | 2.9288e+02 ; | HB2 - CB - HB3 |

```
  6718   6717   6720      1    1.0950e+02    4.1840e+02 ;    HB2 - CB
- CG
  6719   6717   6720      1    1.0950e+02    4.1840e+02 ;    HB3 - CB
- CG
  6720   6723   6724      1    1.0950e+02    4.1840e+02 ;     CG - CD
- HD2
  6720   6723   6725      1    1.0950e+02    4.1840e+02 ;     CG - CD
- HD3
  6720   6723   6726      1    1.1120e+02    6.6944e+02 ;     CG - CD
- NE
  6721   6720   6722      1    1.0950e+02    2.9288e+02 ;    HG2 - CG
- HG3
  6721   6720   6723      1    1.0950e+02    4.1840e+02 ;    HG2 - CG
- CD
  6722   6720   6723      1    1.0950e+02    4.1840e+02 ;    HG3 - CG
- CD
  6723   6726   6727      1    1.1840e+02    4.1840e+02 ;     CD - NE
- HE
  6723   6726   6728      1    1.2320e+02    4.1840e+02 ;     CD - NE
- CZ
  6724   6723   6725      1    1.0950e+02    2.9288e+02 ;    HD2 - CD
- HD3
  6724   6723   6726      1    1.0950e+02    4.1840e+02 ;    HD2 - CD
- NE
  6725   6723   6726      1    1.0950e+02    4.1840e+02 ;    HD3 - CD
- NE
  6726   6728   6729      1    1.2000e+02    5.8576e+02 ;     NE - CZ
- NH1
  6726   6728   6732      1    1.2000e+02    5.8576e+02 ;     NE - CZ
- NH2
  6727   6726   6728      1    1.2000e+02    4.1840e+02 ;     HE - NE
- CZ
  6728   6729   6730      1    1.2000e+02    4.1840e+02 ;     CZ - NH1
- HH11
  6728   6729   6731      1    1.2000e+02    4.1840e+02 ;     CZ - NH1
- HH12
  6728   6732   6733      1    1.2000e+02    4.1840e+02 ;     CZ - NH2
- HH21
  6728   6732   6734      1    1.2000e+02    4.1840e+02 ;     CZ - NH2
- HH22
  6729   6728   6732      1    1.2000e+02    5.8576e+02 ;    NH1 - CZ
- NH2
  6730   6729   6731      1    1.2000e+02    2.9288e+02 ;   HH11 - NH1
- HH12
  6733   6732   6734      1    1.2000e+02    2.9288e+02 ;   HH21 - NH2
- HH22
  6735   6737   6738      1    1.2000e+02    4.1840e+02 ;      C - N
- H
  6735   6737   6739      1    1.2190e+02    4.1840e+02 ;      C - N
- CA
  6736   6735   6737      1    1.2290e+02    6.6944e+02 ;      O - C
- N
```

6737 6739 6740 1 1.0950e+02 4.1840e+02 ; N - CA
- HA
6737 6739 6741 1 1.0970e+02 6.6944e+02 ; N - CA
- CB
6737 6739 6747 1 1.1010e+02 5.2718e+02 ; N - CA
- C
6738 6737 6739 1 1.1804e+02 4.1840e+02 ; H - N
- CA
6739 6741 6742 1 1.0950e+02 4.1840e+02 ; CA - CB
- HB2
6739 6741 6743 1 1.0950e+02 4.1840e+02 ; CA - CB
- HB3
6739 6741 6744 1 1.1110e+02 5.2718e+02 ; CA - CB
- CG
6739 6747 6748 1 1.2040e+02 6.6944e+02 ; CA - C
- O
6739 6747 6749 1 1.1660e+02 5.8576e+02 ; CA - C
- N
6740 6739 6741 1 1.0950e+02 4.1840e+02 ; HA - CA
- CB
6740 6739 6747 1 1.0950e+02 4.1840e+02 ; HA - CA
- C
6741 6739 6747 1 1.1110e+02 5.2718e+02 ; CB - CA
- C
6741 6744 6745 1 1.1700e+02 5.8576e+02 ; CB - CG
- OD1
6741 6744 6746 1 1.1700e+02 5.8576e+02 ; CB - CG
- OD2
6742 6741 6743 1 1.0950e+02 2.9288e+02 ; HB2 - CB
- HB3
6742 6741 6744 1 1.0950e+02 4.1840e+02 ; HB2 - CB
- CG
6743 6741 6744 1 1.0950e+02 4.1840e+02 ; HB3 - CB
- CG
6745 6744 6746 1 1.2600e+02 6.6944e+02 ; OD1 - CG
- OD2
6747 6749 6750 1 1.2000e+02 4.1840e+02 ; C - N
- H
6747 6749 6751 1 1.2190e+02 4.1840e+02 ; C - N
- CA
6748 6747 6749 1 1.2290e+02 6.6944e+02 ; O - C
- N
6749 6751 6752 1 1.0950e+02 4.1840e+02 ; N - CA
- HA
6749 6751 6753 1 1.0970e+02 6.6944e+02 ; N - CA
- CB
6749 6751 6759 1 1.1010e+02 5.2718e+02 ; N - CA
- C
6750 6749 6751 1 1.1804e+02 4.1840e+02 ; H - N
- CA
6751 6753 6754 1 1.0950e+02 4.1840e+02 ; CA - CB
- HB2

| | | | | | | | |
|---|---|---|---|---|---|---|---|
| 6751 | 6753 | 6755 | 1 | 1.0950e+02 | 4.1840e+02 | ; | CA - CB - HB3 |
| 6751 | 6753 | 6756 | 1 | 1.1110e+02 | 5.2718e+02 | ; | CA - CB - CG |
| 6751 | 6759 | 6760 | 1 | 1.2040e+02 | 6.6944e+02 | ; | CA - C - O |
| 6751 | 6759 | 6761 | 1 | 1.1660e+02 | 5.8576e+02 | ; | CA - C - N |
| 6752 | 6751 | 6753 | 1 | 1.0950e+02 | 4.1840e+02 | ; | HA - CA - CB |
| 6752 | 6751 | 6759 | 1 | 1.0950e+02 | 4.1840e+02 | ; | HA - CA - C |
| 6753 | 6751 | 6759 | 1 | 1.1110e+02 | 5.2718e+02 | ; | CB - CA - C |
| 6753 | 6756 | 6757 | 1 | 1.1700e+02 | 5.8576e+02 | ; | CB - CG - OD1 |
| 6753 | 6756 | 6758 | 1 | 1.1700e+02 | 5.8576e+02 | ; | CB - CG - OD2 |
| 6754 | 6753 | 6755 | 1 | 1.0950e+02 | 2.9288e+02 | ; | HB2 - CB - HB3 |
| 6754 | 6753 | 6756 | 1 | 1.0950e+02 | 4.1840e+02 | ; | HB2 - CB - CG |
| 6755 | 6753 | 6756 | 1 | 1.0950e+02 | 4.1840e+02 | ; | HB3 - CB - CG |
| 6757 | 6756 | 6758 | 1 | 1.2600e+02 | 6.6944e+02 | ; | OD1 - CG - OD2 |
| 6759 | 6761 | 6762 | 1 | 1.2000e+02 | 4.1840e+02 | ; | C - N - H |
| 6759 | 6761 | 6763 | 1 | 1.2190e+02 | 4.1840e+02 | ; | C - N - CA |
| 6760 | 6759 | 6761 | 1 | 1.2290e+02 | 6.6944e+02 | ; | O - C - N |
| 6761 | 6763 | 6764 | 1 | 1.0950e+02 | 4.1840e+02 | ; | N - CA - HA |
| 6761 | 6763 | 6765 | 1 | 1.0970e+02 | 6.6944e+02 | ; | N - CA - CB |
| 6761 | 6763 | 6778 | 1 | 1.1010e+02 | 5.2718e+02 | ; | N - CA - C |
| 6762 | 6761 | 6763 | 1 | 1.1804e+02 | 4.1840e+02 | ; | H - N - CA |
| 6763 | 6765 | 6766 | 1 | 1.0950e+02 | 4.1840e+02 | ; | CA - CB - HB |
| 6763 | 6765 | 6767 | 1 | 1.0950e+02 | 3.3472e+02 | ; | CA - CB - CG2 |
| 6763 | 6765 | 6771 | 1 | 1.0950e+02 | 3.3472e+02 | ; | CA - CB - CG1 |
| 6763 | 6778 | 6779 | 1 | 1.2040e+02 | 6.6944e+02 | ; | CA - C - O |
| 6763 | 6778 | 6780 | 1 | 1.1660e+02 | 5.8576e+02 | ; | CA - C - N |
| 6764 | 6763 | 6765 | 1 | 1.0950e+02 | 4.1840e+02 | ; | HA - CA - CB |

```
  6764   6763   6778      1    1.0950e+02    4.1840e+02 ;     HA - CA
- C
  6765   6763   6778      1    1.1110e+02    5.2718e+02 ;     CB - CA
- C
  6765   6767   6768      1    1.0950e+02    4.1840e+02 ;     CB - CG2
- HG21
  6765   6767   6769      1    1.0950e+02    4.1840e+02 ;     CB - CG2
- HG22
  6765   6767   6770      1    1.0950e+02    4.1840e+02 ;     CB - CG2
- HG23
  6765   6771   6772      1    1.0950e+02    4.1840e+02 ;     CB - CG1
- HG12
  6765   6771   6773      1    1.0950e+02    4.1840e+02 ;     CB - CG1
- HG13
  6765   6771   6774      1    1.0950e+02    3.3472e+02 ;     CB - CG1
- CD1
  6766   6765   6767      1    1.0950e+02    4.1840e+02 ;     HB - CB
- CG2
  6766   6765   6771      1    1.0950e+02    4.1840e+02 ;     HB - CB
- CG1
  6767   6765   6771      1    1.0950e+02    3.3472e+02 ;    CG2 - CB
- CG1
  6768   6767   6769      1    1.0950e+02    2.9288e+02 ;   HG21 - CG2
- HG22
  6768   6767   6770      1    1.0950e+02    2.9288e+02 ;   HG21 - CG2
- HG23
  6769   6767   6770      1    1.0950e+02    2.9288e+02 ;   HG22 - CG2
- HG23
  6771   6774   6775      1    1.0950e+02    4.1840e+02 ;    CG1 - CD1
- HD11
  6771   6774   6776      1    1.0950e+02    4.1840e+02 ;    CG1 - CD1
- HD12
  6771   6774   6777      1    1.0950e+02    4.1840e+02 ;    CG1 - CD1
- HD13
  6772   6771   6773      1    1.0950e+02    2.9288e+02 ;   HG12 - CG1
- HG13
  6772   6771   6774      1    1.0950e+02    4.1840e+02 ;   HG12 - CG1
- CD1
  6773   6771   6774      1    1.0950e+02    4.1840e+02 ;   HG13 - CG1
- CD1
  6775   6774   6776      1    1.0950e+02    2.9288e+02 ;   HD11 - CD1
- HD12
  6775   6774   6777      1    1.0950e+02    2.9288e+02 ;   HD11 - CD1
- HD13
  6776   6774   6777      1    1.0950e+02    2.9288e+02 ;   HD12 - CD1
- HD13
  6778   6780   6781      1    1.2000e+02    4.1840e+02 ;      C - N
- H
  6778   6780   6782      1    1.2190e+02    4.1840e+02 ;      C - N
- CA
  6779   6778   6780      1    1.2290e+02    6.6944e+02 ;      O - C
- N
```

```
  6780   6782   6783      1    1.0950e+02    4.1840e+02 ;      N - CA
- HA
  6780   6782   6784      1    1.0970e+02    6.6944e+02 ;      N - CA
- CB
  6780   6782   6793      1    1.1010e+02    5.2718e+02 ;      N - CA
- C
  6781   6780   6782      1    1.1804e+02    4.1840e+02 ;      H - N
- CA
  6782   6784   6785      1    1.0950e+02    4.1840e+02 ;     CA - CB
- HB2
  6782   6784   6786      1    1.0950e+02    4.1840e+02 ;     CA - CB
- HB3
  6782   6784   6787      1    1.0950e+02    3.3472e+02 ;     CA - CB
- CG
  6782   6793   6794      1    1.2040e+02    6.6944e+02 ;     CA - C
- O
  6782   6793   6795      1    1.1660e+02    5.8576e+02 ;     CA - C
- N
  6783   6782   6784      1    1.0950e+02    4.1840e+02 ;     HA - CA
- CB
  6783   6782   6793      1    1.0950e+02    4.1840e+02 ;     HA - CA
- C
  6784   6782   6793      1    1.1110e+02    5.2718e+02 ;     CB - CA
- C
  6784   6787   6788      1    1.0950e+02    4.1840e+02 ;     CB - CG
- HG2
  6784   6787   6789      1    1.0950e+02    4.1840e+02 ;     CB - CG
- HG3
  6784   6787   6790      1    1.1110e+02    5.2718e+02 ;     CB - CG
- CD
  6785   6784   6786      1    1.0950e+02    2.9288e+02 ;    HB2 - CB
- HB3
  6785   6784   6787      1    1.0950e+02    4.1840e+02 ;    HB2 - CB
- CG
  6786   6784   6787      1    1.0950e+02    4.1840e+02 ;    HB3 - CB
- CG
  6787   6790   6791      1    1.1700e+02    5.8576e+02 ;     CG - CD
- OE1
  6787   6790   6792      1    1.1700e+02    5.8576e+02 ;     CG - CD
- OE2
  6788   6787   6789      1    1.0950e+02    2.9288e+02 ;    HG2 - CG
- HG3
  6788   6787   6790      1    1.0950e+02    4.1840e+02 ;    HG2 - CG
- CD
  6789   6787   6790      1    1.0950e+02    4.1840e+02 ;    HG3 - CG
- CD
  6791   6790   6792      1    1.2600e+02    6.6944e+02 ;    OE1 - CD
- OE2
  6793   6795   6796      1    1.2000e+02    4.1840e+02 ;      C - N
- H
  6793   6795   6797      1    1.2190e+02    4.1840e+02 ;      C - N
- CA
```

6794 6793 6795 1 1.2290e+02 6.6944e+02 ; O - C
- N
6795 6797 6798 1 1.0950e+02 4.1840e+02 ; N - CA
- HA
6795 6797 6799 1 1.0970e+02 6.6944e+02 ; N - CA
- CB
6795 6797 6807 1 1.1010e+02 5.2718e+02 ; N - CA
- C
6796 6795 6797 1 1.1804e+02 4.1840e+02 ; H - N
- CA
6797 6799 6800 1 1.0950e+02 4.1840e+02 ; CA - CB
- HB2
6797 6799 6801 1 1.0950e+02 4.1840e+02 ; CA - CB
- HB3
6797 6799 6802 1 1.1110e+02 5.2718e+02 ; CA - CB
- CG
6797 6807 6808 1 1.2040e+02 6.6944e+02 ; CA - C
- O
6797 6807 6809 1 1.1660e+02 5.8576e+02 ; CA - C
- N
6798 6797 6799 1 1.0950e+02 4.1840e+02 ; HA - CA
- CB
6798 6797 6807 1 1.0950e+02 4.1840e+02 ; HA - CA
- C
6799 6797 6807 1 1.1110e+02 5.2718e+02 ; CB - CA
- C
6799 6802 6803 1 1.2040e+02 6.6944e+02 ; CB - CG
- OD1
6799 6802 6804 1 1.1660e+02 5.8576e+02 ; CB - CG
- ND2
6800 6799 6801 1 1.0950e+02 2.9288e+02 ; HB2 - CB
- HB3
6800 6799 6802 1 1.0950e+02 4.1840e+02 ; HB2 - CB
- CG
6801 6799 6802 1 1.0950e+02 4.1840e+02 ; HB3 - CB
- CG
6802 6804 6805 1 1.2000e+02 4.1840e+02 ; CG - ND2
- HD21
6802 6804 6806 1 1.2000e+02 4.1840e+02 ; CG - ND2
- HD22
6803 6802 6804 1 1.2290e+02 6.6944e+02 ; OD1 - CG
- ND2
6805 6804 6806 1 1.2000e+02 2.9288e+02 ; HD21 - ND2
- HD22
6807 6809 6810 1 1.2190e+02 4.1840e+02 ; C - N
- CD
6807 6809 6819 1 1.2190e+02 4.1840e+02 ; C - N
- CA
6808 6807 6809 1 1.2290e+02 6.6944e+02 ; O - C
- N
6809 6810 6811 1 1.0950e+02 4.1840e+02 ; N - CD
- HD2

6809 6810 6812 1 1.0950e+02 4.1840e+02 ; N - CD
- HD3
6809 6810 6813 1 1.0970e+02 6.6944e+02 ; N - CD
- CG
6809 6819 6816 1 1.0970e+02 6.6944e+02 ; N - CA
- CB
6809 6819 6820 1 1.0950e+02 4.1840e+02 ; N - CA
- HA
6809 6819 6821 1 1.1010e+02 5.2718e+02 ; N - CA
- C
6810 6809 6819 1 1.1800e+02 4.1840e+02 ; CD - N
- CA
6810 6813 6814 1 1.0950e+02 4.1840e+02 ; CD - CG
- HG2
6810 6813 6815 1 1.0950e+02 4.1840e+02 ; CD - CG
- HG3
6810 6813 6816 1 1.0950e+02 3.3472e+02 ; CD - CG
- CB
6811 6810 6812 1 1.0950e+02 2.9288e+02 ; HD2 - CD
- HD3
6811 6810 6813 1 1.0950e+02 4.1840e+02 ; HD2 - CD
- CG
6812 6810 6813 1 1.0950e+02 4.1840e+02 ; HD3 - CD
- CG
6813 6816 6817 1 1.0950e+02 4.1840e+02 ; CG - CB
- HB2
6813 6816 6818 1 1.0950e+02 4.1840e+02 ; CG - CB
- HB3
6813 6816 6819 1 1.0950e+02 3.3472e+02 ; CG - CB
- CA
6814 6813 6815 1 1.0950e+02 2.9288e+02 ; HG2 - CG
- HG3
6814 6813 6816 1 1.0950e+02 4.1840e+02 ; HG2 - CG
- CB
6815 6813 6816 1 1.0950e+02 4.1840e+02 ; HG3 - CG
- CB
6816 6819 6820 1 1.0950e+02 4.1840e+02 ; CB - CA
- HA
6816 6819 6821 1 1.1110e+02 5.2718e+02 ; CB - CA
- C
6817 6816 6818 1 1.0950e+02 2.9288e+02 ; HB2 - CB
- HB3
6817 6816 6819 1 1.0950e+02 4.1840e+02 ; HB2 - CB
- CA
6818 6816 6819 1 1.0950e+02 4.1840e+02 ; HB3 - CB
- CA
6819 6821 6822 1 1.2040e+02 6.6944e+02 ; CA - C
- O
6819 6821 6823 1 1.1660e+02 5.8576e+02 ; CA - C
- N
6820 6819 6821 1 1.0950e+02 4.1840e+02 ; HA - CA
- C

```
  6821   6823   6824       1    1.2000e+02    4.1840e+02 ;      C - N
- H
  6821   6823   6825       1    1.2190e+02    4.1840e+02 ;      C - N
- CA
  6822   6821   6823       1    1.2290e+02    6.6944e+02 ;      O - C
- N
  6823   6825   6826       1    1.0950e+02    4.1840e+02 ;      N - CA
- HA
  6823   6825   6827       1    1.0970e+02    6.6944e+02 ;      N - CA
- CB
  6823   6825   6840       1    1.1010e+02    5.2718e+02 ;      N - CA
- C
  6824   6823   6825       1    1.1804e+02    4.1840e+02 ;      H - N
- CA
  6825   6827   6828       1    1.0950e+02    4.1840e+02 ;     CA - CB
- HB2
  6825   6827   6829       1    1.0950e+02    4.1840e+02 ;     CA - CB
- HB3
  6825   6827   6830       1    1.0950e+02    3.3472e+02 ;     CA - CB
- CG
  6825   6840   6841       1    1.2040e+02    6.6944e+02 ;     CA - C
- O
  6825   6840   6842       1    1.1660e+02    5.8576e+02 ;     CA - C
- N
  6826   6825   6827       1    1.0950e+02    4.1840e+02 ;     HA - CA
- CB
  6826   6825   6840       1    1.0950e+02    4.1840e+02 ;     HA - CA
- C
  6827   6825   6840       1    1.1110e+02    5.2718e+02 ;     CB - CA
- C
  6827   6830   6831       1    1.0950e+02    4.1840e+02 ;     CB - CG
- HG
  6827   6830   6832       1    1.0950e+02    3.3472e+02 ;     CB - CG
- CD1
  6827   6830   6836       1    1.0950e+02    3.3472e+02 ;     CB - CG
- CD2
  6828   6827   6829       1    1.0950e+02    2.9288e+02 ;    HB2 - CB
- HB3
  6828   6827   6830       1    1.0950e+02    4.1840e+02 ;    HB2 - CB
- CG
  6829   6827   6830       1    1.0950e+02    4.1840e+02 ;    HB3 - CB
- CG
  6830   6832   6833       1    1.0950e+02    4.1840e+02 ;     CG - CD1
- HD11
  6830   6832   6834       1    1.0950e+02    4.1840e+02 ;     CG - CD1
- HD12
  6830   6832   6835       1    1.0950e+02    4.1840e+02 ;     CG - CD1
- HD13
  6830   6836   6837       1    1.0950e+02    4.1840e+02 ;     CG - CD2
- HD21
  6830   6836   6838       1    1.0950e+02    4.1840e+02 ;     CG - CD2
- HD22
```

6830 6836 6839 1 1.0950e+02 4.1840e+02 ; CG - CD2
- HD23
6831 6830 6832 1 1.0950e+02 4.1840e+02 ; HG - CG
- CD1
6831 6830 6836 1 1.0950e+02 4.1840e+02 ; HG - CG
- CD2
6832 6830 6836 1 1.0950e+02 3.3472e+02 ; CD1 - CG
- CD2
6833 6832 6834 1 1.0950e+02 2.9288e+02 ; HD11 - CD1
- HD12
6833 6832 6835 1 1.0950e+02 2.9288e+02 ; HD11 - CD1
- HD13
6834 6832 6835 1 1.0950e+02 2.9288e+02 ; HD12 - CD1
- HD13
6837 6836 6838 1 1.0950e+02 2.9288e+02 ; HD21 - CD2
- HD22
6837 6836 6839 1 1.0950e+02 2.9288e+02 ; HD21 - CD2
- HD23
6838 6836 6839 1 1.0950e+02 2.9288e+02 ; HD22 - CD2
- HD23
6840 6842 6843 1 1.2000e+02 4.1840e+02 ; C - N
- H
6840 6842 6844 1 1.2190e+02 4.1840e+02 ; C - N
- CA
6841 6840 6842 1 1.2290e+02 6.6944e+02 ; O - C
- N
6842 6844 6845 1 1.0950e+02 4.1840e+02 ; N - CA
- HA
6842 6844 6846 1 1.0970e+02 6.6944e+02 ; N - CA
- CB
6842 6844 6859 1 1.1010e+02 5.2718e+02 ; N - CA
- C
6843 6842 6844 1 1.1804e+02 4.1840e+02 ; H - N
- CA
6844 6846 6847 1 1.0950e+02 4.1840e+02 ; CA - CB
- HB
6844 6846 6848 1 1.0950e+02 3.3472e+02 ; CA - CB
- CG2
6844 6846 6852 1 1.0950e+02 3.3472e+02 ; CA - CB
- CG1
6844 6859 6860 1 1.2040e+02 6.6944e+02 ; CA - C
- O
6844 6859 6861 1 1.1660e+02 5.8576e+02 ; CA - C
- N
6845 6844 6846 1 1.0950e+02 4.1840e+02 ; HA - CA
- CB
6845 6844 6859 1 1.0950e+02 4.1840e+02 ; HA - CA
- C
6846 6844 6859 1 1.1110e+02 5.2718e+02 ; CB - CA
- C
6846 6848 6849 1 1.0950e+02 4.1840e+02 ; CB - CG2
- HG21

6846 6848 6850 1 1.0950e+02 4.1840e+02 ; CB - CG2 - HG22
6846 6848 6851 1 1.0950e+02 4.1840e+02 ; CB - CG2 - HG23
6846 6852 6853 1 1.0950e+02 4.1840e+02 ; CB - CG1 - HG12
6846 6852 6854 1 1.0950e+02 4.1840e+02 ; CB - CG1 - HG13
6846 6852 6855 1 1.0950e+02 3.3472e+02 ; CB - CG1 - CD1
6847 6846 6848 1 1.0950e+02 4.1840e+02 ; HB - CB - CG2
6847 6846 6852 1 1.0950e+02 4.1840e+02 ; HB - CB - CG1
6848 6846 6852 1 1.0950e+02 3.3472e+02 ; CG2 - CB - CG1
6849 6848 6850 1 1.0950e+02 2.9288e+02 ; HG21 - CG2 - HG22
6849 6848 6851 1 1.0950e+02 2.9288e+02 ; HG21 - CG2 - HG23
6850 6848 6851 1 1.0950e+02 2.9288e+02 ; HG22 - CG2 - HG23
6852 6855 6856 1 1.0950e+02 4.1840e+02 ; CG1 - CD1 - HD11
6852 6855 6857 1 1.0950e+02 4.1840e+02 ; CG1 - CD1 - HD12
6852 6855 6858 1 1.0950e+02 4.1840e+02 ; CG1 - CD1 - HD13
6853 6852 6854 1 1.0950e+02 2.9288e+02 ; HG12 - CG1 - HG13
6853 6852 6855 1 1.0950e+02 4.1840e+02 ; HG12 - CG1 - CD1
6854 6852 6855 1 1.0950e+02 4.1840e+02 ; HG13 - CG1 - CD1
6856 6855 6857 1 1.0950e+02 2.9288e+02 ; HD11 - CD1 - HD12
6856 6855 6858 1 1.0950e+02 2.9288e+02 ; HD11 - CD1 - HD13
6857 6855 6858 1 1.0950e+02 2.9288e+02 ; HD12 - CD1 - HD13
6859 6861 6862 1 1.2000e+02 4.1840e+02 ; C - N - H
6859 6861 6863 1 1.2190e+02 4.1840e+02 ; C - N - CA
6860 6859 6861 1 1.2290e+02 6.6944e+02 ; O - C - N
6861 6863 6864 1 1.0950e+02 4.1840e+02 ; N - CA - HA
6861 6863 6865 1 1.0970e+02 6.6944e+02 ; N - CA - CB
6861 6863 6871 1 1.1010e+02 5.2718e+02 ; N - CA - C

6862 6861 6863 1 1.1804e+02 4.1840e+02 ; H - N
- CA
6863 6865 6866 1 1.0950e+02 4.1840e+02 ; CA - CB
- HB2
6863 6865 6867 1 1.0950e+02 4.1840e+02 ; CA - CB
- HB3
6863 6865 6868 1 1.1110e+02 5.2718e+02 ; CA - CB
- CG
6863 6871 6872 1 1.2040e+02 6.6944e+02 ; CA - C
- O
6863 6871 6873 1 1.1660e+02 5.8576e+02 ; CA - C
- N
6864 6863 6865 1 1.0950e+02 4.1840e+02 ; HA - CA
- CB
6864 6863 6871 1 1.0950e+02 4.1840e+02 ; HA - CA
- C
6865 6863 6871 1 1.1110e+02 5.2718e+02 ; CB - CA
- C
6865 6868 6869 1 1.1700e+02 5.8576e+02 ; CB - CG
- OD1
6865 6868 6870 1 1.1700e+02 5.8576e+02 ; CB - CG
- OD2
6866 6865 6867 1 1.0950e+02 2.9288e+02 ; HB2 - CB
- HB3
6866 6865 6868 1 1.0950e+02 4.1840e+02 ; HB2 - CB
- CG
6867 6865 6868 1 1.0950e+02 4.1840e+02 ; HB3 - CB
- CG
6869 6868 6870 1 1.2600e+02 6.6944e+02 ; OD1 - CG
- OD2
6871 6873 6874 1 1.2000e+02 4.1840e+02 ; C - N
- H
6871 6873 6875 1 1.2190e+02 4.1840e+02 ; C - N
- CA
6872 6871 6873 1 1.2290e+02 6.6944e+02 ; O - C
- N
6873 6875 6876 1 1.0950e+02 4.1840e+02 ; N - CA
- HA
6873 6875 6877 1 1.0970e+02 6.6944e+02 ; N - CA
- CB
6873 6875 6893 1 1.1010e+02 5.2718e+02 ; N - CA
- C
6874 6873 6875 1 1.1804e+02 4.1840e+02 ; H - N
- CA
6875 6877 6878 1 1.0950e+02 4.1840e+02 ; CA - CB
- HB2
6875 6877 6879 1 1.0950e+02 4.1840e+02 ; CA - CB
- HB3
6875 6877 6880 1 1.0950e+02 3.3472e+02 ; CA - CB
- CG
6875 6893 6894 1 1.2040e+02 6.6944e+02 ; CA - C
- O

6875 6893 6895 1 1.1660e+02 5.8576e+02 ; CA - C
- N
6876 6875 6877 1 1.0950e+02 4.1840e+02 ; HA - CA
- CB
6876 6875 6893 1 1.0950e+02 4.1840e+02 ; HA - CA
- C
6877 6875 6893 1 1.1110e+02 5.2718e+02 ; CB - CA
- C
6877 6880 6881 1 1.0950e+02 4.1840e+02 ; CB - CG
- HG2
6877 6880 6882 1 1.0950e+02 4.1840e+02 ; CB - CG
- HG3
6877 6880 6883 1 1.0950e+02 3.3472e+02 ; CB - CG
- CD
6878 6877 6879 1 1.0950e+02 2.9288e+02 ; HB2 - CB
- HB3
6878 6877 6880 1 1.0950e+02 4.1840e+02 ; HB2 - CB
- CG
6879 6877 6880 1 1.0950e+02 4.1840e+02 ; HB3 - CB
- CG
6880 6883 6884 1 1.0950e+02 4.1840e+02 ; CG - CD
- HD2
6880 6883 6885 1 1.0950e+02 4.1840e+02 ; CG - CD
- HD3
6880 6883 6886 1 1.0950e+02 3.3472e+02 ; CG - CD
- CE
6881 6880 6882 1 1.0950e+02 2.9288e+02 ; HG2 - CG
- HG3
6881 6880 6883 1 1.0950e+02 4.1840e+02 ; HG2 - CG
- CD
6882 6880 6883 1 1.0950e+02 4.1840e+02 ; HG3 - CG
- CD
6883 6886 6887 1 1.0950e+02 4.1840e+02 ; CD - CE
- HE2
6883 6886 6888 1 1.0950e+02 4.1840e+02 ; CD - CE
- HE3
6883 6886 6889 1 1.1120e+02 6.6944e+02 ; CD - CE
- NZ
6884 6883 6885 1 1.0950e+02 2.9288e+02 ; HD2 - CD
- HD3
6884 6883 6886 1 1.0950e+02 4.1840e+02 ; HD2 - CD
- CE
6885 6883 6886 1 1.0950e+02 4.1840e+02 ; HD3 - CD
- CE
6886 6889 6890 1 1.0950e+02 4.1840e+02 ; CE - NZ
- HZ1
6886 6889 6891 1 1.0950e+02 4.1840e+02 ; CE - NZ
- HZ2
6886 6889 6892 1 1.0950e+02 4.1840e+02 ; CE - NZ
- HZ3
6887 6886 6888 1 1.0950e+02 2.9288e+02 ; HE2 - CE
- HE3

6887 6886 6889 1 1.0950e+02 4.1840e+02 ; HE2 - CE
- NZ
6888 6886 6889 1 1.0950e+02 4.1840e+02 ; HE3 - CE
- NZ
6890 6889 6891 1 1.0950e+02 2.9288e+02 ; HZ1 - NZ
- HZ2
6890 6889 6892 1 1.0950e+02 2.9288e+02 ; HZ1 - NZ
- HZ3
6891 6889 6892 1 1.0950e+02 2.9288e+02 ; HZ2 - NZ
- HZ3
6893 6895 6896 1 1.2000e+02 4.1840e+02 ; C - N
- H
6893 6895 6897 1 1.2190e+02 4.1840e+02 ; C - N
- CA
6894 6893 6895 1 1.2290e+02 6.6944e+02 ; O - C
- N
6895 6897 6898 1 1.0950e+02 4.1840e+02 ; N - CA
- HA
6895 6897 6899 1 1.0970e+02 6.6944e+02 ; N - CA
- CB
6895 6897 6903 1 1.1010e+02 5.2718e+02 ; N - CA
- C
6896 6895 6897 1 1.1804e+02 4.1840e+02 ; H - N
- CA
6897 6899 6900 1 1.0950e+02 4.1840e+02 ; CA - CB
- HB1
6897 6899 6901 1 1.0950e+02 4.1840e+02 ; CA - CB
- HB2
6897 6899 6902 1 1.0950e+02 4.1840e+02 ; CA - CB
- HB3
6897 6903 6904 1 1.2040e+02 6.6944e+02 ; CA - C
- O
6897 6903 6905 1 1.1660e+02 5.8576e+02 ; CA - C
- N
6898 6897 6899 1 1.0950e+02 4.1840e+02 ; HA - CA
- CB
6898 6897 6903 1 1.0950e+02 4.1840e+02 ; HA - CA
- C
6899 6897 6903 1 1.1110e+02 5.2718e+02 ; CB - CA
- C
6900 6899 6901 1 1.0950e+02 2.9288e+02 ; HB1 - CB
- HB2
6900 6899 6902 1 1.0950e+02 2.9288e+02 ; HB1 - CB
- HB3
6901 6899 6902 1 1.0950e+02 2.9288e+02 ; HB2 - CB
- HB3
6903 6905 6906 1 1.2000e+02 4.1840e+02 ; C - N
- H
6903 6905 6907 1 1.2190e+02 4.1840e+02 ; C - N
- CA
6904 6903 6905 1 1.2290e+02 6.6944e+02 ; O - C
- N

6905 6907 6908 1 1.0950e+02 4.1840e+02 ; N - CA
- HA
6905 6907 6909 1 1.0970e+02 6.6944e+02 ; N - CA
- CB
6905 6907 6922 1 1.1010e+02 5.2718e+02 ; N - CA
- C
6906 6905 6907 1 1.1804e+02 4.1840e+02 ; H - N
- CA
6907 6909 6910 1 1.0950e+02 4.1840e+02 ; CA - CB
- HB
6907 6909 6911 1 1.0950e+02 3.3472e+02 ; CA - CB
- CG2
6907 6909 6915 1 1.0950e+02 3.3472e+02 ; CA - CB
- CG1
6907 6922 6923 1 1.2040e+02 6.6944e+02 ; CA - C
- O
6907 6922 6924 1 1.1660e+02 5.8576e+02 ; CA - C
- N
6908 6907 6909 1 1.0950e+02 4.1840e+02 ; HA - CA
- CB
6908 6907 6922 1 1.0950e+02 4.1840e+02 ; HA - CA
- C
6909 6907 6922 1 1.1110e+02 5.2718e+02 ; CB - CA
- C
6909 6911 6912 1 1.0950e+02 4.1840e+02 ; CB - CG2
- HG21
6909 6911 6913 1 1.0950e+02 4.1840e+02 ; CB - CG2
- HG22
6909 6911 6914 1 1.0950e+02 4.1840e+02 ; CB - CG2
- HG23
6909 6915 6916 1 1.0950e+02 4.1840e+02 ; CB - CG1
- HG12
6909 6915 6917 1 1.0950e+02 4.1840e+02 ; CB - CG1
- HG13
6909 6915 6918 1 1.0950e+02 3.3472e+02 ; CB - CG1
- CD1
6910 6909 6911 1 1.0950e+02 4.1840e+02 ; HB - CB
- CG2
6910 6909 6915 1 1.0950e+02 4.1840e+02 ; HB - CB
- CG1
6911 6909 6915 1 1.0950e+02 3.3472e+02 ; CG2 - CB
- CG1
6912 6911 6913 1 1.0950e+02 2.9288e+02 ; HG21 - CG2
- HG22
6912 6911 6914 1 1.0950e+02 2.9288e+02 ; HG21 - CG2
- HG23
6913 6911 6914 1 1.0950e+02 2.9288e+02 ; HG22 - CG2
- HG23
6915 6918 6919 1 1.0950e+02 4.1840e+02 ; CG1 - CD1
- HD11
6915 6918 6920 1 1.0950e+02 4.1840e+02 ; CG1 - CD1
- HD12

```
   6915   6918   6921      1    1.0950e+02    4.1840e+02 ;    CG1 - CD1
 - HD13
   6916   6915   6917      1    1.0950e+02    2.9288e+02 ;   HG12 - CG1
 - HG13
   6916   6915   6918      1    1.0950e+02    4.1840e+02 ;   HG12 - CG1
 - CD1
   6917   6915   6918      1    1.0950e+02    4.1840e+02 ;   HG13 - CG1
 - CD1
   6919   6918   6920      1    1.0950e+02    2.9288e+02 ;   HD11 - CD1
 - HD12
   6919   6918   6921      1    1.0950e+02    2.9288e+02 ;   HD11 - CD1
 - HD13
   6920   6918   6921      1    1.0950e+02    2.9288e+02 ;   HD12 - CD1
 - HD13
   6922   6924   6925      1    1.2000e+02    4.1840e+02 ;      C - N
 - H
   6922   6924   6926      1    1.2190e+02    4.1840e+02 ;      C - N
 - CA
   6923   6922   6924      1    1.2290e+02    6.6944e+02 ;      O - C
 - N
   6924   6926   6927      1    1.0950e+02    4.1840e+02 ;      N - CA
 - HA
   6924   6926   6928      1    1.0970e+02    6.6944e+02 ;      N - CA
 - CB
   6924   6926   6934      1    1.1010e+02    5.2718e+02 ;      N - CA
 - C
   6925   6924   6926      1    1.1804e+02    4.1840e+02 ;      H - N
 - CA
   6926   6928   6929      1    1.0950e+02    4.1840e+02 ;     CA - CB
 - HB2
   6926   6928   6930      1    1.0950e+02    4.1840e+02 ;     CA - CB
 - HB3
   6926   6928   6931      1    1.1110e+02    5.2718e+02 ;     CA - CB
 - CG
   6926   6934   6935      1    1.2040e+02    6.6944e+02 ;     CA - C
 - O
   6926   6934   6936      1    1.1660e+02    5.8576e+02 ;     CA - C
 - N
   6927   6926   6928      1    1.0950e+02    4.1840e+02 ;     HA - CA
 - CB
   6927   6926   6934      1    1.0950e+02    4.1840e+02 ;     HA - CA
 - C
   6928   6926   6934      1    1.1110e+02    5.2718e+02 ;     CB - CA
 - C
   6928   6931   6932      1    1.1700e+02    5.8576e+02 ;     CB - CG
 - OD1
   6928   6931   6933      1    1.1700e+02    5.8576e+02 ;     CB - CG
 - OD2
   6929   6928   6930      1    1.0950e+02    2.9288e+02 ;    HB2 - CB
 - HB3
   6929   6928   6931      1    1.0950e+02    4.1840e+02 ;    HB2 - CB
 - CG
```

```
  6930   6928   6931        1    1.0950e+02    4.1840e+02 ;    HB3 - CB
- CG
  6932   6931   6933        1    1.2600e+02    6.6944e+02 ;    OD1 - CG
- OD2
  6934   6936   6937        1    1.2000e+02    4.1840e+02 ;      C - N
- H
  6934   6936   6938        1    1.2190e+02    4.1840e+02 ;      C - N
- CA
  6935   6934   6936        1    1.2290e+02    6.6944e+02 ;      O - C
- N
  6936   6938   6939        1    1.0950e+02    4.1840e+02 ;      N - CA
- HA
  6936   6938   6940        1    1.0970e+02    6.6944e+02 ;      N - CA
- CB
  6936   6938   6945        1    1.1010e+02    5.2718e+02 ;      N - CA
- C
  6937   6936   6938        1    1.1804e+02    4.1840e+02 ;      H - N
- CA
  6938   6940   6941        1    1.0950e+02    4.1840e+02 ;     CA - CB
- HB2
  6938   6940   6942        1    1.0950e+02    4.1840e+02 ;     CA - CB
- HB3
  6938   6940   6943        1    1.0950e+02    4.1840e+02 ;     CA - CB
- OG
  6938   6945   6946        1    1.2040e+02    6.6944e+02 ;     CA - C
- O
  6938   6945   6947        1    1.1660e+02    5.8576e+02 ;     CA - C
- N
  6939   6938   6940        1    1.0950e+02    4.1840e+02 ;     HA - CA
- CB
  6939   6938   6945        1    1.0950e+02    4.1840e+02 ;     HA - CA
- C
  6940   6938   6945        1    1.1110e+02    5.2718e+02 ;     CB - CA
- C
  6940   6943   6944        1    1.0850e+02    4.6024e+02 ;     CB - OG
- HG
  6941   6940   6942        1    1.0950e+02    2.9288e+02 ;    HB2 - CB
- HB3
  6941   6940   6943        1    1.0950e+02    4.1840e+02 ;    HB2 - CB
- OG
  6942   6940   6943        1    1.0950e+02    4.1840e+02 ;    HB3 - CB
- OG
  6945   6947   6948        1    1.2000e+02    4.1840e+02 ;      C - N
- H
  6945   6947   6949        1    1.2190e+02    4.1840e+02 ;      C - N
- CA
  6946   6945   6947        1    1.2290e+02    6.6944e+02 ;      O - C
- N
  6947   6949   6950        1    1.0950e+02    4.1840e+02 ;      N - CA
- HA
  6947   6949   6951        1    1.0970e+02    6.6944e+02 ;      N - CA
- CB
```

6947 6949 6964 1 1.1010e+02 5.2718e+02 ; N - CA
- C
6948 6947 6949 1 1.1804e+02 4.1840e+02 ; H - N
- CA
6949 6951 6952 1 1.0950e+02 4.1840e+02 ; CA - CB
- HB2
6949 6951 6953 1 1.0950e+02 4.1840e+02 ; CA - CB
- HB3
6949 6951 6954 1 1.0950e+02 3.3472e+02 ; CA - CB
- CG
6949 6964 6965 1 1.2040e+02 6.6944e+02 ; CA - C
- O
6949 6964 6966 1 1.1660e+02 5.8576e+02 ; CA - C
- N
6950 6949 6951 1 1.0950e+02 4.1840e+02 ; HA - CA
- CB
6950 6949 6964 1 1.0950e+02 4.1840e+02 ; HA - CA
- C
6951 6949 6964 1 1.1110e+02 5.2718e+02 ; CB - CA
- C
6951 6954 6955 1 1.0950e+02 4.1840e+02 ; CB - CG
- HG
6951 6954 6956 1 1.0950e+02 3.3472e+02 ; CB - CG
- CD1
6951 6954 6960 1 1.0950e+02 3.3472e+02 ; CB - CG
- CD2
6952 6951 6953 1 1.0950e+02 2.9288e+02 ; HB2 - CB
- HB3
6952 6951 6954 1 1.0950e+02 4.1840e+02 ; HB2 - CB
- CG
6953 6951 6954 1 1.0950e+02 4.1840e+02 ; HB3 - CB
- CG
6954 6956 6957 1 1.0950e+02 4.1840e+02 ; CG - CD1
- HD11
6954 6956 6958 1 1.0950e+02 4.1840e+02 ; CG - CD1
- HD12
6954 6956 6959 1 1.0950e+02 4.1840e+02 ; CG - CD1
- HD13
6954 6960 6961 1 1.0950e+02 4.1840e+02 ; CG - CD2
- HD21
6954 6960 6962 1 1.0950e+02 4.1840e+02 ; CG - CD2
- HD22
6954 6960 6963 1 1.0950e+02 4.1840e+02 ; CG - CD2
- HD23
6955 6954 6956 1 1.0950e+02 4.1840e+02 ; HG - CG
- CD1
6955 6954 6960 1 1.0950e+02 4.1840e+02 ; HG - CG
- CD2
6956 6954 6960 1 1.0950e+02 3.3472e+02 ; CD1 - CG
- CD2
6957 6956 6958 1 1.0950e+02 2.9288e+02 ; HD11 - CD1
- HD12

6957 6956 6959 1 1.0950e+02 2.9288e+02 ; HD11 - CD1
- HD13
6958 6956 6959 1 1.0950e+02 2.9288e+02 ; HD12 - CD1
- HD13
6961 6960 6962 1 1.0950e+02 2.9288e+02 ; HD21 - CD2
- HD22
6961 6960 6963 1 1.0950e+02 2.9288e+02 ; HD21 - CD2
- HD23
6962 6960 6963 1 1.0950e+02 2.9288e+02 ; HD22 - CD2
- HD23
6964 6966 6967 1 1.2000e+02 4.1840e+02 ; C - N
- H
6964 6966 6968 1 1.2190e+02 4.1840e+02 ; C - N
- CA
6965 6964 6966 1 1.2290e+02 6.6944e+02 ; O - C
- N
6966 6968 6969 1 1.0950e+02 4.1840e+02 ; N - CA
- HA
6966 6968 6970 1 1.0970e+02 6.6944e+02 ; N - CA
- CB
6966 6968 6983 1 1.1010e+02 5.2718e+02 ; N - CA
- C
6967 6966 6968 1 1.1804e+02 4.1840e+02 ; H - N
- CA
6968 6970 6971 1 1.0950e+02 4.1840e+02 ; CA - CB
- HB2
6968 6970 6972 1 1.0950e+02 4.1840e+02 ; CA - CB
- HB3
6968 6970 6973 1 1.0950e+02 3.3472e+02 ; CA - CB
- CG
6968 6983 6984 1 1.2040e+02 6.6944e+02 ; CA - C
- O
6968 6983 6985 1 1.1660e+02 5.8576e+02 ; CA - C
- N
6969 6968 6970 1 1.0950e+02 4.1840e+02 ; HA - CA
- CB
6969 6968 6983 1 1.0950e+02 4.1840e+02 ; HA - CA
- C
6970 6968 6983 1 1.1110e+02 5.2718e+02 ; CB - CA
- C
6970 6973 6974 1 1.0950e+02 4.1840e+02 ; CB - CG
- HG
6970 6973 6975 1 1.0950e+02 3.3472e+02 ; CB - CG
- CD1
6970 6973 6979 1 1.0950e+02 3.3472e+02 ; CB - CG
- CD2
6971 6970 6972 1 1.0950e+02 2.9288e+02 ; HB2 - CB
- HB3
6971 6970 6973 1 1.0950e+02 4.1840e+02 ; HB2 - CB
- CG
6972 6970 6973 1 1.0950e+02 4.1840e+02 ; HB3 - CB
- CG

```
  6973   6975   6976       1    1.0950e+02    4.1840e+02 ;     CG - CD1
- HD11
  6973   6975   6977       1    1.0950e+02    4.1840e+02 ;     CG - CD1
- HD12
  6973   6975   6978       1    1.0950e+02    4.1840e+02 ;     CG - CD1
- HD13
  6973   6979   6980       1    1.0950e+02    4.1840e+02 ;     CG - CD2
- HD21
  6973   6979   6981       1    1.0950e+02    4.1840e+02 ;     CG - CD2
- HD22
  6973   6979   6982       1    1.0950e+02    4.1840e+02 ;     CG - CD2
- HD23
  6974   6973   6975       1    1.0950e+02    4.1840e+02 ;     HG - CG
- CD1
  6974   6973   6979       1    1.0950e+02    4.1840e+02 ;     HG - CG
- CD2
  6975   6973   6979       1    1.0950e+02    3.3472e+02 ;    CD1 - CG
- CD2
  6976   6975   6977       1    1.0950e+02    2.9288e+02 ;   HD11 - CD1
- HD12
  6976   6975   6978       1    1.0950e+02    2.9288e+02 ;   HD11 - CD1
- HD13
  6977   6975   6978       1    1.0950e+02    2.9288e+02 ;   HD12 - CD1
- HD13
  6980   6979   6981       1    1.0950e+02    2.9288e+02 ;   HD21 - CD2
- HD22
  6980   6979   6982       1    1.0950e+02    2.9288e+02 ;   HD21 - CD2
- HD23
  6981   6979   6982       1    1.0950e+02    2.9288e+02 ;   HD22 - CD2
- HD23
  6983   6985   6986       1    1.2000e+02    4.1840e+02 ;      C - N
- H
  6983   6985   6987       1    1.2190e+02    4.1840e+02 ;      C - N
- CA
  6984   6983   6985       1    1.2290e+02    6.6944e+02 ;      O - C
- N
  6985   6987   6988       1    1.0950e+02    4.1840e+02 ;      N - CA
- HA
  6985   6987   6989       1    1.0970e+02    6.6944e+02 ;      N - CA
- CB
  6985   6987   7000       1    1.1010e+02    5.2718e+02 ;      N - CA
- C
  6986   6985   6987       1    1.1804e+02    4.1840e+02 ;      H - N
- CA
  6987   6989   6990       1    1.0950e+02    4.1840e+02 ;     CA - CB
- HB2
  6987   6989   6991       1    1.0950e+02    4.1840e+02 ;     CA - CB
- HB3
  6987   6989   6992       1    1.0950e+02    3.3472e+02 ;     CA - CB
- CG
  6987   7000   7001       1    1.2040e+02    6.6944e+02 ;     CA - C
- O
```

6987 7000 7002 1 1.1660e+02 5.8576e+02 ; CA - C
- N
6988 6987 6989 1 1.0950e+02 4.1840e+02 ; HA - CA
- CB
6988 6987 7000 1 1.0950e+02 4.1840e+02 ; HA - CA
- C
6989 6987 7000 1 1.1110e+02 5.2718e+02 ; CB - CA
- C
6989 6992 6993 1 1.0950e+02 4.1840e+02 ; CB - CG
- HG2
6989 6992 6994 1 1.0950e+02 4.1840e+02 ; CB - CG
- HG3
6989 6992 6995 1 1.1110e+02 5.2718e+02 ; CB - CG
- CD
6990 6989 6991 1 1.0950e+02 2.9288e+02 ; HB2 - CB
- HB3
6990 6989 6992 1 1.0950e+02 4.1840e+02 ; HB2 - CB
- CG
6991 6989 6992 1 1.0950e+02 4.1840e+02 ; HB3 - CB
- CG
6992 6995 6996 1 1.2040e+02 6.6944e+02 ; CG - CD
- OE1
6992 6995 6997 1 1.1660e+02 5.8576e+02 ; CG - CD
- NE2
6993 6992 6994 1 1.0950e+02 2.9288e+02 ; HG2 - CG
- HG3
6993 6992 6995 1 1.0950e+02 4.1840e+02 ; HG2 - CG
- CD
6994 6992 6995 1 1.0950e+02 4.1840e+02 ; HG3 - CG
- CD
6995 6997 6998 1 1.2000e+02 4.1840e+02 ; CD - NE2
- HE21
6995 6997 6999 1 1.2000e+02 4.1840e+02 ; CD - NE2
- HE22
6996 6995 6997 1 1.2290e+02 6.6944e+02 ; OE1 - CD
- NE2
6998 6997 6999 1 1.2000e+02 2.9288e+02 ; HE21 - NE2
- HE22
7000 7002 7003 1 1.2000e+02 4.1840e+02 ; C - N
- H
7000 7002 7004 1 1.2190e+02 4.1840e+02 ; C - N
- CA
7001 7000 7002 1 1.2290e+02 6.6944e+02 ; O - C
- N
7002 7004 7005 1 1.0950e+02 4.1840e+02 ; N - CA
- HA
7002 7004 7006 1 1.0970e+02 6.6944e+02 ; N - CA
- CB
7002 7004 7015 1 1.1010e+02 5.2718e+02 ; N - CA
- C
7003 7002 7004 1 1.1804e+02 4.1840e+02 ; H - N
- CA

7004 7006 7007 1 1.0950e+02 4.1840e+02 ; CA - CB
- HB2
7004 7006 7008 1 1.0950e+02 4.1840e+02 ; CA - CB
- HB3
7004 7006 7009 1 1.0950e+02 3.3472e+02 ; CA - CB
- CG
7004 7015 7016 1 1.2040e+02 6.6944e+02 ; CA - C
- O
7004 7015 7017 1 1.1660e+02 5.8576e+02 ; CA - C
- N
7005 7004 7006 1 1.0950e+02 4.1840e+02 ; HA - CA
- CB
7005 7004 7015 1 1.0950e+02 4.1840e+02 ; HA - CA
- C
7006 7004 7015 1 1.1110e+02 5.2718e+02 ; CB - CA
- C
7006 7009 7010 1 1.0950e+02 4.1840e+02 ; CB - CG
- HG2
7006 7009 7011 1 1.0950e+02 4.1840e+02 ; CB - CG
- HG3
7006 7009 7012 1 1.1110e+02 5.2718e+02 ; CB - CG
- CD
7007 7006 7008 1 1.0950e+02 2.9288e+02 ; HB2 - CB
- HB3
7007 7006 7009 1 1.0950e+02 4.1840e+02 ; HB2 - CB
- CG
7008 7006 7009 1 1.0950e+02 4.1840e+02 ; HB3 - CB
- CG
7009 7012 7013 1 1.1700e+02 5.8576e+02 ; CG - CD
- OE1
7009 7012 7014 1 1.1700e+02 5.8576e+02 ; CG - CD
- OE2
7010 7009 7011 1 1.0950e+02 2.9288e+02 ; HG2 - CG
- HG3
7010 7009 7012 1 1.0950e+02 4.1840e+02 ; HG2 - CG
- CD
7011 7009 7012 1 1.0950e+02 4.1840e+02 ; HG3 - CG
- CD
7013 7012 7014 1 1.2600e+02 6.6944e+02 ; OE1 - CD
- OE2
7015 7017 7018 1 1.2000e+02 4.1840e+02 ; C - N
- H
7015 7017 7019 1 1.2190e+02 4.1840e+02 ; C - N
- CA
7016 7015 7017 1 1.2290e+02 6.6944e+02 ; O - C
- N
7017 7019 7020 1 1.0950e+02 4.1840e+02 ; N - CA
- HA
7017 7019 7021 1 1.0970e+02 6.6944e+02 ; N - CA
- CB
7017 7019 7031 1 1.1010e+02 5.2718e+02 ; N - CA
- C

7018 7017 7019 1 1.1804e+02 4.1840e+02 ; H - N
- CA
7019 7021 7022 1 1.0950e+02 4.1840e+02 ; CA - CB
- HB
7019 7021 7023 1 1.0950e+02 3.3472e+02 ; CA - CB
- CG1
7019 7021 7027 1 1.0950e+02 3.3472e+02 ; CA - CB
- CG2
7019 7031 7032 1 1.2040e+02 6.6944e+02 ; CA - C
- O
7019 7031 7033 1 1.1660e+02 5.8576e+02 ; CA - C
- N
7020 7019 7021 1 1.0950e+02 4.1840e+02 ; HA - CA
- CB
7020 7019 7031 1 1.0950e+02 4.1840e+02 ; HA - CA
- C
7021 7019 7031 1 1.1110e+02 5.2718e+02 ; CB - CA
- C
7021 7023 7024 1 1.0950e+02 4.1840e+02 ; CB - CG1
- HG11
7021 7023 7025 1 1.0950e+02 4.1840e+02 ; CB - CG1
- HG12
7021 7023 7026 1 1.0950e+02 4.1840e+02 ; CB - CG1
- HG13
7021 7027 7028 1 1.0950e+02 4.1840e+02 ; CB - CG2
- HG21
7021 7027 7029 1 1.0950e+02 4.1840e+02 ; CB - CG2
- HG22
7021 7027 7030 1 1.0950e+02 4.1840e+02 ; CB - CG2
- HG23
7022 7021 7023 1 1.0950e+02 4.1840e+02 ; HB - CB
- CG1
7022 7021 7027 1 1.0950e+02 4.1840e+02 ; HB - CB
- CG2
7023 7021 7027 1 1.0950e+02 3.3472e+02 ; CG1 - CB
- CG2
7024 7023 7025 1 1.0950e+02 2.9288e+02 ; HG11 - CG1
- HG12
7024 7023 7026 1 1.0950e+02 2.9288e+02 ; HG11 - CG1
- HG13
7025 7023 7026 1 1.0950e+02 2.9288e+02 ; HG12 - CG1
- HG13
7028 7027 7029 1 1.0950e+02 2.9288e+02 ; HG21 - CG2
- HG22
7028 7027 7030 1 1.0950e+02 2.9288e+02 ; HG21 - CG2
- HG23
7029 7027 7030 1 1.0950e+02 2.9288e+02 ; HG22 - CG2
- HG23
7031 7033 7034 1 1.2000e+02 4.1840e+02 ; C - N
- H
7031 7033 7035 1 1.2190e+02 4.1840e+02 ; C - N
- CA

7032 7031 7033 1 1.2290e+02 6.6944e+02 ; O - C
- N
7033 7035 7036 1 1.0950e+02 4.1840e+02 ; N - CA
- HA
7033 7035 7037 1 1.0970e+02 6.6944e+02 ; N - CA
- CB
7033 7035 7043 1 1.1010e+02 5.2718e+02 ; N - CA
- C
7034 7033 7035 1 1.1804e+02 4.1840e+02 ; H - N
- CA
7035 7037 7038 1 1.0950e+02 4.1840e+02 ; CA - CB
- HB2
7035 7037 7039 1 1.0950e+02 4.1840e+02 ; CA - CB
- HB3
7035 7037 7040 1 1.1110e+02 5.2718e+02 ; CA - CB
- CG
7035 7043 7044 1 1.2040e+02 6.6944e+02 ; CA - C
- O
7035 7043 7045 1 1.1660e+02 5.8576e+02 ; CA - C
- N
7036 7035 7037 1 1.0950e+02 4.1840e+02 ; HA - CA
- CB
7036 7035 7043 1 1.0950e+02 4.1840e+02 ; HA - CA
- C
7037 7035 7043 1 1.1110e+02 5.2718e+02 ; CB - CA
- C
7037 7040 7041 1 1.1700e+02 5.8576e+02 ; CB - CG
- OD1
7037 7040 7042 1 1.1700e+02 5.8576e+02 ; CB - CG
- OD2
7038 7037 7039 1 1.0950e+02 2.9288e+02 ; HB2 - CB
- HB3
7038 7037 7040 1 1.0950e+02 4.1840e+02 ; HB2 - CB
- CG
7039 7037 7040 1 1.0950e+02 4.1840e+02 ; HB3 - CB
- CG
7041 7040 7042 1 1.2600e+02 6.6944e+02 ; OD1 - CG
- OD2
7043 7045 7046 1 1.2000e+02 4.1840e+02 ; C - N
- H
7043 7045 7047 1 1.2190e+02 4.1840e+02 ; C - N
- CA
7044 7043 7045 1 1.2290e+02 6.6944e+02 ; O - C
- N
7045 7047 7048 1 1.0950e+02 4.1840e+02 ; N - CA
- HA
7045 7047 7049 1 1.0970e+02 6.6944e+02 ; N - CA
- CB
7045 7047 7062 1 1.1010e+02 5.2718e+02 ; N - CA
- C
7046 7045 7047 1 1.1804e+02 4.1840e+02 ; H - N
- CA

```
  7047   7049   7050      1    1.0950e+02    4.1840e+02 ;     CA - CB
- HB2
  7047   7049   7051      1    1.0950e+02    4.1840e+02 ;     CA - CB
- HB3
  7047   7049   7052      1    1.0950e+02    3.3472e+02 ;     CA - CB
- CG
  7047   7062   7063      1    1.2040e+02    6.6944e+02 ;     CA - C
- O
  7047   7062   7064      1    1.1660e+02    5.8576e+02 ;     CA - C
- N
  7048   7047   7049      1    1.0950e+02    4.1840e+02 ;     HA - CA
- CB
  7048   7047   7062      1    1.0950e+02    4.1840e+02 ;     HA - CA
- C
  7049   7047   7062      1    1.1110e+02    5.2718e+02 ;     CB - CA
- C
  7049   7052   7053      1    1.0950e+02    4.1840e+02 ;     CB - CG
- HG
  7049   7052   7054      1    1.0950e+02    3.3472e+02 ;     CB - CG
- CD1
  7049   7052   7058      1    1.0950e+02    3.3472e+02 ;     CB - CG
- CD2
  7050   7049   7051      1    1.0950e+02    2.9288e+02 ;    HB2 - CB
- HB3
  7050   7049   7052      1    1.0950e+02    4.1840e+02 ;    HB2 - CB
- CG
  7051   7049   7052      1    1.0950e+02    4.1840e+02 ;    HB3 - CB
- CG
  7052   7054   7055      1    1.0950e+02    4.1840e+02 ;     CG - CD1
- HD11
  7052   7054   7056      1    1.0950e+02    4.1840e+02 ;     CG - CD1
- HD12
  7052   7054   7057      1    1.0950e+02    4.1840e+02 ;     CG - CD1
- HD13
  7052   7058   7059      1    1.0950e+02    4.1840e+02 ;     CG - CD2
- HD21
  7052   7058   7060      1    1.0950e+02    4.1840e+02 ;     CG - CD2
- HD22
  7052   7058   7061      1    1.0950e+02    4.1840e+02 ;     CG - CD2
- HD23
  7053   7052   7054      1    1.0950e+02    4.1840e+02 ;     HG - CG
- CD1
  7053   7052   7058      1    1.0950e+02    4.1840e+02 ;     HG - CG
- CD2
  7054   7052   7058      1    1.0950e+02    3.3472e+02 ;    CD1 - CG
- CD2
  7055   7054   7056      1    1.0950e+02    2.9288e+02 ;   HD11 - CD1
- HD12
  7055   7054   7057      1    1.0950e+02    2.9288e+02 ;   HD11 - CD1
- HD13
  7056   7054   7057      1    1.0950e+02    2.9288e+02 ;   HD12 - CD1
- HD13
```

```
  7059   7058   7060      1    1.0950e+02    2.9288e+02 ;   HD21 - CD2
- HD22
  7059   7058   7061      1    1.0950e+02    2.9288e+02 ;   HD21 - CD2
- HD23
  7060   7058   7061      1    1.0950e+02    2.9288e+02 ;   HD22 - CD2
- HD23
  7062   7064   7065      1    1.2000e+02    4.1840e+02 ;      C - N
- H
  7062   7064   7066      1    1.2190e+02    4.1840e+02 ;      C - N
- CA
  7063   7062   7064      1    1.2290e+02    6.6944e+02 ;      O - C
- N
  7064   7066   7067      1    1.0950e+02    4.1840e+02 ;      N - CA
- HA
  7064   7066   7068      1    1.0970e+02    6.6944e+02 ;      N - CA
- CB
  7064   7066   7076      1    1.1010e+02    5.2718e+02 ;      N - CA
- C
  7065   7064   7066      1    1.1804e+02    4.1840e+02 ;      H - N
- CA
  7066   7068   7069      1    1.0950e+02    4.1840e+02 ;     CA - CB
- HB2
  7066   7068   7070      1    1.0950e+02    4.1840e+02 ;     CA - CB
- HB3
  7066   7068   7071      1    1.1110e+02    5.2718e+02 ;     CA - CB
- CG
  7066   7076   7077      1    1.2040e+02    6.6944e+02 ;     CA - C
- O
  7066   7076   7078      1    1.1660e+02    5.8576e+02 ;     CA - C
- N
  7067   7066   7068      1    1.0950e+02    4.1840e+02 ;     HA - CA
- CB
  7067   7066   7076      1    1.0950e+02    4.1840e+02 ;     HA - CA
- C
  7068   7066   7076      1    1.1110e+02    5.2718e+02 ;     CB - CA
- C
  7068   7071   7072      1    1.2040e+02    6.6944e+02 ;     CB - CG
- OD1
  7068   7071   7073      1    1.1660e+02    5.8576e+02 ;     CB - CG
- ND2
  7069   7068   7070      1    1.0950e+02    2.9288e+02 ;    HB2 - CB
- HB3
  7069   7068   7071      1    1.0950e+02    4.1840e+02 ;    HB2 - CB
- CG
  7070   7068   7071      1    1.0950e+02    4.1840e+02 ;    HB3 - CB
- CG
  7071   7073   7074      1    1.2000e+02    4.1840e+02 ;     CG - ND2
- HD21
  7071   7073   7075      1    1.2000e+02    4.1840e+02 ;     CG - ND2
- HD22
  7072   7071   7073      1    1.2290e+02    6.6944e+02 ;    OD1 - CG
- ND2
```

```
  7074   7073   7075       1    1.2000e+02    2.9288e+02 ;   HD21 - ND2
- HD22
  7076   7078   7079       1    1.2000e+02    4.1840e+02 ;      C - N
- H
  7076   7078   7080       1    1.2190e+02    4.1840e+02 ;      C - N
- CA
  7077   7076   7078       1    1.2290e+02    6.6944e+02 ;      O - C
- N
  7078   7080   7081       1    1.0950e+02    4.1840e+02 ;      N - CA
- HA2
  7078   7080   7082       1    1.0950e+02    4.1840e+02 ;      N - CA
- HA3
  7078   7080   7083       1    1.1010e+02    5.2718e+02 ;      N - CA
- C
  7079   7078   7080       1    1.1804e+02    4.1840e+02 ;      H - N
- CA
  7080   7083   7084       1    1.2040e+02    6.6944e+02 ;     CA - C
- O
  7080   7083   7085       1    1.1660e+02    5.8576e+02 ;     CA - C
- N
  7081   7080   7082       1    1.0950e+02    2.9288e+02 ;    HA2 - CA
- HA3
  7081   7080   7083       1    1.0950e+02    4.1840e+02 ;    HA2 - CA
- C
  7082   7080   7083       1    1.0950e+02    4.1840e+02 ;    HA3 - CA
- C
  7083   7085   7086       1    1.2000e+02    4.1840e+02 ;      C - N
- H
  7083   7085   7087       1    1.2190e+02    4.1840e+02 ;      C - N
- CA
  7084   7083   7085       1    1.2290e+02    6.6944e+02 ;      O - C
- N
  7085   7087   7088       1    1.0950e+02    4.1840e+02 ;      N - CA
- HA
  7085   7087   7089       1    1.0970e+02    6.6944e+02 ;      N - CA
- CB
  7085   7087   7095       1    1.1010e+02    5.2718e+02 ;      N - CA
- C
  7086   7085   7087       1    1.1804e+02    4.1840e+02 ;      H - N
- CA
  7087   7089   7090       1    1.0950e+02    4.1840e+02 ;     CA - CB
- HB2
  7087   7089   7091       1    1.0950e+02    4.1840e+02 ;     CA - CB
- HB3
  7087   7089   7092       1    1.1110e+02    5.2718e+02 ;     CA - CB
- CG
  7087   7095   7096       1    1.2040e+02    6.6944e+02 ;     CA - C
- O
  7087   7095   7097       1    1.1660e+02    5.8576e+02 ;     CA - C
- N
  7088   7087   7089       1    1.0950e+02    4.1840e+02 ;     HA - CA
- CB
```

```
  7088   7087   7095      1    1.0950e+02    4.1840e+02 ;     HA - CA
- C
  7089   7087   7095      1    1.1110e+02    5.2718e+02 ;     CB - CA
- C
  7089   7092   7093      1    1.1700e+02    5.8576e+02 ;     CB - CG
- OD1
  7089   7092   7094      1    1.1700e+02    5.8576e+02 ;     CB - CG
- OD2
  7090   7089   7091      1    1.0950e+02    2.9288e+02 ;    HB2 - CB
- HB3
  7090   7089   7092      1    1.0950e+02    4.1840e+02 ;    HB2 - CB
- CG
  7091   7089   7092      1    1.0950e+02    4.1840e+02 ;    HB3 - CB
- CG
  7093   7092   7094      1    1.2600e+02    6.6944e+02 ;    OD1 - CG
- OD2
  7095   7097   7098      1    1.2000e+02    4.1840e+02 ;      C - N
- H
  7095   7097   7099      1    1.2190e+02    4.1840e+02 ;      C - N
- CA
  7096   7095   7097      1    1.2290e+02    6.6944e+02 ;      O - C
- N
  7097   7099   7100      1    1.0950e+02    4.1840e+02 ;      N - CA
- HA2
  7097   7099   7101      1    1.0950e+02    4.1840e+02 ;      N - CA
- HA3
  7097   7099   7102      1    1.1010e+02    5.2718e+02 ;      N - CA
- C
  7098   7097   7099      1    1.1804e+02    4.1840e+02 ;      H - N
- CA
  7099   7102   7103      1    1.2040e+02    6.6944e+02 ;     CA - C
- O
  7099   7102   7104      1    1.1660e+02    5.8576e+02 ;     CA - C
- N
  7100   7099   7101      1    1.0950e+02    2.9288e+02 ;    HA2 - CA
- HA3
  7100   7099   7102      1    1.0950e+02    4.1840e+02 ;    HA2 - CA
- C
  7101   7099   7102      1    1.0950e+02    4.1840e+02 ;    HA3 - CA
- C
  7102   7104   7105      1    1.2000e+02    4.1840e+02 ;      C - N
- H
  7102   7104   7106      1    1.2190e+02    4.1840e+02 ;      C - N
- CA
  7103   7102   7104      1    1.2290e+02    6.6944e+02 ;      O - C
- N
  7104   7106   7107      1    1.0950e+02    4.1840e+02 ;      N - CA
- HA
  7104   7106   7108      1    1.0970e+02    6.6944e+02 ;      N - CA
- CB
  7104   7106   7117      1    1.1010e+02    5.2718e+02 ;      N - CA
- C
```

7105 7104 7106 1 1.1804e+02 4.1840e+02 ; H - N
- CA
7106 7108 7109 1 1.0950e+02 4.1840e+02 ; CA - CB
- HB2
7106 7108 7110 1 1.0950e+02 4.1840e+02 ; CA - CB
- HB3
7106 7108 7111 1 1.0950e+02 3.3472e+02 ; CA - CB
- CG
7106 7117 7118 1 1.2040e+02 6.6944e+02 ; CA - C
- O
7106 7117 7119 1 1.1660e+02 5.8576e+02 ; CA - C
- N
7107 7106 7108 1 1.0950e+02 4.1840e+02 ; HA - CA
- CB
7107 7106 7117 1 1.0950e+02 4.1840e+02 ; HA - CA
- C
7108 7106 7117 1 1.1110e+02 5.2718e+02 ; CB - CA
- C
7108 7111 7112 1 1.0950e+02 4.1840e+02 ; CB - CG
- HG2
7108 7111 7113 1 1.0950e+02 4.1840e+02 ; CB - CG
- HG3
7108 7111 7114 1 1.1110e+02 5.2718e+02 ; CB - CG
- CD
7109 7108 7110 1 1.0950e+02 2.9288e+02 ; HB2 - CB
- HB3
7109 7108 7111 1 1.0950e+02 4.1840e+02 ; HB2 - CB
- CG
7110 7108 7111 1 1.0950e+02 4.1840e+02 ; HB3 - CB
- CG
7111 7114 7115 1 1.1700e+02 5.8576e+02 ; CG - CD
- OE1
7111 7114 7116 1 1.1700e+02 5.8576e+02 ; CG - CD
- OE2
7112 7111 7113 1 1.0950e+02 2.9288e+02 ; HG2 - CG
- HG3
7112 7111 7114 1 1.0950e+02 4.1840e+02 ; HG2 - CG
- CD
7113 7111 7114 1 1.0950e+02 4.1840e+02 ; HG3 - CG
- CD
7115 7114 7116 1 1.2600e+02 6.6944e+02 ; OE1 - CD
- OE2
7117 7119 7120 1 1.2000e+02 4.1840e+02 ; C - N
- H
7117 7119 7121 1 1.2190e+02 4.1840e+02 ; C - N
- CA
7118 7117 7119 1 1.2290e+02 6.6944e+02 ; O - C
- N
7119 7121 7122 1 1.0950e+02 4.1840e+02 ; N - CA
- HA
7119 7121 7123 1 1.0970e+02 6.6944e+02 ; N - CA
- CB

| | | | | | | | |
|---|---|---|---|---|---|---|---|
| 7119 | 7121 | 7136 | 1 | 1.1010e+02 | 5.2718e+02 | ; | N - CA - C |
| 7120 | 7119 | 7121 | 1 | 1.1804e+02 | 4.1840e+02 | ; | H - N - CA |
| 7121 | 7123 | 7124 | 1 | 1.0950e+02 | 4.1840e+02 | ; | CA - CB - HB |
| 7121 | 7123 | 7125 | 1 | 1.0950e+02 | 3.3472e+02 | ; | CA - CB - CG2 |
| 7121 | 7123 | 7129 | 1 | 1.0950e+02 | 3.3472e+02 | ; | CA - CB - CG1 |
| 7121 | 7136 | 7137 | 1 | 1.2040e+02 | 6.6944e+02 | ; | CA - C - O |
| 7121 | 7136 | 7138 | 1 | 1.1660e+02 | 5.8576e+02 | ; | CA - C - N |
| 7122 | 7121 | 7123 | 1 | 1.0950e+02 | 4.1840e+02 | ; | HA - CA - CB |
| 7122 | 7121 | 7136 | 1 | 1.0950e+02 | 4.1840e+02 | ; | HA - CA - C |
| 7123 | 7121 | 7136 | 1 | 1.1110e+02 | 5.2718e+02 | ; | CB - CA - C |
| 7123 | 7125 | 7126 | 1 | 1.0950e+02 | 4.1840e+02 | ; | CB - CG2 - HG21 |
| 7123 | 7125 | 7127 | 1 | 1.0950e+02 | 4.1840e+02 | ; | CB - CG2 - HG22 |
| 7123 | 7125 | 7128 | 1 | 1.0950e+02 | 4.1840e+02 | ; | CB - CG2 - HG23 |
| 7123 | 7129 | 7130 | 1 | 1.0950e+02 | 4.1840e+02 | ; | CB - CG1 - HG12 |
| 7123 | 7129 | 7131 | 1 | 1.0950e+02 | 4.1840e+02 | ; | CB - CG1 - HG13 |
| 7123 | 7129 | 7132 | 1 | 1.0950e+02 | 3.3472e+02 | ; | CB - CG1 - CD1 |
| 7124 | 7123 | 7125 | 1 | 1.0950e+02 | 4.1840e+02 | ; | HB - CB - CG2 |
| 7124 | 7123 | 7129 | 1 | 1.0950e+02 | 4.1840e+02 | ; | HB - CB - CG1 |
| 7125 | 7123 | 7129 | 1 | 1.0950e+02 | 3.3472e+02 | ; | CG2 - CB - CG1 |
| 7126 | 7125 | 7127 | 1 | 1.0950e+02 | 2.9288e+02 | ; | HG21 - CG2 - HG22 |
| 7126 | 7125 | 7128 | 1 | 1.0950e+02 | 2.9288e+02 | ; | HG21 - CG2 - HG23 |
| 7127 | 7125 | 7128 | 1 | 1.0950e+02 | 2.9288e+02 | ; | HG22 - CG2 - HG23 |
| 7129 | 7132 | 7133 | 1 | 1.0950e+02 | 4.1840e+02 | ; | CG1 - CD1 - HD11 |
| 7129 | 7132 | 7134 | 1 | 1.0950e+02 | 4.1840e+02 | ; | CG1 - CD1 - HD12 |
| 7129 | 7132 | 7135 | 1 | 1.0950e+02 | 4.1840e+02 | ; | CG1 - CD1 - HD13 |
| 7130 | 7129 | 7131 | 1 | 1.0950e+02 | 2.9288e+02 | ; | HG12 - CG1 - HG13 |

```
   7130   7129   7132      1    1.0950e+02    4.1840e+02 ;   HG12 - CG1
 - CD1
   7131   7129   7132      1    1.0950e+02    4.1840e+02 ;   HG13 - CG1
 - CD1
   7133   7132   7134      1    1.0950e+02    2.9288e+02 ;   HD11 - CD1
 - HD12
   7133   7132   7135      1    1.0950e+02    2.9288e+02 ;   HD11 - CD1
 - HD13
   7134   7132   7135      1    1.0950e+02    2.9288e+02 ;   HD12 - CD1
 - HD13
   7136   7138   7139      1    1.2000e+02    4.1840e+02 ;      C - N
 - H
   7136   7138   7140      1    1.2190e+02    4.1840e+02 ;      C - N
 - CA
   7137   7136   7138      1    1.2290e+02    6.6944e+02 ;      O - C
 - N
   7138   7140   7141      1    1.0950e+02    4.1840e+02 ;      N - CA
 - HA
   7138   7140   7142      1    1.0970e+02    6.6944e+02 ;      N - CA
 - CB
   7138   7140   7148      1    1.1010e+02    5.2718e+02 ;      N - CA
 - C
   7139   7138   7140      1    1.1804e+02    4.1840e+02 ;      H - N
 - CA
   7140   7142   7143      1    1.0950e+02    4.1840e+02 ;     CA - CB
 - HB2
   7140   7142   7144      1    1.0950e+02    4.1840e+02 ;     CA - CB
 - HB3
   7140   7142   7145      1    1.1110e+02    5.2718e+02 ;     CA - CB
 - CG
   7140   7148   7149      1    1.2040e+02    6.6944e+02 ;     CA - C
 - O
   7140   7148   7150      1    1.1660e+02    5.8576e+02 ;     CA - C
 - N
   7141   7140   7142      1    1.0950e+02    4.1840e+02 ;     HA - CA
 - CB
   7141   7140   7148      1    1.0950e+02    4.1840e+02 ;     HA - CA
 - C
   7142   7140   7148      1    1.1110e+02    5.2718e+02 ;     CB - CA
 - C
   7142   7145   7146      1    1.1700e+02    5.8576e+02 ;     CB - CG
 - OD1
   7142   7145   7147      1    1.1700e+02    5.8576e+02 ;     CB - CG
 - OD2
   7143   7142   7144      1    1.0950e+02    2.9288e+02 ;    HB2 - CB
 - HB3
   7143   7142   7145      1    1.0950e+02    4.1840e+02 ;    HB2 - CB
 - CG
   7144   7142   7145      1    1.0950e+02    4.1840e+02 ;    HB3 - CB
 - CG
   7146   7145   7147      1    1.2600e+02    6.6944e+02 ;    OD1 - CG
 - OD2
```

| | | | | | | | |
|---|---|---|---|---|---|---|---|
| 7148 | 7150 | 7151 | 1 | 1.2000e+02 | 4.1840e+02 | ; | C - N - H |
| 7148 | 7150 | 7152 | 1 | 1.2190e+02 | 4.1840e+02 | ; | C - N - CA |
| 7149 | 7148 | 7150 | 1 | 1.2290e+02 | 6.6944e+02 | ; | O - C - N |
| 7150 | 7152 | 7153 | 1 | 1.0950e+02 | 4.1840e+02 | ; | N - CA - HA |
| 7150 | 7152 | 7154 | 1 | 1.0970e+02 | 6.6944e+02 | ; | N - CA - CB |
| 7150 | 7152 | 7168 | 1 | 1.1010e+02 | 5.2718e+02 | ; | N - CA - C |
| 7151 | 7150 | 7152 | 1 | 1.1804e+02 | 4.1840e+02 | ; | H - N - CA |
| 7152 | 7154 | 7155 | 1 | 1.0950e+02 | 4.1840e+02 | ; | CA - CB - HB2 |
| 7152 | 7154 | 7156 | 1 | 1.0950e+02 | 4.1840e+02 | ; | CA - CB - HB3 |
| 7152 | 7154 | 7157 | 1 | 1.1400e+02 | 5.2718e+02 | ; | CA - CB - CG |
| 7152 | 7168 | 7169 | 1 | 1.2040e+02 | 6.6944e+02 | ; | CA - C - O |
| 7152 | 7168 | 7170 | 1 | 1.1660e+02 | 5.8576e+02 | ; | CA - C - N |
| 7153 | 7152 | 7154 | 1 | 1.0950e+02 | 4.1840e+02 | ; | HA - CA - CB |
| 7153 | 7152 | 7168 | 1 | 1.0950e+02 | 4.1840e+02 | ; | HA - CA - C |
| 7154 | 7152 | 7168 | 1 | 1.1110e+02 | 5.2718e+02 | ; | CB - CA - C |
| 7154 | 7157 | 7158 | 1 | 1.2000e+02 | 5.8576e+02 | ; | CB - CG - CD1 |
| 7154 | 7157 | 7166 | 1 | 1.2000e+02 | 5.8576e+02 | ; | CB - CG - CD2 |
| 7155 | 7154 | 7156 | 1 | 1.0950e+02 | 2.9288e+02 | ; | HB2 - CB - HB3 |
| 7155 | 7154 | 7157 | 1 | 1.0950e+02 | 4.1840e+02 | ; | HB2 - CB - CG |
| 7156 | 7154 | 7157 | 1 | 1.0950e+02 | 4.1840e+02 | ; | HB3 - CB - CG |
| 7157 | 7158 | 7159 | 1 | 1.2000e+02 | 4.1840e+02 | ; | CG - CD1 - HD1 |
| 7157 | 7158 | 7160 | 1 | 1.2000e+02 | 5.2718e+02 | ; | CG - CD1 - CE1 |
| 7157 | 7166 | 7164 | 1 | 1.2000e+02 | 5.2718e+02 | ; | CG - CD2 - CE2 |
| 7157 | 7166 | 7167 | 1 | 1.2000e+02 | 4.1840e+02 | ; | CG - CD2 - HD2 |
| 7158 | 7157 | 7166 | 1 | 1.2000e+02 | 5.2718e+02 | ; | CD1 - CG - CD2 |
| 7158 | 7160 | 7161 | 1 | 1.2000e+02 | 4.1840e+02 | ; | CD1 - CE1 - HE1 |

7158 7160 7162 1 1.2000e+02 5.2718e+02 ; CD1 - CE1
- CZ
7159 7158 7160 1 1.2000e+02 4.1840e+02 ; HD1 - CD1
- CE1
7160 7162 7163 1 1.2000e+02 4.1840e+02 ; CE1 - CZ
- HZ
7160 7162 7164 1 1.2000e+02 5.2718e+02 ; CE1 - CZ
- CE2
7161 7160 7162 1 1.2000e+02 4.1840e+02 ; HE1 - CE1
- CZ
7162 7164 7165 1 1.2000e+02 4.1840e+02 ; CZ - CE2
- HE2
7162 7164 7166 1 1.2000e+02 5.2718e+02 ; CZ - CE2
- CD2
7163 7162 7164 1 1.2000e+02 4.1840e+02 ; HZ - CZ
- CE2
7164 7166 7167 1 1.2000e+02 4.1840e+02 ; CE2 - CD2
- HD2
7165 7164 7166 1 1.2000e+02 4.1840e+02 ; HE2 - CE2
- CD2
7168 7170 7171 1 1.2000e+02 4.1840e+02 ; C - N
- H
7168 7170 7172 1 1.2190e+02 4.1840e+02 ; C - N
- CA
7169 7168 7170 1 1.2290e+02 6.6944e+02 ; O - C
- N
7170 7172 7173 1 1.0950e+02 4.1840e+02 ; N - CA
- HA
7170 7172 7174 1 1.0970e+02 6.6944e+02 ; N - CA
- CB
7170 7172 7186 1 1.1010e+02 5.2718e+02 ; N - CA
- C
7171 7170 7172 1 1.1804e+02 4.1840e+02 ; H - N
- CA
7172 7174 7175 1 1.0950e+02 4.1840e+02 ; CA - CB
- HB2
7172 7174 7176 1 1.0950e+02 4.1840e+02 ; CA - CB
- HB3
7172 7174 7177 1 1.1310e+02 5.2718e+02 ; CA - CB
- CG
7172 7186 7187 1 1.2040e+02 6.6944e+02 ; CA - C
- O
7172 7186 7188 1 1.1660e+02 5.8576e+02 ; CA - C
- N
7173 7172 7174 1 1.0950e+02 4.1840e+02 ; HA - CA
- CB
7173 7172 7186 1 1.0950e+02 4.1840e+02 ; HA - CA
- C
7174 7172 7186 1 1.1110e+02 5.2718e+02 ; CB - CA
- C
7174 7177 7178 1 1.2000e+02 5.8576e+02 ; CB - CG
- ND1

| | | | | | | | |
|---|---|---|---|---|---|---|---|
| 7174 | 7177 | 7184 | 1 | 1.2000e+02 | 5.8576e+02 | ; | CB - CG - CD2 |
| 7175 | 7174 | 7176 | 1 | 1.0950e+02 | 2.9288e+02 | ; | HB2 - CB - HB3 |
| 7175 | 7174 | 7177 | 1 | 1.0950e+02 | 4.1840e+02 | ; | HB2 - CB - CG |
| 7176 | 7174 | 7177 | 1 | 1.0950e+02 | 4.1840e+02 | ; | HB3 - CB - CG |
| 7177 | 7178 | 7179 | 1 | 1.2000e+02 | 4.1840e+02 | ; | CG - ND1 - HD1 |
| 7177 | 7178 | 7180 | 1 | 1.2000e+02 | 5.8576e+02 | ; | CG - ND1 - CE1 |
| 7177 | 7184 | 7182 | 1 | 1.2000e+02 | 5.8576e+02 | ; | CG - CD2 - NE2 |
| 7177 | 7184 | 7185 | 1 | 1.2000e+02 | 4.1840e+02 | ; | CG - CD2 - HD2 |
| 7178 | 7177 | 7184 | 1 | 1.2000e+02 | 5.8576e+02 | ; | ND1 - CG - CD2 |
| 7178 | 7180 | 7181 | 1 | 1.2000e+02 | 4.1840e+02 | ; | ND1 - CE1 - HE1 |
| 7178 | 7180 | 7182 | 1 | 1.2000e+02 | 5.8576e+02 | ; | ND1 - CE1 - NE2 |
| 7179 | 7178 | 7180 | 1 | 1.2000e+02 | 4.1840e+02 | ; | HD1 - ND1 - CE1 |
| 7180 | 7182 | 7183 | 1 | 1.2000e+02 | 4.1840e+02 | ; | CE1 - NE2 - HE2 |
| 7180 | 7182 | 7184 | 1 | 1.2000e+02 | 5.8576e+02 | ; | CE1 - NE2 - CD2 |
| 7181 | 7180 | 7182 | 1 | 1.2000e+02 | 4.1840e+02 | ; | HE1 - CE1 - NE2 |
| 7182 | 7184 | 7185 | 1 | 1.2000e+02 | 4.1840e+02 | ; | NE2 - CD2 - HD2 |
| 7183 | 7182 | 7184 | 1 | 1.2000e+02 | 4.1840e+02 | ; | HE2 - NE2 - CD2 |
| 7186 | 7188 | 7189 | 1 | 1.2000e+02 | 4.1840e+02 | ; | C - N - H |
| 7186 | 7188 | 7190 | 1 | 1.2190e+02 | 4.1840e+02 | ; | C - N - CA |
| 7187 | 7186 | 7188 | 1 | 1.2290e+02 | 6.6944e+02 | ; | O - C - N |
| 7188 | 7190 | 7191 | 1 | 1.0950e+02 | 4.1840e+02 | ; | N - CA - HA |
| 7188 | 7190 | 7192 | 1 | 1.0970e+02 | 6.6944e+02 | ; | N - CA - CB |
| 7188 | 7190 | 7202 | 1 | 1.1010e+02 | 5.2718e+02 | ; | N - CA - C |
| 7189 | 7188 | 7190 | 1 | 1.1804e+02 | 4.1840e+02 | ; | H - N - CA |
| 7190 | 7192 | 7193 | 1 | 1.0950e+02 | 4.1840e+02 | ; | CA - CB - HB2 |
| 7190 | 7192 | 7194 | 1 | 1.0950e+02 | 4.1840e+02 | ; | CA - CB - HB3 |

7190 7192 7195 1 1.0950e+02 3.3472e+02 ; CA - CB
- CG
7190 7202 7203 1 1.2040e+02 6.6944e+02 ; CA - C
- O
7190 7202 7204 1 1.1660e+02 5.8576e+02 ; CA - C
- N
7191 7190 7192 1 1.0950e+02 4.1840e+02 ; HA - CA
- CB
7191 7190 7202 1 1.0950e+02 4.1840e+02 ; HA - CA
- C
7192 7190 7202 1 1.1110e+02 5.2718e+02 ; CB - CA
- C
7192 7195 7196 1 1.0950e+02 4.1840e+02 ; CB - CG
- HG2
7192 7195 7197 1 1.0950e+02 4.1840e+02 ; CB - CG
- HG3
7192 7195 7198 1 1.1110e+02 5.2718e+02 ; CB - CG
- CD
7193 7192 7194 1 1.0950e+02 2.9288e+02 ; HB2 - CB
- HB3
7193 7192 7195 1 1.0950e+02 4.1840e+02 ; HB2 - CB
- CG
7194 7192 7195 1 1.0950e+02 4.1840e+02 ; HB3 - CB
- CG
7195 7198 7199 1 1.2040e+02 6.6944e+02 ; CG - CD
- OE1
7195 7198 7200 1 1.1000e+02 6.6944e+02 ; CG - CD
- OE2
7196 7195 7197 1 1.0950e+02 2.9288e+02 ; HG2 - CG
- HG3
7196 7195 7198 1 1.0950e+02 4.1840e+02 ; HG2 - CG
- CD
7197 7195 7198 1 1.0950e+02 4.1840e+02 ; HG3 - CG
- CD
7198 7200 7201 1 1.1300e+02 4.1840e+02 ; CD - OE2
- HE2
7199 7198 7200 1 1.2000e+02 6.6944e+02 ; OE1 - CD
- OE2
7202 7204 7205 1 1.2000e+02 4.1840e+02 ; C - N
- H
7202 7204 7206 1 1.2190e+02 4.1840e+02 ; C - N
- CA
7203 7202 7204 1 1.2290e+02 6.6944e+02 ; O - C
- N
7204 7206 7207 1 1.0950e+02 4.1840e+02 ; N - CA
- HA
7204 7206 7208 1 1.0970e+02 6.6944e+02 ; N - CA
- CB
7204 7206 7222 1 1.1010e+02 5.2718e+02 ; N - CA
- C
7205 7204 7206 1 1.1804e+02 4.1840e+02 ; H - N
- CA

```
  7206   7208   7209       1    1.0950e+02    4.1840e+02 ;     CA - CB
- HB2
  7206   7208   7210       1    1.0950e+02    4.1840e+02 ;     CA - CB
- HB3
  7206   7208   7211       1    1.1400e+02    5.2718e+02 ;     CA - CB
- CG
  7206   7222   7223       1    1.2040e+02    6.6944e+02 ;     CA - C
- O
  7206   7222   7224       1    1.1660e+02    5.8576e+02 ;     CA - C
- N
  7207   7206   7208       1    1.0950e+02    4.1840e+02 ;     HA - CA
- CB
  7207   7206   7222       1    1.0950e+02    4.1840e+02 ;     HA - CA
- C
  7208   7206   7222       1    1.1110e+02    5.2718e+02 ;     CB - CA
- C
  7208   7211   7212       1    1.2000e+02    5.8576e+02 ;     CB - CG
- CD1
  7208   7211   7220       1    1.2000e+02    5.8576e+02 ;     CB - CG
- CD2
  7209   7208   7210       1    1.0950e+02    2.9288e+02 ;    HB2 - CB
- HB3
  7209   7208   7211       1    1.0950e+02    4.1840e+02 ;    HB2 - CB
- CG
  7210   7208   7211       1    1.0950e+02    4.1840e+02 ;    HB3 - CB
- CG
  7211   7212   7213       1    1.2000e+02    4.1840e+02 ;     CG - CD1
- HD1
  7211   7212   7214       1    1.2000e+02    5.2718e+02 ;     CG - CD1
- CE1
  7211   7220   7218       1    1.2000e+02    5.2718e+02 ;     CG - CD2
- CE2
  7211   7220   7221       1    1.2000e+02    4.1840e+02 ;     CG - CD2
- HD2
  7212   7211   7220       1    1.2000e+02    5.2718e+02 ;    CD1 - CG
- CD2
  7212   7214   7215       1    1.2000e+02    4.1840e+02 ;    CD1 - CE1
- HE1
  7212   7214   7216       1    1.2000e+02    5.2718e+02 ;    CD1 - CE1
- CZ
  7213   7212   7214       1    1.2000e+02    4.1840e+02 ;    HD1 - CD1
- CE1
  7214   7216   7217       1    1.2000e+02    4.1840e+02 ;    CE1 - CZ
- HZ
  7214   7216   7218       1    1.2000e+02    5.2718e+02 ;    CE1 - CZ
- CE2
  7215   7214   7216       1    1.2000e+02    4.1840e+02 ;    HE1 - CE1
- CZ
  7216   7218   7219       1    1.2000e+02    4.1840e+02 ;     CZ - CE2
- HE2
  7216   7218   7220       1    1.2000e+02    5.2718e+02 ;     CZ - CE2
- CD2
```

7217 7216 7218 1 1.2000e+02 4.1840e+02 ; HZ - CZ
- CE2
7218 7220 7221 1 1.2000e+02 4.1840e+02 ; CE2 - CD2
- HD2
7219 7218 7220 1 1.2000e+02 4.1840e+02 ; HE2 - CE2
- CD2
7222 7224 7225 1 1.2000e+02 4.1840e+02 ; C - N
- H
7222 7224 7226 1 1.2190e+02 4.1840e+02 ; C - N
- CA
7223 7222 7224 1 1.2290e+02 6.6944e+02 ; O - C
- N
7224 7226 7227 1 1.0950e+02 4.1840e+02 ; N - CA
- HA
7224 7226 7228 1 1.0970e+02 6.6944e+02 ; N - CA
- CB
7224 7226 7239 1 1.1010e+02 5.2718e+02 ; N - CA
- C
7225 7224 7226 1 1.1804e+02 4.1840e+02 ; H - N
- CA
7226 7228 7229 1 1.0950e+02 4.1840e+02 ; CA - CB
- HB2
7226 7228 7230 1 1.0950e+02 4.1840e+02 ; CA - CB
- HB3
7226 7228 7231 1 1.0950e+02 3.3472e+02 ; CA - CB
- CG
7226 7239 7240 1 1.2040e+02 6.6944e+02 ; CA - C
- O
7226 7239 7241 1 1.1660e+02 5.8576e+02 ; CA - C
- N
7227 7226 7228 1 1.0950e+02 4.1840e+02 ; HA - CA
- CB
7227 7226 7239 1 1.0950e+02 4.1840e+02 ; HA - CA
- C
7228 7226 7239 1 1.1110e+02 5.2718e+02 ; CB - CA
- C
7228 7231 7232 1 1.0950e+02 4.1840e+02 ; CB - CG
- HG2
7228 7231 7233 1 1.0950e+02 4.1840e+02 ; CB - CG
- HG3
7228 7231 7234 1 1.1470e+02 4.1840e+02 ; CB - CG
- SD
7229 7228 7230 1 1.0950e+02 2.9288e+02 ; HB2 - CB
- HB3
7229 7228 7231 1 1.0950e+02 4.1840e+02 ; HB2 - CB
- CG
7230 7228 7231 1 1.0950e+02 4.1840e+02 ; HB3 - CB
- CG
7231 7234 7235 1 9.8900e+01 5.1882e+02 ; CG - SD
- CE
7232 7231 7233 1 1.0950e+02 2.9288e+02 ; HG2 - CG
- HG3

| | | | | | | | | |
|---|---|---|---|---|---|---|---|---|
| 7232 | 7231 | 7234 | 1 | 1.0950e+02 | 4.1840e+02 | ; | HG2 - CG | - SD |
| 7233 | 7231 | 7234 | 1 | 1.0950e+02 | 4.1840e+02 | ; | HG3 - CG | - SD |
| 7234 | 7235 | 7236 | 1 | 1.0950e+02 | 4.1840e+02 | ; | SD - CE | - HE1 |
| 7234 | 7235 | 7237 | 1 | 1.0950e+02 | 4.1840e+02 | ; | SD - CE | - HE2 |
| 7234 | 7235 | 7238 | 1 | 1.0950e+02 | 4.1840e+02 | ; | SD - CE | - HE3 |
| 7236 | 7235 | 7237 | 1 | 1.0950e+02 | 2.9288e+02 | ; | HE1 - CE | - HE2 |
| 7236 | 7235 | 7238 | 1 | 1.0950e+02 | 2.9288e+02 | ; | HE1 - CE | - HE3 |
| 7237 | 7235 | 7238 | 1 | 1.0950e+02 | 2.9288e+02 | ; | HE2 - CE | - HE3 |
| 7239 | 7241 | 7242 | 1 | 1.2000e+02 | 4.1840e+02 | ; | C - N | - H |
| 7239 | 7241 | 7243 | 1 | 1.2190e+02 | 4.1840e+02 | ; | C - N | - CA |
| 7240 | 7239 | 7241 | 1 | 1.2290e+02 | 6.6944e+02 | ; | O - C | - N |
| 7241 | 7243 | 7244 | 1 | 1.0950e+02 | 4.1840e+02 | ; | N - CA | - HA |
| 7241 | 7243 | 7245 | 1 | 1.0970e+02 | 6.6944e+02 | ; | N - CA | - CB |
| 7241 | 7243 | 7258 | 1 | 1.1010e+02 | 5.2718e+02 | ; | N - CA | - C |
| 7242 | 7241 | 7243 | 1 | 1.1804e+02 | 4.1840e+02 | ; | H - N | - CA |
| 7243 | 7245 | 7246 | 1 | 1.0950e+02 | 4.1840e+02 | ; | CA - CB | - HB2 |
| 7243 | 7245 | 7247 | 1 | 1.0950e+02 | 4.1840e+02 | ; | CA - CB | - HB3 |
| 7243 | 7245 | 7248 | 1 | 1.0950e+02 | 3.3472e+02 | ; | CA - CB | - CG |
| 7243 | 7258 | 7259 | 1 | 1.2040e+02 | 6.6944e+02 | ; | CA - C | - O |
| 7243 | 7258 | 7260 | 1 | 1.1660e+02 | 5.8576e+02 | ; | CA - C | - N |
| 7244 | 7243 | 7245 | 1 | 1.0950e+02 | 4.1840e+02 | ; | HA - CA | - CB |
| 7244 | 7243 | 7258 | 1 | 1.0950e+02 | 4.1840e+02 | ; | HA - CA | - C |
| 7245 | 7243 | 7258 | 1 | 1.1110e+02 | 5.2718e+02 | ; | CB - CA | - C |
| 7245 | 7248 | 7249 | 1 | 1.0950e+02 | 4.1840e+02 | ; | CB - CG | - HG |
| 7245 | 7248 | 7250 | 1 | 1.0950e+02 | 3.3472e+02 | ; | CB - CG | - CD1 |
| 7245 | 7248 | 7254 | 1 | 1.0950e+02 | 3.3472e+02 | ; | CB - CG | - CD2 |

7246 7245 7247 1 1.0950e+02 2.9288e+02 ; HB2 - CB
- HB3
7246 7245 7248 1 1.0950e+02 4.1840e+02 ; HB2 - CB
- CG
7247 7245 7248 1 1.0950e+02 4.1840e+02 ; HB3 - CB
- CG
7248 7250 7251 1 1.0950e+02 4.1840e+02 ; CG - CD1
- HD11
7248 7250 7252 1 1.0950e+02 4.1840e+02 ; CG - CD1
- HD12
7248 7250 7253 1 1.0950e+02 4.1840e+02 ; CG - CD1
- HD13
7248 7254 7255 1 1.0950e+02 4.1840e+02 ; CG - CD2
- HD21
7248 7254 7256 1 1.0950e+02 4.1840e+02 ; CG - CD2
- HD22
7248 7254 7257 1 1.0950e+02 4.1840e+02 ; CG - CD2
- HD23
7249 7248 7250 1 1.0950e+02 4.1840e+02 ; HG - CG
- CD1
7249 7248 7254 1 1.0950e+02 4.1840e+02 ; HG - CG
- CD2
7250 7248 7254 1 1.0950e+02 3.3472e+02 ; CD1 - CG
- CD2
7251 7250 7252 1 1.0950e+02 2.9288e+02 ; HD11 - CD1
- HD12
7251 7250 7253 1 1.0950e+02 2.9288e+02 ; HD11 - CD1
- HD13
7252 7250 7253 1 1.0950e+02 2.9288e+02 ; HD12 - CD1
- HD13
7255 7254 7256 1 1.0950e+02 2.9288e+02 ; HD21 - CD2
- HD22
7255 7254 7257 1 1.0950e+02 2.9288e+02 ; HD21 - CD2
- HD23
7256 7254 7257 1 1.0950e+02 2.9288e+02 ; HD22 - CD2
- HD23
7258 7260 7261 1 1.2000e+02 4.1840e+02 ; C - N
- H
7258 7260 7262 1 1.2190e+02 4.1840e+02 ; C - N
- CA
7259 7258 7260 1 1.2290e+02 6.6944e+02 ; O - C
- N
7260 7262 7263 1 1.0950e+02 4.1840e+02 ; N - CA
- HA
7260 7262 7264 1 1.0970e+02 6.6944e+02 ; N - CA
- CB
7260 7262 7275 1 1.1010e+02 5.2718e+02 ; N - CA
- C
7261 7260 7262 1 1.1804e+02 4.1840e+02 ; H - N
- CA
7262 7264 7265 1 1.0950e+02 4.1840e+02 ; CA - CB
- HB2

7262 7264 7266 1 1.0950e+02 4.1840e+02 ; CA - CB
- HB3
7262 7264 7267 1 1.0950e+02 3.3472e+02 ; CA - CB
- CG
7262 7275 7276 1 1.2040e+02 6.6944e+02 ; CA - C
- O
7262 7275 7277 1 1.1660e+02 5.8576e+02 ; CA - C
- N
7263 7262 7264 1 1.0950e+02 4.1840e+02 ; HA - CA
- CB
7263 7262 7275 1 1.0950e+02 4.1840e+02 ; HA - CA
- C
7264 7262 7275 1 1.1110e+02 5.2718e+02 ; CB - CA
- C
7264 7267 7268 1 1.0950e+02 4.1840e+02 ; CB - CG
- HG2
7264 7267 7269 1 1.0950e+02 4.1840e+02 ; CB - CG
- HG3
7264 7267 7270 1 1.1470e+02 4.1840e+02 ; CB - CG
- SD
7265 7264 7266 1 1.0950e+02 2.9288e+02 ; HB2 - CB
- HB3
7265 7264 7267 1 1.0950e+02 4.1840e+02 ; HB2 - CB
- CG
7266 7264 7267 1 1.0950e+02 4.1840e+02 ; HB3 - CB
- CG
7267 7270 7271 1 9.8900e+01 5.1882e+02 ; CG - SD
- CE
7268 7267 7269 1 1.0950e+02 2.9288e+02 ; HG2 - CG
- HG3
7268 7267 7270 1 1.0950e+02 4.1840e+02 ; HG2 - CG
- SD
7269 7267 7270 1 1.0950e+02 4.1840e+02 ; HG3 - CG
- SD
7270 7271 7272 1 1.0950e+02 4.1840e+02 ; SD - CE
- HE1
7270 7271 7273 1 1.0950e+02 4.1840e+02 ; SD - CE
- HE2
7270 7271 7274 1 1.0950e+02 4.1840e+02 ; SD - CE
- HE3
7272 7271 7273 1 1.0950e+02 2.9288e+02 ; HE1 - CE
- HE2
7272 7271 7274 1 1.0950e+02 2.9288e+02 ; HE1 - CE
- HE3
7273 7271 7274 1 1.0950e+02 2.9288e+02 ; HE2 - CE
- HE3
7275 7277 7278 1 1.2000e+02 4.1840e+02 ; C - N
- H
7275 7277 7279 1 1.2190e+02 4.1840e+02 ; C - N
- CA
7276 7275 7277 1 1.2290e+02 6.6944e+02 ; O - C
- N

7277 7279 7280 1 1.0950e+02 4.1840e+02 ; N - CA
- HA
7277 7279 7281 1 1.0970e+02 6.6944e+02 ; N - CA
- CB
7277 7279 7292 1 1.1010e+02 5.2718e+02 ; N - CA
- C
7278 7277 7279 1 1.1804e+02 4.1840e+02 ; H - N
- CA
7279 7281 7282 1 1.0950e+02 4.1840e+02 ; CA - CB
- HB2
7279 7281 7283 1 1.0950e+02 4.1840e+02 ; CA - CB
- HB3
7279 7281 7284 1 1.0950e+02 3.3472e+02 ; CA - CB
- CG
7279 7292 7293 1 1.2040e+02 6.6944e+02 ; CA - C
- O
7279 7292 7294 1 1.1660e+02 5.8576e+02 ; CA - C
- N
7280 7279 7281 1 1.0950e+02 4.1840e+02 ; HA - CA
- CB
7280 7279 7292 1 1.0950e+02 4.1840e+02 ; HA - CA
- C
7281 7279 7292 1 1.1110e+02 5.2718e+02 ; CB - CA
- C
7281 7284 7285 1 1.0950e+02 4.1840e+02 ; CB - CG
- HG2
7281 7284 7286 1 1.0950e+02 4.1840e+02 ; CB - CG
- HG3
7281 7284 7287 1 1.1470e+02 4.1840e+02 ; CB - CG
- SD
7282 7281 7283 1 1.0950e+02 2.9288e+02 ; HB2 - CB
- HB3
7282 7281 7284 1 1.0950e+02 4.1840e+02 ; HB2 - CB
- CG
7283 7281 7284 1 1.0950e+02 4.1840e+02 ; HB3 - CB
- CG
7284 7287 7288 1 9.8900e+01 5.1882e+02 ; CG - SD
- CE
7285 7284 7286 1 1.0950e+02 2.9288e+02 ; HG2 - CG
- HG3
7285 7284 7287 1 1.0950e+02 4.1840e+02 ; HG2 - CG
- SD
7286 7284 7287 1 1.0950e+02 4.1840e+02 ; HG3 - CG
- SD
7287 7288 7289 1 1.0950e+02 4.1840e+02 ; SD - CE
- HE1
7287 7288 7290 1 1.0950e+02 4.1840e+02 ; SD - CE
- HE2
7287 7288 7291 1 1.0950e+02 4.1840e+02 ; SD - CE
- HE3
7289 7288 7290 1 1.0950e+02 2.9288e+02 ; HE1 - CE
- HE2

7289 7288 7291 1 1.0950e+02 2.9288e+02 ; HE1 - CE
- HE3
7290 7288 7291 1 1.0950e+02 2.9288e+02 ; HE2 - CE
- HE3
7292 7294 7295 1 1.2000e+02 4.1840e+02 ; C - N
- H
7292 7294 7296 1 1.2190e+02 4.1840e+02 ; C - N
- CA
7293 7292 7294 1 1.2290e+02 6.6944e+02 ; O - C
- N
7294 7296 7297 1 1.0950e+02 4.1840e+02 ; N - CA
- HA
7294 7296 7298 1 1.0970e+02 6.6944e+02 ; N - CA
- CB
7294 7296 7303 1 1.1010e+02 5.2718e+02 ; N - CA
- C
7295 7294 7296 1 1.1804e+02 4.1840e+02 ; H - N
- CA
7296 7298 7299 1 1.0950e+02 4.1840e+02 ; CA - CB
- HB2
7296 7298 7300 1 1.0950e+02 4.1840e+02 ; CA - CB
- HB3
7296 7298 7301 1 1.0950e+02 4.1840e+02 ; CA - CB
- OG
7296 7303 7304 1 1.2040e+02 6.6944e+02 ; CA - C
- O
7296 7303 7305 1 1.1660e+02 5.8576e+02 ; CA - C
- N
7297 7296 7298 1 1.0950e+02 4.1840e+02 ; HA - CA
- CB
7297 7296 7303 1 1.0950e+02 4.1840e+02 ; HA - CA
- C
7298 7296 7303 1 1.1110e+02 5.2718e+02 ; CB - CA
- C
7298 7301 7302 1 1.0850e+02 4.6024e+02 ; CB - OG
- HG
7299 7298 7300 1 1.0950e+02 2.9288e+02 ; HB2 - CB
- HB3
7299 7298 7301 1 1.0950e+02 4.1840e+02 ; HB2 - CB
- OG
7300 7298 7301 1 1.0950e+02 4.1840e+02 ; HB3 - CB
- OG
7303 7305 7306 1 1.2000e+02 4.1840e+02 ; C - N
- H
7303 7305 7307 1 1.2190e+02 4.1840e+02 ; C - N
- CA
7304 7303 7305 1 1.2290e+02 6.6944e+02 ; O - C
- N
7305 7307 7308 1 1.0950e+02 4.1840e+02 ; N - CA
- HA
7305 7307 7309 1 1.0970e+02 6.6944e+02 ; N - CA
- CB

```
  7305   7307   7325      1    1.1010e+02    5.2718e+02 ;      N - CA
- C
  7306   7305   7307      1    1.1804e+02    4.1840e+02 ;      H - N
- CA
  7307   7309   7310      1    1.0950e+02    4.1840e+02 ;     CA - CB
- HB2
  7307   7309   7311      1    1.0950e+02    4.1840e+02 ;     CA - CB
- HB3
  7307   7309   7312      1    1.0950e+02    3.3472e+02 ;     CA - CB
- CG
  7307   7325   7326      1    1.2040e+02    6.6944e+02 ;     CA - C
- O
  7307   7325   7327      1    1.1660e+02    5.8576e+02 ;     CA - C
- N
  7308   7307   7309      1    1.0950e+02    4.1840e+02 ;     HA - CA
- CB
  7308   7307   7325      1    1.0950e+02    4.1840e+02 ;     HA - CA
- C
  7309   7307   7325      1    1.1110e+02    5.2718e+02 ;     CB - CA
- C
  7309   7312   7313      1    1.0950e+02    4.1840e+02 ;     CB - CG
- HG2
  7309   7312   7314      1    1.0950e+02    4.1840e+02 ;     CB - CG
- HG3
  7309   7312   7315      1    1.0950e+02    3.3472e+02 ;     CB - CG
- CD
  7310   7309   7311      1    1.0950e+02    2.9288e+02 ;    HB2 - CB
- HB3
  7310   7309   7312      1    1.0950e+02    4.1840e+02 ;    HB2 - CB
- CG
  7311   7309   7312      1    1.0950e+02    4.1840e+02 ;    HB3 - CB
- CG
  7312   7315   7316      1    1.0950e+02    4.1840e+02 ;     CG - CD
- HD2
  7312   7315   7317      1    1.0950e+02    4.1840e+02 ;     CG - CD
- HD3
  7312   7315   7318      1    1.0950e+02    3.3472e+02 ;     CG - CD
- CE
  7313   7312   7314      1    1.0950e+02    2.9288e+02 ;    HG2 - CG
- HG3
  7313   7312   7315      1    1.0950e+02    4.1840e+02 ;    HG2 - CG
- CD
  7314   7312   7315      1    1.0950e+02    4.1840e+02 ;    HG3 - CG
- CD
  7315   7318   7319      1    1.0950e+02    4.1840e+02 ;     CD - CE
- HE2
  7315   7318   7320      1    1.0950e+02    4.1840e+02 ;     CD - CE
- HE3
  7315   7318   7321      1    1.1120e+02    6.6944e+02 ;     CD - CE
- NZ
  7316   7315   7317      1    1.0950e+02    2.9288e+02 ;    HD2 - CD
- HD3
```

```
  7316   7315   7318       1    1.0950e+02    4.1840e+02 ;    HD2 - CD
- CE
  7317   7315   7318       1    1.0950e+02    4.1840e+02 ;    HD3 - CD
- CE
  7318   7321   7322       1    1.0950e+02    4.1840e+02 ;     CE - NZ
- HZ1
  7318   7321   7323       1    1.0950e+02    4.1840e+02 ;     CE - NZ
- HZ2
  7318   7321   7324       1    1.0950e+02    4.1840e+02 ;     CE - NZ
- HZ3
  7319   7318   7320       1    1.0950e+02    2.9288e+02 ;    HE2 - CE
- HE3
  7319   7318   7321       1    1.0950e+02    4.1840e+02 ;    HE2 - CE
- NZ
  7320   7318   7321       1    1.0950e+02    4.1840e+02 ;    HE3 - CE
- NZ
  7322   7321   7323       1    1.0950e+02    2.9288e+02 ;    HZ1 - NZ
- HZ2
  7322   7321   7324       1    1.0950e+02    2.9288e+02 ;    HZ1 - NZ
- HZ3
  7323   7321   7324       1    1.0950e+02    2.9288e+02 ;    HZ2 - NZ
- HZ3
  7325   7327   7328       1    1.2000e+02    4.1840e+02 ;      C - N
- H
  7325   7327   7329       1    1.2190e+02    4.1840e+02 ;      C - N
- CA
  7326   7325   7327       1    1.2290e+02    6.6944e+02 ;      O - C
- N
  7327   7329   7330       1    1.0950e+02    4.1840e+02 ;      N - CA
- HA
  7327   7329   7331       1    1.0970e+02    6.6944e+02 ;      N - CA
- CB
  7327   7329   7347       1    1.1010e+02    5.2718e+02 ;      N - CA
- C
  7328   7327   7329       1    1.1804e+02    4.1840e+02 ;      H - N
- CA
  7329   7331   7332       1    1.0950e+02    4.1840e+02 ;     CA - CB
- HB2
  7329   7331   7333       1    1.0950e+02    4.1840e+02 ;     CA - CB
- HB3
  7329   7331   7334       1    1.0950e+02    3.3472e+02 ;     CA - CB
- CG
  7329   7347   7348       1    1.2040e+02    6.6944e+02 ;     CA - C
- O
  7329   7347   7349       1    1.1660e+02    5.8576e+02 ;     CA - C
- N
  7330   7329   7331       1    1.0950e+02    4.1840e+02 ;     HA - CA
- CB
  7330   7329   7347       1    1.0950e+02    4.1840e+02 ;     HA - CA
- C
  7331   7329   7347       1    1.1110e+02    5.2718e+02 ;     CB - CA
- C
```

7331 7334 7335 1 1.0950e+02 4.1840e+02 ; CB - CG
- HG2
7331 7334 7336 1 1.0950e+02 4.1840e+02 ; CB - CG
- HG3
7331 7334 7337 1 1.0950e+02 3.3472e+02 ; CB - CG
- CD
7332 7331 7333 1 1.0950e+02 2.9288e+02 ; HB2 - CB
- HB3
7332 7331 7334 1 1.0950e+02 4.1840e+02 ; HB2 - CB
- CG
7333 7331 7334 1 1.0950e+02 4.1840e+02 ; HB3 - CB
- CG
7334 7337 7338 1 1.0950e+02 4.1840e+02 ; CG - CD
- HD2
7334 7337 7339 1 1.0950e+02 4.1840e+02 ; CG - CD
- HD3
7334 7337 7340 1 1.0950e+02 3.3472e+02 ; CG - CD
- CE
7335 7334 7336 1 1.0950e+02 2.9288e+02 ; HG2 - CG
- HG3
7335 7334 7337 1 1.0950e+02 4.1840e+02 ; HG2 - CG
- CD
7336 7334 7337 1 1.0950e+02 4.1840e+02 ; HG3 - CG
- CD
7337 7340 7341 1 1.0950e+02 4.1840e+02 ; CD - CE
- HE2
7337 7340 7342 1 1.0950e+02 4.1840e+02 ; CD - CE
- HE3
7337 7340 7343 1 1.1120e+02 6.6944e+02 ; CD - CE
- NZ
7338 7337 7339 1 1.0950e+02 2.9288e+02 ; HD2 - CD
- HD3
7338 7337 7340 1 1.0950e+02 4.1840e+02 ; HD2 - CD
- CE
7339 7337 7340 1 1.0950e+02 4.1840e+02 ; HD3 - CD
- CE
7340 7343 7344 1 1.0950e+02 4.1840e+02 ; CE - NZ
- HZ1
7340 7343 7345 1 1.0950e+02 4.1840e+02 ; CE - NZ
- HZ2
7340 7343 7346 1 1.0950e+02 4.1840e+02 ; CE - NZ
- HZ3
7341 7340 7342 1 1.0950e+02 2.9288e+02 ; HE2 - CE
- HE3
7341 7340 7343 1 1.0950e+02 4.1840e+02 ; HE2 - CE
- NZ
7342 7340 7343 1 1.0950e+02 4.1840e+02 ; HE3 - CE
- NZ
7344 7343 7345 1 1.0950e+02 2.9288e+02 ; HZ1 - NZ
- HZ2
7344 7343 7346 1 1.0950e+02 2.9288e+02 ; HZ1 - NZ
- HZ3

7345 7343 7346 1 1.0950e+02 2.9288e+02 ; HZ2 - NZ
- HZ3
7347 7349 7350 1 1.2000e+02 4.1840e+02 ; C - N
- H
7347 7349 7351 1 1.2190e+02 4.1840e+02 ; C - N
- CA
7348 7347 7349 1 1.2290e+02 6.6944e+02 ; O - C
- N
7349 7351 7352 1 1.0950e+02 4.1840e+02 ; N - CA
- HA
7349 7351 7353 1 1.0970e+02 6.6944e+02 ; N - CA
- CB
7349 7351 7369 1 1.1010e+02 5.2718e+02 ; N - CA
- C
7350 7349 7351 1 1.1804e+02 4.1840e+02 ; H - N
- CA
7351 7353 7354 1 1.0950e+02 4.1840e+02 ; CA - CB
- HB2
7351 7353 7355 1 1.0950e+02 4.1840e+02 ; CA - CB
- HB3
7351 7353 7356 1 1.0950e+02 3.3472e+02 ; CA - CB
- CG
7351 7369 7370 1 1.1700e+02 5.8576e+02 ; CA - C
- O
7351 7369 7371 1 1.1700e+02 5.8576e+02 ; CA - C
- OXT
7352 7351 7353 1 1.0950e+02 4.1840e+02 ; HA - CA
- CB
7352 7351 7369 1 1.0950e+02 4.1840e+02 ; HA - CA
- C
7353 7351 7369 1 1.1110e+02 5.2718e+02 ; CB - CA
- C
7353 7356 7357 1 1.0950e+02 4.1840e+02 ; CB - CG
- HG2
7353 7356 7358 1 1.0950e+02 4.1840e+02 ; CB - CG
- HG3
7353 7356 7359 1 1.0950e+02 3.3472e+02 ; CB - CG
- CD
7354 7353 7355 1 1.0950e+02 2.9288e+02 ; HB2 - CB
- HB3
7354 7353 7356 1 1.0950e+02 4.1840e+02 ; HB2 - CB
- CG
7355 7353 7356 1 1.0950e+02 4.1840e+02 ; HB3 - CB
- CG
7356 7359 7360 1 1.0950e+02 4.1840e+02 ; CG - CD
- HD2
7356 7359 7361 1 1.0950e+02 4.1840e+02 ; CG - CD
- HD3
7356 7359 7362 1 1.0950e+02 3.3472e+02 ; CG - CD
- CE
7357 7356 7358 1 1.0950e+02 2.9288e+02 ; HG2 - CG
- HG3

7357 7356 7359 1 1.0950e+02 4.1840e+02 ; HG2 - CG
- CD
7358 7356 7359 1 1.0950e+02 4.1840e+02 ; HG3 - CG
- CD
7359 7362 7363 1 1.0950e+02 4.1840e+02 ; CD - CE
- HE2
7359 7362 7364 1 1.0950e+02 4.1840e+02 ; CD - CE
- HE3
7359 7362 7365 1 1.1120e+02 6.6944e+02 ; CD - CE
- NZ
7360 7359 7361 1 1.0950e+02 2.9288e+02 ; HD2 - CD
- HD3
7360 7359 7362 1 1.0950e+02 4.1840e+02 ; HD2 - CD
- CE
7361 7359 7362 1 1.0950e+02 4.1840e+02 ; HD3 - CD
- CE
7362 7365 7366 1 1.0950e+02 4.1840e+02 ; CE - NZ
- HZ1
7362 7365 7367 1 1.0950e+02 4.1840e+02 ; CE - NZ
- HZ2
7362 7365 7368 1 1.0950e+02 4.1840e+02 ; CE - NZ
- HZ3
7363 7362 7364 1 1.0950e+02 2.9288e+02 ; HE2 - CE
- HE3
7363 7362 7365 1 1.0950e+02 4.1840e+02 ; HE2 - CE
- NZ
7364 7362 7365 1 1.0950e+02 4.1840e+02 ; HE3 - CE
- NZ
7366 7365 7367 1 1.0950e+02 2.9288e+02 ; HZ1 - NZ
- HZ2
7366 7365 7368 1 1.0950e+02 2.9288e+02 ; HZ1 - NZ
- HZ3
7367 7365 7368 1 1.0950e+02 2.9288e+02 ; HZ2 - NZ
- HZ3
7370 7369 7371 1 1.2600e+02 6.6944e+02 ; O - C
- OXT
7372 7373 7374 1 1.2002e+02 5.5731e+02 ; C1 - C2
- C3
7372 7373 7413 1 1.1988e+02 4.0334e+02 ; C1 - C2
- H42
7372 7377 7376 1 1.2069e+02 5.5480e+02 ; C1 - C6
- C5
7372 7377 7415 1 1.1988e+02 4.0334e+02 ; C1 - C6
- H44
7372 7388 7430 1 1.0858e+02 4.1003e+02 ; C1 - O17
- H59
7373 7372 7377 1 1.2002e+02 5.5731e+02 ; C2 - C1
- C6
7373 7372 7388 1 1.1990e+02 5.8158e+02 ; C2 - C1
- O17
7373 7374 7375 1 1.2002e+02 5.5731e+02 ; C2 - C3
- C4

| | | | | | | | |
|---|---|---|---|---|---|---|---|
| 7373 | 7374 | 7414 | 1 | 1.1988e+02 | 4.0334e+02 | ; | C2 - C3 - H43 |
| 7374 | 7373 | 7413 | 1 | 1.1988e+02 | 4.0334e+02 | ; | C3 - C2 - H42 |
| 7374 | 7375 | 7376 | 1 | 1.2069e+02 | 5.5480e+02 | ; | C3 - C4 - C5 |
| 7374 | 7375 | 7387 | 1 | 1.1957e+02 | 5.9078e+02 | ; | C3 - C4 - N16 |
| 7375 | 7374 | 7414 | 1 | 1.1988e+02 | 4.0334e+02 | ; | C4 - C3 - H43 |
| 7375 | 7376 | 7377 | 1 | 1.1838e+02 | 5.5815e+02 | ; | C4 - C5 - C6 |
| 7375 | 7376 | 7379 | 1 | 1.2111e+02 | 5.3555e+02 | ; | C4 - C5 - C8 |
| 7375 | 7387 | 7378 | 1 | 1.1220e+02 | 6.0082e+02 | ; | C4 - N16 - C7 |
| 7375 | 7387 | 7436 | 1 | 1.2691e+02 | 1.2773e+03 | ; | C4 - N16 - RU |
| 7376 | 7375 | 7387 | 1 | 1.1957e+02 | 5.9078e+02 | ; | C5 - C4 - N16 |
| 7376 | 7377 | 7415 | 1 | 1.1986e+02 | 4.0166e+02 | ; | C5 - C6 - H44 |
| 7376 | 7379 | 7378 | 1 | 1.2111e+02 | 5.3555e+02 | ; | C5 - C8 - C7 |
| 7376 | 7379 | 7405 | 1 | 1.2111e+02 | 5.3555e+02 | ; | C5 - C8 - C34 |
| 7377 | 7372 | 7388 | 1 | 1.1990e+02 | 5.8158e+02 | ; | C6 - C1 - O17 |
| 7377 | 7376 | 7379 | 1 | 1.2111e+02 | 5.3555e+02 | ; | C6 - C5 - C8 |
| 7378 | 7379 | 7405 | 1 | 1.1838e+02 | 5.5815e+02 | ; | C7 - C8 - C34 |
| 7378 | 7380 | 7381 | 1 | 1.2002e+02 | 5.5731e+02 | ; | C7 - C9 - C10 |
| 7378 | 7380 | 7386 | 1 | 1.1957e+02 | 5.9078e+02 | ; | C7 - C9 - N15 |
| 7378 | 7387 | 7436 | 1 | 1.2691e+02 | 1.2773e+03 | ; | C7 - N16 - RU |
| 7379 | 7378 | 7380 | 1 | 1.2069e+02 | 5.5480e+02 | ; | C8 - C7 - C9 |
| 7379 | 7378 | 7387 | 1 | 1.1957e+02 | 5.9078e+02 | ; | C8 - C7 - N16 |
| 7379 | 7405 | 7406 | 1 | 1.1152e+02 | 5.7572e+02 | ; | C8 - C34 - C35 |
| 7379 | 7405 | 7412 | 1 | 1.2069e+02 | 5.5480e+02 | ; | C8 - C34 - C41 |
| 7380 | 7378 | 7387 | 1 | 1.1957e+02 | 5.9078e+02 | ; | C9 - C7 - N16 |
| 7380 | 7381 | 7384 | 1 | 1.2060e+02 | 5.5898e+02 | ; | C9 - C10 - C13 |
| 7380 | 7381 | 7412 | 1 | 1.2002e+02 | 5.5731e+02 | ; | C9 - C10 - C41 |

7380 7386 7382 1 1.1994e+02 5.8576e+02 ; C9 - N15 - C11
7380 7386 7436 1 1.1146e+02 1.3327e+03 ; C9 - N15 - RU
7381 7380 7386 1 1.1957e+02 5.9078e+02 ; C10 - C9 - N15
7381 7384 7383 1 1.1700e+02 5.8074e+02 ; C10 - C13 - C12
7381 7384 7417 1 1.2330e+02 4.0083e+02 ; C10 - C13 - H46
7381 7412 7405 1 1.2002e+02 5.5731e+02 ; C10 - C41 - C34
7381 7412 7410 1 1.2060e+02 5.5898e+02 ; C10 - C41 - C39
7382 7383 7384 1 1.2181e+02 5.7990e+02 ; C11 - C12 - C13
7382 7383 7385 1 1.2287e+02 5.6819e+02 ; C11 - C12 - F14
7382 7386 7436 1 1.2985e+02 1.3404e+03 ; C11 - N15 - RU
7383 7382 7386 1 1.2601e+02 5.9580e+02 ; C12 - C11 - N15
7383 7382 7416 1 1.2267e+02 4.1338e+02 ; C12 - C11 - H45
7383 7384 7417 1 1.2043e+02 4.1756e+02 ; C12 - C13 - H46
7384 7381 7412 1 1.2060e+02 5.5898e+02 ; C13 - C10 - C41
7384 7383 7385 1 1.2287e+02 5.6819e+02 ; C13 - C12 - F14
7386 7382 7416 1 1.2099e+02 4.3932e+02 ; N15 - C11 - H45
7386 7436 7387 1 7.9470e+01 1.3390e+03 ; N15 - RU - N16
7386 7436 7389 1 1.4067e+02 0.0000e+00 ; N15 - RU - C18
7386 7436 7390 1 1.0655e+02 1.2648e+03 ; N15 - RU - C19
7386 7436 7391 1 9.9440e+01 1.2369e+03 ; N15 - RU - C20
7386 7436 7392 1 1.2283e+02 4.3179e+01 ; N15 - RU - C21
7386 7436 7393 1 1.5817e+02 9.1504e+02 ; N15 - RU - C22
7386 7436 7394 1 8.9270e+01 1.5543e+03 ; N15 - RU - C23
7387 7436 7389 1 1.3976e+02 5.2886e+01 ; N16 - RU - C18
7387 7436 7390 1 1.5915e+02 7.2308e+02 ; N16 - RU - C19
7387 7436 7391 1 1.2386e+02 6.0919e+01 ; N16 - RU - C20

7387 7436 7392 1 1.0006e+02 9.3730e+02 ; N16 - RU
- C21
7387 7436 7393 1 1.0629e+02 1.2381e+03 ; N16 - RU
- C22
7387 7436 7394 1 8.9820e+01 7.6099e+02 ; N16 - RU
- C23
7389 7390 7391 1 1.2181e+02 5.7990e+02 ; C18 - C19
- C20
7389 7390 7396 1 1.2181e+02 5.7990e+02 ; C18 - C19
- C25
7389 7390 7436 1 7.0640e+01 6.8651e+02 ; C18 - C19
- RU
7389 7393 7392 1 1.2181e+02 5.7990e+02 ; C18 - C22
- C21
7389 7393 7420 1 1.2043e+02 4.1756e+02 ; C18 - C22
- H49
7389 7393 7436 1 7.0410e+01 6.2183e+02 ; C18 - C22
- RU
7389 7436 7390 1 3.6770e+01 5.9664e+02 ; C18 - RU
- C19
7389 7436 7391 1 6.0140e+01 7.8760e+02 ; C18 - RU
- C20
7389 7436 7392 1 5.9990e+01 7.7036e+02 ; C18 - RU
- C21
7389 7436 7393 1 3.6590e+01 7.0977e+02 ; C18 - RU
- C22
7389 7436 7394 1 9.2800e+01 1.3018e+03 ; C18 - RU
- C23
7390 7389 7393 1 1.2181e+02 5.7990e+02 ; C19 - C18
- C22
7390 7389 7418 1 1.2043e+02 4.1756e+02 ; C19 - C18
- H47
7390 7389 7436 1 7.2600e+01 3.7623e+02 ; C19 - C18
- RU
7390 7391 7392 1 1.2181e+02 5.7990e+02 ; C19 - C20
- C21
7390 7391 7435 1 1.2043e+02 4.1756e+02 ; C19 - C20
- H64
7390 7391 7436 1 6.8880e+01 2.8886e+02 ; C19 - C20
- RU
7390 7396 7397 1 1.2367e+02 5.7488e+02 ; C19 - C25
- N26
7390 7396 7398 1 1.3089e+02 5.9915e+02 ; C19 - C25
- O27
7390 7436 7391 1 3.6230e+01 7.4559e+01 ; C19 - RU
- C20
7390 7436 7392 1 5.9660e+01 8.8023e+02 ; C19 - RU
- C21
7390 7436 7393 1 6.1000e+01 1.1206e+03 ; C19 - RU
- C22
7390 7436 7394 1 1.0993e+02 7.2525e+02 ; C19 - RU
- C23

7391 7390 7396 1 1.2181e+02 5.7990e+02 ; C20 - C19 - C25
7391 7390 7436 1 7.4890e+01 5.8819e+02 ; C20 - C19 - RU
7391 7392 7393 1 1.2181e+02 5.7990e+02 ; C20 - C21 - C22
7391 7392 7419 1 1.2043e+02 4.1756e+02 ; C20 - C21 - H48
7391 7392 7436 1 7.2430e+01 3.7288e+02 ; C20 - C21 - RU
7391 7436 7392 1 3.4610e+01 7.5136e+02 ; C20 - RU - C21
7391 7436 7393 1 5.9590e+01 8.8902e+02 ; C20 - RU - C22
7391 7436 7394 1 1.4614e+02 5.1045e+00 ; C20 - RU - C23
7392 7391 7435 1 1.2043e+02 4.1756e+02 ; C21 - C20 - H64
7392 7391 7436 1 7.2960e+01 3.6100e+02 ; C21 - C20 - RU
7392 7393 7420 1 1.2043e+02 4.1756e+02 ; C21 - C22 - H49
7392 7393 7436 1 7.4930e+01 6.4166e+02 ; C21 - C22 - RU
7392 7436 7393 1 3.6050e+01 1.1682e+02 ; C21 - RU - C22
7392 7436 7394 1 1.4753e+02 1.5548e+02 ; C21 - RU - C23
7393 7389 7418 1 1.2043e+02 4.1756e+02 ; C22 - C18 - H47
7393 7389 7436 1 7.3000e+01 3.6108e+02 ; C22 - C18 - RU
7393 7392 7419 1 1.2043e+02 4.1756e+02 ; C22 - C21 - H48
7393 7392 7436 1 6.9020e+01 2.9405e+02 ; C22 - C21 - RU
7393 7436 7394 1 1.1149e+02 7.4634e+02 ; C22 - RU - C23
7395 7394 7436 1 1.7732e+02 5.5497e+02 ; O24 - C23 - RU
7396 7390 7436 1 1.1993e+02 6.1421e+02 ; C25 - C19 - RU
7396 7397 7399 1 1.2010e+02 5.2802e+02 ; C25 - N26 - C28
7396 7397 7434 1 1.1790e+02 3.9832e+02 ; C25 - N26 - H63
7397 7396 7398 1 1.1746e+02 6.2843e+02 ; N26 - C25 - O27
7397 7399 7400 1 1.1161e+02 5.5145e+02 ; N26 - C28 - C29
7397 7399 7401 1 1.1161e+02 5.5145e+02 ; N26 - C28 - C30

7397 7399 7403 1 1.1161e+02 5.5145e+02 ; N26 - C28
- C32
7399 7397 7434 1 1.1768e+02 3.8325e+02 ; C28 - N26
- H63
7399 7400 7431 1 1.0980e+02 3.8744e+02 ; C28 - C29
- H60
7399 7400 7432 1 1.0980e+02 3.8744e+02 ; C28 - C29
- H61
7399 7400 7433 1 1.0980e+02 3.8744e+02 ; C28 - C29
- H62
7399 7401 7402 1 1.1019e+02 5.6484e+02 ; C28 - C30
- O31
7399 7401 7421 1 1.0956e+02 3.8828e+02 ; C28 - C30
- H50
7399 7401 7422 1 1.0956e+02 3.8828e+02 ; C28 - C30
- H51
7399 7403 7404 1 1.1019e+02 5.6484e+02 ; C28 - C32
- O33
7399 7403 7423 1 1.0956e+02 3.8828e+02 ; C28 - C32
- H52
7399 7403 7424 1 1.0956e+02 3.8828e+02 ; C28 - C32
- H53
7400 7399 7401 1 1.1151e+02 5.2635e+02 ; C29 - C28
- C30
7400 7399 7403 1 1.1151e+02 5.2635e+02 ; C29 - C28
- C32
7401 7399 7403 1 1.1151e+02 5.2635e+02 ; C30 - C28
- C32
7401 7402 7425 1 1.0726e+02 3.9664e+02 ; C30 - O31
- H54
7402 7401 7421 1 1.1026e+02 4.2593e+02 ; O31 - C30
- H50
7402 7401 7422 1 1.1026e+02 4.2593e+02 ; O31 - C30
- H51
7403 7404 7429 1 1.0726e+02 3.9664e+02 ; C32 - O33
- H58
7404 7403 7423 1 1.1026e+02 4.2593e+02 ; O33 - C32
- H52
7404 7403 7424 1 1.1026e+02 4.2593e+02 ; O33 - C32
- H53
7405 7406 7407 1 1.2367e+02 5.7488e+02 ; C34 - C35
- N36
7405 7406 7408 1 1.2359e+02 6.0919e+02 ; C34 - C35
- O37
7405 7412 7410 1 1.2060e+02 5.5898e+02 ; C34 - C41
- C39
7406 7405 7412 1 1.2060e+02 5.5898e+02 ; C35 - C34
- C41
7406 7407 7409 1 1.2010e+02 5.2802e+02 ; C35 - N36
- C38
7406 7407 7410 1 1.1675e+02 5.4810e+02 ; C35 - N36
- C39

```
  7407   7406   7408      1    1.1746e+02    6.2843e+02 ;    N36 - C35
- O37
  7407   7409   7426      1    1.0888e+02    4.1673e+02 ;    N36 - C38
- H55
  7407   7409   7427      1    1.0888e+02    4.1673e+02 ;    N36 - C38
- H56
  7407   7409   7428      1    1.0888e+02    4.1673e+02 ;    N36 - C38
- H57
  7407   7410   7411      1    1.1746e+02    6.2843e+02 ;    N36 - C39
- O40
  7407   7410   7412      1    1.2367e+02    5.7488e+02 ;    N36 - C39
- C41
  7409   7407   7410      1    1.2010e+02    5.2802e+02 ;    C38 - N36
- C39
  7411   7410   7412      1    1.2359e+02    6.0919e+02 ;    O40 - C39
- C41
  7418   7389   7436      1    1.2101e+02    2.5790e+02 ;    H47 - C18
- RU
  7419   7392   7436      1    1.2238e+02    2.4368e+02 ;    H48 - C21
- RU
  7420   7393   7436      1    1.2458e+02    2.5372e+02 ;    H49 - C22
- RU
  7421   7401   7422      1    1.0846e+02    3.2803e+02 ;    H50 - C30
- H51
  7423   7403   7424      1    1.0846e+02    3.2803e+02 ;    H52 - C32
- H53
  7426   7409   7427      1    1.0846e+02    3.2803e+02 ;    H55 - C38
- H56
  7426   7409   7428      1    1.0846e+02    3.2803e+02 ;    H55 - C38
- H57
  7427   7409   7428      1    1.0846e+02    3.2803e+02 ;    H56 - C38
- H57
  7431   7400   7432      1    1.0758e+02    3.2970e+02 ;    H60 - C29
- H61
  7431   7400   7433      1    1.0758e+02    3.2970e+02 ;    H60 - C29
- H62
  7432   7400   7433      1    1.0758e+02    3.2970e+02 ;    H61 - C29
- H62
  7435   7391   7436      1    1.2127e+02    2.4878e+02 ;    H64 - C20
- RU

[ dihedrals ] ; propers
; treated as RBs in GROMACS to use combine multiple AMBER torsions per
quartet
;    i      j      k      l   func     C0          C1          C2
C3          C4          C5
     1      5      7      8      3    0.65084    1.95253    0.00000
-2.60338    0.00000    0.00000 ;       N-    CA-    CB-   HB2
     1      5      7      9      3    0.65084    1.95253    0.00000
-2.60338    0.00000    0.00000 ;       N-    CA-    CB-   HB3
     1      5      7     10      3    3.79070    1.39746   -4.61077
```

```
-4.20074    2.51040    0.00000 ;      SG-    CB-    CA-     C
    69      71      73      74      3    0.00000    0.00000    0.00000
0.00000    0.00000    0.00000 ;      C-     N-    CA-    HA
    69      71      73      75      3   11.71520    1.67360   15.06240  -
13.38880    0.00000    0.00000 ;      C-     N-    CA-    CB
    69      71      73      87      3    1.75728    5.27184    2.25936
-7.02912    0.00000    0.00000 ;      C-     N-    CA-     C
    70      69      71      72      3   29.28800   -8.36800  -20.92000
0.00000    0.00000    0.00000 ;      O-     C-     N-     H
    70      69      71      73      3   20.92000    0.00000  -20.92000
0.00000    0.00000    0.00000 ;      O-     C-     N-    CA
    71      73      75      76      3    0.65084    1.95253    0.00000
-2.60338    0.00000    0.00000 ;      N-    CA-    CB-   HB2
    71      73      75      77      3    0.65084    1.95253    0.00000
-2.60338    0.00000    0.00000 ;      N-    CA-    CB-   HB3
    71      73      75      78      3    4.79068    4.02919   -4.82834
-3.66518    2.97901    0.00000 ;      N-    CA-    CB-    CG
    71      73      87      88      3    0.00000    0.00000    0.00000
0.00000    0.00000    0.00000 ;      N-    CA-     C-     O
    71      73      87      89      3   17.40544   -5.02080  -13.22144
9.20480    0.00000    0.00000 ;      N-    CA-     C-     N
    72      71      73      74      3    0.00000    0.00000    0.00000
0.00000    0.00000    0.00000 ;      H-     N-    CA-    HA
    72      71      73      75      3    0.00000    0.00000    0.00000
0.00000    0.00000    0.00000 ;      H-     N-    CA-    CB
    72      71      73      87      3    0.00000    0.00000    0.00000
0.00000    0.00000    0.00000 ;      H-     N-    CA-     C
    73      75      78      79      3    6.81992    9.28011   -2.04179  -
11.48090   -1.23846    0.00000 ;     CA-    CB-    CG-   ND1
    73      75      78      85      3    0.00000    0.00000    0.00000
0.00000    0.00000    0.00000 ;     CA-    CB-    CG-   CD2
    73      87      89      90      3   20.92000    0.00000  -20.92000
0.00000    0.00000    0.00000 ;     CA-     C-     N-     H
    73      87      89      91      3   20.92000    0.00000  -20.92000
0.00000    0.00000    0.00000 ;     CA-     C-     N-    CA
    74      73      75      76      3    0.65084    1.95253    0.00000
-2.60338    0.00000    0.00000 ;     HA-    CA-    CB-   HB2
    74      73      75      77      3    0.65084    1.95253    0.00000
-2.60338    0.00000    0.00000 ;     HA-    CA-    CB-   HB3
    74      73      75      78      3    0.65084    1.95253    0.00000
-2.60338    0.00000    0.00000 ;     HA-    CA-    CB-    CG
    74      73      87      88      3    3.68192   -4.35136    0.00000
1.33888    0.00000    0.00000 ;     HA-    CA-     C-     O
    74      73      87      89      3    0.00000    0.00000    0.00000
0.00000    0.00000    0.00000 ;     HA-    CA-     C-     N
    75      73      87      88      3    0.00000    0.00000    0.00000
0.00000    0.00000    0.00000 ;     CB-    CA-     C-     O
    75      73      87      89      3    2.51040    4.18400    1.67360
-6.69440    0.00000    0.00000 ;     CB-    CA-     C-     N
    75      78      79      80      3   11.71520    0.00000  -11.71520
0.00000    0.00000    0.00000 ;     CB-    CG-   ND1-   HD1
    75      78      79      81      3   11.71520    0.00000  -11.71520
```

0.00000 0.00000 0.00000 ; CB- CG- ND1- CE1
75 78 85 83 3 44.97800 0.00000 -44.97800
0.00000 0.00000 0.00000 ; CB- CG- CD2- NE2
75 78 85 86 3 44.97800 0.00000 -44.97800
0.00000 0.00000 0.00000 ; CB- CG- CD2- HD2
76 75 73 87 3 0.65084 1.95253 0.00000
-2.60338 0.00000 0.00000 ; HB2- CB- CA- C
76 75 78 79 3 0.00000 0.00000 0.00000
0.00000 0.00000 0.00000 ; HB2- CB- CG- ND1
76 75 78 85 3 0.00000 0.00000 0.00000
0.00000 0.00000 0.00000 ; HB2- CB- CG- CD2
77 75 73 87 3 0.65084 1.95253 0.00000
-2.60338 0.00000 0.00000 ; HB3- CB- CA- C
77 75 78 79 3 0.00000 0.00000 0.00000
0.00000 0.00000 0.00000 ; HB3- CB- CG- ND1
77 75 78 85 3 0.00000 0.00000 0.00000
0.00000 0.00000 0.00000 ; HB3- CB- CG- CD2
78 75 73 87 3 3.76560 3.34720 -2.87859
-3.66518 0.83680 0.00000 ; CG- CB- CA- C
78 79 81 82 3 19.45560 0.00000 -19.45560
0.00000 0.00000 0.00000 ; CG- ND1- CE1- HE1
78 79 81 83 3 19.45560 0.00000 -19.45560
0.00000 0.00000 0.00000 ; CG- ND1- CE1- NE2
78 85 83 81 3 12.55200 0.00000 -12.55200
0.00000 0.00000 0.00000 ; CG- CD2- NE2- CE1
78 85 83 84 3 12.55200 0.00000 -12.55200
0.00000 0.00000 0.00000 ; CG- CD2- NE2- HE2
79 78 85 83 3 44.97800 0.00000 -44.97800
0.00000 0.00000 0.00000 ; ND1- CG- CD2- NE2
79 78 85 86 3 44.97800 0.00000 -44.97800
0.00000 0.00000 0.00000 ; ND1- CG- CD2- HD2
79 81 83 84 3 19.45560 0.00000 -19.45560
0.00000 0.00000 0.00000 ; ND1- CE1- NE2- HE2
79 81 83 85 3 19.45560 0.00000 -19.45560
0.00000 0.00000 0.00000 ; ND1- CE1- NE2- CD2
80 79 78 85 3 11.71520 0.00000 -11.71520
0.00000 0.00000 0.00000 ; HD1- ND1- CG- CD2
80 79 81 82 3 19.45560 0.00000 -19.45560
0.00000 0.00000 0.00000 ; HD1- ND1- CE1- HE1
80 79 81 83 3 19.45560 0.00000 -19.45560
0.00000 0.00000 0.00000 ; HD1- ND1- CE1- NE2
81 79 78 85 3 11.71520 0.00000 -11.71520
0.00000 0.00000 0.00000 ; CE1- ND1- CG- CD2
81 83 85 86 3 12.55200 0.00000 -12.55200
0.00000 0.00000 0.00000 ; CE1- NE2- CD2- HD2
82 81 83 84 3 19.45560 0.00000 -19.45560
0.00000 0.00000 0.00000 ; HE1- CE1- NE2- HE2
82 81 83 85 3 19.45560 0.00000 -19.45560
0.00000 0.00000 0.00000 ; HE1- CE1- NE2- CD2
84 83 85 86 3 12.55200 0.00000 -12.55200
0.00000 0.00000 0.00000 ; HE2- NE2- CD2- HD2
87 89 91 92 3 0.00000 0.00000 0.00000

```
0.00000    0.00000    0.00000 ;      C-     N-    CA-    HA
    87     89     91     93      3   11.71520    1.67360   15.06240  -
13.38880    0.00000    0.00000 ;      C-     N-    CA-    CB
    87     89     91     98      3    1.75728    5.27184    2.25936
-7.02912    0.00000    0.00000 ;      C-     N-    CA-     C
    88     87     89     90      3   29.28800   -8.36800  -20.92000
0.00000    0.00000    0.00000 ;      O-     C-     N-     H
    88     87     89     91      3   20.92000    0.00000  -20.92000
0.00000    0.00000    0.00000 ;      O-     C-     N-    CA
    89     91     93     94      3    0.65084    1.95253    0.00000
-2.60338    0.00000    0.00000 ;      N-    CA-    CB-   HB2
    89     91     93     95      3    0.65084    1.95253    0.00000
-2.60338    0.00000    0.00000 ;      N-    CA-    CB-   HB3
    89     91     93     96      3    7.86174    2.24681   -7.41405
-6.71114    5.35552    0.00000 ;      N-    CA-    CB-    OG
    89     91     98     99      3    0.00000    0.00000    0.00000
0.00000    0.00000    0.00000 ;      N-    CA-     C-     O
    89     91     98    100      3   17.40544   -5.02080  -13.22144
9.20480    0.00000    0.00000 ;      N-    CA-     C-     N
    90     89     91     92      3    0.00000    0.00000    0.00000
0.00000    0.00000    0.00000 ;      H-     N-    CA-    HA
    90     89     91     93      3    0.00000    0.00000    0.00000
0.00000    0.00000    0.00000 ;      H-     N-    CA-    CB
    90     89     91     98      3    0.00000    0.00000    0.00000
0.00000    0.00000    0.00000 ;      H-     N-    CA-     C
    91     93     96     97      3    2.05853    2.46856    3.48109
-4.46851    0.23430    0.00000 ;     CA-    CB-    OG-    HG
    91     98    100    101      3   20.92000    0.00000  -20.92000
0.00000    0.00000    0.00000 ;     CA-     C-     N-     H
    91     98    100    102      3   20.92000    0.00000  -20.92000
0.00000    0.00000    0.00000 ;     CA-     C-     N-    CA
    92     91     93     94      3    0.65084    1.95253    0.00000
-2.60338    0.00000    0.00000 ;     HA-    CA-    CB-   HB2
    92     91     93     95      3    0.65084    1.95253    0.00000
-2.60338    0.00000    0.00000 ;     HA-    CA-    CB-   HB3
    92     91     93     96      3    1.04600   -1.04600    0.00000
0.00000    0.00000    0.00000 ;     HA-    CA-    CB-    OG
    92     91     98     99      3    3.68192   -4.35136    0.00000
1.33888    0.00000    0.00000 ;     HA-    CA-     C-     O
    92     91     98    100      3    0.00000    0.00000    0.00000
0.00000    0.00000    0.00000 ;     HA-    CA-     C-     N
    93     91     98     99      3    0.00000    0.00000    0.00000
0.00000    0.00000    0.00000 ;     CB-    CA-     C-     O
    93     91     98    100      3    2.51040    4.18400    1.67360
-6.69440    0.00000    0.00000 ;     CB-    CA-     C-     N
    94     93     91     98      3    0.65084    1.95253    0.00000
-2.60338    0.00000    0.00000 ;    HB2-    CB-    CA-     C
    94     93     96     97      3    0.69733    2.09200    0.00000
-2.78933    0.00000    0.00000 ;    HB2-    CB-    OG-    HG
    95     93     91     98      3    0.65084    1.95253    0.00000
-2.60338    0.00000    0.00000 ;    HB3-    CB-    CA-     C
    95     93     96     97      3    0.69733    2.09200    0.00000
```

-2.78933 0.00000 0.00000 ; HB3- CB- OG- HG
96 93 91 98 3 7.34710 7.79898 -6.14211
-6.71114 4.31789 0.00000 ; OG- CB- CA- C
98 100 102 103 3 0.00000 0.00000 0.00000
0.00000 0.00000 0.00000 ; C- N- CA- HA2
98 100 102 104 3 0.00000 0.00000 0.00000
0.00000 0.00000 0.00000 ; C- N- CA- HA3
98 100 102 105 3 1.75728 5.27184 2.25936
-7.02912 0.00000 0.00000 ; C- N- CA- C
99 98 100 101 3 29.28800 -8.36800 -20.92000
0.00000 0.00000 0.00000 ; O- C- N- H
99 98 100 102 3 20.92000 0.00000 -20.92000
0.00000 0.00000 0.00000 ; O- C- N- CA
100 102 105 106 3 0.00000 0.00000 0.00000
0.00000 0.00000 0.00000 ; N- CA- C- O
100 102 105 107 3 17.40544 -5.02080 -13.22144
9.20480 0.00000 0.00000 ; N- CA- C- N
101 100 102 103 3 0.00000 0.00000 0.00000
0.00000 0.00000 0.00000 ; H- N- CA- HA2
101 100 102 104 3 0.00000 0.00000 0.00000
0.00000 0.00000 0.00000 ; H- N- CA- HA3
101 100 102 105 3 0.00000 0.00000 0.00000
0.00000 0.00000 0.00000 ; H- N- CA- C
102 105 107 108 3 20.92000 0.00000 -20.92000
0.00000 0.00000 0.00000 ; CA- C- N- H
102 105 107 109 3 20.92000 0.00000 -20.92000
0.00000 0.00000 0.00000 ; CA- C- N- CA
103 102 105 106 3 3.68192 -4.35136 0.00000
1.33888 0.00000 0.00000 ; HA2- CA- C- O
103 102 105 107 3 0.00000 0.00000 0.00000
0.00000 0.00000 0.00000 ; HA2- CA- C- N
104 102 105 106 3 3.68192 -4.35136 0.00000
1.33888 0.00000 0.00000 ; HA3- CA- C- O
104 102 105 107 3 0.00000 0.00000 0.00000
0.00000 0.00000 0.00000 ; HA3- CA- C- N
105 107 109 110 3 0.00000 0.00000 0.00000
0.00000 0.00000 0.00000 ; C- N- CA- HA
105 107 109 111 3 11.71520 1.67360 15.06240 -
13.38880 0.00000 0.00000 ; C- N- CA- CB
105 107 109 127 3 1.75728 5.27184 2.25936
-7.02912 0.00000 0.00000 ; C- N- CA- C
106 105 107 108 3 29.28800 -8.36800 -20.92000
0.00000 0.00000 0.00000 ; O- C- N- H
106 105 107 109 3 20.92000 0.00000 -20.92000
0.00000 0.00000 0.00000 ; O- C- N- CA
107 109 111 112 3 0.65084 1.95253 0.00000
-2.60338 0.00000 0.00000 ; N- CA- CB- HB2
107 109 111 113 3 0.65084 1.95253 0.00000
-2.60338 0.00000 0.00000 ; N- CA- CB- HB3
107 109 111 114 3 0.65084 1.95253 0.00000
-2.60338 0.00000 0.00000 ; N- CA- CB- CG
107 109 127 128 3 0.00000 0.00000 0.00000

```
0.00000    0.00000    0.00000 ;       N-    CA-     C-     O
   107     109     127     129       3   17.40544   -5.02080  -13.22144
9.20480    0.00000    0.00000 ;       N-    CA-     C-     N
   108     107     109     110       3    0.00000    0.00000    0.00000
0.00000    0.00000    0.00000 ;       H-     N-    CA-    HA
   108     107     109     111       3    0.00000    0.00000    0.00000
0.00000    0.00000    0.00000 ;       H-     N-    CA-    CB
   108     107     109     127       3    0.00000    0.00000    0.00000
0.00000    0.00000    0.00000 ;       H-     N-    CA-     C
   109     111     114     115       3    0.66944    2.00832    0.00000
-2.67776    0.00000    0.00000 ;      CA-    CB-    CG-   HG2
   109     111     114     116       3    0.66944    2.00832    0.00000
-2.67776    0.00000    0.00000 ;      CA-    CB-    CG-   HG3
   109     111     114     117       3    3.68192    3.09616   -2.09200
-3.01248    0.00000    0.00000 ;      CA-    CB-    CG-    CD
   109     127     129     130       3   20.92000    0.00000  -20.92000
0.00000    0.00000    0.00000 ;      CA-     C-     N-     H
   109     127     129     131       3   20.92000    0.00000  -20.92000
0.00000    0.00000    0.00000 ;      CA-     C-     N-    CA
   110     109     111     112       3    0.65084    1.95253    0.00000
-2.60338    0.00000    0.00000 ;      HA-    CA-    CB-   HB2
   110     109     111     113       3    0.65084    1.95253    0.00000
-2.60338    0.00000    0.00000 ;      HA-    CA-    CB-   HB3
   110     109     111     114       3    0.65084    1.95253    0.00000
-2.60338    0.00000    0.00000 ;      HA-    CA-    CB-    CG
   110     109     127     128       3    3.68192   -4.35136    0.00000
1.33888    0.00000    0.00000 ;      HA-    CA-     C-     O
   110     109     127     129       3    0.00000    0.00000    0.00000
0.00000    0.00000    0.00000 ;      HA-    CA-     C-     N
   111     109     127     128       3    0.00000    0.00000    0.00000
0.00000    0.00000    0.00000 ;      CB-    CA-     C-     O
   111     109     127     129       3    2.51040    4.18400    1.67360
-6.69440    0.00000    0.00000 ;      CB-    CA-     C-     N
   111     114     117     118       3    0.66944    2.00832    0.00000
-2.67776    0.00000    0.00000 ;      CB-    CG-    CD-   HD2
   111     114     117     119       3    0.66944    2.00832    0.00000
-2.67776    0.00000    0.00000 ;      CB-    CG-    CD-   HD3
   111     114     117     120       3    3.68192    3.09616   -2.09200
-3.01248    0.00000    0.00000 ;      CB-    CG-    CD-    CE
   112     111     109     127       3    0.65084    1.95253    0.00000
-2.60338    0.00000    0.00000 ;     HB2-    CB-    CA-     C
   112     111     114     115       3    0.62760    1.88280    0.00000
-2.51040    0.00000    0.00000 ;     HB2-    CB-    CG-   HG2
   112     111     114     116       3    0.62760    1.88280    0.00000
-2.51040    0.00000    0.00000 ;     HB2-    CB-    CG-   HG3
   112     111     114     117       3    0.66944    2.00832    0.00000
-2.67776    0.00000    0.00000 ;     HB2-    CB-    CG-    CD
   113     111     109     127       3    0.65084    1.95253    0.00000
-2.60338    0.00000    0.00000 ;     HB3-    CB-    CA-     C
   113     111     114     115       3    0.62760    1.88280    0.00000
-2.51040    0.00000    0.00000 ;     HB3-    CB-    CG-   HG2
   113     111     114     116       3    0.62760    1.88280    0.00000
```

```
-2.51040    0.00000    0.00000 ;     HB3-    CB-     CG-   HG3
   113    111    114    117      3    0.66944    2.00832    0.00000
-2.67776    0.00000    0.00000 ;     HB3-    CB-     CG-    CD
   114    111    109    127      3    0.65084    1.95253    0.00000
-2.60338    0.00000    0.00000 ;      CG-    CB-     CA-     C
   114    117    120    121      3    0.65084    1.95253    0.00000
-2.60338    0.00000    0.00000 ;      CG-    CD-     CE-   HE2
   114    117    120    122      3    0.65084    1.95253    0.00000
-2.60338    0.00000    0.00000 ;      CG-    CD-     CE-   HE3
   114    117    120    123      3    0.65084    1.95253    0.00000
-2.60338    0.00000    0.00000 ;      CG-    CD-     CE-    NZ
   115    114    117    118      3    0.62760    1.88280    0.00000
-2.51040    0.00000    0.00000 ;     HG2-    CG-     CD-   HD2
   115    114    117    119      3    0.62760    1.88280    0.00000
-2.51040    0.00000    0.00000 ;     HG2-    CG-     CD-   HD3
   115    114    117    120      3    0.66944    2.00832    0.00000
-2.67776    0.00000    0.00000 ;     HG2-    CG-     CD-    CE
   116    114    117    118      3    0.62760    1.88280    0.00000
-2.51040    0.00000    0.00000 ;     HG3-    CG-     CD-   HD2
   116    114    117    119      3    0.62760    1.88280    0.00000
-2.51040    0.00000    0.00000 ;     HG3-    CG-     CD-   HD3
   116    114    117    120      3    0.66944    2.00832    0.00000
-2.67776    0.00000    0.00000 ;     HG3-    CG-     CD-    CE
   117    120    123    124      3    0.65084    1.95253    0.00000
-2.60338    0.00000    0.00000 ;      CD-    CE-     NZ-   HZ1
   117    120    123    125      3    0.65084    1.95253    0.00000
-2.60338    0.00000    0.00000 ;      CD-    CE-     NZ-   HZ2
   117    120    123    126      3    0.65084    1.95253    0.00000
-2.60338    0.00000    0.00000 ;      CD-    CE-     NZ-   HZ3
   118    117    120    121      3    0.65084    1.95253    0.00000
-2.60338    0.00000    0.00000 ;     HD2-    CD-     CE-   HE2
   118    117    120    122      3    0.65084    1.95253    0.00000
-2.60338    0.00000    0.00000 ;     HD2-    CD-     CE-   HE3
   118    117    120    123      3    0.65084    1.95253    0.00000
-2.60338    0.00000    0.00000 ;     HD2-    CD-     CE-    NZ
   119    117    120    121      3    0.65084    1.95253    0.00000
-2.60338    0.00000    0.00000 ;     HD3-    CD-     CE-   HE2
   119    117    120    122      3    0.65084    1.95253    0.00000
-2.60338    0.00000    0.00000 ;     HD3-    CD-     CE-   HE3
   119    117    120    123      3    0.65084    1.95253    0.00000
-2.60338    0.00000    0.00000 ;     HD3-    CD-     CE-    NZ
   121    120    123    124      3    0.65084    1.95253    0.00000
-2.60338    0.00000    0.00000 ;     HE2-    CE-     NZ-   HZ1
   121    120    123    125      3    0.65084    1.95253    0.00000
-2.60338    0.00000    0.00000 ;     HE2-    CE-     NZ-   HZ2
   121    120    123    126      3    0.65084    1.95253    0.00000
-2.60338    0.00000    0.00000 ;     HE2-    CE-     NZ-   HZ3
   122    120    123    124      3    0.65084    1.95253    0.00000
-2.60338    0.00000    0.00000 ;     HE3-    CE-     NZ-   HZ1
   122    120    123    125      3    0.65084    1.95253    0.00000
-2.60338    0.00000    0.00000 ;     HE3-    CE-     NZ-   HZ2
   122    120    123    126      3    0.65084    1.95253    0.00000
```

-2.60338 0.00000 0.00000 ; HE3- CE- NZ- HZ3
127 129 131 132 3 0.00000 0.00000 0.00000
0.00000 0.00000 0.00000 ; C- N- CA- HA
127 129 131 133 3 11.71520 1.67360 15.06240 -
13.38880 0.00000 0.00000 ; C- N- CA- CB
127 129 131 146 3 1.75728 5.27184 2.25936
-7.02912 0.00000 0.00000 ; C- N- CA- C
128 127 129 130 3 29.28800 -8.36800 -20.92000
0.00000 0.00000 0.00000 ; O- C- N- H
128 127 129 131 3 20.92000 0.00000 -20.92000
0.00000 0.00000 0.00000 ; O- C- N- CA
129 131 133 134 3 0.65084 1.95253 0.00000
-2.60338 0.00000 0.00000 ; N- CA- CB- HB2
129 131 133 135 3 0.65084 1.95253 0.00000
-2.60338 0.00000 0.00000 ; N- CA- CB- HB3
129 131 133 136 3 3.79070 1.39746 -4.61077
-2.40998 2.44346 0.00000 ; N- CA- CB- CG
129 131 146 147 3 0.00000 0.00000 0.00000
0.00000 0.00000 0.00000 ; N- CA- C- O
129 131 146 148 3 17.40544 -5.02080 -13.22144
9.20480 0.00000 0.00000 ; N- CA- C- N
130 129 131 132 3 0.00000 0.00000 0.00000
0.00000 0.00000 0.00000 ; H- N- CA- HA
130 129 131 133 3 0.00000 0.00000 0.00000
0.00000 0.00000 0.00000 ; H- N- CA- CB
130 129 131 146 3 0.00000 0.00000 0.00000
0.00000 0.00000 0.00000 ; H- N- CA- C
131 133 136 137 3 0.66944 2.00832 0.00000
-2.67776 0.00000 0.00000 ; CA- CB- CG- HG
131 133 136 138 3 3.90367 0.19665 -6.21742
-2.37651 5.99149 0.00000 ; CA- CB- CG- CD1
131 133 136 142 3 3.90367 0.19665 -6.21742
-2.37651 5.99149 0.00000 ; CA- CB- CG- CD2
131 146 148 149 3 20.92000 0.00000 -20.92000
0.00000 0.00000 0.00000 ; CA- C- N- H
131 146 148 150 3 20.92000 0.00000 -20.92000
0.00000 0.00000 0.00000 ; CA- C- N- CA
132 131 133 134 3 0.65084 1.95253 0.00000
-2.60338 0.00000 0.00000 ; HA- CA- CB- HB2
132 131 133 135 3 0.65084 1.95253 0.00000
-2.60338 0.00000 0.00000 ; HA- CA- CB- HB3
132 131 133 136 3 0.65084 1.95253 0.00000
-2.60338 0.00000 0.00000 ; HA- CA- CB- CG
132 131 146 147 3 3.68192 -4.35136 0.00000
1.33888 0.00000 0.00000 ; HA- CA- C- O
132 131 146 148 3 0.00000 0.00000 0.00000
0.00000 0.00000 0.00000 ; HA- CA- C- N
133 131 146 147 3 0.00000 0.00000 0.00000
0.00000 0.00000 0.00000 ; CB- CA- C- O
133 131 146 148 3 2.51040 4.18400 1.67360
-6.69440 0.00000 0.00000 ; CB- CA- C- N
133 136 138 139 3 0.66944 2.00832 0.00000

```
-2.67776    0.00000    0.00000 ;      CB-    CG-   CD1-  HD11
   133    136    138    140       3    0.66944    2.00832    0.00000
-2.67776    0.00000    0.00000 ;      CB-    CG-   CD1-  HD12
   133    136    138    141       3    0.66944    2.00832    0.00000
-2.67776    0.00000    0.00000 ;      CB-    CG-   CD1-  HD13
   133    136    142    143       3    0.66944    2.00832    0.00000
-2.67776    0.00000    0.00000 ;      CB-    CG-   CD2-  HD21
   133    136    142    144       3    0.66944    2.00832    0.00000
-2.67776    0.00000    0.00000 ;      CB-    CG-   CD2-  HD22
   133    136    142    145       3    0.66944    2.00832    0.00000
-2.67776    0.00000    0.00000 ;      CB-    CG-   CD2-  HD23
   134    133    131    146       3    0.65084    1.95253    0.00000
-2.60338    0.00000    0.00000 ;     HB2-    CB-    CA-     C
   134    133    136    137       3    0.62760    1.88280    0.00000
-2.51040    0.00000    0.00000 ;     HB2-    CB-    CG-    HG
   134    133    136    138       3    0.66944    2.00832    0.00000
-2.67776    0.00000    0.00000 ;     HB2-    CB-    CG-   CD1
   134    133    136    142       3    0.66944    2.00832    0.00000
-2.67776    0.00000    0.00000 ;     HB2-    CB-    CG-   CD2
   135    133    131    146       3    0.65084    1.95253    0.00000
-2.60338    0.00000    0.00000 ;     HB3-    CB-    CA-     C
   135    133    136    137       3    0.62760    1.88280    0.00000
-2.51040    0.00000    0.00000 ;     HB3-    CB-    CG-    HG
   135    133    136    138       3    0.66944    2.00832    0.00000
-2.67776    0.00000    0.00000 ;     HB3-    CB-    CG-   CD1
   135    133    136    142       3    0.66944    2.00832    0.00000
-2.67776    0.00000    0.00000 ;     HB3-    CB-    CG-   CD2
   136    133    131    146       3   10.33448   -1.14642  -11.54784
-2.40998    6.35968    0.00000 ;      CG-    CB-    CA-     C
   137    136    138    139       3    0.62760    1.88280    0.00000
-2.51040    0.00000    0.00000 ;      HG-    CG-   CD1-  HD11
   137    136    138    140       3    0.62760    1.88280    0.00000
-2.51040    0.00000    0.00000 ;      HG-    CG-   CD1-  HD12
   137    136    138    141       3    0.62760    1.88280    0.00000
-2.51040    0.00000    0.00000 ;      HG-    CG-   CD1-  HD13
   137    136    142    143       3    0.62760    1.88280    0.00000
-2.51040    0.00000    0.00000 ;      HG-    CG-   CD2-  HD21
   137    136    142    144       3    0.62760    1.88280    0.00000
-2.51040    0.00000    0.00000 ;      HG-    CG-   CD2-  HD22
   137    136    142    145       3    0.62760    1.88280    0.00000
-2.51040    0.00000    0.00000 ;      HG-    CG-   CD2-  HD23
   138    136    142    143       3    0.66944    2.00832    0.00000
-2.67776    0.00000    0.00000 ;     CD1-    CG-   CD2-  HD21
   138    136    142    144       3    0.66944    2.00832    0.00000
-2.67776    0.00000    0.00000 ;     CD1-    CG-   CD2-  HD22
   138    136    142    145       3    0.66944    2.00832    0.00000
-2.67776    0.00000    0.00000 ;     CD1-    CG-   CD2-  HD23
   139    138    136    142       3    0.66944    2.00832    0.00000
-2.67776    0.00000    0.00000 ;    HD11-   CD1-    CG-   CD2
   140    138    136    142       3    0.66944    2.00832    0.00000
-2.67776    0.00000    0.00000 ;    HD12-   CD1-    CG-   CD2
   141    138    136    142       3    0.66944    2.00832    0.00000
```

```
-2.67776    0.00000    0.00000 ;   HD13-   CD1-    CG-   CD2
   146     148     150     151       3    0.00000    0.00000    0.00000
0.00000    0.00000    0.00000 ;      C-      N-    CA-   HA
   146     148     150     152       3   11.71520    1.67360   15.06240  -
13.38880    0.00000    0.00000 ;      C-      N-    CA-   CB
   146     148     150     161       3    1.75728    5.27184    2.25936
-7.02912    0.00000    0.00000 ;      C-      N-    CA-    C
   147     146     148     149       3   29.28800   -8.36800  -20.92000
0.00000    0.00000    0.00000 ;      O-      C-     N-    H
   147     146     148     150       3   20.92000    0.00000  -20.92000
0.00000    0.00000    0.00000 ;      O-      C-     N-   CA
   148     150     152     153       3    0.65084    1.95253    0.00000
-2.60338    0.00000    0.00000 ;      N-    CA-    CB-   HB2
   148     150     152     154       3    0.65084    1.95253    0.00000
-2.60338    0.00000    0.00000 ;      N-    CA-    CB-   HB3
   148     150     152     155       3    3.21331    2.22589   -4.15053
-2.40998    2.61082    0.00000 ;      N-    CA-    CB-    CG
   148     150     161     162       3    0.00000    0.00000    0.00000
0.00000    0.00000    0.00000 ;      N-    CA-     C-     O
   148     150     161     163       3   17.40544   -5.02080  -13.22144
9.20480    0.00000    0.00000 ;      N-    CA-     C-     N
   149     148     150     151       3    0.00000    0.00000    0.00000
0.00000    0.00000    0.00000 ;      H-      N-    CA-   HA
   149     148     150     152       3    0.00000    0.00000    0.00000
0.00000    0.00000    0.00000 ;      H-      N-    CA-   CB
   149     148     150     161       3    0.00000    0.00000    0.00000
0.00000    0.00000    0.00000 ;      H-      N-    CA-     C
   150     152     155     156       3    0.66944    2.00832    0.00000
-2.67776    0.00000    0.00000 ;     CA-    CB-    CG-   HG2
   150     152     155     157       3    0.66944    2.00832    0.00000
-2.67776    0.00000    0.00000 ;     CA-    CB-    CG-   HG3
   150     152     155     158       3   10.12110   -1.91209    0.01674
10.17549   -1.87443    0.00000 ;     CA-    CB-    CG-    CD
   150     161     163     164       3   20.92000    0.00000  -20.92000
0.00000    0.00000    0.00000 ;     CA-     C-     N-     H
   150     161     163     165       3   20.92000    0.00000  -20.92000
0.00000    0.00000    0.00000 ;     CA-     C-     N-   CA
   151     150     152     153       3    0.65084    1.95253    0.00000
-2.60338    0.00000    0.00000 ;     HA-    CA-    CB-   HB2
   151     150     152     154       3    0.65084    1.95253    0.00000
-2.60338    0.00000    0.00000 ;     HA-    CA-    CB-   HB3
   151     150     152     155       3    0.65084    1.95253    0.00000
-2.60338    0.00000    0.00000 ;     HA-    CA-    CB-    CG
   151     150     161     162       3    3.68192   -4.35136    0.00000
1.33888    0.00000    0.00000 ;     HA-    CA-     C-     O
   151     150     161     163       3    0.00000    0.00000    0.00000
0.00000    0.00000    0.00000 ;     HA-    CA-     C-     N
   152     150     161     162       3    0.00000    0.00000    0.00000
0.00000    0.00000    0.00000 ;     CB-    CA-     C-     O
   152     150     161     163       3    2.51040    4.18400    1.67360
-6.69440    0.00000    0.00000 ;     CB-    CA-     C-     N
   152     155     158     159       3    3.79907    0.00000   -5.40573
```

0.00000    2.14221    0.00000 ;     CB-    CG-    CD-   OE1
152    155    158    160    3    3.79907    0.00000   -5.40573
0.00000    2.14221    0.00000 ;     CB-    CG-    CD-   OE2
153    152    150    161    3    0.65084    1.95253    0.00000
-2.60338    0.00000    0.00000 ;    HB2-    CB-    CA-     C
153    152    155    156    3    0.62760    1.88280    0.00000
-2.51040    0.00000    0.00000 ;    HB2-    CB-    CG-   HG2
153    152    155    157    3    0.62760    1.88280    0.00000
-2.51040    0.00000    0.00000 ;    HB2-    CB-    CG-   HG3
153    152    155    158    3    0.65084    1.95253    0.00000
-2.60338    0.00000    0.00000 ;    HB2-    CB-    CG-    CD
154    152    150    161    3    0.65084    1.95253    0.00000
-2.60338    0.00000    0.00000 ;    HB3-    CB-    CA-     C
154    152    155    156    3    0.62760    1.88280    0.00000
-2.51040    0.00000    0.00000 ;    HB3-    CB-    CG-   HG2
154    152    155    157    3    0.62760    1.88280    0.00000
-2.51040    0.00000    0.00000 ;    HB3-    CB-    CG-   HG3
154    152    155    158    3    0.65084    1.95253    0.00000
-2.60338    0.00000    0.00000 ;    HB3-    CB-    CG-    CD
155    152    150    161    3    6.86594    3.56895   -8.14206
-2.40998    4.85344    0.00000 ;     CG-    CB-    CA-     C
156    155    158    159    3    0.00000    0.00000    0.00000
0.00000    0.00000    0.00000 ;    HG2-    CG-    CD-   OE1
156    155    158    160    3    0.00000    0.00000    0.00000
0.00000    0.00000    0.00000 ;    HG2-    CG-    CD-   OE2
157    155    158    159    3    0.00000    0.00000    0.00000
0.00000    0.00000    0.00000 ;    HG3-    CG-    CD-   OE1
157    155    158    160    3    0.00000    0.00000    0.00000
0.00000    0.00000    0.00000 ;    HG3-    CG-    CD-   OE2
161    163    165    166    3    0.00000    0.00000    0.00000
0.00000    0.00000    0.00000 ;     C-     N-    CA-    HA
161    163    165    167    3   11.71520    1.67360   15.06240  -
13.38880    0.00000    0.00000 ;     C-     N-    CA-    CB
161    163    165    173    3    1.75728    5.27184    2.25936
-7.02912    0.00000    0.00000 ;     C-     N-    CA-     C
162    161    163    164    3   29.28800   -8.36800  -20.92000
0.00000    0.00000    0.00000 ;     O-     C-     N-     H
162    161    163    165    3   20.92000    0.00000  -20.92000
0.00000    0.00000    0.00000 ;     O-     C-     N-    CA
163    165    167    168    3    0.65084    1.95253    0.00000
-2.60338    0.00000    0.00000 ;     N-    CA-    CB-   HB2
163    165    167    169    3    0.65084    1.95253    0.00000
-2.60338    0.00000    0.00000 ;     N-    CA-    CB-   HB3
163    165    167    170    3   15.41386    9.74035   -8.39310
-0.97069    2.97901    0.00000 ;     N-    CA-    CB-    CG
163    165    173    174    3    0.00000    0.00000    0.00000
0.00000    0.00000    0.00000 ;     N-    CA-     C-     O
163    165    173    175    3   17.40544   -5.02080  -13.22144
9.20480    0.00000    0.00000 ;     N-    CA-     C-     N
164    163    165    166    3    0.00000    0.00000    0.00000
0.00000    0.00000    0.00000 ;     H-     N-    CA-    HA
164    163    165    167    3    0.00000    0.00000    0.00000

```
0.00000    0.00000    0.00000 ;       H-     N-     CA-    CB
   164     163     165     173       3    0.00000    0.00000    0.00000
0.00000    0.00000    0.00000 ;       H-     N-     CA-     C
   165     167     170     171       3    6.43499    0.00000   -5.39736
0.00000   -1.03763    0.00000 ;      CA-    CB-     CG-   OD1
   165     167     170     172       3    6.43499    0.00000   -5.39736
0.00000   -1.03763    0.00000 ;      CA-    CB-     CG-   OD2
   165     173     175     176       3   20.92000    0.00000  -20.92000
0.00000    0.00000    0.00000 ;      CA-     C-      N-     H
   165     173     175     177       3   20.92000    0.00000  -20.92000
0.00000    0.00000    0.00000 ;      CA-     C-      N-    CA
   166     165     167     168       3    0.65084    1.95253    0.00000
-2.60338    0.00000    0.00000 ;      HA-    CA-     CB-   HB2
   166     165     167     169       3    0.65084    1.95253    0.00000
-2.60338    0.00000    0.00000 ;      HA-    CA-     CB-   HB3
   166     165     167     170       3    0.65084    1.95253    0.00000
-2.60338    0.00000    0.00000 ;      HA-    CA-     CB-    CG
   166     165     173     174       3    3.68192   -4.35136    0.00000
1.33888    0.00000    0.00000 ;      HA-    CA-      C-     O
   166     165     173     175       3    0.00000    0.00000    0.00000
0.00000    0.00000    0.00000 ;      HA-    CA-      C-     N
   167     165     173     174       3    0.00000    0.00000    0.00000
0.00000    0.00000    0.00000 ;      CB-    CA-      C-     O
   167     165     173     175       3    2.51040    4.18400    1.67360
-6.69440    0.00000    0.00000 ;      CB-    CA-      C-     N
   168     167     165     173       3    0.65084    1.95253    0.00000
-2.60338    0.00000    0.00000 ;     HB2-    CB-     CA-     C
   168     167     170     171       3    0.00000    0.00000    0.00000
0.00000    0.00000    0.00000 ;     HB2-    CB-     CG-   OD1
   168     167     170     172       3    0.00000    0.00000    0.00000
0.00000    0.00000    0.00000 ;     HB2-    CB-     CG-   OD2
   169     167     165     173       3    0.65084    1.95253    0.00000
-2.60338    0.00000    0.00000 ;     HB3-    CB-     CA-     C
   169     167     170     171       3    0.00000    0.00000    0.00000
0.00000    0.00000    0.00000 ;     HB3-    CB-     CG-   OD1
   169     167     170     172       3    0.00000    0.00000    0.00000
0.00000    0.00000    0.00000 ;     HB3-    CB-     CG-   OD2
   170     167     165     173       3    7.14627   -1.04600   -8.99560
-0.97069    5.15469    0.00000 ;      CG-    CB-     CA-     C
   173     175     177     178       3    0.00000    0.00000    0.00000
0.00000    0.00000    0.00000 ;       C-     N-     CA-    HA
   173     175     177     179       3   11.71520    1.67360   15.06240  -
13.38880    0.00000    0.00000 ;       C-     N-     CA-    CB
   173     175     177     195       3    1.75728    5.27184    2.25936
-7.02912    0.00000    0.00000 ;       C-     N-     CA-     C
   174     173     175     176       3   29.28800   -8.36800  -20.92000
0.00000    0.00000    0.00000 ;       O-     C-      N-     H
   174     173     175     177       3   20.92000    0.00000  -20.92000
0.00000    0.00000    0.00000 ;       O-     C-      N-    CA
   175     177     179     180       3    0.65084    1.95253    0.00000
-2.60338    0.00000    0.00000 ;       N-    CA-     CB-   HB2
   175     177     179     181       3    0.65084    1.95253    0.00000
```

```
-2.60338    0.00000    0.00000 ;       N-    CA-    CB-   HB3
   175    177    179    182       3    0.65084    1.95253    0.00000
-2.60338    0.00000    0.00000 ;       N-    CA-    CB-    CG
   175    177    195    196       3    0.00000    0.00000    0.00000
0.00000    0.00000    0.00000 ;       N-    CA-     C-     O
   175    177    195    197       3   17.40544   -5.02080  -13.22144
9.20480    0.00000    0.00000 ;       N-    CA-     C-     N
   176    175    177    178       3    0.00000    0.00000    0.00000
0.00000    0.00000    0.00000 ;       H-     N-    CA-    HA
   176    175    177    179       3    0.00000    0.00000    0.00000
0.00000    0.00000    0.00000 ;       H-     N-    CA-    CB
   176    175    177    195       3    0.00000    0.00000    0.00000
0.00000    0.00000    0.00000 ;       H-     N-    CA-     C
   177    179    182    183       3    0.66944    2.00832    0.00000
-2.67776    0.00000    0.00000 ;      CA-    CB-    CG-   HG2
   177    179    182    184       3    0.66944    2.00832    0.00000
-2.67776    0.00000    0.00000 ;      CA-    CB-    CG-   HG3
   177    179    182    185       3    3.68192    3.09616   -2.09200
-3.01248    0.00000    0.00000 ;      CA-    CB-    CG-    CD
   177    195    197    198       3   20.92000    0.00000  -20.92000
0.00000    0.00000    0.00000 ;      CA-     C-     N-     H
   177    195    197    199       3   20.92000    0.00000  -20.92000
0.00000    0.00000    0.00000 ;      CA-     C-     N-    CA
   178    177    179    180       3    0.65084    1.95253    0.00000
-2.60338    0.00000    0.00000 ;      HA-    CA-    CB-   HB2
   178    177    179    181       3    0.65084    1.95253    0.00000
-2.60338    0.00000    0.00000 ;      HA-    CA-    CB-   HB3
   178    177    179    182       3    0.65084    1.95253    0.00000
-2.60338    0.00000    0.00000 ;      HA-    CA-    CB-    CG
   178    177    195    196       3    3.68192   -4.35136    0.00000
1.33888    0.00000    0.00000 ;      HA-    CA-     C-     O
   178    177    195    197       3    0.00000    0.00000    0.00000
0.00000    0.00000    0.00000 ;      HA-    CA-     C-     N
   179    177    195    196       3    0.00000    0.00000    0.00000
0.00000    0.00000    0.00000 ;      CB-    CA-     C-     O
   179    177    195    197       3    2.51040    4.18400    1.67360
-6.69440    0.00000    0.00000 ;      CB-    CA-     C-     N
   179    182    185    186       3    0.66944    2.00832    0.00000
-2.67776    0.00000    0.00000 ;      CB-    CG-    CD-   HD2
   179    182    185    187       3    0.66944    2.00832    0.00000
-2.67776    0.00000    0.00000 ;      CB-    CG-    CD-   HD3
   179    182    185    188       3    3.68192    3.09616   -2.09200
-3.01248    0.00000    0.00000 ;      CB-    CG-    CD-    CE
   180    179    177    195       3    0.65084    1.95253    0.00000
-2.60338    0.00000    0.00000 ;     HB2-    CB-    CA-     C
   180    179    182    183       3    0.62760    1.88280    0.00000
-2.51040    0.00000    0.00000 ;     HB2-    CB-    CG-   HG2
   180    179    182    184       3    0.62760    1.88280    0.00000
-2.51040    0.00000    0.00000 ;     HB2-    CB-    CG-   HG3
   180    179    182    185       3    0.66944    2.00832    0.00000
-2.67776    0.00000    0.00000 ;     HB2-    CB-    CG-    CD
   181    179    177    195       3    0.65084    1.95253    0.00000
```

```
 -2.60338    0.00000    0.00000 ;     HB3-    CB-    CA-     C
    181    179    182    183      3    0.62760    1.88280    0.00000
 -2.51040    0.00000    0.00000 ;     HB3-    CB-    CG-   HG2
    181    179    182    184      3    0.62760    1.88280    0.00000
 -2.51040    0.00000    0.00000 ;     HB3-    CB-    CG-   HG3
    181    179    182    185      3    0.66944    2.00832    0.00000
 -2.67776    0.00000    0.00000 ;     HB3-    CB-    CG-    CD
    182    179    177    195      3    0.65084    1.95253    0.00000
 -2.60338    0.00000    0.00000 ;      CG-    CB-    CA-     C
    182    185    188    189      3    0.65084    1.95253    0.00000
 -2.60338    0.00000    0.00000 ;      CG-    CD-    CE-   HE2
    182    185    188    190      3    0.65084    1.95253    0.00000
 -2.60338    0.00000    0.00000 ;      CG-    CD-    CE-   HE3
    182    185    188    191      3    0.65084    1.95253    0.00000
 -2.60338    0.00000    0.00000 ;      CG-    CD-    CE-    NZ
    183    182    185    186      3    0.62760    1.88280    0.00000
 -2.51040    0.00000    0.00000 ;     HG2-    CG-    CD-   HD2
    183    182    185    187      3    0.62760    1.88280    0.00000
 -2.51040    0.00000    0.00000 ;     HG2-    CG-    CD-   HD3
    183    182    185    188      3    0.66944    2.00832    0.00000
 -2.67776    0.00000    0.00000 ;     HG2-    CG-    CD-    CE
    184    182    185    186      3    0.62760    1.88280    0.00000
 -2.51040    0.00000    0.00000 ;     HG3-    CG-    CD-   HD2
    184    182    185    187      3    0.62760    1.88280    0.00000
 -2.51040    0.00000    0.00000 ;     HG3-    CG-    CD-   HD3
    184    182    185    188      3    0.66944    2.00832    0.00000
 -2.67776    0.00000    0.00000 ;     HG3-    CG-    CD-    CE
    185    188    191    192      3    0.65084    1.95253    0.00000
 -2.60338    0.00000    0.00000 ;      CD-    CE-    NZ-   HZ1
    185    188    191    193      3    0.65084    1.95253    0.00000
 -2.60338    0.00000    0.00000 ;      CD-    CE-    NZ-   HZ2
    185    188    191    194      3    0.65084    1.95253    0.00000
 -2.60338    0.00000    0.00000 ;      CD-    CE-    NZ-   HZ3
    186    185    188    189      3    0.65084    1.95253    0.00000
 -2.60338    0.00000    0.00000 ;     HD2-    CD-    CE-   HE2
    186    185    188    190      3    0.65084    1.95253    0.00000
 -2.60338    0.00000    0.00000 ;     HD2-    CD-    CE-   HE3
    186    185    188    191      3    0.65084    1.95253    0.00000
 -2.60338    0.00000    0.00000 ;     HD2-    CD-    CE-    NZ
    187    185    188    189      3    0.65084    1.95253    0.00000
 -2.60338    0.00000    0.00000 ;     HD3-    CD-    CE-   HE2
    187    185    188    190      3    0.65084    1.95253    0.00000
 -2.60338    0.00000    0.00000 ;     HD3-    CD-    CE-   HE3
    187    185    188    191      3    0.65084    1.95253    0.00000
 -2.60338    0.00000    0.00000 ;     HD3-    CD-    CE-    NZ
    189    188    191    192      3    0.65084    1.95253    0.00000
 -2.60338    0.00000    0.00000 ;     HE2-    CE-    NZ-   HZ1
    189    188    191    193      3    0.65084    1.95253    0.00000
 -2.60338    0.00000    0.00000 ;     HE2-    CE-    NZ-   HZ2
    189    188    191    194      3    0.65084    1.95253    0.00000
 -2.60338    0.00000    0.00000 ;     HE2-    CE-    NZ-   HZ3
    190    188    191    192      3    0.65084    1.95253    0.00000
```

-2.60338 0.00000 0.00000 ; HE3- CE- NZ- HZ1
190 188 191 193 3 0.65084 1.95253 0.00000
-2.60338 0.00000 0.00000 ; HE3- CE- NZ- HZ2
190 188 191 194 3 0.65084 1.95253 0.00000
-2.60338 0.00000 0.00000 ; HE3- CE- NZ- HZ3
195 197 199 200 3 0.00000 0.00000 0.00000
0.00000 0.00000 0.00000 ; C- N- CA- HA
195 197 199 201 3 11.71520 1.67360 15.06240 -
13.38880 0.00000 0.00000 ; C- N- CA- CB
195 197 199 216 3 1.75728 5.27184 2.25936
-7.02912 0.00000 0.00000 ; C- N- CA- C
196 195 197 198 3 29.28800 -8.36800 -20.92000
0.00000 0.00000 0.00000 ; O- C- N- H
196 195 197 199 3 20.92000 0.00000 -20.92000
0.00000 0.00000 0.00000 ; O- C- N- CA
197 199 201 202 3 0.65084 1.95253 0.00000
-2.60338 0.00000 0.00000 ; N- CA- CB- HB2
197 199 201 203 3 0.65084 1.95253 0.00000
-2.60338 0.00000 0.00000 ; N- CA- CB- HB3
197 199 201 204 3 3.28026 2.46019 -2.19242
-3.21331 -0.23430 0.00000 ; N- CA- CB- CG
197 199 216 217 3 0.00000 0.00000 0.00000
0.00000 0.00000 0.00000 ; N- CA- C- O
197 199 216 218 3 17.40544 -5.02080 -13.22144
9.20480 0.00000 0.00000 ; N- CA- C- N
198 197 199 200 3 0.00000 0.00000 0.00000
0.00000 0.00000 0.00000 ; H- N- CA- HA
198 197 199 201 3 0.00000 0.00000 0.00000
0.00000 0.00000 0.00000 ; H- N- CA- CB
198 197 199 216 3 0.00000 0.00000 0.00000
0.00000 0.00000 0.00000 ; H- N- CA- C
199 201 204 205 3 0.57739 0.00000 1.02926
0.00000 -1.60666 0.00000 ; CA- CB- CG- CD1
199 201 204 214 3 0.57739 0.00000 1.02926
0.00000 -1.60666 0.00000 ; CA- CB- CG- CD2
199 216 218 219 3 20.92000 0.00000 -20.92000
0.00000 0.00000 0.00000 ; CA- C- N- H
199 216 218 220 3 20.92000 0.00000 -20.92000
0.00000 0.00000 0.00000 ; CA- C- N- CA
200 199 201 202 3 0.65084 1.95253 0.00000
-2.60338 0.00000 0.00000 ; HA- CA- CB- HB2
200 199 201 203 3 0.65084 1.95253 0.00000
-2.60338 0.00000 0.00000 ; HA- CA- CB- HB3
200 199 201 204 3 0.65084 1.95253 0.00000
-2.60338 0.00000 0.00000 ; HA- CA- CB- CG
200 199 216 217 3 3.68192 -4.35136 0.00000
1.33888 0.00000 0.00000 ; HA- CA- C- O
200 199 216 218 3 0.00000 0.00000 0.00000
0.00000 0.00000 0.00000 ; HA- CA- C- N
201 199 216 217 3 0.00000 0.00000 0.00000
0.00000 0.00000 0.00000 ; CB- CA- C- O
201 199 216 218 3 2.51040 4.18400 1.67360

-6.69440 0.00000 0.00000 ; CB- CA- C- N
201 204 205 206 3 30.33400 0.00000 -30.33400
0.00000 0.00000 0.00000 ; CB- CG- CD1- HD1
201 204 205 207 3 30.33400 0.00000 -30.33400
0.00000 0.00000 0.00000 ; CB- CG- CD1- CE1
201 204 214 212 3 30.33400 0.00000 -30.33400
0.00000 0.00000 0.00000 ; CB- CG- CD2- CE2
201 204 214 215 3 30.33400 0.00000 -30.33400
0.00000 0.00000 0.00000 ; CB- CG- CD2- HD2
202 201 199 216 3 0.65084 1.95253 0.00000
-2.60338 0.00000 0.00000 ; HB2- CB- CA- C
202 201 204 205 3 0.00000 0.00000 0.00000
0.00000 0.00000 0.00000 ; HB2- CB- CG- CD1
202 201 204 214 3 0.00000 0.00000 0.00000
0.00000 0.00000 0.00000 ; HB2- CB- CG- CD2
203 201 199 216 3 0.65084 1.95253 0.00000
-2.60338 0.00000 0.00000 ; HB3- CB- CA- C
203 201 204 205 3 0.00000 0.00000 0.00000
0.00000 0.00000 0.00000 ; HB3- CB- CG- CD1
203 201 204 214 3 0.00000 0.00000 0.00000
0.00000 0.00000 0.00000 ; HB3- CB- CG- CD2
204 201 199 216 3 4.95804 2.17986 -3.52293
-3.21331 -0.40166 0.00000 ; CG- CB- CA- C
204 205 207 208 3 30.33400 0.00000 -30.33400
0.00000 0.00000 0.00000 ; CG- CD1- CE1- HE1
204 205 207 209 3 30.33400 0.00000 -30.33400
0.00000 0.00000 0.00000 ; CG- CD1- CE1- CZ
204 214 212 209 3 30.33400 0.00000 -30.33400
0.00000 0.00000 0.00000 ; CG- CD2- CE2- CZ
204 214 212 213 3 30.33400 0.00000 -30.33400
0.00000 0.00000 0.00000 ; CG- CD2- CE2- HE2
205 204 214 212 3 30.33400 0.00000 -30.33400
0.00000 0.00000 0.00000 ; CD1- CG- CD2- CE2
205 204 214 215 3 30.33400 0.00000 -30.33400
0.00000 0.00000 0.00000 ; CD1- CG- CD2- HD2
205 207 209 210 3 30.33400 0.00000 -30.33400
0.00000 0.00000 0.00000 ; CD1- CE1- CZ- OH
205 207 209 212 3 30.33400 0.00000 -30.33400
0.00000 0.00000 0.00000 ; CD1- CE1- CZ- CE2
206 205 204 214 3 30.33400 0.00000 -30.33400
0.00000 0.00000 0.00000 ; HD1- CD1- CG- CD2
206 205 207 208 3 30.33400 0.00000 -30.33400
0.00000 0.00000 0.00000 ; HD1- CD1- CE1- HE1
206 205 207 209 3 30.33400 0.00000 -30.33400
0.00000 0.00000 0.00000 ; HD1- CD1- CE1- CZ
207 205 204 214 3 30.33400 0.00000 -30.33400
0.00000 0.00000 0.00000 ; CE1- CD1- CG- CD2
207 209 210 211 3 7.93286 0.00000 -9.56462
0.00000 2.17568 0.00000 ; CE1- CZ- OH- HH
207 209 212 213 3 30.33400 0.00000 -30.33400
0.00000 0.00000 0.00000 ; CE1- CZ- CE2- HE2
207 209 212 214 3 30.33400 0.00000 -30.33400

```
0.00000    0.00000    0.00000 ;    CE1-    CZ-   CE2-   CD2
   208    207    209    210      3   30.33400    0.00000  -30.33400
0.00000    0.00000    0.00000 ;    HE1-   CE1-    CZ-    OH
   208    207    209    212      3   30.33400    0.00000  -30.33400
0.00000    0.00000    0.00000 ;    HE1-   CE1-    CZ-   CE2
   209    212    214    215      3   30.33400    0.00000  -30.33400
0.00000    0.00000    0.00000 ;     CZ-   CE2-   CD2-   HD2
   210    209    212    213      3   30.33400    0.00000  -30.33400
0.00000    0.00000    0.00000 ;     OH-    CZ-   CE2-   HE2
   210    209    212    214      3   30.33400    0.00000  -30.33400
0.00000    0.00000    0.00000 ;     OH-    CZ-   CE2-   CD2
   211    210    209    212      3    7.93286    0.00000   -9.56462
0.00000    2.17568    0.00000 ;     HH-    OH-    CZ-   CE2
   213    212    214    215      3   30.33400    0.00000  -30.33400
0.00000    0.00000    0.00000 ;    HE2-   CE2-   CD2-   HD2
   216    218    220    221      3    0.00000    0.00000    0.00000
0.00000    0.00000    0.00000 ;      C-     N-    CA-    HA
   216    218    220    222      3   11.71520    1.67360   15.06240  -
13.38880    0.00000    0.00000 ;      C-     N-    CA-    CB
   216    218    220    235      3    1.75728    5.27184    2.25936
-7.02912    0.00000    0.00000 ;      C-     N-    CA-     C
   217    216    218    219      3   29.28800   -8.36800  -20.92000
0.00000    0.00000    0.00000 ;      O-     C-     N-     H
   217    216    218    220      3   20.92000    0.00000  -20.92000
0.00000    0.00000    0.00000 ;      O-     C-     N-    CA
   218    220    222    223      3    0.65084    1.95253    0.00000
-2.60338    0.00000    0.00000 ;      N-    CA-    CB-    HB
   218    220    222    224      3    3.83673    0.44769   -1.77402
-2.47693   -0.03347    0.00000 ;      N-    CA-    CB-   CG2
   218    220    222    228      3    2.97482    0.12134    2.04179
-1.89117   -3.24678    0.00000 ;      N-    CA-    CB-   CG1
   218    220    235    236      3    0.00000    0.00000    0.00000
0.00000    0.00000    0.00000 ;      N-    CA-     C-     O
   218    220    235    237      3   17.40544   -5.02080  -13.22144
9.20480    0.00000    0.00000 ;      N-    CA-     C-     N
   219    218    220    221      3    0.00000    0.00000    0.00000
0.00000    0.00000    0.00000 ;      H-     N-    CA-    HA
   219    218    220    222      3    0.00000    0.00000    0.00000
0.00000    0.00000    0.00000 ;      H-     N-    CA-    CB
   219    218    220    235      3    0.00000    0.00000    0.00000
0.00000    0.00000    0.00000 ;      H-     N-    CA-     C
   220    222    224    225      3    0.66944    2.00832    0.00000
-2.67776    0.00000    0.00000 ;     CA-    CB-   CG2-  HG21
   220    222    224    226      3    0.66944    2.00832    0.00000
-2.67776    0.00000    0.00000 ;     CA-    CB-   CG2-  HG22
   220    222    224    227      3    0.66944    2.00832    0.00000
-2.67776    0.00000    0.00000 ;     CA-    CB-   CG2-  HG23
   220    222    228    229      3    0.66944    2.00832    0.00000
-2.67776    0.00000    0.00000 ;     CA-    CB-   CG1-  HG12
   220    222    228    230      3    0.66944    2.00832    0.00000
-2.67776    0.00000    0.00000 ;     CA-    CB-   CG1-  HG13
   220    222    228    231      3    4.24258   -0.52718   -7.25506
```

```
-1.79075    7.69856    0.00000 ;      CA-    CB-   CG1-   CD1
   220    235    237    238      3   20.92000    0.00000  -20.92000
0.00000    0.00000    0.00000 ;      CA-     C-     N-     H
   220    235    237    239      3   20.92000    0.00000  -20.92000
0.00000    0.00000    0.00000 ;      CA-     C-     N-    CA
   221    220    222    223      3    0.65084    1.95253    0.00000
-2.60338    0.00000    0.00000 ;      HA-    CA-    CB-    HB
   221    220    222    224      3    0.65084    1.95253    0.00000
-2.60338    0.00000    0.00000 ;      HA-    CA-    CB-   CG2
   221    220    222    228      3    0.65084    1.95253    0.00000
-2.60338    0.00000    0.00000 ;      HA-    CA-    CB-   CG1
   221    220    235    236      3    3.68192   -4.35136    0.00000
1.33888    0.00000    0.00000 ;      HA-    CA-     C-     O
   221    220    235    237      3    0.00000    0.00000    0.00000
0.00000    0.00000    0.00000 ;      HA-    CA-     C-     N
   222    220    235    236      3    0.00000    0.00000    0.00000
0.00000    0.00000    0.00000 ;      CB-    CA-     C-     O
   222    220    235    237      3    2.51040    4.18400    1.67360
-6.69440    0.00000    0.00000 ;      CB-    CA-     C-     N
   222    228    231    232      3    0.66944    2.00832    0.00000
-2.67776    0.00000    0.00000 ;      CB-   CG1-   CD1-  HD11
   222    228    231    233      3    0.66944    2.00832    0.00000
-2.67776    0.00000    0.00000 ;      CB-   CG1-   CD1-  HD12
   222    228    231    234      3    0.66944    2.00832    0.00000
-2.67776    0.00000    0.00000 ;      CB-   CG1-   CD1-  HD13
   223    222    220    235      3    0.65084    1.95253    0.00000
-2.60338    0.00000    0.00000 ;      HB-    CB-    CA-     C
   223    222    224    225      3    0.62760    1.88280    0.00000
-2.51040    0.00000    0.00000 ;      HB-    CB-   CG2-  HG21
   223    222    224    226      3    0.62760    1.88280    0.00000
-2.51040    0.00000    0.00000 ;      HB-    CB-   CG2-  HG22
   223    222    224    227      3    0.62760    1.88280    0.00000
-2.51040    0.00000    0.00000 ;      HB-    CB-   CG2-  HG23
   223    222    228    229      3    0.62760    1.88280    0.00000
-2.51040    0.00000    0.00000 ;      HB-    CB-   CG1-  HG12
   223    222    228    230      3    0.62760    1.88280    0.00000
-2.51040    0.00000    0.00000 ;      HB-    CB-   CG1-  HG13
   223    222    228    231      3    0.66944    2.00832    0.00000
-2.67776    0.00000    0.00000 ;      HB-    CB-   CG1-   CD1
   224    222    220    235      3    5.67350    3.55640   -6.16722
-2.47693    3.74886    0.00000 ;     CG2-    CB-    CA-     C
   224    222    228    229      3    0.66944    2.00832    0.00000
-2.67776    0.00000    0.00000 ;     CG2-    CB-   CG1-  HG12
   224    222    228    230      3    0.66944    2.00832    0.00000
-2.67776    0.00000    0.00000 ;     CG2-    CB-   CG1-  HG13
   224    222    228    231      3    3.81162    0.49790   -8.14206
-1.79075    7.49773    0.00000 ;     CG2-    CB-   CG1-   CD1
   225    224    222    228      3    0.66944    2.00832    0.00000
-2.67776    0.00000    0.00000 ;    HG21-   CG2-    CB-   CG1
   226    224    222    228      3    0.66944    2.00832    0.00000
-2.67776    0.00000    0.00000 ;    HG22-   CG2-    CB-   CG1
   227    224    222    228      3    0.66944    2.00832    0.00000
```

```
-2.67776    0.00000    0.00000 ;   HG23-   CG2-    CB-   CG1
   228    222    220    235      3    8.26340    0.74057   -9.99976
-1.89117    3.84928    0.00000 ;    CG1-    CB-    CA-     C
   229    228    231    232      3    0.62760    1.88280    0.00000
-2.51040    0.00000    0.00000 ;   HG12-   CG1-   CD1-  HD11
   229    228    231    233      3    0.62760    1.88280    0.00000
-2.51040    0.00000    0.00000 ;   HG12-   CG1-   CD1-  HD12
   229    228    231    234      3    0.62760    1.88280    0.00000
-2.51040    0.00000    0.00000 ;   HG12-   CG1-   CD1-  HD13
   230    228    231    232      3    0.62760    1.88280    0.00000
-2.51040    0.00000    0.00000 ;   HG13-   CG1-   CD1-  HD11
   230    228    231    233      3    0.62760    1.88280    0.00000
-2.51040    0.00000    0.00000 ;   HG13-   CG1-   CD1-  HD12
   230    228    231    234      3    0.62760    1.88280    0.00000
-2.51040    0.00000    0.00000 ;   HG13-   CG1-   CD1-  HD13
   235    237    239    240      3    0.00000    0.00000    0.00000
0.00000    0.00000    0.00000 ;      C-      N-    CA-    HA
   235    237    239    241      3   11.71520    1.67360   15.06240  -
13.38880    0.00000    0.00000 ;      C-      N-    CA-    CB
   235    237    239    254      3    1.75728    5.27184    2.25936
-7.02912    0.00000    0.00000 ;      C-      N-    CA-     C
   236    235    237    238      3   29.28800   -8.36800  -20.92000
0.00000    0.00000    0.00000 ;      O-      C-     N-     H
   236    235    237    239      3   20.92000    0.00000  -20.92000
0.00000    0.00000    0.00000 ;      O-      C-     N-    CA
   237    239    241    242      3    0.65084    1.95253    0.00000
-2.60338    0.00000    0.00000 ;      N-    CA-    CB-    HB
   237    239    241    243      3    3.83673    0.44769   -1.77402
-2.47693   -0.03347    0.00000 ;      N-    CA-    CB-   CG2
   237    239    241    247      3    2.97482    0.12134    2.04179
-1.89117   -3.24678    0.00000 ;      N-    CA-    CB-   CG1
   237    239    254    255      3    0.00000    0.00000    0.00000
0.00000    0.00000    0.00000 ;      N-    CA-     C-     O
   237    239    254    256      3   17.40544   -5.02080  -13.22144
9.20480    0.00000    0.00000 ;      N-    CA-     C-     N
   238    237    239    240      3    0.00000    0.00000    0.00000
0.00000    0.00000    0.00000 ;      H-      N-    CA-    HA
   238    237    239    241      3    0.00000    0.00000    0.00000
0.00000    0.00000    0.00000 ;      H-      N-    CA-    CB
   238    237    239    254      3    0.00000    0.00000    0.00000
0.00000    0.00000    0.00000 ;      H-      N-    CA-     C
   239    241    243    244      3    0.66944    2.00832    0.00000
-2.67776    0.00000    0.00000 ;    CA-    CB-   CG2-  HG21
   239    241    243    245      3    0.66944    2.00832    0.00000
-2.67776    0.00000    0.00000 ;    CA-    CB-   CG2-  HG22
   239    241    243    246      3    0.66944    2.00832    0.00000
-2.67776    0.00000    0.00000 ;    CA-    CB-   CG2-  HG23
   239    241    247    248      3    0.66944    2.00832    0.00000
-2.67776    0.00000    0.00000 ;    CA-    CB-   CG1-  HG12
   239    241    247    249      3    0.66944    2.00832    0.00000
-2.67776    0.00000    0.00000 ;    CA-    CB-   CG1-  HG13
   239    241    247    250      3    4.24258   -0.52718   -7.25506
```

```
-1.79075    7.69856    0.00000 ;      CA-    CB-   CG1-   CD1
   239    254    256    257       3   20.92000    0.00000  -20.92000
0.00000    0.00000    0.00000 ;      CA-     C-     N-     H
   239    254    256    258       3   20.92000    0.00000  -20.92000
0.00000    0.00000    0.00000 ;      CA-     C-     N-    CA
   240    239    241    242       3    0.65084    1.95253    0.00000
-2.60338    0.00000    0.00000 ;      HA-    CA-    CB-    HB
   240    239    241    243       3    0.65084    1.95253    0.00000
-2.60338    0.00000    0.00000 ;      HA-    CA-    CB-   CG2
   240    239    241    247       3    0.65084    1.95253    0.00000
-2.60338    0.00000    0.00000 ;      HA-    CA-    CB-   CG1
   240    239    254    255       3    3.68192   -4.35136    0.00000
1.33888    0.00000    0.00000 ;      HA-    CA-     C-     O
   240    239    254    256       3    0.00000    0.00000    0.00000
0.00000    0.00000    0.00000 ;      HA-    CA-     C-     N
   241    239    254    255       3    0.00000    0.00000    0.00000
0.00000    0.00000    0.00000 ;      CB-    CA-     C-     O
   241    239    254    256       3    2.51040    4.18400    1.67360
-6.69440    0.00000    0.00000 ;      CB-    CA-     C-     N
   241    247    250    251       3    0.66944    2.00832    0.00000
-2.67776    0.00000    0.00000 ;      CB-   CG1-   CD1-  HD11
   241    247    250    252       3    0.66944    2.00832    0.00000
-2.67776    0.00000    0.00000 ;      CB-   CG1-   CD1-  HD12
   241    247    250    253       3    0.66944    2.00832    0.00000
-2.67776    0.00000    0.00000 ;      CB-   CG1-   CD1-  HD13
   242    241    239    254       3    0.65084    1.95253    0.00000
-2.60338    0.00000    0.00000 ;      HB-    CB-    CA-     C
   242    241    243    244       3    0.62760    1.88280    0.00000
-2.51040    0.00000    0.00000 ;      HB-    CB-   CG2-  HG21
   242    241    243    245       3    0.62760    1.88280    0.00000
-2.51040    0.00000    0.00000 ;      HB-    CB-   CG2-  HG22
   242    241    243    246       3    0.62760    1.88280    0.00000
-2.51040    0.00000    0.00000 ;      HB-    CB-   CG2-  HG23
   242    241    247    248       3    0.62760    1.88280    0.00000
-2.51040    0.00000    0.00000 ;      HB-    CB-   CG1-  HG12
   242    241    247    249       3    0.62760    1.88280    0.00000
-2.51040    0.00000    0.00000 ;      HB-    CB-   CG1-  HG13
   242    241    247    250       3    0.66944    2.00832    0.00000
-2.67776    0.00000    0.00000 ;      HB-    CB-   CG1-   CD1
   243    241    239    254       3    5.67350    3.55640   -6.16722
-2.47693    3.74886    0.00000 ;     CG2-    CB-    CA-     C
   243    241    247    248       3    0.66944    2.00832    0.00000
-2.67776    0.00000    0.00000 ;     CG2-    CB-   CG1-  HG12
   243    241    247    249       3    0.66944    2.00832    0.00000
-2.67776    0.00000    0.00000 ;     CG2-    CB-   CG1-  HG13
   243    241    247    250       3    3.81162    0.49790   -8.14206
-1.79075    7.49773    0.00000 ;     CG2-    CB-   CG1-   CD1
   244    243    241    247       3    0.66944    2.00832    0.00000
-2.67776    0.00000    0.00000 ;    HG21-   CG2-    CB-   CG1
   245    243    241    247       3    0.66944    2.00832    0.00000
-2.67776    0.00000    0.00000 ;    HG22-   CG2-    CB-   CG1
   246    243    241    247       3    0.66944    2.00832    0.00000
```

-2.67776 0.00000 0.00000 ; HG23- CG2- CB- CG1
247 241 239 254 3 8.26340 0.74057 -9.99976
-1.89117 3.84928 0.00000 ; CG1- CB- CA- C
248 247 250 251 3 0.62760 1.88280 0.00000
-2.51040 0.00000 0.00000 ; HG12- CG1- CD1- HD11
248 247 250 252 3 0.62760 1.88280 0.00000
-2.51040 0.00000 0.00000 ; HG12- CG1- CD1- HD12
248 247 250 253 3 0.62760 1.88280 0.00000
-2.51040 0.00000 0.00000 ; HG12- CG1- CD1- HD13
249 247 250 251 3 0.62760 1.88280 0.00000
-2.51040 0.00000 0.00000 ; HG13- CG1- CD1- HD11
249 247 250 252 3 0.62760 1.88280 0.00000
-2.51040 0.00000 0.00000 ; HG13- CG1- CD1- HD12
249 247 250 253 3 0.62760 1.88280 0.00000
-2.51040 0.00000 0.00000 ; HG13- CG1- CD1- HD13
254 256 258 259 3 0.00000 0.00000 0.00000
0.00000 0.00000 0.00000 ; C- N- CA- HA
254 256 258 260 3 11.71520 1.67360 15.06240 -
13.38880 0.00000 0.00000 ; C- N- CA- CB
254 256 258 266 3 1.75728 5.27184 2.25936
-7.02912 0.00000 0.00000 ; C- N- CA- C
255 254 256 257 3 29.28800 -8.36800 -20.92000
0.00000 0.00000 0.00000 ; O- C- N- H
255 254 256 258 3 20.92000 0.00000 -20.92000
0.00000 0.00000 0.00000 ; O- C- N- CA
256 258 260 261 3 0.65084 1.95253 0.00000
-2.60338 0.00000 0.00000 ; N- CA- CB- HB2
256 258 260 262 3 0.65084 1.95253 0.00000
-2.60338 0.00000 0.00000 ; N- CA- CB- HB3
256 258 260 263 3 15.41386 9.74035 -8.39310
-0.97069 2.97901 0.00000 ; N- CA- CB- CG
256 258 266 267 3 0.00000 0.00000 0.00000
0.00000 0.00000 0.00000 ; N- CA- C- O
256 258 266 268 3 17.40544 -5.02080 -13.22144
9.20480 0.00000 0.00000 ; N- CA- C- N
257 256 258 259 3 0.00000 0.00000 0.00000
0.00000 0.00000 0.00000 ; H- N- CA- HA
257 256 258 260 3 0.00000 0.00000 0.00000
0.00000 0.00000 0.00000 ; H- N- CA- CB
257 256 258 266 3 0.00000 0.00000 0.00000
0.00000 0.00000 0.00000 ; H- N- CA- C
258 260 263 264 3 6.43499 0.00000 -5.39736
0.00000 -1.03763 0.00000 ; CA- CB- CG- OD1
258 260 263 265 3 6.43499 0.00000 -5.39736
0.00000 -1.03763 0.00000 ; CA- CB- CG- OD2
258 266 268 269 3 20.92000 0.00000 -20.92000
0.00000 0.00000 0.00000 ; CA- C- N- H
258 266 268 270 3 20.92000 0.00000 -20.92000
0.00000 0.00000 0.00000 ; CA- C- N- CA
259 258 260 261 3 0.65084 1.95253 0.00000
-2.60338 0.00000 0.00000 ; HA- CA- CB- HB2
259 258 260 262 3 0.65084 1.95253 0.00000

-2.60338    0.00000    0.00000 ;     HA-    CA-    CB-   HB3
259    258    260    263    3    0.65084    1.95253    0.00000
-2.60338    0.00000    0.00000 ;     HA-    CA-    CB-   CG
259    258    266    267    3    3.68192   -4.35136    0.00000
1.33888    0.00000    0.00000 ;     HA-    CA-    C-    O
259    258    266    268    3    0.00000    0.00000    0.00000
0.00000    0.00000    0.00000 ;     HA-    CA-    C-    N
260    258    266    267    3    0.00000    0.00000    0.00000
0.00000    0.00000    0.00000 ;     CB-    CA-    C-    O
260    258    266    268    3    2.51040    4.18400    1.67360
-6.69440    0.00000    0.00000 ;     CB-    CA-    C-    N
261    260    258    266    3    0.65084    1.95253    0.00000
-2.60338    0.00000    0.00000 ;    HB2-    CB-    CA-    C
261    260    263    264    3    0.00000    0.00000    0.00000
0.00000    0.00000    0.00000 ;    HB2-    CB-    CG-   OD1
261    260    263    265    3    0.00000    0.00000    0.00000
0.00000    0.00000    0.00000 ;    HB2-    CB-    CG-   OD2
262    260    258    266    3    0.65084    1.95253    0.00000
-2.60338    0.00000    0.00000 ;    HB3-    CB-    CA-    C
262    260    263    264    3    0.00000    0.00000    0.00000
0.00000    0.00000    0.00000 ;    HB3-    CB-    CG-   OD1
262    260    263    265    3    0.00000    0.00000    0.00000
0.00000    0.00000    0.00000 ;    HB3-    CB-    CG-   OD2
263    260    258    266    3    7.14627   -1.04600   -8.99560
-0.97069    5.15469    0.00000 ;     CG-    CB-    CA-    C
266    268    270    271    3    0.00000    0.00000    0.00000
0.00000    0.00000    0.00000 ;     C-    N-    CA-    HA
266    268    270    272    3   11.71520    1.67360   15.06240  -
13.38880    0.00000    0.00000 ;     C-    N-    CA-    CB
266    268    270    281    3    1.75728    5.27184    2.25936
-7.02912    0.00000    0.00000 ;     C-    N-    CA-    C
267    266    268    269    3   29.28800   -8.36800  -20.92000
0.00000    0.00000    0.00000 ;     O-    C-    N-    H
267    266    268    270    3   20.92000    0.00000  -20.92000
0.00000    0.00000    0.00000 ;     O-    C-    N-    CA
268    270    272    273    3    0.65084    1.95253    0.00000
-2.60338    0.00000    0.00000 ;     N-    CA-    CB-   HB2
268    270    272    274    3    0.65084    1.95253    0.00000
-2.60338    0.00000    0.00000 ;     N-    CA-    CB-   HB3
268    270    272    275    3    3.21331    2.22589   -4.15053
-2.40998    2.61082    0.00000 ;     N-    CA-    CB-    CG
268    270    281    282    3    0.00000    0.00000    0.00000
0.00000    0.00000    0.00000 ;     N-    CA-    C-    O
268    270    281    283    3   17.40544   -5.02080  -13.22144
9.20480    0.00000    0.00000 ;     N-    CA-    C-    N
269    268    270    271    3    0.00000    0.00000    0.00000
0.00000    0.00000    0.00000 ;     H-    N-    CA-    HA
269    268    270    272    3    0.00000    0.00000    0.00000
0.00000    0.00000    0.00000 ;     H-    N-    CA-    CB
269    268    270    281    3    0.00000    0.00000    0.00000
0.00000    0.00000    0.00000 ;     H-    N-    CA-    C
270    272    275    276    3    0.66944    2.00832    0.00000

```
-2.67776    0.00000    0.00000 ;      CA-    CB-    CG-   HG2
   270    272    275    277      3    0.66944    2.00832    0.00000
-2.67776    0.00000    0.00000 ;      CA-    CB-    CG-   HG3
   270    272    275    278      3   10.12110   -1.91209    0.01674
10.17549   -1.87443    0.00000 ;      CA-    CB-    CG-    CD
   270    281    283    284      3   20.92000    0.00000  -20.92000
0.00000    0.00000    0.00000 ;      CA-     C-     N-     H
   270    281    283    285      3   20.92000    0.00000  -20.92000
0.00000    0.00000    0.00000 ;      CA-     C-     N-    CA
   271    270    272    273      3    0.65084    1.95253    0.00000
-2.60338    0.00000    0.00000 ;      HA-    CA-    CB-   HB2
   271    270    272    274      3    0.65084    1.95253    0.00000
-2.60338    0.00000    0.00000 ;      HA-    CA-    CB-   HB3
   271    270    272    275      3    0.65084    1.95253    0.00000
-2.60338    0.00000    0.00000 ;      HA-    CA-    CB-    CG
   271    270    281    282      3    3.68192   -4.35136    0.00000
1.33888    0.00000    0.00000 ;      HA-    CA-     C-     O
   271    270    281    283      3    0.00000    0.00000    0.00000
0.00000    0.00000    0.00000 ;      HA-    CA-     C-     N
   272    270    281    282      3    0.00000    0.00000    0.00000
0.00000    0.00000    0.00000 ;      CB-    CA-     C-     O
   272    270    281    283      3    2.51040    4.18400    1.67360
-6.69440    0.00000    0.00000 ;      CB-    CA-     C-     N
   272    275    278    279      3    3.79907    0.00000   -5.40573
0.00000    2.14221    0.00000 ;      CB-    CG-    CD-   OE1
   272    275    278    280      3    3.79907    0.00000   -5.40573
0.00000    2.14221    0.00000 ;      CB-    CG-    CD-   OE2
   273    272    270    281      3    0.65084    1.95253    0.00000
-2.60338    0.00000    0.00000 ;     HB2-    CB-    CA-     C
   273    272    275    276      3    0.62760    1.88280    0.00000
-2.51040    0.00000    0.00000 ;     HB2-    CB-    CG-   HG2
   273    272    275    277      3    0.62760    1.88280    0.00000
-2.51040    0.00000    0.00000 ;     HB2-    CB-    CG-   HG3
   273    272    275    278      3    0.65084    1.95253    0.00000
-2.60338    0.00000    0.00000 ;     HB2-    CB-    CG-    CD
   274    272    270    281      3    0.65084    1.95253    0.00000
-2.60338    0.00000    0.00000 ;     HB3-    CB-    CA-     C
   274    272    275    276      3    0.62760    1.88280    0.00000
-2.51040    0.00000    0.00000 ;     HB3-    CB-    CG-   HG2
   274    272    275    277      3    0.62760    1.88280    0.00000
-2.51040    0.00000    0.00000 ;     HB3-    CB-    CG-   HG3
   274    272    275    278      3    0.65084    1.95253    0.00000
-2.60338    0.00000    0.00000 ;     HB3-    CB-    CG-    CD
   275    272    270    281      3    6.86594    3.56895   -8.14206
-2.40998    4.85344    0.00000 ;      CG-    CB-    CA-     C
   276    275    278    279      3    0.00000    0.00000    0.00000
0.00000    0.00000    0.00000 ;     HG2-    CG-    CD-   OE1
   276    275    278    280      3    0.00000    0.00000    0.00000
0.00000    0.00000    0.00000 ;     HG2-    CG-    CD-   OE2
   277    275    278    279      3    0.00000    0.00000    0.00000
0.00000    0.00000    0.00000 ;     HG3-    CG-    CD-   OE1
   277    275    278    280      3    0.00000    0.00000    0.00000
```

0.00000 0.00000 0.00000 ; HG3- CG- CD- OE2
281 283 285 286 3 0.00000 0.00000 0.00000
0.00000 0.00000 0.00000 ; C- N- CA- HA
281 283 285 287 3 11.71520 1.67360 15.06240 -
13.38880 0.00000 0.00000 ; C- N- CA- CB
281 283 285 303 3 1.75728 5.27184 2.25936
-7.02912 0.00000 0.00000 ; C- N- CA- C
282 281 283 284 3 29.28800 -8.36800 -20.92000
0.00000 0.00000 0.00000 ; O- C- N- H
282 281 283 285 3 20.92000 0.00000 -20.92000
0.00000 0.00000 0.00000 ; O- C- N- CA
283 285 287 288 3 0.65084 1.95253 0.00000
-2.60338 0.00000 0.00000 ; N- CA- CB- HB2
283 285 287 289 3 0.65084 1.95253 0.00000
-2.60338 0.00000 0.00000 ; N- CA- CB- HB3
283 285 287 290 3 0.65084 1.95253 0.00000
-2.60338 0.00000 0.00000 ; N- CA- CB- CG
283 285 303 304 3 0.00000 0.00000 0.00000
0.00000 0.00000 0.00000 ; N- CA- C- O
283 285 303 305 3 17.40544 -5.02080 -13.22144
9.20480 0.00000 0.00000 ; N- CA- C- N
284 283 285 286 3 0.00000 0.00000 0.00000
0.00000 0.00000 0.00000 ; H- N- CA- HA
284 283 285 287 3 0.00000 0.00000 0.00000
0.00000 0.00000 0.00000 ; H- N- CA- CB
284 283 285 303 3 0.00000 0.00000 0.00000
0.00000 0.00000 0.00000 ; H- N- CA- C
285 287 290 291 3 0.66944 2.00832 0.00000
-2.67776 0.00000 0.00000 ; CA- CB- CG- HG2
285 287 290 292 3 0.66944 2.00832 0.00000
-2.67776 0.00000 0.00000 ; CA- CB- CG- HG3
285 287 290 293 3 3.68192 3.09616 -2.09200
-3.01248 0.00000 0.00000 ; CA- CB- CG- CD
285 303 305 306 3 20.92000 0.00000 -20.92000
0.00000 0.00000 0.00000 ; CA- C- N- H
285 303 305 307 3 20.92000 0.00000 -20.92000
0.00000 0.00000 0.00000 ; CA- C- N- CA
286 285 287 288 3 0.65084 1.95253 0.00000
-2.60338 0.00000 0.00000 ; HA- CA- CB- HB2
286 285 287 289 3 0.65084 1.95253 0.00000
-2.60338 0.00000 0.00000 ; HA- CA- CB- HB3
286 285 287 290 3 0.65084 1.95253 0.00000
-2.60338 0.00000 0.00000 ; HA- CA- CB- CG
286 285 303 304 3 3.68192 -4.35136 0.00000
1.33888 0.00000 0.00000 ; HA- CA- C- O
286 285 303 305 3 0.00000 0.00000 0.00000
0.00000 0.00000 0.00000 ; HA- CA- C- N
287 285 303 304 3 0.00000 0.00000 0.00000
0.00000 0.00000 0.00000 ; CB- CA- C- O
287 285 303 305 3 2.51040 4.18400 1.67360
-6.69440 0.00000 0.00000 ; CB- CA- C- N
287 290 293 294 3 0.66944 2.00832 0.00000

```
-2.67776    0.00000    0.00000 ;     CB-    CG-    CD-   HD2
   287    290    293    295      3    0.66944    2.00832    0.00000
-2.67776    0.00000    0.00000 ;     CB-    CG-    CD-   HD3
   287    290    293    296      3    3.68192    3.09616   -2.09200
-3.01248    0.00000    0.00000 ;     CB-    CG-    CD-    CE
   288    287    285    303      3    0.65084    1.95253    0.00000
-2.60338    0.00000    0.00000 ;    HB2-    CB-    CA-     C
   288    287    290    291      3    0.62760    1.88280    0.00000
-2.51040    0.00000    0.00000 ;    HB2-    CB-    CG-   HG2
   288    287    290    292      3    0.62760    1.88280    0.00000
-2.51040    0.00000    0.00000 ;    HB2-    CB-    CG-   HG3
   288    287    290    293      3    0.66944    2.00832    0.00000
-2.67776    0.00000    0.00000 ;    HB2-    CB-    CG-    CD
   289    287    285    303      3    0.65084    1.95253    0.00000
-2.60338    0.00000    0.00000 ;    HB3-    CB-    CA-     C
   289    287    290    291      3    0.62760    1.88280    0.00000
-2.51040    0.00000    0.00000 ;    HB3-    CB-    CG-   HG2
   289    287    290    292      3    0.62760    1.88280    0.00000
-2.51040    0.00000    0.00000 ;    HB3-    CB-    CG-   HG3
   289    287    290    293      3    0.66944    2.00832    0.00000
-2.67776    0.00000    0.00000 ;    HB3-    CB-    CG-    CD
   290    287    285    303      3    0.65084    1.95253    0.00000
-2.60338    0.00000    0.00000 ;     CG-    CB-    CA-     C
   290    293    296    297      3    0.65084    1.95253    0.00000
-2.60338    0.00000    0.00000 ;     CG-    CD-    CE-   HE2
   290    293    296    298      3    0.65084    1.95253    0.00000
-2.60338    0.00000    0.00000 ;     CG-    CD-    CE-   HE3
   290    293    296    299      3    0.65084    1.95253    0.00000
-2.60338    0.00000    0.00000 ;     CG-    CD-    CE-    NZ
   291    290    293    294      3    0.62760    1.88280    0.00000
-2.51040    0.00000    0.00000 ;    HG2-    CG-    CD-   HD2
   291    290    293    295      3    0.62760    1.88280    0.00000
-2.51040    0.00000    0.00000 ;    HG2-    CG-    CD-   HD3
   291    290    293    296      3    0.66944    2.00832    0.00000
-2.67776    0.00000    0.00000 ;    HG2-    CG-    CD-    CE
   292    290    293    294      3    0.62760    1.88280    0.00000
-2.51040    0.00000    0.00000 ;    HG3-    CG-    CD-   HD2
   292    290    293    295      3    0.62760    1.88280    0.00000
-2.51040    0.00000    0.00000 ;    HG3-    CG-    CD-   HD3
   292    290    293    296      3    0.66944    2.00832    0.00000
-2.67776    0.00000    0.00000 ;    HG3-    CG-    CD-    CE
   293    296    299    300      3    0.65084    1.95253    0.00000
-2.60338    0.00000    0.00000 ;     CD-    CE-    NZ-   HZ1
   293    296    299    301      3    0.65084    1.95253    0.00000
-2.60338    0.00000    0.00000 ;     CD-    CE-    NZ-   HZ2
   293    296    299    302      3    0.65084    1.95253    0.00000
-2.60338    0.00000    0.00000 ;     CD-    CE-    NZ-   HZ3
   294    293    296    297      3    0.65084    1.95253    0.00000
-2.60338    0.00000    0.00000 ;    HD2-    CD-    CE-   HE2
   294    293    296    298      3    0.65084    1.95253    0.00000
-2.60338    0.00000    0.00000 ;    HD2-    CD-    CE-   HE3
   294    293    296    299      3    0.65084    1.95253    0.00000
```

```
-2.60338    0.00000    0.00000 ;    HD2-    CD-    CE-    NZ
   295    293    296    297      3    0.65084    1.95253    0.00000
-2.60338    0.00000    0.00000 ;    HD3-    CD-    CE-   HE2
   295    293    296    298      3    0.65084    1.95253    0.00000
-2.60338    0.00000    0.00000 ;    HD3-    CD-    CE-   HE3
   295    293    296    299      3    0.65084    1.95253    0.00000
-2.60338    0.00000    0.00000 ;    HD3-    CD-    CE-    NZ
   297    296    299    300      3    0.65084    1.95253    0.00000
-2.60338    0.00000    0.00000 ;    HE2-    CE-    NZ-   HZ1
   297    296    299    301      3    0.65084    1.95253    0.00000
-2.60338    0.00000    0.00000 ;    HE2-    CE-    NZ-   HZ2
   297    296    299    302      3    0.65084    1.95253    0.00000
-2.60338    0.00000    0.00000 ;    HE2-    CE-    NZ-   HZ3
   298    296    299    300      3    0.65084    1.95253    0.00000
-2.60338    0.00000    0.00000 ;    HE3-    CE-    NZ-   HZ1
   298    296    299    301      3    0.65084    1.95253    0.00000
-2.60338    0.00000    0.00000 ;    HE3-    CE-    NZ-   HZ2
   298    296    299    302      3    0.65084    1.95253    0.00000
-2.60338    0.00000    0.00000 ;    HE3-    CE-    NZ-   HZ3
   303    305    307    308      3    0.00000    0.00000    0.00000
0.00000    0.00000    0.00000 ;     C-     N-    CA-    HA
   303    305    307    309      3   11.71520    1.67360   15.06240  -
13.38880    0.00000    0.00000 ;     C-     N-    CA-    CB
   303    305    307    322      3    1.75728    5.27184    2.25936
-7.02912    0.00000    0.00000 ;     C-     N-    CA-     C
   304    303    305    306      3   29.28800   -8.36800  -20.92000
0.00000    0.00000    0.00000 ;     O-     C-     N-     H
   304    303    305    307      3   20.92000    0.00000  -20.92000
0.00000    0.00000    0.00000 ;     O-     C-     N-    CA
   305    307    309    310      3    0.65084    1.95253    0.00000
-2.60338    0.00000    0.00000 ;     N-    CA-    CB-   HB2
   305    307    309    311      3    0.65084    1.95253    0.00000
-2.60338    0.00000    0.00000 ;     N-    CA-    CB-   HB3
   305    307    309    312      3    3.79070    1.39746   -4.61077
-2.40998    2.44346    0.00000 ;     N-    CA-    CB-    CG
   305    307    322    323      3    0.00000    0.00000    0.00000
0.00000    0.00000    0.00000 ;     N-    CA-     C-     O
   305    307    322    324      3   17.40544   -5.02080  -13.22144
9.20480    0.00000    0.00000 ;     N-    CA-     C-     N
   306    305    307    308      3    0.00000    0.00000    0.00000
0.00000    0.00000    0.00000 ;     H-     N-    CA-    HA
   306    305    307    309      3    0.00000    0.00000    0.00000
0.00000    0.00000    0.00000 ;     H-     N-    CA-    CB
   306    305    307    322      3    0.00000    0.00000    0.00000
0.00000    0.00000    0.00000 ;     H-     N-    CA-     C
   307    309    312    313      3    0.66944    2.00832    0.00000
-2.67776    0.00000    0.00000 ;    CA-    CB-    CG-    HG
   307    309    312    314      3    3.90367    0.19665   -6.21742
-2.37651    5.99149    0.00000 ;    CA-    CB-    CG-   CD1
   307    309    312    318      3    3.90367    0.19665   -6.21742
-2.37651    5.99149    0.00000 ;    CA-    CB-    CG-   CD2
   307    322    324    325      3   20.92000    0.00000  -20.92000
```

```
0.00000    0.00000    0.00000 ;      CA-     C-     N-     H
   307    322    324    326      3   20.92000    0.00000  -20.92000
0.00000    0.00000    0.00000 ;      CA-     C-     N-    CA
   308    307    309    310      3    0.65084    1.95253    0.00000
-2.60338    0.00000    0.00000 ;      HA-    CA-    CB-   HB2
   308    307    309    311      3    0.65084    1.95253    0.00000
-2.60338    0.00000    0.00000 ;      HA-    CA-    CB-   HB3
   308    307    309    312      3    0.65084    1.95253    0.00000
-2.60338    0.00000    0.00000 ;      HA-    CA-    CB-    CG
   308    307    322    323      3    3.68192   -4.35136    0.00000
1.33888    0.00000    0.00000 ;      HA-    CA-     C-     O
   308    307    322    324      3    0.00000    0.00000    0.00000
0.00000    0.00000    0.00000 ;      HA-    CA-     C-     N
   309    307    322    323      3    0.00000    0.00000    0.00000
0.00000    0.00000    0.00000 ;      CB-    CA-     C-     O
   309    307    322    324      3    2.51040    4.18400    1.67360
-6.69440    0.00000    0.00000 ;      CB-    CA-     C-     N
   309    312    314    315      3    0.66944    2.00832    0.00000
-2.67776    0.00000    0.00000 ;      CB-    CG-   CD1-  HD11
   309    312    314    316      3    0.66944    2.00832    0.00000
-2.67776    0.00000    0.00000 ;      CB-    CG-   CD1-  HD12
   309    312    314    317      3    0.66944    2.00832    0.00000
-2.67776    0.00000    0.00000 ;      CB-    CG-   CD1-  HD13
   309    312    318    319      3    0.66944    2.00832    0.00000
-2.67776    0.00000    0.00000 ;      CB-    CG-   CD2-  HD21
   309    312    318    320      3    0.66944    2.00832    0.00000
-2.67776    0.00000    0.00000 ;      CB-    CG-   CD2-  HD22
   309    312    318    321      3    0.66944    2.00832    0.00000
-2.67776    0.00000    0.00000 ;      CB-    CG-   CD2-  HD23
   310    309    307    322      3    0.65084    1.95253    0.00000
-2.60338    0.00000    0.00000 ;     HB2-    CB-    CA-     C
   310    309    312    313      3    0.62760    1.88280    0.00000
-2.51040    0.00000    0.00000 ;     HB2-    CB-    CG-    HG
   310    309    312    314      3    0.66944    2.00832    0.00000
-2.67776    0.00000    0.00000 ;     HB2-    CB-    CG-   CD1
   310    309    312    318      3    0.66944    2.00832    0.00000
-2.67776    0.00000    0.00000 ;     HB2-    CB-    CG-   CD2
   311    309    307    322      3    0.65084    1.95253    0.00000
-2.60338    0.00000    0.00000 ;     HB3-    CB-    CA-     C
   311    309    312    313      3    0.62760    1.88280    0.00000
-2.51040    0.00000    0.00000 ;     HB3-    CB-    CG-    HG
   311    309    312    314      3    0.66944    2.00832    0.00000
-2.67776    0.00000    0.00000 ;     HB3-    CB-    CG-   CD1
   311    309    312    318      3    0.66944    2.00832    0.00000
-2.67776    0.00000    0.00000 ;     HB3-    CB-    CG-   CD2
   312    309    307    322      3   10.33448   -1.14642  -11.54784
-2.40998    6.35968    0.00000 ;      CG-    CB-    CA-     C
   313    312    314    315      3    0.62760    1.88280    0.00000
-2.51040    0.00000    0.00000 ;      HG-    CG-   CD1-  HD11
   313    312    314    316      3    0.62760    1.88280    0.00000
-2.51040    0.00000    0.00000 ;      HG-    CG-   CD1-  HD12
   313    312    314    317      3    0.62760    1.88280    0.00000
```

```
-2.51040    0.00000    0.00000 ;      HG-    CG-   CD1-  HD13
   313    312    318    319      3    0.62760    1.88280    0.00000
-2.51040    0.00000    0.00000 ;      HG-    CG-   CD2-  HD21
   313    312    318    320      3    0.62760    1.88280    0.00000
-2.51040    0.00000    0.00000 ;      HG-    CG-   CD2-  HD22
   313    312    318    321      3    0.62760    1.88280    0.00000
-2.51040    0.00000    0.00000 ;      HG-    CG-   CD2-  HD23
   314    312    318    319      3    0.66944    2.00832    0.00000
-2.67776    0.00000    0.00000 ;     CD1-    CG-   CD2-  HD21
   314    312    318    320      3    0.66944    2.00832    0.00000
-2.67776    0.00000    0.00000 ;     CD1-    CG-   CD2-  HD22
   314    312    318    321      3    0.66944    2.00832    0.00000
-2.67776    0.00000    0.00000 ;     CD1-    CG-   CD2-  HD23
   315    314    312    318      3    0.66944    2.00832    0.00000
-2.67776    0.00000    0.00000 ;    HD11-   CD1-    CG-   CD2
   316    314    312    318      3    0.66944    2.00832    0.00000
-2.67776    0.00000    0.00000 ;    HD12-   CD1-    CG-   CD2
   317    314    312    318      3    0.66944    2.00832    0.00000
-2.67776    0.00000    0.00000 ;    HD13-   CD1-    CG-   CD2
   322    324    326    327      3    0.00000    0.00000    0.00000
0.00000    0.00000    0.00000 ;      C-     N-    CA-   HA2
   322    324    326    328      3    0.00000    0.00000    0.00000
0.00000    0.00000    0.00000 ;      C-     N-    CA-   HA3
   322    324    326    329      3    1.75728    5.27184    2.25936
-7.02912    0.00000    0.00000 ;      C-     N-    CA-     C
   323    322    324    325      3   29.28800   -8.36800  -20.92000
0.00000    0.00000    0.00000 ;      O-     C-     N-     H
   323    322    324    326      3   20.92000    0.00000  -20.92000
0.00000    0.00000    0.00000 ;      O-     C-     N-    CA
   324    326    329    330      3    0.00000    0.00000    0.00000
0.00000    0.00000    0.00000 ;      N-    CA-     C-     O
   324    326    329    331      3   17.40544   -5.02080  -13.22144
9.20480    0.00000    0.00000 ;      N-    CA-     C-     N
   325    324    326    327      3    0.00000    0.00000    0.00000
0.00000    0.00000    0.00000 ;      H-     N-    CA-   HA2
   325    324    326    328      3    0.00000    0.00000    0.00000
0.00000    0.00000    0.00000 ;      H-     N-    CA-   HA3
   325    324    326    329      3    0.00000    0.00000    0.00000
0.00000    0.00000    0.00000 ;      H-     N-    CA-     C
   326    329    331    332      3   20.92000    0.00000  -20.92000
0.00000    0.00000    0.00000 ;     CA-     C-     N-     H
   326    329    331    333      3   20.92000    0.00000  -20.92000
0.00000    0.00000    0.00000 ;     CA-     C-     N-    CA
   327    326    329    330      3    3.68192   -4.35136    0.00000
1.33888    0.00000    0.00000 ;    HA2-    CA-     C-     O
   327    326    329    331      3    0.00000    0.00000    0.00000
0.00000    0.00000    0.00000 ;    HA2-    CA-     C-     N
   328    326    329    330      3    3.68192   -4.35136    0.00000
1.33888    0.00000    0.00000 ;    HA3-    CA-     C-     O
   328    326    329    331      3    0.00000    0.00000    0.00000
0.00000    0.00000    0.00000 ;    HA3-    CA-     C-     N
   329    331    333    334      3    0.00000    0.00000    0.00000
```

0.00000 0.00000 0.00000 ; C- N- CA- HA
329 331 333 335 3 11.71520 1.67360 15.06240 -
13.38880 0.00000 0.00000 ; C- N- CA- CB
329 331 333 346 3 1.75728 5.27184 2.25936
-7.02912 0.00000 0.00000 ; C- N- CA- C
330 329 331 332 3 29.28800 -8.36800 -20.92000
0.00000 0.00000 0.00000 ; O- C- N- H
330 329 331 333 3 20.92000 0.00000 -20.92000
0.00000 0.00000 0.00000 ; O- C- N- CA
331 333 335 336 3 0.65084 1.95253 0.00000
-2.60338 0.00000 0.00000 ; N- CA- CB- HB2
331 333 335 337 3 0.65084 1.95253 0.00000
-2.60338 0.00000 0.00000 ; N- CA- CB- HB3
331 333 335 338 3 3.21331 2.22589 -4.15053
-2.40998 2.61082 0.00000 ; N- CA- CB- CG
331 333 346 347 3 0.00000 0.00000 0.00000
0.00000 0.00000 0.00000 ; N- CA- C- O
331 333 346 348 3 17.40544 -5.02080 -13.22144
9.20480 0.00000 0.00000 ; N- CA- C- N
332 331 333 334 3 0.00000 0.00000 0.00000
0.00000 0.00000 0.00000 ; H- N- CA- HA
332 331 333 335 3 0.00000 0.00000 0.00000
0.00000 0.00000 0.00000 ; H- N- CA- CB
332 331 333 346 3 0.00000 0.00000 0.00000
0.00000 0.00000 0.00000 ; H- N- CA- C
333 335 338 339 3 0.66944 2.00832 0.00000
-2.67776 0.00000 0.00000 ; CA- CB- CG- HG2
333 335 338 340 3 0.66944 2.00832 0.00000
-2.67776 0.00000 0.00000 ; CA- CB- CG- HG3
333 335 338 341 3 3.69866 -4.35136 -3.92459
6.89523 4.61914 0.00000 ; CA- CB- CG- CD
333 346 348 349 3 20.92000 0.00000 -20.92000
0.00000 0.00000 0.00000 ; CA- C- N- H
333 346 348 350 3 20.92000 0.00000 -20.92000
0.00000 0.00000 0.00000 ; CA- C- N- CA
334 333 335 336 3 0.65084 1.95253 0.00000
-2.60338 0.00000 0.00000 ; HA- CA- CB- HB2
334 333 335 337 3 0.65084 1.95253 0.00000
-2.60338 0.00000 0.00000 ; HA- CA- CB- HB3
334 333 335 338 3 0.65084 1.95253 0.00000
-2.60338 0.00000 0.00000 ; HA- CA- CB- CG
334 333 346 347 3 3.68192 -4.35136 0.00000
1.33888 0.00000 0.00000 ; HA- CA- C- O
334 333 346 348 3 0.00000 0.00000 0.00000
0.00000 0.00000 0.00000 ; HA- CA- C- N
335 333 346 347 3 0.00000 0.00000 0.00000
0.00000 0.00000 0.00000 ; CB- CA- C- O
335 333 346 348 3 2.51040 4.18400 1.67360
-6.69440 0.00000 0.00000 ; CB- CA- C- N
335 338 341 342 3 0.00000 0.00000 0.00000
0.00000 0.00000 0.00000 ; CB- CG- CD- OE1
335 338 341 343 3 10.32611 1.48114 -8.47678

```
1.42256    1.40582    0.00000 ;      CB-    CG-    CD-   NE2
   336    335    333    346      3    0.65084    1.95253    0.00000
-2.60338    0.00000    0.00000 ;     HB2-    CB-    CA-     C
   336    335    338    339      3    0.62760    1.88280    0.00000
-2.51040    0.00000    0.00000 ;     HB2-    CB-    CG-   HG2
   336    335    338    340      3    0.62760    1.88280    0.00000
-2.51040    0.00000    0.00000 ;     HB2-    CB-    CG-   HG3
   336    335    338    341      3    0.65084    1.95253    0.00000
-2.60338    0.00000    0.00000 ;     HB2-    CB-    CG-    CD
   337    335    333    346      3    0.65084    1.95253    0.00000
-2.60338    0.00000    0.00000 ;     HB3-    CB-    CA-     C
   337    335    338    339      3    0.62760    1.88280    0.00000
-2.51040    0.00000    0.00000 ;     HB3-    CB-    CG-   HG2
   337    335    338    340      3    0.62760    1.88280    0.00000
-2.51040    0.00000    0.00000 ;     HB3-    CB-    CG-   HG3
   337    335    338    341      3    0.65084    1.95253    0.00000
-2.60338    0.00000    0.00000 ;     HB3-    CB-    CG-    CD
   338    335    333    346      3    6.86594    3.56895   -8.14206
-2.40998    4.85344    0.00000 ;      CG-    CB-    CA-     C
   338    341    343    344      3   20.92000    0.00000  -20.92000
0.00000    0.00000    0.00000 ;      CG-    CD-   NE2-  HE21
   338    341    343    345      3   20.92000    0.00000  -20.92000
0.00000    0.00000    0.00000 ;      CG-    CD-   NE2-  HE22
   339    338    341    342      3    3.68192   -4.35136    0.00000
1.33888    0.00000    0.00000 ;     HG2-    CG-    CD-   OE1
   339    338    341    343      3    0.00000    0.00000    0.00000
0.00000    0.00000    0.00000 ;     HG2-    CG-    CD-   NE2
   340    338    341    342      3    3.68192   -4.35136    0.00000
1.33888    0.00000    0.00000 ;     HG3-    CG-    CD-   OE1
   340    338    341    343      3    0.00000    0.00000    0.00000
0.00000    0.00000    0.00000 ;     HG3-    CG-    CD-   NE2
   342    341    343    344      3   29.28800   -8.36800  -20.92000
0.00000    0.00000    0.00000 ;     OE1-    CD-   NE2-  HE21
   342    341    343    345      3   29.28800   -8.36800  -20.92000
0.00000    0.00000    0.00000 ;     OE1-    CD-   NE2-  HE22
   346    348    350    351      3    0.00000    0.00000    0.00000
0.00000    0.00000    0.00000 ;       C-     N-    CA-   HA2
   346    348    350    352      3    0.00000    0.00000    0.00000
0.00000    0.00000    0.00000 ;       C-     N-    CA-   HA3
   346    348    350    353      3    1.75728    5.27184    2.25936
-7.02912    0.00000    0.00000 ;      C-     N-    CA-     C
   347    346    348    349      3   29.28800   -8.36800  -20.92000
0.00000    0.00000    0.00000 ;       O-     C-     N-     H
   347    346    348    350      3   20.92000    0.00000  -20.92000
0.00000    0.00000    0.00000 ;       O-     C-     N-    CA
   348    350    353    354      3    0.00000    0.00000    0.00000
0.00000    0.00000    0.00000 ;       N-    CA-     C-     O
   348    350    353    355      3   17.40544   -5.02080  -13.22144
9.20480    0.00000    0.00000 ;       N-    CA-     C-     N
   349    348    350    351      3    0.00000    0.00000    0.00000
0.00000    0.00000    0.00000 ;       H-     N-    CA-   HA2
   349    348    350    352      3    0.00000    0.00000    0.00000
```

```
0.00000    0.00000    0.00000 ;      H-     N-     CA-   HA3
   349    348    350    353      3    0.00000    0.00000    0.00000
0.00000    0.00000    0.00000 ;      H-     N-     CA-     C
   350    353    355    356      3   20.92000    0.00000  -20.92000
0.00000    0.00000    0.00000 ;     CA-     C-     N-     H
   350    353    355    357      3   20.92000    0.00000  -20.92000
0.00000    0.00000    0.00000 ;     CA-     C-     N-    CA
   351    350    353    354      3    3.68192   -4.35136    0.00000
1.33888    0.00000    0.00000 ;    HA2-    CA-     C-     O
   351    350    353    355      3    0.00000    0.00000    0.00000
0.00000    0.00000    0.00000 ;    HA2-    CA-     C-     N
   352    350    353    354      3    3.68192   -4.35136    0.00000
1.33888    0.00000    0.00000 ;    HA3-    CA-     C-     O
   352    350    353    355      3    0.00000    0.00000    0.00000
0.00000    0.00000    0.00000 ;    HA3-    CA-     C-     N
   353    355    357    358      3    0.00000    0.00000    0.00000
0.00000    0.00000    0.00000 ;      C-     N-     CA-    HA
   353    355    357    359      3   11.71520    1.67360   15.06240  -
13.38880    0.00000    0.00000 ;      C-     N-     CA-    CB
   353    355    357    367      3    1.75728    5.27184    2.25936
-7.02912    0.00000    0.00000 ;      C-     N-     CA-     C
   354    353    355    356      3   29.28800   -8.36800  -20.92000
0.00000    0.00000    0.00000 ;      O-     C-     N-     H
   354    353    355    357      3   20.92000    0.00000  -20.92000
0.00000    0.00000    0.00000 ;      O-     C-     N-    CA
   355    357    359    360      3    0.65084    1.95253    0.00000
-2.60338    0.00000    0.00000 ;      N-    CA-    CB-    HB
   355    357    359    361      3    3.83673    0.44769   -1.77402
-2.47693   -0.03347    0.00000 ;      N-    CA-    CB-   CG2
   355    357    359    365      3    4.93294    1.13386   -3.12963
-5.27184    3.17984    0.00000 ;      N-    CA-    CB-   OG1
   355    357    367    368      3    0.00000    0.00000    0.00000
0.00000    0.00000    0.00000 ;      N-    CA-     C-     O
   355    357    367    369      3   17.40544   -5.02080  -13.22144
9.20480    0.00000    0.00000 ;      N-    CA-     C-     N
   356    355    357    358      3    0.00000    0.00000    0.00000
0.00000    0.00000    0.00000 ;      H-     N-     CA-    HA
   356    355    357    359      3    0.00000    0.00000    0.00000
0.00000    0.00000    0.00000 ;      H-     N-     CA-    CB
   356    355    357    367      3    0.00000    0.00000    0.00000
0.00000    0.00000    0.00000 ;      H-     N-     CA-     C
   357    359    361    362      3    0.66944    2.00832    0.00000
-2.67776    0.00000    0.00000 ;     CA-    CB-   CG2-  HG21
   357    359    361    363      3    0.66944    2.00832    0.00000
-2.67776    0.00000    0.00000 ;     CA-    CB-   CG2-  HG22
   357    359    361    364      3    0.66944    2.00832    0.00000
-2.67776    0.00000    0.00000 ;     CA-    CB-   CG2-  HG23
   357    359    365    366      3    1.12131    2.98738    1.66523
-3.94970    0.43514    0.00000 ;     CA-    CB-   OG1-   HG1
   357    367    369    370      3   20.92000    0.00000  -20.92000
0.00000    0.00000    0.00000 ;     CA-     C-     N-     H
   357    367    369    371      3   20.92000    0.00000  -20.92000
```

0.00000 0.00000 0.00000 ; CA- C- N- CA
358 357 359 360 3 0.65084 1.95253 0.00000
-2.60338 0.00000 0.00000 ; HA- CA- CB- HB
358 357 359 361 3 0.65084 1.95253 0.00000
-2.60338 0.00000 0.00000 ; HA- CA- CB- CG2
358 357 359 365 3 1.04600 -1.04600 0.00000
0.00000 0.00000 0.00000 ; HA- CA- CB- OG1
358 357 367 368 3 3.68192 -4.35136 0.00000
1.33888 0.00000 0.00000 ; HA- CA- C- O
358 357 367 369 3 0.00000 0.00000 0.00000
0.00000 0.00000 0.00000 ; HA- CA- C- N
359 357 367 368 3 0.00000 0.00000 0.00000
0.00000 0.00000 0.00000 ; CB- CA- C- O
359 357 367 369 3 2.51040 4.18400 1.67360
-6.69440 0.00000 0.00000 ; CB- CA- C- N
360 359 357 367 3 0.65084 1.95253 0.00000
-2.60338 0.00000 0.00000 ; HB- CB- CA- C
360 359 361 362 3 0.65084 1.95253 0.00000
-2.60338 0.00000 0.00000 ; HB- CB- CG2- HG21
360 359 361 363 3 0.65084 1.95253 0.00000
-2.60338 0.00000 0.00000 ; HB- CB- CG2- HG22
360 359 361 364 3 0.65084 1.95253 0.00000
-2.60338 0.00000 0.00000 ; HB- CB- CG2- HG23
360 359 365 366 3 0.69733 2.09200 0.00000
-2.78933 0.00000 0.00000 ; HB- CB- OG1- HG1
361 359 357 367 3 5.67350 3.55640 -6.16722
-2.47693 3.74886 0.00000 ; CG2- CB- CA- C
361 359 365 366 3 4.74047 0.27196 -2.26773
-3.94970 1.60666 0.00000 ; CG2- CB- OG1- HG1
362 361 359 365 3 1.04600 -1.04600 0.00000
0.00000 0.00000 0.00000 ; HG21- CG2- CB- OG1
363 361 359 365 3 1.04600 -1.04600 0.00000
0.00000 0.00000 0.00000 ; HG22- CG2- CB- OG1
364 361 359 365 3 1.04600 -1.04600 0.00000
0.00000 0.00000 0.00000 ; HG23- CG2- CB- OG1
365 359 357 367 3 6.53541 6.87013 -6.21742
-5.27184 5.22163 0.00000 ; OG1- CB- CA- C
367 369 371 372 3 0.00000 0.00000 0.00000
0.00000 0.00000 0.00000 ; C- N- CA- HA
367 369 371 373 3 11.71520 1.67360 15.06240 -
13.38880 0.00000 0.00000 ; C- N- CA- CB
367 369 371 388 3 1.75728 5.27184 2.25936
-7.02912 0.00000 0.00000 ; C- N- CA- C
368 367 369 370 3 29.28800 -8.36800 -20.92000
0.00000 0.00000 0.00000 ; O- C- N- H
368 367 369 371 3 20.92000 0.00000 -20.92000
0.00000 0.00000 0.00000 ; O- C- N- CA
369 371 373 374 3 0.65084 1.95253 0.00000
-2.60338 0.00000 0.00000 ; N- CA- CB- HB2
369 371 373 375 3 0.65084 1.95253 0.00000
-2.60338 0.00000 0.00000 ; N- CA- CB- HB3
369 371 373 376 3 3.28026 2.46019 -2.19242

-3.21331   -0.23430    0.00000 ;       N-    CA-    CB-    CG
   369    371    388    389      3    0.00000    0.00000    0.00000
0.00000    0.00000    0.00000 ;      N-    CA-     C-     O
   369    371    388    390      3   17.40544   -5.02080  -13.22144
9.20480    0.00000    0.00000 ;      N-    CA-     C-     N
   370    369    371    372      3    0.00000    0.00000    0.00000
0.00000    0.00000    0.00000 ;      H-     N-    CA-    HA
   370    369    371    373      3    0.00000    0.00000    0.00000
0.00000    0.00000    0.00000 ;      H-     N-    CA-    CB
   370    369    371    388      3    0.00000    0.00000    0.00000
0.00000    0.00000    0.00000 ;      H-     N-    CA-     C
   371    373    376    377      3    0.57739    0.00000    1.02926
0.00000   -1.60666    0.00000 ;     CA-    CB-    CG-   CD1
   371    373    376    386      3    0.57739    0.00000    1.02926
0.00000   -1.60666    0.00000 ;     CA-    CB-    CG-   CD2
   371    388    390    391      3   20.92000    0.00000  -20.92000
0.00000    0.00000    0.00000 ;     CA-     C-     N-     H
   371    388    390    392      3   20.92000    0.00000  -20.92000
0.00000    0.00000    0.00000 ;     CA-     C-     N-    CA
   372    371    373    374      3    0.65084    1.95253    0.00000
-2.60338    0.00000    0.00000 ;     HA-    CA-    CB-   HB2
   372    371    373    375      3    0.65084    1.95253    0.00000
-2.60338    0.00000    0.00000 ;     HA-    CA-    CB-   HB3
   372    371    373    376      3    0.65084    1.95253    0.00000
-2.60338    0.00000    0.00000 ;     HA-    CA-    CB-    CG
   372    371    388    389      3    3.68192   -4.35136    0.00000
1.33888    0.00000    0.00000 ;     HA-    CA-     C-     O
   372    371    388    390      3    0.00000    0.00000    0.00000
0.00000    0.00000    0.00000 ;     HA-    CA-     C-     N
   373    371    388    389      3    0.00000    0.00000    0.00000
0.00000    0.00000    0.00000 ;     CB-    CA-     C-     O
   373    371    388    390      3    2.51040    4.18400    1.67360
-6.69440    0.00000    0.00000 ;     CB-    CA-     C-     N
   373    376    377    378      3   30.33400    0.00000  -30.33400
0.00000    0.00000    0.00000 ;     CB-    CG-   CD1-   HD1
   373    376    377    379      3   30.33400    0.00000  -30.33400
0.00000    0.00000    0.00000 ;     CB-    CG-   CD1-   CE1
   373    376    386    384      3   30.33400    0.00000  -30.33400
0.00000    0.00000    0.00000 ;     CB-    CG-   CD2-   CE2
   373    376    386    387      3   30.33400    0.00000  -30.33400
0.00000    0.00000    0.00000 ;     CB-    CG-   CD2-   HD2
   374    373    371    388      3    0.65084    1.95253    0.00000
-2.60338    0.00000    0.00000 ;    HB2-    CB-    CA-     C
   374    373    376    377      3    0.00000    0.00000    0.00000
0.00000    0.00000    0.00000 ;    HB2-    CB-    CG-   CD1
   374    373    376    386      3    0.00000    0.00000    0.00000
0.00000    0.00000    0.00000 ;    HB2-    CB-    CG-   CD2
   375    373    371    388      3    0.65084    1.95253    0.00000
-2.60338    0.00000    0.00000 ;    HB3-    CB-    CA-     C
   375    373    376    377      3    0.00000    0.00000    0.00000
0.00000    0.00000    0.00000 ;    HB3-    CB-    CG-   CD1
   375    373    376    386      3    0.00000    0.00000    0.00000

```
0.00000    0.00000    0.00000 ;    HB3-    CB-    CG-   CD2
   376    373    371    388      3    4.95804    2.17986   -3.52293
-3.21331   -0.40166    0.00000 ;     CG-    CB-    CA-     C
   376    377    379    380      3   30.33400    0.00000  -30.33400
0.00000    0.00000    0.00000 ;     CG-   CD1-   CE1-   HE1
   376    377    379    381      3   30.33400    0.00000  -30.33400
0.00000    0.00000    0.00000 ;     CG-   CD1-   CE1-    CZ
   376    386    384    381      3   30.33400    0.00000  -30.33400
0.00000    0.00000    0.00000 ;     CG-   CD2-   CE2-    CZ
   376    386    384    385      3   30.33400    0.00000  -30.33400
0.00000    0.00000    0.00000 ;     CG-   CD2-   CE2-   HE2
   377    376    386    384      3   30.33400    0.00000  -30.33400
0.00000    0.00000    0.00000 ;    CD1-    CG-   CD2-   CE2
   377    376    386    387      3   30.33400    0.00000  -30.33400
0.00000    0.00000    0.00000 ;    CD1-    CG-   CD2-   HD2
   377    379    381    382      3   30.33400    0.00000  -30.33400
0.00000    0.00000    0.00000 ;    CD1-   CE1-    CZ-    OH
   377    379    381    384      3   30.33400    0.00000  -30.33400
0.00000    0.00000    0.00000 ;    CD1-   CE1-    CZ-   CE2
   378    377    376    386      3   30.33400    0.00000  -30.33400
0.00000    0.00000    0.00000 ;    HD1-   CD1-    CG-   CD2
   378    377    379    380      3   30.33400    0.00000  -30.33400
0.00000    0.00000    0.00000 ;    HD1-   CD1-   CE1-   HE1
   378    377    379    381      3   30.33400    0.00000  -30.33400
0.00000    0.00000    0.00000 ;    HD1-   CD1-   CE1-    CZ
   379    377    376    386      3   30.33400    0.00000  -30.33400
0.00000    0.00000    0.00000 ;    CE1-   CD1-    CG-   CD2
   379    381    382    383      3    7.93286    0.00000   -9.56462
0.00000    2.17568    0.00000 ;    CE1-    CZ-    OH-    HH
   379    381    384    385      3   30.33400    0.00000  -30.33400
0.00000    0.00000    0.00000 ;    CE1-    CZ-   CE2-   HE2
   379    381    384    386      3   30.33400    0.00000  -30.33400
0.00000    0.00000    0.00000 ;    CE1-    CZ-   CE2-   CD2
   380    379    381    382      3   30.33400    0.00000  -30.33400
0.00000    0.00000    0.00000 ;    HE1-   CE1-    CZ-    OH
   380    379    381    384      3   30.33400    0.00000  -30.33400
0.00000    0.00000    0.00000 ;    HE1-   CE1-    CZ-   CE2
   381    384    386    387      3   30.33400    0.00000  -30.33400
0.00000    0.00000    0.00000 ;     CZ-   CE2-   CD2-   HD2
   382    381    384    385      3   30.33400    0.00000  -30.33400
0.00000    0.00000    0.00000 ;     OH-    CZ-   CE2-   HE2
   382    381    384    386      3   30.33400    0.00000  -30.33400
0.00000    0.00000    0.00000 ;     OH-    CZ-   CE2-   CD2
   383    382    381    384      3    7.93286    0.00000   -9.56462
0.00000    2.17568    0.00000 ;     HH-    OH-    CZ-   CE2
   385    384    386    387      3   30.33400    0.00000  -30.33400
0.00000    0.00000    0.00000 ;    HE2-   CE2-   CD2-   HD2
   388    390    392    393      3    0.00000    0.00000    0.00000
0.00000    0.00000    0.00000 ;      C-     N-    CA-   HA2
   388    390    392    394      3    0.00000    0.00000    0.00000
0.00000    0.00000    0.00000 ;      C-     N-    CA-   HA3
   388    390    392    395      3    1.75728    5.27184    2.25936
```

-7.02912    0.00000    0.00000 ;       C-     N-    CA-     C
   389    388    390    391      3   29.28800   -8.36800  -20.92000
0.00000    0.00000    0.00000 ;       O-     C-     N-     H
   389    388    390    392      3   20.92000    0.00000  -20.92000
0.00000    0.00000    0.00000 ;       O-     C-     N-    CA
   390    392    395    396      3    0.00000    0.00000    0.00000
0.00000    0.00000    0.00000 ;       N-    CA-     C-     O
   390    392    395    397      3   17.40544   -5.02080  -13.22144
9.20480    0.00000    0.00000 ;       N-    CA-     C-     N
   391    390    392    393      3    0.00000    0.00000    0.00000
0.00000    0.00000    0.00000 ;       H-     N-    CA-   HA2
   391    390    392    394      3    0.00000    0.00000    0.00000
0.00000    0.00000    0.00000 ;       H-     N-    CA-   HA3
   391    390    392    395      3    0.00000    0.00000    0.00000
0.00000    0.00000    0.00000 ;       H-     N-    CA-     C
   392    395    397    398      3   20.92000    0.00000  -20.92000
0.00000    0.00000    0.00000 ;      CA-     C-     N-     H
   392    395    397    399      3   20.92000    0.00000  -20.92000
0.00000    0.00000    0.00000 ;      CA-     C-     N-    CA
   393    392    395    396      3    3.68192   -4.35136    0.00000
1.33888    0.00000    0.00000 ;     HA2-    CA-     C-     O
   393    392    395    397      3    0.00000    0.00000    0.00000
0.00000    0.00000    0.00000 ;     HA2-    CA-     C-     N
   394    392    395    396      3    3.68192   -4.35136    0.00000
1.33888    0.00000    0.00000 ;     HA3-    CA-     C-     O
   394    392    395    397      3    0.00000    0.00000    0.00000
0.00000    0.00000    0.00000 ;     HA3-    CA-     C-     N
   395    397    399    400      3    0.00000    0.00000    0.00000
0.00000    0.00000    0.00000 ;       C-     N-    CA-    HA
   395    397    399    401      3   11.71520    1.67360   15.06240  -
13.38880    0.00000    0.00000 ;       C-     N-    CA-    CB
   395    397    399    406      3    1.75728    5.27184    2.25936
-7.02912    0.00000    0.00000 ;       C-     N-    CA-     C
   396    395    397    398      3   29.28800   -8.36800  -20.92000
0.00000    0.00000    0.00000 ;       O-     C-     N-     H
   396    395    397    399      3   20.92000    0.00000  -20.92000
0.00000    0.00000    0.00000 ;       O-     C-     N-    CA
   397    399    401    402      3    0.65084    1.95253    0.00000
-2.60338    0.00000    0.00000 ;       N-    CA-    CB-   HB2
   397    399    401    403      3    0.65084    1.95253    0.00000
-2.60338    0.00000    0.00000 ;       N-    CA-    CB-   HB3
   397    399    401    404      3    6.03751    2.50622   -5.17142
-4.20074    1.10458    0.00000 ;       N-    CA-    CB-    SG
   397    399    406    407      3    0.00000    0.00000    0.00000
0.00000    0.00000    0.00000 ;       N-    CA-     C-     O
   397    399    406    408      3   17.40544   -5.02080  -13.22144
9.20480    0.00000    0.00000 ;       N-    CA-     C-     N
   398    397    399    400      3    0.00000    0.00000    0.00000
0.00000    0.00000    0.00000 ;       H-     N-    CA-    HA
   398    397    399    401      3    0.00000    0.00000    0.00000
0.00000    0.00000    0.00000 ;       H-     N-    CA-    CB
   398    397    399    406      3    0.00000    0.00000    0.00000

0.00000 0.00000 0.00000 ; H- N- CA- C
399 401 404 405 3 1.69034 2.77818 4.11706
-4.21747 1.00416 0.00000 ; CA- CB- SG- HG
399 406 408 409 3 20.92000 0.00000 -20.92000
0.00000 0.00000 0.00000 ; CA- C- N- H
399 406 408 410 3 20.92000 0.00000 -20.92000
0.00000 0.00000 0.00000 ; CA- C- N- CA
400 399 401 402 3 0.65084 1.95253 0.00000
-2.60338 0.00000 0.00000 ; HA- CA- CB- HB2
400 399 401 403 3 0.65084 1.95253 0.00000
-2.60338 0.00000 0.00000 ; HA- CA- CB- HB3
400 399 401 404 3 0.65084 1.95253 0.00000
-2.60338 0.00000 0.00000 ; HA- CA- CB- SG
400 399 406 407 3 3.68192 -4.35136 0.00000
1.33888 0.00000 0.00000 ; HA- CA- C- O
400 399 406 408 3 0.00000 0.00000 0.00000
0.00000 0.00000 0.00000 ; HA- CA- C- N
401 399 406 407 3 0.00000 0.00000 0.00000
0.00000 0.00000 0.00000 ; CB- CA- C- O
401 399 406 408 3 2.51040 4.18400 1.67360
-6.69440 0.00000 0.00000 ; CB- CA- C- N
402 401 399 406 3 0.65084 1.95253 0.00000
-2.60338 0.00000 0.00000 ; HB2- CB- CA- C
402 401 404 405 3 1.04600 3.13800 0.00000
-4.18400 0.00000 0.00000 ; HB2- CB- SG- HG
403 401 399 406 3 0.65084 1.95253 0.00000
-2.60338 0.00000 0.00000 ; HB3- CB- CA- C
403 401 404 405 3 1.04600 3.13800 0.00000
-4.18400 0.00000 0.00000 ; HB3- CB- SG- HG
404 401 399 406 3 5.62330 4.27605 -5.33042
-4.20074 2.51040 0.00000 ; SG- CB- CA- C
406 408 410 411 3 0.00000 0.00000 0.00000
0.00000 0.00000 0.00000 ; C- N- CA- HA
406 408 410 412 3 11.71520 1.67360 15.06240 -
13.38880 0.00000 0.00000 ; C- N- CA- CB
406 408 410 422 3 1.75728 5.27184 2.25936
-7.02912 0.00000 0.00000 ; C- N- CA- C
407 406 408 409 3 29.28800 -8.36800 -20.92000
0.00000 0.00000 0.00000 ; O- C- N- H
407 406 408 410 3 20.92000 0.00000 -20.92000
0.00000 0.00000 0.00000 ; O- C- N- CA
408 410 412 413 3 0.65084 1.95253 0.00000
-2.60338 0.00000 0.00000 ; N- CA- CB- HB
408 410 412 414 3 3.83673 0.44769 -1.77402
-2.47693 -0.03347 0.00000 ; N- CA- CB- CG1
408 410 412 418 3 3.83673 0.44769 -1.77402
-2.47693 -0.03347 0.00000 ; N- CA- CB- CG2
408 410 422 423 3 0.00000 0.00000 0.00000
0.00000 0.00000 0.00000 ; N- CA- C- O
408 410 422 424 3 17.40544 -5.02080 -13.22144
9.20480 0.00000 0.00000 ; N- CA- C- N
409 408 410 411 3 0.00000 0.00000 0.00000

```
0.00000    0.00000    0.00000 ;       H-     N-    CA-    HA
   409    408    410    412       3    0.00000    0.00000    0.00000
0.00000    0.00000    0.00000 ;       H-     N-    CA-    CB
   409    408    410    422       3    0.00000    0.00000    0.00000
0.00000    0.00000    0.00000 ;       H-     N-    CA-     C
   410    412    414    415       3    0.66944    2.00832    0.00000
-2.67776    0.00000    0.00000 ;      CA-    CB-   CG1-  HG11
   410    412    414    416       3    0.66944    2.00832    0.00000
-2.67776    0.00000    0.00000 ;      CA-    CB-   CG1-  HG12
   410    412    414    417       3    0.66944    2.00832    0.00000
-2.67776    0.00000    0.00000 ;      CA-    CB-   CG1-  HG13
   410    412    418    419       3    0.66944    2.00832    0.00000
-2.67776    0.00000    0.00000 ;      CA-    CB-   CG2-  HG21
   410    412    418    420       3    0.66944    2.00832    0.00000
-2.67776    0.00000    0.00000 ;      CA-    CB-   CG2-  HG22
   410    412    418    421       3    0.66944    2.00832    0.00000
-2.67776    0.00000    0.00000 ;      CA-    CB-   CG2-  HG23
   410    422    424    425       3   20.92000    0.00000  -20.92000
0.00000    0.00000    0.00000 ;      CA-     C-     N-     H
   410    422    424    426       3   20.92000    0.00000  -20.92000
0.00000    0.00000    0.00000 ;      CA-     C-     N-    CA
   411    410    412    413       3    0.65084    1.95253    0.00000
-2.60338    0.00000    0.00000 ;      HA-    CA-    CB-    HB
   411    410    412    414       3    0.65084    1.95253    0.00000
-2.60338    0.00000    0.00000 ;      HA-    CA-    CB-   CG1
   411    410    412    418       3    0.65084    1.95253    0.00000
-2.60338    0.00000    0.00000 ;      HA-    CA-    CB-   CG2
   411    410    422    423       3    3.68192   -4.35136    0.00000
1.33888    0.00000    0.00000 ;      HA-    CA-     C-     O
   411    410    422    424       3    0.00000    0.00000    0.00000
0.00000    0.00000    0.00000 ;      HA-    CA-     C-     N
   412    410    422    423       3    0.00000    0.00000    0.00000
0.00000    0.00000    0.00000 ;      CB-    CA-     C-     O
   412    410    422    424       3    2.51040    4.18400    1.67360
-6.69440    0.00000    0.00000 ;      CB-    CA-     C-     N
   413    412    410    422       3    0.65084    1.95253    0.00000
-2.60338    0.00000    0.00000 ;      HB-    CB-    CA-     C
   413    412    414    415       3    0.62760    1.88280    0.00000
-2.51040    0.00000    0.00000 ;      HB-    CB-   CG1-  HG11
   413    412    414    416       3    0.62760    1.88280    0.00000
-2.51040    0.00000    0.00000 ;      HB-    CB-   CG1-  HG12
   413    412    414    417       3    0.62760    1.88280    0.00000
-2.51040    0.00000    0.00000 ;      HB-    CB-   CG1-  HG13
   413    412    418    419       3    0.62760    1.88280    0.00000
-2.51040    0.00000    0.00000 ;      HB-    CB-   CG2-  HG21
   413    412    418    420       3    0.62760    1.88280    0.00000
-2.51040    0.00000    0.00000 ;      HB-    CB-   CG2-  HG22
   413    412    418    421       3    0.62760    1.88280    0.00000
-2.51040    0.00000    0.00000 ;      HB-    CB-   CG2-  HG23
   414    412    410    422       3    5.67350    3.55640   -6.16722
-2.47693    3.74886    0.00000 ;     CG1-    CB-    CA-     C
   414    412    418    419       3    0.66944    2.00832    0.00000
```

```
-2.67776    0.00000    0.00000 ;    CG1-    CB-   CG2-  HG21
   414    412    418    420      3    0.66944    2.00832    0.00000
-2.67776    0.00000    0.00000 ;    CG1-    CB-   CG2-  HG22
   414    412    418    421      3    0.66944    2.00832    0.00000
-2.67776    0.00000    0.00000 ;    CG1-    CB-   CG2-  HG23
   415    414    412    418      3    0.66944    2.00832    0.00000
-2.67776    0.00000    0.00000 ;   HG11-   CG1-    CB-   CG2
   416    414    412    418      3    0.66944    2.00832    0.00000
-2.67776    0.00000    0.00000 ;   HG12-   CG1-    CB-   CG2
   417    414    412    418      3    0.66944    2.00832    0.00000
-2.67776    0.00000    0.00000 ;   HG13-   CG1-    CB-   CG2
   418    412    410    422      3    5.67350    3.55640   -6.16722
-2.47693    3.74886    0.00000 ;    CG2-    CB-    CA-     C
   422    424    426    427      3    0.00000    0.00000    0.00000
0.00000    0.00000    0.00000 ;      C-     N-    CA-    HA
   422    424    426    428      3   11.71520    1.67360   15.06240  -
13.38880    0.00000    0.00000 ;      C-     N-    CA-    CB
   422    424    426    443      3    1.75728    5.27184    2.25936
-7.02912    0.00000    0.00000 ;      C-     N-    CA-     C
   423    422    424    425      3   29.28800   -8.36800  -20.92000
0.00000    0.00000    0.00000 ;      O-     C-     N-     H
   423    422    424    426      3   20.92000    0.00000  -20.92000
0.00000    0.00000    0.00000 ;      O-     C-     N-    CA
   424    426    428    429      3    0.65084    1.95253    0.00000
-2.60338    0.00000    0.00000 ;      N-    CA-    CB-   HB2
   424    426    428    430      3    0.65084    1.95253    0.00000
-2.60338    0.00000    0.00000 ;      N-    CA-    CB-   HB3
   424    426    428    431      3    3.28026    2.46019   -2.19242
-3.21331   -0.23430    0.00000 ;      N-    CA-    CB-    CG
   424    426    443    444      3    0.00000    0.00000    0.00000
0.00000    0.00000    0.00000 ;      N-    CA-     C-     O
   424    426    443    445      3   17.40544   -5.02080  -13.22144
9.20480    0.00000    0.00000 ;      N-    CA-     C-     N
   425    424    426    427      3    0.00000    0.00000    0.00000
0.00000    0.00000    0.00000 ;      H-     N-    CA-    HA
   425    424    426    428      3    0.00000    0.00000    0.00000
0.00000    0.00000    0.00000 ;      H-     N-    CA-    CB
   425    424    426    443      3    0.00000    0.00000    0.00000
0.00000    0.00000    0.00000 ;      H-     N-    CA-     C
   426    428    431    432      3    0.57739    0.00000    1.02926
0.00000   -1.60666    0.00000 ;    CA-    CB-    CG-   CD1
   426    428    431    441      3    0.57739    0.00000    1.02926
0.00000   -1.60666    0.00000 ;    CA-    CB-    CG-   CD2
   426    443    445    446      3   20.92000    0.00000  -20.92000
0.00000    0.00000    0.00000 ;    CA-     C-     N-     H
   426    443    445    447      3   20.92000    0.00000  -20.92000
0.00000    0.00000    0.00000 ;    CA-     C-     N-    CA
   427    426    428    429      3    0.65084    1.95253    0.00000
-2.60338    0.00000    0.00000 ;     HA-    CA-    CB-   HB2
   427    426    428    430      3    0.65084    1.95253    0.00000
-2.60338    0.00000    0.00000 ;     HA-    CA-    CB-   HB3
   427    426    428    431      3    0.65084    1.95253    0.00000
```

```
-2.60338    0.00000    0.00000 ;     HA-    CA-    CB-    CG
   427    426    443    444      3    3.68192   -4.35136    0.00000
1.33888    0.00000    0.00000 ;     HA-    CA-     C-     O
   427    426    443    445      3    0.00000    0.00000    0.00000
0.00000    0.00000    0.00000 ;     HA-    CA-     C-     N
   428    426    443    444      3    0.00000    0.00000    0.00000
0.00000    0.00000    0.00000 ;     CB-    CA-     C-     O
   428    426    443    445      3    2.51040    4.18400    1.67360
-6.69440    0.00000    0.00000 ;     CB-    CA-     C-     N
   428    431    432    433      3   30.33400    0.00000  -30.33400
0.00000    0.00000    0.00000 ;     CB-    CG-   CD1-   HD1
   428    431    432    434      3   30.33400    0.00000  -30.33400
0.00000    0.00000    0.00000 ;     CB-    CG-   CD1-   CE1
   428    431    441    439      3   30.33400    0.00000  -30.33400
0.00000    0.00000    0.00000 ;     CB-    CG-   CD2-   CE2
   428    431    441    442      3   30.33400    0.00000  -30.33400
0.00000    0.00000    0.00000 ;     CB-    CG-   CD2-   HD2
   429    428    426    443      3    0.65084    1.95253    0.00000
-2.60338    0.00000    0.00000 ;    HB2-    CB-    CA-     C
   429    428    431    432      3    0.00000    0.00000    0.00000
0.00000    0.00000    0.00000 ;    HB2-    CB-    CG-   CD1
   429    428    431    441      3    0.00000    0.00000    0.00000
0.00000    0.00000    0.00000 ;    HB2-    CB-    CG-   CD2
   430    428    426    443      3    0.65084    1.95253    0.00000
-2.60338    0.00000    0.00000 ;    HB3-    CB-    CA-     C
   430    428    431    432      3    0.00000    0.00000    0.00000
0.00000    0.00000    0.00000 ;    HB3-    CB-    CG-   CD1
   430    428    431    441      3    0.00000    0.00000    0.00000
0.00000    0.00000    0.00000 ;    HB3-    CB-    CG-   CD2
   431    428    426    443      3    4.95804    2.17986   -3.52293
-3.21331   -0.40166    0.00000 ;     CG-    CB-    CA-     C
   431    432    434    435      3   30.33400    0.00000  -30.33400
0.00000    0.00000    0.00000 ;     CG-   CD1-   CE1-   HE1
   431    432    434    436      3   30.33400    0.00000  -30.33400
0.00000    0.00000    0.00000 ;     CG-   CD1-   CE1-    CZ
   431    441    439    436      3   30.33400    0.00000  -30.33400
0.00000    0.00000    0.00000 ;     CG-   CD2-   CE2-    CZ
   431    441    439    440      3   30.33400    0.00000  -30.33400
0.00000    0.00000    0.00000 ;     CG-   CD2-   CE2-   HE2
   432    431    441    439      3   30.33400    0.00000  -30.33400
0.00000    0.00000    0.00000 ;    CD1-    CG-   CD2-   CE2
   432    431    441    442      3   30.33400    0.00000  -30.33400
0.00000    0.00000    0.00000 ;    CD1-    CG-   CD2-   HD2
   432    434    436    437      3   30.33400    0.00000  -30.33400
0.00000    0.00000    0.00000 ;    CD1-   CE1-    CZ-    OH
   432    434    436    439      3   30.33400    0.00000  -30.33400
0.00000    0.00000    0.00000 ;    CD1-   CE1-    CZ-   CE2
   433    432    431    441      3   30.33400    0.00000  -30.33400
0.00000    0.00000    0.00000 ;    HD1-   CD1-    CG-   CD2
   433    432    434    435      3   30.33400    0.00000  -30.33400
0.00000    0.00000    0.00000 ;    HD1-   CD1-   CE1-   HE1
   433    432    434    436      3   30.33400    0.00000  -30.33400
```

```
0.00000    0.00000    0.00000 ;    HD1-   CD1-   CE1-    CZ
   434    432    431    441      3   30.33400    0.00000  -30.33400
0.00000    0.00000    0.00000 ;    CE1-   CD1-    CG-   CD2
   434    436    437    438      3    7.93286    0.00000   -9.56462
0.00000    2.17568    0.00000 ;    CE1-    CZ-    OH-    HH
   434    436    439    440      3   30.33400    0.00000  -30.33400
0.00000    0.00000    0.00000 ;    CE1-    CZ-   CE2-   HE2
   434    436    439    441      3   30.33400    0.00000  -30.33400
0.00000    0.00000    0.00000 ;    CE1-    CZ-   CE2-   CD2
   435    434    436    437      3   30.33400    0.00000  -30.33400
0.00000    0.00000    0.00000 ;    HE1-   CE1-    CZ-    OH
   435    434    436    439      3   30.33400    0.00000  -30.33400
0.00000    0.00000    0.00000 ;    HE1-   CE1-    CZ-   CE2
   436    439    441    442      3   30.33400    0.00000  -30.33400
0.00000    0.00000    0.00000 ;     CZ-   CE2-   CD2-   HD2
   437    436    439    440      3   30.33400    0.00000  -30.33400
0.00000    0.00000    0.00000 ;     OH-    CZ-   CE2-   HE2
   437    436    439    441      3   30.33400    0.00000  -30.33400
0.00000    0.00000    0.00000 ;     OH-    CZ-   CE2-   CD2
   438    437    436    439      3    7.93286    0.00000   -9.56462
0.00000    2.17568    0.00000 ;     HH-    OH-    CZ-   CE2
   440    439    441    442      3   30.33400    0.00000  -30.33400
0.00000    0.00000    0.00000 ;    HE2-   CE2-   CD2-   HD2
   443    445    447    448      3    0.00000    0.00000    0.00000
0.00000    0.00000    0.00000 ;      C-     N-    CA-    HA
   443    445    447    449      3   11.71520    1.67360   15.06240  -
13.38880    0.00000    0.00000 ;      C-     N-    CA-    CB
   443    445    447    465      3    1.75728    5.27184    2.25936
-7.02912    0.00000    0.00000 ;      C-     N-    CA-     C
   444    443    445    446      3   29.28800   -8.36800  -20.92000
0.00000    0.00000    0.00000 ;      O-     C-     N-     H
   444    443    445    447      3   20.92000    0.00000  -20.92000
0.00000    0.00000    0.00000 ;      O-     C-     N-    CA
   445    447    449    450      3    0.65084    1.95253    0.00000
-2.60338    0.00000    0.00000 ;      N-    CA-    CB-   HB2
   445    447    449    451      3    0.65084    1.95253    0.00000
-2.60338    0.00000    0.00000 ;      N-    CA-    CB-   HB3
   445    447    449    452      3    0.65084    1.95253    0.00000
-2.60338    0.00000    0.00000 ;      N-    CA-    CB-    CG
   445    447    465    466      3    0.00000    0.00000    0.00000
0.00000    0.00000    0.00000 ;      N-    CA-     C-     O
   445    447    465    467      3   17.40544   -5.02080  -13.22144
9.20480    0.00000    0.00000 ;      N-    CA-     C-     N
   446    445    447    448      3    0.00000    0.00000    0.00000
0.00000    0.00000    0.00000 ;      H-     N-    CA-    HA
   446    445    447    449      3    0.00000    0.00000    0.00000
0.00000    0.00000    0.00000 ;      H-     N-    CA-    CB
   446    445    447    465      3    0.00000    0.00000    0.00000
0.00000    0.00000    0.00000 ;      H-     N-    CA-     C
   447    449    452    453      3    0.66944    2.00832    0.00000
-2.67776    0.00000    0.00000 ;     CA-    CB-    CG-   HG2
   447    449    452    454      3    0.66944    2.00832    0.00000
```

-2.67776 0.00000 0.00000 ; CA- CB- CG- HG3
447 449 452 455 3 3.68192 3.09616 -2.09200
-3.01248 0.00000 0.00000 ; CA- CB- CG- CD
447 465 467 468 3 20.92000 0.00000 -20.92000
0.00000 0.00000 0.00000 ; CA- C- N- H
447 465 467 469 3 20.92000 0.00000 -20.92000
0.00000 0.00000 0.00000 ; CA- C- N- CA
448 447 449 450 3 0.65084 1.95253 0.00000
-2.60338 0.00000 0.00000 ; HA- CA- CB- HB2
448 447 449 451 3 0.65084 1.95253 0.00000
-2.60338 0.00000 0.00000 ; HA- CA- CB- HB3
448 447 449 452 3 0.65084 1.95253 0.00000
-2.60338 0.00000 0.00000 ; HA- CA- CB- CG
448 447 465 466 3 3.68192 -4.35136 0.00000
1.33888 0.00000 0.00000 ; HA- CA- C- O
448 447 465 467 3 0.00000 0.00000 0.00000
0.00000 0.00000 0.00000 ; HA- CA- C- N
449 447 465 466 3 0.00000 0.00000 0.00000
0.00000 0.00000 0.00000 ; CB- CA- C- O
449 447 465 467 3 2.51040 4.18400 1.67360
-6.69440 0.00000 0.00000 ; CB- CA- C- N
449 452 455 456 3 0.66944 2.00832 0.00000
-2.67776 0.00000 0.00000 ; CB- CG- CD- HD2
449 452 455 457 3 0.66944 2.00832 0.00000
-2.67776 0.00000 0.00000 ; CB- CG- CD- HD3
449 452 455 458 3 3.68192 3.09616 -2.09200
-3.01248 0.00000 0.00000 ; CB- CG- CD- CE
450 449 447 465 3 0.65084 1.95253 0.00000
-2.60338 0.00000 0.00000 ; HB2- CB- CA- C
450 449 452 453 3 0.62760 1.88280 0.00000
-2.51040 0.00000 0.00000 ; HB2- CB- CG- HG2
450 449 452 454 3 0.62760 1.88280 0.00000
-2.51040 0.00000 0.00000 ; HB2- CB- CG- HG3
450 449 452 455 3 0.66944 2.00832 0.00000
-2.67776 0.00000 0.00000 ; HB2- CB- CG- CD
451 449 447 465 3 0.65084 1.95253 0.00000
-2.60338 0.00000 0.00000 ; HB3- CB- CA- C
451 449 452 453 3 0.62760 1.88280 0.00000
-2.51040 0.00000 0.00000 ; HB3- CB- CG- HG2
451 449 452 454 3 0.62760 1.88280 0.00000
-2.51040 0.00000 0.00000 ; HB3- CB- CG- HG3
451 449 452 455 3 0.66944 2.00832 0.00000
-2.67776 0.00000 0.00000 ; HB3- CB- CG- CD
452 449 447 465 3 0.65084 1.95253 0.00000
-2.60338 0.00000 0.00000 ; CG- CB- CA- C
452 455 458 459 3 0.65084 1.95253 0.00000
-2.60338 0.00000 0.00000 ; CG- CD- CE- HE2
452 455 458 460 3 0.65084 1.95253 0.00000
-2.60338 0.00000 0.00000 ; CG- CD- CE- HE3
452 455 458 461 3 0.65084 1.95253 0.00000
-2.60338 0.00000 0.00000 ; CG- CD- CE- NZ
453 452 455 456 3 0.62760 1.88280 0.00000

```
-2.51040    0.00000    0.00000 ;    HG2-   CG-    CD-   HD2
   453    452    455    457      3    0.62760    1.88280    0.00000
-2.51040    0.00000    0.00000 ;    HG2-   CG-    CD-   HD3
   453    452    455    458      3    0.66944    2.00832    0.00000
-2.67776    0.00000    0.00000 ;    HG2-   CG-    CD-    CE
   454    452    455    456      3    0.62760    1.88280    0.00000
-2.51040    0.00000    0.00000 ;    HG3-   CG-    CD-   HD2
   454    452    455    457      3    0.62760    1.88280    0.00000
-2.51040    0.00000    0.00000 ;    HG3-   CG-    CD-   HD3
   454    452    455    458      3    0.66944    2.00832    0.00000
-2.67776    0.00000    0.00000 ;    HG3-   CG-    CD-    CE
   455    458    461    462      3    0.65084    1.95253    0.00000
-2.60338    0.00000    0.00000 ;     CD-   CE-    NZ-   HZ1
   455    458    461    463      3    0.65084    1.95253    0.00000
-2.60338    0.00000    0.00000 ;     CD-   CE-    NZ-   HZ2
   455    458    461    464      3    0.65084    1.95253    0.00000
-2.60338    0.00000    0.00000 ;     CD-   CE-    NZ-   HZ3
   456    455    458    459      3    0.65084    1.95253    0.00000
-2.60338    0.00000    0.00000 ;    HD2-   CD-    CE-   HE2
   456    455    458    460      3    0.65084    1.95253    0.00000
-2.60338    0.00000    0.00000 ;    HD2-   CD-    CE-   HE3
   456    455    458    461      3    0.65084    1.95253    0.00000
-2.60338    0.00000    0.00000 ;    HD2-   CD-    CE-    NZ
   457    455    458    459      3    0.65084    1.95253    0.00000
-2.60338    0.00000    0.00000 ;    HD3-   CD-    CE-   HE2
   457    455    458    460      3    0.65084    1.95253    0.00000
-2.60338    0.00000    0.00000 ;    HD3-   CD-    CE-   HE3
   457    455    458    461      3    0.65084    1.95253    0.00000
-2.60338    0.00000    0.00000 ;    HD3-   CD-    CE-    NZ
   459    458    461    462      3    0.65084    1.95253    0.00000
-2.60338    0.00000    0.00000 ;    HE2-   CE-    NZ-   HZ1
   459    458    461    463      3    0.65084    1.95253    0.00000
-2.60338    0.00000    0.00000 ;    HE2-   CE-    NZ-   HZ2
   459    458    461    464      3    0.65084    1.95253    0.00000
-2.60338    0.00000    0.00000 ;    HE2-   CE-    NZ-   HZ3
   460    458    461    462      3    0.65084    1.95253    0.00000
-2.60338    0.00000    0.00000 ;    HE3-   CE-    NZ-   HZ1
   460    458    461    463      3    0.65084    1.95253    0.00000
-2.60338    0.00000    0.00000 ;    HE3-   CE-    NZ-   HZ2
   460    458    461    464      3    0.65084    1.95253    0.00000
-2.60338    0.00000    0.00000 ;    HE3-   CE-    NZ-   HZ3
   465    467    469    470      3    0.00000    0.00000    0.00000
0.00000    0.00000    0.00000 ;     C-     N-    CA-   HA2
   465    467    469    471      3    0.00000    0.00000    0.00000
0.00000    0.00000    0.00000 ;     C-     N-    CA-   HA3
   465    467    469    472      3    1.75728    5.27184    2.25936
-7.02912    0.00000    0.00000 ;     C-     N-    CA-     C
   466    465    467    468      3   29.28800   -8.36800  -20.92000
0.00000    0.00000    0.00000 ;     O-     C-     N-     H
   466    465    467    469      3   20.92000    0.00000  -20.92000
0.00000    0.00000    0.00000 ;     O-     C-     N-    CA
   467    469    472    473      3    0.00000    0.00000    0.00000
```

```
0.00000    0.00000    0.00000 ;      N-    CA-     C-     O
   467    469    472    474      3   17.40544   -5.02080  -13.22144
9.20480    0.00000    0.00000 ;      N-    CA-     C-     N
   468    467    469    470      3    0.00000    0.00000    0.00000
0.00000    0.00000    0.00000 ;      H-     N-    CA-   HA2
   468    467    469    471      3    0.00000    0.00000    0.00000
0.00000    0.00000    0.00000 ;      H-     N-    CA-   HA3
   468    467    469    472      3    0.00000    0.00000    0.00000
0.00000    0.00000    0.00000 ;      H-     N-    CA-     C
   469    472    474    475      3   20.92000    0.00000  -20.92000
0.00000    0.00000    0.00000 ;     CA-     C-     N-     H
   469    472    474    476      3   20.92000    0.00000  -20.92000
0.00000    0.00000    0.00000 ;     CA-     C-     N-    CA
   470    469    472    473      3    3.68192   -4.35136    0.00000
1.33888    0.00000    0.00000 ;    HA2-    CA-     C-     O
   470    469    472    474      3    0.00000    0.00000    0.00000
0.00000    0.00000    0.00000 ;    HA2-    CA-     C-     N
   471    469    472    473      3    3.68192   -4.35136    0.00000
1.33888    0.00000    0.00000 ;    HA3-    CA-     C-     O
   471    469    472    474      3    0.00000    0.00000    0.00000
0.00000    0.00000    0.00000 ;    HA3-    CA-     C-     N
   472    474    476    477      3    0.00000    0.00000    0.00000
0.00000    0.00000    0.00000 ;      C-     N-    CA-    HA
   472    474    476    478      3   11.71520    1.67360   15.06240  -
13.38880    0.00000    0.00000 ;      C-     N-    CA-    CB
   472    474    476    491      3    1.75728    5.27184    2.25936
-7.02912    0.00000    0.00000 ;      C-     N-    CA-     C
   473    472    474    475      3   29.28800   -8.36800  -20.92000
0.00000    0.00000    0.00000 ;      O-     C-     N-     H
   473    472    474    476      3   20.92000    0.00000  -20.92000
0.00000    0.00000    0.00000 ;      O-     C-     N-    CA
   474    476    478    479      3    0.65084    1.95253    0.00000
-2.60338    0.00000    0.00000 ;      N-    CA-    CB-    HB
   474    476    478    480      3    3.83673    0.44769   -1.77402
-2.47693   -0.03347    0.00000 ;      N-    CA-    CB-   CG2
   474    476    478    484      3    2.97482    0.12134    2.04179
-1.89117   -3.24678    0.00000 ;      N-    CA-    CB-   CG1
   474    476    491    492      3    0.00000    0.00000    0.00000
0.00000    0.00000    0.00000 ;      N-    CA-     C-     O
   474    476    491    493      3   17.40544   -5.02080  -13.22144
9.20480    0.00000    0.00000 ;      N-    CA-     C-     N
   475    474    476    477      3    0.00000    0.00000    0.00000
0.00000    0.00000    0.00000 ;      H-     N-    CA-    HA
   475    474    476    478      3    0.00000    0.00000    0.00000
0.00000    0.00000    0.00000 ;      H-     N-    CA-    CB
   475    474    476    491      3    0.00000    0.00000    0.00000
0.00000    0.00000    0.00000 ;      H-     N-    CA-     C
   476    478    480    481      3    0.66944    2.00832    0.00000
-2.67776    0.00000    0.00000 ;     CA-    CB-   CG2-  HG21
   476    478    480    482      3    0.66944    2.00832    0.00000
-2.67776    0.00000    0.00000 ;     CA-    CB-   CG2-  HG22
   476    478    480    483      3    0.66944    2.00832    0.00000
```

```
-2.67776    0.00000    0.00000 ;     CA-    CB-   CG2-  HG23
   476    478    484    485      3    0.66944    2.00832    0.00000
-2.67776    0.00000    0.00000 ;     CA-    CB-   CG1-  HG12
   476    478    484    486      3    0.66944    2.00832    0.00000
-2.67776    0.00000    0.00000 ;     CA-    CB-   CG1-  HG13
   476    478    484    487      3    4.24258   -0.52718   -7.25506
-1.79075    7.69856    0.00000 ;     CA-    CB-   CG1-   CD1
   476    491    493    494      3   20.92000    0.00000  -20.92000
0.00000    0.00000    0.00000 ;     CA-     C-     N-     H
   476    491    493    495      3   20.92000    0.00000  -20.92000
0.00000    0.00000    0.00000 ;     CA-     C-     N-    CA
   477    476    478    479      3    0.65084    1.95253    0.00000
-2.60338    0.00000    0.00000 ;     HA-    CA-    CB-    HB
   477    476    478    480      3    0.65084    1.95253    0.00000
-2.60338    0.00000    0.00000 ;     HA-    CA-    CB-   CG2
   477    476    478    484      3    0.65084    1.95253    0.00000
-2.60338    0.00000    0.00000 ;     HA-    CA-    CB-   CG1
   477    476    491    492      3    3.68192   -4.35136    0.00000
1.33888    0.00000    0.00000 ;     HA-    CA-     C-     O
   477    476    491    493      3    0.00000    0.00000    0.00000
0.00000    0.00000    0.00000 ;     HA-    CA-     C-     N
   478    476    491    492      3    0.00000    0.00000    0.00000
0.00000    0.00000    0.00000 ;     CB-    CA-     C-     O
   478    476    491    493      3    2.51040    4.18400    1.67360
-6.69440    0.00000    0.00000 ;     CB-    CA-     C-     N
   478    484    487    488      3    0.66944    2.00832    0.00000
-2.67776    0.00000    0.00000 ;     CB-   CG1-   CD1-  HD11
   478    484    487    489      3    0.66944    2.00832    0.00000
-2.67776    0.00000    0.00000 ;     CB-   CG1-   CD1-  HD12
   478    484    487    490      3    0.66944    2.00832    0.00000
-2.67776    0.00000    0.00000 ;     CB-   CG1-   CD1-  HD13
   479    478    476    491      3    0.65084    1.95253    0.00000
-2.60338    0.00000    0.00000 ;     HB-    CB-    CA-     C
   479    478    480    481      3    0.62760    1.88280    0.00000
-2.51040    0.00000    0.00000 ;     HB-    CB-   CG2-  HG21
   479    478    480    482      3    0.62760    1.88280    0.00000
-2.51040    0.00000    0.00000 ;     HB-    CB-   CG2-  HG22
   479    478    480    483      3    0.62760    1.88280    0.00000
-2.51040    0.00000    0.00000 ;     HB-    CB-   CG2-  HG23
   479    478    484    485      3    0.62760    1.88280    0.00000
-2.51040    0.00000    0.00000 ;     HB-    CB-   CG1-  HG12
   479    478    484    486      3    0.62760    1.88280    0.00000
-2.51040    0.00000    0.00000 ;     HB-    CB-   CG1-  HG13
   479    478    484    487      3    0.66944    2.00832    0.00000
-2.67776    0.00000    0.00000 ;     HB-    CB-   CG1-   CD1
   480    478    476    491      3    5.67350    3.55640   -6.16722
-2.47693    3.74886    0.00000 ;    CG2-    CB-    CA-     C
   480    478    484    485      3    0.66944    2.00832    0.00000
-2.67776    0.00000    0.00000 ;    CG2-    CB-   CG1-  HG12
   480    478    484    486      3    0.66944    2.00832    0.00000
-2.67776    0.00000    0.00000 ;    CG2-    CB-   CG1-  HG13
   480    478    484    487      3    3.81162    0.49790   -8.14206
```

-1.79075 7.49773 0.00000 ; CG2- CB- CG1- CD1
481 480 478 484 3 0.66944 2.00832 0.00000
-2.67776 0.00000 0.00000 ; HG21- CG2- CB- CG1
482 480 478 484 3 0.66944 2.00832 0.00000
-2.67776 0.00000 0.00000 ; HG22- CG2- CB- CG1
483 480 478 484 3 0.66944 2.00832 0.00000
-2.67776 0.00000 0.00000 ; HG23- CG2- CB- CG1
484 478 476 491 3 8.26340 0.74057 -9.99976
-1.89117 3.84928 0.00000 ; CG1- CB- CA- C
485 484 487 488 3 0.62760 1.88280 0.00000
-2.51040 0.00000 0.00000 ; HG12- CG1- CD1- HD11
485 484 487 489 3 0.62760 1.88280 0.00000
-2.51040 0.00000 0.00000 ; HG12- CG1- CD1- HD12
485 484 487 490 3 0.62760 1.88280 0.00000
-2.51040 0.00000 0.00000 ; HG12- CG1- CD1- HD13
486 484 487 488 3 0.62760 1.88280 0.00000
-2.51040 0.00000 0.00000 ; HG13- CG1- CD1- HD11
486 484 487 489 3 0.62760 1.88280 0.00000
-2.51040 0.00000 0.00000 ; HG13- CG1- CD1- HD12
486 484 487 490 3 0.62760 1.88280 0.00000
-2.51040 0.00000 0.00000 ; HG13- CG1- CD1- HD13
491 493 495 496 3 0.00000 0.00000 0.00000
0.00000 0.00000 0.00000 ; C- N- CA- HA
491 493 495 497 3 11.71520 1.67360 15.06240 -
13.38880 0.00000 0.00000 ; C- N- CA- CB
491 493 495 503 3 1.75728 5.27184 2.25936
-7.02912 0.00000 0.00000 ; C- N- CA- C
492 491 493 494 3 29.28800 -8.36800 -20.92000
0.00000 0.00000 0.00000 ; O- C- N- H
492 491 493 495 3 20.92000 0.00000 -20.92000
0.00000 0.00000 0.00000 ; O- C- N- CA
493 495 497 498 3 0.65084 1.95253 0.00000
-2.60338 0.00000 0.00000 ; N- CA- CB- HB2
493 495 497 499 3 0.65084 1.95253 0.00000
-2.60338 0.00000 0.00000 ; N- CA- CB- HB3
493 495 497 500 3 15.41386 9.74035 -8.39310
-0.97069 2.97901 0.00000 ; N- CA- CB- CG
493 495 503 504 3 0.00000 0.00000 0.00000
0.00000 0.00000 0.00000 ; N- CA- C- O
493 495 503 505 3 17.40544 -5.02080 -13.22144
9.20480 0.00000 0.00000 ; N- CA- C- N
494 493 495 496 3 0.00000 0.00000 0.00000
0.00000 0.00000 0.00000 ; H- N- CA- HA
494 493 495 497 3 0.00000 0.00000 0.00000
0.00000 0.00000 0.00000 ; H- N- CA- CB
494 493 495 503 3 0.00000 0.00000 0.00000
0.00000 0.00000 0.00000 ; H- N- CA- C
495 497 500 501 3 6.43499 0.00000 -5.39736
0.00000 -1.03763 0.00000 ; CA- CB- CG- OD1
495 497 500 502 3 6.43499 0.00000 -5.39736
0.00000 -1.03763 0.00000 ; CA- CB- CG- OD2
495 503 505 506 3 20.92000 0.00000 -20.92000

0.00000 0.00000 0.00000 ; CA- C- N- H
495 503 505 507 3 20.92000 0.00000 -20.92000
0.00000 0.00000 0.00000 ; CA- C- N- CA
496 495 497 498 3 0.65084 1.95253 0.00000
-2.60338 0.00000 0.00000 ; HA- CA- CB- HB2
496 495 497 499 3 0.65084 1.95253 0.00000
-2.60338 0.00000 0.00000 ; HA- CA- CB- HB3
496 495 497 500 3 0.65084 1.95253 0.00000
-2.60338 0.00000 0.00000 ; HA- CA- CB- CG
496 495 503 504 3 3.68192 -4.35136 0.00000
1.33888 0.00000 0.00000 ; HA- CA- C- O
496 495 503 505 3 0.00000 0.00000 0.00000
0.00000 0.00000 0.00000 ; HA- CA- C- N
497 495 503 504 3 0.00000 0.00000 0.00000
0.00000 0.00000 0.00000 ; CB- CA- C- O
497 495 503 505 3 2.51040 4.18400 1.67360
-6.69440 0.00000 0.00000 ; CB- CA- C- N
498 497 495 503 3 0.65084 1.95253 0.00000
-2.60338 0.00000 0.00000 ; HB2- CB- CA- C
498 497 500 501 3 0.00000 0.00000 0.00000
0.00000 0.00000 0.00000 ; HB2- CB- CG- OD1
498 497 500 502 3 0.00000 0.00000 0.00000
0.00000 0.00000 0.00000 ; HB2- CB- CG- OD2
499 497 495 503 3 0.65084 1.95253 0.00000
-2.60338 0.00000 0.00000 ; HB3- CB- CA- C
499 497 500 501 3 0.00000 0.00000 0.00000
0.00000 0.00000 0.00000 ; HB3- CB- CG- OD1
499 497 500 502 3 0.00000 0.00000 0.00000
0.00000 0.00000 0.00000 ; HB3- CB- CG- OD2
500 497 495 503 3 7.14627 -1.04600 -8.99560
-0.97069 5.15469 0.00000 ; CG- CB- CA- C
503 505 507 508 3 0.00000 0.00000 0.00000
0.00000 0.00000 0.00000 ; C- N- CA- HA
503 505 507 509 3 11.71520 1.67360 15.06240 -
13.38880 0.00000 0.00000 ; C- N- CA- CB
503 505 507 525 3 1.75728 5.27184 2.25936
-7.02912 0.00000 0.00000 ; C- N- CA- C
504 503 505 506 3 29.28800 -8.36800 -20.92000
0.00000 0.00000 0.00000 ; O- C- N- H
504 503 505 507 3 20.92000 0.00000 -20.92000
0.00000 0.00000 0.00000 ; O- C- N- CA
505 507 509 510 3 0.65084 1.95253 0.00000
-2.60338 0.00000 0.00000 ; N- CA- CB- HB2
505 507 509 511 3 0.65084 1.95253 0.00000
-2.60338 0.00000 0.00000 ; N- CA- CB- HB3
505 507 509 512 3 0.65084 1.95253 0.00000
-2.60338 0.00000 0.00000 ; N- CA- CB- CG
505 507 525 526 3 0.00000 0.00000 0.00000
0.00000 0.00000 0.00000 ; N- CA- C- O
505 507 525 527 3 17.40544 -5.02080 -13.22144
9.20480 0.00000 0.00000 ; N- CA- C- N
506 505 507 508 3 0.00000 0.00000 0.00000

```
0.00000    0.00000    0.00000 ;       H-      N-     CA-    HA
   506     505     507     509       3    0.00000    0.00000    0.00000
0.00000    0.00000    0.00000 ;       H-      N-     CA-    CB
   506     505     507     525       3    0.00000    0.00000    0.00000
0.00000    0.00000    0.00000 ;       H-      N-     CA-     C
   507     509     512     513       3    0.66944    2.00832    0.00000
-2.67776    0.00000    0.00000 ;      CA-    CB-     CG-   HG2
   507     509     512     514       3    0.66944    2.00832    0.00000
-2.67776    0.00000    0.00000 ;      CA-    CB-     CG-   HG3
   507     509     512     515       3    3.68192    3.09616   -2.09200
-3.01248    0.00000    0.00000 ;      CA-    CB-     CG-    CD
   507     525     527     528       3   20.92000    0.00000  -20.92000
0.00000    0.00000    0.00000 ;      CA-      C-      N-     H
   507     525     527     529       3   20.92000    0.00000  -20.92000
0.00000    0.00000    0.00000 ;      CA-      C-      N-    CA
   508     507     509     510       3    0.65084    1.95253    0.00000
-2.60338    0.00000    0.00000 ;      HA-    CA-     CB-   HB2
   508     507     509     511       3    0.65084    1.95253    0.00000
-2.60338    0.00000    0.00000 ;      HA-    CA-     CB-   HB3
   508     507     509     512       3    0.65084    1.95253    0.00000
-2.60338    0.00000    0.00000 ;      HA-    CA-     CB-    CG
   508     507     525     526       3    3.68192   -4.35136    0.00000
1.33888    0.00000    0.00000 ;      HA-    CA-      C-     O
   508     507     525     527       3    0.00000    0.00000    0.00000
0.00000    0.00000    0.00000 ;      HA-    CA-      C-     N
   509     507     525     526       3    0.00000    0.00000    0.00000
0.00000    0.00000    0.00000 ;      CB-    CA-      C-     O
   509     507     525     527       3    2.51040    4.18400    1.67360
-6.69440    0.00000    0.00000 ;      CB-    CA-      C-     N
   509     512     515     516       3    0.66944    2.00832    0.00000
-2.67776    0.00000    0.00000 ;      CB-    CG-     CD-   HD2
   509     512     515     517       3    0.66944    2.00832    0.00000
-2.67776    0.00000    0.00000 ;      CB-    CG-     CD-   HD3
   509     512     515     518       3    3.68192    3.09616   -2.09200
-3.01248    0.00000    0.00000 ;      CB-    CG-     CD-    CE
   510     509     507     525       3    0.65084    1.95253    0.00000
-2.60338    0.00000    0.00000 ;     HB2-    CB-     CA-     C
   510     509     512     513       3    0.62760    1.88280    0.00000
-2.51040    0.00000    0.00000 ;     HB2-    CB-     CG-   HG2
   510     509     512     514       3    0.62760    1.88280    0.00000
-2.51040    0.00000    0.00000 ;     HB2-    CB-     CG-   HG3
   510     509     512     515       3    0.66944    2.00832    0.00000
-2.67776    0.00000    0.00000 ;     HB2-    CB-     CG-    CD
   511     509     507     525       3    0.65084    1.95253    0.00000
-2.60338    0.00000    0.00000 ;     HB3-    CB-     CA-     C
   511     509     512     513       3    0.62760    1.88280    0.00000
-2.51040    0.00000    0.00000 ;     HB3-    CB-     CG-   HG2
   511     509     512     514       3    0.62760    1.88280    0.00000
-2.51040    0.00000    0.00000 ;     HB3-    CB-     CG-   HG3
   511     509     512     515       3    0.66944    2.00832    0.00000
-2.67776    0.00000    0.00000 ;     HB3-    CB-     CG-    CD
   512     509     507     525       3    0.65084    1.95253    0.00000
```

```
-2.60338    0.00000    0.00000 ;      CG-    CB-    CA-     C
   512    515    518    519      3    0.65084    1.95253    0.00000
-2.60338    0.00000    0.00000 ;      CG-    CD-    CE-   HE2
   512    515    518    520      3    0.65084    1.95253    0.00000
-2.60338    0.00000    0.00000 ;      CG-    CD-    CE-   HE3
   512    515    518    521      3    0.65084    1.95253    0.00000
-2.60338    0.00000    0.00000 ;      CG-    CD-    CE-    NZ
   513    512    515    516      3    0.62760    1.88280    0.00000
-2.51040    0.00000    0.00000 ;     HG2-    CG-    CD-   HD2
   513    512    515    517      3    0.62760    1.88280    0.00000
-2.51040    0.00000    0.00000 ;     HG2-    CG-    CD-   HD3
   513    512    515    518      3    0.66944    2.00832    0.00000
-2.67776    0.00000    0.00000 ;     HG2-    CG-    CD-    CE
   514    512    515    516      3    0.62760    1.88280    0.00000
-2.51040    0.00000    0.00000 ;     HG3-    CG-    CD-   HD2
   514    512    515    517      3    0.62760    1.88280    0.00000
-2.51040    0.00000    0.00000 ;     HG3-    CG-    CD-   HD3
   514    512    515    518      3    0.66944    2.00832    0.00000
-2.67776    0.00000    0.00000 ;     HG3-    CG-    CD-    CE
   515    518    521    522      3    0.65084    1.95253    0.00000
-2.60338    0.00000    0.00000 ;      CD-    CE-    NZ-   HZ1
   515    518    521    523      3    0.65084    1.95253    0.00000
-2.60338    0.00000    0.00000 ;      CD-    CE-    NZ-   HZ2
   515    518    521    524      3    0.65084    1.95253    0.00000
-2.60338    0.00000    0.00000 ;      CD-    CE-    NZ-   HZ3
   516    515    518    519      3    0.65084    1.95253    0.00000
-2.60338    0.00000    0.00000 ;     HD2-    CD-    CE-   HE2
   516    515    518    520      3    0.65084    1.95253    0.00000
-2.60338    0.00000    0.00000 ;     HD2-    CD-    CE-   HE3
   516    515    518    521      3    0.65084    1.95253    0.00000
-2.60338    0.00000    0.00000 ;     HD2-    CD-    CE-    NZ
   517    515    518    519      3    0.65084    1.95253    0.00000
-2.60338    0.00000    0.00000 ;     HD3-    CD-    CE-   HE2
   517    515    518    520      3    0.65084    1.95253    0.00000
-2.60338    0.00000    0.00000 ;     HD3-    CD-    CE-   HE3
   517    515    518    521      3    0.65084    1.95253    0.00000
-2.60338    0.00000    0.00000 ;     HD3-    CD-    CE-    NZ
   519    518    521    522      3    0.65084    1.95253    0.00000
-2.60338    0.00000    0.00000 ;     HE2-    CE-    NZ-   HZ1
   519    518    521    523      3    0.65084    1.95253    0.00000
-2.60338    0.00000    0.00000 ;     HE2-    CE-    NZ-   HZ2
   519    518    521    524      3    0.65084    1.95253    0.00000
-2.60338    0.00000    0.00000 ;     HE2-    CE-    NZ-   HZ3
   520    518    521    522      3    0.65084    1.95253    0.00000
-2.60338    0.00000    0.00000 ;     HE3-    CE-    NZ-   HZ1
   520    518    521    523      3    0.65084    1.95253    0.00000
-2.60338    0.00000    0.00000 ;     HE3-    CE-    NZ-   HZ2
   520    518    521    524      3    0.65084    1.95253    0.00000
-2.60338    0.00000    0.00000 ;     HE3-    CE-    NZ-   HZ3
   525    527    529    530      3    0.00000    0.00000    0.00000
0.00000    0.00000    0.00000 ;      C-     N-    CA-    HA
   525    527    529    531      3   11.71520    1.67360   15.06240  -
```

```
13.38880    0.00000    0.00000 ;       C-     N-    CA-    CB
   525    527    529    541      3    1.75728    5.27184    2.25936
-7.02912    0.00000    0.00000 ;       C-     N-    CA-     C
   526    525    527    528      3   29.28800   -8.36800  -20.92000
0.00000    0.00000    0.00000 ;       O-     C-     N-     H
   526    525    527    529      3   20.92000    0.00000  -20.92000
0.00000    0.00000    0.00000 ;       O-     C-     N-    CA
   527    529    531    532      3    0.65084    1.95253    0.00000
-2.60338    0.00000    0.00000 ;       N-    CA-    CB-    HB
   527    529    531    533      3    3.83673    0.44769   -1.77402
-2.47693   -0.03347    0.00000 ;       N-    CA-    CB-   CG1
   527    529    531    537      3    3.83673    0.44769   -1.77402
-2.47693   -0.03347    0.00000 ;       N-    CA-    CB-   CG2
   527    529    541    542      3    0.00000    0.00000    0.00000
0.00000    0.00000    0.00000 ;      N-    CA-     C-     O
   527    529    541    543      3   17.40544   -5.02080  -13.22144
9.20480    0.00000    0.00000 ;      N-    CA-     C-     N
   528    527    529    530      3    0.00000    0.00000    0.00000
0.00000    0.00000    0.00000 ;      H-     N-    CA-    HA
   528    527    529    531      3    0.00000    0.00000    0.00000
0.00000    0.00000    0.00000 ;      H-     N-    CA-    CB
   528    527    529    541      3    0.00000    0.00000    0.00000
0.00000    0.00000    0.00000 ;      H-     N-    CA-     C
   529    531    533    534      3    0.66944    2.00832    0.00000
-2.67776    0.00000    0.00000 ;      CA-    CB-   CG1-  HG11
   529    531    533    535      3    0.66944    2.00832    0.00000
-2.67776    0.00000    0.00000 ;      CA-    CB-   CG1-  HG12
   529    531    533    536      3    0.66944    2.00832    0.00000
-2.67776    0.00000    0.00000 ;      CA-    CB-   CG1-  HG13
   529    531    537    538      3    0.66944    2.00832    0.00000
-2.67776    0.00000    0.00000 ;      CA-    CB-   CG2-  HG21
   529    531    537    539      3    0.66944    2.00832    0.00000
-2.67776    0.00000    0.00000 ;      CA-    CB-   CG2-  HG22
   529    531    537    540      3    0.66944    2.00832    0.00000
-2.67776    0.00000    0.00000 ;      CA-    CB-   CG2-  HG23
   529    541    543    544      3   20.92000    0.00000  -20.92000
0.00000    0.00000    0.00000 ;     CA-     C-     N-     H
   529    541    543    545      3   20.92000    0.00000  -20.92000
0.00000    0.00000    0.00000 ;     CA-     C-     N-    CA
   530    529    531    532      3    0.65084    1.95253    0.00000
-2.60338    0.00000    0.00000 ;      HA-    CA-    CB-    HB
   530    529    531    533      3    0.65084    1.95253    0.00000
-2.60338    0.00000    0.00000 ;      HA-    CA-    CB-   CG1
   530    529    531    537      3    0.65084    1.95253    0.00000
-2.60338    0.00000    0.00000 ;      HA-    CA-    CB-   CG2
   530    529    541    542      3    3.68192   -4.35136    0.00000
1.33888    0.00000    0.00000 ;     HA-    CA-     C-     O
   530    529    541    543      3    0.00000    0.00000    0.00000
0.00000    0.00000    0.00000 ;     HA-    CA-     C-     N
   531    529    541    542      3    0.00000    0.00000    0.00000
0.00000    0.00000    0.00000 ;     CB-    CA-     C-     O
   531    529    541    543      3    2.51040    4.18400    1.67360
```

-6.69440 0.00000 0.00000 ; CB- CA- C- N
532 531 529 541 3 0.65084 1.95253 0.00000
-2.60338 0.00000 0.00000 ; HB- CB- CA- C
532 531 533 534 3 0.62760 1.88280 0.00000
-2.51040 0.00000 0.00000 ; HB- CB- CG1- HG11
532 531 533 535 3 0.62760 1.88280 0.00000
-2.51040 0.00000 0.00000 ; HB- CB- CG1- HG12
532 531 533 536 3 0.62760 1.88280 0.00000
-2.51040 0.00000 0.00000 ; HB- CB- CG1- HG13
532 531 537 538 3 0.62760 1.88280 0.00000
-2.51040 0.00000 0.00000 ; HB- CB- CG2- HG21
532 531 537 539 3 0.62760 1.88280 0.00000
-2.51040 0.00000 0.00000 ; HB- CB- CG2- HG22
532 531 537 540 3 0.62760 1.88280 0.00000
-2.51040 0.00000 0.00000 ; HB- CB- CG2- HG23
533 531 529 541 3 5.67350 3.55640 -6.16722
-2.47693 3.74886 0.00000 ; CG1- CB- CA- C
533 531 537 538 3 0.66944 2.00832 0.00000
-2.67776 0.00000 0.00000 ; CG1- CB- CG2- HG21
533 531 537 539 3 0.66944 2.00832 0.00000
-2.67776 0.00000 0.00000 ; CG1- CB- CG2- HG22
533 531 537 540 3 0.66944 2.00832 0.00000
-2.67776 0.00000 0.00000 ; CG1- CB- CG2- HG23
534 533 531 537 3 0.66944 2.00832 0.00000
-2.67776 0.00000 0.00000 ; HG11- CG1- CB- CG2
535 533 531 537 3 0.66944 2.00832 0.00000
-2.67776 0.00000 0.00000 ; HG12- CG1- CB- CG2
536 533 531 537 3 0.66944 2.00832 0.00000
-2.67776 0.00000 0.00000 ; HG13- CG1- CB- CG2
537 531 529 541 3 5.67350 3.55640 -6.16722
-2.47693 3.74886 0.00000 ; CG2- CB- CA- C
541 543 545 546 3 0.00000 0.00000 0.00000
0.00000 0.00000 0.00000 ; C- N- CA- HA
541 543 545 547 3 11.71520 1.67360 15.06240 -
13.38880 0.00000 0.00000 ; C- N- CA- CB
541 543 545 555 3 1.75728 5.27184 2.25936
-7.02912 0.00000 0.00000 ; C- N- CA- C
542 541 543 544 3 29.28800 -8.36800 -20.92000
0.00000 0.00000 0.00000 ; O- C- N- H
542 541 543 545 3 20.92000 0.00000 -20.92000
0.00000 0.00000 0.00000 ; O- C- N- CA
543 545 547 548 3 0.65084 1.95253 0.00000
-2.60338 0.00000 0.00000 ; N- CA- CB- HB
543 545 547 549 3 3.83673 0.44769 -1.77402
-2.47693 -0.03347 0.00000 ; N- CA- CB- CG2
543 545 547 553 3 4.93294 1.13386 -3.12963
-5.27184 3.17984 0.00000 ; N- CA- CB- OG1
543 545 555 556 3 0.00000 0.00000 0.00000
0.00000 0.00000 0.00000 ; N- CA- C- O
543 545 555 557 3 17.40544 -5.02080 -13.22144
9.20480 0.00000 0.00000 ; N- CA- C- N
544 543 545 546 3 0.00000 0.00000 0.00000

```
0.00000    0.00000    0.00000 ;       H-      N-     CA-    HA
   544     543     545     547       3    0.00000    0.00000    0.00000
0.00000    0.00000    0.00000 ;       H-      N-     CA-    CB
   544     543     545     555       3    0.00000    0.00000    0.00000
0.00000    0.00000    0.00000 ;       H-      N-     CA-     C
   545     547     549     550       3    0.66944    2.00832    0.00000
-2.67776    0.00000    0.00000 ;      CA-    CB-   CG2-  HG21
   545     547     549     551       3    0.66944    2.00832    0.00000
-2.67776    0.00000    0.00000 ;      CA-    CB-   CG2-  HG22
   545     547     549     552       3    0.66944    2.00832    0.00000
-2.67776    0.00000    0.00000 ;      CA-    CB-   CG2-  HG23
   545     547     553     554       3    1.12131    2.98738    1.66523
-3.94970    0.43514    0.00000 ;      CA-    CB-   OG1-   HG1
   545     555     557     558       3   20.92000    0.00000  -20.92000
0.00000    0.00000    0.00000 ;      CA-     C-      N-     H
   545     555     557     559       3   20.92000    0.00000  -20.92000
0.00000    0.00000    0.00000 ;      CA-     C-      N-    CA
   546     545     547     548       3    0.65084    1.95253    0.00000
-2.60338    0.00000    0.00000 ;      HA-    CA-     CB-    HB
   546     545     547     549       3    0.65084    1.95253    0.00000
-2.60338    0.00000    0.00000 ;      HA-    CA-     CB-   CG2
   546     545     547     553       3    1.04600   -1.04600    0.00000
0.00000    0.00000    0.00000 ;      HA-    CA-     CB-   OG1
   546     545     555     556       3    3.68192   -4.35136    0.00000
1.33888    0.00000    0.00000 ;      HA-    CA-      C-     O
   546     545     555     557       3    0.00000    0.00000    0.00000
0.00000    0.00000    0.00000 ;      HA-    CA-      C-     N
   547     545     555     556       3    0.00000    0.00000    0.00000
0.00000    0.00000    0.00000 ;      CB-    CA-      C-     O
   547     545     555     557       3    2.51040    4.18400    1.67360
-6.69440    0.00000    0.00000 ;      CB-    CA-      C-     N
   548     547     545     555       3    0.65084    1.95253    0.00000
-2.60338    0.00000    0.00000 ;      HB-    CB-    CA-     C
   548     547     549     550       3    0.65084    1.95253    0.00000
-2.60338    0.00000    0.00000 ;      HB-    CB-   CG2-  HG21
   548     547     549     551       3    0.65084    1.95253    0.00000
-2.60338    0.00000    0.00000 ;      HB-    CB-   CG2-  HG22
   548     547     549     552       3    0.65084    1.95253    0.00000
-2.60338    0.00000    0.00000 ;      HB-    CB-   CG2-  HG23
   548     547     553     554       3    0.69733    2.09200    0.00000
-2.78933    0.00000    0.00000 ;      HB-    CB-   OG1-   HG1
   549     547     545     555       3    5.67350    3.55640   -6.16722
-2.47693    3.74886    0.00000 ;     CG2-    CB-    CA-     C
   549     547     553     554       3    4.74047    0.27196   -2.26773
-3.94970    1.60666    0.00000 ;     CG2-    CB-   OG1-   HG1
   550     549     547     553       3    1.04600   -1.04600    0.00000
0.00000    0.00000    0.00000 ;   HG21-   CG2-    CB-   OG1
   551     549     547     553       3    1.04600   -1.04600    0.00000
0.00000    0.00000    0.00000 ;   HG22-   CG2-    CB-   OG1
   552     549     547     553       3    1.04600   -1.04600    0.00000
0.00000    0.00000    0.00000 ;   HG23-   CG2-    CB-   OG1
   553     547     545     555       3    6.53541    6.87013   -6.21742
```

-5.27184 5.22163 0.00000 ; OG1- CB- CA- C
555 557 559 560 3 0.00000 0.00000 0.00000
0.00000 0.00000 0.00000 ; C- N- CA- HA
555 557 559 561 3 11.71520 1.67360 15.06240 -
13.38880 0.00000 0.00000 ; C- N- CA- CB
555 557 559 569 3 1.75728 5.27184 2.25936
-7.02912 0.00000 0.00000 ; C- N- CA- C
556 555 557 558 3 29.28800 -8.36800 -20.92000
0.00000 0.00000 0.00000 ; O- C- N- H
556 555 557 559 3 20.92000 0.00000 -20.92000
0.00000 0.00000 0.00000 ; O- C- N- CA
557 559 561 562 3 0.65084 1.95253 0.00000
-2.60338 0.00000 0.00000 ; N- CA- CB- HB2
557 559 561 563 3 0.65084 1.95253 0.00000
-2.60338 0.00000 0.00000 ; N- CA- CB- HB3
557 559 561 564 3 5.99567 3.29281 -4.46014
-0.55229 1.97485 0.00000 ; N- CA- CB- CG
557 559 569 570 3 0.00000 0.00000 0.00000
0.00000 0.00000 0.00000 ; N- CA- C- O
557 559 569 571 3 17.40544 -5.02080 -13.22144
9.20480 0.00000 0.00000 ; N- CA- C- N
558 557 559 560 3 0.00000 0.00000 0.00000
0.00000 0.00000 0.00000 ; H- N- CA- HA
558 557 559 561 3 0.00000 0.00000 0.00000
0.00000 0.00000 0.00000 ; H- N- CA- CB
558 557 559 569 3 0.00000 0.00000 0.00000
0.00000 0.00000 0.00000 ; H- N- CA- C
559 561 564 565 3 0.00000 0.00000 0.00000
0.00000 0.00000 0.00000 ; CA- CB- CG- OD1
559 561 564 566 3 8.84916 -0.31380 -4.32626
5.03754 0.26778 0.00000 ; CA- CB- CG- ND2
559 569 571 572 3 20.92000 0.00000 -20.92000
0.00000 0.00000 0.00000 ; CA- C- N- H
559 569 571 573 3 20.92000 0.00000 -20.92000
0.00000 0.00000 0.00000 ; CA- C- N- CA
560 559 561 562 3 0.65084 1.95253 0.00000
-2.60338 0.00000 0.00000 ; HA- CA- CB- HB2
560 559 561 563 3 0.65084 1.95253 0.00000
-2.60338 0.00000 0.00000 ; HA- CA- CB- HB3
560 559 561 564 3 0.65084 1.95253 0.00000
-2.60338 0.00000 0.00000 ; HA- CA- CB- CG
560 559 569 570 3 3.68192 -4.35136 0.00000
1.33888 0.00000 0.00000 ; HA- CA- C- O
560 559 569 571 3 0.00000 0.00000 0.00000
0.00000 0.00000 0.00000 ; HA- CA- C- N
561 559 569 570 3 0.00000 0.00000 0.00000
0.00000 0.00000 0.00000 ; CB- CA- C- O
561 559 569 571 3 2.51040 4.18400 1.67360
-6.69440 0.00000 0.00000 ; CB- CA- C- N
561 564 566 567 3 20.92000 0.00000 -20.92000
0.00000 0.00000 0.00000 ; CB- CG- ND2- HD21
561 564 566 568 3 20.92000 0.00000 -20.92000

0.00000 0.00000 0.00000 ; CB- CG- ND2- HD22
562 561 559 569 3 0.65084 1.95253 0.00000
-2.60338 0.00000 0.00000 ; HB2- CB- CA- C
562 561 564 565 3 3.68192 -4.35136 0.00000
1.33888 0.00000 0.00000 ; HB2- CB- CG- OD1
562 561 564 566 3 0.00000 0.00000 0.00000
0.00000 0.00000 0.00000 ; HB2- CB- CG- ND2
563 561 559 569 3 0.65084 1.95253 0.00000
-2.60338 0.00000 0.00000 ; HB3- CB- CA- C
563 561 564 565 3 3.68192 -4.35136 0.00000
1.33888 0.00000 0.00000 ; HB3- CB- CG- OD1
563 561 564 566 3 0.00000 0.00000 0.00000
0.00000 0.00000 0.00000 ; HB3- CB- CG- ND2
564 561 559 569 3 7.94542 -3.96225 -6.11701
-0.55229 3.58150 0.00000 ; CG- CB- CA- C
565 564 566 567 3 29.28800 -8.36800 -20.92000
0.00000 0.00000 0.00000 ; OD1- CG- ND2- HD21
565 564 566 568 3 29.28800 -8.36800 -20.92000
0.00000 0.00000 0.00000 ; OD1- CG- ND2- HD22
569 571 573 574 3 0.00000 0.00000 0.00000
0.00000 0.00000 0.00000 ; C- N- CA- HA
569 571 573 575 3 11.71520 1.67360 15.06240 -
13.38880 0.00000 0.00000 ; C- N- CA- CB
569 571 573 586 3 1.75728 5.27184 2.25936
-7.02912 0.00000 0.00000 ; C- N- CA- C
570 569 571 572 3 29.28800 -8.36800 -20.92000
0.00000 0.00000 0.00000 ; O- C- N- H
570 569 571 573 3 20.92000 0.00000 -20.92000
0.00000 0.00000 0.00000 ; O- C- N- CA
571 573 575 576 3 0.65084 1.95253 0.00000
-2.60338 0.00000 0.00000 ; N- CA- CB- HB2
571 573 575 577 3 0.65084 1.95253 0.00000
-2.60338 0.00000 0.00000 ; N- CA- CB- HB3
571 573 575 578 3 3.21331 2.22589 -4.15053
-2.40998 2.61082 0.00000 ; N- CA- CB- CG
571 573 586 587 3 0.00000 0.00000 0.00000
0.00000 0.00000 0.00000 ; N- CA- C- O
571 573 586 588 3 17.40544 -5.02080 -13.22144
9.20480 0.00000 0.00000 ; N- CA- C- N
572 571 573 574 3 0.00000 0.00000 0.00000
0.00000 0.00000 0.00000 ; H- N- CA- HA
572 571 573 575 3 0.00000 0.00000 0.00000
0.00000 0.00000 0.00000 ; H- N- CA- CB
572 571 573 586 3 0.00000 0.00000 0.00000
0.00000 0.00000 0.00000 ; H- N- CA- C
573 575 578 579 3 0.66944 2.00832 0.00000
-2.67776 0.00000 0.00000 ; CA- CB- CG- HG2
573 575 578 580 3 0.66944 2.00832 0.00000
-2.67776 0.00000 0.00000 ; CA- CB- CG- HG3
573 575 578 581 3 3.69866 -4.35136 -3.92459
6.89523 4.61914 0.00000 ; CA- CB- CG- CD
573 586 588 589 3 20.92000 0.00000 -20.92000

```
0.00000    0.00000    0.00000 ;     CA-     C-     N-     H
   573    586    588    590     3   20.92000    0.00000  -20.92000
0.00000    0.00000    0.00000 ;     CA-     C-     N-    CA
   574    573    575    576     3    0.65084    1.95253    0.00000
-2.60338    0.00000    0.00000 ;     HA-    CA-    CB-   HB2
   574    573    575    577     3    0.65084    1.95253    0.00000
-2.60338    0.00000    0.00000 ;     HA-    CA-    CB-   HB3
   574    573    575    578     3    0.65084    1.95253    0.00000
-2.60338    0.00000    0.00000 ;     HA-    CA-    CB-    CG
   574    573    586    587     3    3.68192   -4.35136    0.00000
1.33888    0.00000    0.00000 ;     HA-    CA-     C-     O
   574    573    586    588     3    0.00000    0.00000    0.00000
0.00000    0.00000    0.00000 ;     HA-    CA-     C-     N
   575    573    586    587     3    0.00000    0.00000    0.00000
0.00000    0.00000    0.00000 ;     CB-    CA-     C-     O
   575    573    586    588     3    2.51040    4.18400    1.67360
-6.69440    0.00000    0.00000 ;     CB-    CA-     C-     N
   575    578    581    582     3    0.00000    0.00000    0.00000
0.00000    0.00000    0.00000 ;     CB-    CG-    CD-   OE1
   575    578    581    583     3   10.32611    1.48114   -8.47678
1.42256    1.40582    0.00000 ;     CB-    CG-    CD-   NE2
   576    575    573    586     3    0.65084    1.95253    0.00000
-2.60338    0.00000    0.00000 ;    HB2-    CB-    CA-     C
   576    575    578    579     3    0.62760    1.88280    0.00000
-2.51040    0.00000    0.00000 ;    HB2-    CB-    CG-   HG2
   576    575    578    580     3    0.62760    1.88280    0.00000
-2.51040    0.00000    0.00000 ;    HB2-    CB-    CG-   HG3
   576    575    578    581     3    0.65084    1.95253    0.00000
-2.60338    0.00000    0.00000 ;    HB2-    CB-    CG-    CD
   577    575    573    586     3    0.65084    1.95253    0.00000
-2.60338    0.00000    0.00000 ;    HB3-    CB-    CA-     C
   577    575    578    579     3    0.62760    1.88280    0.00000
-2.51040    0.00000    0.00000 ;    HB3-    CB-    CG-   HG2
   577    575    578    580     3    0.62760    1.88280    0.00000
-2.51040    0.00000    0.00000 ;    HB3-    CB-    CG-   HG3
   577    575    578    581     3    0.65084    1.95253    0.00000
-2.60338    0.00000    0.00000 ;    HB3-    CB-    CG-    CD
   578    575    573    586     3    6.86594    3.56895   -8.14206
-2.40998    4.85344    0.00000 ;     CG-    CB-    CA-     C
   578    581    583    584     3   20.92000    0.00000  -20.92000
0.00000    0.00000    0.00000 ;     CG-    CD-   NE2-  HE21
   578    581    583    585     3   20.92000    0.00000  -20.92000
0.00000    0.00000    0.00000 ;     CG-    CD-   NE2-  HE22
   579    578    581    582     3    3.68192   -4.35136    0.00000
1.33888    0.00000    0.00000 ;    HG2-    CG-    CD-   OE1
   579    578    581    583     3    0.00000    0.00000    0.00000
0.00000    0.00000    0.00000 ;    HG2-    CG-    CD-   NE2
   580    578    581    582     3    3.68192   -4.35136    0.00000
1.33888    0.00000    0.00000 ;    HG3-    CG-    CD-   OE1
   580    578    581    583     3    0.00000    0.00000    0.00000
0.00000    0.00000    0.00000 ;    HG3-    CG-    CD-   NE2
   582    581    583    584     3   29.28800   -8.36800  -20.92000
```

0.00000 0.00000 0.00000 ; OE1- CD- NE2- HE21
582 581 583 585 3 29.28800 -8.36800 -20.92000
0.00000 0.00000 0.00000 ; OE1- CD- NE2- HE22
586 588 590 591 3 0.00000 0.00000 0.00000
0.00000 0.00000 0.00000 ; C- N- CA- HA
586 588 590 592 3 11.71520 1.67360 15.06240 -
13.38880 0.00000 0.00000 ; C- N- CA- CB
586 588 590 605 3 1.75728 5.27184 2.25936
-7.02912 0.00000 0.00000 ; C- N- CA- C
587 586 588 589 3 29.28800 -8.36800 -20.92000
0.00000 0.00000 0.00000 ; O- C- N- H
587 586 588 590 3 20.92000 0.00000 -20.92000
0.00000 0.00000 0.00000 ; O- C- N- CA
588 590 592 593 3 0.65084 1.95253 0.00000
-2.60338 0.00000 0.00000 ; N- CA- CB- HB2
588 590 592 594 3 0.65084 1.95253 0.00000
-2.60338 0.00000 0.00000 ; N- CA- CB- HB3
588 590 592 595 3 3.79070 1.39746 -4.61077
-2.40998 2.44346 0.00000 ; N- CA- CB- CG
588 590 605 606 3 0.00000 0.00000 0.00000
0.00000 0.00000 0.00000 ; N- CA- C- O
588 590 605 607 3 17.40544 -5.02080 -13.22144
9.20480 0.00000 0.00000 ; N- CA- C- N
589 588 590 591 3 0.00000 0.00000 0.00000
0.00000 0.00000 0.00000 ; H- N- CA- HA
589 588 590 592 3 0.00000 0.00000 0.00000
0.00000 0.00000 0.00000 ; H- N- CA- CB
589 588 590 605 3 0.00000 0.00000 0.00000
0.00000 0.00000 0.00000 ; H- N- CA- C
590 592 595 596 3 0.66944 2.00832 0.00000
-2.67776 0.00000 0.00000 ; CA- CB- CG- HG
590 592 595 597 3 3.90367 0.19665 -6.21742
-2.37651 5.99149 0.00000 ; CA- CB- CG- CD1
590 592 595 601 3 3.90367 0.19665 -6.21742
-2.37651 5.99149 0.00000 ; CA- CB- CG- CD2
590 605 607 608 3 20.92000 0.00000 -20.92000
0.00000 0.00000 0.00000 ; CA- C- N- H
590 605 607 609 3 20.92000 0.00000 -20.92000
0.00000 0.00000 0.00000 ; CA- C- N- CA
591 590 592 593 3 0.65084 1.95253 0.00000
-2.60338 0.00000 0.00000 ; HA- CA- CB- HB2
591 590 592 594 3 0.65084 1.95253 0.00000
-2.60338 0.00000 0.00000 ; HA- CA- CB- HB3
591 590 592 595 3 0.65084 1.95253 0.00000
-2.60338 0.00000 0.00000 ; HA- CA- CB- CG
591 590 605 606 3 3.68192 -4.35136 0.00000
1.33888 0.00000 0.00000 ; HA- CA- C- O
591 590 605 607 3 0.00000 0.00000 0.00000
0.00000 0.00000 0.00000 ; HA- CA- C- N
592 590 605 606 3 0.00000 0.00000 0.00000
0.00000 0.00000 0.00000 ; CB- CA- C- O
592 590 605 607 3 2.51040 4.18400 1.67360

```
 -6.69440    0.00000    0.00000 ;      CB-    CA-      C-     N
    592     595     597     598      3    0.66944    2.00832    0.00000
 -2.67776    0.00000    0.00000 ;      CB-    CG-   CD1-  HD11
    592     595     597     599      3    0.66944    2.00832    0.00000
 -2.67776    0.00000    0.00000 ;      CB-    CG-   CD1-  HD12
    592     595     597     600      3    0.66944    2.00832    0.00000
 -2.67776    0.00000    0.00000 ;      CB-    CG-   CD1-  HD13
    592     595     601     602      3    0.66944    2.00832    0.00000
 -2.67776    0.00000    0.00000 ;      CB-    CG-   CD2-  HD21
    592     595     601     603      3    0.66944    2.00832    0.00000
 -2.67776    0.00000    0.00000 ;      CB-    CG-   CD2-  HD22
    592     595     601     604      3    0.66944    2.00832    0.00000
 -2.67776    0.00000    0.00000 ;      CB-    CG-   CD2-  HD23
    593     592     590     605      3    0.65084    1.95253    0.00000
 -2.60338    0.00000    0.00000 ;     HB2-    CB-     CA-      C
    593     592     595     596      3    0.62760    1.88280    0.00000
 -2.51040    0.00000    0.00000 ;     HB2-    CB-     CG-     HG
    593     592     595     597      3    0.66944    2.00832    0.00000
 -2.67776    0.00000    0.00000 ;     HB2-    CB-     CG-    CD1
    593     592     595     601      3    0.66944    2.00832    0.00000
 -2.67776    0.00000    0.00000 ;     HB2-    CB-     CG-    CD2
    594     592     590     605      3    0.65084    1.95253    0.00000
 -2.60338    0.00000    0.00000 ;     HB3-    CB-     CA-      C
    594     592     595     596      3    0.62760    1.88280    0.00000
 -2.51040    0.00000    0.00000 ;     HB3-    CB-     CG-     HG
    594     592     595     597      3    0.66944    2.00832    0.00000
 -2.67776    0.00000    0.00000 ;     HB3-    CB-     CG-    CD1
    594     592     595     601      3    0.66944    2.00832    0.00000
 -2.67776    0.00000    0.00000 ;     HB3-    CB-     CG-    CD2
    595     592     590     605      3   10.33448   -1.14642  -11.54784
 -2.40998    6.35968    0.00000 ;      CG-    CB-     CA-      C
    596     595     597     598      3    0.62760    1.88280    0.00000
 -2.51040    0.00000    0.00000 ;      HG-    CG-   CD1-  HD11
    596     595     597     599      3    0.62760    1.88280    0.00000
 -2.51040    0.00000    0.00000 ;      HG-    CG-   CD1-  HD12
    596     595     597     600      3    0.62760    1.88280    0.00000
 -2.51040    0.00000    0.00000 ;      HG-    CG-   CD1-  HD13
    596     595     601     602      3    0.62760    1.88280    0.00000
 -2.51040    0.00000    0.00000 ;      HG-    CG-   CD2-  HD21
    596     595     601     603      3    0.62760    1.88280    0.00000
 -2.51040    0.00000    0.00000 ;      HG-    CG-   CD2-  HD22
    596     595     601     604      3    0.62760    1.88280    0.00000
 -2.51040    0.00000    0.00000 ;      HG-    CG-   CD2-  HD23
    597     595     601     602      3    0.66944    2.00832    0.00000
 -2.67776    0.00000    0.00000 ;     CD1-    CG-   CD2-  HD21
    597     595     601     603      3    0.66944    2.00832    0.00000
 -2.67776    0.00000    0.00000 ;     CD1-    CG-   CD2-  HD22
    597     595     601     604      3    0.66944    2.00832    0.00000
 -2.67776    0.00000    0.00000 ;     CD1-    CG-   CD2-  HD23
    598     597     595     601      3    0.66944    2.00832    0.00000
 -2.67776    0.00000    0.00000 ;    HD11-   CD1-     CG-    CD2
    599     597     595     601      3    0.66944    2.00832    0.00000
```

```
-2.67776    0.00000    0.00000 ;   HD12-   CD1-    CG-   CD2
   600    597    595    601      3    0.66944    2.00832    0.00000
-2.67776    0.00000    0.00000 ;   HD13-   CD1-    CG-   CD2
   605    607    609    610      3    0.00000    0.00000    0.00000
0.00000    0.00000    0.00000 ;     C-     N-    CA-    HA
   605    607    609    611      3   11.71520    1.67360   15.06240  -
13.38880    0.00000    0.00000 ;     C-     N-    CA-    CB
   605    607    609    626      3    1.75728    5.27184    2.25936
-7.02912    0.00000    0.00000 ;     C-     N-    CA-     C
   606    605    607    608      3   29.28800   -8.36800  -20.92000
0.00000    0.00000    0.00000 ;     O-     C-     N-    H
   606    605    607    609      3   20.92000    0.00000  -20.92000
0.00000    0.00000    0.00000 ;     O-     C-     N-    CA
   607    609    611    612      3    0.65084    1.95253    0.00000
-2.60338    0.00000    0.00000 ;     N-    CA-    CB-   HB2
   607    609    611    613      3    0.65084    1.95253    0.00000
-2.60338    0.00000    0.00000 ;     N-    CA-    CB-   HB3
   607    609    611    614      3    3.28026    2.46019   -2.19242
-3.21331   -0.23430    0.00000 ;     N-    CA-    CB-    CG
   607    609    626    627      3    0.00000    0.00000    0.00000
0.00000    0.00000    0.00000 ;     N-    CA-     C-     O
   607    609    626    628      3   17.40544   -5.02080  -13.22144
9.20480    0.00000    0.00000 ;     N-    CA-     C-     N
   608    607    609    610      3    0.00000    0.00000    0.00000
0.00000    0.00000    0.00000 ;     H-     N-    CA-    HA
   608    607    609    611      3    0.00000    0.00000    0.00000
0.00000    0.00000    0.00000 ;     H-     N-    CA-    CB
   608    607    609    626      3    0.00000    0.00000    0.00000
0.00000    0.00000    0.00000 ;     H-     N-    CA-     C
   609    611    614    615      3    0.57739    0.00000    1.02926
0.00000   -1.60666    0.00000 ;    CA-    CB-    CG-   CD1
   609    611    614    624      3    0.57739    0.00000    1.02926
0.00000   -1.60666    0.00000 ;    CA-    CB-    CG-   CD2
   609    626    628    629      3   20.92000    0.00000  -20.92000
0.00000    0.00000    0.00000 ;    CA-     C-     N-     H
   609    626    628    630      3   20.92000    0.00000  -20.92000
0.00000    0.00000    0.00000 ;    CA-     C-     N-    CA
   610    609    611    612      3    0.65084    1.95253    0.00000
-2.60338    0.00000    0.00000 ;    HA-    CA-    CB-   HB2
   610    609    611    613      3    0.65084    1.95253    0.00000
-2.60338    0.00000    0.00000 ;    HA-    CA-    CB-   HB3
   610    609    611    614      3    0.65084    1.95253    0.00000
-2.60338    0.00000    0.00000 ;    HA-    CA-    CB-    CG
   610    609    626    627      3    3.68192   -4.35136    0.00000
1.33888    0.00000    0.00000 ;    HA-    CA-     C-     O
   610    609    626    628      3    0.00000    0.00000    0.00000
0.00000    0.00000    0.00000 ;    HA-    CA-     C-     N
   611    609    626    627      3    0.00000    0.00000    0.00000
0.00000    0.00000    0.00000 ;    CB-    CA-     C-     O
   611    609    626    628      3    2.51040    4.18400    1.67360
-6.69440    0.00000    0.00000 ;    CB-    CA-     C-     N
   611    614    615    616      3   30.33400    0.00000  -30.33400
```

```
0.00000    0.00000    0.00000 ;     CB-    CG-   CD1-   HD1
   611    614    615    617      3   30.33400    0.00000  -30.33400
0.00000    0.00000    0.00000 ;     CB-    CG-   CD1-   CE1
   611    614    624    622      3   30.33400    0.00000  -30.33400
0.00000    0.00000    0.00000 ;     CB-    CG-   CD2-   CE2
   611    614    624    625      3   30.33400    0.00000  -30.33400
0.00000    0.00000    0.00000 ;     CB-    CG-   CD2-   HD2
   612    611    609    626      3    0.65084    1.95253    0.00000
-2.60338    0.00000    0.00000 ;    HB2-    CB-    CA-     C
   612    611    614    615      3    0.00000    0.00000    0.00000
0.00000    0.00000    0.00000 ;    HB2-    CB-    CG-   CD1
   612    611    614    624      3    0.00000    0.00000    0.00000
0.00000    0.00000    0.00000 ;    HB2-    CB-    CG-   CD2
   613    611    609    626      3    0.65084    1.95253    0.00000
-2.60338    0.00000    0.00000 ;    HB3-    CB-    CA-     C
   613    611    614    615      3    0.00000    0.00000    0.00000
0.00000    0.00000    0.00000 ;    HB3-    CB-    CG-   CD1
   613    611    614    624      3    0.00000    0.00000    0.00000
0.00000    0.00000    0.00000 ;    HB3-    CB-    CG-   CD2
   614    611    609    626      3    4.95804    2.17986   -3.52293
-3.21331   -0.40166    0.00000 ;     CG-    CB-    CA-     C
   614    615    617    618      3   30.33400    0.00000  -30.33400
0.00000    0.00000    0.00000 ;     CG-   CD1-   CE1-   HE1
   614    615    617    619      3   30.33400    0.00000  -30.33400
0.00000    0.00000    0.00000 ;     CG-   CD1-   CE1-    CZ
   614    624    622    619      3   30.33400    0.00000  -30.33400
0.00000    0.00000    0.00000 ;     CG-   CD2-   CE2-    CZ
   614    624    622    623      3   30.33400    0.00000  -30.33400
0.00000    0.00000    0.00000 ;     CG-   CD2-   CE2-   HE2
   615    614    624    622      3   30.33400    0.00000  -30.33400
0.00000    0.00000    0.00000 ;    CD1-    CG-   CD2-   CE2
   615    614    624    625      3   30.33400    0.00000  -30.33400
0.00000    0.00000    0.00000 ;    CD1-    CG-   CD2-   HD2
   615    617    619    620      3   30.33400    0.00000  -30.33400
0.00000    0.00000    0.00000 ;    CD1-   CE1-    CZ-    OH
   615    617    619    622      3   30.33400    0.00000  -30.33400
0.00000    0.00000    0.00000 ;    CD1-   CE1-    CZ-   CE2
   616    615    614    624      3   30.33400    0.00000  -30.33400
0.00000    0.00000    0.00000 ;    HD1-   CD1-    CG-   CD2
   616    615    617    618      3   30.33400    0.00000  -30.33400
0.00000    0.00000    0.00000 ;    HD1-   CD1-   CE1-   HE1
   616    615    617    619      3   30.33400    0.00000  -30.33400
0.00000    0.00000    0.00000 ;    HD1-   CD1-   CE1-    CZ
   617    615    614    624      3   30.33400    0.00000  -30.33400
0.00000    0.00000    0.00000 ;    CE1-   CD1-    CG-   CD2
   617    619    620    621      3    7.93286    0.00000   -9.56462
0.00000    2.17568    0.00000 ;    CE1-    CZ-    OH-    HH
   617    619    622    623      3   30.33400    0.00000  -30.33400
0.00000    0.00000    0.00000 ;    CE1-    CZ-   CE2-   HE2
   617    619    622    624      3   30.33400    0.00000  -30.33400
0.00000    0.00000    0.00000 ;    CE1-    CZ-   CE2-   CD2
   618    617    619    620      3   30.33400    0.00000  -30.33400
```

0.00000 0.00000 0.00000 ; HE1- CE1- CZ- OH
618 617 619 622 3 30.33400 0.00000 -30.33400
0.00000 0.00000 0.00000 ; HE1- CE1- CZ- CE2
619 622 624 625 3 30.33400 0.00000 -30.33400
0.00000 0.00000 0.00000 ; CZ- CE2- CD2- HD2
620 619 622 623 3 30.33400 0.00000 -30.33400
0.00000 0.00000 0.00000 ; OH- CZ- CE2- HE2
620 619 622 624 3 30.33400 0.00000 -30.33400
0.00000 0.00000 0.00000 ; OH- CZ- CE2- CD2
621 620 619 622 3 7.93286 0.00000 -9.56462
0.00000 2.17568 0.00000 ; HH- OH- CZ- CE2
623 622 624 625 3 30.33400 0.00000 -30.33400
0.00000 0.00000 0.00000 ; HE2- CE2- CD2- HD2
626 628 630 631 3 0.00000 0.00000 0.00000
0.00000 0.00000 0.00000 ; C- N- CA- HA
626 628 630 632 3 11.71520 1.67360 15.06240 -
13.38880 0.00000 0.00000 ; C- N- CA- CB
626 628 630 636 3 1.75728 5.27184 2.25936
-7.02912 0.00000 0.00000 ; C- N- CA- C
627 626 628 629 3 29.28800 -8.36800 -20.92000
0.00000 0.00000 0.00000 ; O- C- N- H
627 626 628 630 3 20.92000 0.00000 -20.92000
0.00000 0.00000 0.00000 ; O- C- N- CA
628 630 632 633 3 0.65084 1.95253 0.00000
-2.60338 0.00000 0.00000 ; N- CA- CB- HB1
628 630 632 634 3 0.65084 1.95253 0.00000
-2.60338 0.00000 0.00000 ; N- CA- CB- HB2
628 630 632 635 3 0.65084 1.95253 0.00000
-2.60338 0.00000 0.00000 ; N- CA- CB- HB3
628 630 636 637 3 0.00000 0.00000 0.00000
0.00000 0.00000 0.00000 ; N- CA- C- O
628 630 636 638 3 17.40544 -5.02080 -13.22144
9.20480 0.00000 0.00000 ; N- CA- C- N
629 628 630 631 3 0.00000 0.00000 0.00000
0.00000 0.00000 0.00000 ; H- N- CA- HA
629 628 630 632 3 0.00000 0.00000 0.00000
0.00000 0.00000 0.00000 ; H- N- CA- CB
629 628 630 636 3 0.00000 0.00000 0.00000
0.00000 0.00000 0.00000 ; H- N- CA- C
630 636 638 639 3 20.92000 0.00000 -20.92000
0.00000 0.00000 0.00000 ; CA- C- N- H
630 636 638 640 3 20.92000 0.00000 -20.92000
0.00000 0.00000 0.00000 ; CA- C- N- CA
631 630 632 633 3 0.65084 1.95253 0.00000
-2.60338 0.00000 0.00000 ; HA- CA- CB- HB1
631 630 632 634 3 0.65084 1.95253 0.00000
-2.60338 0.00000 0.00000 ; HA- CA- CB- HB2
631 630 632 635 3 0.65084 1.95253 0.00000
-2.60338 0.00000 0.00000 ; HA- CA- CB- HB3
631 630 636 637 3 3.68192 -4.35136 0.00000
1.33888 0.00000 0.00000 ; HA- CA- C- O
631 630 636 638 3 0.00000 0.00000 0.00000

0.00000 0.00000 0.00000 ; HA- CA- C- N
632 630 636 637 3 0.00000 0.00000 0.00000
0.00000 0.00000 0.00000 ; CB- CA- C- O
632 630 636 638 3 2.51040 4.18400 1.67360
-6.69440 0.00000 0.00000 ; CB- CA- C- N
633 632 630 636 3 0.65084 1.95253 0.00000
-2.60338 0.00000 0.00000 ; HB1- CB- CA- C
634 632 630 636 3 0.65084 1.95253 0.00000
-2.60338 0.00000 0.00000 ; HB2- CB- CA- C
635 632 630 636 3 0.65084 1.95253 0.00000
-2.60338 0.00000 0.00000 ; HB3- CB- CA- C
636 638 640 641 3 0.00000 0.00000 0.00000
0.00000 0.00000 0.00000 ; C- N- CA- HA
636 638 640 642 3 11.71520 1.67360 15.06240 -
13.38880 0.00000 0.00000 ; C- N- CA- CB
636 638 640 655 3 1.75728 5.27184 2.25936
-7.02912 0.00000 0.00000 ; C- N- CA- C
637 636 638 639 3 29.28800 -8.36800 -20.92000
0.00000 0.00000 0.00000 ; O- C- N- H
637 636 638 640 3 20.92000 0.00000 -20.92000
0.00000 0.00000 0.00000 ; O- C- N- CA
638 640 642 643 3 0.65084 1.95253 0.00000
-2.60338 0.00000 0.00000 ; N- CA- CB- HB
638 640 642 644 3 3.83673 0.44769 -1.77402
-2.47693 -0.03347 0.00000 ; N- CA- CB- CG2
638 640 642 648 3 2.97482 0.12134 2.04179
-1.89117 -3.24678 0.00000 ; N- CA- CB- CG1
638 640 655 656 3 0.00000 0.00000 0.00000
0.00000 0.00000 0.00000 ; N- CA- C- O
638 640 655 657 3 17.40544 -5.02080 -13.22144
9.20480 0.00000 0.00000 ; N- CA- C- N
639 638 640 641 3 0.00000 0.00000 0.00000
0.00000 0.00000 0.00000 ; H- N- CA- HA
639 638 640 642 3 0.00000 0.00000 0.00000
0.00000 0.00000 0.00000 ; H- N- CA- CB
639 638 640 655 3 0.00000 0.00000 0.00000
0.00000 0.00000 0.00000 ; H- N- CA- C
640 642 644 645 3 0.66944 2.00832 0.00000
-2.67776 0.00000 0.00000 ; CA- CB- CG2- HG21
640 642 644 646 3 0.66944 2.00832 0.00000
-2.67776 0.00000 0.00000 ; CA- CB- CG2- HG22
640 642 644 647 3 0.66944 2.00832 0.00000
-2.67776 0.00000 0.00000 ; CA- CB- CG2- HG23
640 642 648 649 3 0.66944 2.00832 0.00000
-2.67776 0.00000 0.00000 ; CA- CB- CG1- HG12
640 642 648 650 3 0.66944 2.00832 0.00000
-2.67776 0.00000 0.00000 ; CA- CB- CG1- HG13
640 642 648 651 3 4.24258 -0.52718 -7.25506
-1.79075 7.69856 0.00000 ; CA- CB- CG1- CD1
640 655 657 658 3 20.92000 0.00000 -20.92000
0.00000 0.00000 0.00000 ; CA- C- N- H
640 655 657 659 3 20.92000 0.00000 -20.92000

```
0.00000    0.00000    0.00000 ;      CA-     C-     N-    CA
   641    640    642    643       3    0.65084    1.95253    0.00000
-2.60338    0.00000    0.00000 ;      HA-    CA-    CB-    HB
   641    640    642    644       3    0.65084    1.95253    0.00000
-2.60338    0.00000    0.00000 ;      HA-    CA-    CB-   CG2
   641    640    642    648       3    0.65084    1.95253    0.00000
-2.60338    0.00000    0.00000 ;      HA-    CA-    CB-   CG1
   641    640    655    656       3    3.68192   -4.35136    0.00000
1.33888    0.00000    0.00000 ;      HA-    CA-     C-     O
   641    640    655    657       3    0.00000    0.00000    0.00000
0.00000    0.00000    0.00000 ;      HA-    CA-     C-     N
   642    640    655    656       3    0.00000    0.00000    0.00000
0.00000    0.00000    0.00000 ;      CB-    CA-     C-     O
   642    640    655    657       3    2.51040    4.18400    1.67360
-6.69440    0.00000    0.00000 ;      CB-    CA-     C-     N
   642    648    651    652       3    0.66944    2.00832    0.00000
-2.67776    0.00000    0.00000 ;      CB-   CG1-   CD1-  HD11
   642    648    651    653       3    0.66944    2.00832    0.00000
-2.67776    0.00000    0.00000 ;      CB-   CG1-   CD1-  HD12
   642    648    651    654       3    0.66944    2.00832    0.00000
-2.67776    0.00000    0.00000 ;      CB-   CG1-   CD1-  HD13
   643    642    640    655       3    0.65084    1.95253    0.00000
-2.60338    0.00000    0.00000 ;      HB-    CB-    CA-     C
   643    642    644    645       3    0.62760    1.88280    0.00000
-2.51040    0.00000    0.00000 ;      HB-    CB-   CG2-  HG21
   643    642    644    646       3    0.62760    1.88280    0.00000
-2.51040    0.00000    0.00000 ;      HB-    CB-   CG2-  HG22
   643    642    644    647       3    0.62760    1.88280    0.00000
-2.51040    0.00000    0.00000 ;      HB-    CB-   CG2-  HG23
   643    642    648    649       3    0.62760    1.88280    0.00000
-2.51040    0.00000    0.00000 ;      HB-    CB-   CG1-  HG12
   643    642    648    650       3    0.62760    1.88280    0.00000
-2.51040    0.00000    0.00000 ;      HB-    CB-   CG1-  HG13
   643    642    648    651       3    0.66944    2.00832    0.00000
-2.67776    0.00000    0.00000 ;      HB-    CB-   CG1-   CD1
   644    642    640    655       3    5.67350    3.55640   -6.16722
-2.47693    3.74886    0.00000 ;     CG2-    CB-    CA-     C
   644    642    648    649       3    0.66944    2.00832    0.00000
-2.67776    0.00000    0.00000 ;     CG2-    CB-   CG1-  HG12
   644    642    648    650       3    0.66944    2.00832    0.00000
-2.67776    0.00000    0.00000 ;     CG2-    CB-   CG1-  HG13
   644    642    648    651       3    3.81162    0.49790   -8.14206
-1.79075    7.49773    0.00000 ;     CG2-    CB-   CG1-   CD1
   645    644    642    648       3    0.66944    2.00832    0.00000
-2.67776    0.00000    0.00000 ;    HG21-   CG2-    CB-   CG1
   646    644    642    648       3    0.66944    2.00832    0.00000
-2.67776    0.00000    0.00000 ;    HG22-   CG2-    CB-   CG1
   647    644    642    648       3    0.66944    2.00832    0.00000
-2.67776    0.00000    0.00000 ;    HG23-   CG2-    CB-   CG1
   648    642    640    655       3    8.26340    0.74057   -9.99976
-1.89117    3.84928    0.00000 ;     CG1-    CB-    CA-     C
   649    648    651    652       3    0.62760    1.88280    0.00000
```

```
-2.51040    0.00000    0.00000 ;   HG12-   CG1-   CD1-  HD11
   649    648    651    653      3    0.62760    1.88280    0.00000
-2.51040    0.00000    0.00000 ;   HG12-   CG1-   CD1-  HD12
   649    648    651    654      3    0.62760    1.88280    0.00000
-2.51040    0.00000    0.00000 ;   HG12-   CG1-   CD1-  HD13
   650    648    651    652      3    0.62760    1.88280    0.00000
-2.51040    0.00000    0.00000 ;   HG13-   CG1-   CD1-  HD11
   650    648    651    653      3    0.62760    1.88280    0.00000
-2.51040    0.00000    0.00000 ;   HG13-   CG1-   CD1-  HD12
   650    648    651    654      3    0.62760    1.88280    0.00000
-2.51040    0.00000    0.00000 ;   HG13-   CG1-   CD1-  HD13
   655    657    659    660      3    0.00000    0.00000    0.00000
0.00000    0.00000    0.00000 ;      C-     N-    CA-    HA
   655    657    659    661      3   11.71520    1.67360   15.06240  -
13.38880    0.00000    0.00000 ;      C-     N-    CA-    CB
   655    657    659    677      3    1.75728    5.27184    2.25936
-7.02912    0.00000    0.00000 ;      C-     N-    CA-     C
   656    655    657    658      3   29.28800   -8.36800  -20.92000
0.00000    0.00000    0.00000 ;      O-     C-     N-     H
   656    655    657    659      3   20.92000    0.00000  -20.92000
0.00000    0.00000    0.00000 ;      O-     C-     N-    CA
   657    659    661    662      3    0.65084    1.95253    0.00000
-2.60338    0.00000    0.00000 ;      N-    CA-    CB-   HB2
   657    659    661    663      3    0.65084    1.95253    0.00000
-2.60338    0.00000    0.00000 ;      N-    CA-    CB-   HB3
   657    659    661    664      3    0.65084    1.95253    0.00000
-2.60338    0.00000    0.00000 ;      N-    CA-    CB-    CG
   657    659    677    678      3    0.00000    0.00000    0.00000
0.00000    0.00000    0.00000 ;      N-    CA-     C-     O
   657    659    677    679      3   17.40544   -5.02080  -13.22144
9.20480    0.00000    0.00000 ;      N-    CA-     C-     N
   658    657    659    660      3    0.00000    0.00000    0.00000
0.00000    0.00000    0.00000 ;      H-     N-    CA-    HA
   658    657    659    661      3    0.00000    0.00000    0.00000
0.00000    0.00000    0.00000 ;      H-     N-    CA-    CB
   658    657    659    677      3    0.00000    0.00000    0.00000
0.00000    0.00000    0.00000 ;      H-     N-    CA-     C
   659    661    664    665      3    0.66944    2.00832    0.00000
-2.67776    0.00000    0.00000 ;     CA-    CB-    CG-   HG2
   659    661    664    666      3    0.66944    2.00832    0.00000
-2.67776    0.00000    0.00000 ;     CA-    CB-    CG-   HG3
   659    661    664    667      3    3.68192    3.09616   -2.09200
-3.01248    0.00000    0.00000 ;     CA-    CB-    CG-    CD
   659    677    679    680      3   20.92000    0.00000  -20.92000
0.00000    0.00000    0.00000 ;     CA-     C-     N-     H
   659    677    679    681      3   20.92000    0.00000  -20.92000
0.00000    0.00000    0.00000 ;     CA-     C-     N-    CA
   660    659    661    662      3    0.65084    1.95253    0.00000
-2.60338    0.00000    0.00000 ;     HA-    CA-    CB-   HB2
   660    659    661    663      3    0.65084    1.95253    0.00000
-2.60338    0.00000    0.00000 ;     HA-    CA-    CB-   HB3
   660    659    661    664      3    0.65084    1.95253    0.00000
```

-2.60338 0.00000 0.00000 ; HA- CA- CB- CG
660 659 677 678 3 3.68192 -4.35136 0.00000
1.33888 0.00000 0.00000 ; HA- CA- C- O
660 659 677 679 3 0.00000 0.00000 0.00000
0.00000 0.00000 0.00000 ; HA- CA- C- N
661 659 677 678 3 0.00000 0.00000 0.00000
0.00000 0.00000 0.00000 ; CB- CA- C- O
661 659 677 679 3 2.51040 4.18400 1.67360
-6.69440 0.00000 0.00000 ; CB- CA- C- N
661 664 667 668 3 0.66944 2.00832 0.00000
-2.67776 0.00000 0.00000 ; CB- CG- CD- HD2
661 664 667 669 3 0.66944 2.00832 0.00000
-2.67776 0.00000 0.00000 ; CB- CG- CD- HD3
661 664 667 670 3 3.68192 3.09616 -2.09200
-3.01248 0.00000 0.00000 ; CB- CG- CD- CE
662 661 659 677 3 0.65084 1.95253 0.00000
-2.60338 0.00000 0.00000 ; HB2- CB- CA- C
662 661 664 665 3 0.62760 1.88280 0.00000
-2.51040 0.00000 0.00000 ; HB2- CB- CG- HG2
662 661 664 666 3 0.62760 1.88280 0.00000
-2.51040 0.00000 0.00000 ; HB2- CB- CG- HG3
662 661 664 667 3 0.66944 2.00832 0.00000
-2.67776 0.00000 0.00000 ; HB2- CB- CG- CD
663 661 659 677 3 0.65084 1.95253 0.00000
-2.60338 0.00000 0.00000 ; HB3- CB- CA- C
663 661 664 665 3 0.62760 1.88280 0.00000
-2.51040 0.00000 0.00000 ; HB3- CB- CG- HG2
663 661 664 666 3 0.62760 1.88280 0.00000
-2.51040 0.00000 0.00000 ; HB3- CB- CG- HG3
663 661 664 667 3 0.66944 2.00832 0.00000
-2.67776 0.00000 0.00000 ; HB3- CB- CG- CD
664 661 659 677 3 0.65084 1.95253 0.00000
-2.60338 0.00000 0.00000 ; CG- CB- CA- C
664 667 670 671 3 0.65084 1.95253 0.00000
-2.60338 0.00000 0.00000 ; CG- CD- CE- HE2
664 667 670 672 3 0.65084 1.95253 0.00000
-2.60338 0.00000 0.00000 ; CG- CD- CE- HE3
664 667 670 673 3 0.65084 1.95253 0.00000
-2.60338 0.00000 0.00000 ; CG- CD- CE- NZ
665 664 667 668 3 0.62760 1.88280 0.00000
-2.51040 0.00000 0.00000 ; HG2- CG- CD- HD2
665 664 667 669 3 0.62760 1.88280 0.00000
-2.51040 0.00000 0.00000 ; HG2- CG- CD- HD3
665 664 667 670 3 0.66944 2.00832 0.00000
-2.67776 0.00000 0.00000 ; HG2- CG- CD- CE
666 664 667 668 3 0.62760 1.88280 0.00000
-2.51040 0.00000 0.00000 ; HG3- CG- CD- HD2
666 664 667 669 3 0.62760 1.88280 0.00000
-2.51040 0.00000 0.00000 ; HG3- CG- CD- HD3
666 664 667 670 3 0.66944 2.00832 0.00000
-2.67776 0.00000 0.00000 ; HG3- CG- CD- CE
667 670 673 674 3 0.65084 1.95253 0.00000

-2.60338 0.00000 0.00000 ; CD- CE- NZ- HZ1
667 670 673 675 3 0.65084 1.95253 0.00000
-2.60338 0.00000 0.00000 ; CD- CE- NZ- HZ2
667 670 673 676 3 0.65084 1.95253 0.00000
-2.60338 0.00000 0.00000 ; CD- CE- NZ- HZ3
668 667 670 671 3 0.65084 1.95253 0.00000
-2.60338 0.00000 0.00000 ; HD2- CD- CE- HE2
668 667 670 672 3 0.65084 1.95253 0.00000
-2.60338 0.00000 0.00000 ; HD2- CD- CE- HE3
668 667 670 673 3 0.65084 1.95253 0.00000
-2.60338 0.00000 0.00000 ; HD2- CD- CE- NZ
669 667 670 671 3 0.65084 1.95253 0.00000
-2.60338 0.00000 0.00000 ; HD3- CD- CE- HE2
669 667 670 672 3 0.65084 1.95253 0.00000
-2.60338 0.00000 0.00000 ; HD3- CD- CE- HE3
669 667 670 673 3 0.65084 1.95253 0.00000
-2.60338 0.00000 0.00000 ; HD3- CD- CE- NZ
671 670 673 674 3 0.65084 1.95253 0.00000
-2.60338 0.00000 0.00000 ; HE2- CE- NZ- HZ1
671 670 673 675 3 0.65084 1.95253 0.00000
-2.60338 0.00000 0.00000 ; HE2- CE- NZ- HZ2
671 670 673 676 3 0.65084 1.95253 0.00000
-2.60338 0.00000 0.00000 ; HE2- CE- NZ- HZ3
672 670 673 674 3 0.65084 1.95253 0.00000
-2.60338 0.00000 0.00000 ; HE3- CE- NZ- HZ1
672 670 673 675 3 0.65084 1.95253 0.00000
-2.60338 0.00000 0.00000 ; HE3- CE- NZ- HZ2
672 670 673 676 3 0.65084 1.95253 0.00000
-2.60338 0.00000 0.00000 ; HE3- CE- NZ- HZ3
677 679 681 682 3 0.00000 0.00000 0.00000
0.00000 0.00000 0.00000 ; C- N- CA- HA
677 679 681 683 3 11.71520 1.67360 15.06240 -
13.38880 0.00000 0.00000 ; C- N- CA- CB
677 679 681 692 3 1.75728 5.27184 2.25936
-7.02912 0.00000 0.00000 ; C- N- CA- C
678 677 679 680 3 29.28800 -8.36800 -20.92000
0.00000 0.00000 0.00000 ; O- C- N- H
678 677 679 681 3 20.92000 0.00000 -20.92000
0.00000 0.00000 0.00000 ; O- C- N- CA
679 681 683 684 3 0.65084 1.95253 0.00000
-2.60338 0.00000 0.00000 ; N- CA- CB- HB2
679 681 683 685 3 0.65084 1.95253 0.00000
-2.60338 0.00000 0.00000 ; N- CA- CB- HB3
679 681 683 686 3 3.21331 2.22589 -4.15053
-2.40998 2.61082 0.00000 ; N- CA- CB- CG
679 681 692 693 3 0.00000 0.00000 0.00000
0.00000 0.00000 0.00000 ; N- CA- C- O
679 681 692 694 3 17.40544 -5.02080 -13.22144
9.20480 0.00000 0.00000 ; N- CA- C- N
680 679 681 682 3 0.00000 0.00000 0.00000
0.00000 0.00000 0.00000 ; H- N- CA- HA
680 679 681 683 3 0.00000 0.00000 0.00000

0.00000 0.00000 0.00000 ; H- N- CA- CB
680 679 681 692 3 0.00000 0.00000 0.00000
0.00000 0.00000 0.00000 ; H- N- CA- C
681 683 686 687 3 0.66944 2.00832 0.00000
-2.67776 0.00000 0.00000 ; CA- CB- CG- HG2
681 683 686 688 3 0.66944 2.00832 0.00000
-2.67776 0.00000 0.00000 ; CA- CB- CG- HG3
681 683 686 689 3 10.12110 -1.91209 0.01674
10.17549 -1.87443 0.00000 ; CA- CB- CG- CD
681 692 694 695 3 20.92000 0.00000 -20.92000
0.00000 0.00000 0.00000 ; CA- C- N- H
681 692 694 696 3 20.92000 0.00000 -20.92000
0.00000 0.00000 0.00000 ; CA- C- N- CA
682 681 683 684 3 0.65084 1.95253 0.00000
-2.60338 0.00000 0.00000 ; HA- CA- CB- HB2
682 681 683 685 3 0.65084 1.95253 0.00000
-2.60338 0.00000 0.00000 ; HA- CA- CB- HB3
682 681 683 686 3 0.65084 1.95253 0.00000
-2.60338 0.00000 0.00000 ; HA- CA- CB- CG
682 681 692 693 3 3.68192 -4.35136 0.00000
1.33888 0.00000 0.00000 ; HA- CA- C- O
682 681 692 694 3 0.00000 0.00000 0.00000
0.00000 0.00000 0.00000 ; HA- CA- C- N
683 681 692 693 3 0.00000 0.00000 0.00000
0.00000 0.00000 0.00000 ; CB- CA- C- O
683 681 692 694 3 2.51040 4.18400 1.67360
-6.69440 0.00000 0.00000 ; CB- CA- C- N
683 686 689 690 3 3.79907 0.00000 -5.40573
0.00000 2.14221 0.00000 ; CB- CG- CD- OE1
683 686 689 691 3 3.79907 0.00000 -5.40573
0.00000 2.14221 0.00000 ; CB- CG- CD- OE2
684 683 681 692 3 0.65084 1.95253 0.00000
-2.60338 0.00000 0.00000 ; HB2- CB- CA- C
684 683 686 687 3 0.62760 1.88280 0.00000
-2.51040 0.00000 0.00000 ; HB2- CB- CG- HG2
684 683 686 688 3 0.62760 1.88280 0.00000
-2.51040 0.00000 0.00000 ; HB2- CB- CG- HG3
684 683 686 689 3 0.65084 1.95253 0.00000
-2.60338 0.00000 0.00000 ; HB2- CB- CG- CD
685 683 681 692 3 0.65084 1.95253 0.00000
-2.60338 0.00000 0.00000 ; HB3- CB- CA- C
685 683 686 687 3 0.62760 1.88280 0.00000
-2.51040 0.00000 0.00000 ; HB3- CB- CG- HG2
685 683 686 688 3 0.62760 1.88280 0.00000
-2.51040 0.00000 0.00000 ; HB3- CB- CG- HG3
685 683 686 689 3 0.65084 1.95253 0.00000
-2.60338 0.00000 0.00000 ; HB3- CB- CG- CD
686 683 681 692 3 6.86594 3.56895 -8.14206
-2.40998 4.85344 0.00000 ; CG- CB- CA- C
687 686 689 690 3 0.00000 0.00000 0.00000
0.00000 0.00000 0.00000 ; HG2- CG- CD- OE1
687 686 689 691 3 0.00000 0.00000 0.00000

0.00000 0.00000 0.00000 ; HG2- CG- CD- OE2
688 686 689 690 3 0.00000 0.00000 0.00000
0.00000 0.00000 0.00000 ; HG3- CG- CD- OE1
688 686 689 691 3 0.00000 0.00000 0.00000
0.00000 0.00000 0.00000 ; HG3- CG- CD- OE2
692 694 696 697 3 0.00000 0.00000 0.00000
0.00000 0.00000 0.00000 ; C- N- CA- HA
692 694 696 698 3 11.71520 1.67360 15.06240 -
13.38880 0.00000 0.00000 ; C- N- CA- CB
692 694 696 707 3 1.75728 5.27184 2.25936
-7.02912 0.00000 0.00000 ; C- N- CA- C
693 692 694 695 3 29.28800 -8.36800 -20.92000
0.00000 0.00000 0.00000 ; O- C- N- H
693 692 694 696 3 20.92000 0.00000 -20.92000
0.00000 0.00000 0.00000 ; O- C- N- CA
694 696 698 699 3 0.65084 1.95253 0.00000
-2.60338 0.00000 0.00000 ; N- CA- CB- HB2
694 696 698 700 3 0.65084 1.95253 0.00000
-2.60338 0.00000 0.00000 ; N- CA- CB- HB3
694 696 698 701 3 3.21331 2.22589 -4.15053
-2.40998 2.61082 0.00000 ; N- CA- CB- CG
694 696 707 708 3 0.00000 0.00000 0.00000
0.00000 0.00000 0.00000 ; N- CA- C- O
694 696 707 709 3 17.40544 -5.02080 -13.22144
9.20480 0.00000 0.00000 ; N- CA- C- N
695 694 696 697 3 0.00000 0.00000 0.00000
0.00000 0.00000 0.00000 ; H- N- CA- HA
695 694 696 698 3 0.00000 0.00000 0.00000
0.00000 0.00000 0.00000 ; H- N- CA- CB
695 694 696 707 3 0.00000 0.00000 0.00000
0.00000 0.00000 0.00000 ; H- N- CA- C
696 698 701 702 3 0.66944 2.00832 0.00000
-2.67776 0.00000 0.00000 ; CA- CB- CG- HG2
696 698 701 703 3 0.66944 2.00832 0.00000
-2.67776 0.00000 0.00000 ; CA- CB- CG- HG3
696 698 701 704 3 10.12110 -1.91209 0.01674
10.17549 -1.87443 0.00000 ; CA- CB- CG- CD
696 707 709 710 3 20.92000 0.00000 -20.92000
0.00000 0.00000 0.00000 ; CA- C- N- H
696 707 709 711 3 20.92000 0.00000 -20.92000
0.00000 0.00000 0.00000 ; CA- C- N- CA
697 696 698 699 3 0.65084 1.95253 0.00000
-2.60338 0.00000 0.00000 ; HA- CA- CB- HB2
697 696 698 700 3 0.65084 1.95253 0.00000
-2.60338 0.00000 0.00000 ; HA- CA- CB- HB3
697 696 698 701 3 0.65084 1.95253 0.00000
-2.60338 0.00000 0.00000 ; HA- CA- CB- CG
697 696 707 708 3 3.68192 -4.35136 0.00000
1.33888 0.00000 0.00000 ; HA- CA- C- O
697 696 707 709 3 0.00000 0.00000 0.00000
0.00000 0.00000 0.00000 ; HA- CA- C- N
698 696 707 708 3 0.00000 0.00000 0.00000

0.00000 0.00000 0.00000 ; CB- CA- C- O
698 696 707 709 3 2.51040 4.18400 1.67360
-6.69440 0.00000 0.00000 ; CB- CA- C- N
698 701 704 705 3 3.79907 0.00000 -5.40573
0.00000 2.14221 0.00000 ; CB- CG- CD- OE1
698 701 704 706 3 3.79907 0.00000 -5.40573
0.00000 2.14221 0.00000 ; CB- CG- CD- OE2
699 698 696 707 3 0.65084 1.95253 0.00000
-2.60338 0.00000 0.00000 ; HB2- CB- CA- C
699 698 701 702 3 0.62760 1.88280 0.00000
-2.51040 0.00000 0.00000 ; HB2- CB- CG- HG2
699 698 701 703 3 0.62760 1.88280 0.00000
-2.51040 0.00000 0.00000 ; HB2- CB- CG- HG3
699 698 701 704 3 0.65084 1.95253 0.00000
-2.60338 0.00000 0.00000 ; HB2- CB- CG- CD
700 698 696 707 3 0.65084 1.95253 0.00000
-2.60338 0.00000 0.00000 ; HB3- CB- CA- C
700 698 701 702 3 0.62760 1.88280 0.00000
-2.51040 0.00000 0.00000 ; HB3- CB- CG- HG2
700 698 701 703 3 0.62760 1.88280 0.00000
-2.51040 0.00000 0.00000 ; HB3- CB- CG- HG3
700 698 701 704 3 0.65084 1.95253 0.00000
-2.60338 0.00000 0.00000 ; HB3- CB- CG- CD
701 698 696 707 3 6.86594 3.56895 -8.14206
-2.40998 4.85344 0.00000 ; CG- CB- CA- C
702 701 704 705 3 0.00000 0.00000 0.00000
0.00000 0.00000 0.00000 ; HG2- CG- CD- OE1
702 701 704 706 3 0.00000 0.00000 0.00000
0.00000 0.00000 0.00000 ; HG2- CG- CD- OE2
703 701 704 705 3 0.00000 0.00000 0.00000
0.00000 0.00000 0.00000 ; HG3- CG- CD- OE1
703 701 704 706 3 0.00000 0.00000 0.00000
0.00000 0.00000 0.00000 ; HG3- CG- CD- OE2
707 709 711 712 3 0.00000 0.00000 0.00000
0.00000 0.00000 0.00000 ; C- N- CA- HA
707 709 711 713 3 11.71520 1.67360 15.06240 -
13.38880 0.00000 0.00000 ; C- N- CA- CB
707 709 711 729 3 1.75728 5.27184 2.25936
-7.02912 0.00000 0.00000 ; C- N- CA- C
708 707 709 710 3 29.28800 -8.36800 -20.92000
0.00000 0.00000 0.00000 ; O- C- N- H
708 707 709 711 3 20.92000 0.00000 -20.92000
0.00000 0.00000 0.00000 ; O- C- N- CA
709 711 713 714 3 0.65084 1.95253 0.00000
-2.60338 0.00000 0.00000 ; N- CA- CB- HB2
709 711 713 715 3 0.65084 1.95253 0.00000
-2.60338 0.00000 0.00000 ; N- CA- CB- HB3
709 711 713 716 3 0.65084 1.95253 0.00000
-2.60338 0.00000 0.00000 ; N- CA- CB- CG
709 711 729 730 3 0.00000 0.00000 0.00000
0.00000 0.00000 0.00000 ; N- CA- C- O
709 711 729 731 3 17.40544 -5.02080 -13.22144

-3.66518 2.97901 0.00000 ; N- CA- CB- CG
2063 2065 2078 2079 3 0.00000 0.00000 0.00000
0.00000 0.00000 0.00000 ; N- CA- C- O
2063 2065 2078 2080 3 17.40544 -5.02080 -13.22144
9.20480 0.00000 0.00000 ; N- CA- C- N
2064 2063 2065 2066 3 0.00000 0.00000 0.00000
0.00000 0.00000 0.00000 ; H- N- CA- HA
2064 2063 2065 2067 3 0.00000 0.00000 0.00000
0.00000 0.00000 0.00000 ; H- N- CA- CB
2064 2063 2065 2078 3 0.00000 0.00000 0.00000
0.00000 0.00000 0.00000 ; H- N- CA- C
2065 2067 2070 2071 3 5.98312 6.40152 3.28026 -
12.38464 -1.57318 0.00000 ; CA- CB- CG- ND1
2065 2067 2070 2076 3 0.00000 0.00000 0.00000
0.00000 0.00000 0.00000 ; CA- CB- CG- CD2
2065 2078 2080 2081 3 20.92000 0.00000 -20.92000
0.00000 0.00000 0.00000 ; CA- C- N- H
2065 2078 2080 2082 3 20.92000 0.00000 -20.92000
0.00000 0.00000 0.00000 ; CA- C- N- CA
2066 2065 2067 2068 3 0.65084 1.95253 0.00000
-2.60338 0.00000 0.00000 ; HA- CA- CB- HB2
2066 2065 2067 2069 3 0.65084 1.95253 0.00000
-2.60338 0.00000 0.00000 ; HA- CA- CB- HB3
2066 2065 2067 2070 3 0.65084 1.95253 0.00000
-2.60338 0.00000 0.00000 ; HA- CA- CB- CG
2066 2065 2078 2079 3 3.68192 -4.35136 0.00000
1.33888 0.00000 0.00000 ; HA- CA- C- O
2066 2065 2078 2080 3 0.00000 0.00000 0.00000
0.00000 0.00000 0.00000 ; HA- CA- C- N
2067 2065 2078 2079 3 0.00000 0.00000 0.00000
0.00000 0.00000 0.00000 ; CB- CA- C- O
2067 2065 2078 2080 3 2.51040 4.18400 1.67360
-6.69440 0.00000 0.00000 ; CB- CA- C- N
2067 2070 2071 2072 3 20.08320 0.00000 -20.08320
0.00000 0.00000 0.00000 ; CB- CG- ND1- CE1
2067 2070 2076 2074 3 44.97800 0.00000 -44.97800
0.00000 0.00000 0.00000 ; CB- CG- CD2- NE2
2067 2070 2076 2077 3 44.97800 0.00000 -44.97800
0.00000 0.00000 0.00000 ; CB- CG- CD2- HD2
2068 2067 2065 2078 3 0.65084 1.95253 0.00000
-2.60338 0.00000 0.00000 ; HB2- CB- CA- C
2068 2067 2070 2071 3 0.00000 0.00000 0.00000
0.00000 0.00000 0.00000 ; HB2- CB- CG- ND1
2068 2067 2070 2076 3 0.00000 0.00000 0.00000
0.00000 0.00000 0.00000 ; HB2- CB- CG- CD2
2069 2067 2065 2078 3 0.65084 1.95253 0.00000
-2.60338 0.00000 0.00000 ; HB3- CB- CA- C
2069 2067 2070 2071 3 0.00000 0.00000 0.00000
0.00000 0.00000 0.00000 ; HB3- CB- CG- ND1
2069 2067 2070 2076 3 0.00000 0.00000 0.00000
0.00000 0.00000 0.00000 ; HB3- CB- CG- CD2
2070 2067 2065 2078 3 3.76560 3.34720 -2.87859

-3.66518    0.83680    0.00000 ;      CG-    CB-    CA-     C
  2070   2071   2072   2073      3   41.84000    0.00000  -41.84000
0.00000    0.00000    0.00000 ;     CG-   ND1-   CE1-   HE1
  2070   2071   2072   2074      3   41.84000    0.00000  -41.84000
0.00000    0.00000    0.00000 ;     CG-   ND1-   CE1-   NE2
  2070   2076   2074   2072      3   12.55200    0.00000  -12.55200
0.00000    0.00000    0.00000 ;     CG-   CD2-   NE2-   CE1
  2070   2076   2074   2075      3   12.55200    0.00000  -12.55200
0.00000    0.00000    0.00000 ;     CG-   CD2-   NE2-   HE2
  2071   2070   2076   2074      3   44.97800    0.00000  -44.97800
0.00000    0.00000    0.00000 ;    ND1-    CG-   CD2-   NE2
  2071   2070   2076   2077      3   44.97800    0.00000  -44.97800
0.00000    0.00000    0.00000 ;    ND1-    CG-   CD2-   HD2
  2071   2072   2074   2075      3   19.45560    0.00000  -19.45560
0.00000    0.00000    0.00000 ;    ND1-   CE1-   NE2-   HE2
  2071   2072   2074   2076      3   19.45560    0.00000  -19.45560
0.00000    0.00000    0.00000 ;    ND1-   CE1-   NE2-   CD2
  2072   2071   2070   2076      3   20.08320    0.00000  -20.08320
0.00000    0.00000    0.00000 ;    CE1-   ND1-    CG-   CD2
  2072   2074   2076   2077      3   12.55200    0.00000  -12.55200
0.00000    0.00000    0.00000 ;    CE1-   NE2-   CD2-   HD2
  2073   2072   2074   2075      3   19.45560    0.00000  -19.45560
0.00000    0.00000    0.00000 ;    HE1-   CE1-   NE2-   HE2
  2073   2072   2074   2076      3   19.45560    0.00000  -19.45560
0.00000    0.00000    0.00000 ;    HE1-   CE1-   NE2-   CD2
  2075   2074   2076   2077      3   12.55200    0.00000  -12.55200
0.00000    0.00000    0.00000 ;    HE2-   NE2-   CD2-   HD2
  2078   2080   2082   2083      3    0.00000    0.00000    0.00000
0.00000    0.00000    0.00000 ;      C-     N-    CA-    HA
  2078   2080   2082   2084      3   11.71520    1.67360   15.06240  -
13.38880    0.00000    0.00000 ;      C-     N-    CA-    CB
  2078   2080   2082   2089      3    1.75728    5.27184    2.25936
-7.02912    0.00000    0.00000 ;      C-     N-    CA-     C
  2079   2078   2080   2081      3   29.28800   -8.36800  -20.92000
0.00000    0.00000    0.00000 ;      O-     C-     N-     H
  2079   2078   2080   2082      3   20.92000    0.00000  -20.92000
0.00000    0.00000    0.00000 ;      O-     C-     N-    CA
  2080   2082   2084   2085      3    0.65084    1.95253    0.00000
-2.60338    0.00000    0.00000 ;      N-    CA-    CB-   HB2
  2080   2082   2084   2086      3    0.65084    1.95253    0.00000
-2.60338    0.00000    0.00000 ;      N-    CA-    CB-   HB3
  2080   2082   2084   2087      3    7.86174    2.24681   -7.41405
-6.71114    5.35552    0.00000 ;      N-    CA-    CB-    OG
  2080   2082   2089   2090      3    0.00000    0.00000    0.00000
0.00000    0.00000    0.00000 ;     N-    CA-     C-     O
  2080   2082   2089   2091      3   17.40544   -5.02080  -13.22144
9.20480    0.00000    0.00000 ;     N-    CA-     C-     N
  2081   2080   2082   2083      3    0.00000    0.00000    0.00000
0.00000    0.00000    0.00000 ;     H-     N-    CA-    HA
  2081   2080   2082   2084      3    0.00000    0.00000    0.00000
0.00000    0.00000    0.00000 ;     H-     N-    CA-    CB
  2081   2080   2082   2089      3    0.00000    0.00000    0.00000

0.00000 0.00000 0.00000 ; H- N- CA- C
2082 2084 2087 2088 3 2.05853 2.46856 3.48109
-4.46851 0.23430 0.00000 ; CA- CB- OG- HG
2082 2089 2091 2092 3 20.92000 0.00000 -20.92000
0.00000 0.00000 0.00000 ; CA- C- N- H
2082 2089 2091 2093 3 20.92000 0.00000 -20.92000
0.00000 0.00000 0.00000 ; CA- C- N- CA
2083 2082 2084 2085 3 0.65084 1.95253 0.00000
-2.60338 0.00000 0.00000 ; HA- CA- CB- HB2
2083 2082 2084 2086 3 0.65084 1.95253 0.00000
-2.60338 0.00000 0.00000 ; HA- CA- CB- HB3
2083 2082 2084 2087 3 1.04600 -1.04600 0.00000
0.00000 0.00000 0.00000 ; HA- CA- CB- OG
2083 2082 2089 2090 3 3.68192 -4.35136 0.00000
1.33888 0.00000 0.00000 ; HA- CA- C- O
2083 2082 2089 2091 3 0.00000 0.00000 0.00000
0.00000 0.00000 0.00000 ; HA- CA- C- N
2084 2082 2089 2090 3 0.00000 0.00000 0.00000
0.00000 0.00000 0.00000 ; CB- CA- C- O
2084 2082 2089 2091 3 2.51040 4.18400 1.67360
-6.69440 0.00000 0.00000 ; CB- CA- C- N
2085 2084 2082 2089 3 0.65084 1.95253 0.00000
-2.60338 0.00000 0.00000 ; HB2- CB- CA- C
2085 2084 2087 2088 3 0.69733 2.09200 0.00000
-2.78933 0.00000 0.00000 ; HB2- CB- OG- HG
2086 2084 2082 2089 3 0.65084 1.95253 0.00000
-2.60338 0.00000 0.00000 ; HB3- CB- CA- C
2086 2084 2087 2088 3 0.69733 2.09200 0.00000
-2.78933 0.00000 0.00000 ; HB3- CB- OG- HG
2087 2084 2082 2089 3 7.34710 7.79898 -6.14211
-6.71114 4.31789 0.00000 ; OG- CB- CA- C
2089 2091 2093 2094 3 0.00000 0.00000 0.00000
0.00000 0.00000 0.00000 ; C- N- CA- HA
2089 2091 2093 2095 3 11.71520 1.67360 15.06240 -
13.38880 0.00000 0.00000 ; C- N- CA- CB
2089 2091 2093 2108 3 1.75728 5.27184 2.25936
-7.02912 0.00000 0.00000 ; C- N- CA- C
2090 2089 2091 2092 3 29.28800 -8.36800 -20.92000
0.00000 0.00000 0.00000 ; O- C- N- H
2090 2089 2091 2093 3 20.92000 0.00000 -20.92000
0.00000 0.00000 0.00000 ; O- C- N- CA
2091 2093 2095 2096 3 0.65084 1.95253 0.00000
-2.60338 0.00000 0.00000 ; N- CA- CB- HB2
2091 2093 2095 2097 3 0.65084 1.95253 0.00000
-2.60338 0.00000 0.00000 ; N- CA- CB- HB3
2091 2093 2095 2098 3 3.79070 1.39746 -4.61077
-2.40998 2.44346 0.00000 ; N- CA- CB- CG
2091 2093 2108 2109 3 0.00000 0.00000 0.00000
0.00000 0.00000 0.00000 ; N- CA- C- O
2091 2093 2108 2110 3 17.40544 -5.02080 -13.22144
9.20480 0.00000 0.00000 ; N- CA- C- N
2092 2091 2093 2094 3 0.00000 0.00000 0.00000

```
0.00000    0.00000    0.00000 ;       H-     N-    CA-    HA
  2092   2091   2093   2095      3    0.00000    0.00000    0.00000
0.00000    0.00000    0.00000 ;       H-     N-    CA-    CB
  2092   2091   2093   2108      3    0.00000    0.00000    0.00000
0.00000    0.00000    0.00000 ;       H-     N-    CA-     C
  2093   2095   2098   2099      3    0.66944    2.00832    0.00000
-2.67776    0.00000    0.00000 ;      CA-    CB-    CG-    HG
  2093   2095   2098   2100      3    3.90367    0.19665   -6.21742
-2.37651    5.99149    0.00000 ;      CA-    CB-    CG-   CD1
  2093   2095   2098   2104      3    3.90367    0.19665   -6.21742
-2.37651    5.99149    0.00000 ;      CA-    CB-    CG-   CD2
  2093   2108   2110   2111      3   20.92000    0.00000  -20.92000
0.00000    0.00000    0.00000 ;      CA-     C-     N-     H
  2093   2108   2110   2112      3   20.92000    0.00000  -20.92000
0.00000    0.00000    0.00000 ;      CA-     C-     N-    CA
  2094   2093   2095   2096      3    0.65084    1.95253    0.00000
-2.60338    0.00000    0.00000 ;      HA-    CA-    CB-   HB2
  2094   2093   2095   2097      3    0.65084    1.95253    0.00000
-2.60338    0.00000    0.00000 ;      HA-    CA-    CB-   HB3
  2094   2093   2095   2098      3    0.65084    1.95253    0.00000
-2.60338    0.00000    0.00000 ;      HA-    CA-    CB-    CG
  2094   2093   2108   2109      3    3.68192   -4.35136    0.00000
1.33888    0.00000    0.00000 ;      HA-    CA-     C-     O
  2094   2093   2108   2110      3    0.00000    0.00000    0.00000
0.00000    0.00000    0.00000 ;      HA-    CA-     C-     N
  2095   2093   2108   2109      3    0.00000    0.00000    0.00000
0.00000    0.00000    0.00000 ;      CB-    CA-     C-     O
  2095   2093   2108   2110      3    2.51040    4.18400    1.67360
-6.69440    0.00000    0.00000 ;      CB-    CA-     C-     N
  2095   2098   2100   2101      3    0.66944    2.00832    0.00000
-2.67776    0.00000    0.00000 ;      CB-    CG-   CD1-  HD11
  2095   2098   2100   2102      3    0.66944    2.00832    0.00000
-2.67776    0.00000    0.00000 ;      CB-    CG-   CD1-  HD12
  2095   2098   2100   2103      3    0.66944    2.00832    0.00000
-2.67776    0.00000    0.00000 ;      CB-    CG-   CD1-  HD13
  2095   2098   2104   2105      3    0.66944    2.00832    0.00000
-2.67776    0.00000    0.00000 ;      CB-    CG-   CD2-  HD21
  2095   2098   2104   2106      3    0.66944    2.00832    0.00000
-2.67776    0.00000    0.00000 ;      CB-    CG-   CD2-  HD22
  2095   2098   2104   2107      3    0.66944    2.00832    0.00000
-2.67776    0.00000    0.00000 ;      CB-    CG-   CD2-  HD23
  2096   2095   2093   2108      3    0.65084    1.95253    0.00000
-2.60338    0.00000    0.00000 ;     HB2-    CB-    CA-     C
  2096   2095   2098   2099      3    0.62760    1.88280    0.00000
-2.51040    0.00000    0.00000 ;     HB2-    CB-    CG-    HG
  2096   2095   2098   2100      3    0.66944    2.00832    0.00000
-2.67776    0.00000    0.00000 ;     HB2-    CB-    CG-   CD1
  2096   2095   2098   2104      3    0.66944    2.00832    0.00000
-2.67776    0.00000    0.00000 ;     HB2-    CB-    CG-   CD2
  2097   2095   2093   2108      3    0.65084    1.95253    0.00000
-2.60338    0.00000    0.00000 ;     HB3-    CB-    CA-     C
  2097   2095   2098   2099      3    0.62760    1.88280    0.00000
```

-2.51040 0.00000 0.00000 ; HB3- CB- CG- HG
2097 2095 2098 2100 3 0.66944 2.00832 0.00000
-2.67776 0.00000 0.00000 ; HB3- CB- CG- CD1
2097 2095 2098 2104 3 0.66944 2.00832 0.00000
-2.67776 0.00000 0.00000 ; HB3- CB- CG- CD2
2098 2095 2093 2108 3 10.33448 -1.14642 -11.54784
-2.40998 6.35968 0.00000 ; CG- CB- CA- C
2099 2098 2100 2101 3 0.62760 1.88280 0.00000
-2.51040 0.00000 0.00000 ; HG- CG- CD1- HD11
2099 2098 2100 2102 3 0.62760 1.88280 0.00000
-2.51040 0.00000 0.00000 ; HG- CG- CD1- HD12
2099 2098 2100 2103 3 0.62760 1.88280 0.00000
-2.51040 0.00000 0.00000 ; HG- CG- CD1- HD13
2099 2098 2104 2105 3 0.62760 1.88280 0.00000
-2.51040 0.00000 0.00000 ; HG- CG- CD2- HD21
2099 2098 2104 2106 3 0.62760 1.88280 0.00000
-2.51040 0.00000 0.00000 ; HG- CG- CD2- HD22
2099 2098 2104 2107 3 0.62760 1.88280 0.00000
-2.51040 0.00000 0.00000 ; HG- CG- CD2- HD23
2100 2098 2104 2105 3 0.66944 2.00832 0.00000
-2.67776 0.00000 0.00000 ; CD1- CG- CD2- HD21
2100 2098 2104 2106 3 0.66944 2.00832 0.00000
-2.67776 0.00000 0.00000 ; CD1- CG- CD2- HD22
2100 2098 2104 2107 3 0.66944 2.00832 0.00000
-2.67776 0.00000 0.00000 ; CD1- CG- CD2- HD23
2101 2100 2098 2104 3 0.66944 2.00832 0.00000
-2.67776 0.00000 0.00000 ; HD11- CD1- CG- CD2
2102 2100 2098 2104 3 0.66944 2.00832 0.00000
-2.67776 0.00000 0.00000 ; HD12- CD1- CG- CD2
2103 2100 2098 2104 3 0.66944 2.00832 0.00000
-2.67776 0.00000 0.00000 ; HD13- CD1- CG- CD2
2108 2110 2112 2113 3 0.00000 0.00000 0.00000
0.00000 0.00000 0.00000 ; C- N- CA- HA
2108 2110 2112 2114 3 11.71520 1.67360 15.06240 -
13.38880 0.00000 0.00000 ; C- N- CA- CB
2108 2110 2112 2122 3 1.75728 5.27184 2.25936
-7.02912 0.00000 0.00000 ; C- N- CA- C
2109 2108 2110 2111 3 29.28800 -8.36800 -20.92000
0.00000 0.00000 0.00000 ; O- C- N- H
2109 2108 2110 2112 3 20.92000 0.00000 -20.92000
0.00000 0.00000 0.00000 ; O- C- N- CA
2110 2112 2114 2115 3 0.65084 1.95253 0.00000
-2.60338 0.00000 0.00000 ; N- CA- CB- HB2
2110 2112 2114 2116 3 0.65084 1.95253 0.00000
-2.60338 0.00000 0.00000 ; N- CA- CB- HB3
2110 2112 2114 2117 3 5.99567 3.29281 -4.46014
-0.55229 1.97485 0.00000 ; N- CA- CB- CG
2110 2112 2122 2123 3 0.00000 0.00000 0.00000
0.00000 0.00000 0.00000 ; N- CA- C- O
2110 2112 2122 2124 3 17.40544 -5.02080 -13.22144
9.20480 0.00000 0.00000 ; N- CA- C- N
2111 2110 2112 2113 3 0.00000 0.00000 0.00000

```
0.00000    0.00000    0.00000 ;       H-     N-    CA-    HA
  2111   2110   2112   2114      3    0.00000    0.00000    0.00000
0.00000    0.00000    0.00000 ;       H-     N-    CA-    CB
  2111   2110   2112   2122      3    0.00000    0.00000    0.00000
0.00000    0.00000    0.00000 ;       H-     N-    CA-     C
  2112   2114   2117   2118      3    0.00000    0.00000    0.00000
0.00000    0.00000    0.00000 ;      CA-    CB-    CG-   OD1
  2112   2114   2117   2119      3    8.84916   -0.31380   -4.32626
5.03754    0.26778    0.00000 ;      CA-    CB-    CG-   ND2
  2112   2122   2124   2125      3   20.92000    0.00000  -20.92000
0.00000    0.00000    0.00000 ;      CA-     C-     N-     H
  2112   2122   2124   2126      3   20.92000    0.00000  -20.92000
0.00000    0.00000    0.00000 ;      CA-     C-     N-    CA
  2113   2112   2114   2115      3    0.65084    1.95253    0.00000
-2.60338    0.00000    0.00000 ;      HA-    CA-    CB-   HB2
  2113   2112   2114   2116      3    0.65084    1.95253    0.00000
-2.60338    0.00000    0.00000 ;      HA-    CA-    CB-   HB3
  2113   2112   2114   2117      3    0.65084    1.95253    0.00000
-2.60338    0.00000    0.00000 ;      HA-    CA-    CB-    CG
  2113   2112   2122   2123      3    3.68192   -4.35136    0.00000
1.33888    0.00000    0.00000 ;      HA-    CA-     C-     O
  2113   2112   2122   2124      3    0.00000    0.00000    0.00000
0.00000    0.00000    0.00000 ;      HA-    CA-     C-     N
  2114   2112   2122   2123      3    0.00000    0.00000    0.00000
0.00000    0.00000    0.00000 ;      CB-    CA-     C-     O
  2114   2112   2122   2124      3    2.51040    4.18400    1.67360
-6.69440    0.00000    0.00000 ;      CB-    CA-     C-     N
  2114   2117   2119   2120      3   20.92000    0.00000  -20.92000
0.00000    0.00000    0.00000 ;      CB-    CG-   ND2-  HD21
  2114   2117   2119   2121      3   20.92000    0.00000  -20.92000
0.00000    0.00000    0.00000 ;      CB-    CG-   ND2-  HD22
  2115   2114   2112   2122      3    0.65084    1.95253    0.00000
-2.60338    0.00000    0.00000 ;     HB2-    CB-    CA-     C
  2115   2114   2117   2118      3    3.68192   -4.35136    0.00000
1.33888    0.00000    0.00000 ;     HB2-    CB-    CG-   OD1
  2115   2114   2117   2119      3    0.00000    0.00000    0.00000
0.00000    0.00000    0.00000 ;     HB2-    CB-    CG-   ND2
  2116   2114   2112   2122      3    0.65084    1.95253    0.00000
-2.60338    0.00000    0.00000 ;     HB3-    CB-    CA-     C
  2116   2114   2117   2118      3    3.68192   -4.35136    0.00000
1.33888    0.00000    0.00000 ;     HB3-    CB-    CG-   OD1
  2116   2114   2117   2119      3    0.00000    0.00000    0.00000
0.00000    0.00000    0.00000 ;     HB3-    CB-    CG-   ND2
  2117   2114   2112   2122      3    7.94542   -3.96225   -6.11701
-0.55229    3.58150    0.00000 ;      CG-    CB-    CA-     C
  2118   2117   2119   2120      3   29.28800   -8.36800  -20.92000
0.00000    0.00000    0.00000 ;     OD1-    CG-   ND2-  HD21
  2118   2117   2119   2121      3   29.28800   -8.36800  -20.92000
0.00000    0.00000    0.00000 ;     OD1-    CG-   ND2-  HD22
  2122   2124   2126   2127      3    0.00000    0.00000    0.00000
0.00000    0.00000    0.00000 ;       C-     N-    CA-    HA
  2122   2124   2126   2128      3   11.71520    1.67360   15.06240  -
```

```
13.38880    0.00000    0.00000 ;      C-     N-    CA-    CB
  2122   2124   2126   2141      3    1.75728    5.27184    2.25936
-7.02912    0.00000    0.00000 ;      C-     N-    CA-     C
  2123   2122   2124   2125      3   29.28800   -8.36800  -20.92000
0.00000    0.00000    0.00000 ;     O-     C-     N-     H
  2123   2122   2124   2126      3   20.92000    0.00000  -20.92000
0.00000    0.00000    0.00000 ;     O-     C-     N-    CA
  2124   2126   2128   2129      3    0.65084    1.95253    0.00000
-2.60338    0.00000    0.00000 ;     N-    CA-    CB-    HB
  2124   2126   2128   2130      3    3.83673    0.44769   -1.77402
-2.47693   -0.03347    0.00000 ;     N-    CA-    CB-   CG2
  2124   2126   2128   2134      3    2.97482    0.12134    2.04179
-1.89117   -3.24678    0.00000 ;     N-    CA-    CB-   CG1
  2124   2126   2141   2142      3    0.00000    0.00000    0.00000
0.00000    0.00000    0.00000 ;     N-    CA-     C-     O
  2124   2126   2141   2143      3   17.40544   -5.02080  -13.22144
9.20480    0.00000    0.00000 ;     N-    CA-     C-     N
  2125   2124   2126   2127      3    0.00000    0.00000    0.00000
0.00000    0.00000    0.00000 ;     H-     N-    CA-    HA
  2125   2124   2126   2128      3    0.00000    0.00000    0.00000
0.00000    0.00000    0.00000 ;     H-     N-    CA-    CB
  2125   2124   2126   2141      3    0.00000    0.00000    0.00000
0.00000    0.00000    0.00000 ;     H-     N-    CA-     C
  2126   2128   2130   2131      3    0.66944    2.00832    0.00000
-2.67776    0.00000    0.00000 ;    CA-    CB-   CG2-  HG21
  2126   2128   2130   2132      3    0.66944    2.00832    0.00000
-2.67776    0.00000    0.00000 ;    CA-    CB-   CG2-  HG22
  2126   2128   2130   2133      3    0.66944    2.00832    0.00000
-2.67776    0.00000    0.00000 ;    CA-    CB-   CG2-  HG23
  2126   2128   2134   2135      3    0.66944    2.00832    0.00000
-2.67776    0.00000    0.00000 ;    CA-    CB-   CG1-  HG12
  2126   2128   2134   2136      3    0.66944    2.00832    0.00000
-2.67776    0.00000    0.00000 ;    CA-    CB-   CG1-  HG13
  2126   2128   2134   2137      3    4.24258   -0.52718   -7.25506
-1.79075    7.69856    0.00000 ;    CA-    CB-   CG1-   CD1
  2126   2141   2143   2144      3   20.92000    0.00000  -20.92000
0.00000    0.00000    0.00000 ;    CA-     C-     N-     H
  2126   2141   2143   2145      3   20.92000    0.00000  -20.92000
0.00000    0.00000    0.00000 ;    CA-     C-     N-    CA
  2127   2126   2128   2129      3    0.65084    1.95253    0.00000
-2.60338    0.00000    0.00000 ;    HA-    CA-    CB-    HB
  2127   2126   2128   2130      3    0.65084    1.95253    0.00000
-2.60338    0.00000    0.00000 ;    HA-    CA-    CB-   CG2
  2127   2126   2128   2134      3    0.65084    1.95253    0.00000
-2.60338    0.00000    0.00000 ;    HA-    CA-    CB-   CG1
  2127   2126   2141   2142      3    3.68192   -4.35136    0.00000
1.33888    0.00000    0.00000 ;    HA-    CA-     C-     O
  2127   2126   2141   2143      3    0.00000    0.00000    0.00000
0.00000    0.00000    0.00000 ;    HA-    CA-     C-     N
  2128   2126   2141   2142      3    0.00000    0.00000    0.00000
0.00000    0.00000    0.00000 ;    CB-    CA-     C-     O
  2128   2126   2141   2143      3    2.51040    4.18400    1.67360
```

-6.69440 0.00000 0.00000 ; CB- CA- C- N
2128 2134 2137 2138 3 0.66944 2.00832 0.00000
-2.67776 0.00000 0.00000 ; CB- CG1- CD1- HD11
2128 2134 2137 2139 3 0.66944 2.00832 0.00000
-2.67776 0.00000 0.00000 ; CB- CG1- CD1- HD12
2128 2134 2137 2140 3 0.66944 2.00832 0.00000
-2.67776 0.00000 0.00000 ; CB- CG1- CD1- HD13
2129 2128 2126 2141 3 0.65084 1.95253 0.00000
-2.60338 0.00000 0.00000 ; HB- CB- CA- C
2129 2128 2130 2131 3 0.62760 1.88280 0.00000
-2.51040 0.00000 0.00000 ; HB- CB- CG2- HG21
2129 2128 2130 2132 3 0.62760 1.88280 0.00000
-2.51040 0.00000 0.00000 ; HB- CB- CG2- HG22
2129 2128 2130 2133 3 0.62760 1.88280 0.00000
-2.51040 0.00000 0.00000 ; HB- CB- CG2- HG23
2129 2128 2134 2135 3 0.62760 1.88280 0.00000
-2.51040 0.00000 0.00000 ; HB- CB- CG1- HG12
2129 2128 2134 2136 3 0.62760 1.88280 0.00000
-2.51040 0.00000 0.00000 ; HB- CB- CG1- HG13
2129 2128 2134 2137 3 0.66944 2.00832 0.00000
-2.67776 0.00000 0.00000 ; HB- CB- CG1- CD1
2130 2128 2126 2141 3 5.67350 3.55640 -6.16722
-2.47693 3.74886 0.00000 ; CG2- CB- CA- C
2130 2128 2134 2135 3 0.66944 2.00832 0.00000
-2.67776 0.00000 0.00000 ; CG2- CB- CG1- HG12
2130 2128 2134 2136 3 0.66944 2.00832 0.00000
-2.67776 0.00000 0.00000 ; CG2- CB- CG1- HG13
2130 2128 2134 2137 3 3.81162 0.49790 -8.14206
-1.79075 7.49773 0.00000 ; CG2- CB- CG1- CD1
2131 2130 2128 2134 3 0.66944 2.00832 0.00000
-2.67776 0.00000 0.00000 ; HG21- CG2- CB- CG1
2132 2130 2128 2134 3 0.66944 2.00832 0.00000
-2.67776 0.00000 0.00000 ; HG22- CG2- CB- CG1
2133 2130 2128 2134 3 0.66944 2.00832 0.00000
-2.67776 0.00000 0.00000 ; HG23- CG2- CB- CG1
2134 2128 2126 2141 3 8.26340 0.74057 -9.99976
-1.89117 3.84928 0.00000 ; CG1- CB- CA- C
2135 2134 2137 2138 3 0.62760 1.88280 0.00000
-2.51040 0.00000 0.00000 ; HG12- CG1- CD1- HD11
2135 2134 2137 2139 3 0.62760 1.88280 0.00000
-2.51040 0.00000 0.00000 ; HG12- CG1- CD1- HD12
2135 2134 2137 2140 3 0.62760 1.88280 0.00000
-2.51040 0.00000 0.00000 ; HG12- CG1- CD1- HD13
2136 2134 2137 2138 3 0.62760 1.88280 0.00000
-2.51040 0.00000 0.00000 ; HG13- CG1- CD1- HD11
2136 2134 2137 2139 3 0.62760 1.88280 0.00000
-2.51040 0.00000 0.00000 ; HG13- CG1- CD1- HD12
2136 2134 2137 2140 3 0.62760 1.88280 0.00000
-2.51040 0.00000 0.00000 ; HG13- CG1- CD1- HD13
2141 2143 2145 2146 3 0.00000 0.00000 0.00000
0.00000 0.00000 0.00000 ; C- N- CA- HA
2141 2143 2145 2147 3 11.71520 1.67360 15.06240 -

```
13.38880    0.00000    0.00000 ;       C-     N-    CA-    CB
  2141   2143   2145   2157       3    1.75728    5.27184    2.25936
-7.02912    0.00000    0.00000 ;       C-     N-    CA-     C
  2142   2141   2143   2144       3   29.28800   -8.36800  -20.92000
0.00000    0.00000    0.00000 ;      O-     C-     N-     H
  2142   2141   2143   2145       3   20.92000    0.00000  -20.92000
0.00000    0.00000    0.00000 ;      O-     C-     N-    CA
  2143   2145   2147   2148       3    0.65084    1.95253    0.00000
-2.60338    0.00000    0.00000 ;       N-    CA-    CB-    HB
  2143   2145   2147   2149       3    3.83673    0.44769   -1.77402
-2.47693   -0.03347    0.00000 ;       N-    CA-    CB-   CG1
  2143   2145   2147   2153       3    3.83673    0.44769   -1.77402
-2.47693   -0.03347    0.00000 ;       N-    CA-    CB-   CG2
  2143   2145   2157   2158       3    0.00000    0.00000    0.00000
0.00000    0.00000    0.00000 ;      N-    CA-     C-     O
  2143   2145   2157   2159       3   17.40544   -5.02080  -13.22144
9.20480    0.00000    0.00000 ;      N-    CA-     C-     N
  2144   2143   2145   2146       3    0.00000    0.00000    0.00000
0.00000    0.00000    0.00000 ;      H-     N-    CA-    HA
  2144   2143   2145   2147       3    0.00000    0.00000    0.00000
0.00000    0.00000    0.00000 ;      H-     N-    CA-    CB
  2144   2143   2145   2157       3    0.00000    0.00000    0.00000
0.00000    0.00000    0.00000 ;      H-     N-    CA-     C
  2145   2147   2149   2150       3    0.66944    2.00832    0.00000
-2.67776    0.00000    0.00000 ;      CA-    CB-   CG1-  HG11
  2145   2147   2149   2151       3    0.66944    2.00832    0.00000
-2.67776    0.00000    0.00000 ;      CA-    CB-   CG1-  HG12
  2145   2147   2149   2152       3    0.66944    2.00832    0.00000
-2.67776    0.00000    0.00000 ;      CA-    CB-   CG1-  HG13
  2145   2147   2153   2154       3    0.66944    2.00832    0.00000
-2.67776    0.00000    0.00000 ;      CA-    CB-   CG2-  HG21
  2145   2147   2153   2155       3    0.66944    2.00832    0.00000
-2.67776    0.00000    0.00000 ;      CA-    CB-   CG2-  HG22
  2145   2147   2153   2156       3    0.66944    2.00832    0.00000
-2.67776    0.00000    0.00000 ;      CA-    CB-   CG2-  HG23
  2145   2157   2159   2160       3   20.92000    0.00000  -20.92000
0.00000    0.00000    0.00000 ;     CA-     C-     N-     H
  2145   2157   2159   2161       3   20.92000    0.00000  -20.92000
0.00000    0.00000    0.00000 ;     CA-     C-     N-    CA
  2146   2145   2147   2148       3    0.65084    1.95253    0.00000
-2.60338    0.00000    0.00000 ;      HA-    CA-    CB-    HB
  2146   2145   2147   2149       3    0.65084    1.95253    0.00000
-2.60338    0.00000    0.00000 ;      HA-    CA-    CB-   CG1
  2146   2145   2147   2153       3    0.65084    1.95253    0.00000
-2.60338    0.00000    0.00000 ;      HA-    CA-    CB-   CG2
  2146   2145   2157   2158       3    3.68192   -4.35136    0.00000
1.33888    0.00000    0.00000 ;     HA-    CA-     C-     O
  2146   2145   2157   2159       3    0.00000    0.00000    0.00000
0.00000    0.00000    0.00000 ;     HA-    CA-     C-     N
  2147   2145   2157   2158       3    0.00000    0.00000    0.00000
0.00000    0.00000    0.00000 ;     CB-    CA-     C-     O
  2147   2145   2157   2159       3    2.51040    4.18400    1.67360
```

-6.69440 0.00000 0.00000 ; CB- CA- C- N
2148 2147 2145 2157 3 0.65084 1.95253 0.00000
-2.60338 0.00000 0.00000 ; HB- CB- CA- C
2148 2147 2149 2150 3 0.62760 1.88280 0.00000
-2.51040 0.00000 0.00000 ; HB- CB- CG1- HG11
2148 2147 2149 2151 3 0.62760 1.88280 0.00000
-2.51040 0.00000 0.00000 ; HB- CB- CG1- HG12
2148 2147 2149 2152 3 0.62760 1.88280 0.00000
-2.51040 0.00000 0.00000 ; HB- CB- CG1- HG13
2148 2147 2153 2154 3 0.62760 1.88280 0.00000
-2.51040 0.00000 0.00000 ; HB- CB- CG2- HG21
2148 2147 2153 2155 3 0.62760 1.88280 0.00000
-2.51040 0.00000 0.00000 ; HB- CB- CG2- HG22
2148 2147 2153 2156 3 0.62760 1.88280 0.00000
-2.51040 0.00000 0.00000 ; HB- CB- CG2- HG23
2149 2147 2145 2157 3 5.67350 3.55640 -6.16722
-2.47693 3.74886 0.00000 ; CG1- CB- CA- C
2149 2147 2153 2154 3 0.66944 2.00832 0.00000
-2.67776 0.00000 0.00000 ; CG1- CB- CG2- HG21
2149 2147 2153 2155 3 0.66944 2.00832 0.00000
-2.67776 0.00000 0.00000 ; CG1- CB- CG2- HG22
2149 2147 2153 2156 3 0.66944 2.00832 0.00000
-2.67776 0.00000 0.00000 ; CG1- CB- CG2- HG23
2150 2149 2147 2153 3 0.66944 2.00832 0.00000
-2.67776 0.00000 0.00000 ; HG11- CG1- CB- CG2
2151 2149 2147 2153 3 0.66944 2.00832 0.00000
-2.67776 0.00000 0.00000 ; HG12- CG1- CB- CG2
2152 2149 2147 2153 3 0.66944 2.00832 0.00000
-2.67776 0.00000 0.00000 ; HG13- CG1- CB- CG2
2153 2147 2145 2157 3 5.67350 3.55640 -6.16722
-2.47693 3.74886 0.00000 ; CG2- CB- CA- C
2157 2159 2161 2162 3 0.00000 0.00000 0.00000
0.00000 0.00000 0.00000 ; C- N- CA- HA
2157 2159 2161 2163 3 11.71520 1.67360 15.06240 -
13.38880 0.00000 0.00000 ; C- N- CA- CB
2157 2159 2161 2174 3 1.75728 5.27184 2.25936
-7.02912 0.00000 0.00000 ; C- N- CA- C
2158 2157 2159 2160 3 29.28800 -8.36800 -20.92000
0.00000 0.00000 0.00000 ; O- C- N- H
2158 2157 2159 2161 3 20.92000 0.00000 -20.92000
0.00000 0.00000 0.00000 ; O- C- N- CA
2159 2161 2163 2164 3 0.65084 1.95253 0.00000
-2.60338 0.00000 0.00000 ; N- CA- CB- HB2
2159 2161 2163 2165 3 0.65084 1.95253 0.00000
-2.60338 0.00000 0.00000 ; N- CA- CB- HB3
2159 2161 2163 2166 3 4.79068 4.02919 -4.82834
-3.66518 2.97901 0.00000 ; N- CA- CB- CG
2159 2161 2174 2175 3 0.00000 0.00000 0.00000
0.00000 0.00000 0.00000 ; N- CA- C- O
2159 2161 2174 2176 3 17.40544 -5.02080 -13.22144
9.20480 0.00000 0.00000 ; N- CA- C- N
2160 2159 2161 2162 3 0.00000 0.00000 0.00000

0.00000 0.00000 0.00000 ; H- N- CA- HA
2160 2159 2161 2163 3 0.00000 0.00000 0.00000
0.00000 0.00000 0.00000 ; H- N- CA- CB
2160 2159 2161 2174 3 0.00000 0.00000 0.00000
0.00000 0.00000 0.00000 ; H- N- CA- C
2161 2163 2166 2167 3 5.98312 6.40152 3.28026 -
12.38464 -1.57318 0.00000 ; CA- CB- CG- ND1
2161 2163 2166 2172 3 0.00000 0.00000 0.00000
0.00000 0.00000 0.00000 ; CA- CB- CG- CD2
2161 2174 2176 2177 3 20.92000 0.00000 -20.92000
0.00000 0.00000 0.00000 ; CA- C- N- H
2161 2174 2176 2178 3 20.92000 0.00000 -20.92000
0.00000 0.00000 0.00000 ; CA- C- N- CA
2162 2161 2163 2164 3 0.65084 1.95253 0.00000
-2.60338 0.00000 0.00000 ; HA- CA- CB- HB2
2162 2161 2163 2165 3 0.65084 1.95253 0.00000
-2.60338 0.00000 0.00000 ; HA- CA- CB- HB3
2162 2161 2163 2166 3 0.65084 1.95253 0.00000
-2.60338 0.00000 0.00000 ; HA- CA- CB- CG
2162 2161 2174 2175 3 3.68192 -4.35136 0.00000
1.33888 0.00000 0.00000 ; HA- CA- C- O
2162 2161 2174 2176 3 0.00000 0.00000 0.00000
0.00000 0.00000 0.00000 ; HA- CA- C- N
2163 2161 2174 2175 3 0.00000 0.00000 0.00000
0.00000 0.00000 0.00000 ; CB- CA- C- O
2163 2161 2174 2176 3 2.51040 4.18400 1.67360
-6.69440 0.00000 0.00000 ; CB- CA- C- N
2163 2166 2167 2168 3 20.08320 0.00000 -20.08320
0.00000 0.00000 0.00000 ; CB- CG- ND1- CE1
2163 2166 2172 2170 3 44.97800 0.00000 -44.97800
0.00000 0.00000 0.00000 ; CB- CG- CD2- NE2
2163 2166 2172 2173 3 44.97800 0.00000 -44.97800
0.00000 0.00000 0.00000 ; CB- CG- CD2- HD2
2164 2163 2161 2174 3 0.65084 1.95253 0.00000
-2.60338 0.00000 0.00000 ; HB2- CB- CA- C
2164 2163 2166 2167 3 0.00000 0.00000 0.00000
0.00000 0.00000 0.00000 ; HB2- CB- CG- ND1
2164 2163 2166 2172 3 0.00000 0.00000 0.00000
0.00000 0.00000 0.00000 ; HB2- CB- CG- CD2
2165 2163 2161 2174 3 0.65084 1.95253 0.00000
-2.60338 0.00000 0.00000 ; HB3- CB- CA- C
2165 2163 2166 2167 3 0.00000 0.00000 0.00000
0.00000 0.00000 0.00000 ; HB3- CB- CG- ND1
2165 2163 2166 2172 3 0.00000 0.00000 0.00000
0.00000 0.00000 0.00000 ; HB3- CB- CG- CD2
2166 2163 2161 2174 3 3.76560 3.34720 -2.87859
-3.66518 0.83680 0.00000 ; CG- CB- CA- C
2166 2167 2168 2169 3 41.84000 0.00000 -41.84000
0.00000 0.00000 0.00000 ; CG- ND1- CE1- HE1
2166 2167 2168 2170 3 41.84000 0.00000 -41.84000
0.00000 0.00000 0.00000 ; CG- ND1- CE1- NE2
2166 2172 2170 2168 3 12.55200 0.00000 -12.55200

```
0.00000    0.00000    0.00000 ;      CG-   CD2-   NE2-   CE1
  2166   2172   2170   2171       3   12.55200    0.00000  -12.55200
0.00000    0.00000    0.00000 ;      CG-   CD2-   NE2-   HE2
  2167   2166   2172   2170       3   44.97800    0.00000  -44.97800
0.00000    0.00000    0.00000 ;     ND1-    CG-   CD2-   NE2
  2167   2166   2172   2173       3   44.97800    0.00000  -44.97800
0.00000    0.00000    0.00000 ;     ND1-    CG-   CD2-   HD2
  2167   2168   2170   2171       3   19.45560    0.00000  -19.45560
0.00000    0.00000    0.00000 ;     ND1-   CE1-   NE2-   HE2
  2167   2168   2170   2172       3   19.45560    0.00000  -19.45560
0.00000    0.00000    0.00000 ;     ND1-   CE1-   NE2-   CD2
  2168   2167   2166   2172       3   20.08320    0.00000  -20.08320
0.00000    0.00000    0.00000 ;     CE1-   ND1-    CG-   CD2
  2168   2170   2172   2173       3   12.55200    0.00000  -12.55200
0.00000    0.00000    0.00000 ;     CE1-   NE2-   CD2-   HD2
  2169   2168   2170   2171       3   19.45560    0.00000  -19.45560
0.00000    0.00000    0.00000 ;     HE1-   CE1-   NE2-   HE2
  2169   2168   2170   2172       3   19.45560    0.00000  -19.45560
0.00000    0.00000    0.00000 ;     HE1-   CE1-   NE2-   CD2
  2171   2170   2172   2173       3   12.55200    0.00000  -12.55200
0.00000    0.00000    0.00000 ;     HE2-   NE2-   CD2-   HD2
  2174   2176   2178   2179       3    0.00000    0.00000    0.00000
0.00000    0.00000    0.00000 ;       C-     N-    CA-    HA
  2174   2176   2178   2180       3   11.71520    1.67360   15.06240  -
13.38880    0.00000    0.00000 ;       C-     N-    CA-    CB
  2174   2176   2178   2198       3    1.75728    5.27184    2.25936
-7.02912    0.00000    0.00000 ;       C-     N-    CA-     C
  2175   2174   2176   2177       3   29.28800   -8.36800  -20.92000
0.00000    0.00000    0.00000 ;       O-     C-     N-     H
  2175   2174   2176   2178       3   20.92000    0.00000  -20.92000
0.00000    0.00000    0.00000 ;       O-     C-     N-    CA
  2176   2178   2180   2181       3    0.65084    1.95253    0.00000
-2.60338    0.00000    0.00000 ;       N-    CA-    CB-   HB2
  2176   2178   2180   2182       3    0.65084    1.95253    0.00000
-2.60338    0.00000    0.00000 ;       N-    CA-    CB-   HB3
  2176   2178   2180   2183       3    0.65084    1.95253    0.00000
-2.60338    0.00000    0.00000 ;       N-    CA-    CB-    CG
  2176   2178   2198   2199       3    0.00000    0.00000    0.00000
0.00000    0.00000    0.00000 ;       N-    CA-     C-     O
  2176   2178   2198   2200       3   17.40544   -5.02080  -13.22144
9.20480    0.00000    0.00000 ;       N-    CA-     C-     N
  2177   2176   2178   2179       3    0.00000    0.00000    0.00000
0.00000    0.00000    0.00000 ;       H-     N-    CA-    HA
  2177   2176   2178   2180       3    0.00000    0.00000    0.00000
0.00000    0.00000    0.00000 ;       H-     N-    CA-    CB
  2177   2176   2178   2198       3    0.00000    0.00000    0.00000
0.00000    0.00000    0.00000 ;       H-     N-    CA-     C
  2178   2180   2183   2184       3    0.66944    2.00832    0.00000
-2.67776    0.00000    0.00000 ;      CA-    CB-    CG-   HG2
  2178   2180   2183   2185       3    0.66944    2.00832    0.00000
-2.67776    0.00000    0.00000 ;      CA-    CB-    CG-   HG3
  2178   2180   2183   2186       3    3.68192    3.09616   -2.09200
```

-3.01248    0.00000    0.00000 ;      CA-    CB-    CG-    CD
  2178   2198   2200   2201      3   20.92000    0.00000  -20.92000
0.00000    0.00000    0.00000 ;      CA-    C-     N-     H
  2178   2198   2200   2202      3   20.92000    0.00000  -20.92000
0.00000    0.00000    0.00000 ;      CA-    C-     N-     CA
  2179   2178   2180   2181      3    0.65084    1.95253    0.00000
-2.60338    0.00000    0.00000 ;      HA-    CA-    CB-    HB2
  2179   2178   2180   2182      3    0.65084    1.95253    0.00000
-2.60338    0.00000    0.00000 ;      HA-    CA-    CB-    HB3
  2179   2178   2180   2183      3    0.65084    1.95253    0.00000
-2.60338    0.00000    0.00000 ;      HA-    CA-    CB-    CG
  2179   2178   2198   2199      3    3.68192   -4.35136    0.00000
1.33888    0.00000    0.00000 ;      HA-    CA-    C-     O
  2179   2178   2198   2200      3    0.00000    0.00000    0.00000
0.00000    0.00000    0.00000 ;      HA-    CA-    C-     N
  2180   2178   2198   2199      3    0.00000    0.00000    0.00000
0.00000    0.00000    0.00000 ;      CB-    CA-    C-     O
  2180   2178   2198   2200      3    2.51040    4.18400    1.67360
-6.69440    0.00000    0.00000 ;      CB-    CA-    C-     N
  2180   2183   2186   2187      3    0.65084    1.95253    0.00000
-2.60338    0.00000    0.00000 ;      CB-    CG-    CD-    HD2
  2180   2183   2186   2188      3    0.65084    1.95253    0.00000
-2.60338    0.00000    0.00000 ;      CB-    CG-    CD-    HD3
  2180   2183   2186   2189      3    0.65084    1.95253    0.00000
-2.60338    0.00000    0.00000 ;      CB-    CG-    CD-    NE
  2181   2180   2178   2198      3    0.65084    1.95253    0.00000
-2.60338    0.00000    0.00000 ;      HB2-   CB-    CA-    C
  2181   2180   2183   2184      3    0.62760    1.88280    0.00000
-2.51040    0.00000    0.00000 ;      HB2-   CB-    CG-    HG2
  2181   2180   2183   2185      3    0.62760    1.88280    0.00000
-2.51040    0.00000    0.00000 ;      HB2-   CB-    CG-    HG3
  2181   2180   2183   2186      3    0.66944    2.00832    0.00000
-2.67776    0.00000    0.00000 ;      HB2-   CB-    CG-    CD
  2182   2180   2178   2198      3    0.65084    1.95253    0.00000
-2.60338    0.00000    0.00000 ;      HB3-   CB-    CA-    C
  2182   2180   2183   2184      3    0.62760    1.88280    0.00000
-2.51040    0.00000    0.00000 ;      HB3-   CB-    CG-    HG2
  2182   2180   2183   2185      3    0.62760    1.88280    0.00000
-2.51040    0.00000    0.00000 ;      HB3-   CB-    CG-    HG3
  2182   2180   2183   2186      3    0.66944    2.00832    0.00000
-2.67776    0.00000    0.00000 ;      HB3-   CB-    CG-    CD
  2183   2180   2178   2198      3    0.65084    1.95253    0.00000
-2.60338    0.00000    0.00000 ;      CG-    CB-    CA-    C
  2183   2186   2189   2190      3    0.00000    0.00000    0.00000
0.00000    0.00000    0.00000 ;      CG-    CD-    NE-    HE
  2183   2186   2189   2191      3    0.00000    0.00000    0.00000
0.00000    0.00000    0.00000 ;      CG-    CD-    NE-    CZ
  2184   2183   2186   2187      3    0.65084    1.95253    0.00000
-2.60338    0.00000    0.00000 ;      HG2-   CG-    CD-    HD2
  2184   2183   2186   2188      3    0.65084    1.95253    0.00000
-2.60338    0.00000    0.00000 ;      HG2-   CG-    CD-    HD3
  2184   2183   2186   2189      3    0.65084    1.95253    0.00000

-2.60338 0.00000 0.00000 ; HG2- CG- CD- NE
2185 2183 2186 2187 3 0.65084 1.95253 0.00000
-2.60338 0.00000 0.00000 ; HG3- CG- CD- HD2
2185 2183 2186 2188 3 0.65084 1.95253 0.00000
-2.60338 0.00000 0.00000 ; HG3- CG- CD- HD3
2185 2183 2186 2189 3 0.65084 1.95253 0.00000
-2.60338 0.00000 0.00000 ; HG3- CG- CD- NE
2186 2189 2191 2192 3 20.08320 0.00000 -20.08320
0.00000 0.00000 0.00000 ; CD- NE- CZ- NH1
2186 2189 2191 2195 3 20.08320 0.00000 -20.08320
0.00000 0.00000 0.00000 ; CD- NE- CZ- NH2
2187 2186 2189 2190 3 0.00000 0.00000 0.00000
0.00000 0.00000 0.00000 ; HD2- CD- NE- HE
2187 2186 2189 2191 3 0.00000 0.00000 0.00000
0.00000 0.00000 0.00000 ; HD2- CD- NE- CZ
2188 2186 2189 2190 3 0.00000 0.00000 0.00000
0.00000 0.00000 0.00000 ; HD3- CD- NE- HE
2188 2186 2189 2191 3 0.00000 0.00000 0.00000
0.00000 0.00000 0.00000 ; HD3- CD- NE- CZ
2189 2191 2192 2193 3 20.08320 0.00000 -20.08320
0.00000 0.00000 0.00000 ; NE- CZ- NH1- HH11
2189 2191 2192 2194 3 20.08320 0.00000 -20.08320
0.00000 0.00000 0.00000 ; NE- CZ- NH1- HH12
2189 2191 2195 2196 3 20.08320 0.00000 -20.08320
0.00000 0.00000 0.00000 ; NE- CZ- NH2- HH21
2189 2191 2195 2197 3 20.08320 0.00000 -20.08320
0.00000 0.00000 0.00000 ; NE- CZ- NH2- HH22
2190 2189 2191 2192 3 20.08320 0.00000 -20.08320
0.00000 0.00000 0.00000 ; HE- NE- CZ- NH1
2190 2189 2191 2195 3 20.08320 0.00000 -20.08320
0.00000 0.00000 0.00000 ; HE- NE- CZ- NH2
2192 2191 2195 2196 3 20.08320 0.00000 -20.08320
0.00000 0.00000 0.00000 ; NH1- CZ- NH2- HH21
2192 2191 2195 2197 3 20.08320 0.00000 -20.08320
0.00000 0.00000 0.00000 ; NH1- CZ- NH2- HH22
2193 2192 2191 2195 3 20.08320 0.00000 -20.08320
0.00000 0.00000 0.00000 ; HH11- NH1- CZ- NH2
2194 2192 2191 2195 3 20.08320 0.00000 -20.08320
0.00000 0.00000 0.00000 ; HH12- NH1- CZ- NH2
2198 2200 2202 2203 3 0.00000 0.00000 0.00000
0.00000 0.00000 0.00000 ; C- N- CA- HA
2198 2200 2202 2204 3 11.71520 1.67360 15.06240 -
13.38880 0.00000 0.00000 ; C- N- CA- CB
2198 2200 2202 2210 3 1.75728 5.27184 2.25936
-7.02912 0.00000 0.00000 ; C- N- CA- C
2199 2198 2200 2201 3 29.28800 -8.36800 -20.92000
0.00000 0.00000 0.00000 ; O- C- N- H
2199 2198 2200 2202 3 20.92000 0.00000 -20.92000
0.00000 0.00000 0.00000 ; O- C- N- CA
2200 2202 2204 2205 3 0.65084 1.95253 0.00000
-2.60338 0.00000 0.00000 ; N- CA- CB- HB2
2200 2202 2204 2206 3 0.65084 1.95253 0.00000

-2.60338 0.00000 0.00000 ; N- CA- CB- HB3
2200 2202 2204 2207 3 15.41386 9.74035 -8.39310
-0.97069 2.97901 0.00000 ; N- CA- CB- CG
2200 2202 2210 2211 3 0.00000 0.00000 0.00000
0.00000 0.00000 0.00000 ; N- CA- C- O
2200 2202 2210 2212 3 17.40544 -5.02080 -13.22144
9.20480 0.00000 0.00000 ; N- CA- C- N
2201 2200 2202 2203 3 0.00000 0.00000 0.00000
0.00000 0.00000 0.00000 ; H- N- CA- HA
2201 2200 2202 2204 3 0.00000 0.00000 0.00000
0.00000 0.00000 0.00000 ; H- N- CA- CB
2201 2200 2202 2210 3 0.00000 0.00000 0.00000
0.00000 0.00000 0.00000 ; H- N- CA- C
2202 2204 2207 2208 3 6.43499 0.00000 -5.39736
0.00000 -1.03763 0.00000 ; CA- CB- CG- OD1
2202 2204 2207 2209 3 6.43499 0.00000 -5.39736
0.00000 -1.03763 0.00000 ; CA- CB- CG- OD2
2202 2210 2212 2213 3 20.92000 0.00000 -20.92000
0.00000 0.00000 0.00000 ; CA- C- N- H
2202 2210 2212 2214 3 20.92000 0.00000 -20.92000
0.00000 0.00000 0.00000 ; CA- C- N- CA
2203 2202 2204 2205 3 0.65084 1.95253 0.00000
-2.60338 0.00000 0.00000 ; HA- CA- CB- HB2
2203 2202 2204 2206 3 0.65084 1.95253 0.00000
-2.60338 0.00000 0.00000 ; HA- CA- CB- HB3
2203 2202 2204 2207 3 0.65084 1.95253 0.00000
-2.60338 0.00000 0.00000 ; HA- CA- CB- CG
2203 2202 2210 2211 3 3.68192 -4.35136 0.00000
1.33888 0.00000 0.00000 ; HA- CA- C- O
2203 2202 2210 2212 3 0.00000 0.00000 0.00000
0.00000 0.00000 0.00000 ; HA- CA- C- N
2204 2202 2210 2211 3 0.00000 0.00000 0.00000
0.00000 0.00000 0.00000 ; CB- CA- C- O
2204 2202 2210 2212 3 2.51040 4.18400 1.67360
-6.69440 0.00000 0.00000 ; CB- CA- C- N
2205 2204 2202 2210 3 0.65084 1.95253 0.00000
-2.60338 0.00000 0.00000 ; HB2- CB- CA- C
2205 2204 2207 2208 3 0.00000 0.00000 0.00000
0.00000 0.00000 0.00000 ; HB2- CB- CG- OD1
2205 2204 2207 2209 3 0.00000 0.00000 0.00000
0.00000 0.00000 0.00000 ; HB2- CB- CG- OD2
2206 2204 2202 2210 3 0.65084 1.95253 0.00000
-2.60338 0.00000 0.00000 ; HB3- CB- CA- C
2206 2204 2207 2208 3 0.00000 0.00000 0.00000
0.00000 0.00000 0.00000 ; HB3- CB- CG- OD1
2206 2204 2207 2209 3 0.00000 0.00000 0.00000
0.00000 0.00000 0.00000 ; HB3- CB- CG- OD2
2207 2204 2202 2210 3 7.14627 -1.04600 -8.99560
-0.97069 5.15469 0.00000 ; CG- CB- CA- C
2210 2212 2214 2215 3 0.00000 0.00000 0.00000
0.00000 0.00000 0.00000 ; C- N- CA- HA
2210 2212 2214 2216 3 11.71520 1.67360 15.06240 -

```
13.38880    0.00000    0.00000 ;      C-     N-    CA-    CB
  2210   2212   2214   2229      3    1.75728    5.27184    2.25936
-7.02912    0.00000    0.00000 ;      C-     N-    CA-     C
  2211   2210   2212   2213      3   29.28800   -8.36800  -20.92000
0.00000    0.00000    0.00000 ;      O-     C-     N-     H
  2211   2210   2212   2214      3   20.92000    0.00000  -20.92000
0.00000    0.00000    0.00000 ;      O-     C-     N-    CA
  2212   2214   2216   2217      3    0.65084    1.95253    0.00000
-2.60338    0.00000    0.00000 ;      N-    CA-    CB-   HB2
  2212   2214   2216   2218      3    0.65084    1.95253    0.00000
-2.60338    0.00000    0.00000 ;      N-    CA-    CB-   HB3
  2212   2214   2216   2219      3    3.79070    1.39746   -4.61077
-2.40998    2.44346    0.00000 ;      N-    CA-    CB-    CG
  2212   2214   2229   2230      3    0.00000    0.00000    0.00000
0.00000    0.00000    0.00000 ;      N-    CA-     C-     O
  2212   2214   2229   2231      3   17.40544   -5.02080  -13.22144
9.20480    0.00000    0.00000 ;      N-    CA-     C-     N
  2213   2212   2214   2215      3    0.00000    0.00000    0.00000
0.00000    0.00000    0.00000 ;      H-     N-    CA-    HA
  2213   2212   2214   2216      3    0.00000    0.00000    0.00000
0.00000    0.00000    0.00000 ;      H-     N-    CA-    CB
  2213   2212   2214   2229      3    0.00000    0.00000    0.00000
0.00000    0.00000    0.00000 ;      H-     N-    CA-     C
  2214   2216   2219   2220      3    0.66944    2.00832    0.00000
-2.67776    0.00000    0.00000 ;     CA-    CB-    CG-    HG
  2214   2216   2219   2221      3    3.90367    0.19665   -6.21742
-2.37651    5.99149    0.00000 ;     CA-    CB-    CG-   CD1
  2214   2216   2219   2225      3    3.90367    0.19665   -6.21742
-2.37651    5.99149    0.00000 ;     CA-    CB-    CG-   CD2
  2214   2229   2231   2232      3   20.92000    0.00000  -20.92000
0.00000    0.00000    0.00000 ;     CA-     C-     N-     H
  2214   2229   2231   2233      3   20.92000    0.00000  -20.92000
0.00000    0.00000    0.00000 ;     CA-     C-     N-    CA
  2215   2214   2216   2217      3    0.65084    1.95253    0.00000
-2.60338    0.00000    0.00000 ;     HA-    CA-    CB-   HB2
  2215   2214   2216   2218      3    0.65084    1.95253    0.00000
-2.60338    0.00000    0.00000 ;     HA-    CA-    CB-   HB3
  2215   2214   2216   2219      3    0.65084    1.95253    0.00000
-2.60338    0.00000    0.00000 ;     HA-    CA-    CB-    CG
  2215   2214   2229   2230      3    3.68192   -4.35136    0.00000
1.33888    0.00000    0.00000 ;     HA-    CA-     C-     O
  2215   2214   2229   2231      3    0.00000    0.00000    0.00000
0.00000    0.00000    0.00000 ;     HA-    CA-     C-     N
  2216   2214   2229   2230      3    0.00000    0.00000    0.00000
0.00000    0.00000    0.00000 ;     CB-    CA-     C-     O
  2216   2214   2229   2231      3    2.51040    4.18400    1.67360
-6.69440    0.00000    0.00000 ;     CB-    CA-     C-     N
  2216   2219   2221   2222      3    0.66944    2.00832    0.00000
-2.67776    0.00000    0.00000 ;     CB-    CG-   CD1-  HD11
  2216   2219   2221   2223      3    0.66944    2.00832    0.00000
-2.67776    0.00000    0.00000 ;     CB-    CG-   CD1-  HD12
  2216   2219   2221   2224      3    0.66944    2.00832    0.00000
```

-2.67776 0.00000 0.00000 ; CB- CG- CD1- HD13
2216 2219 2225 2226 3 0.66944 2.00832 0.00000
-2.67776 0.00000 0.00000 ; CB- CG- CD2- HD21
2216 2219 2225 2227 3 0.66944 2.00832 0.00000
-2.67776 0.00000 0.00000 ; CB- CG- CD2- HD22
2216 2219 2225 2228 3 0.66944 2.00832 0.00000
-2.67776 0.00000 0.00000 ; CB- CG- CD2- HD23
2217 2216 2214 2229 3 0.65084 1.95253 0.00000
-2.60338 0.00000 0.00000 ; HB2- CB- CA- C
2217 2216 2219 2220 3 0.62760 1.88280 0.00000
-2.51040 0.00000 0.00000 ; HB2- CB- CG- HG
2217 2216 2219 2221 3 0.66944 2.00832 0.00000
-2.67776 0.00000 0.00000 ; HB2- CB- CG- CD1
2217 2216 2219 2225 3 0.66944 2.00832 0.00000
-2.67776 0.00000 0.00000 ; HB2- CB- CG- CD2
2218 2216 2214 2229 3 0.65084 1.95253 0.00000
-2.60338 0.00000 0.00000 ; HB3- CB- CA- C
2218 2216 2219 2220 3 0.62760 1.88280 0.00000
-2.51040 0.00000 0.00000 ; HB3- CB- CG- HG
2218 2216 2219 2221 3 0.66944 2.00832 0.00000
-2.67776 0.00000 0.00000 ; HB3- CB- CG- CD1
2218 2216 2219 2225 3 0.66944 2.00832 0.00000
-2.67776 0.00000 0.00000 ; HB3- CB- CG- CD2
2219 2216 2214 2229 3 10.33448 -1.14642 -11.54784
-2.40998 6.35968 0.00000 ; CG- CB- CA- C
2220 2219 2221 2222 3 0.62760 1.88280 0.00000
-2.51040 0.00000 0.00000 ; HG- CG- CD1- HD11
2220 2219 2221 2223 3 0.62760 1.88280 0.00000
-2.51040 0.00000 0.00000 ; HG- CG- CD1- HD12
2220 2219 2221 2224 3 0.62760 1.88280 0.00000
-2.51040 0.00000 0.00000 ; HG- CG- CD1- HD13
2220 2219 2225 2226 3 0.62760 1.88280 0.00000
-2.51040 0.00000 0.00000 ; HG- CG- CD2- HD21
2220 2219 2225 2227 3 0.62760 1.88280 0.00000
-2.51040 0.00000 0.00000 ; HG- CG- CD2- HD22
2220 2219 2225 2228 3 0.62760 1.88280 0.00000
-2.51040 0.00000 0.00000 ; HG- CG- CD2- HD23
2221 2219 2225 2226 3 0.66944 2.00832 0.00000
-2.67776 0.00000 0.00000 ; CD1- CG- CD2- HD21
2221 2219 2225 2227 3 0.66944 2.00832 0.00000
-2.67776 0.00000 0.00000 ; CD1- CG- CD2- HD22
2221 2219 2225 2228 3 0.66944 2.00832 0.00000
-2.67776 0.00000 0.00000 ; CD1- CG- CD2- HD23
2222 2221 2219 2225 3 0.66944 2.00832 0.00000
-2.67776 0.00000 0.00000 ; HD11- CD1- CG- CD2
2223 2221 2219 2225 3 0.66944 2.00832 0.00000
-2.67776 0.00000 0.00000 ; HD12- CD1- CG- CD2
2224 2221 2219 2225 3 0.66944 2.00832 0.00000
-2.67776 0.00000 0.00000 ; HD13- CD1- CG- CD2
2229 2231 2233 2234 3 0.00000 0.00000 0.00000
0.00000 0.00000 0.00000 ; C- N- CA- HA
2229 2231 2233 2235 3 11.71520 1.67360 15.06240 -

13.38880    0.00000    0.00000 ;      C-     N-    CA-    CB
  2229   2231   2233   2251      3    1.75728    5.27184    2.25936
-7.02912    0.00000    0.00000 ;      C-     N-    CA-    C
  2230   2229   2231   2232      3   29.28800   -8.36800  -20.92000
0.00000    0.00000    0.00000 ;      O-     C-     N-    H
  2230   2229   2231   2233      3   20.92000    0.00000  -20.92000
0.00000    0.00000    0.00000 ;      O-     C-     N-    CA
  2231   2233   2235   2236      3    0.65084    1.95253    0.00000
-2.60338    0.00000    0.00000 ;      N-    CA-    CB-   HB2
  2231   2233   2235   2237      3    0.65084    1.95253    0.00000
-2.60338    0.00000    0.00000 ;      N-    CA-    CB-   HB3
  2231   2233   2235   2238      3    0.65084    1.95253    0.00000
-2.60338    0.00000    0.00000 ;      N-    CA-    CB-    CG
  2231   2233   2251   2252      3    0.00000    0.00000    0.00000
0.00000    0.00000    0.00000 ;      N-    CA-     C-    O
  2231   2233   2251   2253      3   17.40544   -5.02080  -13.22144
9.20480    0.00000    0.00000 ;      N-    CA-     C-    N
  2232   2231   2233   2234      3    0.00000    0.00000    0.00000
0.00000    0.00000    0.00000 ;      H-     N-    CA-    HA
  2232   2231   2233   2235      3    0.00000    0.00000    0.00000
0.00000    0.00000    0.00000 ;      H-     N-    CA-    CB
  2232   2231   2233   2251      3    0.00000    0.00000    0.00000
0.00000    0.00000    0.00000 ;      H-     N-    CA-    C
  2233   2235   2238   2239      3    0.66944    2.00832    0.00000
-2.67776    0.00000    0.00000 ;     CA-    CB-    CG-   HG2
  2233   2235   2238   2240      3    0.66944    2.00832    0.00000
-2.67776    0.00000    0.00000 ;     CA-    CB-    CG-   HG3
  2233   2235   2238   2241      3    3.68192    3.09616   -2.09200
-3.01248    0.00000    0.00000 ;     CA-    CB-    CG-    CD
  2233   2251   2253   2254      3   20.92000    0.00000  -20.92000
0.00000    0.00000    0.00000 ;     CA-     C-     N-   CD
  2233   2251   2253   2263      3   20.92000    0.00000  -20.92000
0.00000    0.00000    0.00000 ;     CA-     C-     N-   CA
  2234   2233   2235   2236      3    0.65084    1.95253    0.00000
-2.60338    0.00000    0.00000 ;     HA-    CA-    CB-   HB2
  2234   2233   2235   2237      3    0.65084    1.95253    0.00000
-2.60338    0.00000    0.00000 ;     HA-    CA-    CB-   HB3
  2234   2233   2235   2238      3    0.65084    1.95253    0.00000
-2.60338    0.00000    0.00000 ;     HA-    CA-    CB-    CG
  2234   2233   2251   2252      3    3.68192   -4.35136    0.00000
1.33888    0.00000    0.00000 ;     HA-    CA-     C-    O
  2234   2233   2251   2253      3    0.00000    0.00000    0.00000
0.00000    0.00000    0.00000 ;     HA-    CA-     C-    N
  2235   2233   2251   2252      3    0.00000    0.00000    0.00000
0.00000    0.00000    0.00000 ;     CB-    CA-     C-    O
  2235   2233   2251   2253      3    2.51040    4.18400    1.67360
-6.69440    0.00000    0.00000 ;     CB-    CA-     C-     N
  2235   2238   2241   2242      3    0.66944    2.00832    0.00000
-2.67776    0.00000    0.00000 ;     CB-    CG-    CD-   HD2
  2235   2238   2241   2243      3    0.66944    2.00832    0.00000
-2.67776    0.00000    0.00000 ;     CB-    CG-    CD-   HD3
  2235   2238   2241   2244      3    3.68192    3.09616   -2.09200

-3.01248    0.00000    0.00000 ;     CB-    CG-    CD-    CE
 2236   2235   2233   2251      3    0.65084    1.95253    0.00000
-2.60338    0.00000    0.00000 ;    HB2-    CB-    CA-     C
 2236   2235   2238   2239      3    0.62760    1.88280    0.00000
-2.51040    0.00000    0.00000 ;    HB2-    CB-    CG-   HG2
 2236   2235   2238   2240      3    0.62760    1.88280    0.00000
-2.51040    0.00000    0.00000 ;    HB2-    CB-    CG-   HG3
 2236   2235   2238   2241      3    0.66944    2.00832    0.00000
-2.67776    0.00000    0.00000 ;    HB2-    CB-    CG-    CD
 2237   2235   2233   2251      3    0.65084    1.95253    0.00000
-2.60338    0.00000    0.00000 ;    HB3-    CB-    CA-     C
 2237   2235   2238   2239      3    0.62760    1.88280    0.00000
-2.51040    0.00000    0.00000 ;    HB3-    CB-    CG-   HG2
 2237   2235   2238   2240      3    0.62760    1.88280    0.00000
-2.51040    0.00000    0.00000 ;    HB3-    CB-    CG-   HG3
 2237   2235   2238   2241      3    0.66944    2.00832    0.00000
-2.67776    0.00000    0.00000 ;    HB3-    CB-    CG-    CD
 2238   2235   2233   2251      3    0.65084    1.95253    0.00000
-2.60338    0.00000    0.00000 ;     CG-    CB-    CA-     C
 2238   2241   2244   2245      3    0.65084    1.95253    0.00000
-2.60338    0.00000    0.00000 ;     CG-    CD-    CE-   HE2
 2238   2241   2244   2246      3    0.65084    1.95253    0.00000
-2.60338    0.00000    0.00000 ;     CG-    CD-    CE-   HE3
 2238   2241   2244   2247      3    0.65084    1.95253    0.00000
-2.60338    0.00000    0.00000 ;     CG-    CD-    CE-    NZ
 2239   2238   2241   2242      3    0.62760    1.88280    0.00000
-2.51040    0.00000    0.00000 ;    HG2-    CG-    CD-   HD2
 2239   2238   2241   2243      3    0.62760    1.88280    0.00000
-2.51040    0.00000    0.00000 ;    HG2-    CG-    CD-   HD3
 2239   2238   2241   2244      3    0.66944    2.00832    0.00000
-2.67776    0.00000    0.00000 ;    HG2-    CG-    CD-    CE
 2240   2238   2241   2242      3    0.62760    1.88280    0.00000
-2.51040    0.00000    0.00000 ;    HG3-    CG-    CD-   HD2
 2240   2238   2241   2243      3    0.62760    1.88280    0.00000
-2.51040    0.00000    0.00000 ;    HG3-    CG-    CD-   HD3
 2240   2238   2241   2244      3    0.66944    2.00832    0.00000
-2.67776    0.00000    0.00000 ;    HG3-    CG-    CD-    CE
 2241   2244   2247   2248      3    0.65084    1.95253    0.00000
-2.60338    0.00000    0.00000 ;     CD-    CE-    NZ-   HZ1
 2241   2244   2247   2249      3    0.65084    1.95253    0.00000
-2.60338    0.00000    0.00000 ;     CD-    CE-    NZ-   HZ2
 2241   2244   2247   2250      3    0.65084    1.95253    0.00000
-2.60338    0.00000    0.00000 ;     CD-    CE-    NZ-   HZ3
 2242   2241   2244   2245      3    0.65084    1.95253    0.00000
-2.60338    0.00000    0.00000 ;    HD2-    CD-    CE-   HE2
 2242   2241   2244   2246      3    0.65084    1.95253    0.00000
-2.60338    0.00000    0.00000 ;    HD2-    CD-    CE-   HE3
 2242   2241   2244   2247      3    0.65084    1.95253    0.00000
-2.60338    0.00000    0.00000 ;    HD2-    CD-    CE-    NZ
 2243   2241   2244   2245      3    0.65084    1.95253    0.00000
-2.60338    0.00000    0.00000 ;    HD3-    CD-    CE-   HE2
 2243   2241   2244   2246      3    0.65084    1.95253    0.00000

-2.60338 0.00000 0.00000 ; HD3- CD- CE- HE3
2243 2241 2244 2247 3 0.65084 1.95253 0.00000
-2.60338 0.00000 0.00000 ; HD3- CD- CE- NZ
2245 2244 2247 2248 3 0.65084 1.95253 0.00000
-2.60338 0.00000 0.00000 ; HE2- CE- NZ- HZ1
2245 2244 2247 2249 3 0.65084 1.95253 0.00000
-2.60338 0.00000 0.00000 ; HE2- CE- NZ- HZ2
2245 2244 2247 2250 3 0.65084 1.95253 0.00000
-2.60338 0.00000 0.00000 ; HE2- CE- NZ- HZ3
2246 2244 2247 2248 3 0.65084 1.95253 0.00000
-2.60338 0.00000 0.00000 ; HE3- CE- NZ- HZ1
2246 2244 2247 2249 3 0.65084 1.95253 0.00000
-2.60338 0.00000 0.00000 ; HE3- CE- NZ- HZ2
2246 2244 2247 2250 3 0.65084 1.95253 0.00000
-2.60338 0.00000 0.00000 ; HE3- CE- NZ- HZ3
2251 2253 2254 2255 3 0.00000 0.00000 0.00000
0.00000 0.00000 0.00000 ; C- N- CD- HD2
2251 2253 2254 2256 3 0.00000 0.00000 0.00000
0.00000 0.00000 0.00000 ; C- N- CD- HD3
2251 2253 2254 2257 3 10.04160 -3.34720 16.73600
-6.69440 0.00000 0.00000 ; C- N- CD- CG
2251 2253 2263 2260 3 11.71520 1.67360 15.06240 -
13.38880 0.00000 0.00000 ; C- N- CA- CB
2251 2253 2263 2264 3 0.00000 0.00000 0.00000
0.00000 0.00000 0.00000 ; C- N- CA- HA
2251 2253 2263 2265 3 1.75728 5.27184 2.25936
-7.02912 0.00000 0.00000 ; C- N- CA- C
2252 2251 2253 2254 3 20.92000 0.00000 -20.92000
0.00000 0.00000 0.00000 ; O- C- N- CD
2252 2251 2253 2263 3 20.92000 0.00000 -20.92000
0.00000 0.00000 0.00000 ; O- C- N- CA
2253 2254 2257 2258 3 0.65084 1.95253 0.00000
-2.60338 0.00000 0.00000 ; N- CD- CG- HG2
2253 2254 2257 2259 3 0.65084 1.95253 0.00000
-2.60338 0.00000 0.00000 ; N- CD- CG- HG3
2253 2254 2257 2260 3 0.65084 1.95253 0.00000
-2.60338 0.00000 0.00000 ; N- CD- CG- CB
2253 2263 2260 2257 3 0.65084 1.95253 0.00000
-2.60338 0.00000 0.00000 ; N- CA- CB- CG
2253 2263 2260 2261 3 0.65084 1.95253 0.00000
-2.60338 0.00000 0.00000 ; N- CA- CB- HB2
2253 2263 2260 2262 3 0.65084 1.95253 0.00000
-2.60338 0.00000 0.00000 ; N- CA- CB- HB3
2253 2263 2265 2266 3 0.00000 0.00000 0.00000
0.00000 0.00000 0.00000 ; N- CA- C- O
2253 2263 2265 2267 3 17.40544 -5.02080 -13.22144
9.20480 0.00000 0.00000 ; N- CA- C- N
2254 2253 2263 2260 3 0.00000 0.00000 0.00000
0.00000 0.00000 0.00000 ; CD- N- CA- CB
2254 2253 2263 2264 3 0.00000 0.00000 0.00000
0.00000 0.00000 0.00000 ; CD- N- CA- HA
2254 2253 2263 2265 3 0.00000 0.00000 0.00000

0.00000 0.00000 0.00000 ; CD- N- CA- C
2254 2257 2260 2261 3 0.66944 2.00832 0.00000
-2.67776 0.00000 0.00000 ; CD- CG- CB- HB2
2254 2257 2260 2262 3 0.66944 2.00832 0.00000
-2.67776 0.00000 0.00000 ; CD- CG- CB- HB3
2254 2257 2260 2263 3 3.68192 3.09616 -2.09200
-3.01248 0.00000 0.00000 ; CD- CG- CB- CA
2255 2254 2253 2263 3 0.00000 0.00000 0.00000
0.00000 0.00000 0.00000 ; HD2- CD- N- CA
2255 2254 2257 2258 3 0.65084 1.95253 0.00000
-2.60338 0.00000 0.00000 ; HD2- CD- CG- HG2
2255 2254 2257 2259 3 0.65084 1.95253 0.00000
-2.60338 0.00000 0.00000 ; HD2- CD- CG- HG3
2255 2254 2257 2260 3 0.65084 1.95253 0.00000
-2.60338 0.00000 0.00000 ; HD2- CD- CG- CB
2256 2254 2253 2263 3 0.00000 0.00000 0.00000
0.00000 0.00000 0.00000 ; HD3- CD- N- CA
2256 2254 2257 2258 3 0.65084 1.95253 0.00000
-2.60338 0.00000 0.00000 ; HD3- CD- CG- HG2
2256 2254 2257 2259 3 0.65084 1.95253 0.00000
-2.60338 0.00000 0.00000 ; HD3- CD- CG- HG3
2256 2254 2257 2260 3 0.65084 1.95253 0.00000
-2.60338 0.00000 0.00000 ; HD3- CD- CG- CB
2257 2254 2253 2263 3 0.00000 0.00000 0.00000
0.00000 0.00000 0.00000 ; CG- CD- N- CA
2257 2260 2263 2264 3 0.65084 1.95253 0.00000
-2.60338 0.00000 0.00000 ; CG- CB- CA- HA
2257 2260 2263 2265 3 0.65084 1.95253 0.00000
-2.60338 0.00000 0.00000 ; CG- CB- CA- C
2258 2257 2260 2261 3 0.62760 1.88280 0.00000
-2.51040 0.00000 0.00000 ; HG2- CG- CB- HB2
2258 2257 2260 2262 3 0.62760 1.88280 0.00000
-2.51040 0.00000 0.00000 ; HG2- CG- CB- HB3
2258 2257 2260 2263 3 0.66944 2.00832 0.00000
-2.67776 0.00000 0.00000 ; HG2- CG- CB- CA
2259 2257 2260 2261 3 0.62760 1.88280 0.00000
-2.51040 0.00000 0.00000 ; HG3- CG- CB- HB2
2259 2257 2260 2262 3 0.62760 1.88280 0.00000
-2.51040 0.00000 0.00000 ; HG3- CG- CB- HB3
2259 2257 2260 2263 3 0.66944 2.00832 0.00000
-2.67776 0.00000 0.00000 ; HG3- CG- CB- CA
2260 2263 2265 2266 3 0.00000 0.00000 0.00000
0.00000 0.00000 0.00000 ; CB- CA- C- O
2260 2263 2265 2267 3 2.51040 4.18400 1.67360
-6.69440 0.00000 0.00000 ; CB- CA- C- N
2261 2260 2263 2264 3 0.65084 1.95253 0.00000
-2.60338 0.00000 0.00000 ; HB2- CB- CA- HA
2261 2260 2263 2265 3 0.65084 1.95253 0.00000
-2.60338 0.00000 0.00000 ; HB2- CB- CA- C
2262 2260 2263 2264 3 0.65084 1.95253 0.00000
-2.60338 0.00000 0.00000 ; HB3- CB- CA- HA
2262 2260 2263 2265 3 0.65084 1.95253 0.00000

-2.60338 0.00000 0.00000 ; HB3- CB- CA- C
2263 2265 2267 2268 3 20.92000 0.00000 -20.92000
0.00000 0.00000 0.00000 ; CA- C- N- H
2263 2265 2267 2269 3 20.92000 0.00000 -20.92000
0.00000 0.00000 0.00000 ; CA- C- N- CA
2264 2263 2265 2266 3 3.68192 -4.35136 0.00000
1.33888 0.00000 0.00000 ; HA- CA- C- O
2264 2263 2265 2267 3 0.00000 0.00000 0.00000
0.00000 0.00000 0.00000 ; HA- CA- C- N
2265 2267 2269 2270 3 0.00000 0.00000 0.00000
0.00000 0.00000 0.00000 ; C- N- CA- HA
2265 2267 2269 2271 3 11.71520 1.67360 15.06240 -
13.38880 0.00000 0.00000 ; C- N- CA- CB
2265 2267 2269 2280 3 1.75728 5.27184 2.25936
-7.02912 0.00000 0.00000 ; C- N- CA- C
2266 2265 2267 2268 3 29.28800 -8.36800 -20.92000
0.00000 0.00000 0.00000 ; O- C- N- H
2266 2265 2267 2269 3 20.92000 0.00000 -20.92000
0.00000 0.00000 0.00000 ; O- C- N- CA
2267 2269 2271 2272 3 0.65084 1.95253 0.00000
-2.60338 0.00000 0.00000 ; N- CA- CB- HB2
2267 2269 2271 2273 3 0.65084 1.95253 0.00000
-2.60338 0.00000 0.00000 ; N- CA- CB- HB3
2267 2269 2271 2274 3 3.21331 2.22589 -4.15053
-2.40998 2.61082 0.00000 ; N- CA- CB- CG
2267 2269 2280 2281 3 0.00000 0.00000 0.00000
0.00000 0.00000 0.00000 ; N- CA- C- O
2267 2269 2280 2282 3 17.40544 -5.02080 -13.22144
9.20480 0.00000 0.00000 ; N- CA- C- N
2268 2267 2269 2270 3 0.00000 0.00000 0.00000
0.00000 0.00000 0.00000 ; H- N- CA- HA
2268 2267 2269 2271 3 0.00000 0.00000 0.00000
0.00000 0.00000 0.00000 ; H- N- CA- CB
2268 2267 2269 2280 3 0.00000 0.00000 0.00000
0.00000 0.00000 0.00000 ; H- N- CA- C
2269 2271 2274 2275 3 0.66944 2.00832 0.00000
-2.67776 0.00000 0.00000 ; CA- CB- CG- HG2
2269 2271 2274 2276 3 0.66944 2.00832 0.00000
-2.67776 0.00000 0.00000 ; CA- CB- CG- HG3
2269 2271 2274 2277 3 10.12110 -1.91209 0.01674
10.17549 -1.87443 0.00000 ; CA- CB- CG- CD
2269 2280 2282 2283 3 20.92000 0.00000 -20.92000
0.00000 0.00000 0.00000 ; CA- C- N- H
2269 2280 2282 2284 3 20.92000 0.00000 -20.92000
0.00000 0.00000 0.00000 ; CA- C- N- CA
2270 2269 2271 2272 3 0.65084 1.95253 0.00000
-2.60338 0.00000 0.00000 ; HA- CA- CB- HB2
2270 2269 2271 2273 3 0.65084 1.95253 0.00000
-2.60338 0.00000 0.00000 ; HA- CA- CB- HB3
2270 2269 2271 2274 3 0.65084 1.95253 0.00000
-2.60338 0.00000 0.00000 ; HA- CA- CB- CG
2270 2269 2280 2281 3 3.68192 -4.35136 0.00000

1.33888 0.00000 0.00000 ; HA- CA- C- O
2270 2269 2280 2282 3 0.00000 0.00000 0.00000
0.00000 0.00000 0.00000 ; HA- CA- C- N
2271 2269 2280 2281 3 0.00000 0.00000 0.00000
0.00000 0.00000 0.00000 ; CB- CA- C- O
2271 2269 2280 2282 3 2.51040 4.18400 1.67360
-6.69440 0.00000 0.00000 ; CB- CA- C- N
2271 2274 2277 2278 3 3.79907 0.00000 -5.40573
0.00000 2.14221 0.00000 ; CB- CG- CD- OE1
2271 2274 2277 2279 3 3.79907 0.00000 -5.40573
0.00000 2.14221 0.00000 ; CB- CG- CD- OE2
2272 2271 2269 2280 3 0.65084 1.95253 0.00000
-2.60338 0.00000 0.00000 ; HB2- CB- CA- C
2272 2271 2274 2275 3 0.62760 1.88280 0.00000
-2.51040 0.00000 0.00000 ; HB2- CB- CG- HG2
2272 2271 2274 2276 3 0.62760 1.88280 0.00000
-2.51040 0.00000 0.00000 ; HB2- CB- CG- HG3
2272 2271 2274 2277 3 0.65084 1.95253 0.00000
-2.60338 0.00000 0.00000 ; HB2- CB- CG- CD
2273 2271 2269 2280 3 0.65084 1.95253 0.00000
-2.60338 0.00000 0.00000 ; HB3- CB- CA- C
2273 2271 2274 2275 3 0.62760 1.88280 0.00000
-2.51040 0.00000 0.00000 ; HB3- CB- CG- HG2
2273 2271 2274 2276 3 0.62760 1.88280 0.00000
-2.51040 0.00000 0.00000 ; HB3- CB- CG- HG3
2273 2271 2274 2277 3 0.65084 1.95253 0.00000
-2.60338 0.00000 0.00000 ; HB3- CB- CG- CD
2274 2271 2269 2280 3 6.86594 3.56895 -8.14206
-2.40998 4.85344 0.00000 ; CG- CB- CA- C
2275 2274 2277 2278 3 0.00000 0.00000 0.00000
0.00000 0.00000 0.00000 ; HG2- CG- CD- OE1
2275 2274 2277 2279 3 0.00000 0.00000 0.00000
0.00000 0.00000 0.00000 ; HG2- CG- CD- OE2
2276 2274 2277 2278 3 0.00000 0.00000 0.00000
0.00000 0.00000 0.00000 ; HG3- CG- CD- OE1
2276 2274 2277 2279 3 0.00000 0.00000 0.00000
0.00000 0.00000 0.00000 ; HG3- CG- CD- OE2
2280 2282 2284 2285 3 0.00000 0.00000 0.00000
0.00000 0.00000 0.00000 ; C- N- CA- HA
2280 2282 2284 2286 3 11.71520 1.67360 15.06240 -
13.38880 0.00000 0.00000 ; C- N- CA- CB
2280 2282 2284 2294 3 1.75728 5.27184 2.25936
-7.02912 0.00000 0.00000 ; C- N- CA- C
2281 2280 2282 2283 3 29.28800 -8.36800 -20.92000
0.00000 0.00000 0.00000 ; O- C- N- H
2281 2280 2282 2284 3 20.92000 0.00000 -20.92000
0.00000 0.00000 0.00000 ; O- C- N- CA
2282 2284 2286 2287 3 0.65084 1.95253 0.00000
-2.60338 0.00000 0.00000 ; N- CA- CB- HB2
2282 2284 2286 2288 3 0.65084 1.95253 0.00000
-2.60338 0.00000 0.00000 ; N- CA- CB- HB3
2282 2284 2286 2289 3 5.99567 3.29281 -4.46014

-0.55229    1.97485    0.00000 ;       N-    CA-    CB-    CG
  2282   2284   2294   2295      3    0.00000    0.00000    0.00000
0.00000    0.00000    0.00000 ;       N-    CA-     C-     O
  2282   2284   2294   2296      3   17.40544   -5.02080  -13.22144
9.20480    0.00000    0.00000 ;       N-    CA-     C-     N
  2283   2282   2284   2285      3    0.00000    0.00000    0.00000
0.00000    0.00000    0.00000 ;       H-     N-    CA-    HA
  2283   2282   2284   2286      3    0.00000    0.00000    0.00000
0.00000    0.00000    0.00000 ;       H-     N-    CA-    CB
  2283   2282   2284   2294      3    0.00000    0.00000    0.00000
0.00000    0.00000    0.00000 ;       H-     N-    CA-     C
  2284   2286   2289   2290      3    0.00000    0.00000    0.00000
0.00000    0.00000    0.00000 ;      CA-    CB-    CG-   OD1
  2284   2286   2289   2291      3    8.84916   -0.31380   -4.32626
5.03754    0.26778    0.00000 ;      CA-    CB-    CG-   ND2
  2284   2294   2296   2297      3   20.92000    0.00000  -20.92000
0.00000    0.00000    0.00000 ;      CA-     C-     N-     H
  2284   2294   2296   2298      3   20.92000    0.00000  -20.92000
0.00000    0.00000    0.00000 ;      CA-     C-     N-    CA
  2285   2284   2286   2287      3    0.65084    1.95253    0.00000
-2.60338    0.00000    0.00000 ;      HA-    CA-    CB-   HB2
  2285   2284   2286   2288      3    0.65084    1.95253    0.00000
-2.60338    0.00000    0.00000 ;      HA-    CA-    CB-   HB3
  2285   2284   2286   2289      3    0.65084    1.95253    0.00000
-2.60338    0.00000    0.00000 ;      HA-    CA-    CB-    CG
  2285   2284   2294   2295      3    3.68192   -4.35136    0.00000
1.33888    0.00000    0.00000 ;      HA-    CA-     C-     O
  2285   2284   2294   2296      3    0.00000    0.00000    0.00000
0.00000    0.00000    0.00000 ;      HA-    CA-     C-     N
  2286   2284   2294   2295      3    0.00000    0.00000    0.00000
0.00000    0.00000    0.00000 ;      CB-    CA-     C-     O
  2286   2284   2294   2296      3    2.51040    4.18400    1.67360
-6.69440    0.00000    0.00000 ;      CB-    CA-     C-     N
  2286   2289   2291   2292      3   20.92000    0.00000  -20.92000
0.00000    0.00000    0.00000 ;      CB-    CG-   ND2-  HD21
  2286   2289   2291   2293      3   20.92000    0.00000  -20.92000
0.00000    0.00000    0.00000 ;      CB-    CG-   ND2-  HD22
  2287   2286   2284   2294      3    0.65084    1.95253    0.00000
-2.60338    0.00000    0.00000 ;     HB2-    CB-    CA-     C
  2287   2286   2289   2290      3    3.68192   -4.35136    0.00000
1.33888    0.00000    0.00000 ;     HB2-    CB-    CG-   OD1
  2287   2286   2289   2291      3    0.00000    0.00000    0.00000
0.00000    0.00000    0.00000 ;     HB2-    CB-    CG-   ND2
  2288   2286   2284   2294      3    0.65084    1.95253    0.00000
-2.60338    0.00000    0.00000 ;     HB3-    CB-    CA-     C
  2288   2286   2289   2290      3    3.68192   -4.35136    0.00000
1.33888    0.00000    0.00000 ;     HB3-    CB-    CG-   OD1
  2288   2286   2289   2291      3    0.00000    0.00000    0.00000
0.00000    0.00000    0.00000 ;     HB3-    CB-    CG-   ND2
  2289   2286   2284   2294      3    7.94542   -3.96225   -6.11701
-0.55229    3.58150    0.00000 ;      CG-    CB-    CA-     C
  2290   2289   2291   2292      3   29.28800   -8.36800  -20.92000

0.00000 0.00000 0.00000 ; OD1- CG- ND2- HD21
2290 2289 2291 2293 3 29.28800 -8.36800 -20.92000
0.00000 0.00000 0.00000 ; OD1- CG- ND2- HD22
2294 2296 2298 2299 3 0.00000 0.00000 0.00000
0.00000 0.00000 0.00000 ; C- N- CA- HA
2294 2296 2298 2300 3 11.71520 1.67360 15.06240 -
13.38880 0.00000 0.00000 ; C- N- CA- CB
2294 2296 2298 2314 3 1.75728 5.27184 2.25936
-7.02912 0.00000 0.00000 ; C- N- CA- C
2295 2294 2296 2297 3 29.28800 -8.36800 -20.92000
0.00000 0.00000 0.00000 ; O- C- N- H
2295 2294 2296 2298 3 20.92000 0.00000 -20.92000
0.00000 0.00000 0.00000 ; O- C- N- CA
2296 2298 2300 2301 3 0.65084 1.95253 0.00000
-2.60338 0.00000 0.00000 ; N- CA- CB- HB2
2296 2298 2300 2302 3 0.65084 1.95253 0.00000
-2.60338 0.00000 0.00000 ; N- CA- CB- HB3
2296 2298 2300 2303 3 3.28026 2.46019 -2.19242
-3.21331 -0.23430 0.00000 ; N- CA- CB- CG
2296 2298 2314 2315 3 0.00000 0.00000 0.00000
0.00000 0.00000 0.00000 ; N- CA- C- O
2296 2298 2314 2316 3 17.40544 -5.02080 -13.22144
9.20480 0.00000 0.00000 ; N- CA- C- N
2297 2296 2298 2299 3 0.00000 0.00000 0.00000
0.00000 0.00000 0.00000 ; H- N- CA- HA
2297 2296 2298 2300 3 0.00000 0.00000 0.00000
0.00000 0.00000 0.00000 ; H- N- CA- CB
2297 2296 2298 2314 3 0.00000 0.00000 0.00000
0.00000 0.00000 0.00000 ; H- N- CA- C
2298 2300 2303 2304 3 0.57739 0.00000 1.02926
0.00000 -1.60666 0.00000 ; CA- CB- CG- CD1
2298 2300 2303 2312 3 0.57739 0.00000 1.02926
0.00000 -1.60666 0.00000 ; CA- CB- CG- CD2
2298 2314 2316 2317 3 20.92000 0.00000 -20.92000
0.00000 0.00000 0.00000 ; CA- C- N- H
2298 2314 2316 2318 3 20.92000 0.00000 -20.92000
0.00000 0.00000 0.00000 ; CA- C- N- CA
2299 2298 2300 2301 3 0.65084 1.95253 0.00000
-2.60338 0.00000 0.00000 ; HA- CA- CB- HB2
2299 2298 2300 2302 3 0.65084 1.95253 0.00000
-2.60338 0.00000 0.00000 ; HA- CA- CB- HB3
2299 2298 2300 2303 3 0.65084 1.95253 0.00000
-2.60338 0.00000 0.00000 ; HA- CA- CB- CG
2299 2298 2314 2315 3 3.68192 -4.35136 0.00000
1.33888 0.00000 0.00000 ; HA- CA- C- O
2299 2298 2314 2316 3 0.00000 0.00000 0.00000
0.00000 0.00000 0.00000 ; HA- CA- C- N
2300 2298 2314 2315 3 0.00000 0.00000 0.00000
0.00000 0.00000 0.00000 ; CB- CA- C- O
2300 2298 2314 2316 3 2.51040 4.18400 1.67360
-6.69440 0.00000 0.00000 ; CB- CA- C- N
2300 2303 2304 2305 3 30.33400 0.00000 -30.33400

0.00000 0.00000 0.00000 ; CB- CG- CD1- HD1
2300 2303 2304 2306 3 30.33400 0.00000 -30.33400
0.00000 0.00000 0.00000 ; CB- CG- CD1- CE1
2300 2303 2312 2310 3 30.33400 0.00000 -30.33400
0.00000 0.00000 0.00000 ; CB- CG- CD2- CE2
2300 2303 2312 2313 3 30.33400 0.00000 -30.33400
0.00000 0.00000 0.00000 ; CB- CG- CD2- HD2
2301 2300 2298 2314 3 0.65084 1.95253 0.00000
-2.60338 0.00000 0.00000 ; HB2- CB- CA- C
2301 2300 2303 2304 3 0.00000 0.00000 0.00000
0.00000 0.00000 0.00000 ; HB2- CB- CG- CD1
2301 2300 2303 2312 3 0.00000 0.00000 0.00000
0.00000 0.00000 0.00000 ; HB2- CB- CG- CD2
2302 2300 2298 2314 3 0.65084 1.95253 0.00000
-2.60338 0.00000 0.00000 ; HB3- CB- CA- C
2302 2300 2303 2304 3 0.00000 0.00000 0.00000
0.00000 0.00000 0.00000 ; HB3- CB- CG- CD1
2302 2300 2303 2312 3 0.00000 0.00000 0.00000
0.00000 0.00000 0.00000 ; HB3- CB- CG- CD2
2303 2300 2298 2314 3 4.95804 2.17986 -3.52293
-3.21331 -0.40166 0.00000 ; CG- CB- CA- C
2303 2304 2306 2307 3 30.33400 0.00000 -30.33400
0.00000 0.00000 0.00000 ; CG- CD1- CE1- HE1
2303 2304 2306 2308 3 30.33400 0.00000 -30.33400
0.00000 0.00000 0.00000 ; CG- CD1- CE1- CZ
2303 2312 2310 2308 3 30.33400 0.00000 -30.33400
0.00000 0.00000 0.00000 ; CG- CD2- CE2- CZ
2303 2312 2310 2311 3 30.33400 0.00000 -30.33400
0.00000 0.00000 0.00000 ; CG- CD2- CE2- HE2
2304 2303 2312 2310 3 30.33400 0.00000 -30.33400
0.00000 0.00000 0.00000 ; CD1- CG- CD2- CE2
2304 2303 2312 2313 3 30.33400 0.00000 -30.33400
0.00000 0.00000 0.00000 ; CD1- CG- CD2- HD2
2304 2306 2308 2309 3 30.33400 0.00000 -30.33400
0.00000 0.00000 0.00000 ; CD1- CE1- CZ- HZ
2304 2306 2308 2310 3 30.33400 0.00000 -30.33400
0.00000 0.00000 0.00000 ; CD1- CE1- CZ- CE2
2305 2304 2303 2312 3 30.33400 0.00000 -30.33400
0.00000 0.00000 0.00000 ; HD1- CD1- CG- CD2
2305 2304 2306 2307 3 30.33400 0.00000 -30.33400
0.00000 0.00000 0.00000 ; HD1- CD1- CE1- HE1
2305 2304 2306 2308 3 30.33400 0.00000 -30.33400
0.00000 0.00000 0.00000 ; HD1- CD1- CE1- CZ
2306 2304 2303 2312 3 30.33400 0.00000 -30.33400
0.00000 0.00000 0.00000 ; CE1- CD1- CG- CD2
2306 2308 2310 2311 3 30.33400 0.00000 -30.33400
0.00000 0.00000 0.00000 ; CE1- CZ- CE2- HE2
2306 2308 2310 2312 3 30.33400 0.00000 -30.33400
0.00000 0.00000 0.00000 ; CE1- CZ- CE2- CD2
2307 2306 2308 2309 3 30.33400 0.00000 -30.33400
0.00000 0.00000 0.00000 ; HE1- CE1- CZ- HZ
2307 2306 2308 2310 3 30.33400 0.00000 -30.33400

```
0.00000    0.00000    0.00000 ;    HE1-   CE1-    CZ-   CE2
  2308   2310   2312   2313      3   30.33400    0.00000  -30.33400
0.00000    0.00000    0.00000 ;     CZ-   CE2-   CD2-   HD2
  2309   2308   2310   2311      3   30.33400    0.00000  -30.33400
0.00000    0.00000    0.00000 ;     HZ-    CZ-   CE2-   HE2
  2309   2308   2310   2312      3   30.33400    0.00000  -30.33400
0.00000    0.00000    0.00000 ;     HZ-    CZ-   CE2-   CD2
  2311   2310   2312   2313      3   30.33400    0.00000  -30.33400
0.00000    0.00000    0.00000 ;    HE2-   CE2-   CD2-   HD2
  2314   2316   2318   2319      3    0.00000    0.00000    0.00000
0.00000    0.00000    0.00000 ;     C-     N-    CA-    HA
  2314   2316   2318   2320      3   11.71520    1.67360   15.06240  -
13.38880    0.00000    0.00000 ;     C-     N-    CA-    CB
  2314   2316   2318   2333      3    1.75728    5.27184    2.25936
-7.02912    0.00000    0.00000 ;     C-     N-    CA-     C
  2315   2314   2316   2317      3   29.28800   -8.36800  -20.92000
0.00000    0.00000    0.00000 ;     O-     C-     N-     H
  2315   2314   2316   2318      3   20.92000    0.00000  -20.92000
0.00000    0.00000    0.00000 ;     O-     C-     N-    CA
  2316   2318   2320   2321      3    0.65084    1.95253    0.00000
-2.60338    0.00000    0.00000 ;     N-    CA-    CB-   HB2
  2316   2318   2320   2322      3    0.65084    1.95253    0.00000
-2.60338    0.00000    0.00000 ;     N-    CA-    CB-   HB3
  2316   2318   2320   2323      3    3.79070    1.39746   -4.61077
-2.40998    2.44346    0.00000 ;     N-    CA-    CB-    CG
  2316   2318   2333   2334      3    0.00000    0.00000    0.00000
0.00000    0.00000    0.00000 ;     N-    CA-     C-     O
  2316   2318   2333   2335      3   17.40544   -5.02080  -13.22144
9.20480    0.00000    0.00000 ;     N-    CA-     C-     N
  2317   2316   2318   2319      3    0.00000    0.00000    0.00000
0.00000    0.00000    0.00000 ;     H-     N-    CA-    HA
  2317   2316   2318   2320      3    0.00000    0.00000    0.00000
0.00000    0.00000    0.00000 ;     H-     N-    CA-    CB
  2317   2316   2318   2333      3    0.00000    0.00000    0.00000
0.00000    0.00000    0.00000 ;     H-     N-    CA-     C
  2318   2320   2323   2324      3    0.66944    2.00832    0.00000
-2.67776    0.00000    0.00000 ;    CA-    CB-    CG-    HG
  2318   2320   2323   2325      3    3.90367    0.19665   -6.21742
-2.37651    5.99149    0.00000 ;    CA-    CB-    CG-   CD1
  2318   2320   2323   2329      3    3.90367    0.19665   -6.21742
-2.37651    5.99149    0.00000 ;    CA-    CB-    CG-   CD2
  2318   2333   2335   2336      3   20.92000    0.00000  -20.92000
0.00000    0.00000    0.00000 ;    CA-     C-     N-     H
  2318   2333   2335   2337      3   20.92000    0.00000  -20.92000
0.00000    0.00000    0.00000 ;    CA-     C-     N-    CA
  2319   2318   2320   2321      3    0.65084    1.95253    0.00000
-2.60338    0.00000    0.00000 ;    HA-    CA-    CB-   HB2
  2319   2318   2320   2322      3    0.65084    1.95253    0.00000
-2.60338    0.00000    0.00000 ;    HA-    CA-    CB-   HB3
  2319   2318   2320   2323      3    0.65084    1.95253    0.00000
-2.60338    0.00000    0.00000 ;    HA-    CA-    CB-    CG
  2319   2318   2333   2334      3    3.68192   -4.35136    0.00000
```

```
1.33888    0.00000    0.00000 ;     HA-    CA-     C-     O
  2319   2318   2333   2335       3    0.00000    0.00000    0.00000
0.00000    0.00000    0.00000 ;     HA-    CA-     C-     N
  2320   2318   2333   2334       3    0.00000    0.00000    0.00000
0.00000    0.00000    0.00000 ;     CB-    CA-     C-     O
  2320   2318   2333   2335       3    2.51040    4.18400    1.67360
-6.69440    0.00000    0.00000 ;     CB-    CA-     C-     N
  2320   2323   2325   2326       3    0.66944    2.00832    0.00000
-2.67776    0.00000    0.00000 ;     CB-    CG-   CD1-  HD11
  2320   2323   2325   2327       3    0.66944    2.00832    0.00000
-2.67776    0.00000    0.00000 ;     CB-    CG-   CD1-  HD12
  2320   2323   2325   2328       3    0.66944    2.00832    0.00000
-2.67776    0.00000    0.00000 ;     CB-    CG-   CD1-  HD13
  2320   2323   2329   2330       3    0.66944    2.00832    0.00000
-2.67776    0.00000    0.00000 ;     CB-    CG-   CD2-  HD21
  2320   2323   2329   2331       3    0.66944    2.00832    0.00000
-2.67776    0.00000    0.00000 ;     CB-    CG-   CD2-  HD22
  2320   2323   2329   2332       3    0.66944    2.00832    0.00000
-2.67776    0.00000    0.00000 ;     CB-    CG-   CD2-  HD23
  2321   2320   2318   2333       3    0.65084    1.95253    0.00000
-2.60338    0.00000    0.00000 ;    HB2-    CB-    CA-     C
  2321   2320   2323   2324       3    0.62760    1.88280    0.00000
-2.51040    0.00000    0.00000 ;    HB2-    CB-    CG-    HG
  2321   2320   2323   2325       3    0.66944    2.00832    0.00000
-2.67776    0.00000    0.00000 ;    HB2-    CB-    CG-   CD1
  2321   2320   2323   2329       3    0.66944    2.00832    0.00000
-2.67776    0.00000    0.00000 ;    HB2-    CB-    CG-   CD2
  2322   2320   2318   2333       3    0.65084    1.95253    0.00000
-2.60338    0.00000    0.00000 ;    HB3-    CB-    CA-     C
  2322   2320   2323   2324       3    0.62760    1.88280    0.00000
-2.51040    0.00000    0.00000 ;    HB3-    CB-    CG-    HG
  2322   2320   2323   2325       3    0.66944    2.00832    0.00000
-2.67776    0.00000    0.00000 ;    HB3-    CB-    CG-   CD1
  2322   2320   2323   2329       3    0.66944    2.00832    0.00000
-2.67776    0.00000    0.00000 ;    HB3-    CB-    CG-   CD2
  2323   2320   2318   2333       3   10.33448   -1.14642  -11.54784
-2.40998    6.35968    0.00000 ;     CG-    CB-    CA-     C
  2324   2323   2325   2326       3    0.62760    1.88280    0.00000
-2.51040    0.00000    0.00000 ;     HG-    CG-   CD1-  HD11
  2324   2323   2325   2327       3    0.62760    1.88280    0.00000
-2.51040    0.00000    0.00000 ;     HG-    CG-   CD1-  HD12
  2324   2323   2325   2328       3    0.62760    1.88280    0.00000
-2.51040    0.00000    0.00000 ;     HG-    CG-   CD1-  HD13
  2324   2323   2329   2330       3    0.62760    1.88280    0.00000
-2.51040    0.00000    0.00000 ;     HG-    CG-   CD2-  HD21
  2324   2323   2329   2331       3    0.62760    1.88280    0.00000
-2.51040    0.00000    0.00000 ;     HG-    CG-   CD2-  HD22
  2324   2323   2329   2332       3    0.62760    1.88280    0.00000
-2.51040    0.00000    0.00000 ;     HG-    CG-   CD2-  HD23
  2325   2323   2329   2330       3    0.66944    2.00832    0.00000
-2.67776    0.00000    0.00000 ;    CD1-    CG-   CD2-  HD21
  2325   2323   2329   2331       3    0.66944    2.00832    0.00000
```

```
-2.67776    0.00000    0.00000 ;    CD1-    CG-   CD2-  HD22
  2325   2323   2329   2332      3    0.66944    2.00832    0.00000
-2.67776    0.00000    0.00000 ;    CD1-    CG-   CD2-  HD23
  2326   2325   2323   2329      3    0.66944    2.00832    0.00000
-2.67776    0.00000    0.00000 ;   HD11-   CD1-    CG-   CD2
  2327   2325   2323   2329      3    0.66944    2.00832    0.00000
-2.67776    0.00000    0.00000 ;   HD12-   CD1-    CG-   CD2
  2328   2325   2323   2329      3    0.66944    2.00832    0.00000
-2.67776    0.00000    0.00000 ;   HD13-   CD1-    CG-   CD2
  2333   2335   2337   2338      3    0.00000    0.00000    0.00000
0.00000    0.00000    0.00000 ;      C-     N-    CA-    HA
  2333   2335   2337   2339      3   11.71520    1.67360   15.06240  -
13.38880    0.00000    0.00000 ;      C-     N-    CA-    CB
  2333   2335   2337   2353      3    1.75728    5.27184    2.25936
-7.02912    0.00000    0.00000 ;      C-     N-    CA-     C
  2334   2333   2335   2336      3   29.28800   -8.36800  -20.92000
0.00000    0.00000    0.00000 ;      O-     C-     N-     H
  2334   2333   2335   2337      3   20.92000    0.00000  -20.92000
0.00000    0.00000    0.00000 ;      O-     C-     N-    CA
  2335   2337   2339   2340      3    0.65084    1.95253    0.00000
-2.60338    0.00000    0.00000 ;      N-    CA-    CB-   HB2
  2335   2337   2339   2341      3    0.65084    1.95253    0.00000
-2.60338    0.00000    0.00000 ;      N-    CA-    CB-   HB3
  2335   2337   2339   2342      3    3.28026    2.46019   -2.19242
-3.21331   -0.23430    0.00000 ;      N-    CA-    CB-    CG
  2335   2337   2353   2354      3    0.00000    0.00000    0.00000
0.00000    0.00000    0.00000 ;      N-    CA-     C-     O
  2335   2337   2353   2355      3   17.40544   -5.02080  -13.22144
9.20480    0.00000    0.00000 ;      N-    CA-     C-     N
  2336   2335   2337   2338      3    0.00000    0.00000    0.00000
0.00000    0.00000    0.00000 ;      H-     N-    CA-    HA
  2336   2335   2337   2339      3    0.00000    0.00000    0.00000
0.00000    0.00000    0.00000 ;      H-     N-    CA-    CB
  2336   2335   2337   2353      3    0.00000    0.00000    0.00000
0.00000    0.00000    0.00000 ;      H-     N-    CA-     C
  2337   2339   2342   2343      3    0.57739    0.00000    1.02926
0.00000   -1.60666    0.00000 ;    CA-    CB-    CG-   CD1
  2337   2339   2342   2351      3    0.57739    0.00000    1.02926
0.00000   -1.60666    0.00000 ;    CA-    CB-    CG-   CD2
  2337   2353   2355   2356      3   20.92000    0.00000  -20.92000
0.00000    0.00000    0.00000 ;    CA-     C-     N-     H
  2337   2353   2355   2357      3   20.92000    0.00000  -20.92000
0.00000    0.00000    0.00000 ;    CA-     C-     N-    CA
  2338   2337   2339   2340      3    0.65084    1.95253    0.00000
-2.60338    0.00000    0.00000 ;     HA-    CA-    CB-   HB2
  2338   2337   2339   2341      3    0.65084    1.95253    0.00000
-2.60338    0.00000    0.00000 ;     HA-    CA-    CB-   HB3
  2338   2337   2339   2342      3    0.65084    1.95253    0.00000
-2.60338    0.00000    0.00000 ;     HA-    CA-    CB-    CG
  2338   2337   2353   2354      3    3.68192   -4.35136    0.00000
1.33888    0.00000    0.00000 ;    HA-    CA-     C-     O
  2338   2337   2353   2355      3    0.00000    0.00000    0.00000
```

```
0.00000    0.00000    0.00000 ;     HA-    CA-     C-     N
  2339   2337   2353   2354       3    0.00000    0.00000    0.00000
0.00000    0.00000    0.00000 ;     CB-    CA-     C-     O
  2339   2337   2353   2355       3    2.51040    4.18400    1.67360
-6.69440    0.00000    0.00000 ;     CB-    CA-     C-     N
  2339   2342   2343   2344       3   30.33400    0.00000  -30.33400
0.00000    0.00000    0.00000 ;     CB-    CG-   CD1-   HD1
  2339   2342   2343   2345       3   30.33400    0.00000  -30.33400
0.00000    0.00000    0.00000 ;     CB-    CG-   CD1-   CE1
  2339   2342   2351   2349       3   30.33400    0.00000  -30.33400
0.00000    0.00000    0.00000 ;     CB-    CG-   CD2-   CE2
  2339   2342   2351   2352       3   30.33400    0.00000  -30.33400
0.00000    0.00000    0.00000 ;     CB-    CG-   CD2-   HD2
  2340   2339   2337   2353       3    0.65084    1.95253    0.00000
-2.60338    0.00000    0.00000 ;    HB2-    CB-    CA-     C
  2340   2339   2342   2343       3    0.00000    0.00000    0.00000
0.00000    0.00000    0.00000 ;    HB2-    CB-    CG-   CD1
  2340   2339   2342   2351       3    0.00000    0.00000    0.00000
0.00000    0.00000    0.00000 ;    HB2-    CB-    CG-   CD2
  2341   2339   2337   2353       3    0.65084    1.95253    0.00000
-2.60338    0.00000    0.00000 ;    HB3-    CB-    CA-     C
  2341   2339   2342   2343       3    0.00000    0.00000    0.00000
0.00000    0.00000    0.00000 ;    HB3-    CB-    CG-   CD1
  2341   2339   2342   2351       3    0.00000    0.00000    0.00000
0.00000    0.00000    0.00000 ;    HB3-    CB-    CG-   CD2
  2342   2339   2337   2353       3    4.95804    2.17986   -3.52293
-3.21331   -0.40166    0.00000 ;     CG-    CB-    CA-     C
  2342   2343   2345   2346       3   30.33400    0.00000  -30.33400
0.00000    0.00000    0.00000 ;     CG-   CD1-   CE1-   HE1
  2342   2343   2345   2347       3   30.33400    0.00000  -30.33400
0.00000    0.00000    0.00000 ;     CG-   CD1-   CE1-    CZ
  2342   2351   2349   2347       3   30.33400    0.00000  -30.33400
0.00000    0.00000    0.00000 ;     CG-   CD2-   CE2-    CZ
  2342   2351   2349   2350       3   30.33400    0.00000  -30.33400
0.00000    0.00000    0.00000 ;     CG-   CD2-   CE2-   HE2
  2343   2342   2351   2349       3   30.33400    0.00000  -30.33400
0.00000    0.00000    0.00000 ;    CD1-    CG-   CD2-   CE2
  2343   2342   2351   2352       3   30.33400    0.00000  -30.33400
0.00000    0.00000    0.00000 ;    CD1-    CG-   CD2-   HD2
  2343   2345   2347   2348       3   30.33400    0.00000  -30.33400
0.00000    0.00000    0.00000 ;    CD1-   CE1-    CZ-    HZ
  2343   2345   2347   2349       3   30.33400    0.00000  -30.33400
0.00000    0.00000    0.00000 ;    CD1-   CE1-    CZ-   CE2
  2344   2343   2342   2351       3   30.33400    0.00000  -30.33400
0.00000    0.00000    0.00000 ;    HD1-   CD1-    CG-   CD2
  2344   2343   2345   2346       3   30.33400    0.00000  -30.33400
0.00000    0.00000    0.00000 ;    HD1-   CD1-   CE1-   HE1
  2344   2343   2345   2347       3   30.33400    0.00000  -30.33400
0.00000    0.00000    0.00000 ;    HD1-   CD1-   CE1-    CZ
  2345   2343   2342   2351       3   30.33400    0.00000  -30.33400
0.00000    0.00000    0.00000 ;    CE1-   CD1-    CG-   CD2
  2345   2347   2349   2350       3   30.33400    0.00000  -30.33400
```

0.00000 0.00000 0.00000 ; CE1- CZ- CE2- HE2
2345 2347 2349 2351 3 30.33400 0.00000 -30.33400
0.00000 0.00000 0.00000 ; CE1- CZ- CE2- CD2
2346 2345 2347 2348 3 30.33400 0.00000 -30.33400
0.00000 0.00000 0.00000 ; HE1- CE1- CZ- HZ
2346 2345 2347 2349 3 30.33400 0.00000 -30.33400
0.00000 0.00000 0.00000 ; HE1- CE1- CZ- CE2
2347 2349 2351 2352 3 30.33400 0.00000 -30.33400
0.00000 0.00000 0.00000 ; CZ- CE2- CD2- HD2
2348 2347 2349 2350 3 30.33400 0.00000 -30.33400
0.00000 0.00000 0.00000 ; HZ- CZ- CE2- HE2
2348 2347 2349 2351 3 30.33400 0.00000 -30.33400
0.00000 0.00000 0.00000 ; HZ- CZ- CE2- CD2
2350 2349 2351 2352 3 30.33400 0.00000 -30.33400
0.00000 0.00000 0.00000 ; HE2- CE2- CD2- HD2
2353 2355 2357 2358 3 0.00000 0.00000 0.00000
0.00000 0.00000 0.00000 ; C- N- CA- HA
2353 2355 2357 2359 3 11.71520 1.67360 15.06240 -
13.38880 0.00000 0.00000 ; C- N- CA- CB
2353 2355 2357 2370 3 1.75728 5.27184 2.25936
-7.02912 0.00000 0.00000 ; C- N- CA- C
2354 2353 2355 2356 3 29.28800 -8.36800 -20.92000
0.00000 0.00000 0.00000 ; O- C- N- H
2354 2353 2355 2357 3 20.92000 0.00000 -20.92000
0.00000 0.00000 0.00000 ; O- C- N- CA
2355 2357 2359 2360 3 0.65084 1.95253 0.00000
-2.60338 0.00000 0.00000 ; N- CA- CB- HB2
2355 2357 2359 2361 3 0.65084 1.95253 0.00000
-2.60338 0.00000 0.00000 ; N- CA- CB- HB3
2355 2357 2359 2362 3 3.21331 2.22589 -4.15053
-2.40998 2.61082 0.00000 ; N- CA- CB- CG
2355 2357 2370 2371 3 0.00000 0.00000 0.00000
0.00000 0.00000 0.00000 ; N- CA- C- O
2355 2357 2370 2372 3 17.40544 -5.02080 -13.22144
9.20480 0.00000 0.00000 ; N- CA- C- N
2356 2355 2357 2358 3 0.00000 0.00000 0.00000
0.00000 0.00000 0.00000 ; H- N- CA- HA
2356 2355 2357 2359 3 0.00000 0.00000 0.00000
0.00000 0.00000 0.00000 ; H- N- CA- CB
2356 2355 2357 2370 3 0.00000 0.00000 0.00000
0.00000 0.00000 0.00000 ; H- N- CA- C
2357 2359 2362 2363 3 0.66944 2.00832 0.00000
-2.67776 0.00000 0.00000 ; CA- CB- CG- HG2
2357 2359 2362 2364 3 0.66944 2.00832 0.00000
-2.67776 0.00000 0.00000 ; CA- CB- CG- HG3
2357 2359 2362 2365 3 3.69866 -4.35136 -3.92459
6.89523 4.61914 0.00000 ; CA- CB- CG- CD
2357 2370 2372 2373 3 20.92000 0.00000 -20.92000
0.00000 0.00000 0.00000 ; CA- C- N- H
2357 2370 2372 2374 3 20.92000 0.00000 -20.92000
0.00000 0.00000 0.00000 ; CA- C- N- CA
2358 2357 2359 2360 3 0.65084 1.95253 0.00000

```
-2.60338    0.00000    0.00000 ;      HA-    CA-    CB-   HB2
  2358   2357   2359   2361       3    0.65084    1.95253    0.00000
-2.60338    0.00000    0.00000 ;      HA-    CA-    CB-   HB3
  2358   2357   2359   2362       3    0.65084    1.95253    0.00000
-2.60338    0.00000    0.00000 ;      HA-    CA-    CB-    CG
  2358   2357   2370   2371       3    3.68192   -4.35136    0.00000
1.33888    0.00000    0.00000 ;      HA-    CA-     C-     O
  2358   2357   2370   2372       3    0.00000    0.00000    0.00000
0.00000    0.00000    0.00000 ;      HA-    CA-     C-     N
  2359   2357   2370   2371       3    0.00000    0.00000    0.00000
0.00000    0.00000    0.00000 ;      CB-    CA-     C-     O
  2359   2357   2370   2372       3    2.51040    4.18400    1.67360
-6.69440    0.00000    0.00000 ;      CB-    CA-     C-     N
  2359   2362   2365   2366       3    0.00000    0.00000    0.00000
0.00000    0.00000    0.00000 ;      CB-    CG-    CD-   OE1
  2359   2362   2365   2367       3   10.32611    1.48114   -8.47678
1.42256    1.40582    0.00000 ;      CB-    CG-    CD-   NE2
  2360   2359   2357   2370       3    0.65084    1.95253    0.00000
-2.60338    0.00000    0.00000 ;     HB2-    CB-    CA-     C
  2360   2359   2362   2363       3    0.62760    1.88280    0.00000
-2.51040    0.00000    0.00000 ;     HB2-    CB-    CG-   HG2
  2360   2359   2362   2364       3    0.62760    1.88280    0.00000
-2.51040    0.00000    0.00000 ;     HB2-    CB-    CG-   HG3
  2360   2359   2362   2365       3    0.65084    1.95253    0.00000
-2.60338    0.00000    0.00000 ;     HB2-    CB-    CG-    CD
  2361   2359   2357   2370       3    0.65084    1.95253    0.00000
-2.60338    0.00000    0.00000 ;     HB3-    CB-    CA-     C
  2361   2359   2362   2363       3    0.62760    1.88280    0.00000
-2.51040    0.00000    0.00000 ;     HB3-    CB-    CG-   HG2
  2361   2359   2362   2364       3    0.62760    1.88280    0.00000
-2.51040    0.00000    0.00000 ;     HB3-    CB-    CG-   HG3
  2361   2359   2362   2365       3    0.65084    1.95253    0.00000
-2.60338    0.00000    0.00000 ;     HB3-    CB-    CG-    CD
  2362   2359   2357   2370       3    6.86594    3.56895   -8.14206
-2.40998    4.85344    0.00000 ;      CG-    CB-    CA-     C
  2362   2365   2367   2368       3   20.92000    0.00000  -20.92000
0.00000    0.00000    0.00000 ;      CG-    CD-   NE2-  HE21
  2362   2365   2367   2369       3   20.92000    0.00000  -20.92000
0.00000    0.00000    0.00000 ;      CG-    CD-   NE2-  HE22
  2363   2362   2365   2366       3    3.68192   -4.35136    0.00000
1.33888    0.00000    0.00000 ;     HG2-    CG-    CD-   OE1
  2363   2362   2365   2367       3    0.00000    0.00000    0.00000
0.00000    0.00000    0.00000 ;     HG2-    CG-    CD-   NE2
  2364   2362   2365   2366       3    3.68192   -4.35136    0.00000
1.33888    0.00000    0.00000 ;     HG3-    CG-    CD-   OE1
  2364   2362   2365   2367       3    0.00000    0.00000    0.00000
0.00000    0.00000    0.00000 ;     HG3-    CG-    CD-   NE2
  2366   2365   2367   2368       3   29.28800   -8.36800  -20.92000
0.00000    0.00000    0.00000 ;     OE1-    CD-   NE2-  HE21
  2366   2365   2367   2369       3   29.28800   -8.36800  -20.92000
0.00000    0.00000    0.00000 ;     OE1-    CD-   NE2-  HE22
  2370   2372   2374   2375       3    0.00000    0.00000    0.00000
```

0.00000 0.00000 0.00000 ; C- N- CA- HA
2370 2372 2374 2376 3 11.71520 1.67360 15.06240 -
13.38880 0.00000 0.00000 ; C- N- CA- CB
2370 2372 2374 2381 3 1.75728 5.27184 2.25936
-7.02912 0.00000 0.00000 ; C- N- CA- C
2371 2370 2372 2373 3 29.28800 -8.36800 -20.92000
0.00000 0.00000 0.00000 ; O- C- N- H
2371 2370 2372 2374 3 20.92000 0.00000 -20.92000
0.00000 0.00000 0.00000 ; O- C- N- CA
2372 2374 2376 2377 3 0.65084 1.95253 0.00000
-2.60338 0.00000 0.00000 ; N- CA- CB- HB2
2372 2374 2376 2378 3 0.65084 1.95253 0.00000
-2.60338 0.00000 0.00000 ; N- CA- CB- HB3
2372 2374 2376 2379 3 7.86174 2.24681 -7.41405
-6.71114 5.35552 0.00000 ; N- CA- CB- OG
2372 2374 2381 2382 3 0.00000 0.00000 0.00000
0.00000 0.00000 0.00000 ; N- CA- C- O
2372 2374 2381 2383 3 17.40544 -5.02080 -13.22144
9.20480 0.00000 0.00000 ; N- CA- C- N
2373 2372 2374 2375 3 0.00000 0.00000 0.00000
0.00000 0.00000 0.00000 ; H- N- CA- HA
2373 2372 2374 2376 3 0.00000 0.00000 0.00000
0.00000 0.00000 0.00000 ; H- N- CA- CB
2373 2372 2374 2381 3 0.00000 0.00000 0.00000
0.00000 0.00000 0.00000 ; H- N- CA- C
2374 2376 2379 2380 3 2.05853 2.46856 3.48109
-4.46851 0.23430 0.00000 ; CA- CB- OG- HG
2374 2381 2383 2384 3 20.92000 0.00000 -20.92000
0.00000 0.00000 0.00000 ; CA- C- N- H
2374 2381 2383 2385 3 20.92000 0.00000 -20.92000
0.00000 0.00000 0.00000 ; CA- C- N- CA
2375 2374 2376 2377 3 0.65084 1.95253 0.00000
-2.60338 0.00000 0.00000 ; HA- CA- CB- HB2
2375 2374 2376 2378 3 0.65084 1.95253 0.00000
-2.60338 0.00000 0.00000 ; HA- CA- CB- HB3
2375 2374 2376 2379 3 1.04600 -1.04600 0.00000
0.00000 0.00000 0.00000 ; HA- CA- CB- OG
2375 2374 2381 2382 3 3.68192 -4.35136 0.00000
1.33888 0.00000 0.00000 ; HA- CA- C- O
2375 2374 2381 2383 3 0.00000 0.00000 0.00000
0.00000 0.00000 0.00000 ; HA- CA- C- N
2376 2374 2381 2382 3 0.00000 0.00000 0.00000
0.00000 0.00000 0.00000 ; CB- CA- C- O
2376 2374 2381 2383 3 2.51040 4.18400 1.67360
-6.69440 0.00000 0.00000 ; CB- CA- C- N
2377 2376 2374 2381 3 0.65084 1.95253 0.00000
-2.60338 0.00000 0.00000 ; HB2- CB- CA- C
2377 2376 2379 2380 3 0.69733 2.09200 0.00000
-2.78933 0.00000 0.00000 ; HB2- CB- OG- HG
2378 2376 2374 2381 3 0.65084 1.95253 0.00000
-2.60338 0.00000 0.00000 ; HB3- CB- CA- C
2378 2376 2379 2380 3 0.69733 2.09200 0.00000

-2.78933 0.00000 0.00000 ; HB3- CB- OG- HG
2379 2376 2374 2381 3 7.34710 7.79898 -6.14211
-6.71114 4.31789 0.00000 ; OG- CB- CA- C
2381 2383 2385 2386 3 0.00000 0.00000 0.00000
0.00000 0.00000 0.00000 ; C- N- CA- HA
2381 2383 2385 2387 3 11.71520 1.67360 15.06240 -
13.38880 0.00000 0.00000 ; C- N- CA- CB
2381 2383 2385 2396 3 1.75728 5.27184 2.25936
-7.02912 0.00000 0.00000 ; C- N- CA- C
2382 2381 2383 2384 3 29.28800 -8.36800 -20.92000
0.00000 0.00000 0.00000 ; O- C- N- H
2382 2381 2383 2385 3 20.92000 0.00000 -20.92000
0.00000 0.00000 0.00000 ; O- C- N- CA
2383 2385 2387 2388 3 0.65084 1.95253 0.00000
-2.60338 0.00000 0.00000 ; N- CA- CB- HB2
2383 2385 2387 2389 3 0.65084 1.95253 0.00000
-2.60338 0.00000 0.00000 ; N- CA- CB- HB3
2383 2385 2387 2390 3 3.21331 2.22589 -4.15053
-2.40998 2.61082 0.00000 ; N- CA- CB- CG
2383 2385 2396 2397 3 0.00000 0.00000 0.00000
0.00000 0.00000 0.00000 ; N- CA- C- O
2383 2385 2396 2398 3 17.40544 -5.02080 -13.22144
9.20480 0.00000 0.00000 ; N- CA- C- N
2384 2383 2385 2386 3 0.00000 0.00000 0.00000
0.00000 0.00000 0.00000 ; H- N- CA- HA
2384 2383 2385 2387 3 0.00000 0.00000 0.00000
0.00000 0.00000 0.00000 ; H- N- CA- CB
2384 2383 2385 2396 3 0.00000 0.00000 0.00000
0.00000 0.00000 0.00000 ; H- N- CA- C
2385 2387 2390 2391 3 0.66944 2.00832 0.00000
-2.67776 0.00000 0.00000 ; CA- CB- CG- HG2
2385 2387 2390 2392 3 0.66944 2.00832 0.00000
-2.67776 0.00000 0.00000 ; CA- CB- CG- HG3
2385 2387 2390 2393 3 10.12110 -1.91209 0.01674
10.17549 -1.87443 0.00000 ; CA- CB- CG- CD
2385 2396 2398 2399 3 20.92000 0.00000 -20.92000
0.00000 0.00000 0.00000 ; CA- C- N- H
2385 2396 2398 2400 3 20.92000 0.00000 -20.92000
0.00000 0.00000 0.00000 ; CA- C- N- CA
2386 2385 2387 2388 3 0.65084 1.95253 0.00000
-2.60338 0.00000 0.00000 ; HA- CA- CB- HB2
2386 2385 2387 2389 3 0.65084 1.95253 0.00000
-2.60338 0.00000 0.00000 ; HA- CA- CB- HB3
2386 2385 2387 2390 3 0.65084 1.95253 0.00000
-2.60338 0.00000 0.00000 ; HA- CA- CB- CG
2386 2385 2396 2397 3 3.68192 -4.35136 0.00000
1.33888 0.00000 0.00000 ; HA- CA- C- O
2386 2385 2396 2398 3 0.00000 0.00000 0.00000
0.00000 0.00000 0.00000 ; HA- CA- C- N
2387 2385 2396 2397 3 0.00000 0.00000 0.00000
0.00000 0.00000 0.00000 ; CB- CA- C- O
2387 2385 2396 2398 3 2.51040 4.18400 1.67360

-6.69440    0.00000    0.00000 ;     CB-    CA-     C-     N
  2387   2390   2393   2394      3    3.79907    0.00000   -5.40573
0.00000    2.14221    0.00000 ;     CB-    CG-    CD-   OE1
  2387   2390   2393   2395      3    3.79907    0.00000   -5.40573
0.00000    2.14221    0.00000 ;     CB-    CG-    CD-   OE2
  2388   2387   2385   2396      3    0.65084    1.95253    0.00000
-2.60338    0.00000    0.00000 ;    HB2-    CB-    CA-     C
  2388   2387   2390   2391      3    0.62760    1.88280    0.00000
-2.51040    0.00000    0.00000 ;    HB2-    CB-    CG-   HG2
  2388   2387   2390   2392      3    0.62760    1.88280    0.00000
-2.51040    0.00000    0.00000 ;    HB2-    CB-    CG-   HG3
  2388   2387   2390   2393      3    0.65084    1.95253    0.00000
-2.60338    0.00000    0.00000 ;    HB2-    CB-    CG-    CD
  2389   2387   2385   2396      3    0.65084    1.95253    0.00000
-2.60338    0.00000    0.00000 ;    HB3-    CB-    CA-     C
  2389   2387   2390   2391      3    0.62760    1.88280    0.00000
-2.51040    0.00000    0.00000 ;    HB3-    CB-    CG-   HG2
  2389   2387   2390   2392      3    0.62760    1.88280    0.00000
-2.51040    0.00000    0.00000 ;    HB3-    CB-    CG-   HG3
  2389   2387   2390   2393      3    0.65084    1.95253    0.00000
-2.60338    0.00000    0.00000 ;    HB3-    CB-    CG-    CD
  2390   2387   2385   2396      3    6.86594    3.56895   -8.14206
-2.40998    4.85344    0.00000 ;     CG-    CB-    CA-     C
  2391   2390   2393   2394      3    0.00000    0.00000    0.00000
0.00000    0.00000    0.00000 ;    HG2-    CG-    CD-   OE1
  2391   2390   2393   2395      3    0.00000    0.00000    0.00000
0.00000    0.00000    0.00000 ;    HG2-    CG-    CD-   OE2
  2392   2390   2393   2394      3    0.00000    0.00000    0.00000
0.00000    0.00000    0.00000 ;    HG3-    CG-    CD-   OE1
  2392   2390   2393   2395      3    0.00000    0.00000    0.00000
0.00000    0.00000    0.00000 ;    HG3-    CG-    CD-   OE2
  2396   2398   2400   2401      3    0.00000    0.00000    0.00000
0.00000    0.00000    0.00000 ;     C-     N-    CA-    HA
  2396   2398   2400   2402      3   11.71520    1.67360   15.06240  -
13.38880    0.00000    0.00000 ;     C-     N-    CA-    CB
  2396   2398   2400   2410      3    1.75728    5.27184    2.25936
-7.02912    0.00000    0.00000 ;     C-     N-    CA-     C
  2397   2396   2398   2399      3   29.28800   -8.36800  -20.92000
0.00000    0.00000    0.00000 ;     O-     C-     N-     H
  2397   2396   2398   2400      3   20.92000    0.00000  -20.92000
0.00000    0.00000    0.00000 ;     O-     C-     N-    CA
  2398   2400   2402   2403      3    0.65084    1.95253    0.00000
-2.60338    0.00000    0.00000 ;     N-    CA-    CB-   HB2
  2398   2400   2402   2404      3    0.65084    1.95253    0.00000
-2.60338    0.00000    0.00000 ;     N-    CA-    CB-   HB3
  2398   2400   2402   2405      3    5.99567    3.29281   -4.46014
-0.55229    1.97485    0.00000 ;     N-    CA-    CB-    CG
  2398   2400   2410   2411      3    0.00000    0.00000    0.00000
0.00000    0.00000    0.00000 ;     N-    CA-     C-     O
  2398   2400   2410   2412      3   17.40544   -5.02080  -13.22144
9.20480    0.00000    0.00000 ;     N-    CA-     C-     N
  2399   2398   2400   2401      3    0.00000    0.00000    0.00000

```
0.00000    0.00000    0.00000 ;      H-     N-    CA-    HA
  2399   2398   2400   2402      3    0.00000    0.00000    0.00000
0.00000    0.00000    0.00000 ;      H-     N-    CA-    CB
  2399   2398   2400   2410      3    0.00000    0.00000    0.00000
0.00000    0.00000    0.00000 ;      H-     N-    CA-     C
  2400   2402   2405   2406      3    0.00000    0.00000    0.00000
0.00000    0.00000    0.00000 ;     CA-    CB-    CG-   OD1
  2400   2402   2405   2407      3    8.84916   -0.31380   -4.32626
5.03754    0.26778    0.00000 ;     CA-    CB-    CG-   ND2
  2400   2410   2412   2413      3   20.92000    0.00000  -20.92000
0.00000    0.00000    0.00000 ;     CA-     C-     N-     H
  2400   2410   2412   2414      3   20.92000    0.00000  -20.92000
0.00000    0.00000    0.00000 ;     CA-     C-     N-    CA
  2401   2400   2402   2403      3    0.65084    1.95253    0.00000
-2.60338    0.00000    0.00000 ;     HA-    CA-    CB-   HB2
  2401   2400   2402   2404      3    0.65084    1.95253    0.00000
-2.60338    0.00000    0.00000 ;     HA-    CA-    CB-   HB3
  2401   2400   2402   2405      3    0.65084    1.95253    0.00000
-2.60338    0.00000    0.00000 ;     HA-    CA-    CB-    CG
  2401   2400   2410   2411      3    3.68192   -4.35136    0.00000
1.33888    0.00000    0.00000 ;     HA-    CA-     C-     O
  2401   2400   2410   2412      3    0.00000    0.00000    0.00000
0.00000    0.00000    0.00000 ;     HA-    CA-     C-     N
  2402   2400   2410   2411      3    0.00000    0.00000    0.00000
0.00000    0.00000    0.00000 ;     CB-    CA-     C-     O
  2402   2400   2410   2412      3    2.51040    4.18400    1.67360
-6.69440    0.00000    0.00000 ;     CB-    CA-     C-     N
  2402   2405   2407   2408      3   20.92000    0.00000  -20.92000
0.00000    0.00000    0.00000 ;     CB-    CG-   ND2-  HD21
  2402   2405   2407   2409      3   20.92000    0.00000  -20.92000
0.00000    0.00000    0.00000 ;     CB-    CG-   ND2-  HD22
  2403   2402   2400   2410      3    0.65084    1.95253    0.00000
-2.60338    0.00000    0.00000 ;    HB2-    CB-    CA-     C
  2403   2402   2405   2406      3    3.68192   -4.35136    0.00000
1.33888    0.00000    0.00000 ;    HB2-    CB-    CG-   OD1
  2403   2402   2405   2407      3    0.00000    0.00000    0.00000
0.00000    0.00000    0.00000 ;    HB2-    CB-    CG-   ND2
  2404   2402   2400   2410      3    0.65084    1.95253    0.00000
-2.60338    0.00000    0.00000 ;    HB3-    CB-    CA-     C
  2404   2402   2405   2406      3    3.68192   -4.35136    0.00000
1.33888    0.00000    0.00000 ;    HB3-    CB-    CG-   OD1
  2404   2402   2405   2407      3    0.00000    0.00000    0.00000
0.00000    0.00000    0.00000 ;    HB3-    CB-    CG-   ND2
  2405   2402   2400   2410      3    7.94542   -3.96225   -6.11701
-0.55229    3.58150    0.00000 ;     CG-    CB-    CA-     C
  2406   2405   2407   2408      3   29.28800   -8.36800  -20.92000
0.00000    0.00000    0.00000 ;    OD1-    CG-   ND2-  HD21
  2406   2405   2407   2409      3   29.28800   -8.36800  -20.92000
0.00000    0.00000    0.00000 ;    OD1-    CG-   ND2-  HD22
  2410   2412   2414   2415      3    0.00000    0.00000    0.00000
0.00000    0.00000    0.00000 ;      C-     N-    CA-    HA
  2410   2412   2414   2416      3   11.71520    1.67360   15.06240  -
```

```
13.38880    0.00000    0.00000 ;       C-     N-    CA-    CB
  2410   2412   2414   2432       3    1.75728    5.27184    2.25936
-7.02912    0.00000    0.00000 ;       C-     N-    CA-     C
  2411   2410   2412   2413       3   29.28800   -8.36800  -20.92000
0.00000    0.00000    0.00000 ;      O-     C-     N-     H
  2411   2410   2412   2414       3   20.92000    0.00000  -20.92000
0.00000    0.00000    0.00000 ;      O-     C-     N-    CA
  2412   2414   2416   2417       3    0.65084    1.95253    0.00000
-2.60338    0.00000    0.00000 ;       N-    CA-    CB-   HB2
  2412   2414   2416   2418       3    0.65084    1.95253    0.00000
-2.60338    0.00000    0.00000 ;       N-    CA-    CB-   HB3
  2412   2414   2416   2419       3    0.65084    1.95253    0.00000
-2.60338    0.00000    0.00000 ;       N-    CA-    CB-    CG
  2412   2414   2432   2433       3    0.00000    0.00000    0.00000
0.00000    0.00000    0.00000 ;      N-    CA-     C-     O
  2412   2414   2432   2434       3   17.40544   -5.02080  -13.22144
9.20480    0.00000    0.00000 ;      N-    CA-     C-     N
  2413   2412   2414   2415       3    0.00000    0.00000    0.00000
0.00000    0.00000    0.00000 ;      H-     N-    CA-    HA
  2413   2412   2414   2416       3    0.00000    0.00000    0.00000
0.00000    0.00000    0.00000 ;      H-     N-    CA-    CB
  2413   2412   2414   2432       3    0.00000    0.00000    0.00000
0.00000    0.00000    0.00000 ;      H-     N-    CA-     C
  2414   2416   2419   2420       3    0.66944    2.00832    0.00000
-2.67776    0.00000    0.00000 ;      CA-    CB-    CG-   HG2
  2414   2416   2419   2421       3    0.66944    2.00832    0.00000
-2.67776    0.00000    0.00000 ;      CA-    CB-    CG-   HG3
  2414   2416   2419   2422       3    3.68192    3.09616   -2.09200
-3.01248    0.00000    0.00000 ;      CA-    CB-    CG-    CD
  2414   2432   2434   2435       3   20.92000    0.00000  -20.92000
0.00000    0.00000    0.00000 ;     CA-     C-     N-     H
  2414   2432   2434   2436       3   20.92000    0.00000  -20.92000
0.00000    0.00000    0.00000 ;     CA-     C-     N-    CA
  2415   2414   2416   2417       3    0.65084    1.95253    0.00000
-2.60338    0.00000    0.00000 ;      HA-    CA-    CB-   HB2
  2415   2414   2416   2418       3    0.65084    1.95253    0.00000
-2.60338    0.00000    0.00000 ;      HA-    CA-    CB-   HB3
  2415   2414   2416   2419       3    0.65084    1.95253    0.00000
-2.60338    0.00000    0.00000 ;      HA-    CA-    CB-    CG
  2415   2414   2432   2433       3    3.68192   -4.35136    0.00000
1.33888    0.00000    0.00000 ;     HA-    CA-     C-     O
  2415   2414   2432   2434       3    0.00000    0.00000    0.00000
0.00000    0.00000    0.00000 ;     HA-    CA-     C-     N
  2416   2414   2432   2433       3    0.00000    0.00000    0.00000
0.00000    0.00000    0.00000 ;     CB-    CA-     C-     O
  2416   2414   2432   2434       3    2.51040    4.18400    1.67360
-6.69440    0.00000    0.00000 ;      CB-    CA-     C-     N
  2416   2419   2422   2423       3    0.66944    2.00832    0.00000
-2.67776    0.00000    0.00000 ;      CB-    CG-    CD-   HD2
  2416   2419   2422   2424       3    0.66944    2.00832    0.00000
-2.67776    0.00000    0.00000 ;      CB-    CG-    CD-   HD3
  2416   2419   2422   2425       3    3.68192    3.09616   -2.09200
```

```
-3.01248    0.00000    0.00000 ;      CB-    CG-    CD-    CE
  2417   2416   2414   2432      3    0.65084    1.95253    0.00000
-2.60338    0.00000    0.00000 ;     HB2-    CB-    CA-     C
  2417   2416   2419   2420      3    0.62760    1.88280    0.00000
-2.51040    0.00000    0.00000 ;     HB2-    CB-    CG-   HG2
  2417   2416   2419   2421      3    0.62760    1.88280    0.00000
-2.51040    0.00000    0.00000 ;     HB2-    CB-    CG-   HG3
  2417   2416   2419   2422      3    0.66944    2.00832    0.00000
-2.67776    0.00000    0.00000 ;     HB2-    CB-    CG-    CD
  2418   2416   2414   2432      3    0.65084    1.95253    0.00000
-2.60338    0.00000    0.00000 ;     HB3-    CB-    CA-     C
  2418   2416   2419   2420      3    0.62760    1.88280    0.00000
-2.51040    0.00000    0.00000 ;     HB3-    CB-    CG-   HG2
  2418   2416   2419   2421      3    0.62760    1.88280    0.00000
-2.51040    0.00000    0.00000 ;     HB3-    CB-    CG-   HG3
  2418   2416   2419   2422      3    0.66944    2.00832    0.00000
-2.67776    0.00000    0.00000 ;     HB3-    CB-    CG-    CD
  2419   2416   2414   2432      3    0.65084    1.95253    0.00000
-2.60338    0.00000    0.00000 ;      CG-    CB-    CA-     C
  2419   2422   2425   2426      3    0.65084    1.95253    0.00000
-2.60338    0.00000    0.00000 ;      CG-    CD-    CE-   HE2
  2419   2422   2425   2427      3    0.65084    1.95253    0.00000
-2.60338    0.00000    0.00000 ;      CG-    CD-    CE-   HE3
  2419   2422   2425   2428      3    0.65084    1.95253    0.00000
-2.60338    0.00000    0.00000 ;      CG-    CD-    CE-    NZ
  2420   2419   2422   2423      3    0.62760    1.88280    0.00000
-2.51040    0.00000    0.00000 ;     HG2-    CG-    CD-   HD2
  2420   2419   2422   2424      3    0.62760    1.88280    0.00000
-2.51040    0.00000    0.00000 ;     HG2-    CG-    CD-   HD3
  2420   2419   2422   2425      3    0.66944    2.00832    0.00000
-2.67776    0.00000    0.00000 ;     HG2-    CG-    CD-    CE
  2421   2419   2422   2423      3    0.62760    1.88280    0.00000
-2.51040    0.00000    0.00000 ;     HG3-    CG-    CD-   HD2
  2421   2419   2422   2424      3    0.62760    1.88280    0.00000
-2.51040    0.00000    0.00000 ;     HG3-    CG-    CD-   HD3
  2421   2419   2422   2425      3    0.66944    2.00832    0.00000
-2.67776    0.00000    0.00000 ;     HG3-    CG-    CD-    CE
  2422   2425   2428   2429      3    0.65084    1.95253    0.00000
-2.60338    0.00000    0.00000 ;      CD-    CE-    NZ-   HZ1
  2422   2425   2428   2430      3    0.65084    1.95253    0.00000
-2.60338    0.00000    0.00000 ;      CD-    CE-    NZ-   HZ2
  2422   2425   2428   2431      3    0.65084    1.95253    0.00000
-2.60338    0.00000    0.00000 ;      CD-    CE-    NZ-   HZ3
  2423   2422   2425   2426      3    0.65084    1.95253    0.00000
-2.60338    0.00000    0.00000 ;     HD2-    CD-    CE-   HE2
  2423   2422   2425   2427      3    0.65084    1.95253    0.00000
-2.60338    0.00000    0.00000 ;     HD2-    CD-    CE-   HE3
  2423   2422   2425   2428      3    0.65084    1.95253    0.00000
-2.60338    0.00000    0.00000 ;     HD2-    CD-    CE-    NZ
  2424   2422   2425   2426      3    0.65084    1.95253    0.00000
-2.60338    0.00000    0.00000 ;     HD3-    CD-    CE-   HE2
  2424   2422   2425   2427      3    0.65084    1.95253    0.00000
```

-2.60338 0.00000 0.00000 ; HD3- CD- CE- HE3
2424 2422 2425 2428 3 0.65084 1.95253 0.00000
-2.60338 0.00000 0.00000 ; HD3- CD- CE- NZ
2426 2425 2428 2429 3 0.65084 1.95253 0.00000
-2.60338 0.00000 0.00000 ; HE2- CE- NZ- HZ1
2426 2425 2428 2430 3 0.65084 1.95253 0.00000
-2.60338 0.00000 0.00000 ; HE2- CE- NZ- HZ2
2426 2425 2428 2431 3 0.65084 1.95253 0.00000
-2.60338 0.00000 0.00000 ; HE2- CE- NZ- HZ3
2427 2425 2428 2429 3 0.65084 1.95253 0.00000
-2.60338 0.00000 0.00000 ; HE3- CE- NZ- HZ1
2427 2425 2428 2430 3 0.65084 1.95253 0.00000
-2.60338 0.00000 0.00000 ; HE3- CE- NZ- HZ2
2427 2425 2428 2431 3 0.65084 1.95253 0.00000
-2.60338 0.00000 0.00000 ; HE3- CE- NZ- HZ3
2432 2434 2436 2437 3 0.00000 0.00000 0.00000
0.00000 0.00000 0.00000 ; C- N- CA- HA
2432 2434 2436 2438 3 11.71520 1.67360 15.06240 -
13.38880 0.00000 0.00000 ; C- N- CA- CB
2432 2434 2436 2444 3 1.75728 5.27184 2.25936
-7.02912 0.00000 0.00000 ; C- N- CA- C
2433 2432 2434 2435 3 29.28800 -8.36800 -20.92000
0.00000 0.00000 0.00000 ; O- C- N- H
2433 2432 2434 2436 3 20.92000 0.00000 -20.92000
0.00000 0.00000 0.00000 ; O- C- N- CA
2434 2436 2438 2439 3 0.65084 1.95253 0.00000
-2.60338 0.00000 0.00000 ; N- CA- CB- HB2
2434 2436 2438 2440 3 0.65084 1.95253 0.00000
-2.60338 0.00000 0.00000 ; N- CA- CB- HB3
2434 2436 2438 2441 3 15.41386 9.74035 -8.39310
-0.97069 2.97901 0.00000 ; N- CA- CB- CG
2434 2436 2444 2445 3 0.00000 0.00000 0.00000
0.00000 0.00000 0.00000 ; N- CA- C- O
2434 2436 2444 2446 3 17.40544 -5.02080 -13.22144
9.20480 0.00000 0.00000 ; N- CA- C- N
2435 2434 2436 2437 3 0.00000 0.00000 0.00000
0.00000 0.00000 0.00000 ; H- N- CA- HA
2435 2434 2436 2438 3 0.00000 0.00000 0.00000
0.00000 0.00000 0.00000 ; H- N- CA- CB
2435 2434 2436 2444 3 0.00000 0.00000 0.00000
0.00000 0.00000 0.00000 ; H- N- CA- C
2436 2438 2441 2442 3 6.43499 0.00000 -5.39736
0.00000 -1.03763 0.00000 ; CA- CB- CG- OD1
2436 2438 2441 2443 3 6.43499 0.00000 -5.39736
0.00000 -1.03763 0.00000 ; CA- CB- CG- OD2
2436 2444 2446 2447 3 20.92000 0.00000 -20.92000
0.00000 0.00000 0.00000 ; CA- C- N- H
2436 2444 2446 2448 3 20.92000 0.00000 -20.92000
0.00000 0.00000 0.00000 ; CA- C- N- CA
2437 2436 2438 2439 3 0.65084 1.95253 0.00000
-2.60338 0.00000 0.00000 ; HA- CA- CB- HB2
2437 2436 2438 2440 3 0.65084 1.95253 0.00000

```
-2.60338    0.00000    0.00000 ;      HA-    CA-    CB-   HB3
  2437   2436   2438   2441       3    0.65084    1.95253    0.00000
-2.60338    0.00000    0.00000 ;      HA-    CA-    CB-    CG
  2437   2436   2444   2445       3    3.68192   -4.35136    0.00000
1.33888    0.00000    0.00000 ;      HA-    CA-     C-     O
  2437   2436   2444   2446       3    0.00000    0.00000    0.00000
0.00000    0.00000    0.00000 ;      HA-    CA-     C-     N
  2438   2436   2444   2445       3    0.00000    0.00000    0.00000
0.00000    0.00000    0.00000 ;      CB-    CA-     C-     O
  2438   2436   2444   2446       3    2.51040    4.18400    1.67360
-6.69440    0.00000    0.00000 ;      CB-    CA-     C-     N
  2439   2438   2436   2444       3    0.65084    1.95253    0.00000
-2.60338    0.00000    0.00000 ;     HB2-    CB-    CA-     C
  2439   2438   2441   2442       3    0.00000    0.00000    0.00000
0.00000    0.00000    0.00000 ;     HB2-    CB-    CG-   OD1
  2439   2438   2441   2443       3    0.00000    0.00000    0.00000
0.00000    0.00000    0.00000 ;     HB2-    CB-    CG-   OD2
  2440   2438   2436   2444       3    0.65084    1.95253    0.00000
-2.60338    0.00000    0.00000 ;     HB3-    CB-    CA-     C
  2440   2438   2441   2442       3    0.00000    0.00000    0.00000
0.00000    0.00000    0.00000 ;     HB3-    CB-    CG-   OD1
  2440   2438   2441   2443       3    0.00000    0.00000    0.00000
0.00000    0.00000    0.00000 ;     HB3-    CB-    CG-   OD2
  2441   2438   2436   2444       3    7.14627   -1.04600   -8.99560
-0.97069    5.15469    0.00000 ;      CG-    CB-    CA-     C
  2444   2446   2448   2449       3    0.00000    0.00000    0.00000
0.00000    0.00000    0.00000 ;       C-     N-    CA-    HA
  2444   2446   2448   2450       3   11.71520    1.67360   15.06240  -
13.38880    0.00000    0.00000 ;       C-     N-    CA-    CB
  2444   2446   2448   2455       3    1.75728    5.27184    2.25936
-7.02912    0.00000    0.00000 ;       C-     N-    CA-     C
  2445   2444   2446   2447       3   29.28800   -8.36800  -20.92000
0.00000    0.00000    0.00000 ;       O-     C-     N-     H
  2445   2444   2446   2448       3   20.92000    0.00000  -20.92000
0.00000    0.00000    0.00000 ;       O-     C-     N-    CA
  2446   2448   2450   2451       3    0.65084    1.95253    0.00000
-2.60338    0.00000    0.00000 ;       N-    CA-    CB-   HB2
  2446   2448   2450   2452       3    0.65084    1.95253    0.00000
-2.60338    0.00000    0.00000 ;       N-    CA-    CB-   HB3
  2446   2448   2450   2453       3    7.86174    2.24681   -7.41405
-6.71114    5.35552    0.00000 ;       N-    CA-    CB-    OG
  2446   2448   2455   2456       3    0.00000    0.00000    0.00000
0.00000    0.00000    0.00000 ;       N-    CA-     C-     O
  2446   2448   2455   2457       3   17.40544   -5.02080  -13.22144
9.20480    0.00000    0.00000 ;       N-    CA-     C-     N
  2447   2446   2448   2449       3    0.00000    0.00000    0.00000
0.00000    0.00000    0.00000 ;       H-     N-    CA-    HA
  2447   2446   2448   2450       3    0.00000    0.00000    0.00000
0.00000    0.00000    0.00000 ;       H-     N-    CA-    CB
  2447   2446   2448   2455       3    0.00000    0.00000    0.00000
0.00000    0.00000    0.00000 ;       H-     N-    CA-     C
  2448   2450   2453   2454       3    2.05853    2.46856    3.48109
```

-4.46851 0.23430 0.00000 ; CA- CB- OG- HG
2448 2455 2457 2458 3 20.92000 0.00000 -20.92000
0.00000 0.00000 0.00000 ; CA- C- N- H
2448 2455 2457 2459 3 20.92000 0.00000 -20.92000
0.00000 0.00000 0.00000 ; CA- C- N- CA
2449 2448 2450 2451 3 0.65084 1.95253 0.00000
-2.60338 0.00000 0.00000 ; HA- CA- CB- HB2
2449 2448 2450 2452 3 0.65084 1.95253 0.00000
-2.60338 0.00000 0.00000 ; HA- CA- CB- HB3
2449 2448 2450 2453 3 1.04600 -1.04600 0.00000
0.00000 0.00000 0.00000 ; HA- CA- CB- OG
2449 2448 2455 2456 3 3.68192 -4.35136 0.00000
1.33888 0.00000 0.00000 ; HA- CA- C- O
2449 2448 2455 2457 3 0.00000 0.00000 0.00000
0.00000 0.00000 0.00000 ; HA- CA- C- N
2450 2448 2455 2456 3 0.00000 0.00000 0.00000
0.00000 0.00000 0.00000 ; CB- CA- C- O
2450 2448 2455 2457 3 2.51040 4.18400 1.67360
-6.69440 0.00000 0.00000 ; CB- CA- C- N
2451 2450 2448 2455 3 0.65084 1.95253 0.00000
-2.60338 0.00000 0.00000 ; HB2- CB- CA- C
2451 2450 2453 2454 3 0.69733 2.09200 0.00000
-2.78933 0.00000 0.00000 ; HB2- CB- OG- HG
2452 2450 2448 2455 3 0.65084 1.95253 0.00000
-2.60338 0.00000 0.00000 ; HB3- CB- CA- C
2452 2450 2453 2454 3 0.69733 2.09200 0.00000
-2.78933 0.00000 0.00000 ; HB3- CB- OG- HG
2453 2450 2448 2455 3 7.34710 7.79898 -6.14211
-6.71114 4.31789 0.00000 ; OG- CB- CA- C
2455 2457 2459 2460 3 0.00000 0.00000 0.00000
0.00000 0.00000 0.00000 ; C- N- CA- HA
2455 2457 2459 2461 3 11.71520 1.67360 15.06240 -
13.38880 0.00000 0.00000 ; C- N- CA- CB
2455 2457 2459 2474 3 1.75728 5.27184 2.25936
-7.02912 0.00000 0.00000 ; C- N- CA- C
2456 2455 2457 2458 3 29.28800 -8.36800 -20.92000
0.00000 0.00000 0.00000 ; O- C- N- H
2456 2455 2457 2459 3 20.92000 0.00000 -20.92000
0.00000 0.00000 0.00000 ; O- C- N- CA
2457 2459 2461 2462 3 0.65084 1.95253 0.00000
-2.60338 0.00000 0.00000 ; N- CA- CB- HB2
2457 2459 2461 2463 3 0.65084 1.95253 0.00000
-2.60338 0.00000 0.00000 ; N- CA- CB- HB3
2457 2459 2461 2464 3 3.79070 1.39746 -4.61077
-2.40998 2.44346 0.00000 ; N- CA- CB- CG
2457 2459 2474 2475 3 0.00000 0.00000 0.00000
0.00000 0.00000 0.00000 ; N- CA- C- O
2457 2459 2474 2476 3 17.40544 -5.02080 -13.22144
9.20480 0.00000 0.00000 ; N- CA- C- N
2458 2457 2459 2460 3 0.00000 0.00000 0.00000
0.00000 0.00000 0.00000 ; H- N- CA- HA
2458 2457 2459 2461 3 0.00000 0.00000 0.00000

```
0.00000    0.00000    0.00000 ;       H-     N-     CA-    CB
  2458   2457   2459   2474       3    0.00000    0.00000    0.00000
0.00000    0.00000    0.00000 ;       H-     N-     CA-     C
  2459   2461   2464   2465       3    0.66944    2.00832    0.00000
-2.67776    0.00000    0.00000 ;      CA-    CB-    CG-    HG
  2459   2461   2464   2466       3    3.90367    0.19665   -6.21742
-2.37651    5.99149    0.00000 ;      CA-    CB-    CG-   CD1
  2459   2461   2464   2470       3    3.90367    0.19665   -6.21742
-2.37651    5.99149    0.00000 ;      CA-    CB-    CG-   CD2
  2459   2474   2476   2477       3   20.92000    0.00000  -20.92000
0.00000    0.00000    0.00000 ;      CA-     C-     N-     H
  2459   2474   2476   2478       3   20.92000    0.00000  -20.92000
0.00000    0.00000    0.00000 ;      CA-     C-     N-    CA
  2460   2459   2461   2462       3    0.65084    1.95253    0.00000
-2.60338    0.00000    0.00000 ;      HA-    CA-    CB-   HB2
  2460   2459   2461   2463       3    0.65084    1.95253    0.00000
-2.60338    0.00000    0.00000 ;      HA-    CA-    CB-   HB3
  2460   2459   2461   2464       3    0.65084    1.95253    0.00000
-2.60338    0.00000    0.00000 ;      HA-    CA-    CB-    CG
  2460   2459   2474   2475       3    3.68192   -4.35136    0.00000
1.33888    0.00000    0.00000 ;      HA-    CA-     C-     O
  2460   2459   2474   2476       3    0.00000    0.00000    0.00000
0.00000    0.00000    0.00000 ;      HA-    CA-     C-     N
  2461   2459   2474   2475       3    0.00000    0.00000    0.00000
0.00000    0.00000    0.00000 ;      CB-    CA-     C-     O
  2461   2459   2474   2476       3    2.51040    4.18400    1.67360
-6.69440    0.00000    0.00000 ;      CB-    CA-     C-     N
  2461   2464   2466   2467       3    0.66944    2.00832    0.00000
-2.67776    0.00000    0.00000 ;      CB-    CG-   CD1-  HD11
  2461   2464   2466   2468       3    0.66944    2.00832    0.00000
-2.67776    0.00000    0.00000 ;      CB-    CG-   CD1-  HD12
  2461   2464   2466   2469       3    0.66944    2.00832    0.00000
-2.67776    0.00000    0.00000 ;      CB-    CG-   CD1-  HD13
  2461   2464   2470   2471       3    0.66944    2.00832    0.00000
-2.67776    0.00000    0.00000 ;      CB-    CG-   CD2-  HD21
  2461   2464   2470   2472       3    0.66944    2.00832    0.00000
-2.67776    0.00000    0.00000 ;      CB-    CG-   CD2-  HD22
  2461   2464   2470   2473       3    0.66944    2.00832    0.00000
-2.67776    0.00000    0.00000 ;      CB-    CG-   CD2-  HD23
  2462   2461   2459   2474       3    0.65084    1.95253    0.00000
-2.60338    0.00000    0.00000 ;     HB2-    CB-    CA-     C
  2462   2461   2464   2465       3    0.62760    1.88280    0.00000
-2.51040    0.00000    0.00000 ;     HB2-    CB-    CG-    HG
  2462   2461   2464   2466       3    0.66944    2.00832    0.00000
-2.67776    0.00000    0.00000 ;     HB2-    CB-    CG-   CD1
  2462   2461   2464   2470       3    0.66944    2.00832    0.00000
-2.67776    0.00000    0.00000 ;     HB2-    CB-    CG-   CD2
  2463   2461   2459   2474       3    0.65084    1.95253    0.00000
-2.60338    0.00000    0.00000 ;     HB3-    CB-    CA-     C
  2463   2461   2464   2465       3    0.62760    1.88280    0.00000
-2.51040    0.00000    0.00000 ;     HB3-    CB-    CG-    HG
  2463   2461   2464   2466       3    0.66944    2.00832    0.00000
```

-2.67776 0.00000 0.00000 ; HB3- CB- CG- CD1
2463 2461 2464 2470 3 0.66944 2.00832 0.00000
-2.67776 0.00000 0.00000 ; HB3- CB- CG- CD2
2464 2461 2459 2474 3 10.33448 -1.14642 -11.54784
-2.40998 6.35968 0.00000 ; CG- CB- CA- C
2465 2464 2466 2467 3 0.62760 1.88280 0.00000
-2.51040 0.00000 0.00000 ; HG- CG- CD1- HD11
2465 2464 2466 2468 3 0.62760 1.88280 0.00000
-2.51040 0.00000 0.00000 ; HG- CG- CD1- HD12
2465 2464 2466 2469 3 0.62760 1.88280 0.00000
-2.51040 0.00000 0.00000 ; HG- CG- CD1- HD13
2465 2464 2470 2471 3 0.62760 1.88280 0.00000
-2.51040 0.00000 0.00000 ; HG- CG- CD2- HD21
2465 2464 2470 2472 3 0.62760 1.88280 0.00000
-2.51040 0.00000 0.00000 ; HG- CG- CD2- HD22
2465 2464 2470 2473 3 0.62760 1.88280 0.00000
-2.51040 0.00000 0.00000 ; HG- CG- CD2- HD23
2466 2464 2470 2471 3 0.66944 2.00832 0.00000
-2.67776 0.00000 0.00000 ; CD1- CG- CD2- HD21
2466 2464 2470 2472 3 0.66944 2.00832 0.00000
-2.67776 0.00000 0.00000 ; CD1- CG- CD2- HD22
2466 2464 2470 2473 3 0.66944 2.00832 0.00000
-2.67776 0.00000 0.00000 ; CD1- CG- CD2- HD23
2467 2466 2464 2470 3 0.66944 2.00832 0.00000
-2.67776 0.00000 0.00000 ; HD11- CD1- CG- CD2
2468 2466 2464 2470 3 0.66944 2.00832 0.00000
-2.67776 0.00000 0.00000 ; HD12- CD1- CG- CD2
2469 2466 2464 2470 3 0.66944 2.00832 0.00000
-2.67776 0.00000 0.00000 ; HD13- CD1- CG- CD2
2474 2476 2478 2479 3 0.00000 0.00000 0.00000
0.00000 0.00000 0.00000 ; C- N- CA- HA
2474 2476 2478 2480 3 11.71520 1.67360 15.06240 -
13.38880 0.00000 0.00000 ; C- N- CA- CB
2474 2476 2478 2493 3 1.75728 5.27184 2.25936
-7.02912 0.00000 0.00000 ; C- N- CA- C
2475 2474 2476 2477 3 29.28800 -8.36800 -20.92000
0.00000 0.00000 0.00000 ; O- C- N- H
2475 2474 2476 2478 3 20.92000 0.00000 -20.92000
0.00000 0.00000 0.00000 ; O- C- N- CA
2476 2478 2480 2481 3 0.65084 1.95253 0.00000
-2.60338 0.00000 0.00000 ; N- CA- CB- HB2
2476 2478 2480 2482 3 0.65084 1.95253 0.00000
-2.60338 0.00000 0.00000 ; N- CA- CB- HB3
2476 2478 2480 2483 3 3.79070 1.39746 -4.61077
-2.40998 2.44346 0.00000 ; N- CA- CB- CG
2476 2478 2493 2494 3 0.00000 0.00000 0.00000
0.00000 0.00000 0.00000 ; N- CA- C- O
2476 2478 2493 2495 3 17.40544 -5.02080 -13.22144
9.20480 0.00000 0.00000 ; N- CA- C- N
2477 2476 2478 2479 3 0.00000 0.00000 0.00000
0.00000 0.00000 0.00000 ; H- N- CA- HA
2477 2476 2478 2480 3 0.00000 0.00000 0.00000

```
0.00000    0.00000    0.00000 ;       H-     N-     CA-    CB
  2477   2476   2478   2493       3    0.00000    0.00000    0.00000
0.00000    0.00000    0.00000 ;       H-     N-     CA-     C
  2478   2480   2483   2484       3    0.66944    2.00832    0.00000
-2.67776    0.00000    0.00000 ;      CA-    CB-     CG-    HG
  2478   2480   2483   2485       3    3.90367    0.19665   -6.21742
-2.37651    5.99149    0.00000 ;      CA-    CB-     CG-   CD1
  2478   2480   2483   2489       3    3.90367    0.19665   -6.21742
-2.37651    5.99149    0.00000 ;      CA-    CB-     CG-   CD2
  2478   2493   2495   2496       3   20.92000    0.00000  -20.92000
0.00000    0.00000    0.00000 ;      CA-     C-      N-     H
  2478   2493   2495   2497       3   20.92000    0.00000  -20.92000
0.00000    0.00000    0.00000 ;      CA-     C-      N-    CA
  2479   2478   2480   2481       3    0.65084    1.95253    0.00000
-2.60338    0.00000    0.00000 ;      HA-    CA-     CB-   HB2
  2479   2478   2480   2482       3    0.65084    1.95253    0.00000
-2.60338    0.00000    0.00000 ;      HA-    CA-     CB-   HB3
  2479   2478   2480   2483       3    0.65084    1.95253    0.00000
-2.60338    0.00000    0.00000 ;      HA-    CA-     CB-    CG
  2479   2478   2493   2494       3    3.68192   -4.35136    0.00000
1.33888    0.00000    0.00000 ;      HA-    CA-      C-     O
  2479   2478   2493   2495       3    0.00000    0.00000    0.00000
0.00000    0.00000    0.00000 ;      HA-    CA-      C-     N
  2480   2478   2493   2494       3    0.00000    0.00000    0.00000
0.00000    0.00000    0.00000 ;      CB-    CA-      C-     O
  2480   2478   2493   2495       3    2.51040    4.18400    1.67360
-6.69440    0.00000    0.00000 ;      CB-    CA-      C-     N
  2480   2483   2485   2486       3    0.66944    2.00832    0.00000
-2.67776    0.00000    0.00000 ;      CB-    CG-   CD1-  HD11
  2480   2483   2485   2487       3    0.66944    2.00832    0.00000
-2.67776    0.00000    0.00000 ;      CB-    CG-   CD1-  HD12
  2480   2483   2485   2488       3    0.66944    2.00832    0.00000
-2.67776    0.00000    0.00000 ;      CB-    CG-   CD1-  HD13
  2480   2483   2489   2490       3    0.66944    2.00832    0.00000
-2.67776    0.00000    0.00000 ;      CB-    CG-   CD2-  HD21
  2480   2483   2489   2491       3    0.66944    2.00832    0.00000
-2.67776    0.00000    0.00000 ;      CB-    CG-   CD2-  HD22
  2480   2483   2489   2492       3    0.66944    2.00832    0.00000
-2.67776    0.00000    0.00000 ;      CB-    CG-   CD2-  HD23
  2481   2480   2478   2493       3    0.65084    1.95253    0.00000
-2.60338    0.00000    0.00000 ;     HB2-    CB-     CA-     C
  2481   2480   2483   2484       3    0.62760    1.88280    0.00000
-2.51040    0.00000    0.00000 ;     HB2-    CB-     CG-    HG
  2481   2480   2483   2485       3    0.66944    2.00832    0.00000
-2.67776    0.00000    0.00000 ;     HB2-    CB-     CG-   CD1
  2481   2480   2483   2489       3    0.66944    2.00832    0.00000
-2.67776    0.00000    0.00000 ;     HB2-    CB-     CG-   CD2
  2482   2480   2478   2493       3    0.65084    1.95253    0.00000
-2.60338    0.00000    0.00000 ;     HB3-    CB-     CA-     C
  2482   2480   2483   2484       3    0.62760    1.88280    0.00000
-2.51040    0.00000    0.00000 ;     HB3-    CB-     CG-    HG
  2482   2480   2483   2485       3    0.66944    2.00832    0.00000
```

-2.67776 0.00000 0.00000 ; HB3- CB- CG- CD1
2482 2480 2483 2489 3 0.66944 2.00832 0.00000
-2.67776 0.00000 0.00000 ; HB3- CB- CG- CD2
2483 2480 2478 2493 3 10.33448 -1.14642 -11.54784
-2.40998 6.35968 0.00000 ; CG- CB- CA- C
2484 2483 2485 2486 3 0.62760 1.88280 0.00000
-2.51040 0.00000 0.00000 ; HG- CG- CD1- HD11
2484 2483 2485 2487 3 0.62760 1.88280 0.00000
-2.51040 0.00000 0.00000 ; HG- CG- CD1- HD12
2484 2483 2485 2488 3 0.62760 1.88280 0.00000
-2.51040 0.00000 0.00000 ; HG- CG- CD1- HD13
2484 2483 2489 2490 3 0.62760 1.88280 0.00000
-2.51040 0.00000 0.00000 ; HG- CG- CD2- HD21
2484 2483 2489 2491 3 0.62760 1.88280 0.00000
-2.51040 0.00000 0.00000 ; HG- CG- CD2- HD22
2484 2483 2489 2492 3 0.62760 1.88280 0.00000
-2.51040 0.00000 0.00000 ; HG- CG- CD2- HD23
2485 2483 2489 2490 3 0.66944 2.00832 0.00000
-2.67776 0.00000 0.00000 ; CD1- CG- CD2- HD21
2485 2483 2489 2491 3 0.66944 2.00832 0.00000
-2.67776 0.00000 0.00000 ; CD1- CG- CD2- HD22
2485 2483 2489 2492 3 0.66944 2.00832 0.00000
-2.67776 0.00000 0.00000 ; CD1- CG- CD2- HD23
2486 2485 2483 2489 3 0.66944 2.00832 0.00000
-2.67776 0.00000 0.00000 ; HD11- CD1- CG- CD2
2487 2485 2483 2489 3 0.66944 2.00832 0.00000
-2.67776 0.00000 0.00000 ; HD12- CD1- CG- CD2
2488 2485 2483 2489 3 0.66944 2.00832 0.00000
-2.67776 0.00000 0.00000 ; HD13- CD1- CG- CD2
2493 2495 2497 2498 3 0.00000 0.00000 0.00000
0.00000 0.00000 0.00000 ; C- N- CA- HA
2493 2495 2497 2499 3 11.71520 1.67360 15.06240 -
13.38880 0.00000 0.00000 ; C- N- CA- CB
2493 2495 2497 2515 3 1.75728 5.27184 2.25936
-7.02912 0.00000 0.00000 ; C- N- CA- C
2494 2493 2495 2496 3 29.28800 -8.36800 -20.92000
0.00000 0.00000 0.00000 ; O- C- N- H
2494 2493 2495 2497 3 20.92000 0.00000 -20.92000
0.00000 0.00000 0.00000 ; O- C- N- CA
2495 2497 2499 2500 3 0.65084 1.95253 0.00000
-2.60338 0.00000 0.00000 ; N- CA- CB- HB2
2495 2497 2499 2501 3 0.65084 1.95253 0.00000
-2.60338 0.00000 0.00000 ; N- CA- CB- HB3
2495 2497 2499 2502 3 0.65084 1.95253 0.00000
-2.60338 0.00000 0.00000 ; N- CA- CB- CG
2495 2497 2515 2516 3 0.00000 0.00000 0.00000
0.00000 0.00000 0.00000 ; N- CA- C- O
2495 2497 2515 2517 3 17.40544 -5.02080 -13.22144
9.20480 0.00000 0.00000 ; N- CA- C- N
2496 2495 2497 2498 3 0.00000 0.00000 0.00000
0.00000 0.00000 0.00000 ; H- N- CA- HA
2496 2495 2497 2499 3 0.00000 0.00000 0.00000

```
0.00000    0.00000    0.00000 ;      H-    N-    CA-    CB
  2496   2495   2497   2515      3    0.00000    0.00000    0.00000
0.00000    0.00000    0.00000 ;      H-    N-    CA-    C
  2497   2499   2502   2503      3    0.66944    2.00832    0.00000
-2.67776    0.00000    0.00000 ;      CA-    CB-    CG-    HG2
  2497   2499   2502   2504      3    0.66944    2.00832    0.00000
-2.67776    0.00000    0.00000 ;      CA-    CB-    CG-    HG3
  2497   2499   2502   2505      3    3.68192    3.09616   -2.09200
-3.01248    0.00000    0.00000 ;      CA-    CB-    CG-    CD
  2497   2515   2517   2518      3   20.92000    0.00000  -20.92000
0.00000    0.00000    0.00000 ;      CA-    C-    N-    H
  2497   2515   2517   2519      3   20.92000    0.00000  -20.92000
0.00000    0.00000    0.00000 ;      CA-    C-    N-    CA
  2498   2497   2499   2500      3    0.65084    1.95253    0.00000
-2.60338    0.00000    0.00000 ;      HA-    CA-    CB-    HB2
  2498   2497   2499   2501      3    0.65084    1.95253    0.00000
-2.60338    0.00000    0.00000 ;      HA-    CA-    CB-    HB3
  2498   2497   2499   2502      3    0.65084    1.95253    0.00000
-2.60338    0.00000    0.00000 ;      HA-    CA-    CB-    CG
  2498   2497   2515   2516      3    3.68192   -4.35136    0.00000
1.33888    0.00000    0.00000 ;      HA-    CA-    C-    O
  2498   2497   2515   2517      3    0.00000    0.00000    0.00000
0.00000    0.00000    0.00000 ;      HA-    CA-    C-    N
  2499   2497   2515   2516      3    0.00000    0.00000    0.00000
0.00000    0.00000    0.00000 ;      CB-    CA-    C-    O
  2499   2497   2515   2517      3    2.51040    4.18400    1.67360
-6.69440    0.00000    0.00000 ;      CB-    CA-    C-    N
  2499   2502   2505   2506      3    0.66944    2.00832    0.00000
-2.67776    0.00000    0.00000 ;      CB-    CG-    CD-    HD2
  2499   2502   2505   2507      3    0.66944    2.00832    0.00000
-2.67776    0.00000    0.00000 ;      CB-    CG-    CD-    HD3
  2499   2502   2505   2508      3    3.68192    3.09616   -2.09200
-3.01248    0.00000    0.00000 ;      CB-    CG-    CD-    CE
  2500   2499   2497   2515      3    0.65084    1.95253    0.00000
-2.60338    0.00000    0.00000 ;      HB2-    CB-    CA-    C
  2500   2499   2502   2503      3    0.62760    1.88280    0.00000
-2.51040    0.00000    0.00000 ;      HB2-    CB-    CG-    HG2
  2500   2499   2502   2504      3    0.62760    1.88280    0.00000
-2.51040    0.00000    0.00000 ;      HB2-    CB-    CG-    HG3
  2500   2499   2502   2505      3    0.66944    2.00832    0.00000
-2.67776    0.00000    0.00000 ;      HB2-    CB-    CG-    CD
  2501   2499   2497   2515      3    0.65084    1.95253    0.00000
-2.60338    0.00000    0.00000 ;      HB3-    CB-    CA-    C
  2501   2499   2502   2503      3    0.62760    1.88280    0.00000
-2.51040    0.00000    0.00000 ;      HB3-    CB-    CG-    HG2
  2501   2499   2502   2504      3    0.62760    1.88280    0.00000
-2.51040    0.00000    0.00000 ;      HB3-    CB-    CG-    HG3
  2501   2499   2502   2505      3    0.66944    2.00832    0.00000
-2.67776    0.00000    0.00000 ;      HB3-    CB-    CG-    CD
  2502   2499   2497   2515      3    0.65084    1.95253    0.00000
-2.60338    0.00000    0.00000 ;      CG-    CB-    CA-    C
  2502   2505   2508   2509      3    0.65084    1.95253    0.00000
```

-2.60338 0.00000 0.00000 ; CG- CD- CE- HE2
2502 2505 2508 2510 3 0.65084 1.95253 0.00000
-2.60338 0.00000 0.00000 ; CG- CD- CE- HE3
2502 2505 2508 2511 3 0.65084 1.95253 0.00000
-2.60338 0.00000 0.00000 ; CG- CD- CE- NZ
2503 2502 2505 2506 3 0.62760 1.88280 0.00000
-2.51040 0.00000 0.00000 ; HG2- CG- CD- HD2
2503 2502 2505 2507 3 0.62760 1.88280 0.00000
-2.51040 0.00000 0.00000 ; HG2- CG- CD- HD3
2503 2502 2505 2508 3 0.66944 2.00832 0.00000
-2.67776 0.00000 0.00000 ; HG2- CG- CD- CE
2504 2502 2505 2506 3 0.62760 1.88280 0.00000
-2.51040 0.00000 0.00000 ; HG3- CG- CD- HD2
2504 2502 2505 2507 3 0.62760 1.88280 0.00000
-2.51040 0.00000 0.00000 ; HG3- CG- CD- HD3
2504 2502 2505 2508 3 0.66944 2.00832 0.00000
-2.67776 0.00000 0.00000 ; HG3- CG- CD- CE
2505 2508 2511 2512 3 0.65084 1.95253 0.00000
-2.60338 0.00000 0.00000 ; CD- CE- NZ- HZ1
2505 2508 2511 2513 3 0.65084 1.95253 0.00000
-2.60338 0.00000 0.00000 ; CD- CE- NZ- HZ2
2505 2508 2511 2514 3 0.65084 1.95253 0.00000
-2.60338 0.00000 0.00000 ; CD- CE- NZ- HZ3
2506 2505 2508 2509 3 0.65084 1.95253 0.00000
-2.60338 0.00000 0.00000 ; HD2- CD- CE- HE2
2506 2505 2508 2510 3 0.65084 1.95253 0.00000
-2.60338 0.00000 0.00000 ; HD2- CD- CE- HE3
2506 2505 2508 2511 3 0.65084 1.95253 0.00000
-2.60338 0.00000 0.00000 ; HD2- CD- CE- NZ
2507 2505 2508 2509 3 0.65084 1.95253 0.00000
-2.60338 0.00000 0.00000 ; HD3- CD- CE- HE2
2507 2505 2508 2510 3 0.65084 1.95253 0.00000
-2.60338 0.00000 0.00000 ; HD3- CD- CE- HE3
2507 2505 2508 2511 3 0.65084 1.95253 0.00000
-2.60338 0.00000 0.00000 ; HD3- CD- CE- NZ
2509 2508 2511 2512 3 0.65084 1.95253 0.00000
-2.60338 0.00000 0.00000 ; HE2- CE- NZ- HZ1
2509 2508 2511 2513 3 0.65084 1.95253 0.00000
-2.60338 0.00000 0.00000 ; HE2- CE- NZ- HZ2
2509 2508 2511 2514 3 0.65084 1.95253 0.00000
-2.60338 0.00000 0.00000 ; HE2- CE- NZ- HZ3
2510 2508 2511 2512 3 0.65084 1.95253 0.00000
-2.60338 0.00000 0.00000 ; HE3- CE- NZ- HZ1
2510 2508 2511 2513 3 0.65084 1.95253 0.00000
-2.60338 0.00000 0.00000 ; HE3- CE- NZ- HZ2
2510 2508 2511 2514 3 0.65084 1.95253 0.00000
-2.60338 0.00000 0.00000 ; HE3- CE- NZ- HZ3
2515 2517 2519 2520 3 0.00000 0.00000 0.00000
0.00000 0.00000 0.00000 ; C- N- CA- HA
2515 2517 2519 2521 3 11.71520 1.67360 15.06240 -
13.38880 0.00000 0.00000 ; C- N- CA- CB
2515 2517 2519 2534 3 1.75728 5.27184 2.25936

-7.02912 0.00000 0.00000 ; C- N- CA- C
2516 2515 2517 2518 3 29.28800 -8.36800 -20.92000
0.00000 0.00000 0.00000 ; O- C- N- H
2516 2515 2517 2519 3 20.92000 0.00000 -20.92000
0.00000 0.00000 0.00000 ; O- C- N- CA
2517 2519 2521 2522 3 0.65084 1.95253 0.00000
-2.60338 0.00000 0.00000 ; N- CA- CB- HB
2517 2519 2521 2523 3 3.83673 0.44769 -1.77402
-2.47693 -0.03347 0.00000 ; N- CA- CB- CG2
2517 2519 2521 2527 3 2.97482 0.12134 2.04179
-1.89117 -3.24678 0.00000 ; N- CA- CB- CG1
2517 2519 2534 2535 3 0.00000 0.00000 0.00000
0.00000 0.00000 0.00000 ; N- CA- C- O
2517 2519 2534 2536 3 17.40544 -5.02080 -13.22144
9.20480 0.00000 0.00000 ; N- CA- C- N
2518 2517 2519 2520 3 0.00000 0.00000 0.00000
0.00000 0.00000 0.00000 ; H- N- CA- HA
2518 2517 2519 2521 3 0.00000 0.00000 0.00000
0.00000 0.00000 0.00000 ; H- N- CA- CB
2518 2517 2519 2534 3 0.00000 0.00000 0.00000
0.00000 0.00000 0.00000 ; H- N- CA- C
2519 2521 2523 2524 3 0.66944 2.00832 0.00000
-2.67776 0.00000 0.00000 ; CA- CB- CG2- HG21
2519 2521 2523 2525 3 0.66944 2.00832 0.00000
-2.67776 0.00000 0.00000 ; CA- CB- CG2- HG22
2519 2521 2523 2526 3 0.66944 2.00832 0.00000
-2.67776 0.00000 0.00000 ; CA- CB- CG2- HG23
2519 2521 2527 2528 3 0.66944 2.00832 0.00000
-2.67776 0.00000 0.00000 ; CA- CB- CG1- HG12
2519 2521 2527 2529 3 0.66944 2.00832 0.00000
-2.67776 0.00000 0.00000 ; CA- CB- CG1- HG13
2519 2521 2527 2530 3 4.24258 -0.52718 -7.25506
-1.79075 7.69856 0.00000 ; CA- CB- CG1- CD1
2519 2534 2536 2537 3 20.92000 0.00000 -20.92000
0.00000 0.00000 0.00000 ; CA- C- N- H
2519 2534 2536 2538 3 20.92000 0.00000 -20.92000
0.00000 0.00000 0.00000 ; CA- C- N- CA
2520 2519 2521 2522 3 0.65084 1.95253 0.00000
-2.60338 0.00000 0.00000 ; HA- CA- CB- HB
2520 2519 2521 2523 3 0.65084 1.95253 0.00000
-2.60338 0.00000 0.00000 ; HA- CA- CB- CG2
2520 2519 2521 2527 3 0.65084 1.95253 0.00000
-2.60338 0.00000 0.00000 ; HA- CA- CB- CG1
2520 2519 2534 2535 3 3.68192 -4.35136 0.00000
1.33888 0.00000 0.00000 ; HA- CA- C- O
2520 2519 2534 2536 3 0.00000 0.00000 0.00000
0.00000 0.00000 0.00000 ; HA- CA- C- N
2521 2519 2534 2535 3 0.00000 0.00000 0.00000
0.00000 0.00000 0.00000 ; CB- CA- C- O
2521 2519 2534 2536 3 2.51040 4.18400 1.67360
-6.69440 0.00000 0.00000 ; CB- CA- C- N
2521 2527 2530 2531 3 0.66944 2.00832 0.00000

-2.67776 0.00000 0.00000 ; CB- CG1- CD1- HD11
2521 2527 2530 2532 3 0.66944 2.00832 0.00000
-2.67776 0.00000 0.00000 ; CB- CG1- CD1- HD12
2521 2527 2530 2533 3 0.66944 2.00832 0.00000
-2.67776 0.00000 0.00000 ; CB- CG1- CD1- HD13
2522 2521 2519 2534 3 0.65084 1.95253 0.00000
-2.60338 0.00000 0.00000 ; HB- CB- CA- C
2522 2521 2523 2524 3 0.62760 1.88280 0.00000
-2.51040 0.00000 0.00000 ; HB- CB- CG2- HG21
2522 2521 2523 2525 3 0.62760 1.88280 0.00000
-2.51040 0.00000 0.00000 ; HB- CB- CG2- HG22
2522 2521 2523 2526 3 0.62760 1.88280 0.00000
-2.51040 0.00000 0.00000 ; HB- CB- CG2- HG23
2522 2521 2527 2528 3 0.62760 1.88280 0.00000
-2.51040 0.00000 0.00000 ; HB- CB- CG1- HG12
2522 2521 2527 2529 3 0.62760 1.88280 0.00000
-2.51040 0.00000 0.00000 ; HB- CB- CG1- HG13
2522 2521 2527 2530 3 0.66944 2.00832 0.00000
-2.67776 0.00000 0.00000 ; HB- CB- CG1- CD1
2523 2521 2519 2534 3 5.67350 3.55640 -6.16722
-2.47693 3.74886 0.00000 ; CG2- CB- CA- C
2523 2521 2527 2528 3 0.66944 2.00832 0.00000
-2.67776 0.00000 0.00000 ; CG2- CB- CG1- HG12
2523 2521 2527 2529 3 0.66944 2.00832 0.00000
-2.67776 0.00000 0.00000 ; CG2- CB- CG1- HG13
2523 2521 2527 2530 3 3.81162 0.49790 -8.14206
-1.79075 7.49773 0.00000 ; CG2- CB- CG1- CD1
2524 2523 2521 2527 3 0.66944 2.00832 0.00000
-2.67776 0.00000 0.00000 ; HG21- CG2- CB- CG1
2525 2523 2521 2527 3 0.66944 2.00832 0.00000
-2.67776 0.00000 0.00000 ; HG22- CG2- CB- CG1
2526 2523 2521 2527 3 0.66944 2.00832 0.00000
-2.67776 0.00000 0.00000 ; HG23- CG2- CB- CG1
2527 2521 2519 2534 3 8.26340 0.74057 -9.99976
-1.89117 3.84928 0.00000 ; CG1- CB- CA- C
2528 2527 2530 2531 3 0.62760 1.88280 0.00000
-2.51040 0.00000 0.00000 ; HG12- CG1- CD1- HD11
2528 2527 2530 2532 3 0.62760 1.88280 0.00000
-2.51040 0.00000 0.00000 ; HG12- CG1- CD1- HD12
2528 2527 2530 2533 3 0.62760 1.88280 0.00000
-2.51040 0.00000 0.00000 ; HG12- CG1- CD1- HD13
2529 2527 2530 2531 3 0.62760 1.88280 0.00000
-2.51040 0.00000 0.00000 ; HG13- CG1- CD1- HD11
2529 2527 2530 2532 3 0.62760 1.88280 0.00000
-2.51040 0.00000 0.00000 ; HG13- CG1- CD1- HD12
2529 2527 2530 2533 3 0.62760 1.88280 0.00000
-2.51040 0.00000 0.00000 ; HG13- CG1- CD1- HD13
2534 2536 2538 2539 3 0.00000 0.00000 0.00000
0.00000 0.00000 0.00000 ; C- N- CA- HA
2534 2536 2538 2540 3 11.71520 1.67360 15.06240 -
13.38880 0.00000 0.00000 ; C- N- CA- CB
2534 2536 2538 2553 3 1.75728 5.27184 2.25936

```
-7.02912    0.00000    0.00000 ;       C-     N-    CA-     C
  2535   2534   2536   2537       3   29.28800   -8.36800  -20.92000
0.00000    0.00000    0.00000 ;       O-     C-     N-     H
  2535   2534   2536   2538       3   20.92000    0.00000  -20.92000
0.00000    0.00000    0.00000 ;       O-     C-     N-    CA
  2536   2538   2540   2541       3    0.65084    1.95253    0.00000
-2.60338    0.00000    0.00000 ;       N-    CA-    CB-    HB
  2536   2538   2540   2542       3    3.83673    0.44769   -1.77402
-2.47693   -0.03347    0.00000 ;       N-    CA-    CB-   CG2
  2536   2538   2540   2546       3    2.97482    0.12134    2.04179
-1.89117   -3.24678    0.00000 ;       N-    CA-    CB-   CG1
  2536   2538   2553   2554       3    0.00000    0.00000    0.00000
0.00000    0.00000    0.00000 ;       N-    CA-     C-     O
  2536   2538   2553   2555       3   17.40544   -5.02080  -13.22144
9.20480    0.00000    0.00000 ;       N-    CA-     C-     N
  2537   2536   2538   2539       3    0.00000    0.00000    0.00000
0.00000    0.00000    0.00000 ;       H-     N-    CA-    HA
  2537   2536   2538   2540       3    0.00000    0.00000    0.00000
0.00000    0.00000    0.00000 ;       H-     N-    CA-    CB
  2537   2536   2538   2553       3    0.00000    0.00000    0.00000
0.00000    0.00000    0.00000 ;       H-     N-    CA-     C
  2538   2540   2542   2543       3    0.66944    2.00832    0.00000
-2.67776    0.00000    0.00000 ;      CA-    CB-   CG2-  HG21
  2538   2540   2542   2544       3    0.66944    2.00832    0.00000
-2.67776    0.00000    0.00000 ;      CA-    CB-   CG2-  HG22
  2538   2540   2542   2545       3    0.66944    2.00832    0.00000
-2.67776    0.00000    0.00000 ;      CA-    CB-   CG2-  HG23
  2538   2540   2546   2547       3    0.66944    2.00832    0.00000
-2.67776    0.00000    0.00000 ;      CA-    CB-   CG1-  HG12
  2538   2540   2546   2548       3    0.66944    2.00832    0.00000
-2.67776    0.00000    0.00000 ;      CA-    CB-   CG1-  HG13
  2538   2540   2546   2549       3    4.24258   -0.52718   -7.25506
-1.79075    7.69856    0.00000 ;      CA-    CB-   CG1-   CD1
  2538   2553   2555   2556       3   20.92000    0.00000  -20.92000
0.00000    0.00000    0.00000 ;      CA-     C-     N-     H
  2538   2553   2555   2557       3   20.92000    0.00000  -20.92000
0.00000    0.00000    0.00000 ;      CA-     C-     N-    CA
  2539   2538   2540   2541       3    0.65084    1.95253    0.00000
-2.60338    0.00000    0.00000 ;      HA-    CA-    CB-    HB
  2539   2538   2540   2542       3    0.65084    1.95253    0.00000
-2.60338    0.00000    0.00000 ;      HA-    CA-    CB-   CG2
  2539   2538   2540   2546       3    0.65084    1.95253    0.00000
-2.60338    0.00000    0.00000 ;      HA-    CA-    CB-   CG1
  2539   2538   2553   2554       3    3.68192   -4.35136    0.00000
1.33888    0.00000    0.00000 ;      HA-    CA-     C-     O
  2539   2538   2553   2555       3    0.00000    0.00000    0.00000
0.00000    0.00000    0.00000 ;      HA-    CA-     C-     N
  2540   2538   2553   2554       3    0.00000    0.00000    0.00000
0.00000    0.00000    0.00000 ;      CB-    CA-     C-     O
  2540   2538   2553   2555       3    2.51040    4.18400    1.67360
-6.69440    0.00000    0.00000 ;      CB-    CA-     C-     N
  2540   2546   2549   2550       3    0.66944    2.00832    0.00000
```

```
-2.67776    0.00000    0.00000 ;     CB-   CG1-   CD1-  HD11
  2540   2546   2549   2551      3    0.66944    2.00832    0.00000
-2.67776    0.00000    0.00000 ;     CB-   CG1-   CD1-  HD12
  2540   2546   2549   2552      3    0.66944    2.00832    0.00000
-2.67776    0.00000    0.00000 ;     CB-   CG1-   CD1-  HD13
  2541   2540   2538   2553      3    0.65084    1.95253    0.00000
-2.60338    0.00000    0.00000 ;     HB-    CB-    CA-     C
  2541   2540   2542   2543      3    0.62760    1.88280    0.00000
-2.51040    0.00000    0.00000 ;     HB-    CB-   CG2-  HG21
  2541   2540   2542   2544      3    0.62760    1.88280    0.00000
-2.51040    0.00000    0.00000 ;     HB-    CB-   CG2-  HG22
  2541   2540   2542   2545      3    0.62760    1.88280    0.00000
-2.51040    0.00000    0.00000 ;     HB-    CB-   CG2-  HG23
  2541   2540   2546   2547      3    0.62760    1.88280    0.00000
-2.51040    0.00000    0.00000 ;     HB-    CB-   CG1-  HG12
  2541   2540   2546   2548      3    0.62760    1.88280    0.00000
-2.51040    0.00000    0.00000 ;     HB-    CB-   CG1-  HG13
  2541   2540   2546   2549      3    0.66944    2.00832    0.00000
-2.67776    0.00000    0.00000 ;     HB-    CB-   CG1-   CD1
  2542   2540   2538   2553      3    5.67350    3.55640   -6.16722
-2.47693    3.74886    0.00000 ;    CG2-    CB-    CA-     C
  2542   2540   2546   2547      3    0.66944    2.00832    0.00000
-2.67776    0.00000    0.00000 ;    CG2-    CB-   CG1-  HG12
  2542   2540   2546   2548      3    0.66944    2.00832    0.00000
-2.67776    0.00000    0.00000 ;    CG2-    CB-   CG1-  HG13
  2542   2540   2546   2549      3    3.81162    0.49790   -8.14206
-1.79075    7.49773    0.00000 ;    CG2-    CB-   CG1-   CD1
  2543   2542   2540   2546      3    0.66944    2.00832    0.00000
-2.67776    0.00000    0.00000 ;   HG21-   CG2-    CB-   CG1
  2544   2542   2540   2546      3    0.66944    2.00832    0.00000
-2.67776    0.00000    0.00000 ;   HG22-   CG2-    CB-   CG1
  2545   2542   2540   2546      3    0.66944    2.00832    0.00000
-2.67776    0.00000    0.00000 ;   HG23-   CG2-    CB-   CG1
  2546   2540   2538   2553      3    8.26340    0.74057   -9.99976
-1.89117    3.84928    0.00000 ;    CG1-    CB-    CA-     C
  2547   2546   2549   2550      3    0.62760    1.88280    0.00000
-2.51040    0.00000    0.00000 ;   HG12-   CG1-   CD1-  HD11
  2547   2546   2549   2551      3    0.62760    1.88280    0.00000
-2.51040    0.00000    0.00000 ;   HG12-   CG1-   CD1-  HD12
  2547   2546   2549   2552      3    0.62760    1.88280    0.00000
-2.51040    0.00000    0.00000 ;   HG12-   CG1-   CD1-  HD13
  2548   2546   2549   2550      3    0.62760    1.88280    0.00000
-2.51040    0.00000    0.00000 ;   HG13-   CG1-   CD1-  HD11
  2548   2546   2549   2551      3    0.62760    1.88280    0.00000
-2.51040    0.00000    0.00000 ;   HG13-   CG1-   CD1-  HD12
  2548   2546   2549   2552      3    0.62760    1.88280    0.00000
-2.51040    0.00000    0.00000 ;   HG13-   CG1-   CD1-  HD13
  2553   2555   2557   2558      3    0.00000    0.00000    0.00000
0.00000    0.00000    0.00000 ;      C-     N-    CA-    HA
  2553   2555   2557   2559      3   11.71520    1.67360   15.06240  -
13.38880    0.00000    0.00000 ;      C-     N-    CA-    CB
  2553   2555   2557   2565      3    1.75728    5.27184    2.25936
```

```
-7.02912    0.00000    0.00000 ;       C-     N-    CA-     C
  2554   2553   2555   2556       3   29.28800   -8.36800  -20.92000
0.00000    0.00000    0.00000 ;       O-     C-     N-     H
  2554   2553   2555   2557       3   20.92000    0.00000  -20.92000
0.00000    0.00000    0.00000 ;       O-     C-     N-    CA
  2555   2557   2559   2560       3    0.65084    1.95253    0.00000
-2.60338    0.00000    0.00000 ;       N-    CA-    CB-   HB2
  2555   2557   2559   2561       3    0.65084    1.95253    0.00000
-2.60338    0.00000    0.00000 ;       N-    CA-    CB-   HB3
  2555   2557   2559   2562       3   15.41386    9.74035   -8.39310
-0.97069    2.97901    0.00000 ;       N-    CA-    CB-    CG
  2555   2557   2565   2566       3    0.00000    0.00000    0.00000
0.00000    0.00000    0.00000 ;       N-    CA-     C-     O
  2555   2557   2565   2567       3   17.40544   -5.02080  -13.22144
9.20480    0.00000    0.00000 ;       N-    CA-     C-     N
  2556   2555   2557   2558       3    0.00000    0.00000    0.00000
0.00000    0.00000    0.00000 ;       H-     N-    CA-    HA
  2556   2555   2557   2559       3    0.00000    0.00000    0.00000
0.00000    0.00000    0.00000 ;       H-     N-    CA-    CB
  2556   2555   2557   2565       3    0.00000    0.00000    0.00000
0.00000    0.00000    0.00000 ;       H-     N-    CA-     C
  2557   2559   2562   2563       3    6.43499    0.00000   -5.39736
0.00000   -1.03763    0.00000 ;      CA-    CB-    CG-   OD1
  2557   2559   2562   2564       3    6.43499    0.00000   -5.39736
0.00000   -1.03763    0.00000 ;      CA-    CB-    CG-   OD2
  2557   2565   2567   2568       3   20.92000    0.00000  -20.92000
0.00000    0.00000    0.00000 ;      CA-     C-     N-     H
  2557   2565   2567   2569       3   20.92000    0.00000  -20.92000
0.00000    0.00000    0.00000 ;      CA-     C-     N-    CA
  2558   2557   2559   2560       3    0.65084    1.95253    0.00000
-2.60338    0.00000    0.00000 ;      HA-    CA-    CB-   HB2
  2558   2557   2559   2561       3    0.65084    1.95253    0.00000
-2.60338    0.00000    0.00000 ;      HA-    CA-    CB-   HB3
  2558   2557   2559   2562       3    0.65084    1.95253    0.00000
-2.60338    0.00000    0.00000 ;      HA-    CA-    CB-    CG
  2558   2557   2565   2566       3    3.68192   -4.35136    0.00000
1.33888    0.00000    0.00000 ;      HA-    CA-     C-     O
  2558   2557   2565   2567       3    0.00000    0.00000    0.00000
0.00000    0.00000    0.00000 ;      HA-    CA-     C-     N
  2559   2557   2565   2566       3    0.00000    0.00000    0.00000
0.00000    0.00000    0.00000 ;      CB-    CA-     C-     O
  2559   2557   2565   2567       3    2.51040    4.18400    1.67360
-6.69440    0.00000    0.00000 ;      CB-    CA-     C-     N
  2560   2559   2557   2565       3    0.65084    1.95253    0.00000
-2.60338    0.00000    0.00000 ;     HB2-    CB-    CA-     C
  2560   2559   2562   2563       3    0.00000    0.00000    0.00000
0.00000    0.00000    0.00000 ;     HB2-    CB-    CG-   OD1
  2560   2559   2562   2564       3    0.00000    0.00000    0.00000
0.00000    0.00000    0.00000 ;     HB2-    CB-    CG-   OD2
  2561   2559   2557   2565       3    0.65084    1.95253    0.00000
-2.60338    0.00000    0.00000 ;     HB3-    CB-    CA-     C
  2561   2559   2562   2563       3    0.00000    0.00000    0.00000
```

```
0.00000    0.00000    0.00000 ;    HB3-    CB-    CG-   OD1
  2561   2559   2562   2564       3    0.00000    0.00000    0.00000
0.00000    0.00000    0.00000 ;    HB3-    CB-    CG-   OD2
  2562   2559   2557   2565       3    7.14627   -1.04600   -8.99560
-0.97069    5.15469    0.00000 ;     CG-    CB-    CA-     C
  2565   2567   2569   2570       3    0.00000    0.00000    0.00000
0.00000    0.00000    0.00000 ;     C-     N-    CA-    HA
  2565   2567   2569   2571       3   11.71520    1.67360   15.06240  -
13.38880    0.00000    0.00000 ;     C-     N-    CA-    CB
  2565   2567   2569   2585       3    1.75728    5.27184    2.25936
-7.02912    0.00000    0.00000 ;     C-     N-    CA-     C
  2566   2565   2567   2568       3   29.28800   -8.36800  -20.92000
0.00000    0.00000    0.00000 ;     O-     C-     N-     H
  2566   2565   2567   2569       3   20.92000    0.00000  -20.92000
0.00000    0.00000    0.00000 ;     O-     C-     N-    CA
  2567   2569   2571   2572       3    0.65084    1.95253    0.00000
-2.60338    0.00000    0.00000 ;     N-    CA-    CB-   HB2
  2567   2569   2571   2573       3    0.65084    1.95253    0.00000
-2.60338    0.00000    0.00000 ;     N-    CA-    CB-   HB3
  2567   2569   2571   2574       3    3.28026    2.46019   -2.19242
-3.21331   -0.23430    0.00000 ;     N-    CA-    CB-    CG
  2567   2569   2585   2586       3    0.00000    0.00000    0.00000
0.00000    0.00000    0.00000 ;     N-    CA-     C-     O
  2567   2569   2585   2587       3   17.40544   -5.02080  -13.22144
9.20480    0.00000    0.00000 ;     N-    CA-     C-     N
  2568   2567   2569   2570       3    0.00000    0.00000    0.00000
0.00000    0.00000    0.00000 ;     H-     N-    CA-    HA
  2568   2567   2569   2571       3    0.00000    0.00000    0.00000
0.00000    0.00000    0.00000 ;     H-     N-    CA-    CB
  2568   2567   2569   2585       3    0.00000    0.00000    0.00000
0.00000    0.00000    0.00000 ;     H-     N-    CA-     C
  2569   2571   2574   2575       3    0.57739    0.00000    1.02926
0.00000   -1.60666    0.00000 ;    CA-    CB-    CG-   CD1
  2569   2571   2574   2583       3    0.57739    0.00000    1.02926
0.00000   -1.60666    0.00000 ;    CA-    CB-    CG-   CD2
  2569   2585   2587   2588       3   20.92000    0.00000  -20.92000
0.00000    0.00000    0.00000 ;    CA-     C-     N-     H
  2569   2585   2587   2589       3   20.92000    0.00000  -20.92000
0.00000    0.00000    0.00000 ;    CA-     C-     N-    CA
  2570   2569   2571   2572       3    0.65084    1.95253    0.00000
-2.60338    0.00000    0.00000 ;    HA-    CA-    CB-   HB2
  2570   2569   2571   2573       3    0.65084    1.95253    0.00000
-2.60338    0.00000    0.00000 ;    HA-    CA-    CB-   HB3
  2570   2569   2571   2574       3    0.65084    1.95253    0.00000
-2.60338    0.00000    0.00000 ;    HA-    CA-    CB-    CG
  2570   2569   2585   2586       3    3.68192   -4.35136    0.00000
1.33888    0.00000    0.00000 ;    HA-    CA-     C-     O
  2570   2569   2585   2587       3    0.00000    0.00000    0.00000
0.00000    0.00000    0.00000 ;    HA-    CA-     C-     N
  2571   2569   2585   2586       3    0.00000    0.00000    0.00000
0.00000    0.00000    0.00000 ;    CB-    CA-     C-     O
  2571   2569   2585   2587       3    2.51040    4.18400    1.67360
```

-6.69440 0.00000 0.00000 ; CB- CA- C- N
2571 2574 2575 2576 3 30.33400 0.00000 -30.33400
0.00000 0.00000 0.00000 ; CB- CG- CD1- HD1
2571 2574 2575 2577 3 30.33400 0.00000 -30.33400
0.00000 0.00000 0.00000 ; CB- CG- CD1- CE1
2571 2574 2583 2581 3 30.33400 0.00000 -30.33400
0.00000 0.00000 0.00000 ; CB- CG- CD2- CE2
2571 2574 2583 2584 3 30.33400 0.00000 -30.33400
0.00000 0.00000 0.00000 ; CB- CG- CD2- HD2
2572 2571 2569 2585 3 0.65084 1.95253 0.00000
-2.60338 0.00000 0.00000 ; HB2- CB- CA- C
2572 2571 2574 2575 3 0.00000 0.00000 0.00000
0.00000 0.00000 0.00000 ; HB2- CB- CG- CD1
2572 2571 2574 2583 3 0.00000 0.00000 0.00000
0.00000 0.00000 0.00000 ; HB2- CB- CG- CD2
2573 2571 2569 2585 3 0.65084 1.95253 0.00000
-2.60338 0.00000 0.00000 ; HB3- CB- CA- C
2573 2571 2574 2575 3 0.00000 0.00000 0.00000
0.00000 0.00000 0.00000 ; HB3- CB- CG- CD1
2573 2571 2574 2583 3 0.00000 0.00000 0.00000
0.00000 0.00000 0.00000 ; HB3- CB- CG- CD2
2574 2571 2569 2585 3 4.95804 2.17986 -3.52293
-3.21331 -0.40166 0.00000 ; CG- CB- CA- C
2574 2575 2577 2578 3 30.33400 0.00000 -30.33400
0.00000 0.00000 0.00000 ; CG- CD1- CE1- HE1
2574 2575 2577 2579 3 30.33400 0.00000 -30.33400
0.00000 0.00000 0.00000 ; CG- CD1- CE1- CZ
2574 2583 2581 2579 3 30.33400 0.00000 -30.33400
0.00000 0.00000 0.00000 ; CG- CD2- CE2- CZ
2574 2583 2581 2582 3 30.33400 0.00000 -30.33400
0.00000 0.00000 0.00000 ; CG- CD2- CE2- HE2
2575 2574 2583 2581 3 30.33400 0.00000 -30.33400
0.00000 0.00000 0.00000 ; CD1- CG- CD2- CE2
2575 2574 2583 2584 3 30.33400 0.00000 -30.33400
0.00000 0.00000 0.00000 ; CD1- CG- CD2- HD2
2575 2577 2579 2580 3 30.33400 0.00000 -30.33400
0.00000 0.00000 0.00000 ; CD1- CE1- CZ- HZ
2575 2577 2579 2581 3 30.33400 0.00000 -30.33400
0.00000 0.00000 0.00000 ; CD1- CE1- CZ- CE2
2576 2575 2574 2583 3 30.33400 0.00000 -30.33400
0.00000 0.00000 0.00000 ; HD1- CD1- CG- CD2
2576 2575 2577 2578 3 30.33400 0.00000 -30.33400
0.00000 0.00000 0.00000 ; HD1- CD1- CE1- HE1
2576 2575 2577 2579 3 30.33400 0.00000 -30.33400
0.00000 0.00000 0.00000 ; HD1- CD1- CE1- CZ
2577 2575 2574 2583 3 30.33400 0.00000 -30.33400
0.00000 0.00000 0.00000 ; CE1- CD1- CG- CD2
2577 2579 2581 2582 3 30.33400 0.00000 -30.33400
0.00000 0.00000 0.00000 ; CE1- CZ- CE2- HE2
2577 2579 2581 2583 3 30.33400 0.00000 -30.33400
0.00000 0.00000 0.00000 ; CE1- CZ- CE2- CD2
2578 2577 2579 2580 3 30.33400 0.00000 -30.33400

0.00000    0.00000    0.00000 ;    HE1-   CE1-    CZ-    HZ
  2578   2577   2579   2581      3   30.33400    0.00000  -30.33400
0.00000    0.00000    0.00000 ;    HE1-   CE1-    CZ-   CE2
  2579   2581   2583   2584      3   30.33400    0.00000  -30.33400
0.00000    0.00000    0.00000 ;     CZ-   CE2-   CD2-   HD2
  2580   2579   2581   2582      3   30.33400    0.00000  -30.33400
0.00000    0.00000    0.00000 ;     HZ-    CZ-   CE2-   HE2
  2580   2579   2581   2583      3   30.33400    0.00000  -30.33400
0.00000    0.00000    0.00000 ;     HZ-    CZ-   CE2-   CD2
  2582   2581   2583   2584      3   30.33400    0.00000  -30.33400
0.00000    0.00000    0.00000 ;    HE2-   CE2-   CD2-   HD2
  2585   2587   2589   2590      3    0.00000    0.00000    0.00000
0.00000    0.00000    0.00000 ;      C-     N-    CA-   HA2
  2585   2587   2589   2591      3    0.00000    0.00000    0.00000
0.00000    0.00000    0.00000 ;      C-     N-    CA-   HA3
  2585   2587   2589   2592      3    1.75728    5.27184    2.25936
-7.02912    0.00000    0.00000 ;      C-     N-    CA-     C
  2586   2585   2587   2588      3   29.28800   -8.36800  -20.92000
0.00000    0.00000    0.00000 ;      O-     C-     N-     H
  2586   2585   2587   2589      3   20.92000    0.00000  -20.92000
0.00000    0.00000    0.00000 ;      O-     C-     N-    CA
  2587   2589   2592   2593      3    0.00000    0.00000    0.00000
0.00000    0.00000    0.00000 ;      N-    CA-     C-     O
  2587   2589   2592   2594      3   17.40544   -5.02080  -13.22144
9.20480    0.00000    0.00000 ;      N-    CA-     C-     N
  2588   2587   2589   2590      3    0.00000    0.00000    0.00000
0.00000    0.00000    0.00000 ;      H-     N-    CA-   HA2
  2588   2587   2589   2591      3    0.00000    0.00000    0.00000
0.00000    0.00000    0.00000 ;      H-     N-    CA-   HA3
  2588   2587   2589   2592      3    0.00000    0.00000    0.00000
0.00000    0.00000    0.00000 ;      H-     N-    CA-     C
  2589   2592   2594   2595      3   20.92000    0.00000  -20.92000
0.00000    0.00000    0.00000 ;     CA-     C-     N-     H
  2589   2592   2594   2596      3   20.92000    0.00000  -20.92000
0.00000    0.00000    0.00000 ;     CA-     C-     N-    CA
  2590   2589   2592   2593      3    3.68192   -4.35136    0.00000
1.33888    0.00000    0.00000 ;    HA2-    CA-     C-     O
  2590   2589   2592   2594      3    0.00000    0.00000    0.00000
0.00000    0.00000    0.00000 ;    HA2-    CA-     C-     N
  2591   2589   2592   2593      3    3.68192   -4.35136    0.00000
1.33888    0.00000    0.00000 ;    HA3-    CA-     C-     O
  2591   2589   2592   2594      3    0.00000    0.00000    0.00000
0.00000    0.00000    0.00000 ;    HA3-    CA-     C-     N
  2592   2594   2596   2597      3    0.00000    0.00000    0.00000
0.00000    0.00000    0.00000 ;      C-     N-    CA-    HA
  2592   2594   2596   2598      3   11.71520    1.67360   15.06240  -
13.38880    0.00000    0.00000 ;      C-     N-    CA-    CB
  2592   2594   2596   2611      3    1.75728    5.27184    2.25936
-7.02912    0.00000    0.00000 ;      C-     N-    CA-     C
  2593   2592   2594   2595      3   29.28800   -8.36800  -20.92000
0.00000    0.00000    0.00000 ;      O-     C-     N-     H
  2593   2592   2594   2596      3   20.92000    0.00000  -20.92000

```
0.00000    0.00000    0.00000 ;       O-     C-     N-    CA
  2594   2596   2598   2599      3    0.65084    1.95253    0.00000
-2.60338    0.00000    0.00000 ;      N-    CA-    CB-   HB2
  2594   2596   2598   2600      3    0.65084    1.95253    0.00000
-2.60338    0.00000    0.00000 ;      N-    CA-    CB-   HB3
  2594   2596   2598   2601      3    3.79070    1.39746   -4.61077
-2.40998    2.44346    0.00000 ;      N-    CA-    CB-    CG
  2594   2596   2611   2612      3    0.00000    0.00000    0.00000
0.00000    0.00000    0.00000 ;       N-    CA-     C-     O
  2594   2596   2611   2613      3   17.40544   -5.02080  -13.22144
9.20480    0.00000    0.00000 ;       N-    CA-     C-     N
  2595   2594   2596   2597      3    0.00000    0.00000    0.00000
0.00000    0.00000    0.00000 ;       H-     N-    CA-    HA
  2595   2594   2596   2598      3    0.00000    0.00000    0.00000
0.00000    0.00000    0.00000 ;       H-     N-    CA-    CB
  2595   2594   2596   2611      3    0.00000    0.00000    0.00000
0.00000    0.00000    0.00000 ;       H-     N-    CA-     C
  2596   2598   2601   2602      3    0.66944    2.00832    0.00000
-2.67776    0.00000    0.00000 ;     CA-    CB-    CG-    HG
  2596   2598   2601   2603      3    3.90367    0.19665   -6.21742
-2.37651    5.99149    0.00000 ;     CA-    CB-    CG-   CD1
  2596   2598   2601   2607      3    3.90367    0.19665   -6.21742
-2.37651    5.99149    0.00000 ;     CA-    CB-    CG-   CD2
  2596   2611   2613   2614      3   20.92000    0.00000  -20.92000
0.00000    0.00000    0.00000 ;     CA-     C-     N-     H
  2596   2611   2613   2615      3   20.92000    0.00000  -20.92000
0.00000    0.00000    0.00000 ;     CA-     C-     N-    CA
  2597   2596   2598   2599      3    0.65084    1.95253    0.00000
-2.60338    0.00000    0.00000 ;     HA-    CA-    CB-   HB2
  2597   2596   2598   2600      3    0.65084    1.95253    0.00000
-2.60338    0.00000    0.00000 ;     HA-    CA-    CB-   HB3
  2597   2596   2598   2601      3    0.65084    1.95253    0.00000
-2.60338    0.00000    0.00000 ;     HA-    CA-    CB-    CG
  2597   2596   2611   2612      3    3.68192   -4.35136    0.00000
1.33888    0.00000    0.00000 ;     HA-    CA-     C-     O
  2597   2596   2611   2613      3    0.00000    0.00000    0.00000
0.00000    0.00000    0.00000 ;     HA-    CA-     C-     N
  2598   2596   2611   2612      3    0.00000    0.00000    0.00000
0.00000    0.00000    0.00000 ;     CB-    CA-     C-     O
  2598   2596   2611   2613      3    2.51040    4.18400    1.67360
-6.69440    0.00000    0.00000 ;     CB-    CA-     C-     N
  2598   2601   2603   2604      3    0.66944    2.00832    0.00000
-2.67776    0.00000    0.00000 ;     CB-    CG-   CD1-  HD11
  2598   2601   2603   2605      3    0.66944    2.00832    0.00000
-2.67776    0.00000    0.00000 ;     CB-    CG-   CD1-  HD12
  2598   2601   2603   2606      3    0.66944    2.00832    0.00000
-2.67776    0.00000    0.00000 ;     CB-    CG-   CD1-  HD13
  2598   2601   2607   2608      3    0.66944    2.00832    0.00000
-2.67776    0.00000    0.00000 ;     CB-    CG-   CD2-  HD21
  2598   2601   2607   2609      3    0.66944    2.00832    0.00000
-2.67776    0.00000    0.00000 ;     CB-    CG-   CD2-  HD22
  2598   2601   2607   2610      3    0.66944    2.00832    0.00000
```

-2.67776 0.00000 0.00000 ; CB- CG- CD2- HD23
2599 2598 2596 2611 3 0.65084 1.95253 0.00000
-2.60338 0.00000 0.00000 ; HB2- CB- CA- C
2599 2598 2601 2602 3 0.62760 1.88280 0.00000
-2.51040 0.00000 0.00000 ; HB2- CB- CG- HG
2599 2598 2601 2603 3 0.66944 2.00832 0.00000
-2.67776 0.00000 0.00000 ; HB2- CB- CG- CD1
2599 2598 2601 2607 3 0.66944 2.00832 0.00000
-2.67776 0.00000 0.00000 ; HB2- CB- CG- CD2
2600 2598 2596 2611 3 0.65084 1.95253 0.00000
-2.60338 0.00000 0.00000 ; HB3- CB- CA- C
2600 2598 2601 2602 3 0.62760 1.88280 0.00000
-2.51040 0.00000 0.00000 ; HB3- CB- CG- HG
2600 2598 2601 2603 3 0.66944 2.00832 0.00000
-2.67776 0.00000 0.00000 ; HB3- CB- CG- CD1
2600 2598 2601 2607 3 0.66944 2.00832 0.00000
-2.67776 0.00000 0.00000 ; HB3- CB- CG- CD2
2601 2598 2596 2611 3 10.33448 -1.14642 -11.54784
-2.40998 6.35968 0.00000 ; CG- CB- CA- C
2602 2601 2603 2604 3 0.62760 1.88280 0.00000
-2.51040 0.00000 0.00000 ; HG- CG- CD1- HD11
2602 2601 2603 2605 3 0.62760 1.88280 0.00000
-2.51040 0.00000 0.00000 ; HG- CG- CD1- HD12
2602 2601 2603 2606 3 0.62760 1.88280 0.00000
-2.51040 0.00000 0.00000 ; HG- CG- CD1- HD13
2602 2601 2607 2608 3 0.62760 1.88280 0.00000
-2.51040 0.00000 0.00000 ; HG- CG- CD2- HD21
2602 2601 2607 2609 3 0.62760 1.88280 0.00000
-2.51040 0.00000 0.00000 ; HG- CG- CD2- HD22
2602 2601 2607 2610 3 0.62760 1.88280 0.00000
-2.51040 0.00000 0.00000 ; HG- CG- CD2- HD23
2603 2601 2607 2608 3 0.66944 2.00832 0.00000
-2.67776 0.00000 0.00000 ; CD1- CG- CD2- HD21
2603 2601 2607 2609 3 0.66944 2.00832 0.00000
-2.67776 0.00000 0.00000 ; CD1- CG- CD2- HD22
2603 2601 2607 2610 3 0.66944 2.00832 0.00000
-2.67776 0.00000 0.00000 ; CD1- CG- CD2- HD23
2604 2603 2601 2607 3 0.66944 2.00832 0.00000
-2.67776 0.00000 0.00000 ; HD11- CD1- CG- CD2
2605 2603 2601 2607 3 0.66944 2.00832 0.00000
-2.67776 0.00000 0.00000 ; HD12- CD1- CG- CD2
2606 2603 2601 2607 3 0.66944 2.00832 0.00000
-2.67776 0.00000 0.00000 ; HD13- CD1- CG- CD2
2611 2613 2615 2616 3 0.00000 0.00000 0.00000
0.00000 0.00000 0.00000 ; C- N- CA- HA
2611 2613 2615 2617 3 11.71520 1.67360 15.06240 -
13.38880 0.00000 0.00000 ; C- N- CA- CB
2611 2613 2615 2622 3 1.75728 5.27184 2.25936
-7.02912 0.00000 0.00000 ; C- N- CA- C
2612 2611 2613 2614 3 29.28800 -8.36800 -20.92000
0.00000 0.00000 0.00000 ; O- C- N- H
2612 2611 2613 2615 3 20.92000 0.00000 -20.92000

0.00000 0.00000 0.00000 ; O- C- N- CA
2613 2615 2617 2618 3 0.65084 1.95253 0.00000
-2.60338 0.00000 0.00000 ; N- CA- CB- HB2
2613 2615 2617 2619 3 0.65084 1.95253 0.00000
-2.60338 0.00000 0.00000 ; N- CA- CB- HB3
2613 2615 2617 2620 3 7.86174 2.24681 -7.41405
-6.71114 5.35552 0.00000 ; N- CA- CB- OG
2613 2615 2622 2623 3 0.00000 0.00000 0.00000
0.00000 0.00000 0.00000 ; N- CA- C- O
2613 2615 2622 2624 3 17.40544 -5.02080 -13.22144
9.20480 0.00000 0.00000 ; N- CA- C- N
2614 2613 2615 2616 3 0.00000 0.00000 0.00000
0.00000 0.00000 0.00000 ; H- N- CA- HA
2614 2613 2615 2617 3 0.00000 0.00000 0.00000
0.00000 0.00000 0.00000 ; H- N- CA- CB
2614 2613 2615 2622 3 0.00000 0.00000 0.00000
0.00000 0.00000 0.00000 ; H- N- CA- C
2615 2617 2620 2621 3 2.05853 2.46856 3.48109
-4.46851 0.23430 0.00000 ; CA- CB- OG- HG
2615 2622 2624 2625 3 20.92000 0.00000 -20.92000
0.00000 0.00000 0.00000 ; CA- C- N- H
2615 2622 2624 2626 3 20.92000 0.00000 -20.92000
0.00000 0.00000 0.00000 ; CA- C- N- CA
2616 2615 2617 2618 3 0.65084 1.95253 0.00000
-2.60338 0.00000 0.00000 ; HA- CA- CB- HB2
2616 2615 2617 2619 3 0.65084 1.95253 0.00000
-2.60338 0.00000 0.00000 ; HA- CA- CB- HB3
2616 2615 2617 2620 3 1.04600 -1.04600 0.00000
0.00000 0.00000 0.00000 ; HA- CA- CB- OG
2616 2615 2622 2623 3 3.68192 -4.35136 0.00000
1.33888 0.00000 0.00000 ; HA- CA- C- O
2616 2615 2622 2624 3 0.00000 0.00000 0.00000
0.00000 0.00000 0.00000 ; HA- CA- C- N
2617 2615 2622 2623 3 0.00000 0.00000 0.00000
0.00000 0.00000 0.00000 ; CB- CA- C- O
2617 2615 2622 2624 3 2.51040 4.18400 1.67360
-6.69440 0.00000 0.00000 ; CB- CA- C- N
2618 2617 2615 2622 3 0.65084 1.95253 0.00000
-2.60338 0.00000 0.00000 ; HB2- CB- CA- C
2618 2617 2620 2621 3 0.69733 2.09200 0.00000
-2.78933 0.00000 0.00000 ; HB2- CB- OG- HG
2619 2617 2615 2622 3 0.65084 1.95253 0.00000
-2.60338 0.00000 0.00000 ; HB3- CB- CA- C
2619 2617 2620 2621 3 0.69733 2.09200 0.00000
-2.78933 0.00000 0.00000 ; HB3- CB- OG- HG
2620 2617 2615 2622 3 7.34710 7.79898 -6.14211
-6.71114 4.31789 0.00000 ; OG- CB- CA- C
2622 2624 2626 2627 3 0.00000 0.00000 0.00000
0.00000 0.00000 0.00000 ; C- N- CA- HA
2622 2624 2626 2628 3 11.71520 1.67360 15.06240 -
13.38880 0.00000 0.00000 ; C- N- CA- CB
2622 2624 2626 2644 3 1.75728 5.27184 2.25936

-7.02912 0.00000 0.00000 ; C- N- CA- C
2623 2622 2624 2625 3 29.28800 -8.36800 -20.92000
0.00000 0.00000 0.00000 ; O- C- N- H
2623 2622 2624 2626 3 20.92000 0.00000 -20.92000
0.00000 0.00000 0.00000 ; O- C- N- CA
2624 2626 2628 2629 3 0.65084 1.95253 0.00000
-2.60338 0.00000 0.00000 ; N- CA- CB- HB2
2624 2626 2628 2630 3 0.65084 1.95253 0.00000
-2.60338 0.00000 0.00000 ; N- CA- CB- HB3
2624 2626 2628 2631 3 0.65084 1.95253 0.00000
-2.60338 0.00000 0.00000 ; N- CA- CB- CG
2624 2626 2644 2645 3 0.00000 0.00000 0.00000
0.00000 0.00000 0.00000 ; N- CA- C- O
2624 2626 2644 2646 3 17.40544 -5.02080 -13.22144
9.20480 0.00000 0.00000 ; N- CA- C- N
2625 2624 2626 2627 3 0.00000 0.00000 0.00000
0.00000 0.00000 0.00000 ; H- N- CA- HA
2625 2624 2626 2628 3 0.00000 0.00000 0.00000
0.00000 0.00000 0.00000 ; H- N- CA- CB
2625 2624 2626 2644 3 0.00000 0.00000 0.00000
0.00000 0.00000 0.00000 ; H- N- CA- C
2626 2628 2631 2632 3 0.66944 2.00832 0.00000
-2.67776 0.00000 0.00000 ; CA- CB- CG- HG2
2626 2628 2631 2633 3 0.66944 2.00832 0.00000
-2.67776 0.00000 0.00000 ; CA- CB- CG- HG3
2626 2628 2631 2634 3 3.68192 3.09616 -2.09200
-3.01248 0.00000 0.00000 ; CA- CB- CG- CD
2626 2644 2646 2647 3 20.92000 0.00000 -20.92000
0.00000 0.00000 0.00000 ; CA- C- N- H
2626 2644 2646 2648 3 20.92000 0.00000 -20.92000
0.00000 0.00000 0.00000 ; CA- C- N- CA
2627 2626 2628 2629 3 0.65084 1.95253 0.00000
-2.60338 0.00000 0.00000 ; HA- CA- CB- HB2
2627 2626 2628 2630 3 0.65084 1.95253 0.00000
-2.60338 0.00000 0.00000 ; HA- CA- CB- HB3
2627 2626 2628 2631 3 0.65084 1.95253 0.00000
-2.60338 0.00000 0.00000 ; HA- CA- CB- CG
2627 2626 2644 2645 3 3.68192 -4.35136 0.00000
1.33888 0.00000 0.00000 ; HA- CA- C- O
2627 2626 2644 2646 3 0.00000 0.00000 0.00000
0.00000 0.00000 0.00000 ; HA- CA- C- N
2628 2626 2644 2645 3 0.00000 0.00000 0.00000
0.00000 0.00000 0.00000 ; CB- CA- C- O
2628 2626 2644 2646 3 2.51040 4.18400 1.67360
-6.69440 0.00000 0.00000 ; CB- CA- C- N
2628 2631 2634 2635 3 0.66944 2.00832 0.00000
-2.67776 0.00000 0.00000 ; CB- CG- CD- HD2
2628 2631 2634 2636 3 0.66944 2.00832 0.00000
-2.67776 0.00000 0.00000 ; CB- CG- CD- HD3
2628 2631 2634 2637 3 3.68192 3.09616 -2.09200
-3.01248 0.00000 0.00000 ; CB- CG- CD- CE
2629 2628 2626 2644 3 0.65084 1.95253 0.00000

-2.60338 0.00000 0.00000 ; HB2- CB- CA- C
2629 2628 2631 2632 3 0.62760 1.88280 0.00000
-2.51040 0.00000 0.00000 ; HB2- CB- CG- HG2
2629 2628 2631 2633 3 0.62760 1.88280 0.00000
-2.51040 0.00000 0.00000 ; HB2- CB- CG- HG3
2629 2628 2631 2634 3 0.66944 2.00832 0.00000
-2.67776 0.00000 0.00000 ; HB2- CB- CG- CD
2630 2628 2626 2644 3 0.65084 1.95253 0.00000
-2.60338 0.00000 0.00000 ; HB3- CB- CA- C
2630 2628 2631 2632 3 0.62760 1.88280 0.00000
-2.51040 0.00000 0.00000 ; HB3- CB- CG- HG2
2630 2628 2631 2633 3 0.62760 1.88280 0.00000
-2.51040 0.00000 0.00000 ; HB3- CB- CG- HG3
2630 2628 2631 2634 3 0.66944 2.00832 0.00000
-2.67776 0.00000 0.00000 ; HB3- CB- CG- CD
2631 2628 2626 2644 3 0.65084 1.95253 0.00000
-2.60338 0.00000 0.00000 ; CG- CB- CA- C
2631 2634 2637 2638 3 0.65084 1.95253 0.00000
-2.60338 0.00000 0.00000 ; CG- CD- CE- HE2
2631 2634 2637 2639 3 0.65084 1.95253 0.00000
-2.60338 0.00000 0.00000 ; CG- CD- CE- HE3
2631 2634 2637 2640 3 0.65084 1.95253 0.00000
-2.60338 0.00000 0.00000 ; CG- CD- CE- NZ
2632 2631 2634 2635 3 0.62760 1.88280 0.00000
-2.51040 0.00000 0.00000 ; HG2- CG- CD- HD2
2632 2631 2634 2636 3 0.62760 1.88280 0.00000
-2.51040 0.00000 0.00000 ; HG2- CG- CD- HD3
2632 2631 2634 2637 3 0.66944 2.00832 0.00000
-2.67776 0.00000 0.00000 ; HG2- CG- CD- CE
2633 2631 2634 2635 3 0.62760 1.88280 0.00000
-2.51040 0.00000 0.00000 ; HG3- CG- CD- HD2
2633 2631 2634 2636 3 0.62760 1.88280 0.00000
-2.51040 0.00000 0.00000 ; HG3- CG- CD- HD3
2633 2631 2634 2637 3 0.66944 2.00832 0.00000
-2.67776 0.00000 0.00000 ; HG3- CG- CD- CE
2634 2637 2640 2641 3 0.65084 1.95253 0.00000
-2.60338 0.00000 0.00000 ; CD- CE- NZ- HZ1
2634 2637 2640 2642 3 0.65084 1.95253 0.00000
-2.60338 0.00000 0.00000 ; CD- CE- NZ- HZ2
2634 2637 2640 2643 3 0.65084 1.95253 0.00000
-2.60338 0.00000 0.00000 ; CD- CE- NZ- HZ3
2635 2634 2637 2638 3 0.65084 1.95253 0.00000
-2.60338 0.00000 0.00000 ; HD2- CD- CE- HE2
2635 2634 2637 2639 3 0.65084 1.95253 0.00000
-2.60338 0.00000 0.00000 ; HD2- CD- CE- HE3
2635 2634 2637 2640 3 0.65084 1.95253 0.00000
-2.60338 0.00000 0.00000 ; HD2- CD- CE- NZ
2636 2634 2637 2638 3 0.65084 1.95253 0.00000
-2.60338 0.00000 0.00000 ; HD3- CD- CE- HE2
2636 2634 2637 2639 3 0.65084 1.95253 0.00000
-2.60338 0.00000 0.00000 ; HD3- CD- CE- HE3
2636 2634 2637 2640 3 0.65084 1.95253 0.00000

-2.60338 0.00000 0.00000 ; HD3- CD- CE- NZ
2638 2637 2640 2641 3 0.65084 1.95253 0.00000
-2.60338 0.00000 0.00000 ; HE2- CE- NZ- HZ1
2638 2637 2640 2642 3 0.65084 1.95253 0.00000
-2.60338 0.00000 0.00000 ; HE2- CE- NZ- HZ2
2638 2637 2640 2643 3 0.65084 1.95253 0.00000
-2.60338 0.00000 0.00000 ; HE2- CE- NZ- HZ3
2639 2637 2640 2641 3 0.65084 1.95253 0.00000
-2.60338 0.00000 0.00000 ; HE3- CE- NZ- HZ1
2639 2637 2640 2642 3 0.65084 1.95253 0.00000
-2.60338 0.00000 0.00000 ; HE3- CE- NZ- HZ2
2639 2637 2640 2643 3 0.65084 1.95253 0.00000
-2.60338 0.00000 0.00000 ; HE3- CE- NZ- HZ3
2644 2646 2648 2649 3 0.00000 0.00000 0.00000
0.00000 0.00000 0.00000 ; C- N- CA- HA
2644 2646 2648 2650 3 11.71520 1.67360 15.06240 -
13.38880 0.00000 0.00000 ; C- N- CA- CB
2644 2646 2648 2658 3 1.75728 5.27184 2.25936
-7.02912 0.00000 0.00000 ; C- N- CA- C
2645 2644 2646 2647 3 29.28800 -8.36800 -20.92000
0.00000 0.00000 0.00000 ; O- C- N- H
2645 2644 2646 2648 3 20.92000 0.00000 -20.92000
0.00000 0.00000 0.00000 ; O- C- N- CA
2646 2648 2650 2651 3 0.65084 1.95253 0.00000
-2.60338 0.00000 0.00000 ; N- CA- CB- HB2
2646 2648 2650 2652 3 0.65084 1.95253 0.00000
-2.60338 0.00000 0.00000 ; N- CA- CB- HB3
2646 2648 2650 2653 3 5.99567 3.29281 -4.46014
-0.55229 1.97485 0.00000 ; N- CA- CB- CG
2646 2648 2658 2659 3 0.00000 0.00000 0.00000
0.00000 0.00000 0.00000 ; N- CA- C- O
2646 2648 2658 2660 3 17.40544 -5.02080 -13.22144
9.20480 0.00000 0.00000 ; N- CA- C- N
2647 2646 2648 2649 3 0.00000 0.00000 0.00000
0.00000 0.00000 0.00000 ; H- N- CA- HA
2647 2646 2648 2650 3 0.00000 0.00000 0.00000
0.00000 0.00000 0.00000 ; H- N- CA- CB
2647 2646 2648 2658 3 0.00000 0.00000 0.00000
0.00000 0.00000 0.00000 ; H- N- CA- C
2648 2650 2653 2654 3 0.00000 0.00000 0.00000
0.00000 0.00000 0.00000 ; CA- CB- CG- OD1
2648 2650 2653 2655 3 8.84916 -0.31380 -4.32626
5.03754 0.26778 0.00000 ; CA- CB- CG- ND2
2648 2658 2660 2661 3 20.92000 0.00000 -20.92000
0.00000 0.00000 0.00000 ; CA- C- N- H
2648 2658 2660 2662 3 20.92000 0.00000 -20.92000
0.00000 0.00000 0.00000 ; CA- C- N- CA
2649 2648 2650 2651 3 0.65084 1.95253 0.00000
-2.60338 0.00000 0.00000 ; HA- CA- CB- HB2
2649 2648 2650 2652 3 0.65084 1.95253 0.00000
-2.60338 0.00000 0.00000 ; HA- CA- CB- HB3
2649 2648 2650 2653 3 0.65084 1.95253 0.00000

-2.60338 0.00000 0.00000 ; HA- CA- CB- CG
2649 2648 2658 2659 3 3.68192 -4.35136 0.00000
1.33888 0.00000 0.00000 ; HA- CA- C- O
2649 2648 2658 2660 3 0.00000 0.00000 0.00000
0.00000 0.00000 0.00000 ; HA- CA- C- N
2650 2648 2658 2659 3 0.00000 0.00000 0.00000
0.00000 0.00000 0.00000 ; CB- CA- C- O
2650 2648 2658 2660 3 2.51040 4.18400 1.67360
-6.69440 0.00000 0.00000 ; CB- CA- C- N
2650 2653 2655 2656 3 20.92000 0.00000 -20.92000
0.00000 0.00000 0.00000 ; CB- CG- ND2- HD21
2650 2653 2655 2657 3 20.92000 0.00000 -20.92000
0.00000 0.00000 0.00000 ; CB- CG- ND2- HD22
2651 2650 2648 2658 3 0.65084 1.95253 0.00000
-2.60338 0.00000 0.00000 ; HB2- CB- CA- C
2651 2650 2653 2654 3 3.68192 -4.35136 0.00000
1.33888 0.00000 0.00000 ; HB2- CB- CG- OD1
2651 2650 2653 2655 3 0.00000 0.00000 0.00000
0.00000 0.00000 0.00000 ; HB2- CB- CG- ND2
2652 2650 2648 2658 3 0.65084 1.95253 0.00000
-2.60338 0.00000 0.00000 ; HB3- CB- CA- C
2652 2650 2653 2654 3 3.68192 -4.35136 0.00000
1.33888 0.00000 0.00000 ; HB3- CB- CG- OD1
2652 2650 2653 2655 3 0.00000 0.00000 0.00000
0.00000 0.00000 0.00000 ; HB3- CB- CG- ND2
2653 2650 2648 2658 3 7.94542 -3.96225 -6.11701
-0.55229 3.58150 0.00000 ; CG- CB- CA- C
2654 2653 2655 2656 3 29.28800 -8.36800 -20.92000
0.00000 0.00000 0.00000 ; OD1- CG- ND2- HD21
2654 2653 2655 2657 3 29.28800 -8.36800 -20.92000
0.00000 0.00000 0.00000 ; OD1- CG- ND2- HD22
2658 2660 2662 2663 3 0.00000 0.00000 0.00000
0.00000 0.00000 0.00000 ; C- N- CA- HA
2658 2660 2662 2664 3 11.71520 1.67360 15.06240 -
13.38880 0.00000 0.00000 ; C- N- CA- CB
2658 2660 2662 2677 3 1.75728 5.27184 2.25936
-7.02912 0.00000 0.00000 ; C- N- CA- C
2659 2658 2660 2661 3 29.28800 -8.36800 -20.92000
0.00000 0.00000 0.00000 ; O- C- N- H
2659 2658 2660 2662 3 20.92000 0.00000 -20.92000
0.00000 0.00000 0.00000 ; O- C- N- CA
2660 2662 2664 2665 3 0.65084 1.95253 0.00000
-2.60338 0.00000 0.00000 ; N- CA- CB- HB2
2660 2662 2664 2666 3 0.65084 1.95253 0.00000
-2.60338 0.00000 0.00000 ; N- CA- CB- HB3
2660 2662 2664 2667 3 3.79070 1.39746 -4.61077
-2.40998 2.44346 0.00000 ; N- CA- CB- CG
2660 2662 2677 2678 3 0.00000 0.00000 0.00000
0.00000 0.00000 0.00000 ; N- CA- C- O
2660 2662 2677 2679 3 17.40544 -5.02080 -13.22144
9.20480 0.00000 0.00000 ; N- CA- C- N
2661 2660 2662 2663 3 0.00000 0.00000 0.00000

```
0.00000    0.00000    0.00000 ;      H-     N-    CA-    HA
  2661   2660   2662   2664      3    0.00000    0.00000    0.00000
0.00000    0.00000    0.00000 ;      H-     N-    CA-    CB
  2661   2660   2662   2677      3    0.00000    0.00000    0.00000
0.00000    0.00000    0.00000 ;      H-     N-    CA-     C
  2662   2664   2667   2668      3    0.66944    2.00832    0.00000
-2.67776    0.00000    0.00000 ;     CA-    CB-    CG-    HG
  2662   2664   2667   2669      3    3.90367    0.19665   -6.21742
-2.37651    5.99149    0.00000 ;     CA-    CB-    CG-   CD1
  2662   2664   2667   2673      3    3.90367    0.19665   -6.21742
-2.37651    5.99149    0.00000 ;     CA-    CB-    CG-   CD2
  2662   2677   2679   2680      3   20.92000    0.00000  -20.92000
0.00000    0.00000    0.00000 ;     CA-     C-     N-     H
  2662   2677   2679   2681      3   20.92000    0.00000  -20.92000
0.00000    0.00000    0.00000 ;     CA-     C-     N-    CA
  2663   2662   2664   2665      3    0.65084    1.95253    0.00000
-2.60338    0.00000    0.00000 ;     HA-    CA-    CB-   HB2
  2663   2662   2664   2666      3    0.65084    1.95253    0.00000
-2.60338    0.00000    0.00000 ;     HA-    CA-    CB-   HB3
  2663   2662   2664   2667      3    0.65084    1.95253    0.00000
-2.60338    0.00000    0.00000 ;     HA-    CA-    CB-    CG
  2663   2662   2677   2678      3    3.68192   -4.35136    0.00000
1.33888    0.00000    0.00000 ;     HA-    CA-     C-     O
  2663   2662   2677   2679      3    0.00000    0.00000    0.00000
0.00000    0.00000    0.00000 ;     HA-    CA-     C-     N
  2664   2662   2677   2678      3    0.00000    0.00000    0.00000
0.00000    0.00000    0.00000 ;     CB-    CA-     C-     O
  2664   2662   2677   2679      3    2.51040    4.18400    1.67360
-6.69440    0.00000    0.00000 ;     CB-    CA-     C-     N
  2664   2667   2669   2670      3    0.66944    2.00832    0.00000
-2.67776    0.00000    0.00000 ;     CB-    CG-   CD1-  HD11
  2664   2667   2669   2671      3    0.66944    2.00832    0.00000
-2.67776    0.00000    0.00000 ;     CB-    CG-   CD1-  HD12
  2664   2667   2669   2672      3    0.66944    2.00832    0.00000
-2.67776    0.00000    0.00000 ;     CB-    CG-   CD1-  HD13
  2664   2667   2673   2674      3    0.66944    2.00832    0.00000
-2.67776    0.00000    0.00000 ;     CB-    CG-   CD2-  HD21
  2664   2667   2673   2675      3    0.66944    2.00832    0.00000
-2.67776    0.00000    0.00000 ;     CB-    CG-   CD2-  HD22
  2664   2667   2673   2676      3    0.66944    2.00832    0.00000
-2.67776    0.00000    0.00000 ;     CB-    CG-   CD2-  HD23
  2665   2664   2662   2677      3    0.65084    1.95253    0.00000
-2.60338    0.00000    0.00000 ;    HB2-    CB-    CA-     C
  2665   2664   2667   2668      3    0.62760    1.88280    0.00000
-2.51040    0.00000    0.00000 ;    HB2-    CB-    CG-    HG
  2665   2664   2667   2669      3    0.66944    2.00832    0.00000
-2.67776    0.00000    0.00000 ;    HB2-    CB-    CG-   CD1
  2665   2664   2667   2673      3    0.66944    2.00832    0.00000
-2.67776    0.00000    0.00000 ;    HB2-    CB-    CG-   CD2
  2666   2664   2662   2677      3    0.65084    1.95253    0.00000
-2.60338    0.00000    0.00000 ;    HB3-    CB-    CA-     C
  2666   2664   2667   2668      3    0.62760    1.88280    0.00000
```

-2.51040 0.00000 0.00000 ; HB3- CB- CG- HG
2666 2664 2667 2669 3 0.66944 2.00832 0.00000
-2.67776 0.00000 0.00000 ; HB3- CB- CG- CD1
2666 2664 2667 2673 3 0.66944 2.00832 0.00000
-2.67776 0.00000 0.00000 ; HB3- CB- CG- CD2
2667 2664 2662 2677 3 10.33448 -1.14642 -11.54784
-2.40998 6.35968 0.00000 ; CG- CB- CA- C
2668 2667 2669 2670 3 0.62760 1.88280 0.00000
-2.51040 0.00000 0.00000 ; HG- CG- CD1- HD11
2668 2667 2669 2671 3 0.62760 1.88280 0.00000
-2.51040 0.00000 0.00000 ; HG- CG- CD1- HD12
2668 2667 2669 2672 3 0.62760 1.88280 0.00000
-2.51040 0.00000 0.00000 ; HG- CG- CD1- HD13
2668 2667 2673 2674 3 0.62760 1.88280 0.00000
-2.51040 0.00000 0.00000 ; HG- CG- CD2- HD21
2668 2667 2673 2675 3 0.62760 1.88280 0.00000
-2.51040 0.00000 0.00000 ; HG- CG- CD2- HD22
2668 2667 2673 2676 3 0.62760 1.88280 0.00000
-2.51040 0.00000 0.00000 ; HG- CG- CD2- HD23
2669 2667 2673 2674 3 0.66944 2.00832 0.00000
-2.67776 0.00000 0.00000 ; CD1- CG- CD2- HD21
2669 2667 2673 2675 3 0.66944 2.00832 0.00000
-2.67776 0.00000 0.00000 ; CD1- CG- CD2- HD22
2669 2667 2673 2676 3 0.66944 2.00832 0.00000
-2.67776 0.00000 0.00000 ; CD1- CG- CD2- HD23
2670 2669 2667 2673 3 0.66944 2.00832 0.00000
-2.67776 0.00000 0.00000 ; HD11- CD1- CG- CD2
2671 2669 2667 2673 3 0.66944 2.00832 0.00000
-2.67776 0.00000 0.00000 ; HD12- CD1- CG- CD2
2672 2669 2667 2673 3 0.66944 2.00832 0.00000
-2.67776 0.00000 0.00000 ; HD13- CD1- CG- CD2
2677 2679 2681 2682 3 0.00000 0.00000 0.00000
0.00000 0.00000 0.00000 ; C- N- CA- HA2
2677 2679 2681 2683 3 0.00000 0.00000 0.00000
0.00000 0.00000 0.00000 ; C- N- CA- HA3
2677 2679 2681 2684 3 1.75728 5.27184 2.25936
-7.02912 0.00000 0.00000 ; C- N- CA- C
2678 2677 2679 2680 3 29.28800 -8.36800 -20.92000
0.00000 0.00000 0.00000 ; O- C- N- H
2678 2677 2679 2681 3 20.92000 0.00000 -20.92000
0.00000 0.00000 0.00000 ; O- C- N- CA
2679 2681 2684 2685 3 0.00000 0.00000 0.00000
0.00000 0.00000 0.00000 ; N- CA- C- O
2679 2681 2684 2686 3 17.40544 -5.02080 -13.22144
9.20480 0.00000 0.00000 ; N- CA- C- N
2680 2679 2681 2682 3 0.00000 0.00000 0.00000
0.00000 0.00000 0.00000 ; H- N- CA- HA2
2680 2679 2681 2683 3 0.00000 0.00000 0.00000
0.00000 0.00000 0.00000 ; H- N- CA- HA3
2680 2679 2681 2684 3 0.00000 0.00000 0.00000
0.00000 0.00000 0.00000 ; H- N- CA- C
2681 2684 2686 2687 3 20.92000 0.00000 -20.92000

```
0.00000    0.00000    0.00000 ;      CA-     C-     N-     H
  2681   2684   2686   2688      3   20.92000    0.00000  -20.92000
0.00000    0.00000    0.00000 ;      CA-     C-     N-    CA
  2682   2681   2684   2685      3    3.68192   -4.35136    0.00000
1.33888    0.00000    0.00000 ;     HA2-    CA-     C-     O
  2682   2681   2684   2686      3    0.00000    0.00000    0.00000
0.00000    0.00000    0.00000 ;     HA2-    CA-     C-     N
  2683   2681   2684   2685      3    3.68192   -4.35136    0.00000
1.33888    0.00000    0.00000 ;     HA3-    CA-     C-     O
  2683   2681   2684   2686      3    0.00000    0.00000    0.00000
0.00000    0.00000    0.00000 ;     HA3-    CA-     C-     N
  2684   2686   2688   2689      3    0.00000    0.00000    0.00000
0.00000    0.00000    0.00000 ;       C-     N-    CA-    HA
  2684   2686   2688   2690      3   11.71520    1.67360   15.06240  -
13.38880    0.00000    0.00000 ;       C-     N-    CA-    CB
  2684   2686   2688   2698      3    1.75728    5.27184    2.25936
-7.02912    0.00000    0.00000 ;       C-     N-    CA-     C
  2685   2684   2686   2687      3   29.28800   -8.36800  -20.92000
0.00000    0.00000    0.00000 ;       O-     C-     N-     H
  2685   2684   2686   2688      3   20.92000    0.00000  -20.92000
0.00000    0.00000    0.00000 ;       O-     C-     N-    CA
  2686   2688   2690   2691      3    0.65084    1.95253    0.00000
-2.60338    0.00000    0.00000 ;       N-    CA-    CB-    HB
  2686   2688   2690   2692      3    3.83673    0.44769   -1.77402
-2.47693   -0.03347    0.00000 ;       N-    CA-    CB-   CG2
  2686   2688   2690   2696      3    4.93294    1.13386   -3.12963
-5.27184    3.17984    0.00000 ;       N-    CA-    CB-   OG1
  2686   2688   2698   2699      3    0.00000    0.00000    0.00000
0.00000    0.00000    0.00000 ;       N-    CA-     C-     O
  2686   2688   2698   2700      3   17.40544   -5.02080  -13.22144
9.20480    0.00000    0.00000 ;       N-    CA-     C-     N
  2687   2686   2688   2689      3    0.00000    0.00000    0.00000
0.00000    0.00000    0.00000 ;       H-     N-    CA-    HA
  2687   2686   2688   2690      3    0.00000    0.00000    0.00000
0.00000    0.00000    0.00000 ;       H-     N-    CA-    CB
  2687   2686   2688   2698      3    0.00000    0.00000    0.00000
0.00000    0.00000    0.00000 ;       H-     N-    CA-     C
  2688   2690   2692   2693      3    0.66944    2.00832    0.00000
-2.67776    0.00000    0.00000 ;      CA-    CB-   CG2-  HG21
  2688   2690   2692   2694      3    0.66944    2.00832    0.00000
-2.67776    0.00000    0.00000 ;      CA-    CB-   CG2-  HG22
  2688   2690   2692   2695      3    0.66944    2.00832    0.00000
-2.67776    0.00000    0.00000 ;      CA-    CB-   CG2-  HG23
  2688   2690   2696   2697      3    1.12131    2.98738    1.66523
-3.94970    0.43514    0.00000 ;      CA-    CB-   OG1-   HG1
  2688   2698   2700   2701      3   20.92000    0.00000  -20.92000
0.00000    0.00000    0.00000 ;      CA-     C-     N-     H
  2688   2698   2700   2702      3   20.92000    0.00000  -20.92000
0.00000    0.00000    0.00000 ;      CA-     C-     N-    CA
  2689   2688   2690   2691      3    0.65084    1.95253    0.00000
-2.60338    0.00000    0.00000 ;      HA-    CA-    CB-    HB
  2689   2688   2690   2692      3    0.65084    1.95253    0.00000
```

```
-2.60338    0.00000    0.00000 ;      HA-    CA-     CB-   CG2
  2689   2688   2690   2696       3    1.04600   -1.04600    0.00000
0.00000    0.00000    0.00000 ;     HA-    CA-     CB-   OG1
  2689   2688   2698   2699       3    3.68192   -4.35136    0.00000
1.33888    0.00000    0.00000 ;     HA-    CA-      C-     O
  2689   2688   2698   2700       3    0.00000    0.00000    0.00000
0.00000    0.00000    0.00000 ;     HA-    CA-      C-     N
  2690   2688   2698   2699       3    0.00000    0.00000    0.00000
0.00000    0.00000    0.00000 ;     CB-    CA-      C-     O
  2690   2688   2698   2700       3    2.51040    4.18400    1.67360
-6.69440    0.00000    0.00000 ;     CB-    CA-      C-     N
  2691   2690   2688   2698       3    0.65084    1.95253    0.00000
-2.60338    0.00000    0.00000 ;      HB-    CB-     CA-     C
  2691   2690   2692   2693       3    0.65084    1.95253    0.00000
-2.60338    0.00000    0.00000 ;      HB-    CB-   CG2-  HG21
  2691   2690   2692   2694       3    0.65084    1.95253    0.00000
-2.60338    0.00000    0.00000 ;      HB-    CB-   CG2-  HG22
  2691   2690   2692   2695       3    0.65084    1.95253    0.00000
-2.60338    0.00000    0.00000 ;      HB-    CB-   CG2-  HG23
  2691   2690   2696   2697       3    0.69733    2.09200    0.00000
-2.78933    0.00000    0.00000 ;      HB-    CB-   OG1-   HG1
  2692   2690   2688   2698       3    5.67350    3.55640   -6.16722
-2.47693    3.74886    0.00000 ;    CG2-    CB-     CA-     C
  2692   2690   2696   2697       3    4.74047    0.27196   -2.26773
-3.94970    1.60666    0.00000 ;    CG2-    CB-   OG1-   HG1
  2693   2692   2690   2696       3    1.04600   -1.04600    0.00000
0.00000    0.00000    0.00000 ;   HG21-   CG2-     CB-   OG1
  2694   2692   2690   2696       3    1.04600   -1.04600    0.00000
0.00000    0.00000    0.00000 ;   HG22-   CG2-     CB-   OG1
  2695   2692   2690   2696       3    1.04600   -1.04600    0.00000
0.00000    0.00000    0.00000 ;   HG23-   CG2-     CB-   OG1
  2696   2690   2688   2698       3    6.53541    6.87013   -6.21742
-5.27184    5.22163    0.00000 ;    OG1-    CB-     CA-     C
  2698   2700   2702   2703       3    0.00000    0.00000    0.00000
0.00000    0.00000    0.00000 ;      C-     N-     CA-   HA2
  2698   2700   2702   2704       3    0.00000    0.00000    0.00000
0.00000    0.00000    0.00000 ;      C-     N-     CA-   HA3
  2698   2700   2702   2705       3    1.75728    5.27184    2.25936
-7.02912    0.00000    0.00000 ;      C-     N-     CA-     C
  2699   2698   2700   2701       3   29.28800   -8.36800  -20.92000
0.00000    0.00000    0.00000 ;      O-     C-      N-     H
  2699   2698   2700   2702       3   20.92000    0.00000  -20.92000
0.00000    0.00000    0.00000 ;      O-     C-      N-    CA
  2700   2702   2705   2706       3    0.00000    0.00000    0.00000
0.00000    0.00000    0.00000 ;      N-    CA-      C-     O
  2700   2702   2705   2707       3   17.40544   -5.02080  -13.22144
9.20480    0.00000    0.00000 ;      N-    CA-      C-     N
  2701   2700   2702   2703       3    0.00000    0.00000    0.00000
0.00000    0.00000    0.00000 ;      H-     N-     CA-   HA2
  2701   2700   2702   2704       3    0.00000    0.00000    0.00000
0.00000    0.00000    0.00000 ;      H-     N-     CA-   HA3
  2701   2700   2702   2705       3    0.00000    0.00000    0.00000
```

0.00000 0.00000 0.00000 ; H- N- CA- C
2702 2705 2707 2708 3 20.92000 0.00000 -20.92000
0.00000 0.00000 0.00000 ; CA- C- N- H
2702 2705 2707 2709 3 20.92000 0.00000 -20.92000
0.00000 0.00000 0.00000 ; CA- C- N- CA
2703 2702 2705 2706 3 3.68192 -4.35136 0.00000
1.33888 0.00000 0.00000 ; HA2- CA- C- O
2703 2702 2705 2707 3 0.00000 0.00000 0.00000
0.00000 0.00000 0.00000 ; HA2- CA- C- N
2704 2702 2705 2706 3 3.68192 -4.35136 0.00000
1.33888 0.00000 0.00000 ; HA3- CA- C- O
2704 2702 2705 2707 3 0.00000 0.00000 0.00000
0.00000 0.00000 0.00000 ; HA3- CA- C- N
2705 2707 2709 2710 3 0.00000 0.00000 0.00000
0.00000 0.00000 0.00000 ; C- N- CA- HA
2705 2707 2709 2711 3 11.71520 1.67360 15.06240 -
13.38880 0.00000 0.00000 ; C- N- CA- CB
2705 2707 2709 2720 3 1.75728 5.27184 2.25936
-7.02912 0.00000 0.00000 ; C- N- CA- C
2706 2705 2707 2708 3 29.28800 -8.36800 -20.92000
0.00000 0.00000 0.00000 ; O- C- N- H
2706 2705 2707 2709 3 20.92000 0.00000 -20.92000
0.00000 0.00000 0.00000 ; O- C- N- CA
2707 2709 2711 2712 3 0.65084 1.95253 0.00000
-2.60338 0.00000 0.00000 ; N- CA- CB- HB2
2707 2709 2711 2713 3 0.65084 1.95253 0.00000
-2.60338 0.00000 0.00000 ; N- CA- CB- HB3
2707 2709 2711 2714 3 3.21331 2.22589 -4.15053
-2.40998 2.61082 0.00000 ; N- CA- CB- CG
2707 2709 2720 2721 3 0.00000 0.00000 0.00000
0.00000 0.00000 0.00000 ; N- CA- C- O
2707 2709 2720 2722 3 17.40544 -5.02080 -13.22144
9.20480 0.00000 0.00000 ; N- CA- C- N
2708 2707 2709 2710 3 0.00000 0.00000 0.00000
0.00000 0.00000 0.00000 ; H- N- CA- HA
2708 2707 2709 2711 3 0.00000 0.00000 0.00000
0.00000 0.00000 0.00000 ; H- N- CA- CB
2708 2707 2709 2720 3 0.00000 0.00000 0.00000
0.00000 0.00000 0.00000 ; H- N- CA- C
2709 2711 2714 2715 3 0.66944 2.00832 0.00000
-2.67776 0.00000 0.00000 ; CA- CB- CG- HG2
2709 2711 2714 2716 3 0.66944 2.00832 0.00000
-2.67776 0.00000 0.00000 ; CA- CB- CG- HG3
2709 2711 2714 2717 3 10.12110 -1.91209 0.01674
10.17549 -1.87443 0.00000 ; CA- CB- CG- CD
2709 2720 2722 2723 3 20.92000 0.00000 -20.92000
0.00000 0.00000 0.00000 ; CA- C- N- H
2709 2720 2722 2724 3 20.92000 0.00000 -20.92000
0.00000 0.00000 0.00000 ; CA- C- N- CA
2710 2709 2711 2712 3 0.65084 1.95253 0.00000
-2.60338 0.00000 0.00000 ; HA- CA- CB- HB2
2710 2709 2711 2713 3 0.65084 1.95253 0.00000

-2.60338 0.00000 0.00000 ; HA- CA- CB- HB3
2710 2709 2711 2714 3 0.65084 1.95253 0.00000
-2.60338 0.00000 0.00000 ; HA- CA- CB- CG
2710 2709 2720 2721 3 3.68192 -4.35136 0.00000
1.33888 0.00000 0.00000 ; HA- CA- C- O
2710 2709 2720 2722 3 0.00000 0.00000 0.00000
0.00000 0.00000 0.00000 ; HA- CA- C- N
2711 2709 2720 2721 3 0.00000 0.00000 0.00000
0.00000 0.00000 0.00000 ; CB- CA- C- O
2711 2709 2720 2722 3 2.51040 4.18400 1.67360
-6.69440 0.00000 0.00000 ; CB- CA- C- N
2711 2714 2717 2718 3 3.79907 0.00000 -5.40573
0.00000 2.14221 0.00000 ; CB- CG- CD- OE1
2711 2714 2717 2719 3 3.79907 0.00000 -5.40573
0.00000 2.14221 0.00000 ; CB- CG- CD- OE2
2712 2711 2709 2720 3 0.65084 1.95253 0.00000
-2.60338 0.00000 0.00000 ; HB2- CB- CA- C
2712 2711 2714 2715 3 0.62760 1.88280 0.00000
-2.51040 0.00000 0.00000 ; HB2- CB- CG- HG2
2712 2711 2714 2716 3 0.62760 1.88280 0.00000
-2.51040 0.00000 0.00000 ; HB2- CB- CG- HG3
2712 2711 2714 2717 3 0.65084 1.95253 0.00000
-2.60338 0.00000 0.00000 ; HB2- CB- CG- CD
2713 2711 2709 2720 3 0.65084 1.95253 0.00000
-2.60338 0.00000 0.00000 ; HB3- CB- CA- C
2713 2711 2714 2715 3 0.62760 1.88280 0.00000
-2.51040 0.00000 0.00000 ; HB3- CB- CG- HG2
2713 2711 2714 2716 3 0.62760 1.88280 0.00000
-2.51040 0.00000 0.00000 ; HB3- CB- CG- HG3
2713 2711 2714 2717 3 0.65084 1.95253 0.00000
-2.60338 0.00000 0.00000 ; HB3- CB- CG- CD
2714 2711 2709 2720 3 6.86594 3.56895 -8.14206
-2.40998 4.85344 0.00000 ; CG- CB- CA- C
2715 2714 2717 2718 3 0.00000 0.00000 0.00000
0.00000 0.00000 0.00000 ; HG2- CG- CD- OE1
2715 2714 2717 2719 3 0.00000 0.00000 0.00000
0.00000 0.00000 0.00000 ; HG2- CG- CD- OE2
2716 2714 2717 2718 3 0.00000 0.00000 0.00000
0.00000 0.00000 0.00000 ; HG3- CG- CD- OE1
2716 2714 2717 2719 3 0.00000 0.00000 0.00000
0.00000 0.00000 0.00000 ; HG3- CG- CD- OE2
2720 2722 2724 2725 3 0.00000 0.00000 0.00000
0.00000 0.00000 0.00000 ; C- N- CA- HA
2720 2722 2724 2726 3 11.71520 1.67360 15.06240 -
13.38880 0.00000 0.00000 ; C- N- CA- CB
2720 2722 2724 2740 3 1.75728 5.27184 2.25936
-7.02912 0.00000 0.00000 ; C- N- CA- C
2721 2720 2722 2723 3 29.28800 -8.36800 -20.92000
0.00000 0.00000 0.00000 ; O- C- N- H
2721 2720 2722 2724 3 20.92000 0.00000 -20.92000
0.00000 0.00000 0.00000 ; O- C- N- CA
2722 2724 2726 2727 3 0.65084 1.95253 0.00000

```
-2.60338    0.00000    0.00000 ;      N-    CA-    CB-   HB2
  2722   2724   2726   2728      3    0.65084    1.95253    0.00000
-2.60338    0.00000    0.00000 ;      N-    CA-    CB-   HB3
  2722   2724   2726   2729      3    3.28026    2.46019   -2.19242
-3.21331   -0.23430    0.00000 ;      N-    CA-    CB-    CG
  2722   2724   2740   2741      3    0.00000    0.00000    0.00000
0.00000    0.00000    0.00000 ;      N-    CA-     C-     O
  2722   2724   2740   2742      3   17.40544   -5.02080  -13.22144
9.20480    0.00000    0.00000 ;      N-    CA-     C-     N
  2723   2722   2724   2725      3    0.00000    0.00000    0.00000
0.00000    0.00000    0.00000 ;      H-     N-    CA-    HA
  2723   2722   2724   2726      3    0.00000    0.00000    0.00000
0.00000    0.00000    0.00000 ;      H-     N-    CA-    CB
  2723   2722   2724   2740      3    0.00000    0.00000    0.00000
0.00000    0.00000    0.00000 ;      H-     N-    CA-     C
  2724   2726   2729   2730      3    0.57739    0.00000    1.02926
0.00000   -1.60666    0.00000 ;     CA-    CB-    CG-   CD1
  2724   2726   2729   2738      3    0.57739    0.00000    1.02926
0.00000   -1.60666    0.00000 ;     CA-    CB-    CG-   CD2
  2724   2740   2742   2743      3   20.92000    0.00000  -20.92000
0.00000    0.00000    0.00000 ;     CA-     C-     N-     H
  2724   2740   2742   2744      3   20.92000    0.00000  -20.92000
0.00000    0.00000    0.00000 ;     CA-     C-     N-    CA
  2725   2724   2726   2727      3    0.65084    1.95253    0.00000
-2.60338    0.00000    0.00000 ;     HA-    CA-    CB-   HB2
  2725   2724   2726   2728      3    0.65084    1.95253    0.00000
-2.60338    0.00000    0.00000 ;     HA-    CA-    CB-   HB3
  2725   2724   2726   2729      3    0.65084    1.95253    0.00000
-2.60338    0.00000    0.00000 ;     HA-    CA-    CB-    CG
  2725   2724   2740   2741      3    3.68192   -4.35136    0.00000
1.33888    0.00000    0.00000 ;     HA-    CA-     C-     O
  2725   2724   2740   2742      3    0.00000    0.00000    0.00000
0.00000    0.00000    0.00000 ;     HA-    CA-     C-     N
  2726   2724   2740   2741      3    0.00000    0.00000    0.00000
0.00000    0.00000    0.00000 ;     CB-    CA-     C-     O
  2726   2724   2740   2742      3    2.51040    4.18400    1.67360
-6.69440    0.00000    0.00000 ;     CB-    CA-     C-     N
  2726   2729   2730   2731      3   30.33400    0.00000  -30.33400
0.00000    0.00000    0.00000 ;     CB-    CG-   CD1-   HD1
  2726   2729   2730   2732      3   30.33400    0.00000  -30.33400
0.00000    0.00000    0.00000 ;     CB-    CG-   CD1-   CE1
  2726   2729   2738   2736      3   30.33400    0.00000  -30.33400
0.00000    0.00000    0.00000 ;     CB-    CG-   CD2-   CE2
  2726   2729   2738   2739      3   30.33400    0.00000  -30.33400
0.00000    0.00000    0.00000 ;     CB-    CG-   CD2-   HD2
  2727   2726   2724   2740      3    0.65084    1.95253    0.00000
-2.60338    0.00000    0.00000 ;    HB2-    CB-    CA-     C
  2727   2726   2729   2730      3    0.00000    0.00000    0.00000
0.00000    0.00000    0.00000 ;    HB2-    CB-    CG-   CD1
  2727   2726   2729   2738      3    0.00000    0.00000    0.00000
0.00000    0.00000    0.00000 ;    HB2-    CB-    CG-   CD2
  2728   2726   2724   2740      3    0.65084    1.95253    0.00000
```

-2.60338 0.00000 0.00000 ; HB3- CB- CA- C
2728 2726 2729 2730 3 0.00000 0.00000 0.00000
0.00000 0.00000 0.00000 ; HB3- CB- CG- CD1
2728 2726 2729 2738 3 0.00000 0.00000 0.00000
0.00000 0.00000 0.00000 ; HB3- CB- CG- CD2
2729 2726 2724 2740 3 4.95804 2.17986 -3.52293
-3.21331 -0.40166 0.00000 ; CG- CB- CA- C
2729 2730 2732 2733 3 30.33400 0.00000 -30.33400
0.00000 0.00000 0.00000 ; CG- CD1- CE1- HE1
2729 2730 2732 2734 3 30.33400 0.00000 -30.33400
0.00000 0.00000 0.00000 ; CG- CD1- CE1- CZ
2729 2738 2736 2734 3 30.33400 0.00000 -30.33400
0.00000 0.00000 0.00000 ; CG- CD2- CE2- CZ
2729 2738 2736 2737 3 30.33400 0.00000 -30.33400
0.00000 0.00000 0.00000 ; CG- CD2- CE2- HE2
2730 2729 2738 2736 3 30.33400 0.00000 -30.33400
0.00000 0.00000 0.00000 ; CD1- CG- CD2- CE2
2730 2729 2738 2739 3 30.33400 0.00000 -30.33400
0.00000 0.00000 0.00000 ; CD1- CG- CD2- HD2
2730 2732 2734 2735 3 30.33400 0.00000 -30.33400
0.00000 0.00000 0.00000 ; CD1- CE1- CZ- HZ
2730 2732 2734 2736 3 30.33400 0.00000 -30.33400
0.00000 0.00000 0.00000 ; CD1- CE1- CZ- CE2
2731 2730 2729 2738 3 30.33400 0.00000 -30.33400
0.00000 0.00000 0.00000 ; HD1- CD1- CG- CD2
2731 2730 2732 2733 3 30.33400 0.00000 -30.33400
0.00000 0.00000 0.00000 ; HD1- CD1- CE1- HE1
2731 2730 2732 2734 3 30.33400 0.00000 -30.33400
0.00000 0.00000 0.00000 ; HD1- CD1- CE1- CZ
2732 2730 2729 2738 3 30.33400 0.00000 -30.33400
0.00000 0.00000 0.00000 ; CE1- CD1- CG- CD2
2732 2734 2736 2737 3 30.33400 0.00000 -30.33400
0.00000 0.00000 0.00000 ; CE1- CZ- CE2- HE2
2732 2734 2736 2738 3 30.33400 0.00000 -30.33400
0.00000 0.00000 0.00000 ; CE1- CZ- CE2- CD2
2733 2732 2734 2735 3 30.33400 0.00000 -30.33400
0.00000 0.00000 0.00000 ; HE1- CE1- CZ- HZ
2733 2732 2734 2736 3 30.33400 0.00000 -30.33400
0.00000 0.00000 0.00000 ; HE1- CE1- CZ- CE2
2734 2736 2738 2739 3 30.33400 0.00000 -30.33400
0.00000 0.00000 0.00000 ; CZ- CE2- CD2- HD2
2735 2734 2736 2737 3 30.33400 0.00000 -30.33400
0.00000 0.00000 0.00000 ; HZ- CZ- CE2- HE2
2735 2734 2736 2738 3 30.33400 0.00000 -30.33400
0.00000 0.00000 0.00000 ; HZ- CZ- CE2- CD2
2737 2736 2738 2739 3 30.33400 0.00000 -30.33400
0.00000 0.00000 0.00000 ; HE2- CE2- CD2- HD2
2740 2742 2744 2745 3 0.00000 0.00000 0.00000
0.00000 0.00000 0.00000 ; C- N- CA- HA
2740 2742 2744 2746 3 11.71520 1.67360 15.06240 -
13.38880 0.00000 0.00000 ; C- N- CA- CB
2740 2742 2744 2754 3 1.75728 5.27184 2.25936

-7.02912 0.00000 0.00000 ; C- N- CA- C
2741 2740 2742 2743 3 29.28800 -8.36800 -20.92000
0.00000 0.00000 0.00000 ; O- C- N- H
2741 2740 2742 2744 3 20.92000 0.00000 -20.92000
0.00000 0.00000 0.00000 ; O- C- N- CA
2742 2744 2746 2747 3 0.65084 1.95253 0.00000
-2.60338 0.00000 0.00000 ; N- CA- CB- HB
2742 2744 2746 2748 3 3.83673 0.44769 -1.77402
-2.47693 -0.03347 0.00000 ; N- CA- CB- CG2
2742 2744 2746 2752 3 4.93294 1.13386 -3.12963
-5.27184 3.17984 0.00000 ; N- CA- CB- OG1
2742 2744 2754 2755 3 0.00000 0.00000 0.00000
0.00000 0.00000 0.00000 ; N- CA- C- O
2742 2744 2754 2756 3 17.40544 -5.02080 -13.22144
9.20480 0.00000 0.00000 ; N- CA- C- N
2743 2742 2744 2745 3 0.00000 0.00000 0.00000
0.00000 0.00000 0.00000 ; H- N- CA- HA
2743 2742 2744 2746 3 0.00000 0.00000 0.00000
0.00000 0.00000 0.00000 ; H- N- CA- CB
2743 2742 2744 2754 3 0.00000 0.00000 0.00000
0.00000 0.00000 0.00000 ; H- N- CA- C
2744 2746 2748 2749 3 0.66944 2.00832 0.00000
-2.67776 0.00000 0.00000 ; CA- CB- CG2- HG21
2744 2746 2748 2750 3 0.66944 2.00832 0.00000
-2.67776 0.00000 0.00000 ; CA- CB- CG2- HG22
2744 2746 2748 2751 3 0.66944 2.00832 0.00000
-2.67776 0.00000 0.00000 ; CA- CB- CG2- HG23
2744 2746 2752 2753 3 1.12131 2.98738 1.66523
-3.94970 0.43514 0.00000 ; CA- CB- OG1- HG1
2744 2754 2756 2757 3 20.92000 0.00000 -20.92000
0.00000 0.00000 0.00000 ; CA- C- N- H
2744 2754 2756 2758 3 20.92000 0.00000 -20.92000
0.00000 0.00000 0.00000 ; CA- C- N- CA
2745 2744 2746 2747 3 0.65084 1.95253 0.00000
-2.60338 0.00000 0.00000 ; HA- CA- CB- HB
2745 2744 2746 2748 3 0.65084 1.95253 0.00000
-2.60338 0.00000 0.00000 ; HA- CA- CB- CG2
2745 2744 2746 2752 3 1.04600 -1.04600 0.00000
0.00000 0.00000 0.00000 ; HA- CA- CB- OG1
2745 2744 2754 2755 3 3.68192 -4.35136 0.00000
1.33888 0.00000 0.00000 ; HA- CA- C- O
2745 2744 2754 2756 3 0.00000 0.00000 0.00000
0.00000 0.00000 0.00000 ; HA- CA- C- N
2746 2744 2754 2755 3 0.00000 0.00000 0.00000
0.00000 0.00000 0.00000 ; CB- CA- C- O
2746 2744 2754 2756 3 2.51040 4.18400 1.67360
-6.69440 0.00000 0.00000 ; CB- CA- C- N
2747 2746 2744 2754 3 0.65084 1.95253 0.00000
-2.60338 0.00000 0.00000 ; HB- CB- CA- C
2747 2746 2748 2749 3 0.65084 1.95253 0.00000
-2.60338 0.00000 0.00000 ; HB- CB- CG2- HG21
2747 2746 2748 2750 3 0.65084 1.95253 0.00000

-2.60338 0.00000 0.00000 ; HB- CB- CG2- HG22
2747 2746 2748 2751 3 0.65084 1.95253 0.00000
-2.60338 0.00000 0.00000 ; HB- CB- CG2- HG23
2747 2746 2752 2753 3 0.69733 2.09200 0.00000
-2.78933 0.00000 0.00000 ; HB- CB- OG1- HG1
2748 2746 2744 2754 3 5.67350 3.55640 -6.16722
-2.47693 3.74886 0.00000 ; CG2- CB- CA- C
2748 2746 2752 2753 3 4.74047 0.27196 -2.26773
-3.94970 1.60666 0.00000 ; CG2- CB- OG1- HG1
2749 2748 2746 2752 3 1.04600 -1.04600 0.00000
0.00000 0.00000 0.00000 ; HG21- CG2- CB- OG1
2750 2748 2746 2752 3 1.04600 -1.04600 0.00000
0.00000 0.00000 0.00000 ; HG22- CG2- CB- OG1
2751 2748 2746 2752 3 1.04600 -1.04600 0.00000
0.00000 0.00000 0.00000 ; HG23- CG2- CB- OG1
2752 2746 2744 2754 3 6.53541 6.87013 -6.21742
-5.27184 5.22163 0.00000 ; OG1- CB- CA- C
2754 2756 2758 2759 3 0.00000 0.00000 0.00000
0.00000 0.00000 0.00000 ; C- N- CA- HA
2754 2756 2758 2760 3 11.71520 1.67360 15.06240 -
13.38880 0.00000 0.00000 ; C- N- CA- CB
2754 2756 2758 2768 3 1.75728 5.27184 2.25936
-7.02912 0.00000 0.00000 ; C- N- CA- C
2755 2754 2756 2757 3 29.28800 -8.36800 -20.92000
0.00000 0.00000 0.00000 ; O- C- N- H
2755 2754 2756 2758 3 20.92000 0.00000 -20.92000
0.00000 0.00000 0.00000 ; O- C- N- CA
2756 2758 2760 2761 3 0.65084 1.95253 0.00000
-2.60338 0.00000 0.00000 ; N- CA- CB- HB
2756 2758 2760 2762 3 3.83673 0.44769 -1.77402
-2.47693 -0.03347 0.00000 ; N- CA- CB- CG2
2756 2758 2760 2766 3 4.93294 1.13386 -3.12963
-5.27184 3.17984 0.00000 ; N- CA- CB- OG1
2756 2758 2768 2769 3 0.00000 0.00000 0.00000
0.00000 0.00000 0.00000 ; N- CA- C- O
2756 2758 2768 2770 3 17.40544 -5.02080 -13.22144
9.20480 0.00000 0.00000 ; N- CA- C- N
2757 2756 2758 2759 3 0.00000 0.00000 0.00000
0.00000 0.00000 0.00000 ; H- N- CA- HA
2757 2756 2758 2760 3 0.00000 0.00000 0.00000
0.00000 0.00000 0.00000 ; H- N- CA- CB
2757 2756 2758 2768 3 0.00000 0.00000 0.00000
0.00000 0.00000 0.00000 ; H- N- CA- C
2758 2760 2762 2763 3 0.66944 2.00832 0.00000
-2.67776 0.00000 0.00000 ; CA- CB- CG2- HG21
2758 2760 2762 2764 3 0.66944 2.00832 0.00000
-2.67776 0.00000 0.00000 ; CA- CB- CG2- HG22
2758 2760 2762 2765 3 0.66944 2.00832 0.00000
-2.67776 0.00000 0.00000 ; CA- CB- CG2- HG23
2758 2760 2766 2767 3 1.12131 2.98738 1.66523
-3.94970 0.43514 0.00000 ; CA- CB- OG1- HG1
2758 2768 2770 2771 3 20.92000 0.00000 -20.92000

```
0.00000    0.00000    0.00000 ;      CA-     C-      N-      H
  2758   2768   2770   2772       3   20.92000    0.00000  -20.92000
0.00000    0.00000    0.00000 ;      CA-     C-      N-     CA
  2759   2758   2760   2761       3    0.65084    1.95253    0.00000
-2.60338    0.00000    0.00000 ;      HA-    CA-     CB-     HB
  2759   2758   2760   2762       3    0.65084    1.95253    0.00000
-2.60338    0.00000    0.00000 ;      HA-    CA-     CB-    CG2
  2759   2758   2760   2766       3    1.04600   -1.04600    0.00000
0.00000    0.00000    0.00000 ;      HA-    CA-     CB-    OG1
  2759   2758   2768   2769       3    3.68192   -4.35136    0.00000
1.33888    0.00000    0.00000 ;      HA-    CA-      C-      O
  2759   2758   2768   2770       3    0.00000    0.00000    0.00000
0.00000    0.00000    0.00000 ;      HA-    CA-      C-      N
  2760   2758   2768   2769       3    0.00000    0.00000    0.00000
0.00000    0.00000    0.00000 ;      CB-    CA-      C-      O
  2760   2758   2768   2770       3    2.51040    4.18400    1.67360
-6.69440    0.00000    0.00000 ;      CB-    CA-      C-      N
  2761   2760   2758   2768       3    0.65084    1.95253    0.00000
-2.60338    0.00000    0.00000 ;      HB-    CB-     CA-      C
  2761   2760   2762   2763       3    0.65084    1.95253    0.00000
-2.60338    0.00000    0.00000 ;      HB-    CB-    CG2-   HG21
  2761   2760   2762   2764       3    0.65084    1.95253    0.00000
-2.60338    0.00000    0.00000 ;      HB-    CB-    CG2-   HG22
  2761   2760   2762   2765       3    0.65084    1.95253    0.00000
-2.60338    0.00000    0.00000 ;      HB-    CB-    CG2-   HG23
  2761   2760   2766   2767       3    0.69733    2.09200    0.00000
-2.78933    0.00000    0.00000 ;      HB-    CB-    OG1-    HG1
  2762   2760   2758   2768       3    5.67350    3.55640   -6.16722
-2.47693    3.74886    0.00000 ;     CG2-    CB-     CA-      C
  2762   2760   2766   2767       3    4.74047    0.27196   -2.26773
-3.94970    1.60666    0.00000 ;     CG2-    CB-    OG1-    HG1
  2763   2762   2760   2766       3    1.04600   -1.04600    0.00000
0.00000    0.00000    0.00000 ;    HG21-   CG2-     CB-    OG1
  2764   2762   2760   2766       3    1.04600   -1.04600    0.00000
0.00000    0.00000    0.00000 ;    HG22-   CG2-     CB-    OG1
  2765   2762   2760   2766       3    1.04600   -1.04600    0.00000
0.00000    0.00000    0.00000 ;    HG23-   CG2-     CB-    OG1
  2766   2760   2758   2768       3    6.53541    6.87013   -6.21742
-5.27184    5.22163    0.00000 ;     OG1-    CB-     CA-      C
  2768   2770   2772   2773       3    0.00000    0.00000    0.00000
0.00000    0.00000    0.00000 ;       C-     N-     CA-     HA
  2768   2770   2772   2774       3   11.71520    1.67360   15.06240  -
13.38880    0.00000    0.00000 ;       C-     N-     CA-     CB
  2768   2770   2772   2782       3    1.75728    5.27184    2.25936
-7.02912    0.00000    0.00000 ;       C-     N-     CA-      C
  2769   2768   2770   2771       3   29.28800   -8.36800  -20.92000
0.00000    0.00000    0.00000 ;       O-     C-      N-      H
  2769   2768   2770   2772       3   20.92000    0.00000  -20.92000
0.00000    0.00000    0.00000 ;       O-     C-      N-     CA
  2770   2772   2774   2775       3    0.65084    1.95253    0.00000
-2.60338    0.00000    0.00000 ;       N-    CA-     CB-     HB
  2770   2772   2774   2776       3    3.83673    0.44769   -1.77402
```

```
-2.47693   -0.03347    0.00000 ;       N-    CA-    CB-   CG2
  2770   2772   2774   2780       3    4.93294    1.13386   -3.12963
-5.27184    3.17984    0.00000 ;       N-    CA-    CB-   OG1
  2770   2772   2782   2783       3    0.00000    0.00000    0.00000
0.00000    0.00000    0.00000 ;      N-    CA-     C-     O
  2770   2772   2782   2784       3   17.40544   -5.02080  -13.22144
9.20480    0.00000    0.00000 ;      N-    CA-     C-     N
  2771   2770   2772   2773       3    0.00000    0.00000    0.00000
0.00000    0.00000    0.00000 ;      H-     N-    CA-    HA
  2771   2770   2772   2774       3    0.00000    0.00000    0.00000
0.00000    0.00000    0.00000 ;      H-     N-    CA-    CB
  2771   2770   2772   2782       3    0.00000    0.00000    0.00000
0.00000    0.00000    0.00000 ;      H-     N-    CA-     C
  2772   2774   2776   2777       3    0.66944    2.00832    0.00000
-2.67776    0.00000    0.00000 ;      CA-    CB-   CG2-  HG21
  2772   2774   2776   2778       3    0.66944    2.00832    0.00000
-2.67776    0.00000    0.00000 ;      CA-    CB-   CG2-  HG22
  2772   2774   2776   2779       3    0.66944    2.00832    0.00000
-2.67776    0.00000    0.00000 ;      CA-    CB-   CG2-  HG23
  2772   2774   2780   2781       3    1.12131    2.98738    1.66523
-3.94970    0.43514    0.00000 ;      CA-    CB-   OG1-   HG1
  2772   2782   2784   2785       3   20.92000    0.00000  -20.92000
0.00000    0.00000    0.00000 ;     CA-     C-     N-     H
  2772   2782   2784   2786       3   20.92000    0.00000  -20.92000
0.00000    0.00000    0.00000 ;     CA-     C-     N-    CA
  2773   2772   2774   2775       3    0.65084    1.95253    0.00000
-2.60338    0.00000    0.00000 ;      HA-    CA-    CB-    HB
  2773   2772   2774   2776       3    0.65084    1.95253    0.00000
-2.60338    0.00000    0.00000 ;      HA-    CA-    CB-   CG2
  2773   2772   2774   2780       3    1.04600   -1.04600    0.00000
0.00000    0.00000    0.00000 ;     HA-    CA-    CB-   OG1
  2773   2772   2782   2783       3    3.68192   -4.35136    0.00000
1.33888    0.00000    0.00000 ;     HA-    CA-     C-     O
  2773   2772   2782   2784       3    0.00000    0.00000    0.00000
0.00000    0.00000    0.00000 ;     HA-    CA-     C-     N
  2774   2772   2782   2783       3    0.00000    0.00000    0.00000
0.00000    0.00000    0.00000 ;     CB-    CA-     C-     O
  2774   2772   2782   2784       3    2.51040    4.18400    1.67360
-6.69440    0.00000    0.00000 ;      CB-    CA-     C-     N
  2775   2774   2772   2782       3    0.65084    1.95253    0.00000
-2.60338    0.00000    0.00000 ;      HB-    CB-    CA-     C
  2775   2774   2776   2777       3    0.65084    1.95253    0.00000
-2.60338    0.00000    0.00000 ;      HB-    CB-   CG2-  HG21
  2775   2774   2776   2778       3    0.65084    1.95253    0.00000
-2.60338    0.00000    0.00000 ;      HB-    CB-   CG2-  HG22
  2775   2774   2776   2779       3    0.65084    1.95253    0.00000
-2.60338    0.00000    0.00000 ;      HB-    CB-   CG2-  HG23
  2775   2774   2780   2781       3    0.69733    2.09200    0.00000
-2.78933    0.00000    0.00000 ;      HB-    CB-   OG1-   HG1
  2776   2774   2772   2782       3    5.67350    3.55640   -6.16722
-2.47693    3.74886    0.00000 ;     CG2-    CB-    CA-     C
  2776   2774   2780   2781       3    4.74047    0.27196   -2.26773
```

-3.94970 1.60666 0.00000 ; CG2- CB- OG1- HG1
2777 2776 2774 2780 3 1.04600 -1.04600 0.00000
0.00000 0.00000 0.00000 ; HG21- CG2- CB- OG1
2778 2776 2774 2780 3 1.04600 -1.04600 0.00000
0.00000 0.00000 0.00000 ; HG22- CG2- CB- OG1
2779 2776 2774 2780 3 1.04600 -1.04600 0.00000
0.00000 0.00000 0.00000 ; HG23- CG2- CB- OG1
2780 2774 2772 2782 3 6.53541 6.87013 -6.21742
-5.27184 5.22163 0.00000 ; OG1- CB- CA- C
2782 2784 2786 2787 3 0.00000 0.00000 0.00000
0.00000 0.00000 0.00000 ; C- N- CA- HA
2782 2784 2786 2788 3 11.71520 1.67360 15.06240 -
13.38880 0.00000 0.00000 ; C- N- CA- CB
2782 2784 2786 2804 3 1.75728 5.27184 2.25936
-7.02912 0.00000 0.00000 ; C- N- CA- C
2783 2782 2784 2785 3 29.28800 -8.36800 -20.92000
0.00000 0.00000 0.00000 ; O- C- N- H
2783 2782 2784 2786 3 20.92000 0.00000 -20.92000
0.00000 0.00000 0.00000 ; O- C- N- CA
2784 2786 2788 2789 3 0.65084 1.95253 0.00000
-2.60338 0.00000 0.00000 ; N- CA- CB- HB2
2784 2786 2788 2790 3 0.65084 1.95253 0.00000
-2.60338 0.00000 0.00000 ; N- CA- CB- HB3
2784 2786 2788 2791 3 0.65084 1.95253 0.00000
-2.60338 0.00000 0.00000 ; N- CA- CB- CG
2784 2786 2804 2805 3 0.00000 0.00000 0.00000
0.00000 0.00000 0.00000 ; N- CA- C- O
2784 2786 2804 2806 3 17.40544 -5.02080 -13.22144
9.20480 0.00000 0.00000 ; N- CA- C- N
2785 2784 2786 2787 3 0.00000 0.00000 0.00000
0.00000 0.00000 0.00000 ; H- N- CA- HA
2785 2784 2786 2788 3 0.00000 0.00000 0.00000
0.00000 0.00000 0.00000 ; H- N- CA- CB
2785 2784 2786 2804 3 0.00000 0.00000 0.00000
0.00000 0.00000 0.00000 ; H- N- CA- C
2786 2788 2791 2792 3 0.66944 2.00832 0.00000
-2.67776 0.00000 0.00000 ; CA- CB- CG- HG2
2786 2788 2791 2793 3 0.66944 2.00832 0.00000
-2.67776 0.00000 0.00000 ; CA- CB- CG- HG3
2786 2788 2791 2794 3 3.68192 3.09616 -2.09200
-3.01248 0.00000 0.00000 ; CA- CB- CG- CD
2786 2804 2806 2807 3 20.92000 0.00000 -20.92000
0.00000 0.00000 0.00000 ; CA- C- N- H
2786 2804 2806 2808 3 20.92000 0.00000 -20.92000
0.00000 0.00000 0.00000 ; CA- C- N- CA
2787 2786 2788 2789 3 0.65084 1.95253 0.00000
-2.60338 0.00000 0.00000 ; HA- CA- CB- HB2
2787 2786 2788 2790 3 0.65084 1.95253 0.00000
-2.60338 0.00000 0.00000 ; HA- CA- CB- HB3
2787 2786 2788 2791 3 0.65084 1.95253 0.00000
-2.60338 0.00000 0.00000 ; HA- CA- CB- CG
2787 2786 2804 2805 3 3.68192 -4.35136 0.00000

```
1.33888    0.00000    0.00000 ;     HA-    CA-     C-     O
  2787   2786   2804   2806      3    0.00000    0.00000    0.00000
0.00000    0.00000    0.00000 ;     HA-    CA-     C-     N
  2788   2786   2804   2805      3    0.00000    0.00000    0.00000
0.00000    0.00000    0.00000 ;     CB-    CA-     C-     O
  2788   2786   2804   2806      3    2.51040    4.18400    1.67360
-6.69440    0.00000    0.00000 ;     CB-    CA-     C-     N
  2788   2791   2794   2795      3    0.66944    2.00832    0.00000
-2.67776    0.00000    0.00000 ;     CB-    CG-    CD-   HD2
  2788   2791   2794   2796      3    0.66944    2.00832    0.00000
-2.67776    0.00000    0.00000 ;     CB-    CG-    CD-   HD3
  2788   2791   2794   2797      3    3.68192    3.09616   -2.09200
-3.01248    0.00000    0.00000 ;     CB-    CG-    CD-    CE
  2789   2788   2786   2804      3    0.65084    1.95253    0.00000
-2.60338    0.00000    0.00000 ;    HB2-    CB-    CA-     C
  2789   2788   2791   2792      3    0.62760    1.88280    0.00000
-2.51040    0.00000    0.00000 ;    HB2-    CB-    CG-   HG2
  2789   2788   2791   2793      3    0.62760    1.88280    0.00000
-2.51040    0.00000    0.00000 ;    HB2-    CB-    CG-   HG3
  2789   2788   2791   2794      3    0.66944    2.00832    0.00000
-2.67776    0.00000    0.00000 ;    HB2-    CB-    CG-    CD
  2790   2788   2786   2804      3    0.65084    1.95253    0.00000
-2.60338    0.00000    0.00000 ;    HB3-    CB-    CA-     C
  2790   2788   2791   2792      3    0.62760    1.88280    0.00000
-2.51040    0.00000    0.00000 ;    HB3-    CB-    CG-   HG2
  2790   2788   2791   2793      3    0.62760    1.88280    0.00000
-2.51040    0.00000    0.00000 ;    HB3-    CB-    CG-   HG3
  2790   2788   2791   2794      3    0.66944    2.00832    0.00000
-2.67776    0.00000    0.00000 ;    HB3-    CB-    CG-    CD
  2791   2788   2786   2804      3    0.65084    1.95253    0.00000
-2.60338    0.00000    0.00000 ;     CG-    CB-    CA-     C
  2791   2794   2797   2798      3    0.65084    1.95253    0.00000
-2.60338    0.00000    0.00000 ;     CG-    CD-    CE-   HE2
  2791   2794   2797   2799      3    0.65084    1.95253    0.00000
-2.60338    0.00000    0.00000 ;     CG-    CD-    CE-   HE3
  2791   2794   2797   2800      3    0.65084    1.95253    0.00000
-2.60338    0.00000    0.00000 ;     CG-    CD-    CE-    NZ
  2792   2791   2794   2795      3    0.62760    1.88280    0.00000
-2.51040    0.00000    0.00000 ;    HG2-    CG-    CD-   HD2
  2792   2791   2794   2796      3    0.62760    1.88280    0.00000
-2.51040    0.00000    0.00000 ;    HG2-    CG-    CD-   HD3
  2792   2791   2794   2797      3    0.66944    2.00832    0.00000
-2.67776    0.00000    0.00000 ;    HG2-    CG-    CD-    CE
  2793   2791   2794   2795      3    0.62760    1.88280    0.00000
-2.51040    0.00000    0.00000 ;    HG3-    CG-    CD-   HD2
  2793   2791   2794   2796      3    0.62760    1.88280    0.00000
-2.51040    0.00000    0.00000 ;    HG3-    CG-    CD-   HD3
  2793   2791   2794   2797      3    0.66944    2.00832    0.00000
-2.67776    0.00000    0.00000 ;    HG3-    CG-    CD-    CE
  2794   2797   2800   2801      3    0.65084    1.95253    0.00000
-2.60338    0.00000    0.00000 ;     CD-    CE-    NZ-   HZ1
  2794   2797   2800   2802      3    0.65084    1.95253    0.00000
```

-2.60338 0.00000 0.00000 ; CD- CE- NZ- HZ2
2794 2797 2800 2803 3 0.65084 1.95253 0.00000
-2.60338 0.00000 0.00000 ; CD- CE- NZ- HZ3
2795 2794 2797 2798 3 0.65084 1.95253 0.00000
-2.60338 0.00000 0.00000 ; HD2- CD- CE- HE2
2795 2794 2797 2799 3 0.65084 1.95253 0.00000
-2.60338 0.00000 0.00000 ; HD2- CD- CE- HE3
2795 2794 2797 2800 3 0.65084 1.95253 0.00000
-2.60338 0.00000 0.00000 ; HD2- CD- CE- NZ
2796 2794 2797 2798 3 0.65084 1.95253 0.00000
-2.60338 0.00000 0.00000 ; HD3- CD- CE- HE2
2796 2794 2797 2799 3 0.65084 1.95253 0.00000
-2.60338 0.00000 0.00000 ; HD3- CD- CE- HE3
2796 2794 2797 2800 3 0.65084 1.95253 0.00000
-2.60338 0.00000 0.00000 ; HD3- CD- CE- NZ
2798 2797 2800 2801 3 0.65084 1.95253 0.00000
-2.60338 0.00000 0.00000 ; HE2- CE- NZ- HZ1
2798 2797 2800 2802 3 0.65084 1.95253 0.00000
-2.60338 0.00000 0.00000 ; HE2- CE- NZ- HZ2
2798 2797 2800 2803 3 0.65084 1.95253 0.00000
-2.60338 0.00000 0.00000 ; HE2- CE- NZ- HZ3
2799 2797 2800 2801 3 0.65084 1.95253 0.00000
-2.60338 0.00000 0.00000 ; HE3- CE- NZ- HZ1
2799 2797 2800 2802 3 0.65084 1.95253 0.00000
-2.60338 0.00000 0.00000 ; HE3- CE- NZ- HZ2
2799 2797 2800 2803 3 0.65084 1.95253 0.00000
-2.60338 0.00000 0.00000 ; HE3- CE- NZ- HZ3
2804 2806 2808 2809 3 0.00000 0.00000 0.00000
0.00000 0.00000 0.00000 ; C- N- CA- HA
2804 2806 2808 2810 3 11.71520 1.67360 15.06240 -
13.38880 0.00000 0.00000 ; C- N- CA- CB
2804 2806 2808 2814 3 1.75728 5.27184 2.25936
-7.02912 0.00000 0.00000 ; C- N- CA- C
2805 2804 2806 2807 3 29.28800 -8.36800 -20.92000
0.00000 0.00000 0.00000 ; O- C- N- H
2805 2804 2806 2808 3 20.92000 0.00000 -20.92000
0.00000 0.00000 0.00000 ; O- C- N- CA
2806 2808 2810 2811 3 0.65084 1.95253 0.00000
-2.60338 0.00000 0.00000 ; N- CA- CB- HB1
2806 2808 2810 2812 3 0.65084 1.95253 0.00000
-2.60338 0.00000 0.00000 ; N- CA- CB- HB2
2806 2808 2810 2813 3 0.65084 1.95253 0.00000
-2.60338 0.00000 0.00000 ; N- CA- CB- HB3
2806 2808 2814 2815 3 0.00000 0.00000 0.00000
0.00000 0.00000 0.00000 ; N- CA- C- O
2806 2808 2814 2816 3 17.40544 -5.02080 -13.22144
9.20480 0.00000 0.00000 ; N- CA- C- N
2807 2806 2808 2809 3 0.00000 0.00000 0.00000
0.00000 0.00000 0.00000 ; H- N- CA- HA
2807 2806 2808 2810 3 0.00000 0.00000 0.00000
0.00000 0.00000 0.00000 ; H- N- CA- CB
2807 2806 2808 2814 3 0.00000 0.00000 0.00000

0.00000 0.00000 0.00000 ; H- N- CA- C
2808 2814 2816 2817 3 20.92000 0.00000 -20.92000
0.00000 0.00000 0.00000 ; CA- C- N- H
2808 2814 2816 2818 3 20.92000 0.00000 -20.92000
0.00000 0.00000 0.00000 ; CA- C- N- CA
2809 2808 2810 2811 3 0.65084 1.95253 0.00000
-2.60338 0.00000 0.00000 ; HA- CA- CB- HB1
2809 2808 2810 2812 3 0.65084 1.95253 0.00000
-2.60338 0.00000 0.00000 ; HA- CA- CB- HB2
2809 2808 2810 2813 3 0.65084 1.95253 0.00000
-2.60338 0.00000 0.00000 ; HA- CA- CB- HB3
2809 2808 2814 2815 3 3.68192 -4.35136 0.00000
1.33888 0.00000 0.00000 ; HA- CA- C- O
2809 2808 2814 2816 3 0.00000 0.00000 0.00000
0.00000 0.00000 0.00000 ; HA- CA- C- N
2810 2808 2814 2815 3 0.00000 0.00000 0.00000
0.00000 0.00000 0.00000 ; CB- CA- C- O
2810 2808 2814 2816 3 2.51040 4.18400 1.67360
-6.69440 0.00000 0.00000 ; CB- CA- C- N
2811 2810 2808 2814 3 0.65084 1.95253 0.00000
-2.60338 0.00000 0.00000 ; HB1- CB- CA- C
2812 2810 2808 2814 3 0.65084 1.95253 0.00000
-2.60338 0.00000 0.00000 ; HB2- CB- CA- C
2813 2810 2808 2814 3 0.65084 1.95253 0.00000
-2.60338 0.00000 0.00000 ; HB3- CB- CA- C
2814 2816 2818 2819 3 0.00000 0.00000 0.00000
0.00000 0.00000 0.00000 ; C- N- CA- HA2
2814 2816 2818 2820 3 0.00000 0.00000 0.00000
0.00000 0.00000 0.00000 ; C- N- CA- HA3
2814 2816 2818 2821 3 1.75728 5.27184 2.25936
-7.02912 0.00000 0.00000 ; C- N- CA- C
2815 2814 2816 2817 3 29.28800 -8.36800 -20.92000
0.00000 0.00000 0.00000 ; O- C- N- H
2815 2814 2816 2818 3 20.92000 0.00000 -20.92000
0.00000 0.00000 0.00000 ; O- C- N- CA
2816 2818 2821 2822 3 0.00000 0.00000 0.00000
0.00000 0.00000 0.00000 ; N- CA- C- O
2816 2818 2821 2823 3 17.40544 -5.02080 -13.22144
9.20480 0.00000 0.00000 ; N- CA- C- N
2817 2816 2818 2819 3 0.00000 0.00000 0.00000
0.00000 0.00000 0.00000 ; H- N- CA- HA2
2817 2816 2818 2820 3 0.00000 0.00000 0.00000
0.00000 0.00000 0.00000 ; H- N- CA- HA3
2817 2816 2818 2821 3 0.00000 0.00000 0.00000
0.00000 0.00000 0.00000 ; H- N- CA- C
2818 2821 2823 2824 3 20.92000 0.00000 -20.92000
0.00000 0.00000 0.00000 ; CA- C- N- H
2818 2821 2823 2825 3 20.92000 0.00000 -20.92000
0.00000 0.00000 0.00000 ; CA- C- N- CA
2819 2818 2821 2822 3 3.68192 -4.35136 0.00000
1.33888 0.00000 0.00000 ; HA2- CA- C- O
2819 2818 2821 2823 3 0.00000 0.00000 0.00000

0.00000 0.00000 0.00000 ; HA2- CA- C- N
2820 2818 2821 2822 3 3.68192 -4.35136 0.00000
1.33888 0.00000 0.00000 ; HA3- CA- C- O
2820 2818 2821 2823 3 0.00000 0.00000 0.00000
0.00000 0.00000 0.00000 ; HA3- CA- C- N
2821 2823 2825 2826 3 0.00000 0.00000 0.00000
0.00000 0.00000 0.00000 ; C- N- CA- HA
2821 2823 2825 2827 3 11.71520 1.67360 15.06240 -
13.38880 0.00000 0.00000 ; C- N- CA- CB
2821 2823 2825 2835 3 1.75728 5.27184 2.25936
-7.02912 0.00000 0.00000 ; C- N- CA- C
2822 2821 2823 2824 3 29.28800 -8.36800 -20.92000
0.00000 0.00000 0.00000 ; O- C- N- H
2822 2821 2823 2825 3 20.92000 0.00000 -20.92000
0.00000 0.00000 0.00000 ; O- C- N- CA
2823 2825 2827 2828 3 0.65084 1.95253 0.00000
-2.60338 0.00000 0.00000 ; N- CA- CB- HB
2823 2825 2827 2829 3 3.83673 0.44769 -1.77402
-2.47693 -0.03347 0.00000 ; N- CA- CB- CG2
2823 2825 2827 2833 3 4.93294 1.13386 -3.12963
-5.27184 3.17984 0.00000 ; N- CA- CB- OG1
2823 2825 2835 2836 3 0.00000 0.00000 0.00000
0.00000 0.00000 0.00000 ; N- CA- C- O
2823 2825 2835 2837 3 17.40544 -5.02080 -13.22144
9.20480 0.00000 0.00000 ; N- CA- C- N
2824 2823 2825 2826 3 0.00000 0.00000 0.00000
0.00000 0.00000 0.00000 ; H- N- CA- HA
2824 2823 2825 2827 3 0.00000 0.00000 0.00000
0.00000 0.00000 0.00000 ; H- N- CA- CB
2824 2823 2825 2835 3 0.00000 0.00000 0.00000
0.00000 0.00000 0.00000 ; H- N- CA- C
2825 2827 2829 2830 3 0.66944 2.00832 0.00000
-2.67776 0.00000 0.00000 ; CA- CB- CG2- HG21
2825 2827 2829 2831 3 0.66944 2.00832 0.00000
-2.67776 0.00000 0.00000 ; CA- CB- CG2- HG22
2825 2827 2829 2832 3 0.66944 2.00832 0.00000
-2.67776 0.00000 0.00000 ; CA- CB- CG2- HG23
2825 2827 2833 2834 3 1.12131 2.98738 1.66523
-3.94970 0.43514 0.00000 ; CA- CB- OG1- HG1
2825 2835 2837 2838 3 20.92000 0.00000 -20.92000
0.00000 0.00000 0.00000 ; CA- C- N- CD
2825 2835 2837 2847 3 20.92000 0.00000 -20.92000
0.00000 0.00000 0.00000 ; CA- C- N- CA
2826 2825 2827 2828 3 0.65084 1.95253 0.00000
-2.60338 0.00000 0.00000 ; HA- CA- CB- HB
2826 2825 2827 2829 3 0.65084 1.95253 0.00000
-2.60338 0.00000 0.00000 ; HA- CA- CB- CG2
2826 2825 2827 2833 3 1.04600 -1.04600 0.00000
0.00000 0.00000 0.00000 ; HA- CA- CB- OG1
2826 2825 2835 2836 3 3.68192 -4.35136 0.00000
1.33888 0.00000 0.00000 ; HA- CA- C- O
2826 2825 2835 2837 3 0.00000 0.00000 0.00000

0.00000 0.00000 0.00000 ; HA- CA- C- N
2827 2825 2835 2836 3 0.00000 0.00000 0.00000
0.00000 0.00000 0.00000 ; CB- CA- C- O
2827 2825 2835 2837 3 2.51040 4.18400 1.67360
-6.69440 0.00000 0.00000 ; CB- CA- C- N
2828 2827 2825 2835 3 0.65084 1.95253 0.00000
-2.60338 0.00000 0.00000 ; HB- CB- CA- C
2828 2827 2829 2830 3 0.65084 1.95253 0.00000
-2.60338 0.00000 0.00000 ; HB- CB- CG2- HG21
2828 2827 2829 2831 3 0.65084 1.95253 0.00000
-2.60338 0.00000 0.00000 ; HB- CB- CG2- HG22
2828 2827 2829 2832 3 0.65084 1.95253 0.00000
-2.60338 0.00000 0.00000 ; HB- CB- CG2- HG23
2828 2827 2833 2834 3 0.69733 2.09200 0.00000
-2.78933 0.00000 0.00000 ; HB- CB- OG1- HG1
2829 2827 2825 2835 3 5.67350 3.55640 -6.16722
-2.47693 3.74886 0.00000 ; CG2- CB- CA- C
2829 2827 2833 2834 3 4.74047 0.27196 -2.26773
-3.94970 1.60666 0.00000 ; CG2- CB- OG1- HG1
2830 2829 2827 2833 3 1.04600 -1.04600 0.00000
0.00000 0.00000 0.00000 ; HG21- CG2- CB- OG1
2831 2829 2827 2833 3 1.04600 -1.04600 0.00000
0.00000 0.00000 0.00000 ; HG22- CG2- CB- OG1
2832 2829 2827 2833 3 1.04600 -1.04600 0.00000
0.00000 0.00000 0.00000 ; HG23- CG2- CB- OG1
2833 2827 2825 2835 3 6.53541 6.87013 -6.21742
-5.27184 5.22163 0.00000 ; OG1- CB- CA- C
2835 2837 2838 2839 3 0.00000 0.00000 0.00000
0.00000 0.00000 0.00000 ; C- N- CD- HD2
2835 2837 2838 2840 3 0.00000 0.00000 0.00000
0.00000 0.00000 0.00000 ; C- N- CD- HD3
2835 2837 2838 2841 3 10.04160 -3.34720 16.73600
-6.69440 0.00000 0.00000 ; C- N- CD- CG
2835 2837 2847 2844 3 11.71520 1.67360 15.06240 -
13.38880 0.00000 0.00000 ; C- N- CA- CB
2835 2837 2847 2848 3 0.00000 0.00000 0.00000
0.00000 0.00000 0.00000 ; C- N- CA- HA
2835 2837 2847 2849 3 1.75728 5.27184 2.25936
-7.02912 0.00000 0.00000 ; C- N- CA- C
2836 2835 2837 2838 3 20.92000 0.00000 -20.92000
0.00000 0.00000 0.00000 ; O- C- N- CD
2836 2835 2837 2847 3 20.92000 0.00000 -20.92000
0.00000 0.00000 0.00000 ; O- C- N- CA
2837 2838 2841 2842 3 0.65084 1.95253 0.00000
-2.60338 0.00000 0.00000 ; N- CD- CG- HG2
2837 2838 2841 2843 3 0.65084 1.95253 0.00000
-2.60338 0.00000 0.00000 ; N- CD- CG- HG3
2837 2838 2841 2844 3 0.65084 1.95253 0.00000
-2.60338 0.00000 0.00000 ; N- CD- CG- CB
2837 2847 2844 2841 3 0.65084 1.95253 0.00000
-2.60338 0.00000 0.00000 ; N- CA- CB- CG
2837 2847 2844 2845 3 0.65084 1.95253 0.00000

-2.60338 0.00000 0.00000 ; N- CA- CB- HB2
2837 2847 2844 2846 3 0.65084 1.95253 0.00000
-2.60338 0.00000 0.00000 ; N- CA- CB- HB3
2837 2847 2849 2850 3 0.00000 0.00000 0.00000
0.00000 0.00000 0.00000 ; N- CA- C- O
2837 2847 2849 2851 3 17.40544 -5.02080 -13.22144
9.20480 0.00000 0.00000 ; N- CA- C- N
2838 2837 2847 2844 3 0.00000 0.00000 0.00000
0.00000 0.00000 0.00000 ; CD- N- CA- CB
2838 2837 2847 2848 3 0.00000 0.00000 0.00000
0.00000 0.00000 0.00000 ; CD- N- CA- HA
2838 2837 2847 2849 3 0.00000 0.00000 0.00000
0.00000 0.00000 0.00000 ; CD- N- CA- C
2838 2841 2844 2845 3 0.66944 2.00832 0.00000
-2.67776 0.00000 0.00000 ; CD- CG- CB- HB2
2838 2841 2844 2846 3 0.66944 2.00832 0.00000
-2.67776 0.00000 0.00000 ; CD- CG- CB- HB3
2838 2841 2844 2847 3 3.68192 3.09616 -2.09200
-3.01248 0.00000 0.00000 ; CD- CG- CB- CA
2839 2838 2837 2847 3 0.00000 0.00000 0.00000
0.00000 0.00000 0.00000 ; HD2- CD- N- CA
2839 2838 2841 2842 3 0.65084 1.95253 0.00000
-2.60338 0.00000 0.00000 ; HD2- CD- CG- HG2
2839 2838 2841 2843 3 0.65084 1.95253 0.00000
-2.60338 0.00000 0.00000 ; HD2- CD- CG- HG3
2839 2838 2841 2844 3 0.65084 1.95253 0.00000
-2.60338 0.00000 0.00000 ; HD2- CD- CG- CB
2840 2838 2837 2847 3 0.00000 0.00000 0.00000
0.00000 0.00000 0.00000 ; HD3- CD- N- CA
2840 2838 2841 2842 3 0.65084 1.95253 0.00000
-2.60338 0.00000 0.00000 ; HD3- CD- CG- HG2
2840 2838 2841 2843 3 0.65084 1.95253 0.00000
-2.60338 0.00000 0.00000 ; HD3- CD- CG- HG3
2840 2838 2841 2844 3 0.65084 1.95253 0.00000
-2.60338 0.00000 0.00000 ; HD3- CD- CG- CB
2841 2838 2837 2847 3 0.00000 0.00000 0.00000
0.00000 0.00000 0.00000 ; CG- CD- N- CA
2841 2844 2847 2848 3 0.65084 1.95253 0.00000
-2.60338 0.00000 0.00000 ; CG- CB- CA- HA
2841 2844 2847 2849 3 0.65084 1.95253 0.00000
-2.60338 0.00000 0.00000 ; CG- CB- CA- C
2842 2841 2844 2845 3 0.62760 1.88280 0.00000
-2.51040 0.00000 0.00000 ; HG2- CG- CB- HB2
2842 2841 2844 2846 3 0.62760 1.88280 0.00000
-2.51040 0.00000 0.00000 ; HG2- CG- CB- HB3
2842 2841 2844 2847 3 0.66944 2.00832 0.00000
-2.67776 0.00000 0.00000 ; HG2- CG- CB- CA
2843 2841 2844 2845 3 0.62760 1.88280 0.00000
-2.51040 0.00000 0.00000 ; HG3- CG- CB- HB2
2843 2841 2844 2846 3 0.62760 1.88280 0.00000
-2.51040 0.00000 0.00000 ; HG3- CG- CB- HB3
2843 2841 2844 2847 3 0.66944 2.00832 0.00000

-2.67776 0.00000 0.00000 ; HG3- CG- CB- CA
2844 2847 2849 2850 3 0.00000 0.00000 0.00000
0.00000 0.00000 0.00000 ; CB- CA- C- O
2844 2847 2849 2851 3 2.51040 4.18400 1.67360
-6.69440 0.00000 0.00000 ; CB- CA- C- N
2845 2844 2847 2848 3 0.65084 1.95253 0.00000
-2.60338 0.00000 0.00000 ; HB2- CB- CA- HA
2845 2844 2847 2849 3 0.65084 1.95253 0.00000
-2.60338 0.00000 0.00000 ; HB2- CB- CA- C
2846 2844 2847 2848 3 0.65084 1.95253 0.00000
-2.60338 0.00000 0.00000 ; HB3- CB- CA- HA
2846 2844 2847 2849 3 0.65084 1.95253 0.00000
-2.60338 0.00000 0.00000 ; HB3- CB- CA- C
2847 2849 2851 2852 3 20.92000 0.00000 -20.92000
0.00000 0.00000 0.00000 ; CA- C- N- H
2847 2849 2851 2853 3 20.92000 0.00000 -20.92000
0.00000 0.00000 0.00000 ; CA- C- N- CA
2848 2847 2849 2850 3 3.68192 -4.35136 0.00000
1.33888 0.00000 0.00000 ; HA- CA- C- O
2848 2847 2849 2851 3 0.00000 0.00000 0.00000
0.00000 0.00000 0.00000 ; HA- CA- C- N
2849 2851 2853 2854 3 0.00000 0.00000 0.00000
0.00000 0.00000 0.00000 ; C- N- CA- HA
2849 2851 2853 2855 3 11.71520 1.67360 15.06240 -
13.38880 0.00000 0.00000 ; C- N- CA- CB
2849 2851 2853 2870 3 1.75728 5.27184 2.25936
-7.02912 0.00000 0.00000 ; C- N- CA- C
2850 2849 2851 2852 3 29.28800 -8.36800 -20.92000
0.00000 0.00000 0.00000 ; O- C- N- H
2850 2849 2851 2853 3 20.92000 0.00000 -20.92000
0.00000 0.00000 0.00000 ; O- C- N- CA
2851 2853 2855 2856 3 0.65084 1.95253 0.00000
-2.60338 0.00000 0.00000 ; N- CA- CB- HB2
2851 2853 2855 2857 3 0.65084 1.95253 0.00000
-2.60338 0.00000 0.00000 ; N- CA- CB- HB3
2851 2853 2855 2858 3 3.28026 2.46019 -2.19242
-3.21331 -0.23430 0.00000 ; N- CA- CB- CG
2851 2853 2870 2871 3 0.00000 0.00000 0.00000
0.00000 0.00000 0.00000 ; N- CA- C- O
2851 2853 2870 2872 3 17.40544 -5.02080 -13.22144
9.20480 0.00000 0.00000 ; N- CA- C- N
2852 2851 2853 2854 3 0.00000 0.00000 0.00000
0.00000 0.00000 0.00000 ; H- N- CA- HA
2852 2851 2853 2855 3 0.00000 0.00000 0.00000
0.00000 0.00000 0.00000 ; H- N- CA- CB
2852 2851 2853 2870 3 0.00000 0.00000 0.00000
0.00000 0.00000 0.00000 ; H- N- CA- C
2853 2855 2858 2859 3 0.57739 0.00000 1.02926
0.00000 -1.60666 0.00000 ; CA- CB- CG- CD1
2853 2855 2858 2868 3 0.57739 0.00000 1.02926
0.00000 -1.60666 0.00000 ; CA- CB- CG- CD2
2853 2870 2872 2873 3 20.92000 0.00000 -20.92000

```
0.00000    0.00000    0.00000 ;     CA-     C-     N-     H
  2853   2870   2872   2874      3   20.92000    0.00000  -20.92000
0.00000    0.00000    0.00000 ;     CA-     C-     N-    CA
  2854   2853   2855   2856      3    0.65084    1.95253    0.00000
-2.60338    0.00000    0.00000 ;     HA-    CA-    CB-   HB2
  2854   2853   2855   2857      3    0.65084    1.95253    0.00000
-2.60338    0.00000    0.00000 ;     HA-    CA-    CB-   HB3
  2854   2853   2855   2858      3    0.65084    1.95253    0.00000
-2.60338    0.00000    0.00000 ;     HA-    CA-    CB-    CG
  2854   2853   2870   2871      3    3.68192   -4.35136    0.00000
1.33888    0.00000    0.00000 ;     HA-    CA-     C-     O
  2854   2853   2870   2872      3    0.00000    0.00000    0.00000
0.00000    0.00000    0.00000 ;     HA-    CA-     C-     N
  2855   2853   2870   2871      3    0.00000    0.00000    0.00000
0.00000    0.00000    0.00000 ;     CB-    CA-     C-     O
  2855   2853   2870   2872      3    2.51040    4.18400    1.67360
-6.69440    0.00000    0.00000 ;     CB-    CA-     C-     N
  2855   2858   2859   2860      3   30.33400    0.00000  -30.33400
0.00000    0.00000    0.00000 ;     CB-    CG-   CD1-   HD1
  2855   2858   2859   2861      3   30.33400    0.00000  -30.33400
0.00000    0.00000    0.00000 ;     CB-    CG-   CD1-   CE1
  2855   2858   2868   2866      3   30.33400    0.00000  -30.33400
0.00000    0.00000    0.00000 ;     CB-    CG-   CD2-   CE2
  2855   2858   2868   2869      3   30.33400    0.00000  -30.33400
0.00000    0.00000    0.00000 ;     CB-    CG-   CD2-   HD2
  2856   2855   2853   2870      3    0.65084    1.95253    0.00000
-2.60338    0.00000    0.00000 ;    HB2-    CB-    CA-     C
  2856   2855   2858   2859      3    0.00000    0.00000    0.00000
0.00000    0.00000    0.00000 ;    HB2-    CB-    CG-   CD1
  2856   2855   2858   2868      3    0.00000    0.00000    0.00000
0.00000    0.00000    0.00000 ;    HB2-    CB-    CG-   CD2
  2857   2855   2853   2870      3    0.65084    1.95253    0.00000
-2.60338    0.00000    0.00000 ;    HB3-    CB-    CA-     C
  2857   2855   2858   2859      3    0.00000    0.00000    0.00000
0.00000    0.00000    0.00000 ;    HB3-    CB-    CG-   CD1
  2857   2855   2858   2868      3    0.00000    0.00000    0.00000
0.00000    0.00000    0.00000 ;    HB3-    CB-    CG-   CD2
  2858   2855   2853   2870      3    4.95804    2.17986   -3.52293
-3.21331   -0.40166    0.00000 ;     CG-    CB-    CA-     C
  2858   2859   2861   2862      3   30.33400    0.00000  -30.33400
0.00000    0.00000    0.00000 ;     CG-   CD1-   CE1-   HE1
  2858   2859   2861   2863      3   30.33400    0.00000  -30.33400
0.00000    0.00000    0.00000 ;     CG-   CD1-   CE1-    CZ
  2858   2868   2866   2863      3   30.33400    0.00000  -30.33400
0.00000    0.00000    0.00000 ;     CG-   CD2-   CE2-    CZ
  2858   2868   2866   2867      3   30.33400    0.00000  -30.33400
0.00000    0.00000    0.00000 ;     CG-   CD2-   CE2-   HE2
  2859   2858   2868   2866      3   30.33400    0.00000  -30.33400
0.00000    0.00000    0.00000 ;    CD1-    CG-   CD2-   CE2
  2859   2858   2868   2869      3   30.33400    0.00000  -30.33400
0.00000    0.00000    0.00000 ;    CD1-    CG-   CD2-   HD2
  2859   2861   2863   2864      3   30.33400    0.00000  -30.33400
```

```
0.00000    0.00000    0.00000 ;    CD1-   CE1-    CZ-    OH
  2859   2861   2863   2866       3   30.33400    0.00000  -30.33400
0.00000    0.00000    0.00000 ;    CD1-   CE1-    CZ-   CE2
  2860   2859   2858   2868       3   30.33400    0.00000  -30.33400
0.00000    0.00000    0.00000 ;    HD1-   CD1-    CG-   CD2
  2860   2859   2861   2862       3   30.33400    0.00000  -30.33400
0.00000    0.00000    0.00000 ;    HD1-   CD1-   CE1-   HE1
  2860   2859   2861   2863       3   30.33400    0.00000  -30.33400
0.00000    0.00000    0.00000 ;    HD1-   CD1-   CE1-    CZ
  2861   2859   2858   2868       3   30.33400    0.00000  -30.33400
0.00000    0.00000    0.00000 ;    CE1-   CD1-    CG-   CD2
  2861   2863   2864   2865       3    7.93286    0.00000   -9.56462
0.00000    2.17568    0.00000 ;    CE1-    CZ-    OH-    HH
  2861   2863   2866   2867       3   30.33400    0.00000  -30.33400
0.00000    0.00000    0.00000 ;    CE1-    CZ-   CE2-   HE2
  2861   2863   2866   2868       3   30.33400    0.00000  -30.33400
0.00000    0.00000    0.00000 ;    CE1-    CZ-   CE2-   CD2
  2862   2861   2863   2864       3   30.33400    0.00000  -30.33400
0.00000    0.00000    0.00000 ;    HE1-   CE1-    CZ-    OH
  2862   2861   2863   2866       3   30.33400    0.00000  -30.33400
0.00000    0.00000    0.00000 ;    HE1-   CE1-    CZ-   CE2
  2863   2866   2868   2869       3   30.33400    0.00000  -30.33400
0.00000    0.00000    0.00000 ;     CZ-   CE2-   CD2-   HD2
  2864   2863   2866   2867       3   30.33400    0.00000  -30.33400
0.00000    0.00000    0.00000 ;     OH-    CZ-   CE2-   HE2
  2864   2863   2866   2868       3   30.33400    0.00000  -30.33400
0.00000    0.00000    0.00000 ;     OH-    CZ-   CE2-   CD2
  2865   2864   2863   2866       3    7.93286    0.00000   -9.56462
0.00000    2.17568    0.00000 ;     HH-    OH-    CZ-   CE2
  2867   2866   2868   2869       3   30.33400    0.00000  -30.33400
0.00000    0.00000    0.00000 ;    HE2-   CE2-   CD2-   HD2
  2870   2872   2874   2875       3    0.00000    0.00000    0.00000
0.00000    0.00000    0.00000 ;      C-     N-    CA-    HA
  2870   2872   2874   2876       3   11.71520    1.67360   15.06240  -
13.38880    0.00000    0.00000 ;      C-     N-    CA-    CB
  2870   2872   2874   2891       3    1.75728    5.27184    2.25936
-7.02912    0.00000    0.00000 ;      C-     N-    CA-     C
  2871   2870   2872   2873       3   29.28800   -8.36800  -20.92000
0.00000    0.00000    0.00000 ;     O-     C-     N-     H
  2871   2870   2872   2874       3   20.92000    0.00000  -20.92000
0.00000    0.00000    0.00000 ;     O-     C-     N-    CA
  2872   2874   2876   2877       3    0.65084    1.95253    0.00000
-2.60338    0.00000    0.00000 ;      N-    CA-    CB-   HB2
  2872   2874   2876   2878       3    0.65084    1.95253    0.00000
-2.60338    0.00000    0.00000 ;      N-    CA-    CB-   HB3
  2872   2874   2876   2879       3    3.28026    2.46019   -2.19242
-3.21331   -0.23430    0.00000 ;      N-    CA-    CB-    CG
  2872   2874   2891   2892       3    0.00000    0.00000    0.00000
0.00000    0.00000    0.00000 ;     N-    CA-     C-     O
  2872   2874   2891   2893       3   17.40544   -5.02080  -13.22144
9.20480    0.00000    0.00000 ;     N-    CA-     C-     N
  2873   2872   2874   2875       3    0.00000    0.00000    0.00000
```

```
0.00000    0.00000    0.00000 ;       H-      N-     CA-    HA
  2873   2872   2874   2876       3    0.00000    0.00000    0.00000
0.00000    0.00000    0.00000 ;       H-      N-     CA-    CB
  2873   2872   2874   2891       3    0.00000    0.00000    0.00000
0.00000    0.00000    0.00000 ;       H-      N-     CA-     C
  2874   2876   2879   2880       3    0.57739    0.00000    1.02926
0.00000   -1.60666    0.00000 ;      CA-    CB-    CG-   CD1
  2874   2876   2879   2889       3    0.57739    0.00000    1.02926
0.00000   -1.60666    0.00000 ;      CA-    CB-    CG-   CD2
  2874   2891   2893   2894       3   20.92000    0.00000  -20.92000
0.00000    0.00000    0.00000 ;      CA-     C-     N-     H
  2874   2891   2893   2895       3   20.92000    0.00000  -20.92000
0.00000    0.00000    0.00000 ;      CA-     C-     N-    CA
  2875   2874   2876   2877       3    0.65084    1.95253    0.00000
-2.60338    0.00000    0.00000 ;      HA-    CA-    CB-   HB2
  2875   2874   2876   2878       3    0.65084    1.95253    0.00000
-2.60338    0.00000    0.00000 ;      HA-    CA-    CB-   HB3
  2875   2874   2876   2879       3    0.65084    1.95253    0.00000
-2.60338    0.00000    0.00000 ;      HA-    CA-    CB-    CG
  2875   2874   2891   2892       3    3.68192   -4.35136    0.00000
1.33888    0.00000    0.00000 ;      HA-    CA-     C-     O
  2875   2874   2891   2893       3    0.00000    0.00000    0.00000
0.00000    0.00000    0.00000 ;      HA-    CA-     C-     N
  2876   2874   2891   2892       3    0.00000    0.00000    0.00000
0.00000    0.00000    0.00000 ;      CB-    CA-     C-     O
  2876   2874   2891   2893       3    2.51040    4.18400    1.67360
-6.69440    0.00000    0.00000 ;      CB-    CA-     C-     N
  2876   2879   2880   2881       3   30.33400    0.00000  -30.33400
0.00000    0.00000    0.00000 ;      CB-    CG-   CD1-   HD1
  2876   2879   2880   2882       3   30.33400    0.00000  -30.33400
0.00000    0.00000    0.00000 ;      CB-    CG-   CD1-   CE1
  2876   2879   2889   2887       3   30.33400    0.00000  -30.33400
0.00000    0.00000    0.00000 ;      CB-    CG-   CD2-   CE2
  2876   2879   2889   2890       3   30.33400    0.00000  -30.33400
0.00000    0.00000    0.00000 ;      CB-    CG-   CD2-   HD2
  2877   2876   2874   2891       3    0.65084    1.95253    0.00000
-2.60338    0.00000    0.00000 ;     HB2-    CB-    CA-     C
  2877   2876   2879   2880       3    0.00000    0.00000    0.00000
0.00000    0.00000    0.00000 ;     HB2-    CB-    CG-   CD1
  2877   2876   2879   2889       3    0.00000    0.00000    0.00000
0.00000    0.00000    0.00000 ;     HB2-    CB-    CG-   CD2
  2878   2876   2874   2891       3    0.65084    1.95253    0.00000
-2.60338    0.00000    0.00000 ;     HB3-    CB-    CA-     C
  2878   2876   2879   2880       3    0.00000    0.00000    0.00000
0.00000    0.00000    0.00000 ;     HB3-    CB-    CG-   CD1
  2878   2876   2879   2889       3    0.00000    0.00000    0.00000
0.00000    0.00000    0.00000 ;     HB3-    CB-    CG-   CD2
  2879   2876   2874   2891       3    4.95804    2.17986   -3.52293
-3.21331   -0.40166    0.00000 ;      CG-    CB-    CA-     C
  2879   2880   2882   2883       3   30.33400    0.00000  -30.33400
0.00000    0.00000    0.00000 ;      CG-   CD1-   CE1-   HE1
  2879   2880   2882   2884       3   30.33400    0.00000  -30.33400
```

0.00000 0.00000 0.00000 ; CG- CD1- CE1- CZ
2879 2889 2887 2884 3 30.33400 0.00000 -30.33400
0.00000 0.00000 0.00000 ; CG- CD2- CE2- CZ
2879 2889 2887 2888 3 30.33400 0.00000 -30.33400
0.00000 0.00000 0.00000 ; CG- CD2- CE2- HE2
2880 2879 2889 2887 3 30.33400 0.00000 -30.33400
0.00000 0.00000 0.00000 ; CD1- CG- CD2- CE2
2880 2879 2889 2890 3 30.33400 0.00000 -30.33400
0.00000 0.00000 0.00000 ; CD1- CG- CD2- HD2
2880 2882 2884 2885 3 30.33400 0.00000 -30.33400
0.00000 0.00000 0.00000 ; CD1- CE1- CZ- OH
2880 2882 2884 2887 3 30.33400 0.00000 -30.33400
0.00000 0.00000 0.00000 ; CD1- CE1- CZ- CE2
2881 2880 2879 2889 3 30.33400 0.00000 -30.33400
0.00000 0.00000 0.00000 ; HD1- CD1- CG- CD2
2881 2880 2882 2883 3 30.33400 0.00000 -30.33400
0.00000 0.00000 0.00000 ; HD1- CD1- CE1- HE1
2881 2880 2882 2884 3 30.33400 0.00000 -30.33400
0.00000 0.00000 0.00000 ; HD1- CD1- CE1- CZ
2882 2880 2879 2889 3 30.33400 0.00000 -30.33400
0.00000 0.00000 0.00000 ; CE1- CD1- CG- CD2
2882 2884 2885 2886 3 7.93286 0.00000 -9.56462
0.00000 2.17568 0.00000 ; CE1- CZ- OH- HH
2882 2884 2887 2888 3 30.33400 0.00000 -30.33400
0.00000 0.00000 0.00000 ; CE1- CZ- CE2- HE2
2882 2884 2887 2889 3 30.33400 0.00000 -30.33400
0.00000 0.00000 0.00000 ; CE1- CZ- CE2- CD2
2883 2882 2884 2885 3 30.33400 0.00000 -30.33400
0.00000 0.00000 0.00000 ; HE1- CE1- CZ- OH
2883 2882 2884 2887 3 30.33400 0.00000 -30.33400
0.00000 0.00000 0.00000 ; HE1- CE1- CZ- CE2
2884 2887 2889 2890 3 30.33400 0.00000 -30.33400
0.00000 0.00000 0.00000 ; CZ- CE2- CD2- HD2
2885 2884 2887 2888 3 30.33400 0.00000 -30.33400
0.00000 0.00000 0.00000 ; OH- CZ- CE2- HE2
2885 2884 2887 2889 3 30.33400 0.00000 -30.33400
0.00000 0.00000 0.00000 ; OH- CZ- CE2- CD2
2886 2885 2884 2887 3 7.93286 0.00000 -9.56462
0.00000 2.17568 0.00000 ; HH- OH- CZ- CE2
2888 2887 2889 2890 3 30.33400 0.00000 -30.33400
0.00000 0.00000 0.00000 ; HE2- CE2- CD2- HD2
2891 2893 2895 2896 3 0.00000 0.00000 0.00000
0.00000 0.00000 0.00000 ; C- N- CA- HA
2891 2893 2895 2897 3 11.71520 1.67360 15.06240 -
13.38880 0.00000 0.00000 ; C- N- CA- CB
2891 2893 2895 2907 3 1.75728 5.27184 2.25936
-7.02912 0.00000 0.00000 ; C- N- CA- C
2892 2891 2893 2894 3 29.28800 -8.36800 -20.92000
0.00000 0.00000 0.00000 ; O- C- N- H
2892 2891 2893 2895 3 20.92000 0.00000 -20.92000
0.00000 0.00000 0.00000 ; O- C- N- CA
2893 2895 2897 2898 3 0.65084 1.95253 0.00000

```
-2.60338    0.00000    0.00000 ;       N-    CA-    CB-    HB
  2893   2895   2897   2899       3    3.83673    0.44769   -1.77402
-2.47693   -0.03347    0.00000 ;       N-    CA-    CB-   CG1
  2893   2895   2897   2903       3    3.83673    0.44769   -1.77402
-2.47693   -0.03347    0.00000 ;       N-    CA-    CB-   CG2
  2893   2895   2907   2908       3    0.00000    0.00000    0.00000
0.00000    0.00000    0.00000 ;       N-    CA-     C-     O
  2893   2895   2907   2909       3   17.40544   -5.02080  -13.22144
9.20480    0.00000    0.00000 ;       N-    CA-     C-     N
  2894   2893   2895   2896       3    0.00000    0.00000    0.00000
0.00000    0.00000    0.00000 ;       H-     N-    CA-    HA
  2894   2893   2895   2897       3    0.00000    0.00000    0.00000
0.00000    0.00000    0.00000 ;       H-     N-    CA-    CB
  2894   2893   2895   2907       3    0.00000    0.00000    0.00000
0.00000    0.00000    0.00000 ;       H-     N-    CA-     C
  2895   2897   2899   2900       3    0.66944    2.00832    0.00000
-2.67776    0.00000    0.00000 ;      CA-    CB-   CG1-  HG11
  2895   2897   2899   2901       3    0.66944    2.00832    0.00000
-2.67776    0.00000    0.00000 ;      CA-    CB-   CG1-  HG12
  2895   2897   2899   2902       3    0.66944    2.00832    0.00000
-2.67776    0.00000    0.00000 ;      CA-    CB-   CG1-  HG13
  2895   2897   2903   2904       3    0.66944    2.00832    0.00000
-2.67776    0.00000    0.00000 ;      CA-    CB-   CG2-  HG21
  2895   2897   2903   2905       3    0.66944    2.00832    0.00000
-2.67776    0.00000    0.00000 ;      CA-    CB-   CG2-  HG22
  2895   2897   2903   2906       3    0.66944    2.00832    0.00000
-2.67776    0.00000    0.00000 ;      CA-    CB-   CG2-  HG23
  2895   2907   2909   2910       3   20.92000    0.00000  -20.92000
0.00000    0.00000    0.00000 ;      CA-     C-     N-     H
  2895   2907   2909   2911       3   20.92000    0.00000  -20.92000
0.00000    0.00000    0.00000 ;      CA-     C-     N-    CA
  2896   2895   2897   2898       3    0.65084    1.95253    0.00000
-2.60338    0.00000    0.00000 ;      HA-    CA-    CB-    HB
  2896   2895   2897   2899       3    0.65084    1.95253    0.00000
-2.60338    0.00000    0.00000 ;      HA-    CA-    CB-   CG1
  2896   2895   2897   2903       3    0.65084    1.95253    0.00000
-2.60338    0.00000    0.00000 ;      HA-    CA-    CB-   CG2
  2896   2895   2907   2908       3    3.68192   -4.35136    0.00000
1.33888    0.00000    0.00000 ;      HA-    CA-     C-     O
  2896   2895   2907   2909       3    0.00000    0.00000    0.00000
0.00000    0.00000    0.00000 ;      HA-    CA-     C-     N
  2897   2895   2907   2908       3    0.00000    0.00000    0.00000
0.00000    0.00000    0.00000 ;      CB-    CA-     C-     O
  2897   2895   2907   2909       3    2.51040    4.18400    1.67360
-6.69440    0.00000    0.00000 ;      CB-    CA-     C-     N
  2898   2897   2895   2907       3    0.65084    1.95253    0.00000
-2.60338    0.00000    0.00000 ;      HB-    CB-    CA-     C
  2898   2897   2899   2900       3    0.62760    1.88280    0.00000
-2.51040    0.00000    0.00000 ;      HB-    CB-   CG1-  HG11
  2898   2897   2899   2901       3    0.62760    1.88280    0.00000
-2.51040    0.00000    0.00000 ;      HB-    CB-   CG1-  HG12
  2898   2897   2899   2902       3    0.62760    1.88280    0.00000
```

-2.51040    0.00000    0.00000 ;      HB-    CB-   CG1-  HG13
  2898   2897   2903   2904      3    0.62760    1.88280    0.00000
-2.51040    0.00000    0.00000 ;      HB-    CB-   CG2-  HG21
  2898   2897   2903   2905      3    0.62760    1.88280    0.00000
-2.51040    0.00000    0.00000 ;      HB-    CB-   CG2-  HG22
  2898   2897   2903   2906      3    0.62760    1.88280    0.00000
-2.51040    0.00000    0.00000 ;      HB-    CB-   CG2-  HG23
  2899   2897   2895   2907      3    5.67350    3.55640   -6.16722
-2.47693    3.74886    0.00000 ;     CG1-    CB-    CA-     C
  2899   2897   2903   2904      3    0.66944    2.00832    0.00000
-2.67776    0.00000    0.00000 ;     CG1-    CB-   CG2-  HG21
  2899   2897   2903   2905      3    0.66944    2.00832    0.00000
-2.67776    0.00000    0.00000 ;     CG1-    CB-   CG2-  HG22
  2899   2897   2903   2906      3    0.66944    2.00832    0.00000
-2.67776    0.00000    0.00000 ;     CG1-    CB-   CG2-  HG23
  2900   2899   2897   2903      3    0.66944    2.00832    0.00000
-2.67776    0.00000    0.00000 ;    HG11-   CG1-    CB-   CG2
  2901   2899   2897   2903      3    0.66944    2.00832    0.00000
-2.67776    0.00000    0.00000 ;    HG12-   CG1-    CB-   CG2
  2902   2899   2897   2903      3    0.66944    2.00832    0.00000
-2.67776    0.00000    0.00000 ;    HG13-   CG1-    CB-   CG2
  2903   2897   2895   2907      3    5.67350    3.55640   -6.16722
-2.47693    3.74886    0.00000 ;     CG2-    CB-    CA-     C
  2907   2909   2911   2912      3    0.00000    0.00000    0.00000
0.00000    0.00000    0.00000 ;      C-     N-    CA-    HA
  2907   2909   2911   2913      3   11.71520    1.67360   15.06240  -
13.38880    0.00000    0.00000 ;      C-     N-    CA-    CB
  2907   2909   2911   2917      3    1.75728    5.27184    2.25936
-7.02912    0.00000    0.00000 ;      C-     N-    CA-     C
  2908   2907   2909   2910      3   29.28800   -8.36800  -20.92000
0.00000    0.00000    0.00000 ;      O-     C-     N-     H
  2908   2907   2909   2911      3   20.92000    0.00000  -20.92000
0.00000    0.00000    0.00000 ;      O-     C-     N-    CA
  2909   2911   2913   2914      3    0.65084    1.95253    0.00000
-2.60338    0.00000    0.00000 ;      N-    CA-    CB-   HB1
  2909   2911   2913   2915      3    0.65084    1.95253    0.00000
-2.60338    0.00000    0.00000 ;      N-    CA-    CB-   HB2
  2909   2911   2913   2916      3    0.65084    1.95253    0.00000
-2.60338    0.00000    0.00000 ;      N-    CA-    CB-   HB3
  2909   2911   2917   2918      3    0.00000    0.00000    0.00000
0.00000    0.00000    0.00000 ;      N-    CA-     C-     O
  2909   2911   2917   2919      3   17.40544   -5.02080  -13.22144
9.20480    0.00000    0.00000 ;      N-    CA-     C-     N
  2910   2909   2911   2912      3    0.00000    0.00000    0.00000
0.00000    0.00000    0.00000 ;      H-     N-    CA-    HA
  2910   2909   2911   2913      3    0.00000    0.00000    0.00000
0.00000    0.00000    0.00000 ;      H-     N-    CA-    CB
  2910   2909   2911   2917      3    0.00000    0.00000    0.00000
0.00000    0.00000    0.00000 ;      H-     N-    CA-     C
  2911   2917   2919   2920      3   20.92000    0.00000  -20.92000
0.00000    0.00000    0.00000 ;     CA-     C-     N-    CD
  2911   2917   2919   2929      3   20.92000    0.00000  -20.92000

0.00000 0.00000 0.00000 ; CA- C- N- CA
2912 2911 2913 2914 3 0.65084 1.95253 0.00000
-2.60338 0.00000 0.00000 ; HA- CA- CB- HB1
2912 2911 2913 2915 3 0.65084 1.95253 0.00000
-2.60338 0.00000 0.00000 ; HA- CA- CB- HB2
2912 2911 2913 2916 3 0.65084 1.95253 0.00000
-2.60338 0.00000 0.00000 ; HA- CA- CB- HB3
2912 2911 2917 2918 3 3.68192 -4.35136 0.00000
1.33888 0.00000 0.00000 ; HA- CA- C- O
2912 2911 2917 2919 3 0.00000 0.00000 0.00000
0.00000 0.00000 0.00000 ; HA- CA- C- N
2913 2911 2917 2918 3 0.00000 0.00000 0.00000
0.00000 0.00000 0.00000 ; CB- CA- C- O
2913 2911 2917 2919 3 2.51040 4.18400 1.67360
-6.69440 0.00000 0.00000 ; CB- CA- C- N
2914 2913 2911 2917 3 0.65084 1.95253 0.00000
-2.60338 0.00000 0.00000 ; HB1- CB- CA- C
2915 2913 2911 2917 3 0.65084 1.95253 0.00000
-2.60338 0.00000 0.00000 ; HB2- CB- CA- C
2916 2913 2911 2917 3 0.65084 1.95253 0.00000
-2.60338 0.00000 0.00000 ; HB3- CB- CA- C
2917 2919 2920 2921 3 0.00000 0.00000 0.00000
0.00000 0.00000 0.00000 ; C- N- CD- HD2
2917 2919 2920 2922 3 0.00000 0.00000 0.00000
0.00000 0.00000 0.00000 ; C- N- CD- HD3
2917 2919 2920 2923 3 10.04160 -3.34720 16.73600
-6.69440 0.00000 0.00000 ; C- N- CD- CG
2917 2919 2929 2926 3 11.71520 1.67360 15.06240 -
13.38880 0.00000 0.00000 ; C- N- CA- CB
2917 2919 2929 2930 3 0.00000 0.00000 0.00000
0.00000 0.00000 0.00000 ; C- N- CA- HA
2917 2919 2929 2931 3 1.75728 5.27184 2.25936
-7.02912 0.00000 0.00000 ; C- N- CA- C
2918 2917 2919 2920 3 20.92000 0.00000 -20.92000
0.00000 0.00000 0.00000 ; O- C- N- CD
2918 2917 2919 2929 3 20.92000 0.00000 -20.92000
0.00000 0.00000 0.00000 ; O- C- N- CA
2919 2920 2923 2924 3 0.65084 1.95253 0.00000
-2.60338 0.00000 0.00000 ; N- CD- CG- HG2
2919 2920 2923 2925 3 0.65084 1.95253 0.00000
-2.60338 0.00000 0.00000 ; N- CD- CG- HG3
2919 2920 2923 2926 3 0.65084 1.95253 0.00000
-2.60338 0.00000 0.00000 ; N- CD- CG- CB
2919 2929 2926 2923 3 0.65084 1.95253 0.00000
-2.60338 0.00000 0.00000 ; N- CA- CB- CG
2919 2929 2926 2927 3 0.65084 1.95253 0.00000
-2.60338 0.00000 0.00000 ; N- CA- CB- HB2
2919 2929 2926 2928 3 0.65084 1.95253 0.00000
-2.60338 0.00000 0.00000 ; N- CA- CB- HB3
2919 2929 2931 2932 3 0.00000 0.00000 0.00000
0.00000 0.00000 0.00000 ; N- CA- C- O
2919 2929 2931 2933 3 17.40544 -5.02080 -13.22144

9.20480    0.00000    0.00000 ;      N-    CA-     C-     N
  2920   2919   2929   2926      3    0.00000    0.00000    0.00000
0.00000    0.00000    0.00000 ;     CD-     N-    CA-    CB
  2920   2919   2929   2930      3    0.00000    0.00000    0.00000
0.00000    0.00000    0.00000 ;     CD-     N-    CA-    HA
  2920   2919   2929   2931      3    0.00000    0.00000    0.00000
0.00000    0.00000    0.00000 ;     CD-     N-    CA-     C
  2920   2923   2926   2927      3    0.66944    2.00832    0.00000
-2.67776    0.00000    0.00000 ;     CD-    CG-    CB-   HB2
  2920   2923   2926   2928      3    0.66944    2.00832    0.00000
-2.67776    0.00000    0.00000 ;     CD-    CG-    CB-   HB3
  2920   2923   2926   2929      3    3.68192    3.09616   -2.09200
-3.01248    0.00000    0.00000 ;     CD-    CG-    CB-    CA
  2921   2920   2919   2929      3    0.00000    0.00000    0.00000
0.00000    0.00000    0.00000 ;    HD2-    CD-     N-    CA
  2921   2920   2923   2924      3    0.65084    1.95253    0.00000
-2.60338    0.00000    0.00000 ;    HD2-    CD-    CG-   HG2
  2921   2920   2923   2925      3    0.65084    1.95253    0.00000
-2.60338    0.00000    0.00000 ;    HD2-    CD-    CG-   HG3
  2921   2920   2923   2926      3    0.65084    1.95253    0.00000
-2.60338    0.00000    0.00000 ;    HD2-    CD-    CG-    CB
  2922   2920   2919   2929      3    0.00000    0.00000    0.00000
0.00000    0.00000    0.00000 ;    HD3-    CD-     N-    CA
  2922   2920   2923   2924      3    0.65084    1.95253    0.00000
-2.60338    0.00000    0.00000 ;    HD3-    CD-    CG-   HG2
  2922   2920   2923   2925      3    0.65084    1.95253    0.00000
-2.60338    0.00000    0.00000 ;    HD3-    CD-    CG-   HG3
  2922   2920   2923   2926      3    0.65084    1.95253    0.00000
-2.60338    0.00000    0.00000 ;    HD3-    CD-    CG-    CB
  2923   2920   2919   2929      3    0.00000    0.00000    0.00000
0.00000    0.00000    0.00000 ;     CG-    CD-     N-    CA
  2923   2926   2929   2930      3    0.65084    1.95253    0.00000
-2.60338    0.00000    0.00000 ;     CG-    CB-    CA-    HA
  2923   2926   2929   2931      3    0.65084    1.95253    0.00000
-2.60338    0.00000    0.00000 ;     CG-    CB-    CA-     C
  2924   2923   2926   2927      3    0.62760    1.88280    0.00000
-2.51040    0.00000    0.00000 ;    HG2-    CG-    CB-   HB2
  2924   2923   2926   2928      3    0.62760    1.88280    0.00000
-2.51040    0.00000    0.00000 ;    HG2-    CG-    CB-   HB3
  2924   2923   2926   2929      3    0.66944    2.00832    0.00000
-2.67776    0.00000    0.00000 ;    HG2-    CG-    CB-    CA
  2925   2923   2926   2927      3    0.62760    1.88280    0.00000
-2.51040    0.00000    0.00000 ;    HG3-    CG-    CB-   HB2
  2925   2923   2926   2928      3    0.62760    1.88280    0.00000
-2.51040    0.00000    0.00000 ;    HG3-    CG-    CB-   HB3
  2925   2923   2926   2929      3    0.66944    2.00832    0.00000
-2.67776    0.00000    0.00000 ;    HG3-    CG-    CB-    CA
  2926   2929   2931   2932      3    0.00000    0.00000    0.00000
0.00000    0.00000    0.00000 ;     CB-    CA-     C-     O
  2926   2929   2931   2933      3    2.51040    4.18400    1.67360
-6.69440    0.00000    0.00000 ;     CB-    CA-     C-     N
  2927   2926   2929   2930      3    0.65084    1.95253    0.00000

-2.60338 0.00000 0.00000 ; HB2- CB- CA- HA
2927 2926 2929 2931 3 0.65084 1.95253 0.00000
-2.60338 0.00000 0.00000 ; HB2- CB- CA- C
2928 2926 2929 2930 3 0.65084 1.95253 0.00000
-2.60338 0.00000 0.00000 ; HB3- CB- CA- HA
2928 2926 2929 2931 3 0.65084 1.95253 0.00000
-2.60338 0.00000 0.00000 ; HB3- CB- CA- C
2929 2931 2933 2934 3 20.92000 0.00000 -20.92000
0.00000 0.00000 0.00000 ; CA- C- N- H
2929 2931 2933 2935 3 20.92000 0.00000 -20.92000
0.00000 0.00000 0.00000 ; CA- C- N- CA
2930 2929 2931 2932 3 3.68192 -4.35136 0.00000
1.33888 0.00000 0.00000 ; HA- CA- C- O
2930 2929 2931 2933 3 0.00000 0.00000 0.00000
0.00000 0.00000 0.00000 ; HA- CA- C- N
2931 2933 2935 2936 3 0.00000 0.00000 0.00000
0.00000 0.00000 0.00000 ; C- N- CA- HA
2931 2933 2935 2937 3 11.71520 1.67360 15.06240 -
13.38880 0.00000 0.00000 ; C- N- CA- CB
2931 2933 2935 2948 3 1.75728 5.27184 2.25936
-7.02912 0.00000 0.00000 ; C- N- CA- C
2932 2931 2933 2934 3 29.28800 -8.36800 -20.92000
0.00000 0.00000 0.00000 ; O- C- N- H
2932 2931 2933 2935 3 20.92000 0.00000 -20.92000
0.00000 0.00000 0.00000 ; O- C- N- CA
2933 2935 2937 2938 3 0.65084 1.95253 0.00000
-2.60338 0.00000 0.00000 ; N- CA- CB- HB2
2933 2935 2937 2939 3 0.65084 1.95253 0.00000
-2.60338 0.00000 0.00000 ; N- CA- CB- HB3
2933 2935 2937 2940 3 3.21331 2.22589 -4.15053
-2.40998 2.61082 0.00000 ; N- CA- CB- CG
2933 2935 2948 2949 3 0.00000 0.00000 0.00000
0.00000 0.00000 0.00000 ; N- CA- C- O
2933 2935 2948 2950 3 17.40544 -5.02080 -13.22144
9.20480 0.00000 0.00000 ; N- CA- C- N
2934 2933 2935 2936 3 0.00000 0.00000 0.00000
0.00000 0.00000 0.00000 ; H- N- CA- HA
2934 2933 2935 2937 3 0.00000 0.00000 0.00000
0.00000 0.00000 0.00000 ; H- N- CA- CB
2934 2933 2935 2948 3 0.00000 0.00000 0.00000
0.00000 0.00000 0.00000 ; H- N- CA- C
2935 2937 2940 2941 3 0.66944 2.00832 0.00000
-2.67776 0.00000 0.00000 ; CA- CB- CG- HG2
2935 2937 2940 2942 3 0.66944 2.00832 0.00000
-2.67776 0.00000 0.00000 ; CA- CB- CG- HG3
2935 2937 2940 2943 3 3.69866 -4.35136 -3.92459
6.89523 4.61914 0.00000 ; CA- CB- CG- CD
2935 2948 2950 2951 3 20.92000 0.00000 -20.92000
0.00000 0.00000 0.00000 ; CA- C- N- H
2935 2948 2950 2952 3 20.92000 0.00000 -20.92000
0.00000 0.00000 0.00000 ; CA- C- N- CA
2936 2935 2937 2938 3 0.65084 1.95253 0.00000

```
-2.60338    0.00000    0.00000 ;      HA-    CA-    CB-   HB2
  2936   2935   2937   2939       3    0.65084    1.95253    0.00000
-2.60338    0.00000    0.00000 ;      HA-    CA-    CB-   HB3
  2936   2935   2937   2940       3    0.65084    1.95253    0.00000
-2.60338    0.00000    0.00000 ;      HA-    CA-    CB-    CG
  2936   2935   2948   2949       3    3.68192   -4.35136    0.00000
1.33888    0.00000    0.00000 ;      HA-    CA-     C-     O
  2936   2935   2948   2950       3    0.00000    0.00000    0.00000
0.00000    0.00000    0.00000 ;      HA-    CA-     C-     N
  2937   2935   2948   2949       3    0.00000    0.00000    0.00000
0.00000    0.00000    0.00000 ;      CB-    CA-     C-     O
  2937   2935   2948   2950       3    2.51040    4.18400    1.67360
-6.69440    0.00000    0.00000 ;      CB-    CA-     C-     N
  2937   2940   2943   2944       3    0.00000    0.00000    0.00000
0.00000    0.00000    0.00000 ;      CB-    CG-    CD-   OE1
  2937   2940   2943   2945       3   10.32611    1.48114   -8.47678
1.42256    1.40582    0.00000 ;      CB-    CG-    CD-   NE2
  2938   2937   2935   2948       3    0.65084    1.95253    0.00000
-2.60338    0.00000    0.00000 ;     HB2-    CB-    CA-     C
  2938   2937   2940   2941       3    0.62760    1.88280    0.00000
-2.51040    0.00000    0.00000 ;     HB2-    CB-    CG-   HG2
  2938   2937   2940   2942       3    0.62760    1.88280    0.00000
-2.51040    0.00000    0.00000 ;     HB2-    CB-    CG-   HG3
  2938   2937   2940   2943       3    0.65084    1.95253    0.00000
-2.60338    0.00000    0.00000 ;     HB2-    CB-    CG-    CD
  2939   2937   2935   2948       3    0.65084    1.95253    0.00000
-2.60338    0.00000    0.00000 ;     HB3-    CB-    CA-     C
  2939   2937   2940   2941       3    0.62760    1.88280    0.00000
-2.51040    0.00000    0.00000 ;     HB3-    CB-    CG-   HG2
  2939   2937   2940   2942       3    0.62760    1.88280    0.00000
-2.51040    0.00000    0.00000 ;     HB3-    CB-    CG-   HG3
  2939   2937   2940   2943       3    0.65084    1.95253    0.00000
-2.60338    0.00000    0.00000 ;     HB3-    CB-    CG-    CD
  2940   2937   2935   2948       3    6.86594    3.56895   -8.14206
-2.40998    4.85344    0.00000 ;      CG-    CB-    CA-     C
  2940   2943   2945   2946       3   20.92000    0.00000  -20.92000
0.00000    0.00000    0.00000 ;      CG-    CD-   NE2-  HE21
  2940   2943   2945   2947       3   20.92000    0.00000  -20.92000
0.00000    0.00000    0.00000 ;      CG-    CD-   NE2-  HE22
  2941   2940   2943   2944       3    3.68192   -4.35136    0.00000
1.33888    0.00000    0.00000 ;     HG2-    CG-    CD-   OE1
  2941   2940   2943   2945       3    0.00000    0.00000    0.00000
0.00000    0.00000    0.00000 ;     HG2-    CG-    CD-   NE2
  2942   2940   2943   2944       3    3.68192   -4.35136    0.00000
1.33888    0.00000    0.00000 ;     HG3-    CG-    CD-   OE1
  2942   2940   2943   2945       3    0.00000    0.00000    0.00000
0.00000    0.00000    0.00000 ;     HG3-    CG-    CD-   NE2
  2944   2943   2945   2946       3   29.28800   -8.36800  -20.92000
0.00000    0.00000    0.00000 ;     OE1-    CD-   NE2-  HE21
  2944   2943   2945   2947       3   29.28800   -8.36800  -20.92000
0.00000    0.00000    0.00000 ;     OE1-    CD-   NE2-  HE22
  2948   2950   2952   2953       3    0.00000    0.00000    0.00000
```

```
0.00000    0.00000    0.00000 ;       C-     N-     CA-    HA
  2948   2950   2952   2954      3   11.71520    1.67360   15.06240  -
13.38880    0.00000    0.00000 ;       C-     N-     CA-    CB
  2948   2950   2952   2964      3    1.75728    5.27184    2.25936
-7.02912    0.00000    0.00000 ;       C-     N-     CA-     C
  2949   2948   2950   2951      3   29.28800   -8.36800  -20.92000
0.00000    0.00000    0.00000 ;       O-     C-     N-     H
  2949   2948   2950   2952      3   20.92000    0.00000  -20.92000
0.00000    0.00000    0.00000 ;       O-     C-     N-    CA
  2950   2952   2954   2955      3    0.65084    1.95253    0.00000
-2.60338    0.00000    0.00000 ;       N-    CA-    CB-    HB
  2950   2952   2954   2956      3    3.83673    0.44769   -1.77402
-2.47693   -0.03347    0.00000 ;       N-    CA-    CB-   CG1
  2950   2952   2954   2960      3    3.83673    0.44769   -1.77402
-2.47693   -0.03347    0.00000 ;       N-    CA-    CB-   CG2
  2950   2952   2964   2965      3    0.00000    0.00000    0.00000
0.00000    0.00000    0.00000 ;       N-    CA-     C-     O
  2950   2952   2964   2966      3   17.40544   -5.02080  -13.22144
9.20480    0.00000    0.00000 ;       N-    CA-     C-     N
  2951   2950   2952   2953      3    0.00000    0.00000    0.00000
0.00000    0.00000    0.00000 ;       H-     N-    CA-    HA
  2951   2950   2952   2954      3    0.00000    0.00000    0.00000
0.00000    0.00000    0.00000 ;       H-     N-    CA-    CB
  2951   2950   2952   2964      3    0.00000    0.00000    0.00000
0.00000    0.00000    0.00000 ;       H-     N-    CA-     C
  2952   2954   2956   2957      3    0.66944    2.00832    0.00000
-2.67776    0.00000    0.00000 ;      CA-    CB-   CG1-  HG11
  2952   2954   2956   2958      3    0.66944    2.00832    0.00000
-2.67776    0.00000    0.00000 ;      CA-    CB-   CG1-  HG12
  2952   2954   2956   2959      3    0.66944    2.00832    0.00000
-2.67776    0.00000    0.00000 ;      CA-    CB-   CG1-  HG13
  2952   2954   2960   2961      3    0.66944    2.00832    0.00000
-2.67776    0.00000    0.00000 ;      CA-    CB-   CG2-  HG21
  2952   2954   2960   2962      3    0.66944    2.00832    0.00000
-2.67776    0.00000    0.00000 ;      CA-    CB-   CG2-  HG22
  2952   2954   2960   2963      3    0.66944    2.00832    0.00000
-2.67776    0.00000    0.00000 ;      CA-    CB-   CG2-  HG23
  2952   2964   2966   2967      3   20.92000    0.00000  -20.92000
0.00000    0.00000    0.00000 ;      CA-     C-     N-     H
  2952   2964   2966   2968      3   20.92000    0.00000  -20.92000
0.00000    0.00000    0.00000 ;      CA-     C-     N-    CA
  2953   2952   2954   2955      3    0.65084    1.95253    0.00000
-2.60338    0.00000    0.00000 ;      HA-    CA-    CB-    HB
  2953   2952   2954   2956      3    0.65084    1.95253    0.00000
-2.60338    0.00000    0.00000 ;      HA-    CA-    CB-   CG1
  2953   2952   2954   2960      3    0.65084    1.95253    0.00000
-2.60338    0.00000    0.00000 ;      HA-    CA-    CB-   CG2
  2953   2952   2964   2965      3    3.68192   -4.35136    0.00000
1.33888    0.00000    0.00000 ;      HA-    CA-     C-     O
  2953   2952   2964   2966      3    0.00000    0.00000    0.00000
0.00000    0.00000    0.00000 ;      HA-    CA-     C-     N
  2954   2952   2964   2965      3    0.00000    0.00000    0.00000
```

```
0.00000    0.00000    0.00000 ;      CB-    CA-     C-     O
  2954   2952   2964   2966      3    2.51040    4.18400    1.67360
-6.69440    0.00000    0.00000 ;      CB-    CA-     C-     N
  2955   2954   2952   2964      3    0.65084    1.95253    0.00000
-2.60338    0.00000    0.00000 ;      HB-    CB-    CA-     C
  2955   2954   2956   2957      3    0.62760    1.88280    0.00000
-2.51040    0.00000    0.00000 ;      HB-    CB-   CG1-  HG11
  2955   2954   2956   2958      3    0.62760    1.88280    0.00000
-2.51040    0.00000    0.00000 ;      HB-    CB-   CG1-  HG12
  2955   2954   2956   2959      3    0.62760    1.88280    0.00000
-2.51040    0.00000    0.00000 ;      HB-    CB-   CG1-  HG13
  2955   2954   2960   2961      3    0.62760    1.88280    0.00000
-2.51040    0.00000    0.00000 ;      HB-    CB-   CG2-  HG21
  2955   2954   2960   2962      3    0.62760    1.88280    0.00000
-2.51040    0.00000    0.00000 ;      HB-    CB-   CG2-  HG22
  2955   2954   2960   2963      3    0.62760    1.88280    0.00000
-2.51040    0.00000    0.00000 ;      HB-    CB-   CG2-  HG23
  2956   2954   2952   2964      3    5.67350    3.55640   -6.16722
-2.47693    3.74886    0.00000 ;     CG1-    CB-    CA-     C
  2956   2954   2960   2961      3    0.66944    2.00832    0.00000
-2.67776    0.00000    0.00000 ;     CG1-    CB-   CG2-  HG21
  2956   2954   2960   2962      3    0.66944    2.00832    0.00000
-2.67776    0.00000    0.00000 ;     CG1-    CB-   CG2-  HG22
  2956   2954   2960   2963      3    0.66944    2.00832    0.00000
-2.67776    0.00000    0.00000 ;     CG1-    CB-   CG2-  HG23
  2957   2956   2954   2960      3    0.66944    2.00832    0.00000
-2.67776    0.00000    0.00000 ;    HG11-   CG1-     CB-   CG2
  2958   2956   2954   2960      3    0.66944    2.00832    0.00000
-2.67776    0.00000    0.00000 ;    HG12-   CG1-     CB-   CG2
  2959   2956   2954   2960      3    0.66944    2.00832    0.00000
-2.67776    0.00000    0.00000 ;    HG13-   CG1-     CB-   CG2
  2960   2954   2952   2964      3    5.67350    3.55640   -6.16722
-2.47693    3.74886    0.00000 ;     CG2-    CB-    CA-     C
  2964   2966   2968   2969      3    0.00000    0.00000    0.00000
0.00000    0.00000    0.00000 ;      C-     N-    CA-    HA
  2964   2966   2968   2970      3   11.71520    1.67360   15.06240  -
13.38880    0.00000    0.00000 ;      C-     N-    CA-    CB
  2964   2966   2968   2983      3    1.75728    5.27184    2.25936
-7.02912    0.00000    0.00000 ;      C-     N-    CA-     C
  2965   2964   2966   2967      3   29.28800   -8.36800  -20.92000
0.00000    0.00000    0.00000 ;      O-     C-     N-     H
  2965   2964   2966   2968      3   20.92000    0.00000  -20.92000
0.00000    0.00000    0.00000 ;      O-     C-     N-    CA
  2966   2968   2970   2971      3    0.65084    1.95253    0.00000
-2.60338    0.00000    0.00000 ;      N-    CA-    CB-   HB2
  2966   2968   2970   2972      3    0.65084    1.95253    0.00000
-2.60338    0.00000    0.00000 ;      N-    CA-    CB-   HB3
  2966   2968   2970   2973      3    3.79070    1.39746   -4.61077
-2.40998    2.44346    0.00000 ;      N-    CA-    CB-    CG
  2966   2968   2983   2984      3    0.00000    0.00000    0.00000
0.00000    0.00000    0.00000 ;      N-    CA-     C-     O
  2966   2968   2983   2985      3   17.40544   -5.02080  -13.22144
```

```
9.20480    0.00000    0.00000 ;       N-    CA-     C-     N
  2967   2966   2968   2969      3   0.00000    0.00000    0.00000
0.00000    0.00000    0.00000 ;       H-     N-    CA-    HA
  2967   2966   2968   2970      3   0.00000    0.00000    0.00000
0.00000    0.00000    0.00000 ;       H-     N-    CA-    CB
  2967   2966   2968   2983      3   0.00000    0.00000    0.00000
0.00000    0.00000    0.00000 ;       H-     N-    CA-     C
  2968   2970   2973   2974      3   0.66944    2.00832    0.00000
-2.67776    0.00000    0.00000 ;      CA-    CB-    CG-    HG
  2968   2970   2973   2975      3   3.90367    0.19665   -6.21742
-2.37651    5.99149    0.00000 ;      CA-    CB-    CG-   CD1
  2968   2970   2973   2979      3   3.90367    0.19665   -6.21742
-2.37651    5.99149    0.00000 ;      CA-    CB-    CG-   CD2
  2968   2983   2985   2986      3  20.92000    0.00000  -20.92000
0.00000    0.00000    0.00000 ;     CA-     C-     N-     H
  2968   2983   2985   2987      3  20.92000    0.00000  -20.92000
0.00000    0.00000    0.00000 ;     CA-     C-     N-    CA
  2969   2968   2970   2971      3   0.65084    1.95253    0.00000
-2.60338    0.00000    0.00000 ;      HA-    CA-    CB-   HB2
  2969   2968   2970   2972      3   0.65084    1.95253    0.00000
-2.60338    0.00000    0.00000 ;      HA-    CA-    CB-   HB3
  2969   2968   2970   2973      3   0.65084    1.95253    0.00000
-2.60338    0.00000    0.00000 ;      HA-    CA-    CB-    CG
  2969   2968   2983   2984      3   3.68192   -4.35136    0.00000
1.33888    0.00000    0.00000 ;     HA-    CA-     C-     O
  2969   2968   2983   2985      3   0.00000    0.00000    0.00000
0.00000    0.00000    0.00000 ;     HA-    CA-     C-     N
  2970   2968   2983   2984      3   0.00000    0.00000    0.00000
0.00000    0.00000    0.00000 ;     CB-    CA-     C-     O
  2970   2968   2983   2985      3   2.51040    4.18400    1.67360
-6.69440    0.00000    0.00000 ;      CB-    CA-     C-     N
  2970   2973   2975   2976      3   0.66944    2.00832    0.00000
-2.67776    0.00000    0.00000 ;      CB-    CG-   CD1-  HD11
  2970   2973   2975   2977      3   0.66944    2.00832    0.00000
-2.67776    0.00000    0.00000 ;      CB-    CG-   CD1-  HD12
  2970   2973   2975   2978      3   0.66944    2.00832    0.00000
-2.67776    0.00000    0.00000 ;      CB-    CG-   CD1-  HD13
  2970   2973   2979   2980      3   0.66944    2.00832    0.00000
-2.67776    0.00000    0.00000 ;      CB-    CG-   CD2-  HD21
  2970   2973   2979   2981      3   0.66944    2.00832    0.00000
-2.67776    0.00000    0.00000 ;      CB-    CG-   CD2-  HD22
  2970   2973   2979   2982      3   0.66944    2.00832    0.00000
-2.67776    0.00000    0.00000 ;      CB-    CG-   CD2-  HD23
  2971   2970   2968   2983      3   0.65084    1.95253    0.00000
-2.60338    0.00000    0.00000 ;     HB2-    CB-    CA-     C
  2971   2970   2973   2974      3   0.62760    1.88280    0.00000
-2.51040    0.00000    0.00000 ;     HB2-    CB-    CG-    HG
  2971   2970   2973   2975      3   0.66944    2.00832    0.00000
-2.67776    0.00000    0.00000 ;     HB2-    CB-    CG-   CD1
  2971   2970   2973   2979      3   0.66944    2.00832    0.00000
-2.67776    0.00000    0.00000 ;     HB2-    CB-    CG-   CD2
  2972   2970   2968   2983      3   0.65084    1.95253    0.00000
```

-2.60338 0.00000 0.00000 ; HB3- CB- CA- C
2972 2970 2973 2974 3 0.62760 1.88280 0.00000
-2.51040 0.00000 0.00000 ; HB3- CB- CG- HG
2972 2970 2973 2975 3 0.66944 2.00832 0.00000
-2.67776 0.00000 0.00000 ; HB3- CB- CG- CD1
2972 2970 2973 2979 3 0.66944 2.00832 0.00000
-2.67776 0.00000 0.00000 ; HB3- CB- CG- CD2
2973 2970 2968 2983 3 10.33448 -1.14642 -11.54784
-2.40998 6.35968 0.00000 ; CG- CB- CA- C
2974 2973 2975 2976 3 0.62760 1.88280 0.00000
-2.51040 0.00000 0.00000 ; HG- CG- CD1- HD11
2974 2973 2975 2977 3 0.62760 1.88280 0.00000
-2.51040 0.00000 0.00000 ; HG- CG- CD1- HD12
2974 2973 2975 2978 3 0.62760 1.88280 0.00000
-2.51040 0.00000 0.00000 ; HG- CG- CD1- HD13
2974 2973 2979 2980 3 0.62760 1.88280 0.00000
-2.51040 0.00000 0.00000 ; HG- CG- CD2- HD21
2974 2973 2979 2981 3 0.62760 1.88280 0.00000
-2.51040 0.00000 0.00000 ; HG- CG- CD2- HD22
2974 2973 2979 2982 3 0.62760 1.88280 0.00000
-2.51040 0.00000 0.00000 ; HG- CG- CD2- HD23
2975 2973 2979 2980 3 0.66944 2.00832 0.00000
-2.67776 0.00000 0.00000 ; CD1- CG- CD2- HD21
2975 2973 2979 2981 3 0.66944 2.00832 0.00000
-2.67776 0.00000 0.00000 ; CD1- CG- CD2- HD22
2975 2973 2979 2982 3 0.66944 2.00832 0.00000
-2.67776 0.00000 0.00000 ; CD1- CG- CD2- HD23
2976 2975 2973 2979 3 0.66944 2.00832 0.00000
-2.67776 0.00000 0.00000 ; HD11- CD1- CG- CD2
2977 2975 2973 2979 3 0.66944 2.00832 0.00000
-2.67776 0.00000 0.00000 ; HD12- CD1- CG- CD2
2978 2975 2973 2979 3 0.66944 2.00832 0.00000
-2.67776 0.00000 0.00000 ; HD13- CD1- CG- CD2
2983 2985 2987 2988 3 0.00000 0.00000 0.00000
0.00000 0.00000 0.00000 ; C- N- CA- HA
2983 2985 2987 2989 3 11.71520 1.67360 15.06240 -
13.38880 0.00000 0.00000 ; C- N- CA- CB
2983 2985 2987 2995 3 1.75728 5.27184 2.25936
-7.02912 0.00000 0.00000 ; C- N- CA- C
2984 2983 2985 2986 3 29.28800 -8.36800 -20.92000
0.00000 0.00000 0.00000 ; O- C- N- H
2984 2983 2985 2987 3 20.92000 0.00000 -20.92000
0.00000 0.00000 0.00000 ; O- C- N- CA
2985 2987 2989 2990 3 0.65084 1.95253 0.00000
-2.60338 0.00000 0.00000 ; N- CA- CB- HB2
2985 2987 2989 2991 3 0.65084 1.95253 0.00000
-2.60338 0.00000 0.00000 ; N- CA- CB- HB3
2985 2987 2989 2992 3 15.41386 9.74035 -8.39310
-0.97069 2.97901 0.00000 ; N- CA- CB- CG
2985 2987 2995 2996 3 0.00000 0.00000 0.00000
0.00000 0.00000 0.00000 ; N- CA- C- O
2985 2987 2995 2997 3 17.40544 -5.02080 -13.22144

9.20480 0.00000 0.00000 ; N- CA- C- N
2986 2985 2987 2988 3 0.00000 0.00000 0.00000
0.00000 0.00000 0.00000 ; H- N- CA- HA
2986 2985 2987 2989 3 0.00000 0.00000 0.00000
0.00000 0.00000 0.00000 ; H- N- CA- CB
2986 2985 2987 2995 3 0.00000 0.00000 0.00000
0.00000 0.00000 0.00000 ; H- N- CA- C
2987 2989 2992 2993 3 6.43499 0.00000 -5.39736
0.00000 -1.03763 0.00000 ; CA- CB- CG- OD1
2987 2989 2992 2994 3 6.43499 0.00000 -5.39736
0.00000 -1.03763 0.00000 ; CA- CB- CG- OD2
2987 2995 2997 2998 3 20.92000 0.00000 -20.92000
0.00000 0.00000 0.00000 ; CA- C- N- H
2987 2995 2997 2999 3 20.92000 0.00000 -20.92000
0.00000 0.00000 0.00000 ; CA- C- N- CA
2988 2987 2989 2990 3 0.65084 1.95253 0.00000
-2.60338 0.00000 0.00000 ; HA- CA- CB- HB2
2988 2987 2989 2991 3 0.65084 1.95253 0.00000
-2.60338 0.00000 0.00000 ; HA- CA- CB- HB3
2988 2987 2989 2992 3 0.65084 1.95253 0.00000
-2.60338 0.00000 0.00000 ; HA- CA- CB- CG
2988 2987 2995 2996 3 3.68192 -4.35136 0.00000
1.33888 0.00000 0.00000 ; HA- CA- C- O
2988 2987 2995 2997 3 0.00000 0.00000 0.00000
0.00000 0.00000 0.00000 ; HA- CA- C- N
2989 2987 2995 2996 3 0.00000 0.00000 0.00000
0.00000 0.00000 0.00000 ; CB- CA- C- O
2989 2987 2995 2997 3 2.51040 4.18400 1.67360
-6.69440 0.00000 0.00000 ; CB- CA- C- N
2990 2989 2987 2995 3 0.65084 1.95253 0.00000
-2.60338 0.00000 0.00000 ; HB2- CB- CA- C
2990 2989 2992 2993 3 0.00000 0.00000 0.00000
0.00000 0.00000 0.00000 ; HB2- CB- CG- OD1
2990 2989 2992 2994 3 0.00000 0.00000 0.00000
0.00000 0.00000 0.00000 ; HB2- CB- CG- OD2
2991 2989 2987 2995 3 0.65084 1.95253 0.00000
-2.60338 0.00000 0.00000 ; HB3- CB- CA- C
2991 2989 2992 2993 3 0.00000 0.00000 0.00000
0.00000 0.00000 0.00000 ; HB3- CB- CG- OD1
2991 2989 2992 2994 3 0.00000 0.00000 0.00000
0.00000 0.00000 0.00000 ; HB3- CB- CG- OD2
2992 2989 2987 2995 3 7.14627 -1.04600 -8.99560
-0.97069 5.15469 0.00000 ; CG- CB- CA- C
2995 2997 2999 3000 3 0.00000 0.00000 0.00000
0.00000 0.00000 0.00000 ; C- N- CA- HA2
2995 2997 2999 3001 3 0.00000 0.00000 0.00000
0.00000 0.00000 0.00000 ; C- N- CA- HA3
2995 2997 2999 3002 3 1.75728 5.27184 2.25936
-7.02912 0.00000 0.00000 ; C- N- CA- C
2996 2995 2997 2998 3 29.28800 -8.36800 -20.92000
0.00000 0.00000 0.00000 ; O- C- N- H
2996 2995 2997 2999 3 20.92000 0.00000 -20.92000

0.00000 0.00000 0.00000 ; O- C- N- CA
2997 2999 3002 3003 3 0.00000 0.00000 0.00000
0.00000 0.00000 0.00000 ; N- CA- C- O
2997 2999 3002 3004 3 17.40544 -5.02080 -13.22144
9.20480 0.00000 0.00000 ; N- CA- C- N
2998 2997 2999 3000 3 0.00000 0.00000 0.00000
0.00000 0.00000 0.00000 ; H- N- CA- HA2
2998 2997 2999 3001 3 0.00000 0.00000 0.00000
0.00000 0.00000 0.00000 ; H- N- CA- HA3
2998 2997 2999 3002 3 0.00000 0.00000 0.00000
0.00000 0.00000 0.00000 ; H- N- CA- C
2999 3002 3004 3005 3 20.92000 0.00000 -20.92000
0.00000 0.00000 0.00000 ; CA- C- N- H
2999 3002 3004 3006 3 20.92000 0.00000 -20.92000
0.00000 0.00000 0.00000 ; CA- C- N- CA
3000 2999 3002 3003 3 3.68192 -4.35136 0.00000
1.33888 0.00000 0.00000 ; HA2- CA- C- O
3000 2999 3002 3004 3 0.00000 0.00000 0.00000
0.00000 0.00000 0.00000 ; HA2- CA- C- N
3001 2999 3002 3003 3 3.68192 -4.35136 0.00000
1.33888 0.00000 0.00000 ; HA3- CA- C- O
3001 2999 3002 3004 3 0.00000 0.00000 0.00000
0.00000 0.00000 0.00000 ; HA3- CA- C- N
3002 3004 3006 3007 3 0.00000 0.00000 0.00000
0.00000 0.00000 0.00000 ; C- N- CA- HA
3002 3004 3006 3008 3 11.71520 1.67360 15.06240 -
13.38880 0.00000 0.00000 ; C- N- CA- CB
3002 3004 3006 3024 3 1.75728 5.27184 2.25936
-7.02912 0.00000 0.00000 ; C- N- CA- C
3003 3002 3004 3005 3 29.28800 -8.36800 -20.92000
0.00000 0.00000 0.00000 ; O- C- N- H
3003 3002 3004 3006 3 20.92000 0.00000 -20.92000
0.00000 0.00000 0.00000 ; O- C- N- CA
3004 3006 3008 3009 3 0.65084 1.95253 0.00000
-2.60338 0.00000 0.00000 ; N- CA- CB- HB2
3004 3006 3008 3010 3 0.65084 1.95253 0.00000
-2.60338 0.00000 0.00000 ; N- CA- CB- HB3
3004 3006 3008 3011 3 0.65084 1.95253 0.00000
-2.60338 0.00000 0.00000 ; N- CA- CB- CG
3004 3006 3024 3025 3 0.00000 0.00000 0.00000
0.00000 0.00000 0.00000 ; N- CA- C- O
3004 3006 3024 3026 3 17.40544 -5.02080 -13.22144
9.20480 0.00000 0.00000 ; N- CA- C- N
3005 3004 3006 3007 3 0.00000 0.00000 0.00000
0.00000 0.00000 0.00000 ; H- N- CA- HA
3005 3004 3006 3008 3 0.00000 0.00000 0.00000
0.00000 0.00000 0.00000 ; H- N- CA- CB
3005 3004 3006 3024 3 0.00000 0.00000 0.00000
0.00000 0.00000 0.00000 ; H- N- CA- C
3006 3008 3011 3012 3 0.66944 2.00832 0.00000
-2.67776 0.00000 0.00000 ; CA- CB- CG- HG2
3006 3008 3011 3013 3 0.66944 2.00832 0.00000

-2.67776 0.00000 0.00000 ; CA- CB- CG- HG3
3006 3008 3011 3014 3 3.68192 3.09616 -2.09200
-3.01248 0.00000 0.00000 ; CA- CB- CG- CD
3006 3024 3026 3027 3 20.92000 0.00000 -20.92000
0.00000 0.00000 0.00000 ; CA- C- N- H
3006 3024 3026 3028 3 20.92000 0.00000 -20.92000
0.00000 0.00000 0.00000 ; CA- C- N- CA
3007 3006 3008 3009 3 0.65084 1.95253 0.00000
-2.60338 0.00000 0.00000 ; HA- CA- CB- HB2
3007 3006 3008 3010 3 0.65084 1.95253 0.00000
-2.60338 0.00000 0.00000 ; HA- CA- CB- HB3
3007 3006 3008 3011 3 0.65084 1.95253 0.00000
-2.60338 0.00000 0.00000 ; HA- CA- CB- CG
3007 3006 3024 3025 3 3.68192 -4.35136 0.00000
1.33888 0.00000 0.00000 ; HA- CA- C- O
3007 3006 3024 3026 3 0.00000 0.00000 0.00000
0.00000 0.00000 0.00000 ; HA- CA- C- N
3008 3006 3024 3025 3 0.00000 0.00000 0.00000
0.00000 0.00000 0.00000 ; CB- CA- C- O
3008 3006 3024 3026 3 2.51040 4.18400 1.67360
-6.69440 0.00000 0.00000 ; CB- CA- C- N
3008 3011 3014 3015 3 0.66944 2.00832 0.00000
-2.67776 0.00000 0.00000 ; CB- CG- CD- HD2
3008 3011 3014 3016 3 0.66944 2.00832 0.00000
-2.67776 0.00000 0.00000 ; CB- CG- CD- HD3
3008 3011 3014 3017 3 3.68192 3.09616 -2.09200
-3.01248 0.00000 0.00000 ; CB- CG- CD- CE
3009 3008 3006 3024 3 0.65084 1.95253 0.00000
-2.60338 0.00000 0.00000 ; HB2- CB- CA- C
3009 3008 3011 3012 3 0.62760 1.88280 0.00000
-2.51040 0.00000 0.00000 ; HB2- CB- CG- HG2
3009 3008 3011 3013 3 0.62760 1.88280 0.00000
-2.51040 0.00000 0.00000 ; HB2- CB- CG- HG3
3009 3008 3011 3014 3 0.66944 2.00832 0.00000
-2.67776 0.00000 0.00000 ; HB2- CB- CG- CD
3010 3008 3006 3024 3 0.65084 1.95253 0.00000
-2.60338 0.00000 0.00000 ; HB3- CB- CA- C
3010 3008 3011 3012 3 0.62760 1.88280 0.00000
-2.51040 0.00000 0.00000 ; HB3- CB- CG- HG2
3010 3008 3011 3013 3 0.62760 1.88280 0.00000
-2.51040 0.00000 0.00000 ; HB3- CB- CG- HG3
3010 3008 3011 3014 3 0.66944 2.00832 0.00000
-2.67776 0.00000 0.00000 ; HB3- CB- CG- CD
3011 3008 3006 3024 3 0.65084 1.95253 0.00000
-2.60338 0.00000 0.00000 ; CG- CB- CA- C
3011 3014 3017 3018 3 0.65084 1.95253 0.00000
-2.60338 0.00000 0.00000 ; CG- CD- CE- HE2
3011 3014 3017 3019 3 0.65084 1.95253 0.00000
-2.60338 0.00000 0.00000 ; CG- CD- CE- HE3
3011 3014 3017 3020 3 0.65084 1.95253 0.00000
-2.60338 0.00000 0.00000 ; CG- CD- CE- NZ
3012 3011 3014 3015 3 0.62760 1.88280 0.00000

-2.51040 0.00000 0.00000 ; HG2- CG- CD- HD2
3012 3011 3014 3016 3 0.62760 1.88280 0.00000
-2.51040 0.00000 0.00000 ; HG2- CG- CD- HD3
3012 3011 3014 3017 3 0.66944 2.00832 0.00000
-2.67776 0.00000 0.00000 ; HG2- CG- CD- CE
3013 3011 3014 3015 3 0.62760 1.88280 0.00000
-2.51040 0.00000 0.00000 ; HG3- CG- CD- HD2
3013 3011 3014 3016 3 0.62760 1.88280 0.00000
-2.51040 0.00000 0.00000 ; HG3- CG- CD- HD3
3013 3011 3014 3017 3 0.66944 2.00832 0.00000
-2.67776 0.00000 0.00000 ; HG3- CG- CD- CE
3014 3017 3020 3021 3 0.65084 1.95253 0.00000
-2.60338 0.00000 0.00000 ; CD- CE- NZ- HZ1
3014 3017 3020 3022 3 0.65084 1.95253 0.00000
-2.60338 0.00000 0.00000 ; CD- CE- NZ- HZ2
3014 3017 3020 3023 3 0.65084 1.95253 0.00000
-2.60338 0.00000 0.00000 ; CD- CE- NZ- HZ3
3015 3014 3017 3018 3 0.65084 1.95253 0.00000
-2.60338 0.00000 0.00000 ; HD2- CD- CE- HE2
3015 3014 3017 3019 3 0.65084 1.95253 0.00000
-2.60338 0.00000 0.00000 ; HD2- CD- CE- HE3
3015 3014 3017 3020 3 0.65084 1.95253 0.00000
-2.60338 0.00000 0.00000 ; HD2- CD- CE- NZ
3016 3014 3017 3018 3 0.65084 1.95253 0.00000
-2.60338 0.00000 0.00000 ; HD3- CD- CE- HE2
3016 3014 3017 3019 3 0.65084 1.95253 0.00000
-2.60338 0.00000 0.00000 ; HD3- CD- CE- HE3
3016 3014 3017 3020 3 0.65084 1.95253 0.00000
-2.60338 0.00000 0.00000 ; HD3- CD- CE- NZ
3018 3017 3020 3021 3 0.65084 1.95253 0.00000
-2.60338 0.00000 0.00000 ; HE2- CE- NZ- HZ1
3018 3017 3020 3022 3 0.65084 1.95253 0.00000
-2.60338 0.00000 0.00000 ; HE2- CE- NZ- HZ2
3018 3017 3020 3023 3 0.65084 1.95253 0.00000
-2.60338 0.00000 0.00000 ; HE2- CE- NZ- HZ3
3019 3017 3020 3021 3 0.65084 1.95253 0.00000
-2.60338 0.00000 0.00000 ; HE3- CE- NZ- HZ1
3019 3017 3020 3022 3 0.65084 1.95253 0.00000
-2.60338 0.00000 0.00000 ; HE3- CE- NZ- HZ2
3019 3017 3020 3023 3 0.65084 1.95253 0.00000
-2.60338 0.00000 0.00000 ; HE3- CE- NZ- HZ3
3024 3026 3028 3029 3 0.00000 0.00000 0.00000
0.00000 0.00000 0.00000 ; C- N- CA- HA
3024 3026 3028 3030 3 11.71520 1.67360 15.06240 -
13.38880 0.00000 0.00000 ; C- N- CA- CB
3024 3026 3028 3045 3 1.75728 5.27184 2.25936
-7.02912 0.00000 0.00000 ; C- N- CA- C
3025 3024 3026 3027 3 29.28800 -8.36800 -20.92000
0.00000 0.00000 0.00000 ; O- C- N- H
3025 3024 3026 3028 3 20.92000 0.00000 -20.92000
0.00000 0.00000 0.00000 ; O- C- N- CA
3026 3028 3030 3031 3 0.65084 1.95253 0.00000

```
-2.60338    0.00000    0.00000 ;       N-    CA-    CB-   HB2
  3026   3028   3030   3032       3    0.65084    1.95253    0.00000
-2.60338    0.00000    0.00000 ;       N-    CA-    CB-   HB3
  3026   3028   3030   3033       3    3.28026    2.46019   -2.19242
-3.21331   -0.23430    0.00000 ;       N-    CA-    CB-    CG
  3026   3028   3045   3046       3    0.00000    0.00000    0.00000
0.00000    0.00000    0.00000 ;       N-    CA-     C-     O
  3026   3028   3045   3047       3   17.40544   -5.02080  -13.22144
9.20480    0.00000    0.00000 ;       N-    CA-     C-     N
  3027   3026   3028   3029       3    0.00000    0.00000    0.00000
0.00000    0.00000    0.00000 ;       H-     N-    CA-    HA
  3027   3026   3028   3030       3    0.00000    0.00000    0.00000
0.00000    0.00000    0.00000 ;       H-     N-    CA-    CB
  3027   3026   3028   3045       3    0.00000    0.00000    0.00000
0.00000    0.00000    0.00000 ;       H-     N-    CA-     C
  3028   3030   3033   3034       3    0.57739    0.00000    1.02926
0.00000   -1.60666    0.00000 ;      CA-    CB-    CG-   CD1
  3028   3030   3033   3043       3    0.57739    0.00000    1.02926
0.00000   -1.60666    0.00000 ;      CA-    CB-    CG-   CD2
  3028   3045   3047   3048       3   20.92000    0.00000  -20.92000
0.00000    0.00000    0.00000 ;      CA-     C-     N-     H
  3028   3045   3047   3049       3   20.92000    0.00000  -20.92000
0.00000    0.00000    0.00000 ;      CA-     C-     N-    CA
  3029   3028   3030   3031       3    0.65084    1.95253    0.00000
-2.60338    0.00000    0.00000 ;      HA-    CA-    CB-   HB2
  3029   3028   3030   3032       3    0.65084    1.95253    0.00000
-2.60338    0.00000    0.00000 ;      HA-    CA-    CB-   HB3
  3029   3028   3030   3033       3    0.65084    1.95253    0.00000
-2.60338    0.00000    0.00000 ;      HA-    CA-    CB-    CG
  3029   3028   3045   3046       3    3.68192   -4.35136    0.00000
1.33888    0.00000    0.00000 ;      HA-    CA-     C-     O
  3029   3028   3045   3047       3    0.00000    0.00000    0.00000
0.00000    0.00000    0.00000 ;      HA-    CA-     C-     N
  3030   3028   3045   3046       3    0.00000    0.00000    0.00000
0.00000    0.00000    0.00000 ;      CB-    CA-     C-     O
  3030   3028   3045   3047       3    2.51040    4.18400    1.67360
-6.69440    0.00000    0.00000 ;      CB-    CA-     C-     N
  3030   3033   3034   3035       3   30.33400    0.00000  -30.33400
0.00000    0.00000    0.00000 ;      CB-    CG-   CD1-   HD1
  3030   3033   3034   3036       3   30.33400    0.00000  -30.33400
0.00000    0.00000    0.00000 ;      CB-    CG-   CD1-   CE1
  3030   3033   3043   3041       3   30.33400    0.00000  -30.33400
0.00000    0.00000    0.00000 ;      CB-    CG-   CD2-   CE2
  3030   3033   3043   3044       3   30.33400    0.00000  -30.33400
0.00000    0.00000    0.00000 ;      CB-    CG-   CD2-   HD2
  3031   3030   3028   3045       3    0.65084    1.95253    0.00000
-2.60338    0.00000    0.00000 ;     HB2-    CB-    CA-     C
  3031   3030   3033   3034       3    0.00000    0.00000    0.00000
0.00000    0.00000    0.00000 ;     HB2-    CB-    CG-   CD1
  3031   3030   3033   3043       3    0.00000    0.00000    0.00000
0.00000    0.00000    0.00000 ;     HB2-    CB-    CG-   CD2
  3032   3030   3028   3045       3    0.65084    1.95253    0.00000
```

```
-2.60338    0.00000    0.00000 ;    HB3-    CB-    CA-     C
  3032  3030  3033  3034      3    0.00000    0.00000    0.00000
0.00000    0.00000    0.00000 ;    HB3-    CB-    CG-   CD1
  3032  3030  3033  3043      3    0.00000    0.00000    0.00000
0.00000    0.00000    0.00000 ;    HB3-    CB-    CG-   CD2
  3033  3030  3028  3045      3    4.95804    2.17986   -3.52293
-3.21331   -0.40166    0.00000 ;     CG-    CB-    CA-     C
  3033  3034  3036  3037      3   30.33400    0.00000  -30.33400
0.00000    0.00000    0.00000 ;     CG-   CD1-   CE1-   HE1
  3033  3034  3036  3038      3   30.33400    0.00000  -30.33400
0.00000    0.00000    0.00000 ;     CG-   CD1-   CE1-    CZ
  3033  3043  3041  3038      3   30.33400    0.00000  -30.33400
0.00000    0.00000    0.00000 ;     CG-   CD2-   CE2-    CZ
  3033  3043  3041  3042      3   30.33400    0.00000  -30.33400
0.00000    0.00000    0.00000 ;     CG-   CD2-   CE2-   HE2
  3034  3033  3043  3041      3   30.33400    0.00000  -30.33400
0.00000    0.00000    0.00000 ;    CD1-    CG-   CD2-   CE2
  3034  3033  3043  3044      3   30.33400    0.00000  -30.33400
0.00000    0.00000    0.00000 ;    CD1-    CG-   CD2-   HD2
  3034  3036  3038  3039      3   30.33400    0.00000  -30.33400
0.00000    0.00000    0.00000 ;    CD1-   CE1-    CZ-    OH
  3034  3036  3038  3041      3   30.33400    0.00000  -30.33400
0.00000    0.00000    0.00000 ;    CD1-   CE1-    CZ-   CE2
  3035  3034  3033  3043      3   30.33400    0.00000  -30.33400
0.00000    0.00000    0.00000 ;    HD1-   CD1-    CG-   CD2
  3035  3034  3036  3037      3   30.33400    0.00000  -30.33400
0.00000    0.00000    0.00000 ;    HD1-   CD1-   CE1-   HE1
  3035  3034  3036  3038      3   30.33400    0.00000  -30.33400
0.00000    0.00000    0.00000 ;    HD1-   CD1-   CE1-    CZ
  3036  3034  3033  3043      3   30.33400    0.00000  -30.33400
0.00000    0.00000    0.00000 ;    CE1-   CD1-    CG-   CD2
  3036  3038  3039  3040      3    7.93286    0.00000   -9.56462
0.00000    2.17568    0.00000 ;    CE1-    CZ-    OH-    HH
  3036  3038  3041  3042      3   30.33400    0.00000  -30.33400
0.00000    0.00000    0.00000 ;    CE1-    CZ-   CE2-   HE2
  3036  3038  3041  3043      3   30.33400    0.00000  -30.33400
0.00000    0.00000    0.00000 ;    CE1-    CZ-   CE2-   CD2
  3037  3036  3038  3039      3   30.33400    0.00000  -30.33400
0.00000    0.00000    0.00000 ;    HE1-   CE1-    CZ-    OH
  3037  3036  3038  3041      3   30.33400    0.00000  -30.33400
0.00000    0.00000    0.00000 ;    HE1-   CE1-    CZ-   CE2
  3038  3041  3043  3044      3   30.33400    0.00000  -30.33400
0.00000    0.00000    0.00000 ;     CZ-   CE2-   CD2-   HD2
  3039  3038  3041  3042      3   30.33400    0.00000  -30.33400
0.00000    0.00000    0.00000 ;     OH-    CZ-   CE2-   HE2
  3039  3038  3041  3043      3   30.33400    0.00000  -30.33400
0.00000    0.00000    0.00000 ;     OH-    CZ-   CE2-   CD2
  3040  3039  3038  3041      3    7.93286    0.00000   -9.56462
0.00000    2.17568    0.00000 ;     HH-    OH-    CZ-   CE2
  3042  3041  3043  3044      3   30.33400    0.00000  -30.33400
0.00000    0.00000    0.00000 ;    HE2-   CE2-   CD2-   HD2
  3045  3047  3049  3050      3    0.00000    0.00000    0.00000
```

0.00000 0.00000 0.00000 ; C- N- CA- HA
3045 3047 3049 3051 3 11.71520 1.67360 15.06240 -
13.38880 0.00000 0.00000 ; C- N- CA- CB
3045 3047 3049 3057 3 1.75728 5.27184 2.25936
-7.02912 0.00000 0.00000 ; C- N- CA- C
3046 3045 3047 3048 3 29.28800 -8.36800 -20.92000
0.00000 0.00000 0.00000 ; O- C- N- H
3046 3045 3047 3049 3 20.92000 0.00000 -20.92000
0.00000 0.00000 0.00000 ; O- C- N- CA
3047 3049 3051 3052 3 0.65084 1.95253 0.00000
-2.60338 0.00000 0.00000 ; N- CA- CB- HB2
3047 3049 3051 3053 3 0.65084 1.95253 0.00000
-2.60338 0.00000 0.00000 ; N- CA- CB- HB3
3047 3049 3051 3054 3 15.41386 9.74035 -8.39310
-0.97069 2.97901 0.00000 ; N- CA- CB- CG
3047 3049 3057 3058 3 0.00000 0.00000 0.00000
0.00000 0.00000 0.00000 ; N- CA- C- O
3047 3049 3057 3059 3 17.40544 -5.02080 -13.22144
9.20480 0.00000 0.00000 ; N- CA- C- N
3048 3047 3049 3050 3 0.00000 0.00000 0.00000
0.00000 0.00000 0.00000 ; H- N- CA- HA
3048 3047 3049 3051 3 0.00000 0.00000 0.00000
0.00000 0.00000 0.00000 ; H- N- CA- CB
3048 3047 3049 3057 3 0.00000 0.00000 0.00000
0.00000 0.00000 0.00000 ; H- N- CA- C
3049 3051 3054 3055 3 6.43499 0.00000 -5.39736
0.00000 -1.03763 0.00000 ; CA- CB- CG- OD1
3049 3051 3054 3056 3 6.43499 0.00000 -5.39736
0.00000 -1.03763 0.00000 ; CA- CB- CG- OD2
3049 3057 3059 3060 3 20.92000 0.00000 -20.92000
0.00000 0.00000 0.00000 ; CA- C- N- H
3049 3057 3059 3061 3 20.92000 0.00000 -20.92000
0.00000 0.00000 0.00000 ; CA- C- N- CA
3050 3049 3051 3052 3 0.65084 1.95253 0.00000
-2.60338 0.00000 0.00000 ; HA- CA- CB- HB2
3050 3049 3051 3053 3 0.65084 1.95253 0.00000
-2.60338 0.00000 0.00000 ; HA- CA- CB- HB3
3050 3049 3051 3054 3 0.65084 1.95253 0.00000
-2.60338 0.00000 0.00000 ; HA- CA- CB- CG
3050 3049 3057 3058 3 3.68192 -4.35136 0.00000
1.33888 0.00000 0.00000 ; HA- CA- C- O
3050 3049 3057 3059 3 0.00000 0.00000 0.00000
0.00000 0.00000 0.00000 ; HA- CA- C- N
3051 3049 3057 3058 3 0.00000 0.00000 0.00000
0.00000 0.00000 0.00000 ; CB- CA- C- O
3051 3049 3057 3059 3 2.51040 4.18400 1.67360
-6.69440 0.00000 0.00000 ; CB- CA- C- N
3052 3051 3049 3057 3 0.65084 1.95253 0.00000
-2.60338 0.00000 0.00000 ; HB2- CB- CA- C
3052 3051 3054 3055 3 0.00000 0.00000 0.00000
0.00000 0.00000 0.00000 ; HB2- CB- CG- OD1
3052 3051 3054 3056 3 0.00000 0.00000 0.00000

0.00000 0.00000 0.00000 ; HB2- CB- CG- OD2
3053 3051 3049 3057 3 0.65084 1.95253 0.00000
-2.60338 0.00000 0.00000 ; HB3- CB- CA- C
3053 3051 3054 3055 3 0.00000 0.00000 0.00000
0.00000 0.00000 0.00000 ; HB3- CB- CG- OD1
3053 3051 3054 3056 3 0.00000 0.00000 0.00000
0.00000 0.00000 0.00000 ; HB3- CB- CG- OD2
3054 3051 3049 3057 3 7.14627 -1.04600 -8.99560
-0.97069 5.15469 0.00000 ; CG- CB- CA- C
3057 3059 3061 3062 3 0.00000 0.00000 0.00000
0.00000 0.00000 0.00000 ; C- N- CA- HA
3057 3059 3061 3063 3 11.71520 1.67360 15.06240 -
13.38880 0.00000 0.00000 ; C- N- CA- CB
3057 3059 3061 3079 3 1.75728 5.27184 2.25936
-7.02912 0.00000 0.00000 ; C- N- CA- C
3058 3057 3059 3060 3 29.28800 -8.36800 -20.92000
0.00000 0.00000 0.00000 ; O- C- N- H
3058 3057 3059 3061 3 20.92000 0.00000 -20.92000
0.00000 0.00000 0.00000 ; O- C- N- CA
3059 3061 3063 3064 3 0.65084 1.95253 0.00000
-2.60338 0.00000 0.00000 ; N- CA- CB- HB2
3059 3061 3063 3065 3 0.65084 1.95253 0.00000
-2.60338 0.00000 0.00000 ; N- CA- CB- HB3
3059 3061 3063 3066 3 0.65084 1.95253 0.00000
-2.60338 0.00000 0.00000 ; N- CA- CB- CG
3059 3061 3079 3080 3 0.00000 0.00000 0.00000
0.00000 0.00000 0.00000 ; N- CA- C- O
3059 3061 3079 3081 3 17.40544 -5.02080 -13.22144
9.20480 0.00000 0.00000 ; N- CA- C- N
3060 3059 3061 3062 3 0.00000 0.00000 0.00000
0.00000 0.00000 0.00000 ; H- N- CA- HA
3060 3059 3061 3063 3 0.00000 0.00000 0.00000
0.00000 0.00000 0.00000 ; H- N- CA- CB
3060 3059 3061 3079 3 0.00000 0.00000 0.00000
0.00000 0.00000 0.00000 ; H- N- CA- C
3061 3063 3066 3067 3 0.66944 2.00832 0.00000
-2.67776 0.00000 0.00000 ; CA- CB- CG- HG2
3061 3063 3066 3068 3 0.66944 2.00832 0.00000
-2.67776 0.00000 0.00000 ; CA- CB- CG- HG3
3061 3063 3066 3069 3 3.68192 3.09616 -2.09200
-3.01248 0.00000 0.00000 ; CA- CB- CG- CD
3061 3079 3081 3082 3 20.92000 0.00000 -20.92000
0.00000 0.00000 0.00000 ; CA- C- N- H
3061 3079 3081 3083 3 20.92000 0.00000 -20.92000
0.00000 0.00000 0.00000 ; CA- C- N- CA
3062 3061 3063 3064 3 0.65084 1.95253 0.00000
-2.60338 0.00000 0.00000 ; HA- CA- CB- HB2
3062 3061 3063 3065 3 0.65084 1.95253 0.00000
-2.60338 0.00000 0.00000 ; HA- CA- CB- HB3
3062 3061 3063 3066 3 0.65084 1.95253 0.00000
-2.60338 0.00000 0.00000 ; HA- CA- CB- CG
3062 3061 3079 3080 3 3.68192 -4.35136 0.00000

```
1.33888    0.00000    0.00000 ;      HA-    CA-     C-     O
  3062   3061   3079   3081       3    0.00000    0.00000    0.00000
0.00000    0.00000    0.00000 ;      HA-    CA-     C-     N
  3063   3061   3079   3080       3    0.00000    0.00000    0.00000
0.00000    0.00000    0.00000 ;      CB-    CA-     C-     O
  3063   3061   3079   3081       3    2.51040    4.18400    1.67360
-6.69440    0.00000    0.00000 ;      CB-    CA-     C-     N
  3063   3066   3069   3070       3    0.66944    2.00832    0.00000
-2.67776    0.00000    0.00000 ;      CB-    CG-    CD-   HD2
  3063   3066   3069   3071       3    0.66944    2.00832    0.00000
-2.67776    0.00000    0.00000 ;      CB-    CG-    CD-   HD3
  3063   3066   3069   3072       3    3.68192    3.09616   -2.09200
-3.01248    0.00000    0.00000 ;      CB-    CG-    CD-    CE
  3064   3063   3061   3079       3    0.65084    1.95253    0.00000
-2.60338    0.00000    0.00000 ;     HB2-    CB-    CA-     C
  3064   3063   3066   3067       3    0.62760    1.88280    0.00000
-2.51040    0.00000    0.00000 ;     HB2-    CB-    CG-   HG2
  3064   3063   3066   3068       3    0.62760    1.88280    0.00000
-2.51040    0.00000    0.00000 ;     HB2-    CB-    CG-   HG3
  3064   3063   3066   3069       3    0.66944    2.00832    0.00000
-2.67776    0.00000    0.00000 ;     HB2-    CB-    CG-    CD
  3065   3063   3061   3079       3    0.65084    1.95253    0.00000
-2.60338    0.00000    0.00000 ;     HB3-    CB-    CA-     C
  3065   3063   3066   3067       3    0.62760    1.88280    0.00000
-2.51040    0.00000    0.00000 ;     HB3-    CB-    CG-   HG2
  3065   3063   3066   3068       3    0.62760    1.88280    0.00000
-2.51040    0.00000    0.00000 ;     HB3-    CB-    CG-   HG3
  3065   3063   3066   3069       3    0.66944    2.00832    0.00000
-2.67776    0.00000    0.00000 ;     HB3-    CB-    CG-    CD
  3066   3063   3061   3079       3    0.65084    1.95253    0.00000
-2.60338    0.00000    0.00000 ;      CG-    CB-    CA-     C
  3066   3069   3072   3073       3    0.65084    1.95253    0.00000
-2.60338    0.00000    0.00000 ;      CG-    CD-    CE-   HE2
  3066   3069   3072   3074       3    0.65084    1.95253    0.00000
-2.60338    0.00000    0.00000 ;      CG-    CD-    CE-   HE3
  3066   3069   3072   3075       3    0.65084    1.95253    0.00000
-2.60338    0.00000    0.00000 ;      CG-    CD-    CE-    NZ
  3067   3066   3069   3070       3    0.62760    1.88280    0.00000
-2.51040    0.00000    0.00000 ;     HG2-    CG-    CD-   HD2
  3067   3066   3069   3071       3    0.62760    1.88280    0.00000
-2.51040    0.00000    0.00000 ;     HG2-    CG-    CD-   HD3
  3067   3066   3069   3072       3    0.66944    2.00832    0.00000
-2.67776    0.00000    0.00000 ;     HG2-    CG-    CD-    CE
  3068   3066   3069   3070       3    0.62760    1.88280    0.00000
-2.51040    0.00000    0.00000 ;     HG3-    CG-    CD-   HD2
  3068   3066   3069   3071       3    0.62760    1.88280    0.00000
-2.51040    0.00000    0.00000 ;     HG3-    CG-    CD-   HD3
  3068   3066   3069   3072       3    0.66944    2.00832    0.00000
-2.67776    0.00000    0.00000 ;     HG3-    CG-    CD-    CE
  3069   3072   3075   3076       3    0.65084    1.95253    0.00000
-2.60338    0.00000    0.00000 ;      CD-    CE-    NZ-   HZ1
  3069   3072   3075   3077       3    0.65084    1.95253    0.00000
```

-2.60338 0.00000 0.00000 ; CD- CE- NZ- HZ2
3069 3072 3075 3078 3 0.65084 1.95253 0.00000
-2.60338 0.00000 0.00000 ; CD- CE- NZ- HZ3
3070 3069 3072 3073 3 0.65084 1.95253 0.00000
-2.60338 0.00000 0.00000 ; HD2- CD- CE- HE2
3070 3069 3072 3074 3 0.65084 1.95253 0.00000
-2.60338 0.00000 0.00000 ; HD2- CD- CE- HE3
3070 3069 3072 3075 3 0.65084 1.95253 0.00000
-2.60338 0.00000 0.00000 ; HD2- CD- CE- NZ
3071 3069 3072 3073 3 0.65084 1.95253 0.00000
-2.60338 0.00000 0.00000 ; HD3- CD- CE- HE2
3071 3069 3072 3074 3 0.65084 1.95253 0.00000
-2.60338 0.00000 0.00000 ; HD3- CD- CE- HE3
3071 3069 3072 3075 3 0.65084 1.95253 0.00000
-2.60338 0.00000 0.00000 ; HD3- CD- CE- NZ
3073 3072 3075 3076 3 0.65084 1.95253 0.00000
-2.60338 0.00000 0.00000 ; HE2- CE- NZ- HZ1
3073 3072 3075 3077 3 0.65084 1.95253 0.00000
-2.60338 0.00000 0.00000 ; HE2- CE- NZ- HZ2
3073 3072 3075 3078 3 0.65084 1.95253 0.00000
-2.60338 0.00000 0.00000 ; HE2- CE- NZ- HZ3
3074 3072 3075 3076 3 0.65084 1.95253 0.00000
-2.60338 0.00000 0.00000 ; HE3- CE- NZ- HZ1
3074 3072 3075 3077 3 0.65084 1.95253 0.00000
-2.60338 0.00000 0.00000 ; HE3- CE- NZ- HZ2
3074 3072 3075 3078 3 0.65084 1.95253 0.00000
-2.60338 0.00000 0.00000 ; HE3- CE- NZ- HZ3
3079 3081 3083 3084 3 0.00000 0.00000 0.00000
0.00000 0.00000 0.00000 ; C- N- CA- HA
3079 3081 3083 3085 3 11.71520 1.67360 15.06240 -
13.38880 0.00000 0.00000 ; C- N- CA- CB
3079 3081 3083 3100 3 1.75728 5.27184 2.25936
-7.02912 0.00000 0.00000 ; C- N- CA- C
3080 3079 3081 3082 3 29.28800 -8.36800 -20.92000
0.00000 0.00000 0.00000 ; O- C- N- H
3080 3079 3081 3083 3 20.92000 0.00000 -20.92000
0.00000 0.00000 0.00000 ; O- C- N- CA
3081 3083 3085 3086 3 0.65084 1.95253 0.00000
-2.60338 0.00000 0.00000 ; N- CA- CB- HB2
3081 3083 3085 3087 3 0.65084 1.95253 0.00000
-2.60338 0.00000 0.00000 ; N- CA- CB- HB3
3081 3083 3085 3088 3 0.65084 1.95253 0.00000
-2.60338 0.00000 0.00000 ; N- CA- CB- CG
3081 3083 3100 3101 3 0.00000 0.00000 0.00000
0.00000 0.00000 0.00000 ; N- CA- C- O
3081 3083 3100 3102 3 17.40544 -5.02080 -13.22144
9.20480 0.00000 0.00000 ; N- CA- C- N
3082 3081 3083 3084 3 0.00000 0.00000 0.00000
0.00000 0.00000 0.00000 ; H- N- CA- HA
3082 3081 3083 3085 3 0.00000 0.00000 0.00000
0.00000 0.00000 0.00000 ; H- N- CA- CB
3082 3081 3083 3100 3 0.00000 0.00000 0.00000

0.00000    0.00000    0.00000 ;      H-     N-     CA-     C
  3083   3085   3088   3089      3    0.66944    2.00832    0.00000
-2.67776    0.00000    0.00000 ;      CA-    CB-    CG-    HG2
  3083   3085   3088   3090      3    0.66944    2.00832    0.00000
-2.67776    0.00000    0.00000 ;      CA-    CB-    CG-    HG3
  3083   3085   3088   3091      3    3.68192    3.09616   -2.09200
-3.01248    0.00000    0.00000 ;      CA-    CB-    CG-    CD
  3083   3100   3102   3103      3   20.92000    0.00000  -20.92000
0.00000    0.00000    0.00000 ;      CA-     C-     N-     H
  3083   3100   3102   3104      3   20.92000    0.00000  -20.92000
0.00000    0.00000    0.00000 ;      CA-     C-     N-    CA
  3084   3083   3085   3086      3    0.65084    1.95253    0.00000
-2.60338    0.00000    0.00000 ;      HA-    CA-    CB-    HB2
  3084   3083   3085   3087      3    0.65084    1.95253    0.00000
-2.60338    0.00000    0.00000 ;      HA-    CA-    CB-    HB3
  3084   3083   3085   3088      3    0.65084    1.95253    0.00000
-2.60338    0.00000    0.00000 ;      HA-    CA-    CB-    CG
  3084   3083   3100   3101      3    3.68192   -4.35136    0.00000
1.33888    0.00000    0.00000 ;      HA-    CA-     C-     O
  3084   3083   3100   3102      3    0.00000    0.00000    0.00000
0.00000    0.00000    0.00000 ;      HA-    CA-     C-     N
  3085   3083   3100   3101      3    0.00000    0.00000    0.00000
0.00000    0.00000    0.00000 ;      CB-    CA-     C-     O
  3085   3083   3100   3102      3    2.51040    4.18400    1.67360
-6.69440    0.00000    0.00000 ;      CB-    CA-     C-     N
  3085   3088   3091   3092      3    0.66944    2.00832    0.00000
-2.67776    0.00000    0.00000 ;      CB-    CG-    CD-    HD2
  3085   3088   3091   3093      3    0.66944    2.00832    0.00000
-2.67776    0.00000    0.00000 ;      CB-    CG-    CD-    HD3
  3085   3088   3091   3094      3    3.68192    3.09616   -2.09200
-3.01248    0.00000    0.00000 ;      CB-    CG-    CD-    CE
  3086   3085   3083   3100      3    0.65084    1.95253    0.00000
-2.60338    0.00000    0.00000 ;      HB2-    CB-    CA-     C
  3086   3085   3088   3089      3    0.62760    1.88280    0.00000
-2.51040    0.00000    0.00000 ;      HB2-    CB-    CG-    HG2
  3086   3085   3088   3090      3    0.62760    1.88280    0.00000
-2.51040    0.00000    0.00000 ;      HB2-    CB-    CG-    HG3
  3086   3085   3088   3091      3    0.66944    2.00832    0.00000
-2.67776    0.00000    0.00000 ;      HB2-    CB-    CG-    CD
  3087   3085   3083   3100      3    0.65084    1.95253    0.00000
-2.60338    0.00000    0.00000 ;      HB3-    CB-    CA-     C
  3087   3085   3088   3089      3    0.62760    1.88280    0.00000
-2.51040    0.00000    0.00000 ;      HB3-    CB-    CG-    HG2
  3087   3085   3088   3090      3    0.62760    1.88280    0.00000
-2.51040    0.00000    0.00000 ;      HB3-    CB-    CG-    HG3
  3087   3085   3088   3091      3    0.66944    2.00832    0.00000
-2.67776    0.00000    0.00000 ;      HB3-    CB-    CG-    CD
  3088   3085   3083   3100      3    0.65084    1.95253    0.00000
-2.60338    0.00000    0.00000 ;      CG-    CB-    CA-     C
  3088   3091   3094   3095      3    0.65084    1.95253    0.00000
-2.60338    0.00000    0.00000 ;      CG-    CD-    CE-    HE2
  3088   3091   3094   3096      3    0.65084    1.95253    0.00000

```
-2.60338    0.00000    0.00000 ;      CG-    CD-     CE-   HE3
  3088   3091   3094   3097       3    0.65084    1.95253    0.00000
-2.60338    0.00000    0.00000 ;      CG-    CD-     CE-    NZ
  3089   3088   3091   3092       3    0.62760    1.88280    0.00000
-2.51040    0.00000    0.00000 ;     HG2-    CG-     CD-   HD2
  3089   3088   3091   3093       3    0.62760    1.88280    0.00000
-2.51040    0.00000    0.00000 ;     HG2-    CG-     CD-   HD3
  3089   3088   3091   3094       3    0.66944    2.00832    0.00000
-2.67776    0.00000    0.00000 ;     HG2-    CG-     CD-    CE
  3090   3088   3091   3092       3    0.62760    1.88280    0.00000
-2.51040    0.00000    0.00000 ;     HG3-    CG-     CD-   HD2
  3090   3088   3091   3093       3    0.62760    1.88280    0.00000
-2.51040    0.00000    0.00000 ;     HG3-    CG-     CD-   HD3
  3090   3088   3091   3094       3    0.66944    2.00832    0.00000
-2.67776    0.00000    0.00000 ;     HG3-    CG-     CD-    CE
  3091   3094   3097   3098       3    0.65084    1.95253    0.00000
-2.60338    0.00000    0.00000 ;      CD-    CE-     NZ-   HZ2
  3091   3094   3097   3099       3    0.65084    1.95253    0.00000
-2.60338    0.00000    0.00000 ;      CD-    CE-     NZ-   HZ3
  3092   3091   3094   3095       3    0.65084    1.95253    0.00000
-2.60338    0.00000    0.00000 ;     HD2-    CD-     CE-   HE2
  3092   3091   3094   3096       3    0.65084    1.95253    0.00000
-2.60338    0.00000    0.00000 ;     HD2-    CD-     CE-   HE3
  3092   3091   3094   3097       3    0.65084    1.95253    0.00000
-2.60338    0.00000    0.00000 ;     HD2-    CD-     CE-    NZ
  3093   3091   3094   3095       3    0.65084    1.95253    0.00000
-2.60338    0.00000    0.00000 ;     HD3-    CD-     CE-   HE2
  3093   3091   3094   3096       3    0.65084    1.95253    0.00000
-2.60338    0.00000    0.00000 ;     HD3-    CD-     CE-   HE3
  3093   3091   3094   3097       3    0.65084    1.95253    0.00000
-2.60338    0.00000    0.00000 ;     HD3-    CD-     CE-    NZ
  3095   3094   3097   3098       3    0.65084    1.95253    0.00000
-2.60338    0.00000    0.00000 ;     HE2-    CE-     NZ-   HZ2
  3095   3094   3097   3099       3    0.65084    1.95253    0.00000
-2.60338    0.00000    0.00000 ;     HE2-    CE-     NZ-   HZ3
  3096   3094   3097   3098       3    0.65084    1.95253    0.00000
-2.60338    0.00000    0.00000 ;     HE3-    CE-     NZ-   HZ2
  3096   3094   3097   3099       3    0.65084    1.95253    0.00000
-2.60338    0.00000    0.00000 ;     HE3-    CE-     NZ-   HZ3
  3100   3102   3104   3105       3    0.00000    0.00000    0.00000
0.00000    0.00000    0.00000 ;      C-     N-     CA-    HA
  3100   3102   3104   3106       3   11.71520    1.67360   15.06240  -
13.38880    0.00000    0.00000 ;      C-     N-     CA-    CB
  3100   3102   3104   3111       3    1.75728    5.27184    2.25936
-7.02912    0.00000    0.00000 ;      C-     N-     CA-     C
  3101   3100   3102   3103       3   29.28800   -8.36800  -20.92000
0.00000    0.00000    0.00000 ;      O-     C-     N-     H
  3101   3100   3102   3104       3   20.92000    0.00000  -20.92000
0.00000    0.00000    0.00000 ;      O-     C-     N-    CA
  3102   3104   3106   3107       3    0.65084    1.95253    0.00000
-2.60338    0.00000    0.00000 ;      N-    CA-     CB-   HB2
  3102   3104   3106   3108       3    0.65084    1.95253    0.00000
```

-2.60338 0.00000 0.00000 ; N- CA- CB- HB3
3102 3104 3106 3109 3 6.03751 2.50622 -5.17142
-4.20074 1.10458 0.00000 ; N- CA- CB- SG
3102 3104 3111 3112 3 0.00000 0.00000 0.00000
0.00000 0.00000 0.00000 ; N- CA- C- O
3102 3104 3111 3113 3 17.40544 -5.02080 -13.22144
9.20480 0.00000 0.00000 ; N- CA- C- N
3103 3102 3104 3105 3 0.00000 0.00000 0.00000
0.00000 0.00000 0.00000 ; H- N- CA- HA
3103 3102 3104 3106 3 0.00000 0.00000 0.00000
0.00000 0.00000 0.00000 ; H- N- CA- CB
3103 3102 3104 3111 3 0.00000 0.00000 0.00000
0.00000 0.00000 0.00000 ; H- N- CA- C
3104 3106 3109 3110 3 1.69034 2.77818 4.11706
-4.21747 1.00416 0.00000 ; CA- CB- SG- HG
3104 3111 3113 3114 3 20.92000 0.00000 -20.92000
0.00000 0.00000 0.00000 ; CA- C- N- H
3104 3111 3113 3115 3 20.92000 0.00000 -20.92000
0.00000 0.00000 0.00000 ; CA- C- N- CA
3105 3104 3106 3107 3 0.65084 1.95253 0.00000
-2.60338 0.00000 0.00000 ; HA- CA- CB- HB2
3105 3104 3106 3108 3 0.65084 1.95253 0.00000
-2.60338 0.00000 0.00000 ; HA- CA- CB- HB3
3105 3104 3106 3109 3 0.65084 1.95253 0.00000
-2.60338 0.00000 0.00000 ; HA- CA- CB- SG
3105 3104 3111 3112 3 3.68192 -4.35136 0.00000
1.33888 0.00000 0.00000 ; HA- CA- C- O
3105 3104 3111 3113 3 0.00000 0.00000 0.00000
0.00000 0.00000 0.00000 ; HA- CA- C- N
3106 3104 3111 3112 3 0.00000 0.00000 0.00000
0.00000 0.00000 0.00000 ; CB- CA- C- O
3106 3104 3111 3113 3 2.51040 4.18400 1.67360
-6.69440 0.00000 0.00000 ; CB- CA- C- N
3107 3106 3104 3111 3 0.65084 1.95253 0.00000
-2.60338 0.00000 0.00000 ; HB2- CB- CA- C
3107 3106 3109 3110 3 1.04600 3.13800 0.00000
-4.18400 0.00000 0.00000 ; HB2- CB- SG- HG
3108 3106 3104 3111 3 0.65084 1.95253 0.00000
-2.60338 0.00000 0.00000 ; HB3- CB- CA- C
3108 3106 3109 3110 3 1.04600 3.13800 0.00000
-4.18400 0.00000 0.00000 ; HB3- CB- SG- HG
3109 3106 3104 3111 3 5.62330 4.27605 -5.33042
-4.20074 2.51040 0.00000 ; SG- CB- CA- C
3111 3113 3115 3116 3 0.00000 0.00000 0.00000
0.00000 0.00000 0.00000 ; C- N- CA- HA
3111 3113 3115 3117 3 11.71520 1.67360 15.06240 -
13.38880 0.00000 0.00000 ; C- N- CA- CB
3111 3113 3115 3123 3 1.75728 5.27184 2.25936
-7.02912 0.00000 0.00000 ; C- N- CA- C
3112 3111 3113 3114 3 29.28800 -8.36800 -20.92000
0.00000 0.00000 0.00000 ; O- C- N- H
3112 3111 3113 3115 3 20.92000 0.00000 -20.92000

```
0.00000    0.00000    0.00000 ;      O-     C-     N-    CA
  3113   3115   3117   3118      3    0.65084    1.95253    0.00000
-2.60338    0.00000    0.00000 ;      N-    CA-    CB-   HB2
  3113   3115   3117   3119      3    0.65084    1.95253    0.00000
-2.60338    0.00000    0.00000 ;      N-    CA-    CB-   HB3
  3113   3115   3117   3120      3   15.41386    9.74035   -8.39310
-0.97069    2.97901    0.00000 ;      N-    CA-    CB-    CG
  3113   3115   3123   3124      3    0.00000    0.00000    0.00000
0.00000    0.00000    0.00000 ;      N-    CA-     C-     O
  3113   3115   3123   3125      3   17.40544   -5.02080  -13.22144
9.20480    0.00000    0.00000 ;      N-    CA-     C-     N
  3114   3113   3115   3116      3    0.00000    0.00000    0.00000
0.00000    0.00000    0.00000 ;      H-     N-    CA-    HA
  3114   3113   3115   3117      3    0.00000    0.00000    0.00000
0.00000    0.00000    0.00000 ;      H-     N-    CA-    CB
  3114   3113   3115   3123      3    0.00000    0.00000    0.00000
0.00000    0.00000    0.00000 ;      H-     N-    CA-     C
  3115   3117   3120   3121      3    6.43499    0.00000   -5.39736
0.00000   -1.03763    0.00000 ;     CA-    CB-    CG-   OD1
  3115   3117   3120   3122      3    6.43499    0.00000   -5.39736
0.00000   -1.03763    0.00000 ;     CA-    CB-    CG-   OD2
  3115   3123   3125   3126      3   20.92000    0.00000  -20.92000
0.00000    0.00000    0.00000 ;     CA-     C-     N-     H
  3115   3123   3125   3127      3   20.92000    0.00000  -20.92000
0.00000    0.00000    0.00000 ;     CA-     C-     N-    CA
  3116   3115   3117   3118      3    0.65084    1.95253    0.00000
-2.60338    0.00000    0.00000 ;     HA-    CA-    CB-   HB2
  3116   3115   3117   3119      3    0.65084    1.95253    0.00000
-2.60338    0.00000    0.00000 ;     HA-    CA-    CB-   HB3
  3116   3115   3117   3120      3    0.65084    1.95253    0.00000
-2.60338    0.00000    0.00000 ;     HA-    CA-    CB-    CG
  3116   3115   3123   3124      3    3.68192   -4.35136    0.00000
1.33888    0.00000    0.00000 ;     HA-    CA-     C-     O
  3116   3115   3123   3125      3    0.00000    0.00000    0.00000
0.00000    0.00000    0.00000 ;     HA-    CA-     C-     N
  3117   3115   3123   3124      3    0.00000    0.00000    0.00000
0.00000    0.00000    0.00000 ;     CB-    CA-     C-     O
  3117   3115   3123   3125      3    2.51040    4.18400    1.67360
-6.69440    0.00000    0.00000 ;     CB-    CA-     C-     N
  3118   3117   3115   3123      3    0.65084    1.95253    0.00000
-2.60338    0.00000    0.00000 ;    HB2-    CB-    CA-     C
  3118   3117   3120   3121      3    0.00000    0.00000    0.00000
0.00000    0.00000    0.00000 ;    HB2-    CB-    CG-   OD1
  3118   3117   3120   3122      3    0.00000    0.00000    0.00000
0.00000    0.00000    0.00000 ;    HB2-    CB-    CG-   OD2
  3119   3117   3115   3123      3    0.65084    1.95253    0.00000
-2.60338    0.00000    0.00000 ;    HB3-    CB-    CA-     C
  3119   3117   3120   3121      3    0.00000    0.00000    0.00000
0.00000    0.00000    0.00000 ;    HB3-    CB-    CG-   OD1
  3119   3117   3120   3122      3    0.00000    0.00000    0.00000
0.00000    0.00000    0.00000 ;    HB3-    CB-    CG-   OD2
  3120   3117   3115   3123      3    7.14627   -1.04600   -8.99560
```

-0.97069 5.15469 0.00000 ; CG- CB- CA- C
3123 3125 3127 3128 3 0.00000 0.00000 0.00000
0.00000 0.00000 0.00000 ; C- N- CA- HA
3123 3125 3127 3129 3 11.71520 1.67360 15.06240 -
13.38880 0.00000 0.00000 ; C- N- CA- CB
3123 3125 3127 3142 3 1.75728 5.27184 2.25936
-7.02912 0.00000 0.00000 ; C- N- CA- C
3124 3123 3125 3126 3 29.28800 -8.36800 -20.92000
0.00000 0.00000 0.00000 ; O- C- N- H
3124 3123 3125 3127 3 20.92000 0.00000 -20.92000
0.00000 0.00000 0.00000 ; O- C- N- CA
3125 3127 3129 3130 3 0.65084 1.95253 0.00000
-2.60338 0.00000 0.00000 ; N- CA- CB- HB
3125 3127 3129 3131 3 3.83673 0.44769 -1.77402
-2.47693 -0.03347 0.00000 ; N- CA- CB- CG2
3125 3127 3129 3135 3 2.97482 0.12134 2.04179
-1.89117 -3.24678 0.00000 ; N- CA- CB- CG1
3125 3127 3142 3143 3 0.00000 0.00000 0.00000
0.00000 0.00000 0.00000 ; N- CA- C- O
3125 3127 3142 3144 3 17.40544 -5.02080 -13.22144
9.20480 0.00000 0.00000 ; N- CA- C- N
3126 3125 3127 3128 3 0.00000 0.00000 0.00000
0.00000 0.00000 0.00000 ; H- N- CA- HA
3126 3125 3127 3129 3 0.00000 0.00000 0.00000
0.00000 0.00000 0.00000 ; H- N- CA- CB
3126 3125 3127 3142 3 0.00000 0.00000 0.00000
0.00000 0.00000 0.00000 ; H- N- CA- C
3127 3129 3131 3132 3 0.66944 2.00832 0.00000
-2.67776 0.00000 0.00000 ; CA- CB- CG2- HG21
3127 3129 3131 3133 3 0.66944 2.00832 0.00000
-2.67776 0.00000 0.00000 ; CA- CB- CG2- HG22
3127 3129 3131 3134 3 0.66944 2.00832 0.00000
-2.67776 0.00000 0.00000 ; CA- CB- CG2- HG23
3127 3129 3135 3136 3 0.66944 2.00832 0.00000
-2.67776 0.00000 0.00000 ; CA- CB- CG1- HG12
3127 3129 3135 3137 3 0.66944 2.00832 0.00000
-2.67776 0.00000 0.00000 ; CA- CB- CG1- HG13
3127 3129 3135 3138 3 4.24258 -0.52718 -7.25506
-1.79075 7.69856 0.00000 ; CA- CB- CG1- CD1
3127 3142 3144 3145 3 20.92000 0.00000 -20.92000
0.00000 0.00000 0.00000 ; CA- C- N- H
3127 3142 3144 3146 3 20.92000 0.00000 -20.92000
0.00000 0.00000 0.00000 ; CA- C- N- CA
3128 3127 3129 3130 3 0.65084 1.95253 0.00000
-2.60338 0.00000 0.00000 ; HA- CA- CB- HB
3128 3127 3129 3131 3 0.65084 1.95253 0.00000
-2.60338 0.00000 0.00000 ; HA- CA- CB- CG2
3128 3127 3129 3135 3 0.65084 1.95253 0.00000
-2.60338 0.00000 0.00000 ; HA- CA- CB- CG1
3128 3127 3142 3143 3 3.68192 -4.35136 0.00000
1.33888 0.00000 0.00000 ; HA- CA- C- O
3128 3127 3142 3144 3 0.00000 0.00000 0.00000

```
0.00000    0.00000    0.00000 ;     HA-    CA-     C-     N
  3129   3127   3142   3143      3    0.00000    0.00000    0.00000
0.00000    0.00000    0.00000 ;     CB-    CA-     C-     O
  3129   3127   3142   3144      3    2.51040    4.18400    1.67360
-6.69440    0.00000    0.00000 ;     CB-    CA-     C-     N
  3129   3135   3138   3139      3    0.66944    2.00832    0.00000
-2.67776    0.00000    0.00000 ;     CB-   CG1-   CD1-  HD11
  3129   3135   3138   3140      3    0.66944    2.00832    0.00000
-2.67776    0.00000    0.00000 ;     CB-   CG1-   CD1-  HD12
  3129   3135   3138   3141      3    0.66944    2.00832    0.00000
-2.67776    0.00000    0.00000 ;     CB-   CG1-   CD1-  HD13
  3130   3129   3127   3142      3    0.65084    1.95253    0.00000
-2.60338    0.00000    0.00000 ;     HB-    CB-    CA-     C
  3130   3129   3131   3132      3    0.62760    1.88280    0.00000
-2.51040    0.00000    0.00000 ;     HB-    CB-   CG2-  HG21
  3130   3129   3131   3133      3    0.62760    1.88280    0.00000
-2.51040    0.00000    0.00000 ;     HB-    CB-   CG2-  HG22
  3130   3129   3131   3134      3    0.62760    1.88280    0.00000
-2.51040    0.00000    0.00000 ;     HB-    CB-   CG2-  HG23
  3130   3129   3135   3136      3    0.62760    1.88280    0.00000
-2.51040    0.00000    0.00000 ;     HB-    CB-   CG1-  HG12
  3130   3129   3135   3137      3    0.62760    1.88280    0.00000
-2.51040    0.00000    0.00000 ;     HB-    CB-   CG1-  HG13
  3130   3129   3135   3138      3    0.66944    2.00832    0.00000
-2.67776    0.00000    0.00000 ;     HB-    CB-   CG1-   CD1
  3131   3129   3127   3142      3    5.67350    3.55640   -6.16722
-2.47693    3.74886    0.00000 ;    CG2-    CB-    CA-     C
  3131   3129   3135   3136      3    0.66944    2.00832    0.00000
-2.67776    0.00000    0.00000 ;    CG2-    CB-   CG1-  HG12
  3131   3129   3135   3137      3    0.66944    2.00832    0.00000
-2.67776    0.00000    0.00000 ;    CG2-    CB-   CG1-  HG13
  3131   3129   3135   3138      3    3.81162    0.49790   -8.14206
-1.79075    7.49773    0.00000 ;    CG2-    CB-   CG1-   CD1
  3132   3131   3129   3135      3    0.66944    2.00832    0.00000
-2.67776    0.00000    0.00000 ;   HG21-   CG2-     CB-   CG1
  3133   3131   3129   3135      3    0.66944    2.00832    0.00000
-2.67776    0.00000    0.00000 ;   HG22-   CG2-     CB-   CG1
  3134   3131   3129   3135      3    0.66944    2.00832    0.00000
-2.67776    0.00000    0.00000 ;   HG23-   CG2-     CB-   CG1
  3135   3129   3127   3142      3    8.26340    0.74057   -9.99976
-1.89117    3.84928    0.00000 ;    CG1-    CB-    CA-     C
  3136   3135   3138   3139      3    0.62760    1.88280    0.00000
-2.51040    0.00000    0.00000 ;   HG12-   CG1-   CD1-  HD11
  3136   3135   3138   3140      3    0.62760    1.88280    0.00000
-2.51040    0.00000    0.00000 ;   HG12-   CG1-   CD1-  HD12
  3136   3135   3138   3141      3    0.62760    1.88280    0.00000
-2.51040    0.00000    0.00000 ;   HG12-   CG1-   CD1-  HD13
  3137   3135   3138   3139      3    0.62760    1.88280    0.00000
-2.51040    0.00000    0.00000 ;   HG13-   CG1-   CD1-  HD11
  3137   3135   3138   3140      3    0.62760    1.88280    0.00000
-2.51040    0.00000    0.00000 ;   HG13-   CG1-   CD1-  HD12
  3137   3135   3138   3141      3    0.62760    1.88280    0.00000
```

-2.51040 0.00000 0.00000 ; HG13- CG1- CD1- HD13
3142 3144 3146 3147 3 0.00000 0.00000 0.00000
0.00000 0.00000 0.00000 ; C- N- CA- HA
3142 3144 3146 3148 3 11.71520 1.67360 15.06240 -
13.38880 0.00000 0.00000 ; C- N- CA- CB
3142 3144 3146 3166 3 1.75728 5.27184 2.25936
-7.02912 0.00000 0.00000 ; C- N- CA- C
3143 3142 3144 3145 3 29.28800 -8.36800 -20.92000
0.00000 0.00000 0.00000 ; O- C- N- H
3143 3142 3144 3146 3 20.92000 0.00000 -20.92000
0.00000 0.00000 0.00000 ; O- C- N- CA
3144 3146 3148 3149 3 0.65084 1.95253 0.00000
-2.60338 0.00000 0.00000 ; N- CA- CB- HB2
3144 3146 3148 3150 3 0.65084 1.95253 0.00000
-2.60338 0.00000 0.00000 ; N- CA- CB- HB3
3144 3146 3148 3151 3 4.18818 2.60663 -3.65682
-3.91622 1.03763 0.00000 ; N- CA- CB- CG
3144 3146 3166 3167 3 0.00000 0.00000 0.00000
0.00000 0.00000 0.00000 ; N- CA- C- O
3144 3146 3166 3168 3 17.40544 -5.02080 -13.22144
9.20480 0.00000 0.00000 ; N- CA- C- N
3145 3144 3146 3147 3 0.00000 0.00000 0.00000
0.00000 0.00000 0.00000 ; H- N- CA- HA
3145 3144 3146 3148 3 0.00000 0.00000 0.00000
0.00000 0.00000 0.00000 ; H- N- CA- CB
3145 3144 3146 3166 3 0.00000 0.00000 0.00000
0.00000 0.00000 0.00000 ; H- N- CA- C
3146 3148 3151 3152 3 0.00000 0.00000 0.00000
0.00000 0.00000 0.00000 ; CA- CB- CG- CD1
3146 3148 3151 3165 3 4.95386 8.75293 6.59398 -
13.70678 -3.17984 0.00000 ; CA- CB- CG- CD2
3146 3166 3168 3169 3 20.92000 0.00000 -20.92000
0.00000 0.00000 0.00000 ; CA- C- N- H
3146 3166 3168 3170 3 20.92000 0.00000 -20.92000
0.00000 0.00000 0.00000 ; CA- C- N- CA
3147 3146 3148 3149 3 0.65084 1.95253 0.00000
-2.60338 0.00000 0.00000 ; HA- CA- CB- HB2
3147 3146 3148 3150 3 0.65084 1.95253 0.00000
-2.60338 0.00000 0.00000 ; HA- CA- CB- HB3
3147 3146 3148 3151 3 0.65084 1.95253 0.00000
-2.60338 0.00000 0.00000 ; HA- CA- CB- CG
3147 3146 3166 3167 3 3.68192 -4.35136 0.00000
1.33888 0.00000 0.00000 ; HA- CA- C- O
3147 3146 3166 3168 3 0.00000 0.00000 0.00000
0.00000 0.00000 0.00000 ; HA- CA- C- N
3148 3146 3166 3167 3 0.00000 0.00000 0.00000
0.00000 0.00000 0.00000 ; CB- CA- C- O
3148 3146 3166 3168 3 2.51040 4.18400 1.67360
-6.69440 0.00000 0.00000 ; CB- CA- C- N
3148 3151 3152 3153 3 54.60120 0.00000 -54.60120
0.00000 0.00000 0.00000 ; CB- CG- CD1- HD1
3148 3151 3152 3154 3 54.60120 0.00000 -54.60120

0.00000 0.00000 0.00000 ; CB- CG- CD1- NE1
3148 3151 3165 3156 3 14.01640 0.00000 -14.01640
0.00000 0.00000 0.00000 ; CB- CG- CD2- CE2
3148 3151 3165 3163 3 14.01640 0.00000 -14.01640
0.00000 0.00000 0.00000 ; CB- CG- CD2- CE3
3149 3148 3146 3166 3 0.65084 1.95253 0.00000
-2.60338 0.00000 0.00000 ; HB2- CB- CA- C
3149 3148 3151 3152 3 0.00000 0.00000 0.00000
0.00000 0.00000 0.00000 ; HB2- CB- CG- CD1
3149 3148 3151 3165 3 0.00000 0.00000 0.00000
0.00000 0.00000 0.00000 ; HB2- CB- CG- CD2
3150 3148 3146 3166 3 0.65084 1.95253 0.00000
-2.60338 0.00000 0.00000 ; HB3- CB- CA- C
3150 3148 3151 3152 3 0.00000 0.00000 0.00000
0.00000 0.00000 0.00000 ; HB3- CB- CG- CD1
3150 3148 3151 3165 3 0.00000 0.00000 0.00000
0.00000 0.00000 0.00000 ; HB3- CB- CG- CD2
3151 3148 3146 3166 3 4.62332 3.00830 -5.43083
-3.91622 2.47693 0.00000 ; CG- CB- CA- C
3151 3152 3154 3155 3 12.55200 0.00000 -12.55200
0.00000 0.00000 0.00000 ; CG- CD1- NE1- HE1
3151 3152 3154 3156 3 12.55200 0.00000 -12.55200
0.00000 0.00000 0.00000 ; CG- CD1- NE1- CE2
3151 3165 3156 3154 3 25.10400 0.00000 -25.10400
0.00000 0.00000 0.00000 ; CG- CD2- CE2- NE1
3151 3165 3156 3157 3 25.10400 0.00000 -25.10400
0.00000 0.00000 0.00000 ; CG- CD2- CE2- CZ2
3151 3165 3163 3161 3 29.28800 0.00000 -29.28800
0.00000 0.00000 0.00000 ; CG- CD2- CE3- CZ3
3151 3165 3163 3164 3 29.28800 0.00000 -29.28800
0.00000 0.00000 0.00000 ; CG- CD2- CE3- HE3
3152 3151 3165 3156 3 14.01640 0.00000 -14.01640
0.00000 0.00000 0.00000 ; CD1- CG- CD2- CE2
3152 3151 3165 3163 3 14.01640 0.00000 -14.01640
0.00000 0.00000 0.00000 ; CD1- CG- CD2- CE3
3152 3154 3156 3157 3 12.76120 0.00000 -12.76120
0.00000 0.00000 0.00000 ; CD1- NE1- CE2- CZ2
3152 3154 3156 3165 3 12.76120 0.00000 -12.76120
0.00000 0.00000 0.00000 ; CD1- NE1- CE2- CD2
3153 3152 3151 3165 3 54.60120 0.00000 -54.60120
0.00000 0.00000 0.00000 ; HD1- CD1- CG- CD2
3153 3152 3154 3155 3 12.55200 0.00000 -12.55200
0.00000 0.00000 0.00000 ; HD1- CD1- NE1- HE1
3153 3152 3154 3156 3 12.55200 0.00000 -12.55200
0.00000 0.00000 0.00000 ; HD1- CD1- NE1- CE2
3154 3152 3151 3165 3 54.60120 0.00000 -54.60120
0.00000 0.00000 0.00000 ; NE1- CD1- CG- CD2
3154 3156 3157 3158 3 30.33400 0.00000 -30.33400
0.00000 0.00000 0.00000 ; NE1- CE2- CZ2- HZ2
3154 3156 3157 3159 3 30.33400 0.00000 -30.33400
0.00000 0.00000 0.00000 ; NE1- CE2- CZ2- CH2
3154 3156 3165 3163 3 25.10400 0.00000 -25.10400

0.00000 0.00000 0.00000 ; NE1- CE2- CD2- CE3
3155 3154 3156 3157 3 12.76120 0.00000 -12.76120
0.00000 0.00000 0.00000 ; HE1- NE1- CE2- CZ2
3155 3154 3156 3165 3 12.76120 0.00000 -12.76120
0.00000 0.00000 0.00000 ; HE1- NE1- CE2- CD2
3156 3157 3159 3160 3 30.33400 0.00000 -30.33400
0.00000 0.00000 0.00000 ; CE2- CZ2- CH2- HH2
3156 3157 3159 3161 3 30.33400 0.00000 -30.33400
0.00000 0.00000 0.00000 ; CE2- CZ2- CH2- CZ3
3156 3165 3163 3161 3 29.28800 0.00000 -29.28800
0.00000 0.00000 0.00000 ; CE2- CD2- CE3- CZ3
3156 3165 3163 3164 3 29.28800 0.00000 -29.28800
0.00000 0.00000 0.00000 ; CE2- CD2- CE3- HE3
3157 3156 3165 3163 3 25.10400 0.00000 -25.10400
0.00000 0.00000 0.00000 ; CZ2- CE2- CD2- CE3
3157 3159 3161 3162 3 30.33400 0.00000 -30.33400
0.00000 0.00000 0.00000 ; CZ2- CH2- CZ3- HZ3
3157 3159 3161 3163 3 30.33400 0.00000 -30.33400
0.00000 0.00000 0.00000 ; CZ2- CH2- CZ3- CE3
3158 3157 3156 3165 3 30.33400 0.00000 -30.33400
0.00000 0.00000 0.00000 ; HZ2- CZ2- CE2- CD2
3158 3157 3159 3160 3 30.33400 0.00000 -30.33400
0.00000 0.00000 0.00000 ; HZ2- CZ2- CH2- HH2
3158 3157 3159 3161 3 30.33400 0.00000 -30.33400
0.00000 0.00000 0.00000 ; HZ2- CZ2- CH2- CZ3
3159 3157 3156 3165 3 30.33400 0.00000 -30.33400
0.00000 0.00000 0.00000 ; CH2- CZ2- CE2- CD2
3159 3161 3163 3164 3 30.33400 0.00000 -30.33400
0.00000 0.00000 0.00000 ; CH2- CZ3- CE3- HE3
3159 3161 3163 3165 3 30.33400 0.00000 -30.33400
0.00000 0.00000 0.00000 ; CH2- CZ3- CE3- CD2
3160 3159 3161 3162 3 30.33400 0.00000 -30.33400
0.00000 0.00000 0.00000 ; HH2- CH2- CZ3- HZ3
3160 3159 3161 3163 3 30.33400 0.00000 -30.33400
0.00000 0.00000 0.00000 ; HH2- CH2- CZ3- CE3
3162 3161 3163 3164 3 30.33400 0.00000 -30.33400
0.00000 0.00000 0.00000 ; HZ3- CZ3- CE3- HE3
3162 3161 3163 3165 3 30.33400 0.00000 -30.33400
0.00000 0.00000 0.00000 ; HZ3- CZ3- CE3- CD2
3166 3168 3170 3171 3 0.00000 0.00000 0.00000
0.00000 0.00000 0.00000 ; C- N- CA- HA
3166 3168 3170 3172 3 11.71520 1.67360 15.06240 -
13.38880 0.00000 0.00000 ; C- N- CA- CB
3166 3168 3170 3177 3 1.75728 5.27184 2.25936
-7.02912 0.00000 0.00000 ; C- N- CA- C
3167 3166 3168 3169 3 29.28800 -8.36800 -20.92000
0.00000 0.00000 0.00000 ; O- C- N- H
3167 3166 3168 3170 3 20.92000 0.00000 -20.92000
0.00000 0.00000 0.00000 ; O- C- N- CA
3168 3170 3172 3173 3 0.65084 1.95253 0.00000
-2.60338 0.00000 0.00000 ; N- CA- CB- HB2
3168 3170 3172 3174 3 0.65084 1.95253 0.00000

-2.60338 0.00000 0.00000 ; N- CA- CB- HB3
3168 3170 3172 3175 3 7.86174 2.24681 -7.41405
-6.71114 5.35552 0.00000 ; N- CA- CB- OG
3168 3170 3177 3178 3 0.00000 0.00000 0.00000
0.00000 0.00000 0.00000 ; N- CA- C- O
3168 3170 3177 3179 3 17.40544 -5.02080 -13.22144
9.20480 0.00000 0.00000 ; N- CA- C- N
3169 3168 3170 3171 3 0.00000 0.00000 0.00000
0.00000 0.00000 0.00000 ; H- N- CA- HA
3169 3168 3170 3172 3 0.00000 0.00000 0.00000
0.00000 0.00000 0.00000 ; H- N- CA- CB
3169 3168 3170 3177 3 0.00000 0.00000 0.00000
0.00000 0.00000 0.00000 ; H- N- CA- C
3170 3172 3175 3176 3 2.05853 2.46856 3.48109
-4.46851 0.23430 0.00000 ; CA- CB- OG- HG
3170 3177 3179 3180 3 20.92000 0.00000 -20.92000
0.00000 0.00000 0.00000 ; CA- C- N- H
3170 3177 3179 3181 3 20.92000 0.00000 -20.92000
0.00000 0.00000 0.00000 ; CA- C- N- CA
3171 3170 3172 3173 3 0.65084 1.95253 0.00000
-2.60338 0.00000 0.00000 ; HA- CA- CB- HB2
3171 3170 3172 3174 3 0.65084 1.95253 0.00000
-2.60338 0.00000 0.00000 ; HA- CA- CB- HB3
3171 3170 3172 3175 3 1.04600 -1.04600 0.00000
0.00000 0.00000 0.00000 ; HA- CA- CB- OG
3171 3170 3177 3178 3 3.68192 -4.35136 0.00000
1.33888 0.00000 0.00000 ; HA- CA- C- O
3171 3170 3177 3179 3 0.00000 0.00000 0.00000
0.00000 0.00000 0.00000 ; HA- CA- C- N
3172 3170 3177 3178 3 0.00000 0.00000 0.00000
0.00000 0.00000 0.00000 ; CB- CA- C- O
3172 3170 3177 3179 3 2.51040 4.18400 1.67360
-6.69440 0.00000 0.00000 ; CB- CA- C- N
3173 3172 3170 3177 3 0.65084 1.95253 0.00000
-2.60338 0.00000 0.00000 ; HB2- CB- CA- C
3173 3172 3175 3176 3 0.69733 2.09200 0.00000
-2.78933 0.00000 0.00000 ; HB2- CB- OG- HG
3174 3172 3170 3177 3 0.65084 1.95253 0.00000
-2.60338 0.00000 0.00000 ; HB3- CB- CA- C
3174 3172 3175 3176 3 0.69733 2.09200 0.00000
-2.78933 0.00000 0.00000 ; HB3- CB- OG- HG
3175 3172 3170 3177 3 7.34710 7.79898 -6.14211
-6.71114 4.31789 0.00000 ; OG- CB- CA- C
3177 3179 3181 3182 3 0.00000 0.00000 0.00000
0.00000 0.00000 0.00000 ; C- N- CA- HA
3177 3179 3181 3183 3 11.71520 1.67360 15.06240 -
13.38880 0.00000 0.00000 ; C- N- CA- CB
3177 3179 3181 3188 3 1.75728 5.27184 2.25936
-7.02912 0.00000 0.00000 ; C- N- CA- C
3178 3177 3179 3180 3 29.28800 -8.36800 -20.92000
0.00000 0.00000 0.00000 ; O- C- N- H
3178 3177 3179 3181 3 20.92000 0.00000 -20.92000

0.00000 0.00000 0.00000 ; O- C- N- CA
3179 3181 3183 3184 3 0.65084 1.95253 0.00000
-2.60338 0.00000 0.00000 ; N- CA- CB- HB2
3179 3181 3183 3185 3 0.65084 1.95253 0.00000
-2.60338 0.00000 0.00000 ; N- CA- CB- HB3
3179 3181 3183 3186 3 7.86174 2.24681 -7.41405
-6.71114 5.35552 0.00000 ; N- CA- CB- OG
3179 3181 3188 3189 3 0.00000 0.00000 0.00000
0.00000 0.00000 0.00000 ; N- CA- C- O
3179 3181 3188 3190 3 17.40544 -5.02080 -13.22144
9.20480 0.00000 0.00000 ; N- CA- C- N
3180 3179 3181 3182 3 0.00000 0.00000 0.00000
0.00000 0.00000 0.00000 ; H- N- CA- HA
3180 3179 3181 3183 3 0.00000 0.00000 0.00000
0.00000 0.00000 0.00000 ; H- N- CA- CB
3180 3179 3181 3188 3 0.00000 0.00000 0.00000
0.00000 0.00000 0.00000 ; H- N- CA- C
3181 3183 3186 3187 3 2.05853 2.46856 3.48109
-4.46851 0.23430 0.00000 ; CA- CB- OG- HG
3181 3188 3190 3191 3 20.92000 0.00000 -20.92000
0.00000 0.00000 0.00000 ; CA- C- N- H
3181 3188 3190 3192 3 20.92000 0.00000 -20.92000
0.00000 0.00000 0.00000 ; CA- C- N- CA
3182 3181 3183 3184 3 0.65084 1.95253 0.00000
-2.60338 0.00000 0.00000 ; HA- CA- CB- HB2
3182 3181 3183 3185 3 0.65084 1.95253 0.00000
-2.60338 0.00000 0.00000 ; HA- CA- CB- HB3
3182 3181 3183 3186 3 1.04600 -1.04600 0.00000
0.00000 0.00000 0.00000 ; HA- CA- CB- OG
3182 3181 3188 3189 3 3.68192 -4.35136 0.00000
1.33888 0.00000 0.00000 ; HA- CA- C- O
3182 3181 3188 3190 3 0.00000 0.00000 0.00000
0.00000 0.00000 0.00000 ; HA- CA- C- N
3183 3181 3188 3189 3 0.00000 0.00000 0.00000
0.00000 0.00000 0.00000 ; CB- CA- C- O
3183 3181 3188 3190 3 2.51040 4.18400 1.67360
-6.69440 0.00000 0.00000 ; CB- CA- C- N
3184 3183 3181 3188 3 0.65084 1.95253 0.00000
-2.60338 0.00000 0.00000 ; HB2- CB- CA- C
3184 3183 3186 3187 3 0.69733 2.09200 0.00000
-2.78933 0.00000 0.00000 ; HB2- CB- OG- HG
3185 3183 3181 3188 3 0.65084 1.95253 0.00000
-2.60338 0.00000 0.00000 ; HB3- CB- CA- C
3185 3183 3186 3187 3 0.69733 2.09200 0.00000
-2.78933 0.00000 0.00000 ; HB3- CB- OG- HG
3186 3183 3181 3188 3 7.34710 7.79898 -6.14211
-6.71114 4.31789 0.00000 ; OG- CB- CA- C
3188 3190 3192 3193 3 0.00000 0.00000 0.00000
0.00000 0.00000 0.00000 ; C- N- CA- HA2
3188 3190 3192 3194 3 0.00000 0.00000 0.00000
0.00000 0.00000 0.00000 ; C- N- CA- HA3
3188 3190 3192 3195 3 1.75728 5.27184 2.25936

-7.02912 0.00000 0.00000 ; C- N- CA- C
3189 3188 3190 3191 3 29.28800 -8.36800 -20.92000
0.00000 0.00000 0.00000 ; O- C- N- H
3189 3188 3190 3192 3 20.92000 0.00000 -20.92000
0.00000 0.00000 0.00000 ; O- C- N- CA
3190 3192 3195 3196 3 0.00000 0.00000 0.00000
0.00000 0.00000 0.00000 ; N- CA- C- O
3190 3192 3195 3197 3 17.40544 -5.02080 -13.22144
9.20480 0.00000 0.00000 ; N- CA- C- N
3191 3190 3192 3193 3 0.00000 0.00000 0.00000
0.00000 0.00000 0.00000 ; H- N- CA- HA2
3191 3190 3192 3194 3 0.00000 0.00000 0.00000
0.00000 0.00000 0.00000 ; H- N- CA- HA3
3191 3190 3192 3195 3 0.00000 0.00000 0.00000
0.00000 0.00000 0.00000 ; H- N- CA- C
3192 3195 3197 3198 3 20.92000 0.00000 -20.92000
0.00000 0.00000 0.00000 ; CA- C- N- H
3192 3195 3197 3199 3 20.92000 0.00000 -20.92000
0.00000 0.00000 0.00000 ; CA- C- N- CA
3193 3192 3195 3196 3 3.68192 -4.35136 0.00000
1.33888 0.00000 0.00000 ; HA2- CA- C- O
3193 3192 3195 3197 3 0.00000 0.00000 0.00000
0.00000 0.00000 0.00000 ; HA2- CA- C- N
3194 3192 3195 3196 3 3.68192 -4.35136 0.00000
1.33888 0.00000 0.00000 ; HA3- CA- C- O
3194 3192 3195 3197 3 0.00000 0.00000 0.00000
0.00000 0.00000 0.00000 ; HA3- CA- C- N
3195 3197 3199 3200 3 0.00000 0.00000 0.00000
0.00000 0.00000 0.00000 ; C- N- CA- HA
3195 3197 3199 3201 3 11.71520 1.67360 15.06240 -
13.38880 0.00000 0.00000 ; C- N- CA- CB
3195 3197 3199 3211 3 1.75728 5.27184 2.25936
-7.02912 0.00000 0.00000 ; C- N- CA- C
3196 3195 3197 3198 3 29.28800 -8.36800 -20.92000
0.00000 0.00000 0.00000 ; O- C- N- H
3196 3195 3197 3199 3 20.92000 0.00000 -20.92000
0.00000 0.00000 0.00000 ; O- C- N- CA
3197 3199 3201 3202 3 0.65084 1.95253 0.00000
-2.60338 0.00000 0.00000 ; N- CA- CB- HB
3197 3199 3201 3203 3 3.83673 0.44769 -1.77402
-2.47693 -0.03347 0.00000 ; N- CA- CB- CG1
3197 3199 3201 3207 3 3.83673 0.44769 -1.77402
-2.47693 -0.03347 0.00000 ; N- CA- CB- CG2
3197 3199 3211 3212 3 0.00000 0.00000 0.00000
0.00000 0.00000 0.00000 ; N- CA- C- O
3197 3199 3211 3213 3 17.40544 -5.02080 -13.22144
9.20480 0.00000 0.00000 ; N- CA- C- N
3198 3197 3199 3200 3 0.00000 0.00000 0.00000
0.00000 0.00000 0.00000 ; H- N- CA- HA
3198 3197 3199 3201 3 0.00000 0.00000 0.00000
0.00000 0.00000 0.00000 ; H- N- CA- CB
3198 3197 3199 3211 3 0.00000 0.00000 0.00000

```
0.00000    0.00000    0.00000 ;      H-     N-     CA-     C
  3199   3201   3203   3204      3    0.66944    2.00832    0.00000
-2.67776    0.00000    0.00000 ;      CA-    CB-   CG1-  HG11
  3199   3201   3203   3205      3    0.66944    2.00832    0.00000
-2.67776    0.00000    0.00000 ;      CA-    CB-   CG1-  HG12
  3199   3201   3203   3206      3    0.66944    2.00832    0.00000
-2.67776    0.00000    0.00000 ;      CA-    CB-   CG1-  HG13
  3199   3201   3207   3208      3    0.66944    2.00832    0.00000
-2.67776    0.00000    0.00000 ;      CA-    CB-   CG2-  HG21
  3199   3201   3207   3209      3    0.66944    2.00832    0.00000
-2.67776    0.00000    0.00000 ;      CA-    CB-   CG2-  HG22
  3199   3201   3207   3210      3    0.66944    2.00832    0.00000
-2.67776    0.00000    0.00000 ;      CA-    CB-   CG2-  HG23
  3199   3211   3213   3214      3   20.92000    0.00000  -20.92000
0.00000    0.00000    0.00000 ;     CA-     C-     N-     H
  3199   3211   3213   3215      3   20.92000    0.00000  -20.92000
0.00000    0.00000    0.00000 ;     CA-     C-     N-    CA
  3200   3199   3201   3202      3    0.65084    1.95253    0.00000
-2.60338    0.00000    0.00000 ;      HA-    CA-    CB-    HB
  3200   3199   3201   3203      3    0.65084    1.95253    0.00000
-2.60338    0.00000    0.00000 ;      HA-    CA-    CB-   CG1
  3200   3199   3201   3207      3    0.65084    1.95253    0.00000
-2.60338    0.00000    0.00000 ;      HA-    CA-    CB-   CG2
  3200   3199   3211   3212      3    3.68192   -4.35136    0.00000
1.33888    0.00000    0.00000 ;     HA-    CA-     C-     O
  3200   3199   3211   3213      3    0.00000    0.00000    0.00000
0.00000    0.00000    0.00000 ;     HA-    CA-     C-     N
  3201   3199   3211   3212      3    0.00000    0.00000    0.00000
0.00000    0.00000    0.00000 ;     CB-    CA-     C-     O
  3201   3199   3211   3213      3    2.51040    4.18400    1.67360
-6.69440    0.00000    0.00000 ;      CB-    CA-     C-     N
  3202   3201   3199   3211      3    0.65084    1.95253    0.00000
-2.60338    0.00000    0.00000 ;      HB-    CB-    CA-     C
  3202   3201   3203   3204      3    0.62760    1.88280    0.00000
-2.51040    0.00000    0.00000 ;      HB-    CB-   CG1-  HG11
  3202   3201   3203   3205      3    0.62760    1.88280    0.00000
-2.51040    0.00000    0.00000 ;      HB-    CB-   CG1-  HG12
  3202   3201   3203   3206      3    0.62760    1.88280    0.00000
-2.51040    0.00000    0.00000 ;      HB-    CB-   CG1-  HG13
  3202   3201   3207   3208      3    0.62760    1.88280    0.00000
-2.51040    0.00000    0.00000 ;      HB-    CB-   CG2-  HG21
  3202   3201   3207   3209      3    0.62760    1.88280    0.00000
-2.51040    0.00000    0.00000 ;      HB-    CB-   CG2-  HG22
  3202   3201   3207   3210      3    0.62760    1.88280    0.00000
-2.51040    0.00000    0.00000 ;      HB-    CB-   CG2-  HG23
  3203   3201   3199   3211      3    5.67350    3.55640   -6.16722
-2.47693    3.74886    0.00000 ;     CG1-    CB-    CA-     C
  3203   3201   3207   3208      3    0.66944    2.00832    0.00000
-2.67776    0.00000    0.00000 ;     CG1-    CB-   CG2-  HG21
  3203   3201   3207   3209      3    0.66944    2.00832    0.00000
-2.67776    0.00000    0.00000 ;     CG1-    CB-   CG2-  HG22
  3203   3201   3207   3210      3    0.66944    2.00832    0.00000
```

-2.67776    0.00000    0.00000 ;    CG1-    CB-   CG2-  HG23
3204   3203   3201   3207      3    0.66944    2.00832    0.00000
-2.67776    0.00000    0.00000 ;   HG11-   CG1-    CB-   CG2
3205   3203   3201   3207      3    0.66944    2.00832    0.00000
-2.67776    0.00000    0.00000 ;   HG12-   CG1-    CB-   CG2
3206   3203   3201   3207      3    0.66944    2.00832    0.00000
-2.67776    0.00000    0.00000 ;   HG13-   CG1-    CB-   CG2
3207   3201   3199   3211      3    5.67350    3.55640   -6.16722
-2.47693    3.74886    0.00000 ;    CG2-    CB-    CA-     C
3211   3213   3215   3216      3    0.00000    0.00000    0.00000
0.00000    0.00000    0.00000 ;      C-     N-    CA-    HA
3211   3213   3215   3217      3   11.71520    1.67360   15.06240  -
13.38880    0.00000    0.00000 ;      C-     N-    CA-    CB
3211   3213   3215   3230      3    1.75728    5.27184    2.25936
-7.02912    0.00000    0.00000 ;      C-     N-    CA-     C
3212   3211   3213   3214      3   29.28800   -8.36800  -20.92000
0.00000    0.00000    0.00000 ;      O-     C-     N-     H
3212   3211   3213   3215      3   20.92000    0.00000  -20.92000
0.00000    0.00000    0.00000 ;      O-     C-     N-    CA
3213   3215   3217   3218      3    0.65084    1.95253    0.00000
-2.60338    0.00000    0.00000 ;      N-    CA-    CB-    HB
3213   3215   3217   3219      3    3.83673    0.44769   -1.77402
-2.47693   -0.03347    0.00000 ;      N-    CA-    CB-   CG2
3213   3215   3217   3223      3    2.97482    0.12134    2.04179
-1.89117   -3.24678    0.00000 ;      N-    CA-    CB-   CG1
3213   3215   3230   3231      3    0.00000    0.00000    0.00000
0.00000    0.00000    0.00000 ;      N-    CA-     C-     O
3213   3215   3230   3232      3   17.40544   -5.02080  -13.22144
9.20480    0.00000    0.00000 ;      N-    CA-     C-     N
3214   3213   3215   3216      3    0.00000    0.00000    0.00000
0.00000    0.00000    0.00000 ;      H-     N-    CA-    HA
3214   3213   3215   3217      3    0.00000    0.00000    0.00000
0.00000    0.00000    0.00000 ;      H-     N-    CA-    CB
3214   3213   3215   3230      3    0.00000    0.00000    0.00000
0.00000    0.00000    0.00000 ;      H-     N-    CA-     C
3215   3217   3219   3220      3    0.66944    2.00832    0.00000
-2.67776    0.00000    0.00000 ;     CA-    CB-   CG2-  HG21
3215   3217   3219   3221      3    0.66944    2.00832    0.00000
-2.67776    0.00000    0.00000 ;     CA-    CB-   CG2-  HG22
3215   3217   3219   3222      3    0.66944    2.00832    0.00000
-2.67776    0.00000    0.00000 ;     CA-    CB-   CG2-  HG23
3215   3217   3223   3224      3    0.66944    2.00832    0.00000
-2.67776    0.00000    0.00000 ;     CA-    CB-   CG1-  HG12
3215   3217   3223   3225      3    0.66944    2.00832    0.00000
-2.67776    0.00000    0.00000 ;     CA-    CB-   CG1-  HG13
3215   3217   3223   3226      3    4.24258   -0.52718   -7.25506
-1.79075    7.69856    0.00000 ;     CA-    CB-   CG1-   CD1
3215   3230   3232   3233      3   20.92000    0.00000  -20.92000
0.00000    0.00000    0.00000 ;     CA-     C-     N-     H
3215   3230   3232   3234      3   20.92000    0.00000  -20.92000
0.00000    0.00000    0.00000 ;     CA-     C-     N-    CA
3216   3215   3217   3218      3    0.65084    1.95253    0.00000

```
-2.60338    0.00000    0.00000 ;      HA-    CA-     CB-    HB
  3216   3215   3217   3219       3    0.65084    1.95253    0.00000
-2.60338    0.00000    0.00000 ;      HA-    CA-     CB-   CG2
  3216   3215   3217   3223       3    0.65084    1.95253    0.00000
-2.60338    0.00000    0.00000 ;      HA-    CA-     CB-   CG1
  3216   3215   3230   3231       3    3.68192   -4.35136    0.00000
1.33888    0.00000    0.00000 ;      HA-    CA-     C-     O
  3216   3215   3230   3232       3    0.00000    0.00000    0.00000
0.00000    0.00000    0.00000 ;      HA-    CA-     C-     N
  3217   3215   3230   3231       3    0.00000    0.00000    0.00000
0.00000    0.00000    0.00000 ;      CB-    CA-     C-     O
  3217   3215   3230   3232       3    2.51040    4.18400    1.67360
-6.69440    0.00000    0.00000 ;      CB-    CA-     C-     N
  3217   3223   3226   3227       3    0.66944    2.00832    0.00000
-2.67776    0.00000    0.00000 ;      CB-   CG1-   CD1-  HD11
  3217   3223   3226   3228       3    0.66944    2.00832    0.00000
-2.67776    0.00000    0.00000 ;      CB-   CG1-   CD1-  HD12
  3217   3223   3226   3229       3    0.66944    2.00832    0.00000
-2.67776    0.00000    0.00000 ;      CB-   CG1-   CD1-  HD13
  3218   3217   3215   3230       3    0.65084    1.95253    0.00000
-2.60338    0.00000    0.00000 ;      HB-    CB-     CA-     C
  3218   3217   3219   3220       3    0.62760    1.88280    0.00000
-2.51040    0.00000    0.00000 ;      HB-    CB-   CG2-  HG21
  3218   3217   3219   3221       3    0.62760    1.88280    0.00000
-2.51040    0.00000    0.00000 ;      HB-    CB-   CG2-  HG22
  3218   3217   3219   3222       3    0.62760    1.88280    0.00000
-2.51040    0.00000    0.00000 ;      HB-    CB-   CG2-  HG23
  3218   3217   3223   3224       3    0.62760    1.88280    0.00000
-2.51040    0.00000    0.00000 ;      HB-    CB-   CG1-  HG12
  3218   3217   3223   3225       3    0.62760    1.88280    0.00000
-2.51040    0.00000    0.00000 ;      HB-    CB-   CG1-  HG13
  3218   3217   3223   3226       3    0.66944    2.00832    0.00000
-2.67776    0.00000    0.00000 ;      HB-    CB-   CG1-   CD1
  3219   3217   3215   3230       3    5.67350    3.55640   -6.16722
-2.47693    3.74886    0.00000 ;    CG2-    CB-     CA-     C
  3219   3217   3223   3224       3    0.66944    2.00832    0.00000
-2.67776    0.00000    0.00000 ;    CG2-    CB-   CG1-  HG12
  3219   3217   3223   3225       3    0.66944    2.00832    0.00000
-2.67776    0.00000    0.00000 ;    CG2-    CB-   CG1-  HG13
  3219   3217   3223   3226       3    3.81162    0.49790   -8.14206
-1.79075    7.49773    0.00000 ;    CG2-    CB-   CG1-   CD1
  3220   3219   3217   3223       3    0.66944    2.00832    0.00000
-2.67776    0.00000    0.00000 ;   HG21-   CG2-     CB-   CG1
  3221   3219   3217   3223       3    0.66944    2.00832    0.00000
-2.67776    0.00000    0.00000 ;   HG22-   CG2-     CB-   CG1
  3222   3219   3217   3223       3    0.66944    2.00832    0.00000
-2.67776    0.00000    0.00000 ;   HG23-   CG2-     CB-   CG1
  3223   3217   3215   3230       3    8.26340    0.74057   -9.99976
-1.89117    3.84928    0.00000 ;    CG1-    CB-     CA-     C
  3224   3223   3226   3227       3    0.62760    1.88280    0.00000
-2.51040    0.00000    0.00000 ;   HG12-   CG1-   CD1-  HD11
  3224   3223   3226   3228       3    0.62760    1.88280    0.00000
```

-2.51040 0.00000 0.00000 ; HG12- CG1- CD1- HD12
3224 3223 3226 3229 3 0.62760 1.88280 0.00000
-2.51040 0.00000 0.00000 ; HG12- CG1- CD1- HD13
3225 3223 3226 3227 3 0.62760 1.88280 0.00000
-2.51040 0.00000 0.00000 ; HG13- CG1- CD1- HD11
3225 3223 3226 3228 3 0.62760 1.88280 0.00000
-2.51040 0.00000 0.00000 ; HG13- CG1- CD1- HD12
3225 3223 3226 3229 3 0.62760 1.88280 0.00000
-2.51040 0.00000 0.00000 ; HG13- CG1- CD1- HD13
3230 3232 3234 3235 3 0.00000 0.00000 0.00000
0.00000 0.00000 0.00000 ; C- N- CA- HA
3230 3232 3234 3236 3 11.71520 1.67360 15.06240 -
13.38880 0.00000 0.00000 ; C- N- CA- CB
3230 3232 3234 3247 3 1.75728 5.27184 2.25936
-7.02912 0.00000 0.00000 ; C- N- CA- C
3231 3230 3232 3233 3 29.28800 -8.36800 -20.92000
0.00000 0.00000 0.00000 ; O- C- N- H
3231 3230 3232 3234 3 20.92000 0.00000 -20.92000
0.00000 0.00000 0.00000 ; O- C- N- CA
3232 3234 3236 3237 3 0.65084 1.95253 0.00000
-2.60338 0.00000 0.00000 ; N- CA- CB- HB2
3232 3234 3236 3238 3 0.65084 1.95253 0.00000
-2.60338 0.00000 0.00000 ; N- CA- CB- HB3
3232 3234 3236 3239 3 3.21331 2.22589 -4.15053
-2.40998 2.61082 0.00000 ; N- CA- CB- CG
3232 3234 3247 3248 3 0.00000 0.00000 0.00000
0.00000 0.00000 0.00000 ; N- CA- C- O
3232 3234 3247 3249 3 17.40544 -5.02080 -13.22144
9.20480 0.00000 0.00000 ; N- CA- C- N
3233 3232 3234 3235 3 0.00000 0.00000 0.00000
0.00000 0.00000 0.00000 ; H- N- CA- HA
3233 3232 3234 3236 3 0.00000 0.00000 0.00000
0.00000 0.00000 0.00000 ; H- N- CA- CB
3233 3232 3234 3247 3 0.00000 0.00000 0.00000
0.00000 0.00000 0.00000 ; H- N- CA- C
3234 3236 3239 3240 3 0.65084 1.95253 0.00000
-2.60338 0.00000 0.00000 ; CA- CB- CG- HG2
3234 3236 3239 3241 3 0.65084 1.95253 0.00000
-2.60338 0.00000 0.00000 ; CA- CB- CG- HG3
3234 3236 3239 3242 3 2.04598 -1.54390 1.11294
-0.26778 0.93722 0.00000 ; CA- CB- CG- SD
3234 3247 3249 3250 3 20.92000 0.00000 -20.92000
0.00000 0.00000 0.00000 ; CA- C- N- H
3234 3247 3249 3251 3 20.92000 0.00000 -20.92000
0.00000 0.00000 0.00000 ; CA- C- N- CA
3235 3234 3236 3237 3 0.65084 1.95253 0.00000
-2.60338 0.00000 0.00000 ; HA- CA- CB- HB2
3235 3234 3236 3238 3 0.65084 1.95253 0.00000
-2.60338 0.00000 0.00000 ; HA- CA- CB- HB3
3235 3234 3236 3239 3 0.65084 1.95253 0.00000
-2.60338 0.00000 0.00000 ; HA- CA- CB- CG
3235 3234 3247 3248 3 3.68192 -4.35136 0.00000

```
1.33888    0.00000    0.00000 ;     HA-    CA-     C-     O
  3235   3234   3247   3249      3    0.00000    0.00000    0.00000
0.00000    0.00000    0.00000 ;     HA-    CA-     C-     N
  3236   3234   3247   3248      3    0.00000    0.00000    0.00000
0.00000    0.00000    0.00000 ;     CB-    CA-     C-     O
  3236   3234   3247   3249      3    2.51040    4.18400    1.67360
-6.69440    0.00000    0.00000 ;     CB-    CA-     C-     N
  3236   3239   3242   3243      3    3.24260    6.22998    1.79075
-6.92870    1.90790    0.00000 ;     CB-    CG-    SD-    CE
  3237   3236   3234   3247      3    0.65084    1.95253    0.00000
-2.60338    0.00000    0.00000 ;    HB2-    CB-    CA-     C
  3237   3236   3239   3240      3    0.65084    1.95253    0.00000
-2.60338    0.00000    0.00000 ;    HB2-    CB-    CG-   HG2
  3237   3236   3239   3241      3    0.65084    1.95253    0.00000
-2.60338    0.00000    0.00000 ;    HB2-    CB-    CG-   HG3
  3237   3236   3239   3242      3    0.65084    1.95253    0.00000
-2.60338    0.00000    0.00000 ;    HB2-    CB-    CG-    SD
  3238   3236   3234   3247      3    0.65084    1.95253    0.00000
-2.60338    0.00000    0.00000 ;    HB3-    CB-    CA-     C
  3238   3236   3239   3240      3    0.65084    1.95253    0.00000
-2.60338    0.00000    0.00000 ;    HB3-    CB-    CG-   HG2
  3238   3236   3239   3241      3    0.65084    1.95253    0.00000
-2.60338    0.00000    0.00000 ;    HB3-    CB-    CG-   HG3
  3238   3236   3239   3242      3    0.65084    1.95253    0.00000
-2.60338    0.00000    0.00000 ;    HB3-    CB-    CG-    SD
  3239   3236   3234   3247      3    6.86594    3.56895   -8.14206
-2.40998    4.85344    0.00000 ;     CG-    CB-    CA-     C
  3239   3242   3243   3244      3    1.39467    4.18400    0.00000
-5.57867    0.00000    0.00000 ;     CG-    SD-    CE-   HE1
  3239   3242   3243   3245      3    1.39467    4.18400    0.00000
-5.57867    0.00000    0.00000 ;     CG-    SD-    CE-   HE2
  3239   3242   3243   3246      3    1.39467    4.18400    0.00000
-5.57867    0.00000    0.00000 ;     CG-    SD-    CE-   HE3
  3240   3239   3242   3243      3    1.39467    4.18400    0.00000
-5.57867    0.00000    0.00000 ;    HG2-    CG-    SD-    CE
  3241   3239   3242   3243      3    1.39467    4.18400    0.00000
-5.57867    0.00000    0.00000 ;    HG3-    CG-    SD-    CE
  3247   3249   3251   3252      3    0.00000    0.00000    0.00000
0.00000    0.00000    0.00000 ;     C-     N-    CA-    HA
  3247   3249   3251   3253      3   11.71520    1.67360   15.06240  -
13.38880    0.00000    0.00000 ;     C-     N-    CA-    CB
  3247   3249   3251   3268      3    1.75728    5.27184    2.25936
-7.02912    0.00000    0.00000 ;     C-     N-    CA-     C
  3248   3247   3249   3250      3   29.28800   -8.36800  -20.92000
0.00000    0.00000    0.00000 ;     O-     C-     N-     H
  3248   3247   3249   3251      3   20.92000    0.00000  -20.92000
0.00000    0.00000    0.00000 ;     O-     C-     N-    CA
  3249   3251   3253   3254      3    0.65084    1.95253    0.00000
-2.60338    0.00000    0.00000 ;     N-    CA-    CB-   HB2
  3249   3251   3253   3255      3    0.65084    1.95253    0.00000
-2.60338    0.00000    0.00000 ;     N-    CA-    CB-   HB3
  3249   3251   3253   3256      3    3.28026    2.46019   -2.19242
```

-3.21331 -0.23430 0.00000 ; N- CA- CB- CG
3249 3251 3268 3269 3 0.00000 0.00000 0.00000
0.00000 0.00000 0.00000 ; N- CA- C- O
3249 3251 3268 3270 3 17.40544 -5.02080 -13.22144
9.20480 0.00000 0.00000 ; N- CA- C- N
3250 3249 3251 3252 3 0.00000 0.00000 0.00000
0.00000 0.00000 0.00000 ; H- N- CA- HA
3250 3249 3251 3253 3 0.00000 0.00000 0.00000
0.00000 0.00000 0.00000 ; H- N- CA- CB
3250 3249 3251 3268 3 0.00000 0.00000 0.00000
0.00000 0.00000 0.00000 ; H- N- CA- C
3251 3253 3256 3257 3 0.57739 0.00000 1.02926
0.00000 -1.60666 0.00000 ; CA- CB- CG- CD1
3251 3253 3256 3266 3 0.57739 0.00000 1.02926
0.00000 -1.60666 0.00000 ; CA- CB- CG- CD2
3251 3268 3270 3271 3 20.92000 0.00000 -20.92000
0.00000 0.00000 0.00000 ; CA- C- N- H
3251 3268 3270 3272 3 20.92000 0.00000 -20.92000
0.00000 0.00000 0.00000 ; CA- C- N- CA
3252 3251 3253 3254 3 0.65084 1.95253 0.00000
-2.60338 0.00000 0.00000 ; HA- CA- CB- HB2
3252 3251 3253 3255 3 0.65084 1.95253 0.00000
-2.60338 0.00000 0.00000 ; HA- CA- CB- HB3
3252 3251 3253 3256 3 0.65084 1.95253 0.00000
-2.60338 0.00000 0.00000 ; HA- CA- CB- CG
3252 3251 3268 3269 3 3.68192 -4.35136 0.00000
1.33888 0.00000 0.00000 ; HA- CA- C- O
3252 3251 3268 3270 3 0.00000 0.00000 0.00000
0.00000 0.00000 0.00000 ; HA- CA- C- N
3253 3251 3268 3269 3 0.00000 0.00000 0.00000
0.00000 0.00000 0.00000 ; CB- CA- C- O
3253 3251 3268 3270 3 2.51040 4.18400 1.67360
-6.69440 0.00000 0.00000 ; CB- CA- C- N
3253 3256 3257 3258 3 30.33400 0.00000 -30.33400
0.00000 0.00000 0.00000 ; CB- CG- CD1- HD1
3253 3256 3257 3259 3 30.33400 0.00000 -30.33400
0.00000 0.00000 0.00000 ; CB- CG- CD1- CE1
3253 3256 3266 3264 3 30.33400 0.00000 -30.33400
0.00000 0.00000 0.00000 ; CB- CG- CD2- CE2
3253 3256 3266 3267 3 30.33400 0.00000 -30.33400
0.00000 0.00000 0.00000 ; CB- CG- CD2- HD2
3254 3253 3251 3268 3 0.65084 1.95253 0.00000
-2.60338 0.00000 0.00000 ; HB2- CB- CA- C
3254 3253 3256 3257 3 0.00000 0.00000 0.00000
0.00000 0.00000 0.00000 ; HB2- CB- CG- CD1
3254 3253 3256 3266 3 0.00000 0.00000 0.00000
0.00000 0.00000 0.00000 ; HB2- CB- CG- CD2
3255 3253 3251 3268 3 0.65084 1.95253 0.00000
-2.60338 0.00000 0.00000 ; HB3- CB- CA- C
3255 3253 3256 3257 3 0.00000 0.00000 0.00000
0.00000 0.00000 0.00000 ; HB3- CB- CG- CD1
3255 3253 3256 3266 3 0.00000 0.00000 0.00000

0.00000 0.00000 0.00000 ; HB3- CB- CG- CD2
3256 3253 3251 3268 3 4.95804 2.17986 -3.52293
-3.21331 -0.40166 0.00000 ; CG- CB- CA- C
3256 3257 3259 3260 3 30.33400 0.00000 -30.33400
0.00000 0.00000 0.00000 ; CG- CD1- CE1- HE1
3256 3257 3259 3261 3 30.33400 0.00000 -30.33400
0.00000 0.00000 0.00000 ; CG- CD1- CE1- CZ
3256 3266 3264 3261 3 30.33400 0.00000 -30.33400
0.00000 0.00000 0.00000 ; CG- CD2- CE2- CZ
3256 3266 3264 3265 3 30.33400 0.00000 -30.33400
0.00000 0.00000 0.00000 ; CG- CD2- CE2- HE2
3257 3256 3266 3264 3 30.33400 0.00000 -30.33400
0.00000 0.00000 0.00000 ; CD1- CG- CD2- CE2
3257 3256 3266 3267 3 30.33400 0.00000 -30.33400
0.00000 0.00000 0.00000 ; CD1- CG- CD2- HD2
3257 3259 3261 3262 3 30.33400 0.00000 -30.33400
0.00000 0.00000 0.00000 ; CD1- CE1- CZ- OH
3257 3259 3261 3264 3 30.33400 0.00000 -30.33400
0.00000 0.00000 0.00000 ; CD1- CE1- CZ- CE2
3258 3257 3256 3266 3 30.33400 0.00000 -30.33400
0.00000 0.00000 0.00000 ; HD1- CD1- CG- CD2
3258 3257 3259 3260 3 30.33400 0.00000 -30.33400
0.00000 0.00000 0.00000 ; HD1- CD1- CE1- HE1
3258 3257 3259 3261 3 30.33400 0.00000 -30.33400
0.00000 0.00000 0.00000 ; HD1- CD1- CE1- CZ
3259 3257 3256 3266 3 30.33400 0.00000 -30.33400
0.00000 0.00000 0.00000 ; CE1- CD1- CG- CD2
3259 3261 3262 3263 3 7.93286 0.00000 -9.56462
0.00000 2.17568 0.00000 ; CE1- CZ- OH- HH
3259 3261 3264 3265 3 30.33400 0.00000 -30.33400
0.00000 0.00000 0.00000 ; CE1- CZ- CE2- HE2
3259 3261 3264 3266 3 30.33400 0.00000 -30.33400
0.00000 0.00000 0.00000 ; CE1- CZ- CE2- CD2
3260 3259 3261 3262 3 30.33400 0.00000 -30.33400
0.00000 0.00000 0.00000 ; HE1- CE1- CZ- OH
3260 3259 3261 3264 3 30.33400 0.00000 -30.33400
0.00000 0.00000 0.00000 ; HE1- CE1- CZ- CE2
3261 3264 3266 3267 3 30.33400 0.00000 -30.33400
0.00000 0.00000 0.00000 ; CZ- CE2- CD2- HD2
3262 3261 3264 3265 3 30.33400 0.00000 -30.33400
0.00000 0.00000 0.00000 ; OH- CZ- CE2- HE2
3262 3261 3264 3266 3 30.33400 0.00000 -30.33400
0.00000 0.00000 0.00000 ; OH- CZ- CE2- CD2
3263 3262 3261 3264 3 7.93286 0.00000 -9.56462
0.00000 2.17568 0.00000 ; HH- OH- CZ- CE2
3265 3264 3266 3267 3 30.33400 0.00000 -30.33400
0.00000 0.00000 0.00000 ; HE2- CE2- CD2- HD2
3268 3270 3272 3273 3 0.00000 0.00000 0.00000
0.00000 0.00000 0.00000 ; C- N- CA- HA
3268 3270 3272 3274 3 11.71520 1.67360 15.06240 -
13.38880 0.00000 0.00000 ; C- N- CA- CB
3268 3270 3272 3282 3 1.75728 5.27184 2.25936

-7.02912 0.00000 0.00000 ; C- N- CA- C
3269 3268 3270 3271 3 29.28800 -8.36800 -20.92000
0.00000 0.00000 0.00000 ; O- C- N- H
3269 3268 3270 3272 3 20.92000 0.00000 -20.92000
0.00000 0.00000 0.00000 ; O- C- N- CA
3270 3272 3274 3275 3 0.65084 1.95253 0.00000
-2.60338 0.00000 0.00000 ; N- CA- CB- HB
3270 3272 3274 3276 3 3.83673 0.44769 -1.77402
-2.47693 -0.03347 0.00000 ; N- CA- CB- CG2
3270 3272 3274 3280 3 4.93294 1.13386 -3.12963
-5.27184 3.17984 0.00000 ; N- CA- CB- OG1
3270 3272 3282 3283 3 0.00000 0.00000 0.00000
0.00000 0.00000 0.00000 ; N- CA- C- O
3270 3272 3282 3284 3 17.40544 -5.02080 -13.22144
9.20480 0.00000 0.00000 ; N- CA- C- N
3271 3270 3272 3273 3 0.00000 0.00000 0.00000
0.00000 0.00000 0.00000 ; H- N- CA- HA
3271 3270 3272 3274 3 0.00000 0.00000 0.00000
0.00000 0.00000 0.00000 ; H- N- CA- CB
3271 3270 3272 3282 3 0.00000 0.00000 0.00000
0.00000 0.00000 0.00000 ; H- N- CA- C
3272 3274 3276 3277 3 0.66944 2.00832 0.00000
-2.67776 0.00000 0.00000 ; CA- CB- CG2- HG21
3272 3274 3276 3278 3 0.66944 2.00832 0.00000
-2.67776 0.00000 0.00000 ; CA- CB- CG2- HG22
3272 3274 3276 3279 3 0.66944 2.00832 0.00000
-2.67776 0.00000 0.00000 ; CA- CB- CG2- HG23
3272 3274 3280 3281 3 1.12131 2.98738 1.66523
-3.94970 0.43514 0.00000 ; CA- CB- OG1- HG1
3272 3282 3284 3285 3 20.92000 0.00000 -20.92000
0.00000 0.00000 0.00000 ; CA- C- N- H
3272 3282 3284 3286 3 20.92000 0.00000 -20.92000
0.00000 0.00000 0.00000 ; CA- C- N- CA
3273 3272 3274 3275 3 0.65084 1.95253 0.00000
-2.60338 0.00000 0.00000 ; HA- CA- CB- HB
3273 3272 3274 3276 3 0.65084 1.95253 0.00000
-2.60338 0.00000 0.00000 ; HA- CA- CB- CG2
3273 3272 3274 3280 3 1.04600 -1.04600 0.00000
0.00000 0.00000 0.00000 ; HA- CA- CB- OG1
3273 3272 3282 3283 3 3.68192 -4.35136 0.00000
1.33888 0.00000 0.00000 ; HA- CA- C- O
3273 3272 3282 3284 3 0.00000 0.00000 0.00000
0.00000 0.00000 0.00000 ; HA- CA- C- N
3274 3272 3282 3283 3 0.00000 0.00000 0.00000
0.00000 0.00000 0.00000 ; CB- CA- C- O
3274 3272 3282 3284 3 2.51040 4.18400 1.67360
-6.69440 0.00000 0.00000 ; CB- CA- C- N
3275 3274 3272 3282 3 0.65084 1.95253 0.00000
-2.60338 0.00000 0.00000 ; HB- CB- CA- C
3275 3274 3276 3277 3 0.65084 1.95253 0.00000
-2.60338 0.00000 0.00000 ; HB- CB- CG2- HG21
3275 3274 3276 3278 3 0.65084 1.95253 0.00000

-2.60338 0.00000 0.00000 ; HB- CB- CG2- HG22
3275 3274 3276 3279 3 0.65084 1.95253 0.00000
-2.60338 0.00000 0.00000 ; HB- CB- CG2- HG23
3275 3274 3280 3281 3 0.69733 2.09200 0.00000
-2.78933 0.00000 0.00000 ; HB- CB- OG1- HG1
3276 3274 3272 3282 3 5.67350 3.55640 -6.16722
-2.47693 3.74886 0.00000 ; CG2- CB- CA- C
3276 3274 3280 3281 3 4.74047 0.27196 -2.26773
-3.94970 1.60666 0.00000 ; CG2- CB- OG1- HG1
3277 3276 3274 3280 3 1.04600 -1.04600 0.00000
0.00000 0.00000 0.00000 ; HG21- CG2- CB- OG1
3278 3276 3274 3280 3 1.04600 -1.04600 0.00000
0.00000 0.00000 0.00000 ; HG22- CG2- CB- OG1
3279 3276 3274 3280 3 1.04600 -1.04600 0.00000
0.00000 0.00000 0.00000 ; HG23- CG2- CB- OG1
3280 3274 3272 3282 3 6.53541 6.87013 -6.21742
-5.27184 5.22163 0.00000 ; OG1- CB- CA- C
3282 3284 3286 3287 3 0.00000 0.00000 0.00000
0.00000 0.00000 0.00000 ; C- N- CA- HA
3282 3284 3286 3288 3 11.71520 1.67360 15.06240 -
13.38880 0.00000 0.00000 ; C- N- CA- CB
3282 3284 3286 3301 3 1.75728 5.27184 2.25936
-7.02912 0.00000 0.00000 ; C- N- CA- C
3283 3282 3284 3285 3 29.28800 -8.36800 -20.92000
0.00000 0.00000 0.00000 ; O- C- N- H
3283 3282 3284 3286 3 20.92000 0.00000 -20.92000
0.00000 0.00000 0.00000 ; O- C- N- CA
3284 3286 3288 3289 3 0.65084 1.95253 0.00000
-2.60338 0.00000 0.00000 ; N- CA- CB- HB2
3284 3286 3288 3290 3 0.65084 1.95253 0.00000
-2.60338 0.00000 0.00000 ; N- CA- CB- HB3
3284 3286 3288 3291 3 3.79070 1.39746 -4.61077
-2.40998 2.44346 0.00000 ; N- CA- CB- CG
3284 3286 3301 3302 3 0.00000 0.00000 0.00000
0.00000 0.00000 0.00000 ; N- CA- C- O
3284 3286 3301 3303 3 17.40544 -5.02080 -13.22144
9.20480 0.00000 0.00000 ; N- CA- C- N
3285 3284 3286 3287 3 0.00000 0.00000 0.00000
0.00000 0.00000 0.00000 ; H- N- CA- HA
3285 3284 3286 3288 3 0.00000 0.00000 0.00000
0.00000 0.00000 0.00000 ; H- N- CA- CB
3285 3284 3286 3301 3 0.00000 0.00000 0.00000
0.00000 0.00000 0.00000 ; H- N- CA- C
3286 3288 3291 3292 3 0.66944 2.00832 0.00000
-2.67776 0.00000 0.00000 ; CA- CB- CG- HG
3286 3288 3291 3293 3 3.90367 0.19665 -6.21742
-2.37651 5.99149 0.00000 ; CA- CB- CG- CD1
3286 3288 3291 3297 3 3.90367 0.19665 -6.21742
-2.37651 5.99149 0.00000 ; CA- CB- CG- CD2
3286 3301 3303 3304 3 20.92000 0.00000 -20.92000
0.00000 0.00000 0.00000 ; CA- C- N- H
3286 3301 3303 3305 3 20.92000 0.00000 -20.92000

```
0.00000    0.00000    0.00000 ;      CA-     C-      N-    CA
  3287   3286   3288   3289       3    0.65084    1.95253    0.00000
-2.60338    0.00000    0.00000 ;      HA-    CA-     CB-   HB2
  3287   3286   3288   3290       3    0.65084    1.95253    0.00000
-2.60338    0.00000    0.00000 ;      HA-    CA-     CB-   HB3
  3287   3286   3288   3291       3    0.65084    1.95253    0.00000
-2.60338    0.00000    0.00000 ;      HA-    CA-     CB-    CG
  3287   3286   3301   3302       3    3.68192   -4.35136    0.00000
1.33888    0.00000    0.00000 ;      HA-    CA-      C-     O
  3287   3286   3301   3303       3    0.00000    0.00000    0.00000
0.00000    0.00000    0.00000 ;      HA-    CA-      C-     N
  3288   3286   3301   3302       3    0.00000    0.00000    0.00000
0.00000    0.00000    0.00000 ;      CB-    CA-      C-     O
  3288   3286   3301   3303       3    2.51040    4.18400    1.67360
-6.69440    0.00000    0.00000 ;      CB-    CA-      C-     N
  3288   3291   3293   3294       3    0.66944    2.00832    0.00000
-2.67776    0.00000    0.00000 ;      CB-    CG-    CD1-  HD11
  3288   3291   3293   3295       3    0.66944    2.00832    0.00000
-2.67776    0.00000    0.00000 ;      CB-    CG-    CD1-  HD12
  3288   3291   3293   3296       3    0.66944    2.00832    0.00000
-2.67776    0.00000    0.00000 ;      CB-    CG-    CD1-  HD13
  3288   3291   3297   3298       3    0.66944    2.00832    0.00000
-2.67776    0.00000    0.00000 ;      CB-    CG-    CD2-  HD21
  3288   3291   3297   3299       3    0.66944    2.00832    0.00000
-2.67776    0.00000    0.00000 ;      CB-    CG-    CD2-  HD22
  3288   3291   3297   3300       3    0.66944    2.00832    0.00000
-2.67776    0.00000    0.00000 ;      CB-    CG-    CD2-  HD23
  3289   3288   3286   3301       3    0.65084    1.95253    0.00000
-2.60338    0.00000    0.00000 ;     HB2-    CB-     CA-     C
  3289   3288   3291   3292       3    0.62760    1.88280    0.00000
-2.51040    0.00000    0.00000 ;     HB2-    CB-     CG-    HG
  3289   3288   3291   3293       3    0.66944    2.00832    0.00000
-2.67776    0.00000    0.00000 ;     HB2-    CB-     CG-   CD1
  3289   3288   3291   3297       3    0.66944    2.00832    0.00000
-2.67776    0.00000    0.00000 ;     HB2-    CB-     CG-   CD2
  3290   3288   3286   3301       3    0.65084    1.95253    0.00000
-2.60338    0.00000    0.00000 ;     HB3-    CB-     CA-     C
  3290   3288   3291   3292       3    0.62760    1.88280    0.00000
-2.51040    0.00000    0.00000 ;     HB3-    CB-     CG-    HG
  3290   3288   3291   3293       3    0.66944    2.00832    0.00000
-2.67776    0.00000    0.00000 ;     HB3-    CB-     CG-   CD1
  3290   3288   3291   3297       3    0.66944    2.00832    0.00000
-2.67776    0.00000    0.00000 ;     HB3-    CB-     CG-   CD2
  3291   3288   3286   3301       3   10.33448   -1.14642  -11.54784
-2.40998    6.35968    0.00000 ;      CG-    CB-     CA-     C
  3292   3291   3293   3294       3    0.62760    1.88280    0.00000
-2.51040    0.00000    0.00000 ;      HG-    CG-    CD1-  HD11
  3292   3291   3293   3295       3    0.62760    1.88280    0.00000
-2.51040    0.00000    0.00000 ;      HG-    CG-    CD1-  HD12
  3292   3291   3293   3296       3    0.62760    1.88280    0.00000
-2.51040    0.00000    0.00000 ;      HG-    CG-    CD1-  HD13
  3292   3291   3297   3298       3    0.62760    1.88280    0.00000
```

-2.51040 0.00000 0.00000 ; HG- CG- CD2- HD21
3292 3291 3297 3299 3 0.62760 1.88280 0.00000
-2.51040 0.00000 0.00000 ; HG- CG- CD2- HD22
3292 3291 3297 3300 3 0.62760 1.88280 0.00000
-2.51040 0.00000 0.00000 ; HG- CG- CD2- HD23
3293 3291 3297 3298 3 0.66944 2.00832 0.00000
-2.67776 0.00000 0.00000 ; CD1- CG- CD2- HD21
3293 3291 3297 3299 3 0.66944 2.00832 0.00000
-2.67776 0.00000 0.00000 ; CD1- CG- CD2- HD22
3293 3291 3297 3300 3 0.66944 2.00832 0.00000
-2.67776 0.00000 0.00000 ; CD1- CG- CD2- HD23
3294 3293 3291 3297 3 0.66944 2.00832 0.00000
-2.67776 0.00000 0.00000 ; HD11- CD1- CG- CD2
3295 3293 3291 3297 3 0.66944 2.00832 0.00000
-2.67776 0.00000 0.00000 ; HD12- CD1- CG- CD2
3296 3293 3291 3297 3 0.66944 2.00832 0.00000
-2.67776 0.00000 0.00000 ; HD13- CD1- CG- CD2
3301 3303 3305 3306 3 0.00000 0.00000 0.00000
0.00000 0.00000 0.00000 ; C- N- CA- HA
3301 3303 3305 3307 3 11.71520 1.67360 15.06240 -
13.38880 0.00000 0.00000 ; C- N- CA- CB
3301 3303 3305 3320 3 1.75728 5.27184 2.25936
-7.02912 0.00000 0.00000 ; C- N- CA- C
3302 3301 3303 3304 3 29.28800 -8.36800 -20.92000
0.00000 0.00000 0.00000 ; O- C- N- H
3302 3301 3303 3305 3 20.92000 0.00000 -20.92000
0.00000 0.00000 0.00000 ; O- C- N- CA
3303 3305 3307 3308 3 0.65084 1.95253 0.00000
-2.60338 0.00000 0.00000 ; N- CA- CB- HB2
3303 3305 3307 3309 3 0.65084 1.95253 0.00000
-2.60338 0.00000 0.00000 ; N- CA- CB- HB3
3303 3305 3307 3310 3 3.79070 1.39746 -4.61077
-2.40998 2.44346 0.00000 ; N- CA- CB- CG
3303 3305 3320 3321 3 0.00000 0.00000 0.00000
0.00000 0.00000 0.00000 ; N- CA- C- O
3303 3305 3320 3322 3 17.40544 -5.02080 -13.22144
9.20480 0.00000 0.00000 ; N- CA- C- N
3304 3303 3305 3306 3 0.00000 0.00000 0.00000
0.00000 0.00000 0.00000 ; H- N- CA- HA
3304 3303 3305 3307 3 0.00000 0.00000 0.00000
0.00000 0.00000 0.00000 ; H- N- CA- CB
3304 3303 3305 3320 3 0.00000 0.00000 0.00000
0.00000 0.00000 0.00000 ; H- N- CA- C
3305 3307 3310 3311 3 0.66944 2.00832 0.00000
-2.67776 0.00000 0.00000 ; CA- CB- CG- HG
3305 3307 3310 3312 3 3.90367 0.19665 -6.21742
-2.37651 5.99149 0.00000 ; CA- CB- CG- CD1
3305 3307 3310 3316 3 3.90367 0.19665 -6.21742
-2.37651 5.99149 0.00000 ; CA- CB- CG- CD2
3305 3320 3322 3323 3 20.92000 0.00000 -20.92000
0.00000 0.00000 0.00000 ; CA- C- N- H
3305 3320 3322 3324 3 20.92000 0.00000 -20.92000

```
0.00000    0.00000    0.00000 ;      CA-     C-     N-    CA
  3306   3305   3307   3308      3    0.65084    1.95253    0.00000
-2.60338    0.00000    0.00000 ;      HA-    CA-    CB-   HB2
  3306   3305   3307   3309      3    0.65084    1.95253    0.00000
-2.60338    0.00000    0.00000 ;      HA-    CA-    CB-   HB3
  3306   3305   3307   3310      3    0.65084    1.95253    0.00000
-2.60338    0.00000    0.00000 ;      HA-    CA-    CB-    CG
  3306   3305   3320   3321      3    3.68192   -4.35136    0.00000
1.33888    0.00000    0.00000 ;      HA-    CA-     C-     O
  3306   3305   3320   3322      3    0.00000    0.00000    0.00000
0.00000    0.00000    0.00000 ;      HA-    CA-     C-     N
  3307   3305   3320   3321      3    0.00000    0.00000    0.00000
0.00000    0.00000    0.00000 ;      CB-    CA-     C-     O
  3307   3305   3320   3322      3    2.51040    4.18400    1.67360
-6.69440    0.00000    0.00000 ;      CB-    CA-     C-     N
  3307   3310   3312   3313      3    0.66944    2.00832    0.00000
-2.67776    0.00000    0.00000 ;      CB-    CG-   CD1-  HD11
  3307   3310   3312   3314      3    0.66944    2.00832    0.00000
-2.67776    0.00000    0.00000 ;      CB-    CG-   CD1-  HD12
  3307   3310   3312   3315      3    0.66944    2.00832    0.00000
-2.67776    0.00000    0.00000 ;      CB-    CG-   CD1-  HD13
  3307   3310   3316   3317      3    0.66944    2.00832    0.00000
-2.67776    0.00000    0.00000 ;      CB-    CG-   CD2-  HD21
  3307   3310   3316   3318      3    0.66944    2.00832    0.00000
-2.67776    0.00000    0.00000 ;      CB-    CG-   CD2-  HD22
  3307   3310   3316   3319      3    0.66944    2.00832    0.00000
-2.67776    0.00000    0.00000 ;      CB-    CG-   CD2-  HD23
  3308   3307   3305   3320      3    0.65084    1.95253    0.00000
-2.60338    0.00000    0.00000 ;     HB2-    CB-    CA-     C
  3308   3307   3310   3311      3    0.62760    1.88280    0.00000
-2.51040    0.00000    0.00000 ;     HB2-    CB-    CG-    HG
  3308   3307   3310   3312      3    0.66944    2.00832    0.00000
-2.67776    0.00000    0.00000 ;     HB2-    CB-    CG-   CD1
  3308   3307   3310   3316      3    0.66944    2.00832    0.00000
-2.67776    0.00000    0.00000 ;     HB2-    CB-    CG-   CD2
  3309   3307   3305   3320      3    0.65084    1.95253    0.00000
-2.60338    0.00000    0.00000 ;     HB3-    CB-    CA-     C
  3309   3307   3310   3311      3    0.62760    1.88280    0.00000
-2.51040    0.00000    0.00000 ;     HB3-    CB-    CG-    HG
  3309   3307   3310   3312      3    0.66944    2.00832    0.00000
-2.67776    0.00000    0.00000 ;     HB3-    CB-    CG-   CD1
  3309   3307   3310   3316      3    0.66944    2.00832    0.00000
-2.67776    0.00000    0.00000 ;     HB3-    CB-    CG-   CD2
  3310   3307   3305   3320      3   10.33448   -1.14642  -11.54784
-2.40998    6.35968    0.00000 ;      CG-    CB-    CA-     C
  3311   3310   3312   3313      3    0.62760    1.88280    0.00000
-2.51040    0.00000    0.00000 ;      HG-    CG-   CD1-  HD11
  3311   3310   3312   3314      3    0.62760    1.88280    0.00000
-2.51040    0.00000    0.00000 ;      HG-    CG-   CD1-  HD12
  3311   3310   3312   3315      3    0.62760    1.88280    0.00000
-2.51040    0.00000    0.00000 ;      HG-    CG-   CD1-  HD13
  3311   3310   3316   3317      3    0.62760    1.88280    0.00000
```

-2.51040 0.00000 0.00000 ; HG- CG- CD2- HD21
3311 3310 3316 3318 3 0.62760 1.88280 0.00000
-2.51040 0.00000 0.00000 ; HG- CG- CD2- HD22
3311 3310 3316 3319 3 0.62760 1.88280 0.00000
-2.51040 0.00000 0.00000 ; HG- CG- CD2- HD23
3312 3310 3316 3317 3 0.66944 2.00832 0.00000
-2.67776 0.00000 0.00000 ; CD1- CG- CD2- HD21
3312 3310 3316 3318 3 0.66944 2.00832 0.00000
-2.67776 0.00000 0.00000 ; CD1- CG- CD2- HD22
3312 3310 3316 3319 3 0.66944 2.00832 0.00000
-2.67776 0.00000 0.00000 ; CD1- CG- CD2- HD23
3313 3312 3310 3316 3 0.66944 2.00832 0.00000
-2.67776 0.00000 0.00000 ; HD11- CD1- CG- CD2
3314 3312 3310 3316 3 0.66944 2.00832 0.00000
-2.67776 0.00000 0.00000 ; HD12- CD1- CG- CD2
3315 3312 3310 3316 3 0.66944 2.00832 0.00000
-2.67776 0.00000 0.00000 ; HD13- CD1- CG- CD2
3320 3322 3324 3325 3 0.00000 0.00000 0.00000
0.00000 0.00000 0.00000 ; C- N- CA- HA
3320 3322 3324 3326 3 11.71520 1.67360 15.06240 -
13.38880 0.00000 0.00000 ; C- N- CA- CB
3320 3322 3324 3331 3 1.75728 5.27184 2.25936
-7.02912 0.00000 0.00000 ; C- N- CA- C
3321 3320 3322 3323 3 29.28800 -8.36800 -20.92000
0.00000 0.00000 0.00000 ; O- C- N- H
3321 3320 3322 3324 3 20.92000 0.00000 -20.92000
0.00000 0.00000 0.00000 ; O- C- N- CA
3322 3324 3326 3327 3 0.65084 1.95253 0.00000
-2.60338 0.00000 0.00000 ; N- CA- CB- HB2
3322 3324 3326 3328 3 0.65084 1.95253 0.00000
-2.60338 0.00000 0.00000 ; N- CA- CB- HB3
3322 3324 3326 3329 3 6.03751 2.50622 -5.17142
-4.20074 1.10458 0.00000 ; N- CA- CB- SG
3322 3324 3331 3332 3 0.00000 0.00000 0.00000
0.00000 0.00000 0.00000 ; N- CA- C- O
3322 3324 3331 3333 3 17.40544 -5.02080 -13.22144
9.20480 0.00000 0.00000 ; N- CA- C- N
3323 3322 3324 3325 3 0.00000 0.00000 0.00000
0.00000 0.00000 0.00000 ; H- N- CA- HA
3323 3322 3324 3326 3 0.00000 0.00000 0.00000
0.00000 0.00000 0.00000 ; H- N- CA- CB
3323 3322 3324 3331 3 0.00000 0.00000 0.00000
0.00000 0.00000 0.00000 ; H- N- CA- C
3324 3326 3329 3330 3 1.69034 2.77818 4.11706
-4.21747 1.00416 0.00000 ; CA- CB- SG- HG
3324 3331 3333 3334 3 20.92000 0.00000 -20.92000
0.00000 0.00000 0.00000 ; CA- C- N- H
3324 3331 3333 3335 3 20.92000 0.00000 -20.92000
0.00000 0.00000 0.00000 ; CA- C- N- CA
3325 3324 3326 3327 3 0.65084 1.95253 0.00000
-2.60338 0.00000 0.00000 ; HA- CA- CB- HB2
3325 3324 3326 3328 3 0.65084 1.95253 0.00000

```
-2.60338    0.00000    0.00000 ;     HA-    CA-    CB-   HB3
  3325   3324   3326   3329      3    0.65084    1.95253    0.00000
-2.60338    0.00000    0.00000 ;     HA-    CA-    CB-    SG
  3325   3324   3331   3332      3    3.68192   -4.35136    0.00000
1.33888    0.00000    0.00000 ;     HA-    CA-     C-     O
  3325   3324   3331   3333      3    0.00000    0.00000    0.00000
0.00000    0.00000    0.00000 ;     HA-    CA-     C-     N
  3326   3324   3331   3332      3    0.00000    0.00000    0.00000
0.00000    0.00000    0.00000 ;     CB-    CA-     C-     O
  3326   3324   3331   3333      3    2.51040    4.18400    1.67360
-6.69440    0.00000    0.00000 ;     CB-    CA-     C-     N
  3327   3326   3324   3331      3    0.65084    1.95253    0.00000
-2.60338    0.00000    0.00000 ;    HB2-    CB-    CA-     C
  3327   3326   3329   3330      3    1.04600    3.13800    0.00000
-4.18400    0.00000    0.00000 ;    HB2-    CB-    SG-    HG
  3328   3326   3324   3331      3    0.65084    1.95253    0.00000
-2.60338    0.00000    0.00000 ;    HB3-    CB-    CA-     C
  3328   3326   3329   3330      3    1.04600    3.13800    0.00000
-4.18400    0.00000    0.00000 ;    HB3-    CB-    SG-    HG
  3329   3326   3324   3331      3    5.62330    4.27605   -5.33042
-4.20074    2.51040    0.00000 ;     SG-    CB-    CA-     C
  3331   3333   3335   3336      3    0.00000    0.00000    0.00000
0.00000    0.00000    0.00000 ;      C-     N-    CA-   HA2
  3331   3333   3335   3337      3    0.00000    0.00000    0.00000
0.00000    0.00000    0.00000 ;      C-     N-    CA-   HA3
  3331   3333   3335   3338      3    1.75728    5.27184    2.25936
-7.02912    0.00000    0.00000 ;      C-     N-    CA-     C
  3332   3331   3333   3334      3   29.28800   -8.36800  -20.92000
0.00000    0.00000    0.00000 ;      O-     C-     N-     H
  3332   3331   3333   3335      3   20.92000    0.00000  -20.92000
0.00000    0.00000    0.00000 ;      O-     C-     N-    CA
  3333   3335   3338   3339      3    0.00000    0.00000    0.00000
0.00000    0.00000    0.00000 ;      N-    CA-     C-     O
  3333   3335   3338   3340      3   17.40544   -5.02080  -13.22144
9.20480    0.00000    0.00000 ;      N-    CA-     C-     N
  3334   3333   3335   3336      3    0.00000    0.00000    0.00000
0.00000    0.00000    0.00000 ;      H-     N-    CA-   HA2
  3334   3333   3335   3337      3    0.00000    0.00000    0.00000
0.00000    0.00000    0.00000 ;      H-     N-    CA-   HA3
  3334   3333   3335   3338      3    0.00000    0.00000    0.00000
0.00000    0.00000    0.00000 ;      H-     N-    CA-     C
  3335   3338   3340   3341      3   20.92000    0.00000  -20.92000
0.00000    0.00000    0.00000 ;     CA-     C-     N-     H
  3335   3338   3340   3342      3   20.92000    0.00000  -20.92000
0.00000    0.00000    0.00000 ;     CA-     C-     N-    CA
  3336   3335   3338   3339      3    3.68192   -4.35136    0.00000
1.33888    0.00000    0.00000 ;    HA2-    CA-     C-     O
  3336   3335   3338   3340      3    0.00000    0.00000    0.00000
0.00000    0.00000    0.00000 ;    HA2-    CA-     C-     N
  3337   3335   3338   3339      3    3.68192   -4.35136    0.00000
1.33888    0.00000    0.00000 ;    HA3-    CA-     C-     O
  3337   3335   3338   3340      3    0.00000    0.00000    0.00000
```

```
0.00000    0.00000    0.00000 ;    HA3-    CA-     C-     N
  3338   3340   3342   3343      3    0.00000    0.00000    0.00000
0.00000    0.00000    0.00000 ;      C-     N-    CA-    HA
  3338   3340   3342   3344      3   11.71520    1.67360   15.06240  -
13.38880    0.00000    0.00000 ;      C-     N-    CA-    CB
  3338   3340   3342   3359      3    1.75728    5.27184    2.25936
-7.02912    0.00000    0.00000 ;      C-     N-    CA-     C
  3339   3338   3340   3341      3   29.28800   -8.36800  -20.92000
0.00000    0.00000    0.00000 ;      O-     C-     N-     H
  3339   3338   3340   3342      3   20.92000    0.00000  -20.92000
0.00000    0.00000    0.00000 ;      O-     C-     N-    CA
  3340   3342   3344   3345      3    0.65084    1.95253    0.00000
-2.60338    0.00000    0.00000 ;      N-    CA-    CB-   HB2
  3340   3342   3344   3346      3    0.65084    1.95253    0.00000
-2.60338    0.00000    0.00000 ;      N-    CA-    CB-   HB3
  3340   3342   3344   3347      3    3.28026    2.46019   -2.19242
-3.21331   -0.23430    0.00000 ;      N-    CA-    CB-    CG
  3340   3342   3359   3360      3    0.00000    0.00000    0.00000
0.00000    0.00000    0.00000 ;      N-    CA-     C-     O
  3340   3342   3359   3361      3   17.40544   -5.02080  -13.22144
9.20480    0.00000    0.00000 ;      N-    CA-     C-     N
  3341   3340   3342   3343      3    0.00000    0.00000    0.00000
0.00000    0.00000    0.00000 ;      H-     N-    CA-    HA
  3341   3340   3342   3344      3    0.00000    0.00000    0.00000
0.00000    0.00000    0.00000 ;      H-     N-    CA-    CB
  3341   3340   3342   3359      3    0.00000    0.00000    0.00000
0.00000    0.00000    0.00000 ;      H-     N-    CA-     C
  3342   3344   3347   3348      3    0.57739    0.00000    1.02926
0.00000   -1.60666    0.00000 ;     CA-    CB-    CG-   CD1
  3342   3344   3347   3357      3    0.57739    0.00000    1.02926
0.00000   -1.60666    0.00000 ;     CA-    CB-    CG-   CD2
  3342   3359   3361   3362      3   20.92000    0.00000  -20.92000
0.00000    0.00000    0.00000 ;     CA-     C-     N-    CD
  3342   3359   3361   3371      3   20.92000    0.00000  -20.92000
0.00000    0.00000    0.00000 ;     CA-     C-     N-    CA
  3343   3342   3344   3345      3    0.65084    1.95253    0.00000
-2.60338    0.00000    0.00000 ;     HA-    CA-    CB-   HB2
  3343   3342   3344   3346      3    0.65084    1.95253    0.00000
-2.60338    0.00000    0.00000 ;     HA-    CA-    CB-   HB3
  3343   3342   3344   3347      3    0.65084    1.95253    0.00000
-2.60338    0.00000    0.00000 ;     HA-    CA-    CB-    CG
  3343   3342   3359   3360      3    3.68192   -4.35136    0.00000
1.33888    0.00000    0.00000 ;     HA-    CA-     C-     O
  3343   3342   3359   3361      3    0.00000    0.00000    0.00000
0.00000    0.00000    0.00000 ;     HA-    CA-     C-     N
  3344   3342   3359   3360      3    0.00000    0.00000    0.00000
0.00000    0.00000    0.00000 ;     CB-    CA-     C-     O
  3344   3342   3359   3361      3    2.51040    4.18400    1.67360
-6.69440    0.00000    0.00000 ;     CB-    CA-     C-     N
  3344   3347   3348   3349      3   30.33400    0.00000  -30.33400
0.00000    0.00000    0.00000 ;     CB-    CG-   CD1-   HD1
  3344   3347   3348   3350      3   30.33400    0.00000  -30.33400
```

```
0.00000    0.00000    0.00000 ;     CB-    CG-   CD1-   CE1
  3344   3347   3357   3355      3   30.33400    0.00000  -30.33400
0.00000    0.00000    0.00000 ;     CB-    CG-   CD2-   CE2
  3344   3347   3357   3358      3   30.33400    0.00000  -30.33400
0.00000    0.00000    0.00000 ;     CB-    CG-   CD2-   HD2
  3345   3344   3342   3359      3    0.65084    1.95253    0.00000
-2.60338    0.00000    0.00000 ;    HB2-    CB-    CA-     C
  3345   3344   3347   3348      3    0.00000    0.00000    0.00000
0.00000    0.00000    0.00000 ;    HB2-    CB-    CG-   CD1
  3345   3344   3347   3357      3    0.00000    0.00000    0.00000
0.00000    0.00000    0.00000 ;    HB2-    CB-    CG-   CD2
  3346   3344   3342   3359      3    0.65084    1.95253    0.00000
-2.60338    0.00000    0.00000 ;    HB3-    CB-    CA-     C
  3346   3344   3347   3348      3    0.00000    0.00000    0.00000
0.00000    0.00000    0.00000 ;    HB3-    CB-    CG-   CD1
  3346   3344   3347   3357      3    0.00000    0.00000    0.00000
0.00000    0.00000    0.00000 ;    HB3-    CB-    CG-   CD2
  3347   3344   3342   3359      3    4.95804    2.17986   -3.52293
-3.21331   -0.40166    0.00000 ;     CG-    CB-    CA-     C
  3347   3348   3350   3351      3   30.33400    0.00000  -30.33400
0.00000    0.00000    0.00000 ;     CG-   CD1-   CE1-   HE1
  3347   3348   3350   3352      3   30.33400    0.00000  -30.33400
0.00000    0.00000    0.00000 ;     CG-   CD1-   CE1-    CZ
  3347   3357   3355   3352      3   30.33400    0.00000  -30.33400
0.00000    0.00000    0.00000 ;     CG-   CD2-   CE2-    CZ
  3347   3357   3355   3356      3   30.33400    0.00000  -30.33400
0.00000    0.00000    0.00000 ;     CG-   CD2-   CE2-   HE2
  3348   3347   3357   3355      3   30.33400    0.00000  -30.33400
0.00000    0.00000    0.00000 ;    CD1-    CG-   CD2-   CE2
  3348   3347   3357   3358      3   30.33400    0.00000  -30.33400
0.00000    0.00000    0.00000 ;    CD1-    CG-   CD2-   HD2
  3348   3350   3352   3353      3   30.33400    0.00000  -30.33400
0.00000    0.00000    0.00000 ;    CD1-   CE1-    CZ-    OH
  3348   3350   3352   3355      3   30.33400    0.00000  -30.33400
0.00000    0.00000    0.00000 ;    CD1-   CE1-    CZ-   CE2
  3349   3348   3347   3357      3   30.33400    0.00000  -30.33400
0.00000    0.00000    0.00000 ;    HD1-   CD1-    CG-   CD2
  3349   3348   3350   3351      3   30.33400    0.00000  -30.33400
0.00000    0.00000    0.00000 ;    HD1-   CD1-   CE1-   HE1
  3349   3348   3350   3352      3   30.33400    0.00000  -30.33400
0.00000    0.00000    0.00000 ;    HD1-   CD1-   CE1-    CZ
  3350   3348   3347   3357      3   30.33400    0.00000  -30.33400
0.00000    0.00000    0.00000 ;    CE1-   CD1-    CG-   CD2
  3350   3352   3353   3354      3    7.93286    0.00000   -9.56462
0.00000    2.17568    0.00000 ;    CE1-    CZ-    OH-    HH
  3350   3352   3355   3356      3   30.33400    0.00000  -30.33400
0.00000    0.00000    0.00000 ;    CE1-    CZ-   CE2-   HE2
  3350   3352   3355   3357      3   30.33400    0.00000  -30.33400
0.00000    0.00000    0.00000 ;    CE1-    CZ-   CE2-   CD2
  3351   3350   3352   3353      3   30.33400    0.00000  -30.33400
0.00000    0.00000    0.00000 ;    HE1-   CE1-    CZ-    OH
  3351   3350   3352   3355      3   30.33400    0.00000  -30.33400
```

0.00000 0.00000 0.00000 ; HE1- CE1- CZ- CE2
3352 3355 3357 3358 3 30.33400 0.00000 -30.33400
0.00000 0.00000 0.00000 ; CZ- CE2- CD2- HD2
3353 3352 3355 3356 3 30.33400 0.00000 -30.33400
0.00000 0.00000 0.00000 ; OH- CZ- CE2- HE2
3353 3352 3355 3357 3 30.33400 0.00000 -30.33400
0.00000 0.00000 0.00000 ; OH- CZ- CE2- CD2
3354 3353 3352 3355 3 7.93286 0.00000 -9.56462
0.00000 2.17568 0.00000 ; HH- OH- CZ- CE2
3356 3355 3357 3358 3 30.33400 0.00000 -30.33400
0.00000 0.00000 0.00000 ; HE2- CE2- CD2- HD2
3359 3361 3362 3363 3 0.00000 0.00000 0.00000
0.00000 0.00000 0.00000 ; C- N- CD- HD2
3359 3361 3362 3364 3 0.00000 0.00000 0.00000
0.00000 0.00000 0.00000 ; C- N- CD- HD3
3359 3361 3362 3365 3 10.04160 -3.34720 16.73600
-6.69440 0.00000 0.00000 ; C- N- CD- CG
3359 3361 3371 3368 3 11.71520 1.67360 15.06240 -
13.38880 0.00000 0.00000 ; C- N- CA- CB
3359 3361 3371 3372 3 0.00000 0.00000 0.00000
0.00000 0.00000 0.00000 ; C- N- CA- HA
3359 3361 3371 3373 3 1.75728 5.27184 2.25936
-7.02912 0.00000 0.00000 ; C- N- CA- C
3360 3359 3361 3362 3 20.92000 0.00000 -20.92000
0.00000 0.00000 0.00000 ; O- C- N- CD
3360 3359 3361 3371 3 20.92000 0.00000 -20.92000
0.00000 0.00000 0.00000 ; O- C- N- CA
3361 3362 3365 3366 3 0.65084 1.95253 0.00000
-2.60338 0.00000 0.00000 ; N- CD- CG- HG2
3361 3362 3365 3367 3 0.65084 1.95253 0.00000
-2.60338 0.00000 0.00000 ; N- CD- CG- HG3
3361 3362 3365 3368 3 0.65084 1.95253 0.00000
-2.60338 0.00000 0.00000 ; N- CD- CG- CB
3361 3371 3368 3365 3 0.65084 1.95253 0.00000
-2.60338 0.00000 0.00000 ; N- CA- CB- CG
3361 3371 3368 3369 3 0.65084 1.95253 0.00000
-2.60338 0.00000 0.00000 ; N- CA- CB- HB2
3361 3371 3368 3370 3 0.65084 1.95253 0.00000
-2.60338 0.00000 0.00000 ; N- CA- CB- HB3
3361 3371 3373 3374 3 0.00000 0.00000 0.00000
0.00000 0.00000 0.00000 ; N- CA- C- O
3361 3371 3373 3375 3 17.40544 -5.02080 -13.22144
9.20480 0.00000 0.00000 ; N- CA- C- N
3362 3361 3371 3368 3 0.00000 0.00000 0.00000
0.00000 0.00000 0.00000 ; CD- N- CA- CB
3362 3361 3371 3372 3 0.00000 0.00000 0.00000
0.00000 0.00000 0.00000 ; CD- N- CA- HA
3362 3361 3371 3373 3 0.00000 0.00000 0.00000
0.00000 0.00000 0.00000 ; CD- N- CA- C
3362 3365 3368 3369 3 0.66944 2.00832 0.00000
-2.67776 0.00000 0.00000 ; CD- CG- CB- HB2
3362 3365 3368 3370 3 0.66944 2.00832 0.00000

```
-2.67776    0.00000    0.00000 ;      CD-    CG-    CB-   HB3
  3362   3365   3368   3371      3    3.68192    3.09616   -2.09200
-3.01248    0.00000    0.00000 ;      CD-    CG-    CB-    CA
  3363   3362   3361   3371      3    0.00000    0.00000    0.00000
0.00000    0.00000    0.00000 ;     HD2-    CD-     N-    CA
  3363   3362   3365   3366      3    0.65084    1.95253    0.00000
-2.60338    0.00000    0.00000 ;     HD2-    CD-    CG-   HG2
  3363   3362   3365   3367      3    0.65084    1.95253    0.00000
-2.60338    0.00000    0.00000 ;     HD2-    CD-    CG-   HG3
  3363   3362   3365   3368      3    0.65084    1.95253    0.00000
-2.60338    0.00000    0.00000 ;     HD2-    CD-    CG-    CB
  3364   3362   3361   3371      3    0.00000    0.00000    0.00000
0.00000    0.00000    0.00000 ;     HD3-    CD-     N-    CA
  3364   3362   3365   3366      3    0.65084    1.95253    0.00000
-2.60338    0.00000    0.00000 ;     HD3-    CD-    CG-   HG2
  3364   3362   3365   3367      3    0.65084    1.95253    0.00000
-2.60338    0.00000    0.00000 ;     HD3-    CD-    CG-   HG3
  3364   3362   3365   3368      3    0.65084    1.95253    0.00000
-2.60338    0.00000    0.00000 ;     HD3-    CD-    CG-    CB
  3365   3362   3361   3371      3    0.00000    0.00000    0.00000
0.00000    0.00000    0.00000 ;      CG-    CD-     N-    CA
  3365   3368   3371   3372      3    0.65084    1.95253    0.00000
-2.60338    0.00000    0.00000 ;      CG-    CB-    CA-    HA
  3365   3368   3371   3373      3    0.65084    1.95253    0.00000
-2.60338    0.00000    0.00000 ;      CG-    CB-    CA-     C
  3366   3365   3368   3369      3    0.62760    1.88280    0.00000
-2.51040    0.00000    0.00000 ;     HG2-    CG-    CB-   HB2
  3366   3365   3368   3370      3    0.62760    1.88280    0.00000
-2.51040    0.00000    0.00000 ;     HG2-    CG-    CB-   HB3
  3366   3365   3368   3371      3    0.66944    2.00832    0.00000
-2.67776    0.00000    0.00000 ;     HG2-    CG-    CB-    CA
  3367   3365   3368   3369      3    0.62760    1.88280    0.00000
-2.51040    0.00000    0.00000 ;     HG3-    CG-    CB-   HB2
  3367   3365   3368   3370      3    0.62760    1.88280    0.00000
-2.51040    0.00000    0.00000 ;     HG3-    CG-    CB-   HB3
  3367   3365   3368   3371      3    0.66944    2.00832    0.00000
-2.67776    0.00000    0.00000 ;     HG3-    CG-    CB-    CA
  3368   3371   3373   3374      3    0.00000    0.00000    0.00000
0.00000    0.00000    0.00000 ;      CB-    CA-     C-     O
  3368   3371   3373   3375      3    2.51040    4.18400    1.67360
-6.69440    0.00000    0.00000 ;      CB-    CA-     C-     N
  3369   3368   3371   3372      3    0.65084    1.95253    0.00000
-2.60338    0.00000    0.00000 ;     HB2-    CB-    CA-    HA
  3369   3368   3371   3373      3    0.65084    1.95253    0.00000
-2.60338    0.00000    0.00000 ;     HB2-    CB-    CA-     C
  3370   3368   3371   3372      3    0.65084    1.95253    0.00000
-2.60338    0.00000    0.00000 ;     HB3-    CB-    CA-    HA
  3370   3368   3371   3373      3    0.65084    1.95253    0.00000
-2.60338    0.00000    0.00000 ;     HB3-    CB-    CA-     C
  3371   3373   3375   3376      3   20.92000    0.00000  -20.92000
0.00000    0.00000    0.00000 ;      CA-     C-     N-    CD
  3371   3373   3375   3385      3   20.92000    0.00000  -20.92000
```

0.00000 0.00000 0.00000 ; CA- C- N- CA
3372 3371 3373 3374 3 3.68192 -4.35136 0.00000
1.33888 0.00000 0.00000 ; HA- CA- C- O
3372 3371 3373 3375 3 0.00000 0.00000 0.00000
0.00000 0.00000 0.00000 ; HA- CA- C- N
3373 3375 3376 3377 3 0.00000 0.00000 0.00000
0.00000 0.00000 0.00000 ; C- N- CD- HD2
3373 3375 3376 3378 3 0.00000 0.00000 0.00000
0.00000 0.00000 0.00000 ; C- N- CD- HD3
3373 3375 3376 3379 3 10.04160 -3.34720 16.73600
-6.69440 0.00000 0.00000 ; C- N- CD- CG
3373 3375 3385 3382 3 11.71520 1.67360 15.06240 -
13.38880 0.00000 0.00000 ; C- N- CA- CB
3373 3375 3385 3386 3 0.00000 0.00000 0.00000
0.00000 0.00000 0.00000 ; C- N- CA- HA
3373 3375 3385 3387 3 1.75728 5.27184 2.25936
-7.02912 0.00000 0.00000 ; C- N- CA- C
3374 3373 3375 3376 3 20.92000 0.00000 -20.92000
0.00000 0.00000 0.00000 ; O- C- N- CD
3374 3373 3375 3385 3 20.92000 0.00000 -20.92000
0.00000 0.00000 0.00000 ; O- C- N- CA
3375 3376 3379 3380 3 0.65084 1.95253 0.00000
-2.60338 0.00000 0.00000 ; N- CD- CG- HG2
3375 3376 3379 3381 3 0.65084 1.95253 0.00000
-2.60338 0.00000 0.00000 ; N- CD- CG- HG3
3375 3376 3379 3382 3 0.65084 1.95253 0.00000
-2.60338 0.00000 0.00000 ; N- CD- CG- CB
3375 3385 3382 3379 3 0.65084 1.95253 0.00000
-2.60338 0.00000 0.00000 ; N- CA- CB- CG
3375 3385 3382 3383 3 0.65084 1.95253 0.00000
-2.60338 0.00000 0.00000 ; N- CA- CB- HB2
3375 3385 3382 3384 3 0.65084 1.95253 0.00000
-2.60338 0.00000 0.00000 ; N- CA- CB- HB3
3375 3385 3387 3388 3 0.00000 0.00000 0.00000
0.00000 0.00000 0.00000 ; N- CA- C- O
3375 3385 3387 3389 3 17.40544 -5.02080 -13.22144
9.20480 0.00000 0.00000 ; N- CA- C- N
3376 3375 3385 3382 3 0.00000 0.00000 0.00000
0.00000 0.00000 0.00000 ; CD- N- CA- CB
3376 3375 3385 3386 3 0.00000 0.00000 0.00000
0.00000 0.00000 0.00000 ; CD- N- CA- HA
3376 3375 3385 3387 3 0.00000 0.00000 0.00000
0.00000 0.00000 0.00000 ; CD- N- CA- C
3376 3379 3382 3383 3 0.66944 2.00832 0.00000
-2.67776 0.00000 0.00000 ; CD- CG- CB- HB2
3376 3379 3382 3384 3 0.66944 2.00832 0.00000
-2.67776 0.00000 0.00000 ; CD- CG- CB- HB3
3376 3379 3382 3385 3 3.68192 3.09616 -2.09200
-3.01248 0.00000 0.00000 ; CD- CG- CB- CA
3377 3376 3375 3385 3 0.00000 0.00000 0.00000
0.00000 0.00000 0.00000 ; HD2- CD- N- CA
3377 3376 3379 3380 3 0.65084 1.95253 0.00000

```
-2.60338    0.00000    0.00000 ;    HD2-    CD-    CG-   HG2
  3377   3376   3379   3381      3    0.65084    1.95253    0.00000
-2.60338    0.00000    0.00000 ;    HD2-    CD-    CG-   HG3
  3377   3376   3379   3382      3    0.65084    1.95253    0.00000
-2.60338    0.00000    0.00000 ;    HD2-    CD-    CG-    CB
  3378   3376   3375   3385      3    0.00000    0.00000    0.00000
0.00000    0.00000    0.00000 ;    HD3-    CD-     N-    CA
  3378   3376   3379   3380      3    0.65084    1.95253    0.00000
-2.60338    0.00000    0.00000 ;    HD3-    CD-    CG-   HG2
  3378   3376   3379   3381      3    0.65084    1.95253    0.00000
-2.60338    0.00000    0.00000 ;    HD3-    CD-    CG-   HG3
  3378   3376   3379   3382      3    0.65084    1.95253    0.00000
-2.60338    0.00000    0.00000 ;    HD3-    CD-    CG-    CB
  3379   3376   3375   3385      3    0.00000    0.00000    0.00000
0.00000    0.00000    0.00000 ;     CG-    CD-     N-    CA
  3379   3382   3385   3386      3    0.65084    1.95253    0.00000
-2.60338    0.00000    0.00000 ;     CG-    CB-    CA-    HA
  3379   3382   3385   3387      3    0.65084    1.95253    0.00000
-2.60338    0.00000    0.00000 ;     CG-    CB-    CA-     C
  3380   3379   3382   3383      3    0.62760    1.88280    0.00000
-2.51040    0.00000    0.00000 ;    HG2-    CG-    CB-   HB2
  3380   3379   3382   3384      3    0.62760    1.88280    0.00000
-2.51040    0.00000    0.00000 ;    HG2-    CG-    CB-   HB3
  3380   3379   3382   3385      3    0.66944    2.00832    0.00000
-2.67776    0.00000    0.00000 ;    HG2-    CG-    CB-    CA
  3381   3379   3382   3383      3    0.62760    1.88280    0.00000
-2.51040    0.00000    0.00000 ;    HG3-    CG-    CB-   HB2
  3381   3379   3382   3384      3    0.62760    1.88280    0.00000
-2.51040    0.00000    0.00000 ;    HG3-    CG-    CB-   HB3
  3381   3379   3382   3385      3    0.66944    2.00832    0.00000
-2.67776    0.00000    0.00000 ;    HG3-    CG-    CB-    CA
  3382   3385   3387   3388      3    0.00000    0.00000    0.00000
0.00000    0.00000    0.00000 ;     CB-    CA-     C-     O
  3382   3385   3387   3389      3    2.51040    4.18400    1.67360
-6.69440    0.00000    0.00000 ;     CB-    CA-     C-     N
  3383   3382   3385   3386      3    0.65084    1.95253    0.00000
-2.60338    0.00000    0.00000 ;    HB2-    CB-    CA-    HA
  3383   3382   3385   3387      3    0.65084    1.95253    0.00000
-2.60338    0.00000    0.00000 ;    HB2-    CB-    CA-     C
  3384   3382   3385   3386      3    0.65084    1.95253    0.00000
-2.60338    0.00000    0.00000 ;    HB3-    CB-    CA-    HA
  3384   3382   3385   3387      3    0.65084    1.95253    0.00000
-2.60338    0.00000    0.00000 ;    HB3-    CB-    CA-     C
  3385   3387   3389   3390      3   20.92000    0.00000  -20.92000
0.00000    0.00000    0.00000 ;     CA-     C-     N-     H
  3385   3387   3389   3391      3   20.92000    0.00000  -20.92000
0.00000    0.00000    0.00000 ;     CA-     C-     N-    CA
  3386   3385   3387   3388      3    3.68192   -4.35136    0.00000
1.33888    0.00000    0.00000 ;     HA-    CA-     C-     O
  3386   3385   3387   3389      3    0.00000    0.00000    0.00000
0.00000    0.00000    0.00000 ;     HA-    CA-     C-     N
  3387   3389   3391   3392      3    0.00000    0.00000    0.00000
```

0.00000 0.00000 0.00000 ; C- N- CA- HA
3387 3389 3391 3393 3 11.71520 1.67360 15.06240 -
13.38880 0.00000 0.00000 ; C- N- CA- CB
3387 3389 3391 3407 3 1.75728 5.27184 2.25936
-7.02912 0.00000 0.00000 ; C- N- CA- C
3388 3387 3389 3390 3 29.28800 -8.36800 -20.92000
0.00000 0.00000 0.00000 ; O- C- N- H
3388 3387 3389 3391 3 20.92000 0.00000 -20.92000
0.00000 0.00000 0.00000 ; O- C- N- CA
3389 3391 3393 3394 3 0.65084 1.95253 0.00000
-2.60338 0.00000 0.00000 ; N- CA- CB- HB2
3389 3391 3393 3395 3 0.65084 1.95253 0.00000
-2.60338 0.00000 0.00000 ; N- CA- CB- HB3
3389 3391 3393 3396 3 3.28026 2.46019 -2.19242
-3.21331 -0.23430 0.00000 ; N- CA- CB- CG
3389 3391 3407 3408 3 0.00000 0.00000 0.00000
0.00000 0.00000 0.00000 ; N- CA- C- O
3389 3391 3407 3409 3 17.40544 -5.02080 -13.22144
9.20480 0.00000 0.00000 ; N- CA- C- N
3390 3389 3391 3392 3 0.00000 0.00000 0.00000
0.00000 0.00000 0.00000 ; H- N- CA- HA
3390 3389 3391 3393 3 0.00000 0.00000 0.00000
0.00000 0.00000 0.00000 ; H- N- CA- CB
3390 3389 3391 3407 3 0.00000 0.00000 0.00000
0.00000 0.00000 0.00000 ; H- N- CA- C
3391 3393 3396 3397 3 0.57739 0.00000 1.02926
0.00000 -1.60666 0.00000 ; CA- CB- CG- CD1
3391 3393 3396 3405 3 0.57739 0.00000 1.02926
0.00000 -1.60666 0.00000 ; CA- CB- CG- CD2
3391 3407 3409 3410 3 20.92000 0.00000 -20.92000
0.00000 0.00000 0.00000 ; CA- C- N- H
3391 3407 3409 3411 3 20.92000 0.00000 -20.92000
0.00000 0.00000 0.00000 ; CA- C- N- CA
3392 3391 3393 3394 3 0.65084 1.95253 0.00000
-2.60338 0.00000 0.00000 ; HA- CA- CB- HB2
3392 3391 3393 3395 3 0.65084 1.95253 0.00000
-2.60338 0.00000 0.00000 ; HA- CA- CB- HB3
3392 3391 3393 3396 3 0.65084 1.95253 0.00000
-2.60338 0.00000 0.00000 ; HA- CA- CB- CG
3392 3391 3407 3408 3 3.68192 -4.35136 0.00000
1.33888 0.00000 0.00000 ; HA- CA- C- O
3392 3391 3407 3409 3 0.00000 0.00000 0.00000
0.00000 0.00000 0.00000 ; HA- CA- C- N
3393 3391 3407 3408 3 0.00000 0.00000 0.00000
0.00000 0.00000 0.00000 ; CB- CA- C- O
3393 3391 3407 3409 3 2.51040 4.18400 1.67360
-6.69440 0.00000 0.00000 ; CB- CA- C- N
3393 3396 3397 3398 3 30.33400 0.00000 -30.33400
0.00000 0.00000 0.00000 ; CB- CG- CD1- HD1
3393 3396 3397 3399 3 30.33400 0.00000 -30.33400
0.00000 0.00000 0.00000 ; CB- CG- CD1- CE1
3393 3396 3405 3403 3 30.33400 0.00000 -30.33400

0.00000 0.00000 0.00000 ; CB- CG- CD2- CE2
3393 3396 3405 3406 3 30.33400 0.00000 -30.33400
0.00000 0.00000 0.00000 ; CB- CG- CD2- HD2
3394 3393 3391 3407 3 0.65084 1.95253 0.00000
-2.60338 0.00000 0.00000 ; HB2- CB- CA- C
3394 3393 3396 3397 3 0.00000 0.00000 0.00000
0.00000 0.00000 0.00000 ; HB2- CB- CG- CD1
3394 3393 3396 3405 3 0.00000 0.00000 0.00000
0.00000 0.00000 0.00000 ; HB2- CB- CG- CD2
3395 3393 3391 3407 3 0.65084 1.95253 0.00000
-2.60338 0.00000 0.00000 ; HB3- CB- CA- C
3395 3393 3396 3397 3 0.00000 0.00000 0.00000
0.00000 0.00000 0.00000 ; HB3- CB- CG- CD1
3395 3393 3396 3405 3 0.00000 0.00000 0.00000
0.00000 0.00000 0.00000 ; HB3- CB- CG- CD2
3396 3393 3391 3407 3 4.95804 2.17986 -3.52293
-3.21331 -0.40166 0.00000 ; CG- CB- CA- C
3396 3397 3399 3400 3 30.33400 0.00000 -30.33400
0.00000 0.00000 0.00000 ; CG- CD1- CE1- HE1
3396 3397 3399 3401 3 30.33400 0.00000 -30.33400
0.00000 0.00000 0.00000 ; CG- CD1- CE1- CZ
3396 3405 3403 3401 3 30.33400 0.00000 -30.33400
0.00000 0.00000 0.00000 ; CG- CD2- CE2- CZ
3396 3405 3403 3404 3 30.33400 0.00000 -30.33400
0.00000 0.00000 0.00000 ; CG- CD2- CE2- HE2
3397 3396 3405 3403 3 30.33400 0.00000 -30.33400
0.00000 0.00000 0.00000 ; CD1- CG- CD2- CE2
3397 3396 3405 3406 3 30.33400 0.00000 -30.33400
0.00000 0.00000 0.00000 ; CD1- CG- CD2- HD2
3397 3399 3401 3402 3 30.33400 0.00000 -30.33400
0.00000 0.00000 0.00000 ; CD1- CE1- CZ- HZ
3397 3399 3401 3403 3 30.33400 0.00000 -30.33400
0.00000 0.00000 0.00000 ; CD1- CE1- CZ- CE2
3398 3397 3396 3405 3 30.33400 0.00000 -30.33400
0.00000 0.00000 0.00000 ; HD1- CD1- CG- CD2
3398 3397 3399 3400 3 30.33400 0.00000 -30.33400
0.00000 0.00000 0.00000 ; HD1- CD1- CE1- HE1
3398 3397 3399 3401 3 30.33400 0.00000 -30.33400
0.00000 0.00000 0.00000 ; HD1- CD1- CE1- CZ
3399 3397 3396 3405 3 30.33400 0.00000 -30.33400
0.00000 0.00000 0.00000 ; CE1- CD1- CG- CD2
3399 3401 3403 3404 3 30.33400 0.00000 -30.33400
0.00000 0.00000 0.00000 ; CE1- CZ- CE2- HE2
3399 3401 3403 3405 3 30.33400 0.00000 -30.33400
0.00000 0.00000 0.00000 ; CE1- CZ- CE2- CD2
3400 3399 3401 3402 3 30.33400 0.00000 -30.33400
0.00000 0.00000 0.00000 ; HE1- CE1- CZ- HZ
3400 3399 3401 3403 3 30.33400 0.00000 -30.33400
0.00000 0.00000 0.00000 ; HE1- CE1- CZ- CE2
3401 3403 3405 3406 3 30.33400 0.00000 -30.33400
0.00000 0.00000 0.00000 ; CZ- CE2- CD2- HD2
3402 3401 3403 3404 3 30.33400 0.00000 -30.33400

0.00000 0.00000 0.00000 ; HZ- CZ- CE2- HE2
3402 3401 3403 3405 3 30.33400 0.00000 -30.33400
0.00000 0.00000 0.00000 ; HZ- CZ- CE2- CD2
3404 3403 3405 3406 3 30.33400 0.00000 -30.33400
0.00000 0.00000 0.00000 ; HE2- CE2- CD2- HD2
3407 3409 3411 3412 3 0.00000 0.00000 0.00000
0.00000 0.00000 0.00000 ; C- N- CA- HA
3407 3409 3411 3413 3 11.71520 1.67360 15.06240 -
13.38880 0.00000 0.00000 ; C- N- CA- CB
3407 3409 3411 3428 3 1.75728 5.27184 2.25936
-7.02912 0.00000 0.00000 ; C- N- CA- C
3408 3407 3409 3410 3 29.28800 -8.36800 -20.92000
0.00000 0.00000 0.00000 ; O- C- N- H
3408 3407 3409 3411 3 20.92000 0.00000 -20.92000
0.00000 0.00000 0.00000 ; O- C- N- CA
3409 3411 3413 3414 3 0.65084 1.95253 0.00000
-2.60338 0.00000 0.00000 ; N- CA- CB- HB2
3409 3411 3413 3415 3 0.65084 1.95253 0.00000
-2.60338 0.00000 0.00000 ; N- CA- CB- HB3
3409 3411 3413 3416 3 3.28026 2.46019 -2.19242
-3.21331 -0.23430 0.00000 ; N- CA- CB- CG
3409 3411 3428 3429 3 0.00000 0.00000 0.00000
0.00000 0.00000 0.00000 ; N- CA- C- O
3409 3411 3428 3430 3 17.40544 -5.02080 -13.22144
9.20480 0.00000 0.00000 ; N- CA- C- N
3410 3409 3411 3412 3 0.00000 0.00000 0.00000
0.00000 0.00000 0.00000 ; H- N- CA- HA
3410 3409 3411 3413 3 0.00000 0.00000 0.00000
0.00000 0.00000 0.00000 ; H- N- CA- CB
3410 3409 3411 3428 3 0.00000 0.00000 0.00000
0.00000 0.00000 0.00000 ; H- N- CA- C
3411 3413 3416 3417 3 0.57739 0.00000 1.02926
0.00000 -1.60666 0.00000 ; CA- CB- CG- CD1
3411 3413 3416 3426 3 0.57739 0.00000 1.02926
0.00000 -1.60666 0.00000 ; CA- CB- CG- CD2
3411 3428 3430 3431 3 20.92000 0.00000 -20.92000
0.00000 0.00000 0.00000 ; CA- C- N- H
3411 3428 3430 3432 3 20.92000 0.00000 -20.92000
0.00000 0.00000 0.00000 ; CA- C- N- CA
3412 3411 3413 3414 3 0.65084 1.95253 0.00000
-2.60338 0.00000 0.00000 ; HA- CA- CB- HB2
3412 3411 3413 3415 3 0.65084 1.95253 0.00000
-2.60338 0.00000 0.00000 ; HA- CA- CB- HB3
3412 3411 3413 3416 3 0.65084 1.95253 0.00000
-2.60338 0.00000 0.00000 ; HA- CA- CB- CG
3412 3411 3428 3429 3 3.68192 -4.35136 0.00000
1.33888 0.00000 0.00000 ; HA- CA- C- O
3412 3411 3428 3430 3 0.00000 0.00000 0.00000
0.00000 0.00000 0.00000 ; HA- CA- C- N
3413 3411 3428 3429 3 0.00000 0.00000 0.00000
0.00000 0.00000 0.00000 ; CB- CA- C- O
3413 3411 3428 3430 3 2.51040 4.18400 1.67360

-6.69440 0.00000 0.00000 ; CB- CA- C- N
3413 3416 3417 3418 3 30.33400 0.00000 -30.33400
0.00000 0.00000 0.00000 ; CB- CG- CD1- HD1
3413 3416 3417 3419 3 30.33400 0.00000 -30.33400
0.00000 0.00000 0.00000 ; CB- CG- CD1- CE1
3413 3416 3426 3424 3 30.33400 0.00000 -30.33400
0.00000 0.00000 0.00000 ; CB- CG- CD2- CE2
3413 3416 3426 3427 3 30.33400 0.00000 -30.33400
0.00000 0.00000 0.00000 ; CB- CG- CD2- HD2
3414 3413 3411 3428 3 0.65084 1.95253 0.00000
-2.60338 0.00000 0.00000 ; HB2- CB- CA- C
3414 3413 3416 3417 3 0.00000 0.00000 0.00000
0.00000 0.00000 0.00000 ; HB2- CB- CG- CD1
3414 3413 3416 3426 3 0.00000 0.00000 0.00000
0.00000 0.00000 0.00000 ; HB2- CB- CG- CD2
3415 3413 3411 3428 3 0.65084 1.95253 0.00000
-2.60338 0.00000 0.00000 ; HB3- CB- CA- C
3415 3413 3416 3417 3 0.00000 0.00000 0.00000
0.00000 0.00000 0.00000 ; HB3- CB- CG- CD1
3415 3413 3416 3426 3 0.00000 0.00000 0.00000
0.00000 0.00000 0.00000 ; HB3- CB- CG- CD2
3416 3413 3411 3428 3 4.95804 2.17986 -3.52293
-3.21331 -0.40166 0.00000 ; CG- CB- CA- C
3416 3417 3419 3420 3 30.33400 0.00000 -30.33400
0.00000 0.00000 0.00000 ; CG- CD1- CE1- HE1
3416 3417 3419 3421 3 30.33400 0.00000 -30.33400
0.00000 0.00000 0.00000 ; CG- CD1- CE1- CZ
3416 3426 3424 3421 3 30.33400 0.00000 -30.33400
0.00000 0.00000 0.00000 ; CG- CD2- CE2- CZ
3416 3426 3424 3425 3 30.33400 0.00000 -30.33400
0.00000 0.00000 0.00000 ; CG- CD2- CE2- HE2
3417 3416 3426 3424 3 30.33400 0.00000 -30.33400
0.00000 0.00000 0.00000 ; CD1- CG- CD2- CE2
3417 3416 3426 3427 3 30.33400 0.00000 -30.33400
0.00000 0.00000 0.00000 ; CD1- CG- CD2- HD2
3417 3419 3421 3422 3 30.33400 0.00000 -30.33400
0.00000 0.00000 0.00000 ; CD1- CE1- CZ- OH
3417 3419 3421 3424 3 30.33400 0.00000 -30.33400
0.00000 0.00000 0.00000 ; CD1- CE1- CZ- CE2
3418 3417 3416 3426 3 30.33400 0.00000 -30.33400
0.00000 0.00000 0.00000 ; HD1- CD1- CG- CD2
3418 3417 3419 3420 3 30.33400 0.00000 -30.33400
0.00000 0.00000 0.00000 ; HD1- CD1- CE1- HE1
3418 3417 3419 3421 3 30.33400 0.00000 -30.33400
0.00000 0.00000 0.00000 ; HD1- CD1- CE1- CZ
3419 3417 3416 3426 3 30.33400 0.00000 -30.33400
0.00000 0.00000 0.00000 ; CE1- CD1- CG- CD2
3419 3421 3422 3423 3 7.93286 0.00000 -9.56462
0.00000 2.17568 0.00000 ; CE1- CZ- OH- HH
3419 3421 3424 3425 3 30.33400 0.00000 -30.33400
0.00000 0.00000 0.00000 ; CE1- CZ- CE2- HE2
3419 3421 3424 3426 3 30.33400 0.00000 -30.33400

```
0.00000    0.00000    0.00000 ;    CE1-    CZ-   CE2-   CD2
  3420   3419   3421   3422      3   30.33400    0.00000  -30.33400
0.00000    0.00000    0.00000 ;    HE1-   CE1-    CZ-    OH
  3420   3419   3421   3424      3   30.33400    0.00000  -30.33400
0.00000    0.00000    0.00000 ;    HE1-   CE1-    CZ-   CE2
  3421   3424   3426   3427      3   30.33400    0.00000  -30.33400
0.00000    0.00000    0.00000 ;     CZ-   CE2-   CD2-   HD2
  3422   3421   3424   3425      3   30.33400    0.00000  -30.33400
0.00000    0.00000    0.00000 ;     OH-    CZ-   CE2-   HE2
  3422   3421   3424   3426      3   30.33400    0.00000  -30.33400
0.00000    0.00000    0.00000 ;     OH-    CZ-   CE2-   CD2
  3423   3422   3421   3424      3    7.93286    0.00000   -9.56462
0.00000    2.17568    0.00000 ;     HH-    OH-    CZ-   CE2
  3425   3424   3426   3427      3   30.33400    0.00000  -30.33400
0.00000    0.00000    0.00000 ;    HE2-   CE2-   CD2-   HD2
  3428   3430   3432   3433      3    0.00000    0.00000    0.00000
0.00000    0.00000    0.00000 ;      C-     N-    CA-   HA2
  3428   3430   3432   3434      3    0.00000    0.00000    0.00000
0.00000    0.00000    0.00000 ;      C-     N-    CA-   HA3
  3428   3430   3432   3435      3    1.75728    5.27184    2.25936
-7.02912    0.00000    0.00000 ;      C-     N-    CA-     C
  3429   3428   3430   3431      3   29.28800   -8.36800  -20.92000
0.00000    0.00000    0.00000 ;      O-     C-     N-     H
  3429   3428   3430   3432      3   20.92000    0.00000  -20.92000
0.00000    0.00000    0.00000 ;      O-     C-     N-    CA
  3430   3432   3435   3436      3    0.00000    0.00000    0.00000
0.00000    0.00000    0.00000 ;      N-    CA-     C-     O
  3430   3432   3435   3437      3   17.40544   -5.02080  -13.22144
9.20480    0.00000    0.00000 ;      N-    CA-     C-     N
  3431   3430   3432   3433      3    0.00000    0.00000    0.00000
0.00000    0.00000    0.00000 ;      H-     N-    CA-   HA2
  3431   3430   3432   3434      3    0.00000    0.00000    0.00000
0.00000    0.00000    0.00000 ;      H-     N-    CA-   HA3
  3431   3430   3432   3435      3    0.00000    0.00000    0.00000
0.00000    0.00000    0.00000 ;      H-     N-    CA-     C
  3432   3435   3437   3438      3   20.92000    0.00000  -20.92000
0.00000    0.00000    0.00000 ;     CA-     C-     N-     H
  3432   3435   3437   3439      3   20.92000    0.00000  -20.92000
0.00000    0.00000    0.00000 ;     CA-     C-     N-    CA
  3433   3432   3435   3436      3    3.68192   -4.35136    0.00000
1.33888    0.00000    0.00000 ;    HA2-    CA-     C-     O
  3433   3432   3435   3437      3    0.00000    0.00000    0.00000
0.00000    0.00000    0.00000 ;    HA2-    CA-     C-     N
  3434   3432   3435   3436      3    3.68192   -4.35136    0.00000
1.33888    0.00000    0.00000 ;    HA3-    CA-     C-     O
  3434   3432   3435   3437      3    0.00000    0.00000    0.00000
0.00000    0.00000    0.00000 ;    HA3-    CA-     C-     N
  3435   3437   3439   3440      3    0.00000    0.00000    0.00000
0.00000    0.00000    0.00000 ;      C-     N-    CA-    HA
  3435   3437   3439   3441      3   11.71520    1.67360   15.06240  -
13.38880    0.00000    0.00000 ;      C-     N-    CA-    CB
  3435   3437   3439   3447      3    1.75728    5.27184    2.25936
```

-7.02912    0.00000    0.00000 ;      C-     N-    CA-     C
  3436   3435   3437   3438       3   29.28800   -8.36800  -20.92000
0.00000    0.00000    0.00000 ;      O-     C-     N-     H
  3436   3435   3437   3439       3   20.92000    0.00000  -20.92000
0.00000    0.00000    0.00000 ;      O-     C-     N-    CA
  3437   3439   3441   3442       3    0.65084    1.95253    0.00000
-2.60338    0.00000    0.00000 ;      N-    CA-    CB-   HB2
  3437   3439   3441   3443       3    0.65084    1.95253    0.00000
-2.60338    0.00000    0.00000 ;      N-    CA-    CB-   HB3
  3437   3439   3441   3444       3   15.41386    9.74035   -8.39310
-0.97069    2.97901    0.00000 ;      N-    CA-    CB-    CG
  3437   3439   3447   3448       3    0.00000    0.00000    0.00000
0.00000    0.00000    0.00000 ;      N-    CA-     C-     O
  3437   3439   3447   3449       3   17.40544   -5.02080  -13.22144
9.20480    0.00000    0.00000 ;      N-    CA-     C-     N
  3438   3437   3439   3440       3    0.00000    0.00000    0.00000
0.00000    0.00000    0.00000 ;      H-     N-    CA-    HA
  3438   3437   3439   3441       3    0.00000    0.00000    0.00000
0.00000    0.00000    0.00000 ;      H-     N-    CA-    CB
  3438   3437   3439   3447       3    0.00000    0.00000    0.00000
0.00000    0.00000    0.00000 ;      H-     N-    CA-     C
  3439   3441   3444   3445       3    6.43499    0.00000   -5.39736
0.00000   -1.03763    0.00000 ;     CA-    CB-    CG-   OD1
  3439   3441   3444   3446       3    6.43499    0.00000   -5.39736
0.00000   -1.03763    0.00000 ;     CA-    CB-    CG-   OD2
  3439   3447   3449   3450       3   20.92000    0.00000  -20.92000
0.00000    0.00000    0.00000 ;     CA-     C-     N-     H
  3439   3447   3449   3451       3   20.92000    0.00000  -20.92000
0.00000    0.00000    0.00000 ;     CA-     C-     N-    CA
  3440   3439   3441   3442       3    0.65084    1.95253    0.00000
-2.60338    0.00000    0.00000 ;     HA-    CA-    CB-   HB2
  3440   3439   3441   3443       3    0.65084    1.95253    0.00000
-2.60338    0.00000    0.00000 ;     HA-    CA-    CB-   HB3
  3440   3439   3441   3444       3    0.65084    1.95253    0.00000
-2.60338    0.00000    0.00000 ;     HA-    CA-    CB-    CG
  3440   3439   3447   3448       3    3.68192   -4.35136    0.00000
1.33888    0.00000    0.00000 ;     HA-    CA-     C-     O
  3440   3439   3447   3449       3    0.00000    0.00000    0.00000
0.00000    0.00000    0.00000 ;     HA-    CA-     C-     N
  3441   3439   3447   3448       3    0.00000    0.00000    0.00000
0.00000    0.00000    0.00000 ;     CB-    CA-     C-     O
  3441   3439   3447   3449       3    2.51040    4.18400    1.67360
-6.69440    0.00000    0.00000 ;     CB-    CA-     C-     N
  3442   3441   3439   3447       3    0.65084    1.95253    0.00000
-2.60338    0.00000    0.00000 ;    HB2-    CB-    CA-     C
  3442   3441   3444   3445       3    0.00000    0.00000    0.00000
0.00000    0.00000    0.00000 ;    HB2-    CB-    CG-   OD1
  3442   3441   3444   3446       3    0.00000    0.00000    0.00000
0.00000    0.00000    0.00000 ;    HB2-    CB-    CG-   OD2
  3443   3441   3439   3447       3    0.65084    1.95253    0.00000
-2.60338    0.00000    0.00000 ;    HB3-    CB-    CA-     C
  3443   3441   3444   3445       3    0.00000    0.00000    0.00000

0.00000 0.00000 0.00000 ; HB3- CB- CG- OD1
3443 3441 3444 3446 3 0.00000 0.00000 0.00000
0.00000 0.00000 0.00000 ; HB3- CB- CG- OD2
3444 3441 3439 3447 3 7.14627 -1.04600 -8.99560
-0.97069 5.15469 0.00000 ; CG- CB- CA- C
3447 3449 3451 3452 3 0.00000 0.00000 0.00000
0.00000 0.00000 0.00000 ; C- N- CA- HA
3447 3449 3451 3453 3 11.71520 1.67360 15.06240 -
13.38880 0.00000 0.00000 ; C- N- CA- CB
3447 3449 3451 3461 3 1.75728 5.27184 2.25936
-7.02912 0.00000 0.00000 ; C- N- CA- C
3448 3447 3449 3450 3 29.28800 -8.36800 -20.92000
0.00000 0.00000 0.00000 ; O- C- N- H
3448 3447 3449 3451 3 20.92000 0.00000 -20.92000
0.00000 0.00000 0.00000 ; O- C- N- CA
3449 3451 3453 3454 3 0.65084 1.95253 0.00000
-2.60338 0.00000 0.00000 ; N- CA- CB- HB
3449 3451 3453 3455 3 3.83673 0.44769 -1.77402
-2.47693 -0.03347 0.00000 ; N- CA- CB- CG2
3449 3451 3453 3459 3 4.93294 1.13386 -3.12963
-5.27184 3.17984 0.00000 ; N- CA- CB- OG1
3449 3451 3461 3462 3 0.00000 0.00000 0.00000
0.00000 0.00000 0.00000 ; N- CA- C- O
3449 3451 3461 3463 3 17.40544 -5.02080 -13.22144
9.20480 0.00000 0.00000 ; N- CA- C- N
3450 3449 3451 3452 3 0.00000 0.00000 0.00000
0.00000 0.00000 0.00000 ; H- N- CA- HA
3450 3449 3451 3453 3 0.00000 0.00000 0.00000
0.00000 0.00000 0.00000 ; H- N- CA- CB
3450 3449 3451 3461 3 0.00000 0.00000 0.00000
0.00000 0.00000 0.00000 ; H- N- CA- C
3451 3453 3455 3456 3 0.66944 2.00832 0.00000
-2.67776 0.00000 0.00000 ; CA- CB- CG2- HG21
3451 3453 3455 3457 3 0.66944 2.00832 0.00000
-2.67776 0.00000 0.00000 ; CA- CB- CG2- HG22
3451 3453 3455 3458 3 0.66944 2.00832 0.00000
-2.67776 0.00000 0.00000 ; CA- CB- CG2- HG23
3451 3453 3459 3460 3 1.12131 2.98738 1.66523
-3.94970 0.43514 0.00000 ; CA- CB- OG1- HG1
3451 3461 3463 3464 3 20.92000 0.00000 -20.92000
0.00000 0.00000 0.00000 ; CA- C- N- H
3451 3461 3463 3465 3 20.92000 0.00000 -20.92000
0.00000 0.00000 0.00000 ; CA- C- N- CA
3452 3451 3453 3454 3 0.65084 1.95253 0.00000
-2.60338 0.00000 0.00000 ; HA- CA- CB- HB
3452 3451 3453 3455 3 0.65084 1.95253 0.00000
-2.60338 0.00000 0.00000 ; HA- CA- CB- CG2
3452 3451 3453 3459 3 1.04600 -1.04600 0.00000
0.00000 0.00000 0.00000 ; HA- CA- CB- OG1
3452 3451 3461 3462 3 3.68192 -4.35136 0.00000
1.33888 0.00000 0.00000 ; HA- CA- C- O
3452 3451 3461 3463 3 0.00000 0.00000 0.00000

0.00000 0.00000 0.00000 ; HA- CA- C- N
3453 3451 3461 3462 3 0.00000 0.00000 0.00000
0.00000 0.00000 0.00000 ; CB- CA- C- O
3453 3451 3461 3463 3 2.51040 4.18400 1.67360
-6.69440 0.00000 0.00000 ; CB- CA- C- N
3454 3453 3451 3461 3 0.65084 1.95253 0.00000
-2.60338 0.00000 0.00000 ; HB- CB- CA- C
3454 3453 3455 3456 3 0.65084 1.95253 0.00000
-2.60338 0.00000 0.00000 ; HB- CB- CG2- HG21
3454 3453 3455 3457 3 0.65084 1.95253 0.00000
-2.60338 0.00000 0.00000 ; HB- CB- CG2- HG22
3454 3453 3455 3458 3 0.65084 1.95253 0.00000
-2.60338 0.00000 0.00000 ; HB- CB- CG2- HG23
3454 3453 3459 3460 3 0.69733 2.09200 0.00000
-2.78933 0.00000 0.00000 ; HB- CB- OG1- HG1
3455 3453 3451 3461 3 5.67350 3.55640 -6.16722
-2.47693 3.74886 0.00000 ; CG2- CB- CA- C
3455 3453 3459 3460 3 4.74047 0.27196 -2.26773
-3.94970 1.60666 0.00000 ; CG2- CB- OG1- HG1
3456 3455 3453 3459 3 1.04600 -1.04600 0.00000
0.00000 0.00000 0.00000 ; HG21- CG2- CB- OG1
3457 3455 3453 3459 3 1.04600 -1.04600 0.00000
0.00000 0.00000 0.00000 ; HG22- CG2- CB- OG1
3458 3455 3453 3459 3 1.04600 -1.04600 0.00000
0.00000 0.00000 0.00000 ; HG23- CG2- CB- OG1
3459 3453 3451 3461 3 6.53541 6.87013 -6.21742
-5.27184 5.22163 0.00000 ; OG1- CB- CA- C
3461 3463 3465 3466 3 0.00000 0.00000 0.00000
0.00000 0.00000 0.00000 ; C- N- CA- HA
3461 3463 3465 3467 3 11.71520 1.67360 15.06240 -
13.38880 0.00000 0.00000 ; C- N- CA- CB
3461 3463 3465 3473 3 1.75728 5.27184 2.25936
-7.02912 0.00000 0.00000 ; C- N- CA- C
3462 3461 3463 3464 3 29.28800 -8.36800 -20.92000
0.00000 0.00000 0.00000 ; O- C- N- H
3462 3461 3463 3465 3 20.92000 0.00000 -20.92000
0.00000 0.00000 0.00000 ; O- C- N- CA
3463 3465 3467 3468 3 0.65084 1.95253 0.00000
-2.60338 0.00000 0.00000 ; N- CA- CB- HB2
3463 3465 3467 3469 3 0.65084 1.95253 0.00000
-2.60338 0.00000 0.00000 ; N- CA- CB- HB3
3463 3465 3467 3470 3 15.41386 9.74035 -8.39310
-0.97069 2.97901 0.00000 ; N- CA- CB- CG
3463 3465 3473 3474 3 0.00000 0.00000 0.00000
0.00000 0.00000 0.00000 ; N- CA- C- O
3463 3465 3473 3475 3 17.40544 -5.02080 -13.22144
9.20480 0.00000 0.00000 ; N- CA- C- N
3464 3463 3465 3466 3 0.00000 0.00000 0.00000
0.00000 0.00000 0.00000 ; H- N- CA- HA
3464 3463 3465 3467 3 0.00000 0.00000 0.00000
0.00000 0.00000 0.00000 ; H- N- CA- CB
3464 3463 3465 3473 3 0.00000 0.00000 0.00000

0.00000 0.00000 0.00000 ; H- N- CA- C
3465 3467 3470 3471 3 6.43499 0.00000 -5.39736
0.00000 -1.03763 0.00000 ; CA- CB- CG- OD1
3465 3467 3470 3472 3 6.43499 0.00000 -5.39736
0.00000 -1.03763 0.00000 ; CA- CB- CG- OD2
3465 3473 3475 3476 3 20.92000 0.00000 -20.92000
0.00000 0.00000 0.00000 ; CA- C- N- H
3465 3473 3475 3477 3 20.92000 0.00000 -20.92000
0.00000 0.00000 0.00000 ; CA- C- N- CA
3466 3465 3467 3468 3 0.65084 1.95253 0.00000
-2.60338 0.00000 0.00000 ; HA- CA- CB- HB2
3466 3465 3467 3469 3 0.65084 1.95253 0.00000
-2.60338 0.00000 0.00000 ; HA- CA- CB- HB3
3466 3465 3467 3470 3 0.65084 1.95253 0.00000
-2.60338 0.00000 0.00000 ; HA- CA- CB- CG
3466 3465 3473 3474 3 3.68192 -4.35136 0.00000
1.33888 0.00000 0.00000 ; HA- CA- C- O
3466 3465 3473 3475 3 0.00000 0.00000 0.00000
0.00000 0.00000 0.00000 ; HA- CA- C- N
3467 3465 3473 3474 3 0.00000 0.00000 0.00000
0.00000 0.00000 0.00000 ; CB- CA- C- O
3467 3465 3473 3475 3 2.51040 4.18400 1.67360
-6.69440 0.00000 0.00000 ; CB- CA- C- N
3468 3467 3465 3473 3 0.65084 1.95253 0.00000
-2.60338 0.00000 0.00000 ; HB2- CB- CA- C
3468 3467 3470 3471 3 0.00000 0.00000 0.00000
0.00000 0.00000 0.00000 ; HB2- CB- CG- OD1
3468 3467 3470 3472 3 0.00000 0.00000 0.00000
0.00000 0.00000 0.00000 ; HB2- CB- CG- OD2
3469 3467 3465 3473 3 0.65084 1.95253 0.00000
-2.60338 0.00000 0.00000 ; HB3- CB- CA- C
3469 3467 3470 3471 3 0.00000 0.00000 0.00000
0.00000 0.00000 0.00000 ; HB3- CB- CG- OD1
3469 3467 3470 3472 3 0.00000 0.00000 0.00000
0.00000 0.00000 0.00000 ; HB3- CB- CG- OD2
3470 3467 3465 3473 3 7.14627 -1.04600 -8.99560
-0.97069 5.15469 0.00000 ; CG- CB- CA- C
3473 3475 3477 3478 3 0.00000 0.00000 0.00000
0.00000 0.00000 0.00000 ; C- N- CA- HA
3473 3475 3477 3479 3 11.71520 1.67360 15.06240 -
13.38880 0.00000 0.00000 ; C- N- CA- CB
3473 3475 3477 3487 3 1.75728 5.27184 2.25936
-7.02912 0.00000 0.00000 ; C- N- CA- C
3474 3473 3475 3476 3 29.28800 -8.36800 -20.92000
0.00000 0.00000 0.00000 ; O- C- N- H
3474 3473 3475 3477 3 20.92000 0.00000 -20.92000
0.00000 0.00000 0.00000 ; O- C- N- CA
3475 3477 3479 3480 3 0.65084 1.95253 0.00000
-2.60338 0.00000 0.00000 ; N- CA- CB- HB2
3475 3477 3479 3481 3 0.65084 1.95253 0.00000
-2.60338 0.00000 0.00000 ; N- CA- CB- HB3
3475 3477 3479 3482 3 5.99567 3.29281 -4.46014

```
-0.55229    1.97485    0.00000 ;       N-    CA-    CB-    CG
  3475   3477   3487   3488      3    0.00000    0.00000    0.00000
0.00000    0.00000    0.00000 ;      N-    CA-     C-     O
  3475   3477   3487   3489      3   17.40544   -5.02080  -13.22144
9.20480    0.00000    0.00000 ;      N-    CA-     C-     N
  3476   3475   3477   3478      3    0.00000    0.00000    0.00000
0.00000    0.00000    0.00000 ;      H-     N-    CA-    HA
  3476   3475   3477   3479      3    0.00000    0.00000    0.00000
0.00000    0.00000    0.00000 ;      H-     N-    CA-    CB
  3476   3475   3477   3487      3    0.00000    0.00000    0.00000
0.00000    0.00000    0.00000 ;      H-     N-    CA-     C
  3477   3479   3482   3483      3    0.00000    0.00000    0.00000
0.00000    0.00000    0.00000 ;     CA-    CB-    CG-   OD1
  3477   3479   3482   3484      3    8.84916   -0.31380   -4.32626
5.03754    0.26778    0.00000 ;     CA-    CB-    CG-   ND2
  3477   3487   3489   3490      3   20.92000    0.00000  -20.92000
0.00000    0.00000    0.00000 ;     CA-     C-     N-     H
  3477   3487   3489   3491      3   20.92000    0.00000  -20.92000
0.00000    0.00000    0.00000 ;     CA-     C-     N-    CA
  3478   3477   3479   3480      3    0.65084    1.95253    0.00000
-2.60338    0.00000    0.00000 ;     HA-    CA-    CB-   HB2
  3478   3477   3479   3481      3    0.65084    1.95253    0.00000
-2.60338    0.00000    0.00000 ;     HA-    CA-    CB-   HB3
  3478   3477   3479   3482      3    0.65084    1.95253    0.00000
-2.60338    0.00000    0.00000 ;     HA-    CA-    CB-    CG
  3478   3477   3487   3488      3    3.68192   -4.35136    0.00000
1.33888    0.00000    0.00000 ;     HA-    CA-     C-     O
  3478   3477   3487   3489      3    0.00000    0.00000    0.00000
0.00000    0.00000    0.00000 ;     HA-    CA-     C-     N
  3479   3477   3487   3488      3    0.00000    0.00000    0.00000
0.00000    0.00000    0.00000 ;     CB-    CA-     C-     O
  3479   3477   3487   3489      3    2.51040    4.18400    1.67360
-6.69440    0.00000    0.00000 ;     CB-    CA-     C-     N
  3479   3482   3484   3485      3   20.92000    0.00000  -20.92000
0.00000    0.00000    0.00000 ;     CB-    CG-   ND2-  HD21
  3479   3482   3484   3486      3   20.92000    0.00000  -20.92000
0.00000    0.00000    0.00000 ;     CB-    CG-   ND2-  HD22
  3480   3479   3477   3487      3    0.65084    1.95253    0.00000
-2.60338    0.00000    0.00000 ;    HB2-    CB-    CA-     C
  3480   3479   3482   3483      3    3.68192   -4.35136    0.00000
1.33888    0.00000    0.00000 ;    HB2-    CB-    CG-   OD1
  3480   3479   3482   3484      3    0.00000    0.00000    0.00000
0.00000    0.00000    0.00000 ;    HB2-    CB-    CG-   ND2
  3481   3479   3477   3487      3    0.65084    1.95253    0.00000
-2.60338    0.00000    0.00000 ;    HB3-    CB-    CA-     C
  3481   3479   3482   3483      3    3.68192   -4.35136    0.00000
1.33888    0.00000    0.00000 ;    HB3-    CB-    CG-   OD1
  3481   3479   3482   3484      3    0.00000    0.00000    0.00000
0.00000    0.00000    0.00000 ;    HB3-    CB-    CG-   ND2
  3482   3479   3477   3487      3    7.94542   -3.96225   -6.11701
-0.55229    3.58150    0.00000 ;     CG-    CB-    CA-     C
  3483   3482   3484   3485      3   29.28800   -8.36800  -20.92000
```

0.00000 0.00000 0.00000 ; OD1- CG- ND2- HD21
3483 3482 3484 3486 3 29.28800 -8.36800 -20.92000
0.00000 0.00000 0.00000 ; OD1- CG- ND2- HD22
3487 3489 3491 3492 3 0.00000 0.00000 0.00000
0.00000 0.00000 0.00000 ; C- N- CA- HA
3487 3489 3491 3493 3 11.71520 1.67360 15.06240 -
13.38880 0.00000 0.00000 ; C- N- CA- CB
3487 3489 3491 3502 3 1.75728 5.27184 2.25936
-7.02912 0.00000 0.00000 ; C- N- CA- C
3488 3487 3489 3490 3 29.28800 -8.36800 -20.92000
0.00000 0.00000 0.00000 ; O- C- N- H
3488 3487 3489 3491 3 20.92000 0.00000 -20.92000
0.00000 0.00000 0.00000 ; O- C- N- CA
3489 3491 3493 3494 3 0.65084 1.95253 0.00000
-2.60338 0.00000 0.00000 ; N- CA- CB- HB2
3489 3491 3493 3495 3 0.65084 1.95253 0.00000
-2.60338 0.00000 0.00000 ; N- CA- CB- HB3
3489 3491 3493 3496 3 3.21331 2.22589 -4.15053
-2.40998 2.61082 0.00000 ; N- CA- CB- CG
3489 3491 3502 3503 3 0.00000 0.00000 0.00000
0.00000 0.00000 0.00000 ; N- CA- C- O
3489 3491 3502 3504 3 17.40544 -5.02080 -13.22144
9.20480 0.00000 0.00000 ; N- CA- C- N
3490 3489 3491 3492 3 0.00000 0.00000 0.00000
0.00000 0.00000 0.00000 ; H- N- CA- HA
3490 3489 3491 3493 3 0.00000 0.00000 0.00000
0.00000 0.00000 0.00000 ; H- N- CA- CB
3490 3489 3491 3502 3 0.00000 0.00000 0.00000
0.00000 0.00000 0.00000 ; H- N- CA- C
3491 3493 3496 3497 3 0.66944 2.00832 0.00000
-2.67776 0.00000 0.00000 ; CA- CB- CG- HG2
3491 3493 3496 3498 3 0.66944 2.00832 0.00000
-2.67776 0.00000 0.00000 ; CA- CB- CG- HG3
3491 3493 3496 3499 3 10.12110 -1.91209 0.01674
10.17549 -1.87443 0.00000 ; CA- CB- CG- CD
3491 3502 3504 3505 3 20.92000 0.00000 -20.92000
0.00000 0.00000 0.00000 ; CA- C- N- H
3491 3502 3504 3506 3 20.92000 0.00000 -20.92000
0.00000 0.00000 0.00000 ; CA- C- N- CA
3492 3491 3493 3494 3 0.65084 1.95253 0.00000
-2.60338 0.00000 0.00000 ; HA- CA- CB- HB2
3492 3491 3493 3495 3 0.65084 1.95253 0.00000
-2.60338 0.00000 0.00000 ; HA- CA- CB- HB3
3492 3491 3493 3496 3 0.65084 1.95253 0.00000
-2.60338 0.00000 0.00000 ; HA- CA- CB- CG
3492 3491 3502 3503 3 3.68192 -4.35136 0.00000
1.33888 0.00000 0.00000 ; HA- CA- C- O
3492 3491 3502 3504 3 0.00000 0.00000 0.00000
0.00000 0.00000 0.00000 ; HA- CA- C- N
3493 3491 3502 3503 3 0.00000 0.00000 0.00000
0.00000 0.00000 0.00000 ; CB- CA- C- O
3493 3491 3502 3504 3 2.51040 4.18400 1.67360

-6.69440 0.00000 0.00000 ; CB- CA- C- N
3493 3496 3499 3500 3 3.79907 0.00000 -5.40573
0.00000 2.14221 0.00000 ; CB- CG- CD- OE1
3493 3496 3499 3501 3 3.79907 0.00000 -5.40573
0.00000 2.14221 0.00000 ; CB- CG- CD- OE2
3494 3493 3491 3502 3 0.65084 1.95253 0.00000
-2.60338 0.00000 0.00000 ; HB2- CB- CA- C
3494 3493 3496 3497 3 0.62760 1.88280 0.00000
-2.51040 0.00000 0.00000 ; HB2- CB- CG- HG2
3494 3493 3496 3498 3 0.62760 1.88280 0.00000
-2.51040 0.00000 0.00000 ; HB2- CB- CG- HG3
3494 3493 3496 3499 3 0.65084 1.95253 0.00000
-2.60338 0.00000 0.00000 ; HB2- CB- CG- CD
3495 3493 3491 3502 3 0.65084 1.95253 0.00000
-2.60338 0.00000 0.00000 ; HB3- CB- CA- C
3495 3493 3496 3497 3 0.62760 1.88280 0.00000
-2.51040 0.00000 0.00000 ; HB3- CB- CG- HG2
3495 3493 3496 3498 3 0.62760 1.88280 0.00000
-2.51040 0.00000 0.00000 ; HB3- CB- CG- HG3
3495 3493 3496 3499 3 0.65084 1.95253 0.00000
-2.60338 0.00000 0.00000 ; HB3- CB- CG- CD
3496 3493 3491 3502 3 6.86594 3.56895 -8.14206
-2.40998 4.85344 0.00000 ; CG- CB- CA- C
3497 3496 3499 3500 3 0.00000 0.00000 0.00000
0.00000 0.00000 0.00000 ; HG2- CG- CD- OE1
3497 3496 3499 3501 3 0.00000 0.00000 0.00000
0.00000 0.00000 0.00000 ; HG2- CG- CD- OE2
3498 3496 3499 3500 3 0.00000 0.00000 0.00000
0.00000 0.00000 0.00000 ; HG3- CG- CD- OE1
3498 3496 3499 3501 3 0.00000 0.00000 0.00000
0.00000 0.00000 0.00000 ; HG3- CG- CD- OE2
3502 3504 3506 3507 3 0.00000 0.00000 0.00000
0.00000 0.00000 0.00000 ; C- N- CA- HA
3502 3504 3506 3508 3 11.71520 1.67360 15.06240 -
13.38880 0.00000 0.00000 ; C- N- CA- CB
3502 3504 3506 3518 3 1.75728 5.27184 2.25936
-7.02912 0.00000 0.00000 ; C- N- CA- C
3503 3502 3504 3505 3 29.28800 -8.36800 -20.92000
0.00000 0.00000 0.00000 ; O- C- N- H
3503 3502 3504 3506 3 20.92000 0.00000 -20.92000
0.00000 0.00000 0.00000 ; O- C- N- CA
3504 3506 3508 3509 3 0.65084 1.95253 0.00000
-2.60338 0.00000 0.00000 ; N- CA- CB- HB
3504 3506 3508 3510 3 3.83673 0.44769 -1.77402
-2.47693 -0.03347 0.00000 ; N- CA- CB- CG1
3504 3506 3508 3514 3 3.83673 0.44769 -1.77402
-2.47693 -0.03347 0.00000 ; N- CA- CB- CG2
3504 3506 3518 3519 3 0.00000 0.00000 0.00000
0.00000 0.00000 0.00000 ; N- CA- C- O
3504 3506 3518 3520 3 17.40544 -5.02080 -13.22144
9.20480 0.00000 0.00000 ; N- CA- C- N
3505 3504 3506 3507 3 0.00000 0.00000 0.00000

0.00000 0.00000 0.00000 ; H- N- CA- HA
3505 3504 3506 3508 3 0.00000 0.00000 0.00000
0.00000 0.00000 0.00000 ; H- N- CA- CB
3505 3504 3506 3518 3 0.00000 0.00000 0.00000
0.00000 0.00000 0.00000 ; H- N- CA- C
3506 3508 3510 3511 3 0.66944 2.00832 0.00000
-2.67776 0.00000 0.00000 ; CA- CB- CG1- HG11
3506 3508 3510 3512 3 0.66944 2.00832 0.00000
-2.67776 0.00000 0.00000 ; CA- CB- CG1- HG12
3506 3508 3510 3513 3 0.66944 2.00832 0.00000
-2.67776 0.00000 0.00000 ; CA- CB- CG1- HG13
3506 3508 3514 3515 3 0.66944 2.00832 0.00000
-2.67776 0.00000 0.00000 ; CA- CB- CG2- HG21
3506 3508 3514 3516 3 0.66944 2.00832 0.00000
-2.67776 0.00000 0.00000 ; CA- CB- CG2- HG22
3506 3508 3514 3517 3 0.66944 2.00832 0.00000
-2.67776 0.00000 0.00000 ; CA- CB- CG2- HG23
3506 3518 3520 3521 3 20.92000 0.00000 -20.92000
0.00000 0.00000 0.00000 ; CA- C- N- H
3506 3518 3520 3522 3 20.92000 0.00000 -20.92000
0.00000 0.00000 0.00000 ; CA- C- N- CA
3507 3506 3508 3509 3 0.65084 1.95253 0.00000
-2.60338 0.00000 0.00000 ; HA- CA- CB- HB
3507 3506 3508 3510 3 0.65084 1.95253 0.00000
-2.60338 0.00000 0.00000 ; HA- CA- CB- CG1
3507 3506 3508 3514 3 0.65084 1.95253 0.00000
-2.60338 0.00000 0.00000 ; HA- CA- CB- CG2
3507 3506 3518 3519 3 3.68192 -4.35136 0.00000
1.33888 0.00000 0.00000 ; HA- CA- C- O
3507 3506 3518 3520 3 0.00000 0.00000 0.00000
0.00000 0.00000 0.00000 ; HA- CA- C- N
3508 3506 3518 3519 3 0.00000 0.00000 0.00000
0.00000 0.00000 0.00000 ; CB- CA- C- O
3508 3506 3518 3520 3 2.51040 4.18400 1.67360
-6.69440 0.00000 0.00000 ; CB- CA- C- N
3509 3508 3506 3518 3 0.65084 1.95253 0.00000
-2.60338 0.00000 0.00000 ; HB- CB- CA- C
3509 3508 3510 3511 3 0.62760 1.88280 0.00000
-2.51040 0.00000 0.00000 ; HB- CB- CG1- HG11
3509 3508 3510 3512 3 0.62760 1.88280 0.00000
-2.51040 0.00000 0.00000 ; HB- CB- CG1- HG12
3509 3508 3510 3513 3 0.62760 1.88280 0.00000
-2.51040 0.00000 0.00000 ; HB- CB- CG1- HG13
3509 3508 3514 3515 3 0.62760 1.88280 0.00000
-2.51040 0.00000 0.00000 ; HB- CB- CG2- HG21
3509 3508 3514 3516 3 0.62760 1.88280 0.00000
-2.51040 0.00000 0.00000 ; HB- CB- CG2- HG22
3509 3508 3514 3517 3 0.62760 1.88280 0.00000
-2.51040 0.00000 0.00000 ; HB- CB- CG2- HG23
3510 3508 3506 3518 3 5.67350 3.55640 -6.16722
-2.47693 3.74886 0.00000 ; CG1- CB- CA- C
3510 3508 3514 3515 3 0.66944 2.00832 0.00000

-2.67776 0.00000 0.00000 ; CG1- CB- CG2- HG21
3510 3508 3514 3516 3 0.66944 2.00832 0.00000
-2.67776 0.00000 0.00000 ; CG1- CB- CG2- HG22
3510 3508 3514 3517 3 0.66944 2.00832 0.00000
-2.67776 0.00000 0.00000 ; CG1- CB- CG2- HG23
3511 3510 3508 3514 3 0.66944 2.00832 0.00000
-2.67776 0.00000 0.00000 ; HG11- CG1- CB- CG2
3512 3510 3508 3514 3 0.66944 2.00832 0.00000
-2.67776 0.00000 0.00000 ; HG12- CG1- CB- CG2
3513 3510 3508 3514 3 0.66944 2.00832 0.00000
-2.67776 0.00000 0.00000 ; HG13- CG1- CB- CG2
3514 3508 3506 3518 3 5.67350 3.55640 -6.16722
-2.47693 3.74886 0.00000 ; CG2- CB- CA- C
3518 3520 3522 3523 3 0.00000 0.00000 0.00000
0.00000 0.00000 0.00000 ; C- N- CA- HA
3518 3520 3522 3524 3 11.71520 1.67360 15.06240 -
13.38880 0.00000 0.00000 ; C- N- CA- CB
3518 3520 3522 3537 3 1.75728 5.27184 2.25936
-7.02912 0.00000 0.00000 ; C- N- CA- C
3519 3518 3520 3521 3 29.28800 -8.36800 -20.92000
0.00000 0.00000 0.00000 ; O- C- N- H
3519 3518 3520 3522 3 20.92000 0.00000 -20.92000
0.00000 0.00000 0.00000 ; O- C- N- CA
3520 3522 3524 3525 3 0.65084 1.95253 0.00000
-2.60338 0.00000 0.00000 ; N- CA- CB- HB2
3520 3522 3524 3526 3 0.65084 1.95253 0.00000
-2.60338 0.00000 0.00000 ; N- CA- CB- HB3
3520 3522 3524 3527 3 3.79070 1.39746 -4.61077
-2.40998 2.44346 0.00000 ; N- CA- CB- CG
3520 3522 3537 3538 3 0.00000 0.00000 0.00000
0.00000 0.00000 0.00000 ; N- CA- C- O
3520 3522 3537 3539 3 17.40544 -5.02080 -13.22144
9.20480 0.00000 0.00000 ; N- CA- C- N
3521 3520 3522 3523 3 0.00000 0.00000 0.00000
0.00000 0.00000 0.00000 ; H- N- CA- HA
3521 3520 3522 3524 3 0.00000 0.00000 0.00000
0.00000 0.00000 0.00000 ; H- N- CA- CB
3521 3520 3522 3537 3 0.00000 0.00000 0.00000
0.00000 0.00000 0.00000 ; H- N- CA- C
3522 3524 3527 3528 3 0.66944 2.00832 0.00000
-2.67776 0.00000 0.00000 ; CA- CB- CG- HG
3522 3524 3527 3529 3 3.90367 0.19665 -6.21742
-2.37651 5.99149 0.00000 ; CA- CB- CG- CD1
3522 3524 3527 3533 3 3.90367 0.19665 -6.21742
-2.37651 5.99149 0.00000 ; CA- CB- CG- CD2
3522 3537 3539 3540 3 20.92000 0.00000 -20.92000
0.00000 0.00000 0.00000 ; CA- C- N- H
3522 3537 3539 3541 3 20.92000 0.00000 -20.92000
0.00000 0.00000 0.00000 ; CA- C- N- CA
3523 3522 3524 3525 3 0.65084 1.95253 0.00000
-2.60338 0.00000 0.00000 ; HA- CA- CB- HB2
3523 3522 3524 3526 3 0.65084 1.95253 0.00000

-2.60338 0.00000 0.00000 ; HA- CA- CB- HB3
3523 3522 3524 3527 3 0.65084 1.95253 0.00000
-2.60338 0.00000 0.00000 ; HA- CA- CB- CG
3523 3522 3537 3538 3 3.68192 -4.35136 0.00000
1.33888 0.00000 0.00000 ; HA- CA- C- O
3523 3522 3537 3539 3 0.00000 0.00000 0.00000
0.00000 0.00000 0.00000 ; HA- CA- C- N
3524 3522 3537 3538 3 0.00000 0.00000 0.00000
0.00000 0.00000 0.00000 ; CB- CA- C- O
3524 3522 3537 3539 3 2.51040 4.18400 1.67360
-6.69440 0.00000 0.00000 ; CB- CA- C- N
3524 3527 3529 3530 3 0.66944 2.00832 0.00000
-2.67776 0.00000 0.00000 ; CB- CG- CD1- HD11
3524 3527 3529 3531 3 0.66944 2.00832 0.00000
-2.67776 0.00000 0.00000 ; CB- CG- CD1- HD12
3524 3527 3529 3532 3 0.66944 2.00832 0.00000
-2.67776 0.00000 0.00000 ; CB- CG- CD1- HD13
3524 3527 3533 3534 3 0.66944 2.00832 0.00000
-2.67776 0.00000 0.00000 ; CB- CG- CD2- HD21
3524 3527 3533 3535 3 0.66944 2.00832 0.00000
-2.67776 0.00000 0.00000 ; CB- CG- CD2- HD22
3524 3527 3533 3536 3 0.66944 2.00832 0.00000
-2.67776 0.00000 0.00000 ; CB- CG- CD2- HD23
3525 3524 3522 3537 3 0.65084 1.95253 0.00000
-2.60338 0.00000 0.00000 ; HB2- CB- CA- C
3525 3524 3527 3528 3 0.62760 1.88280 0.00000
-2.51040 0.00000 0.00000 ; HB2- CB- CG- HG
3525 3524 3527 3529 3 0.66944 2.00832 0.00000
-2.67776 0.00000 0.00000 ; HB2- CB- CG- CD1
3525 3524 3527 3533 3 0.66944 2.00832 0.00000
-2.67776 0.00000 0.00000 ; HB2- CB- CG- CD2
3526 3524 3522 3537 3 0.65084 1.95253 0.00000
-2.60338 0.00000 0.00000 ; HB3- CB- CA- C
3526 3524 3527 3528 3 0.62760 1.88280 0.00000
-2.51040 0.00000 0.00000 ; HB3- CB- CG- HG
3526 3524 3527 3529 3 0.66944 2.00832 0.00000
-2.67776 0.00000 0.00000 ; HB3- CB- CG- CD1
3526 3524 3527 3533 3 0.66944 2.00832 0.00000
-2.67776 0.00000 0.00000 ; HB3- CB- CG- CD2
3527 3524 3522 3537 3 10.33448 -1.14642 -11.54784
-2.40998 6.35968 0.00000 ; CG- CB- CA- C
3528 3527 3529 3530 3 0.62760 1.88280 0.00000
-2.51040 0.00000 0.00000 ; HG- CG- CD1- HD11
3528 3527 3529 3531 3 0.62760 1.88280 0.00000
-2.51040 0.00000 0.00000 ; HG- CG- CD1- HD12
3528 3527 3529 3532 3 0.62760 1.88280 0.00000
-2.51040 0.00000 0.00000 ; HG- CG- CD1- HD13
3528 3527 3533 3534 3 0.62760 1.88280 0.00000
-2.51040 0.00000 0.00000 ; HG- CG- CD2- HD21
3528 3527 3533 3535 3 0.62760 1.88280 0.00000
-2.51040 0.00000 0.00000 ; HG- CG- CD2- HD22
3528 3527 3533 3536 3 0.62760 1.88280 0.00000

-2.51040    0.00000    0.00000 ;     HG-    CG-    CD2-  HD23
 3529   3527   3533   3534     3    0.66944    2.00832    0.00000
-2.67776    0.00000    0.00000 ;    CD1-    CG-    CD2-  HD21
 3529   3527   3533   3535     3    0.66944    2.00832    0.00000
-2.67776    0.00000    0.00000 ;    CD1-    CG-    CD2-  HD22
 3529   3527   3533   3536     3    0.66944    2.00832    0.00000
-2.67776    0.00000    0.00000 ;    CD1-    CG-    CD2-  HD23
 3530   3529   3527   3533     3    0.66944    2.00832    0.00000
-2.67776    0.00000    0.00000 ;   HD11-   CD1-    CG-   CD2
 3531   3529   3527   3533     3    0.66944    2.00832    0.00000
-2.67776    0.00000    0.00000 ;   HD12-   CD1-    CG-   CD2
 3532   3529   3527   3533     3    0.66944    2.00832    0.00000
-2.67776    0.00000    0.00000 ;   HD13-   CD1-    CG-   CD2
 3537   3539   3541   3542     3    0.00000    0.00000    0.00000
0.00000    0.00000    0.00000 ;     C-     N-    CA-    HA
 3537   3539   3541   3543     3   11.71520    1.67360   15.06240  -
13.38880    0.00000    0.00000 ;     C-     N-    CA-    CB
 3537   3539   3541   3559     3    1.75728    5.27184    2.25936
-7.02912    0.00000    0.00000 ;     C-     N-    CA-     C
 3538   3537   3539   3540     3   29.28800   -8.36800  -20.92000
0.00000    0.00000    0.00000 ;     O-     C-     N-     H
 3538   3537   3539   3541     3   20.92000    0.00000  -20.92000
0.00000    0.00000    0.00000 ;     O-     C-     N-    CA
 3539   3541   3543   3544     3    0.65084    1.95253    0.00000
-2.60338    0.00000    0.00000 ;     N-    CA-    CB-   HB2
 3539   3541   3543   3545     3    0.65084    1.95253    0.00000
-2.60338    0.00000    0.00000 ;     N-    CA-    CB-   HB3
 3539   3541   3543   3546     3    0.65084    1.95253    0.00000
-2.60338    0.00000    0.00000 ;     N-    CA-    CB-    CG
 3539   3541   3559   3560     3    0.00000    0.00000    0.00000
0.00000    0.00000    0.00000 ;     N-    CA-     C-     O
 3539   3541   3559   3561     3   17.40544   -5.02080  -13.22144
9.20480    0.00000    0.00000 ;     N-    CA-     C-     N
 3540   3539   3541   3542     3    0.00000    0.00000    0.00000
0.00000    0.00000    0.00000 ;     H-     N-    CA-    HA
 3540   3539   3541   3543     3    0.00000    0.00000    0.00000
0.00000    0.00000    0.00000 ;     H-     N-    CA-    CB
 3540   3539   3541   3559     3    0.00000    0.00000    0.00000
0.00000    0.00000    0.00000 ;     H-     N-    CA-     C
 3541   3543   3546   3547     3    0.66944    2.00832    0.00000
-2.67776    0.00000    0.00000 ;    CA-    CB-    CG-   HG2
 3541   3543   3546   3548     3    0.66944    2.00832    0.00000
-2.67776    0.00000    0.00000 ;    CA-    CB-    CG-   HG3
 3541   3543   3546   3549     3    3.68192    3.09616   -2.09200
-3.01248    0.00000    0.00000 ;    CA-    CB-    CG-    CD
 3541   3559   3561   3562     3   20.92000    0.00000  -20.92000
0.00000    0.00000    0.00000 ;    CA-     C-     N-     H
 3541   3559   3561   3563     3   20.92000    0.00000  -20.92000
0.00000    0.00000    0.00000 ;    CA-     C-     N-    CA
 3542   3541   3543   3544     3    0.65084    1.95253    0.00000
-2.60338    0.00000    0.00000 ;    HA-    CA-    CB-   HB2
 3542   3541   3543   3545     3    0.65084    1.95253    0.00000

```
-2.60338    0.00000    0.00000 ;      HA-    CA-     CB-   HB3
  3542   3541   3543   3546       3    0.65084    1.95253    0.00000
-2.60338    0.00000    0.00000 ;      HA-    CA-     CB-    CG
  3542   3541   3559   3560       3    3.68192   -4.35136    0.00000
1.33888    0.00000    0.00000 ;      HA-    CA-     C-     O
  3542   3541   3559   3561       3    0.00000    0.00000    0.00000
0.00000    0.00000    0.00000 ;      HA-    CA-     C-     N
  3543   3541   3559   3560       3    0.00000    0.00000    0.00000
0.00000    0.00000    0.00000 ;      CB-    CA-     C-     O
  3543   3541   3559   3561       3    2.51040    4.18400    1.67360
-6.69440    0.00000    0.00000 ;      CB-    CA-     C-     N
  3543   3546   3549   3550       3    0.66944    2.00832    0.00000
-2.67776    0.00000    0.00000 ;      CB-    CG-     CD-   HD2
  3543   3546   3549   3551       3    0.66944    2.00832    0.00000
-2.67776    0.00000    0.00000 ;      CB-    CG-     CD-   HD3
  3543   3546   3549   3552       3    3.68192    3.09616   -2.09200
-3.01248    0.00000    0.00000 ;      CB-    CG-     CD-    CE
  3544   3543   3541   3559       3    0.65084    1.95253    0.00000
-2.60338    0.00000    0.00000 ;     HB2-    CB-     CA-     C
  3544   3543   3546   3547       3    0.62760    1.88280    0.00000
-2.51040    0.00000    0.00000 ;     HB2-    CB-     CG-   HG2
  3544   3543   3546   3548       3    0.62760    1.88280    0.00000
-2.51040    0.00000    0.00000 ;     HB2-    CB-     CG-   HG3
  3544   3543   3546   3549       3    0.66944    2.00832    0.00000
-2.67776    0.00000    0.00000 ;     HB2-    CB-     CG-    CD
  3545   3543   3541   3559       3    0.65084    1.95253    0.00000
-2.60338    0.00000    0.00000 ;     HB3-    CB-     CA-     C
  3545   3543   3546   3547       3    0.62760    1.88280    0.00000
-2.51040    0.00000    0.00000 ;     HB3-    CB-     CG-   HG2
  3545   3543   3546   3548       3    0.62760    1.88280    0.00000
-2.51040    0.00000    0.00000 ;     HB3-    CB-     CG-   HG3
  3545   3543   3546   3549       3    0.66944    2.00832    0.00000
-2.67776    0.00000    0.00000 ;     HB3-    CB-     CG-    CD
  3546   3543   3541   3559       3    0.65084    1.95253    0.00000
-2.60338    0.00000    0.00000 ;      CG-    CB-     CA-     C
  3546   3549   3552   3553       3    0.65084    1.95253    0.00000
-2.60338    0.00000    0.00000 ;      CG-    CD-     CE-   HE2
  3546   3549   3552   3554       3    0.65084    1.95253    0.00000
-2.60338    0.00000    0.00000 ;      CG-    CD-     CE-   HE3
  3546   3549   3552   3555       3    0.65084    1.95253    0.00000
-2.60338    0.00000    0.00000 ;      CG-    CD-     CE-    NZ
  3547   3546   3549   3550       3    0.62760    1.88280    0.00000
-2.51040    0.00000    0.00000 ;     HG2-    CG-     CD-   HD2
  3547   3546   3549   3551       3    0.62760    1.88280    0.00000
-2.51040    0.00000    0.00000 ;     HG2-    CG-     CD-   HD3
  3547   3546   3549   3552       3    0.66944    2.00832    0.00000
-2.67776    0.00000    0.00000 ;     HG2-    CG-     CD-    CE
  3548   3546   3549   3550       3    0.62760    1.88280    0.00000
-2.51040    0.00000    0.00000 ;     HG3-    CG-     CD-   HD2
  3548   3546   3549   3551       3    0.62760    1.88280    0.00000
-2.51040    0.00000    0.00000 ;     HG3-    CG-     CD-   HD3
  3548   3546   3549   3552       3    0.66944    2.00832    0.00000
```

-2.67776 0.00000 0.00000 ; HG3- CG- CD- CE
3549 3552 3555 3556 3 0.65084 1.95253 0.00000
-2.60338 0.00000 0.00000 ; CD- CE- NZ- HZ1
3549 3552 3555 3557 3 0.65084 1.95253 0.00000
-2.60338 0.00000 0.00000 ; CD- CE- NZ- HZ2
3549 3552 3555 3558 3 0.65084 1.95253 0.00000
-2.60338 0.00000 0.00000 ; CD- CE- NZ- HZ3
3550 3549 3552 3553 3 0.65084 1.95253 0.00000
-2.60338 0.00000 0.00000 ; HD2- CD- CE- HE2
3550 3549 3552 3554 3 0.65084 1.95253 0.00000
-2.60338 0.00000 0.00000 ; HD2- CD- CE- HE3
3550 3549 3552 3555 3 0.65084 1.95253 0.00000
-2.60338 0.00000 0.00000 ; HD2- CD- CE- NZ
3551 3549 3552 3553 3 0.65084 1.95253 0.00000
-2.60338 0.00000 0.00000 ; HD3- CD- CE- HE2
3551 3549 3552 3554 3 0.65084 1.95253 0.00000
-2.60338 0.00000 0.00000 ; HD3- CD- CE- HE3
3551 3549 3552 3555 3 0.65084 1.95253 0.00000
-2.60338 0.00000 0.00000 ; HD3- CD- CE- NZ
3553 3552 3555 3556 3 0.65084 1.95253 0.00000
-2.60338 0.00000 0.00000 ; HE2- CE- NZ- HZ1
3553 3552 3555 3557 3 0.65084 1.95253 0.00000
-2.60338 0.00000 0.00000 ; HE2- CE- NZ- HZ2
3553 3552 3555 3558 3 0.65084 1.95253 0.00000
-2.60338 0.00000 0.00000 ; HE2- CE- NZ- HZ3
3554 3552 3555 3556 3 0.65084 1.95253 0.00000
-2.60338 0.00000 0.00000 ; HE3- CE- NZ- HZ1
3554 3552 3555 3557 3 0.65084 1.95253 0.00000
-2.60338 0.00000 0.00000 ; HE3- CE- NZ- HZ2
3554 3552 3555 3558 3 0.65084 1.95253 0.00000
-2.60338 0.00000 0.00000 ; HE3- CE- NZ- HZ3
3559 3561 3563 3564 3 0.00000 0.00000 0.00000
0.00000 0.00000 0.00000 ; C- N- CA- HA
3559 3561 3563 3565 3 11.71520 1.67360 15.06240 -
13.38880 0.00000 0.00000 ; C- N- CA- CB
3559 3561 3563 3581 3 1.75728 5.27184 2.25936
-7.02912 0.00000 0.00000 ; C- N- CA- C
3560 3559 3561 3562 3 29.28800 -8.36800 -20.92000
0.00000 0.00000 0.00000 ; O- C- N- H
3560 3559 3561 3563 3 20.92000 0.00000 -20.92000
0.00000 0.00000 0.00000 ; O- C- N- CA
3561 3563 3565 3566 3 0.65084 1.95253 0.00000
-2.60338 0.00000 0.00000 ; N- CA- CB- HB2
3561 3563 3565 3567 3 0.65084 1.95253 0.00000
-2.60338 0.00000 0.00000 ; N- CA- CB- HB3
3561 3563 3565 3568 3 0.65084 1.95253 0.00000
-2.60338 0.00000 0.00000 ; N- CA- CB- CG
3561 3563 3581 3582 3 0.00000 0.00000 0.00000
0.00000 0.00000 0.00000 ; N- CA- C- O
3561 3563 3581 3583 3 17.40544 -5.02080 -13.22144
9.20480 0.00000 0.00000 ; N- CA- C- N
3562 3561 3563 3564 3 0.00000 0.00000 0.00000

0.00000    0.00000    0.00000 ;       H-     N-     CA-    HA
  3562   3561   3563   3565      3    0.00000    0.00000    0.00000
0.00000    0.00000    0.00000 ;       H-     N-     CA-    CB
  3562   3561   3563   3581      3    0.00000    0.00000    0.00000
0.00000    0.00000    0.00000 ;       H-     N-     CA-     C
  3563   3565   3568   3569      3    0.66944    2.00832    0.00000
-2.67776    0.00000    0.00000 ;      CA-    CB-    CG-   HG2
  3563   3565   3568   3570      3    0.66944    2.00832    0.00000
-2.67776    0.00000    0.00000 ;      CA-    CB-    CG-   HG3
  3563   3565   3568   3571      3    3.68192    3.09616   -2.09200
-3.01248    0.00000    0.00000 ;      CA-    CB-    CG-    CD
  3563   3581   3583   3584      3   20.92000    0.00000  -20.92000
0.00000    0.00000    0.00000 ;      CA-     C-     N-     H
  3563   3581   3583   3585      3   20.92000    0.00000  -20.92000
0.00000    0.00000    0.00000 ;      CA-     C-     N-    CA
  3564   3563   3565   3566      3    0.65084    1.95253    0.00000
-2.60338    0.00000    0.00000 ;      HA-    CA-    CB-   HB2
  3564   3563   3565   3567      3    0.65084    1.95253    0.00000
-2.60338    0.00000    0.00000 ;      HA-    CA-    CB-   HB3
  3564   3563   3565   3568      3    0.65084    1.95253    0.00000
-2.60338    0.00000    0.00000 ;      HA-    CA-    CB-    CG
  3564   3563   3581   3582      3    3.68192   -4.35136    0.00000
1.33888    0.00000    0.00000 ;      HA-    CA-     C-     O
  3564   3563   3581   3583      3    0.00000    0.00000    0.00000
0.00000    0.00000    0.00000 ;      HA-    CA-     C-     N
  3565   3563   3581   3582      3    0.00000    0.00000    0.00000
0.00000    0.00000    0.00000 ;      CB-    CA-     C-     O
  3565   3563   3581   3583      3    2.51040    4.18400    1.67360
-6.69440    0.00000    0.00000 ;      CB-    CA-     C-     N
  3565   3568   3571   3572      3    0.66944    2.00832    0.00000
-2.67776    0.00000    0.00000 ;      CB-    CG-    CD-   HD2
  3565   3568   3571   3573      3    0.66944    2.00832    0.00000
-2.67776    0.00000    0.00000 ;      CB-    CG-    CD-   HD3
  3565   3568   3571   3574      3    3.68192    3.09616   -2.09200
-3.01248    0.00000    0.00000 ;      CB-    CG-    CD-    CE
  3566   3565   3563   3581      3    0.65084    1.95253    0.00000
-2.60338    0.00000    0.00000 ;     HB2-    CB-    CA-     C
  3566   3565   3568   3569      3    0.62760    1.88280    0.00000
-2.51040    0.00000    0.00000 ;     HB2-    CB-    CG-   HG2
  3566   3565   3568   3570      3    0.62760    1.88280    0.00000
-2.51040    0.00000    0.00000 ;     HB2-    CB-    CG-   HG3
  3566   3565   3568   3571      3    0.66944    2.00832    0.00000
-2.67776    0.00000    0.00000 ;     HB2-    CB-    CG-    CD
  3567   3565   3563   3581      3    0.65084    1.95253    0.00000
-2.60338    0.00000    0.00000 ;     HB3-    CB-    CA-     C
  3567   3565   3568   3569      3    0.62760    1.88280    0.00000
-2.51040    0.00000    0.00000 ;     HB3-    CB-    CG-   HG2
  3567   3565   3568   3570      3    0.62760    1.88280    0.00000
-2.51040    0.00000    0.00000 ;     HB3-    CB-    CG-   HG3
  3567   3565   3568   3571      3    0.66944    2.00832    0.00000
-2.67776    0.00000    0.00000 ;     HB3-    CB-    CG-    CD
  3568   3565   3563   3581      3    0.65084    1.95253    0.00000

-2.60338 0.00000 0.00000 ; CG- CB- CA- C
3568 3571 3574 3575 3 0.65084 1.95253 0.00000
-2.60338 0.00000 0.00000 ; CG- CD- CE- HE2
3568 3571 3574 3576 3 0.65084 1.95253 0.00000
-2.60338 0.00000 0.00000 ; CG- CD- CE- HE3
3568 3571 3574 3577 3 0.65084 1.95253 0.00000
-2.60338 0.00000 0.00000 ; CG- CD- CE- NZ
3569 3568 3571 3572 3 0.62760 1.88280 0.00000
-2.51040 0.00000 0.00000 ; HG2- CG- CD- HD2
3569 3568 3571 3573 3 0.62760 1.88280 0.00000
-2.51040 0.00000 0.00000 ; HG2- CG- CD- HD3
3569 3568 3571 3574 3 0.66944 2.00832 0.00000
-2.67776 0.00000 0.00000 ; HG2- CG- CD- CE
3570 3568 3571 3572 3 0.62760 1.88280 0.00000
-2.51040 0.00000 0.00000 ; HG3- CG- CD- HD2
3570 3568 3571 3573 3 0.62760 1.88280 0.00000
-2.51040 0.00000 0.00000 ; HG3- CG- CD- HD3
3570 3568 3571 3574 3 0.66944 2.00832 0.00000
-2.67776 0.00000 0.00000 ; HG3- CG- CD- CE
3571 3574 3577 3578 3 0.65084 1.95253 0.00000
-2.60338 0.00000 0.00000 ; CD- CE- NZ- HZ1
3571 3574 3577 3579 3 0.65084 1.95253 0.00000
-2.60338 0.00000 0.00000 ; CD- CE- NZ- HZ2
3571 3574 3577 3580 3 0.65084 1.95253 0.00000
-2.60338 0.00000 0.00000 ; CD- CE- NZ- HZ3
3572 3571 3574 3575 3 0.65084 1.95253 0.00000
-2.60338 0.00000 0.00000 ; HD2- CD- CE- HE2
3572 3571 3574 3576 3 0.65084 1.95253 0.00000
-2.60338 0.00000 0.00000 ; HD2- CD- CE- HE3
3572 3571 3574 3577 3 0.65084 1.95253 0.00000
-2.60338 0.00000 0.00000 ; HD2- CD- CE- NZ
3573 3571 3574 3575 3 0.65084 1.95253 0.00000
-2.60338 0.00000 0.00000 ; HD3- CD- CE- HE2
3573 3571 3574 3576 3 0.65084 1.95253 0.00000
-2.60338 0.00000 0.00000 ; HD3- CD- CE- HE3
3573 3571 3574 3577 3 0.65084 1.95253 0.00000
-2.60338 0.00000 0.00000 ; HD3- CD- CE- NZ
3575 3574 3577 3578 3 0.65084 1.95253 0.00000
-2.60338 0.00000 0.00000 ; HE2- CE- NZ- HZ1
3575 3574 3577 3579 3 0.65084 1.95253 0.00000
-2.60338 0.00000 0.00000 ; HE2- CE- NZ- HZ2
3575 3574 3577 3580 3 0.65084 1.95253 0.00000
-2.60338 0.00000 0.00000 ; HE2- CE- NZ- HZ3
3576 3574 3577 3578 3 0.65084 1.95253 0.00000
-2.60338 0.00000 0.00000 ; HE3- CE- NZ- HZ1
3576 3574 3577 3579 3 0.65084 1.95253 0.00000
-2.60338 0.00000 0.00000 ; HE3- CE- NZ- HZ2
3576 3574 3577 3580 3 0.65084 1.95253 0.00000
-2.60338 0.00000 0.00000 ; HE3- CE- NZ- HZ3
3581 3583 3585 3586 3 0.00000 0.00000 0.00000
0.00000 0.00000 0.00000 ; C- N- CA- HA
3581 3583 3585 3587 3 11.71520 1.67360 15.06240 -

```
13.38880    0.00000    0.00000 ;       C-     N-    CA-    CB
  3581   3583   3585   3597       3    1.75728    5.27184    2.25936
-7.02912    0.00000    0.00000 ;       C-     N-    CA-     C
  3582   3581   3583   3584       3   29.28800   -8.36800  -20.92000
0.00000    0.00000    0.00000 ;      O-     C-     N-     H
  3582   3581   3583   3585       3   20.92000    0.00000  -20.92000
0.00000    0.00000    0.00000 ;      O-     C-     N-    CA
  3583   3585   3587   3588       3    0.65084    1.95253    0.00000
-2.60338    0.00000    0.00000 ;       N-    CA-    CB-    HB
  3583   3585   3587   3589       3    3.83673    0.44769   -1.77402
-2.47693   -0.03347    0.00000 ;       N-    CA-    CB-   CG1
  3583   3585   3587   3593       3    3.83673    0.44769   -1.77402
-2.47693   -0.03347    0.00000 ;       N-    CA-    CB-   CG2
  3583   3585   3597   3598       3    0.00000    0.00000    0.00000
0.00000    0.00000    0.00000 ;      N-    CA-     C-     O
  3583   3585   3597   3599       3   17.40544   -5.02080  -13.22144
9.20480    0.00000    0.00000 ;      N-    CA-     C-     N
  3584   3583   3585   3586       3    0.00000    0.00000    0.00000
0.00000    0.00000    0.00000 ;      H-     N-    CA-    HA
  3584   3583   3585   3587       3    0.00000    0.00000    0.00000
0.00000    0.00000    0.00000 ;      H-     N-    CA-    CB
  3584   3583   3585   3597       3    0.00000    0.00000    0.00000
0.00000    0.00000    0.00000 ;      H-     N-    CA-     C
  3585   3587   3589   3590       3    0.66944    2.00832    0.00000
-2.67776    0.00000    0.00000 ;      CA-    CB-   CG1-  HG11
  3585   3587   3589   3591       3    0.66944    2.00832    0.00000
-2.67776    0.00000    0.00000 ;      CA-    CB-   CG1-  HG12
  3585   3587   3589   3592       3    0.66944    2.00832    0.00000
-2.67776    0.00000    0.00000 ;      CA-    CB-   CG1-  HG13
  3585   3587   3593   3594       3    0.66944    2.00832    0.00000
-2.67776    0.00000    0.00000 ;      CA-    CB-   CG2-  HG21
  3585   3587   3593   3595       3    0.66944    2.00832    0.00000
-2.67776    0.00000    0.00000 ;      CA-    CB-   CG2-  HG22
  3585   3587   3593   3596       3    0.66944    2.00832    0.00000
-2.67776    0.00000    0.00000 ;      CA-    CB-   CG2-  HG23
  3585   3597   3599   3600       3   20.92000    0.00000  -20.92000
0.00000    0.00000    0.00000 ;     CA-     C-     N-     H
  3585   3597   3599   3601       3   20.92000    0.00000  -20.92000
0.00000    0.00000    0.00000 ;     CA-     C-     N-    CA
  3586   3585   3587   3588       3    0.65084    1.95253    0.00000
-2.60338    0.00000    0.00000 ;      HA-    CA-    CB-    HB
  3586   3585   3587   3589       3    0.65084    1.95253    0.00000
-2.60338    0.00000    0.00000 ;      HA-    CA-    CB-   CG1
  3586   3585   3587   3593       3    0.65084    1.95253    0.00000
-2.60338    0.00000    0.00000 ;      HA-    CA-    CB-   CG2
  3586   3585   3597   3598       3    3.68192   -4.35136    0.00000
1.33888    0.00000    0.00000 ;     HA-    CA-     C-     O
  3586   3585   3597   3599       3    0.00000    0.00000    0.00000
0.00000    0.00000    0.00000 ;     HA-    CA-     C-     N
  3587   3585   3597   3598       3    0.00000    0.00000    0.00000
0.00000    0.00000    0.00000 ;     CB-    CA-     C-     O
  3587   3585   3597   3599       3    2.51040    4.18400    1.67360
```

-6.69440 0.00000 0.00000 ; CB- CA- C- N
3588 3587 3585 3597 3 0.65084 1.95253 0.00000
-2.60338 0.00000 0.00000 ; HB- CB- CA- C
3588 3587 3589 3590 3 0.62760 1.88280 0.00000
-2.51040 0.00000 0.00000 ; HB- CB- CG1- HG11
3588 3587 3589 3591 3 0.62760 1.88280 0.00000
-2.51040 0.00000 0.00000 ; HB- CB- CG1- HG12
3588 3587 3589 3592 3 0.62760 1.88280 0.00000
-2.51040 0.00000 0.00000 ; HB- CB- CG1- HG13
3588 3587 3593 3594 3 0.62760 1.88280 0.00000
-2.51040 0.00000 0.00000 ; HB- CB- CG2- HG21
3588 3587 3593 3595 3 0.62760 1.88280 0.00000
-2.51040 0.00000 0.00000 ; HB- CB- CG2- HG22
3588 3587 3593 3596 3 0.62760 1.88280 0.00000
-2.51040 0.00000 0.00000 ; HB- CB- CG2- HG23
3589 3587 3585 3597 3 5.67350 3.55640 -6.16722
-2.47693 3.74886 0.00000 ; CG1- CB- CA- C
3589 3587 3593 3594 3 0.66944 2.00832 0.00000
-2.67776 0.00000 0.00000 ; CG1- CB- CG2- HG21
3589 3587 3593 3595 3 0.66944 2.00832 0.00000
-2.67776 0.00000 0.00000 ; CG1- CB- CG2- HG22
3589 3587 3593 3596 3 0.66944 2.00832 0.00000
-2.67776 0.00000 0.00000 ; CG1- CB- CG2- HG23
3590 3589 3587 3593 3 0.66944 2.00832 0.00000
-2.67776 0.00000 0.00000 ; HG11- CG1- CB- CG2
3591 3589 3587 3593 3 0.66944 2.00832 0.00000
-2.67776 0.00000 0.00000 ; HG12- CG1- CB- CG2
3592 3589 3587 3593 3 0.66944 2.00832 0.00000
-2.67776 0.00000 0.00000 ; HG13- CG1- CB- CG2
3593 3587 3585 3597 3 5.67350 3.55640 -6.16722
-2.47693 3.74886 0.00000 ; CG2- CB- CA- C
3597 3599 3601 3602 3 0.00000 0.00000 0.00000
0.00000 0.00000 0.00000 ; C- N- CA- HA
3597 3599 3601 3603 3 11.71520 1.67360 15.06240 -
13.38880 0.00000 0.00000 ; C- N- CA- CB
3597 3599 3601 3619 3 1.75728 5.27184 2.25936
-7.02912 0.00000 0.00000 ; C- N- CA- C
3598 3597 3599 3600 3 29.28800 -8.36800 -20.92000
0.00000 0.00000 0.00000 ; O- C- N- H
3598 3597 3599 3601 3 20.92000 0.00000 -20.92000
0.00000 0.00000 0.00000 ; O- C- N- CA
3599 3601 3603 3604 3 0.65084 1.95253 0.00000
-2.60338 0.00000 0.00000 ; N- CA- CB- HB2
3599 3601 3603 3605 3 0.65084 1.95253 0.00000
-2.60338 0.00000 0.00000 ; N- CA- CB- HB3
3599 3601 3603 3606 3 0.65084 1.95253 0.00000
-2.60338 0.00000 0.00000 ; N- CA- CB- CG
3599 3601 3619 3620 3 0.00000 0.00000 0.00000
0.00000 0.00000 0.00000 ; N- CA- C- O
3599 3601 3619 3621 3 17.40544 -5.02080 -13.22144
9.20480 0.00000 0.00000 ; N- CA- C- N
3600 3599 3601 3602 3 0.00000 0.00000 0.00000

```
0.00000    0.00000    0.00000 ;       H-     N-    CA-    HA
  3600   3599   3601   3619      3    0.00000    0.00000    0.00000
0.00000    0.00000    0.00000 ;       H-     N-    CA-    CB
  3600   3599   3601   3603      3    0.00000    0.00000    0.00000
0.00000    0.00000    0.00000 ;       H-     N-    CA-     C
  3601   3603   3606   3607      3    0.66944    2.00832    0.00000
-2.67776    0.00000    0.00000 ;      CA-    CB-    CG-   HG2
  3601   3603   3606   3608      3    0.66944    2.00832    0.00000
-2.67776    0.00000    0.00000 ;      CA-    CB-    CG-   HG3
  3601   3603   3606   3609      3    3.68192    3.09616   -2.09200
-3.01248    0.00000    0.00000 ;      CA-    CB-    CG-    CD
  3601   3619   3621   3622      3   20.92000    0.00000  -20.92000
0.00000    0.00000    0.00000 ;      CA-     C-     N-     H
  3601   3619   3621   3623      3   20.92000    0.00000  -20.92000
0.00000    0.00000    0.00000 ;      CA-     C-     N-    CA
  3602   3601   3603   3604      3    0.65084    1.95253    0.00000
-2.60338    0.00000    0.00000 ;      HA-    CA-    CB-   HB2
  3602   3601   3603   3605      3    0.65084    1.95253    0.00000
-2.60338    0.00000    0.00000 ;      HA-    CA-    CB-   HB3
  3602   3601   3603   3606      3    0.65084    1.95253    0.00000
-2.60338    0.00000    0.00000 ;      HA-    CA-    CB-    CG
  3602   3601   3619   3620      3    3.68192   -4.35136    0.00000
1.33888    0.00000    0.00000 ;      HA-    CA-     C-     O
  3602   3601   3619   3621      3    0.00000    0.00000    0.00000
0.00000    0.00000    0.00000 ;      HA-    CA-     C-     N
  3603   3601   3619   3620      3    0.00000    0.00000    0.00000
0.00000    0.00000    0.00000 ;      CB-    CA-     C-     O
  3603   3601   3619   3621      3    2.51040    4.18400    1.67360
-6.69440    0.00000    0.00000 ;      CB-    CA-     C-     N
  3603   3606   3609   3610      3    0.66944    2.00832    0.00000
-2.67776    0.00000    0.00000 ;      CB-    CG-    CD-   HD2
  3603   3606   3609   3611      3    0.66944    2.00832    0.00000
-2.67776    0.00000    0.00000 ;      CB-    CG-    CD-   HD3
  3603   3606   3609   3612      3    3.68192    3.09616   -2.09200
-3.01248    0.00000    0.00000 ;      CB-    CG-    CD-    CE
  3604   3603   3601   3619      3    0.65084    1.95253    0.00000
-2.60338    0.00000    0.00000 ;     HB2-    CB-    CA-     C
  3604   3603   3606   3607      3    0.62760    1.88280    0.00000
-2.51040    0.00000    0.00000 ;     HB2-    CB-    CG-   HG2
  3604   3603   3606   3608      3    0.62760    1.88280    0.00000
-2.51040    0.00000    0.00000 ;     HB2-    CB-    CG-   HG3
  3604   3603   3606   3609      3    0.66944    2.00832    0.00000
-2.67776    0.00000    0.00000 ;     HB2-    CB-    CG-    CD
  3605   3603   3601   3619      3    0.65084    1.95253    0.00000
-2.60338    0.00000    0.00000 ;     HB3-    CB-    CA-     C
  3605   3603   3606   3607      3    0.62760    1.88280    0.00000
-2.51040    0.00000    0.00000 ;     HB3-    CB-    CG-   HG2
  3605   3603   3606   3608      3    0.62760    1.88280    0.00000
-2.51040    0.00000    0.00000 ;     HB3-    CB-    CG-   HG3
  3605   3603   3606   3609      3    0.66944    2.00832    0.00000
-2.67776    0.00000    0.00000 ;     HB3-    CB-    CG-    CD
  3606   3603   3601   3619      3    0.65084    1.95253    0.00000
```

```
-2.60338    0.00000    0.00000 ;      CG-    CB-    CA-     C
  3606   3609   3612   3614       3    0.65084    1.95253    0.00000
-2.60338    0.00000    0.00000 ;      CG-    CD-    CE-   HE2
  3606   3609   3612   3614       3    0.65084    1.95253    0.00000
-2.60338    0.00000    0.00000 ;      CG-    CD-    CE-   HE3
  3606   3609   3612   3615       3    0.65084    1.95253    0.00000
-2.60338    0.00000    0.00000 ;      CG-    CD-    CE-    NZ
  3607   3606   3609   3610       3    0.62760    1.88280    0.00000
-2.51040    0.00000    0.00000 ;     HG2-    CG-    CD-   HD2
  3607   3606   3609   3611       3    0.62760    1.88280    0.00000
-2.51040    0.00000    0.00000 ;     HG2-    CG-    CD-   HD3
  3607   3606   3609   3612       3    0.66944    2.00832    0.00000
-2.67776    0.00000    0.00000 ;     HG2-    CG-    CD-    CE
  3608   3606   3609   3610       3    0.62760    1.88280    0.00000
-2.51040    0.00000    0.00000 ;     HG3-    CG-    CD-   HD2
  3608   3606   3609   3611       3    0.62760    1.88280    0.00000
-2.51040    0.00000    0.00000 ;     HG3-    CG-    CD-   HD3
  3608   3606   3609   3612       3    0.66944    2.00832    0.00000
-2.67776    0.00000    0.00000 ;     HG3-    CG-    CD-    CE
  3609   3612   3615   3616       3    0.65084    1.95253    0.00000
-2.60338    0.00000    0.00000 ;      CD-    CE-    NZ-   HZ1
  3609   3612   3615   3617       3    0.65084    1.95253    0.00000
-2.60338    0.00000    0.00000 ;      CD-    CE-    NZ-   HZ2
  3609   3612   3615   3618       3    0.65084    1.95253    0.00000
-2.60338    0.00000    0.00000 ;      CD-    CE-    NZ-   HZ3
  3610   3609   3612   3613       3    0.65084    1.95253    0.00000
-2.60338    0.00000    0.00000 ;     HD2-    CD-    CE-   HE2
  3610   3609   3612   3614       3    0.65084    1.95253    0.00000
-2.60338    0.00000    0.00000 ;     HD2-    CD-    CE-   HE3
  3610   3609   3612   3615       3    0.65084    1.95253    0.00000
-2.60338    0.00000    0.00000 ;     HD2-    CD-    CE-    NZ
  3611   3609   3612   3613       3    0.65084    1.95253    0.00000
-2.60338    0.00000    0.00000 ;     HD3-    CD-    CE-   HE2
  3611   3609   3612   3614       3    0.65084    1.95253    0.00000
-2.60338    0.00000    0.00000 ;     HD3-    CD-    CE-   HE3
  3611   3609   3612   3615       3    0.65084    1.95253    0.00000
-2.60338    0.00000    0.00000 ;     HD3-    CD-    CE-    NZ
  3613   3612   3615   3616       3    0.65084    1.95253    0.00000
-2.60338    0.00000    0.00000 ;     HE2-    CE-    NZ-   HZ1
  3613   3612   3615   3617       3    0.65084    1.95253    0.00000
-2.60338    0.00000    0.00000 ;     HE2-    CE-    NZ-   HZ2
  3613   3612   3615   3618       3    0.65084    1.95253    0.00000
-2.60338    0.00000    0.00000 ;     HE2-    CE-    NZ-   HZ3
  3614   3612   3615   3616       3    0.65084    1.95253    0.00000
-2.60338    0.00000    0.00000 ;     HE3-    CE-    NZ-   HZ1
  3614   3612   3615   3617       3    0.65084    1.95253    0.00000
-2.60338    0.00000    0.00000 ;     HE3-    CE-    NZ-   HZ2
  3614   3612   3615   3618       3    0.65084    1.95253    0.00000
-2.60338    0.00000    0.00000 ;     HE3-    CE-    NZ-   HZ3
  3619   3621   3623   3624       3    0.00000    0.00000    0.00000
0.00000    0.00000    0.00000 ;       C-     N-    CA-    HA
  3619   3621   3623   3625       3   11.71520    1.67360   15.06240  -
```

```
13.38880    0.00000    0.00000 ;      C-     N-    CA-    CB
  3619   3621   3623   3641      3    1.75728    5.27184    2.25936
-7.02912    0.00000    0.00000 ;      C-     N-    CA-     C
  3620   3619   3621   3622      3   29.28800   -8.36800  -20.92000
0.00000    0.00000    0.00000 ;      O-     C-     N-     H
  3620   3619   3621   3623      3   20.92000    0.00000  -20.92000
0.00000    0.00000    0.00000 ;      O-     C-     N-    CA
  3621   3623   3625   3626      3    0.65084    1.95253    0.00000
-2.60338    0.00000    0.00000 ;      N-    CA-    CB-   HB2
  3621   3623   3625   3627      3    0.65084    1.95253    0.00000
-2.60338    0.00000    0.00000 ;      N-    CA-    CB-   HB3
  3621   3623   3625   3628      3    0.65084    1.95253    0.00000
-2.60338    0.00000    0.00000 ;      N-    CA-    CB-    CG
  3621   3623   3641   3642      3    0.00000    0.00000    0.00000
0.00000    0.00000    0.00000 ;      N-    CA-     C-     O
  3621   3623   3641   3643      3   17.40544   -5.02080  -13.22144
9.20480    0.00000    0.00000 ;      N-    CA-     C-     N
  3622   3621   3623   3624      3    0.00000    0.00000    0.00000
0.00000    0.00000    0.00000 ;      H-     N-    CA-    HA
  3622   3621   3623   3625      3    0.00000    0.00000    0.00000
0.00000    0.00000    0.00000 ;      H-     N-    CA-    CB
  3622   3621   3623   3641      3    0.00000    0.00000    0.00000
0.00000    0.00000    0.00000 ;      H-     N-    CA-     C
  3623   3625   3628   3629      3    0.66944    2.00832    0.00000
-2.67776    0.00000    0.00000 ;     CA-    CB-    CG-   HG2
  3623   3625   3628   3630      3    0.66944    2.00832    0.00000
-2.67776    0.00000    0.00000 ;     CA-    CB-    CG-   HG3
  3623   3625   3628   3631      3    3.68192    3.09616   -2.09200
-3.01248    0.00000    0.00000 ;     CA-    CB-    CG-    CD
  3623   3641   3643   3644      3   20.92000    0.00000  -20.92000
0.00000    0.00000    0.00000 ;     CA-     C-     N-     H
  3623   3641   3643   3645      3   20.92000    0.00000  -20.92000
0.00000    0.00000    0.00000 ;     CA-     C-     N-    CA
  3624   3623   3625   3626      3    0.65084    1.95253    0.00000
-2.60338    0.00000    0.00000 ;     HA-    CA-    CB-   HB2
  3624   3623   3625   3627      3    0.65084    1.95253    0.00000
-2.60338    0.00000    0.00000 ;     HA-    CA-    CB-   HB3
  3624   3623   3625   3628      3    0.65084    1.95253    0.00000
-2.60338    0.00000    0.00000 ;     HA-    CA-    CB-    CG
  3624   3623   3641   3642      3    3.68192   -4.35136    0.00000
1.33888    0.00000    0.00000 ;     HA-    CA-     C-     O
  3624   3623   3641   3643      3    0.00000    0.00000    0.00000
0.00000    0.00000    0.00000 ;     HA-    CA-     C-     N
  3625   3623   3641   3642      3    0.00000    0.00000    0.00000
0.00000    0.00000    0.00000 ;     CB-    CA-     C-     O
  3625   3623   3641   3643      3    2.51040    4.18400    1.67360
-6.69440    0.00000    0.00000 ;     CB-    CA-     C-     N
  3625   3628   3631   3632      3    0.66944    2.00832    0.00000
-2.67776    0.00000    0.00000 ;     CB-    CG-    CD-   HD2
  3625   3628   3631   3633      3    0.66944    2.00832    0.00000
-2.67776    0.00000    0.00000 ;     CB-    CG-    CD-   HD3
  3625   3628   3631   3634      3    3.68192    3.09616   -2.09200
```

```
-3.01248    0.00000    0.00000 ;     CB-    CG-    CD-    CE
 3626   3625   3623   3641      3    0.65084    1.95253    0.00000
-2.60338    0.00000    0.00000 ;    HB2-    CB-    CA-     C
 3626   3625   3628   3629      3    0.62760    1.88280    0.00000
-2.51040    0.00000    0.00000 ;    HB2-    CB-    CG-   HG2
 3626   3625   3628   3630      3    0.62760    1.88280    0.00000
-2.51040    0.00000    0.00000 ;    HB2-    CB-    CG-   HG3
 3626   3625   3628   3631      3    0.66944    2.00832    0.00000
-2.67776    0.00000    0.00000 ;    HB2-    CB-    CG-    CD
 3627   3625   3623   3641      3    0.65084    1.95253    0.00000
-2.60338    0.00000    0.00000 ;    HB3-    CB-    CA-     C
 3627   3625   3628   3629      3    0.62760    1.88280    0.00000
-2.51040    0.00000    0.00000 ;    HB3-    CB-    CG-   HG2
 3627   3625   3628   3630      3    0.62760    1.88280    0.00000
-2.51040    0.00000    0.00000 ;    HB3-    CB-    CG-   HG3
 3627   3625   3628   3631      3    0.66944    2.00832    0.00000
-2.67776    0.00000    0.00000 ;    HB3-    CB-    CG-    CD
 3628   3625   3623   3641      3    0.65084    1.95253    0.00000
-2.60338    0.00000    0.00000 ;     CG-    CB-    CA-     C
 3628   3631   3634   3635      3    0.65084    1.95253    0.00000
-2.60338    0.00000    0.00000 ;     CG-    CD-    CE-   HE2
 3628   3631   3634   3636      3    0.65084    1.95253    0.00000
-2.60338    0.00000    0.00000 ;     CG-    CD-    CE-   HE3
 3628   3631   3634   3637      3    0.65084    1.95253    0.00000
-2.60338    0.00000    0.00000 ;     CG-    CD-    CE-    NZ
 3629   3628   3631   3632      3    0.62760    1.88280    0.00000
-2.51040    0.00000    0.00000 ;    HG2-    CG-    CD-   HD2
 3629   3628   3631   3633      3    0.62760    1.88280    0.00000
-2.51040    0.00000    0.00000 ;    HG2-    CG-    CD-   HD3
 3629   3628   3631   3634      3    0.66944    2.00832    0.00000
-2.67776    0.00000    0.00000 ;    HG2-    CG-    CD-    CE
 3630   3628   3631   3632      3    0.62760    1.88280    0.00000
-2.51040    0.00000    0.00000 ;    HG3-    CG-    CD-   HD2
 3630   3628   3631   3633      3    0.62760    1.88280    0.00000
-2.51040    0.00000    0.00000 ;    HG3-    CG-    CD-   HD3
 3630   3628   3631   3634      3    0.66944    2.00832    0.00000
-2.67776    0.00000    0.00000 ;    HG3-    CG-    CD-    CE
 3631   3634   3637   3638      3    0.65084    1.95253    0.00000
-2.60338    0.00000    0.00000 ;     CD-    CE-    NZ-   HZ1
 3631   3634   3637   3639      3    0.65084    1.95253    0.00000
-2.60338    0.00000    0.00000 ;     CD-    CE-    NZ-   HZ2
 3631   3634   3637   3640      3    0.65084    1.95253    0.00000
-2.60338    0.00000    0.00000 ;     CD-    CE-    NZ-   HZ3
 3632   3631   3634   3635      3    0.65084    1.95253    0.00000
-2.60338    0.00000    0.00000 ;    HD2-    CD-    CE-   HE2
 3632   3631   3634   3636      3    0.65084    1.95253    0.00000
-2.60338    0.00000    0.00000 ;    HD2-    CD-    CE-   HE3
 3632   3631   3634   3637      3    0.65084    1.95253    0.00000
-2.60338    0.00000    0.00000 ;    HD2-    CD-    CE-    NZ
 3633   3631   3634   3635      3    0.65084    1.95253    0.00000
-2.60338    0.00000    0.00000 ;    HD3-    CD-    CE-   HE2
 3633   3631   3634   3636      3    0.65084    1.95253    0.00000
```

-2.60338 0.00000 0.00000 ; HD3- CD- CE- HE3
3633 3631 3634 3637 3 0.65084 1.95253 0.00000
-2.60338 0.00000 0.00000 ; HD3- CD- CE- NZ
3635 3634 3637 3638 3 0.65084 1.95253 0.00000
-2.60338 0.00000 0.00000 ; HE2- CE- NZ- HZ1
3635 3634 3637 3639 3 0.65084 1.95253 0.00000
-2.60338 0.00000 0.00000 ; HE2- CE- NZ- HZ2
3635 3634 3637 3640 3 0.65084 1.95253 0.00000
-2.60338 0.00000 0.00000 ; HE2- CE- NZ- HZ3
3636 3634 3637 3638 3 0.65084 1.95253 0.00000
-2.60338 0.00000 0.00000 ; HE3- CE- NZ- HZ1
3636 3634 3637 3639 3 0.65084 1.95253 0.00000
-2.60338 0.00000 0.00000 ; HE3- CE- NZ- HZ2
3636 3634 3637 3640 3 0.65084 1.95253 0.00000
-2.60338 0.00000 0.00000 ; HE3- CE- NZ- HZ3
3641 3643 3645 3646 3 0.00000 0.00000 0.00000
0.00000 0.00000 0.00000 ; C- N- CA- HA2
3641 3643 3645 3647 3 0.00000 0.00000 0.00000
0.00000 0.00000 0.00000 ; C- N- CA- HA3
3641 3643 3645 3648 3 1.75728 5.27184 2.25936
-7.02912 0.00000 0.00000 ; C- N- CA- C
3642 3641 3643 3644 3 29.28800 -8.36800 -20.92000
0.00000 0.00000 0.00000 ; O- C- N- H
3642 3641 3643 3645 3 20.92000 0.00000 -20.92000
0.00000 0.00000 0.00000 ; O- C- N- CA
3643 3645 3648 3649 3 0.00000 0.00000 0.00000
0.00000 0.00000 0.00000 ; N- CA- C- O
3643 3645 3648 3650 3 17.40544 -5.02080 -13.22144
9.20480 0.00000 0.00000 ; N- CA- C- N
3644 3643 3645 3646 3 0.00000 0.00000 0.00000
0.00000 0.00000 0.00000 ; H- N- CA- HA2
3644 3643 3645 3647 3 0.00000 0.00000 0.00000
0.00000 0.00000 0.00000 ; H- N- CA- HA3
3644 3643 3645 3648 3 0.00000 0.00000 0.00000
0.00000 0.00000 0.00000 ; H- N- CA- C
3645 3648 3650 3651 3 20.92000 0.00000 -20.92000
0.00000 0.00000 0.00000 ; CA- C- N- H
3645 3648 3650 3652 3 20.92000 0.00000 -20.92000
0.00000 0.00000 0.00000 ; CA- C- N- CA
3646 3645 3648 3649 3 3.68192 -4.35136 0.00000
1.33888 0.00000 0.00000 ; HA2- CA- C- O
3646 3645 3648 3650 3 0.00000 0.00000 0.00000
0.00000 0.00000 0.00000 ; HA2- CA- C- N
3647 3645 3648 3649 3 3.68192 -4.35136 0.00000
1.33888 0.00000 0.00000 ; HA3- CA- C- O
3647 3645 3648 3650 3 0.00000 0.00000 0.00000
0.00000 0.00000 0.00000 ; HA3- CA- C- N
3648 3650 3652 3653 3 0.00000 0.00000 0.00000
0.00000 0.00000 0.00000 ; C- N- CA- HA
3648 3650 3652 3654 3 11.71520 1.67360 15.06240 -
13.38880 0.00000 0.00000 ; C- N- CA- CB
3648 3650 3652 3663 3 1.75728 5.27184 2.25936

-7.02912 0.00000 0.00000 ; C- N- CA- C
3649 3648 3650 3651 3 29.28800 -8.36800 -20.92000
0.00000 0.00000 0.00000 ; O- C- N- H
3649 3648 3650 3652 3 20.92000 0.00000 -20.92000
0.00000 0.00000 0.00000 ; O- C- N- CA
3650 3652 3654 3655 3 0.65084 1.95253 0.00000
-2.60338 0.00000 0.00000 ; N- CA- CB- HB2
3650 3652 3654 3656 3 0.65084 1.95253 0.00000
-2.60338 0.00000 0.00000 ; N- CA- CB- HB3
3650 3652 3654 3657 3 3.21331 2.22589 -4.15053
-2.40998 2.61082 0.00000 ; N- CA- CB- CG
3650 3652 3663 3664 3 0.00000 0.00000 0.00000
0.00000 0.00000 0.00000 ; N- CA- C- O
3650 3652 3663 3665 3 17.40544 -5.02080 -13.22144
9.20480 0.00000 0.00000 ; N- CA- C- N
3651 3650 3652 3653 3 0.00000 0.00000 0.00000
0.00000 0.00000 0.00000 ; H- N- CA- HA
3651 3650 3652 3654 3 0.00000 0.00000 0.00000
0.00000 0.00000 0.00000 ; H- N- CA- CB
3651 3650 3652 3663 3 0.00000 0.00000 0.00000
0.00000 0.00000 0.00000 ; H- N- CA- C
3652 3654 3657 3658 3 0.66944 2.00832 0.00000
-2.67776 0.00000 0.00000 ; CA- CB- CG- HG2
3652 3654 3657 3659 3 0.66944 2.00832 0.00000
-2.67776 0.00000 0.00000 ; CA- CB- CG- HG3
3652 3654 3657 3660 3 10.12110 -1.91209 0.01674
10.17549 -1.87443 0.00000 ; CA- CB- CG- CD
3652 3663 3665 3666 3 20.92000 0.00000 -20.92000
0.00000 0.00000 0.00000 ; CA- C- N- H
3652 3663 3665 3667 3 20.92000 0.00000 -20.92000
0.00000 0.00000 0.00000 ; CA- C- N- CA
3653 3652 3654 3655 3 0.65084 1.95253 0.00000
-2.60338 0.00000 0.00000 ; HA- CA- CB- HB2
3653 3652 3654 3656 3 0.65084 1.95253 0.00000
-2.60338 0.00000 0.00000 ; HA- CA- CB- HB3
3653 3652 3654 3657 3 0.65084 1.95253 0.00000
-2.60338 0.00000 0.00000 ; HA- CA- CB- CG
3653 3652 3663 3664 3 3.68192 -4.35136 0.00000
1.33888 0.00000 0.00000 ; HA- CA- C- O
3653 3652 3663 3665 3 0.00000 0.00000 0.00000
0.00000 0.00000 0.00000 ; HA- CA- C- N
3654 3652 3663 3664 3 0.00000 0.00000 0.00000
0.00000 0.00000 0.00000 ; CB- CA- C- O
3654 3652 3663 3665 3 2.51040 4.18400 1.67360
-6.69440 0.00000 0.00000 ; CB- CA- C- N
3654 3657 3660 3661 3 3.79907 0.00000 -5.40573
0.00000 2.14221 0.00000 ; CB- CG- CD- OE1
3654 3657 3660 3662 3 3.79907 0.00000 -5.40573
0.00000 2.14221 0.00000 ; CB- CG- CD- OE2
3655 3654 3652 3663 3 0.65084 1.95253 0.00000
-2.60338 0.00000 0.00000 ; HB2- CB- CA- C
3655 3654 3657 3658 3 0.62760 1.88280 0.00000

```
-2.51040    0.00000    0.00000 ;    HB2-    CB-    CG-   HG2
  3655   3654   3657   3659      3    0.62760    1.88280    0.00000
-2.51040    0.00000    0.00000 ;    HB2-    CB-    CG-   HG3
  3655   3654   3657   3660      3    0.65084    1.95253    0.00000
-2.60338    0.00000    0.00000 ;    HB2-    CB-    CG-    CD
  3656   3654   3652   3663      3    0.65084    1.95253    0.00000
-2.60338    0.00000    0.00000 ;    HB3-    CB-    CA-     C
  3656   3654   3657   3658      3    0.62760    1.88280    0.00000
-2.51040    0.00000    0.00000 ;    HB3-    CB-    CG-   HG2
  3656   3654   3657   3659      3    0.62760    1.88280    0.00000
-2.51040    0.00000    0.00000 ;    HB3-    CB-    CG-   HG3
  3656   3654   3657   3660      3    0.65084    1.95253    0.00000
-2.60338    0.00000    0.00000 ;    HB3-    CB-    CG-    CD
  3657   3654   3652   3663      3    6.86594    3.56895   -8.14206
-2.40998    4.85344    0.00000 ;     CG-    CB-    CA-     C
  3658   3657   3660   3661      3    0.00000    0.00000    0.00000
0.00000    0.00000    0.00000 ;    HG2-    CG-    CD-   OE1
  3658   3657   3660   3662      3    0.00000    0.00000    0.00000
0.00000    0.00000    0.00000 ;    HG2-    CG-    CD-   OE2
  3659   3657   3660   3661      3    0.00000    0.00000    0.00000
0.00000    0.00000    0.00000 ;    HG3-    CG-    CD-   OE1
  3659   3657   3660   3662      3    0.00000    0.00000    0.00000
0.00000    0.00000    0.00000 ;    HG3-    CG-    CD-   OE2
  3663   3665   3667   3668      3    0.00000    0.00000    0.00000
0.00000    0.00000    0.00000 ;     C-     N-    CA-    HA
  3663   3665   3667   3669      3   11.71520    1.67360   15.06240  -
13.38880    0.00000    0.00000 ;     C-     N-    CA-    CB
  3663   3665   3667   3683      3    1.75728    5.27184    2.25936
-7.02912    0.00000    0.00000 ;     C-     N-    CA-     C
  3664   3663   3665   3666      3   29.28800   -8.36800  -20.92000
0.00000    0.00000    0.00000 ;     O-     C-     N-     H
  3664   3663   3665   3667      3   20.92000    0.00000  -20.92000
0.00000    0.00000    0.00000 ;     O-     C-     N-    CA
  3665   3667   3669   3670      3    0.65084    1.95253    0.00000
-2.60338    0.00000    0.00000 ;     N-    CA-    CB-   HB2
  3665   3667   3669   3671      3    0.65084    1.95253    0.00000
-2.60338    0.00000    0.00000 ;     N-    CA-    CB-   HB3
  3665   3667   3669   3672      3    3.28026    2.46019   -2.19242
-3.21331   -0.23430    0.00000 ;     N-    CA-    CB-    CG
  3665   3667   3683   3684      3    0.00000    0.00000    0.00000
0.00000    0.00000    0.00000 ;     N-    CA-     C-     O
  3665   3667   3683   3685      3   17.40544   -5.02080  -13.22144
9.20480    0.00000    0.00000 ;     N-    CA-     C-     N
  3666   3665   3667   3668      3    0.00000    0.00000    0.00000
0.00000    0.00000    0.00000 ;     H-     N-    CA-    HA
  3666   3665   3667   3669      3    0.00000    0.00000    0.00000
0.00000    0.00000    0.00000 ;     H-     N-    CA-    CB
  3666   3665   3667   3683      3    0.00000    0.00000    0.00000
0.00000    0.00000    0.00000 ;     H-     N-    CA-     C
  3667   3669   3672   3673      3    0.57739    0.00000    1.02926
0.00000   -1.60666    0.00000 ;    CA-    CB-    CG-   CD1
  3667   3669   3672   3681      3    0.57739    0.00000    1.02926
```

0.00000 -1.60666 0.00000 ; CA- CB- CG- CD2
3667 3683 3685 3686 3 20.92000 0.00000 -20.92000
0.00000 0.00000 0.00000 ; CA- C- N- H
3667 3683 3685 3687 3 20.92000 0.00000 -20.92000
0.00000 0.00000 0.00000 ; CA- C- N- CA
3668 3667 3669 3670 3 0.65084 1.95253 0.00000
-2.60338 0.00000 0.00000 ; HA- CA- CB- HB2
3668 3667 3669 3671 3 0.65084 1.95253 0.00000
-2.60338 0.00000 0.00000 ; HA- CA- CB- HB3
3668 3667 3669 3672 3 0.65084 1.95253 0.00000
-2.60338 0.00000 0.00000 ; HA- CA- CB- CG
3668 3667 3683 3684 3 3.68192 -4.35136 0.00000
1.33888 0.00000 0.00000 ; HA- CA- C- O
3668 3667 3683 3685 3 0.00000 0.00000 0.00000
0.00000 0.00000 0.00000 ; HA- CA- C- N
3669 3667 3683 3684 3 0.00000 0.00000 0.00000
0.00000 0.00000 0.00000 ; CB- CA- C- O
3669 3667 3683 3685 3 2.51040 4.18400 1.67360
-6.69440 0.00000 0.00000 ; CB- CA- C- N
3669 3672 3673 3674 3 30.33400 0.00000 -30.33400
0.00000 0.00000 0.00000 ; CB- CG- CD1- HD1
3669 3672 3673 3675 3 30.33400 0.00000 -30.33400
0.00000 0.00000 0.00000 ; CB- CG- CD1- CE1
3669 3672 3681 3679 3 30.33400 0.00000 -30.33400
0.00000 0.00000 0.00000 ; CB- CG- CD2- CE2
3669 3672 3681 3682 3 30.33400 0.00000 -30.33400
0.00000 0.00000 0.00000 ; CB- CG- CD2- HD2
3670 3669 3667 3683 3 0.65084 1.95253 0.00000
-2.60338 0.00000 0.00000 ; HB2- CB- CA- C
3670 3669 3672 3673 3 0.00000 0.00000 0.00000
0.00000 0.00000 0.00000 ; HB2- CB- CG- CD1
3670 3669 3672 3681 3 0.00000 0.00000 0.00000
0.00000 0.00000 0.00000 ; HB2- CB- CG- CD2
3671 3669 3667 3683 3 0.65084 1.95253 0.00000
-2.60338 0.00000 0.00000 ; HB3- CB- CA- C
3671 3669 3672 3673 3 0.00000 0.00000 0.00000
0.00000 0.00000 0.00000 ; HB3- CB- CG- CD1
3671 3669 3672 3681 3 0.00000 0.00000 0.00000
0.00000 0.00000 0.00000 ; HB3- CB- CG- CD2
3672 3669 3667 3683 3 4.95804 2.17986 -3.52293
-3.21331 -0.40166 0.00000 ; CG- CB- CA- C
3672 3673 3675 3676 3 30.33400 0.00000 -30.33400
0.00000 0.00000 0.00000 ; CG- CD1- CE1- HE1
3672 3673 3675 3677 3 30.33400 0.00000 -30.33400
0.00000 0.00000 0.00000 ; CG- CD1- CE1- CZ
3672 3681 3679 3677 3 30.33400 0.00000 -30.33400
0.00000 0.00000 0.00000 ; CG- CD2- CE2- CZ
3672 3681 3679 3680 3 30.33400 0.00000 -30.33400
0.00000 0.00000 0.00000 ; CG- CD2- CE2- HE2
3673 3672 3681 3679 3 30.33400 0.00000 -30.33400
0.00000 0.00000 0.00000 ; CD1- CG- CD2- CE2
3673 3672 3681 3682 3 30.33400 0.00000 -30.33400

0.00000 0.00000 0.00000 ; CD1- CG- CD2- HD2
3673 3675 3677 3678 3 30.33400 0.00000 -30.33400
0.00000 0.00000 0.00000 ; CD1- CE1- CZ- HZ
3673 3675 3677 3679 3 30.33400 0.00000 -30.33400
0.00000 0.00000 0.00000 ; CD1- CE1- CZ- CE2
3674 3673 3672 3681 3 30.33400 0.00000 -30.33400
0.00000 0.00000 0.00000 ; HD1- CD1- CG- CD2
3674 3673 3675 3676 3 30.33400 0.00000 -30.33400
0.00000 0.00000 0.00000 ; HD1- CD1- CE1- HE1
3674 3673 3675 3677 3 30.33400 0.00000 -30.33400
0.00000 0.00000 0.00000 ; HD1- CD1- CE1- CZ
3675 3673 3672 3681 3 30.33400 0.00000 -30.33400
0.00000 0.00000 0.00000 ; CE1- CD1- CG- CD2
3675 3677 3679 3680 3 30.33400 0.00000 -30.33400
0.00000 0.00000 0.00000 ; CE1- CZ- CE2- HE2
3675 3677 3679 3681 3 30.33400 0.00000 -30.33400
0.00000 0.00000 0.00000 ; CE1- CZ- CE2- CD2
3676 3675 3677 3678 3 30.33400 0.00000 -30.33400
0.00000 0.00000 0.00000 ; HE1- CE1- CZ- HZ
3676 3675 3677 3679 3 30.33400 0.00000 -30.33400
0.00000 0.00000 0.00000 ; HE1- CE1- CZ- CE2
3677 3679 3681 3682 3 30.33400 0.00000 -30.33400
0.00000 0.00000 0.00000 ; CZ- CE2- CD2- HD2
3678 3677 3679 3680 3 30.33400 0.00000 -30.33400
0.00000 0.00000 0.00000 ; HZ- CZ- CE2- HE2
3678 3677 3679 3681 3 30.33400 0.00000 -30.33400
0.00000 0.00000 0.00000 ; HZ- CZ- CE2- CD2
3680 3679 3681 3682 3 30.33400 0.00000 -30.33400
0.00000 0.00000 0.00000 ; HE2- CE2- CD2- HD2
3683 3685 3687 3688 3 0.00000 0.00000 0.00000
0.00000 0.00000 0.00000 ; C- N- CA- HA
3683 3685 3687 3689 3 11.71520 1.67360 15.06240 -
13.38880 0.00000 0.00000 ; C- N- CA- CB
3683 3685 3687 3694 3 1.75728 5.27184 2.25936
-7.02912 0.00000 0.00000 ; C- N- CA- C
3684 3683 3685 3686 3 29.28800 -8.36800 -20.92000
0.00000 0.00000 0.00000 ; O- C- N- H
3684 3683 3685 3687 3 20.92000 0.00000 -20.92000
0.00000 0.00000 0.00000 ; O- C- N- CA
3685 3687 3689 3690 3 0.65084 1.95253 0.00000
-2.60338 0.00000 0.00000 ; N- CA- CB- HB2
3685 3687 3689 3691 3 0.65084 1.95253 0.00000
-2.60338 0.00000 0.00000 ; N- CA- CB- HB3
3685 3687 3689 3692 3 6.03751 2.50622 -5.17142
-4.20074 1.10458 0.00000 ; N- CA- CB- SG
3685 3687 3694 3695 3 0.00000 0.00000 0.00000
0.00000 0.00000 0.00000 ; N- CA- C- O
3685 3687 3694 3696 3 17.40544 -5.02080 -13.22144
9.20480 0.00000 0.00000 ; N- CA- C- N
3686 3685 3687 3688 3 0.00000 0.00000 0.00000
0.00000 0.00000 0.00000 ; H- N- CA- HA
3686 3685 3687 3689 3 0.00000 0.00000 0.00000

0.00000 0.00000 0.00000 ; H- N- CA- CB
3686 3685 3687 3694 3 0.00000 0.00000 0.00000
0.00000 0.00000 0.00000 ; H- N- CA- C
3687 3689 3692 3693 3 1.69034 2.77818 4.11706
-4.21747 1.00416 0.00000 ; CA- CB- SG- HG
3687 3694 3696 3697 3 20.92000 0.00000 -20.92000
0.00000 0.00000 0.00000 ; CA- C- N- H
3687 3694 3696 3698 3 20.92000 0.00000 -20.92000
0.00000 0.00000 0.00000 ; CA- C- N- CA
3688 3687 3689 3690 3 0.65084 1.95253 0.00000
-2.60338 0.00000 0.00000 ; HA- CA- CB- HB2
3688 3687 3689 3691 3 0.65084 1.95253 0.00000
-2.60338 0.00000 0.00000 ; HA- CA- CB- HB3
3688 3687 3689 3692 3 0.65084 1.95253 0.00000
-2.60338 0.00000 0.00000 ; HA- CA- CB- SG
3688 3687 3694 3695 3 3.68192 -4.35136 0.00000
1.33888 0.00000 0.00000 ; HA- CA- C- O
3688 3687 3694 3696 3 0.00000 0.00000 0.00000
0.00000 0.00000 0.00000 ; HA- CA- C- N
3689 3687 3694 3695 3 0.00000 0.00000 0.00000
0.00000 0.00000 0.00000 ; CB- CA- C- O
3689 3687 3694 3696 3 2.51040 4.18400 1.67360
-6.69440 0.00000 0.00000 ; CB- CA- C- N
3690 3689 3687 3694 3 0.65084 1.95253 0.00000
-2.60338 0.00000 0.00000 ; HB2- CB- CA- C
3690 3689 3692 3693 3 1.04600 3.13800 0.00000
-4.18400 0.00000 0.00000 ; HB2- CB- SG- HG
3691 3689 3687 3694 3 0.65084 1.95253 0.00000
-2.60338 0.00000 0.00000 ; HB3- CB- CA- C
3691 3689 3692 3693 3 1.04600 3.13800 0.00000
-4.18400 0.00000 0.00000 ; HB3- CB- SG- HG
3692 3689 3687 3694 3 5.62330 4.27605 -5.33042
-4.20074 2.51040 0.00000 ; SG- CB- CA- C
3694 3696 3698 3699 3 0.00000 0.00000 0.00000
0.00000 0.00000 0.00000 ; C- N- CA- HA
3694 3696 3698 3700 3 11.71520 1.67360 15.06240 -
13.38880 0.00000 0.00000 ; C- N- CA- CB
3694 3696 3698 3714 3 1.75728 5.27184 2.25936
-7.02912 0.00000 0.00000 ; C- N- CA- C
3695 3694 3696 3697 3 29.28800 -8.36800 -20.92000
0.00000 0.00000 0.00000 ; O- C- N- H
3695 3694 3696 3698 3 20.92000 0.00000 -20.92000
0.00000 0.00000 0.00000 ; O- C- N- CA
3696 3698 3700 3701 3 0.65084 1.95253 0.00000
-2.60338 0.00000 0.00000 ; N- CA- CB- HB2
3696 3698 3700 3702 3 0.65084 1.95253 0.00000
-2.60338 0.00000 0.00000 ; N- CA- CB- HB3
3696 3698 3700 3703 3 3.28026 2.46019 -2.19242
-3.21331 -0.23430 0.00000 ; N- CA- CB- CG
3696 3698 3714 3715 3 0.00000 0.00000 0.00000
0.00000 0.00000 0.00000 ; N- CA- C- O
3696 3698 3714 3716 3 17.40544 -5.02080 -13.22144

9.20480 0.00000 0.00000 ; N- CA- C- N
3697 3696 3698 3699 3 0.00000 0.00000 0.00000
0.00000 0.00000 0.00000 ; H- N- CA- HA
3697 3696 3698 3700 3 0.00000 0.00000 0.00000
0.00000 0.00000 0.00000 ; H- N- CA- CB
3697 3696 3698 3714 3 0.00000 0.00000 0.00000
0.00000 0.00000 0.00000 ; H- N- CA- C
3698 3700 3703 3704 3 0.57739 0.00000 1.02926
0.00000 -1.60666 0.00000 ; CA- CB- CG- CD1
3698 3700 3703 3712 3 0.57739 0.00000 1.02926
0.00000 -1.60666 0.00000 ; CA- CB- CG- CD2
3698 3714 3716 3717 3 20.92000 0.00000 -20.92000
0.00000 0.00000 0.00000 ; CA- C- N- H
3698 3714 3716 3718 3 20.92000 0.00000 -20.92000
0.00000 0.00000 0.00000 ; CA- C- N- CA
3699 3698 3700 3701 3 0.65084 1.95253 0.00000
-2.60338 0.00000 0.00000 ; HA- CA- CB- HB2
3699 3698 3700 3702 3 0.65084 1.95253 0.00000
-2.60338 0.00000 0.00000 ; HA- CA- CB- HB3
3699 3698 3700 3703 3 0.65084 1.95253 0.00000
-2.60338 0.00000 0.00000 ; HA- CA- CB- CG
3699 3698 3714 3715 3 3.68192 -4.35136 0.00000
1.33888 0.00000 0.00000 ; HA- CA- C- O
3699 3698 3714 3716 3 0.00000 0.00000 0.00000
0.00000 0.00000 0.00000 ; HA- CA- C- N
3700 3698 3714 3715 3 0.00000 0.00000 0.00000
0.00000 0.00000 0.00000 ; CB- CA- C- O
3700 3698 3714 3716 3 2.51040 4.18400 1.67360
-6.69440 0.00000 0.00000 ; CB- CA- C- N
3700 3703 3704 3705 3 30.33400 0.00000 -30.33400
0.00000 0.00000 0.00000 ; CB- CG- CD1- HD1
3700 3703 3704 3706 3 30.33400 0.00000 -30.33400
0.00000 0.00000 0.00000 ; CB- CG- CD1- CE1
3700 3703 3712 3710 3 30.33400 0.00000 -30.33400
0.00000 0.00000 0.00000 ; CB- CG- CD2- CE2
3700 3703 3712 3713 3 30.33400 0.00000 -30.33400
0.00000 0.00000 0.00000 ; CB- CG- CD2- HD2
3701 3700 3698 3714 3 0.65084 1.95253 0.00000
-2.60338 0.00000 0.00000 ; HB2- CB- CA- C
3701 3700 3703 3704 3 0.00000 0.00000 0.00000
0.00000 0.00000 0.00000 ; HB2- CB- CG- CD1
3701 3700 3703 3712 3 0.00000 0.00000 0.00000
0.00000 0.00000 0.00000 ; HB2- CB- CG- CD2
3702 3700 3698 3714 3 0.65084 1.95253 0.00000
-2.60338 0.00000 0.00000 ; HB3- CB- CA- C
3702 3700 3703 3704 3 0.00000 0.00000 0.00000
0.00000 0.00000 0.00000 ; HB3- CB- CG- CD1
3702 3700 3703 3712 3 0.00000 0.00000 0.00000
0.00000 0.00000 0.00000 ; HB3- CB- CG- CD2
3703 3700 3698 3714 3 4.95804 2.17986 -3.52293
-3.21331 -0.40166 0.00000 ; CG- CB- CA- C
3703 3704 3706 3707 3 30.33400 0.00000 -30.33400

```
0.00000    0.00000    0.00000 ;      CG-   CD1-   CE1-   HE1
  3703   3704   3706   3708       3   30.33400    0.00000  -30.33400
0.00000    0.00000    0.00000 ;      CG-   CD1-   CE1-    CZ
  3703   3712   3710   3708       3   30.33400    0.00000  -30.33400
0.00000    0.00000    0.00000 ;      CG-   CD2-   CE2-    CZ
  3703   3712   3710   3711       3   30.33400    0.00000  -30.33400
0.00000    0.00000    0.00000 ;      CG-   CD2-   CE2-   HE2
  3704   3703   3712   3710       3   30.33400    0.00000  -30.33400
0.00000    0.00000    0.00000 ;     CD1-    CG-   CD2-   CE2
  3704   3703   3712   3713       3   30.33400    0.00000  -30.33400
0.00000    0.00000    0.00000 ;     CD1-    CG-   CD2-   HD2
  3704   3706   3708   3709       3   30.33400    0.00000  -30.33400
0.00000    0.00000    0.00000 ;     CD1-   CE1-    CZ-    HZ
  3704   3706   3708   3710       3   30.33400    0.00000  -30.33400
0.00000    0.00000    0.00000 ;     CD1-   CE1-    CZ-   CE2
  3705   3704   3703   3712       3   30.33400    0.00000  -30.33400
0.00000    0.00000    0.00000 ;     HD1-   CD1-    CG-   CD2
  3705   3704   3706   3707       3   30.33400    0.00000  -30.33400
0.00000    0.00000    0.00000 ;     HD1-   CD1-   CE1-   HE1
  3705   3704   3706   3708       3   30.33400    0.00000  -30.33400
0.00000    0.00000    0.00000 ;     HD1-   CD1-   CE1-    CZ
  3706   3704   3703   3712       3   30.33400    0.00000  -30.33400
0.00000    0.00000    0.00000 ;     CE1-   CD1-    CG-   CD2
  3706   3708   3710   3711       3   30.33400    0.00000  -30.33400
0.00000    0.00000    0.00000 ;     CE1-    CZ-   CE2-   HE2
  3706   3708   3710   3712       3   30.33400    0.00000  -30.33400
0.00000    0.00000    0.00000 ;     CE1-    CZ-   CE2-   CD2
  3707   3706   3708   3709       3   30.33400    0.00000  -30.33400
0.00000    0.00000    0.00000 ;     HE1-   CE1-    CZ-    HZ
  3707   3706   3708   3710       3   30.33400    0.00000  -30.33400
0.00000    0.00000    0.00000 ;     HE1-   CE1-    CZ-   CE2
  3708   3710   3712   3713       3   30.33400    0.00000  -30.33400
0.00000    0.00000    0.00000 ;      CZ-   CE2-   CD2-   HD2
  3709   3708   3710   3711       3   30.33400    0.00000  -30.33400
0.00000    0.00000    0.00000 ;      HZ-    CZ-   CE2-   HE2
  3709   3708   3710   3712       3   30.33400    0.00000  -30.33400
0.00000    0.00000    0.00000 ;      HZ-    CZ-   CE2-   CD2
  3711   3710   3712   3713       3   30.33400    0.00000  -30.33400
0.00000    0.00000    0.00000 ;     HE2-   CE2-   CD2-   HD2
  3714   3716   3718   3719       3    0.00000    0.00000    0.00000
0.00000    0.00000    0.00000 ;       C-     N-    CA-    HA
  3714   3716   3718   3720       3   11.71520    1.67360   15.06240  -
13.38880    0.00000    0.00000 ;       C-     N-    CA-    CB
  3714   3716   3718   3735       3    1.75728    5.27184    2.25936
-7.02912    0.00000    0.00000 ;       C-     N-    CA-     C
  3715   3714   3716   3717       3   29.28800   -8.36800  -20.92000
0.00000    0.00000    0.00000 ;       O-     C-     N-     H
  3715   3714   3716   3718       3   20.92000    0.00000  -20.92000
0.00000    0.00000    0.00000 ;       O-     C-     N-    CA
  3716   3718   3720   3721       3    0.65084    1.95253    0.00000
-2.60338    0.00000    0.00000 ;       N-    CA-    CB-   HB2
  3716   3718   3720   3722       3    0.65084    1.95253    0.00000
```

-2.60338 0.00000 0.00000 ; N- CA- CB- HB3
3716 3718 3720 3723 3 3.28026 2.46019 -2.19242
-3.21331 -0.23430 0.00000 ; N- CA- CB- CG
3716 3718 3735 3736 3 0.00000 0.00000 0.00000
0.00000 0.00000 0.00000 ; N- CA- C- O
3716 3718 3735 3737 3 17.40544 -5.02080 -13.22144
9.20480 0.00000 0.00000 ; N- CA- C- N
3717 3716 3718 3719 3 0.00000 0.00000 0.00000
0.00000 0.00000 0.00000 ; H- N- CA- HA
3717 3716 3718 3720 3 0.00000 0.00000 0.00000
0.00000 0.00000 0.00000 ; H- N- CA- CB
3717 3716 3718 3735 3 0.00000 0.00000 0.00000
0.00000 0.00000 0.00000 ; H- N- CA- C
3718 3720 3723 3724 3 0.57739 0.00000 1.02926
0.00000 -1.60666 0.00000 ; CA- CB- CG- CD1
3718 3720 3723 3733 3 0.57739 0.00000 1.02926
0.00000 -1.60666 0.00000 ; CA- CB- CG- CD2
3718 3735 3737 3738 3 20.92000 0.00000 -20.92000
0.00000 0.00000 0.00000 ; CA- C- N- H
3718 3735 3737 3739 3 20.92000 0.00000 -20.92000
0.00000 0.00000 0.00000 ; CA- C- N- CA
3719 3718 3720 3721 3 0.65084 1.95253 0.00000
-2.60338 0.00000 0.00000 ; HA- CA- CB- HB2
3719 3718 3720 3722 3 0.65084 1.95253 0.00000
-2.60338 0.00000 0.00000 ; HA- CA- CB- HB3
3719 3718 3720 3723 3 0.65084 1.95253 0.00000
-2.60338 0.00000 0.00000 ; HA- CA- CB- CG
3719 3718 3735 3736 3 3.68192 -4.35136 0.00000
1.33888 0.00000 0.00000 ; HA- CA- C- O
3719 3718 3735 3737 3 0.00000 0.00000 0.00000
0.00000 0.00000 0.00000 ; HA- CA- C- N
3720 3718 3735 3736 3 0.00000 0.00000 0.00000
0.00000 0.00000 0.00000 ; CB- CA- C- O
3720 3718 3735 3737 3 2.51040 4.18400 1.67360
-6.69440 0.00000 0.00000 ; CB- CA- C- N
3720 3723 3724 3725 3 30.33400 0.00000 -30.33400
0.00000 0.00000 0.00000 ; CB- CG- CD1- HD1
3720 3723 3724 3726 3 30.33400 0.00000 -30.33400
0.00000 0.00000 0.00000 ; CB- CG- CD1- CE1
3720 3723 3733 3731 3 30.33400 0.00000 -30.33400
0.00000 0.00000 0.00000 ; CB- CG- CD2- CE2
3720 3723 3733 3734 3 30.33400 0.00000 -30.33400
0.00000 0.00000 0.00000 ; CB- CG- CD2- HD2
3721 3720 3718 3735 3 0.65084 1.95253 0.00000
-2.60338 0.00000 0.00000 ; HB2- CB- CA- C
3721 3720 3723 3724 3 0.00000 0.00000 0.00000
0.00000 0.00000 0.00000 ; HB2- CB- CG- CD1
3721 3720 3723 3733 3 0.00000 0.00000 0.00000
0.00000 0.00000 0.00000 ; HB2- CB- CG- CD2
3722 3720 3718 3735 3 0.65084 1.95253 0.00000
-2.60338 0.00000 0.00000 ; HB3- CB- CA- C
3722 3720 3723 3724 3 0.00000 0.00000 0.00000

0.00000 0.00000 0.00000 ; HB3- CB- CG- CD1
3722 3720 3723 3733 3 0.00000 0.00000 0.00000
0.00000 0.00000 0.00000 ; HB3- CB- CG- CD2
3723 3720 3718 3735 3 4.95804 2.17986 -3.52293
-3.21331 -0.40166 0.00000 ; CG- CB- CA- C
3723 3724 3726 3727 3 30.33400 0.00000 -30.33400
0.00000 0.00000 0.00000 ; CG- CD1- CE1- HE1
3723 3724 3726 3728 3 30.33400 0.00000 -30.33400
0.00000 0.00000 0.00000 ; CG- CD1- CE1- CZ
3723 3733 3731 3728 3 30.33400 0.00000 -30.33400
0.00000 0.00000 0.00000 ; CG- CD2- CE2- CZ
3723 3733 3731 3732 3 30.33400 0.00000 -30.33400
0.00000 0.00000 0.00000 ; CG- CD2- CE2- HE2
3724 3723 3733 3731 3 30.33400 0.00000 -30.33400
0.00000 0.00000 0.00000 ; CD1- CG- CD2- CE2
3724 3723 3733 3734 3 30.33400 0.00000 -30.33400
0.00000 0.00000 0.00000 ; CD1- CG- CD2- HD2
3724 3726 3728 3729 3 30.33400 0.00000 -30.33400
0.00000 0.00000 0.00000 ; CD1- CE1- CZ- OH
3724 3726 3728 3731 3 30.33400 0.00000 -30.33400
0.00000 0.00000 0.00000 ; CD1- CE1- CZ- CE2
3725 3724 3723 3733 3 30.33400 0.00000 -30.33400
0.00000 0.00000 0.00000 ; HD1- CD1- CG- CD2
3725 3724 3726 3727 3 30.33400 0.00000 -30.33400
0.00000 0.00000 0.00000 ; HD1- CD1- CE1- HE1
3725 3724 3726 3728 3 30.33400 0.00000 -30.33400
0.00000 0.00000 0.00000 ; HD1- CD1- CE1- CZ
3726 3724 3723 3733 3 30.33400 0.00000 -30.33400
0.00000 0.00000 0.00000 ; CE1- CD1- CG- CD2
3726 3728 3729 3730 3 7.93286 0.00000 -9.56462
0.00000 2.17568 0.00000 ; CE1- CZ- OH- HH
3726 3728 3731 3732 3 30.33400 0.00000 -30.33400
0.00000 0.00000 0.00000 ; CE1- CZ- CE2- HE2
3726 3728 3731 3733 3 30.33400 0.00000 -30.33400
0.00000 0.00000 0.00000 ; CE1- CZ- CE2- CD2
3727 3726 3728 3729 3 30.33400 0.00000 -30.33400
0.00000 0.00000 0.00000 ; HE1- CE1- CZ- OH
3727 3726 3728 3731 3 30.33400 0.00000 -30.33400
0.00000 0.00000 0.00000 ; HE1- CE1- CZ- CE2
3728 3731 3733 3734 3 30.33400 0.00000 -30.33400
0.00000 0.00000 0.00000 ; CZ- CE2- CD2- HD2
3729 3728 3731 3732 3 30.33400 0.00000 -30.33400
0.00000 0.00000 0.00000 ; OH- CZ- CE2- HE2
3729 3728 3731 3733 3 30.33400 0.00000 -30.33400
0.00000 0.00000 0.00000 ; OH- CZ- CE2- CD2
3730 3729 3728 3731 3 7.93286 0.00000 -9.56462
0.00000 2.17568 0.00000 ; HH- OH- CZ- CE2
3732 3731 3733 3734 3 30.33400 0.00000 -30.33400
0.00000 0.00000 0.00000 ; HE2- CE2- CD2- HD2
3735 3737 3739 3740 3 0.00000 0.00000 0.00000
0.00000 0.00000 0.00000 ; C- N- CA- HA
3735 3737 3739 3741 3 11.71520 1.67360 15.06240 -

13.38880 0.00000 0.00000 ; C- N- CA- CB
3735 3737 3739 3750 3 1.75728 5.27184 2.25936
-7.02912 0.00000 0.00000 ; C- N- CA- C
3736 3735 3737 3738 3 29.28800 -8.36800 -20.92000
0.00000 0.00000 0.00000 ; O- C- N- H
3736 3735 3737 3739 3 20.92000 0.00000 -20.92000
0.00000 0.00000 0.00000 ; O- C- N- CA
3737 3739 3741 3742 3 0.65084 1.95253 0.00000
-2.60338 0.00000 0.00000 ; N- CA- CB- HB2
3737 3739 3741 3743 3 0.65084 1.95253 0.00000
-2.60338 0.00000 0.00000 ; N- CA- CB- HB3
3737 3739 3741 3744 3 3.21331 2.22589 -4.15053
-2.40998 2.61082 0.00000 ; N- CA- CB- CG
3737 3739 3750 3751 3 0.00000 0.00000 0.00000
0.00000 0.00000 0.00000 ; N- CA- C- O
3737 3739 3750 3752 3 17.40544 -5.02080 -13.22144
9.20480 0.00000 0.00000 ; N- CA- C- N
3738 3737 3739 3740 3 0.00000 0.00000 0.00000
0.00000 0.00000 0.00000 ; H- N- CA- HA
3738 3737 3739 3741 3 0.00000 0.00000 0.00000
0.00000 0.00000 0.00000 ; H- N- CA- CB
3738 3737 3739 3750 3 0.00000 0.00000 0.00000
0.00000 0.00000 0.00000 ; H- N- CA- C
3739 3741 3744 3745 3 0.66944 2.00832 0.00000
-2.67776 0.00000 0.00000 ; CA- CB- CG- HG2
3739 3741 3744 3746 3 0.66944 2.00832 0.00000
-2.67776 0.00000 0.00000 ; CA- CB- CG- HG3
3739 3741 3744 3747 3 10.12110 -1.91209 0.01674
10.17549 -1.87443 0.00000 ; CA- CB- CG- CD
3739 3750 3752 3753 3 20.92000 0.00000 -20.92000
0.00000 0.00000 0.00000 ; CA- C- N- H
3739 3750 3752 3754 3 20.92000 0.00000 -20.92000
0.00000 0.00000 0.00000 ; CA- C- N- CA
3740 3739 3741 3742 3 0.65084 1.95253 0.00000
-2.60338 0.00000 0.00000 ; HA- CA- CB- HB2
3740 3739 3741 3743 3 0.65084 1.95253 0.00000
-2.60338 0.00000 0.00000 ; HA- CA- CB- HB3
3740 3739 3741 3744 3 0.65084 1.95253 0.00000
-2.60338 0.00000 0.00000 ; HA- CA- CB- CG
3740 3739 3750 3751 3 3.68192 -4.35136 0.00000
1.33888 0.00000 0.00000 ; HA- CA- C- O
3740 3739 3750 3752 3 0.00000 0.00000 0.00000
0.00000 0.00000 0.00000 ; HA- CA- C- N
3741 3739 3750 3751 3 0.00000 0.00000 0.00000
0.00000 0.00000 0.00000 ; CB- CA- C- O
3741 3739 3750 3752 3 2.51040 4.18400 1.67360
-6.69440 0.00000 0.00000 ; CB- CA- C- N
3741 3744 3747 3748 3 3.79907 0.00000 -5.40573
0.00000 2.14221 0.00000 ; CB- CG- CD- OE1
3741 3744 3747 3749 3 3.79907 0.00000 -5.40573
0.00000 2.14221 0.00000 ; CB- CG- CD- OE2
3742 3741 3739 3750 3 0.65084 1.95253 0.00000

-2.60338 0.00000 0.00000 ; HB2- CB- CA- C
3742 3741 3744 3745 3 0.62760 1.88280 0.00000
-2.51040 0.00000 0.00000 ; HB2- CB- CG- HG2
3742 3741 3744 3746 3 0.62760 1.88280 0.00000
-2.51040 0.00000 0.00000 ; HB2- CB- CG- HG3
3742 3741 3744 3747 3 0.65084 1.95253 0.00000
-2.60338 0.00000 0.00000 ; HB2- CB- CG- CD
3743 3741 3739 3750 3 0.65084 1.95253 0.00000
-2.60338 0.00000 0.00000 ; HB3- CB- CA- C
3743 3741 3744 3745 3 0.62760 1.88280 0.00000
-2.51040 0.00000 0.00000 ; HB3- CB- CG- HG2
3743 3741 3744 3746 3 0.62760 1.88280 0.00000
-2.51040 0.00000 0.00000 ; HB3- CB- CG- HG3
3743 3741 3744 3747 3 0.65084 1.95253 0.00000
-2.60338 0.00000 0.00000 ; HB3- CB- CG- CD
3744 3741 3739 3750 3 6.86594 3.56895 -8.14206
-2.40998 4.85344 0.00000 ; CG- CB- CA- C
3745 3744 3747 3748 3 0.00000 0.00000 0.00000
0.00000 0.00000 0.00000 ; HG2- CG- CD- OE1
3745 3744 3747 3749 3 0.00000 0.00000 0.00000
0.00000 0.00000 0.00000 ; HG2- CG- CD- OE2
3746 3744 3747 3748 3 0.00000 0.00000 0.00000
0.00000 0.00000 0.00000 ; HG3- CG- CD- OE1
3746 3744 3747 3749 3 0.00000 0.00000 0.00000
0.00000 0.00000 0.00000 ; HG3- CG- CD- OE2
3750 3752 3754 3755 3 0.00000 0.00000 0.00000
0.00000 0.00000 0.00000 ; C- N- CA- HA
3750 3752 3754 3756 3 11.71520 1.67360 15.06240 -
13.38880 0.00000 0.00000 ; C- N- CA- CB
3750 3752 3754 3764 3 1.75728 5.27184 2.25936
-7.02912 0.00000 0.00000 ; C- N- CA- C
3751 3750 3752 3753 3 29.28800 -8.36800 -20.92000
0.00000 0.00000 0.00000 ; O- C- N- H
3751 3750 3752 3754 3 20.92000 0.00000 -20.92000
0.00000 0.00000 0.00000 ; O- C- N- CA
3752 3754 3756 3757 3 0.65084 1.95253 0.00000
-2.60338 0.00000 0.00000 ; N- CA- CB- HB2
3752 3754 3756 3758 3 0.65084 1.95253 0.00000
-2.60338 0.00000 0.00000 ; N- CA- CB- HB3
3752 3754 3756 3759 3 5.99567 3.29281 -4.46014
-0.55229 1.97485 0.00000 ; N- CA- CB- CG
3752 3754 3764 3765 3 0.00000 0.00000 0.00000
0.00000 0.00000 0.00000 ; N- CA- C- O
3752 3754 3764 3766 3 17.40544 -5.02080 -13.22144
9.20480 0.00000 0.00000 ; N- CA- C- N
3753 3752 3754 3755 3 0.00000 0.00000 0.00000
0.00000 0.00000 0.00000 ; H- N- CA- HA
3753 3752 3754 3756 3 0.00000 0.00000 0.00000
0.00000 0.00000 0.00000 ; H- N- CA- CB
3753 3752 3754 3764 3 0.00000 0.00000 0.00000
0.00000 0.00000 0.00000 ; H- N- CA- C
3754 3756 3759 3760 3 0.00000 0.00000 0.00000

```
0.00000    0.00000    0.00000 ;     CA-    CB-    CG-   OD1
  3754   3756   3759   3761       3    8.84916   -0.31380   -4.32626
5.03754    0.26778    0.00000 ;     CA-    CB-    CG-   ND2
  3754   3764   3766   3767       3   20.92000    0.00000  -20.92000
0.00000    0.00000    0.00000 ;     CA-     C-     N-     H
  3754   3764   3766   3768       3   20.92000    0.00000  -20.92000
0.00000    0.00000    0.00000 ;     CA-     C-     N-    CA
  3755   3754   3756   3757       3    0.65084    1.95253    0.00000
-2.60338    0.00000    0.00000 ;     HA-    CA-    CB-   HB2
  3755   3754   3756   3758       3    0.65084    1.95253    0.00000
-2.60338    0.00000    0.00000 ;     HA-    CA-    CB-   HB3
  3755   3754   3756   3759       3    0.65084    1.95253    0.00000
-2.60338    0.00000    0.00000 ;     HA-    CA-    CB-    CG
  3755   3754   3764   3765       3    3.68192   -4.35136    0.00000
1.33888    0.00000    0.00000 ;     HA-    CA-     C-     O
  3755   3754   3764   3766       3    0.00000    0.00000    0.00000
0.00000    0.00000    0.00000 ;     HA-    CA-     C-     N
  3756   3754   3764   3765       3    0.00000    0.00000    0.00000
0.00000    0.00000    0.00000 ;     CB-    CA-     C-     O
  3756   3754   3764   3766       3    2.51040    4.18400    1.67360
-6.69440    0.00000    0.00000 ;     CB-    CA-     C-     N
  3756   3759   3761   3762       3   20.92000    0.00000  -20.92000
0.00000    0.00000    0.00000 ;     CB-    CG-   ND2-  HD21
  3756   3759   3761   3763       3   20.92000    0.00000  -20.92000
0.00000    0.00000    0.00000 ;     CB-    CG-   ND2-  HD22
  3757   3756   3754   3764       3    0.65084    1.95253    0.00000
-2.60338    0.00000    0.00000 ;    HB2-    CB-    CA-     C
  3757   3756   3759   3760       3    3.68192   -4.35136    0.00000
1.33888    0.00000    0.00000 ;    HB2-    CB-    CG-   OD1
  3757   3756   3759   3761       3    0.00000    0.00000    0.00000
0.00000    0.00000    0.00000 ;    HB2-    CB-    CG-   ND2
  3758   3756   3754   3764       3    0.65084    1.95253    0.00000
-2.60338    0.00000    0.00000 ;    HB3-    CB-    CA-     C
  3758   3756   3759   3760       3    3.68192   -4.35136    0.00000
1.33888    0.00000    0.00000 ;    HB3-    CB-    CG-   OD1
  3758   3756   3759   3761       3    0.00000    0.00000    0.00000
0.00000    0.00000    0.00000 ;    HB3-    CB-    CG-   ND2
  3759   3756   3754   3764       3    7.94542   -3.96225   -6.11701
-0.55229    3.58150    0.00000 ;     CG-    CB-    CA-     C
  3760   3759   3761   3762       3   29.28800   -8.36800  -20.92000
0.00000    0.00000    0.00000 ;    OD1-    CG-   ND2-  HD21
  3760   3759   3761   3763       3   29.28800   -8.36800  -20.92000
0.00000    0.00000    0.00000 ;    OD1-    CG-   ND2-  HD22
  3764   3766   3768   3769       3    0.00000    0.00000    0.00000
0.00000    0.00000    0.00000 ;     C-     N-    CA-    HA
  3764   3766   3768   3770       3   11.71520    1.67360   15.06240  -
13.38880    0.00000    0.00000 ;     C-     N-    CA-    CB
  3764   3766   3768   3776       3    1.75728    5.27184    2.25936
-7.02912    0.00000    0.00000 ;     C-     N-    CA-     C
  3765   3764   3766   3767       3   29.28800   -8.36800  -20.92000
0.00000    0.00000    0.00000 ;     O-     C-     N-     H
  3765   3764   3766   3768       3   20.92000    0.00000  -20.92000
```

```
0.00000    0.00000    0.00000 ;       O-     C-     N-    CA
  3766   3768   3770   3771      3    0.65084    1.95253    0.00000
-2.60338    0.00000    0.00000 ;       N-    CA-    CB-   HB2
  3766   3768   3770   3772      3    0.65084    1.95253    0.00000
-2.60338    0.00000    0.00000 ;       N-    CA-    CB-   HB3
  3766   3768   3770   3773      3   15.41386    9.74035   -8.39310
-0.97069    2.97901    0.00000 ;       N-    CA-    CB-    CG
  3766   3768   3776   3777      3    0.00000    0.00000    0.00000
0.00000    0.00000    0.00000 ;       N-    CA-     C-     O
  3766   3768   3776   3778      3   17.40544   -5.02080  -13.22144
9.20480    0.00000    0.00000 ;       N-    CA-     C-     N
  3767   3766   3768   3769      3    0.00000    0.00000    0.00000
0.00000    0.00000    0.00000 ;       H-     N-    CA-    HA
  3767   3766   3768   3770      3    0.00000    0.00000    0.00000
0.00000    0.00000    0.00000 ;       H-     N-    CA-    CB
  3767   3766   3768   3776      3    0.00000    0.00000    0.00000
0.00000    0.00000    0.00000 ;       H-     N-    CA-     C
  3768   3770   3773   3774      3    6.43499    0.00000   -5.39736
0.00000   -1.03763    0.00000 ;      CA-    CB-    CG-   OD1
  3768   3770   3773   3775      3    6.43499    0.00000   -5.39736
0.00000   -1.03763    0.00000 ;      CA-    CB-    CG-   OD2
  3768   3776   3778   3779      3   20.92000    0.00000  -20.92000
0.00000    0.00000    0.00000 ;      CA-     C-     N-     H
  3768   3776   3778   3780      3   20.92000    0.00000  -20.92000
0.00000    0.00000    0.00000 ;      CA-     C-     N-    CA
  3769   3768   3770   3771      3    0.65084    1.95253    0.00000
-2.60338    0.00000    0.00000 ;      HA-    CA-    CB-   HB2
  3769   3768   3770   3772      3    0.65084    1.95253    0.00000
-2.60338    0.00000    0.00000 ;      HA-    CA-    CB-   HB3
  3769   3768   3770   3773      3    0.65084    1.95253    0.00000
-2.60338    0.00000    0.00000 ;      HA-    CA-    CB-    CG
  3769   3768   3776   3777      3    3.68192   -4.35136    0.00000
1.33888    0.00000    0.00000 ;      HA-    CA-     C-     O
  3769   3768   3776   3778      3    0.00000    0.00000    0.00000
0.00000    0.00000    0.00000 ;      HA-    CA-     C-     N
  3770   3768   3776   3777      3    0.00000    0.00000    0.00000
0.00000    0.00000    0.00000 ;      CB-    CA-     C-     O
  3770   3768   3776   3778      3    2.51040    4.18400    1.67360
-6.69440    0.00000    0.00000 ;      CB-    CA-     C-     N
  3771   3770   3768   3776      3    0.65084    1.95253    0.00000
-2.60338    0.00000    0.00000 ;     HB2-    CB-    CA-     C
  3771   3770   3773   3774      3    0.00000    0.00000    0.00000
0.00000    0.00000    0.00000 ;     HB2-    CB-    CG-   OD1
  3771   3770   3773   3775      3    0.00000    0.00000    0.00000
0.00000    0.00000    0.00000 ;     HB2-    CB-    CG-   OD2
  3772   3770   3768   3776      3    0.65084    1.95253    0.00000
-2.60338    0.00000    0.00000 ;     HB3-    CB-    CA-     C
  3772   3770   3773   3774      3    0.00000    0.00000    0.00000
0.00000    0.00000    0.00000 ;     HB3-    CB-    CG-   OD1
  3772   3770   3773   3775      3    0.00000    0.00000    0.00000
0.00000    0.00000    0.00000 ;     HB3-    CB-    CG-   OD2
  3773   3770   3768   3776      3    7.14627   -1.04600   -8.99560
```

-0.97069 5.15469 0.00000 ; CG- CB- CA- C
3776 3778 3780 3781 3 0.00000 0.00000 0.00000
0.00000 0.00000 0.00000 ; C- N- CA- HA
3776 3778 3780 3782 3 11.71520 1.67360 15.06240 -
13.38880 0.00000 0.00000 ; C- N- CA- CB
3776 3778 3780 3800 3 1.75728 5.27184 2.25936
-7.02912 0.00000 0.00000 ; C- N- CA- C
3777 3776 3778 3779 3 29.28800 -8.36800 -20.92000
0.00000 0.00000 0.00000 ; O- C- N- H
3777 3776 3778 3780 3 20.92000 0.00000 -20.92000
0.00000 0.00000 0.00000 ; O- C- N- CA
3778 3780 3782 3783 3 0.65084 1.95253 0.00000
-2.60338 0.00000 0.00000 ; N- CA- CB- HB2
3778 3780 3782 3784 3 0.65084 1.95253 0.00000
-2.60338 0.00000 0.00000 ; N- CA- CB- HB3
3778 3780 3782 3785 3 4.18818 2.60663 -3.65682
-3.91622 1.03763 0.00000 ; N- CA- CB- CG
3778 3780 3800 3801 3 0.00000 0.00000 0.00000
0.00000 0.00000 0.00000 ; N- CA- C- O
3778 3780 3800 3802 3 17.40544 -5.02080 -13.22144
9.20480 0.00000 0.00000 ; N- CA- C- N
3779 3778 3780 3781 3 0.00000 0.00000 0.00000
0.00000 0.00000 0.00000 ; H- N- CA- HA
3779 3778 3780 3782 3 0.00000 0.00000 0.00000
0.00000 0.00000 0.00000 ; H- N- CA- CB
3779 3778 3780 3800 3 0.00000 0.00000 0.00000
0.00000 0.00000 0.00000 ; H- N- CA- C
3780 3782 3785 3786 3 0.00000 0.00000 0.00000
0.00000 0.00000 0.00000 ; CA- CB- CG- CD1
3780 3782 3785 3799 3 4.95386 8.75293 6.59398 -
13.70678 -3.17984 0.00000 ; CA- CB- CG- CD2
3780 3800 3802 3803 3 20.92000 0.00000 -20.92000
0.00000 0.00000 0.00000 ; CA- C- N- H
3780 3800 3802 3804 3 20.92000 0.00000 -20.92000
0.00000 0.00000 0.00000 ; CA- C- N- CA
3781 3780 3782 3783 3 0.65084 1.95253 0.00000
-2.60338 0.00000 0.00000 ; HA- CA- CB- HB2
3781 3780 3782 3784 3 0.65084 1.95253 0.00000
-2.60338 0.00000 0.00000 ; HA- CA- CB- HB3
3781 3780 3782 3785 3 0.65084 1.95253 0.00000
-2.60338 0.00000 0.00000 ; HA- CA- CB- CG
3781 3780 3800 3801 3 3.68192 -4.35136 0.00000
1.33888 0.00000 0.00000 ; HA- CA- C- O
3781 3780 3800 3802 3 0.00000 0.00000 0.00000
0.00000 0.00000 0.00000 ; HA- CA- C- N
3782 3780 3800 3801 3 0.00000 0.00000 0.00000
0.00000 0.00000 0.00000 ; CB- CA- C- O
3782 3780 3800 3802 3 2.51040 4.18400 1.67360
-6.69440 0.00000 0.00000 ; CB- CA- C- N
3782 3785 3786 3787 3 54.60120 0.00000 -54.60120
0.00000 0.00000 0.00000 ; CB- CG- CD1- HD1
3782 3785 3786 3788 3 54.60120 0.00000 -54.60120

```
0.00000    0.00000    0.00000 ;      CB-    CG-    CD1-   NE1
  3782   3785   3799   3790       3   14.01640    0.00000  -14.01640
0.00000    0.00000    0.00000 ;      CB-    CG-    CD2-   CE2
  3782   3785   3799   3797       3   14.01640    0.00000  -14.01640
0.00000    0.00000    0.00000 ;      CB-    CG-    CD2-   CE3
  3783   3782   3780   3800       3    0.65084    1.95253    0.00000
-2.60338    0.00000    0.00000 ;     HB2-    CB-    CA-     C
  3783   3782   3785   3786       3    0.00000    0.00000    0.00000
0.00000    0.00000    0.00000 ;     HB2-    CB-    CG-   CD1
  3783   3782   3785   3799       3    0.00000    0.00000    0.00000
0.00000    0.00000    0.00000 ;     HB2-    CB-    CG-   CD2
  3784   3782   3780   3800       3    0.65084    1.95253    0.00000
-2.60338    0.00000    0.00000 ;     HB3-    CB-    CA-     C
  3784   3782   3785   3786       3    0.00000    0.00000    0.00000
0.00000    0.00000    0.00000 ;     HB3-    CB-    CG-   CD1
  3784   3782   3785   3799       3    0.00000    0.00000    0.00000
0.00000    0.00000    0.00000 ;     HB3-    CB-    CG-   CD2
  3785   3782   3780   3800       3    4.62332    3.00830   -5.43083
-3.91622    2.47693    0.00000 ;      CG-    CB-    CA-     C
  3785   3786   3788   3789       3   12.55200    0.00000  -12.55200
0.00000    0.00000    0.00000 ;      CG-   CD1-   NE1-   HE1
  3785   3786   3788   3790       3   12.55200    0.00000  -12.55200
0.00000    0.00000    0.00000 ;      CG-   CD1-   NE1-   CE2
  3785   3799   3790   3788       3   25.10400    0.00000  -25.10400
0.00000    0.00000    0.00000 ;      CG-   CD2-   CE2-   NE1
  3785   3799   3790   3791       3   25.10400    0.00000  -25.10400
0.00000    0.00000    0.00000 ;      CG-   CD2-   CE2-   CZ2
  3785   3799   3797   3795       3   29.28800    0.00000  -29.28800
0.00000    0.00000    0.00000 ;      CG-   CD2-   CE3-   CZ3
  3785   3799   3797   3798       3   29.28800    0.00000  -29.28800
0.00000    0.00000    0.00000 ;      CG-   CD2-   CE3-   HE3
  3786   3785   3799   3790       3   14.01640    0.00000  -14.01640
0.00000    0.00000    0.00000 ;     CD1-    CG-   CD2-   CE2
  3786   3785   3799   3797       3   14.01640    0.00000  -14.01640
0.00000    0.00000    0.00000 ;     CD1-    CG-   CD2-   CE3
  3786   3788   3790   3791       3   12.76120    0.00000  -12.76120
0.00000    0.00000    0.00000 ;     CD1-   NE1-   CE2-   CZ2
  3786   3788   3790   3799       3   12.76120    0.00000  -12.76120
0.00000    0.00000    0.00000 ;     CD1-   NE1-   CE2-   CD2
  3787   3786   3785   3799       3   54.60120    0.00000  -54.60120
0.00000    0.00000    0.00000 ;     HD1-   CD1-    CG-   CD2
  3787   3786   3788   3789       3   12.55200    0.00000  -12.55200
0.00000    0.00000    0.00000 ;     HD1-   CD1-   NE1-   HE1
  3787   3786   3788   3790       3   12.55200    0.00000  -12.55200
0.00000    0.00000    0.00000 ;     HD1-   CD1-   NE1-   CE2
  3788   3786   3785   3799       3   54.60120    0.00000  -54.60120
0.00000    0.00000    0.00000 ;     NE1-   CD1-    CG-   CD2
  3788   3790   3791   3792       3   30.33400    0.00000  -30.33400
0.00000    0.00000    0.00000 ;     NE1-   CE2-   CZ2-   HZ2
  3788   3790   3791   3793       3   30.33400    0.00000  -30.33400
0.00000    0.00000    0.00000 ;     NE1-   CE2-   CZ2-   CH2
  3788   3790   3799   3797       3   25.10400    0.00000  -25.10400
```

```
0.00000    0.00000    0.00000 ;    NE1-   CE2-   CD2-   CE3
  3789   3788   3790   3791      3   12.76120    0.00000  -12.76120
0.00000    0.00000    0.00000 ;    HE1-   NE1-   CE2-   CZ2
  3789   3788   3790   3799      3   12.76120    0.00000  -12.76120
0.00000    0.00000    0.00000 ;    HE1-   NE1-   CE2-   CD2
  3790   3791   3793   3794      3   30.33400    0.00000  -30.33400
0.00000    0.00000    0.00000 ;    CE2-   CZ2-   CH2-   HH2
  3790   3791   3793   3795      3   30.33400    0.00000  -30.33400
0.00000    0.00000    0.00000 ;    CE2-   CZ2-   CH2-   CZ3
  3790   3799   3797   3795      3   29.28800    0.00000  -29.28800
0.00000    0.00000    0.00000 ;    CE2-   CD2-   CE3-   CZ3
  3790   3799   3797   3798      3   29.28800    0.00000  -29.28800
0.00000    0.00000    0.00000 ;    CE2-   CD2-   CE3-   HE3
  3791   3790   3799   3797      3   25.10400    0.00000  -25.10400
0.00000    0.00000    0.00000 ;    CZ2-   CE2-   CD2-   CE3
  3791   3793   3795   3796      3   30.33400    0.00000  -30.33400
0.00000    0.00000    0.00000 ;    CZ2-   CH2-   CZ3-   HZ3
  3791   3793   3795   3797      3   30.33400    0.00000  -30.33400
0.00000    0.00000    0.00000 ;    CZ2-   CH2-   CZ3-   CE3
  3792   3791   3790   3799      3   30.33400    0.00000  -30.33400
0.00000    0.00000    0.00000 ;    HZ2-   CZ2-   CE2-   CD2
  3792   3791   3793   3794      3   30.33400    0.00000  -30.33400
0.00000    0.00000    0.00000 ;    HZ2-   CZ2-   CH2-   HH2
  3792   3791   3793   3795      3   30.33400    0.00000  -30.33400
0.00000    0.00000    0.00000 ;    HZ2-   CZ2-   CH2-   CZ3
  3793   3791   3790   3799      3   30.33400    0.00000  -30.33400
0.00000    0.00000    0.00000 ;    CH2-   CZ2-   CE2-   CD2
  3793   3795   3797   3798      3   30.33400    0.00000  -30.33400
0.00000    0.00000    0.00000 ;    CH2-   CZ3-   CE3-   HE3
  3793   3795   3797   3799      3   30.33400    0.00000  -30.33400
0.00000    0.00000    0.00000 ;    CH2-   CZ3-   CE3-   CD2
  3794   3793   3795   3796      3   30.33400    0.00000  -30.33400
0.00000    0.00000    0.00000 ;    HH2-   CH2-   CZ3-   HZ3
  3794   3793   3795   3797      3   30.33400    0.00000  -30.33400
0.00000    0.00000    0.00000 ;    HH2-   CH2-   CZ3-   CE3
  3796   3795   3797   3798      3   30.33400    0.00000  -30.33400
0.00000    0.00000    0.00000 ;    HZ3-   CZ3-   CE3-   HE3
  3796   3795   3797   3799      3   30.33400    0.00000  -30.33400
0.00000    0.00000    0.00000 ;    HZ3-   CZ3-   CE3-   CD2
  3800   3802   3804   3805      3    0.00000    0.00000    0.00000
0.00000    0.00000    0.00000 ;      C-     N-    CA-   HA2
  3800   3802   3804   3806      3    0.00000    0.00000    0.00000
0.00000    0.00000    0.00000 ;      C-     N-    CA-   HA3
  3800   3802   3804   3807      3    1.75728    5.27184    2.25936
-7.02912    0.00000    0.00000 ;      C-     N-    CA-     C
  3801   3800   3802   3803      3   29.28800   -8.36800  -20.92000
0.00000    0.00000    0.00000 ;      O-     C-     N-     H
  3801   3800   3802   3804      3   20.92000    0.00000  -20.92000
0.00000    0.00000    0.00000 ;      O-     C-     N-    CA
  3802   3804   3807   3808      3    0.00000    0.00000    0.00000
0.00000    0.00000    0.00000 ;      N-    CA-     C-     O
  3802   3804   3807   3809      3   17.40544   -5.02080  -13.22144
```

9.20480 0.00000 0.00000 ; N- CA- C- N
3803 3802 3804 3805 3 0.00000 0.00000 0.00000
0.00000 0.00000 0.00000 ; H- N- CA- HA2
3803 3802 3804 3806 3 0.00000 0.00000 0.00000
0.00000 0.00000 0.00000 ; H- N- CA- HA3
3803 3802 3804 3807 3 0.00000 0.00000 0.00000
0.00000 0.00000 0.00000 ; H- N- CA- C
3804 3807 3809 3810 3 20.92000 0.00000 -20.92000
0.00000 0.00000 0.00000 ; CA- C- N- H
3804 3807 3809 3811 3 20.92000 0.00000 -20.92000
0.00000 0.00000 0.00000 ; CA- C- N- CA
3805 3804 3807 3808 3 3.68192 -4.35136 0.00000
1.33888 0.00000 0.00000 ; HA2- CA- C- O
3805 3804 3807 3809 3 0.00000 0.00000 0.00000
0.00000 0.00000 0.00000 ; HA2- CA- C- N
3806 3804 3807 3808 3 3.68192 -4.35136 0.00000
1.33888 0.00000 0.00000 ; HA3- CA- C- O
3806 3804 3807 3809 3 0.00000 0.00000 0.00000
0.00000 0.00000 0.00000 ; HA3- CA- C- N
3807 3809 3811 3812 3 0.00000 0.00000 0.00000
0.00000 0.00000 0.00000 ; C- N- CA- HA
3807 3809 3811 3813 3 11.71520 1.67360 15.06240 -
13.38880 0.00000 0.00000 ; C- N- CA- CB
3807 3809 3811 3818 3 1.75728 5.27184 2.25936
-7.02912 0.00000 0.00000 ; C- N- CA- C
3808 3807 3809 3810 3 29.28800 -8.36800 -20.92000
0.00000 0.00000 0.00000 ; O- C- N- H
3808 3807 3809 3811 3 20.92000 0.00000 -20.92000
0.00000 0.00000 0.00000 ; O- C- N- CA
3809 3811 3813 3814 3 0.65084 1.95253 0.00000
-2.60338 0.00000 0.00000 ; N- CA- CB- HB2
3809 3811 3813 3815 3 0.65084 1.95253 0.00000
-2.60338 0.00000 0.00000 ; N- CA- CB- HB3
3809 3811 3813 3816 3 7.86174 2.24681 -7.41405
-6.71114 5.35552 0.00000 ; N- CA- CB- OG
3809 3811 3818 3819 3 0.00000 0.00000 0.00000
0.00000 0.00000 0.00000 ; N- CA- C- O
3809 3811 3818 3820 3 17.40544 -5.02080 -13.22144
9.20480 0.00000 0.00000 ; N- CA- C- N
3810 3809 3811 3812 3 0.00000 0.00000 0.00000
0.00000 0.00000 0.00000 ; H- N- CA- HA
3810 3809 3811 3813 3 0.00000 0.00000 0.00000
0.00000 0.00000 0.00000 ; H- N- CA- CB
3810 3809 3811 3818 3 0.00000 0.00000 0.00000
0.00000 0.00000 0.00000 ; H- N- CA- C
3811 3813 3816 3817 3 2.05853 2.46856 3.48109
-4.46851 0.23430 0.00000 ; CA- CB- OG- HG
3811 3818 3820 3821 3 20.92000 0.00000 -20.92000
0.00000 0.00000 0.00000 ; CA- C- N- H
3811 3818 3820 3822 3 20.92000 0.00000 -20.92000
0.00000 0.00000 0.00000 ; CA- C- N- CA
3812 3811 3813 3814 3 0.65084 1.95253 0.00000

-2.60338 0.00000 0.00000 ; HA- CA- CB- HB2
3812 3811 3813 3815 3 0.65084 1.95253 0.00000
-2.60338 0.00000 0.00000 ; HA- CA- CB- HB3
3812 3811 3813 3816 3 1.04600 -1.04600 0.00000
0.00000 0.00000 0.00000 ; HA- CA- CB- OG
3812 3811 3818 3819 3 3.68192 -4.35136 0.00000
1.33888 0.00000 0.00000 ; HA- CA- C- O
3812 3811 3818 3820 3 0.00000 0.00000 0.00000
0.00000 0.00000 0.00000 ; HA- CA- C- N
3813 3811 3818 3819 3 0.00000 0.00000 0.00000
0.00000 0.00000 0.00000 ; CB- CA- C- O
3813 3811 3818 3820 3 2.51040 4.18400 1.67360
-6.69440 0.00000 0.00000 ; CB- CA- C- N
3814 3813 3811 3818 3 0.65084 1.95253 0.00000
-2.60338 0.00000 0.00000 ; HB2- CB- CA- C
3814 3813 3816 3817 3 0.69733 2.09200 0.00000
-2.78933 0.00000 0.00000 ; HB2- CB- OG- HG
3815 3813 3811 3818 3 0.65084 1.95253 0.00000
-2.60338 0.00000 0.00000 ; HB3- CB- CA- C
3815 3813 3816 3817 3 0.69733 2.09200 0.00000
-2.78933 0.00000 0.00000 ; HB3- CB- OG- HG
3816 3813 3811 3818 3 7.34710 7.79898 -6.14211
-6.71114 4.31789 0.00000 ; OG- CB- CA- C
3818 3820 3822 3823 3 0.00000 0.00000 0.00000
0.00000 0.00000 0.00000 ; C- N- CA- HA
3818 3820 3822 3824 3 11.71520 1.67360 15.06240 -
13.38880 0.00000 0.00000 ; C- N- CA- CB
3818 3820 3822 3837 3 1.75728 5.27184 2.25936
-7.02912 0.00000 0.00000 ; C- N- CA- C
3819 3818 3820 3821 3 29.28800 -8.36800 -20.92000
0.00000 0.00000 0.00000 ; O- C- N- H
3819 3818 3820 3822 3 20.92000 0.00000 -20.92000
0.00000 0.00000 0.00000 ; O- C- N- CA
3820 3822 3824 3825 3 0.65084 1.95253 0.00000
-2.60338 0.00000 0.00000 ; N- CA- CB- HB
3820 3822 3824 3826 3 3.83673 0.44769 -1.77402
-2.47693 -0.03347 0.00000 ; N- CA- CB- CG2
3820 3822 3824 3830 3 2.97482 0.12134 2.04179
-1.89117 -3.24678 0.00000 ; N- CA- CB- CG1
3820 3822 3837 3838 3 0.00000 0.00000 0.00000
0.00000 0.00000 0.00000 ; N- CA- C- O
3820 3822 3837 3839 3 17.40544 -5.02080 -13.22144
9.20480 0.00000 0.00000 ; N- CA- C- N
3821 3820 3822 3823 3 0.00000 0.00000 0.00000
0.00000 0.00000 0.00000 ; H- N- CA- HA
3821 3820 3822 3824 3 0.00000 0.00000 0.00000
0.00000 0.00000 0.00000 ; H- N- CA- CB
3821 3820 3822 3837 3 0.00000 0.00000 0.00000
0.00000 0.00000 0.00000 ; H- N- CA- C
3822 3824 3826 3827 3 0.66944 2.00832 0.00000
-2.67776 0.00000 0.00000 ; CA- CB- CG2- HG21
3822 3824 3826 3828 3 0.66944 2.00832 0.00000

```
-2.67776    0.00000    0.00000 ;      CA-    CB-   CG2-  HG22
  3822   3824   3826   3829      3    0.66944    2.00832    0.00000
-2.67776    0.00000    0.00000 ;      CA-    CB-   CG2-  HG23
  3822   3824   3830   3831      3    0.66944    2.00832    0.00000
-2.67776    0.00000    0.00000 ;      CA-    CB-   CG1-  HG12
  3822   3824   3830   3832      3    0.66944    2.00832    0.00000
-2.67776    0.00000    0.00000 ;      CA-    CB-   CG1-  HG13
  3822   3824   3830   3833      3    4.24258   -0.52718   -7.25506
-1.79075    7.69856    0.00000 ;      CA-    CB-   CG1-   CD1
  3822   3837   3839   3840      3   20.92000    0.00000  -20.92000
0.00000    0.00000    0.00000 ;      CA-     C-     N-     H
  3822   3837   3839   3841      3   20.92000    0.00000  -20.92000
0.00000    0.00000    0.00000 ;      CA-     C-     N-    CA
  3823   3822   3824   3825      3    0.65084    1.95253    0.00000
-2.60338    0.00000    0.00000 ;      HA-    CA-    CB-    HB
  3823   3822   3824   3826      3    0.65084    1.95253    0.00000
-2.60338    0.00000    0.00000 ;      HA-    CA-    CB-   CG2
  3823   3822   3824   3830      3    0.65084    1.95253    0.00000
-2.60338    0.00000    0.00000 ;      HA-    CA-    CB-   CG1
  3823   3822   3837   3838      3    3.68192   -4.35136    0.00000
1.33888    0.00000    0.00000 ;      HA-    CA-     C-     O
  3823   3822   3837   3839      3    0.00000    0.00000    0.00000
0.00000    0.00000    0.00000 ;      HA-    CA-     C-     N
  3824   3822   3837   3838      3    0.00000    0.00000    0.00000
0.00000    0.00000    0.00000 ;      CB-    CA-     C-     O
  3824   3822   3837   3839      3    2.51040    4.18400    1.67360
-6.69440    0.00000    0.00000 ;      CB-    CA-     C-     N
  3824   3830   3833   3834      3    0.66944    2.00832    0.00000
-2.67776    0.00000    0.00000 ;      CB-   CG1-   CD1-  HD11
  3824   3830   3833   3835      3    0.66944    2.00832    0.00000
-2.67776    0.00000    0.00000 ;      CB-   CG1-   CD1-  HD12
  3824   3830   3833   3836      3    0.66944    2.00832    0.00000
-2.67776    0.00000    0.00000 ;      CB-   CG1-   CD1-  HD13
  3825   3824   3822   3837      3    0.65084    1.95253    0.00000
-2.60338    0.00000    0.00000 ;      HB-    CB-    CA-     C
  3825   3824   3826   3827      3    0.62760    1.88280    0.00000
-2.51040    0.00000    0.00000 ;      HB-    CB-   CG2-  HG21
  3825   3824   3826   3828      3    0.62760    1.88280    0.00000
-2.51040    0.00000    0.00000 ;      HB-    CB-   CG2-  HG22
  3825   3824   3826   3829      3    0.62760    1.88280    0.00000
-2.51040    0.00000    0.00000 ;      HB-    CB-   CG2-  HG23
  3825   3824   3830   3831      3    0.62760    1.88280    0.00000
-2.51040    0.00000    0.00000 ;      HB-    CB-   CG1-  HG12
  3825   3824   3830   3832      3    0.62760    1.88280    0.00000
-2.51040    0.00000    0.00000 ;      HB-    CB-   CG1-  HG13
  3825   3824   3830   3833      3    0.66944    2.00832    0.00000
-2.67776    0.00000    0.00000 ;      HB-    CB-   CG1-   CD1
  3826   3824   3822   3837      3    5.67350    3.55640   -6.16722
-2.47693    3.74886    0.00000 ;     CG2-    CB-    CA-     C
  3826   3824   3830   3831      3    0.66944    2.00832    0.00000
-2.67776    0.00000    0.00000 ;     CG2-    CB-   CG1-  HG12
  3826   3824   3830   3832      3    0.66944    2.00832    0.00000
```

-2.67776 0.00000 0.00000 ; CG2- CB- CG1- HG13
3826 3824 3830 3833 3 3.81162 0.49790 -8.14206
-1.79075 7.49773 0.00000 ; CG2- CB- CG1- CD1
3827 3826 3824 3830 3 0.66944 2.00832 0.00000
-2.67776 0.00000 0.00000 ; HG21- CG2- CB- CG1
3828 3826 3824 3830 3 0.66944 2.00832 0.00000
-2.67776 0.00000 0.00000 ; HG22- CG2- CB- CG1
3829 3826 3824 3830 3 0.66944 2.00832 0.00000
-2.67776 0.00000 0.00000 ; HG23- CG2- CB- CG1
3830 3824 3822 3837 3 8.26340 0.74057 -9.99976
-1.89117 3.84928 0.00000 ; CG1- CB- CA- C
3831 3830 3833 3834 3 0.62760 1.88280 0.00000
-2.51040 0.00000 0.00000 ; HG12- CG1- CD1- HD11
3831 3830 3833 3835 3 0.62760 1.88280 0.00000
-2.51040 0.00000 0.00000 ; HG12- CG1- CD1- HD12
3831 3830 3833 3836 3 0.62760 1.88280 0.00000
-2.51040 0.00000 0.00000 ; HG12- CG1- CD1- HD13
3832 3830 3833 3834 3 0.62760 1.88280 0.00000
-2.51040 0.00000 0.00000 ; HG13- CG1- CD1- HD11
3832 3830 3833 3835 3 0.62760 1.88280 0.00000
-2.51040 0.00000 0.00000 ; HG13- CG1- CD1- HD12
3832 3830 3833 3836 3 0.62760 1.88280 0.00000
-2.51040 0.00000 0.00000 ; HG13- CG1- CD1- HD13
3837 3839 3841 3842 3 0.00000 0.00000 0.00000
0.00000 0.00000 0.00000 ; C- N- CA- HA
3837 3839 3841 3843 3 11.71520 1.67360 15.06240 -
13.38880 0.00000 0.00000 ; C- N- CA- CB
3837 3839 3841 3848 3 1.75728 5.27184 2.25936
-7.02912 0.00000 0.00000 ; C- N- CA- C
3838 3837 3839 3840 3 29.28800 -8.36800 -20.92000
0.00000 0.00000 0.00000 ; O- C- N- H
3838 3837 3839 3841 3 20.92000 0.00000 -20.92000
0.00000 0.00000 0.00000 ; O- C- N- CA
3839 3841 3843 3844 3 0.65084 1.95253 0.00000
-2.60338 0.00000 0.00000 ; N- CA- CB- HB2
3839 3841 3843 3845 3 0.65084 1.95253 0.00000
-2.60338 0.00000 0.00000 ; N- CA- CB- HB3
3839 3841 3843 3846 3 7.86174 2.24681 -7.41405
-6.71114 5.35552 0.00000 ; N- CA- CB- OG
3839 3841 3848 3849 3 0.00000 0.00000 0.00000
0.00000 0.00000 0.00000 ; N- CA- C- O
3839 3841 3848 3850 3 17.40544 -5.02080 -13.22144
9.20480 0.00000 0.00000 ; N- CA- C- N
3840 3839 3841 3842 3 0.00000 0.00000 0.00000
0.00000 0.00000 0.00000 ; H- N- CA- HA
3840 3839 3841 3843 3 0.00000 0.00000 0.00000
0.00000 0.00000 0.00000 ; H- N- CA- CB
3840 3839 3841 3848 3 0.00000 0.00000 0.00000
0.00000 0.00000 0.00000 ; H- N- CA- C
3841 3843 3846 3847 3 2.05853 2.46856 3.48109
-4.46851 0.23430 0.00000 ; CA- CB- OG- HG
3841 3848 3850 3851 3 20.92000 0.00000 -20.92000

```
0.00000    0.00000    0.00000 ;      CA-     C-      N-     H
  3841   3848   3850   3852       3   20.92000    0.00000  -20.92000
0.00000    0.00000    0.00000 ;      CA-     C-      N-    CA
  3842   3841   3843   3844       3    0.65084    1.95253    0.00000
-2.60338    0.00000    0.00000 ;      HA-    CA-     CB-   HB2
  3842   3841   3843   3845       3    0.65084    1.95253    0.00000
-2.60338    0.00000    0.00000 ;      HA-    CA-     CB-   HB3
  3842   3841   3843   3846       3    1.04600   -1.04600    0.00000
0.00000    0.00000    0.00000 ;      HA-    CA-     CB-    OG
  3842   3841   3848   3849       3    3.68192   -4.35136    0.00000
1.33888    0.00000    0.00000 ;      HA-    CA-      C-     O
  3842   3841   3848   3850       3    0.00000    0.00000    0.00000
0.00000    0.00000    0.00000 ;      HA-    CA-      C-     N
  3843   3841   3848   3849       3    0.00000    0.00000    0.00000
0.00000    0.00000    0.00000 ;      CB-    CA-      C-     O
  3843   3841   3848   3850       3    2.51040    4.18400    1.67360
-6.69440    0.00000    0.00000 ;      CB-    CA-      C-     N
  3844   3843   3841   3848       3    0.65084    1.95253    0.00000
-2.60338    0.00000    0.00000 ;     HB2-    CB-     CA-     C
  3844   3843   3846   3847       3    0.69733    2.09200    0.00000
-2.78933    0.00000    0.00000 ;     HB2-    CB-     OG-    HG
  3845   3843   3841   3848       3    0.65084    1.95253    0.00000
-2.60338    0.00000    0.00000 ;     HB3-    CB-     CA-     C
  3845   3843   3846   3847       3    0.69733    2.09200    0.00000
-2.78933    0.00000    0.00000 ;     HB3-    CB-     OG-    HG
  3846   3843   3841   3848       3    7.34710    7.79898   -6.14211
-6.71114    4.31789    0.00000 ;      OG-    CB-     CA-     C
  3848   3850   3852   3853       3    0.00000    0.00000    0.00000
0.00000    0.00000    0.00000 ;       C-     N-     CA-    HA
  3848   3850   3852   3854       3   11.71520    1.67360   15.06240  -
13.38880    0.00000    0.00000 ;       C-     N-     CA-    CB
  3848   3850   3852   3859       3    1.75728    5.27184    2.25936
-7.02912    0.00000    0.00000 ;       C-     N-     CA-     C
  3849   3848   3850   3851       3   29.28800   -8.36800  -20.92000
0.00000    0.00000    0.00000 ;       O-     C-      N-     H
  3849   3848   3850   3852       3   20.92000    0.00000  -20.92000
0.00000    0.00000    0.00000 ;       O-     C-      N-    CA
  3850   3852   3854   3855       3    0.65084    1.95253    0.00000
-2.60338    0.00000    0.00000 ;       N-    CA-     CB-   HB2
  3850   3852   3854   3856       3    0.65084    1.95253    0.00000
-2.60338    0.00000    0.00000 ;       N-    CA-     CB-   HB3
  3850   3852   3854   3857       3    7.86174    2.24681   -7.41405
-6.71114    5.35552    0.00000 ;       N-    CA-     CB-    OG
  3850   3852   3859   3860       3    0.00000    0.00000    0.00000
0.00000    0.00000    0.00000 ;       N-    CA-      C-     O
  3850   3852   3859   3861       3   17.40544   -5.02080  -13.22144
9.20480    0.00000    0.00000 ;       N-    CA-      C-     N
  3851   3850   3852   3853       3    0.00000    0.00000    0.00000
0.00000    0.00000    0.00000 ;       H-     N-     CA-    HA
  3851   3850   3852   3854       3    0.00000    0.00000    0.00000
0.00000    0.00000    0.00000 ;       H-     N-     CA-    CB
  3851   3850   3852   3859       3    0.00000    0.00000    0.00000
```

0.00000 0.00000 0.00000 ; H- N- CA- C
3852 3854 3857 3858 3 2.05853 2.46856 3.48109
-4.46851 0.23430 0.00000 ; CA- CB- OG- HG
3852 3859 3861 3862 3 20.92000 0.00000 -20.92000
0.00000 0.00000 0.00000 ; CA- C- N- H
3852 3859 3861 3863 3 20.92000 0.00000 -20.92000
0.00000 0.00000 0.00000 ; CA- C- N- CA
3853 3852 3854 3855 3 0.65084 1.95253 0.00000
-2.60338 0.00000 0.00000 ; HA- CA- CB- HB2
3853 3852 3854 3856 3 0.65084 1.95253 0.00000
-2.60338 0.00000 0.00000 ; HA- CA- CB- HB3
3853 3852 3854 3857 3 1.04600 -1.04600 0.00000
0.00000 0.00000 0.00000 ; HA- CA- CB- OG
3853 3852 3859 3860 3 3.68192 -4.35136 0.00000
1.33888 0.00000 0.00000 ; HA- CA- C- O
3853 3852 3859 3861 3 0.00000 0.00000 0.00000
0.00000 0.00000 0.00000 ; HA- CA- C- N
3854 3852 3859 3860 3 0.00000 0.00000 0.00000
0.00000 0.00000 0.00000 ; CB- CA- C- O
3854 3852 3859 3861 3 2.51040 4.18400 1.67360
-6.69440 0.00000 0.00000 ; CB- CA- C- N
3855 3854 3852 3859 3 0.65084 1.95253 0.00000
-2.60338 0.00000 0.00000 ; HB2- CB- CA- C
3855 3854 3857 3858 3 0.69733 2.09200 0.00000
-2.78933 0.00000 0.00000 ; HB2- CB- OG- HG
3856 3854 3852 3859 3 0.65084 1.95253 0.00000
-2.60338 0.00000 0.00000 ; HB3- CB- CA- C
3856 3854 3857 3858 3 0.69733 2.09200 0.00000
-2.78933 0.00000 0.00000 ; HB3- CB- OG- HG
3857 3854 3852 3859 3 7.34710 7.79898 -6.14211
-6.71114 4.31789 0.00000 ; OG- CB- CA- C
3859 3861 3863 3864 3 0.00000 0.00000 0.00000
0.00000 0.00000 0.00000 ; C- N- CA- HA
3859 3861 3863 3865 3 11.71520 1.67360 15.06240 -
13.38880 0.00000 0.00000 ; C- N- CA- CB
3859 3861 3863 3871 3 1.75728 5.27184 2.25936
-7.02912 0.00000 0.00000 ; C- N- CA- C
3860 3859 3861 3862 3 29.28800 -8.36800 -20.92000
0.00000 0.00000 0.00000 ; O- C- N- H
3860 3859 3861 3863 3 20.92000 0.00000 -20.92000
0.00000 0.00000 0.00000 ; O- C- N- CA
3861 3863 3865 3866 3 0.65084 1.95253 0.00000
-2.60338 0.00000 0.00000 ; N- CA- CB- HB2
3861 3863 3865 3867 3 0.65084 1.95253 0.00000
-2.60338 0.00000 0.00000 ; N- CA- CB- HB3
3861 3863 3865 3868 3 15.41386 9.74035 -8.39310
-0.97069 2.97901 0.00000 ; N- CA- CB- CG
3861 3863 3871 3872 3 0.00000 0.00000 0.00000
0.00000 0.00000 0.00000 ; N- CA- C- O
3861 3863 3871 3873 3 17.40544 -5.02080 -13.22144
9.20480 0.00000 0.00000 ; N- CA- C- N
3862 3861 3863 3864 3 0.00000 0.00000 0.00000

0.00000 0.00000 0.00000 ; H- N- CA- HA
3862 3861 3863 3865 3 0.00000 0.00000 0.00000
0.00000 0.00000 0.00000 ; H- N- CA- CB
3862 3861 3863 3871 3 0.00000 0.00000 0.00000
0.00000 0.00000 0.00000 ; H- N- CA- C
3863 3865 3868 3869 3 6.43499 0.00000 -5.39736
0.00000 -1.03763 0.00000 ; CA- CB- CG- OD1
3863 3865 3868 3870 3 6.43499 0.00000 -5.39736
0.00000 -1.03763 0.00000 ; CA- CB- CG- OD2
3863 3871 3873 3874 3 20.92000 0.00000 -20.92000
0.00000 0.00000 0.00000 ; CA- C- N- H
3863 3871 3873 3875 3 20.92000 0.00000 -20.92000
0.00000 0.00000 0.00000 ; CA- C- N- CA
3864 3863 3865 3866 3 0.65084 1.95253 0.00000
-2.60338 0.00000 0.00000 ; HA- CA- CB- HB2
3864 3863 3865 3867 3 0.65084 1.95253 0.00000
-2.60338 0.00000 0.00000 ; HA- CA- CB- HB3
3864 3863 3865 3868 3 0.65084 1.95253 0.00000
-2.60338 0.00000 0.00000 ; HA- CA- CB- CG
3864 3863 3871 3872 3 3.68192 -4.35136 0.00000
1.33888 0.00000 0.00000 ; HA- CA- C- O
3864 3863 3871 3873 3 0.00000 0.00000 0.00000
0.00000 0.00000 0.00000 ; HA- CA- C- N
3865 3863 3871 3872 3 0.00000 0.00000 0.00000
0.00000 0.00000 0.00000 ; CB- CA- C- O
3865 3863 3871 3873 3 2.51040 4.18400 1.67360
-6.69440 0.00000 0.00000 ; CB- CA- C- N
3866 3865 3863 3871 3 0.65084 1.95253 0.00000
-2.60338 0.00000 0.00000 ; HB2- CB- CA- C
3866 3865 3868 3869 3 0.00000 0.00000 0.00000
0.00000 0.00000 0.00000 ; HB2- CB- CG- OD1
3866 3865 3868 3870 3 0.00000 0.00000 0.00000
0.00000 0.00000 0.00000 ; HB2- CB- CG- OD2
3867 3865 3863 3871 3 0.65084 1.95253 0.00000
-2.60338 0.00000 0.00000 ; HB3- CB- CA- C
3867 3865 3868 3869 3 0.00000 0.00000 0.00000
0.00000 0.00000 0.00000 ; HB3- CB- CG- OD1
3867 3865 3868 3870 3 0.00000 0.00000 0.00000
0.00000 0.00000 0.00000 ; HB3- CB- CG- OD2
3868 3865 3863 3871 3 7.14627 -1.04600 -8.99560
-0.97069 5.15469 0.00000 ; CG- CB- CA- C
3871 3873 3875 3876 3 0.00000 0.00000 0.00000
0.00000 0.00000 0.00000 ; C- N- CA- HA
3871 3873 3875 3877 3 11.71520 1.67360 15.06240 -
13.38880 0.00000 0.00000 ; C- N- CA- CB
3871 3873 3875 3881 3 1.75728 5.27184 2.25936
-7.02912 0.00000 0.00000 ; C- N- CA- C
3872 3871 3873 3874 3 29.28800 -8.36800 -20.92000
0.00000 0.00000 0.00000 ; O- C- N- H
3872 3871 3873 3875 3 20.92000 0.00000 -20.92000
0.00000 0.00000 0.00000 ; O- C- N- CA
3873 3875 3877 3878 3 0.65084 1.95253 0.00000

-2.60338 0.00000 0.00000 ; N- CA- CB- HB1
3873 3875 3877 3879 3 0.65084 1.95253 0.00000
-2.60338 0.00000 0.00000 ; N- CA- CB- HB2
3873 3875 3877 3880 3 0.65084 1.95253 0.00000
-2.60338 0.00000 0.00000 ; N- CA- CB- HB3
3873 3875 3881 3882 3 0.00000 0.00000 0.00000
0.00000 0.00000 0.00000 ; N- CA- C- O
3873 3875 3881 3883 3 17.40544 -5.02080 -13.22144
9.20480 0.00000 0.00000 ; N- CA- C- N
3874 3873 3875 3876 3 0.00000 0.00000 0.00000
0.00000 0.00000 0.00000 ; H- N- CA- HA
3874 3873 3875 3877 3 0.00000 0.00000 0.00000
0.00000 0.00000 0.00000 ; H- N- CA- CB
3874 3873 3875 3881 3 0.00000 0.00000 0.00000
0.00000 0.00000 0.00000 ; H- N- CA- C
3875 3881 3883 3884 3 20.92000 0.00000 -20.92000
0.00000 0.00000 0.00000 ; CA- C- N- H
3875 3881 3883 3885 3 20.92000 0.00000 -20.92000
0.00000 0.00000 0.00000 ; CA- C- N- CA
3876 3875 3877 3878 3 0.65084 1.95253 0.00000
-2.60338 0.00000 0.00000 ; HA- CA- CB- HB1
3876 3875 3877 3879 3 0.65084 1.95253 0.00000
-2.60338 0.00000 0.00000 ; HA- CA- CB- HB2
3876 3875 3877 3880 3 0.65084 1.95253 0.00000
-2.60338 0.00000 0.00000 ; HA- CA- CB- HB3
3876 3875 3881 3882 3 3.68192 -4.35136 0.00000
1.33888 0.00000 0.00000 ; HA- CA- C- O
3876 3875 3881 3883 3 0.00000 0.00000 0.00000
0.00000 0.00000 0.00000 ; HA- CA- C- N
3877 3875 3881 3882 3 0.00000 0.00000 0.00000
0.00000 0.00000 0.00000 ; CB- CA- C- O
3877 3875 3881 3883 3 2.51040 4.18400 1.67360
-6.69440 0.00000 0.00000 ; CB- CA- C- N
3878 3877 3875 3881 3 0.65084 1.95253 0.00000
-2.60338 0.00000 0.00000 ; HB1- CB- CA- C
3879 3877 3875 3881 3 0.65084 1.95253 0.00000
-2.60338 0.00000 0.00000 ; HB2- CB- CA- C
3880 3877 3875 3881 3 0.65084 1.95253 0.00000
-2.60338 0.00000 0.00000 ; HB3- CB- CA- C
3881 3883 3885 3886 3 0.00000 0.00000 0.00000
0.00000 0.00000 0.00000 ; C- N- CA- HA
3881 3883 3885 3887 3 11.71520 1.67360 15.06240 -
13.38880 0.00000 0.00000 ; C- N- CA- CB
3881 3883 3885 3903 3 1.75728 5.27184 2.25936
-7.02912 0.00000 0.00000 ; C- N- CA- C
3882 3881 3883 3884 3 29.28800 -8.36800 -20.92000
0.00000 0.00000 0.00000 ; O- C- N- H
3882 3881 3883 3885 3 20.92000 0.00000 -20.92000
0.00000 0.00000 0.00000 ; O- C- N- CA
3883 3885 3887 3888 3 0.65084 1.95253 0.00000
-2.60338 0.00000 0.00000 ; N- CA- CB- HB2
3883 3885 3887 3889 3 0.65084 1.95253 0.00000

```
-2.60338    0.00000    0.00000 ;      N-    CA-    CB-   HB3
  3883   3885   3887   3890      3    0.65084    1.95253    0.00000
-2.60338    0.00000    0.00000 ;      N-    CA-    CB-    CG
  3883   3885   3903   3904      3    0.00000    0.00000    0.00000
0.00000    0.00000    0.00000 ;      N-    CA-     C-     O
  3883   3885   3903   3905      3   17.40544   -5.02080  -13.22144
9.20480    0.00000    0.00000 ;      N-    CA-     C-     N
  3884   3883   3885   3886      3    0.00000    0.00000    0.00000
0.00000    0.00000    0.00000 ;      H-     N-    CA-    HA
  3884   3883   3885   3887      3    0.00000    0.00000    0.00000
0.00000    0.00000    0.00000 ;      H-     N-    CA-    CB
  3884   3883   3885   3903      3    0.00000    0.00000    0.00000
0.00000    0.00000    0.00000 ;      H-     N-    CA-     C
  3885   3887   3890   3891      3    0.66944    2.00832    0.00000
-2.67776    0.00000    0.00000 ;     CA-    CB-    CG-   HG2
  3885   3887   3890   3892      3    0.66944    2.00832    0.00000
-2.67776    0.00000    0.00000 ;     CA-    CB-    CG-   HG3
  3885   3887   3890   3893      3    3.68192    3.09616   -2.09200
-3.01248    0.00000    0.00000 ;     CA-    CB-    CG-    CD
  3885   3903   3905   3906      3   20.92000    0.00000  -20.92000
0.00000    0.00000    0.00000 ;     CA-     C-     N-     H
  3885   3903   3905   3907      3   20.92000    0.00000  -20.92000
0.00000    0.00000    0.00000 ;     CA-     C-     N-    CA
  3886   3885   3887   3888      3    0.65084    1.95253    0.00000
-2.60338    0.00000    0.00000 ;     HA-    CA-    CB-   HB2
  3886   3885   3887   3889      3    0.65084    1.95253    0.00000
-2.60338    0.00000    0.00000 ;     HA-    CA-    CB-   HB3
  3886   3885   3887   3890      3    0.65084    1.95253    0.00000
-2.60338    0.00000    0.00000 ;     HA-    CA-    CB-    CG
  3886   3885   3903   3904      3    3.68192   -4.35136    0.00000
1.33888    0.00000    0.00000 ;     HA-    CA-     C-     O
  3886   3885   3903   3905      3    0.00000    0.00000    0.00000
0.00000    0.00000    0.00000 ;     HA-    CA-     C-     N
  3887   3885   3903   3904      3    0.00000    0.00000    0.00000
0.00000    0.00000    0.00000 ;     CB-    CA-     C-     O
  3887   3885   3903   3905      3    2.51040    4.18400    1.67360
-6.69440    0.00000    0.00000 ;     CB-    CA-     C-     N
  3887   3890   3893   3894      3    0.66944    2.00832    0.00000
-2.67776    0.00000    0.00000 ;     CB-    CG-    CD-   HD2
  3887   3890   3893   3895      3    0.66944    2.00832    0.00000
-2.67776    0.00000    0.00000 ;     CB-    CG-    CD-   HD3
  3887   3890   3893   3896      3    3.68192    3.09616   -2.09200
-3.01248    0.00000    0.00000 ;     CB-    CG-    CD-    CE
  3888   3887   3885   3903      3    0.65084    1.95253    0.00000
-2.60338    0.00000    0.00000 ;    HB2-    CB-    CA-     C
  3888   3887   3890   3891      3    0.62760    1.88280    0.00000
-2.51040    0.00000    0.00000 ;    HB2-    CB-    CG-   HG2
  3888   3887   3890   3892      3    0.62760    1.88280    0.00000
-2.51040    0.00000    0.00000 ;    HB2-    CB-    CG-   HG3
  3888   3887   3890   3893      3    0.66944    2.00832    0.00000
-2.67776    0.00000    0.00000 ;    HB2-    CB-    CG-    CD
  3889   3887   3885   3903      3    0.65084    1.95253    0.00000
```

```
-2.60338    0.00000    0.00000 ;    HB3-    CB-    CA-     C
  3889   3887   3890   3891      3    0.62760    1.88280    0.00000
-2.51040    0.00000    0.00000 ;    HB3-    CB-    CG-   HG2
  3889   3887   3890   3892      3    0.62760    1.88280    0.00000
-2.51040    0.00000    0.00000 ;    HB3-    CB-    CG-   HG3
  3889   3887   3890   3893      3    0.66944    2.00832    0.00000
-2.67776    0.00000    0.00000 ;    HB3-    CB-    CG-    CD
  3890   3887   3885   3903      3    0.65084    1.95253    0.00000
-2.60338    0.00000    0.00000 ;     CG-    CB-    CA-     C
  3890   3893   3896   3897      3    0.65084    1.95253    0.00000
-2.60338    0.00000    0.00000 ;     CG-    CD-    CE-   HE2
  3890   3893   3896   3898      3    0.65084    1.95253    0.00000
-2.60338    0.00000    0.00000 ;     CG-    CD-    CE-   HE3
  3890   3893   3896   3899      3    0.65084    1.95253    0.00000
-2.60338    0.00000    0.00000 ;     CG-    CD-    CE-    NZ
  3891   3890   3893   3894      3    0.62760    1.88280    0.00000
-2.51040    0.00000    0.00000 ;    HG2-    CG-    CD-   HD2
  3891   3890   3893   3895      3    0.62760    1.88280    0.00000
-2.51040    0.00000    0.00000 ;    HG2-    CG-    CD-   HD3
  3891   3890   3893   3896      3    0.66944    2.00832    0.00000
-2.67776    0.00000    0.00000 ;    HG2-    CG-    CD-    CE
  3892   3890   3893   3894      3    0.62760    1.88280    0.00000
-2.51040    0.00000    0.00000 ;    HG3-    CG-    CD-   HD2
  3892   3890   3893   3895      3    0.62760    1.88280    0.00000
-2.51040    0.00000    0.00000 ;    HG3-    CG-    CD-   HD3
  3892   3890   3893   3896      3    0.66944    2.00832    0.00000
-2.67776    0.00000    0.00000 ;    HG3-    CG-    CD-    CE
  3893   3896   3899   3900      3    0.65084    1.95253    0.00000
-2.60338    0.00000    0.00000 ;     CD-    CE-    NZ-   HZ1
  3893   3896   3899   3901      3    0.65084    1.95253    0.00000
-2.60338    0.00000    0.00000 ;     CD-    CE-    NZ-   HZ2
  3893   3896   3899   3902      3    0.65084    1.95253    0.00000
-2.60338    0.00000    0.00000 ;     CD-    CE-    NZ-   HZ3
  3894   3893   3896   3897      3    0.65084    1.95253    0.00000
-2.60338    0.00000    0.00000 ;    HD2-    CD-    CE-   HE2
  3894   3893   3896   3898      3    0.65084    1.95253    0.00000
-2.60338    0.00000    0.00000 ;    HD2-    CD-    CE-   HE3
  3894   3893   3896   3899      3    0.65084    1.95253    0.00000
-2.60338    0.00000    0.00000 ;    HD2-    CD-    CE-    NZ
  3895   3893   3896   3897      3    0.65084    1.95253    0.00000
-2.60338    0.00000    0.00000 ;    HD3-    CD-    CE-   HE2
  3895   3893   3896   3898      3    0.65084    1.95253    0.00000
-2.60338    0.00000    0.00000 ;    HD3-    CD-    CE-   HE3
  3895   3893   3896   3899      3    0.65084    1.95253    0.00000
-2.60338    0.00000    0.00000 ;    HD3-    CD-    CE-    NZ
  3897   3896   3899   3900      3    0.65084    1.95253    0.00000
-2.60338    0.00000    0.00000 ;    HE2-    CE-    NZ-   HZ1
  3897   3896   3899   3901      3    0.65084    1.95253    0.00000
-2.60338    0.00000    0.00000 ;    HE2-    CE-    NZ-   HZ2
  3897   3896   3899   3902      3    0.65084    1.95253    0.00000
-2.60338    0.00000    0.00000 ;    HE2-    CE-    NZ-   HZ3
  3898   3896   3899   3900      3    0.65084    1.95253    0.00000
```

-2.60338 0.00000 0.00000 ; HE3- CE- NZ- HZ1
3898 3896 3899 3901 3 0.65084 1.95253 0.00000
-2.60338 0.00000 0.00000 ; HE3- CE- NZ- HZ2
3898 3896 3899 3902 3 0.65084 1.95253 0.00000
-2.60338 0.00000 0.00000 ; HE3- CE- NZ- HZ3
3903 3905 3907 3908 3 0.00000 0.00000 0.00000
0.00000 0.00000 0.00000 ; C- N- CA- HA
3903 3905 3907 3909 3 11.71520 1.67360 15.06240 -
13.38880 0.00000 0.00000 ; C- N- CA- CB
3903 3905 3907 3917 3 1.75728 5.27184 2.25936
-7.02912 0.00000 0.00000 ; C- N- CA- C
3904 3903 3905 3906 3 29.28800 -8.36800 -20.92000
0.00000 0.00000 0.00000 ; O- C- N- H
3904 3903 3905 3907 3 20.92000 0.00000 -20.92000
0.00000 0.00000 0.00000 ; O- C- N- CA
3905 3907 3909 3910 3 0.65084 1.95253 0.00000
-2.60338 0.00000 0.00000 ; N- CA- CB- HB2
3905 3907 3909 3911 3 0.65084 1.95253 0.00000
-2.60338 0.00000 0.00000 ; N- CA- CB- HB3
3905 3907 3909 3912 3 5.99567 3.29281 -4.46014
-0.55229 1.97485 0.00000 ; N- CA- CB- CG
3905 3907 3917 3918 3 0.00000 0.00000 0.00000
0.00000 0.00000 0.00000 ; N- CA- C- O
3905 3907 3917 3919 3 17.40544 -5.02080 -13.22144
9.20480 0.00000 0.00000 ; N- CA- C- N
3906 3905 3907 3908 3 0.00000 0.00000 0.00000
0.00000 0.00000 0.00000 ; H- N- CA- HA
3906 3905 3907 3909 3 0.00000 0.00000 0.00000
0.00000 0.00000 0.00000 ; H- N- CA- CB
3906 3905 3907 3917 3 0.00000 0.00000 0.00000
0.00000 0.00000 0.00000 ; H- N- CA- C
3907 3909 3912 3913 3 0.00000 0.00000 0.00000
0.00000 0.00000 0.00000 ; CA- CB- CG- OD1
3907 3909 3912 3914 3 8.84916 -0.31380 -4.32626
5.03754 0.26778 0.00000 ; CA- CB- CG- ND2
3907 3917 3919 3920 3 20.92000 0.00000 -20.92000
0.00000 0.00000 0.00000 ; CA- C- N- H
3907 3917 3919 3921 3 20.92000 0.00000 -20.92000
0.00000 0.00000 0.00000 ; CA- C- N- CA
3908 3907 3909 3910 3 0.65084 1.95253 0.00000
-2.60338 0.00000 0.00000 ; HA- CA- CB- HB2
3908 3907 3909 3911 3 0.65084 1.95253 0.00000
-2.60338 0.00000 0.00000 ; HA- CA- CB- HB3
3908 3907 3909 3912 3 0.65084 1.95253 0.00000
-2.60338 0.00000 0.00000 ; HA- CA- CB- CG
3908 3907 3917 3918 3 3.68192 -4.35136 0.00000
1.33888 0.00000 0.00000 ; HA- CA- C- O
3908 3907 3917 3919 3 0.00000 0.00000 0.00000
0.00000 0.00000 0.00000 ; HA- CA- C- N
3909 3907 3917 3918 3 0.00000 0.00000 0.00000
0.00000 0.00000 0.00000 ; CB- CA- C- O
3909 3907 3917 3919 3 2.51040 4.18400 1.67360

-6.69440 0.00000 0.00000 ; CB- CA- C- N
3909 3912 3914 3915 3 20.92000 0.00000 -20.92000
0.00000 0.00000 0.00000 ; CB- CG- ND2- HD21
3909 3912 3914 3916 3 20.92000 0.00000 -20.92000
0.00000 0.00000 0.00000 ; CB- CG- ND2- HD22
3910 3909 3907 3917 3 0.65084 1.95253 0.00000
-2.60338 0.00000 0.00000 ; HB2- CB- CA- C
3910 3909 3912 3913 3 3.68192 -4.35136 0.00000
1.33888 0.00000 0.00000 ; HB2- CB- CG- OD1
3910 3909 3912 3914 3 0.00000 0.00000 0.00000
0.00000 0.00000 0.00000 ; HB2- CB- CG- ND2
3911 3909 3907 3917 3 0.65084 1.95253 0.00000
-2.60338 0.00000 0.00000 ; HB3- CB- CA- C
3911 3909 3912 3913 3 3.68192 -4.35136 0.00000
1.33888 0.00000 0.00000 ; HB3- CB- CG- OD1
3911 3909 3912 3914 3 0.00000 0.00000 0.00000
0.00000 0.00000 0.00000 ; HB3- CB- CG- ND2
3912 3909 3907 3917 3 7.94542 -3.96225 -6.11701
-0.55229 3.58150 0.00000 ; CG- CB- CA- C
3913 3912 3914 3915 3 29.28800 -8.36800 -20.92000
0.00000 0.00000 0.00000 ; OD1- CG- ND2- HD21
3913 3912 3914 3916 3 29.28800 -8.36800 -20.92000
0.00000 0.00000 0.00000 ; OD1- CG- ND2- HD22
3917 3919 3921 3922 3 0.00000 0.00000 0.00000
0.00000 0.00000 0.00000 ; C- N- CA- HA
3917 3919 3921 3923 3 11.71520 1.67360 15.06240 -
13.38880 0.00000 0.00000 ; C- N- CA- CB
3917 3919 3921 3936 3 1.75728 5.27184 2.25936
-7.02912 0.00000 0.00000 ; C- N- CA- C
3918 3917 3919 3920 3 29.28800 -8.36800 -20.92000
0.00000 0.00000 0.00000 ; O- C- N- H
3918 3917 3919 3921 3 20.92000 0.00000 -20.92000
0.00000 0.00000 0.00000 ; O- C- N- CA
3919 3921 3923 3924 3 0.65084 1.95253 0.00000
-2.60338 0.00000 0.00000 ; N- CA- CB- HB2
3919 3921 3923 3925 3 0.65084 1.95253 0.00000
-2.60338 0.00000 0.00000 ; N- CA- CB- HB3
3919 3921 3923 3926 3 3.79070 1.39746 -4.61077
-2.40998 2.44346 0.00000 ; N- CA- CB- CG
3919 3921 3936 3937 3 0.00000 0.00000 0.00000
0.00000 0.00000 0.00000 ; N- CA- C- O
3919 3921 3936 3938 3 17.40544 -5.02080 -13.22144
9.20480 0.00000 0.00000 ; N- CA- C- N
3920 3919 3921 3922 3 0.00000 0.00000 0.00000
0.00000 0.00000 0.00000 ; H- N- CA- HA
3920 3919 3921 3923 3 0.00000 0.00000 0.00000
0.00000 0.00000 0.00000 ; H- N- CA- CB
3920 3919 3921 3936 3 0.00000 0.00000 0.00000
0.00000 0.00000 0.00000 ; H- N- CA- C
3921 3923 3926 3927 3 0.66944 2.00832 0.00000
-2.67776 0.00000 0.00000 ; CA- CB- CG- HG
3921 3923 3926 3928 3 3.90367 0.19665 -6.21742

```
-2.37651    5.99149    0.00000 ;      CA-    CB-    CG-   CD1
  3921   3923   3926   3932      3    3.90367    0.19665   -6.21742
-2.37651    5.99149    0.00000 ;      CA-    CB-    CG-   CD2
  3921   3936   3938   3939      3   20.92000    0.00000  -20.92000
0.00000    0.00000    0.00000 ;      CA-     C-     N-     H
  3921   3936   3938   3940      3   20.92000    0.00000  -20.92000
0.00000    0.00000    0.00000 ;      CA-     C-     N-    CA
  3922   3921   3923   3924      3    0.65084    1.95253    0.00000
-2.60338    0.00000    0.00000 ;      HA-    CA-    CB-   HB2
  3922   3921   3923   3925      3    0.65084    1.95253    0.00000
-2.60338    0.00000    0.00000 ;      HA-    CA-    CB-   HB3
  3922   3921   3923   3926      3    0.65084    1.95253    0.00000
-2.60338    0.00000    0.00000 ;      HA-    CA-    CB-    CG
  3922   3921   3936   3937      3    3.68192   -4.35136    0.00000
1.33888    0.00000    0.00000 ;      HA-    CA-     C-     O
  3922   3921   3936   3938      3    0.00000    0.00000    0.00000
0.00000    0.00000    0.00000 ;      HA-    CA-     C-     N
  3923   3921   3936   3937      3    0.00000    0.00000    0.00000
0.00000    0.00000    0.00000 ;      CB-    CA-     C-     O
  3923   3921   3936   3938      3    2.51040    4.18400    1.67360
-6.69440    0.00000    0.00000 ;      CB-    CA-     C-     N
  3923   3926   3928   3929      3    0.66944    2.00832    0.00000
-2.67776    0.00000    0.00000 ;      CB-    CG-   CD1-  HD11
  3923   3926   3928   3930      3    0.66944    2.00832    0.00000
-2.67776    0.00000    0.00000 ;      CB-    CG-   CD1-  HD12
  3923   3926   3928   3931      3    0.66944    2.00832    0.00000
-2.67776    0.00000    0.00000 ;      CB-    CG-   CD1-  HD13
  3923   3926   3932   3933      3    0.66944    2.00832    0.00000
-2.67776    0.00000    0.00000 ;      CB-    CG-   CD2-  HD21
  3923   3926   3932   3934      3    0.66944    2.00832    0.00000
-2.67776    0.00000    0.00000 ;      CB-    CG-   CD2-  HD22
  3923   3926   3932   3935      3    0.66944    2.00832    0.00000
-2.67776    0.00000    0.00000 ;      CB-    CG-   CD2-  HD23
  3924   3923   3921   3936      3    0.65084    1.95253    0.00000
-2.60338    0.00000    0.00000 ;     HB2-    CB-    CA-     C
  3924   3923   3926   3927      3    0.62760    1.88280    0.00000
-2.51040    0.00000    0.00000 ;     HB2-    CB-    CG-    HG
  3924   3923   3926   3928      3    0.66944    2.00832    0.00000
-2.67776    0.00000    0.00000 ;     HB2-    CB-    CG-   CD1
  3924   3923   3926   3932      3    0.66944    2.00832    0.00000
-2.67776    0.00000    0.00000 ;     HB2-    CB-    CG-   CD2
  3925   3923   3921   3936      3    0.65084    1.95253    0.00000
-2.60338    0.00000    0.00000 ;     HB3-    CB-    CA-     C
  3925   3923   3926   3927      3    0.62760    1.88280    0.00000
-2.51040    0.00000    0.00000 ;     HB3-    CB-    CG-    HG
  3925   3923   3926   3928      3    0.66944    2.00832    0.00000
-2.67776    0.00000    0.00000 ;     HB3-    CB-    CG-   CD1
  3925   3923   3926   3932      3    0.66944    2.00832    0.00000
-2.67776    0.00000    0.00000 ;     HB3-    CB-    CG-   CD2
  3926   3923   3921   3936      3   10.33448   -1.14642  -11.54784
-2.40998    6.35968    0.00000 ;      CG-    CB-    CA-     C
  3927   3926   3928   3929      3    0.62760    1.88280    0.00000
```

-2.51040 0.00000 0.00000 ; HG- CG- CD1- HD11
3927 3926 3928 3930 3 0.62760 1.88280 0.00000
-2.51040 0.00000 0.00000 ; HG- CG- CD1- HD12
3927 3926 3928 3931 3 0.62760 1.88280 0.00000
-2.51040 0.00000 0.00000 ; HG- CG- CD1- HD13
3927 3926 3932 3933 3 0.62760 1.88280 0.00000
-2.51040 0.00000 0.00000 ; HG- CG- CD2- HD21
3927 3926 3932 3934 3 0.62760 1.88280 0.00000
-2.51040 0.00000 0.00000 ; HG- CG- CD2- HD22
3927 3926 3932 3935 3 0.62760 1.88280 0.00000
-2.51040 0.00000 0.00000 ; HG- CG- CD2- HD23
3928 3926 3932 3933 3 0.66944 2.00832 0.00000
-2.67776 0.00000 0.00000 ; CD1- CG- CD2- HD21
3928 3926 3932 3934 3 0.66944 2.00832 0.00000
-2.67776 0.00000 0.00000 ; CD1- CG- CD2- HD22
3928 3926 3932 3935 3 0.66944 2.00832 0.00000
-2.67776 0.00000 0.00000 ; CD1- CG- CD2- HD23
3929 3928 3926 3932 3 0.66944 2.00832 0.00000
-2.67776 0.00000 0.00000 ; HD11- CD1- CG- CD2
3930 3928 3926 3932 3 0.66944 2.00832 0.00000
-2.67776 0.00000 0.00000 ; HD12- CD1- CG- CD2
3931 3928 3926 3932 3 0.66944 2.00832 0.00000
-2.67776 0.00000 0.00000 ; HD13- CD1- CG- CD2
3936 3938 3940 3941 3 0.00000 0.00000 0.00000
0.00000 0.00000 0.00000 ; C- N- CA- HA
3936 3938 3940 3942 3 11.71520 1.67360 15.06240 -
13.38880 0.00000 0.00000 ; C- N- CA- CB
3936 3938 3940 3955 3 1.75728 5.27184 2.25936
-7.02912 0.00000 0.00000 ; C- N- CA- C
3937 3936 3938 3939 3 29.28800 -8.36800 -20.92000
0.00000 0.00000 0.00000 ; O- C- N- H
3937 3936 3938 3940 3 20.92000 0.00000 -20.92000
0.00000 0.00000 0.00000 ; O- C- N- CA
3938 3940 3942 3943 3 0.65084 1.95253 0.00000
-2.60338 0.00000 0.00000 ; N- CA- CB- HB
3938 3940 3942 3944 3 3.83673 0.44769 -1.77402
-2.47693 -0.03347 0.00000 ; N- CA- CB- CG2
3938 3940 3942 3948 3 2.97482 0.12134 2.04179
-1.89117 -3.24678 0.00000 ; N- CA- CB- CG1
3938 3940 3955 3956 3 0.00000 0.00000 0.00000
0.00000 0.00000 0.00000 ; N- CA- C- O
3938 3940 3955 3957 3 17.40544 -5.02080 -13.22144
9.20480 0.00000 0.00000 ; N- CA- C- N
3939 3938 3940 3941 3 0.00000 0.00000 0.00000
0.00000 0.00000 0.00000 ; H- N- CA- HA
3939 3938 3940 3942 3 0.00000 0.00000 0.00000
0.00000 0.00000 0.00000 ; H- N- CA- CB
3939 3938 3940 3955 3 0.00000 0.00000 0.00000
0.00000 0.00000 0.00000 ; H- N- CA- C
3940 3942 3944 3945 3 0.66944 2.00832 0.00000
-2.67776 0.00000 0.00000 ; CA- CB- CG2- HG21
3940 3942 3944 3946 3 0.66944 2.00832 0.00000

```
-2.67776    0.00000    0.00000 ;      CA-    CB-   CG2-  HG22
  3940   3942   3944   3947       3    0.66944    2.00832    0.00000
-2.67776    0.00000    0.00000 ;      CA-    CB-   CG2-  HG23
  3940   3942   3948   3949       3    0.66944    2.00832    0.00000
-2.67776    0.00000    0.00000 ;      CA-    CB-   CG1-  HG12
  3940   3942   3948   3950       3    0.66944    2.00832    0.00000
-2.67776    0.00000    0.00000 ;      CA-    CB-   CG1-  HG13
  3940   3942   3948   3951       3    4.24258   -0.52718   -7.25506
-1.79075    7.69856    0.00000 ;      CA-    CB-   CG1-   CD1
  3940   3955   3957   3958       3   20.92000    0.00000  -20.92000
0.00000    0.00000    0.00000 ;      CA-     C-     N-     H
  3940   3955   3957   3959       3   20.92000    0.00000  -20.92000
0.00000    0.00000    0.00000 ;      CA-     C-     N-    CA
  3941   3940   3942   3943       3    0.65084    1.95253    0.00000
-2.60338    0.00000    0.00000 ;      HA-    CA-    CB-    HB
  3941   3940   3942   3944       3    0.65084    1.95253    0.00000
-2.60338    0.00000    0.00000 ;      HA-    CA-    CB-   CG2
  3941   3940   3942   3948       3    0.65084    1.95253    0.00000
-2.60338    0.00000    0.00000 ;      HA-    CA-    CB-   CG1
  3941   3940   3955   3956       3    3.68192   -4.35136    0.00000
1.33888    0.00000    0.00000 ;      HA-    CA-     C-     O
  3941   3940   3955   3957       3    0.00000    0.00000    0.00000
0.00000    0.00000    0.00000 ;      HA-    CA-     C-     N
  3942   3940   3955   3956       3    0.00000    0.00000    0.00000
0.00000    0.00000    0.00000 ;      CB-    CA-     C-     O
  3942   3940   3955   3957       3    2.51040    4.18400    1.67360
-6.69440    0.00000    0.00000 ;      CB-    CA-     C-     N
  3942   3948   3951   3952       3    0.66944    2.00832    0.00000
-2.67776    0.00000    0.00000 ;      CB-   CG1-   CD1-  HD11
  3942   3948   3951   3953       3    0.66944    2.00832    0.00000
-2.67776    0.00000    0.00000 ;      CB-   CG1-   CD1-  HD12
  3942   3948   3951   3954       3    0.66944    2.00832    0.00000
-2.67776    0.00000    0.00000 ;      CB-   CG1-   CD1-  HD13
  3943   3942   3940   3955       3    0.65084    1.95253    0.00000
-2.60338    0.00000    0.00000 ;      HB-    CB-    CA-     C
  3943   3942   3944   3945       3    0.62760    1.88280    0.00000
-2.51040    0.00000    0.00000 ;      HB-    CB-   CG2-  HG21
  3943   3942   3944   3946       3    0.62760    1.88280    0.00000
-2.51040    0.00000    0.00000 ;      HB-    CB-   CG2-  HG22
  3943   3942   3944   3947       3    0.62760    1.88280    0.00000
-2.51040    0.00000    0.00000 ;      HB-    CB-   CG2-  HG23
  3943   3942   3948   3949       3    0.62760    1.88280    0.00000
-2.51040    0.00000    0.00000 ;      HB-    CB-   CG1-  HG12
  3943   3942   3948   3950       3    0.62760    1.88280    0.00000
-2.51040    0.00000    0.00000 ;      HB-    CB-   CG1-  HG13
  3943   3942   3948   3951       3    0.66944    2.00832    0.00000
-2.67776    0.00000    0.00000 ;      HB-    CB-   CG1-   CD1
  3944   3942   3940   3955       3    5.67350    3.55640   -6.16722
-2.47693    3.74886    0.00000 ;     CG2-    CB-    CA-     C
  3944   3942   3948   3949       3    0.66944    2.00832    0.00000
-2.67776    0.00000    0.00000 ;     CG2-    CB-   CG1-  HG12
  3944   3942   3948   3950       3    0.66944    2.00832    0.00000
```

-2.67776 0.00000 0.00000 ; CG2- CB- CG1- HG13
3944 3942 3948 3951 3 3.81162 0.49790 -8.14206
-1.79075 7.49773 0.00000 ; CG2- CB- CG1- CD1
3945 3944 3942 3948 3 0.66944 2.00832 0.00000
-2.67776 0.00000 0.00000 ; HG21- CG2- CB- CG1
3946 3944 3942 3948 3 0.66944 2.00832 0.00000
-2.67776 0.00000 0.00000 ; HG22- CG2- CB- CG1
3947 3944 3942 3948 3 0.66944 2.00832 0.00000
-2.67776 0.00000 0.00000 ; HG23- CG2- CB- CG1
3948 3942 3940 3955 3 8.26340 0.74057 -9.99976
-1.89117 3.84928 0.00000 ; CG1- CB- CA- C
3949 3948 3951 3952 3 0.62760 1.88280 0.00000
-2.51040 0.00000 0.00000 ; HG12- CG1- CD1- HD11
3949 3948 3951 3953 3 0.62760 1.88280 0.00000
-2.51040 0.00000 0.00000 ; HG12- CG1- CD1- HD12
3949 3948 3951 3954 3 0.62760 1.88280 0.00000
-2.51040 0.00000 0.00000 ; HG12- CG1- CD1- HD13
3950 3948 3951 3952 3 0.62760 1.88280 0.00000
-2.51040 0.00000 0.00000 ; HG13- CG1- CD1- HD11
3950 3948 3951 3953 3 0.62760 1.88280 0.00000
-2.51040 0.00000 0.00000 ; HG13- CG1- CD1- HD12
3950 3948 3951 3954 3 0.62760 1.88280 0.00000
-2.51040 0.00000 0.00000 ; HG13- CG1- CD1- HD13
3955 3957 3959 3960 3 0.00000 0.00000 0.00000
0.00000 0.00000 0.00000 ; C- N- CA- HA
3955 3957 3959 3961 3 11.71520 1.67360 15.06240 -
13.38880 0.00000 0.00000 ; C- N- CA- CB
3955 3957 3959 3969 3 1.75728 5.27184 2.25936
-7.02912 0.00000 0.00000 ; C- N- CA- C
3956 3955 3957 3958 3 29.28800 -8.36800 -20.92000
0.00000 0.00000 0.00000 ; O- C- N- H
3956 3955 3957 3959 3 20.92000 0.00000 -20.92000
0.00000 0.00000 0.00000 ; O- C- N- CA
3957 3959 3961 3962 3 0.65084 1.95253 0.00000
-2.60338 0.00000 0.00000 ; N- CA- CB- HB
3957 3959 3961 3963 3 3.83673 0.44769 -1.77402
-2.47693 -0.03347 0.00000 ; N- CA- CB- CG2
3957 3959 3961 3967 3 4.93294 1.13386 -3.12963
-5.27184 3.17984 0.00000 ; N- CA- CB- OG1
3957 3959 3969 3970 3 0.00000 0.00000 0.00000
0.00000 0.00000 0.00000 ; N- CA- C- O
3957 3959 3969 3971 3 17.40544 -5.02080 -13.22144
9.20480 0.00000 0.00000 ; N- CA- C- N
3958 3957 3959 3960 3 0.00000 0.00000 0.00000
0.00000 0.00000 0.00000 ; H- N- CA- HA
3958 3957 3959 3961 3 0.00000 0.00000 0.00000
0.00000 0.00000 0.00000 ; H- N- CA- CB
3958 3957 3959 3969 3 0.00000 0.00000 0.00000
0.00000 0.00000 0.00000 ; H- N- CA- C
3959 3961 3963 3964 3 0.66944 2.00832 0.00000
-2.67776 0.00000 0.00000 ; CA- CB- CG2- HG21
3959 3961 3963 3965 3 0.66944 2.00832 0.00000

-2.67776    0.00000    0.00000 ;      CA-    CB-   CG2-  HG22
  3959   3961   3963   3966      3    0.66944    2.00832    0.00000
-2.67776    0.00000    0.00000 ;      CA-    CB-   CG2-  HG23
  3959   3961   3967   3968      3    1.12131    2.98738    1.66523
-3.94970    0.43514    0.00000 ;      CA-    CB-   OG1-   HG1
  3959   3969   3971   3972      3   20.92000    0.00000  -20.92000
0.00000    0.00000    0.00000 ;      CA-     C-     N-     H
  3959   3969   3971   3973      3   20.92000    0.00000  -20.92000
0.00000    0.00000    0.00000 ;      CA-     C-     N-    CA
  3960   3959   3961   3962      3    0.65084    1.95253    0.00000
-2.60338    0.00000    0.00000 ;      HA-    CA-    CB-    HB
  3960   3959   3961   3963      3    0.65084    1.95253    0.00000
-2.60338    0.00000    0.00000 ;      HA-    CA-    CB-   CG2
  3960   3959   3961   3967      3    1.04600   -1.04600    0.00000
0.00000    0.00000    0.00000 ;      HA-    CA-    CB-   OG1
  3960   3959   3969   3970      3    3.68192   -4.35136    0.00000
1.33888    0.00000    0.00000 ;      HA-    CA-     C-     O
  3960   3959   3969   3971      3    0.00000    0.00000    0.00000
0.00000    0.00000    0.00000 ;      HA-    CA-     C-     N
  3961   3959   3969   3970      3    0.00000    0.00000    0.00000
0.00000    0.00000    0.00000 ;      CB-    CA-     C-     O
  3961   3959   3969   3971      3    2.51040    4.18400    1.67360
-6.69440    0.00000    0.00000 ;      CB-    CA-     C-     N
  3962   3961   3959   3969      3    0.65084    1.95253    0.00000
-2.60338    0.00000    0.00000 ;      HB-    CB-    CA-     C
  3962   3961   3963   3964      3    0.65084    1.95253    0.00000
-2.60338    0.00000    0.00000 ;      HB-    CB-   CG2-  HG21
  3962   3961   3963   3965      3    0.65084    1.95253    0.00000
-2.60338    0.00000    0.00000 ;      HB-    CB-   CG2-  HG22
  3962   3961   3963   3966      3    0.65084    1.95253    0.00000
-2.60338    0.00000    0.00000 ;      HB-    CB-   CG2-  HG23
  3962   3961   3967   3968      3    0.69733    2.09200    0.00000
-2.78933    0.00000    0.00000 ;      HB-    CB-   OG1-   HG1
  3963   3961   3959   3969      3    5.67350    3.55640   -6.16722
-2.47693    3.74886    0.00000 ;     CG2-    CB-    CA-     C
  3963   3961   3967   3968      3    4.74047    0.27196   -2.26773
-3.94970    1.60666    0.00000 ;     CG2-    CB-   OG1-   HG1
  3964   3963   3961   3967      3    1.04600   -1.04600    0.00000
0.00000    0.00000    0.00000 ;    HG21-   CG2-    CB-   OG1
  3965   3963   3961   3967      3    1.04600   -1.04600    0.00000
0.00000    0.00000    0.00000 ;    HG22-   CG2-    CB-   OG1
  3966   3963   3961   3967      3    1.04600   -1.04600    0.00000
0.00000    0.00000    0.00000 ;    HG23-   CG2-    CB-   OG1
  3967   3961   3959   3969      3    6.53541    6.87013   -6.21742
-5.27184    5.22163    0.00000 ;     OG1-    CB-    CA-     C
  3969   3971   3973   3974      3    0.00000    0.00000    0.00000
0.00000    0.00000    0.00000 ;       C-     N-    CA-    HA
  3969   3971   3973   3975      3   11.71520    1.67360   15.06240  -
13.38880    0.00000    0.00000 ;       C-     N-    CA-    CB
  3969   3971   3973   3991      3    1.75728    5.27184    2.25936
-7.02912    0.00000    0.00000 ;       C-     N-    CA-     C
  3970   3969   3971   3972      3   29.28800   -8.36800  -20.92000

```
0.00000    0.00000    0.00000 ;      O-     C-     N-     H
  3970   3969   3971   3973      3   20.92000    0.00000  -20.92000
0.00000    0.00000    0.00000 ;      O-     C-     N-    CA
  3971   3973   3975   3976      3    0.65084    1.95253    0.00000
-2.60338    0.00000    0.00000 ;      N-    CA-    CB-   HB2
  3971   3973   3975   3977      3    0.65084    1.95253    0.00000
-2.60338    0.00000    0.00000 ;      N-    CA-    CB-   HB3
  3971   3973   3975   3978      3    0.65084    1.95253    0.00000
-2.60338    0.00000    0.00000 ;      N-    CA-    CB-    CG
  3971   3973   3991   3992      3    0.00000    0.00000    0.00000
0.00000    0.00000    0.00000 ;      N-    CA-     C-     O
  3971   3973   3991   3993      3   17.40544   -5.02080  -13.22144
9.20480    0.00000    0.00000 ;      N-    CA-     C-     N
  3972   3971   3973   3974      3    0.00000    0.00000    0.00000
0.00000    0.00000    0.00000 ;      H-     N-    CA-    HA
  3972   3971   3973   3975      3    0.00000    0.00000    0.00000
0.00000    0.00000    0.00000 ;      H-     N-    CA-    CB
  3972   3971   3973   3991      3    0.00000    0.00000    0.00000
0.00000    0.00000    0.00000 ;      H-     N-    CA-     C
  3973   3975   3978   3979      3    0.66944    2.00832    0.00000
-2.67776    0.00000    0.00000 ;     CA-    CB-    CG-   HG2
  3973   3975   3978   3980      3    0.66944    2.00832    0.00000
-2.67776    0.00000    0.00000 ;     CA-    CB-    CG-   HG3
  3973   3975   3978   3981      3    3.68192    3.09616   -2.09200
-3.01248    0.00000    0.00000 ;     CA-    CB-    CG-    CD
  3973   3991   3993   3994      3   20.92000    0.00000  -20.92000
0.00000    0.00000    0.00000 ;     CA-     C-     N-     H
  3973   3991   3993   3995      3   20.92000    0.00000  -20.92000
0.00000    0.00000    0.00000 ;     CA-     C-     N-    CA
  3974   3973   3975   3976      3    0.65084    1.95253    0.00000
-2.60338    0.00000    0.00000 ;     HA-    CA-    CB-   HB2
  3974   3973   3975   3977      3    0.65084    1.95253    0.00000
-2.60338    0.00000    0.00000 ;     HA-    CA-    CB-   HB3
  3974   3973   3975   3978      3    0.65084    1.95253    0.00000
-2.60338    0.00000    0.00000 ;     HA-    CA-    CB-    CG
  3974   3973   3991   3992      3    3.68192   -4.35136    0.00000
1.33888    0.00000    0.00000 ;     HA-    CA-     C-     O
  3974   3973   3991   3993      3    0.00000    0.00000    0.00000
0.00000    0.00000    0.00000 ;     HA-    CA-     C-     N
  3975   3973   3991   3992      3    0.00000    0.00000    0.00000
0.00000    0.00000    0.00000 ;     CB-    CA-     C-     O
  3975   3973   3991   3993      3    2.51040    4.18400    1.67360
-6.69440    0.00000    0.00000 ;     CB-    CA-     C-     N
  3975   3978   3981   3982      3    0.66944    2.00832    0.00000
-2.67776    0.00000    0.00000 ;     CB-    CG-    CD-   HD2
  3975   3978   3981   3983      3    0.66944    2.00832    0.00000
-2.67776    0.00000    0.00000 ;     CB-    CG-    CD-   HD3
  3975   3978   3981   3984      3    3.68192    3.09616   -2.09200
-3.01248    0.00000    0.00000 ;     CB-    CG-    CD-    CE
  3976   3975   3973   3991      3    0.65084    1.95253    0.00000
-2.60338    0.00000    0.00000 ;    HB2-    CB-    CA-     C
  3976   3975   3978   3979      3    0.62760    1.88280    0.00000
```

```
-2.51040    0.00000    0.00000 ;     HB2-    CB-    CG-   HG2
  3976   3975   3978   3980      3    0.62760    1.88280    0.00000
-2.51040    0.00000    0.00000 ;     HB2-    CB-    CG-   HG3
  3976   3975   3978   3981      3    0.66944    2.00832    0.00000
-2.67776    0.00000    0.00000 ;     HB2-    CB-    CG-    CD
  3977   3975   3973   3991      3    0.65084    1.95253    0.00000
-2.60338    0.00000    0.00000 ;     HB3-    CB-    CA-     C
  3977   3975   3978   3979      3    0.62760    1.88280    0.00000
-2.51040    0.00000    0.00000 ;     HB3-    CB-    CG-   HG2
  3977   3975   3978   3980      3    0.62760    1.88280    0.00000
-2.51040    0.00000    0.00000 ;     HB3-    CB-    CG-   HG3
  3977   3975   3978   3981      3    0.66944    2.00832    0.00000
-2.67776    0.00000    0.00000 ;     HB3-    CB-    CG-    CD
  3978   3975   3973   3991      3    0.65084    1.95253    0.00000
-2.60338    0.00000    0.00000 ;      CG-    CB-    CA-     C
  3978   3981   3984   3985      3    0.65084    1.95253    0.00000
-2.60338    0.00000    0.00000 ;      CG-    CD-    CE-   HE2
  3978   3981   3984   3986      3    0.65084    1.95253    0.00000
-2.60338    0.00000    0.00000 ;      CG-    CD-    CE-   HE3
  3978   3981   3984   3987      3    0.65084    1.95253    0.00000
-2.60338    0.00000    0.00000 ;      CG-    CD-    CE-    NZ
  3979   3978   3981   3982      3    0.62760    1.88280    0.00000
-2.51040    0.00000    0.00000 ;     HG2-    CG-    CD-   HD2
  3979   3978   3981   3983      3    0.62760    1.88280    0.00000
-2.51040    0.00000    0.00000 ;     HG2-    CG-    CD-   HD3
  3979   3978   3981   3984      3    0.66944    2.00832    0.00000
-2.67776    0.00000    0.00000 ;     HG2-    CG-    CD-    CE
  3980   3978   3981   3982      3    0.62760    1.88280    0.00000
-2.51040    0.00000    0.00000 ;     HG3-    CG-    CD-   HD2
  3980   3978   3981   3983      3    0.62760    1.88280    0.00000
-2.51040    0.00000    0.00000 ;     HG3-    CG-    CD-   HD3
  3980   3978   3981   3984      3    0.66944    2.00832    0.00000
-2.67776    0.00000    0.00000 ;     HG3-    CG-    CD-    CE
  3981   3984   3987   3988      3    0.65084    1.95253    0.00000
-2.60338    0.00000    0.00000 ;      CD-    CE-    NZ-   HZ1
  3981   3984   3987   3989      3    0.65084    1.95253    0.00000
-2.60338    0.00000    0.00000 ;      CD-    CE-    NZ-   HZ2
  3981   3984   3987   3990      3    0.65084    1.95253    0.00000
-2.60338    0.00000    0.00000 ;      CD-    CE-    NZ-   HZ3
  3982   3981   3984   3985      3    0.65084    1.95253    0.00000
-2.60338    0.00000    0.00000 ;     HD2-    CD-    CE-   HE2
  3982   3981   3984   3986      3    0.65084    1.95253    0.00000
-2.60338    0.00000    0.00000 ;     HD2-    CD-    CE-   HE3
  3982   3981   3984   3987      3    0.65084    1.95253    0.00000
-2.60338    0.00000    0.00000 ;     HD2-    CD-    CE-    NZ
  3983   3981   3984   3985      3    0.65084    1.95253    0.00000
-2.60338    0.00000    0.00000 ;     HD3-    CD-    CE-   HE2
  3983   3981   3984   3986      3    0.65084    1.95253    0.00000
-2.60338    0.00000    0.00000 ;     HD3-    CD-    CE-   HE3
  3983   3981   3984   3987      3    0.65084    1.95253    0.00000
-2.60338    0.00000    0.00000 ;     HD3-    CD-    CE-    NZ
  3985   3984   3987   3988      3    0.65084    1.95253    0.00000
```

-2.60338 0.00000 0.00000 ; HE2- CE- NZ- HZ1
3985 3984 3987 3989 3 0.65084 1.95253 0.00000
-2.60338 0.00000 0.00000 ; HE2- CE- NZ- HZ2
3985 3984 3987 3990 3 0.65084 1.95253 0.00000
-2.60338 0.00000 0.00000 ; HE2- CE- NZ- HZ3
3986 3984 3987 3988 3 0.65084 1.95253 0.00000
-2.60338 0.00000 0.00000 ; HE3- CE- NZ- HZ1
3986 3984 3987 3989 3 0.65084 1.95253 0.00000
-2.60338 0.00000 0.00000 ; HE3- CE- NZ- HZ2
3986 3984 3987 3990 3 0.65084 1.95253 0.00000
-2.60338 0.00000 0.00000 ; HE3- CE- NZ- HZ3
3991 3993 3995 3996 3 0.00000 0.00000 0.00000
0.00000 0.00000 0.00000 ; C- N- CA- HA
3991 3993 3995 3997 3 11.71520 1.67360 15.06240 -
13.38880 0.00000 0.00000 ; C- N- CA- CB
3991 3993 3995 4010 3 1.75728 5.27184 2.25936
-7.02912 0.00000 0.00000 ; C- N- CA- C
3992 3991 3993 3994 3 29.28800 -8.36800 -20.92000
0.00000 0.00000 0.00000 ; O- C- N- H
3992 3991 3993 3995 3 20.92000 0.00000 -20.92000
0.00000 0.00000 0.00000 ; O- C- N- CA
3993 3995 3997 3998 3 0.65084 1.95253 0.00000
-2.60338 0.00000 0.00000 ; N- CA- CB- HB2
3993 3995 3997 3999 3 0.65084 1.95253 0.00000
-2.60338 0.00000 0.00000 ; N- CA- CB- HB3
3993 3995 3997 4000 3 3.79070 1.39746 -4.61077
-2.40998 2.44346 0.00000 ; N- CA- CB- CG
3993 3995 4010 4011 3 0.00000 0.00000 0.00000
0.00000 0.00000 0.00000 ; N- CA- C- O
3993 3995 4010 4012 3 17.40544 -5.02080 -13.22144
9.20480 0.00000 0.00000 ; N- CA- C- N
3994 3993 3995 3996 3 0.00000 0.00000 0.00000
0.00000 0.00000 0.00000 ; H- N- CA- HA
3994 3993 3995 3997 3 0.00000 0.00000 0.00000
0.00000 0.00000 0.00000 ; H- N- CA- CB
3994 3993 3995 4010 3 0.00000 0.00000 0.00000
0.00000 0.00000 0.00000 ; H- N- CA- C
3995 3997 4000 4001 3 0.66944 2.00832 0.00000
-2.67776 0.00000 0.00000 ; CA- CB- CG- HG
3995 3997 4000 4002 3 3.90367 0.19665 -6.21742
-2.37651 5.99149 0.00000 ; CA- CB- CG- CD1
3995 3997 4000 4006 3 3.90367 0.19665 -6.21742
-2.37651 5.99149 0.00000 ; CA- CB- CG- CD2
3995 4010 4012 4013 3 20.92000 0.00000 -20.92000
0.00000 0.00000 0.00000 ; CA- C- N- H
3995 4010 4012 4014 3 20.92000 0.00000 -20.92000
0.00000 0.00000 0.00000 ; CA- C- N- CA
3996 3995 3997 3998 3 0.65084 1.95253 0.00000
-2.60338 0.00000 0.00000 ; HA- CA- CB- HB2
3996 3995 3997 3999 3 0.65084 1.95253 0.00000
-2.60338 0.00000 0.00000 ; HA- CA- CB- HB3
3996 3995 3997 4000 3 0.65084 1.95253 0.00000

-2.60338 0.00000 0.00000 ; HA- CA- CB- CG
3996 3995 4010 4011 3 3.68192 -4.35136 0.00000
1.33888 0.00000 0.00000 ; HA- CA- C- O
3996 3995 4010 4012 3 0.00000 0.00000 0.00000
0.00000 0.00000 0.00000 ; HA- CA- C- N
3997 3995 4010 4011 3 0.00000 0.00000 0.00000
0.00000 0.00000 0.00000 ; CB- CA- C- O
3997 3995 4010 4012 3 2.51040 4.18400 1.67360
-6.69440 0.00000 0.00000 ; CB- CA- C- N
3997 4000 4002 4003 3 0.66944 2.00832 0.00000
-2.67776 0.00000 0.00000 ; CB- CG- CD1- HD11
3997 4000 4002 4004 3 0.66944 2.00832 0.00000
-2.67776 0.00000 0.00000 ; CB- CG- CD1- HD12
3997 4000 4002 4005 3 0.66944 2.00832 0.00000
-2.67776 0.00000 0.00000 ; CB- CG- CD1- HD13
3997 4000 4006 4007 3 0.66944 2.00832 0.00000
-2.67776 0.00000 0.00000 ; CB- CG- CD2- HD21
3997 4000 4006 4008 3 0.66944 2.00832 0.00000
-2.67776 0.00000 0.00000 ; CB- CG- CD2- HD22
3997 4000 4006 4009 3 0.66944 2.00832 0.00000
-2.67776 0.00000 0.00000 ; CB- CG- CD2- HD23
3998 3997 3995 4010 3 0.65084 1.95253 0.00000
-2.60338 0.00000 0.00000 ; HB2- CB- CA- C
3998 3997 4000 4001 3 0.62760 1.88280 0.00000
-2.51040 0.00000 0.00000 ; HB2- CB- CG- HG
3998 3997 4000 4002 3 0.66944 2.00832 0.00000
-2.67776 0.00000 0.00000 ; HB2- CB- CG- CD1
3998 3997 4000 4006 3 0.66944 2.00832 0.00000
-2.67776 0.00000 0.00000 ; HB2- CB- CG- CD2
3999 3997 3995 4010 3 0.65084 1.95253 0.00000
-2.60338 0.00000 0.00000 ; HB3- CB- CA- C
3999 3997 4000 4001 3 0.62760 1.88280 0.00000
-2.51040 0.00000 0.00000 ; HB3- CB- CG- HG
3999 3997 4000 4002 3 0.66944 2.00832 0.00000
-2.67776 0.00000 0.00000 ; HB3- CB- CG- CD1
3999 3997 4000 4006 3 0.66944 2.00832 0.00000
-2.67776 0.00000 0.00000 ; HB3- CB- CG- CD2
4000 3997 3995 4010 3 10.33448 -1.14642 -11.54784
-2.40998 6.35968 0.00000 ; CG- CB- CA- C
4001 4000 4002 4003 3 0.62760 1.88280 0.00000
-2.51040 0.00000 0.00000 ; HG- CG- CD1- HD11
4001 4000 4002 4004 3 0.62760 1.88280 0.00000
-2.51040 0.00000 0.00000 ; HG- CG- CD1- HD12
4001 4000 4002 4005 3 0.62760 1.88280 0.00000
-2.51040 0.00000 0.00000 ; HG- CG- CD1- HD13
4001 4000 4006 4007 3 0.62760 1.88280 0.00000
-2.51040 0.00000 0.00000 ; HG- CG- CD2- HD21
4001 4000 4006 4008 3 0.62760 1.88280 0.00000
-2.51040 0.00000 0.00000 ; HG- CG- CD2- HD22
4001 4000 4006 4009 3 0.62760 1.88280 0.00000
-2.51040 0.00000 0.00000 ; HG- CG- CD2- HD23
4002 4000 4006 4007 3 0.66944 2.00832 0.00000

```
-2.67776    0.00000    0.00000 ;    CD1-    CG-   CD2-  HD21
  4002   4000   4006   4008      3    0.66944    2.00832    0.00000
-2.67776    0.00000    0.00000 ;    CD1-    CG-   CD2-  HD22
  4002   4000   4006   4009      3    0.66944    2.00832    0.00000
-2.67776    0.00000    0.00000 ;    CD1-    CG-   CD2-  HD23
  4003   4002   4000   4006      3    0.66944    2.00832    0.00000
-2.67776    0.00000    0.00000 ;   HD11-   CD1-    CG-   CD2
  4004   4002   4000   4006      3    0.66944    2.00832    0.00000
-2.67776    0.00000    0.00000 ;   HD12-   CD1-    CG-   CD2
  4005   4002   4000   4006      3    0.66944    2.00832    0.00000
-2.67776    0.00000    0.00000 ;   HD13-   CD1-    CG-   CD2
  4010   4012   4014   4015      3    0.00000    0.00000    0.00000
0.00000    0.00000    0.00000 ;      C-     N-    CA-    HA
  4010   4012   4014   4016      3   11.71520    1.67360   15.06240  -
13.38880    0.00000    0.00000 ;     C-     N-    CA-    CB
  4010   4012   4014   4029      3    1.75728    5.27184    2.25936
-7.02912    0.00000    0.00000 ;     C-     N-    CA-     C
  4011   4010   4012   4013      3   29.28800   -8.36800  -20.92000
0.00000    0.00000    0.00000 ;      O-     C-     N-     H
  4011   4010   4012   4014      3   20.92000    0.00000  -20.92000
0.00000    0.00000    0.00000 ;      O-     C-     N-    CA
  4012   4014   4016   4017      3    0.65084    1.95253    0.00000
-2.60338    0.00000    0.00000 ;     N-    CA-    CB-   HB2
  4012   4014   4016   4018      3    0.65084    1.95253    0.00000
-2.60338    0.00000    0.00000 ;     N-    CA-    CB-   HB3
  4012   4014   4016   4019      3    3.79070    1.39746   -4.61077
-2.40998    2.44346    0.00000 ;     N-    CA-    CB-    CG
  4012   4014   4029   4030      3    0.00000    0.00000    0.00000
0.00000    0.00000    0.00000 ;     N-    CA-     C-     O
  4012   4014   4029   4031      3   17.40544   -5.02080  -13.22144
9.20480    0.00000    0.00000 ;     N-    CA-     C-     N
  4013   4012   4014   4015      3    0.00000    0.00000    0.00000
0.00000    0.00000    0.00000 ;     H-     N-    CA-    HA
  4013   4012   4014   4016      3    0.00000    0.00000    0.00000
0.00000    0.00000    0.00000 ;     H-     N-    CA-    CB
  4013   4012   4014   4029      3    0.00000    0.00000    0.00000
0.00000    0.00000    0.00000 ;     H-     N-    CA-     C
  4014   4016   4019   4020      3    0.66944    2.00832    0.00000
-2.67776    0.00000    0.00000 ;    CA-    CB-    CG-    HG
  4014   4016   4019   4021      3    3.90367    0.19665   -6.21742
-2.37651    5.99149    0.00000 ;    CA-    CB-    CG-   CD1
  4014   4016   4019   4025      3    3.90367    0.19665   -6.21742
-2.37651    5.99149    0.00000 ;    CA-    CB-    CG-   CD2
  4014   4029   4031   4032      3   20.92000    0.00000  -20.92000
0.00000    0.00000    0.00000 ;    CA-     C-     N-     H
  4014   4029   4031   4033      3   20.92000    0.00000  -20.92000
0.00000    0.00000    0.00000 ;    CA-     C-     N-    CA
  4015   4014   4016   4017      3    0.65084    1.95253    0.00000
-2.60338    0.00000    0.00000 ;    HA-    CA-    CB-   HB2
  4015   4014   4016   4018      3    0.65084    1.95253    0.00000
-2.60338    0.00000    0.00000 ;    HA-    CA-    CB-   HB3
  4015   4014   4016   4019      3    0.65084    1.95253    0.00000
```

```
-2.60338    0.00000    0.00000 ;      HA-    CA-    CB-    CG
  4015   4014   4029   4030      3    3.68192   -4.35136    0.00000
1.33888    0.00000    0.00000 ;      HA-    CA-     C-     O
  4015   4014   4029   4031      3    0.00000    0.00000    0.00000
0.00000    0.00000    0.00000 ;      HA-    CA-     C-     N
  4016   4014   4029   4030      3    0.00000    0.00000    0.00000
0.00000    0.00000    0.00000 ;      CB-    CA-     C-     O
  4016   4014   4029   4031      3    2.51040    4.18400    1.67360
-6.69440    0.00000    0.00000 ;      CB-    CA-     C-     N
  4016   4019   4021   4022      3    0.66944    2.00832    0.00000
-2.67776    0.00000    0.00000 ;      CB-    CG-   CD1-  HD11
  4016   4019   4021   4023      3    0.66944    2.00832    0.00000
-2.67776    0.00000    0.00000 ;      CB-    CG-   CD1-  HD12
  4016   4019   4021   4024      3    0.66944    2.00832    0.00000
-2.67776    0.00000    0.00000 ;      CB-    CG-   CD1-  HD13
  4016   4019   4025   4026      3    0.66944    2.00832    0.00000
-2.67776    0.00000    0.00000 ;      CB-    CG-   CD2-  HD21
  4016   4019   4025   4027      3    0.66944    2.00832    0.00000
-2.67776    0.00000    0.00000 ;      CB-    CG-   CD2-  HD22
  4016   4019   4025   4028      3    0.66944    2.00832    0.00000
-2.67776    0.00000    0.00000 ;      CB-    CG-   CD2-  HD23
  4017   4016   4014   4029      3    0.65084    1.95253    0.00000
-2.60338    0.00000    0.00000 ;     HB2-    CB-    CA-     C
  4017   4016   4019   4020      3    0.62760    1.88280    0.00000
-2.51040    0.00000    0.00000 ;     HB2-    CB-    CG-    HG
  4017   4016   4019   4021      3    0.66944    2.00832    0.00000
-2.67776    0.00000    0.00000 ;     HB2-    CB-    CG-   CD1
  4017   4016   4019   4025      3    0.66944    2.00832    0.00000
-2.67776    0.00000    0.00000 ;     HB2-    CB-    CG-   CD2
  4018   4016   4014   4029      3    0.65084    1.95253    0.00000
-2.60338    0.00000    0.00000 ;     HB3-    CB-    CA-     C
  4018   4016   4019   4020      3    0.62760    1.88280    0.00000
-2.51040    0.00000    0.00000 ;     HB3-    CB-    CG-    HG
  4018   4016   4019   4021      3    0.66944    2.00832    0.00000
-2.67776    0.00000    0.00000 ;     HB3-    CB-    CG-   CD1
  4018   4016   4019   4025      3    0.66944    2.00832    0.00000
-2.67776    0.00000    0.00000 ;     HB3-    CB-    CG-   CD2
  4019   4016   4014   4029      3   10.33448   -1.14642  -11.54784
-2.40998    6.35968    0.00000 ;      CG-    CB-    CA-     C
  4020   4019   4021   4022      3    0.62760    1.88280    0.00000
-2.51040    0.00000    0.00000 ;      HG-    CG-   CD1-  HD11
  4020   4019   4021   4023      3    0.62760    1.88280    0.00000
-2.51040    0.00000    0.00000 ;      HG-    CG-   CD1-  HD12
  4020   4019   4021   4024      3    0.62760    1.88280    0.00000
-2.51040    0.00000    0.00000 ;      HG-    CG-   CD1-  HD13
  4020   4019   4025   4026      3    0.62760    1.88280    0.00000
-2.51040    0.00000    0.00000 ;      HG-    CG-   CD2-  HD21
  4020   4019   4025   4027      3    0.62760    1.88280    0.00000
-2.51040    0.00000    0.00000 ;      HG-    CG-   CD2-  HD22
  4020   4019   4025   4028      3    0.62760    1.88280    0.00000
-2.51040    0.00000    0.00000 ;      HG-    CG-   CD2-  HD23
  4021   4019   4025   4026      3    0.66944    2.00832    0.00000
```

```
-2.67776    0.00000    0.00000 ;    CD1-    CG-   CD2-  HD21
  4021   4019   4025   4027      3    0.66944    2.00832    0.00000
-2.67776    0.00000    0.00000 ;    CD1-    CG-   CD2-  HD22
  4021   4019   4025   4028      3    0.66944    2.00832    0.00000
-2.67776    0.00000    0.00000 ;    CD1-    CG-   CD2-  HD23
  4022   4021   4019   4025      3    0.66944    2.00832    0.00000
-2.67776    0.00000    0.00000 ;   HD11-   CD1-    CG-   CD2
  4023   4021   4019   4025      3    0.66944    2.00832    0.00000
-2.67776    0.00000    0.00000 ;   HD12-   CD1-    CG-   CD2
  4024   4021   4019   4025      3    0.66944    2.00832    0.00000
-2.67776    0.00000    0.00000 ;   HD13-   CD1-    CG-   CD2
  4029   4031   4033   4034      3    0.00000    0.00000    0.00000
0.00000    0.00000    0.00000 ;      C-     N-    CA-    HA
  4029   4031   4033   4035      3   11.71520    1.67360   15.06240  -
13.38880    0.00000    0.00000 ;      C-     N-    CA-    CB
  4029   4031   4033   4043      3    1.75728    5.27184    2.25936
-7.02912    0.00000    0.00000 ;      C-     N-    CA-     C
  4030   4029   4031   4032      3   29.28800   -8.36800  -20.92000
0.00000    0.00000    0.00000 ;      O-     C-     N-     H
  4030   4029   4031   4033      3   20.92000    0.00000  -20.92000
0.00000    0.00000    0.00000 ;      O-     C-     N-    CA
  4031   4033   4035   4036      3    0.65084    1.95253    0.00000
-2.60338    0.00000    0.00000 ;      N-    CA-    CB-    HB
  4031   4033   4035   4037      3    3.83673    0.44769   -1.77402
-2.47693   -0.03347    0.00000 ;      N-    CA-    CB-   CG2
  4031   4033   4035   4041      3    4.93294    1.13386   -3.12963
-5.27184    3.17984    0.00000 ;      N-    CA-    CB-   OG1
  4031   4033   4043   4044      3    0.00000    0.00000    0.00000
0.00000    0.00000    0.00000 ;      N-    CA-     C-     O
  4031   4033   4043   4045      3   17.40544   -5.02080  -13.22144
9.20480    0.00000    0.00000 ;      N-    CA-     C-     N
  4032   4031   4033   4034      3    0.00000    0.00000    0.00000
0.00000    0.00000    0.00000 ;      H-     N-    CA-    HA
  4032   4031   4033   4035      3    0.00000    0.00000    0.00000
0.00000    0.00000    0.00000 ;      H-     N-    CA-    CB
  4032   4031   4033   4043      3    0.00000    0.00000    0.00000
0.00000    0.00000    0.00000 ;      H-     N-    CA-     C
  4033   4035   4037   4038      3    0.66944    2.00832    0.00000
-2.67776    0.00000    0.00000 ;     CA-    CB-   CG2-  HG21
  4033   4035   4037   4039      3    0.66944    2.00832    0.00000
-2.67776    0.00000    0.00000 ;     CA-    CB-   CG2-  HG22
  4033   4035   4037   4040      3    0.66944    2.00832    0.00000
-2.67776    0.00000    0.00000 ;     CA-    CB-   CG2-  HG23
  4033   4035   4041   4042      3    1.12131    2.98738    1.66523
-3.94970    0.43514    0.00000 ;     CA-    CB-   OG1-   HG1
  4033   4043   4045   4046      3   20.92000    0.00000  -20.92000
0.00000    0.00000    0.00000 ;     CA-     C-     N-     H
  4033   4043   4045   4047      3   20.92000    0.00000  -20.92000
0.00000    0.00000    0.00000 ;     CA-     C-     N-    CA
  4034   4033   4035   4036      3    0.65084    1.95253    0.00000
-2.60338    0.00000    0.00000 ;     HA-    CA-    CB-    HB
  4034   4033   4035   4037      3    0.65084    1.95253    0.00000
```

-2.60338 0.00000 0.00000 ; HA- CA- CB- CG2
4034 4033 4035 4041 3 1.04600 -1.04600 0.00000
0.00000 0.00000 0.00000 ; HA- CA- CB- OG1
4034 4033 4043 4044 3 3.68192 -4.35136 0.00000
1.33888 0.00000 0.00000 ; HA- CA- C- O
4034 4033 4043 4045 3 0.00000 0.00000 0.00000
0.00000 0.00000 0.00000 ; HA- CA- C- N
4035 4033 4043 4044 3 0.00000 0.00000 0.00000
0.00000 0.00000 0.00000 ; CB- CA- C- O
4035 4033 4043 4045 3 2.51040 4.18400 1.67360
-6.69440 0.00000 0.00000 ; CB- CA- C- N
4036 4035 4033 4043 3 0.65084 1.95253 0.00000
-2.60338 0.00000 0.00000 ; HB- CB- CA- C
4036 4035 4037 4038 3 0.65084 1.95253 0.00000
-2.60338 0.00000 0.00000 ; HB- CB- CG2- HG21
4036 4035 4037 4039 3 0.65084 1.95253 0.00000
-2.60338 0.00000 0.00000 ; HB- CB- CG2- HG22
4036 4035 4037 4040 3 0.65084 1.95253 0.00000
-2.60338 0.00000 0.00000 ; HB- CB- CG2- HG23
4036 4035 4041 4042 3 0.69733 2.09200 0.00000
-2.78933 0.00000 0.00000 ; HB- CB- OG1- HG1
4037 4035 4033 4043 3 5.67350 3.55640 -6.16722
-2.47693 3.74886 0.00000 ; CG2- CB- CA- C
4037 4035 4041 4042 3 4.74047 0.27196 -2.26773
-3.94970 1.60666 0.00000 ; CG2- CB- OG1- HG1
4038 4037 4035 4041 3 1.04600 -1.04600 0.00000
0.00000 0.00000 0.00000 ; HG21- CG2- CB- OG1
4039 4037 4035 4041 3 1.04600 -1.04600 0.00000
0.00000 0.00000 0.00000 ; HG22- CG2- CB- OG1
4040 4037 4035 4041 3 1.04600 -1.04600 0.00000
0.00000 0.00000 0.00000 ; HG23- CG2- CB- OG1
4041 4035 4033 4043 3 6.53541 6.87013 -6.21742
-5.27184 5.22163 0.00000 ; OG1- CB- CA- C
4043 4045 4047 4048 3 0.00000 0.00000 0.00000
0.00000 0.00000 0.00000 ; C- N- CA- HA
4043 4045 4047 4049 3 11.71520 1.67360 15.06240 -
13.38880 0.00000 0.00000 ; C- N- CA- CB
4043 4045 4047 4064 3 1.75728 5.27184 2.25936
-7.02912 0.00000 0.00000 ; C- N- CA- C
4044 4043 4045 4046 3 29.28800 -8.36800 -20.92000
0.00000 0.00000 0.00000 ; O- C- N- H
4044 4043 4045 4047 3 20.92000 0.00000 -20.92000
0.00000 0.00000 0.00000 ; O- C- N- CA
4045 4047 4049 4050 3 0.65084 1.95253 0.00000
-2.60338 0.00000 0.00000 ; N- CA- CB- HB2
4045 4047 4049 4051 3 0.65084 1.95253 0.00000
-2.60338 0.00000 0.00000 ; N- CA- CB- HB3
4045 4047 4049 4052 3 3.28026 2.46019 -2.19242
-3.21331 -0.23430 0.00000 ; N- CA- CB- CG
4045 4047 4064 4065 3 0.00000 0.00000 0.00000
0.00000 0.00000 0.00000 ; N- CA- C- O
4045 4047 4064 4066 3 17.40544 -5.02080 -13.22144

```
9.20480    0.00000    0.00000 ;       N-    CA-     C-     N
  4046   4045   4047   4048      3    0.00000    0.00000    0.00000
0.00000    0.00000    0.00000 ;       H-     N-    CA-    HA
  4046   4045   4047   4049      3    0.00000    0.00000    0.00000
0.00000    0.00000    0.00000 ;       H-     N-    CA-    CB
  4046   4045   4047   4064      3    0.00000    0.00000    0.00000
0.00000    0.00000    0.00000 ;       H-     N-    CA-     C
  4047   4049   4052   4053      3    0.57739    0.00000    1.02926
0.00000   -1.60666    0.00000 ;      CA-    CB-    CG-   CD1
  4047   4049   4052   4062      3    0.57739    0.00000    1.02926
0.00000   -1.60666    0.00000 ;      CA-    CB-    CG-   CD2
  4047   4064   4066   4067      3   20.92000    0.00000  -20.92000
0.00000    0.00000    0.00000 ;      CA-     C-     N-     H
  4047   4064   4066   4068      3   20.92000    0.00000  -20.92000
0.00000    0.00000    0.00000 ;      CA-     C-     N-    CA
  4048   4047   4049   4050      3    0.65084    1.95253    0.00000
-2.60338    0.00000    0.00000 ;      HA-    CA-    CB-   HB2
  4048   4047   4049   4051      3    0.65084    1.95253    0.00000
-2.60338    0.00000    0.00000 ;      HA-    CA-    CB-   HB3
  4048   4047   4049   4052      3    0.65084    1.95253    0.00000
-2.60338    0.00000    0.00000 ;      HA-    CA-    CB-    CG
  4048   4047   4064   4065      3    3.68192   -4.35136    0.00000
1.33888    0.00000    0.00000 ;      HA-    CA-     C-     O
  4048   4047   4064   4066      3    0.00000    0.00000    0.00000
0.00000    0.00000    0.00000 ;      HA-    CA-     C-     N
  4049   4047   4064   4065      3    0.00000    0.00000    0.00000
0.00000    0.00000    0.00000 ;      CB-    CA-     C-     O
  4049   4047   4064   4066      3    2.51040    4.18400    1.67360
-6.69440    0.00000    0.00000 ;      CB-    CA-     C-     N
  4049   4052   4053   4054      3   30.33400    0.00000  -30.33400
0.00000    0.00000    0.00000 ;      CB-    CG-   CD1-   HD1
  4049   4052   4053   4055      3   30.33400    0.00000  -30.33400
0.00000    0.00000    0.00000 ;      CB-    CG-   CD1-   CE1
  4049   4052   4062   4060      3   30.33400    0.00000  -30.33400
0.00000    0.00000    0.00000 ;      CB-    CG-   CD2-   CE2
  4049   4052   4062   4063      3   30.33400    0.00000  -30.33400
0.00000    0.00000    0.00000 ;      CB-    CG-   CD2-   HD2
  4050   4049   4047   4064      3    0.65084    1.95253    0.00000
-2.60338    0.00000    0.00000 ;     HB2-    CB-    CA-     C
  4050   4049   4052   4053      3    0.00000    0.00000    0.00000
0.00000    0.00000    0.00000 ;     HB2-    CB-    CG-   CD1
  4050   4049   4052   4062      3    0.00000    0.00000    0.00000
0.00000    0.00000    0.00000 ;     HB2-    CB-    CG-   CD2
  4051   4049   4047   4064      3    0.65084    1.95253    0.00000
-2.60338    0.00000    0.00000 ;     HB3-    CB-    CA-     C
  4051   4049   4052   4053      3    0.00000    0.00000    0.00000
0.00000    0.00000    0.00000 ;     HB3-    CB-    CG-   CD1
  4051   4049   4052   4062      3    0.00000    0.00000    0.00000
0.00000    0.00000    0.00000 ;     HB3-    CB-    CG-   CD2
  4052   4049   4047   4064      3    4.95804    2.17986   -3.52293
-3.21331   -0.40166    0.00000 ;      CG-    CB-    CA-     C
  4052   4053   4055   4056      3   30.33400    0.00000  -30.33400
```

0.00000 0.00000 0.00000 ; CG- CD1- CE1- HE1
4052 4053 4055 4057 3 30.33400 0.00000 -30.33400
0.00000 0.00000 0.00000 ; CG- CD1- CE1- CZ
4052 4062 4060 4057 3 30.33400 0.00000 -30.33400
0.00000 0.00000 0.00000 ; CG- CD2- CE2- CZ
4052 4062 4060 4061 3 30.33400 0.00000 -30.33400
0.00000 0.00000 0.00000 ; CG- CD2- CE2- HE2
4053 4052 4062 4060 3 30.33400 0.00000 -30.33400
0.00000 0.00000 0.00000 ; CD1- CG- CD2- CE2
4053 4052 4062 4063 3 30.33400 0.00000 -30.33400
0.00000 0.00000 0.00000 ; CD1- CG- CD2- HD2
4053 4055 4057 4058 3 30.33400 0.00000 -30.33400
0.00000 0.00000 0.00000 ; CD1- CE1- CZ- OH
4053 4055 4057 4060 3 30.33400 0.00000 -30.33400
0.00000 0.00000 0.00000 ; CD1- CE1- CZ- CE2
4054 4053 4052 4062 3 30.33400 0.00000 -30.33400
0.00000 0.00000 0.00000 ; HD1- CD1- CG- CD2
4054 4053 4055 4056 3 30.33400 0.00000 -30.33400
0.00000 0.00000 0.00000 ; HD1- CD1- CE1- HE1
4054 4053 4055 4057 3 30.33400 0.00000 -30.33400
0.00000 0.00000 0.00000 ; HD1- CD1- CE1- CZ
4055 4053 4052 4062 3 30.33400 0.00000 -30.33400
0.00000 0.00000 0.00000 ; CE1- CD1- CG- CD2
4055 4057 4058 4059 3 7.93286 0.00000 -9.56462
0.00000 2.17568 0.00000 ; CE1- CZ- OH- HH
4055 4057 4060 4061 3 30.33400 0.00000 -30.33400
0.00000 0.00000 0.00000 ; CE1- CZ- CE2- HE2
4055 4057 4060 4062 3 30.33400 0.00000 -30.33400
0.00000 0.00000 0.00000 ; CE1- CZ- CE2- CD2
4056 4055 4057 4058 3 30.33400 0.00000 -30.33400
0.00000 0.00000 0.00000 ; HE1- CE1- CZ- OH
4056 4055 4057 4060 3 30.33400 0.00000 -30.33400
0.00000 0.00000 0.00000 ; HE1- CE1- CZ- CE2
4057 4060 4062 4063 3 30.33400 0.00000 -30.33400
0.00000 0.00000 0.00000 ; CZ- CE2- CD2- HD2
4058 4057 4060 4061 3 30.33400 0.00000 -30.33400
0.00000 0.00000 0.00000 ; OH- CZ- CE2- HE2
4058 4057 4060 4062 3 30.33400 0.00000 -30.33400
0.00000 0.00000 0.00000 ; OH- CZ- CE2- CD2
4059 4058 4057 4060 3 7.93286 0.00000 -9.56462
0.00000 2.17568 0.00000 ; HH- OH- CZ- CE2
4061 4060 4062 4063 3 30.33400 0.00000 -30.33400
0.00000 0.00000 0.00000 ; HE2- CE2- CD2- HD2
4064 4066 4068 4069 3 0.00000 0.00000 0.00000
0.00000 0.00000 0.00000 ; C- N- CA- HA
4064 4066 4068 4070 3 11.71520 1.67360 15.06240 -
13.38880 0.00000 0.00000 ; C- N- CA- CB
4064 4066 4068 4078 3 1.75728 5.27184 2.25936
-7.02912 0.00000 0.00000 ; C- N- CA- C
4065 4064 4066 4067 3 29.28800 -8.36800 -20.92000
0.00000 0.00000 0.00000 ; O- C- N- H
4065 4064 4066 4068 3 20.92000 0.00000 -20.92000

0.00000 0.00000 0.00000 ; O- C- N- CA
4066 4068 4070 4071 3 0.65084 1.95253 0.00000
-2.60338 0.00000 0.00000 ; N- CA- CB- HB2
4066 4068 4070 4072 3 0.65084 1.95253 0.00000
-2.60338 0.00000 0.00000 ; N- CA- CB- HB3
4066 4068 4070 4073 3 5.99567 3.29281 -4.46014
-0.55229 1.97485 0.00000 ; N- CA- CB- CG
4066 4068 4078 4079 3 0.00000 0.00000 0.00000
0.00000 0.00000 0.00000 ; N- CA- C- O
4066 4068 4078 4080 3 17.40544 -5.02080 -13.22144
9.20480 0.00000 0.00000 ; N- CA- C- N
4067 4066 4068 4069 3 0.00000 0.00000 0.00000
0.00000 0.00000 0.00000 ; H- N- CA- HA
4067 4066 4068 4070 3 0.00000 0.00000 0.00000
0.00000 0.00000 0.00000 ; H- N- CA- CB
4067 4066 4068 4078 3 0.00000 0.00000 0.00000
0.00000 0.00000 0.00000 ; H- N- CA- C
4068 4070 4073 4074 3 0.00000 0.00000 0.00000
0.00000 0.00000 0.00000 ; CA- CB- CG- OD1
4068 4070 4073 4075 3 8.84916 -0.31380 -4.32626
5.03754 0.26778 0.00000 ; CA- CB- CG- ND2
4068 4078 4080 4081 3 20.92000 0.00000 -20.92000
0.00000 0.00000 0.00000 ; CA- C- N- CD
4068 4078 4080 4090 3 20.92000 0.00000 -20.92000
0.00000 0.00000 0.00000 ; CA- C- N- CA
4069 4068 4070 4071 3 0.65084 1.95253 0.00000
-2.60338 0.00000 0.00000 ; HA- CA- CB- HB2
4069 4068 4070 4072 3 0.65084 1.95253 0.00000
-2.60338 0.00000 0.00000 ; HA- CA- CB- HB3
4069 4068 4070 4073 3 0.65084 1.95253 0.00000
-2.60338 0.00000 0.00000 ; HA- CA- CB- CG
4069 4068 4078 4079 3 3.68192 -4.35136 0.00000
1.33888 0.00000 0.00000 ; HA- CA- C- O
4069 4068 4078 4080 3 0.00000 0.00000 0.00000
0.00000 0.00000 0.00000 ; HA- CA- C- N
4070 4068 4078 4079 3 0.00000 0.00000 0.00000
0.00000 0.00000 0.00000 ; CB- CA- C- O
4070 4068 4078 4080 3 2.51040 4.18400 1.67360
-6.69440 0.00000 0.00000 ; CB- CA- C- N
4070 4073 4075 4076 3 20.92000 0.00000 -20.92000
0.00000 0.00000 0.00000 ; CB- CG- ND2- HD21
4070 4073 4075 4077 3 20.92000 0.00000 -20.92000
0.00000 0.00000 0.00000 ; CB- CG- ND2- HD22
4071 4070 4068 4078 3 0.65084 1.95253 0.00000
-2.60338 0.00000 0.00000 ; HB2- CB- CA- C
4071 4070 4073 4074 3 3.68192 -4.35136 0.00000
1.33888 0.00000 0.00000 ; HB2- CB- CG- OD1
4071 4070 4073 4075 3 0.00000 0.00000 0.00000
0.00000 0.00000 0.00000 ; HB2- CB- CG- ND2
4072 4070 4068 4078 3 0.65084 1.95253 0.00000
-2.60338 0.00000 0.00000 ; HB3- CB- CA- C
4072 4070 4073 4074 3 3.68192 -4.35136 0.00000

```
1.33888    0.00000    0.00000 ;    HB3-    CB-    CG-   OD1
  4072   4070   4073   4075      3    0.00000    0.00000    0.00000
0.00000    0.00000    0.00000 ;    HB3-    CB-    CG-   ND2
  4073   4070   4068   4078      3    7.94542   -3.96225   -6.11701
-0.55229    3.58150    0.00000 ;     CG-    CB-    CA-     C
  4074   4073   4075   4076      3   29.28800   -8.36800  -20.92000
0.00000    0.00000    0.00000 ;    OD1-    CG-   ND2-  HD21
  4074   4073   4075   4077      3   29.28800   -8.36800  -20.92000
0.00000    0.00000    0.00000 ;    OD1-    CG-   ND2-  HD22
  4078   4080   4081   4082      3    0.00000    0.00000    0.00000
0.00000    0.00000    0.00000 ;     C-     N-    CD-   HD2
  4078   4080   4081   4083      3    0.00000    0.00000    0.00000
0.00000    0.00000    0.00000 ;     C-     N-    CD-   HD3
  4078   4080   4081   4084      3   10.04160   -3.34720   16.73600
-6.69440    0.00000    0.00000 ;     C-     N-    CD-    CG
  4078   4080   4090   4087      3   11.71520    1.67360   15.06240  -
13.38880    0.00000    0.00000 ;     C-     N-    CA-    CB
  4078   4080   4090   4091      3    0.00000    0.00000    0.00000
0.00000    0.00000    0.00000 ;     C-     N-    CA-    HA
  4078   4080   4090   4092      3    1.75728    5.27184    2.25936
-7.02912    0.00000    0.00000 ;     C-     N-    CA-     C
  4079   4078   4080   4081      3   20.92000    0.00000  -20.92000
0.00000    0.00000    0.00000 ;     O-     C-     N-    CD
  4079   4078   4080   4090      3   20.92000    0.00000  -20.92000
0.00000    0.00000    0.00000 ;     O-     C-     N-    CA
  4080   4081   4084   4085      3    0.65084    1.95253    0.00000
-2.60338    0.00000    0.00000 ;     N-    CD-    CG-   HG2
  4080   4081   4084   4086      3    0.65084    1.95253    0.00000
-2.60338    0.00000    0.00000 ;     N-    CD-    CG-   HG3
  4080   4081   4084   4087      3    0.65084    1.95253    0.00000
-2.60338    0.00000    0.00000 ;     N-    CD-    CG-    CB
  4080   4090   4087   4084      3    0.65084    1.95253    0.00000
-2.60338    0.00000    0.00000 ;     N-    CA-    CB-    CG
  4080   4090   4087   4088      3    0.65084    1.95253    0.00000
-2.60338    0.00000    0.00000 ;     N-    CA-    CB-   HB2
  4080   4090   4087   4089      3    0.65084    1.95253    0.00000
-2.60338    0.00000    0.00000 ;     N-    CA-    CB-   HB3
  4080   4090   4092   4093      3    0.00000    0.00000    0.00000
0.00000    0.00000    0.00000 ;     N-    CA-     C-     O
  4080   4090   4092   4094      3   17.40544   -5.02080  -13.22144
9.20480    0.00000    0.00000 ;     N-    CA-     C-     N
  4081   4080   4090   4087      3    0.00000    0.00000    0.00000
0.00000    0.00000    0.00000 ;    CD-     N-    CA-    CB
  4081   4080   4090   4091      3    0.00000    0.00000    0.00000
0.00000    0.00000    0.00000 ;    CD-     N-    CA-    HA
  4081   4080   4090   4092      3    0.00000    0.00000    0.00000
0.00000    0.00000    0.00000 ;    CD-     N-    CA-     C
  4081   4084   4087   4088      3    0.66944    2.00832    0.00000
-2.67776    0.00000    0.00000 ;    CD-    CG-    CB-   HB2
  4081   4084   4087   4089      3    0.66944    2.00832    0.00000
-2.67776    0.00000    0.00000 ;    CD-    CG-    CB-   HB3
  4081   4084   4087   4090      3    3.68192    3.09616   -2.09200
```

```
-3.01248    0.00000    0.00000 ;     CD-    CG-    CB-    CA
  4082   4081   4080   4090      3    0.00000    0.00000    0.00000
0.00000    0.00000    0.00000 ;    HD2-    CD-     N-    CA
  4082   4081   4084   4085      3    0.65084    1.95253    0.00000
-2.60338    0.00000    0.00000 ;    HD2-    CD-    CG-   HG2
  4082   4081   4084   4086      3    0.65084    1.95253    0.00000
-2.60338    0.00000    0.00000 ;    HD2-    CD-    CG-   HG3
  4082   4081   4084   4087      3    0.65084    1.95253    0.00000
-2.60338    0.00000    0.00000 ;    HD2-    CD-    CG-    CB
  4083   4081   4080   4090      3    0.00000    0.00000    0.00000
0.00000    0.00000    0.00000 ;    HD3-    CD-     N-    CA
  4083   4081   4084   4085      3    0.65084    1.95253    0.00000
-2.60338    0.00000    0.00000 ;    HD3-    CD-    CG-   HG2
  4083   4081   4084   4086      3    0.65084    1.95253    0.00000
-2.60338    0.00000    0.00000 ;    HD3-    CD-    CG-   HG3
  4083   4081   4084   4087      3    0.65084    1.95253    0.00000
-2.60338    0.00000    0.00000 ;    HD3-    CD-    CG-    CB
  4084   4081   4080   4090      3    0.00000    0.00000    0.00000
0.00000    0.00000    0.00000 ;     CG-    CD-     N-    CA
  4084   4087   4090   4091      3    0.65084    1.95253    0.00000
-2.60338    0.00000    0.00000 ;     CG-    CB-    CA-    HA
  4084   4087   4090   4092      3    0.65084    1.95253    0.00000
-2.60338    0.00000    0.00000 ;     CG-    CB-    CA-     C
  4085   4084   4087   4088      3    0.62760    1.88280    0.00000
-2.51040    0.00000    0.00000 ;    HG2-    CG-    CB-   HB2
  4085   4084   4087   4089      3    0.62760    1.88280    0.00000
-2.51040    0.00000    0.00000 ;    HG2-    CG-    CB-   HB3
  4085   4084   4087   4090      3    0.66944    2.00832    0.00000
-2.67776    0.00000    0.00000 ;    HG2-    CG-    CB-    CA
  4086   4084   4087   4088      3    0.62760    1.88280    0.00000
-2.51040    0.00000    0.00000 ;    HG3-    CG-    CB-   HB2
  4086   4084   4087   4089      3    0.62760    1.88280    0.00000
-2.51040    0.00000    0.00000 ;    HG3-    CG-    CB-   HB3
  4086   4084   4087   4090      3    0.66944    2.00832    0.00000
-2.67776    0.00000    0.00000 ;    HG3-    CG-    CB-    CA
  4087   4090   4092   4093      3    0.00000    0.00000    0.00000
0.00000    0.00000    0.00000 ;     CB-    CA-     C-     O
  4087   4090   4092   4094      3    2.51040    4.18400    1.67360
-6.69440    0.00000    0.00000 ;     CB-    CA-     C-     N
  4088   4087   4090   4091      3    0.65084    1.95253    0.00000
-2.60338    0.00000    0.00000 ;    HB2-    CB-    CA-    HA
  4088   4087   4090   4092      3    0.65084    1.95253    0.00000
-2.60338    0.00000    0.00000 ;    HB2-    CB-    CA-     C
  4089   4087   4090   4091      3    0.65084    1.95253    0.00000
-2.60338    0.00000    0.00000 ;    HB3-    CB-    CA-    HA
  4089   4087   4090   4092      3    0.65084    1.95253    0.00000
-2.60338    0.00000    0.00000 ;    HB3-    CB-    CA-     C
  4090   4092   4094   4095      3   20.92000    0.00000  -20.92000
0.00000    0.00000    0.00000 ;     CA-     C-     N-     H
  4090   4092   4094   4096      3   20.92000    0.00000  -20.92000
0.00000    0.00000    0.00000 ;     CA-     C-     N-    CA
  4091   4090   4092   4093      3    3.68192   -4.35136    0.00000
```

1.33888    0.00000    0.00000 ;    HA-    CA-     C-     O
  4091   4090   4092   4094      3    0.00000    0.00000    0.00000
0.00000    0.00000    0.00000 ;    HA-    CA-     C-     N
  4092   4094   4096   4097      3    0.00000    0.00000    0.00000
0.00000    0.00000    0.00000 ;     C-     N-    CA-    HA
  4092   4094   4096   4098      3   11.71520    1.67360   15.06240  -
13.38880    0.00000    0.00000 ;     C-     N-    CA-    CB
  4092   4094   4096   4106      3    1.75728    5.27184    2.25936
-7.02912    0.00000    0.00000 ;     C-     N-    CA-     C
  4093   4092   4094   4095      3   29.28800   -8.36800  -20.92000
0.00000    0.00000    0.00000 ;     O-     C-     N-     H
  4093   4092   4094   4096      3   20.92000    0.00000  -20.92000
0.00000    0.00000    0.00000 ;     O-     C-     N-    CA
  4094   4096   4098   4099      3    0.65084    1.95253    0.00000
-2.60338    0.00000    0.00000 ;     N-    CA-    CB-   HB2
  4094   4096   4098   4100      3    0.65084    1.95253    0.00000
-2.60338    0.00000    0.00000 ;     N-    CA-    CB-   HB3
  4094   4096   4098   4101      3    5.99567    3.29281   -4.46014
-0.55229    1.97485    0.00000 ;     N-    CA-    CB-    CG
  4094   4096   4106   4107      3    0.00000    0.00000    0.00000
0.00000    0.00000    0.00000 ;     N-    CA-     C-     O
  4094   4096   4106   4108      3   17.40544   -5.02080  -13.22144
9.20480    0.00000    0.00000 ;     N-    CA-     C-     N
  4095   4094   4096   4097      3    0.00000    0.00000    0.00000
0.00000    0.00000    0.00000 ;     H-     N-    CA-    HA
  4095   4094   4096   4098      3    0.00000    0.00000    0.00000
0.00000    0.00000    0.00000 ;     H-     N-    CA-    CB
  4095   4094   4096   4106      3    0.00000    0.00000    0.00000
0.00000    0.00000    0.00000 ;     H-     N-    CA-     C
  4096   4098   4101   4102      3    0.00000    0.00000    0.00000
0.00000    0.00000    0.00000 ;    CA-    CB-    CG-   OD1
  4096   4098   4101   4103      3    8.84916   -0.31380   -4.32626
5.03754    0.26778    0.00000 ;    CA-    CB-    CG-   ND2
  4096   4106   4108   4109      3   20.92000    0.00000  -20.92000
0.00000    0.00000    0.00000 ;    CA-     C-     N-     H
  4096   4106   4108   4110      3   20.92000    0.00000  -20.92000
0.00000    0.00000    0.00000 ;    CA-     C-     N-    CA
  4097   4096   4098   4099      3    0.65084    1.95253    0.00000
-2.60338    0.00000    0.00000 ;    HA-    CA-    CB-   HB2
  4097   4096   4098   4100      3    0.65084    1.95253    0.00000
-2.60338    0.00000    0.00000 ;    HA-    CA-    CB-   HB3
  4097   4096   4098   4101      3    0.65084    1.95253    0.00000
-2.60338    0.00000    0.00000 ;    HA-    CA-    CB-    CG
  4097   4096   4106   4107      3    3.68192   -4.35136    0.00000
1.33888    0.00000    0.00000 ;    HA-    CA-     C-     O
  4097   4096   4106   4108      3    0.00000    0.00000    0.00000
0.00000    0.00000    0.00000 ;    HA-    CA-     C-     N
  4098   4096   4106   4107      3    0.00000    0.00000    0.00000
0.00000    0.00000    0.00000 ;    CB-    CA-     C-     O
  4098   4096   4106   4108      3    2.51040    4.18400    1.67360
-6.69440    0.00000    0.00000 ;    CB-    CA-     C-     N
  4098   4101   4103   4104      3   20.92000    0.00000  -20.92000

0.00000 0.00000 0.00000 ; CB- CG- ND2- HD21
4098 4101 4103 4105 3 20.92000 0.00000 -20.92000
0.00000 0.00000 0.00000 ; CB- CG- ND2- HD22
4099 4098 4096 4106 3 0.65084 1.95253 0.00000
-2.60338 0.00000 0.00000 ; HB2- CB- CA- C
4099 4098 4101 4102 3 3.68192 -4.35136 0.00000
1.33888 0.00000 0.00000 ; HB2- CB- CG- OD1
4099 4098 4101 4103 3 0.00000 0.00000 0.00000
0.00000 0.00000 0.00000 ; HB2- CB- CG- ND2
4100 4098 4096 4106 3 0.65084 1.95253 0.00000
-2.60338 0.00000 0.00000 ; HB3- CB- CA- C
4100 4098 4101 4102 3 3.68192 -4.35136 0.00000
1.33888 0.00000 0.00000 ; HB3- CB- CG- OD1
4100 4098 4101 4103 3 0.00000 0.00000 0.00000
0.00000 0.00000 0.00000 ; HB3- CB- CG- ND2
4101 4098 4096 4106 3 7.94542 -3.96225 -6.11701
-0.55229 3.58150 0.00000 ; CG- CB- CA- C
4102 4101 4103 4104 3 29.28800 -8.36800 -20.92000
0.00000 0.00000 0.00000 ; OD1- CG- ND2- HD21
4102 4101 4103 4105 3 29.28800 -8.36800 -20.92000
0.00000 0.00000 0.00000 ; OD1- CG- ND2- HD22
4106 4108 4110 4111 3 0.00000 0.00000 0.00000
0.00000 0.00000 0.00000 ; C- N- CA- HA
4106 4108 4110 4112 3 11.71520 1.67360 15.06240 -
13.38880 0.00000 0.00000 ; C- N- CA- CB
4106 4108 4110 4121 3 1.75728 5.27184 2.25936
-7.02912 0.00000 0.00000 ; C- N- CA- C
4107 4106 4108 4109 3 29.28800 -8.36800 -20.92000
0.00000 0.00000 0.00000 ; O- C- N- H
4107 4106 4108 4110 3 20.92000 0.00000 -20.92000
0.00000 0.00000 0.00000 ; O- C- N- CA
4108 4110 4112 4113 3 0.65084 1.95253 0.00000
-2.60338 0.00000 0.00000 ; N- CA- CB- HB2
4108 4110 4112 4114 3 0.65084 1.95253 0.00000
-2.60338 0.00000 0.00000 ; N- CA- CB- HB3
4108 4110 4112 4115 3 3.21331 2.22589 -4.15053
-2.40998 2.61082 0.00000 ; N- CA- CB- CG
4108 4110 4121 4122 3 0.00000 0.00000 0.00000
0.00000 0.00000 0.00000 ; N- CA- C- O
4108 4110 4121 4123 3 17.40544 -5.02080 -13.22144
9.20480 0.00000 0.00000 ; N- CA- C- N
4109 4108 4110 4111 3 0.00000 0.00000 0.00000
0.00000 0.00000 0.00000 ; H- N- CA- HA
4109 4108 4110 4112 3 0.00000 0.00000 0.00000
0.00000 0.00000 0.00000 ; H- N- CA- CB
4109 4108 4110 4121 3 0.00000 0.00000 0.00000
0.00000 0.00000 0.00000 ; H- N- CA- C
4110 4112 4115 4116 3 0.66944 2.00832 0.00000
-2.67776 0.00000 0.00000 ; CA- CB- CG- HG2
4110 4112 4115 4117 3 0.66944 2.00832 0.00000
-2.67776 0.00000 0.00000 ; CA- CB- CG- HG3
4110 4112 4115 4118 3 10.12110 -1.91209 0.01674

10.17549 -1.87443 0.00000 ; CA- CB- CG- CD
4110 4121 4123 4124 3 20.92000 0.00000 -20.92000
0.00000 0.00000 0.00000 ; CA- C- N- H
4110 4121 4123 4125 3 20.92000 0.00000 -20.92000
0.00000 0.00000 0.00000 ; CA- C- N- CA
4111 4110 4112 4113 3 0.65084 1.95253 0.00000
-2.60338 0.00000 0.00000 ; HA- CA- CB- HB2
4111 4110 4112 4114 3 0.65084 1.95253 0.00000
-2.60338 0.00000 0.00000 ; HA- CA- CB- HB3
4111 4110 4112 4115 3 0.65084 1.95253 0.00000
-2.60338 0.00000 0.00000 ; HA- CA- CB- CG
4111 4110 4121 4122 3 3.68192 -4.35136 0.00000
1.33888 0.00000 0.00000 ; HA- CA- C- O
4111 4110 4121 4123 3 0.00000 0.00000 0.00000
0.00000 0.00000 0.00000 ; HA- CA- C- N
4112 4110 4121 4122 3 0.00000 0.00000 0.00000
0.00000 0.00000 0.00000 ; CB- CA- C- O
4112 4110 4121 4123 3 2.51040 4.18400 1.67360
-6.69440 0.00000 0.00000 ; CB- CA- C- N
4112 4115 4118 4119 3 3.79907 0.00000 -5.40573
0.00000 2.14221 0.00000 ; CB- CG- CD- OE1
4112 4115 4118 4120 3 3.79907 0.00000 -5.40573
0.00000 2.14221 0.00000 ; CB- CG- CD- OE2
4113 4112 4110 4121 3 0.65084 1.95253 0.00000
-2.60338 0.00000 0.00000 ; HB2- CB- CA- C
4113 4112 4115 4116 3 0.62760 1.88280 0.00000
-2.51040 0.00000 0.00000 ; HB2- CB- CG- HG2
4113 4112 4115 4117 3 0.62760 1.88280 0.00000
-2.51040 0.00000 0.00000 ; HB2- CB- CG- HG3
4113 4112 4115 4118 3 0.65084 1.95253 0.00000
-2.60338 0.00000 0.00000 ; HB2- CB- CG- CD
4114 4112 4110 4121 3 0.65084 1.95253 0.00000
-2.60338 0.00000 0.00000 ; HB3- CB- CA- C
4114 4112 4115 4116 3 0.62760 1.88280 0.00000
-2.51040 0.00000 0.00000 ; HB3- CB- CG- HG2
4114 4112 4115 4117 3 0.62760 1.88280 0.00000
-2.51040 0.00000 0.00000 ; HB3- CB- CG- HG3
4114 4112 4115 4118 3 0.65084 1.95253 0.00000
-2.60338 0.00000 0.00000 ; HB3- CB- CG- CD
4115 4112 4110 4121 3 6.86594 3.56895 -8.14206
-2.40998 4.85344 0.00000 ; CG- CB- CA- C
4116 4115 4118 4119 3 0.00000 0.00000 0.00000
0.00000 0.00000 0.00000 ; HG2- CG- CD- OE1
4116 4115 4118 4120 3 0.00000 0.00000 0.00000
0.00000 0.00000 0.00000 ; HG2- CG- CD- OE2
4117 4115 4118 4119 3 0.00000 0.00000 0.00000
0.00000 0.00000 0.00000 ; HG3- CG- CD- OE1
4117 4115 4118 4120 3 0.00000 0.00000 0.00000
0.00000 0.00000 0.00000 ; HG3- CG- CD- OE2
4121 4123 4125 4126 3 0.00000 0.00000 0.00000
0.00000 0.00000 0.00000 ; C- N- CA- HA
4121 4123 4125 4127 3 11.71520 1.67360 15.06240 -

```
13.38880    0.00000    0.00000 ;       C-     N-    CA-    CB
  4121   4123   4125   4145      3    1.75728    5.27184    2.25936
-7.02912    0.00000    0.00000 ;       C-     N-    CA-     C
  4122   4121   4123   4124      3   29.28800   -8.36800  -20.92000
0.00000    0.00000    0.00000 ;      O-     C-     N-     H
  4122   4121   4123   4125      3   20.92000    0.00000  -20.92000
0.00000    0.00000    0.00000 ;      O-     C-     N-    CA
  4123   4125   4127   4128      3    0.65084    1.95253    0.00000
-2.60338    0.00000    0.00000 ;       N-    CA-    CB-   HB2
  4123   4125   4127   4129      3    0.65084    1.95253    0.00000
-2.60338    0.00000    0.00000 ;       N-    CA-    CB-   HB3
  4123   4125   4127   4130      3    0.65084    1.95253    0.00000
-2.60338    0.00000    0.00000 ;       N-    CA-    CB-    CG
  4123   4125   4145   4146      3    0.00000    0.00000    0.00000
0.00000    0.00000    0.00000 ;      N-    CA-     C-     O
  4123   4125   4145   4147      3   17.40544   -5.02080  -13.22144
9.20480    0.00000    0.00000 ;      N-    CA-     C-     N
  4124   4123   4125   4126      3    0.00000    0.00000    0.00000
0.00000    0.00000    0.00000 ;      H-     N-    CA-    HA
  4124   4123   4125   4127      3    0.00000    0.00000    0.00000
0.00000    0.00000    0.00000 ;      H-     N-    CA-    CB
  4124   4123   4125   4145      3    0.00000    0.00000    0.00000
0.00000    0.00000    0.00000 ;      H-     N-    CA-     C
  4125   4127   4130   4131      3    0.66944    2.00832    0.00000
-2.67776    0.00000    0.00000 ;      CA-    CB-    CG-   HG2
  4125   4127   4130   4132      3    0.66944    2.00832    0.00000
-2.67776    0.00000    0.00000 ;      CA-    CB-    CG-   HG3
  4125   4127   4130   4133      3    3.68192    3.09616   -2.09200
-3.01248    0.00000    0.00000 ;      CA-    CB-    CG-    CD
  4125   4145   4147   4148      3   20.92000    0.00000  -20.92000
0.00000    0.00000    0.00000 ;     CA-     C-     N-     H
  4125   4145   4147   4149      3   20.92000    0.00000  -20.92000
0.00000    0.00000    0.00000 ;     CA-     C-     N-    CA
  4126   4125   4127   4128      3    0.65084    1.95253    0.00000
-2.60338    0.00000    0.00000 ;      HA-    CA-    CB-   HB2
  4126   4125   4127   4129      3    0.65084    1.95253    0.00000
-2.60338    0.00000    0.00000 ;      HA-    CA-    CB-   HB3
  4126   4125   4127   4130      3    0.65084    1.95253    0.00000
-2.60338    0.00000    0.00000 ;      HA-    CA-    CB-    CG
  4126   4125   4145   4146      3    3.68192   -4.35136    0.00000
1.33888    0.00000    0.00000 ;     HA-    CA-     C-     O
  4126   4125   4145   4147      3    0.00000    0.00000    0.00000
0.00000    0.00000    0.00000 ;     HA-    CA-     C-     N
  4127   4125   4145   4146      3    0.00000    0.00000    0.00000
0.00000    0.00000    0.00000 ;     CB-    CA-     C-     O
  4127   4125   4145   4147      3    2.51040    4.18400    1.67360
-6.69440    0.00000    0.00000 ;      CB-    CA-     C-     N
  4127   4130   4133   4134      3    0.65084    1.95253    0.00000
-2.60338    0.00000    0.00000 ;      CB-    CG-    CD-   HD2
  4127   4130   4133   4135      3    0.65084    1.95253    0.00000
-2.60338    0.00000    0.00000 ;      CB-    CG-    CD-   HD3
  4127   4130   4133   4136      3    0.65084    1.95253    0.00000
```

```
-2.60338    0.00000    0.00000 ;     CB-    CG-    CD-    NE
  4128   4127   4125   4145      3    0.65084    1.95253    0.00000
-2.60338    0.00000    0.00000 ;    HB2-    CB-    CA-     C
  4128   4127   4130   4131      3    0.62760    1.88280    0.00000
-2.51040    0.00000    0.00000 ;    HB2-    CB-    CG-   HG2
  4128   4127   4130   4132      3    0.62760    1.88280    0.00000
-2.51040    0.00000    0.00000 ;    HB2-    CB-    CG-   HG3
  4128   4127   4130   4133      3    0.66944    2.00832    0.00000
-2.67776    0.00000    0.00000 ;    HB2-    CB-    CG-    CD
  4129   4127   4125   4145      3    0.65084    1.95253    0.00000
-2.60338    0.00000    0.00000 ;    HB3-    CB-    CA-     C
  4129   4127   4130   4131      3    0.62760    1.88280    0.00000
-2.51040    0.00000    0.00000 ;    HB3-    CB-    CG-   HG2
  4129   4127   4130   4132      3    0.62760    1.88280    0.00000
-2.51040    0.00000    0.00000 ;    HB3-    CB-    CG-   HG3
  4129   4127   4130   4133      3    0.66944    2.00832    0.00000
-2.67776    0.00000    0.00000 ;    HB3-    CB-    CG-    CD
  4130   4127   4125   4145      3    0.65084    1.95253    0.00000
-2.60338    0.00000    0.00000 ;     CG-    CB-    CA-     C
  4130   4133   4136   4137      3    0.00000    0.00000    0.00000
0.00000    0.00000    0.00000 ;     CG-    CD-    NE-    HE
  4130   4133   4136   4138      3    0.00000    0.00000    0.00000
0.00000    0.00000    0.00000 ;     CG-    CD-    NE-    CZ
  4131   4130   4133   4134      3    0.65084    1.95253    0.00000
-2.60338    0.00000    0.00000 ;    HG2-    CG-    CD-   HD2
  4131   4130   4133   4135      3    0.65084    1.95253    0.00000
-2.60338    0.00000    0.00000 ;    HG2-    CG-    CD-   HD3
  4131   4130   4133   4136      3    0.65084    1.95253    0.00000
-2.60338    0.00000    0.00000 ;    HG2-    CG-    CD-    NE
  4132   4130   4133   4134      3    0.65084    1.95253    0.00000
-2.60338    0.00000    0.00000 ;    HG3-    CG-    CD-   HD2
  4132   4130   4133   4135      3    0.65084    1.95253    0.00000
-2.60338    0.00000    0.00000 ;    HG3-    CG-    CD-   HD3
  4132   4130   4133   4136      3    0.65084    1.95253    0.00000
-2.60338    0.00000    0.00000 ;    HG3-    CG-    CD-    NE
  4133   4136   4138   4139      3   20.08320    0.00000  -20.08320
0.00000    0.00000    0.00000 ;     CD-    NE-    CZ-   NH1
  4133   4136   4138   4142      3   20.08320    0.00000  -20.08320
0.00000    0.00000    0.00000 ;     CD-    NE-    CZ-   NH2
  4134   4133   4136   4137      3    0.00000    0.00000    0.00000
0.00000    0.00000    0.00000 ;    HD2-    CD-    NE-    HE
  4134   4133   4136   4138      3    0.00000    0.00000    0.00000
0.00000    0.00000    0.00000 ;    HD2-    CD-    NE-    CZ
  4135   4133   4136   4137      3    0.00000    0.00000    0.00000
0.00000    0.00000    0.00000 ;    HD3-    CD-    NE-    HE
  4135   4133   4136   4138      3    0.00000    0.00000    0.00000
0.00000    0.00000    0.00000 ;    HD3-    CD-    NE-    CZ
  4136   4138   4139   4140      3   20.08320    0.00000  -20.08320
0.00000    0.00000    0.00000 ;     NE-    CZ-   NH1-  HH11
  4136   4138   4139   4141      3   20.08320    0.00000  -20.08320
0.00000    0.00000    0.00000 ;     NE-    CZ-   NH1-  HH12
  4136   4138   4142   4143      3   20.08320    0.00000  -20.08320
```

0.00000    0.00000    0.00000 ;      NE-    CZ-   NH2-  HH21
  4136   4138   4142   4144       3   20.08320    0.00000  -20.08320
0.00000    0.00000    0.00000 ;      NE-    CZ-   NH2-  HH22
  4137   4136   4138   4139       3   20.08320    0.00000  -20.08320
0.00000    0.00000    0.00000 ;      HE-    NE-    CZ-   NH1
  4137   4136   4138   4142       3   20.08320    0.00000  -20.08320
0.00000    0.00000    0.00000 ;      HE-    NE-    CZ-   NH2
  4139   4138   4142   4143       3   20.08320    0.00000  -20.08320
0.00000    0.00000    0.00000 ;     NH1-    CZ-   NH2-  HH21
  4139   4138   4142   4144       3   20.08320    0.00000  -20.08320
0.00000    0.00000    0.00000 ;     NH1-    CZ-   NH2-  HH22
  4140   4139   4138   4142       3   20.08320    0.00000  -20.08320
0.00000    0.00000    0.00000 ;    HH11-   NH1-    CZ-   NH2
  4141   4139   4138   4142       3   20.08320    0.00000  -20.08320
0.00000    0.00000    0.00000 ;    HH12-   NH1-    CZ-   NH2
  4145   4147   4149   4150       3    0.00000    0.00000    0.00000
0.00000    0.00000    0.00000 ;       C-     N-    CA-    HA
  4145   4147   4149   4151       3   11.71520    1.67360   15.06240  -
13.38880    0.00000    0.00000 ;       C-     N-    CA-    CB
  4145   4147   4149   4156       3    1.75728    5.27184    2.25936
-7.02912    0.00000    0.00000 ;       C-     N-    CA-     C
  4146   4145   4147   4148       3   29.28800   -8.36800  -20.92000
0.00000    0.00000    0.00000 ;       O-     C-     N-     H
  4146   4145   4147   4149       3   20.92000    0.00000  -20.92000
0.00000    0.00000    0.00000 ;       O-     C-     N-    CA
  4147   4149   4151   4152       3    0.65084    1.95253    0.00000
-2.60338    0.00000    0.00000 ;       N-    CA-    CB-   HB2
  4147   4149   4151   4153       3    0.65084    1.95253    0.00000
-2.60338    0.00000    0.00000 ;       N-    CA-    CB-   HB3
  4147   4149   4151   4154       3    6.03751    2.50622   -5.17142
-4.20074    1.10458    0.00000 ;       N-    CA-    CB-    SG
  4147   4149   4156   4157       3    0.00000    0.00000    0.00000
0.00000    0.00000    0.00000 ;       N-    CA-     C-     O
  4147   4149   4156   4158       3   17.40544   -5.02080  -13.22144
9.20480    0.00000    0.00000 ;       N-    CA-     C-     N
  4148   4147   4149   4150       3    0.00000    0.00000    0.00000
0.00000    0.00000    0.00000 ;       H-     N-    CA-    HA
  4148   4147   4149   4151       3    0.00000    0.00000    0.00000
0.00000    0.00000    0.00000 ;       H-     N-    CA-    CB
  4148   4147   4149   4156       3    0.00000    0.00000    0.00000
0.00000    0.00000    0.00000 ;       H-     N-    CA-     C
  4149   4151   4154   4155       3    1.69034    2.77818    4.11706
-4.21747    1.00416    0.00000 ;      CA-    CB-    SG-    HG
  4149   4156   4158   4159       3   20.92000    0.00000  -20.92000
0.00000    0.00000    0.00000 ;      CA-     C-     N-     H
  4149   4156   4158   4160       3   20.92000    0.00000  -20.92000
0.00000    0.00000    0.00000 ;      CA-     C-     N-    CA
  4150   4149   4151   4152       3    0.65084    1.95253    0.00000
-2.60338    0.00000    0.00000 ;      HA-    CA-    CB-   HB2
  4150   4149   4151   4153       3    0.65084    1.95253    0.00000
-2.60338    0.00000    0.00000 ;      HA-    CA-    CB-   HB3
  4150   4149   4151   4154       3    0.65084    1.95253    0.00000

-2.60338 0.00000 0.00000 ; HA- CA- CB- SG
4150 4149 4156 4157 3 3.68192 -4.35136 0.00000
1.33888 0.00000 0.00000 ; HA- CA- C- O
4150 4149 4156 4158 3 0.00000 0.00000 0.00000
0.00000 0.00000 0.00000 ; HA- CA- C- N
4151 4149 4156 4157 3 0.00000 0.00000 0.00000
0.00000 0.00000 0.00000 ; CB- CA- C- O
4151 4149 4156 4158 3 2.51040 4.18400 1.67360
-6.69440 0.00000 0.00000 ; CB- CA- C- N
4152 4151 4149 4156 3 0.65084 1.95253 0.00000
-2.60338 0.00000 0.00000 ; HB2- CB- CA- C
4152 4151 4154 4155 3 1.04600 3.13800 0.00000
-4.18400 0.00000 0.00000 ; HB2- CB- SG- HG
4153 4151 4149 4156 3 0.65084 1.95253 0.00000
-2.60338 0.00000 0.00000 ; HB3- CB- CA- C
4153 4151 4154 4155 3 1.04600 3.13800 0.00000
-4.18400 0.00000 0.00000 ; HB3- CB- SG- HG
4154 4151 4149 4156 3 5.62330 4.27605 -5.33042
-4.20074 2.51040 0.00000 ; SG- CB- CA- C
4156 4158 4160 4161 3 0.00000 0.00000 0.00000
0.00000 0.00000 0.00000 ; C- N- CA- HA
4156 4158 4160 4162 3 11.71520 1.67360 15.06240 -
13.38880 0.00000 0.00000 ; C- N- CA- CB
4156 4158 4160 4170 3 1.75728 5.27184 2.25936
-7.02912 0.00000 0.00000 ; C- N- CA- C
4157 4156 4158 4159 3 29.28800 -8.36800 -20.92000
0.00000 0.00000 0.00000 ; O- C- N- H
4157 4156 4158 4160 3 20.92000 0.00000 -20.92000
0.00000 0.00000 0.00000 ; O- C- N- CA
4158 4160 4162 4163 3 0.65084 1.95253 0.00000
-2.60338 0.00000 0.00000 ; N- CA- CB- HB
4158 4160 4162 4164 3 3.83673 0.44769 -1.77402
-2.47693 -0.03347 0.00000 ; N- CA- CB- CG2
4158 4160 4162 4168 3 4.93294 1.13386 -3.12963
-5.27184 3.17984 0.00000 ; N- CA- CB- OG1
4158 4160 4170 4171 3 0.00000 0.00000 0.00000
0.00000 0.00000 0.00000 ; N- CA- C- O
4158 4160 4170 4172 3 17.40544 -5.02080 -13.22144
9.20480 0.00000 0.00000 ; N- CA- C- N
4159 4158 4160 4161 3 0.00000 0.00000 0.00000
0.00000 0.00000 0.00000 ; H- N- CA- HA
4159 4158 4160 4162 3 0.00000 0.00000 0.00000
0.00000 0.00000 0.00000 ; H- N- CA- CB
4159 4158 4160 4170 3 0.00000 0.00000 0.00000
0.00000 0.00000 0.00000 ; H- N- CA- C
4160 4162 4164 4165 3 0.66944 2.00832 0.00000
-2.67776 0.00000 0.00000 ; CA- CB- CG2- HG21
4160 4162 4164 4166 3 0.66944 2.00832 0.00000
-2.67776 0.00000 0.00000 ; CA- CB- CG2- HG22
4160 4162 4164 4167 3 0.66944 2.00832 0.00000
-2.67776 0.00000 0.00000 ; CA- CB- CG2- HG23
4160 4162 4168 4169 3 1.12131 2.98738 1.66523

-3.94970    0.43514    0.00000 ;      CA-    CB-   OG1-   HG1
  4160   4170   4172   4173       3   20.92000    0.00000  -20.92000
0.00000    0.00000    0.00000 ;      CA-     C-     N-      H
  4160   4170   4172   4174       3   20.92000    0.00000  -20.92000
0.00000    0.00000    0.00000 ;      CA-     C-     N-     CA
  4161   4160   4162   4163       3    0.65084    1.95253    0.00000
-2.60338    0.00000    0.00000 ;      HA-    CA-    CB-     HB
  4161   4160   4162   4164       3    0.65084    1.95253    0.00000
-2.60338    0.00000    0.00000 ;      HA-    CA-    CB-    CG2
  4161   4160   4162   4168       3    1.04600   -1.04600    0.00000
0.00000    0.00000    0.00000 ;      HA-    CA-    CB-    OG1
  4161   4160   4170   4171       3    3.68192   -4.35136    0.00000
1.33888    0.00000    0.00000 ;      HA-    CA-     C-      O
  4161   4160   4170   4172       3    0.00000    0.00000    0.00000
0.00000    0.00000    0.00000 ;      HA-    CA-     C-      N
  4162   4160   4170   4171       3    0.00000    0.00000    0.00000
0.00000    0.00000    0.00000 ;      CB-    CA-     C-      O
  4162   4160   4170   4172       3    2.51040    4.18400    1.67360
-6.69440    0.00000    0.00000 ;      CB-    CA-     C-      N
  4163   4162   4160   4170       3    0.65084    1.95253    0.00000
-2.60338    0.00000    0.00000 ;      HB-    CB-    CA-      C
  4163   4162   4164   4165       3    0.65084    1.95253    0.00000
-2.60338    0.00000    0.00000 ;      HB-    CB-   CG2-   HG21
  4163   4162   4164   4166       3    0.65084    1.95253    0.00000
-2.60338    0.00000    0.00000 ;      HB-    CB-   CG2-   HG22
  4163   4162   4164   4167       3    0.65084    1.95253    0.00000
-2.60338    0.00000    0.00000 ;      HB-    CB-   CG2-   HG23
  4163   4162   4168   4169       3    0.69733    2.09200    0.00000
-2.78933    0.00000    0.00000 ;      HB-    CB-   OG1-    HG1
  4164   4162   4160   4170       3    5.67350    3.55640   -6.16722
-2.47693    3.74886    0.00000 ;     CG2-    CB-    CA-      C
  4164   4162   4168   4169       3    4.74047    0.27196   -2.26773
-3.94970    1.60666    0.00000 ;     CG2-    CB-   OG1-    HG1
  4165   4164   4162   4168       3    1.04600   -1.04600    0.00000
0.00000    0.00000    0.00000 ;    HG21-   CG2-    CB-    OG1
  4166   4164   4162   4168       3    1.04600   -1.04600    0.00000
0.00000    0.00000    0.00000 ;    HG22-   CG2-    CB-    OG1
  4167   4164   4162   4168       3    1.04600   -1.04600    0.00000
0.00000    0.00000    0.00000 ;    HG23-   CG2-    CB-    OG1
  4168   4162   4160   4170       3    6.53541    6.87013   -6.21742
-5.27184    5.22163    0.00000 ;     OG1-    CB-    CA-      C
  4170   4172   4174   4175       3    0.00000    0.00000    0.00000
0.00000    0.00000    0.00000 ;      C-     N-    CA-     HA
  4170   4172   4174   4176       3   11.71520    1.67360   15.06240  -
13.38880    0.00000    0.00000 ;      C-     N-    CA-     CB
  4170   4172   4174   4189       3    1.75728    5.27184    2.25936
-7.02912    0.00000    0.00000 ;      C-     N-    CA-      C
  4171   4170   4172   4173       3   29.28800   -8.36800  -20.92000
0.00000    0.00000    0.00000 ;      O-     C-     N-      H
  4171   4170   4172   4174       3   20.92000    0.00000  -20.92000
0.00000    0.00000    0.00000 ;      O-     C-     N-     CA
  4172   4174   4176   4177       3    0.65084    1.95253    0.00000

```
-2.60338    0.00000    0.00000 ;       N-    CA-    CB-    HB
  4172   4174   4176   4178       3    3.83673    0.44769   -1.77402
-2.47693   -0.03347    0.00000 ;       N-    CA-    CB-   CG2
  4172   4174   4176   4182       3    2.97482    0.12134    2.04179
-1.89117   -3.24678    0.00000 ;       N-    CA-    CB-   CG1
  4172   4174   4189   4190       3    0.00000    0.00000    0.00000
0.00000    0.00000    0.00000 ;       N-    CA-     C-     O
  4172   4174   4189   4191       3   17.40544   -5.02080  -13.22144
9.20480    0.00000    0.00000 ;       N-    CA-     C-     N
  4173   4172   4174   4175       3    0.00000    0.00000    0.00000
0.00000    0.00000    0.00000 ;       H-     N-    CA-    HA
  4173   4172   4174   4176       3    0.00000    0.00000    0.00000
0.00000    0.00000    0.00000 ;       H-     N-    CA-    CB
  4173   4172   4174   4189       3    0.00000    0.00000    0.00000
0.00000    0.00000    0.00000 ;       H-     N-    CA-     C
  4174   4176   4178   4179       3    0.66944    2.00832    0.00000
-2.67776    0.00000    0.00000 ;      CA-    CB-   CG2-  HG21
  4174   4176   4178   4180       3    0.66944    2.00832    0.00000
-2.67776    0.00000    0.00000 ;      CA-    CB-   CG2-  HG22
  4174   4176   4178   4181       3    0.66944    2.00832    0.00000
-2.67776    0.00000    0.00000 ;      CA-    CB-   CG2-  HG23
  4174   4176   4182   4183       3    0.66944    2.00832    0.00000
-2.67776    0.00000    0.00000 ;      CA-    CB-   CG1-  HG12
  4174   4176   4182   4184       3    0.66944    2.00832    0.00000
-2.67776    0.00000    0.00000 ;      CA-    CB-   CG1-  HG13
  4174   4176   4182   4185       3    4.24258   -0.52718   -7.25506
-1.79075    7.69856    0.00000 ;      CA-    CB-   CG1-   CD1
  4174   4189   4191   4192       3   20.92000    0.00000  -20.92000
0.00000    0.00000    0.00000 ;      CA-     C-     N-     H
  4174   4189   4191   4193       3   20.92000    0.00000  -20.92000
0.00000    0.00000    0.00000 ;      CA-     C-     N-    CA
  4175   4174   4176   4177       3    0.65084    1.95253    0.00000
-2.60338    0.00000    0.00000 ;      HA-    CA-    CB-    HB
  4175   4174   4176   4178       3    0.65084    1.95253    0.00000
-2.60338    0.00000    0.00000 ;      HA-    CA-    CB-   CG2
  4175   4174   4176   4182       3    0.65084    1.95253    0.00000
-2.60338    0.00000    0.00000 ;      HA-    CA-    CB-   CG1
  4175   4174   4189   4190       3    3.68192   -4.35136    0.00000
1.33888    0.00000    0.00000 ;      HA-    CA-     C-     O
  4175   4174   4189   4191       3    0.00000    0.00000    0.00000
0.00000    0.00000    0.00000 ;      HA-    CA-     C-     N
  4176   4174   4189   4190       3    0.00000    0.00000    0.00000
0.00000    0.00000    0.00000 ;      CB-    CA-     C-     O
  4176   4174   4189   4191       3    2.51040    4.18400    1.67360
-6.69440    0.00000    0.00000 ;      CB-    CA-     C-     N
  4176   4182   4185   4186       3    0.66944    2.00832    0.00000
-2.67776    0.00000    0.00000 ;      CB-   CG1-   CD1-  HD11
  4176   4182   4185   4187       3    0.66944    2.00832    0.00000
-2.67776    0.00000    0.00000 ;      CB-   CG1-   CD1-  HD12
  4176   4182   4185   4188       3    0.66944    2.00832    0.00000
-2.67776    0.00000    0.00000 ;      CB-   CG1-   CD1-  HD13
  4177   4176   4174   4189       3    0.65084    1.95253    0.00000
```

-2.60338 0.00000 0.00000 ; HB- CB- CA- C
4177 4176 4178 4179 3 0.62760 1.88280 0.00000
-2.51040 0.00000 0.00000 ; HB- CB- CG2- HG21
4177 4176 4178 4180 3 0.62760 1.88280 0.00000
-2.51040 0.00000 0.00000 ; HB- CB- CG2- HG22
4177 4176 4178 4181 3 0.62760 1.88280 0.00000
-2.51040 0.00000 0.00000 ; HB- CB- CG2- HG23
4177 4176 4182 4183 3 0.62760 1.88280 0.00000
-2.51040 0.00000 0.00000 ; HB- CB- CG1- HG12
4177 4176 4182 4184 3 0.62760 1.88280 0.00000
-2.51040 0.00000 0.00000 ; HB- CB- CG1- HG13
4177 4176 4182 4185 3 0.66944 2.00832 0.00000
-2.67776 0.00000 0.00000 ; HB- CB- CG1- CD1
4178 4176 4174 4189 3 5.67350 3.55640 -6.16722
-2.47693 3.74886 0.00000 ; CG2- CB- CA- C
4178 4176 4182 4183 3 0.66944 2.00832 0.00000
-2.67776 0.00000 0.00000 ; CG2- CB- CG1- HG12
4178 4176 4182 4184 3 0.66944 2.00832 0.00000
-2.67776 0.00000 0.00000 ; CG2- CB- CG1- HG13
4178 4176 4182 4185 3 3.81162 0.49790 -8.14206
-1.79075 7.49773 0.00000 ; CG2- CB- CG1- CD1
4179 4178 4176 4182 3 0.66944 2.00832 0.00000
-2.67776 0.00000 0.00000 ; HG21- CG2- CB- CG1
4180 4178 4176 4182 3 0.66944 2.00832 0.00000
-2.67776 0.00000 0.00000 ; HG22- CG2- CB- CG1
4181 4178 4176 4182 3 0.66944 2.00832 0.00000
-2.67776 0.00000 0.00000 ; HG23- CG2- CB- CG1
4182 4176 4174 4189 3 8.26340 0.74057 -9.99976
-1.89117 3.84928 0.00000 ; CG1- CB- CA- C
4183 4182 4185 4186 3 0.62760 1.88280 0.00000
-2.51040 0.00000 0.00000 ; HG12- CG1- CD1- HD11
4183 4182 4185 4187 3 0.62760 1.88280 0.00000
-2.51040 0.00000 0.00000 ; HG12- CG1- CD1- HD12
4183 4182 4185 4188 3 0.62760 1.88280 0.00000
-2.51040 0.00000 0.00000 ; HG12- CG1- CD1- HD13
4184 4182 4185 4186 3 0.62760 1.88280 0.00000
-2.51040 0.00000 0.00000 ; HG13- CG1- CD1- HD11
4184 4182 4185 4187 3 0.62760 1.88280 0.00000
-2.51040 0.00000 0.00000 ; HG13- CG1- CD1- HD12
4184 4182 4185 4188 3 0.62760 1.88280 0.00000
-2.51040 0.00000 0.00000 ; HG13- CG1- CD1- HD13
4189 4191 4193 4194 3 0.00000 0.00000 0.00000
0.00000 0.00000 0.00000 ; C- N- CA- HA
4189 4191 4193 4195 3 11.71520 1.67360 15.06240 -
13.38880 0.00000 0.00000 ; C- N- CA- CB
4189 4191 4193 4204 3 1.75728 5.27184 2.25936
-7.02912 0.00000 0.00000 ; C- N- CA- C
4190 4189 4191 4192 3 29.28800 -8.36800 -20.92000
0.00000 0.00000 0.00000 ; O- C- N- H
4190 4189 4191 4193 3 20.92000 0.00000 -20.92000
0.00000 0.00000 0.00000 ; O- C- N- CA
4191 4193 4195 4196 3 0.65084 1.95253 0.00000

```
-2.60338    0.00000    0.00000 ;       N-    CA-    CB-   HB2
  4191   4193   4195   4197       3    0.65084    1.95253    0.00000
-2.60338    0.00000    0.00000 ;       N-    CA-    CB-   HB3
  4191   4193   4195   4198       3    3.21331    2.22589   -4.15053
-2.40998    2.61082    0.00000 ;       N-    CA-    CB-    CG
  4191   4193   4204   4205       3    0.00000    0.00000    0.00000
0.00000    0.00000    0.00000 ;       N-    CA-     C-     O
  4191   4193   4204   4206       3   17.40544   -5.02080  -13.22144
9.20480    0.00000    0.00000 ;       N-    CA-     C-     N
  4192   4191   4193   4194       3    0.00000    0.00000    0.00000
0.00000    0.00000    0.00000 ;       H-     N-    CA-    HA
  4192   4191   4193   4195       3    0.00000    0.00000    0.00000
0.00000    0.00000    0.00000 ;       H-     N-    CA-    CB
  4192   4191   4193   4204       3    0.00000    0.00000    0.00000
0.00000    0.00000    0.00000 ;       H-     N-    CA-     C
  4193   4195   4198   4199       3    0.66944    2.00832    0.00000
-2.67776    0.00000    0.00000 ;      CA-    CB-    CG-   HG2
  4193   4195   4198   4200       3    0.66944    2.00832    0.00000
-2.67776    0.00000    0.00000 ;      CA-    CB-    CG-   HG3
  4193   4195   4198   4201       3   10.12110   -1.91209    0.01674
10.17549   -1.87443    0.00000 ;      CA-    CB-    CG-    CD
  4193   4204   4206   4207       3   20.92000    0.00000  -20.92000
0.00000    0.00000    0.00000 ;      CA-     C-     N-     H
  4193   4204   4206   4208       3   20.92000    0.00000  -20.92000
0.00000    0.00000    0.00000 ;      CA-     C-     N-    CA
  4194   4193   4195   4196       3    0.65084    1.95253    0.00000
-2.60338    0.00000    0.00000 ;      HA-    CA-    CB-   HB2
  4194   4193   4195   4197       3    0.65084    1.95253    0.00000
-2.60338    0.00000    0.00000 ;      HA-    CA-    CB-   HB3
  4194   4193   4195   4198       3    0.65084    1.95253    0.00000
-2.60338    0.00000    0.00000 ;      HA-    CA-    CB-    CG
  4194   4193   4204   4205       3    3.68192   -4.35136    0.00000
1.33888    0.00000    0.00000 ;      HA-    CA-     C-     O
  4194   4193   4204   4206       3    0.00000    0.00000    0.00000
0.00000    0.00000    0.00000 ;      HA-    CA-     C-     N
  4195   4193   4204   4205       3    0.00000    0.00000    0.00000
0.00000    0.00000    0.00000 ;      CB-    CA-     C-     O
  4195   4193   4204   4206       3    2.51040    4.18400    1.67360
-6.69440    0.00000    0.00000 ;      CB-    CA-     C-     N
  4195   4198   4201   4202       3    3.79907    0.00000   -5.40573
0.00000    2.14221    0.00000 ;      CB-    CG-    CD-   OE1
  4195   4198   4201   4203       3    3.79907    0.00000   -5.40573
0.00000    2.14221    0.00000 ;      CB-    CG-    CD-   OE2
  4196   4195   4193   4204       3    0.65084    1.95253    0.00000
-2.60338    0.00000    0.00000 ;     HB2-    CB-    CA-     C
  4196   4195   4198   4199       3    0.62760    1.88280    0.00000
-2.51040    0.00000    0.00000 ;     HB2-    CB-    CG-   HG2
  4196   4195   4198   4200       3    0.62760    1.88280    0.00000
-2.51040    0.00000    0.00000 ;     HB2-    CB-    CG-   HG3
  4196   4195   4198   4201       3    0.65084    1.95253    0.00000
-2.60338    0.00000    0.00000 ;     HB2-    CB-    CG-    CD
  4197   4195   4193   4204       3    0.65084    1.95253    0.00000
```

-2.60338 0.00000 0.00000 ; HB3- CB- CA- C
4197 4195 4198 4199 3 0.62760 1.88280 0.00000
-2.51040 0.00000 0.00000 ; HB3- CB- CG- HG2
4197 4195 4198 4200 3 0.62760 1.88280 0.00000
-2.51040 0.00000 0.00000 ; HB3- CB- CG- HG3
4197 4195 4198 4201 3 0.65084 1.95253 0.00000
-2.60338 0.00000 0.00000 ; HB3- CB- CG- CD
4198 4195 4193 4204 3 6.86594 3.56895 -8.14206
-2.40998 4.85344 0.00000 ; CG- CB- CA- C
4199 4198 4201 4202 3 0.00000 0.00000 0.00000
0.00000 0.00000 0.00000 ; HG2- CG- CD- OE1
4199 4198 4201 4203 3 0.00000 0.00000 0.00000
0.00000 0.00000 0.00000 ; HG2- CG- CD- OE2
4200 4198 4201 4202 3 0.00000 0.00000 0.00000
0.00000 0.00000 0.00000 ; HG3- CG- CD- OE1
4200 4198 4201 4203 3 0.00000 0.00000 0.00000
0.00000 0.00000 0.00000 ; HG3- CG- CD- OE2
4204 4206 4208 4209 3 0.00000 0.00000 0.00000
0.00000 0.00000 0.00000 ; C- N- CA- HA
4204 4206 4208 4210 3 11.71520 1.67360 15.06240 -
13.38880 0.00000 0.00000 ; C- N- CA- CB
4204 4206 4208 4219 3 1.75728 5.27184 2.25936
-7.02912 0.00000 0.00000 ; C- N- CA- C
4205 4204 4206 4207 3 29.28800 -8.36800 -20.92000
0.00000 0.00000 0.00000 ; O- C- N- H
4205 4204 4206 4208 3 20.92000 0.00000 -20.92000
0.00000 0.00000 0.00000 ; O- C- N- CA
4206 4208 4210 4211 3 0.65084 1.95253 0.00000
-2.60338 0.00000 0.00000 ; N- CA- CB- HB2
4206 4208 4210 4212 3 0.65084 1.95253 0.00000
-2.60338 0.00000 0.00000 ; N- CA- CB- HB3
4206 4208 4210 4213 3 3.21331 2.22589 -4.15053
-2.40998 2.61082 0.00000 ; N- CA- CB- CG
4206 4208 4219 4220 3 0.00000 0.00000 0.00000
0.00000 0.00000 0.00000 ; N- CA- C- O
4206 4208 4219 4221 3 17.40544 -5.02080 -13.22144
9.20480 0.00000 0.00000 ; N- CA- C- N
4207 4206 4208 4209 3 0.00000 0.00000 0.00000
0.00000 0.00000 0.00000 ; H- N- CA- HA
4207 4206 4208 4210 3 0.00000 0.00000 0.00000
0.00000 0.00000 0.00000 ; H- N- CA- CB
4207 4206 4208 4219 3 0.00000 0.00000 0.00000
0.00000 0.00000 0.00000 ; H- N- CA- C
4208 4210 4213 4214 3 0.66944 2.00832 0.00000
-2.67776 0.00000 0.00000 ; CA- CB- CG- HG2
4208 4210 4213 4215 3 0.66944 2.00832 0.00000
-2.67776 0.00000 0.00000 ; CA- CB- CG- HG3
4208 4210 4213 4216 3 10.12110 -1.91209 0.01674
10.17549 -1.87443 0.00000 ; CA- CB- CG- CD
4208 4219 4221 4222 3 20.92000 0.00000 -20.92000
0.00000 0.00000 0.00000 ; CA- C- N- H
4208 4219 4221 4223 3 20.92000 0.00000 -20.92000

```
0.00000    0.00000    0.00000 ;     CA-     C-     N-    CA
  4209   4208   4210   4211      3    0.65084    1.95253    0.00000
-2.60338    0.00000    0.00000 ;     HA-    CA-    CB-   HB2
  4209   4208   4210   4212      3    0.65084    1.95253    0.00000
-2.60338    0.00000    0.00000 ;     HA-    CA-    CB-   HB3
  4209   4208   4210   4213      3    0.65084    1.95253    0.00000
-2.60338    0.00000    0.00000 ;     HA-    CA-    CB-    CG
  4209   4208   4219   4220      3    3.68192   -4.35136    0.00000
1.33888    0.00000    0.00000 ;     HA-    CA-     C-     O
  4209   4208   4219   4221      3    0.00000    0.00000    0.00000
0.00000    0.00000    0.00000 ;     HA-    CA-     C-     N
  4210   4208   4219   4220      3    0.00000    0.00000    0.00000
0.00000    0.00000    0.00000 ;     CB-    CA-     C-     O
  4210   4208   4219   4221      3    2.51040    4.18400    1.67360
-6.69440    0.00000    0.00000 ;     CB-    CA-     C-     N
  4210   4213   4216   4217      3    3.79907    0.00000   -5.40573
0.00000    2.14221    0.00000 ;     CB-    CG-    CD-   OE1
  4210   4213   4216   4218      3    3.79907    0.00000   -5.40573
0.00000    2.14221    0.00000 ;     CB-    CG-    CD-   OE2
  4211   4210   4208   4219      3    0.65084    1.95253    0.00000
-2.60338    0.00000    0.00000 ;    HB2-    CB-    CA-     C
  4211   4210   4213   4214      3    0.62760    1.88280    0.00000
-2.51040    0.00000    0.00000 ;    HB2-    CB-    CG-   HG2
  4211   4210   4213   4215      3    0.62760    1.88280    0.00000
-2.51040    0.00000    0.00000 ;    HB2-    CB-    CG-   HG3
  4211   4210   4213   4216      3    0.65084    1.95253    0.00000
-2.60338    0.00000    0.00000 ;    HB2-    CB-    CG-    CD
  4212   4210   4208   4219      3    0.65084    1.95253    0.00000
-2.60338    0.00000    0.00000 ;    HB3-    CB-    CA-     C
  4212   4210   4213   4214      3    0.62760    1.88280    0.00000
-2.51040    0.00000    0.00000 ;    HB3-    CB-    CG-   HG2
  4212   4210   4213   4215      3    0.62760    1.88280    0.00000
-2.51040    0.00000    0.00000 ;    HB3-    CB-    CG-   HG3
  4212   4210   4213   4216      3    0.65084    1.95253    0.00000
-2.60338    0.00000    0.00000 ;    HB3-    CB-    CG-    CD
  4213   4210   4208   4219      3    6.86594    3.56895   -8.14206
-2.40998    4.85344    0.00000 ;     CG-    CB-    CA-     C
  4214   4213   4216   4217      3    0.00000    0.00000    0.00000
0.00000    0.00000    0.00000 ;    HG2-    CG-    CD-   OE1
  4214   4213   4216   4218      3    0.00000    0.00000    0.00000
0.00000    0.00000    0.00000 ;    HG2-    CG-    CD-   OE2
  4215   4213   4216   4217      3    0.00000    0.00000    0.00000
0.00000    0.00000    0.00000 ;    HG3-    CG-    CD-   OE1
  4215   4213   4216   4218      3    0.00000    0.00000    0.00000
0.00000    0.00000    0.00000 ;    HG3-    CG-    CD-   OE2
  4219   4221   4223   4224      3    0.00000    0.00000    0.00000
0.00000    0.00000    0.00000 ;     C-     N-    CA-    HA
  4219   4221   4223   4225      3   11.71520    1.67360   15.06240  -
13.38880    0.00000    0.00000 ;     C-     N-    CA-    CB
  4219   4221   4223   4229      3    1.75728    5.27184    2.25936
-7.02912    0.00000    0.00000 ;     C-     N-    CA-     C
  4220   4219   4221   4222      3   29.28800   -8.36800  -20.92000
```

0.00000    0.00000    0.00000 ;     O-    C-    N-    H
  4220   4219   4221   4223      3   20.92000    0.00000  -20.92000
0.00000    0.00000    0.00000 ;     O-    C-    N-    CA
  4221   4223   4225   4226      3    0.65084    1.95253    0.00000
-2.60338    0.00000    0.00000 ;     N-    CA-    CB-    HB1
  4221   4223   4225   4227      3    0.65084    1.95253    0.00000
-2.60338    0.00000    0.00000 ;     N-    CA-    CB-    HB2
  4221   4223   4225   4228      3    0.65084    1.95253    0.00000
-2.60338    0.00000    0.00000 ;     N-    CA-    CB-    HB3
  4221   4223   4229   4230      3    0.00000    0.00000    0.00000
0.00000    0.00000    0.00000 ;     N-    CA-    C-    O
  4221   4223   4229   4231      3   17.40544   -5.02080  -13.22144
9.20480    0.00000    0.00000 ;     N-    CA-    C-    N
  4222   4221   4223   4224      3    0.00000    0.00000    0.00000
0.00000    0.00000    0.00000 ;     H-    N-    CA-    HA
  4222   4221   4223   4225      3    0.00000    0.00000    0.00000
0.00000    0.00000    0.00000 ;     H-    N-    CA-    CB
  4222   4221   4223   4229      3    0.00000    0.00000    0.00000
0.00000    0.00000    0.00000 ;     H-    N-    CA-    C
  4223   4229   4231   4232      3   20.92000    0.00000  -20.92000
0.00000    0.00000    0.00000 ;     CA-    C-    N-    H
  4223   4229   4231   4233      3   20.92000    0.00000  -20.92000
0.00000    0.00000    0.00000 ;     CA-    C-    N-    CA
  4224   4223   4225   4226      3    0.65084    1.95253    0.00000
-2.60338    0.00000    0.00000 ;     HA-    CA-    CB-    HB1
  4224   4223   4225   4227      3    0.65084    1.95253    0.00000
-2.60338    0.00000    0.00000 ;     HA-    CA-    CB-    HB2
  4224   4223   4225   4228      3    0.65084    1.95253    0.00000
-2.60338    0.00000    0.00000 ;     HA-    CA-    CB-    HB3
  4224   4223   4229   4230      3    3.68192   -4.35136    0.00000
1.33888    0.00000    0.00000 ;     HA-    CA-    C-    O
  4224   4223   4229   4231      3    0.00000    0.00000    0.00000
0.00000    0.00000    0.00000 ;     HA-    CA-    C-    N
  4225   4223   4229   4230      3    0.00000    0.00000    0.00000
0.00000    0.00000    0.00000 ;     CB-    CA-    C-    O
  4225   4223   4229   4231      3    2.51040    4.18400    1.67360
-6.69440    0.00000    0.00000 ;     CB-    CA-    C-    N
  4226   4225   4223   4229      3    0.65084    1.95253    0.00000
-2.60338    0.00000    0.00000 ;     HB1-    CB-    CA-    C
  4227   4225   4223   4229      3    0.65084    1.95253    0.00000
-2.60338    0.00000    0.00000 ;     HB2-    CB-    CA-    C
  4228   4225   4223   4229      3    0.65084    1.95253    0.00000
-2.60338    0.00000    0.00000 ;     HB3-    CB-    CA-    C
  4229   4231   4233   4234      3    0.00000    0.00000    0.00000
0.00000    0.00000    0.00000 ;     C-    N-    CA-    HA
  4229   4231   4233   4235      3   11.71520    1.67360   15.06240  -
13.38880    0.00000    0.00000 ;     C-    N-    CA-    CB
  4229   4231   4233   4248      3    1.75728    5.27184    2.25936
-7.02912    0.00000    0.00000 ;     C-    N-    CA-    C
  4230   4229   4231   4232      3   29.28800   -8.36800  -20.92000
0.00000    0.00000    0.00000 ;     O-    C-    N-    H
  4230   4229   4231   4233      3   20.92000    0.00000  -20.92000

```
0.00000    0.00000    0.00000 ;       O-     C-     N-    CA
  4231   4233   4235   4236      3    0.65084    1.95253    0.00000
-2.60338    0.00000    0.00000 ;       N-    CA-    CB-   HB2
  4231   4233   4235   4237      3    0.65084    1.95253    0.00000
-2.60338    0.00000    0.00000 ;       N-    CA-    CB-   HB3
  4231   4233   4235   4238      3    3.79070    1.39746   -4.61077
-2.40998    2.44346    0.00000 ;       N-    CA-    CB-    CG
  4231   4233   4248   4249      3    0.00000    0.00000    0.00000
0.00000    0.00000    0.00000 ;       N-    CA-     C-     O
  4231   4233   4248   4250      3   17.40544   -5.02080  -13.22144
9.20480    0.00000    0.00000 ;       N-    CA-     C-     N
  4232   4231   4233   4234      3    0.00000    0.00000    0.00000
0.00000    0.00000    0.00000 ;       H-     N-    CA-    HA
  4232   4231   4233   4235      3    0.00000    0.00000    0.00000
0.00000    0.00000    0.00000 ;       H-     N-    CA-    CB
  4232   4231   4233   4248      3    0.00000    0.00000    0.00000
0.00000    0.00000    0.00000 ;       H-     N-    CA-     C
  4233   4235   4238   4239      3    0.66944    2.00832    0.00000
-2.67776    0.00000    0.00000 ;      CA-    CB-    CG-    HG
  4233   4235   4238   4240      3    3.90367    0.19665   -6.21742
-2.37651    5.99149    0.00000 ;      CA-    CB-    CG-   CD1
  4233   4235   4238   4244      3    3.90367    0.19665   -6.21742
-2.37651    5.99149    0.00000 ;      CA-    CB-    CG-   CD2
  4233   4248   4250   4251      3   20.92000    0.00000  -20.92000
0.00000    0.00000    0.00000 ;      CA-     C-     N-     H
  4233   4248   4250   4252      3   20.92000    0.00000  -20.92000
0.00000    0.00000    0.00000 ;      CA-     C-     N-    CA
  4234   4233   4235   4236      3    0.65084    1.95253    0.00000
-2.60338    0.00000    0.00000 ;      HA-    CA-    CB-   HB2
  4234   4233   4235   4237      3    0.65084    1.95253    0.00000
-2.60338    0.00000    0.00000 ;      HA-    CA-    CB-   HB3
  4234   4233   4235   4238      3    0.65084    1.95253    0.00000
-2.60338    0.00000    0.00000 ;      HA-    CA-    CB-    CG
  4234   4233   4248   4249      3    3.68192   -4.35136    0.00000
1.33888    0.00000    0.00000 ;      HA-    CA-     C-     O
  4234   4233   4248   4250      3    0.00000    0.00000    0.00000
0.00000    0.00000    0.00000 ;      HA-    CA-     C-     N
  4235   4233   4248   4249      3    0.00000    0.00000    0.00000
0.00000    0.00000    0.00000 ;      CB-    CA-     C-     O
  4235   4233   4248   4250      3    2.51040    4.18400    1.67360
-6.69440    0.00000    0.00000 ;      CB-    CA-     C-     N
  4235   4238   4240   4241      3    0.66944    2.00832    0.00000
-2.67776    0.00000    0.00000 ;      CB-    CG-   CD1-  HD11
  4235   4238   4240   4242      3    0.66944    2.00832    0.00000
-2.67776    0.00000    0.00000 ;      CB-    CG-   CD1-  HD12
  4235   4238   4240   4243      3    0.66944    2.00832    0.00000
-2.67776    0.00000    0.00000 ;      CB-    CG-   CD1-  HD13
  4235   4238   4244   4245      3    0.66944    2.00832    0.00000
-2.67776    0.00000    0.00000 ;      CB-    CG-   CD2-  HD21
  4235   4238   4244   4246      3    0.66944    2.00832    0.00000
-2.67776    0.00000    0.00000 ;      CB-    CG-   CD2-  HD22
  4235   4238   4244   4247      3    0.66944    2.00832    0.00000
```

-2.67776 0.00000 0.00000 ; CB- CG- CD2- HD23
4236 4235 4233 4248 3 0.65084 1.95253 0.00000
-2.60338 0.00000 0.00000 ; HB2- CB- CA- C
4236 4235 4238 4239 3 0.62760 1.88280 0.00000
-2.51040 0.00000 0.00000 ; HB2- CB- CG- HG
4236 4235 4238 4240 3 0.66944 2.00832 0.00000
-2.67776 0.00000 0.00000 ; HB2- CB- CG- CD1
4236 4235 4238 4244 3 0.66944 2.00832 0.00000
-2.67776 0.00000 0.00000 ; HB2- CB- CG- CD2
4237 4235 4233 4248 3 0.65084 1.95253 0.00000
-2.60338 0.00000 0.00000 ; HB3- CB- CA- C
4237 4235 4238 4239 3 0.62760 1.88280 0.00000
-2.51040 0.00000 0.00000 ; HB3- CB- CG- HG
4237 4235 4238 4240 3 0.66944 2.00832 0.00000
-2.67776 0.00000 0.00000 ; HB3- CB- CG- CD1
4237 4235 4238 4244 3 0.66944 2.00832 0.00000
-2.67776 0.00000 0.00000 ; HB3- CB- CG- CD2
4238 4235 4233 4248 3 10.33448 -1.14642 -11.54784
-2.40998 6.35968 0.00000 ; CG- CB- CA- C
4239 4238 4240 4241 3 0.62760 1.88280 0.00000
-2.51040 0.00000 0.00000 ; HG- CG- CD1- HD11
4239 4238 4240 4242 3 0.62760 1.88280 0.00000
-2.51040 0.00000 0.00000 ; HG- CG- CD1- HD12
4239 4238 4240 4243 3 0.62760 1.88280 0.00000
-2.51040 0.00000 0.00000 ; HG- CG- CD1- HD13
4239 4238 4244 4245 3 0.62760 1.88280 0.00000
-2.51040 0.00000 0.00000 ; HG- CG- CD2- HD21
4239 4238 4244 4246 3 0.62760 1.88280 0.00000
-2.51040 0.00000 0.00000 ; HG- CG- CD2- HD22
4239 4238 4244 4247 3 0.62760 1.88280 0.00000
-2.51040 0.00000 0.00000 ; HG- CG- CD2- HD23
4240 4238 4244 4245 3 0.66944 2.00832 0.00000
-2.67776 0.00000 0.00000 ; CD1- CG- CD2- HD21
4240 4238 4244 4246 3 0.66944 2.00832 0.00000
-2.67776 0.00000 0.00000 ; CD1- CG- CD2- HD22
4240 4238 4244 4247 3 0.66944 2.00832 0.00000
-2.67776 0.00000 0.00000 ; CD1- CG- CD2- HD23
4241 4240 4238 4244 3 0.66944 2.00832 0.00000
-2.67776 0.00000 0.00000 ; HD11- CD1- CG- CD2
4242 4240 4238 4244 3 0.66944 2.00832 0.00000
-2.67776 0.00000 0.00000 ; HD12- CD1- CG- CD2
4243 4240 4238 4244 3 0.66944 2.00832 0.00000
-2.67776 0.00000 0.00000 ; HD13- CD1- CG- CD2
4248 4250 4252 4253 3 0.00000 0.00000 0.00000
0.00000 0.00000 0.00000 ; C- N- CA- HA
4248 4250 4252 4254 3 11.71520 1.67360 15.06240 -
13.38880 0.00000 0.00000 ; C- N- CA- CB
4248 4250 4252 4262 3 1.75728 5.27184 2.25936
-7.02912 0.00000 0.00000 ; C- N- CA- C
4249 4248 4250 4251 3 29.28800 -8.36800 -20.92000
0.00000 0.00000 0.00000 ; O- C- N- H
4249 4248 4250 4252 3 20.92000 0.00000 -20.92000

0.00000 0.00000 0.00000 ; O- C- N- CA
4250 4252 4254 4255 3 0.65084 1.95253 0.00000
-2.60338 0.00000 0.00000 ; N- CA- CB- HB2
4250 4252 4254 4256 3 0.65084 1.95253 0.00000
-2.60338 0.00000 0.00000 ; N- CA- CB- HB3
4250 4252 4254 4257 3 5.99567 3.29281 -4.46014
-0.55229 1.97485 0.00000 ; N- CA- CB- CG
4250 4252 4262 4263 3 0.00000 0.00000 0.00000
0.00000 0.00000 0.00000 ; N- CA- C- O
4250 4252 4262 4264 3 17.40544 -5.02080 -13.22144
9.20480 0.00000 0.00000 ; N- CA- C- N
4251 4250 4252 4253 3 0.00000 0.00000 0.00000
0.00000 0.00000 0.00000 ; H- N- CA- HA
4251 4250 4252 4254 3 0.00000 0.00000 0.00000
0.00000 0.00000 0.00000 ; H- N- CA- CB
4251 4250 4252 4262 3 0.00000 0.00000 0.00000
0.00000 0.00000 0.00000 ; H- N- CA- C
4252 4254 4257 4258 3 0.00000 0.00000 0.00000
0.00000 0.00000 0.00000 ; CA- CB- CG- OD1
4252 4254 4257 4259 3 8.84916 -0.31380 -4.32626
5.03754 0.26778 0.00000 ; CA- CB- CG- ND2
4252 4262 4264 4265 3 20.92000 0.00000 -20.92000
0.00000 0.00000 0.00000 ; CA- C- N- H
4252 4262 4264 4266 3 20.92000 0.00000 -20.92000
0.00000 0.00000 0.00000 ; CA- C- N- CA
4253 4252 4254 4255 3 0.65084 1.95253 0.00000
-2.60338 0.00000 0.00000 ; HA- CA- CB- HB2
4253 4252 4254 4256 3 0.65084 1.95253 0.00000
-2.60338 0.00000 0.00000 ; HA- CA- CB- HB3
4253 4252 4254 4257 3 0.65084 1.95253 0.00000
-2.60338 0.00000 0.00000 ; HA- CA- CB- CG
4253 4252 4262 4263 3 3.68192 -4.35136 0.00000
1.33888 0.00000 0.00000 ; HA- CA- C- O
4253 4252 4262 4264 3 0.00000 0.00000 0.00000
0.00000 0.00000 0.00000 ; HA- CA- C- N
4254 4252 4262 4263 3 0.00000 0.00000 0.00000
0.00000 0.00000 0.00000 ; CB- CA- C- O
4254 4252 4262 4264 3 2.51040 4.18400 1.67360
-6.69440 0.00000 0.00000 ; CB- CA- C- N
4254 4257 4259 4260 3 20.92000 0.00000 -20.92000
0.00000 0.00000 0.00000 ; CB- CG- ND2- HD21
4254 4257 4259 4261 3 20.92000 0.00000 -20.92000
0.00000 0.00000 0.00000 ; CB- CG- ND2- HD22
4255 4254 4252 4262 3 0.65084 1.95253 0.00000
-2.60338 0.00000 0.00000 ; HB2- CB- CA- C
4255 4254 4257 4258 3 3.68192 -4.35136 0.00000
1.33888 0.00000 0.00000 ; HB2- CB- CG- OD1
4255 4254 4257 4259 3 0.00000 0.00000 0.00000
0.00000 0.00000 0.00000 ; HB2- CB- CG- ND2
4256 4254 4252 4262 3 0.65084 1.95253 0.00000
-2.60338 0.00000 0.00000 ; HB3- CB- CA- C
4256 4254 4257 4258 3 3.68192 -4.35136 0.00000

```
1.33888    0.00000    0.00000 ;    HB3-    CB-    CG-   OD1
  4256   4254   4257   4259      3    0.00000    0.00000    0.00000
0.00000    0.00000    0.00000 ;    HB3-    CB-    CG-   ND2
  4257   4254   4252   4262      3    7.94542   -3.96225   -6.11701
-0.55229    3.58150    0.00000 ;     CG-    CB-    CA-     C
  4258   4257   4259   4260      3   29.28800   -8.36800  -20.92000
0.00000    0.00000    0.00000 ;    OD1-    CG-   ND2-  HD21
  4258   4257   4259   4261      3   29.28800   -8.36800  -20.92000
0.00000    0.00000    0.00000 ;    OD1-    CG-   ND2-  HD22
  4262   4264   4266   4267      3    0.00000    0.00000    0.00000
0.00000    0.00000    0.00000 ;      C-     N-    CA-    HA
  4262   4264   4266   4268      3   11.71520    1.67360   15.06240  -
13.38880    0.00000    0.00000 ;      C-     N-    CA-    CB
  4262   4264   4266   4280      3    1.75728    5.27184    2.25936
-7.02912    0.00000    0.00000 ;      C-     N-    CA-     C
  4263   4262   4264   4265      3   29.28800   -8.36800  -20.92000
0.00000    0.00000    0.00000 ;      O-     C-     N-     H
  4263   4262   4264   4266      3   20.92000    0.00000  -20.92000
0.00000    0.00000    0.00000 ;      O-     C-     N-    CA
  4264   4266   4268   4269      3    0.65084    1.95253    0.00000
-2.60338    0.00000    0.00000 ;      N-    CA-    CB-   HB2
  4264   4266   4268   4270      3    0.65084    1.95253    0.00000
-2.60338    0.00000    0.00000 ;      N-    CA-    CB-   HB3
  4264   4266   4268   4271      3    4.79068    4.02919   -4.82834
-3.66518    2.97901    0.00000 ;      N-    CA-    CB-    CG
  4264   4266   4280   4281      3    0.00000    0.00000    0.00000
0.00000    0.00000    0.00000 ;      N-    CA-     C-     O
  4264   4266   4280   4282      3   17.40544   -5.02080  -13.22144
9.20480    0.00000    0.00000 ;      N-    CA-     C-     N
  4265   4264   4266   4267      3    0.00000    0.00000    0.00000
0.00000    0.00000    0.00000 ;      H-     N-    CA-    HA
  4265   4264   4266   4268      3    0.00000    0.00000    0.00000
0.00000    0.00000    0.00000 ;      H-     N-    CA-    CB
  4265   4264   4266   4280      3    0.00000    0.00000    0.00000
0.00000    0.00000    0.00000 ;      H-     N-    CA-     C
  4266   4268   4271   4272      3    6.81992    9.28011   -2.04179  -
11.48090   -1.23846    0.00000 ;     CA-    CB-    CG-   ND1
  4266   4268   4271   4278      3    0.00000    0.00000    0.00000
0.00000    0.00000    0.00000 ;     CA-    CB-    CG-   CD2
  4266   4280   4282   4283      3   20.92000    0.00000  -20.92000
0.00000    0.00000    0.00000 ;     CA-     C-     N-    CD
  4266   4280   4282   4292      3   20.92000    0.00000  -20.92000
0.00000    0.00000    0.00000 ;     CA-     C-     N-    CA
  4267   4266   4268   4269      3    0.65084    1.95253    0.00000
-2.60338    0.00000    0.00000 ;     HA-    CA-    CB-   HB2
  4267   4266   4268   4270      3    0.65084    1.95253    0.00000
-2.60338    0.00000    0.00000 ;     HA-    CA-    CB-   HB3
  4267   4266   4268   4271      3    0.65084    1.95253    0.00000
-2.60338    0.00000    0.00000 ;     HA-    CA-    CB-    CG
  4267   4266   4280   4281      3    3.68192   -4.35136    0.00000
1.33888    0.00000    0.00000 ;     HA-    CA-     C-     O
  4267   4266   4280   4282      3    0.00000    0.00000    0.00000
```

0.00000 0.00000 0.00000 ; HA- CA- C- N
4268 4266 4280 4281 3 0.00000 0.00000 0.00000
0.00000 0.00000 0.00000 ; CB- CA- C- O
4268 4266 4280 4282 3 2.51040 4.18400 1.67360
-6.69440 0.00000 0.00000 ; CB- CA- C- N
4268 4271 4272 4273 3 11.71520 0.00000 -11.71520
0.00000 0.00000 0.00000 ; CB- CG- ND1- HD1
4268 4271 4272 4274 3 11.71520 0.00000 -11.71520
0.00000 0.00000 0.00000 ; CB- CG- ND1- CE1
4268 4271 4278 4276 3 44.97800 0.00000 -44.97800
0.00000 0.00000 0.00000 ; CB- CG- CD2- NE2
4268 4271 4278 4279 3 44.97800 0.00000 -44.97800
0.00000 0.00000 0.00000 ; CB- CG- CD2- HD2
4269 4268 4266 4280 3 0.65084 1.95253 0.00000
-2.60338 0.00000 0.00000 ; HB2- CB- CA- C
4269 4268 4271 4272 3 0.00000 0.00000 0.00000
0.00000 0.00000 0.00000 ; HB2- CB- CG- ND1
4269 4268 4271 4278 3 0.00000 0.00000 0.00000
0.00000 0.00000 0.00000 ; HB2- CB- CG- CD2
4270 4268 4266 4280 3 0.65084 1.95253 0.00000
-2.60338 0.00000 0.00000 ; HB3- CB- CA- C
4270 4268 4271 4272 3 0.00000 0.00000 0.00000
0.00000 0.00000 0.00000 ; HB3- CB- CG- ND1
4270 4268 4271 4278 3 0.00000 0.00000 0.00000
0.00000 0.00000 0.00000 ; HB3- CB- CG- CD2
4271 4268 4266 4280 3 3.76560 3.34720 -2.87859
-3.66518 0.83680 0.00000 ; CG- CB- CA- C
4271 4272 4274 4275 3 19.45560 0.00000 -19.45560
0.00000 0.00000 0.00000 ; CG- ND1- CE1- HE1
4271 4272 4274 4276 3 19.45560 0.00000 -19.45560
0.00000 0.00000 0.00000 ; CG- ND1- CE1- NE2
4271 4278 4276 4274 3 12.55200 0.00000 -12.55200
0.00000 0.00000 0.00000 ; CG- CD2- NE2- CE1
4271 4278 4276 4277 3 12.55200 0.00000 -12.55200
0.00000 0.00000 0.00000 ; CG- CD2- NE2- HE2
4272 4271 4278 4276 3 44.97800 0.00000 -44.97800
0.00000 0.00000 0.00000 ; ND1- CG- CD2- NE2
4272 4271 4278 4279 3 44.97800 0.00000 -44.97800
0.00000 0.00000 0.00000 ; ND1- CG- CD2- HD2
4272 4274 4276 4277 3 19.45560 0.00000 -19.45560
0.00000 0.00000 0.00000 ; ND1- CE1- NE2- HE2
4272 4274 4276 4278 3 19.45560 0.00000 -19.45560
0.00000 0.00000 0.00000 ; ND1- CE1- NE2- CD2
4273 4272 4271 4278 3 11.71520 0.00000 -11.71520
0.00000 0.00000 0.00000 ; HD1- ND1- CG- CD2
4273 4272 4274 4275 3 19.45560 0.00000 -19.45560
0.00000 0.00000 0.00000 ; HD1- ND1- CE1- HE1
4273 4272 4274 4276 3 19.45560 0.00000 -19.45560
0.00000 0.00000 0.00000 ; HD1- ND1- CE1- NE2
4274 4272 4271 4278 3 11.71520 0.00000 -11.71520
0.00000 0.00000 0.00000 ; CE1- ND1- CG- CD2
4274 4276 4278 4279 3 12.55200 0.00000 -12.55200

```
0.00000    0.00000    0.00000 ;    CE1-   NE2-   CD2-   HD2
  4275   4274   4276   4277      3   19.45560    0.00000  -19.45560
0.00000    0.00000    0.00000 ;    HE1-   CE1-   NE2-   HE2
  4275   4274   4276   4278      3   19.45560    0.00000  -19.45560
0.00000    0.00000    0.00000 ;    HE1-   CE1-   NE2-   CD2
  4277   4276   4278   4279      3   12.55200    0.00000  -12.55200
0.00000    0.00000    0.00000 ;    HE2-   NE2-   CD2-   HD2
  4280   4282   4283   4284      3    0.00000    0.00000    0.00000
0.00000    0.00000    0.00000 ;     C-     N-    CD-   HD2
  4280   4282   4283   4285      3    0.00000    0.00000    0.00000
0.00000    0.00000    0.00000 ;     C-     N-    CD-   HD3
  4280   4282   4283   4286      3   10.04160   -3.34720   16.73600
-6.69440    0.00000    0.00000 ;     C-     N-    CD-    CG
  4280   4282   4292   4289      3   11.71520    1.67360   15.06240  -
13.38880    0.00000    0.00000 ;     C-     N-    CA-    CB
  4280   4282   4292   4293      3    0.00000    0.00000    0.00000
0.00000    0.00000    0.00000 ;     C-     N-    CA-    HA
  4280   4282   4292   4294      3    1.75728    5.27184    2.25936
-7.02912    0.00000    0.00000 ;     C-     N-    CA-     C
  4281   4280   4282   4283      3   20.92000    0.00000  -20.92000
0.00000    0.00000    0.00000 ;     O-     C-     N-    CD
  4281   4280   4282   4292      3   20.92000    0.00000  -20.92000
0.00000    0.00000    0.00000 ;     O-     C-     N-    CA
  4282   4283   4286   4287      3    0.65084    1.95253    0.00000
-2.60338    0.00000    0.00000 ;     N-    CD-    CG-   HG2
  4282   4283   4286   4288      3    0.65084    1.95253    0.00000
-2.60338    0.00000    0.00000 ;     N-    CD-    CG-   HG3
  4282   4283   4286   4289      3    0.65084    1.95253    0.00000
-2.60338    0.00000    0.00000 ;     N-    CD-    CG-    CB
  4282   4292   4289   4286      3    0.65084    1.95253    0.00000
-2.60338    0.00000    0.00000 ;     N-    CA-    CB-    CG
  4282   4292   4289   4290      3    0.65084    1.95253    0.00000
-2.60338    0.00000    0.00000 ;     N-    CA-    CB-   HB2
  4282   4292   4289   4291      3    0.65084    1.95253    0.00000
-2.60338    0.00000    0.00000 ;     N-    CA-    CB-   HB3
  4282   4292   4294   4295      3    0.00000    0.00000    0.00000
0.00000    0.00000    0.00000 ;     N-    CA-     C-     O
  4282   4292   4294   4296      3   17.40544   -5.02080  -13.22144
9.20480    0.00000    0.00000 ;     N-    CA-     C-     N
  4283   4282   4292   4289      3    0.00000    0.00000    0.00000
0.00000    0.00000    0.00000 ;    CD-     N-    CA-    CB
  4283   4282   4292   4293      3    0.00000    0.00000    0.00000
0.00000    0.00000    0.00000 ;    CD-     N-    CA-    HA
  4283   4282   4292   4294      3    0.00000    0.00000    0.00000
0.00000    0.00000    0.00000 ;    CD-     N-    CA-     C
  4283   4286   4289   4290      3    0.66944    2.00832    0.00000
-2.67776    0.00000    0.00000 ;    CD-    CG-    CB-   HB2
  4283   4286   4289   4291      3    0.66944    2.00832    0.00000
-2.67776    0.00000    0.00000 ;    CD-    CG-    CB-   HB3
  4283   4286   4289   4292      3    3.68192    3.09616   -2.09200
-3.01248    0.00000    0.00000 ;    CD-    CG-    CB-    CA
  4284   4283   4282   4292      3    0.00000    0.00000    0.00000
```

```
0.00000    0.00000    0.00000 ;    HD2-    CD-     N-    CA
  4284   4283   4286   4287      3    0.65084    1.95253    0.00000
-2.60338    0.00000    0.00000 ;    HD2-    CD-    CG-   HG2
  4284   4283   4286   4288      3    0.65084    1.95253    0.00000
-2.60338    0.00000    0.00000 ;    HD2-    CD-    CG-   HG3
  4284   4283   4286   4289      3    0.65084    1.95253    0.00000
-2.60338    0.00000    0.00000 ;    HD2-    CD-    CG-    CB
  4285   4283   4282   4292      3    0.00000    0.00000    0.00000
0.00000    0.00000    0.00000 ;    HD3-    CD-     N-    CA
  4285   4283   4286   4287      3    0.65084    1.95253    0.00000
-2.60338    0.00000    0.00000 ;    HD3-    CD-    CG-   HG2
  4285   4283   4286   4288      3    0.65084    1.95253    0.00000
-2.60338    0.00000    0.00000 ;    HD3-    CD-    CG-   HG3
  4285   4283   4286   4289      3    0.65084    1.95253    0.00000
-2.60338    0.00000    0.00000 ;    HD3-    CD-    CG-    CB
  4286   4283   4282   4292      3    0.00000    0.00000    0.00000
0.00000    0.00000    0.00000 ;     CG-    CD-     N-    CA
  4286   4289   4292   4293      3    0.65084    1.95253    0.00000
-2.60338    0.00000    0.00000 ;     CG-    CB-    CA-    HA
  4286   4289   4292   4294      3    0.65084    1.95253    0.00000
-2.60338    0.00000    0.00000 ;     CG-    CB-    CA-     C
  4287   4286   4289   4290      3    0.62760    1.88280    0.00000
-2.51040    0.00000    0.00000 ;    HG2-    CG-    CB-   HB2
  4287   4286   4289   4291      3    0.62760    1.88280    0.00000
-2.51040    0.00000    0.00000 ;    HG2-    CG-    CB-   HB3
  4287   4286   4289   4292      3    0.66944    2.00832    0.00000
-2.67776    0.00000    0.00000 ;    HG2-    CG-    CB-    CA
  4288   4286   4289   4290      3    0.62760    1.88280    0.00000
-2.51040    0.00000    0.00000 ;    HG3-    CG-    CB-   HB2
  4288   4286   4289   4291      3    0.62760    1.88280    0.00000
-2.51040    0.00000    0.00000 ;    HG3-    CG-    CB-   HB3
  4288   4286   4289   4292      3    0.66944    2.00832    0.00000
-2.67776    0.00000    0.00000 ;    HG3-    CG-    CB-    CA
  4289   4292   4294   4295      3    0.00000    0.00000    0.00000
0.00000    0.00000    0.00000 ;     CB-    CA-     C-     O
  4289   4292   4294   4296      3    2.51040    4.18400    1.67360
-6.69440    0.00000    0.00000 ;     CB-    CA-     C-     N
  4290   4289   4292   4293      3    0.65084    1.95253    0.00000
-2.60338    0.00000    0.00000 ;    HB2-    CB-    CA-    HA
  4290   4289   4292   4294      3    0.65084    1.95253    0.00000
-2.60338    0.00000    0.00000 ;    HB2-    CB-    CA-     C
  4291   4289   4292   4293      3    0.65084    1.95253    0.00000
-2.60338    0.00000    0.00000 ;    HB3-    CB-    CA-    HA
  4291   4289   4292   4294      3    0.65084    1.95253    0.00000
-2.60338    0.00000    0.00000 ;    HB3-    CB-    CA-     C
  4292   4294   4296   4297      3   20.92000    0.00000  -20.92000
0.00000    0.00000    0.00000 ;     CA-     C-     N-     H
  4292   4294   4296   4298      3   20.92000    0.00000  -20.92000
0.00000    0.00000    0.00000 ;     CA-     C-     N-    CA
  4293   4292   4294   4295      3    3.68192   -4.35136    0.00000
1.33888    0.00000    0.00000 ;     HA-    CA-     C-     O
  4293   4292   4294   4296      3    0.00000    0.00000    0.00000
```

0.00000 0.00000 0.00000 ; HA- CA- C- N
4294 4296 4298 4299 3 0.00000 0.00000 0.00000
0.00000 0.00000 0.00000 ; C- N- CA- HA
4294 4296 4298 4300 3 11.71520 1.67360 15.06240 -
13.38880 0.00000 0.00000 ; C- N- CA- CB
4294 4296 4298 4318 3 1.75728 5.27184 2.25936
-7.02912 0.00000 0.00000 ; C- N- CA- C
4295 4294 4296 4297 3 29.28800 -8.36800 -20.92000
0.00000 0.00000 0.00000 ; O- C- N- H
4295 4294 4296 4298 3 20.92000 0.00000 -20.92000
0.00000 0.00000 0.00000 ; O- C- N- CA
4296 4298 4300 4301 3 0.65084 1.95253 0.00000
-2.60338 0.00000 0.00000 ; N- CA- CB- HB2
4296 4298 4300 4302 3 0.65084 1.95253 0.00000
-2.60338 0.00000 0.00000 ; N- CA- CB- HB3
4296 4298 4300 4303 3 4.18818 2.60663 -3.65682
-3.91622 1.03763 0.00000 ; N- CA- CB- CG
4296 4298 4318 4319 3 0.00000 0.00000 0.00000
0.00000 0.00000 0.00000 ; N- CA- C- O
4296 4298 4318 4320 3 17.40544 -5.02080 -13.22144
9.20480 0.00000 0.00000 ; N- CA- C- N
4297 4296 4298 4299 3 0.00000 0.00000 0.00000
0.00000 0.00000 0.00000 ; H- N- CA- HA
4297 4296 4298 4300 3 0.00000 0.00000 0.00000
0.00000 0.00000 0.00000 ; H- N- CA- CB
4297 4296 4298 4318 3 0.00000 0.00000 0.00000
0.00000 0.00000 0.00000 ; H- N- CA- C
4298 4300 4303 4304 3 0.00000 0.00000 0.00000
0.00000 0.00000 0.00000 ; CA- CB- CG- CD1
4298 4300 4303 4317 3 4.95386 8.75293 6.59398 -
13.70678 -3.17984 0.00000 ; CA- CB- CG- CD2
4298 4318 4320 4321 3 20.92000 0.00000 -20.92000
0.00000 0.00000 0.00000 ; CA- C- N- H
4298 4318 4320 4322 3 20.92000 0.00000 -20.92000
0.00000 0.00000 0.00000 ; CA- C- N- CA
4299 4298 4300 4301 3 0.65084 1.95253 0.00000
-2.60338 0.00000 0.00000 ; HA- CA- CB- HB2
4299 4298 4300 4302 3 0.65084 1.95253 0.00000
-2.60338 0.00000 0.00000 ; HA- CA- CB- HB3
4299 4298 4300 4303 3 0.65084 1.95253 0.00000
-2.60338 0.00000 0.00000 ; HA- CA- CB- CG
4299 4298 4318 4319 3 3.68192 -4.35136 0.00000
1.33888 0.00000 0.00000 ; HA- CA- C- O
4299 4298 4318 4320 3 0.00000 0.00000 0.00000
0.00000 0.00000 0.00000 ; HA- CA- C- N
4300 4298 4318 4319 3 0.00000 0.00000 0.00000
0.00000 0.00000 0.00000 ; CB- CA- C- O
4300 4298 4318 4320 3 2.51040 4.18400 1.67360
-6.69440 0.00000 0.00000 ; CB- CA- C- N
4300 4303 4304 4305 3 54.60120 0.00000 -54.60120
0.00000 0.00000 0.00000 ; CB- CG- CD1- HD1
4300 4303 4304 4306 3 54.60120 0.00000 -54.60120

```
0.00000    0.00000    0.00000 ;     CB-    CG-    CD1-   NE1
  4300   4303   4317   4308      3   14.01640    0.00000  -14.01640
0.00000    0.00000    0.00000 ;     CB-    CG-    CD2-   CE2
  4300   4303   4317   4315      3   14.01640    0.00000  -14.01640
0.00000    0.00000    0.00000 ;     CB-    CG-    CD2-   CE3
  4301   4300   4298   4318      3    0.65084    1.95253    0.00000
-2.60338    0.00000    0.00000 ;    HB2-    CB-    CA-     C
  4301   4300   4303   4304      3    0.00000    0.00000    0.00000
0.00000    0.00000    0.00000 ;    HB2-    CB-    CG-   CD1
  4301   4300   4303   4317      3    0.00000    0.00000    0.00000
0.00000    0.00000    0.00000 ;    HB2-    CB-    CG-   CD2
  4302   4300   4298   4318      3    0.65084    1.95253    0.00000
-2.60338    0.00000    0.00000 ;    HB3-    CB-    CA-     C
  4302   4300   4303   4304      3    0.00000    0.00000    0.00000
0.00000    0.00000    0.00000 ;    HB3-    CB-    CG-   CD1
  4302   4300   4303   4317      3    0.00000    0.00000    0.00000
0.00000    0.00000    0.00000 ;    HB3-    CB-    CG-   CD2
  4303   4300   4298   4318      3    4.62332    3.00830   -5.43083
-3.91622    2.47693    0.00000 ;     CG-    CB-    CA-     C
  4303   4304   4306   4307      3   12.55200    0.00000  -12.55200
0.00000    0.00000    0.00000 ;     CG-   CD1-   NE1-   HE1
  4303   4304   4306   4308      3   12.55200    0.00000  -12.55200
0.00000    0.00000    0.00000 ;     CG-   CD1-   NE1-   CE2
  4303   4317   4308   4306      3   25.10400    0.00000  -25.10400
0.00000    0.00000    0.00000 ;     CG-   CD2-   CE2-   NE1
  4303   4317   4308   4309      3   25.10400    0.00000  -25.10400
0.00000    0.00000    0.00000 ;     CG-   CD2-   CE2-   CZ2
  4303   4317   4315   4313      3   29.28800    0.00000  -29.28800
0.00000    0.00000    0.00000 ;     CG-   CD2-   CE3-   CZ3
  4303   4317   4315   4316      3   29.28800    0.00000  -29.28800
0.00000    0.00000    0.00000 ;     CG-   CD2-   CE3-   HE3
  4304   4303   4317   4308      3   14.01640    0.00000  -14.01640
0.00000    0.00000    0.00000 ;    CD1-    CG-   CD2-   CE2
  4304   4303   4317   4315      3   14.01640    0.00000  -14.01640
0.00000    0.00000    0.00000 ;    CD1-    CG-   CD2-   CE3
  4304   4306   4308   4309      3   12.76120    0.00000  -12.76120
0.00000    0.00000    0.00000 ;    CD1-   NE1-   CE2-   CZ2
  4304   4306   4308   4317      3   12.76120    0.00000  -12.76120
0.00000    0.00000    0.00000 ;    CD1-   NE1-   CE2-   CD2
  4305   4304   4303   4317      3   54.60120    0.00000  -54.60120
0.00000    0.00000    0.00000 ;    HD1-   CD1-    CG-   CD2
  4305   4304   4306   4307      3   12.55200    0.00000  -12.55200
0.00000    0.00000    0.00000 ;    HD1-   CD1-   NE1-   HE1
  4305   4304   4306   4308      3   12.55200    0.00000  -12.55200
0.00000    0.00000    0.00000 ;    HD1-   CD1-   NE1-   CE2
  4306   4304   4303   4317      3   54.60120    0.00000  -54.60120
0.00000    0.00000    0.00000 ;    NE1-   CD1-    CG-   CD2
  4306   4308   4309   4310      3   30.33400    0.00000  -30.33400
0.00000    0.00000    0.00000 ;    NE1-   CE2-   CZ2-   HZ2
  4306   4308   4309   4311      3   30.33400    0.00000  -30.33400
0.00000    0.00000    0.00000 ;    NE1-   CE2-   CZ2-   CH2
  4306   4308   4317   4315      3   25.10400    0.00000  -25.10400
```

0.00000 0.00000 0.00000 ; NE1- CE2- CD2- CE3
4307 4306 4308 4309 3 12.76120 0.00000 -12.76120
0.00000 0.00000 0.00000 ; HE1- NE1- CE2- CZ2
4307 4306 4308 4317 3 12.76120 0.00000 -12.76120
0.00000 0.00000 0.00000 ; HE1- NE1- CE2- CD2
4308 4309 4311 4312 3 30.33400 0.00000 -30.33400
0.00000 0.00000 0.00000 ; CE2- CZ2- CH2- HH2
4308 4309 4311 4313 3 30.33400 0.00000 -30.33400
0.00000 0.00000 0.00000 ; CE2- CZ2- CH2- CZ3
4308 4317 4315 4313 3 29.28800 0.00000 -29.28800
0.00000 0.00000 0.00000 ; CE2- CD2- CE3- CZ3
4308 4317 4315 4316 3 29.28800 0.00000 -29.28800
0.00000 0.00000 0.00000 ; CE2- CD2- CE3- HE3
4309 4308 4317 4315 3 25.10400 0.00000 -25.10400
0.00000 0.00000 0.00000 ; CZ2- CE2- CD2- CE3
4309 4311 4313 4314 3 30.33400 0.00000 -30.33400
0.00000 0.00000 0.00000 ; CZ2- CH2- CZ3- HZ3
4309 4311 4313 4315 3 30.33400 0.00000 -30.33400
0.00000 0.00000 0.00000 ; CZ2- CH2- CZ3- CE3
4310 4309 4308 4317 3 30.33400 0.00000 -30.33400
0.00000 0.00000 0.00000 ; HZ2- CZ2- CE2- CD2
4310 4309 4311 4312 3 30.33400 0.00000 -30.33400
0.00000 0.00000 0.00000 ; HZ2- CZ2- CH2- HH2
4310 4309 4311 4313 3 30.33400 0.00000 -30.33400
0.00000 0.00000 0.00000 ; HZ2- CZ2- CH2- CZ3
4311 4309 4308 4317 3 30.33400 0.00000 -30.33400
0.00000 0.00000 0.00000 ; CH2- CZ2- CE2- CD2
4311 4313 4315 4316 3 30.33400 0.00000 -30.33400
0.00000 0.00000 0.00000 ; CH2- CZ3- CE3- HE3
4311 4313 4315 4317 3 30.33400 0.00000 -30.33400
0.00000 0.00000 0.00000 ; CH2- CZ3- CE3- CD2
4312 4311 4313 4314 3 30.33400 0.00000 -30.33400
0.00000 0.00000 0.00000 ; HH2- CH2- CZ3- HZ3
4312 4311 4313 4315 3 30.33400 0.00000 -30.33400
0.00000 0.00000 0.00000 ; HH2- CH2- CZ3- CE3
4314 4313 4315 4316 3 30.33400 0.00000 -30.33400
0.00000 0.00000 0.00000 ; HZ3- CZ3- CE3- HE3
4314 4313 4315 4317 3 30.33400 0.00000 -30.33400
0.00000 0.00000 0.00000 ; HZ3- CZ3- CE3- CD2
4318 4320 4322 4323 3 0.00000 0.00000 0.00000
0.00000 0.00000 0.00000 ; C- N- CA- HA
4318 4320 4322 4324 3 11.71520 1.67360 15.06240 -
13.38880 0.00000 0.00000 ; C- N- CA- CB
4318 4320 4322 4337 3 1.75728 5.27184 2.25936
-7.02912 0.00000 0.00000 ; C- N- CA- C
4319 4318 4320 4321 3 29.28800 -8.36800 -20.92000
0.00000 0.00000 0.00000 ; O- C- N- H
4319 4318 4320 4322 3 20.92000 0.00000 -20.92000
0.00000 0.00000 0.00000 ; O- C- N- CA
4320 4322 4324 4325 3 0.65084 1.95253 0.00000
-2.60338 0.00000 0.00000 ; N- CA- CB- HB
4320 4322 4324 4326 3 3.83673 0.44769 -1.77402

-2.47693 -0.03347 0.00000 ; N- CA- CB- CG2
4320 4322 4324 4330 3 2.97482 0.12134 2.04179
-1.89117 -3.24678 0.00000 ; N- CA- CB- CG1
4320 4322 4337 4338 3 0.00000 0.00000 0.00000
0.00000 0.00000 0.00000 ; N- CA- C- O
4320 4322 4337 4339 3 17.40544 -5.02080 -13.22144
9.20480 0.00000 0.00000 ; N- CA- C- N
4321 4320 4322 4323 3 0.00000 0.00000 0.00000
0.00000 0.00000 0.00000 ; H- N- CA- HA
4321 4320 4322 4324 3 0.00000 0.00000 0.00000
0.00000 0.00000 0.00000 ; H- N- CA- CB
4321 4320 4322 4337 3 0.00000 0.00000 0.00000
0.00000 0.00000 0.00000 ; H- N- CA- C
4322 4324 4326 4327 3 0.66944 2.00832 0.00000
-2.67776 0.00000 0.00000 ; CA- CB- CG2- HG21
4322 4324 4326 4328 3 0.66944 2.00832 0.00000
-2.67776 0.00000 0.00000 ; CA- CB- CG2- HG22
4322 4324 4326 4329 3 0.66944 2.00832 0.00000
-2.67776 0.00000 0.00000 ; CA- CB- CG2- HG23
4322 4324 4330 4331 3 0.66944 2.00832 0.00000
-2.67776 0.00000 0.00000 ; CA- CB- CG1- HG12
4322 4324 4330 4332 3 0.66944 2.00832 0.00000
-2.67776 0.00000 0.00000 ; CA- CB- CG1- HG13
4322 4324 4330 4333 3 4.24258 -0.52718 -7.25506
-1.79075 7.69856 0.00000 ; CA- CB- CG1- CD1
4322 4337 4339 4340 3 20.92000 0.00000 -20.92000
0.00000 0.00000 0.00000 ; CA- C- N- H
4322 4337 4339 4341 3 20.92000 0.00000 -20.92000
0.00000 0.00000 0.00000 ; CA- C- N- CA
4323 4322 4324 4325 3 0.65084 1.95253 0.00000
-2.60338 0.00000 0.00000 ; HA- CA- CB- HB
4323 4322 4324 4326 3 0.65084 1.95253 0.00000
-2.60338 0.00000 0.00000 ; HA- CA- CB- CG2
4323 4322 4324 4330 3 0.65084 1.95253 0.00000
-2.60338 0.00000 0.00000 ; HA- CA- CB- CG1
4323 4322 4337 4338 3 3.68192 -4.35136 0.00000
1.33888 0.00000 0.00000 ; HA- CA- C- O
4323 4322 4337 4339 3 0.00000 0.00000 0.00000
0.00000 0.00000 0.00000 ; HA- CA- C- N
4324 4322 4337 4338 3 0.00000 0.00000 0.00000
0.00000 0.00000 0.00000 ; CB- CA- C- O
4324 4322 4337 4339 3 2.51040 4.18400 1.67360
-6.69440 0.00000 0.00000 ; CB- CA- C- N
4324 4330 4333 4334 3 0.66944 2.00832 0.00000
-2.67776 0.00000 0.00000 ; CB- CG1- CD1- HD11
4324 4330 4333 4335 3 0.66944 2.00832 0.00000
-2.67776 0.00000 0.00000 ; CB- CG1- CD1- HD12
4324 4330 4333 4336 3 0.66944 2.00832 0.00000
-2.67776 0.00000 0.00000 ; CB- CG1- CD1- HD13
4325 4324 4322 4337 3 0.65084 1.95253 0.00000
-2.60338 0.00000 0.00000 ; HB- CB- CA- C
4325 4324 4326 4327 3 0.62760 1.88280 0.00000

-2.51040 0.00000 0.00000 ; HB- CB- CG2- HG21
4325 4324 4326 4328 3 0.62760 1.88280 0.00000
-2.51040 0.00000 0.00000 ; HB- CB- CG2- HG22
4325 4324 4326 4329 3 0.62760 1.88280 0.00000
-2.51040 0.00000 0.00000 ; HB- CB- CG2- HG23
4325 4324 4330 4331 3 0.62760 1.88280 0.00000
-2.51040 0.00000 0.00000 ; HB- CB- CG1- HG12
4325 4324 4330 4332 3 0.62760 1.88280 0.00000
-2.51040 0.00000 0.00000 ; HB- CB- CG1- HG13
4325 4324 4330 4333 3 0.66944 2.00832 0.00000
-2.67776 0.00000 0.00000 ; HB- CB- CG1- CD1
4326 4324 4322 4337 3 5.67350 3.55640 -6.16722
-2.47693 3.74886 0.00000 ; CG2- CB- CA- C
4326 4324 4330 4331 3 0.66944 2.00832 0.00000
-2.67776 0.00000 0.00000 ; CG2- CB- CG1- HG12
4326 4324 4330 4332 3 0.66944 2.00832 0.00000
-2.67776 0.00000 0.00000 ; CG2- CB- CG1- HG13
4326 4324 4330 4333 3 3.81162 0.49790 -8.14206
-1.79075 7.49773 0.00000 ; CG2- CB- CG1- CD1
4327 4326 4324 4330 3 0.66944 2.00832 0.00000
-2.67776 0.00000 0.00000 ; HG21- CG2- CB- CG1
4328 4326 4324 4330 3 0.66944 2.00832 0.00000
-2.67776 0.00000 0.00000 ; HG22- CG2- CB- CG1
4329 4326 4324 4330 3 0.66944 2.00832 0.00000
-2.67776 0.00000 0.00000 ; HG23- CG2- CB- CG1
4330 4324 4322 4337 3 8.26340 0.74057 -9.99976
-1.89117 3.84928 0.00000 ; CG1- CB- CA- C
4331 4330 4333 4334 3 0.62760 1.88280 0.00000
-2.51040 0.00000 0.00000 ; HG12- CG1- CD1- HD11
4331 4330 4333 4335 3 0.62760 1.88280 0.00000
-2.51040 0.00000 0.00000 ; HG12- CG1- CD1- HD12
4331 4330 4333 4336 3 0.62760 1.88280 0.00000
-2.51040 0.00000 0.00000 ; HG12- CG1- CD1- HD13
4332 4330 4333 4334 3 0.62760 1.88280 0.00000
-2.51040 0.00000 0.00000 ; HG13- CG1- CD1- HD11
4332 4330 4333 4335 3 0.62760 1.88280 0.00000
-2.51040 0.00000 0.00000 ; HG13- CG1- CD1- HD12
4332 4330 4333 4336 3 0.62760 1.88280 0.00000
-2.51040 0.00000 0.00000 ; HG13- CG1- CD1- HD13
4337 4339 4341 4342 3 0.00000 0.00000 0.00000
0.00000 0.00000 0.00000 ; C- N- CA- HA
4337 4339 4341 4343 3 11.71520 1.67360 15.06240 -
13.38880 0.00000 0.00000 ; C- N- CA- CB
4337 4339 4341 4351 3 1.75728 5.27184 2.25936
-7.02912 0.00000 0.00000 ; C- N- CA- C
4338 4337 4339 4340 3 29.28800 -8.36800 -20.92000
0.00000 0.00000 0.00000 ; O- C- N- H
4338 4337 4339 4341 3 20.92000 0.00000 -20.92000
0.00000 0.00000 0.00000 ; O- C- N- CA
4339 4341 4343 4344 3 0.65084 1.95253 0.00000
-2.60338 0.00000 0.00000 ; N- CA- CB- HB
4339 4341 4343 4345 3 3.83673 0.44769 -1.77402

-2.47693 -0.03347 0.00000 ; N- CA- CB- CG2
4339 4341 4343 4349 3 4.93294 1.13386 -3.12963
-5.27184 3.17984 0.00000 ; N- CA- CB- OG1
4339 4341 4351 4352 3 0.00000 0.00000 0.00000
0.00000 0.00000 0.00000 ; N- CA- C- O
4339 4341 4351 4353 3 17.40544 -5.02080 -13.22144
9.20480 0.00000 0.00000 ; N- CA- C- N
4340 4339 4341 4342 3 0.00000 0.00000 0.00000
0.00000 0.00000 0.00000 ; H- N- CA- HA
4340 4339 4341 4343 3 0.00000 0.00000 0.00000
0.00000 0.00000 0.00000 ; H- N- CA- CB
4340 4339 4341 4351 3 0.00000 0.00000 0.00000
0.00000 0.00000 0.00000 ; H- N- CA- C
4341 4343 4345 4346 3 0.66944 2.00832 0.00000
-2.67776 0.00000 0.00000 ; CA- CB- CG2- HG21
4341 4343 4345 4347 3 0.66944 2.00832 0.00000
-2.67776 0.00000 0.00000 ; CA- CB- CG2- HG22
4341 4343 4345 4348 3 0.66944 2.00832 0.00000
-2.67776 0.00000 0.00000 ; CA- CB- CG2- HG23
4341 4343 4349 4350 3 1.12131 2.98738 1.66523
-3.94970 0.43514 0.00000 ; CA- CB- OG1- HG1
4341 4351 4353 4354 3 20.92000 0.00000 -20.92000
0.00000 0.00000 0.00000 ; CA- C- N- H
4341 4351 4353 4355 3 20.92000 0.00000 -20.92000
0.00000 0.00000 0.00000 ; CA- C- N- CA
4342 4341 4343 4344 3 0.65084 1.95253 0.00000
-2.60338 0.00000 0.00000 ; HA- CA- CB- HB
4342 4341 4343 4345 3 0.65084 1.95253 0.00000
-2.60338 0.00000 0.00000 ; HA- CA- CB- CG2
4342 4341 4343 4349 3 1.04600 -1.04600 0.00000
0.00000 0.00000 0.00000 ; HA- CA- CB- OG1
4342 4341 4351 4352 3 3.68192 -4.35136 0.00000
1.33888 0.00000 0.00000 ; HA- CA- C- O
4342 4341 4351 4353 3 0.00000 0.00000 0.00000
0.00000 0.00000 0.00000 ; HA- CA- C- N
4343 4341 4351 4352 3 0.00000 0.00000 0.00000
0.00000 0.00000 0.00000 ; CB- CA- C- O
4343 4341 4351 4353 3 2.51040 4.18400 1.67360
-6.69440 0.00000 0.00000 ; CB- CA- C- N
4344 4343 4341 4351 3 0.65084 1.95253 0.00000
-2.60338 0.00000 0.00000 ; HB- CB- CA- C
4344 4343 4345 4346 3 0.65084 1.95253 0.00000
-2.60338 0.00000 0.00000 ; HB- CB- CG2- HG21
4344 4343 4345 4347 3 0.65084 1.95253 0.00000
-2.60338 0.00000 0.00000 ; HB- CB- CG2- HG22
4344 4343 4345 4348 3 0.65084 1.95253 0.00000
-2.60338 0.00000 0.00000 ; HB- CB- CG2- HG23
4344 4343 4349 4350 3 0.69733 2.09200 0.00000
-2.78933 0.00000 0.00000 ; HB- CB- OG1- HG1
4345 4343 4341 4351 3 5.67350 3.55640 -6.16722
-2.47693 3.74886 0.00000 ; CG2- CB- CA- C
4345 4343 4349 4350 3 4.74047 0.27196 -2.26773

-3.94970 1.60666 0.00000 ; CG2- CB- OG1- HG1
4346 4345 4343 4349 3 1.04600 -1.04600 0.00000
0.00000 0.00000 0.00000 ; HG21- CG2- CB- OG1
4347 4345 4343 4349 3 1.04600 -1.04600 0.00000
0.00000 0.00000 0.00000 ; HG22- CG2- CB- OG1
4348 4345 4343 4349 3 1.04600 -1.04600 0.00000
0.00000 0.00000 0.00000 ; HG23- CG2- CB- OG1
4349 4343 4341 4351 3 6.53541 6.87013 -6.21742
-5.27184 5.22163 0.00000 ; OG1- CB- CA- C
4351 4353 4355 4356 3 0.00000 0.00000 0.00000
0.00000 0.00000 0.00000 ; C- N- CA- HA
4351 4353 4355 4357 3 11.71520 1.67360 15.06240 -
13.38880 0.00000 0.00000 ; C- N- CA- CB
4351 4353 4355 4368 3 1.75728 5.27184 2.25936
-7.02912 0.00000 0.00000 ; C- N- CA- C
4352 4351 4353 4354 3 29.28800 -8.36800 -20.92000
0.00000 0.00000 0.00000 ; O- C- N- H
4352 4351 4353 4355 3 20.92000 0.00000 -20.92000
0.00000 0.00000 0.00000 ; O- C- N- CA
4353 4355 4357 4358 3 0.65084 1.95253 0.00000
-2.60338 0.00000 0.00000 ; N- CA- CB- HB2
4353 4355 4357 4359 3 0.65084 1.95253 0.00000
-2.60338 0.00000 0.00000 ; N- CA- CB- HB3
4353 4355 4357 4360 3 3.21331 2.22589 -4.15053
-2.40998 2.61082 0.00000 ; N- CA- CB- CG
4353 4355 4368 4369 3 0.00000 0.00000 0.00000
0.00000 0.00000 0.00000 ; N- CA- C- O
4353 4355 4368 4370 3 17.40544 -5.02080 -13.22144
9.20480 0.00000 0.00000 ; N- CA- C- N
4354 4353 4355 4356 3 0.00000 0.00000 0.00000
0.00000 0.00000 0.00000 ; H- N- CA- HA
4354 4353 4355 4357 3 0.00000 0.00000 0.00000
0.00000 0.00000 0.00000 ; H- N- CA- CB
4354 4353 4355 4368 3 0.00000 0.00000 0.00000
0.00000 0.00000 0.00000 ; H- N- CA- C
4355 4357 4360 4361 3 0.66944 2.00832 0.00000
-2.67776 0.00000 0.00000 ; CA- CB- CG- HG2
4355 4357 4360 4362 3 0.66944 2.00832 0.00000
-2.67776 0.00000 0.00000 ; CA- CB- CG- HG3
4355 4357 4360 4363 3 3.69866 -4.35136 -3.92459
6.89523 4.61914 0.00000 ; CA- CB- CG- CD
4355 4368 4370 4371 3 20.92000 0.00000 -20.92000
0.00000 0.00000 0.00000 ; CA- C- N- H
4355 4368 4370 4372 3 20.92000 0.00000 -20.92000
0.00000 0.00000 0.00000 ; CA- C- N- CA
4356 4355 4357 4358 3 0.65084 1.95253 0.00000
-2.60338 0.00000 0.00000 ; HA- CA- CB- HB2
4356 4355 4357 4359 3 0.65084 1.95253 0.00000
-2.60338 0.00000 0.00000 ; HA- CA- CB- HB3
4356 4355 4357 4360 3 0.65084 1.95253 0.00000
-2.60338 0.00000 0.00000 ; HA- CA- CB- CG
4356 4355 4368 4369 3 3.68192 -4.35136 0.00000

1.33888 0.00000 0.00000 ; HA- CA- C- O
4356 4355 4368 4370 3 0.00000 0.00000 0.00000
0.00000 0.00000 0.00000 ; HA- CA- C- N
4357 4355 4368 4369 3 0.00000 0.00000 0.00000
0.00000 0.00000 0.00000 ; CB- CA- C- O
4357 4355 4368 4370 3 2.51040 4.18400 1.67360
-6.69440 0.00000 0.00000 ; CB- CA- C- N
4357 4360 4363 4364 3 0.00000 0.00000 0.00000
0.00000 0.00000 0.00000 ; CB- CG- CD- OE1
4357 4360 4363 4365 3 10.32611 1.48114 -8.47678
1.42256 1.40582 0.00000 ; CB- CG- CD- NE2
4358 4357 4355 4368 3 0.65084 1.95253 0.00000
-2.60338 0.00000 0.00000 ; HB2- CB- CA- C
4358 4357 4360 4361 3 0.62760 1.88280 0.00000
-2.51040 0.00000 0.00000 ; HB2- CB- CG- HG2
4358 4357 4360 4362 3 0.62760 1.88280 0.00000
-2.51040 0.00000 0.00000 ; HB2- CB- CG- HG3
4358 4357 4360 4363 3 0.65084 1.95253 0.00000
-2.60338 0.00000 0.00000 ; HB2- CB- CG- CD
4359 4357 4355 4368 3 0.65084 1.95253 0.00000
-2.60338 0.00000 0.00000 ; HB3- CB- CA- C
4359 4357 4360 4361 3 0.62760 1.88280 0.00000
-2.51040 0.00000 0.00000 ; HB3- CB- CG- HG2
4359 4357 4360 4362 3 0.62760 1.88280 0.00000
-2.51040 0.00000 0.00000 ; HB3- CB- CG- HG3
4359 4357 4360 4363 3 0.65084 1.95253 0.00000
-2.60338 0.00000 0.00000 ; HB3- CB- CG- CD
4360 4357 4355 4368 3 6.86594 3.56895 -8.14206
-2.40998 4.85344 0.00000 ; CG- CB- CA- C
4360 4363 4365 4366 3 20.92000 0.00000 -20.92000
0.00000 0.00000 0.00000 ; CG- CD- NE2- HE21
4360 4363 4365 4367 3 20.92000 0.00000 -20.92000
0.00000 0.00000 0.00000 ; CG- CD- NE2- HE22
4361 4360 4363 4364 3 3.68192 -4.35136 0.00000
1.33888 0.00000 0.00000 ; HG2- CG- CD- OE1
4361 4360 4363 4365 3 0.00000 0.00000 0.00000
0.00000 0.00000 0.00000 ; HG2- CG- CD- NE2
4362 4360 4363 4364 3 3.68192 -4.35136 0.00000
1.33888 0.00000 0.00000 ; HG3- CG- CD- OE1
4362 4360 4363 4365 3 0.00000 0.00000 0.00000
0.00000 0.00000 0.00000 ; HG3- CG- CD- NE2
4364 4363 4365 4366 3 29.28800 -8.36800 -20.92000
0.00000 0.00000 0.00000 ; OE1- CD- NE2- HE21
4364 4363 4365 4367 3 29.28800 -8.36800 -20.92000
0.00000 0.00000 0.00000 ; OE1- CD- NE2- HE22
4368 4370 4372 4373 3 0.00000 0.00000 0.00000
0.00000 0.00000 0.00000 ; C- N- CA- HA
4368 4370 4372 4374 3 11.71520 1.67360 15.06240 -
13.38880 0.00000 0.00000 ; C- N- CA- CB
4368 4370 4372 4385 3 1.75728 5.27184 2.25936
-7.02912 0.00000 0.00000 ; C- N- CA- C
4369 4368 4370 4371 3 29.28800 -8.36800 -20.92000

0.00000 0.00000 0.00000 ; O- C- N- H
4369 4368 4370 4372 3 20.92000 0.00000 -20.92000
0.00000 0.00000 0.00000 ; O- C- N- CA
4370 4372 4374 4375 3 0.65084 1.95253 0.00000
-2.60338 0.00000 0.00000 ; N- CA- CB- HB2
4370 4372 4374 4376 3 0.65084 1.95253 0.00000
-2.60338 0.00000 0.00000 ; N- CA- CB- HB3
4370 4372 4374 4377 3 3.21331 2.22589 -4.15053
-2.40998 2.61082 0.00000 ; N- CA- CB- CG
4370 4372 4385 4386 3 0.00000 0.00000 0.00000
0.00000 0.00000 0.00000 ; N- CA- C- O
4370 4372 4385 4387 3 17.40544 -5.02080 -13.22144
9.20480 0.00000 0.00000 ; N- CA- C- N
4371 4370 4372 4373 3 0.00000 0.00000 0.00000
0.00000 0.00000 0.00000 ; H- N- CA- HA
4371 4370 4372 4374 3 0.00000 0.00000 0.00000
0.00000 0.00000 0.00000 ; H- N- CA- CB
4371 4370 4372 4385 3 0.00000 0.00000 0.00000
0.00000 0.00000 0.00000 ; H- N- CA- C
4372 4374 4377 4378 3 0.65084 1.95253 0.00000
-2.60338 0.00000 0.00000 ; CA- CB- CG- HG2
4372 4374 4377 4379 3 0.65084 1.95253 0.00000
-2.60338 0.00000 0.00000 ; CA- CB- CG- HG3
4372 4374 4377 4380 3 2.04598 -1.54390 1.11294
-0.26778 0.93722 0.00000 ; CA- CB- CG- SD
4372 4385 4387 4388 3 20.92000 0.00000 -20.92000
0.00000 0.00000 0.00000 ; CA- C- N- H
4372 4385 4387 4389 3 20.92000 0.00000 -20.92000
0.00000 0.00000 0.00000 ; CA- C- N- CA
4373 4372 4374 4375 3 0.65084 1.95253 0.00000
-2.60338 0.00000 0.00000 ; HA- CA- CB- HB2
4373 4372 4374 4376 3 0.65084 1.95253 0.00000
-2.60338 0.00000 0.00000 ; HA- CA- CB- HB3
4373 4372 4374 4377 3 0.65084 1.95253 0.00000
-2.60338 0.00000 0.00000 ; HA- CA- CB- CG
4373 4372 4385 4386 3 3.68192 -4.35136 0.00000
1.33888 0.00000 0.00000 ; HA- CA- C- O
4373 4372 4385 4387 3 0.00000 0.00000 0.00000
0.00000 0.00000 0.00000 ; HA- CA- C- N
4374 4372 4385 4386 3 0.00000 0.00000 0.00000
0.00000 0.00000 0.00000 ; CB- CA- C- O
4374 4372 4385 4387 3 2.51040 4.18400 1.67360
-6.69440 0.00000 0.00000 ; CB- CA- C- N
4374 4377 4380 4381 3 3.24260 6.22998 1.79075
-6.92870 1.90790 0.00000 ; CB- CG- SD- CE
4375 4374 4372 4385 3 0.65084 1.95253 0.00000
-2.60338 0.00000 0.00000 ; HB2- CB- CA- C
4375 4374 4377 4378 3 0.65084 1.95253 0.00000
-2.60338 0.00000 0.00000 ; HB2- CB- CG- HG2
4375 4374 4377 4379 3 0.65084 1.95253 0.00000
-2.60338 0.00000 0.00000 ; HB2- CB- CG- HG3
4375 4374 4377 4380 3 0.65084 1.95253 0.00000

-2.60338 0.00000 0.00000 ; HB2- CB- CG- SD
4376 4374 4372 4385 3 0.65084 1.95253 0.00000
-2.60338 0.00000 0.00000 ; HB3- CB- CA- C
4376 4374 4377 4378 3 0.65084 1.95253 0.00000
-2.60338 0.00000 0.00000 ; HB3- CB- CG- HG2
4376 4374 4377 4379 3 0.65084 1.95253 0.00000
-2.60338 0.00000 0.00000 ; HB3- CB- CG- HG3
4376 4374 4377 4380 3 0.65084 1.95253 0.00000
-2.60338 0.00000 0.00000 ; HB3- CB- CG- SD
4377 4374 4372 4385 3 6.86594 3.56895 -8.14206
-2.40998 4.85344 0.00000 ; CG- CB- CA- C
4377 4380 4381 4382 3 1.39467 4.18400 0.00000
-5.57867 0.00000 0.00000 ; CG- SD- CE- HE1
4377 4380 4381 4383 3 1.39467 4.18400 0.00000
-5.57867 0.00000 0.00000 ; CG- SD- CE- HE2
4377 4380 4381 4384 3 1.39467 4.18400 0.00000
-5.57867 0.00000 0.00000 ; CG- SD- CE- HE3
4378 4377 4380 4381 3 1.39467 4.18400 0.00000
-5.57867 0.00000 0.00000 ; HG2- CG- SD- CE
4379 4377 4380 4381 3 1.39467 4.18400 0.00000
-5.57867 0.00000 0.00000 ; HG3- CG- SD- CE
4385 4387 4389 4390 3 0.00000 0.00000 0.00000
0.00000 0.00000 0.00000 ; C- N- CA- HA
4385 4387 4389 4391 3 11.71520 1.67360 15.06240 -
13.38880 0.00000 0.00000 ; C- N- CA- CB
4385 4387 4389 4399 3 1.75728 5.27184 2.25936
-7.02912 0.00000 0.00000 ; C- N- CA- C
4386 4385 4387 4388 3 29.28800 -8.36800 -20.92000
0.00000 0.00000 0.00000 ; O- C- N- H
4386 4385 4387 4389 3 20.92000 0.00000 -20.92000
0.00000 0.00000 0.00000 ; O- C- N- CA
4387 4389 4391 4392 3 0.65084 1.95253 0.00000
-2.60338 0.00000 0.00000 ; N- CA- CB- HB
4387 4389 4391 4393 3 3.83673 0.44769 -1.77402
-2.47693 -0.03347 0.00000 ; N- CA- CB- CG2
4387 4389 4391 4397 3 4.93294 1.13386 -3.12963
-5.27184 3.17984 0.00000 ; N- CA- CB- OG1
4387 4389 4399 4400 3 0.00000 0.00000 0.00000
0.00000 0.00000 0.00000 ; N- CA- C- O
4387 4389 4399 4401 3 17.40544 -5.02080 -13.22144
9.20480 0.00000 0.00000 ; N- CA- C- N
4388 4387 4389 4390 3 0.00000 0.00000 0.00000
0.00000 0.00000 0.00000 ; H- N- CA- HA
4388 4387 4389 4391 3 0.00000 0.00000 0.00000
0.00000 0.00000 0.00000 ; H- N- CA- CB
4388 4387 4389 4399 3 0.00000 0.00000 0.00000
0.00000 0.00000 0.00000 ; H- N- CA- C
4389 4391 4393 4394 3 0.66944 2.00832 0.00000
-2.67776 0.00000 0.00000 ; CA- CB- CG2- HG21
4389 4391 4393 4395 3 0.66944 2.00832 0.00000
-2.67776 0.00000 0.00000 ; CA- CB- CG2- HG22
4389 4391 4393 4396 3 0.66944 2.00832 0.00000

-2.67776 0.00000 0.00000 ; CA- CB- CG2- HG23
4389 4391 4397 4398 3 1.12131 2.98738 1.66523
-3.94970 0.43514 0.00000 ; CA- CB- OG1- HG1
4389 4399 4401 4402 3 20.92000 0.00000 -20.92000
0.00000 0.00000 0.00000 ; CA- C- N- H
4389 4399 4401 4403 3 20.92000 0.00000 -20.92000
0.00000 0.00000 0.00000 ; CA- C- N- CA
4390 4389 4391 4392 3 0.65084 1.95253 0.00000
-2.60338 0.00000 0.00000 ; HA- CA- CB- HB
4390 4389 4391 4393 3 0.65084 1.95253 0.00000
-2.60338 0.00000 0.00000 ; HA- CA- CB- CG2
4390 4389 4391 4397 3 1.04600 -1.04600 0.00000
0.00000 0.00000 0.00000 ; HA- CA- CB- OG1
4390 4389 4399 4400 3 3.68192 -4.35136 0.00000
1.33888 0.00000 0.00000 ; HA- CA- C- O
4390 4389 4399 4401 3 0.00000 0.00000 0.00000
0.00000 0.00000 0.00000 ; HA- CA- C- N
4391 4389 4399 4400 3 0.00000 0.00000 0.00000
0.00000 0.00000 0.00000 ; CB- CA- C- O
4391 4389 4399 4401 3 2.51040 4.18400 1.67360
-6.69440 0.00000 0.00000 ; CB- CA- C- N
4392 4391 4389 4399 3 0.65084 1.95253 0.00000
-2.60338 0.00000 0.00000 ; HB- CB- CA- C
4392 4391 4393 4394 3 0.65084 1.95253 0.00000
-2.60338 0.00000 0.00000 ; HB- CB- CG2- HG21
4392 4391 4393 4395 3 0.65084 1.95253 0.00000
-2.60338 0.00000 0.00000 ; HB- CB- CG2- HG22
4392 4391 4393 4396 3 0.65084 1.95253 0.00000
-2.60338 0.00000 0.00000 ; HB- CB- CG2- HG23
4392 4391 4397 4398 3 0.69733 2.09200 0.00000
-2.78933 0.00000 0.00000 ; HB- CB- OG1- HG1
4393 4391 4389 4399 3 5.67350 3.55640 -6.16722
-2.47693 3.74886 0.00000 ; CG2- CB- CA- C
4393 4391 4397 4398 3 4.74047 0.27196 -2.26773
-3.94970 1.60666 0.00000 ; CG2- CB- OG1- HG1
4394 4393 4391 4397 3 1.04600 -1.04600 0.00000
0.00000 0.00000 0.00000 ; HG21- CG2- CB- OG1
4395 4393 4391 4397 3 1.04600 -1.04600 0.00000
0.00000 0.00000 0.00000 ; HG22- CG2- CB- OG1
4396 4393 4391 4397 3 1.04600 -1.04600 0.00000
0.00000 0.00000 0.00000 ; HG23- CG2- CB- OG1
4397 4391 4389 4399 3 6.53541 6.87013 -6.21742
-5.27184 5.22163 0.00000 ; OG1- CB- CA- C
4399 4401 4403 4404 3 0.00000 0.00000 0.00000
0.00000 0.00000 0.00000 ; C- N- CA- HA
4399 4401 4403 4405 3 11.71520 1.67360 15.06240 -
13.38880 0.00000 0.00000 ; C- N- CA- CB
4399 4401 4403 4421 3 1.75728 5.27184 2.25936
-7.02912 0.00000 0.00000 ; C- N- CA- C
4400 4399 4401 4402 3 29.28800 -8.36800 -20.92000
0.00000 0.00000 0.00000 ; O- C- N- H
4400 4399 4401 4403 3 20.92000 0.00000 -20.92000

```
0.00000    0.00000    0.00000 ;       O-     C-     N-    CA
  4401   4403   4405   4406       3    0.65084    1.95253    0.00000
-2.60338    0.00000    0.00000 ;       N-    CA-    CB-   HB2
  4401   4403   4405   4407       3    0.65084    1.95253    0.00000
-2.60338    0.00000    0.00000 ;       N-    CA-    CB-   HB3
  4401   4403   4405   4408       3    0.65084    1.95253    0.00000
-2.60338    0.00000    0.00000 ;       N-    CA-    CB-    CG
  4401   4403   4421   4422       3    0.00000    0.00000    0.00000
0.00000    0.00000    0.00000 ;       N-    CA-     C-     O
  4401   4403   4421   4423       3   17.40544   -5.02080  -13.22144
9.20480    0.00000    0.00000 ;       N-    CA-     C-     N
  4402   4401   4403   4404       3    0.00000    0.00000    0.00000
0.00000    0.00000    0.00000 ;       H-     N-    CA-    HA
  4402   4401   4403   4405       3    0.00000    0.00000    0.00000
0.00000    0.00000    0.00000 ;       H-     N-    CA-    CB
  4402   4401   4403   4421       3    0.00000    0.00000    0.00000
0.00000    0.00000    0.00000 ;       H-     N-    CA-     C
  4403   4405   4408   4409       3    0.66944    2.00832    0.00000
-2.67776    0.00000    0.00000 ;      CA-    CB-    CG-   HG2
  4403   4405   4408   4410       3    0.66944    2.00832    0.00000
-2.67776    0.00000    0.00000 ;      CA-    CB-    CG-   HG3
  4403   4405   4408   4411       3    3.68192    3.09616   -2.09200
-3.01248    0.00000    0.00000 ;      CA-    CB-    CG-    CD
  4403   4421   4423   4424       3   20.92000    0.00000  -20.92000
0.00000    0.00000    0.00000 ;      CA-     C-     N-     H
  4403   4421   4423   4425       3   20.92000    0.00000  -20.92000
0.00000    0.00000    0.00000 ;      CA-     C-     N-    CA
  4404   4403   4405   4406       3    0.65084    1.95253    0.00000
-2.60338    0.00000    0.00000 ;      HA-    CA-    CB-   HB2
  4404   4403   4405   4407       3    0.65084    1.95253    0.00000
-2.60338    0.00000    0.00000 ;      HA-    CA-    CB-   HB3
  4404   4403   4405   4408       3    0.65084    1.95253    0.00000
-2.60338    0.00000    0.00000 ;      HA-    CA-    CB-    CG
  4404   4403   4421   4422       3    3.68192   -4.35136    0.00000
1.33888    0.00000    0.00000 ;      HA-    CA-     C-     O
  4404   4403   4421   4423       3    0.00000    0.00000    0.00000
0.00000    0.00000    0.00000 ;      HA-    CA-     C-     N
  4405   4403   4421   4422       3    0.00000    0.00000    0.00000
0.00000    0.00000    0.00000 ;      CB-    CA-     C-     O
  4405   4403   4421   4423       3    2.51040    4.18400    1.67360
-6.69440    0.00000    0.00000 ;      CB-    CA-     C-     N
  4405   4408   4411   4412       3    0.66944    2.00832    0.00000
-2.67776    0.00000    0.00000 ;      CB-    CG-    CD-   HD2
  4405   4408   4411   4413       3    0.66944    2.00832    0.00000
-2.67776    0.00000    0.00000 ;      CB-    CG-    CD-   HD3
  4405   4408   4411   4414       3    3.68192    3.09616   -2.09200
-3.01248    0.00000    0.00000 ;      CB-    CG-    CD-    CE
  4406   4405   4403   4421       3    0.65084    1.95253    0.00000
-2.60338    0.00000    0.00000 ;     HB2-    CB-    CA-     C
  4406   4405   4408   4409       3    0.62760    1.88280    0.00000
-2.51040    0.00000    0.00000 ;     HB2-    CB-    CG-   HG2
  4406   4405   4408   4410       3    0.62760    1.88280    0.00000
```

```
-2.51040    0.00000    0.00000 ;    HB2-    CB-    CG-   HG3
 4406   4405   4408   4411      3    0.66944    2.00832    0.00000
-2.67776    0.00000    0.00000 ;    HB2-    CB-    CG-    CD
 4407   4405   4403   4421      3    0.65084    1.95253    0.00000
-2.60338    0.00000    0.00000 ;    HB3-    CB-    CA-     C
 4407   4405   4408   4409      3    0.62760    1.88280    0.00000
-2.51040    0.00000    0.00000 ;    HB3-    CB-    CG-   HG2
 4407   4405   4408   4410      3    0.62760    1.88280    0.00000
-2.51040    0.00000    0.00000 ;    HB3-    CB-    CG-   HG3
 4407   4405   4408   4411      3    0.66944    2.00832    0.00000
-2.67776    0.00000    0.00000 ;    HB3-    CB-    CG-    CD
 4408   4405   4403   4421      3    0.65084    1.95253    0.00000
-2.60338    0.00000    0.00000 ;     CG-    CB-    CA-     C
 4408   4411   4414   4415      3    0.65084    1.95253    0.00000
-2.60338    0.00000    0.00000 ;     CG-    CD-    CE-   HE2
 4408   4411   4414   4416      3    0.65084    1.95253    0.00000
-2.60338    0.00000    0.00000 ;     CG-    CD-    CE-   HE3
 4408   4411   4414   4417      3    0.65084    1.95253    0.00000
-2.60338    0.00000    0.00000 ;     CG-    CD-    CE-    NZ
 4409   4408   4411   4412      3    0.62760    1.88280    0.00000
-2.51040    0.00000    0.00000 ;    HG2-    CG-    CD-   HD2
 4409   4408   4411   4413      3    0.62760    1.88280    0.00000
-2.51040    0.00000    0.00000 ;    HG2-    CG-    CD-   HD3
 4409   4408   4411   4414      3    0.66944    2.00832    0.00000
-2.67776    0.00000    0.00000 ;    HG2-    CG-    CD-    CE
 4410   4408   4411   4412      3    0.62760    1.88280    0.00000
-2.51040    0.00000    0.00000 ;    HG3-    CG-    CD-   HD2
 4410   4408   4411   4413      3    0.62760    1.88280    0.00000
-2.51040    0.00000    0.00000 ;    HG3-    CG-    CD-   HD3
 4410   4408   4411   4414      3    0.66944    2.00832    0.00000
-2.67776    0.00000    0.00000 ;    HG3-    CG-    CD-    CE
 4411   4414   4417   4418      3    0.65084    1.95253    0.00000
-2.60338    0.00000    0.00000 ;     CD-    CE-    NZ-   HZ1
 4411   4414   4417   4419      3    0.65084    1.95253    0.00000
-2.60338    0.00000    0.00000 ;     CD-    CE-    NZ-   HZ2
 4411   4414   4417   4420      3    0.65084    1.95253    0.00000
-2.60338    0.00000    0.00000 ;     CD-    CE-    NZ-   HZ3
 4412   4411   4414   4415      3    0.65084    1.95253    0.00000
-2.60338    0.00000    0.00000 ;    HD2-    CD-    CE-   HE2
 4412   4411   4414   4416      3    0.65084    1.95253    0.00000
-2.60338    0.00000    0.00000 ;    HD2-    CD-    CE-   HE3
 4412   4411   4414   4417      3    0.65084    1.95253    0.00000
-2.60338    0.00000    0.00000 ;    HD2-    CD-    CE-    NZ
 4413   4411   4414   4415      3    0.65084    1.95253    0.00000
-2.60338    0.00000    0.00000 ;    HD3-    CD-    CE-   HE2
 4413   4411   4414   4416      3    0.65084    1.95253    0.00000
-2.60338    0.00000    0.00000 ;    HD3-    CD-    CE-   HE3
 4413   4411   4414   4417      3    0.65084    1.95253    0.00000
-2.60338    0.00000    0.00000 ;    HD3-    CD-    CE-    NZ
 4415   4414   4417   4418      3    0.65084    1.95253    0.00000
-2.60338    0.00000    0.00000 ;    HE2-    CE-    NZ-   HZ1
 4415   4414   4417   4419      3    0.65084    1.95253    0.00000
```

-2.60338    0.00000    0.00000 ;    HE2-    CE-    NZ-    HZ2
4415    4414    4417    4420    3    0.65084    1.95253    0.00000
-2.60338    0.00000    0.00000 ;    HE2-    CE-    NZ-    HZ3
4416    4414    4417    4418    3    0.65084    1.95253    0.00000
-2.60338    0.00000    0.00000 ;    HE3-    CE-    NZ-    HZ1
4416    4414    4417    4419    3    0.65084    1.95253    0.00000
-2.60338    0.00000    0.00000 ;    HE3-    CE-    NZ-    HZ2
4416    4414    4417    4420    3    0.65084    1.95253    0.00000
-2.60338    0.00000    0.00000 ;    HE3-    CE-    NZ-    HZ3
4421    4423    4425    4426    3    0.00000    0.00000    0.00000
0.00000    0.00000    0.00000 ;    C-    N-    CA-    HA
4421    4423    4425    4427    3    11.71520    1.67360    15.06240    -
13.38880    0.00000    0.00000 ;    C-    N-    CA-    CB
4421    4423    4425    4432    3    1.75728    5.27184    2.25936
-7.02912    0.00000    0.00000 ;    C-    N-    CA-    C
4422    4421    4423    4424    3    29.28800    -8.36800    -20.92000
0.00000    0.00000    0.00000 ;    O-    C-    N-    H
4422    4421    4423    4425    3    20.92000    0.00000    -20.92000
0.00000    0.00000    0.00000 ;    O-    C-    N-    CA
4423    4425    4427    4428    3    0.65084    1.95253    0.00000
-2.60338    0.00000    0.00000 ;    N-    CA-    CB-    HB2
4423    4425    4427    4429    3    0.65084    1.95253    0.00000
-2.60338    0.00000    0.00000 ;    N-    CA-    CB-    HB3
4423    4425    4427    4430    3    7.86174    2.24681    -7.41405
-6.71114    5.35552    0.00000 ;    N-    CA-    CB-    OG
4423    4425    4432    4433    3    0.00000    0.00000    0.00000
0.00000    0.00000    0.00000 ;    N-    CA-    C-    O
4423    4425    4432    4434    3    17.40544    -5.02080    -13.22144
9.20480    0.00000    0.00000 ;    N-    CA-    C-    N
4424    4423    4425    4426    3    0.00000    0.00000    0.00000
0.00000    0.00000    0.00000 ;    H-    N-    CA-    HA
4424    4423    4425    4427    3    0.00000    0.00000    0.00000
0.00000    0.00000    0.00000 ;    H-    N-    CA-    CB
4424    4423    4425    4432    3    0.00000    0.00000    0.00000
0.00000    0.00000    0.00000 ;    H-    N-    CA-    C
4425    4427    4430    4431    3    2.05853    2.46856    3.48109
-4.46851    0.23430    0.00000 ;    CA-    CB-    OG-    HG
4425    4432    4434    4435    3    20.92000    0.00000    -20.92000
0.00000    0.00000    0.00000 ;    CA-    C-    N-    H
4425    4432    4434    4436    3    20.92000    0.00000    -20.92000
0.00000    0.00000    0.00000 ;    CA-    C-    N-    CA
4426    4425    4427    4428    3    0.65084    1.95253    0.00000
-2.60338    0.00000    0.00000 ;    HA-    CA-    CB-    HB2
4426    4425    4427    4429    3    0.65084    1.95253    0.00000
-2.60338    0.00000    0.00000 ;    HA-    CA-    CB-    HB3
4426    4425    4427    4430    3    1.04600    -1.04600    0.00000
0.00000    0.00000    0.00000 ;    HA-    CA-    CB-    OG
4426    4425    4432    4433    3    3.68192    -4.35136    0.00000
1.33888    0.00000    0.00000 ;    HA-    CA-    C-    O
4426    4425    4432    4434    3    0.00000    0.00000    0.00000
0.00000    0.00000    0.00000 ;    HA-    CA-    C-    N
4427    4425    4432    4433    3    0.00000    0.00000    0.00000

0.00000 0.00000 0.00000 ; CB- CA- C- O
4427 4425 4432 4434 3 2.51040 4.18400 1.67360
-6.69440 0.00000 0.00000 ; CB- CA- C- N
4428 4427 4425 4432 3 0.65084 1.95253 0.00000
-2.60338 0.00000 0.00000 ; HB2- CB- CA- C
4428 4427 4430 4431 3 0.69733 2.09200 0.00000
-2.78933 0.00000 0.00000 ; HB2- CB- OG- HG
4429 4427 4425 4432 3 0.65084 1.95253 0.00000
-2.60338 0.00000 0.00000 ; HB3- CB- CA- C
4429 4427 4430 4431 3 0.69733 2.09200 0.00000
-2.78933 0.00000 0.00000 ; HB3- CB- OG- HG
4430 4427 4425 4432 3 7.34710 7.79898 -6.14211
-6.71114 4.31789 0.00000 ; OG- CB- CA- C
4432 4434 4436 4437 3 0.00000 0.00000 0.00000
0.00000 0.00000 0.00000 ; C- N- CA- HA
4432 4434 4436 4438 3 11.71520 1.67360 15.06240 -
13.38880 0.00000 0.00000 ; C- N- CA- CB
4432 4434 4436 4449 3 1.75728 5.27184 2.25936
-7.02912 0.00000 0.00000 ; C- N- CA- C
4433 4432 4434 4435 3 29.28800 -8.36800 -20.92000
0.00000 0.00000 0.00000 ; O- C- N- H
4433 4432 4434 4436 3 20.92000 0.00000 -20.92000
0.00000 0.00000 0.00000 ; O- C- N- CA
4434 4436 4438 4439 3 0.65084 1.95253 0.00000
-2.60338 0.00000 0.00000 ; N- CA- CB- HB2
4434 4436 4438 4440 3 0.65084 1.95253 0.00000
-2.60338 0.00000 0.00000 ; N- CA- CB- HB3
4434 4436 4438 4441 3 4.79068 4.02919 -4.82834
-3.66518 2.97901 0.00000 ; N- CA- CB- CG
4434 4436 4449 4450 3 0.00000 0.00000 0.00000
0.00000 0.00000 0.00000 ; N- CA- C- O
4434 4436 4449 4451 3 17.40544 -5.02080 -13.22144
9.20480 0.00000 0.00000 ; N- CA- C- N
4435 4434 4436 4437 3 0.00000 0.00000 0.00000
0.00000 0.00000 0.00000 ; H- N- CA- HA
4435 4434 4436 4438 3 0.00000 0.00000 0.00000
0.00000 0.00000 0.00000 ; H- N- CA- CB
4435 4434 4436 4449 3 0.00000 0.00000 0.00000
0.00000 0.00000 0.00000 ; H- N- CA- C
4436 4438 4441 4442 3 5.98312 6.40152 3.28026 -
12.38464 -1.57318 0.00000 ; CA- CB- CG- ND1
4436 4438 4441 4447 3 0.00000 0.00000 0.00000
0.00000 0.00000 0.00000 ; CA- CB- CG- CD2
4436 4449 4451 4452 3 20.92000 0.00000 -20.92000
0.00000 0.00000 0.00000 ; CA- C- N- H
4436 4449 4451 4453 3 20.92000 0.00000 -20.92000
0.00000 0.00000 0.00000 ; CA- C- N- CA
4437 4436 4438 4439 3 0.65084 1.95253 0.00000
-2.60338 0.00000 0.00000 ; HA- CA- CB- HB2
4437 4436 4438 4440 3 0.65084 1.95253 0.00000
-2.60338 0.00000 0.00000 ; HA- CA- CB- HB3
4437 4436 4438 4441 3 0.65084 1.95253 0.00000

```
-2.60338    0.00000    0.00000 ;      HA-    CA-    CB-    CG
  4437   4436   4449   4450      3    3.68192   -4.35136    0.00000
1.33888    0.00000    0.00000 ;     HA-    CA-     C-     O
  4437   4436   4449   4451      3    0.00000    0.00000    0.00000
0.00000    0.00000    0.00000 ;     HA-    CA-     C-     N
  4438   4436   4449   4450      3    0.00000    0.00000    0.00000
0.00000    0.00000    0.00000 ;     CB-    CA-     C-     O
  4438   4436   4449   4451      3    2.51040    4.18400    1.67360
-6.69440    0.00000    0.00000 ;      CB-    CA-     C-     N
  4438   4441   4442   4443      3   20.08320    0.00000  -20.08320
0.00000    0.00000    0.00000 ;     CB-    CG-   ND1-   CE1
  4438   4441   4447   4445      3   44.97800    0.00000  -44.97800
0.00000    0.00000    0.00000 ;     CB-    CG-   CD2-   NE2
  4438   4441   4447   4448      3   44.97800    0.00000  -44.97800
0.00000    0.00000    0.00000 ;     CB-    CG-   CD2-   HD2
  4439   4438   4436   4449      3    0.65084    1.95253    0.00000
-2.60338    0.00000    0.00000 ;     HB2-    CB-    CA-     C
  4439   4438   4441   4442      3    0.00000    0.00000    0.00000
0.00000    0.00000    0.00000 ;    HB2-    CB-    CG-   ND1
  4439   4438   4441   4447      3    0.00000    0.00000    0.00000
0.00000    0.00000    0.00000 ;    HB2-    CB-    CG-   CD2
  4440   4438   4436   4449      3    0.65084    1.95253    0.00000
-2.60338    0.00000    0.00000 ;     HB3-    CB-    CA-     C
  4440   4438   4441   4442      3    0.00000    0.00000    0.00000
0.00000    0.00000    0.00000 ;    HB3-    CB-    CG-   ND1
  4440   4438   4441   4447      3    0.00000    0.00000    0.00000
0.00000    0.00000    0.00000 ;    HB3-    CB-    CG-   CD2
  4441   4438   4436   4449      3    3.76560    3.34720   -2.87859
-3.66518    0.83680    0.00000 ;      CG-    CB-    CA-     C
  4441   4442   4443   4444      3   41.84000    0.00000  -41.84000
0.00000    0.00000    0.00000 ;     CG-   ND1-   CE1-   HE1
  4441   4442   4443   4445      3   41.84000    0.00000  -41.84000
0.00000    0.00000    0.00000 ;     CG-   ND1-   CE1-   NE2
  4441   4447   4445   4443      3   12.55200    0.00000  -12.55200
0.00000    0.00000    0.00000 ;     CG-   CD2-   NE2-   CE1
  4441   4447   4445   4446      3   12.55200    0.00000  -12.55200
0.00000    0.00000    0.00000 ;     CG-   CD2-   NE2-   HE2
  4442   4441   4447   4445      3   44.97800    0.00000  -44.97800
0.00000    0.00000    0.00000 ;    ND1-    CG-   CD2-   NE2
  4442   4441   4447   4448      3   44.97800    0.00000  -44.97800
0.00000    0.00000    0.00000 ;    ND1-    CG-   CD2-   HD2
  4442   4443   4445   4446      3   19.45560    0.00000  -19.45560
0.00000    0.00000    0.00000 ;    ND1-   CE1-   NE2-   HE2
  4442   4443   4445   4447      3   19.45560    0.00000  -19.45560
0.00000    0.00000    0.00000 ;    ND1-   CE1-   NE2-   CD2
  4443   4442   4441   4447      3   20.08320    0.00000  -20.08320
0.00000    0.00000    0.00000 ;    CE1-   ND1-    CG-   CD2
  4443   4445   4447   4448      3   12.55200    0.00000  -12.55200
0.00000    0.00000    0.00000 ;    CE1-   NE2-   CD2-   HD2
  4444   4443   4445   4446      3   19.45560    0.00000  -19.45560
0.00000    0.00000    0.00000 ;    HE1-   CE1-   NE2-   HE2
  4444   4443   4445   4447      3   19.45560    0.00000  -19.45560
```

```
0.00000    0.00000    0.00000 ;    HE1-   CE1-   NE2-   CD2
  4446   4445   4447   4448      3   12.55200    0.00000  -12.55200
0.00000    0.00000    0.00000 ;    HE2-   NE2-   CD2-   HD2
  4449   4451   4453   4454      3    0.00000    0.00000    0.00000
0.00000    0.00000    0.00000 ;     C-     N-    CA-    HA
  4449   4451   4453   4455      3   11.71520    1.67360   15.06240  -
13.38880    0.00000    0.00000 ;     C-     N-    CA-    CB
  4449   4451   4453   4464      3    1.75728    5.27184    2.25936
-7.02912    0.00000    0.00000 ;     C-     N-    CA-     C
  4450   4449   4451   4452      3   29.28800   -8.36800  -20.92000
0.00000    0.00000    0.00000 ;     O-     C-     N-     H
  4450   4449   4451   4453      3   20.92000    0.00000  -20.92000
0.00000    0.00000    0.00000 ;     O-     C-     N-    CA
  4451   4453   4455   4456      3    0.65084    1.95253    0.00000
-2.60338    0.00000    0.00000 ;     N-    CA-    CB-   HB2
  4451   4453   4455   4457      3    0.65084    1.95253    0.00000
-2.60338    0.00000    0.00000 ;     N-    CA-    CB-   HB3
  4451   4453   4455   4458      3    3.21331    2.22589   -4.15053
-2.40998    2.61082    0.00000 ;     N-    CA-    CB-    CG
  4451   4453   4464   4465      3    0.00000    0.00000    0.00000
0.00000    0.00000    0.00000 ;     N-    CA-     C-     O
  4451   4453   4464   4466      3   17.40544   -5.02080  -13.22144
9.20480    0.00000    0.00000 ;     N-    CA-     C-     N
  4452   4451   4453   4454      3    0.00000    0.00000    0.00000
0.00000    0.00000    0.00000 ;     H-     N-    CA-    HA
  4452   4451   4453   4455      3    0.00000    0.00000    0.00000
0.00000    0.00000    0.00000 ;     H-     N-    CA-    CB
  4452   4451   4453   4464      3    0.00000    0.00000    0.00000
0.00000    0.00000    0.00000 ;     H-     N-    CA-     C
  4453   4455   4458   4459      3    0.66944    2.00832    0.00000
-2.67776    0.00000    0.00000 ;    CA-    CB-    CG-   HG2
  4453   4455   4458   4460      3    0.66944    2.00832    0.00000
-2.67776    0.00000    0.00000 ;    CA-    CB-    CG-   HG3
  4453   4455   4458   4461      3   10.12110   -1.91209    0.01674
10.17549   -1.87443    0.00000 ;    CA-    CB-    CG-    CD
  4453   4464   4466   4467      3   20.92000    0.00000  -20.92000
0.00000    0.00000    0.00000 ;    CA-     C-     N-     H
  4453   4464   4466   4468      3   20.92000    0.00000  -20.92000
0.00000    0.00000    0.00000 ;    CA-     C-     N-    CA
  4454   4453   4455   4456      3    0.65084    1.95253    0.00000
-2.60338    0.00000    0.00000 ;    HA-    CA-    CB-   HB2
  4454   4453   4455   4457      3    0.65084    1.95253    0.00000
-2.60338    0.00000    0.00000 ;    HA-    CA-    CB-   HB3
  4454   4453   4455   4458      3    0.65084    1.95253    0.00000
-2.60338    0.00000    0.00000 ;    HA-    CA-    CB-    CG
  4454   4453   4464   4465      3    3.68192   -4.35136    0.00000
1.33888    0.00000    0.00000 ;    HA-    CA-     C-     O
  4454   4453   4464   4466      3    0.00000    0.00000    0.00000
0.00000    0.00000    0.00000 ;    HA-    CA-     C-     N
  4455   4453   4464   4465      3    0.00000    0.00000    0.00000
0.00000    0.00000    0.00000 ;    CB-    CA-     C-     O
  4455   4453   4464   4466      3    2.51040    4.18400    1.67360
```

-6.69440 0.00000 0.00000 ; CB- CA- C- N
4455 4458 4461 4462 3 3.79907 0.00000 -5.40573
0.00000 2.14221 0.00000 ; CB- CG- CD- OE1
4455 4458 4461 4463 3 3.79907 0.00000 -5.40573
0.00000 2.14221 0.00000 ; CB- CG- CD- OE2
4456 4455 4453 4464 3 0.65084 1.95253 0.00000
-2.60338 0.00000 0.00000 ; HB2- CB- CA- C
4456 4455 4458 4459 3 0.62760 1.88280 0.00000
-2.51040 0.00000 0.00000 ; HB2- CB- CG- HG2
4456 4455 4458 4460 3 0.62760 1.88280 0.00000
-2.51040 0.00000 0.00000 ; HB2- CB- CG- HG3
4456 4455 4458 4461 3 0.65084 1.95253 0.00000
-2.60338 0.00000 0.00000 ; HB2- CB- CG- CD
4457 4455 4453 4464 3 0.65084 1.95253 0.00000
-2.60338 0.00000 0.00000 ; HB3- CB- CA- C
4457 4455 4458 4459 3 0.62760 1.88280 0.00000
-2.51040 0.00000 0.00000 ; HB3- CB- CG- HG2
4457 4455 4458 4460 3 0.62760 1.88280 0.00000
-2.51040 0.00000 0.00000 ; HB3- CB- CG- HG3
4457 4455 4458 4461 3 0.65084 1.95253 0.00000
-2.60338 0.00000 0.00000 ; HB3- CB- CG- CD
4458 4455 4453 4464 3 6.86594 3.56895 -8.14206
-2.40998 4.85344 0.00000 ; CG- CB- CA- C
4459 4458 4461 4462 3 0.00000 0.00000 0.00000
0.00000 0.00000 0.00000 ; HG2- CG- CD- OE1
4459 4458 4461 4463 3 0.00000 0.00000 0.00000
0.00000 0.00000 0.00000 ; HG2- CG- CD- OE2
4460 4458 4461 4462 3 0.00000 0.00000 0.00000
0.00000 0.00000 0.00000 ; HG3- CG- CD- OE1
4460 4458 4461 4463 3 0.00000 0.00000 0.00000
0.00000 0.00000 0.00000 ; HG3- CG- CD- OE2
4464 4466 4468 4469 3 0.00000 0.00000 0.00000
0.00000 0.00000 0.00000 ; C- N- CA- HA
4464 4466 4468 4470 3 11.71520 1.67360 15.06240 -
13.38880 0.00000 0.00000 ; C- N- CA- CB
4464 4466 4468 4481 3 1.75728 5.27184 2.25936
-7.02912 0.00000 0.00000 ; C- N- CA- C
4465 4464 4466 4467 3 29.28800 -8.36800 -20.92000
0.00000 0.00000 0.00000 ; O- C- N- H
4465 4464 4466 4468 3 20.92000 0.00000 -20.92000
0.00000 0.00000 0.00000 ; O- C- N- CA
4466 4468 4470 4471 3 0.65084 1.95253 0.00000
-2.60338 0.00000 0.00000 ; N- CA- CB- HB2
4466 4468 4470 4472 3 0.65084 1.95253 0.00000
-2.60338 0.00000 0.00000 ; N- CA- CB- HB3
4466 4468 4470 4473 3 4.79068 4.02919 -4.82834
-3.66518 2.97901 0.00000 ; N- CA- CB- CG
4466 4468 4481 4482 3 0.00000 0.00000 0.00000
0.00000 0.00000 0.00000 ; N- CA- C- O
4466 4468 4481 4483 3 17.40544 -5.02080 -13.22144
9.20480 0.00000 0.00000 ; N- CA- C- N
4467 4466 4468 4469 3 0.00000 0.00000 0.00000

0.00000 0.00000 0.00000 ; H- N- CA- HA
4467 4466 4468 4470 3 0.00000 0.00000 0.00000
0.00000 0.00000 0.00000 ; H- N- CA- CB
4467 4466 4468 4481 3 0.00000 0.00000 0.00000
0.00000 0.00000 0.00000 ; H- N- CA- C
4468 4470 4473 4474 3 5.98312 6.40152 3.28026 -
12.38464 -1.57318 0.00000 ; CA- CB- CG- ND1
4468 4470 4473 4479 3 0.00000 0.00000 0.00000
0.00000 0.00000 0.00000 ; CA- CB- CG- CD2
4468 4481 4483 4484 3 20.92000 0.00000 -20.92000
0.00000 0.00000 0.00000 ; CA- C- N- H
4468 4481 4483 4485 3 20.92000 0.00000 -20.92000
0.00000 0.00000 0.00000 ; CA- C- N- CA
4469 4468 4470 4471 3 0.65084 1.95253 0.00000
-2.60338 0.00000 0.00000 ; HA- CA- CB- HB2
4469 4468 4470 4472 3 0.65084 1.95253 0.00000
-2.60338 0.00000 0.00000 ; HA- CA- CB- HB3
4469 4468 4470 4473 3 0.65084 1.95253 0.00000
-2.60338 0.00000 0.00000 ; HA- CA- CB- CG
4469 4468 4481 4482 3 3.68192 -4.35136 0.00000
1.33888 0.00000 0.00000 ; HA- CA- C- O
4469 4468 4481 4483 3 0.00000 0.00000 0.00000
0.00000 0.00000 0.00000 ; HA- CA- C- N
4470 4468 4481 4482 3 0.00000 0.00000 0.00000
0.00000 0.00000 0.00000 ; CB- CA- C- O
4470 4468 4481 4483 3 2.51040 4.18400 1.67360
-6.69440 0.00000 0.00000 ; CB- CA- C- N
4470 4473 4474 4475 3 20.08320 0.00000 -20.08320
0.00000 0.00000 0.00000 ; CB- CG- ND1- CE1
4470 4473 4479 4477 3 44.97800 0.00000 -44.97800
0.00000 0.00000 0.00000 ; CB- CG- CD2- NE2
4470 4473 4479 4480 3 44.97800 0.00000 -44.97800
0.00000 0.00000 0.00000 ; CB- CG- CD2- HD2
4471 4470 4468 4481 3 0.65084 1.95253 0.00000
-2.60338 0.00000 0.00000 ; HB2- CB- CA- C
4471 4470 4473 4474 3 0.00000 0.00000 0.00000
0.00000 0.00000 0.00000 ; HB2- CB- CG- ND1
4471 4470 4473 4479 3 0.00000 0.00000 0.00000
0.00000 0.00000 0.00000 ; HB2- CB- CG- CD2
4472 4470 4468 4481 3 0.65084 1.95253 0.00000
-2.60338 0.00000 0.00000 ; HB3- CB- CA- C
4472 4470 4473 4474 3 0.00000 0.00000 0.00000
0.00000 0.00000 0.00000 ; HB3- CB- CG- ND1
4472 4470 4473 4479 3 0.00000 0.00000 0.00000
0.00000 0.00000 0.00000 ; HB3- CB- CG- CD2
4473 4470 4468 4481 3 3.76560 3.34720 -2.87859
-3.66518 0.83680 0.00000 ; CG- CB- CA- C
4473 4474 4475 4476 3 41.84000 0.00000 -41.84000
0.00000 0.00000 0.00000 ; CG- ND1- CE1- HE1
4473 4474 4475 4477 3 41.84000 0.00000 -41.84000
0.00000 0.00000 0.00000 ; CG- ND1- CE1- NE2
4473 4479 4477 4475 3 12.55200 0.00000 -12.55200

```
0.00000    0.00000    0.00000 ;      CG-   CD2-   NE2-   CE1
  4473   4479   4477   4478       3   12.55200    0.00000  -12.55200
0.00000    0.00000    0.00000 ;      CG-   CD2-   NE2-   HE2
  4474   4473   4479   4477       3   44.97800    0.00000  -44.97800
0.00000    0.00000    0.00000 ;     ND1-    CG-   CD2-   NE2
  4474   4473   4479   4480       3   44.97800    0.00000  -44.97800
0.00000    0.00000    0.00000 ;     ND1-    CG-   CD2-   HD2
  4474   4475   4477   4478       3   19.45560    0.00000  -19.45560
0.00000    0.00000    0.00000 ;     ND1-   CE1-   NE2-   HE2
  4474   4475   4477   4479       3   19.45560    0.00000  -19.45560
0.00000    0.00000    0.00000 ;     ND1-   CE1-   NE2-   CD2
  4475   4474   4473   4479       3   20.08320    0.00000  -20.08320
0.00000    0.00000    0.00000 ;     CE1-   ND1-    CG-   CD2
  4475   4477   4479   4480       3   12.55200    0.00000  -12.55200
0.00000    0.00000    0.00000 ;     CE1-   NE2-   CD2-   HD2
  4476   4475   4477   4478       3   19.45560    0.00000  -19.45560
0.00000    0.00000    0.00000 ;     HE1-   CE1-   NE2-   HE2
  4476   4475   4477   4479       3   19.45560    0.00000  -19.45560
0.00000    0.00000    0.00000 ;     HE1-   CE1-   NE2-   CD2
  4478   4477   4479   4480       3   12.55200    0.00000  -12.55200
0.00000    0.00000    0.00000 ;     HE2-   NE2-   CD2-   HD2
  4481   4483   4485   4486       3    0.00000    0.00000    0.00000
0.00000    0.00000    0.00000 ;       C-     N-    CA-    HA
  4481   4483   4485   4487       3   11.71520    1.67360   15.06240  -
13.38880    0.00000    0.00000 ;       C-     N-    CA-    CB
  4481   4483   4485   4497       3    1.75728    5.27184    2.25936
-7.02912    0.00000    0.00000 ;       C-     N-    CA-     C
  4482   4481   4483   4484       3   29.28800   -8.36800  -20.92000
0.00000    0.00000    0.00000 ;       O-     C-     N-     H
  4482   4481   4483   4485       3   20.92000    0.00000  -20.92000
0.00000    0.00000    0.00000 ;       O-     C-     N-    CA
  4483   4485   4487   4488       3    0.65084    1.95253    0.00000
-2.60338    0.00000    0.00000 ;       N-    CA-    CB-    HB
  4483   4485   4487   4489       3    3.83673    0.44769   -1.77402
-2.47693   -0.03347    0.00000 ;       N-    CA-    CB-   CG1
  4483   4485   4487   4493       3    3.83673    0.44769   -1.77402
-2.47693   -0.03347    0.00000 ;       N-    CA-    CB-   CG2
  4483   4485   4497   4498       3    0.00000    0.00000    0.00000
0.00000    0.00000    0.00000 ;       N-    CA-     C-     O
  4483   4485   4497   4499       3   17.40544   -5.02080  -13.22144
9.20480    0.00000    0.00000 ;       N-    CA-     C-     N
  4484   4483   4485   4486       3    0.00000    0.00000    0.00000
0.00000    0.00000    0.00000 ;       H-     N-    CA-    HA
  4484   4483   4485   4487       3    0.00000    0.00000    0.00000
0.00000    0.00000    0.00000 ;       H-     N-    CA-    CB
  4484   4483   4485   4497       3    0.00000    0.00000    0.00000
0.00000    0.00000    0.00000 ;       H-     N-    CA-     C
  4485   4487   4489   4490       3    0.66944    2.00832    0.00000
-2.67776    0.00000    0.00000 ;      CA-    CB-   CG1-  HG11
  4485   4487   4489   4491       3    0.66944    2.00832    0.00000
-2.67776    0.00000    0.00000 ;      CA-    CB-   CG1-  HG12
  4485   4487   4489   4492       3    0.66944    2.00832    0.00000
```

-2.67776    0.00000    0.00000 ;     CA-    CB-   CG1-  HG13
  4485   4487   4493   4494      3    0.66944    2.00832    0.00000
-2.67776    0.00000    0.00000 ;     CA-    CB-   CG2-  HG21
  4485   4487   4493   4495      3    0.66944    2.00832    0.00000
-2.67776    0.00000    0.00000 ;     CA-    CB-   CG2-  HG22
  4485   4487   4493   4496      3    0.66944    2.00832    0.00000
-2.67776    0.00000    0.00000 ;     CA-    CB-   CG2-  HG23
  4485   4497   4499   4500      3   20.92000    0.00000  -20.92000
0.00000    0.00000    0.00000 ;     CA-     C-     N-     H
  4485   4497   4499   4501      3   20.92000    0.00000  -20.92000
0.00000    0.00000    0.00000 ;     CA-     C-     N-    CA
  4486   4485   4487   4488      3    0.65084    1.95253    0.00000
-2.60338    0.00000    0.00000 ;     HA-    CA-    CB-    HB
  4486   4485   4487   4489      3    0.65084    1.95253    0.00000
-2.60338    0.00000    0.00000 ;     HA-    CA-    CB-   CG1
  4486   4485   4487   4493      3    0.65084    1.95253    0.00000
-2.60338    0.00000    0.00000 ;     HA-    CA-    CB-   CG2
  4486   4485   4497   4498      3    3.68192   -4.35136    0.00000
1.33888    0.00000    0.00000 ;     HA-    CA-     C-     O
  4486   4485   4497   4499      3    0.00000    0.00000    0.00000
0.00000    0.00000    0.00000 ;     HA-    CA-     C-     N
  4487   4485   4497   4498      3    0.00000    0.00000    0.00000
0.00000    0.00000    0.00000 ;     CB-    CA-     C-     O
  4487   4485   4497   4499      3    2.51040    4.18400    1.67360
-6.69440    0.00000    0.00000 ;     CB-    CA-     C-     N
  4488   4487   4485   4497      3    0.65084    1.95253    0.00000
-2.60338    0.00000    0.00000 ;     HB-    CB-    CA-     C
  4488   4487   4489   4490      3    0.62760    1.88280    0.00000
-2.51040    0.00000    0.00000 ;     HB-    CB-   CG1-  HG11
  4488   4487   4489   4491      3    0.62760    1.88280    0.00000
-2.51040    0.00000    0.00000 ;     HB-    CB-   CG1-  HG12
  4488   4487   4489   4492      3    0.62760    1.88280    0.00000
-2.51040    0.00000    0.00000 ;     HB-    CB-   CG1-  HG13
  4488   4487   4493   4494      3    0.62760    1.88280    0.00000
-2.51040    0.00000    0.00000 ;     HB-    CB-   CG2-  HG21
  4488   4487   4493   4495      3    0.62760    1.88280    0.00000
-2.51040    0.00000    0.00000 ;     HB-    CB-   CG2-  HG22
  4488   4487   4493   4496      3    0.62760    1.88280    0.00000
-2.51040    0.00000    0.00000 ;     HB-    CB-   CG2-  HG23
  4489   4487   4485   4497      3    5.67350    3.55640   -6.16722
-2.47693    3.74886    0.00000 ;    CG1-    CB-    CA-     C
  4489   4487   4493   4494      3    0.66944    2.00832    0.00000
-2.67776    0.00000    0.00000 ;    CG1-    CB-   CG2-  HG21
  4489   4487   4493   4495      3    0.66944    2.00832    0.00000
-2.67776    0.00000    0.00000 ;    CG1-    CB-   CG2-  HG22
  4489   4487   4493   4496      3    0.66944    2.00832    0.00000
-2.67776    0.00000    0.00000 ;    CG1-    CB-   CG2-  HG23
  4490   4489   4487   4493      3    0.66944    2.00832    0.00000
-2.67776    0.00000    0.00000 ;   HG11-   CG1-    CB-   CG2
  4491   4489   4487   4493      3    0.66944    2.00832    0.00000
-2.67776    0.00000    0.00000 ;   HG12-   CG1-    CB-   CG2
  4492   4489   4487   4493      3    0.66944    2.00832    0.00000

-2.67776 0.00000 0.00000 ; HG13- CG1- CB- CG2
4493 4487 4485 4497 3 5.67350 3.55640 -6.16722
-2.47693 3.74886 0.00000 ; CG2- CB- CA- C
4497 4499 4501 4502 3 0.00000 0.00000 0.00000
0.00000 0.00000 0.00000 ; C- N- CA- HA
4497 4499 4501 4503 3 11.71520 1.67360 15.06240 -
13.38880 0.00000 0.00000 ; C- N- CA- CB
4497 4499 4501 4512 3 1.75728 5.27184 2.25936
-7.02912 0.00000 0.00000 ; C- N- CA- C
4498 4497 4499 4500 3 29.28800 -8.36800 -20.92000
0.00000 0.00000 0.00000 ; O- C- N- H
4498 4497 4499 4501 3 20.92000 0.00000 -20.92000
0.00000 0.00000 0.00000 ; O- C- N- CA
4499 4501 4503 4504 3 0.65084 1.95253 0.00000
-2.60338 0.00000 0.00000 ; N- CA- CB- HB2
4499 4501 4503 4505 3 0.65084 1.95253 0.00000
-2.60338 0.00000 0.00000 ; N- CA- CB- HB3
4499 4501 4503 4506 3 3.21331 2.22589 -4.15053
-2.40998 2.61082 0.00000 ; N- CA- CB- CG
4499 4501 4512 4513 3 0.00000 0.00000 0.00000
0.00000 0.00000 0.00000 ; N- CA- C- O
4499 4501 4512 4514 3 17.40544 -5.02080 -13.22144
9.20480 0.00000 0.00000 ; N- CA- C- N
4500 4499 4501 4502 3 0.00000 0.00000 0.00000
0.00000 0.00000 0.00000 ; H- N- CA- HA
4500 4499 4501 4503 3 0.00000 0.00000 0.00000
0.00000 0.00000 0.00000 ; H- N- CA- CB
4500 4499 4501 4512 3 0.00000 0.00000 0.00000
0.00000 0.00000 0.00000 ; H- N- CA- C
4501 4503 4506 4507 3 0.66944 2.00832 0.00000
-2.67776 0.00000 0.00000 ; CA- CB- CG- HG2
4501 4503 4506 4508 3 0.66944 2.00832 0.00000
-2.67776 0.00000 0.00000 ; CA- CB- CG- HG3
4501 4503 4506 4509 3 10.12110 -1.91209 0.01674
10.17549 -1.87443 0.00000 ; CA- CB- CG- CD
4501 4512 4514 4515 3 20.92000 0.00000 -20.92000
0.00000 0.00000 0.00000 ; CA- C- N- H
4501 4512 4514 4516 3 20.92000 0.00000 -20.92000
0.00000 0.00000 0.00000 ; CA- C- N- CA
4502 4501 4503 4504 3 0.65084 1.95253 0.00000
-2.60338 0.00000 0.00000 ; HA- CA- CB- HB2
4502 4501 4503 4505 3 0.65084 1.95253 0.00000
-2.60338 0.00000 0.00000 ; HA- CA- CB- HB3
4502 4501 4503 4506 3 0.65084 1.95253 0.00000
-2.60338 0.00000 0.00000 ; HA- CA- CB- CG
4502 4501 4512 4513 3 3.68192 -4.35136 0.00000
1.33888 0.00000 0.00000 ; HA- CA- C- O
4502 4501 4512 4514 3 0.00000 0.00000 0.00000
0.00000 0.00000 0.00000 ; HA- CA- C- N
4503 4501 4512 4513 3 0.00000 0.00000 0.00000
0.00000 0.00000 0.00000 ; CB- CA- C- O
4503 4501 4512 4514 3 2.51040 4.18400 1.67360

-6.69440 0.00000 0.00000 ; CB- CA- C- N
4503 4506 4509 4510 3 3.79907 0.00000 -5.40573
0.00000 2.14221 0.00000 ; CB- CG- CD- OE1
4503 4506 4509 4511 3 3.79907 0.00000 -5.40573
0.00000 2.14221 0.00000 ; CB- CG- CD- OE2
4504 4503 4501 4512 3 0.65084 1.95253 0.00000
-2.60338 0.00000 0.00000 ; HB2- CB- CA- C
4504 4503 4506 4507 3 0.62760 1.88280 0.00000
-2.51040 0.00000 0.00000 ; HB2- CB- CG- HG2
4504 4503 4506 4508 3 0.62760 1.88280 0.00000
-2.51040 0.00000 0.00000 ; HB2- CB- CG- HG3
4504 4503 4506 4509 3 0.65084 1.95253 0.00000
-2.60338 0.00000 0.00000 ; HB2- CB- CG- CD
4505 4503 4501 4512 3 0.65084 1.95253 0.00000
-2.60338 0.00000 0.00000 ; HB3- CB- CA- C
4505 4503 4506 4507 3 0.62760 1.88280 0.00000
-2.51040 0.00000 0.00000 ; HB3- CB- CG- HG2
4505 4503 4506 4508 3 0.62760 1.88280 0.00000
-2.51040 0.00000 0.00000 ; HB3- CB- CG- HG3
4505 4503 4506 4509 3 0.65084 1.95253 0.00000
-2.60338 0.00000 0.00000 ; HB3- CB- CG- CD
4506 4503 4501 4512 3 6.86594 3.56895 -8.14206
-2.40998 4.85344 0.00000 ; CG- CB- CA- C
4507 4506 4509 4510 3 0.00000 0.00000 0.00000
0.00000 0.00000 0.00000 ; HG2- CG- CD- OE1
4507 4506 4509 4511 3 0.00000 0.00000 0.00000
0.00000 0.00000 0.00000 ; HG2- CG- CD- OE2
4508 4506 4509 4510 3 0.00000 0.00000 0.00000
0.00000 0.00000 0.00000 ; HG3- CG- CD- OE1
4508 4506 4509 4511 3 0.00000 0.00000 0.00000
0.00000 0.00000 0.00000 ; HG3- CG- CD- OE2
4512 4514 4516 4517 3 0.00000 0.00000 0.00000
0.00000 0.00000 0.00000 ; C- N- CA- HA
4512 4514 4516 4518 3 11.71520 1.67360 15.06240 -
13.38880 0.00000 0.00000 ; C- N- CA- CB
4512 4514 4516 4531 3 1.75728 5.27184 2.25936
-7.02912 0.00000 0.00000 ; C- N- CA- C
4513 4512 4514 4515 3 29.28800 -8.36800 -20.92000
0.00000 0.00000 0.00000 ; O- C- N- H
4513 4512 4514 4516 3 20.92000 0.00000 -20.92000
0.00000 0.00000 0.00000 ; O- C- N- CA
4514 4516 4518 4519 3 0.65084 1.95253 0.00000
-2.60338 0.00000 0.00000 ; N- CA- CB- HB2
4514 4516 4518 4520 3 0.65084 1.95253 0.00000
-2.60338 0.00000 0.00000 ; N- CA- CB- HB3
4514 4516 4518 4521 3 3.79070 1.39746 -4.61077
-2.40998 2.44346 0.00000 ; N- CA- CB- CG
4514 4516 4531 4532 3 0.00000 0.00000 0.00000
0.00000 0.00000 0.00000 ; N- CA- C- O
4514 4516 4531 4533 3 17.40544 -5.02080 -13.22144
9.20480 0.00000 0.00000 ; N- CA- C- N
4515 4514 4516 4517 3 0.00000 0.00000 0.00000

```
0.00000    0.00000    0.00000 ;      H-     N-    CA-    HA
  4515   4514   4516   4518       3   0.00000    0.00000    0.00000
0.00000    0.00000    0.00000 ;      H-     N-    CA-    CB
  4515   4514   4516   4531       3   0.00000    0.00000    0.00000
0.00000    0.00000    0.00000 ;      H-     N-    CA-     C
  4516   4518   4521   4522       3   0.66944    2.00832    0.00000
-2.67776    0.00000    0.00000 ;     CA-    CB-    CG-    HG
  4516   4518   4521   4523       3   3.90367    0.19665   -6.21742
-2.37651    5.99149    0.00000 ;     CA-    CB-    CG-   CD1
  4516   4518   4521   4527       3   3.90367    0.19665   -6.21742
-2.37651    5.99149    0.00000 ;     CA-    CB-    CG-   CD2
  4516   4531   4533   4534       3  20.92000    0.00000  -20.92000
0.00000    0.00000    0.00000 ;     CA-     C-     N-     H
  4516   4531   4533   4535       3  20.92000    0.00000  -20.92000
0.00000    0.00000    0.00000 ;     CA-     C-     N-    CA
  4517   4516   4518   4519       3   0.65084    1.95253    0.00000
-2.60338    0.00000    0.00000 ;     HA-    CA-    CB-   HB2
  4517   4516   4518   4520       3   0.65084    1.95253    0.00000
-2.60338    0.00000    0.00000 ;     HA-    CA-    CB-   HB3
  4517   4516   4518   4521       3   0.65084    1.95253    0.00000
-2.60338    0.00000    0.00000 ;     HA-    CA-    CB-    CG
  4517   4516   4531   4532       3   3.68192   -4.35136    0.00000
1.33888    0.00000    0.00000 ;     HA-    CA-     C-     O
  4517   4516   4531   4533       3   0.00000    0.00000    0.00000
0.00000    0.00000    0.00000 ;     HA-    CA-     C-     N
  4518   4516   4531   4532       3   0.00000    0.00000    0.00000
0.00000    0.00000    0.00000 ;     CB-    CA-     C-     O
  4518   4516   4531   4533       3   2.51040    4.18400    1.67360
-6.69440    0.00000    0.00000 ;     CB-    CA-     C-     N
  4518   4521   4523   4524       3   0.66944    2.00832    0.00000
-2.67776    0.00000    0.00000 ;     CB-    CG-   CD1-  HD11
  4518   4521   4523   4525       3   0.66944    2.00832    0.00000
-2.67776    0.00000    0.00000 ;     CB-    CG-   CD1-  HD12
  4518   4521   4523   4526       3   0.66944    2.00832    0.00000
-2.67776    0.00000    0.00000 ;     CB-    CG-   CD1-  HD13
  4518   4521   4527   4528       3   0.66944    2.00832    0.00000
-2.67776    0.00000    0.00000 ;     CB-    CG-   CD2-  HD21
  4518   4521   4527   4529       3   0.66944    2.00832    0.00000
-2.67776    0.00000    0.00000 ;     CB-    CG-   CD2-  HD22
  4518   4521   4527   4530       3   0.66944    2.00832    0.00000
-2.67776    0.00000    0.00000 ;     CB-    CG-   CD2-  HD23
  4519   4518   4516   4531       3   0.65084    1.95253    0.00000
-2.60338    0.00000    0.00000 ;    HB2-    CB-    CA-     C
  4519   4518   4521   4522       3   0.62760    1.88280    0.00000
-2.51040    0.00000    0.00000 ;    HB2-    CB-    CG-    HG
  4519   4518   4521   4523       3   0.66944    2.00832    0.00000
-2.67776    0.00000    0.00000 ;    HB2-    CB-    CG-   CD1
  4519   4518   4521   4527       3   0.66944    2.00832    0.00000
-2.67776    0.00000    0.00000 ;    HB2-    CB-    CG-   CD2
  4520   4518   4516   4531       3   0.65084    1.95253    0.00000
-2.60338    0.00000    0.00000 ;    HB3-    CB-    CA-     C
  4520   4518   4521   4522       3   0.62760    1.88280    0.00000
```

-2.51040 0.00000 0.00000 ; HB3- CB- CG- HG
4520 4518 4521 4523 3 0.66944 2.00832 0.00000
-2.67776 0.00000 0.00000 ; HB3- CB- CG- CD1
4520 4518 4521 4527 3 0.66944 2.00832 0.00000
-2.67776 0.00000 0.00000 ; HB3- CB- CG- CD2
4521 4518 4516 4531 3 10.33448 -1.14642 -11.54784
-2.40998 6.35968 0.00000 ; CG- CB- CA- C
4522 4521 4523 4524 3 0.62760 1.88280 0.00000
-2.51040 0.00000 0.00000 ; HG- CG- CD1- HD11
4522 4521 4523 4525 3 0.62760 1.88280 0.00000
-2.51040 0.00000 0.00000 ; HG- CG- CD1- HD12
4522 4521 4523 4526 3 0.62760 1.88280 0.00000
-2.51040 0.00000 0.00000 ; HG- CG- CD1- HD13
4522 4521 4527 4528 3 0.62760 1.88280 0.00000
-2.51040 0.00000 0.00000 ; HG- CG- CD2- HD21
4522 4521 4527 4529 3 0.62760 1.88280 0.00000
-2.51040 0.00000 0.00000 ; HG- CG- CD2- HD22
4522 4521 4527 4530 3 0.62760 1.88280 0.00000
-2.51040 0.00000 0.00000 ; HG- CG- CD2- HD23
4523 4521 4527 4528 3 0.66944 2.00832 0.00000
-2.67776 0.00000 0.00000 ; CD1- CG- CD2- HD21
4523 4521 4527 4529 3 0.66944 2.00832 0.00000
-2.67776 0.00000 0.00000 ; CD1- CG- CD2- HD22
4523 4521 4527 4530 3 0.66944 2.00832 0.00000
-2.67776 0.00000 0.00000 ; CD1- CG- CD2- HD23
4524 4523 4521 4527 3 0.66944 2.00832 0.00000
-2.67776 0.00000 0.00000 ; HD11- CD1- CG- CD2
4525 4523 4521 4527 3 0.66944 2.00832 0.00000
-2.67776 0.00000 0.00000 ; HD12- CD1- CG- CD2
4526 4523 4521 4527 3 0.66944 2.00832 0.00000
-2.67776 0.00000 0.00000 ; HD13- CD1- CG- CD2
4531 4533 4535 4536 3 0.00000 0.00000 0.00000
0.00000 0.00000 0.00000 ; C- N- CA- HA
4531 4533 4535 4537 3 11.71520 1.67360 15.06240 -
13.38880 0.00000 0.00000 ; C- N- CA- CB
4531 4533 4535 4542 3 1.75728 5.27184 2.25936
-7.02912 0.00000 0.00000 ; C- N- CA- C
4532 4531 4533 4534 3 29.28800 -8.36800 -20.92000
0.00000 0.00000 0.00000 ; O- C- N- H
4532 4531 4533 4535 3 20.92000 0.00000 -20.92000
0.00000 0.00000 0.00000 ; O- C- N- CA
4533 4535 4537 4538 3 0.65084 1.95253 0.00000
-2.60338 0.00000 0.00000 ; N- CA- CB- HB2
4533 4535 4537 4539 3 0.65084 1.95253 0.00000
-2.60338 0.00000 0.00000 ; N- CA- CB- HB3
4533 4535 4537 4540 3 7.86174 2.24681 -7.41405
-6.71114 5.35552 0.00000 ; N- CA- CB- OG
4533 4535 4542 4543 3 0.00000 0.00000 0.00000
0.00000 0.00000 0.00000 ; N- CA- C- O
4533 4535 4542 4544 3 17.40544 -5.02080 -13.22144
9.20480 0.00000 0.00000 ; N- CA- C- N
4534 4533 4535 4536 3 0.00000 0.00000 0.00000

```
0.00000    0.00000    0.00000 ;       H-     N-     CA-    HA
  4534   4533   4535   4537      3    0.00000    0.00000    0.00000
0.00000    0.00000    0.00000 ;       H-     N-     CA-    CB
  4534   4533   4535   4542      3    0.00000    0.00000    0.00000
0.00000    0.00000    0.00000 ;       H-     N-     CA-     C
  4535   4537   4540   4541      3    2.05853    2.46856    3.48109
-4.46851    0.23430    0.00000 ;      CA-    CB-    OG-    HG
  4535   4542   4544   4545      3   20.92000    0.00000  -20.92000
0.00000    0.00000    0.00000 ;      CA-     C-     N-     H
  4535   4542   4544   4546      3   20.92000    0.00000  -20.92000
0.00000    0.00000    0.00000 ;      CA-     C-     N-    CA
  4536   4535   4537   4538      3    0.65084    1.95253    0.00000
-2.60338    0.00000    0.00000 ;      HA-    CA-    CB-   HB2
  4536   4535   4537   4539      3    0.65084    1.95253    0.00000
-2.60338    0.00000    0.00000 ;      HA-    CA-    CB-   HB3
  4536   4535   4537   4540      3    1.04600   -1.04600    0.00000
0.00000    0.00000    0.00000 ;      HA-    CA-    CB-   OG
  4536   4535   4542   4543      3    3.68192   -4.35136    0.00000
1.33888    0.00000    0.00000 ;      HA-    CA-     C-     O
  4536   4535   4542   4544      3    0.00000    0.00000    0.00000
0.00000    0.00000    0.00000 ;      HA-    CA-     C-     N
  4537   4535   4542   4543      3    0.00000    0.00000    0.00000
0.00000    0.00000    0.00000 ;      CB-    CA-     C-     O
  4537   4535   4542   4544      3    2.51040    4.18400    1.67360
-6.69440    0.00000    0.00000 ;      CB-    CA-     C-     N
  4538   4537   4535   4542      3    0.65084    1.95253    0.00000
-2.60338    0.00000    0.00000 ;     HB2-    CB-    CA-     C
  4538   4537   4540   4541      3    0.69733    2.09200    0.00000
-2.78933    0.00000    0.00000 ;     HB2-    CB-    OG-    HG
  4539   4537   4535   4542      3    0.65084    1.95253    0.00000
-2.60338    0.00000    0.00000 ;     HB3-    CB-    CA-     C
  4539   4537   4540   4541      3    0.69733    2.09200    0.00000
-2.78933    0.00000    0.00000 ;     HB3-    CB-    OG-    HG
  4540   4537   4535   4542      3    7.34710    7.79898   -6.14211
-6.71114    4.31789    0.00000 ;      OG-    CB-    CA-     C
  4542   4544   4546   4547      3    0.00000    0.00000    0.00000
0.00000    0.00000    0.00000 ;       C-     N-    CA-    HA
  4542   4544   4546   4548      3   11.71520    1.67360   15.06240  -
13.38880    0.00000    0.00000 ;       C-     N-    CA-    CB
  4542   4544   4546   4553      3    1.75728    5.27184    2.25936
-7.02912    0.00000    0.00000 ;       C-     N-    CA-     C
  4543   4542   4544   4545      3   29.28800   -8.36800  -20.92000
0.00000    0.00000    0.00000 ;       O-     C-     N-     H
  4543   4542   4544   4546      3   20.92000    0.00000  -20.92000
0.00000    0.00000    0.00000 ;       O-     C-     N-    CA
  4544   4546   4548   4549      3    0.65084    1.95253    0.00000
-2.60338    0.00000    0.00000 ;       N-    CA-    CB-   HB2
  4544   4546   4548   4550      3    0.65084    1.95253    0.00000
-2.60338    0.00000    0.00000 ;       N-    CA-    CB-   HB3
  4544   4546   4548   4551      3    7.86174    2.24681   -7.41405
-6.71114    5.35552    0.00000 ;       N-    CA-    CB-    OG
  4544   4546   4553   4554      3    0.00000    0.00000    0.00000
```

0.00000 0.00000 0.00000 ; N- CA- C- O
4544 4546 4553 4555 3 17.40544 -5.02080 -13.22144
9.20480 0.00000 0.00000 ; N- CA- C- N
4545 4544 4546 4547 3 0.00000 0.00000 0.00000
0.00000 0.00000 0.00000 ; H- N- CA- HA
4545 4544 4546 4548 3 0.00000 0.00000 0.00000
0.00000 0.00000 0.00000 ; H- N- CA- CB
4545 4544 4546 4553 3 0.00000 0.00000 0.00000
0.00000 0.00000 0.00000 ; H- N- CA- C
4546 4548 4551 4552 3 2.05853 2.46856 3.48109
-4.46851 0.23430 0.00000 ; CA- CB- OG- HG
4546 4553 4555 4556 3 20.92000 0.00000 -20.92000
0.00000 0.00000 0.00000 ; CA- C- N- H
4546 4553 4555 4557 3 20.92000 0.00000 -20.92000
0.00000 0.00000 0.00000 ; CA- C- N- CA
4547 4546 4548 4549 3 0.65084 1.95253 0.00000
-2.60338 0.00000 0.00000 ; HA- CA- CB- HB2
4547 4546 4548 4550 3 0.65084 1.95253 0.00000
-2.60338 0.00000 0.00000 ; HA- CA- CB- HB3
4547 4546 4548 4551 3 1.04600 -1.04600 0.00000
0.00000 0.00000 0.00000 ; HA- CA- CB- OG
4547 4546 4553 4554 3 3.68192 -4.35136 0.00000
1.33888 0.00000 0.00000 ; HA- CA- C- O
4547 4546 4553 4555 3 0.00000 0.00000 0.00000
0.00000 0.00000 0.00000 ; HA- CA- C- N
4548 4546 4553 4554 3 0.00000 0.00000 0.00000
0.00000 0.00000 0.00000 ; CB- CA- C- O
4548 4546 4553 4555 3 2.51040 4.18400 1.67360
-6.69440 0.00000 0.00000 ; CB- CA- C- N
4549 4548 4546 4553 3 0.65084 1.95253 0.00000
-2.60338 0.00000 0.00000 ; HB2- CB- CA- C
4549 4548 4551 4552 3 0.69733 2.09200 0.00000
-2.78933 0.00000 0.00000 ; HB2- CB- OG- HG
4550 4548 4546 4553 3 0.65084 1.95253 0.00000
-2.60338 0.00000 0.00000 ; HB3- CB- CA- C
4550 4548 4551 4552 3 0.69733 2.09200 0.00000
-2.78933 0.00000 0.00000 ; HB3- CB- OG- HG
4551 4548 4546 4553 3 7.34710 7.79898 -6.14211
-6.71114 4.31789 0.00000 ; OG- CB- CA- C
4553 4555 4557 4558 3 0.00000 0.00000 0.00000
0.00000 0.00000 0.00000 ; C- N- CA- HA
4553 4555 4557 4559 3 11.71520 1.67360 15.06240 -
13.38880 0.00000 0.00000 ; C- N- CA- CB
4553 4555 4557 4567 3 1.75728 5.27184 2.25936
-7.02912 0.00000 0.00000 ; C- N- CA- C
4554 4553 4555 4556 3 29.28800 -8.36800 -20.92000
0.00000 0.00000 0.00000 ; O- C- N- H
4554 4553 4555 4557 3 20.92000 0.00000 -20.92000
0.00000 0.00000 0.00000 ; O- C- N- CA
4555 4557 4559 4560 3 0.65084 1.95253 0.00000
-2.60338 0.00000 0.00000 ; N- CA- CB- HB
4555 4557 4559 4561 3 3.83673 0.44769 -1.77402

-2.47693 -0.03347 0.00000 ; N- CA- CB- CG2
4555 4557 4559 4565 3 4.93294 1.13386 -3.12963
-5.27184 3.17984 0.00000 ; N- CA- CB- OG1
4555 4557 4567 4568 3 0.00000 0.00000 0.00000
0.00000 0.00000 0.00000 ; N- CA- C- O
4555 4557 4567 4569 3 17.40544 -5.02080 -13.22144
9.20480 0.00000 0.00000 ; N- CA- C- N
4556 4555 4557 4558 3 0.00000 0.00000 0.00000
0.00000 0.00000 0.00000 ; H- N- CA- HA
4556 4555 4557 4559 3 0.00000 0.00000 0.00000
0.00000 0.00000 0.00000 ; H- N- CA- CB
4556 4555 4557 4567 3 0.00000 0.00000 0.00000
0.00000 0.00000 0.00000 ; H- N- CA- C
4557 4559 4561 4562 3 0.66944 2.00832 0.00000
-2.67776 0.00000 0.00000 ; CA- CB- CG2- HG21
4557 4559 4561 4563 3 0.66944 2.00832 0.00000
-2.67776 0.00000 0.00000 ; CA- CB- CG2- HG22
4557 4559 4561 4564 3 0.66944 2.00832 0.00000
-2.67776 0.00000 0.00000 ; CA- CB- CG2- HG23
4557 4559 4565 4566 3 1.12131 2.98738 1.66523
-3.94970 0.43514 0.00000 ; CA- CB- OG1- HG1
4557 4567 4569 4570 3 20.92000 0.00000 -20.92000
0.00000 0.00000 0.00000 ; CA- C- N- H
4557 4567 4569 4571 3 20.92000 0.00000 -20.92000
0.00000 0.00000 0.00000 ; CA- C- N- CA
4558 4557 4559 4560 3 0.65084 1.95253 0.00000
-2.60338 0.00000 0.00000 ; HA- CA- CB- HB
4558 4557 4559 4561 3 0.65084 1.95253 0.00000
-2.60338 0.00000 0.00000 ; HA- CA- CB- CG2
4558 4557 4559 4565 3 1.04600 -1.04600 0.00000
0.00000 0.00000 0.00000 ; HA- CA- CB- OG1
4558 4557 4567 4568 3 3.68192 -4.35136 0.00000
1.33888 0.00000 0.00000 ; HA- CA- C- O
4558 4557 4567 4569 3 0.00000 0.00000 0.00000
0.00000 0.00000 0.00000 ; HA- CA- C- N
4559 4557 4567 4568 3 0.00000 0.00000 0.00000
0.00000 0.00000 0.00000 ; CB- CA- C- O
4559 4557 4567 4569 3 2.51040 4.18400 1.67360
-6.69440 0.00000 0.00000 ; CB- CA- C- N
4560 4559 4557 4567 3 0.65084 1.95253 0.00000
-2.60338 0.00000 0.00000 ; HB- CB- CA- C
4560 4559 4561 4562 3 0.65084 1.95253 0.00000
-2.60338 0.00000 0.00000 ; HB- CB- CG2- HG21
4560 4559 4561 4563 3 0.65084 1.95253 0.00000
-2.60338 0.00000 0.00000 ; HB- CB- CG2- HG22
4560 4559 4561 4564 3 0.65084 1.95253 0.00000
-2.60338 0.00000 0.00000 ; HB- CB- CG2- HG23
4560 4559 4565 4566 3 0.69733 2.09200 0.00000
-2.78933 0.00000 0.00000 ; HB- CB- OG1- HG1
4561 4559 4557 4567 3 5.67350 3.55640 -6.16722
-2.47693 3.74886 0.00000 ; CG2- CB- CA- C
4561 4559 4565 4566 3 4.74047 0.27196 -2.26773

```
-3.94970    1.60666    0.00000 ;    CG2-    CB-   OG1-   HG1
  4562   4561   4559   4565      3    1.04600   -1.04600    0.00000
0.00000    0.00000    0.00000 ;   HG21-   CG2-    CB-   OG1
  4563   4561   4559   4565      3    1.04600   -1.04600    0.00000
0.00000    0.00000    0.00000 ;   HG22-   CG2-    CB-   OG1
  4564   4561   4559   4565      3    1.04600   -1.04600    0.00000
0.00000    0.00000    0.00000 ;   HG23-   CG2-    CB-   OG1
  4565   4559   4557   4567      3    6.53541    6.87013   -6.21742
-5.27184    5.22163    0.00000 ;    OG1-    CB-    CA-     C
  4567   4569   4571   4572      3    0.00000    0.00000    0.00000
0.00000    0.00000    0.00000 ;      C-     N-    CA-    HA
  4567   4569   4571   4573      3   11.71520    1.67360   15.06240  -
13.38880    0.00000    0.00000 ;      C-     N-    CA-    CB
  4567   4569   4571   4586      3    1.75728    5.27184    2.25936
-7.02912    0.00000    0.00000 ;      C-     N-    CA-     C
  4568   4567   4569   4570      3   29.28800   -8.36800  -20.92000
0.00000    0.00000    0.00000 ;      O-     C-     N-     H
  4568   4567   4569   4571      3   20.92000    0.00000  -20.92000
0.00000    0.00000    0.00000 ;      O-     C-     N-    CA
  4569   4571   4573   4574      3    0.65084    1.95253    0.00000
-2.60338    0.00000    0.00000 ;      N-    CA-    CB-   HB2
  4569   4571   4573   4575      3    0.65084    1.95253    0.00000
-2.60338    0.00000    0.00000 ;      N-    CA-    CB-   HB3
  4569   4571   4573   4576      3    3.79070    1.39746   -4.61077
-2.40998    2.44346    0.00000 ;      N-    CA-    CB-    CG
  4569   4571   4586   4587      3    0.00000    0.00000    0.00000
0.00000    0.00000    0.00000 ;      N-    CA-     C-     O
  4569   4571   4586   4588      3   17.40544   -5.02080  -13.22144
9.20480    0.00000    0.00000 ;      N-    CA-     C-     N
  4570   4569   4571   4572      3    0.00000    0.00000    0.00000
0.00000    0.00000    0.00000 ;      H-     N-    CA-    HA
  4570   4569   4571   4573      3    0.00000    0.00000    0.00000
0.00000    0.00000    0.00000 ;      H-     N-    CA-    CB
  4570   4569   4571   4586      3    0.00000    0.00000    0.00000
0.00000    0.00000    0.00000 ;      H-     N-    CA-     C
  4571   4573   4576   4577      3    0.66944    2.00832    0.00000
-2.67776    0.00000    0.00000 ;     CA-    CB-    CG-    HG
  4571   4573   4576   4578      3    3.90367    0.19665   -6.21742
-2.37651    5.99149    0.00000 ;     CA-    CB-    CG-   CD1
  4571   4573   4576   4582      3    3.90367    0.19665   -6.21742
-2.37651    5.99149    0.00000 ;     CA-    CB-    CG-   CD2
  4571   4586   4588   4589      3   20.92000    0.00000  -20.92000
0.00000    0.00000    0.00000 ;     CA-     C-     N-     H
  4571   4586   4588   4590      3   20.92000    0.00000  -20.92000
0.00000    0.00000    0.00000 ;     CA-     C-     N-    CA
  4572   4571   4573   4574      3    0.65084    1.95253    0.00000
-2.60338    0.00000    0.00000 ;     HA-    CA-    CB-   HB2
  4572   4571   4573   4575      3    0.65084    1.95253    0.00000
-2.60338    0.00000    0.00000 ;     HA-    CA-    CB-   HB3
  4572   4571   4573   4576      3    0.65084    1.95253    0.00000
-2.60338    0.00000    0.00000 ;     HA-    CA-    CB-    CG
  4572   4571   4586   4587      3    3.68192   -4.35136    0.00000
```

```
1.33888    0.00000    0.00000 ;      HA-    CA-     C-     O
  4572   4571   4586   4588       3    0.00000    0.00000    0.00000
0.00000    0.00000    0.00000 ;      HA-    CA-     C-     N
  4573   4571   4586   4587       3    0.00000    0.00000    0.00000
0.00000    0.00000    0.00000 ;      CB-    CA-     C-     O
  4573   4571   4586   4588       3    2.51040    4.18400    1.67360
-6.69440    0.00000    0.00000 ;      CB-    CA-     C-     N
  4573   4576   4578   4579       3    0.66944    2.00832    0.00000
-2.67776    0.00000    0.00000 ;      CB-    CG-   CD1-  HD11
  4573   4576   4578   4580       3    0.66944    2.00832    0.00000
-2.67776    0.00000    0.00000 ;      CB-    CG-   CD1-  HD12
  4573   4576   4578   4581       3    0.66944    2.00832    0.00000
-2.67776    0.00000    0.00000 ;      CB-    CG-   CD1-  HD13
  4573   4576   4582   4583       3    0.66944    2.00832    0.00000
-2.67776    0.00000    0.00000 ;      CB-    CG-   CD2-  HD21
  4573   4576   4582   4584       3    0.66944    2.00832    0.00000
-2.67776    0.00000    0.00000 ;      CB-    CG-   CD2-  HD22
  4573   4576   4582   4585       3    0.66944    2.00832    0.00000
-2.67776    0.00000    0.00000 ;      CB-    CG-   CD2-  HD23
  4574   4573   4571   4586       3    0.65084    1.95253    0.00000
-2.60338    0.00000    0.00000 ;     HB2-    CB-     CA-     C
  4574   4573   4576   4577       3    0.62760    1.88280    0.00000
-2.51040    0.00000    0.00000 ;     HB2-    CB-     CG-    HG
  4574   4573   4576   4578       3    0.66944    2.00832    0.00000
-2.67776    0.00000    0.00000 ;     HB2-    CB-     CG-   CD1
  4574   4573   4576   4582       3    0.66944    2.00832    0.00000
-2.67776    0.00000    0.00000 ;     HB2-    CB-     CG-   CD2
  4575   4573   4571   4586       3    0.65084    1.95253    0.00000
-2.60338    0.00000    0.00000 ;     HB3-    CB-     CA-     C
  4575   4573   4576   4577       3    0.62760    1.88280    0.00000
-2.51040    0.00000    0.00000 ;     HB3-    CB-     CG-    HG
  4575   4573   4576   4578       3    0.66944    2.00832    0.00000
-2.67776    0.00000    0.00000 ;     HB3-    CB-     CG-   CD1
  4575   4573   4576   4582       3    0.66944    2.00832    0.00000
-2.67776    0.00000    0.00000 ;     HB3-    CB-     CG-   CD2
  4576   4573   4571   4586       3   10.33448   -1.14642  -11.54784
-2.40998    6.35968    0.00000 ;      CG-    CB-     CA-     C
  4577   4576   4578   4579       3    0.62760    1.88280    0.00000
-2.51040    0.00000    0.00000 ;      HG-    CG-   CD1-  HD11
  4577   4576   4578   4580       3    0.62760    1.88280    0.00000
-2.51040    0.00000    0.00000 ;      HG-    CG-   CD1-  HD12
  4577   4576   4578   4581       3    0.62760    1.88280    0.00000
-2.51040    0.00000    0.00000 ;      HG-    CG-   CD1-  HD13
  4577   4576   4582   4583       3    0.62760    1.88280    0.00000
-2.51040    0.00000    0.00000 ;      HG-    CG-   CD2-  HD21
  4577   4576   4582   4584       3    0.62760    1.88280    0.00000
-2.51040    0.00000    0.00000 ;      HG-    CG-   CD2-  HD22
  4577   4576   4582   4585       3    0.62760    1.88280    0.00000
-2.51040    0.00000    0.00000 ;      HG-    CG-   CD2-  HD23
  4578   4576   4582   4583       3    0.66944    2.00832    0.00000
-2.67776    0.00000    0.00000 ;     CD1-    CG-   CD2-  HD21
  4578   4576   4582   4584       3    0.66944    2.00832    0.00000
```

-2.67776 0.00000 0.00000 ; CD1- CG- CD2- HD22
4578 4576 4582 4585 3 0.66944 2.00832 0.00000
-2.67776 0.00000 0.00000 ; CD1- CG- CD2- HD23
4579 4578 4576 4582 3 0.66944 2.00832 0.00000
-2.67776 0.00000 0.00000 ; HD11- CD1- CG- CD2
4580 4578 4576 4582 3 0.66944 2.00832 0.00000
-2.67776 0.00000 0.00000 ; HD12- CD1- CG- CD2
4581 4578 4576 4582 3 0.66944 2.00832 0.00000
-2.67776 0.00000 0.00000 ; HD13- CD1- CG- CD2
4586 4588 4590 4591 3 0.00000 0.00000 0.00000
0.00000 0.00000 0.00000 ; C- N- CA- HA
4586 4588 4590 4592 3 11.71520 1.67360 15.06240 -
13.38880 0.00000 0.00000 ; C- N- CA- CB
4586 4588 4590 4605 3 1.75728 5.27184 2.25936
-7.02912 0.00000 0.00000 ; C- N- CA- C
4587 4586 4588 4589 3 29.28800 -8.36800 -20.92000
0.00000 0.00000 0.00000 ; O- C- N- H
4587 4586 4588 4590 3 20.92000 0.00000 -20.92000
0.00000 0.00000 0.00000 ; O- C- N- CA
4588 4590 4592 4593 3 0.65084 1.95253 0.00000
-2.60338 0.00000 0.00000 ; N- CA- CB- HB2
4588 4590 4592 4594 3 0.65084 1.95253 0.00000
-2.60338 0.00000 0.00000 ; N- CA- CB- HB3
4588 4590 4592 4595 3 3.79070 1.39746 -4.61077
-2.40998 2.44346 0.00000 ; N- CA- CB- CG
4588 4590 4605 4606 3 0.00000 0.00000 0.00000
0.00000 0.00000 0.00000 ; N- CA- C- O
4588 4590 4605 4607 3 17.40544 -5.02080 -13.22144
9.20480 0.00000 0.00000 ; N- CA- C- N
4589 4588 4590 4591 3 0.00000 0.00000 0.00000
0.00000 0.00000 0.00000 ; H- N- CA- HA
4589 4588 4590 4592 3 0.00000 0.00000 0.00000
0.00000 0.00000 0.00000 ; H- N- CA- CB
4589 4588 4590 4605 3 0.00000 0.00000 0.00000
0.00000 0.00000 0.00000 ; H- N- CA- C
4590 4592 4595 4596 3 0.66944 2.00832 0.00000
-2.67776 0.00000 0.00000 ; CA- CB- CG- HG
4590 4592 4595 4597 3 3.90367 0.19665 -6.21742
-2.37651 5.99149 0.00000 ; CA- CB- CG- CD1
4590 4592 4595 4601 3 3.90367 0.19665 -6.21742
-2.37651 5.99149 0.00000 ; CA- CB- CG- CD2
4590 4605 4607 4608 3 20.92000 0.00000 -20.92000
0.00000 0.00000 0.00000 ; CA- C- N- H
4590 4605 4607 4609 3 20.92000 0.00000 -20.92000
0.00000 0.00000 0.00000 ; CA- C- N- CA
4591 4590 4592 4593 3 0.65084 1.95253 0.00000
-2.60338 0.00000 0.00000 ; HA- CA- CB- HB2
4591 4590 4592 4594 3 0.65084 1.95253 0.00000
-2.60338 0.00000 0.00000 ; HA- CA- CB- HB3
4591 4590 4592 4595 3 0.65084 1.95253 0.00000
-2.60338 0.00000 0.00000 ; HA- CA- CB- CG
4591 4590 4605 4606 3 3.68192 -4.35136 0.00000

```
1.33888    0.00000    0.00000 ;     HA-    CA-     C-     O
  4591   4590   4605   4607      3   0.00000    0.00000    0.00000
0.00000    0.00000    0.00000 ;     HA-    CA-     C-     N
  4592   4590   4605   4606      3   0.00000    0.00000    0.00000
0.00000    0.00000    0.00000 ;     CB-    CA-     C-     O
  4592   4590   4605   4607      3   2.51040    4.18400    1.67360
-6.69440    0.00000    0.00000 ;     CB-    CA-     C-     N
  4592   4595   4597   4598      3   0.66944    2.00832    0.00000
-2.67776    0.00000    0.00000 ;     CB-    CG-   CD1-  HD11
  4592   4595   4597   4599      3   0.66944    2.00832    0.00000
-2.67776    0.00000    0.00000 ;     CB-    CG-   CD1-  HD12
  4592   4595   4597   4600      3   0.66944    2.00832    0.00000
-2.67776    0.00000    0.00000 ;     CB-    CG-   CD1-  HD13
  4592   4595   4601   4602      3   0.66944    2.00832    0.00000
-2.67776    0.00000    0.00000 ;     CB-    CG-   CD2-  HD21
  4592   4595   4601   4603      3   0.66944    2.00832    0.00000
-2.67776    0.00000    0.00000 ;     CB-    CG-   CD2-  HD22
  4592   4595   4601   4604      3   0.66944    2.00832    0.00000
-2.67776    0.00000    0.00000 ;     CB-    CG-   CD2-  HD23
  4593   4592   4590   4605      3   0.65084    1.95253    0.00000
-2.60338    0.00000    0.00000 ;    HB2-    CB-    CA-     C
  4593   4592   4595   4596      3   0.62760    1.88280    0.00000
-2.51040    0.00000    0.00000 ;    HB2-    CB-    CG-    HG
  4593   4592   4595   4597      3   0.66944    2.00832    0.00000
-2.67776    0.00000    0.00000 ;    HB2-    CB-    CG-   CD1
  4593   4592   4595   4601      3   0.66944    2.00832    0.00000
-2.67776    0.00000    0.00000 ;    HB2-    CB-    CG-   CD2
  4594   4592   4590   4605      3   0.65084    1.95253    0.00000
-2.60338    0.00000    0.00000 ;    HB3-    CB-    CA-     C
  4594   4592   4595   4596      3   0.62760    1.88280    0.00000
-2.51040    0.00000    0.00000 ;    HB3-    CB-    CG-    HG
  4594   4592   4595   4597      3   0.66944    2.00832    0.00000
-2.67776    0.00000    0.00000 ;    HB3-    CB-    CG-   CD1
  4594   4592   4595   4601      3   0.66944    2.00832    0.00000
-2.67776    0.00000    0.00000 ;    HB3-    CB-    CG-   CD2
  4595   4592   4590   4605      3  10.33448   -1.14642  -11.54784
-2.40998    6.35968    0.00000 ;     CG-    CB-    CA-     C
  4596   4595   4597   4598      3   0.62760    1.88280    0.00000
-2.51040    0.00000    0.00000 ;     HG-    CG-   CD1-  HD11
  4596   4595   4597   4599      3   0.62760    1.88280    0.00000
-2.51040    0.00000    0.00000 ;     HG-    CG-   CD1-  HD12
  4596   4595   4597   4600      3   0.62760    1.88280    0.00000
-2.51040    0.00000    0.00000 ;     HG-    CG-   CD1-  HD13
  4596   4595   4601   4602      3   0.62760    1.88280    0.00000
-2.51040    0.00000    0.00000 ;     HG-    CG-   CD2-  HD21
  4596   4595   4601   4603      3   0.62760    1.88280    0.00000
-2.51040    0.00000    0.00000 ;     HG-    CG-   CD2-  HD22
  4596   4595   4601   4604      3   0.62760    1.88280    0.00000
-2.51040    0.00000    0.00000 ;     HG-    CG-   CD2-  HD23
  4597   4595   4601   4602      3   0.66944    2.00832    0.00000
-2.67776    0.00000    0.00000 ;    CD1-    CG-   CD2-  HD21
  4597   4595   4601   4603      3   0.66944    2.00832    0.00000
```

-2.67776 0.00000 0.00000 ; CD1- CG- CD2- HD22
4597 4595 4601 4604 3 0.66944 2.00832 0.00000
-2.67776 0.00000 0.00000 ; CD1- CG- CD2- HD23
4598 4597 4595 4601 3 0.66944 2.00832 0.00000
-2.67776 0.00000 0.00000 ; HD11- CD1- CG- CD2
4599 4597 4595 4601 3 0.66944 2.00832 0.00000
-2.67776 0.00000 0.00000 ; HD12- CD1- CG- CD2
4600 4597 4595 4601 3 0.66944 2.00832 0.00000
-2.67776 0.00000 0.00000 ; HD13- CD1- CG- CD2
4605 4607 4609 4610 3 0.00000 0.00000 0.00000
0.00000 0.00000 0.00000 ; C- N- CA- HA
4605 4607 4609 4611 3 11.71520 1.67360 15.06240 -
13.38880 0.00000 0.00000 ; C- N- CA- CB
4605 4607 4609 4627 3 1.75728 5.27184 2.25936
-7.02912 0.00000 0.00000 ; C- N- CA- C
4606 4605 4607 4608 3 29.28800 -8.36800 -20.92000
0.00000 0.00000 0.00000 ; O- C- N- H
4606 4605 4607 4609 3 20.92000 0.00000 -20.92000
0.00000 0.00000 0.00000 ; O- C- N- CA
4607 4609 4611 4612 3 0.65084 1.95253 0.00000
-2.60338 0.00000 0.00000 ; N- CA- CB- HB2
4607 4609 4611 4613 3 0.65084 1.95253 0.00000
-2.60338 0.00000 0.00000 ; N- CA- CB- HB3
4607 4609 4611 4614 3 0.65084 1.95253 0.00000
-2.60338 0.00000 0.00000 ; N- CA- CB- CG
4607 4609 4627 4628 3 0.00000 0.00000 0.00000
0.00000 0.00000 0.00000 ; N- CA- C- O
4607 4609 4627 4629 3 17.40544 -5.02080 -13.22144
9.20480 0.00000 0.00000 ; N- CA- C- N
4608 4607 4609 4610 3 0.00000 0.00000 0.00000
0.00000 0.00000 0.00000 ; H- N- CA- HA
4608 4607 4609 4611 3 0.00000 0.00000 0.00000
0.00000 0.00000 0.00000 ; H- N- CA- CB
4608 4607 4609 4627 3 0.00000 0.00000 0.00000
0.00000 0.00000 0.00000 ; H- N- CA- C
4609 4611 4614 4615 3 0.66944 2.00832 0.00000
-2.67776 0.00000 0.00000 ; CA- CB- CG- HG2
4609 4611 4614 4616 3 0.66944 2.00832 0.00000
-2.67776 0.00000 0.00000 ; CA- CB- CG- HG3
4609 4611 4614 4617 3 3.68192 3.09616 -2.09200
-3.01248 0.00000 0.00000 ; CA- CB- CG- CD
4609 4627 4629 4630 3 20.92000 0.00000 -20.92000
0.00000 0.00000 0.00000 ; CA- C- N- H
4609 4627 4629 4631 3 20.92000 0.00000 -20.92000
0.00000 0.00000 0.00000 ; CA- C- N- CA
4610 4609 4611 4612 3 0.65084 1.95253 0.00000
-2.60338 0.00000 0.00000 ; HA- CA- CB- HB2
4610 4609 4611 4613 3 0.65084 1.95253 0.00000
-2.60338 0.00000 0.00000 ; HA- CA- CB- HB3
4610 4609 4611 4614 3 0.65084 1.95253 0.00000
-2.60338 0.00000 0.00000 ; HA- CA- CB- CG
4610 4609 4627 4628 3 3.68192 -4.35136 0.00000

1.33888 0.00000 0.00000 ; HA- CA- C- O
4610 4609 4627 4629 3 0.00000 0.00000 0.00000
0.00000 0.00000 0.00000 ; HA- CA- C- N
4611 4609 4627 4628 3 0.00000 0.00000 0.00000
0.00000 0.00000 0.00000 ; CB- CA- C- O
4611 4609 4627 4629 3 2.51040 4.18400 1.67360
-6.69440 0.00000 0.00000 ; CB- CA- C- N
4611 4614 4617 4618 3 0.66944 2.00832 0.00000
-2.67776 0.00000 0.00000 ; CB- CG- CD- HD2
4611 4614 4617 4619 3 0.66944 2.00832 0.00000
-2.67776 0.00000 0.00000 ; CB- CG- CD- HD3
4611 4614 4617 4620 3 3.68192 3.09616 -2.09200
-3.01248 0.00000 0.00000 ; CB- CG- CD- CE
4612 4611 4609 4627 3 0.65084 1.95253 0.00000
-2.60338 0.00000 0.00000 ; HB2- CB- CA- C
4612 4611 4614 4615 3 0.62760 1.88280 0.00000
-2.51040 0.00000 0.00000 ; HB2- CB- CG- HG2
4612 4611 4614 4616 3 0.62760 1.88280 0.00000
-2.51040 0.00000 0.00000 ; HB2- CB- CG- HG3
4612 4611 4614 4617 3 0.66944 2.00832 0.00000
-2.67776 0.00000 0.00000 ; HB2- CB- CG- CD
4613 4611 4609 4627 3 0.65084 1.95253 0.00000
-2.60338 0.00000 0.00000 ; HB3- CB- CA- C
4613 4611 4614 4615 3 0.62760 1.88280 0.00000
-2.51040 0.00000 0.00000 ; HB3- CB- CG- HG2
4613 4611 4614 4616 3 0.62760 1.88280 0.00000
-2.51040 0.00000 0.00000 ; HB3- CB- CG- HG3
4613 4611 4614 4617 3 0.66944 2.00832 0.00000
-2.67776 0.00000 0.00000 ; HB3- CB- CG- CD
4614 4611 4609 4627 3 0.65084 1.95253 0.00000
-2.60338 0.00000 0.00000 ; CG- CB- CA- C
4614 4617 4620 4621 3 0.65084 1.95253 0.00000
-2.60338 0.00000 0.00000 ; CG- CD- CE- HE2
4614 4617 4620 4622 3 0.65084 1.95253 0.00000
-2.60338 0.00000 0.00000 ; CG- CD- CE- HE3
4614 4617 4620 4623 3 0.65084 1.95253 0.00000
-2.60338 0.00000 0.00000 ; CG- CD- CE- NZ
4615 4614 4617 4618 3 0.62760 1.88280 0.00000
-2.51040 0.00000 0.00000 ; HG2- CG- CD- HD2
4615 4614 4617 4619 3 0.62760 1.88280 0.00000
-2.51040 0.00000 0.00000 ; HG2- CG- CD- HD3
4615 4614 4617 4620 3 0.66944 2.00832 0.00000
-2.67776 0.00000 0.00000 ; HG2- CG- CD- CE
4616 4614 4617 4618 3 0.62760 1.88280 0.00000
-2.51040 0.00000 0.00000 ; HG3- CG- CD- HD2
4616 4614 4617 4619 3 0.62760 1.88280 0.00000
-2.51040 0.00000 0.00000 ; HG3- CG- CD- HD3
4616 4614 4617 4620 3 0.66944 2.00832 0.00000
-2.67776 0.00000 0.00000 ; HG3- CG- CD- CE
4617 4620 4623 4624 3 0.65084 1.95253 0.00000
-2.60338 0.00000 0.00000 ; CD- CE- NZ- HZ1
4617 4620 4623 4625 3 0.65084 1.95253 0.00000

-2.60338 0.00000 0.00000 ; CD- CE- NZ- HZ2
4617 4620 4623 4626 3 0.65084 1.95253 0.00000
-2.60338 0.00000 0.00000 ; CD- CE- NZ- HZ3
4618 4617 4620 4621 3 0.65084 1.95253 0.00000
-2.60338 0.00000 0.00000 ; HD2- CD- CE- HE2
4618 4617 4620 4622 3 0.65084 1.95253 0.00000
-2.60338 0.00000 0.00000 ; HD2- CD- CE- HE3
4618 4617 4620 4623 3 0.65084 1.95253 0.00000
-2.60338 0.00000 0.00000 ; HD2- CD- CE- NZ
4619 4617 4620 4621 3 0.65084 1.95253 0.00000
-2.60338 0.00000 0.00000 ; HD3- CD- CE- HE2
4619 4617 4620 4622 3 0.65084 1.95253 0.00000
-2.60338 0.00000 0.00000 ; HD3- CD- CE- HE3
4619 4617 4620 4623 3 0.65084 1.95253 0.00000
-2.60338 0.00000 0.00000 ; HD3- CD- CE- NZ
4621 4620 4623 4624 3 0.65084 1.95253 0.00000
-2.60338 0.00000 0.00000 ; HE2- CE- NZ- HZ1
4621 4620 4623 4625 3 0.65084 1.95253 0.00000
-2.60338 0.00000 0.00000 ; HE2- CE- NZ- HZ2
4621 4620 4623 4626 3 0.65084 1.95253 0.00000
-2.60338 0.00000 0.00000 ; HE2- CE- NZ- HZ3
4622 4620 4623 4624 3 0.65084 1.95253 0.00000
-2.60338 0.00000 0.00000 ; HE3- CE- NZ- HZ1
4622 4620 4623 4625 3 0.65084 1.95253 0.00000
-2.60338 0.00000 0.00000 ; HE3- CE- NZ- HZ2
4622 4620 4623 4626 3 0.65084 1.95253 0.00000
-2.60338 0.00000 0.00000 ; HE3- CE- NZ- HZ3
4627 4629 4631 4632 3 0.00000 0.00000 0.00000
0.00000 0.00000 0.00000 ; C- N- CA- HA
4627 4629 4631 4633 3 11.71520 1.67360 15.06240 -
13.38880 0.00000 0.00000 ; C- N- CA- CB
4627 4629 4631 4641 3 1.75728 5.27184 2.25936
-7.02912 0.00000 0.00000 ; C- N- CA- C
4628 4627 4629 4630 3 29.28800 -8.36800 -20.92000
0.00000 0.00000 0.00000 ; O- C- N- H
4628 4627 4629 4631 3 20.92000 0.00000 -20.92000
0.00000 0.00000 0.00000 ; O- C- N- CA
4629 4631 4633 4634 3 0.65084 1.95253 0.00000
-2.60338 0.00000 0.00000 ; N- CA- CB- HB2
4629 4631 4633 4635 3 0.65084 1.95253 0.00000
-2.60338 0.00000 0.00000 ; N- CA- CB- HB3
4629 4631 4633 4636 3 5.99567 3.29281 -4.46014
-0.55229 1.97485 0.00000 ; N- CA- CB- CG
4629 4631 4641 4642 3 0.00000 0.00000 0.00000
0.00000 0.00000 0.00000 ; N- CA- C- O
4629 4631 4641 4643 3 17.40544 -5.02080 -13.22144
9.20480 0.00000 0.00000 ; N- CA- C- N
4630 4629 4631 4632 3 0.00000 0.00000 0.00000
0.00000 0.00000 0.00000 ; H- N- CA- HA
4630 4629 4631 4633 3 0.00000 0.00000 0.00000
0.00000 0.00000 0.00000 ; H- N- CA- CB
4630 4629 4631 4641 3 0.00000 0.00000 0.00000

```
0.00000    0.00000    0.00000 ;      H-     N-    CA-     C
  4631   4633   4636   4637       3    0.00000    0.00000    0.00000
0.00000    0.00000    0.00000 ;     CA-    CB-    CG-   OD1
  4631   4633   4636   4638       3    8.84916   -0.31380   -4.32626
5.03754    0.26778    0.00000 ;     CA-    CB-    CG-   ND2
  4631   4641   4643   4644       3   20.92000    0.00000  -20.92000
0.00000    0.00000    0.00000 ;     CA-     C-     N-     H
  4631   4641   4643   4645       3   20.92000    0.00000  -20.92000
0.00000    0.00000    0.00000 ;     CA-     C-     N-    CA
  4632   4631   4633   4634       3    0.65084    1.95253    0.00000
-2.60338    0.00000    0.00000 ;     HA-    CA-    CB-   HB2
  4632   4631   4633   4635       3    0.65084    1.95253    0.00000
-2.60338    0.00000    0.00000 ;     HA-    CA-    CB-   HB3
  4632   4631   4633   4636       3    0.65084    1.95253    0.00000
-2.60338    0.00000    0.00000 ;     HA-    CA-    CB-    CG
  4632   4631   4641   4642       3    3.68192   -4.35136    0.00000
1.33888    0.00000    0.00000 ;     HA-    CA-     C-     O
  4632   4631   4641   4643       3    0.00000    0.00000    0.00000
0.00000    0.00000    0.00000 ;     HA-    CA-     C-     N
  4633   4631   4641   4642       3    0.00000    0.00000    0.00000
0.00000    0.00000    0.00000 ;     CB-    CA-     C-     O
  4633   4631   4641   4643       3    2.51040    4.18400    1.67360
-6.69440    0.00000    0.00000 ;     CB-    CA-     C-     N
  4633   4636   4638   4639       3   20.92000    0.00000  -20.92000
0.00000    0.00000    0.00000 ;     CB-    CG-   ND2-  HD21
  4633   4636   4638   4640       3   20.92000    0.00000  -20.92000
0.00000    0.00000    0.00000 ;     CB-    CG-   ND2-  HD22
  4634   4633   4631   4641       3    0.65084    1.95253    0.00000
-2.60338    0.00000    0.00000 ;    HB2-    CB-    CA-     C
  4634   4633   4636   4637       3    3.68192   -4.35136    0.00000
1.33888    0.00000    0.00000 ;    HB2-    CB-    CG-   OD1
  4634   4633   4636   4638       3    0.00000    0.00000    0.00000
0.00000    0.00000    0.00000 ;    HB2-    CB-    CG-   ND2
  4635   4633   4631   4641       3    0.65084    1.95253    0.00000
-2.60338    0.00000    0.00000 ;    HB3-    CB-    CA-     C
  4635   4633   4636   4637       3    3.68192   -4.35136    0.00000
1.33888    0.00000    0.00000 ;    HB3-    CB-    CG-   OD1
  4635   4633   4636   4638       3    0.00000    0.00000    0.00000
0.00000    0.00000    0.00000 ;    HB3-    CB-    CG-   ND2
  4636   4633   4631   4641       3    7.94542   -3.96225   -6.11701
-0.55229    3.58150    0.00000 ;     CG-    CB-    CA-     C
  4637   4636   4638   4639       3   29.28800   -8.36800  -20.92000
0.00000    0.00000    0.00000 ;    OD1-    CG-   ND2-  HD21
  4637   4636   4638   4640       3   29.28800   -8.36800  -20.92000
0.00000    0.00000    0.00000 ;    OD1-    CG-   ND2-  HD22
  4641   4643   4645   4646       3    0.00000    0.00000    0.00000
0.00000    0.00000    0.00000 ;      C-     N-    CA-    HA
  4641   4643   4645   4647       3   11.71520    1.67360   15.06240  -
13.38880    0.00000    0.00000 ;      C-     N-    CA-    CB
  4641   4643   4645   4660       3    1.75728    5.27184    2.25936
-7.02912    0.00000    0.00000 ;      C-     N-    CA-     C
  4642   4641   4643   4644       3   29.28800   -8.36800  -20.92000
```

0.00000 0.00000 0.00000 ; O- C- N- H
4642 4641 4643 4645 3 20.92000 0.00000 -20.92000
0.00000 0.00000 0.00000 ; O- C- N- CA
4643 4645 4647 4648 3 0.65084 1.95253 0.00000
-2.60338 0.00000 0.00000 ; N- CA- CB- HB2
4643 4645 4647 4649 3 0.65084 1.95253 0.00000
-2.60338 0.00000 0.00000 ; N- CA- CB- HB3
4643 4645 4647 4650 3 3.79070 1.39746 -4.61077
-2.40998 2.44346 0.00000 ; N- CA- CB- CG
4643 4645 4660 4661 3 0.00000 0.00000 0.00000
0.00000 0.00000 0.00000 ; N- CA- C- O
4643 4645 4660 4662 3 17.40544 -5.02080 -13.22144
9.20480 0.00000 0.00000 ; N- CA- C- N
4644 4643 4645 4646 3 0.00000 0.00000 0.00000
0.00000 0.00000 0.00000 ; H- N- CA- HA
4644 4643 4645 4647 3 0.00000 0.00000 0.00000
0.00000 0.00000 0.00000 ; H- N- CA- CB
4644 4643 4645 4660 3 0.00000 0.00000 0.00000
0.00000 0.00000 0.00000 ; H- N- CA- C
4645 4647 4650 4651 3 0.66944 2.00832 0.00000
-2.67776 0.00000 0.00000 ; CA- CB- CG- HG
4645 4647 4650 4652 3 3.90367 0.19665 -6.21742
-2.37651 5.99149 0.00000 ; CA- CB- CG- CD1
4645 4647 4650 4656 3 3.90367 0.19665 -6.21742
-2.37651 5.99149 0.00000 ; CA- CB- CG- CD2
4645 4660 4662 4663 3 20.92000 0.00000 -20.92000
0.00000 0.00000 0.00000 ; CA- C- N- H
4645 4660 4662 4664 3 20.92000 0.00000 -20.92000
0.00000 0.00000 0.00000 ; CA- C- N- CA
4646 4645 4647 4648 3 0.65084 1.95253 0.00000
-2.60338 0.00000 0.00000 ; HA- CA- CB- HB2
4646 4645 4647 4649 3 0.65084 1.95253 0.00000
-2.60338 0.00000 0.00000 ; HA- CA- CB- HB3
4646 4645 4647 4650 3 0.65084 1.95253 0.00000
-2.60338 0.00000 0.00000 ; HA- CA- CB- CG
4646 4645 4660 4661 3 3.68192 -4.35136 0.00000
1.33888 0.00000 0.00000 ; HA- CA- C- O
4646 4645 4660 4662 3 0.00000 0.00000 0.00000
0.00000 0.00000 0.00000 ; HA- CA- C- N
4647 4645 4660 4661 3 0.00000 0.00000 0.00000
0.00000 0.00000 0.00000 ; CB- CA- C- O
4647 4645 4660 4662 3 2.51040 4.18400 1.67360
-6.69440 0.00000 0.00000 ; CB- CA- C- N
4647 4650 4652 4653 3 0.66944 2.00832 0.00000
-2.67776 0.00000 0.00000 ; CB- CG- CD1- HD11
4647 4650 4652 4654 3 0.66944 2.00832 0.00000
-2.67776 0.00000 0.00000 ; CB- CG- CD1- HD12
4647 4650 4652 4655 3 0.66944 2.00832 0.00000
-2.67776 0.00000 0.00000 ; CB- CG- CD1- HD13
4647 4650 4656 4657 3 0.66944 2.00832 0.00000
-2.67776 0.00000 0.00000 ; CB- CG- CD2- HD21
4647 4650 4656 4658 3 0.66944 2.00832 0.00000

-2.67776    0.00000    0.00000 ;     CB-    CG-   CD2-  HD22
4647   4650   4656   4659      3    0.66944    2.00832    0.00000
-2.67776    0.00000    0.00000 ;     CB-    CG-   CD2-  HD23
4648   4647   4645   4660      3    0.65084    1.95253    0.00000
-2.60338    0.00000    0.00000 ;    HB2-    CB-    CA-     C
4648   4647   4650   4651      3    0.62760    1.88280    0.00000
-2.51040    0.00000    0.00000 ;    HB2-    CB-    CG-    HG
4648   4647   4650   4652      3    0.66944    2.00832    0.00000
-2.67776    0.00000    0.00000 ;    HB2-    CB-    CG-   CD1
4648   4647   4650   4656      3    0.66944    2.00832    0.00000
-2.67776    0.00000    0.00000 ;    HB2-    CB-    CG-   CD2
4649   4647   4645   4660      3    0.65084    1.95253    0.00000
-2.60338    0.00000    0.00000 ;    HB3-    CB-    CA-     C
4649   4647   4650   4651      3    0.62760    1.88280    0.00000
-2.51040    0.00000    0.00000 ;    HB3-    CB-    CG-    HG
4649   4647   4650   4652      3    0.66944    2.00832    0.00000
-2.67776    0.00000    0.00000 ;    HB3-    CB-    CG-   CD1
4649   4647   4650   4656      3    0.66944    2.00832    0.00000
-2.67776    0.00000    0.00000 ;    HB3-    CB-    CG-   CD2
4650   4647   4645   4660      3   10.33448   -1.14642  -11.54784
-2.40998    6.35968    0.00000 ;     CG-    CB-    CA-     C
4651   4650   4652   4653      3    0.62760    1.88280    0.00000
-2.51040    0.00000    0.00000 ;     HG-    CG-   CD1-  HD11
4651   4650   4652   4654      3    0.62760    1.88280    0.00000
-2.51040    0.00000    0.00000 ;     HG-    CG-   CD1-  HD12
4651   4650   4652   4655      3    0.62760    1.88280    0.00000
-2.51040    0.00000    0.00000 ;     HG-    CG-   CD1-  HD13
4651   4650   4656   4657      3    0.62760    1.88280    0.00000
-2.51040    0.00000    0.00000 ;     HG-    CG-   CD2-  HD21
4651   4650   4656   4658      3    0.62760    1.88280    0.00000
-2.51040    0.00000    0.00000 ;     HG-    CG-   CD2-  HD22
4651   4650   4656   4659      3    0.62760    1.88280    0.00000
-2.51040    0.00000    0.00000 ;     HG-    CG-   CD2-  HD23
4652   4650   4656   4657      3    0.66944    2.00832    0.00000
-2.67776    0.00000    0.00000 ;    CD1-    CG-   CD2-  HD21
4652   4650   4656   4658      3    0.66944    2.00832    0.00000
-2.67776    0.00000    0.00000 ;    CD1-    CG-   CD2-  HD22
4652   4650   4656   4659      3    0.66944    2.00832    0.00000
-2.67776    0.00000    0.00000 ;    CD1-    CG-   CD2-  HD23
4653   4652   4650   4656      3    0.66944    2.00832    0.00000
-2.67776    0.00000    0.00000 ;   HD11-   CD1-    CG-   CD2
4654   4652   4650   4656      3    0.66944    2.00832    0.00000
-2.67776    0.00000    0.00000 ;   HD12-   CD1-    CG-   CD2
4655   4652   4650   4656      3    0.66944    2.00832    0.00000
-2.67776    0.00000    0.00000 ;   HD13-   CD1-    CG-   CD2
4660   4662   4664   4665      3    0.00000    0.00000    0.00000
0.00000    0.00000    0.00000 ;      C-     N-    CA-    HA
4660   4662   4664   4666      3   11.71520    1.67360   15.06240  -
13.38880    0.00000    0.00000 ;      C-     N-    CA-    CB
4660   4662   4664   4682      3    1.75728    5.27184    2.25936
-7.02912    0.00000    0.00000 ;      C-     N-    CA-     C
4661   4660   4662   4663      3   29.28800   -8.36800  -20.92000

```
0.00000    0.00000    0.00000 ;       O-     C-     N-     H
  4661   4660   4662   4664      3   20.92000    0.00000  -20.92000
0.00000    0.00000    0.00000 ;       O-     C-     N-    CA
  4662   4664   4666   4667      3    0.65084    1.95253    0.00000
-2.60338    0.00000    0.00000 ;      N-    CA-    CB-   HB2
  4662   4664   4666   4668      3    0.65084    1.95253    0.00000
-2.60338    0.00000    0.00000 ;      N-    CA-    CB-   HB3
  4662   4664   4666   4669      3    0.65084    1.95253    0.00000
-2.60338    0.00000    0.00000 ;      N-    CA-    CB-    CG
  4662   4664   4682   4683      3    0.00000    0.00000    0.00000
0.00000    0.00000    0.00000 ;       N-    CA-     C-     O
  4662   4664   4682   4684      3   17.40544   -5.02080  -13.22144
9.20480    0.00000    0.00000 ;       N-    CA-     C-     N
  4663   4662   4664   4665      3    0.00000    0.00000    0.00000
0.00000    0.00000    0.00000 ;       H-     N-    CA-    HA
  4663   4662   4664   4666      3    0.00000    0.00000    0.00000
0.00000    0.00000    0.00000 ;       H-     N-    CA-    CB
  4663   4662   4664   4682      3    0.00000    0.00000    0.00000
0.00000    0.00000    0.00000 ;       H-     N-    CA-     C
  4664   4666   4669   4670      3    0.66944    2.00832    0.00000
-2.67776    0.00000    0.00000 ;     CA-    CB-    CG-   HG2
  4664   4666   4669   4671      3    0.66944    2.00832    0.00000
-2.67776    0.00000    0.00000 ;     CA-    CB-    CG-   HG3
  4664   4666   4669   4672      3    3.68192    3.09616   -2.09200
-3.01248    0.00000    0.00000 ;     CA-    CB-    CG-    CD
  4664   4682   4684   4685      3   20.92000    0.00000  -20.92000
0.00000    0.00000    0.00000 ;      CA-     C-     N-     H
  4664   4682   4684   4686      3   20.92000    0.00000  -20.92000
0.00000    0.00000    0.00000 ;      CA-     C-     N-    CA
  4665   4664   4666   4667      3    0.65084    1.95253    0.00000
-2.60338    0.00000    0.00000 ;     HA-    CA-    CB-   HB2
  4665   4664   4666   4668      3    0.65084    1.95253    0.00000
-2.60338    0.00000    0.00000 ;     HA-    CA-    CB-   HB3
  4665   4664   4666   4669      3    0.65084    1.95253    0.00000
-2.60338    0.00000    0.00000 ;     HA-    CA-    CB-    CG
  4665   4664   4682   4683      3    3.68192   -4.35136    0.00000
1.33888    0.00000    0.00000 ;      HA-    CA-     C-     O
  4665   4664   4682   4684      3    0.00000    0.00000    0.00000
0.00000    0.00000    0.00000 ;      HA-    CA-     C-     N
  4666   4664   4682   4683      3    0.00000    0.00000    0.00000
0.00000    0.00000    0.00000 ;      CB-    CA-     C-     O
  4666   4664   4682   4684      3    2.51040    4.18400    1.67360
-6.69440    0.00000    0.00000 ;     CB-    CA-     C-     N
  4666   4669   4672   4673      3    0.66944    2.00832    0.00000
-2.67776    0.00000    0.00000 ;     CB-    CG-    CD-   HD2
  4666   4669   4672   4674      3    0.66944    2.00832    0.00000
-2.67776    0.00000    0.00000 ;     CB-    CG-    CD-   HD3
  4666   4669   4672   4675      3    3.68192    3.09616   -2.09200
-3.01248    0.00000    0.00000 ;     CB-    CG-    CD-    CE
  4667   4666   4664   4682      3    0.65084    1.95253    0.00000
-2.60338    0.00000    0.00000 ;    HB2-    CB-    CA-     C
  4667   4666   4669   4670      3    0.62760    1.88280    0.00000
```

```
-2.51040    0.00000    0.00000 ;    HB2-   CB-    CG-   HG2
 4667   4666   4669   4671      3    0.62760    1.88280    0.00000
-2.51040    0.00000    0.00000 ;    HB2-   CB-    CG-   HG3
 4667   4666   4669   4672      3    0.66944    2.00832    0.00000
-2.67776    0.00000    0.00000 ;    HB2-   CB-    CG-    CD
 4668   4666   4664   4682      3    0.65084    1.95253    0.00000
-2.60338    0.00000    0.00000 ;    HB3-   CB-    CA-     C
 4668   4666   4669   4670      3    0.62760    1.88280    0.00000
-2.51040    0.00000    0.00000 ;    HB3-   CB-    CG-   HG2
 4668   4666   4669   4671      3    0.62760    1.88280    0.00000
-2.51040    0.00000    0.00000 ;    HB3-   CB-    CG-   HG3
 4668   4666   4669   4672      3    0.66944    2.00832    0.00000
-2.67776    0.00000    0.00000 ;    HB3-   CB-    CG-    CD
 4669   4666   4664   4682      3    0.65084    1.95253    0.00000
-2.60338    0.00000    0.00000 ;     CG-   CB-    CA-     C
 4669   4672   4675   4676      3    0.65084    1.95253    0.00000
-2.60338    0.00000    0.00000 ;     CG-   CD-    CE-   HE2
 4669   4672   4675   4677      3    0.65084    1.95253    0.00000
-2.60338    0.00000    0.00000 ;     CG-   CD-    CE-   HE3
 4669   4672   4675   4678      3    0.65084    1.95253    0.00000
-2.60338    0.00000    0.00000 ;     CG-   CD-    CE-    NZ
 4670   4669   4672   4673      3    0.62760    1.88280    0.00000
-2.51040    0.00000    0.00000 ;    HG2-   CG-    CD-   HD2
 4670   4669   4672   4674      3    0.62760    1.88280    0.00000
-2.51040    0.00000    0.00000 ;    HG2-   CG-    CD-   HD3
 4670   4669   4672   4675      3    0.66944    2.00832    0.00000
-2.67776    0.00000    0.00000 ;    HG2-   CG-    CD-    CE
 4671   4669   4672   4673      3    0.62760    1.88280    0.00000
-2.51040    0.00000    0.00000 ;    HG3-   CG-    CD-   HD2
 4671   4669   4672   4674      3    0.62760    1.88280    0.00000
-2.51040    0.00000    0.00000 ;    HG3-   CG-    CD-   HD3
 4671   4669   4672   4675      3    0.66944    2.00832    0.00000
-2.67776    0.00000    0.00000 ;    HG3-   CG-    CD-    CE
 4672   4675   4678   4679      3    0.65084    1.95253    0.00000
-2.60338    0.00000    0.00000 ;     CD-   CE-    NZ-   HZ1
 4672   4675   4678   4680      3    0.65084    1.95253    0.00000
-2.60338    0.00000    0.00000 ;     CD-   CE-    NZ-   HZ2
 4672   4675   4678   4681      3    0.65084    1.95253    0.00000
-2.60338    0.00000    0.00000 ;     CD-   CE-    NZ-   HZ3
 4673   4672   4675   4676      3    0.65084    1.95253    0.00000
-2.60338    0.00000    0.00000 ;    HD2-   CD-    CE-   HE2
 4673   4672   4675   4677      3    0.65084    1.95253    0.00000
-2.60338    0.00000    0.00000 ;    HD2-   CD-    CE-   HE3
 4673   4672   4675   4678      3    0.65084    1.95253    0.00000
-2.60338    0.00000    0.00000 ;    HD2-   CD-    CE-    NZ
 4674   4672   4675   4676      3    0.65084    1.95253    0.00000
-2.60338    0.00000    0.00000 ;    HD3-   CD-    CE-   HE2
 4674   4672   4675   4677      3    0.65084    1.95253    0.00000
-2.60338    0.00000    0.00000 ;    HD3-   CD-    CE-   HE3
 4674   4672   4675   4678      3    0.65084    1.95253    0.00000
-2.60338    0.00000    0.00000 ;    HD3-   CD-    CE-    NZ
 4676   4675   4678   4679      3    0.65084    1.95253    0.00000
```

```
-2.60338    0.00000    0.00000 ;    HE2-    CE-    NZ-   HZ1
  4676   4675   4678   4680       3    0.65084    1.95253    0.00000
-2.60338    0.00000    0.00000 ;    HE2-    CE-    NZ-   HZ2
  4676   4675   4678   4681       3    0.65084    1.95253    0.00000
-2.60338    0.00000    0.00000 ;    HE2-    CE-    NZ-   HZ3
  4677   4675   4678   4679       3    0.65084    1.95253    0.00000
-2.60338    0.00000    0.00000 ;    HE3-    CE-    NZ-   HZ1
  4677   4675   4678   4680       3    0.65084    1.95253    0.00000
-2.60338    0.00000    0.00000 ;    HE3-    CE-    NZ-   HZ2
  4677   4675   4678   4681       3    0.65084    1.95253    0.00000
-2.60338    0.00000    0.00000 ;    HE3-    CE-    NZ-   HZ3
  4682   4684   4686   4687       3    0.00000    0.00000    0.00000
0.00000    0.00000    0.00000 ;    C-    N-    CA-    HA
  4682   4684   4686   4688       3   11.71520    1.67360   15.06240  -
13.38880    0.00000    0.00000 ;    C-    N-    CA-    CB
  4682   4684   4686   4696       3    1.75728    5.27184    2.25936
-7.02912    0.00000    0.00000 ;    C-    N-    CA-    C
  4683   4682   4684   4685       3   29.28800   -8.36800  -20.92000
0.00000    0.00000    0.00000 ;    O-    C-    N-    H
  4683   4682   4684   4686       3   20.92000    0.00000  -20.92000
0.00000    0.00000    0.00000 ;    O-    C-    N-   CA
  4684   4686   4688   4689       3    0.65084    1.95253    0.00000
-2.60338    0.00000    0.00000 ;    N-    CA-    CB-   HB2
  4684   4686   4688   4690       3    0.65084    1.95253    0.00000
-2.60338    0.00000    0.00000 ;    N-    CA-    CB-   HB3
  4684   4686   4688   4691       3    5.99567    3.29281   -4.46014
-0.55229    1.97485    0.00000 ;    N-    CA-    CB-    CG
  4684   4686   4696   4697       3    0.00000    0.00000    0.00000
0.00000    0.00000    0.00000 ;    N-    CA-    C-    O
  4684   4686   4696   4698       3   17.40544   -5.02080  -13.22144
9.20480    0.00000    0.00000 ;    N-    CA-    C-    N
  4685   4684   4686   4687       3    0.00000    0.00000    0.00000
0.00000    0.00000    0.00000 ;    H-    N-    CA-   HA
  4685   4684   4686   4688       3    0.00000    0.00000    0.00000
0.00000    0.00000    0.00000 ;    H-    N-    CA-   CB
  4685   4684   4686   4696       3    0.00000    0.00000    0.00000
0.00000    0.00000    0.00000 ;    H-    N-    CA-    C
  4686   4688   4691   4692       3    0.00000    0.00000    0.00000
0.00000    0.00000    0.00000 ;    CA-    CB-    CG-   OD1
  4686   4688   4691   4693       3    8.84916   -0.31380   -4.32626
5.03754    0.26778    0.00000 ;    CA-    CB-    CG-   ND2
  4686   4696   4698   4699       3   20.92000    0.00000  -20.92000
0.00000    0.00000    0.00000 ;    CA-    C-    N-    H
  4686   4696   4698   4700       3   20.92000    0.00000  -20.92000
0.00000    0.00000    0.00000 ;    CA-    C-    N-   CA
  4687   4686   4688   4689       3    0.65084    1.95253    0.00000
-2.60338    0.00000    0.00000 ;    HA-    CA-    CB-   HB2
  4687   4686   4688   4690       3    0.65084    1.95253    0.00000
-2.60338    0.00000    0.00000 ;    HA-    CA-    CB-   HB3
  4687   4686   4688   4691       3    0.65084    1.95253    0.00000
-2.60338    0.00000    0.00000 ;    HA-    CA-    CB-    CG
  4687   4686   4696   4697       3    3.68192   -4.35136    0.00000
```

1.33888 0.00000 0.00000 ; HA- CA- C- O
4687 4686 4696 4698 3 0.00000 0.00000 0.00000
0.00000 0.00000 0.00000 ; HA- CA- C- N
4688 4686 4696 4697 3 0.00000 0.00000 0.00000
0.00000 0.00000 0.00000 ; CB- CA- C- O
4688 4686 4696 4698 3 2.51040 4.18400 1.67360
-6.69440 0.00000 0.00000 ; CB- CA- C- N
4688 4691 4693 4694 3 20.92000 0.00000 -20.92000
0.00000 0.00000 0.00000 ; CB- CG- ND2- HD21
4688 4691 4693 4695 3 20.92000 0.00000 -20.92000
0.00000 0.00000 0.00000 ; CB- CG- ND2- HD22
4689 4688 4686 4696 3 0.65084 1.95253 0.00000
-2.60338 0.00000 0.00000 ; HB2- CB- CA- C
4689 4688 4691 4692 3 3.68192 -4.35136 0.00000
1.33888 0.00000 0.00000 ; HB2- CB- CG- OD1
4689 4688 4691 4693 3 0.00000 0.00000 0.00000
0.00000 0.00000 0.00000 ; HB2- CB- CG- ND2
4690 4688 4686 4696 3 0.65084 1.95253 0.00000
-2.60338 0.00000 0.00000 ; HB3- CB- CA- C
4690 4688 4691 4692 3 3.68192 -4.35136 0.00000
1.33888 0.00000 0.00000 ; HB3- CB- CG- OD1
4690 4688 4691 4693 3 0.00000 0.00000 0.00000
0.00000 0.00000 0.00000 ; HB3- CB- CG- ND2
4691 4688 4686 4696 3 7.94542 -3.96225 -6.11701
-0.55229 3.58150 0.00000 ; CG- CB- CA- C
4692 4691 4693 4694 3 29.28800 -8.36800 -20.92000
0.00000 0.00000 0.00000 ; OD1- CG- ND2- HD21
4692 4691 4693 4695 3 29.28800 -8.36800 -20.92000
0.00000 0.00000 0.00000 ; OD1- CG- ND2- HD22
4696 4698 4700 4701 3 0.00000 0.00000 0.00000
0.00000 0.00000 0.00000 ; C- N- CA- HA
4696 4698 4700 4702 3 11.71520 1.67360 15.06240 -
13.38880 0.00000 0.00000 ; C- N- CA- CB
4696 4698 4700 4716 3 1.75728 5.27184 2.25936
-7.02912 0.00000 0.00000 ; C- N- CA- C
4697 4696 4698 4699 3 29.28800 -8.36800 -20.92000
0.00000 0.00000 0.00000 ; O- C- N- H
4697 4696 4698 4700 3 20.92000 0.00000 -20.92000
0.00000 0.00000 0.00000 ; O- C- N- CA
4698 4700 4702 4703 3 0.65084 1.95253 0.00000
-2.60338 0.00000 0.00000 ; N- CA- CB- HB2
4698 4700 4702 4704 3 0.65084 1.95253 0.00000
-2.60338 0.00000 0.00000 ; N- CA- CB- HB3
4698 4700 4702 4705 3 3.28026 2.46019 -2.19242
-3.21331 -0.23430 0.00000 ; N- CA- CB- CG
4698 4700 4716 4717 3 0.00000 0.00000 0.00000
0.00000 0.00000 0.00000 ; N- CA- C- O
4698 4700 4716 4718 3 17.40544 -5.02080 -13.22144
9.20480 0.00000 0.00000 ; N- CA- C- N
4699 4698 4700 4701 3 0.00000 0.00000 0.00000
0.00000 0.00000 0.00000 ; H- N- CA- HA
4699 4698 4700 4702 3 0.00000 0.00000 0.00000

```
0.00000    0.00000    0.00000 ;      H-     N-    CA-    CB
  4699   4698   4700   4716      3    0.00000    0.00000    0.00000
0.00000    0.00000    0.00000 ;      H-     N-    CA-     C
  4700   4702   4705   4706      3    0.57739    0.00000    1.02926
0.00000   -1.60666    0.00000 ;     CA-    CB-    CG-   CD1
  4700   4702   4705   4714      3    0.57739    0.00000    1.02926
0.00000   -1.60666    0.00000 ;     CA-    CB-    CG-   CD2
  4700   4716   4718   4719      3   20.92000    0.00000  -20.92000
0.00000    0.00000    0.00000 ;     CA-     C-     N-     H
  4700   4716   4718   4720      3   20.92000    0.00000  -20.92000
0.00000    0.00000    0.00000 ;     CA-     C-     N-    CA
  4701   4700   4702   4703      3    0.65084    1.95253    0.00000
-2.60338    0.00000    0.00000 ;     HA-    CA-    CB-   HB2
  4701   4700   4702   4704      3    0.65084    1.95253    0.00000
-2.60338    0.00000    0.00000 ;     HA-    CA-    CB-   HB3
  4701   4700   4702   4705      3    0.65084    1.95253    0.00000
-2.60338    0.00000    0.00000 ;     HA-    CA-    CB-    CG
  4701   4700   4716   4717      3    3.68192   -4.35136    0.00000
1.33888    0.00000    0.00000 ;     HA-    CA-     C-     O
  4701   4700   4716   4718      3    0.00000    0.00000    0.00000
0.00000    0.00000    0.00000 ;     HA-    CA-     C-     N
  4702   4700   4716   4717      3    0.00000    0.00000    0.00000
0.00000    0.00000    0.00000 ;     CB-    CA-     C-     O
  4702   4700   4716   4718      3    2.51040    4.18400    1.67360
-6.69440    0.00000    0.00000 ;     CB-    CA-     C-     N
  4702   4705   4706   4707      3   30.33400    0.00000  -30.33400
0.00000    0.00000    0.00000 ;     CB-    CG-   CD1-   HD1
  4702   4705   4706   4708      3   30.33400    0.00000  -30.33400
0.00000    0.00000    0.00000 ;     CB-    CG-   CD1-   CE1
  4702   4705   4714   4712      3   30.33400    0.00000  -30.33400
0.00000    0.00000    0.00000 ;     CB-    CG-   CD2-   CE2
  4702   4705   4714   4715      3   30.33400    0.00000  -30.33400
0.00000    0.00000    0.00000 ;     CB-    CG-   CD2-   HD2
  4703   4702   4700   4716      3    0.65084    1.95253    0.00000
-2.60338    0.00000    0.00000 ;    HB2-    CB-    CA-     C
  4703   4702   4705   4706      3    0.00000    0.00000    0.00000
0.00000    0.00000    0.00000 ;    HB2-    CB-    CG-   CD1
  4703   4702   4705   4714      3    0.00000    0.00000    0.00000
0.00000    0.00000    0.00000 ;    HB2-    CB-    CG-   CD2
  4704   4702   4700   4716      3    0.65084    1.95253    0.00000
-2.60338    0.00000    0.00000 ;    HB3-    CB-    CA-     C
  4704   4702   4705   4706      3    0.00000    0.00000    0.00000
0.00000    0.00000    0.00000 ;    HB3-    CB-    CG-   CD1
  4704   4702   4705   4714      3    0.00000    0.00000    0.00000
0.00000    0.00000    0.00000 ;    HB3-    CB-    CG-   CD2
  4705   4702   4700   4716      3    4.95804    2.17986   -3.52293
-3.21331   -0.40166    0.00000 ;     CG-    CB-    CA-     C
  4705   4706   4708   4709      3   30.33400    0.00000  -30.33400
0.00000    0.00000    0.00000 ;     CG-   CD1-   CE1-   HE1
  4705   4706   4708   4710      3   30.33400    0.00000  -30.33400
0.00000    0.00000    0.00000 ;     CG-   CD1-   CE1-    CZ
  4705   4714   4712   4710      3   30.33400    0.00000  -30.33400
```

```
0.00000    0.00000    0.00000 ;     CG-   CD2-   CE2-    CZ
  4705   4714   4712   4713      3   30.33400    0.00000  -30.33400
0.00000    0.00000    0.00000 ;     CG-   CD2-   CE2-   HE2
  4706   4705   4714   4712      3   30.33400    0.00000  -30.33400
0.00000    0.00000    0.00000 ;    CD1-    CG-   CD2-   CE2
  4706   4705   4714   4715      3   30.33400    0.00000  -30.33400
0.00000    0.00000    0.00000 ;    CD1-    CG-   CD2-   HD2
  4706   4708   4710   4711      3   30.33400    0.00000  -30.33400
0.00000    0.00000    0.00000 ;    CD1-   CE1-    CZ-    HZ
  4706   4708   4710   4712      3   30.33400    0.00000  -30.33400
0.00000    0.00000    0.00000 ;    CD1-   CE1-    CZ-   CE2
  4707   4706   4705   4714      3   30.33400    0.00000  -30.33400
0.00000    0.00000    0.00000 ;    HD1-   CD1-    CG-   CD2
  4707   4706   4708   4709      3   30.33400    0.00000  -30.33400
0.00000    0.00000    0.00000 ;    HD1-   CD1-   CE1-   HE1
  4707   4706   4708   4710      3   30.33400    0.00000  -30.33400
0.00000    0.00000    0.00000 ;    HD1-   CD1-   CE1-    CZ
  4708   4706   4705   4714      3   30.33400    0.00000  -30.33400
0.00000    0.00000    0.00000 ;    CE1-   CD1-    CG-   CD2
  4708   4710   4712   4713      3   30.33400    0.00000  -30.33400
0.00000    0.00000    0.00000 ;    CE1-    CZ-   CE2-   HE2
  4708   4710   4712   4714      3   30.33400    0.00000  -30.33400
0.00000    0.00000    0.00000 ;    CE1-    CZ-   CE2-   CD2
  4709   4708   4710   4711      3   30.33400    0.00000  -30.33400
0.00000    0.00000    0.00000 ;    HE1-   CE1-    CZ-    HZ
  4709   4708   4710   4712      3   30.33400    0.00000  -30.33400
0.00000    0.00000    0.00000 ;    HE1-   CE1-    CZ-   CE2
  4710   4712   4714   4715      3   30.33400    0.00000  -30.33400
0.00000    0.00000    0.00000 ;     CZ-   CE2-   CD2-   HD2
  4711   4710   4712   4713      3   30.33400    0.00000  -30.33400
0.00000    0.00000    0.00000 ;     HZ-    CZ-   CE2-   HE2
  4711   4710   4712   4714      3   30.33400    0.00000  -30.33400
0.00000    0.00000    0.00000 ;     HZ-    CZ-   CE2-   CD2
  4713   4712   4714   4715      3   30.33400    0.00000  -30.33400
0.00000    0.00000    0.00000 ;    HE2-   CE2-   CD2-   HD2
  4716   4718   4720   4721      3    0.00000    0.00000    0.00000
0.00000    0.00000    0.00000 ;      C-     N-    CA-    HA
  4716   4718   4720   4722      3   11.71520    1.67360   15.06240  -
13.38880    0.00000    0.00000 ;      C-     N-    CA-    CB
  4716   4718   4720   4738      3    1.75728    5.27184    2.25936
-7.02912    0.00000    0.00000 ;      C-     N-    CA-     C
  4717   4716   4718   4719      3   29.28800   -8.36800  -20.92000
0.00000    0.00000    0.00000 ;      O-     C-     N-     H
  4717   4716   4718   4720      3   20.92000    0.00000  -20.92000
0.00000    0.00000    0.00000 ;      O-     C-     N-    CA
  4718   4720   4722   4723      3    0.65084    1.95253    0.00000
-2.60338    0.00000    0.00000 ;      N-    CA-    CB-   HB2
  4718   4720   4722   4724      3    0.65084    1.95253    0.00000
-2.60338    0.00000    0.00000 ;      N-    CA-    CB-   HB3
  4718   4720   4722   4725      3    0.65084    1.95253    0.00000
-2.60338    0.00000    0.00000 ;      N-    CA-    CB-    CG
  4718   4720   4738   4739      3    0.00000    0.00000    0.00000
```

```
0.00000    0.00000    0.00000 ;       N-    CA-     C-     O
  4718   4720   4738   4740       3   17.40544   -5.02080  -13.22144
9.20480    0.00000    0.00000 ;       N-    CA-     C-     N
  4719   4718   4720   4721       3    0.00000    0.00000    0.00000
0.00000    0.00000    0.00000 ;       H-     N-    CA-    HA
  4719   4718   4720   4722       3    0.00000    0.00000    0.00000
0.00000    0.00000    0.00000 ;       H-     N-    CA-    CB
  4719   4718   4720   4738       3    0.00000    0.00000    0.00000
0.00000    0.00000    0.00000 ;       H-     N-    CA-     C
  4720   4722   4725   4726       3    0.66944    2.00832    0.00000
-2.67776    0.00000    0.00000 ;      CA-    CB-    CG-   HG2
  4720   4722   4725   4727       3    0.66944    2.00832    0.00000
-2.67776    0.00000    0.00000 ;      CA-    CB-    CG-   HG3
  4720   4722   4725   4728       3    3.68192    3.09616   -2.09200
-3.01248    0.00000    0.00000 ;      CA-    CB-    CG-    CD
  4720   4738   4740   4741       3   20.92000    0.00000  -20.92000
0.00000    0.00000    0.00000 ;      CA-     C-     N-     H
  4720   4738   4740   4742       3   20.92000    0.00000  -20.92000
0.00000    0.00000    0.00000 ;      CA-     C-     N-    CA
  4721   4720   4722   4723       3    0.65084    1.95253    0.00000
-2.60338    0.00000    0.00000 ;      HA-    CA-    CB-   HB2
  4721   4720   4722   4724       3    0.65084    1.95253    0.00000
-2.60338    0.00000    0.00000 ;      HA-    CA-    CB-   HB3
  4721   4720   4722   4725       3    0.65084    1.95253    0.00000
-2.60338    0.00000    0.00000 ;      HA-    CA-    CB-    CG
  4721   4720   4738   4739       3    3.68192   -4.35136    0.00000
1.33888    0.00000    0.00000 ;      HA-    CA-     C-     O
  4721   4720   4738   4740       3    0.00000    0.00000    0.00000
0.00000    0.00000    0.00000 ;      HA-    CA-     C-     N
  4722   4720   4738   4739       3    0.00000    0.00000    0.00000
0.00000    0.00000    0.00000 ;      CB-    CA-     C-     O
  4722   4720   4738   4740       3    2.51040    4.18400    1.67360
-6.69440    0.00000    0.00000 ;      CB-    CA-     C-     N
  4722   4725   4728   4729       3    0.66944    2.00832    0.00000
-2.67776    0.00000    0.00000 ;      CB-    CG-    CD-   HD2
  4722   4725   4728   4730       3    0.66944    2.00832    0.00000
-2.67776    0.00000    0.00000 ;      CB-    CG-    CD-   HD3
  4722   4725   4728   4731       3    3.68192    3.09616   -2.09200
-3.01248    0.00000    0.00000 ;      CB-    CG-    CD-    CE
  4723   4722   4720   4738       3    0.65084    1.95253    0.00000
-2.60338    0.00000    0.00000 ;     HB2-    CB-    CA-     C
  4723   4722   4725   4726       3    0.62760    1.88280    0.00000
-2.51040    0.00000    0.00000 ;     HB2-    CB-    CG-   HG2
  4723   4722   4725   4727       3    0.62760    1.88280    0.00000
-2.51040    0.00000    0.00000 ;     HB2-    CB-    CG-   HG3
  4723   4722   4725   4728       3    0.66944    2.00832    0.00000
-2.67776    0.00000    0.00000 ;     HB2-    CB-    CG-    CD
  4724   4722   4720   4738       3    0.65084    1.95253    0.00000
-2.60338    0.00000    0.00000 ;     HB3-    CB-    CA-     C
  4724   4722   4725   4726       3    0.62760    1.88280    0.00000
-2.51040    0.00000    0.00000 ;     HB3-    CB-    CG-   HG2
  4724   4722   4725   4727       3    0.62760    1.88280    0.00000
```

```
-2.51040    0.00000    0.00000 ;    HB3-    CB-    CG-   HG3
  4724   4722   4725   4728      3    0.66944    2.00832    0.00000
-2.67776    0.00000    0.00000 ;    HB3-    CB-    CG-    CD
  4725   4722   4720   4738      3    0.65084    1.95253    0.00000
-2.60338    0.00000    0.00000 ;     CG-    CB-    CA-     C
  4725   4728   4731   4732      3    0.65084    1.95253    0.00000
-2.60338    0.00000    0.00000 ;     CG-    CD-    CE-   HE2
  4725   4728   4731   4733      3    0.65084    1.95253    0.00000
-2.60338    0.00000    0.00000 ;     CG-    CD-    CE-   HE3
  4725   4728   4731   4734      3    0.65084    1.95253    0.00000
-2.60338    0.00000    0.00000 ;     CG-    CD-    CE-    NZ
  4726   4725   4728   4729      3    0.62760    1.88280    0.00000
-2.51040    0.00000    0.00000 ;    HG2-    CG-    CD-   HD2
  4726   4725   4728   4730      3    0.62760    1.88280    0.00000
-2.51040    0.00000    0.00000 ;    HG2-    CG-    CD-   HD3
  4726   4725   4728   4731      3    0.66944    2.00832    0.00000
-2.67776    0.00000    0.00000 ;    HG2-    CG-    CD-    CE
  4727   4725   4728   4729      3    0.62760    1.88280    0.00000
-2.51040    0.00000    0.00000 ;    HG3-    CG-    CD-   HD2
  4727   4725   4728   4730      3    0.62760    1.88280    0.00000
-2.51040    0.00000    0.00000 ;    HG3-    CG-    CD-   HD3
  4727   4725   4728   4731      3    0.66944    2.00832    0.00000
-2.67776    0.00000    0.00000 ;    HG3-    CG-    CD-    CE
  4728   4731   4734   4735      3    0.65084    1.95253    0.00000
-2.60338    0.00000    0.00000 ;     CD-    CE-    NZ-   HZ1
  4728   4731   4734   4736      3    0.65084    1.95253    0.00000
-2.60338    0.00000    0.00000 ;     CD-    CE-    NZ-   HZ2
  4728   4731   4734   4737      3    0.65084    1.95253    0.00000
-2.60338    0.00000    0.00000 ;     CD-    CE-    NZ-   HZ3
  4729   4728   4731   4732      3    0.65084    1.95253    0.00000
-2.60338    0.00000    0.00000 ;    HD2-    CD-    CE-   HE2
  4729   4728   4731   4733      3    0.65084    1.95253    0.00000
-2.60338    0.00000    0.00000 ;    HD2-    CD-    CE-   HE3
  4729   4728   4731   4734      3    0.65084    1.95253    0.00000
-2.60338    0.00000    0.00000 ;    HD2-    CD-    CE-    NZ
  4730   4728   4731   4732      3    0.65084    1.95253    0.00000
-2.60338    0.00000    0.00000 ;    HD3-    CD-    CE-   HE2
  4730   4728   4731   4733      3    0.65084    1.95253    0.00000
-2.60338    0.00000    0.00000 ;    HD3-    CD-    CE-   HE3
  4730   4728   4731   4734      3    0.65084    1.95253    0.00000
-2.60338    0.00000    0.00000 ;    HD3-    CD-    CE-    NZ
  4732   4731   4734   4735      3    0.65084    1.95253    0.00000
-2.60338    0.00000    0.00000 ;    HE2-    CE-    NZ-   HZ1
  4732   4731   4734   4736      3    0.65084    1.95253    0.00000
-2.60338    0.00000    0.00000 ;    HE2-    CE-    NZ-   HZ2
  4732   4731   4734   4737      3    0.65084    1.95253    0.00000
-2.60338    0.00000    0.00000 ;    HE2-    CE-    NZ-   HZ3
  4733   4731   4734   4735      3    0.65084    1.95253    0.00000
-2.60338    0.00000    0.00000 ;    HE3-    CE-    NZ-   HZ1
  4733   4731   4734   4736      3    0.65084    1.95253    0.00000
-2.60338    0.00000    0.00000 ;    HE3-    CE-    NZ-   HZ2
  4733   4731   4734   4737      3    0.65084    1.95253    0.00000
```

-2.60338 0.00000 0.00000 ; HE3- CE- NZ- HZ3
4738 4740 4742 4743 3 0.00000 0.00000 0.00000
0.00000 0.00000 0.00000 ; C- N- CA- HA
4738 4740 4742 4744 3 11.71520 1.67360 15.06240 -
13.38880 0.00000 0.00000 ; C- N- CA- CB
4738 4740 4742 4760 3 1.75728 5.27184 2.25936
-7.02912 0.00000 0.00000 ; C- N- CA- C
4739 4738 4740 4741 3 29.28800 -8.36800 -20.92000
0.00000 0.00000 0.00000 ; O- C- N- H
4739 4738 4740 4742 3 20.92000 0.00000 -20.92000
0.00000 0.00000 0.00000 ; O- C- N- CA
4740 4742 4744 4745 3 0.65084 1.95253 0.00000
-2.60338 0.00000 0.00000 ; N- CA- CB- HB2
4740 4742 4744 4746 3 0.65084 1.95253 0.00000
-2.60338 0.00000 0.00000 ; N- CA- CB- HB3
4740 4742 4744 4747 3 0.65084 1.95253 0.00000
-2.60338 0.00000 0.00000 ; N- CA- CB- CG
4740 4742 4760 4761 3 0.00000 0.00000 0.00000
0.00000 0.00000 0.00000 ; N- CA- C- O
4740 4742 4760 4762 3 17.40544 -5.02080 -13.22144
9.20480 0.00000 0.00000 ; N- CA- C- N
4741 4740 4742 4743 3 0.00000 0.00000 0.00000
0.00000 0.00000 0.00000 ; H- N- CA- HA
4741 4740 4742 4744 3 0.00000 0.00000 0.00000
0.00000 0.00000 0.00000 ; H- N- CA- CB
4741 4740 4742 4760 3 0.00000 0.00000 0.00000
0.00000 0.00000 0.00000 ; H- N- CA- C
4742 4744 4747 4748 3 0.66944 2.00832 0.00000
-2.67776 0.00000 0.00000 ; CA- CB- CG- HG2
4742 4744 4747 4749 3 0.66944 2.00832 0.00000
-2.67776 0.00000 0.00000 ; CA- CB- CG- HG3
4742 4744 4747 4750 3 3.68192 3.09616 -2.09200
-3.01248 0.00000 0.00000 ; CA- CB- CG- CD
4742 4760 4762 4763 3 20.92000 0.00000 -20.92000
0.00000 0.00000 0.00000 ; CA- C- N- H
4742 4760 4762 4764 3 20.92000 0.00000 -20.92000
0.00000 0.00000 0.00000 ; CA- C- N- CA
4743 4742 4744 4745 3 0.65084 1.95253 0.00000
-2.60338 0.00000 0.00000 ; HA- CA- CB- HB2
4743 4742 4744 4746 3 0.65084 1.95253 0.00000
-2.60338 0.00000 0.00000 ; HA- CA- CB- HB3
4743 4742 4744 4747 3 0.65084 1.95253 0.00000
-2.60338 0.00000 0.00000 ; HA- CA- CB- CG
4743 4742 4760 4761 3 3.68192 -4.35136 0.00000
1.33888 0.00000 0.00000 ; HA- CA- C- O
4743 4742 4760 4762 3 0.00000 0.00000 0.00000
0.00000 0.00000 0.00000 ; HA- CA- C- N
4744 4742 4760 4761 3 0.00000 0.00000 0.00000
0.00000 0.00000 0.00000 ; CB- CA- C- O
4744 4742 4760 4762 3 2.51040 4.18400 1.67360
-6.69440 0.00000 0.00000 ; CB- CA- C- N
4744 4747 4750 4751 3 0.66944 2.00832 0.00000

-2.67776    0.00000    0.00000 ;      CB-    CG-    CD-   HD2
  4744   4747   4750   4752      3    0.66944    2.00832    0.00000
-2.67776    0.00000    0.00000 ;      CB-    CG-    CD-   HD3
  4744   4747   4750   4753      3    3.68192    3.09616   -2.09200
-3.01248    0.00000    0.00000 ;      CB-    CG-    CD-    CE
  4745   4744   4742   4760      3    0.65084    1.95253    0.00000
-2.60338    0.00000    0.00000 ;     HB2-    CB-    CA-     C
  4745   4744   4747   4748      3    0.62760    1.88280    0.00000
-2.51040    0.00000    0.00000 ;     HB2-    CB-    CG-   HG2
  4745   4744   4747   4749      3    0.62760    1.88280    0.00000
-2.51040    0.00000    0.00000 ;     HB2-    CB-    CG-   HG3
  4745   4744   4747   4750      3    0.66944    2.00832    0.00000
-2.67776    0.00000    0.00000 ;     HB2-    CB-    CG-    CD
  4746   4744   4742   4760      3    0.65084    1.95253    0.00000
-2.60338    0.00000    0.00000 ;     HB3-    CB-    CA-     C
  4746   4744   4747   4748      3    0.62760    1.88280    0.00000
-2.51040    0.00000    0.00000 ;     HB3-    CB-    CG-   HG2
  4746   4744   4747   4749      3    0.62760    1.88280    0.00000
-2.51040    0.00000    0.00000 ;     HB3-    CB-    CG-   HG3
  4746   4744   4747   4750      3    0.66944    2.00832    0.00000
-2.67776    0.00000    0.00000 ;     HB3-    CB-    CG-    CD
  4747   4744   4742   4760      3    0.65084    1.95253    0.00000
-2.60338    0.00000    0.00000 ;      CG-    CB-    CA-     C
  4747   4750   4753   4754      3    0.65084    1.95253    0.00000
-2.60338    0.00000    0.00000 ;      CG-    CD-    CE-   HE2
  4747   4750   4753   4755      3    0.65084    1.95253    0.00000
-2.60338    0.00000    0.00000 ;      CG-    CD-    CE-   HE3
  4747   4750   4753   4756      3    0.65084    1.95253    0.00000
-2.60338    0.00000    0.00000 ;      CG-    CD-    CE-    NZ
  4748   4747   4750   4751      3    0.62760    1.88280    0.00000
-2.51040    0.00000    0.00000 ;     HG2-    CG-    CD-   HD2
  4748   4747   4750   4752      3    0.62760    1.88280    0.00000
-2.51040    0.00000    0.00000 ;     HG2-    CG-    CD-   HD3
  4748   4747   4750   4753      3    0.66944    2.00832    0.00000
-2.67776    0.00000    0.00000 ;     HG2-    CG-    CD-    CE
  4749   4747   4750   4751      3    0.62760    1.88280    0.00000
-2.51040    0.00000    0.00000 ;     HG3-    CG-    CD-   HD2
  4749   4747   4750   4752      3    0.62760    1.88280    0.00000
-2.51040    0.00000    0.00000 ;     HG3-    CG-    CD-   HD3
  4749   4747   4750   4753      3    0.66944    2.00832    0.00000
-2.67776    0.00000    0.00000 ;     HG3-    CG-    CD-    CE
  4750   4753   4756   4757      3    0.65084    1.95253    0.00000
-2.60338    0.00000    0.00000 ;      CD-    CE-    NZ-   HZ1
  4750   4753   4756   4758      3    0.65084    1.95253    0.00000
-2.60338    0.00000    0.00000 ;      CD-    CE-    NZ-   HZ2
  4750   4753   4756   4759      3    0.65084    1.95253    0.00000
-2.60338    0.00000    0.00000 ;      CD-    CE-    NZ-   HZ3
  4751   4750   4753   4754      3    0.65084    1.95253    0.00000
-2.60338    0.00000    0.00000 ;     HD2-    CD-    CE-   HE2
  4751   4750   4753   4755      3    0.65084    1.95253    0.00000
-2.60338    0.00000    0.00000 ;     HD2-    CD-    CE-   HE3
  4751   4750   4753   4756      3    0.65084    1.95253    0.00000

```
-2.60338    0.00000    0.00000 ;    HD2-    CD-    CE-    NZ
  4752   4750   4753   4754      3    0.65084    1.95253    0.00000
-2.60338    0.00000    0.00000 ;    HD3-    CD-    CE-   HE2
  4752   4750   4753   4755      3    0.65084    1.95253    0.00000
-2.60338    0.00000    0.00000 ;    HD3-    CD-    CE-   HE3
  4752   4750   4753   4756      3    0.65084    1.95253    0.00000
-2.60338    0.00000    0.00000 ;    HD3-    CD-    CE-    NZ
  4754   4753   4756   4757      3    0.65084    1.95253    0.00000
-2.60338    0.00000    0.00000 ;    HE2-    CE-    NZ-   HZ1
  4754   4753   4756   4758      3    0.65084    1.95253    0.00000
-2.60338    0.00000    0.00000 ;    HE2-    CE-    NZ-   HZ2
  4754   4753   4756   4759      3    0.65084    1.95253    0.00000
-2.60338    0.00000    0.00000 ;    HE2-    CE-    NZ-   HZ3
  4755   4753   4756   4757      3    0.65084    1.95253    0.00000
-2.60338    0.00000    0.00000 ;    HE3-    CE-    NZ-   HZ1
  4755   4753   4756   4758      3    0.65084    1.95253    0.00000
-2.60338    0.00000    0.00000 ;    HE3-    CE-    NZ-   HZ2
  4755   4753   4756   4759      3    0.65084    1.95253    0.00000
-2.60338    0.00000    0.00000 ;    HE3-    CE-    NZ-   HZ3
  4760   4762   4764   4765      3    0.00000    0.00000    0.00000
0.00000    0.00000    0.00000 ;     C-    N-    CA-    HA
  4760   4762   4764   4766      3   11.71520    1.67360   15.06240  -
13.38880    0.00000    0.00000 ;     C-    N-    CA-    CB
  4760   4762   4764   4775      3    1.75728    5.27184    2.25936
-7.02912    0.00000    0.00000 ;     C-    N-    CA-     C
  4761   4760   4762   4763      3   29.28800   -8.36800  -20.92000
0.00000    0.00000    0.00000 ;     O-     C-    N-     H
  4761   4760   4762   4764      3   20.92000    0.00000  -20.92000
0.00000    0.00000    0.00000 ;     O-     C-    N-    CA
  4762   4764   4766   4767      3    0.65084    1.95253    0.00000
-2.60338    0.00000    0.00000 ;     N-    CA-    CB-   HB2
  4762   4764   4766   4768      3    0.65084    1.95253    0.00000
-2.60338    0.00000    0.00000 ;     N-    CA-    CB-   HB3
  4762   4764   4766   4769      3    3.21331    2.22589   -4.15053
-2.40998    2.61082    0.00000 ;     N-    CA-    CB-    CG
  4762   4764   4775   4776      3    0.00000    0.00000    0.00000
0.00000    0.00000    0.00000 ;     N-    CA-     C-     O
  4762   4764   4775   4777      3   17.40544   -5.02080  -13.22144
9.20480    0.00000    0.00000 ;     N-    CA-     C-     N
  4763   4762   4764   4765      3    0.00000    0.00000    0.00000
0.00000    0.00000    0.00000 ;     H-     N-    CA-    HA
  4763   4762   4764   4766      3    0.00000    0.00000    0.00000
0.00000    0.00000    0.00000 ;     H-     N-    CA-    CB
  4763   4762   4764   4775      3    0.00000    0.00000    0.00000
0.00000    0.00000    0.00000 ;     H-     N-    CA-     C
  4764   4766   4769   4770      3    0.66944    2.00832    0.00000
-2.67776    0.00000    0.00000 ;    CA-    CB-    CG-   HG2
  4764   4766   4769   4771      3    0.66944    2.00832    0.00000
-2.67776    0.00000    0.00000 ;    CA-    CB-    CG-   HG3
  4764   4766   4769   4772      3   10.12110   -1.91209    0.01674
10.17549   -1.87443    0.00000 ;    CA-    CB-    CG-    CD
  4764   4775   4777   4778      3   20.92000    0.00000  -20.92000
```

0.00000 0.00000 0.00000 ; CA- C- N- H
4764 4775 4777 4779 3 20.92000 0.00000 -20.92000
0.00000 0.00000 0.00000 ; CA- C- N- CA
4765 4764 4766 4767 3 0.65084 1.95253 0.00000
-2.60338 0.00000 0.00000 ; HA- CA- CB- HB2
4765 4764 4766 4768 3 0.65084 1.95253 0.00000
-2.60338 0.00000 0.00000 ; HA- CA- CB- HB3
4765 4764 4766 4769 3 0.65084 1.95253 0.00000
-2.60338 0.00000 0.00000 ; HA- CA- CB- CG
4765 4764 4775 4776 3 3.68192 -4.35136 0.00000
1.33888 0.00000 0.00000 ; HA- CA- C- O
4765 4764 4775 4777 3 0.00000 0.00000 0.00000
0.00000 0.00000 0.00000 ; HA- CA- C- N
4766 4764 4775 4776 3 0.00000 0.00000 0.00000
0.00000 0.00000 0.00000 ; CB- CA- C- O
4766 4764 4775 4777 3 2.51040 4.18400 1.67360
-6.69440 0.00000 0.00000 ; CB- CA- C- N
4766 4769 4772 4773 3 3.79907 0.00000 -5.40573
0.00000 2.14221 0.00000 ; CB- CG- CD- OE1
4766 4769 4772 4774 3 3.79907 0.00000 -5.40573
0.00000 2.14221 0.00000 ; CB- CG- CD- OE2
4767 4766 4764 4775 3 0.65084 1.95253 0.00000
-2.60338 0.00000 0.00000 ; HB2- CB- CA- C
4767 4766 4769 4770 3 0.62760 1.88280 0.00000
-2.51040 0.00000 0.00000 ; HB2- CB- CG- HG2
4767 4766 4769 4771 3 0.62760 1.88280 0.00000
-2.51040 0.00000 0.00000 ; HB2- CB- CG- HG3
4767 4766 4769 4772 3 0.65084 1.95253 0.00000
-2.60338 0.00000 0.00000 ; HB2- CB- CG- CD
4768 4766 4764 4775 3 0.65084 1.95253 0.00000
-2.60338 0.00000 0.00000 ; HB3- CB- CA- C
4768 4766 4769 4770 3 0.62760 1.88280 0.00000
-2.51040 0.00000 0.00000 ; HB3- CB- CG- HG2
4768 4766 4769 4771 3 0.62760 1.88280 0.00000
-2.51040 0.00000 0.00000 ; HB3- CB- CG- HG3
4768 4766 4769 4772 3 0.65084 1.95253 0.00000
-2.60338 0.00000 0.00000 ; HB3- CB- CG- CD
4769 4766 4764 4775 3 6.86594 3.56895 -8.14206
-2.40998 4.85344 0.00000 ; CG- CB- CA- C
4770 4769 4772 4773 3 0.00000 0.00000 0.00000
0.00000 0.00000 0.00000 ; HG2- CG- CD- OE1
4770 4769 4772 4774 3 0.00000 0.00000 0.00000
0.00000 0.00000 0.00000 ; HG2- CG- CD- OE2
4771 4769 4772 4773 3 0.00000 0.00000 0.00000
0.00000 0.00000 0.00000 ; HG3- CG- CD- OE1
4771 4769 4772 4774 3 0.00000 0.00000 0.00000
0.00000 0.00000 0.00000 ; HG3- CG- CD- OE2
4775 4777 4779 4780 3 0.00000 0.00000 0.00000
0.00000 0.00000 0.00000 ; C- N- CA- HA
4775 4777 4779 4781 3 11.71520 1.67360 15.06240 -
13.38880 0.00000 0.00000 ; C- N- CA- CB
4775 4777 4779 4789 3 1.75728 5.27184 2.25936

-7.02912 0.00000 0.00000 ; C- N- CA- C
4776 4775 4777 4778 3 29.28800 -8.36800 -20.92000
0.00000 0.00000 0.00000 ; O- C- N- H
4776 4775 4777 4779 3 20.92000 0.00000 -20.92000
0.00000 0.00000 0.00000 ; O- C- N- CA
4777 4779 4781 4782 3 0.65084 1.95253 0.00000
-2.60338 0.00000 0.00000 ; N- CA- CB- HB2
4777 4779 4781 4783 3 0.65084 1.95253 0.00000
-2.60338 0.00000 0.00000 ; N- CA- CB- HB3
4777 4779 4781 4784 3 5.99567 3.29281 -4.46014
-0.55229 1.97485 0.00000 ; N- CA- CB- CG
4777 4779 4789 4790 3 0.00000 0.00000 0.00000
0.00000 0.00000 0.00000 ; N- CA- C- O
4777 4779 4789 4791 3 17.40544 -5.02080 -13.22144
9.20480 0.00000 0.00000 ; N- CA- C- N
4778 4777 4779 4780 3 0.00000 0.00000 0.00000
0.00000 0.00000 0.00000 ; H- N- CA- HA
4778 4777 4779 4781 3 0.00000 0.00000 0.00000
0.00000 0.00000 0.00000 ; H- N- CA- CB
4778 4777 4779 4789 3 0.00000 0.00000 0.00000
0.00000 0.00000 0.00000 ; H- N- CA- C
4779 4781 4784 4785 3 0.00000 0.00000 0.00000
0.00000 0.00000 0.00000 ; CA- CB- CG- OD1
4779 4781 4784 4786 3 8.84916 -0.31380 -4.32626
5.03754 0.26778 0.00000 ; CA- CB- CG- ND2
4779 4789 4791 4792 3 20.92000 0.00000 -20.92000
0.00000 0.00000 0.00000 ; CA- C- N- H
4779 4789 4791 4793 3 20.92000 0.00000 -20.92000
0.00000 0.00000 0.00000 ; CA- C- N- CA
4780 4779 4781 4782 3 0.65084 1.95253 0.00000
-2.60338 0.00000 0.00000 ; HA- CA- CB- HB2
4780 4779 4781 4783 3 0.65084 1.95253 0.00000
-2.60338 0.00000 0.00000 ; HA- CA- CB- HB3
4780 4779 4781 4784 3 0.65084 1.95253 0.00000
-2.60338 0.00000 0.00000 ; HA- CA- CB- CG
4780 4779 4789 4790 3 3.68192 -4.35136 0.00000
1.33888 0.00000 0.00000 ; HA- CA- C- O
4780 4779 4789 4791 3 0.00000 0.00000 0.00000
0.00000 0.00000 0.00000 ; HA- CA- C- N
4781 4779 4789 4790 3 0.00000 0.00000 0.00000
0.00000 0.00000 0.00000 ; CB- CA- C- O
4781 4779 4789 4791 3 2.51040 4.18400 1.67360
-6.69440 0.00000 0.00000 ; CB- CA- C- N
4781 4784 4786 4787 3 20.92000 0.00000 -20.92000
0.00000 0.00000 0.00000 ; CB- CG- ND2- HD21
4781 4784 4786 4788 3 20.92000 0.00000 -20.92000
0.00000 0.00000 0.00000 ; CB- CG- ND2- HD22
4782 4781 4779 4789 3 0.65084 1.95253 0.00000
-2.60338 0.00000 0.00000 ; HB2- CB- CA- C
4782 4781 4784 4785 3 3.68192 -4.35136 0.00000
1.33888 0.00000 0.00000 ; HB2- CB- CG- OD1
4782 4781 4784 4786 3 0.00000 0.00000 0.00000

0.00000    0.00000    0.00000 ;    HB2-    CB-    CG-   ND2
  4783   4781   4779   4789      3    0.65084    1.95253    0.00000
-2.60338    0.00000    0.00000 ;    HB3-    CB-    CA-     C
  4783   4781   4784   4785      3    3.68192   -4.35136    0.00000
1.33888    0.00000    0.00000 ;    HB3-    CB-    CG-   OD1
  4783   4781   4784   4786      3    0.00000    0.00000    0.00000
0.00000    0.00000    0.00000 ;    HB3-    CB-    CG-   ND2
  4784   4781   4779   4789      3    7.94542   -3.96225   -6.11701
-0.55229    3.58150    0.00000 ;     CG-    CB-    CA-     C
  4785   4784   4786   4787      3   29.28800   -8.36800  -20.92000
0.00000    0.00000    0.00000 ;    OD1-    CG-   ND2-  HD21
  4785   4784   4786   4788      3   29.28800   -8.36800  -20.92000
0.00000    0.00000    0.00000 ;    OD1-    CG-   ND2-  HD22
  4789   4791   4793   4794      3    0.00000    0.00000    0.00000
0.00000    0.00000    0.00000 ;     C-     N-    CA-    HA
  4789   4791   4793   4795      3   11.71520    1.67360   15.06240  -
13.38880    0.00000    0.00000 ;     C-     N-    CA-    CB
  4789   4791   4793   4804      3    1.75728    5.27184    2.25936
-7.02912    0.00000    0.00000 ;     C-     N-    CA-     C
  4790   4789   4791   4792      3   29.28800   -8.36800  -20.92000
0.00000    0.00000    0.00000 ;     O-     C-     N-     H
  4790   4789   4791   4793      3   20.92000    0.00000  -20.92000
0.00000    0.00000    0.00000 ;     O-     C-     N-    CA
  4791   4793   4795   4796      3    0.65084    1.95253    0.00000
-2.60338    0.00000    0.00000 ;     N-    CA-    CB-   HB2
  4791   4793   4795   4797      3    0.65084    1.95253    0.00000
-2.60338    0.00000    0.00000 ;     N-    CA-    CB-   HB3
  4791   4793   4795   4798      3    3.21331    2.22589   -4.15053
-2.40998    2.61082    0.00000 ;     N-    CA-    CB-    CG
  4791   4793   4804   4805      3    0.00000    0.00000    0.00000
0.00000    0.00000    0.00000 ;     N-    CA-     C-     O
  4791   4793   4804   4806      3   17.40544   -5.02080  -13.22144
9.20480    0.00000    0.00000 ;     N-    CA-     C-     N
  4792   4791   4793   4794      3    0.00000    0.00000    0.00000
0.00000    0.00000    0.00000 ;     H-     N-    CA-    HA
  4792   4791   4793   4795      3    0.00000    0.00000    0.00000
0.00000    0.00000    0.00000 ;     H-     N-    CA-    CB
  4792   4791   4793   4804      3    0.00000    0.00000    0.00000
0.00000    0.00000    0.00000 ;     H-     N-    CA-     C
  4793   4795   4798   4799      3    0.66944    2.00832    0.00000
-2.67776    0.00000    0.00000 ;    CA-    CB-    CG-   HG2
  4793   4795   4798   4800      3    0.66944    2.00832    0.00000
-2.67776    0.00000    0.00000 ;    CA-    CB-    CG-   HG3
  4793   4795   4798   4801      3   10.12110   -1.91209    0.01674
10.17549   -1.87443    0.00000 ;    CA-    CB-    CG-    CD
  4793   4804   4806   4807      3   20.92000    0.00000  -20.92000
0.00000    0.00000    0.00000 ;    CA-     C-     N-     H
  4793   4804   4806   4808      3   20.92000    0.00000  -20.92000
0.00000    0.00000    0.00000 ;    CA-     C-     N-    CA
  4794   4793   4795   4796      3    0.65084    1.95253    0.00000
-2.60338    0.00000    0.00000 ;    HA-    CA-    CB-   HB2
  4794   4793   4795   4797      3    0.65084    1.95253    0.00000

-2.60338 0.00000 0.00000 ; HA- CA- CB- HB3
4794 4793 4795 4798 3 0.65084 1.95253 0.00000
-2.60338 0.00000 0.00000 ; HA- CA- CB- CG
4794 4793 4804 4805 3 3.68192 -4.35136 0.00000
1.33888 0.00000 0.00000 ; HA- CA- C- O
4794 4793 4804 4806 3 0.00000 0.00000 0.00000
0.00000 0.00000 0.00000 ; HA- CA- C- N
4795 4793 4804 4805 3 0.00000 0.00000 0.00000
0.00000 0.00000 0.00000 ; CB- CA- C- O
4795 4793 4804 4806 3 2.51040 4.18400 1.67360
-6.69440 0.00000 0.00000 ; CB- CA- C- N
4795 4798 4801 4802 3 3.79907 0.00000 -5.40573
0.00000 2.14221 0.00000 ; CB- CG- CD- OE1
4795 4798 4801 4803 3 3.79907 0.00000 -5.40573
0.00000 2.14221 0.00000 ; CB- CG- CD- OE2
4796 4795 4793 4804 3 0.65084 1.95253 0.00000
-2.60338 0.00000 0.00000 ; HB2- CB- CA- C
4796 4795 4798 4799 3 0.62760 1.88280 0.00000
-2.51040 0.00000 0.00000 ; HB2- CB- CG- HG2
4796 4795 4798 4800 3 0.62760 1.88280 0.00000
-2.51040 0.00000 0.00000 ; HB2- CB- CG- HG3
4796 4795 4798 4801 3 0.65084 1.95253 0.00000
-2.60338 0.00000 0.00000 ; HB2- CB- CG- CD
4797 4795 4793 4804 3 0.65084 1.95253 0.00000
-2.60338 0.00000 0.00000 ; HB3- CB- CA- C
4797 4795 4798 4799 3 0.62760 1.88280 0.00000
-2.51040 0.00000 0.00000 ; HB3- CB- CG- HG2
4797 4795 4798 4800 3 0.62760 1.88280 0.00000
-2.51040 0.00000 0.00000 ; HB3- CB- CG- HG3
4797 4795 4798 4801 3 0.65084 1.95253 0.00000
-2.60338 0.00000 0.00000 ; HB3- CB- CG- CD
4798 4795 4793 4804 3 6.86594 3.56895 -8.14206
-2.40998 4.85344 0.00000 ; CG- CB- CA- C
4799 4798 4801 4802 3 0.00000 0.00000 0.00000
0.00000 0.00000 0.00000 ; HG2- CG- CD- OE1
4799 4798 4801 4803 3 0.00000 0.00000 0.00000
0.00000 0.00000 0.00000 ; HG2- CG- CD- OE2
4800 4798 4801 4802 3 0.00000 0.00000 0.00000
0.00000 0.00000 0.00000 ; HG3- CG- CD- OE1
4800 4798 4801 4803 3 0.00000 0.00000 0.00000
0.00000 0.00000 0.00000 ; HG3- CG- CD- OE2
4804 4806 4808 4809 3 0.00000 0.00000 0.00000
0.00000 0.00000 0.00000 ; C- N- CA- HA
4804 4806 4808 4810 3 11.71520 1.67360 15.06240 -
13.38880 0.00000 0.00000 ; C- N- CA- CB
4804 4806 4808 4823 3 1.75728 5.27184 2.25936
-7.02912 0.00000 0.00000 ; C- N- CA- C
4805 4804 4806 4807 3 29.28800 -8.36800 -20.92000
0.00000 0.00000 0.00000 ; O- C- N- H
4805 4804 4806 4808 3 20.92000 0.00000 -20.92000
0.00000 0.00000 0.00000 ; O- C- N- CA
4806 4808 4810 4811 3 0.65084 1.95253 0.00000

-2.60338    0.00000    0.00000 ;       N-    CA-    CB-   HB2
  4806   4808   4810   4812      3    0.65084    1.95253    0.00000
-2.60338    0.00000    0.00000 ;       N-    CA-    CB-   HB3
  4806   4808   4810   4813      3    3.79070    1.39746   -4.61077
-2.40998    2.44346    0.00000 ;       N-    CA-    CB-    CG
  4806   4808   4823   4824      3    0.00000    0.00000    0.00000
0.00000    0.00000    0.00000 ;       N-    CA-     C-     O
  4806   4808   4823   4825      3   17.40544   -5.02080  -13.22144
9.20480    0.00000    0.00000 ;       N-    CA-     C-     N
  4807   4806   4808   4809      3    0.00000    0.00000    0.00000
0.00000    0.00000    0.00000 ;       H-     N-    CA-    HA
  4807   4806   4808   4810      3    0.00000    0.00000    0.00000
0.00000    0.00000    0.00000 ;       H-     N-    CA-    CB
  4807   4806   4808   4823      3    0.00000    0.00000    0.00000
0.00000    0.00000    0.00000 ;       H-     N-    CA-     C
  4808   4810   4813   4814      3    0.66944    2.00832    0.00000
-2.67776    0.00000    0.00000 ;      CA-    CB-    CG-    HG
  4808   4810   4813   4815      3    3.90367    0.19665   -6.21742
-2.37651    5.99149    0.00000 ;      CA-    CB-    CG-   CD1
  4808   4810   4813   4819      3    3.90367    0.19665   -6.21742
-2.37651    5.99149    0.00000 ;      CA-    CB-    CG-   CD2
  4808   4823   4825   4826      3   20.92000    0.00000  -20.92000
0.00000    0.00000    0.00000 ;      CA-     C-     N-     H
  4808   4823   4825   4827      3   20.92000    0.00000  -20.92000
0.00000    0.00000    0.00000 ;      CA-     C-     N-    CA
  4809   4808   4810   4811      3    0.65084    1.95253    0.00000
-2.60338    0.00000    0.00000 ;      HA-    CA-    CB-   HB2
  4809   4808   4810   4812      3    0.65084    1.95253    0.00000
-2.60338    0.00000    0.00000 ;      HA-    CA-    CB-   HB3
  4809   4808   4810   4813      3    0.65084    1.95253    0.00000
-2.60338    0.00000    0.00000 ;      HA-    CA-    CB-    CG
  4809   4808   4823   4824      3    3.68192   -4.35136    0.00000
1.33888    0.00000    0.00000 ;      HA-    CA-     C-     O
  4809   4808   4823   4825      3    0.00000    0.00000    0.00000
0.00000    0.00000    0.00000 ;      HA-    CA-     C-     N
  4810   4808   4823   4824      3    0.00000    0.00000    0.00000
0.00000    0.00000    0.00000 ;      CB-    CA-     C-     O
  4810   4808   4823   4825      3    2.51040    4.18400    1.67360
-6.69440    0.00000    0.00000 ;      CB-    CA-     C-     N
  4810   4813   4815   4816      3    0.66944    2.00832    0.00000
-2.67776    0.00000    0.00000 ;      CB-    CG-   CD1-  HD11
  4810   4813   4815   4817      3    0.66944    2.00832    0.00000
-2.67776    0.00000    0.00000 ;      CB-    CG-   CD1-  HD12
  4810   4813   4815   4818      3    0.66944    2.00832    0.00000
-2.67776    0.00000    0.00000 ;      CB-    CG-   CD1-  HD13
  4810   4813   4819   4820      3    0.66944    2.00832    0.00000
-2.67776    0.00000    0.00000 ;      CB-    CG-   CD2-  HD21
  4810   4813   4819   4821      3    0.66944    2.00832    0.00000
-2.67776    0.00000    0.00000 ;      CB-    CG-   CD2-  HD22
  4810   4813   4819   4822      3    0.66944    2.00832    0.00000
-2.67776    0.00000    0.00000 ;      CB-    CG-   CD2-  HD23
  4811   4810   4808   4823      3    0.65084    1.95253    0.00000

-2.60338 0.00000 0.00000 ; HB2- CB- CA- C
4811 4810 4813 4814 3 0.62760 1.88280 0.00000
-2.51040 0.00000 0.00000 ; HB2- CB- CG- HG
4811 4810 4813 4815 3 0.66944 2.00832 0.00000
-2.67776 0.00000 0.00000 ; HB2- CB- CG- CD1
4811 4810 4813 4819 3 0.66944 2.00832 0.00000
-2.67776 0.00000 0.00000 ; HB2- CB- CG- CD2
4812 4810 4808 4823 3 0.65084 1.95253 0.00000
-2.60338 0.00000 0.00000 ; HB3- CB- CA- C
4812 4810 4813 4814 3 0.62760 1.88280 0.00000
-2.51040 0.00000 0.00000 ; HB3- CB- CG- HG
4812 4810 4813 4815 3 0.66944 2.00832 0.00000
-2.67776 0.00000 0.00000 ; HB3- CB- CG- CD1
4812 4810 4813 4819 3 0.66944 2.00832 0.00000
-2.67776 0.00000 0.00000 ; HB3- CB- CG- CD2
4813 4810 4808 4823 3 10.33448 -1.14642 -11.54784
-2.40998 6.35968 0.00000 ; CG- CB- CA- C
4814 4813 4815 4816 3 0.62760 1.88280 0.00000
-2.51040 0.00000 0.00000 ; HG- CG- CD1- HD11
4814 4813 4815 4817 3 0.62760 1.88280 0.00000
-2.51040 0.00000 0.00000 ; HG- CG- CD1- HD12
4814 4813 4815 4818 3 0.62760 1.88280 0.00000
-2.51040 0.00000 0.00000 ; HG- CG- CD1- HD13
4814 4813 4819 4820 3 0.62760 1.88280 0.00000
-2.51040 0.00000 0.00000 ; HG- CG- CD2- HD21
4814 4813 4819 4821 3 0.62760 1.88280 0.00000
-2.51040 0.00000 0.00000 ; HG- CG- CD2- HD22
4814 4813 4819 4822 3 0.62760 1.88280 0.00000
-2.51040 0.00000 0.00000 ; HG- CG- CD2- HD23
4815 4813 4819 4820 3 0.66944 2.00832 0.00000
-2.67776 0.00000 0.00000 ; CD1- CG- CD2- HD21
4815 4813 4819 4821 3 0.66944 2.00832 0.00000
-2.67776 0.00000 0.00000 ; CD1- CG- CD2- HD22
4815 4813 4819 4822 3 0.66944 2.00832 0.00000
-2.67776 0.00000 0.00000 ; CD1- CG- CD2- HD23
4816 4815 4813 4819 3 0.66944 2.00832 0.00000
-2.67776 0.00000 0.00000 ; HD11- CD1- CG- CD2
4817 4815 4813 4819 3 0.66944 2.00832 0.00000
-2.67776 0.00000 0.00000 ; HD12- CD1- CG- CD2
4818 4815 4813 4819 3 0.66944 2.00832 0.00000
-2.67776 0.00000 0.00000 ; HD13- CD1- CG- CD2
4823 4825 4827 4828 3 0.00000 0.00000 0.00000
0.00000 0.00000 0.00000 ; C- N- CA- HA
4823 4825 4827 4829 3 11.71520 1.67360 15.06240 -
13.38880 0.00000 0.00000 ; C- N- CA- CB
4823 4825 4827 4845 3 1.75728 5.27184 2.25936
-7.02912 0.00000 0.00000 ; C- N- CA- C
4824 4823 4825 4826 3 29.28800 -8.36800 -20.92000
0.00000 0.00000 0.00000 ; O- C- N- H
4824 4823 4825 4827 3 20.92000 0.00000 -20.92000
0.00000 0.00000 0.00000 ; O- C- N- CA
4825 4827 4829 4830 3 0.65084 1.95253 0.00000

```
-2.60338    0.00000    0.00000 ;       N-    CA-    CB-   HB2
  4825   4827   4829   4831      3    0.65084    1.95253    0.00000
-2.60338    0.00000    0.00000 ;       N-    CA-    CB-   HB3
  4825   4827   4829   4832      3    0.65084    1.95253    0.00000
-2.60338    0.00000    0.00000 ;       N-    CA-    CB-    CG
  4825   4827   4845   4846      3    0.00000    0.00000    0.00000
0.00000    0.00000    0.00000 ;       N-    CA-     C-     O
  4825   4827   4845   4847      3   17.40544   -5.02080  -13.22144
9.20480    0.00000    0.00000 ;       N-    CA-     C-     N
  4826   4825   4827   4828      3    0.00000    0.00000    0.00000
0.00000    0.00000    0.00000 ;       H-     N-    CA-    HA
  4826   4825   4827   4829      3    0.00000    0.00000    0.00000
0.00000    0.00000    0.00000 ;       H-     N-    CA-    CB
  4826   4825   4827   4845      3    0.00000    0.00000    0.00000
0.00000    0.00000    0.00000 ;       H-     N-    CA-     C
  4827   4829   4832   4833      3    0.66944    2.00832    0.00000
-2.67776    0.00000    0.00000 ;      CA-    CB-    CG-   HG2
  4827   4829   4832   4834      3    0.66944    2.00832    0.00000
-2.67776    0.00000    0.00000 ;      CA-    CB-    CG-   HG3
  4827   4829   4832   4835      3    3.68192    3.09616   -2.09200
-3.01248    0.00000    0.00000 ;      CA-    CB-    CG-    CD
  4827   4845   4847   4848      3   20.92000    0.00000  -20.92000
0.00000    0.00000    0.00000 ;      CA-     C-     N-     H
  4827   4845   4847   4849      3   20.92000    0.00000  -20.92000
0.00000    0.00000    0.00000 ;      CA-     C-     N-    CA
  4828   4827   4829   4830      3    0.65084    1.95253    0.00000
-2.60338    0.00000    0.00000 ;      HA-    CA-    CB-   HB2
  4828   4827   4829   4831      3    0.65084    1.95253    0.00000
-2.60338    0.00000    0.00000 ;      HA-    CA-    CB-   HB3
  4828   4827   4829   4832      3    0.65084    1.95253    0.00000
-2.60338    0.00000    0.00000 ;      HA-    CA-    CB-    CG
  4828   4827   4845   4846      3    3.68192   -4.35136    0.00000
1.33888    0.00000    0.00000 ;      HA-    CA-     C-     O
  4828   4827   4845   4847      3    0.00000    0.00000    0.00000
0.00000    0.00000    0.00000 ;      HA-    CA-     C-     N
  4829   4827   4845   4846      3    0.00000    0.00000    0.00000
0.00000    0.00000    0.00000 ;      CB-    CA-     C-     O
  4829   4827   4845   4847      3    2.51040    4.18400    1.67360
-6.69440    0.00000    0.00000 ;      CB-    CA-     C-     N
  4829   4832   4835   4836      3    0.66944    2.00832    0.00000
-2.67776    0.00000    0.00000 ;      CB-    CG-    CD-   HD2
  4829   4832   4835   4837      3    0.66944    2.00832    0.00000
-2.67776    0.00000    0.00000 ;      CB-    CG-    CD-   HD3
  4829   4832   4835   4838      3    3.68192    3.09616   -2.09200
-3.01248    0.00000    0.00000 ;      CB-    CG-    CD-    CE
  4830   4829   4827   4845      3    0.65084    1.95253    0.00000
-2.60338    0.00000    0.00000 ;     HB2-    CB-    CA-     C
  4830   4829   4832   4833      3    0.62760    1.88280    0.00000
-2.51040    0.00000    0.00000 ;     HB2-    CB-    CG-   HG2
  4830   4829   4832   4834      3    0.62760    1.88280    0.00000
-2.51040    0.00000    0.00000 ;     HB2-    CB-    CG-   HG3
  4830   4829   4832   4835      3    0.66944    2.00832    0.00000
```

```
-2.67776    0.00000    0.00000 ;     HB2-    CB-    CG-    CD
  4831   4829   4827   4845      3    0.65084    1.95253    0.00000
-2.60338    0.00000    0.00000 ;     HB3-    CB-    CA-     C
  4831   4829   4832   4833      3    0.62760    1.88280    0.00000
-2.51040    0.00000    0.00000 ;     HB3-    CB-    CG-   HG2
  4831   4829   4832   4834      3    0.62760    1.88280    0.00000
-2.51040    0.00000    0.00000 ;     HB3-    CB-    CG-   HG3
  4831   4829   4832   4835      3    0.66944    2.00832    0.00000
-2.67776    0.00000    0.00000 ;     HB3-    CB-    CG-    CD
  4832   4829   4827   4845      3    0.65084    1.95253    0.00000
-2.60338    0.00000    0.00000 ;      CG-    CB-    CA-     C
  4832   4835   4838   4839      3    0.65084    1.95253    0.00000
-2.60338    0.00000    0.00000 ;      CG-    CD-    CE-   HE2
  4832   4835   4838   4840      3    0.65084    1.95253    0.00000
-2.60338    0.00000    0.00000 ;      CG-    CD-    CE-   HE3
  4832   4835   4838   4841      3    0.65084    1.95253    0.00000
-2.60338    0.00000    0.00000 ;      CG-    CD-    CE-    NZ
  4833   4832   4835   4836      3    0.62760    1.88280    0.00000
-2.51040    0.00000    0.00000 ;     HG2-    CG-    CD-   HD2
  4833   4832   4835   4837      3    0.62760    1.88280    0.00000
-2.51040    0.00000    0.00000 ;     HG2-    CG-    CD-   HD3
  4833   4832   4835   4838      3    0.66944    2.00832    0.00000
-2.67776    0.00000    0.00000 ;     HG2-    CG-    CD-    CE
  4834   4832   4835   4836      3    0.62760    1.88280    0.00000
-2.51040    0.00000    0.00000 ;     HG3-    CG-    CD-   HD2
  4834   4832   4835   4837      3    0.62760    1.88280    0.00000
-2.51040    0.00000    0.00000 ;     HG3-    CG-    CD-   HD3
  4834   4832   4835   4838      3    0.66944    2.00832    0.00000
-2.67776    0.00000    0.00000 ;     HG3-    CG-    CD-    CE
  4835   4838   4841   4842      3    0.65084    1.95253    0.00000
-2.60338    0.00000    0.00000 ;      CD-    CE-    NZ-   HZ1
  4835   4838   4841   4843      3    0.65084    1.95253    0.00000
-2.60338    0.00000    0.00000 ;      CD-    CE-    NZ-   HZ2
  4835   4838   4841   4844      3    0.65084    1.95253    0.00000
-2.60338    0.00000    0.00000 ;      CD-    CE-    NZ-   HZ3
  4836   4835   4838   4839      3    0.65084    1.95253    0.00000
-2.60338    0.00000    0.00000 ;     HD2-    CD-    CE-   HE2
  4836   4835   4838   4840      3    0.65084    1.95253    0.00000
-2.60338    0.00000    0.00000 ;     HD2-    CD-    CE-   HE3
  4836   4835   4838   4841      3    0.65084    1.95253    0.00000
-2.60338    0.00000    0.00000 ;     HD2-    CD-    CE-    NZ
  4837   4835   4838   4839      3    0.65084    1.95253    0.00000
-2.60338    0.00000    0.00000 ;     HD3-    CD-    CE-   HE2
  4837   4835   4838   4840      3    0.65084    1.95253    0.00000
-2.60338    0.00000    0.00000 ;     HD3-    CD-    CE-   HE3
  4837   4835   4838   4841      3    0.65084    1.95253    0.00000
-2.60338    0.00000    0.00000 ;     HD3-    CD-    CE-    NZ
  4839   4838   4841   4842      3    0.65084    1.95253    0.00000
-2.60338    0.00000    0.00000 ;     HE2-    CE-    NZ-   HZ1
  4839   4838   4841   4843      3    0.65084    1.95253    0.00000
-2.60338    0.00000    0.00000 ;     HE2-    CE-    NZ-   HZ2
  4839   4838   4841   4844      3    0.65084    1.95253    0.00000
```

-2.60338 0.00000 0.00000 ; HE2- CE- NZ- HZ3
4840 4838 4841 4842 3 0.65084 1.95253 0.00000
-2.60338 0.00000 0.00000 ; HE3- CE- NZ- HZ1
4840 4838 4841 4843 3 0.65084 1.95253 0.00000
-2.60338 0.00000 0.00000 ; HE3- CE- NZ- HZ2
4840 4838 4841 4844 3 0.65084 1.95253 0.00000
-2.60338 0.00000 0.00000 ; HE3- CE- NZ- HZ3
4845 4847 4849 4850 3 0.00000 0.00000 0.00000
0.00000 0.00000 0.00000 ; C- N- CA- HA
4845 4847 4849 4851 3 11.71520 1.67360 15.06240 -
13.38880 0.00000 0.00000 ; C- N- CA- CB
4845 4847 4849 4867 3 1.75728 5.27184 2.25936
-7.02912 0.00000 0.00000 ; C- N- CA- C
4846 4845 4847 4848 3 29.28800 -8.36800 -20.92000
0.00000 0.00000 0.00000 ; O- C- N- H
4846 4845 4847 4849 3 20.92000 0.00000 -20.92000
0.00000 0.00000 0.00000 ; O- C- N- CA
4847 4849 4851 4852 3 0.65084 1.95253 0.00000
-2.60338 0.00000 0.00000 ; N- CA- CB- HB2
4847 4849 4851 4853 3 0.65084 1.95253 0.00000
-2.60338 0.00000 0.00000 ; N- CA- CB- HB3
4847 4849 4851 4854 3 0.65084 1.95253 0.00000
-2.60338 0.00000 0.00000 ; N- CA- CB- CG
4847 4849 4867 4868 3 0.00000 0.00000 0.00000
0.00000 0.00000 0.00000 ; N- CA- C- O
4847 4849 4867 4869 3 17.40544 -5.02080 -13.22144
9.20480 0.00000 0.00000 ; N- CA- C- N
4848 4847 4849 4850 3 0.00000 0.00000 0.00000
0.00000 0.00000 0.00000 ; H- N- CA- HA
4848 4847 4849 4851 3 0.00000 0.00000 0.00000
0.00000 0.00000 0.00000 ; H- N- CA- CB
4848 4847 4849 4867 3 0.00000 0.00000 0.00000
0.00000 0.00000 0.00000 ; H- N- CA- C
4849 4851 4854 4855 3 0.66944 2.00832 0.00000
-2.67776 0.00000 0.00000 ; CA- CB- CG- HG2
4849 4851 4854 4856 3 0.66944 2.00832 0.00000
-2.67776 0.00000 0.00000 ; CA- CB- CG- HG3
4849 4851 4854 4857 3 3.68192 3.09616 -2.09200
-3.01248 0.00000 0.00000 ; CA- CB- CG- CD
4849 4867 4869 4870 3 20.92000 0.00000 -20.92000
0.00000 0.00000 0.00000 ; CA- C- N- H
4849 4867 4869 4871 3 20.92000 0.00000 -20.92000
0.00000 0.00000 0.00000 ; CA- C- N- CA
4850 4849 4851 4852 3 0.65084 1.95253 0.00000
-2.60338 0.00000 0.00000 ; HA- CA- CB- HB2
4850 4849 4851 4853 3 0.65084 1.95253 0.00000
-2.60338 0.00000 0.00000 ; HA- CA- CB- HB3
4850 4849 4851 4854 3 0.65084 1.95253 0.00000
-2.60338 0.00000 0.00000 ; HA- CA- CB- CG
4850 4849 4867 4868 3 3.68192 -4.35136 0.00000
1.33888 0.00000 0.00000 ; HA- CA- C- O
4850 4849 4867 4869 3 0.00000 0.00000 0.00000

```
0.00000    0.00000    0.00000 ;      HA-    CA-     C-     N
  4851   4849   4867   4868       3    0.00000    0.00000    0.00000
0.00000    0.00000    0.00000 ;      CB-    CA-     C-     O
  4851   4849   4867   4869       3    2.51040    4.18400    1.67360
-6.69440    0.00000    0.00000 ;      CB-    CA-     C-     N
  4851   4854   4857   4858       3    0.66944    2.00832    0.00000
-2.67776    0.00000    0.00000 ;      CB-    CG-    CD-   HD2
  4851   4854   4857   4859       3    0.66944    2.00832    0.00000
-2.67776    0.00000    0.00000 ;      CB-    CG-    CD-   HD3
  4851   4854   4857   4860       3    3.68192    3.09616   -2.09200
-3.01248    0.00000    0.00000 ;      CB-    CG-    CD-    CE
  4852   4851   4849   4867       3    0.65084    1.95253    0.00000
-2.60338    0.00000    0.00000 ;     HB2-    CB-    CA-     C
  4852   4851   4854   4855       3    0.62760    1.88280    0.00000
-2.51040    0.00000    0.00000 ;     HB2-    CB-    CG-   HG2
  4852   4851   4854   4856       3    0.62760    1.88280    0.00000
-2.51040    0.00000    0.00000 ;     HB2-    CB-    CG-   HG3
  4852   4851   4854   4857       3    0.66944    2.00832    0.00000
-2.67776    0.00000    0.00000 ;     HB2-    CB-    CG-    CD
  4853   4851   4849   4867       3    0.65084    1.95253    0.00000
-2.60338    0.00000    0.00000 ;     HB3-    CB-    CA-     C
  4853   4851   4854   4855       3    0.62760    1.88280    0.00000
-2.51040    0.00000    0.00000 ;     HB3-    CB-    CG-   HG2
  4853   4851   4854   4856       3    0.62760    1.88280    0.00000
-2.51040    0.00000    0.00000 ;     HB3-    CB-    CG-   HG3
  4853   4851   4854   4857       3    0.66944    2.00832    0.00000
-2.67776    0.00000    0.00000 ;     HB3-    CB-    CG-    CD
  4854   4851   4849   4867       3    0.65084    1.95253    0.00000
-2.60338    0.00000    0.00000 ;      CG-    CB-    CA-     C
  4854   4857   4860   4861       3    0.65084    1.95253    0.00000
-2.60338    0.00000    0.00000 ;      CG-    CD-    CE-   HE2
  4854   4857   4860   4862       3    0.65084    1.95253    0.00000
-2.60338    0.00000    0.00000 ;      CG-    CD-    CE-   HE3
  4854   4857   4860   4863       3    0.65084    1.95253    0.00000
-2.60338    0.00000    0.00000 ;      CG-    CD-    CE-    NZ
  4855   4854   4857   4858       3    0.62760    1.88280    0.00000
-2.51040    0.00000    0.00000 ;     HG2-    CG-    CD-   HD2
  4855   4854   4857   4859       3    0.62760    1.88280    0.00000
-2.51040    0.00000    0.00000 ;     HG2-    CG-    CD-   HD3
  4855   4854   4857   4860       3    0.66944    2.00832    0.00000
-2.67776    0.00000    0.00000 ;     HG2-    CG-    CD-    CE
  4856   4854   4857   4858       3    0.62760    1.88280    0.00000
-2.51040    0.00000    0.00000 ;     HG3-    CG-    CD-   HD2
  4856   4854   4857   4859       3    0.62760    1.88280    0.00000
-2.51040    0.00000    0.00000 ;     HG3-    CG-    CD-   HD3
  4856   4854   4857   4860       3    0.66944    2.00832    0.00000
-2.67776    0.00000    0.00000 ;     HG3-    CG-    CD-    CE
  4857   4860   4863   4864       3    0.65084    1.95253    0.00000
-2.60338    0.00000    0.00000 ;      CD-    CE-    NZ-   HZ1
  4857   4860   4863   4865       3    0.65084    1.95253    0.00000
-2.60338    0.00000    0.00000 ;      CD-    CE-    NZ-   HZ2
  4857   4860   4863   4866       3    0.65084    1.95253    0.00000
```

-2.60338 0.00000 0.00000 ; CD- CE- NZ- HZ3
4858 4857 4860 4861 3 0.65084 1.95253 0.00000
-2.60338 0.00000 0.00000 ; HD2- CD- CE- HE2
4858 4857 4860 4862 3 0.65084 1.95253 0.00000
-2.60338 0.00000 0.00000 ; HD2- CD- CE- HE3
4858 4857 4860 4863 3 0.65084 1.95253 0.00000
-2.60338 0.00000 0.00000 ; HD2- CD- CE- NZ
4859 4857 4860 4861 3 0.65084 1.95253 0.00000
-2.60338 0.00000 0.00000 ; HD3- CD- CE- HE2
4859 4857 4860 4862 3 0.65084 1.95253 0.00000
-2.60338 0.00000 0.00000 ; HD3- CD- CE- HE3
4859 4857 4860 4863 3 0.65084 1.95253 0.00000
-2.60338 0.00000 0.00000 ; HD3- CD- CE- NZ
4861 4860 4863 4864 3 0.65084 1.95253 0.00000
-2.60338 0.00000 0.00000 ; HE2- CE- NZ- HZ1
4861 4860 4863 4865 3 0.65084 1.95253 0.00000
-2.60338 0.00000 0.00000 ; HE2- CE- NZ- HZ2
4861 4860 4863 4866 3 0.65084 1.95253 0.00000
-2.60338 0.00000 0.00000 ; HE2- CE- NZ- HZ3
4862 4860 4863 4864 3 0.65084 1.95253 0.00000
-2.60338 0.00000 0.00000 ; HE3- CE- NZ- HZ1
4862 4860 4863 4865 3 0.65084 1.95253 0.00000
-2.60338 0.00000 0.00000 ; HE3- CE- NZ- HZ2
4862 4860 4863 4866 3 0.65084 1.95253 0.00000
-2.60338 0.00000 0.00000 ; HE3- CE- NZ- HZ3
4867 4869 4871 4872 3 0.00000 0.00000 0.00000
0.00000 0.00000 0.00000 ; C- N- CA- HA
4867 4869 4871 4873 3 11.71520 1.67360 15.06240 -
13.38880 0.00000 0.00000 ; C- N- CA- CB
4867 4869 4871 4886 3 1.75728 5.27184 2.25936
-7.02912 0.00000 0.00000 ; C- N- CA- C
4868 4867 4869 4870 3 29.28800 -8.36800 -20.92000
0.00000 0.00000 0.00000 ; O- C- N- H
4868 4867 4869 4871 3 20.92000 0.00000 -20.92000
0.00000 0.00000 0.00000 ; O- C- N- CA
4869 4871 4873 4874 3 0.65084 1.95253 0.00000
-2.60338 0.00000 0.00000 ; N- CA- CB- HB
4869 4871 4873 4875 3 3.83673 0.44769 -1.77402
-2.47693 -0.03347 0.00000 ; N- CA- CB- CG2
4869 4871 4873 4879 3 2.97482 0.12134 2.04179
-1.89117 -3.24678 0.00000 ; N- CA- CB- CG1
4869 4871 4886 4887 3 0.00000 0.00000 0.00000
0.00000 0.00000 0.00000 ; N- CA- C- O
4869 4871 4886 4888 3 17.40544 -5.02080 -13.22144
9.20480 0.00000 0.00000 ; N- CA- C- N
4870 4869 4871 4872 3 0.00000 0.00000 0.00000
0.00000 0.00000 0.00000 ; H- N- CA- HA
4870 4869 4871 4873 3 0.00000 0.00000 0.00000
0.00000 0.00000 0.00000 ; H- N- CA- CB
4870 4869 4871 4886 3 0.00000 0.00000 0.00000
0.00000 0.00000 0.00000 ; H- N- CA- C
4871 4873 4875 4876 3 0.66944 2.00832 0.00000

-2.67776 0.00000 0.00000 ; CA- CB- CG2- HG21
4871 4873 4875 4877 3 0.66944 2.00832 0.00000
-2.67776 0.00000 0.00000 ; CA- CB- CG2- HG22
4871 4873 4875 4878 3 0.66944 2.00832 0.00000
-2.67776 0.00000 0.00000 ; CA- CB- CG2- HG23
4871 4873 4879 4880 3 0.66944 2.00832 0.00000
-2.67776 0.00000 0.00000 ; CA- CB- CG1- HG12
4871 4873 4879 4881 3 0.66944 2.00832 0.00000
-2.67776 0.00000 0.00000 ; CA- CB- CG1- HG13
4871 4873 4879 4882 3 4.24258 -0.52718 -7.25506
-1.79075 7.69856 0.00000 ; CA- CB- CG1- CD1
4871 4886 4888 4889 3 20.92000 0.00000 -20.92000
0.00000 0.00000 0.00000 ; CA- C- N- H
4871 4886 4888 4890 3 20.92000 0.00000 -20.92000
0.00000 0.00000 0.00000 ; CA- C- N- CA
4872 4871 4873 4874 3 0.65084 1.95253 0.00000
-2.60338 0.00000 0.00000 ; HA- CA- CB- HB
4872 4871 4873 4875 3 0.65084 1.95253 0.00000
-2.60338 0.00000 0.00000 ; HA- CA- CB- CG2
4872 4871 4873 4879 3 0.65084 1.95253 0.00000
-2.60338 0.00000 0.00000 ; HA- CA- CB- CG1
4872 4871 4886 4887 3 3.68192 -4.35136 0.00000
1.33888 0.00000 0.00000 ; HA- CA- C- O
4872 4871 4886 4888 3 0.00000 0.00000 0.00000
0.00000 0.00000 0.00000 ; HA- CA- C- N
4873 4871 4886 4887 3 0.00000 0.00000 0.00000
0.00000 0.00000 0.00000 ; CB- CA- C- O
4873 4871 4886 4888 3 2.51040 4.18400 1.67360
-6.69440 0.00000 0.00000 ; CB- CA- C- N
4873 4879 4882 4883 3 0.66944 2.00832 0.00000
-2.67776 0.00000 0.00000 ; CB- CG1- CD1- HD11
4873 4879 4882 4884 3 0.66944 2.00832 0.00000
-2.67776 0.00000 0.00000 ; CB- CG1- CD1- HD12
4873 4879 4882 4885 3 0.66944 2.00832 0.00000
-2.67776 0.00000 0.00000 ; CB- CG1- CD1- HD13
4874 4873 4871 4886 3 0.65084 1.95253 0.00000
-2.60338 0.00000 0.00000 ; HB- CB- CA- C
4874 4873 4875 4876 3 0.62760 1.88280 0.00000
-2.51040 0.00000 0.00000 ; HB- CB- CG2- HG21
4874 4873 4875 4877 3 0.62760 1.88280 0.00000
-2.51040 0.00000 0.00000 ; HB- CB- CG2- HG22
4874 4873 4875 4878 3 0.62760 1.88280 0.00000
-2.51040 0.00000 0.00000 ; HB- CB- CG2- HG23
4874 4873 4879 4880 3 0.62760 1.88280 0.00000
-2.51040 0.00000 0.00000 ; HB- CB- CG1- HG12
4874 4873 4879 4881 3 0.62760 1.88280 0.00000
-2.51040 0.00000 0.00000 ; HB- CB- CG1- HG13
4874 4873 4879 4882 3 0.66944 2.00832 0.00000
-2.67776 0.00000 0.00000 ; HB- CB- CG1- CD1
4875 4873 4871 4886 3 5.67350 3.55640 -6.16722
-2.47693 3.74886 0.00000 ; CG2- CB- CA- C
4875 4873 4879 4880 3 0.66944 2.00832 0.00000

-2.67776 0.00000 0.00000 ; CG2- CB- CG1- HG12
4875 4873 4879 4881 3 0.66944 2.00832 0.00000
-2.67776 0.00000 0.00000 ; CG2- CB- CG1- HG13
4875 4873 4879 4882 3 3.81162 0.49790 -8.14206
-1.79075 7.49773 0.00000 ; CG2- CB- CG1- CD1
4876 4875 4873 4879 3 0.66944 2.00832 0.00000
-2.67776 0.00000 0.00000 ; HG21- CG2- CB- CG1
4877 4875 4873 4879 3 0.66944 2.00832 0.00000
-2.67776 0.00000 0.00000 ; HG22- CG2- CB- CG1
4878 4875 4873 4879 3 0.66944 2.00832 0.00000
-2.67776 0.00000 0.00000 ; HG23- CG2- CB- CG1
4879 4873 4871 4886 3 8.26340 0.74057 -9.99976
-1.89117 3.84928 0.00000 ; CG1- CB- CA- C
4880 4879 4882 4883 3 0.62760 1.88280 0.00000
-2.51040 0.00000 0.00000 ; HG12- CG1- CD1- HD11
4880 4879 4882 4884 3 0.62760 1.88280 0.00000
-2.51040 0.00000 0.00000 ; HG12- CG1- CD1- HD12
4880 4879 4882 4885 3 0.62760 1.88280 0.00000
-2.51040 0.00000 0.00000 ; HG12- CG1- CD1- HD13
4881 4879 4882 4883 3 0.62760 1.88280 0.00000
-2.51040 0.00000 0.00000 ; HG13- CG1- CD1- HD11
4881 4879 4882 4884 3 0.62760 1.88280 0.00000
-2.51040 0.00000 0.00000 ; HG13- CG1- CD1- HD12
4881 4879 4882 4885 3 0.62760 1.88280 0.00000
-2.51040 0.00000 0.00000 ; HG13- CG1- CD1- HD13
4886 4888 4890 4891 3 0.00000 0.00000 0.00000
0.00000 0.00000 0.00000 ; C- N- CA- HA
4886 4888 4890 4892 3 11.71520 1.67360 15.06240 -
13.38880 0.00000 0.00000 ; C- N- CA- CB
4886 4888 4890 4896 3 1.75728 5.27184 2.25936
-7.02912 0.00000 0.00000 ; C- N- CA- C
4887 4886 4888 4889 3 29.28800 -8.36800 -20.92000
0.00000 0.00000 0.00000 ; O- C- N- H
4887 4886 4888 4890 3 20.92000 0.00000 -20.92000
0.00000 0.00000 0.00000 ; O- C- N- CA
4888 4890 4892 4893 3 0.65084 1.95253 0.00000
-2.60338 0.00000 0.00000 ; N- CA- CB- HB1
4888 4890 4892 4894 3 0.65084 1.95253 0.00000
-2.60338 0.00000 0.00000 ; N- CA- CB- HB2
4888 4890 4892 4895 3 0.65084 1.95253 0.00000
-2.60338 0.00000 0.00000 ; N- CA- CB- HB3
4888 4890 4896 4897 3 0.00000 0.00000 0.00000
0.00000 0.00000 0.00000 ; N- CA- C- O
4888 4890 4896 4898 3 17.40544 -5.02080 -13.22144
9.20480 0.00000 0.00000 ; N- CA- C- N
4889 4888 4890 4891 3 0.00000 0.00000 0.00000
0.00000 0.00000 0.00000 ; H- N- CA- HA
4889 4888 4890 4892 3 0.00000 0.00000 0.00000
0.00000 0.00000 0.00000 ; H- N- CA- CB
4889 4888 4890 4896 3 0.00000 0.00000 0.00000
0.00000 0.00000 0.00000 ; H- N- CA- C
4890 4896 4898 4899 3 20.92000 0.00000 -20.92000

```
0.00000    0.00000    0.00000 ;      CA-     C-     N-     H
  4890   4896   4898   4900      3   20.92000    0.00000  -20.92000
0.00000    0.00000    0.00000 ;      CA-     C-     N-    CA
  4891   4890   4892   4893      3    0.65084    1.95253    0.00000
-2.60338    0.00000    0.00000 ;      HA-    CA-    CB-   HB1
  4891   4890   4892   4894      3    0.65084    1.95253    0.00000
-2.60338    0.00000    0.00000 ;      HA-    CA-    CB-   HB2
  4891   4890   4892   4895      3    0.65084    1.95253    0.00000
-2.60338    0.00000    0.00000 ;      HA-    CA-    CB-   HB3
  4891   4890   4896   4897      3    3.68192   -4.35136    0.00000
1.33888    0.00000    0.00000 ;      HA-    CA-     C-     O
  4891   4890   4896   4898      3    0.00000    0.00000    0.00000
0.00000    0.00000    0.00000 ;      HA-    CA-     C-     N
  4892   4890   4896   4897      3    0.00000    0.00000    0.00000
0.00000    0.00000    0.00000 ;      CB-    CA-     C-     O
  4892   4890   4896   4898      3    2.51040    4.18400    1.67360
-6.69440    0.00000    0.00000 ;      CB-    CA-     C-     N
  4893   4892   4890   4896      3    0.65084    1.95253    0.00000
-2.60338    0.00000    0.00000 ;     HB1-    CB-    CA-     C
  4894   4892   4890   4896      3    0.65084    1.95253    0.00000
-2.60338    0.00000    0.00000 ;     HB2-    CB-    CA-     C
  4895   4892   4890   4896      3    0.65084    1.95253    0.00000
-2.60338    0.00000    0.00000 ;     HB3-    CB-    CA-     C
  4896   4898   4900   4901      3    0.00000    0.00000    0.00000
0.00000    0.00000    0.00000 ;       C-     N-    CA-    HA
  4896   4898   4900   4902      3   11.71520    1.67360   15.06240  -
13.38880    0.00000    0.00000 ;       C-     N-    CA-    CB
  4896   4898   4900   4915      3    1.75728    5.27184    2.25936
-7.02912    0.00000    0.00000 ;       C-     N-    CA-     C
  4897   4896   4898   4899      3   29.28800   -8.36800  -20.92000
0.00000    0.00000    0.00000 ;       O-     C-     N-     H
  4897   4896   4898   4900      3   20.92000    0.00000  -20.92000
0.00000    0.00000    0.00000 ;       O-     C-     N-    CA
  4898   4900   4902   4903      3    0.65084    1.95253    0.00000
-2.60338    0.00000    0.00000 ;       N-    CA-    CB-   HB2
  4898   4900   4902   4904      3    0.65084    1.95253    0.00000
-2.60338    0.00000    0.00000 ;       N-    CA-    CB-   HB3
  4898   4900   4902   4905      3    3.79070    1.39746   -4.61077
-2.40998    2.44346    0.00000 ;       N-    CA-    CB-    CG
  4898   4900   4915   4916      3    0.00000    0.00000    0.00000
0.00000    0.00000    0.00000 ;       N-    CA-     C-     O
  4898   4900   4915   4917      3   17.40544   -5.02080  -13.22144
9.20480    0.00000    0.00000 ;       N-    CA-     C-     N
  4899   4898   4900   4901      3    0.00000    0.00000    0.00000
0.00000    0.00000    0.00000 ;       H-     N-    CA-    HA
  4899   4898   4900   4902      3    0.00000    0.00000    0.00000
0.00000    0.00000    0.00000 ;       H-     N-    CA-    CB
  4899   4898   4900   4915      3    0.00000    0.00000    0.00000
0.00000    0.00000    0.00000 ;       H-     N-    CA-     C
  4900   4902   4905   4906      3    0.66944    2.00832    0.00000
-2.67776    0.00000    0.00000 ;      CA-    CB-    CG-    HG
  4900   4902   4905   4907      3    3.90367    0.19665   -6.21742
```

-2.37651 5.99149 0.00000 ; CA- CB- CG- CD1
4900 4902 4905 4911 3 3.90367 0.19665 -6.21742
-2.37651 5.99149 0.00000 ; CA- CB- CG- CD2
4900 4915 4917 4918 3 20.92000 0.00000 -20.92000
0.00000 0.00000 0.00000 ; CA- C- N- H
4900 4915 4917 4919 3 20.92000 0.00000 -20.92000
0.00000 0.00000 0.00000 ; CA- C- N- CA
4901 4900 4902 4903 3 0.65084 1.95253 0.00000
-2.60338 0.00000 0.00000 ; HA- CA- CB- HB2
4901 4900 4902 4904 3 0.65084 1.95253 0.00000
-2.60338 0.00000 0.00000 ; HA- CA- CB- HB3
4901 4900 4902 4905 3 0.65084 1.95253 0.00000
-2.60338 0.00000 0.00000 ; HA- CA- CB- CG
4901 4900 4915 4916 3 3.68192 -4.35136 0.00000
1.33888 0.00000 0.00000 ; HA- CA- C- O
4901 4900 4915 4917 3 0.00000 0.00000 0.00000
0.00000 0.00000 0.00000 ; HA- CA- C- N
4902 4900 4915 4916 3 0.00000 0.00000 0.00000
0.00000 0.00000 0.00000 ; CB- CA- C- O
4902 4900 4915 4917 3 2.51040 4.18400 1.67360
-6.69440 0.00000 0.00000 ; CB- CA- C- N
4902 4905 4907 4908 3 0.66944 2.00832 0.00000
-2.67776 0.00000 0.00000 ; CB- CG- CD1- HD11
4902 4905 4907 4909 3 0.66944 2.00832 0.00000
-2.67776 0.00000 0.00000 ; CB- CG- CD1- HD12
4902 4905 4907 4910 3 0.66944 2.00832 0.00000
-2.67776 0.00000 0.00000 ; CB- CG- CD1- HD13
4902 4905 4911 4912 3 0.66944 2.00832 0.00000
-2.67776 0.00000 0.00000 ; CB- CG- CD2- HD21
4902 4905 4911 4913 3 0.66944 2.00832 0.00000
-2.67776 0.00000 0.00000 ; CB- CG- CD2- HD22
4902 4905 4911 4914 3 0.66944 2.00832 0.00000
-2.67776 0.00000 0.00000 ; CB- CG- CD2- HD23
4903 4902 4900 4915 3 0.65084 1.95253 0.00000
-2.60338 0.00000 0.00000 ; HB2- CB- CA- C
4903 4902 4905 4906 3 0.62760 1.88280 0.00000
-2.51040 0.00000 0.00000 ; HB2- CB- CG- HG
4903 4902 4905 4907 3 0.66944 2.00832 0.00000
-2.67776 0.00000 0.00000 ; HB2- CB- CG- CD1
4903 4902 4905 4911 3 0.66944 2.00832 0.00000
-2.67776 0.00000 0.00000 ; HB2- CB- CG- CD2
4904 4902 4900 4915 3 0.65084 1.95253 0.00000
-2.60338 0.00000 0.00000 ; HB3- CB- CA- C
4904 4902 4905 4906 3 0.62760 1.88280 0.00000
-2.51040 0.00000 0.00000 ; HB3- CB- CG- HG
4904 4902 4905 4907 3 0.66944 2.00832 0.00000
-2.67776 0.00000 0.00000 ; HB3- CB- CG- CD1
4904 4902 4905 4911 3 0.66944 2.00832 0.00000
-2.67776 0.00000 0.00000 ; HB3- CB- CG- CD2
4905 4902 4900 4915 3 10.33448 -1.14642 -11.54784
-2.40998 6.35968 0.00000 ; CG- CB- CA- C
4906 4905 4907 4908 3 0.62760 1.88280 0.00000

```
-2.51040    0.00000    0.00000 ;      HG-    CG-   CD1-  HD11
  4906   4905   4907   4909       3    0.62760    1.88280    0.00000
-2.51040    0.00000    0.00000 ;      HG-    CG-   CD1-  HD12
  4906   4905   4907   4910       3    0.62760    1.88280    0.00000
-2.51040    0.00000    0.00000 ;      HG-    CG-   CD1-  HD13
  4906   4905   4911   4912       3    0.62760    1.88280    0.00000
-2.51040    0.00000    0.00000 ;      HG-    CG-   CD2-  HD21
  4906   4905   4911   4913       3    0.62760    1.88280    0.00000
-2.51040    0.00000    0.00000 ;      HG-    CG-   CD2-  HD22
  4906   4905   4911   4914       3    0.62760    1.88280    0.00000
-2.51040    0.00000    0.00000 ;      HG-    CG-   CD2-  HD23
  4907   4905   4911   4912       3    0.66944    2.00832    0.00000
-2.67776    0.00000    0.00000 ;     CD1-    CG-   CD2-  HD21
  4907   4905   4911   4913       3    0.66944    2.00832    0.00000
-2.67776    0.00000    0.00000 ;     CD1-    CG-   CD2-  HD22
  4907   4905   4911   4914       3    0.66944    2.00832    0.00000
-2.67776    0.00000    0.00000 ;     CD1-    CG-   CD2-  HD23
  4908   4907   4905   4911       3    0.66944    2.00832    0.00000
-2.67776    0.00000    0.00000 ;   HD11-   CD1-    CG-   CD2
  4909   4907   4905   4911       3    0.66944    2.00832    0.00000
-2.67776    0.00000    0.00000 ;   HD12-   CD1-    CG-   CD2
  4910   4907   4905   4911       3    0.66944    2.00832    0.00000
-2.67776    0.00000    0.00000 ;   HD13-   CD1-    CG-   CD2
  4915   4917   4919   4920       3    0.00000    0.00000    0.00000
0.00000    0.00000    0.00000 ;       C-     N-    CA-    HA
  4915   4917   4919   4921       3   11.71520    1.67360   15.06240  -
13.38880    0.00000    0.00000 ;      C-     N-    CA-    CB
  4915   4917   4919   4929       3    1.75728    5.27184    2.25936
-7.02912    0.00000    0.00000 ;      C-     N-    CA-     C
  4916   4915   4917   4918       3   29.28800   -8.36800  -20.92000
0.00000    0.00000    0.00000 ;       O-     C-     N-     H
  4916   4915   4917   4919       3   20.92000    0.00000  -20.92000
0.00000    0.00000    0.00000 ;       O-     C-     N-    CA
  4917   4919   4921   4922       3    0.65084    1.95253    0.00000
-2.60338    0.00000    0.00000 ;      N-    CA-    CB-    HB
  4917   4919   4921   4923       3    3.83673    0.44769   -1.77402
-2.47693   -0.03347    0.00000 ;      N-    CA-    CB-   CG2
  4917   4919   4921   4927       3    4.93294    1.13386   -3.12963
-5.27184    3.17984    0.00000 ;      N-    CA-    CB-   OG1
  4917   4919   4929   4930       3    0.00000    0.00000    0.00000
0.00000    0.00000    0.00000 ;       N-    CA-     C-     O
  4917   4919   4929   4931       3   17.40544   -5.02080  -13.22144
9.20480    0.00000    0.00000 ;       N-    CA-     C-     N
  4918   4917   4919   4920       3    0.00000    0.00000    0.00000
0.00000    0.00000    0.00000 ;       H-     N-    CA-    HA
  4918   4917   4919   4921       3    0.00000    0.00000    0.00000
0.00000    0.00000    0.00000 ;       H-     N-    CA-    CB
  4918   4917   4919   4929       3    0.00000    0.00000    0.00000
0.00000    0.00000    0.00000 ;       H-     N-    CA-     C
  4919   4921   4923   4924       3    0.66944    2.00832    0.00000
-2.67776    0.00000    0.00000 ;      CA-    CB-   CG2-  HG21
  4919   4921   4923   4925       3    0.66944    2.00832    0.00000
```

-2.67776    0.00000    0.00000 ;     CA-    CB-   CG2-  HG22
  4919   4921   4923   4926      3    0.66944    2.00832    0.00000
-2.67776    0.00000    0.00000 ;     CA-    CB-   CG2-  HG23
  4919   4921   4927   4928      3    1.12131    2.98738    1.66523
-3.94970    0.43514    0.00000 ;     CA-    CB-   OG1-   HG1
  4919   4929   4931   4932      3   20.92000    0.00000  -20.92000
0.00000    0.00000    0.00000 ;     CA-     C-     N-     H
  4919   4929   4931   4933      3   20.92000    0.00000  -20.92000
0.00000    0.00000    0.00000 ;     CA-     C-     N-    CA
  4920   4919   4921   4922      3    0.65084    1.95253    0.00000
-2.60338    0.00000    0.00000 ;     HA-    CA-    CB-    HB
  4920   4919   4921   4923      3    0.65084    1.95253    0.00000
-2.60338    0.00000    0.00000 ;     HA-    CA-    CB-   CG2
  4920   4919   4921   4927      3    1.04600   -1.04600    0.00000
0.00000    0.00000    0.00000 ;     HA-    CA-    CB-   OG1
  4920   4919   4929   4930      3    3.68192   -4.35136    0.00000
1.33888    0.00000    0.00000 ;     HA-    CA-     C-     O
  4920   4919   4929   4931      3    0.00000    0.00000    0.00000
0.00000    0.00000    0.00000 ;     HA-    CA-     C-     N
  4921   4919   4929   4930      3    0.00000    0.00000    0.00000
0.00000    0.00000    0.00000 ;     CB-    CA-     C-     O
  4921   4919   4929   4931      3    2.51040    4.18400    1.67360
-6.69440    0.00000    0.00000 ;     CB-    CA-     C-     N
  4922   4921   4919   4929      3    0.65084    1.95253    0.00000
-2.60338    0.00000    0.00000 ;     HB-    CB-    CA-     C
  4922   4921   4923   4924      3    0.65084    1.95253    0.00000
-2.60338    0.00000    0.00000 ;     HB-    CB-   CG2-  HG21
  4922   4921   4923   4925      3    0.65084    1.95253    0.00000
-2.60338    0.00000    0.00000 ;     HB-    CB-   CG2-  HG22
  4922   4921   4923   4926      3    0.65084    1.95253    0.00000
-2.60338    0.00000    0.00000 ;     HB-    CB-   CG2-  HG23
  4922   4921   4927   4928      3    0.69733    2.09200    0.00000
-2.78933    0.00000    0.00000 ;     HB-    CB-   OG1-   HG1
  4923   4921   4919   4929      3    5.67350    3.55640   -6.16722
-2.47693    3.74886    0.00000 ;    CG2-    CB-    CA-     C
  4923   4921   4927   4928      3    4.74047    0.27196   -2.26773
-3.94970    1.60666    0.00000 ;    CG2-    CB-   OG1-   HG1
  4924   4923   4921   4927      3    1.04600   -1.04600    0.00000
0.00000    0.00000    0.00000 ;   HG21-   CG2-    CB-   OG1
  4925   4923   4921   4927      3    1.04600   -1.04600    0.00000
0.00000    0.00000    0.00000 ;   HG22-   CG2-    CB-   OG1
  4926   4923   4921   4927      3    1.04600   -1.04600    0.00000
0.00000    0.00000    0.00000 ;   HG23-   CG2-    CB-   OG1
  4927   4921   4919   4929      3    6.53541    6.87013   -6.21742
-5.27184    5.22163    0.00000 ;    OG1-    CB-    CA-     C
  4929   4931   4933   4934      3    0.00000    0.00000    0.00000
0.00000    0.00000    0.00000 ;      C-     N-    CA-    HA
  4929   4931   4933   4935      3   11.71520    1.67360   15.06240  -
13.38880    0.00000    0.00000 ;      C-     N-    CA-    CB
  4929   4931   4933   4948      3    1.75728    5.27184    2.25936
-7.02912    0.00000    0.00000 ;      C-     N-    CA-     C
  4930   4929   4931   4932      3   29.28800   -8.36800  -20.92000

```
0.00000    0.00000    0.00000 ;       O-     C-     N-     H
  4930   4929   4931   4933       3   20.92000    0.00000  -20.92000
0.00000    0.00000    0.00000 ;       O-     C-     N-    CA
  4931   4933   4935   4936       3    0.65084    1.95253    0.00000
-2.60338    0.00000    0.00000 ;      N-    CA-    CB-    HB
  4931   4933   4935   4937       3    3.83673    0.44769   -1.77402
-2.47693   -0.03347    0.00000 ;      N-    CA-    CB-   CG2
  4931   4933   4935   4941       3    2.97482    0.12134    2.04179
-1.89117   -3.24678    0.00000 ;      N-    CA-    CB-   CG1
  4931   4933   4948   4949       3    0.00000    0.00000    0.00000
0.00000    0.00000    0.00000 ;      N-    CA-     C-     O
  4931   4933   4948   4950       3   17.40544   -5.02080  -13.22144
9.20480    0.00000    0.00000 ;      N-    CA-     C-     N
  4932   4931   4933   4934       3    0.00000    0.00000    0.00000
0.00000    0.00000    0.00000 ;      H-     N-    CA-    HA
  4932   4931   4933   4935       3    0.00000    0.00000    0.00000
0.00000    0.00000    0.00000 ;      H-     N-    CA-    CB
  4932   4931   4933   4948       3    0.00000    0.00000    0.00000
0.00000    0.00000    0.00000 ;      H-     N-    CA-     C
  4933   4935   4937   4938       3    0.66944    2.00832    0.00000
-2.67776    0.00000    0.00000 ;     CA-    CB-   CG2-  HG21
  4933   4935   4937   4939       3    0.66944    2.00832    0.00000
-2.67776    0.00000    0.00000 ;     CA-    CB-   CG2-  HG22
  4933   4935   4937   4940       3    0.66944    2.00832    0.00000
-2.67776    0.00000    0.00000 ;     CA-    CB-   CG2-  HG23
  4933   4935   4941   4942       3    0.66944    2.00832    0.00000
-2.67776    0.00000    0.00000 ;     CA-    CB-   CG1-  HG12
  4933   4935   4941   4943       3    0.66944    2.00832    0.00000
-2.67776    0.00000    0.00000 ;     CA-    CB-   CG1-  HG13
  4933   4935   4941   4944       3    4.24258   -0.52718   -7.25506
-1.79075    7.69856    0.00000 ;     CA-    CB-   CG1-   CD1
  4933   4948   4950   4951       3   20.92000    0.00000  -20.92000
0.00000    0.00000    0.00000 ;     CA-     C-     N-     H
  4933   4948   4950   4952       3   20.92000    0.00000  -20.92000
0.00000    0.00000    0.00000 ;     CA-     C-     N-    CA
  4934   4933   4935   4936       3    0.65084    1.95253    0.00000
-2.60338    0.00000    0.00000 ;     HA-    CA-    CB-    HB
  4934   4933   4935   4937       3    0.65084    1.95253    0.00000
-2.60338    0.00000    0.00000 ;     HA-    CA-    CB-   CG2
  4934   4933   4935   4941       3    0.65084    1.95253    0.00000
-2.60338    0.00000    0.00000 ;     HA-    CA-    CB-   CG1
  4934   4933   4948   4949       3    3.68192   -4.35136    0.00000
1.33888    0.00000    0.00000 ;     HA-    CA-     C-     O
  4934   4933   4948   4950       3    0.00000    0.00000    0.00000
0.00000    0.00000    0.00000 ;     HA-    CA-     C-     N
  4935   4933   4948   4949       3    0.00000    0.00000    0.00000
0.00000    0.00000    0.00000 ;     CB-    CA-     C-     O
  4935   4933   4948   4950       3    2.51040    4.18400    1.67360
-6.69440    0.00000    0.00000 ;     CB-    CA-     C-     N
  4935   4941   4944   4945       3    0.66944    2.00832    0.00000
-2.67776    0.00000    0.00000 ;     CB-   CG1-   CD1-  HD11
  4935   4941   4944   4946       3    0.66944    2.00832    0.00000
```

-2.67776 0.00000 0.00000 ; CB- CG1- CD1- HD12
4935 4941 4944 4947 3 0.66944 2.00832 0.00000
-2.67776 0.00000 0.00000 ; CB- CG1- CD1- HD13
4936 4935 4933 4948 3 0.65084 1.95253 0.00000
-2.60338 0.00000 0.00000 ; HB- CB- CA- C
4936 4935 4937 4938 3 0.62760 1.88280 0.00000
-2.51040 0.00000 0.00000 ; HB- CB- CG2- HG21
4936 4935 4937 4939 3 0.62760 1.88280 0.00000
-2.51040 0.00000 0.00000 ; HB- CB- CG2- HG22
4936 4935 4937 4940 3 0.62760 1.88280 0.00000
-2.51040 0.00000 0.00000 ; HB- CB- CG2- HG23
4936 4935 4941 4942 3 0.62760 1.88280 0.00000
-2.51040 0.00000 0.00000 ; HB- CB- CG1- HG12
4936 4935 4941 4943 3 0.62760 1.88280 0.00000
-2.51040 0.00000 0.00000 ; HB- CB- CG1- HG13
4936 4935 4941 4944 3 0.66944 2.00832 0.00000
-2.67776 0.00000 0.00000 ; HB- CB- CG1- CD1
4937 4935 4933 4948 3 5.67350 3.55640 -6.16722
-2.47693 3.74886 0.00000 ; CG2- CB- CA- C
4937 4935 4941 4942 3 0.66944 2.00832 0.00000
-2.67776 0.00000 0.00000 ; CG2- CB- CG1- HG12
4937 4935 4941 4943 3 0.66944 2.00832 0.00000
-2.67776 0.00000 0.00000 ; CG2- CB- CG1- HG13
4937 4935 4941 4944 3 3.81162 0.49790 -8.14206
-1.79075 7.49773 0.00000 ; CG2- CB- CG1- CD1
4938 4937 4935 4941 3 0.66944 2.00832 0.00000
-2.67776 0.00000 0.00000 ; HG21- CG2- CB- CG1
4939 4937 4935 4941 3 0.66944 2.00832 0.00000
-2.67776 0.00000 0.00000 ; HG22- CG2- CB- CG1
4940 4937 4935 4941 3 0.66944 2.00832 0.00000
-2.67776 0.00000 0.00000 ; HG23- CG2- CB- CG1
4941 4935 4933 4948 3 8.26340 0.74057 -9.99976
-1.89117 3.84928 0.00000 ; CG1- CB- CA- C
4942 4941 4944 4945 3 0.62760 1.88280 0.00000
-2.51040 0.00000 0.00000 ; HG12- CG1- CD1- HD11
4942 4941 4944 4946 3 0.62760 1.88280 0.00000
-2.51040 0.00000 0.00000 ; HG12- CG1- CD1- HD12
4942 4941 4944 4947 3 0.62760 1.88280 0.00000
-2.51040 0.00000 0.00000 ; HG12- CG1- CD1- HD13
4943 4941 4944 4945 3 0.62760 1.88280 0.00000
-2.51040 0.00000 0.00000 ; HG13- CG1- CD1- HD11
4943 4941 4944 4946 3 0.62760 1.88280 0.00000
-2.51040 0.00000 0.00000 ; HG13- CG1- CD1- HD12
4943 4941 4944 4947 3 0.62760 1.88280 0.00000
-2.51040 0.00000 0.00000 ; HG13- CG1- CD1- HD13
4948 4950 4952 4953 3 0.00000 0.00000 0.00000
0.00000 0.00000 0.00000 ; C- N- CA- HA
4948 4950 4952 4954 3 11.71520 1.67360 15.06240 -
13.38880 0.00000 0.00000 ; C- N- CA- CB
4948 4950 4952 4967 3 1.75728 5.27184 2.25936
-7.02912 0.00000 0.00000 ; C- N- CA- C
4949 4948 4950 4951 3 29.28800 -8.36800 -20.92000

```
0.00000    0.00000    0.00000 ;       O-     C-     N-     H
  4949   4948   4950   4952       3   20.92000    0.00000  -20.92000
0.00000    0.00000    0.00000 ;       O-     C-     N-    CA
  4950   4952   4954   4955       3    0.65084    1.95253    0.00000
-2.60338    0.00000    0.00000 ;      N-    CA-    CB-    HB
  4950   4952   4954   4956       3    3.83673    0.44769   -1.77402
-2.47693   -0.03347    0.00000 ;      N-    CA-    CB-   CG2
  4950   4952   4954   4960       3    2.97482    0.12134    2.04179
-1.89117   -3.24678    0.00000 ;      N-    CA-    CB-   CG1
  4950   4952   4967   4968       3    0.00000    0.00000    0.00000
0.00000    0.00000    0.00000 ;      N-    CA-     C-     O
  4950   4952   4967   4969       3   17.40544   -5.02080  -13.22144
9.20480    0.00000    0.00000 ;      N-    CA-     C-     N
  4951   4950   4952   4953       3    0.00000    0.00000    0.00000
0.00000    0.00000    0.00000 ;      H-     N-    CA-    HA
  4951   4950   4952   4954       3    0.00000    0.00000    0.00000
0.00000    0.00000    0.00000 ;      H-     N-    CA-    CB
  4951   4950   4952   4967       3    0.00000    0.00000    0.00000
0.00000    0.00000    0.00000 ;      H-     N-    CA-     C
  4952   4954   4956   4957       3    0.66944    2.00832    0.00000
-2.67776    0.00000    0.00000 ;     CA-    CB-   CG2-  HG21
  4952   4954   4956   4958       3    0.66944    2.00832    0.00000
-2.67776    0.00000    0.00000 ;     CA-    CB-   CG2-  HG22
  4952   4954   4956   4959       3    0.66944    2.00832    0.00000
-2.67776    0.00000    0.00000 ;     CA-    CB-   CG2-  HG23
  4952   4954   4960   4961       3    0.66944    2.00832    0.00000
-2.67776    0.00000    0.00000 ;     CA-    CB-   CG1-  HG12
  4952   4954   4960   4962       3    0.66944    2.00832    0.00000
-2.67776    0.00000    0.00000 ;     CA-    CB-   CG1-  HG13
  4952   4954   4960   4963       3    4.24258   -0.52718   -7.25506
-1.79075    7.69856    0.00000 ;     CA-    CB-   CG1-   CD1
  4952   4967   4969   4970       3   20.92000    0.00000  -20.92000
0.00000    0.00000    0.00000 ;     CA-     C-     N-     H
  4952   4967   4969   4971       3   20.92000    0.00000  -20.92000
0.00000    0.00000    0.00000 ;     CA-     C-     N-    CA
  4953   4952   4954   4955       3    0.65084    1.95253    0.00000
-2.60338    0.00000    0.00000 ;     HA-    CA-    CB-    HB
  4953   4952   4954   4956       3    0.65084    1.95253    0.00000
-2.60338    0.00000    0.00000 ;     HA-    CA-    CB-   CG2
  4953   4952   4954   4960       3    0.65084    1.95253    0.00000
-2.60338    0.00000    0.00000 ;     HA-    CA-    CB-   CG1
  4953   4952   4967   4968       3    3.68192   -4.35136    0.00000
1.33888    0.00000    0.00000 ;     HA-    CA-     C-     O
  4953   4952   4967   4969       3    0.00000    0.00000    0.00000
0.00000    0.00000    0.00000 ;     HA-    CA-     C-     N
  4954   4952   4967   4968       3    0.00000    0.00000    0.00000
0.00000    0.00000    0.00000 ;     CB-    CA-     C-     O
  4954   4952   4967   4969       3    2.51040    4.18400    1.67360
-6.69440    0.00000    0.00000 ;     CB-    CA-     C-     N
  4954   4960   4963   4964       3    0.66944    2.00832    0.00000
-2.67776    0.00000    0.00000 ;     CB-   CG1-   CD1-  HD11
  4954   4960   4963   4965       3    0.66944    2.00832    0.00000
```

-2.67776 0.00000 0.00000 ; CB- CG1- CD1- HD12
4954 4960 4963 4966 3 0.66944 2.00832 0.00000
-2.67776 0.00000 0.00000 ; CB- CG1- CD1- HD13
4955 4954 4952 4967 3 0.65084 1.95253 0.00000
-2.60338 0.00000 0.00000 ; HB- CB- CA- C
4955 4954 4956 4957 3 0.62760 1.88280 0.00000
-2.51040 0.00000 0.00000 ; HB- CB- CG2- HG21
4955 4954 4956 4958 3 0.62760 1.88280 0.00000
-2.51040 0.00000 0.00000 ; HB- CB- CG2- HG22
4955 4954 4956 4959 3 0.62760 1.88280 0.00000
-2.51040 0.00000 0.00000 ; HB- CB- CG2- HG23
4955 4954 4960 4961 3 0.62760 1.88280 0.00000
-2.51040 0.00000 0.00000 ; HB- CB- CG1- HG12
4955 4954 4960 4962 3 0.62760 1.88280 0.00000
-2.51040 0.00000 0.00000 ; HB- CB- CG1- HG13
4955 4954 4960 4963 3 0.66944 2.00832 0.00000
-2.67776 0.00000 0.00000 ; HB- CB- CG1- CD1
4956 4954 4952 4967 3 5.67350 3.55640 -6.16722
-2.47693 3.74886 0.00000 ; CG2- CB- CA- C
4956 4954 4960 4961 3 0.66944 2.00832 0.00000
-2.67776 0.00000 0.00000 ; CG2- CB- CG1- HG12
4956 4954 4960 4962 3 0.66944 2.00832 0.00000
-2.67776 0.00000 0.00000 ; CG2- CB- CG1- HG13
4956 4954 4960 4963 3 3.81162 0.49790 -8.14206
-1.79075 7.49773 0.00000 ; CG2- CB- CG1- CD1
4957 4956 4954 4960 3 0.66944 2.00832 0.00000
-2.67776 0.00000 0.00000 ; HG21- CG2- CB- CG1
4958 4956 4954 4960 3 0.66944 2.00832 0.00000
-2.67776 0.00000 0.00000 ; HG22- CG2- CB- CG1
4959 4956 4954 4960 3 0.66944 2.00832 0.00000
-2.67776 0.00000 0.00000 ; HG23- CG2- CB- CG1
4960 4954 4952 4967 3 8.26340 0.74057 -9.99976
-1.89117 3.84928 0.00000 ; CG1- CB- CA- C
4961 4960 4963 4964 3 0.62760 1.88280 0.00000
-2.51040 0.00000 0.00000 ; HG12- CG1- CD1- HD11
4961 4960 4963 4965 3 0.62760 1.88280 0.00000
-2.51040 0.00000 0.00000 ; HG12- CG1- CD1- HD12
4961 4960 4963 4966 3 0.62760 1.88280 0.00000
-2.51040 0.00000 0.00000 ; HG12- CG1- CD1- HD13
4962 4960 4963 4964 3 0.62760 1.88280 0.00000
-2.51040 0.00000 0.00000 ; HG13- CG1- CD1- HD11
4962 4960 4963 4965 3 0.62760 1.88280 0.00000
-2.51040 0.00000 0.00000 ; HG13- CG1- CD1- HD12
4962 4960 4963 4966 3 0.62760 1.88280 0.00000
-2.51040 0.00000 0.00000 ; HG13- CG1- CD1- HD13
4967 4969 4971 4972 3 0.00000 0.00000 0.00000
0.00000 0.00000 0.00000 ; C- N- CA- HA
4967 4969 4971 4973 3 11.71520 1.67360 15.06240 -
13.38880 0.00000 0.00000 ; C- N- CA- CB
4967 4969 4971 4977 3 1.75728 5.27184 2.25936
-7.02912 0.00000 0.00000 ; C- N- CA- C
4968 4967 4969 4970 3 29.28800 -8.36800 -20.92000

0.00000 0.00000 0.00000 ; O- C- N- H
4968 4967 4969 4971 3 20.92000 0.00000 -20.92000
0.00000 0.00000 0.00000 ; O- C- N- CA
4969 4971 4973 4974 3 0.65084 1.95253 0.00000
-2.60338 0.00000 0.00000 ; N- CA- CB- HB1
4969 4971 4973 4975 3 0.65084 1.95253 0.00000
-2.60338 0.00000 0.00000 ; N- CA- CB- HB2
4969 4971 4973 4976 3 0.65084 1.95253 0.00000
-2.60338 0.00000 0.00000 ; N- CA- CB- HB3
4969 4971 4977 4978 3 0.00000 0.00000 0.00000
0.00000 0.00000 0.00000 ; N- CA- C- O
4969 4971 4977 4979 3 17.40544 -5.02080 -13.22144
9.20480 0.00000 0.00000 ; N- CA- C- N
4970 4969 4971 4972 3 0.00000 0.00000 0.00000
0.00000 0.00000 0.00000 ; H- N- CA- HA
4970 4969 4971 4973 3 0.00000 0.00000 0.00000
0.00000 0.00000 0.00000 ; H- N- CA- CB
4970 4969 4971 4977 3 0.00000 0.00000 0.00000
0.00000 0.00000 0.00000 ; H- N- CA- C
4971 4977 4979 4980 3 20.92000 0.00000 -20.92000
0.00000 0.00000 0.00000 ; CA- C- N- H
4971 4977 4979 4981 3 20.92000 0.00000 -20.92000
0.00000 0.00000 0.00000 ; CA- C- N- CA
4972 4971 4973 4974 3 0.65084 1.95253 0.00000
-2.60338 0.00000 0.00000 ; HA- CA- CB- HB1
4972 4971 4973 4975 3 0.65084 1.95253 0.00000
-2.60338 0.00000 0.00000 ; HA- CA- CB- HB2
4972 4971 4973 4976 3 0.65084 1.95253 0.00000
-2.60338 0.00000 0.00000 ; HA- CA- CB- HB3
4972 4971 4977 4978 3 3.68192 -4.35136 0.00000
1.33888 0.00000 0.00000 ; HA- CA- C- O
4972 4971 4977 4979 3 0.00000 0.00000 0.00000
0.00000 0.00000 0.00000 ; HA- CA- C- N
4973 4971 4977 4978 3 0.00000 0.00000 0.00000
0.00000 0.00000 0.00000 ; CB- CA- C- O
4973 4971 4977 4979 3 2.51040 4.18400 1.67360
-6.69440 0.00000 0.00000 ; CB- CA- C- N
4974 4973 4971 4977 3 0.65084 1.95253 0.00000
-2.60338 0.00000 0.00000 ; HB1- CB- CA- C
4975 4973 4971 4977 3 0.65084 1.95253 0.00000
-2.60338 0.00000 0.00000 ; HB2- CB- CA- C
4976 4973 4971 4977 3 0.65084 1.95253 0.00000
-2.60338 0.00000 0.00000 ; HB3- CB- CA- C
4977 4979 4981 4982 3 0.00000 0.00000 0.00000
0.00000 0.00000 0.00000 ; C- N- CA- HA
4977 4979 4981 4983 3 11.71520 1.67360 15.06240 -
13.38880 0.00000 0.00000 ; C- N- CA- CB
4977 4979 4981 4999 3 1.75728 5.27184 2.25936
-7.02912 0.00000 0.00000 ; C- N- CA- C
4978 4977 4979 4980 3 29.28800 -8.36800 -20.92000
0.00000 0.00000 0.00000 ; O- C- N- H
4978 4977 4979 4981 3 20.92000 0.00000 -20.92000

```
0.00000    0.00000    0.00000 ;      O-     C-     N-    CA
  4979   4981   4983   4984      3    0.65084    1.95253    0.00000
-2.60338    0.00000    0.00000 ;      N-    CA-    CB-   HB2
  4979   4981   4983   4985      3    0.65084    1.95253    0.00000
-2.60338    0.00000    0.00000 ;      N-    CA-    CB-   HB3
  4979   4981   4983   4986      3    0.65084    1.95253    0.00000
-2.60338    0.00000    0.00000 ;      N-    CA-    CB-    CG
  4979   4981   4999   5000      3    0.00000    0.00000    0.00000
0.00000    0.00000    0.00000 ;      N-    CA-     C-     O
  4979   4981   4999   5001      3   17.40544   -5.02080  -13.22144
9.20480    0.00000    0.00000 ;      N-    CA-     C-     N
  4980   4979   4981   4982      3    0.00000    0.00000    0.00000
0.00000    0.00000    0.00000 ;      H-     N-    CA-    HA
  4980   4979   4981   4983      3    0.00000    0.00000    0.00000
0.00000    0.00000    0.00000 ;      H-     N-    CA-    CB
  4980   4979   4981   4999      3    0.00000    0.00000    0.00000
0.00000    0.00000    0.00000 ;      H-     N-    CA-     C
  4981   4983   4986   4987      3    0.66944    2.00832    0.00000
-2.67776    0.00000    0.00000 ;     CA-    CB-    CG-   HG2
  4981   4983   4986   4988      3    0.66944    2.00832    0.00000
-2.67776    0.00000    0.00000 ;     CA-    CB-    CG-   HG3
  4981   4983   4986   4989      3    3.68192    3.09616   -2.09200
-3.01248    0.00000    0.00000 ;     CA-    CB-    CG-    CD
  4981   4999   5001   5002      3   20.92000    0.00000  -20.92000
0.00000    0.00000    0.00000 ;     CA-     C-     N-     H
  4981   4999   5001   5003      3   20.92000    0.00000  -20.92000
0.00000    0.00000    0.00000 ;     CA-     C-     N-    CA
  4982   4981   4983   4984      3    0.65084    1.95253    0.00000
-2.60338    0.00000    0.00000 ;     HA-    CA-    CB-   HB2
  4982   4981   4983   4985      3    0.65084    1.95253    0.00000
-2.60338    0.00000    0.00000 ;     HA-    CA-    CB-   HB3
  4982   4981   4983   4986      3    0.65084    1.95253    0.00000
-2.60338    0.00000    0.00000 ;     HA-    CA-    CB-    CG
  4982   4981   4999   5000      3    3.68192   -4.35136    0.00000
1.33888    0.00000    0.00000 ;     HA-    CA-     C-     O
  4982   4981   4999   5001      3    0.00000    0.00000    0.00000
0.00000    0.00000    0.00000 ;     HA-    CA-     C-     N
  4983   4981   4999   5000      3    0.00000    0.00000    0.00000
0.00000    0.00000    0.00000 ;     CB-    CA-     C-     O
  4983   4981   4999   5001      3    2.51040    4.18400    1.67360
-6.69440    0.00000    0.00000 ;     CB-    CA-     C-     N
  4983   4986   4989   4990      3    0.66944    2.00832    0.00000
-2.67776    0.00000    0.00000 ;     CB-    CG-    CD-   HD2
  4983   4986   4989   4991      3    0.66944    2.00832    0.00000
-2.67776    0.00000    0.00000 ;     CB-    CG-    CD-   HD3
  4983   4986   4989   4992      3    3.68192    3.09616   -2.09200
-3.01248    0.00000    0.00000 ;     CB-    CG-    CD-    CE
  4984   4983   4981   4999      3    0.65084    1.95253    0.00000
-2.60338    0.00000    0.00000 ;    HB2-    CB-    CA-     C
  4984   4983   4986   4987      3    0.62760    1.88280    0.00000
-2.51040    0.00000    0.00000 ;    HB2-    CB-    CG-   HG2
  4984   4983   4986   4988      3    0.62760    1.88280    0.00000
```

-2.51040 0.00000 0.00000 ; HB2- CB- CG- HG3
4984 4983 4986 4989 3 0.66944 2.00832 0.00000
-2.67776 0.00000 0.00000 ; HB2- CB- CG- CD
4985 4983 4981 4999 3 0.65084 1.95253 0.00000
-2.60338 0.00000 0.00000 ; HB3- CB- CA- C
4985 4983 4986 4987 3 0.62760 1.88280 0.00000
-2.51040 0.00000 0.00000 ; HB3- CB- CG- HG2
4985 4983 4986 4988 3 0.62760 1.88280 0.00000
-2.51040 0.00000 0.00000 ; HB3- CB- CG- HG3
4985 4983 4986 4989 3 0.66944 2.00832 0.00000
-2.67776 0.00000 0.00000 ; HB3- CB- CG- CD
4986 4983 4981 4999 3 0.65084 1.95253 0.00000
-2.60338 0.00000 0.00000 ; CG- CB- CA- C
4986 4989 4992 4993 3 0.65084 1.95253 0.00000
-2.60338 0.00000 0.00000 ; CG- CD- CE- HE2
4986 4989 4992 4994 3 0.65084 1.95253 0.00000
-2.60338 0.00000 0.00000 ; CG- CD- CE- HE3
4986 4989 4992 4995 3 0.65084 1.95253 0.00000
-2.60338 0.00000 0.00000 ; CG- CD- CE- NZ
4987 4986 4989 4990 3 0.62760 1.88280 0.00000
-2.51040 0.00000 0.00000 ; HG2- CG- CD- HD2
4987 4986 4989 4991 3 0.62760 1.88280 0.00000
-2.51040 0.00000 0.00000 ; HG2- CG- CD- HD3
4987 4986 4989 4992 3 0.66944 2.00832 0.00000
-2.67776 0.00000 0.00000 ; HG2- CG- CD- CE
4988 4986 4989 4990 3 0.62760 1.88280 0.00000
-2.51040 0.00000 0.00000 ; HG3- CG- CD- HD2
4988 4986 4989 4991 3 0.62760 1.88280 0.00000
-2.51040 0.00000 0.00000 ; HG3- CG- CD- HD3
4988 4986 4989 4992 3 0.66944 2.00832 0.00000
-2.67776 0.00000 0.00000 ; HG3- CG- CD- CE
4989 4992 4995 4996 3 0.65084 1.95253 0.00000
-2.60338 0.00000 0.00000 ; CD- CE- NZ- HZ1
4989 4992 4995 4997 3 0.65084 1.95253 0.00000
-2.60338 0.00000 0.00000 ; CD- CE- NZ- HZ2
4989 4992 4995 4998 3 0.65084 1.95253 0.00000
-2.60338 0.00000 0.00000 ; CD- CE- NZ- HZ3
4990 4989 4992 4993 3 0.65084 1.95253 0.00000
-2.60338 0.00000 0.00000 ; HD2- CD- CE- HE2
4990 4989 4992 4994 3 0.65084 1.95253 0.00000
-2.60338 0.00000 0.00000 ; HD2- CD- CE- HE3
4990 4989 4992 4995 3 0.65084 1.95253 0.00000
-2.60338 0.00000 0.00000 ; HD2- CD- CE- NZ
4991 4989 4992 4993 3 0.65084 1.95253 0.00000
-2.60338 0.00000 0.00000 ; HD3- CD- CE- HE2
4991 4989 4992 4994 3 0.65084 1.95253 0.00000
-2.60338 0.00000 0.00000 ; HD3- CD- CE- HE3
4991 4989 4992 4995 3 0.65084 1.95253 0.00000
-2.60338 0.00000 0.00000 ; HD3- CD- CE- NZ
4993 4992 4995 4996 3 0.65084 1.95253 0.00000
-2.60338 0.00000 0.00000 ; HE2- CE- NZ- HZ1
4993 4992 4995 4997 3 0.65084 1.95253 0.00000

-2.60338 0.00000 0.00000 ; HE2- CE- NZ- HZ2
4993 4992 4995 4998 3 0.65084 1.95253 0.00000
-2.60338 0.00000 0.00000 ; HE2- CE- NZ- HZ3
4994 4992 4995 4996 3 0.65084 1.95253 0.00000
-2.60338 0.00000 0.00000 ; HE3- CE- NZ- HZ1
4994 4992 4995 4997 3 0.65084 1.95253 0.00000
-2.60338 0.00000 0.00000 ; HE3- CE- NZ- HZ2
4994 4992 4995 4998 3 0.65084 1.95253 0.00000
-2.60338 0.00000 0.00000 ; HE3- CE- NZ- HZ3
4999 5001 5003 5004 3 0.00000 0.00000 0.00000
0.00000 0.00000 0.00000 ; C- N- CA- HA
4999 5001 5003 5005 3 11.71520 1.67360 15.06240 -
13.38880 0.00000 0.00000 ; C- N- CA- CB
4999 5001 5003 5016 3 1.75728 5.27184 2.25936
-7.02912 0.00000 0.00000 ; C- N- CA- C
5000 4999 5001 5002 3 29.28800 -8.36800 -20.92000
0.00000 0.00000 0.00000 ; O- C- N- H
5000 4999 5001 5003 3 20.92000 0.00000 -20.92000
0.00000 0.00000 0.00000 ; O- C- N- CA
5001 5003 5005 5006 3 0.65084 1.95253 0.00000
-2.60338 0.00000 0.00000 ; N- CA- CB- HB2
5001 5003 5005 5007 3 0.65084 1.95253 0.00000
-2.60338 0.00000 0.00000 ; N- CA- CB- HB3
5001 5003 5005 5008 3 4.79068 4.02919 -4.82834
-3.66518 2.97901 0.00000 ; N- CA- CB- CG
5001 5003 5016 5017 3 0.00000 0.00000 0.00000
0.00000 0.00000 0.00000 ; N- CA- C- O
5001 5003 5016 5018 3 17.40544 -5.02080 -13.22144
9.20480 0.00000 0.00000 ; N- CA- C- N
5002 5001 5003 5004 3 0.00000 0.00000 0.00000
0.00000 0.00000 0.00000 ; H- N- CA- HA
5002 5001 5003 5005 3 0.00000 0.00000 0.00000
0.00000 0.00000 0.00000 ; H- N- CA- CB
5002 5001 5003 5016 3 0.00000 0.00000 0.00000
0.00000 0.00000 0.00000 ; H- N- CA- C
5003 5005 5008 5009 3 5.98312 6.40152 3.28026 -
12.38464 -1.57318 0.00000 ; CA- CB- CG- ND1
5003 5005 5008 5014 3 0.00000 0.00000 0.00000
0.00000 0.00000 0.00000 ; CA- CB- CG- CD2
5003 5016 5018 5019 3 20.92000 0.00000 -20.92000
0.00000 0.00000 0.00000 ; CA- C- N- H
5003 5016 5018 5020 3 20.92000 0.00000 -20.92000
0.00000 0.00000 0.00000 ; CA- C- N- CA
5004 5003 5005 5006 3 0.65084 1.95253 0.00000
-2.60338 0.00000 0.00000 ; HA- CA- CB- HB2
5004 5003 5005 5007 3 0.65084 1.95253 0.00000
-2.60338 0.00000 0.00000 ; HA- CA- CB- HB3
5004 5003 5005 5008 3 0.65084 1.95253 0.00000
-2.60338 0.00000 0.00000 ; HA- CA- CB- CG
5004 5003 5016 5017 3 3.68192 -4.35136 0.00000
1.33888 0.00000 0.00000 ; HA- CA- C- O
5004 5003 5016 5018 3 0.00000 0.00000 0.00000

```
0.00000    0.00000    0.00000 ;     HA-    CA-     C-     N
  5005   5003   5016   5017      3    0.00000    0.00000    0.00000
0.00000    0.00000    0.00000 ;     CB-    CA-     C-     O
  5005   5003   5016   5018      3    2.51040    4.18400    1.67360
-6.69440    0.00000    0.00000 ;     CB-    CA-     C-     N
  5005   5008   5009   5010      3   20.08320    0.00000  -20.08320
0.00000    0.00000    0.00000 ;     CB-    CG-   ND1-   CE1
  5005   5008   5014   5012      3   44.97800    0.00000  -44.97800
0.00000    0.00000    0.00000 ;     CB-    CG-   CD2-   NE2
  5005   5008   5014   5015      3   44.97800    0.00000  -44.97800
0.00000    0.00000    0.00000 ;     CB-    CG-   CD2-   HD2
  5006   5005   5003   5016      3    0.65084    1.95253    0.00000
-2.60338    0.00000    0.00000 ;    HB2-    CB-    CA-     C
  5006   5005   5008   5009      3    0.00000    0.00000    0.00000
0.00000    0.00000    0.00000 ;    HB2-    CB-    CG-   ND1
  5006   5005   5008   5014      3    0.00000    0.00000    0.00000
0.00000    0.00000    0.00000 ;    HB2-    CB-    CG-   CD2
  5007   5005   5003   5016      3    0.65084    1.95253    0.00000
-2.60338    0.00000    0.00000 ;    HB3-    CB-    CA-     C
  5007   5005   5008   5009      3    0.00000    0.00000    0.00000
0.00000    0.00000    0.00000 ;    HB3-    CB-    CG-   ND1
  5007   5005   5008   5014      3    0.00000    0.00000    0.00000
0.00000    0.00000    0.00000 ;    HB3-    CB-    CG-   CD2
  5008   5005   5003   5016      3    3.76560    3.34720   -2.87859
-3.66518    0.83680    0.00000 ;     CG-    CB-    CA-     C
  5008   5009   5010   5011      3   41.84000    0.00000  -41.84000
0.00000    0.00000    0.00000 ;     CG-   ND1-   CE1-   HE1
  5008   5009   5010   5012      3   41.84000    0.00000  -41.84000
0.00000    0.00000    0.00000 ;     CG-   ND1-   CE1-   NE2
  5008   5014   5012   5010      3   12.55200    0.00000  -12.55200
0.00000    0.00000    0.00000 ;     CG-   CD2-   NE2-   CE1
  5008   5014   5012   5013      3   12.55200    0.00000  -12.55200
0.00000    0.00000    0.00000 ;     CG-   CD2-   NE2-   HE2
  5009   5008   5014   5012      3   44.97800    0.00000  -44.97800
0.00000    0.00000    0.00000 ;    ND1-    CG-   CD2-   NE2
  5009   5008   5014   5015      3   44.97800    0.00000  -44.97800
0.00000    0.00000    0.00000 ;    ND1-    CG-   CD2-   HD2
  5009   5010   5012   5013      3   19.45560    0.00000  -19.45560
0.00000    0.00000    0.00000 ;    ND1-   CE1-   NE2-   HE2
  5009   5010   5012   5014      3   19.45560    0.00000  -19.45560
0.00000    0.00000    0.00000 ;    ND1-   CE1-   NE2-   CD2
  5010   5009   5008   5014      3   20.08320    0.00000  -20.08320
0.00000    0.00000    0.00000 ;    CE1-   ND1-    CG-   CD2
  5010   5012   5014   5015      3   12.55200    0.00000  -12.55200
0.00000    0.00000    0.00000 ;    CE1-   NE2-   CD2-   HD2
  5011   5010   5012   5013      3   19.45560    0.00000  -19.45560
0.00000    0.00000    0.00000 ;    HE1-   CE1-   NE2-   HE2
  5011   5010   5012   5014      3   19.45560    0.00000  -19.45560
0.00000    0.00000    0.00000 ;    HE1-   CE1-   NE2-   CD2
  5013   5012   5014   5015      3   12.55200    0.00000  -12.55200
0.00000    0.00000    0.00000 ;    HE2-   NE2-   CD2-   HD2
  5016   5018   5020   5021      3    0.00000    0.00000    0.00000
```

```
0.00000    0.00000    0.00000 ;      C-     N-    CA-    HA
  5016   5018   5020   5022      3   11.71520    1.67360   15.06240  -
13.38880    0.00000    0.00000 ;      C-     N-    CA-    CB
  5016   5018   5020   5035      3    1.75728    5.27184    2.25936
-7.02912    0.00000    0.00000 ;      C-     N-    CA-     C
  5017   5016   5018   5019      3   29.28800   -8.36800  -20.92000
0.00000    0.00000    0.00000 ;      O-     C-     N-     H
  5017   5016   5018   5020      3   20.92000    0.00000  -20.92000
0.00000    0.00000    0.00000 ;      O-     C-     N-    CA
  5018   5020   5022   5023      3    0.65084    1.95253    0.00000
-2.60338    0.00000    0.00000 ;      N-    CA-    CB-   HB2
  5018   5020   5022   5024      3    0.65084    1.95253    0.00000
-2.60338    0.00000    0.00000 ;      N-    CA-    CB-   HB3
  5018   5020   5022   5025      3    3.79070    1.39746   -4.61077
-2.40998    2.44346    0.00000 ;      N-    CA-    CB-    CG
  5018   5020   5035   5036      3    0.00000    0.00000    0.00000
0.00000    0.00000    0.00000 ;      N-    CA-     C-     O
  5018   5020   5035   5037      3   17.40544   -5.02080  -13.22144
9.20480    0.00000    0.00000 ;      N-    CA-     C-     N
  5019   5018   5020   5021      3    0.00000    0.00000    0.00000
0.00000    0.00000    0.00000 ;      H-     N-    CA-    HA
  5019   5018   5020   5022      3    0.00000    0.00000    0.00000
0.00000    0.00000    0.00000 ;      H-     N-    CA-    CB
  5019   5018   5020   5035      3    0.00000    0.00000    0.00000
0.00000    0.00000    0.00000 ;      H-     N-    CA-     C
  5020   5022   5025   5026      3    0.66944    2.00832    0.00000
-2.67776    0.00000    0.00000 ;     CA-    CB-    CG-    HG
  5020   5022   5025   5027      3    3.90367    0.19665   -6.21742
-2.37651    5.99149    0.00000 ;     CA-    CB-    CG-   CD1
  5020   5022   5025   5031      3    3.90367    0.19665   -6.21742
-2.37651    5.99149    0.00000 ;     CA-    CB-    CG-   CD2
  5020   5035   5037   5038      3   20.92000    0.00000  -20.92000
0.00000    0.00000    0.00000 ;     CA-     C-     N-     H
  5020   5035   5037   5039      3   20.92000    0.00000  -20.92000
0.00000    0.00000    0.00000 ;     CA-     C-     N-    CA
  5021   5020   5022   5023      3    0.65084    1.95253    0.00000
-2.60338    0.00000    0.00000 ;     HA-    CA-    CB-   HB2
  5021   5020   5022   5024      3    0.65084    1.95253    0.00000
-2.60338    0.00000    0.00000 ;     HA-    CA-    CB-   HB3
  5021   5020   5022   5025      3    0.65084    1.95253    0.00000
-2.60338    0.00000    0.00000 ;     HA-    CA-    CB-    CG
  5021   5020   5035   5036      3    3.68192   -4.35136    0.00000
1.33888    0.00000    0.00000 ;     HA-    CA-     C-     O
  5021   5020   5035   5037      3    0.00000    0.00000    0.00000
0.00000    0.00000    0.00000 ;     HA-    CA-     C-     N
  5022   5020   5035   5036      3    0.00000    0.00000    0.00000
0.00000    0.00000    0.00000 ;     CB-    CA-     C-     O
  5022   5020   5035   5037      3    2.51040    4.18400    1.67360
-6.69440    0.00000    0.00000 ;     CB-    CA-     C-     N
  5022   5025   5027   5028      3    0.66944    2.00832    0.00000
-2.67776    0.00000    0.00000 ;     CB-    CG-   CD1-  HD11
  5022   5025   5027   5029      3    0.66944    2.00832    0.00000
```

```
-2.67776    0.00000    0.00000 ;     CB-    CG-   CD1-  HD12
  5022   5025   5027   5030      3    0.66944    2.00832    0.00000
-2.67776    0.00000    0.00000 ;     CB-    CG-   CD1-  HD13
  5022   5025   5031   5032      3    0.66944    2.00832    0.00000
-2.67776    0.00000    0.00000 ;     CB-    CG-   CD2-  HD21
  5022   5025   5031   5033      3    0.66944    2.00832    0.00000
-2.67776    0.00000    0.00000 ;     CB-    CG-   CD2-  HD22
  5022   5025   5031   5034      3    0.66944    2.00832    0.00000
-2.67776    0.00000    0.00000 ;     CB-    CG-   CD2-  HD23
  5023   5022   5020   5035      3    0.65084    1.95253    0.00000
-2.60338    0.00000    0.00000 ;    HB2-    CB-    CA-     C
  5023   5022   5025   5026      3    0.62760    1.88280    0.00000
-2.51040    0.00000    0.00000 ;    HB2-    CB-    CG-    HG
  5023   5022   5025   5027      3    0.66944    2.00832    0.00000
-2.67776    0.00000    0.00000 ;    HB2-    CB-    CG-   CD1
  5023   5022   5025   5031      3    0.66944    2.00832    0.00000
-2.67776    0.00000    0.00000 ;    HB2-    CB-    CG-   CD2
  5024   5022   5020   5035      3    0.65084    1.95253    0.00000
-2.60338    0.00000    0.00000 ;    HB3-    CB-    CA-     C
  5024   5022   5025   5026      3    0.62760    1.88280    0.00000
-2.51040    0.00000    0.00000 ;    HB3-    CB-    CG-    HG
  5024   5022   5025   5027      3    0.66944    2.00832    0.00000
-2.67776    0.00000    0.00000 ;    HB3-    CB-    CG-   CD1
  5024   5022   5025   5031      3    0.66944    2.00832    0.00000
-2.67776    0.00000    0.00000 ;    HB3-    CB-    CG-   CD2
  5025   5022   5020   5035      3   10.33448   -1.14642  -11.54784
-2.40998    6.35968    0.00000 ;     CG-    CB-    CA-     C
  5026   5025   5027   5028      3    0.62760    1.88280    0.00000
-2.51040    0.00000    0.00000 ;     HG-    CG-   CD1-  HD11
  5026   5025   5027   5029      3    0.62760    1.88280    0.00000
-2.51040    0.00000    0.00000 ;     HG-    CG-   CD1-  HD12
  5026   5025   5027   5030      3    0.62760    1.88280    0.00000
-2.51040    0.00000    0.00000 ;     HG-    CG-   CD1-  HD13
  5026   5025   5031   5032      3    0.62760    1.88280    0.00000
-2.51040    0.00000    0.00000 ;     HG-    CG-   CD2-  HD21
  5026   5025   5031   5033      3    0.62760    1.88280    0.00000
-2.51040    0.00000    0.00000 ;     HG-    CG-   CD2-  HD22
  5026   5025   5031   5034      3    0.62760    1.88280    0.00000
-2.51040    0.00000    0.00000 ;     HG-    CG-   CD2-  HD23
  5027   5025   5031   5032      3    0.66944    2.00832    0.00000
-2.67776    0.00000    0.00000 ;    CD1-    CG-   CD2-  HD21
  5027   5025   5031   5033      3    0.66944    2.00832    0.00000
-2.67776    0.00000    0.00000 ;    CD1-    CG-   CD2-  HD22
  5027   5025   5031   5034      3    0.66944    2.00832    0.00000
-2.67776    0.00000    0.00000 ;    CD1-    CG-   CD2-  HD23
  5028   5027   5025   5031      3    0.66944    2.00832    0.00000
-2.67776    0.00000    0.00000 ;   HD11-   CD1-    CG-   CD2
  5029   5027   5025   5031      3    0.66944    2.00832    0.00000
-2.67776    0.00000    0.00000 ;   HD12-   CD1-    CG-   CD2
  5030   5027   5025   5031      3    0.66944    2.00832    0.00000
-2.67776    0.00000    0.00000 ;   HD13-   CD1-    CG-   CD2
  5035   5037   5039   5040      3    0.00000    0.00000    0.00000
```

0.00000 0.00000 0.00000 ; C- N- CA- HA
5035 5037 5039 5041 3 11.71520 1.67360 15.06240 -
13.38880 0.00000 0.00000 ; C- N- CA- CB
5035 5037 5039 5046 3 1.75728 5.27184 2.25936
-7.02912 0.00000 0.00000 ; C- N- CA- C
5036 5035 5037 5038 3 29.28800 -8.36800 -20.92000
0.00000 0.00000 0.00000 ; O- C- N- H
5036 5035 5037 5039 3 20.92000 0.00000 -20.92000
0.00000 0.00000 0.00000 ; O- C- N- CA
5037 5039 5041 5042 3 0.65084 1.95253 0.00000
-2.60338 0.00000 0.00000 ; N- CA- CB- HB2
5037 5039 5041 5043 3 0.65084 1.95253 0.00000
-2.60338 0.00000 0.00000 ; N- CA- CB- HB3
5037 5039 5041 5044 3 6.03751 2.50622 -5.17142
-4.20074 1.10458 0.00000 ; N- CA- CB- SG
5037 5039 5046 5047 3 0.00000 0.00000 0.00000
0.00000 0.00000 0.00000 ; N- CA- C- O
5037 5039 5046 5048 3 17.40544 -5.02080 -13.22144
9.20480 0.00000 0.00000 ; N- CA- C- N
5038 5037 5039 5040 3 0.00000 0.00000 0.00000
0.00000 0.00000 0.00000 ; H- N- CA- HA
5038 5037 5039 5041 3 0.00000 0.00000 0.00000
0.00000 0.00000 0.00000 ; H- N- CA- CB
5038 5037 5039 5046 3 0.00000 0.00000 0.00000
0.00000 0.00000 0.00000 ; H- N- CA- C
5039 5041 5044 5045 3 1.69034 2.77818 4.11706
-4.21747 1.00416 0.00000 ; CA- CB- SG- HG
5039 5046 5048 5049 3 20.92000 0.00000 -20.92000
0.00000 0.00000 0.00000 ; CA- C- N- H
5039 5046 5048 5050 3 20.92000 0.00000 -20.92000
0.00000 0.00000 0.00000 ; CA- C- N- CA
5040 5039 5041 5042 3 0.65084 1.95253 0.00000
-2.60338 0.00000 0.00000 ; HA- CA- CB- HB2
5040 5039 5041 5043 3 0.65084 1.95253 0.00000
-2.60338 0.00000 0.00000 ; HA- CA- CB- HB3
5040 5039 5041 5044 3 0.65084 1.95253 0.00000
-2.60338 0.00000 0.00000 ; HA- CA- CB- SG
5040 5039 5046 5047 3 3.68192 -4.35136 0.00000
1.33888 0.00000 0.00000 ; HA- CA- C- O
5040 5039 5046 5048 3 0.00000 0.00000 0.00000
0.00000 0.00000 0.00000 ; HA- CA- C- N
5041 5039 5046 5047 3 0.00000 0.00000 0.00000
0.00000 0.00000 0.00000 ; CB- CA- C- O
5041 5039 5046 5048 3 2.51040 4.18400 1.67360
-6.69440 0.00000 0.00000 ; CB- CA- C- N
5042 5041 5039 5046 3 0.65084 1.95253 0.00000
-2.60338 0.00000 0.00000 ; HB2- CB- CA- C
5042 5041 5044 5045 3 1.04600 3.13800 0.00000
-4.18400 0.00000 0.00000 ; HB2- CB- SG- HG
5043 5041 5039 5046 3 0.65084 1.95253 0.00000
-2.60338 0.00000 0.00000 ; HB3- CB- CA- C
5043 5041 5044 5045 3 1.04600 3.13800 0.00000

-4.18400 0.00000 0.00000 ; HB3- CB- SG- HG
5044 5041 5039 5046 3 5.62330 4.27605 -5.33042
-4.20074 2.51040 0.00000 ; SG- CB- CA- C
5046 5048 5050 5051 3 0.00000 0.00000 0.00000
0.00000 0.00000 0.00000 ; C- N- CA- HA
5046 5048 5050 5052 3 11.71520 1.67360 15.06240 -
13.38880 0.00000 0.00000 ; C- N- CA- CB
5046 5048 5050 5058 3 1.75728 5.27184 2.25936
-7.02912 0.00000 0.00000 ; C- N- CA- C
5047 5046 5048 5049 3 29.28800 -8.36800 -20.92000
0.00000 0.00000 0.00000 ; O- C- N- H
5047 5046 5048 5050 3 20.92000 0.00000 -20.92000
0.00000 0.00000 0.00000 ; O- C- N- CA
5048 5050 5052 5053 3 0.65084 1.95253 0.00000
-2.60338 0.00000 0.00000 ; N- CA- CB- HB2
5048 5050 5052 5054 3 0.65084 1.95253 0.00000
-2.60338 0.00000 0.00000 ; N- CA- CB- HB3
5048 5050 5052 5055 3 15.41386 9.74035 -8.39310
-0.97069 2.97901 0.00000 ; N- CA- CB- CG
5048 5050 5058 5059 3 0.00000 0.00000 0.00000
0.00000 0.00000 0.00000 ; N- CA- C- O
5048 5050 5058 5060 3 17.40544 -5.02080 -13.22144
9.20480 0.00000 0.00000 ; N- CA- C- N
5049 5048 5050 5051 3 0.00000 0.00000 0.00000
0.00000 0.00000 0.00000 ; H- N- CA- HA
5049 5048 5050 5052 3 0.00000 0.00000 0.00000
0.00000 0.00000 0.00000 ; H- N- CA- CB
5049 5048 5050 5058 3 0.00000 0.00000 0.00000
0.00000 0.00000 0.00000 ; H- N- CA- C
5050 5052 5055 5056 3 6.43499 0.00000 -5.39736
0.00000 -1.03763 0.00000 ; CA- CB- CG- OD1
5050 5052 5055 5057 3 6.43499 0.00000 -5.39736
0.00000 -1.03763 0.00000 ; CA- CB- CG- OD2
5050 5058 5060 5061 3 20.92000 0.00000 -20.92000
0.00000 0.00000 0.00000 ; CA- C- N- H
5050 5058 5060 5062 3 20.92000 0.00000 -20.92000
0.00000 0.00000 0.00000 ; CA- C- N- CA
5051 5050 5052 5053 3 0.65084 1.95253 0.00000
-2.60338 0.00000 0.00000 ; HA- CA- CB- HB2
5051 5050 5052 5054 3 0.65084 1.95253 0.00000
-2.60338 0.00000 0.00000 ; HA- CA- CB- HB3
5051 5050 5052 5055 3 0.65084 1.95253 0.00000
-2.60338 0.00000 0.00000 ; HA- CA- CB- CG
5051 5050 5058 5059 3 3.68192 -4.35136 0.00000
1.33888 0.00000 0.00000 ; HA- CA- C- O
5051 5050 5058 5060 3 0.00000 0.00000 0.00000
0.00000 0.00000 0.00000 ; HA- CA- C- N
5052 5050 5058 5059 3 0.00000 0.00000 0.00000
0.00000 0.00000 0.00000 ; CB- CA- C- O
5052 5050 5058 5060 3 2.51040 4.18400 1.67360
-6.69440 0.00000 0.00000 ; CB- CA- C- N
5053 5052 5050 5058 3 0.65084 1.95253 0.00000

```
-2.60338    0.00000    0.00000 ;    HB2-    CB-    CA-     C
  5053   5052   5055   5056      3    0.00000    0.00000    0.00000
0.00000    0.00000    0.00000 ;    HB2-    CB-    CG-   OD1
  5053   5052   5055   5057      3    0.00000    0.00000    0.00000
0.00000    0.00000    0.00000 ;    HB2-    CB-    CG-   OD2
  5054   5052   5050   5058      3    0.65084    1.95253    0.00000
-2.60338    0.00000    0.00000 ;    HB3-    CB-    CA-     C
  5054   5052   5055   5056      3    0.00000    0.00000    0.00000
0.00000    0.00000    0.00000 ;    HB3-    CB-    CG-   OD1
  5054   5052   5055   5057      3    0.00000    0.00000    0.00000
0.00000    0.00000    0.00000 ;    HB3-    CB-    CG-   OD2
  5055   5052   5050   5058      3    7.14627   -1.04600   -8.99560
-0.97069    5.15469    0.00000 ;     CG-    CB-    CA-     C
  5058   5060   5062   5063      3    0.00000    0.00000    0.00000
0.00000    0.00000    0.00000 ;     C-     N-    CA-    HA
  5058   5060   5062   5064      3   11.71520    1.67360   15.06240  -
13.38880    0.00000    0.00000 ;     C-     N-    CA-    CB
  5058   5060   5062   5074      3    1.75728    5.27184    2.25936
-7.02912    0.00000    0.00000 ;     C-     N-    CA-     C
  5059   5058   5060   5061      3   29.28800   -8.36800  -20.92000
0.00000    0.00000    0.00000 ;     O-     C-     N-     H
  5059   5058   5060   5062      3   20.92000    0.00000  -20.92000
0.00000    0.00000    0.00000 ;     O-     C-     N-    CA
  5060   5062   5064   5065      3    0.65084    1.95253    0.00000
-2.60338    0.00000    0.00000 ;     N-    CA-    CB-    HB
  5060   5062   5064   5066      3    3.83673    0.44769   -1.77402
-2.47693   -0.03347    0.00000 ;     N-    CA-    CB-   CG1
  5060   5062   5064   5070      3    3.83673    0.44769   -1.77402
-2.47693   -0.03347    0.00000 ;     N-    CA-    CB-   CG2
  5060   5062   5074   5075      3    0.00000    0.00000    0.00000
0.00000    0.00000    0.00000 ;     N-    CA-     C-     O
  5060   5062   5074   5076      3   17.40544   -5.02080  -13.22144
9.20480    0.00000    0.00000 ;     N-    CA-     C-     N
  5061   5060   5062   5063      3    0.00000    0.00000    0.00000
0.00000    0.00000    0.00000 ;     H-     N-    CA-    HA
  5061   5060   5062   5064      3    0.00000    0.00000    0.00000
0.00000    0.00000    0.00000 ;     H-     N-    CA-    CB
  5061   5060   5062   5074      3    0.00000    0.00000    0.00000
0.00000    0.00000    0.00000 ;     H-     N-    CA-     C
  5062   5064   5066   5067      3    0.66944    2.00832    0.00000
-2.67776    0.00000    0.00000 ;    CA-    CB-   CG1-  HG11
  5062   5064   5066   5068      3    0.66944    2.00832    0.00000
-2.67776    0.00000    0.00000 ;    CA-    CB-   CG1-  HG12
  5062   5064   5066   5069      3    0.66944    2.00832    0.00000
-2.67776    0.00000    0.00000 ;    CA-    CB-   CG1-  HG13
  5062   5064   5070   5071      3    0.66944    2.00832    0.00000
-2.67776    0.00000    0.00000 ;    CA-    CB-   CG2-  HG21
  5062   5064   5070   5072      3    0.66944    2.00832    0.00000
-2.67776    0.00000    0.00000 ;    CA-    CB-   CG2-  HG22
  5062   5064   5070   5073      3    0.66944    2.00832    0.00000
-2.67776    0.00000    0.00000 ;    CA-    CB-   CG2-  HG23
  5062   5074   5076   5077      3   20.92000    0.00000  -20.92000
```

```
0.00000    0.00000    0.00000 ;     CA-    C-     N-    H
  5062   5074   5076   5078       3   20.92000    0.00000  -20.92000
0.00000    0.00000    0.00000 ;     CA-    C-     N-   CA
  5063   5062   5064   5065       3    0.65084    1.95253    0.00000
-2.60338    0.00000    0.00000 ;     HA-   CA-    CB-   HB
  5063   5062   5064   5066       3    0.65084    1.95253    0.00000
-2.60338    0.00000    0.00000 ;     HA-   CA-    CB-  CG1
  5063   5062   5064   5070       3    0.65084    1.95253    0.00000
-2.60338    0.00000    0.00000 ;     HA-   CA-    CB-  CG2
  5063   5062   5074   5075       3    3.68192   -4.35136    0.00000
1.33888    0.00000    0.00000 ;     HA-   CA-     C-    O
  5063   5062   5074   5076       3    0.00000    0.00000    0.00000
0.00000    0.00000    0.00000 ;     HA-   CA-     C-    N
  5064   5062   5074   5075       3    0.00000    0.00000    0.00000
0.00000    0.00000    0.00000 ;     CB-   CA-     C-    O
  5064   5062   5074   5076       3    2.51040    4.18400    1.67360
-6.69440    0.00000    0.00000 ;     CB-   CA-     C-    N
  5065   5064   5062   5074       3    0.65084    1.95253    0.00000
-2.60338    0.00000    0.00000 ;     HB-   CB-    CA-    C
  5065   5064   5066   5067       3    0.62760    1.88280    0.00000
-2.51040    0.00000    0.00000 ;     HB-   CB-   CG1-  HG11
  5065   5064   5066   5068       3    0.62760    1.88280    0.00000
-2.51040    0.00000    0.00000 ;     HB-   CB-   CG1-  HG12
  5065   5064   5066   5069       3    0.62760    1.88280    0.00000
-2.51040    0.00000    0.00000 ;     HB-   CB-   CG1-  HG13
  5065   5064   5070   5071       3    0.62760    1.88280    0.00000
-2.51040    0.00000    0.00000 ;     HB-   CB-   CG2-  HG21
  5065   5064   5070   5072       3    0.62760    1.88280    0.00000
-2.51040    0.00000    0.00000 ;     HB-   CB-   CG2-  HG22
  5065   5064   5070   5073       3    0.62760    1.88280    0.00000
-2.51040    0.00000    0.00000 ;     HB-   CB-   CG2-  HG23
  5066   5064   5062   5074       3    5.67350    3.55640   -6.16722
-2.47693    3.74886    0.00000 ;    CG1-   CB-    CA-    C
  5066   5064   5070   5071       3    0.66944    2.00832    0.00000
-2.67776    0.00000    0.00000 ;    CG1-   CB-   CG2-  HG21
  5066   5064   5070   5072       3    0.66944    2.00832    0.00000
-2.67776    0.00000    0.00000 ;    CG1-   CB-   CG2-  HG22
  5066   5064   5070   5073       3    0.66944    2.00832    0.00000
-2.67776    0.00000    0.00000 ;    CG1-   CB-   CG2-  HG23
  5067   5066   5064   5070       3    0.66944    2.00832    0.00000
-2.67776    0.00000    0.00000 ;   HG11-  CG1-    CB-  CG2
  5068   5066   5064   5070       3    0.66944    2.00832    0.00000
-2.67776    0.00000    0.00000 ;   HG12-  CG1-    CB-  CG2
  5069   5066   5064   5070       3    0.66944    2.00832    0.00000
-2.67776    0.00000    0.00000 ;   HG13-  CG1-    CB-  CG2
  5070   5064   5062   5074       3    5.67350    3.55640   -6.16722
-2.47693    3.74886    0.00000 ;    CG2-   CB-    CA-    C
  5074   5076   5078   5079       3    0.00000    0.00000    0.00000
0.00000    0.00000    0.00000 ;      C-    N-   CA-   HA
  5074   5076   5078   5080       3   11.71520    1.67360   15.06240  -
13.38880    0.00000    0.00000 ;      C-    N-   CA-   CB
  5074   5076   5078   5089       3    1.75728    5.27184    2.25936
```

-7.02912 0.00000 0.00000 ; C- N- CA- C
5075 5074 5076 5077 3 29.28800 -8.36800 -20.92000
0.00000 0.00000 0.00000 ; O- C- N- H
5075 5074 5076 5078 3 20.92000 0.00000 -20.92000
0.00000 0.00000 0.00000 ; O- C- N- CA
5076 5078 5080 5081 3 0.65084 1.95253 0.00000
-2.60338 0.00000 0.00000 ; N- CA- CB- HB2
5076 5078 5080 5082 3 0.65084 1.95253 0.00000
-2.60338 0.00000 0.00000 ; N- CA- CB- HB3
5076 5078 5080 5083 3 3.21331 2.22589 -4.15053
-2.40998 2.61082 0.00000 ; N- CA- CB- CG
5076 5078 5089 5090 3 0.00000 0.00000 0.00000
0.00000 0.00000 0.00000 ; N- CA- C- O
5076 5078 5089 5091 3 17.40544 -5.02080 -13.22144
9.20480 0.00000 0.00000 ; N- CA- C- N
5077 5076 5078 5079 3 0.00000 0.00000 0.00000
0.00000 0.00000 0.00000 ; H- N- CA- HA
5077 5076 5078 5080 3 0.00000 0.00000 0.00000
0.00000 0.00000 0.00000 ; H- N- CA- CB
5077 5076 5078 5089 3 0.00000 0.00000 0.00000
0.00000 0.00000 0.00000 ; H- N- CA- C
5078 5080 5083 5084 3 0.66944 2.00832 0.00000
-2.67776 0.00000 0.00000 ; CA- CB- CG- HG2
5078 5080 5083 5085 3 0.66944 2.00832 0.00000
-2.67776 0.00000 0.00000 ; CA- CB- CG- HG3
5078 5080 5083 5086 3 10.12110 -1.91209 0.01674
10.17549 -1.87443 0.00000 ; CA- CB- CG- CD
5078 5089 5091 5092 3 20.92000 0.00000 -20.92000
0.00000 0.00000 0.00000 ; CA- C- N- H
5078 5089 5091 5093 3 20.92000 0.00000 -20.92000
0.00000 0.00000 0.00000 ; CA- C- N- CA
5079 5078 5080 5081 3 0.65084 1.95253 0.00000
-2.60338 0.00000 0.00000 ; HA- CA- CB- HB2
5079 5078 5080 5082 3 0.65084 1.95253 0.00000
-2.60338 0.00000 0.00000 ; HA- CA- CB- HB3
5079 5078 5080 5083 3 0.65084 1.95253 0.00000
-2.60338 0.00000 0.00000 ; HA- CA- CB- CG
5079 5078 5089 5090 3 3.68192 -4.35136 0.00000
1.33888 0.00000 0.00000 ; HA- CA- C- O
5079 5078 5089 5091 3 0.00000 0.00000 0.00000
0.00000 0.00000 0.00000 ; HA- CA- C- N
5080 5078 5089 5090 3 0.00000 0.00000 0.00000
0.00000 0.00000 0.00000 ; CB- CA- C- O
5080 5078 5089 5091 3 2.51040 4.18400 1.67360
-6.69440 0.00000 0.00000 ; CB- CA- C- N
5080 5083 5086 5087 3 3.79907 0.00000 -5.40573
0.00000 2.14221 0.00000 ; CB- CG- CD- OE1
5080 5083 5086 5088 3 3.79907 0.00000 -5.40573
0.00000 2.14221 0.00000 ; CB- CG- CD- OE2
5081 5080 5078 5089 3 0.65084 1.95253 0.00000
-2.60338 0.00000 0.00000 ; HB2- CB- CA- C
5081 5080 5083 5084 3 0.62760 1.88280 0.00000

-2.51040 0.00000 0.00000 ; HB2- CB- CG- HG2
5081 5080 5083 5085 3 0.62760 1.88280 0.00000
-2.51040 0.00000 0.00000 ; HB2- CB- CG- HG3
5081 5080 5083 5086 3 0.65084 1.95253 0.00000
-2.60338 0.00000 0.00000 ; HB2- CB- CG- CD
5082 5080 5078 5089 3 0.65084 1.95253 0.00000
-2.60338 0.00000 0.00000 ; HB3- CB- CA- C
5082 5080 5083 5084 3 0.62760 1.88280 0.00000
-2.51040 0.00000 0.00000 ; HB3- CB- CG- HG2
5082 5080 5083 5085 3 0.62760 1.88280 0.00000
-2.51040 0.00000 0.00000 ; HB3- CB- CG- HG3
5082 5080 5083 5086 3 0.65084 1.95253 0.00000
-2.60338 0.00000 0.00000 ; HB3- CB- CG- CD
5083 5080 5078 5089 3 6.86594 3.56895 -8.14206
-2.40998 4.85344 0.00000 ; CG- CB- CA- C
5084 5083 5086 5087 3 0.00000 0.00000 0.00000
0.00000 0.00000 0.00000 ; HG2- CG- CD- OE1
5084 5083 5086 5088 3 0.00000 0.00000 0.00000
0.00000 0.00000 0.00000 ; HG2- CG- CD- OE2
5085 5083 5086 5087 3 0.00000 0.00000 0.00000
0.00000 0.00000 0.00000 ; HG3- CG- CD- OE1
5085 5083 5086 5088 3 0.00000 0.00000 0.00000
0.00000 0.00000 0.00000 ; HG3- CG- CD- OE2
5089 5091 5093 5094 3 0.00000 0.00000 0.00000
0.00000 0.00000 0.00000 ; C- N- CA- HA
5089 5091 5093 5095 3 11.71520 1.67360 15.06240 -
13.38880 0.00000 0.00000 ; C- N- CA- CB
5089 5091 5093 5108 3 1.75728 5.27184 2.25936
-7.02912 0.00000 0.00000 ; C- N- CA- C
5090 5089 5091 5092 3 29.28800 -8.36800 -20.92000
0.00000 0.00000 0.00000 ; O- C- N- H
5090 5089 5091 5093 3 20.92000 0.00000 -20.92000
0.00000 0.00000 0.00000 ; O- C- N- CA
5091 5093 5095 5096 3 0.65084 1.95253 0.00000
-2.60338 0.00000 0.00000 ; N- CA- CB- HB
5091 5093 5095 5097 3 3.83673 0.44769 -1.77402
-2.47693 -0.03347 0.00000 ; N- CA- CB- CG2
5091 5093 5095 5101 3 2.97482 0.12134 2.04179
-1.89117 -3.24678 0.00000 ; N- CA- CB- CG1
5091 5093 5108 5109 3 0.00000 0.00000 0.00000
0.00000 0.00000 0.00000 ; N- CA- C- O
5091 5093 5108 5110 3 17.40544 -5.02080 -13.22144
9.20480 0.00000 0.00000 ; N- CA- C- N
5092 5091 5093 5094 3 0.00000 0.00000 0.00000
0.00000 0.00000 0.00000 ; H- N- CA- HA
5092 5091 5093 5095 3 0.00000 0.00000 0.00000
0.00000 0.00000 0.00000 ; H- N- CA- CB
5092 5091 5093 5108 3 0.00000 0.00000 0.00000
0.00000 0.00000 0.00000 ; H- N- CA- C
5093 5095 5097 5098 3 0.66944 2.00832 0.00000
-2.67776 0.00000 0.00000 ; CA- CB- CG2- HG21
5093 5095 5097 5099 3 0.66944 2.00832 0.00000

```
-2.67776    0.00000    0.00000 ;     CA-    CB-   CG2-  HG22
  5093   5095   5097   5100      3    0.66944    2.00832    0.00000
-2.67776    0.00000    0.00000 ;     CA-    CB-   CG2-  HG23
  5093   5095   5101   5102      3    0.66944    2.00832    0.00000
-2.67776    0.00000    0.00000 ;     CA-    CB-   CG1-  HG12
  5093   5095   5101   5103      3    0.66944    2.00832    0.00000
-2.67776    0.00000    0.00000 ;     CA-    CB-   CG1-  HG13
  5093   5095   5101   5104      3    4.24258   -0.52718   -7.25506
-1.79075    7.69856    0.00000 ;     CA-    CB-   CG1-   CD1
  5093   5108   5110   5111      3   20.92000    0.00000  -20.92000
0.00000    0.00000    0.00000 ;     CA-     C-     N-     H
  5093   5108   5110   5112      3   20.92000    0.00000  -20.92000
0.00000    0.00000    0.00000 ;     CA-     C-     N-    CA
  5094   5093   5095   5096      3    0.65084    1.95253    0.00000
-2.60338    0.00000    0.00000 ;     HA-    CA-    CB-    HB
  5094   5093   5095   5097      3    0.65084    1.95253    0.00000
-2.60338    0.00000    0.00000 ;     HA-    CA-    CB-   CG2
  5094   5093   5095   5101      3    0.65084    1.95253    0.00000
-2.60338    0.00000    0.00000 ;     HA-    CA-    CB-   CG1
  5094   5093   5108   5109      3    3.68192   -4.35136    0.00000
1.33888    0.00000    0.00000 ;     HA-    CA-     C-     O
  5094   5093   5108   5110      3    0.00000    0.00000    0.00000
0.00000    0.00000    0.00000 ;     HA-    CA-     C-     N
  5095   5093   5108   5109      3    0.00000    0.00000    0.00000
0.00000    0.00000    0.00000 ;     CB-    CA-     C-     O
  5095   5093   5108   5110      3    2.51040    4.18400    1.67360
-6.69440    0.00000    0.00000 ;     CB-    CA-     C-     N
  5095   5101   5104   5105      3    0.66944    2.00832    0.00000
-2.67776    0.00000    0.00000 ;     CB-   CG1-   CD1-  HD11
  5095   5101   5104   5106      3    0.66944    2.00832    0.00000
-2.67776    0.00000    0.00000 ;     CB-   CG1-   CD1-  HD12
  5095   5101   5104   5107      3    0.66944    2.00832    0.00000
-2.67776    0.00000    0.00000 ;     CB-   CG1-   CD1-  HD13
  5096   5095   5093   5108      3    0.65084    1.95253    0.00000
-2.60338    0.00000    0.00000 ;     HB-    CB-    CA-     C
  5096   5095   5097   5098      3    0.62760    1.88280    0.00000
-2.51040    0.00000    0.00000 ;     HB-    CB-   CG2-  HG21
  5096   5095   5097   5099      3    0.62760    1.88280    0.00000
-2.51040    0.00000    0.00000 ;     HB-    CB-   CG2-  HG22
  5096   5095   5097   5100      3    0.62760    1.88280    0.00000
-2.51040    0.00000    0.00000 ;     HB-    CB-   CG2-  HG23
  5096   5095   5101   5102      3    0.62760    1.88280    0.00000
-2.51040    0.00000    0.00000 ;     HB-    CB-   CG1-  HG12
  5096   5095   5101   5103      3    0.62760    1.88280    0.00000
-2.51040    0.00000    0.00000 ;     HB-    CB-   CG1-  HG13
  5096   5095   5101   5104      3    0.66944    2.00832    0.00000
-2.67776    0.00000    0.00000 ;     HB-    CB-   CG1-   CD1
  5097   5095   5093   5108      3    5.67350    3.55640   -6.16722
-2.47693    3.74886    0.00000 ;    CG2-    CB-    CA-     C
  5097   5095   5101   5102      3    0.66944    2.00832    0.00000
-2.67776    0.00000    0.00000 ;    CG2-    CB-   CG1-  HG12
  5097   5095   5101   5103      3    0.66944    2.00832    0.00000
```

-2.67776 0.00000 0.00000 ; CG2- CB- CG1- HG13
5097 5095 5101 5104 3 3.81162 0.49790 -8.14206
-1.79075 7.49773 0.00000 ; CG2- CB- CG1- CD1
5098 5097 5095 5101 3 0.66944 2.00832 0.00000
-2.67776 0.00000 0.00000 ; HG21- CG2- CB- CG1
5099 5097 5095 5101 3 0.66944 2.00832 0.00000
-2.67776 0.00000 0.00000 ; HG22- CG2- CB- CG1
5100 5097 5095 5101 3 0.66944 2.00832 0.00000
-2.67776 0.00000 0.00000 ; HG23- CG2- CB- CG1
5101 5095 5093 5108 3 8.26340 0.74057 -9.99976
-1.89117 3.84928 0.00000 ; CG1- CB- CA- C
5102 5101 5104 5105 3 0.62760 1.88280 0.00000
-2.51040 0.00000 0.00000 ; HG12- CG1- CD1- HD11
5102 5101 5104 5106 3 0.62760 1.88280 0.00000
-2.51040 0.00000 0.00000 ; HG12- CG1- CD1- HD12
5102 5101 5104 5107 3 0.62760 1.88280 0.00000
-2.51040 0.00000 0.00000 ; HG12- CG1- CD1- HD13
5103 5101 5104 5105 3 0.62760 1.88280 0.00000
-2.51040 0.00000 0.00000 ; HG13- CG1- CD1- HD11
5103 5101 5104 5106 3 0.62760 1.88280 0.00000
-2.51040 0.00000 0.00000 ; HG13- CG1- CD1- HD12
5103 5101 5104 5107 3 0.62760 1.88280 0.00000
-2.51040 0.00000 0.00000 ; HG13- CG1- CD1- HD13
5108 5110 5112 5113 3 0.00000 0.00000 0.00000
0.00000 0.00000 0.00000 ; C- N- CA- HA
5108 5110 5112 5114 3 11.71520 1.67360 15.06240 -
13.38880 0.00000 0.00000 ; C- N- CA- CB
5108 5110 5112 5122 3 1.75728 5.27184 2.25936
-7.02912 0.00000 0.00000 ; C- N- CA- C
5109 5108 5110 5111 3 29.28800 -8.36800 -20.92000
0.00000 0.00000 0.00000 ; O- C- N- H
5109 5108 5110 5112 3 20.92000 0.00000 -20.92000
0.00000 0.00000 0.00000 ; O- C- N- CA
5110 5112 5114 5115 3 0.65084 1.95253 0.00000
-2.60338 0.00000 0.00000 ; N- CA- CB- HB2
5110 5112 5114 5116 3 0.65084 1.95253 0.00000
-2.60338 0.00000 0.00000 ; N- CA- CB- HB3
5110 5112 5114 5117 3 5.99567 3.29281 -4.46014
-0.55229 1.97485 0.00000 ; N- CA- CB- CG
5110 5112 5122 5123 3 0.00000 0.00000 0.00000
0.00000 0.00000 0.00000 ; N- CA- C- O
5110 5112 5122 5124 3 17.40544 -5.02080 -13.22144
9.20480 0.00000 0.00000 ; N- CA- C- N
5111 5110 5112 5113 3 0.00000 0.00000 0.00000
0.00000 0.00000 0.00000 ; H- N- CA- HA
5111 5110 5112 5114 3 0.00000 0.00000 0.00000
0.00000 0.00000 0.00000 ; H- N- CA- CB
5111 5110 5112 5122 3 0.00000 0.00000 0.00000
0.00000 0.00000 0.00000 ; H- N- CA- C
5112 5114 5117 5118 3 0.00000 0.00000 0.00000
0.00000 0.00000 0.00000 ; CA- CB- CG- OD1
5112 5114 5117 5119 3 8.84916 -0.31380 -4.32626

```
5.03754    0.26778    0.00000 ;      CA-    CB-    CG-   ND2
  5112   5122   5124   5125      3   20.92000    0.00000  -20.92000
0.00000    0.00000    0.00000 ;      CA-     C-     N-     H
  5112   5122   5124   5126      3   20.92000    0.00000  -20.92000
0.00000    0.00000    0.00000 ;      CA-     C-     N-    CA
  5113   5112   5114   5115      3    0.65084    1.95253    0.00000
-2.60338    0.00000    0.00000 ;     HA-    CA-    CB-   HB2
  5113   5112   5114   5116      3    0.65084    1.95253    0.00000
-2.60338    0.00000    0.00000 ;     HA-    CA-    CB-   HB3
  5113   5112   5114   5117      3    0.65084    1.95253    0.00000
-2.60338    0.00000    0.00000 ;     HA-    CA-    CB-    CG
  5113   5112   5122   5123      3    3.68192   -4.35136    0.00000
1.33888    0.00000    0.00000 ;      HA-    CA-     C-     O
  5113   5112   5122   5124      3    0.00000    0.00000    0.00000
0.00000    0.00000    0.00000 ;      HA-    CA-     C-     N
  5114   5112   5122   5123      3    0.00000    0.00000    0.00000
0.00000    0.00000    0.00000 ;      CB-    CA-     C-     O
  5114   5112   5122   5124      3    2.51040    4.18400    1.67360
-6.69440    0.00000    0.00000 ;     CB-    CA-     C-     N
  5114   5117   5119   5120      3   20.92000    0.00000  -20.92000
0.00000    0.00000    0.00000 ;      CB-    CG-   ND2-  HD21
  5114   5117   5119   5121      3   20.92000    0.00000  -20.92000
0.00000    0.00000    0.00000 ;      CB-    CG-   ND2-  HD22
  5115   5114   5112   5122      3    0.65084    1.95253    0.00000
-2.60338    0.00000    0.00000 ;    HB2-    CB-    CA-     C
  5115   5114   5117   5118      3    3.68192   -4.35136    0.00000
1.33888    0.00000    0.00000 ;     HB2-    CB-    CG-   OD1
  5115   5114   5117   5119      3    0.00000    0.00000    0.00000
0.00000    0.00000    0.00000 ;     HB2-    CB-    CG-   ND2
  5116   5114   5112   5122      3    0.65084    1.95253    0.00000
-2.60338    0.00000    0.00000 ;    HB3-    CB-    CA-     C
  5116   5114   5117   5118      3    3.68192   -4.35136    0.00000
1.33888    0.00000    0.00000 ;     HB3-    CB-    CG-   OD1
  5116   5114   5117   5119      3    0.00000    0.00000    0.00000
0.00000    0.00000    0.00000 ;     HB3-    CB-    CG-   ND2
  5117   5114   5112   5122      3    7.94542   -3.96225   -6.11701
-0.55229    3.58150    0.00000 ;     CG-    CB-    CA-     C
  5118   5117   5119   5120      3   29.28800   -8.36800  -20.92000
0.00000    0.00000    0.00000 ;     OD1-    CG-   ND2-  HD21
  5118   5117   5119   5121      3   29.28800   -8.36800  -20.92000
0.00000    0.00000    0.00000 ;     OD1-    CG-   ND2-  HD22
  5122   5124   5126   5127      3    0.00000    0.00000    0.00000
0.00000    0.00000    0.00000 ;       C-     N-    CA-    HA
  5122   5124   5126   5128      3   11.71520    1.67360   15.06240  -
13.38880    0.00000    0.00000 ;      C-     N-    CA-    CB
  5122   5124   5126   5136      3    1.75728    5.27184    2.25936
-7.02912    0.00000    0.00000 ;      C-     N-    CA-     C
  5123   5122   5124   5125      3   29.28800   -8.36800  -20.92000
0.00000    0.00000    0.00000 ;       O-     C-     N-     H
  5123   5122   5124   5126      3   20.92000    0.00000  -20.92000
0.00000    0.00000    0.00000 ;       O-     C-     N-    CA
  5124   5126   5128   5129      3    0.65084    1.95253    0.00000
```

```
-2.60338    0.00000    0.00000 ;       N-    CA-    CB-   HB2
  5124   5126   5128   5130      3    0.65084    1.95253    0.00000
-2.60338    0.00000    0.00000 ;       N-    CA-    CB-   HB3
  5124   5126   5128   5131      3    5.99567    3.29281   -4.46014
-0.55229    1.97485    0.00000 ;       N-    CA-    CB-    CG
  5124   5126   5136   5137      3    0.00000    0.00000    0.00000
0.00000    0.00000    0.00000 ;       N-    CA-     C-     O
  5124   5126   5136   5138      3   17.40544   -5.02080  -13.22144
9.20480    0.00000    0.00000 ;       N-    CA-     C-     N
  5125   5124   5126   5127      3    0.00000    0.00000    0.00000
0.00000    0.00000    0.00000 ;       H-     N-    CA-    HA
  5125   5124   5126   5128      3    0.00000    0.00000    0.00000
0.00000    0.00000    0.00000 ;       H-     N-    CA-    CB
  5125   5124   5126   5136      3    0.00000    0.00000    0.00000
0.00000    0.00000    0.00000 ;       H-     N-    CA-     C
  5126   5128   5131   5132      3    0.00000    0.00000    0.00000
0.00000    0.00000    0.00000 ;      CA-    CB-    CG-   OD1
  5126   5128   5131   5133      3    8.84916   -0.31380   -4.32626
5.03754    0.26778    0.00000 ;      CA-    CB-    CG-   ND2
  5126   5136   5138   5139      3   20.92000    0.00000  -20.92000
0.00000    0.00000    0.00000 ;      CA-     C-     N-     H
  5126   5136   5138   5140      3   20.92000    0.00000  -20.92000
0.00000    0.00000    0.00000 ;      CA-     C-     N-    CA
  5127   5126   5128   5129      3    0.65084    1.95253    0.00000
-2.60338    0.00000    0.00000 ;      HA-    CA-    CB-   HB2
  5127   5126   5128   5130      3    0.65084    1.95253    0.00000
-2.60338    0.00000    0.00000 ;      HA-    CA-    CB-   HB3
  5127   5126   5128   5131      3    0.65084    1.95253    0.00000
-2.60338    0.00000    0.00000 ;      HA-    CA-    CB-    CG
  5127   5126   5136   5137      3    3.68192   -4.35136    0.00000
1.33888    0.00000    0.00000 ;      HA-    CA-     C-     O
  5127   5126   5136   5138      3    0.00000    0.00000    0.00000
0.00000    0.00000    0.00000 ;      HA-    CA-     C-     N
  5128   5126   5136   5137      3    0.00000    0.00000    0.00000
0.00000    0.00000    0.00000 ;      CB-    CA-     C-     O
  5128   5126   5136   5138      3    2.51040    4.18400    1.67360
-6.69440    0.00000    0.00000 ;      CB-    CA-     C-     N
  5128   5131   5133   5134      3   20.92000    0.00000  -20.92000
0.00000    0.00000    0.00000 ;      CB-    CG-   ND2-  HD21
  5128   5131   5133   5135      3   20.92000    0.00000  -20.92000
0.00000    0.00000    0.00000 ;      CB-    CG-   ND2-  HD22
  5129   5128   5126   5136      3    0.65084    1.95253    0.00000
-2.60338    0.00000    0.00000 ;     HB2-    CB-    CA-     C
  5129   5128   5131   5132      3    3.68192   -4.35136    0.00000
1.33888    0.00000    0.00000 ;     HB2-    CB-    CG-   OD1
  5129   5128   5131   5133      3    0.00000    0.00000    0.00000
0.00000    0.00000    0.00000 ;     HB2-    CB-    CG-   ND2
  5130   5128   5126   5136      3    0.65084    1.95253    0.00000
-2.60338    0.00000    0.00000 ;     HB3-    CB-    CA-     C
  5130   5128   5131   5132      3    3.68192   -4.35136    0.00000
1.33888    0.00000    0.00000 ;     HB3-    CB-    CG-   OD1
  5130   5128   5131   5133      3    0.00000    0.00000    0.00000
```

0.00000 0.00000 0.00000 ; HB3- CB- CG- ND2
5131 5128 5126 5136 3 7.94542 -3.96225 -6.11701
-0.55229 3.58150 0.00000 ; CG- CB- CA- C
5132 5131 5133 5134 3 29.28800 -8.36800 -20.92000
0.00000 0.00000 0.00000 ; OD1- CG- ND2- HD21
5132 5131 5133 5135 3 29.28800 -8.36800 -20.92000
0.00000 0.00000 0.00000 ; OD1- CG- ND2- HD22
5136 5138 5140 5141 3 0.00000 0.00000 0.00000
0.00000 0.00000 0.00000 ; C- N- CA- HA
5136 5138 5140 5142 3 11.71520 1.67360 15.06240 -
13.38880 0.00000 0.00000 ; C- N- CA- CB
5136 5138 5140 5155 3 1.75728 5.27184 2.25936
-7.02912 0.00000 0.00000 ; C- N- CA- C
5137 5136 5138 5139 3 29.28800 -8.36800 -20.92000
0.00000 0.00000 0.00000 ; O- C- N- H
5137 5136 5138 5140 3 20.92000 0.00000 -20.92000
0.00000 0.00000 0.00000 ; O- C- N- CA
5138 5140 5142 5143 3 0.65084 1.95253 0.00000
-2.60338 0.00000 0.00000 ; N- CA- CB- HB2
5138 5140 5142 5144 3 0.65084 1.95253 0.00000
-2.60338 0.00000 0.00000 ; N- CA- CB- HB3
5138 5140 5142 5145 3 3.79070 1.39746 -4.61077
-2.40998 2.44346 0.00000 ; N- CA- CB- CG
5138 5140 5155 5156 3 0.00000 0.00000 0.00000
0.00000 0.00000 0.00000 ; N- CA- C- O
5138 5140 5155 5157 3 17.40544 -5.02080 -13.22144
9.20480 0.00000 0.00000 ; N- CA- C- N
5139 5138 5140 5141 3 0.00000 0.00000 0.00000
0.00000 0.00000 0.00000 ; H- N- CA- HA
5139 5138 5140 5142 3 0.00000 0.00000 0.00000
0.00000 0.00000 0.00000 ; H- N- CA- CB
5139 5138 5140 5155 3 0.00000 0.00000 0.00000
0.00000 0.00000 0.00000 ; H- N- CA- C
5140 5142 5145 5146 3 0.66944 2.00832 0.00000
-2.67776 0.00000 0.00000 ; CA- CB- CG- HG
5140 5142 5145 5147 3 3.90367 0.19665 -6.21742
-2.37651 5.99149 0.00000 ; CA- CB- CG- CD1
5140 5142 5145 5151 3 3.90367 0.19665 -6.21742
-2.37651 5.99149 0.00000 ; CA- CB- CG- CD2
5140 5155 5157 5158 3 20.92000 0.00000 -20.92000
0.00000 0.00000 0.00000 ; CA- C- N- H
5140 5155 5157 5159 3 20.92000 0.00000 -20.92000
0.00000 0.00000 0.00000 ; CA- C- N- CA
5141 5140 5142 5143 3 0.65084 1.95253 0.00000
-2.60338 0.00000 0.00000 ; HA- CA- CB- HB2
5141 5140 5142 5144 3 0.65084 1.95253 0.00000
-2.60338 0.00000 0.00000 ; HA- CA- CB- HB3
5141 5140 5142 5145 3 0.65084 1.95253 0.00000
-2.60338 0.00000 0.00000 ; HA- CA- CB- CG
5141 5140 5155 5156 3 3.68192 -4.35136 0.00000
1.33888 0.00000 0.00000 ; HA- CA- C- O
5141 5140 5155 5157 3 0.00000 0.00000 0.00000

0.00000 0.00000 0.00000 ; HA- CA- C- N
5142 5140 5155 5156 3 0.00000 0.00000 0.00000
0.00000 0.00000 0.00000 ; CB- CA- C- O
5142 5140 5155 5157 3 2.51040 4.18400 1.67360
-6.69440 0.00000 0.00000 ; CB- CA- C- N
5142 5145 5147 5148 3 0.66944 2.00832 0.00000
-2.67776 0.00000 0.00000 ; CB- CG- CD1- HD11
5142 5145 5147 5149 3 0.66944 2.00832 0.00000
-2.67776 0.00000 0.00000 ; CB- CG- CD1- HD12
5142 5145 5147 5150 3 0.66944 2.00832 0.00000
-2.67776 0.00000 0.00000 ; CB- CG- CD1- HD13
5142 5145 5151 5152 3 0.66944 2.00832 0.00000
-2.67776 0.00000 0.00000 ; CB- CG- CD2- HD21
5142 5145 5151 5153 3 0.66944 2.00832 0.00000
-2.67776 0.00000 0.00000 ; CB- CG- CD2- HD22
5142 5145 5151 5154 3 0.66944 2.00832 0.00000
-2.67776 0.00000 0.00000 ; CB- CG- CD2- HD23
5143 5142 5140 5155 3 0.65084 1.95253 0.00000
-2.60338 0.00000 0.00000 ; HB2- CB- CA- C
5143 5142 5145 5146 3 0.62760 1.88280 0.00000
-2.51040 0.00000 0.00000 ; HB2- CB- CG- HG
5143 5142 5145 5147 3 0.66944 2.00832 0.00000
-2.67776 0.00000 0.00000 ; HB2- CB- CG- CD1
5143 5142 5145 5151 3 0.66944 2.00832 0.00000
-2.67776 0.00000 0.00000 ; HB2- CB- CG- CD2
5144 5142 5140 5155 3 0.65084 1.95253 0.00000
-2.60338 0.00000 0.00000 ; HB3- CB- CA- C
5144 5142 5145 5146 3 0.62760 1.88280 0.00000
-2.51040 0.00000 0.00000 ; HB3- CB- CG- HG
5144 5142 5145 5147 3 0.66944 2.00832 0.00000
-2.67776 0.00000 0.00000 ; HB3- CB- CG- CD1
5144 5142 5145 5151 3 0.66944 2.00832 0.00000
-2.67776 0.00000 0.00000 ; HB3- CB- CG- CD2
5145 5142 5140 5155 3 10.33448 -1.14642 -11.54784
-2.40998 6.35968 0.00000 ; CG- CB- CA- C
5146 5145 5147 5148 3 0.62760 1.88280 0.00000
-2.51040 0.00000 0.00000 ; HG- CG- CD1- HD11
5146 5145 5147 5149 3 0.62760 1.88280 0.00000
-2.51040 0.00000 0.00000 ; HG- CG- CD1- HD12
5146 5145 5147 5150 3 0.62760 1.88280 0.00000
-2.51040 0.00000 0.00000 ; HG- CG- CD1- HD13
5146 5145 5151 5152 3 0.62760 1.88280 0.00000
-2.51040 0.00000 0.00000 ; HG- CG- CD2- HD21
5146 5145 5151 5153 3 0.62760 1.88280 0.00000
-2.51040 0.00000 0.00000 ; HG- CG- CD2- HD22
5146 5145 5151 5154 3 0.62760 1.88280 0.00000
-2.51040 0.00000 0.00000 ; HG- CG- CD2- HD23
5147 5145 5151 5152 3 0.66944 2.00832 0.00000
-2.67776 0.00000 0.00000 ; CD1- CG- CD2- HD21
5147 5145 5151 5153 3 0.66944 2.00832 0.00000
-2.67776 0.00000 0.00000 ; CD1- CG- CD2- HD22
5147 5145 5151 5154 3 0.66944 2.00832 0.00000

-2.67776 0.00000 0.00000 ; CD1- CG- CD2- HD23
5148 5147 5145 5151 3 0.66944 2.00832 0.00000
-2.67776 0.00000 0.00000 ; HD11- CD1- CG- CD2
5149 5147 5145 5151 3 0.66944 2.00832 0.00000
-2.67776 0.00000 0.00000 ; HD12- CD1- CG- CD2
5150 5147 5145 5151 3 0.66944 2.00832 0.00000
-2.67776 0.00000 0.00000 ; HD13- CD1- CG- CD2
5155 5157 5159 5160 3 0.00000 0.00000 0.00000
0.00000 0.00000 0.00000 ; C- N- CA- HA
5155 5157 5159 5161 3 11.71520 1.67360 15.06240 -
13.38880 0.00000 0.00000 ; C- N- CA- CB
5155 5157 5159 5179 3 1.75728 5.27184 2.25936
-7.02912 0.00000 0.00000 ; C- N- CA- C
5156 5155 5157 5158 3 29.28800 -8.36800 -20.92000
0.00000 0.00000 0.00000 ; O- C- N- H
5156 5155 5157 5159 3 20.92000 0.00000 -20.92000
0.00000 0.00000 0.00000 ; O- C- N- CA
5157 5159 5161 5162 3 0.65084 1.95253 0.00000
-2.60338 0.00000 0.00000 ; N- CA- CB- HB2
5157 5159 5161 5163 3 0.65084 1.95253 0.00000
-2.60338 0.00000 0.00000 ; N- CA- CB- HB3
5157 5159 5161 5164 3 0.65084 1.95253 0.00000
-2.60338 0.00000 0.00000 ; N- CA- CB- CG
5157 5159 5179 5180 3 0.00000 0.00000 0.00000
0.00000 0.00000 0.00000 ; N- CA- C- O
5157 5159 5179 5181 3 17.40544 -5.02080 -13.22144
9.20480 0.00000 0.00000 ; N- CA- C- N
5158 5157 5159 5160 3 0.00000 0.00000 0.00000
0.00000 0.00000 0.00000 ; H- N- CA- HA
5158 5157 5159 5161 3 0.00000 0.00000 0.00000
0.00000 0.00000 0.00000 ; H- N- CA- CB
5158 5157 5159 5179 3 0.00000 0.00000 0.00000
0.00000 0.00000 0.00000 ; H- N- CA- C
5159 5161 5164 5165 3 0.66944 2.00832 0.00000
-2.67776 0.00000 0.00000 ; CA- CB- CG- HG2
5159 5161 5164 5166 3 0.66944 2.00832 0.00000
-2.67776 0.00000 0.00000 ; CA- CB- CG- HG3
5159 5161 5164 5167 3 3.68192 3.09616 -2.09200
-3.01248 0.00000 0.00000 ; CA- CB- CG- CD
5159 5179 5181 5182 3 20.92000 0.00000 -20.92000
0.00000 0.00000 0.00000 ; CA- C- N- H
5159 5179 5181 5183 3 20.92000 0.00000 -20.92000
0.00000 0.00000 0.00000 ; CA- C- N- CA
5160 5159 5161 5162 3 0.65084 1.95253 0.00000
-2.60338 0.00000 0.00000 ; HA- CA- CB- HB2
5160 5159 5161 5163 3 0.65084 1.95253 0.00000
-2.60338 0.00000 0.00000 ; HA- CA- CB- HB3
5160 5159 5161 5164 3 0.65084 1.95253 0.00000
-2.60338 0.00000 0.00000 ; HA- CA- CB- CG
5160 5159 5179 5180 3 3.68192 -4.35136 0.00000
1.33888 0.00000 0.00000 ; HA- CA- C- O
5160 5159 5179 5181 3 0.00000 0.00000 0.00000

```
0.00000    0.00000    0.00000 ;     HA-    CA-     C-     N
  5161   5159   5179   5180      3    0.00000    0.00000    0.00000
0.00000    0.00000    0.00000 ;     CB-    CA-     C-     O
  5161   5159   5179   5181      3    2.51040    4.18400    1.67360
-6.69440    0.00000    0.00000 ;     CB-    CA-     C-     N
  5161   5164   5167   5168      3    0.65084    1.95253    0.00000
-2.60338    0.00000    0.00000 ;     CB-    CG-    CD-   HD2
  5161   5164   5167   5169      3    0.65084    1.95253    0.00000
-2.60338    0.00000    0.00000 ;     CB-    CG-    CD-   HD3
  5161   5164   5167   5170      3    0.65084    1.95253    0.00000
-2.60338    0.00000    0.00000 ;     CB-    CG-    CD-    NE
  5162   5161   5159   5179      3    0.65084    1.95253    0.00000
-2.60338    0.00000    0.00000 ;    HB2-    CB-    CA-     C
  5162   5161   5164   5165      3    0.62760    1.88280    0.00000
-2.51040    0.00000    0.00000 ;    HB2-    CB-    CG-   HG2
  5162   5161   5164   5166      3    0.62760    1.88280    0.00000
-2.51040    0.00000    0.00000 ;    HB2-    CB-    CG-   HG3
  5162   5161   5164   5167      3    0.66944    2.00832    0.00000
-2.67776    0.00000    0.00000 ;    HB2-    CB-    CG-    CD
  5163   5161   5159   5179      3    0.65084    1.95253    0.00000
-2.60338    0.00000    0.00000 ;    HB3-    CB-    CA-     C
  5163   5161   5164   5165      3    0.62760    1.88280    0.00000
-2.51040    0.00000    0.00000 ;    HB3-    CB-    CG-   HG2
  5163   5161   5164   5166      3    0.62760    1.88280    0.00000
-2.51040    0.00000    0.00000 ;    HB3-    CB-    CG-   HG3
  5163   5161   5164   5167      3    0.66944    2.00832    0.00000
-2.67776    0.00000    0.00000 ;    HB3-    CB-    CG-    CD
  5164   5161   5159   5179      3    0.65084    1.95253    0.00000
-2.60338    0.00000    0.00000 ;     CG-    CB-    CA-     C
  5164   5167   5170   5171      3    0.00000    0.00000    0.00000
0.00000    0.00000    0.00000 ;     CG-    CD-    NE-    HE
  5164   5167   5170   5172      3    0.00000    0.00000    0.00000
0.00000    0.00000    0.00000 ;     CG-    CD-    NE-    CZ
  5165   5164   5167   5168      3    0.65084    1.95253    0.00000
-2.60338    0.00000    0.00000 ;    HG2-    CG-    CD-   HD2
  5165   5164   5167   5169      3    0.65084    1.95253    0.00000
-2.60338    0.00000    0.00000 ;    HG2-    CG-    CD-   HD3
  5165   5164   5167   5170      3    0.65084    1.95253    0.00000
-2.60338    0.00000    0.00000 ;    HG2-    CG-    CD-    NE
  5166   5164   5167   5168      3    0.65084    1.95253    0.00000
-2.60338    0.00000    0.00000 ;    HG3-    CG-    CD-   HD2
  5166   5164   5167   5169      3    0.65084    1.95253    0.00000
-2.60338    0.00000    0.00000 ;    HG3-    CG-    CD-   HD3
  5166   5164   5167   5170      3    0.65084    1.95253    0.00000
-2.60338    0.00000    0.00000 ;    HG3-    CG-    CD-    NE
  5167   5170   5172   5173      3   20.08320    0.00000  -20.08320
0.00000    0.00000    0.00000 ;     CD-    NE-    CZ-   NH1
  5167   5170   5172   5176      3   20.08320    0.00000  -20.08320
0.00000    0.00000    0.00000 ;     CD-    NE-    CZ-   NH2
  5168   5167   5170   5171      3    0.00000    0.00000    0.00000
0.00000    0.00000    0.00000 ;    HD2-    CD-    NE-    HE
  5168   5167   5170   5172      3    0.00000    0.00000    0.00000
```

0.00000    0.00000    0.00000 ;    HD2-    CD-    NE-    CZ
  5169   5167   5170   5171      3    0.00000    0.00000    0.00000
0.00000    0.00000    0.00000 ;    HD3-    CD-    NE-    HE
  5169   5167   5170   5172      3    0.00000    0.00000    0.00000
0.00000    0.00000    0.00000 ;    HD3-    CD-    NE-    CZ
  5170   5172   5173   5174      3   20.08320    0.00000  -20.08320
0.00000    0.00000    0.00000 ;     NE-    CZ-   NH1-  HH11
  5170   5172   5173   5175      3   20.08320    0.00000  -20.08320
0.00000    0.00000    0.00000 ;     NE-    CZ-   NH1-  HH12
  5170   5172   5176   5177      3   20.08320    0.00000  -20.08320
0.00000    0.00000    0.00000 ;     NE-    CZ-   NH2-  HH21
  5170   5172   5176   5178      3   20.08320    0.00000  -20.08320
0.00000    0.00000    0.00000 ;     NE-    CZ-   NH2-  HH22
  5171   5170   5172   5173      3   20.08320    0.00000  -20.08320
0.00000    0.00000    0.00000 ;     HE-    NE-    CZ-   NH1
  5171   5170   5172   5176      3   20.08320    0.00000  -20.08320
0.00000    0.00000    0.00000 ;     HE-    NE-    CZ-   NH2
  5173   5172   5176   5177      3   20.08320    0.00000  -20.08320
0.00000    0.00000    0.00000 ;    NH1-    CZ-   NH2-  HH21
  5173   5172   5176   5178      3   20.08320    0.00000  -20.08320
0.00000    0.00000    0.00000 ;    NH1-    CZ-   NH2-  HH22
  5174   5173   5172   5176      3   20.08320    0.00000  -20.08320
0.00000    0.00000    0.00000 ;   HH11-   NH1-    CZ-   NH2
  5175   5173   5172   5176      3   20.08320    0.00000  -20.08320
0.00000    0.00000    0.00000 ;   HH12-   NH1-    CZ-   NH2
  5179   5181   5183   5184      3    0.00000    0.00000    0.00000
0.00000    0.00000    0.00000 ;      C-     N-    CA-    HA
  5179   5181   5183   5185      3   11.71520    1.67360   15.06240  -
13.38880    0.00000    0.00000 ;      C-     N-    CA-    CB
  5179   5181   5183   5193      3    1.75728    5.27184    2.25936
-7.02912    0.00000    0.00000 ;      C-     N-    CA-     C
  5180   5179   5181   5182      3   29.28800   -8.36800  -20.92000
0.00000    0.00000    0.00000 ;      O-     C-     N-     H
  5180   5179   5181   5183      3   20.92000    0.00000  -20.92000
0.00000    0.00000    0.00000 ;      O-     C-     N-    CA
  5181   5183   5185   5186      3    0.65084    1.95253    0.00000
-2.60338    0.00000    0.00000 ;      N-    CA-    CB-   HB2
  5181   5183   5185   5187      3    0.65084    1.95253    0.00000
-2.60338    0.00000    0.00000 ;      N-    CA-    CB-   HB3
  5181   5183   5185   5188      3    5.99567    3.29281   -4.46014
-0.55229    1.97485    0.00000 ;      N-    CA-    CB-    CG
  5181   5183   5193   5194      3    0.00000    0.00000    0.00000
0.00000    0.00000    0.00000 ;      N-    CA-     C-     O
  5181   5183   5193   5195      3   17.40544   -5.02080  -13.22144
9.20480    0.00000    0.00000 ;      N-    CA-     C-     N
  5182   5181   5183   5184      3    0.00000    0.00000    0.00000
0.00000    0.00000    0.00000 ;      H-     N-    CA-    HA
  5182   5181   5183   5185      3    0.00000    0.00000    0.00000
0.00000    0.00000    0.00000 ;      H-     N-    CA-    CB
  5182   5181   5183   5193      3    0.00000    0.00000    0.00000
0.00000    0.00000    0.00000 ;      H-     N-    CA-     C
  5183   5185   5188   5189      3    0.00000    0.00000    0.00000

0.00000 0.00000 0.00000 ; CA- CB- CG- OD1
5183 5185 5188 5190 3 8.84916 -0.31380 -4.32626
5.03754 0.26778 0.00000 ; CA- CB- CG- ND2
5183 5193 5195 5196 3 20.92000 0.00000 -20.92000
0.00000 0.00000 0.00000 ; CA- C- N- H
5183 5193 5195 5197 3 20.92000 0.00000 -20.92000
0.00000 0.00000 0.00000 ; CA- C- N- CA
5184 5183 5185 5186 3 0.65084 1.95253 0.00000
-2.60338 0.00000 0.00000 ; HA- CA- CB- HB2
5184 5183 5185 5187 3 0.65084 1.95253 0.00000
-2.60338 0.00000 0.00000 ; HA- CA- CB- HB3
5184 5183 5185 5188 3 0.65084 1.95253 0.00000
-2.60338 0.00000 0.00000 ; HA- CA- CB- CG
5184 5183 5193 5194 3 3.68192 -4.35136 0.00000
1.33888 0.00000 0.00000 ; HA- CA- C- O
5184 5183 5193 5195 3 0.00000 0.00000 0.00000
0.00000 0.00000 0.00000 ; HA- CA- C- N
5185 5183 5193 5194 3 0.00000 0.00000 0.00000
0.00000 0.00000 0.00000 ; CB- CA- C- O
5185 5183 5193 5195 3 2.51040 4.18400 1.67360
-6.69440 0.00000 0.00000 ; CB- CA- C- N
5185 5188 5190 5191 3 20.92000 0.00000 -20.92000
0.00000 0.00000 0.00000 ; CB- CG- ND2- HD21
5185 5188 5190 5192 3 20.92000 0.00000 -20.92000
0.00000 0.00000 0.00000 ; CB- CG- ND2- HD22
5186 5185 5183 5193 3 0.65084 1.95253 0.00000
-2.60338 0.00000 0.00000 ; HB2- CB- CA- C
5186 5185 5188 5189 3 3.68192 -4.35136 0.00000
1.33888 0.00000 0.00000 ; HB2- CB- CG- OD1
5186 5185 5188 5190 3 0.00000 0.00000 0.00000
0.00000 0.00000 0.00000 ; HB2- CB- CG- ND2
5187 5185 5183 5193 3 0.65084 1.95253 0.00000
-2.60338 0.00000 0.00000 ; HB3- CB- CA- C
5187 5185 5188 5189 3 3.68192 -4.35136 0.00000
1.33888 0.00000 0.00000 ; HB3- CB- CG- OD1
5187 5185 5188 5190 3 0.00000 0.00000 0.00000
0.00000 0.00000 0.00000 ; HB3- CB- CG- ND2
5188 5185 5183 5193 3 7.94542 -3.96225 -6.11701
-0.55229 3.58150 0.00000 ; CG- CB- CA- C
5189 5188 5190 5191 3 29.28800 -8.36800 -20.92000
0.00000 0.00000 0.00000 ; OD1- CG- ND2- HD21
5189 5188 5190 5192 3 29.28800 -8.36800 -20.92000
0.00000 0.00000 0.00000 ; OD1- CG- ND2- HD22
5193 5195 5197 5198 3 0.00000 0.00000 0.00000
0.00000 0.00000 0.00000 ; C- N- CA- HA
5193 5195 5197 5199 3 11.71520 1.67360 15.06240 -
13.38880 0.00000 0.00000 ; C- N- CA- CB
5193 5195 5197 5212 3 1.75728 5.27184 2.25936
-7.02912 0.00000 0.00000 ; C- N- CA- C
5194 5193 5195 5196 3 29.28800 -8.36800 -20.92000
0.00000 0.00000 0.00000 ; O- C- N- H
5194 5193 5195 5197 3 20.92000 0.00000 -20.92000

0.00000    0.00000    0.00000 ;      O-     C-     N-    CA
 5195   5197   5199   5200       3    0.65084    1.95253    0.00000
-2.60338    0.00000    0.00000 ;      N-    CA-    CB-    HB
 5195   5197   5199   5201       3    3.83673    0.44769   -1.77402
-2.47693   -0.03347    0.00000 ;      N-    CA-    CB-   CG2
 5195   5197   5199   5205       3    2.97482    0.12134    2.04179
-1.89117   -3.24678    0.00000 ;      N-    CA-    CB-   CG1
 5195   5197   5212   5213       3    0.00000    0.00000    0.00000
0.00000    0.00000    0.00000 ;      N-    CA-     C-     O
 5195   5197   5212   5214       3   17.40544   -5.02080  -13.22144
9.20480    0.00000    0.00000 ;      N-    CA-     C-     N
 5196   5195   5197   5198       3    0.00000    0.00000    0.00000
0.00000    0.00000    0.00000 ;      H-     N-    CA-    HA
 5196   5195   5197   5199       3    0.00000    0.00000    0.00000
0.00000    0.00000    0.00000 ;      H-     N-    CA-    CB
 5196   5195   5197   5212       3    0.00000    0.00000    0.00000
0.00000    0.00000    0.00000 ;      H-     N-    CA-     C
 5197   5199   5201   5202       3    0.66944    2.00832    0.00000
-2.67776    0.00000    0.00000 ;     CA-    CB-   CG2-  HG21
 5197   5199   5201   5203       3    0.66944    2.00832    0.00000
-2.67776    0.00000    0.00000 ;     CA-    CB-   CG2-  HG22
 5197   5199   5201   5204       3    0.66944    2.00832    0.00000
-2.67776    0.00000    0.00000 ;     CA-    CB-   CG2-  HG23
 5197   5199   5205   5206       3    0.66944    2.00832    0.00000
-2.67776    0.00000    0.00000 ;     CA-    CB-   CG1-  HG12
 5197   5199   5205   5207       3    0.66944    2.00832    0.00000
-2.67776    0.00000    0.00000 ;     CA-    CB-   CG1-  HG13
 5197   5199   5205   5208       3    4.24258   -0.52718   -7.25506
-1.79075    7.69856    0.00000 ;     CA-    CB-   CG1-   CD1
 5197   5212   5214   5215       3   20.92000    0.00000  -20.92000
0.00000    0.00000    0.00000 ;     CA-     C-     N-     H
 5197   5212   5214   5216       3   20.92000    0.00000  -20.92000
0.00000    0.00000    0.00000 ;     CA-     C-     N-    CA
 5198   5197   5199   5200       3    0.65084    1.95253    0.00000
-2.60338    0.00000    0.00000 ;     HA-    CA-    CB-    HB
 5198   5197   5199   5201       3    0.65084    1.95253    0.00000
-2.60338    0.00000    0.00000 ;     HA-    CA-    CB-   CG2
 5198   5197   5199   5205       3    0.65084    1.95253    0.00000
-2.60338    0.00000    0.00000 ;     HA-    CA-    CB-   CG1
 5198   5197   5212   5213       3    3.68192   -4.35136    0.00000
1.33888    0.00000    0.00000 ;     HA-    CA-     C-     O
 5198   5197   5212   5214       3    0.00000    0.00000    0.00000
0.00000    0.00000    0.00000 ;     HA-    CA-     C-     N
 5199   5197   5212   5213       3    0.00000    0.00000    0.00000
0.00000    0.00000    0.00000 ;     CB-    CA-     C-     O
 5199   5197   5212   5214       3    2.51040    4.18400    1.67360
-6.69440    0.00000    0.00000 ;     CB-    CA-     C-     N
 5199   5205   5208   5209       3    0.66944    2.00832    0.00000
-2.67776    0.00000    0.00000 ;     CB-   CG1-   CD1-  HD11
 5199   5205   5208   5210       3    0.66944    2.00832    0.00000
-2.67776    0.00000    0.00000 ;     CB-   CG1-   CD1-  HD12
 5199   5205   5208   5211       3    0.66944    2.00832    0.00000

-2.67776 0.00000 0.00000 ; CB- CG1- CD1- HD13
5200 5199 5197 5212 3 0.65084 1.95253 0.00000
-2.60338 0.00000 0.00000 ; HB- CB- CA- C
5200 5199 5201 5202 3 0.62760 1.88280 0.00000
-2.51040 0.00000 0.00000 ; HB- CB- CG2- HG21
5200 5199 5201 5203 3 0.62760 1.88280 0.00000
-2.51040 0.00000 0.00000 ; HB- CB- CG2- HG22
5200 5199 5201 5204 3 0.62760 1.88280 0.00000
-2.51040 0.00000 0.00000 ; HB- CB- CG2- HG23
5200 5199 5205 5206 3 0.62760 1.88280 0.00000
-2.51040 0.00000 0.00000 ; HB- CB- CG1- HG12
5200 5199 5205 5207 3 0.62760 1.88280 0.00000
-2.51040 0.00000 0.00000 ; HB- CB- CG1- HG13
5200 5199 5205 5208 3 0.66944 2.00832 0.00000
-2.67776 0.00000 0.00000 ; HB- CB- CG1- CD1
5201 5199 5197 5212 3 5.67350 3.55640 -6.16722
-2.47693 3.74886 0.00000 ; CG2- CB- CA- C
5201 5199 5205 5206 3 0.66944 2.00832 0.00000
-2.67776 0.00000 0.00000 ; CG2- CB- CG1- HG12
5201 5199 5205 5207 3 0.66944 2.00832 0.00000
-2.67776 0.00000 0.00000 ; CG2- CB- CG1- HG13
5201 5199 5205 5208 3 3.81162 0.49790 -8.14206
-1.79075 7.49773 0.00000 ; CG2- CB- CG1- CD1
5202 5201 5199 5205 3 0.66944 2.00832 0.00000
-2.67776 0.00000 0.00000 ; HG21- CG2- CB- CG1
5203 5201 5199 5205 3 0.66944 2.00832 0.00000
-2.67776 0.00000 0.00000 ; HG22- CG2- CB- CG1
5204 5201 5199 5205 3 0.66944 2.00832 0.00000
-2.67776 0.00000 0.00000 ; HG23- CG2- CB- CG1
5205 5199 5197 5212 3 8.26340 0.74057 -9.99976
-1.89117 3.84928 0.00000 ; CG1- CB- CA- C
5206 5205 5208 5209 3 0.62760 1.88280 0.00000
-2.51040 0.00000 0.00000 ; HG12- CG1- CD1- HD11
5206 5205 5208 5210 3 0.62760 1.88280 0.00000
-2.51040 0.00000 0.00000 ; HG12- CG1- CD1- HD12
5206 5205 5208 5211 3 0.62760 1.88280 0.00000
-2.51040 0.00000 0.00000 ; HG12- CG1- CD1- HD13
5207 5205 5208 5209 3 0.62760 1.88280 0.00000
-2.51040 0.00000 0.00000 ; HG13- CG1- CD1- HD11
5207 5205 5208 5210 3 0.62760 1.88280 0.00000
-2.51040 0.00000 0.00000 ; HG13- CG1- CD1- HD12
5207 5205 5208 5211 3 0.62760 1.88280 0.00000
-2.51040 0.00000 0.00000 ; HG13- CG1- CD1- HD13
5212 5214 5216 5217 3 0.00000 0.00000 0.00000
0.00000 0.00000 0.00000 ; C- N- CA- HA
5212 5214 5216 5218 3 11.71520 1.67360 15.06240 -
13.38880 0.00000 0.00000 ; C- N- CA- CB
5212 5214 5216 5232 3 1.75728 5.27184 2.25936
-7.02912 0.00000 0.00000 ; C- N- CA- C
5213 5212 5214 5215 3 29.28800 -8.36800 -20.92000
0.00000 0.00000 0.00000 ; O- C- N- H
5213 5212 5214 5216 3 20.92000 0.00000 -20.92000

```
0.00000    0.00000    0.00000 ;       O-     C-     N-    CA
  5214   5216   5218   5219      3    0.65084    1.95253    0.00000
-2.60338    0.00000    0.00000 ;       N-    CA-    CB-   HB2
  5214   5216   5218   5220      3    0.65084    1.95253    0.00000
-2.60338    0.00000    0.00000 ;       N-    CA-    CB-   HB3
  5214   5216   5218   5221      3    3.28026    2.46019   -2.19242
-3.21331   -0.23430    0.00000 ;       N-    CA-    CB-    CG
  5214   5216   5232   5233      3    0.00000    0.00000    0.00000
0.00000    0.00000    0.00000 ;       N-    CA-     C-     O
  5214   5216   5232   5234      3   17.40544   -5.02080  -13.22144
9.20480    0.00000    0.00000 ;       N-    CA-     C-     N
  5215   5214   5216   5217      3    0.00000    0.00000    0.00000
0.00000    0.00000    0.00000 ;       H-     N-    CA-    HA
  5215   5214   5216   5218      3    0.00000    0.00000    0.00000
0.00000    0.00000    0.00000 ;       H-     N-    CA-    CB
  5215   5214   5216   5232      3    0.00000    0.00000    0.00000
0.00000    0.00000    0.00000 ;       H-     N-    CA-     C
  5216   5218   5221   5222      3    0.57739    0.00000    1.02926
0.00000   -1.60666    0.00000 ;     CA-    CB-    CG-   CD1
  5216   5218   5221   5230      3    0.57739    0.00000    1.02926
0.00000   -1.60666    0.00000 ;     CA-    CB-    CG-   CD2
  5216   5232   5234   5235      3   20.92000    0.00000  -20.92000
0.00000    0.00000    0.00000 ;     CA-     C-     N-     H
  5216   5232   5234   5236      3   20.92000    0.00000  -20.92000
0.00000    0.00000    0.00000 ;     CA-     C-     N-    CA
  5217   5216   5218   5219      3    0.65084    1.95253    0.00000
-2.60338    0.00000    0.00000 ;      HA-    CA-    CB-   HB2
  5217   5216   5218   5220      3    0.65084    1.95253    0.00000
-2.60338    0.00000    0.00000 ;      HA-    CA-    CB-   HB3
  5217   5216   5218   5221      3    0.65084    1.95253    0.00000
-2.60338    0.00000    0.00000 ;      HA-    CA-    CB-    CG
  5217   5216   5232   5233      3    3.68192   -4.35136    0.00000
1.33888    0.00000    0.00000 ;     HA-    CA-     C-     O
  5217   5216   5232   5234      3    0.00000    0.00000    0.00000
0.00000    0.00000    0.00000 ;     HA-    CA-     C-     N
  5218   5216   5232   5233      3    0.00000    0.00000    0.00000
0.00000    0.00000    0.00000 ;     CB-    CA-     C-     O
  5218   5216   5232   5234      3    2.51040    4.18400    1.67360
-6.69440    0.00000    0.00000 ;      CB-    CA-     C-     N
  5218   5221   5222   5223      3   30.33400    0.00000  -30.33400
0.00000    0.00000    0.00000 ;     CB-    CG-   CD1-   HD1
  5218   5221   5222   5224      3   30.33400    0.00000  -30.33400
0.00000    0.00000    0.00000 ;     CB-    CG-   CD1-   CE1
  5218   5221   5230   5228      3   30.33400    0.00000  -30.33400
0.00000    0.00000    0.00000 ;     CB-    CG-   CD2-   CE2
  5218   5221   5230   5231      3   30.33400    0.00000  -30.33400
0.00000    0.00000    0.00000 ;     CB-    CG-   CD2-   HD2
  5219   5218   5216   5232      3    0.65084    1.95253    0.00000
-2.60338    0.00000    0.00000 ;     HB2-    CB-    CA-     C
  5219   5218   5221   5222      3    0.00000    0.00000    0.00000
0.00000    0.00000    0.00000 ;    HB2-    CB-    CG-   CD1
  5219   5218   5221   5230      3    0.00000    0.00000    0.00000
```

```
0.00000    0.00000    0.00000 ;    HB2-    CB-    CG-   CD2
  5220   5218   5216   5232      3    0.65084    1.95253    0.00000
-2.60338    0.00000    0.00000 ;    HB3-    CB-    CA-     C
  5220   5218   5221   5222      3    0.00000    0.00000    0.00000
0.00000    0.00000    0.00000 ;    HB3-    CB-    CG-   CD1
  5220   5218   5221   5230      3    0.00000    0.00000    0.00000
0.00000    0.00000    0.00000 ;    HB3-    CB-    CG-   CD2
  5221   5218   5216   5232      3    4.95804    2.17986   -3.52293
-3.21331   -0.40166    0.00000 ;    CG-    CB-    CA-     C
  5221   5222   5224   5225      3   30.33400    0.00000  -30.33400
0.00000    0.00000    0.00000 ;    CG-   CD1-   CE1-   HE1
  5221   5222   5224   5226      3   30.33400    0.00000  -30.33400
0.00000    0.00000    0.00000 ;    CG-   CD1-   CE1-    CZ
  5221   5230   5228   5226      3   30.33400    0.00000  -30.33400
0.00000    0.00000    0.00000 ;    CG-   CD2-   CE2-    CZ
  5221   5230   5228   5229      3   30.33400    0.00000  -30.33400
0.00000    0.00000    0.00000 ;    CG-   CD2-   CE2-   HE2
  5222   5221   5230   5228      3   30.33400    0.00000  -30.33400
0.00000    0.00000    0.00000 ;   CD1-    CG-   CD2-   CE2
  5222   5221   5230   5231      3   30.33400    0.00000  -30.33400
0.00000    0.00000    0.00000 ;   CD1-    CG-   CD2-   HD2
  5222   5224   5226   5227      3   30.33400    0.00000  -30.33400
0.00000    0.00000    0.00000 ;   CD1-   CE1-    CZ-    HZ
  5222   5224   5226   5228      3   30.33400    0.00000  -30.33400
0.00000    0.00000    0.00000 ;   CD1-   CE1-    CZ-   CE2
  5223   5222   5221   5230      3   30.33400    0.00000  -30.33400
0.00000    0.00000    0.00000 ;   HD1-   CD1-    CG-   CD2
  5223   5222   5224   5225      3   30.33400    0.00000  -30.33400
0.00000    0.00000    0.00000 ;   HD1-   CD1-   CE1-   HE1
  5223   5222   5224   5226      3   30.33400    0.00000  -30.33400
0.00000    0.00000    0.00000 ;   HD1-   CD1-   CE1-    CZ
  5224   5222   5221   5230      3   30.33400    0.00000  -30.33400
0.00000    0.00000    0.00000 ;   CE1-   CD1-    CG-   CD2
  5224   5226   5228   5229      3   30.33400    0.00000  -30.33400
0.00000    0.00000    0.00000 ;   CE1-    CZ-   CE2-   HE2
  5224   5226   5228   5230      3   30.33400    0.00000  -30.33400
0.00000    0.00000    0.00000 ;   CE1-    CZ-   CE2-   CD2
  5225   5224   5226   5227      3   30.33400    0.00000  -30.33400
0.00000    0.00000    0.00000 ;   HE1-   CE1-    CZ-    HZ
  5225   5224   5226   5228      3   30.33400    0.00000  -30.33400
0.00000    0.00000    0.00000 ;   HE1-   CE1-    CZ-   CE2
  5226   5228   5230   5231      3   30.33400    0.00000  -30.33400
0.00000    0.00000    0.00000 ;    CZ-   CE2-   CD2-   HD2
  5227   5226   5228   5229      3   30.33400    0.00000  -30.33400
0.00000    0.00000    0.00000 ;    HZ-    CZ-   CE2-   HE2
  5227   5226   5228   5230      3   30.33400    0.00000  -30.33400
0.00000    0.00000    0.00000 ;    HZ-    CZ-   CE2-   CD2
  5229   5228   5230   5231      3   30.33400    0.00000  -30.33400
0.00000    0.00000    0.00000 ;   HE2-   CE2-   CD2-   HD2
  5232   5234   5236   5237      3    0.00000    0.00000    0.00000
0.00000    0.00000    0.00000 ;     C-     N-    CA-    HA
  5232   5234   5236   5238      3   11.71520    1.67360   15.06240  -
```

```
13.38880    0.00000    0.00000 ;       C-     N-    CA-    CB
  5232   5234   5236   5251       3    1.75728    5.27184    2.25936
-7.02912    0.00000    0.00000 ;       C-     N-    CA-     C
  5233   5232   5234   5235       3   29.28800   -8.36800  -20.92000
0.00000    0.00000    0.00000 ;       O-     C-     N-     H
  5233   5232   5234   5236       3   20.92000    0.00000  -20.92000
0.00000    0.00000    0.00000 ;       O-     C-     N-    CA
  5234   5236   5238   5239       3    0.65084    1.95253    0.00000
-2.60338    0.00000    0.00000 ;       N-    CA-    CB-    HB
  5234   5236   5238   5240       3    3.83673    0.44769   -1.77402
-2.47693   -0.03347    0.00000 ;       N-    CA-    CB-   CG2
  5234   5236   5238   5244       3    2.97482    0.12134    2.04179
-1.89117   -3.24678    0.00000 ;       N-    CA-    CB-   CG1
  5234   5236   5251   5252       3    0.00000    0.00000    0.00000
0.00000    0.00000    0.00000 ;       N-    CA-     C-     O
  5234   5236   5251   5253       3   17.40544   -5.02080  -13.22144
9.20480    0.00000    0.00000 ;       N-    CA-     C-     N
  5235   5234   5236   5237       3    0.00000    0.00000    0.00000
0.00000    0.00000    0.00000 ;       H-     N-    CA-    HA
  5235   5234   5236   5238       3    0.00000    0.00000    0.00000
0.00000    0.00000    0.00000 ;       H-     N-    CA-    CB
  5235   5234   5236   5251       3    0.00000    0.00000    0.00000
0.00000    0.00000    0.00000 ;       H-     N-    CA-     C
  5236   5238   5240   5241       3    0.66944    2.00832    0.00000
-2.67776    0.00000    0.00000 ;      CA-    CB-   CG2-  HG21
  5236   5238   5240   5242       3    0.66944    2.00832    0.00000
-2.67776    0.00000    0.00000 ;      CA-    CB-   CG2-  HG22
  5236   5238   5240   5243       3    0.66944    2.00832    0.00000
-2.67776    0.00000    0.00000 ;      CA-    CB-   CG2-  HG23
  5236   5238   5244   5245       3    0.66944    2.00832    0.00000
-2.67776    0.00000    0.00000 ;      CA-    CB-   CG1-  HG12
  5236   5238   5244   5246       3    0.66944    2.00832    0.00000
-2.67776    0.00000    0.00000 ;      CA-    CB-   CG1-  HG13
  5236   5238   5244   5247       3    4.24258   -0.52718   -7.25506
-1.79075    7.69856    0.00000 ;      CA-    CB-   CG1-   CD1
  5236   5251   5253   5254       3   20.92000    0.00000  -20.92000
0.00000    0.00000    0.00000 ;      CA-     C-     N-     H
  5236   5251   5253   5255       3   20.92000    0.00000  -20.92000
0.00000    0.00000    0.00000 ;      CA-     C-     N-    CA
  5237   5236   5238   5239       3    0.65084    1.95253    0.00000
-2.60338    0.00000    0.00000 ;      HA-    CA-    CB-    HB
  5237   5236   5238   5240       3    0.65084    1.95253    0.00000
-2.60338    0.00000    0.00000 ;      HA-    CA-    CB-   CG2
  5237   5236   5238   5244       3    0.65084    1.95253    0.00000
-2.60338    0.00000    0.00000 ;      HA-    CA-    CB-   CG1
  5237   5236   5251   5252       3    3.68192   -4.35136    0.00000
1.33888    0.00000    0.00000 ;      HA-    CA-     C-     O
  5237   5236   5251   5253       3    0.00000    0.00000    0.00000
0.00000    0.00000    0.00000 ;      HA-    CA-     C-     N
  5238   5236   5251   5252       3    0.00000    0.00000    0.00000
0.00000    0.00000    0.00000 ;      CB-    CA-     C-     O
  5238   5236   5251   5253       3    2.51040    4.18400    1.67360
```

-6.69440 0.00000 0.00000 ; CB- CA- C- N
5238 5244 5247 5248 3 0.66944 2.00832 0.00000
-2.67776 0.00000 0.00000 ; CB- CG1- CD1- HD11
5238 5244 5247 5249 3 0.66944 2.00832 0.00000
-2.67776 0.00000 0.00000 ; CB- CG1- CD1- HD12
5238 5244 5247 5250 3 0.66944 2.00832 0.00000
-2.67776 0.00000 0.00000 ; CB- CG1- CD1- HD13
5239 5238 5236 5251 3 0.65084 1.95253 0.00000
-2.60338 0.00000 0.00000 ; HB- CB- CA- C
5239 5238 5240 5241 3 0.62760 1.88280 0.00000
-2.51040 0.00000 0.00000 ; HB- CB- CG2- HG21
5239 5238 5240 5242 3 0.62760 1.88280 0.00000
-2.51040 0.00000 0.00000 ; HB- CB- CG2- HG22
5239 5238 5240 5243 3 0.62760 1.88280 0.00000
-2.51040 0.00000 0.00000 ; HB- CB- CG2- HG23
5239 5238 5244 5245 3 0.62760 1.88280 0.00000
-2.51040 0.00000 0.00000 ; HB- CB- CG1- HG12
5239 5238 5244 5246 3 0.62760 1.88280 0.00000
-2.51040 0.00000 0.00000 ; HB- CB- CG1- HG13
5239 5238 5244 5247 3 0.66944 2.00832 0.00000
-2.67776 0.00000 0.00000 ; HB- CB- CG1- CD1
5240 5238 5236 5251 3 5.67350 3.55640 -6.16722
-2.47693 3.74886 0.00000 ; CG2- CB- CA- C
5240 5238 5244 5245 3 0.66944 2.00832 0.00000
-2.67776 0.00000 0.00000 ; CG2- CB- CG1- HG12
5240 5238 5244 5246 3 0.66944 2.00832 0.00000
-2.67776 0.00000 0.00000 ; CG2- CB- CG1- HG13
5240 5238 5244 5247 3 3.81162 0.49790 -8.14206
-1.79075 7.49773 0.00000 ; CG2- CB- CG1- CD1
5241 5240 5238 5244 3 0.66944 2.00832 0.00000
-2.67776 0.00000 0.00000 ; HG21- CG2- CB- CG1
5242 5240 5238 5244 3 0.66944 2.00832 0.00000
-2.67776 0.00000 0.00000 ; HG22- CG2- CB- CG1
5243 5240 5238 5244 3 0.66944 2.00832 0.00000
-2.67776 0.00000 0.00000 ; HG23- CG2- CB- CG1
5244 5238 5236 5251 3 8.26340 0.74057 -9.99976
-1.89117 3.84928 0.00000 ; CG1- CB- CA- C
5245 5244 5247 5248 3 0.62760 1.88280 0.00000
-2.51040 0.00000 0.00000 ; HG12- CG1- CD1- HD11
5245 5244 5247 5249 3 0.62760 1.88280 0.00000
-2.51040 0.00000 0.00000 ; HG12- CG1- CD1- HD12
5245 5244 5247 5250 3 0.62760 1.88280 0.00000
-2.51040 0.00000 0.00000 ; HG12- CG1- CD1- HD13
5246 5244 5247 5248 3 0.62760 1.88280 0.00000
-2.51040 0.00000 0.00000 ; HG13- CG1- CD1- HD11
5246 5244 5247 5249 3 0.62760 1.88280 0.00000
-2.51040 0.00000 0.00000 ; HG13- CG1- CD1- HD12
5246 5244 5247 5250 3 0.62760 1.88280 0.00000
-2.51040 0.00000 0.00000 ; HG13- CG1- CD1- HD13
5251 5253 5255 5256 3 0.00000 0.00000 0.00000
0.00000 0.00000 0.00000 ; C- N- CA- HA
5251 5253 5255 5257 3 11.71520 1.67360 15.06240 -

```
13.38880    0.00000    0.00000 ;       C-     N-    CA-    CB
  5251  5253  5255  5261      3    1.75728    5.27184    2.25936
-7.02912    0.00000    0.00000 ;       C-     N-    CA-     C
  5252  5251  5253  5254      3   29.28800   -8.36800  -20.92000
0.00000    0.00000    0.00000 ;      O-     C-     N-     H
  5252  5251  5253  5255      3   20.92000    0.00000  -20.92000
0.00000    0.00000    0.00000 ;      O-     C-     N-    CA
  5253  5255  5257  5258      3    0.65084    1.95253    0.00000
-2.60338    0.00000    0.00000 ;      N-    CA-    CB-   HB1
  5253  5255  5257  5259      3    0.65084    1.95253    0.00000
-2.60338    0.00000    0.00000 ;      N-    CA-    CB-   HB2
  5253  5255  5257  5260      3    0.65084    1.95253    0.00000
-2.60338    0.00000    0.00000 ;      N-    CA-    CB-   HB3
  5253  5255  5261  5262      3    0.00000    0.00000    0.00000
0.00000    0.00000    0.00000 ;      N-    CA-     C-     O
  5253  5255  5261  5263      3   17.40544   -5.02080  -13.22144
9.20480    0.00000    0.00000 ;      N-    CA-     C-     N
  5254  5253  5255  5256      3    0.00000    0.00000    0.00000
0.00000    0.00000    0.00000 ;      H-     N-    CA-    HA
  5254  5253  5255  5257      3    0.00000    0.00000    0.00000
0.00000    0.00000    0.00000 ;      H-     N-    CA-    CB
  5254  5253  5255  5261      3    0.00000    0.00000    0.00000
0.00000    0.00000    0.00000 ;      H-     N-    CA-     C
  5255  5261  5263  5264      3   20.92000    0.00000  -20.92000
0.00000    0.00000    0.00000 ;     CA-     C-     N-     H
  5255  5261  5263  5265      3   20.92000    0.00000  -20.92000
0.00000    0.00000    0.00000 ;     CA-     C-     N-    CA
  5256  5255  5257  5258      3    0.65084    1.95253    0.00000
-2.60338    0.00000    0.00000 ;      HA-    CA-    CB-   HB1
  5256  5255  5257  5259      3    0.65084    1.95253    0.00000
-2.60338    0.00000    0.00000 ;      HA-    CA-    CB-   HB2
  5256  5255  5257  5260      3    0.65084    1.95253    0.00000
-2.60338    0.00000    0.00000 ;      HA-    CA-    CB-   HB3
  5256  5255  5261  5262      3    3.68192   -4.35136    0.00000
1.33888    0.00000    0.00000 ;     HA-    CA-     C-     O
  5256  5255  5261  5263      3    0.00000    0.00000    0.00000
0.00000    0.00000    0.00000 ;     HA-    CA-     C-     N
  5257  5255  5261  5262      3    0.00000    0.00000    0.00000
0.00000    0.00000    0.00000 ;     CB-    CA-     C-     O
  5257  5255  5261  5263      3    2.51040    4.18400    1.67360
-6.69440    0.00000    0.00000 ;     CB-    CA-     C-     N
  5258  5257  5255  5261      3    0.65084    1.95253    0.00000
-2.60338    0.00000    0.00000 ;    HB1-    CB-    CA-     C
  5259  5257  5255  5261      3    0.65084    1.95253    0.00000
-2.60338    0.00000    0.00000 ;    HB2-    CB-    CA-     C
  5260  5257  5255  5261      3    0.65084    1.95253    0.00000
-2.60338    0.00000    0.00000 ;    HB3-    CB-    CA-     C
  5261  5263  5265  5266      3    0.00000    0.00000    0.00000
0.00000    0.00000    0.00000 ;      C-     N-    CA-    HA
  5261  5263  5265  5267      3   11.71520    1.67360   15.06240  -
13.38880    0.00000    0.00000 ;       C-     N-    CA-    CB
  5261  5263  5265  5280      3    1.75728    5.27184    2.25936
```

-7.02912 0.00000 0.00000 ; C- N- CA- C
5262 5261 5263 5264 3 29.28800 -8.36800 -20.92000
0.00000 0.00000 0.00000 ; O- C- N- H
5262 5261 5263 5265 3 20.92000 0.00000 -20.92000
0.00000 0.00000 0.00000 ; O- C- N- CA
5263 5265 5267 5268 3 0.65084 1.95253 0.00000
-2.60338 0.00000 0.00000 ; N- CA- CB- HB2
5263 5265 5267 5269 3 0.65084 1.95253 0.00000
-2.60338 0.00000 0.00000 ; N- CA- CB- HB3
5263 5265 5267 5270 3 3.79070 1.39746 -4.61077
-2.40998 2.44346 0.00000 ; N- CA- CB- CG
5263 5265 5280 5281 3 0.00000 0.00000 0.00000
0.00000 0.00000 0.00000 ; N- CA- C- O
5263 5265 5280 5282 3 17.40544 -5.02080 -13.22144
9.20480 0.00000 0.00000 ; N- CA- C- N
5264 5263 5265 5266 3 0.00000 0.00000 0.00000
0.00000 0.00000 0.00000 ; H- N- CA- HA
5264 5263 5265 5267 3 0.00000 0.00000 0.00000
0.00000 0.00000 0.00000 ; H- N- CA- CB
5264 5263 5265 5280 3 0.00000 0.00000 0.00000
0.00000 0.00000 0.00000 ; H- N- CA- C
5265 5267 5270 5271 3 0.66944 2.00832 0.00000
-2.67776 0.00000 0.00000 ; CA- CB- CG- HG
5265 5267 5270 5272 3 3.90367 0.19665 -6.21742
-2.37651 5.99149 0.00000 ; CA- CB- CG- CD1
5265 5267 5270 5276 3 3.90367 0.19665 -6.21742
-2.37651 5.99149 0.00000 ; CA- CB- CG- CD2
5265 5280 5282 5283 3 20.92000 0.00000 -20.92000
0.00000 0.00000 0.00000 ; CA- C- N- H
5265 5280 5282 5284 3 20.92000 0.00000 -20.92000
0.00000 0.00000 0.00000 ; CA- C- N- CA
5266 5265 5267 5268 3 0.65084 1.95253 0.00000
-2.60338 0.00000 0.00000 ; HA- CA- CB- HB2
5266 5265 5267 5269 3 0.65084 1.95253 0.00000
-2.60338 0.00000 0.00000 ; HA- CA- CB- HB3
5266 5265 5267 5270 3 0.65084 1.95253 0.00000
-2.60338 0.00000 0.00000 ; HA- CA- CB- CG
5266 5265 5280 5281 3 3.68192 -4.35136 0.00000
1.33888 0.00000 0.00000 ; HA- CA- C- O
5266 5265 5280 5282 3 0.00000 0.00000 0.00000
0.00000 0.00000 0.00000 ; HA- CA- C- N
5267 5265 5280 5281 3 0.00000 0.00000 0.00000
0.00000 0.00000 0.00000 ; CB- CA- C- O
5267 5265 5280 5282 3 2.51040 4.18400 1.67360
-6.69440 0.00000 0.00000 ; CB- CA- C- N
5267 5270 5272 5273 3 0.66944 2.00832 0.00000
-2.67776 0.00000 0.00000 ; CB- CG- CD1- HD11
5267 5270 5272 5274 3 0.66944 2.00832 0.00000
-2.67776 0.00000 0.00000 ; CB- CG- CD1- HD12
5267 5270 5272 5275 3 0.66944 2.00832 0.00000
-2.67776 0.00000 0.00000 ; CB- CG- CD1- HD13
5267 5270 5276 5277 3 0.66944 2.00832 0.00000

-2.67776 0.00000 0.00000 ; CB- CG- CD2- HD21
5267 5270 5276 5278 3 0.66944 2.00832 0.00000
-2.67776 0.00000 0.00000 ; CB- CG- CD2- HD22
5267 5270 5276 5279 3 0.66944 2.00832 0.00000
-2.67776 0.00000 0.00000 ; CB- CG- CD2- HD23
5268 5267 5265 5280 3 0.65084 1.95253 0.00000
-2.60338 0.00000 0.00000 ; HB2- CB- CA- C
5268 5267 5270 5271 3 0.62760 1.88280 0.00000
-2.51040 0.00000 0.00000 ; HB2- CB- CG- HG
5268 5267 5270 5272 3 0.66944 2.00832 0.00000
-2.67776 0.00000 0.00000 ; HB2- CB- CG- CD1
5268 5267 5270 5276 3 0.66944 2.00832 0.00000
-2.67776 0.00000 0.00000 ; HB2- CB- CG- CD2
5269 5267 5265 5280 3 0.65084 1.95253 0.00000
-2.60338 0.00000 0.00000 ; HB3- CB- CA- C
5269 5267 5270 5271 3 0.62760 1.88280 0.00000
-2.51040 0.00000 0.00000 ; HB3- CB- CG- HG
5269 5267 5270 5272 3 0.66944 2.00832 0.00000
-2.67776 0.00000 0.00000 ; HB3- CB- CG- CD1
5269 5267 5270 5276 3 0.66944 2.00832 0.00000
-2.67776 0.00000 0.00000 ; HB3- CB- CG- CD2
5270 5267 5265 5280 3 10.33448 -1.14642 -11.54784
-2.40998 6.35968 0.00000 ; CG- CB- CA- C
5271 5270 5272 5273 3 0.62760 1.88280 0.00000
-2.51040 0.00000 0.00000 ; HG- CG- CD1- HD11
5271 5270 5272 5274 3 0.62760 1.88280 0.00000
-2.51040 0.00000 0.00000 ; HG- CG- CD1- HD12
5271 5270 5272 5275 3 0.62760 1.88280 0.00000
-2.51040 0.00000 0.00000 ; HG- CG- CD1- HD13
5271 5270 5276 5277 3 0.62760 1.88280 0.00000
-2.51040 0.00000 0.00000 ; HG- CG- CD2- HD21
5271 5270 5276 5278 3 0.62760 1.88280 0.00000
-2.51040 0.00000 0.00000 ; HG- CG- CD2- HD22
5271 5270 5276 5279 3 0.62760 1.88280 0.00000
-2.51040 0.00000 0.00000 ; HG- CG- CD2- HD23
5272 5270 5276 5277 3 0.66944 2.00832 0.00000
-2.67776 0.00000 0.00000 ; CD1- CG- CD2- HD21
5272 5270 5276 5278 3 0.66944 2.00832 0.00000
-2.67776 0.00000 0.00000 ; CD1- CG- CD2- HD22
5272 5270 5276 5279 3 0.66944 2.00832 0.00000
-2.67776 0.00000 0.00000 ; CD1- CG- CD2- HD23
5273 5272 5270 5276 3 0.66944 2.00832 0.00000
-2.67776 0.00000 0.00000 ; HD11- CD1- CG- CD2
5274 5272 5270 5276 3 0.66944 2.00832 0.00000
-2.67776 0.00000 0.00000 ; HD12- CD1- CG- CD2
5275 5272 5270 5276 3 0.66944 2.00832 0.00000
-2.67776 0.00000 0.00000 ; HD13- CD1- CG- CD2
5280 5282 5284 5285 3 0.00000 0.00000 0.00000
0.00000 0.00000 0.00000 ; C- N- CA- HA
5280 5282 5284 5286 3 11.71520 1.67360 15.06240 -
13.38880 0.00000 0.00000 ; C- N- CA- CB
5280 5282 5284 5292 3 1.75728 5.27184 2.25936

-7.02912 0.00000 0.00000 ; C- N- CA- C
5281 5280 5282 5283 3 29.28800 -8.36800 -20.92000
0.00000 0.00000 0.00000 ; O- C- N- H
5281 5280 5282 5284 3 20.92000 0.00000 -20.92000
0.00000 0.00000 0.00000 ; O- C- N- CA
5282 5284 5286 5287 3 0.65084 1.95253 0.00000
-2.60338 0.00000 0.00000 ; N- CA- CB- HB2
5282 5284 5286 5288 3 0.65084 1.95253 0.00000
-2.60338 0.00000 0.00000 ; N- CA- CB- HB3
5282 5284 5286 5289 3 15.41386 9.74035 -8.39310
-0.97069 2.97901 0.00000 ; N- CA- CB- CG
5282 5284 5292 5293 3 0.00000 0.00000 0.00000
0.00000 0.00000 0.00000 ; N- CA- C- O
5282 5284 5292 5294 3 17.40544 -5.02080 -13.22144
9.20480 0.00000 0.00000 ; N- CA- C- N
5283 5282 5284 5285 3 0.00000 0.00000 0.00000
0.00000 0.00000 0.00000 ; H- N- CA- HA
5283 5282 5284 5286 3 0.00000 0.00000 0.00000
0.00000 0.00000 0.00000 ; H- N- CA- CB
5283 5282 5284 5292 3 0.00000 0.00000 0.00000
0.00000 0.00000 0.00000 ; H- N- CA- C
5284 5286 5289 5290 3 6.43499 0.00000 -5.39736
0.00000 -1.03763 0.00000 ; CA- CB- CG- OD1
5284 5286 5289 5291 3 6.43499 0.00000 -5.39736
0.00000 -1.03763 0.00000 ; CA- CB- CG- OD2
5284 5292 5294 5295 3 20.92000 0.00000 -20.92000
0.00000 0.00000 0.00000 ; CA- C- N- H
5284 5292 5294 5296 3 20.92000 0.00000 -20.92000
0.00000 0.00000 0.00000 ; CA- C- N- CA
5285 5284 5286 5287 3 0.65084 1.95253 0.00000
-2.60338 0.00000 0.00000 ; HA- CA- CB- HB2
5285 5284 5286 5288 3 0.65084 1.95253 0.00000
-2.60338 0.00000 0.00000 ; HA- CA- CB- HB3
5285 5284 5286 5289 3 0.65084 1.95253 0.00000
-2.60338 0.00000 0.00000 ; HA- CA- CB- CG
5285 5284 5292 5293 3 3.68192 -4.35136 0.00000
1.33888 0.00000 0.00000 ; HA- CA- C- O
5285 5284 5292 5294 3 0.00000 0.00000 0.00000
0.00000 0.00000 0.00000 ; HA- CA- C- N
5286 5284 5292 5293 3 0.00000 0.00000 0.00000
0.00000 0.00000 0.00000 ; CB- CA- C- O
5286 5284 5292 5294 3 2.51040 4.18400 1.67360
-6.69440 0.00000 0.00000 ; CB- CA- C- N
5287 5286 5284 5292 3 0.65084 1.95253 0.00000
-2.60338 0.00000 0.00000 ; HB2- CB- CA- C
5287 5286 5289 5290 3 0.00000 0.00000 0.00000
0.00000 0.00000 0.00000 ; HB2- CB- CG- OD1
5287 5286 5289 5291 3 0.00000 0.00000 0.00000
0.00000 0.00000 0.00000 ; HB2- CB- CG- OD2
5288 5286 5284 5292 3 0.65084 1.95253 0.00000
-2.60338 0.00000 0.00000 ; HB3- CB- CA- C
5288 5286 5289 5290 3 0.00000 0.00000 0.00000

0.00000 0.00000 0.00000 ; HB3- CB- CG- OD1
5288 5286 5289 5291 3 0.00000 0.00000 0.00000
0.00000 0.00000 0.00000 ; HB3- CB- CG- OD2
5289 5286 5284 5292 3 7.14627 -1.04600 -8.99560
-0.97069 5.15469 0.00000 ; CG- CB- CA- C
5292 5294 5296 5297 3 0.00000 0.00000 0.00000
0.00000 0.00000 0.00000 ; C- N- CA- HA
5292 5294 5296 5298 3 11.71520 1.67360 15.06240 -
13.38880 0.00000 0.00000 ; C- N- CA- CB
5292 5294 5296 5308 3 1.75728 5.27184 2.25936
-7.02912 0.00000 0.00000 ; C- N- CA- C
5293 5292 5294 5295 3 29.28800 -8.36800 -20.92000
0.00000 0.00000 0.00000 ; O- C- N- H
5293 5292 5294 5296 3 20.92000 0.00000 -20.92000
0.00000 0.00000 0.00000 ; O- C- N- CA
5294 5296 5298 5299 3 0.65084 1.95253 0.00000
-2.60338 0.00000 0.00000 ; N- CA- CB- HB
5294 5296 5298 5300 3 3.83673 0.44769 -1.77402
-2.47693 -0.03347 0.00000 ; N- CA- CB- CG1
5294 5296 5298 5304 3 3.83673 0.44769 -1.77402
-2.47693 -0.03347 0.00000 ; N- CA- CB- CG2
5294 5296 5308 5309 3 0.00000 0.00000 0.00000
0.00000 0.00000 0.00000 ; N- CA- C- O
5294 5296 5308 5310 3 17.40544 -5.02080 -13.22144
9.20480 0.00000 0.00000 ; N- CA- C- N
5295 5294 5296 5297 3 0.00000 0.00000 0.00000
0.00000 0.00000 0.00000 ; H- N- CA- HA
5295 5294 5296 5298 3 0.00000 0.00000 0.00000
0.00000 0.00000 0.00000 ; H- N- CA- CB
5295 5294 5296 5308 3 0.00000 0.00000 0.00000
0.00000 0.00000 0.00000 ; H- N- CA- C
5296 5298 5300 5301 3 0.66944 2.00832 0.00000
-2.67776 0.00000 0.00000 ; CA- CB- CG1- HG11
5296 5298 5300 5302 3 0.66944 2.00832 0.00000
-2.67776 0.00000 0.00000 ; CA- CB- CG1- HG12
5296 5298 5300 5303 3 0.66944 2.00832 0.00000
-2.67776 0.00000 0.00000 ; CA- CB- CG1- HG13
5296 5298 5304 5305 3 0.66944 2.00832 0.00000
-2.67776 0.00000 0.00000 ; CA- CB- CG2- HG21
5296 5298 5304 5306 3 0.66944 2.00832 0.00000
-2.67776 0.00000 0.00000 ; CA- CB- CG2- HG22
5296 5298 5304 5307 3 0.66944 2.00832 0.00000
-2.67776 0.00000 0.00000 ; CA- CB- CG2- HG23
5296 5308 5310 5311 3 20.92000 0.00000 -20.92000
0.00000 0.00000 0.00000 ; CA- C- N- H
5296 5308 5310 5312 3 20.92000 0.00000 -20.92000
0.00000 0.00000 0.00000 ; CA- C- N- CA
5297 5296 5298 5299 3 0.65084 1.95253 0.00000
-2.60338 0.00000 0.00000 ; HA- CA- CB- HB
5297 5296 5298 5300 3 0.65084 1.95253 0.00000
-2.60338 0.00000 0.00000 ; HA- CA- CB- CG1
5297 5296 5298 5304 3 0.65084 1.95253 0.00000

```
-2.60338    0.00000    0.00000 ;      HA-    CA-     CB-   CG2
  5297   5296   5308   5309       3    3.68192   -4.35136    0.00000
1.33888    0.00000    0.00000 ;      HA-    CA-     C-     O
  5297   5296   5308   5310       3    0.00000    0.00000    0.00000
0.00000    0.00000    0.00000 ;      HA-    CA-     C-     N
  5298   5296   5308   5309       3    0.00000    0.00000    0.00000
0.00000    0.00000    0.00000 ;      CB-    CA-     C-     O
  5298   5296   5308   5310       3    2.51040    4.18400    1.67360
-6.69440    0.00000    0.00000 ;      CB-    CA-     C-     N
  5299   5298   5296   5308       3    0.65084    1.95253    0.00000
-2.60338    0.00000    0.00000 ;      HB-    CB-     CA-     C
  5299   5298   5300   5301       3    0.62760    1.88280    0.00000
-2.51040    0.00000    0.00000 ;      HB-    CB-    CG1-  HG11
  5299   5298   5300   5302       3    0.62760    1.88280    0.00000
-2.51040    0.00000    0.00000 ;      HB-    CB-    CG1-  HG12
  5299   5298   5300   5303       3    0.62760    1.88280    0.00000
-2.51040    0.00000    0.00000 ;      HB-    CB-    CG1-  HG13
  5299   5298   5304   5305       3    0.62760    1.88280    0.00000
-2.51040    0.00000    0.00000 ;      HB-    CB-    CG2-  HG21
  5299   5298   5304   5306       3    0.62760    1.88280    0.00000
-2.51040    0.00000    0.00000 ;      HB-    CB-    CG2-  HG22
  5299   5298   5304   5307       3    0.62760    1.88280    0.00000
-2.51040    0.00000    0.00000 ;      HB-    CB-    CG2-  HG23
  5300   5298   5296   5308       3    5.67350    3.55640   -6.16722
-2.47693    3.74886    0.00000 ;     CG1-    CB-     CA-     C
  5300   5298   5304   5305       3    0.66944    2.00832    0.00000
-2.67776    0.00000    0.00000 ;     CG1-    CB-    CG2-  HG21
  5300   5298   5304   5306       3    0.66944    2.00832    0.00000
-2.67776    0.00000    0.00000 ;     CG1-    CB-    CG2-  HG22
  5300   5298   5304   5307       3    0.66944    2.00832    0.00000
-2.67776    0.00000    0.00000 ;     CG1-    CB-    CG2-  HG23
  5301   5300   5298   5304       3    0.66944    2.00832    0.00000
-2.67776    0.00000    0.00000 ;    HG11-   CG1-     CB-   CG2
  5302   5300   5298   5304       3    0.66944    2.00832    0.00000
-2.67776    0.00000    0.00000 ;    HG12-   CG1-     CB-   CG2
  5303   5300   5298   5304       3    0.66944    2.00832    0.00000
-2.67776    0.00000    0.00000 ;    HG13-   CG1-     CB-   CG2
  5304   5298   5296   5308       3    5.67350    3.55640   -6.16722
-2.47693    3.74886    0.00000 ;     CG2-    CB-     CA-     C
  5308   5310   5312   5313       3    0.00000    0.00000    0.00000
0.00000    0.00000    0.00000 ;       C-     N-     CA-    HA
  5308   5310   5312   5314       3   11.71520    1.67360   15.06240  -
13.38880    0.00000    0.00000 ;       C-     N-     CA-    CB
  5308   5310   5312   5320       3    1.75728    5.27184    2.25936
-7.02912    0.00000    0.00000 ;       C-     N-     CA-     C
  5309   5308   5310   5311       3   29.28800   -8.36800  -20.92000
0.00000    0.00000    0.00000 ;      O-     C-      N-     H
  5309   5308   5310   5312       3   20.92000    0.00000  -20.92000
0.00000    0.00000    0.00000 ;      O-     C-      N-    CA
  5310   5312   5314   5315       3    0.65084    1.95253    0.00000
-2.60338    0.00000    0.00000 ;       N-    CA-     CB-   HB2
  5310   5312   5314   5316       3    0.65084    1.95253    0.00000
```

-2.60338    0.00000    0.00000 ;     N-    CA-    CB-   HB3
 5310   5312   5314   5317      3   15.41386    9.74035   -8.39310
-0.97069    2.97901    0.00000 ;     N-    CA-    CB-    CG
 5310   5312   5320   5321      3    0.00000    0.00000    0.00000
0.00000    0.00000    0.00000 ;     N-    CA-     C-     O
 5310   5312   5320   5322      3   17.40544   -5.02080  -13.22144
9.20480    0.00000    0.00000 ;     N-    CA-     C-     N
 5311   5310   5312   5313      3    0.00000    0.00000    0.00000
0.00000    0.00000    0.00000 ;     H-     N-    CA-    HA
 5311   5310   5312   5314      3    0.00000    0.00000    0.00000
0.00000    0.00000    0.00000 ;     H-     N-    CA-    CB
 5311   5310   5312   5320      3    0.00000    0.00000    0.00000
0.00000    0.00000    0.00000 ;     H-     N-    CA-     C
 5312   5314   5317   5318      3    6.43499    0.00000   -5.39736
0.00000   -1.03763    0.00000 ;    CA-    CB-    CG-   OD1
 5312   5314   5317   5319      3    6.43499    0.00000   -5.39736
0.00000   -1.03763    0.00000 ;    CA-    CB-    CG-   OD2
 5312   5320   5322   5323      3   20.92000    0.00000  -20.92000
0.00000    0.00000    0.00000 ;    CA-     C-     N-     H
 5312   5320   5322   5324      3   20.92000    0.00000  -20.92000
0.00000    0.00000    0.00000 ;    CA-     C-     N-    CA
 5313   5312   5314   5315      3    0.65084    1.95253    0.00000
-2.60338    0.00000    0.00000 ;    HA-    CA-    CB-   HB2
 5313   5312   5314   5316      3    0.65084    1.95253    0.00000
-2.60338    0.00000    0.00000 ;    HA-    CA-    CB-   HB3
 5313   5312   5314   5317      3    0.65084    1.95253    0.00000
-2.60338    0.00000    0.00000 ;    HA-    CA-    CB-    CG
 5313   5312   5320   5321      3    3.68192   -4.35136    0.00000
1.33888    0.00000    0.00000 ;    HA-    CA-     C-     O
 5313   5312   5320   5322      3    0.00000    0.00000    0.00000
0.00000    0.00000    0.00000 ;    HA-    CA-     C-     N
 5314   5312   5320   5321      3    0.00000    0.00000    0.00000
0.00000    0.00000    0.00000 ;    CB-    CA-     C-     O
 5314   5312   5320   5322      3    2.51040    4.18400    1.67360
-6.69440    0.00000    0.00000 ;    CB-    CA-     C-     N
 5315   5314   5312   5320      3    0.65084    1.95253    0.00000
-2.60338    0.00000    0.00000 ;   HB2-    CB-    CA-     C
 5315   5314   5317   5318      3    0.00000    0.00000    0.00000
0.00000    0.00000    0.00000 ;   HB2-    CB-    CG-   OD1
 5315   5314   5317   5319      3    0.00000    0.00000    0.00000
0.00000    0.00000    0.00000 ;   HB2-    CB-    CG-   OD2
 5316   5314   5312   5320      3    0.65084    1.95253    0.00000
-2.60338    0.00000    0.00000 ;   HB3-    CB-    CA-     C
 5316   5314   5317   5318      3    0.00000    0.00000    0.00000
0.00000    0.00000    0.00000 ;   HB3-    CB-    CG-   OD1
 5316   5314   5317   5319      3    0.00000    0.00000    0.00000
0.00000    0.00000    0.00000 ;   HB3-    CB-    CG-   OD2
 5317   5314   5312   5320      3    7.14627   -1.04600   -8.99560
-0.97069    5.15469    0.00000 ;    CG-    CB-    CA-     C
 5320   5322   5324   5325      3    0.00000    0.00000    0.00000
0.00000    0.00000    0.00000 ;     C-     N-    CA-    HA
 5320   5322   5324   5326      3   11.71520    1.67360   15.06240  -

```
13.38880    0.00000    0.00000 ;       C-     N-    CA-    CB
  5320   5322   5324   5334      3    1.75728    5.27184    2.25936
-7.02912    0.00000    0.00000 ;       C-     N-    CA-     C
  5321   5320   5322   5323      3   29.28800   -8.36800  -20.92000
0.00000    0.00000    0.00000 ;      O-     C-     N-     H
  5321   5320   5322   5324      3   20.92000    0.00000  -20.92000
0.00000    0.00000    0.00000 ;      O-     C-     N-    CA
  5322   5324   5326   5327      3    0.65084    1.95253    0.00000
-2.60338    0.00000    0.00000 ;      N-    CA-    CB-   HB2
  5322   5324   5326   5328      3    0.65084    1.95253    0.00000
-2.60338    0.00000    0.00000 ;      N-    CA-    CB-   HB3
  5322   5324   5326   5329      3    5.99567    3.29281   -4.46014
-0.55229    1.97485    0.00000 ;      N-    CA-    CB-    CG
  5322   5324   5334   5335      3    0.00000    0.00000    0.00000
0.00000    0.00000    0.00000 ;      N-    CA-     C-     O
  5322   5324   5334   5336      3   17.40544   -5.02080  -13.22144
9.20480    0.00000    0.00000 ;      N-    CA-     C-     N
  5323   5322   5324   5325      3    0.00000    0.00000    0.00000
0.00000    0.00000    0.00000 ;      H-     N-    CA-    HA
  5323   5322   5324   5326      3    0.00000    0.00000    0.00000
0.00000    0.00000    0.00000 ;      H-     N-    CA-    CB
  5323   5322   5324   5334      3    0.00000    0.00000    0.00000
0.00000    0.00000    0.00000 ;      H-     N-    CA-     C
  5324   5326   5329   5330      3    0.00000    0.00000    0.00000
0.00000    0.00000    0.00000 ;     CA-    CB-    CG-   OD1
  5324   5326   5329   5331      3    8.84916   -0.31380   -4.32626
5.03754    0.26778    0.00000 ;     CA-    CB-    CG-   ND2
  5324   5334   5336   5337      3   20.92000    0.00000  -20.92000
0.00000    0.00000    0.00000 ;     CA-     C-     N-     H
  5324   5334   5336   5338      3   20.92000    0.00000  -20.92000
0.00000    0.00000    0.00000 ;     CA-     C-     N-    CA
  5325   5324   5326   5327      3    0.65084    1.95253    0.00000
-2.60338    0.00000    0.00000 ;     HA-    CA-    CB-   HB2
  5325   5324   5326   5328      3    0.65084    1.95253    0.00000
-2.60338    0.00000    0.00000 ;     HA-    CA-    CB-   HB3
  5325   5324   5326   5329      3    0.65084    1.95253    0.00000
-2.60338    0.00000    0.00000 ;     HA-    CA-    CB-    CG
  5325   5324   5334   5335      3    3.68192   -4.35136    0.00000
1.33888    0.00000    0.00000 ;     HA-    CA-     C-     O
  5325   5324   5334   5336      3    0.00000    0.00000    0.00000
0.00000    0.00000    0.00000 ;     HA-    CA-     C-     N
  5326   5324   5334   5335      3    0.00000    0.00000    0.00000
0.00000    0.00000    0.00000 ;     CB-    CA-     C-     O
  5326   5324   5334   5336      3    2.51040    4.18400    1.67360
-6.69440    0.00000    0.00000 ;     CB-    CA-     C-     N
  5326   5329   5331   5332      3   20.92000    0.00000  -20.92000
0.00000    0.00000    0.00000 ;     CB-    CG-   ND2-  HD21
  5326   5329   5331   5333      3   20.92000    0.00000  -20.92000
0.00000    0.00000    0.00000 ;     CB-    CG-   ND2-  HD22
  5327   5326   5324   5334      3    0.65084    1.95253    0.00000
-2.60338    0.00000    0.00000 ;     HB2-    CB-    CA-     C
  5327   5326   5329   5330      3    3.68192   -4.35136    0.00000
```

1.33888 0.00000 0.00000 ; HB2- CB- CG- OD1
5327 5326 5329 5331 3 0.00000 0.00000 0.00000
0.00000 0.00000 0.00000 ; HB2- CB- CG- ND2
5328 5326 5324 5334 3 0.65084 1.95253 0.00000
-2.60338 0.00000 0.00000 ; HB3- CB- CA- C
5328 5326 5329 5330 3 3.68192 -4.35136 0.00000
1.33888 0.00000 0.00000 ; HB3- CB- CG- OD1
5328 5326 5329 5331 3 0.00000 0.00000 0.00000
0.00000 0.00000 0.00000 ; HB3- CB- CG- ND2
5329 5326 5324 5334 3 7.94542 -3.96225 -6.11701
-0.55229 3.58150 0.00000 ; CG- CB- CA- C
5330 5329 5331 5332 3 29.28800 -8.36800 -20.92000
0.00000 0.00000 0.00000 ; OD1- CG- ND2- HD21
5330 5329 5331 5333 3 29.28800 -8.36800 -20.92000
0.00000 0.00000 0.00000 ; OD1- CG- ND2- HD22
5334 5336 5338 5339 3 0.00000 0.00000 0.00000
0.00000 0.00000 0.00000 ; C- N- CA- HA
5334 5336 5338 5340 3 11.71520 1.67360 15.06240 -
13.38880 0.00000 0.00000 ; C- N- CA- CB
5334 5336 5338 5345 3 1.75728 5.27184 2.25936
-7.02912 0.00000 0.00000 ; C- N- CA- C
5335 5334 5336 5337 3 29.28800 -8.36800 -20.92000
0.00000 0.00000 0.00000 ; O- C- N- H
5335 5334 5336 5338 3 20.92000 0.00000 -20.92000
0.00000 0.00000 0.00000 ; O- C- N- CA
5336 5338 5340 5341 3 0.65084 1.95253 0.00000
-2.60338 0.00000 0.00000 ; N- CA- CB- HB2
5336 5338 5340 5342 3 0.65084 1.95253 0.00000
-2.60338 0.00000 0.00000 ; N- CA- CB- HB3
5336 5338 5340 5343 3 7.86174 2.24681 -7.41405
-6.71114 5.35552 0.00000 ; N- CA- CB- OG
5336 5338 5345 5346 3 0.00000 0.00000 0.00000
0.00000 0.00000 0.00000 ; N- CA- C- O
5336 5338 5345 5347 3 17.40544 -5.02080 -13.22144
9.20480 0.00000 0.00000 ; N- CA- C- N
5337 5336 5338 5339 3 0.00000 0.00000 0.00000
0.00000 0.00000 0.00000 ; H- N- CA- HA
5337 5336 5338 5340 3 0.00000 0.00000 0.00000
0.00000 0.00000 0.00000 ; H- N- CA- CB
5337 5336 5338 5345 3 0.00000 0.00000 0.00000
0.00000 0.00000 0.00000 ; H- N- CA- C
5338 5340 5343 5344 3 2.05853 2.46856 3.48109
-4.46851 0.23430 0.00000 ; CA- CB- OG- HG
5338 5345 5347 5348 3 20.92000 0.00000 -20.92000
0.00000 0.00000 0.00000 ; CA- C- N- H
5338 5345 5347 5349 3 20.92000 0.00000 -20.92000
0.00000 0.00000 0.00000 ; CA- C- N- CA
5339 5338 5340 5341 3 0.65084 1.95253 0.00000
-2.60338 0.00000 0.00000 ; HA- CA- CB- HB2
5339 5338 5340 5342 3 0.65084 1.95253 0.00000
-2.60338 0.00000 0.00000 ; HA- CA- CB- HB3
5339 5338 5340 5343 3 1.04600 -1.04600 0.00000

0.00000 0.00000 0.00000 ; HA- CA- CB- OG
5339 5338 5345 5346 3 3.68192 -4.35136 0.00000
1.33888 0.00000 0.00000 ; HA- CA- C- O
5339 5338 5345 5347 3 0.00000 0.00000 0.00000
0.00000 0.00000 0.00000 ; HA- CA- C- N
5340 5338 5345 5346 3 0.00000 0.00000 0.00000
0.00000 0.00000 0.00000 ; CB- CA- C- O
5340 5338 5345 5347 3 2.51040 4.18400 1.67360
-6.69440 0.00000 0.00000 ; CB- CA- C- N
5341 5340 5338 5345 3 0.65084 1.95253 0.00000
-2.60338 0.00000 0.00000 ; HB2- CB- CA- C
5341 5340 5343 5344 3 0.69733 2.09200 0.00000
-2.78933 0.00000 0.00000 ; HB2- CB- OG- HG
5342 5340 5338 5345 3 0.65084 1.95253 0.00000
-2.60338 0.00000 0.00000 ; HB3- CB- CA- C
5342 5340 5343 5344 3 0.69733 2.09200 0.00000
-2.78933 0.00000 0.00000 ; HB3- CB- OG- HG
5343 5340 5338 5345 3 7.34710 7.79898 -6.14211
-6.71114 4.31789 0.00000 ; OG- CB- CA- C
5345 5347 5349 5350 3 0.00000 0.00000 0.00000
0.00000 0.00000 0.00000 ; C- N- CA- HA2
5345 5347 5349 5351 3 0.00000 0.00000 0.00000
0.00000 0.00000 0.00000 ; C- N- CA- HA3
5345 5347 5349 5352 3 1.75728 5.27184 2.25936
-7.02912 0.00000 0.00000 ; C- N- CA- C
5346 5345 5347 5348 3 29.28800 -8.36800 -20.92000
0.00000 0.00000 0.00000 ; O- C- N- H
5346 5345 5347 5349 3 20.92000 0.00000 -20.92000
0.00000 0.00000 0.00000 ; O- C- N- CA
5347 5349 5352 5353 3 0.00000 0.00000 0.00000
0.00000 0.00000 0.00000 ; N- CA- C- O
5347 5349 5352 5354 3 17.40544 -5.02080 -13.22144
9.20480 0.00000 0.00000 ; N- CA- C- N
5348 5347 5349 5350 3 0.00000 0.00000 0.00000
0.00000 0.00000 0.00000 ; H- N- CA- HA2
5348 5347 5349 5351 3 0.00000 0.00000 0.00000
0.00000 0.00000 0.00000 ; H- N- CA- HA3
5348 5347 5349 5352 3 0.00000 0.00000 0.00000
0.00000 0.00000 0.00000 ; H- N- CA- C
5349 5352 5354 5355 3 20.92000 0.00000 -20.92000
0.00000 0.00000 0.00000 ; CA- C- N- H
5349 5352 5354 5356 3 20.92000 0.00000 -20.92000
0.00000 0.00000 0.00000 ; CA- C- N- CA
5350 5349 5352 5353 3 3.68192 -4.35136 0.00000
1.33888 0.00000 0.00000 ; HA2- CA- C- O
5350 5349 5352 5354 3 0.00000 0.00000 0.00000
0.00000 0.00000 0.00000 ; HA2- CA- C- N
5351 5349 5352 5353 3 3.68192 -4.35136 0.00000
1.33888 0.00000 0.00000 ; HA3- CA- C- O
5351 5349 5352 5354 3 0.00000 0.00000 0.00000
0.00000 0.00000 0.00000 ; HA3- CA- C- N
5352 5354 5356 5357 3 0.00000 0.00000 0.00000

0.00000 0.00000 0.00000 ; C- N- CA- HA
5352 5354 5356 5358 3 11.71520 1.67360 15.06240 -
13.38880 0.00000 0.00000 ; C- N- CA- CB
5352 5354 5356 5366 3 1.75728 5.27184 2.25936
-7.02912 0.00000 0.00000 ; C- N- CA- C
5353 5352 5354 5355 3 29.28800 -8.36800 -20.92000
0.00000 0.00000 0.00000 ; O- C- N- H
5353 5352 5354 5356 3 20.92000 0.00000 -20.92000
0.00000 0.00000 0.00000 ; O- C- N- CA
5354 5356 5358 5359 3 0.65084 1.95253 0.00000
-2.60338 0.00000 0.00000 ; N- CA- CB- HB
5354 5356 5358 5360 3 3.83673 0.44769 -1.77402
-2.47693 -0.03347 0.00000 ; N- CA- CB- CG2
5354 5356 5358 5364 3 4.93294 1.13386 -3.12963
-5.27184 3.17984 0.00000 ; N- CA- CB- OG1
5354 5356 5366 5367 3 0.00000 0.00000 0.00000
0.00000 0.00000 0.00000 ; N- CA- C- O
5354 5356 5366 5368 3 17.40544 -5.02080 -13.22144
9.20480 0.00000 0.00000 ; N- CA- C- N
5355 5354 5356 5357 3 0.00000 0.00000 0.00000
0.00000 0.00000 0.00000 ; H- N- CA- HA
5355 5354 5356 5358 3 0.00000 0.00000 0.00000
0.00000 0.00000 0.00000 ; H- N- CA- CB
5355 5354 5356 5366 3 0.00000 0.00000 0.00000
0.00000 0.00000 0.00000 ; H- N- CA- C
5356 5358 5360 5361 3 0.66944 2.00832 0.00000
-2.67776 0.00000 0.00000 ; CA- CB- CG2- HG21
5356 5358 5360 5362 3 0.66944 2.00832 0.00000
-2.67776 0.00000 0.00000 ; CA- CB- CG2- HG22
5356 5358 5360 5363 3 0.66944 2.00832 0.00000
-2.67776 0.00000 0.00000 ; CA- CB- CG2- HG23
5356 5358 5364 5365 3 1.12131 2.98738 1.66523
-3.94970 0.43514 0.00000 ; CA- CB- OG1- HG1
5356 5366 5368 5369 3 20.92000 0.00000 -20.92000
0.00000 0.00000 0.00000 ; CA- C- N- H
5356 5366 5368 5370 3 20.92000 0.00000 -20.92000
0.00000 0.00000 0.00000 ; CA- C- N- CA
5357 5356 5358 5359 3 0.65084 1.95253 0.00000
-2.60338 0.00000 0.00000 ; HA- CA- CB- HB
5357 5356 5358 5360 3 0.65084 1.95253 0.00000
-2.60338 0.00000 0.00000 ; HA- CA- CB- CG2
5357 5356 5358 5364 3 1.04600 -1.04600 0.00000
0.00000 0.00000 0.00000 ; HA- CA- CB- OG1
5357 5356 5366 5367 3 3.68192 -4.35136 0.00000
1.33888 0.00000 0.00000 ; HA- CA- C- O
5357 5356 5366 5368 3 0.00000 0.00000 0.00000
0.00000 0.00000 0.00000 ; HA- CA- C- N
5358 5356 5366 5367 3 0.00000 0.00000 0.00000
0.00000 0.00000 0.00000 ; CB- CA- C- O
5358 5356 5366 5368 3 2.51040 4.18400 1.67360
-6.69440 0.00000 0.00000 ; CB- CA- C- N
5359 5358 5356 5366 3 0.65084 1.95253 0.00000

-2.60338 0.00000 0.00000 ; HB- CB- CA- C
5359 5358 5360 5361 3 0.65084 1.95253 0.00000
-2.60338 0.00000 0.00000 ; HB- CB- CG2- HG21
5359 5358 5360 5362 3 0.65084 1.95253 0.00000
-2.60338 0.00000 0.00000 ; HB- CB- CG2- HG22
5359 5358 5360 5363 3 0.65084 1.95253 0.00000
-2.60338 0.00000 0.00000 ; HB- CB- CG2- HG23
5359 5358 5364 5365 3 0.69733 2.09200 0.00000
-2.78933 0.00000 0.00000 ; HB- CB- OG1- HG1
5360 5358 5356 5366 3 5.67350 3.55640 -6.16722
-2.47693 3.74886 0.00000 ; CG2- CB- CA- C
5360 5358 5364 5365 3 4.74047 0.27196 -2.26773
-3.94970 1.60666 0.00000 ; CG2- CB- OG1- HG1
5361 5360 5358 5364 3 1.04600 -1.04600 0.00000
0.00000 0.00000 0.00000 ; HG21- CG2- CB- OG1
5362 5360 5358 5364 3 1.04600 -1.04600 0.00000
0.00000 0.00000 0.00000 ; HG22- CG2- CB- OG1
5363 5360 5358 5364 3 1.04600 -1.04600 0.00000
0.00000 0.00000 0.00000 ; HG23- CG2- CB- OG1
5364 5358 5356 5366 3 6.53541 6.87013 -6.21742
-5.27184 5.22163 0.00000 ; OG1- CB- CA- C
5366 5368 5370 5371 3 0.00000 0.00000 0.00000
0.00000 0.00000 0.00000 ; C- N- CA- HA
5366 5368 5370 5372 3 11.71520 1.67360 15.06240 -
13.38880 0.00000 0.00000 ; C- N- CA- CB
5366 5368 5370 5385 3 1.75728 5.27184 2.25936
-7.02912 0.00000 0.00000 ; C- N- CA- C
5367 5366 5368 5369 3 29.28800 -8.36800 -20.92000
0.00000 0.00000 0.00000 ; O- C- N- H
5367 5366 5368 5370 3 20.92000 0.00000 -20.92000
0.00000 0.00000 0.00000 ; O- C- N- CA
5368 5370 5372 5373 3 0.65084 1.95253 0.00000
-2.60338 0.00000 0.00000 ; N- CA- CB- HB2
5368 5370 5372 5374 3 0.65084 1.95253 0.00000
-2.60338 0.00000 0.00000 ; N- CA- CB- HB3
5368 5370 5372 5375 3 3.79070 1.39746 -4.61077
-2.40998 2.44346 0.00000 ; N- CA- CB- CG
5368 5370 5385 5386 3 0.00000 0.00000 0.00000
0.00000 0.00000 0.00000 ; N- CA- C- O
5368 5370 5385 5387 3 17.40544 -5.02080 -13.22144
9.20480 0.00000 0.00000 ; N- CA- C- N
5369 5368 5370 5371 3 0.00000 0.00000 0.00000
0.00000 0.00000 0.00000 ; H- N- CA- HA
5369 5368 5370 5372 3 0.00000 0.00000 0.00000
0.00000 0.00000 0.00000 ; H- N- CA- CB
5369 5368 5370 5385 3 0.00000 0.00000 0.00000
0.00000 0.00000 0.00000 ; H- N- CA- C
5370 5372 5375 5376 3 0.66944 2.00832 0.00000
-2.67776 0.00000 0.00000 ; CA- CB- CG- HG
5370 5372 5375 5377 3 3.90367 0.19665 -6.21742
-2.37651 5.99149 0.00000 ; CA- CB- CG- CD1
5370 5372 5375 5381 3 3.90367 0.19665 -6.21742

```
-2.37651    5.99149    0.00000 ;      CA-    CB-    CG-   CD2
  5370   5385   5387   5388       3   20.92000    0.00000  -20.92000
0.00000    0.00000    0.00000 ;     CA-     C-     N-     H
  5370   5385   5387   5389       3   20.92000    0.00000  -20.92000
0.00000    0.00000    0.00000 ;     CA-     C-     N-    CA
  5371   5370   5372   5373       3    0.65084    1.95253    0.00000
-2.60338    0.00000    0.00000 ;      HA-    CA-    CB-   HB2
  5371   5370   5372   5374       3    0.65084    1.95253    0.00000
-2.60338    0.00000    0.00000 ;      HA-    CA-    CB-   HB3
  5371   5370   5372   5375       3    0.65084    1.95253    0.00000
-2.60338    0.00000    0.00000 ;      HA-    CA-    CB-    CG
  5371   5370   5385   5386       3    3.68192   -4.35136    0.00000
1.33888    0.00000    0.00000 ;     HA-    CA-     C-     O
  5371   5370   5385   5387       3    0.00000    0.00000    0.00000
0.00000    0.00000    0.00000 ;     HA-    CA-     C-     N
  5372   5370   5385   5386       3    0.00000    0.00000    0.00000
0.00000    0.00000    0.00000 ;     CB-    CA-     C-     O
  5372   5370   5385   5387       3    2.51040    4.18400    1.67360
-6.69440    0.00000    0.00000 ;      CB-    CA-     C-     N
  5372   5375   5377   5378       3    0.66944    2.00832    0.00000
-2.67776    0.00000    0.00000 ;      CB-    CG-   CD1-  HD11
  5372   5375   5377   5379       3    0.66944    2.00832    0.00000
-2.67776    0.00000    0.00000 ;      CB-    CG-   CD1-  HD12
  5372   5375   5377   5380       3    0.66944    2.00832    0.00000
-2.67776    0.00000    0.00000 ;      CB-    CG-   CD1-  HD13
  5372   5375   5381   5382       3    0.66944    2.00832    0.00000
-2.67776    0.00000    0.00000 ;      CB-    CG-   CD2-  HD21
  5372   5375   5381   5383       3    0.66944    2.00832    0.00000
-2.67776    0.00000    0.00000 ;      CB-    CG-   CD2-  HD22
  5372   5375   5381   5384       3    0.66944    2.00832    0.00000
-2.67776    0.00000    0.00000 ;      CB-    CG-   CD2-  HD23
  5373   5372   5370   5385       3    0.65084    1.95253    0.00000
-2.60338    0.00000    0.00000 ;     HB2-    CB-    CA-     C
  5373   5372   5375   5376       3    0.62760    1.88280    0.00000
-2.51040    0.00000    0.00000 ;     HB2-    CB-    CG-    HG
  5373   5372   5375   5377       3    0.66944    2.00832    0.00000
-2.67776    0.00000    0.00000 ;     HB2-    CB-    CG-   CD1
  5373   5372   5375   5381       3    0.66944    2.00832    0.00000
-2.67776    0.00000    0.00000 ;     HB2-    CB-    CG-   CD2
  5374   5372   5370   5385       3    0.65084    1.95253    0.00000
-2.60338    0.00000    0.00000 ;     HB3-    CB-    CA-     C
  5374   5372   5375   5376       3    0.62760    1.88280    0.00000
-2.51040    0.00000    0.00000 ;     HB3-    CB-    CG-    HG
  5374   5372   5375   5377       3    0.66944    2.00832    0.00000
-2.67776    0.00000    0.00000 ;     HB3-    CB-    CG-   CD1
  5374   5372   5375   5381       3    0.66944    2.00832    0.00000
-2.67776    0.00000    0.00000 ;     HB3-    CB-    CG-   CD2
  5375   5372   5370   5385       3   10.33448   -1.14642  -11.54784
-2.40998    6.35968    0.00000 ;      CG-    CB-    CA-     C
  5376   5375   5377   5378       3    0.62760    1.88280    0.00000
-2.51040    0.00000    0.00000 ;      HG-    CG-   CD1-  HD11
  5376   5375   5377   5379       3    0.62760    1.88280    0.00000
```

-2.51040 0.00000 0.00000 ; HG- CG- CD1- HD12
5376 5375 5377 5380 3 0.62760 1.88280 0.00000
-2.51040 0.00000 0.00000 ; HG- CG- CD1- HD13
5376 5375 5381 5382 3 0.62760 1.88280 0.00000
-2.51040 0.00000 0.00000 ; HG- CG- CD2- HD21
5376 5375 5381 5383 3 0.62760 1.88280 0.00000
-2.51040 0.00000 0.00000 ; HG- CG- CD2- HD22
5376 5375 5381 5384 3 0.62760 1.88280 0.00000
-2.51040 0.00000 0.00000 ; HG- CG- CD2- HD23
5377 5375 5381 5382 3 0.66944 2.00832 0.00000
-2.67776 0.00000 0.00000 ; CD1- CG- CD2- HD21
5377 5375 5381 5383 3 0.66944 2.00832 0.00000
-2.67776 0.00000 0.00000 ; CD1- CG- CD2- HD22
5377 5375 5381 5384 3 0.66944 2.00832 0.00000
-2.67776 0.00000 0.00000 ; CD1- CG- CD2- HD23
5378 5377 5375 5381 3 0.66944 2.00832 0.00000
-2.67776 0.00000 0.00000 ; HD11- CD1- CG- CD2
5379 5377 5375 5381 3 0.66944 2.00832 0.00000
-2.67776 0.00000 0.00000 ; HD12- CD1- CG- CD2
5380 5377 5375 5381 3 0.66944 2.00832 0.00000
-2.67776 0.00000 0.00000 ; HD13- CD1- CG- CD2
5385 5387 5389 5390 3 0.00000 0.00000 0.00000
0.00000 0.00000 0.00000 ; C- N- CA- HA
5385 5387 5389 5391 3 11.71520 1.67360 15.06240 -
13.38880 0.00000 0.00000 ; C- N- CA- CB
5385 5387 5389 5396 3 1.75728 5.27184 2.25936
-7.02912 0.00000 0.00000 ; C- N- CA- C
5386 5385 5387 5388 3 29.28800 -8.36800 -20.92000
0.00000 0.00000 0.00000 ; O- C- N- H
5386 5385 5387 5389 3 20.92000 0.00000 -20.92000
0.00000 0.00000 0.00000 ; O- C- N- CA
5387 5389 5391 5392 3 0.65084 1.95253 0.00000
-2.60338 0.00000 0.00000 ; N- CA- CB- HB2
5387 5389 5391 5393 3 0.65084 1.95253 0.00000
-2.60338 0.00000 0.00000 ; N- CA- CB- HB3
5387 5389 5391 5394 3 7.86174 2.24681 -7.41405
-6.71114 5.35552 0.00000 ; N- CA- CB- OG
5387 5389 5396 5397 3 0.00000 0.00000 0.00000
0.00000 0.00000 0.00000 ; N- CA- C- O
5387 5389 5396 5398 3 17.40544 -5.02080 -13.22144
9.20480 0.00000 0.00000 ; N- CA- C- N
5388 5387 5389 5390 3 0.00000 0.00000 0.00000
0.00000 0.00000 0.00000 ; H- N- CA- HA
5388 5387 5389 5391 3 0.00000 0.00000 0.00000
0.00000 0.00000 0.00000 ; H- N- CA- CB
5388 5387 5389 5396 3 0.00000 0.00000 0.00000
0.00000 0.00000 0.00000 ; H- N- CA- C
5389 5391 5394 5395 3 2.05853 2.46856 3.48109
-4.46851 0.23430 0.00000 ; CA- CB- OG- HG
5389 5396 5398 5399 3 20.92000 0.00000 -20.92000
0.00000 0.00000 0.00000 ; CA- C- N- H
5389 5396 5398 5400 3 20.92000 0.00000 -20.92000

0.00000    0.00000    0.00000 ;     CA-    C-    N-    CA
5390   5389   5391   5392     3   0.65084   1.95253   0.00000
-2.60338    0.00000    0.00000 ;     HA-    CA-    CB-    HB2
5390   5389   5391   5393     3   0.65084   1.95253   0.00000
-2.60338    0.00000    0.00000 ;     HA-    CA-    CB-    HB3
5390   5389   5391   5394     3   1.04600   -1.04600   0.00000
0.00000    0.00000    0.00000 ;     HA-    CA-    CB-    OG
5390   5389   5396   5397     3   3.68192   -4.35136   0.00000
1.33888    0.00000    0.00000 ;     HA-    CA-    C-    O
5390   5389   5396   5398     3   0.00000   0.00000   0.00000
0.00000    0.00000    0.00000 ;     HA-    CA-    C-    N
5391   5389   5396   5397     3   0.00000   0.00000   0.00000
0.00000    0.00000    0.00000 ;     CB-    CA-    C-    O
5391   5389   5396   5398     3   2.51040   4.18400   1.67360
-6.69440    0.00000    0.00000 ;     CB-    CA-    C-    N
5392   5391   5389   5396     3   0.65084   1.95253   0.00000
-2.60338    0.00000    0.00000 ;     HB2-    CB-    CA-    C
5392   5391   5394   5395     3   0.69733   2.09200   0.00000
-2.78933    0.00000    0.00000 ;     HB2-    CB-    OG-    HG
5393   5391   5389   5396     3   0.65084   1.95253   0.00000
-2.60338    0.00000    0.00000 ;     HB3-    CB-    CA-    C
5393   5391   5394   5395     3   0.69733   2.09200   0.00000
-2.78933    0.00000    0.00000 ;     HB3-    CB-    OG-    HG
5394   5391   5389   5396     3   7.34710   7.79898   -6.14211
-6.71114    4.31789    0.00000 ;     OG-    CB-    CA-    C
5396   5398   5400   5401     3   0.00000   0.00000   0.00000
0.00000    0.00000    0.00000 ;     C-    N-    CA-    HA
5396   5398   5400   5402     3   11.71520   1.67360   15.06240   -
13.38880    0.00000    0.00000 ;     C-    N-    CA-    CB
5396   5398   5400   5407     3   1.75728   5.27184   2.25936
-7.02912    0.00000    0.00000 ;     C-    N-    CA-    C
5397   5396   5398   5399     3   29.28800   -8.36800   -20.92000
0.00000    0.00000    0.00000 ;     O-    C-    N-    H
5397   5396   5398   5400     3   20.92000   0.00000   -20.92000
0.00000    0.00000    0.00000 ;     O-    C-    N-    CA
5398   5400   5402   5403     3   0.65084   1.95253   0.00000
-2.60338    0.00000    0.00000 ;     N-    CA-    CB-    HB2
5398   5400   5402   5404     3   0.65084   1.95253   0.00000
-2.60338    0.00000    0.00000 ;     N-    CA-    CB-    HB3
5398   5400   5402   5405     3   7.86174   2.24681   -7.41405
-6.71114    5.35552    0.00000 ;     N-    CA-    CB-    OG
5398   5400   5407   5408     3   0.00000   0.00000   0.00000
0.00000    0.00000    0.00000 ;     N-    CA-    C-    O
5398   5400   5407   5409     3   17.40544   -5.02080   -13.22144
9.20480    0.00000    0.00000 ;     N-    CA-    C-    N
5399   5398   5400   5401     3   0.00000   0.00000   0.00000
0.00000    0.00000    0.00000 ;     H-    N-    CA-    HA
5399   5398   5400   5402     3   0.00000   0.00000   0.00000
0.00000    0.00000    0.00000 ;     H-    N-    CA-    CB
5399   5398   5400   5407     3   0.00000   0.00000   0.00000
0.00000    0.00000    0.00000 ;     H-    N-    CA-    C
5400   5402   5405   5406     3   2.05853   2.46856   3.48109

-4.46851 0.23430 0.00000 ; CA- CB- OG- HG
5400 5407 5409 5410 3 20.92000 0.00000 -20.92000
0.00000 0.00000 0.00000 ; CA- C- N- H
5400 5407 5409 5411 3 20.92000 0.00000 -20.92000
0.00000 0.00000 0.00000 ; CA- C- N- CA
5401 5400 5402 5403 3 0.65084 1.95253 0.00000
-2.60338 0.00000 0.00000 ; HA- CA- CB- HB2
5401 5400 5402 5404 3 0.65084 1.95253 0.00000
-2.60338 0.00000 0.00000 ; HA- CA- CB- HB3
5401 5400 5402 5405 3 1.04600 -1.04600 0.00000
0.00000 0.00000 0.00000 ; HA- CA- CB- OG
5401 5400 5407 5408 3 3.68192 -4.35136 0.00000
1.33888 0.00000 0.00000 ; HA- CA- C- O
5401 5400 5407 5409 3 0.00000 0.00000 0.00000
0.00000 0.00000 0.00000 ; HA- CA- C- N
5402 5400 5407 5408 3 0.00000 0.00000 0.00000
0.00000 0.00000 0.00000 ; CB- CA- C- O
5402 5400 5407 5409 3 2.51040 4.18400 1.67360
-6.69440 0.00000 0.00000 ; CB- CA- C- N
5403 5402 5400 5407 3 0.65084 1.95253 0.00000
-2.60338 0.00000 0.00000 ; HB2- CB- CA- C
5403 5402 5405 5406 3 0.69733 2.09200 0.00000
-2.78933 0.00000 0.00000 ; HB2- CB- OG- HG
5404 5402 5400 5407 3 0.65084 1.95253 0.00000
-2.60338 0.00000 0.00000 ; HB3- CB- CA- C
5404 5402 5405 5406 3 0.69733 2.09200 0.00000
-2.78933 0.00000 0.00000 ; HB3- CB- OG- HG
5405 5402 5400 5407 3 7.34710 7.79898 -6.14211
-6.71114 4.31789 0.00000 ; OG- CB- CA- C
5407 5409 5411 5412 3 0.00000 0.00000 0.00000
0.00000 0.00000 0.00000 ; C- N- CA- HA
5407 5409 5411 5413 3 11.71520 1.67360 15.06240 -
13.38880 0.00000 0.00000 ; C- N- CA- CB
5407 5409 5411 5424 3 1.75728 5.27184 2.25936
-7.02912 0.00000 0.00000 ; C- N- CA- C
5408 5407 5409 5410 3 29.28800 -8.36800 -20.92000
0.00000 0.00000 0.00000 ; O- C- N- H
5408 5407 5409 5411 3 20.92000 0.00000 -20.92000
0.00000 0.00000 0.00000 ; O- C- N- CA
5409 5411 5413 5414 3 0.65084 1.95253 0.00000
-2.60338 0.00000 0.00000 ; N- CA- CB- HB2
5409 5411 5413 5415 3 0.65084 1.95253 0.00000
-2.60338 0.00000 0.00000 ; N- CA- CB- HB3
5409 5411 5413 5416 3 3.21331 2.22589 -4.15053
-2.40998 2.61082 0.00000 ; N- CA- CB- CG
5409 5411 5424 5425 3 0.00000 0.00000 0.00000
0.00000 0.00000 0.00000 ; N- CA- C- O
5409 5411 5424 5426 3 17.40544 -5.02080 -13.22144
9.20480 0.00000 0.00000 ; N- CA- C- N
5410 5409 5411 5412 3 0.00000 0.00000 0.00000
0.00000 0.00000 0.00000 ; H- N- CA- HA
5410 5409 5411 5413 3 0.00000 0.00000 0.00000

```
0.00000    0.00000    0.00000 ;      H-     N-    CA-    CB
  5410   5409   5411   5424      3    0.00000    0.00000    0.00000
0.00000    0.00000    0.00000 ;      H-     N-    CA-     C
  5411   5413   5416   5417      3    0.66944    2.00832    0.00000
-2.67776    0.00000    0.00000 ;     CA-    CB-    CG-   HG2
  5411   5413   5416   5418      3    0.66944    2.00832    0.00000
-2.67776    0.00000    0.00000 ;     CA-    CB-    CG-   HG3
  5411   5413   5416   5419      3    3.69866   -4.35136   -3.92459
6.89523    4.61914    0.00000 ;     CA-    CB-    CG-    CD
  5411   5424   5426   5427      3   20.92000    0.00000  -20.92000
0.00000    0.00000    0.00000 ;     CA-     C-     N-     H
  5411   5424   5426   5428      3   20.92000    0.00000  -20.92000
0.00000    0.00000    0.00000 ;     CA-     C-     N-    CA
  5412   5411   5413   5414      3    0.65084    1.95253    0.00000
-2.60338    0.00000    0.00000 ;     HA-    CA-    CB-   HB2
  5412   5411   5413   5415      3    0.65084    1.95253    0.00000
-2.60338    0.00000    0.00000 ;     HA-    CA-    CB-   HB3
  5412   5411   5413   5416      3    0.65084    1.95253    0.00000
-2.60338    0.00000    0.00000 ;     HA-    CA-    CB-    CG
  5412   5411   5424   5425      3    3.68192   -4.35136    0.00000
1.33888    0.00000    0.00000 ;     HA-    CA-     C-     O
  5412   5411   5424   5426      3    0.00000    0.00000    0.00000
0.00000    0.00000    0.00000 ;     HA-    CA-     C-     N
  5413   5411   5424   5425      3    0.00000    0.00000    0.00000
0.00000    0.00000    0.00000 ;     CB-    CA-     C-     O
  5413   5411   5424   5426      3    2.51040    4.18400    1.67360
-6.69440    0.00000    0.00000 ;     CB-    CA-     C-     N
  5413   5416   5419   5420      3    0.00000    0.00000    0.00000
0.00000    0.00000    0.00000 ;     CB-    CG-    CD-   OE1
  5413   5416   5419   5421      3   10.32611    1.48114   -8.47678
1.42256    1.40582    0.00000 ;     CB-    CG-    CD-   NE2
  5414   5413   5411   5424      3    0.65084    1.95253    0.00000
-2.60338    0.00000    0.00000 ;    HB2-    CB-    CA-     C
  5414   5413   5416   5417      3    0.62760    1.88280    0.00000
-2.51040    0.00000    0.00000 ;    HB2-    CB-    CG-   HG2
  5414   5413   5416   5418      3    0.62760    1.88280    0.00000
-2.51040    0.00000    0.00000 ;    HB2-    CB-    CG-   HG3
  5414   5413   5416   5419      3    0.65084    1.95253    0.00000
-2.60338    0.00000    0.00000 ;    HB2-    CB-    CG-    CD
  5415   5413   5411   5424      3    0.65084    1.95253    0.00000
-2.60338    0.00000    0.00000 ;    HB3-    CB-    CA-     C
  5415   5413   5416   5417      3    0.62760    1.88280    0.00000
-2.51040    0.00000    0.00000 ;    HB3-    CB-    CG-   HG2
  5415   5413   5416   5418      3    0.62760    1.88280    0.00000
-2.51040    0.00000    0.00000 ;    HB3-    CB-    CG-   HG3
  5415   5413   5416   5419      3    0.65084    1.95253    0.00000
-2.60338    0.00000    0.00000 ;    HB3-    CB-    CG-    CD
  5416   5413   5411   5424      3    6.86594    3.56895   -8.14206
-2.40998    4.85344    0.00000 ;     CG-    CB-    CA-     C
  5416   5419   5421   5422      3   20.92000    0.00000  -20.92000
0.00000    0.00000    0.00000 ;     CG-    CD-   NE2-  HE21
  5416   5419   5421   5423      3   20.92000    0.00000  -20.92000
```

0.00000 0.00000 0.00000 ; CG- CD- NE2- HE22
5417 5416 5419 5420 3 3.68192 -4.35136 0.00000
1.33888 0.00000 0.00000 ; HG2- CG- CD- OE1
5417 5416 5419 5421 3 0.00000 0.00000 0.00000
0.00000 0.00000 0.00000 ; HG2- CG- CD- NE2
5418 5416 5419 5420 3 3.68192 -4.35136 0.00000
1.33888 0.00000 0.00000 ; HG3- CG- CD- OE1
5418 5416 5419 5421 3 0.00000 0.00000 0.00000
0.00000 0.00000 0.00000 ; HG3- CG- CD- NE2
5420 5419 5421 5422 3 29.28800 -8.36800 -20.92000
0.00000 0.00000 0.00000 ; OE1- CD- NE2- HE21
5420 5419 5421 5423 3 29.28800 -8.36800 -20.92000
0.00000 0.00000 0.00000 ; OE1- CD- NE2- HE22
5424 5426 5428 5429 3 0.00000 0.00000 0.00000
0.00000 0.00000 0.00000 ; C- N- CA- HA
5424 5426 5428 5430 3 11.71520 1.67360 15.06240 -
13.38880 0.00000 0.00000 ; C- N- CA- CB
5424 5426 5428 5440 3 1.75728 5.27184 2.25936
-7.02912 0.00000 0.00000 ; C- N- CA- C
5425 5424 5426 5427 3 29.28800 -8.36800 -20.92000
0.00000 0.00000 0.00000 ; O- C- N- H
5425 5424 5426 5428 3 20.92000 0.00000 -20.92000
0.00000 0.00000 0.00000 ; O- C- N- CA
5426 5428 5430 5431 3 0.65084 1.95253 0.00000
-2.60338 0.00000 0.00000 ; N- CA- CB- HB2
5426 5428 5430 5432 3 0.65084 1.95253 0.00000
-2.60338 0.00000 0.00000 ; N- CA- CB- HB3
5426 5428 5430 5433 3 3.21331 2.22589 -4.15053
-2.40998 2.61082 0.00000 ; N- CA- CB- CG
5426 5428 5440 5441 3 0.00000 0.00000 0.00000
0.00000 0.00000 0.00000 ; N- CA- C- O
5426 5428 5440 5442 3 17.40544 -5.02080 -13.22144
9.20480 0.00000 0.00000 ; N- CA- C- N
5427 5426 5428 5429 3 0.00000 0.00000 0.00000
0.00000 0.00000 0.00000 ; H- N- CA- HA
5427 5426 5428 5430 3 0.00000 0.00000 0.00000
0.00000 0.00000 0.00000 ; H- N- CA- CB
5427 5426 5428 5440 3 0.00000 0.00000 0.00000
0.00000 0.00000 0.00000 ; H- N- CA- C
5428 5430 5433 5434 3 0.66944 2.00832 0.00000
-2.67776 0.00000 0.00000 ; CA- CB- CG- HG2
5428 5430 5433 5435 3 0.66944 2.00832 0.00000
-2.67776 0.00000 0.00000 ; CA- CB- CG- HG3
5428 5430 5433 5436 3 3.69866 -4.35136 -3.92459
6.89523 4.61914 0.00000 ; CA- CB- CG- CD
5428 5440 5442 5443 3 20.92000 0.00000 -20.92000
0.00000 0.00000 0.00000 ; CA- C- N- H
5428 5440 5442 5444 3 20.92000 0.00000 -20.92000
0.00000 0.00000 0.00000 ; CA- C- N- CA
5429 5428 5430 5431 3 0.65084 1.95253 0.00000
-2.60338 0.00000 0.00000 ; HA- CA- CB- HB2
5429 5428 5430 5432 3 0.65084 1.95253 0.00000

```
-2.60338    0.00000    0.00000 ;      HA-    CA-    CB-   HB3
  5429   5428   5430   5433      3    0.65084    1.95253    0.00000
-2.60338    0.00000    0.00000 ;      HA-    CA-    CB-    CG
  5429   5428   5440   5441      3    3.68192   -4.35136    0.00000
1.33888    0.00000    0.00000 ;      HA-    CA-     C-     O
  5429   5428   5440   5442      3    0.00000    0.00000    0.00000
0.00000    0.00000    0.00000 ;      HA-    CA-     C-     N
  5430   5428   5440   5441      3    0.00000    0.00000    0.00000
0.00000    0.00000    0.00000 ;      CB-    CA-     C-     O
  5430   5428   5440   5442      3    2.51040    4.18400    1.67360
-6.69440    0.00000    0.00000 ;      CB-    CA-     C-     N
  5430   5433   5436   5437      3    0.00000    0.00000    0.00000
0.00000    0.00000    0.00000 ;      CB-    CG-    CD-   OE1
  5430   5433   5436   5438      3   12.79049    3.13382   -7.02912
0.41840   -2.20915    0.00000 ;      CB-    CG-    CD-   OE2
  5431   5430   5428   5440      3    0.65084    1.95253    0.00000
-2.60338    0.00000    0.00000 ;     HB2-    CB-    CA-     C
  5431   5430   5433   5434      3    0.62760    1.88280    0.00000
-2.51040    0.00000    0.00000 ;     HB2-    CB-    CG-   HG2
  5431   5430   5433   5435      3    0.62760    1.88280    0.00000
-2.51040    0.00000    0.00000 ;     HB2-    CB-    CG-   HG3
  5431   5430   5433   5436      3    0.65084    1.95253    0.00000
-2.60338    0.00000    0.00000 ;     HB2-    CB-    CG-    CD
  5432   5430   5428   5440      3    0.65084    1.95253    0.00000
-2.60338    0.00000    0.00000 ;     HB3-    CB-    CA-     C
  5432   5430   5433   5434      3    0.62760    1.88280    0.00000
-2.51040    0.00000    0.00000 ;     HB3-    CB-    CG-   HG2
  5432   5430   5433   5435      3    0.62760    1.88280    0.00000
-2.51040    0.00000    0.00000 ;     HB3-    CB-    CG-   HG3
  5432   5430   5433   5436      3    0.65084    1.95253    0.00000
-2.60338    0.00000    0.00000 ;     HB3-    CB-    CG-    CD
  5433   5430   5428   5440      3    6.86594    3.56895   -8.14206
-2.40998    4.85344    0.00000 ;      CG-    CB-    CA-     C
  5433   5436   5438   5439      3   27.46796    7.88684  -26.42614
-8.01654    3.78234    0.00000 ;      CG-    CD-   OE2-   HE2
  5434   5433   5436   5437      3    3.68192   -4.35136    0.00000
1.33888    0.00000    0.00000 ;     HG2-    CG-    CD-   OE1
  5434   5433   5436   5438      3    0.00000    0.00000    0.00000
0.00000    0.00000    0.00000 ;     HG2-    CG-    CD-   OE2
  5435   5433   5436   5437      3    3.68192   -4.35136    0.00000
1.33888    0.00000    0.00000 ;     HG3-    CG-    CD-   OE1
  5435   5433   5436   5438      3    0.00000    0.00000    0.00000
0.00000    0.00000    0.00000 ;     HG3-    CG-    CD-   OE2
  5437   5436   5438   5439      3   27.19600   -7.94960  -19.24640
0.00000    0.00000    0.00000 ;     OE1-    CD-   OE2-   HE2
  5440   5442   5444   5445      3    0.00000    0.00000    0.00000
0.00000    0.00000    0.00000 ;      C-     N-    CA-    HA
  5440   5442   5444   5446      3   11.71520    1.67360   15.06240  -
13.38880    0.00000    0.00000 ;      C-     N-    CA-    CB
  5440   5442   5444   5459      3    1.75728    5.27184    2.25936
-7.02912    0.00000    0.00000 ;      C-     N-    CA-     C
  5441   5440   5442   5443      3   29.28800   -8.36800  -20.92000
```

```
0.00000    0.00000    0.00000 ;      O-     C-     N-     H
  5441   5440   5442   5444       3   20.92000    0.00000  -20.92000
0.00000    0.00000    0.00000 ;      O-     C-     N-    CA
  5442   5444   5446   5447       3    0.65084    1.95253    0.00000
-2.60338    0.00000    0.00000 ;      N-    CA-    CB-    HB
  5442   5444   5446   5448       3    3.83673    0.44769   -1.77402
-2.47693   -0.03347    0.00000 ;      N-    CA-    CB-   CG2
  5442   5444   5446   5452       3    2.97482    0.12134    2.04179
-1.89117   -3.24678    0.00000 ;      N-    CA-    CB-   CG1
  5442   5444   5459   5460       3    0.00000    0.00000    0.00000
0.00000    0.00000    0.00000 ;      N-    CA-     C-     O
  5442   5444   5459   5461       3   17.40544   -5.02080  -13.22144
9.20480    0.00000    0.00000 ;      N-    CA-     C-     N
  5443   5442   5444   5445       3    0.00000    0.00000    0.00000
0.00000    0.00000    0.00000 ;      H-     N-    CA-    HA
  5443   5442   5444   5446       3    0.00000    0.00000    0.00000
0.00000    0.00000    0.00000 ;      H-     N-    CA-    CB
  5443   5442   5444   5459       3    0.00000    0.00000    0.00000
0.00000    0.00000    0.00000 ;      H-     N-    CA-     C
  5444   5446   5448   5449       3    0.66944    2.00832    0.00000
-2.67776    0.00000    0.00000 ;     CA-    CB-   CG2-  HG21
  5444   5446   5448   5450       3    0.66944    2.00832    0.00000
-2.67776    0.00000    0.00000 ;     CA-    CB-   CG2-  HG22
  5444   5446   5448   5451       3    0.66944    2.00832    0.00000
-2.67776    0.00000    0.00000 ;     CA-    CB-   CG2-  HG23
  5444   5446   5452   5453       3    0.66944    2.00832    0.00000
-2.67776    0.00000    0.00000 ;     CA-    CB-   CG1-  HG12
  5444   5446   5452   5454       3    0.66944    2.00832    0.00000
-2.67776    0.00000    0.00000 ;     CA-    CB-   CG1-  HG13
  5444   5446   5452   5455       3    4.24258   -0.52718   -7.25506
-1.79075    7.69856    0.00000 ;     CA-    CB-   CG1-   CD1
  5444   5459   5461   5462       3   20.92000    0.00000  -20.92000
0.00000    0.00000    0.00000 ;     CA-     C-     N-     H
  5444   5459   5461   5463       3   20.92000    0.00000  -20.92000
0.00000    0.00000    0.00000 ;     CA-     C-     N-    CA
  5445   5444   5446   5447       3    0.65084    1.95253    0.00000
-2.60338    0.00000    0.00000 ;     HA-    CA-    CB-    HB
  5445   5444   5446   5448       3    0.65084    1.95253    0.00000
-2.60338    0.00000    0.00000 ;     HA-    CA-    CB-   CG2
  5445   5444   5446   5452       3    0.65084    1.95253    0.00000
-2.60338    0.00000    0.00000 ;     HA-    CA-    CB-   CG1
  5445   5444   5459   5460       3    3.68192   -4.35136    0.00000
1.33888    0.00000    0.00000 ;     HA-    CA-     C-     O
  5445   5444   5459   5461       3    0.00000    0.00000    0.00000
0.00000    0.00000    0.00000 ;     HA-    CA-     C-     N
  5446   5444   5459   5460       3    0.00000    0.00000    0.00000
0.00000    0.00000    0.00000 ;     CB-    CA-     C-     O
  5446   5444   5459   5461       3    2.51040    4.18400    1.67360
-6.69440    0.00000    0.00000 ;     CB-    CA-     C-     N
  5446   5452   5455   5456       3    0.66944    2.00832    0.00000
-2.67776    0.00000    0.00000 ;     CB-   CG1-   CD1-  HD11
  5446   5452   5455   5457       3    0.66944    2.00832    0.00000
```

-2.67776 0.00000 0.00000 ; CB- CG1- CD1- HD12
5446 5452 5455 5458 3 0.66944 2.00832 0.00000
-2.67776 0.00000 0.00000 ; CB- CG1- CD1- HD13
5447 5446 5444 5459 3 0.65084 1.95253 0.00000
-2.60338 0.00000 0.00000 ; HB- CB- CA- C
5447 5446 5448 5449 3 0.62760 1.88280 0.00000
-2.51040 0.00000 0.00000 ; HB- CB- CG2- HG21
5447 5446 5448 5450 3 0.62760 1.88280 0.00000
-2.51040 0.00000 0.00000 ; HB- CB- CG2- HG22
5447 5446 5448 5451 3 0.62760 1.88280 0.00000
-2.51040 0.00000 0.00000 ; HB- CB- CG2- HG23
5447 5446 5452 5453 3 0.62760 1.88280 0.00000
-2.51040 0.00000 0.00000 ; HB- CB- CG1- HG12
5447 5446 5452 5454 3 0.62760 1.88280 0.00000
-2.51040 0.00000 0.00000 ; HB- CB- CG1- HG13
5447 5446 5452 5455 3 0.66944 2.00832 0.00000
-2.67776 0.00000 0.00000 ; HB- CB- CG1- CD1
5448 5446 5444 5459 3 5.67350 3.55640 -6.16722
-2.47693 3.74886 0.00000 ; CG2- CB- CA- C
5448 5446 5452 5453 3 0.66944 2.00832 0.00000
-2.67776 0.00000 0.00000 ; CG2- CB- CG1- HG12
5448 5446 5452 5454 3 0.66944 2.00832 0.00000
-2.67776 0.00000 0.00000 ; CG2- CB- CG1- HG13
5448 5446 5452 5455 3 3.81162 0.49790 -8.14206
-1.79075 7.49773 0.00000 ; CG2- CB- CG1- CD1
5449 5448 5446 5452 3 0.66944 2.00832 0.00000
-2.67776 0.00000 0.00000 ; HG21- CG2- CB- CG1
5450 5448 5446 5452 3 0.66944 2.00832 0.00000
-2.67776 0.00000 0.00000 ; HG22- CG2- CB- CG1
5451 5448 5446 5452 3 0.66944 2.00832 0.00000
-2.67776 0.00000 0.00000 ; HG23- CG2- CB- CG1
5452 5446 5444 5459 3 8.26340 0.74057 -9.99976
-1.89117 3.84928 0.00000 ; CG1- CB- CA- C
5453 5452 5455 5456 3 0.62760 1.88280 0.00000
-2.51040 0.00000 0.00000 ; HG12- CG1- CD1- HD11
5453 5452 5455 5457 3 0.62760 1.88280 0.00000
-2.51040 0.00000 0.00000 ; HG12- CG1- CD1- HD12
5453 5452 5455 5458 3 0.62760 1.88280 0.00000
-2.51040 0.00000 0.00000 ; HG12- CG1- CD1- HD13
5454 5452 5455 5456 3 0.62760 1.88280 0.00000
-2.51040 0.00000 0.00000 ; HG13- CG1- CD1- HD11
5454 5452 5455 5457 3 0.62760 1.88280 0.00000
-2.51040 0.00000 0.00000 ; HG13- CG1- CD1- HD12
5454 5452 5455 5458 3 0.62760 1.88280 0.00000
-2.51040 0.00000 0.00000 ; HG13- CG1- CD1- HD13
5459 5461 5463 5464 3 0.00000 0.00000 0.00000
0.00000 0.00000 0.00000 ; C- N- CA- HA
5459 5461 5463 5465 3 11.71520 1.67360 15.06240 -
13.38880 0.00000 0.00000 ; C- N- CA- CB
5459 5461 5463 5478 3 1.75728 5.27184 2.25936
-7.02912 0.00000 0.00000 ; C- N- CA- C
5460 5459 5461 5462 3 29.28800 -8.36800 -20.92000

```
0.00000    0.00000    0.00000 ;       O-     C-     N-     H
  5460   5459   5461   5463      3   20.92000    0.00000  -20.92000
0.00000    0.00000    0.00000 ;       O-     C-     N-    CA
  5461   5463   5465   5466      3    0.65084    1.95253    0.00000
-2.60338    0.00000    0.00000 ;      N-    CA-    CB-   HB2
  5461   5463   5465   5467      3    0.65084    1.95253    0.00000
-2.60338    0.00000    0.00000 ;      N-    CA-    CB-   HB3
  5461   5463   5465   5468      3    3.79070    1.39746   -4.61077
-2.40998    2.44346    0.00000 ;      N-    CA-    CB-    CG
  5461   5463   5478   5479      3    0.00000    0.00000    0.00000
0.00000    0.00000    0.00000 ;      N-    CA-     C-     O
  5461   5463   5478   5480      3   17.40544   -5.02080  -13.22144
9.20480    0.00000    0.00000 ;      N-    CA-     C-     N
  5462   5461   5463   5464      3    0.00000    0.00000    0.00000
0.00000    0.00000    0.00000 ;      H-     N-    CA-    HA
  5462   5461   5463   5465      3    0.00000    0.00000    0.00000
0.00000    0.00000    0.00000 ;      H-     N-    CA-    CB
  5462   5461   5463   5478      3    0.00000    0.00000    0.00000
0.00000    0.00000    0.00000 ;      H-     N-    CA-     C
  5463   5465   5468   5469      3    0.66944    2.00832    0.00000
-2.67776    0.00000    0.00000 ;     CA-    CB-    CG-    HG
  5463   5465   5468   5470      3    3.90367    0.19665   -6.21742
-2.37651    5.99149    0.00000 ;     CA-    CB-    CG-   CD1
  5463   5465   5468   5474      3    3.90367    0.19665   -6.21742
-2.37651    5.99149    0.00000 ;     CA-    CB-    CG-   CD2
  5463   5478   5480   5481      3   20.92000    0.00000  -20.92000
0.00000    0.00000    0.00000 ;     CA-     C-     N-     H
  5463   5478   5480   5482      3   20.92000    0.00000  -20.92000
0.00000    0.00000    0.00000 ;     CA-     C-     N-    CA
  5464   5463   5465   5466      3    0.65084    1.95253    0.00000
-2.60338    0.00000    0.00000 ;     HA-    CA-    CB-   HB2
  5464   5463   5465   5467      3    0.65084    1.95253    0.00000
-2.60338    0.00000    0.00000 ;     HA-    CA-    CB-   HB3
  5464   5463   5465   5468      3    0.65084    1.95253    0.00000
-2.60338    0.00000    0.00000 ;     HA-    CA-    CB-    CG
  5464   5463   5478   5479      3    3.68192   -4.35136    0.00000
1.33888    0.00000    0.00000 ;     HA-    CA-     C-     O
  5464   5463   5478   5480      3    0.00000    0.00000    0.00000
0.00000    0.00000    0.00000 ;     HA-    CA-     C-     N
  5465   5463   5478   5479      3    0.00000    0.00000    0.00000
0.00000    0.00000    0.00000 ;     CB-    CA-     C-     O
  5465   5463   5478   5480      3    2.51040    4.18400    1.67360
-6.69440    0.00000    0.00000 ;     CB-    CA-     C-     N
  5465   5468   5470   5471      3    0.66944    2.00832    0.00000
-2.67776    0.00000    0.00000 ;     CB-    CG-   CD1-  HD11
  5465   5468   5470   5472      3    0.66944    2.00832    0.00000
-2.67776    0.00000    0.00000 ;     CB-    CG-   CD1-  HD12
  5465   5468   5470   5473      3    0.66944    2.00832    0.00000
-2.67776    0.00000    0.00000 ;     CB-    CG-   CD1-  HD13
  5465   5468   5474   5475      3    0.66944    2.00832    0.00000
-2.67776    0.00000    0.00000 ;     CB-    CG-   CD2-  HD21
  5465   5468   5474   5476      3    0.66944    2.00832    0.00000
```

-2.67776 0.00000 0.00000 ; CB- CG- CD2- HD22
5465 5468 5474 5477 3 0.66944 2.00832 0.00000
-2.67776 0.00000 0.00000 ; CB- CG- CD2- HD23
5466 5465 5463 5478 3 0.65084 1.95253 0.00000
-2.60338 0.00000 0.00000 ; HB2- CB- CA- C
5466 5465 5468 5469 3 0.62760 1.88280 0.00000
-2.51040 0.00000 0.00000 ; HB2- CB- CG- HG
5466 5465 5468 5470 3 0.66944 2.00832 0.00000
-2.67776 0.00000 0.00000 ; HB2- CB- CG- CD1
5466 5465 5468 5474 3 0.66944 2.00832 0.00000
-2.67776 0.00000 0.00000 ; HB2- CB- CG- CD2
5467 5465 5463 5478 3 0.65084 1.95253 0.00000
-2.60338 0.00000 0.00000 ; HB3- CB- CA- C
5467 5465 5468 5469 3 0.62760 1.88280 0.00000
-2.51040 0.00000 0.00000 ; HB3- CB- CG- HG
5467 5465 5468 5470 3 0.66944 2.00832 0.00000
-2.67776 0.00000 0.00000 ; HB3- CB- CG- CD1
5467 5465 5468 5474 3 0.66944 2.00832 0.00000
-2.67776 0.00000 0.00000 ; HB3- CB- CG- CD2
5468 5465 5463 5478 3 10.33448 -1.14642 -11.54784
-2.40998 6.35968 0.00000 ; CG- CB- CA- C
5469 5468 5470 5471 3 0.62760 1.88280 0.00000
-2.51040 0.00000 0.00000 ; HG- CG- CD1- HD11
5469 5468 5470 5472 3 0.62760 1.88280 0.00000
-2.51040 0.00000 0.00000 ; HG- CG- CD1- HD12
5469 5468 5470 5473 3 0.62760 1.88280 0.00000
-2.51040 0.00000 0.00000 ; HG- CG- CD1- HD13
5469 5468 5474 5475 3 0.62760 1.88280 0.00000
-2.51040 0.00000 0.00000 ; HG- CG- CD2- HD21
5469 5468 5474 5476 3 0.62760 1.88280 0.00000
-2.51040 0.00000 0.00000 ; HG- CG- CD2- HD22
5469 5468 5474 5477 3 0.62760 1.88280 0.00000
-2.51040 0.00000 0.00000 ; HG- CG- CD2- HD23
5470 5468 5474 5475 3 0.66944 2.00832 0.00000
-2.67776 0.00000 0.00000 ; CD1- CG- CD2- HD21
5470 5468 5474 5476 3 0.66944 2.00832 0.00000
-2.67776 0.00000 0.00000 ; CD1- CG- CD2- HD22
5470 5468 5474 5477 3 0.66944 2.00832 0.00000
-2.67776 0.00000 0.00000 ; CD1- CG- CD2- HD23
5471 5470 5468 5474 3 0.66944 2.00832 0.00000
-2.67776 0.00000 0.00000 ; HD11- CD1- CG- CD2
5472 5470 5468 5474 3 0.66944 2.00832 0.00000
-2.67776 0.00000 0.00000 ; HD12- CD1- CG- CD2
5473 5470 5468 5474 3 0.66944 2.00832 0.00000
-2.67776 0.00000 0.00000 ; HD13- CD1- CG- CD2
5478 5480 5482 5483 3 0.00000 0.00000 0.00000
0.00000 0.00000 0.00000 ; C- N- CA- HA
5478 5480 5482 5484 3 11.71520 1.67360 15.06240 -
13.38880 0.00000 0.00000 ; C- N- CA- CB
5478 5480 5482 5490 3 1.75728 5.27184 2.25936
-7.02912 0.00000 0.00000 ; C- N- CA- C
5479 5478 5480 5481 3 29.28800 -8.36800 -20.92000

```
0.00000    0.00000    0.00000 ;       O-      C-      N-      H
  5479   5478   5480   5482       3   20.92000    0.00000  -20.92000
0.00000    0.00000    0.00000 ;       O-      C-      N-     CA
  5480   5482   5484   5485       3    0.65084    1.95253    0.00000
-2.60338    0.00000    0.00000 ;      N-     CA-     CB-    HB2
  5480   5482   5484   5486       3    0.65084    1.95253    0.00000
-2.60338    0.00000    0.00000 ;      N-     CA-     CB-    HB3
  5480   5482   5484   5487       3   15.41386    9.74035   -8.39310
-0.97069    2.97901    0.00000 ;      N-     CA-     CB-     CG
  5480   5482   5490   5491       3    0.00000    0.00000    0.00000
0.00000    0.00000    0.00000 ;       N-     CA-      C-      O
  5480   5482   5490   5492       3   17.40544   -5.02080  -13.22144
9.20480    0.00000    0.00000 ;       N-     CA-      C-      N
  5481   5480   5482   5483       3    0.00000    0.00000    0.00000
0.00000    0.00000    0.00000 ;       H-      N-     CA-     HA
  5481   5480   5482   5484       3    0.00000    0.00000    0.00000
0.00000    0.00000    0.00000 ;       H-      N-     CA-     CB
  5481   5480   5482   5490       3    0.00000    0.00000    0.00000
0.00000    0.00000    0.00000 ;       H-      N-     CA-      C
  5482   5484   5487   5488       3    6.43499    0.00000   -5.39736
0.00000   -1.03763    0.00000 ;      CA-     CB-     CG-    OD1
  5482   5484   5487   5489       3    6.43499    0.00000   -5.39736
0.00000   -1.03763    0.00000 ;      CA-     CB-     CG-    OD2
  5482   5490   5492   5493       3   20.92000    0.00000  -20.92000
0.00000    0.00000    0.00000 ;      CA-      C-      N-      H
  5482   5490   5492   5494       3   20.92000    0.00000  -20.92000
0.00000    0.00000    0.00000 ;      CA-      C-      N-     CA
  5483   5482   5484   5485       3    0.65084    1.95253    0.00000
-2.60338    0.00000    0.00000 ;     HA-     CA-     CB-    HB2
  5483   5482   5484   5486       3    0.65084    1.95253    0.00000
-2.60338    0.00000    0.00000 ;     HA-     CA-     CB-    HB3
  5483   5482   5484   5487       3    0.65084    1.95253    0.00000
-2.60338    0.00000    0.00000 ;     HA-     CA-     CB-     CG
  5483   5482   5490   5491       3    3.68192   -4.35136    0.00000
1.33888    0.00000    0.00000 ;      HA-     CA-      C-      O
  5483   5482   5490   5492       3    0.00000    0.00000    0.00000
0.00000    0.00000    0.00000 ;      HA-     CA-      C-      N
  5484   5482   5490   5491       3    0.00000    0.00000    0.00000
0.00000    0.00000    0.00000 ;      CB-     CA-      C-      O
  5484   5482   5490   5492       3    2.51040    4.18400    1.67360
-6.69440    0.00000    0.00000 ;     CB-     CA-      C-      N
  5485   5484   5482   5490       3    0.65084    1.95253    0.00000
-2.60338    0.00000    0.00000 ;    HB2-     CB-     CA-      C
  5485   5484   5487   5488       3    0.00000    0.00000    0.00000
0.00000    0.00000    0.00000 ;     HB2-     CB-     CG-    OD1
  5485   5484   5487   5489       3    0.00000    0.00000    0.00000
0.00000    0.00000    0.00000 ;     HB2-     CB-     CG-    OD2
  5486   5484   5482   5490       3    0.65084    1.95253    0.00000
-2.60338    0.00000    0.00000 ;    HB3-     CB-     CA-      C
  5486   5484   5487   5488       3    0.00000    0.00000    0.00000
0.00000    0.00000    0.00000 ;     HB3-     CB-     CG-    OD1
  5486   5484   5487   5489       3    0.00000    0.00000    0.00000
```

0.00000 0.00000 0.00000 ; HB3- CB- CG- OD2
5487 5484 5482 5490 3 7.14627 -1.04600 -8.99560
-0.97069 5.15469 0.00000 ; CG- CB- CA- C
5490 5492 5494 5495 3 0.00000 0.00000 0.00000
0.00000 0.00000 0.00000 ; C- N- CA- HA2
5490 5492 5494 5496 3 0.00000 0.00000 0.00000
0.00000 0.00000 0.00000 ; C- N- CA- HA3
5490 5492 5494 5497 3 1.75728 5.27184 2.25936
-7.02912 0.00000 0.00000 ; C- N- CA- C
5491 5490 5492 5493 3 29.28800 -8.36800 -20.92000
0.00000 0.00000 0.00000 ; O- C- N- H
5491 5490 5492 5494 3 20.92000 0.00000 -20.92000
0.00000 0.00000 0.00000 ; O- C- N- CA
5492 5494 5497 5498 3 0.00000 0.00000 0.00000
0.00000 0.00000 0.00000 ; N- CA- C- O
5492 5494 5497 5499 3 17.40544 -5.02080 -13.22144
9.20480 0.00000 0.00000 ; N- CA- C- N
5493 5492 5494 5495 3 0.00000 0.00000 0.00000
0.00000 0.00000 0.00000 ; H- N- CA- HA2
5493 5492 5494 5496 3 0.00000 0.00000 0.00000
0.00000 0.00000 0.00000 ; H- N- CA- HA3
5493 5492 5494 5497 3 0.00000 0.00000 0.00000
0.00000 0.00000 0.00000 ; H- N- CA- C
5494 5497 5499 5500 3 20.92000 0.00000 -20.92000
0.00000 0.00000 0.00000 ; CA- C- N- H
5494 5497 5499 5501 3 20.92000 0.00000 -20.92000
0.00000 0.00000 0.00000 ; CA- C- N- CA
5495 5494 5497 5498 3 3.68192 -4.35136 0.00000
1.33888 0.00000 0.00000 ; HA2- CA- C- O
5495 5494 5497 5499 3 0.00000 0.00000 0.00000
0.00000 0.00000 0.00000 ; HA2- CA- C- N
5496 5494 5497 5498 3 3.68192 -4.35136 0.00000
1.33888 0.00000 0.00000 ; HA3- CA- C- O
5496 5494 5497 5499 3 0.00000 0.00000 0.00000
0.00000 0.00000 0.00000 ; HA3- CA- C- N
5497 5499 5501 5502 3 0.00000 0.00000 0.00000
0.00000 0.00000 0.00000 ; C- N- CA- HA
5497 5499 5501 5503 3 11.71520 1.67360 15.06240 -
13.38880 0.00000 0.00000 ; C- N- CA- CB
5497 5499 5501 5516 3 1.75728 5.27184 2.25936
-7.02912 0.00000 0.00000 ; C- N- CA- C
5498 5497 5499 5500 3 29.28800 -8.36800 -20.92000
0.00000 0.00000 0.00000 ; O- C- N- H
5498 5497 5499 5501 3 20.92000 0.00000 -20.92000
0.00000 0.00000 0.00000 ; O- C- N- CA
5499 5501 5503 5504 3 0.65084 1.95253 0.00000
-2.60338 0.00000 0.00000 ; N- CA- CB- HB2
5499 5501 5503 5505 3 0.65084 1.95253 0.00000
-2.60338 0.00000 0.00000 ; N- CA- CB- HB3
5499 5501 5503 5506 3 3.79070 1.39746 -4.61077
-2.40998 2.44346 0.00000 ; N- CA- CB- CG
5499 5501 5516 5517 3 0.00000 0.00000 0.00000

```
0.00000    0.00000    0.00000 ;       N-    CA-     C-     O
  5499   5501   5516   5518       3   17.40544   -5.02080  -13.22144
9.20480    0.00000    0.00000 ;       N-    CA-     C-     N
  5500   5499   5501   5502       3    0.00000    0.00000    0.00000
0.00000    0.00000    0.00000 ;       H-     N-    CA-    HA
  5500   5499   5501   5503       3    0.00000    0.00000    0.00000
0.00000    0.00000    0.00000 ;       H-     N-    CA-    CB
  5500   5499   5501   5516       3    0.00000    0.00000    0.00000
0.00000    0.00000    0.00000 ;       H-     N-    CA-     C
  5501   5503   5506   5507       3    0.66944    2.00832    0.00000
-2.67776    0.00000    0.00000 ;      CA-    CB-    CG-    HG
  5501   5503   5506   5508       3    3.90367    0.19665   -6.21742
-2.37651    5.99149    0.00000 ;      CA-    CB-    CG-   CD1
  5501   5503   5506   5512       3    3.90367    0.19665   -6.21742
-2.37651    5.99149    0.00000 ;      CA-    CB-    CG-   CD2
  5501   5516   5518   5519       3   20.92000    0.00000  -20.92000
0.00000    0.00000    0.00000 ;      CA-     C-     N-     H
  5501   5516   5518   5520       3   20.92000    0.00000  -20.92000
0.00000    0.00000    0.00000 ;      CA-     C-     N-    CA
  5502   5501   5503   5504       3    0.65084    1.95253    0.00000
-2.60338    0.00000    0.00000 ;      HA-    CA-    CB-   HB2
  5502   5501   5503   5505       3    0.65084    1.95253    0.00000
-2.60338    0.00000    0.00000 ;      HA-    CA-    CB-   HB3
  5502   5501   5503   5506       3    0.65084    1.95253    0.00000
-2.60338    0.00000    0.00000 ;      HA-    CA-    CB-    CG
  5502   5501   5516   5517       3    3.68192   -4.35136    0.00000
1.33888    0.00000    0.00000 ;      HA-    CA-     C-     O
  5502   5501   5516   5518       3    0.00000    0.00000    0.00000
0.00000    0.00000    0.00000 ;      HA-    CA-     C-     N
  5503   5501   5516   5517       3    0.00000    0.00000    0.00000
0.00000    0.00000    0.00000 ;      CB-    CA-     C-     O
  5503   5501   5516   5518       3    2.51040    4.18400    1.67360
-6.69440    0.00000    0.00000 ;      CB-    CA-     C-     N
  5503   5506   5508   5509       3    0.66944    2.00832    0.00000
-2.67776    0.00000    0.00000 ;      CB-    CG-   CD1-  HD11
  5503   5506   5508   5510       3    0.66944    2.00832    0.00000
-2.67776    0.00000    0.00000 ;      CB-    CG-   CD1-  HD12
  5503   5506   5508   5511       3    0.66944    2.00832    0.00000
-2.67776    0.00000    0.00000 ;      CB-    CG-   CD1-  HD13
  5503   5506   5512   5513       3    0.66944    2.00832    0.00000
-2.67776    0.00000    0.00000 ;      CB-    CG-   CD2-  HD21
  5503   5506   5512   5514       3    0.66944    2.00832    0.00000
-2.67776    0.00000    0.00000 ;      CB-    CG-   CD2-  HD22
  5503   5506   5512   5515       3    0.66944    2.00832    0.00000
-2.67776    0.00000    0.00000 ;      CB-    CG-   CD2-  HD23
  5504   5503   5501   5516       3    0.65084    1.95253    0.00000
-2.60338    0.00000    0.00000 ;     HB2-    CB-    CA-     C
  5504   5503   5506   5507       3    0.62760    1.88280    0.00000
-2.51040    0.00000    0.00000 ;     HB2-    CB-    CG-    HG
  5504   5503   5506   5508       3    0.66944    2.00832    0.00000
-2.67776    0.00000    0.00000 ;     HB2-    CB-    CG-   CD1
  5504   5503   5506   5512       3    0.66944    2.00832    0.00000
```

-2.67776 0.00000 0.00000 ; HB2- CB- CG- CD2
5505 5503 5501 5516 3 0.65084 1.95253 0.00000
-2.60338 0.00000 0.00000 ; HB3- CB- CA- C
5505 5503 5506 5507 3 0.62760 1.88280 0.00000
-2.51040 0.00000 0.00000 ; HB3- CB- CG- HG
5505 5503 5506 5508 3 0.66944 2.00832 0.00000
-2.67776 0.00000 0.00000 ; HB3- CB- CG- CD1
5505 5503 5506 5512 3 0.66944 2.00832 0.00000
-2.67776 0.00000 0.00000 ; HB3- CB- CG- CD2
5506 5503 5501 5516 3 10.33448 -1.14642 -11.54784
-2.40998 6.35968 0.00000 ; CG- CB- CA- C
5507 5506 5508 5509 3 0.62760 1.88280 0.00000
-2.51040 0.00000 0.00000 ; HG- CG- CD1- HD11
5507 5506 5508 5510 3 0.62760 1.88280 0.00000
-2.51040 0.00000 0.00000 ; HG- CG- CD1- HD12
5507 5506 5508 5511 3 0.62760 1.88280 0.00000
-2.51040 0.00000 0.00000 ; HG- CG- CD1- HD13
5507 5506 5512 5513 3 0.62760 1.88280 0.00000
-2.51040 0.00000 0.00000 ; HG- CG- CD2- HD21
5507 5506 5512 5514 3 0.62760 1.88280 0.00000
-2.51040 0.00000 0.00000 ; HG- CG- CD2- HD22
5507 5506 5512 5515 3 0.62760 1.88280 0.00000
-2.51040 0.00000 0.00000 ; HG- CG- CD2- HD23
5508 5506 5512 5513 3 0.66944 2.00832 0.00000
-2.67776 0.00000 0.00000 ; CD1- CG- CD2- HD21
5508 5506 5512 5514 3 0.66944 2.00832 0.00000
-2.67776 0.00000 0.00000 ; CD1- CG- CD2- HD22
5508 5506 5512 5515 3 0.66944 2.00832 0.00000
-2.67776 0.00000 0.00000 ; CD1- CG- CD2- HD23
5509 5508 5506 5512 3 0.66944 2.00832 0.00000
-2.67776 0.00000 0.00000 ; HD11- CD1- CG- CD2
5510 5508 5506 5512 3 0.66944 2.00832 0.00000
-2.67776 0.00000 0.00000 ; HD12- CD1- CG- CD2
5511 5508 5506 5512 3 0.66944 2.00832 0.00000
-2.67776 0.00000 0.00000 ; HD13- CD1- CG- CD2
5516 5518 5520 5521 3 0.00000 0.00000 0.00000
0.00000 0.00000 0.00000 ; C- N- CA- HA
5516 5518 5520 5522 3 11.71520 1.67360 15.06240 -
13.38880 0.00000 0.00000 ; C- N- CA- CB
5516 5518 5520 5538 3 1.75728 5.27184 2.25936
-7.02912 0.00000 0.00000 ; C- N- CA- C
5517 5516 5518 5519 3 29.28800 -8.36800 -20.92000
0.00000 0.00000 0.00000 ; O- C- N- H
5517 5516 5518 5520 3 20.92000 0.00000 -20.92000
0.00000 0.00000 0.00000 ; O- C- N- CA
5518 5520 5522 5523 3 0.65084 1.95253 0.00000
-2.60338 0.00000 0.00000 ; N- CA- CB- HB2
5518 5520 5522 5524 3 0.65084 1.95253 0.00000
-2.60338 0.00000 0.00000 ; N- CA- CB- HB3
5518 5520 5522 5525 3 0.65084 1.95253 0.00000
-2.60338 0.00000 0.00000 ; N- CA- CB- CG
5518 5520 5538 5539 3 0.00000 0.00000 0.00000

0.00000    0.00000    0.00000 ;      N-    CA-     C-     O
  5518   5520   5538   5540      3   17.40544   -5.02080  -13.22144
9.20480    0.00000    0.00000 ;      N-    CA-     C-     N
  5519   5518   5520   5521      3    0.00000    0.00000    0.00000
0.00000    0.00000    0.00000 ;      H-     N-    CA-    HA
  5519   5518   5520   5522      3    0.00000    0.00000    0.00000
0.00000    0.00000    0.00000 ;      H-     N-    CA-    CB
  5519   5518   5520   5538      3    0.00000    0.00000    0.00000
0.00000    0.00000    0.00000 ;      H-     N-    CA-     C
  5520   5522   5525   5526      3    0.66944    2.00832    0.00000
-2.67776    0.00000    0.00000 ;     CA-    CB-    CG-   HG2
  5520   5522   5525   5527      3    0.66944    2.00832    0.00000
-2.67776    0.00000    0.00000 ;     CA-    CB-    CG-   HG3
  5520   5522   5525   5528      3    3.68192    3.09616   -2.09200
-3.01248    0.00000    0.00000 ;     CA-    CB-    CG-    CD
  5520   5538   5540   5541      3   20.92000    0.00000  -20.92000
0.00000    0.00000    0.00000 ;     CA-     C-     N-     H
  5520   5538   5540   5542      3   20.92000    0.00000  -20.92000
0.00000    0.00000    0.00000 ;     CA-     C-     N-    CA
  5521   5520   5522   5523      3    0.65084    1.95253    0.00000
-2.60338    0.00000    0.00000 ;     HA-    CA-    CB-   HB2
  5521   5520   5522   5524      3    0.65084    1.95253    0.00000
-2.60338    0.00000    0.00000 ;     HA-    CA-    CB-   HB3
  5521   5520   5522   5525      3    0.65084    1.95253    0.00000
-2.60338    0.00000    0.00000 ;     HA-    CA-    CB-    CG
  5521   5520   5538   5539      3    3.68192   -4.35136    0.00000
1.33888    0.00000    0.00000 ;     HA-    CA-     C-     O
  5521   5520   5538   5540      3    0.00000    0.00000    0.00000
0.00000    0.00000    0.00000 ;     HA-    CA-     C-     N
  5522   5520   5538   5539      3    0.00000    0.00000    0.00000
0.00000    0.00000    0.00000 ;     CB-    CA-     C-     O
  5522   5520   5538   5540      3    2.51040    4.18400    1.67360
-6.69440    0.00000    0.00000 ;     CB-    CA-     C-     N
  5522   5525   5528   5529      3    0.66944    2.00832    0.00000
-2.67776    0.00000    0.00000 ;     CB-    CG-    CD-   HD2
  5522   5525   5528   5530      3    0.66944    2.00832    0.00000
-2.67776    0.00000    0.00000 ;     CB-    CG-    CD-   HD3
  5522   5525   5528   5531      3    3.68192    3.09616   -2.09200
-3.01248    0.00000    0.00000 ;     CB-    CG-    CD-    CE
  5523   5522   5520   5538      3    0.65084    1.95253    0.00000
-2.60338    0.00000    0.00000 ;    HB2-    CB-    CA-     C
  5523   5522   5525   5526      3    0.62760    1.88280    0.00000
-2.51040    0.00000    0.00000 ;    HB2-    CB-    CG-   HG2
  5523   5522   5525   5527      3    0.62760    1.88280    0.00000
-2.51040    0.00000    0.00000 ;    HB2-    CB-    CG-   HG3
  5523   5522   5525   5528      3    0.66944    2.00832    0.00000
-2.67776    0.00000    0.00000 ;    HB2-    CB-    CG-    CD
  5524   5522   5520   5538      3    0.65084    1.95253    0.00000
-2.60338    0.00000    0.00000 ;    HB3-    CB-    CA-     C
  5524   5522   5525   5526      3    0.62760    1.88280    0.00000
-2.51040    0.00000    0.00000 ;    HB3-    CB-    CG-   HG2
  5524   5522   5525   5527      3    0.62760    1.88280    0.00000

```
-2.51040    0.00000    0.00000 ;    HB3-    CB-    CG-   HG3
  5524   5522   5525   5528      3    0.66944    2.00832    0.00000
-2.67776    0.00000    0.00000 ;    HB3-    CB-    CG-    CD
  5525   5522   5520   5538      3    0.65084    1.95253    0.00000
-2.60338    0.00000    0.00000 ;     CG-    CB-    CA-     C
  5525   5528   5531   5532      3    0.65084    1.95253    0.00000
-2.60338    0.00000    0.00000 ;     CG-    CD-    CE-   HE2
  5525   5528   5531   5533      3    0.65084    1.95253    0.00000
-2.60338    0.00000    0.00000 ;     CG-    CD-    CE-   HE3
  5525   5528   5531   5534      3    0.65084    1.95253    0.00000
-2.60338    0.00000    0.00000 ;     CG-    CD-    CE-    NZ
  5526   5525   5528   5529      3    0.62760    1.88280    0.00000
-2.51040    0.00000    0.00000 ;    HG2-    CG-    CD-   HD2
  5526   5525   5528   5530      3    0.62760    1.88280    0.00000
-2.51040    0.00000    0.00000 ;    HG2-    CG-    CD-   HD3
  5526   5525   5528   5531      3    0.66944    2.00832    0.00000
-2.67776    0.00000    0.00000 ;    HG2-    CG-    CD-    CE
  5527   5525   5528   5529      3    0.62760    1.88280    0.00000
-2.51040    0.00000    0.00000 ;    HG3-    CG-    CD-   HD2
  5527   5525   5528   5530      3    0.62760    1.88280    0.00000
-2.51040    0.00000    0.00000 ;    HG3-    CG-    CD-   HD3
  5527   5525   5528   5531      3    0.66944    2.00832    0.00000
-2.67776    0.00000    0.00000 ;    HG3-    CG-    CD-    CE
  5528   5531   5534   5535      3    0.65084    1.95253    0.00000
-2.60338    0.00000    0.00000 ;     CD-    CE-    NZ-   HZ1
  5528   5531   5534   5536      3    0.65084    1.95253    0.00000
-2.60338    0.00000    0.00000 ;     CD-    CE-    NZ-   HZ2
  5528   5531   5534   5537      3    0.65084    1.95253    0.00000
-2.60338    0.00000    0.00000 ;     CD-    CE-    NZ-   HZ3
  5529   5528   5531   5532      3    0.65084    1.95253    0.00000
-2.60338    0.00000    0.00000 ;    HD2-    CD-    CE-   HE2
  5529   5528   5531   5533      3    0.65084    1.95253    0.00000
-2.60338    0.00000    0.00000 ;    HD2-    CD-    CE-   HE3
  5529   5528   5531   5534      3    0.65084    1.95253    0.00000
-2.60338    0.00000    0.00000 ;    HD2-    CD-    CE-    NZ
  5530   5528   5531   5532      3    0.65084    1.95253    0.00000
-2.60338    0.00000    0.00000 ;    HD3-    CD-    CE-   HE2
  5530   5528   5531   5533      3    0.65084    1.95253    0.00000
-2.60338    0.00000    0.00000 ;    HD3-    CD-    CE-   HE3
  5530   5528   5531   5534      3    0.65084    1.95253    0.00000
-2.60338    0.00000    0.00000 ;    HD3-    CD-    CE-    NZ
  5532   5531   5534   5535      3    0.65084    1.95253    0.00000
-2.60338    0.00000    0.00000 ;    HE2-    CE-    NZ-   HZ1
  5532   5531   5534   5536      3    0.65084    1.95253    0.00000
-2.60338    0.00000    0.00000 ;    HE2-    CE-    NZ-   HZ2
  5532   5531   5534   5537      3    0.65084    1.95253    0.00000
-2.60338    0.00000    0.00000 ;    HE2-    CE-    NZ-   HZ3
  5533   5531   5534   5535      3    0.65084    1.95253    0.00000
-2.60338    0.00000    0.00000 ;    HE3-    CE-    NZ-   HZ1
  5533   5531   5534   5536      3    0.65084    1.95253    0.00000
-2.60338    0.00000    0.00000 ;    HE3-    CE-    NZ-   HZ2
  5533   5531   5534   5537      3    0.65084    1.95253    0.00000
```

-2.60338 0.00000 0.00000 ; HE3- CE- NZ- HZ3
5538 5540 5542 5543 3 0.00000 0.00000 0.00000
0.00000 0.00000 0.00000 ; C- N- CA- HA
5538 5540 5542 5544 3 11.71520 1.67360 15.06240 -
13.38880 0.00000 0.00000 ; C- N- CA- CB
5538 5540 5542 5560 3 1.75728 5.27184 2.25936
-7.02912 0.00000 0.00000 ; C- N- CA- C
5539 5538 5540 5541 3 29.28800 -8.36800 -20.92000
0.00000 0.00000 0.00000 ; O- C- N- H
5539 5538 5540 5542 3 20.92000 0.00000 -20.92000
0.00000 0.00000 0.00000 ; O- C- N- CA
5540 5542 5544 5545 3 0.65084 1.95253 0.00000
-2.60338 0.00000 0.00000 ; N- CA- CB- HB2
5540 5542 5544 5546 3 0.65084 1.95253 0.00000
-2.60338 0.00000 0.00000 ; N- CA- CB- HB3
5540 5542 5544 5547 3 0.65084 1.95253 0.00000
-2.60338 0.00000 0.00000 ; N- CA- CB- CG
5540 5542 5560 5561 3 0.00000 0.00000 0.00000
0.00000 0.00000 0.00000 ; N- CA- C- O
5540 5542 5560 5562 3 17.40544 -5.02080 -13.22144
9.20480 0.00000 0.00000 ; N- CA- C- N
5541 5540 5542 5543 3 0.00000 0.00000 0.00000
0.00000 0.00000 0.00000 ; H- N- CA- HA
5541 5540 5542 5544 3 0.00000 0.00000 0.00000
0.00000 0.00000 0.00000 ; H- N- CA- CB
5541 5540 5542 5560 3 0.00000 0.00000 0.00000
0.00000 0.00000 0.00000 ; H- N- CA- C
5542 5544 5547 5548 3 0.66944 2.00832 0.00000
-2.67776 0.00000 0.00000 ; CA- CB- CG- HG2
5542 5544 5547 5549 3 0.66944 2.00832 0.00000
-2.67776 0.00000 0.00000 ; CA- CB- CG- HG3
5542 5544 5547 5550 3 3.68192 3.09616 -2.09200
-3.01248 0.00000 0.00000 ; CA- CB- CG- CD
5542 5560 5562 5563 3 20.92000 0.00000 -20.92000
0.00000 0.00000 0.00000 ; CA- C- N- H
5542 5560 5562 5564 3 20.92000 0.00000 -20.92000
0.00000 0.00000 0.00000 ; CA- C- N- CA
5543 5542 5544 5545 3 0.65084 1.95253 0.00000
-2.60338 0.00000 0.00000 ; HA- CA- CB- HB2
5543 5542 5544 5546 3 0.65084 1.95253 0.00000
-2.60338 0.00000 0.00000 ; HA- CA- CB- HB3
5543 5542 5544 5547 3 0.65084 1.95253 0.00000
-2.60338 0.00000 0.00000 ; HA- CA- CB- CG
5543 5542 5560 5561 3 3.68192 -4.35136 0.00000
1.33888 0.00000 0.00000 ; HA- CA- C- O
5543 5542 5560 5562 3 0.00000 0.00000 0.00000
0.00000 0.00000 0.00000 ; HA- CA- C- N
5544 5542 5560 5561 3 0.00000 0.00000 0.00000
0.00000 0.00000 0.00000 ; CB- CA- C- O
5544 5542 5560 5562 3 2.51040 4.18400 1.67360
-6.69440 0.00000 0.00000 ; CB- CA- C- N
5544 5547 5550 5551 3 0.66944 2.00832 0.00000

```
-2.67776    0.00000    0.00000 ;      CB-    CG-    CD-   HD2
  5544   5547   5550   5552      3    0.66944    2.00832    0.00000
-2.67776    0.00000    0.00000 ;      CB-    CG-    CD-   HD3
  5544   5547   5550   5553      3    3.68192    3.09616   -2.09200
-3.01248    0.00000    0.00000 ;      CB-    CG-    CD-    CE
  5545   5544   5542   5560      3    0.65084    1.95253    0.00000
-2.60338    0.00000    0.00000 ;     HB2-    CB-    CA-     C
  5545   5544   5547   5548      3    0.62760    1.88280    0.00000
-2.51040    0.00000    0.00000 ;     HB2-    CB-    CG-   HG2
  5545   5544   5547   5549      3    0.62760    1.88280    0.00000
-2.51040    0.00000    0.00000 ;     HB2-    CB-    CG-   HG3
  5545   5544   5547   5550      3    0.66944    2.00832    0.00000
-2.67776    0.00000    0.00000 ;     HB2-    CB-    CG-    CD
  5546   5544   5542   5560      3    0.65084    1.95253    0.00000
-2.60338    0.00000    0.00000 ;     HB3-    CB-    CA-     C
  5546   5544   5547   5548      3    0.62760    1.88280    0.00000
-2.51040    0.00000    0.00000 ;     HB3-    CB-    CG-   HG2
  5546   5544   5547   5549      3    0.62760    1.88280    0.00000
-2.51040    0.00000    0.00000 ;     HB3-    CB-    CG-   HG3
  5546   5544   5547   5550      3    0.66944    2.00832    0.00000
-2.67776    0.00000    0.00000 ;     HB3-    CB-    CG-    CD
  5547   5544   5542   5560      3    0.65084    1.95253    0.00000
-2.60338    0.00000    0.00000 ;      CG-    CB-    CA-     C
  5547   5550   5553   5554      3    0.65084    1.95253    0.00000
-2.60338    0.00000    0.00000 ;      CG-    CD-    CE-   HE2
  5547   5550   5553   5555      3    0.65084    1.95253    0.00000
-2.60338    0.00000    0.00000 ;      CG-    CD-    CE-   HE3
  5547   5550   5553   5556      3    0.65084    1.95253    0.00000
-2.60338    0.00000    0.00000 ;      CG-    CD-    CE-    NZ
  5548   5547   5550   5551      3    0.62760    1.88280    0.00000
-2.51040    0.00000    0.00000 ;     HG2-    CG-    CD-   HD2
  5548   5547   5550   5552      3    0.62760    1.88280    0.00000
-2.51040    0.00000    0.00000 ;     HG2-    CG-    CD-   HD3
  5548   5547   5550   5553      3    0.66944    2.00832    0.00000
-2.67776    0.00000    0.00000 ;     HG2-    CG-    CD-    CE
  5549   5547   5550   5551      3    0.62760    1.88280    0.00000
-2.51040    0.00000    0.00000 ;     HG3-    CG-    CD-   HD2
  5549   5547   5550   5552      3    0.62760    1.88280    0.00000
-2.51040    0.00000    0.00000 ;     HG3-    CG-    CD-   HD3
  5549   5547   5550   5553      3    0.66944    2.00832    0.00000
-2.67776    0.00000    0.00000 ;     HG3-    CG-    CD-    CE
  5550   5553   5556   5557      3    0.65084    1.95253    0.00000
-2.60338    0.00000    0.00000 ;      CD-    CE-    NZ-   HZ1
  5550   5553   5556   5558      3    0.65084    1.95253    0.00000
-2.60338    0.00000    0.00000 ;      CD-    CE-    NZ-   HZ2
  5550   5553   5556   5559      3    0.65084    1.95253    0.00000
-2.60338    0.00000    0.00000 ;      CD-    CE-    NZ-   HZ3
  5551   5550   5553   5554      3    0.65084    1.95253    0.00000
-2.60338    0.00000    0.00000 ;     HD2-    CD-    CE-   HE2
  5551   5550   5553   5555      3    0.65084    1.95253    0.00000
-2.60338    0.00000    0.00000 ;     HD2-    CD-    CE-   HE3
  5551   5550   5553   5556      3    0.65084    1.95253    0.00000
```

-2.60338 0.00000 0.00000 ; HD2- CD- CE- NZ
5552 5550 5553 5554 3 0.65084 1.95253 0.00000
-2.60338 0.00000 0.00000 ; HD3- CD- CE- HE2
5552 5550 5553 5555 3 0.65084 1.95253 0.00000
-2.60338 0.00000 0.00000 ; HD3- CD- CE- HE3
5552 5550 5553 5556 3 0.65084 1.95253 0.00000
-2.60338 0.00000 0.00000 ; HD3- CD- CE- NZ
5554 5553 5556 5557 3 0.65084 1.95253 0.00000
-2.60338 0.00000 0.00000 ; HE2- CE- NZ- HZ1
5554 5553 5556 5558 3 0.65084 1.95253 0.00000
-2.60338 0.00000 0.00000 ; HE2- CE- NZ- HZ2
5554 5553 5556 5559 3 0.65084 1.95253 0.00000
-2.60338 0.00000 0.00000 ; HE2- CE- NZ- HZ3
5555 5553 5556 5557 3 0.65084 1.95253 0.00000
-2.60338 0.00000 0.00000 ; HE3- CE- NZ- HZ1
5555 5553 5556 5558 3 0.65084 1.95253 0.00000
-2.60338 0.00000 0.00000 ; HE3- CE- NZ- HZ2
5555 5553 5556 5559 3 0.65084 1.95253 0.00000
-2.60338 0.00000 0.00000 ; HE3- CE- NZ- HZ3
5560 5562 5564 5565 3 0.00000 0.00000 0.00000
0.00000 0.00000 0.00000 ; C- N- CA- HA
5560 5562 5564 5566 3 11.71520 1.67360 15.06240 -
13.38880 0.00000 0.00000 ; C- N- CA- CB
5560 5562 5564 5579 3 1.75728 5.27184 2.25936
-7.02912 0.00000 0.00000 ; C- N- CA- C
5561 5560 5562 5563 3 29.28800 -8.36800 -20.92000
0.00000 0.00000 0.00000 ; O- C- N- H
5561 5560 5562 5564 3 20.92000 0.00000 -20.92000
0.00000 0.00000 0.00000 ; O- C- N- CA
5562 5564 5566 5567 3 0.65084 1.95253 0.00000
-2.60338 0.00000 0.00000 ; N- CA- CB- HB
5562 5564 5566 5568 3 3.83673 0.44769 -1.77402
-2.47693 -0.03347 0.00000 ; N- CA- CB- CG2
5562 5564 5566 5572 3 2.97482 0.12134 2.04179
-1.89117 -3.24678 0.00000 ; N- CA- CB- CG1
5562 5564 5579 5580 3 0.00000 0.00000 0.00000
0.00000 0.00000 0.00000 ; N- CA- C- O
5562 5564 5579 5581 3 17.40544 -5.02080 -13.22144
9.20480 0.00000 0.00000 ; N- CA- C- N
5563 5562 5564 5565 3 0.00000 0.00000 0.00000
0.00000 0.00000 0.00000 ; H- N- CA- HA
5563 5562 5564 5566 3 0.00000 0.00000 0.00000
0.00000 0.00000 0.00000 ; H- N- CA- CB
5563 5562 5564 5579 3 0.00000 0.00000 0.00000
0.00000 0.00000 0.00000 ; H- N- CA- C
5564 5566 5568 5569 3 0.66944 2.00832 0.00000
-2.67776 0.00000 0.00000 ; CA- CB- CG2- HG21
5564 5566 5568 5570 3 0.66944 2.00832 0.00000
-2.67776 0.00000 0.00000 ; CA- CB- CG2- HG22
5564 5566 5568 5571 3 0.66944 2.00832 0.00000
-2.67776 0.00000 0.00000 ; CA- CB- CG2- HG23
5564 5566 5572 5573 3 0.66944 2.00832 0.00000

```
-2.67776    0.00000    0.00000 ;      CA-    CB-   CG1-  HG12
  5564   5566   5572   5574       3    0.66944    2.00832    0.00000
-2.67776    0.00000    0.00000 ;      CA-    CB-   CG1-  HG13
  5564   5566   5572   5575       3    4.24258   -0.52718   -7.25506
-1.79075    7.69856    0.00000 ;      CA-    CB-   CG1-   CD1
  5564   5579   5581   5582       3   20.92000    0.00000  -20.92000
0.00000    0.00000    0.00000 ;      CA-     C-     N-     H
  5564   5579   5581   5583       3   20.92000    0.00000  -20.92000
0.00000    0.00000    0.00000 ;      CA-     C-     N-    CA
  5565   5564   5566   5567       3    0.65084    1.95253    0.00000
-2.60338    0.00000    0.00000 ;      HA-    CA-    CB-    HB
  5565   5564   5566   5568       3    0.65084    1.95253    0.00000
-2.60338    0.00000    0.00000 ;      HA-    CA-    CB-   CG2
  5565   5564   5566   5572       3    0.65084    1.95253    0.00000
-2.60338    0.00000    0.00000 ;      HA-    CA-    CB-   CG1
  5565   5564   5579   5580       3    3.68192   -4.35136    0.00000
1.33888    0.00000    0.00000 ;      HA-    CA-     C-     O
  5565   5564   5579   5581       3    0.00000    0.00000    0.00000
0.00000    0.00000    0.00000 ;      HA-    CA-     C-     N
  5566   5564   5579   5580       3    0.00000    0.00000    0.00000
0.00000    0.00000    0.00000 ;      CB-    CA-     C-     O
  5566   5564   5579   5581       3    2.51040    4.18400    1.67360
-6.69440    0.00000    0.00000 ;      CB-    CA-     C-     N
  5566   5572   5575   5576       3    0.66944    2.00832    0.00000
-2.67776    0.00000    0.00000 ;      CB-   CG1-   CD1-  HD11
  5566   5572   5575   5577       3    0.66944    2.00832    0.00000
-2.67776    0.00000    0.00000 ;      CB-   CG1-   CD1-  HD12
  5566   5572   5575   5578       3    0.66944    2.00832    0.00000
-2.67776    0.00000    0.00000 ;      CB-   CG1-   CD1-  HD13
  5567   5566   5564   5579       3    0.65084    1.95253    0.00000
-2.60338    0.00000    0.00000 ;      HB-    CB-    CA-     C
  5567   5566   5568   5569       3    0.62760    1.88280    0.00000
-2.51040    0.00000    0.00000 ;      HB-    CB-   CG2-  HG21
  5567   5566   5568   5570       3    0.62760    1.88280    0.00000
-2.51040    0.00000    0.00000 ;      HB-    CB-   CG2-  HG22
  5567   5566   5568   5571       3    0.62760    1.88280    0.00000
-2.51040    0.00000    0.00000 ;      HB-    CB-   CG2-  HG23
  5567   5566   5572   5573       3    0.62760    1.88280    0.00000
-2.51040    0.00000    0.00000 ;      HB-    CB-   CG1-  HG12
  5567   5566   5572   5574       3    0.62760    1.88280    0.00000
-2.51040    0.00000    0.00000 ;      HB-    CB-   CG1-  HG13
  5567   5566   5572   5575       3    0.66944    2.00832    0.00000
-2.67776    0.00000    0.00000 ;      HB-    CB-   CG1-   CD1
  5568   5566   5564   5579       3    5.67350    3.55640   -6.16722
-2.47693    3.74886    0.00000 ;     CG2-    CB-    CA-     C
  5568   5566   5572   5573       3    0.66944    2.00832    0.00000
-2.67776    0.00000    0.00000 ;     CG2-    CB-   CG1-  HG12
  5568   5566   5572   5574       3    0.66944    2.00832    0.00000
-2.67776    0.00000    0.00000 ;     CG2-    CB-   CG1-  HG13
  5568   5566   5572   5575       3    3.81162    0.49790   -8.14206
-1.79075    7.49773    0.00000 ;     CG2-    CB-   CG1-   CD1
  5569   5568   5566   5572       3    0.66944    2.00832    0.00000
```

-2.67776 0.00000 0.00000 ; HG21- CG2- CB- CG1
5570 5568 5566 5572 3 0.66944 2.00832 0.00000
-2.67776 0.00000 0.00000 ; HG22- CG2- CB- CG1
5571 5568 5566 5572 3 0.66944 2.00832 0.00000
-2.67776 0.00000 0.00000 ; HG23- CG2- CB- CG1
5572 5566 5564 5579 3 8.26340 0.74057 -9.99976
-1.89117 3.84928 0.00000 ; CG1- CB- CA- C
5573 5572 5575 5576 3 0.62760 1.88280 0.00000
-2.51040 0.00000 0.00000 ; HG12- CG1- CD1- HD11
5573 5572 5575 5577 3 0.62760 1.88280 0.00000
-2.51040 0.00000 0.00000 ; HG12- CG1- CD1- HD12
5573 5572 5575 5578 3 0.62760 1.88280 0.00000
-2.51040 0.00000 0.00000 ; HG12- CG1- CD1- HD13
5574 5572 5575 5576 3 0.62760 1.88280 0.00000
-2.51040 0.00000 0.00000 ; HG13- CG1- CD1- HD11
5574 5572 5575 5577 3 0.62760 1.88280 0.00000
-2.51040 0.00000 0.00000 ; HG13- CG1- CD1- HD12
5574 5572 5575 5578 3 0.62760 1.88280 0.00000
-2.51040 0.00000 0.00000 ; HG13- CG1- CD1- HD13
5579 5581 5583 5584 3 0.00000 0.00000 0.00000
0.00000 0.00000 0.00000 ; C- N- CA- HA2
5579 5581 5583 5585 3 0.00000 0.00000 0.00000
0.00000 0.00000 0.00000 ; C- N- CA- HA3
5579 5581 5583 5586 3 1.75728 5.27184 2.25936
-7.02912 0.00000 0.00000 ; C- N- CA- C
5580 5579 5581 5582 3 29.28800 -8.36800 -20.92000
0.00000 0.00000 0.00000 ; O- C- N- H
5580 5579 5581 5583 3 20.92000 0.00000 -20.92000
0.00000 0.00000 0.00000 ; O- C- N- CA
5581 5583 5586 5587 3 0.00000 0.00000 0.00000
0.00000 0.00000 0.00000 ; N- CA- C- O
5581 5583 5586 5588 3 17.40544 -5.02080 -13.22144
9.20480 0.00000 0.00000 ; N- CA- C- N
5582 5581 5583 5584 3 0.00000 0.00000 0.00000
0.00000 0.00000 0.00000 ; H- N- CA- HA2
5582 5581 5583 5585 3 0.00000 0.00000 0.00000
0.00000 0.00000 0.00000 ; H- N- CA- HA3
5582 5581 5583 5586 3 0.00000 0.00000 0.00000
0.00000 0.00000 0.00000 ; H- N- CA- C
5583 5586 5588 5589 3 20.92000 0.00000 -20.92000
0.00000 0.00000 0.00000 ; CA- C- N- H
5583 5586 5588 5590 3 20.92000 0.00000 -20.92000
0.00000 0.00000 0.00000 ; CA- C- N- CA
5584 5583 5586 5587 3 3.68192 -4.35136 0.00000
1.33888 0.00000 0.00000 ; HA2- CA- C- O
5584 5583 5586 5588 3 0.00000 0.00000 0.00000
0.00000 0.00000 0.00000 ; HA2- CA- C- N
5585 5583 5586 5587 3 3.68192 -4.35136 0.00000
1.33888 0.00000 0.00000 ; HA3- CA- C- O
5585 5583 5586 5588 3 0.00000 0.00000 0.00000
0.00000 0.00000 0.00000 ; HA3- CA- C- N
5586 5588 5590 5591 3 0.00000 0.00000 0.00000

0.00000 0.00000 0.00000 ; C- N- CA- HA
5586 5588 5590 5592 3 11.71520 1.67360 15.06240 -
13.38880 0.00000 0.00000 ; C- N- CA- CB
5586 5588 5590 5607 3 1.75728 5.27184 2.25936
-7.02912 0.00000 0.00000 ; C- N- CA- C
5587 5586 5588 5589 3 29.28800 -8.36800 -20.92000
0.00000 0.00000 0.00000 ; O- C- N- H
5587 5586 5588 5590 3 20.92000 0.00000 -20.92000
0.00000 0.00000 0.00000 ; O- C- N- CA
5588 5590 5592 5593 3 0.65084 1.95253 0.00000
-2.60338 0.00000 0.00000 ; N- CA- CB- HB2
5588 5590 5592 5594 3 0.65084 1.95253 0.00000
-2.60338 0.00000 0.00000 ; N- CA- CB- HB3
5588 5590 5592 5595 3 3.28026 2.46019 -2.19242
-3.21331 -0.23430 0.00000 ; N- CA- CB- CG
5588 5590 5607 5608 3 0.00000 0.00000 0.00000
0.00000 0.00000 0.00000 ; N- CA- C- O
5588 5590 5607 5609 3 17.40544 -5.02080 -13.22144
9.20480 0.00000 0.00000 ; N- CA- C- N
5589 5588 5590 5591 3 0.00000 0.00000 0.00000
0.00000 0.00000 0.00000 ; H- N- CA- HA
5589 5588 5590 5592 3 0.00000 0.00000 0.00000
0.00000 0.00000 0.00000 ; H- N- CA- CB
5589 5588 5590 5607 3 0.00000 0.00000 0.00000
0.00000 0.00000 0.00000 ; H- N- CA- C
5590 5592 5595 5596 3 0.57739 0.00000 1.02926
0.00000 -1.60666 0.00000 ; CA- CB- CG- CD1
5590 5592 5595 5605 3 0.57739 0.00000 1.02926
0.00000 -1.60666 0.00000 ; CA- CB- CG- CD2
5590 5607 5609 5610 3 20.92000 0.00000 -20.92000
0.00000 0.00000 0.00000 ; CA- C- N- H
5590 5607 5609 5611 3 20.92000 0.00000 -20.92000
0.00000 0.00000 0.00000 ; CA- C- N- CA
5591 5590 5592 5593 3 0.65084 1.95253 0.00000
-2.60338 0.00000 0.00000 ; HA- CA- CB- HB2
5591 5590 5592 5594 3 0.65084 1.95253 0.00000
-2.60338 0.00000 0.00000 ; HA- CA- CB- HB3
5591 5590 5592 5595 3 0.65084 1.95253 0.00000
-2.60338 0.00000 0.00000 ; HA- CA- CB- CG
5591 5590 5607 5608 3 3.68192 -4.35136 0.00000
1.33888 0.00000 0.00000 ; HA- CA- C- O
5591 5590 5607 5609 3 0.00000 0.00000 0.00000
0.00000 0.00000 0.00000 ; HA- CA- C- N
5592 5590 5607 5608 3 0.00000 0.00000 0.00000
0.00000 0.00000 0.00000 ; CB- CA- C- O
5592 5590 5607 5609 3 2.51040 4.18400 1.67360
-6.69440 0.00000 0.00000 ; CB- CA- C- N
5592 5595 5596 5597 3 30.33400 0.00000 -30.33400
0.00000 0.00000 0.00000 ; CB- CG- CD1- HD1
5592 5595 5596 5598 3 30.33400 0.00000 -30.33400
0.00000 0.00000 0.00000 ; CB- CG- CD1- CE1
5592 5595 5605 5603 3 30.33400 0.00000 -30.33400

```
0.00000    0.00000    0.00000 ;      CB-    CG-   CD2-   CE2
  5592   5595   5605   5606      3   30.33400    0.00000  -30.33400
0.00000    0.00000    0.00000 ;      CB-    CG-   CD2-   HD2
  5593   5592   5590   5607      3    0.65084    1.95253    0.00000
-2.60338    0.00000    0.00000 ;    HB2-    CB-    CA-     C
  5593   5592   5595   5596      3    0.00000    0.00000    0.00000
0.00000    0.00000    0.00000 ;    HB2-    CB-    CG-   CD1
  5593   5592   5595   5605      3    0.00000    0.00000    0.00000
0.00000    0.00000    0.00000 ;    HB2-    CB-    CG-   CD2
  5594   5592   5590   5607      3    0.65084    1.95253    0.00000
-2.60338    0.00000    0.00000 ;    HB3-    CB-    CA-     C
  5594   5592   5595   5596      3    0.00000    0.00000    0.00000
0.00000    0.00000    0.00000 ;    HB3-    CB-    CG-   CD1
  5594   5592   5595   5605      3    0.00000    0.00000    0.00000
0.00000    0.00000    0.00000 ;    HB3-    CB-    CG-   CD2
  5595   5592   5590   5607      3    4.95804    2.17986   -3.52293
-3.21331   -0.40166    0.00000 ;     CG-    CB-    CA-     C
  5595   5596   5598   5599      3   30.33400    0.00000  -30.33400
0.00000    0.00000    0.00000 ;      CG-   CD1-   CE1-   HE1
  5595   5596   5598   5600      3   30.33400    0.00000  -30.33400
0.00000    0.00000    0.00000 ;      CG-   CD1-   CE1-    CZ
  5595   5605   5603   5600      3   30.33400    0.00000  -30.33400
0.00000    0.00000    0.00000 ;      CG-   CD2-   CE2-    CZ
  5595   5605   5603   5604      3   30.33400    0.00000  -30.33400
0.00000    0.00000    0.00000 ;      CG-   CD2-   CE2-   HE2
  5596   5595   5605   5603      3   30.33400    0.00000  -30.33400
0.00000    0.00000    0.00000 ;    CD1-    CG-   CD2-   CE2
  5596   5595   5605   5606      3   30.33400    0.00000  -30.33400
0.00000    0.00000    0.00000 ;    CD1-    CG-   CD2-   HD2
  5596   5598   5600   5601      3   30.33400    0.00000  -30.33400
0.00000    0.00000    0.00000 ;    CD1-   CE1-    CZ-    OH
  5596   5598   5600   5603      3   30.33400    0.00000  -30.33400
0.00000    0.00000    0.00000 ;    CD1-   CE1-    CZ-   CE2
  5597   5596   5595   5605      3   30.33400    0.00000  -30.33400
0.00000    0.00000    0.00000 ;    HD1-   CD1-    CG-   CD2
  5597   5596   5598   5599      3   30.33400    0.00000  -30.33400
0.00000    0.00000    0.00000 ;    HD1-   CD1-   CE1-   HE1
  5597   5596   5598   5600      3   30.33400    0.00000  -30.33400
0.00000    0.00000    0.00000 ;    HD1-   CD1-   CE1-    CZ
  5598   5596   5595   5605      3   30.33400    0.00000  -30.33400
0.00000    0.00000    0.00000 ;    CE1-   CD1-    CG-   CD2
  5598   5600   5601   5602      3    7.93286    0.00000   -9.56462
0.00000    2.17568    0.00000 ;    CE1-    CZ-    OH-    HH
  5598   5600   5603   5604      3   30.33400    0.00000  -30.33400
0.00000    0.00000    0.00000 ;    CE1-    CZ-   CE2-   HE2
  5598   5600   5603   5605      3   30.33400    0.00000  -30.33400
0.00000    0.00000    0.00000 ;    CE1-    CZ-   CE2-   CD2
  5599   5598   5600   5601      3   30.33400    0.00000  -30.33400
0.00000    0.00000    0.00000 ;    HE1-   CE1-    CZ-    OH
  5599   5598   5600   5603      3   30.33400    0.00000  -30.33400
0.00000    0.00000    0.00000 ;    HE1-   CE1-    CZ-   CE2
  5600   5603   5605   5606      3   30.33400    0.00000  -30.33400
```

0.00000 0.00000 0.00000 ; CZ- CE2- CD2- HD2
5601 5600 5603 5604 3 30.33400 0.00000 -30.33400
0.00000 0.00000 0.00000 ; OH- CZ- CE2- HE2
5601 5600 5603 5605 3 30.33400 0.00000 -30.33400
0.00000 0.00000 0.00000 ; OH- CZ- CE2- CD2
5602 5601 5600 5603 3 7.93286 0.00000 -9.56462
0.00000 2.17568 0.00000 ; HH- OH- CZ- CE2
5604 5603 5605 5606 3 30.33400 0.00000 -30.33400
0.00000 0.00000 0.00000 ; HE2- CE2- CD2- HD2
5607 5609 5611 5612 3 0.00000 0.00000 0.00000
0.00000 0.00000 0.00000 ; C- N- CA- HA
5607 5609 5611 5613 3 11.71520 1.67360 15.06240 -
13.38880 0.00000 0.00000 ; C- N- CA- CB
5607 5609 5611 5624 3 1.75728 5.27184 2.25936
-7.02912 0.00000 0.00000 ; C- N- CA- C
5608 5607 5609 5610 3 29.28800 -8.36800 -20.92000
0.00000 0.00000 0.00000 ; O- C- N- H
5608 5607 5609 5611 3 20.92000 0.00000 -20.92000
0.00000 0.00000 0.00000 ; O- C- N- CA
5609 5611 5613 5614 3 0.65084 1.95253 0.00000
-2.60338 0.00000 0.00000 ; N- CA- CB- HB2
5609 5611 5613 5615 3 0.65084 1.95253 0.00000
-2.60338 0.00000 0.00000 ; N- CA- CB- HB3
5609 5611 5613 5616 3 3.21331 2.22589 -4.15053
-2.40998 2.61082 0.00000 ; N- CA- CB- CG
5609 5611 5624 5625 3 0.00000 0.00000 0.00000
0.00000 0.00000 0.00000 ; N- CA- C- O
5609 5611 5624 5626 3 17.40544 -5.02080 -13.22144
9.20480 0.00000 0.00000 ; N- CA- C- N
5610 5609 5611 5612 3 0.00000 0.00000 0.00000
0.00000 0.00000 0.00000 ; H- N- CA- HA
5610 5609 5611 5613 3 0.00000 0.00000 0.00000
0.00000 0.00000 0.00000 ; H- N- CA- CB
5610 5609 5611 5624 3 0.00000 0.00000 0.00000
0.00000 0.00000 0.00000 ; H- N- CA- C
5611 5613 5616 5617 3 0.66944 2.00832 0.00000
-2.67776 0.00000 0.00000 ; CA- CB- CG- HG2
5611 5613 5616 5618 3 0.66944 2.00832 0.00000
-2.67776 0.00000 0.00000 ; CA- CB- CG- HG3
5611 5613 5616 5619 3 3.69866 -4.35136 -3.92459
6.89523 4.61914 0.00000 ; CA- CB- CG- CD
5611 5624 5626 5627 3 20.92000 0.00000 -20.92000
0.00000 0.00000 0.00000 ; CA- C- N- H
5611 5624 5626 5628 3 20.92000 0.00000 -20.92000
0.00000 0.00000 0.00000 ; CA- C- N- CA
5612 5611 5613 5614 3 0.65084 1.95253 0.00000
-2.60338 0.00000 0.00000 ; HA- CA- CB- HB2
5612 5611 5613 5615 3 0.65084 1.95253 0.00000
-2.60338 0.00000 0.00000 ; HA- CA- CB- HB3
5612 5611 5613 5616 3 0.65084 1.95253 0.00000
-2.60338 0.00000 0.00000 ; HA- CA- CB- CG
5612 5611 5624 5625 3 3.68192 -4.35136 0.00000

1.33888 0.00000 0.00000 ; HA- CA- C- O
5612 5611 5624 5626 3 0.00000 0.00000 0.00000
0.00000 0.00000 0.00000 ; HA- CA- C- N
5613 5611 5624 5625 3 0.00000 0.00000 0.00000
0.00000 0.00000 0.00000 ; CB- CA- C- O
5613 5611 5624 5626 3 2.51040 4.18400 1.67360
-6.69440 0.00000 0.00000 ; CB- CA- C- N
5613 5616 5619 5620 3 0.00000 0.00000 0.00000
0.00000 0.00000 0.00000 ; CB- CG- CD- OE1
5613 5616 5619 5621 3 10.32611 1.48114 -8.47678
1.42256 1.40582 0.00000 ; CB- CG- CD- NE2
5614 5613 5611 5624 3 0.65084 1.95253 0.00000
-2.60338 0.00000 0.00000 ; HB2- CB- CA- C
5614 5613 5616 5617 3 0.62760 1.88280 0.00000
-2.51040 0.00000 0.00000 ; HB2- CB- CG- HG2
5614 5613 5616 5618 3 0.62760 1.88280 0.00000
-2.51040 0.00000 0.00000 ; HB2- CB- CG- HG3
5614 5613 5616 5619 3 0.65084 1.95253 0.00000
-2.60338 0.00000 0.00000 ; HB2- CB- CG- CD
5615 5613 5611 5624 3 0.65084 1.95253 0.00000
-2.60338 0.00000 0.00000 ; HB3- CB- CA- C
5615 5613 5616 5617 3 0.62760 1.88280 0.00000
-2.51040 0.00000 0.00000 ; HB3- CB- CG- HG2
5615 5613 5616 5618 3 0.62760 1.88280 0.00000
-2.51040 0.00000 0.00000 ; HB3- CB- CG- HG3
5615 5613 5616 5619 3 0.65084 1.95253 0.00000
-2.60338 0.00000 0.00000 ; HB3- CB- CG- CD
5616 5613 5611 5624 3 6.86594 3.56895 -8.14206
-2.40998 4.85344 0.00000 ; CG- CB- CA- C
5616 5619 5621 5622 3 20.92000 0.00000 -20.92000
0.00000 0.00000 0.00000 ; CG- CD- NE2- HE21
5616 5619 5621 5623 3 20.92000 0.00000 -20.92000
0.00000 0.00000 0.00000 ; CG- CD- NE2- HE22
5617 5616 5619 5620 3 3.68192 -4.35136 0.00000
1.33888 0.00000 0.00000 ; HG2- CG- CD- OE1
5617 5616 5619 5621 3 0.00000 0.00000 0.00000
0.00000 0.00000 0.00000 ; HG2- CG- CD- NE2
5618 5616 5619 5620 3 3.68192 -4.35136 0.00000
1.33888 0.00000 0.00000 ; HG3- CG- CD- OE1
5618 5616 5619 5621 3 0.00000 0.00000 0.00000
0.00000 0.00000 0.00000 ; HG3- CG- CD- NE2
5620 5619 5621 5622 3 29.28800 -8.36800 -20.92000
0.00000 0.00000 0.00000 ; OE1- CD- NE2- HE21
5620 5619 5621 5623 3 29.28800 -8.36800 -20.92000
0.00000 0.00000 0.00000 ; OE1- CD- NE2- HE22
5624 5626 5628 5629 3 0.00000 0.00000 0.00000
0.00000 0.00000 0.00000 ; C- N- CA- HA
5624 5626 5628 5630 3 11.71520 1.67360 15.06240 -
13.38880 0.00000 0.00000 ; C- N- CA- CB
5624 5626 5628 5646 3 1.75728 5.27184 2.25936
-7.02912 0.00000 0.00000 ; C- N- CA- C
5625 5624 5626 5627 3 29.28800 -8.36800 -20.92000

```
0.00000    0.00000    0.00000 ;       O-     C-     N-     H
  5625   5624   5626   5628      3   20.92000    0.00000  -20.92000
0.00000    0.00000    0.00000 ;       O-     C-     N-    CA
  5626   5628   5630   5631      3    0.65084    1.95253    0.00000
-2.60338    0.00000    0.00000 ;      N-    CA-    CB-   HB2
  5626   5628   5630   5632      3    0.65084    1.95253    0.00000
-2.60338    0.00000    0.00000 ;      N-    CA-    CB-   HB3
  5626   5628   5630   5633      3    0.65084    1.95253    0.00000
-2.60338    0.00000    0.00000 ;      N-    CA-    CB-    CG
  5626   5628   5646   5647      3    0.00000    0.00000    0.00000
0.00000    0.00000    0.00000 ;      N-    CA-     C-     O
  5626   5628   5646   5648      3   17.40544   -5.02080  -13.22144
9.20480    0.00000    0.00000 ;      N-    CA-     C-     N
  5627   5626   5628   5629      3    0.00000    0.00000    0.00000
0.00000    0.00000    0.00000 ;      H-     N-    CA-    HA
  5627   5626   5628   5630      3    0.00000    0.00000    0.00000
0.00000    0.00000    0.00000 ;      H-     N-    CA-    CB
  5627   5626   5628   5646      3    0.00000    0.00000    0.00000
0.00000    0.00000    0.00000 ;      H-     N-    CA-     C
  5628   5630   5633   5634      3    0.66944    2.00832    0.00000
-2.67776    0.00000    0.00000 ;     CA-    CB-    CG-   HG2
  5628   5630   5633   5635      3    0.66944    2.00832    0.00000
-2.67776    0.00000    0.00000 ;     CA-    CB-    CG-   HG3
  5628   5630   5633   5636      3    3.68192    3.09616   -2.09200
-3.01248    0.00000    0.00000 ;     CA-    CB-    CG-    CD
  5628   5646   5648   5649      3   20.92000    0.00000  -20.92000
0.00000    0.00000    0.00000 ;     CA-     C-     N-     H
  5628   5646   5648   5650      3   20.92000    0.00000  -20.92000
0.00000    0.00000    0.00000 ;     CA-     C-     N-    CA
  5629   5628   5630   5631      3    0.65084    1.95253    0.00000
-2.60338    0.00000    0.00000 ;     HA-    CA-    CB-   HB2
  5629   5628   5630   5632      3    0.65084    1.95253    0.00000
-2.60338    0.00000    0.00000 ;     HA-    CA-    CB-   HB3
  5629   5628   5630   5633      3    0.65084    1.95253    0.00000
-2.60338    0.00000    0.00000 ;     HA-    CA-    CB-    CG
  5629   5628   5646   5647      3    3.68192   -4.35136    0.00000
1.33888    0.00000    0.00000 ;     HA-    CA-     C-     O
  5629   5628   5646   5648      3    0.00000    0.00000    0.00000
0.00000    0.00000    0.00000 ;     HA-    CA-     C-     N
  5630   5628   5646   5647      3    0.00000    0.00000    0.00000
0.00000    0.00000    0.00000 ;     CB-    CA-     C-     O
  5630   5628   5646   5648      3    2.51040    4.18400    1.67360
-6.69440    0.00000    0.00000 ;     CB-    CA-     C-     N
  5630   5633   5636   5637      3    0.66944    2.00832    0.00000
-2.67776    0.00000    0.00000 ;     CB-    CG-    CD-   HD2
  5630   5633   5636   5638      3    0.66944    2.00832    0.00000
-2.67776    0.00000    0.00000 ;     CB-    CG-    CD-   HD3
  5630   5633   5636   5639      3    3.68192    3.09616   -2.09200
-3.01248    0.00000    0.00000 ;     CB-    CG-    CD-    CE
  5631   5630   5628   5646      3    0.65084    1.95253    0.00000
-2.60338    0.00000    0.00000 ;    HB2-    CB-    CA-     C
  5631   5630   5633   5634      3    0.62760    1.88280    0.00000
```

```
-2.51040    0.00000    0.00000 ;    HB2-    CB-    CG-   HG2
  5631  5630  5633  5635     3   0.62760   1.88280   0.00000
-2.51040    0.00000    0.00000 ;    HB2-    CB-    CG-   HG3
  5631  5630  5633  5636     3   0.66944   2.00832   0.00000
-2.67776    0.00000    0.00000 ;    HB2-    CB-    CG-    CD
  5632  5630  5628  5646     3   0.65084   1.95253   0.00000
-2.60338    0.00000    0.00000 ;    HB3-    CB-    CA-     C
  5632  5630  5633  5634     3   0.62760   1.88280   0.00000
-2.51040    0.00000    0.00000 ;    HB3-    CB-    CG-   HG2
  5632  5630  5633  5635     3   0.62760   1.88280   0.00000
-2.51040    0.00000    0.00000 ;    HB3-    CB-    CG-   HG3
  5632  5630  5633  5636     3   0.66944   2.00832   0.00000
-2.67776    0.00000    0.00000 ;    HB3-    CB-    CG-    CD
  5633  5630  5628  5646     3   0.65084   1.95253   0.00000
-2.60338    0.00000    0.00000 ;     CG-    CB-    CA-     C
  5633  5636  5639  5640     3   0.65084   1.95253   0.00000
-2.60338    0.00000    0.00000 ;     CG-    CD-    CE-   HE2
  5633  5636  5639  5641     3   0.65084   1.95253   0.00000
-2.60338    0.00000    0.00000 ;     CG-    CD-    CE-   HE3
  5633  5636  5639  5642     3   0.65084   1.95253   0.00000
-2.60338    0.00000    0.00000 ;     CG-    CD-    CE-    NZ
  5634  5633  5636  5637     3   0.62760   1.88280   0.00000
-2.51040    0.00000    0.00000 ;    HG2-    CG-    CD-   HD2
  5634  5633  5636  5638     3   0.62760   1.88280   0.00000
-2.51040    0.00000    0.00000 ;    HG2-    CG-    CD-   HD3
  5634  5633  5636  5639     3   0.66944   2.00832   0.00000
-2.67776    0.00000    0.00000 ;    HG2-    CG-    CD-    CE
  5635  5633  5636  5637     3   0.62760   1.88280   0.00000
-2.51040    0.00000    0.00000 ;    HG3-    CG-    CD-   HD2
  5635  5633  5636  5638     3   0.62760   1.88280   0.00000
-2.51040    0.00000    0.00000 ;    HG3-    CG-    CD-   HD3
  5635  5633  5636  5639     3   0.66944   2.00832   0.00000
-2.67776    0.00000    0.00000 ;    HG3-    CG-    CD-    CE
  5636  5639  5642  5643     3   0.65084   1.95253   0.00000
-2.60338    0.00000    0.00000 ;     CD-    CE-    NZ-   HZ1
  5636  5639  5642  5644     3   0.65084   1.95253   0.00000
-2.60338    0.00000    0.00000 ;     CD-    CE-    NZ-   HZ2
  5636  5639  5642  5645     3   0.65084   1.95253   0.00000
-2.60338    0.00000    0.00000 ;     CD-    CE-    NZ-   HZ3
  5637  5636  5639  5640     3   0.65084   1.95253   0.00000
-2.60338    0.00000    0.00000 ;    HD2-    CD-    CE-   HE2
  5637  5636  5639  5641     3   0.65084   1.95253   0.00000
-2.60338    0.00000    0.00000 ;    HD2-    CD-    CE-   HE3
  5637  5636  5639  5642     3   0.65084   1.95253   0.00000
-2.60338    0.00000    0.00000 ;    HD2-    CD-    CE-    NZ
  5638  5636  5639  5640     3   0.65084   1.95253   0.00000
-2.60338    0.00000    0.00000 ;    HD3-    CD-    CE-   HE2
  5638  5636  5639  5641     3   0.65084   1.95253   0.00000
-2.60338    0.00000    0.00000 ;    HD3-    CD-    CE-   HE3
  5638  5636  5639  5642     3   0.65084   1.95253   0.00000
-2.60338    0.00000    0.00000 ;    HD3-    CD-    CE-    NZ
  5640  5639  5642  5643     3   0.65084   1.95253   0.00000
```

-2.60338 0.00000 0.00000 ; HE2- CE- NZ- HZ1
5640 5639 5642 5644 3 0.65084 1.95253 0.00000
-2.60338 0.00000 0.00000 ; HE2- CE- NZ- HZ2
5640 5639 5642 5645 3 0.65084 1.95253 0.00000
-2.60338 0.00000 0.00000 ; HE2- CE- NZ- HZ3
5641 5639 5642 5643 3 0.65084 1.95253 0.00000
-2.60338 0.00000 0.00000 ; HE3- CE- NZ- HZ1
5641 5639 5642 5644 3 0.65084 1.95253 0.00000
-2.60338 0.00000 0.00000 ; HE3- CE- NZ- HZ2
5641 5639 5642 5645 3 0.65084 1.95253 0.00000
-2.60338 0.00000 0.00000 ; HE3- CE- NZ- HZ3
5646 5648 5650 5651 3 0.00000 0.00000 0.00000
0.00000 0.00000 0.00000 ; C- N- CA- HA
5646 5648 5650 5652 3 11.71520 1.67360 15.06240 -
13.38880 0.00000 0.00000 ; C- N- CA- CB
5646 5648 5650 5665 3 1.75728 5.27184 2.25936
-7.02912 0.00000 0.00000 ; C- N- CA- C
5647 5646 5648 5649 3 29.28800 -8.36800 -20.92000
0.00000 0.00000 0.00000 ; O- C- N- H
5647 5646 5648 5650 3 20.92000 0.00000 -20.92000
0.00000 0.00000 0.00000 ; O- C- N- CA
5648 5650 5652 5653 3 0.65084 1.95253 0.00000
-2.60338 0.00000 0.00000 ; N- CA- CB- HB
5648 5650 5652 5654 3 3.83673 0.44769 -1.77402
-2.47693 -0.03347 0.00000 ; N- CA- CB- CG2
5648 5650 5652 5658 3 2.97482 0.12134 2.04179
-1.89117 -3.24678 0.00000 ; N- CA- CB- CG1
5648 5650 5665 5666 3 0.00000 0.00000 0.00000
0.00000 0.00000 0.00000 ; N- CA- C- O
5648 5650 5665 5667 3 17.40544 -5.02080 -13.22144
9.20480 0.00000 0.00000 ; N- CA- C- N
5649 5648 5650 5651 3 0.00000 0.00000 0.00000
0.00000 0.00000 0.00000 ; H- N- CA- HA
5649 5648 5650 5652 3 0.00000 0.00000 0.00000
0.00000 0.00000 0.00000 ; H- N- CA- CB
5649 5648 5650 5665 3 0.00000 0.00000 0.00000
0.00000 0.00000 0.00000 ; H- N- CA- C
5650 5652 5654 5655 3 0.66944 2.00832 0.00000
-2.67776 0.00000 0.00000 ; CA- CB- CG2- HG21
5650 5652 5654 5656 3 0.66944 2.00832 0.00000
-2.67776 0.00000 0.00000 ; CA- CB- CG2- HG22
5650 5652 5654 5657 3 0.66944 2.00832 0.00000
-2.67776 0.00000 0.00000 ; CA- CB- CG2- HG23
5650 5652 5658 5659 3 0.66944 2.00832 0.00000
-2.67776 0.00000 0.00000 ; CA- CB- CG1- HG12
5650 5652 5658 5660 3 0.66944 2.00832 0.00000
-2.67776 0.00000 0.00000 ; CA- CB- CG1- HG13
5650 5652 5658 5661 3 4.24258 -0.52718 -7.25506
-1.79075 7.69856 0.00000 ; CA- CB- CG1- CD1
5650 5665 5667 5668 3 20.92000 0.00000 -20.92000
0.00000 0.00000 0.00000 ; CA- C- N- CD
5650 5665 5667 5677 3 20.92000 0.00000 -20.92000

0.00000 0.00000 0.00000 ; CA- C- N- CA
5651 5650 5652 5653 3 0.65084 1.95253 0.00000
-2.60338 0.00000 0.00000 ; HA- CA- CB- HB
5651 5650 5652 5654 3 0.65084 1.95253 0.00000
-2.60338 0.00000 0.00000 ; HA- CA- CB- CG2
5651 5650 5652 5658 3 0.65084 1.95253 0.00000
-2.60338 0.00000 0.00000 ; HA- CA- CB- CG1
5651 5650 5665 5666 3 3.68192 -4.35136 0.00000
1.33888 0.00000 0.00000 ; HA- CA- C- O
5651 5650 5665 5667 3 0.00000 0.00000 0.00000
0.00000 0.00000 0.00000 ; HA- CA- C- N
5652 5650 5665 5666 3 0.00000 0.00000 0.00000
0.00000 0.00000 0.00000 ; CB- CA- C- O
5652 5650 5665 5667 3 2.51040 4.18400 1.67360
-6.69440 0.00000 0.00000 ; CB- CA- C- N
5652 5658 5661 5662 3 0.66944 2.00832 0.00000
-2.67776 0.00000 0.00000 ; CB- CG1- CD1- HD11
5652 5658 5661 5663 3 0.66944 2.00832 0.00000
-2.67776 0.00000 0.00000 ; CB- CG1- CD1- HD12
5652 5658 5661 5664 3 0.66944 2.00832 0.00000
-2.67776 0.00000 0.00000 ; CB- CG1- CD1- HD13
5653 5652 5650 5665 3 0.65084 1.95253 0.00000
-2.60338 0.00000 0.00000 ; HB- CB- CA- C
5653 5652 5654 5655 3 0.62760 1.88280 0.00000
-2.51040 0.00000 0.00000 ; HB- CB- CG2- HG21
5653 5652 5654 5656 3 0.62760 1.88280 0.00000
-2.51040 0.00000 0.00000 ; HB- CB- CG2- HG22
5653 5652 5654 5657 3 0.62760 1.88280 0.00000
-2.51040 0.00000 0.00000 ; HB- CB- CG2- HG23
5653 5652 5658 5659 3 0.62760 1.88280 0.00000
-2.51040 0.00000 0.00000 ; HB- CB- CG1- HG12
5653 5652 5658 5660 3 0.62760 1.88280 0.00000
-2.51040 0.00000 0.00000 ; HB- CB- CG1- HG13
5653 5652 5658 5661 3 0.66944 2.00832 0.00000
-2.67776 0.00000 0.00000 ; HB- CB- CG1- CD1
5654 5652 5650 5665 3 5.67350 3.55640 -6.16722
-2.47693 3.74886 0.00000 ; CG2- CB- CA- C
5654 5652 5658 5659 3 0.66944 2.00832 0.00000
-2.67776 0.00000 0.00000 ; CG2- CB- CG1- HG12
5654 5652 5658 5660 3 0.66944 2.00832 0.00000
-2.67776 0.00000 0.00000 ; CG2- CB- CG1- HG13
5654 5652 5658 5661 3 3.81162 0.49790 -8.14206
-1.79075 7.49773 0.00000 ; CG2- CB- CG1- CD1
5655 5654 5652 5658 3 0.66944 2.00832 0.00000
-2.67776 0.00000 0.00000 ; HG21- CG2- CB- CG1
5656 5654 5652 5658 3 0.66944 2.00832 0.00000
-2.67776 0.00000 0.00000 ; HG22- CG2- CB- CG1
5657 5654 5652 5658 3 0.66944 2.00832 0.00000
-2.67776 0.00000 0.00000 ; HG23- CG2- CB- CG1
5658 5652 5650 5665 3 8.26340 0.74057 -9.99976
-1.89117 3.84928 0.00000 ; CG1- CB- CA- C
5659 5658 5661 5662 3 0.62760 1.88280 0.00000

-2.51040 0.00000 0.00000 ; HG12- CG1- CD1- HD11
5659 5658 5661 5663 3 0.62760 1.88280 0.00000
-2.51040 0.00000 0.00000 ; HG12- CG1- CD1- HD12
5659 5658 5661 5664 3 0.62760 1.88280 0.00000
-2.51040 0.00000 0.00000 ; HG12- CG1- CD1- HD13
5660 5658 5661 5662 3 0.62760 1.88280 0.00000
-2.51040 0.00000 0.00000 ; HG13- CG1- CD1- HD11
5660 5658 5661 5663 3 0.62760 1.88280 0.00000
-2.51040 0.00000 0.00000 ; HG13- CG1- CD1- HD12
5660 5658 5661 5664 3 0.62760 1.88280 0.00000
-2.51040 0.00000 0.00000 ; HG13- CG1- CD1- HD13
5665 5667 5668 5669 3 0.00000 0.00000 0.00000
0.00000 0.00000 0.00000 ; C- N- CD- HD2
5665 5667 5668 5670 3 0.00000 0.00000 0.00000
0.00000 0.00000 0.00000 ; C- N- CD- HD3
5665 5667 5668 5671 3 10.04160 -3.34720 16.73600
-6.69440 0.00000 0.00000 ; C- N- CD- CG
5665 5667 5677 5674 3 11.71520 1.67360 15.06240 -
13.38880 0.00000 0.00000 ; C- N- CA- CB
5665 5667 5677 5678 3 0.00000 0.00000 0.00000
0.00000 0.00000 0.00000 ; C- N- CA- HA
5665 5667 5677 5679 3 1.75728 5.27184 2.25936
-7.02912 0.00000 0.00000 ; C- N- CA- C
5666 5665 5667 5668 3 20.92000 0.00000 -20.92000
0.00000 0.00000 0.00000 ; O- C- N- CD
5666 5665 5667 5677 3 20.92000 0.00000 -20.92000
0.00000 0.00000 0.00000 ; O- C- N- CA
5667 5668 5671 5672 3 0.65084 1.95253 0.00000
-2.60338 0.00000 0.00000 ; N- CD- CG- HG2
5667 5668 5671 5673 3 0.65084 1.95253 0.00000
-2.60338 0.00000 0.00000 ; N- CD- CG- HG3
5667 5668 5671 5674 3 0.65084 1.95253 0.00000
-2.60338 0.00000 0.00000 ; N- CD- CG- CB
5667 5677 5674 5671 3 0.65084 1.95253 0.00000
-2.60338 0.00000 0.00000 ; N- CA- CB- CG
5667 5677 5674 5675 3 0.65084 1.95253 0.00000
-2.60338 0.00000 0.00000 ; N- CA- CB- HB2
5667 5677 5674 5676 3 0.65084 1.95253 0.00000
-2.60338 0.00000 0.00000 ; N- CA- CB- HB3
5667 5677 5679 5680 3 0.00000 0.00000 0.00000
0.00000 0.00000 0.00000 ; N- CA- C- O
5667 5677 5679 5681 3 17.40544 -5.02080 -13.22144
9.20480 0.00000 0.00000 ; N- CA- C- N
5668 5667 5677 5674 3 0.00000 0.00000 0.00000
0.00000 0.00000 0.00000 ; CD- N- CA- CB
5668 5667 5677 5678 3 0.00000 0.00000 0.00000
0.00000 0.00000 0.00000 ; CD- N- CA- HA
5668 5667 5677 5679 3 0.00000 0.00000 0.00000
0.00000 0.00000 0.00000 ; CD- N- CA- C
5668 5671 5674 5675 3 0.66944 2.00832 0.00000
-2.67776 0.00000 0.00000 ; CD- CG- CB- HB2
5668 5671 5674 5676 3 0.66944 2.00832 0.00000

-2.67776    0.00000    0.00000 ;     CD-    CG-    CB-   HB3
  5668   5671   5674   5677      3    3.68192    3.09616   -2.09200
-3.01248    0.00000    0.00000 ;     CD-    CG-    CB-    CA
  5669   5668   5667   5677      3    0.00000    0.00000    0.00000
0.00000    0.00000    0.00000 ;    HD2-    CD-     N-    CA
  5669   5668   5671   5672      3    0.65084    1.95253    0.00000
-2.60338    0.00000    0.00000 ;    HD2-    CD-    CG-   HG2
  5669   5668   5671   5673      3    0.65084    1.95253    0.00000
-2.60338    0.00000    0.00000 ;    HD2-    CD-    CG-   HG3
  5669   5668   5671   5674      3    0.65084    1.95253    0.00000
-2.60338    0.00000    0.00000 ;    HD2-    CD-    CG-    CB
  5670   5668   5667   5677      3    0.00000    0.00000    0.00000
0.00000    0.00000    0.00000 ;    HD3-    CD-     N-    CA
  5670   5668   5671   5672      3    0.65084    1.95253    0.00000
-2.60338    0.00000    0.00000 ;    HD3-    CD-    CG-   HG2
  5670   5668   5671   5673      3    0.65084    1.95253    0.00000
-2.60338    0.00000    0.00000 ;    HD3-    CD-    CG-   HG3
  5670   5668   5671   5674      3    0.65084    1.95253    0.00000
-2.60338    0.00000    0.00000 ;    HD3-    CD-    CG-    CB
  5671   5668   5667   5677      3    0.00000    0.00000    0.00000
0.00000    0.00000    0.00000 ;     CG-    CD-     N-    CA
  5671   5674   5677   5678      3    0.65084    1.95253    0.00000
-2.60338    0.00000    0.00000 ;     CG-    CB-    CA-    HA
  5671   5674   5677   5679      3    0.65084    1.95253    0.00000
-2.60338    0.00000    0.00000 ;     CG-    CB-    CA-     C
  5672   5671   5674   5675      3    0.62760    1.88280    0.00000
-2.51040    0.00000    0.00000 ;    HG2-    CG-    CB-   HB2
  5672   5671   5674   5676      3    0.62760    1.88280    0.00000
-2.51040    0.00000    0.00000 ;    HG2-    CG-    CB-   HB3
  5672   5671   5674   5677      3    0.66944    2.00832    0.00000
-2.67776    0.00000    0.00000 ;    HG2-    CG-    CB-    CA
  5673   5671   5674   5675      3    0.62760    1.88280    0.00000
-2.51040    0.00000    0.00000 ;    HG3-    CG-    CB-   HB2
  5673   5671   5674   5676      3    0.62760    1.88280    0.00000
-2.51040    0.00000    0.00000 ;    HG3-    CG-    CB-   HB3
  5673   5671   5674   5677      3    0.66944    2.00832    0.00000
-2.67776    0.00000    0.00000 ;    HG3-    CG-    CB-    CA
  5674   5677   5679   5680      3    0.00000    0.00000    0.00000
0.00000    0.00000    0.00000 ;     CB-    CA-     C-     O
  5674   5677   5679   5681      3    2.51040    4.18400    1.67360
-6.69440    0.00000    0.00000 ;     CB-    CA-     C-     N
  5675   5674   5677   5678      3    0.65084    1.95253    0.00000
-2.60338    0.00000    0.00000 ;    HB2-    CB-    CA-    HA
  5675   5674   5677   5679      3    0.65084    1.95253    0.00000
-2.60338    0.00000    0.00000 ;    HB2-    CB-    CA-     C
  5676   5674   5677   5678      3    0.65084    1.95253    0.00000
-2.60338    0.00000    0.00000 ;    HB3-    CB-    CA-    HA
  5676   5674   5677   5679      3    0.65084    1.95253    0.00000
-2.60338    0.00000    0.00000 ;    HB3-    CB-    CA-     C
  5677   5679   5681   5682      3   20.92000    0.00000  -20.92000
0.00000    0.00000    0.00000 ;     CA-     C-     N-    CD
  5677   5679   5681   5691      3   20.92000    0.00000  -20.92000

0.00000 0.00000 0.00000 ; CA- C- N- CA
5678 5677 5679 5680 3 3.68192 -4.35136 0.00000
1.33888 0.00000 0.00000 ; HA- CA- C- O
5678 5677 5679 5681 3 0.00000 0.00000 0.00000
0.00000 0.00000 0.00000 ; HA- CA- C- N
5679 5681 5682 5683 3 0.00000 0.00000 0.00000
0.00000 0.00000 0.00000 ; C- N- CD- HD2
5679 5681 5682 5684 3 0.00000 0.00000 0.00000
0.00000 0.00000 0.00000 ; C- N- CD- HD3
5679 5681 5682 5685 3 10.04160 -3.34720 16.73600
-6.69440 0.00000 0.00000 ; C- N- CD- CG
5679 5681 5691 5688 3 11.71520 1.67360 15.06240 -
13.38880 0.00000 0.00000 ; C- N- CA- CB
5679 5681 5691 5692 3 0.00000 0.00000 0.00000
0.00000 0.00000 0.00000 ; C- N- CA- HA
5679 5681 5691 5693 3 1.75728 5.27184 2.25936
-7.02912 0.00000 0.00000 ; C- N- CA- C
5680 5679 5681 5682 3 20.92000 0.00000 -20.92000
0.00000 0.00000 0.00000 ; O- C- N- CD
5680 5679 5681 5691 3 20.92000 0.00000 -20.92000
0.00000 0.00000 0.00000 ; O- C- N- CA
5681 5682 5685 5686 3 0.65084 1.95253 0.00000
-2.60338 0.00000 0.00000 ; N- CD- CG- HG2
5681 5682 5685 5687 3 0.65084 1.95253 0.00000
-2.60338 0.00000 0.00000 ; N- CD- CG- HG3
5681 5682 5685 5688 3 0.65084 1.95253 0.00000
-2.60338 0.00000 0.00000 ; N- CD- CG- CB
5681 5691 5688 5685 3 0.65084 1.95253 0.00000
-2.60338 0.00000 0.00000 ; N- CA- CB- CG
5681 5691 5688 5689 3 0.65084 1.95253 0.00000
-2.60338 0.00000 0.00000 ; N- CA- CB- HB2
5681 5691 5688 5690 3 0.65084 1.95253 0.00000
-2.60338 0.00000 0.00000 ; N- CA- CB- HB3
5681 5691 5693 5694 3 0.00000 0.00000 0.00000
0.00000 0.00000 0.00000 ; N- CA- C- O
5681 5691 5693 5695 3 17.40544 -5.02080 -13.22144
9.20480 0.00000 0.00000 ; N- CA- C- N
5682 5681 5691 5688 3 0.00000 0.00000 0.00000
0.00000 0.00000 0.00000 ; CD- N- CA- CB
5682 5681 5691 5692 3 0.00000 0.00000 0.00000
0.00000 0.00000 0.00000 ; CD- N- CA- HA
5682 5681 5691 5693 3 0.00000 0.00000 0.00000
0.00000 0.00000 0.00000 ; CD- N- CA- C
5682 5685 5688 5689 3 0.66944 2.00832 0.00000
-2.67776 0.00000 0.00000 ; CD- CG- CB- HB2
5682 5685 5688 5690 3 0.66944 2.00832 0.00000
-2.67776 0.00000 0.00000 ; CD- CG- CB- HB3
5682 5685 5688 5691 3 3.68192 3.09616 -2.09200
-3.01248 0.00000 0.00000 ; CD- CG- CB- CA
5683 5682 5681 5691 3 0.00000 0.00000 0.00000
0.00000 0.00000 0.00000 ; HD2- CD- N- CA
5683 5682 5685 5686 3 0.65084 1.95253 0.00000

```
-2.60338    0.00000    0.00000 ;    HD2-    CD-    CG-   HG2
  5683   5682   5685   5687      3    0.65084    1.95253    0.00000
-2.60338    0.00000    0.00000 ;    HD2-    CD-    CG-   HG3
  5683   5682   5685   5688      3    0.65084    1.95253    0.00000
-2.60338    0.00000    0.00000 ;    HD2-    CD-    CG-    CB
  5684   5682   5681   5691      3    0.00000    0.00000    0.00000
0.00000    0.00000    0.00000 ;    HD3-    CD-     N-    CA
  5684   5682   5685   5686      3    0.65084    1.95253    0.00000
-2.60338    0.00000    0.00000 ;    HD3-    CD-    CG-   HG2
  5684   5682   5685   5687      3    0.65084    1.95253    0.00000
-2.60338    0.00000    0.00000 ;    HD3-    CD-    CG-   HG3
  5684   5682   5685   5688      3    0.65084    1.95253    0.00000
-2.60338    0.00000    0.00000 ;    HD3-    CD-    CG-    CB
  5685   5682   5681   5691      3    0.00000    0.00000    0.00000
0.00000    0.00000    0.00000 ;     CG-    CD-     N-    CA
  5685   5688   5691   5692      3    0.65084    1.95253    0.00000
-2.60338    0.00000    0.00000 ;     CG-    CB-    CA-    HA
  5685   5688   5691   5693      3    0.65084    1.95253    0.00000
-2.60338    0.00000    0.00000 ;     CG-    CB-    CA-     C
  5686   5685   5688   5689      3    0.62760    1.88280    0.00000
-2.51040    0.00000    0.00000 ;    HG2-    CG-    CB-   HB2
  5686   5685   5688   5690      3    0.62760    1.88280    0.00000
-2.51040    0.00000    0.00000 ;    HG2-    CG-    CB-   HB3
  5686   5685   5688   5691      3    0.66944    2.00832    0.00000
-2.67776    0.00000    0.00000 ;    HG2-    CG-    CB-    CA
  5687   5685   5688   5689      3    0.62760    1.88280    0.00000
-2.51040    0.00000    0.00000 ;    HG3-    CG-    CB-   HB2
  5687   5685   5688   5690      3    0.62760    1.88280    0.00000
-2.51040    0.00000    0.00000 ;    HG3-    CG-    CB-   HB3
  5687   5685   5688   5691      3    0.66944    2.00832    0.00000
-2.67776    0.00000    0.00000 ;    HG3-    CG-    CB-    CA
  5688   5691   5693   5694      3    0.00000    0.00000    0.00000
0.00000    0.00000    0.00000 ;     CB-    CA-     C-     O
  5688   5691   5693   5695      3    2.51040    4.18400    1.67360
-6.69440    0.00000    0.00000 ;     CB-    CA-     C-     N
  5689   5688   5691   5692      3    0.65084    1.95253    0.00000
-2.60338    0.00000    0.00000 ;    HB2-    CB-    CA-    HA
  5689   5688   5691   5693      3    0.65084    1.95253    0.00000
-2.60338    0.00000    0.00000 ;    HB2-    CB-    CA-     C
  5690   5688   5691   5692      3    0.65084    1.95253    0.00000
-2.60338    0.00000    0.00000 ;    HB3-    CB-    CA-    HA
  5690   5688   5691   5693      3    0.65084    1.95253    0.00000
-2.60338    0.00000    0.00000 ;    HB3-    CB-    CA-     C
  5691   5693   5695   5696      3   20.92000    0.00000  -20.92000
0.00000    0.00000    0.00000 ;     CA-     C-     N-     H
  5691   5693   5695   5697      3   20.92000    0.00000  -20.92000
0.00000    0.00000    0.00000 ;     CA-     C-     N-    CA
  5692   5691   5693   5694      3    3.68192   -4.35136    0.00000
1.33888    0.00000    0.00000 ;     HA-    CA-     C-     O
  5692   5691   5693   5695      3    0.00000    0.00000    0.00000
0.00000    0.00000    0.00000 ;     HA-    CA-     C-     N
  5693   5695   5697   5698      3    0.00000    0.00000    0.00000
```

0.00000 0.00000 0.00000 ; C- N- CA- HA
5693 5695 5697 5699 3 11.71520 1.67360 15.06240 -
13.38880 0.00000 0.00000 ; C- N- CA- CB
5693 5695 5697 5705 3 1.75728 5.27184 2.25936
-7.02912 0.00000 0.00000 ; C- N- CA- C
5694 5693 5695 5696 3 29.28800 -8.36800 -20.92000
0.00000 0.00000 0.00000 ; O- C- N- H
5694 5693 5695 5697 3 20.92000 0.00000 -20.92000
0.00000 0.00000 0.00000 ; O- C- N- CA
5695 5697 5699 5700 3 0.65084 1.95253 0.00000
-2.60338 0.00000 0.00000 ; N- CA- CB- HB2
5695 5697 5699 5701 3 0.65084 1.95253 0.00000
-2.60338 0.00000 0.00000 ; N- CA- CB- HB3
5695 5697 5699 5702 3 15.41386 9.74035 -8.39310
-0.97069 2.97901 0.00000 ; N- CA- CB- CG
5695 5697 5705 5706 3 0.00000 0.00000 0.00000
0.00000 0.00000 0.00000 ; N- CA- C- O
5695 5697 5705 5707 3 17.40544 -5.02080 -13.22144
9.20480 0.00000 0.00000 ; N- CA- C- N
5696 5695 5697 5698 3 0.00000 0.00000 0.00000
0.00000 0.00000 0.00000 ; H- N- CA- HA
5696 5695 5697 5699 3 0.00000 0.00000 0.00000
0.00000 0.00000 0.00000 ; H- N- CA- CB
5696 5695 5697 5705 3 0.00000 0.00000 0.00000
0.00000 0.00000 0.00000 ; H- N- CA- C
5697 5699 5702 5703 3 6.43499 0.00000 -5.39736
0.00000 -1.03763 0.00000 ; CA- CB- CG- OD1
5697 5699 5702 5704 3 6.43499 0.00000 -5.39736
0.00000 -1.03763 0.00000 ; CA- CB- CG- OD2
5697 5705 5707 5708 3 20.92000 0.00000 -20.92000
0.00000 0.00000 0.00000 ; CA- C- N- H
5697 5705 5707 5709 3 20.92000 0.00000 -20.92000
0.00000 0.00000 0.00000 ; CA- C- N- CA
5698 5697 5699 5700 3 0.65084 1.95253 0.00000
-2.60338 0.00000 0.00000 ; HA- CA- CB- HB2
5698 5697 5699 5701 3 0.65084 1.95253 0.00000
-2.60338 0.00000 0.00000 ; HA- CA- CB- HB3
5698 5697 5699 5702 3 0.65084 1.95253 0.00000
-2.60338 0.00000 0.00000 ; HA- CA- CB- CG
5698 5697 5705 5706 3 3.68192 -4.35136 0.00000
1.33888 0.00000 0.00000 ; HA- CA- C- O
5698 5697 5705 5707 3 0.00000 0.00000 0.00000
0.00000 0.00000 0.00000 ; HA- CA- C- N
5699 5697 5705 5706 3 0.00000 0.00000 0.00000
0.00000 0.00000 0.00000 ; CB- CA- C- O
5699 5697 5705 5707 3 2.51040 4.18400 1.67360
-6.69440 0.00000 0.00000 ; CB- CA- C- N
5700 5699 5697 5705 3 0.65084 1.95253 0.00000
-2.60338 0.00000 0.00000 ; HB2- CB- CA- C
5700 5699 5702 5703 3 0.00000 0.00000 0.00000
0.00000 0.00000 0.00000 ; HB2- CB- CG- OD1
5700 5699 5702 5704 3 0.00000 0.00000 0.00000

0.00000 0.00000 0.00000 ; HB2- CB- CG- OD2
5701 5699 5697 5705 3 0.65084 1.95253 0.00000
-2.60338 0.00000 0.00000 ; HB3- CB- CA- C
5701 5699 5702 5703 3 0.00000 0.00000 0.00000
0.00000 0.00000 0.00000 ; HB3- CB- CG- OD1
5701 5699 5702 5704 3 0.00000 0.00000 0.00000
0.00000 0.00000 0.00000 ; HB3- CB- CG- OD2
5702 5699 5697 5705 3 7.14627 -1.04600 -8.99560
-0.97069 5.15469 0.00000 ; CG- CB- CA- C
5705 5707 5709 5710 3 0.00000 0.00000 0.00000
0.00000 0.00000 0.00000 ; C- N- CA- HA
5705 5707 5709 5711 3 11.71520 1.67360 15.06240 -
13.38880 0.00000 0.00000 ; C- N- CA- CB
5705 5707 5709 5724 3 1.75728 5.27184 2.25936
-7.02912 0.00000 0.00000 ; C- N- CA- C
5706 5705 5707 5708 3 29.28800 -8.36800 -20.92000
0.00000 0.00000 0.00000 ; O- C- N- H
5706 5705 5707 5709 3 20.92000 0.00000 -20.92000
0.00000 0.00000 0.00000 ; O- C- N- CA
5707 5709 5711 5712 3 0.65084 1.95253 0.00000
-2.60338 0.00000 0.00000 ; N- CA- CB- HB
5707 5709 5711 5713 3 3.83673 0.44769 -1.77402
-2.47693 -0.03347 0.00000 ; N- CA- CB- CG2
5707 5709 5711 5717 3 2.97482 0.12134 2.04179
-1.89117 -3.24678 0.00000 ; N- CA- CB- CG1
5707 5709 5724 5725 3 0.00000 0.00000 0.00000
0.00000 0.00000 0.00000 ; N- CA- C- O
5707 5709 5724 5726 3 17.40544 -5.02080 -13.22144
9.20480 0.00000 0.00000 ; N- CA- C- N
5708 5707 5709 5710 3 0.00000 0.00000 0.00000
0.00000 0.00000 0.00000 ; H- N- CA- HA
5708 5707 5709 5711 3 0.00000 0.00000 0.00000
0.00000 0.00000 0.00000 ; H- N- CA- CB
5708 5707 5709 5724 3 0.00000 0.00000 0.00000
0.00000 0.00000 0.00000 ; H- N- CA- C
5709 5711 5713 5714 3 0.66944 2.00832 0.00000
-2.67776 0.00000 0.00000 ; CA- CB- CG2- HG21
5709 5711 5713 5715 3 0.66944 2.00832 0.00000
-2.67776 0.00000 0.00000 ; CA- CB- CG2- HG22
5709 5711 5713 5716 3 0.66944 2.00832 0.00000
-2.67776 0.00000 0.00000 ; CA- CB- CG2- HG23
5709 5711 5717 5718 3 0.66944 2.00832 0.00000
-2.67776 0.00000 0.00000 ; CA- CB- CG1- HG12
5709 5711 5717 5719 3 0.66944 2.00832 0.00000
-2.67776 0.00000 0.00000 ; CA- CB- CG1- HG13
5709 5711 5717 5720 3 4.24258 -0.52718 -7.25506
-1.79075 7.69856 0.00000 ; CA- CB- CG1- CD1
5709 5724 5726 5727 3 20.92000 0.00000 -20.92000
0.00000 0.00000 0.00000 ; CA- C- N- H
5709 5724 5726 5728 3 20.92000 0.00000 -20.92000
0.00000 0.00000 0.00000 ; CA- C- N- CA
5710 5709 5711 5712 3 0.65084 1.95253 0.00000

```
-2.60338    0.00000    0.00000 ;      HA-    CA-    CB-    HB
  5710   5709   5711   5713       3    0.65084    1.95253    0.00000
-2.60338    0.00000    0.00000 ;      HA-    CA-    CB-   CG2
  5710   5709   5711   5717       3    0.65084    1.95253    0.00000
-2.60338    0.00000    0.00000 ;      HA-    CA-    CB-   CG1
  5710   5709   5724   5725       3    3.68192   -4.35136    0.00000
1.33888    0.00000    0.00000 ;      HA-    CA-     C-     O
  5710   5709   5724   5726       3    0.00000    0.00000    0.00000
0.00000    0.00000    0.00000 ;      HA-    CA-     C-     N
  5711   5709   5724   5725       3    0.00000    0.00000    0.00000
0.00000    0.00000    0.00000 ;      CB-    CA-     C-     O
  5711   5709   5724   5726       3    2.51040    4.18400    1.67360
-6.69440    0.00000    0.00000 ;      CB-    CA-     C-     N
  5711   5717   5720   5721       3    0.66944    2.00832    0.00000
-2.67776    0.00000    0.00000 ;      CB-   CG1-   CD1-  HD11
  5711   5717   5720   5722       3    0.66944    2.00832    0.00000
-2.67776    0.00000    0.00000 ;      CB-   CG1-   CD1-  HD12
  5711   5717   5720   5723       3    0.66944    2.00832    0.00000
-2.67776    0.00000    0.00000 ;      CB-   CG1-   CD1-  HD13
  5712   5711   5709   5724       3    0.65084    1.95253    0.00000
-2.60338    0.00000    0.00000 ;      HB-    CB-    CA-     C
  5712   5711   5713   5714       3    0.62760    1.88280    0.00000
-2.51040    0.00000    0.00000 ;      HB-    CB-   CG2-  HG21
  5712   5711   5713   5715       3    0.62760    1.88280    0.00000
-2.51040    0.00000    0.00000 ;      HB-    CB-   CG2-  HG22
  5712   5711   5713   5716       3    0.62760    1.88280    0.00000
-2.51040    0.00000    0.00000 ;      HB-    CB-   CG2-  HG23
  5712   5711   5717   5718       3    0.62760    1.88280    0.00000
-2.51040    0.00000    0.00000 ;      HB-    CB-   CG1-  HG12
  5712   5711   5717   5719       3    0.62760    1.88280    0.00000
-2.51040    0.00000    0.00000 ;      HB-    CB-   CG1-  HG13
  5712   5711   5717   5720       3    0.66944    2.00832    0.00000
-2.67776    0.00000    0.00000 ;      HB-    CB-   CG1-   CD1
  5713   5711   5709   5724       3    5.67350    3.55640   -6.16722
-2.47693    3.74886    0.00000 ;     CG2-    CB-    CA-     C
  5713   5711   5717   5718       3    0.66944    2.00832    0.00000
-2.67776    0.00000    0.00000 ;     CG2-    CB-   CG1-  HG12
  5713   5711   5717   5719       3    0.66944    2.00832    0.00000
-2.67776    0.00000    0.00000 ;     CG2-    CB-   CG1-  HG13
  5713   5711   5717   5720       3    3.81162    0.49790   -8.14206
-1.79075    7.49773    0.00000 ;     CG2-    CB-   CG1-   CD1
  5714   5713   5711   5717       3    0.66944    2.00832    0.00000
-2.67776    0.00000    0.00000 ;    HG21-   CG2-    CB-   CG1
  5715   5713   5711   5717       3    0.66944    2.00832    0.00000
-2.67776    0.00000    0.00000 ;    HG22-   CG2-    CB-   CG1
  5716   5713   5711   5717       3    0.66944    2.00832    0.00000
-2.67776    0.00000    0.00000 ;    HG23-   CG2-    CB-   CG1
  5717   5711   5709   5724       3    8.26340    0.74057   -9.99976
-1.89117    3.84928    0.00000 ;     CG1-    CB-    CA-     C
  5718   5717   5720   5721       3    0.62760    1.88280    0.00000
-2.51040    0.00000    0.00000 ;    HG12-   CG1-   CD1-  HD11
  5718   5717   5720   5722       3    0.62760    1.88280    0.00000
```

-2.51040 0.00000 0.00000 ; HG12- CG1- CD1- HD12
5718 5717 5720 5723 3 0.62760 1.88280 0.00000
-2.51040 0.00000 0.00000 ; HG12- CG1- CD1- HD13
5719 5717 5720 5721 3 0.62760 1.88280 0.00000
-2.51040 0.00000 0.00000 ; HG13- CG1- CD1- HD11
5719 5717 5720 5722 3 0.62760 1.88280 0.00000
-2.51040 0.00000 0.00000 ; HG13- CG1- CD1- HD12
5719 5717 5720 5723 3 0.62760 1.88280 0.00000
-2.51040 0.00000 0.00000 ; HG13- CG1- CD1- HD13
5724 5726 5728 5729 3 0.00000 0.00000 0.00000
0.00000 0.00000 0.00000 ; C- N- CA- HA
5724 5726 5728 5730 3 11.71520 1.67360 15.06240 -
13.38880 0.00000 0.00000 ; C- N- CA- CB
5724 5726 5728 5741 3 1.75728 5.27184 2.25936
-7.02912 0.00000 0.00000 ; C- N- CA- C
5725 5724 5726 5727 3 29.28800 -8.36800 -20.92000
0.00000 0.00000 0.00000 ; O- C- N- H
5725 5724 5726 5728 3 20.92000 0.00000 -20.92000
0.00000 0.00000 0.00000 ; O- C- N- CA
5726 5728 5730 5731 3 0.65084 1.95253 0.00000
-2.60338 0.00000 0.00000 ; N- CA- CB- HB2
5726 5728 5730 5732 3 0.65084 1.95253 0.00000
-2.60338 0.00000 0.00000 ; N- CA- CB- HB3
5726 5728 5730 5733 3 4.79068 4.02919 -4.82834
-3.66518 2.97901 0.00000 ; N- CA- CB- CG
5726 5728 5741 5742 3 0.00000 0.00000 0.00000
0.00000 0.00000 0.00000 ; N- CA- C- O
5726 5728 5741 5743 3 17.40544 -5.02080 -13.22144
9.20480 0.00000 0.00000 ; N- CA- C- N
5727 5726 5728 5729 3 0.00000 0.00000 0.00000
0.00000 0.00000 0.00000 ; H- N- CA- HA
5727 5726 5728 5730 3 0.00000 0.00000 0.00000
0.00000 0.00000 0.00000 ; H- N- CA- CB
5727 5726 5728 5741 3 0.00000 0.00000 0.00000
0.00000 0.00000 0.00000 ; H- N- CA- C
5728 5730 5733 5734 3 5.98312 6.40152 3.28026 -
12.38464 -1.57318 0.00000 ; CA- CB- CG- ND1
5728 5730 5733 5739 3 0.00000 0.00000 0.00000
0.00000 0.00000 0.00000 ; CA- CB- CG- CD2
5728 5741 5743 5744 3 20.92000 0.00000 -20.92000
0.00000 0.00000 0.00000 ; CA- C- N- H
5728 5741 5743 5745 3 20.92000 0.00000 -20.92000
0.00000 0.00000 0.00000 ; CA- C- N- CA
5729 5728 5730 5731 3 0.65084 1.95253 0.00000
-2.60338 0.00000 0.00000 ; HA- CA- CB- HB2
5729 5728 5730 5732 3 0.65084 1.95253 0.00000
-2.60338 0.00000 0.00000 ; HA- CA- CB- HB3
5729 5728 5730 5733 3 0.65084 1.95253 0.00000
-2.60338 0.00000 0.00000 ; HA- CA- CB- CG
5729 5728 5741 5742 3 3.68192 -4.35136 0.00000
1.33888 0.00000 0.00000 ; HA- CA- C- O
5729 5728 5741 5743 3 0.00000 0.00000 0.00000

```
0.00000    0.00000    0.00000 ;     HA-    CA-     C-     N
  5730   5728   5741   5742      3    0.00000    0.00000    0.00000
0.00000    0.00000    0.00000 ;     CB-    CA-     C-     O
  5730   5728   5741   5743      3    2.51040    4.18400    1.67360
-6.69440    0.00000    0.00000 ;     CB-    CA-     C-     N
  5730   5733   5734   5735      3   20.08320    0.00000  -20.08320
0.00000    0.00000    0.00000 ;     CB-    CG-   ND1-   CE1
  5730   5733   5739   5737      3   44.97800    0.00000  -44.97800
0.00000    0.00000    0.00000 ;     CB-    CG-   CD2-   NE2
  5730   5733   5739   5740      3   44.97800    0.00000  -44.97800
0.00000    0.00000    0.00000 ;     CB-    CG-   CD2-   HD2
  5731   5730   5728   5741      3    0.65084    1.95253    0.00000
-2.60338    0.00000    0.00000 ;    HB2-    CB-    CA-     C
  5731   5730   5733   5734      3    0.00000    0.00000    0.00000
0.00000    0.00000    0.00000 ;    HB2-    CB-    CG-   ND1
  5731   5730   5733   5739      3    0.00000    0.00000    0.00000
0.00000    0.00000    0.00000 ;    HB2-    CB-    CG-   CD2
  5732   5730   5728   5741      3    0.65084    1.95253    0.00000
-2.60338    0.00000    0.00000 ;    HB3-    CB-    CA-     C
  5732   5730   5733   5734      3    0.00000    0.00000    0.00000
0.00000    0.00000    0.00000 ;    HB3-    CB-    CG-   ND1
  5732   5730   5733   5739      3    0.00000    0.00000    0.00000
0.00000    0.00000    0.00000 ;    HB3-    CB-    CG-   CD2
  5733   5730   5728   5741      3    3.76560    3.34720   -2.87859
-3.66518    0.83680    0.00000 ;     CG-    CB-    CA-     C
  5733   5734   5735   5736      3   41.84000    0.00000  -41.84000
0.00000    0.00000    0.00000 ;     CG-   ND1-   CE1-   HE1
  5733   5734   5735   5737      3   41.84000    0.00000  -41.84000
0.00000    0.00000    0.00000 ;     CG-   ND1-   CE1-   NE2
  5733   5739   5737   5735      3   12.55200    0.00000  -12.55200
0.00000    0.00000    0.00000 ;     CG-   CD2-   NE2-   CE1
  5733   5739   5737   5738      3   12.55200    0.00000  -12.55200
0.00000    0.00000    0.00000 ;     CG-   CD2-   NE2-   HE2
  5734   5733   5739   5737      3   44.97800    0.00000  -44.97800
0.00000    0.00000    0.00000 ;    ND1-    CG-   CD2-   NE2
  5734   5733   5739   5740      3   44.97800    0.00000  -44.97800
0.00000    0.00000    0.00000 ;    ND1-    CG-   CD2-   HD2
  5734   5735   5737   5738      3   19.45560    0.00000  -19.45560
0.00000    0.00000    0.00000 ;    ND1-   CE1-   NE2-   HE2
  5734   5735   5737   5739      3   19.45560    0.00000  -19.45560
0.00000    0.00000    0.00000 ;    ND1-   CE1-   NE2-   CD2
  5735   5734   5733   5739      3   20.08320    0.00000  -20.08320
0.00000    0.00000    0.00000 ;    CE1-   ND1-    CG-   CD2
  5735   5737   5739   5740      3   12.55200    0.00000  -12.55200
0.00000    0.00000    0.00000 ;    CE1-   NE2-   CD2-   HD2
  5736   5735   5737   5738      3   19.45560    0.00000  -19.45560
0.00000    0.00000    0.00000 ;    HE1-   CE1-   NE2-   HE2
  5736   5735   5737   5739      3   19.45560    0.00000  -19.45560
0.00000    0.00000    0.00000 ;    HE1-   CE1-   NE2-   CD2
  5738   5737   5739   5740      3   12.55200    0.00000  -12.55200
0.00000    0.00000    0.00000 ;    HE2-   NE2-   CD2-   HD2
  5741   5743   5745   5746      3    0.00000    0.00000    0.00000
```

0.00000 0.00000 0.00000 ; C- N- CA- HA
5741 5743 5745 5747 3 11.71520 1.67360 15.06240 -
13.38880 0.00000 0.00000 ; C- N- CA- CB
5741 5743 5745 5758 3 1.75728 5.27184 2.25936
-7.02912 0.00000 0.00000 ; C- N- CA- C
5742 5741 5743 5744 3 29.28800 -8.36800 -20.92000
0.00000 0.00000 0.00000 ; O- C- N- H
5742 5741 5743 5745 3 20.92000 0.00000 -20.92000
0.00000 0.00000 0.00000 ; O- C- N- CA
5743 5745 5747 5748 3 0.65084 1.95253 0.00000
-2.60338 0.00000 0.00000 ; N- CA- CB- HB2
5743 5745 5747 5749 3 0.65084 1.95253 0.00000
-2.60338 0.00000 0.00000 ; N- CA- CB- HB3
5743 5745 5747 5750 3 3.21331 2.22589 -4.15053
-2.40998 2.61082 0.00000 ; N- CA- CB- CG
5743 5745 5758 5759 3 0.00000 0.00000 0.00000
0.00000 0.00000 0.00000 ; N- CA- C- O
5743 5745 5758 5760 3 17.40544 -5.02080 -13.22144
9.20480 0.00000 0.00000 ; N- CA- C- N
5744 5743 5745 5746 3 0.00000 0.00000 0.00000
0.00000 0.00000 0.00000 ; H- N- CA- HA
5744 5743 5745 5747 3 0.00000 0.00000 0.00000
0.00000 0.00000 0.00000 ; H- N- CA- CB
5744 5743 5745 5758 3 0.00000 0.00000 0.00000
0.00000 0.00000 0.00000 ; H- N- CA- C
5745 5747 5750 5751 3 0.66944 2.00832 0.00000
-2.67776 0.00000 0.00000 ; CA- CB- CG- HG2
5745 5747 5750 5752 3 0.66944 2.00832 0.00000
-2.67776 0.00000 0.00000 ; CA- CB- CG- HG3
5745 5747 5750 5753 3 3.69866 -4.35136 -3.92459
6.89523 4.61914 0.00000 ; CA- CB- CG- CD
5745 5758 5760 5761 3 20.92000 0.00000 -20.92000
0.00000 0.00000 0.00000 ; CA- C- N- H
5745 5758 5760 5762 3 20.92000 0.00000 -20.92000
0.00000 0.00000 0.00000 ; CA- C- N- CA
5746 5745 5747 5748 3 0.65084 1.95253 0.00000
-2.60338 0.00000 0.00000 ; HA- CA- CB- HB2
5746 5745 5747 5749 3 0.65084 1.95253 0.00000
-2.60338 0.00000 0.00000 ; HA- CA- CB- HB3
5746 5745 5747 5750 3 0.65084 1.95253 0.00000
-2.60338 0.00000 0.00000 ; HA- CA- CB- CG
5746 5745 5758 5759 3 3.68192 -4.35136 0.00000
1.33888 0.00000 0.00000 ; HA- CA- C- O
5746 5745 5758 5760 3 0.00000 0.00000 0.00000
0.00000 0.00000 0.00000 ; HA- CA- C- N
5747 5745 5758 5759 3 0.00000 0.00000 0.00000
0.00000 0.00000 0.00000 ; CB- CA- C- O
5747 5745 5758 5760 3 2.51040 4.18400 1.67360
-6.69440 0.00000 0.00000 ; CB- CA- C- N
5747 5750 5753 5754 3 0.00000 0.00000 0.00000
0.00000 0.00000 0.00000 ; CB- CG- CD- OE1
5747 5750 5753 5755 3 10.32611 1.48114 -8.47678

```
1.42256    1.40582    0.00000 ;      CB-    CG-    CD-   NE2
  5748   5747   5745   5758       3    0.65084    1.95253    0.00000
-2.60338    0.00000    0.00000 ;     HB2-    CB-    CA-     C
  5748   5747   5750   5751       3    0.62760    1.88280    0.00000
-2.51040    0.00000    0.00000 ;     HB2-    CB-    CG-   HG2
  5748   5747   5750   5752       3    0.62760    1.88280    0.00000
-2.51040    0.00000    0.00000 ;     HB2-    CB-    CG-   HG3
  5748   5747   5750   5753       3    0.65084    1.95253    0.00000
-2.60338    0.00000    0.00000 ;     HB2-    CB-    CG-    CD
  5749   5747   5745   5758       3    0.65084    1.95253    0.00000
-2.60338    0.00000    0.00000 ;     HB3-    CB-    CA-     C
  5749   5747   5750   5751       3    0.62760    1.88280    0.00000
-2.51040    0.00000    0.00000 ;     HB3-    CB-    CG-   HG2
  5749   5747   5750   5752       3    0.62760    1.88280    0.00000
-2.51040    0.00000    0.00000 ;     HB3-    CB-    CG-   HG3
  5749   5747   5750   5753       3    0.65084    1.95253    0.00000
-2.60338    0.00000    0.00000 ;     HB3-    CB-    CG-    CD
  5750   5747   5745   5758       3    6.86594    3.56895   -8.14206
-2.40998    4.85344    0.00000 ;      CG-    CB-    CA-     C
  5750   5753   5755   5756       3   20.92000    0.00000  -20.92000
0.00000    0.00000    0.00000 ;      CG-    CD-   NE2-  HE21
  5750   5753   5755   5757       3   20.92000    0.00000  -20.92000
0.00000    0.00000    0.00000 ;      CG-    CD-   NE2-  HE22
  5751   5750   5753   5754       3    3.68192   -4.35136    0.00000
1.33888    0.00000    0.00000 ;     HG2-    CG-    CD-   OE1
  5751   5750   5753   5755       3    0.00000    0.00000    0.00000
0.00000    0.00000    0.00000 ;     HG2-    CG-    CD-   NE2
  5752   5750   5753   5754       3    3.68192   -4.35136    0.00000
1.33888    0.00000    0.00000 ;     HG3-    CG-    CD-   OE1
  5752   5750   5753   5755       3    0.00000    0.00000    0.00000
0.00000    0.00000    0.00000 ;     HG3-    CG-    CD-   NE2
  5754   5753   5755   5756       3   29.28800   -8.36800  -20.92000
0.00000    0.00000    0.00000 ;     OE1-    CD-   NE2-  HE21
  5754   5753   5755   5757       3   29.28800   -8.36800  -20.92000
0.00000    0.00000    0.00000 ;     OE1-    CD-   NE2-  HE22
  5758   5760   5762   5763       3    0.00000    0.00000    0.00000
0.00000    0.00000    0.00000 ;       C-     N-    CA-    HA
  5758   5760   5762   5764       3   11.71520    1.67360   15.06240  -
13.38880    0.00000    0.00000 ;       C-     N-    CA-    CB
  5758   5760   5762   5774       3    1.75728    5.27184    2.25936
-7.02912    0.00000    0.00000 ;       C-     N-    CA-     C
  5759   5758   5760   5761       3   29.28800   -8.36800  -20.92000
0.00000    0.00000    0.00000 ;       O-     C-     N-     H
  5759   5758   5760   5762       3   20.92000    0.00000  -20.92000
0.00000    0.00000    0.00000 ;       O-     C-     N-    CA
  5760   5762   5764   5765       3    0.65084    1.95253    0.00000
-2.60338    0.00000    0.00000 ;       N-    CA-    CB-    HB
  5760   5762   5764   5766       3    3.83673    0.44769   -1.77402
-2.47693   -0.03347    0.00000 ;       N-    CA-    CB-   CG1
  5760   5762   5764   5770       3    3.83673    0.44769   -1.77402
-2.47693   -0.03347    0.00000 ;       N-    CA-    CB-   CG2
  5760   5762   5774   5775       3    0.00000    0.00000    0.00000
```

```
0.00000    0.00000    0.00000 ;       N-    CA-     C-     O
  5760   5762   5774   5776      3   17.40544   -5.02080  -13.22144
9.20480    0.00000    0.00000 ;       N-    CA-     C-     N
  5761   5760   5762   5763      3    0.00000    0.00000    0.00000
0.00000    0.00000    0.00000 ;       H-     N-    CA-    HA
  5761   5760   5762   5764      3    0.00000    0.00000    0.00000
0.00000    0.00000    0.00000 ;       H-     N-    CA-    CB
  5761   5760   5762   5774      3    0.00000    0.00000    0.00000
0.00000    0.00000    0.00000 ;       H-     N-    CA-     C
  5762   5764   5766   5767      3    0.66944    2.00832    0.00000
-2.67776    0.00000    0.00000 ;      CA-    CB-   CG1-  HG11
  5762   5764   5766   5768      3    0.66944    2.00832    0.00000
-2.67776    0.00000    0.00000 ;      CA-    CB-   CG1-  HG12
  5762   5764   5766   5769      3    0.66944    2.00832    0.00000
-2.67776    0.00000    0.00000 ;      CA-    CB-   CG1-  HG13
  5762   5764   5770   5771      3    0.66944    2.00832    0.00000
-2.67776    0.00000    0.00000 ;      CA-    CB-   CG2-  HG21
  5762   5764   5770   5772      3    0.66944    2.00832    0.00000
-2.67776    0.00000    0.00000 ;      CA-    CB-   CG2-  HG22
  5762   5764   5770   5773      3    0.66944    2.00832    0.00000
-2.67776    0.00000    0.00000 ;      CA-    CB-   CG2-  HG23
  5762   5774   5776   5777      3   20.92000    0.00000  -20.92000
0.00000    0.00000    0.00000 ;      CA-     C-     N-     H
  5762   5774   5776   5778      3   20.92000    0.00000  -20.92000
0.00000    0.00000    0.00000 ;      CA-     C-     N-    CA
  5763   5762   5764   5765      3    0.65084    1.95253    0.00000
-2.60338    0.00000    0.00000 ;      HA-    CA-    CB-    HB
  5763   5762   5764   5766      3    0.65084    1.95253    0.00000
-2.60338    0.00000    0.00000 ;      HA-    CA-    CB-   CG1
  5763   5762   5764   5770      3    0.65084    1.95253    0.00000
-2.60338    0.00000    0.00000 ;      HA-    CA-    CB-   CG2
  5763   5762   5774   5775      3    3.68192   -4.35136    0.00000
1.33888    0.00000    0.00000 ;      HA-    CA-     C-     O
  5763   5762   5774   5776      3    0.00000    0.00000    0.00000
0.00000    0.00000    0.00000 ;      HA-    CA-     C-     N
  5764   5762   5774   5775      3    0.00000    0.00000    0.00000
0.00000    0.00000    0.00000 ;      CB-    CA-     C-     O
  5764   5762   5774   5776      3    2.51040    4.18400    1.67360
-6.69440    0.00000    0.00000 ;      CB-    CA-     C-     N
  5765   5764   5762   5774      3    0.65084    1.95253    0.00000
-2.60338    0.00000    0.00000 ;      HB-    CB-    CA-     C
  5765   5764   5766   5767      3    0.62760    1.88280    0.00000
-2.51040    0.00000    0.00000 ;      HB-    CB-   CG1-  HG11
  5765   5764   5766   5768      3    0.62760    1.88280    0.00000
-2.51040    0.00000    0.00000 ;      HB-    CB-   CG1-  HG12
  5765   5764   5766   5769      3    0.62760    1.88280    0.00000
-2.51040    0.00000    0.00000 ;      HB-    CB-   CG1-  HG13
  5765   5764   5770   5771      3    0.62760    1.88280    0.00000
-2.51040    0.00000    0.00000 ;      HB-    CB-   CG2-  HG21
  5765   5764   5770   5772      3    0.62760    1.88280    0.00000
-2.51040    0.00000    0.00000 ;      HB-    CB-   CG2-  HG22
  5765   5764   5770   5773      3    0.62760    1.88280    0.00000
```

-2.51040 0.00000 0.00000 ; HB- CB- CG2- HG23
5766 5764 5762 5774 3 5.67350 3.55640 -6.16722
-2.47693 3.74886 0.00000 ; CG1- CB- CA- C
5766 5764 5770 5771 3 0.66944 2.00832 0.00000
-2.67776 0.00000 0.00000 ; CG1- CB- CG2- HG21
5766 5764 5770 5772 3 0.66944 2.00832 0.00000
-2.67776 0.00000 0.00000 ; CG1- CB- CG2- HG22
5766 5764 5770 5773 3 0.66944 2.00832 0.00000
-2.67776 0.00000 0.00000 ; CG1- CB- CG2- HG23
5767 5766 5764 5770 3 0.66944 2.00832 0.00000
-2.67776 0.00000 0.00000 ; HG11- CG1- CB- CG2
5768 5766 5764 5770 3 0.66944 2.00832 0.00000
-2.67776 0.00000 0.00000 ; HG12- CG1- CB- CG2
5769 5766 5764 5770 3 0.66944 2.00832 0.00000
-2.67776 0.00000 0.00000 ; HG13- CG1- CB- CG2
5770 5764 5762 5774 3 5.67350 3.55640 -6.16722
-2.47693 3.74886 0.00000 ; CG2- CB- CA- C
5774 5776 5778 5779 3 0.00000 0.00000 0.00000
0.00000 0.00000 0.00000 ; C- N- CA- HA
5774 5776 5778 5780 3 11.71520 1.67360 15.06240 -
13.38880 0.00000 0.00000 ; C- N- CA- CB
5774 5776 5778 5793 3 1.75728 5.27184 2.25936
-7.02912 0.00000 0.00000 ; C- N- CA- C
5775 5774 5776 5777 3 29.28800 -8.36800 -20.92000
0.00000 0.00000 0.00000 ; O- C- N- H
5775 5774 5776 5778 3 20.92000 0.00000 -20.92000
0.00000 0.00000 0.00000 ; O- C- N- CA
5776 5778 5780 5781 3 0.65084 1.95253 0.00000
-2.60338 0.00000 0.00000 ; N- CA- CB- HB2
5776 5778 5780 5782 3 0.65084 1.95253 0.00000
-2.60338 0.00000 0.00000 ; N- CA- CB- HB3
5776 5778 5780 5783 3 3.79070 1.39746 -4.61077
-2.40998 2.44346 0.00000 ; N- CA- CB- CG
5776 5778 5793 5794 3 0.00000 0.00000 0.00000
0.00000 0.00000 0.00000 ; N- CA- C- O
5776 5778 5793 5795 3 17.40544 -5.02080 -13.22144
9.20480 0.00000 0.00000 ; N- CA- C- N
5777 5776 5778 5779 3 0.00000 0.00000 0.00000
0.00000 0.00000 0.00000 ; H- N- CA- HA
5777 5776 5778 5780 3 0.00000 0.00000 0.00000
0.00000 0.00000 0.00000 ; H- N- CA- CB
5777 5776 5778 5793 3 0.00000 0.00000 0.00000
0.00000 0.00000 0.00000 ; H- N- CA- C
5778 5780 5783 5784 3 0.66944 2.00832 0.00000
-2.67776 0.00000 0.00000 ; CA- CB- CG- HG
5778 5780 5783 5785 3 3.90367 0.19665 -6.21742
-2.37651 5.99149 0.00000 ; CA- CB- CG- CD1
5778 5780 5783 5789 3 3.90367 0.19665 -6.21742
-2.37651 5.99149 0.00000 ; CA- CB- CG- CD2
5778 5793 5795 5796 3 20.92000 0.00000 -20.92000
0.00000 0.00000 0.00000 ; CA- C- N- H
5778 5793 5795 5797 3 20.92000 0.00000 -20.92000

```
0.00000    0.00000    0.00000 ;      CA-     C-     N-    CA
  5779   5778   5780   5781       3    0.65084    1.95253    0.00000
-2.60338    0.00000    0.00000 ;     HA-    CA-    CB-   HB2
  5779   5778   5780   5782       3    0.65084    1.95253    0.00000
-2.60338    0.00000    0.00000 ;     HA-    CA-    CB-   HB3
  5779   5778   5780   5783       3    0.65084    1.95253    0.00000
-2.60338    0.00000    0.00000 ;     HA-    CA-    CB-    CG
  5779   5778   5793   5794       3    3.68192   -4.35136    0.00000
1.33888    0.00000    0.00000 ;      HA-    CA-     C-     O
  5779   5778   5793   5795       3    0.00000    0.00000    0.00000
0.00000    0.00000    0.00000 ;      HA-    CA-     C-     N
  5780   5778   5793   5794       3    0.00000    0.00000    0.00000
0.00000    0.00000    0.00000 ;      CB-    CA-     C-     O
  5780   5778   5793   5795       3    2.51040    4.18400    1.67360
-6.69440    0.00000    0.00000 ;     CB-    CA-     C-     N
  5780   5783   5785   5786       3    0.66944    2.00832    0.00000
-2.67776    0.00000    0.00000 ;     CB-    CG-   CD1-  HD11
  5780   5783   5785   5787       3    0.66944    2.00832    0.00000
-2.67776    0.00000    0.00000 ;     CB-    CG-   CD1-  HD12
  5780   5783   5785   5788       3    0.66944    2.00832    0.00000
-2.67776    0.00000    0.00000 ;     CB-    CG-   CD1-  HD13
  5780   5783   5789   5790       3    0.66944    2.00832    0.00000
-2.67776    0.00000    0.00000 ;     CB-    CG-   CD2-  HD21
  5780   5783   5789   5791       3    0.66944    2.00832    0.00000
-2.67776    0.00000    0.00000 ;     CB-    CG-   CD2-  HD22
  5780   5783   5789   5792       3    0.66944    2.00832    0.00000
-2.67776    0.00000    0.00000 ;     CB-    CG-   CD2-  HD23
  5781   5780   5778   5793       3    0.65084    1.95253    0.00000
-2.60338    0.00000    0.00000 ;    HB2-    CB-    CA-     C
  5781   5780   5783   5784       3    0.62760    1.88280    0.00000
-2.51040    0.00000    0.00000 ;    HB2-    CB-    CG-    HG
  5781   5780   5783   5785       3    0.66944    2.00832    0.00000
-2.67776    0.00000    0.00000 ;    HB2-    CB-    CG-   CD1
  5781   5780   5783   5789       3    0.66944    2.00832    0.00000
-2.67776    0.00000    0.00000 ;    HB2-    CB-    CG-   CD2
  5782   5780   5778   5793       3    0.65084    1.95253    0.00000
-2.60338    0.00000    0.00000 ;    HB3-    CB-    CA-     C
  5782   5780   5783   5784       3    0.62760    1.88280    0.00000
-2.51040    0.00000    0.00000 ;    HB3-    CB-    CG-    HG
  5782   5780   5783   5785       3    0.66944    2.00832    0.00000
-2.67776    0.00000    0.00000 ;    HB3-    CB-    CG-   CD1
  5782   5780   5783   5789       3    0.66944    2.00832    0.00000
-2.67776    0.00000    0.00000 ;    HB3-    CB-    CG-   CD2
  5783   5780   5778   5793       3   10.33448   -1.14642  -11.54784
-2.40998    6.35968    0.00000 ;     CG-    CB-    CA-     C
  5784   5783   5785   5786       3    0.62760    1.88280    0.00000
-2.51040    0.00000    0.00000 ;     HG-    CG-   CD1-  HD11
  5784   5783   5785   5787       3    0.62760    1.88280    0.00000
-2.51040    0.00000    0.00000 ;     HG-    CG-   CD1-  HD12
  5784   5783   5785   5788       3    0.62760    1.88280    0.00000
-2.51040    0.00000    0.00000 ;     HG-    CG-   CD1-  HD13
  5784   5783   5789   5790       3    0.62760    1.88280    0.00000
```

-2.51040 0.00000 0.00000 ; HG- CG- CD2- HD21
5784 5783 5789 5791 3 0.62760 1.88280 0.00000
-2.51040 0.00000 0.00000 ; HG- CG- CD2- HD22
5784 5783 5789 5792 3 0.62760 1.88280 0.00000
-2.51040 0.00000 0.00000 ; HG- CG- CD2- HD23
5785 5783 5789 5790 3 0.66944 2.00832 0.00000
-2.67776 0.00000 0.00000 ; CD1- CG- CD2- HD21
5785 5783 5789 5791 3 0.66944 2.00832 0.00000
-2.67776 0.00000 0.00000 ; CD1- CG- CD2- HD22
5785 5783 5789 5792 3 0.66944 2.00832 0.00000
-2.67776 0.00000 0.00000 ; CD1- CG- CD2- HD23
5786 5785 5783 5789 3 0.66944 2.00832 0.00000
-2.67776 0.00000 0.00000 ; HD11- CD1- CG- CD2
5787 5785 5783 5789 3 0.66944 2.00832 0.00000
-2.67776 0.00000 0.00000 ; HD12- CD1- CG- CD2
5788 5785 5783 5789 3 0.66944 2.00832 0.00000
-2.67776 0.00000 0.00000 ; HD13- CD1- CG- CD2
5793 5795 5797 5798 3 0.00000 0.00000 0.00000
0.00000 0.00000 0.00000 ; C- N- CA- HA
5793 5795 5797 5799 3 11.71520 1.67360 15.06240 -
13.38880 0.00000 0.00000 ; C- N- CA- CB
5793 5795 5797 5817 3 1.75728 5.27184 2.25936
-7.02912 0.00000 0.00000 ; C- N- CA- C
5794 5793 5795 5796 3 29.28800 -8.36800 -20.92000
0.00000 0.00000 0.00000 ; O- C- N- H
5794 5793 5795 5797 3 20.92000 0.00000 -20.92000
0.00000 0.00000 0.00000 ; O- C- N- CA
5795 5797 5799 5800 3 0.65084 1.95253 0.00000
-2.60338 0.00000 0.00000 ; N- CA- CB- HB2
5795 5797 5799 5801 3 0.65084 1.95253 0.00000
-2.60338 0.00000 0.00000 ; N- CA- CB- HB3
5795 5797 5799 5802 3 0.65084 1.95253 0.00000
-2.60338 0.00000 0.00000 ; N- CA- CB- CG
5795 5797 5817 5818 3 0.00000 0.00000 0.00000
0.00000 0.00000 0.00000 ; N- CA- C- O
5795 5797 5817 5819 3 17.40544 -5.02080 -13.22144
9.20480 0.00000 0.00000 ; N- CA- C- N
5796 5795 5797 5798 3 0.00000 0.00000 0.00000
0.00000 0.00000 0.00000 ; H- N- CA- HA
5796 5795 5797 5799 3 0.00000 0.00000 0.00000
0.00000 0.00000 0.00000 ; H- N- CA- CB
5796 5795 5797 5817 3 0.00000 0.00000 0.00000
0.00000 0.00000 0.00000 ; H- N- CA- C
5797 5799 5802 5803 3 0.66944 2.00832 0.00000
-2.67776 0.00000 0.00000 ; CA- CB- CG- HG2
5797 5799 5802 5804 3 0.66944 2.00832 0.00000
-2.67776 0.00000 0.00000 ; CA- CB- CG- HG3
5797 5799 5802 5805 3 3.68192 3.09616 -2.09200
-3.01248 0.00000 0.00000 ; CA- CB- CG- CD
5797 5817 5819 5820 3 20.92000 0.00000 -20.92000
0.00000 0.00000 0.00000 ; CA- C- N- H
5797 5817 5819 5821 3 20.92000 0.00000 -20.92000

```
0.00000    0.00000    0.00000 ;      CA-     C-     N-    CA
  5798   5797   5799   5800       3    0.65084    1.95253    0.00000
-2.60338    0.00000    0.00000 ;      HA-    CA-    CB-   HB2
  5798   5797   5799   5801       3    0.65084    1.95253    0.00000
-2.60338    0.00000    0.00000 ;      HA-    CA-    CB-   HB3
  5798   5797   5799   5802       3    0.65084    1.95253    0.00000
-2.60338    0.00000    0.00000 ;      HA-    CA-    CB-    CG
  5798   5797   5817   5818       3    3.68192   -4.35136    0.00000
1.33888    0.00000    0.00000 ;      HA-    CA-     C-     O
  5798   5797   5817   5819       3    0.00000    0.00000    0.00000
0.00000    0.00000    0.00000 ;      HA-    CA-     C-     N
  5799   5797   5817   5818       3    0.00000    0.00000    0.00000
0.00000    0.00000    0.00000 ;      CB-    CA-     C-     O
  5799   5797   5817   5819       3    2.51040    4.18400    1.67360
-6.69440    0.00000    0.00000 ;      CB-    CA-     C-     N
  5799   5802   5805   5806       3    0.65084    1.95253    0.00000
-2.60338    0.00000    0.00000 ;      CB-    CG-    CD-   HD2
  5799   5802   5805   5807       3    0.65084    1.95253    0.00000
-2.60338    0.00000    0.00000 ;      CB-    CG-    CD-   HD3
  5799   5802   5805   5808       3    0.65084    1.95253    0.00000
-2.60338    0.00000    0.00000 ;      CB-    CG-    CD-    NE
  5800   5799   5797   5817       3    0.65084    1.95253    0.00000
-2.60338    0.00000    0.00000 ;     HB2-    CB-    CA-     C
  5800   5799   5802   5803       3    0.62760    1.88280    0.00000
-2.51040    0.00000    0.00000 ;     HB2-    CB-    CG-   HG2
  5800   5799   5802   5804       3    0.62760    1.88280    0.00000
-2.51040    0.00000    0.00000 ;     HB2-    CB-    CG-   HG3
  5800   5799   5802   5805       3    0.66944    2.00832    0.00000
-2.67776    0.00000    0.00000 ;     HB2-    CB-    CG-    CD
  5801   5799   5797   5817       3    0.65084    1.95253    0.00000
-2.60338    0.00000    0.00000 ;     HB3-    CB-    CA-     C
  5801   5799   5802   5803       3    0.62760    1.88280    0.00000
-2.51040    0.00000    0.00000 ;     HB3-    CB-    CG-   HG2
  5801   5799   5802   5804       3    0.62760    1.88280    0.00000
-2.51040    0.00000    0.00000 ;     HB3-    CB-    CG-   HG3
  5801   5799   5802   5805       3    0.66944    2.00832    0.00000
-2.67776    0.00000    0.00000 ;     HB3-    CB-    CG-    CD
  5802   5799   5797   5817       3    0.65084    1.95253    0.00000
-2.60338    0.00000    0.00000 ;      CG-    CB-    CA-     C
  5802   5805   5808   5809       3    0.00000    0.00000    0.00000
0.00000    0.00000    0.00000 ;      CG-    CD-    NE-    HE
  5802   5805   5808   5810       3    0.00000    0.00000    0.00000
0.00000    0.00000    0.00000 ;      CG-    CD-    NE-    CZ
  5803   5802   5805   5806       3    0.65084    1.95253    0.00000
-2.60338    0.00000    0.00000 ;     HG2-    CG-    CD-   HD2
  5803   5802   5805   5807       3    0.65084    1.95253    0.00000
-2.60338    0.00000    0.00000 ;     HG2-    CG-    CD-   HD3
  5803   5802   5805   5808       3    0.65084    1.95253    0.00000
-2.60338    0.00000    0.00000 ;     HG2-    CG-    CD-    NE
  5804   5802   5805   5806       3    0.65084    1.95253    0.00000
-2.60338    0.00000    0.00000 ;     HG3-    CG-    CD-   HD2
  5804   5802   5805   5807       3    0.65084    1.95253    0.00000
```

-2.60338    0.00000    0.00000 ;    HG3-    CG-    CD-   HD3
  5804   5802   5805   5808      3    0.65084    1.95253    0.00000
-2.60338    0.00000    0.00000 ;    HG3-    CG-    CD-    NE
  5805   5808   5810   5811      3   20.08320    0.00000  -20.08320
0.00000    0.00000    0.00000 ;     CD-    NE-    CZ-   NH1
  5805   5808   5810   5814      3   20.08320    0.00000  -20.08320
0.00000    0.00000    0.00000 ;     CD-    NE-    CZ-   NH2
  5806   5805   5808   5809      3    0.00000    0.00000    0.00000
0.00000    0.00000    0.00000 ;    HD2-    CD-    NE-    HE
  5806   5805   5808   5810      3    0.00000    0.00000    0.00000
0.00000    0.00000    0.00000 ;    HD2-    CD-    NE-    CZ
  5807   5805   5808   5809      3    0.00000    0.00000    0.00000
0.00000    0.00000    0.00000 ;    HD3-    CD-    NE-    HE
  5807   5805   5808   5810      3    0.00000    0.00000    0.00000
0.00000    0.00000    0.00000 ;    HD3-    CD-    NE-    CZ
  5808   5810   5811   5812      3   20.08320    0.00000  -20.08320
0.00000    0.00000    0.00000 ;     NE-    CZ-   NH1-  HH11
  5808   5810   5811   5813      3   20.08320    0.00000  -20.08320
0.00000    0.00000    0.00000 ;     NE-    CZ-   NH1-  HH12
  5808   5810   5814   5815      3   20.08320    0.00000  -20.08320
0.00000    0.00000    0.00000 ;     NE-    CZ-   NH2-  HH21
  5808   5810   5814   5816      3   20.08320    0.00000  -20.08320
0.00000    0.00000    0.00000 ;     NE-    CZ-   NH2-  HH22
  5809   5808   5810   5811      3   20.08320    0.00000  -20.08320
0.00000    0.00000    0.00000 ;     HE-    NE-    CZ-   NH1
  5809   5808   5810   5814      3   20.08320    0.00000  -20.08320
0.00000    0.00000    0.00000 ;     HE-    NE-    CZ-   NH2
  5811   5810   5814   5815      3   20.08320    0.00000  -20.08320
0.00000    0.00000    0.00000 ;    NH1-    CZ-   NH2-  HH21
  5811   5810   5814   5816      3   20.08320    0.00000  -20.08320
0.00000    0.00000    0.00000 ;    NH1-    CZ-   NH2-  HH22
  5812   5811   5810   5814      3   20.08320    0.00000  -20.08320
0.00000    0.00000    0.00000 ;   HH11-   NH1-    CZ-   NH2
  5813   5811   5810   5814      3   20.08320    0.00000  -20.08320
0.00000    0.00000    0.00000 ;   HH12-   NH1-    CZ-   NH2
  5817   5819   5821   5822      3    0.00000    0.00000    0.00000
0.00000    0.00000    0.00000 ;      C-     N-    CA-    HA
  5817   5819   5821   5823      3   11.71520    1.67360   15.06240  -
13.38880    0.00000    0.00000 ;      C-     N-    CA-    CB
  5817   5819   5821   5829      3    1.75728    5.27184    2.25936
-7.02912    0.00000    0.00000 ;      C-     N-    CA-     C
  5818   5817   5819   5820      3   29.28800   -8.36800  -20.92000
0.00000    0.00000    0.00000 ;      O-     C-     N-     H
  5818   5817   5819   5821      3   20.92000    0.00000  -20.92000
0.00000    0.00000    0.00000 ;      O-     C-     N-    CA
  5819   5821   5823   5824      3    0.65084    1.95253    0.00000
-2.60338    0.00000    0.00000 ;      N-    CA-    CB-   HB2
  5819   5821   5823   5825      3    0.65084    1.95253    0.00000
-2.60338    0.00000    0.00000 ;      N-    CA-    CB-   HB3
  5819   5821   5823   5826      3   15.41386    9.74035   -8.39310
-0.97069    2.97901    0.00000 ;      N-    CA-    CB-    CG
  5819   5821   5829   5830      3    0.00000    0.00000    0.00000

0.00000    0.00000    0.00000 ;     N-    CA-     C-     O
  5819   5821   5829   5831      3   17.40544   -5.02080  -13.22144
9.20480    0.00000    0.00000 ;     N-    CA-     C-     N
  5820   5819   5821   5822      3    0.00000    0.00000    0.00000
0.00000    0.00000    0.00000 ;     H-     N-    CA-    HA
  5820   5819   5821   5823      3    0.00000    0.00000    0.00000
0.00000    0.00000    0.00000 ;     H-     N-    CA-    CB
  5820   5819   5821   5829      3    0.00000    0.00000    0.00000
0.00000    0.00000    0.00000 ;     H-     N-    CA-     C
  5821   5823   5826   5827      3    6.43499    0.00000   -5.39736
0.00000   -1.03763    0.00000 ;    CA-    CB-    CG-   OD1
  5821   5823   5826   5828      3    6.43499    0.00000   -5.39736
0.00000   -1.03763    0.00000 ;    CA-    CB-    CG-   OD2
  5821   5829   5831   5832      3   20.92000    0.00000  -20.92000
0.00000    0.00000    0.00000 ;    CA-     C-     N-     H
  5821   5829   5831   5833      3   20.92000    0.00000  -20.92000
0.00000    0.00000    0.00000 ;    CA-     C-     N-    CA
  5822   5821   5823   5824      3    0.65084    1.95253    0.00000
-2.60338    0.00000    0.00000 ;    HA-    CA-    CB-   HB2
  5822   5821   5823   5825      3    0.65084    1.95253    0.00000
-2.60338    0.00000    0.00000 ;    HA-    CA-    CB-   HB3
  5822   5821   5823   5826      3    0.65084    1.95253    0.00000
-2.60338    0.00000    0.00000 ;    HA-    CA-    CB-    CG
  5822   5821   5829   5830      3    3.68192   -4.35136    0.00000
1.33888    0.00000    0.00000 ;    HA-    CA-     C-     O
  5822   5821   5829   5831      3    0.00000    0.00000    0.00000
0.00000    0.00000    0.00000 ;    HA-    CA-     C-     N
  5823   5821   5829   5830      3    0.00000    0.00000    0.00000
0.00000    0.00000    0.00000 ;    CB-    CA-     C-     O
  5823   5821   5829   5831      3    2.51040    4.18400    1.67360
-6.69440    0.00000    0.00000 ;    CB-    CA-     C-     N
  5824   5823   5821   5829      3    0.65084    1.95253    0.00000
-2.60338    0.00000    0.00000 ;   HB2-    CB-    CA-     C
  5824   5823   5826   5827      3    0.00000    0.00000    0.00000
0.00000    0.00000    0.00000 ;   HB2-    CB-    CG-   OD1
  5824   5823   5826   5828      3    0.00000    0.00000    0.00000
0.00000    0.00000    0.00000 ;   HB2-    CB-    CG-   OD2
  5825   5823   5821   5829      3    0.65084    1.95253    0.00000
-2.60338    0.00000    0.00000 ;   HB3-    CB-    CA-     C
  5825   5823   5826   5827      3    0.00000    0.00000    0.00000
0.00000    0.00000    0.00000 ;   HB3-    CB-    CG-   OD1
  5825   5823   5826   5828      3    0.00000    0.00000    0.00000
0.00000    0.00000    0.00000 ;   HB3-    CB-    CG-   OD2
  5826   5823   5821   5829      3    7.14627   -1.04600   -8.99560
-0.97069    5.15469    0.00000 ;    CG-    CB-    CA-     C
  5829   5831   5833   5834      3    0.00000    0.00000    0.00000
0.00000    0.00000    0.00000 ;     C-     N-    CA-    HA
  5829   5831   5833   5835      3   11.71520    1.67360   15.06240  -
13.38880    0.00000    0.00000 ;     C-     N-    CA-    CB
  5829   5831   5833   5848      3    1.75728    5.27184    2.25936
-7.02912    0.00000    0.00000 ;     C-     N-    CA-     C
  5830   5829   5831   5832      3   29.28800   -8.36800  -20.92000

```
0.00000    0.00000    0.00000 ;       O-     C-     N-     H
  5830   5829   5831   5833       3   20.92000    0.00000  -20.92000
0.00000    0.00000    0.00000 ;       O-     C-     N-    CA
  5831   5833   5835   5836       3    0.65084    1.95253    0.00000
-2.60338    0.00000    0.00000 ;       N-    CA-    CB-    HB
  5831   5833   5835   5837       3    3.83673    0.44769   -1.77402
-2.47693   -0.03347    0.00000 ;       N-    CA-    CB-   CG2
  5831   5833   5835   5841       3    2.97482    0.12134    2.04179
-1.89117   -3.24678    0.00000 ;       N-    CA-    CB-   CG1
  5831   5833   5848   5849       3    0.00000    0.00000    0.00000
0.00000    0.00000    0.00000 ;       N-    CA-     C-     O
  5831   5833   5848   5850       3   17.40544   -5.02080  -13.22144
9.20480    0.00000    0.00000 ;       N-    CA-     C-     N
  5832   5831   5833   5834       3    0.00000    0.00000    0.00000
0.00000    0.00000    0.00000 ;       H-     N-    CA-    HA
  5832   5831   5833   5835       3    0.00000    0.00000    0.00000
0.00000    0.00000    0.00000 ;       H-     N-    CA-    CB
  5832   5831   5833   5848       3    0.00000    0.00000    0.00000
0.00000    0.00000    0.00000 ;       H-     N-    CA-     C
  5833   5835   5837   5838       3    0.66944    2.00832    0.00000
-2.67776    0.00000    0.00000 ;      CA-    CB-   CG2-  HG21
  5833   5835   5837   5839       3    0.66944    2.00832    0.00000
-2.67776    0.00000    0.00000 ;      CA-    CB-   CG2-  HG22
  5833   5835   5837   5840       3    0.66944    2.00832    0.00000
-2.67776    0.00000    0.00000 ;      CA-    CB-   CG2-  HG23
  5833   5835   5841   5842       3    0.66944    2.00832    0.00000
-2.67776    0.00000    0.00000 ;      CA-    CB-   CG1-  HG12
  5833   5835   5841   5843       3    0.66944    2.00832    0.00000
-2.67776    0.00000    0.00000 ;      CA-    CB-   CG1-  HG13
  5833   5835   5841   5844       3    4.24258   -0.52718   -7.25506
-1.79075    7.69856    0.00000 ;      CA-    CB-   CG1-   CD1
  5833   5848   5850   5851       3   20.92000    0.00000  -20.92000
0.00000    0.00000    0.00000 ;      CA-     C-     N-     H
  5833   5848   5850   5852       3   20.92000    0.00000  -20.92000
0.00000    0.00000    0.00000 ;      CA-     C-     N-    CA
  5834   5833   5835   5836       3    0.65084    1.95253    0.00000
-2.60338    0.00000    0.00000 ;      HA-    CA-    CB-    HB
  5834   5833   5835   5837       3    0.65084    1.95253    0.00000
-2.60338    0.00000    0.00000 ;      HA-    CA-    CB-   CG2
  5834   5833   5835   5841       3    0.65084    1.95253    0.00000
-2.60338    0.00000    0.00000 ;      HA-    CA-    CB-   CG1
  5834   5833   5848   5849       3    3.68192   -4.35136    0.00000
1.33888    0.00000    0.00000 ;      HA-    CA-     C-     O
  5834   5833   5848   5850       3    0.00000    0.00000    0.00000
0.00000    0.00000    0.00000 ;      HA-    CA-     C-     N
  5835   5833   5848   5849       3    0.00000    0.00000    0.00000
0.00000    0.00000    0.00000 ;      CB-    CA-     C-     O
  5835   5833   5848   5850       3    2.51040    4.18400    1.67360
-6.69440    0.00000    0.00000 ;      CB-    CA-     C-     N
  5835   5841   5844   5845       3    0.66944    2.00832    0.00000
-2.67776    0.00000    0.00000 ;      CB-   CG1-   CD1-  HD11
  5835   5841   5844   5846       3    0.66944    2.00832    0.00000
```

-2.67776    0.00000    0.00000 ;      CB-   CG1-   CD1-  HD12
  5835   5841   5844   5847      3    0.66944    2.00832    0.00000
-2.67776    0.00000    0.00000 ;      CB-   CG1-   CD1-  HD13
  5836   5835   5833   5848      3    0.65084    1.95253    0.00000
-2.60338    0.00000    0.00000 ;      HB-    CB-    CA-     C
  5836   5835   5837   5838      3    0.62760    1.88280    0.00000
-2.51040    0.00000    0.00000 ;      HB-    CB-   CG2-  HG21
  5836   5835   5837   5839      3    0.62760    1.88280    0.00000
-2.51040    0.00000    0.00000 ;      HB-    CB-   CG2-  HG22
  5836   5835   5837   5840      3    0.62760    1.88280    0.00000
-2.51040    0.00000    0.00000 ;      HB-    CB-   CG2-  HG23
  5836   5835   5841   5842      3    0.62760    1.88280    0.00000
-2.51040    0.00000    0.00000 ;      HB-    CB-   CG1-  HG12
  5836   5835   5841   5843      3    0.62760    1.88280    0.00000
-2.51040    0.00000    0.00000 ;      HB-    CB-   CG1-  HG13
  5836   5835   5841   5844      3    0.66944    2.00832    0.00000
-2.67776    0.00000    0.00000 ;      HB-    CB-   CG1-   CD1
  5837   5835   5833   5848      3    5.67350    3.55640   -6.16722
-2.47693    3.74886    0.00000 ;     CG2-    CB-    CA-     C
  5837   5835   5841   5842      3    0.66944    2.00832    0.00000
-2.67776    0.00000    0.00000 ;     CG2-    CB-   CG1-  HG12
  5837   5835   5841   5843      3    0.66944    2.00832    0.00000
-2.67776    0.00000    0.00000 ;     CG2-    CB-   CG1-  HG13
  5837   5835   5841   5844      3    3.81162    0.49790   -8.14206
-1.79075    7.49773    0.00000 ;     CG2-    CB-   CG1-   CD1
  5838   5837   5835   5841      3    0.66944    2.00832    0.00000
-2.67776    0.00000    0.00000 ;    HG21-   CG2-    CB-   CG1
  5839   5837   5835   5841      3    0.66944    2.00832    0.00000
-2.67776    0.00000    0.00000 ;    HG22-   CG2-    CB-   CG1
  5840   5837   5835   5841      3    0.66944    2.00832    0.00000
-2.67776    0.00000    0.00000 ;    HG23-   CG2-    CB-   CG1
  5841   5835   5833   5848      3    8.26340    0.74057   -9.99976
-1.89117    3.84928    0.00000 ;     CG1-    CB-    CA-     C
  5842   5841   5844   5845      3    0.62760    1.88280    0.00000
-2.51040    0.00000    0.00000 ;    HG12-   CG1-   CD1-  HD11
  5842   5841   5844   5846      3    0.62760    1.88280    0.00000
-2.51040    0.00000    0.00000 ;    HG12-   CG1-   CD1-  HD12
  5842   5841   5844   5847      3    0.62760    1.88280    0.00000
-2.51040    0.00000    0.00000 ;    HG12-   CG1-   CD1-  HD13
  5843   5841   5844   5845      3    0.62760    1.88280    0.00000
-2.51040    0.00000    0.00000 ;    HG13-   CG1-   CD1-  HD11
  5843   5841   5844   5846      3    0.62760    1.88280    0.00000
-2.51040    0.00000    0.00000 ;    HG13-   CG1-   CD1-  HD12
  5843   5841   5844   5847      3    0.62760    1.88280    0.00000
-2.51040    0.00000    0.00000 ;    HG13-   CG1-   CD1-  HD13
  5848   5850   5852   5853      3    0.00000    0.00000    0.00000
0.00000    0.00000    0.00000 ;       C-     N-    CA-    HA
  5848   5850   5852   5854      3   11.71520    1.67360   15.06240  -
13.38880    0.00000    0.00000 ;       C-     N-    CA-    CB
  5848   5850   5852   5860      3    1.75728    5.27184    2.25936
-7.02912    0.00000    0.00000 ;       C-     N-    CA-     C
  5849   5848   5850   5851      3   29.28800   -8.36800  -20.92000

```
0.00000    0.00000    0.00000 ;       O-      C-      N-      H
  5849   5848   5850   5852       3   20.92000    0.00000  -20.92000
0.00000    0.00000    0.00000 ;       O-      C-      N-     CA
  5850   5852   5854   5855       3    0.65084    1.95253    0.00000
-2.60338    0.00000    0.00000 ;       N-     CA-     CB-    HB2
  5850   5852   5854   5856       3    0.65084    1.95253    0.00000
-2.60338    0.00000    0.00000 ;       N-     CA-     CB-    HB3
  5850   5852   5854   5857       3   15.41386    9.74035   -8.39310
-0.97069    2.97901    0.00000 ;       N-     CA-     CB-     CG
  5850   5852   5860   5861       3    0.00000    0.00000    0.00000
0.00000    0.00000    0.00000 ;       N-     CA-      C-      O
  5850   5852   5860   5862       3   17.40544   -5.02080  -13.22144
9.20480    0.00000    0.00000 ;       N-     CA-      C-      N
  5851   5850   5852   5853       3    0.00000    0.00000    0.00000
0.00000    0.00000    0.00000 ;       H-      N-     CA-     HA
  5851   5850   5852   5854       3    0.00000    0.00000    0.00000
0.00000    0.00000    0.00000 ;       H-      N-     CA-     CB
  5851   5850   5852   5860       3    0.00000    0.00000    0.00000
0.00000    0.00000    0.00000 ;       H-      N-     CA-      C
  5852   5854   5857   5858       3    6.43499    0.00000   -5.39736
0.00000   -1.03763    0.00000 ;      CA-     CB-     CG-    OD1
  5852   5854   5857   5859       3    6.43499    0.00000   -5.39736
0.00000   -1.03763    0.00000 ;      CA-     CB-     CG-    OD2
  5852   5860   5862   5863       3   20.92000    0.00000  -20.92000
0.00000    0.00000    0.00000 ;      CA-      C-      N-      H
  5852   5860   5862   5864       3   20.92000    0.00000  -20.92000
0.00000    0.00000    0.00000 ;      CA-      C-      N-     CA
  5853   5852   5854   5855       3    0.65084    1.95253    0.00000
-2.60338    0.00000    0.00000 ;      HA-     CA-     CB-    HB2
  5853   5852   5854   5856       3    0.65084    1.95253    0.00000
-2.60338    0.00000    0.00000 ;      HA-     CA-     CB-    HB3
  5853   5852   5854   5857       3    0.65084    1.95253    0.00000
-2.60338    0.00000    0.00000 ;      HA-     CA-     CB-     CG
  5853   5852   5860   5861       3    3.68192   -4.35136    0.00000
1.33888    0.00000    0.00000 ;      HA-     CA-      C-      O
  5853   5852   5860   5862       3    0.00000    0.00000    0.00000
0.00000    0.00000    0.00000 ;      HA-     CA-      C-      N
  5854   5852   5860   5861       3    0.00000    0.00000    0.00000
0.00000    0.00000    0.00000 ;      CB-     CA-      C-      O
  5854   5852   5860   5862       3    2.51040    4.18400    1.67360
-6.69440    0.00000    0.00000 ;      CB-     CA-      C-      N
  5855   5854   5852   5860       3    0.65084    1.95253    0.00000
-2.60338    0.00000    0.00000 ;     HB2-     CB-     CA-      C
  5855   5854   5857   5858       3    0.00000    0.00000    0.00000
0.00000    0.00000    0.00000 ;     HB2-     CB-     CG-    OD1
  5855   5854   5857   5859       3    0.00000    0.00000    0.00000
0.00000    0.00000    0.00000 ;     HB2-     CB-     CG-    OD2
  5856   5854   5852   5860       3    0.65084    1.95253    0.00000
-2.60338    0.00000    0.00000 ;     HB3-     CB-     CA-      C
  5856   5854   5857   5858       3    0.00000    0.00000    0.00000
0.00000    0.00000    0.00000 ;     HB3-     CB-     CG-    OD1
  5856   5854   5857   5859       3    0.00000    0.00000    0.00000
```

0.00000 0.00000 0.00000 ; HB3- CB- CG- OD2
5857 5854 5852 5860 3 7.14627 -1.04600 -8.99560
-0.97069 5.15469 0.00000 ; CG- CB- CA- C
5860 5862 5864 5865 3 0.00000 0.00000 0.00000
0.00000 0.00000 0.00000 ; C- N- CA- HA
5860 5862 5864 5866 3 11.71520 1.67360 15.06240 -
13.38880 0.00000 0.00000 ; C- N- CA- CB
5860 5862 5864 5871 3 1.75728 5.27184 2.25936
-7.02912 0.00000 0.00000 ; C- N- CA- C
5861 5860 5862 5863 3 29.28800 -8.36800 -20.92000
0.00000 0.00000 0.00000 ; O- C- N- H
5861 5860 5862 5864 3 20.92000 0.00000 -20.92000
0.00000 0.00000 0.00000 ; O- C- N- CA
5862 5864 5866 5867 3 0.65084 1.95253 0.00000
-2.60338 0.00000 0.00000 ; N- CA- CB- HB2
5862 5864 5866 5868 3 0.65084 1.95253 0.00000
-2.60338 0.00000 0.00000 ; N- CA- CB- HB3
5862 5864 5866 5869 3 7.86174 2.24681 -7.41405
-6.71114 5.35552 0.00000 ; N- CA- CB- OG
5862 5864 5871 5872 3 0.00000 0.00000 0.00000
0.00000 0.00000 0.00000 ; N- CA- C- O
5862 5864 5871 5873 3 17.40544 -5.02080 -13.22144
9.20480 0.00000 0.00000 ; N- CA- C- N
5863 5862 5864 5865 3 0.00000 0.00000 0.00000
0.00000 0.00000 0.00000 ; H- N- CA- HA
5863 5862 5864 5866 3 0.00000 0.00000 0.00000
0.00000 0.00000 0.00000 ; H- N- CA- CB
5863 5862 5864 5871 3 0.00000 0.00000 0.00000
0.00000 0.00000 0.00000 ; H- N- CA- C
5864 5866 5869 5870 3 2.05853 2.46856 3.48109
-4.46851 0.23430 0.00000 ; CA- CB- OG- HG
5864 5871 5873 5874 3 20.92000 0.00000 -20.92000
0.00000 0.00000 0.00000 ; CA- C- N- H
5864 5871 5873 5875 3 20.92000 0.00000 -20.92000
0.00000 0.00000 0.00000 ; CA- C- N- CA
5865 5864 5866 5867 3 0.65084 1.95253 0.00000
-2.60338 0.00000 0.00000 ; HA- CA- CB- HB2
5865 5864 5866 5868 3 0.65084 1.95253 0.00000
-2.60338 0.00000 0.00000 ; HA- CA- CB- HB3
5865 5864 5866 5869 3 1.04600 -1.04600 0.00000
0.00000 0.00000 0.00000 ; HA- CA- CB- OG
5865 5864 5871 5872 3 3.68192 -4.35136 0.00000
1.33888 0.00000 0.00000 ; HA- CA- C- O
5865 5864 5871 5873 3 0.00000 0.00000 0.00000
0.00000 0.00000 0.00000 ; HA- CA- C- N
5866 5864 5871 5872 3 0.00000 0.00000 0.00000
0.00000 0.00000 0.00000 ; CB- CA- C- O
5866 5864 5871 5873 3 2.51040 4.18400 1.67360
-6.69440 0.00000 0.00000 ; CB- CA- C- N
5867 5866 5864 5871 3 0.65084 1.95253 0.00000
-2.60338 0.00000 0.00000 ; HB2- CB- CA- C
5867 5866 5869 5870 3 0.69733 2.09200 0.00000

```
-2.78933    0.00000    0.00000 ;    HB2-    CB-    OG-    HG
  5868   5866   5864   5871      3    0.65084    1.95253    0.00000
-2.60338    0.00000    0.00000 ;    HB3-    CB-    CA-    C
  5868   5866   5869   5870      3    0.69733    2.09200    0.00000
-2.78933    0.00000    0.00000 ;    HB3-    CB-    OG-    HG
  5869   5866   5864   5871      3    7.34710    7.79898   -6.14211
-6.71114    4.31789    0.00000 ;    OG-    CB-    CA-    C
  5871   5873   5875   5876      3    0.00000    0.00000    0.00000
0.00000    0.00000    0.00000 ;    C-    N-    CA-    HA
  5871   5873   5875   5877      3   11.71520    1.67360   15.06240  -
13.38880    0.00000    0.00000 ;    C-    N-    CA-    CB
  5871   5873   5875   5885      3    1.75728    5.27184    2.25936
-7.02912    0.00000    0.00000 ;    C-    N-    CA-    C
  5872   5871   5873   5874      3   29.28800   -8.36800  -20.92000
0.00000    0.00000    0.00000 ;    O-    C-    N-    H
  5872   5871   5873   5875      3   20.92000    0.00000  -20.92000
0.00000    0.00000    0.00000 ;    O-    C-    N-    CA
  5873   5875   5877   5878      3    0.65084    1.95253    0.00000
-2.60338    0.00000    0.00000 ;    N-    CA-    CB-    HB2
  5873   5875   5877   5879      3    0.65084    1.95253    0.00000
-2.60338    0.00000    0.00000 ;    N-    CA-    CB-    HB3
  5873   5875   5877   5880      3    5.99567    3.29281   -4.46014
-0.55229    1.97485    0.00000 ;    N-    CA-    CB-    CG
  5873   5875   5885   5886      3    0.00000    0.00000    0.00000
0.00000    0.00000    0.00000 ;    N-    CA-    C-    O
  5873   5875   5885   5887      3   17.40544   -5.02080  -13.22144
9.20480    0.00000    0.00000 ;    N-    CA-    C-    N
  5874   5873   5875   5876      3    0.00000    0.00000    0.00000
0.00000    0.00000    0.00000 ;    H-    N-    CA-    HA
  5874   5873   5875   5877      3    0.00000    0.00000    0.00000
0.00000    0.00000    0.00000 ;    H-    N-    CA-    CB
  5874   5873   5875   5885      3    0.00000    0.00000    0.00000
0.00000    0.00000    0.00000 ;    H-    N-    CA-    C
  5875   5877   5880   5881      3    0.00000    0.00000    0.00000
0.00000    0.00000    0.00000 ;    CA-    CB-    CG-    OD1
  5875   5877   5880   5882      3    8.84916   -0.31380   -4.32626
5.03754    0.26778    0.00000 ;    CA-    CB-    CG-    ND2
  5875   5885   5887   5888      3   20.92000    0.00000  -20.92000
0.00000    0.00000    0.00000 ;    CA-    C-    N-    H
  5875   5885   5887   5889      3   20.92000    0.00000  -20.92000
0.00000    0.00000    0.00000 ;    CA-    C-    N-    CA
  5876   5875   5877   5878      3    0.65084    1.95253    0.00000
-2.60338    0.00000    0.00000 ;    HA-    CA-    CB-    HB2
  5876   5875   5877   5879      3    0.65084    1.95253    0.00000
-2.60338    0.00000    0.00000 ;    HA-    CA-    CB-    HB3
  5876   5875   5877   5880      3    0.65084    1.95253    0.00000
-2.60338    0.00000    0.00000 ;    HA-    CA-    CB-    CG
  5876   5875   5885   5886      3    3.68192   -4.35136    0.00000
1.33888    0.00000    0.00000 ;    HA-    CA-    C-    O
  5876   5875   5885   5887      3    0.00000    0.00000    0.00000
0.00000    0.00000    0.00000 ;    HA-    CA-    C-    N
  5877   5875   5885   5886      3    0.00000    0.00000    0.00000
```

0.00000 0.00000 0.00000 ; CB- CA- C- O
5877 5875 5885 5887 3 2.51040 4.18400 1.67360
-6.69440 0.00000 0.00000 ; CB- CA- C- N
5877 5880 5882 5883 3 20.92000 0.00000 -20.92000
0.00000 0.00000 0.00000 ; CB- CG- ND2- HD21
5877 5880 5882 5884 3 20.92000 0.00000 -20.92000
0.00000 0.00000 0.00000 ; CB- CG- ND2- HD22
5878 5877 5875 5885 3 0.65084 1.95253 0.00000
-2.60338 0.00000 0.00000 ; HB2- CB- CA- C
5878 5877 5880 5881 3 3.68192 -4.35136 0.00000
1.33888 0.00000 0.00000 ; HB2- CB- CG- OD1
5878 5877 5880 5882 3 0.00000 0.00000 0.00000
0.00000 0.00000 0.00000 ; HB2- CB- CG- ND2
5879 5877 5875 5885 3 0.65084 1.95253 0.00000
-2.60338 0.00000 0.00000 ; HB3- CB- CA- C
5879 5877 5880 5881 3 3.68192 -4.35136 0.00000
1.33888 0.00000 0.00000 ; HB3- CB- CG- OD1
5879 5877 5880 5882 3 0.00000 0.00000 0.00000
0.00000 0.00000 0.00000 ; HB3- CB- CG- ND2
5880 5877 5875 5885 3 7.94542 -3.96225 -6.11701
-0.55229 3.58150 0.00000 ; CG- CB- CA- C
5881 5880 5882 5883 3 29.28800 -8.36800 -20.92000
0.00000 0.00000 0.00000 ; OD1- CG- ND2- HD21
5881 5880 5882 5884 3 29.28800 -8.36800 -20.92000
0.00000 0.00000 0.00000 ; OD1- CG- ND2- HD22
5885 5887 5889 5890 3 0.00000 0.00000 0.00000
0.00000 0.00000 0.00000 ; C- N- CA- HA
5885 5887 5889 5891 3 11.71520 1.67360 15.06240 -
13.38880 0.00000 0.00000 ; C- N- CA- CB
5885 5887 5889 5895 3 1.75728 5.27184 2.25936
-7.02912 0.00000 0.00000 ; C- N- CA- C
5886 5885 5887 5888 3 29.28800 -8.36800 -20.92000
0.00000 0.00000 0.00000 ; O- C- N- H
5886 5885 5887 5889 3 20.92000 0.00000 -20.92000
0.00000 0.00000 0.00000 ; O- C- N- CA
5887 5889 5891 5892 3 0.65084 1.95253 0.00000
-2.60338 0.00000 0.00000 ; N- CA- CB- HB1
5887 5889 5891 5893 3 0.65084 1.95253 0.00000
-2.60338 0.00000 0.00000 ; N- CA- CB- HB2
5887 5889 5891 5894 3 0.65084 1.95253 0.00000
-2.60338 0.00000 0.00000 ; N- CA- CB- HB3
5887 5889 5895 5896 3 0.00000 0.00000 0.00000
0.00000 0.00000 0.00000 ; N- CA- C- O
5887 5889 5895 5897 3 17.40544 -5.02080 -13.22144
9.20480 0.00000 0.00000 ; N- CA- C- N
5888 5887 5889 5890 3 0.00000 0.00000 0.00000
0.00000 0.00000 0.00000 ; H- N- CA- HA
5888 5887 5889 5891 3 0.00000 0.00000 0.00000
0.00000 0.00000 0.00000 ; H- N- CA- CB
5888 5887 5889 5895 3 0.00000 0.00000 0.00000
0.00000 0.00000 0.00000 ; H- N- CA- C
5889 5895 5897 5898 3 20.92000 0.00000 -20.92000

0.00000 0.00000 0.00000 ; CA- C- N- H
5889 5895 5897 5899 3 20.92000 0.00000 -20.92000
0.00000 0.00000 0.00000 ; CA- C- N- CA
5890 5889 5891 5892 3 0.65084 1.95253 0.00000
-2.60338 0.00000 0.00000 ; HA- CA- CB- HB1
5890 5889 5891 5893 3 0.65084 1.95253 0.00000
-2.60338 0.00000 0.00000 ; HA- CA- CB- HB2
5890 5889 5891 5894 3 0.65084 1.95253 0.00000
-2.60338 0.00000 0.00000 ; HA- CA- CB- HB3
5890 5889 5895 5896 3 3.68192 -4.35136 0.00000
1.33888 0.00000 0.00000 ; HA- CA- C- O
5890 5889 5895 5897 3 0.00000 0.00000 0.00000
0.00000 0.00000 0.00000 ; HA- CA- C- N
5891 5889 5895 5896 3 0.00000 0.00000 0.00000
0.00000 0.00000 0.00000 ; CB- CA- C- O
5891 5889 5895 5897 3 2.51040 4.18400 1.67360
-6.69440 0.00000 0.00000 ; CB- CA- C- N
5892 5891 5889 5895 3 0.65084 1.95253 0.00000
-2.60338 0.00000 0.00000 ; HB1- CB- CA- C
5893 5891 5889 5895 3 0.65084 1.95253 0.00000
-2.60338 0.00000 0.00000 ; HB2- CB- CA- C
5894 5891 5889 5895 3 0.65084 1.95253 0.00000
-2.60338 0.00000 0.00000 ; HB3- CB- CA- C
5895 5897 5899 5900 3 0.00000 0.00000 0.00000
0.00000 0.00000 0.00000 ; C- N- CA- HA
5895 5897 5899 5901 3 11.71520 1.67360 15.06240 -
13.38880 0.00000 0.00000 ; C- N- CA- CB
5895 5897 5899 5906 3 1.75728 5.27184 2.25936
-7.02912 0.00000 0.00000 ; C- N- CA- C
5896 5895 5897 5898 3 29.28800 -8.36800 -20.92000
0.00000 0.00000 0.00000 ; O- C- N- H
5896 5895 5897 5899 3 20.92000 0.00000 -20.92000
0.00000 0.00000 0.00000 ; O- C- N- CA
5897 5899 5901 5902 3 0.65084 1.95253 0.00000
-2.60338 0.00000 0.00000 ; N- CA- CB- HB2
5897 5899 5901 5903 3 0.65084 1.95253 0.00000
-2.60338 0.00000 0.00000 ; N- CA- CB- HB3
5897 5899 5901 5904 3 7.86174 2.24681 -7.41405
-6.71114 5.35552 0.00000 ; N- CA- CB- OG
5897 5899 5906 5907 3 0.00000 0.00000 0.00000
0.00000 0.00000 0.00000 ; N- CA- C- O
5897 5899 5906 5908 3 17.40544 -5.02080 -13.22144
9.20480 0.00000 0.00000 ; N- CA- C- N
5898 5897 5899 5900 3 0.00000 0.00000 0.00000
0.00000 0.00000 0.00000 ; H- N- CA- HA
5898 5897 5899 5901 3 0.00000 0.00000 0.00000
0.00000 0.00000 0.00000 ; H- N- CA- CB
5898 5897 5899 5906 3 0.00000 0.00000 0.00000
0.00000 0.00000 0.00000 ; H- N- CA- C
5899 5901 5904 5905 3 2.05853 2.46856 3.48109
-4.46851 0.23430 0.00000 ; CA- CB- OG- HG
5899 5906 5908 5909 3 20.92000 0.00000 -20.92000

```
0.00000    0.00000    0.00000 ;     CA-     C-     N-     H
  5899   5906   5908   5910      3   20.92000    0.00000  -20.92000
0.00000    0.00000    0.00000 ;     CA-     C-     N-    CA
  5900   5899   5901   5902      3    0.65084    1.95253    0.00000
-2.60338    0.00000    0.00000 ;     HA-    CA-    CB-   HB2
  5900   5899   5901   5903      3    0.65084    1.95253    0.00000
-2.60338    0.00000    0.00000 ;     HA-    CA-    CB-   HB3
  5900   5899   5901   5904      3    1.04600   -1.04600    0.00000
0.00000    0.00000    0.00000 ;     HA-    CA-    CB-    OG
  5900   5899   5906   5907      3    3.68192   -4.35136    0.00000
1.33888    0.00000    0.00000 ;     HA-    CA-     C-     O
  5900   5899   5906   5908      3    0.00000    0.00000    0.00000
0.00000    0.00000    0.00000 ;     HA-    CA-     C-     N
  5901   5899   5906   5907      3    0.00000    0.00000    0.00000
0.00000    0.00000    0.00000 ;     CB-    CA-     C-     O
  5901   5899   5906   5908      3    2.51040    4.18400    1.67360
-6.69440    0.00000    0.00000 ;     CB-    CA-     C-     N
  5902   5901   5899   5906      3    0.65084    1.95253    0.00000
-2.60338    0.00000    0.00000 ;    HB2-    CB-    CA-     C
  5902   5901   5904   5905      3    0.69733    2.09200    0.00000
-2.78933    0.00000    0.00000 ;    HB2-    CB-    OG-    HG
  5903   5901   5899   5906      3    0.65084    1.95253    0.00000
-2.60338    0.00000    0.00000 ;    HB3-    CB-    CA-     C
  5903   5901   5904   5905      3    0.69733    2.09200    0.00000
-2.78933    0.00000    0.00000 ;    HB3-    CB-    OG-    HG
  5904   5901   5899   5906      3    7.34710    7.79898   -6.14211
-6.71114    4.31789    0.00000 ;     OG-    CB-    CA-     C
  5906   5908   5910   5911      3    0.00000    0.00000    0.00000
0.00000    0.00000    0.00000 ;      C-     N-    CA-   HA2
  5906   5908   5910   5912      3    0.00000    0.00000    0.00000
0.00000    0.00000    0.00000 ;      C-     N-    CA-   HA3
  5906   5908   5910   5913      3    1.75728    5.27184    2.25936
-7.02912    0.00000    0.00000 ;      C-     N-    CA-     C
  5907   5906   5908   5909      3   29.28800   -8.36800  -20.92000
0.00000    0.00000    0.00000 ;      O-     C-     N-     H
  5907   5906   5908   5910      3   20.92000    0.00000  -20.92000
0.00000    0.00000    0.00000 ;      O-     C-     N-    CA
  5908   5910   5913   5914      3    0.00000    0.00000    0.00000
0.00000    0.00000    0.00000 ;      N-    CA-     C-     O
  5908   5910   5913   5915      3   17.40544   -5.02080  -13.22144
9.20480    0.00000    0.00000 ;      N-    CA-     C-     N
  5909   5908   5910   5911      3    0.00000    0.00000    0.00000
0.00000    0.00000    0.00000 ;      H-     N-    CA-   HA2
  5909   5908   5910   5912      3    0.00000    0.00000    0.00000
0.00000    0.00000    0.00000 ;      H-     N-    CA-   HA3
  5909   5908   5910   5913      3    0.00000    0.00000    0.00000
0.00000    0.00000    0.00000 ;      H-     N-    CA-     C
  5910   5913   5915   5916      3   20.92000    0.00000  -20.92000
0.00000    0.00000    0.00000 ;     CA-     C-     N-     H
  5910   5913   5915   5917      3   20.92000    0.00000  -20.92000
0.00000    0.00000    0.00000 ;     CA-     C-     N-    CA
  5911   5910   5913   5914      3    3.68192   -4.35136    0.00000
```

```
1.33888    0.00000    0.00000 ;    HA2-    CA-     C-     O
  5911   5910   5913   5915      3    0.00000    0.00000    0.00000
0.00000    0.00000    0.00000 ;    HA2-    CA-     C-     N
  5912   5910   5913   5914      3    3.68192   -4.35136    0.00000
1.33888    0.00000    0.00000 ;    HA3-    CA-     C-     O
  5912   5910   5913   5915      3    0.00000    0.00000    0.00000
0.00000    0.00000    0.00000 ;    HA3-    CA-     C-     N
  5913   5915   5917   5918      3    0.00000    0.00000    0.00000
0.00000    0.00000    0.00000 ;      C-     N-    CA-    HA
  5913   5915   5917   5919      3   11.71520    1.67360   15.06240  -
13.38880    0.00000    0.00000 ;      C-     N-    CA-    CB
  5913   5915   5917   5930      3    1.75728    5.27184    2.25936
-7.02912    0.00000    0.00000 ;      C-     N-    CA-     C
  5914   5913   5915   5916      3   29.28800   -8.36800  -20.92000
0.00000    0.00000    0.00000 ;      O-     C-     N-     H
  5914   5913   5915   5917      3   20.92000    0.00000  -20.92000
0.00000    0.00000    0.00000 ;      O-     C-     N-    CA
  5915   5917   5919   5920      3    0.65084    1.95253    0.00000
-2.60338    0.00000    0.00000 ;      N-    CA-    CB-   HB2
  5915   5917   5919   5921      3    0.65084    1.95253    0.00000
-2.60338    0.00000    0.00000 ;      N-    CA-    CB-   HB3
  5915   5917   5919   5922      3    3.21331    2.22589   -4.15053
-2.40998    2.61082    0.00000 ;      N-    CA-    CB-    CG
  5915   5917   5930   5931      3    0.00000    0.00000    0.00000
0.00000    0.00000    0.00000 ;      N-    CA-     C-     O
  5915   5917   5930   5932      3   17.40544   -5.02080  -13.22144
9.20480    0.00000    0.00000 ;      N-    CA-     C-     N
  5916   5915   5917   5918      3    0.00000    0.00000    0.00000
0.00000    0.00000    0.00000 ;      H-     N-    CA-    HA
  5916   5915   5917   5919      3    0.00000    0.00000    0.00000
0.00000    0.00000    0.00000 ;      H-     N-    CA-    CB
  5916   5915   5917   5930      3    0.00000    0.00000    0.00000
0.00000    0.00000    0.00000 ;      H-     N-    CA-     C
  5917   5919   5922   5923      3    0.66944    2.00832    0.00000
-2.67776    0.00000    0.00000 ;     CA-    CB-    CG-   HG2
  5917   5919   5922   5924      3    0.66944    2.00832    0.00000
-2.67776    0.00000    0.00000 ;     CA-    CB-    CG-   HG3
  5917   5919   5922   5925      3    3.69866   -4.35136   -3.92459
6.89523    4.61914    0.00000 ;     CA-    CB-    CG-    CD
  5917   5930   5932   5933      3   20.92000    0.00000  -20.92000
0.00000    0.00000    0.00000 ;     CA-     C-     N-     H
  5917   5930   5932   5934      3   20.92000    0.00000  -20.92000
0.00000    0.00000    0.00000 ;     CA-     C-     N-    CA
  5918   5917   5919   5920      3    0.65084    1.95253    0.00000
-2.60338    0.00000    0.00000 ;     HA-    CA-    CB-   HB2
  5918   5917   5919   5921      3    0.65084    1.95253    0.00000
-2.60338    0.00000    0.00000 ;     HA-    CA-    CB-   HB3
  5918   5917   5919   5922      3    0.65084    1.95253    0.00000
-2.60338    0.00000    0.00000 ;     HA-    CA-    CB-    CG
  5918   5917   5930   5931      3    3.68192   -4.35136    0.00000
1.33888    0.00000    0.00000 ;     HA-    CA-     C-     O
  5918   5917   5930   5932      3    0.00000    0.00000    0.00000
```

0.00000 0.00000 0.00000 ; HA- CA- C- N
5919 5917 5930 5931 3 0.00000 0.00000 0.00000
0.00000 0.00000 0.00000 ; CB- CA- C- O
5919 5917 5930 5932 3 2.51040 4.18400 1.67360
-6.69440 0.00000 0.00000 ; CB- CA- C- N
5919 5922 5925 5926 3 0.00000 0.00000 0.00000
0.00000 0.00000 0.00000 ; CB- CG- CD- OE1
5919 5922 5925 5927 3 10.32611 1.48114 -8.47678
1.42256 1.40582 0.00000 ; CB- CG- CD- NE2
5920 5919 5917 5930 3 0.65084 1.95253 0.00000
-2.60338 0.00000 0.00000 ; HB2- CB- CA- C
5920 5919 5922 5923 3 0.62760 1.88280 0.00000
-2.51040 0.00000 0.00000 ; HB2- CB- CG- HG2
5920 5919 5922 5924 3 0.62760 1.88280 0.00000
-2.51040 0.00000 0.00000 ; HB2- CB- CG- HG3
5920 5919 5922 5925 3 0.65084 1.95253 0.00000
-2.60338 0.00000 0.00000 ; HB2- CB- CG- CD
5921 5919 5917 5930 3 0.65084 1.95253 0.00000
-2.60338 0.00000 0.00000 ; HB3- CB- CA- C
5921 5919 5922 5923 3 0.62760 1.88280 0.00000
-2.51040 0.00000 0.00000 ; HB3- CB- CG- HG2
5921 5919 5922 5924 3 0.62760 1.88280 0.00000
-2.51040 0.00000 0.00000 ; HB3- CB- CG- HG3
5921 5919 5922 5925 3 0.65084 1.95253 0.00000
-2.60338 0.00000 0.00000 ; HB3- CB- CG- CD
5922 5919 5917 5930 3 6.86594 3.56895 -8.14206
-2.40998 4.85344 0.00000 ; CG- CB- CA- C
5922 5925 5927 5928 3 20.92000 0.00000 -20.92000
0.00000 0.00000 0.00000 ; CG- CD- NE2- HE21
5922 5925 5927 5929 3 20.92000 0.00000 -20.92000
0.00000 0.00000 0.00000 ; CG- CD- NE2- HE22
5923 5922 5925 5926 3 3.68192 -4.35136 0.00000
1.33888 0.00000 0.00000 ; HG2- CG- CD- OE1
5923 5922 5925 5927 3 0.00000 0.00000 0.00000
0.00000 0.00000 0.00000 ; HG2- CG- CD- NE2
5924 5922 5925 5926 3 3.68192 -4.35136 0.00000
1.33888 0.00000 0.00000 ; HG3- CG- CD- OE1
5924 5922 5925 5927 3 0.00000 0.00000 0.00000
0.00000 0.00000 0.00000 ; HG3- CG- CD- NE2
5926 5925 5927 5928 3 29.28800 -8.36800 -20.92000
0.00000 0.00000 0.00000 ; OE1- CD- NE2- HE21
5926 5925 5927 5929 3 29.28800 -8.36800 -20.92000
0.00000 0.00000 0.00000 ; OE1- CD- NE2- HE22
5930 5932 5934 5935 3 0.00000 0.00000 0.00000
0.00000 0.00000 0.00000 ; C- N- CA- HA
5930 5932 5934 5936 3 11.71520 1.67360 15.06240 -
13.38880 0.00000 0.00000 ; C- N- CA- CB
5930 5932 5934 5949 3 1.75728 5.27184 2.25936
-7.02912 0.00000 0.00000 ; C- N- CA- C
5931 5930 5932 5933 3 29.28800 -8.36800 -20.92000
0.00000 0.00000 0.00000 ; O- C- N- H
5931 5930 5932 5934 3 20.92000 0.00000 -20.92000

```
0.00000    0.00000    0.00000 ;       O-     C-     N-    CA
  5932   5934   5936   5937      3    0.65084    1.95253    0.00000
-2.60338    0.00000    0.00000 ;       N-    CA-    CB-    HB
  5932   5934   5936   5938      3    3.83673    0.44769   -1.77402
-2.47693   -0.03347    0.00000 ;       N-    CA-    CB-   CG2
  5932   5934   5936   5942      3    2.97482    0.12134    2.04179
-1.89117   -3.24678    0.00000 ;       N-    CA-    CB-   CG1
  5932   5934   5949   5950      3    0.00000    0.00000    0.00000
0.00000    0.00000    0.00000 ;       N-    CA-     C-     O
  5932   5934   5949   5951      3   17.40544   -5.02080  -13.22144
9.20480    0.00000    0.00000 ;       N-    CA-     C-     N
  5933   5932   5934   5935      3    0.00000    0.00000    0.00000
0.00000    0.00000    0.00000 ;       H-     N-    CA-    HA
  5933   5932   5934   5936      3    0.00000    0.00000    0.00000
0.00000    0.00000    0.00000 ;       H-     N-    CA-    CB
  5933   5932   5934   5949      3    0.00000    0.00000    0.00000
0.00000    0.00000    0.00000 ;       H-     N-    CA-     C
  5934   5936   5938   5939      3    0.66944    2.00832    0.00000
-2.67776    0.00000    0.00000 ;      CA-    CB-   CG2-  HG21
  5934   5936   5938   5940      3    0.66944    2.00832    0.00000
-2.67776    0.00000    0.00000 ;      CA-    CB-   CG2-  HG22
  5934   5936   5938   5941      3    0.66944    2.00832    0.00000
-2.67776    0.00000    0.00000 ;      CA-    CB-   CG2-  HG23
  5934   5936   5942   5943      3    0.66944    2.00832    0.00000
-2.67776    0.00000    0.00000 ;      CA-    CB-   CG1-  HG12
  5934   5936   5942   5944      3    0.66944    2.00832    0.00000
-2.67776    0.00000    0.00000 ;      CA-    CB-   CG1-  HG13
  5934   5936   5942   5945      3    4.24258   -0.52718   -7.25506
-1.79075    7.69856    0.00000 ;      CA-    CB-   CG1-   CD1
  5934   5949   5951   5952      3   20.92000    0.00000  -20.92000
0.00000    0.00000    0.00000 ;      CA-     C-     N-     H
  5934   5949   5951   5953      3   20.92000    0.00000  -20.92000
0.00000    0.00000    0.00000 ;      CA-     C-     N-    CA
  5935   5934   5936   5937      3    0.65084    1.95253    0.00000
-2.60338    0.00000    0.00000 ;      HA-    CA-    CB-    HB
  5935   5934   5936   5938      3    0.65084    1.95253    0.00000
-2.60338    0.00000    0.00000 ;      HA-    CA-    CB-   CG2
  5935   5934   5936   5942      3    0.65084    1.95253    0.00000
-2.60338    0.00000    0.00000 ;      HA-    CA-    CB-   CG1
  5935   5934   5949   5950      3    3.68192   -4.35136    0.00000
1.33888    0.00000    0.00000 ;      HA-    CA-     C-     O
  5935   5934   5949   5951      3    0.00000    0.00000    0.00000
0.00000    0.00000    0.00000 ;      HA-    CA-     C-     N
  5936   5934   5949   5950      3    0.00000    0.00000    0.00000
0.00000    0.00000    0.00000 ;      CB-    CA-     C-     O
  5936   5934   5949   5951      3    2.51040    4.18400    1.67360
-6.69440    0.00000    0.00000 ;      CB-    CA-     C-     N
  5936   5942   5945   5946      3    0.66944    2.00832    0.00000
-2.67776    0.00000    0.00000 ;      CB-   CG1-   CD1-  HD11
  5936   5942   5945   5947      3    0.66944    2.00832    0.00000
-2.67776    0.00000    0.00000 ;      CB-   CG1-   CD1-  HD12
  5936   5942   5945   5948      3    0.66944    2.00832    0.00000
```

-2.67776 0.00000 0.00000 ; CB- CG1- CD1- HD13
5937 5936 5934 5949 3 0.65084 1.95253 0.00000
-2.60338 0.00000 0.00000 ; HB- CB- CA- C
5937 5936 5938 5939 3 0.62760 1.88280 0.00000
-2.51040 0.00000 0.00000 ; HB- CB- CG2- HG21
5937 5936 5938 5940 3 0.62760 1.88280 0.00000
-2.51040 0.00000 0.00000 ; HB- CB- CG2- HG22
5937 5936 5938 5941 3 0.62760 1.88280 0.00000
-2.51040 0.00000 0.00000 ; HB- CB- CG2- HG23
5937 5936 5942 5943 3 0.62760 1.88280 0.00000
-2.51040 0.00000 0.00000 ; HB- CB- CG1- HG12
5937 5936 5942 5944 3 0.62760 1.88280 0.00000
-2.51040 0.00000 0.00000 ; HB- CB- CG1- HG13
5937 5936 5942 5945 3 0.66944 2.00832 0.00000
-2.67776 0.00000 0.00000 ; HB- CB- CG1- CD1
5938 5936 5934 5949 3 5.67350 3.55640 -6.16722
-2.47693 3.74886 0.00000 ; CG2- CB- CA- C
5938 5936 5942 5943 3 0.66944 2.00832 0.00000
-2.67776 0.00000 0.00000 ; CG2- CB- CG1- HG12
5938 5936 5942 5944 3 0.66944 2.00832 0.00000
-2.67776 0.00000 0.00000 ; CG2- CB- CG1- HG13
5938 5936 5942 5945 3 3.81162 0.49790 -8.14206
-1.79075 7.49773 0.00000 ; CG2- CB- CG1- CD1
5939 5938 5936 5942 3 0.66944 2.00832 0.00000
-2.67776 0.00000 0.00000 ; HG21- CG2- CB- CG1
5940 5938 5936 5942 3 0.66944 2.00832 0.00000
-2.67776 0.00000 0.00000 ; HG22- CG2- CB- CG1
5941 5938 5936 5942 3 0.66944 2.00832 0.00000
-2.67776 0.00000 0.00000 ; HG23- CG2- CB- CG1
5942 5936 5934 5949 3 8.26340 0.74057 -9.99976
-1.89117 3.84928 0.00000 ; CG1- CB- CA- C
5943 5942 5945 5946 3 0.62760 1.88280 0.00000
-2.51040 0.00000 0.00000 ; HG12- CG1- CD1- HD11
5943 5942 5945 5947 3 0.62760 1.88280 0.00000
-2.51040 0.00000 0.00000 ; HG12- CG1- CD1- HD12
5943 5942 5945 5948 3 0.62760 1.88280 0.00000
-2.51040 0.00000 0.00000 ; HG12- CG1- CD1- HD13
5944 5942 5945 5946 3 0.62760 1.88280 0.00000
-2.51040 0.00000 0.00000 ; HG13- CG1- CD1- HD11
5944 5942 5945 5947 3 0.62760 1.88280 0.00000
-2.51040 0.00000 0.00000 ; HG13- CG1- CD1- HD12
5944 5942 5945 5948 3 0.62760 1.88280 0.00000
-2.51040 0.00000 0.00000 ; HG13- CG1- CD1- HD13
5949 5951 5953 5954 3 0.00000 0.00000 0.00000
0.00000 0.00000 0.00000 ; C- N- CA- HA
5949 5951 5953 5955 3 11.71520 1.67360 15.06240 -
13.38880 0.00000 0.00000 ; C- N- CA- CB
5949 5951 5953 5966 3 1.75728 5.27184 2.25936
-7.02912 0.00000 0.00000 ; C- N- CA- C
5950 5949 5951 5952 3 29.28800 -8.36800 -20.92000
0.00000 0.00000 0.00000 ; O- C- N- H
5950 5949 5951 5953 3 20.92000 0.00000 -20.92000

0.00000 0.00000 0.00000 ; O- C- N- CA
5951 5953 5955 5956 3 0.65084 1.95253 0.00000
-2.60338 0.00000 0.00000 ; N- CA- CB- HB2
5951 5953 5955 5957 3 0.65084 1.95253 0.00000
-2.60338 0.00000 0.00000 ; N- CA- CB- HB3
5951 5953 5955 5958 3 4.79068 4.02919 -4.82834
-3.66518 2.97901 0.00000 ; N- CA- CB- CG
5951 5953 5966 5967 3 0.00000 0.00000 0.00000
0.00000 0.00000 0.00000 ; N- CA- C- O
5951 5953 5966 5968 3 17.40544 -5.02080 -13.22144
9.20480 0.00000 0.00000 ; N- CA- C- N
5952 5951 5953 5954 3 0.00000 0.00000 0.00000
0.00000 0.00000 0.00000 ; H- N- CA- HA
5952 5951 5953 5955 3 0.00000 0.00000 0.00000
0.00000 0.00000 0.00000 ; H- N- CA- CB
5952 5951 5953 5966 3 0.00000 0.00000 0.00000
0.00000 0.00000 0.00000 ; H- N- CA- C
5953 5955 5958 5959 3 5.98312 6.40152 3.28026 -
12.38464 -1.57318 0.00000 ; CA- CB- CG- ND1
5953 5955 5958 5964 3 0.00000 0.00000 0.00000
0.00000 0.00000 0.00000 ; CA- CB- CG- CD2
5953 5966 5968 5969 3 20.92000 0.00000 -20.92000
0.00000 0.00000 0.00000 ; CA- C- N- H
5953 5966 5968 5970 3 20.92000 0.00000 -20.92000
0.00000 0.00000 0.00000 ; CA- C- N- CA
5954 5953 5955 5956 3 0.65084 1.95253 0.00000
-2.60338 0.00000 0.00000 ; HA- CA- CB- HB2
5954 5953 5955 5957 3 0.65084 1.95253 0.00000
-2.60338 0.00000 0.00000 ; HA- CA- CB- HB3
5954 5953 5955 5958 3 0.65084 1.95253 0.00000
-2.60338 0.00000 0.00000 ; HA- CA- CB- CG
5954 5953 5966 5967 3 3.68192 -4.35136 0.00000
1.33888 0.00000 0.00000 ; HA- CA- C- O
5954 5953 5966 5968 3 0.00000 0.00000 0.00000
0.00000 0.00000 0.00000 ; HA- CA- C- N
5955 5953 5966 5967 3 0.00000 0.00000 0.00000
0.00000 0.00000 0.00000 ; CB- CA- C- O
5955 5953 5966 5968 3 2.51040 4.18400 1.67360
-6.69440 0.00000 0.00000 ; CB- CA- C- N
5955 5958 5959 5960 3 20.08320 0.00000 -20.08320
0.00000 0.00000 0.00000 ; CB- CG- ND1- CE1
5955 5958 5964 5962 3 44.97800 0.00000 -44.97800
0.00000 0.00000 0.00000 ; CB- CG- CD2- NE2
5955 5958 5964 5965 3 44.97800 0.00000 -44.97800
0.00000 0.00000 0.00000 ; CB- CG- CD2- HD2
5956 5955 5953 5966 3 0.65084 1.95253 0.00000
-2.60338 0.00000 0.00000 ; HB2- CB- CA- C
5956 5955 5958 5959 3 0.00000 0.00000 0.00000
0.00000 0.00000 0.00000 ; HB2- CB- CG- ND1
5956 5955 5958 5964 3 0.00000 0.00000 0.00000
0.00000 0.00000 0.00000 ; HB2- CB- CG- CD2
5957 5955 5953 5966 3 0.65084 1.95253 0.00000

-2.60338 0.00000 0.00000 ; HB3- CB- CA- C
5957 5955 5958 5959 3 0.00000 0.00000 0.00000
0.00000 0.00000 0.00000 ; HB3- CB- CG- ND1
5957 5955 5958 5964 3 0.00000 0.00000 0.00000
0.00000 0.00000 0.00000 ; HB3- CB- CG- CD2
5958 5955 5953 5966 3 3.76560 3.34720 -2.87859
-3.66518 0.83680 0.00000 ; CG- CB- CA- C
5958 5959 5960 5961 3 41.84000 0.00000 -41.84000
0.00000 0.00000 0.00000 ; CG- ND1- CE1- HE1
5958 5959 5960 5962 3 41.84000 0.00000 -41.84000
0.00000 0.00000 0.00000 ; CG- ND1- CE1- NE2
5958 5964 5962 5960 3 12.55200 0.00000 -12.55200
0.00000 0.00000 0.00000 ; CG- CD2- NE2- CE1
5958 5964 5962 5963 3 12.55200 0.00000 -12.55200
0.00000 0.00000 0.00000 ; CG- CD2- NE2- HE2
5959 5958 5964 5962 3 44.97800 0.00000 -44.97800
0.00000 0.00000 0.00000 ; ND1- CG- CD2- NE2
5959 5958 5964 5965 3 44.97800 0.00000 -44.97800
0.00000 0.00000 0.00000 ; ND1- CG- CD2- HD2
5959 5960 5962 5963 3 19.45560 0.00000 -19.45560
0.00000 0.00000 0.00000 ; ND1- CE1- NE2- HE2
5959 5960 5962 5964 3 19.45560 0.00000 -19.45560
0.00000 0.00000 0.00000 ; ND1- CE1- NE2- CD2
5960 5959 5958 5964 3 20.08320 0.00000 -20.08320
0.00000 0.00000 0.00000 ; CE1- ND1- CG- CD2
5960 5962 5964 5965 3 12.55200 0.00000 -12.55200
0.00000 0.00000 0.00000 ; CE1- NE2- CD2- HD2
5961 5960 5962 5963 3 19.45560 0.00000 -19.45560
0.00000 0.00000 0.00000 ; HE1- CE1- NE2- HE2
5961 5960 5962 5964 3 19.45560 0.00000 -19.45560
0.00000 0.00000 0.00000 ; HE1- CE1- NE2- CD2
5963 5962 5964 5965 3 12.55200 0.00000 -12.55200
0.00000 0.00000 0.00000 ; HE2- NE2- CD2- HD2
5966 5968 5970 5971 3 0.00000 0.00000 0.00000
0.00000 0.00000 0.00000 ; C- N- CA- HA
5966 5968 5970 5972 3 11.71520 1.67360 15.06240 -
13.38880 0.00000 0.00000 ; C- N- CA- CB
5966 5968 5970 5987 3 1.75728 5.27184 2.25936
-7.02912 0.00000 0.00000 ; C- N- CA- C
5967 5966 5968 5969 3 29.28800 -8.36800 -20.92000
0.00000 0.00000 0.00000 ; O- C- N- H
5967 5966 5968 5970 3 20.92000 0.00000 -20.92000
0.00000 0.00000 0.00000 ; O- C- N- CA
5968 5970 5972 5973 3 0.65084 1.95253 0.00000
-2.60338 0.00000 0.00000 ; N- CA- CB- HB2
5968 5970 5972 5974 3 0.65084 1.95253 0.00000
-2.60338 0.00000 0.00000 ; N- CA- CB- HB3
5968 5970 5972 5975 3 3.28026 2.46019 -2.19242
-3.21331 -0.23430 0.00000 ; N- CA- CB- CG
5968 5970 5987 5988 3 0.00000 0.00000 0.00000
0.00000 0.00000 0.00000 ; N- CA- C- O
5968 5970 5987 5989 3 17.40544 -5.02080 -13.22144

```
9.20480    0.00000    0.00000 ;      N-    CA-     C-     N
  5969   5968   5970   5971      3    0.00000    0.00000    0.00000
0.00000    0.00000    0.00000 ;      H-     N-    CA-    HA
  5969   5968   5970   5972      3    0.00000    0.00000    0.00000
0.00000    0.00000    0.00000 ;      H-     N-    CA-    CB
  5969   5968   5970   5987      3    0.00000    0.00000    0.00000
0.00000    0.00000    0.00000 ;      H-     N-    CA-     C
  5970   5972   5975   5976      3    0.57739    0.00000    1.02926
0.00000   -1.60666    0.00000 ;     CA-    CB-    CG-   CD1
  5970   5972   5975   5985      3    0.57739    0.00000    1.02926
0.00000   -1.60666    0.00000 ;     CA-    CB-    CG-   CD2
  5970   5987   5989   5990      3   20.92000    0.00000  -20.92000
0.00000    0.00000    0.00000 ;     CA-     C-     N-     H
  5970   5987   5989   5991      3   20.92000    0.00000  -20.92000
0.00000    0.00000    0.00000 ;     CA-     C-     N-    CA
  5971   5970   5972   5973      3    0.65084    1.95253    0.00000
-2.60338    0.00000    0.00000 ;     HA-    CA-    CB-   HB2
  5971   5970   5972   5974      3    0.65084    1.95253    0.00000
-2.60338    0.00000    0.00000 ;     HA-    CA-    CB-   HB3
  5971   5970   5972   5975      3    0.65084    1.95253    0.00000
-2.60338    0.00000    0.00000 ;     HA-    CA-    CB-    CG
  5971   5970   5987   5988      3    3.68192   -4.35136    0.00000
1.33888    0.00000    0.00000 ;     HA-    CA-     C-     O
  5971   5970   5987   5989      3    0.00000    0.00000    0.00000
0.00000    0.00000    0.00000 ;     HA-    CA-     C-     N
  5972   5970   5987   5988      3    0.00000    0.00000    0.00000
0.00000    0.00000    0.00000 ;     CB-    CA-     C-     O
  5972   5970   5987   5989      3    2.51040    4.18400    1.67360
-6.69440    0.00000    0.00000 ;     CB-    CA-     C-     N
  5972   5975   5976   5977      3   30.33400    0.00000  -30.33400
0.00000    0.00000    0.00000 ;     CB-    CG-   CD1-   HD1
  5972   5975   5976   5978      3   30.33400    0.00000  -30.33400
0.00000    0.00000    0.00000 ;     CB-    CG-   CD1-   CE1
  5972   5975   5985   5983      3   30.33400    0.00000  -30.33400
0.00000    0.00000    0.00000 ;     CB-    CG-   CD2-   CE2
  5972   5975   5985   5986      3   30.33400    0.00000  -30.33400
0.00000    0.00000    0.00000 ;     CB-    CG-   CD2-   HD2
  5973   5972   5970   5987      3    0.65084    1.95253    0.00000
-2.60338    0.00000    0.00000 ;    HB2-    CB-    CA-     C
  5973   5972   5975   5976      3    0.00000    0.00000    0.00000
0.00000    0.00000    0.00000 ;    HB2-    CB-    CG-   CD1
  5973   5972   5975   5985      3    0.00000    0.00000    0.00000
0.00000    0.00000    0.00000 ;    HB2-    CB-    CG-   CD2
  5974   5972   5970   5987      3    0.65084    1.95253    0.00000
-2.60338    0.00000    0.00000 ;    HB3-    CB-    CA-     C
  5974   5972   5975   5976      3    0.00000    0.00000    0.00000
0.00000    0.00000    0.00000 ;    HB3-    CB-    CG-   CD1
  5974   5972   5975   5985      3    0.00000    0.00000    0.00000
0.00000    0.00000    0.00000 ;    HB3-    CB-    CG-   CD2
  5975   5972   5970   5987      3    4.95804    2.17986   -3.52293
-3.21331   -0.40166    0.00000 ;     CG-    CB-    CA-     C
  5975   5976   5978   5979      3   30.33400    0.00000  -30.33400
```

0.00000 0.00000 0.00000 ; CG- CD1- CE1- HE1
5975 5976 5978 5980 3 30.33400 0.00000 -30.33400
0.00000 0.00000 0.00000 ; CG- CD1- CE1- CZ
5975 5985 5983 5980 3 30.33400 0.00000 -30.33400
0.00000 0.00000 0.00000 ; CG- CD2- CE2- CZ
5975 5985 5983 5984 3 30.33400 0.00000 -30.33400
0.00000 0.00000 0.00000 ; CG- CD2- CE2- HE2
5976 5975 5985 5983 3 30.33400 0.00000 -30.33400
0.00000 0.00000 0.00000 ; CD1- CG- CD2- CE2
5976 5975 5985 5986 3 30.33400 0.00000 -30.33400
0.00000 0.00000 0.00000 ; CD1- CG- CD2- HD2
5976 5978 5980 5981 3 30.33400 0.00000 -30.33400
0.00000 0.00000 0.00000 ; CD1- CE1- CZ- OH
5976 5978 5980 5983 3 30.33400 0.00000 -30.33400
0.00000 0.00000 0.00000 ; CD1- CE1- CZ- CE2
5977 5976 5975 5985 3 30.33400 0.00000 -30.33400
0.00000 0.00000 0.00000 ; HD1- CD1- CG- CD2
5977 5976 5978 5979 3 30.33400 0.00000 -30.33400
0.00000 0.00000 0.00000 ; HD1- CD1- CE1- HE1
5977 5976 5978 5980 3 30.33400 0.00000 -30.33400
0.00000 0.00000 0.00000 ; HD1- CD1- CE1- CZ
5978 5976 5975 5985 3 30.33400 0.00000 -30.33400
0.00000 0.00000 0.00000 ; CE1- CD1- CG- CD2
5978 5980 5981 5982 3 7.93286 0.00000 -9.56462
0.00000 2.17568 0.00000 ; CE1- CZ- OH- HH
5978 5980 5983 5984 3 30.33400 0.00000 -30.33400
0.00000 0.00000 0.00000 ; CE1- CZ- CE2- HE2
5978 5980 5983 5985 3 30.33400 0.00000 -30.33400
0.00000 0.00000 0.00000 ; CE1- CZ- CE2- CD2
5979 5978 5980 5981 3 30.33400 0.00000 -30.33400
0.00000 0.00000 0.00000 ; HE1- CE1- CZ- OH
5979 5978 5980 5983 3 30.33400 0.00000 -30.33400
0.00000 0.00000 0.00000 ; HE1- CE1- CZ- CE2
5980 5983 5985 5986 3 30.33400 0.00000 -30.33400
0.00000 0.00000 0.00000 ; CZ- CE2- CD2- HD2
5981 5980 5983 5984 3 30.33400 0.00000 -30.33400
0.00000 0.00000 0.00000 ; OH- CZ- CE2- HE2
5981 5980 5983 5985 3 30.33400 0.00000 -30.33400
0.00000 0.00000 0.00000 ; OH- CZ- CE2- CD2
5982 5981 5980 5983 3 7.93286 0.00000 -9.56462
0.00000 2.17568 0.00000 ; HH- OH- CZ- CE2
5984 5983 5985 5986 3 30.33400 0.00000 -30.33400
0.00000 0.00000 0.00000 ; HE2- CE2- CD2- HD2
5987 5989 5991 5992 3 0.00000 0.00000 0.00000
0.00000 0.00000 0.00000 ; C- N- CA- HA
5987 5989 5991 5993 3 11.71520 1.67360 15.06240 -
13.38880 0.00000 0.00000 ; C- N- CA- CB
5987 5989 5991 6001 3 1.75728 5.27184 2.25936
-7.02912 0.00000 0.00000 ; C- N- CA- C
5988 5987 5989 5990 3 29.28800 -8.36800 -20.92000
0.00000 0.00000 0.00000 ; O- C- N- H
5988 5987 5989 5991 3 20.92000 0.00000 -20.92000

```
0.00000    0.00000    0.00000 ;       O-     C-     N-    CA
  5989   5991   5993   5994       3    0.65084    1.95253    0.00000
-2.60338    0.00000    0.00000 ;       N-    CA-    CB-    HB
  5989   5991   5993   5995       3    3.83673    0.44769   -1.77402
-2.47693   -0.03347    0.00000 ;       N-    CA-    CB-   CG2
  5989   5991   5993   5999       3    4.93294    1.13386   -3.12963
-5.27184    3.17984    0.00000 ;       N-    CA-    CB-   OG1
  5989   5991   6001   6002       3    0.00000    0.00000    0.00000
0.00000    0.00000    0.00000 ;       N-    CA-     C-     O
  5989   5991   6001   6003       3   17.40544   -5.02080  -13.22144
9.20480    0.00000    0.00000 ;       N-    CA-     C-     N
  5990   5989   5991   5992       3    0.00000    0.00000    0.00000
0.00000    0.00000    0.00000 ;       H-     N-    CA-    HA
  5990   5989   5991   5993       3    0.00000    0.00000    0.00000
0.00000    0.00000    0.00000 ;       H-     N-    CA-    CB
  5990   5989   5991   6001       3    0.00000    0.00000    0.00000
0.00000    0.00000    0.00000 ;       H-     N-    CA-     C
  5991   5993   5995   5996       3    0.66944    2.00832    0.00000
-2.67776    0.00000    0.00000 ;      CA-    CB-   CG2-  HG21
  5991   5993   5995   5997       3    0.66944    2.00832    0.00000
-2.67776    0.00000    0.00000 ;      CA-    CB-   CG2-  HG22
  5991   5993   5995   5998       3    0.66944    2.00832    0.00000
-2.67776    0.00000    0.00000 ;      CA-    CB-   CG2-  HG23
  5991   5993   5999   6000       3    1.12131    2.98738    1.66523
-3.94970    0.43514    0.00000 ;      CA-    CB-   OG1-   HG1
  5991   6001   6003   6004       3   20.92000    0.00000  -20.92000
0.00000    0.00000    0.00000 ;      CA-     C-     N-     H
  5991   6001   6003   6005       3   20.92000    0.00000  -20.92000
0.00000    0.00000    0.00000 ;      CA-     C-     N-    CA
  5992   5991   5993   5994       3    0.65084    1.95253    0.00000
-2.60338    0.00000    0.00000 ;      HA-    CA-    CB-    HB
  5992   5991   5993   5995       3    0.65084    1.95253    0.00000
-2.60338    0.00000    0.00000 ;      HA-    CA-    CB-   CG2
  5992   5991   5993   5999       3    1.04600   -1.04600    0.00000
0.00000    0.00000    0.00000 ;      HA-    CA-    CB-   OG1
  5992   5991   6001   6002       3    3.68192   -4.35136    0.00000
1.33888    0.00000    0.00000 ;      HA-    CA-     C-     O
  5992   5991   6001   6003       3    0.00000    0.00000    0.00000
0.00000    0.00000    0.00000 ;      HA-    CA-     C-     N
  5993   5991   6001   6002       3    0.00000    0.00000    0.00000
0.00000    0.00000    0.00000 ;      CB-    CA-     C-     O
  5993   5991   6001   6003       3    2.51040    4.18400    1.67360
-6.69440    0.00000    0.00000 ;      CB-    CA-     C-     N
  5994   5993   5991   6001       3    0.65084    1.95253    0.00000
-2.60338    0.00000    0.00000 ;      HB-    CB-    CA-     C
  5994   5993   5995   5996       3    0.65084    1.95253    0.00000
-2.60338    0.00000    0.00000 ;      HB-    CB-   CG2-  HG21
  5994   5993   5995   5997       3    0.65084    1.95253    0.00000
-2.60338    0.00000    0.00000 ;      HB-    CB-   CG2-  HG22
  5994   5993   5995   5998       3    0.65084    1.95253    0.00000
-2.60338    0.00000    0.00000 ;      HB-    CB-   CG2-  HG23
  5994   5993   5999   6000       3    0.69733    2.09200    0.00000
```

-2.78933 0.00000 0.00000 ; HB- CB- OG1- HG1
5995 5993 5991 6001 3 5.67350 3.55640 -6.16722
-2.47693 3.74886 0.00000 ; CG2- CB- CA- C
5995 5993 5999 6000 3 4.74047 0.27196 -2.26773
-3.94970 1.60666 0.00000 ; CG2- CB- OG1- HG1
5996 5995 5993 5999 3 1.04600 -1.04600 0.00000
0.00000 0.00000 0.00000 ; HG21- CG2- CB- OG1
5997 5995 5993 5999 3 1.04600 -1.04600 0.00000
0.00000 0.00000 0.00000 ; HG22- CG2- CB- OG1
5998 5995 5993 5999 3 1.04600 -1.04600 0.00000
0.00000 0.00000 0.00000 ; HG23- CG2- CB- OG1
5999 5993 5991 6001 3 6.53541 6.87013 -6.21742
-5.27184 5.22163 0.00000 ; OG1- CB- CA- C
6001 6003 6005 6006 3 0.00000 0.00000 0.00000
0.00000 0.00000 0.00000 ; C- N- CA- HA
6001 6003 6005 6007 3 11.71520 1.67360 15.06240 -
13.38880 0.00000 0.00000 ; C- N- CA- CB
6001 6003 6005 6013 3 1.75728 5.27184 2.25936
-7.02912 0.00000 0.00000 ; C- N- CA- C
6002 6001 6003 6004 3 29.28800 -8.36800 -20.92000
0.00000 0.00000 0.00000 ; O- C- N- H
6002 6001 6003 6005 3 20.92000 0.00000 -20.92000
0.00000 0.00000 0.00000 ; O- C- N- CA
6003 6005 6007 6008 3 0.65084 1.95253 0.00000
-2.60338 0.00000 0.00000 ; N- CA- CB- HB2
6003 6005 6007 6009 3 0.65084 1.95253 0.00000
-2.60338 0.00000 0.00000 ; N- CA- CB- HB3
6003 6005 6007 6010 3 15.41386 9.74035 -8.39310
-0.97069 2.97901 0.00000 ; N- CA- CB- CG
6003 6005 6013 6014 3 0.00000 0.00000 0.00000
0.00000 0.00000 0.00000 ; N- CA- C- O
6003 6005 6013 6015 3 17.40544 -5.02080 -13.22144
9.20480 0.00000 0.00000 ; N- CA- C- N
6004 6003 6005 6006 3 0.00000 0.00000 0.00000
0.00000 0.00000 0.00000 ; H- N- CA- HA
6004 6003 6005 6007 3 0.00000 0.00000 0.00000
0.00000 0.00000 0.00000 ; H- N- CA- CB
6004 6003 6005 6013 3 0.00000 0.00000 0.00000
0.00000 0.00000 0.00000 ; H- N- CA- C
6005 6007 6010 6011 3 6.43499 0.00000 -5.39736
0.00000 -1.03763 0.00000 ; CA- CB- CG- OD1
6005 6007 6010 6012 3 6.43499 0.00000 -5.39736
0.00000 -1.03763 0.00000 ; CA- CB- CG- OD2
6005 6013 6015 6016 3 20.92000 0.00000 -20.92000
0.00000 0.00000 0.00000 ; CA- C- N- H
6005 6013 6015 6017 3 20.92000 0.00000 -20.92000
0.00000 0.00000 0.00000 ; CA- C- N- CA
6006 6005 6007 6008 3 0.65084 1.95253 0.00000
-2.60338 0.00000 0.00000 ; HA- CA- CB- HB2
6006 6005 6007 6009 3 0.65084 1.95253 0.00000
-2.60338 0.00000 0.00000 ; HA- CA- CB- HB3
6006 6005 6007 6010 3 0.65084 1.95253 0.00000

-2.60338    0.00000    0.00000 ;      HA-    CA-     CB-     CG
  6006   6005   6013   6014      3    3.68192   -4.35136    0.00000
1.33888    0.00000    0.00000 ;      HA-    CA-     C-     O
  6006   6005   6013   6015      3    0.00000    0.00000    0.00000
0.00000    0.00000    0.00000 ;      HA-    CA-     C-     N
  6007   6005   6013   6014      3    0.00000    0.00000    0.00000
0.00000    0.00000    0.00000 ;      CB-    CA-     C-     O
  6007   6005   6013   6015      3    2.51040    4.18400    1.67360
-6.69440    0.00000    0.00000 ;      CB-    CA-     C-     N
  6008   6007   6005   6013      3    0.65084    1.95253    0.00000
-2.60338    0.00000    0.00000 ;     HB2-    CB-     CA-     C
  6008   6007   6010   6011      3    0.00000    0.00000    0.00000
0.00000    0.00000    0.00000 ;     HB2-    CB-     CG-    OD1
  6008   6007   6010   6012      3    0.00000    0.00000    0.00000
0.00000    0.00000    0.00000 ;     HB2-    CB-     CG-    OD2
  6009   6007   6005   6013      3    0.65084    1.95253    0.00000
-2.60338    0.00000    0.00000 ;     HB3-    CB-     CA-     C
  6009   6007   6010   6011      3    0.00000    0.00000    0.00000
0.00000    0.00000    0.00000 ;     HB3-    CB-     CG-    OD1
  6009   6007   6010   6012      3    0.00000    0.00000    0.00000
0.00000    0.00000    0.00000 ;     HB3-    CB-     CG-    OD2
  6010   6007   6005   6013      3    7.14627   -1.04600   -8.99560
-0.97069    5.15469    0.00000 ;      CG-    CB-     CA-     C
  6013   6015   6017   6018      3    0.00000    0.00000    0.00000
0.00000    0.00000    0.00000 ;      C-     N-     CA-    HA
  6013   6015   6017   6019      3   11.71520    1.67360   15.06240  -
13.38880    0.00000    0.00000 ;      C-     N-     CA-    CB
  6013   6015   6017   6033      3    1.75728    5.27184    2.25936
-7.02912    0.00000    0.00000 ;      C-     N-     CA-     C
  6014   6013   6015   6016      3   29.28800   -8.36800  -20.92000
0.00000    0.00000    0.00000 ;      O-     C-     N-     H
  6014   6013   6015   6017      3   20.92000    0.00000  -20.92000
0.00000    0.00000    0.00000 ;      O-     C-     N-    CA
  6015   6017   6019   6020      3    0.65084    1.95253    0.00000
-2.60338    0.00000    0.00000 ;      N-    CA-     CB-    HB2
  6015   6017   6019   6021      3    0.65084    1.95253    0.00000
-2.60338    0.00000    0.00000 ;      N-    CA-     CB-    HB3
  6015   6017   6019   6022      3    3.28026    2.46019   -2.19242
-3.21331   -0.23430    0.00000 ;      N-    CA-     CB-    CG
  6015   6017   6033   6034      3    0.00000    0.00000    0.00000
0.00000    0.00000    0.00000 ;      N-    CA-     C-     O
  6015   6017   6033   6035      3   17.40544   -5.02080  -13.22144
9.20480    0.00000    0.00000 ;      N-    CA-     C-     N
  6016   6015   6017   6018      3    0.00000    0.00000    0.00000
0.00000    0.00000    0.00000 ;      H-     N-     CA-    HA
  6016   6015   6017   6019      3    0.00000    0.00000    0.00000
0.00000    0.00000    0.00000 ;      H-     N-     CA-    CB
  6016   6015   6017   6033      3    0.00000    0.00000    0.00000
0.00000    0.00000    0.00000 ;      H-     N-     CA-     C
  6017   6019   6022   6023      3    0.57739    0.00000    1.02926
0.00000   -1.60666    0.00000 ;     CA-    CB-     CG-    CD1
  6017   6019   6022   6031      3    0.57739    0.00000    1.02926

0.00000 -1.60666 0.00000 ; CA- CB- CG- CD2
6017 6033 6035 6036 3 20.92000 0.00000 -20.92000
0.00000 0.00000 0.00000 ; CA- C- N- H
6017 6033 6035 6037 3 20.92000 0.00000 -20.92000
0.00000 0.00000 0.00000 ; CA- C- N- CA
6018 6017 6019 6020 3 0.65084 1.95253 0.00000
-2.60338 0.00000 0.00000 ; HA- CA- CB- HB2
6018 6017 6019 6021 3 0.65084 1.95253 0.00000
-2.60338 0.00000 0.00000 ; HA- CA- CB- HB3
6018 6017 6019 6022 3 0.65084 1.95253 0.00000
-2.60338 0.00000 0.00000 ; HA- CA- CB- CG
6018 6017 6033 6034 3 3.68192 -4.35136 0.00000
1.33888 0.00000 0.00000 ; HA- CA- C- O
6018 6017 6033 6035 3 0.00000 0.00000 0.00000
0.00000 0.00000 0.00000 ; HA- CA- C- N
6019 6017 6033 6034 3 0.00000 0.00000 0.00000
0.00000 0.00000 0.00000 ; CB- CA- C- O
6019 6017 6033 6035 3 2.51040 4.18400 1.67360
-6.69440 0.00000 0.00000 ; CB- CA- C- N
6019 6022 6023 6024 3 30.33400 0.00000 -30.33400
0.00000 0.00000 0.00000 ; CB- CG- CD1- HD1
6019 6022 6023 6025 3 30.33400 0.00000 -30.33400
0.00000 0.00000 0.00000 ; CB- CG- CD1- CE1
6019 6022 6031 6029 3 30.33400 0.00000 -30.33400
0.00000 0.00000 0.00000 ; CB- CG- CD2- CE2
6019 6022 6031 6032 3 30.33400 0.00000 -30.33400
0.00000 0.00000 0.00000 ; CB- CG- CD2- HD2
6020 6019 6017 6033 3 0.65084 1.95253 0.00000
-2.60338 0.00000 0.00000 ; HB2- CB- CA- C
6020 6019 6022 6023 3 0.00000 0.00000 0.00000
0.00000 0.00000 0.00000 ; HB2- CB- CG- CD1
6020 6019 6022 6031 3 0.00000 0.00000 0.00000
0.00000 0.00000 0.00000 ; HB2- CB- CG- CD2
6021 6019 6017 6033 3 0.65084 1.95253 0.00000
-2.60338 0.00000 0.00000 ; HB3- CB- CA- C
6021 6019 6022 6023 3 0.00000 0.00000 0.00000
0.00000 0.00000 0.00000 ; HB3- CB- CG- CD1
6021 6019 6022 6031 3 0.00000 0.00000 0.00000
0.00000 0.00000 0.00000 ; HB3- CB- CG- CD2
6022 6019 6017 6033 3 4.95804 2.17986 -3.52293
-3.21331 -0.40166 0.00000 ; CG- CB- CA- C
6022 6023 6025 6026 3 30.33400 0.00000 -30.33400
0.00000 0.00000 0.00000 ; CG- CD1- CE1- HE1
6022 6023 6025 6027 3 30.33400 0.00000 -30.33400
0.00000 0.00000 0.00000 ; CG- CD1- CE1- CZ
6022 6031 6029 6027 3 30.33400 0.00000 -30.33400
0.00000 0.00000 0.00000 ; CG- CD2- CE2- CZ
6022 6031 6029 6030 3 30.33400 0.00000 -30.33400
0.00000 0.00000 0.00000 ; CG- CD2- CE2- HE2
6023 6022 6031 6029 3 30.33400 0.00000 -30.33400
0.00000 0.00000 0.00000 ; CD1- CG- CD2- CE2
6023 6022 6031 6032 3 30.33400 0.00000 -30.33400

```
0.00000    0.00000    0.00000 ;    CD1-    CG-   CD2-   HD2
  6023   6025   6027   6028       3   30.33400    0.00000  -30.33400
0.00000    0.00000    0.00000 ;    CD1-   CE1-    CZ-    HZ
  6023   6025   6027   6029       3   30.33400    0.00000  -30.33400
0.00000    0.00000    0.00000 ;    CD1-   CE1-    CZ-   CE2
  6024   6023   6022   6031       3   30.33400    0.00000  -30.33400
0.00000    0.00000    0.00000 ;    HD1-   CD1-    CG-   CD2
  6024   6023   6025   6026       3   30.33400    0.00000  -30.33400
0.00000    0.00000    0.00000 ;    HD1-   CD1-   CE1-   HE1
  6024   6023   6025   6027       3   30.33400    0.00000  -30.33400
0.00000    0.00000    0.00000 ;    HD1-   CD1-   CE1-    CZ
  6025   6023   6022   6031       3   30.33400    0.00000  -30.33400
0.00000    0.00000    0.00000 ;    CE1-   CD1-    CG-   CD2
  6025   6027   6029   6030       3   30.33400    0.00000  -30.33400
0.00000    0.00000    0.00000 ;    CE1-    CZ-   CE2-   HE2
  6025   6027   6029   6031       3   30.33400    0.00000  -30.33400
0.00000    0.00000    0.00000 ;    CE1-    CZ-   CE2-   CD2
  6026   6025   6027   6028       3   30.33400    0.00000  -30.33400
0.00000    0.00000    0.00000 ;    HE1-   CE1-    CZ-    HZ
  6026   6025   6027   6029       3   30.33400    0.00000  -30.33400
0.00000    0.00000    0.00000 ;    HE1-   CE1-    CZ-   CE2
  6027   6029   6031   6032       3   30.33400    0.00000  -30.33400
0.00000    0.00000    0.00000 ;     CZ-   CE2-   CD2-   HD2
  6028   6027   6029   6030       3   30.33400    0.00000  -30.33400
0.00000    0.00000    0.00000 ;     HZ-    CZ-   CE2-   HE2
  6028   6027   6029   6031       3   30.33400    0.00000  -30.33400
0.00000    0.00000    0.00000 ;     HZ-    CZ-   CE2-   CD2
  6030   6029   6031   6032       3   30.33400    0.00000  -30.33400
0.00000    0.00000    0.00000 ;    HE2-   CE2-   CD2-   HD2
  6033   6035   6037   6038       3    0.00000    0.00000    0.00000
0.00000    0.00000    0.00000 ;      C-     N-    CA-    HA
  6033   6035   6037   6039       3   11.71520    1.67360   15.06240  -
13.38880    0.00000    0.00000 ;      C-     N-    CA-    CB
  6033   6035   6037   6052       3    1.75728    5.27184    2.25936
-7.02912    0.00000    0.00000 ;      C-     N-    CA-     C
  6034   6033   6035   6036       3   29.28800   -8.36800  -20.92000
0.00000    0.00000    0.00000 ;      O-     C-     N-     H
  6034   6033   6035   6037       3   20.92000    0.00000  -20.92000
0.00000    0.00000    0.00000 ;      O-     C-     N-    CA
  6035   6037   6039   6040       3    0.65084    1.95253    0.00000
-2.60338    0.00000    0.00000 ;      N-    CA-    CB-   HB2
  6035   6037   6039   6041       3    0.65084    1.95253    0.00000
-2.60338    0.00000    0.00000 ;      N-    CA-    CB-   HB3
  6035   6037   6039   6042       3    3.79070    1.39746   -4.61077
-2.40998    2.44346    0.00000 ;      N-    CA-    CB-    CG
  6035   6037   6052   6053       3    0.00000    0.00000    0.00000
0.00000    0.00000    0.00000 ;      N-    CA-     C-     O
  6035   6037   6052   6054       3   17.40544   -5.02080  -13.22144
9.20480    0.00000    0.00000 ;      N-    CA-     C-     N
  6036   6035   6037   6038       3    0.00000    0.00000    0.00000
0.00000    0.00000    0.00000 ;      H-     N-    CA-    HA
  6036   6035   6037   6039       3    0.00000    0.00000    0.00000
```

0.00000 0.00000 0.00000 ; H- N- CA- CB
6036 6035 6037 6052 3 0.00000 0.00000 0.00000
0.00000 0.00000 0.00000 ; H- N- CA- C
6037 6039 6042 6043 3 0.66944 2.00832 0.00000
-2.67776 0.00000 0.00000 ; CA- CB- CG- HG
6037 6039 6042 6044 3 3.90367 0.19665 -6.21742
-2.37651 5.99149 0.00000 ; CA- CB- CG- CD1
6037 6039 6042 6048 3 3.90367 0.19665 -6.21742
-2.37651 5.99149 0.00000 ; CA- CB- CG- CD2
6037 6052 6054 6055 3 20.92000 0.00000 -20.92000
0.00000 0.00000 0.00000 ; CA- C- N- H
6037 6052 6054 6056 3 20.92000 0.00000 -20.92000
0.00000 0.00000 0.00000 ; CA- C- N- CA
6038 6037 6039 6040 3 0.65084 1.95253 0.00000
-2.60338 0.00000 0.00000 ; HA- CA- CB- HB2
6038 6037 6039 6041 3 0.65084 1.95253 0.00000
-2.60338 0.00000 0.00000 ; HA- CA- CB- HB3
6038 6037 6039 6042 3 0.65084 1.95253 0.00000
-2.60338 0.00000 0.00000 ; HA- CA- CB- CG
6038 6037 6052 6053 3 3.68192 -4.35136 0.00000
1.33888 0.00000 0.00000 ; HA- CA- C- O
6038 6037 6052 6054 3 0.00000 0.00000 0.00000
0.00000 0.00000 0.00000 ; HA- CA- C- N
6039 6037 6052 6053 3 0.00000 0.00000 0.00000
0.00000 0.00000 0.00000 ; CB- CA- C- O
6039 6037 6052 6054 3 2.51040 4.18400 1.67360
-6.69440 0.00000 0.00000 ; CB- CA- C- N
6039 6042 6044 6045 3 0.66944 2.00832 0.00000
-2.67776 0.00000 0.00000 ; CB- CG- CD1- HD11
6039 6042 6044 6046 3 0.66944 2.00832 0.00000
-2.67776 0.00000 0.00000 ; CB- CG- CD1- HD12
6039 6042 6044 6047 3 0.66944 2.00832 0.00000
-2.67776 0.00000 0.00000 ; CB- CG- CD1- HD13
6039 6042 6048 6049 3 0.66944 2.00832 0.00000
-2.67776 0.00000 0.00000 ; CB- CG- CD2- HD21
6039 6042 6048 6050 3 0.66944 2.00832 0.00000
-2.67776 0.00000 0.00000 ; CB- CG- CD2- HD22
6039 6042 6048 6051 3 0.66944 2.00832 0.00000
-2.67776 0.00000 0.00000 ; CB- CG- CD2- HD23
6040 6039 6037 6052 3 0.65084 1.95253 0.00000
-2.60338 0.00000 0.00000 ; HB2- CB- CA- C
6040 6039 6042 6043 3 0.62760 1.88280 0.00000
-2.51040 0.00000 0.00000 ; HB2- CB- CG- HG
6040 6039 6042 6044 3 0.66944 2.00832 0.00000
-2.67776 0.00000 0.00000 ; HB2- CB- CG- CD1
6040 6039 6042 6048 3 0.66944 2.00832 0.00000
-2.67776 0.00000 0.00000 ; HB2- CB- CG- CD2
6041 6039 6037 6052 3 0.65084 1.95253 0.00000
-2.60338 0.00000 0.00000 ; HB3- CB- CA- C
6041 6039 6042 6043 3 0.62760 1.88280 0.00000
-2.51040 0.00000 0.00000 ; HB3- CB- CG- HG
6041 6039 6042 6044 3 0.66944 2.00832 0.00000

```
-2.67776    0.00000    0.00000 ;     HB3-    CB-     CG-   CD1
  6041   6039   6042   6048       3    0.66944    2.00832    0.00000
-2.67776    0.00000    0.00000 ;     HB3-    CB-     CG-   CD2
  6042   6039   6037   6052       3   10.33448   -1.14642  -11.54784
-2.40998    6.35968    0.00000 ;      CG-    CB-     CA-     C
  6043   6042   6044   6045       3    0.62760    1.88280    0.00000
-2.51040    0.00000    0.00000 ;      HG-    CG-    CD1-  HD11
  6043   6042   6044   6046       3    0.62760    1.88280    0.00000
-2.51040    0.00000    0.00000 ;      HG-    CG-    CD1-  HD12
  6043   6042   6044   6047       3    0.62760    1.88280    0.00000
-2.51040    0.00000    0.00000 ;      HG-    CG-    CD1-  HD13
  6043   6042   6048   6049       3    0.62760    1.88280    0.00000
-2.51040    0.00000    0.00000 ;      HG-    CG-    CD2-  HD21
  6043   6042   6048   6050       3    0.62760    1.88280    0.00000
-2.51040    0.00000    0.00000 ;      HG-    CG-    CD2-  HD22
  6043   6042   6048   6051       3    0.62760    1.88280    0.00000
-2.51040    0.00000    0.00000 ;      HG-    CG-    CD2-  HD23
  6044   6042   6048   6049       3    0.66944    2.00832    0.00000
-2.67776    0.00000    0.00000 ;     CD1-    CG-    CD2-  HD21
  6044   6042   6048   6050       3    0.66944    2.00832    0.00000
-2.67776    0.00000    0.00000 ;     CD1-    CG-    CD2-  HD22
  6044   6042   6048   6051       3    0.66944    2.00832    0.00000
-2.67776    0.00000    0.00000 ;     CD1-    CG-    CD2-  HD23
  6045   6044   6042   6048       3    0.66944    2.00832    0.00000
-2.67776    0.00000    0.00000 ;    HD11-   CD1-     CG-   CD2
  6046   6044   6042   6048       3    0.66944    2.00832    0.00000
-2.67776    0.00000    0.00000 ;    HD12-   CD1-     CG-   CD2
  6047   6044   6042   6048       3    0.66944    2.00832    0.00000
-2.67776    0.00000    0.00000 ;    HD13-   CD1-     CG-   CD2
  6052   6054   6056   6057       3    0.00000    0.00000    0.00000
0.00000    0.00000    0.00000 ;       C-      N-     CA-    HA
  6052   6054   6056   6058       3   11.71520    1.67360   15.06240  -
13.38880    0.00000    0.00000 ;       C-      N-     CA-    CB
  6052   6054   6056   6062       3    1.75728    5.27184    2.25936
-7.02912    0.00000    0.00000 ;       C-      N-     CA-     C
  6053   6052   6054   6055       3   29.28800   -8.36800  -20.92000
0.00000    0.00000    0.00000 ;       O-      C-      N-     H
  6053   6052   6054   6056       3   20.92000    0.00000  -20.92000
0.00000    0.00000    0.00000 ;       O-      C-      N-    CA
  6054   6056   6058   6059       3    0.65084    1.95253    0.00000
-2.60338    0.00000    0.00000 ;       N-    CA-     CB-   HB1
  6054   6056   6058   6060       3    0.65084    1.95253    0.00000
-2.60338    0.00000    0.00000 ;       N-    CA-     CB-   HB2
  6054   6056   6058   6061       3    0.65084    1.95253    0.00000
-2.60338    0.00000    0.00000 ;       N-    CA-     CB-   HB3
  6054   6056   6062   6063       3    0.00000    0.00000    0.00000
0.00000    0.00000    0.00000 ;       N-    CA-      C-     O
  6054   6056   6062   6064       3   17.40544   -5.02080  -13.22144
9.20480    0.00000    0.00000 ;       N-    CA-      C-     N
  6055   6054   6056   6057       3    0.00000    0.00000    0.00000
0.00000    0.00000    0.00000 ;       H-      N-     CA-    HA
  6055   6054   6056   6058       3    0.00000    0.00000    0.00000
```

```
0.00000    0.00000    0.00000 ;      H-     N-    CA-    CB
  6055   6054   6056   6062      3    0.00000    0.00000    0.00000
0.00000    0.00000    0.00000 ;      H-     N-    CA-     C
  6056   6062   6064   6065      3   20.92000    0.00000  -20.92000
0.00000    0.00000    0.00000 ;     CA-     C-     N-     H
  6056   6062   6064   6066      3   20.92000    0.00000  -20.92000
0.00000    0.00000    0.00000 ;     CA-     C-     N-    CA
  6057   6056   6058   6059      3    0.65084    1.95253    0.00000
-2.60338    0.00000    0.00000 ;     HA-    CA-    CB-   HB1
  6057   6056   6058   6060      3    0.65084    1.95253    0.00000
-2.60338    0.00000    0.00000 ;     HA-    CA-    CB-   HB2
  6057   6056   6058   6061      3    0.65084    1.95253    0.00000
-2.60338    0.00000    0.00000 ;     HA-    CA-    CB-   HB3
  6057   6056   6062   6063      3    3.68192   -4.35136    0.00000
1.33888    0.00000    0.00000 ;     HA-    CA-     C-     O
  6057   6056   6062   6064      3    0.00000    0.00000    0.00000
0.00000    0.00000    0.00000 ;     HA-    CA-     C-     N
  6058   6056   6062   6063      3    0.00000    0.00000    0.00000
0.00000    0.00000    0.00000 ;     CB-    CA-     C-     O
  6058   6056   6062   6064      3    2.51040    4.18400    1.67360
-6.69440    0.00000    0.00000 ;     CB-    CA-     C-     N
  6059   6058   6056   6062      3    0.65084    1.95253    0.00000
-2.60338    0.00000    0.00000 ;    HB1-    CB-    CA-     C
  6060   6058   6056   6062      3    0.65084    1.95253    0.00000
-2.60338    0.00000    0.00000 ;    HB2-    CB-    CA-     C
  6061   6058   6056   6062      3    0.65084    1.95253    0.00000
-2.60338    0.00000    0.00000 ;    HB3-    CB-    CA-     C
  6062   6064   6066   6067      3    0.00000    0.00000    0.00000
0.00000    0.00000    0.00000 ;      C-     N-    CA-    HA
  6062   6064   6066   6068      3   11.71520    1.67360   15.06240  -
13.38880    0.00000    0.00000 ;      C-     N-    CA-    CB
  6062   6064   6066   6072      3    1.75728    5.27184    2.25936
-7.02912    0.00000    0.00000 ;      C-     N-    CA-     C
  6063   6062   6064   6065      3   29.28800   -8.36800  -20.92000
0.00000    0.00000    0.00000 ;      O-     C-     N-     H
  6063   6062   6064   6066      3   20.92000    0.00000  -20.92000
0.00000    0.00000    0.00000 ;      O-     C-     N-    CA
  6064   6066   6068   6069      3    0.65084    1.95253    0.00000
-2.60338    0.00000    0.00000 ;      N-    CA-    CB-   HB1
  6064   6066   6068   6070      3    0.65084    1.95253    0.00000
-2.60338    0.00000    0.00000 ;      N-    CA-    CB-   HB2
  6064   6066   6068   6071      3    0.65084    1.95253    0.00000
-2.60338    0.00000    0.00000 ;      N-    CA-    CB-   HB3
  6064   6066   6072   6073      3    0.00000    0.00000    0.00000
0.00000    0.00000    0.00000 ;      N-    CA-     C-     O
  6064   6066   6072   6074      3   17.40544   -5.02080  -13.22144
9.20480    0.00000    0.00000 ;      N-    CA-     C-     N
  6065   6064   6066   6067      3    0.00000    0.00000    0.00000
0.00000    0.00000    0.00000 ;      H-     N-    CA-    HA
  6065   6064   6066   6068      3    0.00000    0.00000    0.00000
0.00000    0.00000    0.00000 ;      H-     N-    CA-    CB
  6065   6064   6066   6072      3    0.00000    0.00000    0.00000
```

0.00000 0.00000 0.00000 ; H- N- CA- C
6066 6072 6074 6075 3 20.92000 0.00000 -20.92000
0.00000 0.00000 0.00000 ; CA- C- N- H
6066 6072 6074 6076 3 20.92000 0.00000 -20.92000
0.00000 0.00000 0.00000 ; CA- C- N- CA
6067 6066 6068 6069 3 0.65084 1.95253 0.00000
-2.60338 0.00000 0.00000 ; HA- CA- CB- HB1
6067 6066 6068 6070 3 0.65084 1.95253 0.00000
-2.60338 0.00000 0.00000 ; HA- CA- CB- HB2
6067 6066 6068 6071 3 0.65084 1.95253 0.00000
-2.60338 0.00000 0.00000 ; HA- CA- CB- HB3
6067 6066 6072 6073 3 3.68192 -4.35136 0.00000
1.33888 0.00000 0.00000 ; HA- CA- C- O
6067 6066 6072 6074 3 0.00000 0.00000 0.00000
0.00000 0.00000 0.00000 ; HA- CA- C- N
6068 6066 6072 6073 3 0.00000 0.00000 0.00000
0.00000 0.00000 0.00000 ; CB- CA- C- O
6068 6066 6072 6074 3 2.51040 4.18400 1.67360
-6.69440 0.00000 0.00000 ; CB- CA- C- N
6069 6068 6066 6072 3 0.65084 1.95253 0.00000
-2.60338 0.00000 0.00000 ; HB1- CB- CA- C
6070 6068 6066 6072 3 0.65084 1.95253 0.00000
-2.60338 0.00000 0.00000 ; HB2- CB- CA- C
6071 6068 6066 6072 3 0.65084 1.95253 0.00000
-2.60338 0.00000 0.00000 ; HB3- CB- CA- C
6072 6074 6076 6077 3 0.00000 0.00000 0.00000
0.00000 0.00000 0.00000 ; C- N- CA- HA
6072 6074 6076 6078 3 11.71520 1.67360 15.06240 -
13.38880 0.00000 0.00000 ; C- N- CA- CB
6072 6074 6076 6086 3 1.75728 5.27184 2.25936
-7.02912 0.00000 0.00000 ; C- N- CA- C
6073 6072 6074 6075 3 29.28800 -8.36800 -20.92000
0.00000 0.00000 0.00000 ; O- C- N- H
6073 6072 6074 6076 3 20.92000 0.00000 -20.92000
0.00000 0.00000 0.00000 ; O- C- N- CA
6074 6076 6078 6079 3 0.65084 1.95253 0.00000
-2.60338 0.00000 0.00000 ; N- CA- CB- HB
6074 6076 6078 6080 3 3.83673 0.44769 -1.77402
-2.47693 -0.03347 0.00000 ; N- CA- CB- CG2
6074 6076 6078 6084 3 4.93294 1.13386 -3.12963
-5.27184 3.17984 0.00000 ; N- CA- CB- OG1
6074 6076 6086 6087 3 0.00000 0.00000 0.00000
0.00000 0.00000 0.00000 ; N- CA- C- O
6074 6076 6086 6088 3 17.40544 -5.02080 -13.22144
9.20480 0.00000 0.00000 ; N- CA- C- N
6075 6074 6076 6077 3 0.00000 0.00000 0.00000
0.00000 0.00000 0.00000 ; H- N- CA- HA
6075 6074 6076 6078 3 0.00000 0.00000 0.00000
0.00000 0.00000 0.00000 ; H- N- CA- CB
6075 6074 6076 6086 3 0.00000 0.00000 0.00000
0.00000 0.00000 0.00000 ; H- N- CA- C
6076 6078 6080 6081 3 0.66944 2.00832 0.00000

-2.67776 0.00000 0.00000 ; CA- CB- CG2- HG21
6076 6078 6080 6082 3 0.66944 2.00832 0.00000
-2.67776 0.00000 0.00000 ; CA- CB- CG2- HG22
6076 6078 6080 6083 3 0.66944 2.00832 0.00000
-2.67776 0.00000 0.00000 ; CA- CB- CG2- HG23
6076 6078 6084 6085 3 1.12131 2.98738 1.66523
-3.94970 0.43514 0.00000 ; CA- CB- OG1- HG1
6076 6086 6088 6089 3 20.92000 0.00000 -20.92000
0.00000 0.00000 0.00000 ; CA- C- N- H
6076 6086 6088 6090 3 20.92000 0.00000 -20.92000
0.00000 0.00000 0.00000 ; CA- C- N- CA
6077 6076 6078 6079 3 0.65084 1.95253 0.00000
-2.60338 0.00000 0.00000 ; HA- CA- CB- HB
6077 6076 6078 6080 3 0.65084 1.95253 0.00000
-2.60338 0.00000 0.00000 ; HA- CA- CB- CG2
6077 6076 6078 6084 3 1.04600 -1.04600 0.00000
0.00000 0.00000 0.00000 ; HA- CA- CB- OG1
6077 6076 6086 6087 3 3.68192 -4.35136 0.00000
1.33888 0.00000 0.00000 ; HA- CA- C- O
6077 6076 6086 6088 3 0.00000 0.00000 0.00000
0.00000 0.00000 0.00000 ; HA- CA- C- N
6078 6076 6086 6087 3 0.00000 0.00000 0.00000
0.00000 0.00000 0.00000 ; CB- CA- C- O
6078 6076 6086 6088 3 2.51040 4.18400 1.67360
-6.69440 0.00000 0.00000 ; CB- CA- C- N
6079 6078 6076 6086 3 0.65084 1.95253 0.00000
-2.60338 0.00000 0.00000 ; HB- CB- CA- C
6079 6078 6080 6081 3 0.65084 1.95253 0.00000
-2.60338 0.00000 0.00000 ; HB- CB- CG2- HG21
6079 6078 6080 6082 3 0.65084 1.95253 0.00000
-2.60338 0.00000 0.00000 ; HB- CB- CG2- HG22
6079 6078 6080 6083 3 0.65084 1.95253 0.00000
-2.60338 0.00000 0.00000 ; HB- CB- CG2- HG23
6079 6078 6084 6085 3 0.69733 2.09200 0.00000
-2.78933 0.00000 0.00000 ; HB- CB- OG1- HG1
6080 6078 6076 6086 3 5.67350 3.55640 -6.16722
-2.47693 3.74886 0.00000 ; CG2- CB- CA- C
6080 6078 6084 6085 3 4.74047 0.27196 -2.26773
-3.94970 1.60666 0.00000 ; CG2- CB- OG1- HG1
6081 6080 6078 6084 3 1.04600 -1.04600 0.00000
0.00000 0.00000 0.00000 ; HG21- CG2- CB- OG1
6082 6080 6078 6084 3 1.04600 -1.04600 0.00000
0.00000 0.00000 0.00000 ; HG22- CG2- CB- OG1
6083 6080 6078 6084 3 1.04600 -1.04600 0.00000
0.00000 0.00000 0.00000 ; HG23- CG2- CB- OG1
6084 6078 6076 6086 3 6.53541 6.87013 -6.21742
-5.27184 5.22163 0.00000 ; OG1- CB- CA- C
6086 6088 6090 6091 3 0.00000 0.00000 0.00000
0.00000 0.00000 0.00000 ; C- N- CA- HA
6086 6088 6090 6092 3 11.71520 1.67360 15.06240 -
13.38880 0.00000 0.00000 ; C- N- CA- CB
6086 6088 6090 6105 3 1.75728 5.27184 2.25936

-7.02912 0.00000 0.00000 ; C- N- CA- C
6087 6086 6088 6089 3 29.28800 -8.36800 -20.92000
0.00000 0.00000 0.00000 ; O- C- N- H
6087 6086 6088 6090 3 20.92000 0.00000 -20.92000
0.00000 0.00000 0.00000 ; O- C- N- CA
6088 6090 6092 6093 3 0.65084 1.95253 0.00000
-2.60338 0.00000 0.00000 ; N- CA- CB- HB
6088 6090 6092 6094 3 3.83673 0.44769 -1.77402
-2.47693 -0.03347 0.00000 ; N- CA- CB- CG2
6088 6090 6092 6098 3 2.97482 0.12134 2.04179
-1.89117 -3.24678 0.00000 ; N- CA- CB- CG1
6088 6090 6105 6106 3 0.00000 0.00000 0.00000
0.00000 0.00000 0.00000 ; N- CA- C- O
6088 6090 6105 6107 3 17.40544 -5.02080 -13.22144
9.20480 0.00000 0.00000 ; N- CA- C- N
6089 6088 6090 6091 3 0.00000 0.00000 0.00000
0.00000 0.00000 0.00000 ; H- N- CA- HA
6089 6088 6090 6092 3 0.00000 0.00000 0.00000
0.00000 0.00000 0.00000 ; H- N- CA- CB
6089 6088 6090 6105 3 0.00000 0.00000 0.00000
0.00000 0.00000 0.00000 ; H- N- CA- C
6090 6092 6094 6095 3 0.66944 2.00832 0.00000
-2.67776 0.00000 0.00000 ; CA- CB- CG2- HG21
6090 6092 6094 6096 3 0.66944 2.00832 0.00000
-2.67776 0.00000 0.00000 ; CA- CB- CG2- HG22
6090 6092 6094 6097 3 0.66944 2.00832 0.00000
-2.67776 0.00000 0.00000 ; CA- CB- CG2- HG23
6090 6092 6098 6099 3 0.66944 2.00832 0.00000
-2.67776 0.00000 0.00000 ; CA- CB- CG1- HG12
6090 6092 6098 6100 3 0.66944 2.00832 0.00000
-2.67776 0.00000 0.00000 ; CA- CB- CG1- HG13
6090 6092 6098 6101 3 4.24258 -0.52718 -7.25506
-1.79075 7.69856 0.00000 ; CA- CB- CG1- CD1
6090 6105 6107 6108 3 20.92000 0.00000 -20.92000
0.00000 0.00000 0.00000 ; CA- C- N- H
6090 6105 6107 6109 3 20.92000 0.00000 -20.92000
0.00000 0.00000 0.00000 ; CA- C- N- CA
6091 6090 6092 6093 3 0.65084 1.95253 0.00000
-2.60338 0.00000 0.00000 ; HA- CA- CB- HB
6091 6090 6092 6094 3 0.65084 1.95253 0.00000
-2.60338 0.00000 0.00000 ; HA- CA- CB- CG2
6091 6090 6092 6098 3 0.65084 1.95253 0.00000
-2.60338 0.00000 0.00000 ; HA- CA- CB- CG1
6091 6090 6105 6106 3 3.68192 -4.35136 0.00000
1.33888 0.00000 0.00000 ; HA- CA- C- O
6091 6090 6105 6107 3 0.00000 0.00000 0.00000
0.00000 0.00000 0.00000 ; HA- CA- C- N
6092 6090 6105 6106 3 0.00000 0.00000 0.00000
0.00000 0.00000 0.00000 ; CB- CA- C- O
6092 6090 6105 6107 3 2.51040 4.18400 1.67360
-6.69440 0.00000 0.00000 ; CB- CA- C- N
6092 6098 6101 6102 3 0.66944 2.00832 0.00000

-2.67776 0.00000 0.00000 ; CB- CG1- CD1- HD11
6092 6098 6101 6103 3 0.66944 2.00832 0.00000
-2.67776 0.00000 0.00000 ; CB- CG1- CD1- HD12
6092 6098 6101 6104 3 0.66944 2.00832 0.00000
-2.67776 0.00000 0.00000 ; CB- CG1- CD1- HD13
6093 6092 6090 6105 3 0.65084 1.95253 0.00000
-2.60338 0.00000 0.00000 ; HB- CB- CA- C
6093 6092 6094 6095 3 0.62760 1.88280 0.00000
-2.51040 0.00000 0.00000 ; HB- CB- CG2- HG21
6093 6092 6094 6096 3 0.62760 1.88280 0.00000
-2.51040 0.00000 0.00000 ; HB- CB- CG2- HG22
6093 6092 6094 6097 3 0.62760 1.88280 0.00000
-2.51040 0.00000 0.00000 ; HB- CB- CG2- HG23
6093 6092 6098 6099 3 0.62760 1.88280 0.00000
-2.51040 0.00000 0.00000 ; HB- CB- CG1- HG12
6093 6092 6098 6100 3 0.62760 1.88280 0.00000
-2.51040 0.00000 0.00000 ; HB- CB- CG1- HG13
6093 6092 6098 6101 3 0.66944 2.00832 0.00000
-2.67776 0.00000 0.00000 ; HB- CB- CG1- CD1
6094 6092 6090 6105 3 5.67350 3.55640 -6.16722
-2.47693 3.74886 0.00000 ; CG2- CB- CA- C
6094 6092 6098 6099 3 0.66944 2.00832 0.00000
-2.67776 0.00000 0.00000 ; CG2- CB- CG1- HG12
6094 6092 6098 6100 3 0.66944 2.00832 0.00000
-2.67776 0.00000 0.00000 ; CG2- CB- CG1- HG13
6094 6092 6098 6101 3 3.81162 0.49790 -8.14206
-1.79075 7.49773 0.00000 ; CG2- CB- CG1- CD1
6095 6094 6092 6098 3 0.66944 2.00832 0.00000
-2.67776 0.00000 0.00000 ; HG21- CG2- CB- CG1
6096 6094 6092 6098 3 0.66944 2.00832 0.00000
-2.67776 0.00000 0.00000 ; HG22- CG2- CB- CG1
6097 6094 6092 6098 3 0.66944 2.00832 0.00000
-2.67776 0.00000 0.00000 ; HG23- CG2- CB- CG1
6098 6092 6090 6105 3 8.26340 0.74057 -9.99976
-1.89117 3.84928 0.00000 ; CG1- CB- CA- C
6099 6098 6101 6102 3 0.62760 1.88280 0.00000
-2.51040 0.00000 0.00000 ; HG12- CG1- CD1- HD11
6099 6098 6101 6103 3 0.62760 1.88280 0.00000
-2.51040 0.00000 0.00000 ; HG12- CG1- CD1- HD12
6099 6098 6101 6104 3 0.62760 1.88280 0.00000
-2.51040 0.00000 0.00000 ; HG12- CG1- CD1- HD13
6100 6098 6101 6102 3 0.62760 1.88280 0.00000
-2.51040 0.00000 0.00000 ; HG13- CG1- CD1- HD11
6100 6098 6101 6103 3 0.62760 1.88280 0.00000
-2.51040 0.00000 0.00000 ; HG13- CG1- CD1- HD12
6100 6098 6101 6104 3 0.62760 1.88280 0.00000
-2.51040 0.00000 0.00000 ; HG13- CG1- CD1- HD13
6105 6107 6109 6110 3 0.00000 0.00000 0.00000
0.00000 0.00000 0.00000 ; C- N- CA- HA
6105 6107 6109 6111 3 11.71520 1.67360 15.06240 -
13.38880 0.00000 0.00000 ; C- N- CA- CB
6105 6107 6109 6117 3 1.75728 5.27184 2.25936

-7.02912 0.00000 0.00000 ; C- N- CA- C
6106 6105 6107 6108 3 29.28800 -8.36800 -20.92000
0.00000 0.00000 0.00000 ; O- C- N- H
6106 6105 6107 6109 3 20.92000 0.00000 -20.92000
0.00000 0.00000 0.00000 ; O- C- N- CA
6107 6109 6111 6112 3 0.65084 1.95253 0.00000
-2.60338 0.00000 0.00000 ; N- CA- CB- HB2
6107 6109 6111 6113 3 0.65084 1.95253 0.00000
-2.60338 0.00000 0.00000 ; N- CA- CB- HB3
6107 6109 6111 6114 3 15.41386 9.74035 -8.39310
-0.97069 2.97901 0.00000 ; N- CA- CB- CG
6107 6109 6117 6118 3 0.00000 0.00000 0.00000
0.00000 0.00000 0.00000 ; N- CA- C- O
6107 6109 6117 6119 3 17.40544 -5.02080 -13.22144
9.20480 0.00000 0.00000 ; N- CA- C- N
6108 6107 6109 6110 3 0.00000 0.00000 0.00000
0.00000 0.00000 0.00000 ; H- N- CA- HA
6108 6107 6109 6111 3 0.00000 0.00000 0.00000
0.00000 0.00000 0.00000 ; H- N- CA- CB
6108 6107 6109 6117 3 0.00000 0.00000 0.00000
0.00000 0.00000 0.00000 ; H- N- CA- C
6109 6111 6114 6115 3 6.43499 0.00000 -5.39736
0.00000 -1.03763 0.00000 ; CA- CB- CG- OD1
6109 6111 6114 6116 3 6.43499 0.00000 -5.39736
0.00000 -1.03763 0.00000 ; CA- CB- CG- OD2
6109 6117 6119 6120 3 20.92000 0.00000 -20.92000
0.00000 0.00000 0.00000 ; CA- C- N- H
6109 6117 6119 6121 3 20.92000 0.00000 -20.92000
0.00000 0.00000 0.00000 ; CA- C- N- CA
6110 6109 6111 6112 3 0.65084 1.95253 0.00000
-2.60338 0.00000 0.00000 ; HA- CA- CB- HB2
6110 6109 6111 6113 3 0.65084 1.95253 0.00000
-2.60338 0.00000 0.00000 ; HA- CA- CB- HB3
6110 6109 6111 6114 3 0.65084 1.95253 0.00000
-2.60338 0.00000 0.00000 ; HA- CA- CB- CG
6110 6109 6117 6118 3 3.68192 -4.35136 0.00000
1.33888 0.00000 0.00000 ; HA- CA- C- O
6110 6109 6117 6119 3 0.00000 0.00000 0.00000
0.00000 0.00000 0.00000 ; HA- CA- C- N
6111 6109 6117 6118 3 0.00000 0.00000 0.00000
0.00000 0.00000 0.00000 ; CB- CA- C- O
6111 6109 6117 6119 3 2.51040 4.18400 1.67360
-6.69440 0.00000 0.00000 ; CB- CA- C- N
6112 6111 6109 6117 3 0.65084 1.95253 0.00000
-2.60338 0.00000 0.00000 ; HB2- CB- CA- C
6112 6111 6114 6115 3 0.00000 0.00000 0.00000
0.00000 0.00000 0.00000 ; HB2- CB- CG- OD1
6112 6111 6114 6116 3 0.00000 0.00000 0.00000
0.00000 0.00000 0.00000 ; HB2- CB- CG- OD2
6113 6111 6109 6117 3 0.65084 1.95253 0.00000
-2.60338 0.00000 0.00000 ; HB3- CB- CA- C
6113 6111 6114 6115 3 0.00000 0.00000 0.00000

0.00000 0.00000 0.00000 ; HB3- CB- CG- OD1
6113 6111 6114 6116 3 0.00000 0.00000 0.00000
0.00000 0.00000 0.00000 ; HB3- CB- CG- OD2
6114 6111 6109 6117 3 7.14627 -1.04600 -8.99560
-0.97069 5.15469 0.00000 ; CG- CB- CA- C
6117 6119 6121 6122 3 0.00000 0.00000 0.00000
0.00000 0.00000 0.00000 ; C- N- CA- HA
6117 6119 6121 6123 3 11.71520 1.67360 15.06240 -
13.38880 0.00000 0.00000 ; C- N- CA- CB
6117 6119 6121 6139 3 1.75728 5.27184 2.25936
-7.02912 0.00000 0.00000 ; C- N- CA- C
6118 6117 6119 6120 3 29.28800 -8.36800 -20.92000
0.00000 0.00000 0.00000 ; O- C- N- H
6118 6117 6119 6121 3 20.92000 0.00000 -20.92000
0.00000 0.00000 0.00000 ; O- C- N- CA
6119 6121 6123 6124 3 0.65084 1.95253 0.00000
-2.60338 0.00000 0.00000 ; N- CA- CB- HB2
6119 6121 6123 6125 3 0.65084 1.95253 0.00000
-2.60338 0.00000 0.00000 ; N- CA- CB- HB3
6119 6121 6123 6126 3 0.65084 1.95253 0.00000
-2.60338 0.00000 0.00000 ; N- CA- CB- CG
6119 6121 6139 6140 3 0.00000 0.00000 0.00000
0.00000 0.00000 0.00000 ; N- CA- C- O
6119 6121 6139 6141 3 17.40544 -5.02080 -13.22144
9.20480 0.00000 0.00000 ; N- CA- C- N
6120 6119 6121 6122 3 0.00000 0.00000 0.00000
0.00000 0.00000 0.00000 ; H- N- CA- HA
6120 6119 6121 6123 3 0.00000 0.00000 0.00000
0.00000 0.00000 0.00000 ; H- N- CA- CB
6120 6119 6121 6139 3 0.00000 0.00000 0.00000
0.00000 0.00000 0.00000 ; H- N- CA- C
6121 6123 6126 6127 3 0.66944 2.00832 0.00000
-2.67776 0.00000 0.00000 ; CA- CB- CG- HG2
6121 6123 6126 6128 3 0.66944 2.00832 0.00000
-2.67776 0.00000 0.00000 ; CA- CB- CG- HG3
6121 6123 6126 6129 3 3.68192 3.09616 -2.09200
-3.01248 0.00000 0.00000 ; CA- CB- CG- CD
6121 6139 6141 6142 3 20.92000 0.00000 -20.92000
0.00000 0.00000 0.00000 ; CA- C- N- H
6121 6139 6141 6143 3 20.92000 0.00000 -20.92000
0.00000 0.00000 0.00000 ; CA- C- N- CA
6122 6121 6123 6124 3 0.65084 1.95253 0.00000
-2.60338 0.00000 0.00000 ; HA- CA- CB- HB2
6122 6121 6123 6125 3 0.65084 1.95253 0.00000
-2.60338 0.00000 0.00000 ; HA- CA- CB- HB3
6122 6121 6123 6126 3 0.65084 1.95253 0.00000
-2.60338 0.00000 0.00000 ; HA- CA- CB- CG
6122 6121 6139 6140 3 3.68192 -4.35136 0.00000
1.33888 0.00000 0.00000 ; HA- CA- C- O
6122 6121 6139 6141 3 0.00000 0.00000 0.00000
0.00000 0.00000 0.00000 ; HA- CA- C- N
6123 6121 6139 6140 3 0.00000 0.00000 0.00000

```
0.00000    0.00000    0.00000 ;      CB-    CA-     C-     O
  6123   6121   6139   6141      3    2.51040    4.18400    1.67360
-6.69440    0.00000    0.00000 ;      CB-    CA-     C-     N
  6123   6126   6129   6130      3    0.66944    2.00832    0.00000
-2.67776    0.00000    0.00000 ;      CB-    CG-     CD-   HD2
  6123   6126   6129   6131      3    0.66944    2.00832    0.00000
-2.67776    0.00000    0.00000 ;      CB-    CG-     CD-   HD3
  6123   6126   6129   6132      3    3.68192    3.09616   -2.09200
-3.01248    0.00000    0.00000 ;      CB-    CG-     CD-    CE
  6124   6123   6121   6139      3    0.65084    1.95253    0.00000
-2.60338    0.00000    0.00000 ;     HB2-    CB-     CA-     C
  6124   6123   6126   6127      3    0.62760    1.88280    0.00000
-2.51040    0.00000    0.00000 ;     HB2-    CB-     CG-   HG2
  6124   6123   6126   6128      3    0.62760    1.88280    0.00000
-2.51040    0.00000    0.00000 ;     HB2-    CB-     CG-   HG3
  6124   6123   6126   6129      3    0.66944    2.00832    0.00000
-2.67776    0.00000    0.00000 ;     HB2-    CB-     CG-    CD
  6125   6123   6121   6139      3    0.65084    1.95253    0.00000
-2.60338    0.00000    0.00000 ;     HB3-    CB-     CA-     C
  6125   6123   6126   6127      3    0.62760    1.88280    0.00000
-2.51040    0.00000    0.00000 ;     HB3-    CB-     CG-   HG2
  6125   6123   6126   6128      3    0.62760    1.88280    0.00000
-2.51040    0.00000    0.00000 ;     HB3-    CB-     CG-   HG3
  6125   6123   6126   6129      3    0.66944    2.00832    0.00000
-2.67776    0.00000    0.00000 ;     HB3-    CB-     CG-    CD
  6126   6123   6121   6139      3    0.65084    1.95253    0.00000
-2.60338    0.00000    0.00000 ;      CG-    CB-     CA-     C
  6126   6129   6132   6133      3    0.65084    1.95253    0.00000
-2.60338    0.00000    0.00000 ;      CG-    CD-     CE-   HE2
  6126   6129   6132   6134      3    0.65084    1.95253    0.00000
-2.60338    0.00000    0.00000 ;      CG-    CD-     CE-   HE3
  6126   6129   6132   6135      3    0.65084    1.95253    0.00000
-2.60338    0.00000    0.00000 ;      CG-    CD-     CE-    NZ
  6127   6126   6129   6130      3    0.62760    1.88280    0.00000
-2.51040    0.00000    0.00000 ;     HG2-    CG-     CD-   HD2
  6127   6126   6129   6131      3    0.62760    1.88280    0.00000
-2.51040    0.00000    0.00000 ;     HG2-    CG-     CD-   HD3
  6127   6126   6129   6132      3    0.66944    2.00832    0.00000
-2.67776    0.00000    0.00000 ;     HG2-    CG-     CD-    CE
  6128   6126   6129   6130      3    0.62760    1.88280    0.00000
-2.51040    0.00000    0.00000 ;     HG3-    CG-     CD-   HD2
  6128   6126   6129   6131      3    0.62760    1.88280    0.00000
-2.51040    0.00000    0.00000 ;     HG3-    CG-     CD-   HD3
  6128   6126   6129   6132      3    0.66944    2.00832    0.00000
-2.67776    0.00000    0.00000 ;     HG3-    CG-     CD-    CE
  6129   6132   6135   6136      3    0.65084    1.95253    0.00000
-2.60338    0.00000    0.00000 ;      CD-    CE-     NZ-   HZ1
  6129   6132   6135   6137      3    0.65084    1.95253    0.00000
-2.60338    0.00000    0.00000 ;      CD-    CE-     NZ-   HZ2
  6129   6132   6135   6138      3    0.65084    1.95253    0.00000
-2.60338    0.00000    0.00000 ;      CD-    CE-     NZ-   HZ3
  6130   6129   6132   6133      3    0.65084    1.95253    0.00000
```

-2.60338 0.00000 0.00000 ; HD2- CD- CE- HE2
6130 6129 6132 6134 3 0.65084 1.95253 0.00000
-2.60338 0.00000 0.00000 ; HD2- CD- CE- HE3
6130 6129 6132 6135 3 0.65084 1.95253 0.00000
-2.60338 0.00000 0.00000 ; HD2- CD- CE- NZ
6131 6129 6132 6133 3 0.65084 1.95253 0.00000
-2.60338 0.00000 0.00000 ; HD3- CD- CE- HE2
6131 6129 6132 6134 3 0.65084 1.95253 0.00000
-2.60338 0.00000 0.00000 ; HD3- CD- CE- HE3
6131 6129 6132 6135 3 0.65084 1.95253 0.00000
-2.60338 0.00000 0.00000 ; HD3- CD- CE- NZ
6133 6132 6135 6136 3 0.65084 1.95253 0.00000
-2.60338 0.00000 0.00000 ; HE2- CE- NZ- HZ1
6133 6132 6135 6137 3 0.65084 1.95253 0.00000
-2.60338 0.00000 0.00000 ; HE2- CE- NZ- HZ2
6133 6132 6135 6138 3 0.65084 1.95253 0.00000
-2.60338 0.00000 0.00000 ; HE2- CE- NZ- HZ3
6134 6132 6135 6136 3 0.65084 1.95253 0.00000
-2.60338 0.00000 0.00000 ; HE3- CE- NZ- HZ1
6134 6132 6135 6137 3 0.65084 1.95253 0.00000
-2.60338 0.00000 0.00000 ; HE3- CE- NZ- HZ2
6134 6132 6135 6138 3 0.65084 1.95253 0.00000
-2.60338 0.00000 0.00000 ; HE3- CE- NZ- HZ3
6139 6141 6143 6144 3 0.00000 0.00000 0.00000
0.00000 0.00000 0.00000 ; C- N- CA- HA
6139 6141 6143 6145 3 11.71520 1.67360 15.06240 -
13.38880 0.00000 0.00000 ; C- N- CA- CB
6139 6141 6143 6156 3 1.75728 5.27184 2.25936
-7.02912 0.00000 0.00000 ; C- N- CA- C
6140 6139 6141 6142 3 29.28800 -8.36800 -20.92000
0.00000 0.00000 0.00000 ; O- C- N- H
6140 6139 6141 6143 3 20.92000 0.00000 -20.92000
0.00000 0.00000 0.00000 ; O- C- N- CA
6141 6143 6145 6146 3 0.65084 1.95253 0.00000
-2.60338 0.00000 0.00000 ; N- CA- CB- HB2
6141 6143 6145 6147 3 0.65084 1.95253 0.00000
-2.60338 0.00000 0.00000 ; N- CA- CB- HB3
6141 6143 6145 6148 3 3.21331 2.22589 -4.15053
-2.40998 2.61082 0.00000 ; N- CA- CB- CG
6141 6143 6156 6157 3 0.00000 0.00000 0.00000
0.00000 0.00000 0.00000 ; N- CA- C- O
6141 6143 6156 6158 3 17.40544 -5.02080 -13.22144
9.20480 0.00000 0.00000 ; N- CA- C- N
6142 6141 6143 6144 3 0.00000 0.00000 0.00000
0.00000 0.00000 0.00000 ; H- N- CA- HA
6142 6141 6143 6145 3 0.00000 0.00000 0.00000
0.00000 0.00000 0.00000 ; H- N- CA- CB
6142 6141 6143 6156 3 0.00000 0.00000 0.00000
0.00000 0.00000 0.00000 ; H- N- CA- C
6143 6145 6148 6149 3 0.66944 2.00832 0.00000
-2.67776 0.00000 0.00000 ; CA- CB- CG- HG2
6143 6145 6148 6150 3 0.66944 2.00832 0.00000

-2.67776    0.00000    0.00000 ;    CA-    CB-    CG-   HG3
  6143   6145   6148   6151      3    3.69866   -4.35136   -3.92459
6.89523    4.61914    0.00000 ;    CA-    CB-    CG-    CD
  6143   6156   6158   6159      3   20.92000    0.00000  -20.92000
0.00000    0.00000    0.00000 ;    CA-     C-     N-     H
  6143   6156   6158   6160      3   20.92000    0.00000  -20.92000
0.00000    0.00000    0.00000 ;    CA-     C-     N-    CA
  6144   6143   6145   6146      3    0.65084    1.95253    0.00000
-2.60338    0.00000    0.00000 ;    HA-    CA-    CB-   HB2
  6144   6143   6145   6147      3    0.65084    1.95253    0.00000
-2.60338    0.00000    0.00000 ;    HA-    CA-    CB-   HB3
  6144   6143   6145   6148      3    0.65084    1.95253    0.00000
-2.60338    0.00000    0.00000 ;    HA-    CA-    CB-    CG
  6144   6143   6156   6157      3    3.68192   -4.35136    0.00000
1.33888    0.00000    0.00000 ;    HA-    CA-     C-     O
  6144   6143   6156   6158      3    0.00000    0.00000    0.00000
0.00000    0.00000    0.00000 ;    HA-    CA-     C-     N
  6145   6143   6156   6157      3    0.00000    0.00000    0.00000
0.00000    0.00000    0.00000 ;    CB-    CA-     C-     O
  6145   6143   6156   6158      3    2.51040    4.18400    1.67360
-6.69440    0.00000    0.00000 ;    CB-    CA-     C-     N
  6145   6148   6151   6152      3    0.00000    0.00000    0.00000
0.00000    0.00000    0.00000 ;    CB-    CG-    CD-   OE1
  6145   6148   6151   6153      3   10.32611    1.48114   -8.47678
1.42256    1.40582    0.00000 ;    CB-    CG-    CD-   NE2
  6146   6145   6143   6156      3    0.65084    1.95253    0.00000
-2.60338    0.00000    0.00000 ;   HB2-    CB-    CA-     C
  6146   6145   6148   6149      3    0.62760    1.88280    0.00000
-2.51040    0.00000    0.00000 ;   HB2-    CB-    CG-   HG2
  6146   6145   6148   6150      3    0.62760    1.88280    0.00000
-2.51040    0.00000    0.00000 ;   HB2-    CB-    CG-   HG3
  6146   6145   6148   6151      3    0.65084    1.95253    0.00000
-2.60338    0.00000    0.00000 ;   HB2-    CB-    CG-    CD
  6147   6145   6143   6156      3    0.65084    1.95253    0.00000
-2.60338    0.00000    0.00000 ;   HB3-    CB-    CA-     C
  6147   6145   6148   6149      3    0.62760    1.88280    0.00000
-2.51040    0.00000    0.00000 ;   HB3-    CB-    CG-   HG2
  6147   6145   6148   6150      3    0.62760    1.88280    0.00000
-2.51040    0.00000    0.00000 ;   HB3-    CB-    CG-   HG3
  6147   6145   6148   6151      3    0.65084    1.95253    0.00000
-2.60338    0.00000    0.00000 ;   HB3-    CB-    CG-    CD
  6148   6145   6143   6156      3    6.86594    3.56895   -8.14206
-2.40998    4.85344    0.00000 ;    CG-    CB-    CA-     C
  6148   6151   6153   6154      3   20.92000    0.00000  -20.92000
0.00000    0.00000    0.00000 ;    CG-    CD-   NE2-  HE21
  6148   6151   6153   6155      3   20.92000    0.00000  -20.92000
0.00000    0.00000    0.00000 ;    CG-    CD-   NE2-  HE22
  6149   6148   6151   6152      3    3.68192   -4.35136    0.00000
1.33888    0.00000    0.00000 ;   HG2-    CG-    CD-   OE1
  6149   6148   6151   6153      3    0.00000    0.00000    0.00000
0.00000    0.00000    0.00000 ;   HG2-    CG-    CD-   NE2
  6150   6148   6151   6152      3    3.68192   -4.35136    0.00000

1.33888 0.00000 0.00000 ; HG3- CG- CD- OE1
6150 6148 6151 6153 3 0.00000 0.00000 0.00000
0.00000 0.00000 0.00000 ; HG3- CG- CD- NE2
6152 6151 6153 6154 3 29.28800 -8.36800 -20.92000
0.00000 0.00000 0.00000 ; OE1- CD- NE2- HE21
6152 6151 6153 6155 3 29.28800 -8.36800 -20.92000
0.00000 0.00000 0.00000 ; OE1- CD- NE2- HE22
6156 6158 6160 6161 3 0.00000 0.00000 0.00000
0.00000 0.00000 0.00000 ; C- N- CA- HA
6156 6158 6160 6162 3 11.71520 1.67360 15.06240 -
13.38880 0.00000 0.00000 ; C- N- CA- CB
6156 6158 6160 6170 3 1.75728 5.27184 2.25936
-7.02912 0.00000 0.00000 ; C- N- CA- C
6157 6156 6158 6159 3 29.28800 -8.36800 -20.92000
0.00000 0.00000 0.00000 ; O- C- N- H
6157 6156 6158 6160 3 20.92000 0.00000 -20.92000
0.00000 0.00000 0.00000 ; O- C- N- CA
6158 6160 6162 6163 3 0.65084 1.95253 0.00000
-2.60338 0.00000 0.00000 ; N- CA- CB- HB
6158 6160 6162 6164 3 3.83673 0.44769 -1.77402
-2.47693 -0.03347 0.00000 ; N- CA- CB- CG2
6158 6160 6162 6168 3 4.93294 1.13386 -3.12963
-5.27184 3.17984 0.00000 ; N- CA- CB- OG1
6158 6160 6170 6171 3 0.00000 0.00000 0.00000
0.00000 0.00000 0.00000 ; N- CA- C- O
6158 6160 6170 6172 3 17.40544 -5.02080 -13.22144
9.20480 0.00000 0.00000 ; N- CA- C- N
6159 6158 6160 6161 3 0.00000 0.00000 0.00000
0.00000 0.00000 0.00000 ; H- N- CA- HA
6159 6158 6160 6162 3 0.00000 0.00000 0.00000
0.00000 0.00000 0.00000 ; H- N- CA- CB
6159 6158 6160 6170 3 0.00000 0.00000 0.00000
0.00000 0.00000 0.00000 ; H- N- CA- C
6160 6162 6164 6165 3 0.66944 2.00832 0.00000
-2.67776 0.00000 0.00000 ; CA- CB- CG2- HG21
6160 6162 6164 6166 3 0.66944 2.00832 0.00000
-2.67776 0.00000 0.00000 ; CA- CB- CG2- HG22
6160 6162 6164 6167 3 0.66944 2.00832 0.00000
-2.67776 0.00000 0.00000 ; CA- CB- CG2- HG23
6160 6162 6168 6169 3 1.12131 2.98738 1.66523
-3.94970 0.43514 0.00000 ; CA- CB- OG1- HG1
6160 6170 6172 6173 3 20.92000 0.00000 -20.92000
0.00000 0.00000 0.00000 ; CA- C- N- H
6160 6170 6172 6174 3 20.92000 0.00000 -20.92000
0.00000 0.00000 0.00000 ; CA- C- N- CA
6161 6160 6162 6163 3 0.65084 1.95253 0.00000
-2.60338 0.00000 0.00000 ; HA- CA- CB- HB
6161 6160 6162 6164 3 0.65084 1.95253 0.00000
-2.60338 0.00000 0.00000 ; HA- CA- CB- CG2
6161 6160 6162 6168 3 1.04600 -1.04600 0.00000
0.00000 0.00000 0.00000 ; HA- CA- CB- OG1
6161 6160 6170 6171 3 3.68192 -4.35136 0.00000

1.33888 0.00000 0.00000 ; HA- CA- C- O
6161 6160 6170 6172 3 0.00000 0.00000 0.00000
0.00000 0.00000 0.00000 ; HA- CA- C- N
6162 6160 6170 6171 3 0.00000 0.00000 0.00000
0.00000 0.00000 0.00000 ; CB- CA- C- O
6162 6160 6170 6172 3 2.51040 4.18400 1.67360
-6.69440 0.00000 0.00000 ; CB- CA- C- N
6163 6162 6160 6170 3 0.65084 1.95253 0.00000
-2.60338 0.00000 0.00000 ; HB- CB- CA- C
6163 6162 6164 6165 3 0.65084 1.95253 0.00000
-2.60338 0.00000 0.00000 ; HB- CB- CG2- HG21
6163 6162 6164 6166 3 0.65084 1.95253 0.00000
-2.60338 0.00000 0.00000 ; HB- CB- CG2- HG22
6163 6162 6164 6167 3 0.65084 1.95253 0.00000
-2.60338 0.00000 0.00000 ; HB- CB- CG2- HG23
6163 6162 6168 6169 3 0.69733 2.09200 0.00000
-2.78933 0.00000 0.00000 ; HB- CB- OG1- HG1
6164 6162 6160 6170 3 5.67350 3.55640 -6.16722
-2.47693 3.74886 0.00000 ; CG2- CB- CA- C
6164 6162 6168 6169 3 4.74047 0.27196 -2.26773
-3.94970 1.60666 0.00000 ; CG2- CB- OG1- HG1
6165 6164 6162 6168 3 1.04600 -1.04600 0.00000
0.00000 0.00000 0.00000 ; HG21- CG2- CB- OG1
6166 6164 6162 6168 3 1.04600 -1.04600 0.00000
0.00000 0.00000 0.00000 ; HG22- CG2- CB- OG1
6167 6164 6162 6168 3 1.04600 -1.04600 0.00000
0.00000 0.00000 0.00000 ; HG23- CG2- CB- OG1
6168 6162 6160 6170 3 6.53541 6.87013 -6.21742
-5.27184 5.22163 0.00000 ; OG1- CB- CA- C
6170 6172 6174 6175 3 0.00000 0.00000 0.00000
0.00000 0.00000 0.00000 ; C- N- CA- HA
6170 6172 6174 6176 3 11.71520 1.67360 15.06240 -
13.38880 0.00000 0.00000 ; C- N- CA- CB
6170 6172 6174 6191 3 1.75728 5.27184 2.25936
-7.02912 0.00000 0.00000 ; C- N- CA- C
6171 6170 6172 6173 3 29.28800 -8.36800 -20.92000
0.00000 0.00000 0.00000 ; O- C- N- H
6171 6170 6172 6174 3 20.92000 0.00000 -20.92000
0.00000 0.00000 0.00000 ; O- C- N- CA
6172 6174 6176 6177 3 0.65084 1.95253 0.00000
-2.60338 0.00000 0.00000 ; N- CA- CB- HB2
6172 6174 6176 6178 3 0.65084 1.95253 0.00000
-2.60338 0.00000 0.00000 ; N- CA- CB- HB3
6172 6174 6176 6179 3 3.28026 2.46019 -2.19242
-3.21331 -0.23430 0.00000 ; N- CA- CB- CG
6172 6174 6191 6192 3 0.00000 0.00000 0.00000
0.00000 0.00000 0.00000 ; N- CA- C- O
6172 6174 6191 6193 3 17.40544 -5.02080 -13.22144
9.20480 0.00000 0.00000 ; N- CA- C- N
6173 6172 6174 6175 3 0.00000 0.00000 0.00000
0.00000 0.00000 0.00000 ; H- N- CA- HA
6173 6172 6174 6176 3 0.00000 0.00000 0.00000

0.00000 0.00000 0.00000 ; H- N- CA- CB
6173 6172 6174 6191 3 0.00000 0.00000 0.00000
0.00000 0.00000 0.00000 ; H- N- CA- C
6174 6176 6179 6180 3 0.57739 0.00000 1.02926
0.00000 -1.60666 0.00000 ; CA- CB- CG- CD1
6174 6176 6179 6189 3 0.57739 0.00000 1.02926
0.00000 -1.60666 0.00000 ; CA- CB- CG- CD2
6174 6191 6193 6194 3 20.92000 0.00000 -20.92000
0.00000 0.00000 0.00000 ; CA- C- N- H
6174 6191 6193 6195 3 20.92000 0.00000 -20.92000
0.00000 0.00000 0.00000 ; CA- C- N- CA
6175 6174 6176 6177 3 0.65084 1.95253 0.00000
-2.60338 0.00000 0.00000 ; HA- CA- CB- HB2
6175 6174 6176 6178 3 0.65084 1.95253 0.00000
-2.60338 0.00000 0.00000 ; HA- CA- CB- HB3
6175 6174 6176 6179 3 0.65084 1.95253 0.00000
-2.60338 0.00000 0.00000 ; HA- CA- CB- CG
6175 6174 6191 6192 3 3.68192 -4.35136 0.00000
1.33888 0.00000 0.00000 ; HA- CA- C- O
6175 6174 6191 6193 3 0.00000 0.00000 0.00000
0.00000 0.00000 0.00000 ; HA- CA- C- N
6176 6174 6191 6192 3 0.00000 0.00000 0.00000
0.00000 0.00000 0.00000 ; CB- CA- C- O
6176 6174 6191 6193 3 2.51040 4.18400 1.67360
-6.69440 0.00000 0.00000 ; CB- CA- C- N
6176 6179 6180 6181 3 30.33400 0.00000 -30.33400
0.00000 0.00000 0.00000 ; CB- CG- CD1- HD1
6176 6179 6180 6182 3 30.33400 0.00000 -30.33400
0.00000 0.00000 0.00000 ; CB- CG- CD1- CE1
6176 6179 6189 6187 3 30.33400 0.00000 -30.33400
0.00000 0.00000 0.00000 ; CB- CG- CD2- CE2
6176 6179 6189 6190 3 30.33400 0.00000 -30.33400
0.00000 0.00000 0.00000 ; CB- CG- CD2- HD2
6177 6176 6174 6191 3 0.65084 1.95253 0.00000
-2.60338 0.00000 0.00000 ; HB2- CB- CA- C
6177 6176 6179 6180 3 0.00000 0.00000 0.00000
0.00000 0.00000 0.00000 ; HB2- CB- CG- CD1
6177 6176 6179 6189 3 0.00000 0.00000 0.00000
0.00000 0.00000 0.00000 ; HB2- CB- CG- CD2
6178 6176 6174 6191 3 0.65084 1.95253 0.00000
-2.60338 0.00000 0.00000 ; HB3- CB- CA- C
6178 6176 6179 6180 3 0.00000 0.00000 0.00000
0.00000 0.00000 0.00000 ; HB3- CB- CG- CD1
6178 6176 6179 6189 3 0.00000 0.00000 0.00000
0.00000 0.00000 0.00000 ; HB3- CB- CG- CD2
6179 6176 6174 6191 3 4.95804 2.17986 -3.52293
-3.21331 -0.40166 0.00000 ; CG- CB- CA- C
6179 6180 6182 6183 3 30.33400 0.00000 -30.33400
0.00000 0.00000 0.00000 ; CG- CD1- CE1- HE1
6179 6180 6182 6184 3 30.33400 0.00000 -30.33400
0.00000 0.00000 0.00000 ; CG- CD1- CE1- CZ
6179 6189 6187 6184 3 30.33400 0.00000 -30.33400

0.00000 0.00000 0.00000 ; CG- CD2- CE2- CZ
6179 6189 6187 6188 3 30.33400 0.00000 -30.33400
0.00000 0.00000 0.00000 ; CG- CD2- CE2- HE2
6180 6179 6189 6187 3 30.33400 0.00000 -30.33400
0.00000 0.00000 0.00000 ; CD1- CG- CD2- CE2
6180 6179 6189 6190 3 30.33400 0.00000 -30.33400
0.00000 0.00000 0.00000 ; CD1- CG- CD2- HD2
6180 6182 6184 6185 3 30.33400 0.00000 -30.33400
0.00000 0.00000 0.00000 ; CD1- CE1- CZ- OH
6180 6182 6184 6187 3 30.33400 0.00000 -30.33400
0.00000 0.00000 0.00000 ; CD1- CE1- CZ- CE2
6181 6180 6179 6189 3 30.33400 0.00000 -30.33400
0.00000 0.00000 0.00000 ; HD1- CD1- CG- CD2
6181 6180 6182 6183 3 30.33400 0.00000 -30.33400
0.00000 0.00000 0.00000 ; HD1- CD1- CE1- HE1
6181 6180 6182 6184 3 30.33400 0.00000 -30.33400
0.00000 0.00000 0.00000 ; HD1- CD1- CE1- CZ
6182 6180 6179 6189 3 30.33400 0.00000 -30.33400
0.00000 0.00000 0.00000 ; CE1- CD1- CG- CD2
6182 6184 6185 6186 3 7.93286 0.00000 -9.56462
0.00000 2.17568 0.00000 ; CE1- CZ- OH- HH
6182 6184 6187 6188 3 30.33400 0.00000 -30.33400
0.00000 0.00000 0.00000 ; CE1- CZ- CE2- HE2
6182 6184 6187 6189 3 30.33400 0.00000 -30.33400
0.00000 0.00000 0.00000 ; CE1- CZ- CE2- CD2
6183 6182 6184 6185 3 30.33400 0.00000 -30.33400
0.00000 0.00000 0.00000 ; HE1- CE1- CZ- OH
6183 6182 6184 6187 3 30.33400 0.00000 -30.33400
0.00000 0.00000 0.00000 ; HE1- CE1- CZ- CE2
6184 6187 6189 6190 3 30.33400 0.00000 -30.33400
0.00000 0.00000 0.00000 ; CZ- CE2- CD2- HD2
6185 6184 6187 6188 3 30.33400 0.00000 -30.33400
0.00000 0.00000 0.00000 ; OH- CZ- CE2- HE2
6185 6184 6187 6189 3 30.33400 0.00000 -30.33400
0.00000 0.00000 0.00000 ; OH- CZ- CE2- CD2
6186 6185 6184 6187 3 7.93286 0.00000 -9.56462
0.00000 2.17568 0.00000 ; HH- OH- CZ- CE2
6188 6187 6189 6190 3 30.33400 0.00000 -30.33400
0.00000 0.00000 0.00000 ; HE2- CE2- CD2- HD2
6191 6193 6195 6196 3 0.00000 0.00000 0.00000
0.00000 0.00000 0.00000 ; C- N- CA- HA
6191 6193 6195 6197 3 11.71520 1.67360 15.06240 -
13.38880 0.00000 0.00000 ; C- N- CA- CB
6191 6193 6195 6210 3 1.75728 5.27184 2.25936
-7.02912 0.00000 0.00000 ; C- N- CA- C
6192 6191 6193 6194 3 29.28800 -8.36800 -20.92000
0.00000 0.00000 0.00000 ; O- C- N- H
6192 6191 6193 6195 3 20.92000 0.00000 -20.92000
0.00000 0.00000 0.00000 ; O- C- N- CA
6193 6195 6197 6198 3 0.65084 1.95253 0.00000
-2.60338 0.00000 0.00000 ; N- CA- CB- HB2
6193 6195 6197 6199 3 0.65084 1.95253 0.00000

```
-2.60338    0.00000    0.00000 ;       N-    CA-    CB-   HB3
  6193   6195   6197   6200      3    3.79070    1.39746   -4.61077
-2.40998    2.44346    0.00000 ;       N-    CA-    CB-    CG
  6193   6195   6210   6211      3    0.00000    0.00000    0.00000
0.00000    0.00000    0.00000 ;       N-    CA-     C-     O
  6193   6195   6210   6212      3   17.40544   -5.02080  -13.22144
9.20480    0.00000    0.00000 ;       N-    CA-     C-     N
  6194   6193   6195   6196      3    0.00000    0.00000    0.00000
0.00000    0.00000    0.00000 ;       H-     N-    CA-    HA
  6194   6193   6195   6197      3    0.00000    0.00000    0.00000
0.00000    0.00000    0.00000 ;       H-     N-    CA-    CB
  6194   6193   6195   6210      3    0.00000    0.00000    0.00000
0.00000    0.00000    0.00000 ;       H-     N-    CA-     C
  6195   6197   6200   6201      3    0.66944    2.00832    0.00000
-2.67776    0.00000    0.00000 ;      CA-    CB-    CG-    HG
  6195   6197   6200   6202      3    3.90367    0.19665   -6.21742
-2.37651    5.99149    0.00000 ;      CA-    CB-    CG-   CD1
  6195   6197   6200   6206      3    3.90367    0.19665   -6.21742
-2.37651    5.99149    0.00000 ;      CA-    CB-    CG-   CD2
  6195   6210   6212   6213      3   20.92000    0.00000  -20.92000
0.00000    0.00000    0.00000 ;      CA-     C-     N-     H
  6195   6210   6212   6214      3   20.92000    0.00000  -20.92000
0.00000    0.00000    0.00000 ;      CA-     C-     N-    CA
  6196   6195   6197   6198      3    0.65084    1.95253    0.00000
-2.60338    0.00000    0.00000 ;      HA-    CA-    CB-   HB2
  6196   6195   6197   6199      3    0.65084    1.95253    0.00000
-2.60338    0.00000    0.00000 ;      HA-    CA-    CB-   HB3
  6196   6195   6197   6200      3    0.65084    1.95253    0.00000
-2.60338    0.00000    0.00000 ;      HA-    CA-    CB-    CG
  6196   6195   6210   6211      3    3.68192   -4.35136    0.00000
1.33888    0.00000    0.00000 ;      HA-    CA-     C-     O
  6196   6195   6210   6212      3    0.00000    0.00000    0.00000
0.00000    0.00000    0.00000 ;      HA-    CA-     C-     N
  6197   6195   6210   6211      3    0.00000    0.00000    0.00000
0.00000    0.00000    0.00000 ;      CB-    CA-     C-     O
  6197   6195   6210   6212      3    2.51040    4.18400    1.67360
-6.69440    0.00000    0.00000 ;      CB-    CA-     C-     N
  6197   6200   6202   6203      3    0.66944    2.00832    0.00000
-2.67776    0.00000    0.00000 ;      CB-    CG-   CD1-  HD11
  6197   6200   6202   6204      3    0.66944    2.00832    0.00000
-2.67776    0.00000    0.00000 ;      CB-    CG-   CD1-  HD12
  6197   6200   6202   6205      3    0.66944    2.00832    0.00000
-2.67776    0.00000    0.00000 ;      CB-    CG-   CD1-  HD13
  6197   6200   6206   6207      3    0.66944    2.00832    0.00000
-2.67776    0.00000    0.00000 ;      CB-    CG-   CD2-  HD21
  6197   6200   6206   6208      3    0.66944    2.00832    0.00000
-2.67776    0.00000    0.00000 ;      CB-    CG-   CD2-  HD22
  6197   6200   6206   6209      3    0.66944    2.00832    0.00000
-2.67776    0.00000    0.00000 ;      CB-    CG-   CD2-  HD23
  6198   6197   6195   6210      3    0.65084    1.95253    0.00000
-2.60338    0.00000    0.00000 ;     HB2-    CB-    CA-     C
  6198   6197   6200   6201      3    0.62760    1.88280    0.00000
```

```
-2.51040    0.00000    0.00000 ;    HB2-    CB-    CG-    HG
  6198   6197   6200   6202      3    0.66944    2.00832    0.00000
-2.67776    0.00000    0.00000 ;    HB2-    CB-    CG-   CD1
  6198   6197   6200   6206      3    0.66944    2.00832    0.00000
-2.67776    0.00000    0.00000 ;    HB2-    CB-    CG-   CD2
  6199   6197   6195   6210      3    0.65084    1.95253    0.00000
-2.60338    0.00000    0.00000 ;    HB3-    CB-    CA-     C
  6199   6197   6200   6201      3    0.62760    1.88280    0.00000
-2.51040    0.00000    0.00000 ;    HB3-    CB-    CG-    HG
  6199   6197   6200   6202      3    0.66944    2.00832    0.00000
-2.67776    0.00000    0.00000 ;    HB3-    CB-    CG-   CD1
  6199   6197   6200   6206      3    0.66944    2.00832    0.00000
-2.67776    0.00000    0.00000 ;    HB3-    CB-    CG-   CD2
  6200   6197   6195   6210      3   10.33448   -1.14642  -11.54784
-2.40998    6.35968    0.00000 ;     CG-    CB-    CA-     C
  6201   6200   6202   6203      3    0.62760    1.88280    0.00000
-2.51040    0.00000    0.00000 ;     HG-    CG-   CD1-  HD11
  6201   6200   6202   6204      3    0.62760    1.88280    0.00000
-2.51040    0.00000    0.00000 ;     HG-    CG-   CD1-  HD12
  6201   6200   6202   6205      3    0.62760    1.88280    0.00000
-2.51040    0.00000    0.00000 ;     HG-    CG-   CD1-  HD13
  6201   6200   6206   6207      3    0.62760    1.88280    0.00000
-2.51040    0.00000    0.00000 ;     HG-    CG-   CD2-  HD21
  6201   6200   6206   6208      3    0.62760    1.88280    0.00000
-2.51040    0.00000    0.00000 ;     HG-    CG-   CD2-  HD22
  6201   6200   6206   6209      3    0.62760    1.88280    0.00000
-2.51040    0.00000    0.00000 ;     HG-    CG-   CD2-  HD23
  6202   6200   6206   6207      3    0.66944    2.00832    0.00000
-2.67776    0.00000    0.00000 ;    CD1-    CG-   CD2-  HD21
  6202   6200   6206   6208      3    0.66944    2.00832    0.00000
-2.67776    0.00000    0.00000 ;    CD1-    CG-   CD2-  HD22
  6202   6200   6206   6209      3    0.66944    2.00832    0.00000
-2.67776    0.00000    0.00000 ;    CD1-    CG-   CD2-  HD23
  6203   6202   6200   6206      3    0.66944    2.00832    0.00000
-2.67776    0.00000    0.00000 ;   HD11-   CD1-    CG-   CD2
  6204   6202   6200   6206      3    0.66944    2.00832    0.00000
-2.67776    0.00000    0.00000 ;   HD12-   CD1-    CG-   CD2
  6205   6202   6200   6206      3    0.66944    2.00832    0.00000
-2.67776    0.00000    0.00000 ;   HD13-   CD1-    CG-   CD2
  6210   6212   6214   6215      3    0.00000    0.00000    0.00000
0.00000    0.00000    0.00000 ;     C-     N-    CA-    HA
  6210   6212   6214   6216      3   11.71520    1.67360   15.06240  -
13.38880    0.00000    0.00000 ;     C-     N-    CA-    CB
  6210   6212   6214   6232      3    1.75728    5.27184    2.25936
-7.02912    0.00000    0.00000 ;     C-     N-    CA-     C
  6211   6210   6212   6213      3   29.28800   -8.36800  -20.92000
0.00000    0.00000    0.00000 ;     O-     C-     N-     H
  6211   6210   6212   6214      3   20.92000    0.00000  -20.92000
0.00000    0.00000    0.00000 ;     O-     C-     N-    CA
  6212   6214   6216   6217      3    0.65084    1.95253    0.00000
-2.60338    0.00000    0.00000 ;     N-    CA-    CB-   HB2
  6212   6214   6216   6218      3    0.65084    1.95253    0.00000
```

```
-2.60338    0.00000    0.00000 ;      N-    CA-    CB-   HB3
  6212   6214   6216   6219      3    0.65084    1.95253    0.00000
-2.60338    0.00000    0.00000 ;      N-    CA-    CB-    CG
  6212   6214   6232   6233      3    0.00000    0.00000    0.00000
0.00000    0.00000    0.00000 ;      N-    CA-     C-     O
  6212   6214   6232   6234      3   17.40544   -5.02080  -13.22144
9.20480    0.00000    0.00000 ;      N-    CA-     C-     N
  6213   6212   6214   6215      3    0.00000    0.00000    0.00000
0.00000    0.00000    0.00000 ;      H-     N-    CA-    HA
  6213   6212   6214   6216      3    0.00000    0.00000    0.00000
0.00000    0.00000    0.00000 ;      H-     N-    CA-    CB
  6213   6212   6214   6232      3    0.00000    0.00000    0.00000
0.00000    0.00000    0.00000 ;      H-     N-    CA-     C
  6214   6216   6219   6220      3    0.66944    2.00832    0.00000
-2.67776    0.00000    0.00000 ;     CA-    CB-    CG-   HG2
  6214   6216   6219   6221      3    0.66944    2.00832    0.00000
-2.67776    0.00000    0.00000 ;     CA-    CB-    CG-   HG3
  6214   6216   6219   6222      3    3.68192    3.09616   -2.09200
-3.01248    0.00000    0.00000 ;     CA-    CB-    CG-    CD
  6214   6232   6234   6235      3   20.92000    0.00000  -20.92000
0.00000    0.00000    0.00000 ;     CA-     C-     N-     H
  6214   6232   6234   6236      3   20.92000    0.00000  -20.92000
0.00000    0.00000    0.00000 ;     CA-     C-     N-    CA
  6215   6214   6216   6217      3    0.65084    1.95253    0.00000
-2.60338    0.00000    0.00000 ;     HA-    CA-    CB-   HB2
  6215   6214   6216   6218      3    0.65084    1.95253    0.00000
-2.60338    0.00000    0.00000 ;     HA-    CA-    CB-   HB3
  6215   6214   6216   6219      3    0.65084    1.95253    0.00000
-2.60338    0.00000    0.00000 ;     HA-    CA-    CB-    CG
  6215   6214   6232   6233      3    3.68192   -4.35136    0.00000
1.33888    0.00000    0.00000 ;     HA-    CA-     C-     O
  6215   6214   6232   6234      3    0.00000    0.00000    0.00000
0.00000    0.00000    0.00000 ;     HA-    CA-     C-     N
  6216   6214   6232   6233      3    0.00000    0.00000    0.00000
0.00000    0.00000    0.00000 ;     CB-    CA-     C-     O
  6216   6214   6232   6234      3    2.51040    4.18400    1.67360
-6.69440    0.00000    0.00000 ;     CB-    CA-     C-     N
  6216   6219   6222   6223      3    0.66944    2.00832    0.00000
-2.67776    0.00000    0.00000 ;     CB-    CG-    CD-   HD2
  6216   6219   6222   6224      3    0.66944    2.00832    0.00000
-2.67776    0.00000    0.00000 ;     CB-    CG-    CD-   HD3
  6216   6219   6222   6225      3    3.68192    3.09616   -2.09200
-3.01248    0.00000    0.00000 ;     CB-    CG-    CD-    CE
  6217   6216   6214   6232      3    0.65084    1.95253    0.00000
-2.60338    0.00000    0.00000 ;    HB2-    CB-    CA-     C
  6217   6216   6219   6220      3    0.62760    1.88280    0.00000
-2.51040    0.00000    0.00000 ;    HB2-    CB-    CG-   HG2
  6217   6216   6219   6221      3    0.62760    1.88280    0.00000
-2.51040    0.00000    0.00000 ;    HB2-    CB-    CG-   HG3
  6217   6216   6219   6222      3    0.66944    2.00832    0.00000
-2.67776    0.00000    0.00000 ;    HB2-    CB-    CG-    CD
  6218   6216   6214   6232      3    0.65084    1.95253    0.00000
```

```
-2.60338    0.00000    0.00000 ;    HB3-    CB-    CA-     C
 6218   6216   6219   6220      3    0.62760    1.88280    0.00000
-2.51040    0.00000    0.00000 ;    HB3-    CB-    CG-   HG2
 6218   6216   6219   6221      3    0.62760    1.88280    0.00000
-2.51040    0.00000    0.00000 ;    HB3-    CB-    CG-   HG3
 6218   6216   6219   6222      3    0.66944    2.00832    0.00000
-2.67776    0.00000    0.00000 ;    HB3-    CB-    CG-    CD
 6219   6216   6214   6232      3    0.65084    1.95253    0.00000
-2.60338    0.00000    0.00000 ;     CG-    CB-    CA-     C
 6219   6222   6225   6226      3    0.65084    1.95253    0.00000
-2.60338    0.00000    0.00000 ;     CG-    CD-    CE-   HE2
 6219   6222   6225   6227      3    0.65084    1.95253    0.00000
-2.60338    0.00000    0.00000 ;     CG-    CD-    CE-   HE3
 6219   6222   6225   6228      3    0.65084    1.95253    0.00000
-2.60338    0.00000    0.00000 ;     CG-    CD-    CE-    NZ
 6220   6219   6222   6223      3    0.62760    1.88280    0.00000
-2.51040    0.00000    0.00000 ;    HG2-    CG-    CD-   HD2
 6220   6219   6222   6224      3    0.62760    1.88280    0.00000
-2.51040    0.00000    0.00000 ;    HG2-    CG-    CD-   HD3
 6220   6219   6222   6225      3    0.66944    2.00832    0.00000
-2.67776    0.00000    0.00000 ;    HG2-    CG-    CD-    CE
 6221   6219   6222   6223      3    0.62760    1.88280    0.00000
-2.51040    0.00000    0.00000 ;    HG3-    CG-    CD-   HD2
 6221   6219   6222   6224      3    0.62760    1.88280    0.00000
-2.51040    0.00000    0.00000 ;    HG3-    CG-    CD-   HD3
 6221   6219   6222   6225      3    0.66944    2.00832    0.00000
-2.67776    0.00000    0.00000 ;    HG3-    CG-    CD-    CE
 6222   6225   6228   6229      3    0.65084    1.95253    0.00000
-2.60338    0.00000    0.00000 ;     CD-    CE-    NZ-   HZ1
 6222   6225   6228   6230      3    0.65084    1.95253    0.00000
-2.60338    0.00000    0.00000 ;     CD-    CE-    NZ-   HZ2
 6222   6225   6228   6231      3    0.65084    1.95253    0.00000
-2.60338    0.00000    0.00000 ;     CD-    CE-    NZ-   HZ3
 6223   6222   6225   6226      3    0.65084    1.95253    0.00000
-2.60338    0.00000    0.00000 ;    HD2-    CD-    CE-   HE2
 6223   6222   6225   6227      3    0.65084    1.95253    0.00000
-2.60338    0.00000    0.00000 ;    HD2-    CD-    CE-   HE3
 6223   6222   6225   6228      3    0.65084    1.95253    0.00000
-2.60338    0.00000    0.00000 ;    HD2-    CD-    CE-    NZ
 6224   6222   6225   6226      3    0.65084    1.95253    0.00000
-2.60338    0.00000    0.00000 ;    HD3-    CD-    CE-   HE2
 6224   6222   6225   6227      3    0.65084    1.95253    0.00000
-2.60338    0.00000    0.00000 ;    HD3-    CD-    CE-   HE3
 6224   6222   6225   6228      3    0.65084    1.95253    0.00000
-2.60338    0.00000    0.00000 ;    HD3-    CD-    CE-    NZ
 6226   6225   6228   6229      3    0.65084    1.95253    0.00000
-2.60338    0.00000    0.00000 ;    HE2-    CE-    NZ-   HZ1
 6226   6225   6228   6230      3    0.65084    1.95253    0.00000
-2.60338    0.00000    0.00000 ;    HE2-    CE-    NZ-   HZ2
 6226   6225   6228   6231      3    0.65084    1.95253    0.00000
-2.60338    0.00000    0.00000 ;    HE2-    CE-    NZ-   HZ3
 6227   6225   6228   6229      3    0.65084    1.95253    0.00000
```

-2.60338 0.00000 0.00000 ; HE3- CE- NZ- HZ1
6227 6225 6228 6230 3 0.65084 1.95253 0.00000
-2.60338 0.00000 0.00000 ; HE3- CE- NZ- HZ2
6227 6225 6228 6231 3 0.65084 1.95253 0.00000
-2.60338 0.00000 0.00000 ; HE3- CE- NZ- HZ3
6232 6234 6236 6237 3 0.00000 0.00000 0.00000
0.00000 0.00000 0.00000 ; C- N- CA- HA
6232 6234 6236 6238 3 11.71520 1.67360 15.06240 -
13.38880 0.00000 0.00000 ; C- N- CA- CB
6232 6234 6236 6254 3 1.75728 5.27184 2.25936
-7.02912 0.00000 0.00000 ; C- N- CA- C
6233 6232 6234 6235 3 29.28800 -8.36800 -20.92000
0.00000 0.00000 0.00000 ; O- C- N- H
6233 6232 6234 6236 3 20.92000 0.00000 -20.92000
0.00000 0.00000 0.00000 ; O- C- N- CA
6234 6236 6238 6239 3 0.65084 1.95253 0.00000
-2.60338 0.00000 0.00000 ; N- CA- CB- HB2
6234 6236 6238 6240 3 0.65084 1.95253 0.00000
-2.60338 0.00000 0.00000 ; N- CA- CB- HB3
6234 6236 6238 6241 3 0.65084 1.95253 0.00000
-2.60338 0.00000 0.00000 ; N- CA- CB- CG
6234 6236 6254 6255 3 0.00000 0.00000 0.00000
0.00000 0.00000 0.00000 ; N- CA- C- O
6234 6236 6254 6256 3 17.40544 -5.02080 -13.22144
9.20480 0.00000 0.00000 ; N- CA- C- N
6235 6234 6236 6237 3 0.00000 0.00000 0.00000
0.00000 0.00000 0.00000 ; H- N- CA- HA
6235 6234 6236 6238 3 0.00000 0.00000 0.00000
0.00000 0.00000 0.00000 ; H- N- CA- CB
6235 6234 6236 6254 3 0.00000 0.00000 0.00000
0.00000 0.00000 0.00000 ; H- N- CA- C
6236 6238 6241 6242 3 0.66944 2.00832 0.00000
-2.67776 0.00000 0.00000 ; CA- CB- CG- HG2
6236 6238 6241 6243 3 0.66944 2.00832 0.00000
-2.67776 0.00000 0.00000 ; CA- CB- CG- HG3
6236 6238 6241 6244 3 3.68192 3.09616 -2.09200
-3.01248 0.00000 0.00000 ; CA- CB- CG- CD
6236 6254 6256 6257 3 20.92000 0.00000 -20.92000
0.00000 0.00000 0.00000 ; CA- C- N- H
6236 6254 6256 6258 3 20.92000 0.00000 -20.92000
0.00000 0.00000 0.00000 ; CA- C- N- CA
6237 6236 6238 6239 3 0.65084 1.95253 0.00000
-2.60338 0.00000 0.00000 ; HA- CA- CB- HB2
6237 6236 6238 6240 3 0.65084 1.95253 0.00000
-2.60338 0.00000 0.00000 ; HA- CA- CB- HB3
6237 6236 6238 6241 3 0.65084 1.95253 0.00000
-2.60338 0.00000 0.00000 ; HA- CA- CB- CG
6237 6236 6254 6255 3 3.68192 -4.35136 0.00000
1.33888 0.00000 0.00000 ; HA- CA- C- O
6237 6236 6254 6256 3 0.00000 0.00000 0.00000
0.00000 0.00000 0.00000 ; HA- CA- C- N
6238 6236 6254 6255 3 0.00000 0.00000 0.00000

```
0.00000    0.00000    0.00000 ;     CB-    CA-     C-     O
  6238   6236   6254   6256      3    2.51040    4.18400    1.67360
-6.69440    0.00000    0.00000 ;     CB-    CA-     C-     N
  6238   6241   6244   6245      3    0.66944    2.00832    0.00000
-2.67776    0.00000    0.00000 ;     CB-    CG-     CD-    HD2
  6238   6241   6244   6246      3    0.66944    2.00832    0.00000
-2.67776    0.00000    0.00000 ;     CB-    CG-     CD-    HD3
  6238   6241   6244   6247      3    3.68192    3.09616   -2.09200
-3.01248    0.00000    0.00000 ;     CB-    CG-     CD-     CE
  6239   6238   6236   6254      3    0.65084    1.95253    0.00000
-2.60338    0.00000    0.00000 ;    HB2-    CB-     CA-     C
  6239   6238   6241   6242      3    0.62760    1.88280    0.00000
-2.51040    0.00000    0.00000 ;    HB2-    CB-     CG-    HG2
  6239   6238   6241   6243      3    0.62760    1.88280    0.00000
-2.51040    0.00000    0.00000 ;    HB2-    CB-     CG-    HG3
  6239   6238   6241   6244      3    0.66944    2.00832    0.00000
-2.67776    0.00000    0.00000 ;    HB2-    CB-     CG-     CD
  6240   6238   6236   6254      3    0.65084    1.95253    0.00000
-2.60338    0.00000    0.00000 ;    HB3-    CB-     CA-     C
  6240   6238   6241   6242      3    0.62760    1.88280    0.00000
-2.51040    0.00000    0.00000 ;    HB3-    CB-     CG-    HG2
  6240   6238   6241   6243      3    0.62760    1.88280    0.00000
-2.51040    0.00000    0.00000 ;    HB3-    CB-     CG-    HG3
  6240   6238   6241   6244      3    0.66944    2.00832    0.00000
-2.67776    0.00000    0.00000 ;    HB3-    CB-     CG-     CD
  6241   6238   6236   6254      3    0.65084    1.95253    0.00000
-2.60338    0.00000    0.00000 ;     CG-    CB-     CA-     C
  6241   6244   6247   6248      3    0.65084    1.95253    0.00000
-2.60338    0.00000    0.00000 ;     CG-    CD-     CE-    HE2
  6241   6244   6247   6249      3    0.65084    1.95253    0.00000
-2.60338    0.00000    0.00000 ;     CG-    CD-     CE-    HE3
  6241   6244   6247   6250      3    0.65084    1.95253    0.00000
-2.60338    0.00000    0.00000 ;     CG-    CD-     CE-     NZ
  6242   6241   6244   6245      3    0.62760    1.88280    0.00000
-2.51040    0.00000    0.00000 ;    HG2-    CG-     CD-    HD2
  6242   6241   6244   6246      3    0.62760    1.88280    0.00000
-2.51040    0.00000    0.00000 ;    HG2-    CG-     CD-    HD3
  6242   6241   6244   6247      3    0.66944    2.00832    0.00000
-2.67776    0.00000    0.00000 ;    HG2-    CG-     CD-     CE
  6243   6241   6244   6245      3    0.62760    1.88280    0.00000
-2.51040    0.00000    0.00000 ;    HG3-    CG-     CD-    HD2
  6243   6241   6244   6246      3    0.62760    1.88280    0.00000
-2.51040    0.00000    0.00000 ;    HG3-    CG-     CD-    HD3
  6243   6241   6244   6247      3    0.66944    2.00832    0.00000
-2.67776    0.00000    0.00000 ;    HG3-    CG-     CD-     CE
  6244   6247   6250   6251      3    0.65084    1.95253    0.00000
-2.60338    0.00000    0.00000 ;     CD-    CE-     NZ-    HZ1
  6244   6247   6250   6252      3    0.65084    1.95253    0.00000
-2.60338    0.00000    0.00000 ;     CD-    CE-     NZ-    HZ2
  6244   6247   6250   6253      3    0.65084    1.95253    0.00000
-2.60338    0.00000    0.00000 ;     CD-    CE-     NZ-    HZ3
  6245   6244   6247   6248      3    0.65084    1.95253    0.00000
```

-2.60338 0.00000 0.00000 ; HD2- CD- CE- HE2
6245 6244 6247 6249 3 0.65084 1.95253 0.00000
-2.60338 0.00000 0.00000 ; HD2- CD- CE- HE3
6245 6244 6247 6250 3 0.65084 1.95253 0.00000
-2.60338 0.00000 0.00000 ; HD2- CD- CE- NZ
6246 6244 6247 6248 3 0.65084 1.95253 0.00000
-2.60338 0.00000 0.00000 ; HD3- CD- CE- HE2
6246 6244 6247 6249 3 0.65084 1.95253 0.00000
-2.60338 0.00000 0.00000 ; HD3- CD- CE- HE3
6246 6244 6247 6250 3 0.65084 1.95253 0.00000
-2.60338 0.00000 0.00000 ; HD3- CD- CE- NZ
6248 6247 6250 6251 3 0.65084 1.95253 0.00000
-2.60338 0.00000 0.00000 ; HE2- CE- NZ- HZ1
6248 6247 6250 6252 3 0.65084 1.95253 0.00000
-2.60338 0.00000 0.00000 ; HE2- CE- NZ- HZ2
6248 6247 6250 6253 3 0.65084 1.95253 0.00000
-2.60338 0.00000 0.00000 ; HE2- CE- NZ- HZ3
6249 6247 6250 6251 3 0.65084 1.95253 0.00000
-2.60338 0.00000 0.00000 ; HE3- CE- NZ- HZ1
6249 6247 6250 6252 3 0.65084 1.95253 0.00000
-2.60338 0.00000 0.00000 ; HE3- CE- NZ- HZ2
6249 6247 6250 6253 3 0.65084 1.95253 0.00000
-2.60338 0.00000 0.00000 ; HE3- CE- NZ- HZ3
6254 6256 6258 6259 3 0.00000 0.00000 0.00000
0.00000 0.00000 0.00000 ; C- N- CA- HA
6254 6256 6258 6260 3 11.71520 1.67360 15.06240 -
13.38880 0.00000 0.00000 ; C- N- CA- CB
6254 6256 6258 6269 3 1.75728 5.27184 2.25936
-7.02912 0.00000 0.00000 ; C- N- CA- C
6255 6254 6256 6257 3 29.28800 -8.36800 -20.92000
0.00000 0.00000 0.00000 ; O- C- N- H
6255 6254 6256 6258 3 20.92000 0.00000 -20.92000
0.00000 0.00000 0.00000 ; O- C- N- CA
6256 6258 6260 6261 3 0.65084 1.95253 0.00000
-2.60338 0.00000 0.00000 ; N- CA- CB- HB2
6256 6258 6260 6262 3 0.65084 1.95253 0.00000
-2.60338 0.00000 0.00000 ; N- CA- CB- HB3
6256 6258 6260 6263 3 3.21331 2.22589 -4.15053
-2.40998 2.61082 0.00000 ; N- CA- CB- CG
6256 6258 6269 6270 3 0.00000 0.00000 0.00000
0.00000 0.00000 0.00000 ; N- CA- C- O
6256 6258 6269 6271 3 17.40544 -5.02080 -13.22144
9.20480 0.00000 0.00000 ; N- CA- C- N
6257 6256 6258 6259 3 0.00000 0.00000 0.00000
0.00000 0.00000 0.00000 ; H- N- CA- HA
6257 6256 6258 6260 3 0.00000 0.00000 0.00000
0.00000 0.00000 0.00000 ; H- N- CA- CB
6257 6256 6258 6269 3 0.00000 0.00000 0.00000
0.00000 0.00000 0.00000 ; H- N- CA- C
6258 6260 6263 6264 3 0.66944 2.00832 0.00000
-2.67776 0.00000 0.00000 ; CA- CB- CG- HG2
6258 6260 6263 6265 3 0.66944 2.00832 0.00000

```
-2.67776    0.00000    0.00000 ;      CA-    CB-    CG-   HG3
  6258   6260   6263   6266       3   10.12110   -1.91209    0.01674
10.17549   -1.87443    0.00000 ;      CA-    CB-    CG-    CD
  6258   6269   6271   6272       3   20.92000    0.00000  -20.92000
0.00000    0.00000    0.00000 ;     CA-     C-     N-     H
  6258   6269   6271   6273       3   20.92000    0.00000  -20.92000
0.00000    0.00000    0.00000 ;     CA-     C-     N-    CA
  6259   6258   6260   6261       3    0.65084    1.95253    0.00000
-2.60338    0.00000    0.00000 ;      HA-    CA-    CB-   HB2
  6259   6258   6260   6262       3    0.65084    1.95253    0.00000
-2.60338    0.00000    0.00000 ;      HA-    CA-    CB-   HB3
  6259   6258   6260   6263       3    0.65084    1.95253    0.00000
-2.60338    0.00000    0.00000 ;      HA-    CA-    CB-    CG
  6259   6258   6269   6270       3    3.68192   -4.35136    0.00000
1.33888    0.00000    0.00000 ;     HA-    CA-     C-     O
  6259   6258   6269   6271       3    0.00000    0.00000    0.00000
0.00000    0.00000    0.00000 ;     HA-    CA-     C-     N
  6260   6258   6269   6270       3    0.00000    0.00000    0.00000
0.00000    0.00000    0.00000 ;     CB-    CA-     C-     O
  6260   6258   6269   6271       3    2.51040    4.18400    1.67360
-6.69440    0.00000    0.00000 ;      CB-    CA-     C-     N
  6260   6263   6266   6267       3    3.79907    0.00000   -5.40573
0.00000    2.14221    0.00000 ;     CB-    CG-    CD-   OE1
  6260   6263   6266   6268       3    3.79907    0.00000   -5.40573
0.00000    2.14221    0.00000 ;     CB-    CG-    CD-   OE2
  6261   6260   6258   6269       3    0.65084    1.95253    0.00000
-2.60338    0.00000    0.00000 ;     HB2-    CB-    CA-     C
  6261   6260   6263   6264       3    0.62760    1.88280    0.00000
-2.51040    0.00000    0.00000 ;     HB2-    CB-    CG-   HG2
  6261   6260   6263   6265       3    0.62760    1.88280    0.00000
-2.51040    0.00000    0.00000 ;     HB2-    CB-    CG-   HG3
  6261   6260   6263   6266       3    0.65084    1.95253    0.00000
-2.60338    0.00000    0.00000 ;     HB2-    CB-    CG-    CD
  6262   6260   6258   6269       3    0.65084    1.95253    0.00000
-2.60338    0.00000    0.00000 ;     HB3-    CB-    CA-     C
  6262   6260   6263   6264       3    0.62760    1.88280    0.00000
-2.51040    0.00000    0.00000 ;     HB3-    CB-    CG-   HG2
  6262   6260   6263   6265       3    0.62760    1.88280    0.00000
-2.51040    0.00000    0.00000 ;     HB3-    CB-    CG-   HG3
  6262   6260   6263   6266       3    0.65084    1.95253    0.00000
-2.60338    0.00000    0.00000 ;     HB3-    CB-    CG-    CD
  6263   6260   6258   6269       3    6.86594    3.56895   -8.14206
-2.40998    4.85344    0.00000 ;      CG-    CB-    CA-     C
  6264   6263   6266   6267       3    0.00000    0.00000    0.00000
0.00000    0.00000    0.00000 ;    HG2-    CG-    CD-   OE1
  6264   6263   6266   6268       3    0.00000    0.00000    0.00000
0.00000    0.00000    0.00000 ;    HG2-    CG-    CD-   OE2
  6265   6263   6266   6267       3    0.00000    0.00000    0.00000
0.00000    0.00000    0.00000 ;    HG3-    CG-    CD-   OE1
  6265   6263   6266   6268       3    0.00000    0.00000    0.00000
0.00000    0.00000    0.00000 ;    HG3-    CG-    CD-   OE2
  6269   6271   6273   6274       3    0.00000    0.00000    0.00000
```

0.00000    0.00000    0.00000 ;       C-     N-     CA-    HA
  6269   6271   6273   6275      3   11.71520    1.67360   15.06240  -
13.38880    0.00000    0.00000 ;       C-     N-     CA-    CB
  6269   6271   6273   6285      3    1.75728    5.27184    2.25936
-7.02912    0.00000    0.00000 ;       C-     N-     CA-     C
  6270   6269   6271   6272      3   29.28800   -8.36800  -20.92000
0.00000    0.00000    0.00000 ;       O-     C-     N-     H
  6270   6269   6271   6273      3   20.92000    0.00000  -20.92000
0.00000    0.00000    0.00000 ;       O-     C-     N-    CA
  6271   6273   6275   6276      3    0.65084    1.95253    0.00000
-2.60338    0.00000    0.00000 ;       N-    CA-    CB-    HB
  6271   6273   6275   6277      3    3.83673    0.44769   -1.77402
-2.47693   -0.03347    0.00000 ;       N-    CA-    CB-   CG1
  6271   6273   6275   6281      3    3.83673    0.44769   -1.77402
-2.47693   -0.03347    0.00000 ;       N-    CA-    CB-   CG2
  6271   6273   6285   6286      3    0.00000    0.00000    0.00000
0.00000    0.00000    0.00000 ;       N-    CA-     C-     O
  6271   6273   6285   6287      3   17.40544   -5.02080  -13.22144
9.20480    0.00000    0.00000 ;       N-    CA-     C-     N
  6272   6271   6273   6274      3    0.00000    0.00000    0.00000
0.00000    0.00000    0.00000 ;       H-     N-    CA-    HA
  6272   6271   6273   6275      3    0.00000    0.00000    0.00000
0.00000    0.00000    0.00000 ;       H-     N-    CA-    CB
  6272   6271   6273   6285      3    0.00000    0.00000    0.00000
0.00000    0.00000    0.00000 ;       H-     N-    CA-     C
  6273   6275   6277   6278      3    0.66944    2.00832    0.00000
-2.67776    0.00000    0.00000 ;      CA-    CB-   CG1-  HG11
  6273   6275   6277   6279      3    0.66944    2.00832    0.00000
-2.67776    0.00000    0.00000 ;      CA-    CB-   CG1-  HG12
  6273   6275   6277   6280      3    0.66944    2.00832    0.00000
-2.67776    0.00000    0.00000 ;      CA-    CB-   CG1-  HG13
  6273   6275   6281   6282      3    0.66944    2.00832    0.00000
-2.67776    0.00000    0.00000 ;      CA-    CB-   CG2-  HG21
  6273   6275   6281   6283      3    0.66944    2.00832    0.00000
-2.67776    0.00000    0.00000 ;      CA-    CB-   CG2-  HG22
  6273   6275   6281   6284      3    0.66944    2.00832    0.00000
-2.67776    0.00000    0.00000 ;      CA-    CB-   CG2-  HG23
  6273   6285   6287   6288      3   20.92000    0.00000  -20.92000
0.00000    0.00000    0.00000 ;      CA-     C-     N-     H
  6273   6285   6287   6289      3   20.92000    0.00000  -20.92000
0.00000    0.00000    0.00000 ;      CA-     C-     N-    CA
  6274   6273   6275   6276      3    0.65084    1.95253    0.00000
-2.60338    0.00000    0.00000 ;      HA-    CA-    CB-    HB
  6274   6273   6275   6277      3    0.65084    1.95253    0.00000
-2.60338    0.00000    0.00000 ;      HA-    CA-    CB-   CG1
  6274   6273   6275   6281      3    0.65084    1.95253    0.00000
-2.60338    0.00000    0.00000 ;      HA-    CA-    CB-   CG2
  6274   6273   6285   6286      3    3.68192   -4.35136    0.00000
1.33888    0.00000    0.00000 ;      HA-    CA-     C-     O
  6274   6273   6285   6287      3    0.00000    0.00000    0.00000
0.00000    0.00000    0.00000 ;      HA-    CA-     C-     N
  6275   6273   6285   6286      3    0.00000    0.00000    0.00000

0.00000 0.00000 0.00000 ; CB- CA- C- O
6275 6273 6285 6287 3 2.51040 4.18400 1.67360
-6.69440 0.00000 0.00000 ; CB- CA- C- N
6276 6275 6273 6285 3 0.65084 1.95253 0.00000
-2.60338 0.00000 0.00000 ; HB- CB- CA- C
6276 6275 6277 6278 3 0.62760 1.88280 0.00000
-2.51040 0.00000 0.00000 ; HB- CB- CG1- HG11
6276 6275 6277 6279 3 0.62760 1.88280 0.00000
-2.51040 0.00000 0.00000 ; HB- CB- CG1- HG12
6276 6275 6277 6280 3 0.62760 1.88280 0.00000
-2.51040 0.00000 0.00000 ; HB- CB- CG1- HG13
6276 6275 6281 6282 3 0.62760 1.88280 0.00000
-2.51040 0.00000 0.00000 ; HB- CB- CG2- HG21
6276 6275 6281 6283 3 0.62760 1.88280 0.00000
-2.51040 0.00000 0.00000 ; HB- CB- CG2- HG22
6276 6275 6281 6284 3 0.62760 1.88280 0.00000
-2.51040 0.00000 0.00000 ; HB- CB- CG2- HG23
6277 6275 6273 6285 3 5.67350 3.55640 -6.16722
-2.47693 3.74886 0.00000 ; CG1- CB- CA- C
6277 6275 6281 6282 3 0.66944 2.00832 0.00000
-2.67776 0.00000 0.00000 ; CG1- CB- CG2- HG21
6277 6275 6281 6283 3 0.66944 2.00832 0.00000
-2.67776 0.00000 0.00000 ; CG1- CB- CG2- HG22
6277 6275 6281 6284 3 0.66944 2.00832 0.00000
-2.67776 0.00000 0.00000 ; CG1- CB- CG2- HG23
6278 6277 6275 6281 3 0.66944 2.00832 0.00000
-2.67776 0.00000 0.00000 ; HG11- CG1- CB- CG2
6279 6277 6275 6281 3 0.66944 2.00832 0.00000
-2.67776 0.00000 0.00000 ; HG12- CG1- CB- CG2
6280 6277 6275 6281 3 0.66944 2.00832 0.00000
-2.67776 0.00000 0.00000 ; HG13- CG1- CB- CG2
6281 6275 6273 6285 3 5.67350 3.55640 -6.16722
-2.47693 3.74886 0.00000 ; CG2- CB- CA- C
6285 6287 6289 6290 3 0.00000 0.00000 0.00000
0.00000 0.00000 0.00000 ; C- N- CA- HA
6285 6287 6289 6291 3 11.71520 1.67360 15.06240 -
13.38880 0.00000 0.00000 ; C- N- CA- CB
6285 6287 6289 6296 3 1.75728 5.27184 2.25936
-7.02912 0.00000 0.00000 ; C- N- CA- C
6286 6285 6287 6288 3 29.28800 -8.36800 -20.92000
0.00000 0.00000 0.00000 ; O- C- N- H
6286 6285 6287 6289 3 20.92000 0.00000 -20.92000
0.00000 0.00000 0.00000 ; O- C- N- CA
6287 6289 6291 6292 3 0.65084 1.95253 0.00000
-2.60338 0.00000 0.00000 ; N- CA- CB- HB2
6287 6289 6291 6293 3 0.65084 1.95253 0.00000
-2.60338 0.00000 0.00000 ; N- CA- CB- HB3
6287 6289 6291 6294 3 6.03751 2.50622 -5.17142
-4.20074 1.10458 0.00000 ; N- CA- CB- SG
6287 6289 6296 6297 3 0.00000 0.00000 0.00000
0.00000 0.00000 0.00000 ; N- CA- C- O
6287 6289 6296 6298 3 17.40544 -5.02080 -13.22144

```
9.20480    0.00000    0.00000 ;       N-    CA-     C-     N
  6288   6287   6289   6290       3    0.00000    0.00000    0.00000
0.00000    0.00000    0.00000 ;       H-     N-    CA-    HA
  6288   6287   6289   6291       3    0.00000    0.00000    0.00000
0.00000    0.00000    0.00000 ;       H-     N-    CA-    CB
  6288   6287   6289   6296       3    0.00000    0.00000    0.00000
0.00000    0.00000    0.00000 ;       H-     N-    CA-     C
  6289   6291   6294   6295       3    1.69034    2.77818    4.11706
-4.21747    1.00416    0.00000 ;      CA-    CB-    SG-    HG
  6289   6296   6298   6299       3   20.92000    0.00000  -20.92000
0.00000    0.00000    0.00000 ;      CA-     C-     N-     H
  6289   6296   6298   6300       3   20.92000    0.00000  -20.92000
0.00000    0.00000    0.00000 ;      CA-     C-     N-    CA
  6290   6289   6291   6292       3    0.65084    1.95253    0.00000
-2.60338    0.00000    0.00000 ;      HA-    CA-    CB-   HB2
  6290   6289   6291   6293       3    0.65084    1.95253    0.00000
-2.60338    0.00000    0.00000 ;      HA-    CA-    CB-   HB3
  6290   6289   6291   6294       3    0.65084    1.95253    0.00000
-2.60338    0.00000    0.00000 ;      HA-    CA-    CB-    SG
  6290   6289   6296   6297       3    3.68192   -4.35136    0.00000
1.33888    0.00000    0.00000 ;      HA-    CA-     C-     O
  6290   6289   6296   6298       3    0.00000    0.00000    0.00000
0.00000    0.00000    0.00000 ;      HA-    CA-     C-     N
  6291   6289   6296   6297       3    0.00000    0.00000    0.00000
0.00000    0.00000    0.00000 ;      CB-    CA-     C-     O
  6291   6289   6296   6298       3    2.51040    4.18400    1.67360
-6.69440    0.00000    0.00000 ;      CB-    CA-     C-     N
  6292   6291   6289   6296       3    0.65084    1.95253    0.00000
-2.60338    0.00000    0.00000 ;     HB2-    CB-    CA-     C
  6292   6291   6294   6295       3    1.04600    3.13800    0.00000
-4.18400    0.00000    0.00000 ;     HB2-    CB-    SG-    HG
  6293   6291   6289   6296       3    0.65084    1.95253    0.00000
-2.60338    0.00000    0.00000 ;     HB3-    CB-    CA-     C
  6293   6291   6294   6295       3    1.04600    3.13800    0.00000
-4.18400    0.00000    0.00000 ;     HB3-    CB-    SG-    HG
  6294   6291   6289   6296       3    5.62330    4.27605   -5.33042
-4.20074    2.51040    0.00000 ;      SG-    CB-    CA-     C
  6296   6298   6300   6301       3    0.00000    0.00000    0.00000
0.00000    0.00000    0.00000 ;       C-     N-    CA-    HA
  6296   6298   6300   6302       3   11.71520    1.67360   15.06240  -
13.38880    0.00000    0.00000 ;      C-     N-    CA-    CB
  6296   6298   6300   6315       3    1.75728    5.27184    2.25936
-7.02912    0.00000    0.00000 ;      C-     N-    CA-     C
  6297   6296   6298   6299       3   29.28800   -8.36800  -20.92000
0.00000    0.00000    0.00000 ;       O-     C-     N-     H
  6297   6296   6298   6300       3   20.92000    0.00000  -20.92000
0.00000    0.00000    0.00000 ;       O-     C-     N-    CA
  6298   6300   6302   6303       3    0.65084    1.95253    0.00000
-2.60338    0.00000    0.00000 ;      N-    CA-    CB-   HB2
  6298   6300   6302   6304       3    0.65084    1.95253    0.00000
-2.60338    0.00000    0.00000 ;      N-    CA-    CB-   HB3
  6298   6300   6302   6305       3    3.79070    1.39746   -4.61077
```

-2.40998 2.44346 0.00000 ; N- CA- CB- CG
6298 6300 6315 6316 3 0.00000 0.00000 0.00000
0.00000 0.00000 0.00000 ; N- CA- C- O
6298 6300 6315 6317 3 17.40544 -5.02080 -13.22144
9.20480 0.00000 0.00000 ; N- CA- C- N
6299 6298 6300 6301 3 0.00000 0.00000 0.00000
0.00000 0.00000 0.00000 ; H- N- CA- HA
6299 6298 6300 6302 3 0.00000 0.00000 0.00000
0.00000 0.00000 0.00000 ; H- N- CA- CB
6299 6298 6300 6315 3 0.00000 0.00000 0.00000
0.00000 0.00000 0.00000 ; H- N- CA- C
6300 6302 6305 6306 3 0.66944 2.00832 0.00000
-2.67776 0.00000 0.00000 ; CA- CB- CG- HG
6300 6302 6305 6307 3 3.90367 0.19665 -6.21742
-2.37651 5.99149 0.00000 ; CA- CB- CG- CD1
6300 6302 6305 6311 3 3.90367 0.19665 -6.21742
-2.37651 5.99149 0.00000 ; CA- CB- CG- CD2
6300 6315 6317 6318 3 20.92000 0.00000 -20.92000
0.00000 0.00000 0.00000 ; CA- C- N- H
6300 6315 6317 6319 3 20.92000 0.00000 -20.92000
0.00000 0.00000 0.00000 ; CA- C- N- CA
6301 6300 6302 6303 3 0.65084 1.95253 0.00000
-2.60338 0.00000 0.00000 ; HA- CA- CB- HB2
6301 6300 6302 6304 3 0.65084 1.95253 0.00000
-2.60338 0.00000 0.00000 ; HA- CA- CB- HB3
6301 6300 6302 6305 3 0.65084 1.95253 0.00000
-2.60338 0.00000 0.00000 ; HA- CA- CB- CG
6301 6300 6315 6316 3 3.68192 -4.35136 0.00000
1.33888 0.00000 0.00000 ; HA- CA- C- O
6301 6300 6315 6317 3 0.00000 0.00000 0.00000
0.00000 0.00000 0.00000 ; HA- CA- C- N
6302 6300 6315 6316 3 0.00000 0.00000 0.00000
0.00000 0.00000 0.00000 ; CB- CA- C- O
6302 6300 6315 6317 3 2.51040 4.18400 1.67360
-6.69440 0.00000 0.00000 ; CB- CA- C- N
6302 6305 6307 6308 3 0.66944 2.00832 0.00000
-2.67776 0.00000 0.00000 ; CB- CG- CD1- HD11
6302 6305 6307 6309 3 0.66944 2.00832 0.00000
-2.67776 0.00000 0.00000 ; CB- CG- CD1- HD12
6302 6305 6307 6310 3 0.66944 2.00832 0.00000
-2.67776 0.00000 0.00000 ; CB- CG- CD1- HD13
6302 6305 6311 6312 3 0.66944 2.00832 0.00000
-2.67776 0.00000 0.00000 ; CB- CG- CD2- HD21
6302 6305 6311 6313 3 0.66944 2.00832 0.00000
-2.67776 0.00000 0.00000 ; CB- CG- CD2- HD22
6302 6305 6311 6314 3 0.66944 2.00832 0.00000
-2.67776 0.00000 0.00000 ; CB- CG- CD2- HD23
6303 6302 6300 6315 3 0.65084 1.95253 0.00000
-2.60338 0.00000 0.00000 ; HB2- CB- CA- C
6303 6302 6305 6306 3 0.62760 1.88280 0.00000
-2.51040 0.00000 0.00000 ; HB2- CB- CG- HG
6303 6302 6305 6307 3 0.66944 2.00832 0.00000

-2.67776 0.00000 0.00000 ; HB2- CB- CG- CD1
6303 6302 6305 6311 3 0.66944 2.00832 0.00000
-2.67776 0.00000 0.00000 ; HB2- CB- CG- CD2
6304 6302 6300 6315 3 0.65084 1.95253 0.00000
-2.60338 0.00000 0.00000 ; HB3- CB- CA- C
6304 6302 6305 6306 3 0.62760 1.88280 0.00000
-2.51040 0.00000 0.00000 ; HB3- CB- CG- HG
6304 6302 6305 6307 3 0.66944 2.00832 0.00000
-2.67776 0.00000 0.00000 ; HB3- CB- CG- CD1
6304 6302 6305 6311 3 0.66944 2.00832 0.00000
-2.67776 0.00000 0.00000 ; HB3- CB- CG- CD2
6305 6302 6300 6315 3 10.33448 -1.14642 -11.54784
-2.40998 6.35968 0.00000 ; CG- CB- CA- C
6306 6305 6307 6308 3 0.62760 1.88280 0.00000
-2.51040 0.00000 0.00000 ; HG- CG- CD1- HD11
6306 6305 6307 6309 3 0.62760 1.88280 0.00000
-2.51040 0.00000 0.00000 ; HG- CG- CD1- HD12
6306 6305 6307 6310 3 0.62760 1.88280 0.00000
-2.51040 0.00000 0.00000 ; HG- CG- CD1- HD13
6306 6305 6311 6312 3 0.62760 1.88280 0.00000
-2.51040 0.00000 0.00000 ; HG- CG- CD2- HD21
6306 6305 6311 6313 3 0.62760 1.88280 0.00000
-2.51040 0.00000 0.00000 ; HG- CG- CD2- HD22
6306 6305 6311 6314 3 0.62760 1.88280 0.00000
-2.51040 0.00000 0.00000 ; HG- CG- CD2- HD23
6307 6305 6311 6312 3 0.66944 2.00832 0.00000
-2.67776 0.00000 0.00000 ; CD1- CG- CD2- HD21
6307 6305 6311 6313 3 0.66944 2.00832 0.00000
-2.67776 0.00000 0.00000 ; CD1- CG- CD2- HD22
6307 6305 6311 6314 3 0.66944 2.00832 0.00000
-2.67776 0.00000 0.00000 ; CD1- CG- CD2- HD23
6308 6307 6305 6311 3 0.66944 2.00832 0.00000
-2.67776 0.00000 0.00000 ; HD11- CD1- CG- CD2
6309 6307 6305 6311 3 0.66944 2.00832 0.00000
-2.67776 0.00000 0.00000 ; HD12- CD1- CG- CD2
6310 6307 6305 6311 3 0.66944 2.00832 0.00000
-2.67776 0.00000 0.00000 ; HD13- CD1- CG- CD2
6315 6317 6319 6320 3 0.00000 0.00000 0.00000
0.00000 0.00000 0.00000 ; C- N- CA- HA
6315 6317 6319 6321 3 11.71520 1.67360 15.06240 -
13.38880 0.00000 0.00000 ; C- N- CA- CB
6315 6317 6319 6334 3 1.75728 5.27184 2.25936
-7.02912 0.00000 0.00000 ; C- N- CA- C
6316 6315 6317 6318 3 29.28800 -8.36800 -20.92000
0.00000 0.00000 0.00000 ; O- C- N- H
6316 6315 6317 6319 3 20.92000 0.00000 -20.92000
0.00000 0.00000 0.00000 ; O- C- N- CA
6317 6319 6321 6322 3 0.65084 1.95253 0.00000
-2.60338 0.00000 0.00000 ; N- CA- CB- HB
6317 6319 6321 6323 3 3.83673 0.44769 -1.77402
-2.47693 -0.03347 0.00000 ; N- CA- CB- CG2
6317 6319 6321 6327 3 2.97482 0.12134 2.04179

-1.89117 -3.24678 0.00000 ; N- CA- CB- CG1
6317 6319 6334 6335 3 0.00000 0.00000 0.00000
0.00000 0.00000 0.00000 ; N- CA- C- O
6317 6319 6334 6336 3 17.40544 -5.02080 -13.22144
9.20480 0.00000 0.00000 ; N- CA- C- N
6318 6317 6319 6320 3 0.00000 0.00000 0.00000
0.00000 0.00000 0.00000 ; H- N- CA- HA
6318 6317 6319 6321 3 0.00000 0.00000 0.00000
0.00000 0.00000 0.00000 ; H- N- CA- CB
6318 6317 6319 6334 3 0.00000 0.00000 0.00000
0.00000 0.00000 0.00000 ; H- N- CA- C
6319 6321 6323 6324 3 0.66944 2.00832 0.00000
-2.67776 0.00000 0.00000 ; CA- CB- CG2- HG21
6319 6321 6323 6325 3 0.66944 2.00832 0.00000
-2.67776 0.00000 0.00000 ; CA- CB- CG2- HG22
6319 6321 6323 6326 3 0.66944 2.00832 0.00000
-2.67776 0.00000 0.00000 ; CA- CB- CG2- HG23
6319 6321 6327 6328 3 0.66944 2.00832 0.00000
-2.67776 0.00000 0.00000 ; CA- CB- CG1- HG12
6319 6321 6327 6329 3 0.66944 2.00832 0.00000
-2.67776 0.00000 0.00000 ; CA- CB- CG1- HG13
6319 6321 6327 6330 3 4.24258 -0.52718 -7.25506
-1.79075 7.69856 0.00000 ; CA- CB- CG1- CD1
6319 6334 6336 6337 3 20.92000 0.00000 -20.92000
0.00000 0.00000 0.00000 ; CA- C- N- CD
6319 6334 6336 6346 3 20.92000 0.00000 -20.92000
0.00000 0.00000 0.00000 ; CA- C- N- CA
6320 6319 6321 6322 3 0.65084 1.95253 0.00000
-2.60338 0.00000 0.00000 ; HA- CA- CB- HB
6320 6319 6321 6323 3 0.65084 1.95253 0.00000
-2.60338 0.00000 0.00000 ; HA- CA- CB- CG2
6320 6319 6321 6327 3 0.65084 1.95253 0.00000
-2.60338 0.00000 0.00000 ; HA- CA- CB- CG1
6320 6319 6334 6335 3 3.68192 -4.35136 0.00000
1.33888 0.00000 0.00000 ; HA- CA- C- O
6320 6319 6334 6336 3 0.00000 0.00000 0.00000
0.00000 0.00000 0.00000 ; HA- CA- C- N
6321 6319 6334 6335 3 0.00000 0.00000 0.00000
0.00000 0.00000 0.00000 ; CB- CA- C- O
6321 6319 6334 6336 3 2.51040 4.18400 1.67360
-6.69440 0.00000 0.00000 ; CB- CA- C- N
6321 6327 6330 6331 3 0.66944 2.00832 0.00000
-2.67776 0.00000 0.00000 ; CB- CG1- CD1- HD11
6321 6327 6330 6332 3 0.66944 2.00832 0.00000
-2.67776 0.00000 0.00000 ; CB- CG1- CD1- HD12
6321 6327 6330 6333 3 0.66944 2.00832 0.00000
-2.67776 0.00000 0.00000 ; CB- CG1- CD1- HD13
6322 6321 6319 6334 3 0.65084 1.95253 0.00000
-2.60338 0.00000 0.00000 ; HB- CB- CA- C
6322 6321 6323 6324 3 0.62760 1.88280 0.00000
-2.51040 0.00000 0.00000 ; HB- CB- CG2- HG21
6322 6321 6323 6325 3 0.62760 1.88280 0.00000

-2.51040 0.00000 0.00000 ; HB- CB- CG2- HG22
6322 6321 6323 6326 3 0.62760 1.88280 0.00000
-2.51040 0.00000 0.00000 ; HB- CB- CG2- HG23
6322 6321 6327 6328 3 0.62760 1.88280 0.00000
-2.51040 0.00000 0.00000 ; HB- CB- CG1- HG12
6322 6321 6327 6329 3 0.62760 1.88280 0.00000
-2.51040 0.00000 0.00000 ; HB- CB- CG1- HG13
6322 6321 6327 6330 3 0.66944 2.00832 0.00000
-2.67776 0.00000 0.00000 ; HB- CB- CG1- CD1
6323 6321 6319 6334 3 5.67350 3.55640 -6.16722
-2.47693 3.74886 0.00000 ; CG2- CB- CA- C
6323 6321 6327 6328 3 0.66944 2.00832 0.00000
-2.67776 0.00000 0.00000 ; CG2- CB- CG1- HG12
6323 6321 6327 6329 3 0.66944 2.00832 0.00000
-2.67776 0.00000 0.00000 ; CG2- CB- CG1- HG13
6323 6321 6327 6330 3 3.81162 0.49790 -8.14206
-1.79075 7.49773 0.00000 ; CG2- CB- CG1- CD1
6324 6323 6321 6327 3 0.66944 2.00832 0.00000
-2.67776 0.00000 0.00000 ; HG21- CG2- CB- CG1
6325 6323 6321 6327 3 0.66944 2.00832 0.00000
-2.67776 0.00000 0.00000 ; HG22- CG2- CB- CG1
6326 6323 6321 6327 3 0.66944 2.00832 0.00000
-2.67776 0.00000 0.00000 ; HG23- CG2- CB- CG1
6327 6321 6319 6334 3 8.26340 0.74057 -9.99976
-1.89117 3.84928 0.00000 ; CG1- CB- CA- C
6328 6327 6330 6331 3 0.62760 1.88280 0.00000
-2.51040 0.00000 0.00000 ; HG12- CG1- CD1- HD11
6328 6327 6330 6332 3 0.62760 1.88280 0.00000
-2.51040 0.00000 0.00000 ; HG12- CG1- CD1- HD12
6328 6327 6330 6333 3 0.62760 1.88280 0.00000
-2.51040 0.00000 0.00000 ; HG12- CG1- CD1- HD13
6329 6327 6330 6331 3 0.62760 1.88280 0.00000
-2.51040 0.00000 0.00000 ; HG13- CG1- CD1- HD11
6329 6327 6330 6332 3 0.62760 1.88280 0.00000
-2.51040 0.00000 0.00000 ; HG13- CG1- CD1- HD12
6329 6327 6330 6333 3 0.62760 1.88280 0.00000
-2.51040 0.00000 0.00000 ; HG13- CG1- CD1- HD13
6334 6336 6337 6338 3 0.00000 0.00000 0.00000
0.00000 0.00000 0.00000 ; C- N- CD- HD2
6334 6336 6337 6339 3 0.00000 0.00000 0.00000
0.00000 0.00000 0.00000 ; C- N- CD- HD3
6334 6336 6337 6340 3 10.04160 -3.34720 16.73600
-6.69440 0.00000 0.00000 ; C- N- CD- CG
6334 6336 6346 6343 3 11.71520 1.67360 15.06240 -
13.38880 0.00000 0.00000 ; C- N- CA- CB
6334 6336 6346 6347 3 0.00000 0.00000 0.00000
0.00000 0.00000 0.00000 ; C- N- CA- HA
6334 6336 6346 6348 3 1.75728 5.27184 2.25936
-7.02912 0.00000 0.00000 ; C- N- CA- C
6335 6334 6336 6337 3 20.92000 0.00000 -20.92000
0.00000 0.00000 0.00000 ; O- C- N- CD
6335 6334 6336 6346 3 20.92000 0.00000 -20.92000

```
0.00000    0.00000    0.00000 ;       O-     C-     N-    CA
  6336   6337   6340   6341      3    0.65084    1.95253    0.00000
-2.60338    0.00000    0.00000 ;       N-    CD-    CG-   HG2
  6336   6337   6340   6342      3    0.65084    1.95253    0.00000
-2.60338    0.00000    0.00000 ;       N-    CD-    CG-   HG3
  6336   6337   6340   6343      3    0.65084    1.95253    0.00000
-2.60338    0.00000    0.00000 ;       N-    CD-    CG-    CB
  6336   6346   6343   6340      3    0.65084    1.95253    0.00000
-2.60338    0.00000    0.00000 ;       N-    CA-    CB-    CG
  6336   6346   6343   6344      3    0.65084    1.95253    0.00000
-2.60338    0.00000    0.00000 ;       N-    CA-    CB-   HB2
  6336   6346   6343   6345      3    0.65084    1.95253    0.00000
-2.60338    0.00000    0.00000 ;       N-    CA-    CB-   HB3
  6336   6346   6348   6349      3    0.00000    0.00000    0.00000
0.00000    0.00000    0.00000 ;       N-    CA-     C-     O
  6336   6346   6348   6350      3   17.40544   -5.02080  -13.22144
9.20480    0.00000    0.00000 ;       N-    CA-     C-     N
  6337   6336   6346   6343      3    0.00000    0.00000    0.00000
0.00000    0.00000    0.00000 ;      CD-     N-    CA-    CB
  6337   6336   6346   6347      3    0.00000    0.00000    0.00000
0.00000    0.00000    0.00000 ;      CD-     N-    CA-    HA
  6337   6336   6346   6348      3    0.00000    0.00000    0.00000
0.00000    0.00000    0.00000 ;      CD-     N-    CA-     C
  6337   6340   6343   6344      3    0.66944    2.00832    0.00000
-2.67776    0.00000    0.00000 ;      CD-    CG-    CB-   HB2
  6337   6340   6343   6345      3    0.66944    2.00832    0.00000
-2.67776    0.00000    0.00000 ;      CD-    CG-    CB-   HB3
  6337   6340   6343   6346      3    3.68192    3.09616   -2.09200
-3.01248    0.00000    0.00000 ;      CD-    CG-    CB-    CA
  6338   6337   6336   6346      3    0.00000    0.00000    0.00000
0.00000    0.00000    0.00000 ;     HD2-    CD-     N-    CA
  6338   6337   6340   6341      3    0.65084    1.95253    0.00000
-2.60338    0.00000    0.00000 ;     HD2-    CD-    CG-   HG2
  6338   6337   6340   6342      3    0.65084    1.95253    0.00000
-2.60338    0.00000    0.00000 ;     HD2-    CD-    CG-   HG3
  6338   6337   6340   6343      3    0.65084    1.95253    0.00000
-2.60338    0.00000    0.00000 ;     HD2-    CD-    CG-    CB
  6339   6337   6336   6346      3    0.00000    0.00000    0.00000
0.00000    0.00000    0.00000 ;     HD3-    CD-     N-    CA
  6339   6337   6340   6341      3    0.65084    1.95253    0.00000
-2.60338    0.00000    0.00000 ;     HD3-    CD-    CG-   HG2
  6339   6337   6340   6342      3    0.65084    1.95253    0.00000
-2.60338    0.00000    0.00000 ;     HD3-    CD-    CG-   HG3
  6339   6337   6340   6343      3    0.65084    1.95253    0.00000
-2.60338    0.00000    0.00000 ;     HD3-    CD-    CG-    CB
  6340   6337   6336   6346      3    0.00000    0.00000    0.00000
0.00000    0.00000    0.00000 ;      CG-    CD-     N-    CA
  6340   6343   6346   6347      3    0.65084    1.95253    0.00000
-2.60338    0.00000    0.00000 ;      CG-    CB-    CA-    HA
  6340   6343   6346   6348      3    0.65084    1.95253    0.00000
-2.60338    0.00000    0.00000 ;      CG-    CB-    CA-     C
  6341   6340   6343   6344      3    0.62760    1.88280    0.00000
```

-2.51040 0.00000 0.00000 ; HG2- CG- CB- HB2
6341 6340 6343 6345 3 0.62760 1.88280 0.00000
-2.51040 0.00000 0.00000 ; HG2- CG- CB- HB3
6341 6340 6343 6346 3 0.66944 2.00832 0.00000
-2.67776 0.00000 0.00000 ; HG2- CG- CB- CA
6342 6340 6343 6344 3 0.62760 1.88280 0.00000
-2.51040 0.00000 0.00000 ; HG3- CG- CB- HB2
6342 6340 6343 6345 3 0.62760 1.88280 0.00000
-2.51040 0.00000 0.00000 ; HG3- CG- CB- HB3
6342 6340 6343 6346 3 0.66944 2.00832 0.00000
-2.67776 0.00000 0.00000 ; HG3- CG- CB- CA
6343 6346 6348 6349 3 0.00000 0.00000 0.00000
0.00000 0.00000 0.00000 ; CB- CA- C- O
6343 6346 6348 6350 3 2.51040 4.18400 1.67360
-6.69440 0.00000 0.00000 ; CB- CA- C- N
6344 6343 6346 6347 3 0.65084 1.95253 0.00000
-2.60338 0.00000 0.00000 ; HB2- CB- CA- HA
6344 6343 6346 6348 3 0.65084 1.95253 0.00000
-2.60338 0.00000 0.00000 ; HB2- CB- CA- C
6345 6343 6346 6347 3 0.65084 1.95253 0.00000
-2.60338 0.00000 0.00000 ; HB3- CB- CA- HA
6345 6343 6346 6348 3 0.65084 1.95253 0.00000
-2.60338 0.00000 0.00000 ; HB3- CB- CA- C
6346 6348 6350 6351 3 20.92000 0.00000 -20.92000
0.00000 0.00000 0.00000 ; CA- C- N- H
6346 6348 6350 6352 3 20.92000 0.00000 -20.92000
0.00000 0.00000 0.00000 ; CA- C- N- CA
6347 6346 6348 6349 3 3.68192 -4.35136 0.00000
1.33888 0.00000 0.00000 ; HA- CA- C- O
6347 6346 6348 6350 3 0.00000 0.00000 0.00000
0.00000 0.00000 0.00000 ; HA- CA- C- N
6348 6350 6352 6353 3 0.00000 0.00000 0.00000
0.00000 0.00000 0.00000 ; C- N- CA- HA
6348 6350 6352 6354 3 11.71520 1.67360 15.06240 -
13.38880 0.00000 0.00000 ; C- N- CA- CB
6348 6350 6352 6368 3 1.75728 5.27184 2.25936
-7.02912 0.00000 0.00000 ; C- N- CA- C
6349 6348 6350 6351 3 29.28800 -8.36800 -20.92000
0.00000 0.00000 0.00000 ; O- C- N- H
6349 6348 6350 6352 3 20.92000 0.00000 -20.92000
0.00000 0.00000 0.00000 ; O- C- N- CA
6350 6352 6354 6355 3 0.65084 1.95253 0.00000
-2.60338 0.00000 0.00000 ; N- CA- CB- HB2
6350 6352 6354 6356 3 0.65084 1.95253 0.00000
-2.60338 0.00000 0.00000 ; N- CA- CB- HB3
6350 6352 6354 6357 3 3.28026 2.46019 -2.19242
-3.21331 -0.23430 0.00000 ; N- CA- CB- CG
6350 6352 6368 6369 3 0.00000 0.00000 0.00000
0.00000 0.00000 0.00000 ; N- CA- C- O
6350 6352 6368 6370 3 17.40544 -5.02080 -13.22144
9.20480 0.00000 0.00000 ; N- CA- C- N
6351 6350 6352 6353 3 0.00000 0.00000 0.00000

0.00000 0.00000 0.00000 ; H- N- CA- HA
6351 6350 6352 6354 3 0.00000 0.00000 0.00000
0.00000 0.00000 0.00000 ; H- N- CA- CB
6351 6350 6352 6368 3 0.00000 0.00000 0.00000
0.00000 0.00000 0.00000 ; H- N- CA- C
6352 6354 6357 6358 3 0.57739 0.00000 1.02926
0.00000 -1.60666 0.00000 ; CA- CB- CG- CD1
6352 6354 6357 6366 3 0.57739 0.00000 1.02926
0.00000 -1.60666 0.00000 ; CA- CB- CG- CD2
6352 6368 6370 6371 3 20.92000 0.00000 -20.92000
0.00000 0.00000 0.00000 ; CA- C- N- H
6352 6368 6370 6372 3 20.92000 0.00000 -20.92000
0.00000 0.00000 0.00000 ; CA- C- N- CA
6353 6352 6354 6355 3 0.65084 1.95253 0.00000
-2.60338 0.00000 0.00000 ; HA- CA- CB- HB2
6353 6352 6354 6356 3 0.65084 1.95253 0.00000
-2.60338 0.00000 0.00000 ; HA- CA- CB- HB3
6353 6352 6354 6357 3 0.65084 1.95253 0.00000
-2.60338 0.00000 0.00000 ; HA- CA- CB- CG
6353 6352 6368 6369 3 3.68192 -4.35136 0.00000
1.33888 0.00000 0.00000 ; HA- CA- C- O
6353 6352 6368 6370 3 0.00000 0.00000 0.00000
0.00000 0.00000 0.00000 ; HA- CA- C- N
6354 6352 6368 6369 3 0.00000 0.00000 0.00000
0.00000 0.00000 0.00000 ; CB- CA- C- O
6354 6352 6368 6370 3 2.51040 4.18400 1.67360
-6.69440 0.00000 0.00000 ; CB- CA- C- N
6354 6357 6358 6359 3 30.33400 0.00000 -30.33400
0.00000 0.00000 0.00000 ; CB- CG- CD1- HD1
6354 6357 6358 6360 3 30.33400 0.00000 -30.33400
0.00000 0.00000 0.00000 ; CB- CG- CD1- CE1
6354 6357 6366 6364 3 30.33400 0.00000 -30.33400
0.00000 0.00000 0.00000 ; CB- CG- CD2- CE2
6354 6357 6366 6367 3 30.33400 0.00000 -30.33400
0.00000 0.00000 0.00000 ; CB- CG- CD2- HD2
6355 6354 6352 6368 3 0.65084 1.95253 0.00000
-2.60338 0.00000 0.00000 ; HB2- CB- CA- C
6355 6354 6357 6358 3 0.00000 0.00000 0.00000
0.00000 0.00000 0.00000 ; HB2- CB- CG- CD1
6355 6354 6357 6366 3 0.00000 0.00000 0.00000
0.00000 0.00000 0.00000 ; HB2- CB- CG- CD2
6356 6354 6352 6368 3 0.65084 1.95253 0.00000
-2.60338 0.00000 0.00000 ; HB3- CB- CA- C
6356 6354 6357 6358 3 0.00000 0.00000 0.00000
0.00000 0.00000 0.00000 ; HB3- CB- CG- CD1
6356 6354 6357 6366 3 0.00000 0.00000 0.00000
0.00000 0.00000 0.00000 ; HB3- CB- CG- CD2
6357 6354 6352 6368 3 4.95804 2.17986 -3.52293
-3.21331 -0.40166 0.00000 ; CG- CB- CA- C
6357 6358 6360 6361 3 30.33400 0.00000 -30.33400
0.00000 0.00000 0.00000 ; CG- CD1- CE1- HE1
6357 6358 6360 6362 3 30.33400 0.00000 -30.33400

```
0.00000    0.00000    0.00000 ;     CG-   CD1-   CE1-    CZ
  6357   6366   6364   6362      3   30.33400    0.00000  -30.33400
0.00000    0.00000    0.00000 ;     CG-   CD2-   CE2-    CZ
  6357   6366   6364   6365      3   30.33400    0.00000  -30.33400
0.00000    0.00000    0.00000 ;     CG-   CD2-   CE2-   HE2
  6358   6357   6366   6364      3   30.33400    0.00000  -30.33400
0.00000    0.00000    0.00000 ;    CD1-    CG-   CD2-   CE2
  6358   6357   6366   6367      3   30.33400    0.00000  -30.33400
0.00000    0.00000    0.00000 ;    CD1-    CG-   CD2-   HD2
  6358   6360   6362   6363      3   30.33400    0.00000  -30.33400
0.00000    0.00000    0.00000 ;    CD1-   CE1-    CZ-    HZ
  6358   6360   6362   6364      3   30.33400    0.00000  -30.33400
0.00000    0.00000    0.00000 ;    CD1-   CE1-    CZ-   CE2
  6359   6358   6357   6366      3   30.33400    0.00000  -30.33400
0.00000    0.00000    0.00000 ;    HD1-   CD1-    CG-   CD2
  6359   6358   6360   6361      3   30.33400    0.00000  -30.33400
0.00000    0.00000    0.00000 ;    HD1-   CD1-   CE1-   HE1
  6359   6358   6360   6362      3   30.33400    0.00000  -30.33400
0.00000    0.00000    0.00000 ;    HD1-   CD1-   CE1-    CZ
  6360   6358   6357   6366      3   30.33400    0.00000  -30.33400
0.00000    0.00000    0.00000 ;    CE1-   CD1-    CG-   CD2
  6360   6362   6364   6365      3   30.33400    0.00000  -30.33400
0.00000    0.00000    0.00000 ;    CE1-    CZ-   CE2-   HE2
  6360   6362   6364   6366      3   30.33400    0.00000  -30.33400
0.00000    0.00000    0.00000 ;    CE1-    CZ-   CE2-   CD2
  6361   6360   6362   6363      3   30.33400    0.00000  -30.33400
0.00000    0.00000    0.00000 ;    HE1-   CE1-    CZ-    HZ
  6361   6360   6362   6364      3   30.33400    0.00000  -30.33400
0.00000    0.00000    0.00000 ;    HE1-   CE1-    CZ-   CE2
  6362   6364   6366   6367      3   30.33400    0.00000  -30.33400
0.00000    0.00000    0.00000 ;     CZ-   CE2-   CD2-   HD2
  6363   6362   6364   6365      3   30.33400    0.00000  -30.33400
0.00000    0.00000    0.00000 ;     HZ-    CZ-   CE2-   HE2
  6363   6362   6364   6366      3   30.33400    0.00000  -30.33400
0.00000    0.00000    0.00000 ;     HZ-    CZ-   CE2-   CD2
  6365   6364   6366   6367      3   30.33400    0.00000  -30.33400
0.00000    0.00000    0.00000 ;    HE2-   CE2-   CD2-   HD2
  6368   6370   6372   6373      3    0.00000    0.00000    0.00000
0.00000    0.00000    0.00000 ;      C-     N-    CA-    HA
  6368   6370   6372   6374      3   11.71520    1.67360   15.06240  -
13.38880    0.00000    0.00000 ;      C-     N-    CA-    CB
  6368   6370   6372   6390      3    1.75728    5.27184    2.25936
-7.02912    0.00000    0.00000 ;      C-     N-    CA-     C
  6369   6368   6370   6371      3   29.28800   -8.36800  -20.92000
0.00000    0.00000    0.00000 ;      O-     C-     N-     H
  6369   6368   6370   6372      3   20.92000    0.00000  -20.92000
0.00000    0.00000    0.00000 ;      O-     C-     N-    CA
  6370   6372   6374   6375      3    0.65084    1.95253    0.00000
-2.60338    0.00000    0.00000 ;      N-    CA-    CB-   HB2
  6370   6372   6374   6376      3    0.65084    1.95253    0.00000
-2.60338    0.00000    0.00000 ;      N-    CA-    CB-   HB3
  6370   6372   6374   6377      3    0.65084    1.95253    0.00000
```

-2.60338 0.00000 0.00000 ; N- CA- CB- CG
6370 6372 6390 6391 3 0.00000 0.00000 0.00000
0.00000 0.00000 0.00000 ; N- CA- C- O
6370 6372 6390 6392 3 17.40544 -5.02080 -13.22144
9.20480 0.00000 0.00000 ; N- CA- C- N
6371 6370 6372 6373 3 0.00000 0.00000 0.00000
0.00000 0.00000 0.00000 ; H- N- CA- HA
6371 6370 6372 6374 3 0.00000 0.00000 0.00000
0.00000 0.00000 0.00000 ; H- N- CA- CB
6371 6370 6372 6390 3 0.00000 0.00000 0.00000
0.00000 0.00000 0.00000 ; H- N- CA- C
6372 6374 6377 6378 3 0.66944 2.00832 0.00000
-2.67776 0.00000 0.00000 ; CA- CB- CG- HG2
6372 6374 6377 6379 3 0.66944 2.00832 0.00000
-2.67776 0.00000 0.00000 ; CA- CB- CG- HG3
6372 6374 6377 6380 3 3.68192 3.09616 -2.09200
-3.01248 0.00000 0.00000 ; CA- CB- CG- CD
6372 6390 6392 6393 3 20.92000 0.00000 -20.92000
0.00000 0.00000 0.00000 ; CA- C- N- H
6372 6390 6392 6394 3 20.92000 0.00000 -20.92000
0.00000 0.00000 0.00000 ; CA- C- N- CA
6373 6372 6374 6375 3 0.65084 1.95253 0.00000
-2.60338 0.00000 0.00000 ; HA- CA- CB- HB2
6373 6372 6374 6376 3 0.65084 1.95253 0.00000
-2.60338 0.00000 0.00000 ; HA- CA- CB- HB3
6373 6372 6374 6377 3 0.65084 1.95253 0.00000
-2.60338 0.00000 0.00000 ; HA- CA- CB- CG
6373 6372 6390 6391 3 3.68192 -4.35136 0.00000
1.33888 0.00000 0.00000 ; HA- CA- C- O
6373 6372 6390 6392 3 0.00000 0.00000 0.00000
0.00000 0.00000 0.00000 ; HA- CA- C- N
6374 6372 6390 6391 3 0.00000 0.00000 0.00000
0.00000 0.00000 0.00000 ; CB- CA- C- O
6374 6372 6390 6392 3 2.51040 4.18400 1.67360
-6.69440 0.00000 0.00000 ; CB- CA- C- N
6374 6377 6380 6381 3 0.66944 2.00832 0.00000
-2.67776 0.00000 0.00000 ; CB- CG- CD- HD2
6374 6377 6380 6382 3 0.66944 2.00832 0.00000
-2.67776 0.00000 0.00000 ; CB- CG- CD- HD3
6374 6377 6380 6383 3 3.68192 3.09616 -2.09200
-3.01248 0.00000 0.00000 ; CB- CG- CD- CE
6375 6374 6372 6390 3 0.65084 1.95253 0.00000
-2.60338 0.00000 0.00000 ; HB2- CB- CA- C
6375 6374 6377 6378 3 0.62760 1.88280 0.00000
-2.51040 0.00000 0.00000 ; HB2- CB- CG- HG2
6375 6374 6377 6379 3 0.62760 1.88280 0.00000
-2.51040 0.00000 0.00000 ; HB2- CB- CG- HG3
6375 6374 6377 6380 3 0.66944 2.00832 0.00000
-2.67776 0.00000 0.00000 ; HB2- CB- CG- CD
6376 6374 6372 6390 3 0.65084 1.95253 0.00000
-2.60338 0.00000 0.00000 ; HB3- CB- CA- C
6376 6374 6377 6378 3 0.62760 1.88280 0.00000

```
-2.51040    0.00000    0.00000 ;     HB3-    CB-     CG-   HG2
 6376   6374   6377   6379       3    0.62760    1.88280    0.00000
-2.51040    0.00000    0.00000 ;     HB3-    CB-     CG-   HG3
 6376   6374   6377   6380       3    0.66944    2.00832    0.00000
-2.67776    0.00000    0.00000 ;     HB3-    CB-     CG-    CD
 6377   6374   6372   6390       3    0.65084    1.95253    0.00000
-2.60338    0.00000    0.00000 ;      CG-    CB-     CA-     C
 6377   6380   6383   6384       3    0.65084    1.95253    0.00000
-2.60338    0.00000    0.00000 ;      CG-    CD-     CE-   HE2
 6377   6380   6383   6385       3    0.65084    1.95253    0.00000
-2.60338    0.00000    0.00000 ;      CG-    CD-     CE-   HE3
 6377   6380   6383   6386       3    0.65084    1.95253    0.00000
-2.60338    0.00000    0.00000 ;      CG-    CD-     CE-    NZ
 6378   6377   6380   6381       3    0.62760    1.88280    0.00000
-2.51040    0.00000    0.00000 ;     HG2-    CG-     CD-   HD2
 6378   6377   6380   6382       3    0.62760    1.88280    0.00000
-2.51040    0.00000    0.00000 ;     HG2-    CG-     CD-   HD3
 6378   6377   6380   6383       3    0.66944    2.00832    0.00000
-2.67776    0.00000    0.00000 ;     HG2-    CG-     CD-    CE
 6379   6377   6380   6381       3    0.62760    1.88280    0.00000
-2.51040    0.00000    0.00000 ;     HG3-    CG-     CD-   HD2
 6379   6377   6380   6382       3    0.62760    1.88280    0.00000
-2.51040    0.00000    0.00000 ;     HG3-    CG-     CD-   HD3
 6379   6377   6380   6383       3    0.66944    2.00832    0.00000
-2.67776    0.00000    0.00000 ;     HG3-    CG-     CD-    CE
 6380   6383   6386   6387       3    0.65084    1.95253    0.00000
-2.60338    0.00000    0.00000 ;      CD-    CE-     NZ-   HZ1
 6380   6383   6386   6388       3    0.65084    1.95253    0.00000
-2.60338    0.00000    0.00000 ;      CD-    CE-     NZ-   HZ2
 6380   6383   6386   6389       3    0.65084    1.95253    0.00000
-2.60338    0.00000    0.00000 ;      CD-    CE-     NZ-   HZ3
 6381   6380   6383   6384       3    0.65084    1.95253    0.00000
-2.60338    0.00000    0.00000 ;     HD2-    CD-     CE-   HE2
 6381   6380   6383   6385       3    0.65084    1.95253    0.00000
-2.60338    0.00000    0.00000 ;     HD2-    CD-     CE-   HE3
 6381   6380   6383   6386       3    0.65084    1.95253    0.00000
-2.60338    0.00000    0.00000 ;     HD2-    CD-     CE-    NZ
 6382   6380   6383   6384       3    0.65084    1.95253    0.00000
-2.60338    0.00000    0.00000 ;     HD3-    CD-     CE-   HE2
 6382   6380   6383   6385       3    0.65084    1.95253    0.00000
-2.60338    0.00000    0.00000 ;     HD3-    CD-     CE-   HE3
 6382   6380   6383   6386       3    0.65084    1.95253    0.00000
-2.60338    0.00000    0.00000 ;     HD3-    CD-     CE-    NZ
 6384   6383   6386   6387       3    0.65084    1.95253    0.00000
-2.60338    0.00000    0.00000 ;     HE2-    CE-     NZ-   HZ1
 6384   6383   6386   6388       3    0.65084    1.95253    0.00000
-2.60338    0.00000    0.00000 ;     HE2-    CE-     NZ-   HZ2
 6384   6383   6386   6389       3    0.65084    1.95253    0.00000
-2.60338    0.00000    0.00000 ;     HE2-    CE-     NZ-   HZ3
 6385   6383   6386   6387       3    0.65084    1.95253    0.00000
-2.60338    0.00000    0.00000 ;     HE3-    CE-     NZ-   HZ1
 6385   6383   6386   6388       3    0.65084    1.95253    0.00000
```

-2.60338 0.00000 0.00000 ; HE3- CE- NZ- HZ2
6385 6383 6386 6389 3 0.65084 1.95253 0.00000
-2.60338 0.00000 0.00000 ; HE3- CE- NZ- HZ3
6390 6392 6394 6395 3 0.00000 0.00000 0.00000
0.00000 0.00000 0.00000 ; C- N- CA- HA
6390 6392 6394 6396 3 11.71520 1.67360 15.06240 -
13.38880 0.00000 0.00000 ; C- N- CA- CB
6390 6392 6394 6410 3 1.75728 5.27184 2.25936
-7.02912 0.00000 0.00000 ; C- N- CA- C
6391 6390 6392 6393 3 29.28800 -8.36800 -20.92000
0.00000 0.00000 0.00000 ; O- C- N- H
6391 6390 6392 6394 3 20.92000 0.00000 -20.92000
0.00000 0.00000 0.00000 ; O- C- N- CA
6392 6394 6396 6397 3 0.65084 1.95253 0.00000
-2.60338 0.00000 0.00000 ; N- CA- CB- HB2
6392 6394 6396 6398 3 0.65084 1.95253 0.00000
-2.60338 0.00000 0.00000 ; N- CA- CB- HB3
6392 6394 6396 6399 3 3.28026 2.46019 -2.19242
-3.21331 -0.23430 0.00000 ; N- CA- CB- CG
6392 6394 6410 6411 3 0.00000 0.00000 0.00000
0.00000 0.00000 0.00000 ; N- CA- C- O
6392 6394 6410 6412 3 17.40544 -5.02080 -13.22144
9.20480 0.00000 0.00000 ; N- CA- C- N
6393 6392 6394 6395 3 0.00000 0.00000 0.00000
0.00000 0.00000 0.00000 ; H- N- CA- HA
6393 6392 6394 6396 3 0.00000 0.00000 0.00000
0.00000 0.00000 0.00000 ; H- N- CA- CB
6393 6392 6394 6410 3 0.00000 0.00000 0.00000
0.00000 0.00000 0.00000 ; H- N- CA- C
6394 6396 6399 6400 3 0.57739 0.00000 1.02926
0.00000 -1.60666 0.00000 ; CA- CB- CG- CD1
6394 6396 6399 6408 3 0.57739 0.00000 1.02926
0.00000 -1.60666 0.00000 ; CA- CB- CG- CD2
6394 6410 6412 6413 3 20.92000 0.00000 -20.92000
0.00000 0.00000 0.00000 ; CA- C- N- H
6394 6410 6412 6414 3 20.92000 0.00000 -20.92000
0.00000 0.00000 0.00000 ; CA- C- N- CA
6395 6394 6396 6397 3 0.65084 1.95253 0.00000
-2.60338 0.00000 0.00000 ; HA- CA- CB- HB2
6395 6394 6396 6398 3 0.65084 1.95253 0.00000
-2.60338 0.00000 0.00000 ; HA- CA- CB- HB3
6395 6394 6396 6399 3 0.65084 1.95253 0.00000
-2.60338 0.00000 0.00000 ; HA- CA- CB- CG
6395 6394 6410 6411 3 3.68192 -4.35136 0.00000
1.33888 0.00000 0.00000 ; HA- CA- C- O
6395 6394 6410 6412 3 0.00000 0.00000 0.00000
0.00000 0.00000 0.00000 ; HA- CA- C- N
6396 6394 6410 6411 3 0.00000 0.00000 0.00000
0.00000 0.00000 0.00000 ; CB- CA- C- O
6396 6394 6410 6412 3 2.51040 4.18400 1.67360
-6.69440 0.00000 0.00000 ; CB- CA- C- N
6396 6399 6400 6401 3 30.33400 0.00000 -30.33400

```
0.00000    0.00000    0.00000 ;     CB-    CG-   CD1-   HD1
  6396   6399   6400   6402      3   30.33400    0.00000  -30.33400
0.00000    0.00000    0.00000 ;     CB-    CG-   CD1-   CE1
  6396   6399   6408   6406      3   30.33400    0.00000  -30.33400
0.00000    0.00000    0.00000 ;     CB-    CG-   CD2-   CE2
  6396   6399   6408   6409      3   30.33400    0.00000  -30.33400
0.00000    0.00000    0.00000 ;     CB-    CG-   CD2-   HD2
  6397   6396   6394   6410      3    0.65084    1.95253    0.00000
-2.60338    0.00000    0.00000 ;    HB2-    CB-    CA-     C
  6397   6396   6399   6400      3    0.00000    0.00000    0.00000
0.00000    0.00000    0.00000 ;    HB2-    CB-    CG-   CD1
  6397   6396   6399   6408      3    0.00000    0.00000    0.00000
0.00000    0.00000    0.00000 ;    HB2-    CB-    CG-   CD2
  6398   6396   6394   6410      3    0.65084    1.95253    0.00000
-2.60338    0.00000    0.00000 ;    HB3-    CB-    CA-     C
  6398   6396   6399   6400      3    0.00000    0.00000    0.00000
0.00000    0.00000    0.00000 ;    HB3-    CB-    CG-   CD1
  6398   6396   6399   6408      3    0.00000    0.00000    0.00000
0.00000    0.00000    0.00000 ;    HB3-    CB-    CG-   CD2
  6399   6396   6394   6410      3    4.95804    2.17986   -3.52293
-3.21331   -0.40166    0.00000 ;     CG-    CB-    CA-     C
  6399   6400   6402   6403      3   30.33400    0.00000  -30.33400
0.00000    0.00000    0.00000 ;     CG-   CD1-   CE1-   HE1
  6399   6400   6402   6404      3   30.33400    0.00000  -30.33400
0.00000    0.00000    0.00000 ;     CG-   CD1-   CE1-    CZ
  6399   6408   6406   6404      3   30.33400    0.00000  -30.33400
0.00000    0.00000    0.00000 ;     CG-   CD2-   CE2-    CZ
  6399   6408   6406   6407      3   30.33400    0.00000  -30.33400
0.00000    0.00000    0.00000 ;     CG-   CD2-   CE2-   HE2
  6400   6399   6408   6406      3   30.33400    0.00000  -30.33400
0.00000    0.00000    0.00000 ;    CD1-    CG-   CD2-   CE2
  6400   6399   6408   6409      3   30.33400    0.00000  -30.33400
0.00000    0.00000    0.00000 ;    CD1-    CG-   CD2-   HD2
  6400   6402   6404   6405      3   30.33400    0.00000  -30.33400
0.00000    0.00000    0.00000 ;    CD1-   CE1-    CZ-    HZ
  6400   6402   6404   6406      3   30.33400    0.00000  -30.33400
0.00000    0.00000    0.00000 ;    CD1-   CE1-    CZ-   CE2
  6401   6400   6399   6408      3   30.33400    0.00000  -30.33400
0.00000    0.00000    0.00000 ;    HD1-   CD1-    CG-   CD2
  6401   6400   6402   6403      3   30.33400    0.00000  -30.33400
0.00000    0.00000    0.00000 ;    HD1-   CD1-   CE1-   HE1
  6401   6400   6402   6404      3   30.33400    0.00000  -30.33400
0.00000    0.00000    0.00000 ;    HD1-   CD1-   CE1-    CZ
  6402   6400   6399   6408      3   30.33400    0.00000  -30.33400
0.00000    0.00000    0.00000 ;    CE1-   CD1-    CG-   CD2
  6402   6404   6406   6407      3   30.33400    0.00000  -30.33400
0.00000    0.00000    0.00000 ;    CE1-    CZ-   CE2-   HE2
  6402   6404   6406   6408      3   30.33400    0.00000  -30.33400
0.00000    0.00000    0.00000 ;    CE1-    CZ-   CE2-   CD2
  6403   6402   6404   6405      3   30.33400    0.00000  -30.33400
0.00000    0.00000    0.00000 ;    HE1-   CE1-    CZ-    HZ
  6403   6402   6404   6406      3   30.33400    0.00000  -30.33400
```

0.00000 0.00000 0.00000 ; HE1- CE1- CZ- CE2
6404 6406 6408 6409 3 30.33400 0.00000 -30.33400
0.00000 0.00000 0.00000 ; CZ- CE2- CD2- HD2
6405 6404 6406 6407 3 30.33400 0.00000 -30.33400
0.00000 0.00000 0.00000 ; HZ- CZ- CE2- HE2
6405 6404 6406 6408 3 30.33400 0.00000 -30.33400
0.00000 0.00000 0.00000 ; HZ- CZ- CE2- CD2
6407 6406 6408 6409 3 30.33400 0.00000 -30.33400
0.00000 0.00000 0.00000 ; HE2- CE2- CD2- HD2
6410 6412 6414 6415 3 0.00000 0.00000 0.00000
0.00000 0.00000 0.00000 ; C- N- CA- HA
6410 6412 6414 6416 3 11.71520 1.67360 15.06240 -
13.38880 0.00000 0.00000 ; C- N- CA- CB
6410 6412 6414 6430 3 1.75728 5.27184 2.25936
-7.02912 0.00000 0.00000 ; C- N- CA- C
6411 6410 6412 6413 3 29.28800 -8.36800 -20.92000
0.00000 0.00000 0.00000 ; O- C- N- H
6411 6410 6412 6414 3 20.92000 0.00000 -20.92000
0.00000 0.00000 0.00000 ; O- C- N- CA
6412 6414 6416 6417 3 0.65084 1.95253 0.00000
-2.60338 0.00000 0.00000 ; N- CA- CB- HB2
6412 6414 6416 6418 3 0.65084 1.95253 0.00000
-2.60338 0.00000 0.00000 ; N- CA- CB- HB3
6412 6414 6416 6419 3 3.28026 2.46019 -2.19242
-3.21331 -0.23430 0.00000 ; N- CA- CB- CG
6412 6414 6430 6431 3 0.00000 0.00000 0.00000
0.00000 0.00000 0.00000 ; N- CA- C- O
6412 6414 6430 6432 3 17.40544 -5.02080 -13.22144
9.20480 0.00000 0.00000 ; N- CA- C- N
6413 6412 6414 6415 3 0.00000 0.00000 0.00000
0.00000 0.00000 0.00000 ; H- N- CA- HA
6413 6412 6414 6416 3 0.00000 0.00000 0.00000
0.00000 0.00000 0.00000 ; H- N- CA- CB
6413 6412 6414 6430 3 0.00000 0.00000 0.00000
0.00000 0.00000 0.00000 ; H- N- CA- C
6414 6416 6419 6420 3 0.57739 0.00000 1.02926
0.00000 -1.60666 0.00000 ; CA- CB- CG- CD1
6414 6416 6419 6428 3 0.57739 0.00000 1.02926
0.00000 -1.60666 0.00000 ; CA- CB- CG- CD2
6414 6430 6432 6433 3 20.92000 0.00000 -20.92000
0.00000 0.00000 0.00000 ; CA- C- N- H
6414 6430 6432 6434 3 20.92000 0.00000 -20.92000
0.00000 0.00000 0.00000 ; CA- C- N- CA
6415 6414 6416 6417 3 0.65084 1.95253 0.00000
-2.60338 0.00000 0.00000 ; HA- CA- CB- HB2
6415 6414 6416 6418 3 0.65084 1.95253 0.00000
-2.60338 0.00000 0.00000 ; HA- CA- CB- HB3
6415 6414 6416 6419 3 0.65084 1.95253 0.00000
-2.60338 0.00000 0.00000 ; HA- CA- CB- CG
6415 6414 6430 6431 3 3.68192 -4.35136 0.00000
1.33888 0.00000 0.00000 ; HA- CA- C- O
6415 6414 6430 6432 3 0.00000 0.00000 0.00000

```
0.00000    0.00000    0.00000 ;     HA-    CA-     C-     N
  6416   6414   6430   6431      3    0.00000    0.00000    0.00000
0.00000    0.00000    0.00000 ;     CB-    CA-     C-     O
  6416   6414   6430   6432      3    2.51040    4.18400    1.67360
-6.69440    0.00000    0.00000 ;     CB-    CA-     C-     N
  6416   6419   6420   6421      3   30.33400    0.00000  -30.33400
0.00000    0.00000    0.00000 ;     CB-    CG-   CD1-   HD1
  6416   6419   6420   6422      3   30.33400    0.00000  -30.33400
0.00000    0.00000    0.00000 ;     CB-    CG-   CD1-   CE1
  6416   6419   6428   6426      3   30.33400    0.00000  -30.33400
0.00000    0.00000    0.00000 ;     CB-    CG-   CD2-   CE2
  6416   6419   6428   6429      3   30.33400    0.00000  -30.33400
0.00000    0.00000    0.00000 ;     CB-    CG-   CD2-   HD2
  6417   6416   6414   6430      3    0.65084    1.95253    0.00000
-2.60338    0.00000    0.00000 ;    HB2-    CB-    CA-     C
  6417   6416   6419   6420      3    0.00000    0.00000    0.00000
0.00000    0.00000    0.00000 ;    HB2-    CB-    CG-   CD1
  6417   6416   6419   6428      3    0.00000    0.00000    0.00000
0.00000    0.00000    0.00000 ;    HB2-    CB-    CG-   CD2
  6418   6416   6414   6430      3    0.65084    1.95253    0.00000
-2.60338    0.00000    0.00000 ;    HB3-    CB-    CA-     C
  6418   6416   6419   6420      3    0.00000    0.00000    0.00000
0.00000    0.00000    0.00000 ;    HB3-    CB-    CG-   CD1
  6418   6416   6419   6428      3    0.00000    0.00000    0.00000
0.00000    0.00000    0.00000 ;    HB3-    CB-    CG-   CD2
  6419   6416   6414   6430      3    4.95804    2.17986   -3.52293
-3.21331   -0.40166    0.00000 ;     CG-    CB-    CA-     C
  6419   6420   6422   6423      3   30.33400    0.00000  -30.33400
0.00000    0.00000    0.00000 ;     CG-   CD1-   CE1-   HE1
  6419   6420   6422   6424      3   30.33400    0.00000  -30.33400
0.00000    0.00000    0.00000 ;     CG-   CD1-   CE1-    CZ
  6419   6428   6426   6424      3   30.33400    0.00000  -30.33400
0.00000    0.00000    0.00000 ;     CG-   CD2-   CE2-    CZ
  6419   6428   6426   6427      3   30.33400    0.00000  -30.33400
0.00000    0.00000    0.00000 ;     CG-   CD2-   CE2-   HE2
  6420   6419   6428   6426      3   30.33400    0.00000  -30.33400
0.00000    0.00000    0.00000 ;    CD1-    CG-   CD2-   CE2
  6420   6419   6428   6429      3   30.33400    0.00000  -30.33400
0.00000    0.00000    0.00000 ;    CD1-    CG-   CD2-   HD2
  6420   6422   6424   6425      3   30.33400    0.00000  -30.33400
0.00000    0.00000    0.00000 ;    CD1-   CE1-    CZ-    HZ
  6420   6422   6424   6426      3   30.33400    0.00000  -30.33400
0.00000    0.00000    0.00000 ;    CD1-   CE1-    CZ-   CE2
  6421   6420   6419   6428      3   30.33400    0.00000  -30.33400
0.00000    0.00000    0.00000 ;    HD1-   CD1-    CG-   CD2
  6421   6420   6422   6423      3   30.33400    0.00000  -30.33400
0.00000    0.00000    0.00000 ;    HD1-   CD1-   CE1-   HE1
  6421   6420   6422   6424      3   30.33400    0.00000  -30.33400
0.00000    0.00000    0.00000 ;    HD1-   CD1-   CE1-    CZ
  6422   6420   6419   6428      3   30.33400    0.00000  -30.33400
0.00000    0.00000    0.00000 ;    CE1-   CD1-    CG-   CD2
  6422   6424   6426   6427      3   30.33400    0.00000  -30.33400
```

0.00000 0.00000 0.00000 ; CE1- CZ- CE2- HE2
6422 6424 6426 6428 3 30.33400 0.00000 -30.33400
0.00000 0.00000 0.00000 ; CE1- CZ- CE2- CD2
6423 6422 6424 6425 3 30.33400 0.00000 -30.33400
0.00000 0.00000 0.00000 ; HE1- CE1- CZ- HZ
6423 6422 6424 6426 3 30.33400 0.00000 -30.33400
0.00000 0.00000 0.00000 ; HE1- CE1- CZ- CE2
6424 6426 6428 6429 3 30.33400 0.00000 -30.33400
0.00000 0.00000 0.00000 ; CZ- CE2- CD2- HD2
6425 6424 6426 6427 3 30.33400 0.00000 -30.33400
0.00000 0.00000 0.00000 ; HZ- CZ- CE2- HE2
6425 6424 6426 6428 3 30.33400 0.00000 -30.33400
0.00000 0.00000 0.00000 ; HZ- CZ- CE2- CD2
6427 6426 6428 6429 3 30.33400 0.00000 -30.33400
0.00000 0.00000 0.00000 ; HE2- CE2- CD2- HD2
6430 6432 6434 6435 3 0.00000 0.00000 0.00000
0.00000 0.00000 0.00000 ; C- N- CA- HA
6430 6432 6434 6436 3 11.71520 1.67360 15.06240 -
13.38880 0.00000 0.00000 ; C- N- CA- CB
6430 6432 6434 6442 3 1.75728 5.27184 2.25936
-7.02912 0.00000 0.00000 ; C- N- CA- C
6431 6430 6432 6433 3 29.28800 -8.36800 -20.92000
0.00000 0.00000 0.00000 ; O- C- N- H
6431 6430 6432 6434 3 20.92000 0.00000 -20.92000
0.00000 0.00000 0.00000 ; O- C- N- CA
6432 6434 6436 6437 3 0.65084 1.95253 0.00000
-2.60338 0.00000 0.00000 ; N- CA- CB- HB2
6432 6434 6436 6438 3 0.65084 1.95253 0.00000
-2.60338 0.00000 0.00000 ; N- CA- CB- HB3
6432 6434 6436 6439 3 15.41386 9.74035 -8.39310
-0.97069 2.97901 0.00000 ; N- CA- CB- CG
6432 6434 6442 6443 3 0.00000 0.00000 0.00000
0.00000 0.00000 0.00000 ; N- CA- C- O
6432 6434 6442 6444 3 17.40544 -5.02080 -13.22144
9.20480 0.00000 0.00000 ; N- CA- C- N
6433 6432 6434 6435 3 0.00000 0.00000 0.00000
0.00000 0.00000 0.00000 ; H- N- CA- HA
6433 6432 6434 6436 3 0.00000 0.00000 0.00000
0.00000 0.00000 0.00000 ; H- N- CA- CB
6433 6432 6434 6442 3 0.00000 0.00000 0.00000
0.00000 0.00000 0.00000 ; H- N- CA- C
6434 6436 6439 6440 3 6.43499 0.00000 -5.39736
0.00000 -1.03763 0.00000 ; CA- CB- CG- OD1
6434 6436 6439 6441 3 6.43499 0.00000 -5.39736
0.00000 -1.03763 0.00000 ; CA- CB- CG- OD2
6434 6442 6444 6445 3 20.92000 0.00000 -20.92000
0.00000 0.00000 0.00000 ; CA- C- N- H
6434 6442 6444 6446 3 20.92000 0.00000 -20.92000
0.00000 0.00000 0.00000 ; CA- C- N- CA
6435 6434 6436 6437 3 0.65084 1.95253 0.00000
-2.60338 0.00000 0.00000 ; HA- CA- CB- HB2
6435 6434 6436 6438 3 0.65084 1.95253 0.00000

```
-2.60338    0.00000    0.00000 ;      HA-    CA-    CB-   HB3
  6435   6434   6436   6439      3    0.65084    1.95253    0.00000
-2.60338    0.00000    0.00000 ;      HA-    CA-    CB-   CG
  6435   6434   6442   6443      3    3.68192   -4.35136    0.00000
1.33888    0.00000    0.00000 ;      HA-    CA-    C-    O
  6435   6434   6442   6444      3    0.00000    0.00000    0.00000
0.00000    0.00000    0.00000 ;      HA-    CA-    C-    N
  6436   6434   6442   6443      3    0.00000    0.00000    0.00000
0.00000    0.00000    0.00000 ;      CB-    CA-    C-    O
  6436   6434   6442   6444      3    2.51040    4.18400    1.67360
-6.69440    0.00000    0.00000 ;      CB-    CA-    C-    N
  6437   6436   6434   6442      3    0.65084    1.95253    0.00000
-2.60338    0.00000    0.00000 ;     HB2-    CB-    CA-    C
  6437   6436   6439   6440      3    0.00000    0.00000    0.00000
0.00000    0.00000    0.00000 ;     HB2-    CB-    CG-   OD1
  6437   6436   6439   6441      3    0.00000    0.00000    0.00000
0.00000    0.00000    0.00000 ;     HB2-    CB-    CG-   OD2
  6438   6436   6434   6442      3    0.65084    1.95253    0.00000
-2.60338    0.00000    0.00000 ;     HB3-    CB-    CA-    C
  6438   6436   6439   6440      3    0.00000    0.00000    0.00000
0.00000    0.00000    0.00000 ;     HB3-    CB-    CG-   OD1
  6438   6436   6439   6441      3    0.00000    0.00000    0.00000
0.00000    0.00000    0.00000 ;     HB3-    CB-    CG-   OD2
  6439   6436   6434   6442      3    7.14627   -1.04600   -8.99560
-0.97069    5.15469    0.00000 ;      CG-    CB-    CA-    C
  6442   6444   6446   6447      3    0.00000    0.00000    0.00000
0.00000    0.00000    0.00000 ;      C-     N-    CA-    HA
  6442   6444   6446   6448      3   11.71520    1.67360   15.06240  -
13.38880    0.00000    0.00000 ;      C-     N-    CA-    CB
  6442   6444   6446   6461      3    1.75728    5.27184    2.25936
-7.02912    0.00000    0.00000 ;      C-     N-    CA-    C
  6443   6442   6444   6445      3   29.28800   -8.36800  -20.92000
0.00000    0.00000    0.00000 ;      O-     C-     N-    H
  6443   6442   6444   6446      3   20.92000    0.00000  -20.92000
0.00000    0.00000    0.00000 ;      O-     C-     N-    CA
  6444   6446   6448   6449      3    0.65084    1.95253    0.00000
-2.60338    0.00000    0.00000 ;      N-    CA-    CB-    HB
  6444   6446   6448   6450      3    3.83673    0.44769   -1.77402
-2.47693   -0.03347    0.00000 ;      N-    CA-    CB-   CG2
  6444   6446   6448   6454      3    2.97482    0.12134    2.04179
-1.89117   -3.24678    0.00000 ;      N-    CA-    CB-   CG1
  6444   6446   6461   6462      3    0.00000    0.00000    0.00000
0.00000    0.00000    0.00000 ;      N-    CA-    C-    O
  6444   6446   6461   6463      3   17.40544   -5.02080  -13.22144
9.20480    0.00000    0.00000 ;      N-    CA-    C-    N
  6445   6444   6446   6447      3    0.00000    0.00000    0.00000
0.00000    0.00000    0.00000 ;      H-     N-    CA-    HA
  6445   6444   6446   6448      3    0.00000    0.00000    0.00000
0.00000    0.00000    0.00000 ;      H-     N-    CA-    CB
  6445   6444   6446   6461      3    0.00000    0.00000    0.00000
0.00000    0.00000    0.00000 ;      H-     N-    CA-    C
  6446   6448   6450   6451      3    0.66944    2.00832    0.00000
```

```
-2.67776    0.00000    0.00000 ;      CA-    CB-   CG2-  HG21
  6446   6448   6450   6452      3    0.66944    2.00832    0.00000
-2.67776    0.00000    0.00000 ;      CA-    CB-   CG2-  HG22
  6446   6448   6450   6453      3    0.66944    2.00832    0.00000
-2.67776    0.00000    0.00000 ;      CA-    CB-   CG2-  HG23
  6446   6448   6454   6455      3    0.66944    2.00832    0.00000
-2.67776    0.00000    0.00000 ;      CA-    CB-   CG1-  HG12
  6446   6448   6454   6456      3    0.66944    2.00832    0.00000
-2.67776    0.00000    0.00000 ;      CA-    CB-   CG1-  HG13
  6446   6448   6454   6457      3    4.24258   -0.52718   -7.25506
-1.79075    7.69856    0.00000 ;      CA-    CB-   CG1-   CD1
  6446   6461   6463   6464      3   20.92000    0.00000  -20.92000
0.00000    0.00000    0.00000 ;      CA-     C-     N-     H
  6446   6461   6463   6465      3   20.92000    0.00000  -20.92000
0.00000    0.00000    0.00000 ;      CA-     C-     N-    CA
  6447   6446   6448   6449      3    0.65084    1.95253    0.00000
-2.60338    0.00000    0.00000 ;      HA-    CA-    CB-    HB
  6447   6446   6448   6450      3    0.65084    1.95253    0.00000
-2.60338    0.00000    0.00000 ;      HA-    CA-    CB-   CG2
  6447   6446   6448   6454      3    0.65084    1.95253    0.00000
-2.60338    0.00000    0.00000 ;      HA-    CA-    CB-   CG1
  6447   6446   6461   6462      3    3.68192   -4.35136    0.00000
1.33888    0.00000    0.00000 ;      HA-    CA-     C-     O
  6447   6446   6461   6463      3    0.00000    0.00000    0.00000
0.00000    0.00000    0.00000 ;      HA-    CA-     C-     N
  6448   6446   6461   6462      3    0.00000    0.00000    0.00000
0.00000    0.00000    0.00000 ;      CB-    CA-     C-     O
  6448   6446   6461   6463      3    2.51040    4.18400    1.67360
-6.69440    0.00000    0.00000 ;      CB-    CA-     C-     N
  6448   6454   6457   6458      3    0.66944    2.00832    0.00000
-2.67776    0.00000    0.00000 ;      CB-   CG1-   CD1-  HD11
  6448   6454   6457   6459      3    0.66944    2.00832    0.00000
-2.67776    0.00000    0.00000 ;      CB-   CG1-   CD1-  HD12
  6448   6454   6457   6460      3    0.66944    2.00832    0.00000
-2.67776    0.00000    0.00000 ;      CB-   CG1-   CD1-  HD13
  6449   6448   6446   6461      3    0.65084    1.95253    0.00000
-2.60338    0.00000    0.00000 ;      HB-    CB-    CA-     C
  6449   6448   6450   6451      3    0.62760    1.88280    0.00000
-2.51040    0.00000    0.00000 ;      HB-    CB-   CG2-  HG21
  6449   6448   6450   6452      3    0.62760    1.88280    0.00000
-2.51040    0.00000    0.00000 ;      HB-    CB-   CG2-  HG22
  6449   6448   6450   6453      3    0.62760    1.88280    0.00000
-2.51040    0.00000    0.00000 ;      HB-    CB-   CG2-  HG23
  6449   6448   6454   6455      3    0.62760    1.88280    0.00000
-2.51040    0.00000    0.00000 ;      HB-    CB-   CG1-  HG12
  6449   6448   6454   6456      3    0.62760    1.88280    0.00000
-2.51040    0.00000    0.00000 ;      HB-    CB-   CG1-  HG13
  6449   6448   6454   6457      3    0.66944    2.00832    0.00000
-2.67776    0.00000    0.00000 ;      HB-    CB-   CG1-   CD1
  6450   6448   6446   6461      3    5.67350    3.55640   -6.16722
-2.47693    3.74886    0.00000 ;     CG2-    CB-    CA-     C
  6450   6448   6454   6455      3    0.66944    2.00832    0.00000
```

-2.67776 0.00000 0.00000 ; CG2- CB- CG1- HG12
6450 6448 6454 6456 3 0.66944 2.00832 0.00000
-2.67776 0.00000 0.00000 ; CG2- CB- CG1- HG13
6450 6448 6454 6457 3 3.81162 0.49790 -8.14206
-1.79075 7.49773 0.00000 ; CG2- CB- CG1- CD1
6451 6450 6448 6454 3 0.66944 2.00832 0.00000
-2.67776 0.00000 0.00000 ; HG21- CG2- CB- CG1
6452 6450 6448 6454 3 0.66944 2.00832 0.00000
-2.67776 0.00000 0.00000 ; HG22- CG2- CB- CG1
6453 6450 6448 6454 3 0.66944 2.00832 0.00000
-2.67776 0.00000 0.00000 ; HG23- CG2- CB- CG1
6454 6448 6446 6461 3 8.26340 0.74057 -9.99976
-1.89117 3.84928 0.00000 ; CG1- CB- CA- C
6455 6454 6457 6458 3 0.62760 1.88280 0.00000
-2.51040 0.00000 0.00000 ; HG12- CG1- CD1- HD11
6455 6454 6457 6459 3 0.62760 1.88280 0.00000
-2.51040 0.00000 0.00000 ; HG12- CG1- CD1- HD12
6455 6454 6457 6460 3 0.62760 1.88280 0.00000
-2.51040 0.00000 0.00000 ; HG12- CG1- CD1- HD13
6456 6454 6457 6458 3 0.62760 1.88280 0.00000
-2.51040 0.00000 0.00000 ; HG13- CG1- CD1- HD11
6456 6454 6457 6459 3 0.62760 1.88280 0.00000
-2.51040 0.00000 0.00000 ; HG13- CG1- CD1- HD12
6456 6454 6457 6460 3 0.62760 1.88280 0.00000
-2.51040 0.00000 0.00000 ; HG13- CG1- CD1- HD13
6461 6463 6465 6466 3 0.00000 0.00000 0.00000
0.00000 0.00000 0.00000 ; C- N- CA- HA
6461 6463 6465 6467 3 11.71520 1.67360 15.06240 -
13.38880 0.00000 0.00000 ; C- N- CA- CB
6461 6463 6465 6473 3 1.75728 5.27184 2.25936
-7.02912 0.00000 0.00000 ; C- N- CA- C
6462 6461 6463 6464 3 29.28800 -8.36800 -20.92000
0.00000 0.00000 0.00000 ; O- C- N- H
6462 6461 6463 6465 3 20.92000 0.00000 -20.92000
0.00000 0.00000 0.00000 ; O- C- N- CA
6463 6465 6467 6468 3 0.65084 1.95253 0.00000
-2.60338 0.00000 0.00000 ; N- CA- CB- HB2
6463 6465 6467 6469 3 0.65084 1.95253 0.00000
-2.60338 0.00000 0.00000 ; N- CA- CB- HB3
6463 6465 6467 6470 3 15.41386 9.74035 -8.39310
-0.97069 2.97901 0.00000 ; N- CA- CB- CG
6463 6465 6473 6474 3 0.00000 0.00000 0.00000
0.00000 0.00000 0.00000 ; N- CA- C- O
6463 6465 6473 6475 3 17.40544 -5.02080 -13.22144
9.20480 0.00000 0.00000 ; N- CA- C- N
6464 6463 6465 6466 3 0.00000 0.00000 0.00000
0.00000 0.00000 0.00000 ; H- N- CA- HA
6464 6463 6465 6467 3 0.00000 0.00000 0.00000
0.00000 0.00000 0.00000 ; H- N- CA- CB
6464 6463 6465 6473 3 0.00000 0.00000 0.00000
0.00000 0.00000 0.00000 ; H- N- CA- C
6465 6467 6470 6471 3 6.43499 0.00000 -5.39736

```
0.00000   -1.03763    0.00000 ;      CA-     CB-     CG-   OD1
  6465   6467   6470   6472       3    6.43499    0.00000   -5.39736
0.00000   -1.03763    0.00000 ;      CA-     CB-     CG-   OD2
  6465   6473   6475   6476       3   20.92000    0.00000  -20.92000
0.00000    0.00000    0.00000 ;      CA-      C-      N-     H
  6465   6473   6475   6477       3   20.92000    0.00000  -20.92000
0.00000    0.00000    0.00000 ;      CA-      C-      N-    CA
  6466   6465   6467   6468       3    0.65084    1.95253    0.00000
-2.60338    0.00000    0.00000 ;      HA-     CA-     CB-   HB2
  6466   6465   6467   6469       3    0.65084    1.95253    0.00000
-2.60338    0.00000    0.00000 ;      HA-     CA-     CB-   HB3
  6466   6465   6467   6470       3    0.65084    1.95253    0.00000
-2.60338    0.00000    0.00000 ;      HA-     CA-     CB-    CG
  6466   6465   6473   6474       3    3.68192   -4.35136    0.00000
1.33888    0.00000    0.00000 ;      HA-     CA-      C-     O
  6466   6465   6473   6475       3    0.00000    0.00000    0.00000
0.00000    0.00000    0.00000 ;      HA-     CA-      C-     N
  6467   6465   6473   6474       3    0.00000    0.00000    0.00000
0.00000    0.00000    0.00000 ;      CB-     CA-      C-     O
  6467   6465   6473   6475       3    2.51040    4.18400    1.67360
-6.69440    0.00000    0.00000 ;      CB-     CA-      C-     N
  6468   6467   6465   6473       3    0.65084    1.95253    0.00000
-2.60338    0.00000    0.00000 ;     HB2-     CB-     CA-     C
  6468   6467   6470   6471       3    0.00000    0.00000    0.00000
0.00000    0.00000    0.00000 ;     HB2-     CB-     CG-   OD1
  6468   6467   6470   6472       3    0.00000    0.00000    0.00000
0.00000    0.00000    0.00000 ;     HB2-     CB-     CG-   OD2
  6469   6467   6465   6473       3    0.65084    1.95253    0.00000
-2.60338    0.00000    0.00000 ;     HB3-     CB-     CA-     C
  6469   6467   6470   6471       3    0.00000    0.00000    0.00000
0.00000    0.00000    0.00000 ;     HB3-     CB-     CG-   OD1
  6469   6467   6470   6472       3    0.00000    0.00000    0.00000
0.00000    0.00000    0.00000 ;     HB3-     CB-     CG-   OD2
  6470   6467   6465   6473       3    7.14627   -1.04600   -8.99560
-0.97069    5.15469    0.00000 ;      CG-     CB-     CA-     C
  6473   6475   6477   6478       3    0.00000    0.00000    0.00000
0.00000    0.00000    0.00000 ;       C-      N-     CA-   HA2
  6473   6475   6477   6479       3    0.00000    0.00000    0.00000
0.00000    0.00000    0.00000 ;       C-      N-     CA-   HA3
  6473   6475   6477   6480       3    1.75728    5.27184    2.25936
-7.02912    0.00000    0.00000 ;       C-      N-     CA-     C
  6474   6473   6475   6476       3   29.28800   -8.36800  -20.92000
0.00000    0.00000    0.00000 ;       O-      C-      N-     H
  6474   6473   6475   6477       3   20.92000    0.00000  -20.92000
0.00000    0.00000    0.00000 ;       O-      C-      N-    CA
  6475   6477   6480   6481       3    0.00000    0.00000    0.00000
0.00000    0.00000    0.00000 ;       N-     CA-      C-     O
  6475   6477   6480   6482       3   17.40544   -5.02080  -13.22144
9.20480    0.00000    0.00000 ;       N-     CA-      C-     N
  6476   6475   6477   6478       3    0.00000    0.00000    0.00000
0.00000    0.00000    0.00000 ;       H-      N-     CA-   HA2
  6476   6475   6477   6479       3    0.00000    0.00000    0.00000
```

0.00000 0.00000 0.00000 ; H- N- CA- HA3
6476 6475 6477 6480 3 0.00000 0.00000 0.00000
0.00000 0.00000 0.00000 ; H- N- CA- C
6477 6480 6482 6483 3 20.92000 0.00000 -20.92000
0.00000 0.00000 0.00000 ; CA- C- N- H
6477 6480 6482 6484 3 20.92000 0.00000 -20.92000
0.00000 0.00000 0.00000 ; CA- C- N- CA
6478 6477 6480 6481 3 3.68192 -4.35136 0.00000
1.33888 0.00000 0.00000 ; HA2- CA- C- O
6478 6477 6480 6482 3 0.00000 0.00000 0.00000
0.00000 0.00000 0.00000 ; HA2- CA- C- N
6479 6477 6480 6481 3 3.68192 -4.35136 0.00000
1.33888 0.00000 0.00000 ; HA3- CA- C- O
6479 6477 6480 6482 3 0.00000 0.00000 0.00000
0.00000 0.00000 0.00000 ; HA3- CA- C- N
6480 6482 6484 6485 3 0.00000 0.00000 0.00000
0.00000 0.00000 0.00000 ; C- N- CA- HA
6480 6482 6484 6486 3 11.71520 1.67360 15.06240 -
13.38880 0.00000 0.00000 ; C- N- CA- CB
6480 6482 6484 6494 3 1.75728 5.27184 2.25936
-7.02912 0.00000 0.00000 ; C- N- CA- C
6481 6480 6482 6483 3 29.28800 -8.36800 -20.92000
0.00000 0.00000 0.00000 ; O- C- N- H
6481 6480 6482 6484 3 20.92000 0.00000 -20.92000
0.00000 0.00000 0.00000 ; O- C- N- CA
6482 6484 6486 6487 3 0.65084 1.95253 0.00000
-2.60338 0.00000 0.00000 ; N- CA- CB- HB2
6482 6484 6486 6488 3 0.65084 1.95253 0.00000
-2.60338 0.00000 0.00000 ; N- CA- CB- HB3
6482 6484 6486 6489 3 5.99567 3.29281 -4.46014
-0.55229 1.97485 0.00000 ; N- CA- CB- CG
6482 6484 6494 6495 3 0.00000 0.00000 0.00000
0.00000 0.00000 0.00000 ; N- CA- C- O
6482 6484 6494 6496 3 17.40544 -5.02080 -13.22144
9.20480 0.00000 0.00000 ; N- CA- C- N
6483 6482 6484 6485 3 0.00000 0.00000 0.00000
0.00000 0.00000 0.00000 ; H- N- CA- HA
6483 6482 6484 6486 3 0.00000 0.00000 0.00000
0.00000 0.00000 0.00000 ; H- N- CA- CB
6483 6482 6484 6494 3 0.00000 0.00000 0.00000
0.00000 0.00000 0.00000 ; H- N- CA- C
6484 6486 6489 6490 3 0.00000 0.00000 0.00000
0.00000 0.00000 0.00000 ; CA- CB- CG- OD1
6484 6486 6489 6491 3 8.84916 -0.31380 -4.32626
5.03754 0.26778 0.00000 ; CA- CB- CG- ND2
6484 6494 6496 6497 3 20.92000 0.00000 -20.92000
0.00000 0.00000 0.00000 ; CA- C- N- H
6484 6494 6496 6498 3 20.92000 0.00000 -20.92000
0.00000 0.00000 0.00000 ; CA- C- N- CA
6485 6484 6486 6487 3 0.65084 1.95253 0.00000
-2.60338 0.00000 0.00000 ; HA- CA- CB- HB2
6485 6484 6486 6488 3 0.65084 1.95253 0.00000

```
-2.60338    0.00000    0.00000 ;      HA-    CA-    CB-   HB3
  6485   6484   6486   6489      3    0.65084    1.95253    0.00000
-2.60338    0.00000    0.00000 ;      HA-    CA-    CB-    CG
  6485   6484   6494   6495      3    3.68192   -4.35136    0.00000
1.33888    0.00000    0.00000 ;      HA-    CA-     C-     O
  6485   6484   6494   6496      3    0.00000    0.00000    0.00000
0.00000    0.00000    0.00000 ;      HA-    CA-     C-     N
  6486   6484   6494   6495      3    0.00000    0.00000    0.00000
0.00000    0.00000    0.00000 ;      CB-    CA-     C-     O
  6486   6484   6494   6496      3    2.51040    4.18400    1.67360
-6.69440    0.00000    0.00000 ;      CB-    CA-     C-     N
  6486   6489   6491   6492      3   20.92000    0.00000  -20.92000
0.00000    0.00000    0.00000 ;      CB-    CG-   ND2-  HD21
  6486   6489   6491   6493      3   20.92000    0.00000  -20.92000
0.00000    0.00000    0.00000 ;      CB-    CG-   ND2-  HD22
  6487   6486   6484   6494      3    0.65084    1.95253    0.00000
-2.60338    0.00000    0.00000 ;     HB2-    CB-    CA-     C
  6487   6486   6489   6490      3    3.68192   -4.35136    0.00000
1.33888    0.00000    0.00000 ;     HB2-    CB-    CG-   OD1
  6487   6486   6489   6491      3    0.00000    0.00000    0.00000
0.00000    0.00000    0.00000 ;     HB2-    CB-    CG-   ND2
  6488   6486   6484   6494      3    0.65084    1.95253    0.00000
-2.60338    0.00000    0.00000 ;     HB3-    CB-    CA-     C
  6488   6486   6489   6490      3    3.68192   -4.35136    0.00000
1.33888    0.00000    0.00000 ;     HB3-    CB-    CG-   OD1
  6488   6486   6489   6491      3    0.00000    0.00000    0.00000
0.00000    0.00000    0.00000 ;     HB3-    CB-    CG-   ND2
  6489   6486   6484   6494      3    7.94542   -3.96225   -6.11701
-0.55229    3.58150    0.00000 ;      CG-    CB-    CA-     C
  6490   6489   6491   6492      3   29.28800   -8.36800  -20.92000
0.00000    0.00000    0.00000 ;     OD1-    CG-   ND2-  HD21
  6490   6489   6491   6493      3   29.28800   -8.36800  -20.92000
0.00000    0.00000    0.00000 ;     OD1-    CG-   ND2-  HD22
  6494   6496   6498   6499      3    0.00000    0.00000    0.00000
0.00000    0.00000    0.00000 ;       C-     N-    CA-   HA2
  6494   6496   6498   6500      3    0.00000    0.00000    0.00000
0.00000    0.00000    0.00000 ;       C-     N-    CA-   HA3
  6494   6496   6498   6501      3    1.75728    5.27184    2.25936
-7.02912    0.00000    0.00000 ;       C-     N-    CA-     C
  6495   6494   6496   6497      3   29.28800   -8.36800  -20.92000
0.00000    0.00000    0.00000 ;       O-     C-     N-     H
  6495   6494   6496   6498      3   20.92000    0.00000  -20.92000
0.00000    0.00000    0.00000 ;       O-     C-     N-    CA
  6496   6498   6501   6502      3    0.00000    0.00000    0.00000
0.00000    0.00000    0.00000 ;       N-    CA-     C-     O
  6496   6498   6501   6503      3   17.40544   -5.02080  -13.22144
9.20480    0.00000    0.00000 ;       N-    CA-     C-     N
  6497   6496   6498   6499      3    0.00000    0.00000    0.00000
0.00000    0.00000    0.00000 ;       H-     N-    CA-   HA2
  6497   6496   6498   6500      3    0.00000    0.00000    0.00000
0.00000    0.00000    0.00000 ;       H-     N-    CA-   HA3
  6497   6496   6498   6501      3    0.00000    0.00000    0.00000
```

0.00000 0.00000 0.00000 ; H- N- CA- C
6498 6501 6503 6504 3 20.92000 0.00000 -20.92000
0.00000 0.00000 0.00000 ; CA- C- N- H
6498 6501 6503 6505 3 20.92000 0.00000 -20.92000
0.00000 0.00000 0.00000 ; CA- C- N- CA
6499 6498 6501 6502 3 3.68192 -4.35136 0.00000
1.33888 0.00000 0.00000 ; HA2- CA- C- O
6499 6498 6501 6503 3 0.00000 0.00000 0.00000
0.00000 0.00000 0.00000 ; HA2- CA- C- N
6500 6498 6501 6502 3 3.68192 -4.35136 0.00000
1.33888 0.00000 0.00000 ; HA3- CA- C- O
6500 6498 6501 6503 3 0.00000 0.00000 0.00000
0.00000 0.00000 0.00000 ; HA3- CA- C- N
6501 6503 6505 6506 3 0.00000 0.00000 0.00000
0.00000 0.00000 0.00000 ; C- N- CA- HA
6501 6503 6505 6507 3 11.71520 1.67360 15.06240 -
13.38880 0.00000 0.00000 ; C- N- CA- CB
6501 6503 6505 6523 3 1.75728 5.27184 2.25936
-7.02912 0.00000 0.00000 ; C- N- CA- C
6502 6501 6503 6504 3 29.28800 -8.36800 -20.92000
0.00000 0.00000 0.00000 ; O- C- N- H
6502 6501 6503 6505 3 20.92000 0.00000 -20.92000
0.00000 0.00000 0.00000 ; O- C- N- CA
6503 6505 6507 6508 3 0.65084 1.95253 0.00000
-2.60338 0.00000 0.00000 ; N- CA- CB- HB2
6503 6505 6507 6509 3 0.65084 1.95253 0.00000
-2.60338 0.00000 0.00000 ; N- CA- CB- HB3
6503 6505 6507 6510 3 0.65084 1.95253 0.00000
-2.60338 0.00000 0.00000 ; N- CA- CB- CG
6503 6505 6523 6524 3 0.00000 0.00000 0.00000
0.00000 0.00000 0.00000 ; N- CA- C- O
6503 6505 6523 6525 3 17.40544 -5.02080 -13.22144
9.20480 0.00000 0.00000 ; N- CA- C- N
6504 6503 6505 6506 3 0.00000 0.00000 0.00000
0.00000 0.00000 0.00000 ; H- N- CA- HA
6504 6503 6505 6507 3 0.00000 0.00000 0.00000
0.00000 0.00000 0.00000 ; H- N- CA- CB
6504 6503 6505 6523 3 0.00000 0.00000 0.00000
0.00000 0.00000 0.00000 ; H- N- CA- C
6505 6507 6510 6511 3 0.66944 2.00832 0.00000
-2.67776 0.00000 0.00000 ; CA- CB- CG- HG2
6505 6507 6510 6512 3 0.66944 2.00832 0.00000
-2.67776 0.00000 0.00000 ; CA- CB- CG- HG3
6505 6507 6510 6513 3 3.68192 3.09616 -2.09200
-3.01248 0.00000 0.00000 ; CA- CB- CG- CD
6505 6523 6525 6526 3 20.92000 0.00000 -20.92000
0.00000 0.00000 0.00000 ; CA- C- N- H
6505 6523 6525 6527 3 20.92000 0.00000 -20.92000
0.00000 0.00000 0.00000 ; CA- C- N- CA
6506 6505 6507 6508 3 0.65084 1.95253 0.00000
-2.60338 0.00000 0.00000 ; HA- CA- CB- HB2
6506 6505 6507 6509 3 0.65084 1.95253 0.00000

```
-2.60338    0.00000    0.00000 ;      HA-    CA-     CB-   HB3
  6506   6505   6507   6510       3    0.65084    1.95253    0.00000
-2.60338    0.00000    0.00000 ;      HA-    CA-     CB-    CG
  6506   6505   6523   6524       3    3.68192   -4.35136    0.00000
1.33888    0.00000    0.00000 ;      HA-    CA-     C-     O
  6506   6505   6523   6525       3    0.00000    0.00000    0.00000
0.00000    0.00000    0.00000 ;      HA-    CA-     C-     N
  6507   6505   6523   6524       3    0.00000    0.00000    0.00000
0.00000    0.00000    0.00000 ;      CB-    CA-     C-     O
  6507   6505   6523   6525       3    2.51040    4.18400    1.67360
-6.69440    0.00000    0.00000 ;      CB-    CA-     C-     N
  6507   6510   6513   6514       3    0.66944    2.00832    0.00000
-2.67776    0.00000    0.00000 ;      CB-    CG-     CD-   HD2
  6507   6510   6513   6515       3    0.66944    2.00832    0.00000
-2.67776    0.00000    0.00000 ;      CB-    CG-     CD-   HD3
  6507   6510   6513   6516       3    3.68192    3.09616   -2.09200
-3.01248    0.00000    0.00000 ;      CB-    CG-     CD-    CE
  6508   6507   6505   6523       3    0.65084    1.95253    0.00000
-2.60338    0.00000    0.00000 ;     HB2-    CB-     CA-     C
  6508   6507   6510   6511       3    0.62760    1.88280    0.00000
-2.51040    0.00000    0.00000 ;     HB2-    CB-     CG-   HG2
  6508   6507   6510   6512       3    0.62760    1.88280    0.00000
-2.51040    0.00000    0.00000 ;     HB2-    CB-     CG-   HG3
  6508   6507   6510   6513       3    0.66944    2.00832    0.00000
-2.67776    0.00000    0.00000 ;     HB2-    CB-     CG-    CD
  6509   6507   6505   6523       3    0.65084    1.95253    0.00000
-2.60338    0.00000    0.00000 ;     HB3-    CB-     CA-     C
  6509   6507   6510   6511       3    0.62760    1.88280    0.00000
-2.51040    0.00000    0.00000 ;     HB3-    CB-     CG-   HG2
  6509   6507   6510   6512       3    0.62760    1.88280    0.00000
-2.51040    0.00000    0.00000 ;     HB3-    CB-     CG-   HG3
  6509   6507   6510   6513       3    0.66944    2.00832    0.00000
-2.67776    0.00000    0.00000 ;     HB3-    CB-     CG-    CD
  6510   6507   6505   6523       3    0.65084    1.95253    0.00000
-2.60338    0.00000    0.00000 ;      CG-    CB-     CA-     C
  6510   6513   6516   6517       3    0.65084    1.95253    0.00000
-2.60338    0.00000    0.00000 ;      CG-    CD-     CE-   HE2
  6510   6513   6516   6518       3    0.65084    1.95253    0.00000
-2.60338    0.00000    0.00000 ;      CG-    CD-     CE-   HE3
  6510   6513   6516   6519       3    0.65084    1.95253    0.00000
-2.60338    0.00000    0.00000 ;      CG-    CD-     CE-    NZ
  6511   6510   6513   6514       3    0.62760    1.88280    0.00000
-2.51040    0.00000    0.00000 ;     HG2-    CG-     CD-   HD2
  6511   6510   6513   6515       3    0.62760    1.88280    0.00000
-2.51040    0.00000    0.00000 ;     HG2-    CG-     CD-   HD3
  6511   6510   6513   6516       3    0.66944    2.00832    0.00000
-2.67776    0.00000    0.00000 ;     HG2-    CG-     CD-    CE
  6512   6510   6513   6514       3    0.62760    1.88280    0.00000
-2.51040    0.00000    0.00000 ;     HG3-    CG-     CD-   HD2
  6512   6510   6513   6515       3    0.62760    1.88280    0.00000
-2.51040    0.00000    0.00000 ;     HG3-    CG-     CD-   HD3
  6512   6510   6513   6516       3    0.66944    2.00832    0.00000
```

-2.67776    0.00000    0.00000 ;    HG3-    CG-    CD-    CE
6513   6516   6519   6520      3    0.65084    1.95253    0.00000
-2.60338    0.00000    0.00000 ;    CD-    CE-    NZ-   HZ1
6513   6516   6519   6521      3    0.65084    1.95253    0.00000
-2.60338    0.00000    0.00000 ;    CD-    CE-    NZ-   HZ2
6513   6516   6519   6522      3    0.65084    1.95253    0.00000
-2.60338    0.00000    0.00000 ;    CD-    CE-    NZ-   HZ3
6514   6513   6516   6517      3    0.65084    1.95253    0.00000
-2.60338    0.00000    0.00000 ;    HD2-    CD-    CE-   HE2
6514   6513   6516   6518      3    0.65084    1.95253    0.00000
-2.60338    0.00000    0.00000 ;    HD2-    CD-    CE-   HE3
6514   6513   6516   6519      3    0.65084    1.95253    0.00000
-2.60338    0.00000    0.00000 ;    HD2-    CD-    CE-    NZ
6515   6513   6516   6517      3    0.65084    1.95253    0.00000
-2.60338    0.00000    0.00000 ;    HD3-    CD-    CE-   HE2
6515   6513   6516   6518      3    0.65084    1.95253    0.00000
-2.60338    0.00000    0.00000 ;    HD3-    CD-    CE-   HE3
6515   6513   6516   6519      3    0.65084    1.95253    0.00000
-2.60338    0.00000    0.00000 ;    HD3-    CD-    CE-    NZ
6517   6516   6519   6520      3    0.65084    1.95253    0.00000
-2.60338    0.00000    0.00000 ;    HE2-    CE-    NZ-   HZ1
6517   6516   6519   6521      3    0.65084    1.95253    0.00000
-2.60338    0.00000    0.00000 ;    HE2-    CE-    NZ-   HZ2
6517   6516   6519   6522      3    0.65084    1.95253    0.00000
-2.60338    0.00000    0.00000 ;    HE2-    CE-    NZ-   HZ3
6518   6516   6519   6520      3    0.65084    1.95253    0.00000
-2.60338    0.00000    0.00000 ;    HE3-    CE-    NZ-   HZ1
6518   6516   6519   6521      3    0.65084    1.95253    0.00000
-2.60338    0.00000    0.00000 ;    HE3-    CE-    NZ-   HZ2
6518   6516   6519   6522      3    0.65084    1.95253    0.00000
-2.60338    0.00000    0.00000 ;    HE3-    CE-    NZ-   HZ3
6523   6525   6527   6528      3    0.00000    0.00000    0.00000
0.00000    0.00000    0.00000 ;    C-    N-    CA-   HA
6523   6525   6527   6529      3   11.71520    1.67360   15.06240  -
13.38880    0.00000    0.00000 ;    C-    N-    CA-   CB
6523   6525   6527   6542      3    1.75728    5.27184    2.25936
-7.02912    0.00000    0.00000 ;    C-    N-    CA-    C
6524   6523   6525   6526      3   29.28800   -8.36800  -20.92000
0.00000    0.00000    0.00000 ;    O-    C-    N-    H
6524   6523   6525   6527      3   20.92000    0.00000  -20.92000
0.00000    0.00000    0.00000 ;    O-    C-    N-   CA
6525   6527   6529   6530      3    0.65084    1.95253    0.00000
-2.60338    0.00000    0.00000 ;    N-    CA-    CB-    HB
6525   6527   6529   6531      3    3.83673    0.44769   -1.77402
-2.47693   -0.03347    0.00000 ;    N-    CA-    CB-   CG2
6525   6527   6529   6535      3    2.97482    0.12134    2.04179
-1.89117   -3.24678    0.00000 ;    N-    CA-    CB-   CG1
6525   6527   6542   6543      3    0.00000    0.00000    0.00000
0.00000    0.00000    0.00000 ;    N-    CA-    C-    O
6525   6527   6542   6544      3   17.40544   -5.02080  -13.22144
9.20480    0.00000    0.00000 ;    N-    CA-    C-    N
6526   6525   6527   6528      3    0.00000    0.00000    0.00000

0.00000    0.00000    0.00000 ;       H-     N-     CA-    HA
  6526   6525   6527   6529       3    0.00000    0.00000    0.00000
0.00000    0.00000    0.00000 ;       H-     N-     CA-    CB
  6526   6525   6527   6542       3    0.00000    0.00000    0.00000
0.00000    0.00000    0.00000 ;       H-     N-     CA-     C
  6527   6529   6531   6532       3    0.66944    2.00832    0.00000
-2.67776    0.00000    0.00000 ;      CA-    CB-   CG2-  HG21
  6527   6529   6531   6533       3    0.66944    2.00832    0.00000
-2.67776    0.00000    0.00000 ;      CA-    CB-   CG2-  HG22
  6527   6529   6531   6534       3    0.66944    2.00832    0.00000
-2.67776    0.00000    0.00000 ;      CA-    CB-   CG2-  HG23
  6527   6529   6535   6536       3    0.66944    2.00832    0.00000
-2.67776    0.00000    0.00000 ;      CA-    CB-   CG1-  HG12
  6527   6529   6535   6537       3    0.66944    2.00832    0.00000
-2.67776    0.00000    0.00000 ;      CA-    CB-   CG1-  HG13
  6527   6529   6535   6538       3    4.24258   -0.52718   -7.25506
-1.79075    7.69856    0.00000 ;      CA-    CB-   CG1-   CD1
  6527   6542   6544   6545       3   20.92000    0.00000  -20.92000
0.00000    0.00000    0.00000 ;      CA-     C-     N-     H
  6527   6542   6544   6546       3   20.92000    0.00000  -20.92000
0.00000    0.00000    0.00000 ;      CA-     C-     N-    CA
  6528   6527   6529   6530       3    0.65084    1.95253    0.00000
-2.60338    0.00000    0.00000 ;      HA-    CA-    CB-    HB
  6528   6527   6529   6531       3    0.65084    1.95253    0.00000
-2.60338    0.00000    0.00000 ;      HA-    CA-    CB-   CG2
  6528   6527   6529   6535       3    0.65084    1.95253    0.00000
-2.60338    0.00000    0.00000 ;      HA-    CA-    CB-   CG1
  6528   6527   6542   6543       3    3.68192   -4.35136    0.00000
1.33888    0.00000    0.00000 ;      HA-    CA-     C-     O
  6528   6527   6542   6544       3    0.00000    0.00000    0.00000
0.00000    0.00000    0.00000 ;      HA-    CA-     C-     N
  6529   6527   6542   6543       3    0.00000    0.00000    0.00000
0.00000    0.00000    0.00000 ;      CB-    CA-     C-     O
  6529   6527   6542   6544       3    2.51040    4.18400    1.67360
-6.69440    0.00000    0.00000 ;      CB-    CA-     C-     N
  6529   6535   6538   6539       3    0.66944    2.00832    0.00000
-2.67776    0.00000    0.00000 ;      CB-   CG1-   CD1-  HD11
  6529   6535   6538   6540       3    0.66944    2.00832    0.00000
-2.67776    0.00000    0.00000 ;      CB-   CG1-   CD1-  HD12
  6529   6535   6538   6541       3    0.66944    2.00832    0.00000
-2.67776    0.00000    0.00000 ;      CB-   CG1-   CD1-  HD13
  6530   6529   6527   6542       3    0.65084    1.95253    0.00000
-2.60338    0.00000    0.00000 ;      HB-    CB-    CA-     C
  6530   6529   6531   6532       3    0.62760    1.88280    0.00000
-2.51040    0.00000    0.00000 ;      HB-    CB-   CG2-  HG21
  6530   6529   6531   6533       3    0.62760    1.88280    0.00000
-2.51040    0.00000    0.00000 ;      HB-    CB-   CG2-  HG22
  6530   6529   6531   6534       3    0.62760    1.88280    0.00000
-2.51040    0.00000    0.00000 ;      HB-    CB-   CG2-  HG23
  6530   6529   6535   6536       3    0.62760    1.88280    0.00000
-2.51040    0.00000    0.00000 ;      HB-    CB-   CG1-  HG12
  6530   6529   6535   6537       3    0.62760    1.88280    0.00000

```
-2.51040    0.00000    0.00000 ;      HB-    CB-   CG1-  HG13
  6530   6529   6535   6538      3    0.66944    2.00832    0.00000
-2.67776    0.00000    0.00000 ;      HB-    CB-   CG1-   CD1
  6531   6529   6527   6542      3    5.67350    3.55640   -6.16722
-2.47693    3.74886    0.00000 ;     CG2-    CB-    CA-     C
  6531   6529   6535   6536      3    0.66944    2.00832    0.00000
-2.67776    0.00000    0.00000 ;     CG2-    CB-   CG1-  HG12
  6531   6529   6535   6537      3    0.66944    2.00832    0.00000
-2.67776    0.00000    0.00000 ;     CG2-    CB-   CG1-  HG13
  6531   6529   6535   6538      3    3.81162    0.49790   -8.14206
-1.79075    7.49773    0.00000 ;     CG2-    CB-   CG1-   CD1
  6532   6531   6529   6535      3    0.66944    2.00832    0.00000
-2.67776    0.00000    0.00000 ;    HG21-   CG2-    CB-   CG1
  6533   6531   6529   6535      3    0.66944    2.00832    0.00000
-2.67776    0.00000    0.00000 ;    HG22-   CG2-    CB-   CG1
  6534   6531   6529   6535      3    0.66944    2.00832    0.00000
-2.67776    0.00000    0.00000 ;    HG23-   CG2-    CB-   CG1
  6535   6529   6527   6542      3    8.26340    0.74057   -9.99976
-1.89117    3.84928    0.00000 ;     CG1-    CB-    CA-     C
  6536   6535   6538   6539      3    0.62760    1.88280    0.00000
-2.51040    0.00000    0.00000 ;    HG12-   CG1-   CD1-  HD11
  6536   6535   6538   6540      3    0.62760    1.88280    0.00000
-2.51040    0.00000    0.00000 ;    HG12-   CG1-   CD1-  HD12
  6536   6535   6538   6541      3    0.62760    1.88280    0.00000
-2.51040    0.00000    0.00000 ;    HG12-   CG1-   CD1-  HD13
  6537   6535   6538   6539      3    0.62760    1.88280    0.00000
-2.51040    0.00000    0.00000 ;    HG13-   CG1-   CD1-  HD11
  6537   6535   6538   6540      3    0.62760    1.88280    0.00000
-2.51040    0.00000    0.00000 ;    HG13-   CG1-   CD1-  HD12
  6537   6535   6538   6541      3    0.62760    1.88280    0.00000
-2.51040    0.00000    0.00000 ;    HG13-   CG1-   CD1-  HD13
  6542   6544   6546   6547      3    0.00000    0.00000    0.00000
0.00000    0.00000    0.00000 ;      C-     N-    CA-    HA
  6542   6544   6546   6548      3   11.71520    1.67360   15.06240  -
13.38880    0.00000    0.00000 ;      C-     N-    CA-    CB
  6542   6544   6546   6553      3    1.75728    5.27184    2.25936
-7.02912    0.00000    0.00000 ;      C-     N-    CA-     C
  6543   6542   6544   6545      3   29.28800   -8.36800  -20.92000
0.00000    0.00000    0.00000 ;      O-     C-     N-     H
  6543   6542   6544   6546      3   20.92000    0.00000  -20.92000
0.00000    0.00000    0.00000 ;      O-     C-     N-    CA
  6544   6546   6548   6549      3    0.65084    1.95253    0.00000
-2.60338    0.00000    0.00000 ;      N-    CA-    CB-   HB2
  6544   6546   6548   6550      3    0.65084    1.95253    0.00000
-2.60338    0.00000    0.00000 ;      N-    CA-    CB-   HB3
  6544   6546   6548   6551      3    7.86174    2.24681   -7.41405
-6.71114    5.35552    0.00000 ;      N-    CA-    CB-    OG
  6544   6546   6553   6554      3    0.00000    0.00000    0.00000
0.00000    0.00000    0.00000 ;      N-    CA-     C-     O
  6544   6546   6553   6555      3   17.40544   -5.02080  -13.22144
9.20480    0.00000    0.00000 ;      N-    CA-     C-     N
  6545   6544   6546   6547      3    0.00000    0.00000    0.00000
```

0.00000 0.00000 0.00000 ; H- N- CA- HA
6545 6544 6546 6548 3 0.00000 0.00000 0.00000
0.00000 0.00000 0.00000 ; H- N- CA- CB
6545 6544 6546 6553 3 0.00000 0.00000 0.00000
0.00000 0.00000 0.00000 ; H- N- CA- C
6546 6548 6551 6552 3 2.05853 2.46856 3.48109
-4.46851 0.23430 0.00000 ; CA- CB- OG- HG
6546 6553 6555 6556 3 20.92000 0.00000 -20.92000
0.00000 0.00000 0.00000 ; CA- C- N- H
6546 6553 6555 6557 3 20.92000 0.00000 -20.92000
0.00000 0.00000 0.00000 ; CA- C- N- CA
6547 6546 6548 6549 3 0.65084 1.95253 0.00000
-2.60338 0.00000 0.00000 ; HA- CA- CB- HB2
6547 6546 6548 6550 3 0.65084 1.95253 0.00000
-2.60338 0.00000 0.00000 ; HA- CA- CB- HB3
6547 6546 6548 6551 3 1.04600 -1.04600 0.00000
0.00000 0.00000 0.00000 ; HA- CA- CB- OG
6547 6546 6553 6554 3 3.68192 -4.35136 0.00000
1.33888 0.00000 0.00000 ; HA- CA- C- O
6547 6546 6553 6555 3 0.00000 0.00000 0.00000
0.00000 0.00000 0.00000 ; HA- CA- C- N
6548 6546 6553 6554 3 0.00000 0.00000 0.00000
0.00000 0.00000 0.00000 ; CB- CA- C- O
6548 6546 6553 6555 3 2.51040 4.18400 1.67360
-6.69440 0.00000 0.00000 ; CB- CA- C- N
6549 6548 6546 6553 3 0.65084 1.95253 0.00000
-2.60338 0.00000 0.00000 ; HB2- CB- CA- C
6549 6548 6551 6552 3 0.69733 2.09200 0.00000
-2.78933 0.00000 0.00000 ; HB2- CB- OG- HG
6550 6548 6546 6553 3 0.65084 1.95253 0.00000
-2.60338 0.00000 0.00000 ; HB3- CB- CA- C
6550 6548 6551 6552 3 0.69733 2.09200 0.00000
-2.78933 0.00000 0.00000 ; HB3- CB- OG- HG
6551 6548 6546 6553 3 7.34710 7.79898 -6.14211
-6.71114 4.31789 0.00000 ; OG- CB- CA- C
6553 6555 6557 6558 3 0.00000 0.00000 0.00000
0.00000 0.00000 0.00000 ; C- N- CA- HA
6553 6555 6557 6559 3 11.71520 1.67360 15.06240 -
13.38880 0.00000 0.00000 ; C- N- CA- CB
6553 6555 6557 6569 3 1.75728 5.27184 2.25936
-7.02912 0.00000 0.00000 ; C- N- CA- C
6554 6553 6555 6556 3 29.28800 -8.36800 -20.92000
0.00000 0.00000 0.00000 ; O- C- N- H
6554 6553 6555 6557 3 20.92000 0.00000 -20.92000
0.00000 0.00000 0.00000 ; O- C- N- CA
6555 6557 6559 6560 3 0.65084 1.95253 0.00000
-2.60338 0.00000 0.00000 ; N- CA- CB- HB
6555 6557 6559 6561 3 3.83673 0.44769 -1.77402
-2.47693 -0.03347 0.00000 ; N- CA- CB- CG1
6555 6557 6559 6565 3 3.83673 0.44769 -1.77402
-2.47693 -0.03347 0.00000 ; N- CA- CB- CG2
6555 6557 6569 6570 3 0.00000 0.00000 0.00000

```
0.00000    0.00000    0.00000 ;       N-    CA-     C-     O
  6555   6557   6569   6571      3   17.40544   -5.02080  -13.22144
9.20480    0.00000    0.00000 ;       N-    CA-     C-     N
  6556   6555   6557   6558      3    0.00000    0.00000    0.00000
0.00000    0.00000    0.00000 ;       H-     N-    CA-    HA
  6556   6555   6557   6559      3    0.00000    0.00000    0.00000
0.00000    0.00000    0.00000 ;       H-     N-    CA-    CB
  6556   6555   6557   6569      3    0.00000    0.00000    0.00000
0.00000    0.00000    0.00000 ;       H-     N-    CA-     C
  6557   6559   6561   6562      3    0.66944    2.00832    0.00000
-2.67776    0.00000    0.00000 ;      CA-    CB-   CG1-  HG11
  6557   6559   6561   6563      3    0.66944    2.00832    0.00000
-2.67776    0.00000    0.00000 ;      CA-    CB-   CG1-  HG12
  6557   6559   6561   6564      3    0.66944    2.00832    0.00000
-2.67776    0.00000    0.00000 ;      CA-    CB-   CG1-  HG13
  6557   6559   6565   6566      3    0.66944    2.00832    0.00000
-2.67776    0.00000    0.00000 ;      CA-    CB-   CG2-  HG21
  6557   6559   6565   6567      3    0.66944    2.00832    0.00000
-2.67776    0.00000    0.00000 ;      CA-    CB-   CG2-  HG22
  6557   6559   6565   6568      3    0.66944    2.00832    0.00000
-2.67776    0.00000    0.00000 ;      CA-    CB-   CG2-  HG23
  6557   6569   6571   6572      3   20.92000    0.00000  -20.92000
0.00000    0.00000    0.00000 ;      CA-     C-     N-     H
  6557   6569   6571   6573      3   20.92000    0.00000  -20.92000
0.00000    0.00000    0.00000 ;      CA-     C-     N-    CA
  6558   6557   6559   6560      3    0.65084    1.95253    0.00000
-2.60338    0.00000    0.00000 ;      HA-    CA-    CB-    HB
  6558   6557   6559   6561      3    0.65084    1.95253    0.00000
-2.60338    0.00000    0.00000 ;      HA-    CA-    CB-   CG1
  6558   6557   6559   6565      3    0.65084    1.95253    0.00000
-2.60338    0.00000    0.00000 ;      HA-    CA-    CB-   CG2
  6558   6557   6569   6570      3    3.68192   -4.35136    0.00000
1.33888    0.00000    0.00000 ;      HA-    CA-     C-     O
  6558   6557   6569   6571      3    0.00000    0.00000    0.00000
0.00000    0.00000    0.00000 ;      HA-    CA-     C-     N
  6559   6557   6569   6570      3    0.00000    0.00000    0.00000
0.00000    0.00000    0.00000 ;      CB-    CA-     C-     O
  6559   6557   6569   6571      3    2.51040    4.18400    1.67360
-6.69440    0.00000    0.00000 ;      CB-    CA-     C-     N
  6560   6559   6557   6569      3    0.65084    1.95253    0.00000
-2.60338    0.00000    0.00000 ;      HB-    CB-    CA-     C
  6560   6559   6561   6562      3    0.62760    1.88280    0.00000
-2.51040    0.00000    0.00000 ;      HB-    CB-   CG1-  HG11
  6560   6559   6561   6563      3    0.62760    1.88280    0.00000
-2.51040    0.00000    0.00000 ;      HB-    CB-   CG1-  HG12
  6560   6559   6561   6564      3    0.62760    1.88280    0.00000
-2.51040    0.00000    0.00000 ;      HB-    CB-   CG1-  HG13
  6560   6559   6565   6566      3    0.62760    1.88280    0.00000
-2.51040    0.00000    0.00000 ;      HB-    CB-   CG2-  HG21
  6560   6559   6565   6567      3    0.62760    1.88280    0.00000
-2.51040    0.00000    0.00000 ;      HB-    CB-   CG2-  HG22
  6560   6559   6565   6568      3    0.62760    1.88280    0.00000
```

-2.51040    0.00000    0.00000 ;      HB-    CB-   CG2-  HG23
  6561  6559  6557  6569      3    5.67350    3.55640   -6.16722
-2.47693    3.74886    0.00000 ;     CG1-    CB-    CA-     C
  6561  6559  6565  6566      3    0.66944    2.00832    0.00000
-2.67776    0.00000    0.00000 ;     CG1-    CB-   CG2-  HG21
  6561  6559  6565  6567      3    0.66944    2.00832    0.00000
-2.67776    0.00000    0.00000 ;     CG1-    CB-   CG2-  HG22
  6561  6559  6565  6568      3    0.66944    2.00832    0.00000
-2.67776    0.00000    0.00000 ;     CG1-    CB-   CG2-  HG23
  6562  6561  6559  6565      3    0.66944    2.00832    0.00000
-2.67776    0.00000    0.00000 ;    HG11-   CG1-    CB-   CG2
  6563  6561  6559  6565      3    0.66944    2.00832    0.00000
-2.67776    0.00000    0.00000 ;    HG12-   CG1-    CB-   CG2
  6564  6561  6559  6565      3    0.66944    2.00832    0.00000
-2.67776    0.00000    0.00000 ;    HG13-   CG1-    CB-   CG2
  6565  6559  6557  6569      3    5.67350    3.55640   -6.16722
-2.47693    3.74886    0.00000 ;     CG2-    CB-    CA-     C
  6569  6571  6573  6574      3    0.00000    0.00000    0.00000
0.00000    0.00000    0.00000 ;      C-     N-    CA-    HA
  6569  6571  6573  6575      3   11.71520    1.67360   15.06240  -
13.38880    0.00000    0.00000 ;      C-     N-    CA-    CB
  6569  6571  6573  6584      3    1.75728    5.27184    2.25936
-7.02912    0.00000    0.00000 ;      C-     N-    CA-     C
  6570  6569  6571  6572      3   29.28800   -8.36800  -20.92000
0.00000    0.00000    0.00000 ;      O-     C-     N-     H
  6570  6569  6571  6573      3   20.92000    0.00000  -20.92000
0.00000    0.00000    0.00000 ;      O-     C-     N-    CA
  6571  6573  6575  6576      3    0.65084    1.95253    0.00000
-2.60338    0.00000    0.00000 ;      N-    CA-    CB-   HB2
  6571  6573  6575  6577      3    0.65084    1.95253    0.00000
-2.60338    0.00000    0.00000 ;      N-    CA-    CB-   HB3
  6571  6573  6575  6578      3    3.21331    2.22589   -4.15053
-2.40998    2.61082    0.00000 ;      N-    CA-    CB-    CG
  6571  6573  6584  6585      3    0.00000    0.00000    0.00000
0.00000    0.00000    0.00000 ;      N-    CA-     C-     O
  6571  6573  6584  6586      3   17.40544   -5.02080  -13.22144
9.20480    0.00000    0.00000 ;      N-    CA-     C-     N
  6572  6571  6573  6574      3    0.00000    0.00000    0.00000
0.00000    0.00000    0.00000 ;      H-     N-    CA-    HA
  6572  6571  6573  6575      3    0.00000    0.00000    0.00000
0.00000    0.00000    0.00000 ;      H-     N-    CA-    CB
  6572  6571  6573  6584      3    0.00000    0.00000    0.00000
0.00000    0.00000    0.00000 ;      H-     N-    CA-     C
  6573  6575  6578  6579      3    0.66944    2.00832    0.00000
-2.67776    0.00000    0.00000 ;     CA-    CB-    CG-   HG2
  6573  6575  6578  6580      3    0.66944    2.00832    0.00000
-2.67776    0.00000    0.00000 ;     CA-    CB-    CG-   HG3
  6573  6575  6578  6581      3   10.12110   -1.91209    0.01674
10.17549   -1.87443    0.00000 ;     CA-    CB-    CG-    CD
  6573  6584  6586  6587      3   20.92000    0.00000  -20.92000
0.00000    0.00000    0.00000 ;     CA-     C-     N-     H
  6573  6584  6586  6588      3   20.92000    0.00000  -20.92000

0.00000 0.00000 0.00000 ; CA- C- N- CA
6574 6573 6575 6576 3 0.65084 1.95253 0.00000
-2.60338 0.00000 0.00000 ; HA- CA- CB- HB2
6574 6573 6575 6577 3 0.65084 1.95253 0.00000
-2.60338 0.00000 0.00000 ; HA- CA- CB- HB3
6574 6573 6575 6578 3 0.65084 1.95253 0.00000
-2.60338 0.00000 0.00000 ; HA- CA- CB- CG
6574 6573 6584 6585 3 3.68192 -4.35136 0.00000
1.33888 0.00000 0.00000 ; HA- CA- C- O
6574 6573 6584 6586 3 0.00000 0.00000 0.00000
0.00000 0.00000 0.00000 ; HA- CA- C- N
6575 6573 6584 6585 3 0.00000 0.00000 0.00000
0.00000 0.00000 0.00000 ; CB- CA- C- O
6575 6573 6584 6586 3 2.51040 4.18400 1.67360
-6.69440 0.00000 0.00000 ; CB- CA- C- N
6575 6578 6581 6582 3 3.79907 0.00000 -5.40573
0.00000 2.14221 0.00000 ; CB- CG- CD- OE1
6575 6578 6581 6583 3 3.79907 0.00000 -5.40573
0.00000 2.14221 0.00000 ; CB- CG- CD- OE2
6576 6575 6573 6584 3 0.65084 1.95253 0.00000
-2.60338 0.00000 0.00000 ; HB2- CB- CA- C
6576 6575 6578 6579 3 0.62760 1.88280 0.00000
-2.51040 0.00000 0.00000 ; HB2- CB- CG- HG2
6576 6575 6578 6580 3 0.62760 1.88280 0.00000
-2.51040 0.00000 0.00000 ; HB2- CB- CG- HG3
6576 6575 6578 6581 3 0.65084 1.95253 0.00000
-2.60338 0.00000 0.00000 ; HB2- CB- CG- CD
6577 6575 6573 6584 3 0.65084 1.95253 0.00000
-2.60338 0.00000 0.00000 ; HB3- CB- CA- C
6577 6575 6578 6579 3 0.62760 1.88280 0.00000
-2.51040 0.00000 0.00000 ; HB3- CB- CG- HG2
6577 6575 6578 6580 3 0.62760 1.88280 0.00000
-2.51040 0.00000 0.00000 ; HB3- CB- CG- HG3
6577 6575 6578 6581 3 0.65084 1.95253 0.00000
-2.60338 0.00000 0.00000 ; HB3- CB- CG- CD
6578 6575 6573 6584 3 6.86594 3.56895 -8.14206
-2.40998 4.85344 0.00000 ; CG- CB- CA- C
6579 6578 6581 6582 3 0.00000 0.00000 0.00000
0.00000 0.00000 0.00000 ; HG2- CG- CD- OE1
6579 6578 6581 6583 3 0.00000 0.00000 0.00000
0.00000 0.00000 0.00000 ; HG2- CG- CD- OE2
6580 6578 6581 6582 3 0.00000 0.00000 0.00000
0.00000 0.00000 0.00000 ; HG3- CG- CD- OE1
6580 6578 6581 6583 3 0.00000 0.00000 0.00000
0.00000 0.00000 0.00000 ; HG3- CG- CD- OE2
6584 6586 6588 6589 3 0.00000 0.00000 0.00000
0.00000 0.00000 0.00000 ; C- N- CA- HA
6584 6586 6588 6590 3 11.71520 1.67360 15.06240 -
13.38880 0.00000 0.00000 ; C- N- CA- CB
6584 6586 6588 6600 3 1.75728 5.27184 2.25936
-7.02912 0.00000 0.00000 ; C- N- CA- C
6585 6584 6586 6587 3 29.28800 -8.36800 -20.92000

0.00000 0.00000 0.00000 ; O- C- N- H
6585 6584 6586 6588 3 20.92000 0.00000 -20.92000
0.00000 0.00000 0.00000 ; O- C- N- CA
6586 6588 6590 6591 3 0.65084 1.95253 0.00000
-2.60338 0.00000 0.00000 ; N- CA- CB- HB2
6586 6588 6590 6592 3 0.65084 1.95253 0.00000
-2.60338 0.00000 0.00000 ; N- CA- CB- HB3
6586 6588 6590 6593 3 3.21331 2.22589 -4.15053
-2.40998 2.61082 0.00000 ; N- CA- CB- CG
6586 6588 6600 6601 3 0.00000 0.00000 0.00000
0.00000 0.00000 0.00000 ; N- CA- C- O
6586 6588 6600 6602 3 17.40544 -5.02080 -13.22144
9.20480 0.00000 0.00000 ; N- CA- C- N
6587 6586 6588 6589 3 0.00000 0.00000 0.00000
0.00000 0.00000 0.00000 ; H- N- CA- HA
6587 6586 6588 6590 3 0.00000 0.00000 0.00000
0.00000 0.00000 0.00000 ; H- N- CA- CB
6587 6586 6588 6600 3 0.00000 0.00000 0.00000
0.00000 0.00000 0.00000 ; H- N- CA- C
6588 6590 6593 6594 3 0.66944 2.00832 0.00000
-2.67776 0.00000 0.00000 ; CA- CB- CG- HG2
6588 6590 6593 6595 3 0.66944 2.00832 0.00000
-2.67776 0.00000 0.00000 ; CA- CB- CG- HG3
6588 6590 6593 6596 3 3.69866 -4.35136 -3.92459
6.89523 4.61914 0.00000 ; CA- CB- CG- CD
6588 6600 6602 6603 3 20.92000 0.00000 -20.92000
0.00000 0.00000 0.00000 ; CA- C- N- H
6588 6600 6602 6604 3 20.92000 0.00000 -20.92000
0.00000 0.00000 0.00000 ; CA- C- N- CA
6589 6588 6590 6591 3 0.65084 1.95253 0.00000
-2.60338 0.00000 0.00000 ; HA- CA- CB- HB2
6589 6588 6590 6592 3 0.65084 1.95253 0.00000
-2.60338 0.00000 0.00000 ; HA- CA- CB- HB3
6589 6588 6590 6593 3 0.65084 1.95253 0.00000
-2.60338 0.00000 0.00000 ; HA- CA- CB- CG
6589 6588 6600 6601 3 3.68192 -4.35136 0.00000
1.33888 0.00000 0.00000 ; HA- CA- C- O
6589 6588 6600 6602 3 0.00000 0.00000 0.00000
0.00000 0.00000 0.00000 ; HA- CA- C- N
6590 6588 6600 6601 3 0.00000 0.00000 0.00000
0.00000 0.00000 0.00000 ; CB- CA- C- O
6590 6588 6600 6602 3 2.51040 4.18400 1.67360
-6.69440 0.00000 0.00000 ; CB- CA- C- N
6590 6593 6596 6597 3 0.00000 0.00000 0.00000
0.00000 0.00000 0.00000 ; CB- CG- CD- OE1
6590 6593 6596 6598 3 12.79049 3.13382 -7.02912
0.41840 -2.20915 0.00000 ; CB- CG- CD- OE2
6591 6590 6588 6600 3 0.65084 1.95253 0.00000
-2.60338 0.00000 0.00000 ; HB2- CB- CA- C
6591 6590 6593 6594 3 0.62760 1.88280 0.00000
-2.51040 0.00000 0.00000 ; HB2- CB- CG- HG2
6591 6590 6593 6595 3 0.62760 1.88280 0.00000

-2.51040 0.00000 0.00000 ; HB2- CB- CG- HG3
6591 6590 6593 6596 3 0.65084 1.95253 0.00000
-2.60338 0.00000 0.00000 ; HB2- CB- CG- CD
6592 6590 6588 6600 3 0.65084 1.95253 0.00000
-2.60338 0.00000 0.00000 ; HB3- CB- CA- C
6592 6590 6593 6594 3 0.62760 1.88280 0.00000
-2.51040 0.00000 0.00000 ; HB3- CB- CG- HG2
6592 6590 6593 6595 3 0.62760 1.88280 0.00000
-2.51040 0.00000 0.00000 ; HB3- CB- CG- HG3
6592 6590 6593 6596 3 0.65084 1.95253 0.00000
-2.60338 0.00000 0.00000 ; HB3- CB- CG- CD
6593 6590 6588 6600 3 6.86594 3.56895 -8.14206
-2.40998 4.85344 0.00000 ; CG- CB- CA- C
6593 6596 6598 6599 3 27.46796 7.88684 -26.42614
-8.01654 3.78234 0.00000 ; CG- CD- OE2- HE2
6594 6593 6596 6597 3 3.68192 -4.35136 0.00000
1.33888 0.00000 0.00000 ; HG2- CG- CD- OE1
6594 6593 6596 6598 3 0.00000 0.00000 0.00000
0.00000 0.00000 0.00000 ; HG2- CG- CD- OE2
6595 6593 6596 6597 3 3.68192 -4.35136 0.00000
1.33888 0.00000 0.00000 ; HG3- CG- CD- OE1
6595 6593 6596 6598 3 0.00000 0.00000 0.00000
0.00000 0.00000 0.00000 ; HG3- CG- CD- OE2
6597 6596 6598 6599 3 27.19600 -7.94960 -19.24640
0.00000 0.00000 0.00000 ; OE1- CD- OE2- HE2
6600 6602 6604 6605 3 0.00000 0.00000 0.00000
0.00000 0.00000 0.00000 ; C- N- CA- HA
6600 6602 6604 6606 3 11.71520 1.67360 15.06240 -
13.38880 0.00000 0.00000 ; C- N- CA- CB
6600 6602 6604 6619 3 1.75728 5.27184 2.25936
-7.02912 0.00000 0.00000 ; C- N- CA- C
6601 6600 6602 6603 3 29.28800 -8.36800 -20.92000
0.00000 0.00000 0.00000 ; O- C- N- H
6601 6600 6602 6604 3 20.92000 0.00000 -20.92000
0.00000 0.00000 0.00000 ; O- C- N- CA
6602 6604 6606 6607 3 0.65084 1.95253 0.00000
-2.60338 0.00000 0.00000 ; N- CA- CB- HB2
6602 6604 6606 6608 3 0.65084 1.95253 0.00000
-2.60338 0.00000 0.00000 ; N- CA- CB- HB3
6602 6604 6606 6609 3 3.79070 1.39746 -4.61077
-2.40998 2.44346 0.00000 ; N- CA- CB- CG
6602 6604 6619 6620 3 0.00000 0.00000 0.00000
0.00000 0.00000 0.00000 ; N- CA- C- O
6602 6604 6619 6621 3 17.40544 -5.02080 -13.22144
9.20480 0.00000 0.00000 ; N- CA- C- N
6603 6602 6604 6605 3 0.00000 0.00000 0.00000
0.00000 0.00000 0.00000 ; H- N- CA- HA
6603 6602 6604 6606 3 0.00000 0.00000 0.00000
0.00000 0.00000 0.00000 ; H- N- CA- CB
6603 6602 6604 6619 3 0.00000 0.00000 0.00000
0.00000 0.00000 0.00000 ; H- N- CA- C
6604 6606 6609 6610 3 0.66944 2.00832 0.00000

```
-2.67776    0.00000    0.00000 ;      CA-    CB-    CG-    HG
  6604   6606   6609   6611       3    3.90367    0.19665   -6.21742
-2.37651    5.99149    0.00000 ;      CA-    CB-    CG-   CD1
  6604   6606   6609   6615       3    3.90367    0.19665   -6.21742
-2.37651    5.99149    0.00000 ;      CA-    CB-    CG-   CD2
  6604   6619   6621   6622       3   20.92000    0.00000  -20.92000
0.00000    0.00000    0.00000 ;      CA-     C-     N-     H
  6604   6619   6621   6623       3   20.92000    0.00000  -20.92000
0.00000    0.00000    0.00000 ;      CA-     C-     N-    CA
  6605   6604   6606   6607       3    0.65084    1.95253    0.00000
-2.60338    0.00000    0.00000 ;      HA-    CA-    CB-   HB2
  6605   6604   6606   6608       3    0.65084    1.95253    0.00000
-2.60338    0.00000    0.00000 ;      HA-    CA-    CB-   HB3
  6605   6604   6606   6609       3    0.65084    1.95253    0.00000
-2.60338    0.00000    0.00000 ;      HA-    CA-    CB-    CG
  6605   6604   6619   6620       3    3.68192   -4.35136    0.00000
1.33888    0.00000    0.00000 ;      HA-    CA-     C-     O
  6605   6604   6619   6621       3    0.00000    0.00000    0.00000
0.00000    0.00000    0.00000 ;      HA-    CA-     C-     N
  6606   6604   6619   6620       3    0.00000    0.00000    0.00000
0.00000    0.00000    0.00000 ;      CB-    CA-     C-     O
  6606   6604   6619   6621       3    2.51040    4.18400    1.67360
-6.69440    0.00000    0.00000 ;      CB-    CA-     C-     N
  6606   6609   6611   6612       3    0.66944    2.00832    0.00000
-2.67776    0.00000    0.00000 ;      CB-    CG-   CD1-  HD11
  6606   6609   6611   6613       3    0.66944    2.00832    0.00000
-2.67776    0.00000    0.00000 ;      CB-    CG-   CD1-  HD12
  6606   6609   6611   6614       3    0.66944    2.00832    0.00000
-2.67776    0.00000    0.00000 ;      CB-    CG-   CD1-  HD13
  6606   6609   6615   6616       3    0.66944    2.00832    0.00000
-2.67776    0.00000    0.00000 ;      CB-    CG-   CD2-  HD21
  6606   6609   6615   6617       3    0.66944    2.00832    0.00000
-2.67776    0.00000    0.00000 ;      CB-    CG-   CD2-  HD22
  6606   6609   6615   6618       3    0.66944    2.00832    0.00000
-2.67776    0.00000    0.00000 ;      CB-    CG-   CD2-  HD23
  6607   6606   6604   6619       3    0.65084    1.95253    0.00000
-2.60338    0.00000    0.00000 ;     HB2-    CB-    CA-     C
  6607   6606   6609   6610       3    0.62760    1.88280    0.00000
-2.51040    0.00000    0.00000 ;     HB2-    CB-    CG-    HG
  6607   6606   6609   6611       3    0.66944    2.00832    0.00000
-2.67776    0.00000    0.00000 ;     HB2-    CB-    CG-   CD1
  6607   6606   6609   6615       3    0.66944    2.00832    0.00000
-2.67776    0.00000    0.00000 ;     HB2-    CB-    CG-   CD2
  6608   6606   6604   6619       3    0.65084    1.95253    0.00000
-2.60338    0.00000    0.00000 ;     HB3-    CB-    CA-     C
  6608   6606   6609   6610       3    0.62760    1.88280    0.00000
-2.51040    0.00000    0.00000 ;     HB3-    CB-    CG-    HG
  6608   6606   6609   6611       3    0.66944    2.00832    0.00000
-2.67776    0.00000    0.00000 ;     HB3-    CB-    CG-   CD1
  6608   6606   6609   6615       3    0.66944    2.00832    0.00000
-2.67776    0.00000    0.00000 ;     HB3-    CB-    CG-   CD2
  6609   6606   6604   6619       3   10.33448   -1.14642  -11.54784
```

-2.40998 6.35968 0.00000 ; CG- CB- CA- C
6610 6609 6611 6612 3 0.62760 1.88280 0.00000
-2.51040 0.00000 0.00000 ; HG- CG- CD1- HD11
6610 6609 6611 6613 3 0.62760 1.88280 0.00000
-2.51040 0.00000 0.00000 ; HG- CG- CD1- HD12
6610 6609 6611 6614 3 0.62760 1.88280 0.00000
-2.51040 0.00000 0.00000 ; HG- CG- CD1- HD13
6610 6609 6615 6616 3 0.62760 1.88280 0.00000
-2.51040 0.00000 0.00000 ; HG- CG- CD2- HD21
6610 6609 6615 6617 3 0.62760 1.88280 0.00000
-2.51040 0.00000 0.00000 ; HG- CG- CD2- HD22
6610 6609 6615 6618 3 0.62760 1.88280 0.00000
-2.51040 0.00000 0.00000 ; HG- CG- CD2- HD23
6611 6609 6615 6616 3 0.66944 2.00832 0.00000
-2.67776 0.00000 0.00000 ; CD1- CG- CD2- HD21
6611 6609 6615 6617 3 0.66944 2.00832 0.00000
-2.67776 0.00000 0.00000 ; CD1- CG- CD2- HD22
6611 6609 6615 6618 3 0.66944 2.00832 0.00000
-2.67776 0.00000 0.00000 ; CD1- CG- CD2- HD23
6612 6611 6609 6615 3 0.66944 2.00832 0.00000
-2.67776 0.00000 0.00000 ; HD11- CD1- CG- CD2
6613 6611 6609 6615 3 0.66944 2.00832 0.00000
-2.67776 0.00000 0.00000 ; HD12- CD1- CG- CD2
6614 6611 6609 6615 3 0.66944 2.00832 0.00000
-2.67776 0.00000 0.00000 ; HD13- CD1- CG- CD2
6619 6621 6623 6624 3 0.00000 0.00000 0.00000
0.00000 0.00000 0.00000 ; C- N- CA- HA
6619 6621 6623 6625 3 11.71520 1.67360 15.06240 -
13.38880 0.00000 0.00000 ; C- N- CA- CB
6619 6621 6623 6641 3 1.75728 5.27184 2.25936
-7.02912 0.00000 0.00000 ; C- N- CA- C
6620 6619 6621 6622 3 29.28800 -8.36800 -20.92000
0.00000 0.00000 0.00000 ; O- C- N- H
6620 6619 6621 6623 3 20.92000 0.00000 -20.92000
0.00000 0.00000 0.00000 ; O- C- N- CA
6621 6623 6625 6626 3 0.65084 1.95253 0.00000
-2.60338 0.00000 0.00000 ; N- CA- CB- HB2
6621 6623 6625 6627 3 0.65084 1.95253 0.00000
-2.60338 0.00000 0.00000 ; N- CA- CB- HB3
6621 6623 6625 6628 3 0.65084 1.95253 0.00000
-2.60338 0.00000 0.00000 ; N- CA- CB- CG
6621 6623 6641 6642 3 0.00000 0.00000 0.00000
0.00000 0.00000 0.00000 ; N- CA- C- O
6621 6623 6641 6643 3 17.40544 -5.02080 -13.22144
9.20480 0.00000 0.00000 ; N- CA- C- N
6622 6621 6623 6624 3 0.00000 0.00000 0.00000
0.00000 0.00000 0.00000 ; H- N- CA- HA
6622 6621 6623 6625 3 0.00000 0.00000 0.00000
0.00000 0.00000 0.00000 ; H- N- CA- CB
6622 6621 6623 6641 3 0.00000 0.00000 0.00000
0.00000 0.00000 0.00000 ; H- N- CA- C
6623 6625 6628 6629 3 0.66944 2.00832 0.00000

```
-2.67776    0.00000    0.00000 ;      CA-    CB-    CG-   HG2
  6623   6625   6628   6630       3    0.66944    2.00832    0.00000
-2.67776    0.00000    0.00000 ;      CA-    CB-    CG-   HG3
  6623   6625   6628   6631       3    3.68192    3.09616   -2.09200
-3.01248    0.00000    0.00000 ;      CA-    CB-    CG-    CD
  6623   6641   6643   6644       3   20.92000    0.00000  -20.92000
0.00000    0.00000    0.00000 ;      CA-     C-     N-     H
  6623   6641   6643   6645       3   20.92000    0.00000  -20.92000
0.00000    0.00000    0.00000 ;      CA-     C-     N-    CA
  6624   6623   6625   6626       3    0.65084    1.95253    0.00000
-2.60338    0.00000    0.00000 ;      HA-    CA-    CB-   HB2
  6624   6623   6625   6627       3    0.65084    1.95253    0.00000
-2.60338    0.00000    0.00000 ;      HA-    CA-    CB-   HB3
  6624   6623   6625   6628       3    0.65084    1.95253    0.00000
-2.60338    0.00000    0.00000 ;      HA-    CA-    CB-    CG
  6624   6623   6641   6642       3    3.68192   -4.35136    0.00000
1.33888    0.00000    0.00000 ;      HA-    CA-     C-     O
  6624   6623   6641   6643       3    0.00000    0.00000    0.00000
0.00000    0.00000    0.00000 ;      HA-    CA-     C-     N
  6625   6623   6641   6642       3    0.00000    0.00000    0.00000
0.00000    0.00000    0.00000 ;      CB-    CA-     C-     O
  6625   6623   6641   6643       3    2.51040    4.18400    1.67360
-6.69440    0.00000    0.00000 ;      CB-    CA-     C-     N
  6625   6628   6631   6632       3    0.66944    2.00832    0.00000
-2.67776    0.00000    0.00000 ;      CB-    CG-    CD-   HD2
  6625   6628   6631   6633       3    0.66944    2.00832    0.00000
-2.67776    0.00000    0.00000 ;      CB-    CG-    CD-   HD3
  6625   6628   6631   6634       3    3.68192    3.09616   -2.09200
-3.01248    0.00000    0.00000 ;      CB-    CG-    CD-    CE
  6626   6625   6623   6641       3    0.65084    1.95253    0.00000
-2.60338    0.00000    0.00000 ;     HB2-    CB-    CA-     C
  6626   6625   6628   6629       3    0.62760    1.88280    0.00000
-2.51040    0.00000    0.00000 ;     HB2-    CB-    CG-   HG2
  6626   6625   6628   6630       3    0.62760    1.88280    0.00000
-2.51040    0.00000    0.00000 ;     HB2-    CB-    CG-   HG3
  6626   6625   6628   6631       3    0.66944    2.00832    0.00000
-2.67776    0.00000    0.00000 ;     HB2-    CB-    CG-    CD
  6627   6625   6623   6641       3    0.65084    1.95253    0.00000
-2.60338    0.00000    0.00000 ;     HB3-    CB-    CA-     C
  6627   6625   6628   6629       3    0.62760    1.88280    0.00000
-2.51040    0.00000    0.00000 ;     HB3-    CB-    CG-   HG2
  6627   6625   6628   6630       3    0.62760    1.88280    0.00000
-2.51040    0.00000    0.00000 ;     HB3-    CB-    CG-   HG3
  6627   6625   6628   6631       3    0.66944    2.00832    0.00000
-2.67776    0.00000    0.00000 ;     HB3-    CB-    CG-    CD
  6628   6625   6623   6641       3    0.65084    1.95253    0.00000
-2.60338    0.00000    0.00000 ;      CG-    CB-    CA-     C
  6628   6631   6634   6635       3    0.65084    1.95253    0.00000
-2.60338    0.00000    0.00000 ;      CG-    CD-    CE-   HE2
  6628   6631   6634   6636       3    0.65084    1.95253    0.00000
-2.60338    0.00000    0.00000 ;      CG-    CD-    CE-   HE3
  6628   6631   6634   6637       3    0.65084    1.95253    0.00000
```

-2.60338    0.00000    0.00000 ;     CG-    CD-    CE-    NZ
6629   6628   6631   6632      3    0.62760    1.88280    0.00000
-2.51040    0.00000    0.00000 ;    HG2-    CG-    CD-   HD2
6629   6628   6631   6633      3    0.62760    1.88280    0.00000
-2.51040    0.00000    0.00000 ;    HG2-    CG-    CD-   HD3
6629   6628   6631   6634      3    0.66944    2.00832    0.00000
-2.67776    0.00000    0.00000 ;    HG2-    CG-    CD-    CE
6630   6628   6631   6632      3    0.62760    1.88280    0.00000
-2.51040    0.00000    0.00000 ;    HG3-    CG-    CD-   HD2
6630   6628   6631   6633      3    0.62760    1.88280    0.00000
-2.51040    0.00000    0.00000 ;    HG3-    CG-    CD-   HD3
6630   6628   6631   6634      3    0.66944    2.00832    0.00000
-2.67776    0.00000    0.00000 ;    HG3-    CG-    CD-    CE
6631   6634   6637   6638      3    0.65084    1.95253    0.00000
-2.60338    0.00000    0.00000 ;     CD-    CE-    NZ-   HZ1
6631   6634   6637   6639      3    0.65084    1.95253    0.00000
-2.60338    0.00000    0.00000 ;     CD-    CE-    NZ-   HZ2
6631   6634   6637   6640      3    0.65084    1.95253    0.00000
-2.60338    0.00000    0.00000 ;     CD-    CE-    NZ-   HZ3
6632   6631   6634   6635      3    0.65084    1.95253    0.00000
-2.60338    0.00000    0.00000 ;    HD2-    CD-    CE-   HE2
6632   6631   6634   6636      3    0.65084    1.95253    0.00000
-2.60338    0.00000    0.00000 ;    HD2-    CD-    CE-   HE3
6632   6631   6634   6637      3    0.65084    1.95253    0.00000
-2.60338    0.00000    0.00000 ;    HD2-    CD-    CE-    NZ
6633   6631   6634   6635      3    0.65084    1.95253    0.00000
-2.60338    0.00000    0.00000 ;    HD3-    CD-    CE-   HE2
6633   6631   6634   6636      3    0.65084    1.95253    0.00000
-2.60338    0.00000    0.00000 ;    HD3-    CD-    CE-   HE3
6633   6631   6634   6637      3    0.65084    1.95253    0.00000
-2.60338    0.00000    0.00000 ;    HD3-    CD-    CE-    NZ
6635   6634   6637   6638      3    0.65084    1.95253    0.00000
-2.60338    0.00000    0.00000 ;    HE2-    CE-    NZ-   HZ1
6635   6634   6637   6639      3    0.65084    1.95253    0.00000
-2.60338    0.00000    0.00000 ;    HE2-    CE-    NZ-   HZ2
6635   6634   6637   6640      3    0.65084    1.95253    0.00000
-2.60338    0.00000    0.00000 ;    HE2-    CE-    NZ-   HZ3
6636   6634   6637   6638      3    0.65084    1.95253    0.00000
-2.60338    0.00000    0.00000 ;    HE3-    CE-    NZ-   HZ1
6636   6634   6637   6639      3    0.65084    1.95253    0.00000
-2.60338    0.00000    0.00000 ;    HE3-    CE-    NZ-   HZ2
6636   6634   6637   6640      3    0.65084    1.95253    0.00000
-2.60338    0.00000    0.00000 ;    HE3-    CE-    NZ-   HZ3
6641   6643   6645   6646      3    0.00000    0.00000    0.00000
0.00000    0.00000    0.00000 ;     C-     N-    CA-    HA
6641   6643   6645   6647      3   11.71520    1.67360   15.06240  -
13.38880    0.00000    0.00000 ;     C-     N-    CA-    CB
6641   6643   6645   6665      3    1.75728    5.27184    2.25936
-7.02912    0.00000    0.00000 ;     C-     N-    CA-     C
6642   6641   6643   6644      3   29.28800   -8.36800  -20.92000
0.00000    0.00000    0.00000 ;     O-     C-     N-     H
6642   6641   6643   6645      3   20.92000    0.00000  -20.92000

```
0.00000    0.00000    0.00000 ;       O-     C-     N-    CA
  6643   6645   6647   6648       3    0.65084    1.95253    0.00000
-2.60338    0.00000    0.00000 ;       N-    CA-    CB-   HB2
  6643   6645   6647   6649       3    0.65084    1.95253    0.00000
-2.60338    0.00000    0.00000 ;       N-    CA-    CB-   HB3
  6643   6645   6647   6650       3    0.65084    1.95253    0.00000
-2.60338    0.00000    0.00000 ;       N-    CA-    CB-    CG
  6643   6645   6665   6666       3    0.00000    0.00000    0.00000
0.00000    0.00000    0.00000 ;       N-    CA-     C-     O
  6643   6645   6665   6667       3   17.40544   -5.02080  -13.22144
9.20480    0.00000    0.00000 ;       N-    CA-     C-     N
  6644   6643   6645   6646       3    0.00000    0.00000    0.00000
0.00000    0.00000    0.00000 ;       H-     N-    CA-    HA
  6644   6643   6645   6647       3    0.00000    0.00000    0.00000
0.00000    0.00000    0.00000 ;       H-     N-    CA-    CB
  6644   6643   6645   6665       3    0.00000    0.00000    0.00000
0.00000    0.00000    0.00000 ;       H-     N-    CA-     C
  6645   6647   6650   6651       3    0.66944    2.00832    0.00000
-2.67776    0.00000    0.00000 ;      CA-    CB-    CG-   HG2
  6645   6647   6650   6652       3    0.66944    2.00832    0.00000
-2.67776    0.00000    0.00000 ;      CA-    CB-    CG-   HG3
  6645   6647   6650   6653       3    3.68192    3.09616   -2.09200
-3.01248    0.00000    0.00000 ;      CA-    CB-    CG-    CD
  6645   6665   6667   6668       3   20.92000    0.00000  -20.92000
0.00000    0.00000    0.00000 ;      CA-     C-     N-     H
  6645   6665   6667   6669       3   20.92000    0.00000  -20.92000
0.00000    0.00000    0.00000 ;      CA-     C-     N-    CA
  6646   6645   6647   6648       3    0.65084    1.95253    0.00000
-2.60338    0.00000    0.00000 ;      HA-    CA-    CB-   HB2
  6646   6645   6647   6649       3    0.65084    1.95253    0.00000
-2.60338    0.00000    0.00000 ;      HA-    CA-    CB-   HB3
  6646   6645   6647   6650       3    0.65084    1.95253    0.00000
-2.60338    0.00000    0.00000 ;      HA-    CA-    CB-    CG
  6646   6645   6665   6666       3    3.68192   -4.35136    0.00000
1.33888    0.00000    0.00000 ;      HA-    CA-     C-     O
  6646   6645   6665   6667       3    0.00000    0.00000    0.00000
0.00000    0.00000    0.00000 ;      HA-    CA-     C-     N
  6647   6645   6665   6666       3    0.00000    0.00000    0.00000
0.00000    0.00000    0.00000 ;      CB-    CA-     C-     O
  6647   6645   6665   6667       3    2.51040    4.18400    1.67360
-6.69440    0.00000    0.00000 ;      CB-    CA-     C-     N
  6647   6650   6653   6654       3    0.65084    1.95253    0.00000
-2.60338    0.00000    0.00000 ;      CB-    CG-    CD-   HD2
  6647   6650   6653   6655       3    0.65084    1.95253    0.00000
-2.60338    0.00000    0.00000 ;      CB-    CG-    CD-   HD3
  6647   6650   6653   6656       3    0.65084    1.95253    0.00000
-2.60338    0.00000    0.00000 ;      CB-    CG-    CD-    NE
  6648   6647   6645   6665       3    0.65084    1.95253    0.00000
-2.60338    0.00000    0.00000 ;     HB2-    CB-    CA-     C
  6648   6647   6650   6651       3    0.62760    1.88280    0.00000
-2.51040    0.00000    0.00000 ;     HB2-    CB-    CG-   HG2
  6648   6647   6650   6652       3    0.62760    1.88280    0.00000
```

```
-2.51040    0.00000    0.00000 ;    HB2-   CB-    CG-   HG3
  6648   6647   6650   6653      3    0.66944    2.00832    0.00000
-2.67776    0.00000    0.00000 ;    HB2-   CB-    CG-    CD
  6649   6647   6645   6665      3    0.65084    1.95253    0.00000
-2.60338    0.00000    0.00000 ;    HB3-   CB-    CA-     C
  6649   6647   6650   6651      3    0.62760    1.88280    0.00000
-2.51040    0.00000    0.00000 ;    HB3-   CB-    CG-   HG2
  6649   6647   6650   6652      3    0.62760    1.88280    0.00000
-2.51040    0.00000    0.00000 ;    HB3-   CB-    CG-   HG3
  6649   6647   6650   6653      3    0.66944    2.00832    0.00000
-2.67776    0.00000    0.00000 ;    HB3-   CB-    CG-    CD
  6650   6647   6645   6665      3    0.65084    1.95253    0.00000
-2.60338    0.00000    0.00000 ;     CG-   CB-    CA-     C
  6650   6653   6656   6657      3    0.00000    0.00000    0.00000
0.00000    0.00000    0.00000 ;     CG-   CD-    NE-    HE
  6650   6653   6656   6658      3    0.00000    0.00000    0.00000
0.00000    0.00000    0.00000 ;     CG-   CD-    NE-    CZ
  6651   6650   6653   6654      3    0.65084    1.95253    0.00000
-2.60338    0.00000    0.00000 ;    HG2-   CG-    CD-   HD2
  6651   6650   6653   6655      3    0.65084    1.95253    0.00000
-2.60338    0.00000    0.00000 ;    HG2-   CG-    CD-   HD3
  6651   6650   6653   6656      3    0.65084    1.95253    0.00000
-2.60338    0.00000    0.00000 ;    HG2-   CG-    CD-    NE
  6652   6650   6653   6654      3    0.65084    1.95253    0.00000
-2.60338    0.00000    0.00000 ;    HG3-   CG-    CD-   HD2
  6652   6650   6653   6655      3    0.65084    1.95253    0.00000
-2.60338    0.00000    0.00000 ;    HG3-   CG-    CD-   HD3
  6652   6650   6653   6656      3    0.65084    1.95253    0.00000
-2.60338    0.00000    0.00000 ;    HG3-   CG-    CD-    NE
  6653   6656   6658   6659      3   20.08320    0.00000  -20.08320
0.00000    0.00000    0.00000 ;     CD-   NE-    CZ-   NH1
  6653   6656   6658   6662      3   20.08320    0.00000  -20.08320
0.00000    0.00000    0.00000 ;     CD-   NE-    CZ-   NH2
  6654   6653   6656   6657      3    0.00000    0.00000    0.00000
0.00000    0.00000    0.00000 ;    HD2-   CD-    NE-    HE
  6654   6653   6656   6658      3    0.00000    0.00000    0.00000
0.00000    0.00000    0.00000 ;    HD2-   CD-    NE-    CZ
  6655   6653   6656   6657      3    0.00000    0.00000    0.00000
0.00000    0.00000    0.00000 ;    HD3-   CD-    NE-    HE
  6655   6653   6656   6658      3    0.00000    0.00000    0.00000
0.00000    0.00000    0.00000 ;    HD3-   CD-    NE-    CZ
  6656   6658   6659   6660      3   20.08320    0.00000  -20.08320
0.00000    0.00000    0.00000 ;     NE-   CZ-   NH1-  HH11
  6656   6658   6659   6661      3   20.08320    0.00000  -20.08320
0.00000    0.00000    0.00000 ;     NE-   CZ-   NH1-  HH12
  6656   6658   6662   6663      3   20.08320    0.00000  -20.08320
0.00000    0.00000    0.00000 ;     NE-   CZ-   NH2-  HH21
  6656   6658   6662   6664      3   20.08320    0.00000  -20.08320
0.00000    0.00000    0.00000 ;     NE-   CZ-   NH2-  HH22
  6657   6656   6658   6659      3   20.08320    0.00000  -20.08320
0.00000    0.00000    0.00000 ;     HE-   NE-    CZ-   NH1
  6657   6656   6658   6662      3   20.08320    0.00000  -20.08320
```

0.00000 0.00000 0.00000 ; HE- NE- CZ- NH2
6659 6658 6662 6663 3 20.08320 0.00000 -20.08320
0.00000 0.00000 0.00000 ; NH1- CZ- NH2- HH21
6659 6658 6662 6664 3 20.08320 0.00000 -20.08320
0.00000 0.00000 0.00000 ; NH1- CZ- NH2- HH22
6660 6659 6658 6662 3 20.08320 0.00000 -20.08320
0.00000 0.00000 0.00000 ; HH11- NH1- CZ- NH2
6661 6659 6658 6662 3 20.08320 0.00000 -20.08320
0.00000 0.00000 0.00000 ; HH12- NH1- CZ- NH2
6665 6667 6669 6670 3 0.00000 0.00000 0.00000
0.00000 0.00000 0.00000 ; C- N- CA- HA
6665 6667 6669 6671 3 11.71520 1.67360 15.06240 -
13.38880 0.00000 0.00000 ; C- N- CA- CB
6665 6667 6669 6684 3 1.75728 5.27184 2.25936
-7.02912 0.00000 0.00000 ; C- N- CA- C
6666 6665 6667 6668 3 29.28800 -8.36800 -20.92000
0.00000 0.00000 0.00000 ; O- C- N- H
6666 6665 6667 6669 3 20.92000 0.00000 -20.92000
0.00000 0.00000 0.00000 ; O- C- N- CA
6667 6669 6671 6672 3 0.65084 1.95253 0.00000
-2.60338 0.00000 0.00000 ; N- CA- CB- HB
6667 6669 6671 6673 3 3.83673 0.44769 -1.77402
-2.47693 -0.03347 0.00000 ; N- CA- CB- CG2
6667 6669 6671 6677 3 2.97482 0.12134 2.04179
-1.89117 -3.24678 0.00000 ; N- CA- CB- CG1
6667 6669 6684 6685 3 0.00000 0.00000 0.00000
0.00000 0.00000 0.00000 ; N- CA- C- O
6667 6669 6684 6686 3 17.40544 -5.02080 -13.22144
9.20480 0.00000 0.00000 ; N- CA- C- N
6668 6667 6669 6670 3 0.00000 0.00000 0.00000
0.00000 0.00000 0.00000 ; H- N- CA- HA
6668 6667 6669 6671 3 0.00000 0.00000 0.00000
0.00000 0.00000 0.00000 ; H- N- CA- CB
6668 6667 6669 6684 3 0.00000 0.00000 0.00000
0.00000 0.00000 0.00000 ; H- N- CA- C
6669 6671 6673 6674 3 0.66944 2.00832 0.00000
-2.67776 0.00000 0.00000 ; CA- CB- CG2- HG21
6669 6671 6673 6675 3 0.66944 2.00832 0.00000
-2.67776 0.00000 0.00000 ; CA- CB- CG2- HG22
6669 6671 6673 6676 3 0.66944 2.00832 0.00000
-2.67776 0.00000 0.00000 ; CA- CB- CG2- HG23
6669 6671 6677 6678 3 0.66944 2.00832 0.00000
-2.67776 0.00000 0.00000 ; CA- CB- CG1- HG12
6669 6671 6677 6679 3 0.66944 2.00832 0.00000
-2.67776 0.00000 0.00000 ; CA- CB- CG1- HG13
6669 6671 6677 6680 3 4.24258 -0.52718 -7.25506
-1.79075 7.69856 0.00000 ; CA- CB- CG1- CD1
6669 6684 6686 6687 3 20.92000 0.00000 -20.92000
0.00000 0.00000 0.00000 ; CA- C- N- H
6669 6684 6686 6688 3 20.92000 0.00000 -20.92000
0.00000 0.00000 0.00000 ; CA- C- N- CA
6670 6669 6671 6672 3 0.65084 1.95253 0.00000

```
-2.60338    0.00000    0.00000 ;      HA-    CA-    CB-    HB
  6670   6669   6671   6673      3    0.65084    1.95253    0.00000
-2.60338    0.00000    0.00000 ;      HA-    CA-    CB-   CG2
  6670   6669   6671   6677      3    0.65084    1.95253    0.00000
-2.60338    0.00000    0.00000 ;      HA-    CA-    CB-   CG1
  6670   6669   6684   6685      3    3.68192   -4.35136    0.00000
1.33888    0.00000    0.00000 ;      HA-    CA-     C-     O
  6670   6669   6684   6686      3    0.00000    0.00000    0.00000
0.00000    0.00000    0.00000 ;      HA-    CA-     C-     N
  6671   6669   6684   6685      3    0.00000    0.00000    0.00000
0.00000    0.00000    0.00000 ;      CB-    CA-     C-     O
  6671   6669   6684   6686      3    2.51040    4.18400    1.67360
-6.69440    0.00000    0.00000 ;      CB-    CA-     C-     N
  6671   6677   6680   6681      3    0.66944    2.00832    0.00000
-2.67776    0.00000    0.00000 ;      CB-   CG1-   CD1-  HD11
  6671   6677   6680   6682      3    0.66944    2.00832    0.00000
-2.67776    0.00000    0.00000 ;      CB-   CG1-   CD1-  HD12
  6671   6677   6680   6683      3    0.66944    2.00832    0.00000
-2.67776    0.00000    0.00000 ;      CB-   CG1-   CD1-  HD13
  6672   6671   6669   6684      3    0.65084    1.95253    0.00000
-2.60338    0.00000    0.00000 ;      HB-    CB-    CA-     C
  6672   6671   6673   6674      3    0.62760    1.88280    0.00000
-2.51040    0.00000    0.00000 ;      HB-    CB-   CG2-  HG21
  6672   6671   6673   6675      3    0.62760    1.88280    0.00000
-2.51040    0.00000    0.00000 ;      HB-    CB-   CG2-  HG22
  6672   6671   6673   6676      3    0.62760    1.88280    0.00000
-2.51040    0.00000    0.00000 ;      HB-    CB-   CG2-  HG23
  6672   6671   6677   6678      3    0.62760    1.88280    0.00000
-2.51040    0.00000    0.00000 ;      HB-    CB-   CG1-  HG12
  6672   6671   6677   6679      3    0.62760    1.88280    0.00000
-2.51040    0.00000    0.00000 ;      HB-    CB-   CG1-  HG13
  6672   6671   6677   6680      3    0.66944    2.00832    0.00000
-2.67776    0.00000    0.00000 ;      HB-    CB-   CG1-   CD1
  6673   6671   6669   6684      3    5.67350    3.55640   -6.16722
-2.47693    3.74886    0.00000 ;     CG2-    CB-    CA-     C
  6673   6671   6677   6678      3    0.66944    2.00832    0.00000
-2.67776    0.00000    0.00000 ;     CG2-    CB-   CG1-  HG12
  6673   6671   6677   6679      3    0.66944    2.00832    0.00000
-2.67776    0.00000    0.00000 ;     CG2-    CB-   CG1-  HG13
  6673   6671   6677   6680      3    3.81162    0.49790   -8.14206
-1.79075    7.49773    0.00000 ;     CG2-    CB-   CG1-   CD1
  6674   6673   6671   6677      3    0.66944    2.00832    0.00000
-2.67776    0.00000    0.00000 ;    HG21-   CG2-    CB-   CG1
  6675   6673   6671   6677      3    0.66944    2.00832    0.00000
-2.67776    0.00000    0.00000 ;    HG22-   CG2-    CB-   CG1
  6676   6673   6671   6677      3    0.66944    2.00832    0.00000
-2.67776    0.00000    0.00000 ;    HG23-   CG2-    CB-   CG1
  6677   6671   6669   6684      3    8.26340    0.74057   -9.99976
-1.89117    3.84928    0.00000 ;     CG1-    CB-    CA-     C
  6678   6677   6680   6681      3    0.62760    1.88280    0.00000
-2.51040    0.00000    0.00000 ;    HG12-   CG1-   CD1-  HD11
  6678   6677   6680   6682      3    0.62760    1.88280    0.00000
```

```
-2.51040    0.00000    0.00000 ;   HG12-   CG1-   CD1-  HD12
  6678   6677   6680   6683      3    0.62760    1.88280    0.00000
-2.51040    0.00000    0.00000 ;   HG12-   CG1-   CD1-  HD13
  6679   6677   6680   6681      3    0.62760    1.88280    0.00000
-2.51040    0.00000    0.00000 ;   HG13-   CG1-   CD1-  HD11
  6679   6677   6680   6682      3    0.62760    1.88280    0.00000
-2.51040    0.00000    0.00000 ;   HG13-   CG1-   CD1-  HD12
  6679   6677   6680   6683      3    0.62760    1.88280    0.00000
-2.51040    0.00000    0.00000 ;   HG13-   CG1-   CD1-  HD13
  6684   6686   6688   6689      3    0.00000    0.00000    0.00000
0.00000    0.00000    0.00000 ;      C-     N-    CA-    HA
  6684   6686   6688   6690      3   11.71520    1.67360   15.06240  -
13.38880    0.00000    0.00000 ;      C-     N-    CA-    CB
  6684   6686   6688   6704      3    1.75728    5.27184    2.25936
-7.02912    0.00000    0.00000 ;      C-     N-    CA-     C
  6685   6684   6686   6687      3   29.28800   -8.36800  -20.92000
0.00000    0.00000    0.00000 ;      O-     C-     N-     H
  6685   6684   6686   6688      3   20.92000    0.00000  -20.92000
0.00000    0.00000    0.00000 ;      O-     C-     N-    CA
  6686   6688   6690   6691      3    0.65084    1.95253    0.00000
-2.60338    0.00000    0.00000 ;      N-    CA-    CB-   HB2
  6686   6688   6690   6692      3    0.65084    1.95253    0.00000
-2.60338    0.00000    0.00000 ;      N-    CA-    CB-   HB3
  6686   6688   6690   6693      3    3.28026    2.46019   -2.19242
-3.21331   -0.23430    0.00000 ;      N-    CA-    CB-    CG
  6686   6688   6704   6705      3    0.00000    0.00000    0.00000
0.00000    0.00000    0.00000 ;     N-    CA-     C-     O
  6686   6688   6704   6706      3   17.40544   -5.02080  -13.22144
9.20480    0.00000    0.00000 ;     N-    CA-     C-     N
  6687   6686   6688   6689      3    0.00000    0.00000    0.00000
0.00000    0.00000    0.00000 ;     H-     N-    CA-    HA
  6687   6686   6688   6690      3    0.00000    0.00000    0.00000
0.00000    0.00000    0.00000 ;     H-     N-    CA-    CB
  6687   6686   6688   6704      3    0.00000    0.00000    0.00000
0.00000    0.00000    0.00000 ;     H-     N-    CA-     C
  6688   6690   6693   6694      3    0.57739    0.00000    1.02926
0.00000   -1.60666    0.00000 ;    CA-    CB-    CG-   CD1
  6688   6690   6693   6702      3    0.57739    0.00000    1.02926
0.00000   -1.60666    0.00000 ;    CA-    CB-    CG-   CD2
  6688   6704   6706   6707      3   20.92000    0.00000  -20.92000
0.00000    0.00000    0.00000 ;    CA-     C-     N-     H
  6688   6704   6706   6708      3   20.92000    0.00000  -20.92000
0.00000    0.00000    0.00000 ;    CA-     C-     N-    CA
  6689   6688   6690   6691      3    0.65084    1.95253    0.00000
-2.60338    0.00000    0.00000 ;     HA-    CA-    CB-   HB2
  6689   6688   6690   6692      3    0.65084    1.95253    0.00000
-2.60338    0.00000    0.00000 ;     HA-    CA-    CB-   HB3
  6689   6688   6690   6693      3    0.65084    1.95253    0.00000
-2.60338    0.00000    0.00000 ;     HA-    CA-    CB-    CG
  6689   6688   6704   6705      3    3.68192   -4.35136    0.00000
1.33888    0.00000    0.00000 ;    HA-    CA-     C-     O
  6689   6688   6704   6706      3    0.00000    0.00000    0.00000
```

```
0.00000    0.00000    0.00000 ;     HA-    CA-     C-     N
  6690   6688   6704   6705      3    0.00000    0.00000    0.00000
0.00000    0.00000    0.00000 ;     CB-    CA-     C-     O
  6690   6688   6704   6706      3    2.51040    4.18400    1.67360
-6.69440    0.00000    0.00000 ;     CB-    CA-     C-     N
  6690   6693   6694   6695      3   30.33400    0.00000  -30.33400
0.00000    0.00000    0.00000 ;     CB-    CG-   CD1-   HD1
  6690   6693   6694   6696      3   30.33400    0.00000  -30.33400
0.00000    0.00000    0.00000 ;     CB-    CG-   CD1-   CE1
  6690   6693   6702   6700      3   30.33400    0.00000  -30.33400
0.00000    0.00000    0.00000 ;     CB-    CG-   CD2-   CE2
  6690   6693   6702   6703      3   30.33400    0.00000  -30.33400
0.00000    0.00000    0.00000 ;     CB-    CG-   CD2-   HD2
  6691   6690   6688   6704      3    0.65084    1.95253    0.00000
-2.60338    0.00000    0.00000 ;    HB2-    CB-    CA-     C
  6691   6690   6693   6694      3    0.00000    0.00000    0.00000
0.00000    0.00000    0.00000 ;    HB2-    CB-    CG-   CD1
  6691   6690   6693   6702      3    0.00000    0.00000    0.00000
0.00000    0.00000    0.00000 ;    HB2-    CB-    CG-   CD2
  6692   6690   6688   6704      3    0.65084    1.95253    0.00000
-2.60338    0.00000    0.00000 ;    HB3-    CB-    CA-     C
  6692   6690   6693   6694      3    0.00000    0.00000    0.00000
0.00000    0.00000    0.00000 ;    HB3-    CB-    CG-   CD1
  6692   6690   6693   6702      3    0.00000    0.00000    0.00000
0.00000    0.00000    0.00000 ;    HB3-    CB-    CG-   CD2
  6693   6690   6688   6704      3    4.95804    2.17986   -3.52293
-3.21331   -0.40166    0.00000 ;     CG-    CB-    CA-     C
  6693   6694   6696   6697      3   30.33400    0.00000  -30.33400
0.00000    0.00000    0.00000 ;     CG-   CD1-   CE1-   HE1
  6693   6694   6696   6698      3   30.33400    0.00000  -30.33400
0.00000    0.00000    0.00000 ;     CG-   CD1-   CE1-    CZ
  6693   6702   6700   6698      3   30.33400    0.00000  -30.33400
0.00000    0.00000    0.00000 ;     CG-   CD2-   CE2-    CZ
  6693   6702   6700   6701      3   30.33400    0.00000  -30.33400
0.00000    0.00000    0.00000 ;     CG-   CD2-   CE2-   HE2
  6694   6693   6702   6700      3   30.33400    0.00000  -30.33400
0.00000    0.00000    0.00000 ;    CD1-    CG-   CD2-   CE2
  6694   6693   6702   6703      3   30.33400    0.00000  -30.33400
0.00000    0.00000    0.00000 ;    CD1-    CG-   CD2-   HD2
  6694   6696   6698   6699      3   30.33400    0.00000  -30.33400
0.00000    0.00000    0.00000 ;    CD1-   CE1-    CZ-    HZ
  6694   6696   6698   6700      3   30.33400    0.00000  -30.33400
0.00000    0.00000    0.00000 ;    CD1-   CE1-    CZ-   CE2
  6695   6694   6693   6702      3   30.33400    0.00000  -30.33400
0.00000    0.00000    0.00000 ;    HD1-   CD1-    CG-   CD2
  6695   6694   6696   6697      3   30.33400    0.00000  -30.33400
0.00000    0.00000    0.00000 ;    HD1-   CD1-   CE1-   HE1
  6695   6694   6696   6698      3   30.33400    0.00000  -30.33400
0.00000    0.00000    0.00000 ;    HD1-   CD1-   CE1-    CZ
  6696   6694   6693   6702      3   30.33400    0.00000  -30.33400
0.00000    0.00000    0.00000 ;    CE1-   CD1-    CG-   CD2
  6696   6698   6700   6701      3   30.33400    0.00000  -30.33400
```

0.00000 0.00000 0.00000 ; CE1- CZ- CE2- HE2
6696 6698 6700 6702 3 30.33400 0.00000 -30.33400
0.00000 0.00000 0.00000 ; CE1- CZ- CE2- CD2
6697 6696 6698 6699 3 30.33400 0.00000 -30.33400
0.00000 0.00000 0.00000 ; HE1- CE1- CZ- HZ
6697 6696 6698 6700 3 30.33400 0.00000 -30.33400
0.00000 0.00000 0.00000 ; HE1- CE1- CZ- CE2
6698 6700 6702 6703 3 30.33400 0.00000 -30.33400
0.00000 0.00000 0.00000 ; CZ- CE2- CD2- HD2
6699 6698 6700 6701 3 30.33400 0.00000 -30.33400
0.00000 0.00000 0.00000 ; HZ- CZ- CE2- HE2
6699 6698 6700 6702 3 30.33400 0.00000 -30.33400
0.00000 0.00000 0.00000 ; HZ- CZ- CE2- CD2
6701 6700 6702 6703 3 30.33400 0.00000 -30.33400
0.00000 0.00000 0.00000 ; HE2- CE2- CD2- HD2
6704 6706 6708 6709 3 0.00000 0.00000 0.00000
0.00000 0.00000 0.00000 ; C- N- CA- HA2
6704 6706 6708 6710 3 0.00000 0.00000 0.00000
0.00000 0.00000 0.00000 ; C- N- CA- HA3
6704 6706 6708 6711 3 1.75728 5.27184 2.25936
-7.02912 0.00000 0.00000 ; C- N- CA- C
6705 6704 6706 6707 3 29.28800 -8.36800 -20.92000
0.00000 0.00000 0.00000 ; O- C- N- H
6705 6704 6706 6708 3 20.92000 0.00000 -20.92000
0.00000 0.00000 0.00000 ; O- C- N- CA
6706 6708 6711 6712 3 0.00000 0.00000 0.00000
0.00000 0.00000 0.00000 ; N- CA- C- O
6706 6708 6711 6713 3 17.40544 -5.02080 -13.22144
9.20480 0.00000 0.00000 ; N- CA- C- N
6707 6706 6708 6709 3 0.00000 0.00000 0.00000
0.00000 0.00000 0.00000 ; H- N- CA- HA2
6707 6706 6708 6710 3 0.00000 0.00000 0.00000
0.00000 0.00000 0.00000 ; H- N- CA- HA3
6707 6706 6708 6711 3 0.00000 0.00000 0.00000
0.00000 0.00000 0.00000 ; H- N- CA- C
6708 6711 6713 6714 3 20.92000 0.00000 -20.92000
0.00000 0.00000 0.00000 ; CA- C- N- H
6708 6711 6713 6715 3 20.92000 0.00000 -20.92000
0.00000 0.00000 0.00000 ; CA- C- N- CA
6709 6708 6711 6712 3 3.68192 -4.35136 0.00000
1.33888 0.00000 0.00000 ; HA2- CA- C- O
6709 6708 6711 6713 3 0.00000 0.00000 0.00000
0.00000 0.00000 0.00000 ; HA2- CA- C- N
6710 6708 6711 6712 3 3.68192 -4.35136 0.00000
1.33888 0.00000 0.00000 ; HA3- CA- C- O
6710 6708 6711 6713 3 0.00000 0.00000 0.00000
0.00000 0.00000 0.00000 ; HA3- CA- C- N
6711 6713 6715 6716 3 0.00000 0.00000 0.00000
0.00000 0.00000 0.00000 ; C- N- CA- HA
6711 6713 6715 6717 3 11.71520 1.67360 15.06240 -
13.38880 0.00000 0.00000 ; C- N- CA- CB
6711 6713 6715 6735 3 1.75728 5.27184 2.25936

-7.02912 0.00000 0.00000 ; C- N- CA- C
6712 6711 6713 6714 3 29.28800 -8.36800 -20.92000
0.00000 0.00000 0.00000 ; O- C- N- H
6712 6711 6713 6715 3 20.92000 0.00000 -20.92000
0.00000 0.00000 0.00000 ; O- C- N- CA
6713 6715 6717 6718 3 0.65084 1.95253 0.00000
-2.60338 0.00000 0.00000 ; N- CA- CB- HB2
6713 6715 6717 6719 3 0.65084 1.95253 0.00000
-2.60338 0.00000 0.00000 ; N- CA- CB- HB3
6713 6715 6717 6720 3 0.65084 1.95253 0.00000
-2.60338 0.00000 0.00000 ; N- CA- CB- CG
6713 6715 6735 6736 3 0.00000 0.00000 0.00000
0.00000 0.00000 0.00000 ; N- CA- C- O
6713 6715 6735 6737 3 17.40544 -5.02080 -13.22144
9.20480 0.00000 0.00000 ; N- CA- C- N
6714 6713 6715 6716 3 0.00000 0.00000 0.00000
0.00000 0.00000 0.00000 ; H- N- CA- HA
6714 6713 6715 6717 3 0.00000 0.00000 0.00000
0.00000 0.00000 0.00000 ; H- N- CA- CB
6714 6713 6715 6735 3 0.00000 0.00000 0.00000
0.00000 0.00000 0.00000 ; H- N- CA- C
6715 6717 6720 6721 3 0.66944 2.00832 0.00000
-2.67776 0.00000 0.00000 ; CA- CB- CG- HG2
6715 6717 6720 6722 3 0.66944 2.00832 0.00000
-2.67776 0.00000 0.00000 ; CA- CB- CG- HG3
6715 6717 6720 6723 3 3.68192 3.09616 -2.09200
-3.01248 0.00000 0.00000 ; CA- CB- CG- CD
6715 6735 6737 6738 3 20.92000 0.00000 -20.92000
0.00000 0.00000 0.00000 ; CA- C- N- H
6715 6735 6737 6739 3 20.92000 0.00000 -20.92000
0.00000 0.00000 0.00000 ; CA- C- N- CA
6716 6715 6717 6718 3 0.65084 1.95253 0.00000
-2.60338 0.00000 0.00000 ; HA- CA- CB- HB2
6716 6715 6717 6719 3 0.65084 1.95253 0.00000
-2.60338 0.00000 0.00000 ; HA- CA- CB- HB3
6716 6715 6717 6720 3 0.65084 1.95253 0.00000
-2.60338 0.00000 0.00000 ; HA- CA- CB- CG
6716 6715 6735 6736 3 3.68192 -4.35136 0.00000
1.33888 0.00000 0.00000 ; HA- CA- C- O
6716 6715 6735 6737 3 0.00000 0.00000 0.00000
0.00000 0.00000 0.00000 ; HA- CA- C- N
6717 6715 6735 6736 3 0.00000 0.00000 0.00000
0.00000 0.00000 0.00000 ; CB- CA- C- O
6717 6715 6735 6737 3 2.51040 4.18400 1.67360
-6.69440 0.00000 0.00000 ; CB- CA- C- N
6717 6720 6723 6724 3 0.65084 1.95253 0.00000
-2.60338 0.00000 0.00000 ; CB- CG- CD- HD2
6717 6720 6723 6725 3 0.65084 1.95253 0.00000
-2.60338 0.00000 0.00000 ; CB- CG- CD- HD3
6717 6720 6723 6726 3 0.65084 1.95253 0.00000
-2.60338 0.00000 0.00000 ; CB- CG- CD- NE
6718 6717 6715 6735 3 0.65084 1.95253 0.00000

```
-2.60338    0.00000    0.00000 ;    HB2-    CB-    CA-     C
  6718   6717   6720   6721      3    0.62760    1.88280    0.00000
-2.51040    0.00000    0.00000 ;    HB2-    CB-    CG-   HG2
  6718   6717   6720   6722      3    0.62760    1.88280    0.00000
-2.51040    0.00000    0.00000 ;    HB2-    CB-    CG-   HG3
  6718   6717   6720   6723      3    0.66944    2.00832    0.00000
-2.67776    0.00000    0.00000 ;    HB2-    CB-    CG-    CD
  6719   6717   6715   6735      3    0.65084    1.95253    0.00000
-2.60338    0.00000    0.00000 ;    HB3-    CB-    CA-     C
  6719   6717   6720   6721      3    0.62760    1.88280    0.00000
-2.51040    0.00000    0.00000 ;    HB3-    CB-    CG-   HG2
  6719   6717   6720   6722      3    0.62760    1.88280    0.00000
-2.51040    0.00000    0.00000 ;    HB3-    CB-    CG-   HG3
  6719   6717   6720   6723      3    0.66944    2.00832    0.00000
-2.67776    0.00000    0.00000 ;    HB3-    CB-    CG-    CD
  6720   6717   6715   6735      3    0.65084    1.95253    0.00000
-2.60338    0.00000    0.00000 ;     CG-    CB-    CA-     C
  6720   6723   6726   6727      3    0.00000    0.00000    0.00000
0.00000    0.00000    0.00000 ;     CG-    CD-    NE-    HE
  6720   6723   6726   6728      3    0.00000    0.00000    0.00000
0.00000    0.00000    0.00000 ;     CG-    CD-    NE-    CZ
  6721   6720   6723   6724      3    0.65084    1.95253    0.00000
-2.60338    0.00000    0.00000 ;    HG2-    CG-    CD-   HD2
  6721   6720   6723   6725      3    0.65084    1.95253    0.00000
-2.60338    0.00000    0.00000 ;    HG2-    CG-    CD-   HD3
  6721   6720   6723   6726      3    0.65084    1.95253    0.00000
-2.60338    0.00000    0.00000 ;    HG2-    CG-    CD-    NE
  6722   6720   6723   6724      3    0.65084    1.95253    0.00000
-2.60338    0.00000    0.00000 ;    HG3-    CG-    CD-   HD2
  6722   6720   6723   6725      3    0.65084    1.95253    0.00000
-2.60338    0.00000    0.00000 ;    HG3-    CG-    CD-   HD3
  6722   6720   6723   6726      3    0.65084    1.95253    0.00000
-2.60338    0.00000    0.00000 ;    HG3-    CG-    CD-    NE
  6723   6726   6728   6729      3   20.08320    0.00000  -20.08320
0.00000    0.00000    0.00000 ;     CD-    NE-    CZ-   NH1
  6723   6726   6728   6732      3   20.08320    0.00000  -20.08320
0.00000    0.00000    0.00000 ;     CD-    NE-    CZ-   NH2
  6724   6723   6726   6727      3    0.00000    0.00000    0.00000
0.00000    0.00000    0.00000 ;    HD2-    CD-    NE-    HE
  6724   6723   6726   6728      3    0.00000    0.00000    0.00000
0.00000    0.00000    0.00000 ;    HD2-    CD-    NE-    CZ
  6725   6723   6726   6727      3    0.00000    0.00000    0.00000
0.00000    0.00000    0.00000 ;    HD3-    CD-    NE-    HE
  6725   6723   6726   6728      3    0.00000    0.00000    0.00000
0.00000    0.00000    0.00000 ;    HD3-    CD-    NE-    CZ
  6726   6728   6729   6730      3   20.08320    0.00000  -20.08320
0.00000    0.00000    0.00000 ;     NE-    CZ-   NH1-  HH11
  6726   6728   6729   6731      3   20.08320    0.00000  -20.08320
0.00000    0.00000    0.00000 ;     NE-    CZ-   NH1-  HH12
  6726   6728   6732   6733      3   20.08320    0.00000  -20.08320
0.00000    0.00000    0.00000 ;     NE-    CZ-   NH2-  HH21
  6726   6728   6732   6734      3   20.08320    0.00000  -20.08320
```

```
0.00000    0.00000    0.00000 ;     NE-    CZ-   NH2-  HH22
  6727   6726   6728   6729      3   20.08320    0.00000  -20.08320
0.00000    0.00000    0.00000 ;     HE-    NE-    CZ-   NH1
  6727   6726   6728   6732      3   20.08320    0.00000  -20.08320
0.00000    0.00000    0.00000 ;     HE-    NE-    CZ-   NH2
  6729   6728   6732   6733      3   20.08320    0.00000  -20.08320
0.00000    0.00000    0.00000 ;    NH1-    CZ-   NH2-  HH21
  6729   6728   6732   6734      3   20.08320    0.00000  -20.08320
0.00000    0.00000    0.00000 ;    NH1-    CZ-   NH2-  HH22
  6730   6729   6728   6732      3   20.08320    0.00000  -20.08320
0.00000    0.00000    0.00000 ;   HH11-   NH1-    CZ-   NH2
  6731   6729   6728   6732      3   20.08320    0.00000  -20.08320
0.00000    0.00000    0.00000 ;   HH12-   NH1-    CZ-   NH2
  6735   6737   6739   6740      3    0.00000    0.00000    0.00000
0.00000    0.00000    0.00000 ;      C-     N-    CA-    HA
  6735   6737   6739   6741      3   11.71520    1.67360   15.06240  -
13.38880    0.00000    0.00000 ;      C-     N-    CA-    CB
  6735   6737   6739   6747      3    1.75728    5.27184    2.25936
-7.02912    0.00000    0.00000 ;      C-     N-    CA-     C
  6736   6735   6737   6738      3   29.28800   -8.36800  -20.92000
0.00000    0.00000    0.00000 ;      O-     C-     N-     H
  6736   6735   6737   6739      3   20.92000    0.00000  -20.92000
0.00000    0.00000    0.00000 ;      O-     C-     N-    CA
  6737   6739   6741   6742      3    0.65084    1.95253    0.00000
-2.60338    0.00000    0.00000 ;      N-    CA-    CB-   HB2
  6737   6739   6741   6743      3    0.65084    1.95253    0.00000
-2.60338    0.00000    0.00000 ;      N-    CA-    CB-   HB3
  6737   6739   6741   6744      3   15.41386    9.74035   -8.39310
-0.97069    2.97901    0.00000 ;      N-    CA-    CB-    CG
  6737   6739   6747   6748      3    0.00000    0.00000    0.00000
0.00000    0.00000    0.00000 ;      N-    CA-     C-     O
  6737   6739   6747   6749      3   17.40544   -5.02080  -13.22144
9.20480    0.00000    0.00000 ;      N-    CA-     C-     N
  6738   6737   6739   6740      3    0.00000    0.00000    0.00000
0.00000    0.00000    0.00000 ;      H-     N-    CA-    HA
  6738   6737   6739   6741      3    0.00000    0.00000    0.00000
0.00000    0.00000    0.00000 ;      H-     N-    CA-    CB
  6738   6737   6739   6747      3    0.00000    0.00000    0.00000
0.00000    0.00000    0.00000 ;      H-     N-    CA-     C
  6739   6741   6744   6745      3    6.43499    0.00000   -5.39736
0.00000   -1.03763    0.00000 ;     CA-    CB-    CG-   OD1
  6739   6741   6744   6746      3    6.43499    0.00000   -5.39736
0.00000   -1.03763    0.00000 ;     CA-    CB-    CG-   OD2
  6739   6747   6749   6750      3   20.92000    0.00000  -20.92000
0.00000    0.00000    0.00000 ;     CA-     C-     N-     H
  6739   6747   6749   6751      3   20.92000    0.00000  -20.92000
0.00000    0.00000    0.00000 ;     CA-     C-     N-    CA
  6740   6739   6741   6742      3    0.65084    1.95253    0.00000
-2.60338    0.00000    0.00000 ;     HA-    CA-    CB-   HB2
  6740   6739   6741   6743      3    0.65084    1.95253    0.00000
-2.60338    0.00000    0.00000 ;     HA-    CA-    CB-   HB3
  6740   6739   6741   6744      3    0.65084    1.95253    0.00000
```

-2.60338 0.00000 0.00000 ; HA- CA- CB- CG
6740 6739 6747 6748 3 3.68192 -4.35136 0.00000
1.33888 0.00000 0.00000 ; HA- CA- C- O
6740 6739 6747 6749 3 0.00000 0.00000 0.00000
0.00000 0.00000 0.00000 ; HA- CA- C- N
6741 6739 6747 6748 3 0.00000 0.00000 0.00000
0.00000 0.00000 0.00000 ; CB- CA- C- O
6741 6739 6747 6749 3 2.51040 4.18400 1.67360
-6.69440 0.00000 0.00000 ; CB- CA- C- N
6742 6741 6739 6747 3 0.65084 1.95253 0.00000
-2.60338 0.00000 0.00000 ; HB2- CB- CA- C
6742 6741 6744 6745 3 0.00000 0.00000 0.00000
0.00000 0.00000 0.00000 ; HB2- CB- CG- OD1
6742 6741 6744 6746 3 0.00000 0.00000 0.00000
0.00000 0.00000 0.00000 ; HB2- CB- CG- OD2
6743 6741 6739 6747 3 0.65084 1.95253 0.00000
-2.60338 0.00000 0.00000 ; HB3- CB- CA- C
6743 6741 6744 6745 3 0.00000 0.00000 0.00000
0.00000 0.00000 0.00000 ; HB3- CB- CG- OD1
6743 6741 6744 6746 3 0.00000 0.00000 0.00000
0.00000 0.00000 0.00000 ; HB3- CB- CG- OD2
6744 6741 6739 6747 3 7.14627 -1.04600 -8.99560
-0.97069 5.15469 0.00000 ; CG- CB- CA- C
6747 6749 6751 6752 3 0.00000 0.00000 0.00000
0.00000 0.00000 0.00000 ; C- N- CA- HA
6747 6749 6751 6753 3 11.71520 1.67360 15.06240 -
13.38880 0.00000 0.00000 ; C- N- CA- CB
6747 6749 6751 6759 3 1.75728 5.27184 2.25936
-7.02912 0.00000 0.00000 ; C- N- CA- C
6748 6747 6749 6750 3 29.28800 -8.36800 -20.92000
0.00000 0.00000 0.00000 ; O- C- N- H
6748 6747 6749 6751 3 20.92000 0.00000 -20.92000
0.00000 0.00000 0.00000 ; O- C- N- CA
6749 6751 6753 6754 3 0.65084 1.95253 0.00000
-2.60338 0.00000 0.00000 ; N- CA- CB- HB2
6749 6751 6753 6755 3 0.65084 1.95253 0.00000
-2.60338 0.00000 0.00000 ; N- CA- CB- HB3
6749 6751 6753 6756 3 15.41386 9.74035 -8.39310
-0.97069 2.97901 0.00000 ; N- CA- CB- CG
6749 6751 6759 6760 3 0.00000 0.00000 0.00000
0.00000 0.00000 0.00000 ; N- CA- C- O
6749 6751 6759 6761 3 17.40544 -5.02080 -13.22144
9.20480 0.00000 0.00000 ; N- CA- C- N
6750 6749 6751 6752 3 0.00000 0.00000 0.00000
0.00000 0.00000 0.00000 ; H- N- CA- HA
6750 6749 6751 6753 3 0.00000 0.00000 0.00000
0.00000 0.00000 0.00000 ; H- N- CA- CB
6750 6749 6751 6759 3 0.00000 0.00000 0.00000
0.00000 0.00000 0.00000 ; H- N- CA- C
6751 6753 6756 6757 3 6.43499 0.00000 -5.39736
0.00000 -1.03763 0.00000 ; CA- CB- CG- OD1
6751 6753 6756 6758 3 6.43499 0.00000 -5.39736

0.00000   -1.03763    0.00000 ;      CA-    CB-    CG-   OD2
  6751   6759   6761   6762       3   20.92000    0.00000  -20.92000
0.00000    0.00000    0.00000 ;      CA-     C-     N-     H
  6751   6759   6761   6763       3   20.92000    0.00000  -20.92000
0.00000    0.00000    0.00000 ;      CA-     C-     N-    CA
  6752   6751   6753   6754       3    0.65084    1.95253    0.00000
-2.60338    0.00000    0.00000 ;      HA-    CA-    CB-   HB2
  6752   6751   6753   6755       3    0.65084    1.95253    0.00000
-2.60338    0.00000    0.00000 ;      HA-    CA-    CB-   HB3
  6752   6751   6753   6756       3    0.65084    1.95253    0.00000
-2.60338    0.00000    0.00000 ;      HA-    CA-    CB-    CG
  6752   6751   6759   6760       3    3.68192   -4.35136    0.00000
1.33888    0.00000    0.00000 ;      HA-    CA-     C-     O
  6752   6751   6759   6761       3    0.00000    0.00000    0.00000
0.00000    0.00000    0.00000 ;      HA-    CA-     C-     N
  6753   6751   6759   6760       3    0.00000    0.00000    0.00000
0.00000    0.00000    0.00000 ;      CB-    CA-     C-     O
  6753   6751   6759   6761       3    2.51040    4.18400    1.67360
-6.69440    0.00000    0.00000 ;      CB-    CA-     C-     N
  6754   6753   6751   6759       3    0.65084    1.95253    0.00000
-2.60338    0.00000    0.00000 ;     HB2-    CB-    CA-     C
  6754   6753   6756   6757       3    0.00000    0.00000    0.00000
0.00000    0.00000    0.00000 ;     HB2-    CB-    CG-   OD1
  6754   6753   6756   6758       3    0.00000    0.00000    0.00000
0.00000    0.00000    0.00000 ;     HB2-    CB-    CG-   OD2
  6755   6753   6751   6759       3    0.65084    1.95253    0.00000
-2.60338    0.00000    0.00000 ;     HB3-    CB-    CA-     C
  6755   6753   6756   6757       3    0.00000    0.00000    0.00000
0.00000    0.00000    0.00000 ;     HB3-    CB-    CG-   OD1
  6755   6753   6756   6758       3    0.00000    0.00000    0.00000
0.00000    0.00000    0.00000 ;     HB3-    CB-    CG-   OD2
  6756   6753   6751   6759       3    7.14627   -1.04600   -8.99560
-0.97069    5.15469    0.00000 ;      CG-    CB-    CA-     C
  6759   6761   6763   6764       3    0.00000    0.00000    0.00000
0.00000    0.00000    0.00000 ;       C-     N-    CA-    HA
  6759   6761   6763   6765       3   11.71520    1.67360   15.06240  -
13.38880    0.00000    0.00000 ;       C-     N-    CA-    CB
  6759   6761   6763   6778       3    1.75728    5.27184    2.25936
-7.02912    0.00000    0.00000 ;       C-     N-    CA-     C
  6760   6759   6761   6762       3   29.28800   -8.36800  -20.92000
0.00000    0.00000    0.00000 ;      O-     C-     N-     H
  6760   6759   6761   6763       3   20.92000    0.00000  -20.92000
0.00000    0.00000    0.00000 ;      O-     C-     N-    CA
  6761   6763   6765   6766       3    0.65084    1.95253    0.00000
-2.60338    0.00000    0.00000 ;       N-    CA-    CB-    HB
  6761   6763   6765   6767       3    3.83673    0.44769   -1.77402
-2.47693   -0.03347    0.00000 ;       N-    CA-    CB-   CG2
  6761   6763   6765   6771       3    2.97482    0.12134    2.04179
-1.89117   -3.24678    0.00000 ;       N-    CA-    CB-   CG1
  6761   6763   6778   6779       3    0.00000    0.00000    0.00000
0.00000    0.00000    0.00000 ;      N-    CA-     C-     O
  6761   6763   6778   6780       3   17.40544   -5.02080  -13.22144

```
9.20480    0.00000    0.00000 ;      N-    CA-     C-     N
  6762   6761   6763   6764      3    0.00000    0.00000    0.00000
0.00000    0.00000    0.00000 ;      H-     N-    CA-    HA
  6762   6761   6763   6765      3    0.00000    0.00000    0.00000
0.00000    0.00000    0.00000 ;      H-     N-    CA-    CB
  6762   6761   6763   6778      3    0.00000    0.00000    0.00000
0.00000    0.00000    0.00000 ;      H-     N-    CA-     C
  6763   6765   6767   6768      3    0.66944    2.00832    0.00000
-2.67776    0.00000    0.00000 ;     CA-    CB-   CG2-  HG21
  6763   6765   6767   6769      3    0.66944    2.00832    0.00000
-2.67776    0.00000    0.00000 ;     CA-    CB-   CG2-  HG22
  6763   6765   6767   6770      3    0.66944    2.00832    0.00000
-2.67776    0.00000    0.00000 ;     CA-    CB-   CG2-  HG23
  6763   6765   6771   6772      3    0.66944    2.00832    0.00000
-2.67776    0.00000    0.00000 ;     CA-    CB-   CG1-  HG12
  6763   6765   6771   6773      3    0.66944    2.00832    0.00000
-2.67776    0.00000    0.00000 ;     CA-    CB-   CG1-  HG13
  6763   6765   6771   6774      3    4.24258   -0.52718   -7.25506
-1.79075    7.69856    0.00000 ;     CA-    CB-   CG1-   CD1
  6763   6778   6780   6781      3   20.92000    0.00000  -20.92000
0.00000    0.00000    0.00000 ;     CA-     C-     N-     H
  6763   6778   6780   6782      3   20.92000    0.00000  -20.92000
0.00000    0.00000    0.00000 ;     CA-     C-     N-    CA
  6764   6763   6765   6766      3    0.65084    1.95253    0.00000
-2.60338    0.00000    0.00000 ;     HA-    CA-    CB-    HB
  6764   6763   6765   6767      3    0.65084    1.95253    0.00000
-2.60338    0.00000    0.00000 ;     HA-    CA-    CB-   CG2
  6764   6763   6765   6771      3    0.65084    1.95253    0.00000
-2.60338    0.00000    0.00000 ;     HA-    CA-    CB-   CG1
  6764   6763   6778   6779      3    3.68192   -4.35136    0.00000
1.33888    0.00000    0.00000 ;     HA-    CA-     C-     O
  6764   6763   6778   6780      3    0.00000    0.00000    0.00000
0.00000    0.00000    0.00000 ;     HA-    CA-     C-     N
  6765   6763   6778   6779      3    0.00000    0.00000    0.00000
0.00000    0.00000    0.00000 ;     CB-    CA-     C-     O
  6765   6763   6778   6780      3    2.51040    4.18400    1.67360
-6.69440    0.00000    0.00000 ;     CB-    CA-     C-     N
  6765   6771   6774   6775      3    0.66944    2.00832    0.00000
-2.67776    0.00000    0.00000 ;     CB-   CG1-   CD1-  HD11
  6765   6771   6774   6776      3    0.66944    2.00832    0.00000
-2.67776    0.00000    0.00000 ;     CB-   CG1-   CD1-  HD12
  6765   6771   6774   6777      3    0.66944    2.00832    0.00000
-2.67776    0.00000    0.00000 ;     CB-   CG1-   CD1-  HD13
  6766   6765   6763   6778      3    0.65084    1.95253    0.00000
-2.60338    0.00000    0.00000 ;     HB-    CB-    CA-     C
  6766   6765   6767   6768      3    0.62760    1.88280    0.00000
-2.51040    0.00000    0.00000 ;     HB-    CB-   CG2-  HG21
  6766   6765   6767   6769      3    0.62760    1.88280    0.00000
-2.51040    0.00000    0.00000 ;     HB-    CB-   CG2-  HG22
  6766   6765   6767   6770      3    0.62760    1.88280    0.00000
-2.51040    0.00000    0.00000 ;     HB-    CB-   CG2-  HG23
  6766   6765   6771   6772      3    0.62760    1.88280    0.00000
```

-2.51040 0.00000 0.00000 ; HB- CB- CG1- HG12
6766 6765 6771 6773 3 0.62760 1.88280 0.00000
-2.51040 0.00000 0.00000 ; HB- CB- CG1- HG13
6766 6765 6771 6774 3 0.66944 2.00832 0.00000
-2.67776 0.00000 0.00000 ; HB- CB- CG1- CD1
6767 6765 6763 6778 3 5.67350 3.55640 -6.16722
-2.47693 3.74886 0.00000 ; CG2- CB- CA- C
6767 6765 6771 6772 3 0.66944 2.00832 0.00000
-2.67776 0.00000 0.00000 ; CG2- CB- CG1- HG12
6767 6765 6771 6773 3 0.66944 2.00832 0.00000
-2.67776 0.00000 0.00000 ; CG2- CB- CG1- HG13
6767 6765 6771 6774 3 3.81162 0.49790 -8.14206
-1.79075 7.49773 0.00000 ; CG2- CB- CG1- CD1
6768 6767 6765 6771 3 0.66944 2.00832 0.00000
-2.67776 0.00000 0.00000 ; HG21- CG2- CB- CG1
6769 6767 6765 6771 3 0.66944 2.00832 0.00000
-2.67776 0.00000 0.00000 ; HG22- CG2- CB- CG1
6770 6767 6765 6771 3 0.66944 2.00832 0.00000
-2.67776 0.00000 0.00000 ; HG23- CG2- CB- CG1
6771 6765 6763 6778 3 8.26340 0.74057 -9.99976
-1.89117 3.84928 0.00000 ; CG1- CB- CA- C
6772 6771 6774 6775 3 0.62760 1.88280 0.00000
-2.51040 0.00000 0.00000 ; HG12- CG1- CD1- HD11
6772 6771 6774 6776 3 0.62760 1.88280 0.00000
-2.51040 0.00000 0.00000 ; HG12- CG1- CD1- HD12
6772 6771 6774 6777 3 0.62760 1.88280 0.00000
-2.51040 0.00000 0.00000 ; HG12- CG1- CD1- HD13
6773 6771 6774 6775 3 0.62760 1.88280 0.00000
-2.51040 0.00000 0.00000 ; HG13- CG1- CD1- HD11
6773 6771 6774 6776 3 0.62760 1.88280 0.00000
-2.51040 0.00000 0.00000 ; HG13- CG1- CD1- HD12
6773 6771 6774 6777 3 0.62760 1.88280 0.00000
-2.51040 0.00000 0.00000 ; HG13- CG1- CD1- HD13
6778 6780 6782 6783 3 0.00000 0.00000 0.00000
0.00000 0.00000 0.00000 ; C- N- CA- HA
6778 6780 6782 6784 3 11.71520 1.67360 15.06240 -
13.38880 0.00000 0.00000 ; C- N- CA- CB
6778 6780 6782 6793 3 1.75728 5.27184 2.25936
-7.02912 0.00000 0.00000 ; C- N- CA- C
6779 6778 6780 6781 3 29.28800 -8.36800 -20.92000
0.00000 0.00000 0.00000 ; O- C- N- H
6779 6778 6780 6782 3 20.92000 0.00000 -20.92000
0.00000 0.00000 0.00000 ; O- C- N- CA
6780 6782 6784 6785 3 0.65084 1.95253 0.00000
-2.60338 0.00000 0.00000 ; N- CA- CB- HB2
6780 6782 6784 6786 3 0.65084 1.95253 0.00000
-2.60338 0.00000 0.00000 ; N- CA- CB- HB3
6780 6782 6784 6787 3 3.21331 2.22589 -4.15053
-2.40998 2.61082 0.00000 ; N- CA- CB- CG
6780 6782 6793 6794 3 0.00000 0.00000 0.00000
0.00000 0.00000 0.00000 ; N- CA- C- O
6780 6782 6793 6795 3 17.40544 -5.02080 -13.22144

```
9.20480    0.00000    0.00000 ;      N-    CA-     C-     N
  6781   6780   6782   6783       3    0.00000    0.00000    0.00000
0.00000    0.00000    0.00000 ;      H-     N-    CA-    HA
  6781   6780   6782   6784       3    0.00000    0.00000    0.00000
0.00000    0.00000    0.00000 ;      H-     N-    CA-    CB
  6781   6780   6782   6793       3    0.00000    0.00000    0.00000
0.00000    0.00000    0.00000 ;      H-     N-    CA-     C
  6782   6784   6787   6788       3    0.66944    2.00832    0.00000
-2.67776    0.00000    0.00000 ;     CA-    CB-    CG-   HG2
  6782   6784   6787   6789       3    0.66944    2.00832    0.00000
-2.67776    0.00000    0.00000 ;     CA-    CB-    CG-   HG3
  6782   6784   6787   6790       3   10.12110   -1.91209    0.01674
10.17549   -1.87443    0.00000 ;     CA-    CB-    CG-    CD
  6782   6793   6795   6796       3   20.92000    0.00000  -20.92000
0.00000    0.00000    0.00000 ;     CA-     C-     N-     H
  6782   6793   6795   6797       3   20.92000    0.00000  -20.92000
0.00000    0.00000    0.00000 ;     CA-     C-     N-    CA
  6783   6782   6784   6785       3    0.65084    1.95253    0.00000
-2.60338    0.00000    0.00000 ;     HA-    CA-    CB-   HB2
  6783   6782   6784   6786       3    0.65084    1.95253    0.00000
-2.60338    0.00000    0.00000 ;     HA-    CA-    CB-   HB3
  6783   6782   6784   6787       3    0.65084    1.95253    0.00000
-2.60338    0.00000    0.00000 ;     HA-    CA-    CB-    CG
  6783   6782   6793   6794       3    3.68192   -4.35136    0.00000
1.33888    0.00000    0.00000 ;     HA-    CA-     C-     O
  6783   6782   6793   6795       3    0.00000    0.00000    0.00000
0.00000    0.00000    0.00000 ;     HA-    CA-     C-     N
  6784   6782   6793   6794       3    0.00000    0.00000    0.00000
0.00000    0.00000    0.00000 ;     CB-    CA-     C-     O
  6784   6782   6793   6795       3    2.51040    4.18400    1.67360
-6.69440    0.00000    0.00000 ;     CB-    CA-     C-     N
  6784   6787   6790   6791       3    3.79907    0.00000   -5.40573
0.00000    2.14221    0.00000 ;     CB-    CG-    CD-   OE1
  6784   6787   6790   6792       3    3.79907    0.00000   -5.40573
0.00000    2.14221    0.00000 ;     CB-    CG-    CD-   OE2
  6785   6784   6782   6793       3    0.65084    1.95253    0.00000
-2.60338    0.00000    0.00000 ;    HB2-    CB-    CA-     C
  6785   6784   6787   6788       3    0.62760    1.88280    0.00000
-2.51040    0.00000    0.00000 ;    HB2-    CB-    CG-   HG2
  6785   6784   6787   6789       3    0.62760    1.88280    0.00000
-2.51040    0.00000    0.00000 ;    HB2-    CB-    CG-   HG3
  6785   6784   6787   6790       3    0.65084    1.95253    0.00000
-2.60338    0.00000    0.00000 ;    HB2-    CB-    CG-    CD
  6786   6784   6782   6793       3    0.65084    1.95253    0.00000
-2.60338    0.00000    0.00000 ;    HB3-    CB-    CA-     C
  6786   6784   6787   6788       3    0.62760    1.88280    0.00000
-2.51040    0.00000    0.00000 ;    HB3-    CB-    CG-   HG2
  6786   6784   6787   6789       3    0.62760    1.88280    0.00000
-2.51040    0.00000    0.00000 ;    HB3-    CB-    CG-   HG3
  6786   6784   6787   6790       3    0.65084    1.95253    0.00000
-2.60338    0.00000    0.00000 ;    HB3-    CB-    CG-    CD
  6787   6784   6782   6793       3    6.86594    3.56895   -8.14206
```

-2.40998 4.85344 0.00000 ; CG- CB- CA- C
6788 6787 6790 6791 3 0.00000 0.00000 0.00000
0.00000 0.00000 0.00000 ; HG2- CG- CD- OE1
6788 6787 6790 6792 3 0.00000 0.00000 0.00000
0.00000 0.00000 0.00000 ; HG2- CG- CD- OE2
6789 6787 6790 6791 3 0.00000 0.00000 0.00000
0.00000 0.00000 0.00000 ; HG3- CG- CD- OE1
6789 6787 6790 6792 3 0.00000 0.00000 0.00000
0.00000 0.00000 0.00000 ; HG3- CG- CD- OE2
6793 6795 6797 6798 3 0.00000 0.00000 0.00000
0.00000 0.00000 0.00000 ; C- N- CA- HA
6793 6795 6797 6799 3 11.71520 1.67360 15.06240 -
13.38880 0.00000 0.00000 ; C- N- CA- CB
6793 6795 6797 6807 3 1.75728 5.27184 2.25936
-7.02912 0.00000 0.00000 ; C- N- CA- C
6794 6793 6795 6796 3 29.28800 -8.36800 -20.92000
0.00000 0.00000 0.00000 ; O- C- N- H
6794 6793 6795 6797 3 20.92000 0.00000 -20.92000
0.00000 0.00000 0.00000 ; O- C- N- CA
6795 6797 6799 6800 3 0.65084 1.95253 0.00000
-2.60338 0.00000 0.00000 ; N- CA- CB- HB2
6795 6797 6799 6801 3 0.65084 1.95253 0.00000
-2.60338 0.00000 0.00000 ; N- CA- CB- HB3
6795 6797 6799 6802 3 5.99567 3.29281 -4.46014
-0.55229 1.97485 0.00000 ; N- CA- CB- CG
6795 6797 6807 6808 3 0.00000 0.00000 0.00000
0.00000 0.00000 0.00000 ; N- CA- C- O
6795 6797 6807 6809 3 17.40544 -5.02080 -13.22144
9.20480 0.00000 0.00000 ; N- CA- C- N
6796 6795 6797 6798 3 0.00000 0.00000 0.00000
0.00000 0.00000 0.00000 ; H- N- CA- HA
6796 6795 6797 6799 3 0.00000 0.00000 0.00000
0.00000 0.00000 0.00000 ; H- N- CA- CB
6796 6795 6797 6807 3 0.00000 0.00000 0.00000
0.00000 0.00000 0.00000 ; H- N- CA- C
6797 6799 6802 6803 3 0.00000 0.00000 0.00000
0.00000 0.00000 0.00000 ; CA- CB- CG- OD1
6797 6799 6802 6804 3 8.84916 -0.31380 -4.32626
5.03754 0.26778 0.00000 ; CA- CB- CG- ND2
6797 6807 6809 6810 3 20.92000 0.00000 -20.92000
0.00000 0.00000 0.00000 ; CA- C- N- CD
6797 6807 6809 6819 3 20.92000 0.00000 -20.92000
0.00000 0.00000 0.00000 ; CA- C- N- CA
6798 6797 6799 6800 3 0.65084 1.95253 0.00000
-2.60338 0.00000 0.00000 ; HA- CA- CB- HB2
6798 6797 6799 6801 3 0.65084 1.95253 0.00000
-2.60338 0.00000 0.00000 ; HA- CA- CB- HB3
6798 6797 6799 6802 3 0.65084 1.95253 0.00000
-2.60338 0.00000 0.00000 ; HA- CA- CB- CG
6798 6797 6807 6808 3 3.68192 -4.35136 0.00000
1.33888 0.00000 0.00000 ; HA- CA- C- O
6798 6797 6807 6809 3 0.00000 0.00000 0.00000

0.00000    0.00000    0.00000 ;     HA-    CA-     C-     N
  6799   6797   6807   6808      3    0.00000    0.00000    0.00000
0.00000    0.00000    0.00000 ;     CB-    CA-     C-     O
  6799   6797   6807   6809      3    2.51040    4.18400    1.67360
-6.69440    0.00000    0.00000 ;     CB-    CA-     C-     N
  6799   6802   6804   6805      3   20.92000    0.00000  -20.92000
0.00000    0.00000    0.00000 ;     CB-    CG-   ND2-  HD21
  6799   6802   6804   6806      3   20.92000    0.00000  -20.92000
0.00000    0.00000    0.00000 ;     CB-    CG-   ND2-  HD22
  6800   6799   6797   6807      3    0.65084    1.95253    0.00000
-2.60338    0.00000    0.00000 ;    HB2-    CB-    CA-     C
  6800   6799   6802   6803      3    3.68192   -4.35136    0.00000
1.33888    0.00000    0.00000 ;    HB2-    CB-    CG-   OD1
  6800   6799   6802   6804      3    0.00000    0.00000    0.00000
0.00000    0.00000    0.00000 ;    HB2-    CB-    CG-   ND2
  6801   6799   6797   6807      3    0.65084    1.95253    0.00000
-2.60338    0.00000    0.00000 ;    HB3-    CB-    CA-     C
  6801   6799   6802   6803      3    3.68192   -4.35136    0.00000
1.33888    0.00000    0.00000 ;    HB3-    CB-    CG-   OD1
  6801   6799   6802   6804      3    0.00000    0.00000    0.00000
0.00000    0.00000    0.00000 ;    HB3-    CB-    CG-   ND2
  6802   6799   6797   6807      3    7.94542   -3.96225   -6.11701
-0.55229    3.58150    0.00000 ;     CG-    CB-    CA-     C
  6803   6802   6804   6805      3   29.28800   -8.36800  -20.92000
0.00000    0.00000    0.00000 ;    OD1-    CG-   ND2-  HD21
  6803   6802   6804   6806      3   29.28800   -8.36800  -20.92000
0.00000    0.00000    0.00000 ;    OD1-    CG-   ND2-  HD22
  6807   6809   6810   6811      3    0.00000    0.00000    0.00000
0.00000    0.00000    0.00000 ;     C-     N-    CD-   HD2
  6807   6809   6810   6812      3    0.00000    0.00000    0.00000
0.00000    0.00000    0.00000 ;     C-     N-    CD-   HD3
  6807   6809   6810   6813      3   10.04160   -3.34720   16.73600
-6.69440    0.00000    0.00000 ;     C-     N-    CD-    CG
  6807   6809   6819   6816      3   11.71520    1.67360   15.06240  -
13.38880    0.00000    0.00000 ;     C-     N-    CA-    CB
  6807   6809   6819   6820      3    0.00000    0.00000    0.00000
0.00000    0.00000    0.00000 ;     C-     N-    CA-    HA
  6807   6809   6819   6821      3    1.75728    5.27184    2.25936
-7.02912    0.00000    0.00000 ;     C-     N-    CA-     C
  6808   6807   6809   6810      3   20.92000    0.00000  -20.92000
0.00000    0.00000    0.00000 ;     O-     C-     N-    CD
  6808   6807   6809   6819      3   20.92000    0.00000  -20.92000
0.00000    0.00000    0.00000 ;     O-     C-     N-    CA
  6809   6810   6813   6814      3    0.65084    1.95253    0.00000
-2.60338    0.00000    0.00000 ;     N-    CD-    CG-   HG2
  6809   6810   6813   6815      3    0.65084    1.95253    0.00000
-2.60338    0.00000    0.00000 ;     N-    CD-    CG-   HG3
  6809   6810   6813   6816      3    0.65084    1.95253    0.00000
-2.60338    0.00000    0.00000 ;     N-    CD-    CG-    CB
  6809   6819   6816   6813      3    0.65084    1.95253    0.00000
-2.60338    0.00000    0.00000 ;     N-    CA-    CB-    CG
  6809   6819   6816   6817      3    0.65084    1.95253    0.00000

```
-2.60338    0.00000    0.00000 ;       N-    CA-    CB-   HB2
  6809   6819   6816   6818      3    0.65084    1.95253    0.00000
-2.60338    0.00000    0.00000 ;       N-    CA-    CB-   HB3
  6809   6819   6821   6822      3    0.00000    0.00000    0.00000
0.00000    0.00000    0.00000 ;       N-    CA-     C-     O
  6809   6819   6821   6823      3   17.40544   -5.02080  -13.22144
9.20480    0.00000    0.00000 ;       N-    CA-     C-     N
  6810   6809   6819   6816      3    0.00000    0.00000    0.00000
0.00000    0.00000    0.00000 ;      CD-     N-    CA-    CB
  6810   6809   6819   6820      3    0.00000    0.00000    0.00000
0.00000    0.00000    0.00000 ;      CD-     N-    CA-    HA
  6810   6809   6819   6821      3    0.00000    0.00000    0.00000
0.00000    0.00000    0.00000 ;      CD-     N-    CA-     C
  6810   6813   6816   6817      3    0.66944    2.00832    0.00000
-2.67776    0.00000    0.00000 ;      CD-    CG-    CB-   HB2
  6810   6813   6816   6818      3    0.66944    2.00832    0.00000
-2.67776    0.00000    0.00000 ;      CD-    CG-    CB-   HB3
  6810   6813   6816   6819      3    3.68192    3.09616   -2.09200
-3.01248    0.00000    0.00000 ;      CD-    CG-    CB-    CA
  6811   6810   6809   6819      3    0.00000    0.00000    0.00000
0.00000    0.00000    0.00000 ;     HD2-    CD-     N-    CA
  6811   6810   6813   6814      3    0.65084    1.95253    0.00000
-2.60338    0.00000    0.00000 ;     HD2-    CD-    CG-   HG2
  6811   6810   6813   6815      3    0.65084    1.95253    0.00000
-2.60338    0.00000    0.00000 ;     HD2-    CD-    CG-   HG3
  6811   6810   6813   6816      3    0.65084    1.95253    0.00000
-2.60338    0.00000    0.00000 ;     HD2-    CD-    CG-    CB
  6812   6810   6809   6819      3    0.00000    0.00000    0.00000
0.00000    0.00000    0.00000 ;     HD3-    CD-     N-    CA
  6812   6810   6813   6814      3    0.65084    1.95253    0.00000
-2.60338    0.00000    0.00000 ;     HD3-    CD-    CG-   HG2
  6812   6810   6813   6815      3    0.65084    1.95253    0.00000
-2.60338    0.00000    0.00000 ;     HD3-    CD-    CG-   HG3
  6812   6810   6813   6816      3    0.65084    1.95253    0.00000
-2.60338    0.00000    0.00000 ;     HD3-    CD-    CG-    CB
  6813   6810   6809   6819      3    0.00000    0.00000    0.00000
0.00000    0.00000    0.00000 ;      CG-    CD-     N-    CA
  6813   6816   6819   6820      3    0.65084    1.95253    0.00000
-2.60338    0.00000    0.00000 ;      CG-    CB-    CA-    HA
  6813   6816   6819   6821      3    0.65084    1.95253    0.00000
-2.60338    0.00000    0.00000 ;      CG-    CB-    CA-     C
  6814   6813   6816   6817      3    0.62760    1.88280    0.00000
-2.51040    0.00000    0.00000 ;     HG2-    CG-    CB-   HB2
  6814   6813   6816   6818      3    0.62760    1.88280    0.00000
-2.51040    0.00000    0.00000 ;     HG2-    CG-    CB-   HB3
  6814   6813   6816   6819      3    0.66944    2.00832    0.00000
-2.67776    0.00000    0.00000 ;     HG2-    CG-    CB-    CA
  6815   6813   6816   6817      3    0.62760    1.88280    0.00000
-2.51040    0.00000    0.00000 ;     HG3-    CG-    CB-   HB2
  6815   6813   6816   6818      3    0.62760    1.88280    0.00000
-2.51040    0.00000    0.00000 ;     HG3-    CG-    CB-   HB3
  6815   6813   6816   6819      3    0.66944    2.00832    0.00000
```

-2.67776    0.00000    0.00000 ;    HG3-    CG-    CB-    CA
  6816   6819   6821   6822     3    0.00000    0.00000    0.00000
0.00000    0.00000    0.00000 ;    CB-    CA-    C-    O
  6816   6819   6821   6823     3    2.51040    4.18400    1.67360
-6.69440    0.00000    0.00000 ;    CB-    CA-    C-    N
  6817   6816   6819   6820     3    0.65084    1.95253    0.00000
-2.60338    0.00000    0.00000 ;    HB2-    CB-    CA-    HA
  6817   6816   6819   6821     3    0.65084    1.95253    0.00000
-2.60338    0.00000    0.00000 ;    HB2-    CB-    CA-    C
  6818   6816   6819   6820     3    0.65084    1.95253    0.00000
-2.60338    0.00000    0.00000 ;    HB3-    CB-    CA-    HA
  6818   6816   6819   6821     3    0.65084    1.95253    0.00000
-2.60338    0.00000    0.00000 ;    HB3-    CB-    CA-    C
  6819   6821   6823   6824     3   20.92000    0.00000  -20.92000
0.00000    0.00000    0.00000 ;    CA-    C-    N-    H
  6819   6821   6823   6825     3   20.92000    0.00000  -20.92000
0.00000    0.00000    0.00000 ;    CA-    C-    N-    CA
  6820   6819   6821   6822     3    3.68192   -4.35136    0.00000
1.33888    0.00000    0.00000 ;    HA-    CA-    C-    O
  6820   6819   6821   6823     3    0.00000    0.00000    0.00000
0.00000    0.00000    0.00000 ;    HA-    CA-    C-    N
  6821   6823   6825   6826     3    0.00000    0.00000    0.00000
0.00000    0.00000    0.00000 ;    C-    N-    CA-    HA
  6821   6823   6825   6827     3   11.71520    1.67360   15.06240  -
13.38880    0.00000    0.00000 ;    C-    N-    CA-    CB
  6821   6823   6825   6840     3    1.75728    5.27184    2.25936
-7.02912    0.00000    0.00000 ;    C-    N-    CA-    C
  6822   6821   6823   6824     3   29.28800   -8.36800  -20.92000
0.00000    0.00000    0.00000 ;    O-    C-    N-    H
  6822   6821   6823   6825     3   20.92000    0.00000  -20.92000
0.00000    0.00000    0.00000 ;    O-    C-    N-    CA
  6823   6825   6827   6828     3    0.65084    1.95253    0.00000
-2.60338    0.00000    0.00000 ;    N-    CA-    CB-    HB2
  6823   6825   6827   6829     3    0.65084    1.95253    0.00000
-2.60338    0.00000    0.00000 ;    N-    CA-    CB-    HB3
  6823   6825   6827   6830     3    3.79070    1.39746   -4.61077
-2.40998    2.44346    0.00000 ;    N-    CA-    CB-    CG
  6823   6825   6840   6841     3    0.00000    0.00000    0.00000
0.00000    0.00000    0.00000 ;    N-    CA-    C-    O
  6823   6825   6840   6842     3   17.40544   -5.02080  -13.22144
9.20480    0.00000    0.00000 ;    N-    CA-    C-    N
  6824   6823   6825   6826     3    0.00000    0.00000    0.00000
0.00000    0.00000    0.00000 ;    H-    N-    CA-    HA
  6824   6823   6825   6827     3    0.00000    0.00000    0.00000
0.00000    0.00000    0.00000 ;    H-    N-    CA-    CB
  6824   6823   6825   6840     3    0.00000    0.00000    0.00000
0.00000    0.00000    0.00000 ;    H-    N-    CA-    C
  6825   6827   6830   6831     3    0.66944    2.00832    0.00000
-2.67776    0.00000    0.00000 ;    CA-    CB-    CG-    HG
  6825   6827   6830   6832     3    3.90367    0.19665   -6.21742
-2.37651    5.99149    0.00000 ;    CA-    CB-    CG-    CD1
  6825   6827   6830   6836     3    3.90367    0.19665   -6.21742

```
-2.37651    5.99149    0.00000 ;     CA-    CB-    CG-   CD2
  6825   6840   6842   6843      3   20.92000    0.00000  -20.92000
0.00000    0.00000    0.00000 ;     CA-    C-     N-     H
  6825   6840   6842   6844      3   20.92000    0.00000  -20.92000
0.00000    0.00000    0.00000 ;     CA-    C-     N-    CA
  6826   6825   6827   6828      3    0.65084    1.95253    0.00000
-2.60338    0.00000    0.00000 ;     HA-    CA-    CB-   HB2
  6826   6825   6827   6829      3    0.65084    1.95253    0.00000
-2.60338    0.00000    0.00000 ;     HA-    CA-    CB-   HB3
  6826   6825   6827   6830      3    0.65084    1.95253    0.00000
-2.60338    0.00000    0.00000 ;     HA-    CA-    CB-    CG
  6826   6825   6840   6841      3    3.68192   -4.35136    0.00000
1.33888    0.00000    0.00000 ;     HA-    CA-     C-     O
  6826   6825   6840   6842      3    0.00000    0.00000    0.00000
0.00000    0.00000    0.00000 ;     HA-    CA-     C-     N
  6827   6825   6840   6841      3    0.00000    0.00000    0.00000
0.00000    0.00000    0.00000 ;     CB-    CA-     C-     O
  6827   6825   6840   6842      3    2.51040    4.18400    1.67360
-6.69440    0.00000    0.00000 ;     CB-    CA-     C-     N
  6827   6830   6832   6833      3    0.66944    2.00832    0.00000
-2.67776    0.00000    0.00000 ;     CB-    CG-   CD1-  HD11
  6827   6830   6832   6834      3    0.66944    2.00832    0.00000
-2.67776    0.00000    0.00000 ;     CB-    CG-   CD1-  HD12
  6827   6830   6832   6835      3    0.66944    2.00832    0.00000
-2.67776    0.00000    0.00000 ;     CB-    CG-   CD1-  HD13
  6827   6830   6836   6837      3    0.66944    2.00832    0.00000
-2.67776    0.00000    0.00000 ;     CB-    CG-   CD2-  HD21
  6827   6830   6836   6838      3    0.66944    2.00832    0.00000
-2.67776    0.00000    0.00000 ;     CB-    CG-   CD2-  HD22
  6827   6830   6836   6839      3    0.66944    2.00832    0.00000
-2.67776    0.00000    0.00000 ;     CB-    CG-   CD2-  HD23
  6828   6827   6825   6840      3    0.65084    1.95253    0.00000
-2.60338    0.00000    0.00000 ;    HB2-    CB-    CA-     C
  6828   6827   6830   6831      3    0.62760    1.88280    0.00000
-2.51040    0.00000    0.00000 ;    HB2-    CB-    CG-    HG
  6828   6827   6830   6832      3    0.66944    2.00832    0.00000
-2.67776    0.00000    0.00000 ;    HB2-    CB-    CG-   CD1
  6828   6827   6830   6836      3    0.66944    2.00832    0.00000
-2.67776    0.00000    0.00000 ;    HB2-    CB-    CG-   CD2
  6829   6827   6825   6840      3    0.65084    1.95253    0.00000
-2.60338    0.00000    0.00000 ;    HB3-    CB-    CA-     C
  6829   6827   6830   6831      3    0.62760    1.88280    0.00000
-2.51040    0.00000    0.00000 ;    HB3-    CB-    CG-    HG
  6829   6827   6830   6832      3    0.66944    2.00832    0.00000
-2.67776    0.00000    0.00000 ;    HB3-    CB-    CG-   CD1
  6829   6827   6830   6836      3    0.66944    2.00832    0.00000
-2.67776    0.00000    0.00000 ;    HB3-    CB-    CG-   CD2
  6830   6827   6825   6840      3   10.33448   -1.14642  -11.54784
-2.40998    6.35968    0.00000 ;     CG-    CB-    CA-     C
  6831   6830   6832   6833      3    0.62760    1.88280    0.00000
-2.51040    0.00000    0.00000 ;     HG-    CG-   CD1-  HD11
  6831   6830   6832   6834      3    0.62760    1.88280    0.00000
```

```
-2.51040    0.00000    0.00000 ;     HG-    CG-   CD1-  HD12
  6831   6830   6832   6835       3    0.62760    1.88280    0.00000
-2.51040    0.00000    0.00000 ;     HG-    CG-   CD1-  HD13
  6831   6830   6836   6837       3    0.62760    1.88280    0.00000
-2.51040    0.00000    0.00000 ;     HG-    CG-   CD2-  HD21
  6831   6830   6836   6838       3    0.62760    1.88280    0.00000
-2.51040    0.00000    0.00000 ;     HG-    CG-   CD2-  HD22
  6831   6830   6836   6839       3    0.62760    1.88280    0.00000
-2.51040    0.00000    0.00000 ;     HG-    CG-   CD2-  HD23
  6832   6830   6836   6837       3    0.66944    2.00832    0.00000
-2.67776    0.00000    0.00000 ;    CD1-    CG-   CD2-  HD21
  6832   6830   6836   6838       3    0.66944    2.00832    0.00000
-2.67776    0.00000    0.00000 ;    CD1-    CG-   CD2-  HD22
  6832   6830   6836   6839       3    0.66944    2.00832    0.00000
-2.67776    0.00000    0.00000 ;    CD1-    CG-   CD2-  HD23
  6833   6832   6830   6836       3    0.66944    2.00832    0.00000
-2.67776    0.00000    0.00000 ;   HD11-   CD1-    CG-   CD2
  6834   6832   6830   6836       3    0.66944    2.00832    0.00000
-2.67776    0.00000    0.00000 ;   HD12-   CD1-    CG-   CD2
  6835   6832   6830   6836       3    0.66944    2.00832    0.00000
-2.67776    0.00000    0.00000 ;   HD13-   CD1-    CG-   CD2
  6840   6842   6844   6845       3    0.00000    0.00000    0.00000
0.00000    0.00000    0.00000 ;      C-     N-    CA-    HA
  6840   6842   6844   6846       3   11.71520    1.67360   15.06240  -
13.38880    0.00000    0.00000 ;      C-     N-    CA-    CB
  6840   6842   6844   6859       3    1.75728    5.27184    2.25936
-7.02912    0.00000    0.00000 ;      C-     N-    CA-     C
  6841   6840   6842   6843       3   29.28800   -8.36800  -20.92000
0.00000    0.00000    0.00000 ;      O-     C-     N-     H
  6841   6840   6842   6844       3   20.92000    0.00000  -20.92000
0.00000    0.00000    0.00000 ;      O-     C-     N-    CA
  6842   6844   6846   6847       3    0.65084    1.95253    0.00000
-2.60338    0.00000    0.00000 ;      N-    CA-    CB-    HB
  6842   6844   6846   6848       3    3.83673    0.44769   -1.77402
-2.47693   -0.03347    0.00000 ;      N-    CA-    CB-   CG2
  6842   6844   6846   6852       3    2.97482    0.12134    2.04179
-1.89117   -3.24678    0.00000 ;      N-    CA-    CB-   CG1
  6842   6844   6859   6860       3    0.00000    0.00000    0.00000
0.00000    0.00000    0.00000 ;      N-    CA-     C-     O
  6842   6844   6859   6861       3   17.40544   -5.02080  -13.22144
9.20480    0.00000    0.00000 ;      N-    CA-     C-     N
  6843   6842   6844   6845       3    0.00000    0.00000    0.00000
0.00000    0.00000    0.00000 ;      H-     N-    CA-    HA
  6843   6842   6844   6846       3    0.00000    0.00000    0.00000
0.00000    0.00000    0.00000 ;      H-     N-    CA-    CB
  6843   6842   6844   6859       3    0.00000    0.00000    0.00000
0.00000    0.00000    0.00000 ;      H-     N-    CA-     C
  6844   6846   6848   6849       3    0.66944    2.00832    0.00000
-2.67776    0.00000    0.00000 ;     CA-    CB-   CG2-  HG21
  6844   6846   6848   6850       3    0.66944    2.00832    0.00000
-2.67776    0.00000    0.00000 ;     CA-    CB-   CG2-  HG22
  6844   6846   6848   6851       3    0.66944    2.00832    0.00000
```

```
-2.67776    0.00000    0.00000 ;      CA-    CB-   CG2-  HG23
  6844   6846   6852   6853       3    0.66944    2.00832    0.00000
-2.67776    0.00000    0.00000 ;      CA-    CB-   CG1-  HG12
  6844   6846   6852   6854       3    0.66944    2.00832    0.00000
-2.67776    0.00000    0.00000 ;      CA-    CB-   CG1-  HG13
  6844   6846   6852   6855       3    4.24258   -0.52718   -7.25506
-1.79075    7.69856    0.00000 ;      CA-    CB-   CG1-   CD1
  6844   6859   6861   6862       3   20.92000    0.00000  -20.92000
0.00000    0.00000    0.00000 ;      CA-     C-     N-     H
  6844   6859   6861   6863       3   20.92000    0.00000  -20.92000
0.00000    0.00000    0.00000 ;      CA-     C-     N-    CA
  6845   6844   6846   6847       3    0.65084    1.95253    0.00000
-2.60338    0.00000    0.00000 ;      HA-    CA-    CB-    HB
  6845   6844   6846   6848       3    0.65084    1.95253    0.00000
-2.60338    0.00000    0.00000 ;      HA-    CA-    CB-   CG2
  6845   6844   6846   6852       3    0.65084    1.95253    0.00000
-2.60338    0.00000    0.00000 ;      HA-    CA-    CB-   CG1
  6845   6844   6859   6860       3    3.68192   -4.35136    0.00000
1.33888    0.00000    0.00000 ;      HA-    CA-     C-     O
  6845   6844   6859   6861       3    0.00000    0.00000    0.00000
0.00000    0.00000    0.00000 ;      HA-    CA-     C-     N
  6846   6844   6859   6860       3    0.00000    0.00000    0.00000
0.00000    0.00000    0.00000 ;      CB-    CA-     C-     O
  6846   6844   6859   6861       3    2.51040    4.18400    1.67360
-6.69440    0.00000    0.00000 ;      CB-    CA-     C-     N
  6846   6852   6855   6856       3    0.66944    2.00832    0.00000
-2.67776    0.00000    0.00000 ;      CB-   CG1-   CD1-  HD11
  6846   6852   6855   6857       3    0.66944    2.00832    0.00000
-2.67776    0.00000    0.00000 ;      CB-   CG1-   CD1-  HD12
  6846   6852   6855   6858       3    0.66944    2.00832    0.00000
-2.67776    0.00000    0.00000 ;      CB-   CG1-   CD1-  HD13
  6847   6846   6844   6859       3    0.65084    1.95253    0.00000
-2.60338    0.00000    0.00000 ;      HB-    CB-    CA-     C
  6847   6846   6848   6849       3    0.62760    1.88280    0.00000
-2.51040    0.00000    0.00000 ;      HB-    CB-   CG2-  HG21
  6847   6846   6848   6850       3    0.62760    1.88280    0.00000
-2.51040    0.00000    0.00000 ;      HB-    CB-   CG2-  HG22
  6847   6846   6848   6851       3    0.62760    1.88280    0.00000
-2.51040    0.00000    0.00000 ;      HB-    CB-   CG2-  HG23
  6847   6846   6852   6853       3    0.62760    1.88280    0.00000
-2.51040    0.00000    0.00000 ;      HB-    CB-   CG1-  HG12
  6847   6846   6852   6854       3    0.62760    1.88280    0.00000
-2.51040    0.00000    0.00000 ;      HB-    CB-   CG1-  HG13
  6847   6846   6852   6855       3    0.66944    2.00832    0.00000
-2.67776    0.00000    0.00000 ;      HB-    CB-   CG1-   CD1
  6848   6846   6844   6859       3    5.67350    3.55640   -6.16722
-2.47693    3.74886    0.00000 ;     CG2-    CB-    CA-     C
  6848   6846   6852   6853       3    0.66944    2.00832    0.00000
-2.67776    0.00000    0.00000 ;     CG2-    CB-   CG1-  HG12
  6848   6846   6852   6854       3    0.66944    2.00832    0.00000
-2.67776    0.00000    0.00000 ;     CG2-    CB-   CG1-  HG13
  6848   6846   6852   6855       3    3.81162    0.49790   -8.14206
```

-1.79075 7.49773 0.00000 ; CG2- CB- CG1- CD1
6849 6848 6846 6852 3 0.66944 2.00832 0.00000
-2.67776 0.00000 0.00000 ; HG21- CG2- CB- CG1
6850 6848 6846 6852 3 0.66944 2.00832 0.00000
-2.67776 0.00000 0.00000 ; HG22- CG2- CB- CG1
6851 6848 6846 6852 3 0.66944 2.00832 0.00000
-2.67776 0.00000 0.00000 ; HG23- CG2- CB- CG1
6852 6846 6844 6859 3 8.26340 0.74057 -9.99976
-1.89117 3.84928 0.00000 ; CG1- CB- CA- C
6853 6852 6855 6856 3 0.62760 1.88280 0.00000
-2.51040 0.00000 0.00000 ; HG12- CG1- CD1- HD11
6853 6852 6855 6857 3 0.62760 1.88280 0.00000
-2.51040 0.00000 0.00000 ; HG12- CG1- CD1- HD12
6853 6852 6855 6858 3 0.62760 1.88280 0.00000
-2.51040 0.00000 0.00000 ; HG12- CG1- CD1- HD13
6854 6852 6855 6856 3 0.62760 1.88280 0.00000
-2.51040 0.00000 0.00000 ; HG13- CG1- CD1- HD11
6854 6852 6855 6857 3 0.62760 1.88280 0.00000
-2.51040 0.00000 0.00000 ; HG13- CG1- CD1- HD12
6854 6852 6855 6858 3 0.62760 1.88280 0.00000
-2.51040 0.00000 0.00000 ; HG13- CG1- CD1- HD13
6859 6861 6863 6864 3 0.00000 0.00000 0.00000
0.00000 0.00000 0.00000 ; C- N- CA- HA
6859 6861 6863 6865 3 11.71520 1.67360 15.06240 -
13.38880 0.00000 0.00000 ; C- N- CA- CB
6859 6861 6863 6871 3 1.75728 5.27184 2.25936
-7.02912 0.00000 0.00000 ; C- N- CA- C
6860 6859 6861 6862 3 29.28800 -8.36800 -20.92000
0.00000 0.00000 0.00000 ; O- C- N- H
6860 6859 6861 6863 3 20.92000 0.00000 -20.92000
0.00000 0.00000 0.00000 ; O- C- N- CA
6861 6863 6865 6866 3 0.65084 1.95253 0.00000
-2.60338 0.00000 0.00000 ; N- CA- CB- HB2
6861 6863 6865 6867 3 0.65084 1.95253 0.00000
-2.60338 0.00000 0.00000 ; N- CA- CB- HB3
6861 6863 6865 6868 3 15.41386 9.74035 -8.39310
-0.97069 2.97901 0.00000 ; N- CA- CB- CG
6861 6863 6871 6872 3 0.00000 0.00000 0.00000
0.00000 0.00000 0.00000 ; N- CA- C- O
6861 6863 6871 6873 3 17.40544 -5.02080 -13.22144
9.20480 0.00000 0.00000 ; N- CA- C- N
6862 6861 6863 6864 3 0.00000 0.00000 0.00000
0.00000 0.00000 0.00000 ; H- N- CA- HA
6862 6861 6863 6865 3 0.00000 0.00000 0.00000
0.00000 0.00000 0.00000 ; H- N- CA- CB
6862 6861 6863 6871 3 0.00000 0.00000 0.00000
0.00000 0.00000 0.00000 ; H- N- CA- C
6863 6865 6868 6869 3 6.43499 0.00000 -5.39736
0.00000 -1.03763 0.00000 ; CA- CB- CG- OD1
6863 6865 6868 6870 3 6.43499 0.00000 -5.39736
0.00000 -1.03763 0.00000 ; CA- CB- CG- OD2
6863 6871 6873 6874 3 20.92000 0.00000 -20.92000

```
0.00000    0.00000    0.00000 ;      CA-     C-     N-     H
  6863   6871   6873   6875       3   20.92000    0.00000  -20.92000
0.00000    0.00000    0.00000 ;      CA-     C-     N-    CA
  6864   6863   6865   6866       3    0.65084    1.95253    0.00000
-2.60338    0.00000    0.00000 ;      HA-    CA-    CB-   HB2
  6864   6863   6865   6867       3    0.65084    1.95253    0.00000
-2.60338    0.00000    0.00000 ;      HA-    CA-    CB-   HB3
  6864   6863   6865   6868       3    0.65084    1.95253    0.00000
-2.60338    0.00000    0.00000 ;      HA-    CA-    CB-    CG
  6864   6863   6871   6872       3    3.68192   -4.35136    0.00000
1.33888    0.00000    0.00000 ;      HA-    CA-     C-     O
  6864   6863   6871   6873       3    0.00000    0.00000    0.00000
0.00000    0.00000    0.00000 ;      HA-    CA-     C-     N
  6865   6863   6871   6872       3    0.00000    0.00000    0.00000
0.00000    0.00000    0.00000 ;      CB-    CA-     C-     O
  6865   6863   6871   6873       3    2.51040    4.18400    1.67360
-6.69440    0.00000    0.00000 ;      CB-    CA-     C-     N
  6866   6865   6863   6871       3    0.65084    1.95253    0.00000
-2.60338    0.00000    0.00000 ;     HB2-    CB-    CA-     C
  6866   6865   6868   6869       3    0.00000    0.00000    0.00000
0.00000    0.00000    0.00000 ;     HB2-    CB-    CG-   OD1
  6866   6865   6868   6870       3    0.00000    0.00000    0.00000
0.00000    0.00000    0.00000 ;     HB2-    CB-    CG-   OD2
  6867   6865   6863   6871       3    0.65084    1.95253    0.00000
-2.60338    0.00000    0.00000 ;     HB3-    CB-    CA-     C
  6867   6865   6868   6869       3    0.00000    0.00000    0.00000
0.00000    0.00000    0.00000 ;     HB3-    CB-    CG-   OD1
  6867   6865   6868   6870       3    0.00000    0.00000    0.00000
0.00000    0.00000    0.00000 ;     HB3-    CB-    CG-   OD2
  6868   6865   6863   6871       3    7.14627   -1.04600   -8.99560
-0.97069    5.15469    0.00000 ;      CG-    CB-    CA-     C
  6871   6873   6875   6876       3    0.00000    0.00000    0.00000
0.00000    0.00000    0.00000 ;       C-     N-    CA-    HA
  6871   6873   6875   6877       3   11.71520    1.67360   15.06240  -
13.38880    0.00000    0.00000 ;       C-     N-    CA-    CB
  6871   6873   6875   6893       3    1.75728    5.27184    2.25936
-7.02912    0.00000    0.00000 ;       C-     N-    CA-     C
  6872   6871   6873   6874       3   29.28800   -8.36800  -20.92000
0.00000    0.00000    0.00000 ;       O-     C-     N-     H
  6872   6871   6873   6875       3   20.92000    0.00000  -20.92000
0.00000    0.00000    0.00000 ;       O-     C-     N-    CA
  6873   6875   6877   6878       3    0.65084    1.95253    0.00000
-2.60338    0.00000    0.00000 ;       N-    CA-    CB-   HB2
  6873   6875   6877   6879       3    0.65084    1.95253    0.00000
-2.60338    0.00000    0.00000 ;       N-    CA-    CB-   HB3
  6873   6875   6877   6880       3    0.65084    1.95253    0.00000
-2.60338    0.00000    0.00000 ;       N-    CA-    CB-    CG
  6873   6875   6893   6894       3    0.00000    0.00000    0.00000
0.00000    0.00000    0.00000 ;       N-    CA-     C-     O
  6873   6875   6893   6895       3   17.40544   -5.02080  -13.22144
9.20480    0.00000    0.00000 ;       N-    CA-     C-     N
  6874   6873   6875   6876       3    0.00000    0.00000    0.00000
```

```
0.00000    0.00000    0.00000 ;       H-      N-     CA-    HA
  6874   6873   6875   6877       3    0.00000    0.00000    0.00000
0.00000    0.00000    0.00000 ;       H-      N-     CA-    CB
  6874   6873   6875   6893       3    0.00000    0.00000    0.00000
0.00000    0.00000    0.00000 ;       H-      N-     CA-     C
  6875   6877   6880   6881       3    0.66944    2.00832    0.00000
-2.67776    0.00000    0.00000 ;      CA-    CB-    CG-   HG2
  6875   6877   6880   6882       3    0.66944    2.00832    0.00000
-2.67776    0.00000    0.00000 ;      CA-    CB-    CG-   HG3
  6875   6877   6880   6883       3    3.68192    3.09616   -2.09200
-3.01248    0.00000    0.00000 ;      CA-    CB-    CG-    CD
  6875   6893   6895   6896       3   20.92000    0.00000  -20.92000
0.00000    0.00000    0.00000 ;      CA-     C-     N-     H
  6875   6893   6895   6897       3   20.92000    0.00000  -20.92000
0.00000    0.00000    0.00000 ;      CA-     C-     N-    CA
  6876   6875   6877   6878       3    0.65084    1.95253    0.00000
-2.60338    0.00000    0.00000 ;      HA-    CA-    CB-   HB2
  6876   6875   6877   6879       3    0.65084    1.95253    0.00000
-2.60338    0.00000    0.00000 ;      HA-    CA-    CB-   HB3
  6876   6875   6877   6880       3    0.65084    1.95253    0.00000
-2.60338    0.00000    0.00000 ;      HA-    CA-    CB-    CG
  6876   6875   6893   6894       3    3.68192   -4.35136    0.00000
1.33888    0.00000    0.00000 ;      HA-    CA-     C-     O
  6876   6875   6893   6895       3    0.00000    0.00000    0.00000
0.00000    0.00000    0.00000 ;      HA-    CA-     C-     N
  6877   6875   6893   6894       3    0.00000    0.00000    0.00000
0.00000    0.00000    0.00000 ;      CB-    CA-     C-     O
  6877   6875   6893   6895       3    2.51040    4.18400    1.67360
-6.69440    0.00000    0.00000 ;      CB-    CA-     C-     N
  6877   6880   6883   6884       3    0.66944    2.00832    0.00000
-2.67776    0.00000    0.00000 ;      CB-    CG-    CD-   HD2
  6877   6880   6883   6885       3    0.66944    2.00832    0.00000
-2.67776    0.00000    0.00000 ;      CB-    CG-    CD-   HD3
  6877   6880   6883   6886       3    3.68192    3.09616   -2.09200
-3.01248    0.00000    0.00000 ;      CB-    CG-    CD-    CE
  6878   6877   6875   6893       3    0.65084    1.95253    0.00000
-2.60338    0.00000    0.00000 ;     HB2-    CB-    CA-     C
  6878   6877   6880   6881       3    0.62760    1.88280    0.00000
-2.51040    0.00000    0.00000 ;     HB2-    CB-    CG-   HG2
  6878   6877   6880   6882       3    0.62760    1.88280    0.00000
-2.51040    0.00000    0.00000 ;     HB2-    CB-    CG-   HG3
  6878   6877   6880   6883       3    0.66944    2.00832    0.00000
-2.67776    0.00000    0.00000 ;     HB2-    CB-    CG-    CD
  6879   6877   6875   6893       3    0.65084    1.95253    0.00000
-2.60338    0.00000    0.00000 ;     HB3-    CB-    CA-     C
  6879   6877   6880   6881       3    0.62760    1.88280    0.00000
-2.51040    0.00000    0.00000 ;     HB3-    CB-    CG-   HG2
  6879   6877   6880   6882       3    0.62760    1.88280    0.00000
-2.51040    0.00000    0.00000 ;     HB3-    CB-    CG-   HG3
  6879   6877   6880   6883       3    0.66944    2.00832    0.00000
-2.67776    0.00000    0.00000 ;     HB3-    CB-    CG-    CD
  6880   6877   6875   6893       3    0.65084    1.95253    0.00000
```

-2.60338 0.00000 0.00000 ; CG- CB- CA- C
6880 6883 6886 6887 3 0.65084 1.95253 0.00000
-2.60338 0.00000 0.00000 ; CG- CD- CE- HE2
6880 6883 6886 6888 3 0.65084 1.95253 0.00000
-2.60338 0.00000 0.00000 ; CG- CD- CE- HE3
6880 6883 6886 6889 3 0.65084 1.95253 0.00000
-2.60338 0.00000 0.00000 ; CG- CD- CE- NZ
6881 6880 6883 6884 3 0.62760 1.88280 0.00000
-2.51040 0.00000 0.00000 ; HG2- CG- CD- HD2
6881 6880 6883 6885 3 0.62760 1.88280 0.00000
-2.51040 0.00000 0.00000 ; HG2- CG- CD- HD3
6881 6880 6883 6886 3 0.66944 2.00832 0.00000
-2.67776 0.00000 0.00000 ; HG2- CG- CD- CE
6882 6880 6883 6884 3 0.62760 1.88280 0.00000
-2.51040 0.00000 0.00000 ; HG3- CG- CD- HD2
6882 6880 6883 6885 3 0.62760 1.88280 0.00000
-2.51040 0.00000 0.00000 ; HG3- CG- CD- HD3
6882 6880 6883 6886 3 0.66944 2.00832 0.00000
-2.67776 0.00000 0.00000 ; HG3- CG- CD- CE
6883 6886 6889 6890 3 0.65084 1.95253 0.00000
-2.60338 0.00000 0.00000 ; CD- CE- NZ- HZ1
6883 6886 6889 6891 3 0.65084 1.95253 0.00000
-2.60338 0.00000 0.00000 ; CD- CE- NZ- HZ2
6883 6886 6889 6892 3 0.65084 1.95253 0.00000
-2.60338 0.00000 0.00000 ; CD- CE- NZ- HZ3
6884 6883 6886 6887 3 0.65084 1.95253 0.00000
-2.60338 0.00000 0.00000 ; HD2- CD- CE- HE2
6884 6883 6886 6888 3 0.65084 1.95253 0.00000
-2.60338 0.00000 0.00000 ; HD2- CD- CE- HE3
6884 6883 6886 6889 3 0.65084 1.95253 0.00000
-2.60338 0.00000 0.00000 ; HD2- CD- CE- NZ
6885 6883 6886 6887 3 0.65084 1.95253 0.00000
-2.60338 0.00000 0.00000 ; HD3- CD- CE- HE2
6885 6883 6886 6888 3 0.65084 1.95253 0.00000
-2.60338 0.00000 0.00000 ; HD3- CD- CE- HE3
6885 6883 6886 6889 3 0.65084 1.95253 0.00000
-2.60338 0.00000 0.00000 ; HD3- CD- CE- NZ
6887 6886 6889 6890 3 0.65084 1.95253 0.00000
-2.60338 0.00000 0.00000 ; HE2- CE- NZ- HZ1
6887 6886 6889 6891 3 0.65084 1.95253 0.00000
-2.60338 0.00000 0.00000 ; HE2- CE- NZ- HZ2
6887 6886 6889 6892 3 0.65084 1.95253 0.00000
-2.60338 0.00000 0.00000 ; HE2- CE- NZ- HZ3
6888 6886 6889 6890 3 0.65084 1.95253 0.00000
-2.60338 0.00000 0.00000 ; HE3- CE- NZ- HZ1
6888 6886 6889 6891 3 0.65084 1.95253 0.00000
-2.60338 0.00000 0.00000 ; HE3- CE- NZ- HZ2
6888 6886 6889 6892 3 0.65084 1.95253 0.00000
-2.60338 0.00000 0.00000 ; HE3- CE- NZ- HZ3
6893 6895 6897 6898 3 0.00000 0.00000 0.00000
0.00000 0.00000 0.00000 ; C- N- CA- HA
6893 6895 6897 6899 3 11.71520 1.67360 15.06240 -

13.38880 0.00000 0.00000 ; C- N- CA- CB
6893 6895 6897 6903 3 1.75728 5.27184 2.25936
-7.02912 0.00000 0.00000 ; C- N- CA- C
6894 6893 6895 6896 3 29.28800 -8.36800 -20.92000
0.00000 0.00000 0.00000 ; O- C- N- H
6894 6893 6895 6897 3 20.92000 0.00000 -20.92000
0.00000 0.00000 0.00000 ; O- C- N- CA
6895 6897 6899 6900 3 0.65084 1.95253 0.00000
-2.60338 0.00000 0.00000 ; N- CA- CB- HB1
6895 6897 6899 6901 3 0.65084 1.95253 0.00000
-2.60338 0.00000 0.00000 ; N- CA- CB- HB2
6895 6897 6899 6902 3 0.65084 1.95253 0.00000
-2.60338 0.00000 0.00000 ; N- CA- CB- HB3
6895 6897 6903 6904 3 0.00000 0.00000 0.00000
0.00000 0.00000 0.00000 ; N- CA- C- O
6895 6897 6903 6905 3 17.40544 -5.02080 -13.22144
9.20480 0.00000 0.00000 ; N- CA- C- N
6896 6895 6897 6898 3 0.00000 0.00000 0.00000
0.00000 0.00000 0.00000 ; H- N- CA- HA
6896 6895 6897 6899 3 0.00000 0.00000 0.00000
0.00000 0.00000 0.00000 ; H- N- CA- CB
6896 6895 6897 6903 3 0.00000 0.00000 0.00000
0.00000 0.00000 0.00000 ; H- N- CA- C
6897 6903 6905 6906 3 20.92000 0.00000 -20.92000
0.00000 0.00000 0.00000 ; CA- C- N- H
6897 6903 6905 6907 3 20.92000 0.00000 -20.92000
0.00000 0.00000 0.00000 ; CA- C- N- CA
6898 6897 6899 6900 3 0.65084 1.95253 0.00000
-2.60338 0.00000 0.00000 ; HA- CA- CB- HB1
6898 6897 6899 6901 3 0.65084 1.95253 0.00000
-2.60338 0.00000 0.00000 ; HA- CA- CB- HB2
6898 6897 6899 6902 3 0.65084 1.95253 0.00000
-2.60338 0.00000 0.00000 ; HA- CA- CB- HB3
6898 6897 6903 6904 3 3.68192 -4.35136 0.00000
1.33888 0.00000 0.00000 ; HA- CA- C- O
6898 6897 6903 6905 3 0.00000 0.00000 0.00000
0.00000 0.00000 0.00000 ; HA- CA- C- N
6899 6897 6903 6904 3 0.00000 0.00000 0.00000
0.00000 0.00000 0.00000 ; CB- CA- C- O
6899 6897 6903 6905 3 2.51040 4.18400 1.67360
-6.69440 0.00000 0.00000 ; CB- CA- C- N
6900 6899 6897 6903 3 0.65084 1.95253 0.00000
-2.60338 0.00000 0.00000 ; HB1- CB- CA- C
6901 6899 6897 6903 3 0.65084 1.95253 0.00000
-2.60338 0.00000 0.00000 ; HB2- CB- CA- C
6902 6899 6897 6903 3 0.65084 1.95253 0.00000
-2.60338 0.00000 0.00000 ; HB3- CB- CA- C
6903 6905 6907 6908 3 0.00000 0.00000 0.00000
0.00000 0.00000 0.00000 ; C- N- CA- HA
6903 6905 6907 6909 3 11.71520 1.67360 15.06240 -
13.38880 0.00000 0.00000 ; C- N- CA- CB
6903 6905 6907 6922 3 1.75728 5.27184 2.25936

```
-7.02912    0.00000    0.00000 ;       C-     N-    CA-     C
  6904   6903   6905   6906      3   29.28800   -8.36800  -20.92000
0.00000    0.00000    0.00000 ;      O-     C-     N-     H
  6904   6903   6905   6907      3   20.92000    0.00000  -20.92000
0.00000    0.00000    0.00000 ;      O-     C-     N-    CA
  6905   6907   6909   6910      3    0.65084    1.95253    0.00000
-2.60338    0.00000    0.00000 ;      N-    CA-    CB-    HB
  6905   6907   6909   6911      3    3.83673    0.44769   -1.77402
-2.47693   -0.03347    0.00000 ;      N-    CA-    CB-   CG2
  6905   6907   6909   6915      3    2.97482    0.12134    2.04179
-1.89117   -3.24678    0.00000 ;      N-    CA-    CB-   CG1
  6905   6907   6922   6923      3    0.00000    0.00000    0.00000
0.00000    0.00000    0.00000 ;      N-    CA-     C-     O
  6905   6907   6922   6924      3   17.40544   -5.02080  -13.22144
9.20480    0.00000    0.00000 ;      N-    CA-     C-     N
  6906   6905   6907   6908      3    0.00000    0.00000    0.00000
0.00000    0.00000    0.00000 ;      H-     N-    CA-    HA
  6906   6905   6907   6909      3    0.00000    0.00000    0.00000
0.00000    0.00000    0.00000 ;      H-     N-    CA-    CB
  6906   6905   6907   6922      3    0.00000    0.00000    0.00000
0.00000    0.00000    0.00000 ;      H-     N-    CA-     C
  6907   6909   6911   6912      3    0.66944    2.00832    0.00000
-2.67776    0.00000    0.00000 ;     CA-    CB-   CG2-  HG21
  6907   6909   6911   6913      3    0.66944    2.00832    0.00000
-2.67776    0.00000    0.00000 ;     CA-    CB-   CG2-  HG22
  6907   6909   6911   6914      3    0.66944    2.00832    0.00000
-2.67776    0.00000    0.00000 ;     CA-    CB-   CG2-  HG23
  6907   6909   6915   6916      3    0.66944    2.00832    0.00000
-2.67776    0.00000    0.00000 ;     CA-    CB-   CG1-  HG12
  6907   6909   6915   6917      3    0.66944    2.00832    0.00000
-2.67776    0.00000    0.00000 ;     CA-    CB-   CG1-  HG13
  6907   6909   6915   6918      3    4.24258   -0.52718   -7.25506
-1.79075    7.69856    0.00000 ;     CA-    CB-   CG1-   CD1
  6907   6922   6924   6925      3   20.92000    0.00000  -20.92000
0.00000    0.00000    0.00000 ;     CA-     C-     N-     H
  6907   6922   6924   6926      3   20.92000    0.00000  -20.92000
0.00000    0.00000    0.00000 ;     CA-     C-     N-    CA
  6908   6907   6909   6910      3    0.65084    1.95253    0.00000
-2.60338    0.00000    0.00000 ;     HA-    CA-    CB-    HB
  6908   6907   6909   6911      3    0.65084    1.95253    0.00000
-2.60338    0.00000    0.00000 ;     HA-    CA-    CB-   CG2
  6908   6907   6909   6915      3    0.65084    1.95253    0.00000
-2.60338    0.00000    0.00000 ;     HA-    CA-    CB-   CG1
  6908   6907   6922   6923      3    3.68192   -4.35136    0.00000
1.33888    0.00000    0.00000 ;     HA-    CA-     C-     O
  6908   6907   6922   6924      3    0.00000    0.00000    0.00000
0.00000    0.00000    0.00000 ;     HA-    CA-     C-     N
  6909   6907   6922   6923      3    0.00000    0.00000    0.00000
0.00000    0.00000    0.00000 ;     CB-    CA-     C-     O
  6909   6907   6922   6924      3    2.51040    4.18400    1.67360
-6.69440    0.00000    0.00000 ;     CB-    CA-     C-     N
  6909   6915   6918   6919      3    0.66944    2.00832    0.00000
```

-2.67776 0.00000 0.00000 ; CB- CG1- CD1- HD11
6909 6915 6918 6920 3 0.66944 2.00832 0.00000
-2.67776 0.00000 0.00000 ; CB- CG1- CD1- HD12
6909 6915 6918 6921 3 0.66944 2.00832 0.00000
-2.67776 0.00000 0.00000 ; CB- CG1- CD1- HD13
6910 6909 6907 6922 3 0.65084 1.95253 0.00000
-2.60338 0.00000 0.00000 ; HB- CB- CA- C
6910 6909 6911 6912 3 0.62760 1.88280 0.00000
-2.51040 0.00000 0.00000 ; HB- CB- CG2- HG21
6910 6909 6911 6913 3 0.62760 1.88280 0.00000
-2.51040 0.00000 0.00000 ; HB- CB- CG2- HG22
6910 6909 6911 6914 3 0.62760 1.88280 0.00000
-2.51040 0.00000 0.00000 ; HB- CB- CG2- HG23
6910 6909 6915 6916 3 0.62760 1.88280 0.00000
-2.51040 0.00000 0.00000 ; HB- CB- CG1- HG12
6910 6909 6915 6917 3 0.62760 1.88280 0.00000
-2.51040 0.00000 0.00000 ; HB- CB- CG1- HG13
6910 6909 6915 6918 3 0.66944 2.00832 0.00000
-2.67776 0.00000 0.00000 ; HB- CB- CG1- CD1
6911 6909 6907 6922 3 5.67350 3.55640 -6.16722
-2.47693 3.74886 0.00000 ; CG2- CB- CA- C
6911 6909 6915 6916 3 0.66944 2.00832 0.00000
-2.67776 0.00000 0.00000 ; CG2- CB- CG1- HG12
6911 6909 6915 6917 3 0.66944 2.00832 0.00000
-2.67776 0.00000 0.00000 ; CG2- CB- CG1- HG13
6911 6909 6915 6918 3 3.81162 0.49790 -8.14206
-1.79075 7.49773 0.00000 ; CG2- CB- CG1- CD1
6912 6911 6909 6915 3 0.66944 2.00832 0.00000
-2.67776 0.00000 0.00000 ; HG21- CG2- CB- CG1
6913 6911 6909 6915 3 0.66944 2.00832 0.00000
-2.67776 0.00000 0.00000 ; HG22- CG2- CB- CG1
6914 6911 6909 6915 3 0.66944 2.00832 0.00000
-2.67776 0.00000 0.00000 ; HG23- CG2- CB- CG1
6915 6909 6907 6922 3 8.26340 0.74057 -9.99976
-1.89117 3.84928 0.00000 ; CG1- CB- CA- C
6916 6915 6918 6919 3 0.62760 1.88280 0.00000
-2.51040 0.00000 0.00000 ; HG12- CG1- CD1- HD11
6916 6915 6918 6920 3 0.62760 1.88280 0.00000
-2.51040 0.00000 0.00000 ; HG12- CG1- CD1- HD12
6916 6915 6918 6921 3 0.62760 1.88280 0.00000
-2.51040 0.00000 0.00000 ; HG12- CG1- CD1- HD13
6917 6915 6918 6919 3 0.62760 1.88280 0.00000
-2.51040 0.00000 0.00000 ; HG13- CG1- CD1- HD11
6917 6915 6918 6920 3 0.62760 1.88280 0.00000
-2.51040 0.00000 0.00000 ; HG13- CG1- CD1- HD12
6917 6915 6918 6921 3 0.62760 1.88280 0.00000
-2.51040 0.00000 0.00000 ; HG13- CG1- CD1- HD13
6922 6924 6926 6927 3 0.00000 0.00000 0.00000
0.00000 0.00000 0.00000 ; C- N- CA- HA
6922 6924 6926 6928 3 11.71520 1.67360 15.06240 -
13.38880 0.00000 0.00000 ; C- N- CA- CB
6922 6924 6926 6934 3 1.75728 5.27184 2.25936

```
-7.02912    0.00000    0.00000 ;       C-     N-    CA-     C
  6923   6922   6924   6925      3   29.28800   -8.36800  -20.92000
0.00000    0.00000    0.00000 ;       O-     C-     N-     H
  6923   6922   6924   6926      3   20.92000    0.00000  -20.92000
0.00000    0.00000    0.00000 ;       O-     C-     N-    CA
  6924   6926   6928   6929      3    0.65084    1.95253    0.00000
-2.60338    0.00000    0.00000 ;       N-    CA-    CB-   HB2
  6924   6926   6928   6930      3    0.65084    1.95253    0.00000
-2.60338    0.00000    0.00000 ;       N-    CA-    CB-   HB3
  6924   6926   6928   6931      3   15.41386    9.74035   -8.39310
-0.97069    2.97901    0.00000 ;       N-    CA-    CB-    CG
  6924   6926   6934   6935      3    0.00000    0.00000    0.00000
0.00000    0.00000    0.00000 ;       N-    CA-     C-     O
  6924   6926   6934   6936      3   17.40544   -5.02080  -13.22144
9.20480    0.00000    0.00000 ;       N-    CA-     C-     N
  6925   6924   6926   6927      3    0.00000    0.00000    0.00000
0.00000    0.00000    0.00000 ;       H-     N-    CA-    HA
  6925   6924   6926   6928      3    0.00000    0.00000    0.00000
0.00000    0.00000    0.00000 ;       H-     N-    CA-    CB
  6925   6924   6926   6934      3    0.00000    0.00000    0.00000
0.00000    0.00000    0.00000 ;       H-     N-    CA-     C
  6926   6928   6931   6932      3    6.43499    0.00000   -5.39736
0.00000   -1.03763    0.00000 ;      CA-    CB-    CG-   OD1
  6926   6928   6931   6933      3    6.43499    0.00000   -5.39736
0.00000   -1.03763    0.00000 ;      CA-    CB-    CG-   OD2
  6926   6934   6936   6937      3   20.92000    0.00000  -20.92000
0.00000    0.00000    0.00000 ;      CA-     C-     N-     H
  6926   6934   6936   6938      3   20.92000    0.00000  -20.92000
0.00000    0.00000    0.00000 ;      CA-     C-     N-    CA
  6927   6926   6928   6929      3    0.65084    1.95253    0.00000
-2.60338    0.00000    0.00000 ;      HA-    CA-    CB-   HB2
  6927   6926   6928   6930      3    0.65084    1.95253    0.00000
-2.60338    0.00000    0.00000 ;      HA-    CA-    CB-   HB3
  6927   6926   6928   6931      3    0.65084    1.95253    0.00000
-2.60338    0.00000    0.00000 ;      HA-    CA-    CB-    CG
  6927   6926   6934   6935      3    3.68192   -4.35136    0.00000
1.33888    0.00000    0.00000 ;      HA-    CA-     C-     O
  6927   6926   6934   6936      3    0.00000    0.00000    0.00000
0.00000    0.00000    0.00000 ;      HA-    CA-     C-     N
  6928   6926   6934   6935      3    0.00000    0.00000    0.00000
0.00000    0.00000    0.00000 ;      CB-    CA-     C-     O
  6928   6926   6934   6936      3    2.51040    4.18400    1.67360
-6.69440    0.00000    0.00000 ;      CB-    CA-     C-     N
  6929   6928   6926   6934      3    0.65084    1.95253    0.00000
-2.60338    0.00000    0.00000 ;     HB2-    CB-    CA-     C
  6929   6928   6931   6932      3    0.00000    0.00000    0.00000
0.00000    0.00000    0.00000 ;     HB2-    CB-    CG-   OD1
  6929   6928   6931   6933      3    0.00000    0.00000    0.00000
0.00000    0.00000    0.00000 ;     HB2-    CB-    CG-   OD2
  6930   6928   6926   6934      3    0.65084    1.95253    0.00000
-2.60338    0.00000    0.00000 ;     HB3-    CB-    CA-     C
  6930   6928   6931   6932      3    0.00000    0.00000    0.00000
```

```
0.00000    0.00000    0.00000 ;    HB3-    CB-    CG-   OD1
  6930   6928   6931   6933      3    0.00000    0.00000    0.00000
0.00000    0.00000    0.00000 ;    HB3-    CB-    CG-   OD2
  6931   6928   6926   6934      3    7.14627   -1.04600   -8.99560
-0.97069    5.15469    0.00000 ;     CG-    CB-    CA-     C
  6934   6936   6938   6939      3    0.00000    0.00000    0.00000
0.00000    0.00000    0.00000 ;     C-     N-    CA-    HA
  6934   6936   6938   6940      3   11.71520    1.67360   15.06240  -
13.38880    0.00000    0.00000 ;     C-     N-    CA-    CB
  6934   6936   6938   6945      3    1.75728    5.27184    2.25936
-7.02912    0.00000    0.00000 ;     C-     N-    CA-     C
  6935   6934   6936   6937      3   29.28800   -8.36800  -20.92000
0.00000    0.00000    0.00000 ;     O-     C-     N-     H
  6935   6934   6936   6938      3   20.92000    0.00000  -20.92000
0.00000    0.00000    0.00000 ;     O-     C-     N-    CA
  6936   6938   6940   6941      3    0.65084    1.95253    0.00000
-2.60338    0.00000    0.00000 ;     N-    CA-    CB-   HB2
  6936   6938   6940   6942      3    0.65084    1.95253    0.00000
-2.60338    0.00000    0.00000 ;     N-    CA-    CB-   HB3
  6936   6938   6940   6943      3    7.86174    2.24681   -7.41405
-6.71114    5.35552    0.00000 ;     N-    CA-    CB-    OG
  6936   6938   6945   6946      3    0.00000    0.00000    0.00000
0.00000    0.00000    0.00000 ;     N-    CA-     C-     O
  6936   6938   6945   6947      3   17.40544   -5.02080  -13.22144
9.20480    0.00000    0.00000 ;     N-    CA-     C-     N
  6937   6936   6938   6939      3    0.00000    0.00000    0.00000
0.00000    0.00000    0.00000 ;     H-     N-    CA-    HA
  6937   6936   6938   6940      3    0.00000    0.00000    0.00000
0.00000    0.00000    0.00000 ;     H-     N-    CA-    CB
  6937   6936   6938   6945      3    0.00000    0.00000    0.00000
0.00000    0.00000    0.00000 ;     H-     N-    CA-     C
  6938   6940   6943   6944      3    2.05853    2.46856    3.48109
-4.46851    0.23430    0.00000 ;    CA-    CB-    OG-    HG
  6938   6945   6947   6948      3   20.92000    0.00000  -20.92000
0.00000    0.00000    0.00000 ;    CA-     C-     N-     H
  6938   6945   6947   6949      3   20.92000    0.00000  -20.92000
0.00000    0.00000    0.00000 ;    CA-     C-     N-    CA
  6939   6938   6940   6941      3    0.65084    1.95253    0.00000
-2.60338    0.00000    0.00000 ;    HA-    CA-    CB-   HB2
  6939   6938   6940   6942      3    0.65084    1.95253    0.00000
-2.60338    0.00000    0.00000 ;    HA-    CA-    CB-   HB3
  6939   6938   6940   6943      3    1.04600   -1.04600    0.00000
0.00000    0.00000    0.00000 ;    HA-    CA-    CB-    OG
  6939   6938   6945   6946      3    3.68192   -4.35136    0.00000
1.33888    0.00000    0.00000 ;    HA-    CA-     C-     O
  6939   6938   6945   6947      3    0.00000    0.00000    0.00000
0.00000    0.00000    0.00000 ;    HA-    CA-     C-     N
  6940   6938   6945   6946      3    0.00000    0.00000    0.00000
0.00000    0.00000    0.00000 ;    CB-    CA-     C-     O
  6940   6938   6945   6947      3    2.51040    4.18400    1.67360
-6.69440    0.00000    0.00000 ;    CB-    CA-     C-     N
  6941   6940   6938   6945      3    0.65084    1.95253    0.00000
```

```
-2.60338    0.00000    0.00000 ;    HB2-   CB-    CA-     C
  6941   6940   6943   6944      3    0.69733    2.09200    0.00000
-2.78933    0.00000    0.00000 ;    HB2-   CB-    OG-    HG
  6942   6940   6938   6945      3    0.65084    1.95253    0.00000
-2.60338    0.00000    0.00000 ;    HB3-   CB-    CA-     C
  6942   6940   6943   6944      3    0.69733    2.09200    0.00000
-2.78933    0.00000    0.00000 ;    HB3-   CB-    OG-    HG
  6943   6940   6938   6945      3    7.34710    7.79898   -6.14211
-6.71114    4.31789    0.00000 ;     OG-   CB-    CA-     C
  6945   6947   6949   6950      3    0.00000    0.00000    0.00000
0.00000    0.00000    0.00000 ;     C-     N-    CA-    HA
  6945   6947   6949   6951      3   11.71520    1.67360   15.06240  -
13.38880    0.00000    0.00000 ;     C-     N-    CA-    CB
  6945   6947   6949   6964      3    1.75728    5.27184    2.25936
-7.02912    0.00000    0.00000 ;     C-     N-    CA-     C
  6946   6945   6947   6948      3   29.28800   -8.36800  -20.92000
0.00000    0.00000    0.00000 ;     O-     C-     N-     H
  6946   6945   6947   6949      3   20.92000    0.00000  -20.92000
0.00000    0.00000    0.00000 ;     O-     C-     N-    CA
  6947   6949   6951   6952      3    0.65084    1.95253    0.00000
-2.60338    0.00000    0.00000 ;     N-    CA-    CB-   HB2
  6947   6949   6951   6953      3    0.65084    1.95253    0.00000
-2.60338    0.00000    0.00000 ;     N-    CA-    CB-   HB3
  6947   6949   6951   6954      3    3.79070    1.39746   -4.61077
-2.40998    2.44346    0.00000 ;     N-    CA-    CB-    CG
  6947   6949   6964   6965      3    0.00000    0.00000    0.00000
0.00000    0.00000    0.00000 ;     N-    CA-     C-     O
  6947   6949   6964   6966      3   17.40544   -5.02080  -13.22144
9.20480    0.00000    0.00000 ;     N-    CA-     C-     N
  6948   6947   6949   6950      3    0.00000    0.00000    0.00000
0.00000    0.00000    0.00000 ;     H-     N-    CA-    HA
  6948   6947   6949   6951      3    0.00000    0.00000    0.00000
0.00000    0.00000    0.00000 ;     H-     N-    CA-    CB
  6948   6947   6949   6964      3    0.00000    0.00000    0.00000
0.00000    0.00000    0.00000 ;     H-     N-    CA-     C
  6949   6951   6954   6955      3    0.66944    2.00832    0.00000
-2.67776    0.00000    0.00000 ;    CA-    CB-    CG-    HG
  6949   6951   6954   6956      3    3.90367    0.19665   -6.21742
-2.37651    5.99149    0.00000 ;    CA-    CB-    CG-   CD1
  6949   6951   6954   6960      3    3.90367    0.19665   -6.21742
-2.37651    5.99149    0.00000 ;    CA-    CB-    CG-   CD2
  6949   6964   6966   6967      3   20.92000    0.00000  -20.92000
0.00000    0.00000    0.00000 ;    CA-     C-     N-     H
  6949   6964   6966   6968      3   20.92000    0.00000  -20.92000
0.00000    0.00000    0.00000 ;    CA-     C-     N-    CA
  6950   6949   6951   6952      3    0.65084    1.95253    0.00000
-2.60338    0.00000    0.00000 ;    HA-    CA-    CB-   HB2
  6950   6949   6951   6953      3    0.65084    1.95253    0.00000
-2.60338    0.00000    0.00000 ;    HA-    CA-    CB-   HB3
  6950   6949   6951   6954      3    0.65084    1.95253    0.00000
-2.60338    0.00000    0.00000 ;    HA-    CA-    CB-    CG
  6950   6949   6964   6965      3    3.68192   -4.35136    0.00000
```

```
1.33888    0.00000    0.00000 ;      HA-    CA-     C-     O
  6950   6949   6964   6966       3    0.00000    0.00000    0.00000
0.00000    0.00000    0.00000 ;      HA-    CA-     C-     N
  6951   6949   6964   6965       3    0.00000    0.00000    0.00000
0.00000    0.00000    0.00000 ;      CB-    CA-     C-     O
  6951   6949   6964   6966       3    2.51040    4.18400    1.67360
-6.69440    0.00000    0.00000 ;     CB-    CA-     C-     N
  6951   6954   6956   6957       3    0.66944    2.00832    0.00000
-2.67776    0.00000    0.00000 ;     CB-    CG-   CD1-  HD11
  6951   6954   6956   6958       3    0.66944    2.00832    0.00000
-2.67776    0.00000    0.00000 ;     CB-    CG-   CD1-  HD12
  6951   6954   6956   6959       3    0.66944    2.00832    0.00000
-2.67776    0.00000    0.00000 ;     CB-    CG-   CD1-  HD13
  6951   6954   6960   6961       3    0.66944    2.00832    0.00000
-2.67776    0.00000    0.00000 ;     CB-    CG-   CD2-  HD21
  6951   6954   6960   6962       3    0.66944    2.00832    0.00000
-2.67776    0.00000    0.00000 ;     CB-    CG-   CD2-  HD22
  6951   6954   6960   6963       3    0.66944    2.00832    0.00000
-2.67776    0.00000    0.00000 ;     CB-    CG-   CD2-  HD23
  6952   6951   6949   6964       3    0.65084    1.95253    0.00000
-2.60338    0.00000    0.00000 ;    HB2-    CB-     CA-     C
  6952   6951   6954   6955       3    0.62760    1.88280    0.00000
-2.51040    0.00000    0.00000 ;    HB2-    CB-     CG-    HG
  6952   6951   6954   6956       3    0.66944    2.00832    0.00000
-2.67776    0.00000    0.00000 ;    HB2-    CB-     CG-   CD1
  6952   6951   6954   6960       3    0.66944    2.00832    0.00000
-2.67776    0.00000    0.00000 ;    HB2-    CB-     CG-   CD2
  6953   6951   6949   6964       3    0.65084    1.95253    0.00000
-2.60338    0.00000    0.00000 ;    HB3-    CB-     CA-     C
  6953   6951   6954   6955       3    0.62760    1.88280    0.00000
-2.51040    0.00000    0.00000 ;    HB3-    CB-     CG-    HG
  6953   6951   6954   6956       3    0.66944    2.00832    0.00000
-2.67776    0.00000    0.00000 ;    HB3-    CB-     CG-   CD1
  6953   6951   6954   6960       3    0.66944    2.00832    0.00000
-2.67776    0.00000    0.00000 ;    HB3-    CB-     CG-   CD2
  6954   6951   6949   6964       3   10.33448   -1.14642  -11.54784
-2.40998    6.35968    0.00000 ;     CG-    CB-     CA-     C
  6955   6954   6956   6957       3    0.62760    1.88280    0.00000
-2.51040    0.00000    0.00000 ;     HG-    CG-   CD1-  HD11
  6955   6954   6956   6958       3    0.62760    1.88280    0.00000
-2.51040    0.00000    0.00000 ;     HG-    CG-   CD1-  HD12
  6955   6954   6956   6959       3    0.62760    1.88280    0.00000
-2.51040    0.00000    0.00000 ;     HG-    CG-   CD1-  HD13
  6955   6954   6960   6961       3    0.62760    1.88280    0.00000
-2.51040    0.00000    0.00000 ;     HG-    CG-   CD2-  HD21
  6955   6954   6960   6962       3    0.62760    1.88280    0.00000
-2.51040    0.00000    0.00000 ;     HG-    CG-   CD2-  HD22
  6955   6954   6960   6963       3    0.62760    1.88280    0.00000
-2.51040    0.00000    0.00000 ;     HG-    CG-   CD2-  HD23
  6956   6954   6960   6961       3    0.66944    2.00832    0.00000
-2.67776    0.00000    0.00000 ;    CD1-    CG-   CD2-  HD21
  6956   6954   6960   6962       3    0.66944    2.00832    0.00000
```

```
-2.67776    0.00000    0.00000 ;    CD1-    CG-   CD2-  HD22
  6956   6954   6960   6963       3    0.66944    2.00832    0.00000
-2.67776    0.00000    0.00000 ;    CD1-    CG-   CD2-  HD23
  6957   6956   6954   6960       3    0.66944    2.00832    0.00000
-2.67776    0.00000    0.00000 ;   HD11-   CD1-    CG-   CD2
  6958   6956   6954   6960       3    0.66944    2.00832    0.00000
-2.67776    0.00000    0.00000 ;   HD12-   CD1-    CG-   CD2
  6959   6956   6954   6960       3    0.66944    2.00832    0.00000
-2.67776    0.00000    0.00000 ;   HD13-   CD1-    CG-   CD2
  6964   6966   6968   6969       3    0.00000    0.00000    0.00000
0.00000    0.00000    0.00000 ;      C-     N-    CA-    HA
  6964   6966   6968   6970       3   11.71520    1.67360   15.06240  -
13.38880    0.00000    0.00000 ;      C-     N-    CA-    CB
  6964   6966   6968   6983       3    1.75728    5.27184    2.25936
-7.02912    0.00000    0.00000 ;      C-     N-    CA-     C
  6965   6964   6966   6967       3   29.28800   -8.36800  -20.92000
0.00000    0.00000    0.00000 ;      O-     C-     N-     H
  6965   6964   6966   6968       3   20.92000    0.00000  -20.92000
0.00000    0.00000    0.00000 ;      O-     C-     N-    CA
  6966   6968   6970   6971       3    0.65084    1.95253    0.00000
-2.60338    0.00000    0.00000 ;      N-    CA-    CB-   HB2
  6966   6968   6970   6972       3    0.65084    1.95253    0.00000
-2.60338    0.00000    0.00000 ;      N-    CA-    CB-   HB3
  6966   6968   6970   6973       3    3.79070    1.39746   -4.61077
-2.40998    2.44346    0.00000 ;      N-    CA-    CB-    CG
  6966   6968   6983   6984       3    0.00000    0.00000    0.00000
0.00000    0.00000    0.00000 ;      N-    CA-     C-     O
  6966   6968   6983   6985       3   17.40544   -5.02080  -13.22144
9.20480    0.00000    0.00000 ;      N-    CA-     C-     N
  6967   6966   6968   6969       3    0.00000    0.00000    0.00000
0.00000    0.00000    0.00000 ;      H-     N-    CA-    HA
  6967   6966   6968   6970       3    0.00000    0.00000    0.00000
0.00000    0.00000    0.00000 ;      H-     N-    CA-    CB
  6967   6966   6968   6983       3    0.00000    0.00000    0.00000
0.00000    0.00000    0.00000 ;      H-     N-    CA-     C
  6968   6970   6973   6974       3    0.66944    2.00832    0.00000
-2.67776    0.00000    0.00000 ;     CA-    CB-    CG-    HG
  6968   6970   6973   6975       3    3.90367    0.19665   -6.21742
-2.37651    5.99149    0.00000 ;     CA-    CB-    CG-   CD1
  6968   6970   6973   6979       3    3.90367    0.19665   -6.21742
-2.37651    5.99149    0.00000 ;     CA-    CB-    CG-   CD2
  6968   6983   6985   6986       3   20.92000    0.00000  -20.92000
0.00000    0.00000    0.00000 ;     CA-     C-     N-     H
  6968   6983   6985   6987       3   20.92000    0.00000  -20.92000
0.00000    0.00000    0.00000 ;     CA-     C-     N-    CA
  6969   6968   6970   6971       3    0.65084    1.95253    0.00000
-2.60338    0.00000    0.00000 ;     HA-    CA-    CB-   HB2
  6969   6968   6970   6972       3    0.65084    1.95253    0.00000
-2.60338    0.00000    0.00000 ;     HA-    CA-    CB-   HB3
  6969   6968   6970   6973       3    0.65084    1.95253    0.00000
-2.60338    0.00000    0.00000 ;     HA-    CA-    CB-    CG
  6969   6968   6983   6984       3    3.68192   -4.35136    0.00000
```

```
1.33888    0.00000    0.00000 ;     HA-    CA-     C-     O
  6969   6968   6983   6985      3   0.00000    0.00000    0.00000
0.00000    0.00000    0.00000 ;     HA-    CA-     C-     N
  6970   6968   6983   6984      3   0.00000    0.00000    0.00000
0.00000    0.00000    0.00000 ;     CB-    CA-     C-     O
  6970   6968   6983   6985      3   2.51040    4.18400    1.67360
-6.69440    0.00000    0.00000 ;     CB-    CA-     C-     N
  6970   6973   6975   6976      3   0.66944    2.00832    0.00000
-2.67776    0.00000    0.00000 ;     CB-    CG-   CD1-  HD11
  6970   6973   6975   6977      3   0.66944    2.00832    0.00000
-2.67776    0.00000    0.00000 ;     CB-    CG-   CD1-  HD12
  6970   6973   6975   6978      3   0.66944    2.00832    0.00000
-2.67776    0.00000    0.00000 ;     CB-    CG-   CD1-  HD13
  6970   6973   6979   6980      3   0.66944    2.00832    0.00000
-2.67776    0.00000    0.00000 ;     CB-    CG-   CD2-  HD21
  6970   6973   6979   6981      3   0.66944    2.00832    0.00000
-2.67776    0.00000    0.00000 ;     CB-    CG-   CD2-  HD22
  6970   6973   6979   6982      3   0.66944    2.00832    0.00000
-2.67776    0.00000    0.00000 ;     CB-    CG-   CD2-  HD23
  6971   6970   6968   6983      3   0.65084    1.95253    0.00000
-2.60338    0.00000    0.00000 ;    HB2-    CB-     CA-     C
  6971   6970   6973   6974      3   0.62760    1.88280    0.00000
-2.51040    0.00000    0.00000 ;    HB2-    CB-     CG-    HG
  6971   6970   6973   6975      3   0.66944    2.00832    0.00000
-2.67776    0.00000    0.00000 ;    HB2-    CB-     CG-   CD1
  6971   6970   6973   6979      3   0.66944    2.00832    0.00000
-2.67776    0.00000    0.00000 ;    HB2-    CB-     CG-   CD2
  6972   6970   6968   6983      3   0.65084    1.95253    0.00000
-2.60338    0.00000    0.00000 ;    HB3-    CB-     CA-     C
  6972   6970   6973   6974      3   0.62760    1.88280    0.00000
-2.51040    0.00000    0.00000 ;    HB3-    CB-     CG-    HG
  6972   6970   6973   6975      3   0.66944    2.00832    0.00000
-2.67776    0.00000    0.00000 ;    HB3-    CB-     CG-   CD1
  6972   6970   6973   6979      3   0.66944    2.00832    0.00000
-2.67776    0.00000    0.00000 ;    HB3-    CB-     CG-   CD2
  6973   6970   6968   6983      3  10.33448   -1.14642  -11.54784
-2.40998    6.35968    0.00000 ;     CG-    CB-     CA-     C
  6974   6973   6975   6976      3   0.62760    1.88280    0.00000
-2.51040    0.00000    0.00000 ;     HG-    CG-   CD1-  HD11
  6974   6973   6975   6977      3   0.62760    1.88280    0.00000
-2.51040    0.00000    0.00000 ;     HG-    CG-   CD1-  HD12
  6974   6973   6975   6978      3   0.62760    1.88280    0.00000
-2.51040    0.00000    0.00000 ;     HG-    CG-   CD1-  HD13
  6974   6973   6979   6980      3   0.62760    1.88280    0.00000
-2.51040    0.00000    0.00000 ;     HG-    CG-   CD2-  HD21
  6974   6973   6979   6981      3   0.62760    1.88280    0.00000
-2.51040    0.00000    0.00000 ;     HG-    CG-   CD2-  HD22
  6974   6973   6979   6982      3   0.62760    1.88280    0.00000
-2.51040    0.00000    0.00000 ;     HG-    CG-   CD2-  HD23
  6975   6973   6979   6980      3   0.66944    2.00832    0.00000
-2.67776    0.00000    0.00000 ;    CD1-    CG-   CD2-  HD21
  6975   6973   6979   6981      3   0.66944    2.00832    0.00000
```

-2.67776 0.00000 0.00000 ; CD1- CG- CD2- HD22
6975 6973 6979 6982 3 0.66944 2.00832 0.00000
-2.67776 0.00000 0.00000 ; CD1- CG- CD2- HD23
6976 6975 6973 6979 3 0.66944 2.00832 0.00000
-2.67776 0.00000 0.00000 ; HD11- CD1- CG- CD2
6977 6975 6973 6979 3 0.66944 2.00832 0.00000
-2.67776 0.00000 0.00000 ; HD12- CD1- CG- CD2
6978 6975 6973 6979 3 0.66944 2.00832 0.00000
-2.67776 0.00000 0.00000 ; HD13- CD1- CG- CD2
6983 6985 6987 6988 3 0.00000 0.00000 0.00000
0.00000 0.00000 0.00000 ; C- N- CA- HA
6983 6985 6987 6989 3 11.71520 1.67360 15.06240 -
13.38880 0.00000 0.00000 ; C- N- CA- CB
6983 6985 6987 7000 3 1.75728 5.27184 2.25936
-7.02912 0.00000 0.00000 ; C- N- CA- C
6984 6983 6985 6986 3 29.28800 -8.36800 -20.92000
0.00000 0.00000 0.00000 ; O- C- N- H
6984 6983 6985 6987 3 20.92000 0.00000 -20.92000
0.00000 0.00000 0.00000 ; O- C- N- CA
6985 6987 6989 6990 3 0.65084 1.95253 0.00000
-2.60338 0.00000 0.00000 ; N- CA- CB- HB2
6985 6987 6989 6991 3 0.65084 1.95253 0.00000
-2.60338 0.00000 0.00000 ; N- CA- CB- HB3
6985 6987 6989 6992 3 3.21331 2.22589 -4.15053
-2.40998 2.61082 0.00000 ; N- CA- CB- CG
6985 6987 7000 7001 3 0.00000 0.00000 0.00000
0.00000 0.00000 0.00000 ; N- CA- C- O
6985 6987 7000 7002 3 17.40544 -5.02080 -13.22144
9.20480 0.00000 0.00000 ; N- CA- C- N
6986 6985 6987 6988 3 0.00000 0.00000 0.00000
0.00000 0.00000 0.00000 ; H- N- CA- HA
6986 6985 6987 6989 3 0.00000 0.00000 0.00000
0.00000 0.00000 0.00000 ; H- N- CA- CB
6986 6985 6987 7000 3 0.00000 0.00000 0.00000
0.00000 0.00000 0.00000 ; H- N- CA- C
6987 6989 6992 6993 3 0.66944 2.00832 0.00000
-2.67776 0.00000 0.00000 ; CA- CB- CG- HG2
6987 6989 6992 6994 3 0.66944 2.00832 0.00000
-2.67776 0.00000 0.00000 ; CA- CB- CG- HG3
6987 6989 6992 6995 3 3.69866 -4.35136 -3.92459
6.89523 4.61914 0.00000 ; CA- CB- CG- CD
6987 7000 7002 7003 3 20.92000 0.00000 -20.92000
0.00000 0.00000 0.00000 ; CA- C- N- H
6987 7000 7002 7004 3 20.92000 0.00000 -20.92000
0.00000 0.00000 0.00000 ; CA- C- N- CA
6988 6987 6989 6990 3 0.65084 1.95253 0.00000
-2.60338 0.00000 0.00000 ; HA- CA- CB- HB2
6988 6987 6989 6991 3 0.65084 1.95253 0.00000
-2.60338 0.00000 0.00000 ; HA- CA- CB- HB3
6988 6987 6989 6992 3 0.65084 1.95253 0.00000
-2.60338 0.00000 0.00000 ; HA- CA- CB- CG
6988 6987 7000 7001 3 3.68192 -4.35136 0.00000

1.33888 0.00000 0.00000 ; HA- CA- C- O
6988 6987 7000 7002 3 0.00000 0.00000 0.00000
0.00000 0.00000 0.00000 ; HA- CA- C- N
6989 6987 7000 7001 3 0.00000 0.00000 0.00000
0.00000 0.00000 0.00000 ; CB- CA- C- O
6989 6987 7000 7002 3 2.51040 4.18400 1.67360
-6.69440 0.00000 0.00000 ; CB- CA- C- N
6989 6992 6995 6996 3 0.00000 0.00000 0.00000
0.00000 0.00000 0.00000 ; CB- CG- CD- OE1
6989 6992 6995 6997 3 10.32611 1.48114 -8.47678
1.42256 1.40582 0.00000 ; CB- CG- CD- NE2
6990 6989 6987 7000 3 0.65084 1.95253 0.00000
-2.60338 0.00000 0.00000 ; HB2- CB- CA- C
6990 6989 6992 6993 3 0.62760 1.88280 0.00000
-2.51040 0.00000 0.00000 ; HB2- CB- CG- HG2
6990 6989 6992 6994 3 0.62760 1.88280 0.00000
-2.51040 0.00000 0.00000 ; HB2- CB- CG- HG3
6990 6989 6992 6995 3 0.65084 1.95253 0.00000
-2.60338 0.00000 0.00000 ; HB2- CB- CG- CD
6991 6989 6987 7000 3 0.65084 1.95253 0.00000
-2.60338 0.00000 0.00000 ; HB3- CB- CA- C
6991 6989 6992 6993 3 0.62760 1.88280 0.00000
-2.51040 0.00000 0.00000 ; HB3- CB- CG- HG2
6991 6989 6992 6994 3 0.62760 1.88280 0.00000
-2.51040 0.00000 0.00000 ; HB3- CB- CG- HG3
6991 6989 6992 6995 3 0.65084 1.95253 0.00000
-2.60338 0.00000 0.00000 ; HB3- CB- CG- CD
6992 6989 6987 7000 3 6.86594 3.56895 -8.14206
-2.40998 4.85344 0.00000 ; CG- CB- CA- C
6992 6995 6997 6998 3 20.92000 0.00000 -20.92000
0.00000 0.00000 0.00000 ; CG- CD- NE2- HE21
6992 6995 6997 6999 3 20.92000 0.00000 -20.92000
0.00000 0.00000 0.00000 ; CG- CD- NE2- HE22
6993 6992 6995 6996 3 3.68192 -4.35136 0.00000
1.33888 0.00000 0.00000 ; HG2- CG- CD- OE1
6993 6992 6995 6997 3 0.00000 0.00000 0.00000
0.00000 0.00000 0.00000 ; HG2- CG- CD- NE2
6994 6992 6995 6996 3 3.68192 -4.35136 0.00000
1.33888 0.00000 0.00000 ; HG3- CG- CD- OE1
6994 6992 6995 6997 3 0.00000 0.00000 0.00000
0.00000 0.00000 0.00000 ; HG3- CG- CD- NE2
6996 6995 6997 6998 3 29.28800 -8.36800 -20.92000
0.00000 0.00000 0.00000 ; OE1- CD- NE2- HE21
6996 6995 6997 6999 3 29.28800 -8.36800 -20.92000
0.00000 0.00000 0.00000 ; OE1- CD- NE2- HE22
7000 7002 7004 7005 3 0.00000 0.00000 0.00000
0.00000 0.00000 0.00000 ; C- N- CA- HA
7000 7002 7004 7006 3 11.71520 1.67360 15.06240 -
13.38880 0.00000 0.00000 ; C- N- CA- CB
7000 7002 7004 7015 3 1.75728 5.27184 2.25936
-7.02912 0.00000 0.00000 ; C- N- CA- C
7001 7000 7002 7003 3 29.28800 -8.36800 -20.92000

```
0.00000    0.00000    0.00000 ;       O-     C-     N-     H
  7001   7000   7002   7004       3   20.92000    0.00000  -20.92000
0.00000    0.00000    0.00000 ;       O-     C-     N-    CA
  7002   7004   7006   7007       3    0.65084    1.95253    0.00000
-2.60338    0.00000    0.00000 ;       N-    CA-    CB-   HB2
  7002   7004   7006   7008       3    0.65084    1.95253    0.00000
-2.60338    0.00000    0.00000 ;       N-    CA-    CB-   HB3
  7002   7004   7006   7009       3    3.21331    2.22589   -4.15053
-2.40998    2.61082    0.00000 ;       N-    CA-    CB-    CG
  7002   7004   7015   7016       3    0.00000    0.00000    0.00000
0.00000    0.00000    0.00000 ;       N-    CA-     C-     O
  7002   7004   7015   7017       3   17.40544   -5.02080  -13.22144
9.20480    0.00000    0.00000 ;       N-    CA-     C-     N
  7003   7002   7004   7005       3    0.00000    0.00000    0.00000
0.00000    0.00000    0.00000 ;       H-     N-    CA-    HA
  7003   7002   7004   7006       3    0.00000    0.00000    0.00000
0.00000    0.00000    0.00000 ;       H-     N-    CA-    CB
  7003   7002   7004   7015       3    0.00000    0.00000    0.00000
0.00000    0.00000    0.00000 ;       H-     N-    CA-     C
  7004   7006   7009   7010       3    0.66944    2.00832    0.00000
-2.67776    0.00000    0.00000 ;      CA-    CB-    CG-   HG2
  7004   7006   7009   7011       3    0.66944    2.00832    0.00000
-2.67776    0.00000    0.00000 ;      CA-    CB-    CG-   HG3
  7004   7006   7009   7012       3   10.12110   -1.91209    0.01674
10.17549   -1.87443    0.00000 ;      CA-    CB-    CG-    CD
  7004   7015   7017   7018       3   20.92000    0.00000  -20.92000
0.00000    0.00000    0.00000 ;      CA-     C-     N-     H
  7004   7015   7017   7019       3   20.92000    0.00000  -20.92000
0.00000    0.00000    0.00000 ;      CA-     C-     N-    CA
  7005   7004   7006   7007       3    0.65084    1.95253    0.00000
-2.60338    0.00000    0.00000 ;      HA-    CA-    CB-   HB2
  7005   7004   7006   7008       3    0.65084    1.95253    0.00000
-2.60338    0.00000    0.00000 ;      HA-    CA-    CB-   HB3
  7005   7004   7006   7009       3    0.65084    1.95253    0.00000
-2.60338    0.00000    0.00000 ;      HA-    CA-    CB-    CG
  7005   7004   7015   7016       3    3.68192   -4.35136    0.00000
1.33888    0.00000    0.00000 ;      HA-    CA-     C-     O
  7005   7004   7015   7017       3    0.00000    0.00000    0.00000
0.00000    0.00000    0.00000 ;      HA-    CA-     C-     N
  7006   7004   7015   7016       3    0.00000    0.00000    0.00000
0.00000    0.00000    0.00000 ;      CB-    CA-     C-     O
  7006   7004   7015   7017       3    2.51040    4.18400    1.67360
-6.69440    0.00000    0.00000 ;      CB-    CA-     C-     N
  7006   7009   7012   7013       3    3.79907    0.00000   -5.40573
0.00000    2.14221    0.00000 ;      CB-    CG-    CD-   OE1
  7006   7009   7012   7014       3    3.79907    0.00000   -5.40573
0.00000    2.14221    0.00000 ;      CB-    CG-    CD-   OE2
  7007   7006   7004   7015       3    0.65084    1.95253    0.00000
-2.60338    0.00000    0.00000 ;     HB2-    CB-    CA-     C
  7007   7006   7009   7010       3    0.62760    1.88280    0.00000
-2.51040    0.00000    0.00000 ;     HB2-    CB-    CG-   HG2
  7007   7006   7009   7011       3    0.62760    1.88280    0.00000
```

-2.51040 0.00000 0.00000 ; HB2- CB- CG- HG3
7007 7006 7009 7012 3 0.65084 1.95253 0.00000
-2.60338 0.00000 0.00000 ; HB2- CB- CG- CD
7008 7006 7004 7015 3 0.65084 1.95253 0.00000
-2.60338 0.00000 0.00000 ; HB3- CB- CA- C
7008 7006 7009 7010 3 0.62760 1.88280 0.00000
-2.51040 0.00000 0.00000 ; HB3- CB- CG- HG2
7008 7006 7009 7011 3 0.62760 1.88280 0.00000
-2.51040 0.00000 0.00000 ; HB3- CB- CG- HG3
7008 7006 7009 7012 3 0.65084 1.95253 0.00000
-2.60338 0.00000 0.00000 ; HB3- CB- CG- CD
7009 7006 7004 7015 3 6.86594 3.56895 -8.14206
-2.40998 4.85344 0.00000 ; CG- CB- CA- C
7010 7009 7012 7013 3 0.00000 0.00000 0.00000
0.00000 0.00000 0.00000 ; HG2- CG- CD- OE1
7010 7009 7012 7014 3 0.00000 0.00000 0.00000
0.00000 0.00000 0.00000 ; HG2- CG- CD- OE2
7011 7009 7012 7013 3 0.00000 0.00000 0.00000
0.00000 0.00000 0.00000 ; HG3- CG- CD- OE1
7011 7009 7012 7014 3 0.00000 0.00000 0.00000
0.00000 0.00000 0.00000 ; HG3- CG- CD- OE2
7015 7017 7019 7020 3 0.00000 0.00000 0.00000
0.00000 0.00000 0.00000 ; C- N- CA- HA
7015 7017 7019 7021 3 11.71520 1.67360 15.06240 -
13.38880 0.00000 0.00000 ; C- N- CA- CB
7015 7017 7019 7031 3 1.75728 5.27184 2.25936
-7.02912 0.00000 0.00000 ; C- N- CA- C
7016 7015 7017 7018 3 29.28800 -8.36800 -20.92000
0.00000 0.00000 0.00000 ; O- C- N- H
7016 7015 7017 7019 3 20.92000 0.00000 -20.92000
0.00000 0.00000 0.00000 ; O- C- N- CA
7017 7019 7021 7022 3 0.65084 1.95253 0.00000
-2.60338 0.00000 0.00000 ; N- CA- CB- HB
7017 7019 7021 7023 3 3.83673 0.44769 -1.77402
-2.47693 -0.03347 0.00000 ; N- CA- CB- CG1
7017 7019 7021 7027 3 3.83673 0.44769 -1.77402
-2.47693 -0.03347 0.00000 ; N- CA- CB- CG2
7017 7019 7031 7032 3 0.00000 0.00000 0.00000
0.00000 0.00000 0.00000 ; N- CA- C- O
7017 7019 7031 7033 3 17.40544 -5.02080 -13.22144
9.20480 0.00000 0.00000 ; N- CA- C- N
7018 7017 7019 7020 3 0.00000 0.00000 0.00000
0.00000 0.00000 0.00000 ; H- N- CA- HA
7018 7017 7019 7021 3 0.00000 0.00000 0.00000
0.00000 0.00000 0.00000 ; H- N- CA- CB
7018 7017 7019 7031 3 0.00000 0.00000 0.00000
0.00000 0.00000 0.00000 ; H- N- CA- C
7019 7021 7023 7024 3 0.66944 2.00832 0.00000
-2.67776 0.00000 0.00000 ; CA- CB- CG1- HG11
7019 7021 7023 7025 3 0.66944 2.00832 0.00000
-2.67776 0.00000 0.00000 ; CA- CB- CG1- HG12
7019 7021 7023 7026 3 0.66944 2.00832 0.00000

```
-2.67776    0.00000    0.00000 ;     CA-    CB-   CG1-  HG13
  7019   7021   7027   7028      3    0.66944    2.00832    0.00000
-2.67776    0.00000    0.00000 ;     CA-    CB-   CG2-  HG21
  7019   7021   7027   7029      3    0.66944    2.00832    0.00000
-2.67776    0.00000    0.00000 ;     CA-    CB-   CG2-  HG22
  7019   7021   7027   7030      3    0.66944    2.00832    0.00000
-2.67776    0.00000    0.00000 ;     CA-    CB-   CG2-  HG23
  7019   7031   7033   7034      3   20.92000    0.00000  -20.92000
0.00000    0.00000    0.00000 ;     CA-     C-     N-     H
  7019   7031   7033   7035      3   20.92000    0.00000  -20.92000
0.00000    0.00000    0.00000 ;     CA-     C-     N-    CA
  7020   7019   7021   7022      3    0.65084    1.95253    0.00000
-2.60338    0.00000    0.00000 ;     HA-    CA-    CB-    HB
  7020   7019   7021   7023      3    0.65084    1.95253    0.00000
-2.60338    0.00000    0.00000 ;     HA-    CA-    CB-   CG1
  7020   7019   7021   7027      3    0.65084    1.95253    0.00000
-2.60338    0.00000    0.00000 ;     HA-    CA-    CB-   CG2
  7020   7019   7031   7032      3    3.68192   -4.35136    0.00000
1.33888    0.00000    0.00000 ;     HA-    CA-     C-     O
  7020   7019   7031   7033      3    0.00000    0.00000    0.00000
0.00000    0.00000    0.00000 ;     HA-    CA-     C-     N
  7021   7019   7031   7032      3    0.00000    0.00000    0.00000
0.00000    0.00000    0.00000 ;     CB-    CA-     C-     O
  7021   7019   7031   7033      3    2.51040    4.18400    1.67360
-6.69440    0.00000    0.00000 ;     CB-    CA-     C-     N
  7022   7021   7019   7031      3    0.65084    1.95253    0.00000
-2.60338    0.00000    0.00000 ;     HB-    CB-    CA-     C
  7022   7021   7023   7024      3    0.62760    1.88280    0.00000
-2.51040    0.00000    0.00000 ;     HB-    CB-   CG1-  HG11
  7022   7021   7023   7025      3    0.62760    1.88280    0.00000
-2.51040    0.00000    0.00000 ;     HB-    CB-   CG1-  HG12
  7022   7021   7023   7026      3    0.62760    1.88280    0.00000
-2.51040    0.00000    0.00000 ;     HB-    CB-   CG1-  HG13
  7022   7021   7027   7028      3    0.62760    1.88280    0.00000
-2.51040    0.00000    0.00000 ;     HB-    CB-   CG2-  HG21
  7022   7021   7027   7029      3    0.62760    1.88280    0.00000
-2.51040    0.00000    0.00000 ;     HB-    CB-   CG2-  HG22
  7022   7021   7027   7030      3    0.62760    1.88280    0.00000
-2.51040    0.00000    0.00000 ;     HB-    CB-   CG2-  HG23
  7023   7021   7019   7031      3    5.67350    3.55640   -6.16722
-2.47693    3.74886    0.00000 ;    CG1-    CB-    CA-     C
  7023   7021   7027   7028      3    0.66944    2.00832    0.00000
-2.67776    0.00000    0.00000 ;    CG1-    CB-   CG2-  HG21
  7023   7021   7027   7029      3    0.66944    2.00832    0.00000
-2.67776    0.00000    0.00000 ;    CG1-    CB-   CG2-  HG22
  7023   7021   7027   7030      3    0.66944    2.00832    0.00000
-2.67776    0.00000    0.00000 ;    CG1-    CB-   CG2-  HG23
  7024   7023   7021   7027      3    0.66944    2.00832    0.00000
-2.67776    0.00000    0.00000 ;   HG11-   CG1-    CB-   CG2
  7025   7023   7021   7027      3    0.66944    2.00832    0.00000
-2.67776    0.00000    0.00000 ;   HG12-   CG1-    CB-   CG2
  7026   7023   7021   7027      3    0.66944    2.00832    0.00000
```

-2.67776    0.00000    0.00000 ;   HG13-   CG1-    CB-   CG2
 7027   7021   7019   7031      3    5.67350    3.55640   -6.16722
-2.47693    3.74886    0.00000 ;    CG2-    CB-    CA-     C
 7031   7033   7035   7036      3    0.00000    0.00000    0.00000
0.00000    0.00000    0.00000 ;      C-     N-    CA-    HA
 7031   7033   7035   7037      3   11.71520    1.67360   15.06240  -
13.38880    0.00000    0.00000 ;     C-     N-    CA-    CB
 7031   7033   7035   7043      3    1.75728    5.27184    2.25936
-7.02912    0.00000    0.00000 ;     C-     N-    CA-     C
 7032   7031   7033   7034      3   29.28800   -8.36800  -20.92000
0.00000    0.00000    0.00000 ;      O-     C-     N-    H
 7032   7031   7033   7035      3   20.92000    0.00000  -20.92000
0.00000    0.00000    0.00000 ;      O-     C-     N-    CA
 7033   7035   7037   7038      3    0.65084    1.95253    0.00000
-2.60338    0.00000    0.00000 ;     N-    CA-    CB-   HB2
 7033   7035   7037   7039      3    0.65084    1.95253    0.00000
-2.60338    0.00000    0.00000 ;     N-    CA-    CB-   HB3
 7033   7035   7037   7040      3   15.41386    9.74035   -8.39310
-0.97069    2.97901    0.00000 ;     N-    CA-    CB-    CG
 7033   7035   7043   7044      3    0.00000    0.00000    0.00000
0.00000    0.00000    0.00000 ;      N-    CA-     C-     O
 7033   7035   7043   7045      3   17.40544   -5.02080  -13.22144
9.20480    0.00000    0.00000 ;      N-    CA-     C-     N
 7034   7033   7035   7036      3    0.00000    0.00000    0.00000
0.00000    0.00000    0.00000 ;      H-     N-    CA-    HA
 7034   7033   7035   7037      3    0.00000    0.00000    0.00000
0.00000    0.00000    0.00000 ;      H-     N-    CA-    CB
 7034   7033   7035   7043      3    0.00000    0.00000    0.00000
0.00000    0.00000    0.00000 ;      H-     N-    CA-     C
 7035   7037   7040   7041      3    6.43499    0.00000   -5.39736
0.00000   -1.03763    0.00000 ;     CA-    CB-    CG-   OD1
 7035   7037   7040   7042      3    6.43499    0.00000   -5.39736
0.00000   -1.03763    0.00000 ;     CA-    CB-    CG-   OD2
 7035   7043   7045   7046      3   20.92000    0.00000  -20.92000
0.00000    0.00000    0.00000 ;     CA-     C-     N-     H
 7035   7043   7045   7047      3   20.92000    0.00000  -20.92000
0.00000    0.00000    0.00000 ;     CA-     C-     N-    CA
 7036   7035   7037   7038      3    0.65084    1.95253    0.00000
-2.60338    0.00000    0.00000 ;     HA-    CA-    CB-   HB2
 7036   7035   7037   7039      3    0.65084    1.95253    0.00000
-2.60338    0.00000    0.00000 ;     HA-    CA-    CB-   HB3
 7036   7035   7037   7040      3    0.65084    1.95253    0.00000
-2.60338    0.00000    0.00000 ;     HA-    CA-    CB-    CG
 7036   7035   7043   7044      3    3.68192   -4.35136    0.00000
1.33888    0.00000    0.00000 ;     HA-    CA-     C-     O
 7036   7035   7043   7045      3    0.00000    0.00000    0.00000
0.00000    0.00000    0.00000 ;     HA-    CA-     C-     N
 7037   7035   7043   7044      3    0.00000    0.00000    0.00000
0.00000    0.00000    0.00000 ;     CB-    CA-     C-     O
 7037   7035   7043   7045      3    2.51040    4.18400    1.67360
-6.69440    0.00000    0.00000 ;     CB-    CA-     C-     N
 7038   7037   7035   7043      3    0.65084    1.95253    0.00000

```
-2.60338    0.00000    0.00000 ;     HB2-    CB-     CA-     C
  7038   7037   7040   7041       3    0.00000    0.00000    0.00000
0.00000    0.00000    0.00000 ;    HB2-    CB-    CG-   OD1
  7038   7037   7040   7042       3    0.00000    0.00000    0.00000
0.00000    0.00000    0.00000 ;    HB2-    CB-    CG-   OD2
  7039   7037   7035   7043       3    0.65084    1.95253    0.00000
-2.60338    0.00000    0.00000 ;     HB3-    CB-     CA-     C
  7039   7037   7040   7041       3    0.00000    0.00000    0.00000
0.00000    0.00000    0.00000 ;    HB3-    CB-    CG-   OD1
  7039   7037   7040   7042       3    0.00000    0.00000    0.00000
0.00000    0.00000    0.00000 ;    HB3-    CB-    CG-   OD2
  7040   7037   7035   7043       3    7.14627   -1.04600   -8.99560
-0.97069    5.15469    0.00000 ;      CG-    CB-     CA-     C
  7043   7045   7047   7048       3    0.00000    0.00000    0.00000
0.00000    0.00000    0.00000 ;      C-     N-    CA-    HA
  7043   7045   7047   7049       3   11.71520    1.67360   15.06240  -
13.38880    0.00000    0.00000 ;      C-     N-    CA-    CB
  7043   7045   7047   7062       3    1.75728    5.27184    2.25936
-7.02912    0.00000    0.00000 ;      C-     N-    CA-     C
  7044   7043   7045   7046       3   29.28800   -8.36800  -20.92000
0.00000    0.00000    0.00000 ;      O-    C-     N-    H
  7044   7043   7045   7047       3   20.92000    0.00000  -20.92000
0.00000    0.00000    0.00000 ;      O-     C-     N-    CA
  7045   7047   7049   7050       3    0.65084    1.95253    0.00000
-2.60338    0.00000    0.00000 ;      N-    CA-    CB-   HB2
  7045   7047   7049   7051       3    0.65084    1.95253    0.00000
-2.60338    0.00000    0.00000 ;      N-    CA-    CB-   HB3
  7045   7047   7049   7052       3    3.79070    1.39746   -4.61077
-2.40998    2.44346    0.00000 ;      N-    CA-    CB-    CG
  7045   7047   7062   7063       3    0.00000    0.00000    0.00000
0.00000    0.00000    0.00000 ;      N-    CA-     C-     O
  7045   7047   7062   7064       3   17.40544   -5.02080  -13.22144
9.20480    0.00000    0.00000 ;      N-    CA-     C-     N
  7046   7045   7047   7048       3    0.00000    0.00000    0.00000
0.00000    0.00000    0.00000 ;      H-     N-    CA-    HA
  7046   7045   7047   7049       3    0.00000    0.00000    0.00000
0.00000    0.00000    0.00000 ;      H-     N-    CA-    CB
  7046   7045   7047   7062       3    0.00000    0.00000    0.00000
0.00000    0.00000    0.00000 ;      H-     N-    CA-     C
  7047   7049   7052   7053       3    0.66944    2.00832    0.00000
-2.67776    0.00000    0.00000 ;     CA-    CB-    CG-    HG
  7047   7049   7052   7054       3    3.90367    0.19665   -6.21742
-2.37651    5.99149    0.00000 ;     CA-    CB-    CG-   CD1
  7047   7049   7052   7058       3    3.90367    0.19665   -6.21742
-2.37651    5.99149    0.00000 ;     CA-    CB-    CG-   CD2
  7047   7062   7064   7065       3   20.92000    0.00000  -20.92000
0.00000    0.00000    0.00000 ;     CA-     C-     N-     H
  7047   7062   7064   7066       3   20.92000    0.00000  -20.92000
0.00000    0.00000    0.00000 ;     CA-     C-     N-    CA
  7048   7047   7049   7050       3    0.65084    1.95253    0.00000
-2.60338    0.00000    0.00000 ;     HA-    CA-    CB-   HB2
  7048   7047   7049   7051       3    0.65084    1.95253    0.00000
```

```
-2.60338    0.00000    0.00000 ;      HA-    CA-    CB-   HB3
  7048   7047   7049   7052       3    0.65084    1.95253    0.00000
-2.60338    0.00000    0.00000 ;      HA-    CA-    CB-    CG
  7048   7047   7062   7063       3    3.68192   -4.35136    0.00000
1.33888    0.00000    0.00000 ;      HA-    CA-     C-     O
  7048   7047   7062   7064       3    0.00000    0.00000    0.00000
0.00000    0.00000    0.00000 ;      HA-    CA-     C-     N
  7049   7047   7062   7063       3    0.00000    0.00000    0.00000
0.00000    0.00000    0.00000 ;      CB-    CA-     C-     O
  7049   7047   7062   7064       3    2.51040    4.18400    1.67360
-6.69440    0.00000    0.00000 ;      CB-    CA-     C-     N
  7049   7052   7054   7055       3    0.66944    2.00832    0.00000
-2.67776    0.00000    0.00000 ;      CB-    CG-   CD1-  HD11
  7049   7052   7054   7056       3    0.66944    2.00832    0.00000
-2.67776    0.00000    0.00000 ;      CB-    CG-   CD1-  HD12
  7049   7052   7054   7057       3    0.66944    2.00832    0.00000
-2.67776    0.00000    0.00000 ;      CB-    CG-   CD1-  HD13
  7049   7052   7058   7059       3    0.66944    2.00832    0.00000
-2.67776    0.00000    0.00000 ;      CB-    CG-   CD2-  HD21
  7049   7052   7058   7060       3    0.66944    2.00832    0.00000
-2.67776    0.00000    0.00000 ;      CB-    CG-   CD2-  HD22
  7049   7052   7058   7061       3    0.66944    2.00832    0.00000
-2.67776    0.00000    0.00000 ;      CB-    CG-   CD2-  HD23
  7050   7049   7047   7062       3    0.65084    1.95253    0.00000
-2.60338    0.00000    0.00000 ;     HB2-    CB-    CA-     C
  7050   7049   7052   7053       3    0.62760    1.88280    0.00000
-2.51040    0.00000    0.00000 ;     HB2-    CB-    CG-    HG
  7050   7049   7052   7054       3    0.66944    2.00832    0.00000
-2.67776    0.00000    0.00000 ;     HB2-    CB-    CG-   CD1
  7050   7049   7052   7058       3    0.66944    2.00832    0.00000
-2.67776    0.00000    0.00000 ;     HB2-    CB-    CG-   CD2
  7051   7049   7047   7062       3    0.65084    1.95253    0.00000
-2.60338    0.00000    0.00000 ;     HB3-    CB-    CA-     C
  7051   7049   7052   7053       3    0.62760    1.88280    0.00000
-2.51040    0.00000    0.00000 ;     HB3-    CB-    CG-    HG
  7051   7049   7052   7054       3    0.66944    2.00832    0.00000
-2.67776    0.00000    0.00000 ;     HB3-    CB-    CG-   CD1
  7051   7049   7052   7058       3    0.66944    2.00832    0.00000
-2.67776    0.00000    0.00000 ;     HB3-    CB-    CG-   CD2
  7052   7049   7047   7062       3   10.33448   -1.14642  -11.54784
-2.40998    6.35968    0.00000 ;      CG-    CB-    CA-     C
  7053   7052   7054   7055       3    0.62760    1.88280    0.00000
-2.51040    0.00000    0.00000 ;      HG-    CG-   CD1-  HD11
  7053   7052   7054   7056       3    0.62760    1.88280    0.00000
-2.51040    0.00000    0.00000 ;      HG-    CG-   CD1-  HD12
  7053   7052   7054   7057       3    0.62760    1.88280    0.00000
-2.51040    0.00000    0.00000 ;      HG-    CG-   CD1-  HD13
  7053   7052   7058   7059       3    0.62760    1.88280    0.00000
-2.51040    0.00000    0.00000 ;      HG-    CG-   CD2-  HD21
  7053   7052   7058   7060       3    0.62760    1.88280    0.00000
-2.51040    0.00000    0.00000 ;      HG-    CG-   CD2-  HD22
  7053   7052   7058   7061       3    0.62760    1.88280    0.00000
```

-2.51040 0.00000 0.00000 ; HG- CG- CD2- HD23
7054 7052 7058 7059 3 0.66944 2.00832 0.00000
-2.67776 0.00000 0.00000 ; CD1- CG- CD2- HD21
7054 7052 7058 7060 3 0.66944 2.00832 0.00000
-2.67776 0.00000 0.00000 ; CD1- CG- CD2- HD22
7054 7052 7058 7061 3 0.66944 2.00832 0.00000
-2.67776 0.00000 0.00000 ; CD1- CG- CD2- HD23
7055 7054 7052 7058 3 0.66944 2.00832 0.00000
-2.67776 0.00000 0.00000 ; HD11- CD1- CG- CD2
7056 7054 7052 7058 3 0.66944 2.00832 0.00000
-2.67776 0.00000 0.00000 ; HD12- CD1- CG- CD2
7057 7054 7052 7058 3 0.66944 2.00832 0.00000
-2.67776 0.00000 0.00000 ; HD13- CD1- CG- CD2
7062 7064 7066 7067 3 0.00000 0.00000 0.00000
0.00000 0.00000 0.00000 ; C- N- CA- HA
7062 7064 7066 7068 3 11.71520 1.67360 15.06240 -
13.38880 0.00000 0.00000 ; C- N- CA- CB
7062 7064 7066 7076 3 1.75728 5.27184 2.25936
-7.02912 0.00000 0.00000 ; C- N- CA- C
7063 7062 7064 7065 3 29.28800 -8.36800 -20.92000
0.00000 0.00000 0.00000 ; O- C- N- H
7063 7062 7064 7066 3 20.92000 0.00000 -20.92000
0.00000 0.00000 0.00000 ; O- C- N- CA
7064 7066 7068 7069 3 0.65084 1.95253 0.00000
-2.60338 0.00000 0.00000 ; N- CA- CB- HB2
7064 7066 7068 7070 3 0.65084 1.95253 0.00000
-2.60338 0.00000 0.00000 ; N- CA- CB- HB3
7064 7066 7068 7071 3 5.99567 3.29281 -4.46014
-0.55229 1.97485 0.00000 ; N- CA- CB- CG
7064 7066 7076 7077 3 0.00000 0.00000 0.00000
0.00000 0.00000 0.00000 ; N- CA- C- O
7064 7066 7076 7078 3 17.40544 -5.02080 -13.22144
9.20480 0.00000 0.00000 ; N- CA- C- N
7065 7064 7066 7067 3 0.00000 0.00000 0.00000
0.00000 0.00000 0.00000 ; H- N- CA- HA
7065 7064 7066 7068 3 0.00000 0.00000 0.00000
0.00000 0.00000 0.00000 ; H- N- CA- CB
7065 7064 7066 7076 3 0.00000 0.00000 0.00000
0.00000 0.00000 0.00000 ; H- N- CA- C
7066 7068 7071 7072 3 0.00000 0.00000 0.00000
0.00000 0.00000 0.00000 ; CA- CB- CG- OD1
7066 7068 7071 7073 3 8.84916 -0.31380 -4.32626
5.03754 0.26778 0.00000 ; CA- CB- CG- ND2
7066 7076 7078 7079 3 20.92000 0.00000 -20.92000
0.00000 0.00000 0.00000 ; CA- C- N- H
7066 7076 7078 7080 3 20.92000 0.00000 -20.92000
0.00000 0.00000 0.00000 ; CA- C- N- CA
7067 7066 7068 7069 3 0.65084 1.95253 0.00000
-2.60338 0.00000 0.00000 ; HA- CA- CB- HB2
7067 7066 7068 7070 3 0.65084 1.95253 0.00000
-2.60338 0.00000 0.00000 ; HA- CA- CB- HB3
7067 7066 7068 7071 3 0.65084 1.95253 0.00000

```
-2.60338    0.00000    0.00000 ;     HA-    CA-    CB-    CG
  7067   7066   7076   7077      3    3.68192   -4.35136    0.00000
1.33888    0.00000    0.00000 ;     HA-    CA-     C-     O
  7067   7066   7076   7078      3    0.00000    0.00000    0.00000
0.00000    0.00000    0.00000 ;     HA-    CA-     C-     N
  7068   7066   7076   7077      3    0.00000    0.00000    0.00000
0.00000    0.00000    0.00000 ;     CB-    CA-     C-     O
  7068   7066   7076   7078      3    2.51040    4.18400    1.67360
-6.69440    0.00000    0.00000 ;     CB-    CA-     C-     N
  7068   7071   7073   7074      3   20.92000    0.00000  -20.92000
0.00000    0.00000    0.00000 ;     CB-    CG-   ND2-  HD21
  7068   7071   7073   7075      3   20.92000    0.00000  -20.92000
0.00000    0.00000    0.00000 ;     CB-    CG-   ND2-  HD22
  7069   7068   7066   7076      3    0.65084    1.95253    0.00000
-2.60338    0.00000    0.00000 ;    HB2-    CB-    CA-     C
  7069   7068   7071   7072      3    3.68192   -4.35136    0.00000
1.33888    0.00000    0.00000 ;    HB2-    CB-    CG-   OD1
  7069   7068   7071   7073      3    0.00000    0.00000    0.00000
0.00000    0.00000    0.00000 ;    HB2-    CB-    CG-   ND2
  7070   7068   7066   7076      3    0.65084    1.95253    0.00000
-2.60338    0.00000    0.00000 ;    HB3-    CB-    CA-     C
  7070   7068   7071   7072      3    3.68192   -4.35136    0.00000
1.33888    0.00000    0.00000 ;    HB3-    CB-    CG-   OD1
  7070   7068   7071   7073      3    0.00000    0.00000    0.00000
0.00000    0.00000    0.00000 ;    HB3-    CB-    CG-   ND2
  7071   7068   7066   7076      3    7.94542   -3.96225   -6.11701
-0.55229    3.58150    0.00000 ;     CG-    CB-    CA-     C
  7072   7071   7073   7074      3   29.28800   -8.36800  -20.92000
0.00000    0.00000    0.00000 ;    OD1-    CG-   ND2-  HD21
  7072   7071   7073   7075      3   29.28800   -8.36800  -20.92000
0.00000    0.00000    0.00000 ;    OD1-    CG-   ND2-  HD22
  7076   7078   7080   7081      3    0.00000    0.00000    0.00000
0.00000    0.00000    0.00000 ;     C-     N-    CA-   HA2
  7076   7078   7080   7082      3    0.00000    0.00000    0.00000
0.00000    0.00000    0.00000 ;     C-     N-    CA-   HA3
  7076   7078   7080   7083      3    1.75728    5.27184    2.25936
-7.02912    0.00000    0.00000 ;     C-     N-    CA-     C
  7077   7076   7078   7079      3   29.28800   -8.36800  -20.92000
0.00000    0.00000    0.00000 ;     O-     C-     N-     H
  7077   7076   7078   7080      3   20.92000    0.00000  -20.92000
0.00000    0.00000    0.00000 ;     O-     C-     N-    CA
  7078   7080   7083   7084      3    0.00000    0.00000    0.00000
0.00000    0.00000    0.00000 ;     N-    CA-     C-     O
  7078   7080   7083   7085      3   17.40544   -5.02080  -13.22144
9.20480    0.00000    0.00000 ;     N-    CA-     C-     N
  7079   7078   7080   7081      3    0.00000    0.00000    0.00000
0.00000    0.00000    0.00000 ;     H-     N-    CA-   HA2
  7079   7078   7080   7082      3    0.00000    0.00000    0.00000
0.00000    0.00000    0.00000 ;     H-     N-    CA-   HA3
  7079   7078   7080   7083      3    0.00000    0.00000    0.00000
0.00000    0.00000    0.00000 ;     H-     N-    CA-     C
  7080   7083   7085   7086      3   20.92000    0.00000  -20.92000
```

0.00000 0.00000 0.00000 ; CA- C- N- H
7080 7083 7085 7087 3 20.92000 0.00000 -20.92000
0.00000 0.00000 0.00000 ; CA- C- N- CA
7081 7080 7083 7084 3 3.68192 -4.35136 0.00000
1.33888 0.00000 0.00000 ; HA2- CA- C- O
7081 7080 7083 7085 3 0.00000 0.00000 0.00000
0.00000 0.00000 0.00000 ; HA2- CA- C- N
7082 7080 7083 7084 3 3.68192 -4.35136 0.00000
1.33888 0.00000 0.00000 ; HA3- CA- C- O
7082 7080 7083 7085 3 0.00000 0.00000 0.00000
0.00000 0.00000 0.00000 ; HA3- CA- C- N
7083 7085 7087 7088 3 0.00000 0.00000 0.00000
0.00000 0.00000 0.00000 ; C- N- CA- HA
7083 7085 7087 7089 3 11.71520 1.67360 15.06240 -
13.38880 0.00000 0.00000 ; C- N- CA- CB
7083 7085 7087 7095 3 1.75728 5.27184 2.25936
-7.02912 0.00000 0.00000 ; C- N- CA- C
7084 7083 7085 7086 3 29.28800 -8.36800 -20.92000
0.00000 0.00000 0.00000 ; O- C- N- H
7084 7083 7085 7087 3 20.92000 0.00000 -20.92000
0.00000 0.00000 0.00000 ; O- C- N- CA
7085 7087 7089 7090 3 0.65084 1.95253 0.00000
-2.60338 0.00000 0.00000 ; N- CA- CB- HB2
7085 7087 7089 7091 3 0.65084 1.95253 0.00000
-2.60338 0.00000 0.00000 ; N- CA- CB- HB3
7085 7087 7089 7092 3 15.41386 9.74035 -8.39310
-0.97069 2.97901 0.00000 ; N- CA- CB- CG
7085 7087 7095 7096 3 0.00000 0.00000 0.00000
0.00000 0.00000 0.00000 ; N- CA- C- O
7085 7087 7095 7097 3 17.40544 -5.02080 -13.22144
9.20480 0.00000 0.00000 ; N- CA- C- N
7086 7085 7087 7088 3 0.00000 0.00000 0.00000
0.00000 0.00000 0.00000 ; H- N- CA- HA
7086 7085 7087 7089 3 0.00000 0.00000 0.00000
0.00000 0.00000 0.00000 ; H- N- CA- CB
7086 7085 7087 7095 3 0.00000 0.00000 0.00000
0.00000 0.00000 0.00000 ; H- N- CA- C
7087 7089 7092 7093 3 6.43499 0.00000 -5.39736
0.00000 -1.03763 0.00000 ; CA- CB- CG- OD1
7087 7089 7092 7094 3 6.43499 0.00000 -5.39736
0.00000 -1.03763 0.00000 ; CA- CB- CG- OD2
7087 7095 7097 7098 3 20.92000 0.00000 -20.92000
0.00000 0.00000 0.00000 ; CA- C- N- H
7087 7095 7097 7099 3 20.92000 0.00000 -20.92000
0.00000 0.00000 0.00000 ; CA- C- N- CA
7088 7087 7089 7090 3 0.65084 1.95253 0.00000
-2.60338 0.00000 0.00000 ; HA- CA- CB- HB2
7088 7087 7089 7091 3 0.65084 1.95253 0.00000
-2.60338 0.00000 0.00000 ; HA- CA- CB- HB3
7088 7087 7089 7092 3 0.65084 1.95253 0.00000
-2.60338 0.00000 0.00000 ; HA- CA- CB- CG
7088 7087 7095 7096 3 3.68192 -4.35136 0.00000

```
1.33888    0.00000    0.00000 ;     HA-    CA-     C-     O
  7088   7087   7095   7097       3    0.00000    0.00000    0.00000
0.00000    0.00000    0.00000 ;     HA-    CA-     C-     N
  7089   7087   7095   7096       3    0.00000    0.00000    0.00000
0.00000    0.00000    0.00000 ;     CB-    CA-     C-     O
  7089   7087   7095   7097       3    2.51040    4.18400    1.67360
-6.69440    0.00000    0.00000 ;     CB-    CA-     C-     N
  7090   7089   7087   7095       3    0.65084    1.95253    0.00000
-2.60338    0.00000    0.00000 ;    HB2-    CB-    CA-     C
  7090   7089   7092   7093       3    0.00000    0.00000    0.00000
0.00000    0.00000    0.00000 ;    HB2-    CB-    CG-   OD1
  7090   7089   7092   7094       3    0.00000    0.00000    0.00000
0.00000    0.00000    0.00000 ;    HB2-    CB-    CG-   OD2
  7091   7089   7087   7095       3    0.65084    1.95253    0.00000
-2.60338    0.00000    0.00000 ;    HB3-    CB-    CA-     C
  7091   7089   7092   7093       3    0.00000    0.00000    0.00000
0.00000    0.00000    0.00000 ;    HB3-    CB-    CG-   OD1
  7091   7089   7092   7094       3    0.00000    0.00000    0.00000
0.00000    0.00000    0.00000 ;    HB3-    CB-    CG-   OD2
  7092   7089   7087   7095       3    7.14627   -1.04600   -8.99560
-0.97069    5.15469    0.00000 ;     CG-    CB-    CA-     C
  7095   7097   7099   7100       3    0.00000    0.00000    0.00000
0.00000    0.00000    0.00000 ;      C-     N-    CA-   HA2
  7095   7097   7099   7101       3    0.00000    0.00000    0.00000
0.00000    0.00000    0.00000 ;      C-     N-    CA-   HA3
  7095   7097   7099   7102       3    1.75728    5.27184    2.25936
-7.02912    0.00000    0.00000 ;      C-     N-    CA-     C
  7096   7095   7097   7098       3   29.28800   -8.36800  -20.92000
0.00000    0.00000    0.00000 ;      O-     C-     N-     H
  7096   7095   7097   7099       3   20.92000    0.00000  -20.92000
0.00000    0.00000    0.00000 ;      O-     C-     N-    CA
  7097   7099   7102   7103       3    0.00000    0.00000    0.00000
0.00000    0.00000    0.00000 ;      N-    CA-     C-     O
  7097   7099   7102   7104       3   17.40544   -5.02080  -13.22144
9.20480    0.00000    0.00000 ;      N-    CA-     C-     N
  7098   7097   7099   7100       3    0.00000    0.00000    0.00000
0.00000    0.00000    0.00000 ;      H-     N-    CA-   HA2
  7098   7097   7099   7101       3    0.00000    0.00000    0.00000
0.00000    0.00000    0.00000 ;      H-     N-    CA-   HA3
  7098   7097   7099   7102       3    0.00000    0.00000    0.00000
0.00000    0.00000    0.00000 ;      H-     N-    CA-     C
  7099   7102   7104   7105       3   20.92000    0.00000  -20.92000
0.00000    0.00000    0.00000 ;     CA-     C-     N-     H
  7099   7102   7104   7106       3   20.92000    0.00000  -20.92000
0.00000    0.00000    0.00000 ;     CA-     C-     N-    CA
  7100   7099   7102   7103       3    3.68192   -4.35136    0.00000
1.33888    0.00000    0.00000 ;    HA2-    CA-     C-     O
  7100   7099   7102   7104       3    0.00000    0.00000    0.00000
0.00000    0.00000    0.00000 ;    HA2-    CA-     C-     N
  7101   7099   7102   7103       3    3.68192   -4.35136    0.00000
1.33888    0.00000    0.00000 ;    HA3-    CA-     C-     O
  7101   7099   7102   7104       3    0.00000    0.00000    0.00000
```

0.00000 0.00000 0.00000 ; HA3- CA- C- N
7102 7104 7106 7107 3 0.00000 0.00000 0.00000
0.00000 0.00000 0.00000 ; C- N- CA- HA
7102 7104 7106 7108 3 11.71520 1.67360 15.06240 -
13.38880 0.00000 0.00000 ; C- N- CA- CB
7102 7104 7106 7117 3 1.75728 5.27184 2.25936
-7.02912 0.00000 0.00000 ; C- N- CA- C
7103 7102 7104 7105 3 29.28800 -8.36800 -20.92000
0.00000 0.00000 0.00000 ; O- C- N- H
7103 7102 7104 7106 3 20.92000 0.00000 -20.92000
0.00000 0.00000 0.00000 ; O- C- N- CA
7104 7106 7108 7109 3 0.65084 1.95253 0.00000
-2.60338 0.00000 0.00000 ; N- CA- CB- HB2
7104 7106 7108 7110 3 0.65084 1.95253 0.00000
-2.60338 0.00000 0.00000 ; N- CA- CB- HB3
7104 7106 7108 7111 3 3.21331 2.22589 -4.15053
-2.40998 2.61082 0.00000 ; N- CA- CB- CG
7104 7106 7117 7118 3 0.00000 0.00000 0.00000
0.00000 0.00000 0.00000 ; N- CA- C- O
7104 7106 7117 7119 3 17.40544 -5.02080 -13.22144
9.20480 0.00000 0.00000 ; N- CA- C- N
7105 7104 7106 7107 3 0.00000 0.00000 0.00000
0.00000 0.00000 0.00000 ; H- N- CA- HA
7105 7104 7106 7108 3 0.00000 0.00000 0.00000
0.00000 0.00000 0.00000 ; H- N- CA- CB
7105 7104 7106 7117 3 0.00000 0.00000 0.00000
0.00000 0.00000 0.00000 ; H- N- CA- C
7106 7108 7111 7112 3 0.66944 2.00832 0.00000
-2.67776 0.00000 0.00000 ; CA- CB- CG- HG2
7106 7108 7111 7113 3 0.66944 2.00832 0.00000
-2.67776 0.00000 0.00000 ; CA- CB- CG- HG3
7106 7108 7111 7114 3 10.12110 -1.91209 0.01674
10.17549 -1.87443 0.00000 ; CA- CB- CG- CD
7106 7117 7119 7120 3 20.92000 0.00000 -20.92000
0.00000 0.00000 0.00000 ; CA- C- N- H
7106 7117 7119 7121 3 20.92000 0.00000 -20.92000
0.00000 0.00000 0.00000 ; CA- C- N- CA
7107 7106 7108 7109 3 0.65084 1.95253 0.00000
-2.60338 0.00000 0.00000 ; HA- CA- CB- HB2
7107 7106 7108 7110 3 0.65084 1.95253 0.00000
-2.60338 0.00000 0.00000 ; HA- CA- CB- HB3
7107 7106 7108 7111 3 0.65084 1.95253 0.00000
-2.60338 0.00000 0.00000 ; HA- CA- CB- CG
7107 7106 7117 7118 3 3.68192 -4.35136 0.00000
1.33888 0.00000 0.00000 ; HA- CA- C- O
7107 7106 7117 7119 3 0.00000 0.00000 0.00000
0.00000 0.00000 0.00000 ; HA- CA- C- N
7108 7106 7117 7118 3 0.00000 0.00000 0.00000
0.00000 0.00000 0.00000 ; CB- CA- C- O
7108 7106 7117 7119 3 2.51040 4.18400 1.67360
-6.69440 0.00000 0.00000 ; CB- CA- C- N
7108 7111 7114 7115 3 3.79907 0.00000 -5.40573

0.00000 2.14221 0.00000 ; CB- CG- CD- OE1
7108 7111 7114 7116 3 3.79907 0.00000 -5.40573
0.00000 2.14221 0.00000 ; CB- CG- CD- OE2
7109 7108 7106 7117 3 0.65084 1.95253 0.00000
-2.60338 0.00000 0.00000 ; HB2- CB- CA- C
7109 7108 7111 7112 3 0.62760 1.88280 0.00000
-2.51040 0.00000 0.00000 ; HB2- CB- CG- HG2
7109 7108 7111 7113 3 0.62760 1.88280 0.00000
-2.51040 0.00000 0.00000 ; HB2- CB- CG- HG3
7109 7108 7111 7114 3 0.65084 1.95253 0.00000
-2.60338 0.00000 0.00000 ; HB2- CB- CG- CD
7110 7108 7106 7117 3 0.65084 1.95253 0.00000
-2.60338 0.00000 0.00000 ; HB3- CB- CA- C
7110 7108 7111 7112 3 0.62760 1.88280 0.00000
-2.51040 0.00000 0.00000 ; HB3- CB- CG- HG2
7110 7108 7111 7113 3 0.62760 1.88280 0.00000
-2.51040 0.00000 0.00000 ; HB3- CB- CG- HG3
7110 7108 7111 7114 3 0.65084 1.95253 0.00000
-2.60338 0.00000 0.00000 ; HB3- CB- CG- CD
7111 7108 7106 7117 3 6.86594 3.56895 -8.14206
-2.40998 4.85344 0.00000 ; CG- CB- CA- C
7112 7111 7114 7115 3 0.00000 0.00000 0.00000
0.00000 0.00000 0.00000 ; HG2- CG- CD- OE1
7112 7111 7114 7116 3 0.00000 0.00000 0.00000
0.00000 0.00000 0.00000 ; HG2- CG- CD- OE2
7113 7111 7114 7115 3 0.00000 0.00000 0.00000
0.00000 0.00000 0.00000 ; HG3- CG- CD- OE1
7113 7111 7114 7116 3 0.00000 0.00000 0.00000
0.00000 0.00000 0.00000 ; HG3- CG- CD- OE2
7117 7119 7121 7122 3 0.00000 0.00000 0.00000
0.00000 0.00000 0.00000 ; C- N- CA- HA
7117 7119 7121 7123 3 11.71520 1.67360 15.06240 -
13.38880 0.00000 0.00000 ; C- N- CA- CB
7117 7119 7121 7136 3 1.75728 5.27184 2.25936
-7.02912 0.00000 0.00000 ; C- N- CA- C
7118 7117 7119 7120 3 29.28800 -8.36800 -20.92000
0.00000 0.00000 0.00000 ; O- C- N- H
7118 7117 7119 7121 3 20.92000 0.00000 -20.92000
0.00000 0.00000 0.00000 ; O- C- N- CA
7119 7121 7123 7124 3 0.65084 1.95253 0.00000
-2.60338 0.00000 0.00000 ; N- CA- CB- HB
7119 7121 7123 7125 3 3.83673 0.44769 -1.77402
-2.47693 -0.03347 0.00000 ; N- CA- CB- CG2
7119 7121 7123 7129 3 2.97482 0.12134 2.04179
-1.89117 -3.24678 0.00000 ; N- CA- CB- CG1
7119 7121 7136 7137 3 0.00000 0.00000 0.00000
0.00000 0.00000 0.00000 ; N- CA- C- O
7119 7121 7136 7138 3 17.40544 -5.02080 -13.22144
9.20480 0.00000 0.00000 ; N- CA- C- N
7120 7119 7121 7122 3 0.00000 0.00000 0.00000
0.00000 0.00000 0.00000 ; H- N- CA- HA
7120 7119 7121 7123 3 0.00000 0.00000 0.00000

```
0.00000    0.00000    0.00000 ;      H-     N-    CA-    CB
  7120   7119   7121   7136      3    0.00000    0.00000    0.00000
0.00000    0.00000    0.00000 ;      H-     N-    CA-     C
  7121   7123   7125   7126      3    0.66944    2.00832    0.00000
-2.67776    0.00000    0.00000 ;      CA-    CB-   CG2-  HG21
  7121   7123   7125   7127      3    0.66944    2.00832    0.00000
-2.67776    0.00000    0.00000 ;      CA-    CB-   CG2-  HG22
  7121   7123   7125   7128      3    0.66944    2.00832    0.00000
-2.67776    0.00000    0.00000 ;      CA-    CB-   CG2-  HG23
  7121   7123   7129   7130      3    0.66944    2.00832    0.00000
-2.67776    0.00000    0.00000 ;      CA-    CB-   CG1-  HG12
  7121   7123   7129   7131      3    0.66944    2.00832    0.00000
-2.67776    0.00000    0.00000 ;      CA-    CB-   CG1-  HG13
  7121   7123   7129   7132      3    4.24258   -0.52718   -7.25506
-1.79075    7.69856    0.00000 ;      CA-    CB-   CG1-   CD1
  7121   7136   7138   7139      3   20.92000    0.00000  -20.92000
0.00000    0.00000    0.00000 ;     CA-     C-     N-     H
  7121   7136   7138   7140      3   20.92000    0.00000  -20.92000
0.00000    0.00000    0.00000 ;     CA-     C-     N-    CA
  7122   7121   7123   7124      3    0.65084    1.95253    0.00000
-2.60338    0.00000    0.00000 ;      HA-    CA-    CB-    HB
  7122   7121   7123   7125      3    0.65084    1.95253    0.00000
-2.60338    0.00000    0.00000 ;      HA-    CA-    CB-   CG2
  7122   7121   7123   7129      3    0.65084    1.95253    0.00000
-2.60338    0.00000    0.00000 ;      HA-    CA-    CB-   CG1
  7122   7121   7136   7137      3    3.68192   -4.35136    0.00000
1.33888    0.00000    0.00000 ;     HA-    CA-     C-     O
  7122   7121   7136   7138      3    0.00000    0.00000    0.00000
0.00000    0.00000    0.00000 ;     HA-    CA-     C-     N
  7123   7121   7136   7137      3    0.00000    0.00000    0.00000
0.00000    0.00000    0.00000 ;     CB-    CA-     C-     O
  7123   7121   7136   7138      3    2.51040    4.18400    1.67360
-6.69440    0.00000    0.00000 ;      CB-    CA-     C-     N
  7123   7129   7132   7133      3    0.66944    2.00832    0.00000
-2.67776    0.00000    0.00000 ;      CB-   CG1-   CD1-  HD11
  7123   7129   7132   7134      3    0.66944    2.00832    0.00000
-2.67776    0.00000    0.00000 ;      CB-   CG1-   CD1-  HD12
  7123   7129   7132   7135      3    0.66944    2.00832    0.00000
-2.67776    0.00000    0.00000 ;      CB-   CG1-   CD1-  HD13
  7124   7123   7121   7136      3    0.65084    1.95253    0.00000
-2.60338    0.00000    0.00000 ;      HB-    CB-    CA-     C
  7124   7123   7125   7126      3    0.62760    1.88280    0.00000
-2.51040    0.00000    0.00000 ;      HB-    CB-   CG2-  HG21
  7124   7123   7125   7127      3    0.62760    1.88280    0.00000
-2.51040    0.00000    0.00000 ;      HB-    CB-   CG2-  HG22
  7124   7123   7125   7128      3    0.62760    1.88280    0.00000
-2.51040    0.00000    0.00000 ;      HB-    CB-   CG2-  HG23
  7124   7123   7129   7130      3    0.62760    1.88280    0.00000
-2.51040    0.00000    0.00000 ;      HB-    CB-   CG1-  HG12
  7124   7123   7129   7131      3    0.62760    1.88280    0.00000
-2.51040    0.00000    0.00000 ;      HB-    CB-   CG1-  HG13
  7124   7123   7129   7132      3    0.66944    2.00832    0.00000
```

```
-2.67776    0.00000    0.00000 ;     HB-    CB-   CG1-   CD1
  7125   7123   7121   7136       3    5.67350    3.55640   -6.16722
-2.47693    3.74886    0.00000 ;    CG2-    CB-    CA-     C
  7125   7123   7129   7130       3    0.66944    2.00832    0.00000
-2.67776    0.00000    0.00000 ;    CG2-    CB-   CG1-  HG12
  7125   7123   7129   7131       3    0.66944    2.00832    0.00000
-2.67776    0.00000    0.00000 ;    CG2-    CB-   CG1-  HG13
  7125   7123   7129   7132       3    3.81162    0.49790   -8.14206
-1.79075    7.49773    0.00000 ;    CG2-    CB-   CG1-   CD1
  7126   7125   7123   7129       3    0.66944    2.00832    0.00000
-2.67776    0.00000    0.00000 ;   HG21-   CG2-    CB-   CG1
  7127   7125   7123   7129       3    0.66944    2.00832    0.00000
-2.67776    0.00000    0.00000 ;   HG22-   CG2-    CB-   CG1
  7128   7125   7123   7129       3    0.66944    2.00832    0.00000
-2.67776    0.00000    0.00000 ;   HG23-   CG2-    CB-   CG1
  7129   7123   7121   7136       3    8.26340    0.74057   -9.99976
-1.89117    3.84928    0.00000 ;    CG1-    CB-    CA-     C
  7130   7129   7132   7133       3    0.62760    1.88280    0.00000
-2.51040    0.00000    0.00000 ;   HG12-   CG1-   CD1-  HD11
  7130   7129   7132   7134       3    0.62760    1.88280    0.00000
-2.51040    0.00000    0.00000 ;   HG12-   CG1-   CD1-  HD12
  7130   7129   7132   7135       3    0.62760    1.88280    0.00000
-2.51040    0.00000    0.00000 ;   HG12-   CG1-   CD1-  HD13
  7131   7129   7132   7133       3    0.62760    1.88280    0.00000
-2.51040    0.00000    0.00000 ;   HG13-   CG1-   CD1-  HD11
  7131   7129   7132   7134       3    0.62760    1.88280    0.00000
-2.51040    0.00000    0.00000 ;   HG13-   CG1-   CD1-  HD12
  7131   7129   7132   7135       3    0.62760    1.88280    0.00000
-2.51040    0.00000    0.00000 ;   HG13-   CG1-   CD1-  HD13
  7136   7138   7140   7141       3    0.00000    0.00000    0.00000
0.00000    0.00000    0.00000 ;      C-     N-    CA-    HA
  7136   7138   7140   7142       3   11.71520    1.67360   15.06240  -
13.38880    0.00000    0.00000 ;      C-     N-    CA-    CB
  7136   7138   7140   7148       3    1.75728    5.27184    2.25936
-7.02912    0.00000    0.00000 ;      C-     N-    CA-     C
  7137   7136   7138   7139       3   29.28800   -8.36800  -20.92000
0.00000    0.00000    0.00000 ;      O-     C-     N-     H
  7137   7136   7138   7140       3   20.92000    0.00000  -20.92000
0.00000    0.00000    0.00000 ;      O-     C-     N-    CA
  7138   7140   7142   7143       3    0.65084    1.95253    0.00000
-2.60338    0.00000    0.00000 ;      N-    CA-    CB-   HB2
  7138   7140   7142   7144       3    0.65084    1.95253    0.00000
-2.60338    0.00000    0.00000 ;      N-    CA-    CB-   HB3
  7138   7140   7142   7145       3   15.41386    9.74035   -8.39310
-0.97069    2.97901    0.00000 ;      N-    CA-    CB-    CG
  7138   7140   7148   7149       3    0.00000    0.00000    0.00000
0.00000    0.00000    0.00000 ;      N-    CA-     C-     O
  7138   7140   7148   7150       3   17.40544   -5.02080  -13.22144
9.20480    0.00000    0.00000 ;      N-    CA-     C-     N
  7139   7138   7140   7141       3    0.00000    0.00000    0.00000
0.00000    0.00000    0.00000 ;      H-     N-    CA-    HA
  7139   7138   7140   7142       3    0.00000    0.00000    0.00000
```

```
0.00000    0.00000    0.00000 ;      H-     N-    CA-    CB
  7139   7138   7140   7148       3    0.00000    0.00000    0.00000
0.00000    0.00000    0.00000 ;      H-     N-    CA-     C
  7140   7142   7145   7146       3    6.43499    0.00000   -5.39736
0.00000   -1.03763    0.00000 ;     CA-    CB-    CG-   OD1
  7140   7142   7145   7147       3    6.43499    0.00000   -5.39736
0.00000   -1.03763    0.00000 ;     CA-    CB-    CG-   OD2
  7140   7148   7150   7151       3   20.92000    0.00000  -20.92000
0.00000    0.00000    0.00000 ;     CA-     C-     N-     H
  7140   7148   7150   7152       3   20.92000    0.00000  -20.92000
0.00000    0.00000    0.00000 ;     CA-     C-     N-    CA
  7141   7140   7142   7143       3    0.65084    1.95253    0.00000
-2.60338    0.00000    0.00000 ;     HA-    CA-    CB-   HB2
  7141   7140   7142   7144       3    0.65084    1.95253    0.00000
-2.60338    0.00000    0.00000 ;     HA-    CA-    CB-   HB3
  7141   7140   7142   7145       3    0.65084    1.95253    0.00000
-2.60338    0.00000    0.00000 ;     HA-    CA-    CB-    CG
  7141   7140   7148   7149       3    3.68192   -4.35136    0.00000
1.33888    0.00000    0.00000 ;     HA-    CA-     C-     O
  7141   7140   7148   7150       3    0.00000    0.00000    0.00000
0.00000    0.00000    0.00000 ;     HA-    CA-     C-     N
  7142   7140   7148   7149       3    0.00000    0.00000    0.00000
0.00000    0.00000    0.00000 ;     CB-    CA-     C-     O
  7142   7140   7148   7150       3    2.51040    4.18400    1.67360
-6.69440    0.00000    0.00000 ;     CB-    CA-     C-     N
  7143   7142   7140   7148       3    0.65084    1.95253    0.00000
-2.60338    0.00000    0.00000 ;    HB2-    CB-    CA-     C
  7143   7142   7145   7146       3    0.00000    0.00000    0.00000
0.00000    0.00000    0.00000 ;    HB2-    CB-    CG-   OD1
  7143   7142   7145   7147       3    0.00000    0.00000    0.00000
0.00000    0.00000    0.00000 ;    HB2-    CB-    CG-   OD2
  7144   7142   7140   7148       3    0.65084    1.95253    0.00000
-2.60338    0.00000    0.00000 ;    HB3-    CB-    CA-     C
  7144   7142   7145   7146       3    0.00000    0.00000    0.00000
0.00000    0.00000    0.00000 ;    HB3-    CB-    CG-   OD1
  7144   7142   7145   7147       3    0.00000    0.00000    0.00000
0.00000    0.00000    0.00000 ;    HB3-    CB-    CG-   OD2
  7145   7142   7140   7148       3    7.14627   -1.04600   -8.99560
-0.97069    5.15469    0.00000 ;     CG-    CB-    CA-     C
  7148   7150   7152   7153       3    0.00000    0.00000    0.00000
0.00000    0.00000    0.00000 ;      C-     N-    CA-    HA
  7148   7150   7152   7154       3   11.71520    1.67360   15.06240  -
13.38880    0.00000    0.00000 ;      C-     N-    CA-    CB
  7148   7150   7152   7168       3    1.75728    5.27184    2.25936
-7.02912    0.00000    0.00000 ;      C-     N-    CA-     C
  7149   7148   7150   7151       3   29.28800   -8.36800  -20.92000
0.00000    0.00000    0.00000 ;      O-     C-     N-     H
  7149   7148   7150   7152       3   20.92000    0.00000  -20.92000
0.00000    0.00000    0.00000 ;      O-     C-     N-    CA
  7150   7152   7154   7155       3    0.65084    1.95253    0.00000
-2.60338    0.00000    0.00000 ;      N-    CA-    CB-   HB2
  7150   7152   7154   7156       3    0.65084    1.95253    0.00000
```

-2.60338 0.00000 0.00000 ; N- CA- CB- HB3
7150 7152 7154 7157 3 3.28026 2.46019 -2.19242
-3.21331 -0.23430 0.00000 ; N- CA- CB- CG
7150 7152 7168 7169 3 0.00000 0.00000 0.00000
0.00000 0.00000 0.00000 ; N- CA- C- O
7150 7152 7168 7170 3 17.40544 -5.02080 -13.22144
9.20480 0.00000 0.00000 ; N- CA- C- N
7151 7150 7152 7153 3 0.00000 0.00000 0.00000
0.00000 0.00000 0.00000 ; H- N- CA- HA
7151 7150 7152 7154 3 0.00000 0.00000 0.00000
0.00000 0.00000 0.00000 ; H- N- CA- CB
7151 7150 7152 7168 3 0.00000 0.00000 0.00000
0.00000 0.00000 0.00000 ; H- N- CA- C
7152 7154 7157 7158 3 0.57739 0.00000 1.02926
0.00000 -1.60666 0.00000 ; CA- CB- CG- CD1
7152 7154 7157 7166 3 0.57739 0.00000 1.02926
0.00000 -1.60666 0.00000 ; CA- CB- CG- CD2
7152 7168 7170 7171 3 20.92000 0.00000 -20.92000
0.00000 0.00000 0.00000 ; CA- C- N- H
7152 7168 7170 7172 3 20.92000 0.00000 -20.92000
0.00000 0.00000 0.00000 ; CA- C- N- CA
7153 7152 7154 7155 3 0.65084 1.95253 0.00000
-2.60338 0.00000 0.00000 ; HA- CA- CB- HB2
7153 7152 7154 7156 3 0.65084 1.95253 0.00000
-2.60338 0.00000 0.00000 ; HA- CA- CB- HB3
7153 7152 7154 7157 3 0.65084 1.95253 0.00000
-2.60338 0.00000 0.00000 ; HA- CA- CB- CG
7153 7152 7168 7169 3 3.68192 -4.35136 0.00000
1.33888 0.00000 0.00000 ; HA- CA- C- O
7153 7152 7168 7170 3 0.00000 0.00000 0.00000
0.00000 0.00000 0.00000 ; HA- CA- C- N
7154 7152 7168 7169 3 0.00000 0.00000 0.00000
0.00000 0.00000 0.00000 ; CB- CA- C- O
7154 7152 7168 7170 3 2.51040 4.18400 1.67360
-6.69440 0.00000 0.00000 ; CB- CA- C- N
7154 7157 7158 7159 3 30.33400 0.00000 -30.33400
0.00000 0.00000 0.00000 ; CB- CG- CD1- HD1
7154 7157 7158 7160 3 30.33400 0.00000 -30.33400
0.00000 0.00000 0.00000 ; CB- CG- CD1- CE1
7154 7157 7166 7164 3 30.33400 0.00000 -30.33400
0.00000 0.00000 0.00000 ; CB- CG- CD2- CE2
7154 7157 7166 7167 3 30.33400 0.00000 -30.33400
0.00000 0.00000 0.00000 ; CB- CG- CD2- HD2
7155 7154 7152 7168 3 0.65084 1.95253 0.00000
-2.60338 0.00000 0.00000 ; HB2- CB- CA- C
7155 7154 7157 7158 3 0.00000 0.00000 0.00000
0.00000 0.00000 0.00000 ; HB2- CB- CG- CD1
7155 7154 7157 7166 3 0.00000 0.00000 0.00000
0.00000 0.00000 0.00000 ; HB2- CB- CG- CD2
7156 7154 7152 7168 3 0.65084 1.95253 0.00000
-2.60338 0.00000 0.00000 ; HB3- CB- CA- C
7156 7154 7157 7158 3 0.00000 0.00000 0.00000

```
0.00000    0.00000    0.00000 ;    HB3-    CB-    CG-   CD1
  7156   7154   7157   7166       3    0.00000    0.00000    0.00000
0.00000    0.00000    0.00000 ;    HB3-    CB-    CG-   CD2
  7157   7154   7152   7168       3    4.95804    2.17986   -3.52293
-3.21331   -0.40166    0.00000 ;     CG-    CB-    CA-     C
  7157   7158   7160   7161       3   30.33400    0.00000  -30.33400
0.00000    0.00000    0.00000 ;     CG-   CD1-   CE1-   HE1
  7157   7158   7160   7162       3   30.33400    0.00000  -30.33400
0.00000    0.00000    0.00000 ;     CG-   CD1-   CE1-    CZ
  7157   7166   7164   7162       3   30.33400    0.00000  -30.33400
0.00000    0.00000    0.00000 ;     CG-   CD2-   CE2-    CZ
  7157   7166   7164   7165       3   30.33400    0.00000  -30.33400
0.00000    0.00000    0.00000 ;     CG-   CD2-   CE2-   HE2
  7158   7157   7166   7164       3   30.33400    0.00000  -30.33400
0.00000    0.00000    0.00000 ;    CD1-    CG-   CD2-   CE2
  7158   7157   7166   7167       3   30.33400    0.00000  -30.33400
0.00000    0.00000    0.00000 ;    CD1-    CG-   CD2-   HD2
  7158   7160   7162   7163       3   30.33400    0.00000  -30.33400
0.00000    0.00000    0.00000 ;    CD1-   CE1-    CZ-    HZ
  7158   7160   7162   7164       3   30.33400    0.00000  -30.33400
0.00000    0.00000    0.00000 ;    CD1-   CE1-    CZ-   CE2
  7159   7158   7157   7166       3   30.33400    0.00000  -30.33400
0.00000    0.00000    0.00000 ;    HD1-   CD1-    CG-   CD2
  7159   7158   7160   7161       3   30.33400    0.00000  -30.33400
0.00000    0.00000    0.00000 ;    HD1-   CD1-   CE1-   HE1
  7159   7158   7160   7162       3   30.33400    0.00000  -30.33400
0.00000    0.00000    0.00000 ;    HD1-   CD1-   CE1-    CZ
  7160   7158   7157   7166       3   30.33400    0.00000  -30.33400
0.00000    0.00000    0.00000 ;    CE1-   CD1-    CG-   CD2
  7160   7162   7164   7165       3   30.33400    0.00000  -30.33400
0.00000    0.00000    0.00000 ;    CE1-    CZ-   CE2-   HE2
  7160   7162   7164   7166       3   30.33400    0.00000  -30.33400
0.00000    0.00000    0.00000 ;    CE1-    CZ-   CE2-   CD2
  7161   7160   7162   7163       3   30.33400    0.00000  -30.33400
0.00000    0.00000    0.00000 ;    HE1-   CE1-    CZ-    HZ
  7161   7160   7162   7164       3   30.33400    0.00000  -30.33400
0.00000    0.00000    0.00000 ;    HE1-   CE1-    CZ-   CE2
  7162   7164   7166   7167       3   30.33400    0.00000  -30.33400
0.00000    0.00000    0.00000 ;     CZ-   CE2-   CD2-   HD2
  7163   7162   7164   7165       3   30.33400    0.00000  -30.33400
0.00000    0.00000    0.00000 ;     HZ-    CZ-   CE2-   HE2
  7163   7162   7164   7166       3   30.33400    0.00000  -30.33400
0.00000    0.00000    0.00000 ;     HZ-    CZ-   CE2-   CD2
  7165   7164   7166   7167       3   30.33400    0.00000  -30.33400
0.00000    0.00000    0.00000 ;    HE2-   CE2-   CD2-   HD2
  7168   7170   7172   7173       3    0.00000    0.00000    0.00000
0.00000    0.00000    0.00000 ;      C-     N-    CA-    HA
  7168   7170   7172   7174       3   11.71520    1.67360   15.06240  -
13.38880    0.00000    0.00000 ;      C-     N-    CA-    CB
  7168   7170   7172   7186       3    1.75728    5.27184    2.25936
-7.02912    0.00000    0.00000 ;      C-     N-    CA-     C
  7169   7168   7170   7171       3   29.28800   -8.36800  -20.92000
```

0.00000 0.00000 0.00000 ; O- C- N- H
7169 7168 7170 7172 3 20.92000 0.00000 -20.92000
0.00000 0.00000 0.00000 ; O- C- N- CA
7170 7172 7174 7175 3 0.65084 1.95253 0.00000
-2.60338 0.00000 0.00000 ; N- CA- CB- HB2
7170 7172 7174 7176 3 0.65084 1.95253 0.00000
-2.60338 0.00000 0.00000 ; N- CA- CB- HB3
7170 7172 7174 7177 3 4.79068 4.02919 -4.82834
-3.66518 2.97901 0.00000 ; N- CA- CB- CG
7170 7172 7186 7187 3 0.00000 0.00000 0.00000
0.00000 0.00000 0.00000 ; N- CA- C- O
7170 7172 7186 7188 3 17.40544 -5.02080 -13.22144
9.20480 0.00000 0.00000 ; N- CA- C- N
7171 7170 7172 7173 3 0.00000 0.00000 0.00000
0.00000 0.00000 0.00000 ; H- N- CA- HA
7171 7170 7172 7174 3 0.00000 0.00000 0.00000
0.00000 0.00000 0.00000 ; H- N- CA- CB
7171 7170 7172 7186 3 0.00000 0.00000 0.00000
0.00000 0.00000 0.00000 ; H- N- CA- C
7172 7174 7177 7178 3 6.81992 9.28011 -2.04179 -
11.48090 -1.23846 0.00000 ; CA- CB- CG- ND1
7172 7174 7177 7184 3 0.00000 0.00000 0.00000
0.00000 0.00000 0.00000 ; CA- CB- CG- CD2
7172 7186 7188 7189 3 20.92000 0.00000 -20.92000
0.00000 0.00000 0.00000 ; CA- C- N- H
7172 7186 7188 7190 3 20.92000 0.00000 -20.92000
0.00000 0.00000 0.00000 ; CA- C- N- CA
7173 7172 7174 7175 3 0.65084 1.95253 0.00000
-2.60338 0.00000 0.00000 ; HA- CA- CB- HB2
7173 7172 7174 7176 3 0.65084 1.95253 0.00000
-2.60338 0.00000 0.00000 ; HA- CA- CB- HB3
7173 7172 7174 7177 3 0.65084 1.95253 0.00000
-2.60338 0.00000 0.00000 ; HA- CA- CB- CG
7173 7172 7186 7187 3 3.68192 -4.35136 0.00000
1.33888 0.00000 0.00000 ; HA- CA- C- O
7173 7172 7186 7188 3 0.00000 0.00000 0.00000
0.00000 0.00000 0.00000 ; HA- CA- C- N
7174 7172 7186 7187 3 0.00000 0.00000 0.00000
0.00000 0.00000 0.00000 ; CB- CA- C- O
7174 7172 7186 7188 3 2.51040 4.18400 1.67360
-6.69440 0.00000 0.00000 ; CB- CA- C- N
7174 7177 7178 7179 3 11.71520 0.00000 -11.71520
0.00000 0.00000 0.00000 ; CB- CG- ND1- HD1
7174 7177 7178 7180 3 11.71520 0.00000 -11.71520
0.00000 0.00000 0.00000 ; CB- CG- ND1- CE1
7174 7177 7184 7182 3 44.97800 0.00000 -44.97800
0.00000 0.00000 0.00000 ; CB- CG- CD2- NE2
7174 7177 7184 7185 3 44.97800 0.00000 -44.97800
0.00000 0.00000 0.00000 ; CB- CG- CD2- HD2
7175 7174 7172 7186 3 0.65084 1.95253 0.00000
-2.60338 0.00000 0.00000 ; HB2- CB- CA- C
7175 7174 7177 7178 3 0.00000 0.00000 0.00000

0.00000 0.00000 0.00000 ; HB2- CB- CG- ND1
7175 7174 7177 7184 3 0.00000 0.00000 0.00000
0.00000 0.00000 0.00000 ; HB2- CB- CG- CD2
7176 7174 7172 7186 3 0.65084 1.95253 0.00000
-2.60338 0.00000 0.00000 ; HB3- CB- CA- C
7176 7174 7177 7178 3 0.00000 0.00000 0.00000
0.00000 0.00000 0.00000 ; HB3- CB- CG- ND1
7176 7174 7177 7184 3 0.00000 0.00000 0.00000
0.00000 0.00000 0.00000 ; HB3- CB- CG- CD2
7177 7174 7172 7186 3 3.76560 3.34720 -2.87859
-3.66518 0.83680 0.00000 ; CG- CB- CA- C
7177 7178 7180 7181 3 19.45560 0.00000 -19.45560
0.00000 0.00000 0.00000 ; CG- ND1- CE1- HE1
7177 7178 7180 7182 3 19.45560 0.00000 -19.45560
0.00000 0.00000 0.00000 ; CG- ND1- CE1- NE2
7177 7184 7182 7180 3 12.55200 0.00000 -12.55200
0.00000 0.00000 0.00000 ; CG- CD2- NE2- CE1
7177 7184 7182 7183 3 12.55200 0.00000 -12.55200
0.00000 0.00000 0.00000 ; CG- CD2- NE2- HE2
7178 7177 7184 7182 3 44.97800 0.00000 -44.97800
0.00000 0.00000 0.00000 ; ND1- CG- CD2- NE2
7178 7177 7184 7185 3 44.97800 0.00000 -44.97800
0.00000 0.00000 0.00000 ; ND1- CG- CD2- HD2
7178 7180 7182 7183 3 19.45560 0.00000 -19.45560
0.00000 0.00000 0.00000 ; ND1- CE1- NE2- HE2
7178 7180 7182 7184 3 19.45560 0.00000 -19.45560
0.00000 0.00000 0.00000 ; ND1- CE1- NE2- CD2
7179 7178 7177 7184 3 11.71520 0.00000 -11.71520
0.00000 0.00000 0.00000 ; HD1- ND1- CG- CD2
7179 7178 7180 7181 3 19.45560 0.00000 -19.45560
0.00000 0.00000 0.00000 ; HD1- ND1- CE1- HE1
7179 7178 7180 7182 3 19.45560 0.00000 -19.45560
0.00000 0.00000 0.00000 ; HD1- ND1- CE1- NE2
7180 7178 7177 7184 3 11.71520 0.00000 -11.71520
0.00000 0.00000 0.00000 ; CE1- ND1- CG- CD2
7180 7182 7184 7185 3 12.55200 0.00000 -12.55200
0.00000 0.00000 0.00000 ; CE1- NE2- CD2- HD2
7181 7180 7182 7183 3 19.45560 0.00000 -19.45560
0.00000 0.00000 0.00000 ; HE1- CE1- NE2- HE2
7181 7180 7182 7184 3 19.45560 0.00000 -19.45560
0.00000 0.00000 0.00000 ; HE1- CE1- NE2- CD2
7183 7182 7184 7185 3 12.55200 0.00000 -12.55200
0.00000 0.00000 0.00000 ; HE2- NE2- CD2- HD2
7186 7188 7190 7191 3 0.00000 0.00000 0.00000
0.00000 0.00000 0.00000 ; C- N- CA- HA
7186 7188 7190 7192 3 11.71520 1.67360 15.06240 -
13.38880 0.00000 0.00000 ; C- N- CA- CB
7186 7188 7190 7202 3 1.75728 5.27184 2.25936
-7.02912 0.00000 0.00000 ; C- N- CA- C
7187 7186 7188 7189 3 29.28800 -8.36800 -20.92000
0.00000 0.00000 0.00000 ; O- C- N- H
7187 7186 7188 7190 3 20.92000 0.00000 -20.92000

```
0.00000    0.00000    0.00000 ;       O-     C-     N-    CA
  7188   7190   7192   7193      3    0.65084    1.95253    0.00000
-2.60338    0.00000    0.00000 ;       N-    CA-    CB-   HB2
  7188   7190   7192   7194      3    0.65084    1.95253    0.00000
-2.60338    0.00000    0.00000 ;       N-    CA-    CB-   HB3
  7188   7190   7192   7195      3    3.21331    2.22589   -4.15053
-2.40998    2.61082    0.00000 ;       N-    CA-    CB-    CG
  7188   7190   7202   7203      3    0.00000    0.00000    0.00000
0.00000    0.00000    0.00000 ;       N-    CA-     C-     O
  7188   7190   7202   7204      3   17.40544   -5.02080  -13.22144
9.20480    0.00000    0.00000 ;       N-    CA-     C-     N
  7189   7188   7190   7191      3    0.00000    0.00000    0.00000
0.00000    0.00000    0.00000 ;       H-     N-    CA-    HA
  7189   7188   7190   7192      3    0.00000    0.00000    0.00000
0.00000    0.00000    0.00000 ;       H-     N-    CA-    CB
  7189   7188   7190   7202      3    0.00000    0.00000    0.00000
0.00000    0.00000    0.00000 ;       H-     N-    CA-     C
  7190   7192   7195   7196      3    0.66944    2.00832    0.00000
-2.67776    0.00000    0.00000 ;      CA-    CB-    CG-   HG2
  7190   7192   7195   7197      3    0.66944    2.00832    0.00000
-2.67776    0.00000    0.00000 ;      CA-    CB-    CG-   HG3
  7190   7192   7195   7198      3    3.69866   -4.35136   -3.92459
6.89523    4.61914    0.00000 ;      CA-    CB-    CG-    CD
  7190   7202   7204   7205      3   20.92000    0.00000  -20.92000
0.00000    0.00000    0.00000 ;      CA-     C-     N-     H
  7190   7202   7204   7206      3   20.92000    0.00000  -20.92000
0.00000    0.00000    0.00000 ;      CA-     C-     N-    CA
  7191   7190   7192   7193      3    0.65084    1.95253    0.00000
-2.60338    0.00000    0.00000 ;      HA-    CA-    CB-   HB2
  7191   7190   7192   7194      3    0.65084    1.95253    0.00000
-2.60338    0.00000    0.00000 ;      HA-    CA-    CB-   HB3
  7191   7190   7192   7195      3    0.65084    1.95253    0.00000
-2.60338    0.00000    0.00000 ;      HA-    CA-    CB-    CG
  7191   7190   7202   7203      3    3.68192   -4.35136    0.00000
1.33888    0.00000    0.00000 ;      HA-    CA-     C-     O
  7191   7190   7202   7204      3    0.00000    0.00000    0.00000
0.00000    0.00000    0.00000 ;      HA-    CA-     C-     N
  7192   7190   7202   7203      3    0.00000    0.00000    0.00000
0.00000    0.00000    0.00000 ;      CB-    CA-     C-     O
  7192   7190   7202   7204      3    2.51040    4.18400    1.67360
-6.69440    0.00000    0.00000 ;      CB-    CA-     C-     N
  7192   7195   7198   7199      3    0.00000    0.00000    0.00000
0.00000    0.00000    0.00000 ;      CB-    CG-    CD-   OE1
  7192   7195   7198   7200      3   12.79049    3.13382   -7.02912
0.41840   -2.20915    0.00000 ;      CB-    CG-    CD-   OE2
  7193   7192   7190   7202      3    0.65084    1.95253    0.00000
-2.60338    0.00000    0.00000 ;     HB2-    CB-    CA-     C
  7193   7192   7195   7196      3    0.62760    1.88280    0.00000
-2.51040    0.00000    0.00000 ;     HB2-    CB-    CG-   HG2
  7193   7192   7195   7197      3    0.62760    1.88280    0.00000
-2.51040    0.00000    0.00000 ;     HB2-    CB-    CG-   HG3
  7193   7192   7195   7198      3    0.65084    1.95253    0.00000
```

-2.60338    0.00000    0.00000 ;    HB2-    CB-    CG-    CD
  7194   7192   7190   7202       3    0.65084    1.95253    0.00000
-2.60338    0.00000    0.00000 ;    HB3-    CB-    CA-    C
  7194   7192   7195   7196       3    0.62760    1.88280    0.00000
-2.51040    0.00000    0.00000 ;    HB3-    CB-    CG-   HG2
  7194   7192   7195   7197       3    0.62760    1.88280    0.00000
-2.51040    0.00000    0.00000 ;    HB3-    CB-    CG-   HG3
  7194   7192   7195   7198       3    0.65084    1.95253    0.00000
-2.60338    0.00000    0.00000 ;    HB3-    CB-    CG-    CD
  7195   7192   7190   7202       3    6.86594    3.56895   -8.14206
-2.40998    4.85344    0.00000 ;    CG-    CB-    CA-    C
  7195   7198   7200   7201       3   27.46796    7.88684  -26.42614
-8.01654    3.78234    0.00000 ;    CG-    CD-   OE2-   HE2
  7196   7195   7198   7199       3    3.68192   -4.35136    0.00000
1.33888    0.00000    0.00000 ;    HG2-    CG-    CD-   OE1
  7196   7195   7198   7200       3    0.00000    0.00000    0.00000
0.00000    0.00000    0.00000 ;    HG2-    CG-    CD-   OE2
  7197   7195   7198   7199       3    3.68192   -4.35136    0.00000
1.33888    0.00000    0.00000 ;    HG3-    CG-    CD-   OE1
  7197   7195   7198   7200       3    0.00000    0.00000    0.00000
0.00000    0.00000    0.00000 ;    HG3-    CG-    CD-   OE2
  7199   7198   7200   7201       3   27.19600   -7.94960  -19.24640
0.00000    0.00000    0.00000 ;    OE1-    CD-   OE2-   HE2
  7202   7204   7206   7207       3    0.00000    0.00000    0.00000
0.00000    0.00000    0.00000 ;    C-    N-    CA-    HA
  7202   7204   7206   7208       3   11.71520    1.67360   15.06240  -
13.38880    0.00000    0.00000 ;    C-    N-    CA-    CB
  7202   7204   7206   7222       3    1.75728    5.27184    2.25936
-7.02912    0.00000    0.00000 ;    C-    N-    CA-    C
  7203   7202   7204   7205       3   29.28800   -8.36800  -20.92000
0.00000    0.00000    0.00000 ;    O-    C-    N-    H
  7203   7202   7204   7206       3   20.92000    0.00000  -20.92000
0.00000    0.00000    0.00000 ;    O-    C-    N-    CA
  7204   7206   7208   7209       3    0.65084    1.95253    0.00000
-2.60338    0.00000    0.00000 ;    N-    CA-    CB-   HB2
  7204   7206   7208   7210       3    0.65084    1.95253    0.00000
-2.60338    0.00000    0.00000 ;    N-    CA-    CB-   HB3
  7204   7206   7208   7211       3    3.28026    2.46019   -2.19242
-3.21331   -0.23430    0.00000 ;    N-    CA-    CB-    CG
  7204   7206   7222   7223       3    0.00000    0.00000    0.00000
0.00000    0.00000    0.00000 ;    N-    CA-    C-    O
  7204   7206   7222   7224       3   17.40544   -5.02080  -13.22144
9.20480    0.00000    0.00000 ;    N-    CA-    C-    N
  7205   7204   7206   7207       3    0.00000    0.00000    0.00000
0.00000    0.00000    0.00000 ;    H-    N-    CA-    HA
  7205   7204   7206   7208       3    0.00000    0.00000    0.00000
0.00000    0.00000    0.00000 ;    H-    N-    CA-    CB
  7205   7204   7206   7222       3    0.00000    0.00000    0.00000
0.00000    0.00000    0.00000 ;    H-    N-    CA-    C
  7206   7208   7211   7212       3    0.57739    0.00000    1.02926
0.00000   -1.60666    0.00000 ;    CA-    CB-    CG-   CD1
  7206   7208   7211   7220       3    0.57739    0.00000    1.02926

0.00000   -1.60666    0.00000 ;      CA-     CB-     CG-   CD2
  7206   7222   7224   7225       3   20.92000    0.00000  -20.92000
0.00000    0.00000    0.00000 ;      CA-      C-      N-     H
  7206   7222   7224   7226       3   20.92000    0.00000  -20.92000
0.00000    0.00000    0.00000 ;      CA-      C-      N-    CA
  7207   7206   7208   7209       3    0.65084    1.95253    0.00000
-2.60338    0.00000    0.00000 ;      HA-     CA-     CB-   HB2
  7207   7206   7208   7210       3    0.65084    1.95253    0.00000
-2.60338    0.00000    0.00000 ;      HA-     CA-     CB-   HB3
  7207   7206   7208   7211       3    0.65084    1.95253    0.00000
-2.60338    0.00000    0.00000 ;      HA-     CA-     CB-    CG
  7207   7206   7222   7223       3    3.68192   -4.35136    0.00000
1.33888    0.00000    0.00000 ;      HA-     CA-      C-     O
  7207   7206   7222   7224       3    0.00000    0.00000    0.00000
0.00000    0.00000    0.00000 ;      HA-     CA-      C-     N
  7208   7206   7222   7223       3    0.00000    0.00000    0.00000
0.00000    0.00000    0.00000 ;      CB-     CA-      C-     O
  7208   7206   7222   7224       3    2.51040    4.18400    1.67360
-6.69440    0.00000    0.00000 ;      CB-     CA-      C-     N
  7208   7211   7212   7213       3   30.33400    0.00000  -30.33400
0.00000    0.00000    0.00000 ;      CB-     CG-    CD1-   HD1
  7208   7211   7212   7214       3   30.33400    0.00000  -30.33400
0.00000    0.00000    0.00000 ;      CB-     CG-    CD1-   CE1
  7208   7211   7220   7218       3   30.33400    0.00000  -30.33400
0.00000    0.00000    0.00000 ;      CB-     CG-    CD2-   CE2
  7208   7211   7220   7221       3   30.33400    0.00000  -30.33400
0.00000    0.00000    0.00000 ;      CB-     CG-    CD2-   HD2
  7209   7208   7206   7222       3    0.65084    1.95253    0.00000
-2.60338    0.00000    0.00000 ;     HB2-     CB-     CA-     C
  7209   7208   7211   7212       3    0.00000    0.00000    0.00000
0.00000    0.00000    0.00000 ;     HB2-     CB-     CG-   CD1
  7209   7208   7211   7220       3    0.00000    0.00000    0.00000
0.00000    0.00000    0.00000 ;     HB2-     CB-     CG-   CD2
  7210   7208   7206   7222       3    0.65084    1.95253    0.00000
-2.60338    0.00000    0.00000 ;     HB3-     CB-     CA-     C
  7210   7208   7211   7212       3    0.00000    0.00000    0.00000
0.00000    0.00000    0.00000 ;     HB3-     CB-     CG-   CD1
  7210   7208   7211   7220       3    0.00000    0.00000    0.00000
0.00000    0.00000    0.00000 ;     HB3-     CB-     CG-   CD2
  7211   7208   7206   7222       3    4.95804    2.17986   -3.52293
-3.21331   -0.40166    0.00000 ;      CG-     CB-     CA-     C
  7211   7212   7214   7215       3   30.33400    0.00000  -30.33400
0.00000    0.00000    0.00000 ;      CG-    CD1-    CE1-   HE1
  7211   7212   7214   7216       3   30.33400    0.00000  -30.33400
0.00000    0.00000    0.00000 ;      CG-    CD1-    CE1-    CZ
  7211   7220   7218   7216       3   30.33400    0.00000  -30.33400
0.00000    0.00000    0.00000 ;      CG-    CD2-    CE2-    CZ
  7211   7220   7218   7219       3   30.33400    0.00000  -30.33400
0.00000    0.00000    0.00000 ;      CG-    CD2-    CE2-   HE2
  7212   7211   7220   7218       3   30.33400    0.00000  -30.33400
0.00000    0.00000    0.00000 ;     CD1-     CG-    CD2-   CE2
  7212   7211   7220   7221       3   30.33400    0.00000  -30.33400

```
0.00000    0.00000    0.00000 ;    CD1-    CG-   CD2-   HD2
  7212   7214   7216   7217       3   30.33400    0.00000  -30.33400
0.00000    0.00000    0.00000 ;    CD1-   CE1-    CZ-    HZ
  7212   7214   7216   7218       3   30.33400    0.00000  -30.33400
0.00000    0.00000    0.00000 ;    CD1-   CE1-    CZ-   CE2
  7213   7212   7211   7220       3   30.33400    0.00000  -30.33400
0.00000    0.00000    0.00000 ;    HD1-   CD1-    CG-   CD2
  7213   7212   7214   7215       3   30.33400    0.00000  -30.33400
0.00000    0.00000    0.00000 ;    HD1-   CD1-   CE1-   HE1
  7213   7212   7214   7216       3   30.33400    0.00000  -30.33400
0.00000    0.00000    0.00000 ;    HD1-   CD1-   CE1-    CZ
  7214   7212   7211   7220       3   30.33400    0.00000  -30.33400
0.00000    0.00000    0.00000 ;    CE1-   CD1-    CG-   CD2
  7214   7216   7218   7219       3   30.33400    0.00000  -30.33400
0.00000    0.00000    0.00000 ;    CE1-    CZ-   CE2-   HE2
  7214   7216   7218   7220       3   30.33400    0.00000  -30.33400
0.00000    0.00000    0.00000 ;    CE1-    CZ-   CE2-   CD2
  7215   7214   7216   7217       3   30.33400    0.00000  -30.33400
0.00000    0.00000    0.00000 ;    HE1-   CE1-    CZ-    HZ
  7215   7214   7216   7218       3   30.33400    0.00000  -30.33400
0.00000    0.00000    0.00000 ;    HE1-   CE1-    CZ-   CE2
  7216   7218   7220   7221       3   30.33400    0.00000  -30.33400
0.00000    0.00000    0.00000 ;     CZ-   CE2-   CD2-   HD2
  7217   7216   7218   7219       3   30.33400    0.00000  -30.33400
0.00000    0.00000    0.00000 ;     HZ-    CZ-   CE2-   HE2
  7217   7216   7218   7220       3   30.33400    0.00000  -30.33400
0.00000    0.00000    0.00000 ;     HZ-    CZ-   CE2-   CD2
  7219   7218   7220   7221       3   30.33400    0.00000  -30.33400
0.00000    0.00000    0.00000 ;    HE2-   CE2-   CD2-   HD2
  7222   7224   7226   7227       3    0.00000    0.00000    0.00000
0.00000    0.00000    0.00000 ;      C-     N-    CA-    HA
  7222   7224   7226   7228       3   11.71520    1.67360   15.06240  -
13.38880    0.00000    0.00000 ;      C-     N-    CA-    CB
  7222   7224   7226   7239       3    1.75728    5.27184    2.25936
-7.02912    0.00000    0.00000 ;      C-     N-    CA-     C
  7223   7222   7224   7225       3   29.28800   -8.36800  -20.92000
0.00000    0.00000    0.00000 ;      O-     C-     N-     H
  7223   7222   7224   7226       3   20.92000    0.00000  -20.92000
0.00000    0.00000    0.00000 ;      O-     C-     N-    CA
  7224   7226   7228   7229       3    0.65084    1.95253    0.00000
-2.60338    0.00000    0.00000 ;      N-    CA-    CB-   HB2
  7224   7226   7228   7230       3    0.65084    1.95253    0.00000
-2.60338    0.00000    0.00000 ;      N-    CA-    CB-   HB3
  7224   7226   7228   7231       3    3.21331    2.22589   -4.15053
-2.40998    2.61082    0.00000 ;      N-    CA-    CB-    CG
  7224   7226   7239   7240       3    0.00000    0.00000    0.00000
0.00000    0.00000    0.00000 ;      N-    CA-     C-     O
  7224   7226   7239   7241       3   17.40544   -5.02080  -13.22144
9.20480    0.00000    0.00000 ;      N-    CA-     C-     N
  7225   7224   7226   7227       3    0.00000    0.00000    0.00000
0.00000    0.00000    0.00000 ;      H-     N-    CA-    HA
  7225   7224   7226   7228       3    0.00000    0.00000    0.00000
```

```
0.00000    0.00000    0.00000 ;      H-     N-    CA-    CB
  7225   7224   7226   7239      3    0.00000    0.00000    0.00000
0.00000    0.00000    0.00000 ;      H-     N-    CA-     C
  7226   7228   7231   7232      3    0.65084    1.95253    0.00000
-2.60338    0.00000    0.00000 ;     CA-    CB-    CG-   HG2
  7226   7228   7231   7233      3    0.65084    1.95253    0.00000
-2.60338    0.00000    0.00000 ;     CA-    CB-    CG-   HG3
  7226   7228   7231   7234      3    2.04598   -1.54390    1.11294
-0.26778    0.93722    0.00000 ;     CA-    CB-    CG-    SD
  7226   7239   7241   7242      3   20.92000    0.00000  -20.92000
0.00000    0.00000    0.00000 ;     CA-     C-     N-     H
  7226   7239   7241   7243      3   20.92000    0.00000  -20.92000
0.00000    0.00000    0.00000 ;     CA-     C-     N-    CA
  7227   7226   7228   7229      3    0.65084    1.95253    0.00000
-2.60338    0.00000    0.00000 ;     HA-    CA-    CB-   HB2
  7227   7226   7228   7230      3    0.65084    1.95253    0.00000
-2.60338    0.00000    0.00000 ;     HA-    CA-    CB-   HB3
  7227   7226   7228   7231      3    0.65084    1.95253    0.00000
-2.60338    0.00000    0.00000 ;     HA-    CA-    CB-    CG
  7227   7226   7239   7240      3    3.68192   -4.35136    0.00000
1.33888    0.00000    0.00000 ;     HA-    CA-     C-     O
  7227   7226   7239   7241      3    0.00000    0.00000    0.00000
0.00000    0.00000    0.00000 ;     HA-    CA-     C-     N
  7228   7226   7239   7240      3    0.00000    0.00000    0.00000
0.00000    0.00000    0.00000 ;     CB-    CA-     C-     O
  7228   7226   7239   7241      3    2.51040    4.18400    1.67360
-6.69440    0.00000    0.00000 ;     CB-    CA-     C-     N
  7228   7231   7234   7235      3    3.24260    6.22998    1.79075
-6.92870    1.90790    0.00000 ;     CB-    CG-    SD-    CE
  7229   7228   7226   7239      3    0.65084    1.95253    0.00000
-2.60338    0.00000    0.00000 ;    HB2-    CB-    CA-     C
  7229   7228   7231   7232      3    0.65084    1.95253    0.00000
-2.60338    0.00000    0.00000 ;    HB2-    CB-    CG-   HG2
  7229   7228   7231   7233      3    0.65084    1.95253    0.00000
-2.60338    0.00000    0.00000 ;    HB2-    CB-    CG-   HG3
  7229   7228   7231   7234      3    0.65084    1.95253    0.00000
-2.60338    0.00000    0.00000 ;    HB2-    CB-    CG-    SD
  7230   7228   7226   7239      3    0.65084    1.95253    0.00000
-2.60338    0.00000    0.00000 ;    HB3-    CB-    CA-     C
  7230   7228   7231   7232      3    0.65084    1.95253    0.00000
-2.60338    0.00000    0.00000 ;    HB3-    CB-    CG-   HG2
  7230   7228   7231   7233      3    0.65084    1.95253    0.00000
-2.60338    0.00000    0.00000 ;    HB3-    CB-    CG-   HG3
  7230   7228   7231   7234      3    0.65084    1.95253    0.00000
-2.60338    0.00000    0.00000 ;    HB3-    CB-    CG-    SD
  7231   7228   7226   7239      3    6.86594    3.56895   -8.14206
-2.40998    4.85344    0.00000 ;     CG-    CB-    CA-     C
  7231   7234   7235   7236      3    1.39467    4.18400    0.00000
-5.57867    0.00000    0.00000 ;     CG-    SD-    CE-   HE1
  7231   7234   7235   7237      3    1.39467    4.18400    0.00000
-5.57867    0.00000    0.00000 ;     CG-    SD-    CE-   HE2
  7231   7234   7235   7238      3    1.39467    4.18400    0.00000
```

-5.57867 0.00000 0.00000 ; CG- SD- CE- HE3
7232 7231 7234 7235 3 1.39467 4.18400 0.00000
-5.57867 0.00000 0.00000 ; HG2- CG- SD- CE
7233 7231 7234 7235 3 1.39467 4.18400 0.00000
-5.57867 0.00000 0.00000 ; HG3- CG- SD- CE
7239 7241 7243 7244 3 0.00000 0.00000 0.00000
0.00000 0.00000 0.00000 ; C- N- CA- HA
7239 7241 7243 7245 3 11.71520 1.67360 15.06240 -
13.38880 0.00000 0.00000 ; C- N- CA- CB
7239 7241 7243 7258 3 1.75728 5.27184 2.25936
-7.02912 0.00000 0.00000 ; C- N- CA- C
7240 7239 7241 7242 3 29.28800 -8.36800 -20.92000
0.00000 0.00000 0.00000 ; O- C- N- H
7240 7239 7241 7243 3 20.92000 0.00000 -20.92000
0.00000 0.00000 0.00000 ; O- C- N- CA
7241 7243 7245 7246 3 0.65084 1.95253 0.00000
-2.60338 0.00000 0.00000 ; N- CA- CB- HB2
7241 7243 7245 7247 3 0.65084 1.95253 0.00000
-2.60338 0.00000 0.00000 ; N- CA- CB- HB3
7241 7243 7245 7248 3 3.79070 1.39746 -4.61077
-2.40998 2.44346 0.00000 ; N- CA- CB- CG
7241 7243 7258 7259 3 0.00000 0.00000 0.00000
0.00000 0.00000 0.00000 ; N- CA- C- O
7241 7243 7258 7260 3 17.40544 -5.02080 -13.22144
9.20480 0.00000 0.00000 ; N- CA- C- N
7242 7241 7243 7244 3 0.00000 0.00000 0.00000
0.00000 0.00000 0.00000 ; H- N- CA- HA
7242 7241 7243 7245 3 0.00000 0.00000 0.00000
0.00000 0.00000 0.00000 ; H- N- CA- CB
7242 7241 7243 7258 3 0.00000 0.00000 0.00000
0.00000 0.00000 0.00000 ; H- N- CA- C
7243 7245 7248 7249 3 0.66944 2.00832 0.00000
-2.67776 0.00000 0.00000 ; CA- CB- CG- HG
7243 7245 7248 7250 3 3.90367 0.19665 -6.21742
-2.37651 5.99149 0.00000 ; CA- CB- CG- CD1
7243 7245 7248 7254 3 3.90367 0.19665 -6.21742
-2.37651 5.99149 0.00000 ; CA- CB- CG- CD2
7243 7258 7260 7261 3 20.92000 0.00000 -20.92000
0.00000 0.00000 0.00000 ; CA- C- N- H
7243 7258 7260 7262 3 20.92000 0.00000 -20.92000
0.00000 0.00000 0.00000 ; CA- C- N- CA
7244 7243 7245 7246 3 0.65084 1.95253 0.00000
-2.60338 0.00000 0.00000 ; HA- CA- CB- HB2
7244 7243 7245 7247 3 0.65084 1.95253 0.00000
-2.60338 0.00000 0.00000 ; HA- CA- CB- HB3
7244 7243 7245 7248 3 0.65084 1.95253 0.00000
-2.60338 0.00000 0.00000 ; HA- CA- CB- CG
7244 7243 7258 7259 3 3.68192 -4.35136 0.00000
1.33888 0.00000 0.00000 ; HA- CA- C- O
7244 7243 7258 7260 3 0.00000 0.00000 0.00000
0.00000 0.00000 0.00000 ; HA- CA- C- N
7245 7243 7258 7259 3 0.00000 0.00000 0.00000

```
0.00000    0.00000    0.00000 ;      CB-    CA-     C-     O
  7245   7243   7258   7260      3    2.51040    4.18400    1.67360
-6.69440    0.00000    0.00000 ;     CB-    CA-     C-     N
  7245   7248   7250   7251      3    0.66944    2.00832    0.00000
-2.67776    0.00000    0.00000 ;     CB-    CG-   CD1-  HD11
  7245   7248   7250   7252      3    0.66944    2.00832    0.00000
-2.67776    0.00000    0.00000 ;     CB-    CG-   CD1-  HD12
  7245   7248   7250   7253      3    0.66944    2.00832    0.00000
-2.67776    0.00000    0.00000 ;     CB-    CG-   CD1-  HD13
  7245   7248   7254   7255      3    0.66944    2.00832    0.00000
-2.67776    0.00000    0.00000 ;     CB-    CG-   CD2-  HD21
  7245   7248   7254   7256      3    0.66944    2.00832    0.00000
-2.67776    0.00000    0.00000 ;     CB-    CG-   CD2-  HD22
  7245   7248   7254   7257      3    0.66944    2.00832    0.00000
-2.67776    0.00000    0.00000 ;     CB-    CG-   CD2-  HD23
  7246   7245   7243   7258      3    0.65084    1.95253    0.00000
-2.60338    0.00000    0.00000 ;    HB2-    CB-    CA-     C
  7246   7245   7248   7249      3    0.62760    1.88280    0.00000
-2.51040    0.00000    0.00000 ;    HB2-    CB-    CG-    HG
  7246   7245   7248   7250      3    0.66944    2.00832    0.00000
-2.67776    0.00000    0.00000 ;    HB2-    CB-    CG-   CD1
  7246   7245   7248   7254      3    0.66944    2.00832    0.00000
-2.67776    0.00000    0.00000 ;    HB2-    CB-    CG-   CD2
  7247   7245   7243   7258      3    0.65084    1.95253    0.00000
-2.60338    0.00000    0.00000 ;    HB3-    CB-    CA-     C
  7247   7245   7248   7249      3    0.62760    1.88280    0.00000
-2.51040    0.00000    0.00000 ;    HB3-    CB-    CG-    HG
  7247   7245   7248   7250      3    0.66944    2.00832    0.00000
-2.67776    0.00000    0.00000 ;    HB3-    CB-    CG-   CD1
  7247   7245   7248   7254      3    0.66944    2.00832    0.00000
-2.67776    0.00000    0.00000 ;    HB3-    CB-    CG-   CD2
  7248   7245   7243   7258      3   10.33448   -1.14642  -11.54784
-2.40998    6.35968    0.00000 ;     CG-    CB-    CA-     C
  7249   7248   7250   7251      3    0.62760    1.88280    0.00000
-2.51040    0.00000    0.00000 ;     HG-    CG-   CD1-  HD11
  7249   7248   7250   7252      3    0.62760    1.88280    0.00000
-2.51040    0.00000    0.00000 ;     HG-    CG-   CD1-  HD12
  7249   7248   7250   7253      3    0.62760    1.88280    0.00000
-2.51040    0.00000    0.00000 ;     HG-    CG-   CD1-  HD13
  7249   7248   7254   7255      3    0.62760    1.88280    0.00000
-2.51040    0.00000    0.00000 ;     HG-    CG-   CD2-  HD21
  7249   7248   7254   7256      3    0.62760    1.88280    0.00000
-2.51040    0.00000    0.00000 ;     HG-    CG-   CD2-  HD22
  7249   7248   7254   7257      3    0.62760    1.88280    0.00000
-2.51040    0.00000    0.00000 ;     HG-    CG-   CD2-  HD23
  7250   7248   7254   7255      3    0.66944    2.00832    0.00000
-2.67776    0.00000    0.00000 ;    CD1-    CG-   CD2-  HD21
  7250   7248   7254   7256      3    0.66944    2.00832    0.00000
-2.67776    0.00000    0.00000 ;    CD1-    CG-   CD2-  HD22
  7250   7248   7254   7257      3    0.66944    2.00832    0.00000
-2.67776    0.00000    0.00000 ;    CD1-    CG-   CD2-  HD23
  7251   7250   7248   7254      3    0.66944    2.00832    0.00000
```

```
-2.67776    0.00000    0.00000 ;   HD11-   CD1-    CG-   CD2
  7252   7250   7248   7254      3    0.66944    2.00832    0.00000
-2.67776    0.00000    0.00000 ;   HD12-   CD1-    CG-   CD2
  7253   7250   7248   7254      3    0.66944    2.00832    0.00000
-2.67776    0.00000    0.00000 ;   HD13-   CD1-    CG-   CD2
  7258   7260   7262   7263      3    0.00000    0.00000    0.00000
0.00000    0.00000    0.00000 ;      C-     N-    CA-   HA
  7258   7260   7262   7264      3   11.71520    1.67360   15.06240  -
13.38880    0.00000    0.00000 ;      C-     N-    CA-   CB
  7258   7260   7262   7275      3    1.75728    5.27184    2.25936
-7.02912    0.00000    0.00000 ;      C-     N-    CA-    C
  7259   7258   7260   7261      3   29.28800   -8.36800  -20.92000
0.00000    0.00000    0.00000 ;      O-     C-     N-    H
  7259   7258   7260   7262      3   20.92000    0.00000  -20.92000
0.00000    0.00000    0.00000 ;      O-     C-     N-   CA
  7260   7262   7264   7265      3    0.65084    1.95253    0.00000
-2.60338    0.00000    0.00000 ;      N-    CA-    CB-   HB2
  7260   7262   7264   7266      3    0.65084    1.95253    0.00000
-2.60338    0.00000    0.00000 ;      N-    CA-    CB-   HB3
  7260   7262   7264   7267      3    3.21331    2.22589   -4.15053
-2.40998    2.61082    0.00000 ;      N-    CA-    CB-    CG
  7260   7262   7275   7276      3    0.00000    0.00000    0.00000
0.00000    0.00000    0.00000 ;      N-    CA-     C-    O
  7260   7262   7275   7277      3   17.40544   -5.02080  -13.22144
9.20480    0.00000    0.00000 ;      N-    CA-     C-    N
  7261   7260   7262   7263      3    0.00000    0.00000    0.00000
0.00000    0.00000    0.00000 ;      H-     N-    CA-   HA
  7261   7260   7262   7264      3    0.00000    0.00000    0.00000
0.00000    0.00000    0.00000 ;      H-     N-    CA-   CB
  7261   7260   7262   7275      3    0.00000    0.00000    0.00000
0.00000    0.00000    0.00000 ;      H-     N-    CA-    C
  7262   7264   7267   7268      3    0.65084    1.95253    0.00000
-2.60338    0.00000    0.00000 ;     CA-    CB-    CG-   HG2
  7262   7264   7267   7269      3    0.65084    1.95253    0.00000
-2.60338    0.00000    0.00000 ;     CA-    CB-    CG-   HG3
  7262   7264   7267   7270      3    2.04598   -1.54390    1.11294
-0.26778    0.93722    0.00000 ;     CA-    CB-    CG-    SD
  7262   7275   7277   7278      3   20.92000    0.00000  -20.92000
0.00000    0.00000    0.00000 ;     CA-     C-     N-    H
  7262   7275   7277   7279      3   20.92000    0.00000  -20.92000
0.00000    0.00000    0.00000 ;     CA-     C-     N-   CA
  7263   7262   7264   7265      3    0.65084    1.95253    0.00000
-2.60338    0.00000    0.00000 ;     HA-    CA-    CB-   HB2
  7263   7262   7264   7266      3    0.65084    1.95253    0.00000
-2.60338    0.00000    0.00000 ;     HA-    CA-    CB-   HB3
  7263   7262   7264   7267      3    0.65084    1.95253    0.00000
-2.60338    0.00000    0.00000 ;     HA-    CA-    CB-    CG
  7263   7262   7275   7276      3    3.68192   -4.35136    0.00000
1.33888    0.00000    0.00000 ;     HA-    CA-     C-    O
  7263   7262   7275   7277      3    0.00000    0.00000    0.00000
0.00000    0.00000    0.00000 ;     HA-    CA-     C-    N
  7264   7262   7275   7276      3    0.00000    0.00000    0.00000
```

```
0.00000    0.00000    0.00000 ;      CB-    CA-     C-     O
  7264   7262   7275   7277      3    2.51040    4.18400    1.67360
-6.69440    0.00000    0.00000 ;      CB-    CA-     C-     N
  7264   7267   7270   7271      3    3.24260    6.22998    1.79075
-6.92870    1.90790    0.00000 ;      CB-    CG-    SD-    CE
  7265   7264   7262   7275      3    0.65084    1.95253    0.00000
-2.60338    0.00000    0.00000 ;     HB2-    CB-    CA-     C
  7265   7264   7267   7268      3    0.65084    1.95253    0.00000
-2.60338    0.00000    0.00000 ;     HB2-    CB-    CG-   HG2
  7265   7264   7267   7269      3    0.65084    1.95253    0.00000
-2.60338    0.00000    0.00000 ;     HB2-    CB-    CG-   HG3
  7265   7264   7267   7270      3    0.65084    1.95253    0.00000
-2.60338    0.00000    0.00000 ;     HB2-    CB-    CG-    SD
  7266   7264   7262   7275      3    0.65084    1.95253    0.00000
-2.60338    0.00000    0.00000 ;     HB3-    CB-    CA-     C
  7266   7264   7267   7268      3    0.65084    1.95253    0.00000
-2.60338    0.00000    0.00000 ;     HB3-    CB-    CG-   HG2
  7266   7264   7267   7269      3    0.65084    1.95253    0.00000
-2.60338    0.00000    0.00000 ;     HB3-    CB-    CG-   HG3
  7266   7264   7267   7270      3    0.65084    1.95253    0.00000
-2.60338    0.00000    0.00000 ;     HB3-    CB-    CG-    SD
  7267   7264   7262   7275      3    6.86594    3.56895   -8.14206
-2.40998    4.85344    0.00000 ;      CG-    CB-    CA-     C
  7267   7270   7271   7272      3    1.39467    4.18400    0.00000
-5.57867    0.00000    0.00000 ;      CG-    SD-    CE-   HE1
  7267   7270   7271   7273      3    1.39467    4.18400    0.00000
-5.57867    0.00000    0.00000 ;      CG-    SD-    CE-   HE2
  7267   7270   7271   7274      3    1.39467    4.18400    0.00000
-5.57867    0.00000    0.00000 ;      CG-    SD-    CE-   HE3
  7268   7267   7270   7271      3    1.39467    4.18400    0.00000
-5.57867    0.00000    0.00000 ;     HG2-    CG-    SD-    CE
  7269   7267   7270   7271      3    1.39467    4.18400    0.00000
-5.57867    0.00000    0.00000 ;     HG3-    CG-    SD-    CE
  7275   7277   7279   7280      3    0.00000    0.00000    0.00000
0.00000    0.00000    0.00000 ;      C-     N-    CA-    HA
  7275   7277   7279   7281      3   11.71520    1.67360   15.06240  -
13.38880    0.00000    0.00000 ;      C-     N-    CA-    CB
  7275   7277   7279   7292      3    1.75728    5.27184    2.25936
-7.02912    0.00000    0.00000 ;      C-     N-    CA-     C
  7276   7275   7277   7278      3   29.28800   -8.36800  -20.92000
0.00000    0.00000    0.00000 ;      O-     C-     N-     H
  7276   7275   7277   7279      3   20.92000    0.00000  -20.92000
0.00000    0.00000    0.00000 ;      O-     C-     N-    CA
  7277   7279   7281   7282      3    0.65084    1.95253    0.00000
-2.60338    0.00000    0.00000 ;      N-    CA-    CB-   HB2
  7277   7279   7281   7283      3    0.65084    1.95253    0.00000
-2.60338    0.00000    0.00000 ;      N-    CA-    CB-   HB3
  7277   7279   7281   7284      3    3.21331    2.22589   -4.15053
-2.40998    2.61082    0.00000 ;      N-    CA-    CB-    CG
  7277   7279   7292   7293      3    0.00000    0.00000    0.00000
0.00000    0.00000    0.00000 ;      N-    CA-     C-     O
  7277   7279   7292   7294      3   17.40544   -5.02080  -13.22144
```

```
9.20480    0.00000    0.00000 ;      N-    CA-     C-     N
  7278   7277   7279   7280      3    0.00000    0.00000    0.00000
0.00000    0.00000    0.00000 ;      H-     N-    CA-    HA
  7278   7277   7279   7281      3    0.00000    0.00000    0.00000
0.00000    0.00000    0.00000 ;      H-     N-    CA-    CB
  7278   7277   7279   7292      3    0.00000    0.00000    0.00000
0.00000    0.00000    0.00000 ;      H-     N-    CA-     C
  7279   7281   7284   7285      3    0.65084    1.95253    0.00000
-2.60338    0.00000    0.00000 ;     CA-    CB-    CG-   HG2
  7279   7281   7284   7286      3    0.65084    1.95253    0.00000
-2.60338    0.00000    0.00000 ;     CA-    CB-    CG-   HG3
  7279   7281   7284   7287      3    2.04598   -1.54390    1.11294
-0.26778    0.93722    0.00000 ;     CA-    CB-    CG-    SD
  7279   7292   7294   7295      3   20.92000    0.00000  -20.92000
0.00000    0.00000    0.00000 ;     CA-     C-     N-     H
  7279   7292   7294   7296      3   20.92000    0.00000  -20.92000
0.00000    0.00000    0.00000 ;     CA-     C-     N-    CA
  7280   7279   7281   7282      3    0.65084    1.95253    0.00000
-2.60338    0.00000    0.00000 ;     HA-    CA-    CB-   HB2
  7280   7279   7281   7283      3    0.65084    1.95253    0.00000
-2.60338    0.00000    0.00000 ;     HA-    CA-    CB-   HB3
  7280   7279   7281   7284      3    0.65084    1.95253    0.00000
-2.60338    0.00000    0.00000 ;     HA-    CA-    CB-    CG
  7280   7279   7292   7293      3    3.68192   -4.35136    0.00000
1.33888    0.00000    0.00000 ;     HA-    CA-     C-     O
  7280   7279   7292   7294      3    0.00000    0.00000    0.00000
0.00000    0.00000    0.00000 ;     HA-    CA-     C-     N
  7281   7279   7292   7293      3    0.00000    0.00000    0.00000
0.00000    0.00000    0.00000 ;     CB-    CA-     C-     O
  7281   7279   7292   7294      3    2.51040    4.18400    1.67360
-6.69440    0.00000    0.00000 ;     CB-    CA-     C-     N
  7281   7284   7287   7288      3    3.24260    6.22998    1.79075
-6.92870    1.90790    0.00000 ;     CB-    CG-    SD-    CE
  7282   7281   7279   7292      3    0.65084    1.95253    0.00000
-2.60338    0.00000    0.00000 ;    HB2-    CB-    CA-     C
  7282   7281   7284   7285      3    0.65084    1.95253    0.00000
-2.60338    0.00000    0.00000 ;    HB2-    CB-    CG-   HG2
  7282   7281   7284   7286      3    0.65084    1.95253    0.00000
-2.60338    0.00000    0.00000 ;    HB2-    CB-    CG-   HG3
  7282   7281   7284   7287      3    0.65084    1.95253    0.00000
-2.60338    0.00000    0.00000 ;    HB2-    CB-    CG-    SD
  7283   7281   7279   7292      3    0.65084    1.95253    0.00000
-2.60338    0.00000    0.00000 ;    HB3-    CB-    CA-     C
  7283   7281   7284   7285      3    0.65084    1.95253    0.00000
-2.60338    0.00000    0.00000 ;    HB3-    CB-    CG-   HG2
  7283   7281   7284   7286      3    0.65084    1.95253    0.00000
-2.60338    0.00000    0.00000 ;    HB3-    CB-    CG-   HG3
  7283   7281   7284   7287      3    0.65084    1.95253    0.00000
-2.60338    0.00000    0.00000 ;    HB3-    CB-    CG-    SD
  7284   7281   7279   7292      3    6.86594    3.56895   -8.14206
-2.40998    4.85344    0.00000 ;     CG-    CB-    CA-     C
  7284   7287   7288   7289      3    1.39467    4.18400    0.00000
```

-5.57867 0.00000 0.00000 ; CG- SD- CE- HE1
7284 7287 7288 7290 3 1.39467 4.18400 0.00000
-5.57867 0.00000 0.00000 ; CG- SD- CE- HE2
7284 7287 7288 7291 3 1.39467 4.18400 0.00000
-5.57867 0.00000 0.00000 ; CG- SD- CE- HE3
7285 7284 7287 7288 3 1.39467 4.18400 0.00000
-5.57867 0.00000 0.00000 ; HG2- CG- SD- CE
7286 7284 7287 7288 3 1.39467 4.18400 0.00000
-5.57867 0.00000 0.00000 ; HG3- CG- SD- CE
7292 7294 7296 7297 3 0.00000 0.00000 0.00000
0.00000 0.00000 0.00000 ; C- N- CA- HA
7292 7294 7296 7298 3 11.71520 1.67360 15.06240 -
13.38880 0.00000 0.00000 ; C- N- CA- CB
7292 7294 7296 7303 3 1.75728 5.27184 2.25936
-7.02912 0.00000 0.00000 ; C- N- CA- C
7293 7292 7294 7295 3 29.28800 -8.36800 -20.92000
0.00000 0.00000 0.00000 ; O- C- N- H
7293 7292 7294 7296 3 20.92000 0.00000 -20.92000
0.00000 0.00000 0.00000 ; O- C- N- CA
7294 7296 7298 7299 3 0.65084 1.95253 0.00000
-2.60338 0.00000 0.00000 ; N- CA- CB- HB2
7294 7296 7298 7300 3 0.65084 1.95253 0.00000
-2.60338 0.00000 0.00000 ; N- CA- CB- HB3
7294 7296 7298 7301 3 7.86174 2.24681 -7.41405
-6.71114 5.35552 0.00000 ; N- CA- CB- OG
7294 7296 7303 7304 3 0.00000 0.00000 0.00000
0.00000 0.00000 0.00000 ; N- CA- C- O
7294 7296 7303 7305 3 17.40544 -5.02080 -13.22144
9.20480 0.00000 0.00000 ; N- CA- C- N
7295 7294 7296 7297 3 0.00000 0.00000 0.00000
0.00000 0.00000 0.00000 ; H- N- CA- HA
7295 7294 7296 7298 3 0.00000 0.00000 0.00000
0.00000 0.00000 0.00000 ; H- N- CA- CB
7295 7294 7296 7303 3 0.00000 0.00000 0.00000
0.00000 0.00000 0.00000 ; H- N- CA- C
7296 7298 7301 7302 3 2.05853 2.46856 3.48109
-4.46851 0.23430 0.00000 ; CA- CB- OG- HG
7296 7303 7305 7306 3 20.92000 0.00000 -20.92000
0.00000 0.00000 0.00000 ; CA- C- N- H
7296 7303 7305 7307 3 20.92000 0.00000 -20.92000
0.00000 0.00000 0.00000 ; CA- C- N- CA
7297 7296 7298 7299 3 0.65084 1.95253 0.00000
-2.60338 0.00000 0.00000 ; HA- CA- CB- HB2
7297 7296 7298 7300 3 0.65084 1.95253 0.00000
-2.60338 0.00000 0.00000 ; HA- CA- CB- HB3
7297 7296 7298 7301 3 1.04600 -1.04600 0.00000
0.00000 0.00000 0.00000 ; HA- CA- CB- OG
7297 7296 7303 7304 3 3.68192 -4.35136 0.00000
1.33888 0.00000 0.00000 ; HA- CA- C- O
7297 7296 7303 7305 3 0.00000 0.00000 0.00000
0.00000 0.00000 0.00000 ; HA- CA- C- N
7298 7296 7303 7304 3 0.00000 0.00000 0.00000

0.00000    0.00000    0.00000 ;      CB-    CA-     C-     O
  7298   7296   7303   7305      3    2.51040    4.18400    1.67360
-6.69440    0.00000    0.00000 ;     CB-    CA-     C-     N
  7299   7298   7296   7303      3    0.65084    1.95253    0.00000
-2.60338    0.00000    0.00000 ;    HB2-    CB-    CA-     C
  7299   7298   7301   7302      3    0.69733    2.09200    0.00000
-2.78933    0.00000    0.00000 ;    HB2-    CB-    OG-    HG
  7300   7298   7296   7303      3    0.65084    1.95253    0.00000
-2.60338    0.00000    0.00000 ;    HB3-    CB-    CA-     C
  7300   7298   7301   7302      3    0.69733    2.09200    0.00000
-2.78933    0.00000    0.00000 ;    HB3-    CB-    OG-    HG
  7301   7298   7296   7303      3    7.34710    7.79898   -6.14211
-6.71114    4.31789    0.00000 ;     OG-    CB-    CA-     C
  7303   7305   7307   7308      3    0.00000    0.00000    0.00000
0.00000    0.00000    0.00000 ;      C-     N-    CA-    HA
  7303   7305   7307   7309      3   11.71520    1.67360   15.06240  -
13.38880    0.00000    0.00000 ;      C-     N-    CA-    CB
  7303   7305   7307   7325      3    1.75728    5.27184    2.25936
-7.02912    0.00000    0.00000 ;      C-     N-    CA-     C
  7304   7303   7305   7306      3   29.28800   -8.36800  -20.92000
0.00000    0.00000    0.00000 ;      O-     C-     N-     H
  7304   7303   7305   7307      3   20.92000    0.00000  -20.92000
0.00000    0.00000    0.00000 ;      O-     C-     N-    CA
  7305   7307   7309   7310      3    0.65084    1.95253    0.00000
-2.60338    0.00000    0.00000 ;      N-    CA-    CB-   HB2
  7305   7307   7309   7311      3    0.65084    1.95253    0.00000
-2.60338    0.00000    0.00000 ;      N-    CA-    CB-   HB3
  7305   7307   7309   7312      3    0.65084    1.95253    0.00000
-2.60338    0.00000    0.00000 ;      N-    CA-    CB-    CG
  7305   7307   7325   7326      3    0.00000    0.00000    0.00000
0.00000    0.00000    0.00000 ;      N-    CA-     C-     O
  7305   7307   7325   7327      3   17.40544   -5.02080  -13.22144
9.20480    0.00000    0.00000 ;      N-    CA-     C-     N
  7306   7305   7307   7308      3    0.00000    0.00000    0.00000
0.00000    0.00000    0.00000 ;      H-     N-    CA-    HA
  7306   7305   7307   7309      3    0.00000    0.00000    0.00000
0.00000    0.00000    0.00000 ;      H-     N-    CA-    CB
  7306   7305   7307   7325      3    0.00000    0.00000    0.00000
0.00000    0.00000    0.00000 ;      H-     N-    CA-     C
  7307   7309   7312   7313      3    0.66944    2.00832    0.00000
-2.67776    0.00000    0.00000 ;     CA-    CB-    CG-   HG2
  7307   7309   7312   7314      3    0.66944    2.00832    0.00000
-2.67776    0.00000    0.00000 ;     CA-    CB-    CG-   HG3
  7307   7309   7312   7315      3    3.68192    3.09616   -2.09200
-3.01248    0.00000    0.00000 ;     CA-    CB-    CG-    CD
  7307   7325   7327   7328      3   20.92000    0.00000  -20.92000
0.00000    0.00000    0.00000 ;     CA-     C-     N-     H
  7307   7325   7327   7329      3   20.92000    0.00000  -20.92000
0.00000    0.00000    0.00000 ;     CA-     C-     N-    CA
  7308   7307   7309   7310      3    0.65084    1.95253    0.00000
-2.60338    0.00000    0.00000 ;     HA-    CA-    CB-   HB2
  7308   7307   7309   7311      3    0.65084    1.95253    0.00000

```
-2.60338    0.00000    0.00000 ;      HA-    CA-     CB-   HB3
  7308   7307   7309   7312       3    0.65084    1.95253    0.00000
-2.60338    0.00000    0.00000 ;      HA-    CA-     CB-    CG
  7308   7307   7325   7326       3    3.68192   -4.35136    0.00000
1.33888    0.00000    0.00000 ;      HA-    CA-     C-     O
  7308   7307   7325   7327       3    0.00000    0.00000    0.00000
0.00000    0.00000    0.00000 ;      HA-    CA-     C-     N
  7309   7307   7325   7326       3    0.00000    0.00000    0.00000
0.00000    0.00000    0.00000 ;      CB-    CA-     C-     O
  7309   7307   7325   7327       3    2.51040    4.18400    1.67360
-6.69440    0.00000    0.00000 ;      CB-    CA-     C-     N
  7309   7312   7315   7316       3    0.66944    2.00832    0.00000
-2.67776    0.00000    0.00000 ;      CB-    CG-     CD-   HD2
  7309   7312   7315   7317       3    0.66944    2.00832    0.00000
-2.67776    0.00000    0.00000 ;      CB-    CG-     CD-   HD3
  7309   7312   7315   7318       3    3.68192    3.09616   -2.09200
-3.01248    0.00000    0.00000 ;      CB-    CG-     CD-    CE
  7310   7309   7307   7325       3    0.65084    1.95253    0.00000
-2.60338    0.00000    0.00000 ;     HB2-    CB-     CA-     C
  7310   7309   7312   7313       3    0.62760    1.88280    0.00000
-2.51040    0.00000    0.00000 ;     HB2-    CB-     CG-   HG2
  7310   7309   7312   7314       3    0.62760    1.88280    0.00000
-2.51040    0.00000    0.00000 ;     HB2-    CB-     CG-   HG3
  7310   7309   7312   7315       3    0.66944    2.00832    0.00000
-2.67776    0.00000    0.00000 ;     HB2-    CB-     CG-    CD
  7311   7309   7307   7325       3    0.65084    1.95253    0.00000
-2.60338    0.00000    0.00000 ;     HB3-    CB-     CA-     C
  7311   7309   7312   7313       3    0.62760    1.88280    0.00000
-2.51040    0.00000    0.00000 ;     HB3-    CB-     CG-   HG2
  7311   7309   7312   7314       3    0.62760    1.88280    0.00000
-2.51040    0.00000    0.00000 ;     HB3-    CB-     CG-   HG3
  7311   7309   7312   7315       3    0.66944    2.00832    0.00000
-2.67776    0.00000    0.00000 ;     HB3-    CB-     CG-    CD
  7312   7309   7307   7325       3    0.65084    1.95253    0.00000
-2.60338    0.00000    0.00000 ;      CG-    CB-     CA-     C
  7312   7315   7318   7319       3    0.65084    1.95253    0.00000
-2.60338    0.00000    0.00000 ;      CG-    CD-     CE-   HE2
  7312   7315   7318   7320       3    0.65084    1.95253    0.00000
-2.60338    0.00000    0.00000 ;      CG-    CD-     CE-   HE3
  7312   7315   7318   7321       3    0.65084    1.95253    0.00000
-2.60338    0.00000    0.00000 ;      CG-    CD-     CE-    NZ
  7313   7312   7315   7316       3    0.62760    1.88280    0.00000
-2.51040    0.00000    0.00000 ;     HG2-    CG-     CD-   HD2
  7313   7312   7315   7317       3    0.62760    1.88280    0.00000
-2.51040    0.00000    0.00000 ;     HG2-    CG-     CD-   HD3
  7313   7312   7315   7318       3    0.66944    2.00832    0.00000
-2.67776    0.00000    0.00000 ;     HG2-    CG-     CD-    CE
  7314   7312   7315   7316       3    0.62760    1.88280    0.00000
-2.51040    0.00000    0.00000 ;     HG3-    CG-     CD-   HD2
  7314   7312   7315   7317       3    0.62760    1.88280    0.00000
-2.51040    0.00000    0.00000 ;     HG3-    CG-     CD-   HD3
  7314   7312   7315   7318       3    0.66944    2.00832    0.00000
```

-2.67776    0.00000    0.00000 ;    HG3-    CG-    CD-    CE
 7315   7318   7321   7322      3    0.65084    1.95253    0.00000
-2.60338    0.00000    0.00000 ;    CD-    CE-    NZ-   HZ1
 7315   7318   7321   7323      3    0.65084    1.95253    0.00000
-2.60338    0.00000    0.00000 ;    CD-    CE-    NZ-   HZ2
 7315   7318   7321   7324      3    0.65084    1.95253    0.00000
-2.60338    0.00000    0.00000 ;    CD-    CE-    NZ-   HZ3
 7316   7315   7318   7319      3    0.65084    1.95253    0.00000
-2.60338    0.00000    0.00000 ;    HD2-    CD-    CE-   HE2
 7316   7315   7318   7320      3    0.65084    1.95253    0.00000
-2.60338    0.00000    0.00000 ;    HD2-    CD-    CE-   HE3
 7316   7315   7318   7321      3    0.65084    1.95253    0.00000
-2.60338    0.00000    0.00000 ;    HD2-    CD-    CE-    NZ
 7317   7315   7318   7319      3    0.65084    1.95253    0.00000
-2.60338    0.00000    0.00000 ;    HD3-    CD-    CE-   HE2
 7317   7315   7318   7320      3    0.65084    1.95253    0.00000
-2.60338    0.00000    0.00000 ;    HD3-    CD-    CE-   HE3
 7317   7315   7318   7321      3    0.65084    1.95253    0.00000
-2.60338    0.00000    0.00000 ;    HD3-    CD-    CE-    NZ
 7319   7318   7321   7322      3    0.65084    1.95253    0.00000
-2.60338    0.00000    0.00000 ;    HE2-    CE-    NZ-   HZ1
 7319   7318   7321   7323      3    0.65084    1.95253    0.00000
-2.60338    0.00000    0.00000 ;    HE2-    CE-    NZ-   HZ2
 7319   7318   7321   7324      3    0.65084    1.95253    0.00000
-2.60338    0.00000    0.00000 ;    HE2-    CE-    NZ-   HZ3
 7320   7318   7321   7322      3    0.65084    1.95253    0.00000
-2.60338    0.00000    0.00000 ;    HE3-    CE-    NZ-   HZ1
 7320   7318   7321   7323      3    0.65084    1.95253    0.00000
-2.60338    0.00000    0.00000 ;    HE3-    CE-    NZ-   HZ2
 7320   7318   7321   7324      3    0.65084    1.95253    0.00000
-2.60338    0.00000    0.00000 ;    HE3-    CE-    NZ-   HZ3
 7325   7327   7329   7330      3    0.00000    0.00000    0.00000
0.00000    0.00000    0.00000 ;    C-    N-    CA-    HA
 7325   7327   7329   7331      3   11.71520    1.67360   15.06240  -
13.38880    0.00000    0.00000 ;    C-    N-    CA-    CB
 7325   7327   7329   7347      3    1.75728    5.27184    2.25936
-7.02912    0.00000    0.00000 ;    C-    N-    CA-    C
 7326   7325   7327   7328      3   29.28800   -8.36800  -20.92000
0.00000    0.00000    0.00000 ;    O-    C-    N-    H
 7326   7325   7327   7329      3   20.92000    0.00000  -20.92000
0.00000    0.00000    0.00000 ;    O-    C-    N-    CA
 7327   7329   7331   7332      3    0.65084    1.95253    0.00000
-2.60338    0.00000    0.00000 ;    N-    CA-    CB-   HB2
 7327   7329   7331   7333      3    0.65084    1.95253    0.00000
-2.60338    0.00000    0.00000 ;    N-    CA-    CB-   HB3
 7327   7329   7331   7334      3    0.65084    1.95253    0.00000
-2.60338    0.00000    0.00000 ;    N-    CA-    CB-    CG
 7327   7329   7347   7348      3    0.00000    0.00000    0.00000
0.00000    0.00000    0.00000 ;    N-    CA-    C-    O
 7327   7329   7347   7349      3   17.40544   -5.02080  -13.22144
9.20480    0.00000    0.00000 ;    N-    CA-    C-    N
 7328   7327   7329   7330      3    0.00000    0.00000    0.00000

```
0.00000    0.00000    0.00000 ;      H-     N-    CA-    HA
  7328   7327   7329   7331       3    0.00000    0.00000    0.00000
0.00000    0.00000    0.00000 ;      H-     N-    CA-    CB
  7328   7327   7329   7347       3    0.00000    0.00000    0.00000
0.00000    0.00000    0.00000 ;      H-     N-    CA-     C
  7329   7331   7334   7335       3    0.66944    2.00832    0.00000
-2.67776    0.00000    0.00000 ;     CA-    CB-    CG-   HG2
  7329   7331   7334   7336       3    0.66944    2.00832    0.00000
-2.67776    0.00000    0.00000 ;     CA-    CB-    CG-   HG3
  7329   7331   7334   7337       3    3.68192    3.09616   -2.09200
-3.01248    0.00000    0.00000 ;     CA-    CB-    CG-    CD
  7329   7347   7349   7350       3   20.92000    0.00000  -20.92000
0.00000    0.00000    0.00000 ;     CA-     C-     N-     H
  7329   7347   7349   7351       3   20.92000    0.00000  -20.92000
0.00000    0.00000    0.00000 ;     CA-     C-     N-    CA
  7330   7329   7331   7332       3    0.65084    1.95253    0.00000
-2.60338    0.00000    0.00000 ;     HA-    CA-    CB-   HB2
  7330   7329   7331   7333       3    0.65084    1.95253    0.00000
-2.60338    0.00000    0.00000 ;     HA-    CA-    CB-   HB3
  7330   7329   7331   7334       3    0.65084    1.95253    0.00000
-2.60338    0.00000    0.00000 ;     HA-    CA-    CB-    CG
  7330   7329   7347   7348       3    3.68192   -4.35136    0.00000
1.33888    0.00000    0.00000 ;     HA-    CA-     C-     O
  7330   7329   7347   7349       3    0.00000    0.00000    0.00000
0.00000    0.00000    0.00000 ;     HA-    CA-     C-     N
  7331   7329   7347   7348       3    0.00000    0.00000    0.00000
0.00000    0.00000    0.00000 ;     CB-    CA-     C-     O
  7331   7329   7347   7349       3    2.51040    4.18400    1.67360
-6.69440    0.00000    0.00000 ;     CB-    CA-     C-     N
  7331   7334   7337   7338       3    0.66944    2.00832    0.00000
-2.67776    0.00000    0.00000 ;     CB-    CG-    CD-   HD2
  7331   7334   7337   7339       3    0.66944    2.00832    0.00000
-2.67776    0.00000    0.00000 ;     CB-    CG-    CD-   HD3
  7331   7334   7337   7340       3    3.68192    3.09616   -2.09200
-3.01248    0.00000    0.00000 ;     CB-    CG-    CD-    CE
  7332   7331   7329   7347       3    0.65084    1.95253    0.00000
-2.60338    0.00000    0.00000 ;    HB2-    CB-    CA-     C
  7332   7331   7334   7335       3    0.62760    1.88280    0.00000
-2.51040    0.00000    0.00000 ;    HB2-    CB-    CG-   HG2
  7332   7331   7334   7336       3    0.62760    1.88280    0.00000
-2.51040    0.00000    0.00000 ;    HB2-    CB-    CG-   HG3
  7332   7331   7334   7337       3    0.66944    2.00832    0.00000
-2.67776    0.00000    0.00000 ;    HB2-    CB-    CG-    CD
  7333   7331   7329   7347       3    0.65084    1.95253    0.00000
-2.60338    0.00000    0.00000 ;    HB3-    CB-    CA-     C
  7333   7331   7334   7335       3    0.62760    1.88280    0.00000
-2.51040    0.00000    0.00000 ;    HB3-    CB-    CG-   HG2
  7333   7331   7334   7336       3    0.62760    1.88280    0.00000
-2.51040    0.00000    0.00000 ;    HB3-    CB-    CG-   HG3
  7333   7331   7334   7337       3    0.66944    2.00832    0.00000
-2.67776    0.00000    0.00000 ;    HB3-    CB-    CG-    CD
  7334   7331   7329   7347       3    0.65084    1.95253    0.00000
```

-2.60338 0.00000 0.00000 ; CG- CB- CA- C
7334 7337 7340 7341 3 0.65084 1.95253 0.00000
-2.60338 0.00000 0.00000 ; CG- CD- CE- HE2
7334 7337 7340 7342 3 0.65084 1.95253 0.00000
-2.60338 0.00000 0.00000 ; CG- CD- CE- HE3
7334 7337 7340 7343 3 0.65084 1.95253 0.00000
-2.60338 0.00000 0.00000 ; CG- CD- CE- NZ
7335 7334 7337 7338 3 0.62760 1.88280 0.00000
-2.51040 0.00000 0.00000 ; HG2- CG- CD- HD2
7335 7334 7337 7339 3 0.62760 1.88280 0.00000
-2.51040 0.00000 0.00000 ; HG2- CG- CD- HD3
7335 7334 7337 7340 3 0.66944 2.00832 0.00000
-2.67776 0.00000 0.00000 ; HG2- CG- CD- CE
7336 7334 7337 7338 3 0.62760 1.88280 0.00000
-2.51040 0.00000 0.00000 ; HG3- CG- CD- HD2
7336 7334 7337 7339 3 0.62760 1.88280 0.00000
-2.51040 0.00000 0.00000 ; HG3- CG- CD- HD3
7336 7334 7337 7340 3 0.66944 2.00832 0.00000
-2.67776 0.00000 0.00000 ; HG3- CG- CD- CE
7337 7340 7343 7344 3 0.65084 1.95253 0.00000
-2.60338 0.00000 0.00000 ; CD- CE- NZ- HZ1
7337 7340 7343 7345 3 0.65084 1.95253 0.00000
-2.60338 0.00000 0.00000 ; CD- CE- NZ- HZ2
7337 7340 7343 7346 3 0.65084 1.95253 0.00000
-2.60338 0.00000 0.00000 ; CD- CE- NZ- HZ3
7338 7337 7340 7341 3 0.65084 1.95253 0.00000
-2.60338 0.00000 0.00000 ; HD2- CD- CE- HE2
7338 7337 7340 7342 3 0.65084 1.95253 0.00000
-2.60338 0.00000 0.00000 ; HD2- CD- CE- HE3
7338 7337 7340 7343 3 0.65084 1.95253 0.00000
-2.60338 0.00000 0.00000 ; HD2- CD- CE- NZ
7339 7337 7340 7341 3 0.65084 1.95253 0.00000
-2.60338 0.00000 0.00000 ; HD3- CD- CE- HE2
7339 7337 7340 7342 3 0.65084 1.95253 0.00000
-2.60338 0.00000 0.00000 ; HD3- CD- CE- HE3
7339 7337 7340 7343 3 0.65084 1.95253 0.00000
-2.60338 0.00000 0.00000 ; HD3- CD- CE- NZ
7341 7340 7343 7344 3 0.65084 1.95253 0.00000
-2.60338 0.00000 0.00000 ; HE2- CE- NZ- HZ1
7341 7340 7343 7345 3 0.65084 1.95253 0.00000
-2.60338 0.00000 0.00000 ; HE2- CE- NZ- HZ2
7341 7340 7343 7346 3 0.65084 1.95253 0.00000
-2.60338 0.00000 0.00000 ; HE2- CE- NZ- HZ3
7342 7340 7343 7344 3 0.65084 1.95253 0.00000
-2.60338 0.00000 0.00000 ; HE3- CE- NZ- HZ1
7342 7340 7343 7345 3 0.65084 1.95253 0.00000
-2.60338 0.00000 0.00000 ; HE3- CE- NZ- HZ2
7342 7340 7343 7346 3 0.65084 1.95253 0.00000
-2.60338 0.00000 0.00000 ; HE3- CE- NZ- HZ3
7347 7349 7351 7352 3 0.00000 0.00000 0.00000
0.00000 0.00000 0.00000 ; C- N- CA- HA
7347 7349 7351 7353 3 11.71520 1.67360 15.06240 -

```
13.38880    0.00000    0.00000 ;      C-     N-    CA-    CB
  7347   7349   7351   7369      3    1.75728    5.27184    2.25936
-7.02912    0.00000    0.00000 ;      C-     N-    CA-     C
  7348   7347   7349   7350      3   29.28800   -8.36800  -20.92000
0.00000    0.00000    0.00000 ;      O-     C-     N-     H
  7348   7347   7349   7351      3   20.92000    0.00000  -20.92000
0.00000    0.00000    0.00000 ;      O-     C-     N-    CA
  7349   7351   7353   7354      3    0.65084    1.95253    0.00000
-2.60338    0.00000    0.00000 ;      N-    CA-    CB-   HB2
  7349   7351   7353   7355      3    0.65084    1.95253    0.00000
-2.60338    0.00000    0.00000 ;      N-    CA-    CB-   HB3
  7349   7351   7353   7356      3    0.65084    1.95253    0.00000
-2.60338    0.00000    0.00000 ;      N-    CA-    CB-    CG
  7349   7351   7369   7370      3    0.00000    0.00000    0.00000
0.00000    0.00000    0.00000 ;      N-    CA-     C-     O
  7349   7351   7369   7371      3    0.00000    0.00000    0.00000
0.00000    0.00000    0.00000 ;      N-    CA-     C-   OXT
  7350   7349   7351   7352      3    0.00000    0.00000    0.00000
0.00000    0.00000    0.00000 ;      H-     N-    CA-    HA
  7350   7349   7351   7353      3    0.00000    0.00000    0.00000
0.00000    0.00000    0.00000 ;      H-     N-    CA-    CB
  7350   7349   7351   7369      3    0.00000    0.00000    0.00000
0.00000    0.00000    0.00000 ;      H-     N-    CA-     C
  7351   7353   7356   7357      3    0.66944    2.00832    0.00000
-2.67776    0.00000    0.00000 ;     CA-    CB-    CG-   HG2
  7351   7353   7356   7358      3    0.66944    2.00832    0.00000
-2.67776    0.00000    0.00000 ;     CA-    CB-    CG-   HG3
  7351   7353   7356   7359      3    3.68192    3.09616   -2.09200
-3.01248    0.00000    0.00000 ;     CA-    CB-    CG-    CD
  7352   7351   7353   7354      3    0.65084    1.95253    0.00000
-2.60338    0.00000    0.00000 ;     HA-    CA-    CB-   HB2
  7352   7351   7353   7355      3    0.65084    1.95253    0.00000
-2.60338    0.00000    0.00000 ;     HA-    CA-    CB-   HB3
  7352   7351   7353   7356      3    0.65084    1.95253    0.00000
-2.60338    0.00000    0.00000 ;     HA-    CA-    CB-    CG
  7352   7351   7369   7370      3    0.00000    0.00000    0.00000
0.00000    0.00000    0.00000 ;     HA-    CA-     C-     O
  7352   7351   7369   7371      3    0.00000    0.00000    0.00000
0.00000    0.00000    0.00000 ;     HA-    CA-     C-   OXT
  7353   7351   7369   7370      3    0.00000    0.00000    0.00000
0.00000    0.00000    0.00000 ;     CB-    CA-     C-     O
  7353   7351   7369   7371      3    0.00000    0.00000    0.00000
0.00000    0.00000    0.00000 ;     CB-    CA-     C-   OXT
  7353   7356   7359   7360      3    0.66944    2.00832    0.00000
-2.67776    0.00000    0.00000 ;     CB-    CG-    CD-   HD2
  7353   7356   7359   7361      3    0.66944    2.00832    0.00000
-2.67776    0.00000    0.00000 ;     CB-    CG-    CD-   HD3
  7353   7356   7359   7362      3    3.68192    3.09616   -2.09200
-3.01248    0.00000    0.00000 ;     CB-    CG-    CD-    CE
  7354   7353   7351   7369      3    0.65084    1.95253    0.00000
-2.60338    0.00000    0.00000 ;    HB2-    CB-    CA-     C
  7354   7353   7356   7357      3    0.62760    1.88280    0.00000
```

```
 -2.51040    0.00000    0.00000 ;     HB2-    CB-     CG-   HG2
  7354   7353   7356   7358      3    0.62760    1.88280    0.00000
 -2.51040    0.00000    0.00000 ;     HB2-    CB-     CG-   HG3
  7354   7353   7356   7359      3    0.66944    2.00832    0.00000
 -2.67776    0.00000    0.00000 ;     HB2-    CB-     CG-    CD
  7355   7353   7351   7369      3    0.65084    1.95253    0.00000
 -2.60338    0.00000    0.00000 ;     HB3-    CB-     CA-     C
  7355   7353   7356   7357      3    0.62760    1.88280    0.00000
 -2.51040    0.00000    0.00000 ;     HB3-    CB-     CG-   HG2
  7355   7353   7356   7358      3    0.62760    1.88280    0.00000
 -2.51040    0.00000    0.00000 ;     HB3-    CB-     CG-   HG3
  7355   7353   7356   7359      3    0.66944    2.00832    0.00000
 -2.67776    0.00000    0.00000 ;     HB3-    CB-     CG-    CD
  7356   7353   7351   7369      3    0.65084    1.95253    0.00000
 -2.60338    0.00000    0.00000 ;      CG-    CB-     CA-     C
  7356   7359   7362   7363      3    0.65084    1.95253    0.00000
 -2.60338    0.00000    0.00000 ;      CG-    CD-     CE-   HE2
  7356   7359   7362   7364      3    0.65084    1.95253    0.00000
 -2.60338    0.00000    0.00000 ;      CG-    CD-     CE-   HE3
  7356   7359   7362   7365      3    0.65084    1.95253    0.00000
 -2.60338    0.00000    0.00000 ;      CG-    CD-     CE-    NZ
  7357   7356   7359   7360      3    0.62760    1.88280    0.00000
 -2.51040    0.00000    0.00000 ;     HG2-    CG-     CD-   HD2
  7357   7356   7359   7361      3    0.62760    1.88280    0.00000
 -2.51040    0.00000    0.00000 ;     HG2-    CG-     CD-   HD3
  7357   7356   7359   7362      3    0.66944    2.00832    0.00000
 -2.67776    0.00000    0.00000 ;     HG2-    CG-     CD-    CE
  7358   7356   7359   7360      3    0.62760    1.88280    0.00000
 -2.51040    0.00000    0.00000 ;     HG3-    CG-     CD-   HD2
  7358   7356   7359   7361      3    0.62760    1.88280    0.00000
 -2.51040    0.00000    0.00000 ;     HG3-    CG-     CD-   HD3
  7358   7356   7359   7362      3    0.66944    2.00832    0.00000
 -2.67776    0.00000    0.00000 ;     HG3-    CG-     CD-    CE
  7359   7362   7365   7366      3    0.65084    1.95253    0.00000
 -2.60338    0.00000    0.00000 ;      CD-    CE-     NZ-   HZ1
  7359   7362   7365   7367      3    0.65084    1.95253    0.00000
 -2.60338    0.00000    0.00000 ;      CD-    CE-     NZ-   HZ2
  7359   7362   7365   7368      3    0.65084    1.95253    0.00000
 -2.60338    0.00000    0.00000 ;      CD-    CE-     NZ-   HZ3
  7360   7359   7362   7363      3    0.65084    1.95253    0.00000
 -2.60338    0.00000    0.00000 ;     HD2-    CD-     CE-   HE2
  7360   7359   7362   7364      3    0.65084    1.95253    0.00000
 -2.60338    0.00000    0.00000 ;     HD2-    CD-     CE-   HE3
  7360   7359   7362   7365      3    0.65084    1.95253    0.00000
 -2.60338    0.00000    0.00000 ;     HD2-    CD-     CE-    NZ
  7361   7359   7362   7363      3    0.65084    1.95253    0.00000
 -2.60338    0.00000    0.00000 ;     HD3-    CD-     CE-   HE2
  7361   7359   7362   7364      3    0.65084    1.95253    0.00000
 -2.60338    0.00000    0.00000 ;     HD3-    CD-     CE-   HE3
  7361   7359   7362   7365      3    0.65084    1.95253    0.00000
 -2.60338    0.00000    0.00000 ;     HD3-    CD-     CE-    NZ
  7363   7362   7365   7366      3    0.65084    1.95253    0.00000
```

```
-2.60338    0.00000    0.00000 ;     HE2-    CE-     NZ-   HZ1
  7363   7362   7365   7367       3    0.65084    1.95253    0.00000
-2.60338    0.00000    0.00000 ;     HE2-    CE-     NZ-   HZ2
  7363   7362   7365   7368       3    0.65084    1.95253    0.00000
-2.60338    0.00000    0.00000 ;     HE2-    CE-     NZ-   HZ3
  7364   7362   7365   7366       3    0.65084    1.95253    0.00000
-2.60338    0.00000    0.00000 ;     HE3-    CE-     NZ-   HZ1
  7364   7362   7365   7367       3    0.65084    1.95253    0.00000
-2.60338    0.00000    0.00000 ;     HE3-    CE-     NZ-   HZ2
  7364   7362   7365   7368       3    0.65084    1.95253    0.00000
-2.60338    0.00000    0.00000 ;     HE3-    CE-     NZ-   HZ3
  7372   7373   7374   7375       3   30.33400    0.00000  -30.33400
0.00000    0.00000    0.00000 ;      C1-    C2-     C3-    C4
  7372   7373   7374   7414       3   30.33400    0.00000  -30.33400
0.00000    0.00000    0.00000 ;      C1-    C2-     C3-   H43
  7372   7377   7376   7375       3   30.33400    0.00000  -30.33400
0.00000    0.00000    0.00000 ;      C1-    C6-     C5-    C4
  7372   7377   7376   7379       3   30.33400    0.00000  -30.33400
0.00000    0.00000    0.00000 ;      C1-    C6-     C5-    C8
  7373   7372   7377   7376       3   30.33400    0.00000  -30.33400
0.00000    0.00000    0.00000 ;      C2-    C1-     C6-    C5
  7373   7372   7377   7415       3   30.33400    0.00000  -30.33400
0.00000    0.00000    0.00000 ;      C2-    C1-     C6-   H44
  7373   7372   7388   7430       3    7.53120    0.00000   -7.53120
0.00000    0.00000    0.00000 ;      C2-    C1-    O17-   H59
  7373   7374   7375   7376       3   30.33400    0.00000  -30.33400
0.00000    0.00000    0.00000 ;      C2-    C3-     C4-    C5
  7373   7374   7375   7387       3   30.33400    0.00000  -30.33400
0.00000    0.00000    0.00000 ;      C2-    C3-     C4-   N16
  7374   7373   7372   7377       3   30.33400    0.00000  -30.33400
0.00000    0.00000    0.00000 ;      C3-    C2-     C1-    C6
  7374   7373   7372   7388       3   30.33400    0.00000  -30.33400
0.00000    0.00000    0.00000 ;      C3-    C2-     C1-   O17
  7374   7375   7376   7377       3   30.33400    0.00000  -30.33400
0.00000    0.00000    0.00000 ;      C3-    C4-     C5-    C6
  7374   7375   7376   7379       3   30.33400    0.00000  -30.33400
0.00000    0.00000    0.00000 ;      C3-    C4-     C5-    C8
  7374   7375   7387   7378       3    0.00000    0.00000    0.00000
0.00000    0.00000    0.00000 ;      C3-    C4-    N16-    C7
  7374   7375   7387   7436       3    0.00000    0.00000    0.00000
0.00000    0.00000    0.00000 ;      C3-    C4-    N16-    RU
  7375   7374   7373   7413       3   30.33400    0.00000  -30.33400
0.00000    0.00000    0.00000 ;      C4-    C3-     C2-   H42
  7375   7376   7377   7415       3   30.33400    0.00000  -30.33400
0.00000    0.00000    0.00000 ;      C4-    C5-     C6-   H44
  7375   7376   7379   7378       3    8.36800    0.00000   -8.36800
0.00000    0.00000    0.00000 ;      C4-    C5-     C8-    C7
  7375   7376   7379   7405       3    8.36800    0.00000   -8.36800
0.00000    0.00000    0.00000 ;      C4-    C5-     C8-   C34
  7375   7387   7378   7379       3    0.00000    0.00000    0.00000
0.00000    0.00000    0.00000 ;      C4-   N16-     C7-    C8
  7375   7387   7378   7380       3    0.00000    0.00000    0.00000
```

0.00000 0.00000 0.00000 ; C4- N16- C7- C9
7375 7387 7436 7386 3 0.00000 0.00000 0.00000
0.00000 0.00000 0.00000 ; C4- N16- RU- N15
7375 7387 7436 7389 3 0.00000 0.00000 0.00000
0.00000 0.00000 0.00000 ; C4- N16- RU- C18
7375 7387 7436 7390 3 0.00000 0.00000 0.00000
0.00000 0.00000 0.00000 ; C4- N16- RU- C19
7375 7387 7436 7391 3 0.00000 0.00000 0.00000
0.00000 0.00000 0.00000 ; C4- N16- RU- C20
7375 7387 7436 7392 3 0.00000 0.00000 0.00000
0.00000 0.00000 0.00000 ; C4- N16- RU- C21
7375 7387 7436 7393 3 0.00000 0.00000 0.00000
0.00000 0.00000 0.00000 ; C4- N16- RU- C22
7375 7387 7436 7394 3 0.00000 0.00000 0.00000
0.00000 0.00000 0.00000 ; C4- N16- RU- C23
7376 7375 7374 7414 3 30.33400 0.00000 -30.33400
0.00000 0.00000 0.00000 ; C5- C4- C3- H43
7376 7375 7387 7378 3 0.00000 0.00000 0.00000
0.00000 0.00000 0.00000 ; C5- C4- N16- C7
7376 7375 7387 7436 3 0.00000 0.00000 0.00000
0.00000 0.00000 0.00000 ; C5- C4- N16- RU
7376 7377 7372 7388 3 30.33400 0.00000 -30.33400
0.00000 0.00000 0.00000 ; C5- C6- C1- O17
7376 7379 7378 7380 3 30.33400 0.00000 -30.33400
0.00000 0.00000 0.00000 ; C5- C8- C7- C9
7376 7379 7378 7387 3 30.33400 0.00000 -30.33400
0.00000 0.00000 0.00000 ; C5- C8- C7- N16
7376 7379 7405 7406 3 30.33400 0.00000 -30.33400
0.00000 0.00000 0.00000 ; C5- C8- C34- C35
7376 7379 7405 7412 3 30.33400 0.00000 -30.33400
0.00000 0.00000 0.00000 ; C5- C8- C34- C41
7377 7372 7373 7413 3 30.33400 0.00000 -30.33400
0.00000 0.00000 0.00000 ; C6- C1- C2- H42
7377 7372 7388 7430 3 7.53120 0.00000 -7.53120
0.00000 0.00000 0.00000 ; C6- C1- O17- H59
7377 7376 7375 7387 3 30.33400 0.00000 -30.33400
0.00000 0.00000 0.00000 ; C6- C5- C4- N16
7377 7376 7379 7378 3 8.36800 0.00000 -8.36800
0.00000 0.00000 0.00000 ; C6- C5- C8- C7
7377 7376 7379 7405 3 8.36800 0.00000 -8.36800
0.00000 0.00000 0.00000 ; C6- C5- C8- C34
7378 7379 7405 7406 3 30.33400 0.00000 -30.33400
0.00000 0.00000 0.00000 ; C7- C8- C34- C35
7378 7379 7405 7412 3 30.33400 0.00000 -30.33400
0.00000 0.00000 0.00000 ; C7- C8- C34- C41
7378 7380 7381 7384 3 30.33400 0.00000 -30.33400
0.00000 0.00000 0.00000 ; C7- C9- C10- C13
7378 7380 7381 7412 3 30.33400 0.00000 -30.33400
0.00000 0.00000 0.00000 ; C7- C9- C10- C41
7378 7380 7386 7382 3 0.00000 0.00000 0.00000
0.00000 0.00000 0.00000 ; C7- C9- N15- C11
7378 7380 7386 7436 3 0.00000 0.00000 0.00000

```
0.00000    0.00000    0.00000 ;     C7-    C9-   N15-    RU
  7378   7387   7436   7386      3    0.00000    0.00000    0.00000
0.00000    0.00000    0.00000 ;     C7-   N16-    RU-   N15
  7378   7387   7436   7389      3    0.00000    0.00000    0.00000
0.00000    0.00000    0.00000 ;     C7-   N16-    RU-   C18
  7378   7387   7436   7390      3    0.00000    0.00000    0.00000
0.00000    0.00000    0.00000 ;     C7-   N16-    RU-   C19
  7378   7387   7436   7391      3    0.00000    0.00000    0.00000
0.00000    0.00000    0.00000 ;     C7-   N16-    RU-   C20
  7378   7387   7436   7392      3    0.00000    0.00000    0.00000
0.00000    0.00000    0.00000 ;     C7-   N16-    RU-   C21
  7378   7387   7436   7393      3    0.00000    0.00000    0.00000
0.00000    0.00000    0.00000 ;     C7-   N16-    RU-   C22
  7378   7387   7436   7394      3    0.00000    0.00000    0.00000
0.00000    0.00000    0.00000 ;     C7-   N16-    RU-   C23
  7379   7376   7375   7387      3   30.33400    0.00000  -30.33400
0.00000    0.00000    0.00000 ;     C8-    C5-    C4-   N16
  7379   7376   7377   7415      3   30.33400    0.00000  -30.33400
0.00000    0.00000    0.00000 ;     C8-    C5-    C6-   H44
  7379   7378   7380   7381      3   30.33400    0.00000  -30.33400
0.00000    0.00000    0.00000 ;     C8-    C7-    C9-   C10
  7379   7378   7380   7386      3   30.33400    0.00000  -30.33400
0.00000    0.00000    0.00000 ;     C8-    C7-    C9-   N15
  7379   7378   7387   7436      3    0.00000    0.00000    0.00000
0.00000    0.00000    0.00000 ;     C8-    C7-   N16-    RU
  7379   7405   7406   7407      3    5.85760    0.00000   -5.85760
0.00000    0.00000    0.00000 ;     C8-   C34-   C35-   N36
  7379   7405   7406   7408      3    5.85760    0.00000   -5.85760
0.00000    0.00000    0.00000 ;     C8-   C34-   C35-   O37
  7379   7405   7412   7381      3   30.33400    0.00000  -30.33400
0.00000    0.00000    0.00000 ;     C8-   C34-   C41-   C10
  7379   7405   7412   7410      3   30.33400    0.00000  -30.33400
0.00000    0.00000    0.00000 ;     C8-   C34-   C41-   C39
  7380   7378   7379   7405      3   30.33400    0.00000  -30.33400
0.00000    0.00000    0.00000 ;     C9-    C7-    C8-   C34
  7380   7378   7387   7436      3    0.00000    0.00000    0.00000
0.00000    0.00000    0.00000 ;     C9-    C7-   N16-    RU
  7380   7381   7384   7383      3    5.85760    0.00000   -5.85760
0.00000    0.00000    0.00000 ;     C9-   C10-   C13-   C12
  7380   7381   7384   7417      3    5.85760    0.00000   -5.85760
0.00000    0.00000    0.00000 ;     C9-   C10-   C13-   H46
  7380   7381   7412   7405      3   30.33400    0.00000  -30.33400
0.00000    0.00000    0.00000 ;     C9-   C10-   C41-   C34
  7380   7381   7412   7410      3   30.33400    0.00000  -30.33400
0.00000    0.00000    0.00000 ;     C9-   C10-   C41-   C39
  7380   7386   7382   7383      3   34.72720    0.00000  -34.72720
0.00000    0.00000    0.00000 ;     C9-   N15-   C11-   C12
  7380   7386   7382   7416      3   34.72720    0.00000  -34.72720
0.00000    0.00000    0.00000 ;     C9-   N15-   C11-   H45
  7380   7386   7436   7387      3    0.00000    0.00000    0.00000
0.00000    0.00000    0.00000 ;     C9-   N15-    RU-   N16
  7380   7386   7436   7389      3    0.00000    0.00000    0.00000
```

```
0.00000    0.00000    0.00000 ;     C9-   N15-    RU-   C18
  7380   7386   7436   7390      3    0.00000    0.00000    0.00000
0.00000    0.00000    0.00000 ;     C9-   N15-    RU-   C19
  7380   7386   7436   7391      3    0.00000    0.00000    0.00000
0.00000    0.00000    0.00000 ;     C9-   N15-    RU-   C20
  7380   7386   7436   7392      3    0.00000    0.00000    0.00000
0.00000    0.00000    0.00000 ;     C9-   N15-    RU-   C21
  7380   7386   7436   7393      3    0.00000    0.00000    0.00000
0.00000    0.00000    0.00000 ;     C9-   N15-    RU-   C22
  7380   7386   7436   7394      3    0.00000    0.00000    0.00000
0.00000    0.00000    0.00000 ;     C9-   N15-    RU-   C23
  7381   7380   7378   7387      3   30.33400    0.00000  -30.33400
0.00000    0.00000    0.00000 ;    C10-    C9-    C7-   N16
  7381   7380   7386   7382      3    0.00000    0.00000    0.00000
0.00000    0.00000    0.00000 ;    C10-    C9-   N15-   C11
  7381   7380   7386   7436      3    0.00000    0.00000    0.00000
0.00000    0.00000    0.00000 ;    C10-    C9-   N15-    RU
  7381   7384   7383   7382      3   55.64720    0.00000  -55.64720
0.00000    0.00000    0.00000 ;    C10-   C13-   C12-   C11
  7381   7384   7383   7385      3   55.64720    0.00000  -55.64720
0.00000    0.00000    0.00000 ;    C10-   C13-   C12-   F14
  7381   7412   7405   7406      3   30.33400    0.00000  -30.33400
0.00000    0.00000    0.00000 ;    C10-   C41-   C34-   C35
  7381   7412   7410   7407      3    5.85760    0.00000   -5.85760
0.00000    0.00000    0.00000 ;    C10-   C41-   C39-   N36
  7381   7412   7410   7411      3    5.85760    0.00000   -5.85760
0.00000    0.00000    0.00000 ;    C10-   C41-   C39-   O40
  7382   7383   7384   7417      3   55.64720    0.00000  -55.64720
0.00000    0.00000    0.00000 ;    C11-   C12-   C13-   H46
  7382   7386   7436   7387      3    0.00000    0.00000    0.00000
0.00000    0.00000    0.00000 ;    C11-   N15-    RU-   N16
  7382   7386   7436   7389      3    0.00000    0.00000    0.00000
0.00000    0.00000    0.00000 ;    C11-   N15-    RU-   C18
  7382   7386   7436   7390      3    0.00000    0.00000    0.00000
0.00000    0.00000    0.00000 ;    C11-   N15-    RU-   C19
  7382   7386   7436   7391      3    0.00000    0.00000    0.00000
0.00000    0.00000    0.00000 ;    C11-   N15-    RU-   C20
  7382   7386   7436   7392      3    0.00000    0.00000    0.00000
0.00000    0.00000    0.00000 ;    C11-   N15-    RU-   C21
  7382   7386   7436   7393      3    0.00000    0.00000    0.00000
0.00000    0.00000    0.00000 ;    C11-   N15-    RU-   C22
  7382   7386   7436   7394      3    0.00000    0.00000    0.00000
0.00000    0.00000    0.00000 ;    C11-   N15-    RU-   C23
  7383   7382   7386   7436      3    0.00000    0.00000    0.00000
0.00000    0.00000    0.00000 ;    C12-   C11-   N15-    RU
  7383   7384   7381   7412      3    5.85760    0.00000   -5.85760
0.00000    0.00000    0.00000 ;    C12-   C13-   C10-   C41
  7384   7381   7380   7386      3   30.33400    0.00000  -30.33400
0.00000    0.00000    0.00000 ;    C13-   C10-    C9-   N15
  7384   7381   7412   7405      3   30.33400    0.00000  -30.33400
0.00000    0.00000    0.00000 ;    C13-   C10-   C41-   C34
  7384   7381   7412   7410      3   30.33400    0.00000  -30.33400
```

```
0.00000    0.00000    0.00000 ;    C13-   C10-   C41-   C39
  7384   7383   7382   7386      3   55.64720    0.00000  -55.64720
0.00000    0.00000    0.00000 ;    C13-   C12-   C11-   N15
  7384   7383   7382   7416      3   55.64720    0.00000  -55.64720
0.00000    0.00000    0.00000 ;    C13-   C12-   C11-   H45
  7385   7383   7382   7386      3   55.64720    0.00000  -55.64720
0.00000    0.00000    0.00000 ;    F14-   C12-   C11-   N15
  7385   7383   7382   7416      3   55.64720    0.00000  -55.64720
0.00000    0.00000    0.00000 ;    F14-   C12-   C11-   H45
  7385   7383   7384   7417      3   55.64720    0.00000  -55.64720
0.00000    0.00000    0.00000 ;    F14-   C12-   C13-   H46
  7386   7380   7378   7387      3   30.33400    0.00000  -30.33400
0.00000    0.00000    0.00000 ;    N15-    C9-    C7-   N16
  7386   7380   7381   7412      3   30.33400    0.00000  -30.33400
0.00000    0.00000    0.00000 ;    N15-    C9-   C10-   C41
  7386   7436   7389   7390      3    0.00000    0.00000    0.00000
0.00000    0.00000    0.00000 ;    N15-    RU-   C18-   C19
  7386   7436   7389   7393      3    0.00000    0.00000    0.00000
0.00000    0.00000    0.00000 ;    N15-    RU-   C18-   C22
  7386   7436   7389   7418      3    0.00000    0.00000    0.00000
0.00000    0.00000    0.00000 ;    N15-    RU-   C18-   H47
  7386   7436   7390   7389      3    0.00000    0.00000    0.00000
0.00000    0.00000    0.00000 ;    N15-    RU-   C19-   C18
  7386   7436   7390   7391      3    0.00000    0.00000    0.00000
0.00000    0.00000    0.00000 ;    N15-    RU-   C19-   C20
  7386   7436   7390   7396      3    0.00000    0.00000    0.00000
0.00000    0.00000    0.00000 ;    N15-    RU-   C19-   C25
  7386   7436   7391   7390      3    0.00000    0.00000    0.00000
0.00000    0.00000    0.00000 ;    N15-    RU-   C20-   C19
  7386   7436   7391   7392      3    0.00000    0.00000    0.00000
0.00000    0.00000    0.00000 ;    N15-    RU-   C20-   C21
  7386   7436   7391   7435      3    0.00000    0.00000    0.00000
0.00000    0.00000    0.00000 ;    N15-    RU-   C20-   H64
  7386   7436   7392   7391      3    0.00000    0.00000    0.00000
0.00000    0.00000    0.00000 ;    N15-    RU-   C21-   C20
  7386   7436   7392   7393      3    0.00000    0.00000    0.00000
0.00000    0.00000    0.00000 ;    N15-    RU-   C21-   C22
  7386   7436   7392   7419      3    0.00000    0.00000    0.00000
0.00000    0.00000    0.00000 ;    N15-    RU-   C21-   H48
  7386   7436   7393   7389      3    0.00000    0.00000    0.00000
0.00000    0.00000    0.00000 ;    N15-    RU-   C22-   C18
  7386   7436   7393   7392      3    0.00000    0.00000    0.00000
0.00000    0.00000    0.00000 ;    N15-    RU-   C22-   C21
  7386   7436   7393   7420      3    0.00000    0.00000    0.00000
0.00000    0.00000    0.00000 ;    N15-    RU-   C22-   H49
  7386   7436   7394   7395      3    0.00000    0.00000    0.00000
0.00000    0.00000    0.00000 ;    N15-    RU-   C23-   O24
  7387   7375   7374   7414      3   30.33400    0.00000  -30.33400
0.00000    0.00000    0.00000 ;    N16-    C4-    C3-   H43
  7387   7378   7379   7405      3   30.33400    0.00000  -30.33400
0.00000    0.00000    0.00000 ;    N16-    C7-    C8-   C34
  7387   7436   7389   7390      3    0.00000    0.00000    0.00000
```

0.00000 0.00000 0.00000 ; N16- RU- C18- C19
7387 7436 7389 7393 3 0.00000 0.00000 0.00000
0.00000 0.00000 0.00000 ; N16- RU- C18- C22
7387 7436 7389 7418 3 0.00000 0.00000 0.00000
0.00000 0.00000 0.00000 ; N16- RU- C18- H47
7387 7436 7390 7389 3 0.00000 0.00000 0.00000
0.00000 0.00000 0.00000 ; N16- RU- C19- C18
7387 7436 7390 7391 3 0.00000 0.00000 0.00000
0.00000 0.00000 0.00000 ; N16- RU- C19- C20
7387 7436 7390 7396 3 0.00000 0.00000 0.00000
0.00000 0.00000 0.00000 ; N16- RU- C19- C25
7387 7436 7391 7390 3 0.00000 0.00000 0.00000
0.00000 0.00000 0.00000 ; N16- RU- C20- C19
7387 7436 7391 7392 3 0.00000 0.00000 0.00000
0.00000 0.00000 0.00000 ; N16- RU- C20- C21
7387 7436 7391 7435 3 0.00000 0.00000 0.00000
0.00000 0.00000 0.00000 ; N16- RU- C20- H64
7387 7436 7392 7391 3 0.00000 0.00000 0.00000
0.00000 0.00000 0.00000 ; N16- RU- C21- C20
7387 7436 7392 7393 3 0.00000 0.00000 0.00000
0.00000 0.00000 0.00000 ; N16- RU- C21- C22
7387 7436 7392 7419 3 0.00000 0.00000 0.00000
0.00000 0.00000 0.00000 ; N16- RU- C21- H48
7387 7436 7393 7389 3 0.00000 0.00000 0.00000
0.00000 0.00000 0.00000 ; N16- RU- C22- C18
7387 7436 7393 7392 3 0.00000 0.00000 0.00000
0.00000 0.00000 0.00000 ; N16- RU- C22- C21
7387 7436 7393 7420 3 0.00000 0.00000 0.00000
0.00000 0.00000 0.00000 ; N16- RU- C22- H49
7387 7436 7394 7395 3 0.00000 0.00000 0.00000
0.00000 0.00000 0.00000 ; N16- RU- C23- O24
7388 7372 7373 7413 3 30.33400 0.00000 -30.33400
0.00000 0.00000 0.00000 ; O17- C1- C2- H42
7388 7372 7377 7415 3 30.33400 0.00000 -30.33400
0.00000 0.00000 0.00000 ; O17- C1- C6- H44
7389 7390 7391 7392 3 0.00000 0.00000 0.00000
0.00000 0.00000 0.00000 ; C18- C19- C20- C21
7389 7390 7391 7435 3 0.00000 0.00000 0.00000
0.00000 0.00000 0.00000 ; C18- C19- C20- H64
7389 7390 7391 7436 3 0.00000 0.00000 0.00000
0.00000 0.00000 0.00000 ; C18- C19- C20- RU
7389 7390 7396 7397 3 0.00000 0.00000 0.00000
0.00000 0.00000 0.00000 ; C18- C19- C25- N26
7389 7390 7396 7398 3 0.00000 0.00000 0.00000
0.00000 0.00000 0.00000 ; C18- C19- C25- O27
7389 7390 7436 7391 3 0.00000 0.00000 0.00000
0.00000 0.00000 0.00000 ; C18- C19- RU- C20
7389 7390 7436 7392 3 0.00000 0.00000 0.00000
0.00000 0.00000 0.00000 ; C18- C19- RU- C21
7389 7390 7436 7393 3 0.00000 0.00000 0.00000
0.00000 0.00000 0.00000 ; C18- C19- RU- C22
7389 7390 7436 7394 3 0.00000 0.00000 0.00000

```
0.00000    0.00000    0.00000 ;    C18-   C19-    RU-   C23
  7389   7393   7392   7391      3    0.00000    0.00000    0.00000
0.00000    0.00000    0.00000 ;    C18-   C22-   C21-   C20
  7389   7393   7392   7419      3    0.00000    0.00000    0.00000
0.00000    0.00000    0.00000 ;    C18-   C22-   C21-   H48
  7389   7393   7392   7436      3    0.00000    0.00000    0.00000
0.00000    0.00000    0.00000 ;    C18-   C22-   C21-    RU
  7389   7393   7436   7390      3    0.00000    0.00000    0.00000
0.00000    0.00000    0.00000 ;    C18-   C22-    RU-   C19
  7389   7393   7436   7391      3    0.00000    0.00000    0.00000
0.00000    0.00000    0.00000 ;    C18-   C22-    RU-   C20
  7389   7393   7436   7392      3    0.00000    0.00000    0.00000
0.00000    0.00000    0.00000 ;    C18-   C22-    RU-   C21
  7389   7393   7436   7394      3    0.00000    0.00000    0.00000
0.00000    0.00000    0.00000 ;    C18-   C22-    RU-   C23
  7389   7436   7390   7391      3    0.00000    0.00000    0.00000
0.00000    0.00000    0.00000 ;    C18-    RU-   C19-   C20
  7389   7436   7390   7396      3    0.00000    0.00000    0.00000
0.00000    0.00000    0.00000 ;    C18-    RU-   C19-   C25
  7389   7436   7391   7390      3    0.00000    0.00000    0.00000
0.00000    0.00000    0.00000 ;    C18-    RU-   C20-   C19
  7389   7436   7391   7392      3    0.00000    0.00000    0.00000
0.00000    0.00000    0.00000 ;    C18-    RU-   C20-   C21
  7389   7436   7391   7435      3    0.00000    0.00000    0.00000
0.00000    0.00000    0.00000 ;    C18-    RU-   C20-   H64
  7389   7436   7392   7391      3    0.00000    0.00000    0.00000
0.00000    0.00000    0.00000 ;    C18-    RU-   C21-   C20
  7389   7436   7392   7393      3    0.00000    0.00000    0.00000
0.00000    0.00000    0.00000 ;    C18-    RU-   C21-   C22
  7389   7436   7392   7419      3    0.00000    0.00000    0.00000
0.00000    0.00000    0.00000 ;    C18-    RU-   C21-   H48
  7389   7436   7393   7392      3    0.00000    0.00000    0.00000
0.00000    0.00000    0.00000 ;    C18-    RU-   C22-   C21
  7389   7436   7393   7420      3    0.00000    0.00000    0.00000
0.00000    0.00000    0.00000 ;    C18-    RU-   C22-   H49
  7389   7436   7394   7395      3    0.00000    0.00000    0.00000
0.00000    0.00000    0.00000 ;    C18-    RU-   C23-   O24
  7390   7389   7393   7392      3    0.00000    0.00000    0.00000
0.00000    0.00000    0.00000 ;    C19-   C18-   C22-   C21
  7390   7389   7393   7420      3    0.00000    0.00000    0.00000
0.00000    0.00000    0.00000 ;    C19-   C18-   C22-   H49
  7390   7389   7393   7436      3    0.00000    0.00000    0.00000
0.00000    0.00000    0.00000 ;    C19-   C18-   C22-    RU
  7390   7389   7436   7391      3    0.00000    0.00000    0.00000
0.00000    0.00000    0.00000 ;    C19-   C18-    RU-   C20
  7390   7389   7436   7392      3    0.00000    0.00000    0.00000
0.00000    0.00000    0.00000 ;    C19-   C18-    RU-   C21
  7390   7389   7436   7393      3    0.00000    0.00000    0.00000
0.00000    0.00000    0.00000 ;    C19-   C18-    RU-   C22
  7390   7389   7436   7394      3    0.00000    0.00000    0.00000
0.00000    0.00000    0.00000 ;    C19-   C18-    RU-   C23
  7390   7391   7392   7393      3    0.00000    0.00000    0.00000
```

```
0.00000    0.00000    0.00000 ;    C19-   C20-   C21-   C22
  7390   7391   7392   7419      3    0.00000    0.00000    0.00000
0.00000    0.00000    0.00000 ;    C19-   C20-   C21-   H48
  7390   7391   7392   7436      3    0.00000    0.00000    0.00000
0.00000    0.00000    0.00000 ;    C19-   C20-   C21-    RU
  7390   7391   7436   7392      3    0.00000    0.00000    0.00000
0.00000    0.00000    0.00000 ;    C19-   C20-    RU-   C21
  7390   7391   7436   7393      3    0.00000    0.00000    0.00000
0.00000    0.00000    0.00000 ;    C19-   C20-    RU-   C22
  7390   7391   7436   7394      3    0.00000    0.00000    0.00000
0.00000    0.00000    0.00000 ;    C19-   C20-    RU-   C23
  7390   7396   7397   7399      3    5.43920    0.00000   -5.43920
0.00000    0.00000    0.00000 ;    C19-   C25-   N26-   C28
  7390   7396   7397   7434      3    5.43920    0.00000   -5.43920
0.00000    0.00000    0.00000 ;    C19-   C25-   N26-   H63
  7390   7436   7389   7393      3    0.00000    0.00000    0.00000
0.00000    0.00000    0.00000 ;    C19-    RU-   C18-   C22
  7390   7436   7389   7418      3    0.00000    0.00000    0.00000
0.00000    0.00000    0.00000 ;    C19-    RU-   C18-   H47
  7390   7436   7391   7392      3    0.00000    0.00000    0.00000
0.00000    0.00000    0.00000 ;    C19-    RU-   C20-   C21
  7390   7436   7391   7435      3    0.00000    0.00000    0.00000
0.00000    0.00000    0.00000 ;    C19-    RU-   C20-   H64
  7390   7436   7392   7391      3    0.00000    0.00000    0.00000
0.00000    0.00000    0.00000 ;    C19-    RU-   C21-   C20
  7390   7436   7392   7393      3    0.00000    0.00000    0.00000
0.00000    0.00000    0.00000 ;    C19-    RU-   C21-   C22
  7390   7436   7392   7419      3    0.00000    0.00000    0.00000
0.00000    0.00000    0.00000 ;    C19-    RU-   C21-   H48
  7390   7436   7393   7392      3    0.00000    0.00000    0.00000
0.00000    0.00000    0.00000 ;    C19-    RU-   C22-   C21
  7390   7436   7393   7420      3    0.00000    0.00000    0.00000
0.00000    0.00000    0.00000 ;    C19-    RU-   C22-   H49
  7390   7436   7394   7395      3    0.00000    0.00000    0.00000
0.00000    0.00000    0.00000 ;    C19-    RU-   C23-   O24
  7391   7390   7389   7393      3    0.00000    0.00000    0.00000
0.00000    0.00000    0.00000 ;    C20-   C19-   C18-   C22
  7391   7390   7389   7418      3    0.00000    0.00000    0.00000
0.00000    0.00000    0.00000 ;    C20-   C19-   C18-   H47
  7391   7390   7389   7436      3    0.00000    0.00000    0.00000
0.00000    0.00000    0.00000 ;    C20-   C19-   C18-    RU
  7391   7390   7396   7397      3    0.00000    0.00000    0.00000
0.00000    0.00000    0.00000 ;    C20-   C19-   C25-   N26
  7391   7390   7396   7398      3    0.00000    0.00000    0.00000
0.00000    0.00000    0.00000 ;    C20-   C19-   C25-   O27
  7391   7390   7436   7392      3    0.00000    0.00000    0.00000
0.00000    0.00000    0.00000 ;    C20-   C19-    RU-   C21
  7391   7390   7436   7393      3    0.00000    0.00000    0.00000
0.00000    0.00000    0.00000 ;    C20-   C19-    RU-   C22
  7391   7390   7436   7394      3    0.00000    0.00000    0.00000
0.00000    0.00000    0.00000 ;    C20-   C19-    RU-   C23
  7391   7392   7393   7420      3    0.00000    0.00000    0.00000
```

```
0.00000    0.00000    0.00000 ;    C20-   C21-   C22-   H49
  7391   7392   7393   7436      3    0.00000    0.00000    0.00000
0.00000    0.00000    0.00000 ;    C20-   C21-   C22-    RU
  7391   7392   7436   7393      3    0.00000    0.00000    0.00000
0.00000    0.00000    0.00000 ;    C20-   C21-    RU-   C22
  7391   7392   7436   7394      3    0.00000    0.00000    0.00000
0.00000    0.00000    0.00000 ;    C20-   C21-    RU-   C23
  7391   7436   7389   7393      3    0.00000    0.00000    0.00000
0.00000    0.00000    0.00000 ;    C20-    RU-   C18-   C22
  7391   7436   7389   7418      3    0.00000    0.00000    0.00000
0.00000    0.00000    0.00000 ;    C20-    RU-   C18-   H47
  7391   7436   7390   7396      3    0.00000    0.00000    0.00000
0.00000    0.00000    0.00000 ;    C20-    RU-   C19-   C25
  7391   7436   7392   7393      3    0.00000    0.00000    0.00000
0.00000    0.00000    0.00000 ;    C20-    RU-   C21-   C22
  7391   7436   7392   7419      3    0.00000    0.00000    0.00000
0.00000    0.00000    0.00000 ;    C20-    RU-   C21-   H48
  7391   7436   7393   7392      3    0.00000    0.00000    0.00000
0.00000    0.00000    0.00000 ;    C20-    RU-   C22-   C21
  7391   7436   7393   7420      3    0.00000    0.00000    0.00000
0.00000    0.00000    0.00000 ;    C20-    RU-   C22-   H49
  7391   7436   7394   7395      3    0.00000    0.00000    0.00000
0.00000    0.00000    0.00000 ;    C20-    RU-   C23-   O24
  7392   7391   7390   7396      3    0.00000    0.00000    0.00000
0.00000    0.00000    0.00000 ;    C21-   C20-   C19-   C25
  7392   7391   7390   7436      3    0.00000    0.00000    0.00000
0.00000    0.00000    0.00000 ;    C21-   C20-   C19-    RU
  7392   7391   7436   7393      3    0.00000    0.00000    0.00000
0.00000    0.00000    0.00000 ;    C21-   C20-    RU-   C22
  7392   7391   7436   7394      3    0.00000    0.00000    0.00000
0.00000    0.00000    0.00000 ;    C21-   C20-    RU-   C23
  7392   7393   7389   7418      3    0.00000    0.00000    0.00000
0.00000    0.00000    0.00000 ;    C21-   C22-   C18-   H47
  7392   7393   7389   7436      3    0.00000    0.00000    0.00000
0.00000    0.00000    0.00000 ;    C21-   C22-   C18-    RU
  7392   7393   7436   7394      3    0.00000    0.00000    0.00000
0.00000    0.00000    0.00000 ;    C21-   C22-    RU-   C23
  7392   7436   7389   7393      3    0.00000    0.00000    0.00000
0.00000    0.00000    0.00000 ;    C21-    RU-   C18-   C22
  7392   7436   7389   7418      3    0.00000    0.00000    0.00000
0.00000    0.00000    0.00000 ;    C21-    RU-   C18-   H47
  7392   7436   7390   7396      3    0.00000    0.00000    0.00000
0.00000    0.00000    0.00000 ;    C21-    RU-   C19-   C25
  7392   7436   7391   7435      3    0.00000    0.00000    0.00000
0.00000    0.00000    0.00000 ;    C21-    RU-   C20-   H64
  7392   7436   7393   7420      3    0.00000    0.00000    0.00000
0.00000    0.00000    0.00000 ;    C21-    RU-   C22-   H49
  7392   7436   7394   7395      3    0.00000    0.00000    0.00000
0.00000    0.00000    0.00000 ;    C21-    RU-   C23-   O24
  7393   7389   7390   7396      3    0.00000    0.00000    0.00000
0.00000    0.00000    0.00000 ;    C22-   C18-   C19-   C25
  7393   7389   7390   7436      3    0.00000    0.00000    0.00000
```

```
0.00000    0.00000    0.00000 ;    C22-   C18-   C19-    RU
  7393   7389   7436   7394      3    0.00000    0.00000    0.00000
0.00000    0.00000    0.00000 ;    C22-   C18-    RU-   C23
  7393   7392   7391   7435      3    0.00000    0.00000    0.00000
0.00000    0.00000    0.00000 ;    C22-   C21-   C20-   H64
  7393   7392   7391   7436      3    0.00000    0.00000    0.00000
0.00000    0.00000    0.00000 ;    C22-   C21-   C20-    RU
  7393   7392   7436   7394      3    0.00000    0.00000    0.00000
0.00000    0.00000    0.00000 ;    C22-   C21-    RU-   C23
  7393   7436   7389   7418      3    0.00000    0.00000    0.00000
0.00000    0.00000    0.00000 ;    C22-    RU-   C18-   H47
  7393   7436   7390   7396      3    0.00000    0.00000    0.00000
0.00000    0.00000    0.00000 ;    C22-    RU-   C19-   C25
  7393   7436   7391   7435      3    0.00000    0.00000    0.00000
0.00000    0.00000    0.00000 ;    C22-    RU-   C20-   H64
  7393   7436   7392   7419      3    0.00000    0.00000    0.00000
0.00000    0.00000    0.00000 ;    C22-    RU-   C21-   H48
  7393   7436   7394   7395      3    0.00000    0.00000    0.00000
0.00000    0.00000    0.00000 ;    C22-    RU-   C23-   O24
  7394   7436   7389   7418      3    0.00000    0.00000    0.00000
0.00000    0.00000    0.00000 ;    C23-    RU-   C18-   H47
  7394   7436   7390   7396      3    0.00000    0.00000    0.00000
0.00000    0.00000    0.00000 ;    C23-    RU-   C19-   C25
  7394   7436   7391   7435      3    0.00000    0.00000    0.00000
0.00000    0.00000    0.00000 ;    C23-    RU-   C20-   H64
  7394   7436   7392   7419      3    0.00000    0.00000    0.00000
0.00000    0.00000    0.00000 ;    C23-    RU-   C21-   H48
  7394   7436   7393   7420      3    0.00000    0.00000    0.00000
0.00000    0.00000    0.00000 ;    C23-    RU-   C22-   H49
  7396   7390   7389   7418      3   55.64720    0.00000  -55.64720
0.00000    0.00000    0.00000 ;    C25-   C19-   C18-   H47
  7396   7390   7389   7436      3    0.00000    0.00000    0.00000
0.00000    0.00000    0.00000 ;    C25-   C19-   C18-    RU
  7396   7390   7391   7435      3   55.64720    0.00000  -55.64720
0.00000    0.00000    0.00000 ;    C25-   C19-   C20-   H64
  7396   7390   7391   7436      3    0.00000    0.00000    0.00000
0.00000    0.00000    0.00000 ;    C25-   C19-   C20-    RU
  7396   7397   7399   7400      3    0.00000    0.00000    0.00000
0.00000    0.00000    0.00000 ;    C25-   N26-   C28-   C29
  7396   7397   7399   7401      3    0.00000    0.00000    0.00000
0.00000    0.00000    0.00000 ;    C25-   N26-   C28-   C30
  7396   7397   7399   7403      3    0.00000    0.00000    0.00000
0.00000    0.00000    0.00000 ;    C25-   N26-   C28-   C32
  7397   7396   7390   7436      3    0.00000    0.00000    0.00000
0.00000    0.00000    0.00000 ;    N26-   C25-   C19-    RU
  7397   7399   7400   7431      3    0.65084    1.95253    0.00000
-2.60338    0.00000    0.00000 ;    N26-   C28-   C29-   H60
  7397   7399   7400   7432      3    0.65084    1.95253    0.00000
-2.60338    0.00000    0.00000 ;    N26-   C28-   C29-   H61
  7397   7399   7400   7433      3    0.65084    1.95253    0.00000
-2.60338    0.00000    0.00000 ;    N26-   C28-   C29-   H62
  7397   7399   7401   7402      3    0.65084    1.95253    0.00000
```

```
-2.60338    0.00000    0.00000 ;    N26-   C28-   C30-   O31
  7397   7399   7401   7421       3    0.65084    1.95253    0.00000
-2.60338    0.00000    0.00000 ;    N26-   C28-   C30-   H50
  7397   7399   7401   7422       3    0.65084    1.95253    0.00000
-2.60338    0.00000    0.00000 ;    N26-   C28-   C30-   H51
  7397   7399   7403   7404       3    0.65084    1.95253    0.00000
-2.60338    0.00000    0.00000 ;    N26-   C28-   C32-   O33
  7397   7399   7403   7423       3    0.65084    1.95253    0.00000
-2.60338    0.00000    0.00000 ;    N26-   C28-   C32-   H52
  7397   7399   7403   7424       3    0.65084    1.95253    0.00000
-2.60338    0.00000    0.00000 ;    N26-   C28-   C32-   H53
  7398   7396   7390   7436       3    0.00000    0.00000    0.00000
0.00000    0.00000    0.00000 ;    O27-   C25-   C19-   RU
  7398   7396   7397   7399       3    5.43920    0.00000   -5.43920
0.00000    0.00000    0.00000 ;    O27-   C25-   N26-   C28
  7398   7396   7397   7434       3    5.43920    0.00000   -5.43920
0.00000    0.00000    0.00000 ;    O27-   C25-   N26-   H63
  7399   7401   7402   7425       3    1.71544    0.96232    0.00000
-2.67776    0.00000    0.00000 ;    C28-   C30-   O31-   H54
  7399   7403   7404   7429       3    1.71544    0.96232    0.00000
-2.67776    0.00000    0.00000 ;    C28-   C32-   O33-   H58
  7400   7399   7397   7434       3    0.00000    0.00000    0.00000
0.00000    0.00000    0.00000 ;    C29-   C28-   N26-   H63
  7400   7399   7401   7402       3    0.65084    1.95253    0.00000
-2.60338    0.00000    0.00000 ;    C29-   C28-   C30-   O31
  7400   7399   7401   7421       3    0.65084    1.95253    0.00000
-2.60338    0.00000    0.00000 ;    C29-   C28-   C30-   H50
  7400   7399   7401   7422       3    0.65084    1.95253    0.00000
-2.60338    0.00000    0.00000 ;    C29-   C28-   C30-   H51
  7400   7399   7403   7404       3    0.65084    1.95253    0.00000
-2.60338    0.00000    0.00000 ;    C29-   C28-   C32-   O33
  7400   7399   7403   7423       3    0.65084    1.95253    0.00000
-2.60338    0.00000    0.00000 ;    C29-   C28-   C32-   H52
  7400   7399   7403   7424       3    0.65084    1.95253    0.00000
-2.60338    0.00000    0.00000 ;    C29-   C28-   C32-   H53
  7401   7399   7397   7434       3    0.00000    0.00000    0.00000
0.00000    0.00000    0.00000 ;    C30-   C28-   N26-   H63
  7401   7399   7400   7431       3    0.66944    2.00832    0.00000
-2.67776    0.00000    0.00000 ;    C30-   C28-   C29-   H60
  7401   7399   7400   7432       3    0.66944    2.00832    0.00000
-2.67776    0.00000    0.00000 ;    C30-   C28-   C29-   H61
  7401   7399   7400   7433       3    0.66944    2.00832    0.00000
-2.67776    0.00000    0.00000 ;    C30-   C28-   C29-   H62
  7401   7399   7403   7404       3    0.65084    1.95253    0.00000
-2.60338    0.00000    0.00000 ;    C30-   C28-   C32-   O33
  7401   7399   7403   7423       3    0.65084    1.95253    0.00000
-2.60338    0.00000    0.00000 ;    C30-   C28-   C32-   H52
  7401   7399   7403   7424       3    0.65084    1.95253    0.00000
-2.60338    0.00000    0.00000 ;    C30-   C28-   C32-   H53
  7402   7401   7399   7403       3    0.65084    1.95253    0.00000
-2.60338    0.00000    0.00000 ;    O31-   C30-   C28-   C32
  7403   7399   7397   7434       3    0.00000    0.00000    0.00000
```

0.00000 0.00000 0.00000 ; C32- C28- N26- H63
7403 7399 7400 7431 3 0.66944 2.00832 0.00000
-2.67776 0.00000 0.00000 ; C32- C28- C29- H60
7403 7399 7400 7432 3 0.66944 2.00832 0.00000
-2.67776 0.00000 0.00000 ; C32- C28- C29- H61
7403 7399 7400 7433 3 0.66944 2.00832 0.00000
-2.67776 0.00000 0.00000 ; C32- C28- C29- H62
7403 7399 7401 7421 3 0.65084 1.95253 0.00000
-2.60338 0.00000 0.00000 ; C32- C28- C30- H50
7403 7399 7401 7422 3 0.65084 1.95253 0.00000
-2.60338 0.00000 0.00000 ; C32- C28- C30- H51
7405 7406 7407 7409 3 5.43920 0.00000 -5.43920
0.00000 0.00000 0.00000 ; C34- C35- N36- C38
7405 7406 7407 7410 3 5.43920 0.00000 -5.43920
0.00000 0.00000 0.00000 ; C34- C35- N36- C39
7405 7412 7410 7407 3 5.85760 0.00000 -5.85760
0.00000 0.00000 0.00000 ; C34- C41- C39- N36
7405 7412 7410 7411 3 5.85760 0.00000 -5.85760
0.00000 0.00000 0.00000 ; C34- C41- C39- O40
7406 7405 7412 7410 3 30.33400 0.00000 -30.33400
0.00000 0.00000 0.00000 ; C35- C34- C41- C39
7406 7407 7409 7426 3 0.00000 0.00000 0.00000
0.00000 0.00000 0.00000 ; C35- N36- C38- H55
7406 7407 7409 7427 3 0.00000 0.00000 0.00000
0.00000 0.00000 0.00000 ; C35- N36- C38- H56
7406 7407 7409 7428 3 0.00000 0.00000 0.00000
0.00000 0.00000 0.00000 ; C35- N36- C38- H57
7406 7407 7410 7411 3 5.43920 0.00000 -5.43920
0.00000 0.00000 0.00000 ; C35- N36- C39- O40
7406 7407 7410 7412 3 5.43920 0.00000 -5.43920
0.00000 0.00000 0.00000 ; C35- N36- C39- C41
7407 7406 7405 7412 3 5.85760 0.00000 -5.85760
0.00000 0.00000 0.00000 ; N36- C35- C34- C41
7408 7406 7405 7412 3 5.85760 0.00000 -5.85760
0.00000 0.00000 0.00000 ; O37- C35- C34- C41
7408 7406 7407 7409 3 5.43920 0.00000 -5.43920
0.00000 0.00000 0.00000 ; O37- C35- N36- C38
7408 7406 7407 7410 3 5.43920 0.00000 -5.43920
0.00000 0.00000 0.00000 ; O37- C35- N36- C39
7409 7407 7410 7411 3 5.43920 0.00000 -5.43920
0.00000 0.00000 0.00000 ; C38- N36- C39- O40
7409 7407 7410 7412 3 5.43920 0.00000 -5.43920
0.00000 0.00000 0.00000 ; C38- N36- C39- C41
7410 7407 7409 7426 3 0.00000 0.00000 0.00000
0.00000 0.00000 0.00000 ; C39- N36- C38- H55
7410 7407 7409 7427 3 0.00000 0.00000 0.00000
0.00000 0.00000 0.00000 ; C39- N36- C38- H56
7410 7407 7409 7428 3 0.00000 0.00000 0.00000
0.00000 0.00000 0.00000 ; C39- N36- C38- H57
7412 7381 7384 7417 3 5.85760 0.00000 -5.85760
0.00000 0.00000 0.00000 ; C41- C10- C13- H46
7413 7373 7374 7414 3 30.33400 0.00000 -30.33400

```
0.00000    0.00000    0.00000 ;    H42-    C2-    C3-   H43
  7416   7382   7386   7436       3    0.00000    0.00000    0.00000
0.00000    0.00000    0.00000 ;    H45-   C11-   N15-    RU
  7418   7389   7390   7436       3    0.00000    0.00000    0.00000
0.00000    0.00000    0.00000 ;    H47-   C18-   C19-    RU
  7418   7389   7393   7420       3   55.64720    0.00000  -55.64720
0.00000    0.00000    0.00000 ;    H47-   C18-   C22-   H49
  7418   7389   7393   7436       3    0.00000    0.00000    0.00000
0.00000    0.00000    0.00000 ;    H47-   C18-   C22-    RU
  7419   7392   7391   7435       3   55.64720    0.00000  -55.64720
0.00000    0.00000    0.00000 ;    H48-   C21-   C20-   H64
  7419   7392   7391   7436       3    0.00000    0.00000    0.00000
0.00000    0.00000    0.00000 ;    H48-   C21-   C20-    RU
  7419   7392   7393   7420       3   55.64720    0.00000  -55.64720
0.00000    0.00000    0.00000 ;    H48-   C21-   C22-   H49
  7419   7392   7393   7436       3    0.00000    0.00000    0.00000
0.00000    0.00000    0.00000 ;    H48-   C21-   C22-    RU
  7420   7393   7389   7436       3    0.00000    0.00000    0.00000
0.00000    0.00000    0.00000 ;    H49-   C22-   C18-    RU
  7420   7393   7392   7436       3    0.00000    0.00000    0.00000
0.00000    0.00000    0.00000 ;    H49-   C22-   C21-    RU
  7421   7401   7402   7425       3    0.69733    2.09200    0.00000
-2.78933    0.00000    0.00000 ;    H50-   C30-   O31-   H54
  7422   7401   7402   7425       3    0.69733    2.09200    0.00000
-2.78933    0.00000    0.00000 ;    H51-   C30-   O31-   H54
  7423   7403   7404   7429       3    0.69733    2.09200    0.00000
-2.78933    0.00000    0.00000 ;    H52-   C32-   O33-   H58
  7424   7403   7404   7429       3    0.69733    2.09200    0.00000
-2.78933    0.00000    0.00000 ;    H53-   C32-   O33-   H58
  7435   7391   7390   7436       3    0.00000    0.00000    0.00000
0.00000    0.00000    0.00000 ;    H64-   C20-   C19-    RU
  7435   7391   7392   7436       3    0.00000    0.00000    0.00000
0.00000    0.00000    0.00000 ;    H64-   C20-   C21-    RU

[ dihedrals ] ; impropers
; treated as propers in GROMACS to use correct AMBER analytical
function
;    i      j      k      l   func   phase     kd      pn
     5     22     20     21      1   180.00  43.93200   2 ;     CA-
N-     C-     O
    20     24     22     23      1   180.00   4.60240   2 ;      C-
CA-     N-     H
    24     41     39     40      1   180.00  43.93200   2 ;     CA-
N-     C-     O
    39     43     41     42      1   180.00   4.60240   2 ;      C-
CA-     N-     H
    43     60     58     59      1   180.00  43.93200   2 ;     CA-
N-     C-     O
    58     62     60     61      1   180.00   4.60240   2 ;      C-
CA-     N-     H
    62     71     69     70      1   180.00  43.93200   2 ;     CA-
```

```
N-      C-      O
    69      73      71      72       1   180.00   4.60240   2 ;      C-
CA-     N-      H
    73      89      87      88       1   180.00  43.93200   2 ;     CA-
N-      C-      O
    75      85      78      79       1   180.00   4.60240   2 ;     CB-
CD2-    CG-   ND1
    78      81      79      80       1   180.00   4.18400   2 ;     CG-
CE1-   ND1-   HD1
    78      83      85      86       1   180.00   4.60240   2 ;     CG-
NE2-   CD2-   HD2
    79      83      81      82       1   180.00   4.60240   2 ;    ND1-
NE2-   CE1-   HE1
    81      85      83      84       1   180.00   4.18400   2 ;    CE1-
CD2-   NE2-   HE2
    87      91      89      90       1   180.00   4.60240   2 ;      C-
CA-     N-      H
    91     100      98      99       1   180.00  43.93200   2 ;     CA-
N-      C-      O
    98     102     100     101       1   180.00   4.60240   2 ;      C-
CA-     N-      H
   102     107     105     106       1   180.00  43.93200   2 ;     CA-
N-      C-      O
   105     109     107     108       1   180.00   4.60240   2 ;      C-
CA-     N-      H
   109     129     127     128       1   180.00  43.93200   2 ;     CA-
N-      C-      O
   127     131     129     130       1   180.00   4.60240   2 ;      C-
CA-     N-      H
   131     148     146     147       1   180.00  43.93200   2 ;     CA-
N-      C-      O
   146     150     148     149       1   180.00   4.60240   2 ;      C-
CA-     N-      H
   150     163     161     162       1   180.00  43.93200   2 ;     CA-
N-      C-      O
   155     159     158     160       1   180.00  43.93200   2 ;     CG-
OE1-    CD-   OE2
   161     165     163     164       1   180.00   4.60240   2 ;      C-
CA-     N-      H
   165     175     173     174       1   180.00  43.93200   2 ;     CA-
N-      C-      O
   167     171     170     172       1   180.00  43.93200   2 ;     CB-
OD1-    CG-   OD2
   173     177     175     176       1   180.00   4.60240   2 ;      C-
CA-     N-      H
   177     197     195     196       1   180.00  43.93200   2 ;     CA-
N-      C-      O
   195     199     197     198       1   180.00   4.60240   2 ;      C-
CA-     N-      H
   199     218     216     217       1   180.00  43.93200   2 ;     CA-
N-      C-      O
   204     207     205     206       1   180.00   4.60240   2 ;     CG-
```

```
CE1-   CD1-   HD1
   204    212    214    215       1   180.00   4.60240   2 ;     CG-
CE2-   CD2-   HD2
   205    209    207    208       1   180.00   4.60240   2 ;    CD1-
CZ-   CE1-   HE1
   205    214    204    201       1   180.00   4.60240   2 ;    CD1-
CD2-    CG-    CB
   207    212    209    210       1   180.00   4.60240   2 ;    CE1-
CE2-    CZ-    OH
   209    214    212    213       1   180.00   4.60240   2 ;     CZ-
CD2-   CE2-   HE2
   216    220    218    219       1   180.00   4.60240   2 ;      C-
CA-     N-     H
   220    237    235    236       1   180.00  43.93200   2 ;     CA-
N-     C-     O
   235    239    237    238       1   180.00   4.60240   2 ;      C-
CA-     N-     H
   239    256    254    255       1   180.00  43.93200   2 ;     CA-
N-     C-     O
   254    258    256    257       1   180.00   4.60240   2 ;      C-
CA-     N-     H
   258    268    266    267       1   180.00  43.93200   2 ;     CA-
N-     C-     O
   260    264    263    265       1   180.00  43.93200   2 ;     CB-
OD1-    CG-   OD2
   266    270    268    269       1   180.00   4.60240   2 ;      C-
CA-     N-     H
   270    283    281    282       1   180.00  43.93200   2 ;     CA-
N-     C-     O
   275    279    278    280       1   180.00  43.93200   2 ;     CG-
OE1-    CD-   OE2
   281    285    283    284       1   180.00   4.60240   2 ;      C-
CA-     N-     H
   285    305    303    304       1   180.00  43.93200   2 ;     CA-
N-     C-     O
   303    307    305    306       1   180.00   4.60240   2 ;      C-
CA-     N-     H
   307    324    322    323       1   180.00  43.93200   2 ;     CA-
N-     C-     O
   322    326    324    325       1   180.00   4.60240   2 ;      C-
CA-     N-     H
   326    331    329    330       1   180.00  43.93200   2 ;     CA-
N-     C-     O
   329    333    331    332       1   180.00   4.60240   2 ;      C-
CA-     N-     H
   333    348    346    347       1   180.00  43.93200   2 ;     CA-
N-     C-     O
   338    343    341    342       1   180.00  43.93200   2 ;     CG-
NE2-    CD-   OE1
   341    344    343    345       1   180.00   4.18400   2 ;     CD-
HE21-   NE2-  HE22
   346    350    348    349       1   180.00   4.60240   2 ;      C-
```

CA- N- H
350 355 353 354 1 180.00 43.93200 2 ; CA-
N- C- O
353 357 355 356 1 180.00 4.60240 2 ; C-
CA- N- H
357 369 367 368 1 180.00 43.93200 2 ; CA-
N- C- O
367 371 369 370 1 180.00 4.60240 2 ; C-
CA- N- H
371 390 388 389 1 180.00 43.93200 2 ; CA-
N- C- O
376 379 377 378 1 180.00 4.60240 2 ; CG-
CE1- CD1- HD1
376 384 386 387 1 180.00 4.60240 2 ; CG-
CE2- CD2- HD2
377 381 379 380 1 180.00 4.60240 2 ; CD1-
CZ- CE1- HE1
377 386 376 373 1 180.00 4.60240 2 ; CD1-
CD2- CG- CB
379 384 381 382 1 180.00 4.60240 2 ; CE1-
CE2- CZ- OH
381 386 384 385 1 180.00 4.60240 2 ; CZ-
CD2- CE2- HE2
388 392 390 391 1 180.00 4.60240 2 ; C-
CA- N- H
392 397 395 396 1 180.00 43.93200 2 ; CA-
N- C- O
395 399 397 398 1 180.00 4.60240 2 ; C-
CA- N- H
399 408 406 407 1 180.00 43.93200 2 ; CA-
N- C- O
406 410 408 409 1 180.00 4.60240 2 ; C-
CA- N- H
410 424 422 423 1 180.00 43.93200 2 ; CA-
N- C- O
422 426 424 425 1 180.00 4.60240 2 ; C-
CA- N- H
426 445 443 444 1 180.00 43.93200 2 ; CA-
N- C- O
431 434 432 433 1 180.00 4.60240 2 ; CG-
CE1- CD1- HD1
431 439 441 442 1 180.00 4.60240 2 ; CG-
CE2- CD2- HD2
432 436 434 435 1 180.00 4.60240 2 ; CD1-
CZ- CE1- HE1
432 441 431 428 1 180.00 4.60240 2 ; CD1-
CD2- CG- CB
434 439 436 437 1 180.00 4.60240 2 ; CE1-
CE2- CZ- OH
436 441 439 440 1 180.00 4.60240 2 ; CZ-
CD2- CE2- HE2
443 447 445 446 1 180.00 4.60240 2 ; C-

```
CA-     N-     H
   447    467    465    466       1   180.00  43.93200   2 ;     CA-
N-     C-     O
   465    469    467    468       1   180.00   4.60240   2 ;      C-
CA-     N-     H
   469    474    472    473       1   180.00  43.93200   2 ;     CA-
N-     C-     O
   472    476    474    475       1   180.00   4.60240   2 ;      C-
CA-     N-     H
   476    493    491    492       1   180.00  43.93200   2 ;     CA-
N-     C-     O
   491    495    493    494       1   180.00   4.60240   2 ;      C-
CA-     N-     H
   495    505    503    504       1   180.00  43.93200   2 ;     CA-
N-     C-     O
   497    501    500    502       1   180.00  43.93200   2 ;     CB-
OD1-    CG-   OD2
   503    507    505    506       1   180.00   4.60240   2 ;      C-
CA-     N-     H
   507    527    525    526       1   180.00  43.93200   2 ;     CA-
N-     C-     O
   525    529    527    528       1   180.00   4.60240   2 ;      C-
CA-     N-     H
   529    543    541    542       1   180.00  43.93200   2 ;     CA-
N-     C-     O
   541    545    543    544       1   180.00   4.60240   2 ;      C-
CA-     N-     H
   545    557    555    556       1   180.00  43.93200   2 ;     CA-
N-     C-     O
   555    559    557    558       1   180.00   4.60240   2 ;      C-
CA-     N-     H
   559    571    569    570       1   180.00  43.93200   2 ;     CA-
N-     C-     O
   561    566    564    565       1   180.00  43.93200   2 ;     CB-
ND2-    CG-   OD1
   564    567    566    568       1   180.00   4.18400   2 ;     CG-
HD21-   ND2-  HD22
   569    573    571    572       1   180.00   4.60240   2 ;      C-
CA-     N-     H
   573    588    586    587       1   180.00  43.93200   2 ;     CA-
N-     C-     O
   578    583    581    582       1   180.00  43.93200   2 ;     CG-
NE2-    CD-   OE1
   581    584    583    585       1   180.00   4.18400   2 ;     CD-
HE21-   NE2-  HE22
   586    590    588    589       1   180.00   4.60240   2 ;      C-
CA-     N-     H
   590    607    605    606       1   180.00  43.93200   2 ;     CA-
N-     C-     O
   605    609    607    608       1   180.00   4.60240   2 ;      C-
CA-     N-     H
   609    628    626    627       1   180.00  43.93200   2 ;     CA-
```

```
N-     C-     O
   614    617    615    616       1   180.00   4.60240   2 ;     CG-
CE1-   CD1-   HD1
   614    622    624    625       1   180.00   4.60240   2 ;     CG-
CE2-   CD2-   HD2
   615    619    617    618       1   180.00   4.60240   2 ;    CD1-
CZ-   CE1-   HE1
   615    624    614    611       1   180.00   4.60240   2 ;    CD1-
CD2-    CG-    CB
   617    622    619    620       1   180.00   4.60240   2 ;    CE1-
CE2-    CZ-    OH
   619    624    622    623       1   180.00   4.60240   2 ;     CZ-
CD2-   CE2-   HE2
   626    630    628    629       1   180.00   4.60240   2 ;      C-
CA-     N-     H
   630    638    636    637       1   180.00  43.93200   2 ;     CA-
N-     C-     O
   636    640    638    639       1   180.00   4.60240   2 ;      C-
CA-     N-     H
   640    657    655    656       1   180.00  43.93200   2 ;     CA-
N-     C-     O
   655    659    657    658       1   180.00   4.60240   2 ;      C-
CA-     N-     H
   659    679    677    678       1   180.00  43.93200   2 ;     CA-
N-     C-     O
   677    681    679    680       1   180.00   4.60240   2 ;      C-
CA-     N-     H
   681    694    692    693       1   180.00  43.93200   2 ;     CA-
N-     C-     O
   686    690    689    691       1   180.00  43.93200   2 ;     CG-
OE1-    CD-   OE2
   692    696    694    695       1   180.00   4.60240   2 ;      C-
CA-     N-     H
   696    709    707    708       1   180.00  43.93200   2 ;     CA-
N-     C-     O
   701    705    704    706       1   180.00  43.93200   2 ;     CG-
OE1-    CD-   OE2
   707    711    709    710       1   180.00   4.60240   2 ;      C-
CA-     N-     H
   711    731    729    730       1   180.00  43.93200   2 ;     CA-
N-     C-     O
   729    733    731    732       1   180.00   4.60240   2 ;      C-
CA-     N-     H
   733    753    751    752       1   180.00  43.93200   2 ;     CA-
N-     C-     O
   751    755    753    754       1   180.00   4.60240   2 ;      C-
CA-     N-     H
   755    765    763    764       1   180.00  43.93200   2 ;     CA-
N-     C-     O
   757    761    760    762       1   180.00  43.93200   2 ;     CB-
OD1-    CG-   OD2
   763    767    765    766       1   180.00   4.60240   2 ;      C-
```

CD2-    CG-    CB
  2033   2038   2035   2036      1   180.00   4.60240   2 ;    CE1-
CE2-    CZ-    OH
  2035   2040   2038   2039      1   180.00   4.60240   2 ;     CZ-
CD2-   CE2-   HE2
  2042   2046   2044   2045      1   180.00   4.60240   2 ;      C-
CA-     N-     H
  2046   2063   2061   2062      1   180.00  43.93200   2 ;     CA-
N-     C-     O
  2061   2065   2063   2064      1   180.00   4.60240   2 ;      C-
CA-     N-     H
  2065   2080   2078   2079      1   180.00  43.93200   2 ;     CA-
N-     C-     O
  2067   2076   2070   2071      1   180.00   4.60240   2 ;     CB-
CD2-    CG-   ND1
  2070   2074   2076   2077      1   180.00   4.60240   2 ;     CG-
NE2-   CD2-   HD2
  2071   2074   2072   2073      1   180.00   4.60240   2 ;    ND1-
NE2-   CE1-   HE1
  2072   2076   2074   2075      1   180.00   4.18400   2 ;    CE1-
CD2-   NE2-   HE2
  2078   2082   2080   2081      1   180.00   4.60240   2 ;      C-
CA-     N-     H
  2082   2091   2089   2090      1   180.00  43.93200   2 ;     CA-
N-     C-     O
  2089   2093   2091   2092      1   180.00   4.60240   2 ;      C-
CA-     N-     H
  2093   2110   2108   2109      1   180.00  43.93200   2 ;     CA-
N-     C-     O
  2108   2112   2110   2111      1   180.00   4.60240   2 ;      C-
CA-     N-     H
  2112   2124   2122   2123      1   180.00  43.93200   2 ;     CA-
N-     C-     O
  2114   2119   2117   2118      1   180.00  43.93200   2 ;     CB-
ND2-    CG-   OD1
  2117   2120   2119   2121      1   180.00   4.18400   2 ;     CG-
HD21-   ND2-  HD22
  2122   2126   2124   2125      1   180.00   4.60240   2 ;      C-
CA-     N-     H
  2126   2143   2141   2142      1   180.00  43.93200   2 ;     CA-
N-     C-     O
  2141   2145   2143   2144      1   180.00   4.60240   2 ;      C-
CA-     N-     H
  2145   2159   2157   2158      1   180.00  43.93200   2 ;     CA-
N-     C-     O
  2157   2161   2159   2160      1   180.00   4.60240   2 ;      C-
CA-     N-     H
  2161   2176   2174   2175      1   180.00  43.93200   2 ;     CA-
N-     C-     O
  2163   2172   2166   2167      1   180.00   4.60240   2 ;     CB-
CD2-    CG-   ND1
  2166   2170   2172   2173      1   180.00   4.60240   2 ;     CG-

NE2-   CD2-   HD2
  2167   2170   2168   2169       1   180.00   4.60240   2 ;    ND1-
NE2-   CE1-   HE1
  2168   2172   2170   2171       1   180.00   4.18400   2 ;    CE1-
CD2-   NE2-   HE2
  2174   2178   2176   2177       1   180.00   4.60240   2 ;      C-
CA-    N-     H
  2178   2200   2198   2199       1   180.00  43.93200   2 ;     CA-
N-     C-     O
  2186   2191   2189   2190       1   180.00   4.18400   2 ;     CD-
CZ-    NE-    HE
  2189   2192   2191   2195       1   180.00  43.93200   2 ;     NE-
NH1-    CZ-   NH2
  2191   2193   2192   2194       1   180.00   4.18400   2 ;     CZ-
HH11-   NH1-  HH12
  2191   2196   2195   2197       1   180.00   4.18400   2 ;     CZ-
HH21-   NH2-  HH22
  2198   2202   2200   2201       1   180.00   4.60240   2 ;      C-
CA-    N-     H
  2202   2212   2210   2211       1   180.00  43.93200   2 ;     CA-
N-     C-     O
  2204   2208   2207   2209       1   180.00  43.93200   2 ;     CB-
OD1-    CG-   OD2
  2210   2214   2212   2213       1   180.00   4.60240   2 ;      C-
CA-    N-     H
  2214   2231   2229   2230       1   180.00  43.93200   2 ;     CA-
N-     C-     O
  2229   2233   2231   2232       1   180.00   4.60240   2 ;      C-
CA-    N-     H
  2233   2253   2251   2252       1   180.00  43.93200   2 ;     CA-
N-     C-     O
  2251   2254   2253   2263       1   180.00   4.18400   2 ;      C-
CD-    N-     CA
  2263   2267   2265   2266       1   180.00  43.93200   2 ;     CA-
N-     C-     O
  2265   2269   2267   2268       1   180.00   4.60240   2 ;      C-
CA-    N-     H
  2269   2282   2280   2281       1   180.00  43.93200   2 ;     CA-
N-     C-     O
  2274   2278   2277   2279       1   180.00  43.93200   2 ;     CG-
OE1-    CD-   OE2
  2280   2284   2282   2283       1   180.00   4.60240   2 ;      C-
CA-    N-     H
  2284   2296   2294   2295       1   180.00  43.93200   2 ;     CA-
N-     C-     O
  2286   2291   2289   2290       1   180.00  43.93200   2 ;     CB-
ND2-    CG-   OD1
  2289   2292   2291   2293       1   180.00   4.18400   2 ;     CG-
HD21-   ND2-  HD22
  2294   2298   2296   2297       1   180.00   4.60240   2 ;      C-
CA-    N-     H
  2298   2316   2314   2315       1   180.00  43.93200   2 ;     CA-

N-     C-     O
  2303   2306   2304   2305       1   180.00   4.60240   2 ;     CG-
CE1-   CD1-   HD1
  2303   2310   2312   2313       1   180.00   4.60240   2 ;     CG-
CE2-   CD2-   HD2
  2304   2308   2306   2307       1   180.00   4.60240   2 ;    CD1-
CZ-   CE1-   HE1
  2304   2312   2303   2300       1   180.00   4.60240   2 ;    CD1-
CD2-    CG-    CB
  2306   2310   2308   2309       1   180.00   4.60240   2 ;    CE1-
CE2-    CZ-    HZ
  2308   2312   2310   2311       1   180.00   4.60240   2 ;     CZ-
CD2-   CE2-   HE2
  2314   2318   2316   2317       1   180.00   4.60240   2 ;      C-
CA-     N-     H
  2318   2335   2333   2334       1   180.00  43.93200   2 ;     CA-
N-     C-     O
  2333   2337   2335   2336       1   180.00   4.60240   2 ;      C-
CA-     N-     H
  2337   2355   2353   2354       1   180.00  43.93200   2 ;     CA-
N-     C-     O
  2342   2345   2343   2344       1   180.00   4.60240   2 ;     CG-
CE1-   CD1-   HD1
  2342   2349   2351   2352       1   180.00   4.60240   2 ;     CG-
CE2-   CD2-   HD2
  2343   2347   2345   2346       1   180.00   4.60240   2 ;    CD1-
CZ-   CE1-   HE1
  2343   2351   2342   2339       1   180.00   4.60240   2 ;    CD1-
CD2-    CG-    CB
  2345   2349   2347   2348       1   180.00   4.60240   2 ;    CE1-
CE2-    CZ-    HZ
  2347   2351   2349   2350       1   180.00   4.60240   2 ;     CZ-
CD2-   CE2-   HE2
  2353   2357   2355   2356       1   180.00   4.60240   2 ;      C-
CA-     N-     H
  2357   2372   2370   2371       1   180.00  43.93200   2 ;     CA-
N-     C-     O
  2362   2367   2365   2366       1   180.00  43.93200   2 ;     CG-
NE2-    CD-   OE1
  2365   2368   2367   2369       1   180.00   4.18400   2 ;     CD-
HE21-   NE2-  HE22
  2370   2374   2372   2373       1   180.00   4.60240   2 ;      C-
CA-     N-     H
  2374   2383   2381   2382       1   180.00  43.93200   2 ;     CA-
N-     C-     O
  2381   2385   2383   2384       1   180.00   4.60240   2 ;      C-
CA-     N-     H
  2385   2398   2396   2397       1   180.00  43.93200   2 ;     CA-
N-     C-     O
  2390   2394   2393   2395       1   180.00  43.93200   2 ;     CG-
OE1-    CD-   OE2
  2396   2400   2398   2399       1   180.00   4.60240   2 ;      C-

CA-     N-     H
  2400   2412   2410   2411       1   180.00  43.93200   2 ;     CA-
N-     C-     O
  2402   2407   2405   2406       1   180.00  43.93200   2 ;     CB-
ND2-    CG-   OD1
  2405   2408   2407   2409       1   180.00   4.18400   2 ;     CG-
HD21-   ND2-  HD22
  2410   2414   2412   2413       1   180.00   4.60240   2 ;      C-
CA-     N-     H
  2414   2434   2432   2433       1   180.00  43.93200   2 ;     CA-
N-     C-     O
  2432   2436   2434   2435       1   180.00   4.60240   2 ;      C-
CA-     N-     H
  2436   2446   2444   2445       1   180.00  43.93200   2 ;     CA-
N-     C-     O
  2438   2442   2441   2443       1   180.00  43.93200   2 ;     CB-
OD1-    CG-   OD2
  2444   2448   2446   2447       1   180.00   4.60240   2 ;      C-
CA-     N-     H
  2448   2457   2455   2456       1   180.00  43.93200   2 ;     CA-
N-     C-     O
  2455   2459   2457   2458       1   180.00   4.60240   2 ;      C-
CA-     N-     H
  2459   2476   2474   2475       1   180.00  43.93200   2 ;     CA-
N-     C-     O
  2474   2478   2476   2477       1   180.00   4.60240   2 ;      C-
CA-     N-     H
  2478   2495   2493   2494       1   180.00  43.93200   2 ;     CA-
N-     C-     O
  2493   2497   2495   2496       1   180.00   4.60240   2 ;      C-
CA-     N-     H
  2497   2517   2515   2516       1   180.00  43.93200   2 ;     CA-
N-     C-     O
  2515   2519   2517   2518       1   180.00   4.60240   2 ;      C-
CA-     N-     H
  2519   2536   2534   2535       1   180.00  43.93200   2 ;     CA-
N-     C-     O
  2534   2538   2536   2537       1   180.00   4.60240   2 ;      C-
CA-     N-     H
  2538   2555   2553   2554       1   180.00  43.93200   2 ;     CA-
N-     C-     O
  2553   2557   2555   2556       1   180.00   4.60240   2 ;      C-
CA-     N-     H
  2557   2567   2565   2566       1   180.00  43.93200   2 ;     CA-
N-     C-     O
  2559   2563   2562   2564       1   180.00  43.93200   2 ;     CB-
OD1-    CG-   OD2
  2565   2569   2567   2568       1   180.00   4.60240   2 ;      C-
CA-     N-     H
  2569   2587   2585   2586       1   180.00  43.93200   2 ;     CA-
N-     C-     O
  2574   2577   2575   2576       1   180.00   4.60240   2 ;     CG-

CE1- CD1- HD1
2574 2581 2583 2584 1 180.00 4.60240 2 ; CG-
CE2- CD2- HD2
2575 2579 2577 2578 1 180.00 4.60240 2 ; CD1-
CZ- CE1- HE1
2575 2583 2574 2571 1 180.00 4.60240 2 ; CD1-
CD2- CG- CB
2577 2581 2579 2580 1 180.00 4.60240 2 ; CE1-
CE2- CZ- HZ
2579 2583 2581 2582 1 180.00 4.60240 2 ; CZ-
CD2- CE2- HE2
2585 2589 2587 2588 1 180.00 4.60240 2 ; C-
CA- N- H
2589 2594 2592 2593 1 180.00 43.93200 2 ; CA-
N- C- O
2592 2596 2594 2595 1 180.00 4.60240 2 ; C-
CA- N- H
2596 2613 2611 2612 1 180.00 43.93200 2 ; CA-
N- C- O
2611 2615 2613 2614 1 180.00 4.60240 2 ; C-
CA- N- H
2615 2624 2622 2623 1 180.00 43.93200 2 ; CA-
N- C- O
2622 2626 2624 2625 1 180.00 4.60240 2 ; C-
CA- N- H
2626 2646 2644 2645 1 180.00 43.93200 2 ; CA-
N- C- O
2644 2648 2646 2647 1 180.00 4.60240 2 ; C-
CA- N- H
2648 2660 2658 2659 1 180.00 43.93200 2 ; CA-
N- C- O
2650 2655 2653 2654 1 180.00 43.93200 2 ; CB-
ND2- CG- OD1
2653 2656 2655 2657 1 180.00 4.18400 2 ; CG-
HD21- ND2- HD22
2658 2662 2660 2661 1 180.00 4.60240 2 ; C-
CA- N- H
2662 2679 2677 2678 1 180.00 43.93200 2 ; CA-
N- C- O
2677 2681 2679 2680 1 180.00 4.60240 2 ; C-
CA- N- H
2681 2686 2684 2685 1 180.00 43.93200 2 ; CA-
N- C- O
2684 2688 2686 2687 1 180.00 4.60240 2 ; C-
CA- N- H
2688 2700 2698 2699 1 180.00 43.93200 2 ; CA-
N- C- O
2698 2702 2700 2701 1 180.00 4.60240 2 ; C-
CA- N- H
2702 2707 2705 2706 1 180.00 43.93200 2 ; CA-
N- C- O
2705 2709 2707 2708 1 180.00 4.60240 2 ; C-

CA- N- H
2709 2722 2720 2721 1 180.00 43.93200 2 ; CA-
N- C- O
2714 2718 2717 2719 1 180.00 43.93200 2 ; CG-
OE1- CD- OE2
2720 2724 2722 2723 1 180.00 4.60240 2 ; C-
CA- N- H
2724 2742 2740 2741 1 180.00 43.93200 2 ; CA-
N- C- O
2729 2732 2730 2731 1 180.00 4.60240 2 ; CG-
CE1- CD1- HD1
2729 2736 2738 2739 1 180.00 4.60240 2 ; CG-
CE2- CD2- HD2
2730 2734 2732 2733 1 180.00 4.60240 2 ; CD1-
CZ- CE1- HE1
2730 2738 2729 2726 1 180.00 4.60240 2 ; CD1-
CD2- CG- CB
2732 2736 2734 2735 1 180.00 4.60240 2 ; CE1-
CE2- CZ- HZ
2734 2738 2736 2737 1 180.00 4.60240 2 ; CZ-
CD2- CE2- HE2
2740 2744 2742 2743 1 180.00 4.60240 2 ; C-
CA- N- H
2744 2756 2754 2755 1 180.00 43.93200 2 ; CA-
N- C- O
2754 2758 2756 2757 1 180.00 4.60240 2 ; C-
CA- N- H
2758 2770 2768 2769 1 180.00 43.93200 2 ; CA-
N- C- O
2768 2772 2770 2771 1 180.00 4.60240 2 ; C-
CA- N- H
2772 2784 2782 2783 1 180.00 43.93200 2 ; CA-
N- C- O
2782 2786 2784 2785 1 180.00 4.60240 2 ; C-
CA- N- H
2786 2806 2804 2805 1 180.00 43.93200 2 ; CA-
N- C- O
2804 2808 2806 2807 1 180.00 4.60240 2 ; C-
CA- N- H
2808 2816 2814 2815 1 180.00 43.93200 2 ; CA-
N- C- O
2814 2818 2816 2817 1 180.00 4.60240 2 ; C-
CA- N- H
2818 2823 2821 2822 1 180.00 43.93200 2 ; CA-
N- C- O
2821 2825 2823 2824 1 180.00 4.60240 2 ; C-
CA- N- H
2825 2837 2835 2836 1 180.00 43.93200 2 ; CA-
N- C- O
2835 2838 2837 2847 1 180.00 4.18400 2 ; C-
CD- N- CA
2847 2851 2849 2850 1 180.00 43.93200 2 ; CA-

N- C- O
2849 2853 2851 2852 1 180.00 4.60240 2 ; C-
CA- N- H
2853 2872 2870 2871 1 180.00 43.93200 2 ; CA-
N- C- O
2858 2861 2859 2860 1 180.00 4.60240 2 ; CG-
CE1- CD1- HD1
2858 2866 2868 2869 1 180.00 4.60240 2 ; CG-
CE2- CD2- HD2
2859 2863 2861 2862 1 180.00 4.60240 2 ; CD1-
CZ- CE1- HE1
2859 2868 2858 2855 1 180.00 4.60240 2 ; CD1-
CD2- CG- CB
2861 2866 2863 2864 1 180.00 4.60240 2 ; CE1-
CE2- CZ- OH
2863 2868 2866 2867 1 180.00 4.60240 2 ; CZ-
CD2- CE2- HE2
2870 2874 2872 2873 1 180.00 4.60240 2 ; C-
CA- N- H
2874 2893 2891 2892 1 180.00 43.93200 2 ; CA-
N- C- O
2879 2882 2880 2881 1 180.00 4.60240 2 ; CG-
CE1- CD1- HD1
2879 2887 2889 2890 1 180.00 4.60240 2 ; CG-
CE2- CD2- HD2
2880 2884 2882 2883 1 180.00 4.60240 2 ; CD1-
CZ- CE1- HE1
2880 2889 2879 2876 1 180.00 4.60240 2 ; CD1-
CD2- CG- CB
2882 2887 2884 2885 1 180.00 4.60240 2 ; CE1-
CE2- CZ- OH
2884 2889 2887 2888 1 180.00 4.60240 2 ; CZ-
CD2- CE2- HE2
2891 2895 2893 2894 1 180.00 4.60240 2 ; C-
CA- N- H
2895 2909 2907 2908 1 180.00 43.93200 2 ; CA-
N- C- O
2907 2911 2909 2910 1 180.00 4.60240 2 ; C-
CA- N- H
2911 2919 2917 2918 1 180.00 43.93200 2 ; CA-
N- C- O
2917 2920 2919 2929 1 180.00 4.18400 2 ; C-
CD- N- CA
2929 2933 2931 2932 1 180.00 43.93200 2 ; CA-
N- C- O
2931 2935 2933 2934 1 180.00 4.60240 2 ; C-
CA- N- H
2935 2950 2948 2949 1 180.00 43.93200 2 ; CA-
N- C- O
2940 2945 2943 2944 1 180.00 43.93200 2 ; CG-
NE2- CD- OE1
2943 2946 2945 2947 1 180.00 4.18400 2 ; CD-

HE21- NE2- HE22
2948 2952 2950 2951 1 180.00 4.60240 2 ; C-
CA- N- H
2952 2966 2964 2965 1 180.00 43.93200 2 ; CA-
N- C- O
2964 2968 2966 2967 1 180.00 4.60240 2 ; C-
CA- N- H
2968 2985 2983 2984 1 180.00 43.93200 2 ; CA-
N- C- O
2983 2987 2985 2986 1 180.00 4.60240 2 ; C-
CA- N- H
2987 2997 2995 2996 1 180.00 43.93200 2 ; CA-
N- C- O
2989 2993 2992 2994 1 180.00 43.93200 2 ; CB-
OD1- CG- OD2
2995 2999 2997 2998 1 180.00 4.60240 2 ; C-
CA- N- H
2999 3004 3002 3003 1 180.00 43.93200 2 ; CA-
N- C- O
3002 3006 3004 3005 1 180.00 4.60240 2 ; C-
CA- N- H
3006 3026 3024 3025 1 180.00 43.93200 2 ; CA-
N- C- O
3024 3028 3026 3027 1 180.00 4.60240 2 ; C-
CA- N- H
3028 3047 3045 3046 1 180.00 43.93200 2 ; CA-
N- C- O
3033 3036 3034 3035 1 180.00 4.60240 2 ; CG-
CE1- CD1- HD1
3033 3041 3043 3044 1 180.00 4.60240 2 ; CG-
CE2- CD2- HD2
3034 3038 3036 3037 1 180.00 4.60240 2 ; CD1-
CZ- CE1- HE1
3034 3043 3033 3030 1 180.00 4.60240 2 ; CD1-
CD2- CG- CB
3036 3041 3038 3039 1 180.00 4.60240 2 ; CE1-
CE2- CZ- OH
3038 3043 3041 3042 1 180.00 4.60240 2 ; CZ-
CD2- CE2- HE2
3045 3049 3047 3048 1 180.00 4.60240 2 ; C-
CA- N- H
3049 3059 3057 3058 1 180.00 43.93200 2 ; CA-
N- C- O
3051 3055 3054 3056 1 180.00 43.93200 2 ; CB-
OD1- CG- OD2
3057 3061 3059 3060 1 180.00 4.60240 2 ; C-
CA- N- H
3061 3081 3079 3080 1 180.00 43.93200 2 ; CA-
N- C- O
3079 3083 3081 3082 1 180.00 4.60240 2 ; C-
CA- N- H
3083 3102 3100 3101 1 180.00 43.93200 2 ; CA-

```
N-      C-      O
  3100   3104   3102   3103       1   180.00   4.60240   2 ;      C-
CA-     N-      H
  3104   3113   3111   3112       1   180.00  43.93200   2 ;     CA-
N-      C-      O
  3111   3115   3113   3114       1   180.00   4.60240   2 ;      C-
CA-     N-      H
  3115   3125   3123   3124       1   180.00  43.93200   2 ;     CA-
N-      C-      O
  3117   3121   3120   3122       1   180.00  43.93200   2 ;     CB-
OD1-    CG-    OD2
  3123   3127   3125   3126       1   180.00   4.60240   2 ;      C-
CA-     N-      H
  3127   3144   3142   3143       1   180.00  43.93200   2 ;     CA-
N-      C-      O
  3142   3146   3144   3145       1   180.00   4.60240   2 ;      C-
CA-     N-      H
  3146   3168   3166   3167       1   180.00  43.93200   2 ;     CA-
N-      C-      O
  3151   3154   3152   3153       1   180.00   4.60240   2 ;     CG-
NE1-   CD1-    HD1
  3152   3156   3154   3155       1   180.00   4.18400   2 ;    CD1-
CE2-   NE1-    HE1
  3156   3159   3157   3158       1   180.00   4.60240   2 ;    CE2-
CH2-   CZ2-    HZ2
  3157   3161   3159   3160       1   180.00   4.60240   2 ;    CZ2-
CZ3-   CH2-    HH2
  3159   3163   3161   3162       1   180.00   4.60240   2 ;    CH2-
CE3-   CZ3-    HZ3
  3161   3165   3163   3164       1   180.00   4.60240   2 ;    CZ3-
CD2-   CE3-    HE3
  3165   3148   3151   3152       1   180.00   4.60240   2 ;    CD2-
CB-     CG-   CD1
  3166   3170   3168   3169       1   180.00   4.60240   2 ;      C-
CA-     N-      H
  3170   3179   3177   3178       1   180.00  43.93200   2 ;     CA-
N-      C-      O
  3177   3181   3179   3180       1   180.00   4.60240   2 ;      C-
CA-     N-      H
  3181   3190   3188   3189       1   180.00  43.93200   2 ;     CA-
N-      C-      O
  3188   3192   3190   3191       1   180.00   4.60240   2 ;      C-
CA-     N-      H
  3192   3197   3195   3196       1   180.00  43.93200   2 ;     CA-
N-      C-      O
  3195   3199   3197   3198       1   180.00   4.60240   2 ;      C-
CA-     N-      H
  3199   3213   3211   3212       1   180.00  43.93200   2 ;     CA-
N-      C-      O
  3211   3215   3213   3214       1   180.00   4.60240   2 ;      C-
CA-     N-      H
  3215   3232   3230   3231       1   180.00  43.93200   2 ;     CA-
```

N- C- O
3230 3234 3232 3233 1 180.00 4.60240 2 ; C-
CA- N- H
3234 3249 3247 3248 1 180.00 43.93200 2 ; CA-
N- C- O
3247 3251 3249 3250 1 180.00 4.60240 2 ; C-
CA- N- H
3251 3270 3268 3269 1 180.00 43.93200 2 ; CA-
N- C- O
3256 3259 3257 3258 1 180.00 4.60240 2 ; CG-
CE1- CD1- HD1
3256 3264 3266 3267 1 180.00 4.60240 2 ; CG-
CE2- CD2- HD2
3257 3261 3259 3260 1 180.00 4.60240 2 ; CD1-
CZ- CE1- HE1
3257 3266 3256 3253 1 180.00 4.60240 2 ; CD1-
CD2- CG- CB
3259 3264 3261 3262 1 180.00 4.60240 2 ; CE1-
CE2- CZ- OH
3261 3266 3264 3265 1 180.00 4.60240 2 ; CZ-
CD2- CE2- HE2
3268 3272 3270 3271 1 180.00 4.60240 2 ; C-
CA- N- H
3272 3284 3282 3283 1 180.00 43.93200 2 ; CA-
N- C- O
3282 3286 3284 3285 1 180.00 4.60240 2 ; C-
CA- N- H
3286 3303 3301 3302 1 180.00 43.93200 2 ; CA-
N- C- O
3301 3305 3303 3304 1 180.00 4.60240 2 ; C-
CA- N- H
3305 3322 3320 3321 1 180.00 43.93200 2 ; CA-
N- C- O
3320 3324 3322 3323 1 180.00 4.60240 2 ; C-
CA- N- H
3324 3333 3331 3332 1 180.00 43.93200 2 ; CA-
N- C- O
3331 3335 3333 3334 1 180.00 4.60240 2 ; C-
CA- N- H
3335 3340 3338 3339 1 180.00 43.93200 2 ; CA-
N- C- O
3338 3342 3340 3341 1 180.00 4.60240 2 ; C-
CA- N- H
3342 3361 3359 3360 1 180.00 43.93200 2 ; CA-
N- C- O
3347 3350 3348 3349 1 180.00 4.60240 2 ; CG-
CE1- CD1- HD1
3347 3355 3357 3358 1 180.00 4.60240 2 ; CG-
CE2- CD2- HD2
3348 3352 3350 3351 1 180.00 4.60240 2 ; CD1-
CZ- CE1- HE1
3348 3357 3347 3344 1 180.00 4.60240 2 ; CD1-

```
CD2-    CG-    CB
  3350   3355   3352   3353       1   180.00   4.60240   2 ;    CE1-
CE2-    CZ-    OH
  3352   3357   3355   3356       1   180.00   4.60240   2 ;     CZ-
CD2-   CE2-   HE2
  3359   3362   3361   3371       1   180.00   4.18400   2 ;      C-
CD-     N-    CA
  3371   3375   3373   3374       1   180.00  43.93200   2 ;     CA-
N-     C-     O
  3373   3376   3375   3385       1   180.00   4.18400   2 ;      C-
CD-     N-    CA
  3385   3389   3387   3388       1   180.00  43.93200   2 ;     CA-
N-     C-     O
  3387   3391   3389   3390       1   180.00   4.60240   2 ;      C-
CA-     N-     H
  3391   3409   3407   3408       1   180.00  43.93200   2 ;     CA-
N-     C-     O
  3396   3399   3397   3398       1   180.00   4.60240   2 ;     CG-
CE1-   CD1-   HD1
  3396   3403   3405   3406       1   180.00   4.60240   2 ;     CG-
CE2-   CD2-   HD2
  3397   3401   3399   3400       1   180.00   4.60240   2 ;    CD1-
CZ-   CE1-   HE1
  3397   3405   3396   3393       1   180.00   4.60240   2 ;    CD1-
CD2-    CG-    CB
  3399   3403   3401   3402       1   180.00   4.60240   2 ;    CE1-
CE2-    CZ-    HZ
  3401   3405   3403   3404       1   180.00   4.60240   2 ;     CZ-
CD2-   CE2-   HE2
  3407   3411   3409   3410       1   180.00   4.60240   2 ;      C-
CA-     N-     H
  3411   3430   3428   3429       1   180.00  43.93200   2 ;     CA-
N-     C-     O
  3416   3419   3417   3418       1   180.00   4.60240   2 ;     CG-
CE1-   CD1-   HD1
  3416   3424   3426   3427       1   180.00   4.60240   2 ;     CG-
CE2-   CD2-   HD2
  3417   3421   3419   3420       1   180.00   4.60240   2 ;    CD1-
CZ-   CE1-   HE1
  3417   3426   3416   3413       1   180.00   4.60240   2 ;    CD1-
CD2-    CG-    CB
  3419   3424   3421   3422       1   180.00   4.60240   2 ;    CE1-
CE2-    CZ-    OH
  3421   3426   3424   3425       1   180.00   4.60240   2 ;     CZ-
CD2-   CE2-   HE2
  3428   3432   3430   3431       1   180.00   4.60240   2 ;      C-
CA-     N-     H
  3432   3437   3435   3436       1   180.00  43.93200   2 ;     CA-
N-     C-     O
  3435   3439   3437   3438       1   180.00   4.60240   2 ;      C-
CA-     N-     H
  3439   3449   3447   3448       1   180.00  43.93200   2 ;     CA-
```

```
N-      C-      O
  3441   3445   3444   3446       1   180.00  43.93200   2 ;     CB-
OD1-    CG-   OD2
  3447   3451   3449   3450       1   180.00   4.60240   2 ;      C-
CA-     N-     H
  3451   3463   3461   3462       1   180.00  43.93200   2 ;     CA-
N-      C-      O
  3461   3465   3463   3464       1   180.00   4.60240   2 ;      C-
CA-     N-     H
  3465   3475   3473   3474       1   180.00  43.93200   2 ;     CA-
N-      C-      O
  3467   3471   3470   3472       1   180.00  43.93200   2 ;     CB-
OD1-    CG-   OD2
  3473   3477   3475   3476       1   180.00   4.60240   2 ;      C-
CA-     N-     H
  3477   3489   3487   3488       1   180.00  43.93200   2 ;     CA-
N-      C-      O
  3479   3484   3482   3483       1   180.00  43.93200   2 ;     CB-
ND2-    CG-   OD1
  3482   3485   3484   3486       1   180.00   4.18400   2 ;     CG-
HD21-   ND2-  HD22
  3487   3491   3489   3490       1   180.00   4.60240   2 ;      C-
CA-     N-     H
  3491   3504   3502   3503       1   180.00  43.93200   2 ;     CA-
N-      C-      O
  3496   3500   3499   3501       1   180.00  43.93200   2 ;     CG-
OE1-    CD-   OE2
  3502   3506   3504   3505       1   180.00   4.60240   2 ;      C-
CA-     N-     H
  3506   3520   3518   3519       1   180.00  43.93200   2 ;     CA-
N-      C-      O
  3518   3522   3520   3521       1   180.00   4.60240   2 ;      C-
CA-     N-     H
  3522   3539   3537   3538       1   180.00  43.93200   2 ;     CA-
N-      C-      O
  3537   3541   3539   3540       1   180.00   4.60240   2 ;      C-
CA-     N-     H
  3541   3561   3559   3560       1   180.00  43.93200   2 ;     CA-
N-      C-      O
  3559   3563   3561   3562       1   180.00   4.60240   2 ;      C-
CA-     N-     H
  3563   3583   3581   3582       1   180.00  43.93200   2 ;     CA-
N-      C-      O
  3581   3585   3583   3584       1   180.00   4.60240   2 ;      C-
CA-     N-     H
  3585   3599   3597   3598       1   180.00  43.93200   2 ;     CA-
N-      C-      O
  3597   3601   3599   3600       1   180.00   4.60240   2 ;      C-
CA-     N-     H
  3601   3621   3619   3620       1   180.00  43.93200   2 ;     CA-
N-      C-      O
  3619   3623   3621   3622       1   180.00   4.60240   2 ;      C-
```

CA-     N-     H
  3623   3643   3641   3642       1   180.00  43.93200   2 ;     CA-
N-     C-     O
  3641   3645   3643   3644       1   180.00   4.60240   2 ;      C-
CA-     N-     H
  3645   3650   3648   3649       1   180.00  43.93200   2 ;     CA-
N-     C-     O
  3648   3652   3650   3651       1   180.00   4.60240   2 ;      C-
CA-     N-     H
  3652   3665   3663   3664       1   180.00  43.93200   2 ;     CA-
N-     C-     O
  3657   3661   3660   3662       1   180.00  43.93200   2 ;     CG-
OE1-    CD-   OE2
  3663   3667   3665   3666       1   180.00   4.60240   2 ;      C-
CA-     N-     H
  3667   3685   3683   3684       1   180.00  43.93200   2 ;     CA-
N-     C-     O
  3672   3675   3673   3674       1   180.00   4.60240   2 ;     CG-
CE1-   CD1-   HD1
  3672   3679   3681   3682       1   180.00   4.60240   2 ;     CG-
CE2-   CD2-   HD2
  3673   3677   3675   3676       1   180.00   4.60240   2 ;    CD1-
CZ-   CE1-   HE1
  3673   3681   3672   3669       1   180.00   4.60240   2 ;    CD1-
CD2-    CG-    CB
  3675   3679   3677   3678       1   180.00   4.60240   2 ;    CE1-
CE2-    CZ-    HZ
  3677   3681   3679   3680       1   180.00   4.60240   2 ;     CZ-
CD2-   CE2-   HE2
  3683   3687   3685   3686       1   180.00   4.60240   2 ;      C-
CA-     N-     H
  3687   3696   3694   3695       1   180.00  43.93200   2 ;     CA-
N-     C-     O
  3694   3698   3696   3697       1   180.00   4.60240   2 ;      C-
CA-     N-     H
  3698   3716   3714   3715       1   180.00  43.93200   2 ;     CA-
N-     C-     O
  3703   3706   3704   3705       1   180.00   4.60240   2 ;     CG-
CE1-   CD1-   HD1
  3703   3710   3712   3713       1   180.00   4.60240   2 ;     CG-
CE2-   CD2-   HD2
  3704   3708   3706   3707       1   180.00   4.60240   2 ;    CD1-
CZ-   CE1-   HE1
  3704   3712   3703   3700       1   180.00   4.60240   2 ;    CD1-
CD2-    CG-    CB
  3706   3710   3708   3709       1   180.00   4.60240   2 ;    CE1-
CE2-    CZ-    HZ
  3708   3712   3710   3711       1   180.00   4.60240   2 ;     CZ-
CD2-   CE2-   HE2
  3714   3718   3716   3717       1   180.00   4.60240   2 ;      C-
CA-     N-     H
  3718   3737   3735   3736       1   180.00  43.93200   2 ;     CA-

N- C- O
3723 3726 3724 3725 1 180.00 4.60240 2 ; CG-
CE1- CD1- HD1
3723 3731 3733 3734 1 180.00 4.60240 2 ; CG-
CE2- CD2- HD2
3724 3728 3726 3727 1 180.00 4.60240 2 ; CD1-
CZ- CE1- HE1
3724 3733 3723 3720 1 180.00 4.60240 2 ; CD1-
CD2- CG- CB
3726 3731 3728 3729 1 180.00 4.60240 2 ; CE1-
CE2- CZ- OH
3728 3733 3731 3732 1 180.00 4.60240 2 ; CZ-
CD2- CE2- HE2
3735 3739 3737 3738 1 180.00 4.60240 2 ; C-
CA- N- H
3739 3752 3750 3751 1 180.00 43.93200 2 ; CA-
N- C- O
3744 3748 3747 3749 1 180.00 43.93200 2 ; CG-
OE1- CD- OE2
3750 3754 3752 3753 1 180.00 4.60240 2 ; C-
CA- N- H
3754 3766 3764 3765 1 180.00 43.93200 2 ; CA-
N- C- O
3756 3761 3759 3760 1 180.00 43.93200 2 ; CB-
ND2- CG- OD1
3759 3762 3761 3763 1 180.00 4.18400 2 ; CG-
HD21- ND2- HD22
3764 3768 3766 3767 1 180.00 4.60240 2 ; C-
CA- N- H
3768 3778 3776 3777 1 180.00 43.93200 2 ; CA-
N- C- O
3770 3774 3773 3775 1 180.00 43.93200 2 ; CB-
OD1- CG- OD2
3776 3780 3778 3779 1 180.00 4.60240 2 ; C-
CA- N- H
3780 3802 3800 3801 1 180.00 43.93200 2 ; CA-
N- C- O
3785 3788 3786 3787 1 180.00 4.60240 2 ; CG-
NE1- CD1- HD1
3786 3790 3788 3789 1 180.00 4.18400 2 ; CD1-
CE2- NE1- HE1
3790 3793 3791 3792 1 180.00 4.60240 2 ; CE2-
CH2- CZ2- HZ2
3791 3795 3793 3794 1 180.00 4.60240 2 ; CZ2-
CZ3- CH2- HH2
3793 3797 3795 3796 1 180.00 4.60240 2 ; CH2-
CE3- CZ3- HZ3
3795 3799 3797 3798 1 180.00 4.60240 2 ; CZ3-
CD2- CE3- HE3
3799 3782 3785 3786 1 180.00 4.60240 2 ; CD2-
CB- CG- CD1
3800 3804 3802 3803 1 180.00 4.60240 2 ; C-

CA- N- H
3804 3809 3807 3808 1 180.00 43.93200 2 ; CA-
N- C- O
3807 3811 3809 3810 1 180.00 4.60240 2 ; C-
CA- N- H
3811 3820 3818 3819 1 180.00 43.93200 2 ; CA-
N- C- O
3818 3822 3820 3821 1 180.00 4.60240 2 ; C-
CA- N- H
3822 3839 3837 3838 1 180.00 43.93200 2 ; CA-
N- C- O
3837 3841 3839 3840 1 180.00 4.60240 2 ; C-
CA- N- H
3841 3850 3848 3849 1 180.00 43.93200 2 ; CA-
N- C- O
3848 3852 3850 3851 1 180.00 4.60240 2 ; C-
CA- N- H
3852 3861 3859 3860 1 180.00 43.93200 2 ; CA-
N- C- O
3859 3863 3861 3862 1 180.00 4.60240 2 ; C-
CA- N- H
3863 3873 3871 3872 1 180.00 43.93200 2 ; CA-
N- C- O
3865 3869 3868 3870 1 180.00 43.93200 2 ; CB-
OD1- CG- OD2
3871 3875 3873 3874 1 180.00 4.60240 2 ; C-
CA- N- H
3875 3883 3881 3882 1 180.00 43.93200 2 ; CA-
N- C- O
3881 3885 3883 3884 1 180.00 4.60240 2 ; C-
CA- N- H
3885 3905 3903 3904 1 180.00 43.93200 2 ; CA-
N- C- O
3903 3907 3905 3906 1 180.00 4.60240 2 ; C-
CA- N- H
3907 3919 3917 3918 1 180.00 43.93200 2 ; CA-
N- C- O
3909 3914 3912 3913 1 180.00 43.93200 2 ; CB-
ND2- CG- OD1
3912 3915 3914 3916 1 180.00 4.18400 2 ; CG-
HD21- ND2- HD22
3917 3921 3919 3920 1 180.00 4.60240 2 ; C-
CA- N- H
3921 3938 3936 3937 1 180.00 43.93200 2 ; CA-
N- C- O
3936 3940 3938 3939 1 180.00 4.60240 2 ; C-
CA- N- H
3940 3957 3955 3956 1 180.00 43.93200 2 ; CA-
N- C- O
3955 3959 3957 3958 1 180.00 4.60240 2 ; C-
CA- N- H
3959 3971 3969 3970 1 180.00 43.93200 2 ; CA-

N- C- O
3969 3973 3971 3972 1 180.00 4.60240 2 ; C-
CA- N- H
3973 3993 3991 3992 1 180.00 43.93200 2 ; CA-
N- C- O
3991 3995 3993 3994 1 180.00 4.60240 2 ; C-
CA- N- H
3995 4012 4010 4011 1 180.00 43.93200 2 ; CA-
N- C- O
4010 4014 4012 4013 1 180.00 4.60240 2 ; C-
CA- N- H
4014 4031 4029 4030 1 180.00 43.93200 2 ; CA-
N- C- O
4029 4033 4031 4032 1 180.00 4.60240 2 ; C-
CA- N- H
4033 4045 4043 4044 1 180.00 43.93200 2 ; CA-
N- C- O
4043 4047 4045 4046 1 180.00 4.60240 2 ; C-
CA- N- H
4047 4066 4064 4065 1 180.00 43.93200 2 ; CA-
N- C- O
4052 4055 4053 4054 1 180.00 4.60240 2 ; CG-
CE1- CD1- HD1
4052 4060 4062 4063 1 180.00 4.60240 2 ; CG-
CE2- CD2- HD2
4053 4057 4055 4056 1 180.00 4.60240 2 ; CD1-
CZ- CE1- HE1
4053 4062 4052 4049 1 180.00 4.60240 2 ; CD1-
CD2- CG- CB
4055 4060 4057 4058 1 180.00 4.60240 2 ; CE1-
CE2- CZ- OH
4057 4062 4060 4061 1 180.00 4.60240 2 ; CZ-
CD2- CE2- HE2
4064 4068 4066 4067 1 180.00 4.60240 2 ; C-
CA- N- H
4068 4080 4078 4079 1 180.00 43.93200 2 ; CA-
N- C- O
4070 4075 4073 4074 1 180.00 43.93200 2 ; CB-
ND2- CG- OD1
4073 4076 4075 4077 1 180.00 4.18400 2 ; CG-
HD21- ND2- HD22
4078 4081 4080 4090 1 180.00 4.18400 2 ; C-
CD- N- CA
4090 4094 4092 4093 1 180.00 43.93200 2 ; CA-
N- C- O
4092 4096 4094 4095 1 180.00 4.60240 2 ; C-
CA- N- H
4096 4108 4106 4107 1 180.00 43.93200 2 ; CA-
N- C- O
4098 4103 4101 4102 1 180.00 43.93200 2 ; CB-
ND2- CG- OD1
4101 4104 4103 4105 1 180.00 4.18400 2 ; CG-

HD21- ND2- HD22
4106 4110 4108 4109 1 180.00 4.60240 2 ; C-
CA- N- H
4110 4123 4121 4122 1 180.00 43.93200 2 ; CA-
N- C- O
4115 4119 4118 4120 1 180.00 43.93200 2 ; CG-
OE1- CD- OE2
4121 4125 4123 4124 1 180.00 4.60240 2 ; C-
CA- N- H
4125 4147 4145 4146 1 180.00 43.93200 2 ; CA-
N- C- O
4133 4138 4136 4137 1 180.00 4.18400 2 ; CD-
CZ- NE- HE
4136 4139 4138 4142 1 180.00 43.93200 2 ; NE-
NH1- CZ- NH2
4138 4140 4139 4141 1 180.00 4.18400 2 ; CZ-
HH11- NH1- HH12
4138 4143 4142 4144 1 180.00 4.18400 2 ; CZ-
HH21- NH2- HH22
4145 4149 4147 4148 1 180.00 4.60240 2 ; C-
CA- N- H
4149 4158 4156 4157 1 180.00 43.93200 2 ; CA-
N- C- O
4156 4160 4158 4159 1 180.00 4.60240 2 ; C-
CA- N- H
4160 4172 4170 4171 1 180.00 43.93200 2 ; CA-
N- C- O
4170 4174 4172 4173 1 180.00 4.60240 2 ; C-
CA- N- H
4174 4191 4189 4190 1 180.00 43.93200 2 ; CA-
N- C- O
4189 4193 4191 4192 1 180.00 4.60240 2 ; C-
CA- N- H
4193 4206 4204 4205 1 180.00 43.93200 2 ; CA-
N- C- O
4198 4202 4201 4203 1 180.00 43.93200 2 ; CG-
OE1- CD- OE2
4204 4208 4206 4207 1 180.00 4.60240 2 ; C-
CA- N- H
4208 4221 4219 4220 1 180.00 43.93200 2 ; CA-
N- C- O
4213 4217 4216 4218 1 180.00 43.93200 2 ; CG-
OE1- CD- OE2
4219 4223 4221 4222 1 180.00 4.60240 2 ; C-
CA- N- H
4223 4231 4229 4230 1 180.00 43.93200 2 ; CA-
N- C- O
4229 4233 4231 4232 1 180.00 4.60240 2 ; C-
CA- N- H
4233 4250 4248 4249 1 180.00 43.93200 2 ; CA-
N- C- O
4248 4252 4250 4251 1 180.00 4.60240 2 ; C-

CA-     N-     H
  4252  4264  4262  4263      1  180.00  43.93200  2 ;     CA-
N-     C-     O
  4254  4259  4257  4258      1  180.00  43.93200  2 ;     CB-
ND2-   CG-   OD1
  4257  4260  4259  4261      1  180.00   4.18400  2 ;     CG-
HD21-  ND2-  HD22
  4262  4266  4264  4265      1  180.00   4.60240  2 ;      C-
CA-     N-     H
  4266  4282  4280  4281      1  180.00  43.93200  2 ;     CA-
N-     C-     O
  4268  4278  4271  4272      1  180.00   4.60240  2 ;     CB-
CD2-   CG-   ND1
  4271  4274  4272  4273      1  180.00   4.18400  2 ;     CG-
CE1-   ND1-   HD1
  4271  4276  4278  4279      1  180.00   4.60240  2 ;     CG-
NE2-   CD2-   HD2
  4272  4276  4274  4275      1  180.00   4.60240  2 ;    ND1-
NE2-   CE1-   HE1
  4274  4278  4276  4277      1  180.00   4.18400  2 ;    CE1-
CD2-   NE2-   HE2
  4280  4283  4282  4292      1  180.00   4.18400  2 ;      C-
CD-     N-    CA
  4292  4296  4294  4295      1  180.00  43.93200  2 ;     CA-
N-     C-     O
  4294  4298  4296  4297      1  180.00   4.60240  2 ;      C-
CA-     N-     H
  4298  4320  4318  4319      1  180.00  43.93200  2 ;     CA-
N-     C-     O
  4303  4306  4304  4305      1  180.00   4.60240  2 ;     CG-
NE1-   CD1-   HD1
  4304  4308  4306  4307      1  180.00   4.18400  2 ;    CD1-
CE2-   NE1-   HE1
  4308  4311  4309  4310      1  180.00   4.60240  2 ;    CE2-
CH2-   CZ2-   HZ2
  4309  4313  4311  4312      1  180.00   4.60240  2 ;    CZ2-
CZ3-   CH2-   HH2
  4311  4315  4313  4314      1  180.00   4.60240  2 ;    CH2-
CE3-   CZ3-   HZ3
  4313  4317  4315  4316      1  180.00   4.60240  2 ;    CZ3-
CD2-   CE3-   HE3
  4317  4300  4303  4304      1  180.00   4.60240  2 ;    CD2-
CB-    CG-   CD1
  4318  4322  4320  4321      1  180.00   4.60240  2 ;      C-
CA-     N-     H
  4322  4339  4337  4338      1  180.00  43.93200  2 ;     CA-
N-     C-     O
  4337  4341  4339  4340      1  180.00   4.60240  2 ;      C-
CA-     N-     H
  4341  4353  4351  4352      1  180.00  43.93200  2 ;     CA-
N-     C-     O
  4351  4355  4353  4354      1  180.00   4.60240  2 ;      C-

CA-     N-     H
  4355   4370   4368   4369       1   180.00  43.93200   2 ;     CA-
N-     C-     O
  4360   4365   4363   4364       1   180.00  43.93200   2 ;     CG-
NE2-    CD-   OE1
  4363   4366   4365   4367       1   180.00   4.18400   2 ;     CD-
HE21-   NE2-  HE22
  4368   4372   4370   4371       1   180.00   4.60240   2 ;      C-
CA-     N-     H
  4372   4387   4385   4386       1   180.00  43.93200   2 ;     CA-
N-     C-     O
  4385   4389   4387   4388       1   180.00   4.60240   2 ;      C-
CA-     N-     H
  4389   4401   4399   4400       1   180.00  43.93200   2 ;     CA-
N-     C-     O
  4399   4403   4401   4402       1   180.00   4.60240   2 ;      C-
CA-     N-     H
  4403   4423   4421   4422       1   180.00  43.93200   2 ;     CA-
N-     C-     O
  4421   4425   4423   4424       1   180.00   4.60240   2 ;      C-
CA-     N-     H
  4425   4434   4432   4433       1   180.00  43.93200   2 ;     CA-
N-     C-     O
  4432   4436   4434   4435       1   180.00   4.60240   2 ;      C-
CA-     N-     H
  4436   4451   4449   4450       1   180.00  43.93200   2 ;     CA-
N-     C-     O
  4438   4447   4441   4442       1   180.00   4.60240   2 ;     CB-
CD2-    CG-   ND1
  4441   4445   4447   4448       1   180.00   4.60240   2 ;     CG-
NE2-   CD2-   HD2
  4442   4445   4443   4444       1   180.00   4.60240   2 ;    ND1-
NE2-   CE1-   HE1
  4443   4447   4445   4446       1   180.00   4.18400   2 ;    CE1-
CD2-   NE2-   HE2
  4449   4453   4451   4452       1   180.00   4.60240   2 ;      C-
CA-     N-     H
  4453   4466   4464   4465       1   180.00  43.93200   2 ;     CA-
N-     C-     O
  4458   4462   4461   4463       1   180.00  43.93200   2 ;     CG-
OE1-    CD-   OE2
  4464   4468   4466   4467       1   180.00   4.60240   2 ;      C-
CA-     N-     H
  4468   4483   4481   4482       1   180.00  43.93200   2 ;     CA-
N-     C-     O
  4470   4479   4473   4474       1   180.00   4.60240   2 ;     CB-
CD2-    CG-   ND1
  4473   4477   4479   4480       1   180.00   4.60240   2 ;     CG-
NE2-   CD2-   HD2
  4474   4477   4475   4476       1   180.00   4.60240   2 ;    ND1-
NE2-   CE1-   HE1
  4475   4479   4477   4478       1   180.00   4.18400   2 ;    CE1-

```
CD2-   NE2-   HE2
  4481  4485  4483  4484      1  180.00   4.60240  2 ;     C-
CA-    N-    H
  4485  4499  4497  4498      1  180.00  43.93200  2 ;    CA-
N-    C-    O
  4497  4501  4499  4500      1  180.00   4.60240  2 ;     C-
CA-    N-    H
  4501  4514  4512  4513      1  180.00  43.93200  2 ;    CA-
N-    C-    O
  4506  4510  4509  4511      1  180.00  43.93200  2 ;    CG-
OE1-   CD-   OE2
  4512  4516  4514  4515      1  180.00   4.60240  2 ;     C-
CA-    N-    H
  4516  4533  4531  4532      1  180.00  43.93200  2 ;    CA-
N-    C-    O
  4531  4535  4533  4534      1  180.00   4.60240  2 ;     C-
CA-    N-    H
  4535  4544  4542  4543      1  180.00  43.93200  2 ;    CA-
N-    C-    O
  4542  4546  4544  4545      1  180.00   4.60240  2 ;     C-
CA-    N-    H
  4546  4555  4553  4554      1  180.00  43.93200  2 ;    CA-
N-    C-    O
  4553  4557  4555  4556      1  180.00   4.60240  2 ;     C-
CA-    N-    H
  4557  4569  4567  4568      1  180.00  43.93200  2 ;    CA-
N-    C-    O
  4567  4571  4569  4570      1  180.00   4.60240  2 ;     C-
CA-    N-    H
  4571  4588  4586  4587      1  180.00  43.93200  2 ;    CA-
N-    C-    O
  4586  4590  4588  4589      1  180.00   4.60240  2 ;     C-
CA-    N-    H
  4590  4607  4605  4606      1  180.00  43.93200  2 ;    CA-
N-    C-    O
  4605  4609  4607  4608      1  180.00   4.60240  2 ;     C-
CA-    N-    H
  4609  4629  4627  4628      1  180.00  43.93200  2 ;    CA-
N-    C-    O
  4627  4631  4629  4630      1  180.00   4.60240  2 ;     C-
CA-    N-    H
  4631  4643  4641  4642      1  180.00  43.93200  2 ;    CA-
N-    C-    O
  4633  4638  4636  4637      1  180.00  43.93200  2 ;    CB-
ND2-   CG-   OD1
  4636  4639  4638  4640      1  180.00   4.18400  2 ;    CG-
HD21-   ND2-  HD22
  4641  4645  4643  4644      1  180.00   4.60240  2 ;     C-
CA-    N-    H
  4645  4662  4660  4661      1  180.00  43.93200  2 ;    CA-
N-    C-    O
  4660  4664  4662  4663      1  180.00   4.60240  2 ;     C-
```

CA- N- H
4664 4684 4682 4683 1 180.00 43.93200 2 ; CA-
N- C- O
4682 4686 4684 4685 1 180.00 4.60240 2 ; C-
CA- N- H
4686 4698 4696 4697 1 180.00 43.93200 2 ; CA-
N- C- O
4688 4693 4691 4692 1 180.00 43.93200 2 ; CB-
ND2- CG- OD1
4691 4694 4693 4695 1 180.00 4.18400 2 ; CG-
HD21- ND2- HD22
4696 4700 4698 4699 1 180.00 4.60240 2 ; C-
CA- N- H
4700 4718 4716 4717 1 180.00 43.93200 2 ; CA-
N- C- O
4705 4708 4706 4707 1 180.00 4.60240 2 ; CG-
CE1- CD1- HD1
4705 4712 4714 4715 1 180.00 4.60240 2 ; CG-
CE2- CD2- HD2
4706 4710 4708 4709 1 180.00 4.60240 2 ; CD1-
CZ- CE1- HE1
4706 4714 4705 4702 1 180.00 4.60240 2 ; CD1-
CD2- CG- CB
4708 4712 4710 4711 1 180.00 4.60240 2 ; CE1-
CE2- CZ- HZ
4710 4714 4712 4713 1 180.00 4.60240 2 ; CZ-
CD2- CE2- HE2
4716 4720 4718 4719 1 180.00 4.60240 2 ; C-
CA- N- H
4720 4740 4738 4739 1 180.00 43.93200 2 ; CA-
N- C- O
4738 4742 4740 4741 1 180.00 4.60240 2 ; C-
CA- N- H
4742 4762 4760 4761 1 180.00 43.93200 2 ; CA-
N- C- O
4760 4764 4762 4763 1 180.00 4.60240 2 ; C-
CA- N- H
4764 4777 4775 4776 1 180.00 43.93200 2 ; CA-
N- C- O
4769 4773 4772 4774 1 180.00 43.93200 2 ; CG-
OE1- CD- OE2
4775 4779 4777 4778 1 180.00 4.60240 2 ; C-
CA- N- H
4779 4791 4789 4790 1 180.00 43.93200 2 ; CA-
N- C- O
4781 4786 4784 4785 1 180.00 43.93200 2 ; CB-
ND2- CG- OD1
4784 4787 4786 4788 1 180.00 4.18400 2 ; CG-
HD21- ND2- HD22
4789 4793 4791 4792 1 180.00 4.60240 2 ; C-
CA- N- H
4793 4806 4804 4805 1 180.00 43.93200 2 ; CA-

N-      C-      O
  4798   4802   4801   4803       1   180.00  43.93200   2 ;     CG-
OE1-    CD-   OE2
  4804   4808   4806   4807       1   180.00   4.60240   2 ;      C-
CA-     N-     H
  4808   4825   4823   4824       1   180.00  43.93200   2 ;     CA-
N-      C-      O
  4823   4827   4825   4826       1   180.00   4.60240   2 ;      C-
CA-     N-     H
  4827   4847   4845   4846       1   180.00  43.93200   2 ;     CA-
N-      C-      O
  4845   4849   4847   4848       1   180.00   4.60240   2 ;      C-
CA-     N-     H
  4849   4869   4867   4868       1   180.00  43.93200   2 ;     CA-
N-      C-      O
  4867   4871   4869   4870       1   180.00   4.60240   2 ;      C-
CA-     N-     H
  4871   4888   4886   4887       1   180.00  43.93200   2 ;     CA-
N-      C-      O
  4886   4890   4888   4889       1   180.00   4.60240   2 ;      C-
CA-     N-     H
  4890   4898   4896   4897       1   180.00  43.93200   2 ;     CA-
N-      C-      O
  4896   4900   4898   4899       1   180.00   4.60240   2 ;      C-
CA-     N-     H
  4900   4917   4915   4916       1   180.00  43.93200   2 ;     CA-
N-      C-      O
  4915   4919   4917   4918       1   180.00   4.60240   2 ;      C-
CA-     N-     H
  4919   4931   4929   4930       1   180.00  43.93200   2 ;     CA-
N-      C-      O
  4929   4933   4931   4932       1   180.00   4.60240   2 ;      C-
CA-     N-     H
  4933   4950   4948   4949       1   180.00  43.93200   2 ;     CA-
N-      C-      O
  4948   4952   4950   4951       1   180.00   4.60240   2 ;      C-
CA-     N-     H
  4952   4969   4967   4968       1   180.00  43.93200   2 ;     CA-
N-      C-      O
  4967   4971   4969   4970       1   180.00   4.60240   2 ;      C-
CA-     N-     H
  4971   4979   4977   4978       1   180.00  43.93200   2 ;     CA-
N-      C-      O
  4977   4981   4979   4980       1   180.00   4.60240   2 ;      C-
CA-     N-     H
  4981   5001   4999   5000       1   180.00  43.93200   2 ;     CA-
N-      C-      O
  4999   5003   5001   5002       1   180.00   4.60240   2 ;      C-
CA-     N-     H
  5003   5018   5016   5017       1   180.00  43.93200   2 ;     CA-
N-      C-      O
  5005   5014   5008   5009       1   180.00   4.60240   2 ;     CB-

```
CD2-    CG-   ND1
  5008   5012   5014   5015       1   180.00   4.60240   2 ;     CG-
NE2-   CD2-   HD2
  5009   5012   5010   5011       1   180.00   4.60240   2 ;    ND1-
NE2-   CE1-   HE1
  5010   5014   5012   5013       1   180.00   4.18400   2 ;    CE1-
CD2-   NE2-   HE2
  5016   5020   5018   5019       1   180.00   4.60240   2 ;      C-
CA-     N-     H
  5020   5037   5035   5036       1   180.00  43.93200   2 ;     CA-
N-     C-     O
  5035   5039   5037   5038       1   180.00   4.60240   2 ;      C-
CA-     N-     H
  5039   5048   5046   5047       1   180.00  43.93200   2 ;     CA-
N-     C-     O
  5046   5050   5048   5049       1   180.00   4.60240   2 ;      C-
CA-     N-     H
  5050   5060   5058   5059       1   180.00  43.93200   2 ;     CA-
N-     C-     O
  5052   5056   5055   5057       1   180.00  43.93200   2 ;     CB-
OD1-    CG-   OD2
  5058   5062   5060   5061       1   180.00   4.60240   2 ;      C-
CA-     N-     H
  5062   5076   5074   5075       1   180.00  43.93200   2 ;     CA-
N-     C-     O
  5074   5078   5076   5077       1   180.00   4.60240   2 ;      C-
CA-     N-     H
  5078   5091   5089   5090       1   180.00  43.93200   2 ;     CA-
N-     C-     O
  5083   5087   5086   5088       1   180.00  43.93200   2 ;     CG-
OE1-    CD-   OE2
  5089   5093   5091   5092       1   180.00   4.60240   2 ;      C-
CA-     N-     H
  5093   5110   5108   5109       1   180.00  43.93200   2 ;     CA-
N-     C-     O
  5108   5112   5110   5111       1   180.00   4.60240   2 ;      C-
CA-     N-     H
  5112   5124   5122   5123       1   180.00  43.93200   2 ;     CA-
N-     C-     O
  5114   5119   5117   5118       1   180.00  43.93200   2 ;     CB-
ND2-    CG-   OD1
  5117   5120   5119   5121       1   180.00   4.18400   2 ;     CG-
HD21-   ND2-  HD22
  5122   5126   5124   5125       1   180.00   4.60240   2 ;      C-
CA-     N-     H
  5126   5138   5136   5137       1   180.00  43.93200   2 ;     CA-
N-     C-     O
  5128   5133   5131   5132       1   180.00  43.93200   2 ;     CB-
ND2-    CG-   OD1
  5131   5134   5133   5135       1   180.00   4.18400   2 ;     CG-
HD21-   ND2-  HD22
  5136   5140   5138   5139       1   180.00   4.60240   2 ;      C-
```

CA- N- H
5140 5157 5155 5156 1 180.00 43.93200 2 ; CA-
N- C- O
5155 5159 5157 5158 1 180.00 4.60240 2 ; C-
CA- N- H
5159 5181 5179 5180 1 180.00 43.93200 2 ; CA-
N- C- O
5167 5172 5170 5171 1 180.00 4.18400 2 ; CD-
CZ- NE- HE
5170 5173 5172 5176 1 180.00 43.93200 2 ; NE-
NH1- CZ- NH2
5172 5174 5173 5175 1 180.00 4.18400 2 ; CZ-
HH11- NH1- HH12
5172 5177 5176 5178 1 180.00 4.18400 2 ; CZ-
HH21- NH2- HH22
5179 5183 5181 5182 1 180.00 4.60240 2 ; C-
CA- N- H
5183 5195 5193 5194 1 180.00 43.93200 2 ; CA-
N- C- O
5185 5190 5188 5189 1 180.00 43.93200 2 ; CB-
ND2- CG- OD1
5188 5191 5190 5192 1 180.00 4.18400 2 ; CG-
HD21- ND2- HD22
5193 5197 5195 5196 1 180.00 4.60240 2 ; C-
CA- N- H
5197 5214 5212 5213 1 180.00 43.93200 2 ; CA-
N- C- O
5212 5216 5214 5215 1 180.00 4.60240 2 ; C-
CA- N- H
5216 5234 5232 5233 1 180.00 43.93200 2 ; CA-
N- C- O
5221 5224 5222 5223 1 180.00 4.60240 2 ; CG-
CE1- CD1- HD1
5221 5228 5230 5231 1 180.00 4.60240 2 ; CG-
CE2- CD2- HD2
5222 5226 5224 5225 1 180.00 4.60240 2 ; CD1-
CZ- CE1- HE1
5222 5230 5221 5218 1 180.00 4.60240 2 ; CD1-
CD2- CG- CB
5224 5228 5226 5227 1 180.00 4.60240 2 ; CE1-
CE2- CZ- HZ
5226 5230 5228 5229 1 180.00 4.60240 2 ; CZ-
CD2- CE2- HE2
5232 5236 5234 5235 1 180.00 4.60240 2 ; C-
CA- N- H
5236 5253 5251 5252 1 180.00 43.93200 2 ; CA-
N- C- O
5251 5255 5253 5254 1 180.00 4.60240 2 ; C-
CA- N- H
5255 5263 5261 5262 1 180.00 43.93200 2 ; CA-
N- C- O
5261 5265 5263 5264 1 180.00 4.60240 2 ; C-

```
CA-     N-     H
  5265  5282  5280  5281      1   180.00  43.93200   2 ;     CA-
N-     C-     O
  5280  5284  5282  5283      1   180.00   4.60240   2 ;      C-
CA-     N-     H
  5284  5294  5292  5293      1   180.00  43.93200   2 ;     CA-
N-     C-     O
  5286  5290  5289  5291      1   180.00  43.93200   2 ;     CB-
OD1-    CG-   OD2
  5292  5296  5294  5295      1   180.00   4.60240   2 ;      C-
CA-     N-     H
  5296  5310  5308  5309      1   180.00  43.93200   2 ;     CA-
N-     C-     O
  5308  5312  5310  5311      1   180.00   4.60240   2 ;      C-
CA-     N-     H
  5312  5322  5320  5321      1   180.00  43.93200   2 ;     CA-
N-     C-     O
  5314  5318  5317  5319      1   180.00  43.93200   2 ;     CB-
OD1-    CG-   OD2
  5320  5324  5322  5323      1   180.00   4.60240   2 ;      C-
CA-     N-     H
  5324  5336  5334  5335      1   180.00  43.93200   2 ;     CA-
N-     C-     O
  5326  5331  5329  5330      1   180.00  43.93200   2 ;     CB-
ND2-    CG-   OD1
  5329  5332  5331  5333      1   180.00   4.18400   2 ;     CG-
HD21-   ND2-  HD22
  5334  5338  5336  5337      1   180.00   4.60240   2 ;      C-
CA-     N-     H
  5338  5347  5345  5346      1   180.00  43.93200   2 ;     CA-
N-     C-     O
  5345  5349  5347  5348      1   180.00   4.60240   2 ;      C-
CA-     N-     H
  5349  5354  5352  5353      1   180.00  43.93200   2 ;     CA-
N-     C-     O
  5352  5356  5354  5355      1   180.00   4.60240   2 ;      C-
CA-     N-     H
  5356  5368  5366  5367      1   180.00  43.93200   2 ;     CA-
N-     C-     O
  5366  5370  5368  5369      1   180.00   4.60240   2 ;      C-
CA-     N-     H
  5370  5387  5385  5386      1   180.00  43.93200   2 ;     CA-
N-     C-     O
  5385  5389  5387  5388      1   180.00   4.60240   2 ;      C-
CA-     N-     H
  5389  5398  5396  5397      1   180.00  43.93200   2 ;     CA-
N-     C-     O
  5396  5400  5398  5399      1   180.00   4.60240   2 ;      C-
CA-     N-     H
  5400  5409  5407  5408      1   180.00  43.93200   2 ;     CA-
N-     C-     O
  5407  5411  5409  5410      1   180.00   4.60240   2 ;      C-
```

CA- N- H
5411 5426 5424 5425 1 180.00 43.93200 2 ; CA-
N- C- O
5416 5421 5419 5420 1 180.00 43.93200 2 ; CG-
NE2- CD- OE1
5419 5422 5421 5423 1 180.00 4.18400 2 ; CD-
HE21- NE2- HE22
5424 5428 5426 5427 1 180.00 4.60240 2 ; C-
CA- N- H
5428 5442 5440 5441 1 180.00 43.93200 2 ; CA-
N- C- O
5433 5437 5436 5438 1 180.00 43.93200 2 ; CG-
OE1- CD- OE2
5440 5444 5442 5443 1 180.00 4.60240 2 ; C-
CA- N- H
5444 5461 5459 5460 1 180.00 43.93200 2 ; CA-
N- C- O
5459 5463 5461 5462 1 180.00 4.60240 2 ; C-
CA- N- H
5463 5480 5478 5479 1 180.00 43.93200 2 ; CA-
N- C- O
5478 5482 5480 5481 1 180.00 4.60240 2 ; C-
CA- N- H
5482 5492 5490 5491 1 180.00 43.93200 2 ; CA-
N- C- O
5484 5488 5487 5489 1 180.00 43.93200 2 ; CB-
OD1- CG- OD2
5490 5494 5492 5493 1 180.00 4.60240 2 ; C-
CA- N- H
5494 5499 5497 5498 1 180.00 43.93200 2 ; CA-
N- C- O
5497 5501 5499 5500 1 180.00 4.60240 2 ; C-
CA- N- H
5501 5518 5516 5517 1 180.00 43.93200 2 ; CA-
N- C- O
5516 5520 5518 5519 1 180.00 4.60240 2 ; C-
CA- N- H
5520 5540 5538 5539 1 180.00 43.93200 2 ; CA-
N- C- O
5538 5542 5540 5541 1 180.00 4.60240 2 ; C-
CA- N- H
5542 5562 5560 5561 1 180.00 43.93200 2 ; CA-
N- C- O
5560 5564 5562 5563 1 180.00 4.60240 2 ; C-
CA- N- H
5564 5581 5579 5580 1 180.00 43.93200 2 ; CA-
N- C- O
5579 5583 5581 5582 1 180.00 4.60240 2 ; C-
CA- N- H
5583 5588 5586 5587 1 180.00 43.93200 2 ; CA-
N- C- O
5586 5590 5588 5589 1 180.00 4.60240 2 ; C-

CA-     N-     H
  5590   5609   5607   5608      1   180.00  43.93200   2 ;     CA-
N-     C-     O
  5595   5598   5596   5597      1   180.00   4.60240   2 ;     CG-
CE1-   CD1-   HD1
  5595   5603   5605   5606      1   180.00   4.60240   2 ;     CG-
CE2-   CD2-   HD2
  5596   5600   5598   5599      1   180.00   4.60240   2 ;    CD1-
CZ-   CE1-   HE1
  5596   5605   5595   5592      1   180.00   4.60240   2 ;    CD1-
CD2-    CG-    CB
  5598   5603   5600   5601      1   180.00   4.60240   2 ;    CE1-
CE2-    CZ-    OH
  5600   5605   5603   5604      1   180.00   4.60240   2 ;     CZ-
CD2-   CE2-   HE2
  5607   5611   5609   5610      1   180.00   4.60240   2 ;      C-
CA-     N-     H
  5611   5626   5624   5625      1   180.00  43.93200   2 ;     CA-
N-     C-     O
  5616   5621   5619   5620      1   180.00  43.93200   2 ;     CG-
NE2-    CD-   OE1
  5619   5622   5621   5623      1   180.00   4.18400   2 ;     CD-
HE21-   NE2-  HE22
  5624   5628   5626   5627      1   180.00   4.60240   2 ;      C-
CA-     N-     H
  5628   5648   5646   5647      1   180.00  43.93200   2 ;     CA-
N-     C-     O
  5646   5650   5648   5649      1   180.00   4.60240   2 ;      C-
CA-     N-     H
  5650   5667   5665   5666      1   180.00  43.93200   2 ;     CA-
N-     C-     O
  5665   5668   5667   5677      1   180.00   4.18400   2 ;      C-
CD-     N-    CA
  5677   5681   5679   5680      1   180.00  43.93200   2 ;     CA-
N-     C-     O
  5679   5682   5681   5691      1   180.00   4.18400   2 ;      C-
CD-     N-    CA
  5691   5695   5693   5694      1   180.00  43.93200   2 ;     CA-
N-     C-     O
  5693   5697   5695   5696      1   180.00   4.60240   2 ;      C-
CA-     N-     H
  5697   5707   5705   5706      1   180.00  43.93200   2 ;     CA-
N-     C-     O
  5699   5703   5702   5704      1   180.00  43.93200   2 ;     CB-
OD1-    CG-   OD2
  5705   5709   5707   5708      1   180.00   4.60240   2 ;      C-
CA-     N-     H
  5709   5726   5724   5725      1   180.00  43.93200   2 ;     CA-
N-     C-     O
  5724   5728   5726   5727      1   180.00   4.60240   2 ;      C-
CA-     N-     H
  5728   5743   5741   5742      1   180.00  43.93200   2 ;     CA-

N- C- O
5730 5739 5733 5734 1 180.00 4.60240 2 ; CB-
CD2- CG- ND1
5733 5737 5739 5740 1 180.00 4.60240 2 ; CG-
NE2- CD2- HD2
5734 5737 5735 5736 1 180.00 4.60240 2 ; ND1-
NE2- CE1- HE1
5735 5739 5737 5738 1 180.00 4.18400 2 ; CE1-
CD2- NE2- HE2
5741 5745 5743 5744 1 180.00 4.60240 2 ; C-
CA- N- H
5745 5760 5758 5759 1 180.00 43.93200 2 ; CA-
N- C- O
5750 5755 5753 5754 1 180.00 43.93200 2 ; CG-
NE2- CD- OE1
5753 5756 5755 5757 1 180.00 4.18400 2 ; CD-
HE21- NE2- HE22
5758 5762 5760 5761 1 180.00 4.60240 2 ; C-
CA- N- H
5762 5776 5774 5775 1 180.00 43.93200 2 ; CA-
N- C- O
5774 5778 5776 5777 1 180.00 4.60240 2 ; C-
CA- N- H
5778 5795 5793 5794 1 180.00 43.93200 2 ; CA-
N- C- O
5793 5797 5795 5796 1 180.00 4.60240 2 ; C-
CA- N- H
5797 5819 5817 5818 1 180.00 43.93200 2 ; CA-
N- C- O
5805 5810 5808 5809 1 180.00 4.18400 2 ; CD-
CZ- NE- HE
5808 5811 5810 5814 1 180.00 43.93200 2 ; NE-
NH1- CZ- NH2
5810 5812 5811 5813 1 180.00 4.18400 2 ; CZ-
HH11- NH1- HH12
5810 5815 5814 5816 1 180.00 4.18400 2 ; CZ-
HH21- NH2- HH22
5817 5821 5819 5820 1 180.00 4.60240 2 ; C-
CA- N- H
5821 5831 5829 5830 1 180.00 43.93200 2 ; CA-
N- C- O
5823 5827 5826 5828 1 180.00 43.93200 2 ; CB-
OD1- CG- OD2
5829 5833 5831 5832 1 180.00 4.60240 2 ; C-
CA- N- H
5833 5850 5848 5849 1 180.00 43.93200 2 ; CA-
N- C- O
5848 5852 5850 5851 1 180.00 4.60240 2 ; C-
CA- N- H
5852 5862 5860 5861 1 180.00 43.93200 2 ; CA-
N- C- O
5854 5858 5857 5859 1 180.00 43.93200 2 ; CB-

OD1- CG- OD2
5860 5864 5862 5863 1 180.00 4.60240 2 ; C-
CA- N- H
5864 5873 5871 5872 1 180.00 43.93200 2 ; CA-
N- C- O
5871 5875 5873 5874 1 180.00 4.60240 2 ; C-
CA- N- H
5875 5887 5885 5886 1 180.00 43.93200 2 ; CA-
N- C- O
5877 5882 5880 5881 1 180.00 43.93200 2 ; CB-
ND2- CG- OD1
5880 5883 5882 5884 1 180.00 4.18400 2 ; CG-
HD21- ND2- HD22
5885 5889 5887 5888 1 180.00 4.60240 2 ; C-
CA- N- H
5889 5897 5895 5896 1 180.00 43.93200 2 ; CA-
N- C- O
5895 5899 5897 5898 1 180.00 4.60240 2 ; C-
CA- N- H
5899 5908 5906 5907 1 180.00 43.93200 2 ; CA-
N- C- O
5906 5910 5908 5909 1 180.00 4.60240 2 ; C-
CA- N- H
5910 5915 5913 5914 1 180.00 43.93200 2 ; CA-
N- C- O
5913 5917 5915 5916 1 180.00 4.60240 2 ; C-
CA- N- H
5917 5932 5930 5931 1 180.00 43.93200 2 ; CA-
N- C- O
5922 5927 5925 5926 1 180.00 43.93200 2 ; CG-
NE2- CD- OE1
5925 5928 5927 5929 1 180.00 4.18400 2 ; CD-
HE21- NE2- HE22
5930 5934 5932 5933 1 180.00 4.60240 2 ; C-
CA- N- H
5934 5951 5949 5950 1 180.00 43.93200 2 ; CA-
N- C- O
5949 5953 5951 5952 1 180.00 4.60240 2 ; C-
CA- N- H
5953 5968 5966 5967 1 180.00 43.93200 2 ; CA-
N- C- O
5955 5964 5958 5959 1 180.00 4.60240 2 ; CB-
CD2- CG- ND1
5958 5962 5964 5965 1 180.00 4.60240 2 ; CG-
NE2- CD2- HD2
5959 5962 5960 5961 1 180.00 4.60240 2 ; ND1-
NE2- CE1- HE1
5960 5964 5962 5963 1 180.00 4.18400 2 ; CE1-
CD2- NE2- HE2
5966 5970 5968 5969 1 180.00 4.60240 2 ; C-
CA- N- H
5970 5989 5987 5988 1 180.00 43.93200 2 ; CA-

```
N-     C-     O
  5975   5978   5976   5977       1   180.00   4.60240   2 ;     CG-
CE1-   CD1-   HD1
  5975   5983   5985   5986       1   180.00   4.60240   2 ;     CG-
CE2-   CD2-   HD2
  5976   5980   5978   5979       1   180.00   4.60240   2 ;    CD1-
CZ-   CE1-   HE1
  5976   5985   5975   5972       1   180.00   4.60240   2 ;    CD1-
CD2-    CG-    CB
  5978   5983   5980   5981       1   180.00   4.60240   2 ;    CE1-
CE2-    CZ-    OH
  5980   5985   5983   5984       1   180.00   4.60240   2 ;     CZ-
CD2-   CE2-   HE2
  5987   5991   5989   5990       1   180.00   4.60240   2 ;      C-
CA-     N-     H
  5991   6003   6001   6002       1   180.00  43.93200   2 ;     CA-
N-     C-     O
  6001   6005   6003   6004       1   180.00   4.60240   2 ;      C-
CA-     N-     H
  6005   6015   6013   6014       1   180.00  43.93200   2 ;     CA-
N-     C-     O
  6007   6011   6010   6012       1   180.00  43.93200   2 ;     CB-
OD1-    CG-   OD2
  6013   6017   6015   6016       1   180.00   4.60240   2 ;      C-
CA-     N-     H
  6017   6035   6033   6034       1   180.00  43.93200   2 ;     CA-
N-     C-     O
  6022   6025   6023   6024       1   180.00   4.60240   2 ;     CG-
CE1-   CD1-   HD1
  6022   6029   6031   6032       1   180.00   4.60240   2 ;     CG-
CE2-   CD2-   HD2
  6023   6027   6025   6026       1   180.00   4.60240   2 ;    CD1-
CZ-   CE1-   HE1
  6023   6031   6022   6019       1   180.00   4.60240   2 ;    CD1-
CD2-    CG-    CB
  6025   6029   6027   6028       1   180.00   4.60240   2 ;    CE1-
CE2-    CZ-    HZ
  6027   6031   6029   6030       1   180.00   4.60240   2 ;     CZ-
CD2-   CE2-   HE2
  6033   6037   6035   6036       1   180.00   4.60240   2 ;      C-
CA-     N-     H
  6037   6054   6052   6053       1   180.00  43.93200   2 ;     CA-
N-     C-     O
  6052   6056   6054   6055       1   180.00   4.60240   2 ;      C-
CA-     N-     H
  6056   6064   6062   6063       1   180.00  43.93200   2 ;     CA-
N-     C-     O
  6062   6066   6064   6065       1   180.00   4.60240   2 ;      C-
CA-     N-     H
  6066   6074   6072   6073       1   180.00  43.93200   2 ;     CA-
N-     C-     O
  6072   6076   6074   6075       1   180.00   4.60240   2 ;      C-
```

```
CA-     N-     H
  6076   6088   6086   6087       1   180.00  43.93200   2 ;     CA-
N-     C-     O
  6086   6090   6088   6089       1   180.00   4.60240   2 ;      C-
CA-     N-     H
  6090   6107   6105   6106       1   180.00  43.93200   2 ;     CA-
N-     C-     O
  6105   6109   6107   6108       1   180.00   4.60240   2 ;      C-
CA-     N-     H
  6109   6119   6117   6118       1   180.00  43.93200   2 ;     CA-
N-     C-     O
  6111   6115   6114   6116       1   180.00  43.93200   2 ;     CB-
OD1-    CG-   OD2
  6117   6121   6119   6120       1   180.00   4.60240   2 ;      C-
CA-     N-     H
  6121   6141   6139   6140       1   180.00  43.93200   2 ;     CA-
N-     C-     O
  6139   6143   6141   6142       1   180.00   4.60240   2 ;      C-
CA-     N-     H
  6143   6158   6156   6157       1   180.00  43.93200   2 ;     CA-
N-     C-     O
  6148   6153   6151   6152       1   180.00  43.93200   2 ;     CG-
NE2-    CD-   OE1
  6151   6154   6153   6155       1   180.00   4.18400   2 ;     CD-
HE21-   NE2-  HE22
  6156   6160   6158   6159       1   180.00   4.60240   2 ;      C-
CA-     N-     H
  6160   6172   6170   6171       1   180.00  43.93200   2 ;     CA-
N-     C-     O
  6170   6174   6172   6173       1   180.00   4.60240   2 ;      C-
CA-     N-     H
  6174   6193   6191   6192       1   180.00  43.93200   2 ;     CA-
N-     C-     O
  6179   6182   6180   6181       1   180.00   4.60240   2 ;     CG-
CE1-   CD1-   HD1
  6179   6187   6189   6190       1   180.00   4.60240   2 ;     CG-
CE2-   CD2-   HD2
  6180   6184   6182   6183       1   180.00   4.60240   2 ;    CD1-
CZ-   CE1-   HE1
  6180   6189   6179   6176       1   180.00   4.60240   2 ;    CD1-
CD2-    CG-    CB
  6182   6187   6184   6185       1   180.00   4.60240   2 ;    CE1-
CE2-    CZ-    OH
  6184   6189   6187   6188       1   180.00   4.60240   2 ;     CZ-
CD2-   CE2-   HE2
  6191   6195   6193   6194       1   180.00   4.60240   2 ;      C-
CA-     N-     H
  6195   6212   6210   6211       1   180.00  43.93200   2 ;     CA-
N-     C-     O
  6210   6214   6212   6213       1   180.00   4.60240   2 ;      C-
CA-     N-     H
  6214   6234   6232   6233       1   180.00  43.93200   2 ;     CA-
```

```
N-     C-     O
  6232   6236   6234   6235       1   180.00   4.60240   2 ;      C-
CA-     N-     H
  6236   6256   6254   6255       1   180.00  43.93200   2 ;     CA-
N-     C-     O
  6254   6258   6256   6257       1   180.00   4.60240   2 ;      C-
CA-     N-     H
  6258   6271   6269   6270       1   180.00  43.93200   2 ;     CA-
N-     C-     O
  6263   6267   6266   6268       1   180.00  43.93200   2 ;     CG-
OE1-    CD-   OE2
  6269   6273   6271   6272       1   180.00   4.60240   2 ;      C-
CA-     N-     H
  6273   6287   6285   6286       1   180.00  43.93200   2 ;     CA-
N-     C-     O
  6285   6289   6287   6288       1   180.00   4.60240   2 ;      C-
CA-     N-     H
  6289   6298   6296   6297       1   180.00  43.93200   2 ;     CA-
N-     C-     O
  6296   6300   6298   6299       1   180.00   4.60240   2 ;      C-
CA-     N-     H
  6300   6317   6315   6316       1   180.00  43.93200   2 ;     CA-
N-     C-     O
  6315   6319   6317   6318       1   180.00   4.60240   2 ;      C-
CA-     N-     H
  6319   6336   6334   6335       1   180.00  43.93200   2 ;     CA-
N-     C-     O
  6334   6337   6336   6346       1   180.00   4.18400   2 ;      C-
CD-     N-    CA
  6346   6350   6348   6349       1   180.00  43.93200   2 ;     CA-
N-     C-     O
  6348   6352   6350   6351       1   180.00   4.60240   2 ;      C-
CA-     N-     H
  6352   6370   6368   6369       1   180.00  43.93200   2 ;     CA-
N-     C-     O
  6357   6360   6358   6359       1   180.00   4.60240   2 ;     CG-
CE1-   CD1-   HD1
  6357   6364   6366   6367       1   180.00   4.60240   2 ;     CG-
CE2-   CD2-   HD2
  6358   6362   6360   6361       1   180.00   4.60240   2 ;    CD1-
CZ-   CE1-   HE1
  6358   6366   6357   6354       1   180.00   4.60240   2 ;    CD1-
CD2-    CG-    CB
  6360   6364   6362   6363       1   180.00   4.60240   2 ;    CE1-
CE2-    CZ-    HZ
  6362   6366   6364   6365       1   180.00   4.60240   2 ;     CZ-
CD2-   CE2-   HE2
  6368   6372   6370   6371       1   180.00   4.60240   2 ;      C-
CA-     N-     H
  6372   6392   6390   6391       1   180.00  43.93200   2 ;     CA-
N-     C-     O
  6390   6394   6392   6393       1   180.00   4.60240   2 ;      C-
```

```
CA-     N-     H
  6394   6412   6410   6411       1   180.00  43.93200   2 ;     CA-
N-     C-     O
  6399   6402   6400   6401       1   180.00   4.60240   2 ;     CG-
CE1-   CD1-   HD1
  6399   6406   6408   6409       1   180.00   4.60240   2 ;     CG-
CE2-   CD2-   HD2
  6400   6404   6402   6403       1   180.00   4.60240   2 ;    CD1-
CZ-   CE1-   HE1
  6400   6408   6399   6396       1   180.00   4.60240   2 ;    CD1-
CD2-    CG-    CB
  6402   6406   6404   6405       1   180.00   4.60240   2 ;    CE1-
CE2-    CZ-    HZ
  6404   6408   6406   6407       1   180.00   4.60240   2 ;     CZ-
CD2-   CE2-   HE2
  6410   6414   6412   6413       1   180.00   4.60240   2 ;      C-
CA-     N-     H
  6414   6432   6430   6431       1   180.00  43.93200   2 ;     CA-
N-     C-     O
  6419   6422   6420   6421       1   180.00   4.60240   2 ;     CG-
CE1-   CD1-   HD1
  6419   6426   6428   6429       1   180.00   4.60240   2 ;     CG-
CE2-   CD2-   HD2
  6420   6424   6422   6423       1   180.00   4.60240   2 ;    CD1-
CZ-   CE1-   HE1
  6420   6428   6419   6416       1   180.00   4.60240   2 ;    CD1-
CD2-    CG-    CB
  6422   6426   6424   6425       1   180.00   4.60240   2 ;    CE1-
CE2-    CZ-    HZ
  6424   6428   6426   6427       1   180.00   4.60240   2 ;     CZ-
CD2-   CE2-   HE2
  6430   6434   6432   6433       1   180.00   4.60240   2 ;      C-
CA-     N-     H
  6434   6444   6442   6443       1   180.00  43.93200   2 ;     CA-
N-     C-     O
  6436   6440   6439   6441       1   180.00  43.93200   2 ;     CB-
OD1-    CG-   OD2
  6442   6446   6444   6445       1   180.00   4.60240   2 ;      C-
CA-     N-     H
  6446   6463   6461   6462       1   180.00  43.93200   2 ;     CA-
N-     C-     O
  6461   6465   6463   6464       1   180.00   4.60240   2 ;      C-
CA-     N-     H
  6465   6475   6473   6474       1   180.00  43.93200   2 ;     CA-
N-     C-     O
  6467   6471   6470   6472       1   180.00  43.93200   2 ;     CB-
OD1-    CG-   OD2
  6473   6477   6475   6476       1   180.00   4.60240   2 ;      C-
CA-     N-     H
  6477   6482   6480   6481       1   180.00  43.93200   2 ;     CA-
N-     C-     O
  6480   6484   6482   6483       1   180.00   4.60240   2 ;      C-
```

```
CA-     N-     H
  6484  6496  6494  6495      1  180.00  43.93200   2 ;     CA-
N-     C-     O
  6486  6491  6489  6490      1  180.00  43.93200   2 ;     CB-
ND2-    CG-   OD1
  6489  6492  6491  6493      1  180.00   4.18400   2 ;     CG-
HD21-   ND2-  HD22
  6494  6498  6496  6497      1  180.00   4.60240   2 ;      C-
CA-     N-     H
  6498  6503  6501  6502      1  180.00  43.93200   2 ;     CA-
N-     C-     O
  6501  6505  6503  6504      1  180.00   4.60240   2 ;      C-
CA-     N-     H
  6505  6525  6523  6524      1  180.00  43.93200   2 ;     CA-
N-     C-     O
  6523  6527  6525  6526      1  180.00   4.60240   2 ;      C-
CA-     N-     H
  6527  6544  6542  6543      1  180.00  43.93200   2 ;     CA-
N-     C-     O
  6542  6546  6544  6545      1  180.00   4.60240   2 ;      C-
CA-     N-     H
  6546  6555  6553  6554      1  180.00  43.93200   2 ;     CA-
N-     C-     O
  6553  6557  6555  6556      1  180.00   4.60240   2 ;      C-
CA-     N-     H
  6557  6571  6569  6570      1  180.00  43.93200   2 ;     CA-
N-     C-     O
  6569  6573  6571  6572      1  180.00   4.60240   2 ;      C-
CA-     N-     H
  6573  6586  6584  6585      1  180.00  43.93200   2 ;     CA-
N-     C-     O
  6578  6582  6581  6583      1  180.00  43.93200   2 ;     CG-
OE1-    CD-   OE2
  6584  6588  6586  6587      1  180.00   4.60240   2 ;      C-
CA-     N-     H
  6588  6602  6600  6601      1  180.00  43.93200   2 ;     CA-
N-     C-     O
  6593  6597  6596  6598      1  180.00  43.93200   2 ;     CG-
OE1-    CD-   OE2
  6600  6604  6602  6603      1  180.00   4.60240   2 ;      C-
CA-     N-     H
  6604  6621  6619  6620      1  180.00  43.93200   2 ;     CA-
N-     C-     O
  6619  6623  6621  6622      1  180.00   4.60240   2 ;      C-
CA-     N-     H
  6623  6643  6641  6642      1  180.00  43.93200   2 ;     CA-
N-     C-     O
  6641  6645  6643  6644      1  180.00   4.60240   2 ;      C-
CA-     N-     H
  6645  6667  6665  6666      1  180.00  43.93200   2 ;     CA-
N-     C-     O
  6653  6658  6656  6657      1  180.00   4.18400   2 ;     CD-
```

CZ- NE- HE
6656 6659 6658 6662 1 180.00 43.93200 2 ; NE-
NH1- CZ- NH2
6658 6660 6659 6661 1 180.00 4.18400 2 ; CZ-
HH11- NH1- HH12
6658 6663 6662 6664 1 180.00 4.18400 2 ; CZ-
HH21- NH2- HH22
6665 6669 6667 6668 1 180.00 4.60240 2 ; C-
CA- N- H
6669 6686 6684 6685 1 180.00 43.93200 2 ; CA-
N- C- O
6684 6688 6686 6687 1 180.00 4.60240 2 ; C-
CA- N- H
6688 6706 6704 6705 1 180.00 43.93200 2 ; CA-
N- C- O
6693 6696 6694 6695 1 180.00 4.60240 2 ; CG-
CE1- CD1- HD1
6693 6700 6702 6703 1 180.00 4.60240 2 ; CG-
CE2- CD2- HD2
6694 6698 6696 6697 1 180.00 4.60240 2 ; CD1-
CZ- CE1- HE1
6694 6702 6693 6690 1 180.00 4.60240 2 ; CD1-
CD2- CG- CB
6696 6700 6698 6699 1 180.00 4.60240 2 ; CE1-
CE2- CZ- HZ
6698 6702 6700 6701 1 180.00 4.60240 2 ; CZ-
CD2- CE2- HE2
6704 6708 6706 6707 1 180.00 4.60240 2 ; C-
CA- N- H
6708 6713 6711 6712 1 180.00 43.93200 2 ; CA-
N- C- O
6711 6715 6713 6714 1 180.00 4.60240 2 ; C-
CA- N- H
6715 6737 6735 6736 1 180.00 43.93200 2 ; CA-
N- C- O
6723 6728 6726 6727 1 180.00 4.18400 2 ; CD-
CZ- NE- HE
6726 6729 6728 6732 1 180.00 43.93200 2 ; NE-
NH1- CZ- NH2
6728 6730 6729 6731 1 180.00 4.18400 2 ; CZ-
HH11- NH1- HH12
6728 6733 6732 6734 1 180.00 4.18400 2 ; CZ-
HH21- NH2- HH22
6735 6739 6737 6738 1 180.00 4.60240 2 ; C-
CA- N- H
6739 6749 6747 6748 1 180.00 43.93200 2 ; CA-
N- C- O
6741 6745 6744 6746 1 180.00 43.93200 2 ; CB-
OD1- CG- OD2
6747 6751 6749 6750 1 180.00 4.60240 2 ; C-
CA- N- H
6751 6761 6759 6760 1 180.00 43.93200 2 ; CA-

N-      C-      O
  6753   6757   6756   6758       1   180.00  43.93200   2 ;     CB-
OD1-    CG-   OD2
  6759   6763   6761   6762       1   180.00   4.60240   2 ;      C-
CA-     N-     H
  6763   6780   6778   6779       1   180.00  43.93200   2 ;     CA-
N-      C-      O
  6778   6782   6780   6781       1   180.00   4.60240   2 ;      C-
CA-     N-     H
  6782   6795   6793   6794       1   180.00  43.93200   2 ;     CA-
N-      C-      O
  6787   6791   6790   6792       1   180.00  43.93200   2 ;     CG-
OE1-    CD-   OE2
  6793   6797   6795   6796       1   180.00   4.60240   2 ;      C-
CA-     N-     H
  6797   6809   6807   6808       1   180.00  43.93200   2 ;     CA-
N-      C-      O
  6799   6804   6802   6803       1   180.00  43.93200   2 ;     CB-
ND2-    CG-   OD1
  6802   6805   6804   6806       1   180.00   4.18400   2 ;     CG-
HD21-   ND2-  HD22
  6807   6810   6809   6819       1   180.00   4.18400   2 ;      C-
CD-     N-     CA
  6819   6823   6821   6822       1   180.00  43.93200   2 ;     CA-
N-      C-      O
  6821   6825   6823   6824       1   180.00   4.60240   2 ;      C-
CA-     N-     H
  6825   6842   6840   6841       1   180.00  43.93200   2 ;     CA-
N-      C-      O
  6840   6844   6842   6843       1   180.00   4.60240   2 ;      C-
CA-     N-     H
  6844   6861   6859   6860       1   180.00  43.93200   2 ;     CA-
N-      C-      O
  6859   6863   6861   6862       1   180.00   4.60240   2 ;      C-
CA-     N-     H
  6863   6873   6871   6872       1   180.00  43.93200   2 ;     CA-
N-      C-      O
  6865   6869   6868   6870       1   180.00  43.93200   2 ;     CB-
OD1-    CG-   OD2
  6871   6875   6873   6874       1   180.00   4.60240   2 ;      C-
CA-     N-     H
  6875   6895   6893   6894       1   180.00  43.93200   2 ;     CA-
N-      C-      O
  6893   6897   6895   6896       1   180.00   4.60240   2 ;      C-
CA-     N-     H
  6897   6905   6903   6904       1   180.00  43.93200   2 ;     CA-
N-      C-      O
  6903   6907   6905   6906       1   180.00   4.60240   2 ;      C-
CA-     N-     H
  6907   6924   6922   6923       1   180.00  43.93200   2 ;     CA-
N-      C-      O
  6922   6926   6924   6925       1   180.00   4.60240   2 ;      C-

CA-     N-     H
  6926   6936   6934   6935       1   180.00  43.93200   2 ;     CA-
N-     C-     O
  6928   6932   6931   6933       1   180.00  43.93200   2 ;     CB-
OD1-    CG-   OD2
  6934   6938   6936   6937       1   180.00   4.60240   2 ;      C-
CA-     N-     H
  6938   6947   6945   6946       1   180.00  43.93200   2 ;     CA-
N-     C-     O
  6945   6949   6947   6948       1   180.00   4.60240   2 ;      C-
CA-     N-     H
  6949   6966   6964   6965       1   180.00  43.93200   2 ;     CA-
N-     C-     O
  6964   6968   6966   6967       1   180.00   4.60240   2 ;      C-
CA-     N-     H
  6968   6985   6983   6984       1   180.00  43.93200   2 ;     CA-
N-     C-     O
  6983   6987   6985   6986       1   180.00   4.60240   2 ;      C-
CA-     N-     H
  6987   7002   7000   7001       1   180.00  43.93200   2 ;     CA-
N-     C-     O
  6992   6997   6995   6996       1   180.00  43.93200   2 ;     CG-
NE2-    CD-   OE1
  6995   6998   6997   6999       1   180.00   4.18400   2 ;     CD-
HE21-   NE2-  HE22
  7000   7004   7002   7003       1   180.00   4.60240   2 ;      C-
CA-     N-     H
  7004   7017   7015   7016       1   180.00  43.93200   2 ;     CA-
N-     C-     O
  7009   7013   7012   7014       1   180.00  43.93200   2 ;     CG-
OE1-    CD-   OE2
  7015   7019   7017   7018       1   180.00   4.60240   2 ;      C-
CA-     N-     H
  7019   7033   7031   7032       1   180.00  43.93200   2 ;     CA-
N-     C-     O
  7031   7035   7033   7034       1   180.00   4.60240   2 ;      C-
CA-     N-     H
  7035   7045   7043   7044       1   180.00  43.93200   2 ;     CA-
N-     C-     O
  7037   7041   7040   7042       1   180.00  43.93200   2 ;     CB-
OD1-    CG-   OD2
  7043   7047   7045   7046       1   180.00   4.60240   2 ;      C-
CA-     N-     H
  7047   7064   7062   7063       1   180.00  43.93200   2 ;     CA-
N-     C-     O
  7062   7066   7064   7065       1   180.00   4.60240   2 ;      C-
CA-     N-     H
  7066   7078   7076   7077       1   180.00  43.93200   2 ;     CA-
N-     C-     O
  7068   7073   7071   7072       1   180.00  43.93200   2 ;     CB-
ND2-    CG-   OD1
  7071   7074   7073   7075       1   180.00   4.18400   2 ;     CG-

```
HD21-   ND2-  HD22
  7076   7080   7078   7079       1   180.00   4.60240   2 ;      C-
CA-     N-     H
  7080   7085   7083   7084       1   180.00  43.93200   2 ;     CA-
N-     C-     O
  7083   7087   7085   7086       1   180.00   4.60240   2 ;      C-
CA-     N-     H
  7087   7097   7095   7096       1   180.00  43.93200   2 ;     CA-
N-     C-     O
  7089   7093   7092   7094       1   180.00  43.93200   2 ;     CB-
OD1-    CG-   OD2
  7095   7099   7097   7098       1   180.00   4.60240   2 ;      C-
CA-     N-     H
  7099   7104   7102   7103       1   180.00  43.93200   2 ;     CA-
N-     C-     O
  7102   7106   7104   7105       1   180.00   4.60240   2 ;      C-
CA-     N-     H
  7106   7119   7117   7118       1   180.00  43.93200   2 ;     CA-
N-     C-     O
  7111   7115   7114   7116       1   180.00  43.93200   2 ;     CG-
OE1-    CD-   OE2
  7117   7121   7119   7120       1   180.00   4.60240   2 ;      C-
CA-     N-     H
  7121   7138   7136   7137       1   180.00  43.93200   2 ;     CA-
N-     C-     O
  7136   7140   7138   7139       1   180.00   4.60240   2 ;      C-
CA-     N-     H
  7140   7150   7148   7149       1   180.00  43.93200   2 ;     CA-
N-     C-     O
  7142   7146   7145   7147       1   180.00  43.93200   2 ;     CB-
OD1-    CG-   OD2
  7148   7152   7150   7151       1   180.00   4.60240   2 ;      C-
CA-     N-     H
  7152   7170   7168   7169       1   180.00  43.93200   2 ;     CA-
N-     C-     O
  7157   7160   7158   7159       1   180.00   4.60240   2 ;     CG-
CE1-   CD1-   HD1
  7157   7164   7166   7167       1   180.00   4.60240   2 ;     CG-
CE2-   CD2-   HD2
  7158   7162   7160   7161       1   180.00   4.60240   2 ;    CD1-
CZ-   CE1-   HE1
  7158   7166   7157   7154       1   180.00   4.60240   2 ;    CD1-
CD2-    CG-    CB
  7160   7164   7162   7163       1   180.00   4.60240   2 ;    CE1-
CE2-    CZ-    HZ
  7162   7166   7164   7165       1   180.00   4.60240   2 ;     CZ-
CD2-   CE2-   HE2
  7168   7172   7170   7171       1   180.00   4.60240   2 ;      C-
CA-     N-     H
  7172   7188   7186   7187       1   180.00  43.93200   2 ;     CA-
N-     C-     O
  7174   7184   7177   7178       1   180.00   4.60240   2 ;     CB-
```

```
CD2-    CG-   ND1
  7177   7180   7178   7179       1   180.00   4.18400   2 ;     CG-
CE1-   ND1-   HD1
  7177   7182   7184   7185       1   180.00   4.60240   2 ;     CG-
NE2-   CD2-   HD2
  7178   7182   7180   7181       1   180.00   4.60240   2 ;    ND1-
NE2-   CE1-   HE1
  7180   7184   7182   7183       1   180.00   4.18400   2 ;    CE1-
CD2-   NE2-   HE2
  7186   7190   7188   7189       1   180.00   4.60240   2 ;      C-
CA-     N-     H
  7190   7204   7202   7203       1   180.00  43.93200   2 ;     CA-
N-     C-     O
  7195   7199   7198   7200       1   180.00  43.93200   2 ;     CG-
OE1-    CD-   OE2
  7202   7206   7204   7205       1   180.00   4.60240   2 ;      C-
CA-     N-     H
  7206   7224   7222   7223       1   180.00  43.93200   2 ;     CA-
N-     C-     O
  7211   7214   7212   7213       1   180.00   4.60240   2 ;     CG-
CE1-   CD1-   HD1
  7211   7218   7220   7221       1   180.00   4.60240   2 ;     CG-
CE2-   CD2-   HD2
  7212   7216   7214   7215       1   180.00   4.60240   2 ;    CD1-
CZ-   CE1-   HE1
  7212   7220   7211   7208       1   180.00   4.60240   2 ;    CD1-
CD2-    CG-    CB
  7214   7218   7216   7217       1   180.00   4.60240   2 ;    CE1-
CE2-    CZ-    HZ
  7216   7220   7218   7219       1   180.00   4.60240   2 ;     CZ-
CD2-   CE2-   HE2
  7222   7226   7224   7225       1   180.00   4.60240   2 ;      C-
CA-     N-     H
  7226   7241   7239   7240       1   180.00  43.93200   2 ;     CA-
N-     C-     O
  7239   7243   7241   7242       1   180.00   4.60240   2 ;      C-
CA-     N-     H
  7243   7260   7258   7259       1   180.00  43.93200   2 ;     CA-
N-     C-     O
  7258   7262   7260   7261       1   180.00   4.60240   2 ;      C-
CA-     N-     H
  7262   7277   7275   7276       1   180.00  43.93200   2 ;     CA-
N-     C-     O
  7275   7279   7277   7278       1   180.00   4.60240   2 ;      C-
CA-     N-     H
  7279   7294   7292   7293       1   180.00  43.93200   2 ;     CA-
N-     C-     O
  7292   7296   7294   7295       1   180.00   4.60240   2 ;      C-
CA-     N-     H
  7296   7305   7303   7304       1   180.00  43.93200   2 ;     CA-
N-     C-     O
  7303   7307   7305   7306       1   180.00   4.60240   2 ;      C-
```

```
CA-     N-     H
  7307   7327   7325   7326       1   180.00  43.93200   2 ;     CA-
N-     C-     O
  7325   7329   7327   7328       1   180.00   4.60240   2 ;      C-
CA-     N-     H
  7329   7349   7347   7348       1   180.00  43.93200   2 ;     CA-
N-     C-     O
  7347   7351   7349   7350       1   180.00   4.60240   2 ;      C-
CA-     N-     H
  7351   7370   7369   7371       1   180.00  43.93200   2 ;     CA-
O-     C-   OXT
  7372   7374   7373   7413       1   180.00   4.60240   2 ;     C1-
C3-    C2-   H42
  7372   7376   7377   7415       1   180.00   4.60240   2 ;     C1-
C5-    C6-   H44
  7373   7375   7374   7414       1   180.00   4.60240   2 ;     C2-
C4-    C3-   H43
  7373   7377   7372   7388       1   180.00   4.60240   2 ;     C2-
C6-    C1-   O17
  7375   7377   7376   7379       1   180.00   4.60240   2 ;     C4-
C6-    C5-    C8
  7378   7405   7379   7376       1   180.00   4.60240   2 ;     C7-
C34-    C8-    C5
  7381   7405   7412   7410       1   180.00   4.60240   2 ;    C10-
C34-   C41-   C39
  7382   7384   7383   7385       1   180.00   4.60240   2 ;    C11-
C13-   C12-   F14
  7383   7381   7384   7417       1   180.00   4.60240   2 ;    C12-
C10-   C13-   H46
  7384   7380   7381   7412       1   180.00   4.60240   2 ;    C13-
C9-   C10-   C41
  7386   7378   7380   7381       1   180.00   4.60240   2 ;    N15-
C7-    C9-   C10
  7386   7383   7382   7416       1   180.00   4.60240   2 ;    N15-
C12-   C11-   H45
  7387   7374   7375   7376       1   180.00   4.60240   2 ;    N16-
C3-    C4-    C5
  7387   7380   7378   7379       1   180.00   4.60240   2 ;    N16-
C9-    C7-    C8
  7389   7391   7390   7396       1   180.00   4.60240   2 ;    C18-
C20-   C19-   C25
  7389   7392   7393   7420       1   180.00   4.60240   2 ;    C18-
C21-   C22-   H49
  7390   7392   7391   7435       1   180.00   4.60240   2 ;    C19-
C21-   C20-   H64
  7390   7393   7389   7418       1   180.00   4.60240   2 ;    C19-
C22-   C18-   H47
  7390   7397   7396   7398       1   180.00   4.60240   2 ;    C19-
N26-   C25-   O27
  7391   7393   7392   7419       1   180.00   4.60240   2 ;    C20-
C22-   C21-   H48
  7396   7399   7397   7434       1   180.00   4.60240   2 ;    C25-
```

```
C28-   N26-   H63
  7405   7407   7406   7408      1   180.00   4.60240   2 ;    C34-
N36-   C35-   O37
  7406   7410   7407   7409      1   180.00   4.60240   2 ;    C35-
C39-   N36-   C38
  7406   7412   7405   7379      1   180.00   4.60240   2 ;    C35-
C41-   C34-    C8
  7412   7407   7410   7411      1   180.00   4.60240   2 ;    C41-
N36-   C39-   O40

[ system ]
 4MVF_E52_dry

[ molecules ]
; Compound        nmols
 4MVF_E52_dry     1
```
